# Systems-level proteomic models of cotton fiber development: A high-resolution resource to analyze cell dynamics and trait engineering

**DOI:** 10.1101/2025.02.05.636703

**Authors:** Youngwoo Lee, Pengcheng Yang, Heena Rani, Gideon Miller, Corrinne E Grover, Sivakumar Swaminathan, Olga A Zabotina, Jonathan F Wendel, Jun Xie, Daniel B Szymanski

## Abstract

The shapes and material properties of cotton seed coat trichoblasts are the basis of a multibillion-dollar natural fiber industry. As such, these highly specialized cells are low-hanging fruit for intentional trait engineering. However, broad successes will require more mechanistic knowledge about their systems-level cellular controls. This time-series study integrates daily measurements of purified fiber transcriptomes and proteomes with multiscale fiber phenotyping datasets that span the same developmental interval. Abundance profiles of the subcellular proteomes are the foundation of the analyses. This resource article provides direct information concerning which homoeologs operate and informative depictions of how compartmentalized cellular systems change during developmental transitions. Prediction accuracy was partially validated by analysis of the protein expression group 11, which contained multiple known secondary cell wall cellulose synthases and dozens of unknown proteins and an averaged profile that was strongly correlated with a sharp state transition in cellulose microfibril alignment and increased cellulose content. The dataset as a whole can serve as a hypothesis-generating machine to guide future experiments that relate to cell shape and growth rate control, reversible tissue formation, and cell wall remodeling. Integration of mRNA and protein abundance revealed widespread evidence for post-transcriptional control. In addition, there were hundreds of transcriptionally controlled genes with differing timepoints of transition. This latter gene set can be used to more reliably analyze transcriptional control networks and to generate collections of gene expression drivers for cotton fiber research. The protein and transcript data are organized into user-friendly tables and a web interface that can be searched using any plant ortholog of interest based on developmental time, abundance, annotations, or phenotypic association.

## Introduction

Cotton (*Gossypium* spp.) is the world’s most important fiber crop and the foundation of a multibillion-dollar renewable textile industry (Wilkins et al., 2010; Huang et al., 2021). Tens of thousands of unicellular seed coat trichoblasts synchronously differentiate and transform their length-to-width aspect ratios to ∼3,000:1 during development (Stewart, 1975). At maturity, each fiber contains a helically patterned cellulose-rich wall that desiccates after programmed cell death to generate harvestable thin, strong, spinnable fibers (Seagull, 1992; Kim and Triplett, 2001; Butterworth et al., 2009). Fiber size, spinnability, and strength are the output of a morphogenetic program of anisotropic diffuse growth (Ryser, 1977; Yanagisawa et al., 2022) with an underestimated degree of complexity (Fig. 1). Phenotyping daily reveals an early tapering phase that is completed by 2 days post anthesis (DPA), during which the hemispherical fiber initial is converted to a thin elongated trichoblast (Graham and Haigler, 2021; Yanagisawa et al., 2022). After tapering, fiber cells form a tissue and growth rate decelerates (Singh et al., 2009; Wilson et al., 2024), and eventually the cell wall (CW) thickens toward the end of a protracted phase of decelerating anisotropic diffuse growth (Schubert et al., 1973; Wilson et al., 2024). During growth, the cells are also simultaneously modulating plasmodesmata functions and turgor pressure (Ruan et al., 2001). Given progress in understanding the molecular and biomechanical control of trichoblast morphogenesis (Yanagisawa et al., 2015; Yu et al., 2019; Yanagisawa et al., 2022), there are emerging opportunities to use biotechnological interventions to engineer fiber traits (Qin et al., 2022; Liu et al., 2025; Sun et al., 2025). The goal of this research is to integrate high-temporal resolution quantitative proteomics, RNA-seq, and phenotyping to provide a database that broadly predicts cellular networks that mediate key developmental transitions. This study also introduces several methodological advances that enhance the precision and interpretability of large-scale proteomic resources. These include a novel daily sampling strategy that captures fine-scale developmental transitions, label-free quantification optimized for subcellular fractions that enables reproducible abundance profiling across six orders of magnitude, and integrated RNA-protein-phenotype cross-correlation analyses that together provide unprecedented temporal resolution to predict functional modules and their developmental roles.

**Figure 1.**
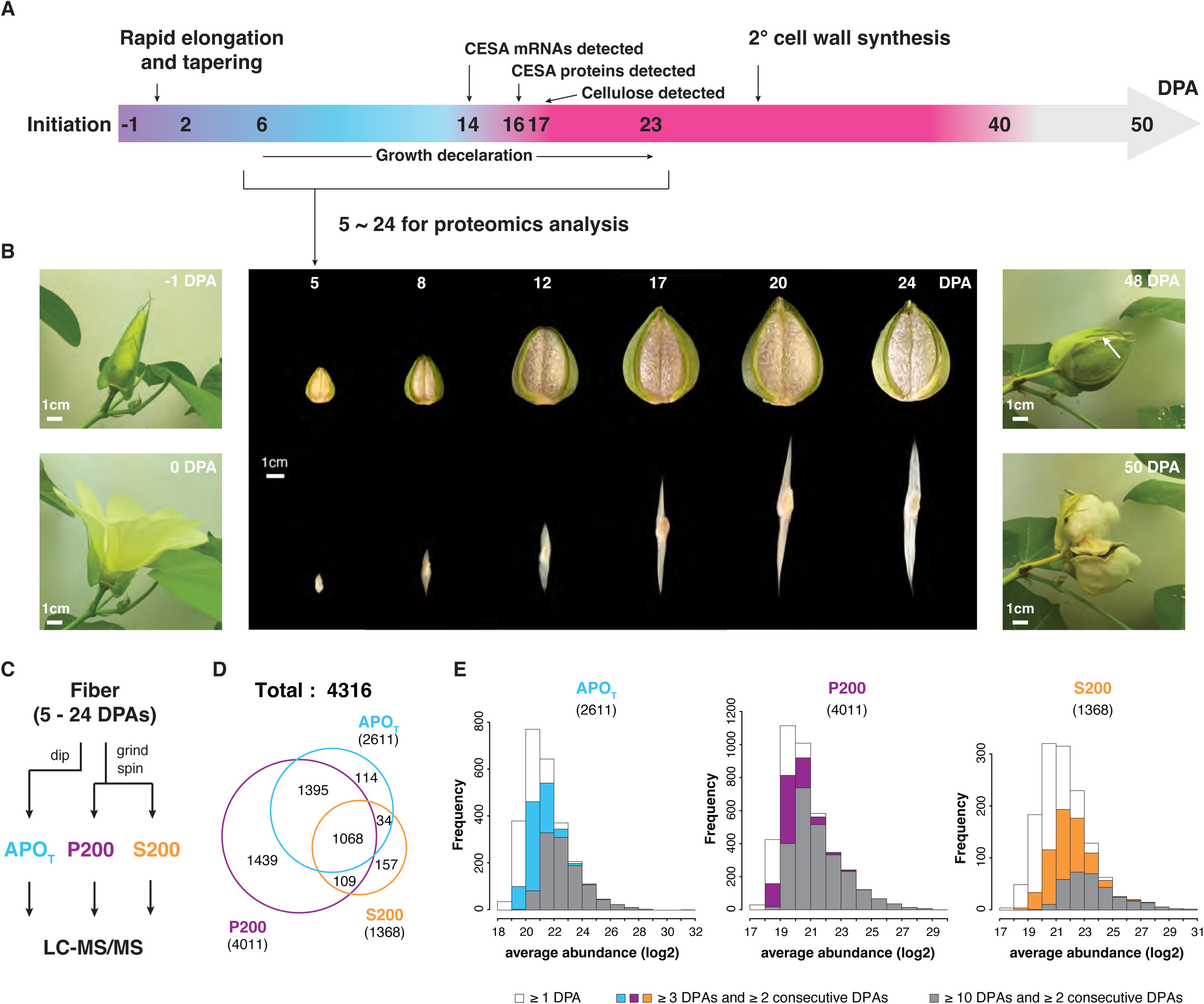
Deep proteome profiling across fiber development in cotton (*G. hirsutum*) plants: cell fractionation and quantitative proteomics. (A) Key developmental stages of cotton fibers, adapted and re-illustrated based on published descriptions (Swaminathan et al., 2024; Wilson et al., 2024; Grover et al., 2025). Flower opening day is defined as 0 days post anthesis (DPA). (B) Phenotypes of flowers, dissected bolls, and combed fibers at selected time points for proteomics analysis. The white arrow indicates the opening of the boll at 48 DPA. One sepal was removed from the flower for visualization. The entire developmental progression, from anthesis at 0 DPA to boll opening at 48 DPA, is captured in Supplementary Video S1, which provides a continuous visualization of fiber growth dynamics shown in Fig. 1, A and B. (C) Cell fractionation of fiber samples collected daily from 5 to 24 DPA in triplicate for quantitative proteomics analysis (Lee et al., 2025). (D) Proteome coverage across the three different subcellular compartments. (E) Distributions of average protein abundances across DPAs showing missing values. Abbreviations: APO_T_, apoplast; P200, crude microsome; S200, soluble fractions. Parentheses indicate the number of proteins in each cellular proteome.

Fiber development is coordinated with the organ-scale growth and physiology of the enclosing capsule (colloquially termed a “boll”) chamber (Fig. 1, A and B); the process also includes reversible transitions between cell-autonomous growth and tissue morphogenesis during which the cells are tightly adhered (Singh et al., 2009). Cotton has long served as a useful model for CW assembly and morphogenesis (Meinert and Delmer, 1977; Seagull, 1993; Tiwari and Wilkins, 1995), and components of the molecular control continue to emerge (Huang et al., 2021). A highly conserved cellulose patterning module underlies polarized diffuse growth in the plant kingdom (Baskin, 2005), and the details of how interactions among cortical microtubules, patterned cellulose synthase (CESA) activity, and CW stress-sensing operate to control cell shape changes are beginning to emerge (Coen and Cosgrove, 2023; Li et al., 2023). Strong lateral interactions between microfibrils (Yanagisawa et al., 2015; Zhang et al., 2021; Yanagisawa et al., 2022) and interactions between fiber and matrix components generate domains of the CW that deform asymmetrically in response to turgor-generated mechanical stress in the CW (Yanagisawa et al., 2015; Zhang et al., 2021; Yanagisawa et al., 2022). Spatially patterned cortical microtubules influence microfibril deposition throughout development to maintain cell diameter and pattern a helical secondary CW (SCW) (Seagull, 1992; Tiwari and Wilkins, 1995; Yanagisawa et al., 2022; Wilson et al., 2024). Important cotton microtubule-associated proteins are being discovered using forward and reverse genetic approaches (Rajangam et al., 2008; Zang et al., 2021; Chen et al., 2024). The microtubule-CESA module does not function in isolation; actin-based intracellular transport, vesicle trafficking, metabolism and redox control, and hormone signaling function in concert during fiber development (Huang et al., 2021).

Proteomic profiling provides a way to obtain direct measurements of the protein machineries and post-translational controls that enable effective morphogenetic work and coordinate diverse functions, some of which reside in different subcellular compartments. Historically, proteomic analyses of cotton fibers have identified proteins with altered abundances as a function of domestication (Bao et al., 2011; Hu et al., 2013; Qin et al., 2017), hormone or inhibitor responses (Wang et al., 2020; Yang et al., 2023), and developmental stages (Mujahid et al., 2016; Zhou et al., 2019; Jiang et al., 2022). In general, the number of discovered proteins is low, and temporal sampling of fiber development is limited to 5-day intervals at best. Gel-free workflows and increasingly sensitive instrumentation enable the quantification of thousands, rather than dozens, of proteins in a typical experiment (Hu et al., 2015; Lee and Szymanski, 2021; Yang et al., 2023). This presents an opportunity to gain broad insights into the protein dynamics that underlie the numerous developmental transitions during fiber development that are not known.

Here we combined daily sampling and clean cell fractionation procedures to analyze how protein abundances change in distinct subcellular compartments at a daily resolution. Our unique analytical pipeline allowed us to generate thousands of reliable protein abundance profiles spanning a broad period of fiber development that encompasses primary cell wall synthesis and the transition into secondary wall synthesis. Clustering methods identified unique protein expression (PE) groups that reflected specialized functions of the apoplast, cytosol, and membrane-associated proteins. Multiple groups had clear temporal transitions and broad memberships across the cell fractions, providing specific predictions of when, which, and how cellular systems operate. Time-series correlation analysis with a wide range of phenotypes quantified at the same daily resolution provided a way to predict suites of genes that coordinate the unidirectional progression of fiber development. Integration of RNA-seq and proteomic data can provide large-scale predictions about general modes of gene expression control (Nie et al., 2006; Manzoni et al., 2018). The proteomic data and their integration were partially validated by the accurate predictions of groups of known proteins with known localizations and stage-specific functions. The protein groups contain more than just the knowns and have a broad relevance to the field of cell wall assembly (Hoffmann et al., 2021) because they broadly predict enzymes, cytoskeletal, and vesicle trafficking machineries that mediate the transition to secondary wall synthesis. Combinations of cross-correlation analysis and machine learning methods globally predict modes of transcriptional and post-transcriptional control, providing large-scale mechanistic predictions. Collectively, our resource article is broadly relevant to fiber growth control and provides valuable data to fuel synthetic biology, comprehensive analyses of cellular processes controlling morphology, and eventually trait engineering. The data are organized within searchable and user-friendly supplemental tables. All protein abundance profiles and associated RNA-seq data are publicly accessible via a dedicated web portal (plant-proteomics.shinyapps.io/cotton-fiber-omics-atlas/).

## Results

### Protein profiling of developing cotton fiber by proteomics

Fiber development is reproducible across a wide range of environmental conditions (Schubert et al., 1973; Jan et al., 2022), including fibers that are cultured in vitro (Meinert and Delmer, 1977). A central goal of this study was to generate reliable methods to collect time-series data of protein abundance in multiple subcellular fractions from purified fiber tissue (Fig. 1).

Methods to isolate highly purified fiber samples and generate high-quality cell fractions with limited cross-contamination with broken cells were developed (Lee et al., 2025). The method enables precursor ion-based quantification of a diverse population of proteins within and between cell fractions. Here we scaled up the procedure to increase coverage of the crude cytosolic (S200) fraction. Each replicate included multiple locules from a single boll and included three replicates daily from 5-24 DPA (Fig. 1C). Protein yields in the different cell fractions are included in Supplementary Data Set S1. The earliest timepoint was chosen based on the minimum requirements for adequate protein precipitation from the soluble fractions, and the final time point was selected in order to capture a complete transition to SCW synthesis.

A crude apoplast (APO_T_) fraction was generated after careful dissection and brief immersion of the intact locules in the isolation buffer. This protocol minimizes contamination from broken cells and has been shown to generate a distinct apoplast fraction with unique protein multimerization behaviors and composition compared to cytosol and microsome fractions (Lee et al., 2025). After immersion, fibers were manually dissected, homogenized, and subjected to differential centrifugation to generate a crude microsomal (P200) and a soluble (S200) fraction (Fig. 1C). Across the 5 to 24 DPA window, 4316 proteins were reproducibly identified and quantified in 2 of 3 biological replicates of at least one cell fraction, providing the most comprehensive cotton fiber proteome coverage to date (Fig. 1D; Supplementary Data Set S3).

Cotton (*G. hirsutum*) is an allopolyploid with two subgenomes (A and D), and as such the duplicated genes (homoeologs) are encoded by the respective subgenomes, such that they potentially can be distinguished based on diverged peptide sequences (Hu et al., 2015; Lee and Szymanski, 2021). Homoeologs present challenges for accurate identifications and were defined as present if a homoeolog displayed at least 1/4 of the unique peptides compared to the leading protein. Diagnostic homoeolog-specific peptides above this threshold were detected for ∼55% of the protein groups, and these proteins are annotated with “A” and “D” suffixes following the Locus ID (Supplementary Data Set S3). The mRNA and protein search databases (Supplementary Files S1 & S2) were generated based on aggregated A- and D-subgenome SNPs relative to a base diploid assembly (Grover et al., 2025). These databases enabled high coverage matches with the MS spectra and accurate comparisons of transcript and protein abundance profiles for specific homoeologs (see below). The search databases and locus ID conventions used here include cross-referenced locus IDs across several commonly used cotton genome assemblies to facilitate gene matching (Supplementary Data Set S2D).

The P200 fraction consistently had the most diverse protein population over the time course (Supplementary Data Set S3C), and, as observed previously (Lee et al., 2025), was more similar to the APO_T_ than to the S200. The APO_T_ fraction primarily consists of membrane-enclosed particles with a highly heterogeneous composition (Lee et al., 2025). Widespread overlap among the different cell fractions can occur because many proteins are dually localized (McBride et al., 2017). Our pipeline generated protein quantification across over 6 orders of magnitude (Fig. 1E). The protein types amenable to proteomic analyses are extremely diverse (Supplementary Data Set S3). The coverage of metabolism in this dataset is deep (Supplementary Fig. S1) and provides broad experimental evidence for the specific gene products that operate in central metabolism, phenylpropanoid biosynthesis (Tuttle et al., 2015; Yang et al., 2023), fatty acid synthesis (Qin et al., 2007), and cell wall assembly (Pettolino et al., 2022).

A protein was retained for abundance profiling across the time course if it had non-zero values in three or more fractions with at least two of them on 2 consecutive days. The protein population composed of those detected in at least 2 of 3 consecutive DPAs and the subset detected in 10 or more DPAs had similar intensity distributions (Fig. 1E; Supplementary Fig. S5E), indicating that signal intensity is not the primary determinant of expression profile variability. The raw and normalized intensity data and plotted distributions are provided in Supplementary Data Set S3 and Supplementary Figs. S2-S4. Our deep proteome profiling provided a broad depiction of compartmentalized protein behaviors during development, identifying diverse groups of proteins that coordinate fiber development. However, the widespread occurrence of missing values between biological replicates is a widely recognized shortcoming of label-free quantitative proteomics (Jörnsten et al., 2005; Wang et al., 2020; Hamid et al., 2022). We developed a highly stringent data filtering procedure to minimize the effects of artifactual missing values between DPA, eliminate potentially unreliable profiles, and eliminate unreliable group members with low signal to generate a restricted dataset containing informative expression profiles from three subcellular locales (see Materials and Methods).

To identify expression groups, the informative profiles were analyzed using a machine learning-based clustering method termed Self-Organizing Map (SOM) analysis (Kohonen, 1990; Wehrens and Kruisselbrink, 2018). In an SOM, the input data are mapped onto a discrete topological grid formed by a set of groups. The groups are typically organized in a 2-dimensional grid, with rows and columns. As the model is trained on input data, the groups gradually adjust their weights to become more similar to the pattern of the input data in their neighborhood, resulting in clusters of the input profile data. The number of SOM groups was determined by evaluating cluster stability, ensuring that proteins consistently grouped with the same or neighboring clusters as the number of groups increased. This strategy minimized empty or single-member clusters, providing meaningful resolution and interpretability. Each SOM group has a “code” that reflects a central expression pattern of the members (Fig. 2A; Supplementary Data Set S4). A group was scored as unreliable if: 1) the code profile was punctuated by missing values across the timeline; 2) the profile pattern reflected variability in protein coverage in specific DPAs (Supplementary Data Set S3D); or 3) the pattern was defined by strings of three adjacent zeros that could simply reflect random gaps in the data that were not filled by interpolation. Following this filtering, 39 groups were classified as unreliable, and the remaining 44 potentially informative groups were subjected to further filtering to remove unreliable members with discordant features in their expression.

**Figure 2.**
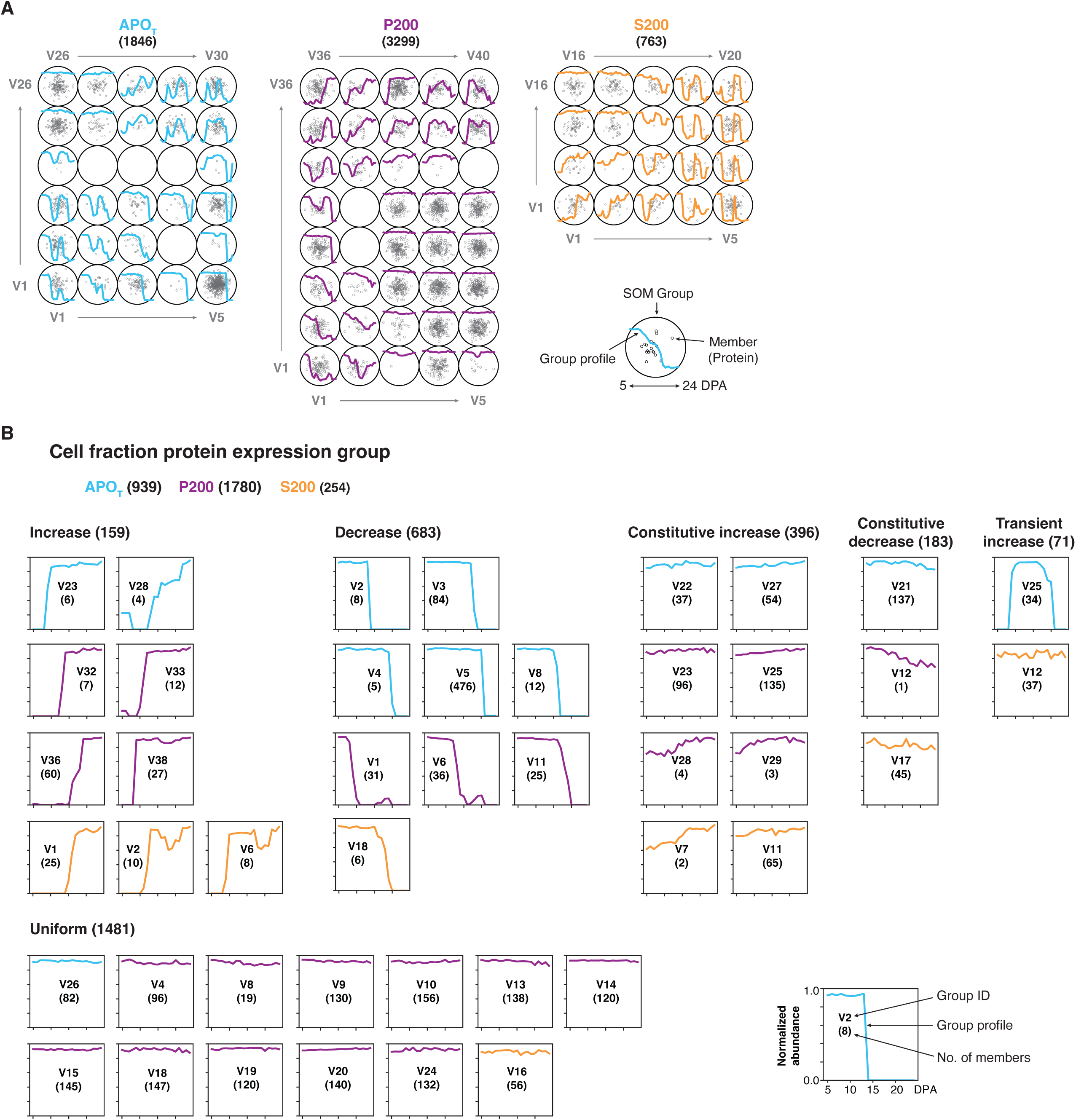
Generating a core set of protein expression modules during cotton fiber development. (A) Grouping of similar protein profiles in the three subcellular proteomes using a machine learning-based self-organizing map (SOM) approach. Each dot represents an individual protein member, positioned according to the similarity of its developmental abundance profile relative to neighboring proteins in the SOM grid. The number of SOM groups was selected based on group density and distribution to minimize empty groups or multiple groups with similar patterns. (B) A set of informative protein expression groups across fiber development in the three different cellular fractions. Supplementary Data Set S3 aggregates and organizes the resulting data into user-friendly tables containing raw, normalized, and interpolated protein abundance values across subcellular fractions and developmental time points. Abbreviations: APO_T_, apoplast; P200, crude microsome, S200, soluble fractions. Parentheses indicate the number of proteins in each cellular proteome or group.

A multi-step quality control procedure was then applied to each SOM cluster to exclude outliers, unreliable expression patterns, and low-abundance proteins, yielding high-confidence protein expression profiles (Supplementary Figs. S2-S5). For each group, member intensities were normalized to a 0-1 range based on their maximum values and averaged to generate representative expression patterns on a consistent scale (Fig. 2B). This process resulted in 44 distinct cell fraction protein expression group patterns (12 in APO_T_, 23 in P200, and 9 in S200), where a 10% change in the normalized scale corresponds to approximately a 4-fold difference in abundance (or 2-fold on the log2 scale) due to the high signal intensity of the included profiles. In the end, these 44 expression groups provide large-scale functional data on gene sets and cellular activities (Fig. 2B; Supplementary Data Sets S4 and S5). Supplementary Data Sets S4 and S5 provide user-friendly searchable tables of normalized and interpolated reliable protein abundance values for all reproducibly quantified proteins across subcellular fractions and developmental time points. The dataset can be searched or filtered by locus ID, functional annotation, subcellular localization, or developmental trend. The protein data provide clues about gene functions that may operate during specific developmental windows in particular subcellular locales.

Many of the profiles were non-uniform, with sharp abundance transitions often reflecting a shift to or from consecutive days of non-detection. Those classified as “increasing” had a clearly defined timepoint at which the protein transitioned from below to above the detection limit (Fig. 2B). The inverse was true for the profiles classified as “decreasing.” We hypothesize that these clearly varying expression patterns reflect rearrangements of suites of cellular machineries as the fiber population undergoes developmental transitions. Some non-uniform expression groups were defined by a specific subcellular compartment. Gene Ontology (GO) enrichment analysis shows that P200 protein expression groups with an increasing pattern have hydrolase activity, guanyl nucleotide binding, and catalytic activity terms, whereas S200 was enriched in the response to stress term (Supplementary Data Set S6). Constitutively expressed proteins classified as non-uniform were flagged if there was at least a 4-fold increase or decrease in the mean protein abundance of the group members during development (Supplementary Data Set S5). These “constitutive increase/decrease” groups may reflect broad reprogramming of gene expression relative to the transition to SCW synthesis (Grover et al., 2025) and/or proteins with unequal activities in the elongation and SCW phases. A large number of proteins had flat relative abundances and were classified as uniform. These proteins may define part of the “constitutivome” that includes essential functions that operate across the developmental phases.

### Integration of protein expression groups with fiber phenotypes: validation of PE11 with known secondary cell wall synthesis proteins

We next wanted to correlate protein expression profiles with phenotypic transitions. To generate a merged set of profiles aggregating protein abundance dynamics across the cell fractions, we turned again to SOM analysis. After evaluating SOM configurations with 9, 20, 30, and 40 groups, we selected the 20 groups as it minimized empty or single-member clusters while maintaining stable group memberships (Supplementary Data Set S7A; Supplementary Figs. S6-S9). This configuration yielded 15 distinct merged protein expression groups (PE1-PE15) (Fig. 3A). The merged groups are designated as protein expression (PE) modules, each representing a distinct pattern of protein abundance dynamics across development. Eleven PE groups contained clearly distinct and non-uniform profiles (Fig. 3A; Supplementary Data Set S7). Daily sampling enabled these expression groups to be resolved, and they may have interdependent subcellular functions. However, additional work with daily or even diurnal sampling is needed to define more clearly the temporal sequence and functions of PE module members. Our current data do not enable us to conclude that all PE members function at the same time to achieve a single morphological transition.

**Figure 3.**
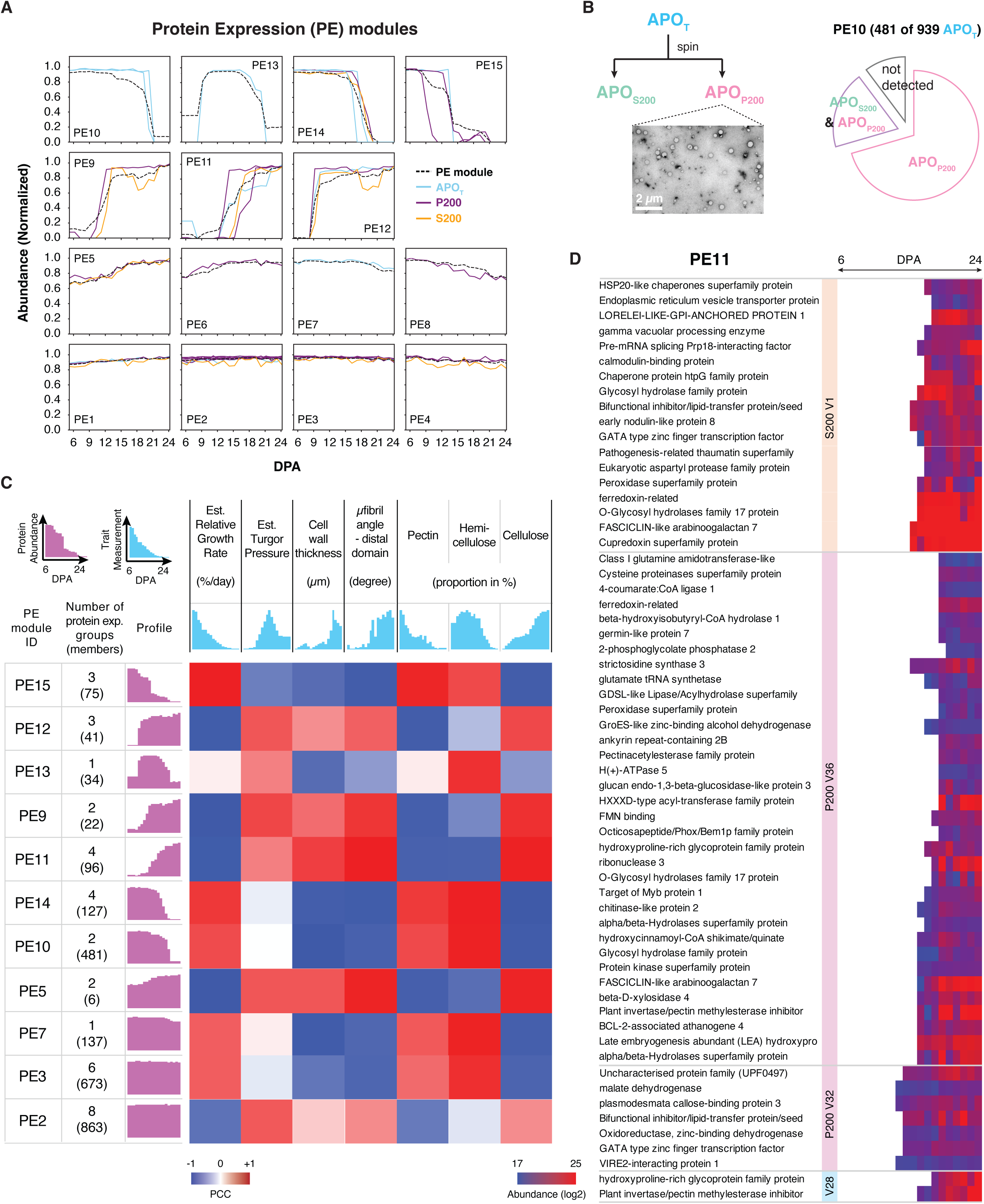
Integration of protein expression (PE) module dynamics with fiber phenotypes. (A) Protein expression modules merged from the APO_T_, P200, and S200 cellular fraction protein expression groups. The merged clusters are designated as PE modules, each representing a distinct pattern of abundance dynamics across development. (B) PE10 module reflects developmentally regulated extracellular vesicles. The APO_P200_ and APO_S200_ fractions were isolated using the pipeline (Lee et al., 2025), which also provided the proteomic data for the comparison analysis. The TEM image shows vesicles present in APO_P200_ fraction. Overlap of APO_T_ proteins in PE10 with APO_P200_ proteins is indicated (Supplementary Data Set S7D). (C) Integration analysis of fiber phenotypes and protein network modules. Pearson correlation coefficient (PCC) between protein patterns and phenotypes (Swaminathan et al., 2024; Wilson et al., 2024) was calculated. Module profiles correspond to the representative PE module profiles shown in (A). Supplementary Data Set S8 compiles the results of correlation analyses between PE modules and developmental phenotypes. (D) PE11 protein expression module showing similar and unique compartment-specific profiles associated with increasing cellulose content. Supplementary Data Set S9 provides a comprehensive list of proteins in PE11 and their subcellular localizations, enabling exploration of the molecular composition of the cellulose synthesis machinery and related pathways. Abbreviations: APO_T_, apoplast; P200, crude microsome; S200, soluble fractions. Parentheses indicate the number of proteins in each cellular proteome or group.

The mean normalized intensities of the PE members were used to estimate the group behavior and to test for correlations with phenotypes. Aggregation of seven developmentally variable fiber phenotypes from the same or similar developmental windows (Swaminathan et al., 2024; Wilson et al., 2024) provided a way to test for significant associations between PE groups and phenotypic transitions using correlation analyses (Fig. 3C; Supplementary Data Set S8). The results of these analyses are compiled in Supplementary Data Set S8, which integrates quantitative phenotypic profiles with PE modules. This dataset enables users to identify protein expressions that are significantly associated with specific developmental or biochemical traits, such as growth rate, wall composition, or the onset of SCW synthesis, providing a broad functional framework for linking molecular networks to measurable fiber phenotypes. As a key validating example, the PE11 group was most strongly positively correlated with the transition to SCW synthesis, which is characterized by transitions in cellulose microfibril angle, increases in cell wall thickness, and an increasing proportion of cellulose in the cell wall (Fig. 3C; Supplementary Data Set S8B). PE11 contained proteins present in all three subcellular compartments and included many known proteins involved in CW biosynthesis, as well as many previously uncharacterized proteins (Fig. 3D; Supplementary Data Sets S8 and S9). PE11 also contained *G. hirsutum* orthologs of each of the SCW-associated CESAs (Kim et al., 2019; Grover et al., 2025) that were induced at ∼16 DPA to similar levels (Fig. 4D; Supplementary Data Set S9A). PE14 was negatively correlated with SCW transition and may contain proteins that are specifically involved in the fiber elongation and remodeling of the primary CW (Fig. 3C; Supplementary Data Sets S7C and S8B). PE10 was also negatively correlated with SCW transition, but this group was composed solely of APO_T_ members. By comparing these data with those in a purified cotton EV dataset (Lee et al., 2025), we found that PE10 members were either specific to the particulate fraction or dual-localized in the soluble and particulate fractions of the apoplast (Supplementary Data Set S7D). PE10 contained more than half of the total APO_T_ proteome. The particulate fraction was enriched in spherical stain-excluding compartments that could be visualized using negative staining and transmission electron microscopy (Fig. 3B). To determine if the apoplastic particles (APO_p200_) had a unique polysaccharide composition, their glycome profiles were compared to those of the P200 and APO_s200_ fractions. Several epitopes related to xylan and xyloglucan were specifically enriched in the APO_p200_ particles compared to the other cell fractions and many other epitopes were selectively depleted (Supplementary Data Set S10A). These data provide further evidence for their unique composition and may reflect their mode of generation. PE7 and PE13 were also APO_T_-specific expression groups with similar profiles (Fig. 3C). These profiles may reflect the developmentally regulated biogenesis of extracellular vesicles.

**Figure 4.**
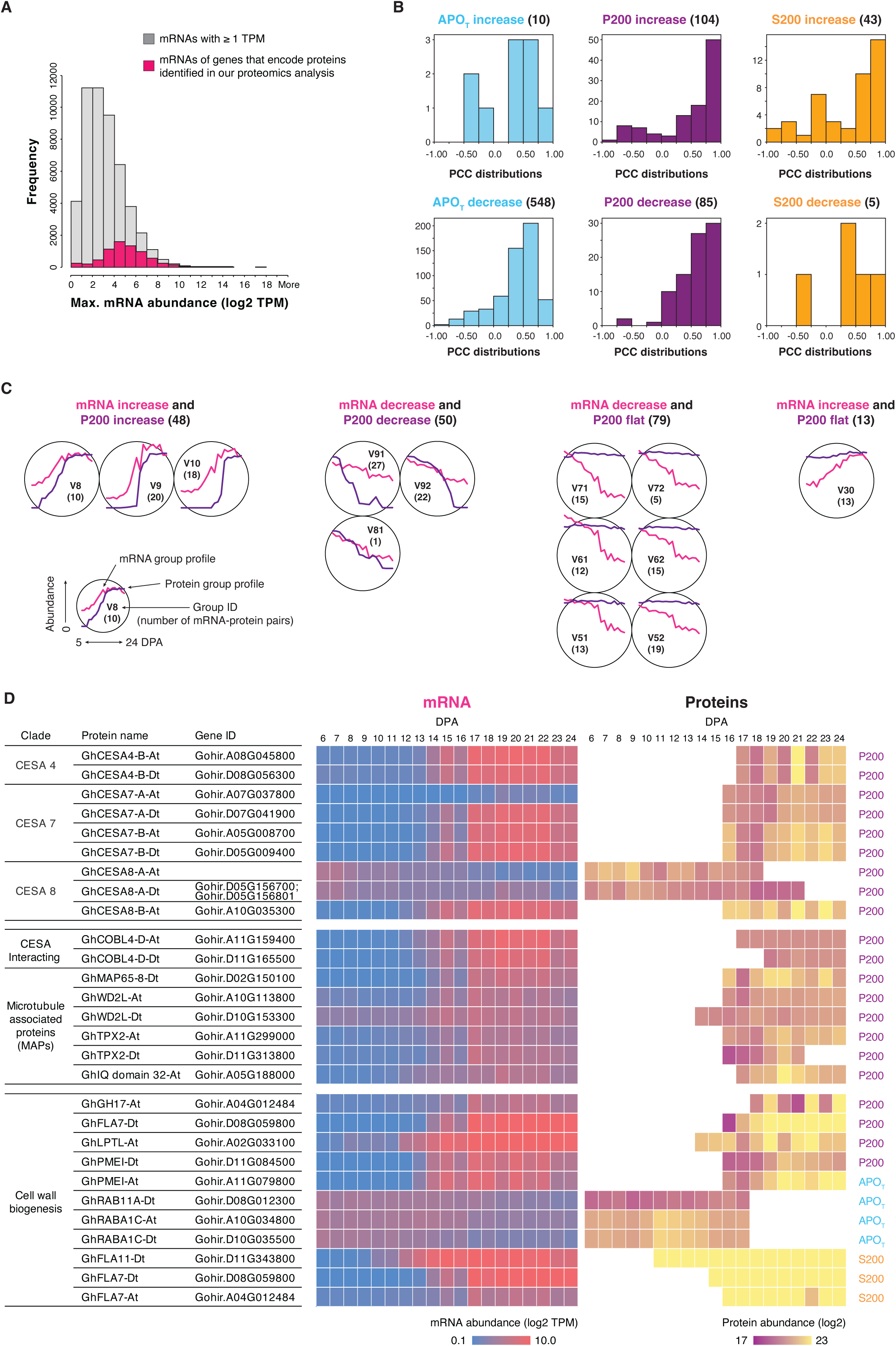
Global analysis of transcript and protein profiles across fiber development. (A) Coverage of proteins present in the cell fraction protein expression groups over the cotton fiber transcriptome data (Grover et al., 2025). Maximum mRNA and protein abundances across fiber development were visualized. The filtered transcriptome dataset is available in Supplementary Data Set S11. (B) Pearson correlation coefficient (PCC) between mRNAs and proteins for Case 1 proteins (see Materials and Methods). These proteins were present within the cell fraction protein expression groups that show “increase” or “decrease” profiles as defined in Fig. 2B. Parentheses indicate the number of proteins. (C) SOM-based weighted cross-correlation of paired mRNAs and proteins that have flat profiles across development. Protein patterns were defined in Fig. 2B, and mRNA patterns were defined using a fold change threshold (8-fold abundance change). The complete SOM result can be found in Supplementary Data Set S13 and Supplementary Fig. S13. (D) Transcriptionally controlled, known protein markers that function in SCW synthesis. Example mRNA and protein abundance profiles are shown for transcriptionally regulated genes present both in the PE11 module and assigned to the V8, V9, V10, and V92 groups of the weighted cross-correlation SOM result in Fig. 4C. A broader list of predicted transcriptionally regulated cell-wall proteins, including their abundance profiles and PCC values, is provided in Supplementary Data Set S14. At and Dt indicate homoeologs for the A- and D-subgenomes, respectively. Abbreviations: APO_T_, apoplast; P200, crude microsome; S200, soluble fractions.

PE15 module declined from maximal levels at 5 DPA and was most strongly correlated with the decelerating relative growth rate and a reduced proportion of pectin (Fig. 3C). PE15 may contain groups of proteins associated with a fiber state with a faster relative growth rate. PE9 and PE12 were anti-correlated with the relative growth rate and included auxin-response factors, ABA receptors, and vesicle trafficking machineries (Supplementary Data Set S7C, column T, PE9 module) that enable persistent axial growth and/or early components that prime the cell for the transition to tissue-based morphogenesis (Singh et al., 2009) or SCW synthesis (Pettolino et al., 2022). Many of the merged protein groups correlated with phenotypic profiles, and these analyses provide models of how diverse types of gene products function in multiple cellular locations to coordinate developmental transitions.

### Integration of mRNA and protein dynamics: distinct modes of gene expression control

To globally analyze potential modes of gene expression control, we integrated the proteomic profile data with fiber transcript data obtained using the same conditions and developmental time points (Grover et al., 2025). Pearson correlation coefficients (PCC) were calculated for 6809 genes in which protein profiles could be matched to a single homoeolog (see Materials and Methods, case 1) or a pair of homoeologs (Fig. 4A; Supplementary Figs. S10-S12, case 2). We initially focused on proteins with a non-uniform expression pattern, i.e., the mRNA/protein correlations in the “increasing” or “decreasing” categories that were stronger in the P200 and S200 fractions than the APO_T_ fraction (Fig. 4B; Supplementary Data Set S12). ∼48% of mRNA/protein pairs in P200 and S200 had PCC values > 0.75, and protein abundance could be explained by transcriptional control. In general, the APO_T_ proteins in both expression groups were weakly correlated with their mRNAs because this cell fraction was dominated by putative extracellular vesicle proteins that may have reduced abundance after ∼20 DPA (Fig. 3A). In addition to the APO_T_ groups, those showing a negative correlation could be explained by the spatial and functional compartmentalization of gene expression regulation, highlighting the importance of subcellular localization information to understand mRNA-protein correlations. However, there was evidence for transcriptionally regulated proteins in the apoplast. A cotton ortholog of the secreted signaling molecule STOMAGEN, usually associated with non-cell-autonomous cell proliferation (Sugano et al., 2010; Cui et al., 2014), was downregulated at ∼14 DPA and correlated with transcript levels (Supplementary Data Set S7).

To conduct a more open-ended test for commonly observed or non-linear patterns of gene expression control, a weighted SOM clustering was conducted by combining the mRNA and protein data for 1,228 different P200 proteins with informative profiles (Fig. 4C; Supplementary Data Set S13). The full clustering result is shown graphically in Supplementary Fig. S13. The analysis produced three groups (V8-V10) with a total of 48 members with coherently increasing protein and mRNA expression, and three groups (V81,V91,V92) with a total of 50 members with coherently decreasing expression (Fig. 4C), of which 48 of 98 were captured in the PCC analysis described in Fig. 4B. Among the large population of proteins with a more uniform expression, there were 79 genes in which mRNA levels fell by at least 8-fold over the time course (Fig. 4C). These examples could reflect post-transcriptional controls in which the mRNA translation efficiency increases or protein stability increases during development. In V30, transcript levels increased 8-fold, but protein levels were stable (Fig. 3C), indicating that translation of the mRNA may be regulated or the steady state of the protein may be the result of physical interactions with a stabilizing factor. The SOM-weighted cross-correlation analyses provide quantitative information about how diverse types and timings of gene expression control operate during fiber development.

To partially validate our analyses with knowns, we queried our list of genes predicted to be subject to temporal transcriptional control with those known to be involved in CW assembly or the transition to SCW synthesis (Supplementary Data Set S14). For example, among the 67 proteins present in the PE11 expression group, 44 were predicted to be transcriptionally regulated. Several orthologs of documented players in SCW synthesis (e.g., cell wall CESAs, COBRAs, and WAVE-DAMPENED-LIKE2 proteins) were reliably detected in the P200 fraction from fiber cell extracts (Fig. 4D) (Wen et al., 2022; Chen et al., 2024; He et al., 2024). GhCESA4-B, 7-A/B, and 8-B proteins were first detected with a ∼2-day lag compared to when the corresponding mRNAs first increased. A database of cotton orthologs to predicted Arabidopsis cell wall genes (Cell Wall Genomics database; https://cellwall.genomics.purdue.edu/) was created, and a subset of those was present in the set of predicted transcriptionally controlled genes (Supplementary Data Set S14). Supplementary Data Set S14 offers a systematically organized reference linking these regulated, cell wall-related gene families with their mRNA and protein profiles, providing a practical guide for selecting candidate loci for functional analyses and for identifying stage-specific promoters as tools to engineer fiber traits. These data summarized in this resource article provide a new view of the regulatory complexities of fiber development and a source of promoters and gene products that appear to be subject to transcriptional control and more easily modulated in transformed lines.

## Discussion

In this resource article, we combine sensitive and accurate quantitative proteomics with the synchronized process of cotton fiber morphogenesis to broadly predict protein networks that orchestrate fiber development. The data are derived from a single-cell type that develops in the insulated environment of the developing capsule. Daily sampling reveals an unexpected complexity: an extended phase of growth deceleration in early development, followed by multiple waves of gene activation and suppression that precede the transition to SCW synthesis and multiple phases of cellulose microfibril and cell wall matrix remodeling (Swaminathan et al., 2024; Wilson et al., 2024; Grover et al., 2025; Lee et al., 2025). These transitions are not all anchored to a single general transition to SCW synthesis. Our easily searchable proteomic datasets provide a new conceptual framework to analyze how networks of proteins in distinct subcellular locations coordinate multiple sequential cellular transitions that define fiber traits. All datasets generated and analyzed in this study—including normalized, interpolated protein abundances (Supplementary Data Set S4), integrated protein expression (PE) modules (S7), correlation analyses linking PE modules to phenotypes (S8), subcellular composition and functional enrichment data (S9), and transcriptionally regulated gene lists (S14)—are publicly accessible in a simple, searchable interface, providing a comprehensive framework for hypothesis generation and functional discoveries across laboratories.

The proteomic resources generated here provide experimental support for over 4,300 fiber proteins, and reliable abundance profiles over a wide window of development for more than 2,900. This dataset provides useful time-series data on the localization and abundance of a functionally diverse population of enzymes, hormone signaling proteins like auxin response factors and ABA receptors, as well as proteins that span the full breadth of molecular functions in the cell. There are many enzymes that are specifically associated with fiber elongation, including those involved in flavonoid, anthocyanin, and wogonin metabolism (Supplementary Fig. S1; Supplementary Data Sets S5 and S6). The roles of these small molecules in fiber morphogenesis merit further analyses. Additionally, these protein and RNA-seq data are publicly available in a search interface (plant-proteomics.shinyapps.io/cotton-fiber-omics-atlas/).

The SOM clustering method and cross-correlation with time-resolved phenotypes were used to generate models of how protein groups in multiple cellular compartments are linked to specific developmental transitions (Fig. 3). The merged profiles captured constitutively expressed proteins with uniform (PE1,PE2,PE3,PE4), increasing (PE5,PE6), and decreasing (PE7,PE8) abundances across the timeline. These proteins may provide functions that are more likely to be subject to post-translational controls. Many proteins were assigned to groups with clear time points of abundance change (PE9-PE12,PE14-PE15). This approach enabled us to identify groups of up- and down-trending proteins that transition at a similar time point. There were expression group pairs that were most strongly associated with growth rate variability (PE15,PE12) and others with the transition to SCW synthesis (PE11,PE14). These data provide systems-level views into how proteins in multiple subcellular locations might work together to mediate developmental transitions. These data can be used to guide future experiments that aim to analyze a wide range of cellular processes and developmental transitions. These pathways and players have a broad relevance to plant biology because they are likely to be conserved across many plant species and cell types that rely on cellulose-based morphogenetic systems (Hoffmann et al., 2021).

In the case of the transition to SCW synthesis, many key players are conserved across species and cell types with a thickened SCW (Kumar et al., 2016). Our dataset contains orthologs of each of the three CESA clades (Wen et al., 2022) and COBRA-like (He et al., 2024) proteins that are known to influence SCW synthesis in cotton, and many of them appear to be transcriptionally regulated (Figs. 3D and 4D). SCW patterning is a complex process that includes a radical reorganization of the cytoskeletal, vesicle trafficking, and cell wall assembly systems (Hoffmann et al., 2021). This proteomics analysis provides new insights into the particular CESAs and systems of proteins that operate prior to (PE14) and after (PE11) the transition to SCW synthesis. Interestingly, CESA8A proteins, classified as SCW CESAs based on their sequence, accumulated at high levels in growing fibers (Fig. 4D). Homomeric CESA8 complexes can synthesize cellulose (Purushotham et al., 2016; Purushotham et al., 2020), and it seems likely that GhCESA8A is involved in primary cell wall assembly.

Some of the expression groups may reflect reversible transitions in cell-cell adhesion as fibers interconvert between cell-autonomous and tissue-like morphogenesis at ∼5 and ∼20 DPA (Singh et al., 2009). These different modes of growth may partially explain relative growth rate deceleration and the functional roles of protein expression groups correlated with growth rate control (Fig. 3C). A prior publication on the apoplast proteome (Lee et al., 2025) and the time-series analyses here point to the existence of heterogeneously sized membrane-enclosed compartments in the extracellular space (Fig. 3B). The merged expression group profiles PE13 and PE10 reflect time points of transition to and from tissue-based morphogenesis (Fig. 3A). We speculate that these vesicle-enclosed compartments cannot simply diffuse from the plasma membrane to the extracellular space but are generated when cell-to-cell contact structures that include cytoplasmic components experience nanoscale mechanical failure as bundled fibers grow at different rates or directions.

Integration of proteomic and RNA-seq data can reveal distinct modes of gene expression control and define key temporal relationships between transcriptional upregulation, protein detection, and associated phenotypes (e.g., increases in cellulose content at 17 DPA). We identified a large set of transcriptionally activated and repressed genes at different stages of development (Fig. 4B and 4C). These gene classifications are likely reliable as they identified specific *G. hirsutum* loci encoding orthologs of known players in trichoblast development (Wen et al., 2022; Chen et al., 2024; He et al., 2024) as well as a broader set of unknown genes that have been annotated to have CW-related functions (Fig. 4D). These gene sets provide a potential toolkit of promoters and genes that may be effectively used to modulate fiber traits. Other potentially useful gene sets with constitutive expression or discordant transcript and protein profiles likely reflect unknown post-translational control mechanisms and will require further research to develop methods to alter their activities.

### Conclusions and future challenges

The true complexity of cotton fiber development is underappreciated; nonetheless, it is an experimentally tractable, single-cell type system that has enormous potential to drive basic plant biology research and the architectural engineering of crop traits. Our proteome-centric dataset advances the field greatly by directly linking reliable homoeolog localization and abundance measurements to a high temporal resolution developmental timeline and a wide array of known fiber phenotypes. Our data-dependent LC-MS/MS workflow predicted functional associations for thousands of proteins and can broadly drive hypothesis testing. The barriers to cotton fiber transformation are diminished, so it is becoming possible to validate these predictions at a reasonable scale (Kangben et al., 2024). The data also have broad utility to broadly guide experiments in contexts that include cell wall remodeling and morphogenesis control. There are clear paths to increased proteome coverage. New instrumentation and methods for data-independent acquisition/DIA can easily generate 3 to 5-fold increases in coverage and fewer missing values (Meier et al., 2020). Improved coverage and deeper insights into endomembrane system functions will require new methods for organelle separation and parallel LC-MS/MS analyses.

This data resource can serve to bridge gaps between key genes and the cell wall material properties that determine architectural traits. Finite element biomechanical models of growing cells and tissue predict key CW material properties and CW geometry/force-sensing mechanisms that are central to morphogenesis (Yanagisawa et al., 2022; Li et al., 2023). Finding the functional linkages among key glycome epitopes, polysaccharide-modifying protein systems, and cell wall properties provides a path to trait engineering. We anticipate that growing proteomic datasets and additional high-dimensional transcriptomic and metabolomic data (Tuttle et al., 2015; Qin et al., 2022; Sun et al., 2025) will be fully aggregated in public databases (Yu et al., 2021; Yang et al., 2023) to fuel new machine learning and AI-driven approaches that predict functional relationships among genetic maps, molecular features, cellular functions, and architectural traits.

## Materials and methods

### Cotton growth conditions

*Gossypium hirsutum* cv. TM1 plants were cultivated in Conviron^®^ E15 growth chambers (Conviron, Pembina, ND, USA) at Purdue University Plant Growth Center as previously described (Grover et al., 2025; Lee et al., 2025). Seeds were sown in 3-gallon pots containing a soil mixture composed of soil, perlite, bark, and chicken grit in a 4:2:2:1 ratio. Growth chambers were maintained at 50-70% relative humidity with a day/night temperature of 28/23°C and 16/8-hour photoperiod at a light intensity of 500 μmol m ² s ¹. The light sources included 28 Sylvania F72T12/CW/VHO 100 W and 4 Sylvania F24T12/CW/HO 35 W fluorescent lamps, as well as 12 GE 60 W 130 V A19 incandescent bulbs. Thirty-minute, two-step ramp-up and ramp-down periods (15 minutes at 166 μmol photons m ² s ¹ followed by 15 minutes at 333 μmol photons m ² s ¹) were programmed at the beginning and end of the daytime period, respectively. At anthesis, flowers were marked as 0 days post anthesis (0 DPA). Three bolls were collected daily from 5 to 24 DPA, yielding a total of 60 samples (20 days × 3 replicates).

Harvested cotton bolls were immediately kept on ice and carefully dissected to obtain four intact ovules from each boll.

### Cell fractionation into apoplast (APO_T_), microsome (P200), and cytosolic (S200) fractions

APO_T_, P200, and S200 fractions were prepared as previously described (Lee et al., 2025). One intact ovule (700 mg) from each locule was dipped in 5 mL of pre-chilled microsome isolation buffer (MIB) [50 mM HEPES/KOH (pH 7.5), 250 mM sorbitol, 50 mM KOAc, 2 mM Mg(OAc)_2_, 1 mM EDTA, 1 mM EGTA, 1 mM dithiothreitol (DTT), 2 mM PMSF, and 1% (v/v) protease inhibitor cocktail (160 mg/mL benzamidine-HCl, 100 mg/mL leupeptin, 12 mg/mL phenanthroline, 0.1 mg/mL aprotinin, and 0.1 mg/mL pepstatin A)]. After 10 minutes of incubation under gentle shaking, the resulting solution was filtered through two layers of cheesecloth. This entire APO_T_ fraction was then subjected to protein precipitation using a cold acetone method.

Cotton fibers, single-celled trichomes that develop on the ovule epidermis, are referred to here as “fibers” for clarity. For P200 and S200 fractions, fibers were directly isolated from the ovule material used for the APO_T_ preparation described above (Lee et al., 2025). Purified fresh fiber tissue was homogenized in cold MIB (sample-to-MIB ratio of 1:4) using a Polytron homogenizer (Brinkmann Instruments, New York, NY, USA) with a chilled blade tip for 10 seconds of grinding, followed by 1 minute on ice and an additional 10 seconds of grinding. The homogenate was filtered through four layers of cheesecloth pre-soaked in cold MIB, and the cheesecloth was further squeezed with chilled forceps to recover the remaining extract. The filtered homogenate was centrifuged at 1,000 × g for 10 minutes at 4°C using an Allegra X-30R centrifuge (Beckman Coulter Life Sciences, Indianapolis, IN, USA) to remove debris. The supernatant was then ultracentrifuged at 200,000 × g for 20 minutes at 4°C using a Beckman Optima Ultracentrifuge with a TLA110 rotor (Beckman Coulter Life Sciences, Indianapolis, IN, USA) to obtain a cytosolic (S200) fraction and a pellet. The pellet was washed with cold MIB, incubated on ice for 10 minutes, and ultracentrifuged again under the same conditions as above; this washing step was repeated once. The final pellet, defined as a microsome fraction (P200), was directly subjected to protein precipitation using a cold acetone method. Protein yield information for the three cell fractions is provided in Supplementary Data Set S1.

### Determination of protein concentration

For the APO_T_ and S200 cellular fractions, dried pellets from acetone precipitation were dissolved and denatured in 100 μL of 8 M urea for 1 hour at room temperature. For the P200 fraction, 200 μL of 8 M urea was directly added to the final pellet and incubated for 1 hour at room temperature to denature membrane-associated proteins. Undissolved debris was removed by centrifugation at 12,000 × g for 15 minutes using an Allegra X-30R centrifuge (Beckman Coulter Life Sciences, Indianapolis, IN, USA). Protein concentration was determined using a BCA assay kit according to the manufacturer’s protocol (Thermo Fisher Scientific, Waltham, MA, USA).

### LC-MS/MS sample preparation

For LC-MS/MS analysis, proteins were digested with trypsin as described previously (McBride et al., 2019; Lee et al., 2025). All protein samples obtained from the APO_T_ and S200 fractions, as well as 50 μg of protein from the P200 fraction, were used for digestion, and MS/MS data were acquired as previously described (Lee et al., 2025).

### Peptide identification and quantification

Andromeda search engine in MaxQuant (version 1.6.14.0) was used for protein identification and relative protein abundance quantification (Tyanova et al., 2016). Searches were conducted as described previously (Lee et al., 2021; Lee et al., 2025). To minimize locus identity (Locus ID) assignment errors between cell fractions, all LC-MS/MS raw files were searched in a single MaxQuant job, and signal intensities were normalized across all time points (Cox et al., 2014; Tyanova et al., 2016). Separate parameter groups were defined for each of the three cellular fractions to allow fraction-specific settings (Supplementary Data Set S2). The search parameters were as follows: match between runs was set to the default setting; 1% protein and peptide false discovery rate (FDR) was set; 1 was chosen for the minimum unique peptides; label-free quantification was selected within each parameter group; all other parameters were set to default. A species-specific and homoeolog-diagnostic cotton reference proteome was built using *G. raimondii* genome annotation (Grover et al., 2025) to facilitate proteome and transcriptome data integration (Grover et al., 2025). This reference proteome (Supplementary File S2) includes species- and homoeolog-specific single nucleotide polymorphisms (SNPs) and had been validated to perform well in *G. hirsutum* (Hu et al., 2021). To promote consistent locus ID assignments across annotated *Gossypium hirsutum* genome versions, a locus ID conversion table is provided. Identified peptides, proteins, and a locus ID conversion table are available in Supplementary Data Set S2.

### Data filtering for protein identification in each cellular fraction (APO_T_, P200, S200)

Clear criteria were developed to define protein presence in each cellular fraction, as previously described (Lee et al., 2025; Lee et al., 2025). First, proteins with MS/MS spectral counts ≥ 1 in at least two out of three biological replicates were chosen as identified proteins in each cellular fraction. Second, among these, proteins with valid label-free quantification (LFQ) values in at least two out of three replicates were retained as reproducibly quantified proteins. The reproducible protein sets defined by MS/MS counts and LFQ criteria are provided in Supplementary Data Set S3, A and B, respectively.

### Homoeolog and ortholog determination

Cultivated cotton (*G. hirsutum*) is an allotetraploid formed through the hybridization of two diploid progenitor species that contributed the A and D subgenomes (Wendel, 1989).

Standard protein grouping algorithms, such as MaxQuant leading protein output, are often inadequate for resolving the unique complexities of this genome, particularly, the high sequence identity shared between homoeologs. To address this, we utilized unique peptide information as diagnostic, homoeolog-specific peptide evidence (Hu et al., 2013; Lee and Szymanski, 2021) to define protein identifications.

### Step 1: Isoform consolidation

To eliminate redundancy and streamline the analysis, all isoforms belonging to the same gene locus (identified by their shared Locus ID prefix, e.g., Gorai.001G012200) were consolidated into a single representative protein entry. This step ensured that each locus was represented by a single high-confidence identifier, eliminating redundant entries for the subsequent subgenome determination in Step 2.

### Step 2: Peptide evidence filtering and thresholding

Homoeolog discrimination was based exclusively on unique peptide evidence reported in the “Peptide counts (unique)” column of the MaxQuant proteinGroups.txt file. For each protein group, we implemented a 25% (1:4) ratio threshold of unique peptide counts, calculated as the count for any other protein in the group relative to the leading protein count (defined as the first protein entry with the highest count). Any protein in the group was defined as identified if its unique peptide count was at least 25% of the leading protein’s count. This served as a useful method to distinguish between subgenome-specific homoeologs and lower-resolution identifications that could not reliably distinguish multiple orthologous group members.

### Step 3: Homoeolog classification

Filtered protein assignments were categorized into three distinct cases across the cotton annotation columns in Supplementary Data Sets:

- **Case 1:** Unambiguous homoeologs, where all unique peptides mapped exclusively to either the A or D subgenome (3,189 of 5,360 total proteins).
- **Case 2:** Homoeologous pair assignments, where unique peptides mapped to both subgenomes with proportional support for each protein meeting or exceeding the 25% threshold relative to the leading protein (2171 of 5,360 total proteins).
- **Case 3:** Ambiguous ortholog assignments, where unique peptides mapped to multiple homoeologs from multiple loci; these uncertain assignments were retained for completeness but flagged for caution (398 of 5,360 total proteins).

Protein abundances were maintained as discrete MaxQuant protein groups and were not duplicated across individual homoeologs for the SOM analyses (Supplementary Data Set S3B).

### Profile interpolation

The dataset contains many missing values for protein abundance at a subset of DPAs. Low abundance proteins often had profiles riddled with three or more missing adjacent values and were deemed unreliable. Profiles with a gap of three missing values were therefore not interpolated. Two missing adjacent values, represented as zeros, at either end or within the middle of a protein abundance profile were interpolated. For a single zero occurring in the middle of a profile, the missing value was replaced by the average of its neighboring non-zero values. When two consecutive zeros occurred, linear interpolation was applied. At the two ends of a profile, a single zero was replaced by the adjacent non-zero value, whereas two consecutive zeros were replaced by the nearest non-zero value. These interpolated protein abundance profiles were used for downstream analyses and are available in Supplementary Data Set S3C.

### Self-organizing maps (SOM) for protein expression patterns

Self-organizing maps (SOM), a robust unsupervised machine learning method (Kohonen, 1990), were used to analyze protein expression patterns. We applied a SOM algorithm (Wehrens and Kruisselbrink, 2018; Yang et al., 2025) based on an inner product distance metric to cluster the interpolated protein abundance profiles from each cell fraction. We performed SOM with various numbers of clusters (e.g., 9, 20, 30, and 40) and qualitatively evaluated clustering stability. We selected the number of clusters that consistently grouped proteins within the same or neighboring clusters while avoiding many empty or very small clusters. The resulting SOM clusters defined representative protein expression profile patterns (Fig. 2A). Clustering results for all protein expression groups are available in Supplementary Data Set S4 and Supplementary Figs. S2-S4. Furthermore, resulting clusters were subjected to rigorous quality control filtering. Cluster profiles were flagged as unreliable if they mirrored the overall distribution pattern of protein coverage (Supplementary Data Set S3D) or were dominated by patterns that can be explained by the interpolation, e.g., a gap of three consecutive zeros. After this quality assessment, 12 groups in APO_T_, 23 groups in P200, and 9 groups in S200 fractions were retained as groups containing informative protein profiles (Supplementary Data Set S4).

### Quality control for reliable protein expression patterns

To ensure reliable protein expression patterns, a multi-step quality control procedure was applied to each SOM cluster to remove outliers, unreliable profiles, and low abundance profiles (Supplementary Figs. S2-S4).

### Step 1: Outlier detection

Outliers were identified by calculating all pairwise similarities defined by the inner product of two profiles in a cluster:

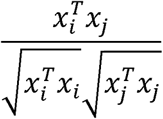

If the similarity between a protein profile within a group and the group centroid was less than either the group’s median similarity or 0.8, the protein was treated as an outlier and removed (Supplementary Figs. S2-S4, Step 1).

### Step 2: Removal of unreliable profiles

Unreliable profiles were defined as those with isolated non-zeros, including profiles with a single non-zero value flanked by zeros or three consecutive zeros in the middle of the expression pattern. These profiles were removed from the clusters (Supplementary Figs. S2-S4, Step 2).

### Step 3: Filtering based on abundance

To further validate the clustering and filtering steps, we conducted a detailed analysis to assess the relationship between protein expression classes, including non-reproducible proteins, outliers (Step 1), unreliable profiles (Step 2), and reliable profiles, and signal intensity (Supplementary Fig. S5). Proteins with below median abundance values across DPAs were more likely to exhibit missing values and contain more zeros in their profiles. When the median abundance dropped below 20 (log2-transformed) for APO_T_, 19 for P200, and 20 for S200, these proteins were considered low abundance and removed. These cutoffs were derived from the median abundance values of proteins excluded during the earlier steps (Supplementary Figs. S2-S4, Step 3).

### Informative cell fraction protein expression group patterns

To generate representative expression patterns for each protein group, member intensities were normalized to a 0-1 range based on their maximum values within the group (Fig. 2B). The normalized profiles were then averaged to produce a consistent scale across all patterns. This procedure yielded 44 distinct cell fraction protein expression group patterns (Fig. 2B), including 12 in APO_T_, 23 in P200, and 9 in S200, as detailed in Supplementary Data Set S5. In this dataset, a 10% absolute change (increase or decrease) in the normalized profile corresponds approximately to a 4-fold change in actual protein abundance, equivalent to a 2-fold change on the log2 scale used in the figures and data. This scaling relationship arises because the reliable profiles used in these protein expression groups have high baseline abundances, with average log2-transformed abundances of at least 20 (Supplementary Fig. S5).

### Defining Protein Expression (PE) modules for integrative analyses

Similar protein expression patterns were merged into modules through an additional SOM clustering analysis applied to the 44 previously generated protein expression group patterns. SOM outputs with 9, 20, 30, and 40 groups were evaluated to determine the optimal cluster resolution (Supplementary Data Set 7A; Supplementary Figs. S6-S9). As the number of groups increased, the SOM produced a higher proportion of empty clusters or clusters containing only a single group pattern, typically with flat profiles, while the overall cluster memberships remained largely unchanged. Consequently, we selected the 20-cluster SOM, which defined 15 distinct merged protein expression modules (Fig. 3A). For clarity, these merged protein expression groups are hereafter designated as protein expression (PE) modules. The profiles of the PE modules, together with their constituent proteins, were subsequently used as protein phenotypes for integrative analyses with phenotypic and transcriptomic data, available in Supplementary Data Set S7, B and C.

### Glycome profiling

APO_T_ fraction was further separated into APO_S200_ and APO_P200_ subfractions by ultracentrifugation as described previously (Lee et al., 2025). The supernatant (APO_S200_ fraction) was completely lyophilized. The pellet containing apoplastic vesicles (APO_P200_ fraction) was lysed by sonication and subjected to an additional ultracentrifugation step to collect the soluble fraction containing released polysaccharides (Okekeogbu et al., 2019). Lyophilized APO_S200_ and APO_P200_ fractions were redissolved in water, and their sugar contents were quantified (Supplementary Data Set S10A). Two micrograms of polysaccharides from each fraction were spotted into the wells of a 96-well plate and probed with 71 different antibodies (Supplementary Data Set S10B), as previously described (Swaminathan et al., 2024). Crude microsomes obtained from lysed fiber cells (P200) were processed identically and used as an additional control.

### Fiber Phenotype Data Acquisition

Phenotypic data for correlation analysis were collected from cotton fibers grown under identical conditions over the same developmental timeline as the proteomic samples. Data for seven key fiber traits, such as estimated relative growth rate, turgor pressure, cell wall thickness, microfibril angle, and proportional contents of pectin, hemicellulose, and cellulose, were aggregated from previous studies. For complete details on the methodologies used to acquire these measurements, please refer to Swaminathan et al. (2024) and Wilson et al. (2024).

### Transcriptome Data Source and Processing

The transcriptomic data used for mRNA-protein correlation analyses were obtained from a previously published study (Grover et al., 2025), which profiled cotton fibers across the same developmental time points (5-24 DPA) and under identical growth conditions as described here. We downloaded the Supplementary Table 1, which contains transcript profiles for 74,776 genes, and filtered out low abundance transcripts with fewer than one transcript per kilobase million (TPM) to retain high-quality, reliable transcriptome information. The resulting dataset, comprising 50,512 protein-coding genes, is provided in Supplementary Data Set S11 and was used for subsequent comparative analyses.

### Correlation analysis and classification of mRNA and protein pairs

To evaluate the relationship between transcript and protein levels, we performed a paired analysis of abundance profiles. Utilizing the homoeolog and ortholog assignments established previously, we linked each protein’s developmental abundance pattern (5-24 DPA) to its corresponding mRNA data from the filtered transcriptome (Supplementary Data Set S11).

Pearson correlation coefficients (PCC) were calculated for these mRNA-protein pairs across each cell fraction. Where a protein group was associated with multiple homoeologs (Cases 2 and 3), the protein abundance profile was duplicated for each homoeolog in the group during RNA-protein correlation testing. For these instances, we retained the mRNA-protein pair that yielded the highest PCC for all downstream analyses. The resulting PCCs for each mRNA-protein pair within each cell fraction are provided in Supplementary Data Set S12, with corresponding profile visualizations in Supplementary Figs. S10-S12.

### RNA-protein relationship patterns via SOM

To identify relationships between mRNAs and proteins that exhibit constitutively expressed profile patterns, Case 1 P200 proteins were paired with their corresponding transcripts. We then applied SOM clustering to these mRNA-protein pairs, using a 100-group arrangement (10 rows by 10 columns; Supplementary Fig. S13). The resulting SOM clusters defined the relationship patterns between mRNAs and proteins across cotton development. Details of the clustering results and the profiles of paired mRNAs and proteins are provided in Supplementary Data Set S13.

### Electron and fluorescence microscopy of isolated extracellular vesicles (APO_p200_)

Apoplastic particulate subfraction (APO_p200_) was enriched from a freshly prepared APO_T_ fraction obtained from four intact ovules of a 9 DPA boll by ultracentrifugation at 200,000 × g for 20 minutes at 4°C using a Beckman Optima Ultracentrifuge. The APO_p200_ fraction was diluted 100-fold in phosphate buffered saline (pH 7.0). One microliter of the resuspended APO_p200_ vesicles was negatively stained with 1% uranyl acetate on plasma-treated copper grids. Imaging was performed at an accelerating voltage of 40 to 100 kV using a Philips CM-100 Transmission Electron Microscope (Philips/FEI Corporation, Eindhoven, Holland).

### Statistical tests and data analysis

Statistical analysis was performed using R version 4.2.0 (R Core Team, 2018) on RStudio 2022.07.1 (RStudio Team, 2018). The Flexible Self-Organizing Maps in Kohonen 3.0 package for R (Wehrens and Kruisselbrink, 2018). Microsoft Excel on Office 365 for Mac was used to organize and display the analyzed data.

## Supporting information

Supplementary Data Set S1

Supplementary Data Set S2

Supplementary Data Set S3

Supplementary Data Set S4

Supplementary Data Set S5

Supplementary Data Set S6

Supplementary Data Set S7

Supplementary Data Set S8

Supplementary Data Set S9

Supplementary Data Set S10

Supplementary Data Set S11

Supplementary Data Set S12

Supplementary Data Set S13

Supplementary Data Set S14

## Code availability

The source codes for the protein SOM clustering analyses and the weighted cross-correlation SOM clustering analysis are publicly available on GitHub (https://github.com/yangpengchengstat/Proteomics-based-models-of-cotton-fiber-protein-dynamics-and-gene-expression-control).

## Data availability

The protein abundance profiles and associated RNA-seq data are accessible via a searchable online atlas at plant-proteomics.shinyapps.io/cotton-fiber-omics-atlas/. The mass spectrometry proteomics data have been deposited to the ProteomeXchange Consortium via the PRIDE partner repository (Perez-Riverol et al., 2021) with the dataset identifier PXD051704.

## Acknowledgements

This material is based upon work supported by the National Science Foundation under Grant No. 1951819 to D.B.S. We would like to thank Dr. Uma Aryal at the Purdue Proteomics Facility for running the LC-MS/MS samples. We are grateful to Eileen L. Mallery for growing cotton plants and to Dr. Pragya Barua for helping with sample preparation.

## Author contributions

Y.L.: Conceptualization, Methodology, Data Curation, Formal analysis, Investigation, Validation, Software, Visualization, Writing - Original Draft, Writing - Review & Editing

P.Y.: Methodology, Data Curation, Formal analysis, Software, Visualization, Writing - Review & Editing

H.R.: Methodology, Data Curation, Investigation, Visualization

G.M.: Data Curation, Software

C.E.G.: Methodology, Resources, Writing - Review & Editing

S.S.: Methodology, Resources, Investigation, Writing - Review & Editing

O.A.Z.: Conceptualization, Methodology, Resources, Writing - Review & Editing, Supervision, Project administration, Funding acquisition

J.F.W.: Conceptualization, Methodology, Resources, Writing - Review & Editing, Supervision, Project administration, Funding acquisition

J.X.: Conceptualization, Methodology, Formal analysis, Writing - Original Draft, Writing - Review & Editing, Supervision, Project administration, Funding acquisition

D.B.S.: Conceptualization, Methodology, Formal analysis, Investigation, Validation, Writing - Original Draft, Writing - Review & Editing, Supervision, Project administration, Funding acquisition

## Supplementary information

### Supplementary Figures

Supplementary Figure S1. Enzyme coverage of metabolic pathways.

Supplementary Figure S2. SOM grouping and profile filtering (APO_T_).

Supplementary Figure S3. SOM grouping and profile filtering (P200).

Supplementary Figure S4. SOM grouping and profile filtering (S200).

Supplementary Figure S5. Analysis of proteins removed at each data filtering step during the identification of reliable members.

Supplementary Figure S6. Merged SOM protein expression modules (9-cluster model).

Supplementary Figure S7. Merged SOM protein expression modules (20-cluster model).

Supplementary Figure S8. Merged SOM protein expression modules (30-cluster model).

Supplementary Figure S9. Merged SOM protein expression modules (40-cluster model).

Supplementary Figure S10. Profiles of paired mRNAs and proteins (APO_T_).

Supplementary Figure S11. Profiles of paired mRNAs and proteins (P200).

Supplementary Figure S12. Profiles of paired mRNAs and proteins (S200).

Supplementary Figure S13. Weighted cross-correlation SOM of paired mRNAs and proteins (P200).

### Supplementary Video

Supplementary Video S1. A movie of cotton flower and boll development (−5 to 53 DPA).

### Supplementary Files

Supplementary File S1. The mRNA search database (AD1_AtDt.transcripts.fa).

Supplementary File S2. The protein reference proteome (AD1_AtDt.pep.fasta).

### Supplementary Data Sets

Supplementary Data Set S1. Fiber protein yields (APO_T_, P200, and S200).

Supplementary Data Set S2. Raw abundances of peptides and proteins (APO_T_, P200, and S200).

- Raw peptide and protein abundance information obtained from a MaxQuant search.
- Locus ID conversion table from the Gorai to the Gohir ID systems.

Supplementary Data Set S3. Raw and interpolated protein profiles across fiber development (APO_T_, P200, and S200).

- Average abundances of proteins identified by MS/MS count(s) > 0 in at least 2 out of the 3 replicates and quantified by LFQ > 0 in at least 2 out of the 3 replicates.
- **Interpolated protein profiles (S3C).**
- Number of proteins at each filtering step (unfiltered raw data, MS/MS, and LFQ).

Supplementary Data Set S4. SOM clustering results and filtering used to generate protein expression group profiles (APO_T_, P200, and S200).

- SOM group profiles in each cell fraction.
- **Profiles of members in each SOM group and a summary of the data filtering.**

Supplementary Data Set S5. SOM protein expression group profiles (APO_T_, P200, and S200).

- **SOM groups and their profiles in the three cell fractions (visualized in Fig. 2B).**

Supplementary Data Set S6. Gene Ontology enrichment analysis of P200 and S200 protein expression groups with increasing patterns.

Supplementary Data Set S7. Merged SOM protein expression (PE) modules.

- Optimization of the number of groups defined for SOM-based clustering.
- **Informative PE modules provided with representative expression profiles and individual member protein profiles.** Reliable profiles from Supplementary Data Set S4 were pooled; localization and cell fraction group information are provided in column O.
- PE10 APO_T_ analysis result.

Supplementary Data Set S8. Integrative analyses of fiber phenotypes and merged protein expression (PE) modules.

- **Phenotype estimation data (profiles of each phenotype from 6 to 24 DPA).**
- PCCs and p-values for correlations between phenotypes and merged PE modules (visualized in Fig. 3C).

Supplementary Data Set S9. Gene Ontology enrichment analysis of the PE11 module (visualized in Fig. 3D).

Supplementary Data Set S10. Glycome profiling of polysaccharide epitopes (APO_p200_ vs. APO_s200_ vs. P200).

Supplementary Data Set S11. Abundance profiles of transcripts in cotton fiber across development (Grover et al., 2025).

- Filtered mRNA dataset downloaded from the Grover et al., 2025 study.

Supplementary Data Set S12. Pearson correlation coefficient (PCC) analyses between mRNAs and proteins present in cell fraction protein expression groups with trend-up or trend-down patterns.

Supplementary Data Set S13. Weighted cross-correlation SOM of paired mRNAs and proteins (P200).

- SOM group profiles with fold changes in their expression profiles (visualized in Fig. 4C).
- mRNA and protein profiles assigned to each SOM group.

Supplementary Data Set S14. Integrative analysis of mRNAs and proteins annotated as cell wall-related gene families.

- The list of Cell Wall Related Gene Families was downloaded from https://cellwall.genomics.purdue.edu/.
- Data integration and correlation analysis results for cell wall-related gene families with their mRNA and protein profiles.

