## Supplementary Figure S1 for "Systems-level proteomic models of cotton fiber development: A high-resolution resource to analyze cell dynamics and trait engineering"

**Supplementary Figure S1. Enzyme coverage of metabolic pathways.**

Cotton gene IDs were mapped onto metabolic pathways using Cellular Overview/Omics viewer at Plant Metabolic Network GraimondiiCyc v4.0.0 (https://pmn.plantcyc.org/overviewsWeb/celOv.shtml?orgid=GRAIMONDII).

1A. Pathway map of reproducible detected $APO_T$ proteins (present in Figure 1C).

1B. Pathway map of reproducible detected P200 proteins (present in Figure 1C).
Because Plant Metabolic Network does not allow a search with a large number (>900) of gene IDs in a query, the two pathway mappings (1/2 and 2/2) were generated in the two separated searches.

1C. Pathway map of reproducible detected S200 proteins (present in Figure 1C).

1D. Pathway map of cytoplasmic proteins (P200 and S200) with informative profiles (present in Figure 2B).

1E. Pathway map of extracellular proteins ($APO_T$) with informative profiles (present in Figure 2B).

1A. Pathway map of reproducible detected $APO_T$ proteins (present in Figure 1C).

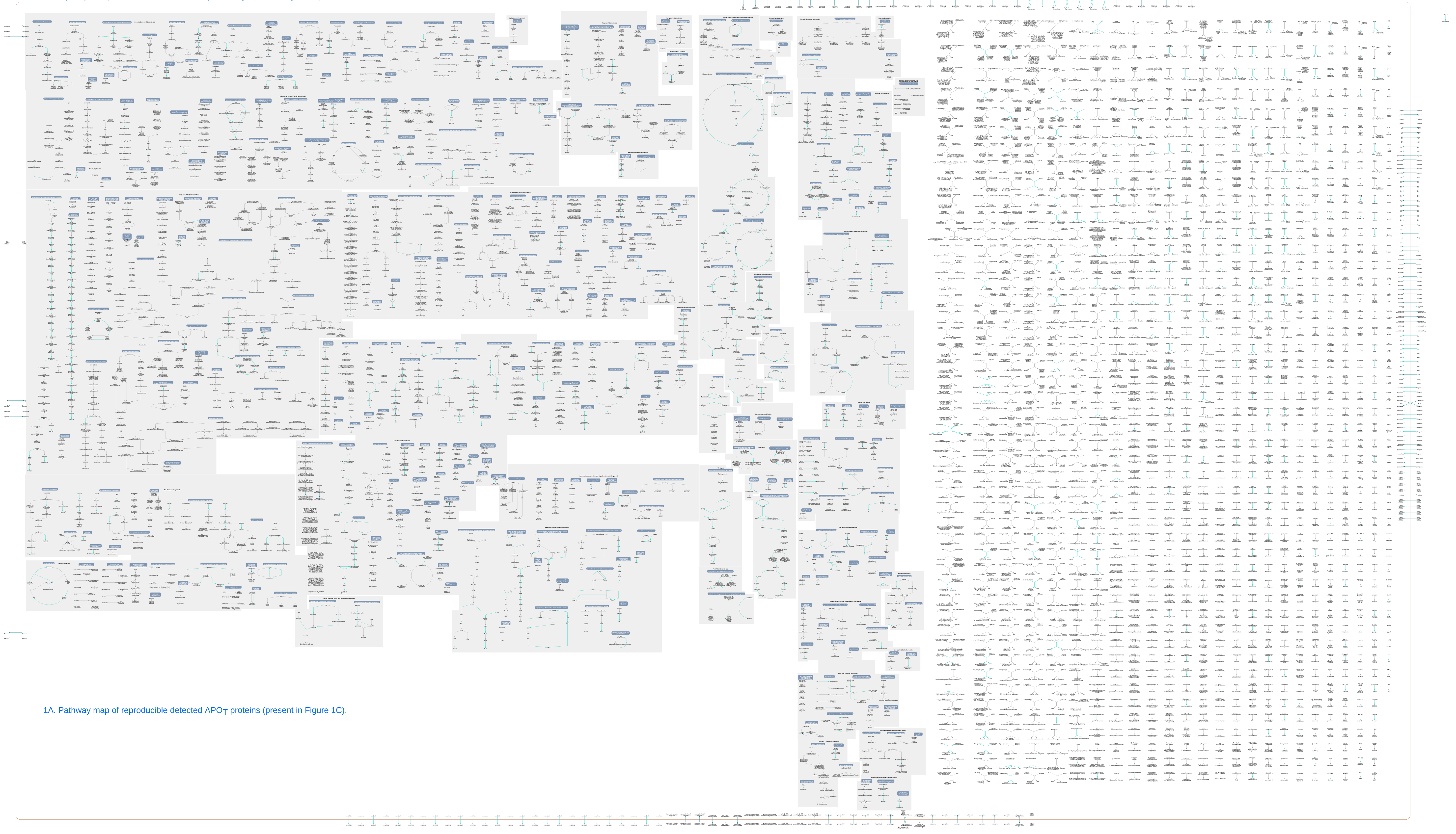

1A. Pathway map of reproducible detected $APO_T$ proteins (present in Figure 1C).

1B. 1/2 Pathway map of reproducible detected P200 proteins (present in Figure 1C).

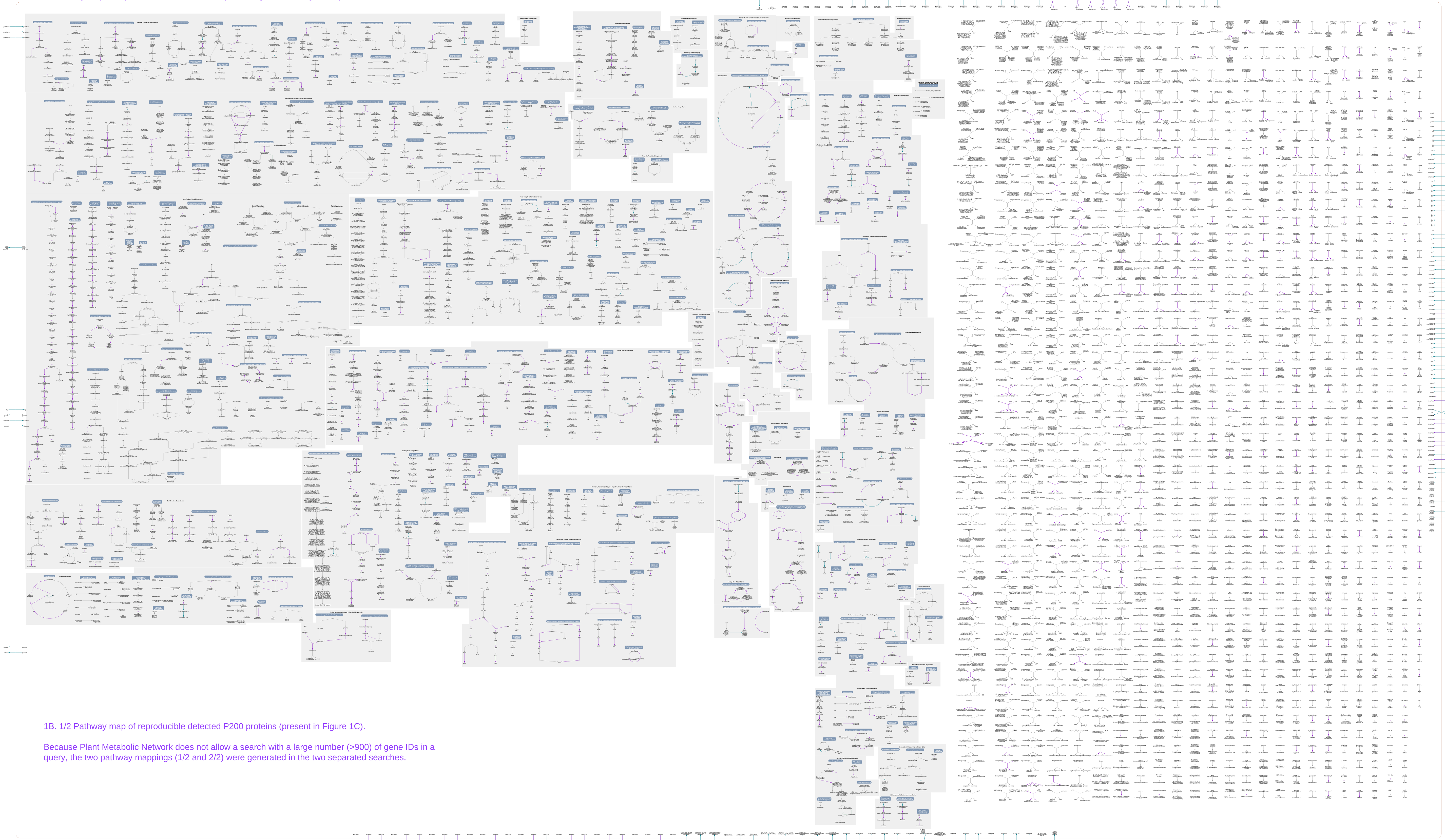

1B. 1/2 Pathway map of reproducible detected P200 proteins (present in Figure 1C).

Because Plant Metabolic Network does not allow a search with a large number (>900) of gene IDs in a query, the two pathway mappings (1/2 and 2/2) were generated in the two separated searches.

1B. 2/2 Pathway map of reproducible detected P200 proteins (present in Figure 1C).

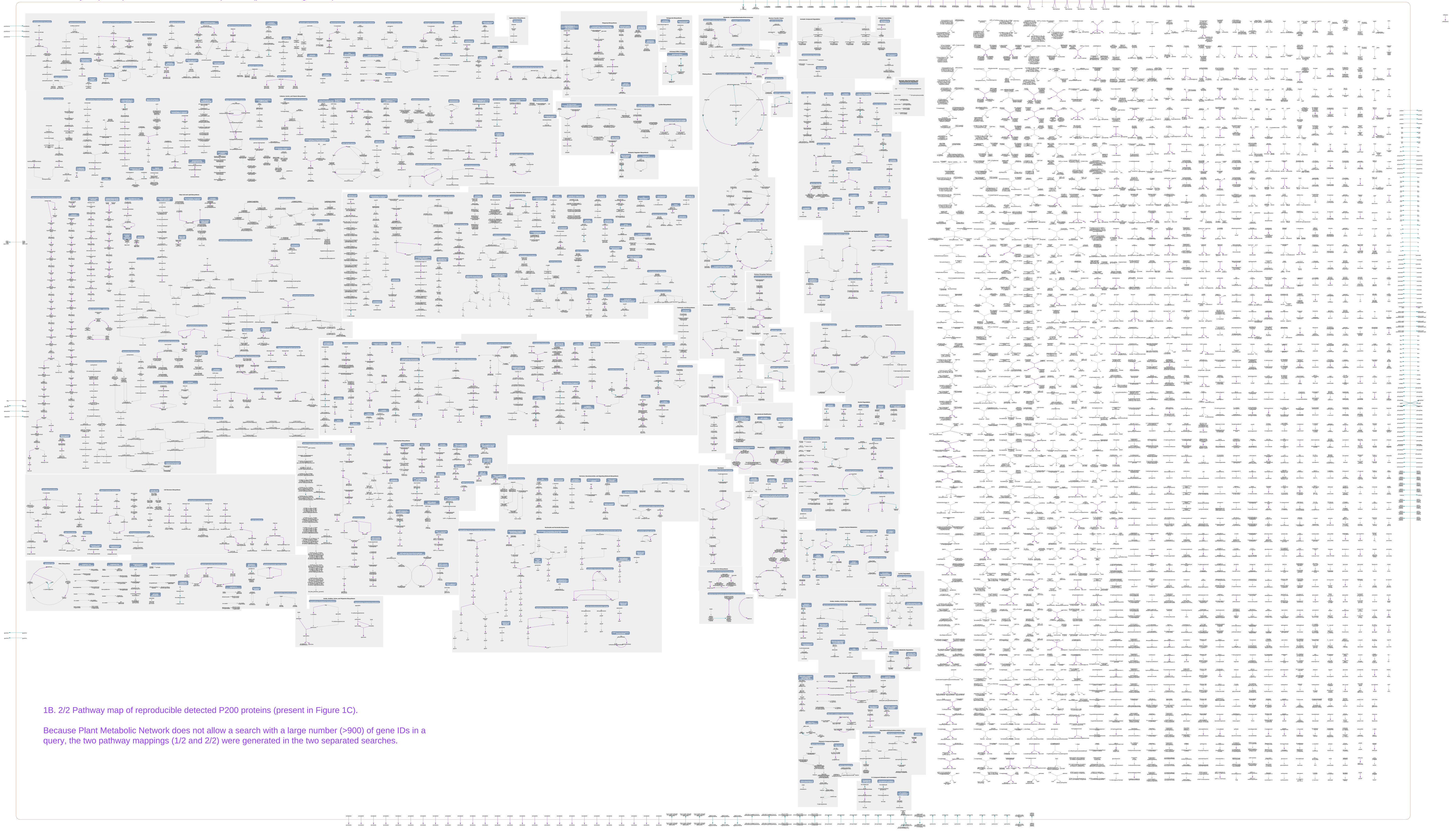

1B. 2/2 Pathway map of reproducible detected P200 proteins (present in Figure 1C).

Because Plant Metabolic Network does not allow a search with a large number (>900) of gene IDs in a query, the two pathway mappings (1/2 and 2/2) were generated in the two separated searches.

1C. Pathway map of reproducible detected S200 proteins (present in Figure 1C).

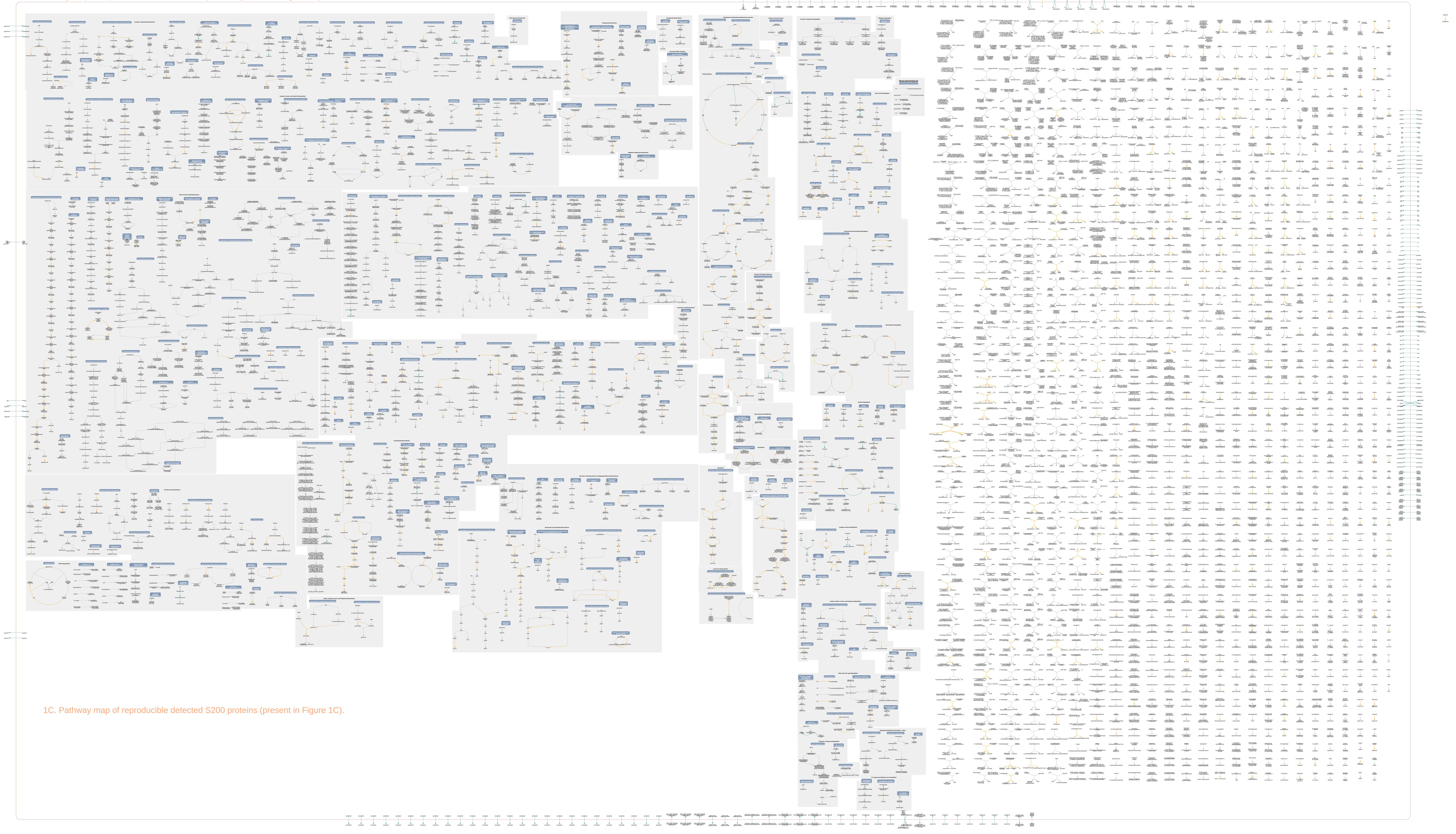

1C. Pathway map of reproducible detected S200 proteins (present in Figure 1C).

1D. Pathway map of cytoplasmic proteins (P200 and S200) with informative profiles (present in Figure 2B).

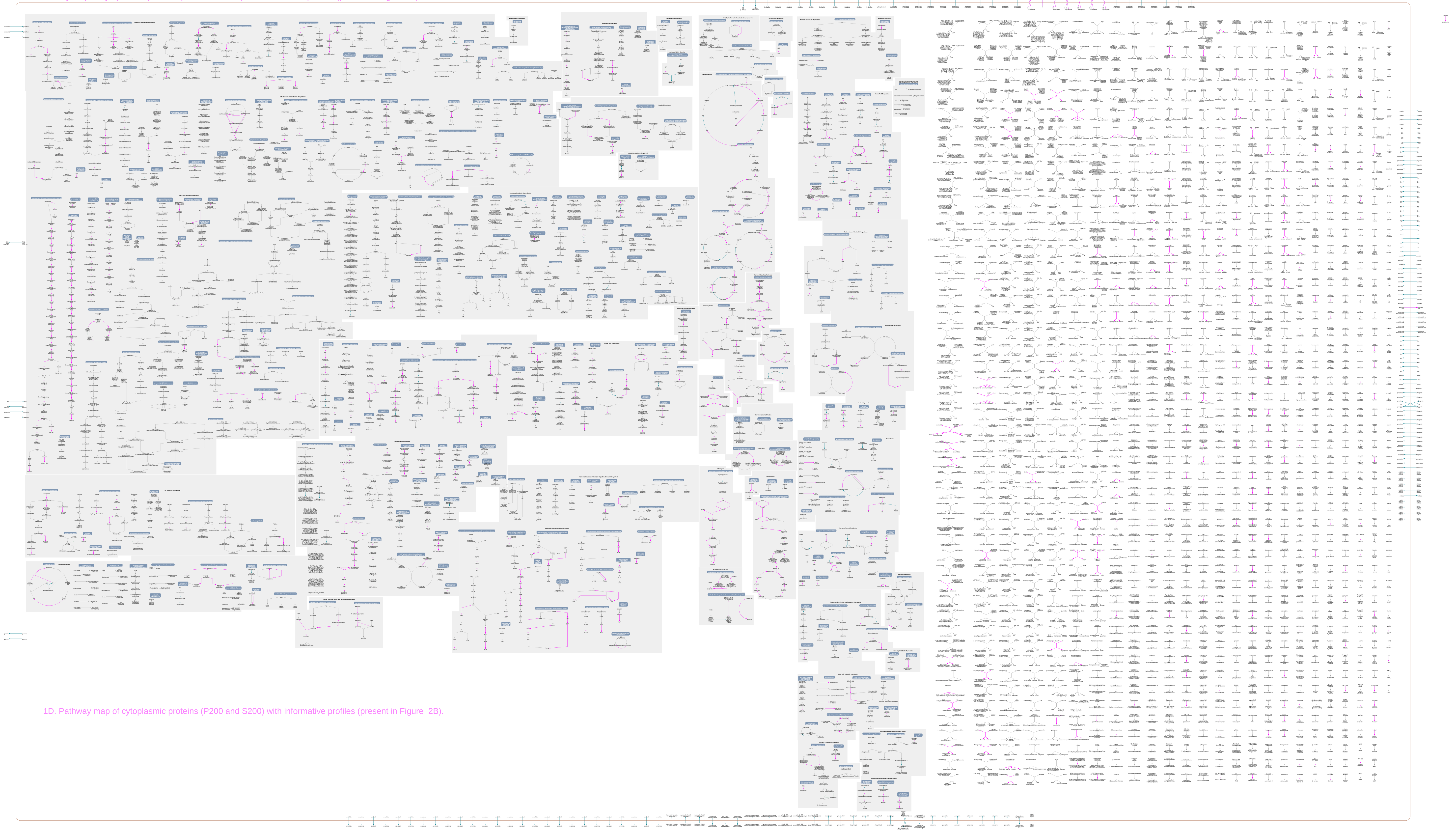

1D. Pathway map of cytoplasmic proteins (P200 and S200) with informative profiles (present in Figure 2B).

### 1E. Pathway map of extracellular proteins (APO$_T$) with informative profiles (present in Figure 2B).

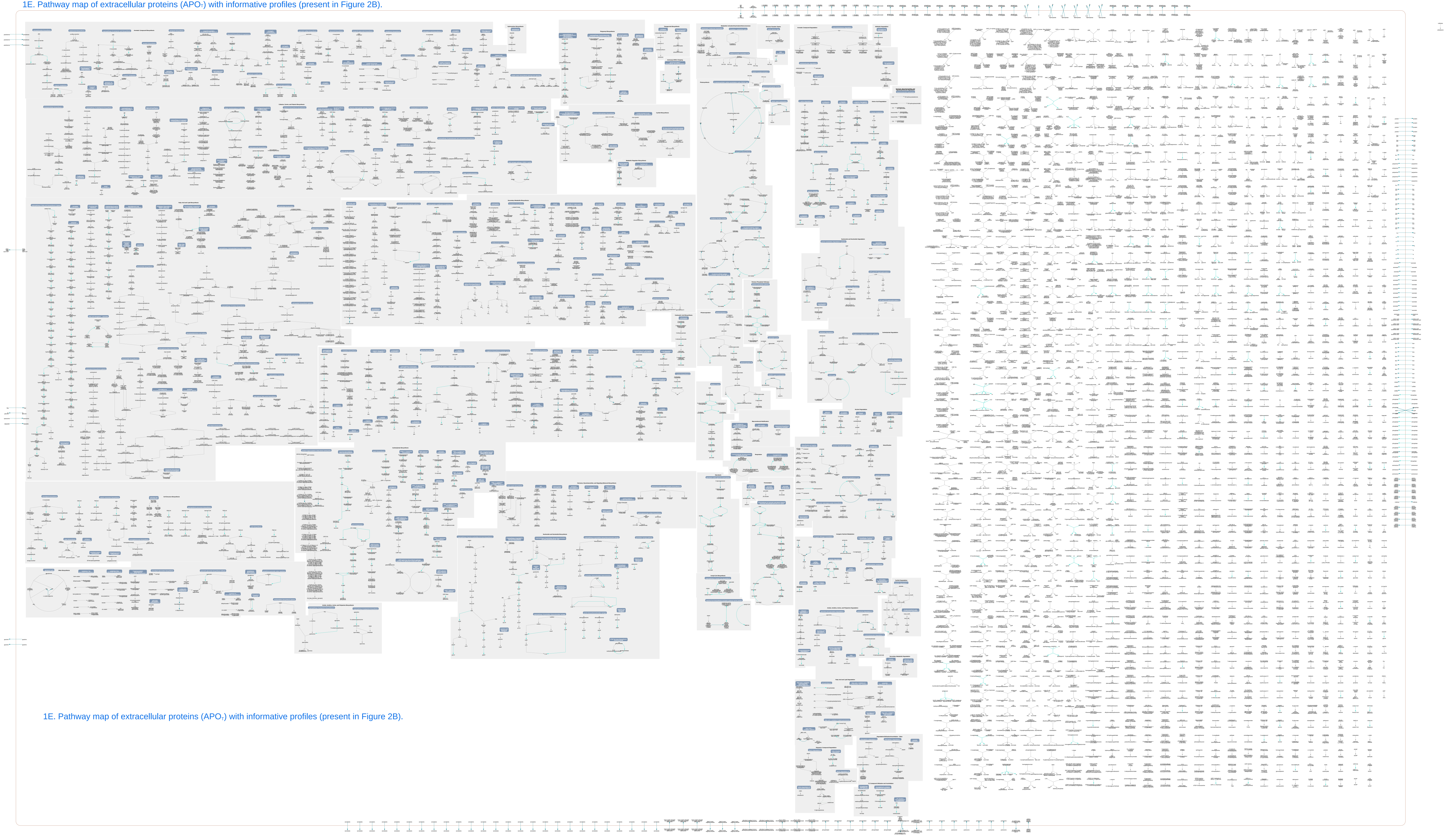

1E. Pathway map of extracellular proteins (APO$_T$) with informative profiles (present in Figure 2B).
