## Supplementary Figure S2 for "Systems-level proteomic models of cotton fiber development: A high-resolution resource to analyze cell dynamics and trait engineering"

**Histogram of the average abundances of non−zero values**

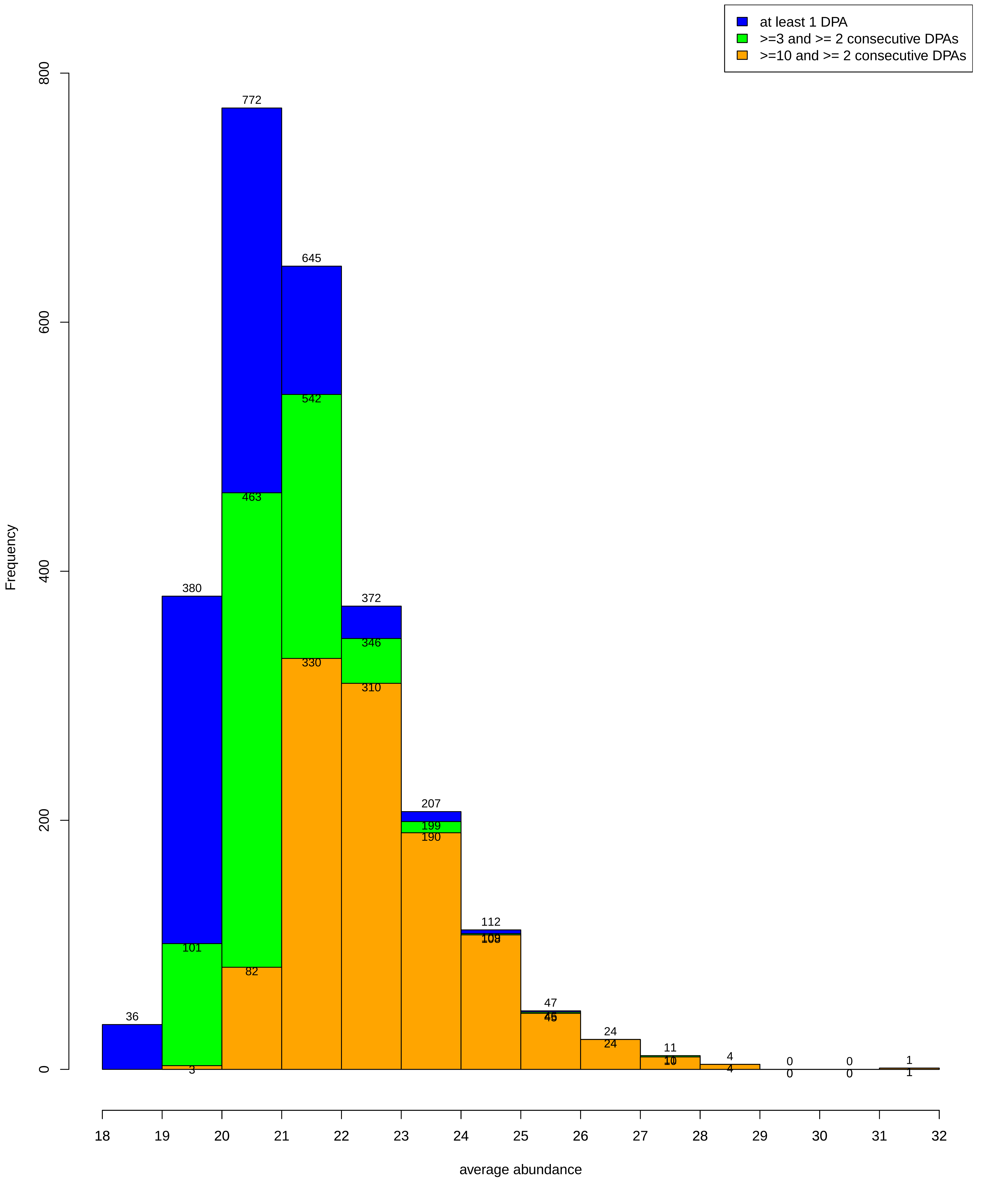

Codes

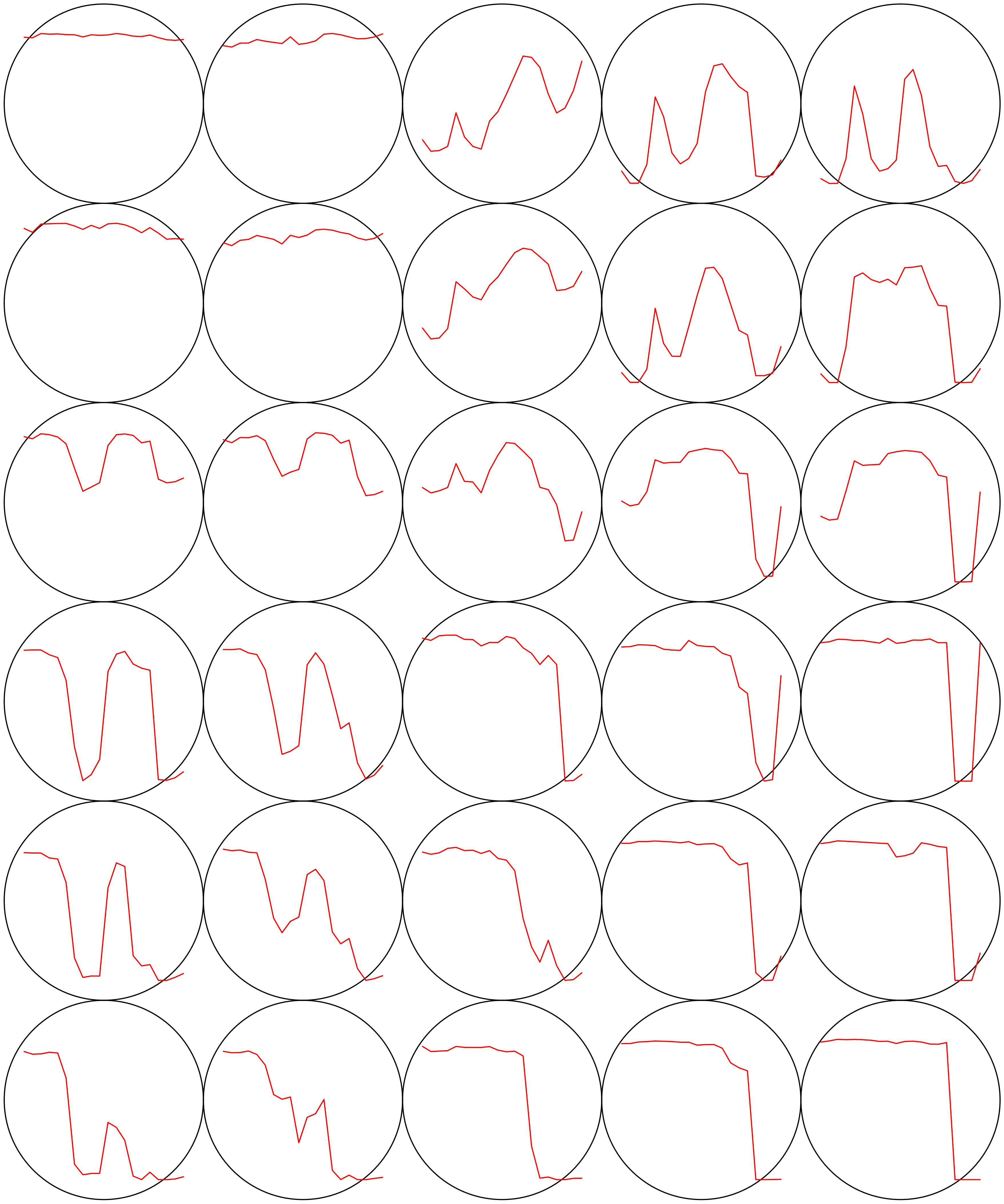

### Count plot

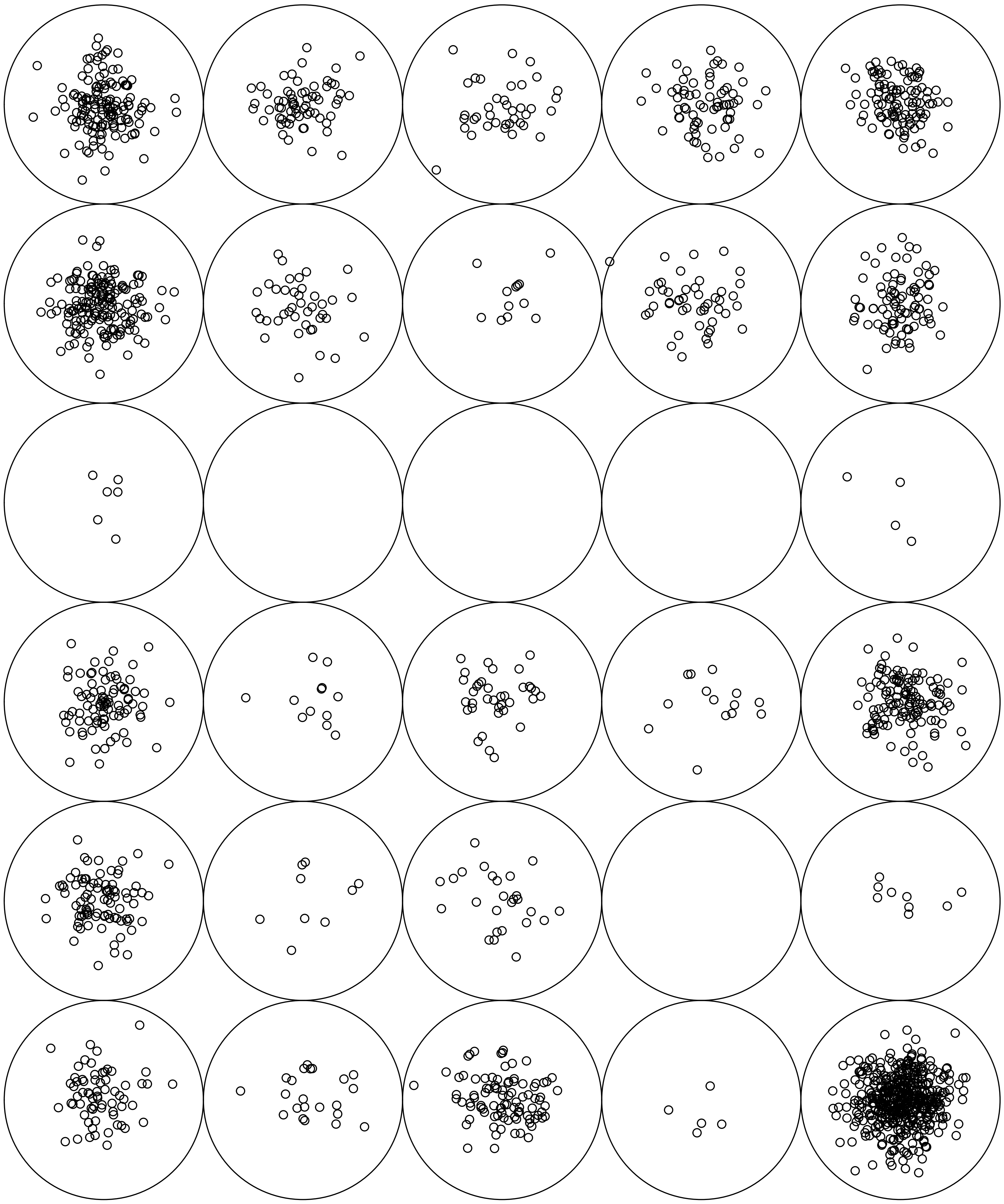

V1

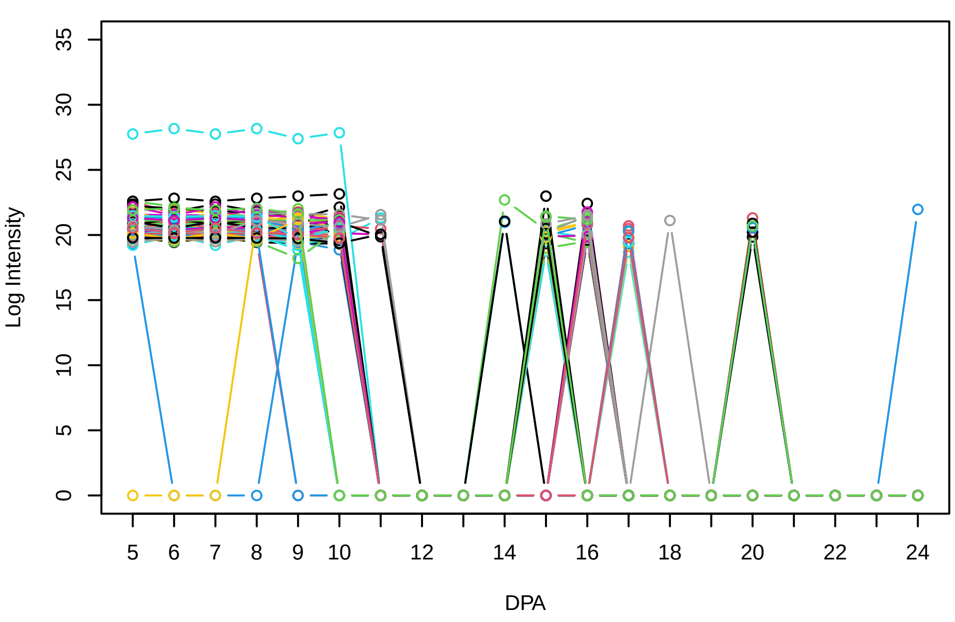


Histogram of the average abundances of non−zero values

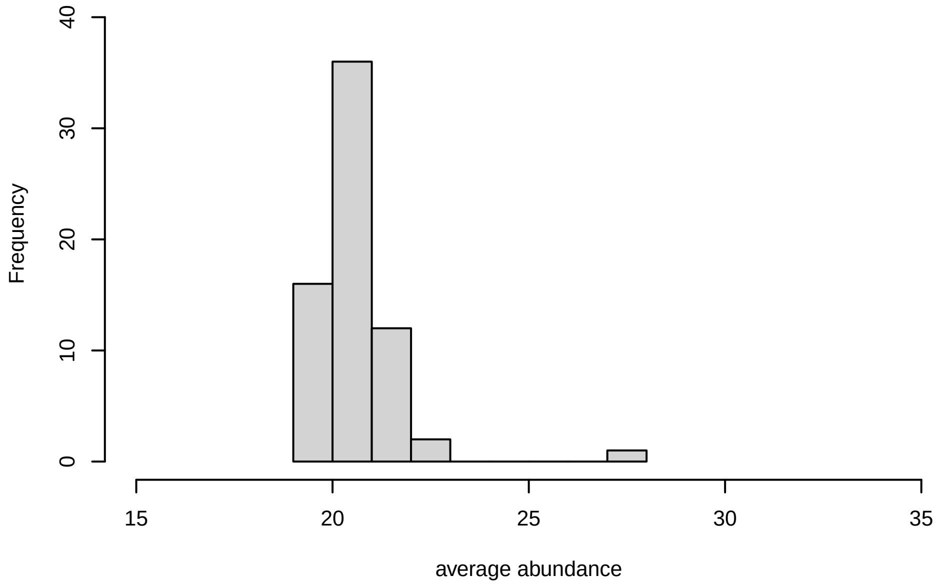


V1, after remove outlier and cutoff is 0.8

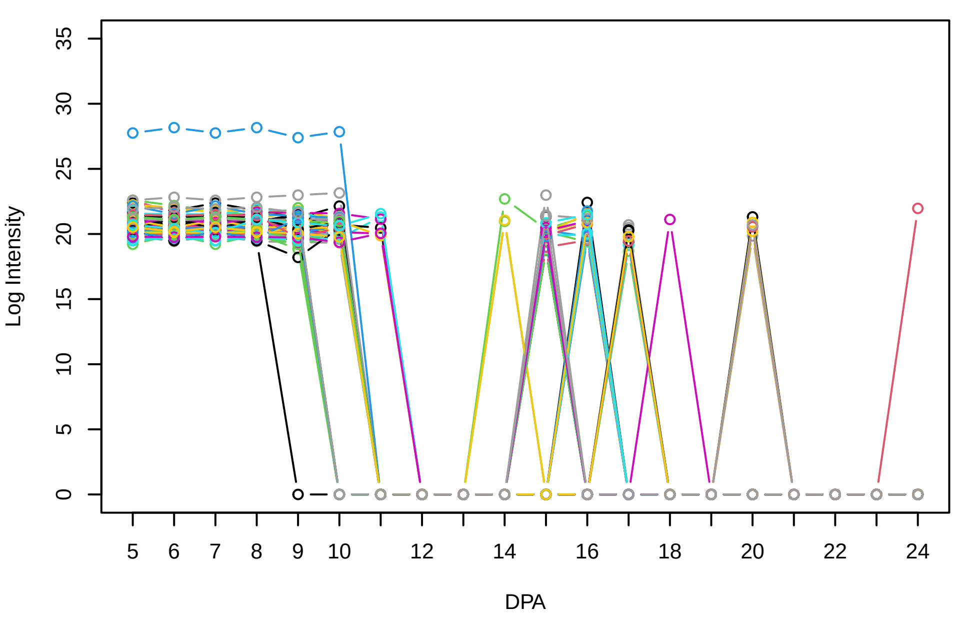


Histogram of the average abundances of non−outlier proteins

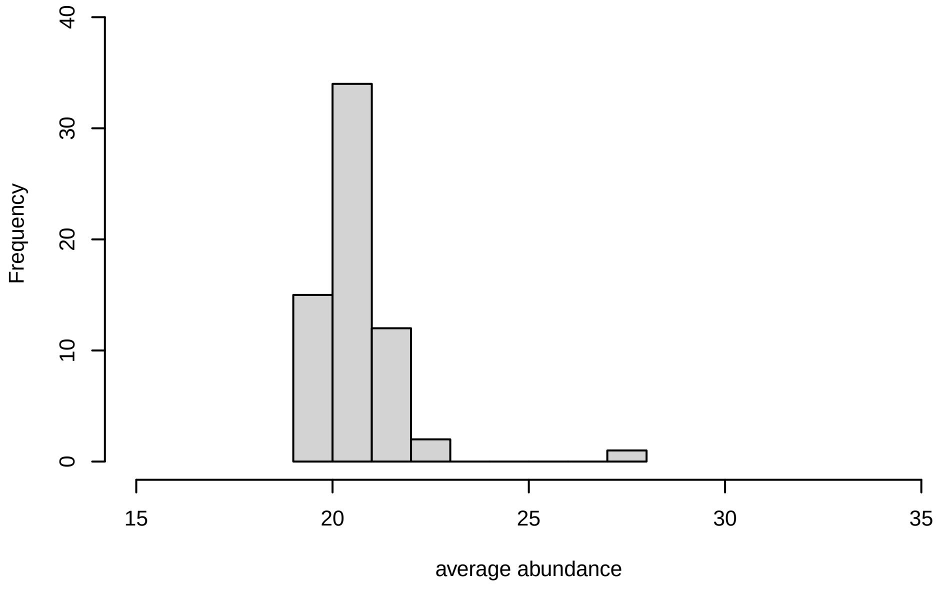


V1, after remove unreliable

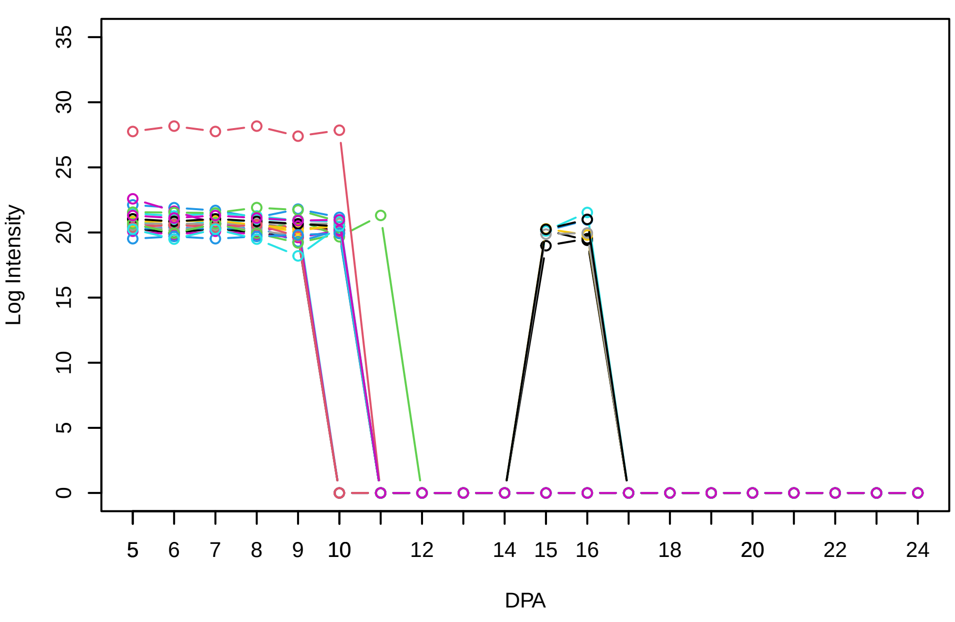


Histogram of the average abundances of reliable proteins

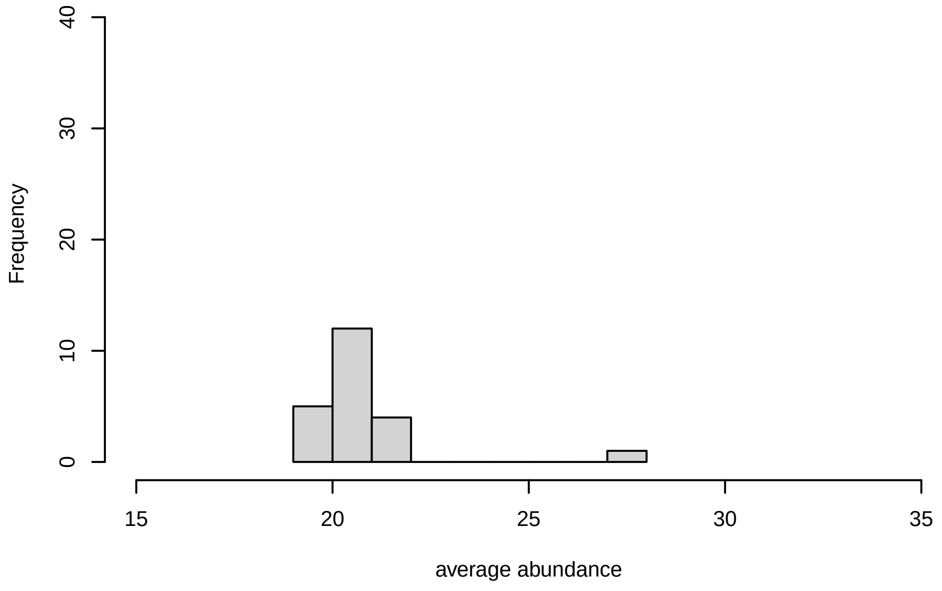


V1, after remove low abundance

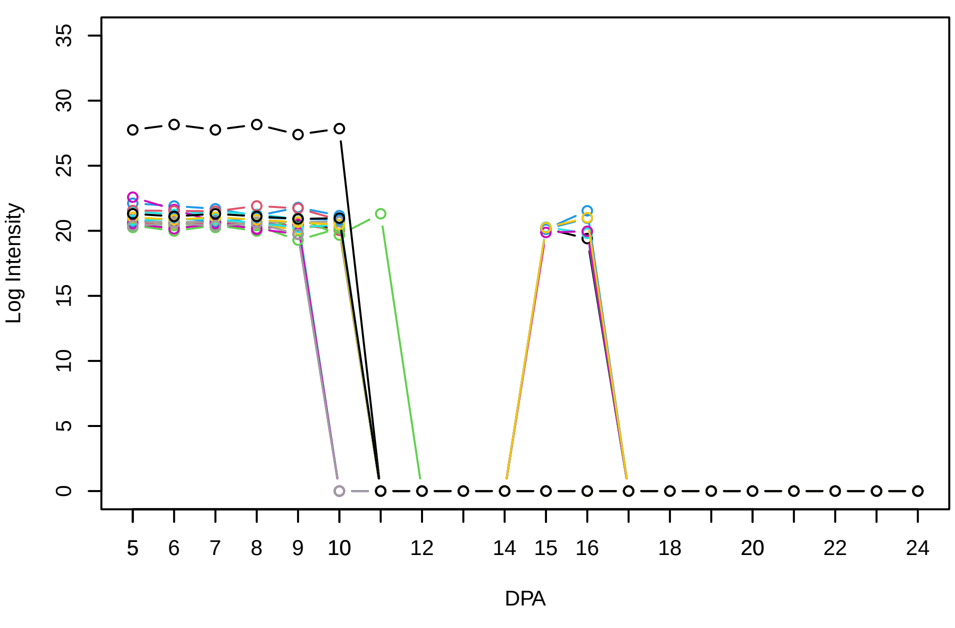


Histogram of the average abundances of high abundance proteins

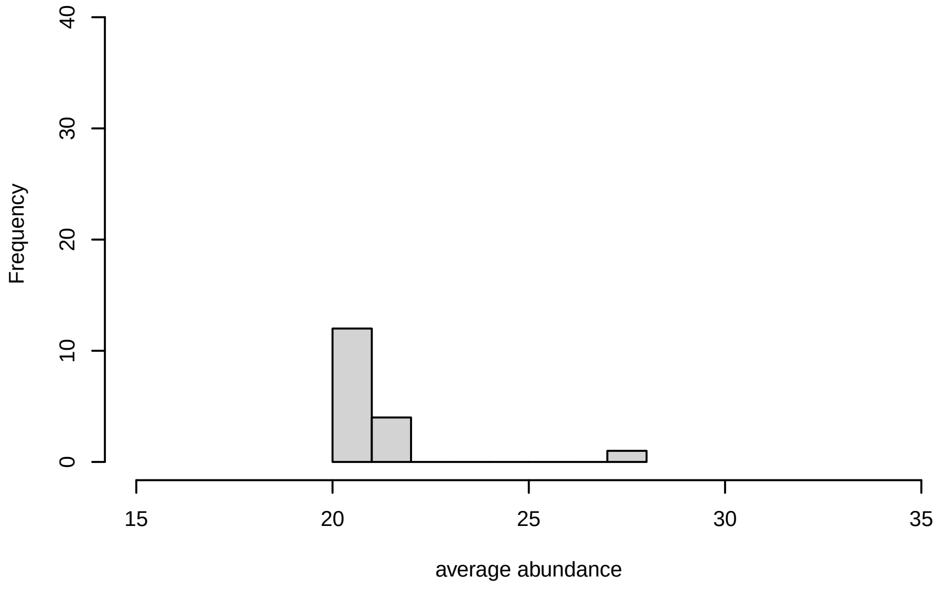

V2

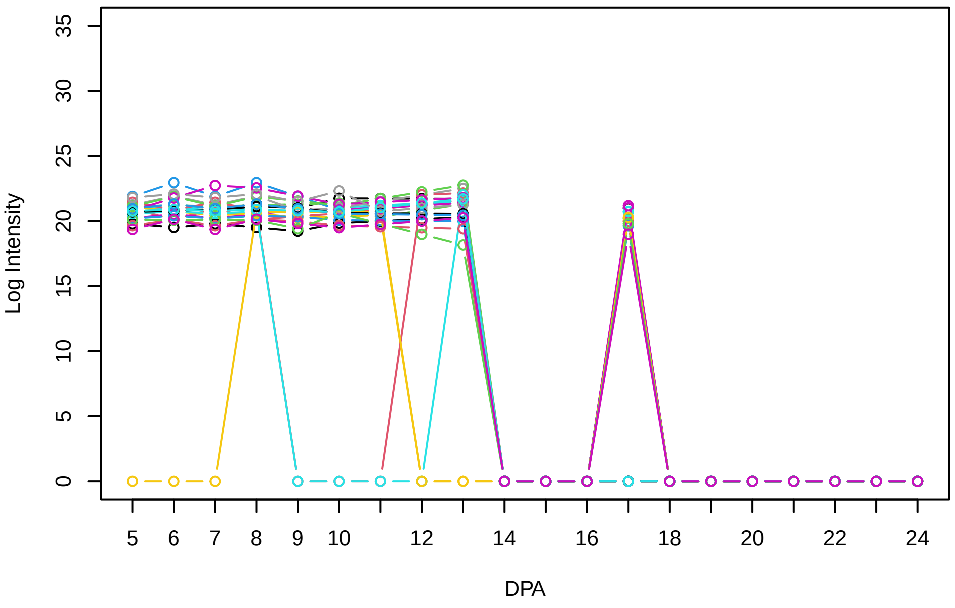


Histogram of the average abundances of non−zero values

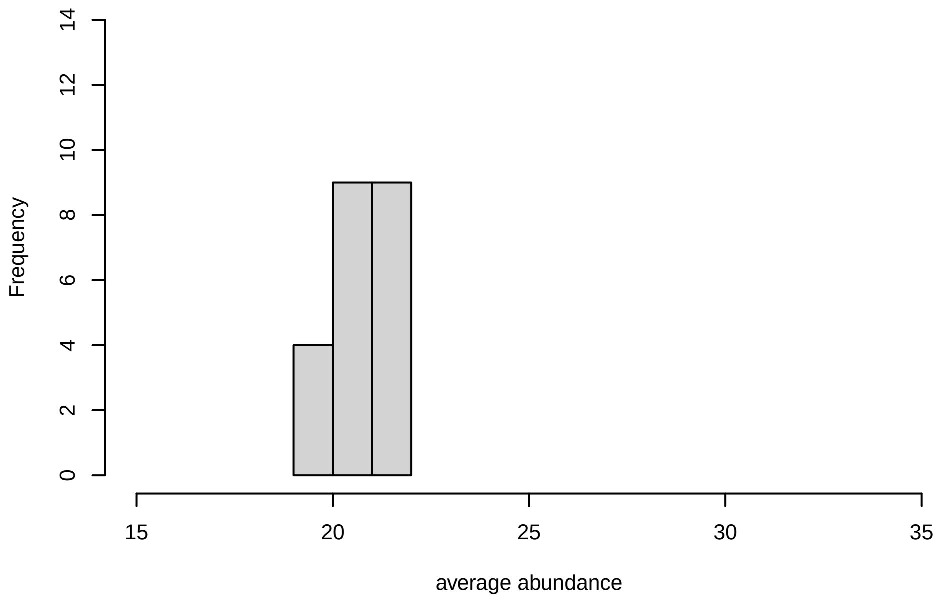


V2, after remove outlier and cutoff is 0.8

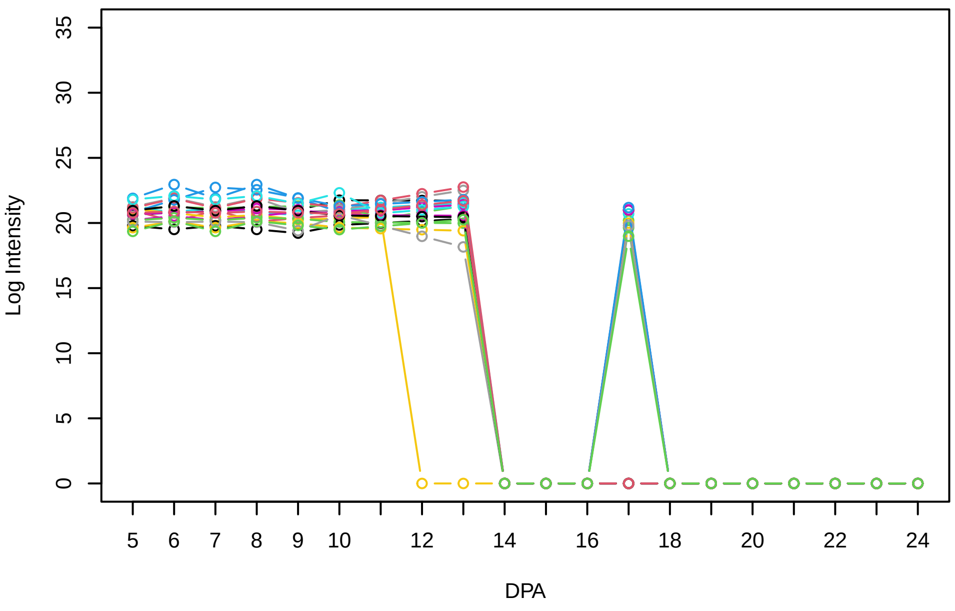


Histogram of the average abundances of non−outlier proteins

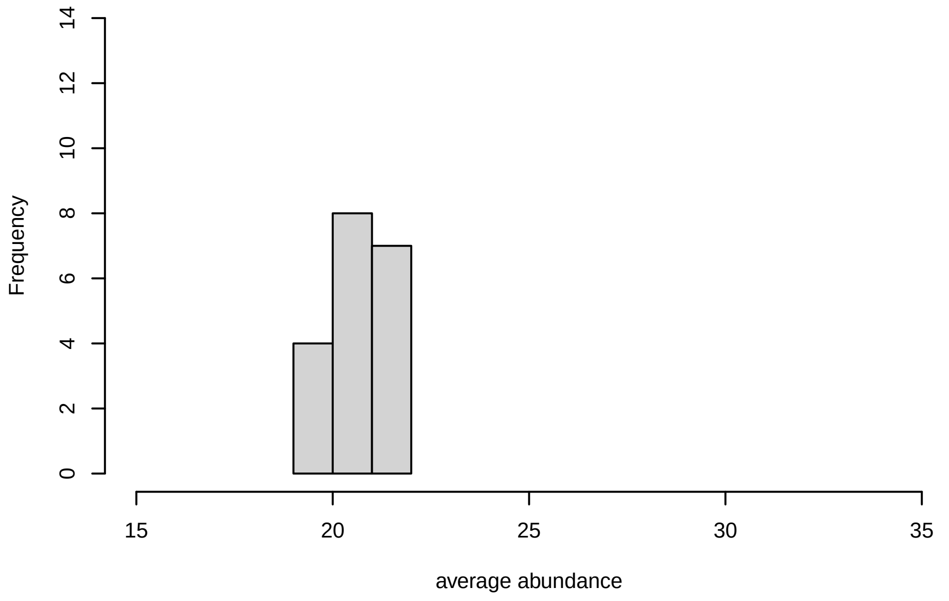


V2, after remove unreliable

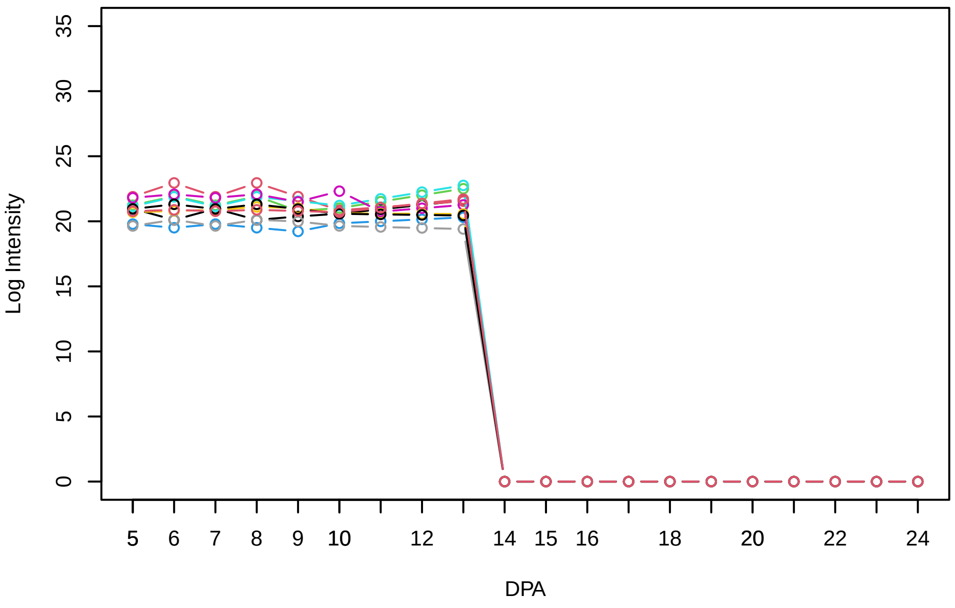


Histogram of the average abundances of reliable proteins

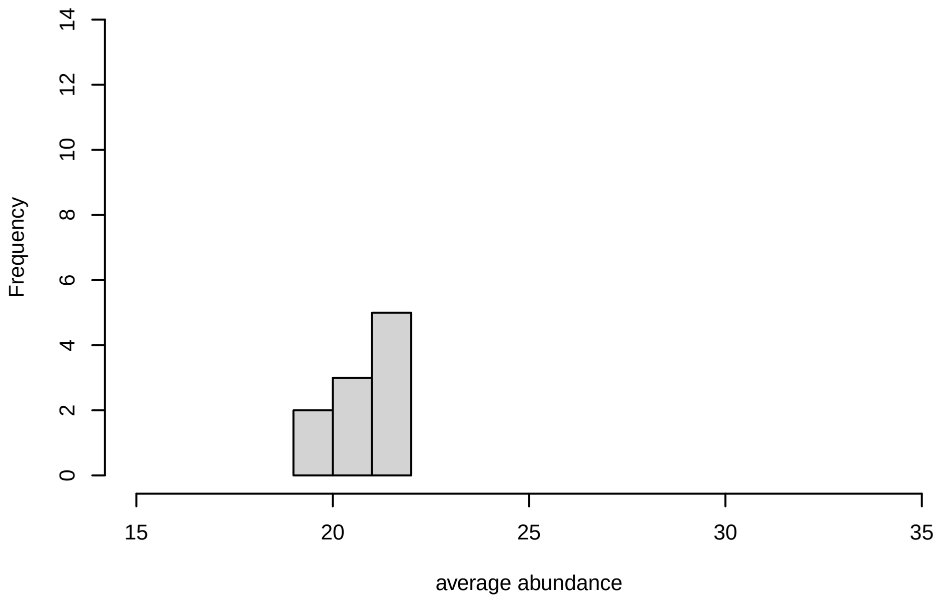


V2, after remove low abundance

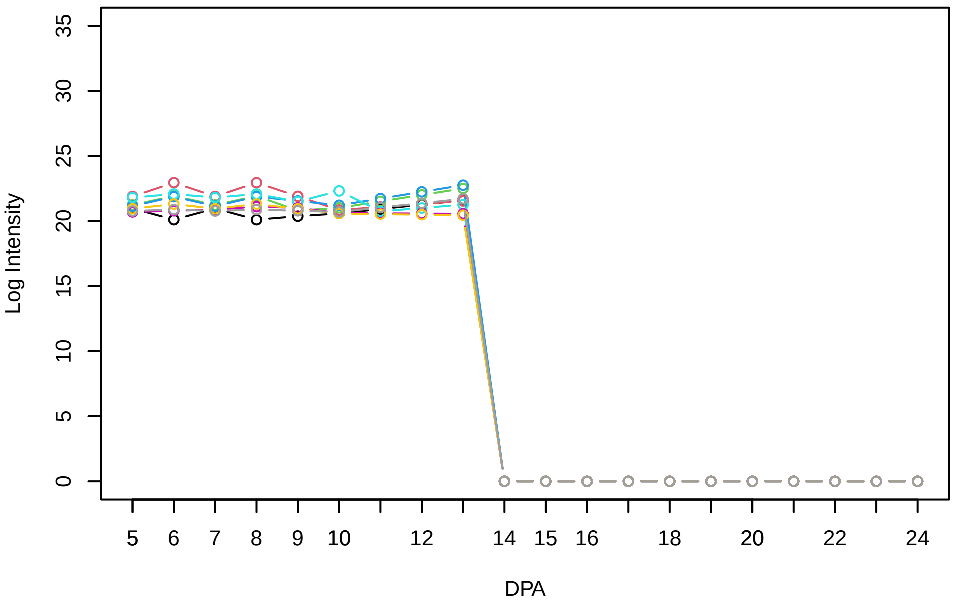


Histogram of the average abundances of high abundance proteins

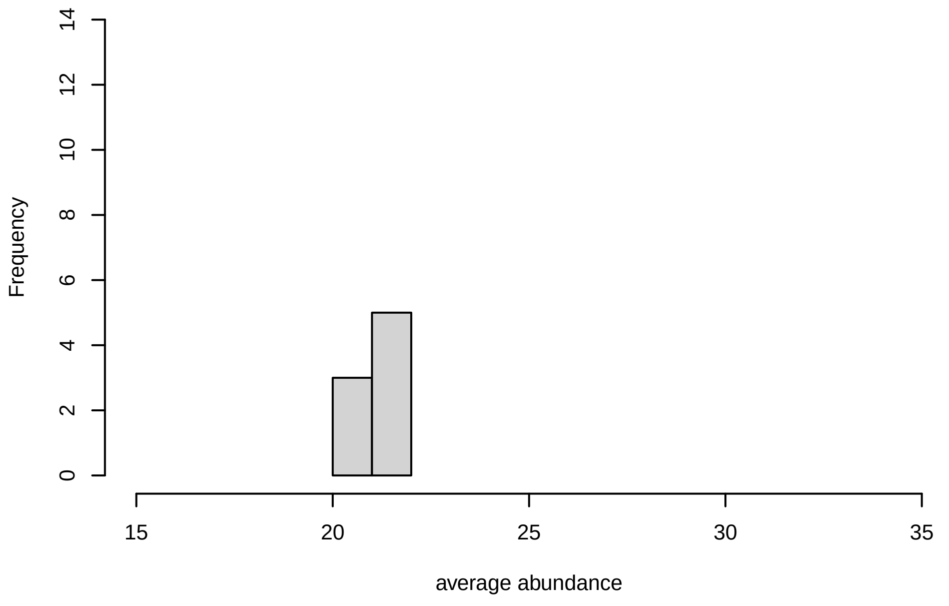

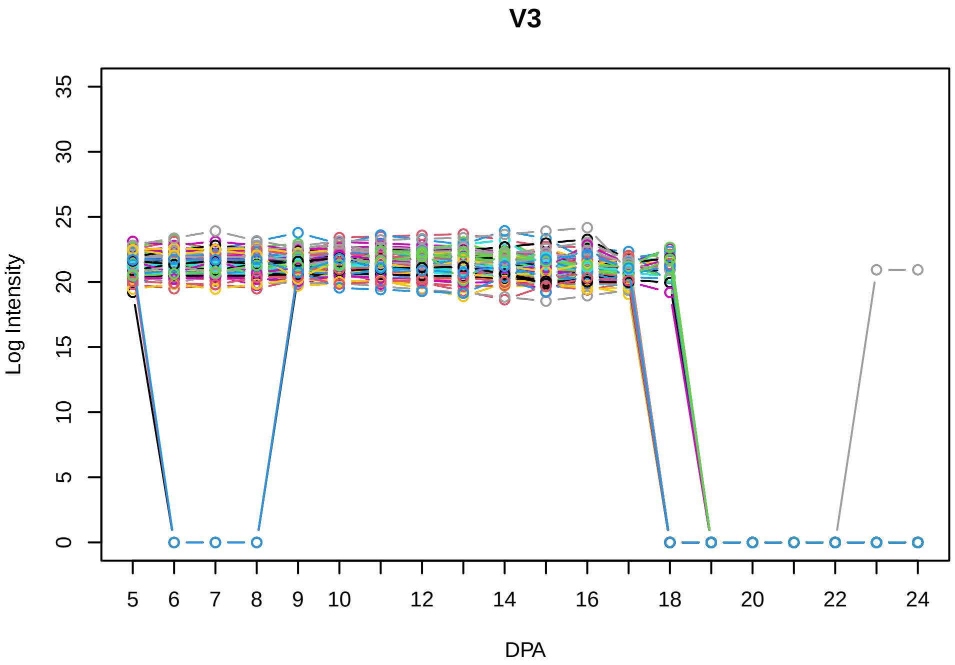
V3
Log Intensity
DPA

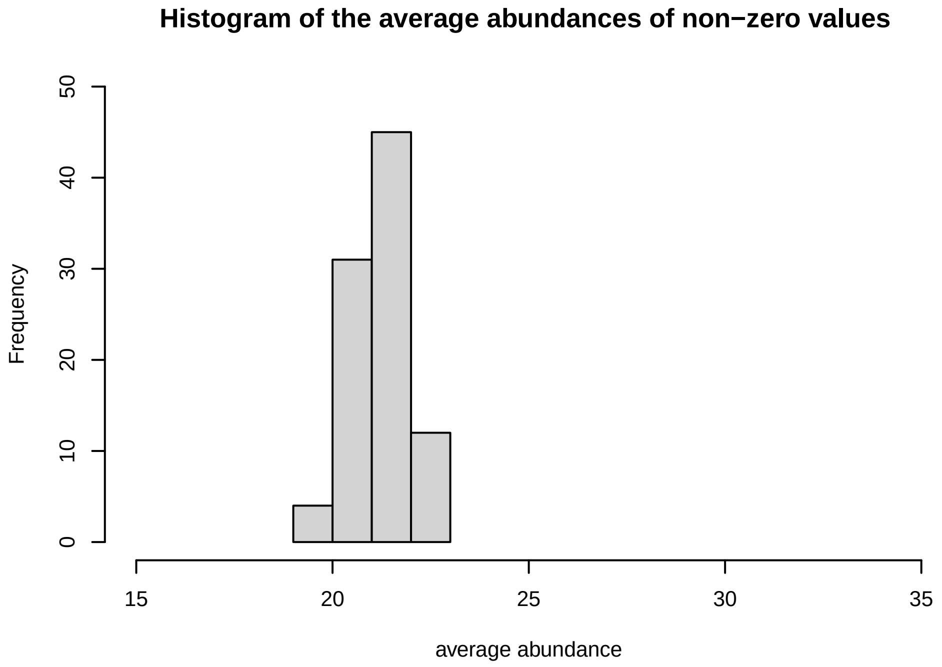
Histogram of the average abundances of non−zero values
Frequency
average abundance

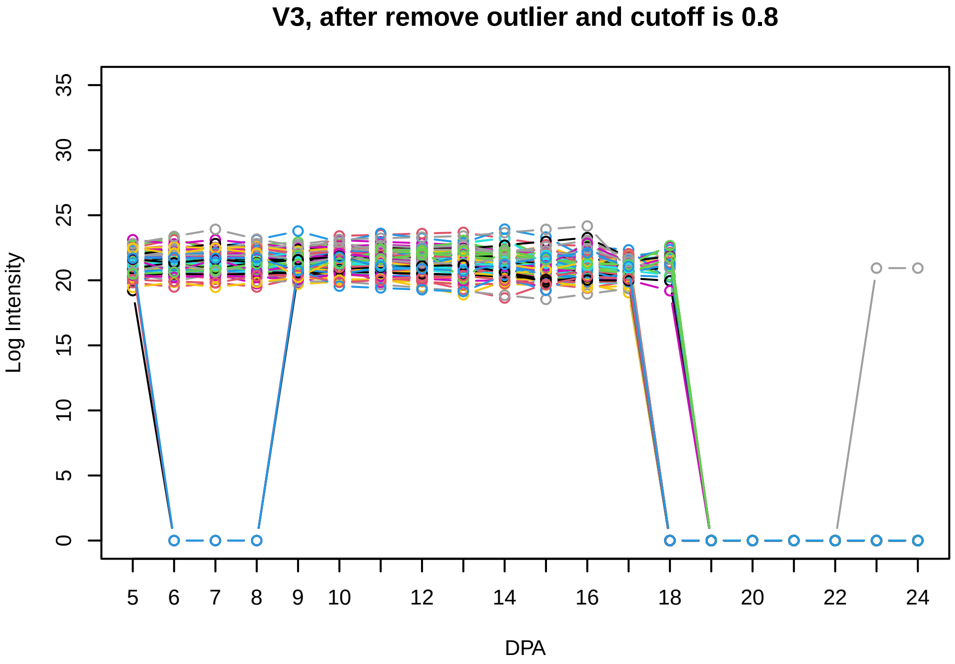
V3, after remove outlier and cutoff is 0.8
Log Intensity
DPA

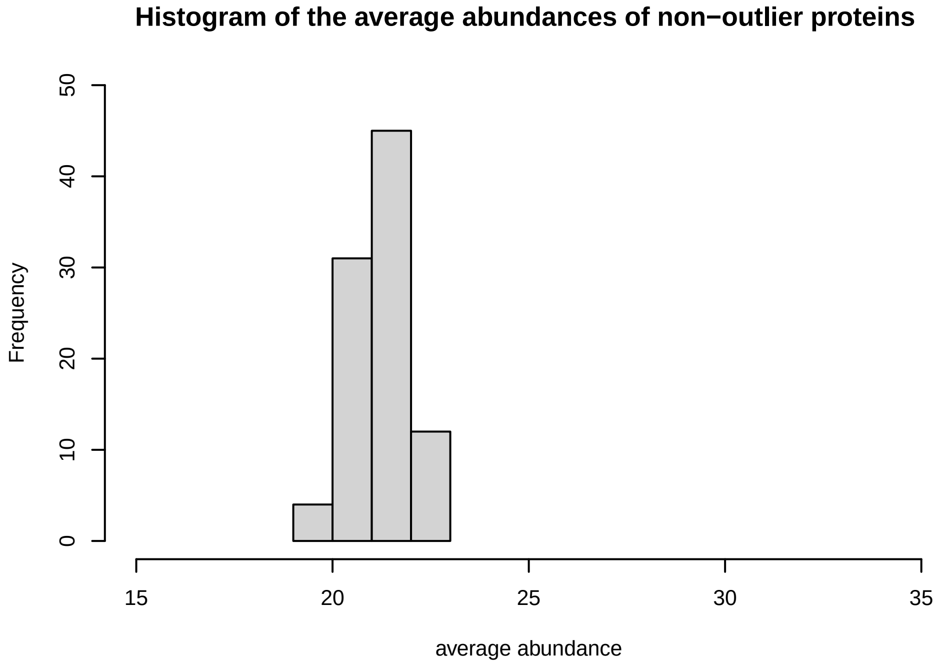
Histogram of the average abundances of non−outlier proteins
Frequency
average abundance

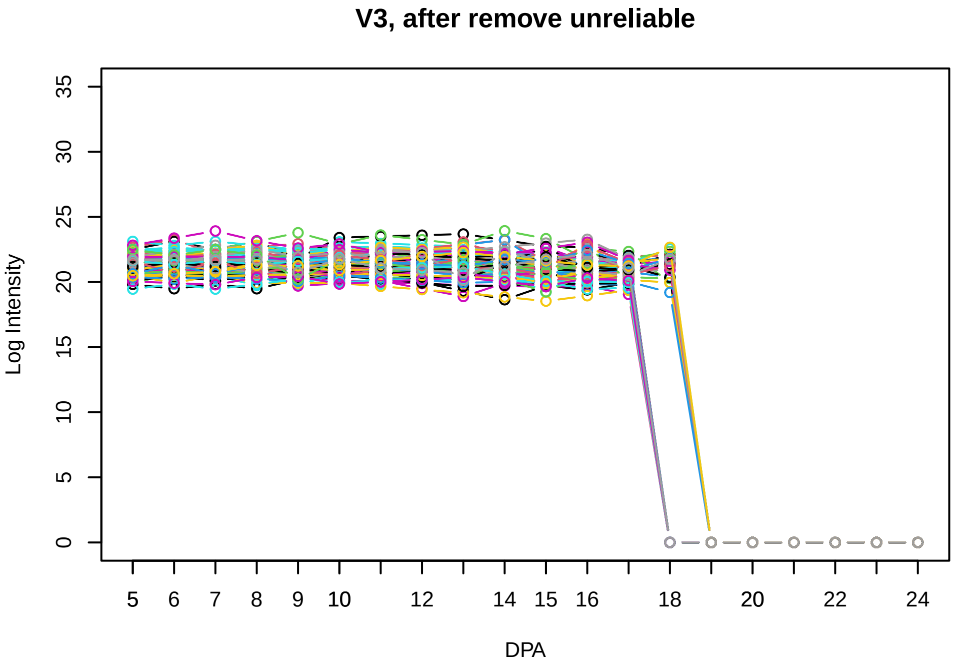
V3, after remove unreliable
Log Intensity
DPA

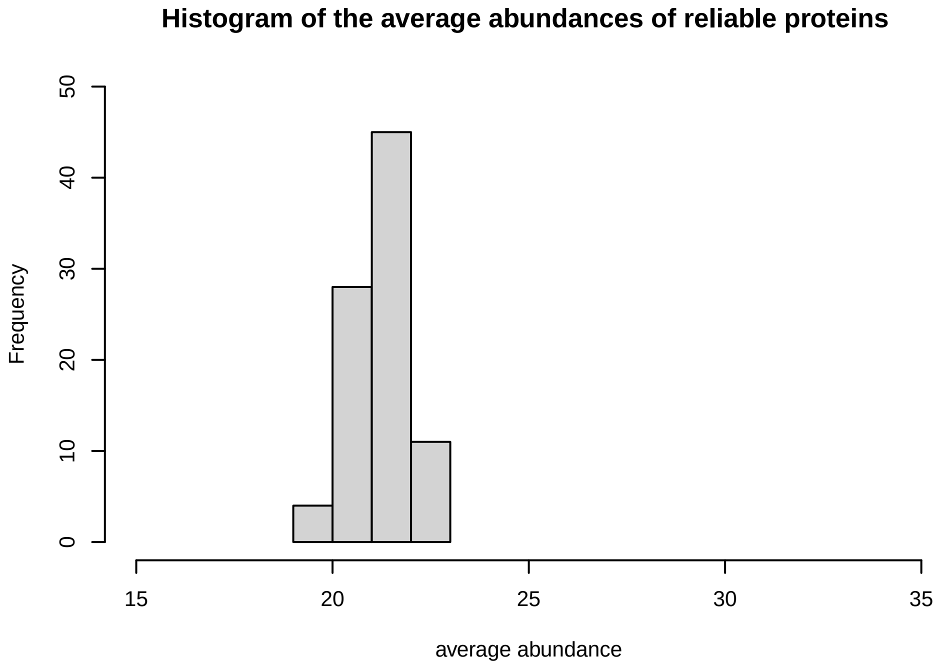
Histogram of the average abundances of reliable proteins
Frequency
average abundance

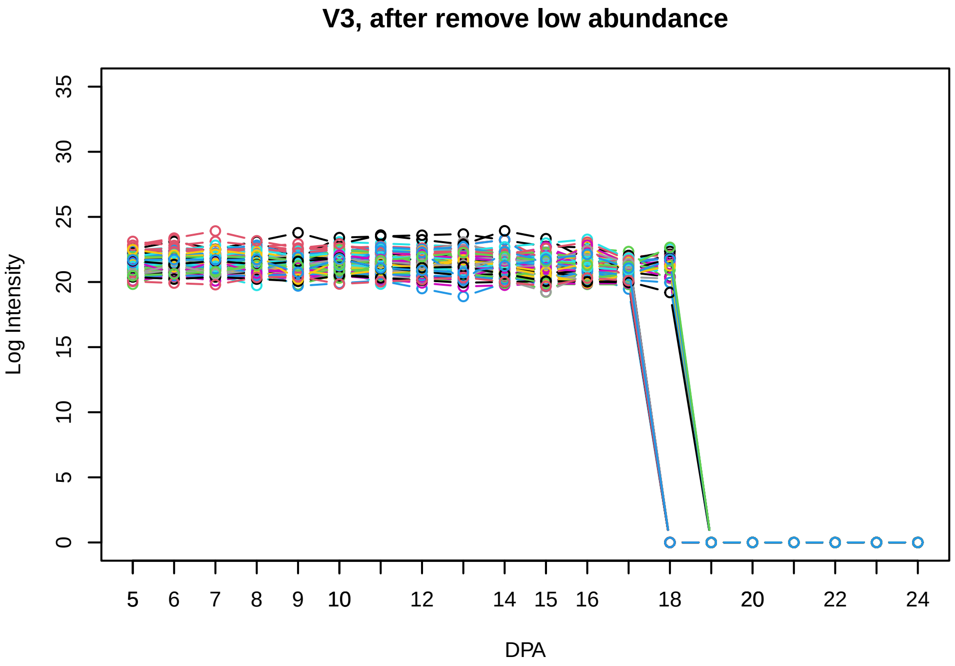
V3, after remove low abundance
Log Intensity
DPA

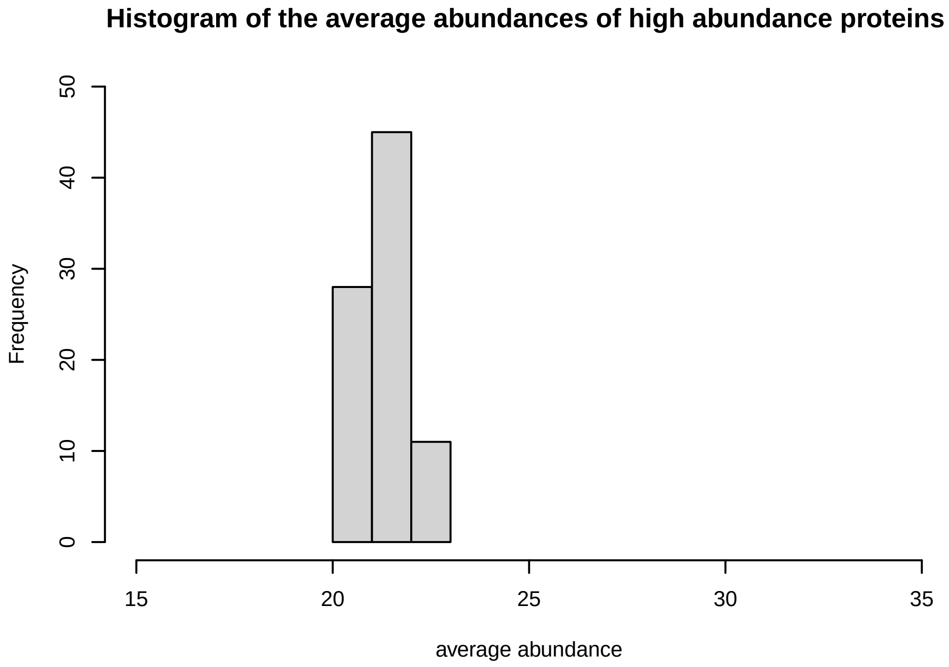
Histogram of the average abundances of high abundance proteins
Frequency
average abundance

## V4

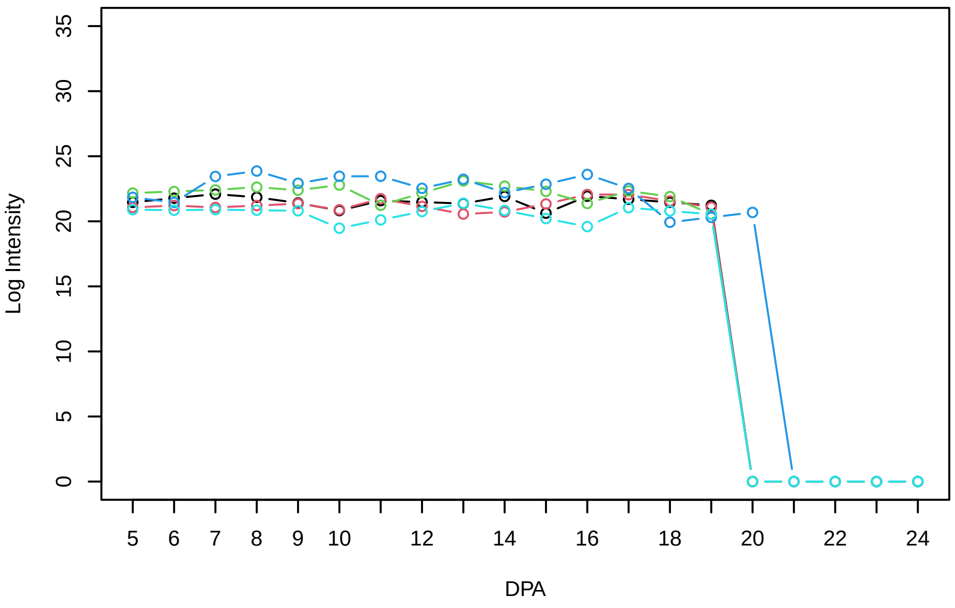


### Histogram of the average abundances of non−zero values

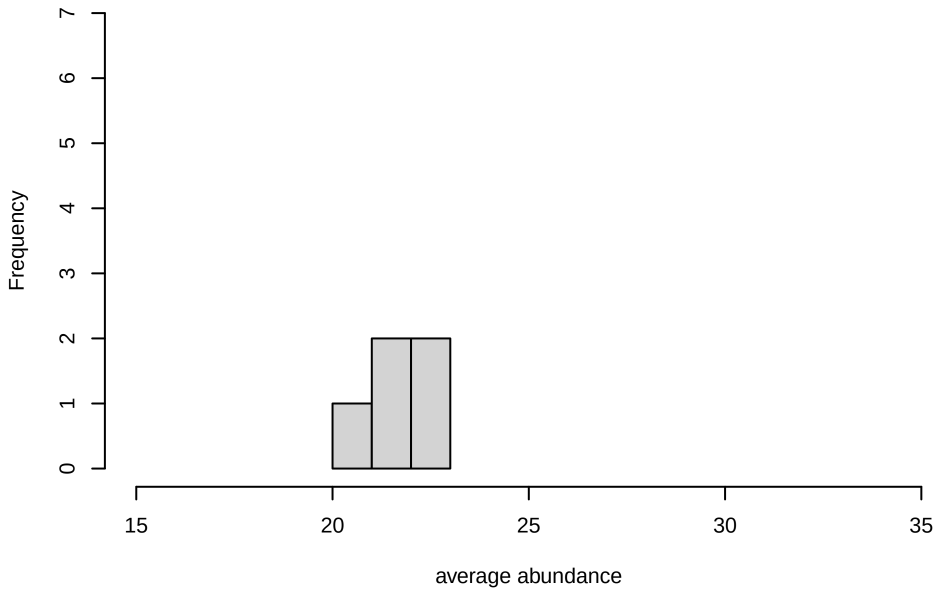


### V4, after remove outlier and cutoff is 0.8

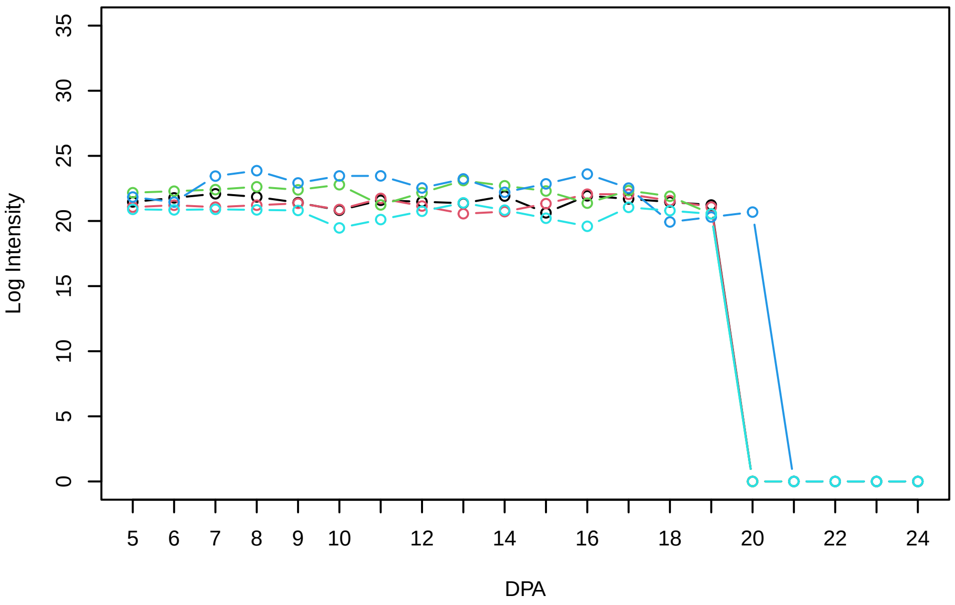


### Histogram of the average abundances of non−outlier proteins

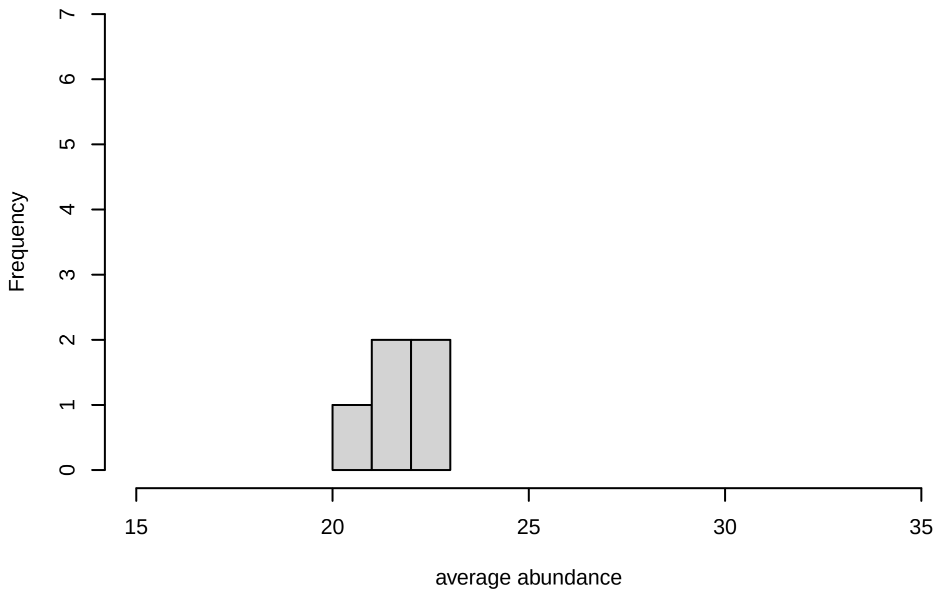


### V4, after remove unreliable

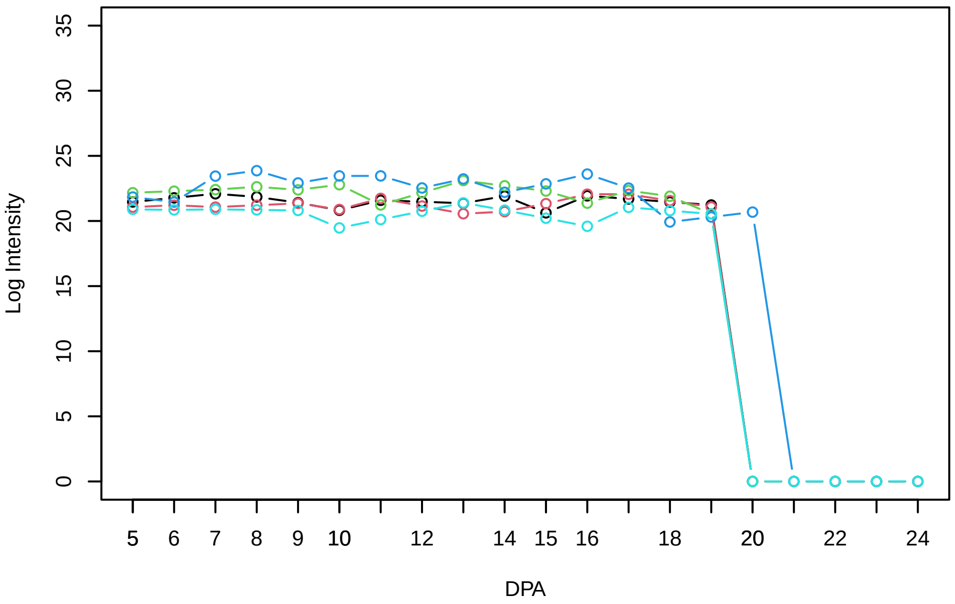


### Histogram of the average abundances of reliable proteins

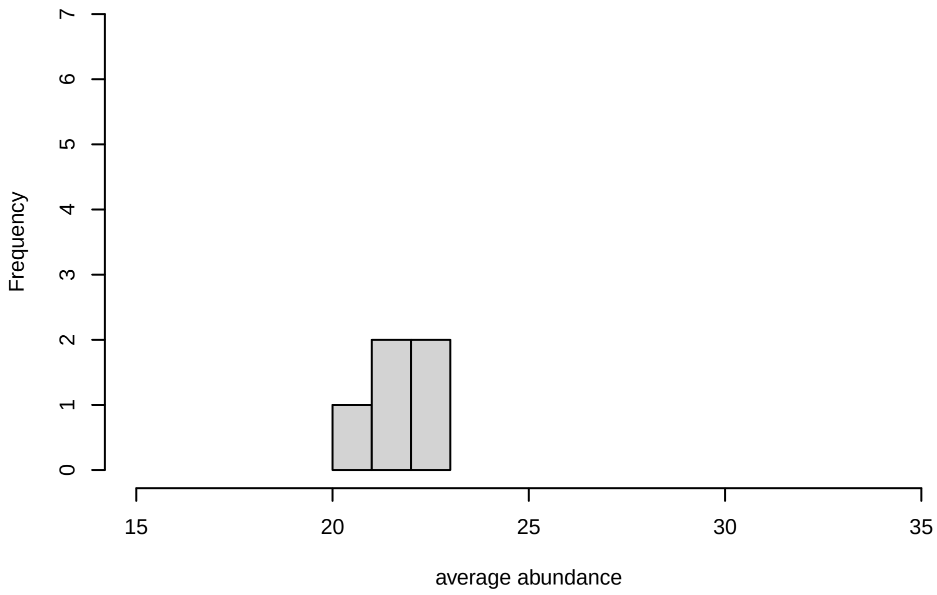


### V4, after remove low abundance

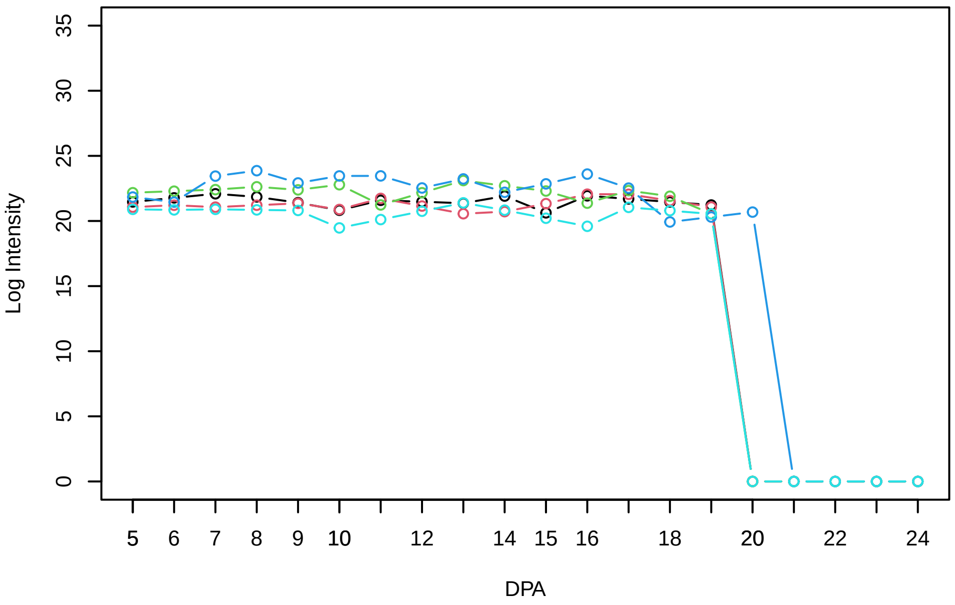


### Histogram of the average abundances of high abundance proteins

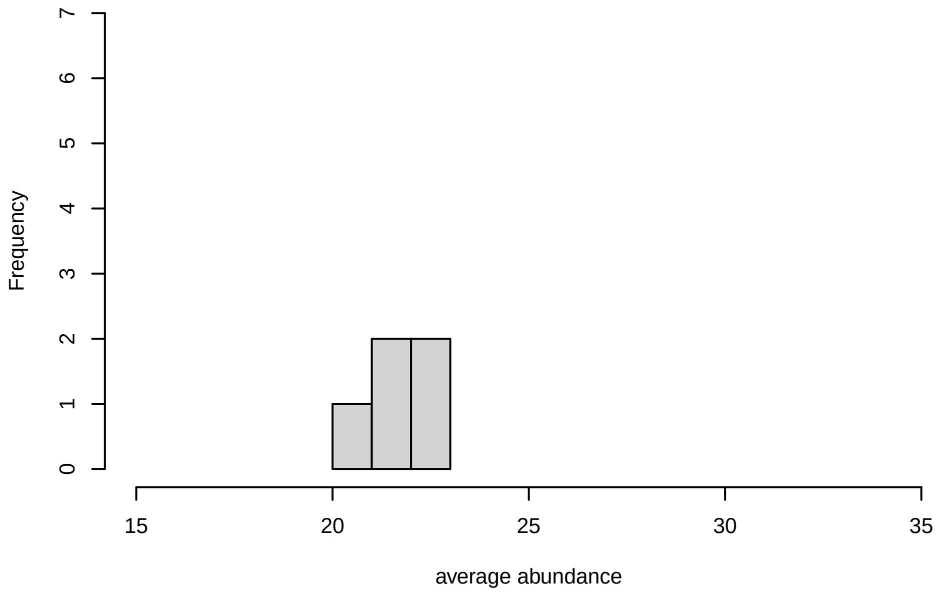

V5

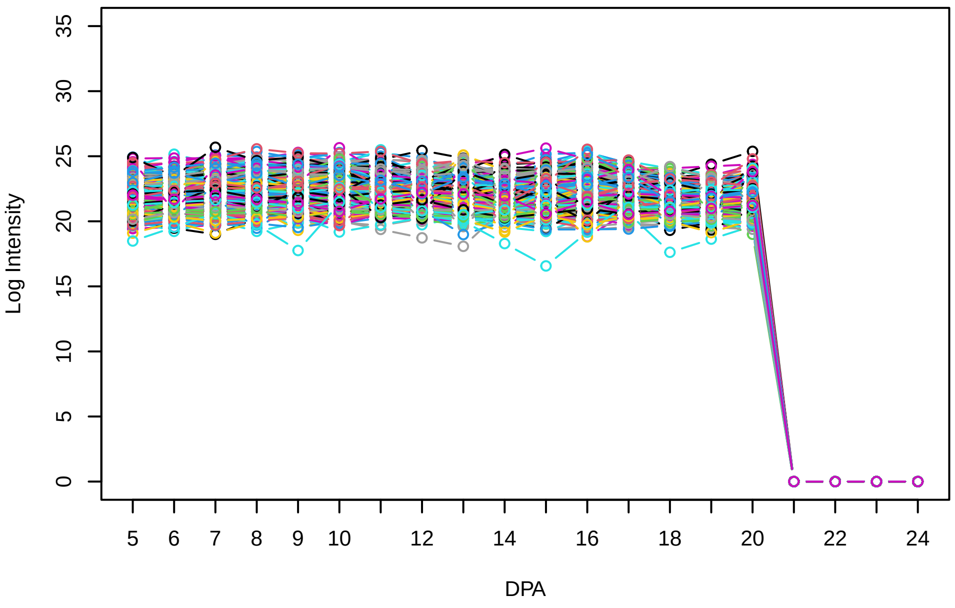


Histogram of the average abundances of non−zero values

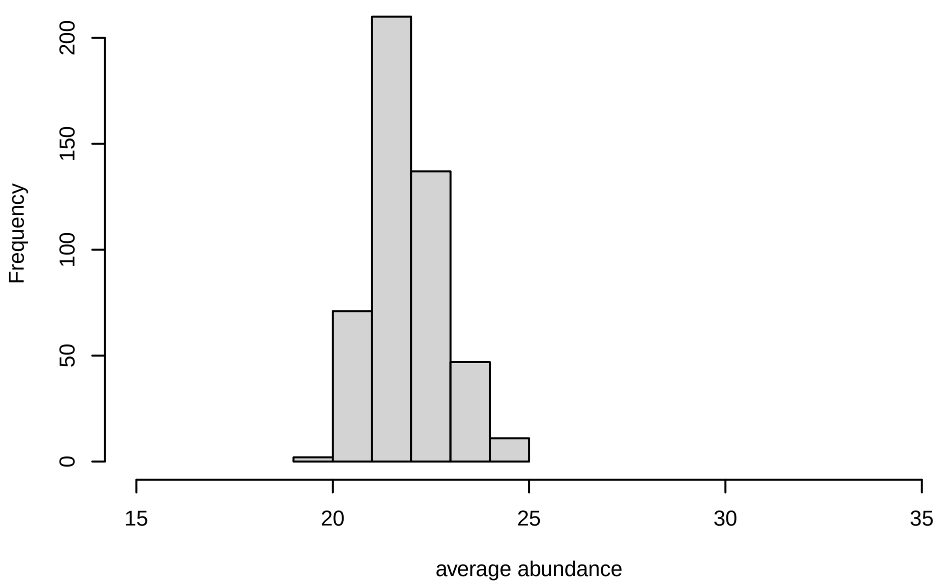


V5, after remove outlier and cutoff is 0.8

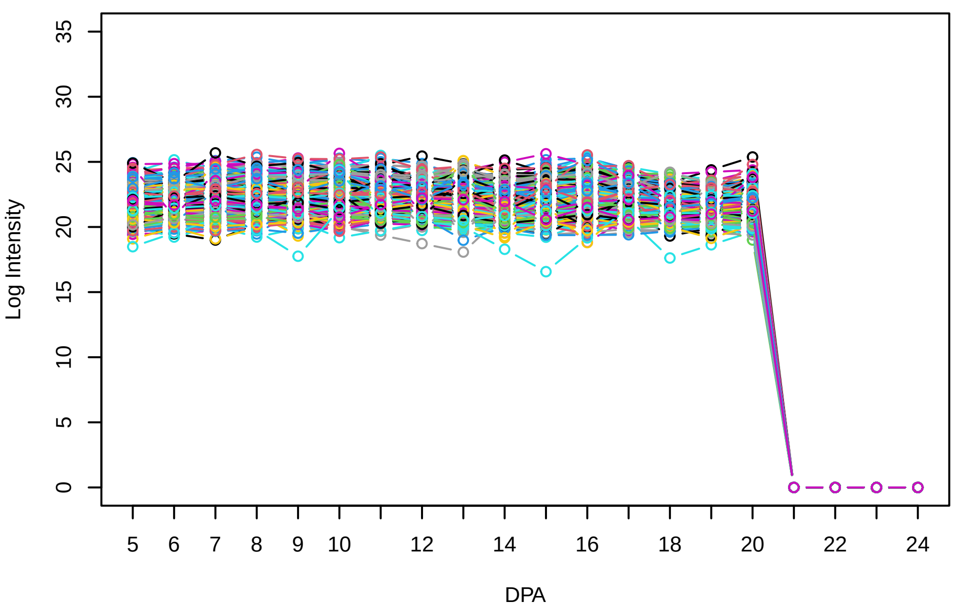


Histogram of the average abundances of non−outlier proteins

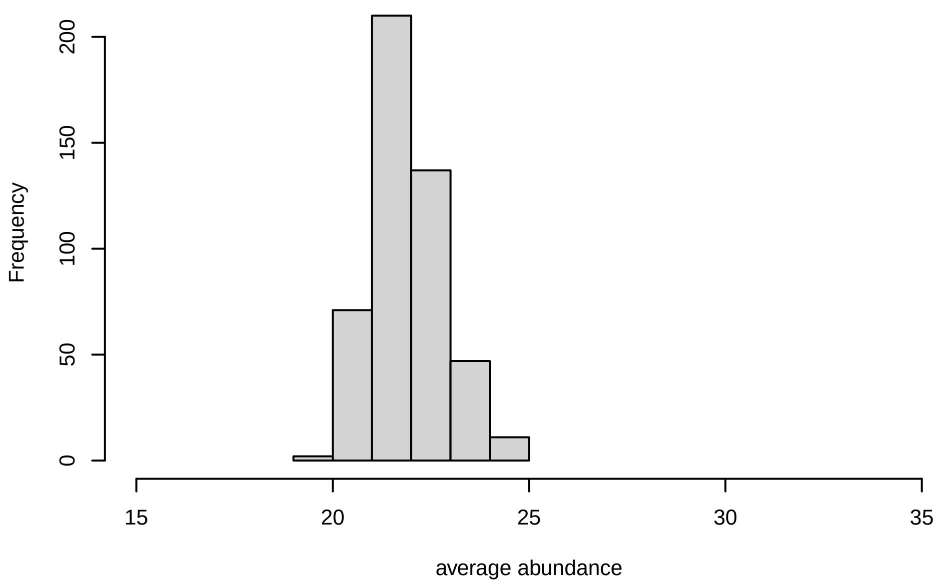


V5, after remove unreliable

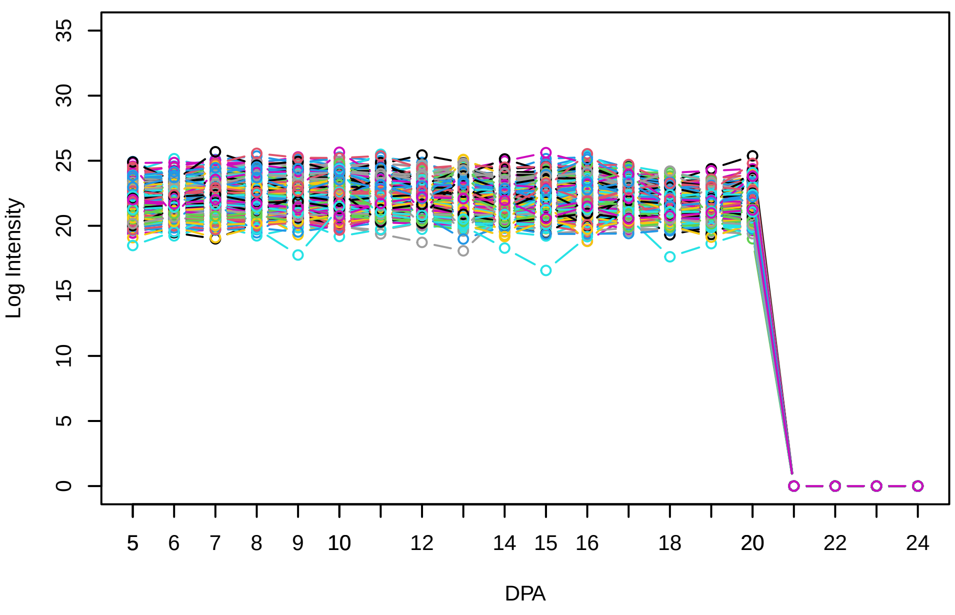


Histogram of the average abundances of reliable proteins

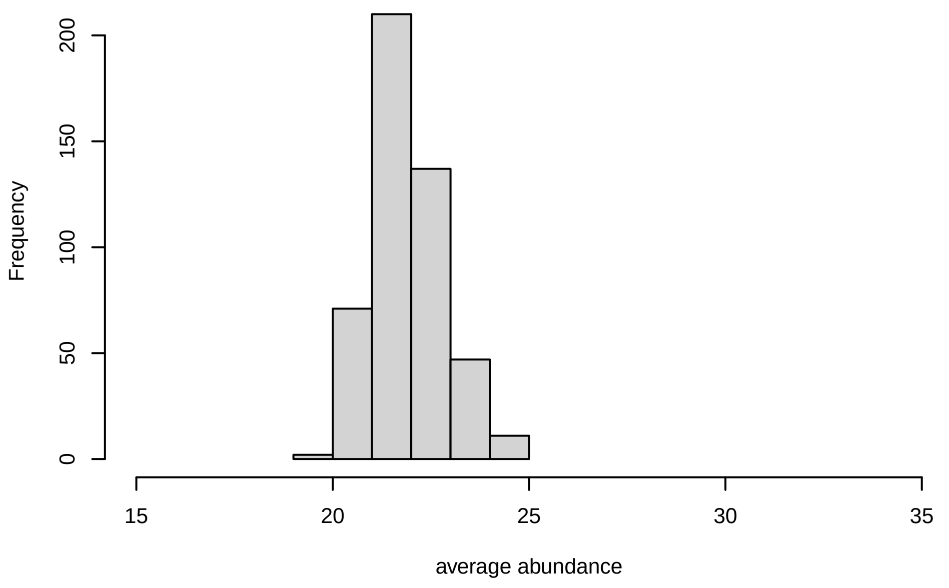


V5, after remove low abundance

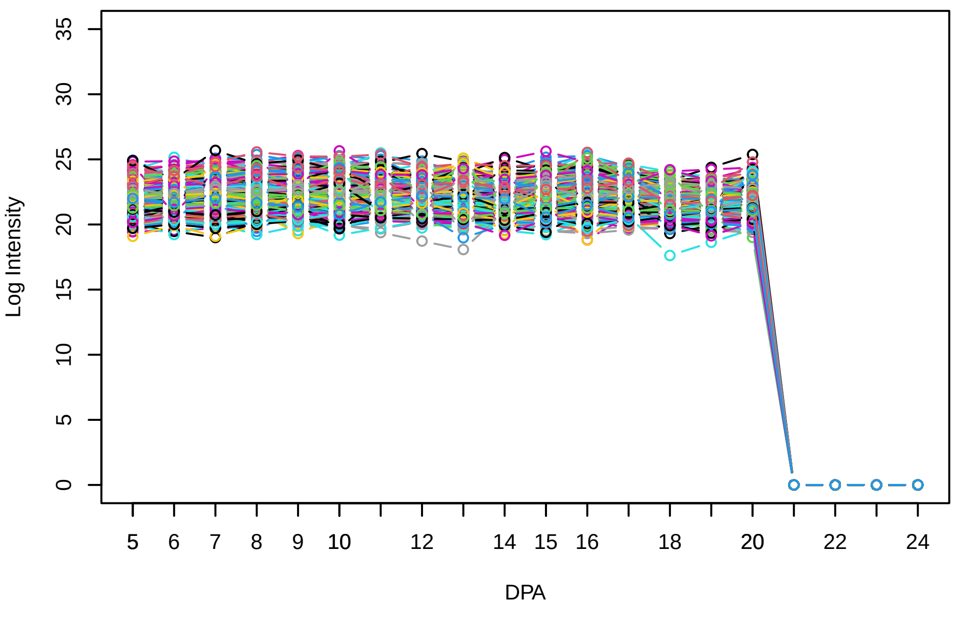


Histogram of the average abundances of high abundance proteins

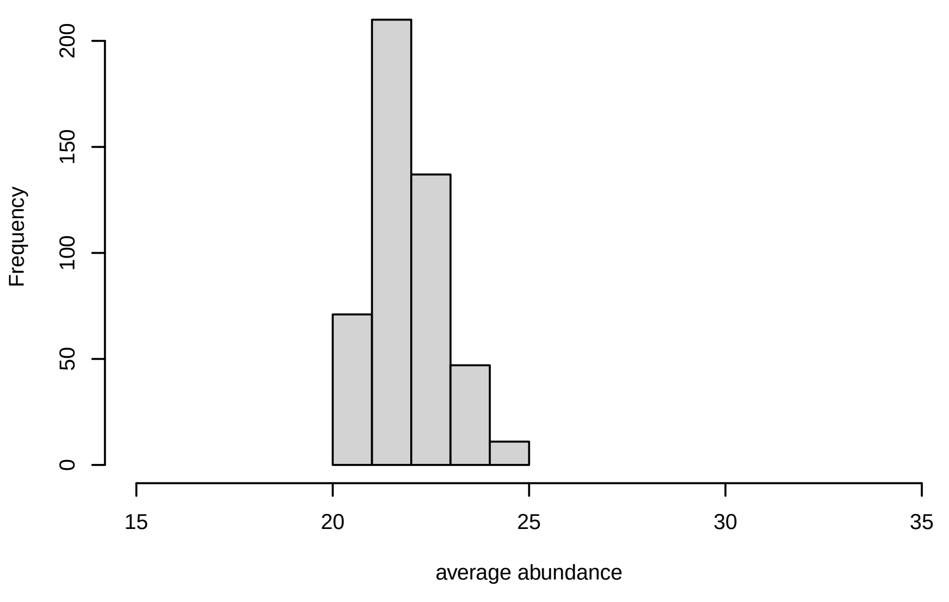

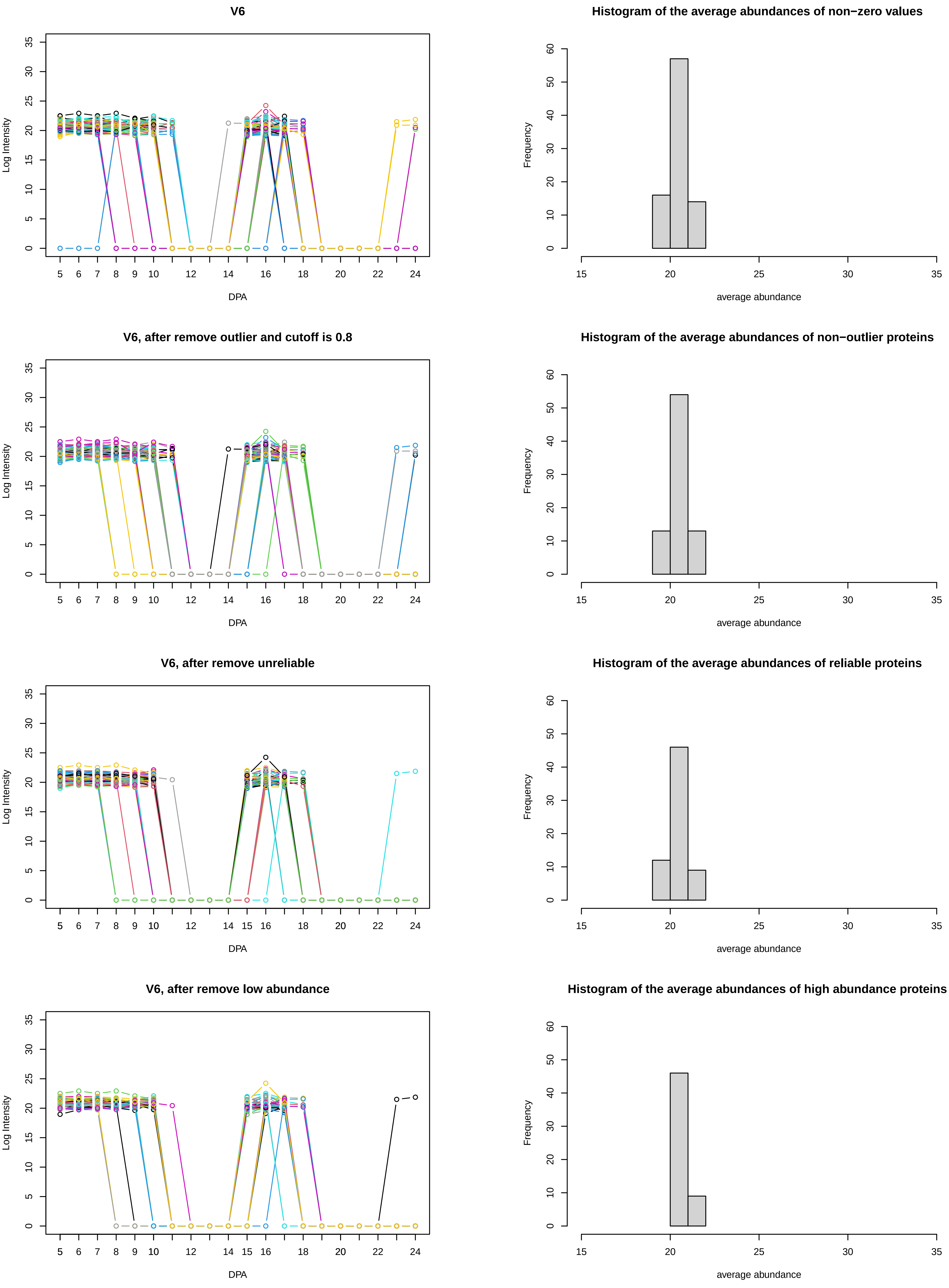
V6
Histogram of the average abundances of non−zero values
V6, after remove outlier and cutoff is 0.8
Histogram of the average abundances of non−outlier proteins
V6, after remove unreliable
Histogram of the average abundances of reliable proteins
V6, after remove low abundance
Histogram of the average abundances of high abundance proteins
Log Intensity
DPA
Frequency
average abundance

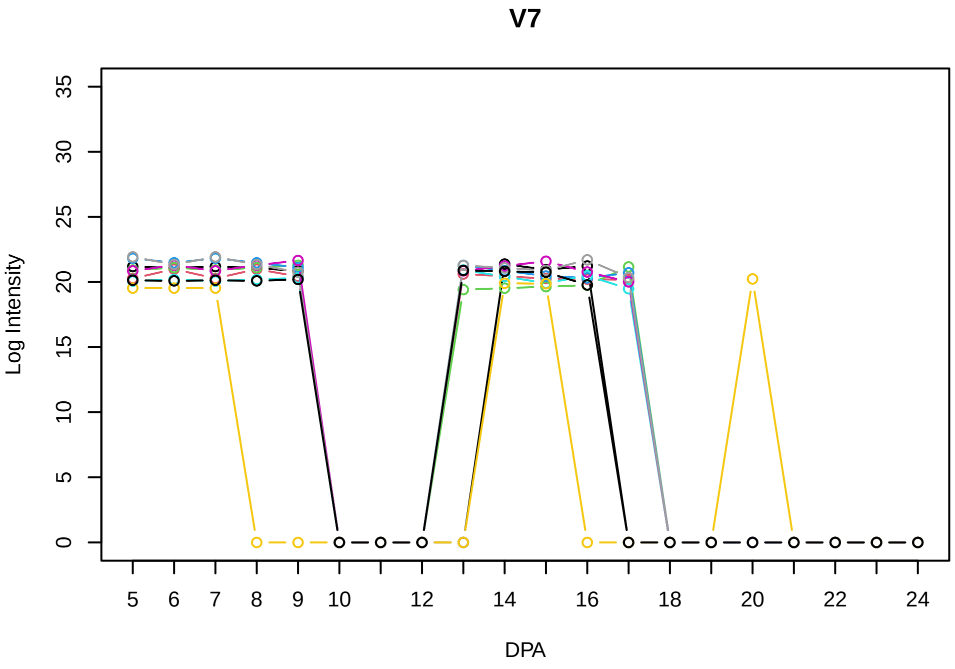
V7
Log Intensity
DPA

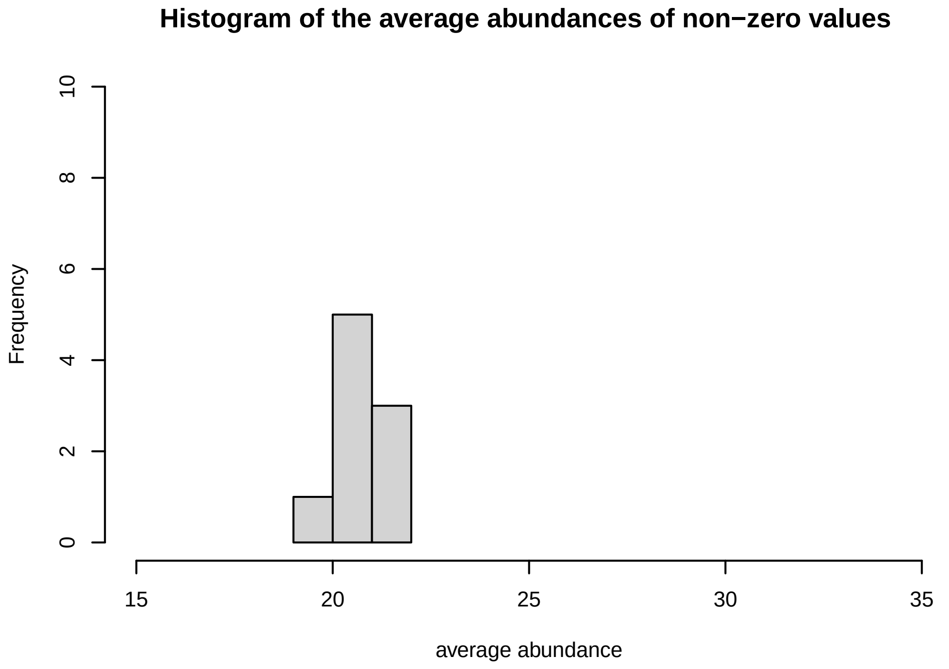
Histogram of the average abundances of non−zero values
Frequency
average abundance

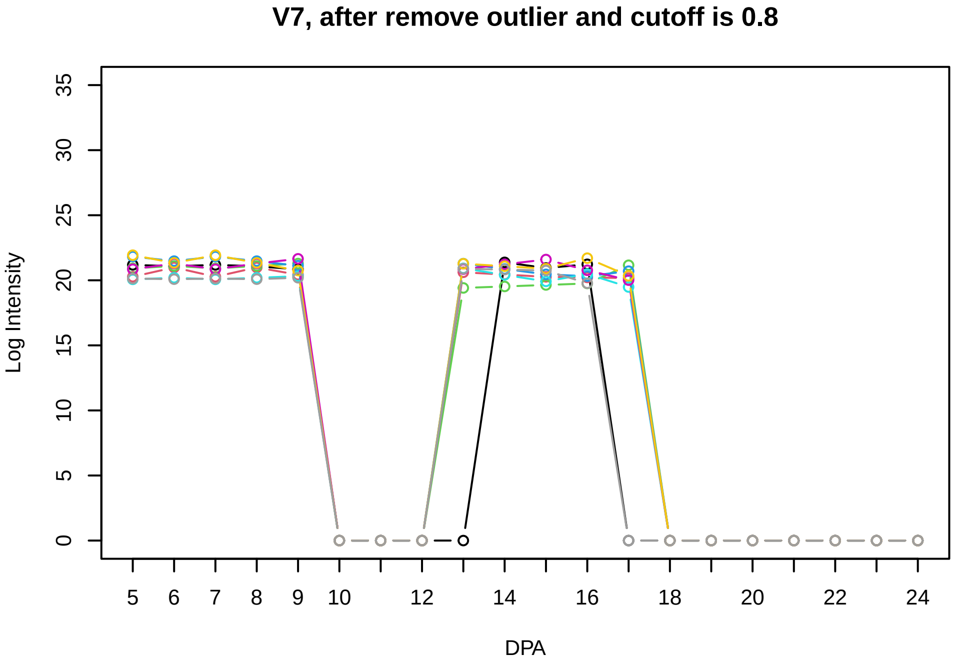
V7, after remove outlier and cutoff is 0.8
Log Intensity
DPA

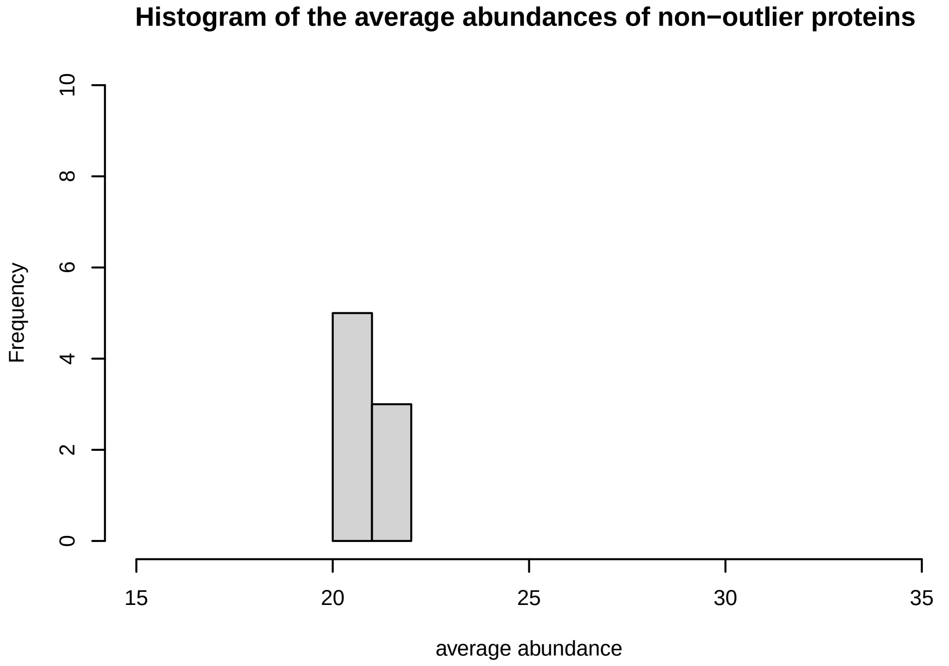
Histogram of the average abundances of non−outlier proteins
Frequency
average abundance

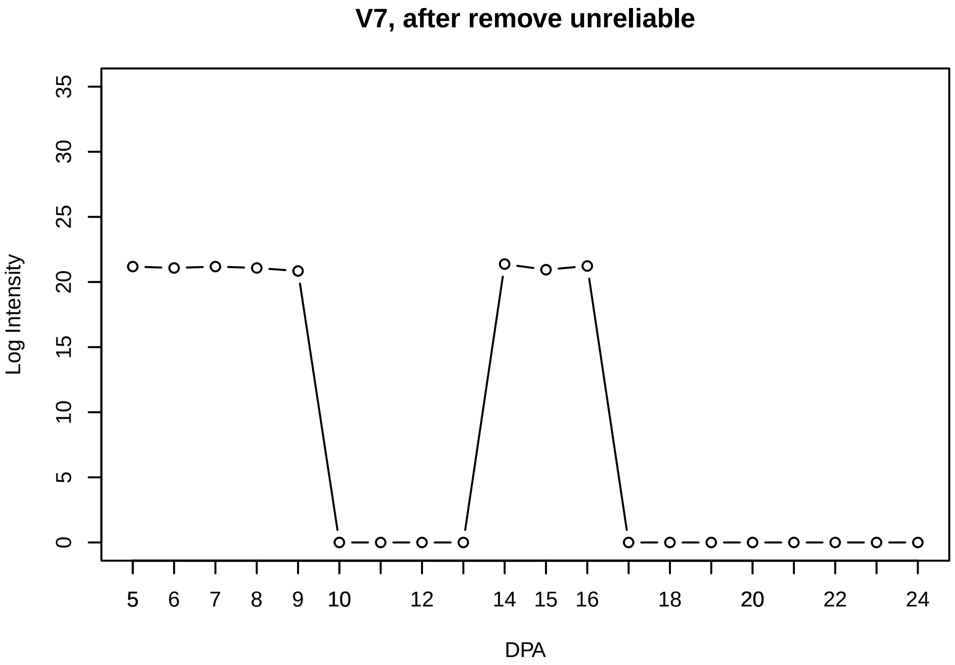
V7, after remove unreliable
Log Intensity
DPA

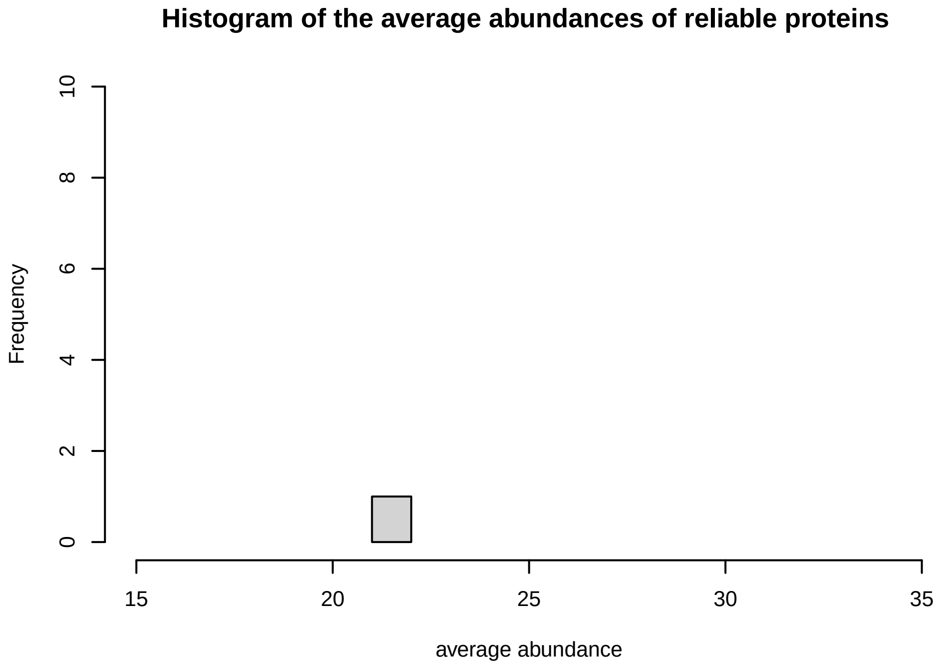
Histogram of the average abundances of reliable proteins
Frequency
average abundance

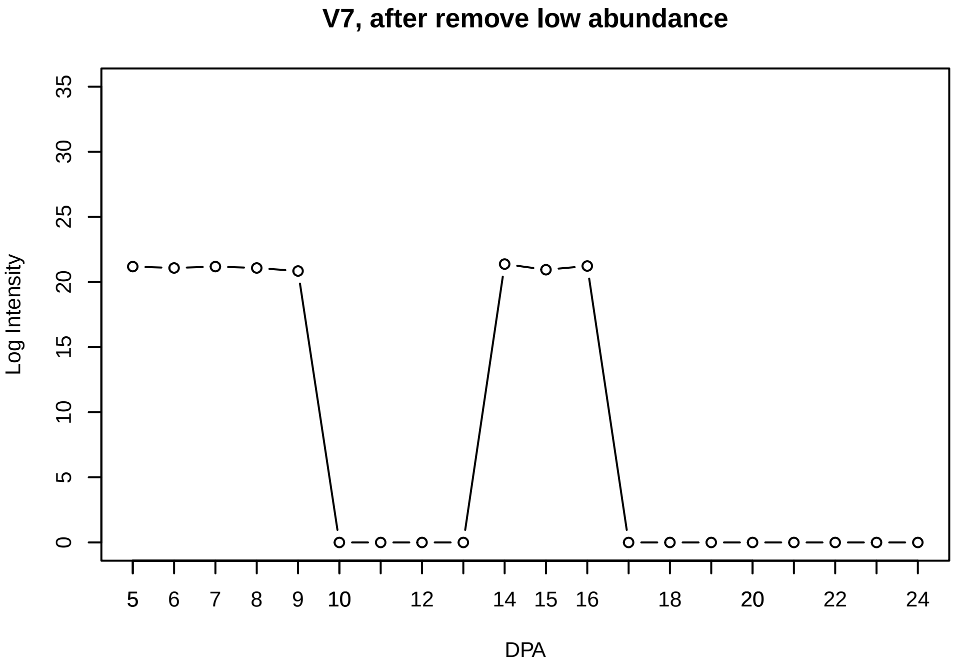
V7, after remove low abundance
Log Intensity
DPA

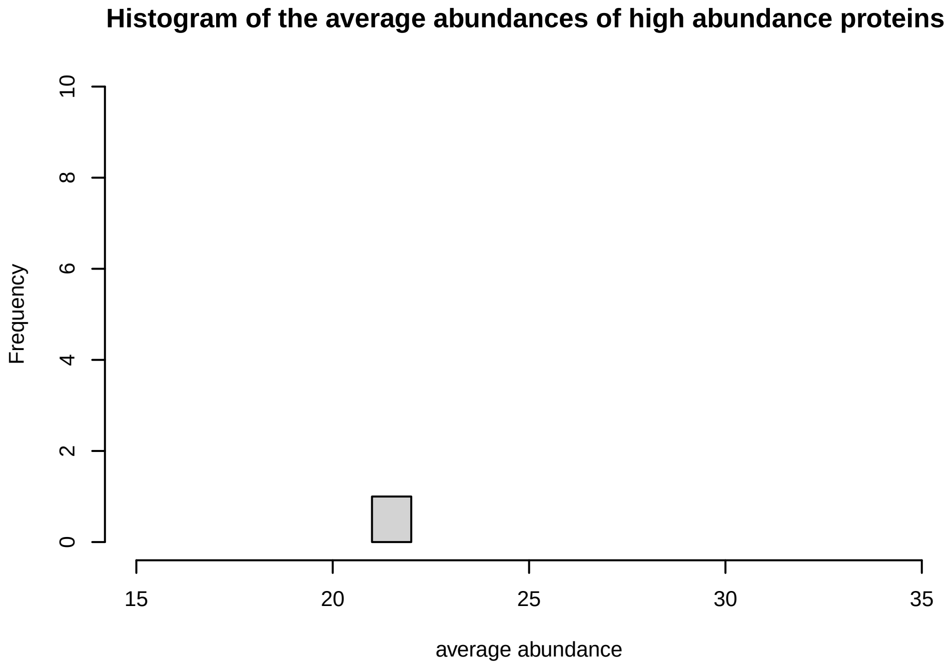
Histogram of the average abundances of high abundance proteins
Frequency
average abundance

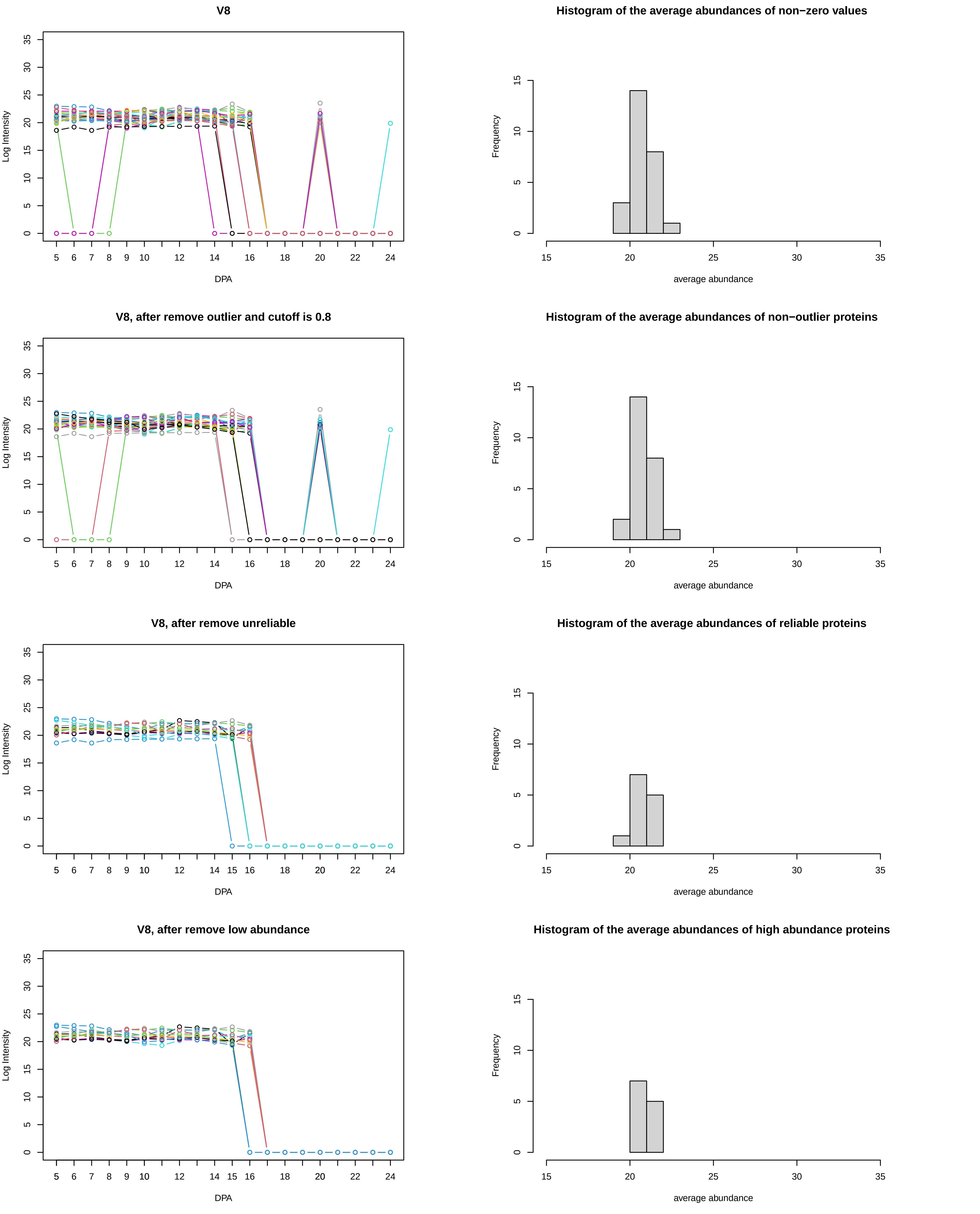

V8
Log Intensity
DPA
Histogram of the average abundances of non−zero values
Frequency
average abundance
V8, after remove outlier and cutoff is 0.8
Log Intensity
DPA
Histogram of the average abundances of non−outlier proteins
Frequency
average abundance
V8, after remove unreliable
Log Intensity
DPA
Histogram of the average abundances of reliable proteins
Frequency
average abundance
V8, after remove low abundance
Log Intensity
DPA
Histogram of the average abundances of high abundance proteins
Frequency
average abundance

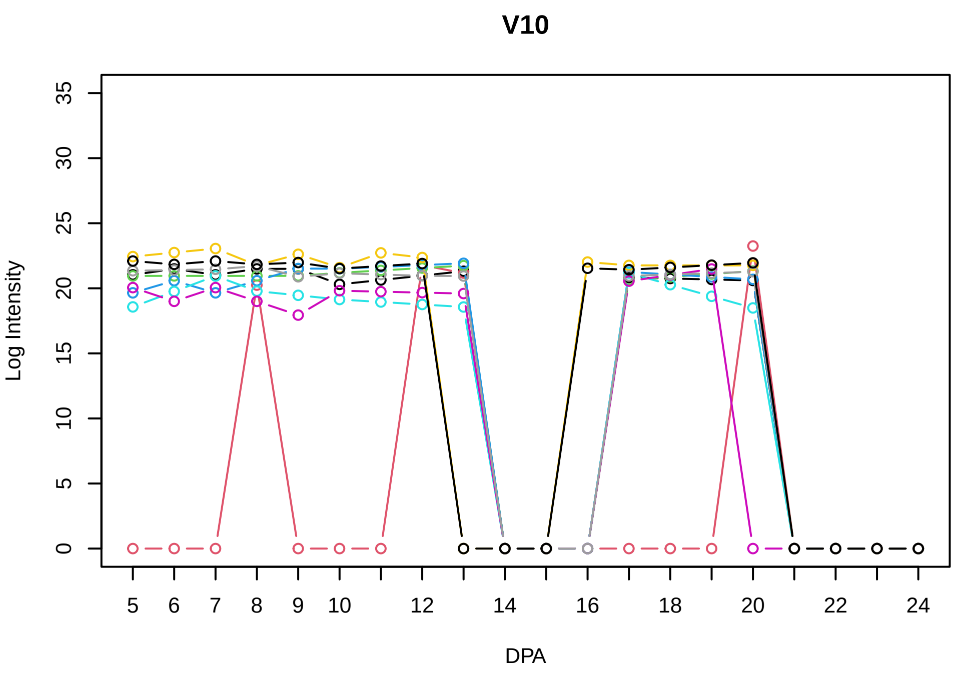
V10
Log Intensity
DPA
0
5
10
15
20
25
30
35
5
6
7
8
9
10
12
14
16
18
20
22
24

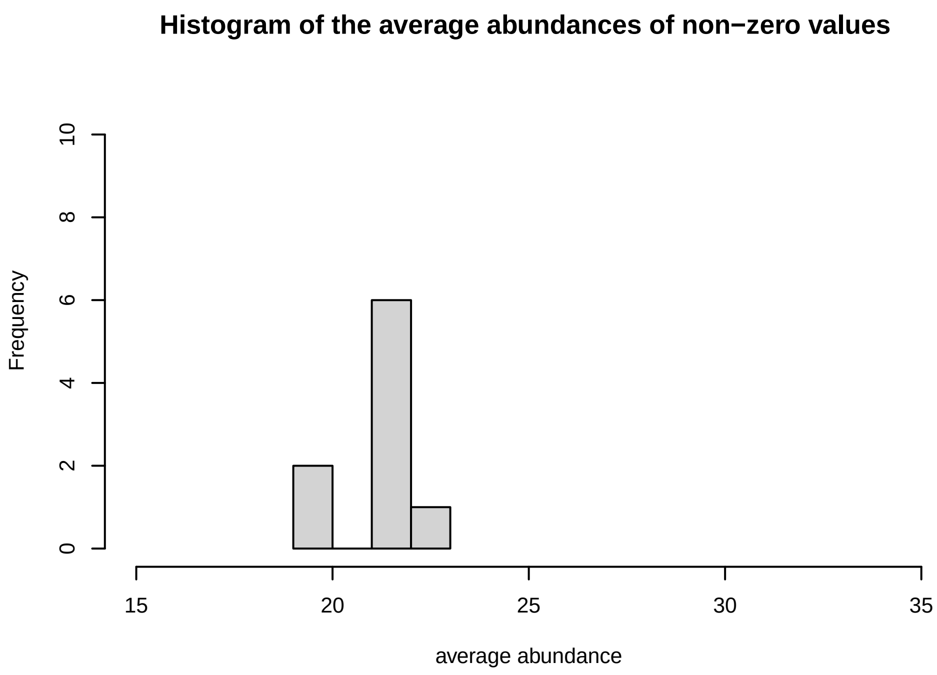
Histogram of the average abundances of non−zero values
Frequency
average abundance
0
2
4
6
8
10
15
20
25
30
35

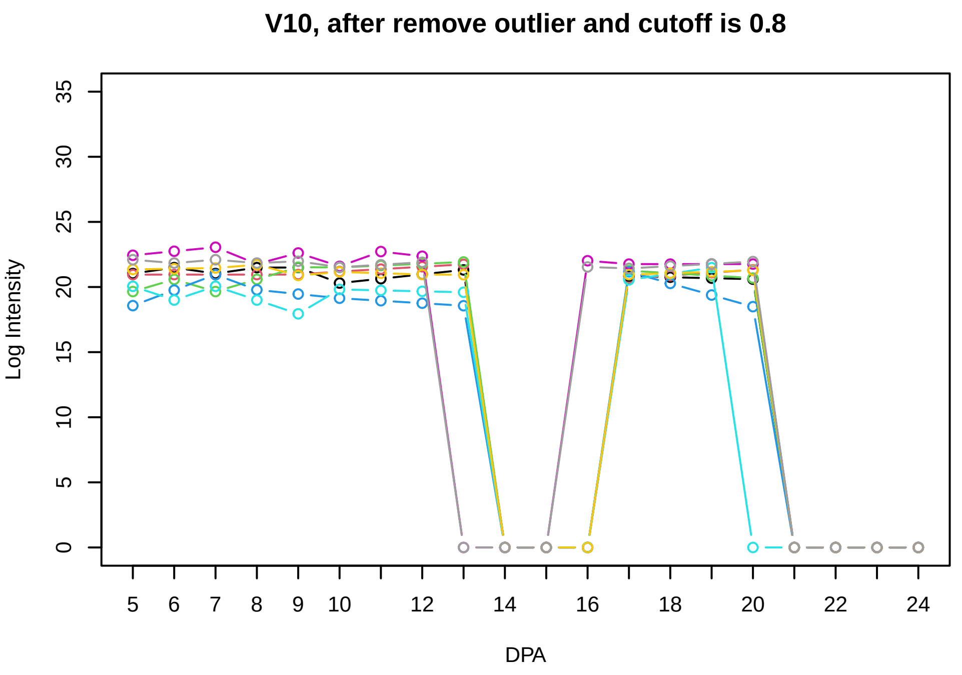
V10, after remove outlier and cutoff is 0.8
Log Intensity
DPA
0
5
10
15
20
25
30
35
5
6
7
8
9
10
12
14
16
18
20
22
24

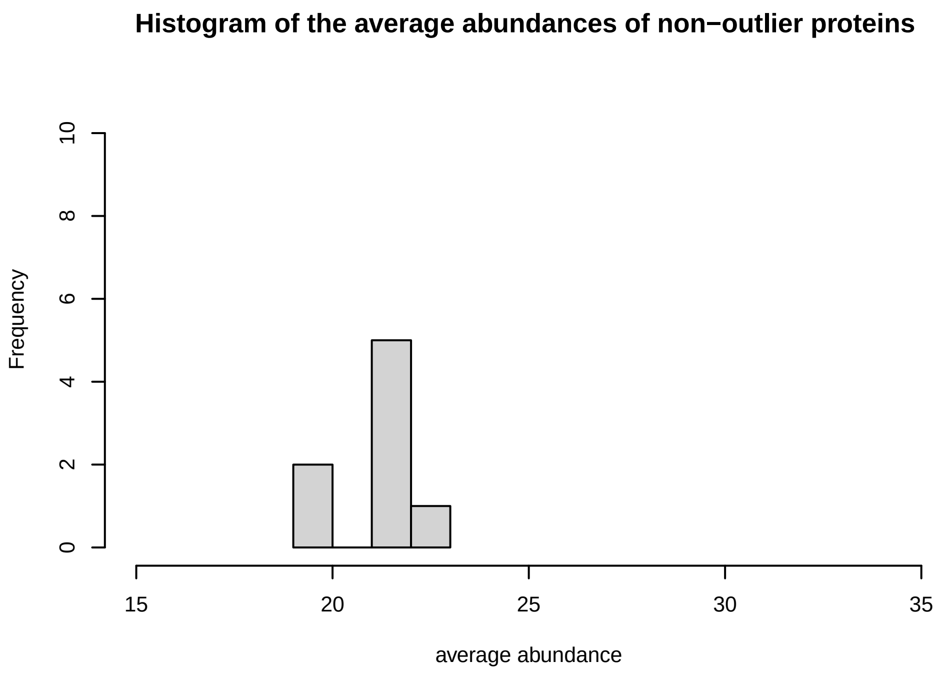
Histogram of the average abundances of non−outlier proteins
Frequency
average abundance
0
2
4
6
8
10
15
20
25
30
35

**V11**

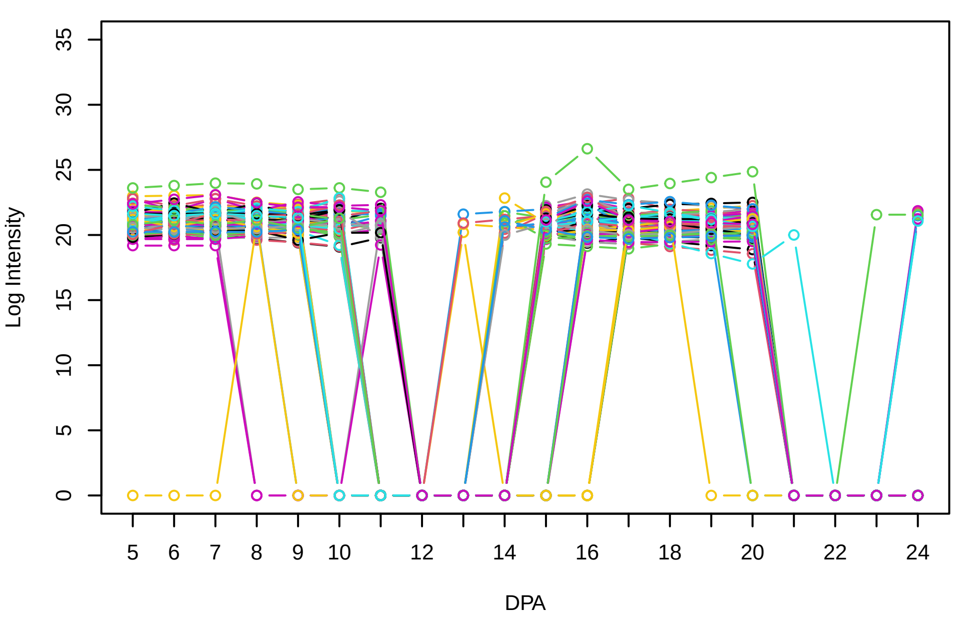


**Histogram of the average abundances of non−zero values**

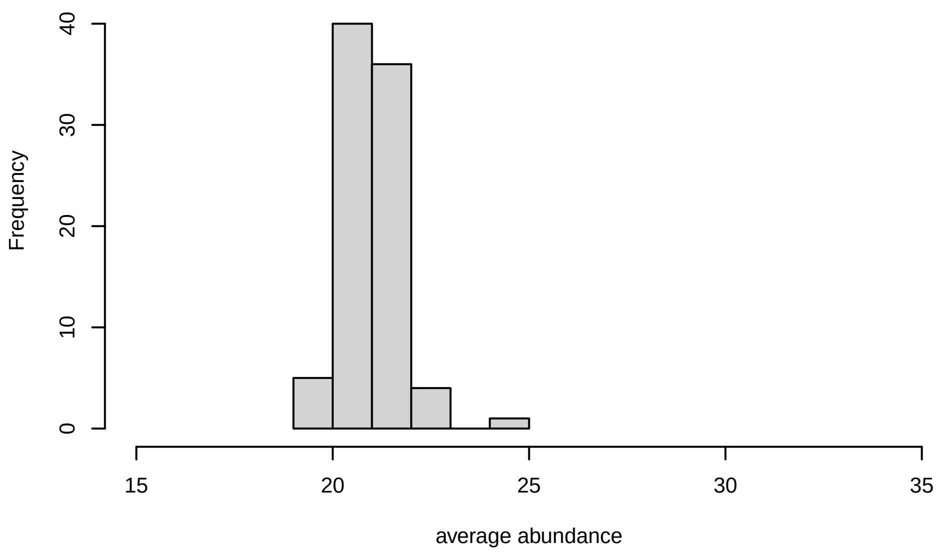


**V11, after remove outlier and cutoff is 0.8**

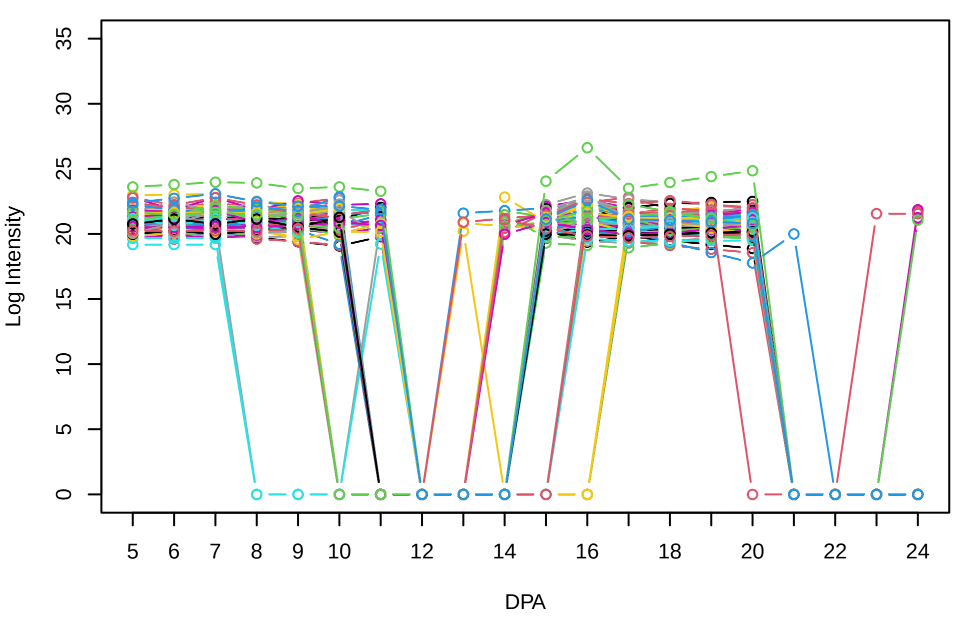


**Histogram of the average abundances of non−outlier proteins**

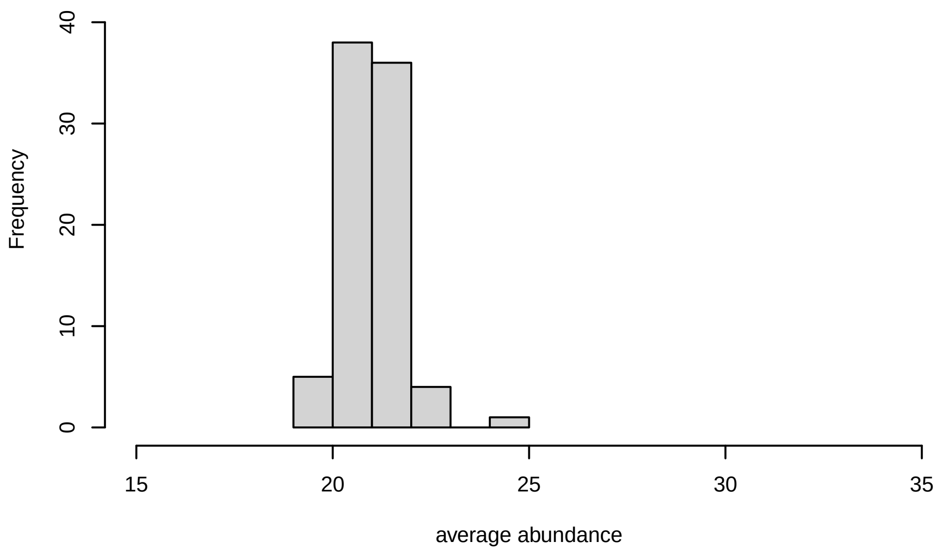


**V11, after remove unreliable**

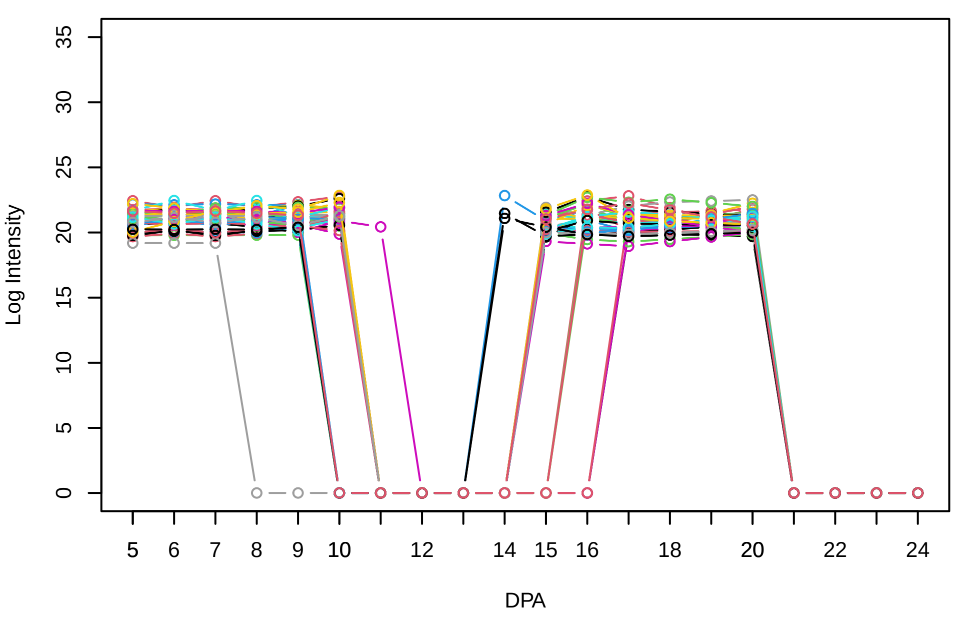


**Histogram of the average abundances of reliable proteins**

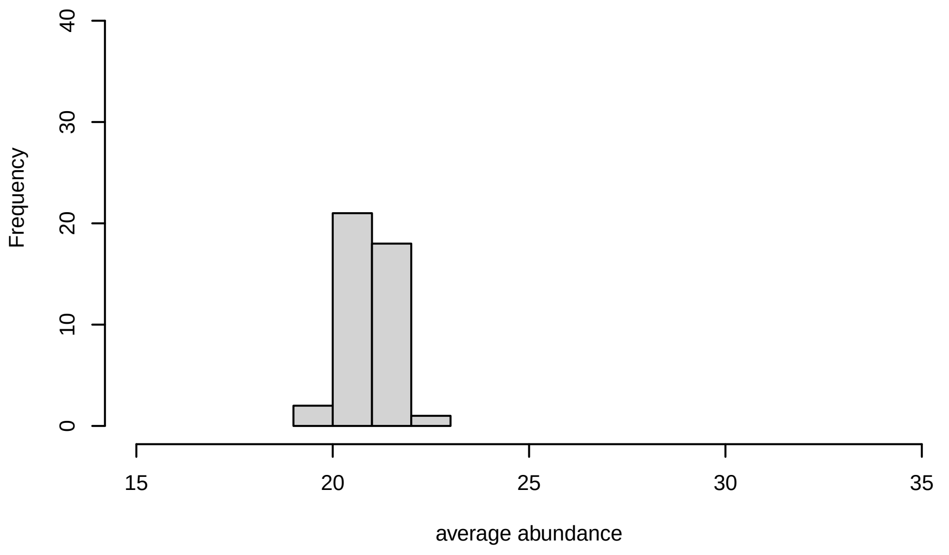


**V11, after remove low abundance**

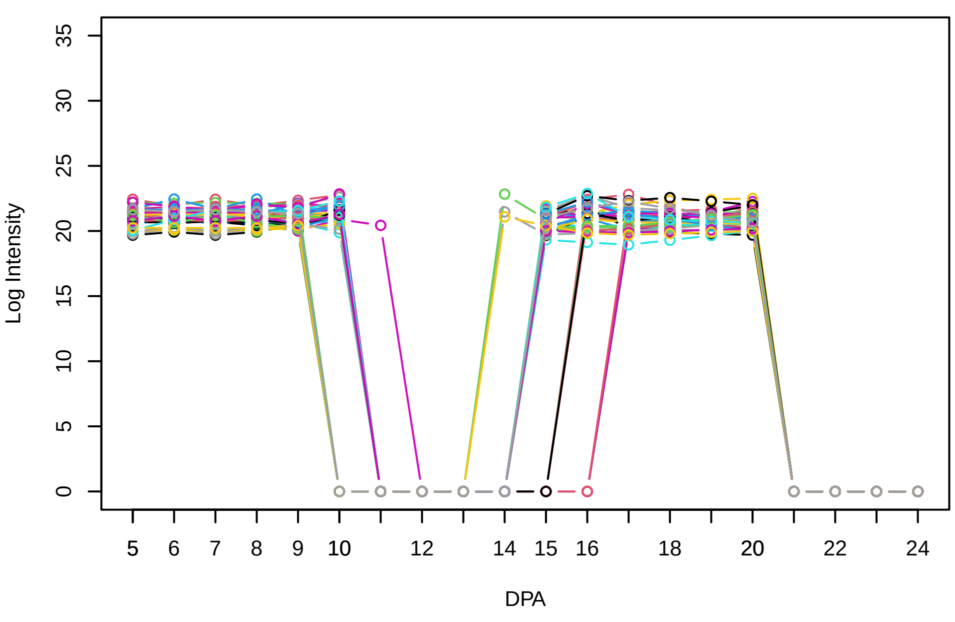


**Histogram of the average abundances of high abundance proteins**

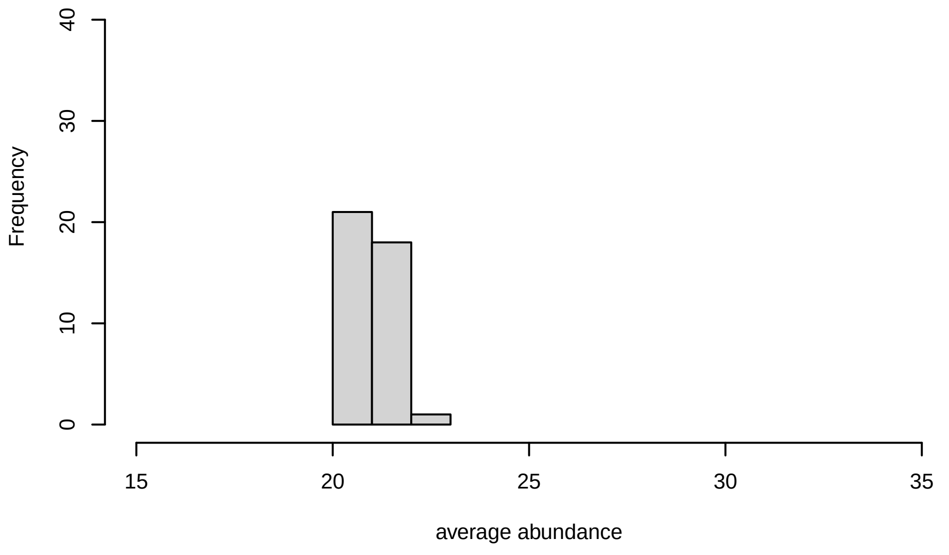

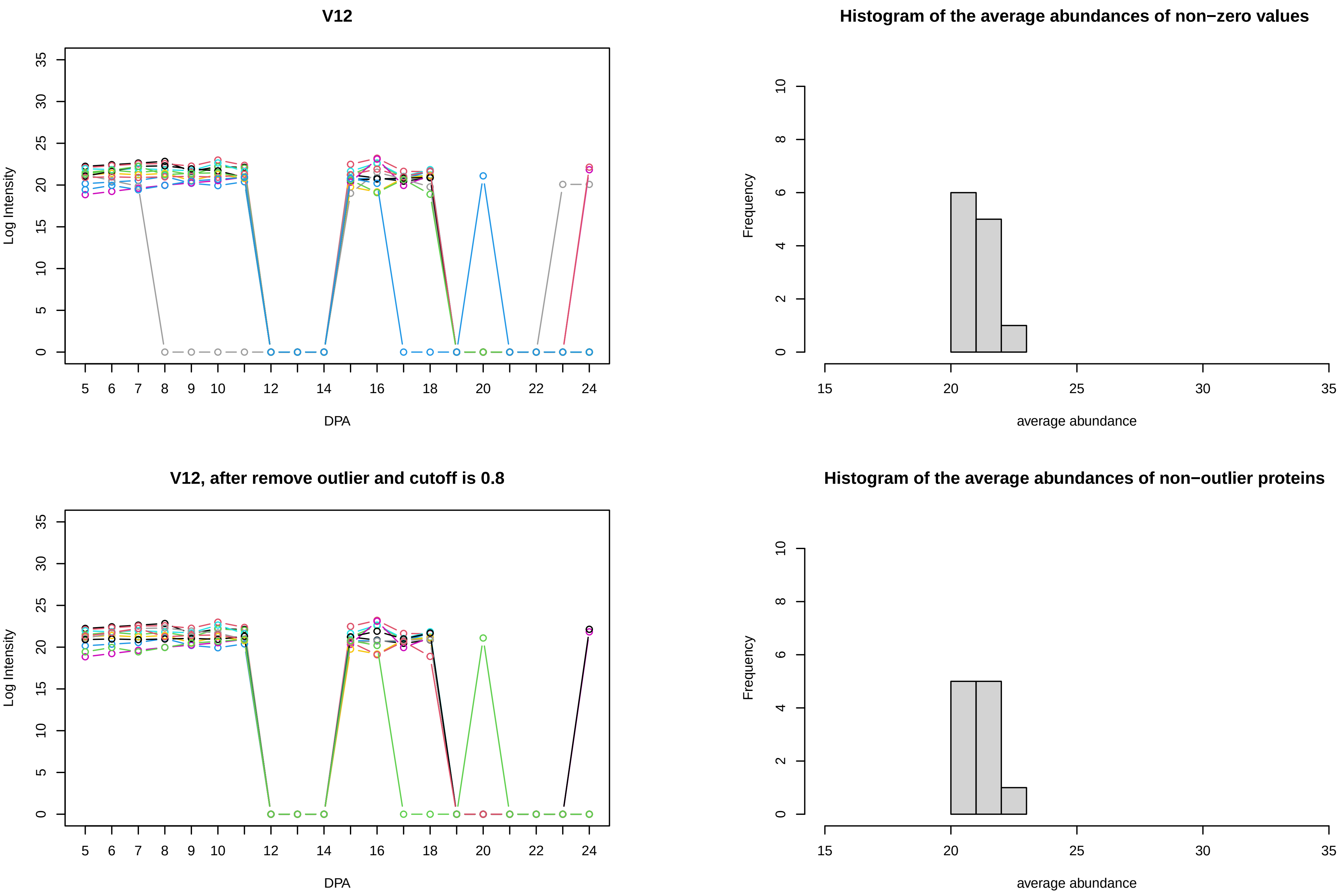
V12
Log Intensity
DPA
Histogram of the average abundances of non−zero values
Frequency
average abundance
V12, after remove outlier and cutoff is 0.8
Log Intensity
DPA
Histogram of the average abundances of non−outlier proteins
Frequency
average abundance

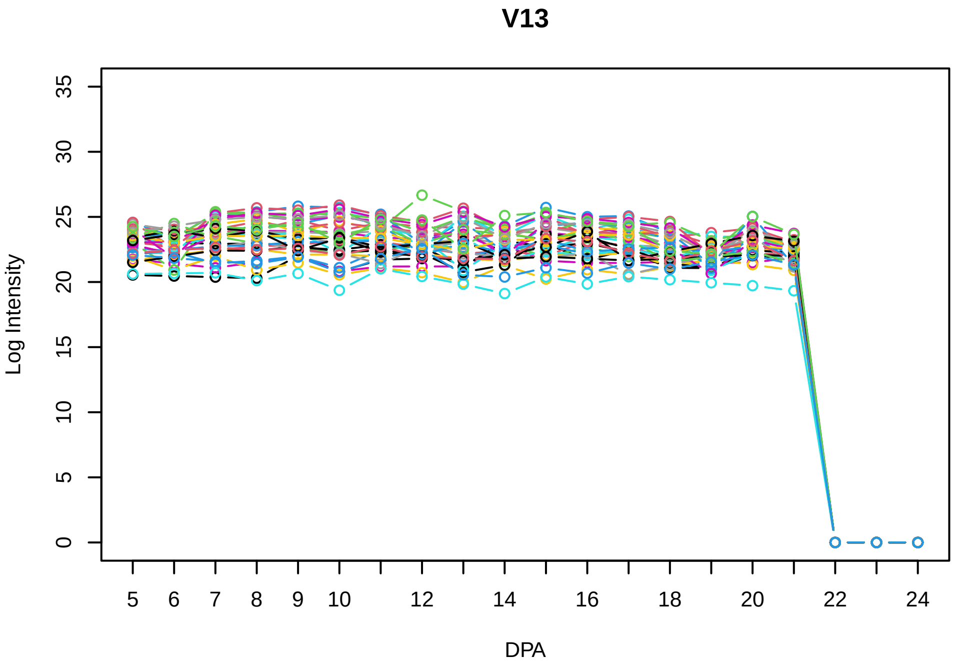
V13
Log Intensity
DPA
0
5
10
15
20
25
30
35
5
6
7
8
9
10
12
14
16
18
20
22
24

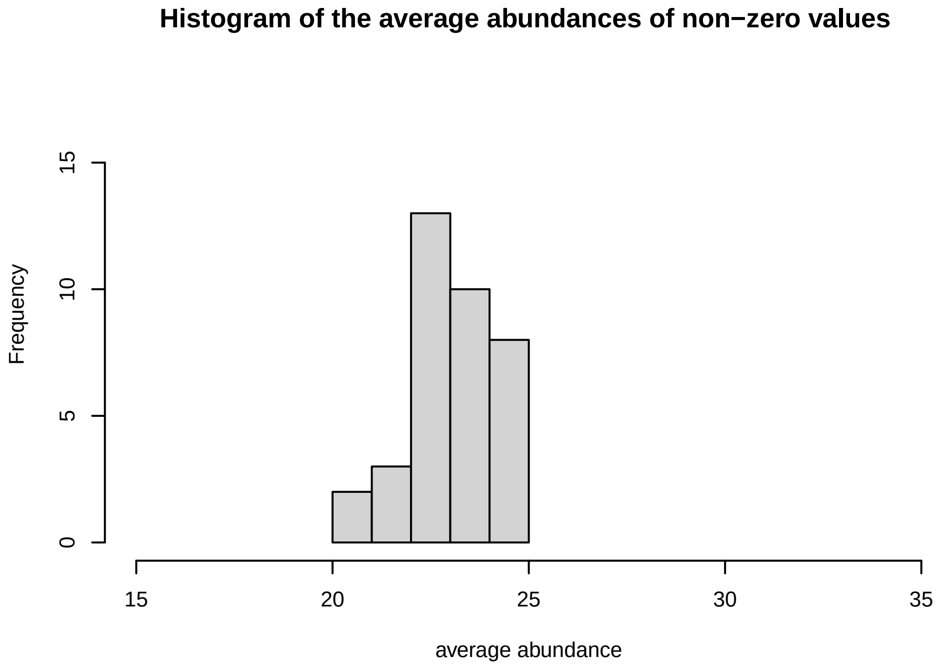
Histogram of the average abundances of non−zero values
Frequency
average abundance
0
5
10
15
15
20
25
30
35

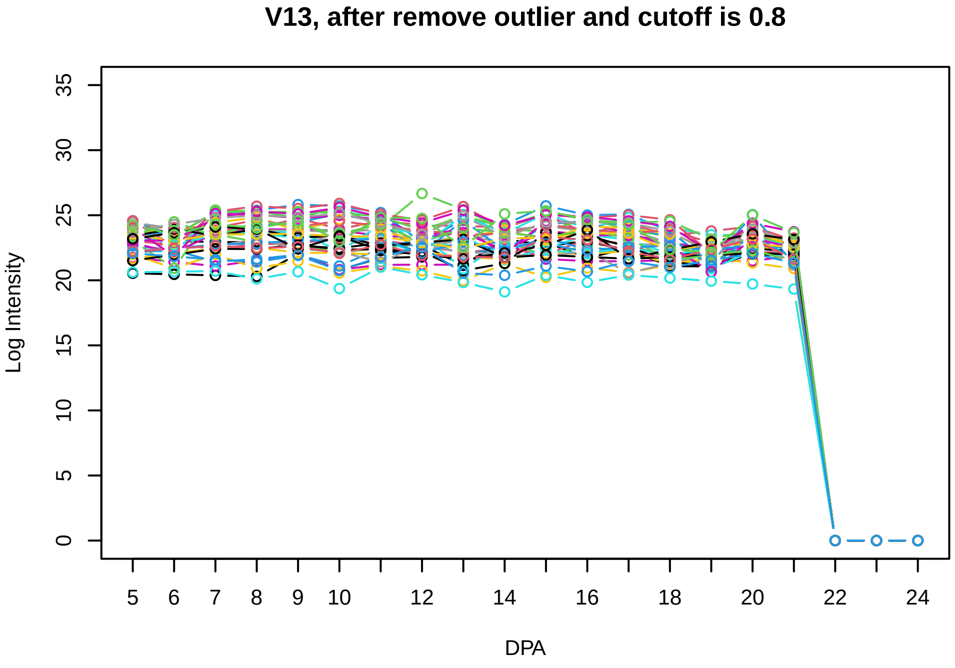
V13, after remove outlier and cutoff is 0.8
Log Intensity
DPA
0
5
10
15
20
25
30
35
5
6
7
8
9
10
12
14
16
18
20
22
24

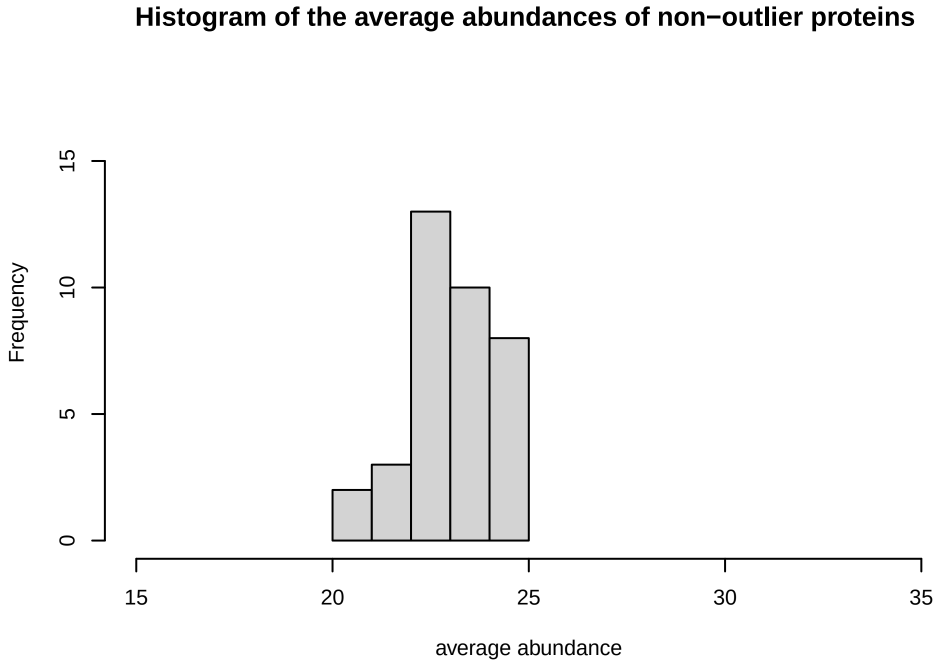
Histogram of the average abundances of non−outlier proteins
Frequency
average abundance
0
5
10
15
15
20
25
30
35

**V14**

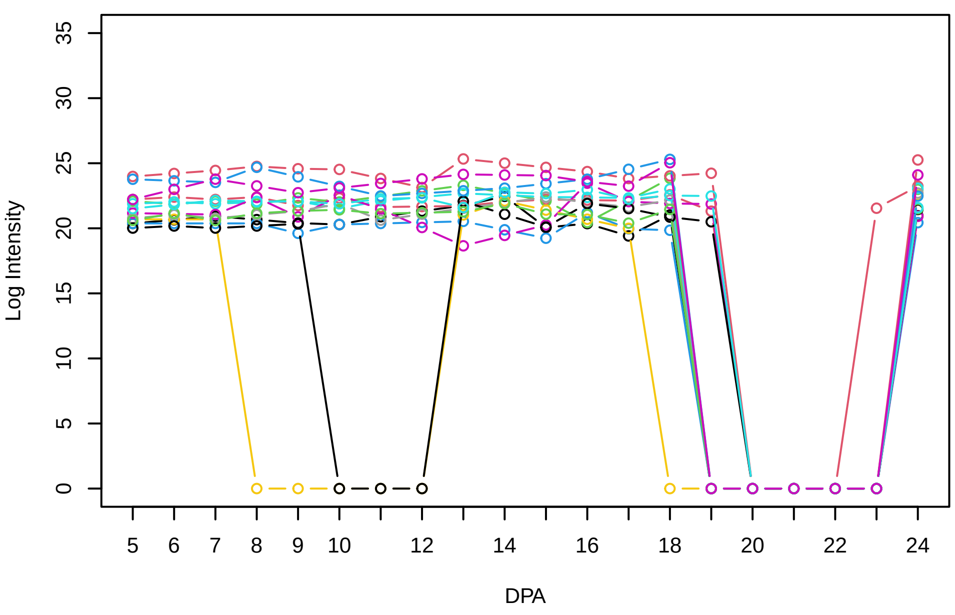


**Histogram of the average abundances of non−zero values**

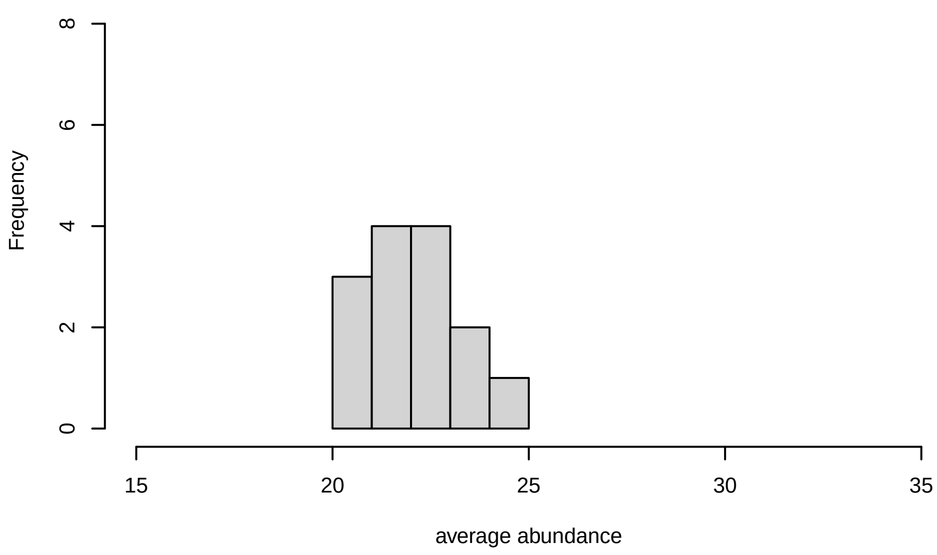


**V14, after remove outlier and cutoff is 0.8**

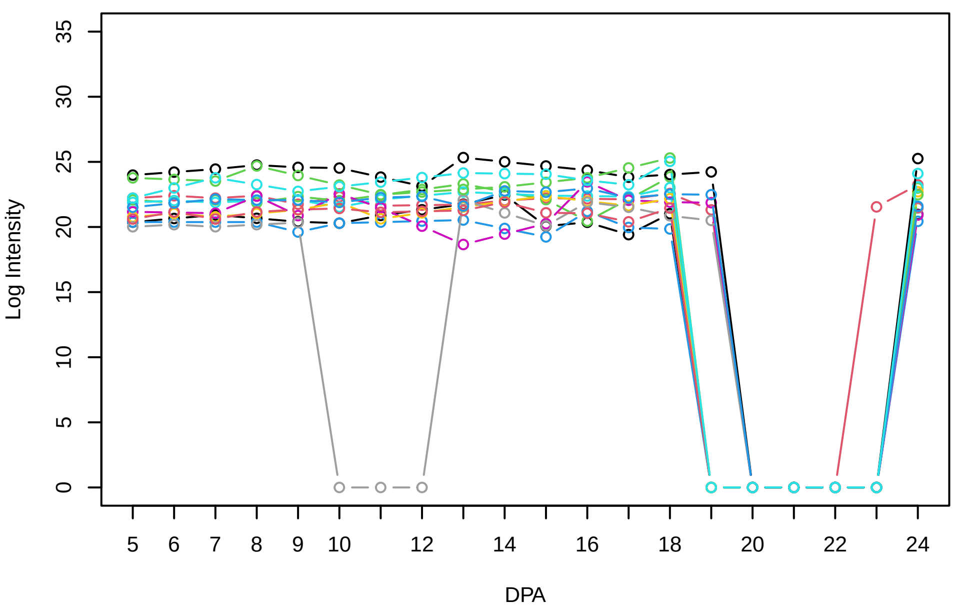


**Histogram of the average abundances of non−outlier proteins**

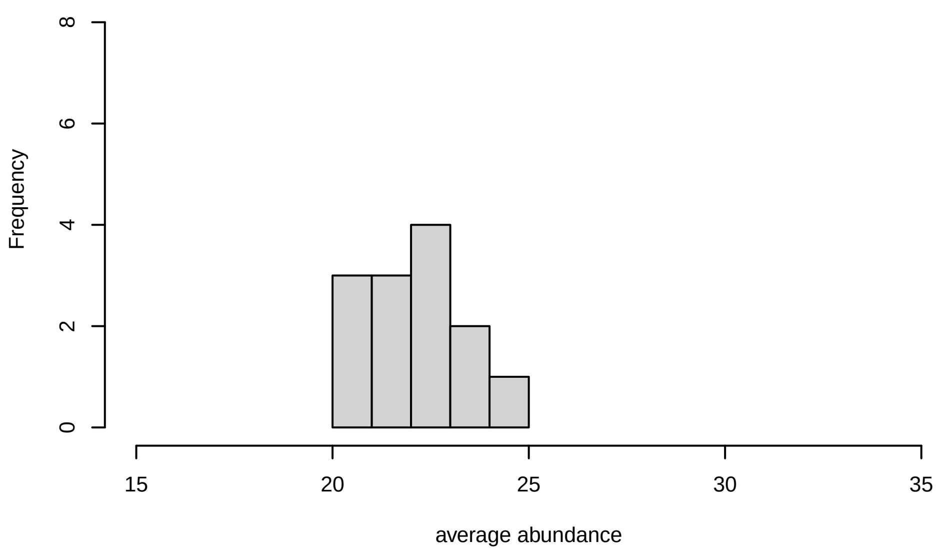

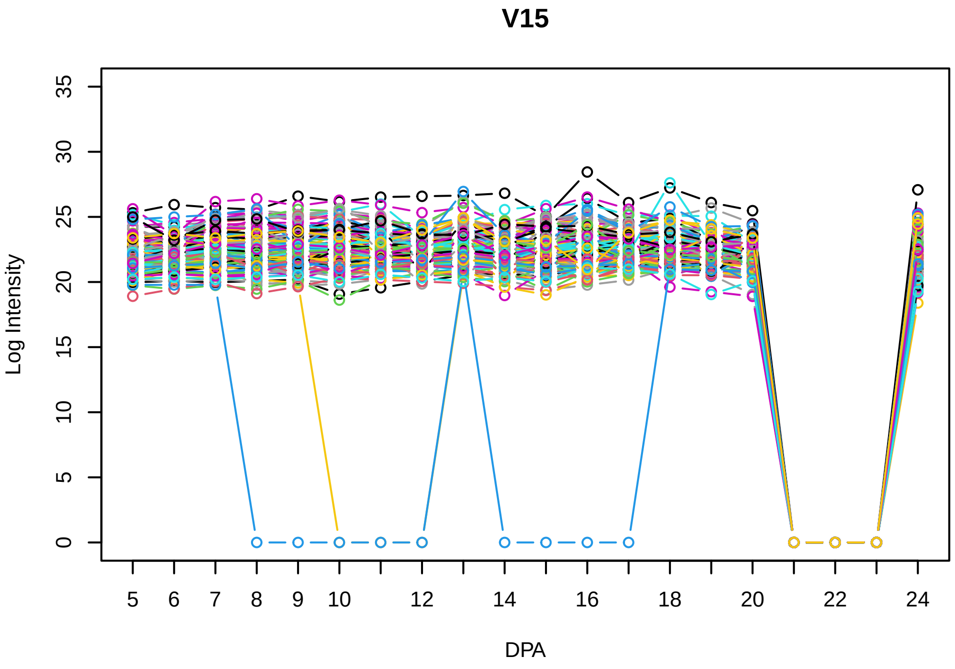
V15
Log Intensity
DPA
0
5
10
15
20
25
30
35
5
6
7
8
9
10
12
14
16
18
20
22
24

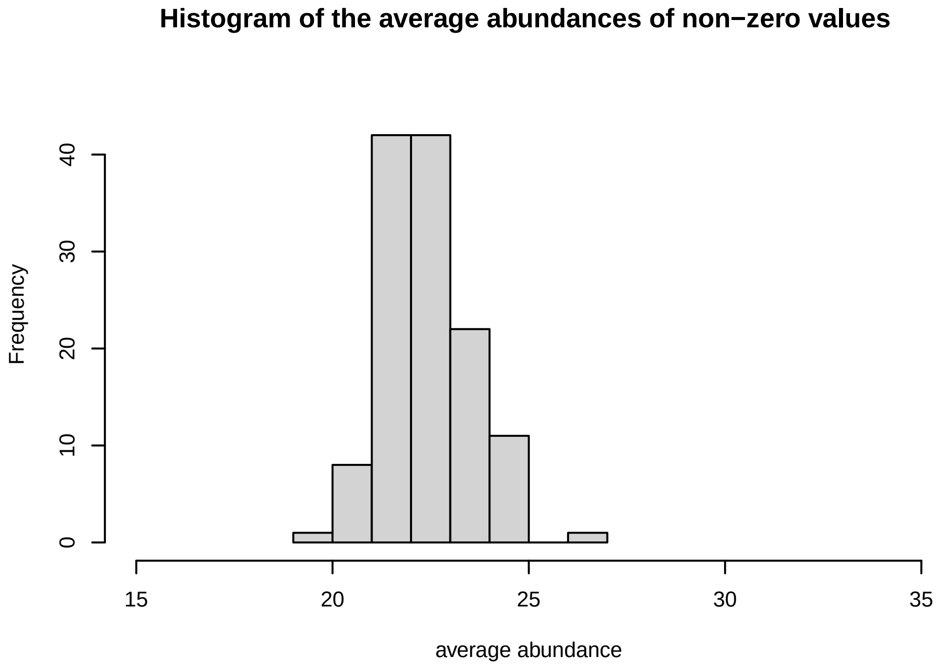
Histogram of the average abundances of non−zero values
Frequency
average abundance
0
10
20
30
40
15
20
25
30
35

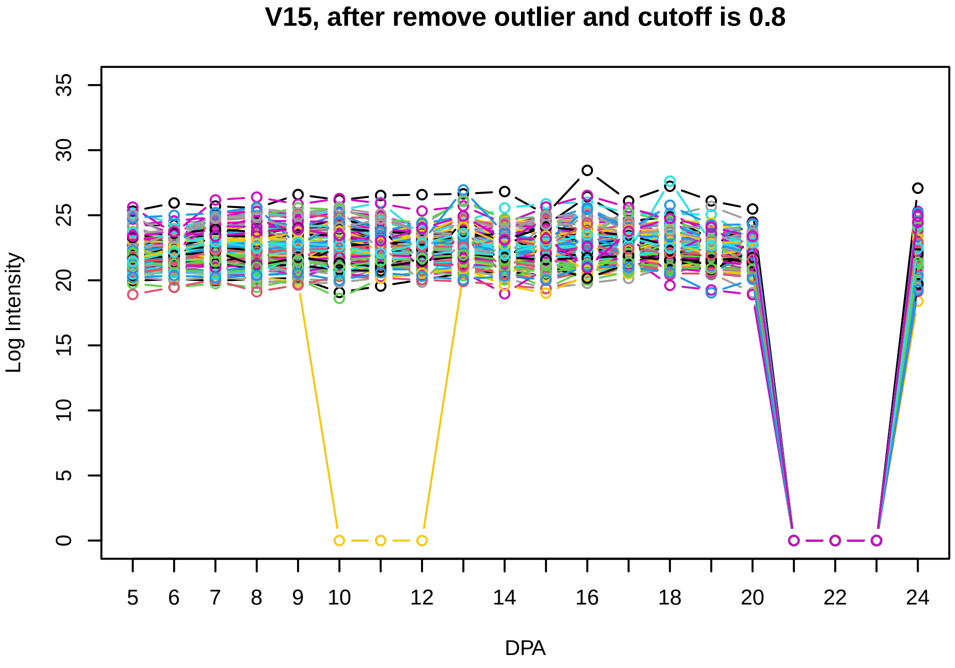
V15, after remove outlier and cutoff is 0.8
Log Intensity
DPA
0
5
10
15
20
25
30
35
5
6
7
8
9
10
12
14
16
18
20
22
24

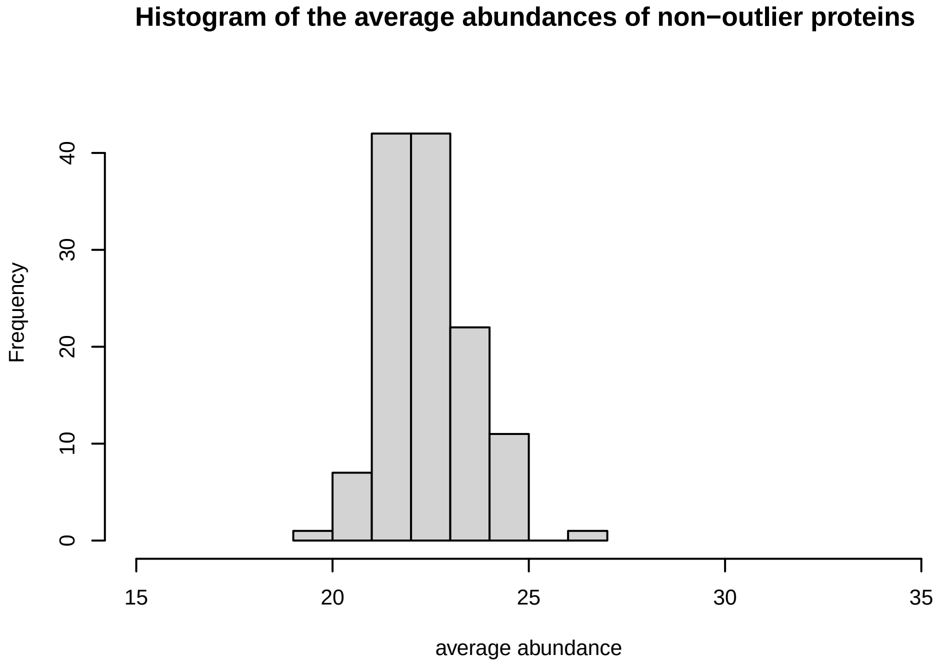
Histogram of the average abundances of non−outlier proteins
Frequency
average abundance
0
10
20
30
40
15
20
25
30
35

V16

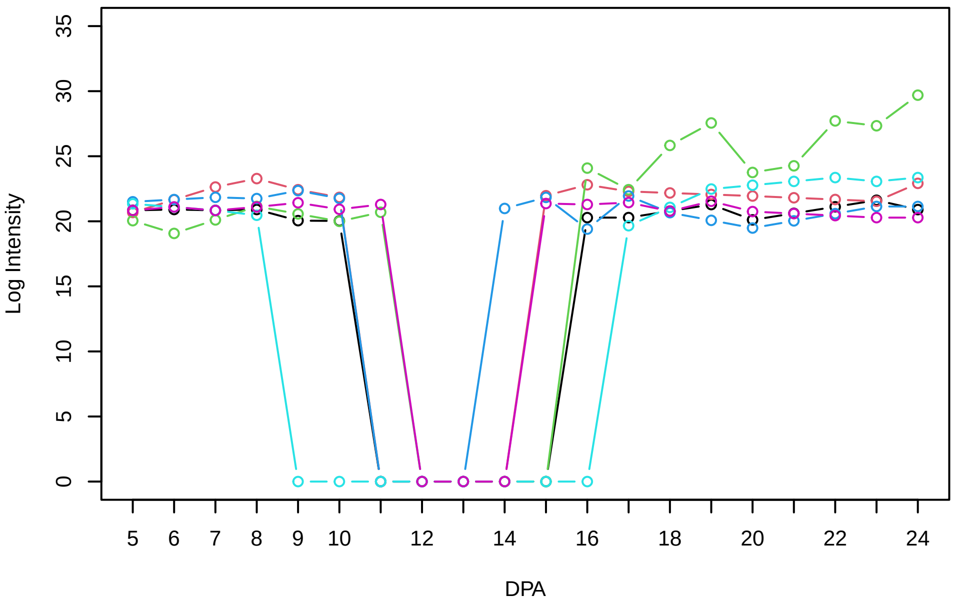


Histogram of the average abundances of non−zero values

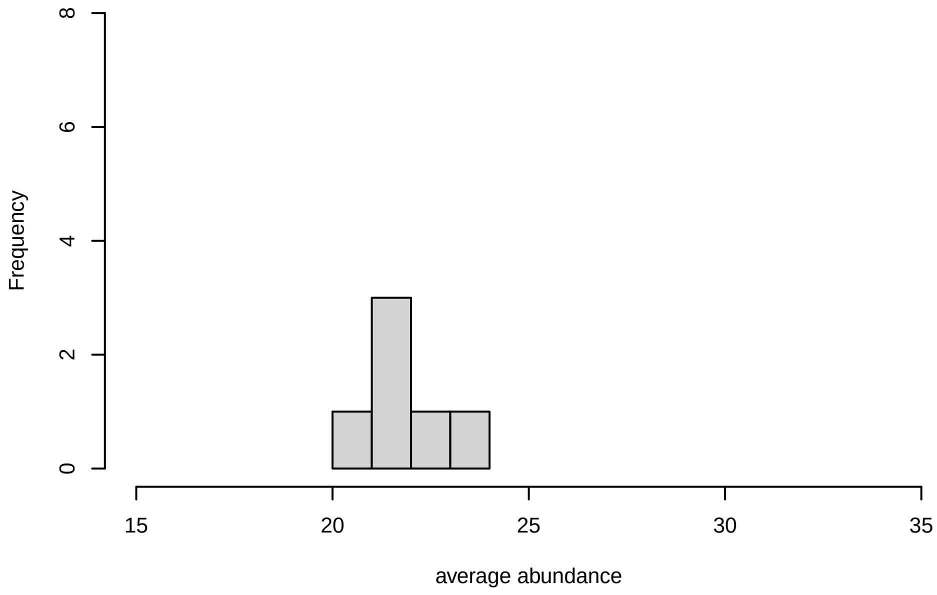


V16, after remove outlier and cutoff is 0.8

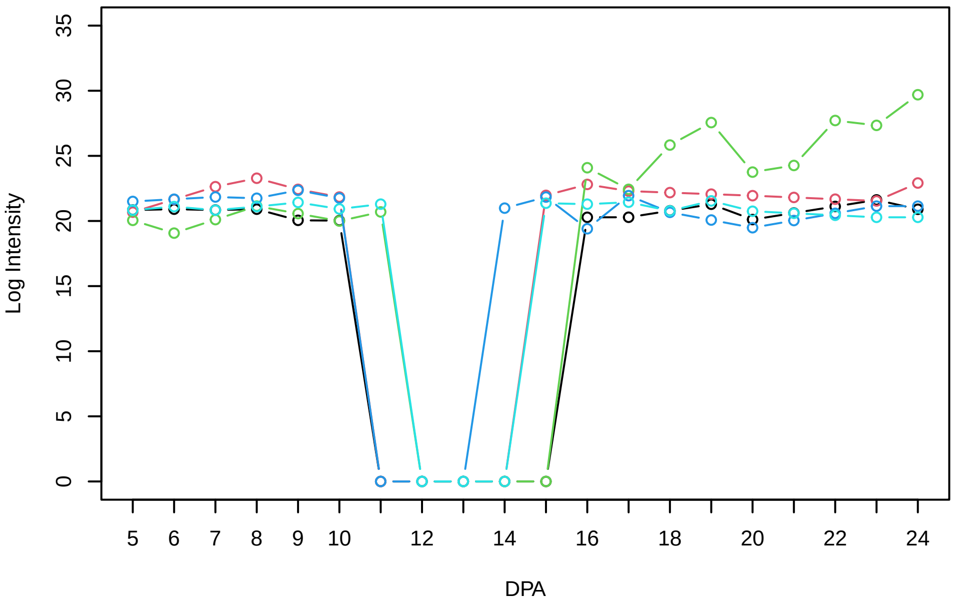


Histogram of the average abundances of non−outlier proteins

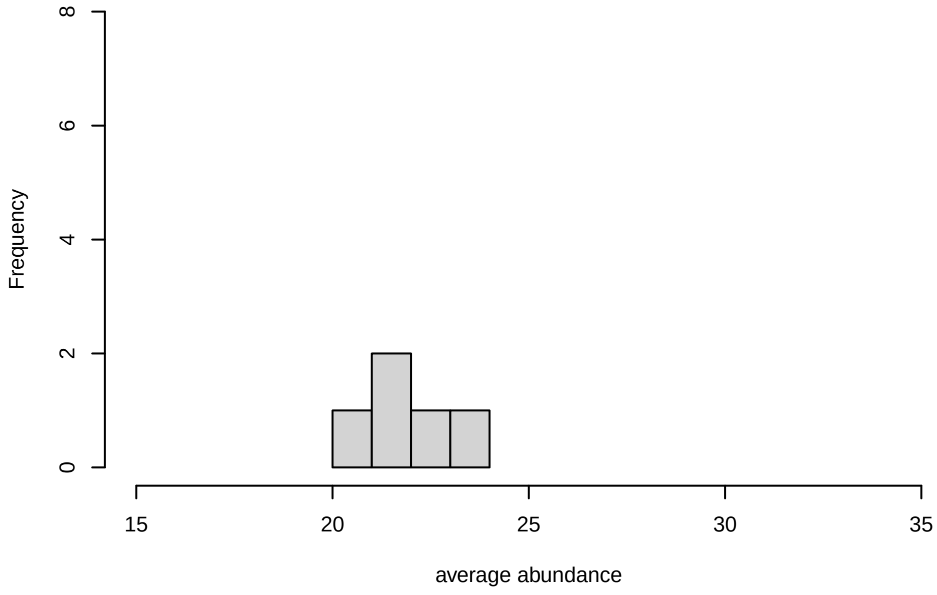


V16, after remove unreliable

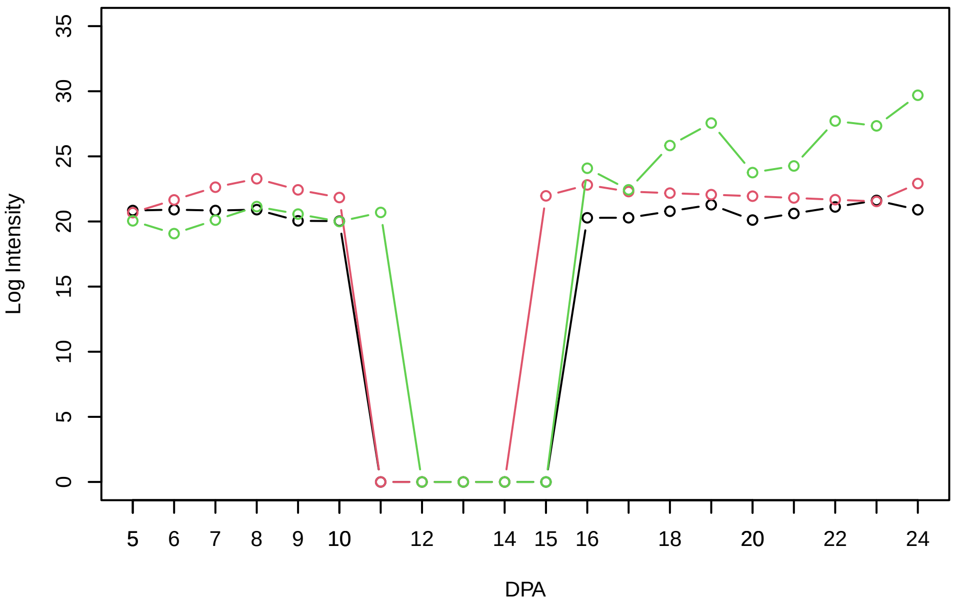


Histogram of the average abundances of reliable proteins

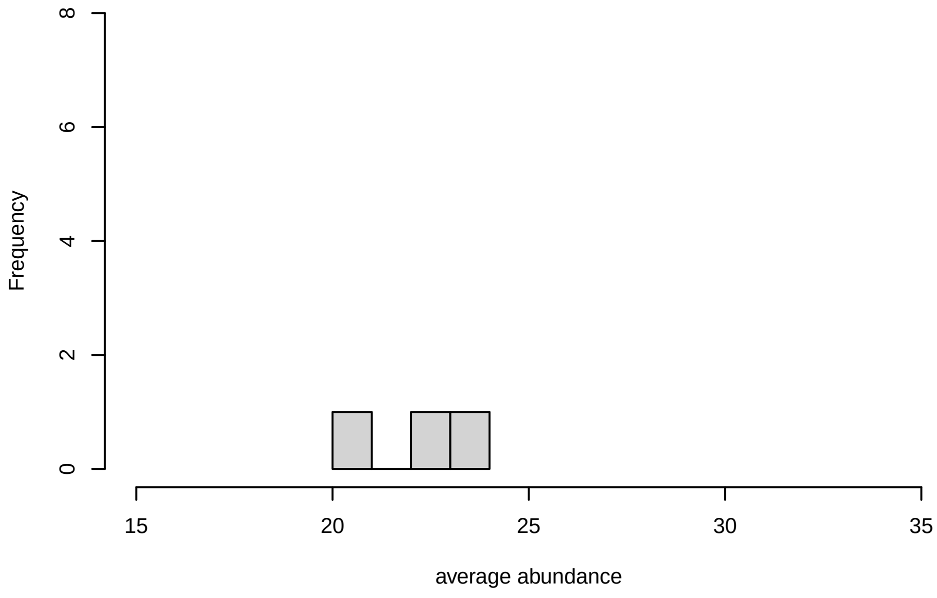


V16, after remove low abundance

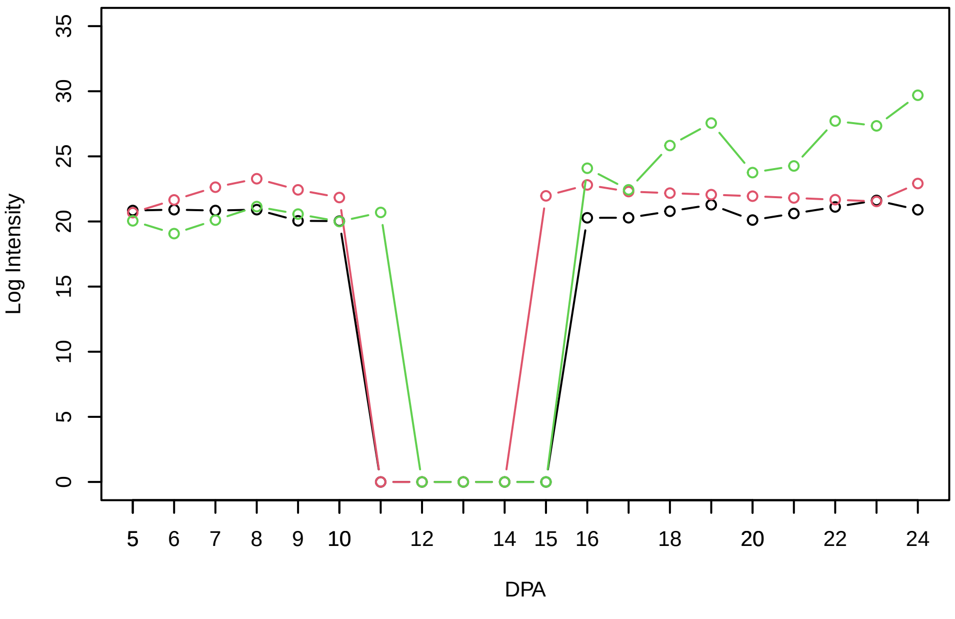


Histogram of the average abundances of high abundance proteins

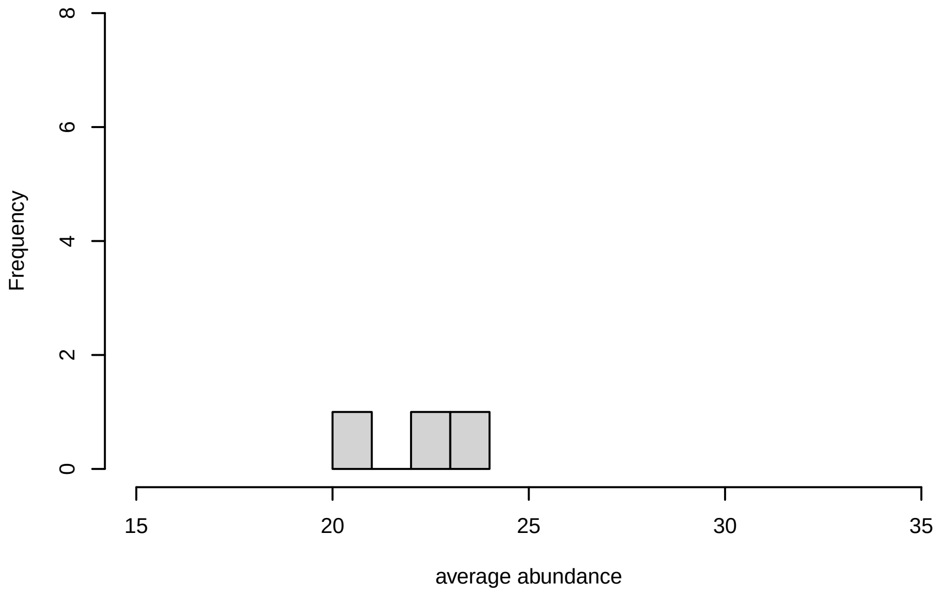

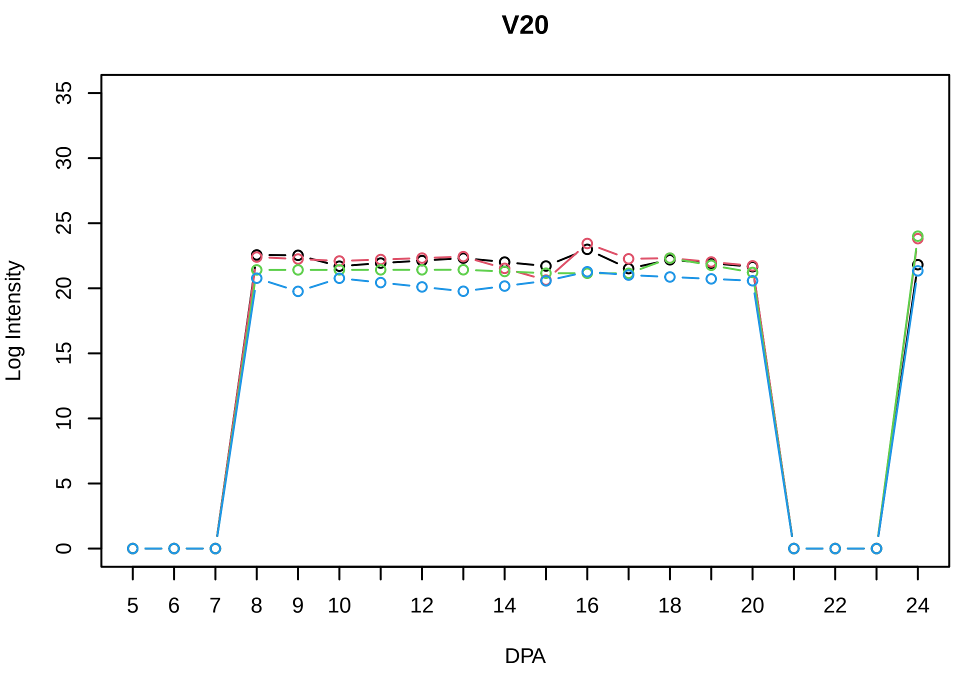
V20
Log Intensity
DPA

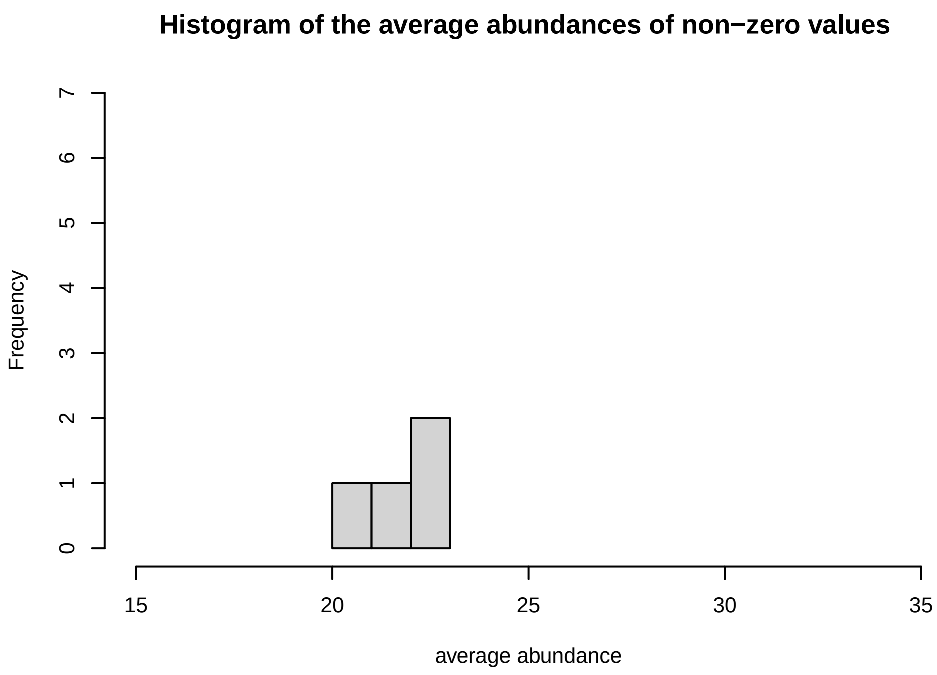
Histogram of the average abundances of non−zero values
Frequency
average abundance

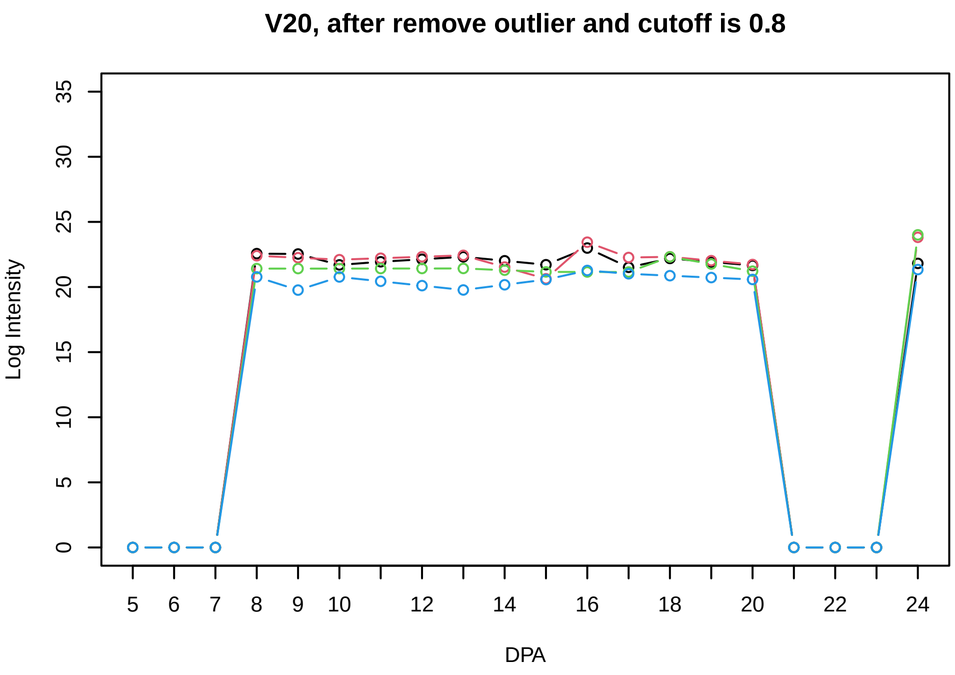
V20, after remove outlier and cutoff is 0.8
Log Intensity
DPA

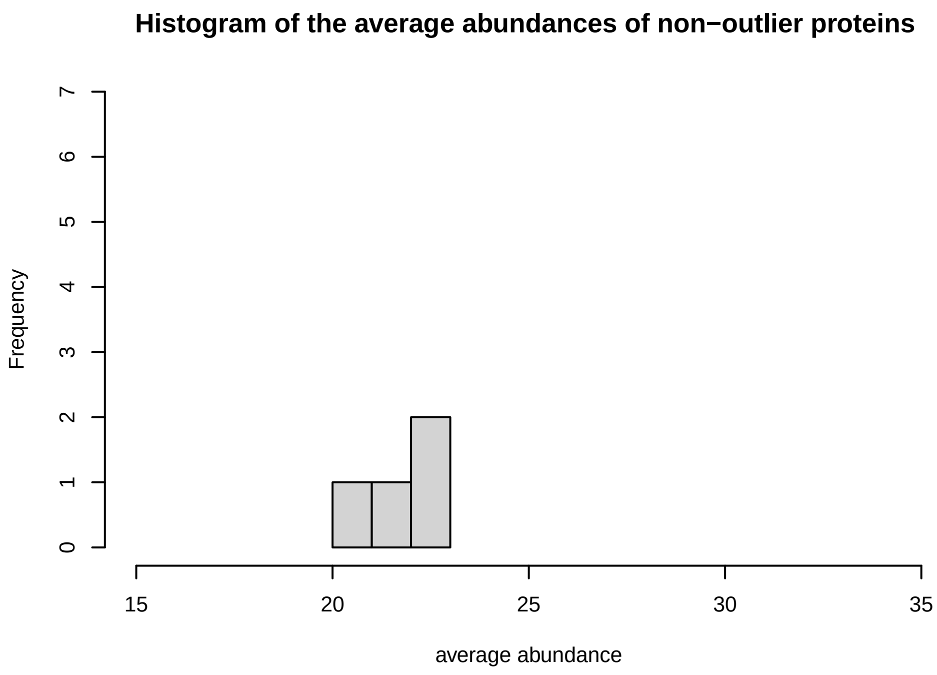
Histogram of the average abundances of non−outlier proteins
Frequency
average abundance

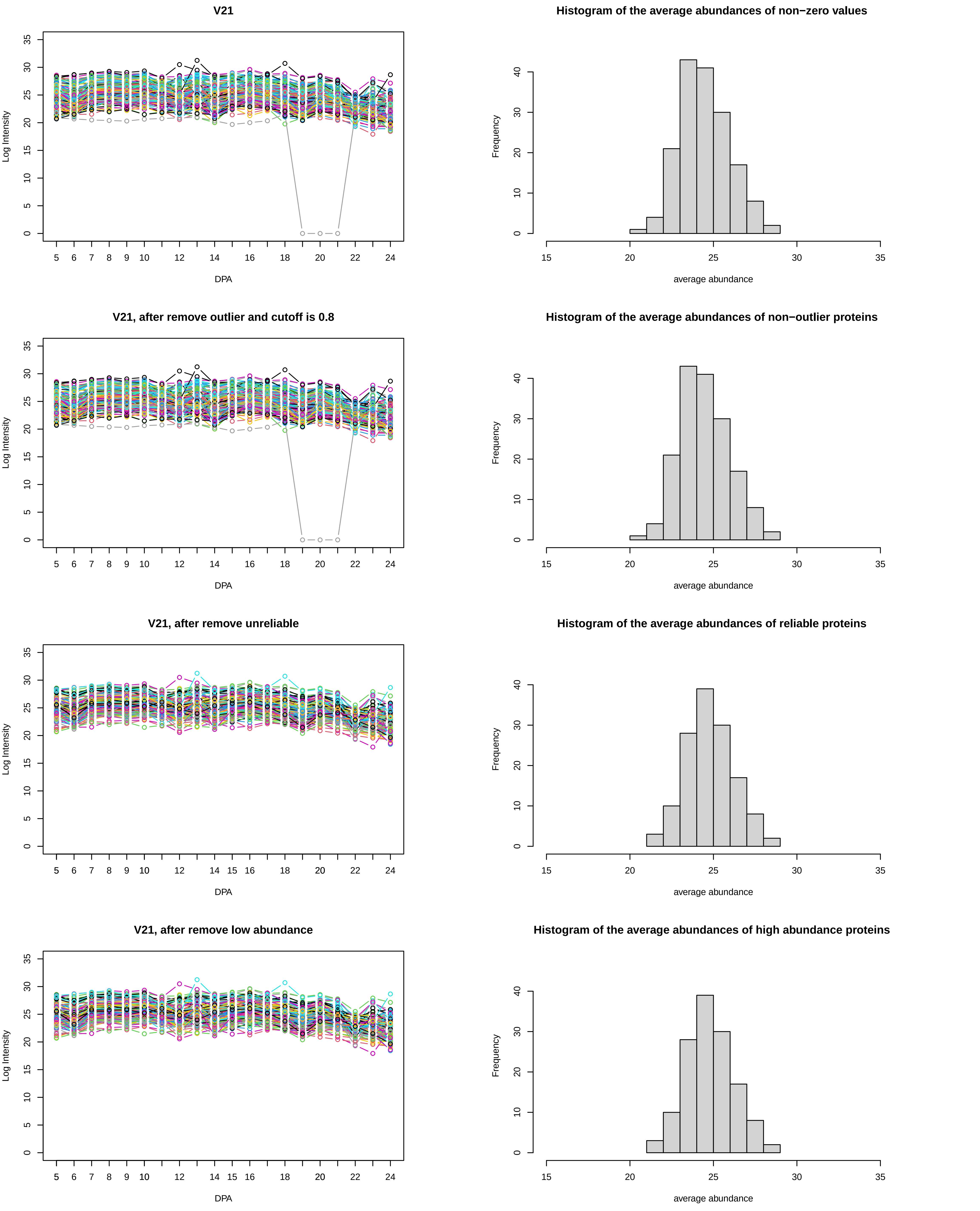
V21
Log Intensity
DPA
Histogram of the average abundances of non−zero values
Frequency
average abundance
V21, after remove outlier and cutoff is 0.8
Log Intensity
DPA
Histogram of the average abundances of non−outlier proteins
Frequency
average abundance
V21, after remove unreliable
Log Intensity
DPA
Histogram of the average abundances of reliable proteins
Frequency
average abundance
V21, after remove low abundance
Log Intensity
DPA
Histogram of the average abundances of high abundance proteins
Frequency
average abundance

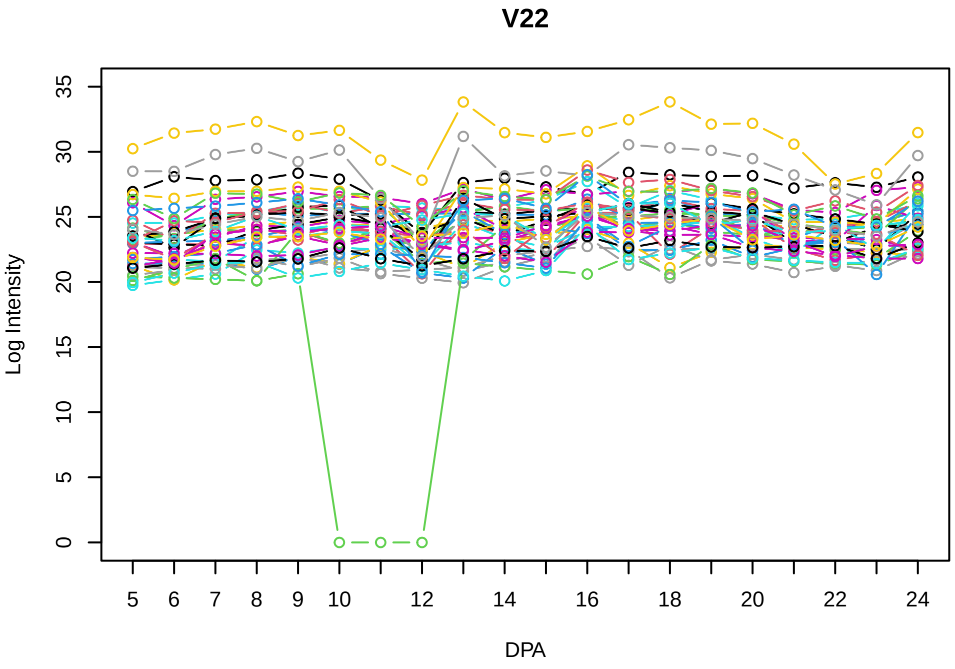
V22
Log Intensity
DPA
5 6 7 8 9 10 12 14 16 18 20 22 24
0 5 10 15 20 25 30 35

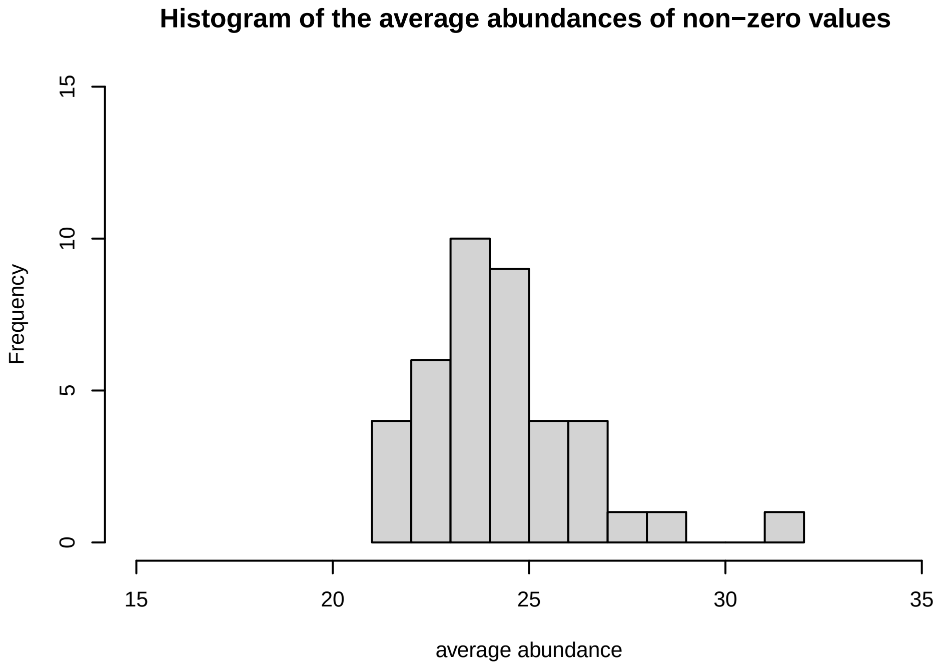
Histogram of the average abundances of non−zero values
Frequency
average abundance
15 20 25 30 35
0 5 10 15

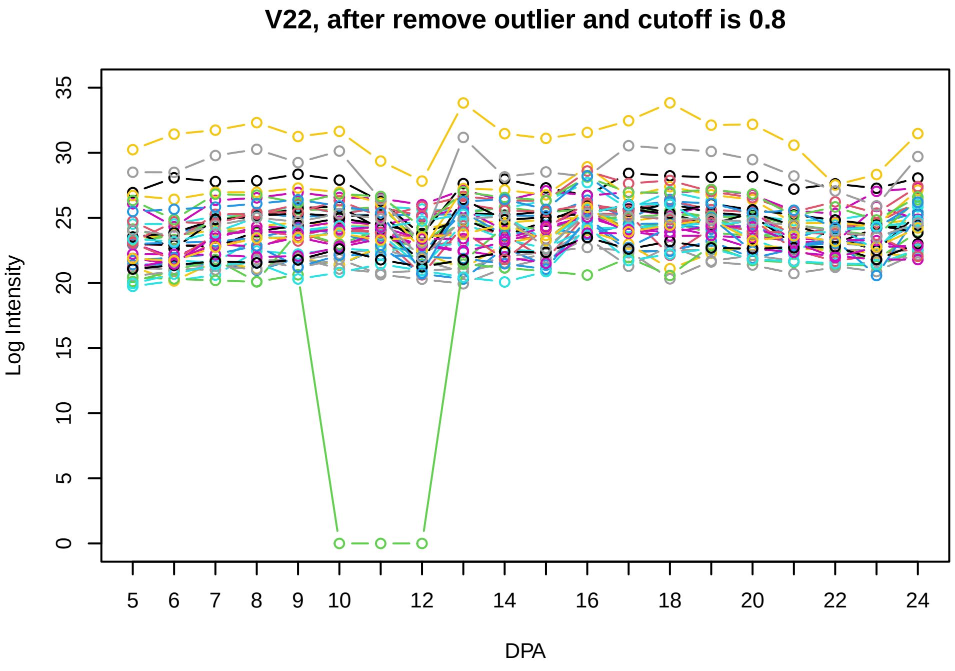
V22, after remove outlier and cutoff is 0.8
Log Intensity
DPA
5 6 7 8 9 10 12 14 16 18 20 22 24
0 5 10 15 20 25 30 35

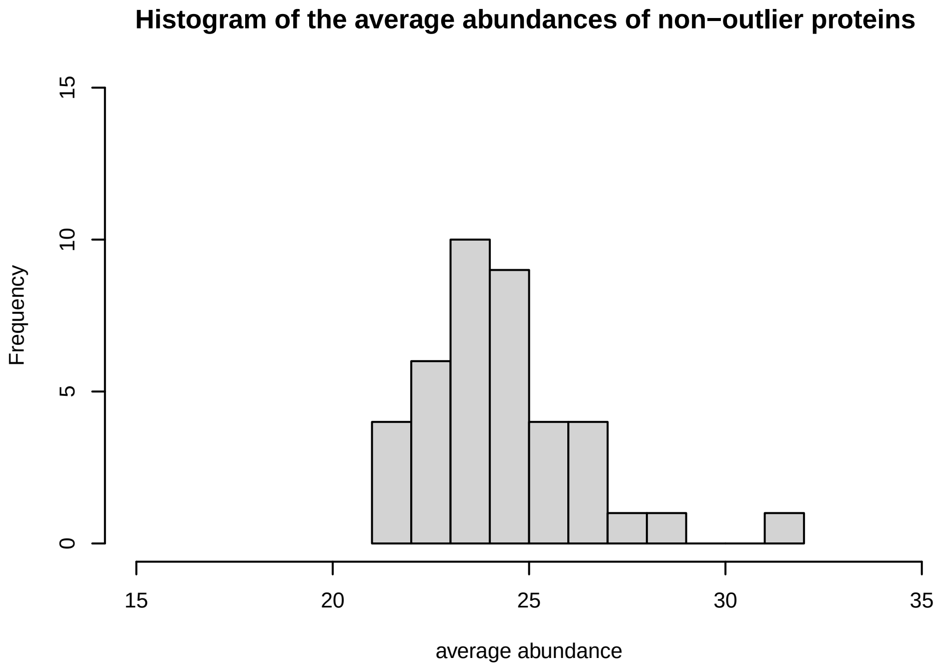
Histogram of the average abundances of non−outlier proteins
Frequency
average abundance
15 20 25 30 35
0 5 10 15

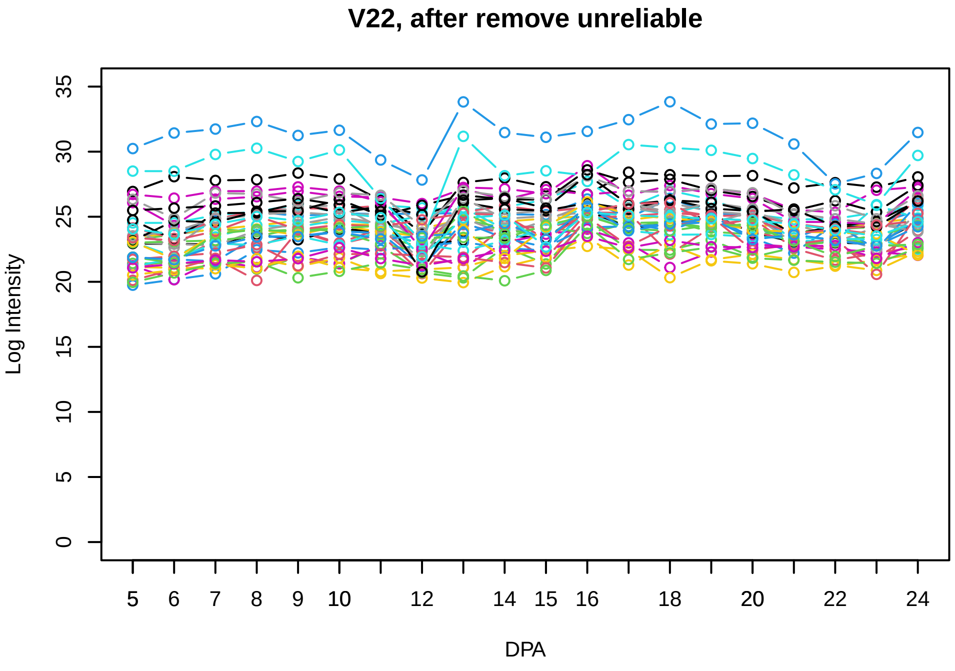
V22, after remove unreliable
Log Intensity
DPA
5 6 7 8 9 10 12 14 15 16 18 20 22 24
0 5 10 15 20 25 30 35

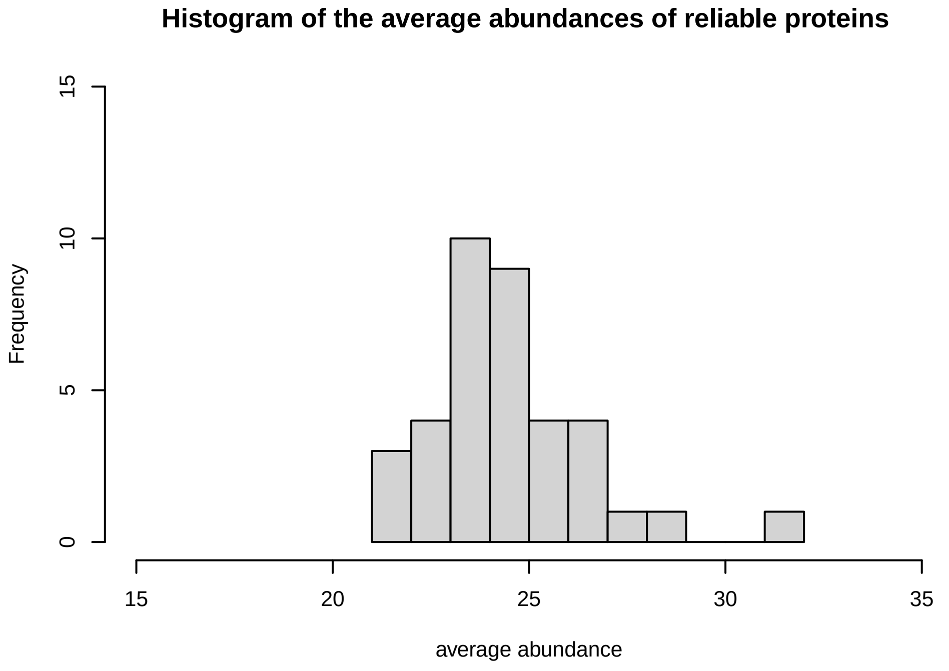
Histogram of the average abundances of reliable proteins
Frequency
average abundance
15 20 25 30 35
0 5 10 15

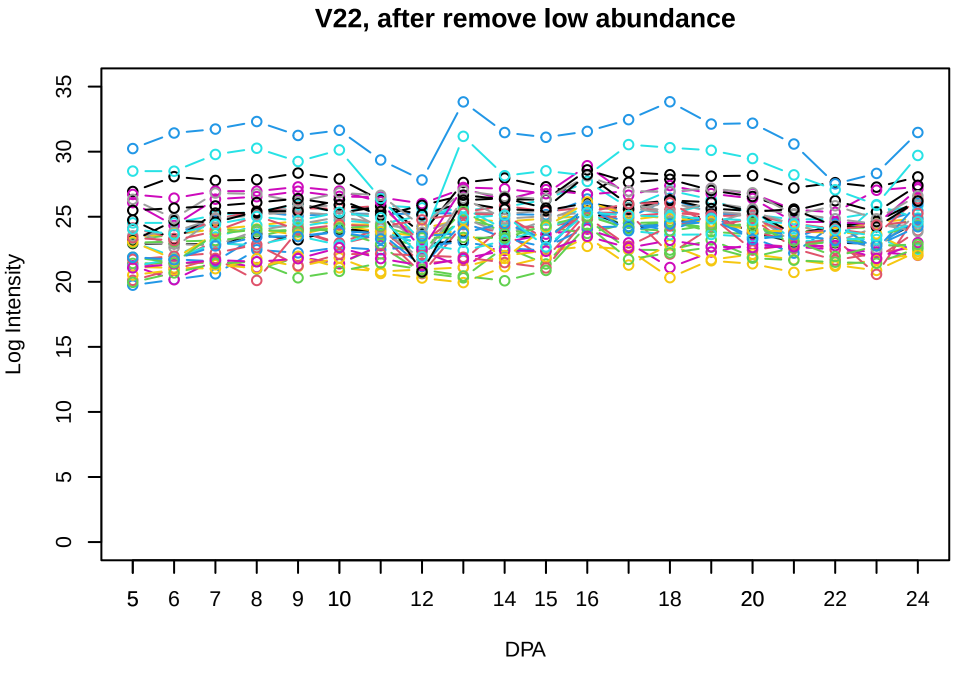
V22, after remove low abundance
Log Intensity
DPA
5 6 7 8 9 10 12 14 15 16 18 20 22 24
0 5 10 15 20 25 30 35

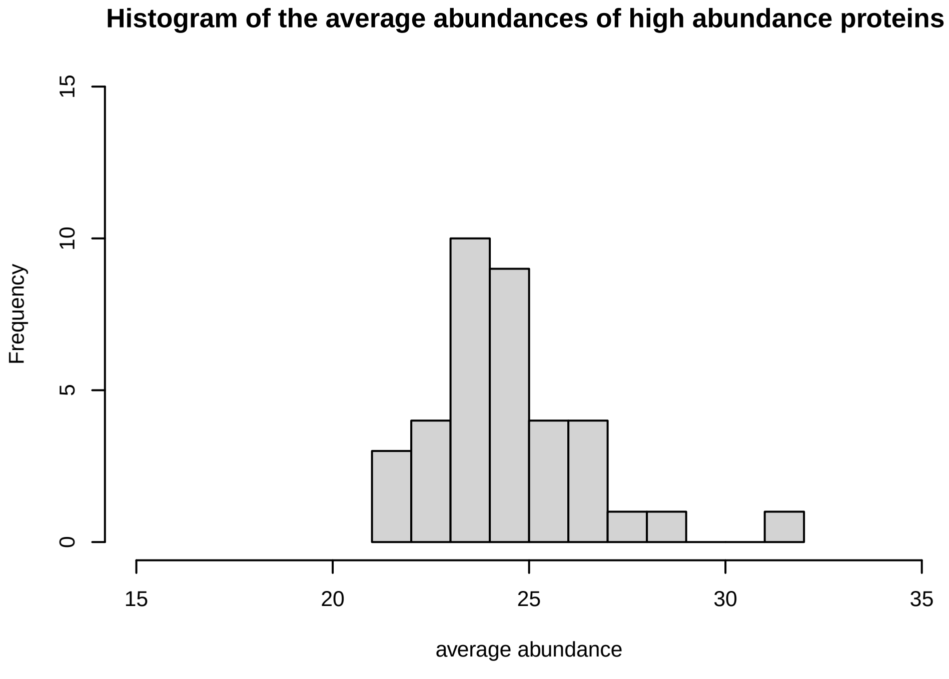
Histogram of the average abundances of high abundance proteins
Frequency
average abundance
15 20 25 30 35
0 5 10 15

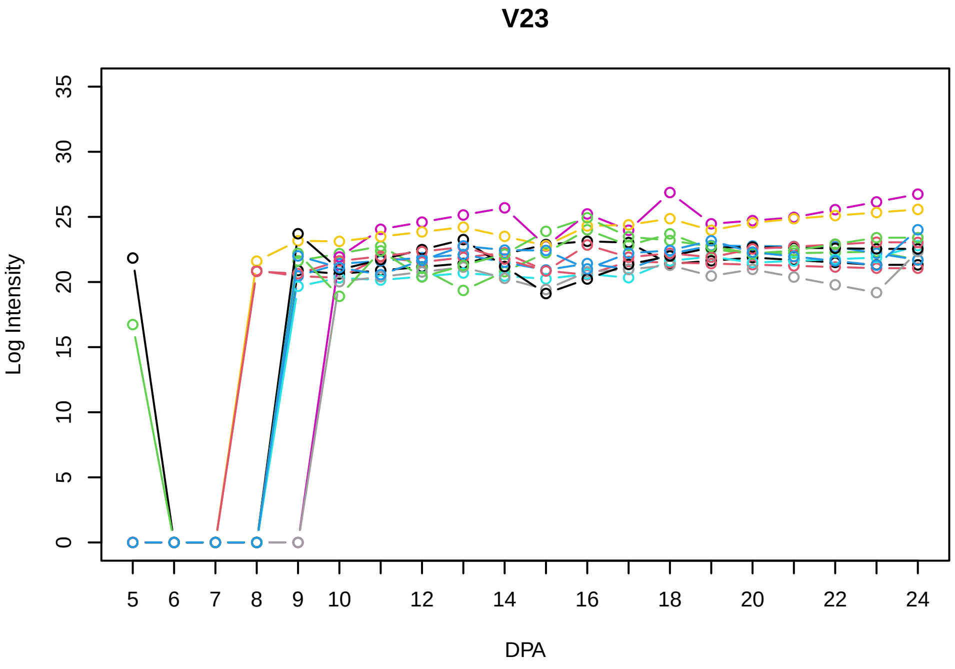
V23
Log Intensity
DPA

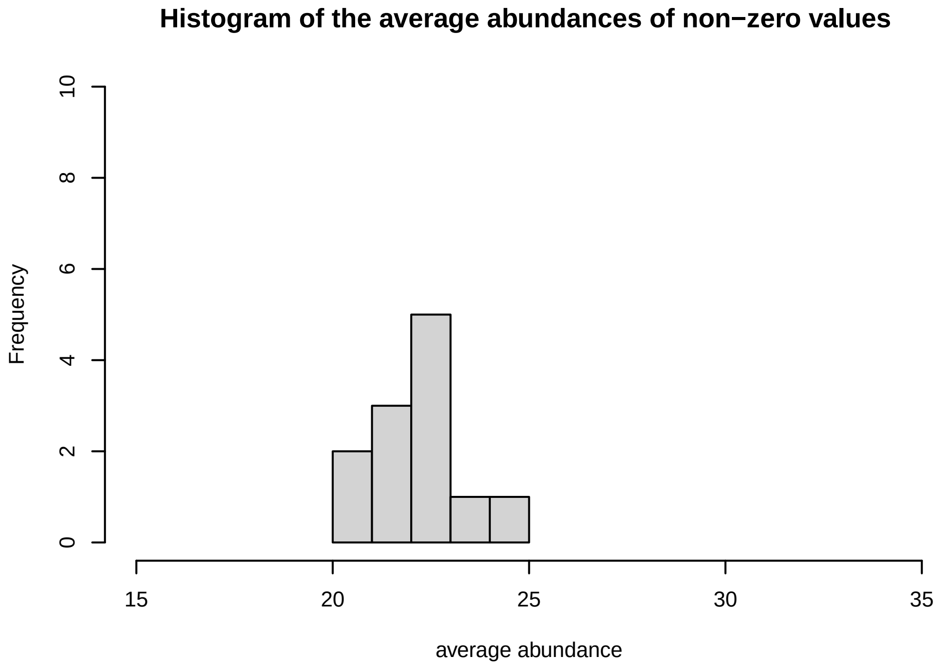
Histogram of the average abundances of non−zero values
Frequency
average abundance

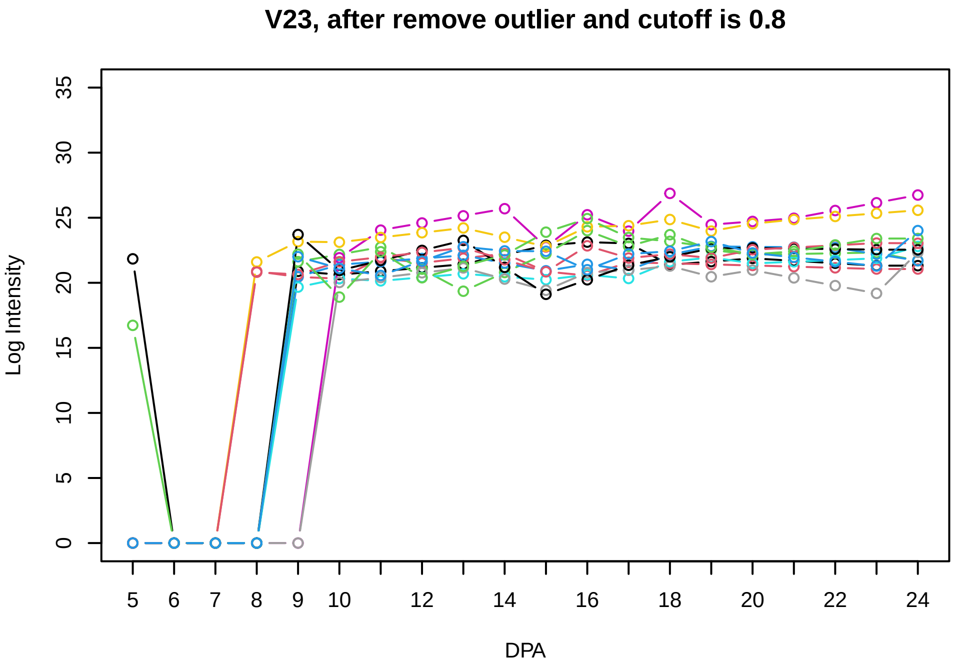
V23, after remove outlier and cutoff is 0.8
Log Intensity
DPA

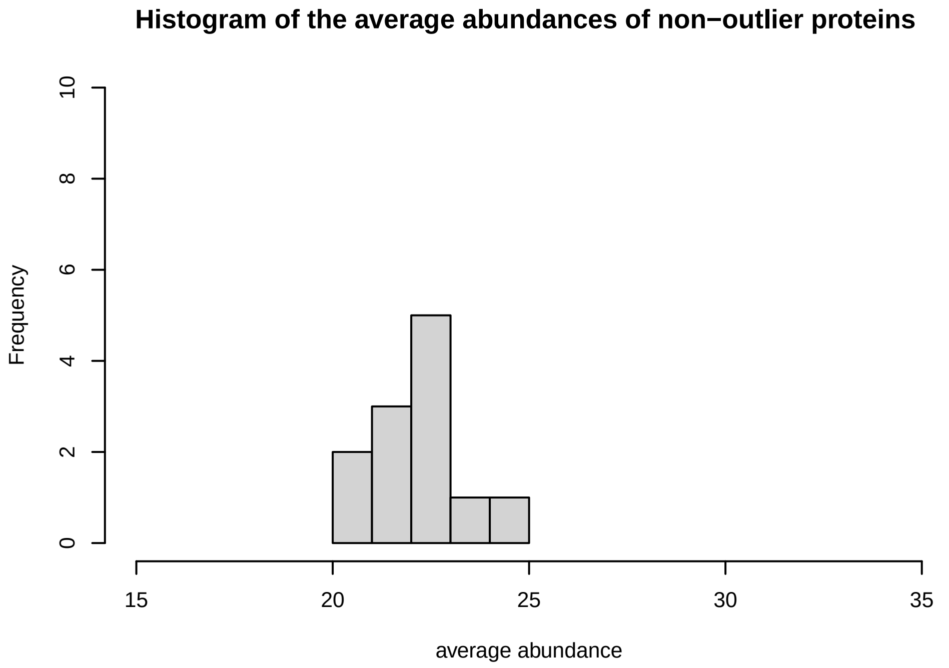
Histogram of the average abundances of non−outlier proteins
Frequency
average abundance

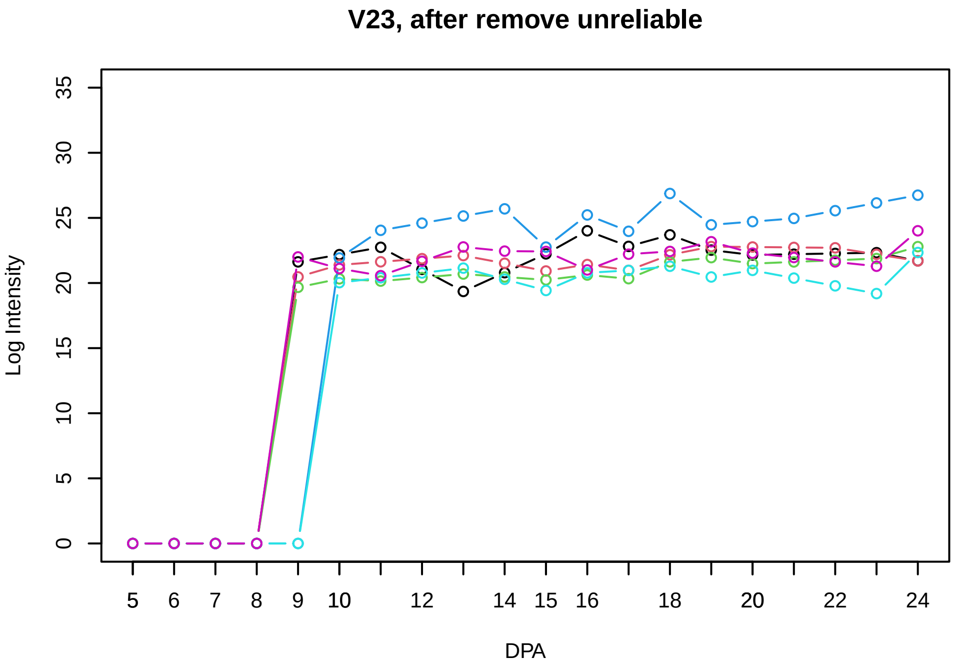
V23, after remove unreliable
Log Intensity
DPA

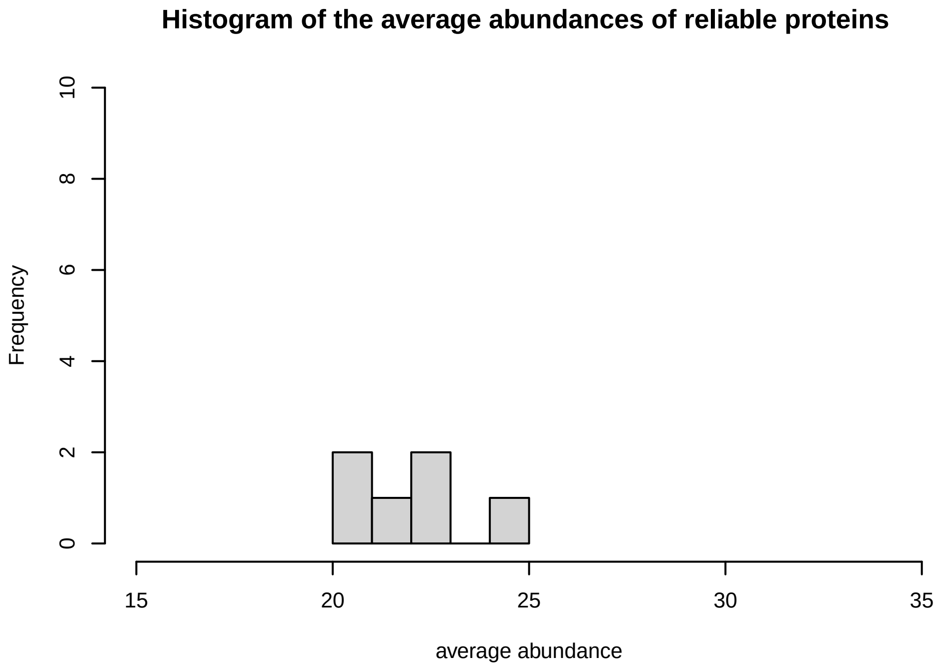
Histogram of the average abundances of reliable proteins
Frequency
average abundance

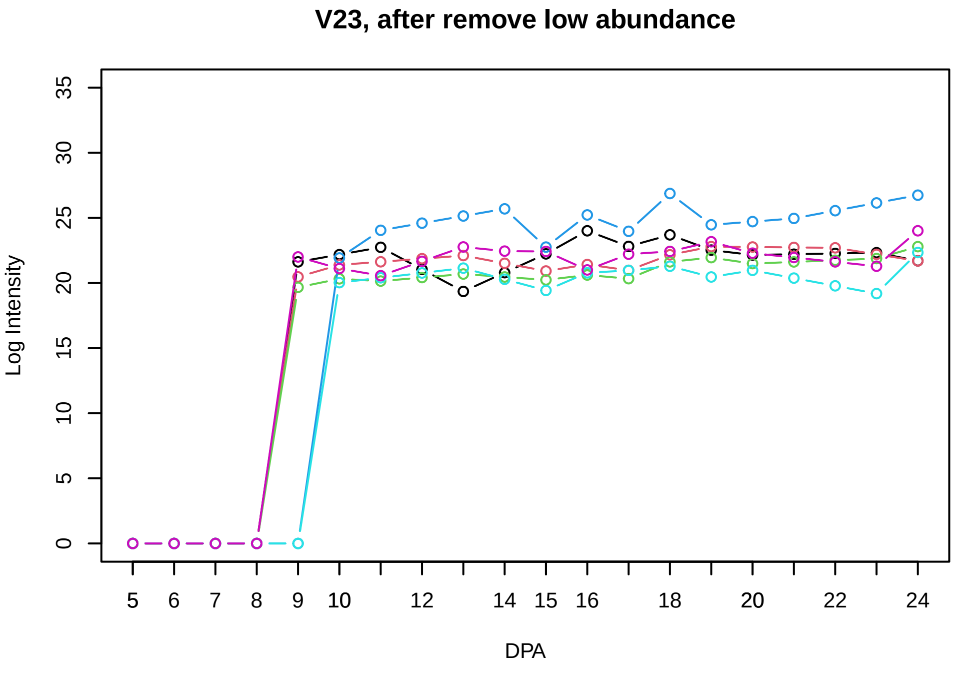
V23, after remove low abundance
Log Intensity
DPA

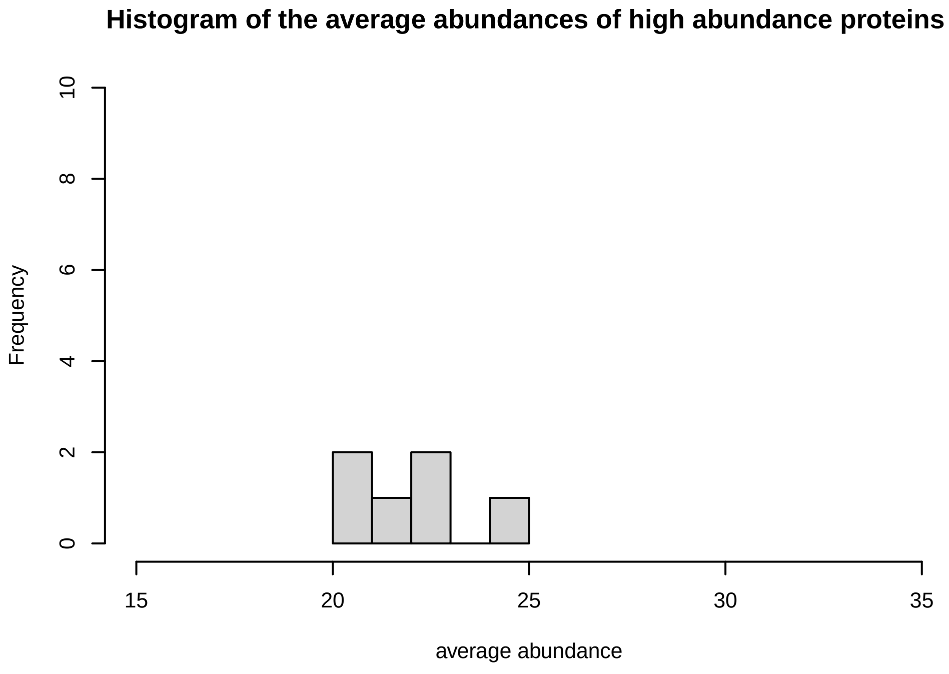
Histogram of the average abundances of high abundance proteins
Frequency
average abundance

**V24**

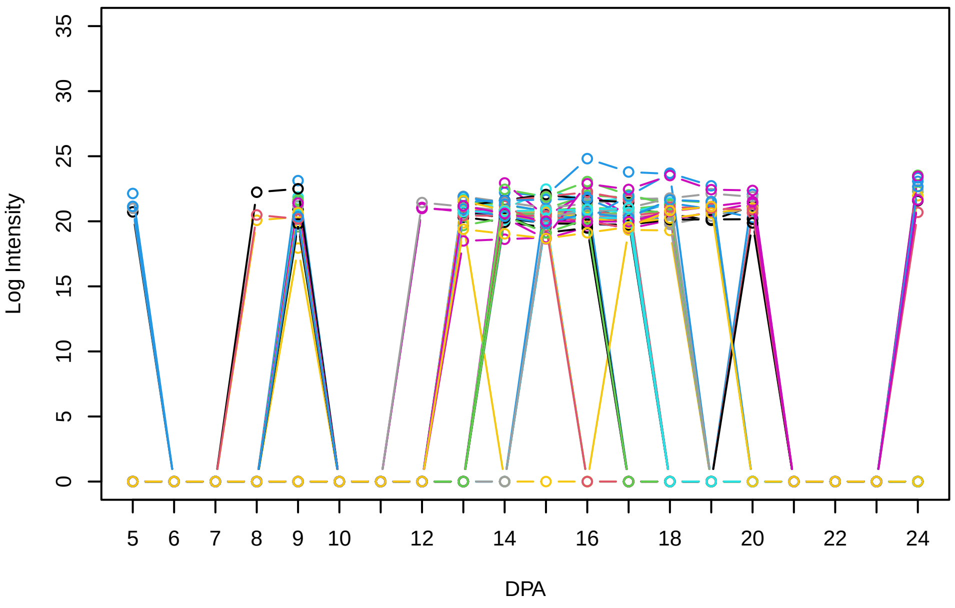


**Histogram of the average abundances of non−zero values**

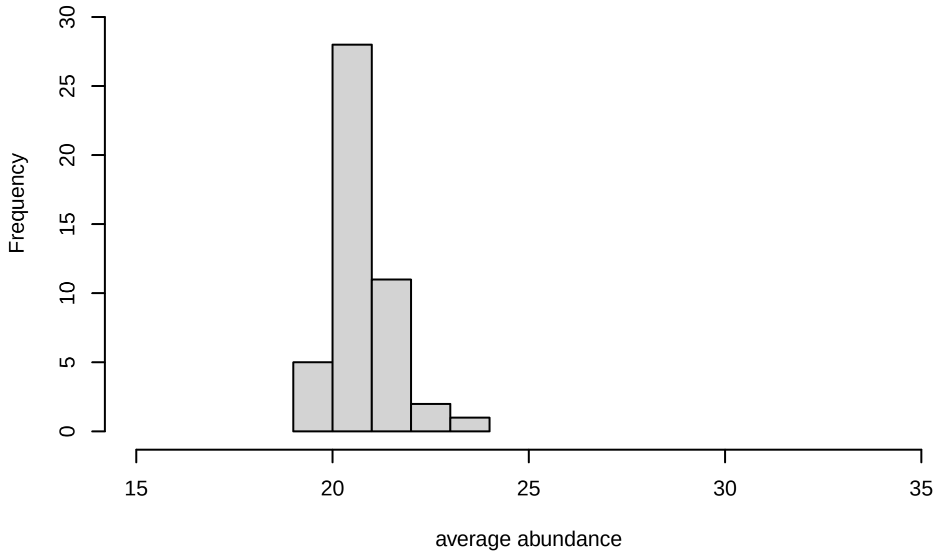


**V24, after remove outlier and cutoff is 0.673441816473663**

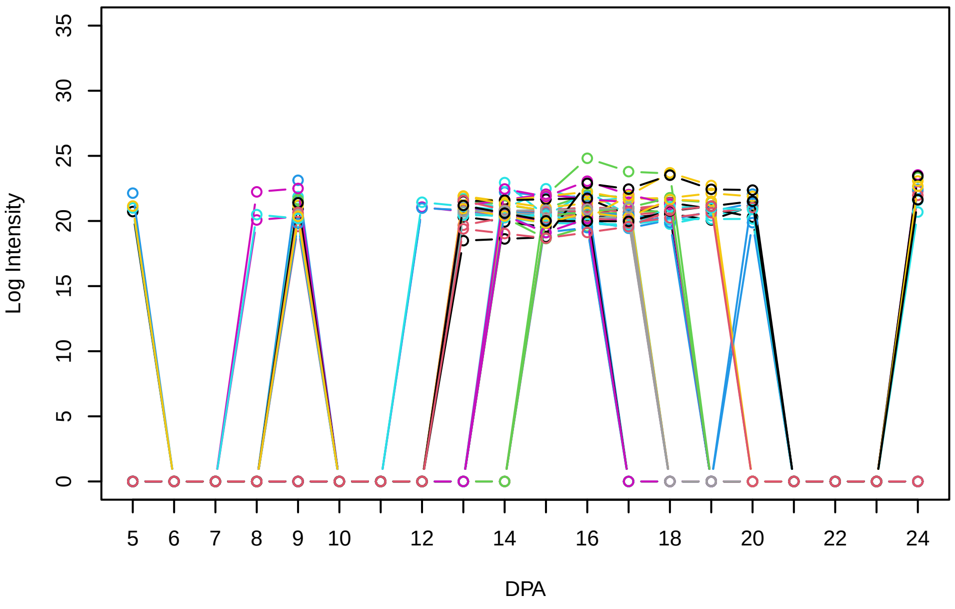


**Histogram of the average abundances of non−outlier proteins**

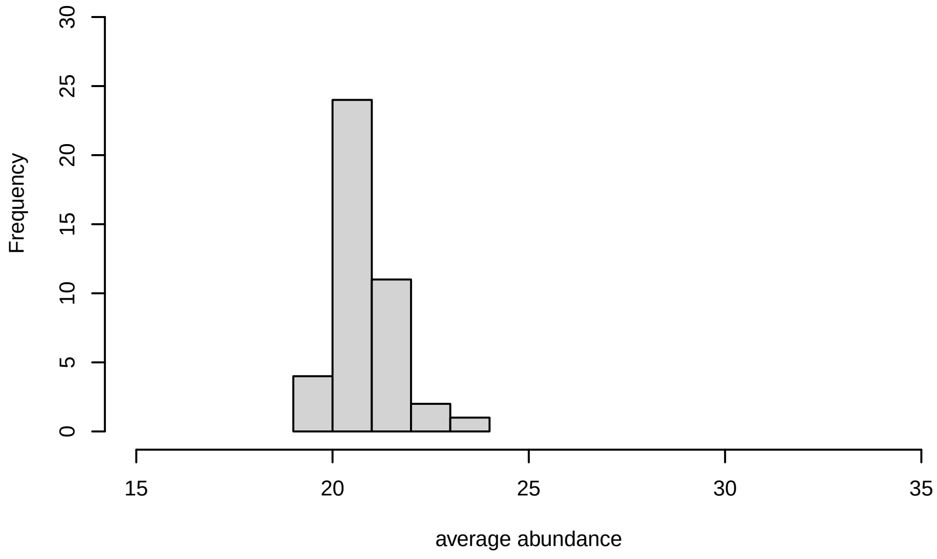


**V24, after remove unreliable**

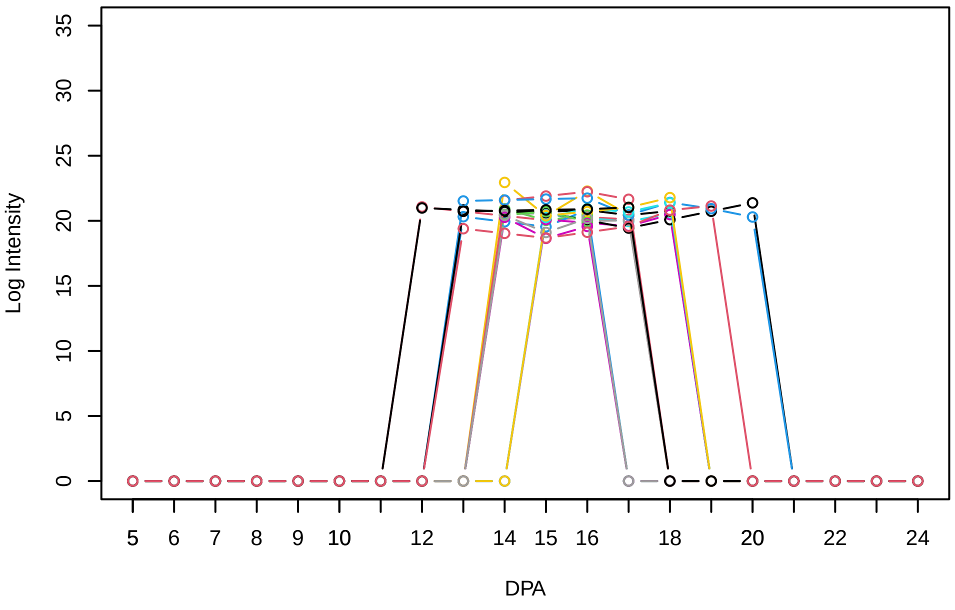


**Histogram of the average abundances of reliable proteins**

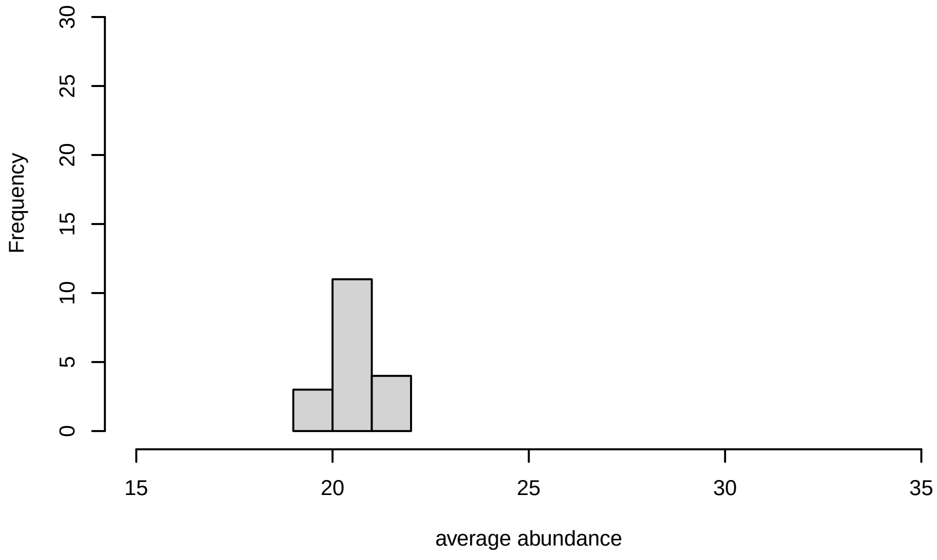


**V24, after remove low abundance**

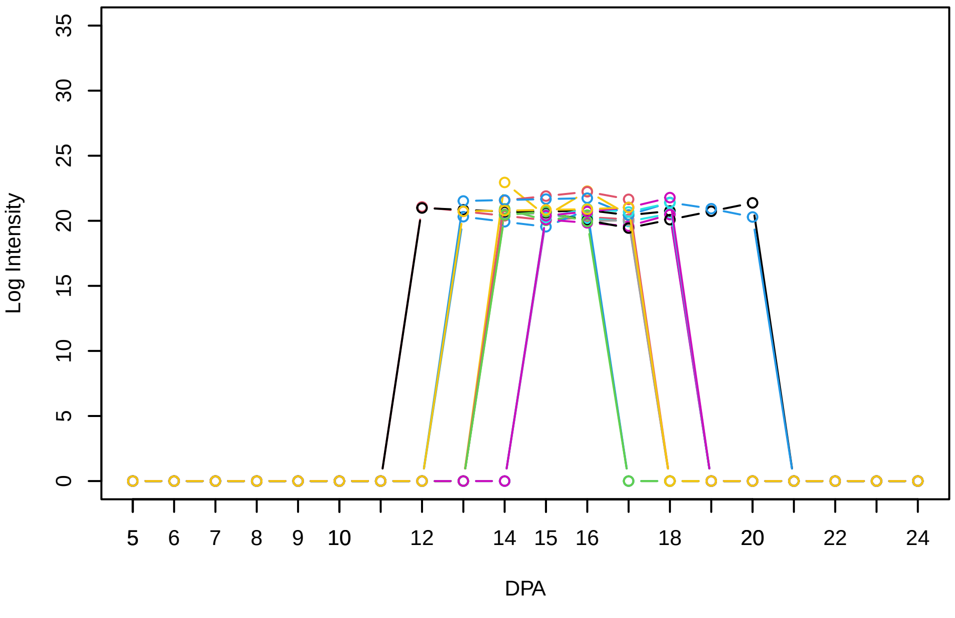


**Histogram of the average abundances of high abundance proteins**

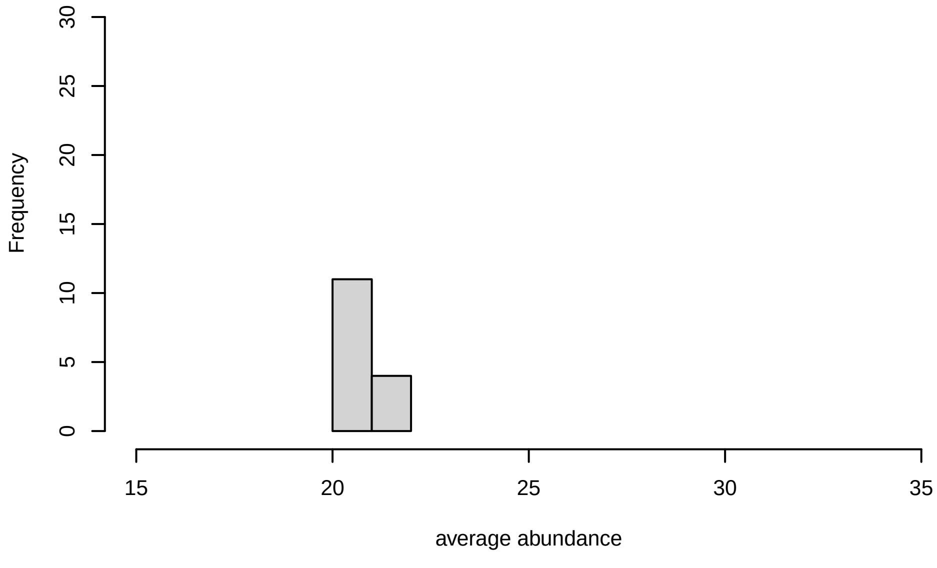

V25

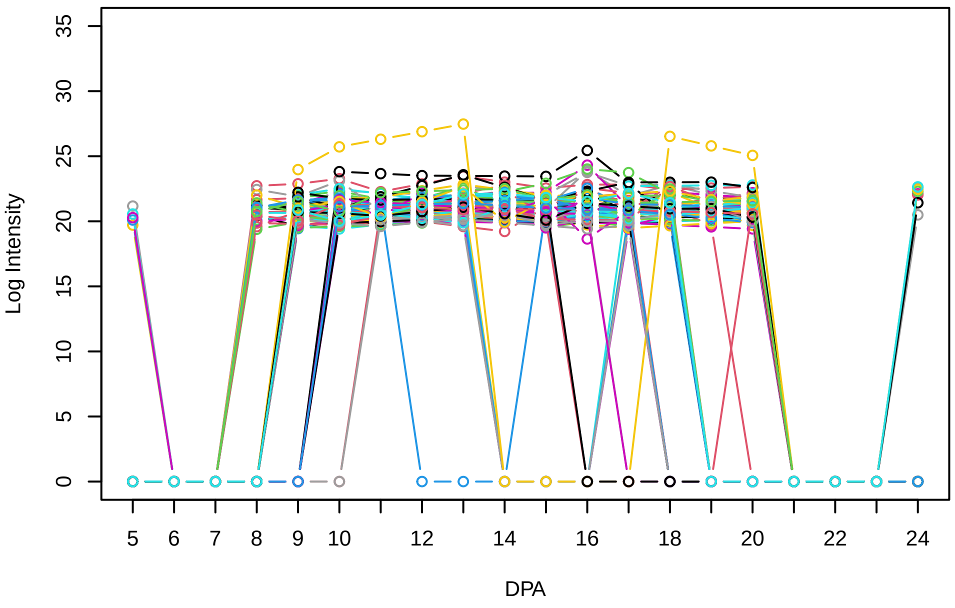


Histogram of the average abundances of non−zero values

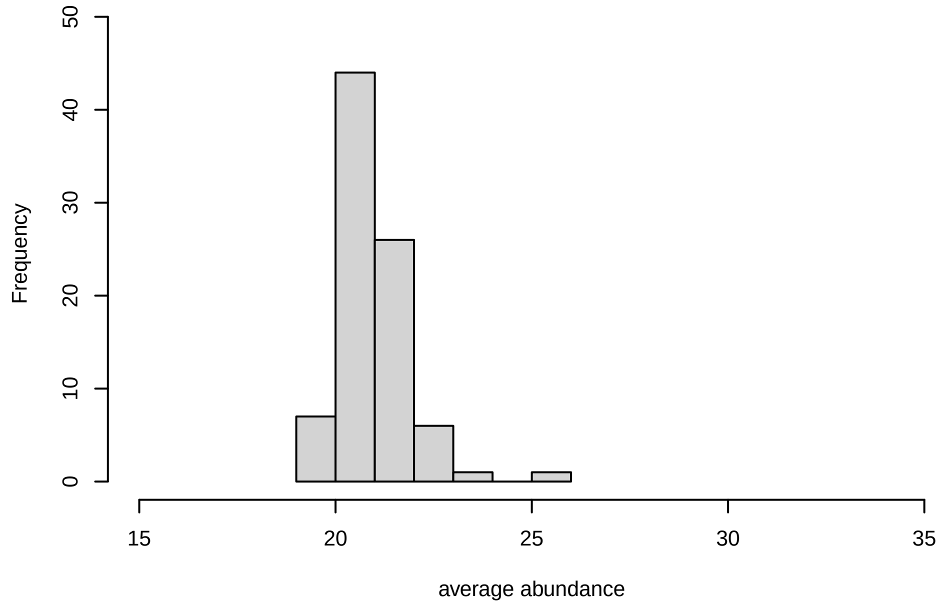


V25, after remove outlier and cutoff is 0.8

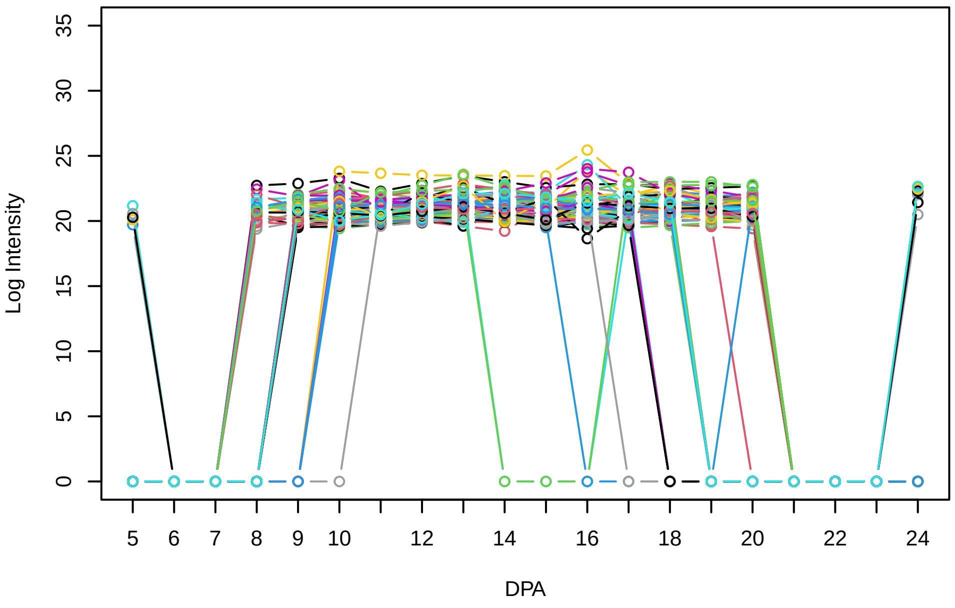


Histogram of the average abundances of non−outlier proteins

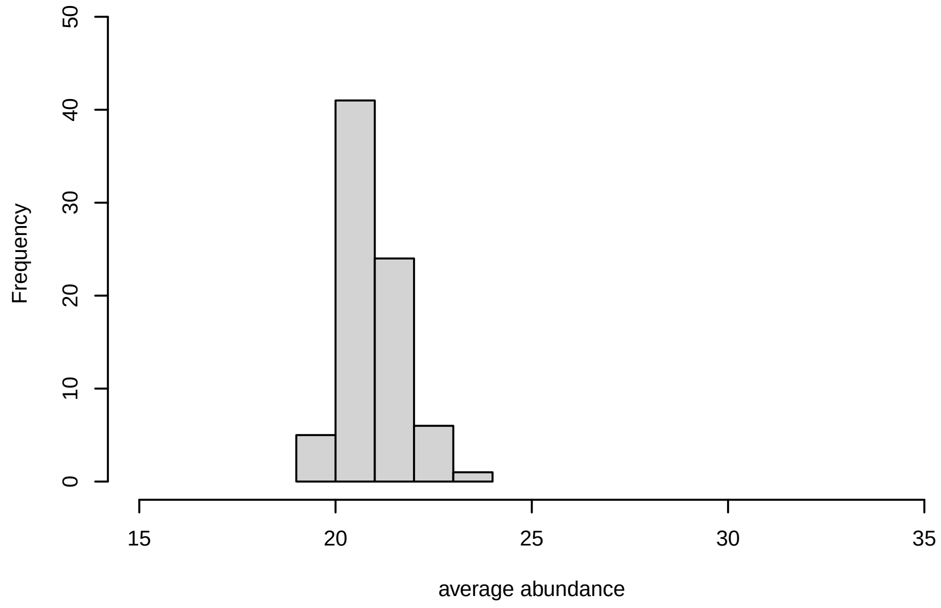


V25, after remove unreliable

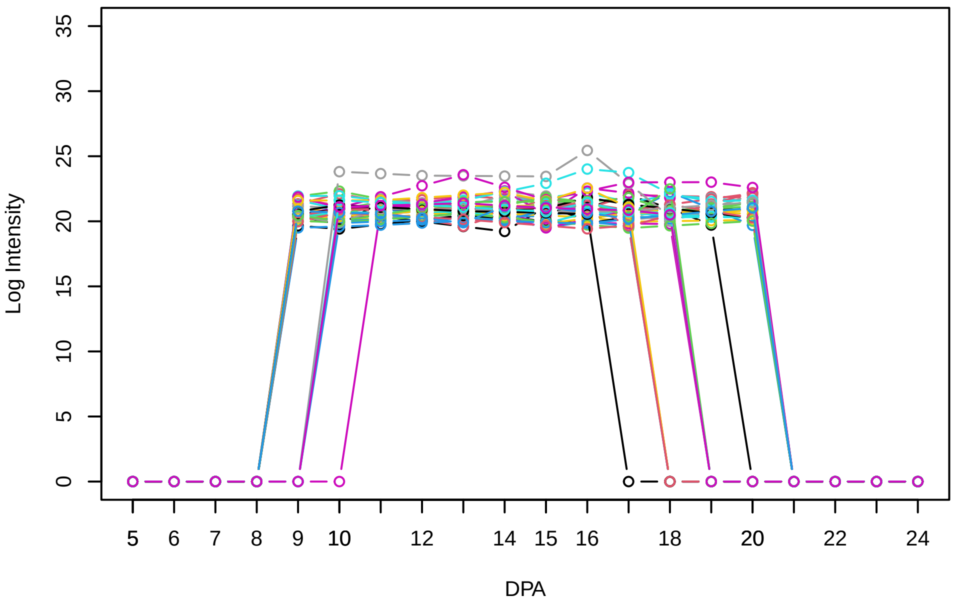


Histogram of the average abundances of reliable proteins

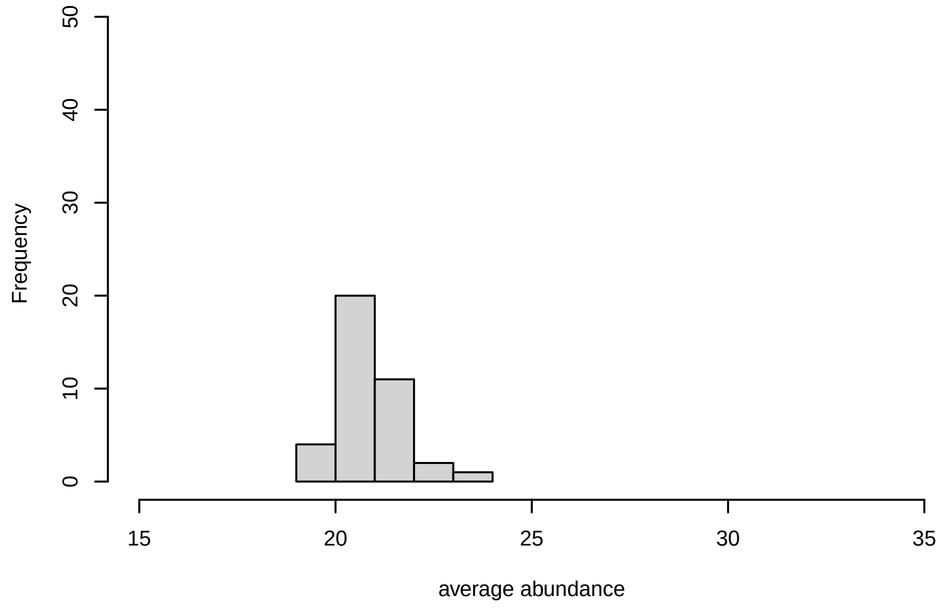


V25, after remove low abundance

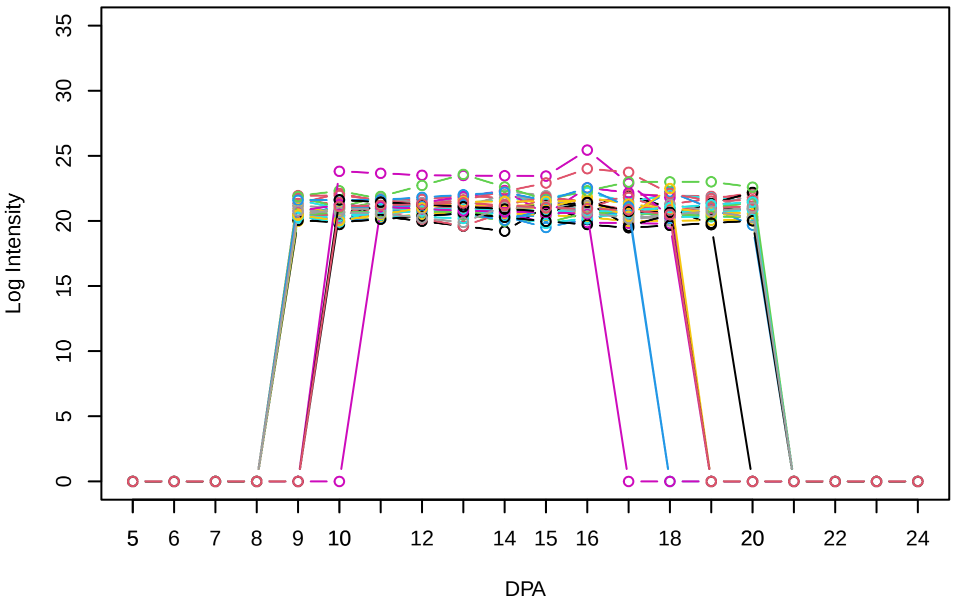


Histogram of the average abundances of high abundance proteins

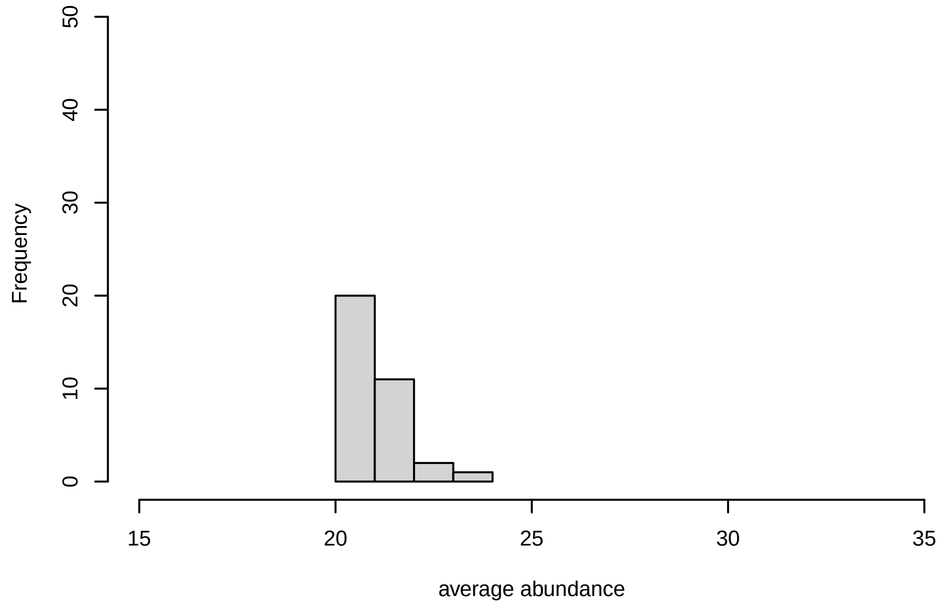

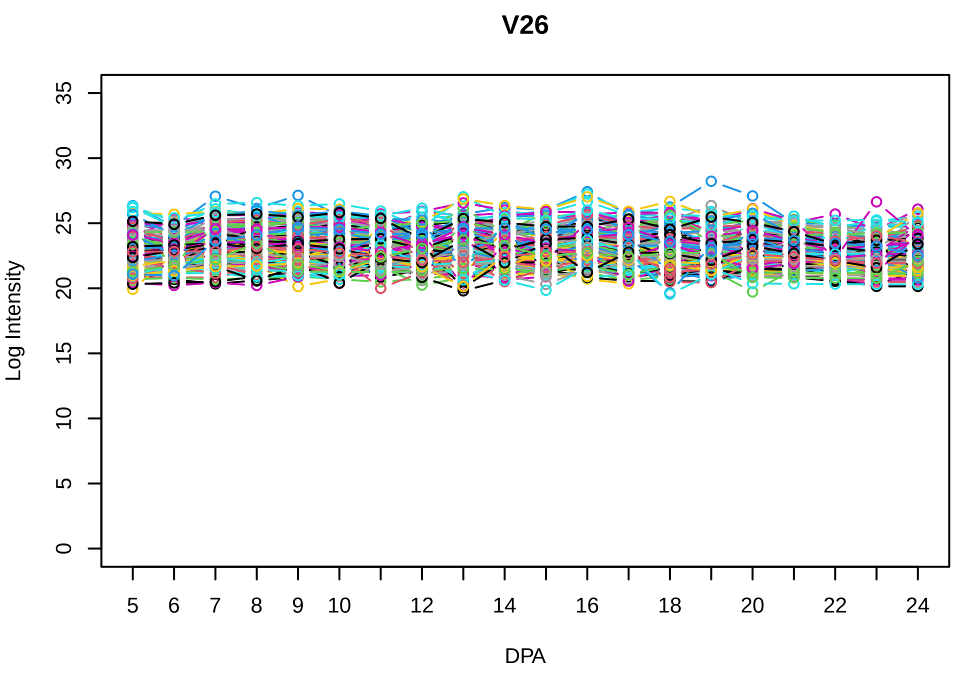
V26
Log Intensity
DPA

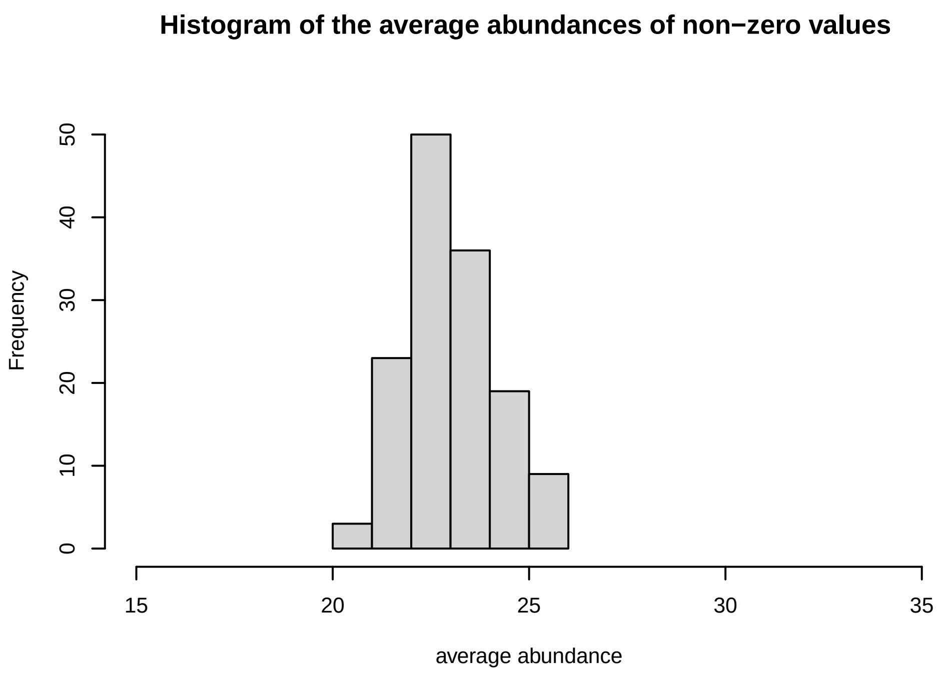
Histogram of the average abundances of non−zero values
Frequency
average abundance

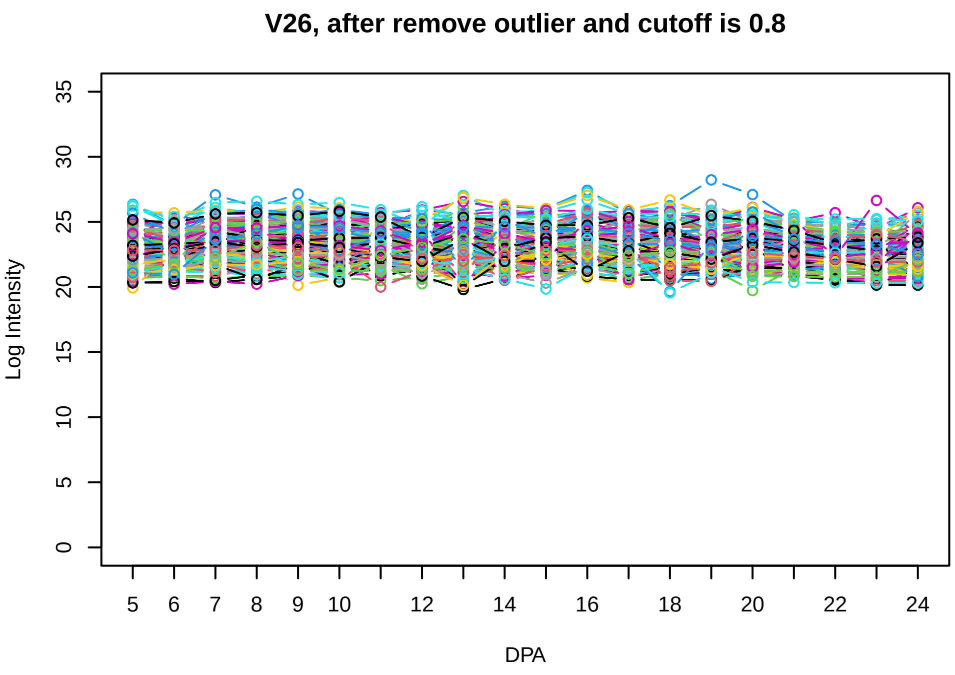
V26, after remove outlier and cutoff is 0.8
Log Intensity
DPA

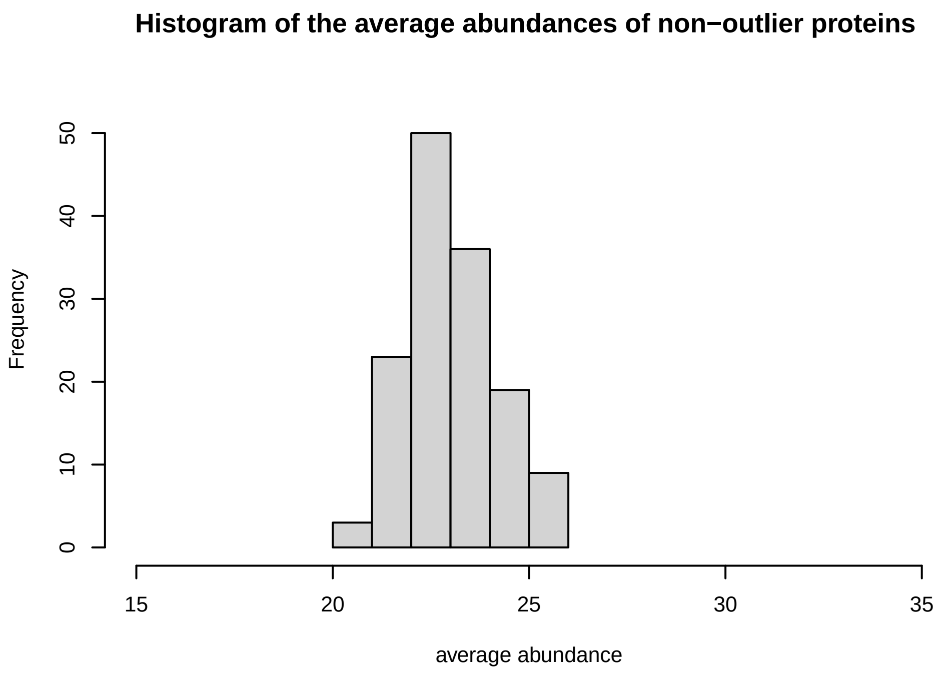
Histogram of the average abundances of non−outlier proteins
Frequency
average abundance

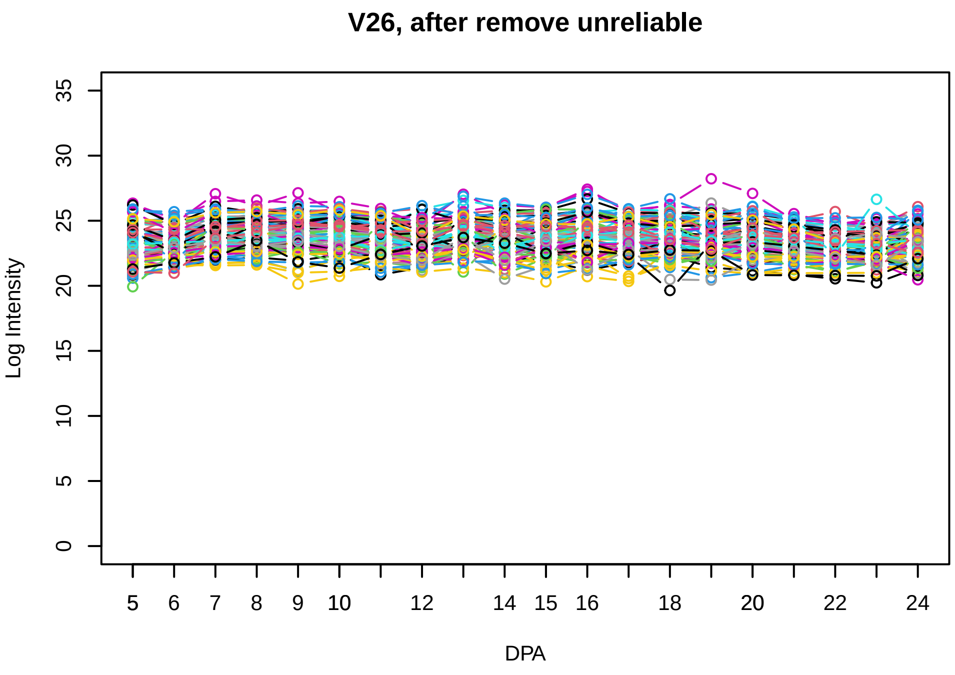
V26, after remove unreliable
Log Intensity
DPA

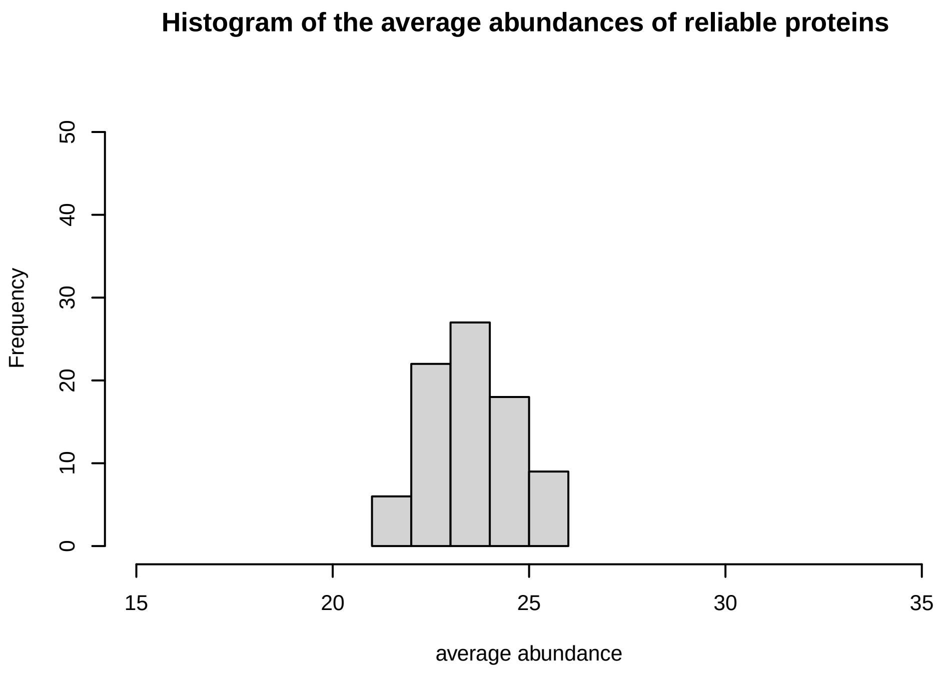
Histogram of the average abundances of reliable proteins
Frequency
average abundance

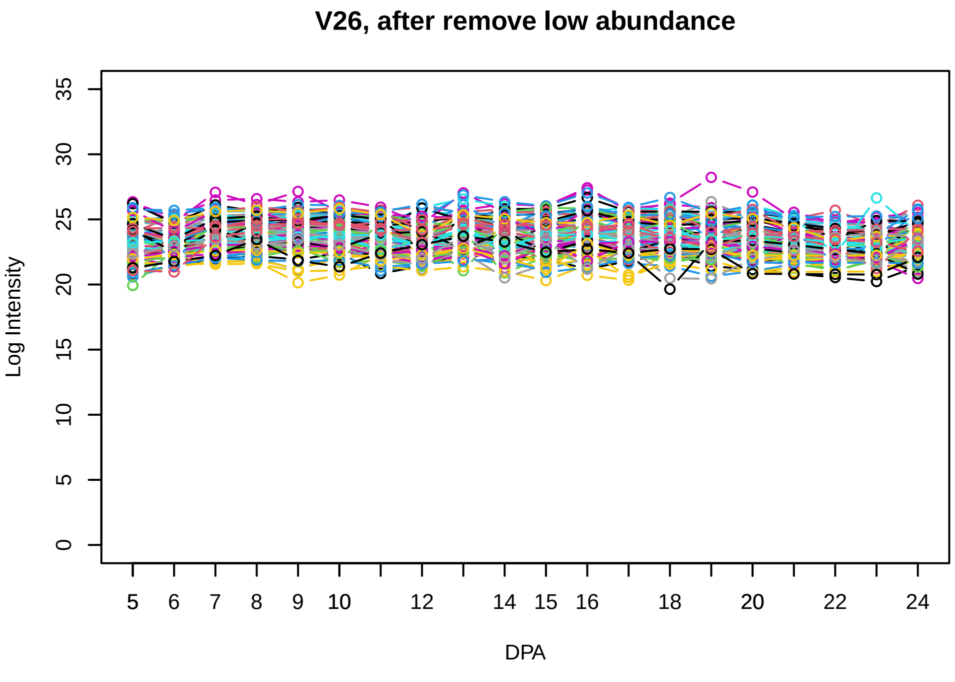
V26, after remove low abundance
Log Intensity
DPA

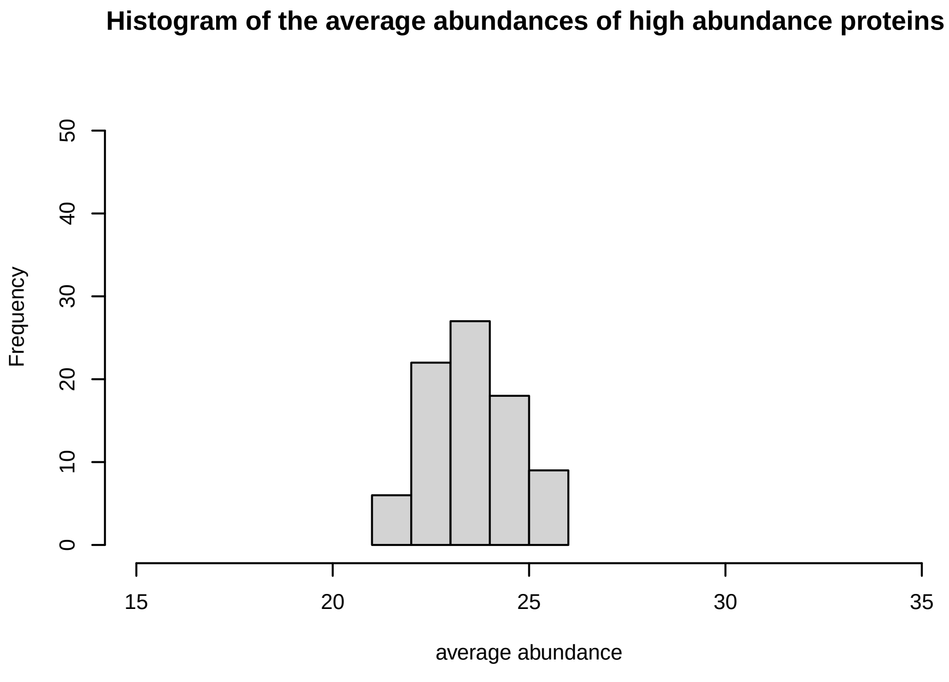
Histogram of the average abundances of high abundance proteins
Frequency
average abundance

**V27**

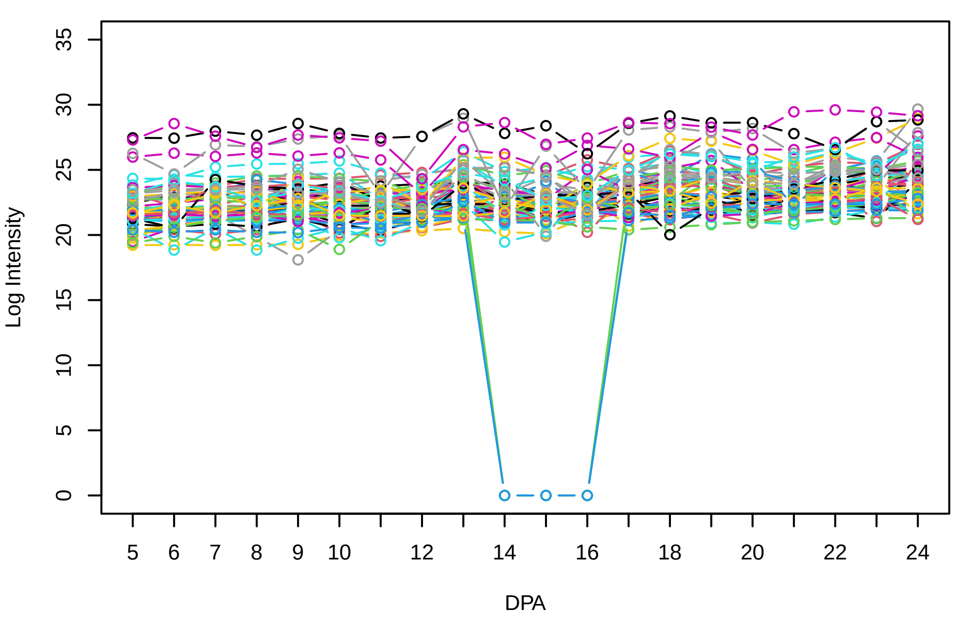


**Histogram of the average abundances of non−zero values**

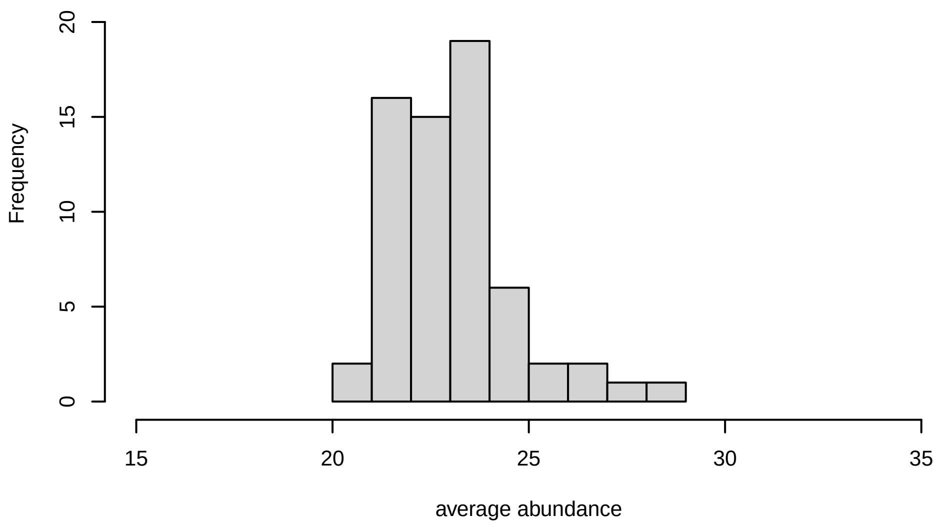


**V27, after remove outlier and cutoff is 0.8**

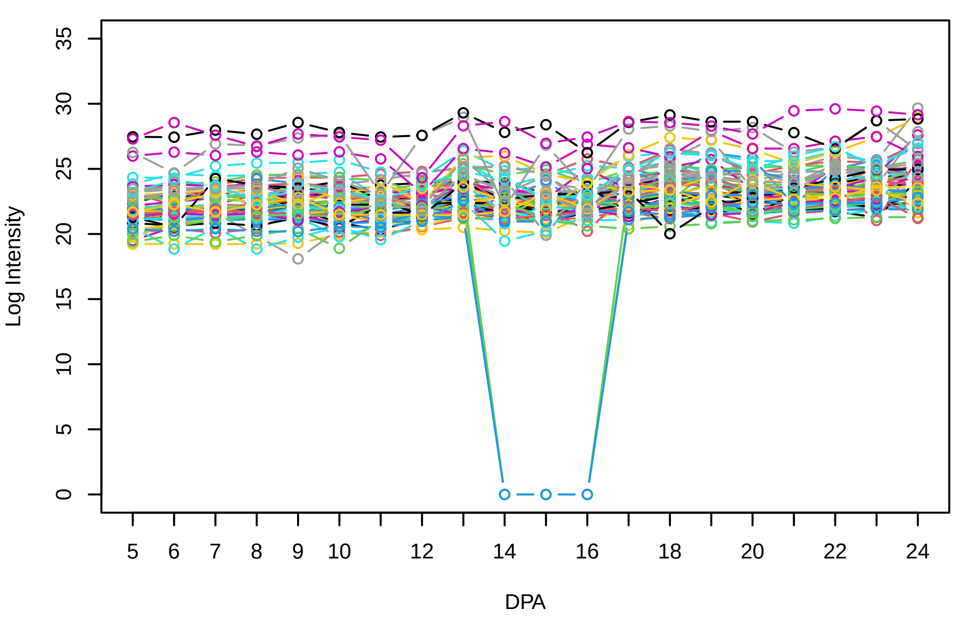


**Histogram of the average abundances of non−outlier proteins**

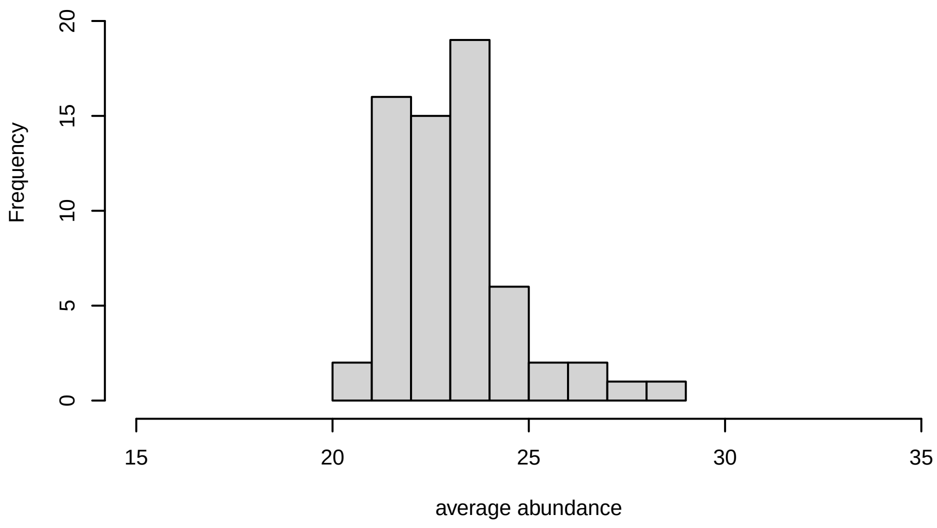


**V27, after remove unreliable**

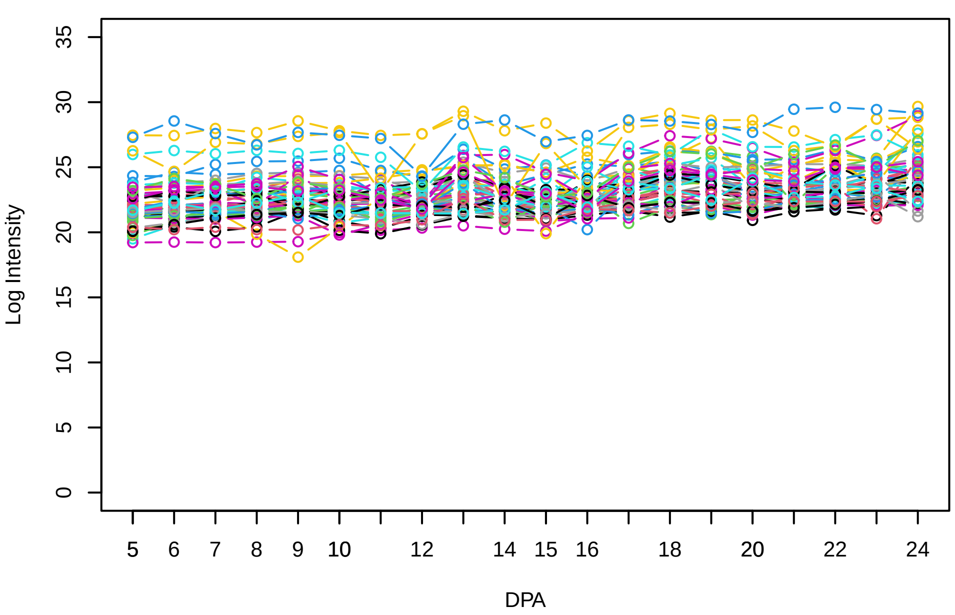


**Histogram of the average abundances of reliable proteins**

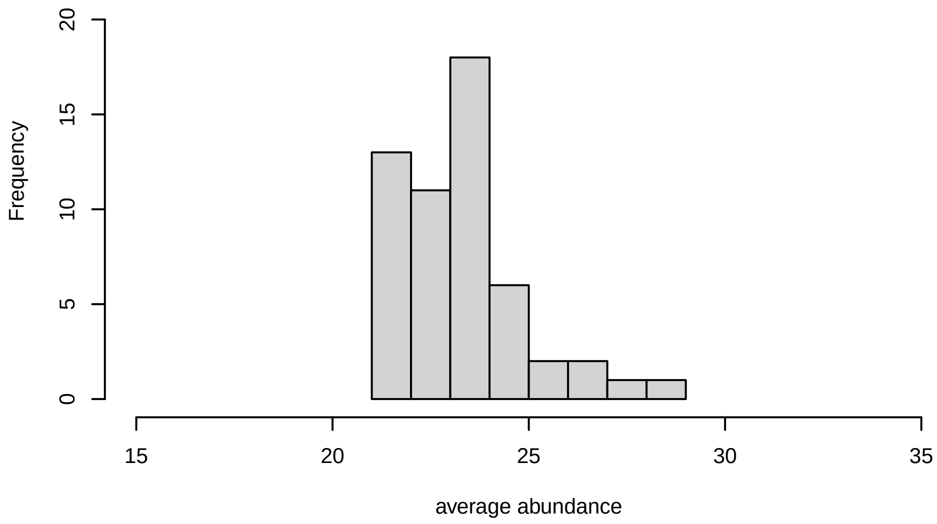


**V27, after remove low abundance**

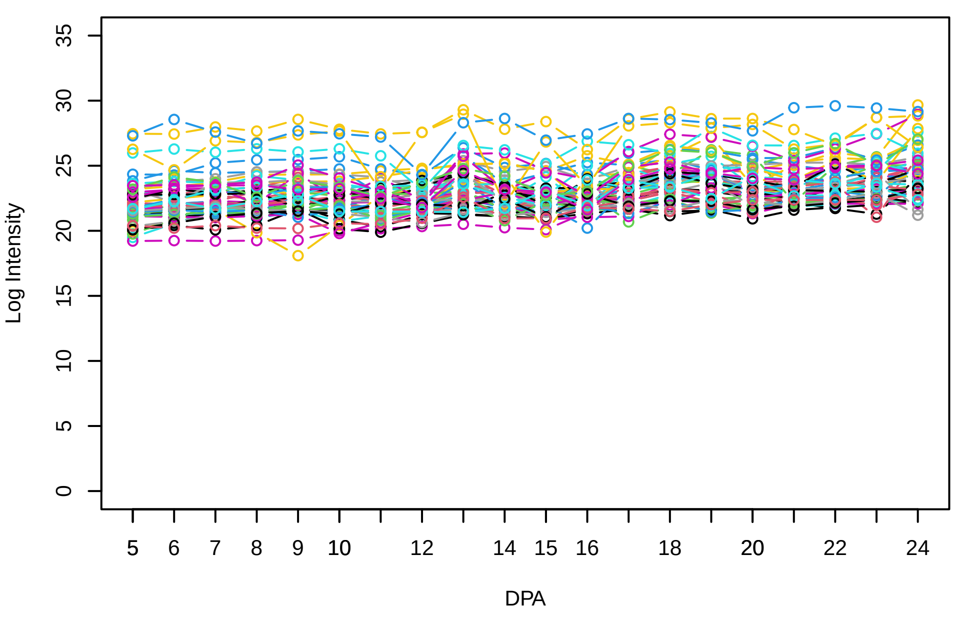


**Histogram of the average abundances of high abundance proteins**

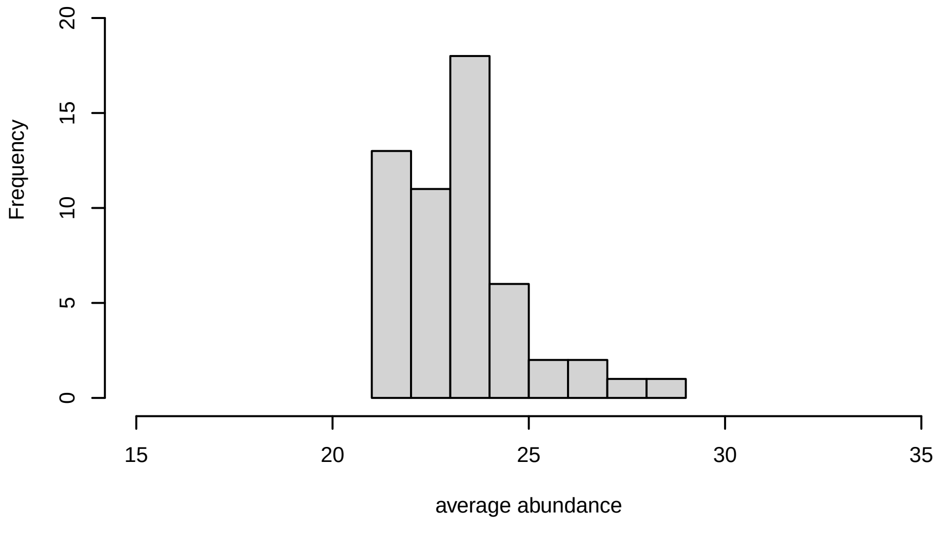

**V28**

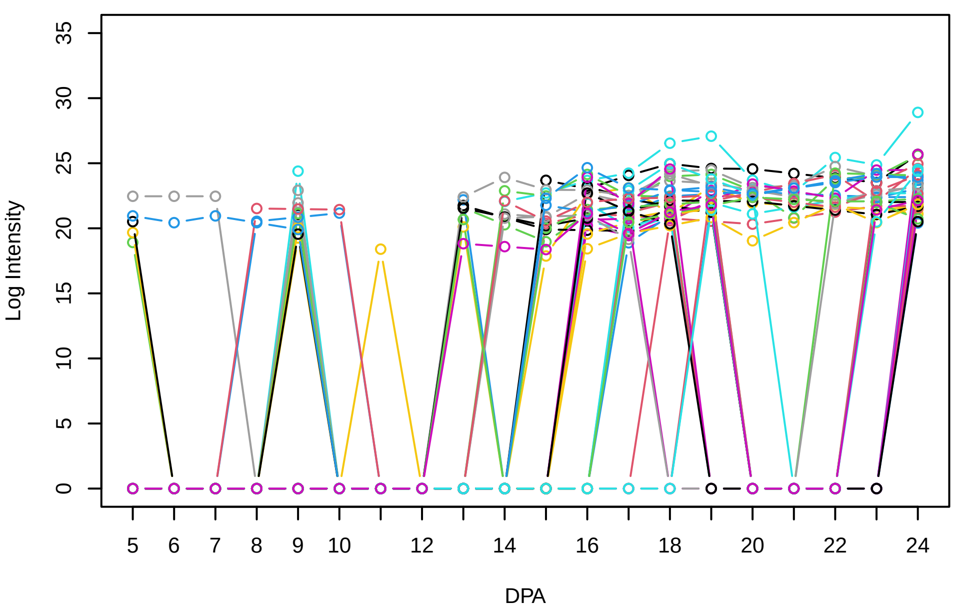


**Histogram of the average abundances of non−zero values**

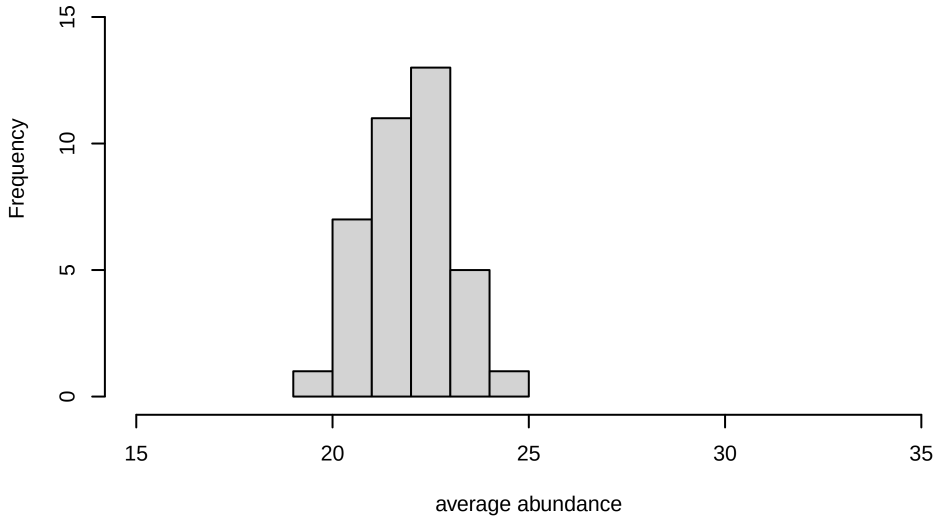


**V28, after remove outlier and cutoff is 0.656944625404285**

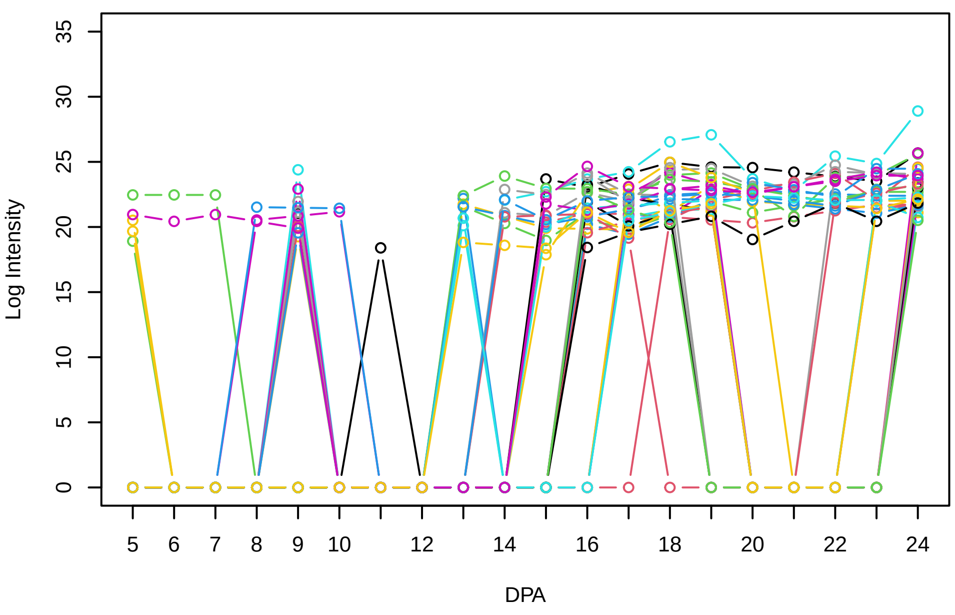


**Histogram of the average abundances of non−outlier proteins**

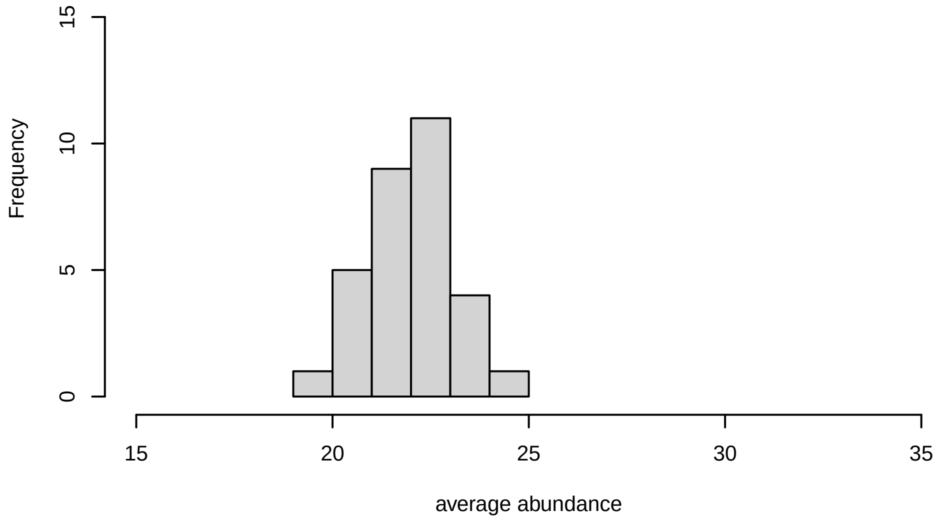


**V28, after remove unreliable**

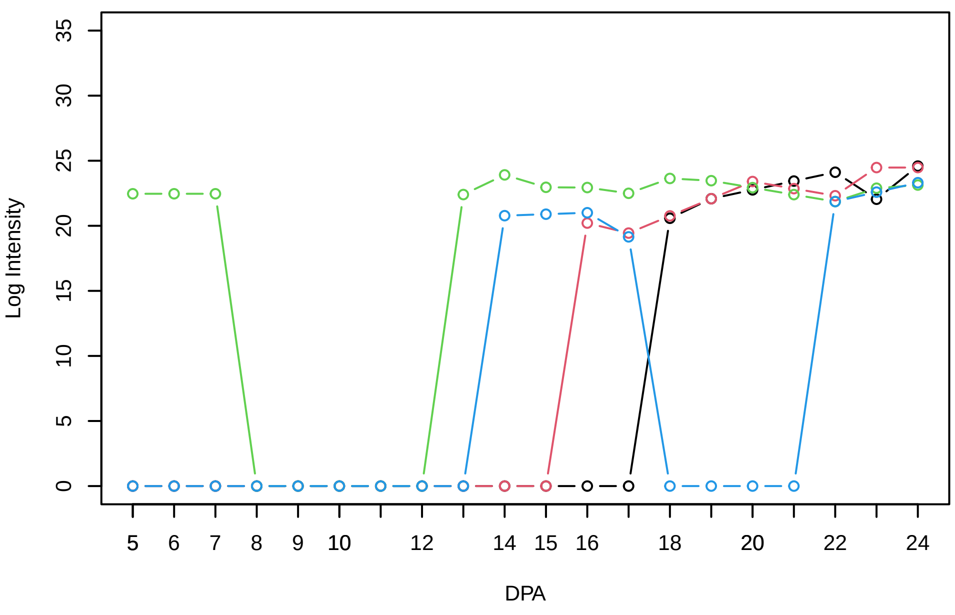


**Histogram of the average abundances of reliable proteins**

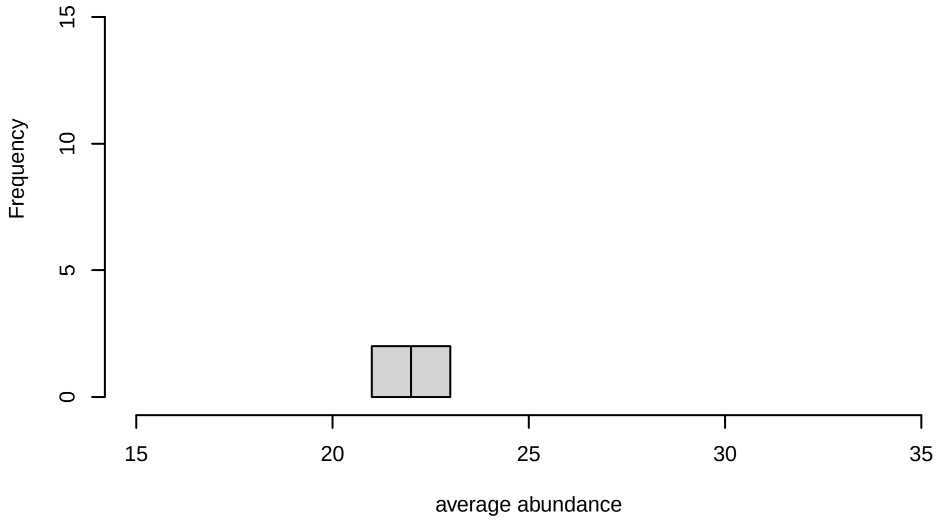


**V28, after remove low abundance**

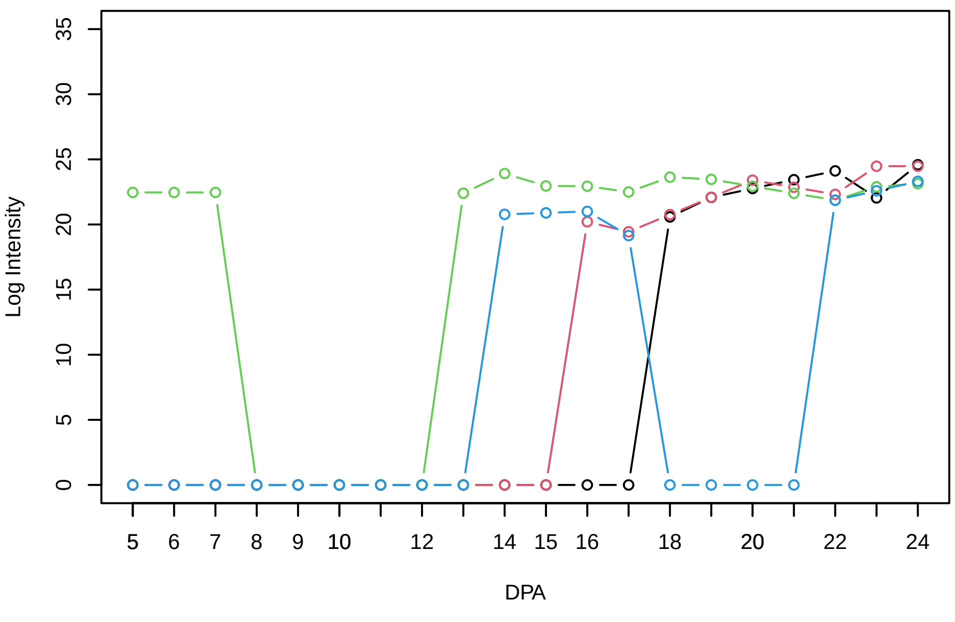


**Histogram of the average abundances of high abundance proteins**

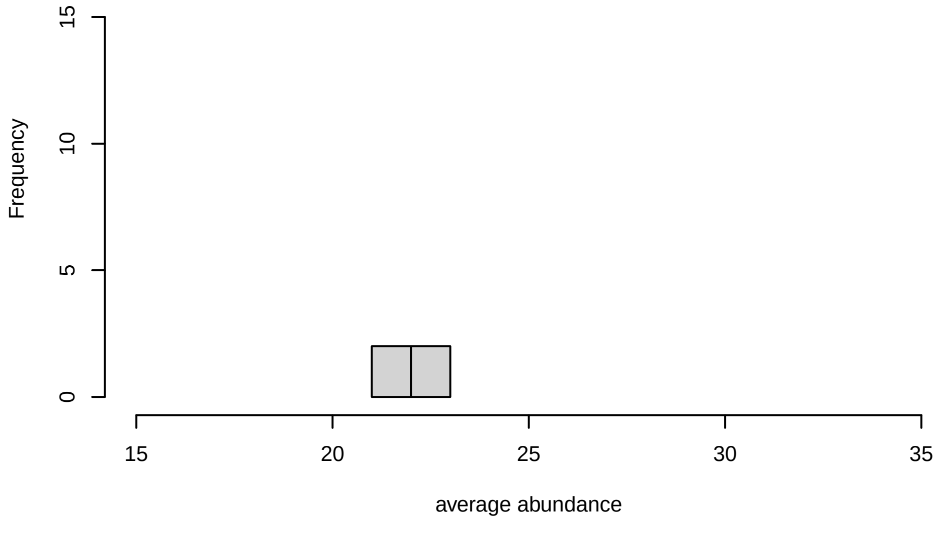

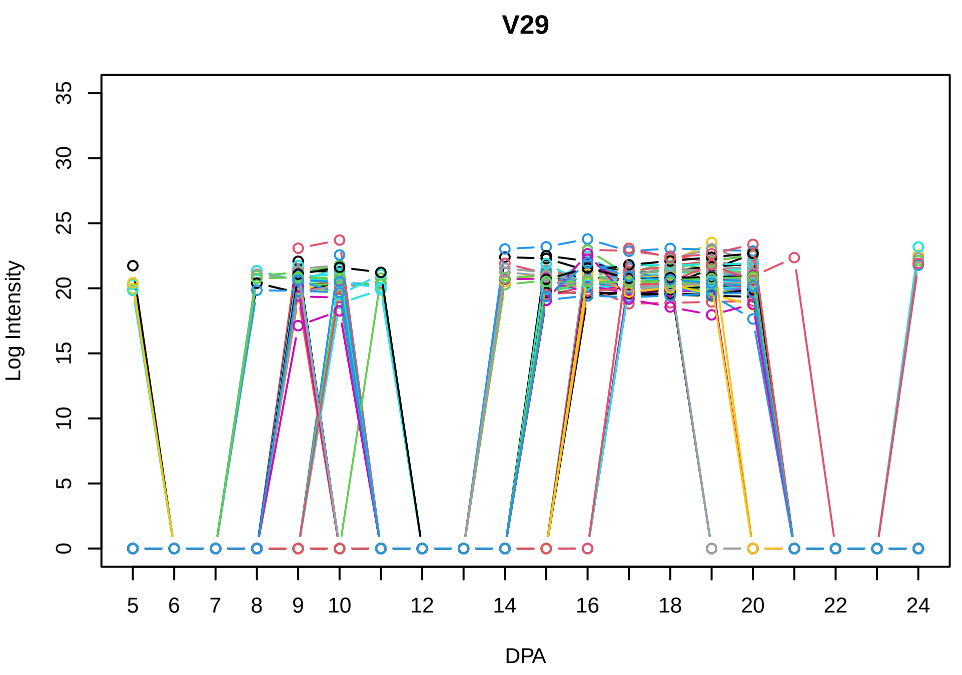
V29
Log Intensity
DPA

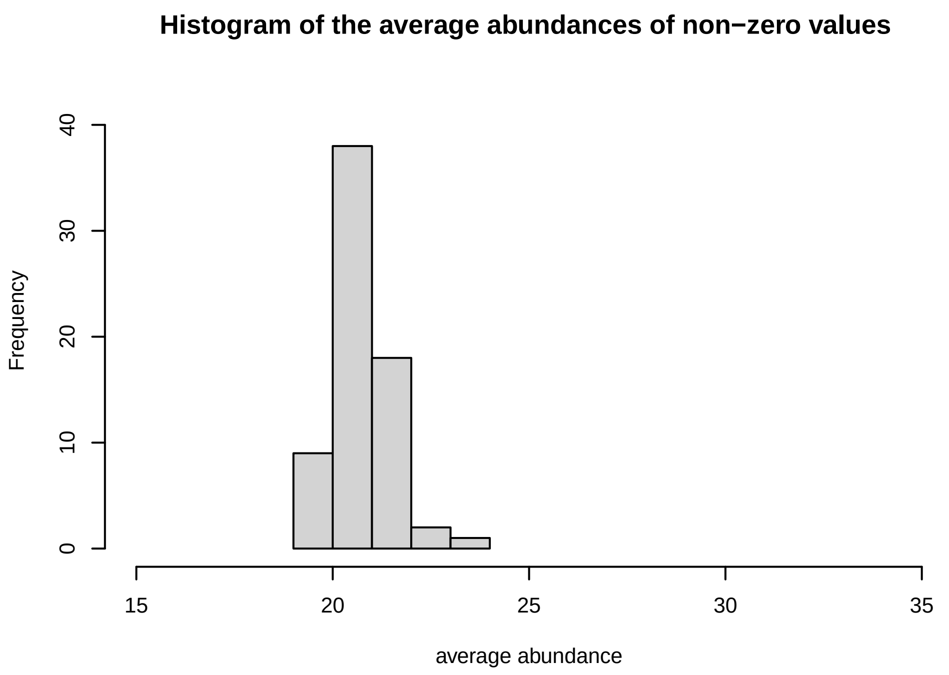
Histogram of the average abundances of non−zero values
Frequency
average abundance

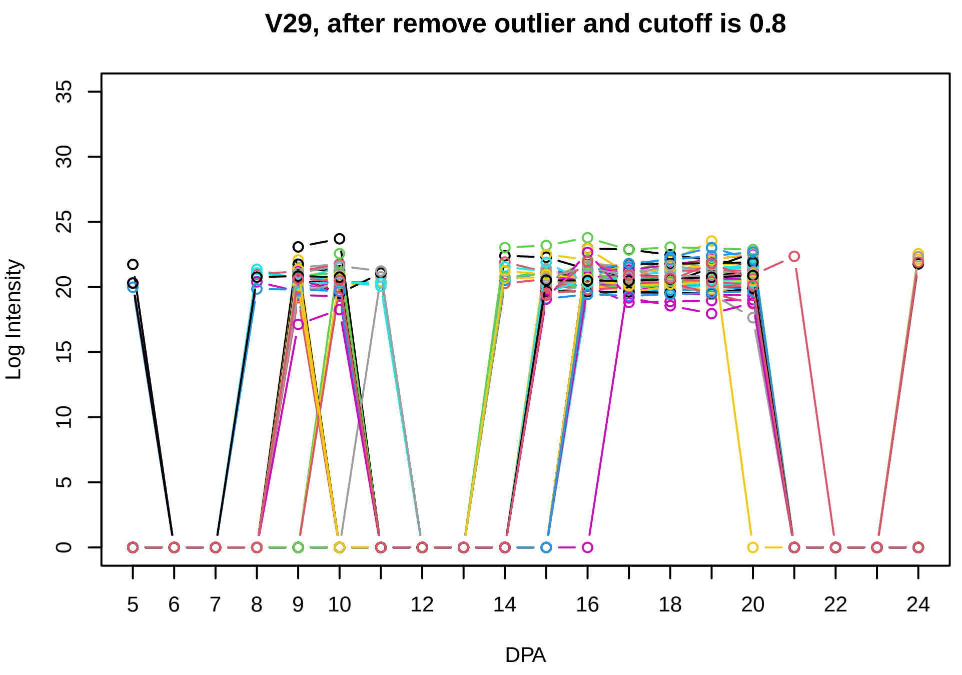
V29, after remove outlier and cutoff is 0.8
Log Intensity
DPA

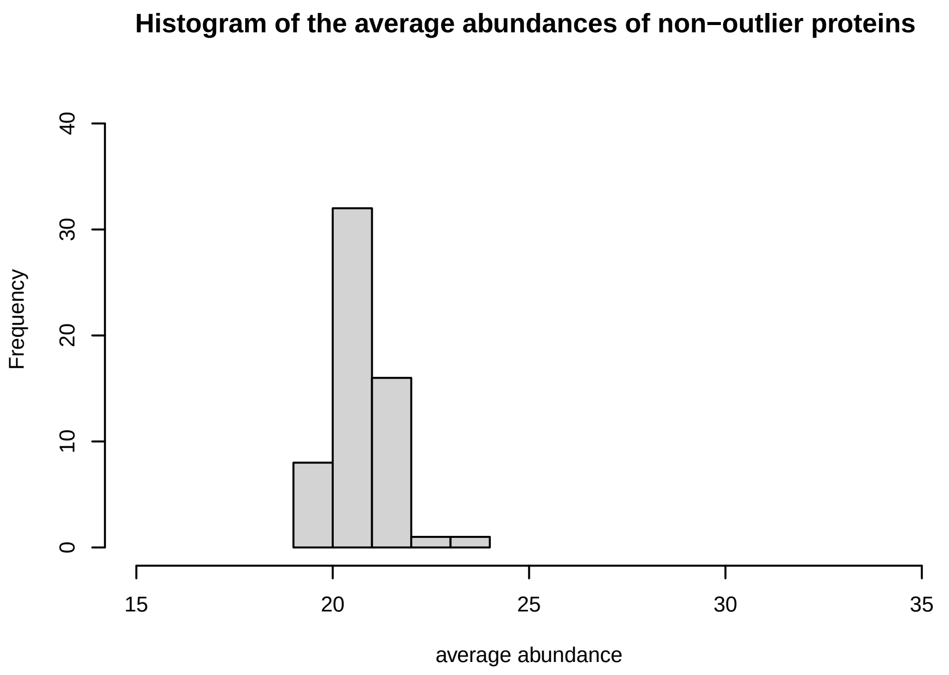
Histogram of the average abundances of non−outlier proteins
Frequency
average abundance

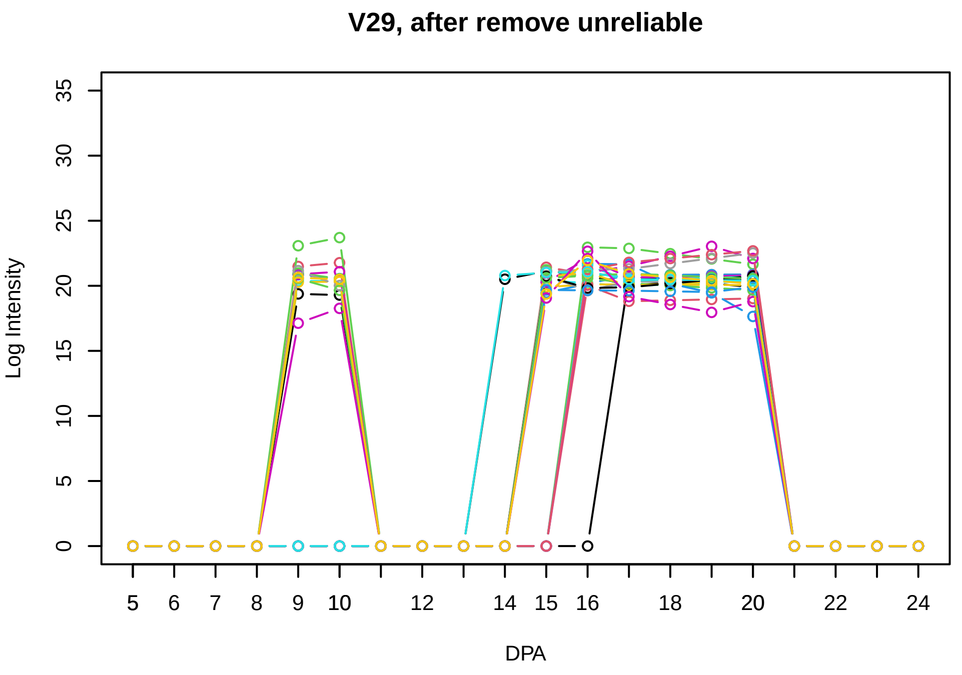
V29, after remove unreliable
Log Intensity
DPA

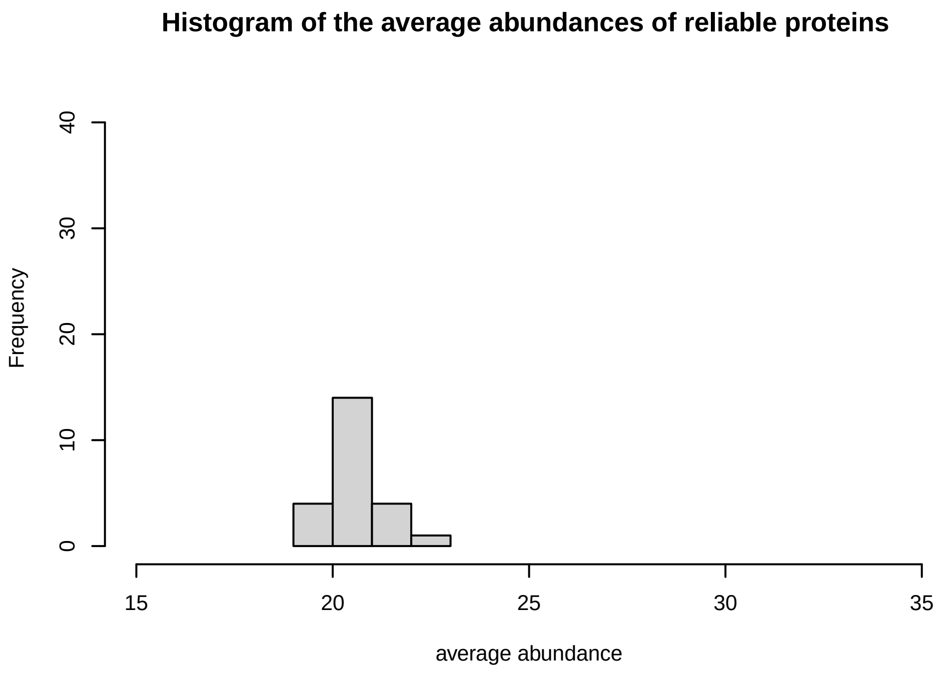
Histogram of the average abundances of reliable proteins
Frequency
average abundance

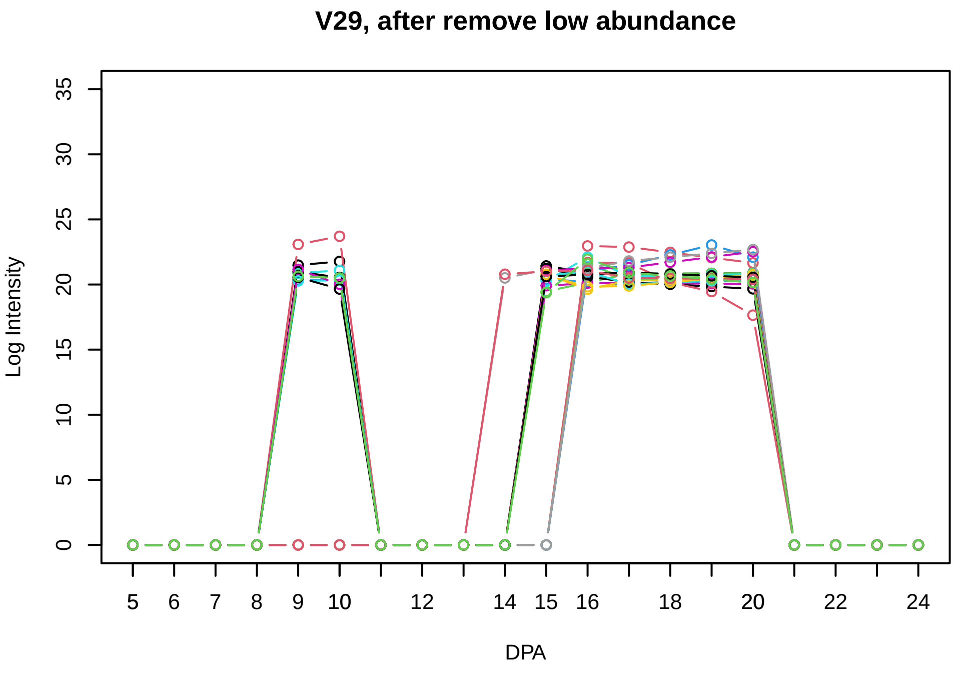
V29, after remove low abundance
Log Intensity
DPA

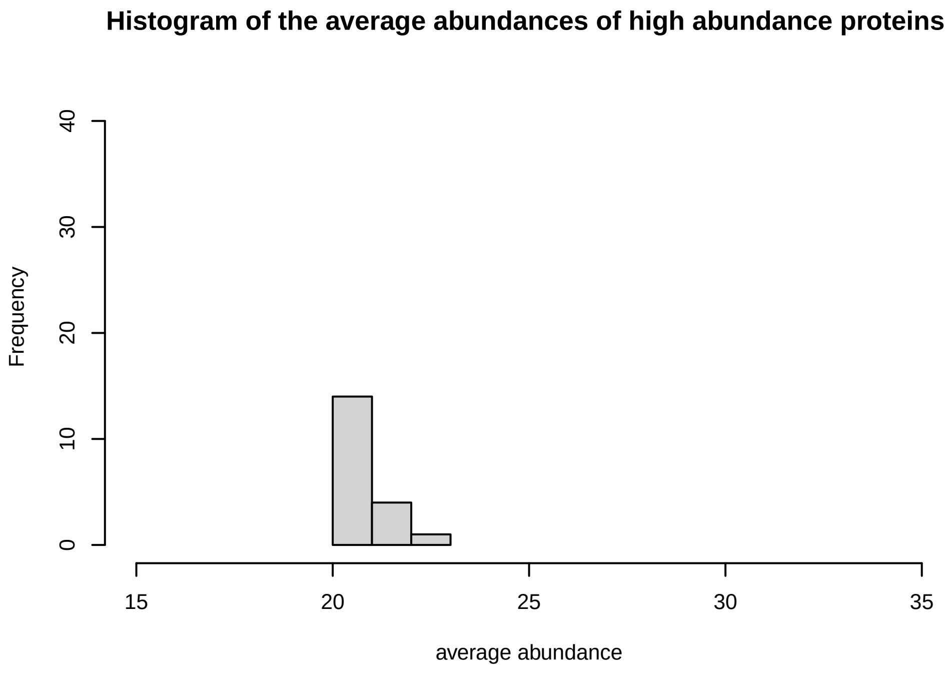
Histogram of the average abundances of high abundance proteins
Frequency
average abundance

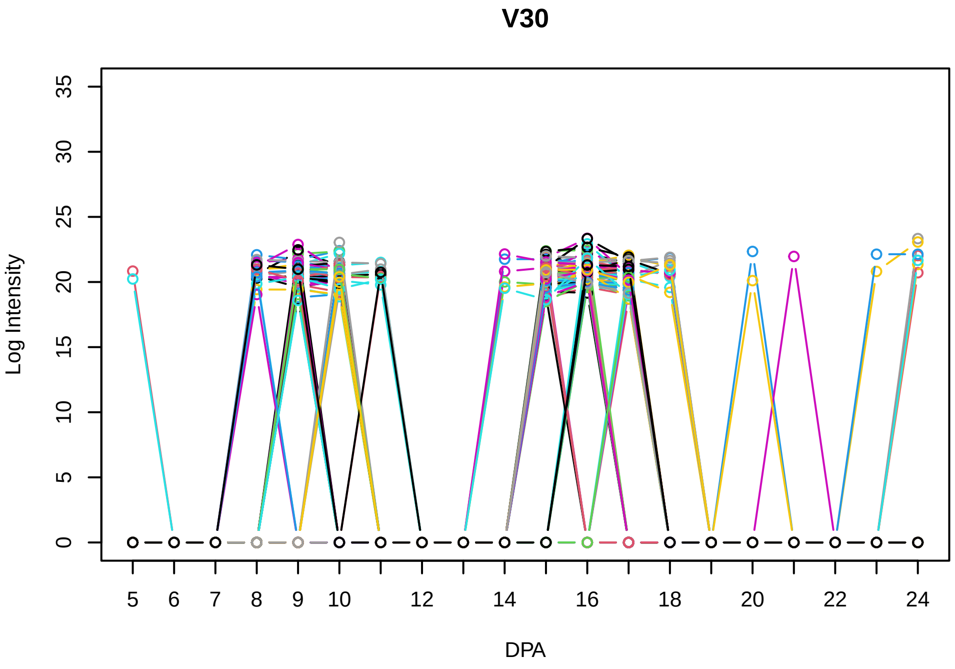
V30
Log Intensity
DPA

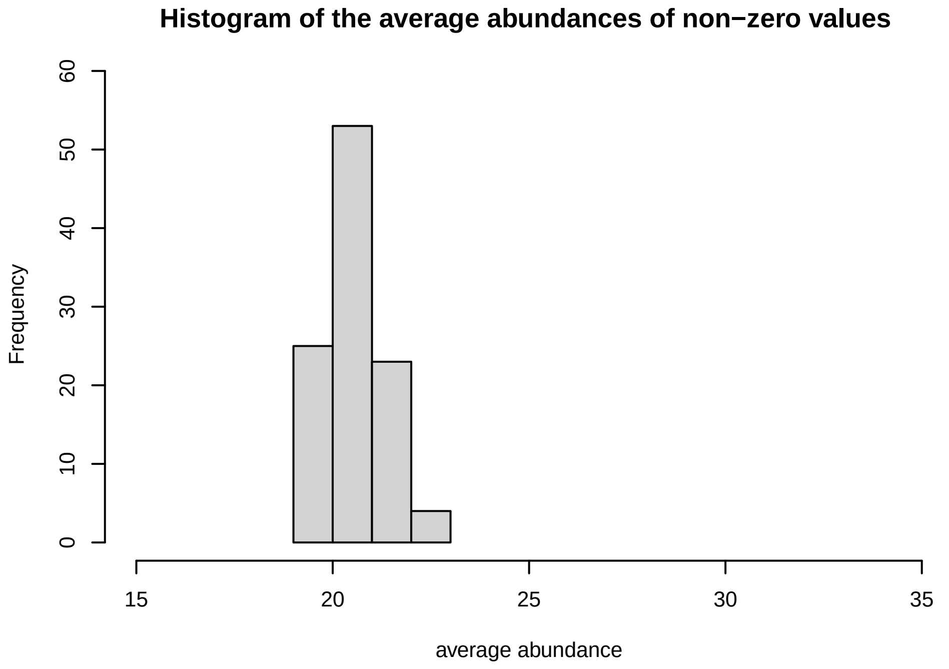
Histogram of the average abundances of non−zero values
Frequency
average abundance

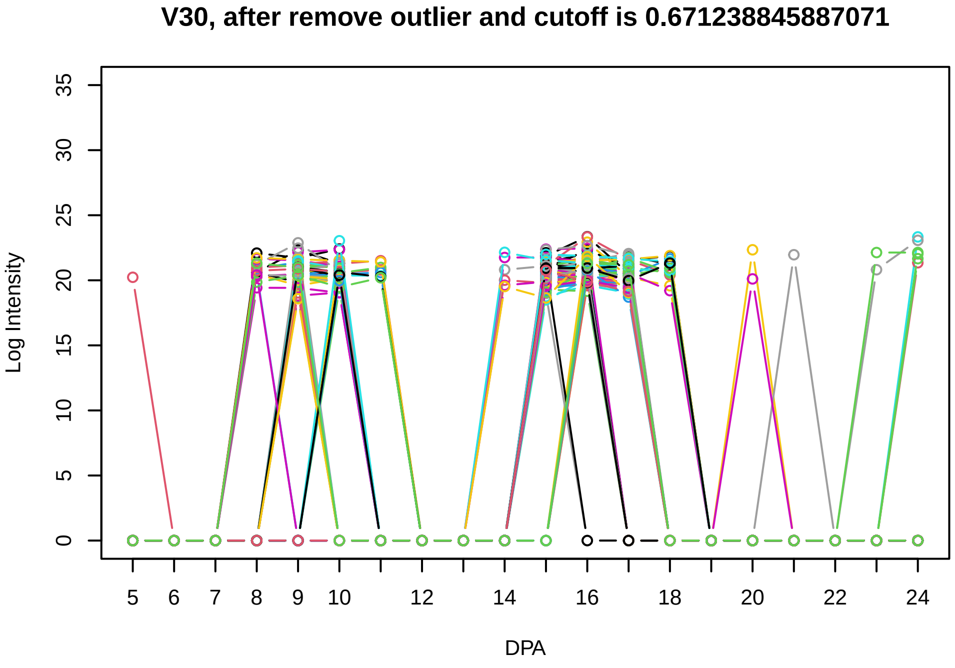
V30, after remove outlier and cutoff is 0.671238845887071
Log Intensity
DPA

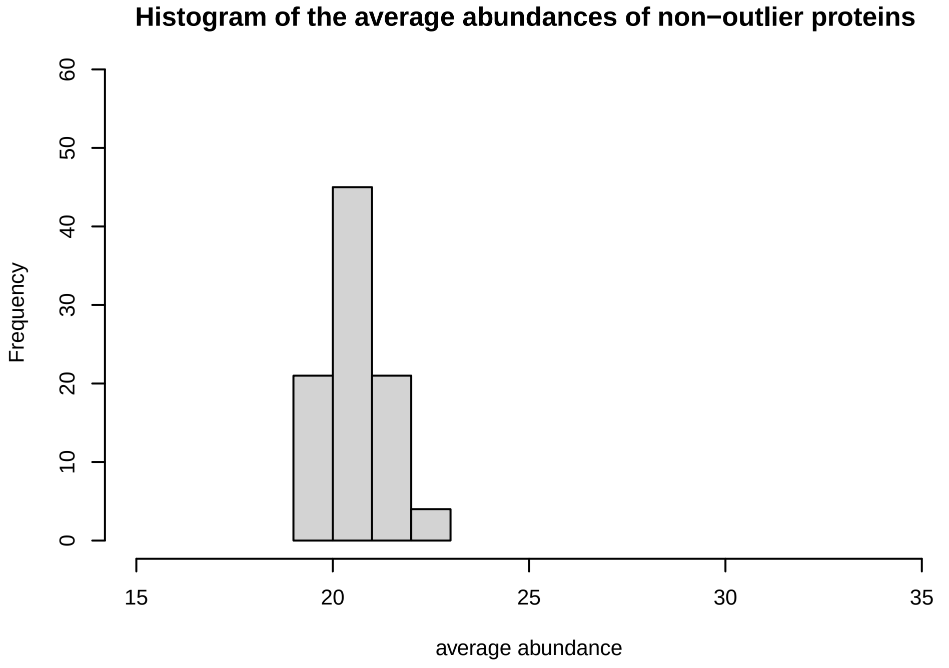
Histogram of the average abundances of non−outlier proteins
Frequency
average abundance

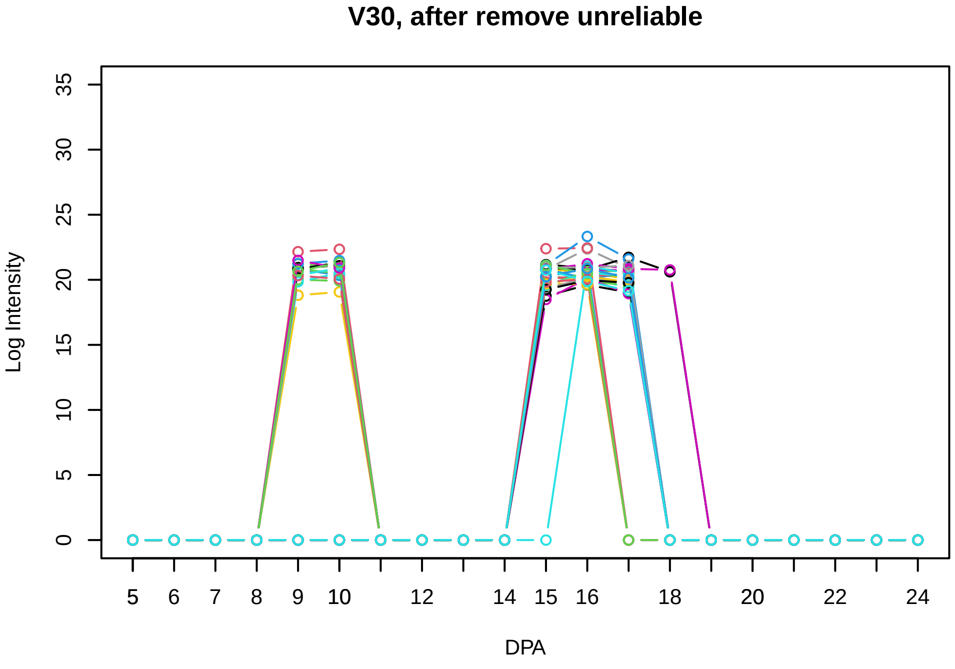
V30, after remove unreliable
Log Intensity
DPA

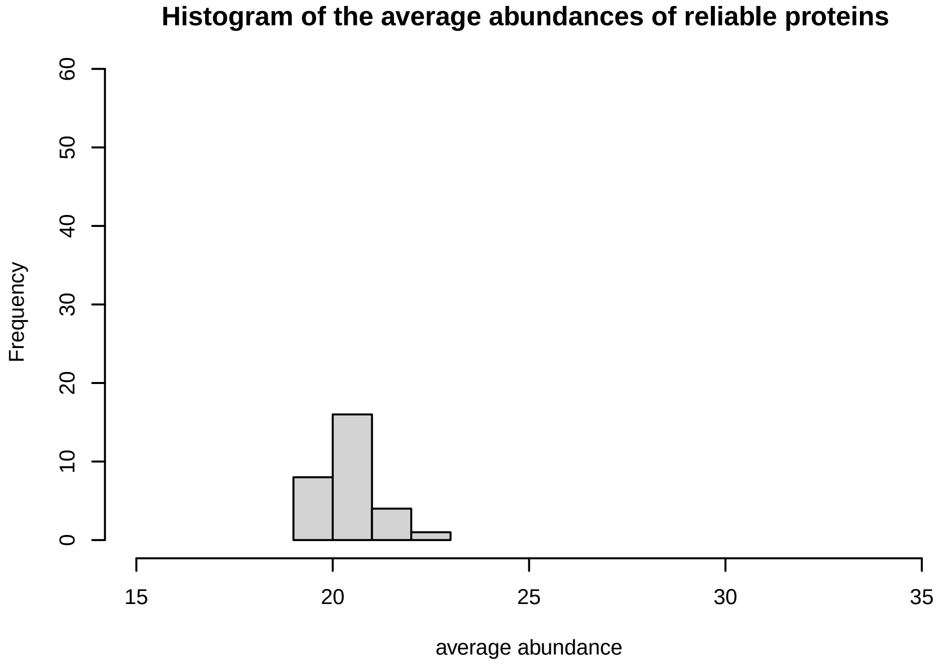
Histogram of the average abundances of reliable proteins
Frequency
average abundance

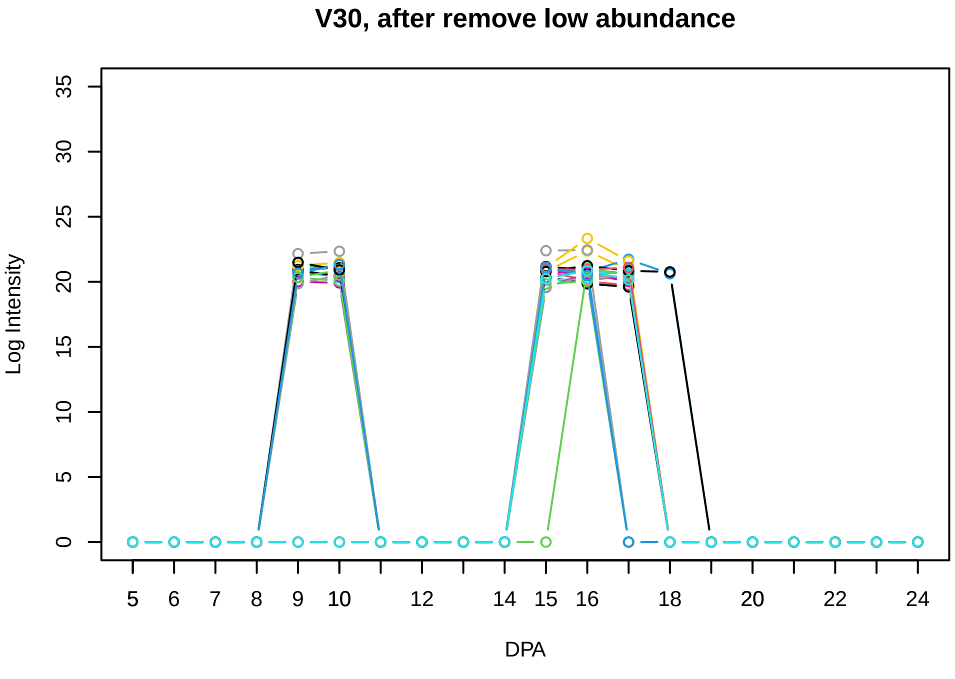
V30, after remove low abundance
Log Intensity
DPA

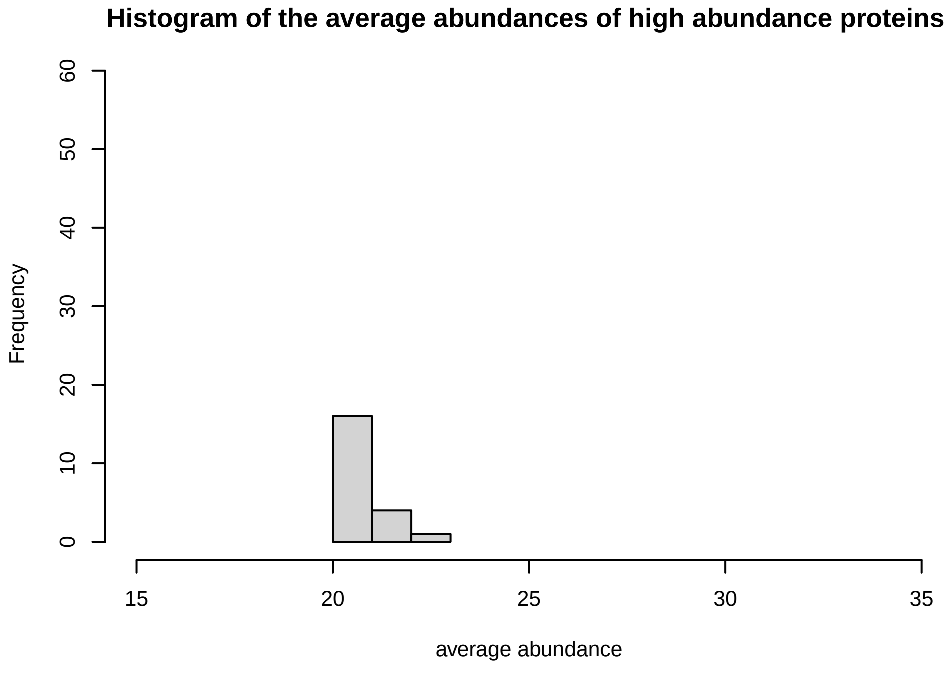
Histogram of the average abundances of high abundance proteins
Frequency
average abundance
