## Supplementary Figure S3 for "Systems-level proteomic models of cotton fiber development: A high-resolution resource to analyze cell dynamics and trait engineering"

**Histogram of the average abundances of non−zero values**

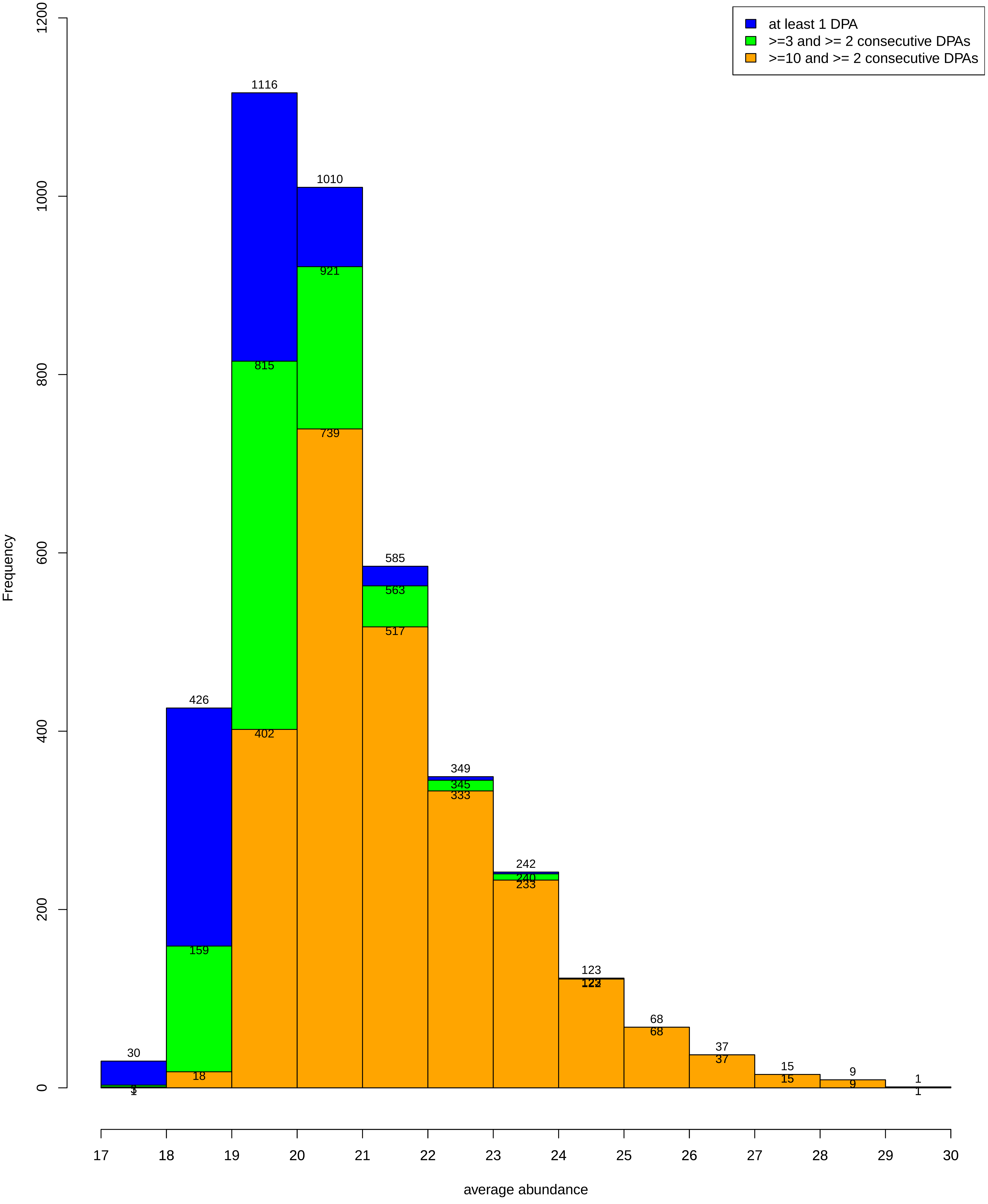

**Codes**

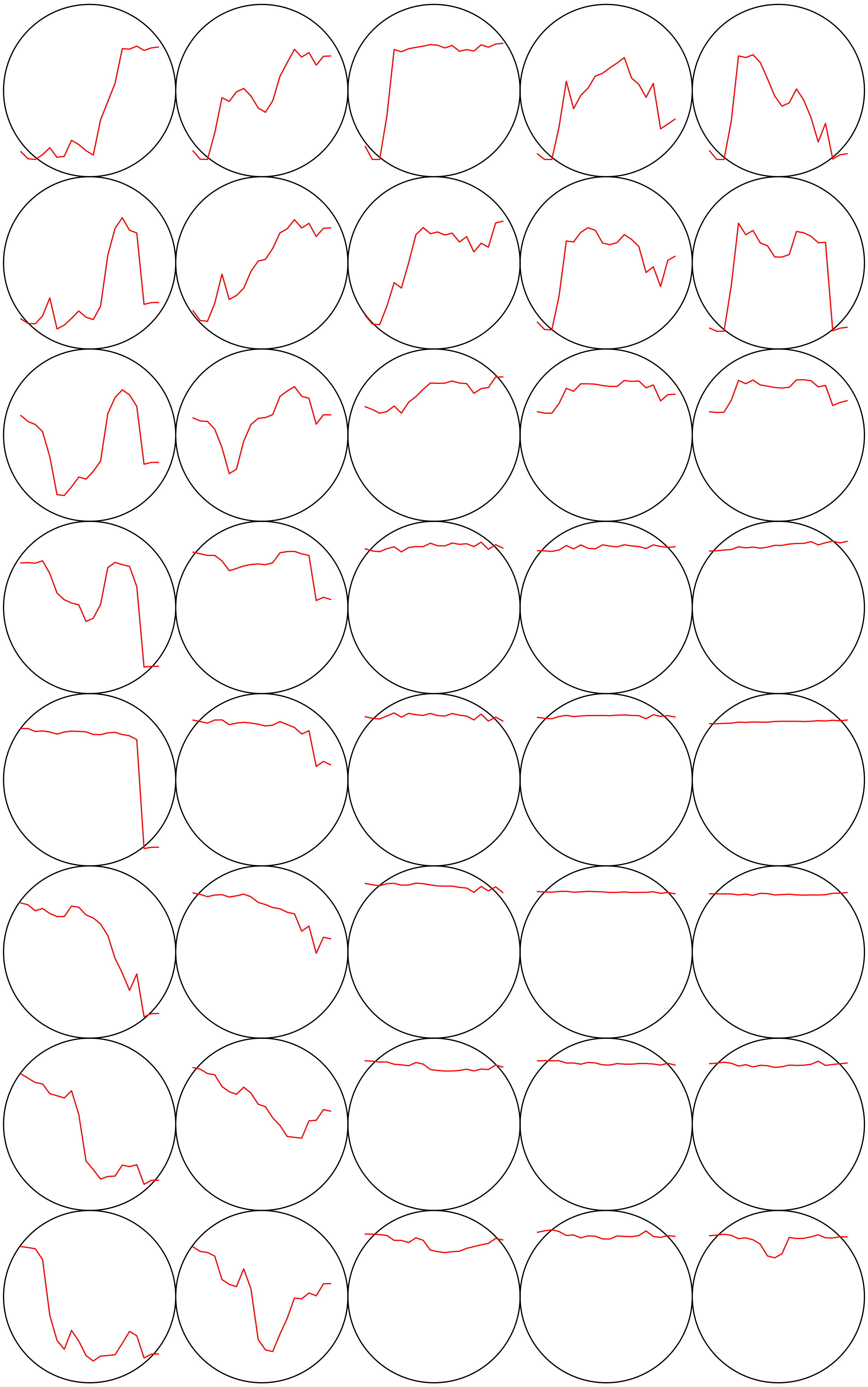

**Count plot**

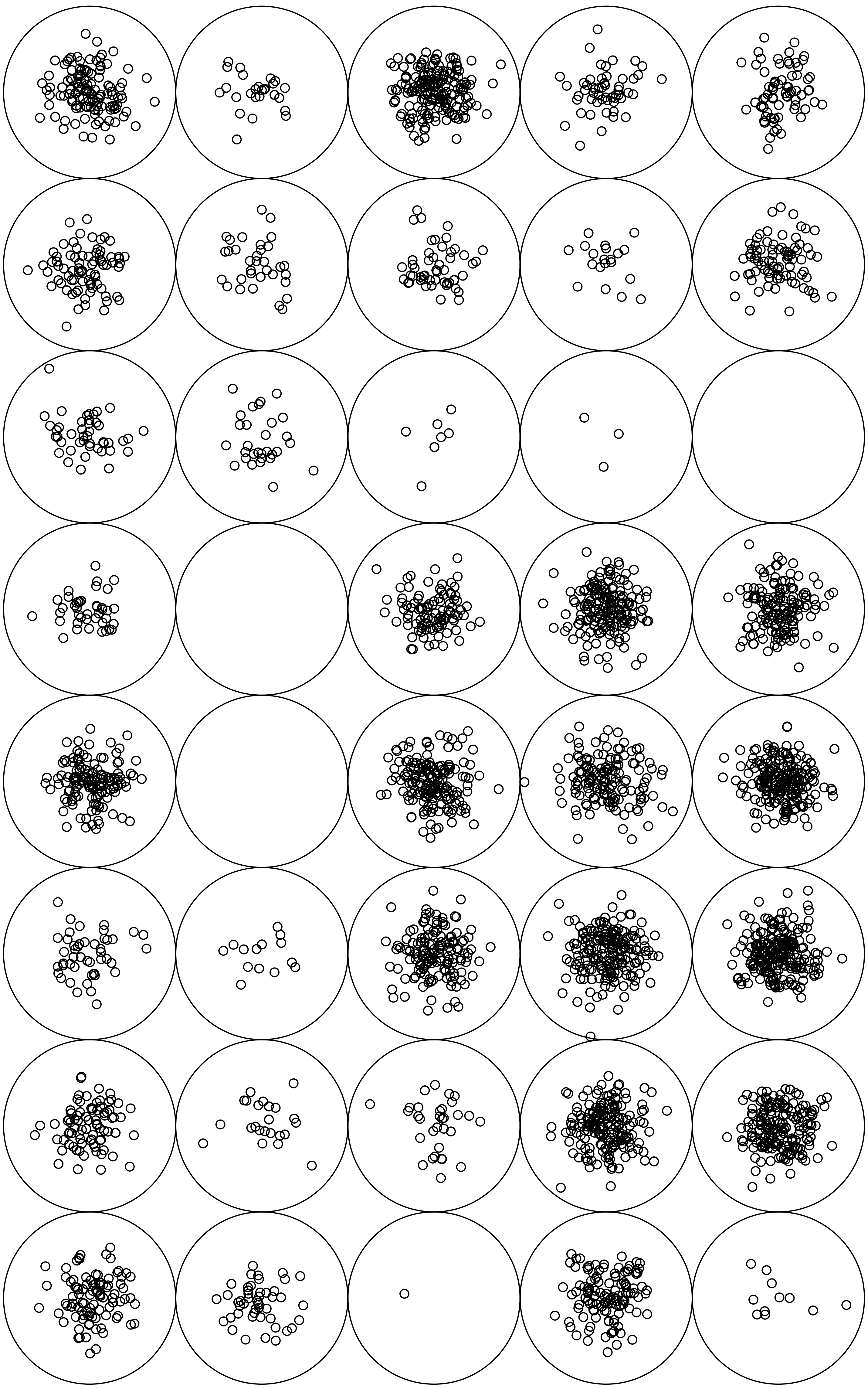

**V1**

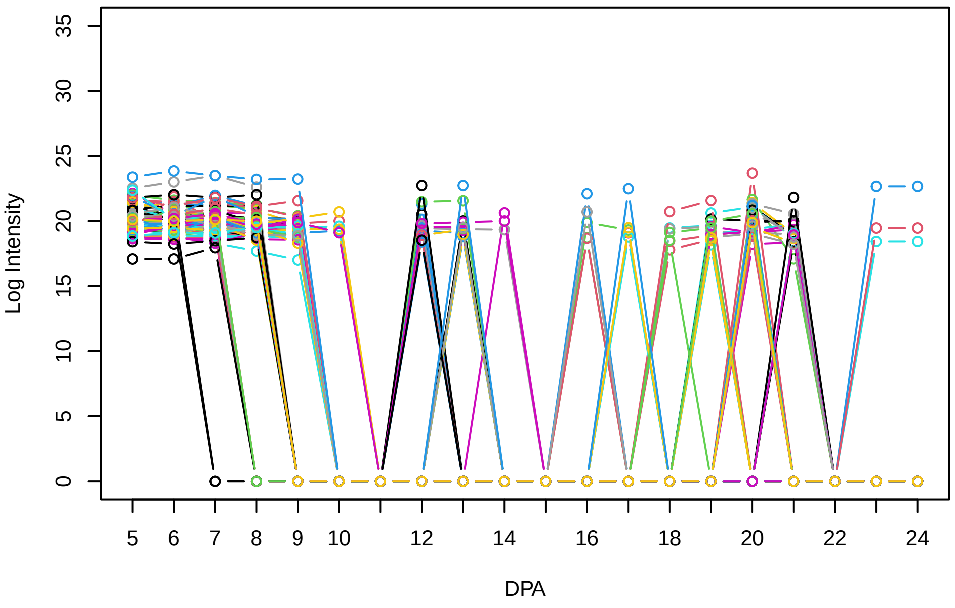


**Histogram of the average abundances of non−zero values**

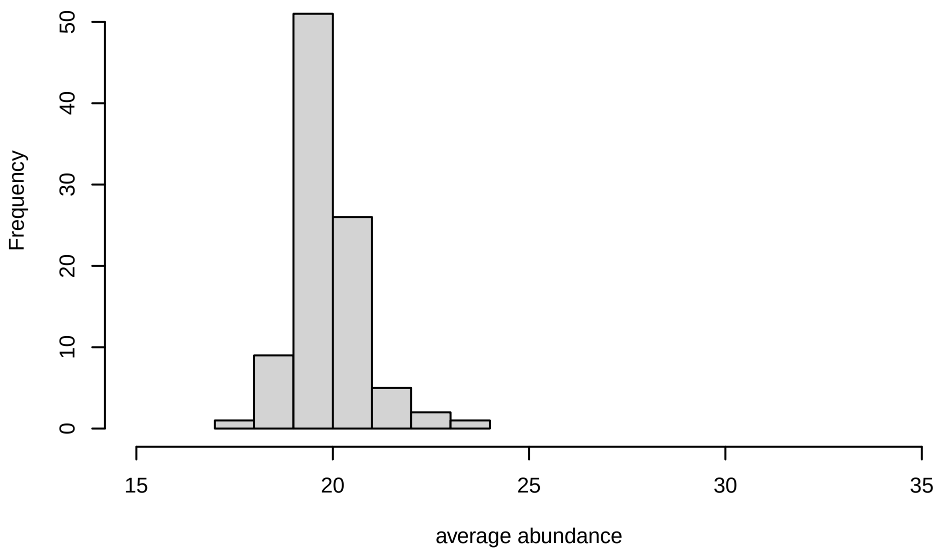


**V1, after remove outlier and cutoff is 0.751856023633583**

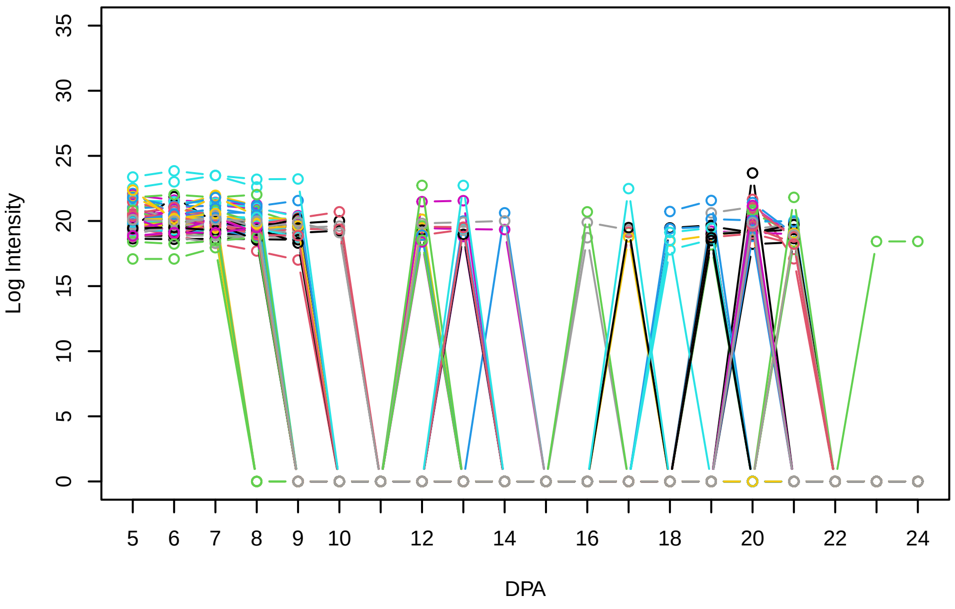


**Histogram of the average abundances of non−outlier proteins**

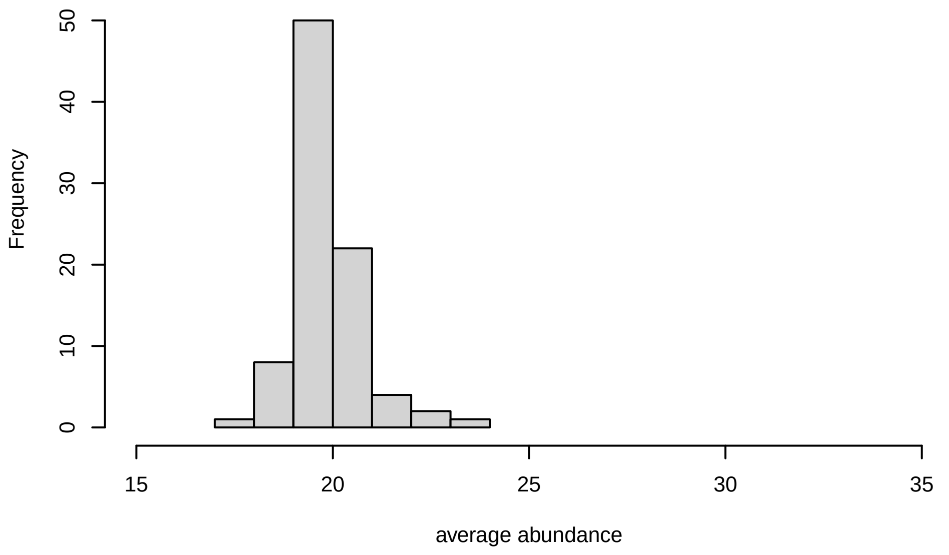


**V1, after remove unreliable**

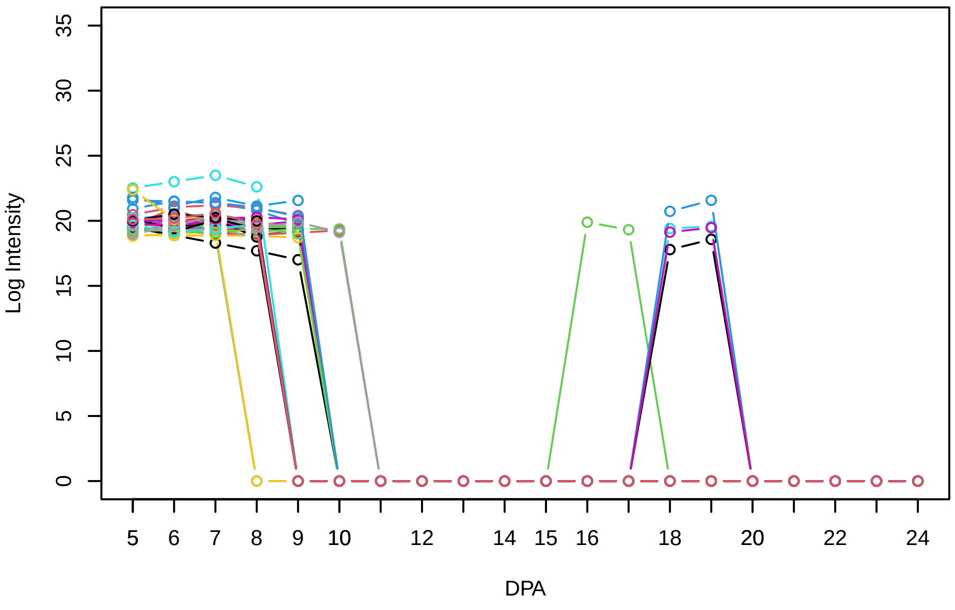


**Histogram of the average abundances of reliable proteins**

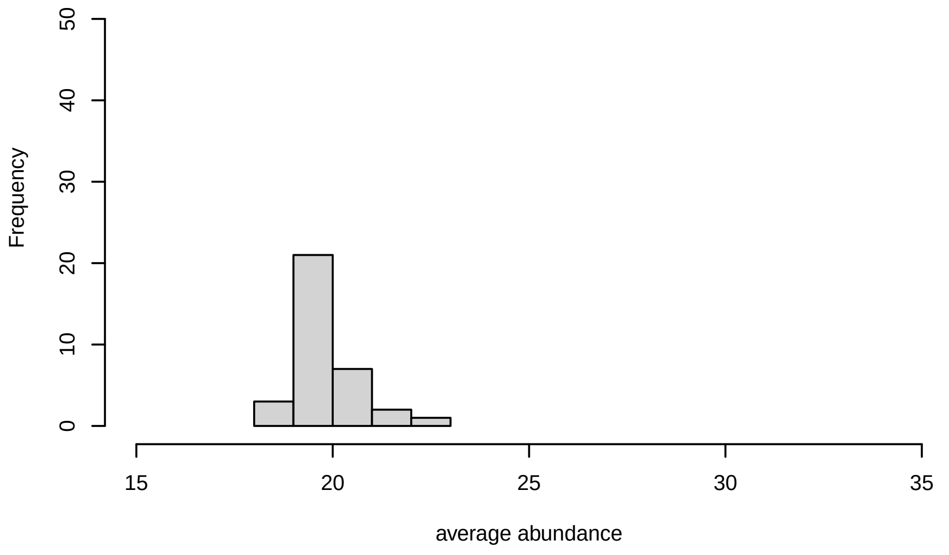


**V1, after remove low abundance**

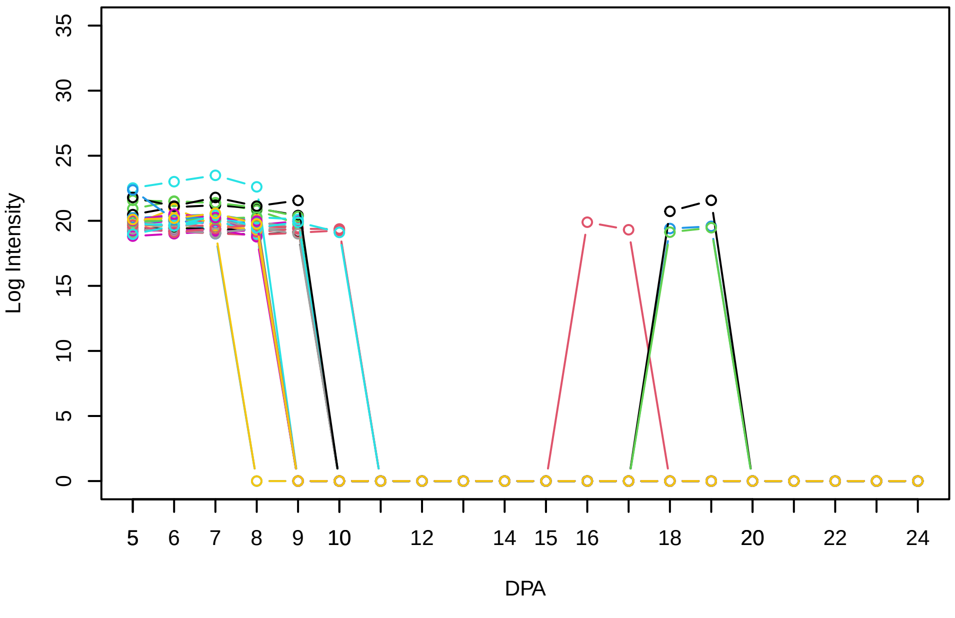


**Histogram of the average abundances of high abundance proteins**

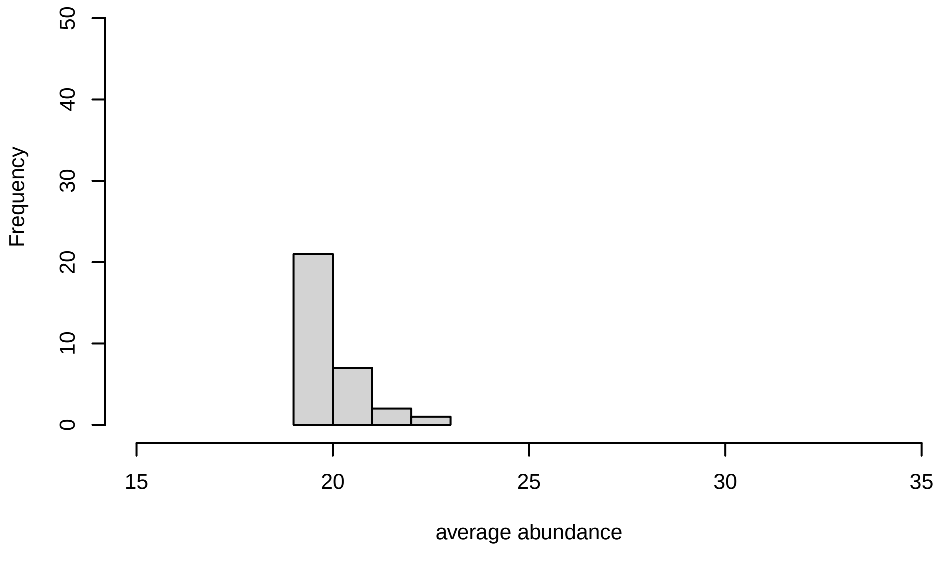

V2

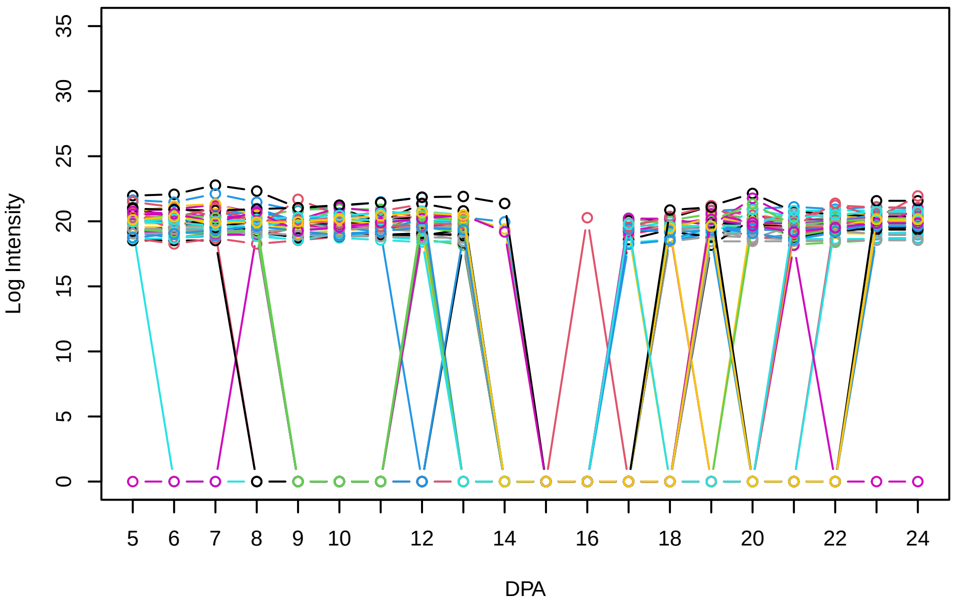


Histogram of the average abundances of non−zero values

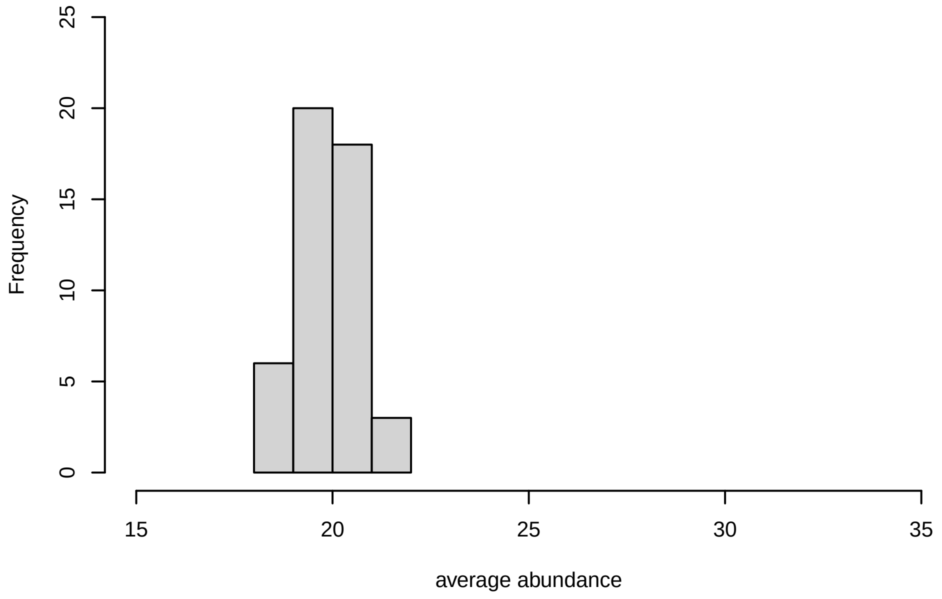


V2, after remove outlier and cutoff is 0.8

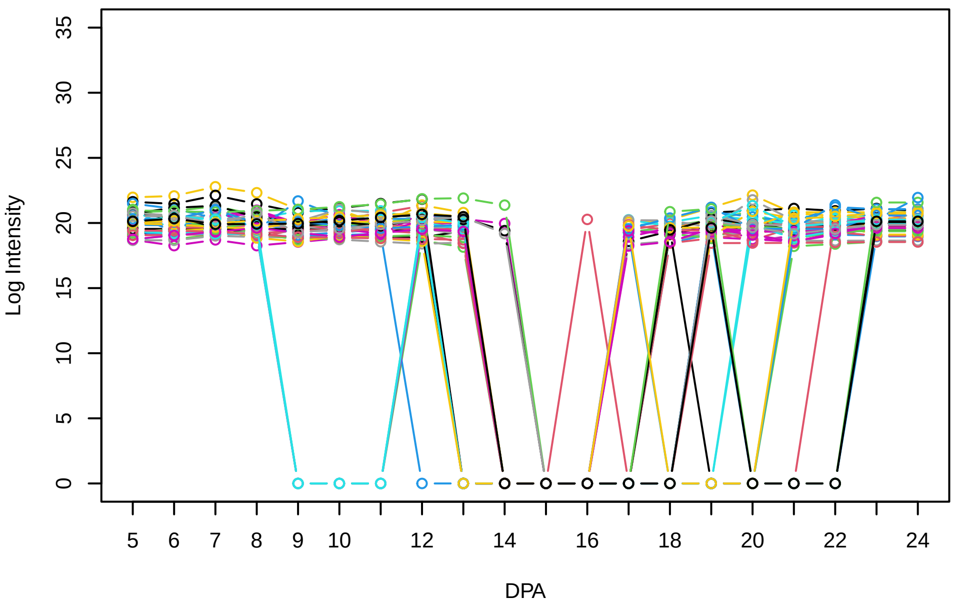


Histogram of the average abundances of non−outlier proteins

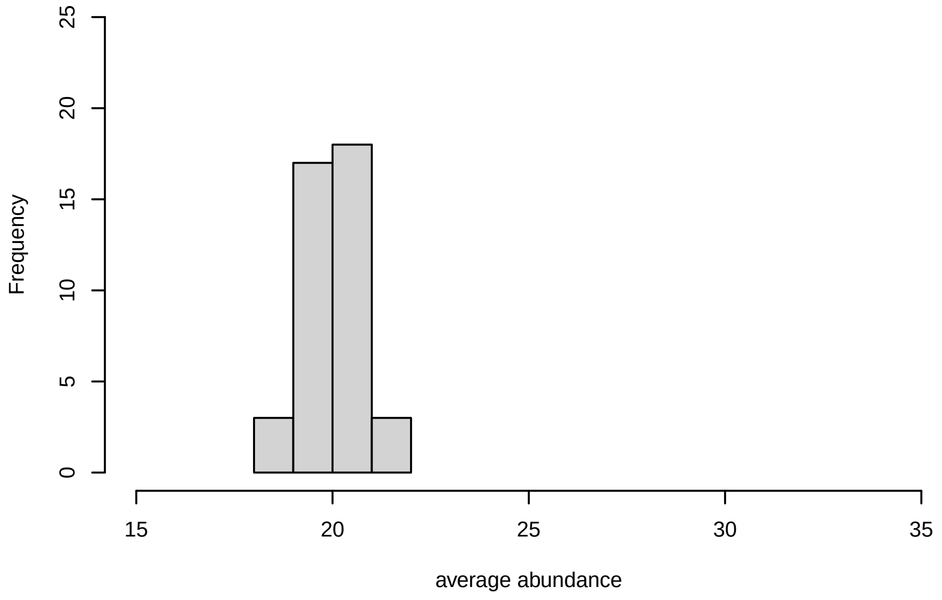


V2, after remove unreliable

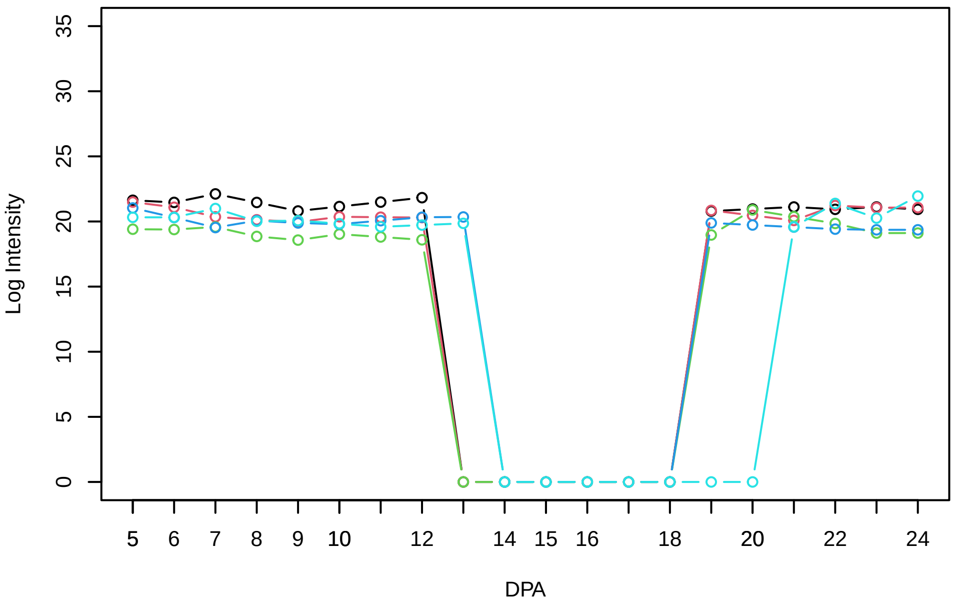


Histogram of the average abundances of reliable proteins

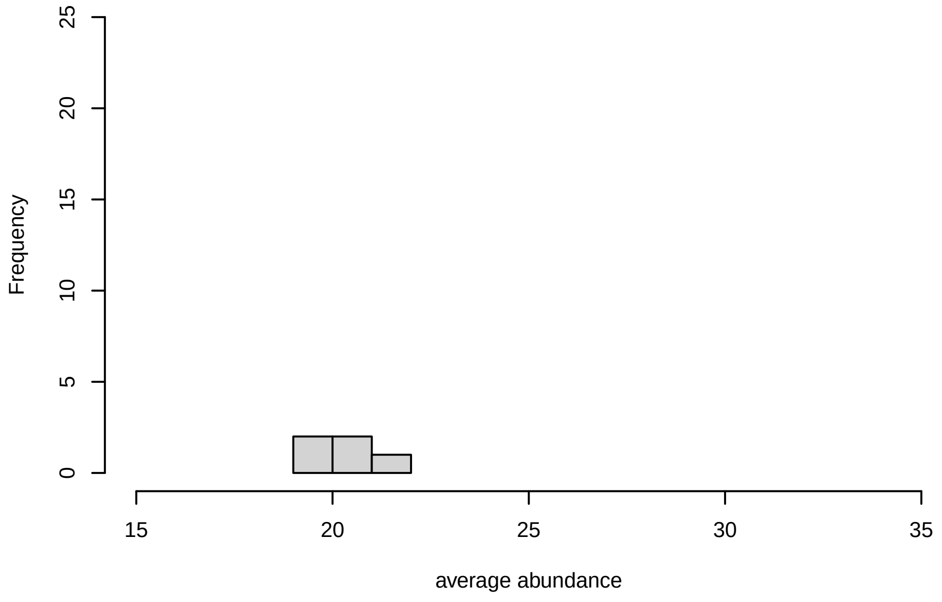


V2, after remove low abundance

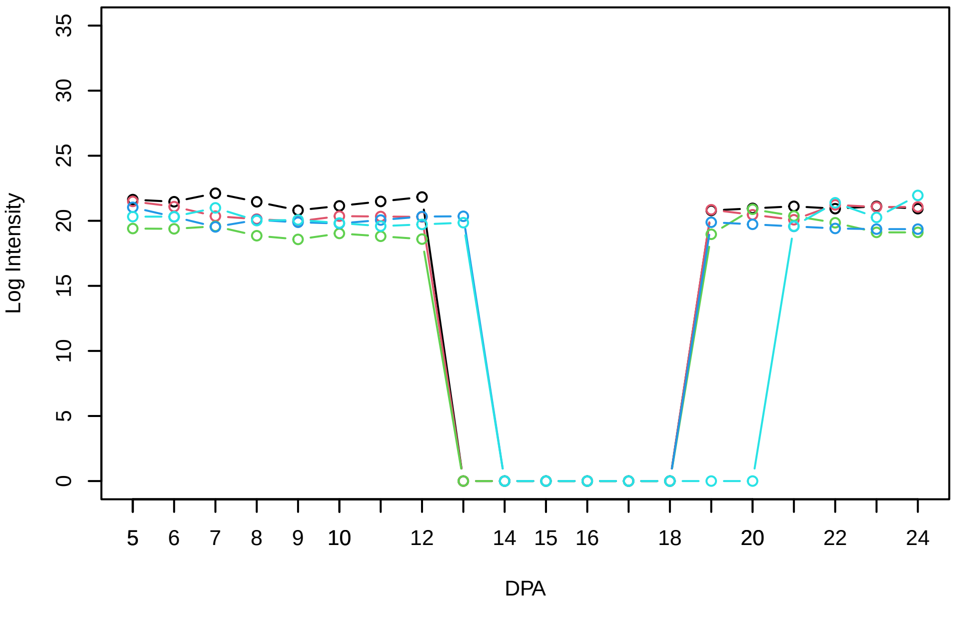


Histogram of the average abundances of high abundance proteins

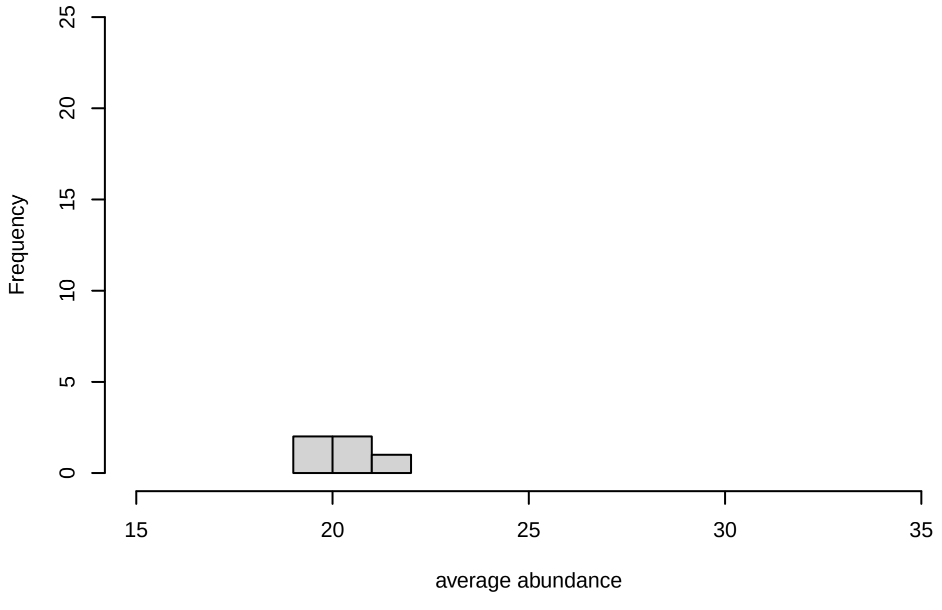

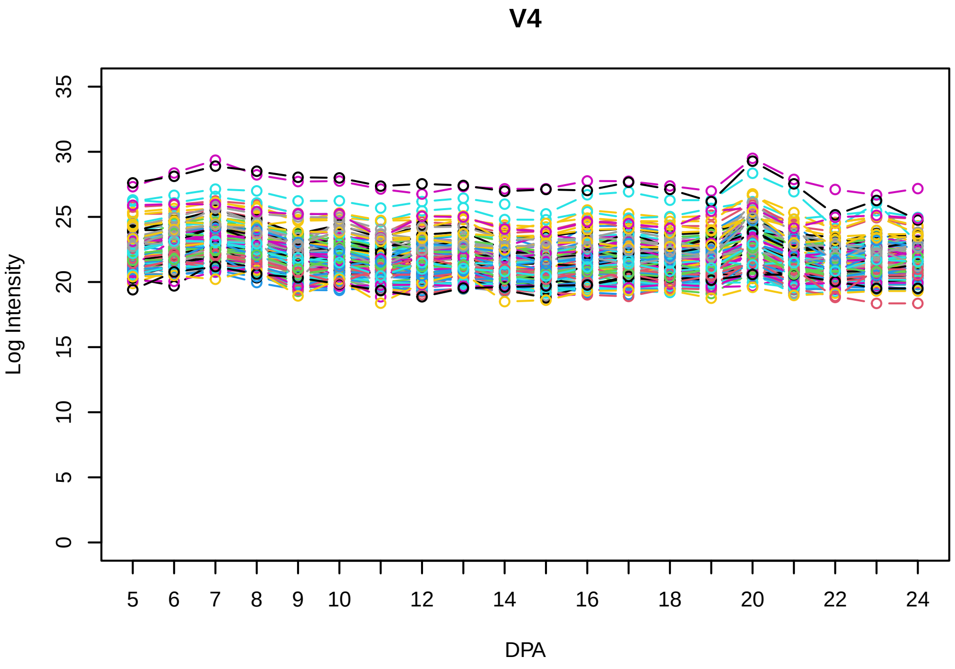
V4
Log Intensity
DPA

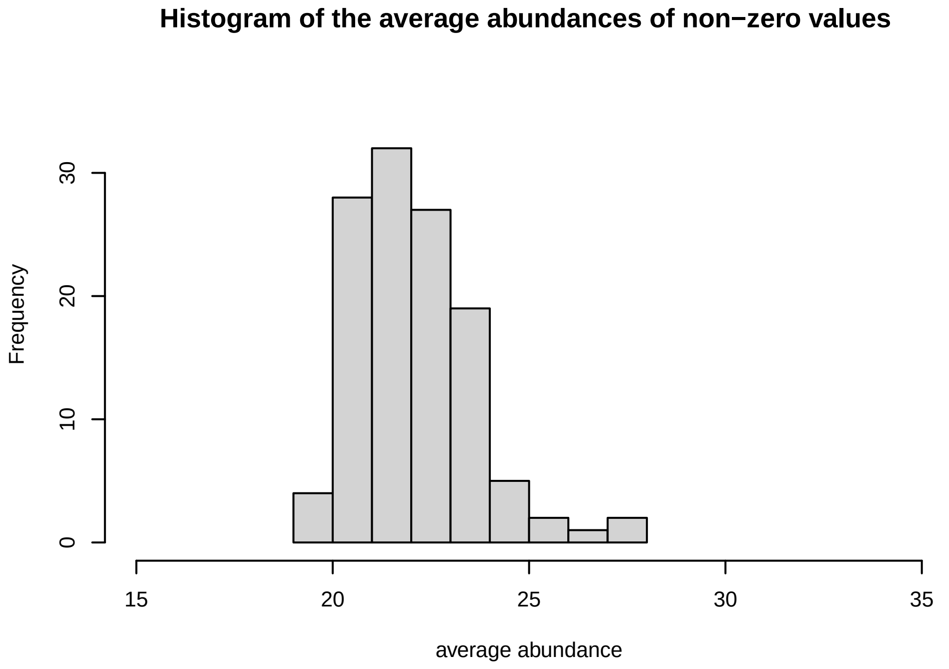
Histogram of the average abundances of non−zero values
Frequency
average abundance

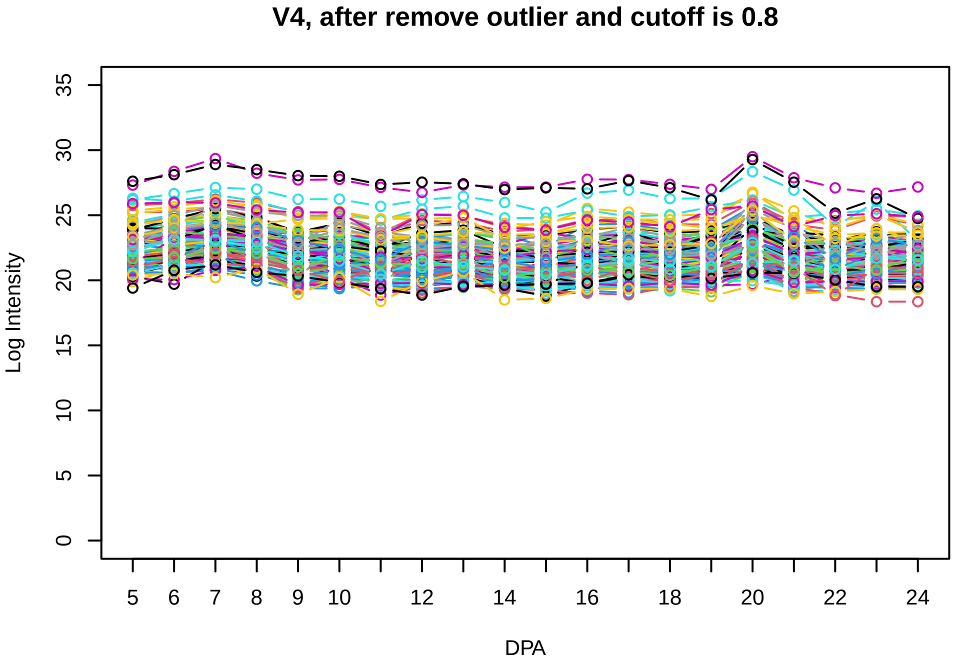
V4, after remove outlier and cutoff is 0.8
Log Intensity
DPA

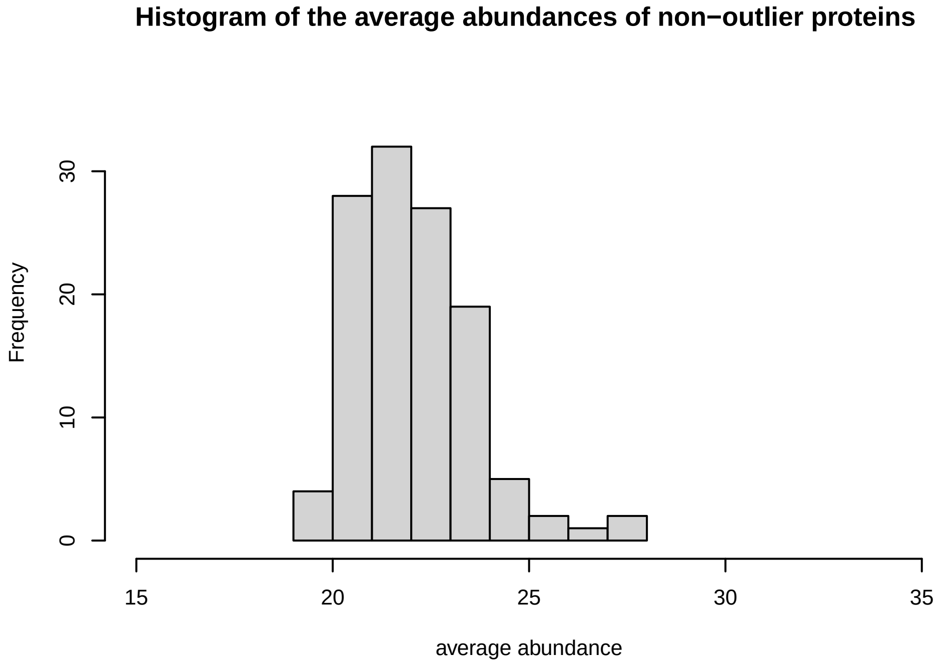
Histogram of the average abundances of non−outlier proteins
Frequency
average abundance

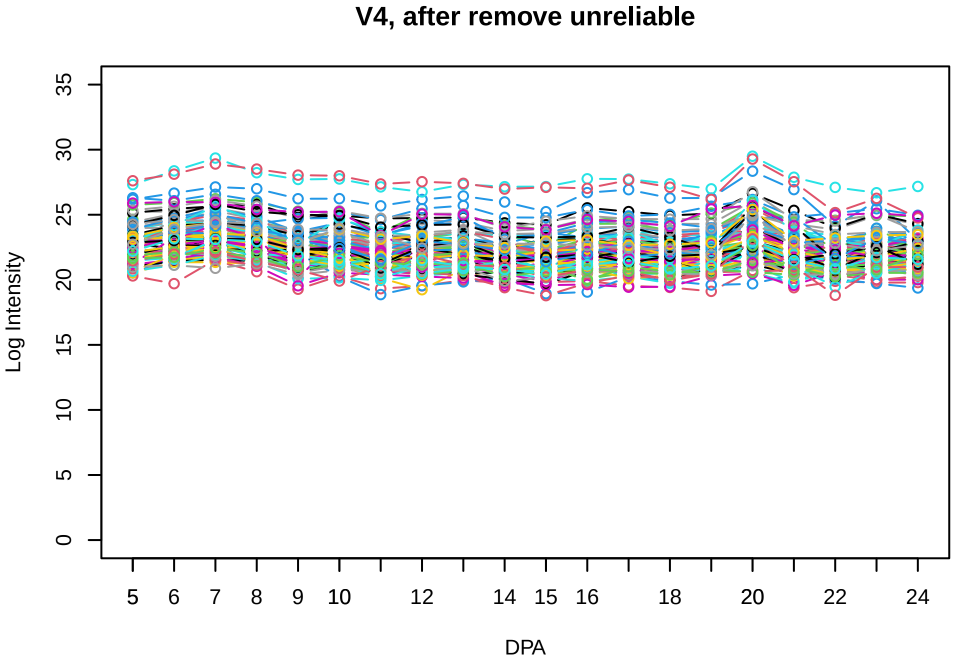
V4, after remove unreliable
Log Intensity
DPA

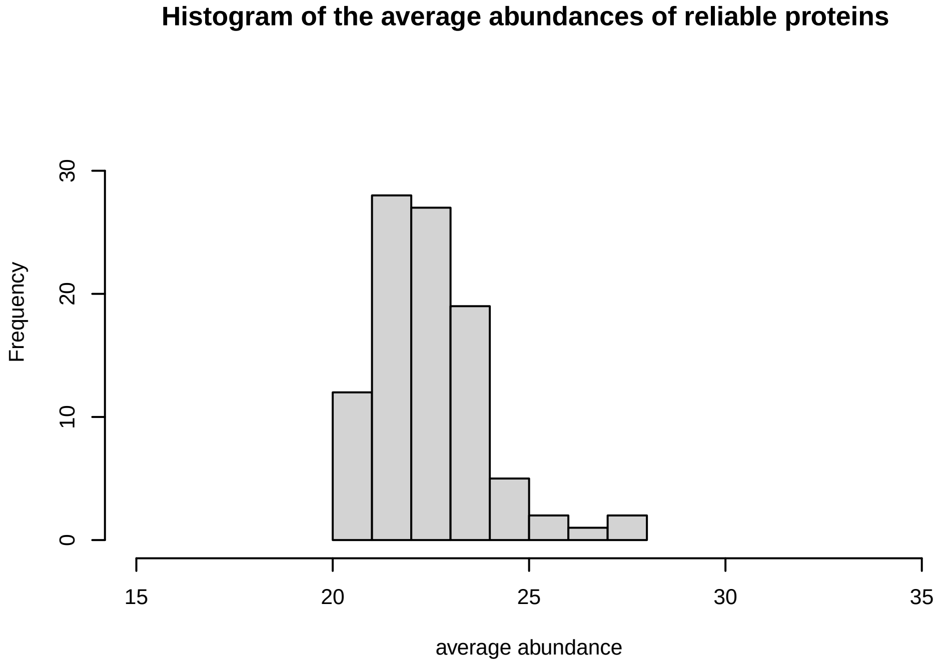
Histogram of the average abundances of reliable proteins
Frequency
average abundance

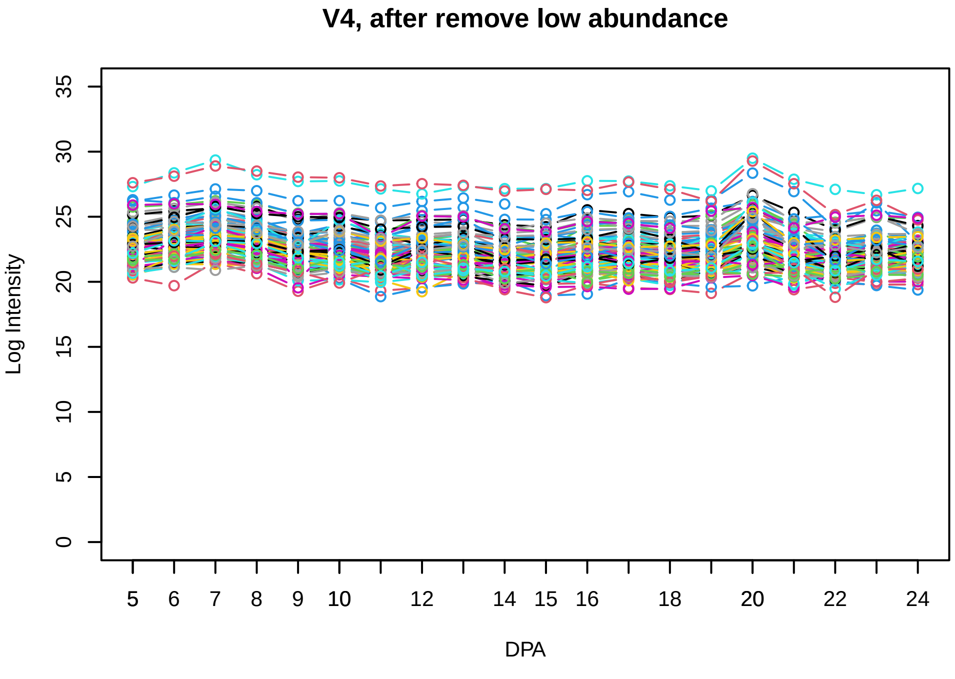
V4, after remove low abundance
Log Intensity
DPA

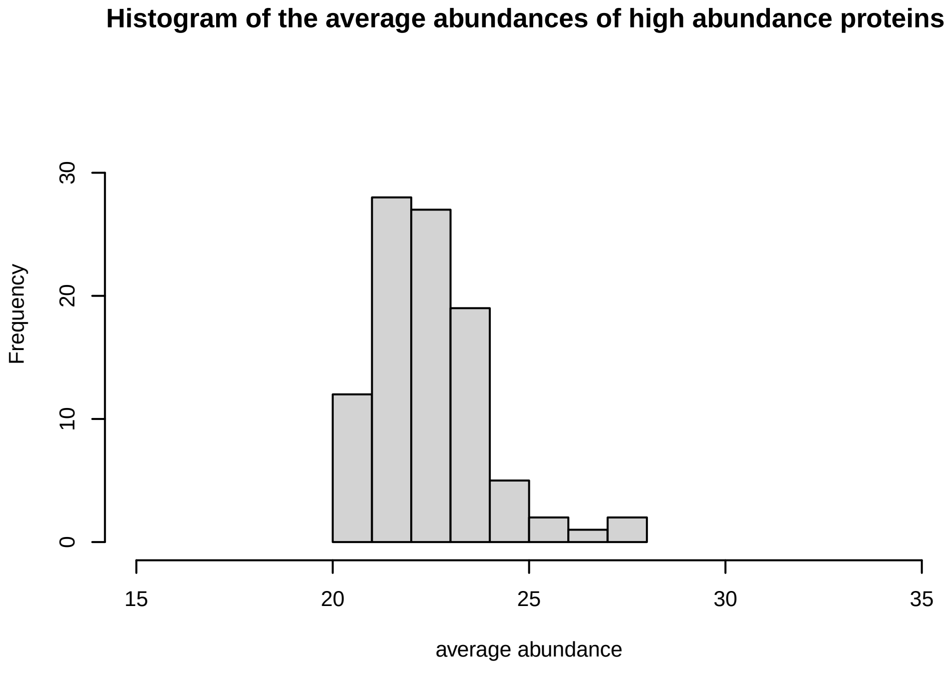
Histogram of the average abundances of high abundance proteins
Frequency
average abundance

**V5**

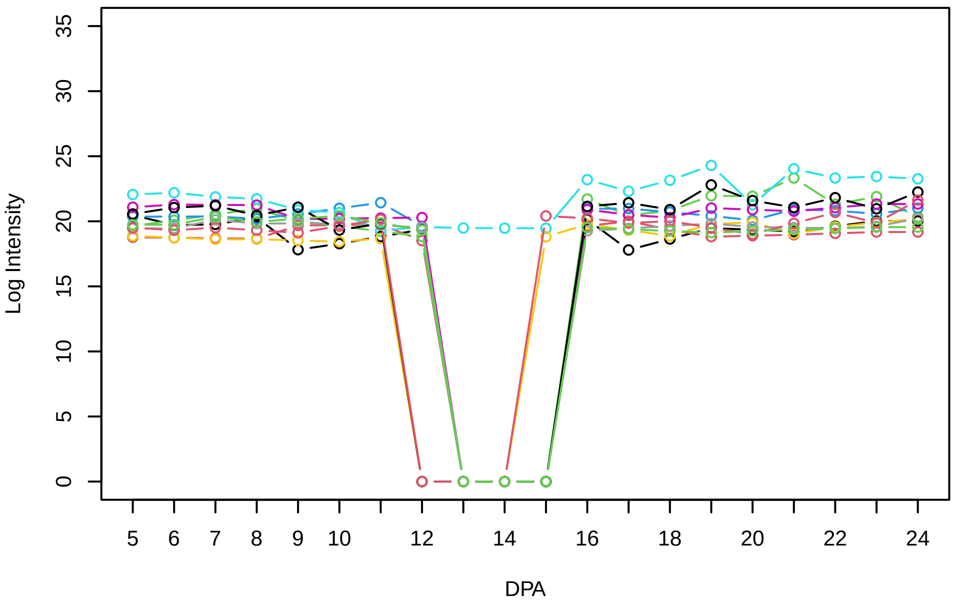


**Histogram of the average abundances of non−zero values**

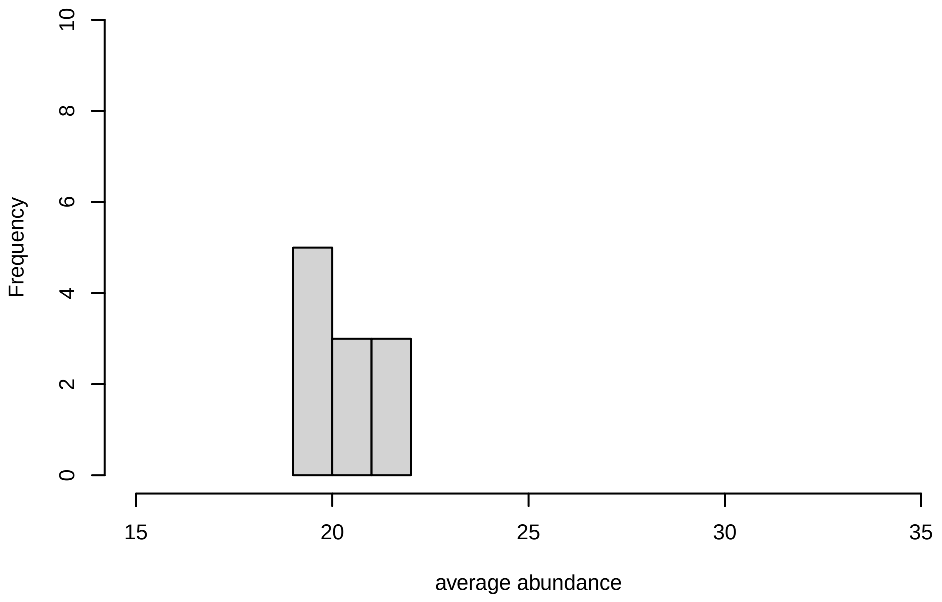


**V5, after remove outlier and cutoff is 0.8**

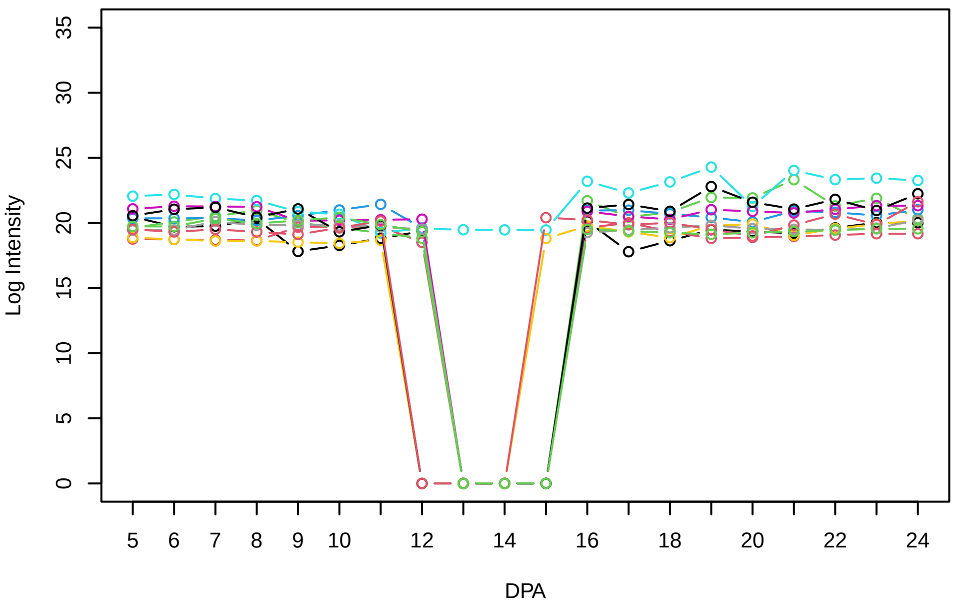


**Histogram of the average abundances of non−outlier proteins**

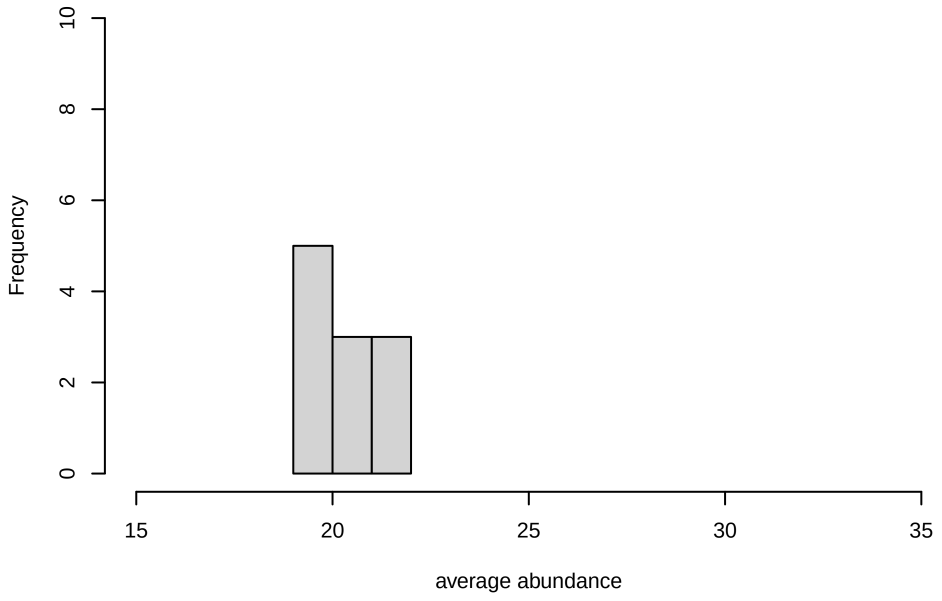


**V5, after remove unreliable**

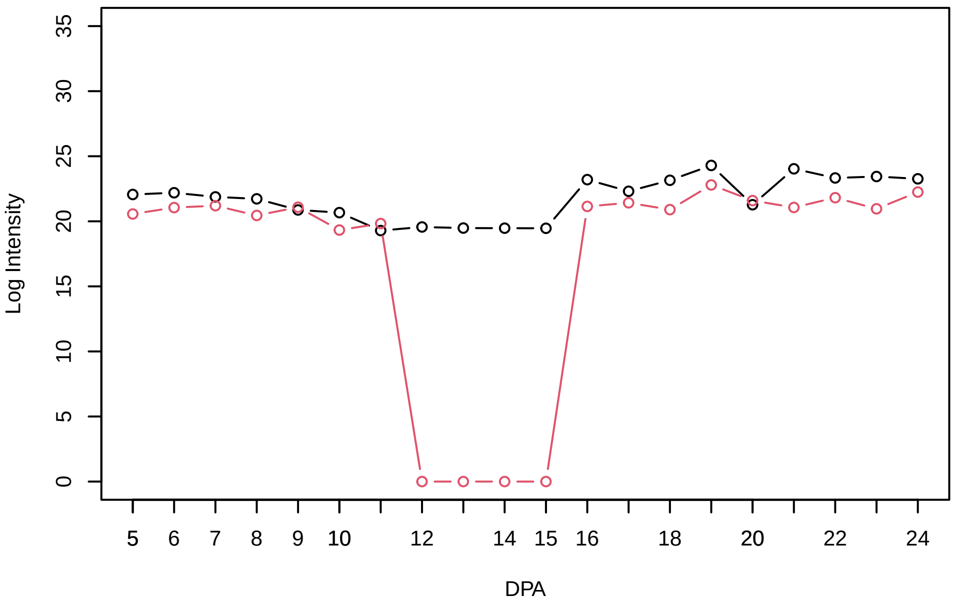


**Histogram of the average abundances of reliable proteins**

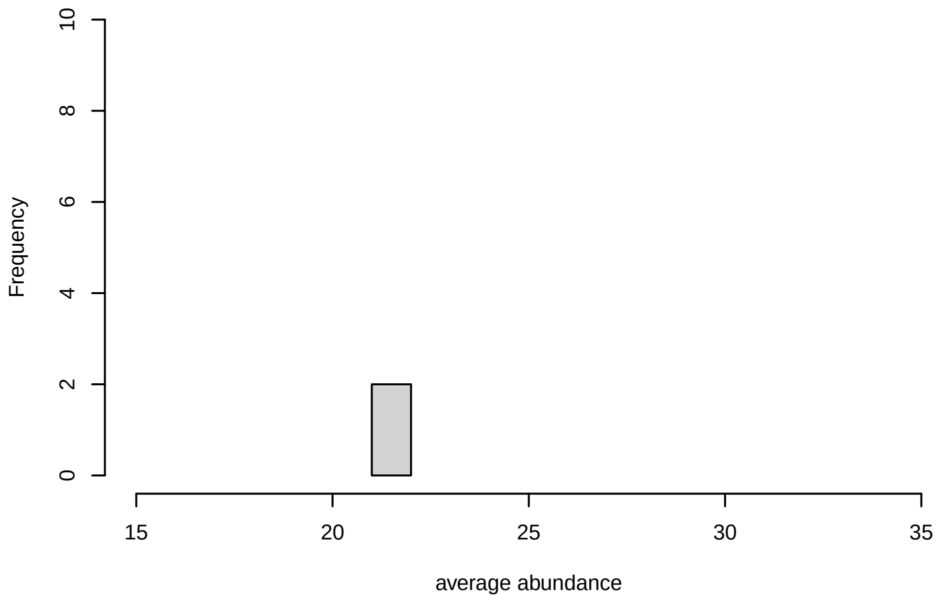


**V5, after remove low abundance**

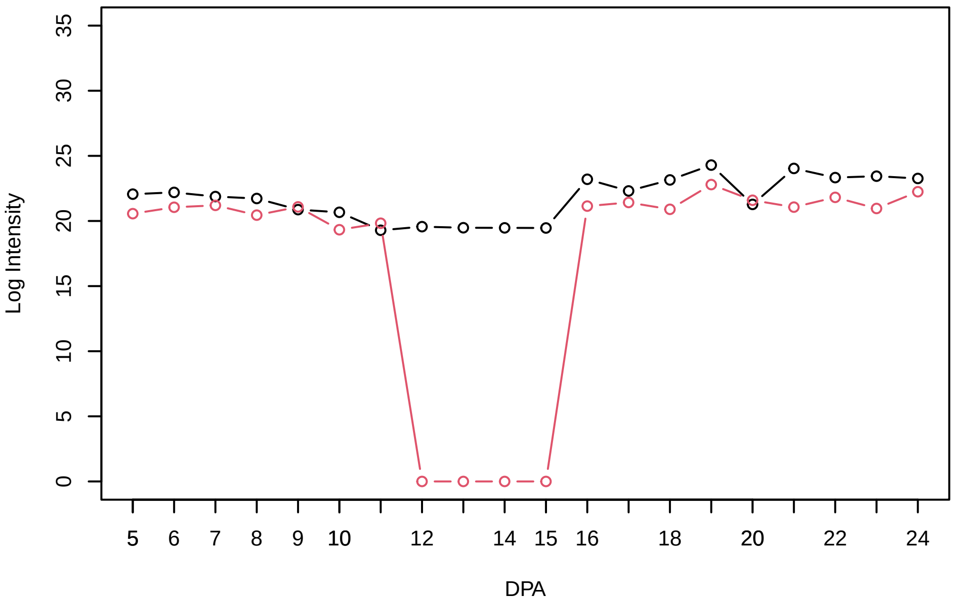


**Histogram of the average abundances of high abundance proteins**

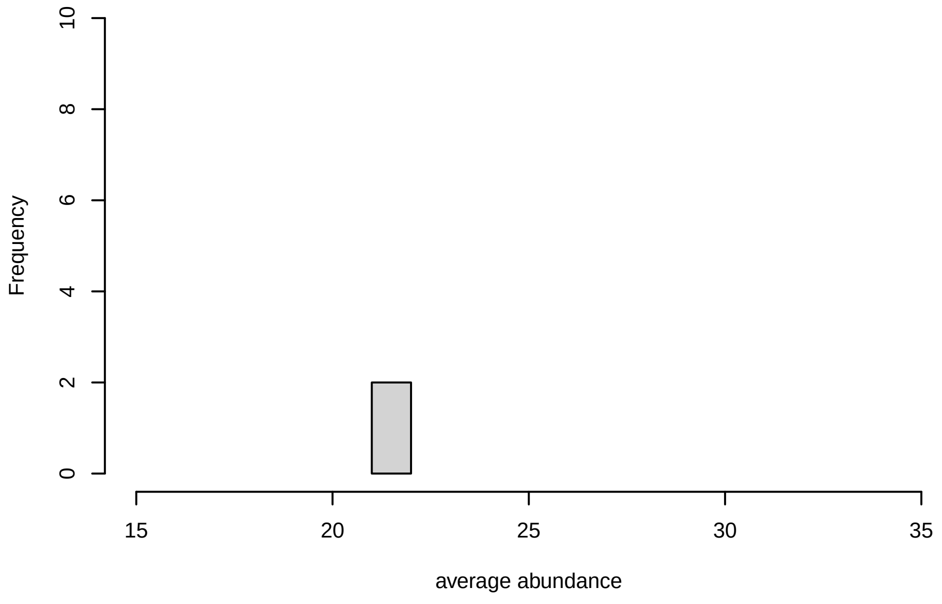

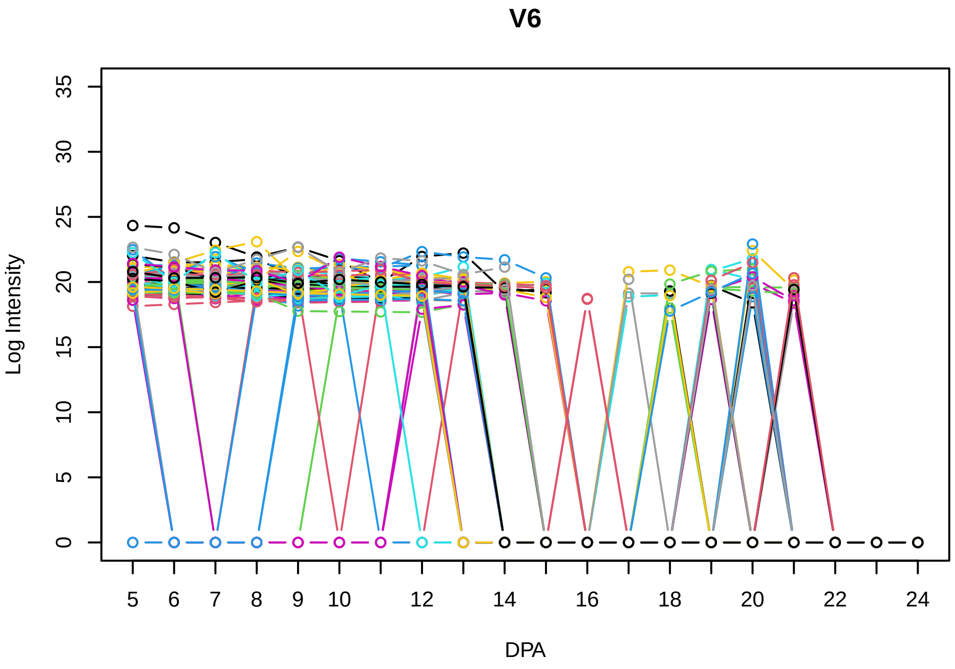
V6
Log Intensity
DPA

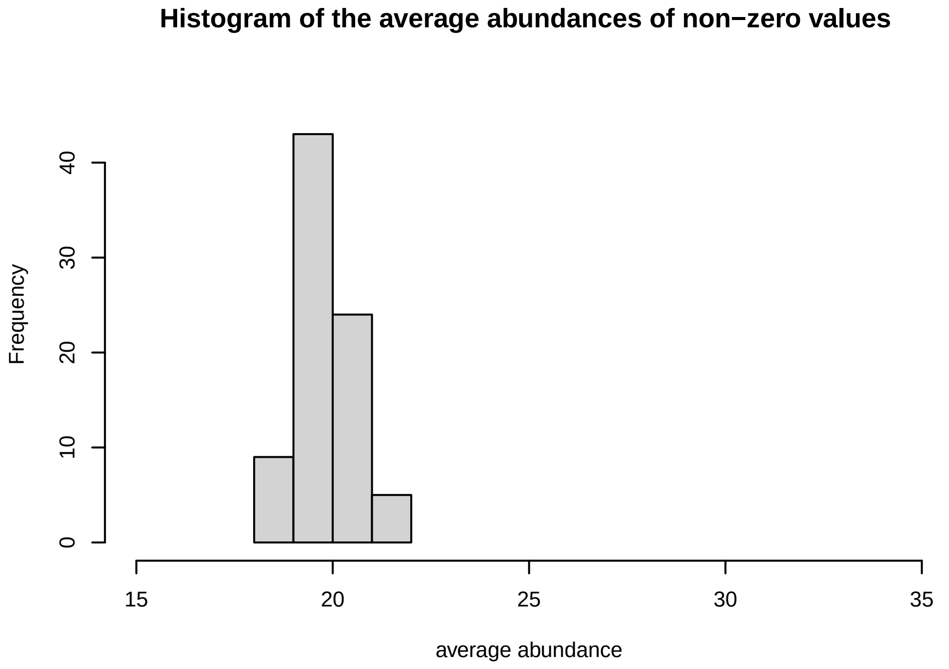
Histogram of the average abundances of non−zero values
Frequency
average abundance

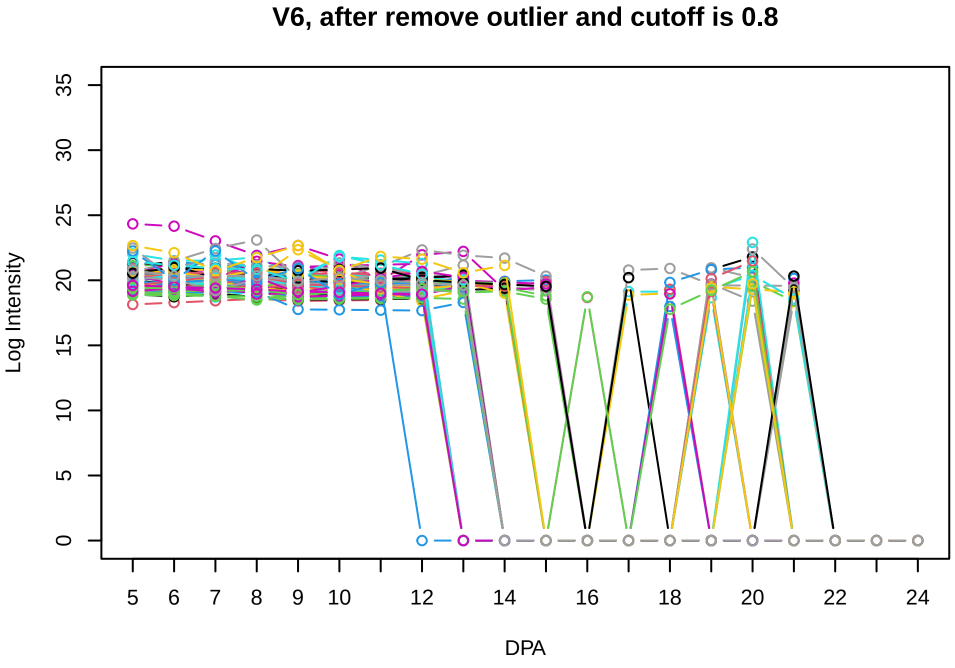
V6, after remove outlier and cutoff is 0.8
Log Intensity
DPA

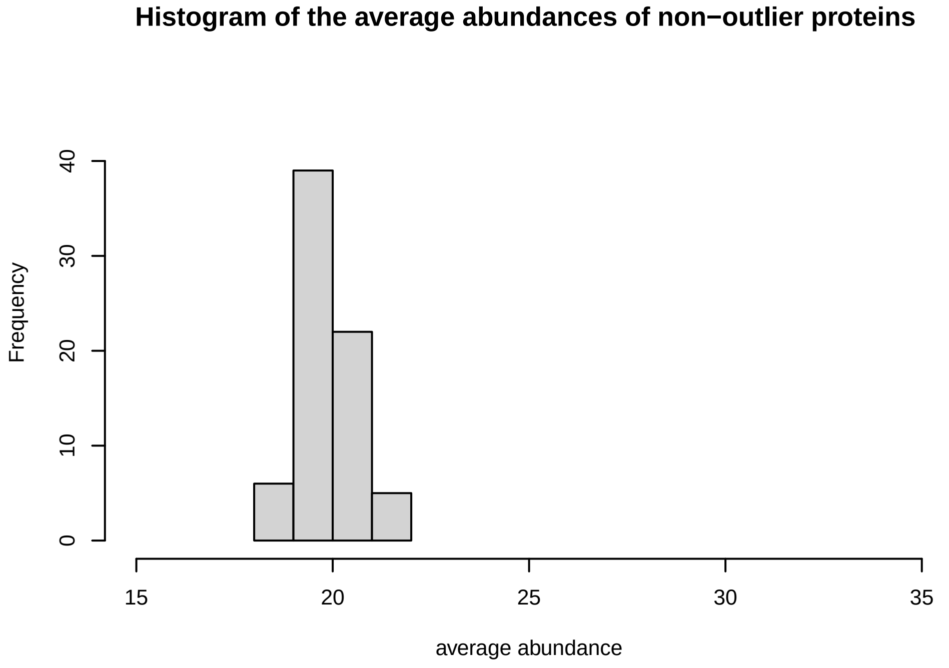
Histogram of the average abundances of non−outlier proteins
Frequency
average abundance

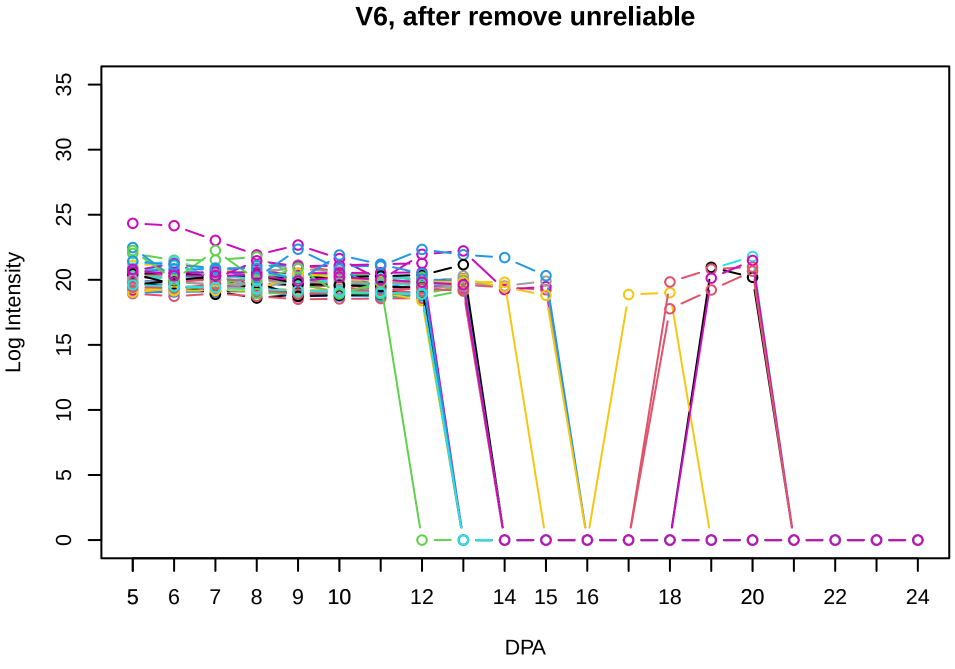
V6, after remove unreliable
Log Intensity
DPA

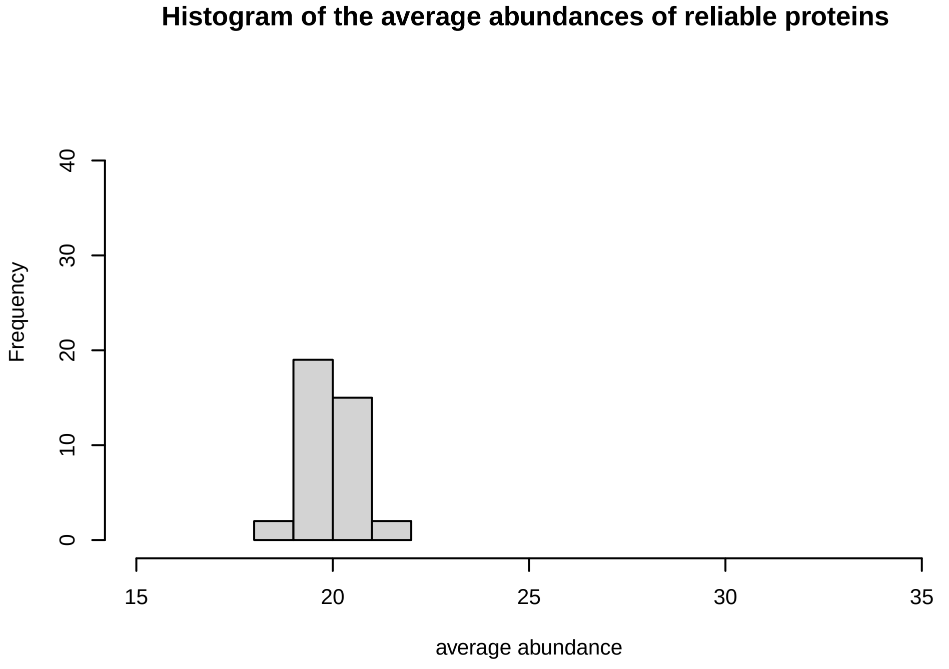
Histogram of the average abundances of reliable proteins
Frequency
average abundance

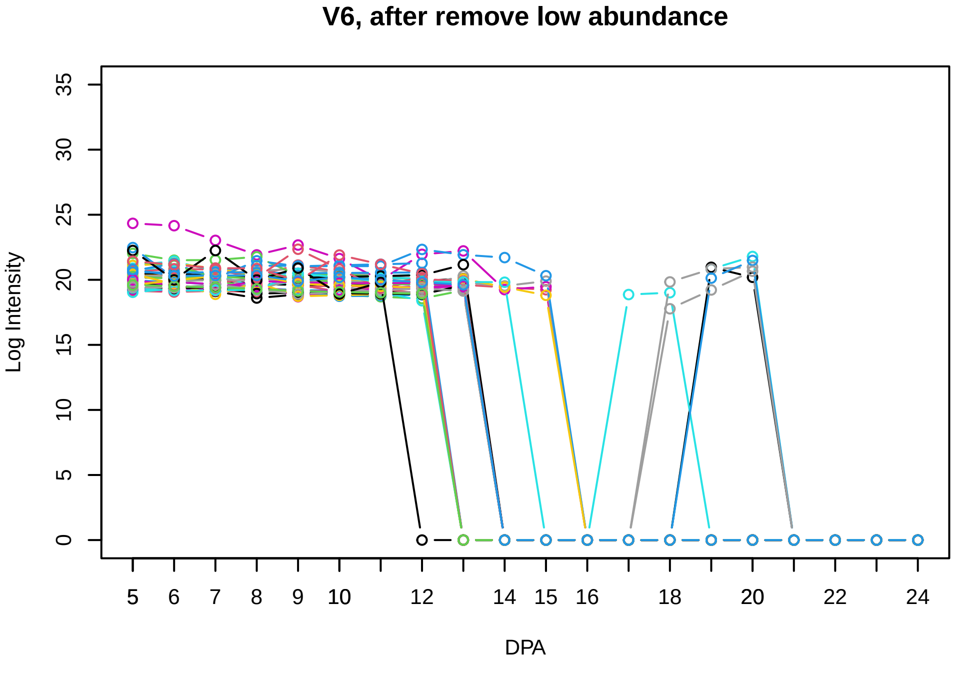
V6, after remove low abundance
Log Intensity
DPA

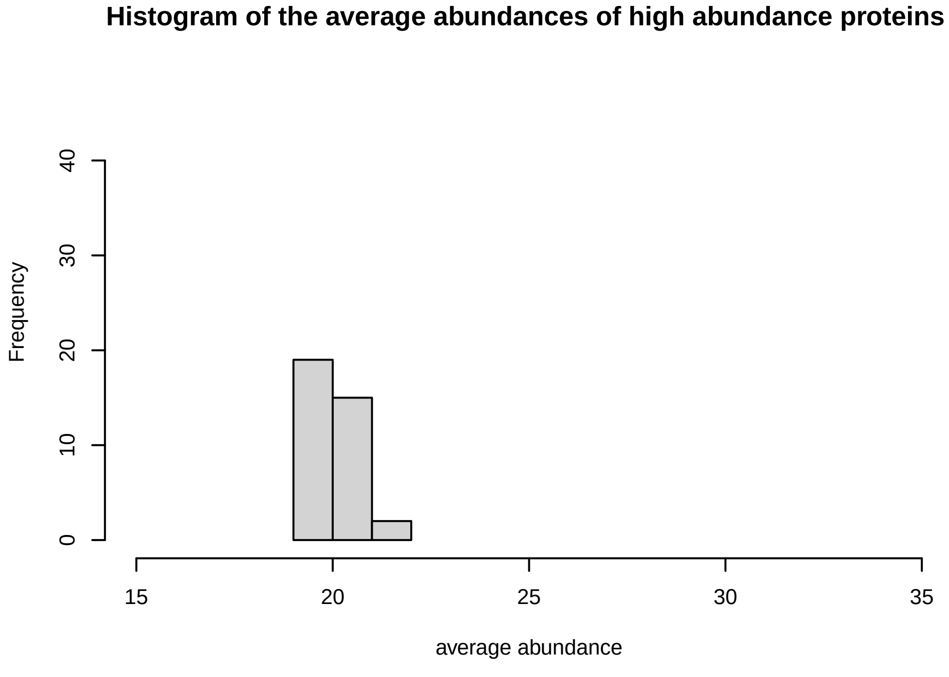
Histogram of the average abundances of high abundance proteins
Frequency
average abundance

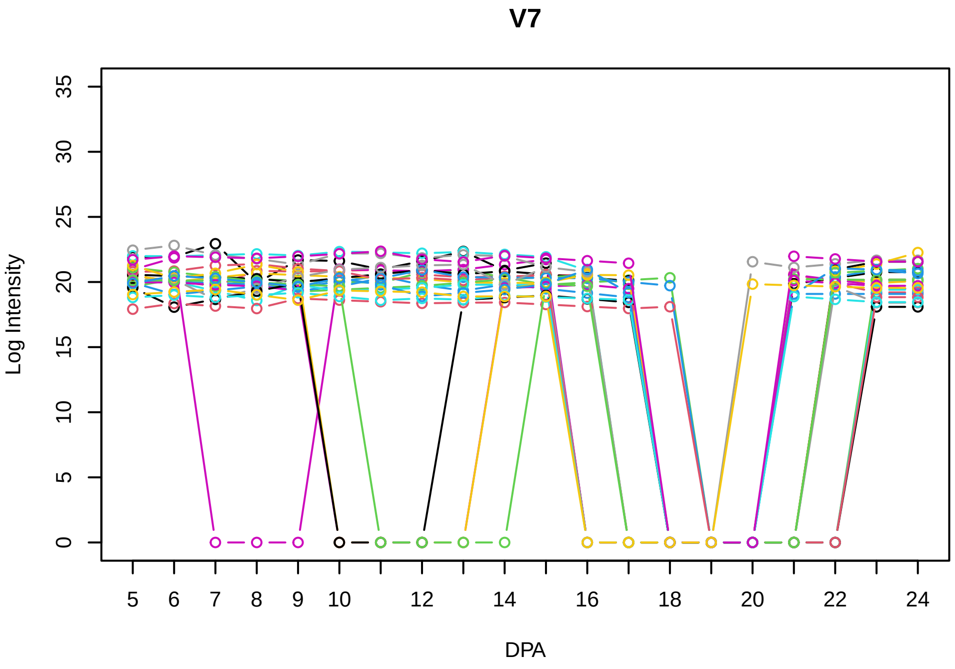
V7
Log Intensity
DPA

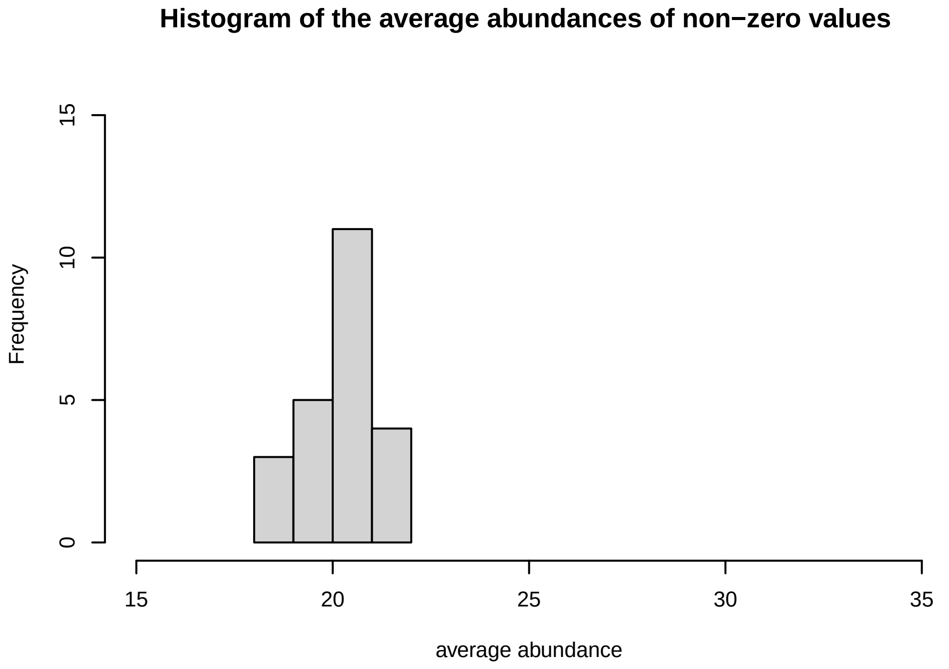
Histogram of the average abundances of non−zero values
Frequency
average abundance

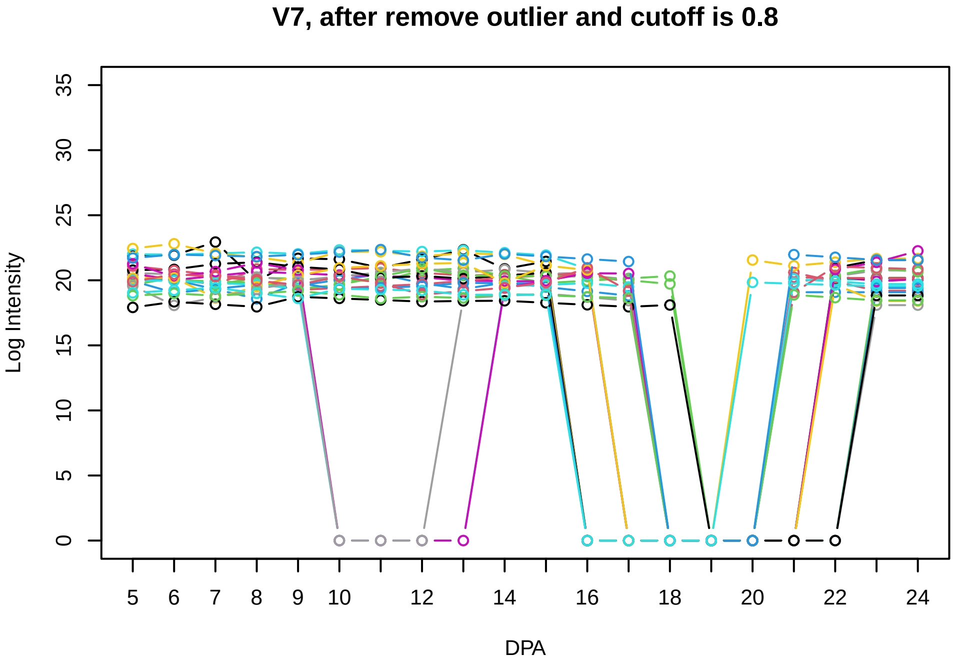
V7, after remove outlier and cutoff is 0.8
Log Intensity
DPA

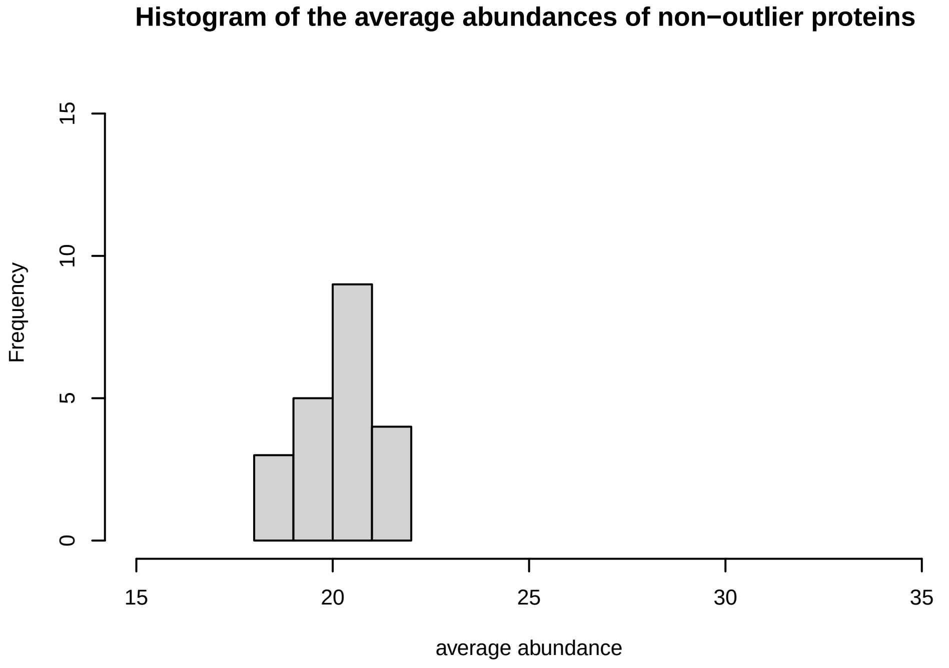
Histogram of the average abundances of non−outlier proteins
Frequency
average abundance

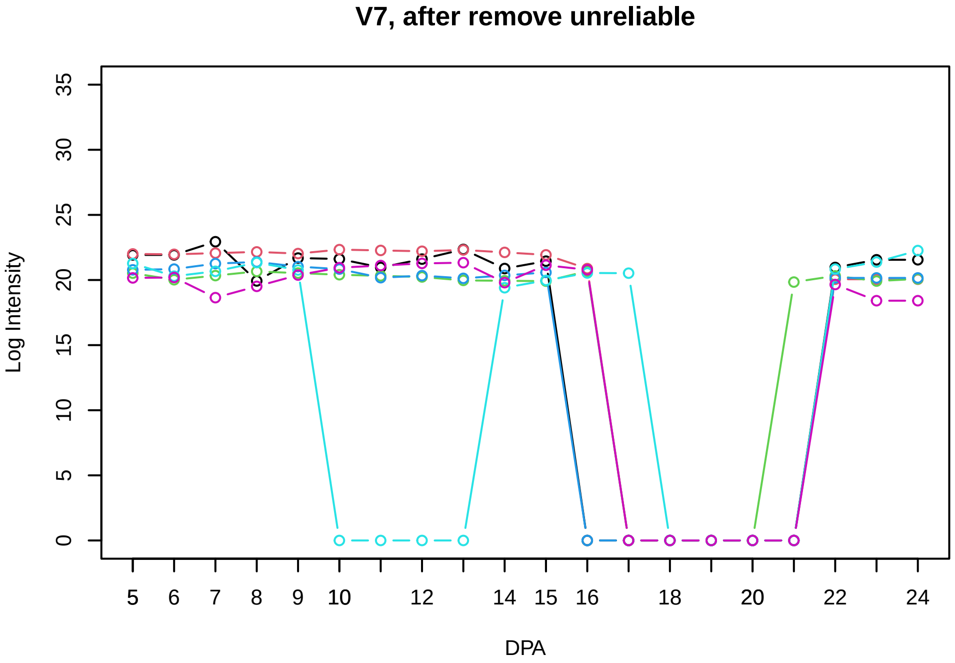
V7, after remove unreliable
Log Intensity
DPA

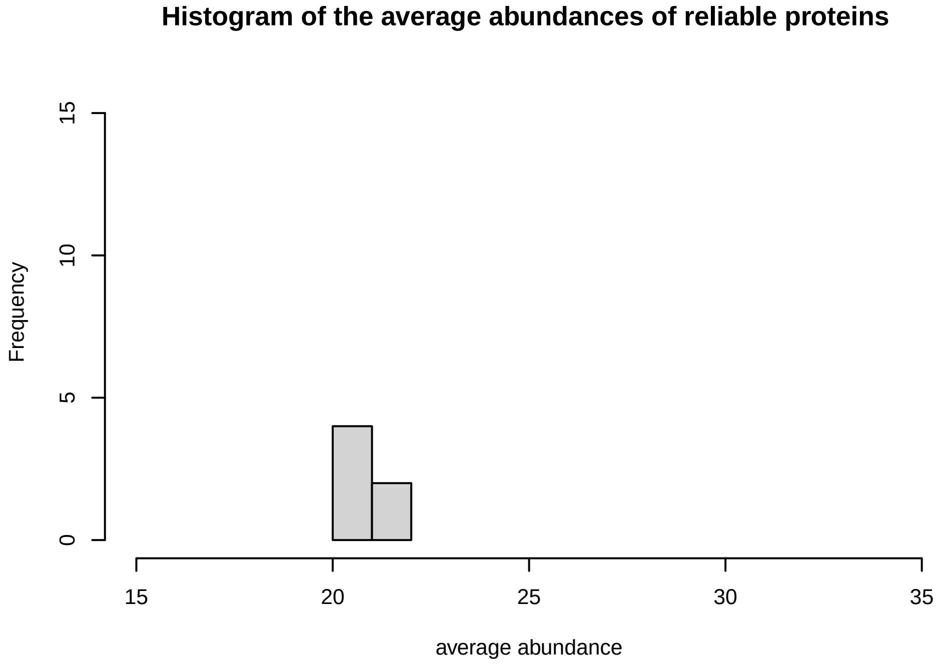
Histogram of the average abundances of reliable proteins
Frequency
average abundance

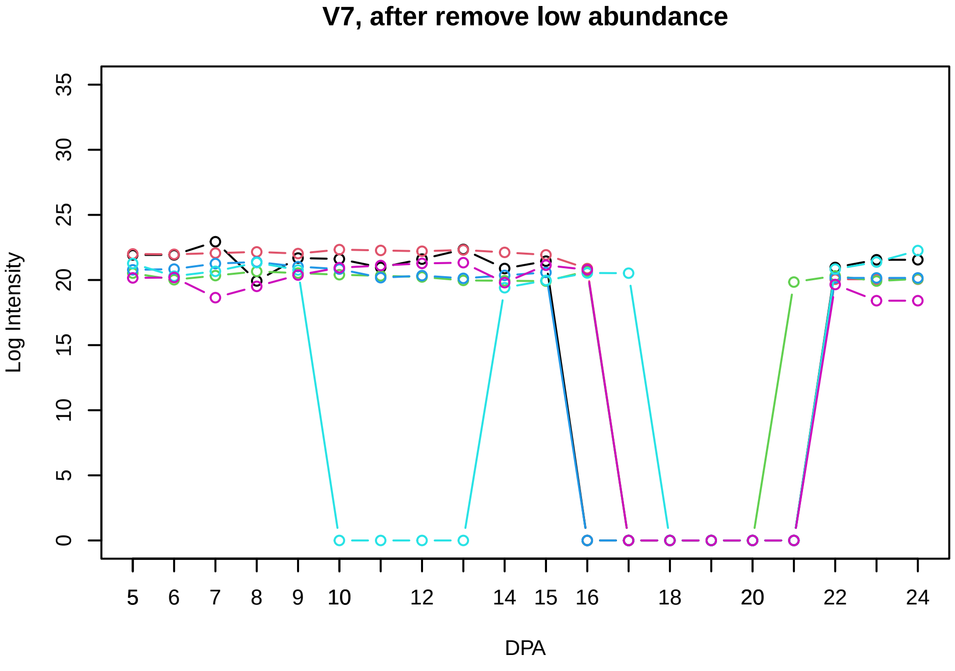
V7, after remove low abundance
Log Intensity
DPA

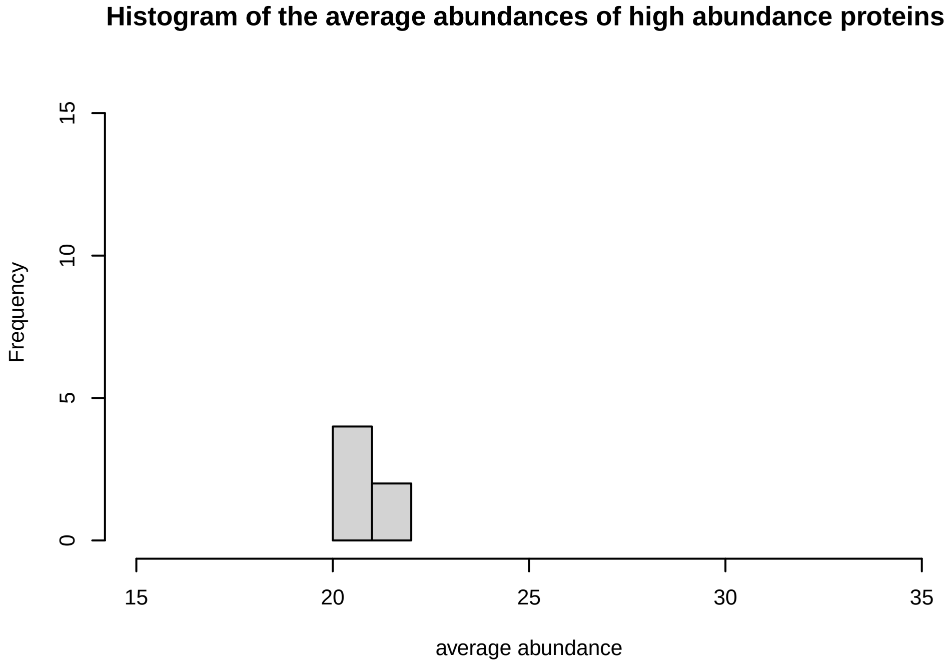
Histogram of the average abundances of high abundance proteins
Frequency
average abundance

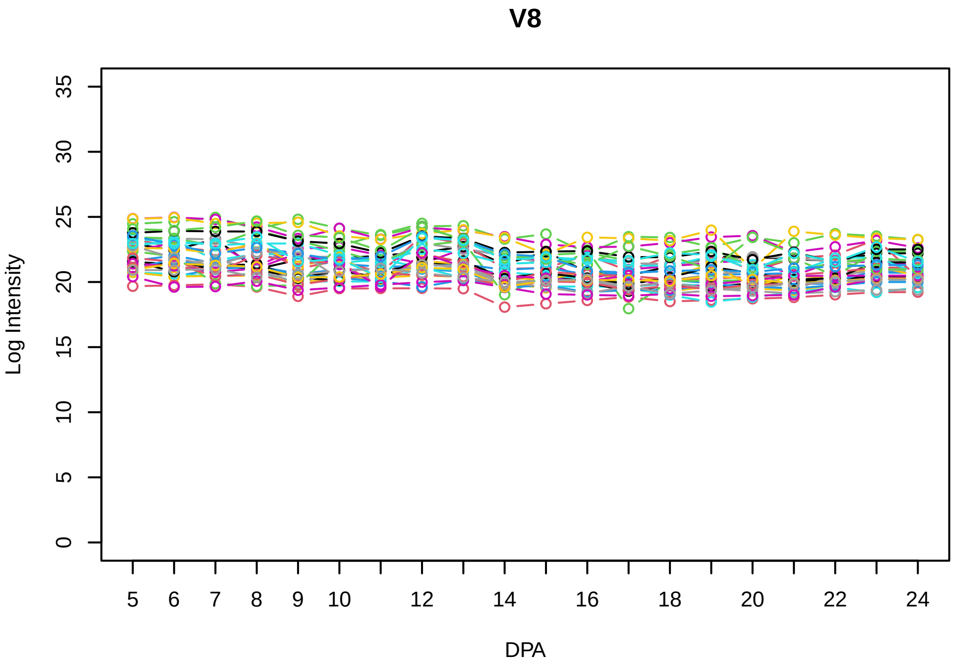
V8
Log Intensity
DPA

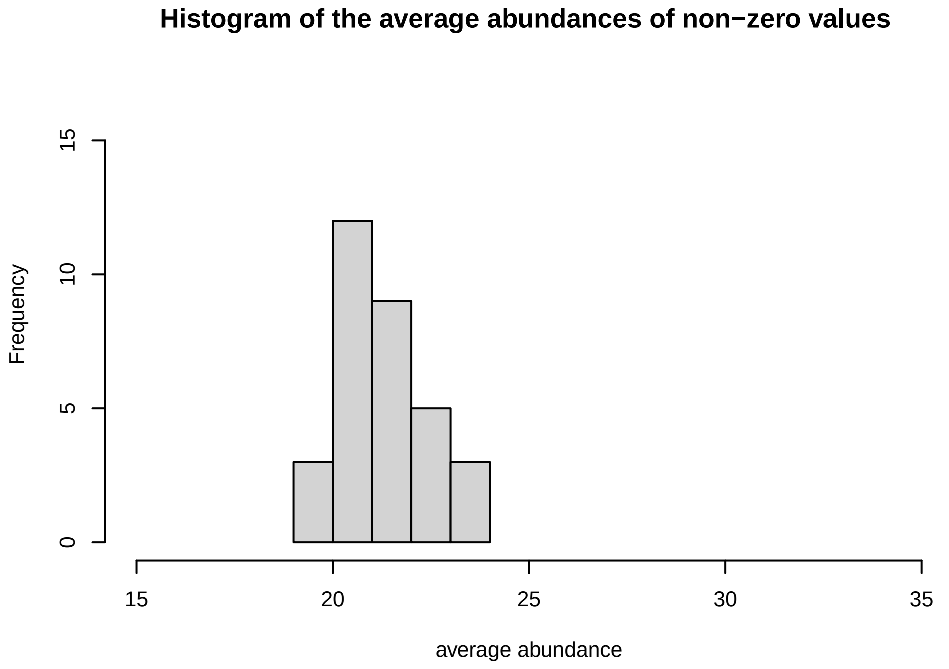
Histogram of the average abundances of non−zero values
Frequency
average abundance

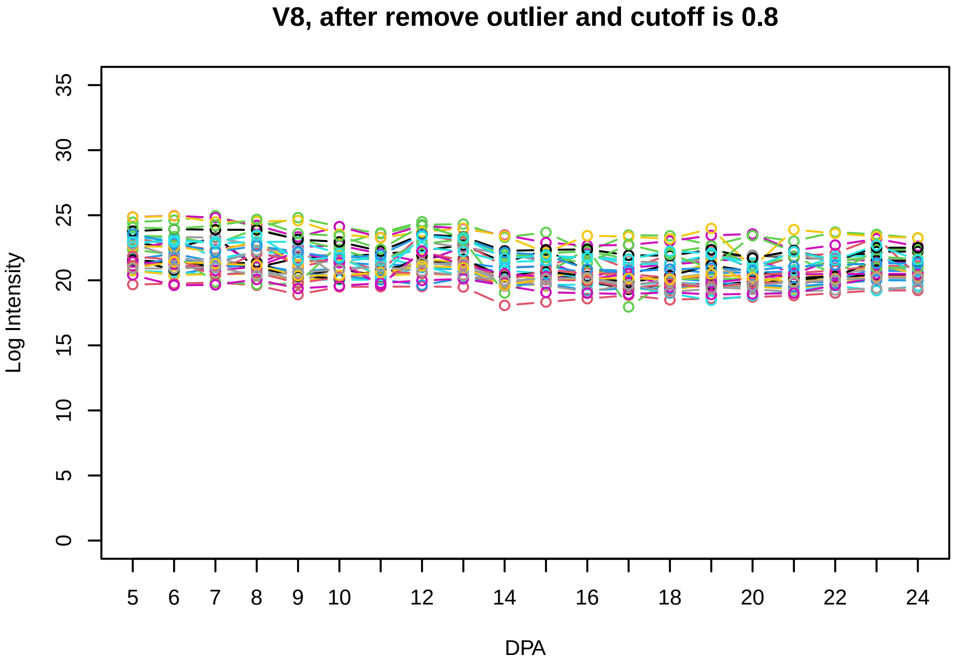
V8, after remove outlier and cutoff is 0.8
Log Intensity
DPA

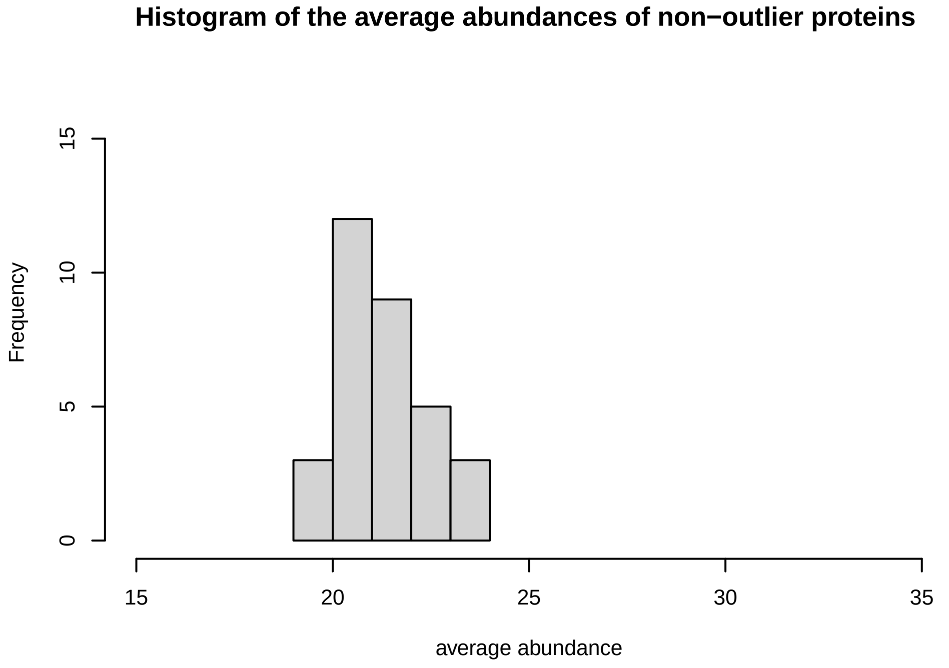
Histogram of the average abundances of non−outlier proteins
Frequency
average abundance

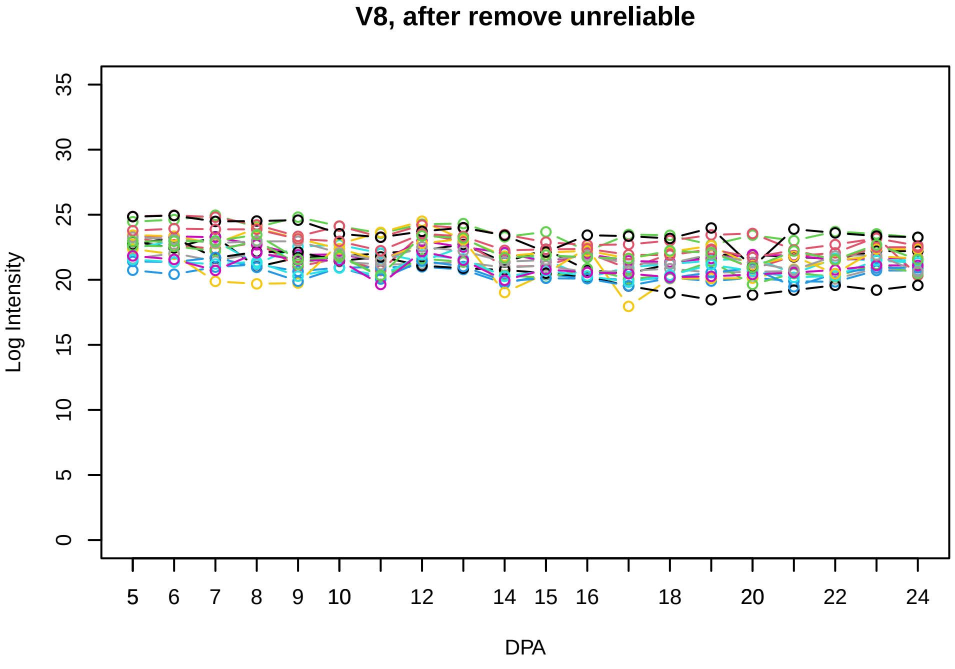
V8, after remove unreliable
Log Intensity
DPA

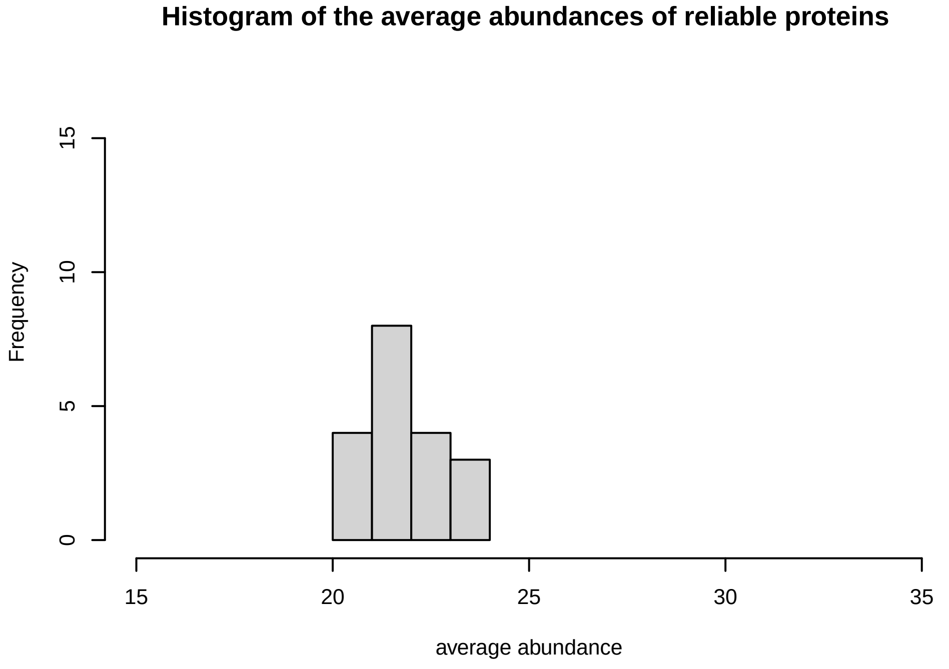
Histogram of the average abundances of reliable proteins
Frequency
average abundance

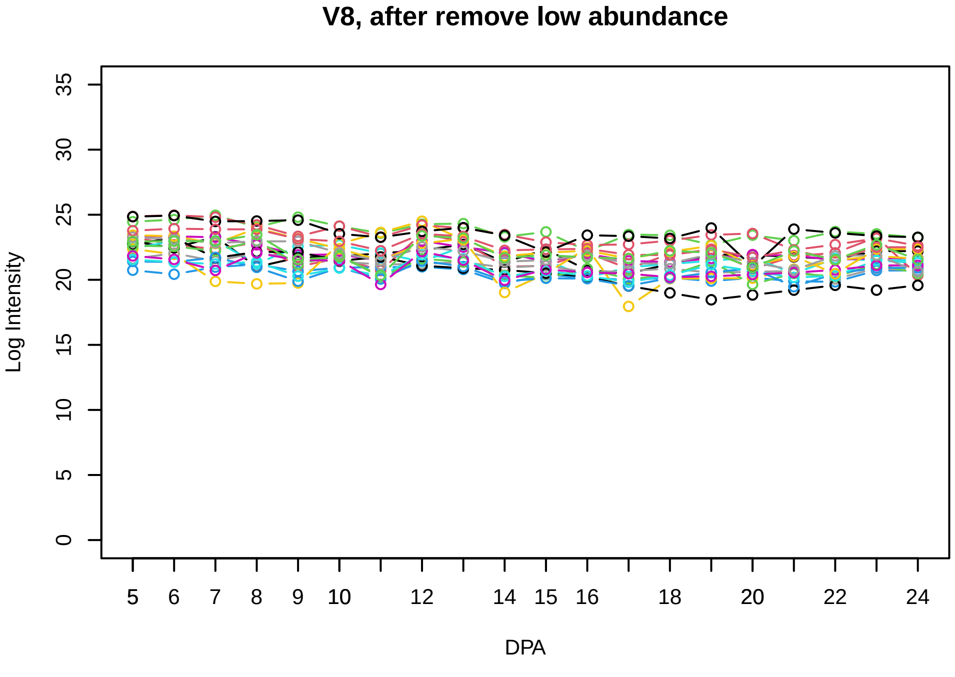
V8, after remove low abundance
Log Intensity
DPA

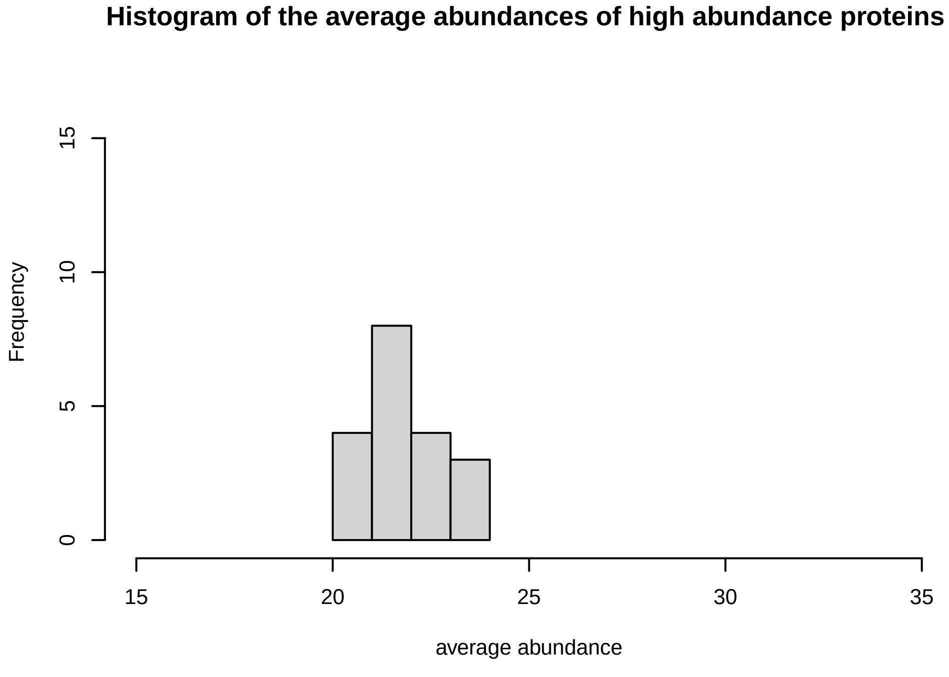
Histogram of the average abundances of high abundance proteins
Frequency
average abundance

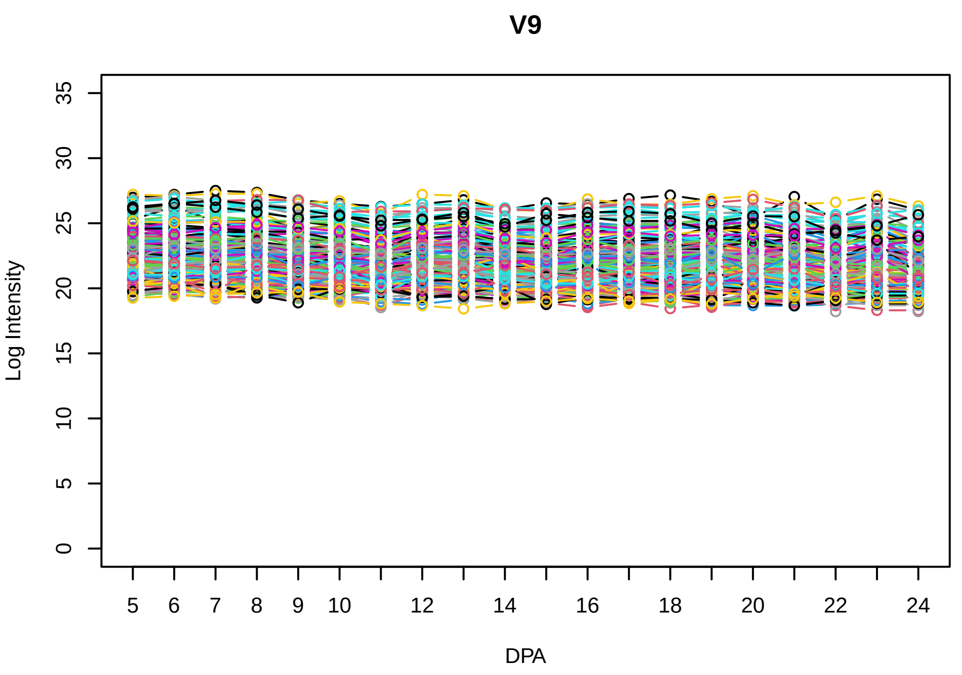
V9
Log Intensity
DPA

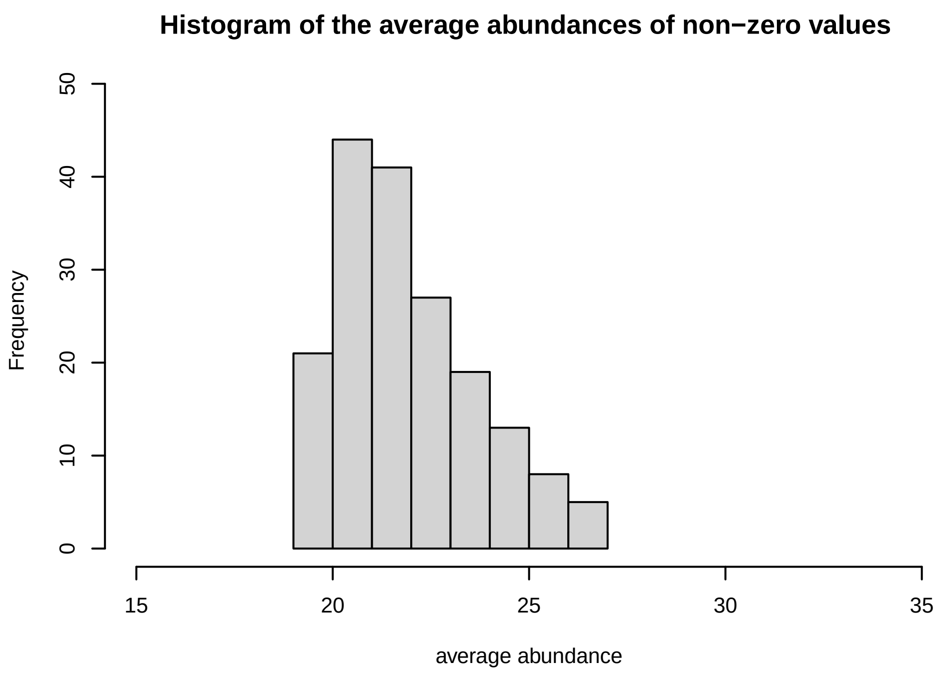
Histogram of the average abundances of non−zero values
Frequency
average abundance

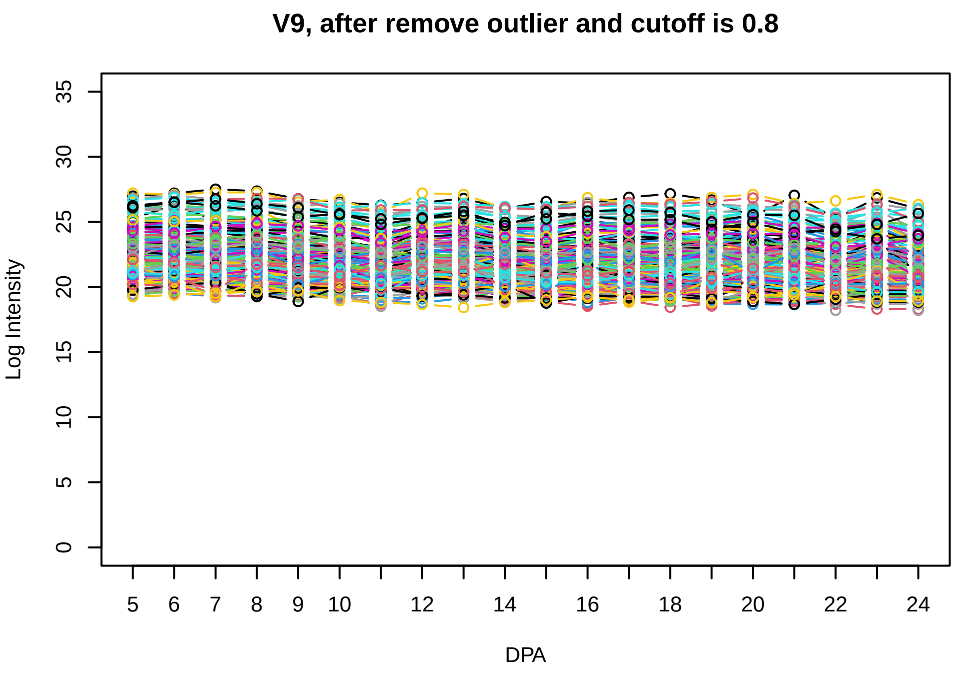
V9, after remove outlier and cutoff is 0.8
Log Intensity
DPA

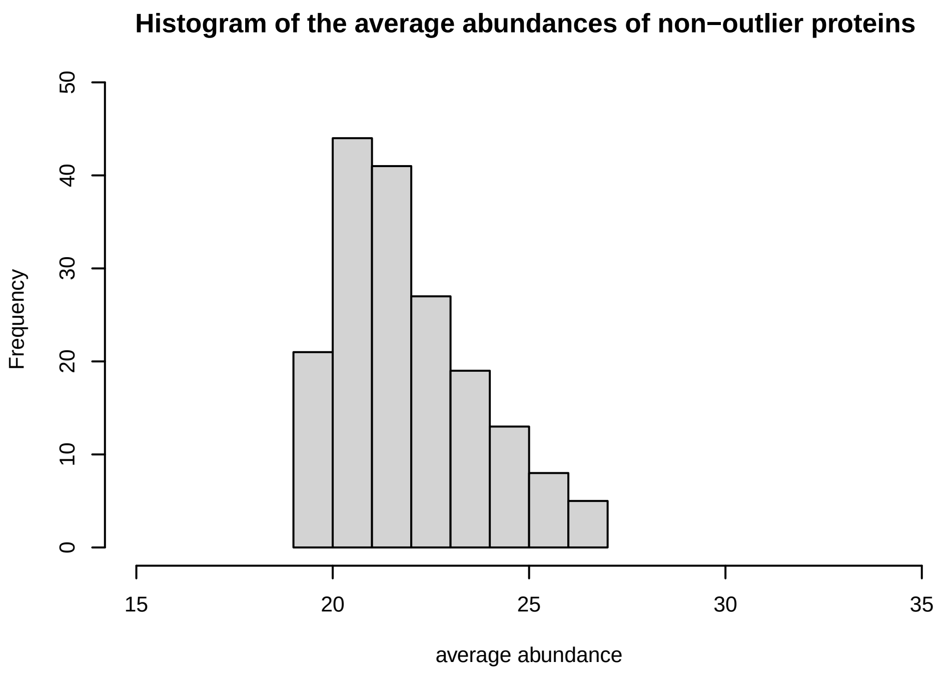
Histogram of the average abundances of non−outlier proteins
Frequency
average abundance

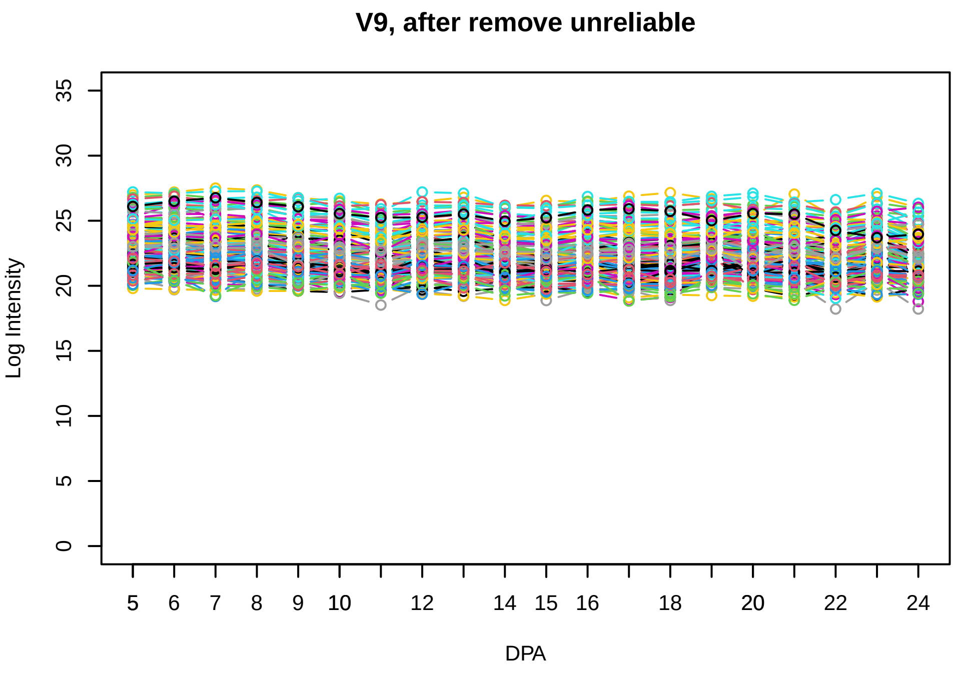
V9, after remove unreliable
Log Intensity
DPA

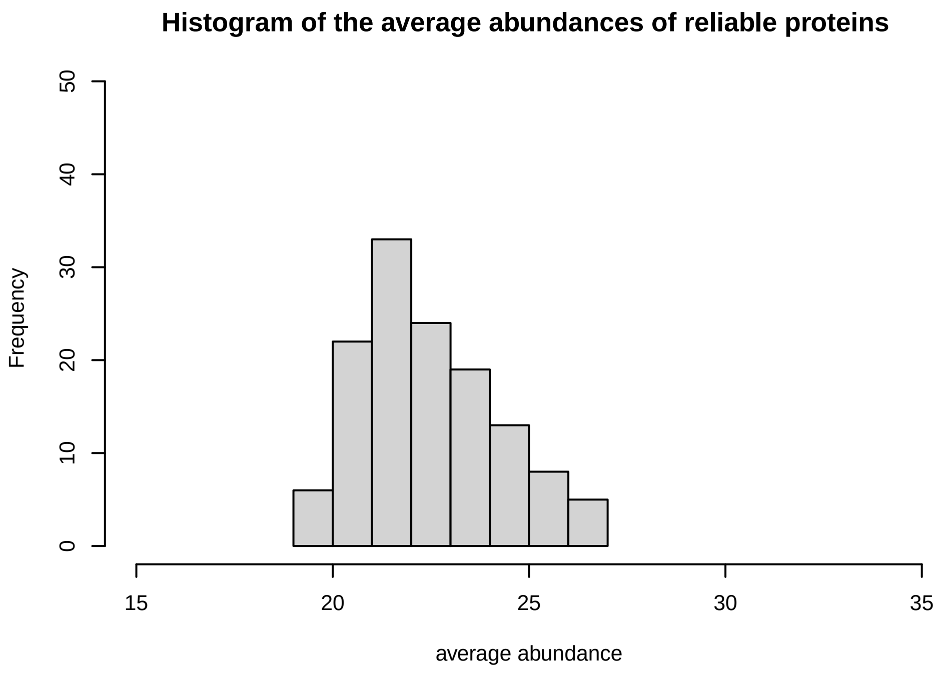
Histogram of the average abundances of reliable proteins
Frequency
average abundance

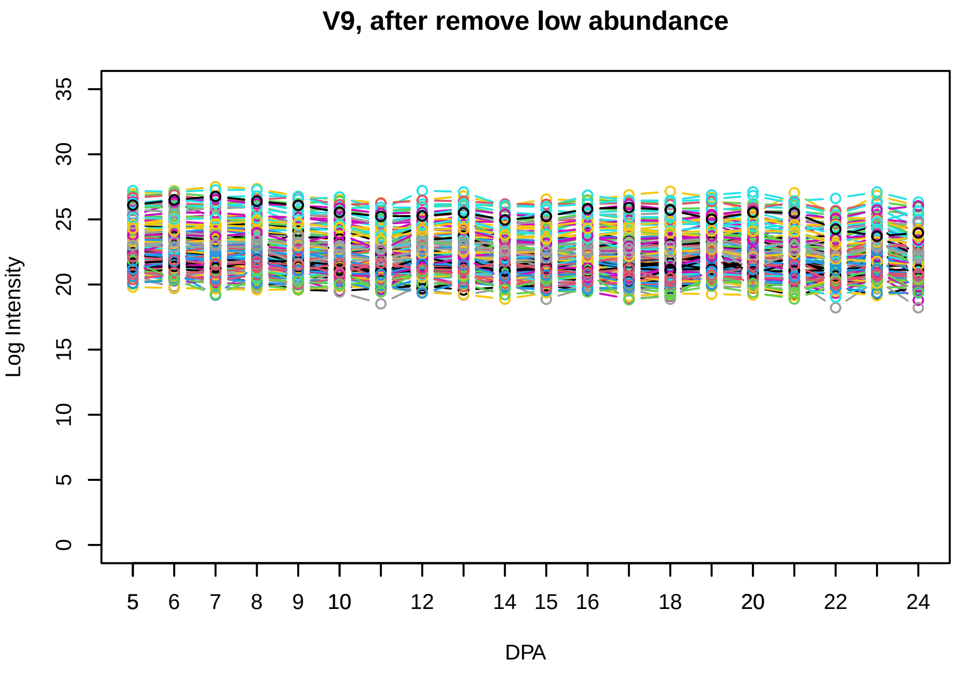
V9, after remove low abundance
Log Intensity
DPA

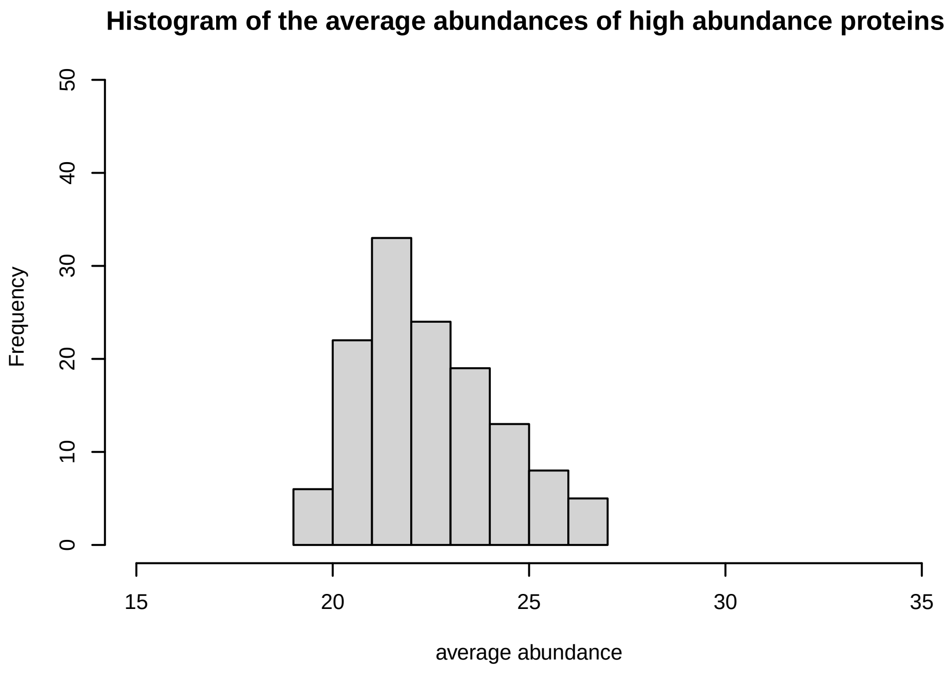
Histogram of the average abundances of high abundance proteins
Frequency
average abundance

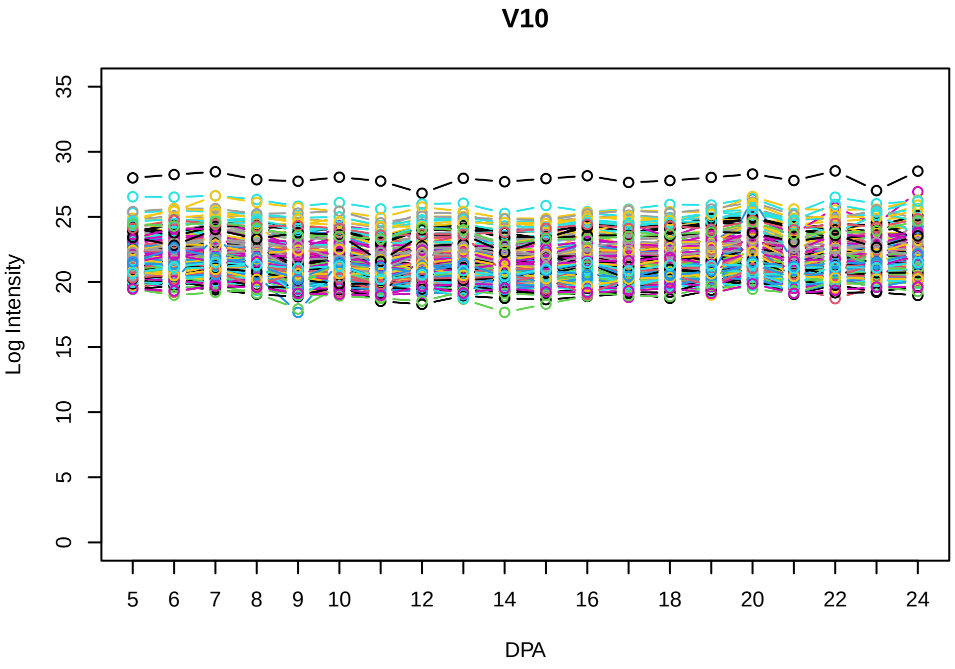
V10
Log Intensity
DPA

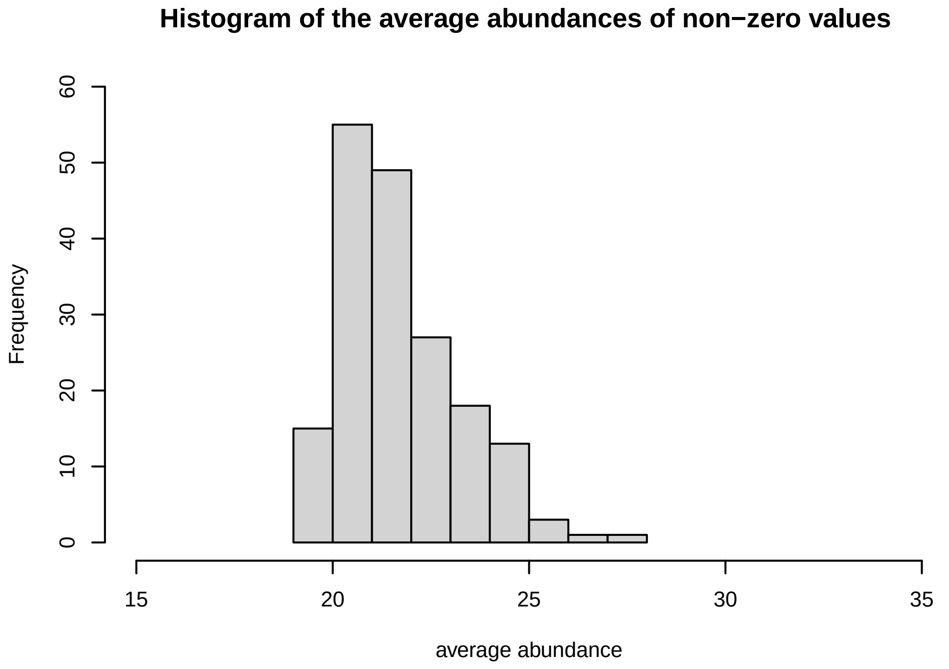
Histogram of the average abundances of non−zero values
Frequency
average abundance

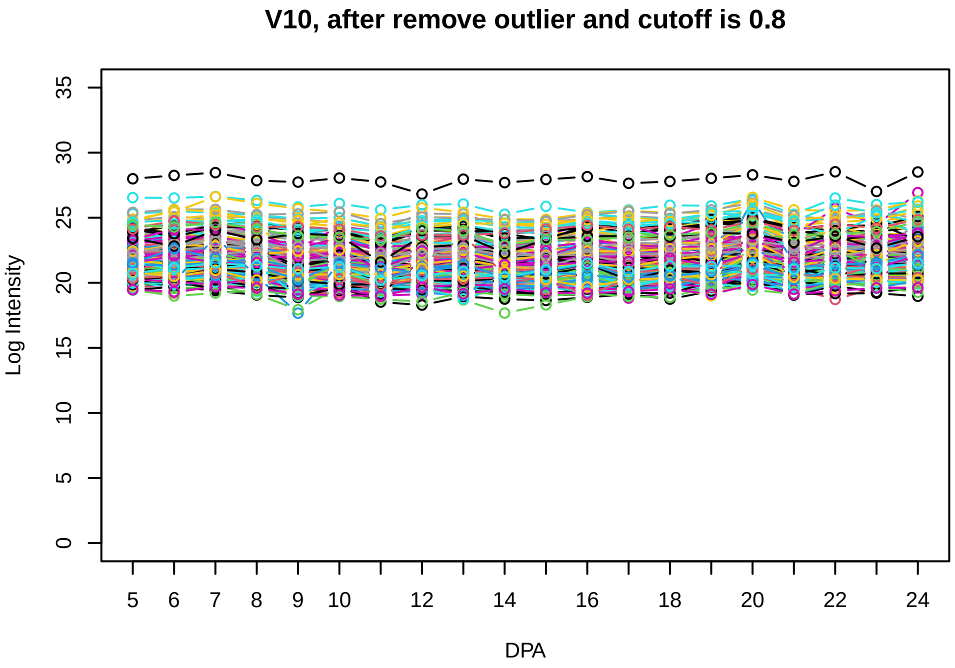
V10, after remove outlier and cutoff is 0.8
Log Intensity
DPA

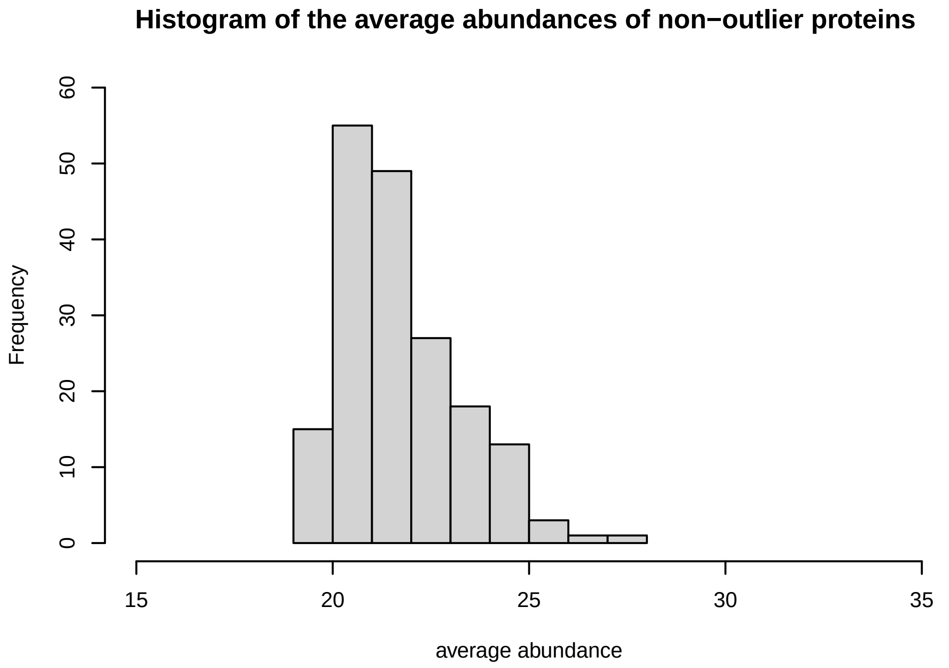
Histogram of the average abundances of non−outlier proteins
Frequency
average abundance

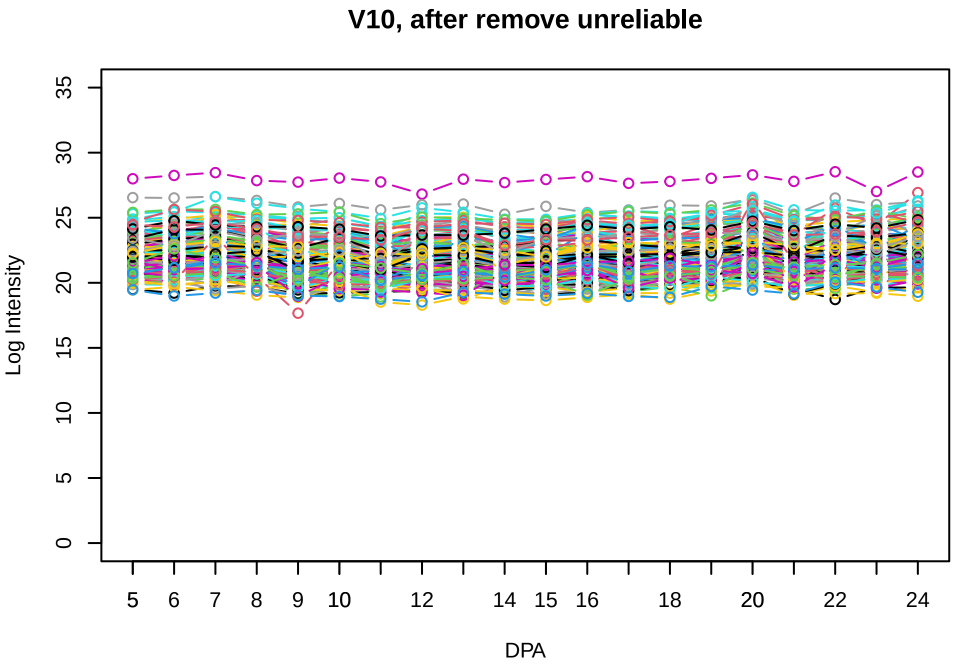
V10, after remove unreliable
Log Intensity
DPA

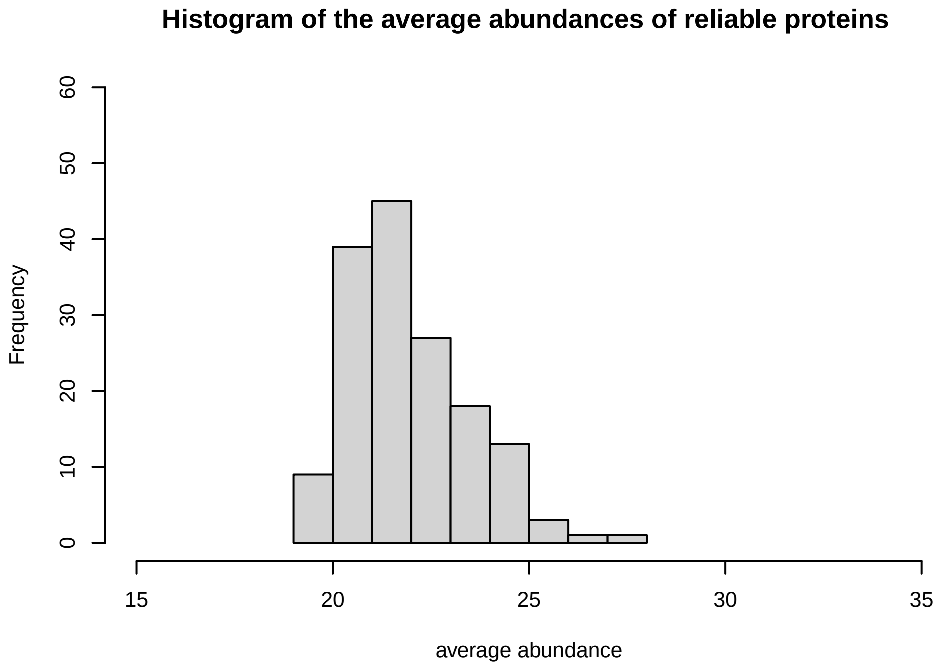
Histogram of the average abundances of reliable proteins
Frequency
average abundance

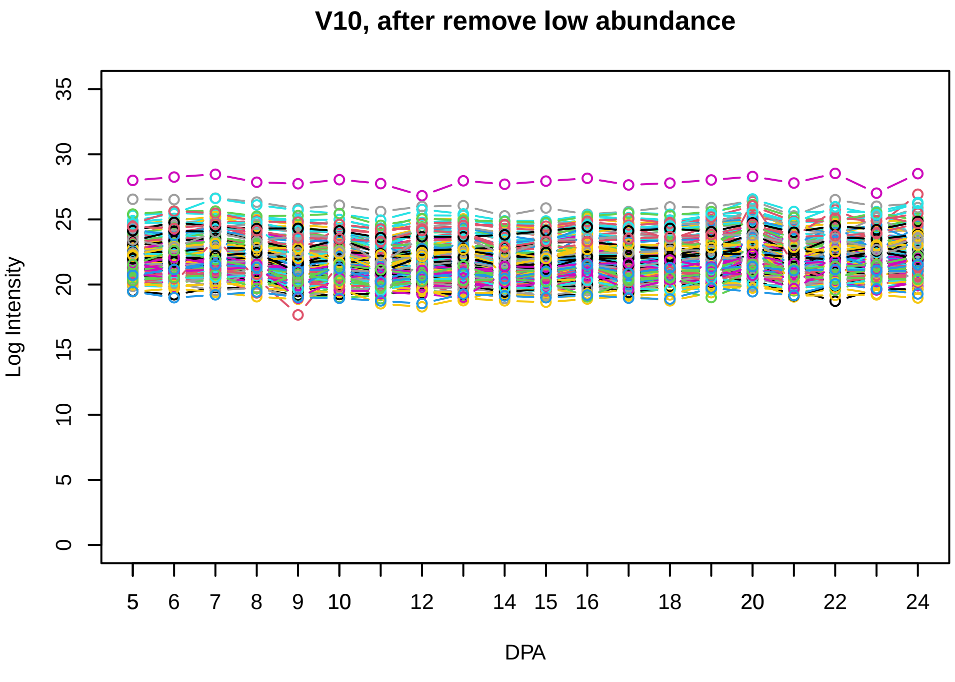
V10, after remove low abundance
Log Intensity
DPA

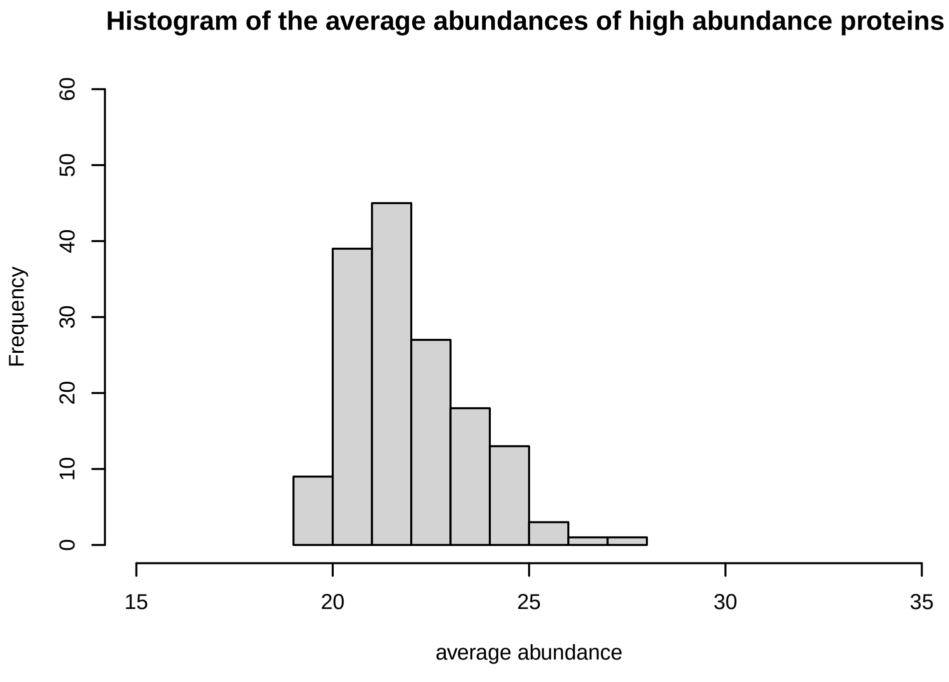
Histogram of the average abundances of high abundance proteins
Frequency
average abundance

V11

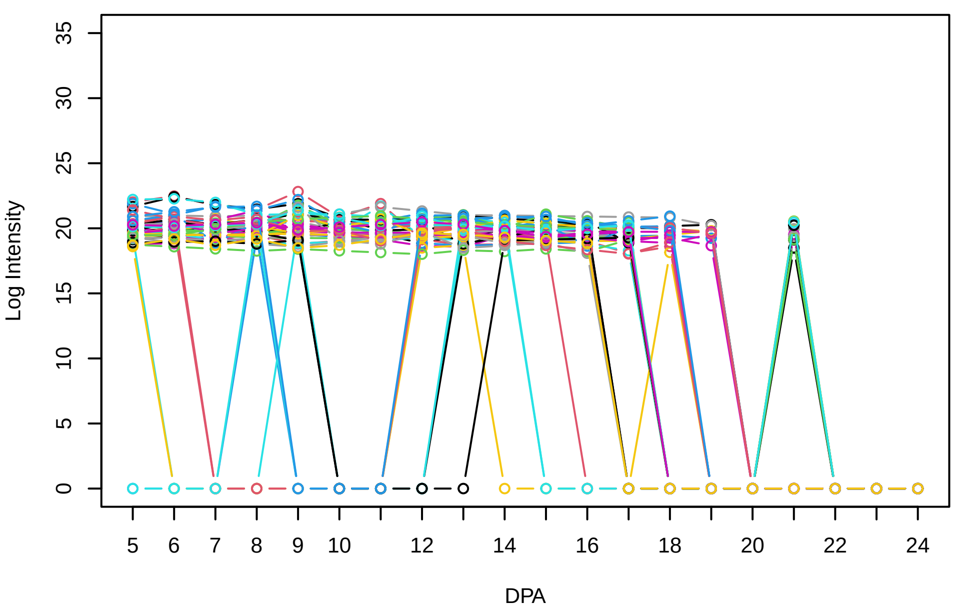


Histogram of the average abundances of non−zero values

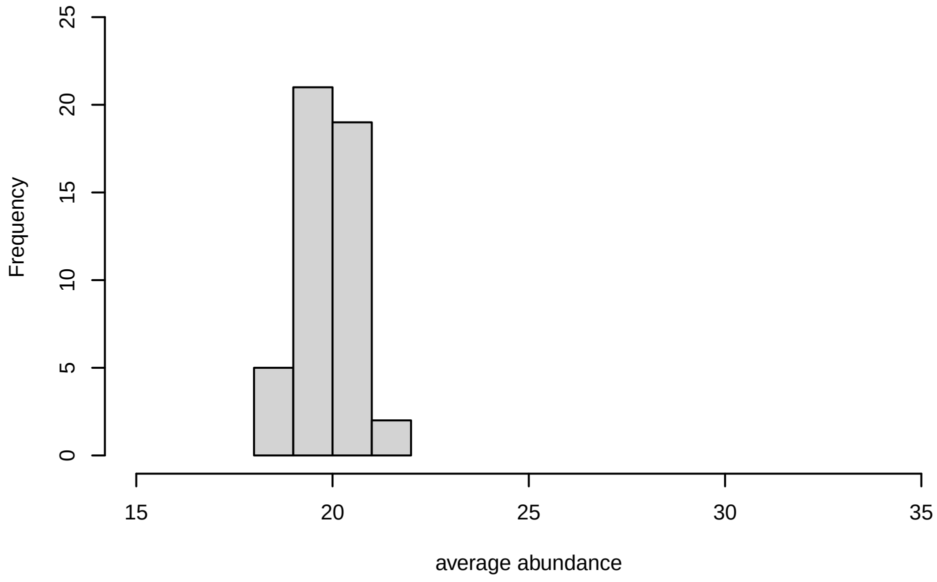


V11, after remove outlier and cutoff is 0.8

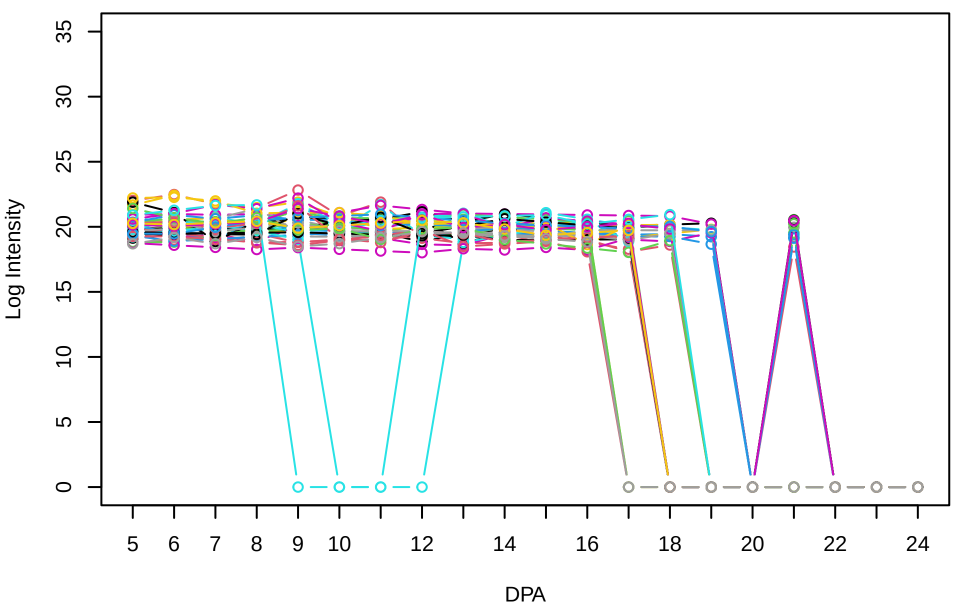


Histogram of the average abundances of non−outlier proteins

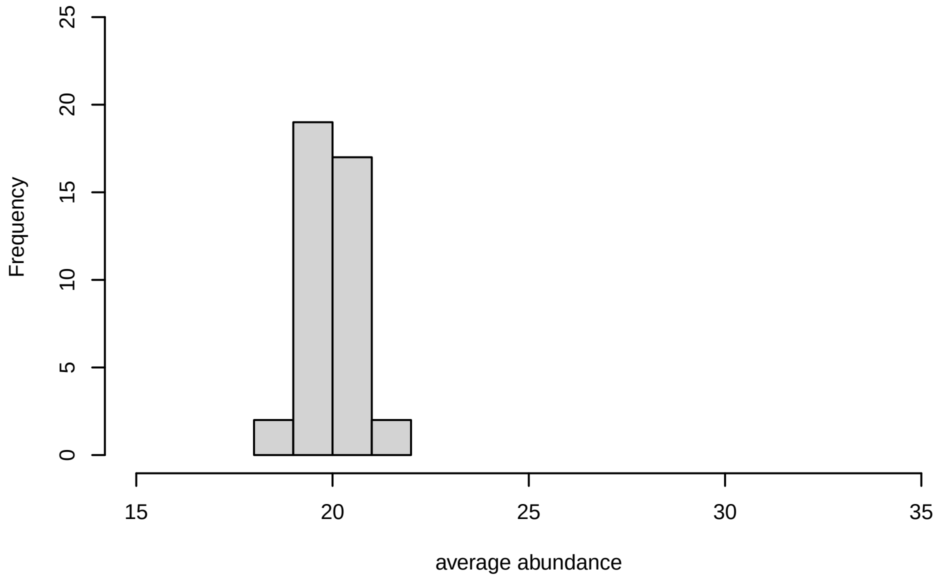


V11, after remove unreliable

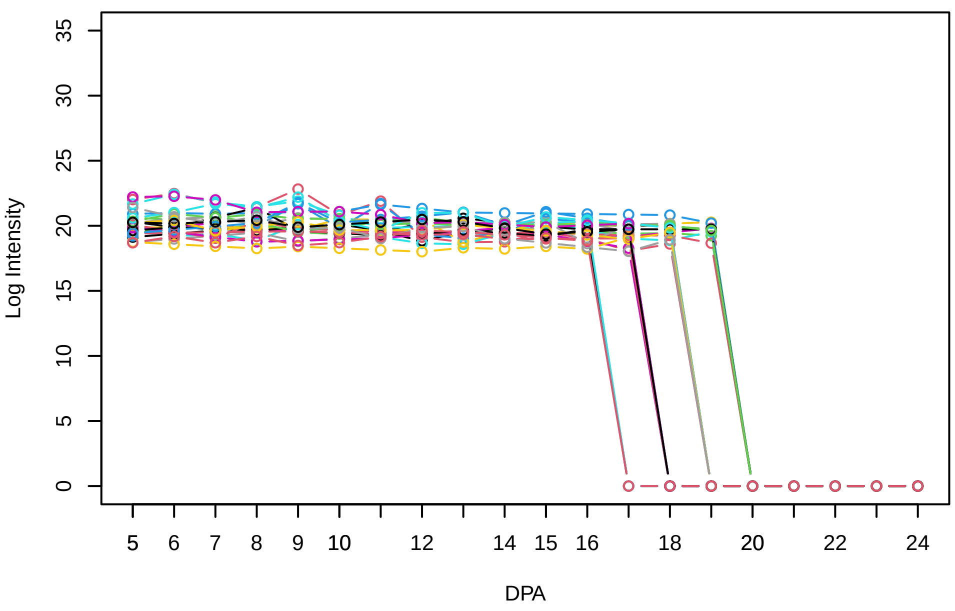


Histogram of the average abundances of reliable proteins

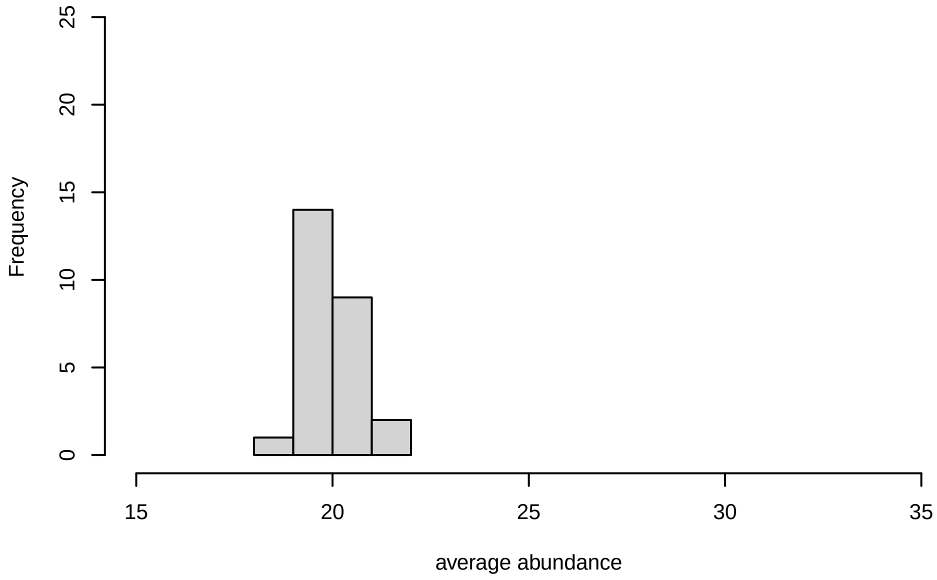


V11, after remove low abundance

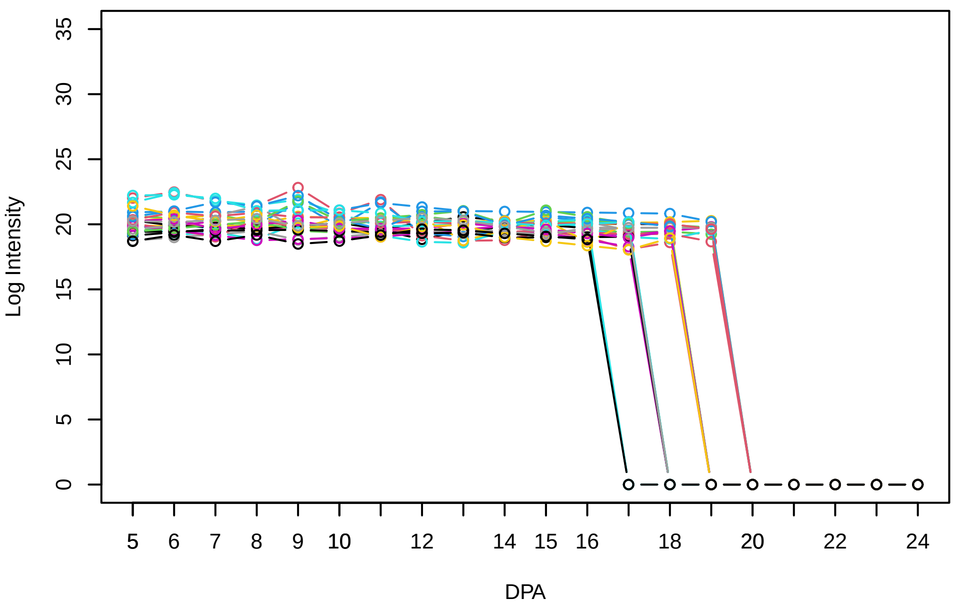


Histogram of the average abundances of high abundance proteins

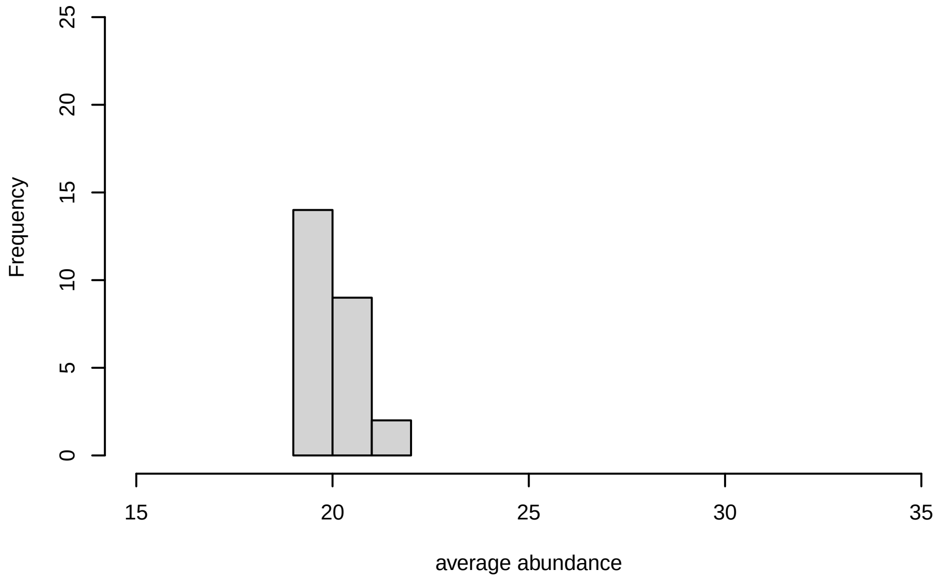

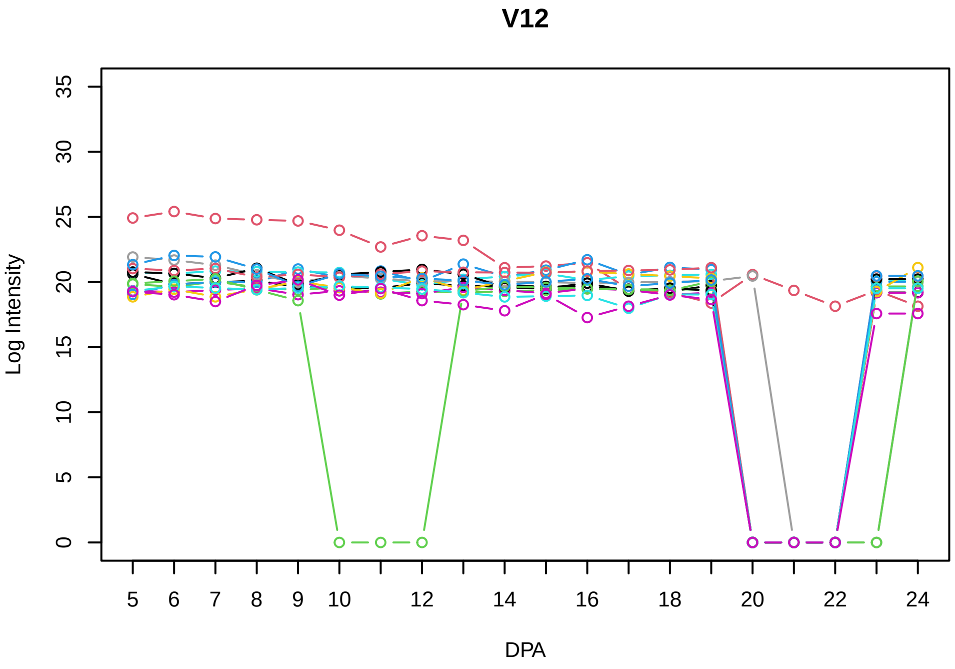
V12
Log Intensity
DPA

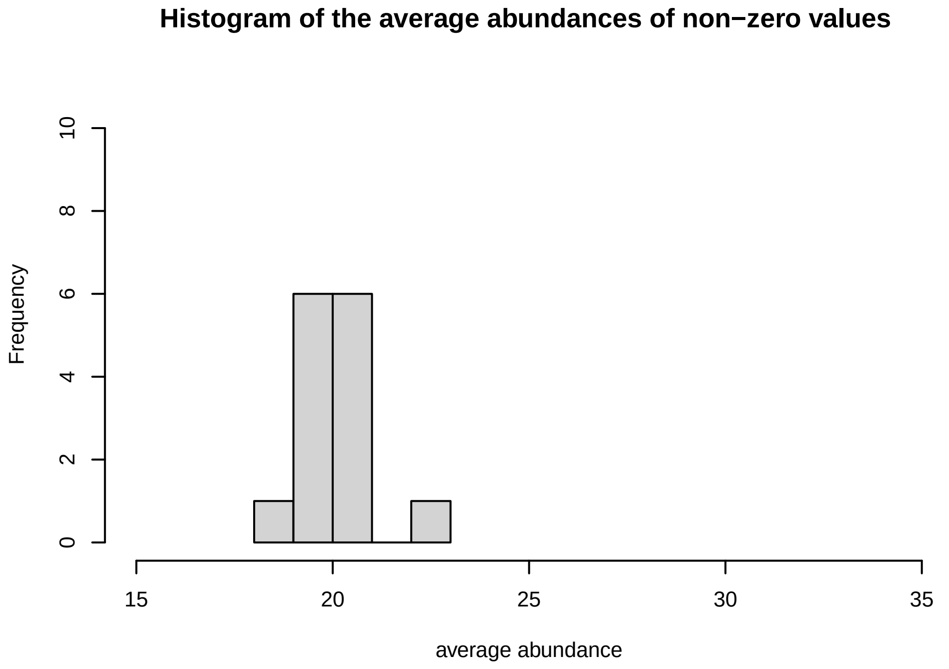
Histogram of the average abundances of non−zero values
Frequency
average abundance

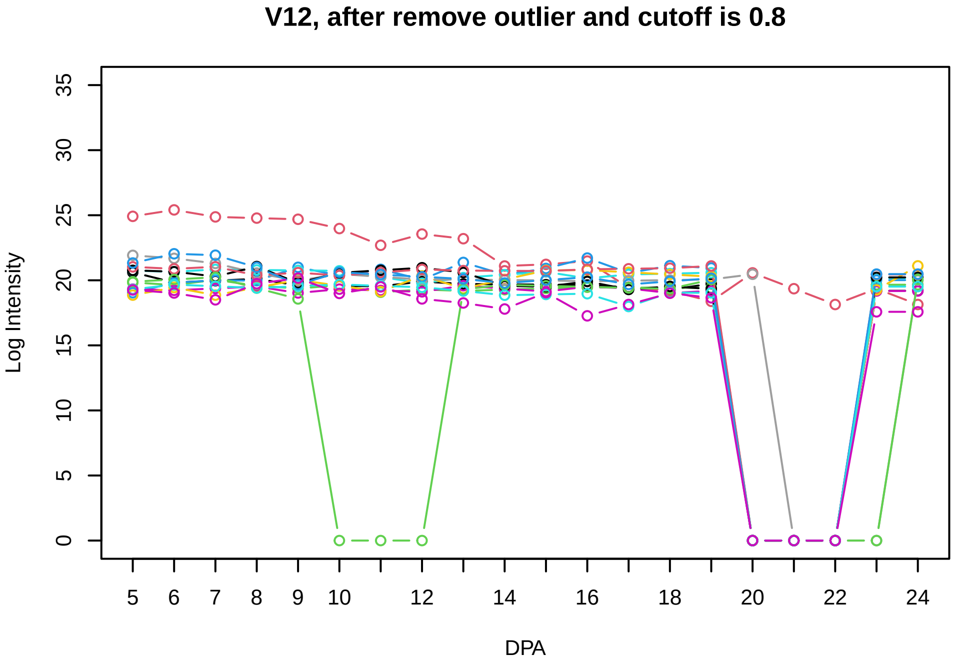
V12, after remove outlier and cutoff is 0.8
Log Intensity
DPA

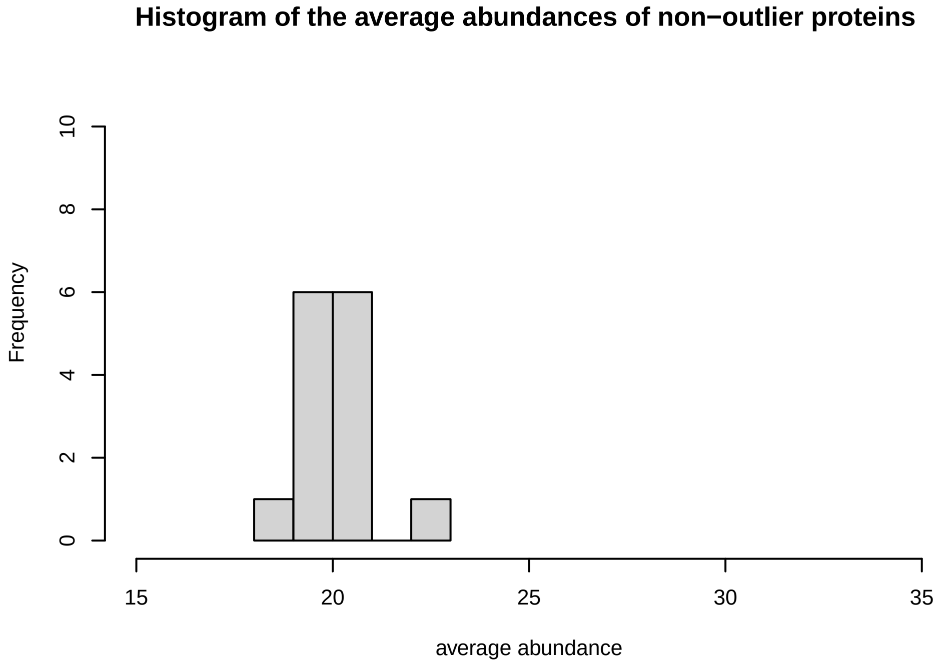
Histogram of the average abundances of non−outlier proteins
Frequency
average abundance

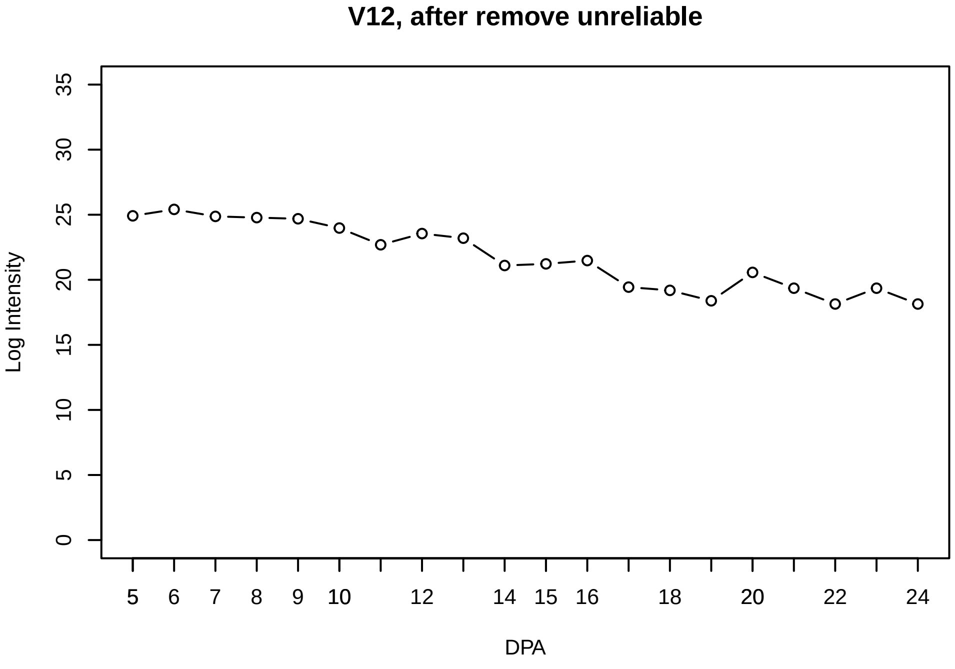
V12, after remove unreliable
Log Intensity
DPA

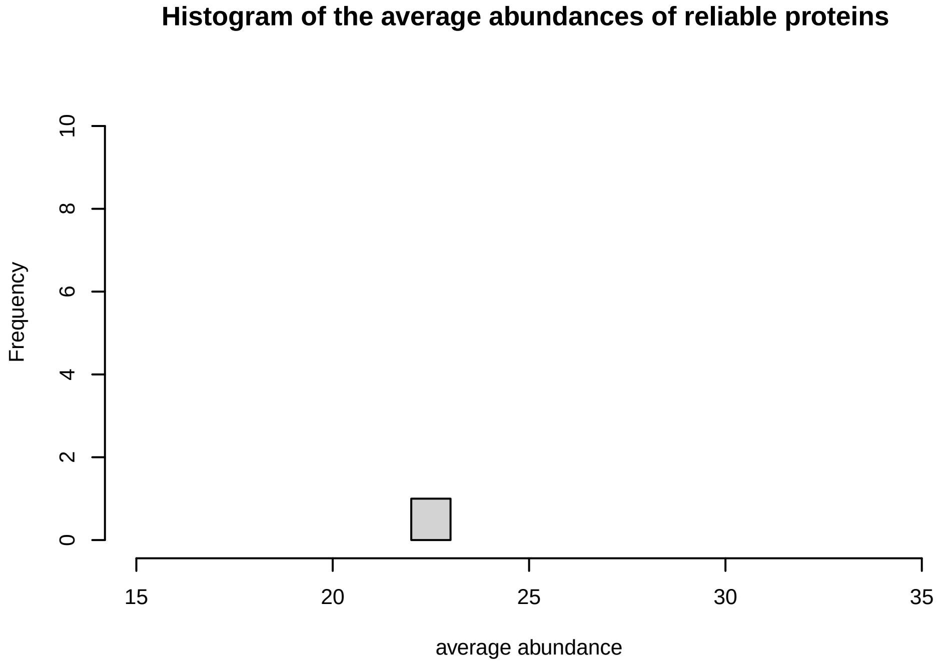
Histogram of the average abundances of reliable proteins
Frequency
average abundance

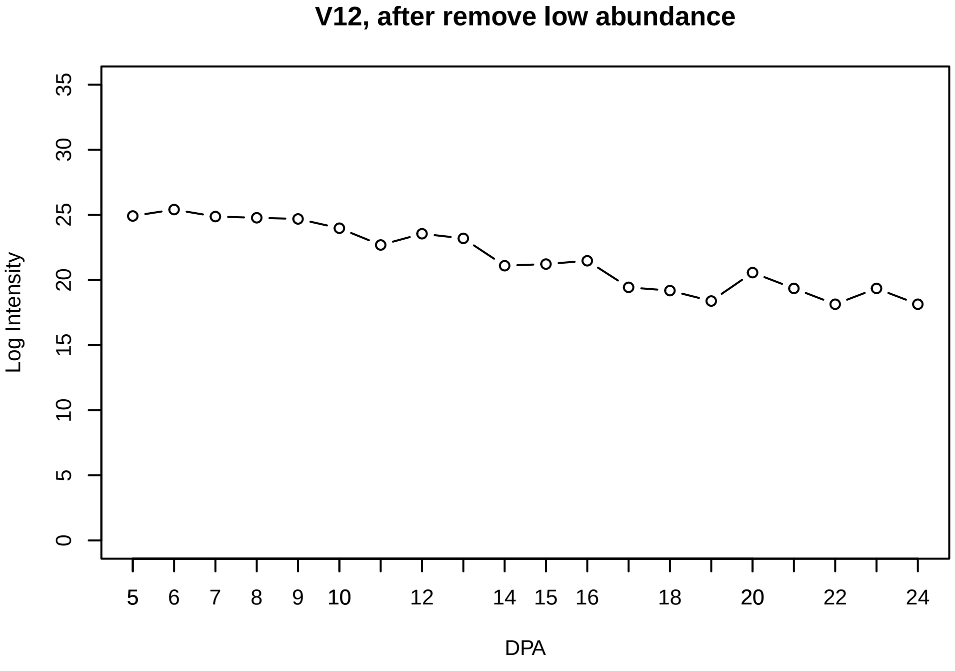
V12, after remove low abundance
Log Intensity
DPA

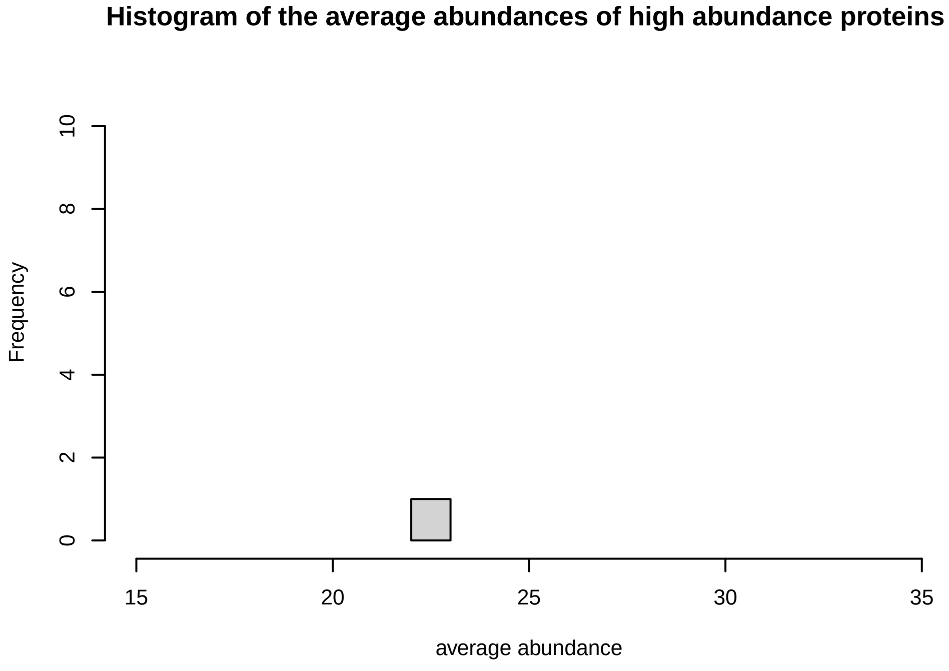
Histogram of the average abundances of high abundance proteins
Frequency
average abundance

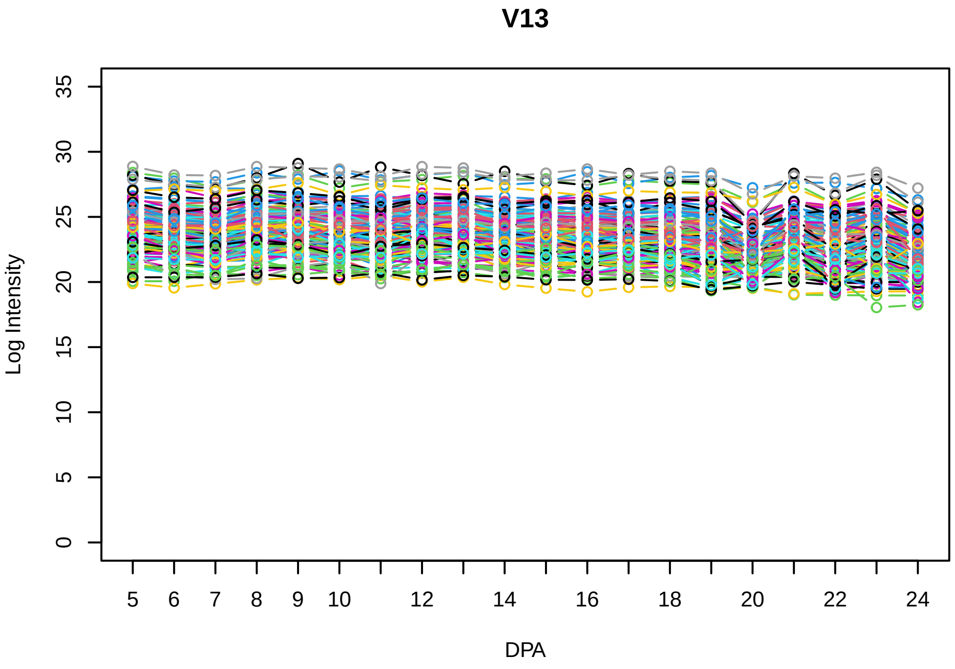
V13
Log Intensity
DPA

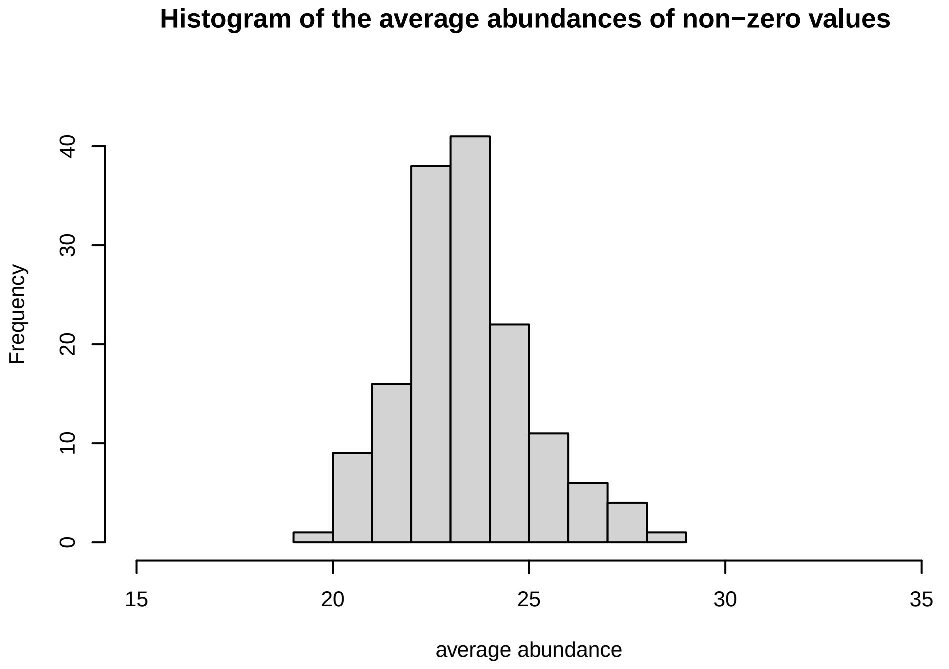
Histogram of the average abundances of non−zero values
Frequency
average abundance

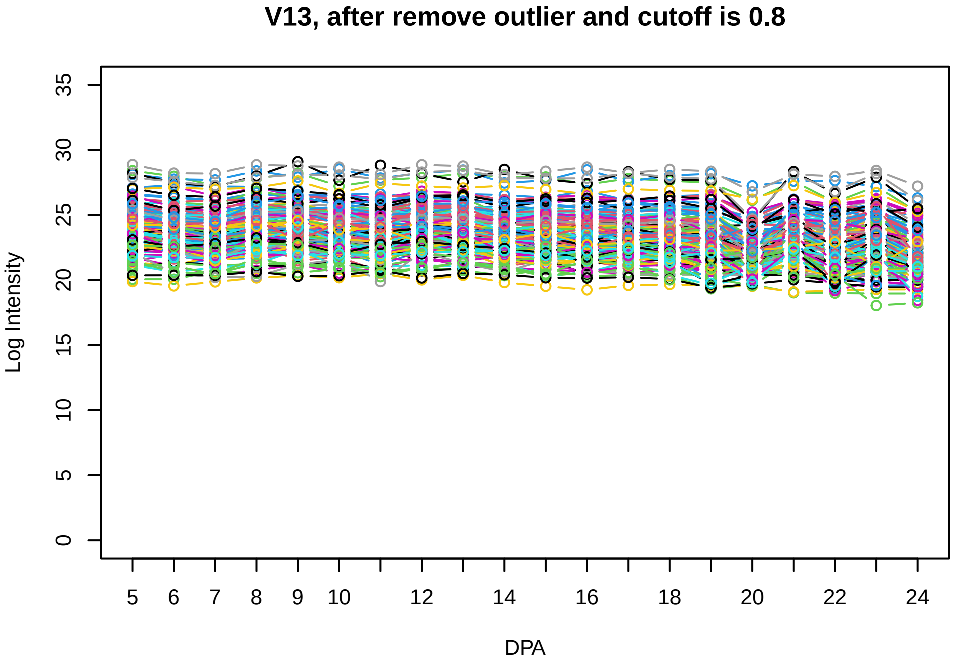
V13, after remove outlier and cutoff is 0.8
Log Intensity
DPA

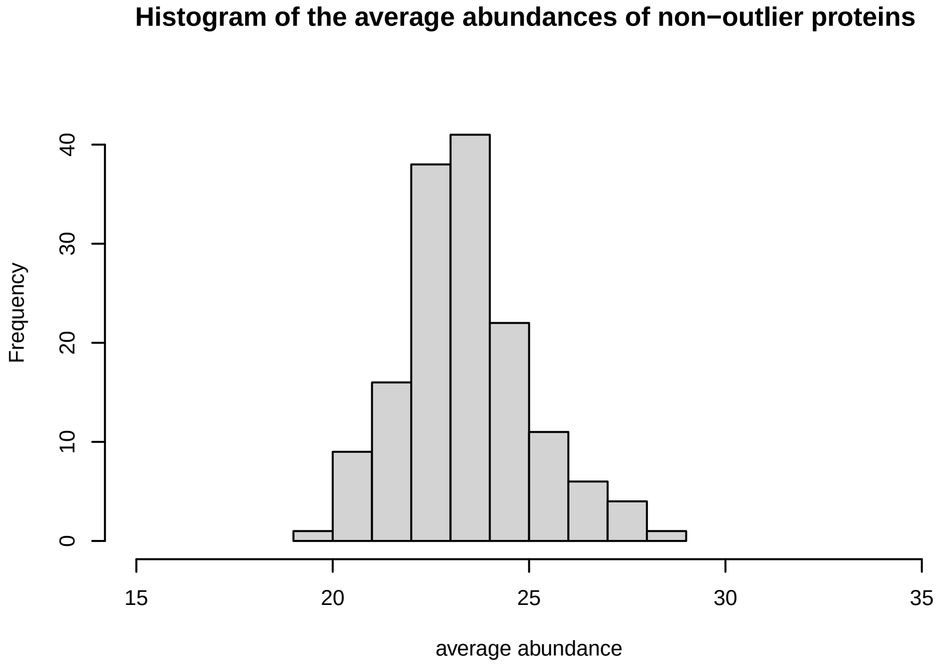
Histogram of the average abundances of non−outlier proteins
Frequency
average abundance

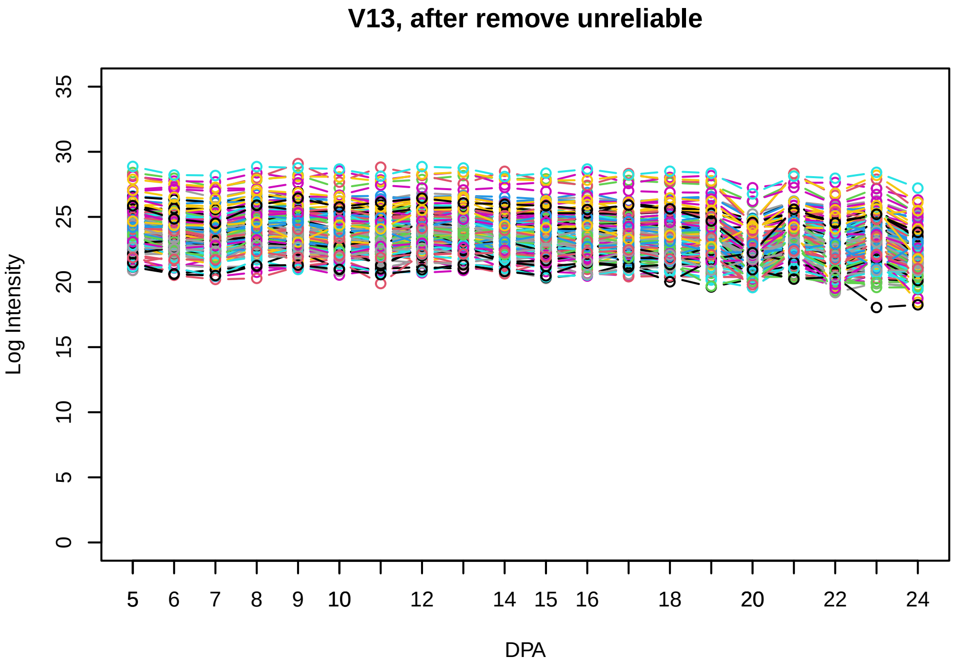
V13, after remove unreliable
Log Intensity
DPA

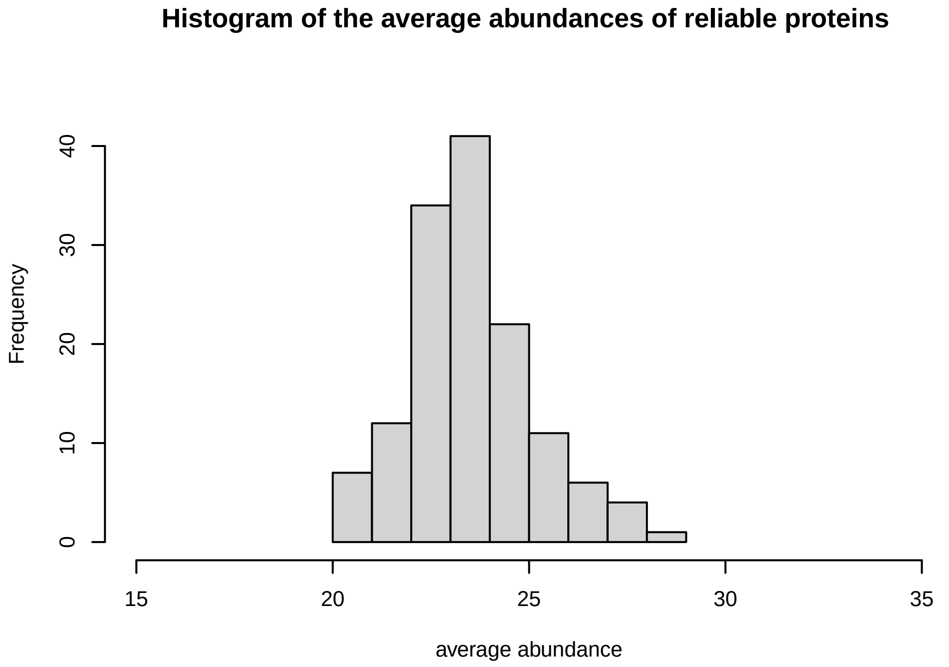
Histogram of the average abundances of reliable proteins
Frequency
average abundance

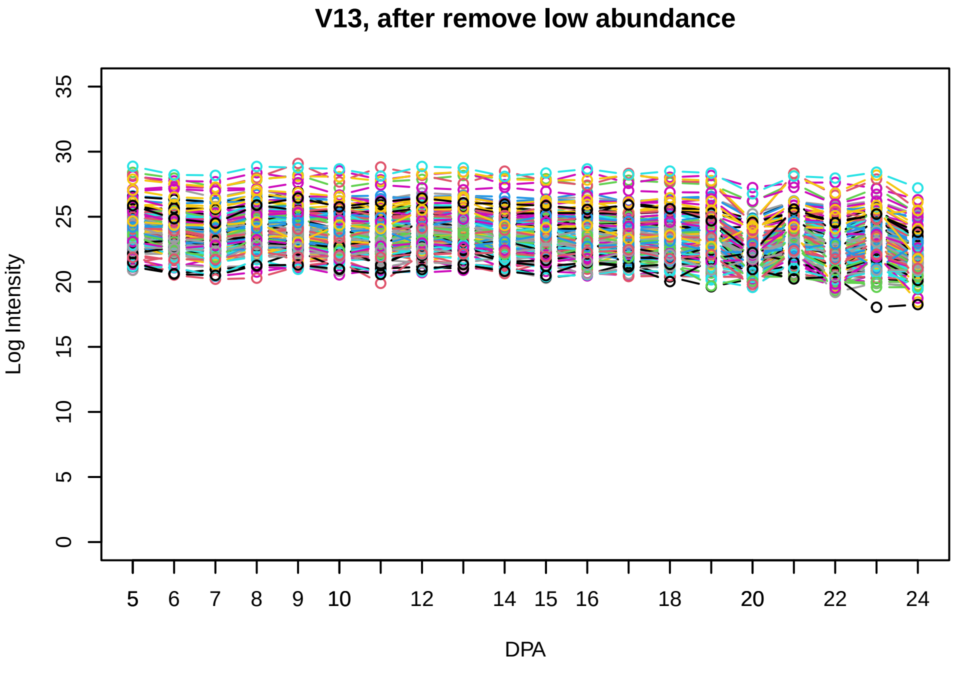
V13, after remove low abundance
Log Intensity
DPA

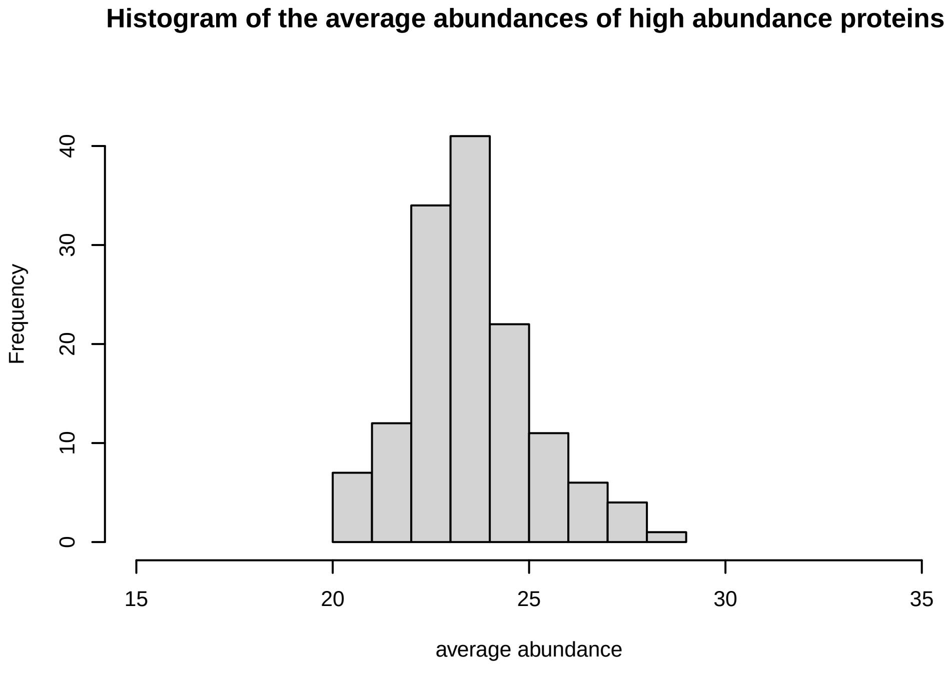
Histogram of the average abundances of high abundance proteins
Frequency
average abundance

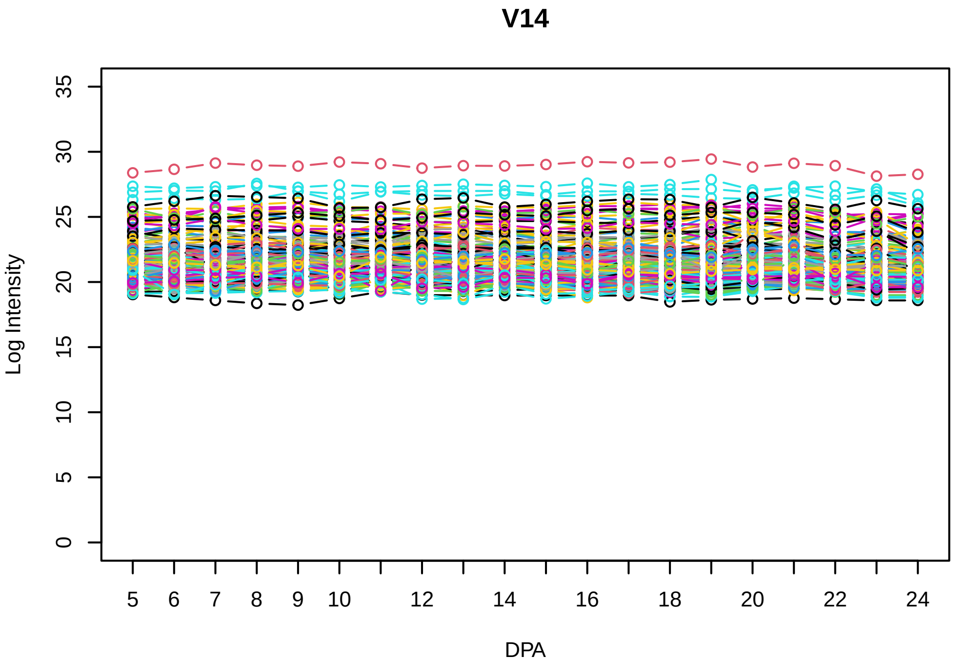
V14
Log Intensity
DPA

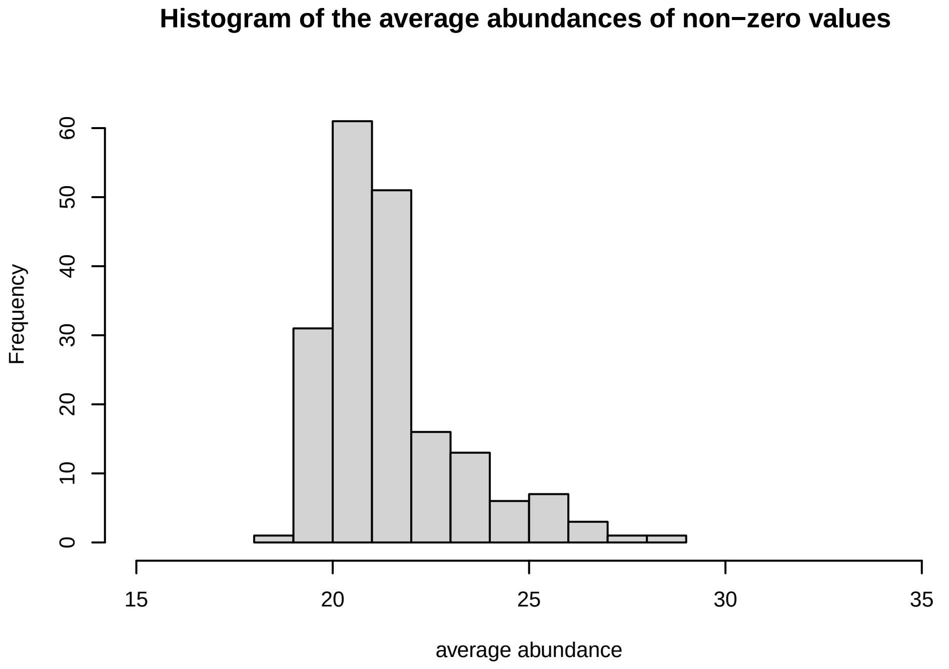
Histogram of the average abundances of non−zero values
Frequency
average abundance

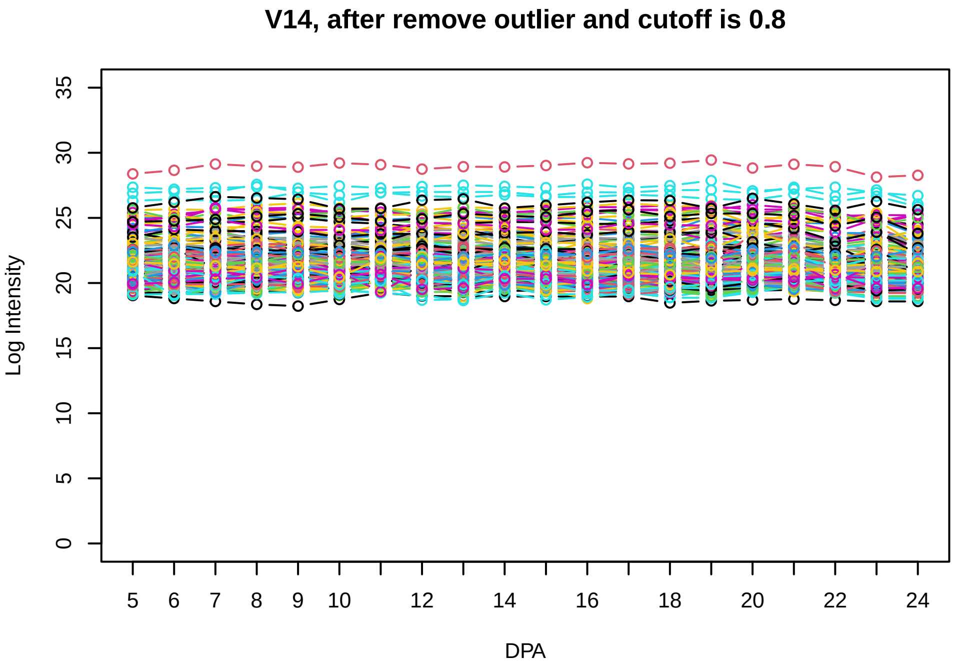
V14, after remove outlier and cutoff is 0.8
Log Intensity
DPA

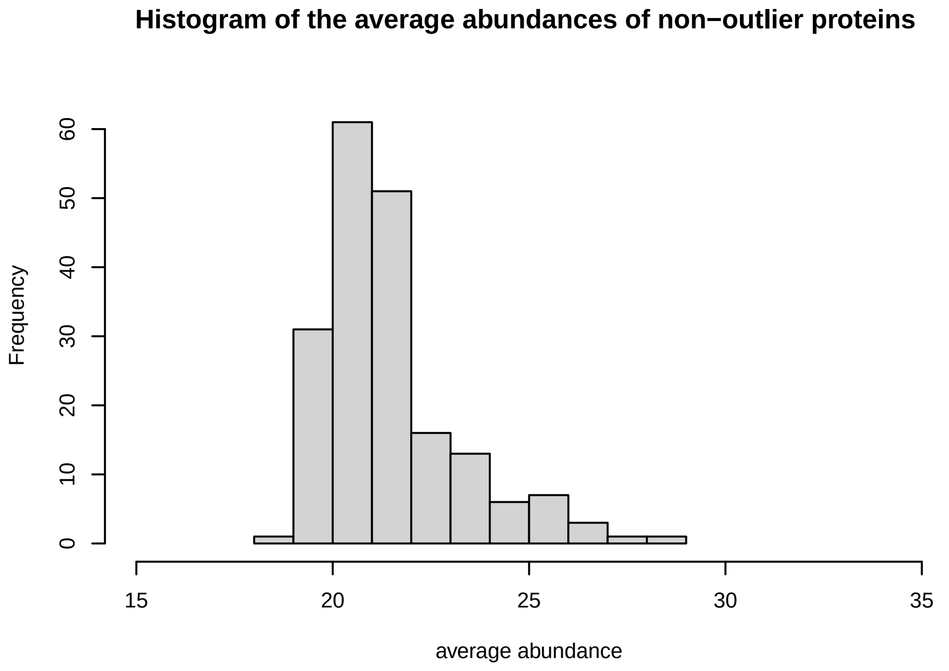
Histogram of the average abundances of non−outlier proteins
Frequency
average abundance

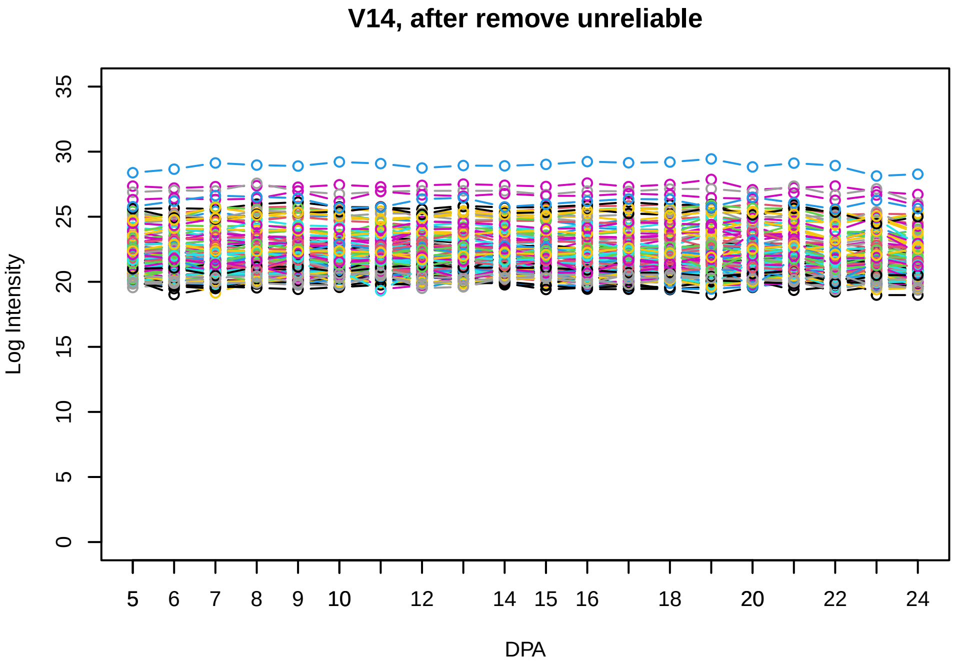
V14, after remove unreliable
Log Intensity
DPA

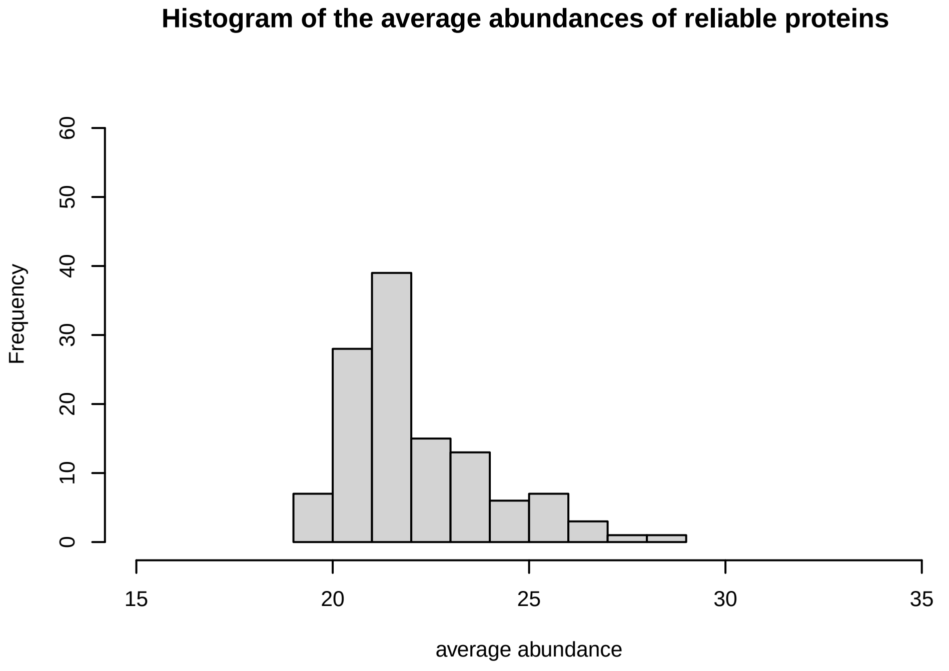
Histogram of the average abundances of reliable proteins
Frequency
average abundance

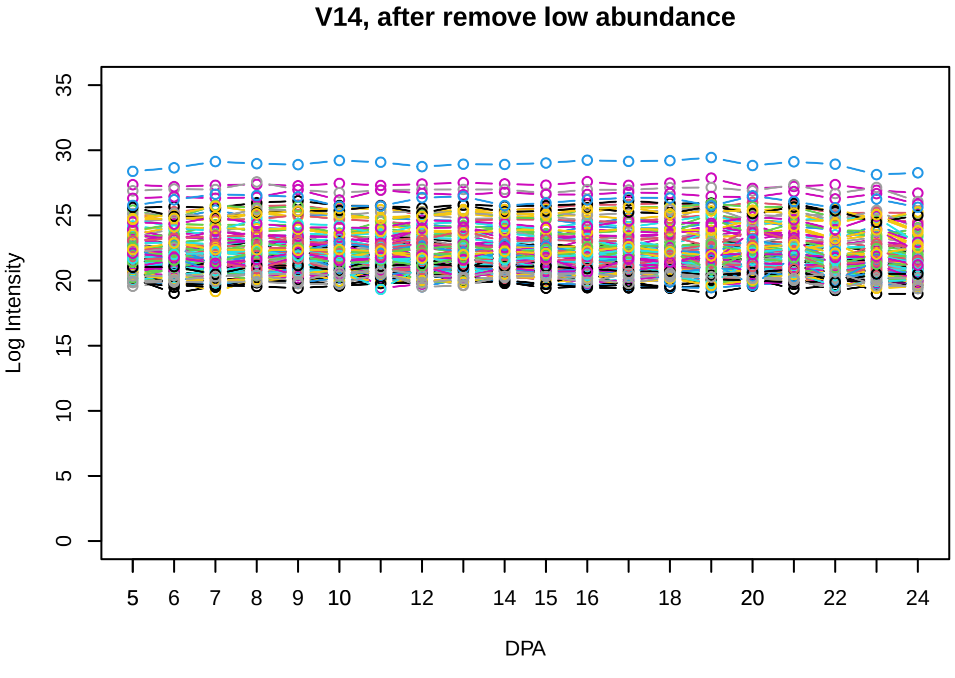
V14, after remove low abundance
Log Intensity
DPA

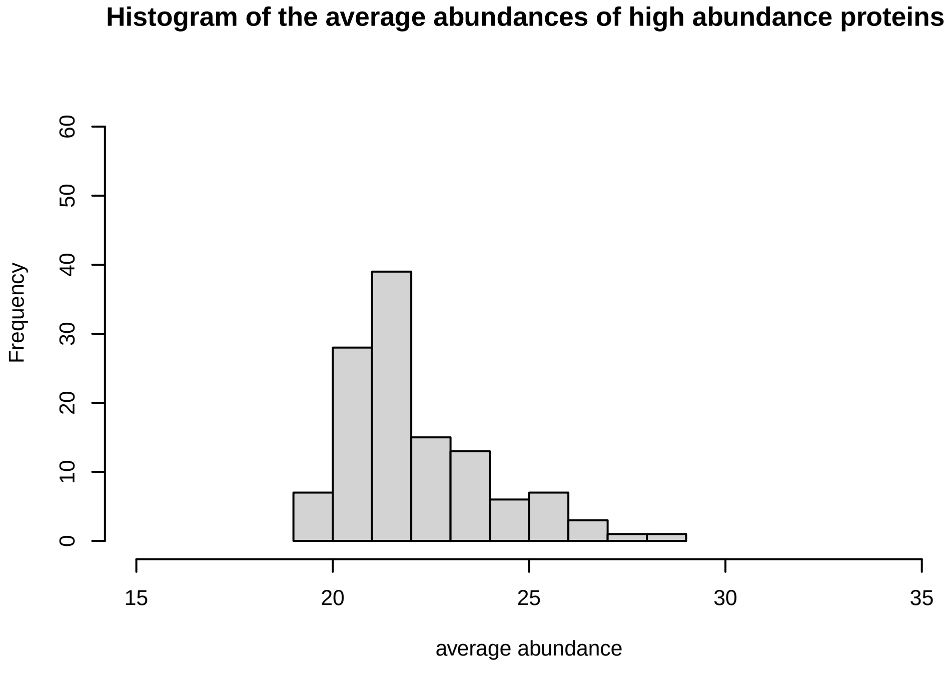
Histogram of the average abundances of high abundance proteins
Frequency
average abundance

**V15**

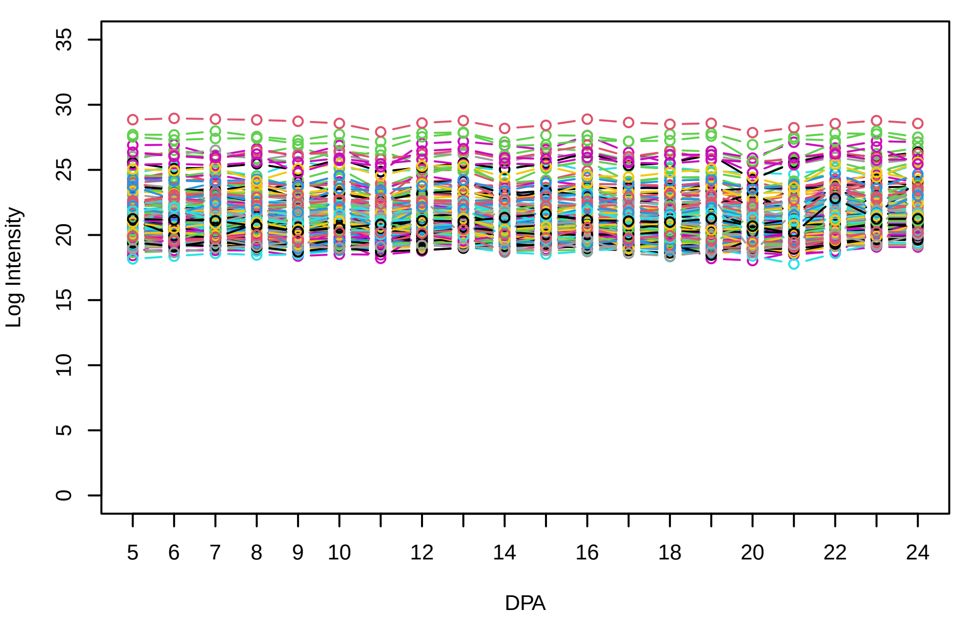


**Histogram of the average abundances of non−zero values**

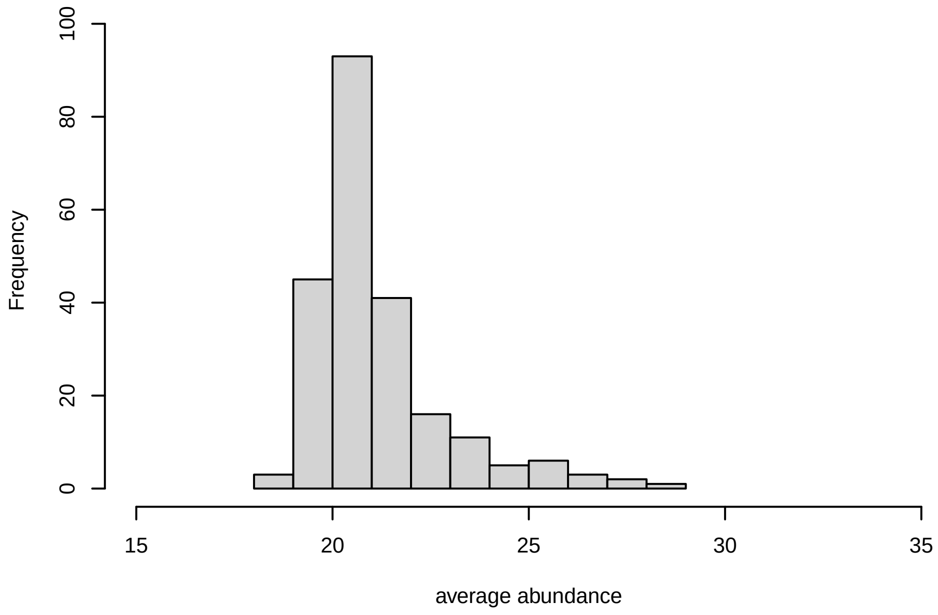


**V15, after remove outlier and cutoff is 0.8**

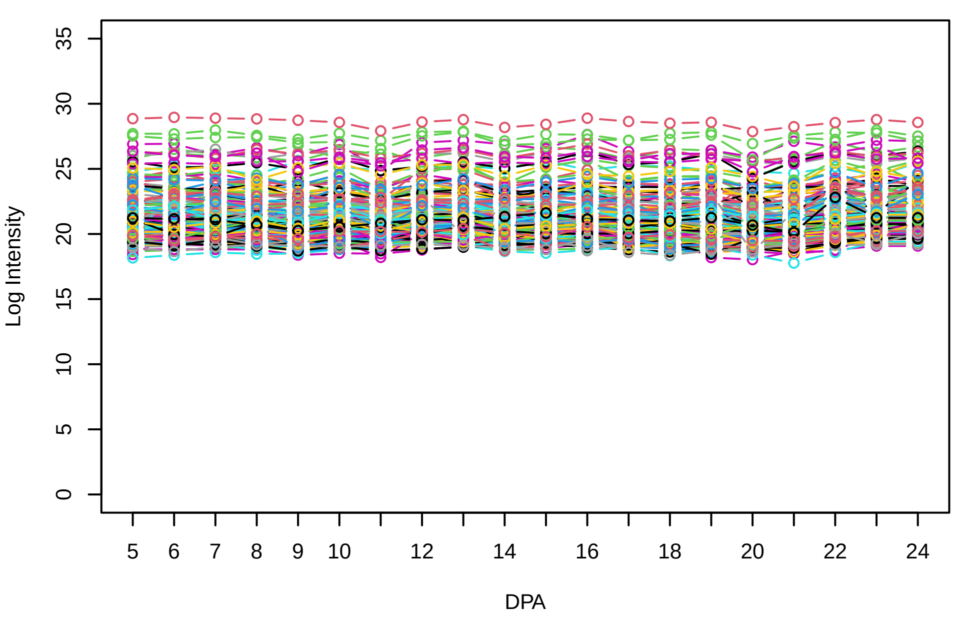


**Histogram of the average abundances of non−outlier proteins**

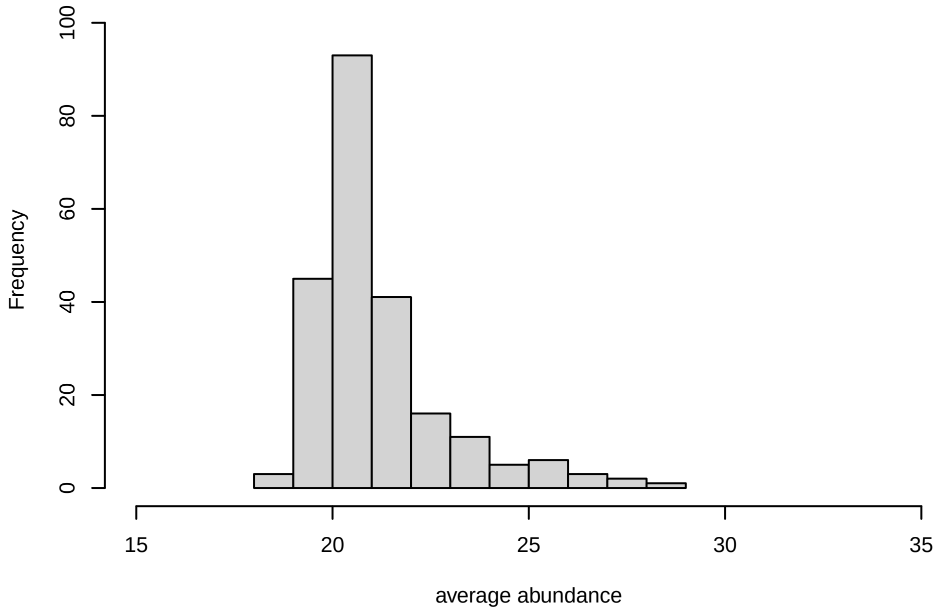


**V15, after remove unreliable**

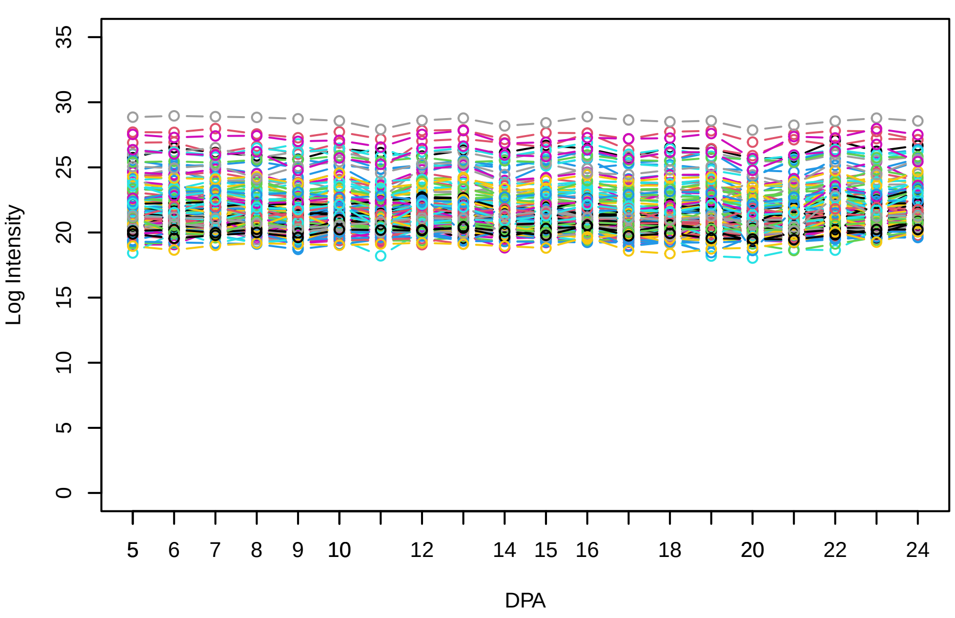


**Histogram of the average abundances of reliable proteins**

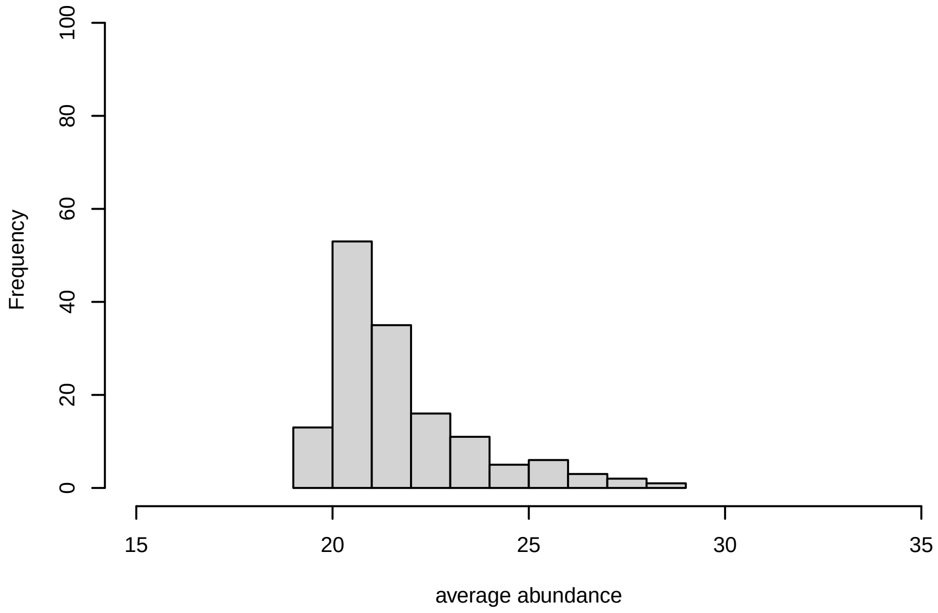


**V15, after remove low abundance**

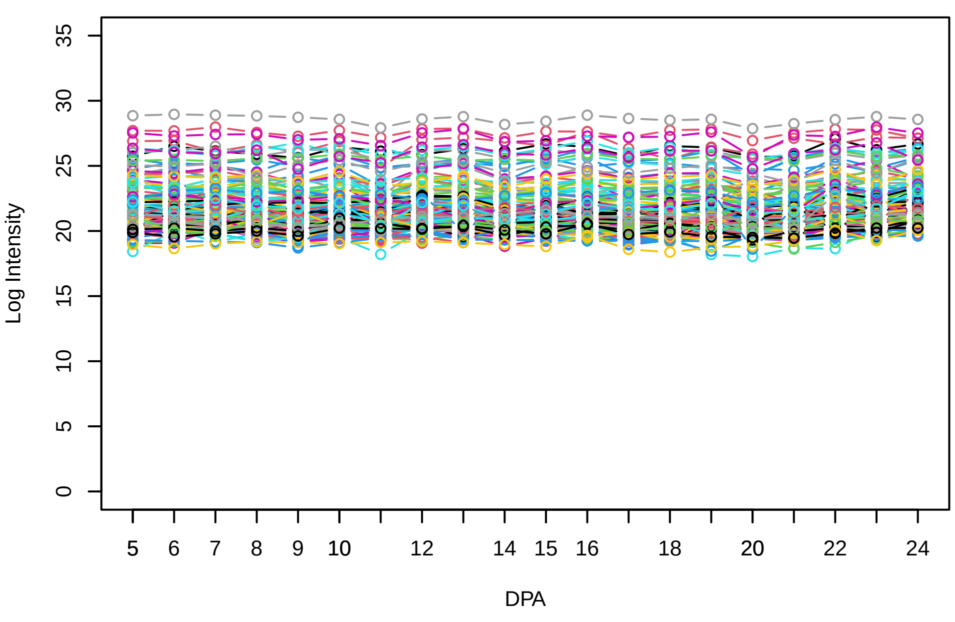


**Histogram of the average abundances of high abundance proteins**

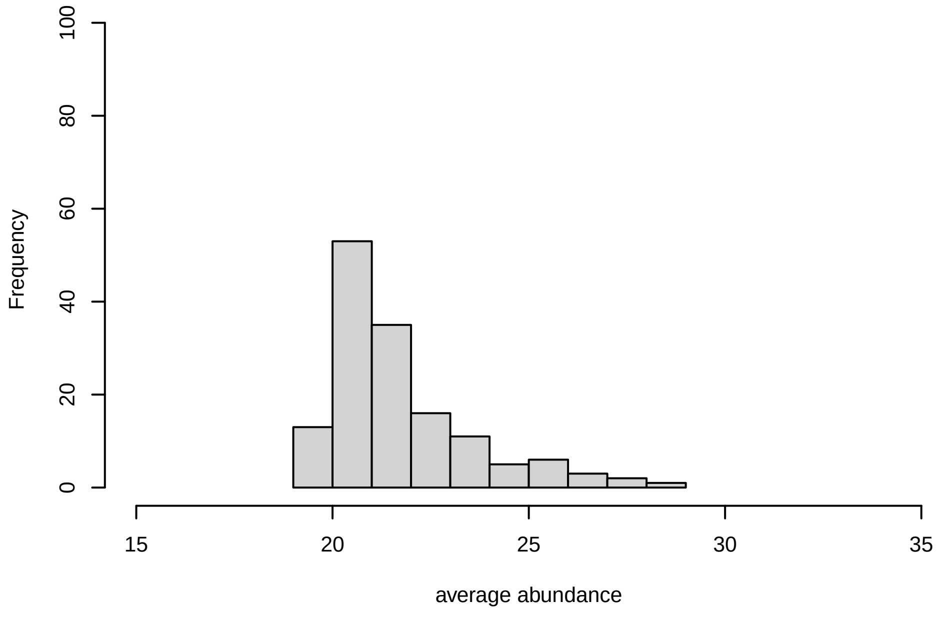

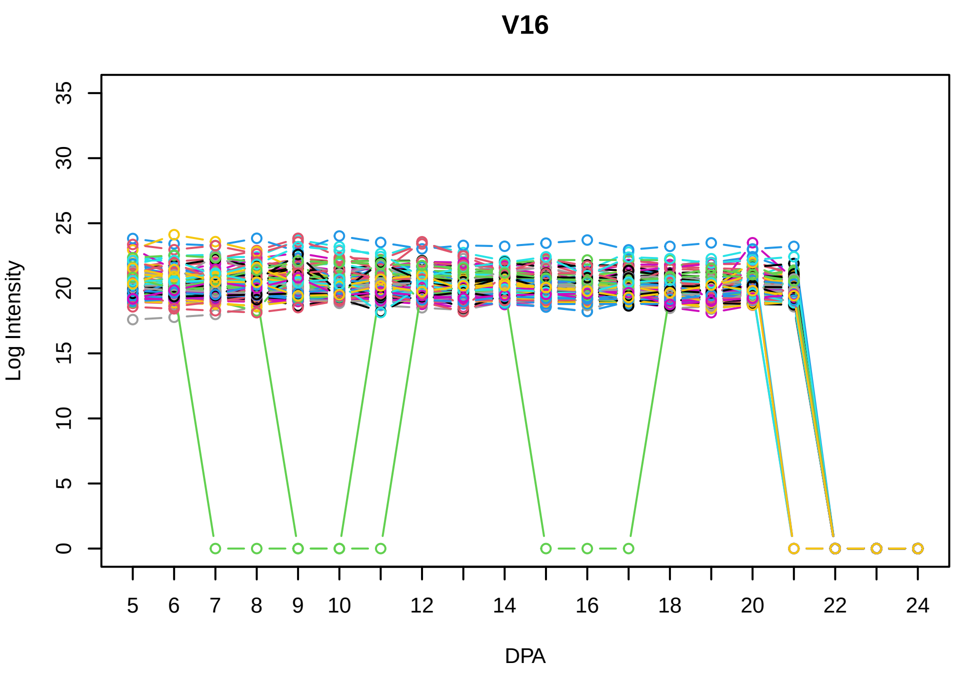
V16
Log Intensity
DPA

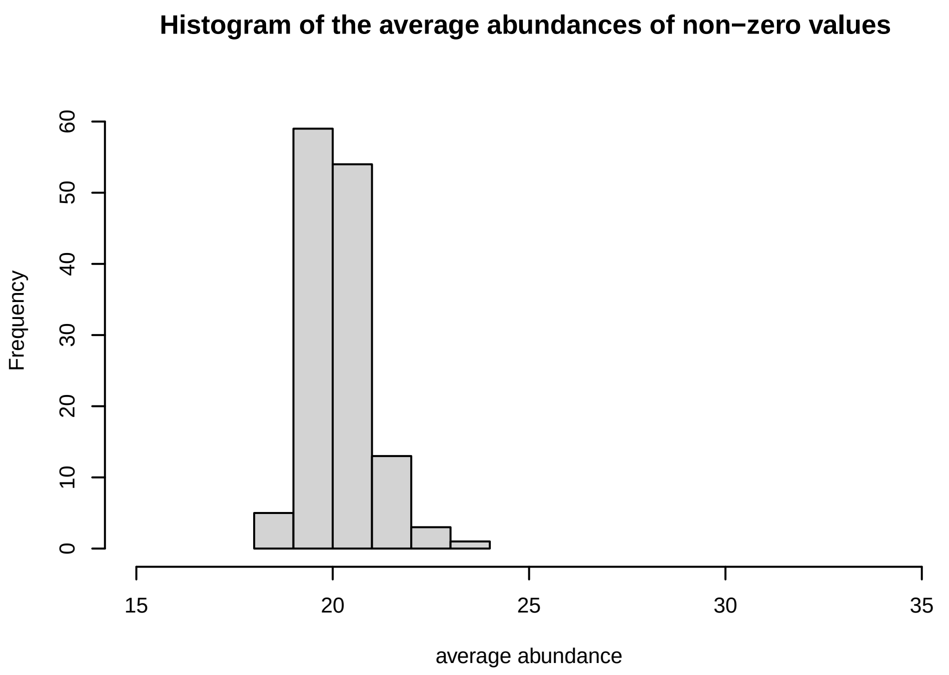
Histogram of the average abundances of non−zero values
Frequency
average abundance

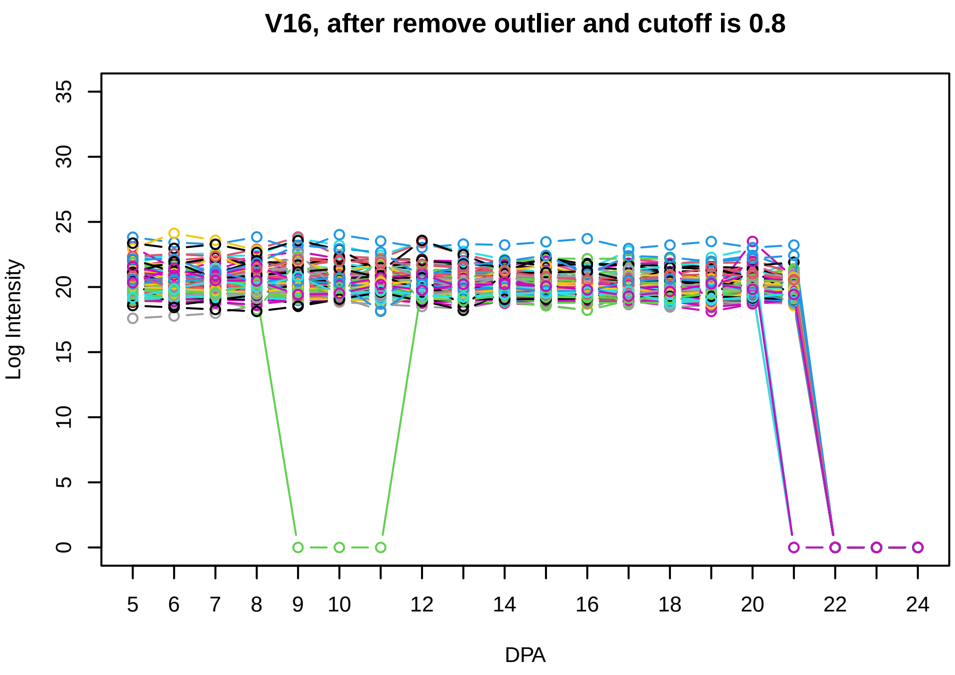
V16, after remove outlier and cutoff is 0.8
Log Intensity
DPA

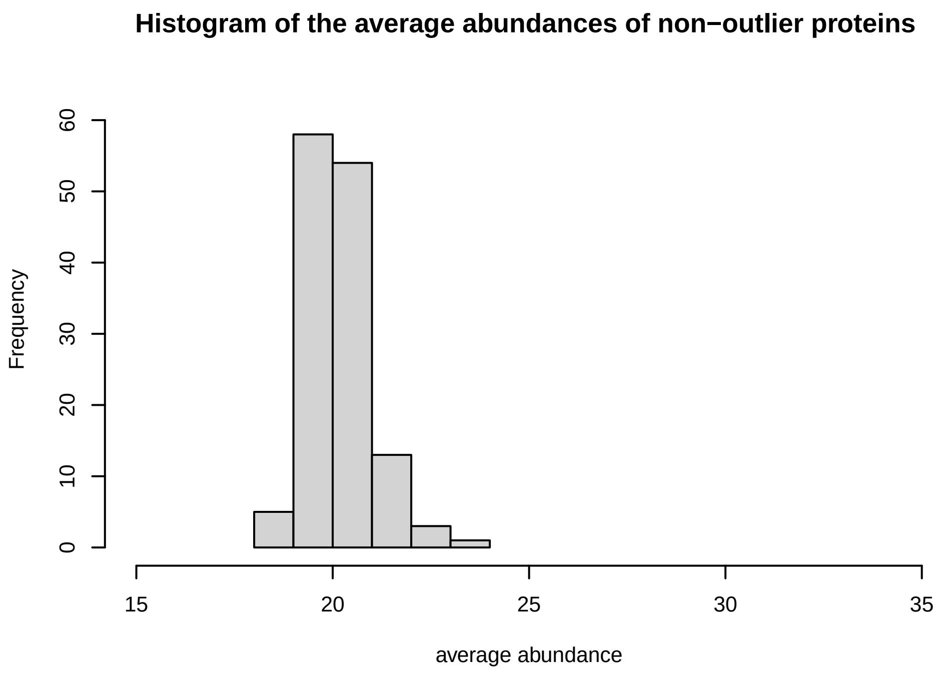
Histogram of the average abundances of non−outlier proteins
Frequency
average abundance

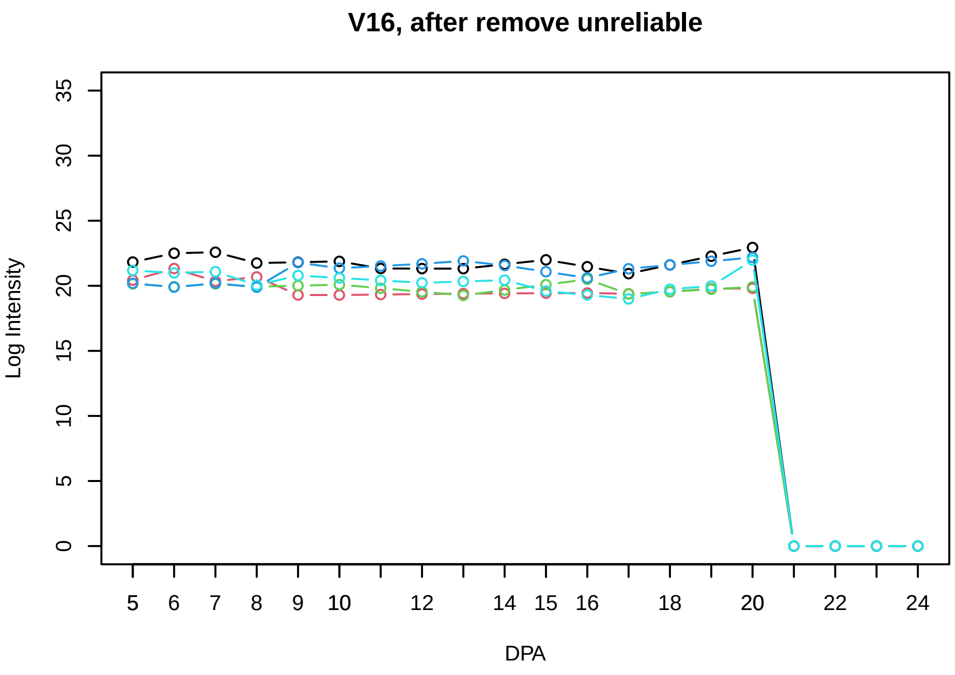
V16, after remove unreliable
Log Intensity
DPA

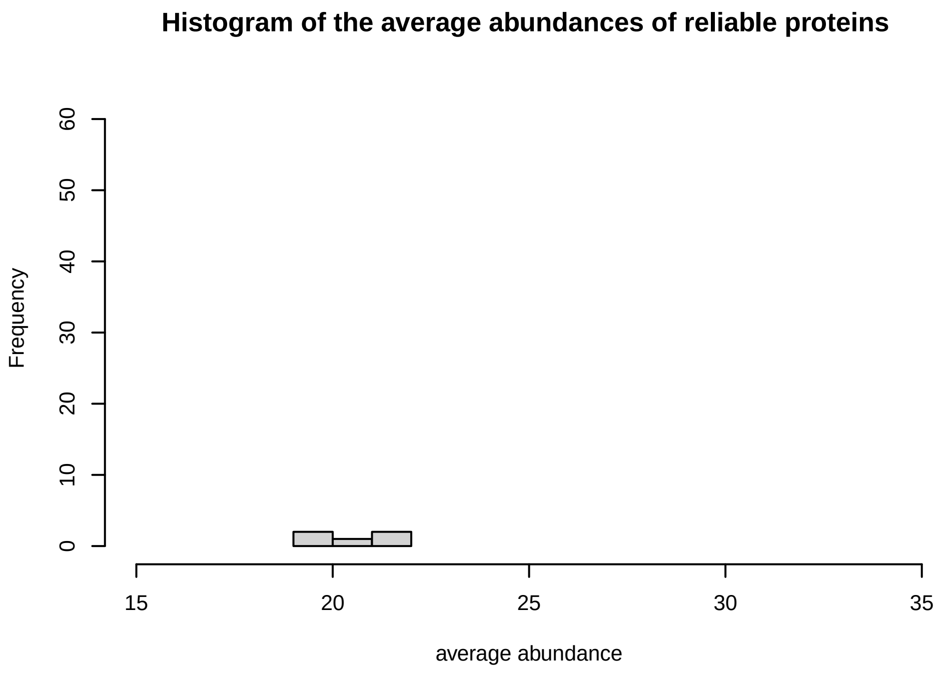
Histogram of the average abundances of reliable proteins
Frequency
average abundance

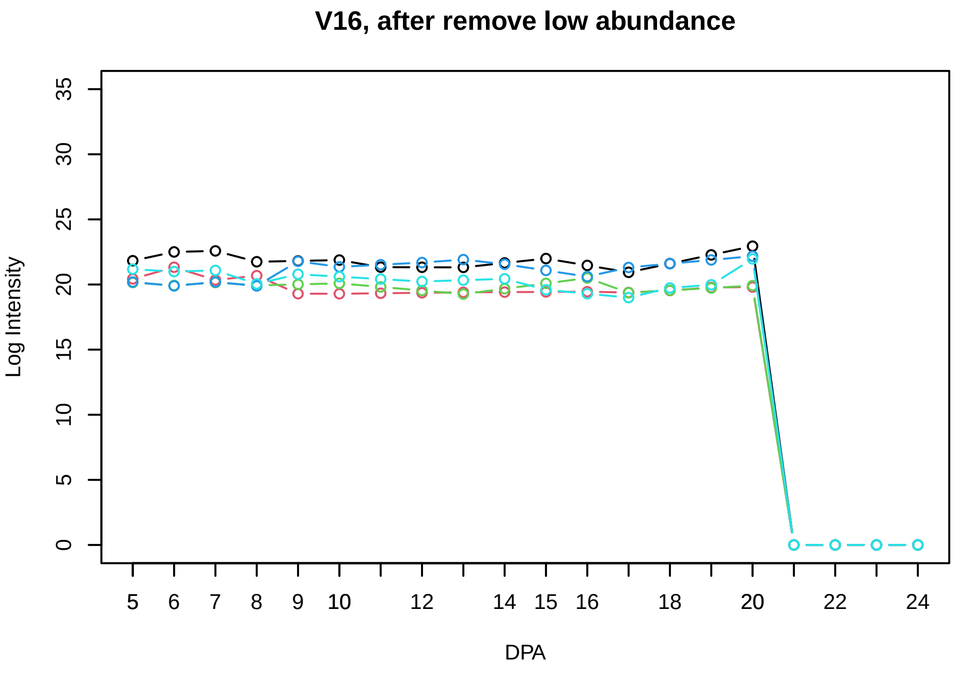
V16, after remove low abundance
Log Intensity
DPA

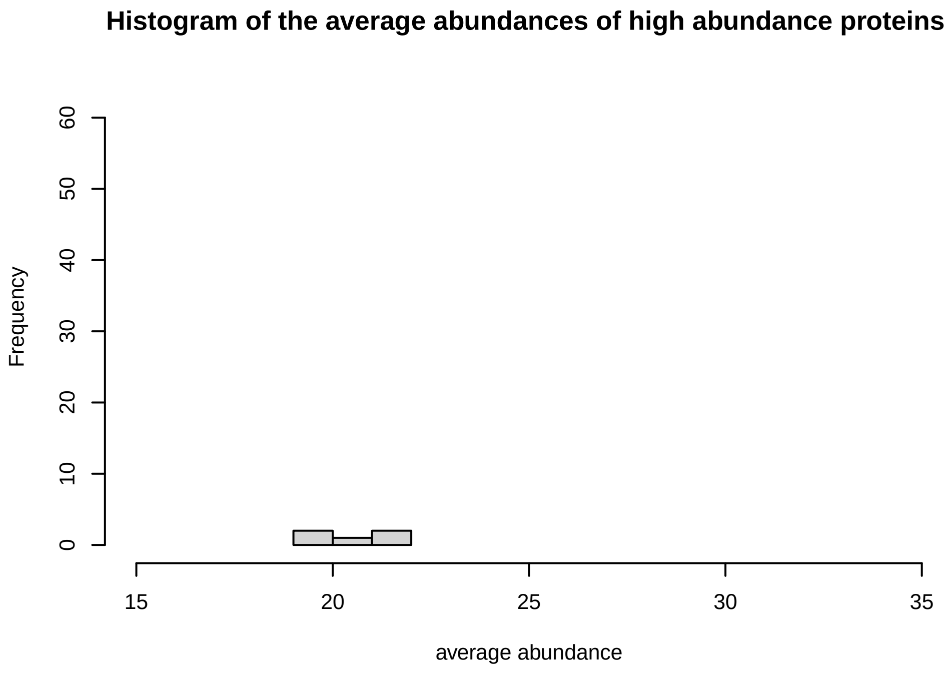
Histogram of the average abundances of high abundance proteins
Frequency
average abundance

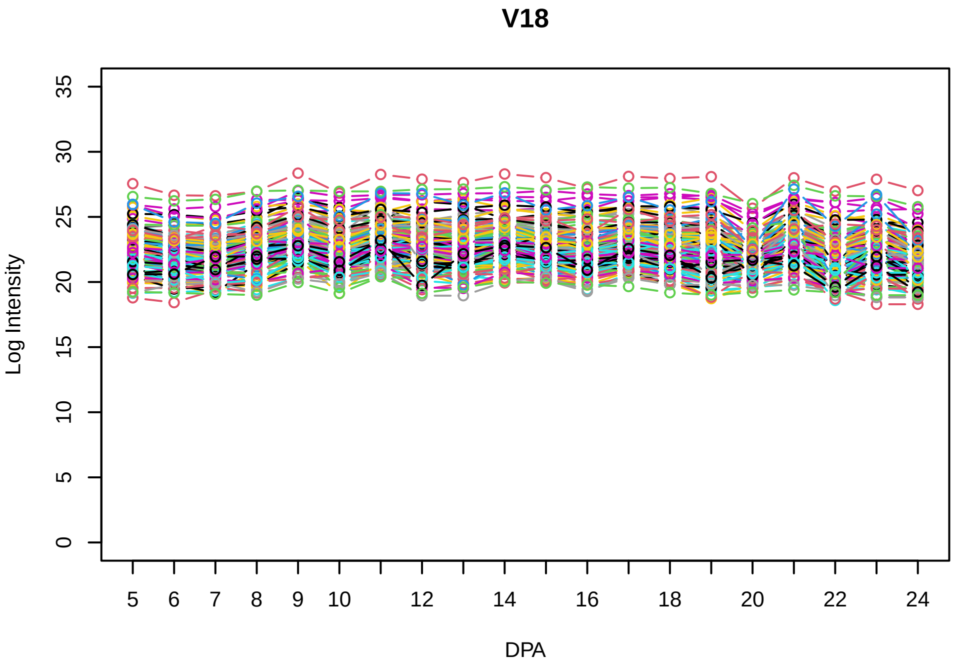
V18
Log Intensity
DPA
0 5 10 15 20 25 30 35
5 6 7 8 9 10 12 14 16 18 20 22 24

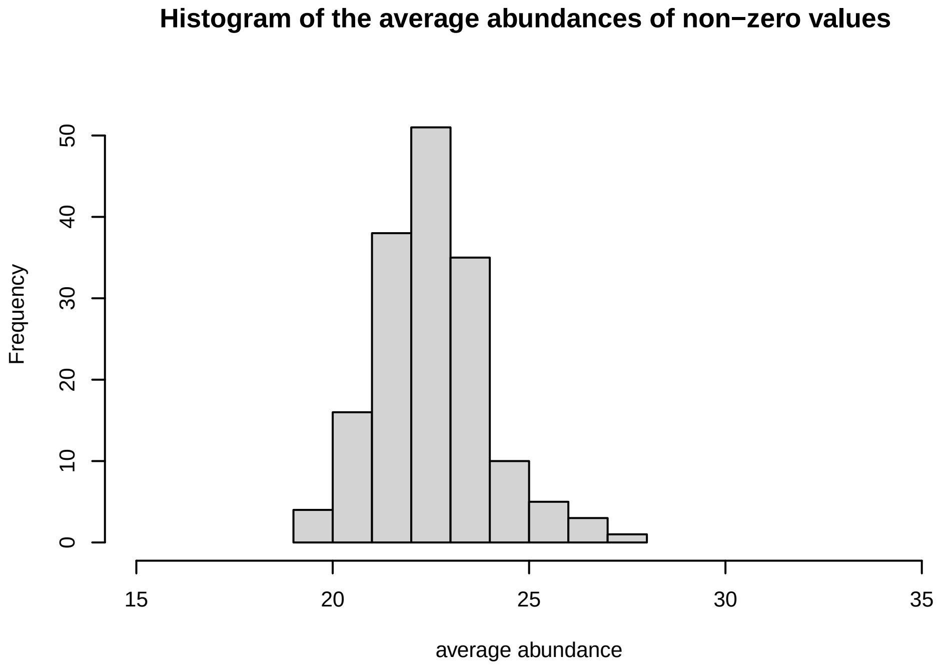
Histogram of the average abundances of non−zero values
Frequency
average abundance
0 10 20 30 40 50
15 20 25 30 35

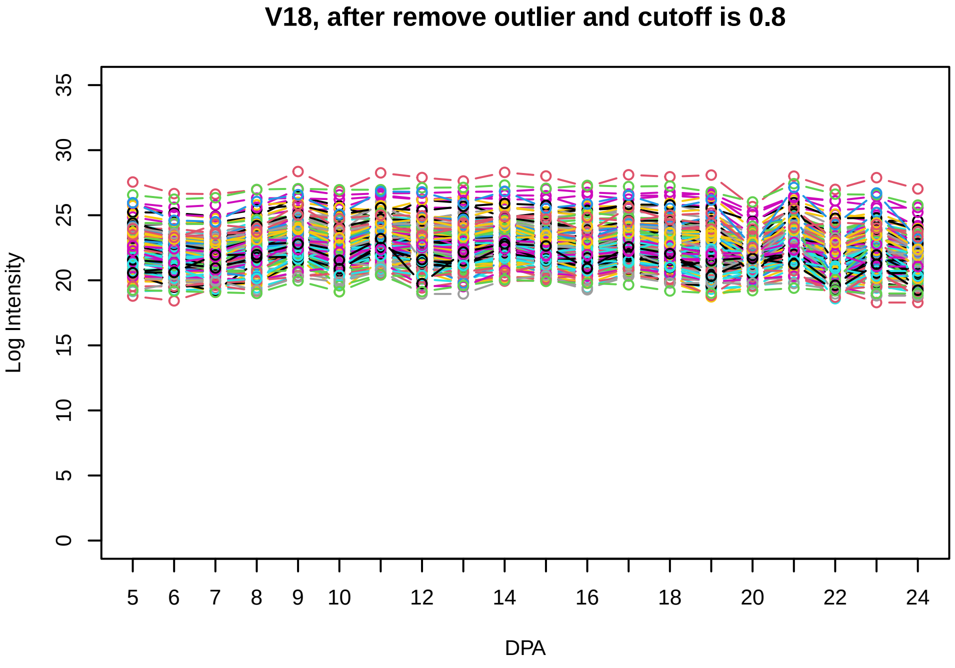
V18, after remove outlier and cutoff is 0.8
Log Intensity
DPA
0 5 10 15 20 25 30 35
5 6 7 8 9 10 12 14 16 18 20 22 24

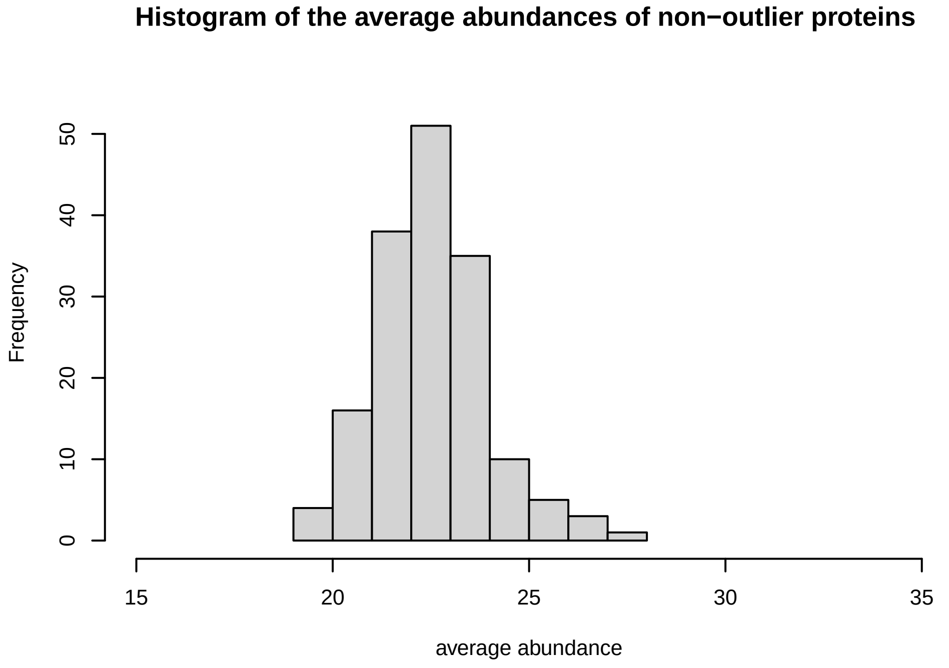
Histogram of the average abundances of non−outlier proteins
Frequency
average abundance
0 10 20 30 40 50
15 20 25 30 35

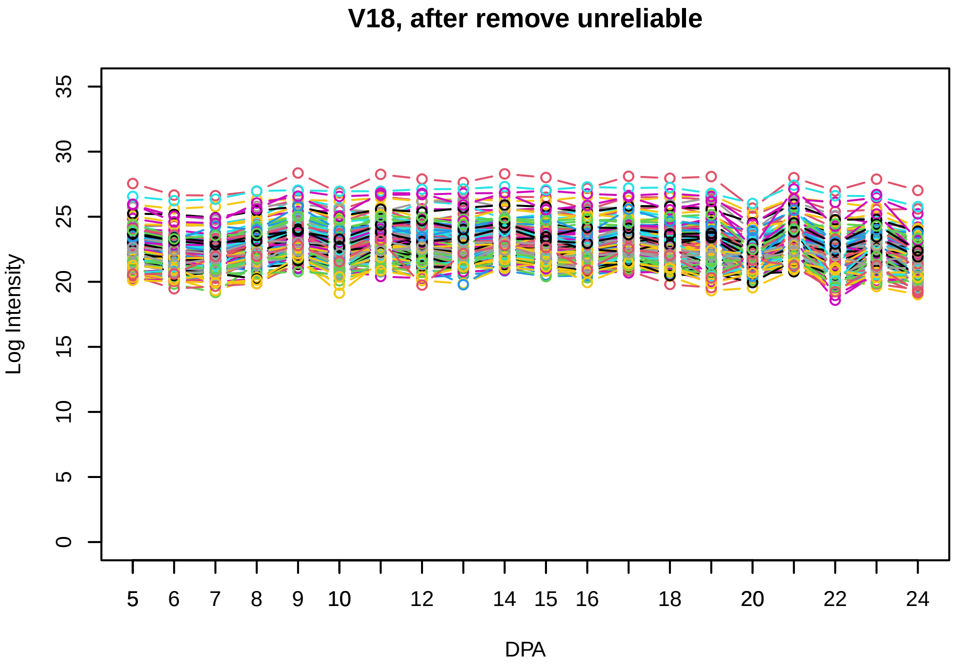
V18, after remove unreliable
Log Intensity
DPA
0 5 10 15 20 25 30 35
5 6 7 8 9 10 12 14 15 16 18 20 22 24

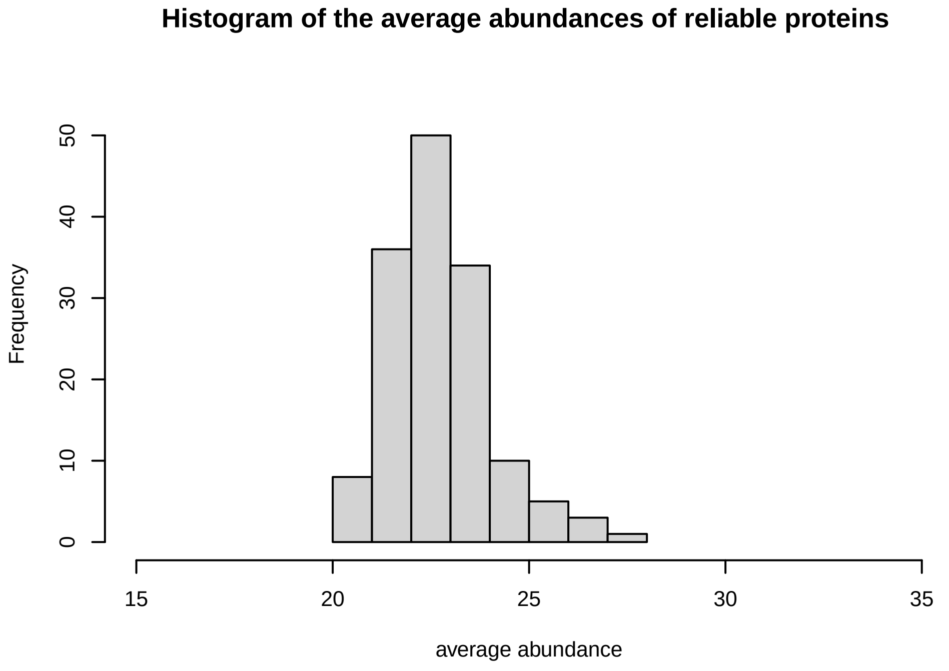
Histogram of the average abundances of reliable proteins
Frequency
average abundance
0 10 20 30 40 50
15 20 25 30 35

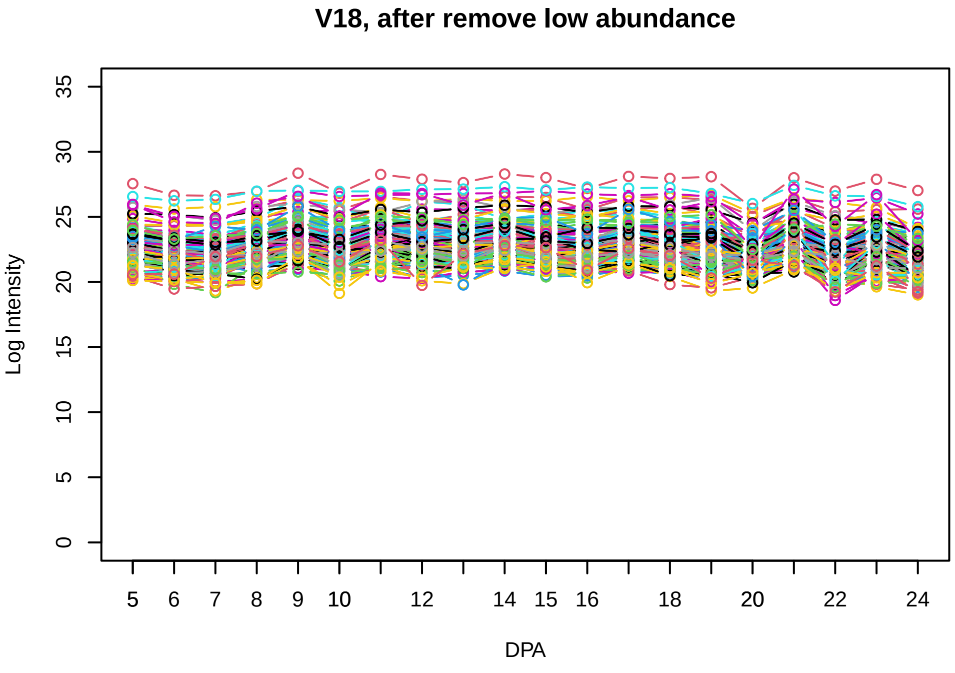
V18, after remove low abundance
Log Intensity
DPA
0 5 10 15 20 25 30 35
5 6 7 8 9 10 12 14 15 16 18 20 22 24

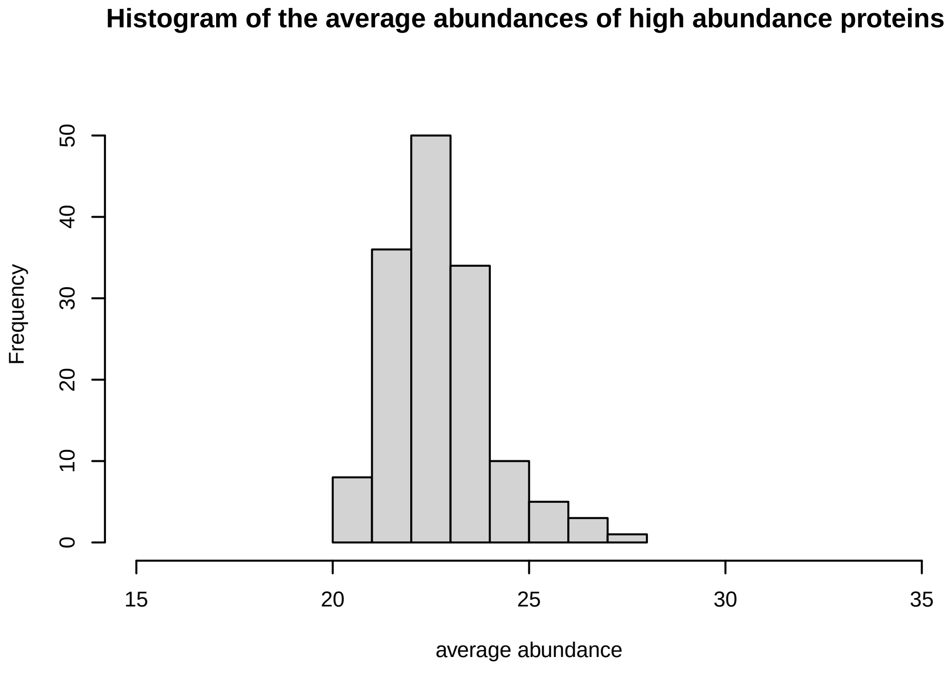
Histogram of the average abundances of high abundance proteins
Frequency
average abundance
0 10 20 30 40 50
15 20 25 30 35

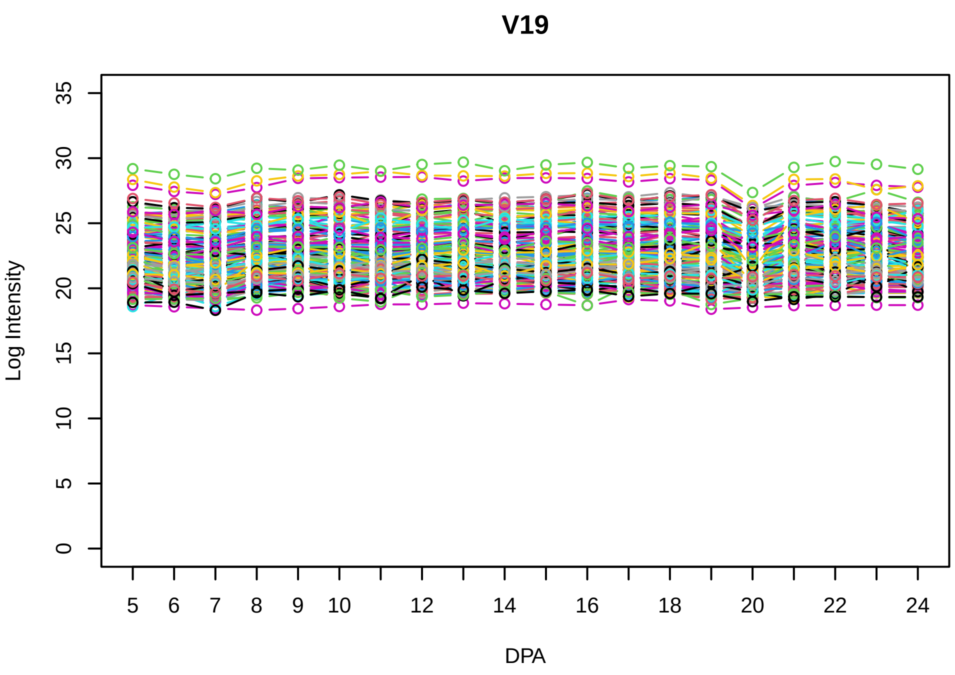
V19
Log Intensity
DPA

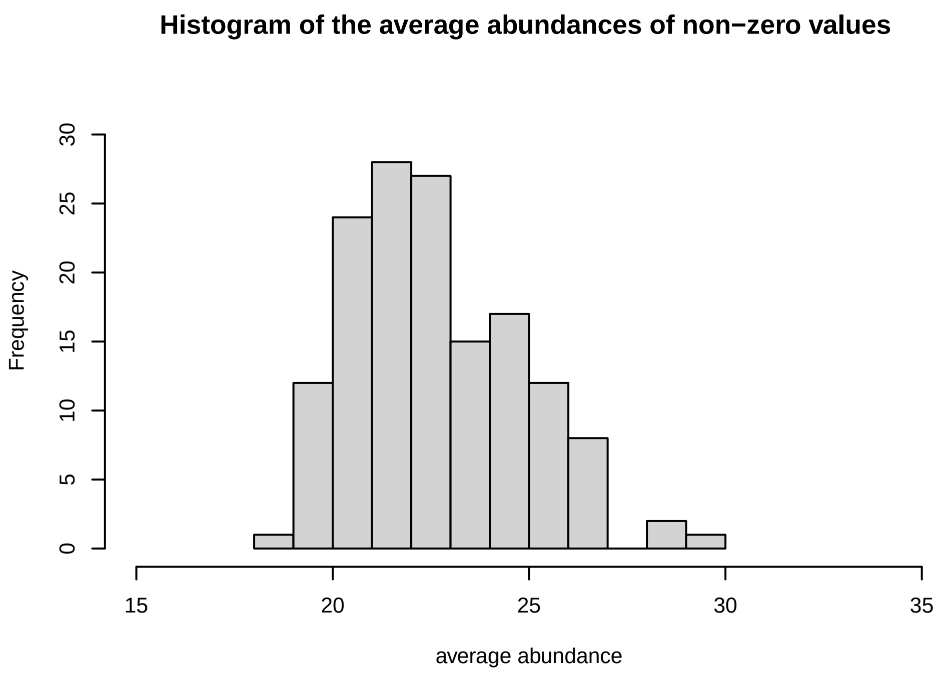
Histogram of the average abundances of non−zero values
Frequency
average abundance

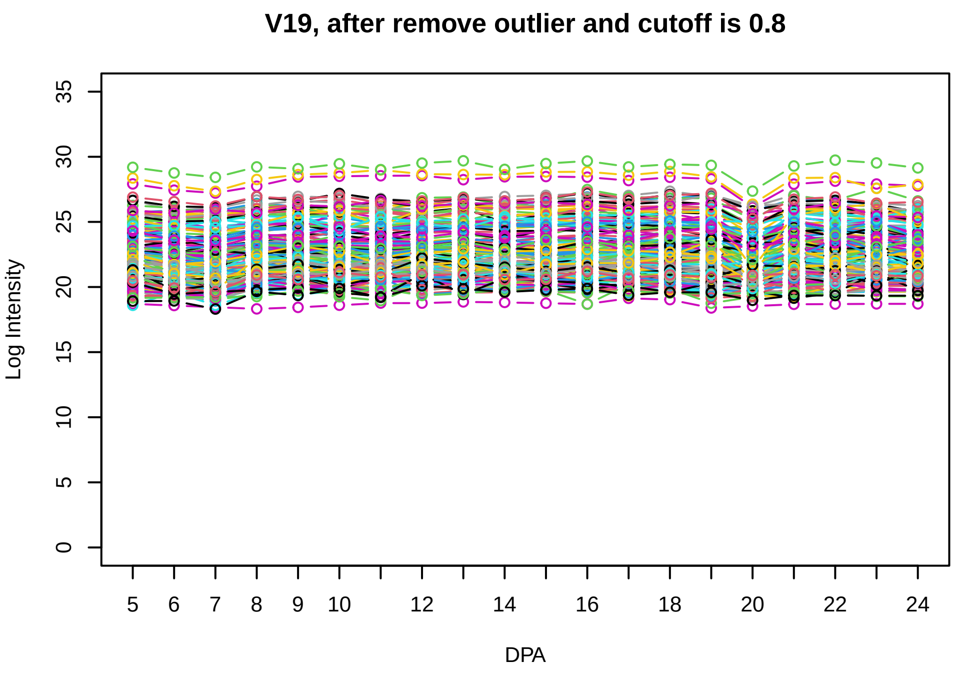
V19, after remove outlier and cutoff is 0.8
Log Intensity
DPA

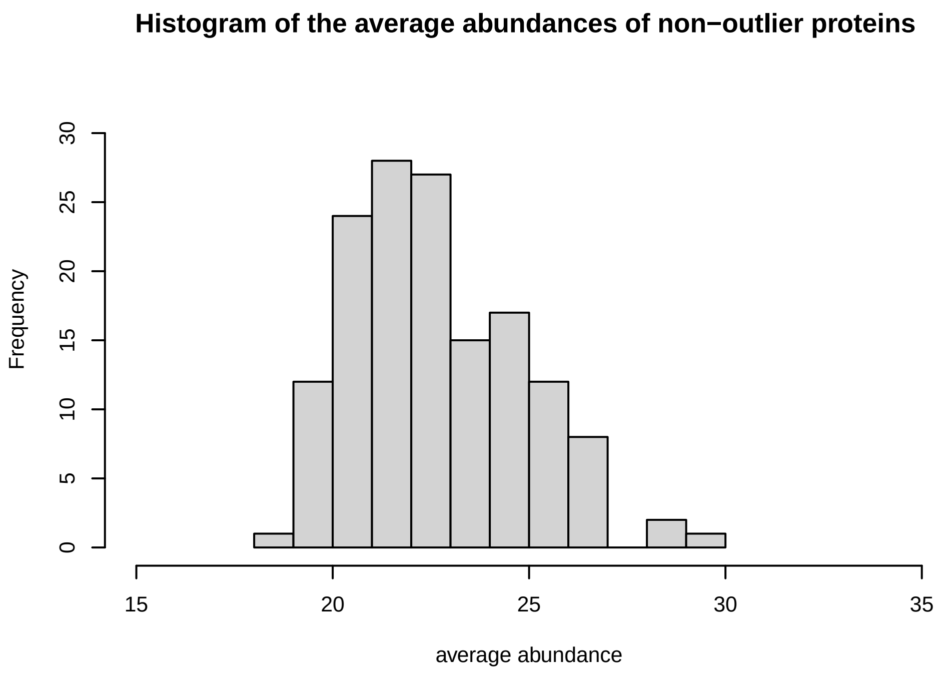
Histogram of the average abundances of non−outlier proteins
Frequency
average abundance

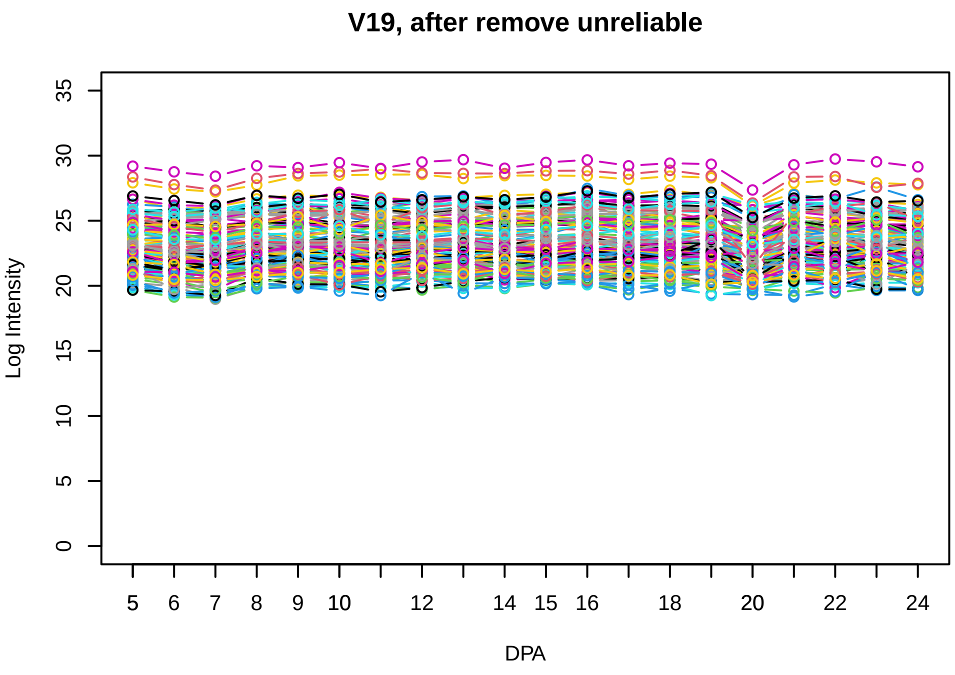
V19, after remove unreliable
Log Intensity
DPA

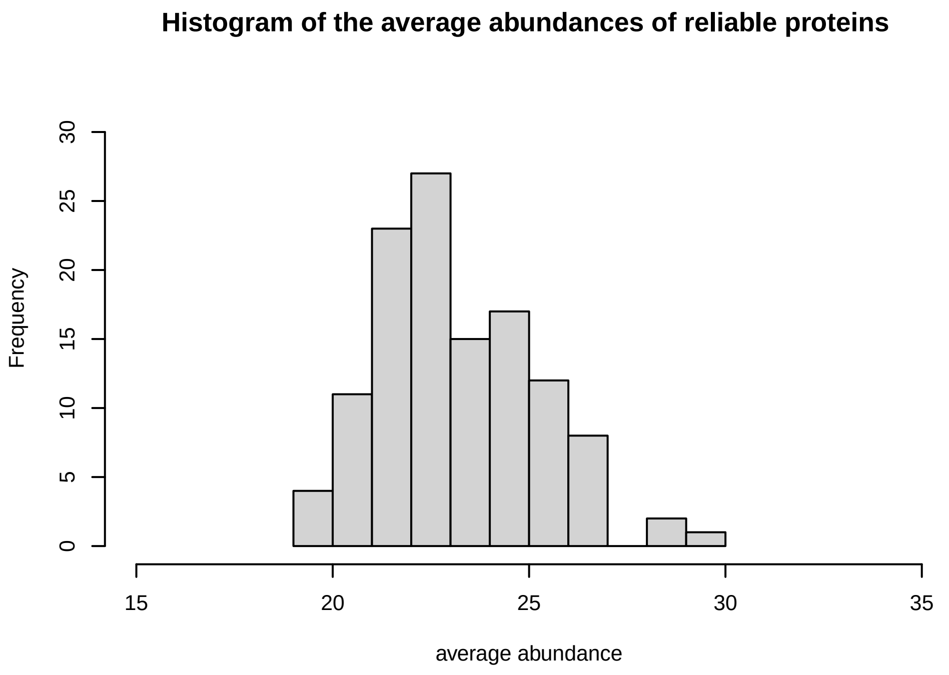
Histogram of the average abundances of reliable proteins
Frequency
average abundance

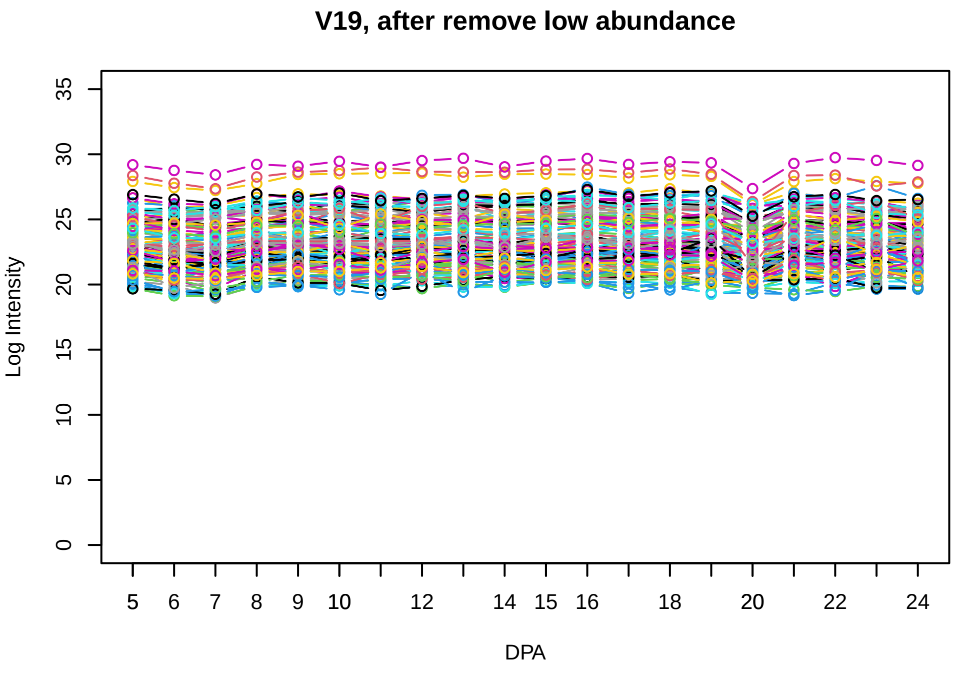
V19, after remove low abundance
Log Intensity
DPA

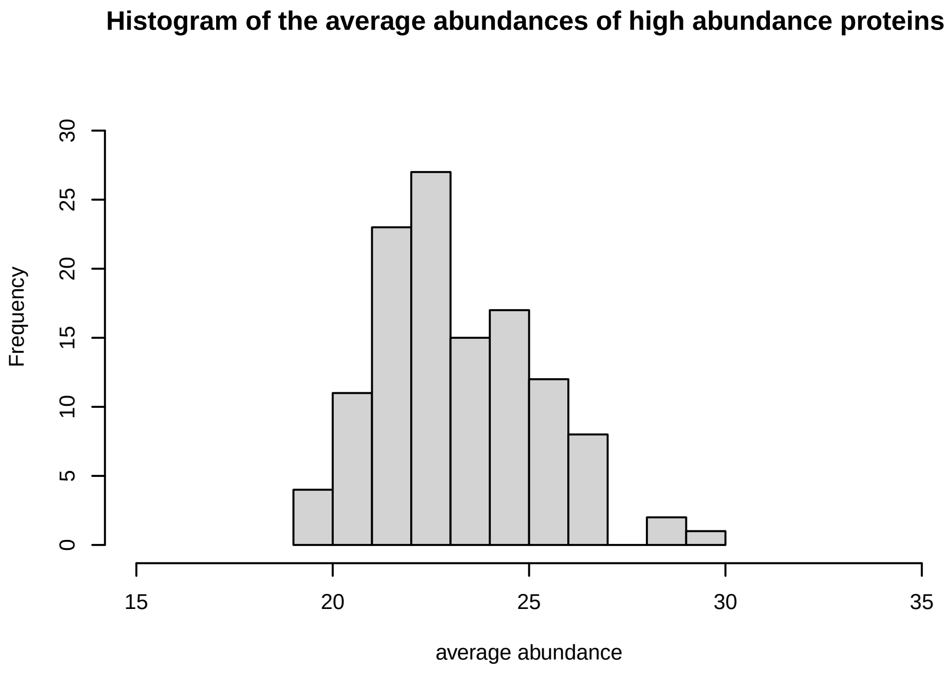
Histogram of the average abundances of high abundance proteins
Frequency
average abundance

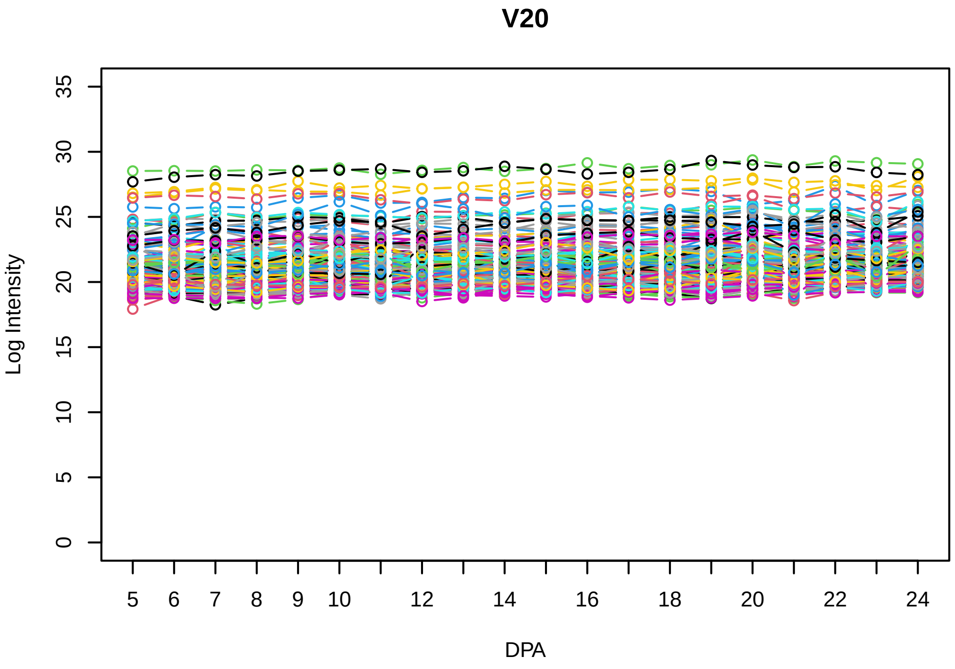
V20
Log Intensity
DPA

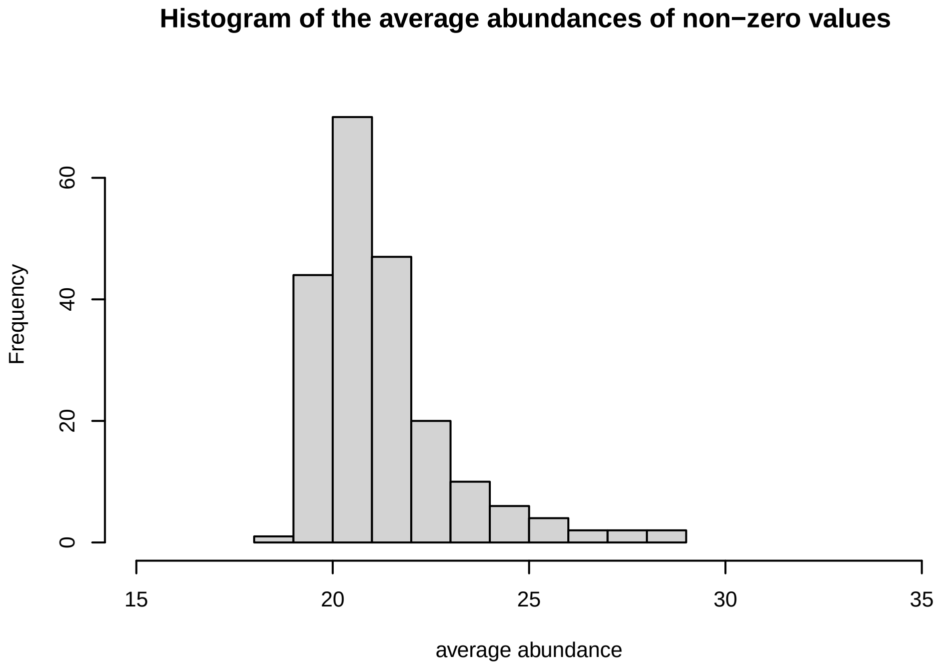
Histogram of the average abundances of non−zero values
Frequency
average abundance

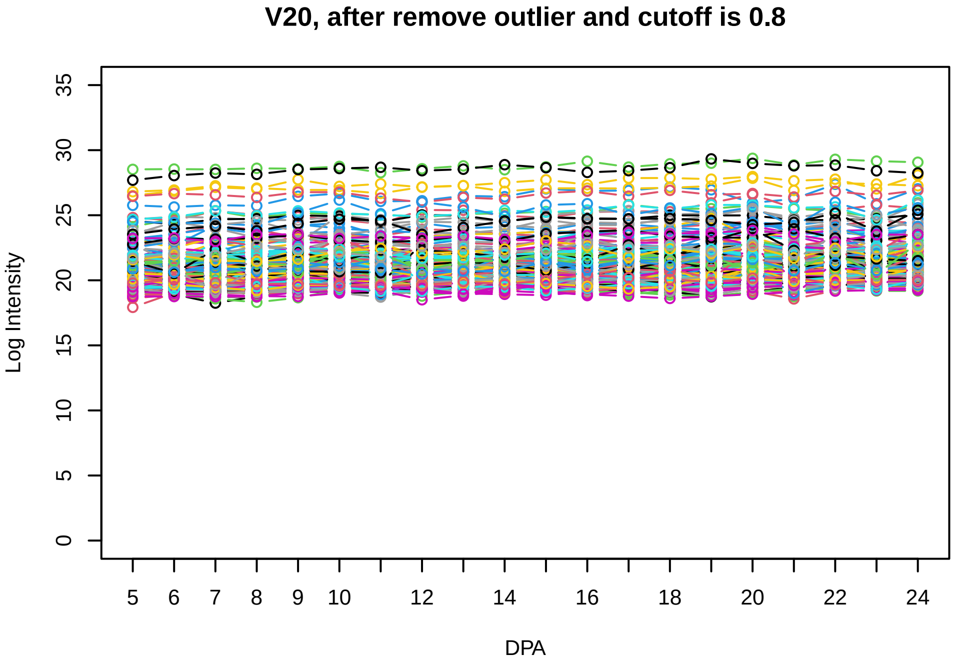
V20, after remove outlier and cutoff is 0.8
Log Intensity
DPA

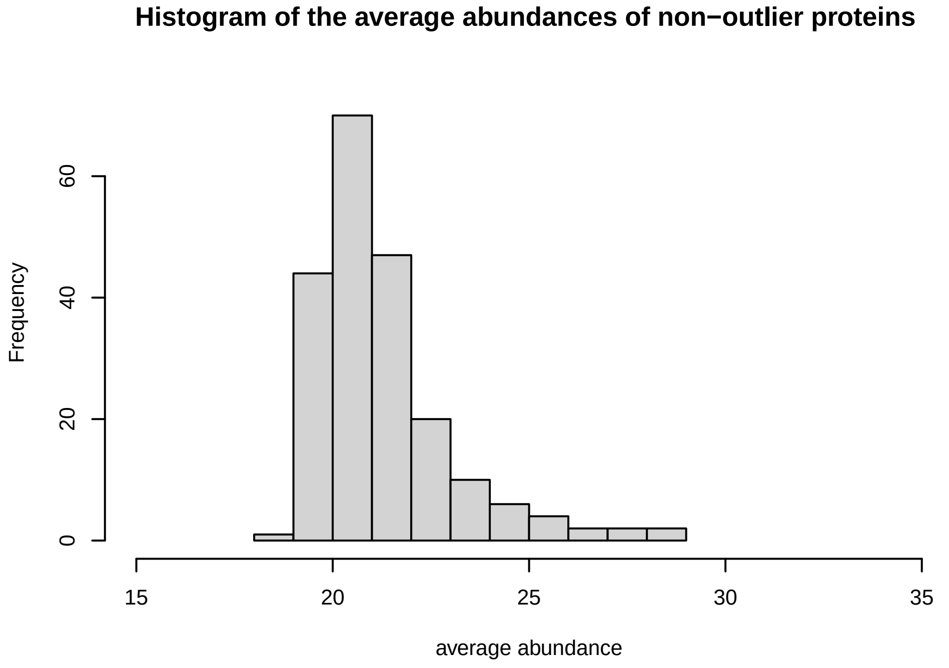
Histogram of the average abundances of non−outlier proteins
Frequency
average abundance

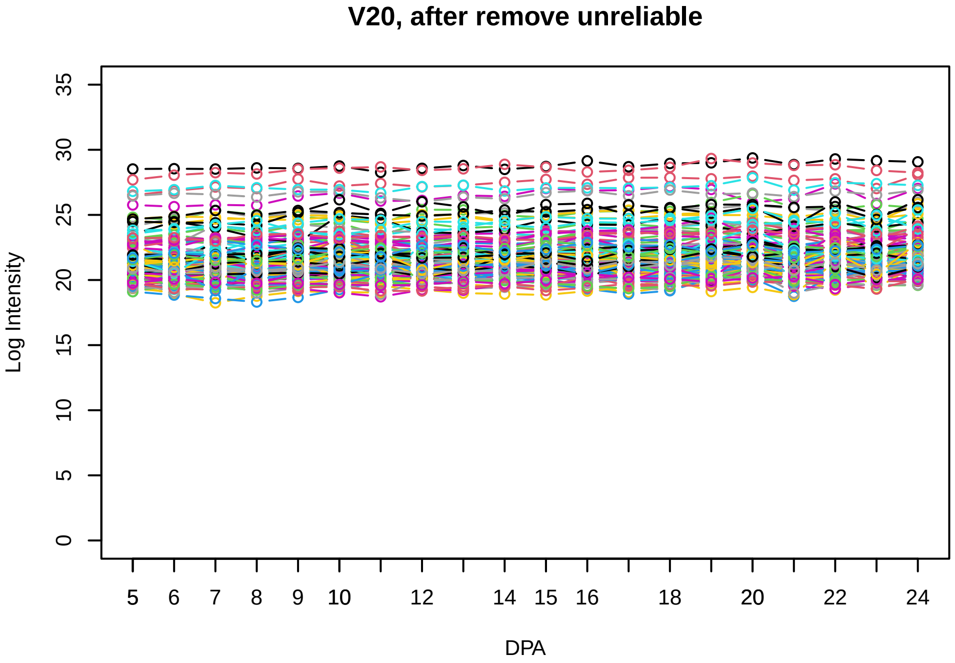
V20, after remove unreliable
Log Intensity
DPA

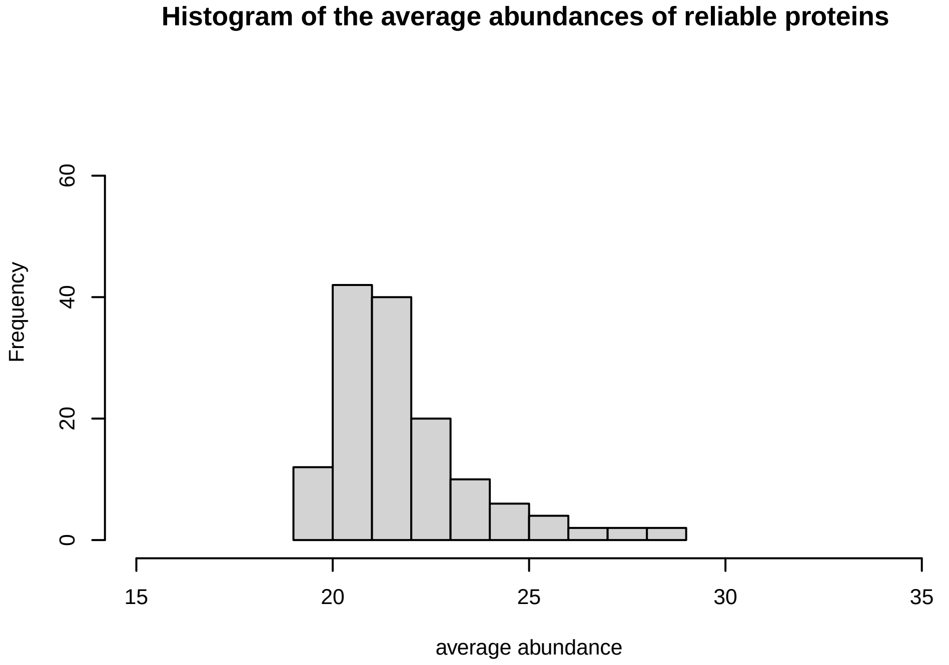
Histogram of the average abundances of reliable proteins
Frequency
average abundance

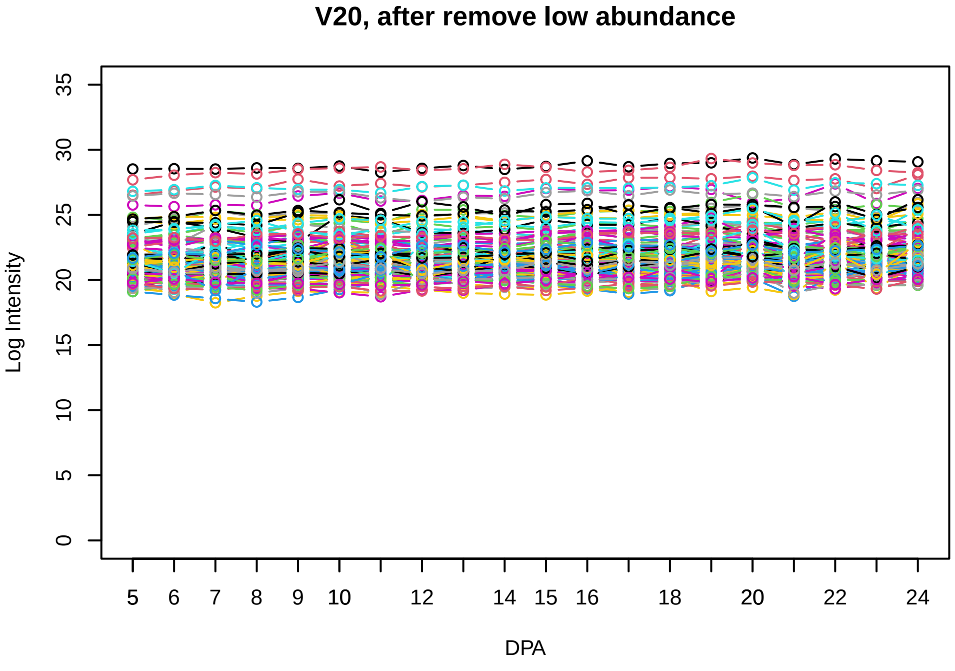
V20, after remove low abundance
Log Intensity
DPA

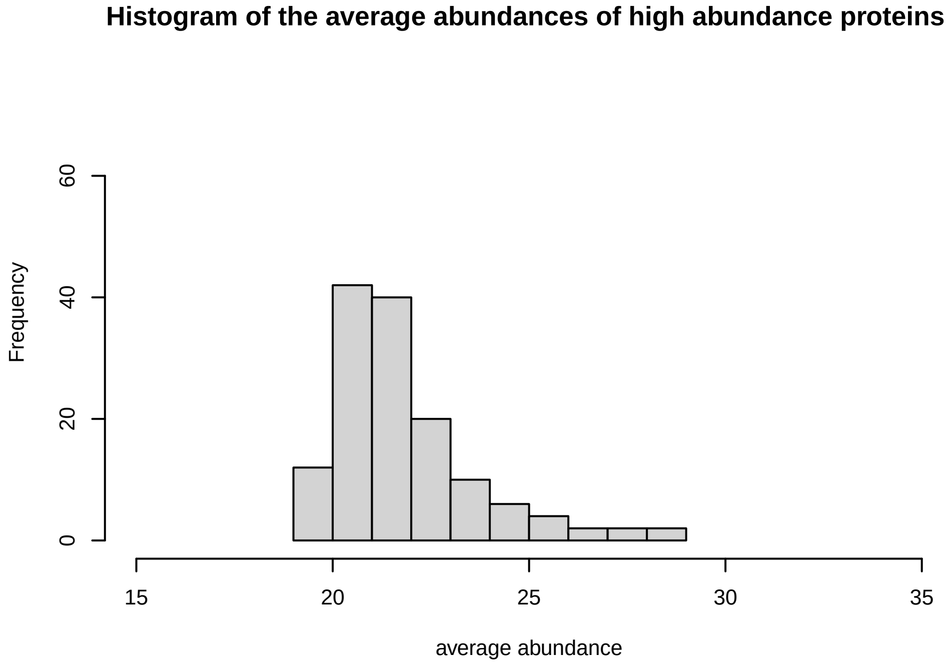
Histogram of the average abundances of high abundance proteins
Frequency
average abundance

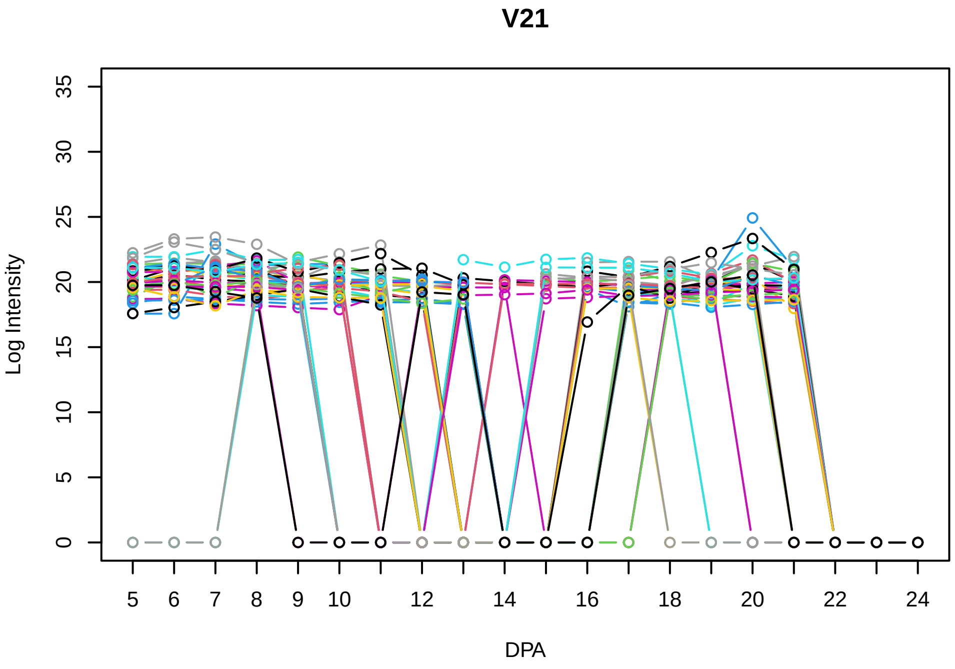
V21
Log Intensity
DPA

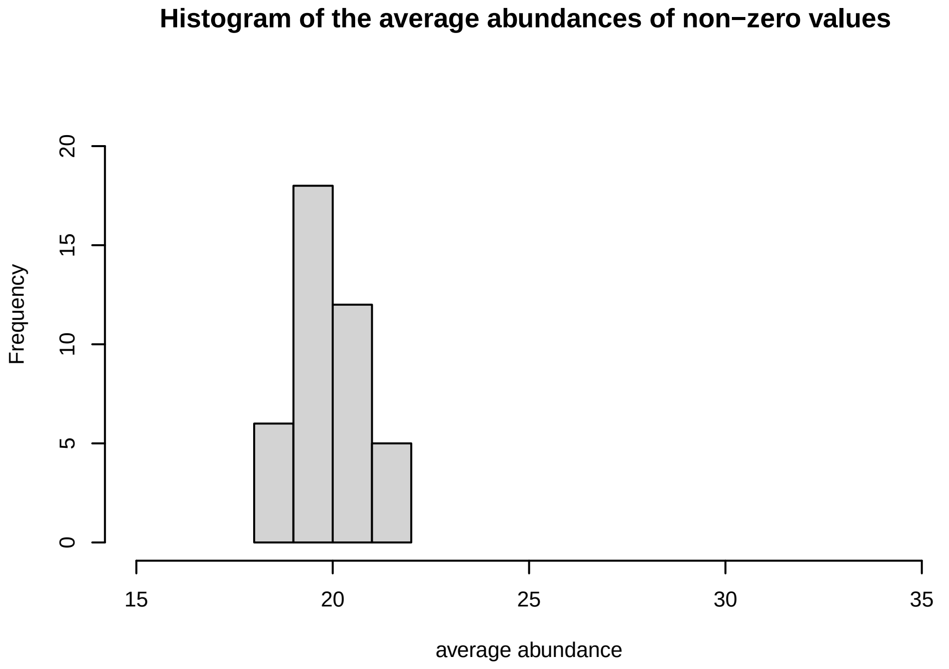
Histogram of the average abundances of non−zero values
Frequency
average abundance

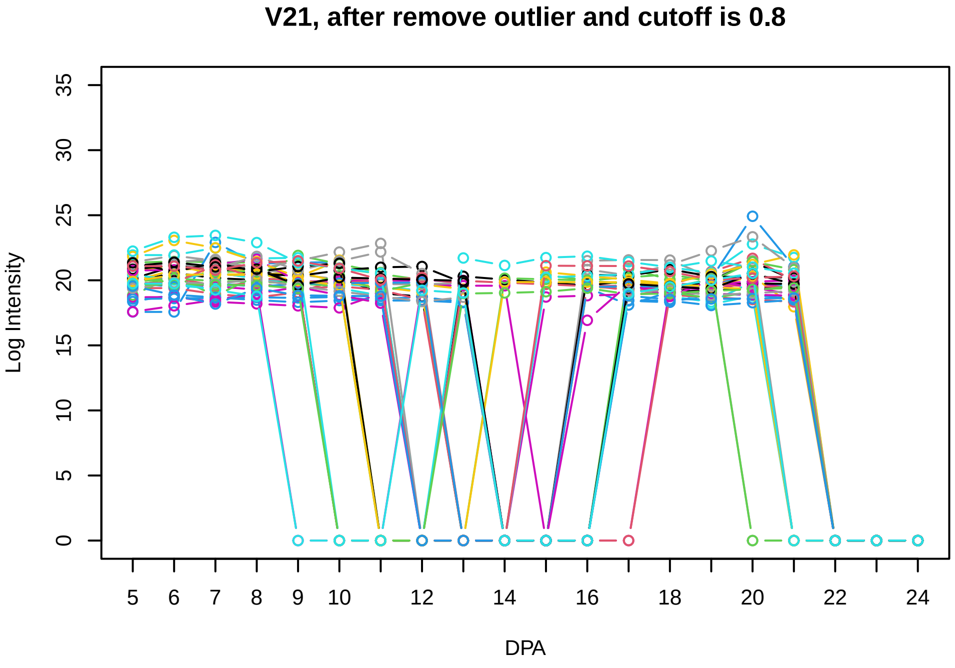
V21, after remove outlier and cutoff is 0.8
Log Intensity
DPA

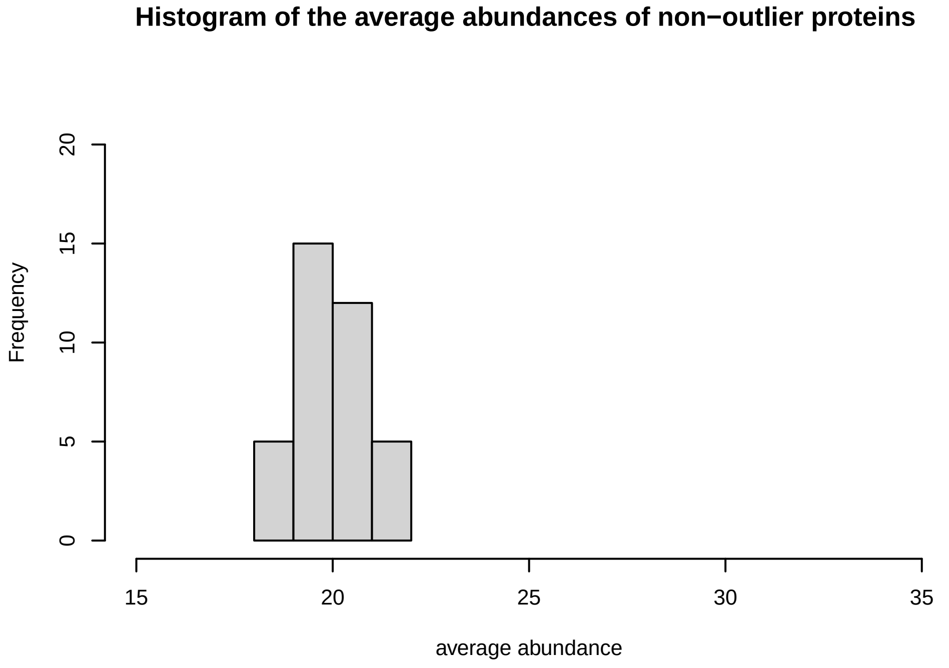
Histogram of the average abundances of non−outlier proteins
Frequency
average abundance

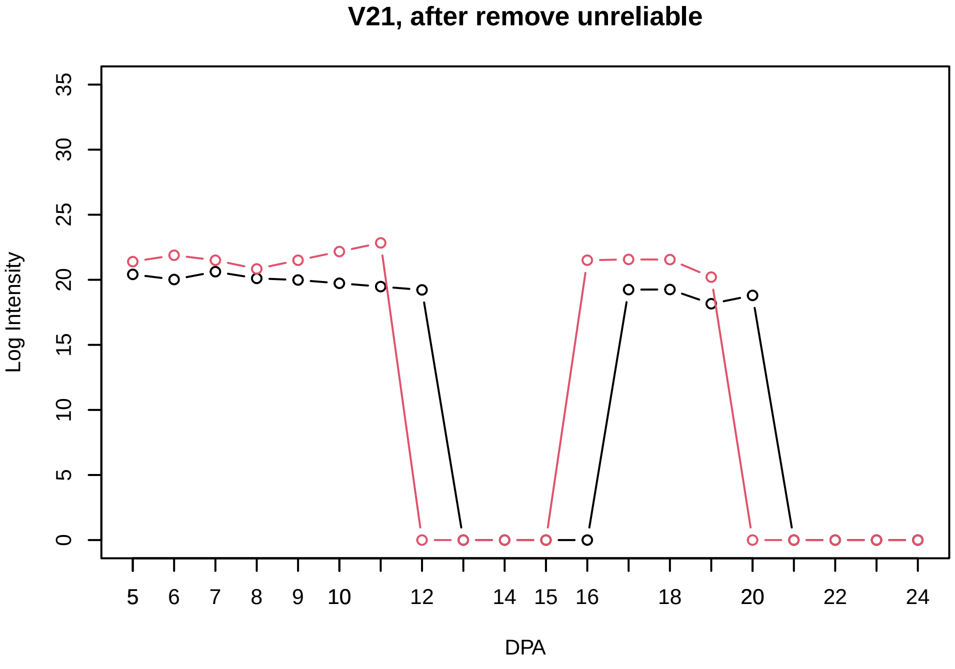
V21, after remove unreliable
Log Intensity
DPA

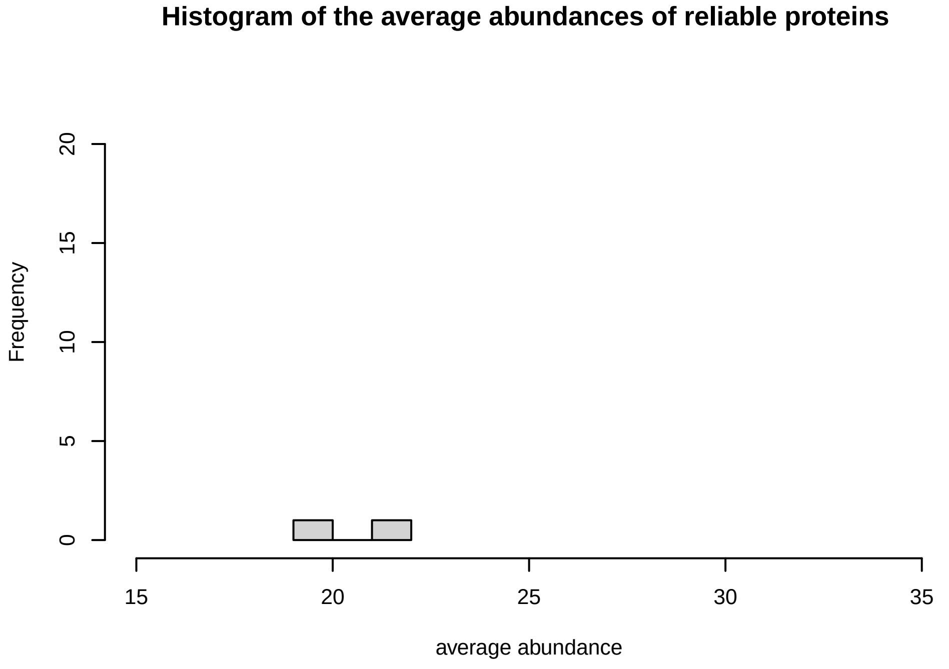
Histogram of the average abundances of reliable proteins
Frequency
average abundance

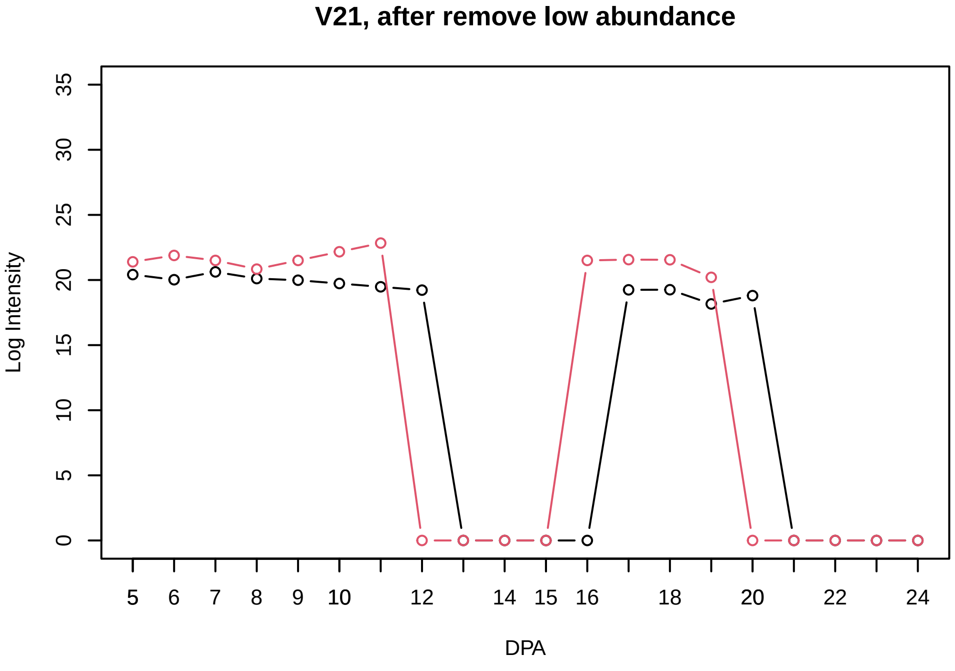
V21, after remove low abundance
Log Intensity
DPA

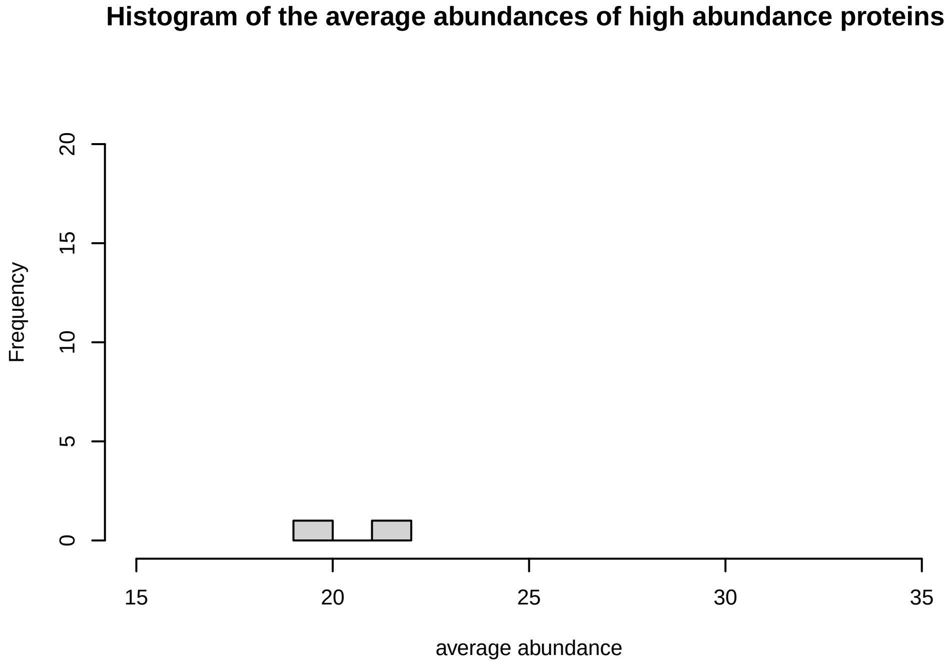
Histogram of the average abundances of high abundance proteins
Frequency
average abundance

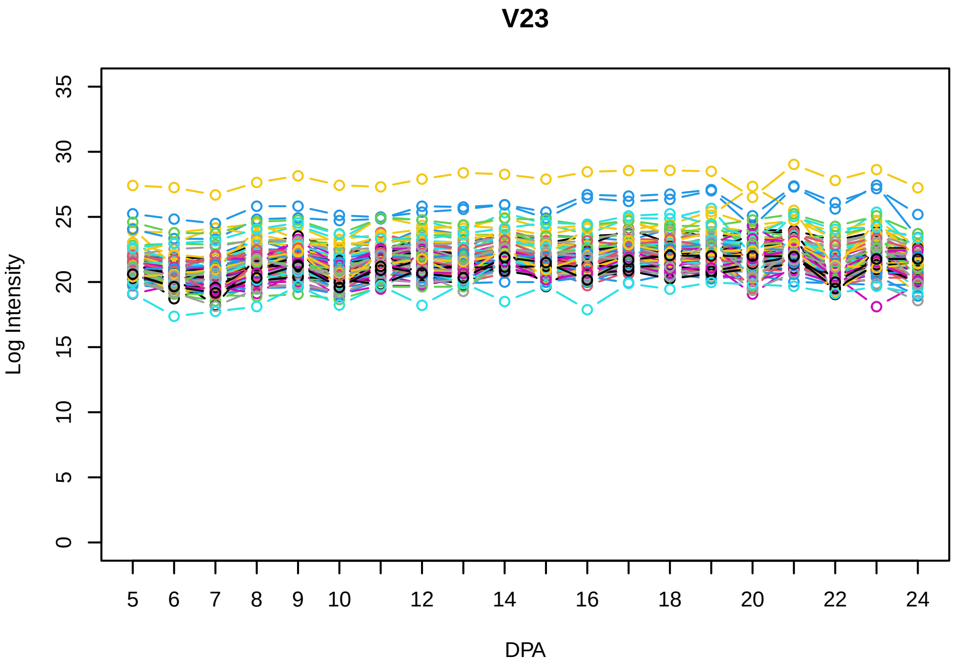

V23
Log Intensity
DPA
0 5 10 15 20 25 30 35
5 6 7 8 9 10 12 14 16 18 20 22 24


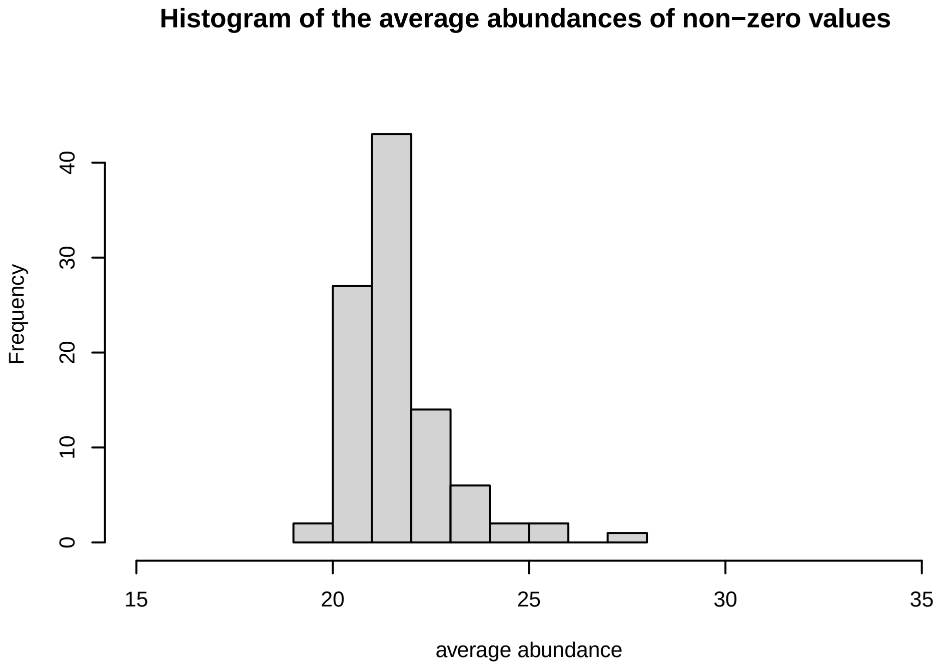

Histogram of the average abundances of non−zero values
Frequency
average abundance
0 10 20 30 40
15 20 25 30 35


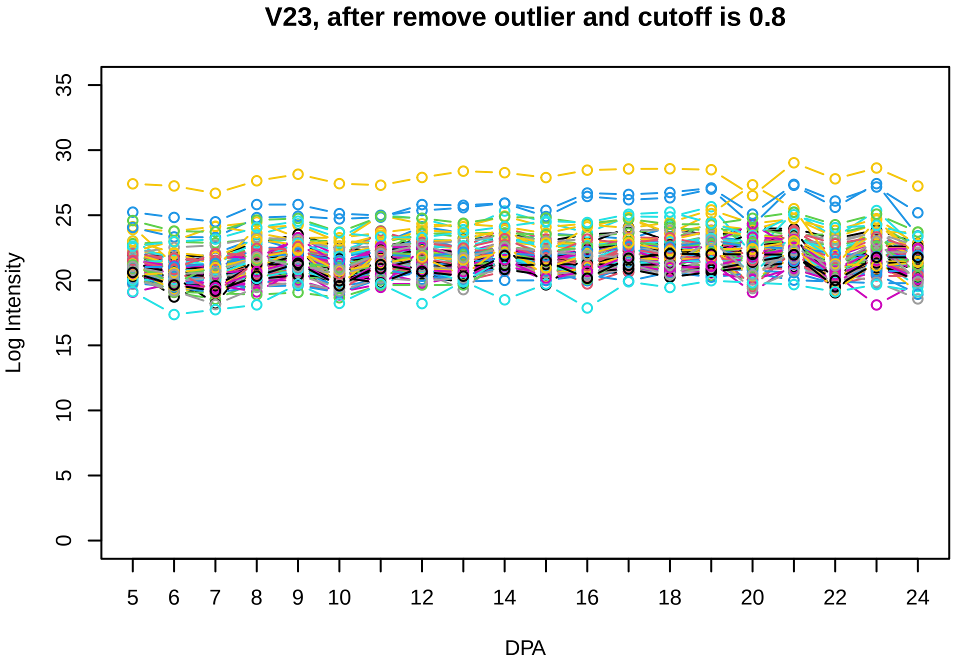

V23, after remove outlier and cutoff is 0.8
Log Intensity
DPA
0 5 10 15 20 25 30 35
5 6 7 8 9 10 12 14 16 18 20 22 24


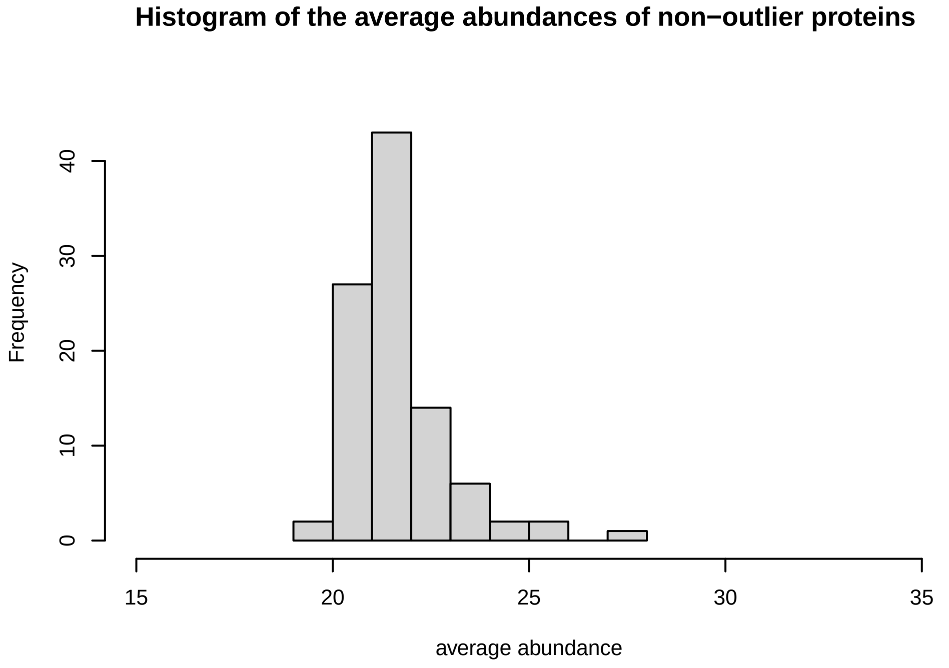

Histogram of the average abundances of non−outlier proteins
Frequency
average abundance
0 10 20 30 40
15 20 25 30 35


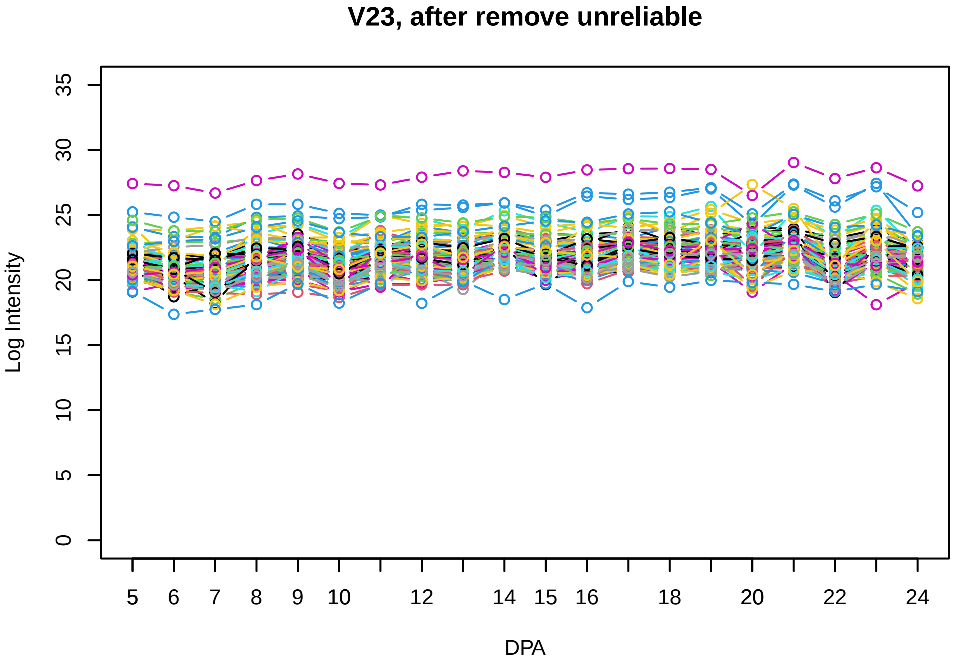

V23, after remove unreliable
Log Intensity
DPA
0 5 10 15 20 25 30 35
5 6 7 8 9 10 12 14 15 16 18 20 22 24


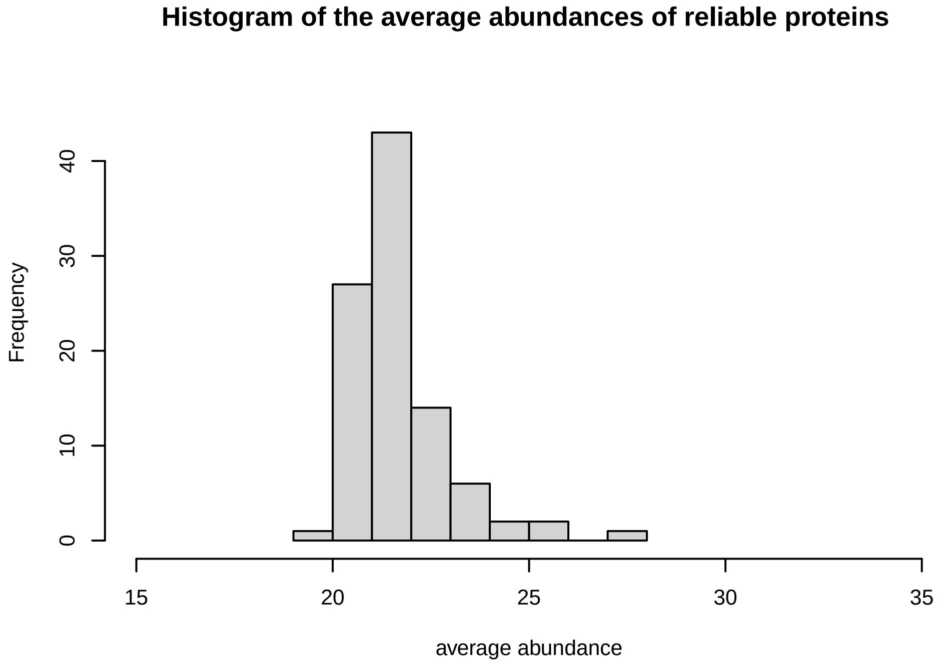

Histogram of the average abundances of reliable proteins
Frequency
average abundance
0 10 20 30 40
15 20 25 30 35


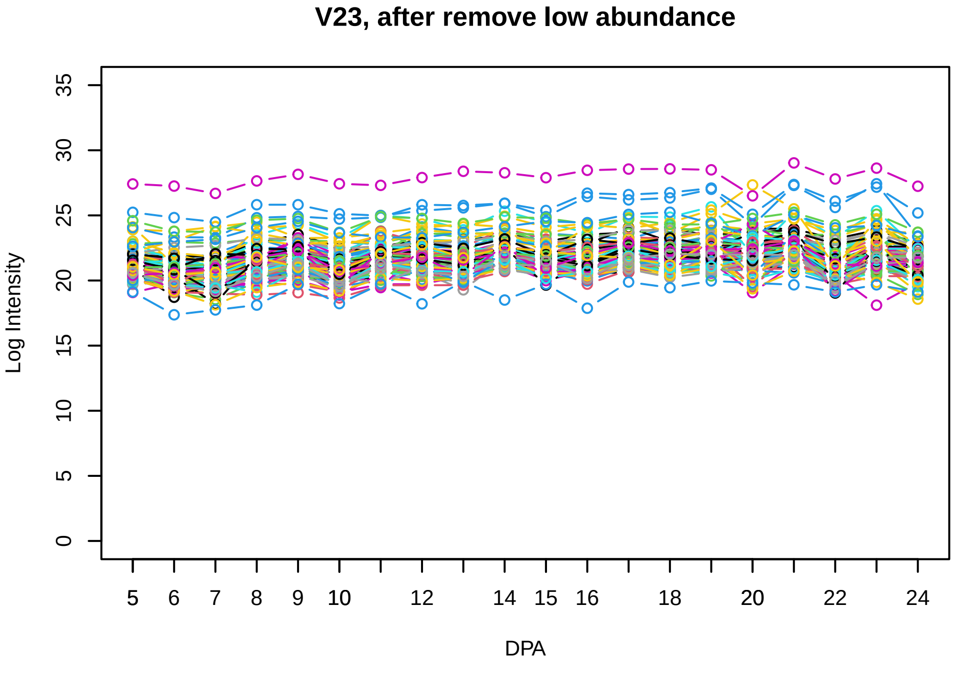

V23, after remove low abundance
Log Intensity
DPA
0 5 10 15 20 25 30 35
5 6 7 8 9 10 12 14 15 16 18 20 22 24


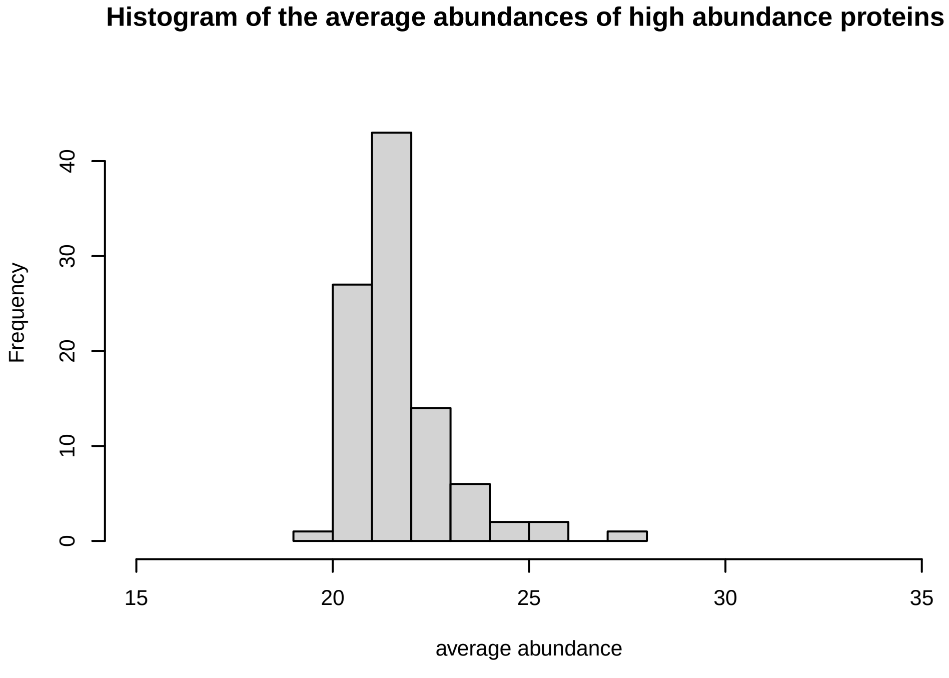

Histogram of the average abundances of high abundance proteins
Frequency
average abundance
0 10 20 30 40
15 20 25 30 35

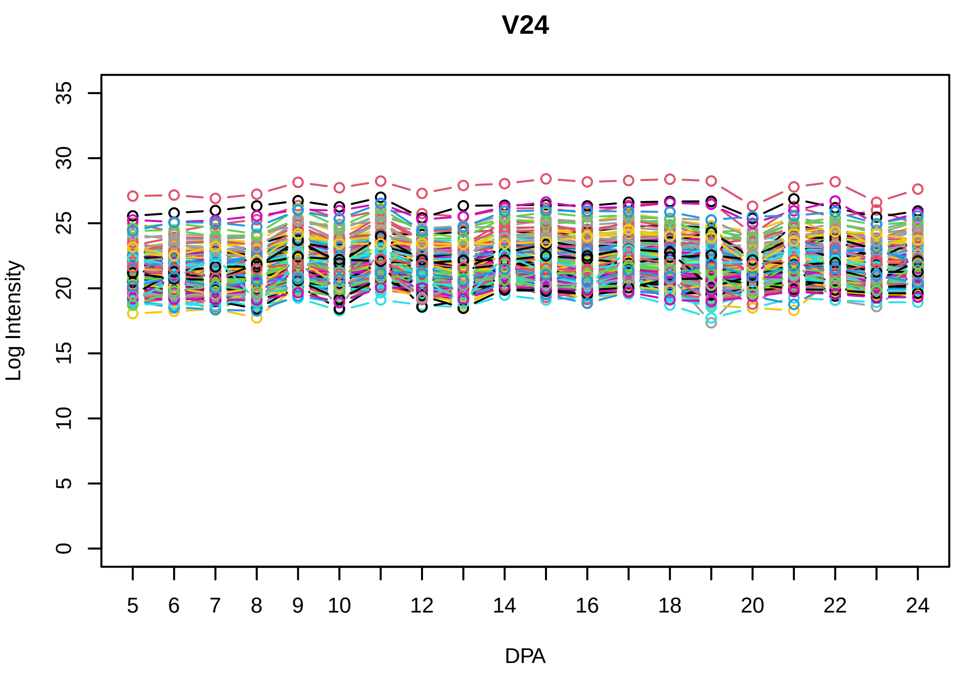
V24
Log Intensity
DPA

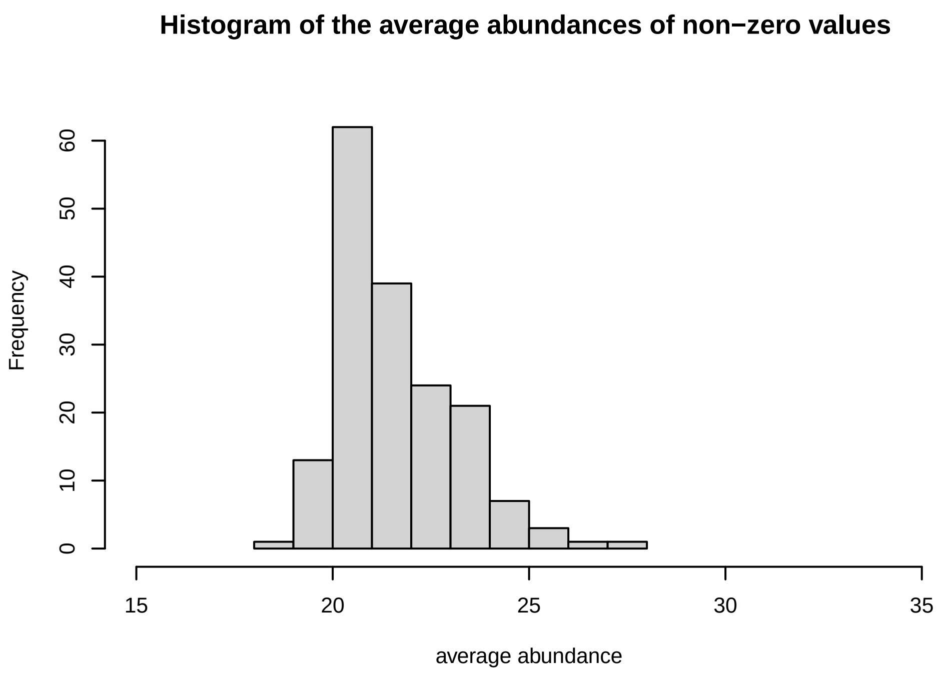
Histogram of the average abundances of non−zero values
Frequency
average abundance

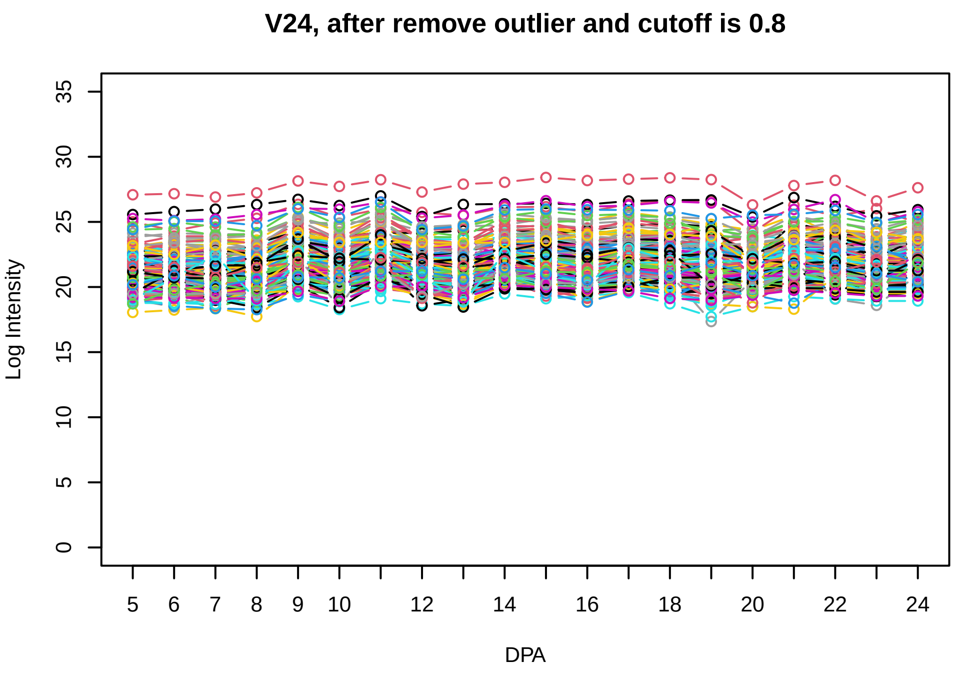
V24, after remove outlier and cutoff is 0.8
Log Intensity
DPA

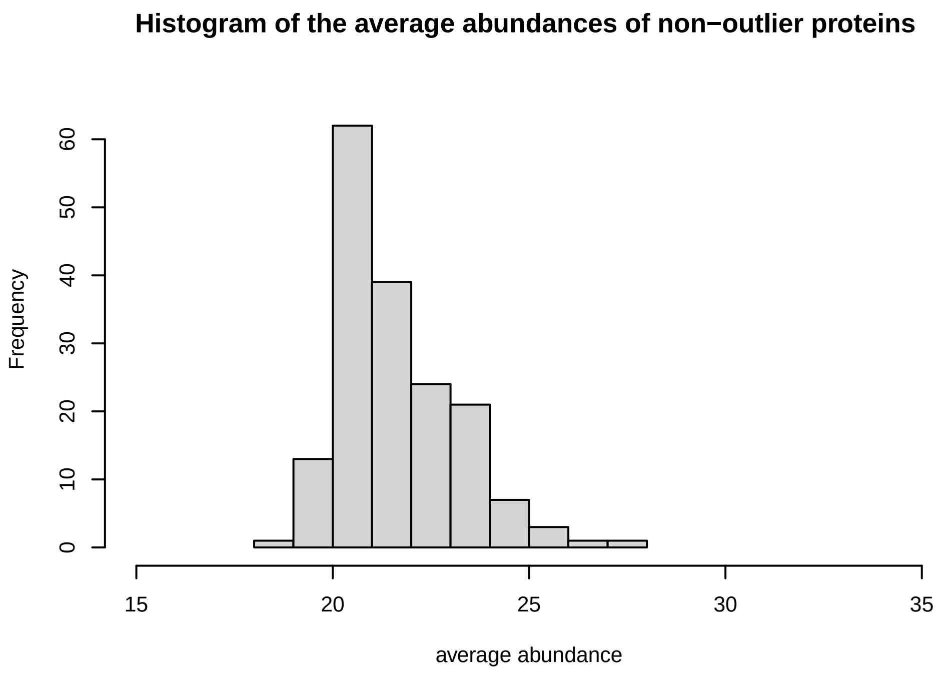
Histogram of the average abundances of non−outlier proteins
Frequency
average abundance

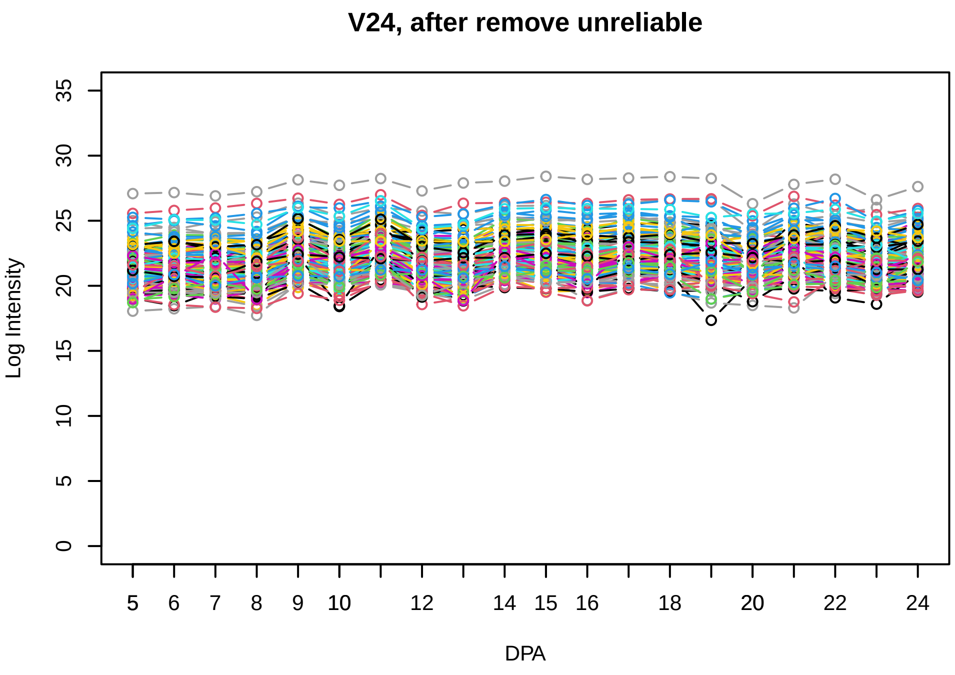
V24, after remove unreliable
Log Intensity
DPA

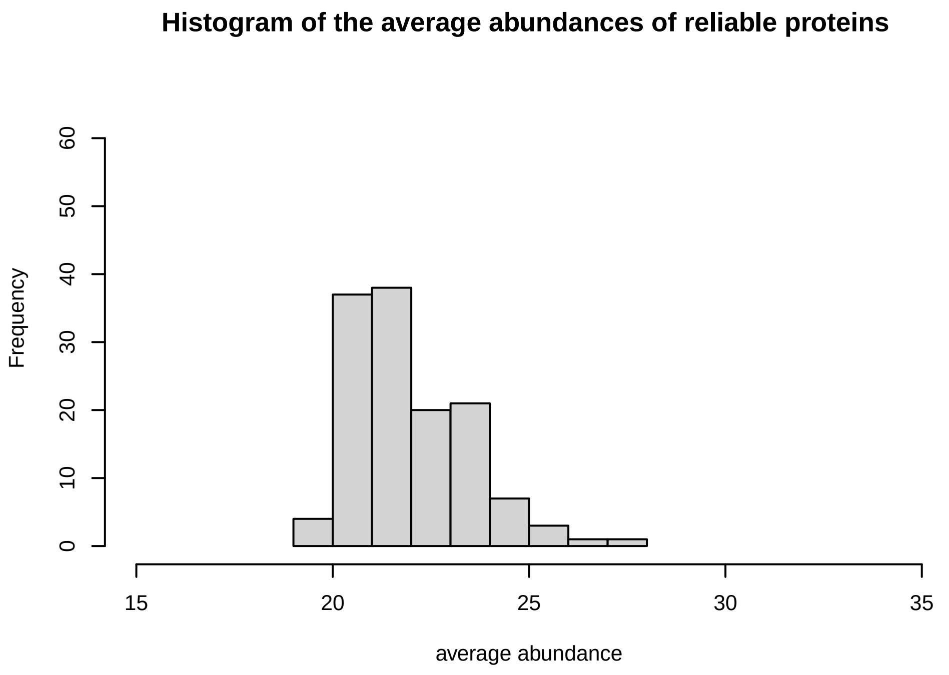
Histogram of the average abundances of reliable proteins
Frequency
average abundance

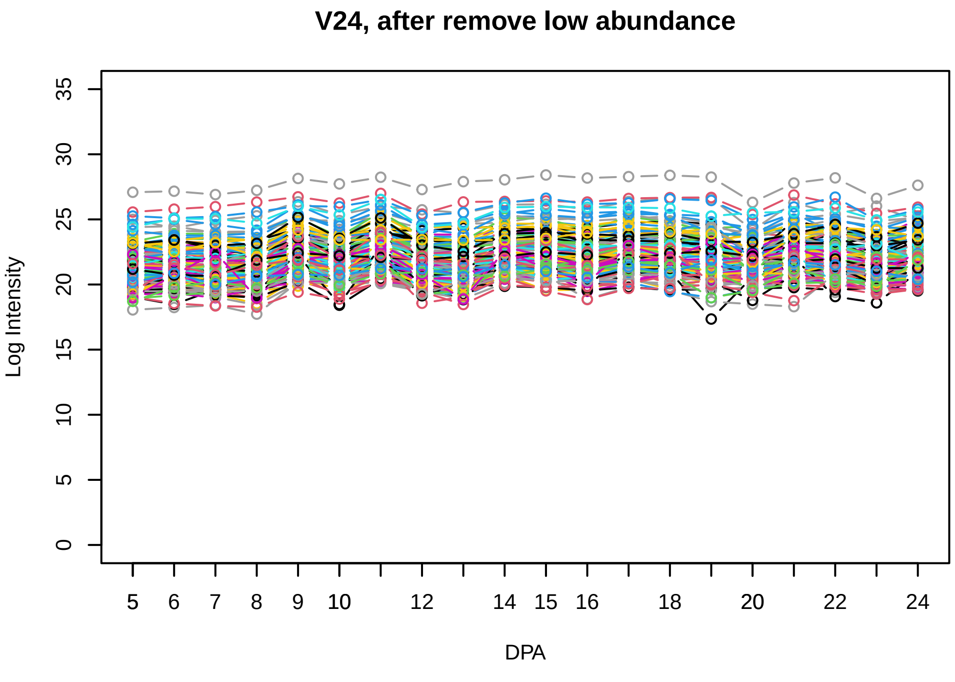
V24, after remove low abundance
Log Intensity
DPA

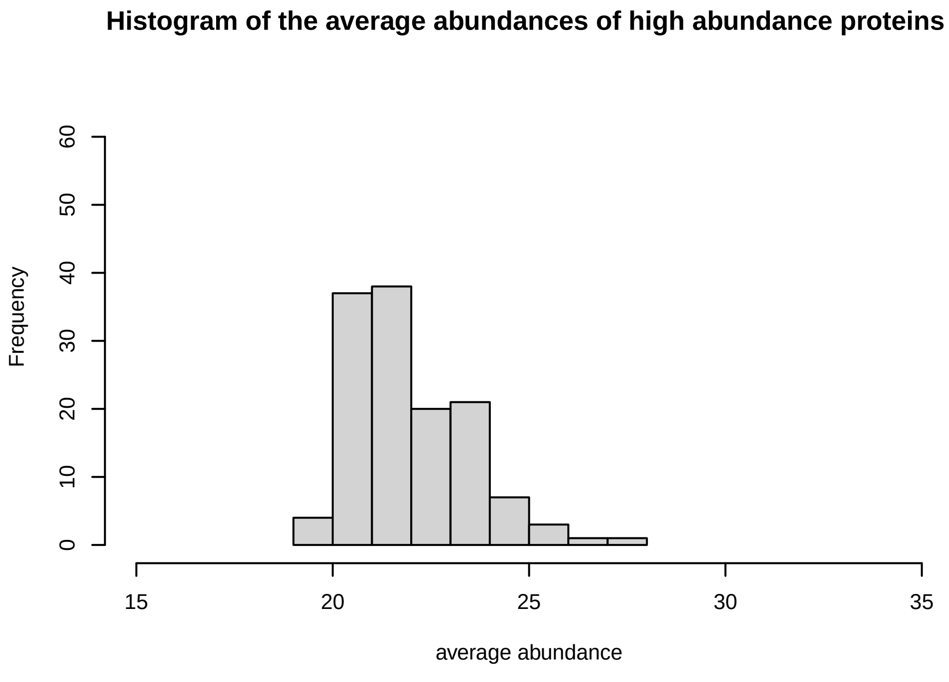
Histogram of the average abundances of high abundance proteins
Frequency
average abundance

**V25**

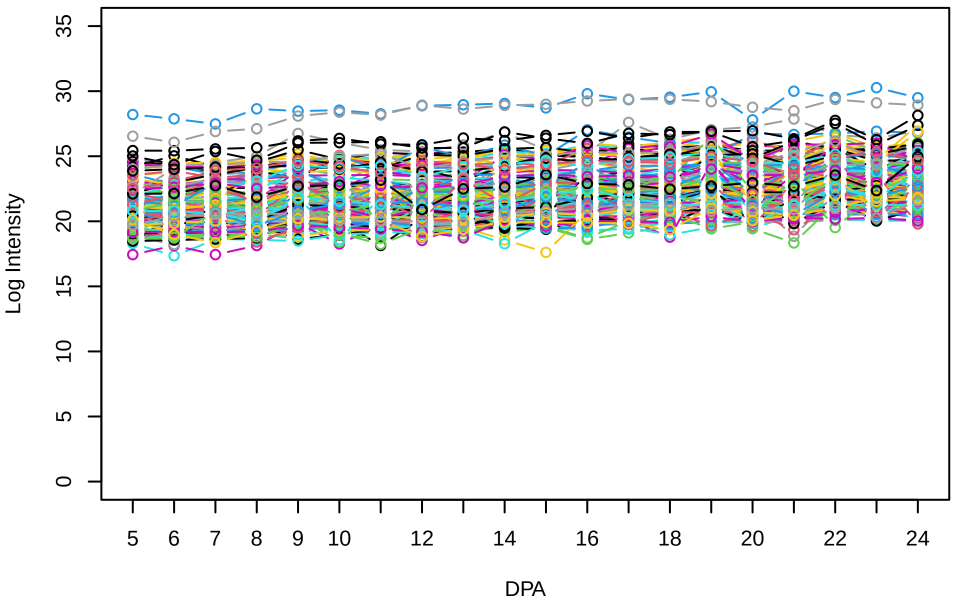


**Histogram of the average abundances of non−zero values**

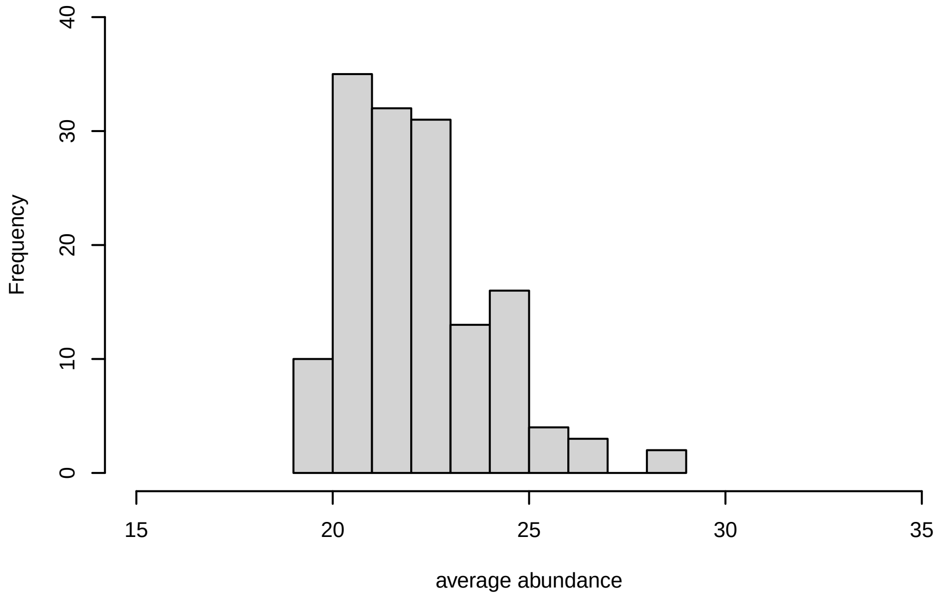


**V25, after remove outlier and cutoff is 0.8**

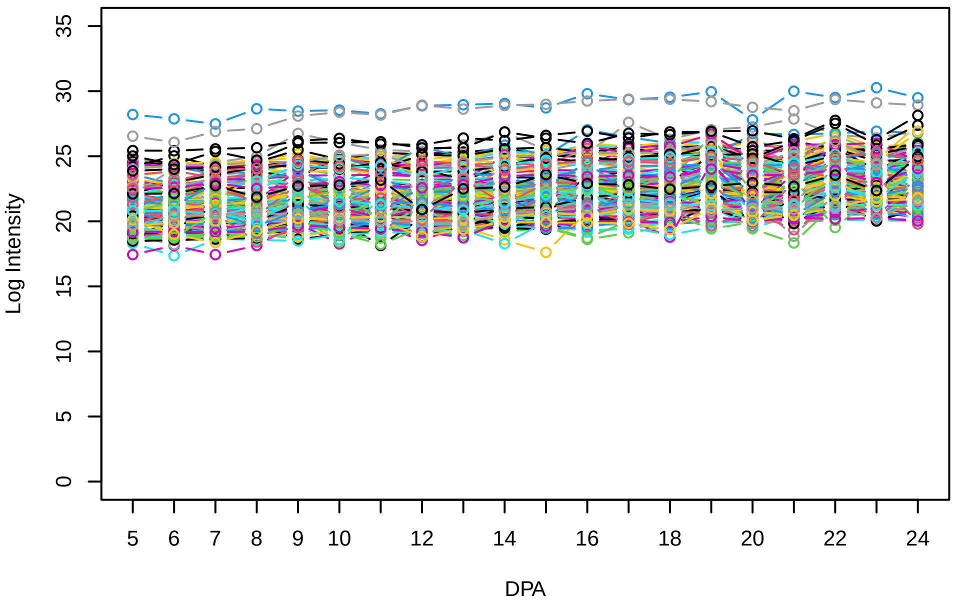


**Histogram of the average abundances of non−outlier proteins**

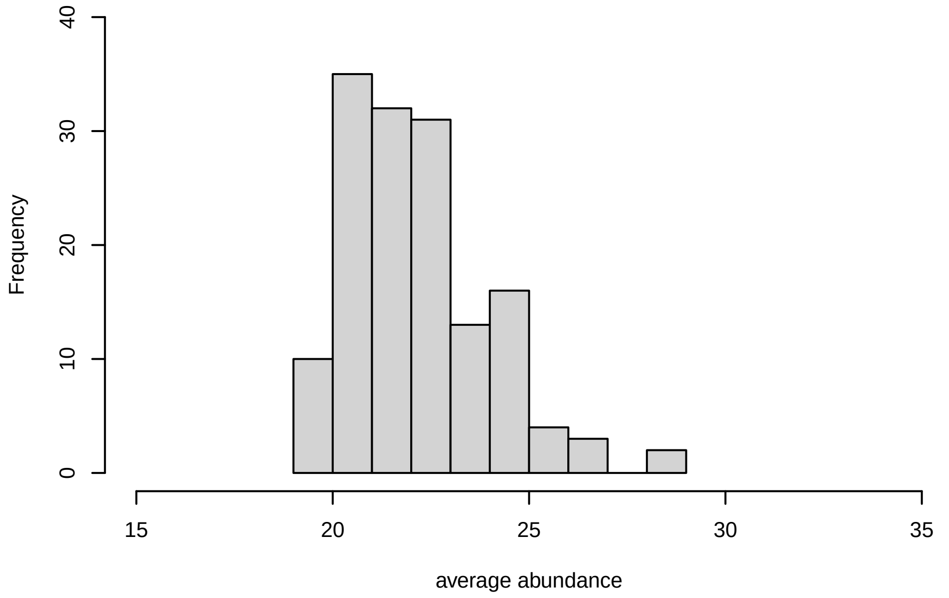


**V25, after remove unreliable**

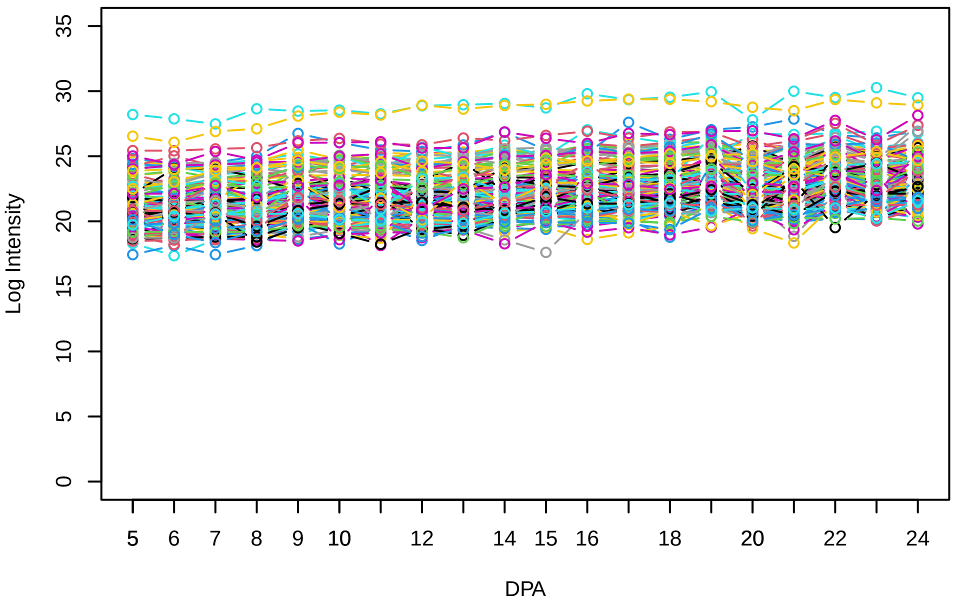


**Histogram of the average abundances of reliable proteins**

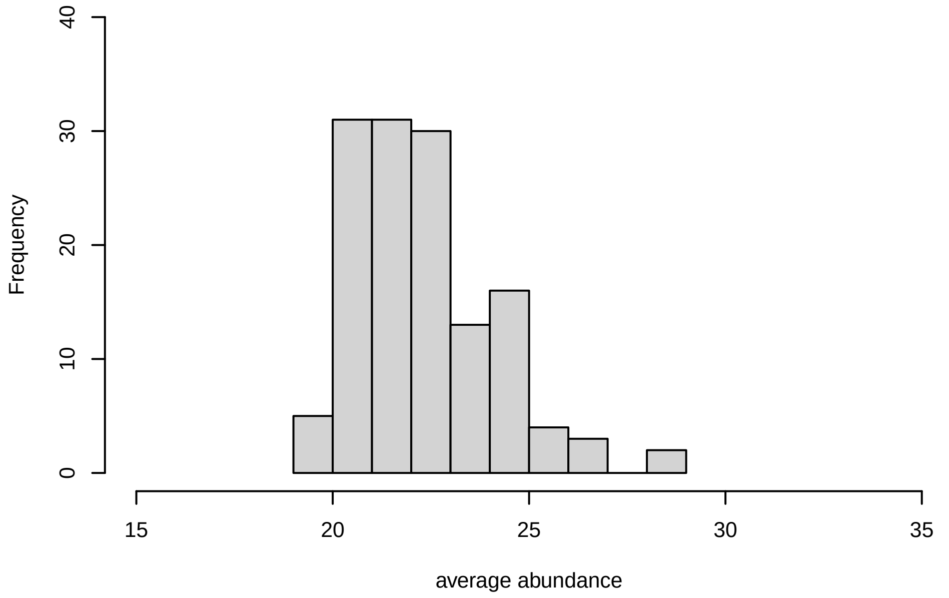


**V25, after remove low abundance**

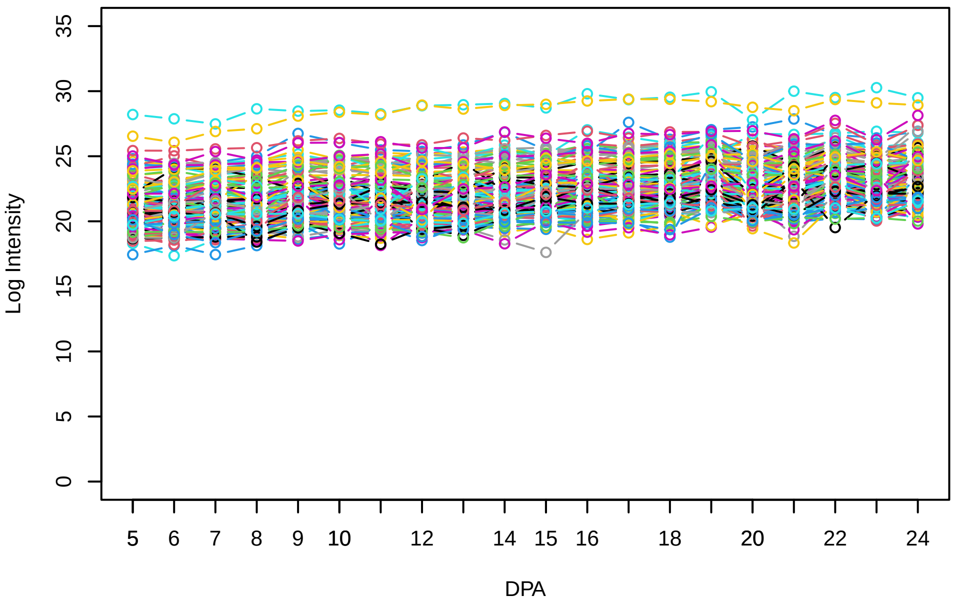


**Histogram of the average abundances of high abundance proteins**

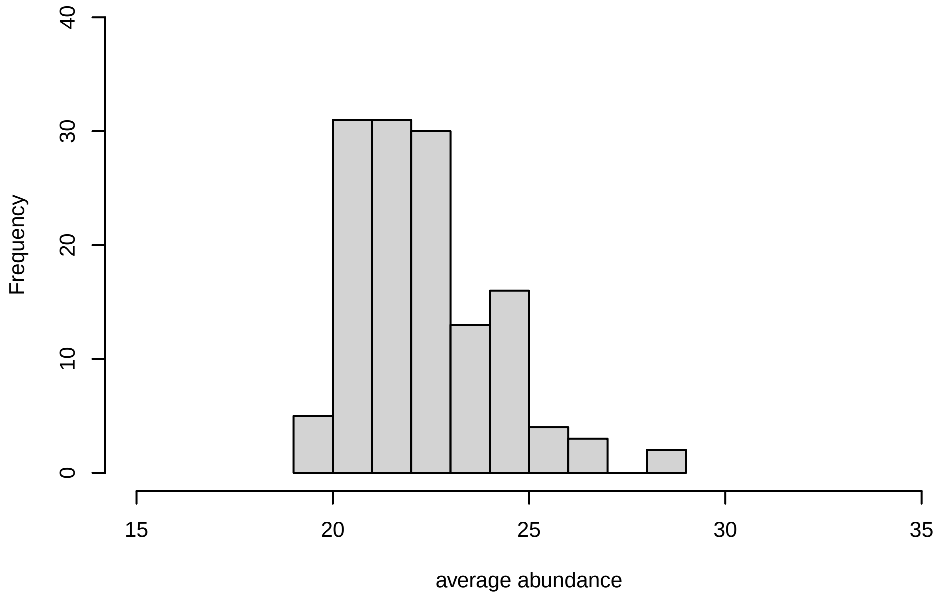

V26

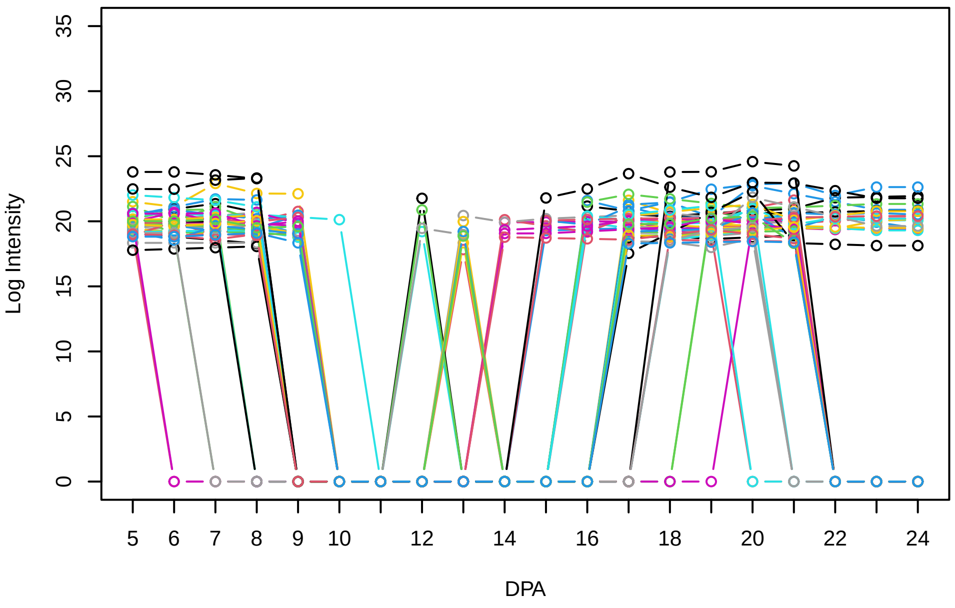


Histogram of the average abundances of non−zero values

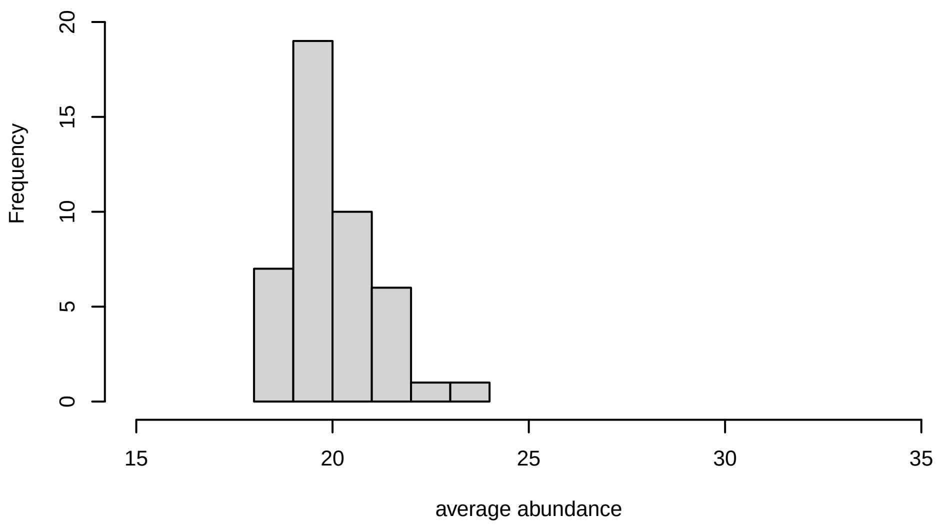


V26, after remove outlier and cutoff is 0.78944379457436

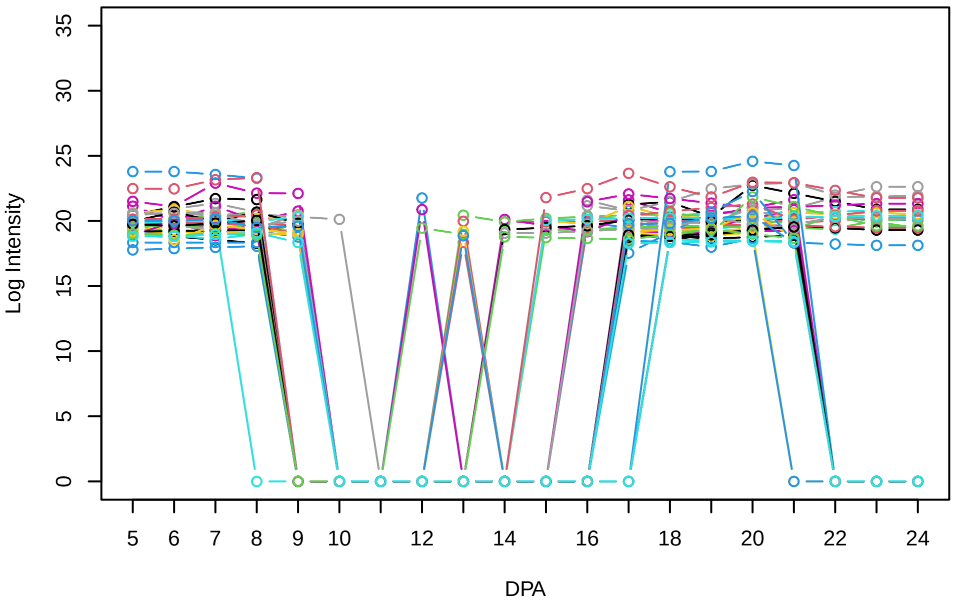


Histogram of the average abundances of non−outlier proteins

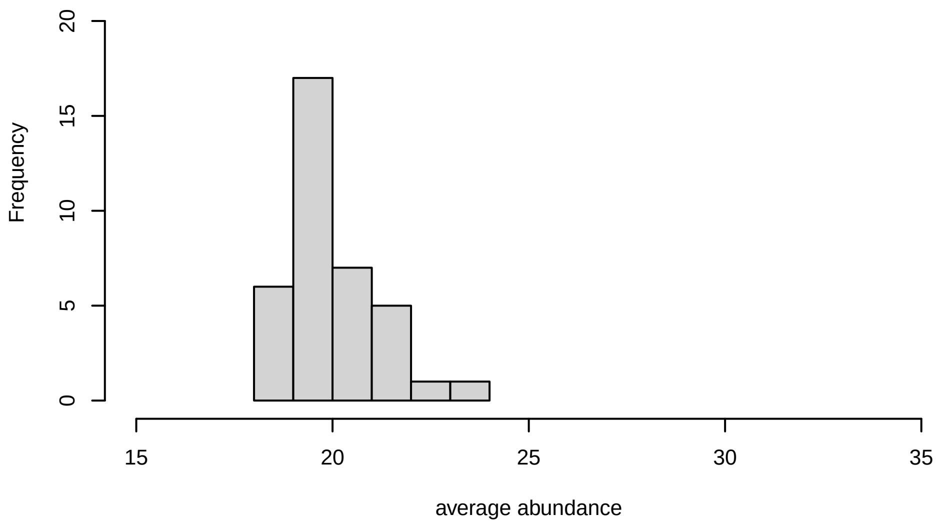


V26, after remove unreliable

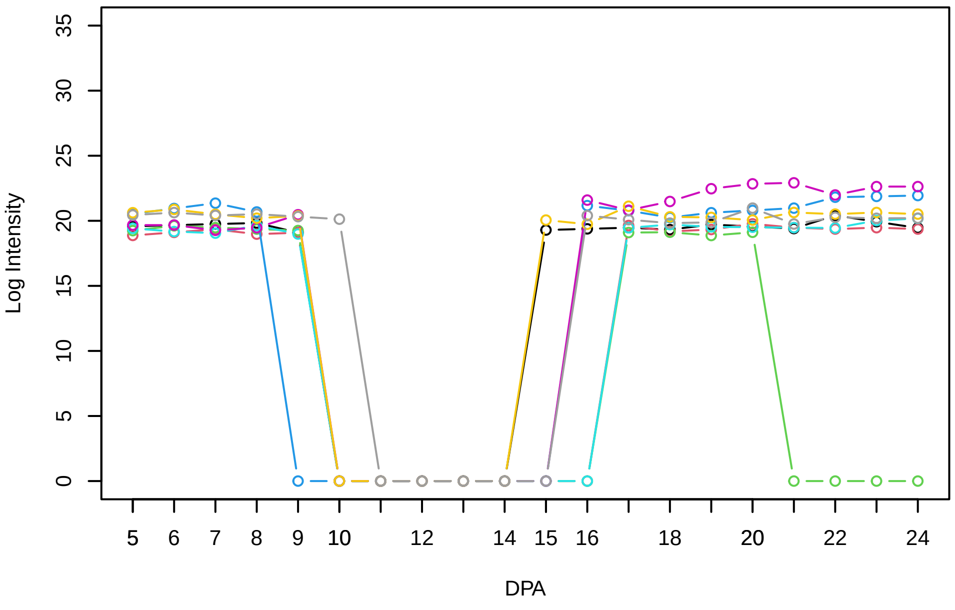


Histogram of the average abundances of reliable proteins

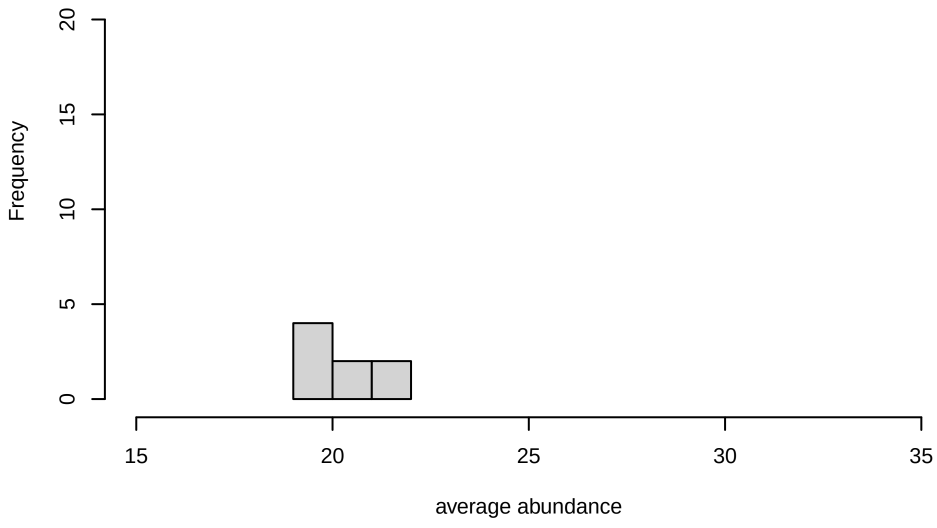


V26, after remove low abundance

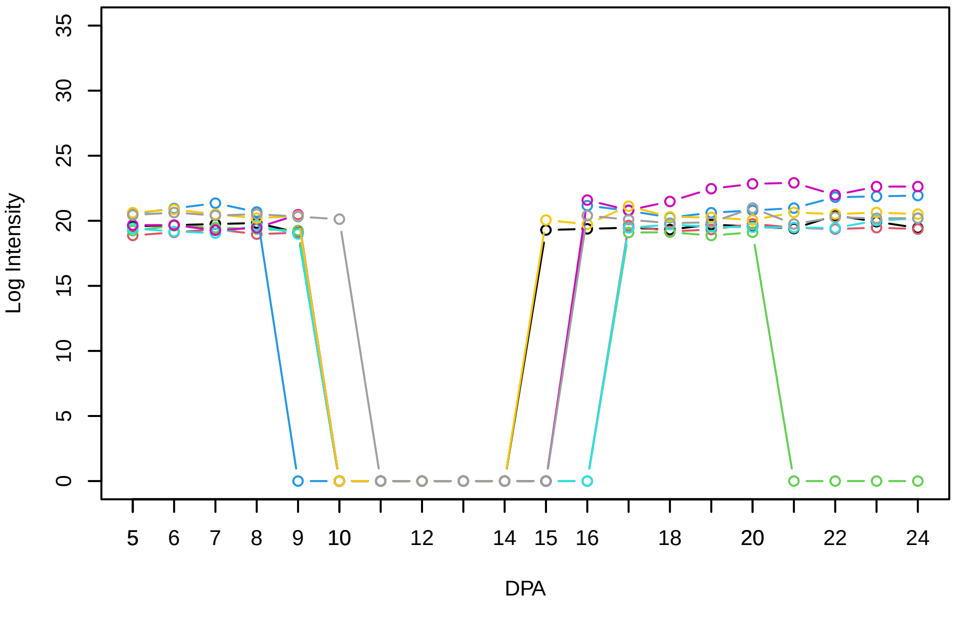


Histogram of the average abundances of high abundance proteins

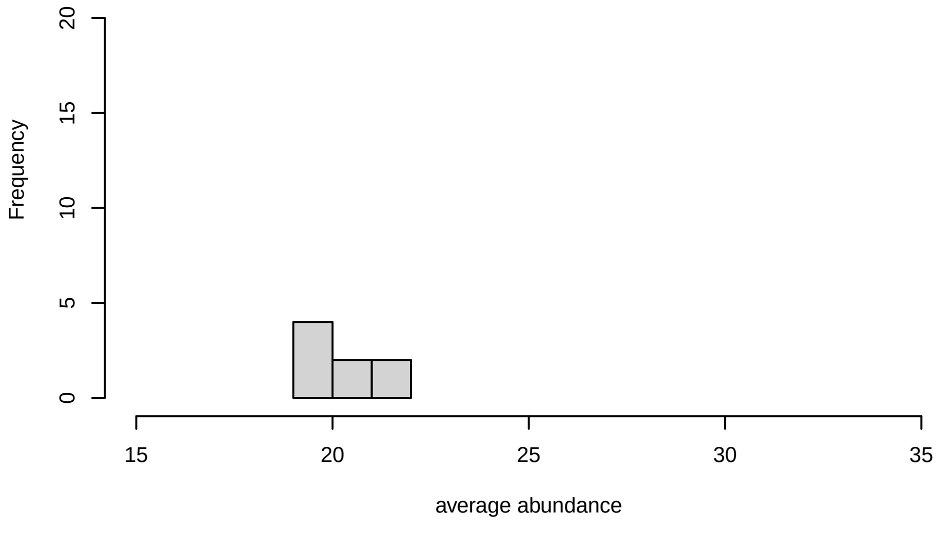

V27

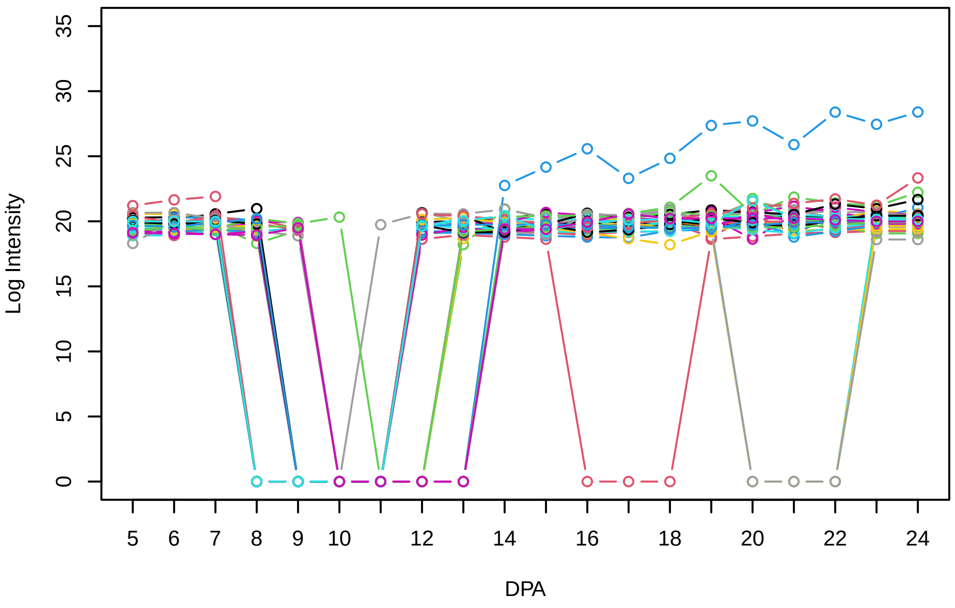


Histogram of the average abundances of non−zero values

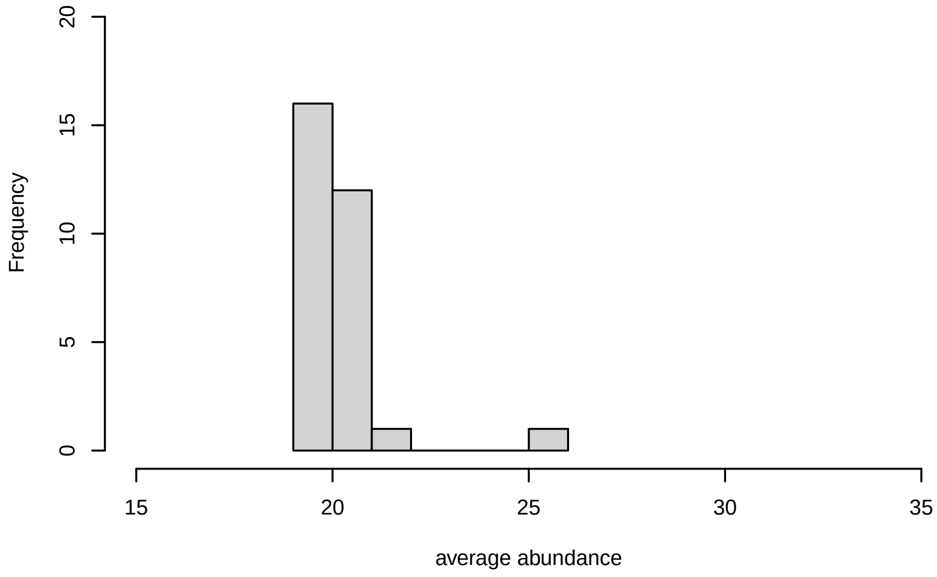


V27, after remove outlier and cutoff is 0.8

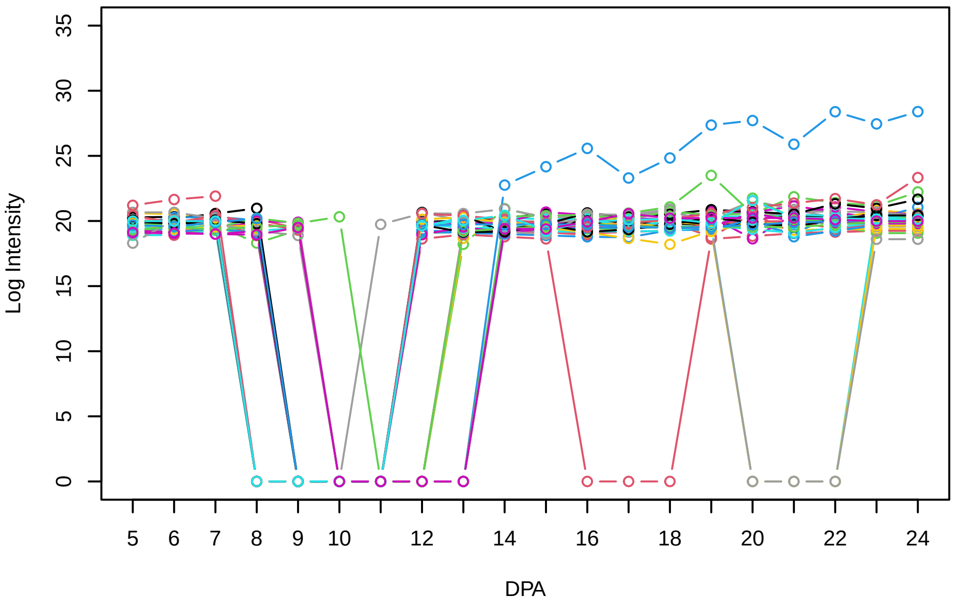


Histogram of the average abundances of non−outlier proteins

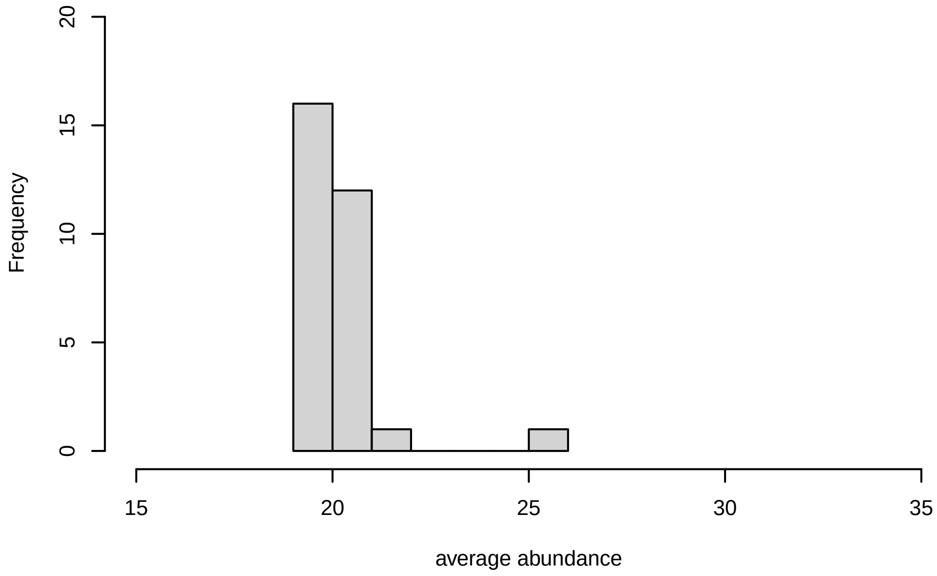


V27, after remove unreliable

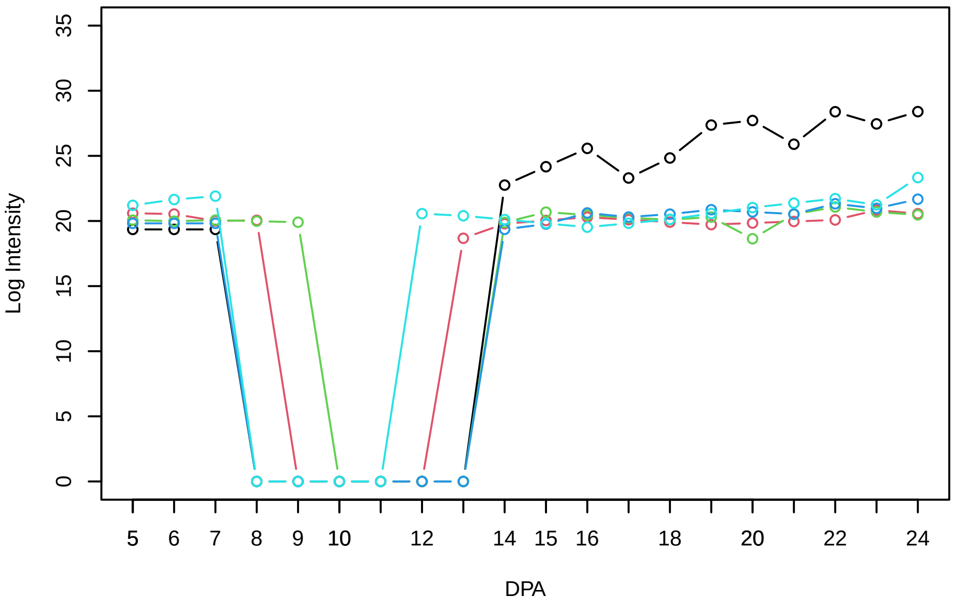


Histogram of the average abundances of reliable proteins

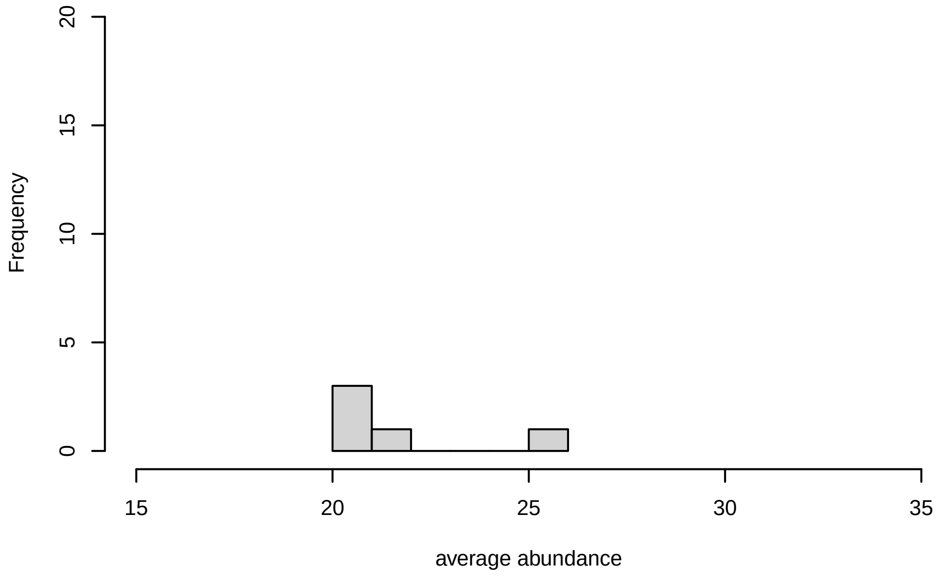


V27, after remove low abundance

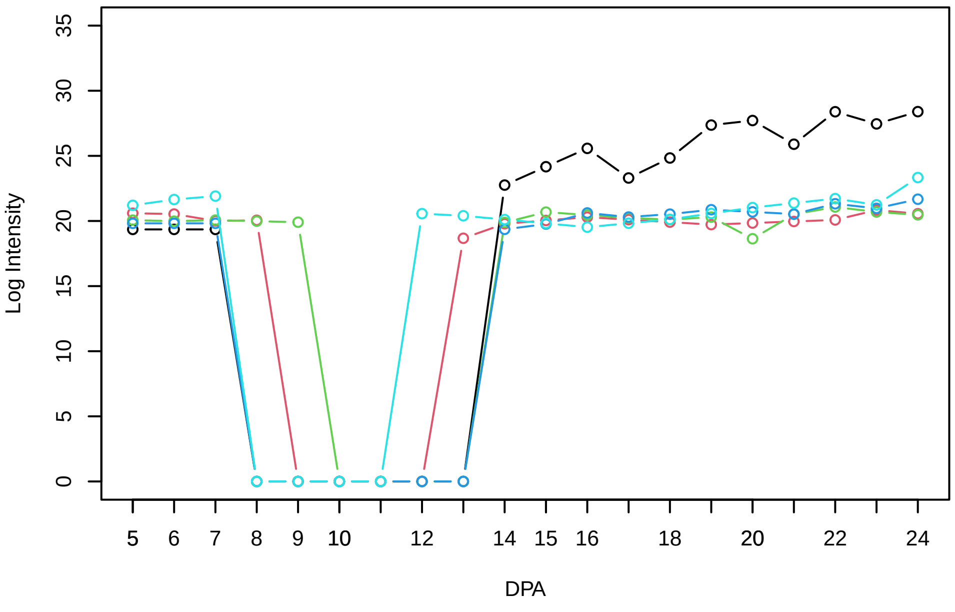


Histogram of the average abundances of high abundance proteins

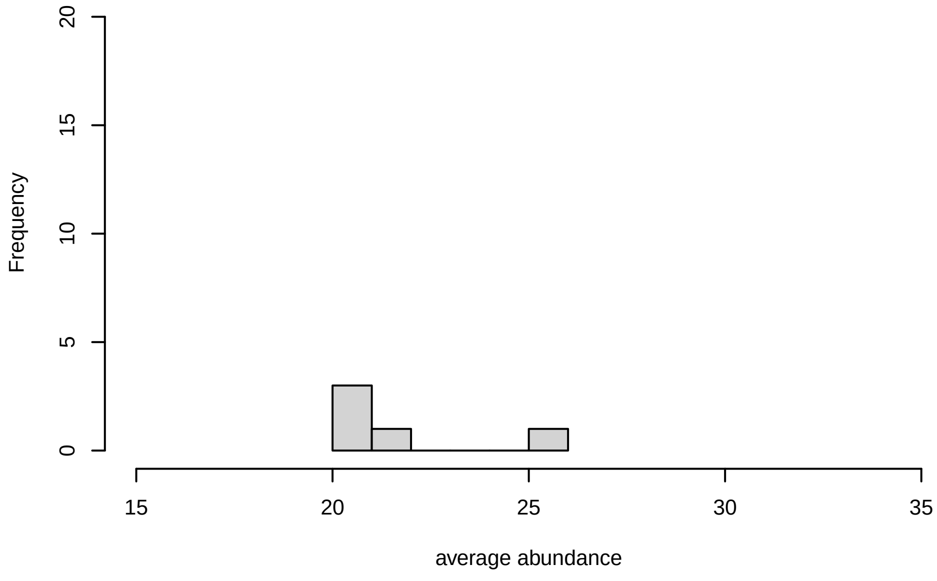

V28

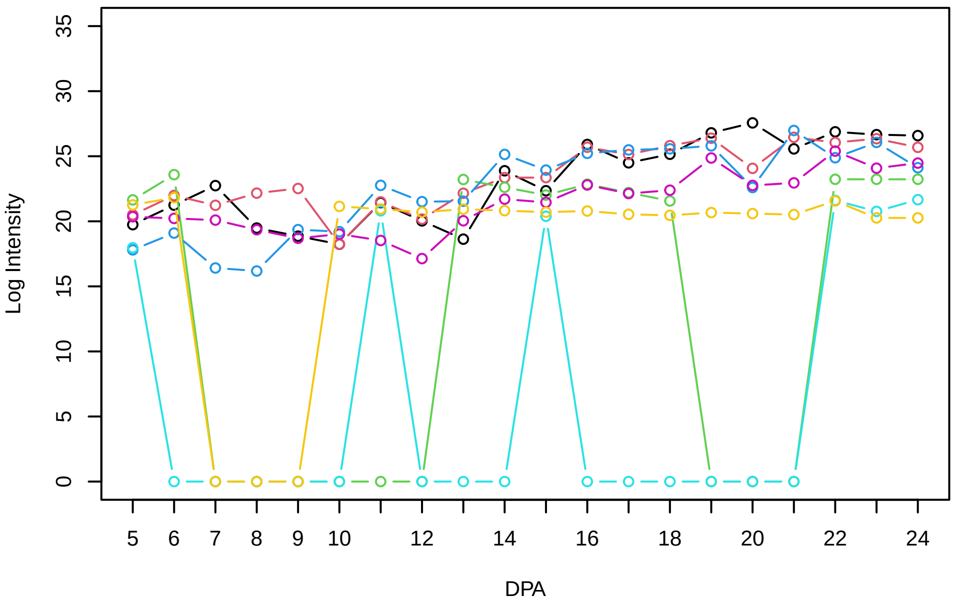


Histogram of the average abundances of non−zero values

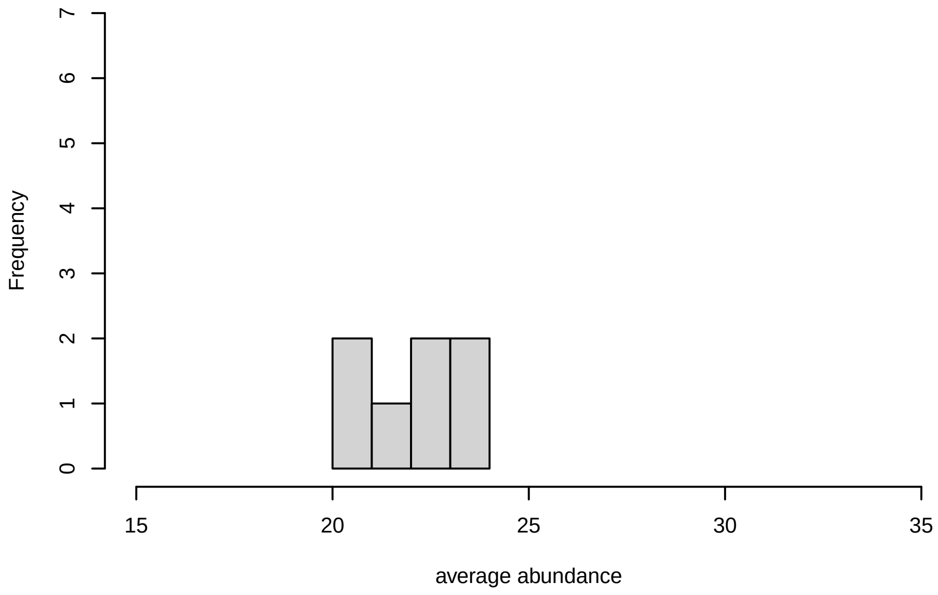


V28, after remove outlier and cutoff is 0.8

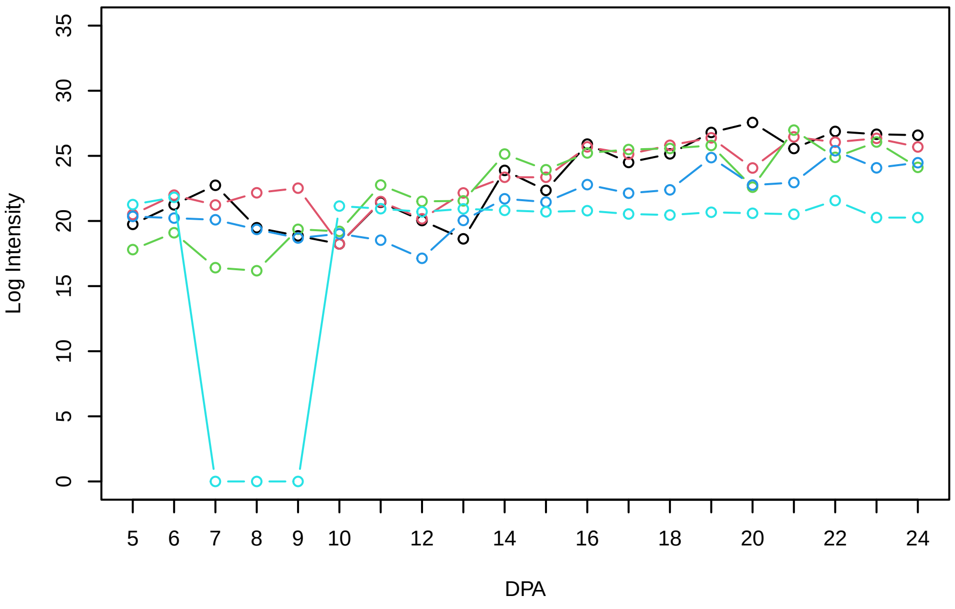


Histogram of the average abundances of non−outlier proteins

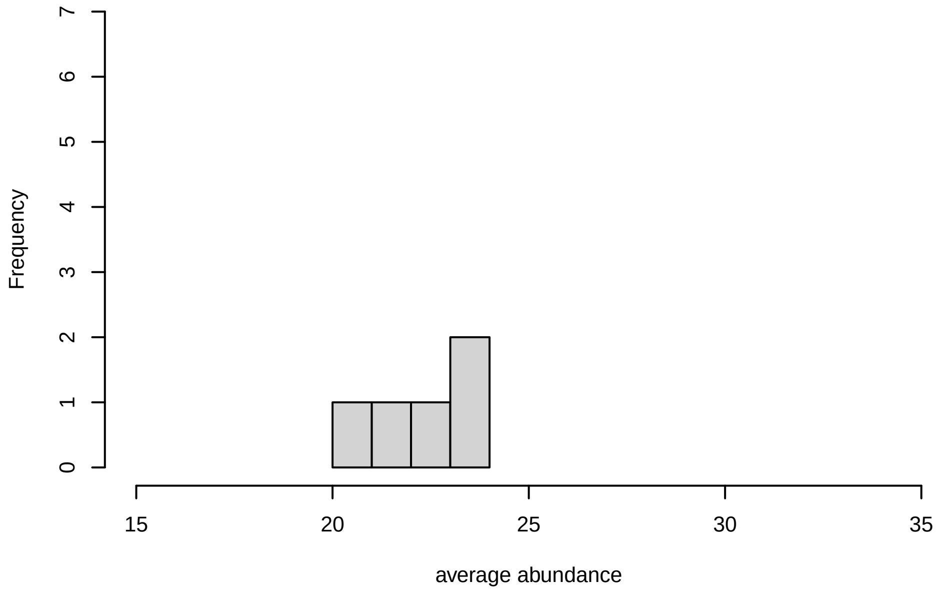


V28, after remove unreliable

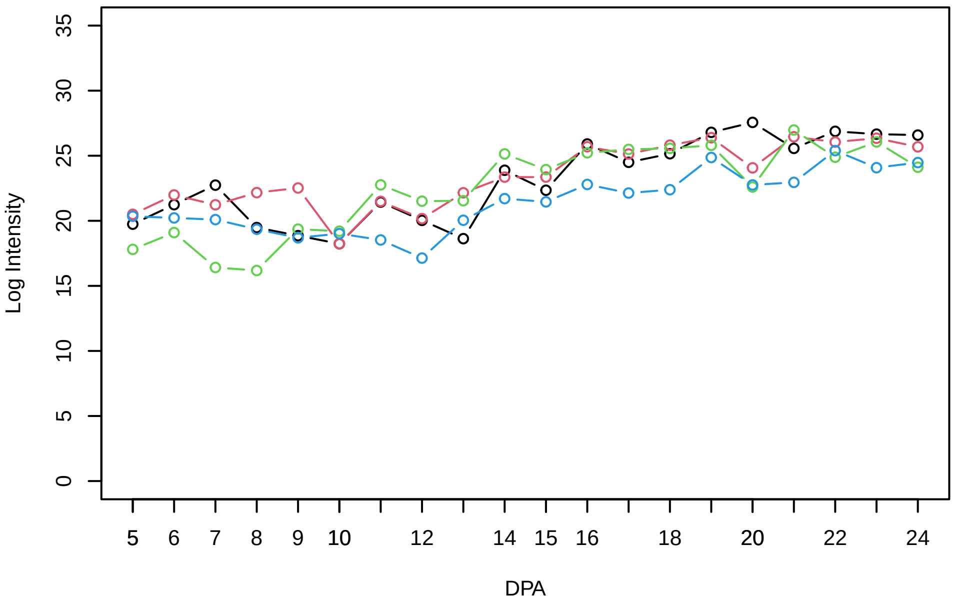


Histogram of the average abundances of reliable proteins

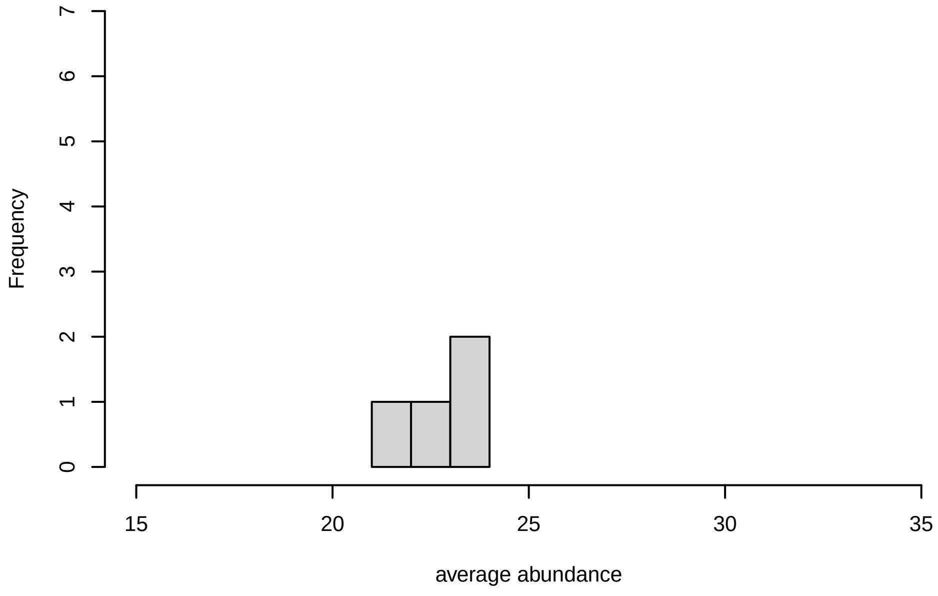


V28, after remove low abundance

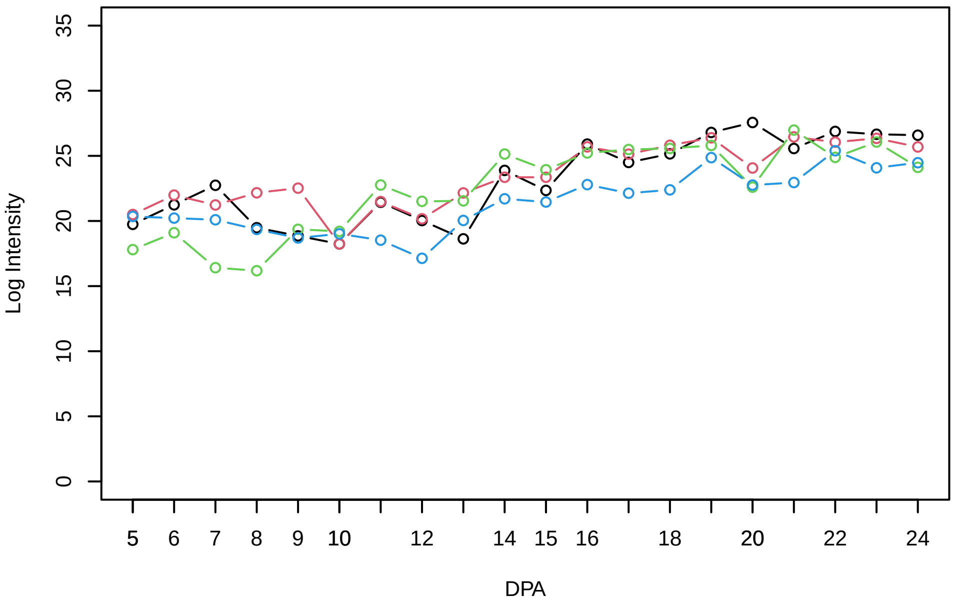


Histogram of the average abundances of high abundance proteins

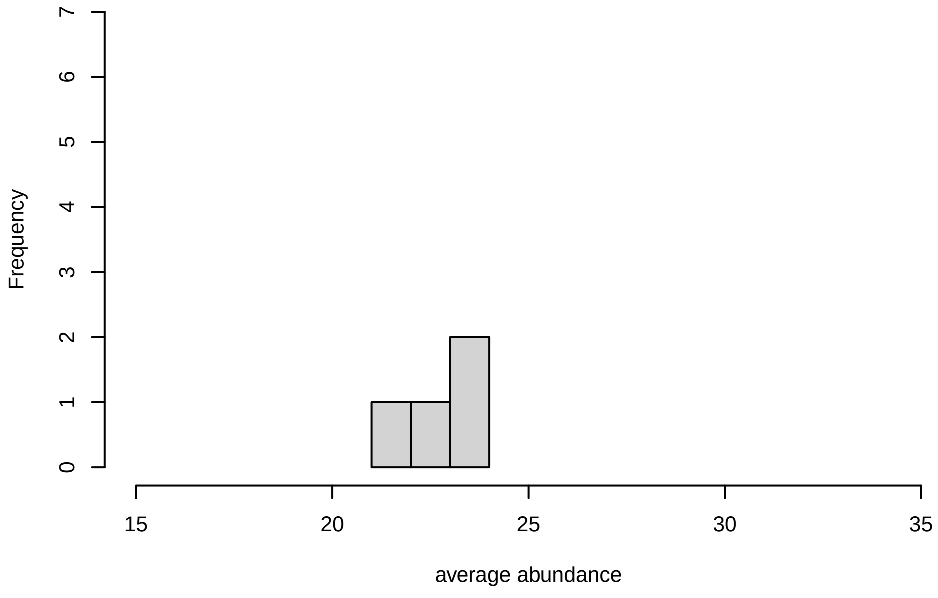

**V29**

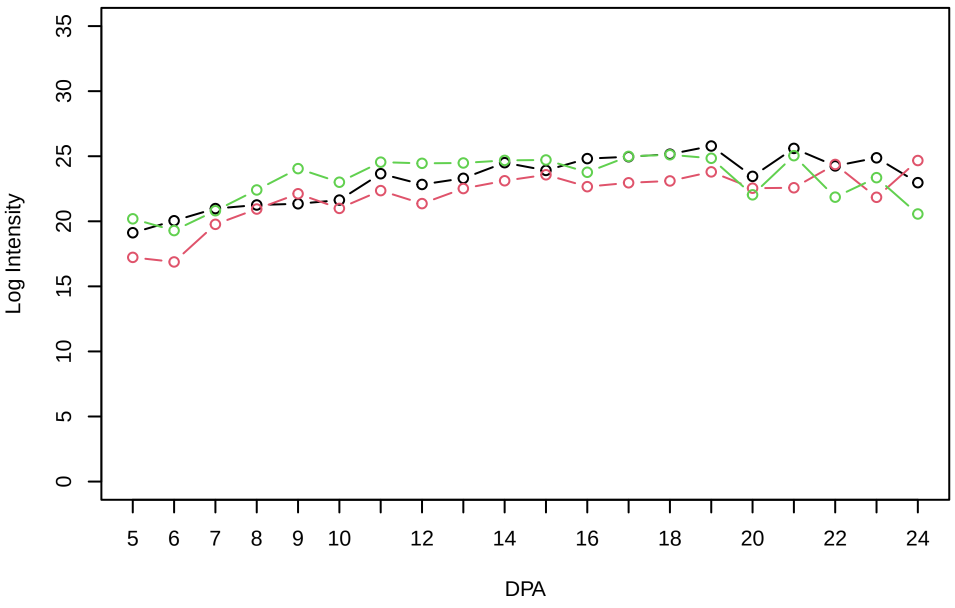


**Histogram of the average abundances of non−zero values**

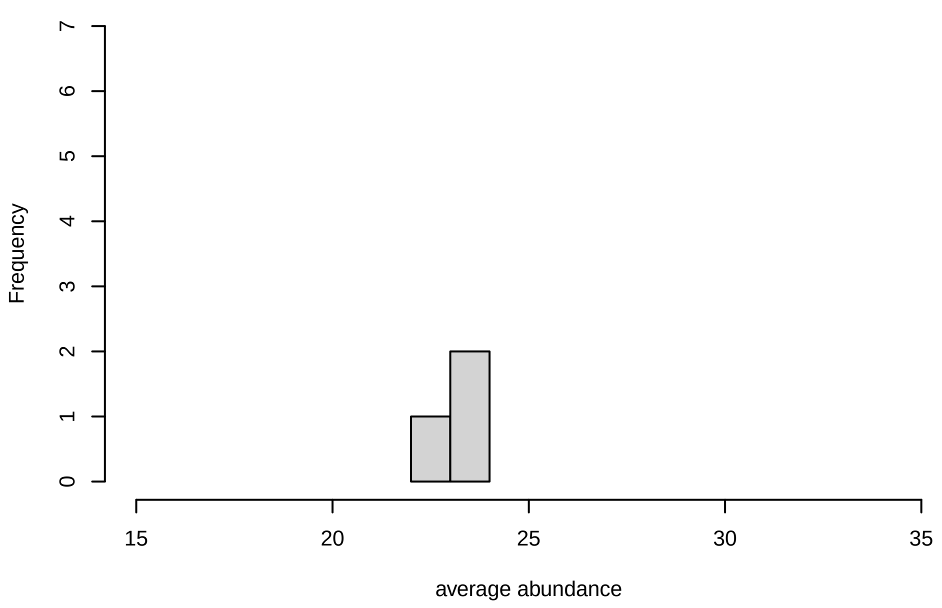


**V29, after remove outlier and cutoff is 0.8**

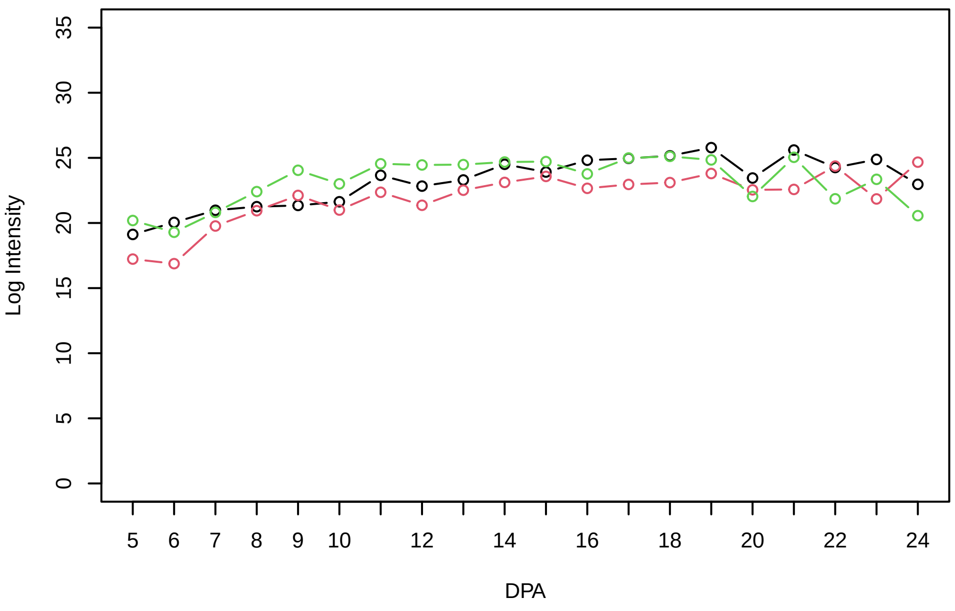


**Histogram of the average abundances of non−outlier proteins**

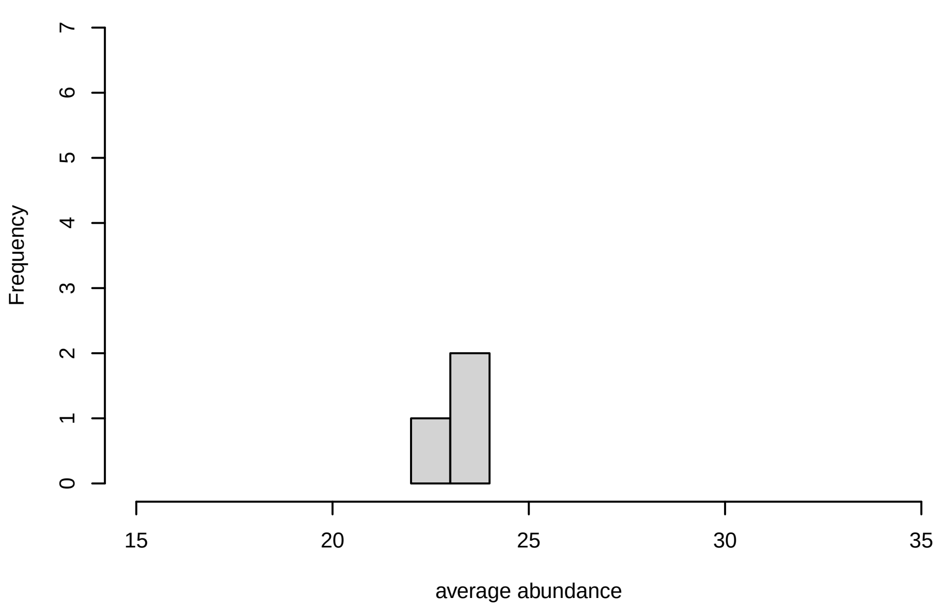


**V29, after remove unreliable**

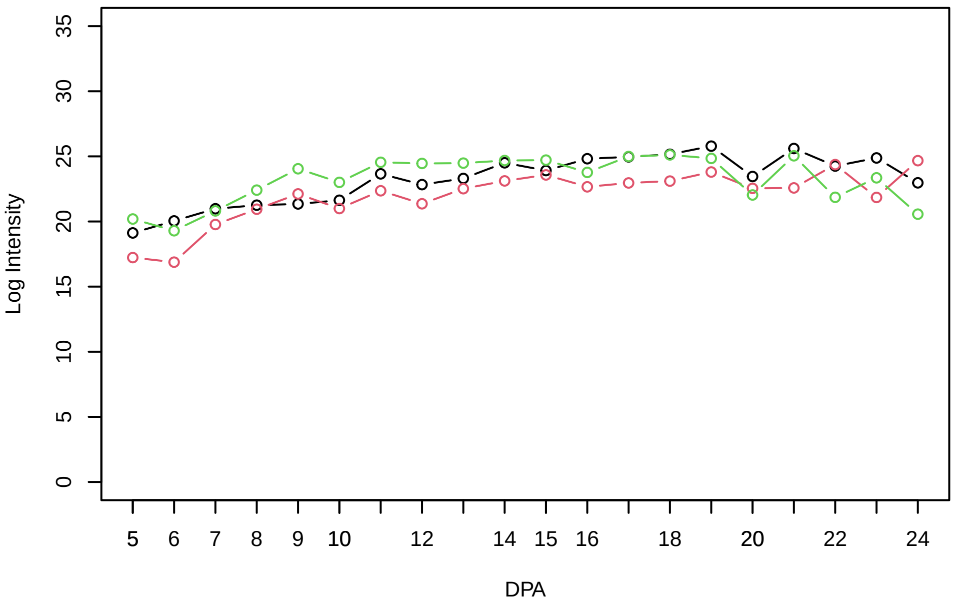


**Histogram of the average abundances of reliable proteins**

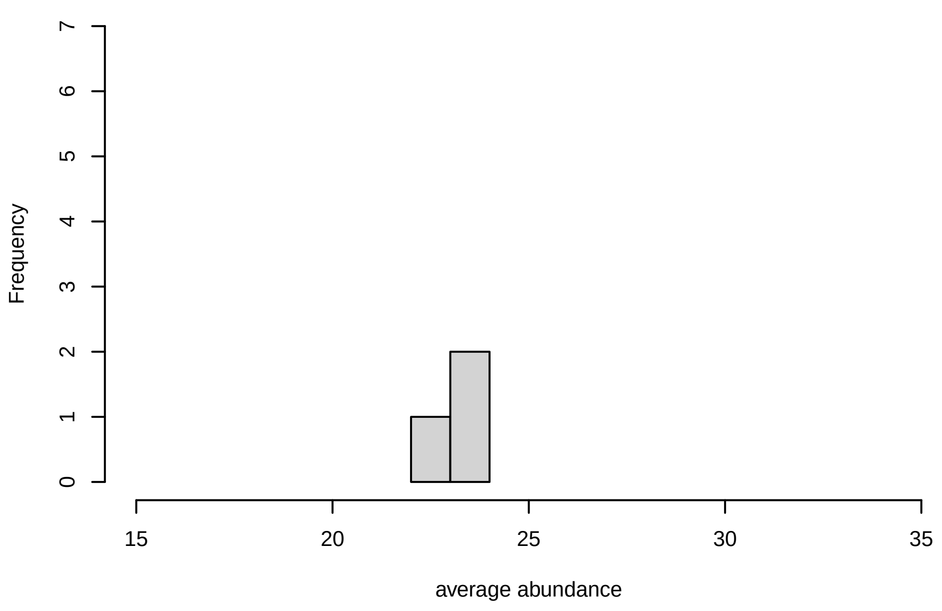


**V29, after remove low abundance**

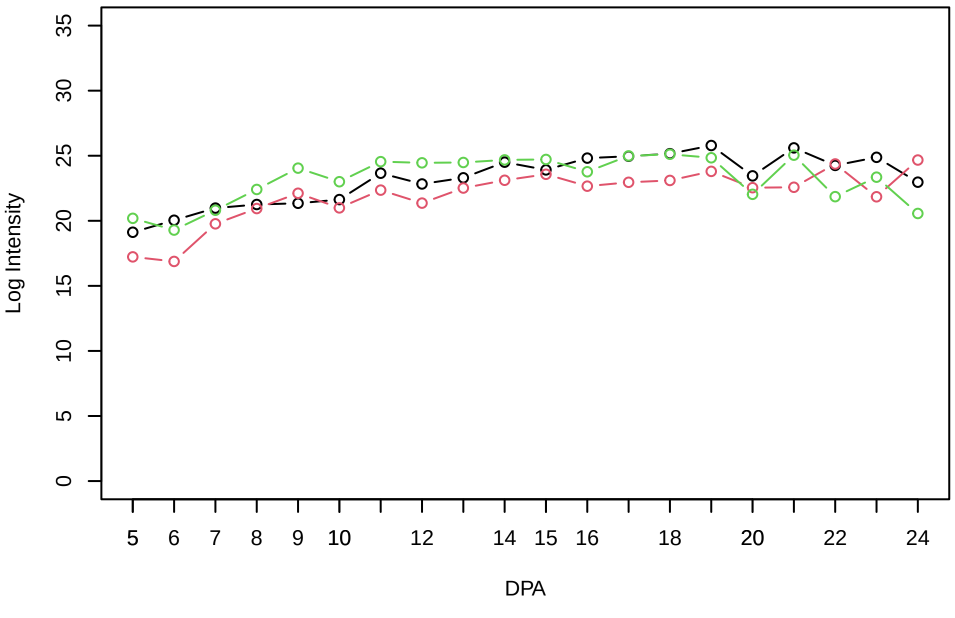


**Histogram of the average abundances of high abundance proteins**

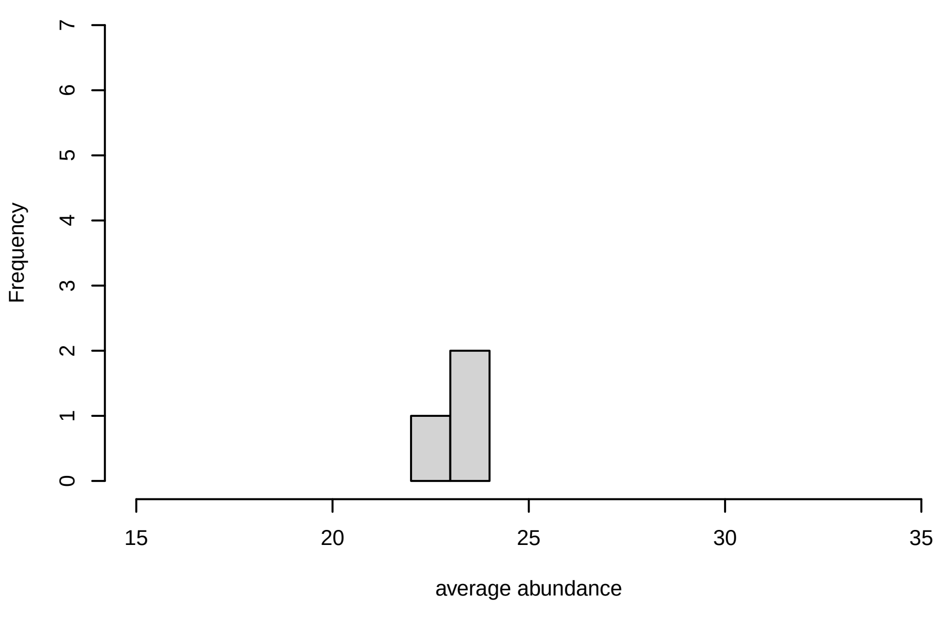

**V31**

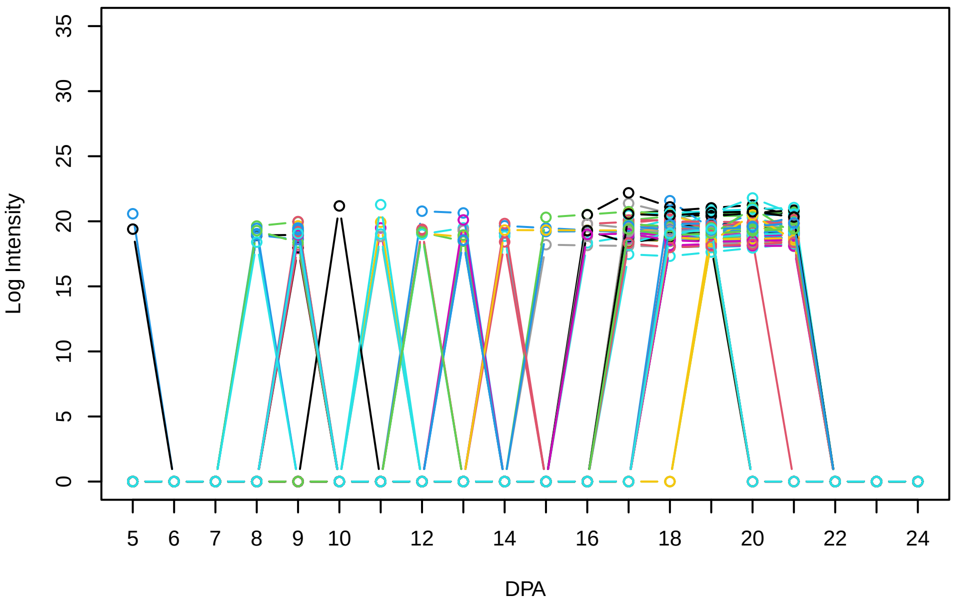


**Histogram of the average abundances of non−zero values**

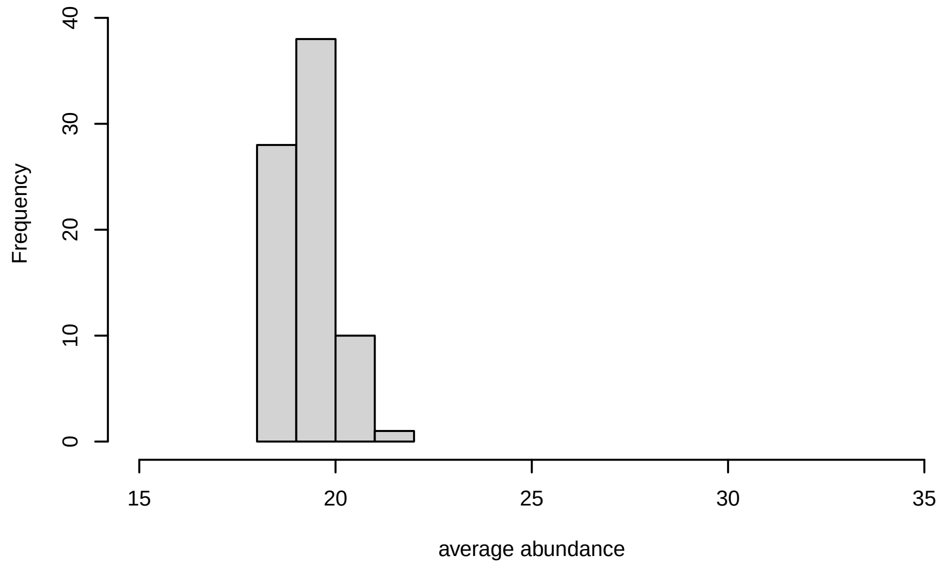


**V31, after remove outlier and cutoff is 0.774357732358406**

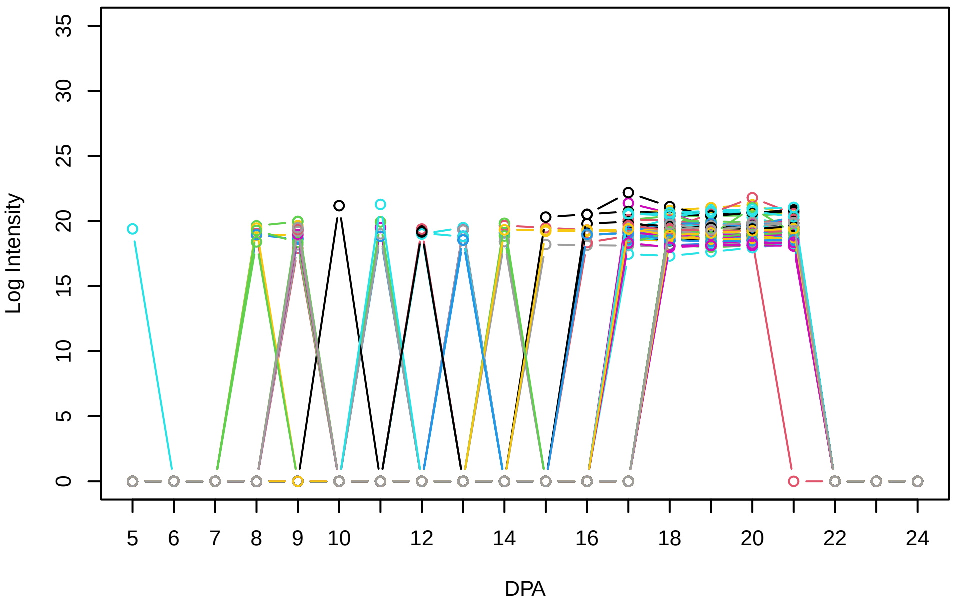


**Histogram of the average abundances of non−outlier proteins**

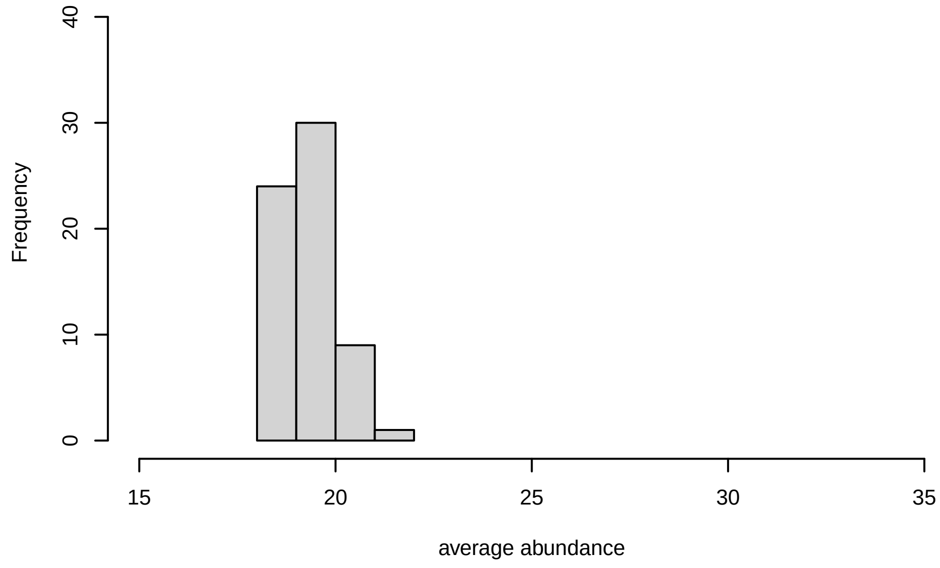

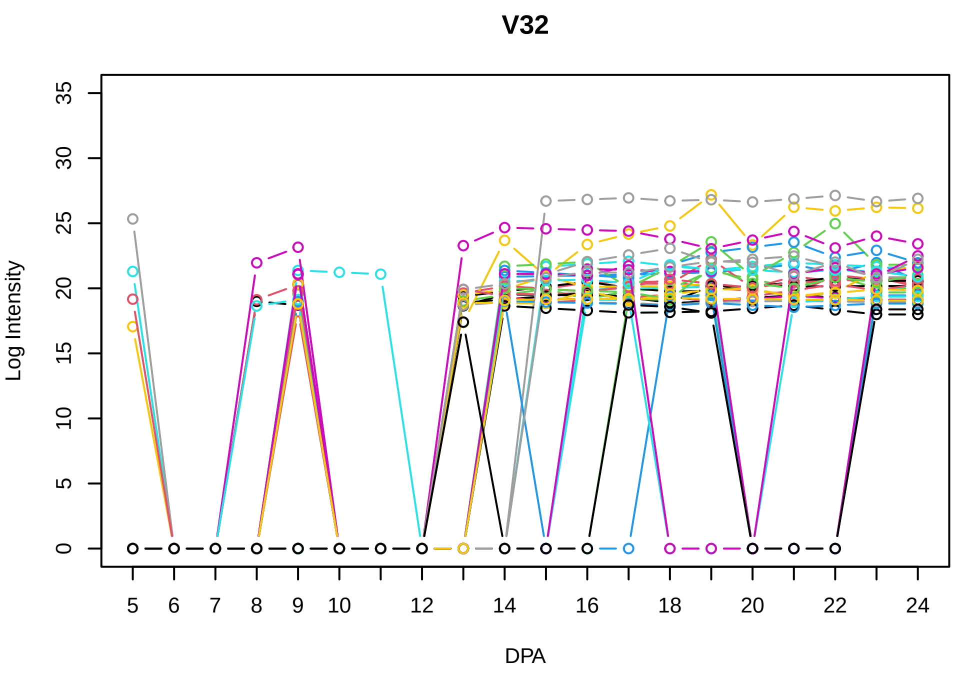
V32
Log Intensity
DPA

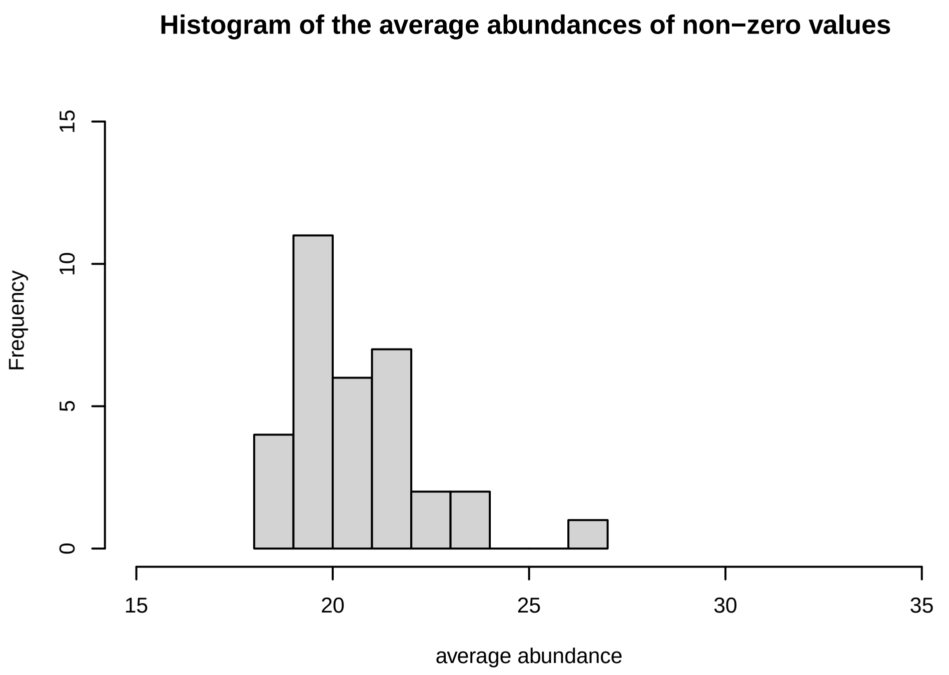
Histogram of the average abundances of non−zero values
Frequency
average abundance

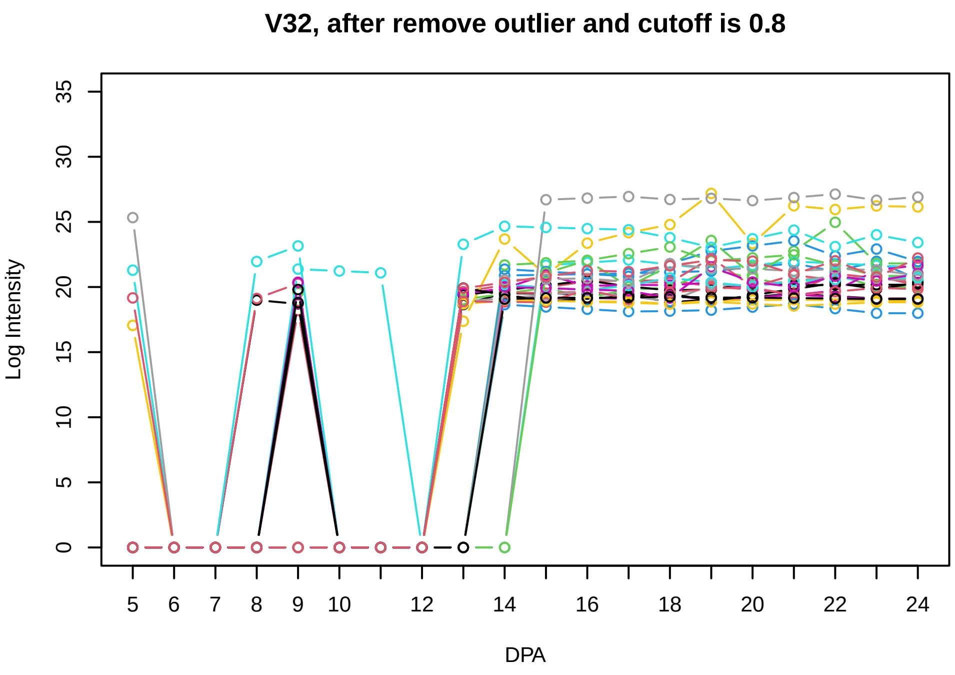
V32, after remove outlier and cutoff is 0.8
Log Intensity
DPA

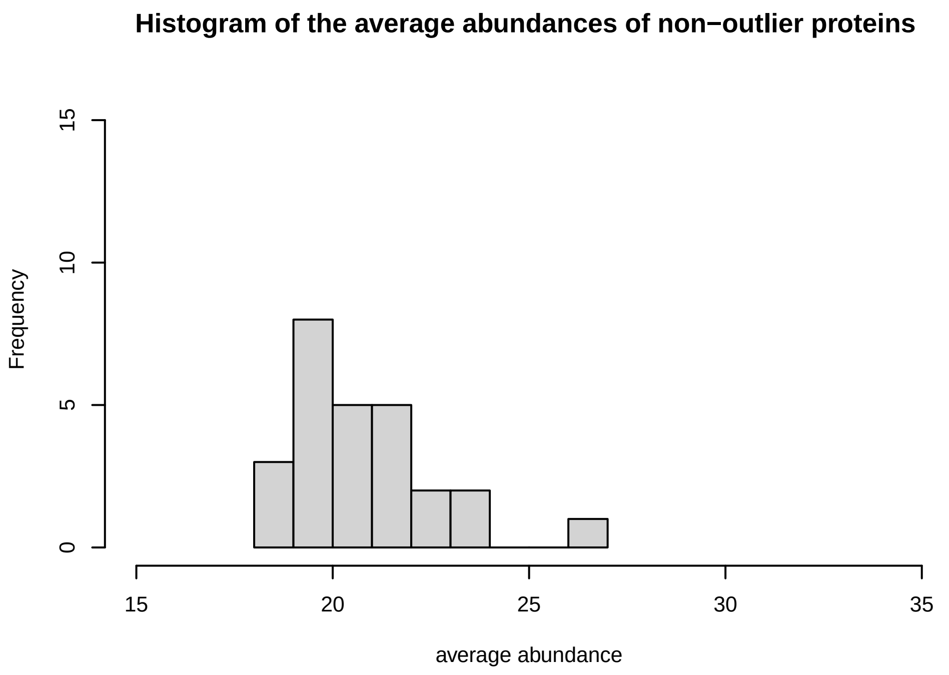
Histogram of the average abundances of non−outlier proteins
Frequency
average abundance

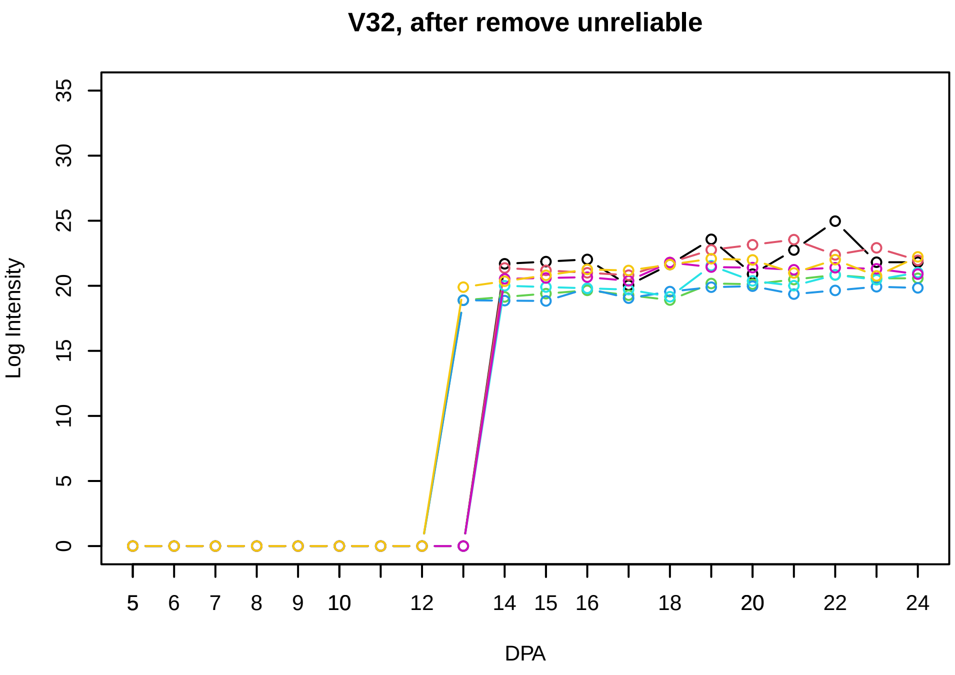
V32, after remove unreliable
Log Intensity
DPA

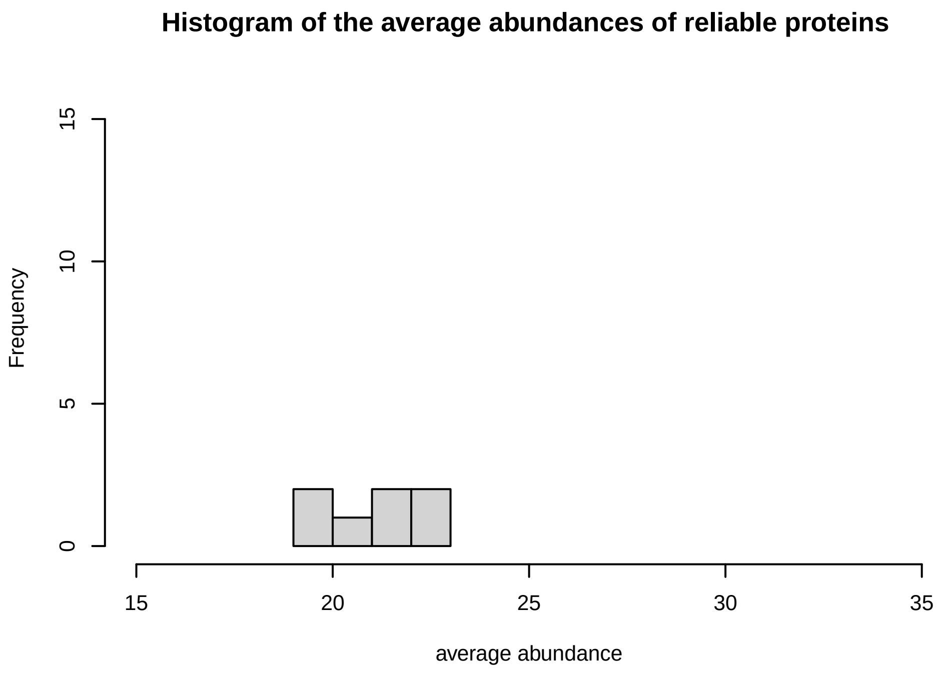
Histogram of the average abundances of reliable proteins
Frequency
average abundance

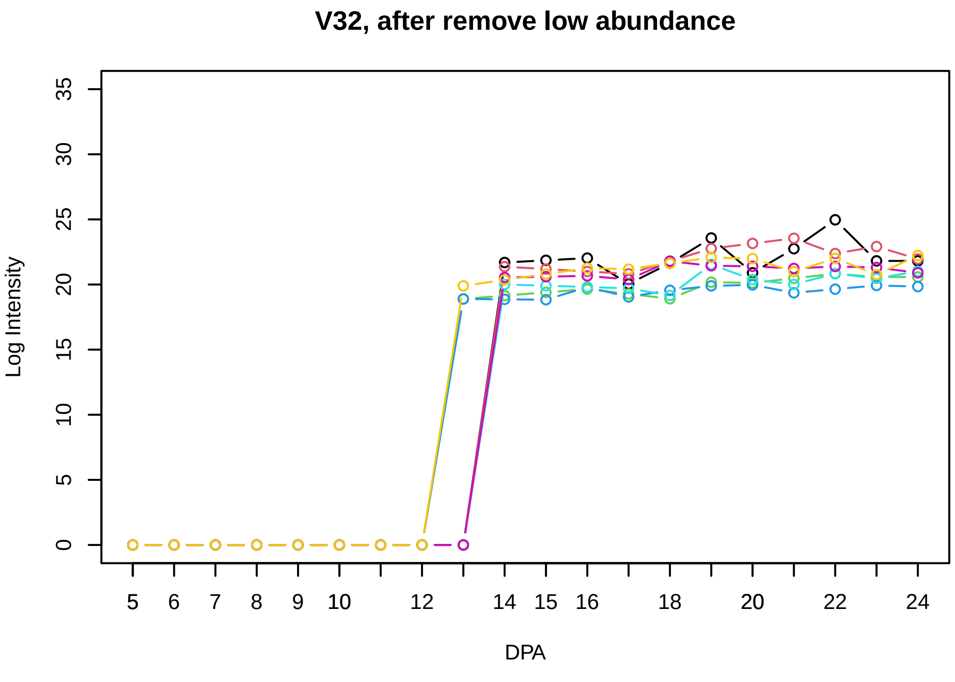
V32, after remove low abundance
Log Intensity
DPA

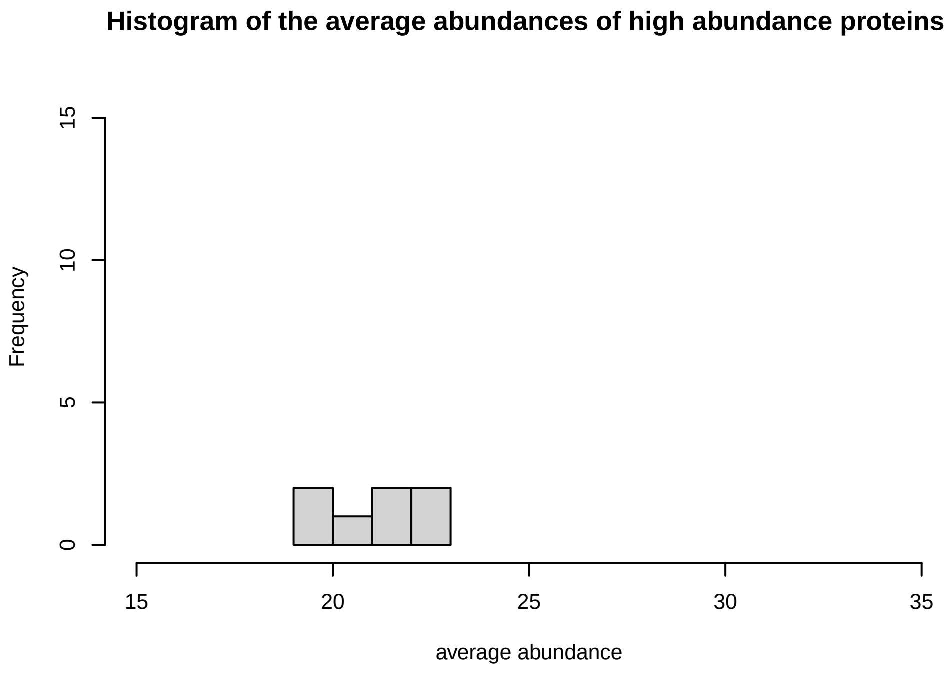
Histogram of the average abundances of high abundance proteins
Frequency
average abundance

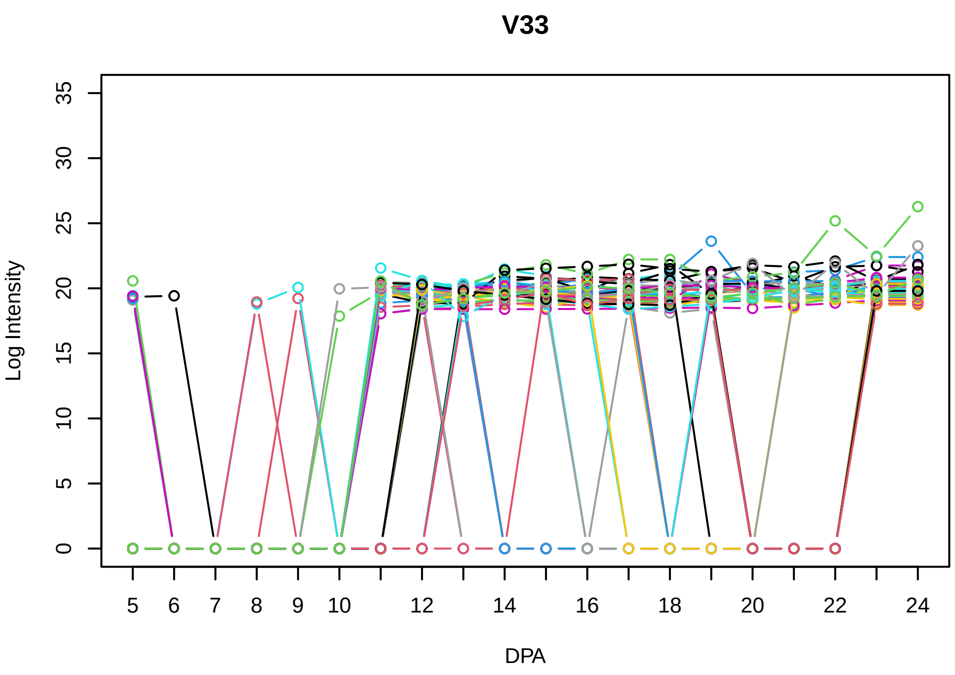
V33
Log Intensity
DPA
0 5 10 15 20 25 30 35
5 6 7 8 9 10 12 14 16 18 20 22 24

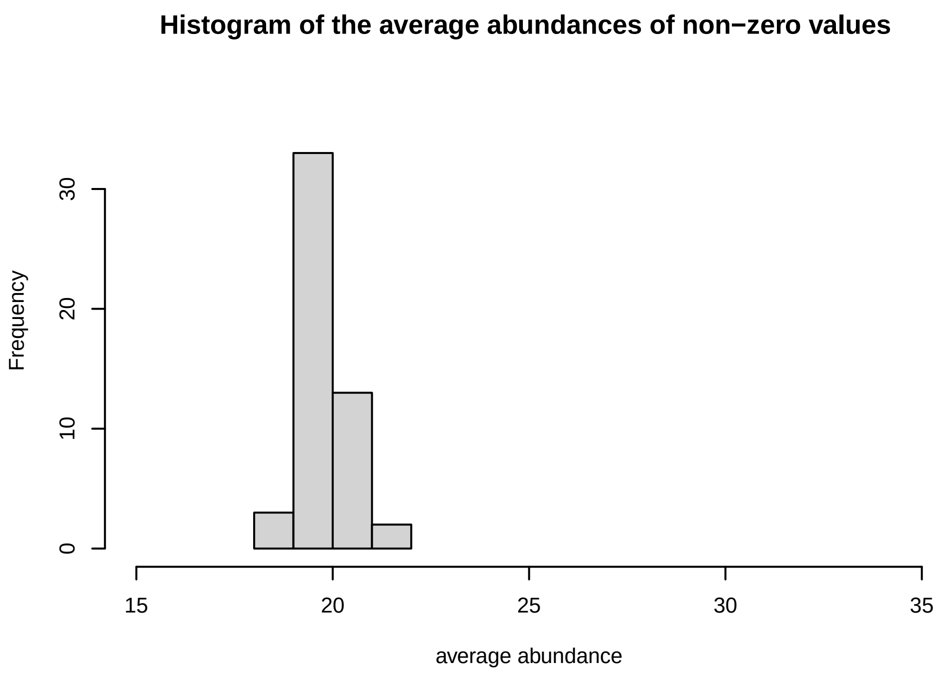
Histogram of the average abundances of non−zero values
Frequency
average abundance
0 10 20 30
15 20 25 30 35

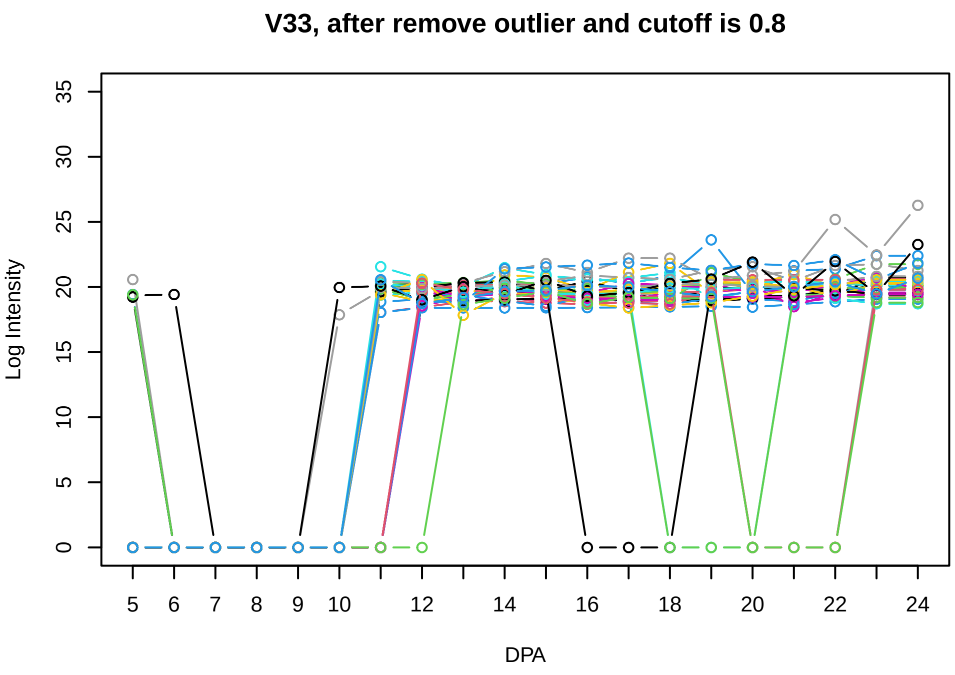
V33, after remove outlier and cutoff is 0.8
Log Intensity
DPA
0 5 10 15 20 25 30 35
5 6 7 8 9 10 12 14 16 18 20 22 24

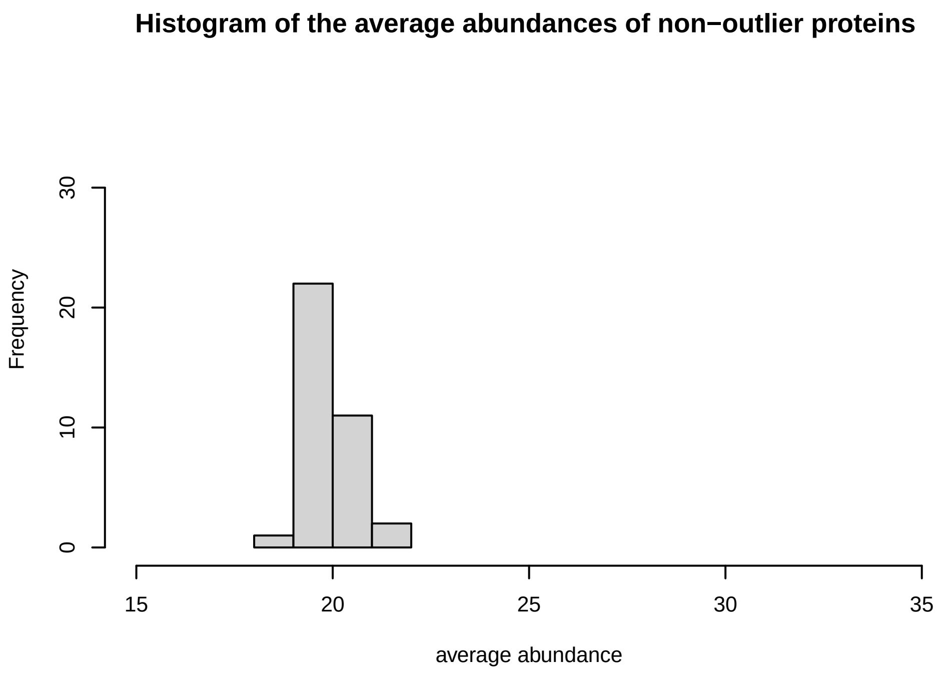
Histogram of the average abundances of non−outlier proteins
Frequency
average abundance
0 10 20 30
15 20 25 30 35

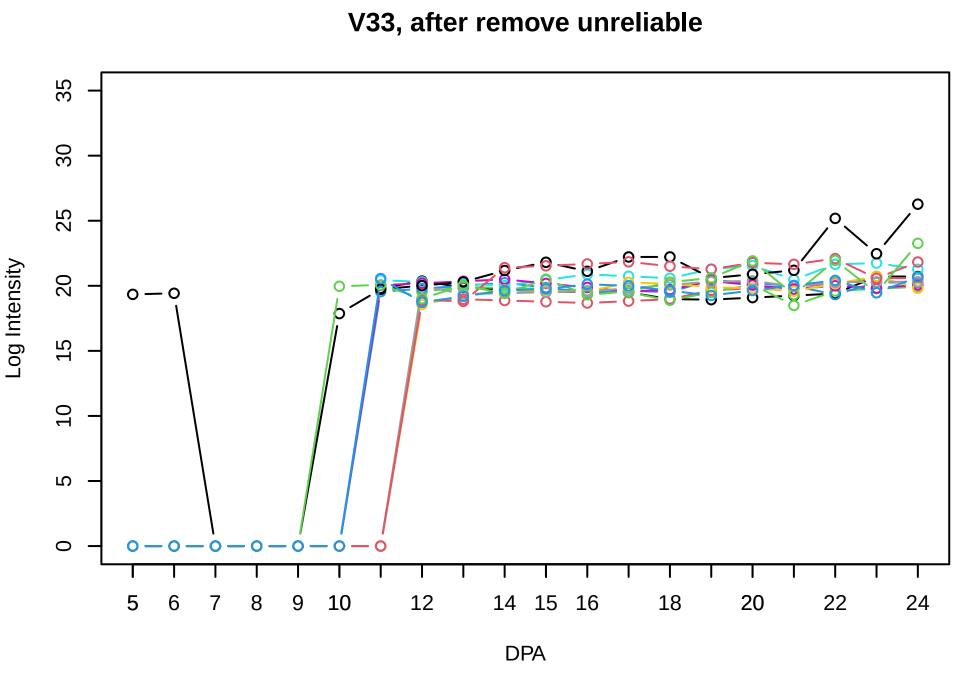
V33, after remove unreliable
Log Intensity
DPA
0 5 10 15 20 25 30 35
5 6 7 8 9 10 12 14 15 16 18 20 22 24

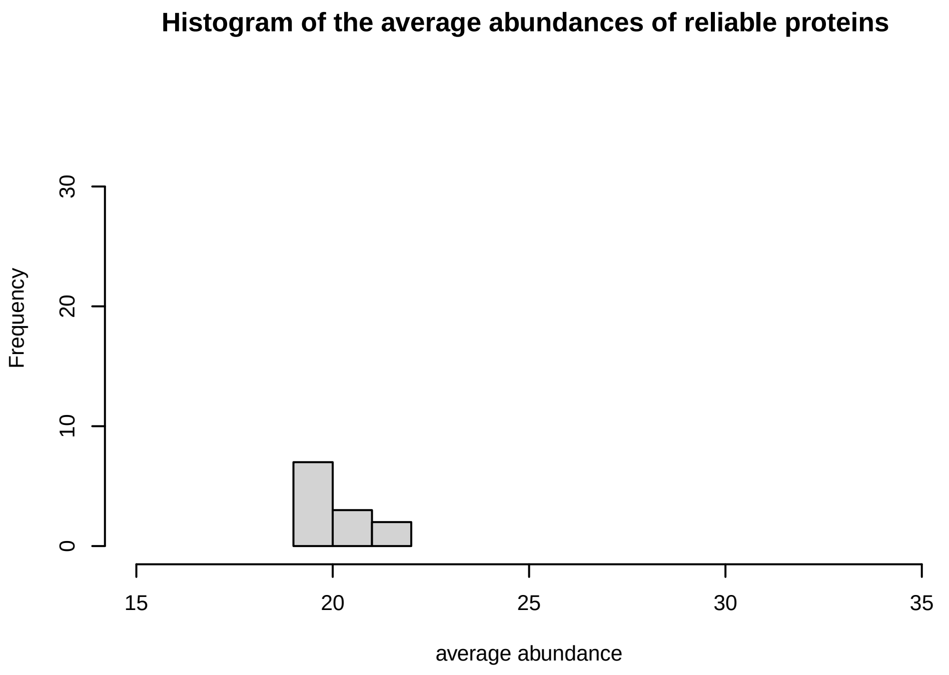
Histogram of the average abundances of reliable proteins
Frequency
average abundance
0 10 20 30
15 20 25 30 35

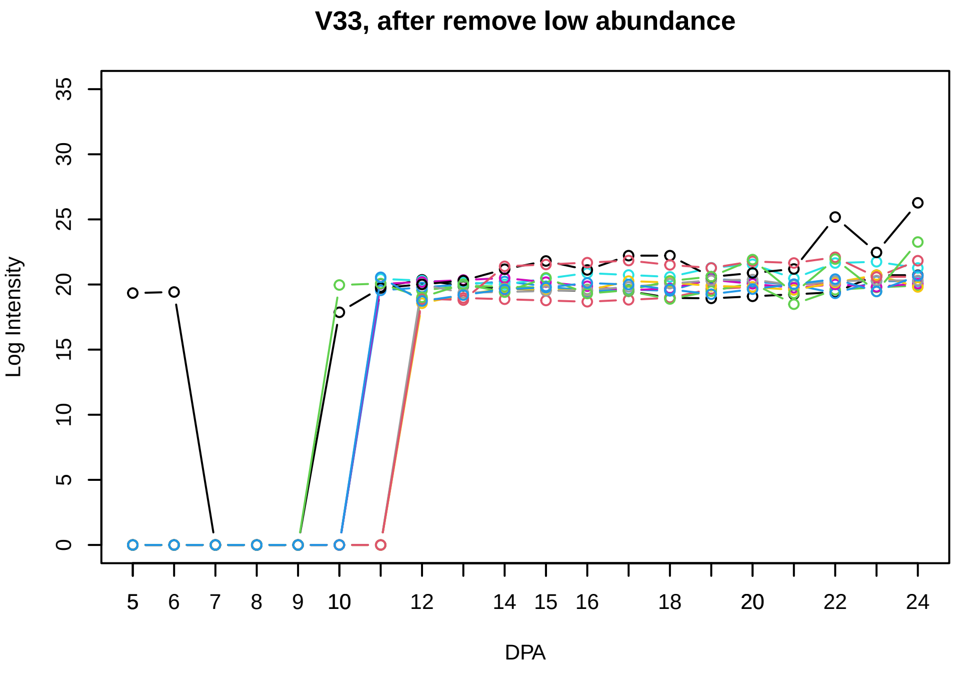
V33, after remove low abundance
Log Intensity
DPA
0 5 10 15 20 25 30 35
5 6 7 8 9 10 12 14 15 16 18 20 22 24

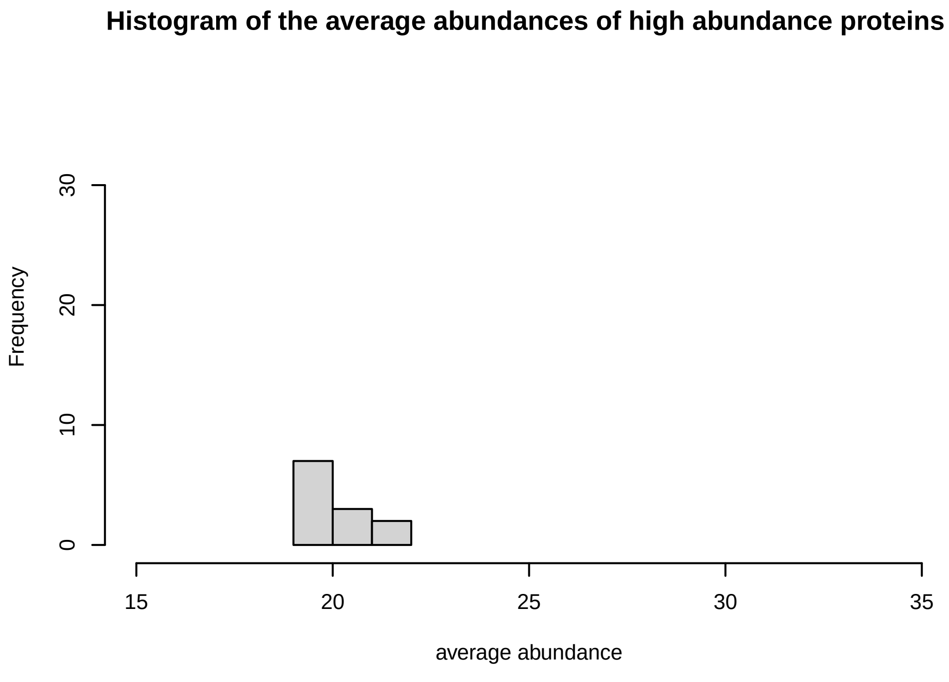
Histogram of the average abundances of high abundance proteins
Frequency
average abundance
0 10 20 30
15 20 25 30 35

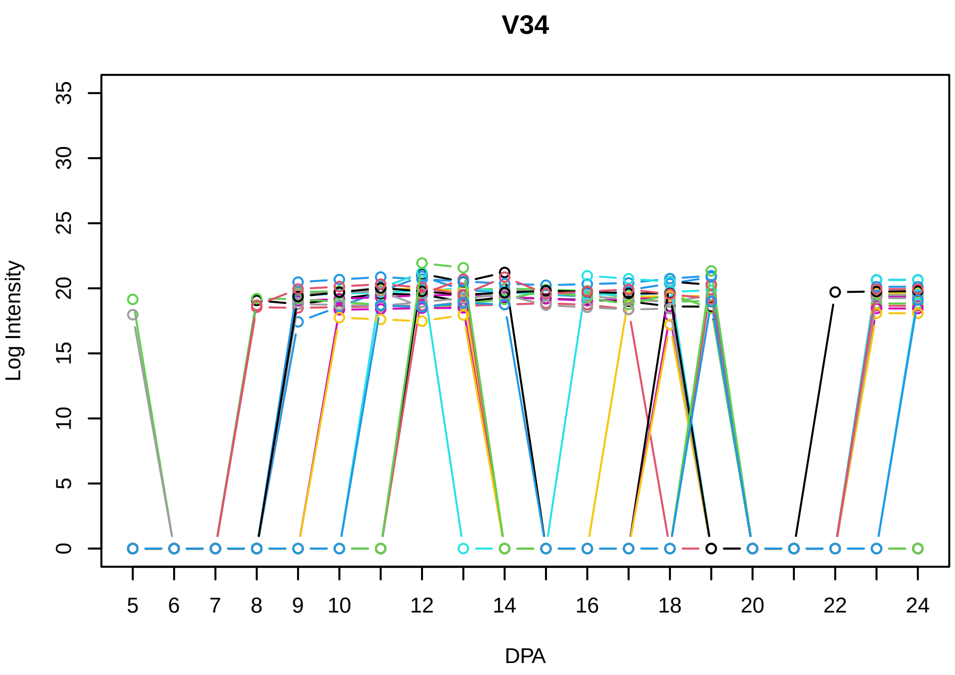
V34
Log Intensity
DPA
0
5
10
15
20
25
30
35
5
6
7
8
9
10
12
14
16
18
20
22
24

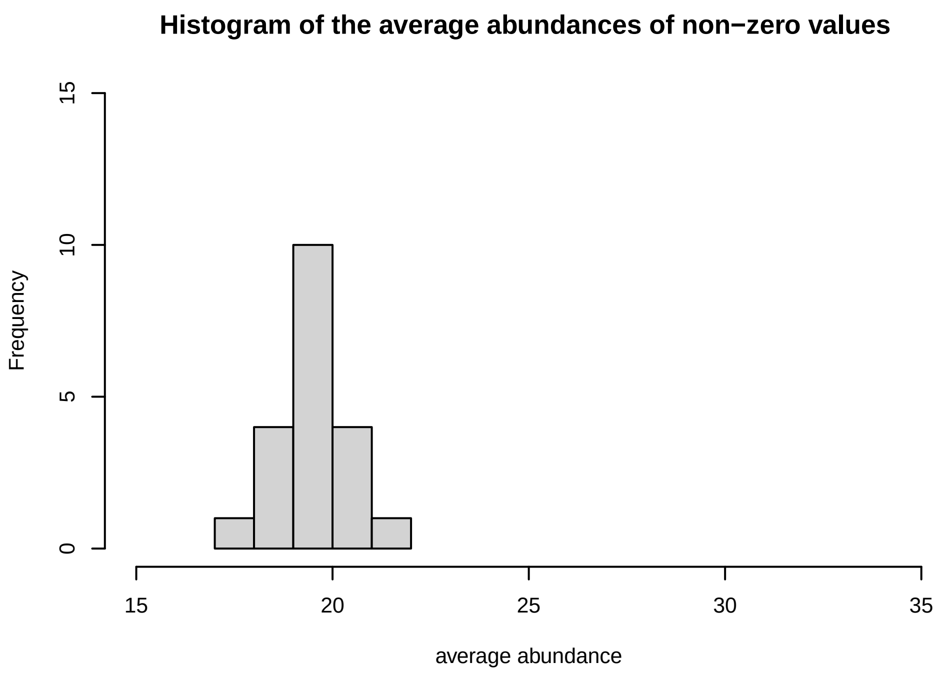
Histogram of the average abundances of non−zero values
Frequency
average abundance
0
5
10
15
15
20
25
30
35

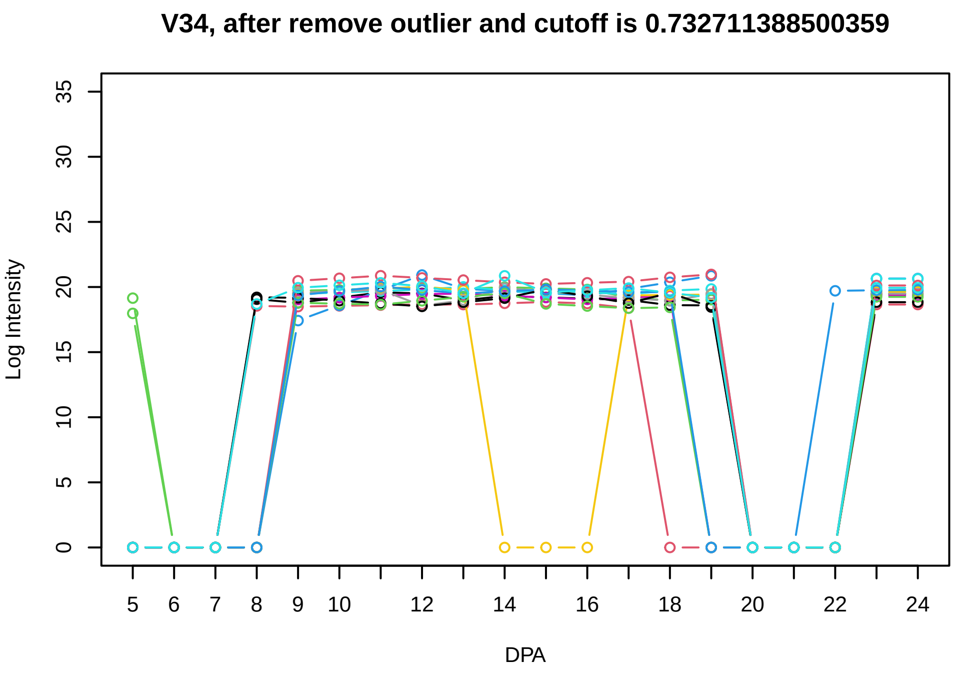
V34, after remove outlier and cutoff is 0.732711388500359
Log Intensity
DPA
0
5
10
15
20
25
30
35
5
6
7
8
9
10
12
14
16
18
20
22
24

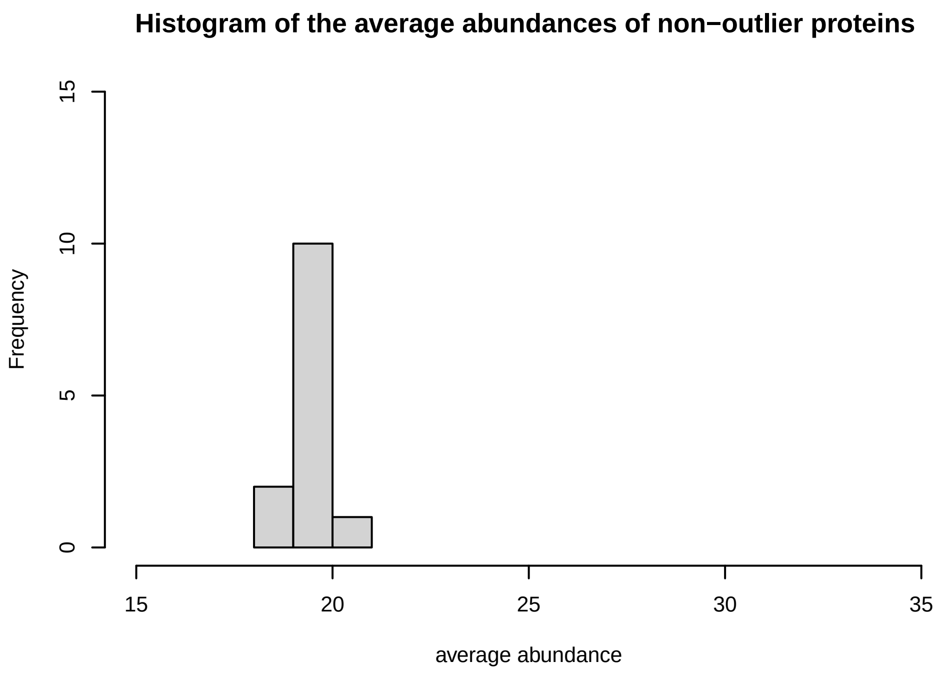
Histogram of the average abundances of non−outlier proteins
Frequency
average abundance
0
5
10
15
15
20
25
30
35

V35

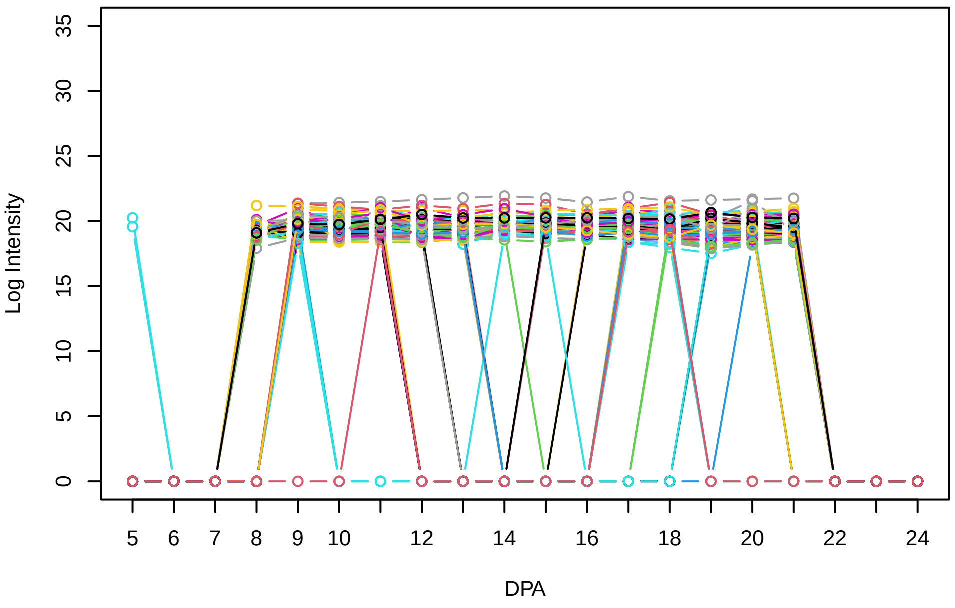


Histogram of the average abundances of non−zero values

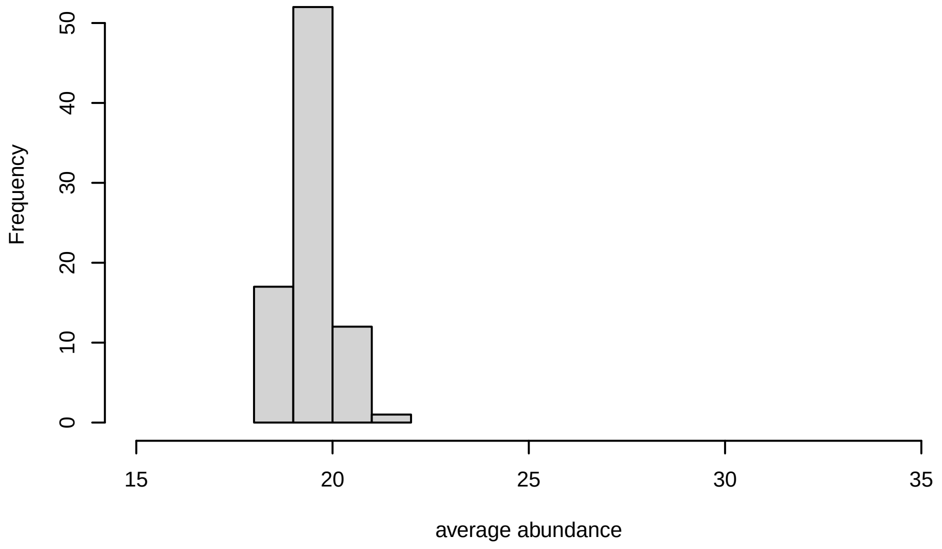


V35, after remove outlier and cutoff is 0.8

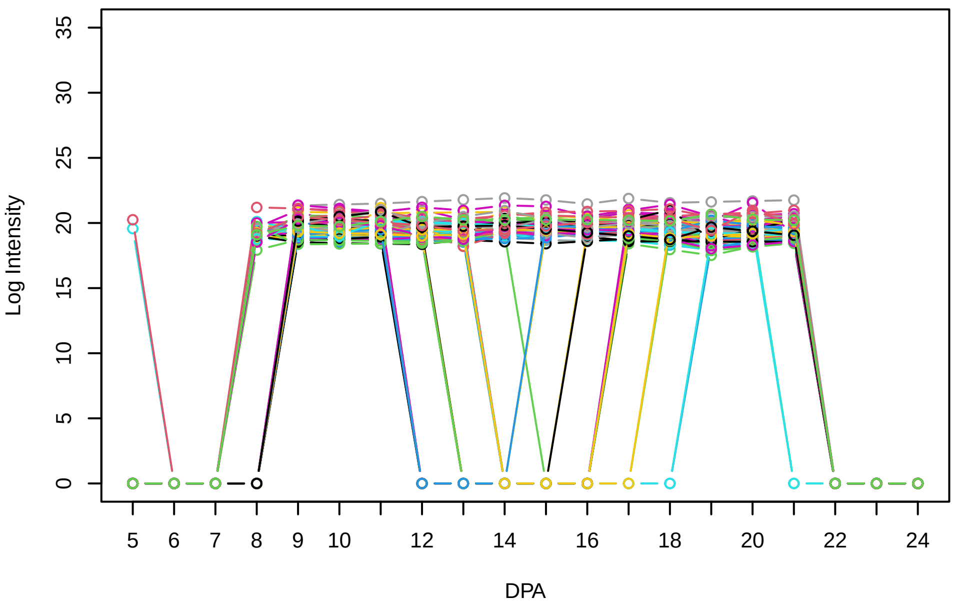


Histogram of the average abundances of non−outlier proteins

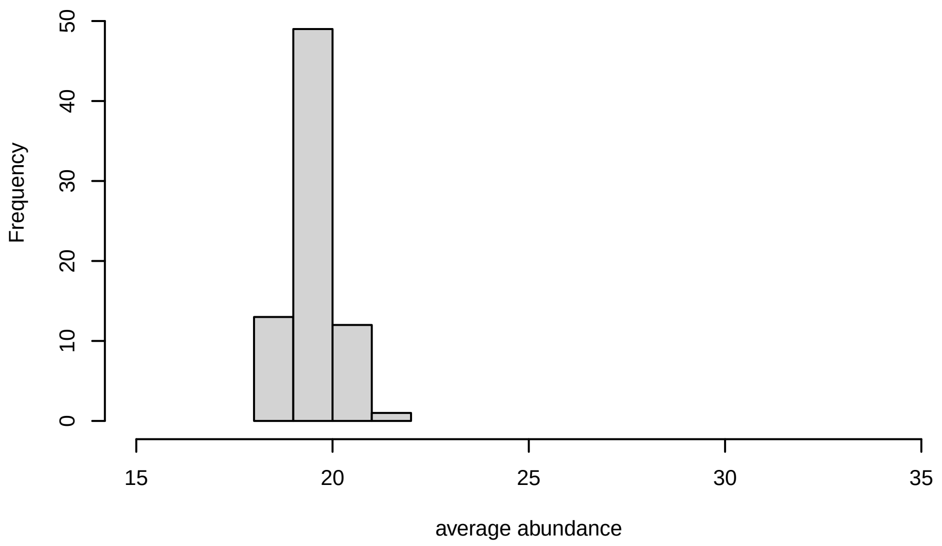


V35, after remove unreliable

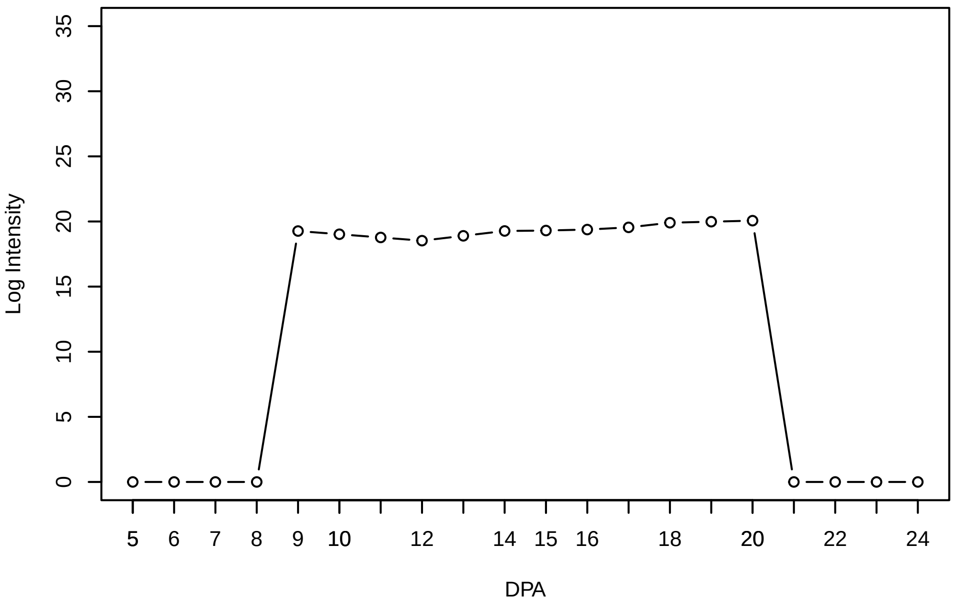


Histogram of the average abundances of reliable proteins

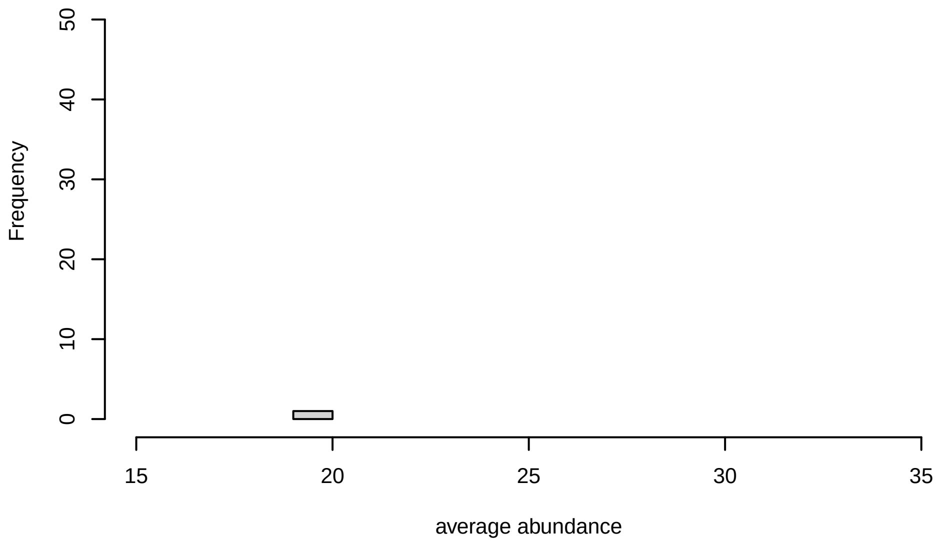


V35, after remove low abundance

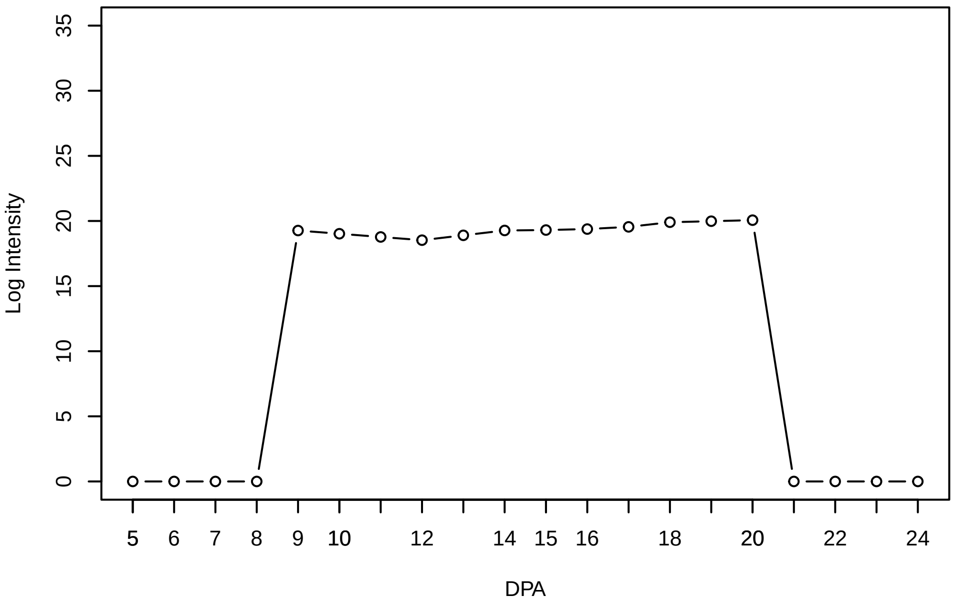


Histogram of the average abundances of high abundance proteins

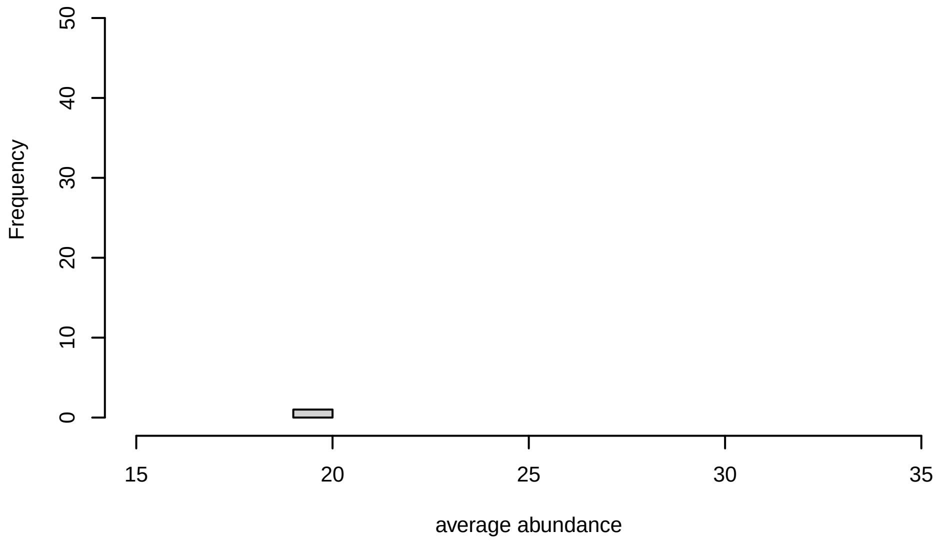

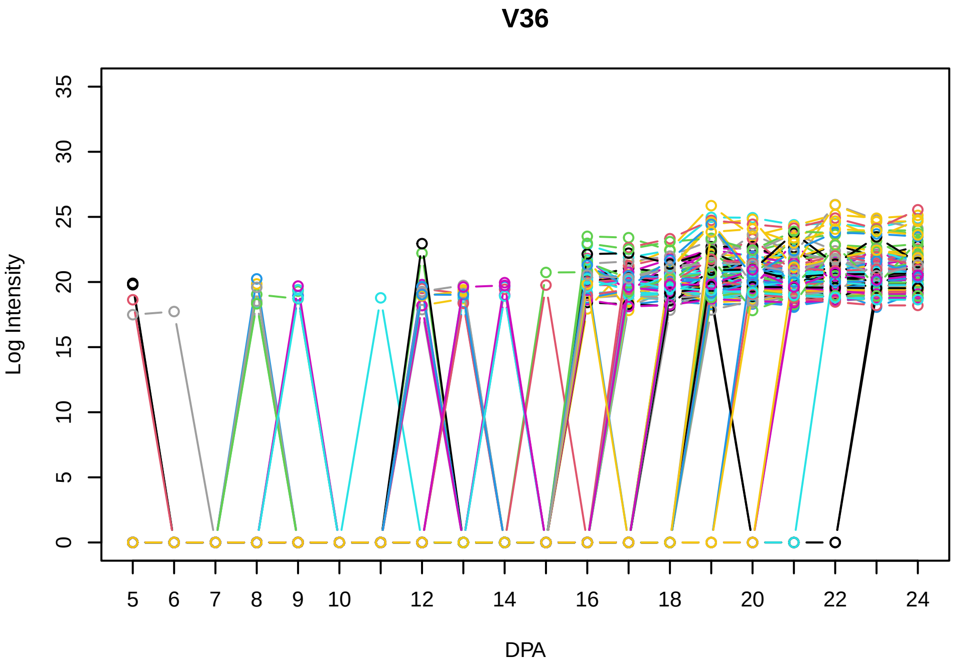
V36
Log Intensity
DPA

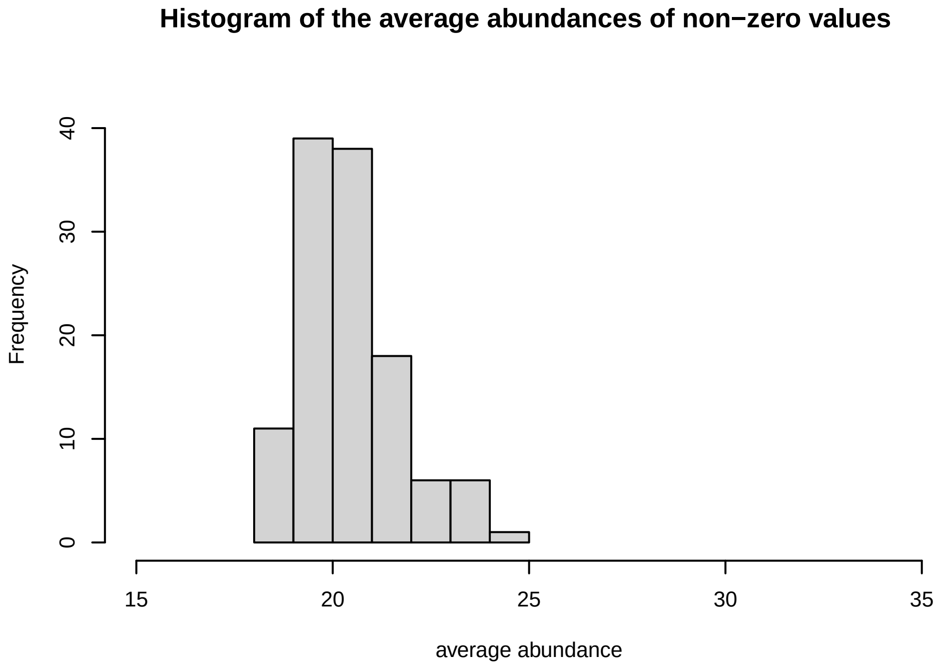
Histogram of the average abundances of non−zero values
Frequency
average abundance

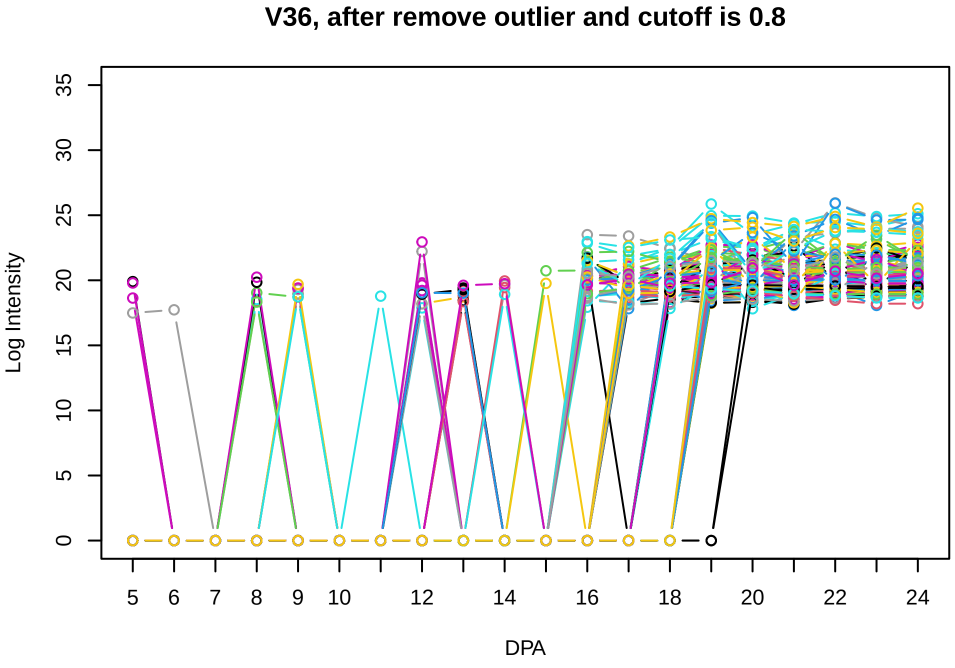
V36, after remove outlier and cutoff is 0.8
Log Intensity
DPA

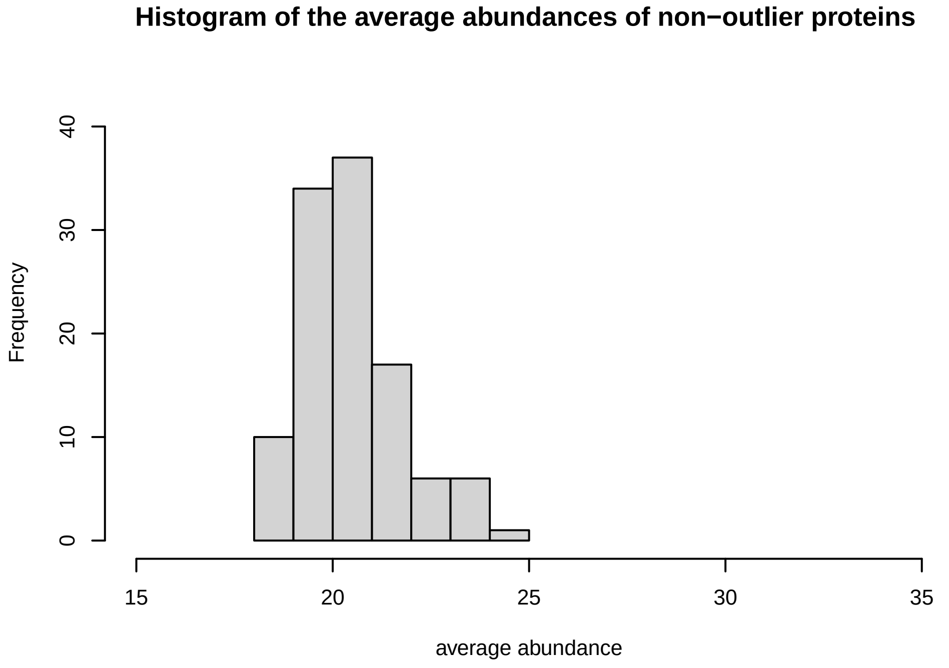
Histogram of the average abundances of non−outlier proteins
Frequency
average abundance

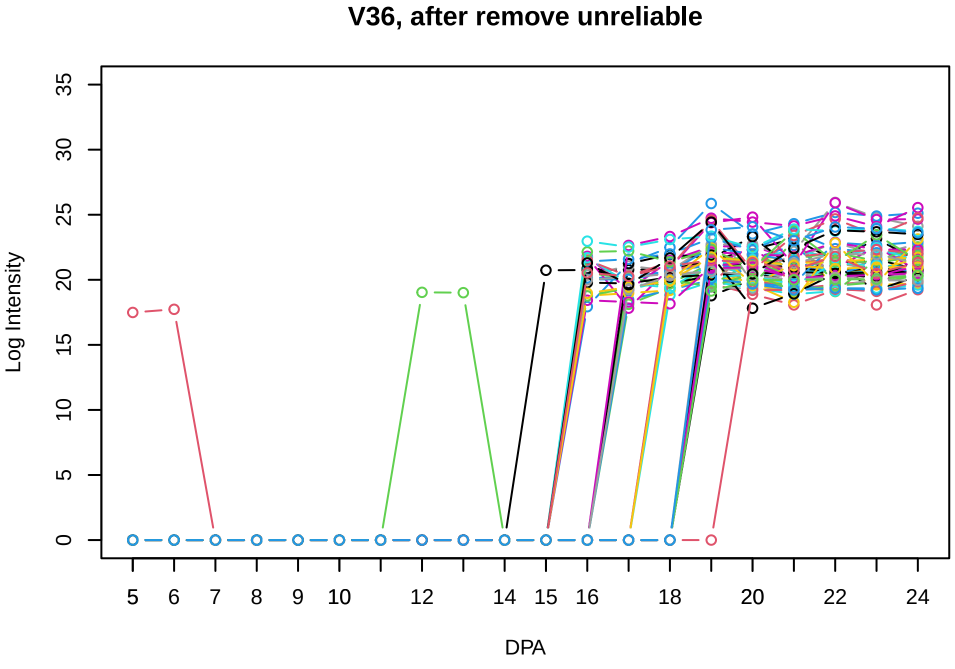
V36, after remove unreliable
Log Intensity
DPA

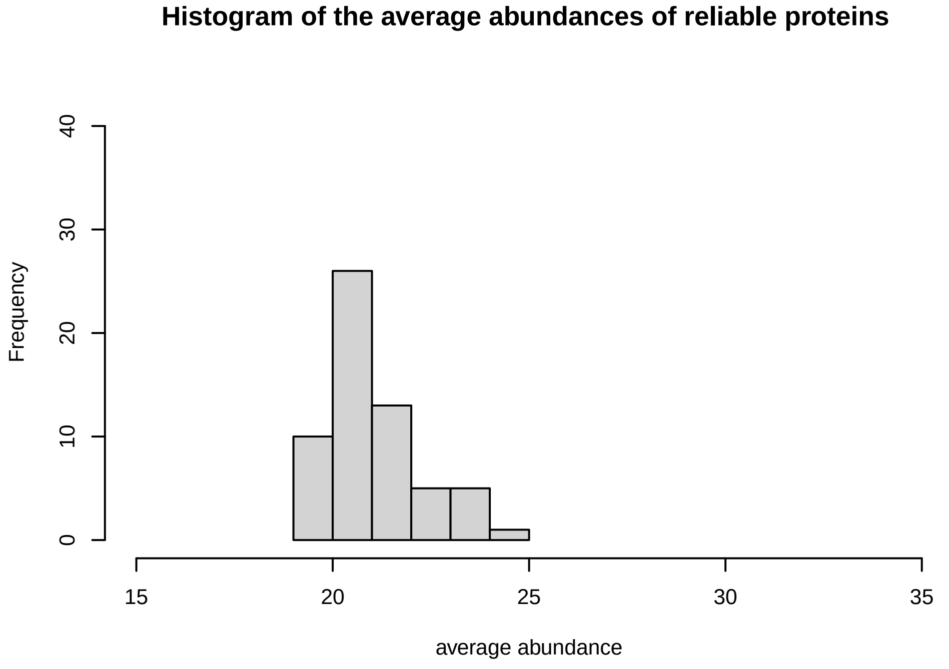
Histogram of the average abundances of reliable proteins
Frequency
average abundance

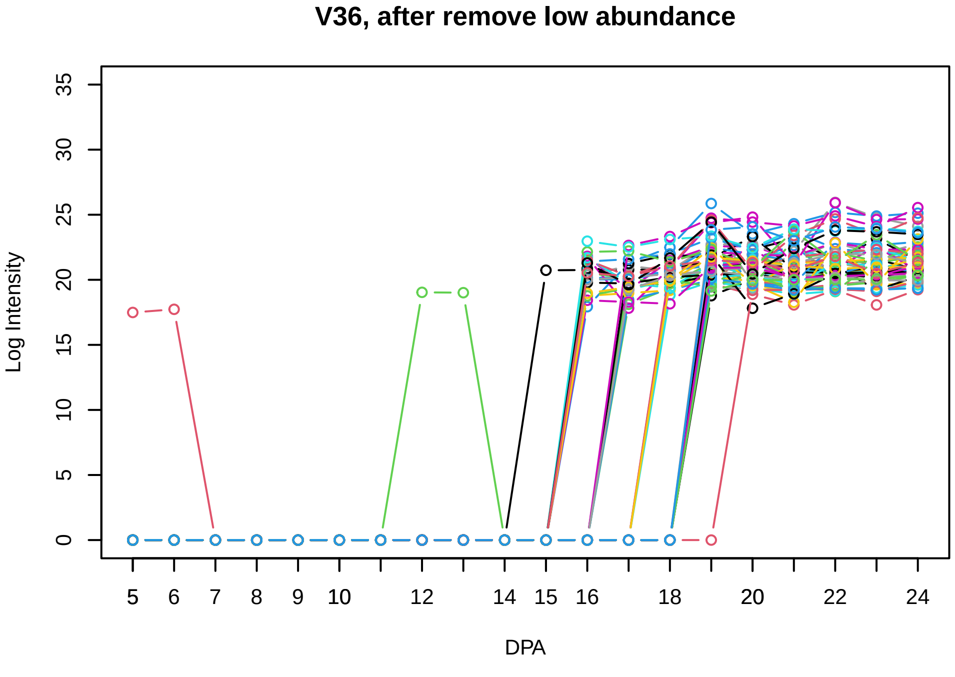
V36, after remove low abundance
Log Intensity
DPA

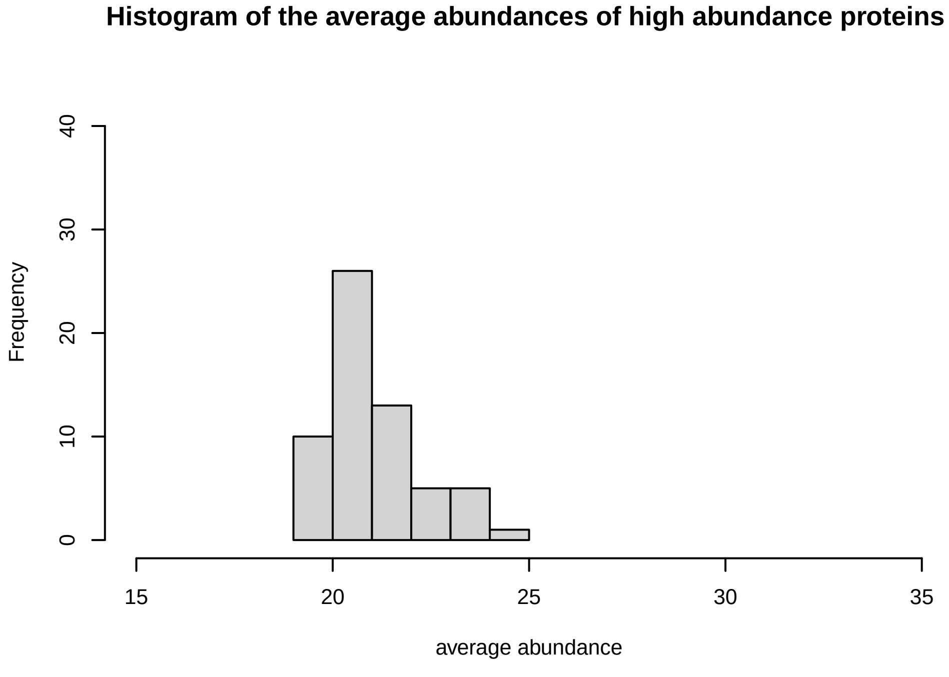
Histogram of the average abundances of high abundance proteins
Frequency
average abundance

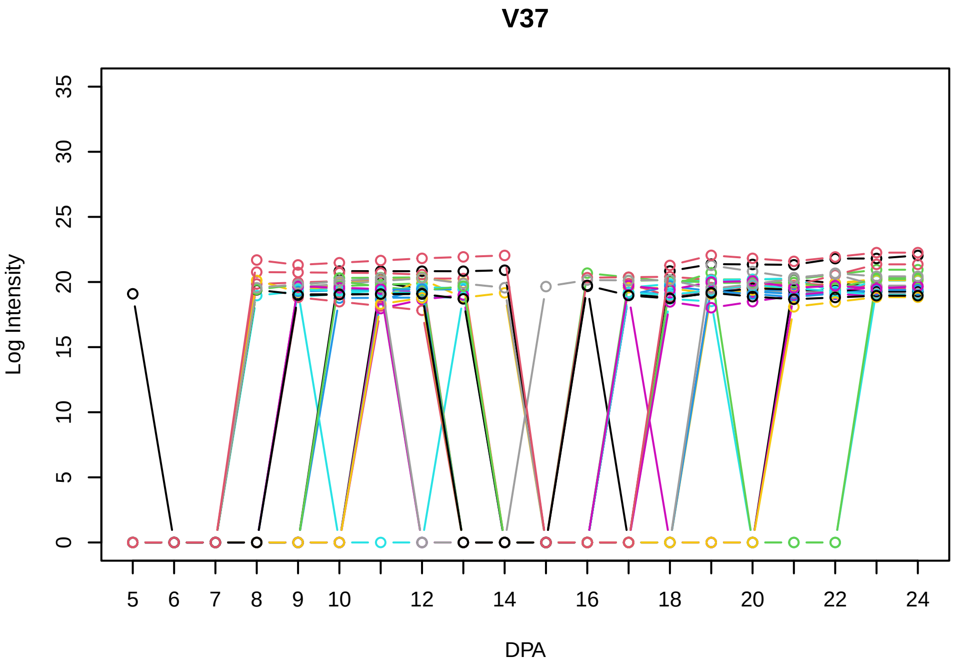
V37
Log Intensity
DPA

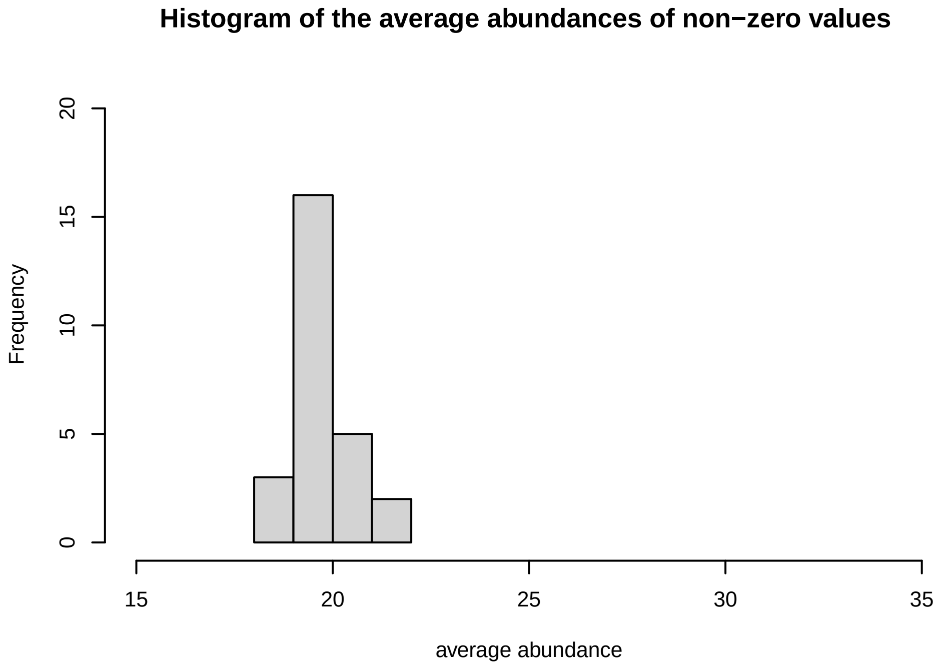
Histogram of the average abundances of non−zero values
Frequency
average abundance

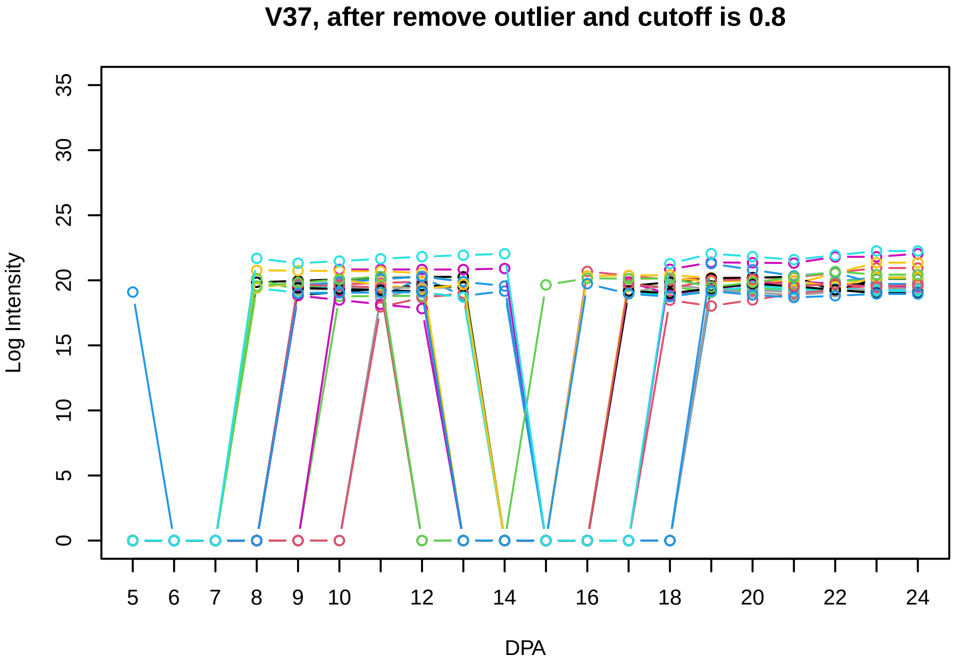
V37, after remove outlier and cutoff is 0.8
Log Intensity
DPA

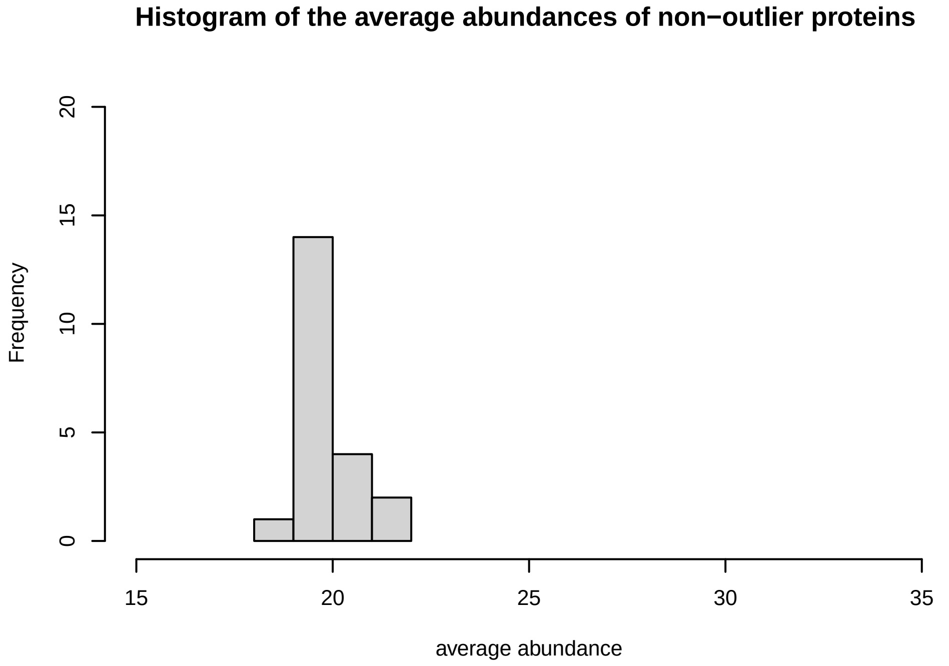
Histogram of the average abundances of non−outlier proteins
Frequency
average abundance

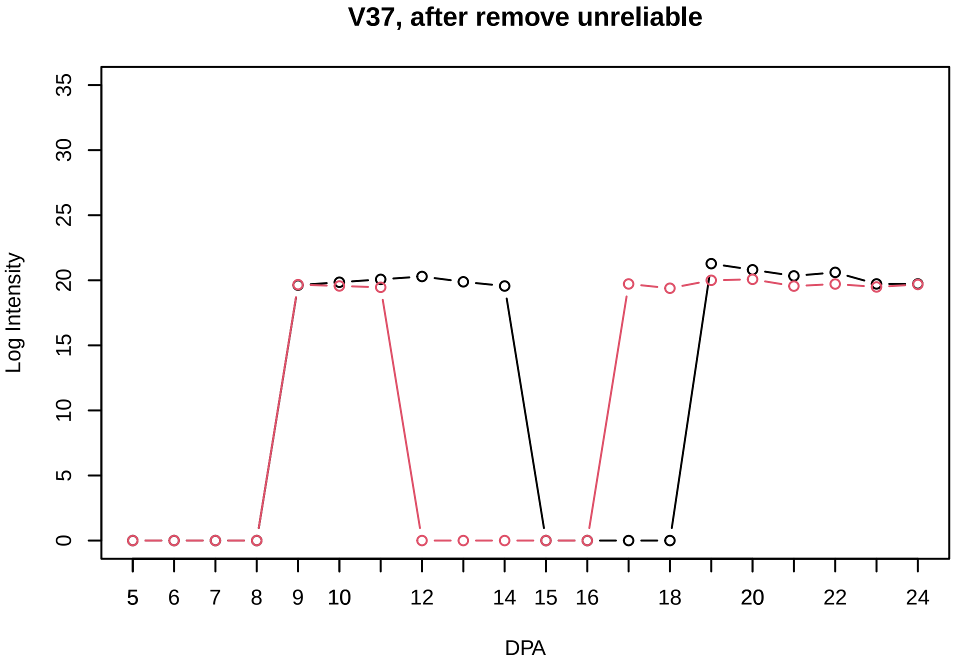
V37, after remove unreliable
Log Intensity
DPA

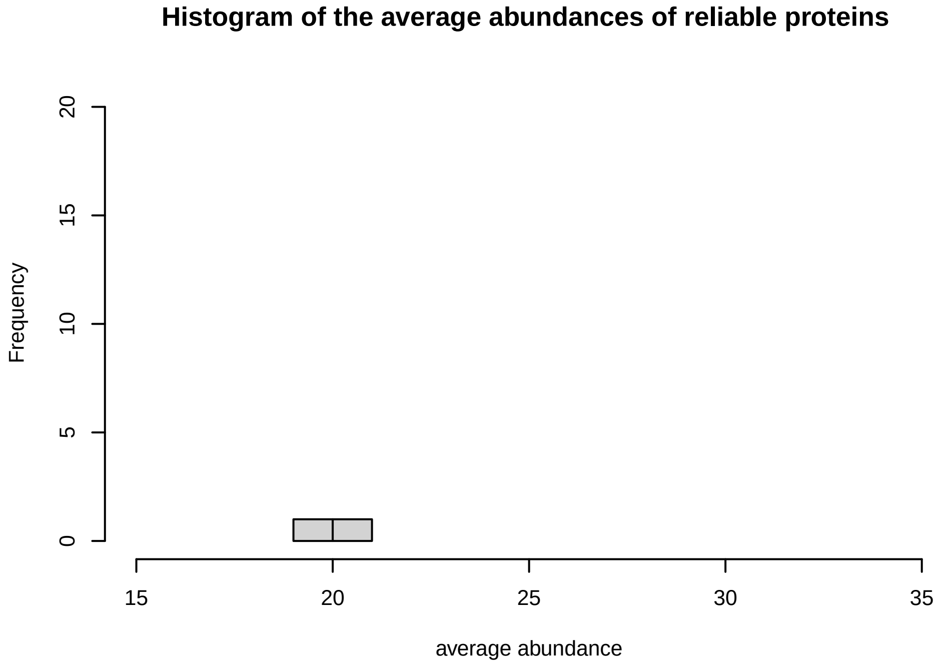
Histogram of the average abundances of reliable proteins
Frequency
average abundance

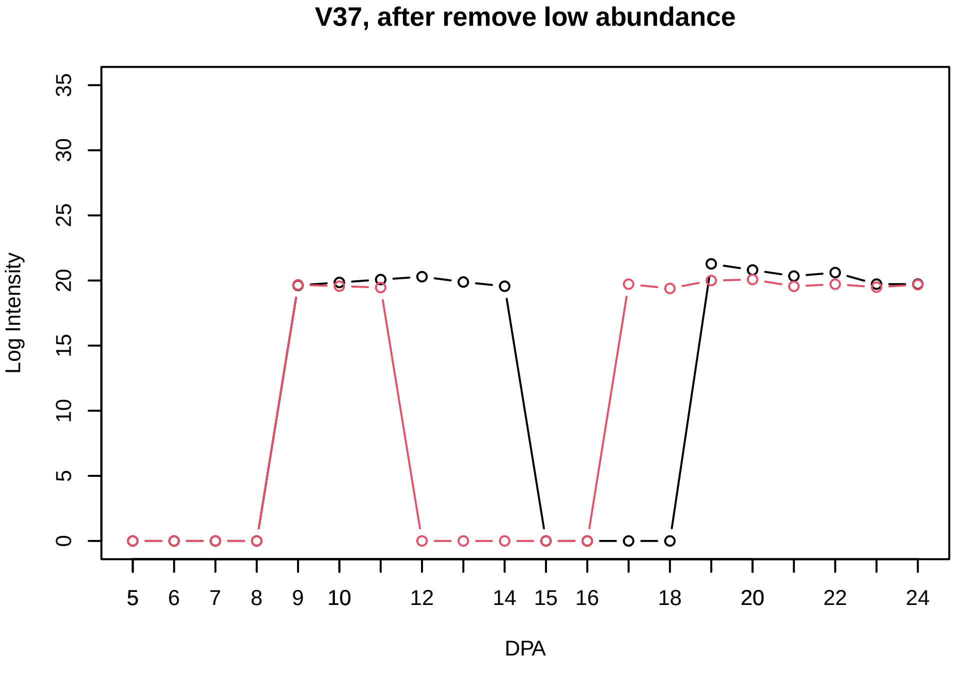
V37, after remove low abundance
Log Intensity
DPA

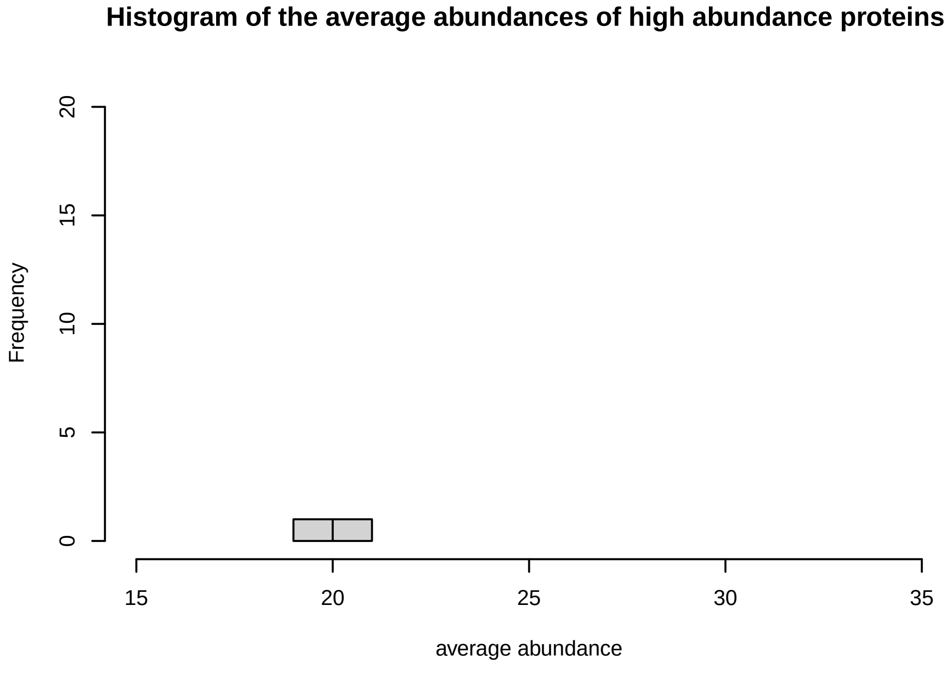
Histogram of the average abundances of high abundance proteins
Frequency
average abundance

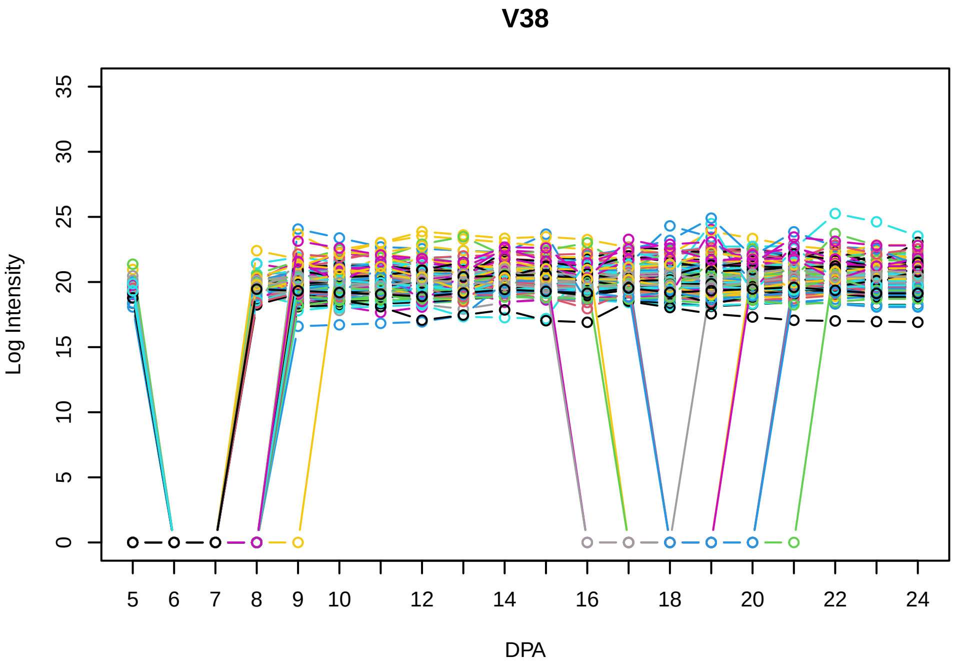
V38
Log Intensity
DPA

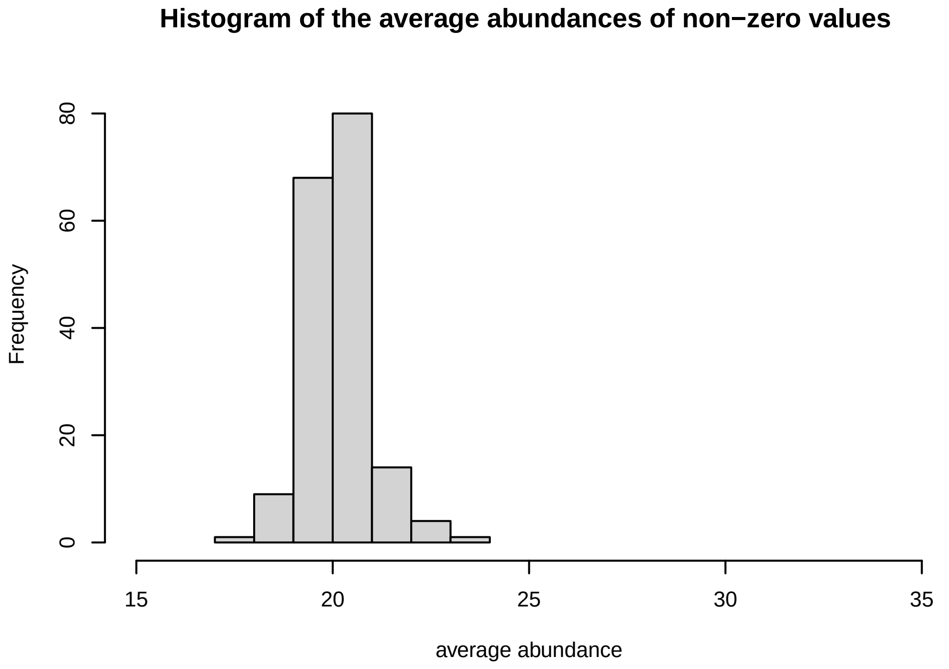
Histogram of the average abundances of non−zero values
Frequency
average abundance

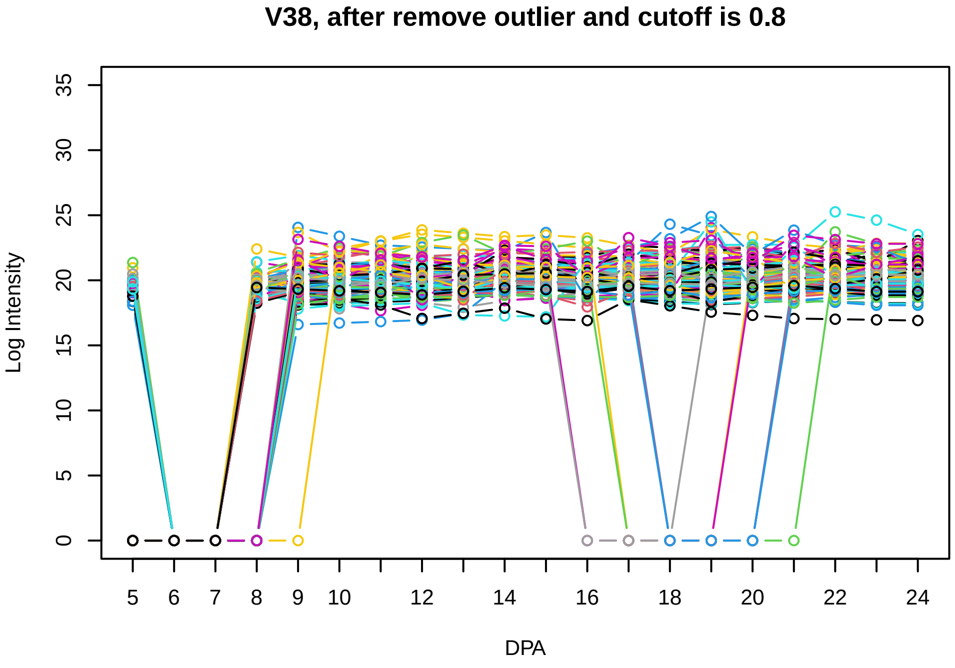
V38, after remove outlier and cutoff is 0.8
Log Intensity
DPA

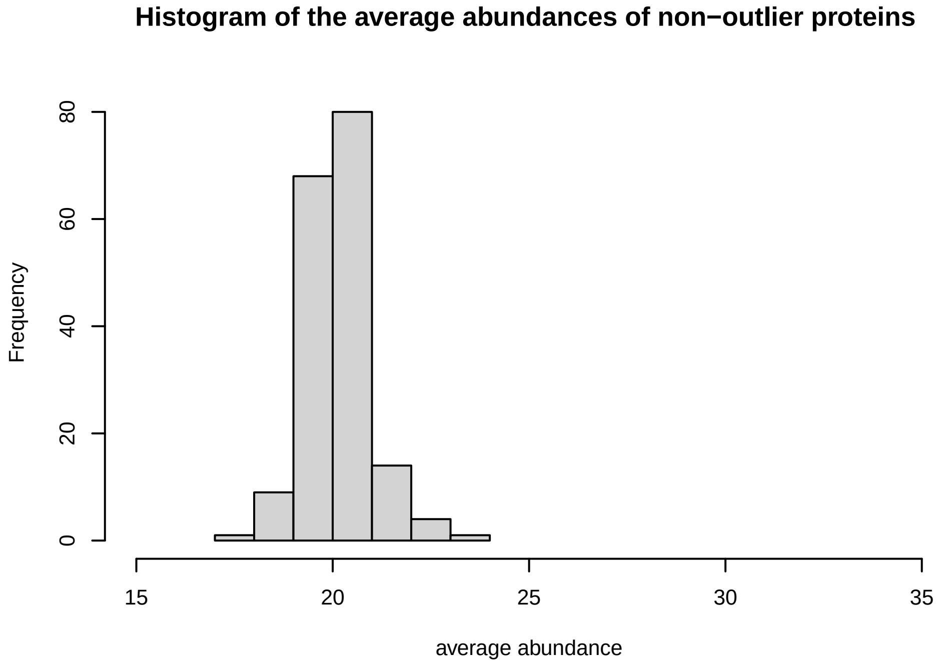
Histogram of the average abundances of non−outlier proteins
Frequency
average abundance

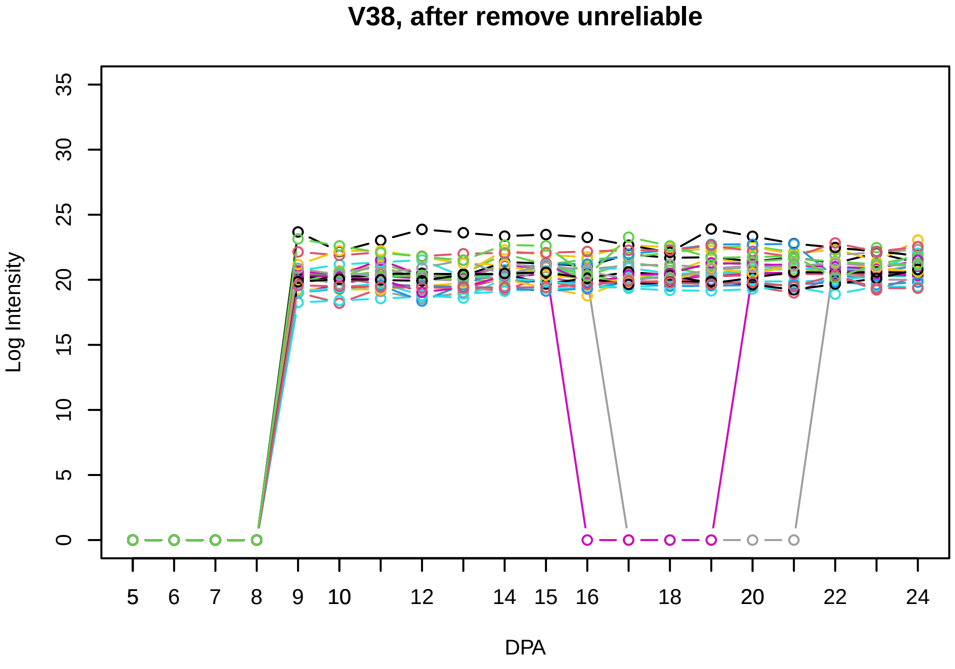
V38, after remove unreliable
Log Intensity
DPA

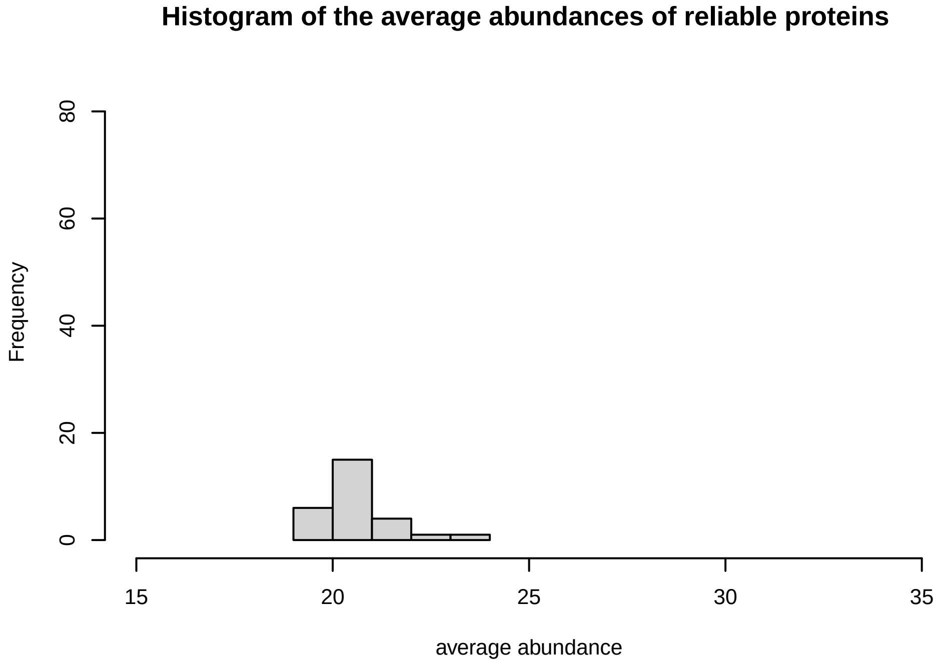
Histogram of the average abundances of reliable proteins
Frequency
average abundance

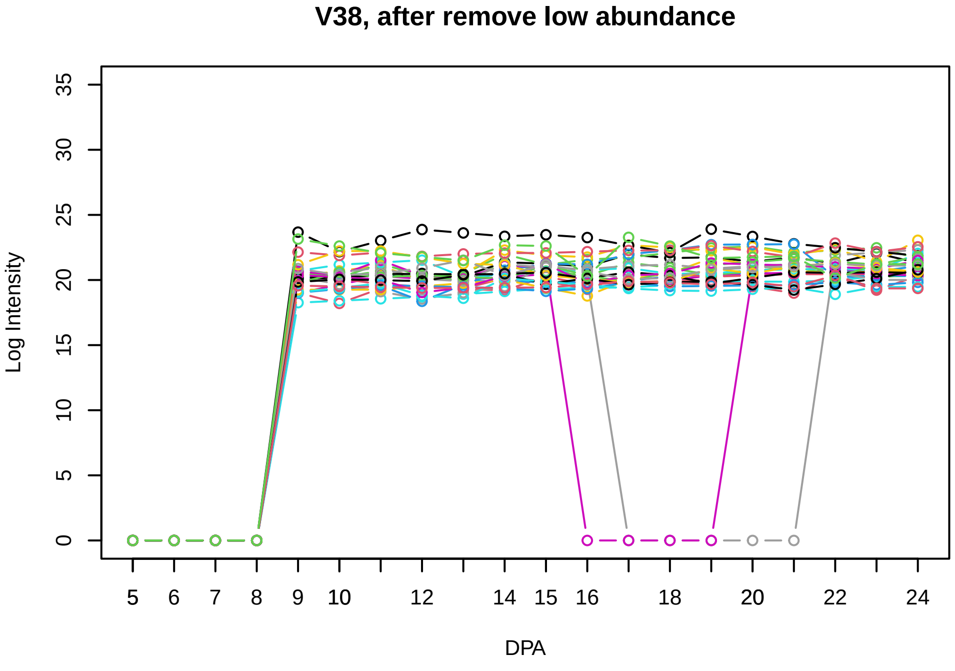
V38, after remove low abundance
Log Intensity
DPA

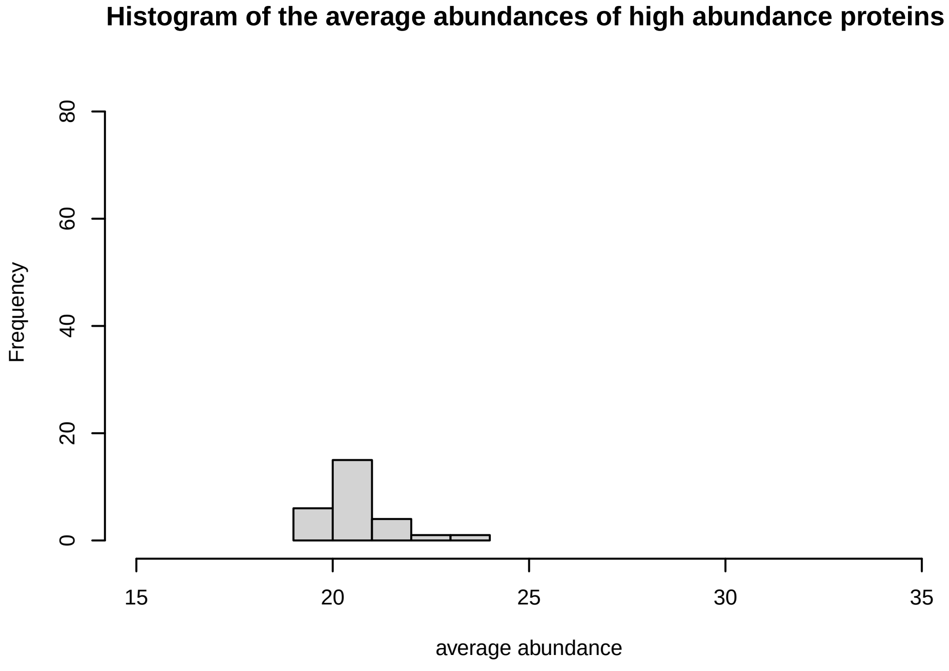
Histogram of the average abundances of high abundance proteins
Frequency
average abundance

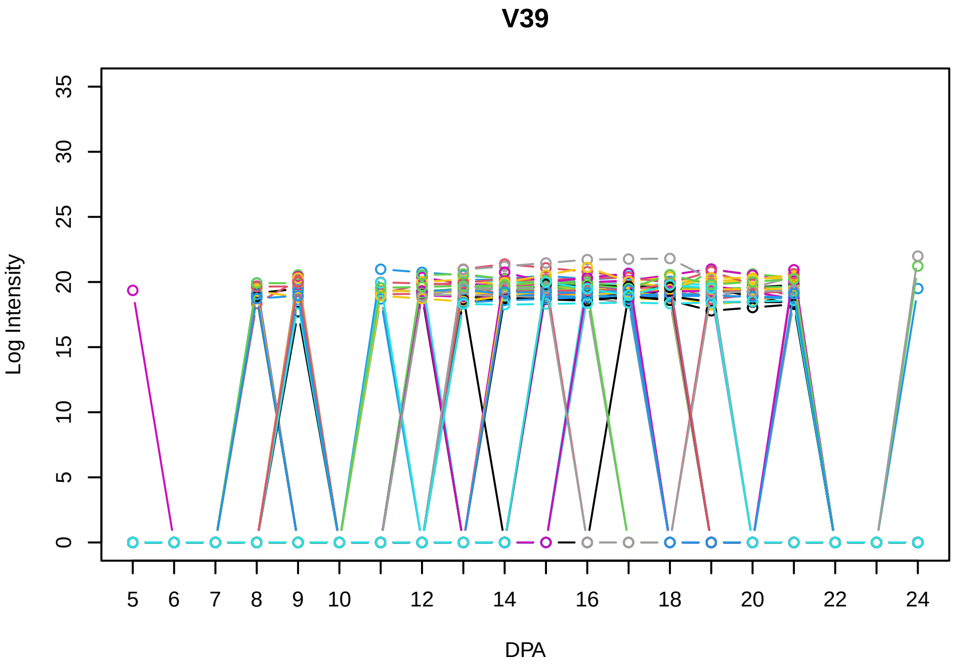
V39
Log Intensity
DPA

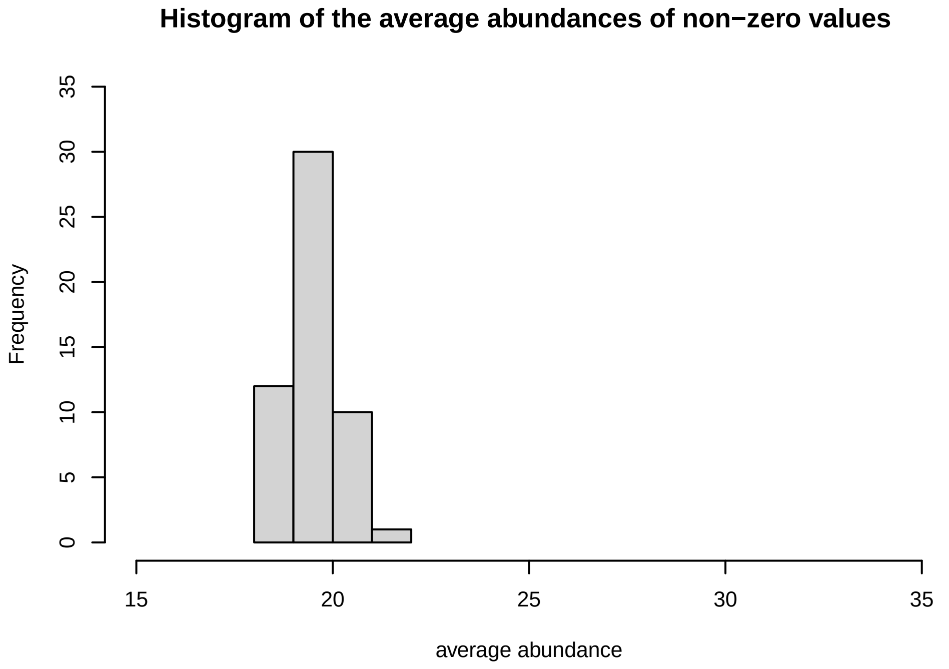
Histogram of the average abundances of non−zero values
Frequency
average abundance

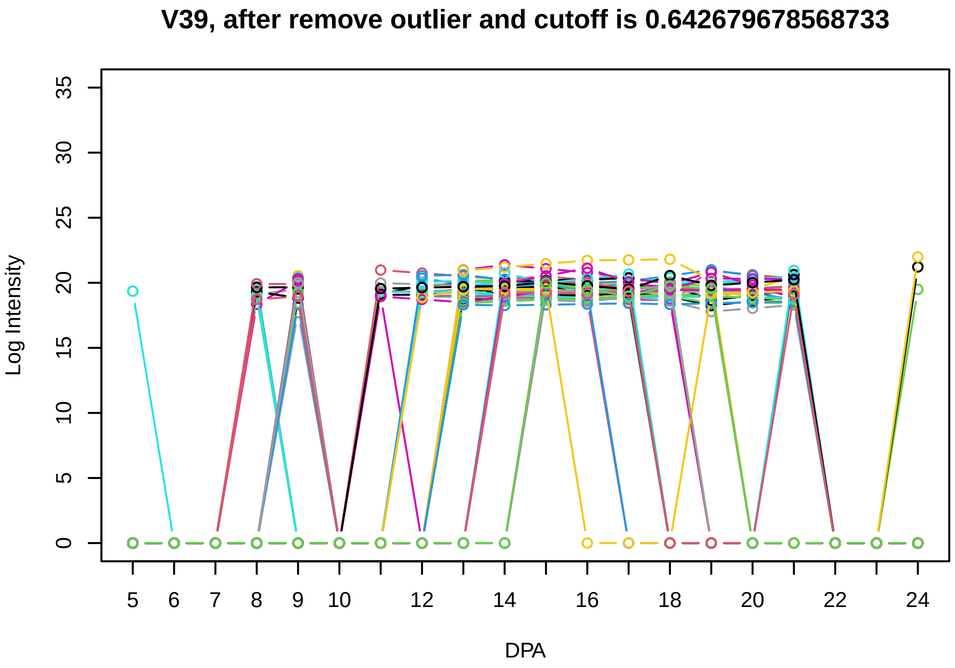
V39, after remove outlier and cutoff is 0.642679678568733
Log Intensity
DPA

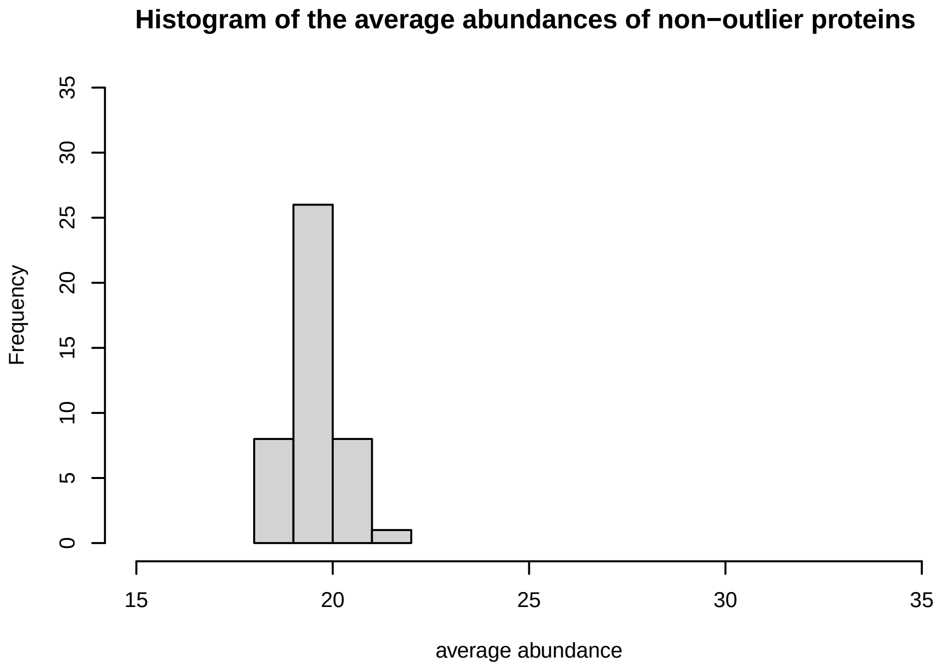
Histogram of the average abundances of non−outlier proteins
Frequency
average abundance

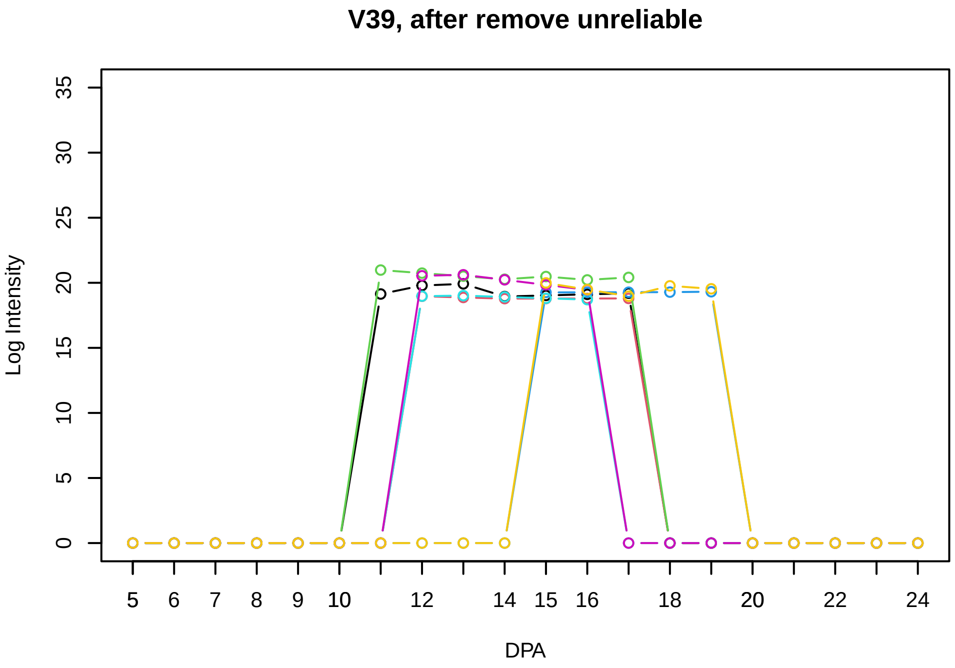
V39, after remove unreliable
Log Intensity
DPA

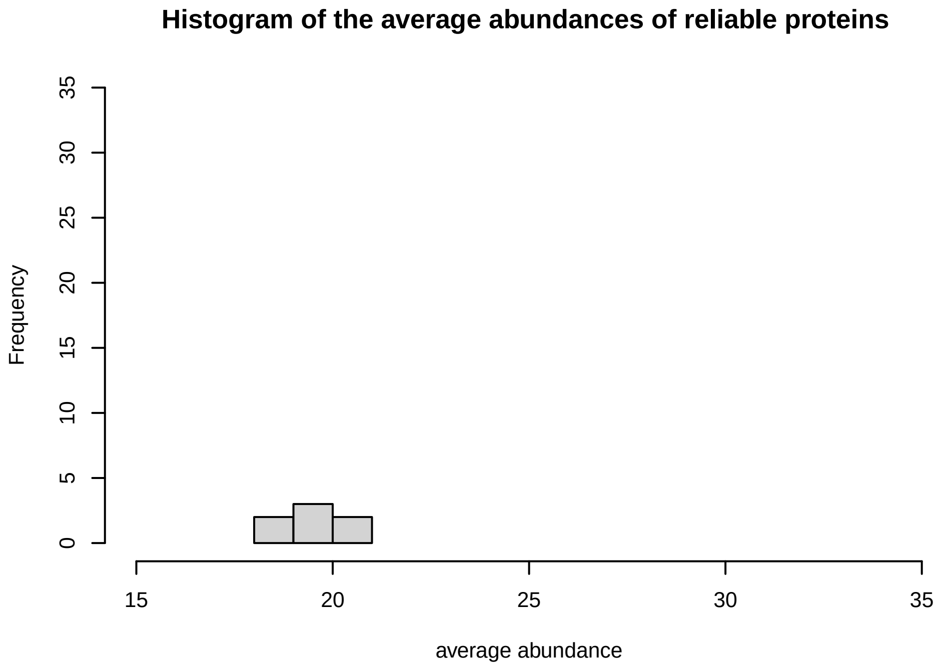
Histogram of the average abundances of reliable proteins
Frequency
average abundance

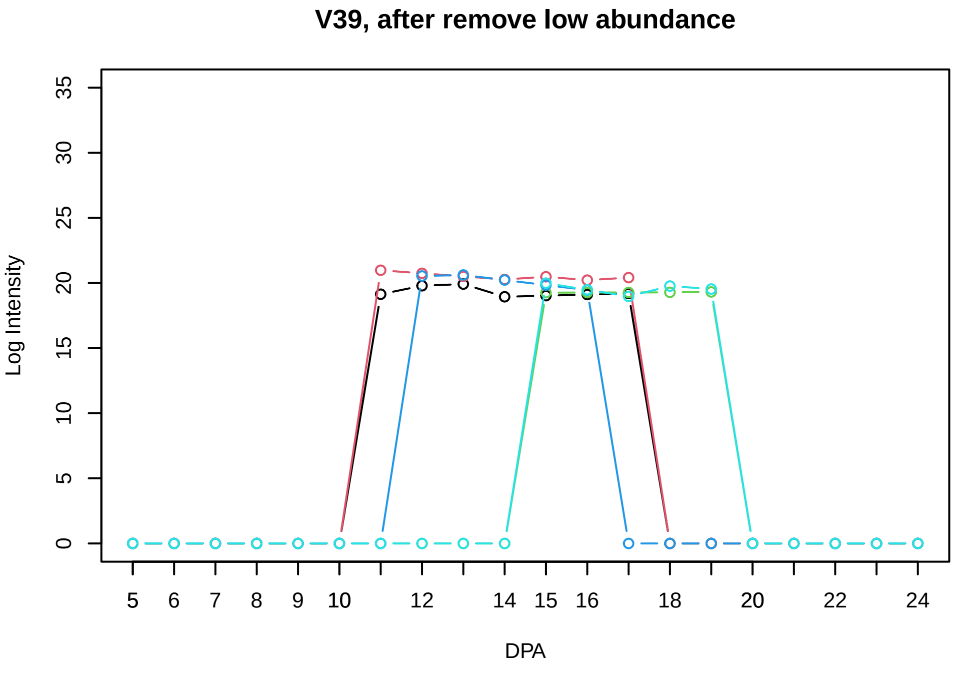
V39, after remove low abundance
Log Intensity
DPA

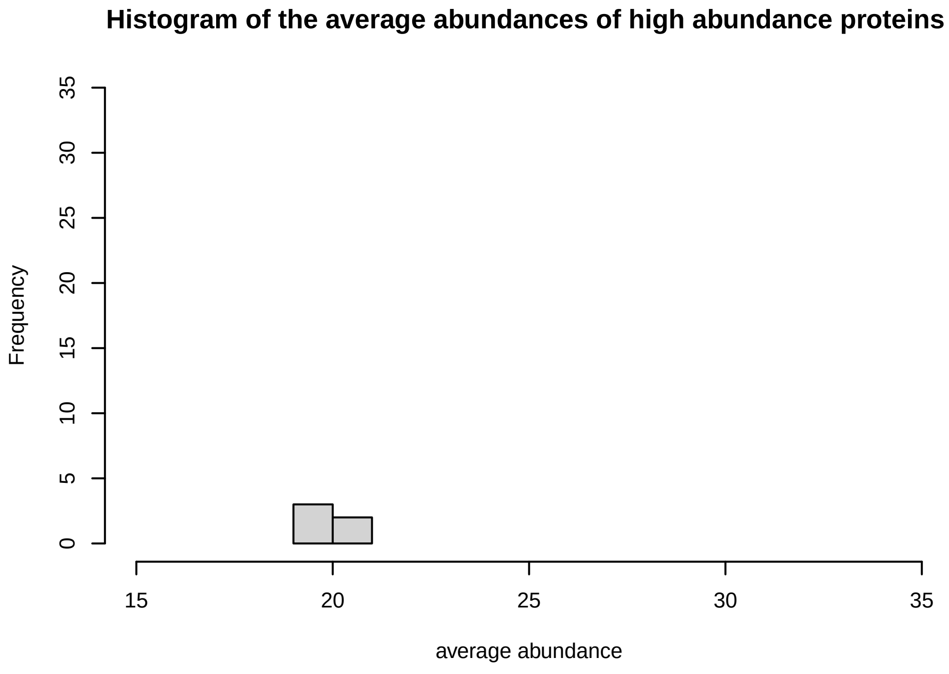
Histogram of the average abundances of high abundance proteins
Frequency
average abundance

**V40**

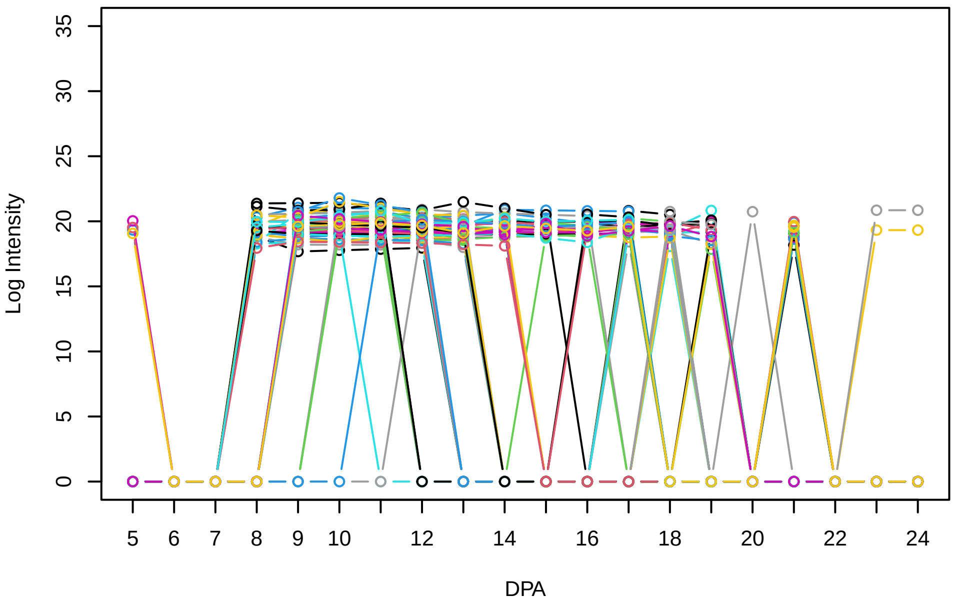


**Histogram of the average abundances of non−zero values**

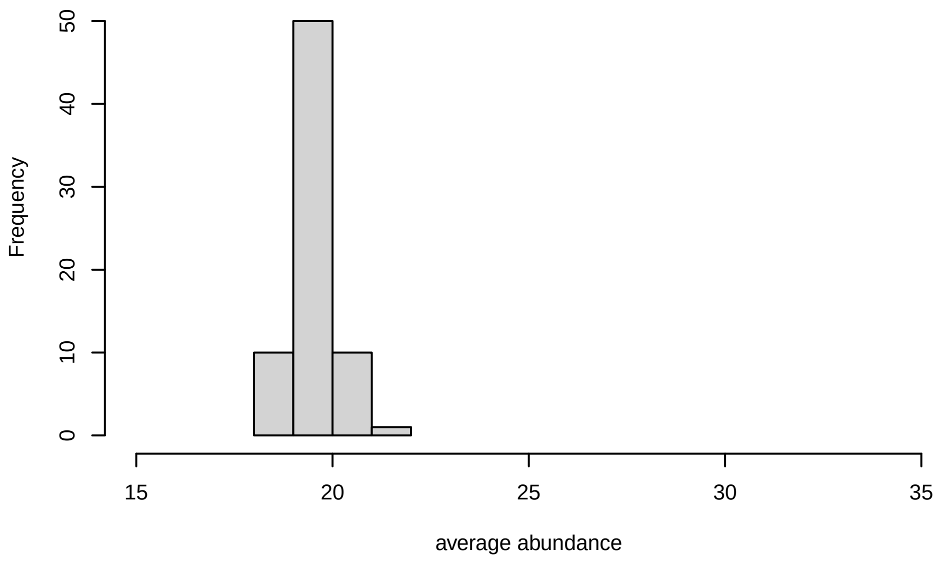


**V40, after remove outlier and cutoff is 0.687304641725951**

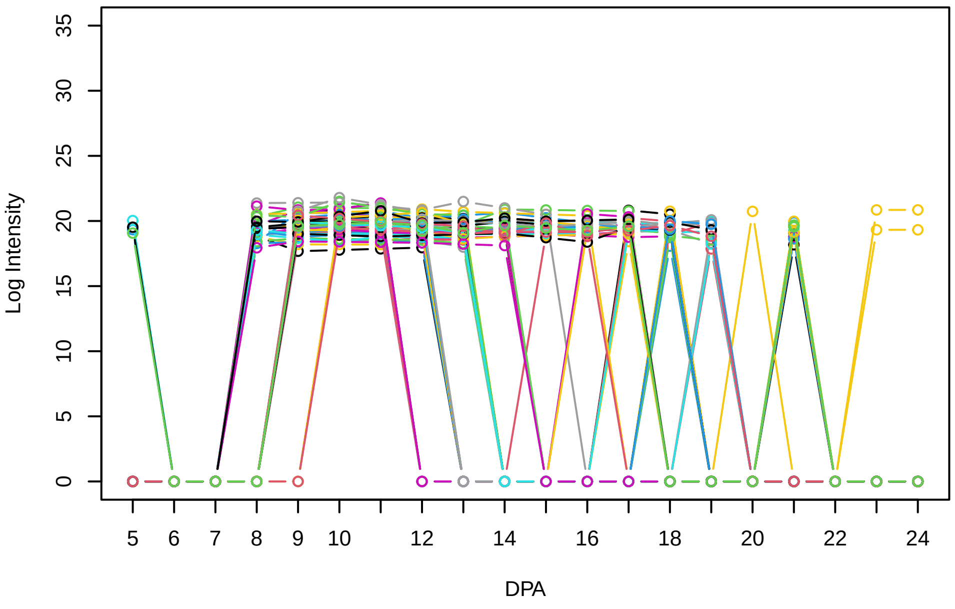


**Histogram of the average abundances of non−outlier proteins**

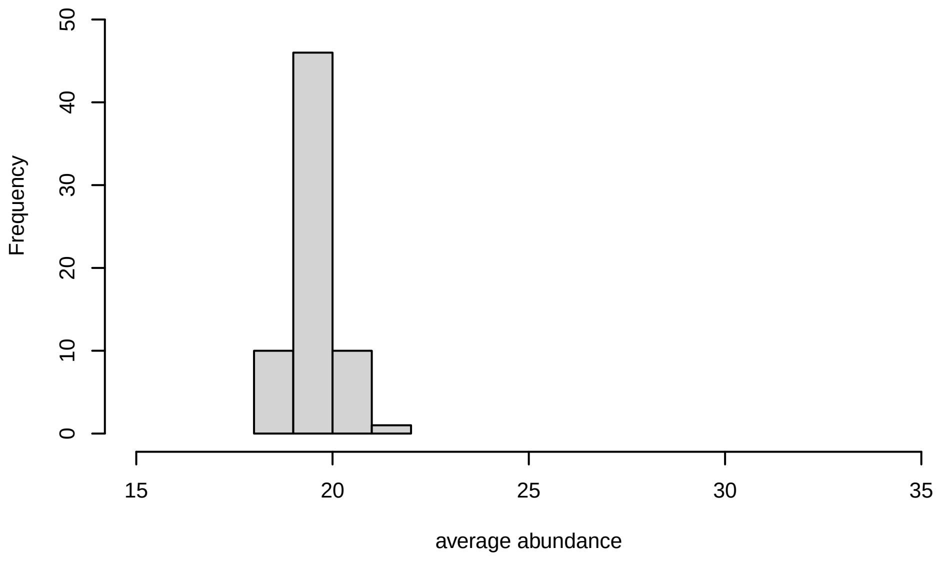


**V40, after remove unreliable**

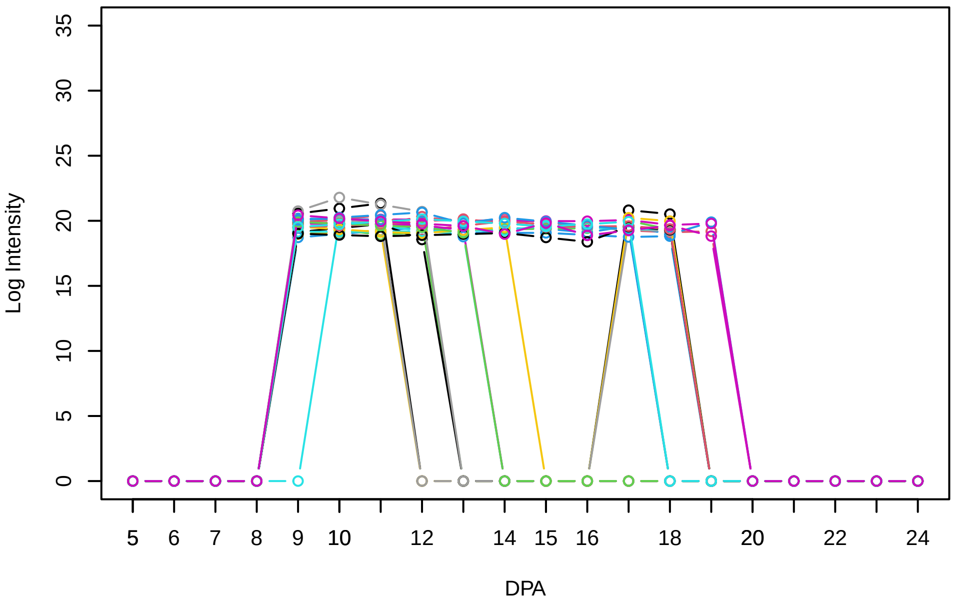


**Histogram of the average abundances of reliable proteins**

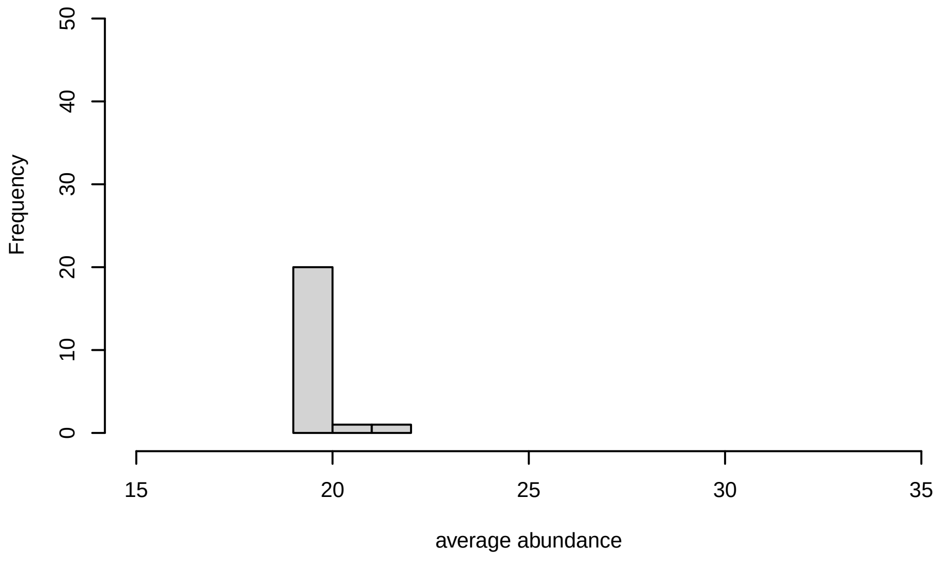


**V40, after remove low abundance**

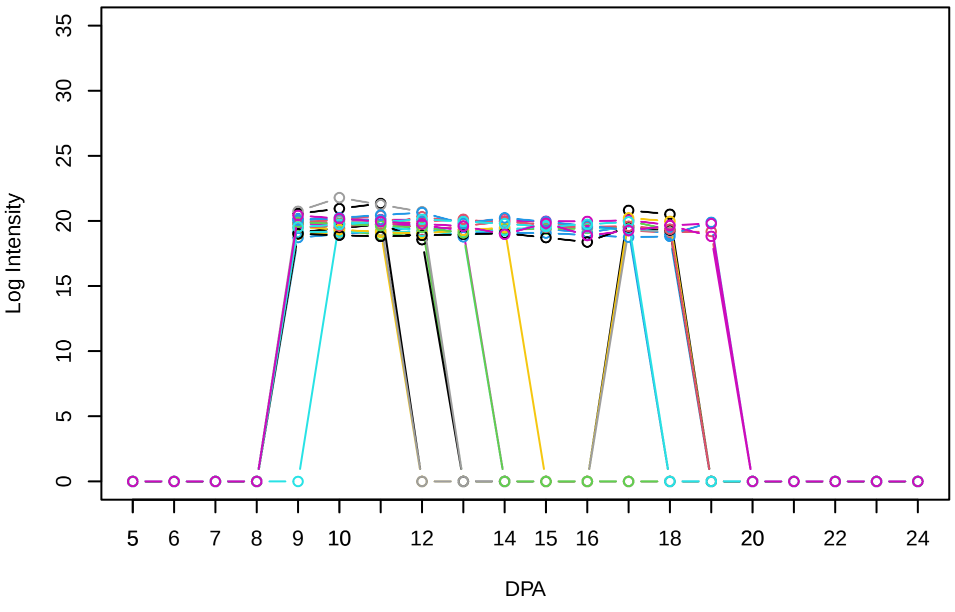


**Histogram of the average abundances of high abundance proteins**

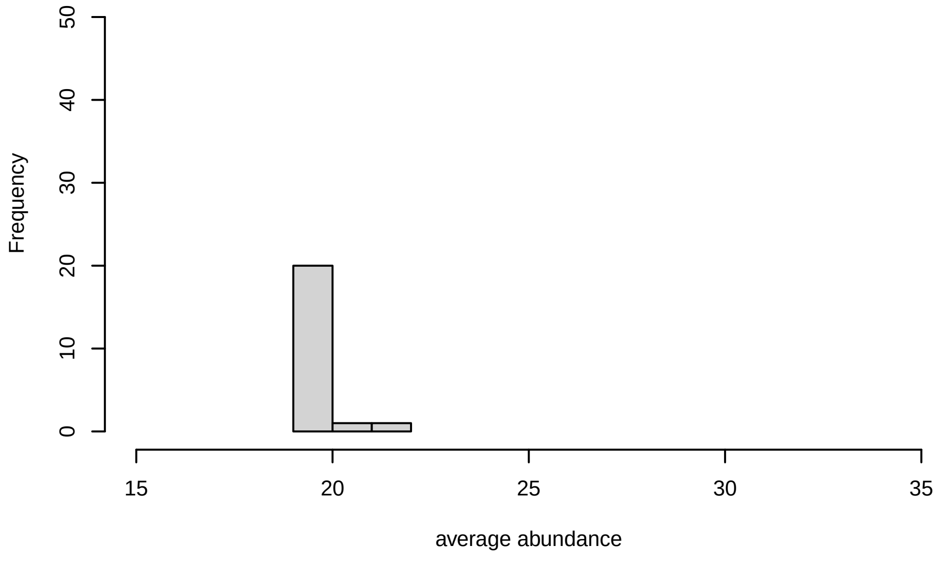
