## Supplementary Figure S4 for "Systems-level proteomic models of cotton fiber development: A high-resolution resource to analyze cell dynamics and trait engineering"

**Histogram of the average abundances of non−zero values**

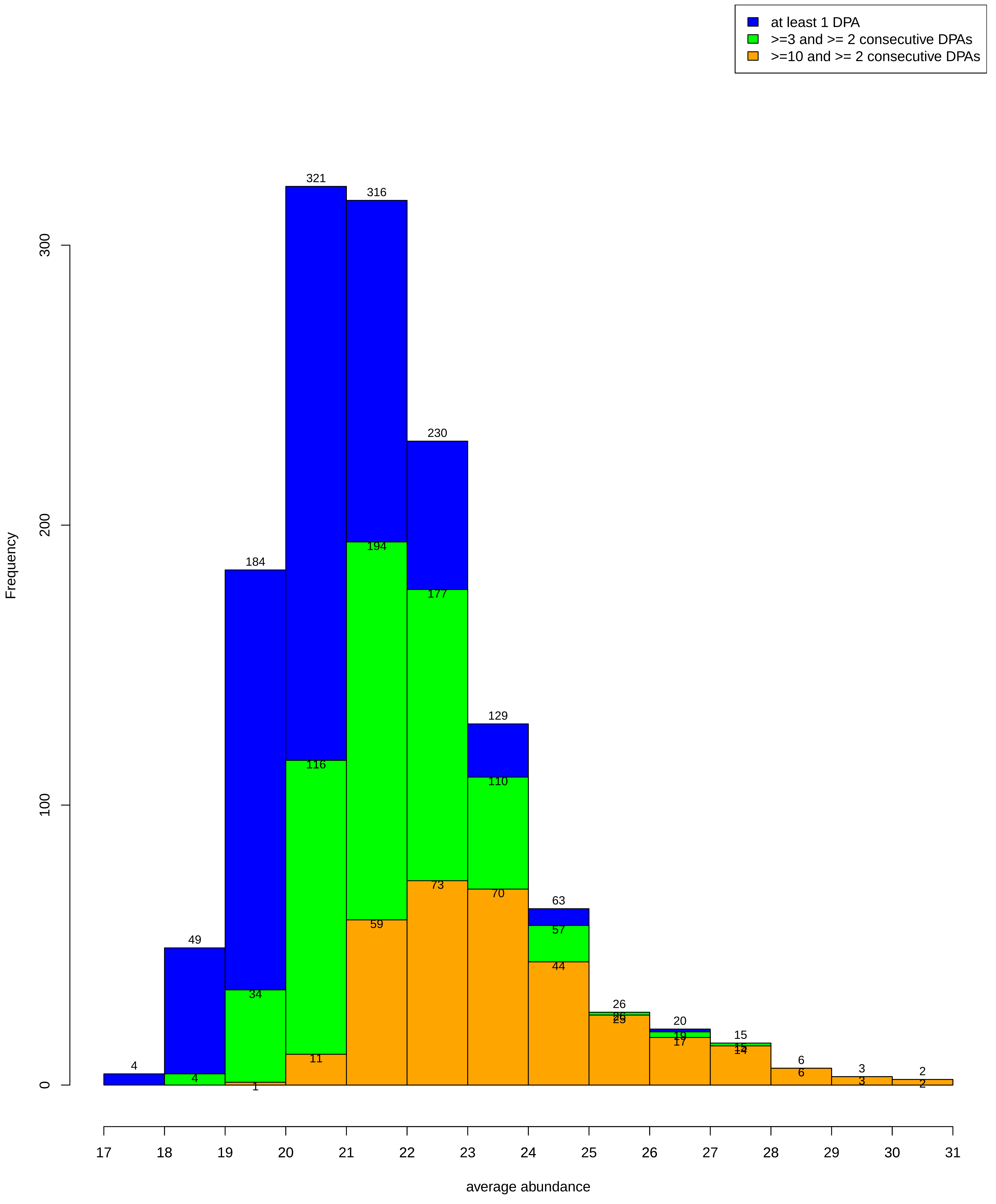

**Codes**

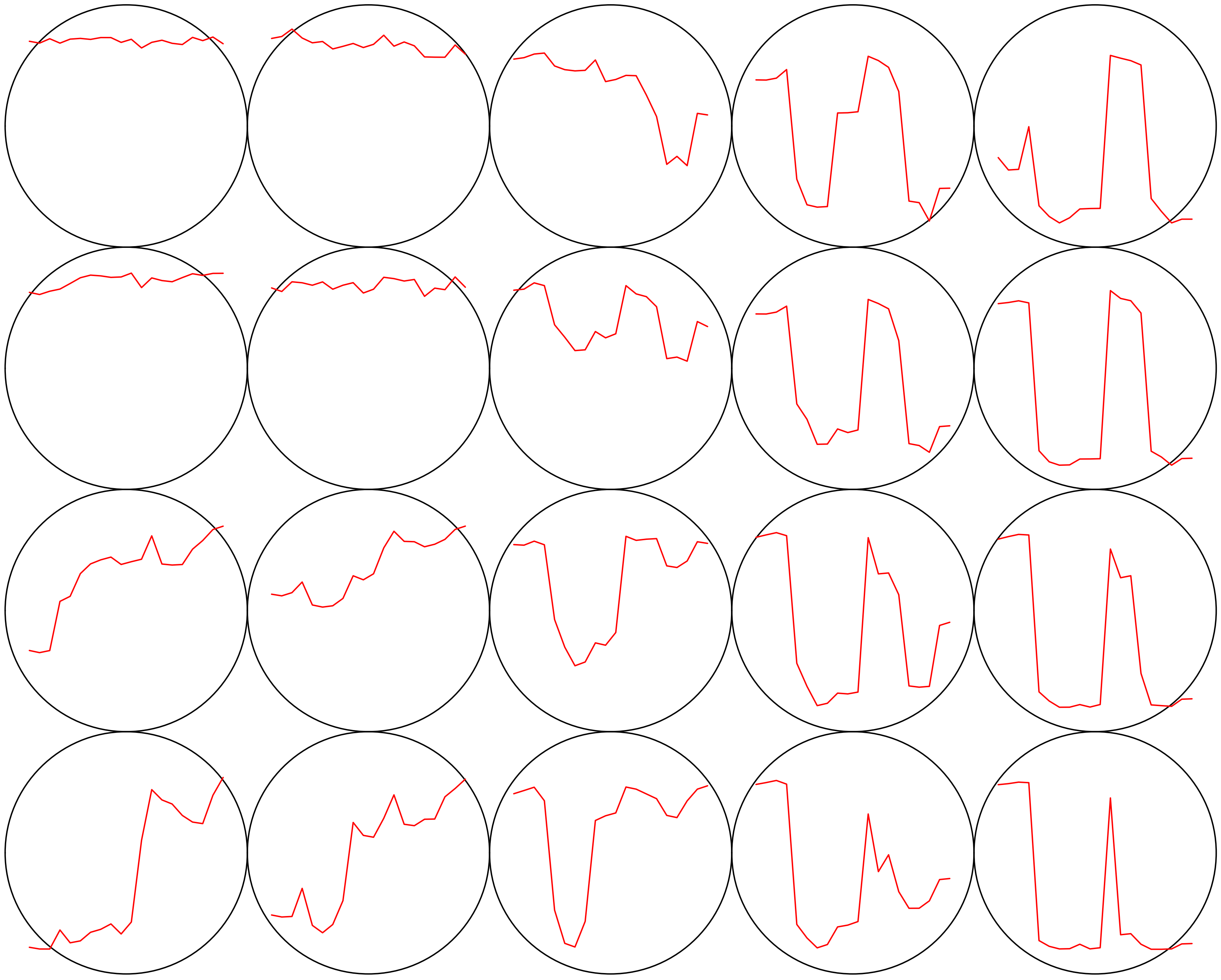

**Count plot**

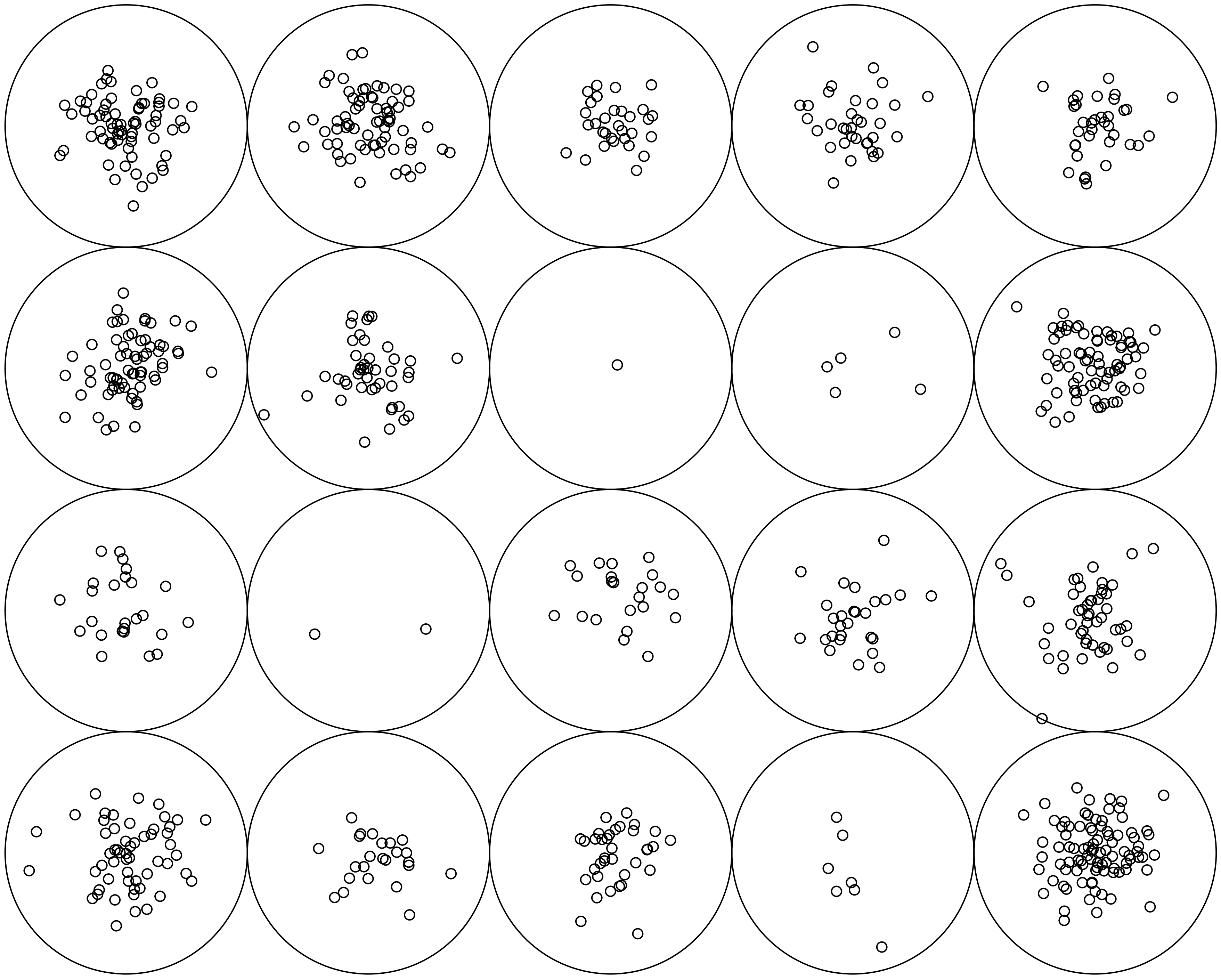

V1

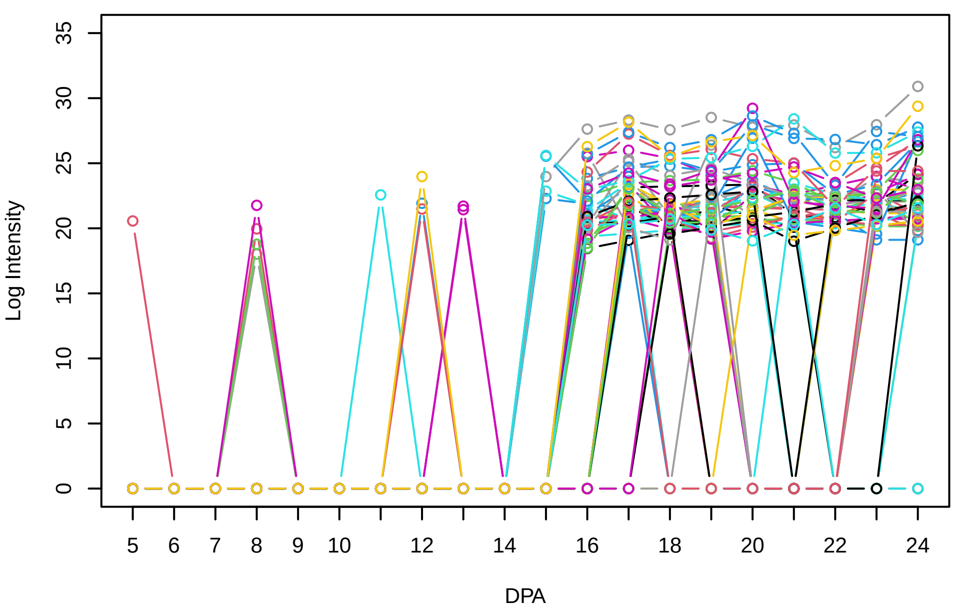


Histogram of the average abundances of non−zero values

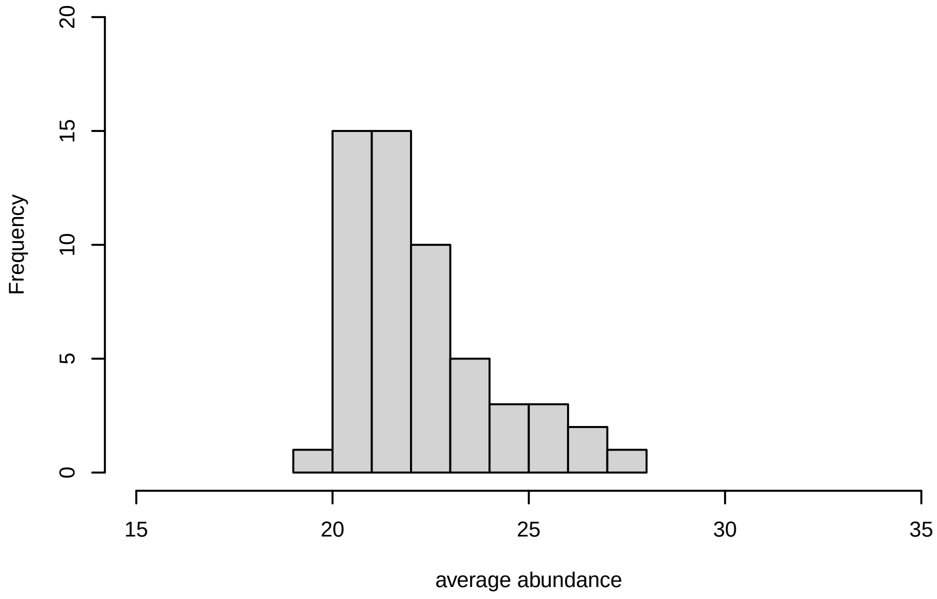


V1, after remove outlier and cutoff is 0.8

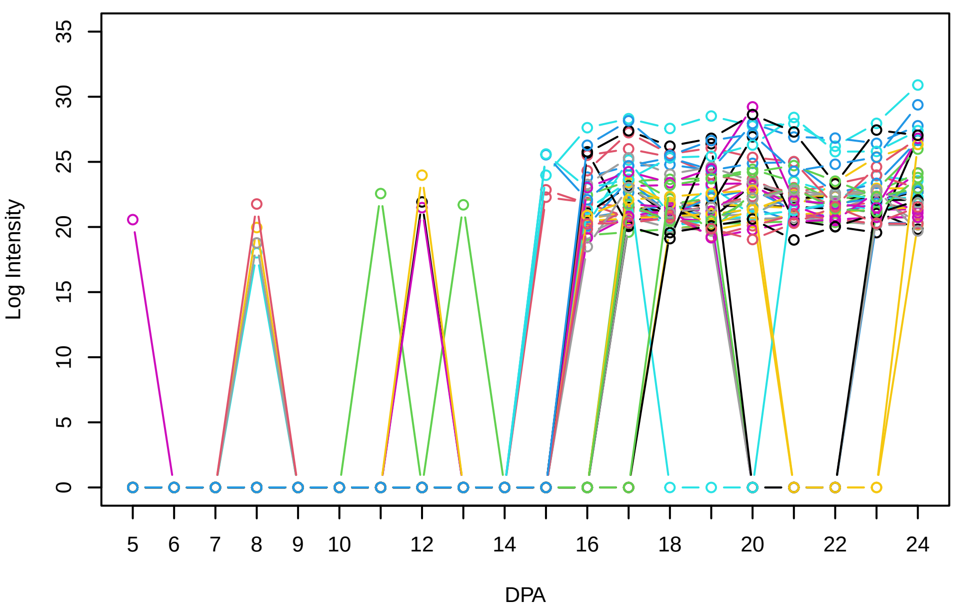


Histogram of the average abundances of non−outlier proteins

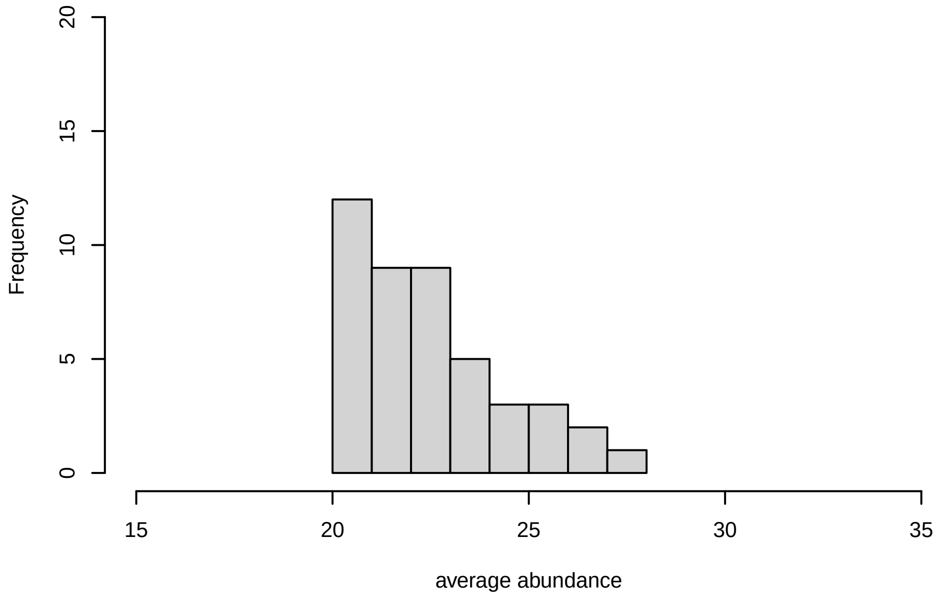


V1, after remove unreliable

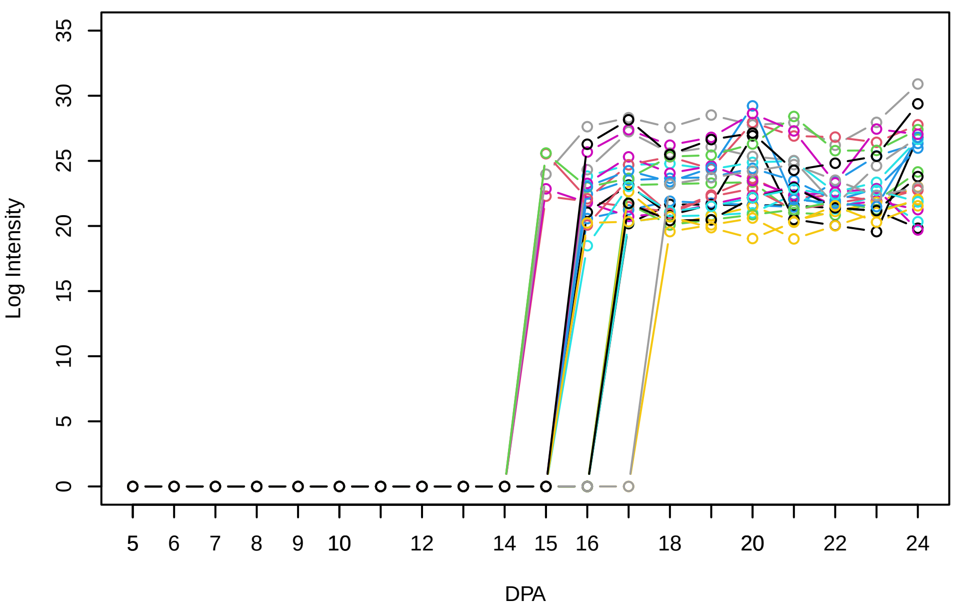


Histogram of the average abundances of reliable proteins

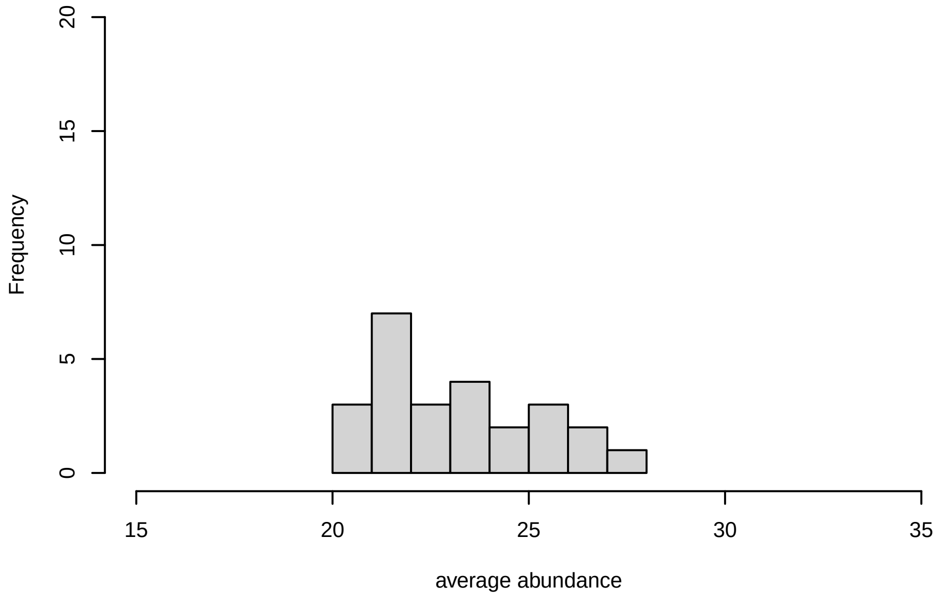


V1, after remove low abundance

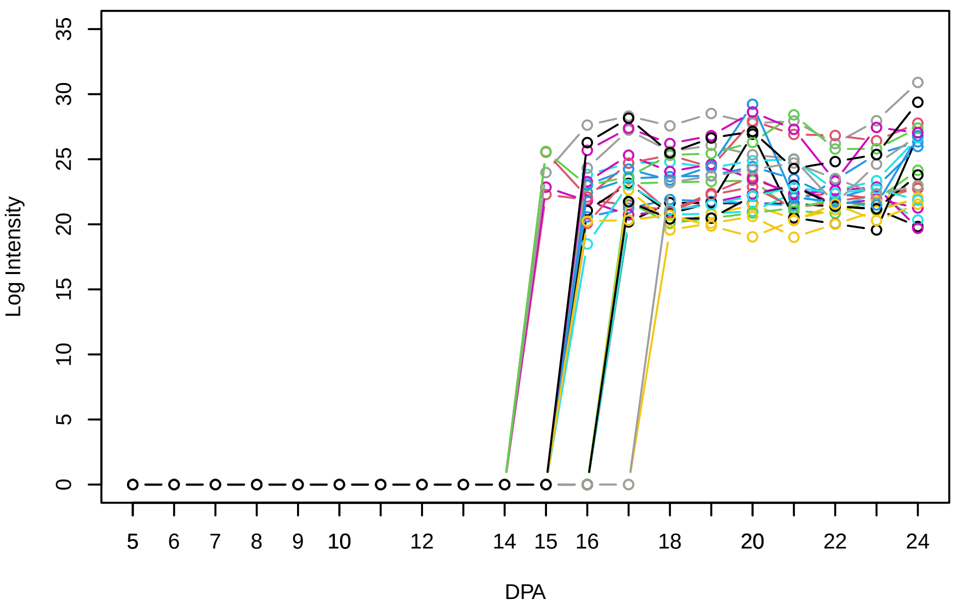


Histogram of the average abundances of high abundance proteins

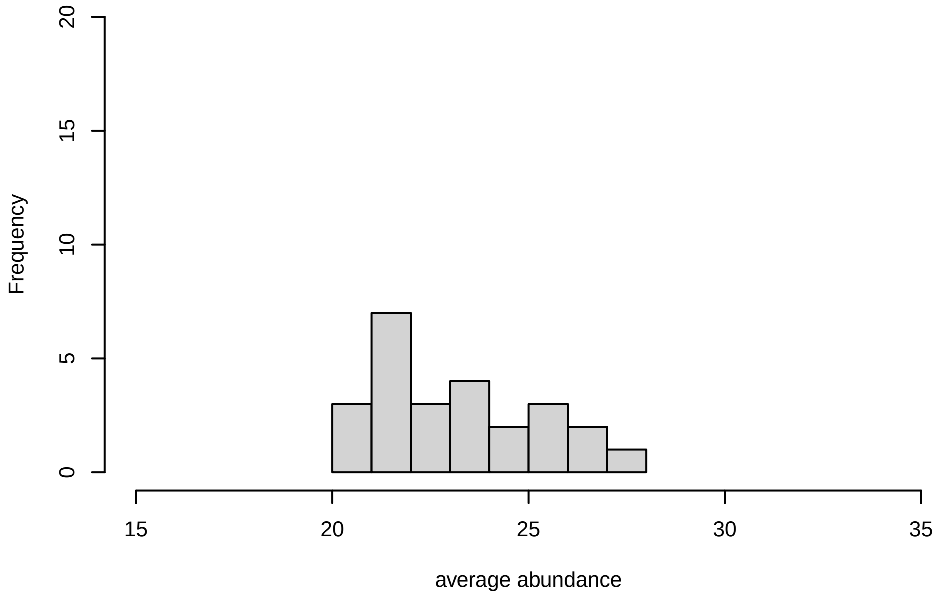

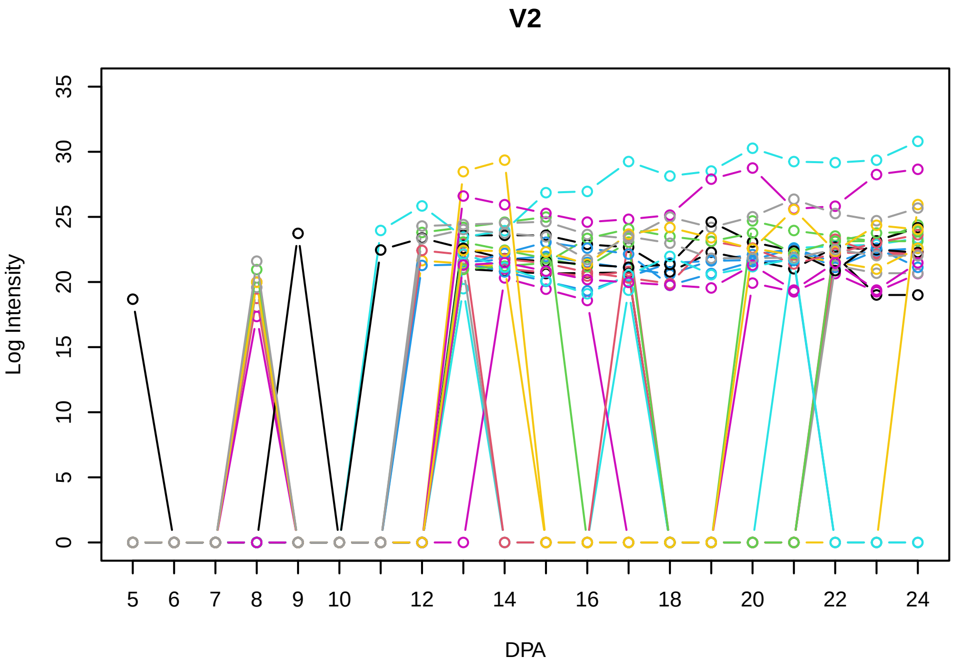
V2
Log Intensity
DPA

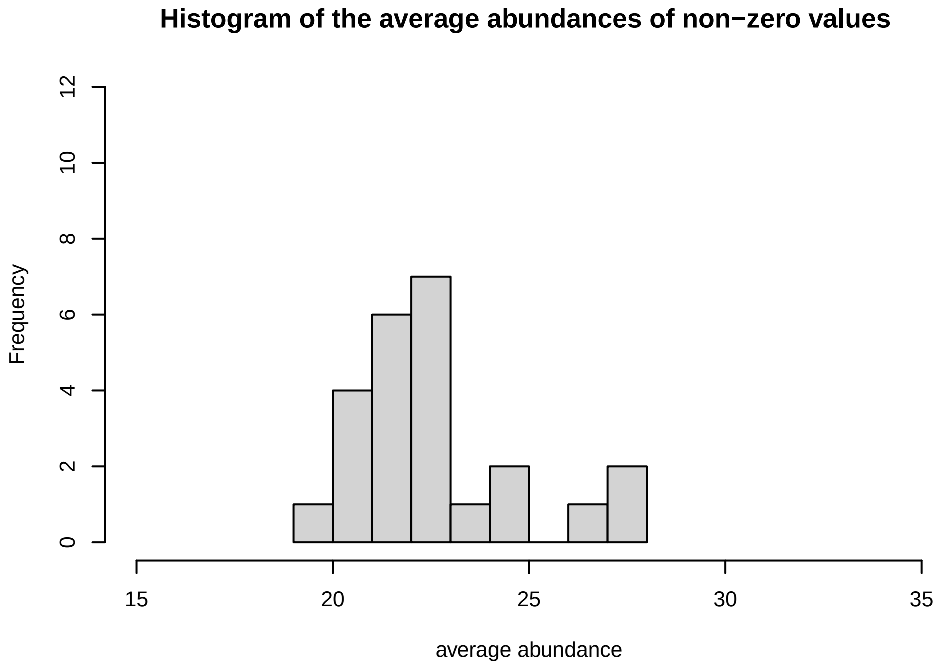
Histogram of the average abundances of non−zero values
Frequency
average abundance

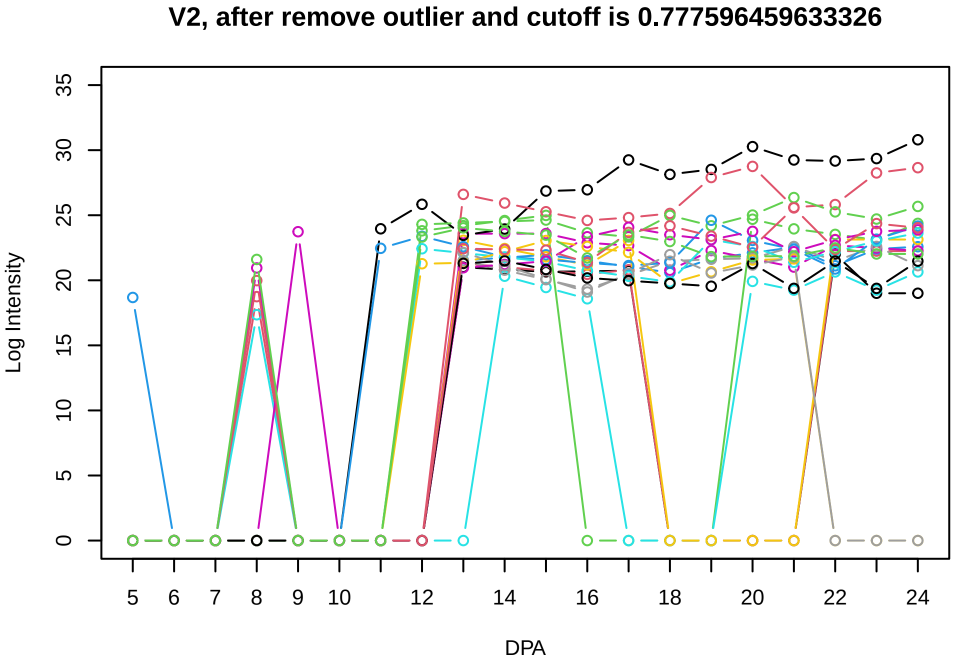
V2, after remove outlier and cutoff is 0.777596459633326
Log Intensity
DPA

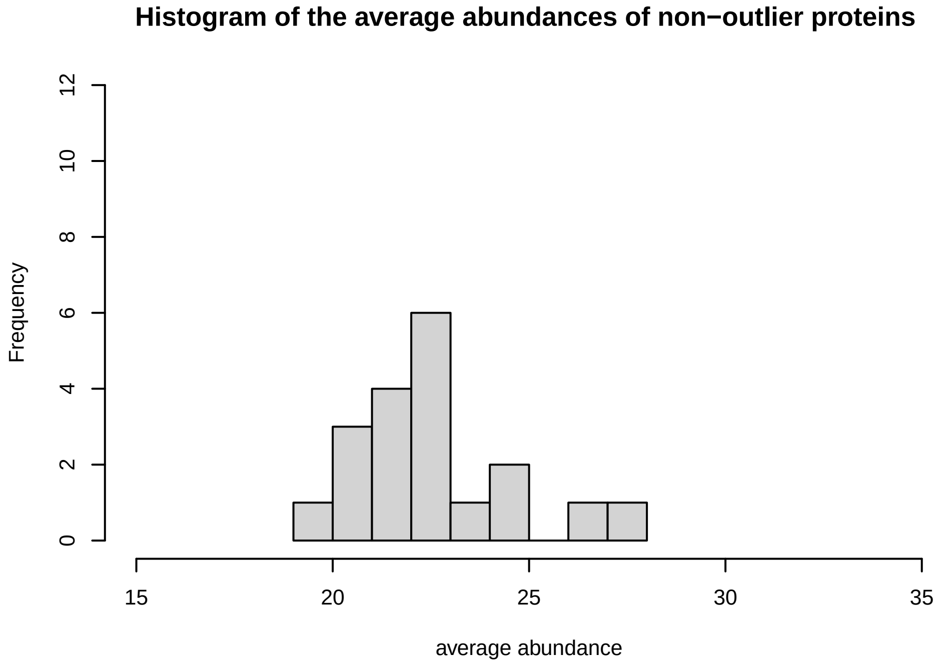
Histogram of the average abundances of non−outlier proteins
Frequency
average abundance

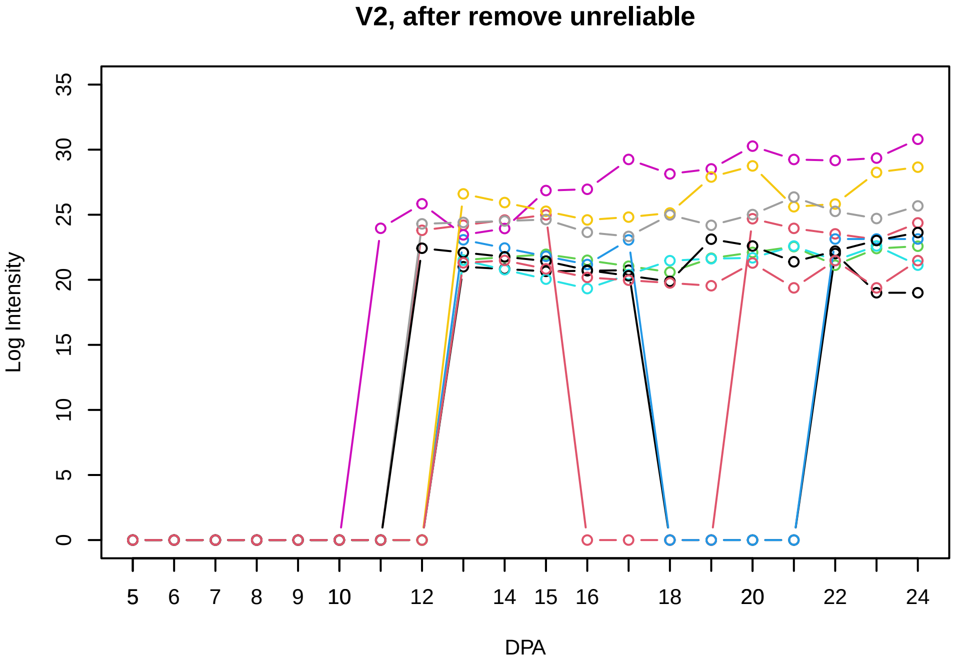
V2, after remove unreliable
Log Intensity
DPA

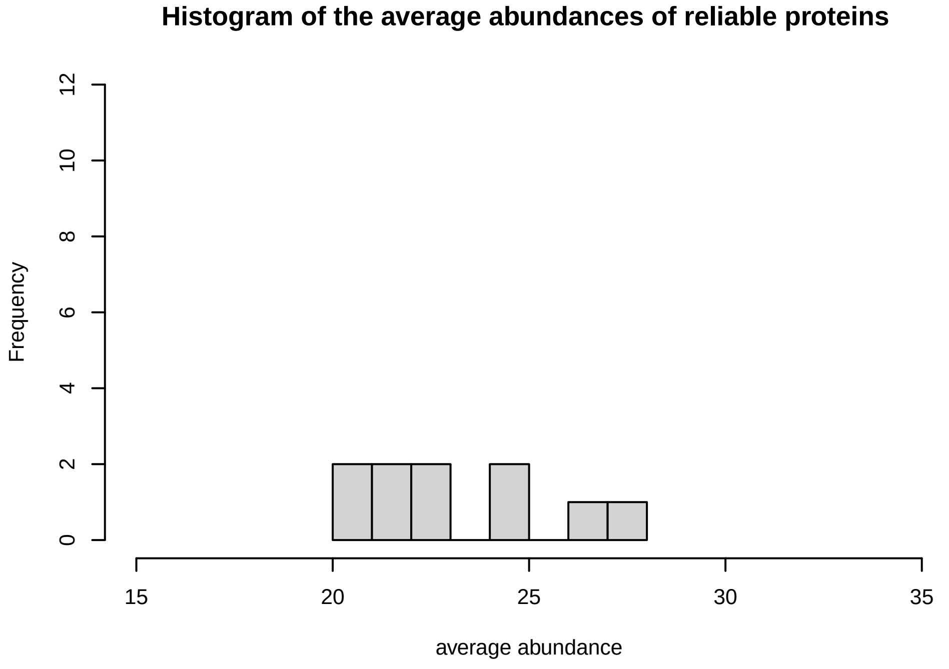
Histogram of the average abundances of reliable proteins
Frequency
average abundance

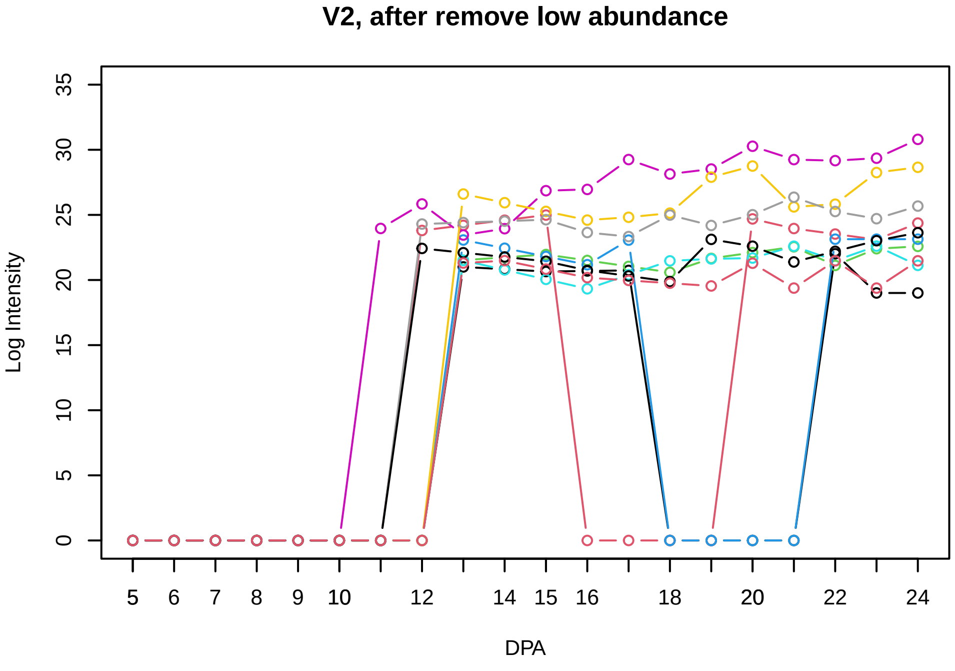
V2, after remove low abundance
Log Intensity
DPA

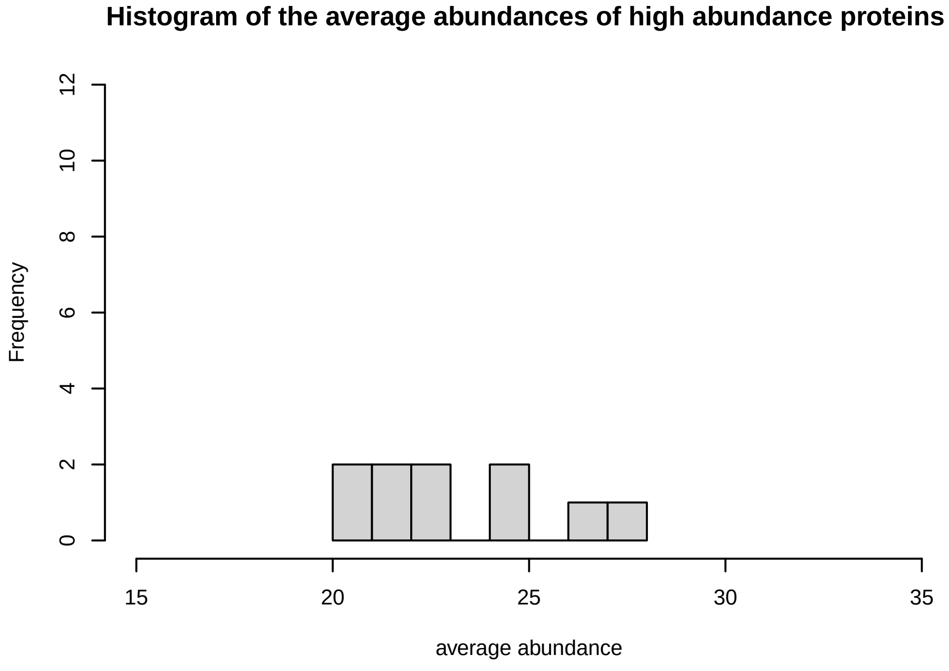
Histogram of the average abundances of high abundance proteins
Frequency
average abundance

V3

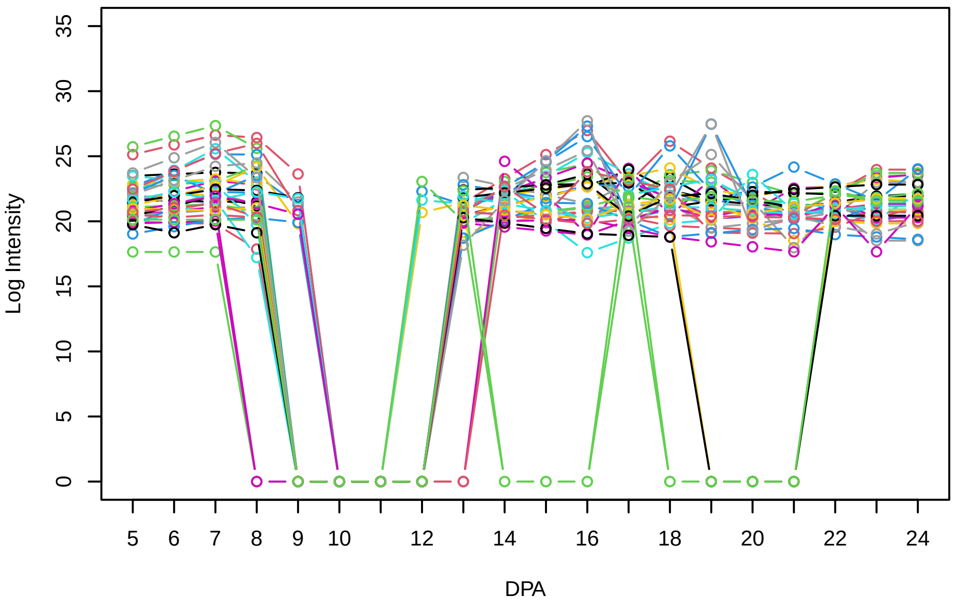


Histogram of the average abundances of non−zero values

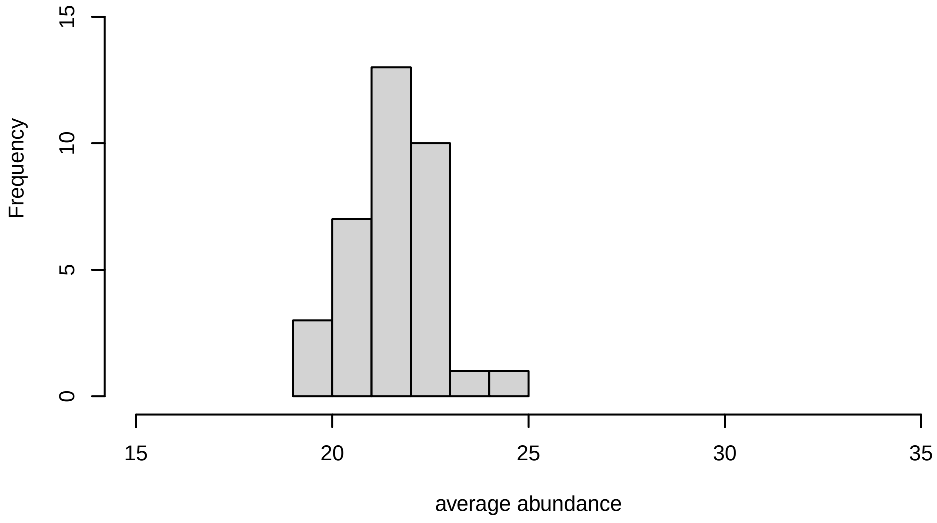


V3, after remove outlier and cutoff is 0.8

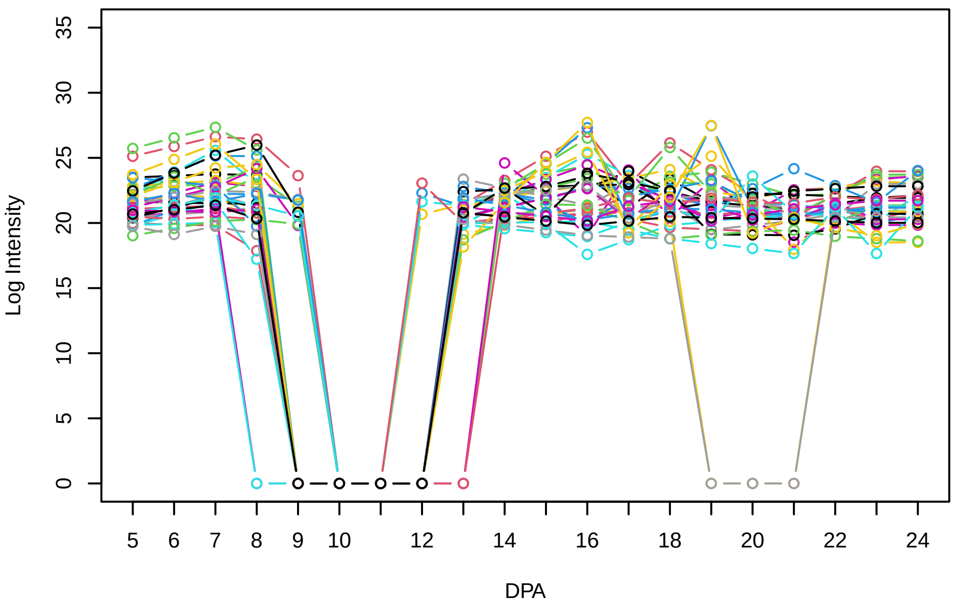


Histogram of the average abundances of non−outlier proteins

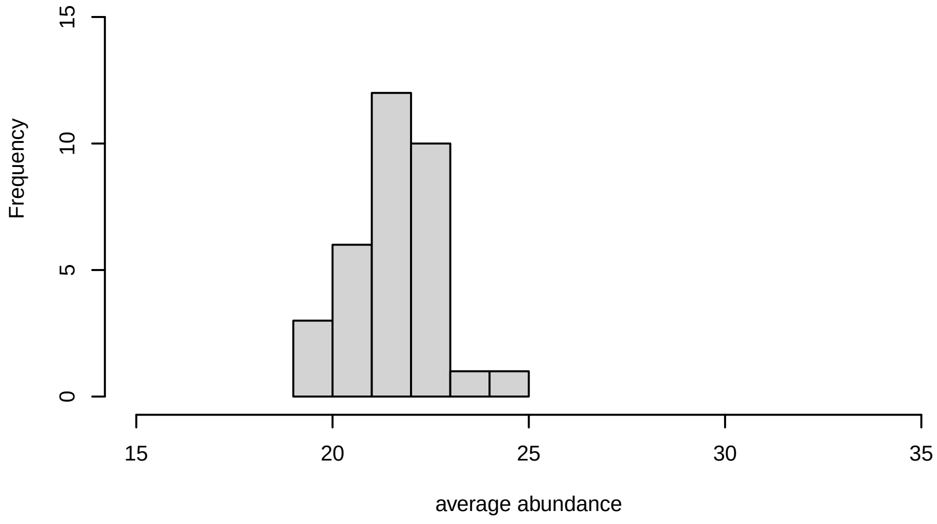


V3, after remove unreliable

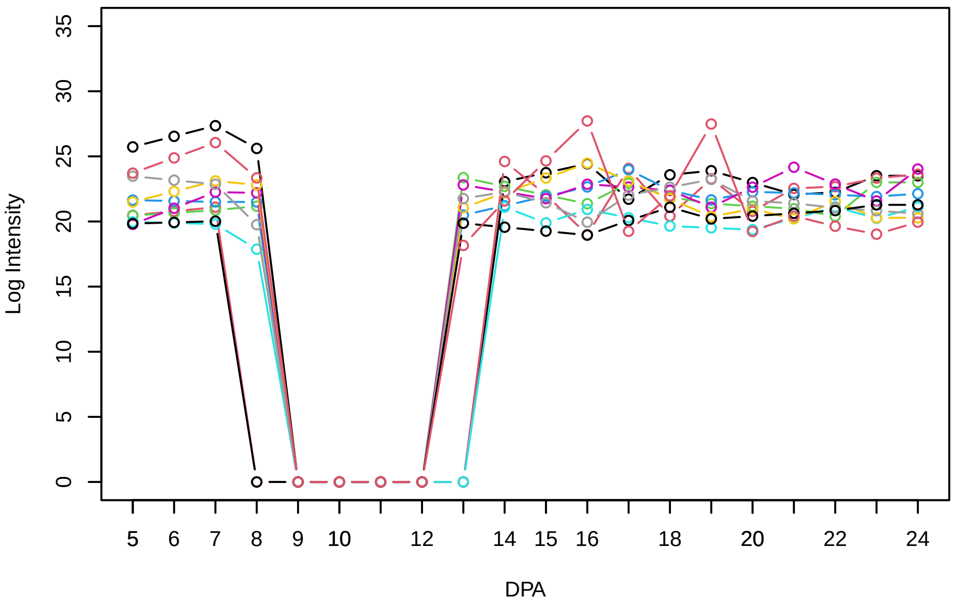


Histogram of the average abundances of reliable proteins

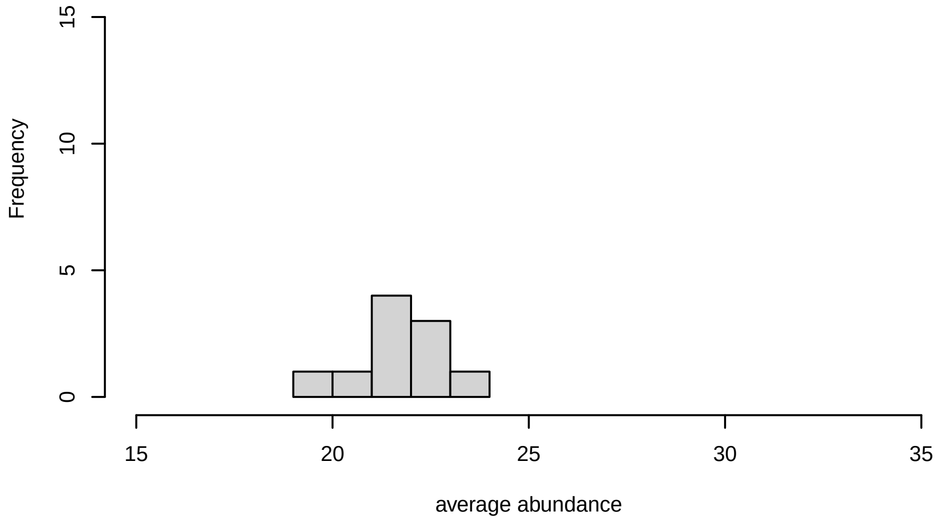


V3, after remove low abundance

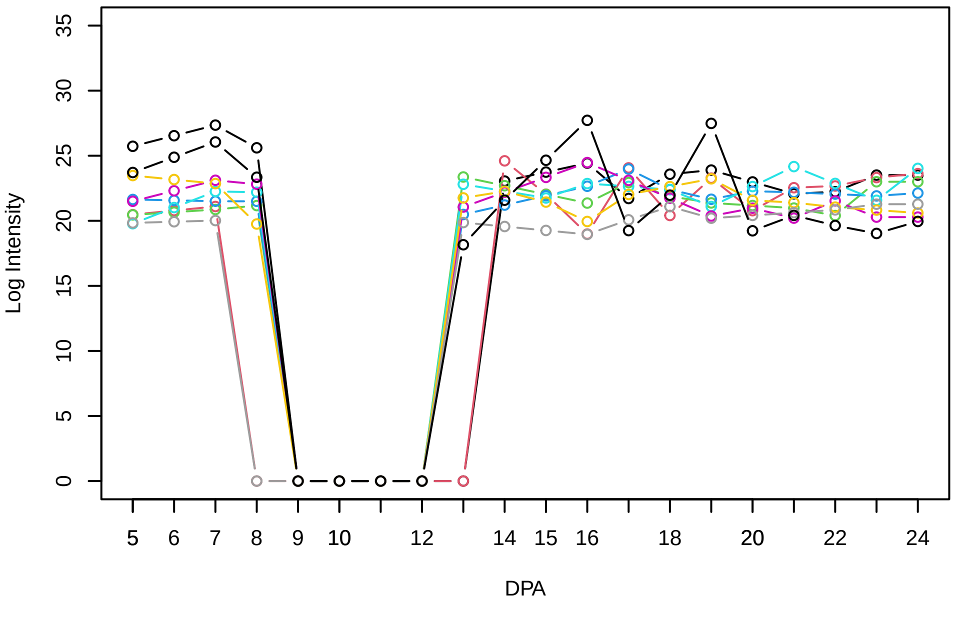


Histogram of the average abundances of high abundance proteins

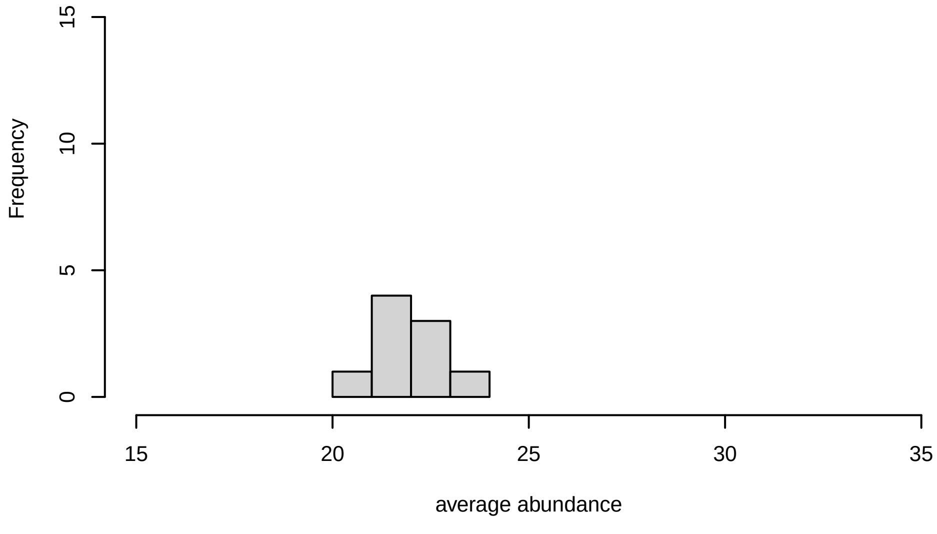

V4

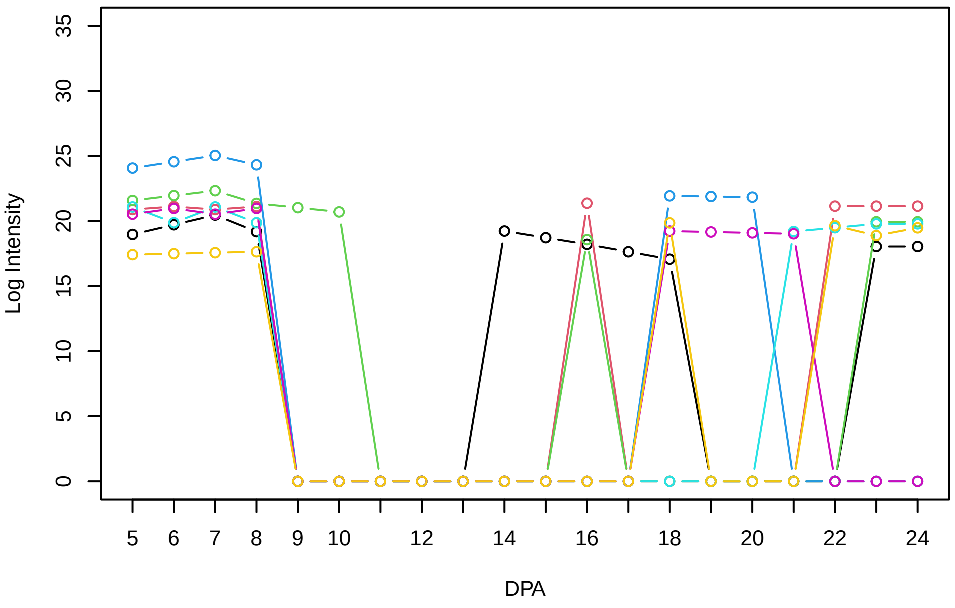


Histogram of the average abundances of non−zero values

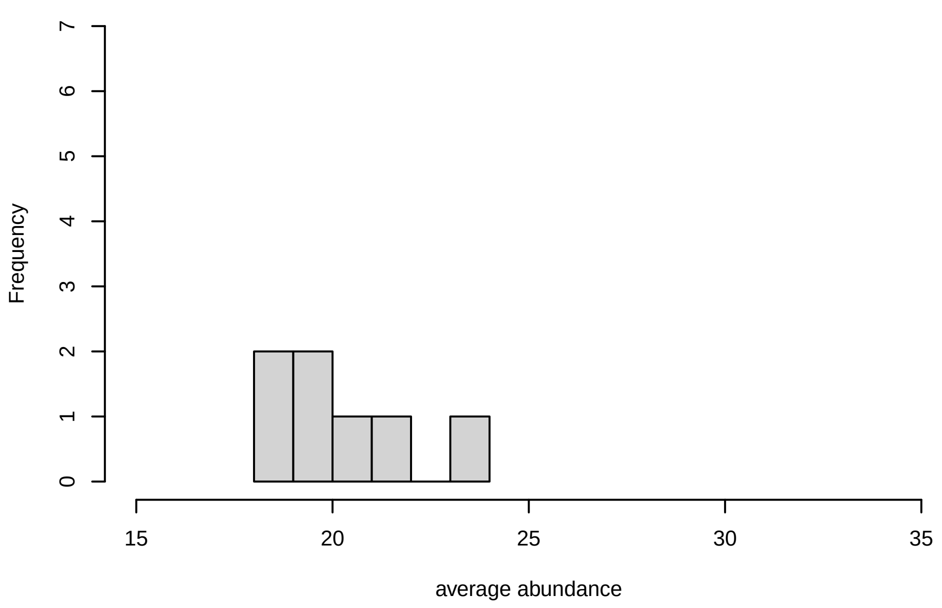


V4, after remove outlier and cutoff is 0.663396977903535

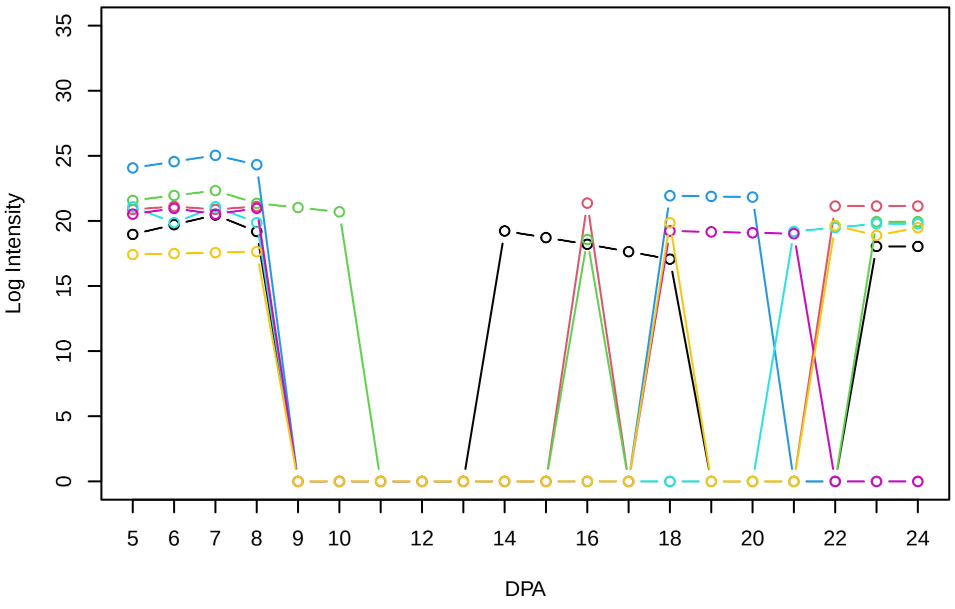


Histogram of the average abundances of non−outlier proteins

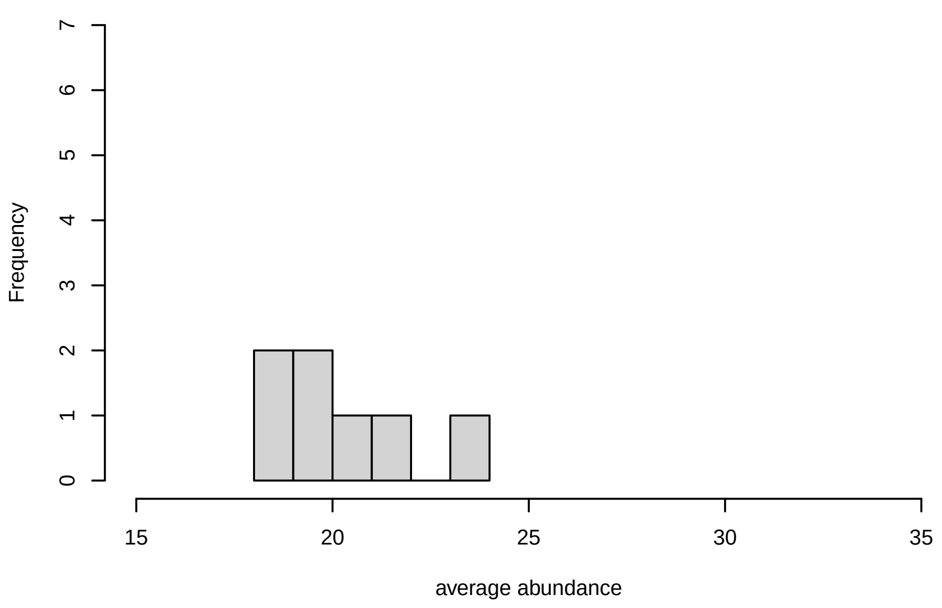


V4, after remove unreliable

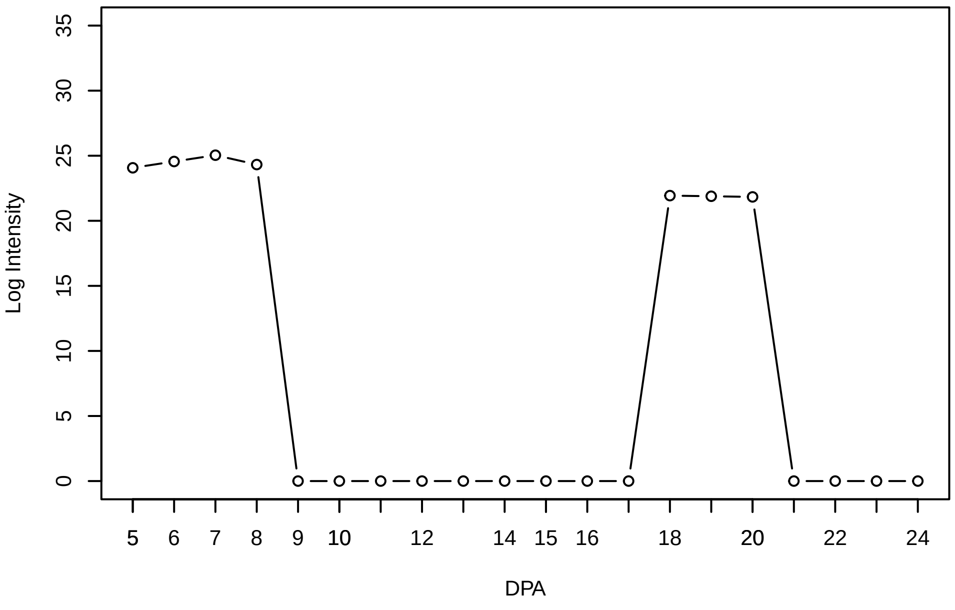


Histogram of the average abundances of reliable proteins

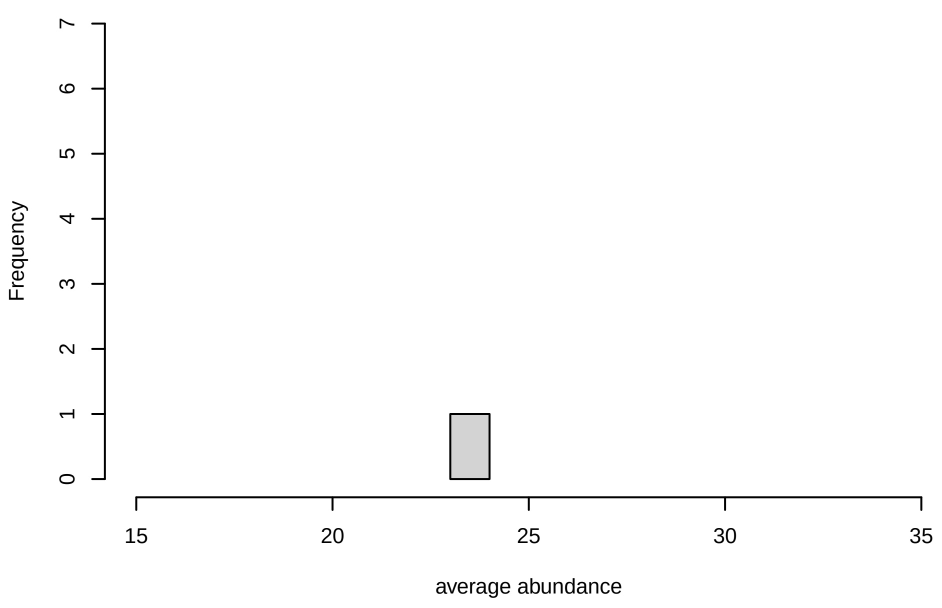


V4, after remove low abundance

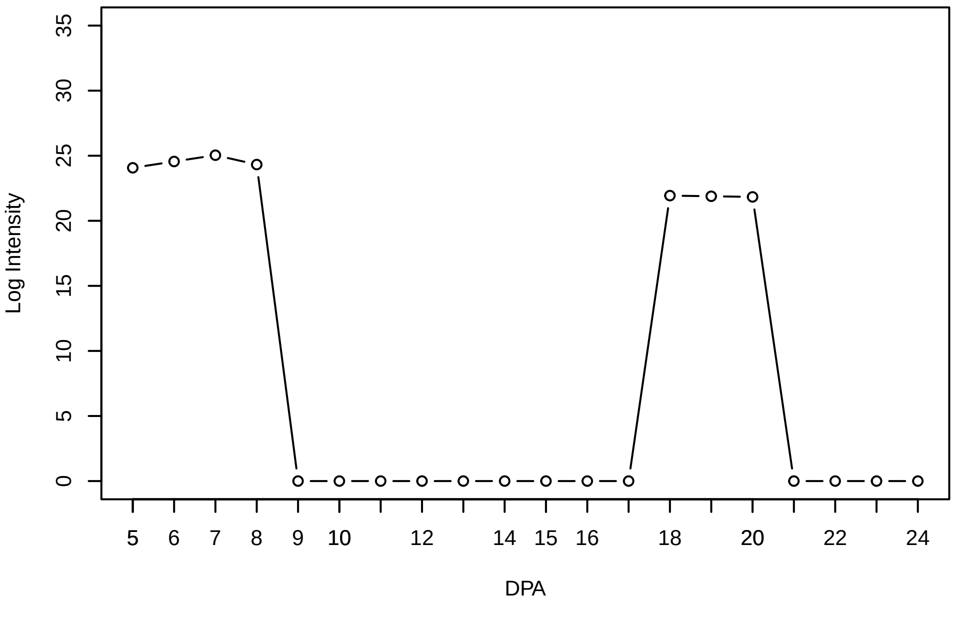


Histogram of the average abundances of high abundance proteins

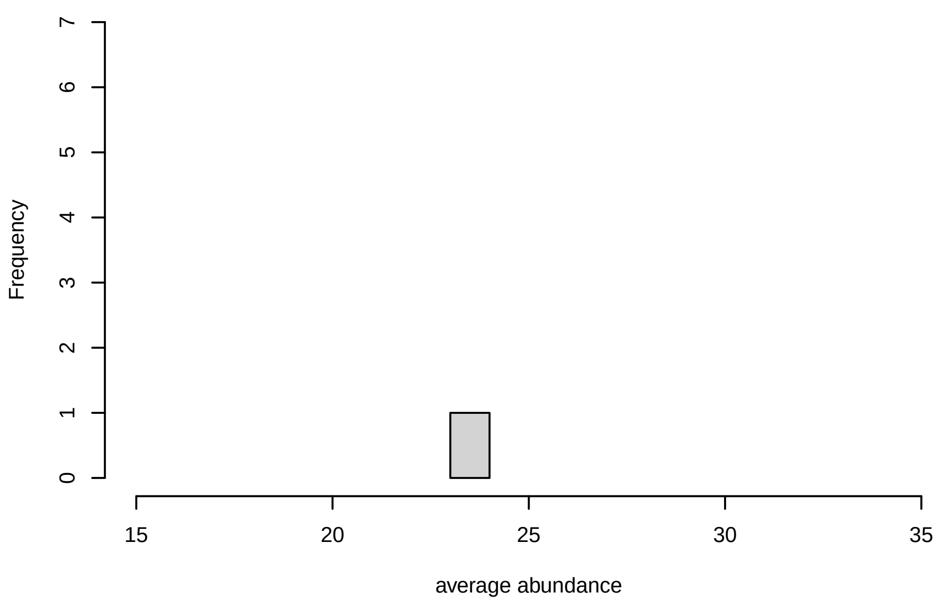

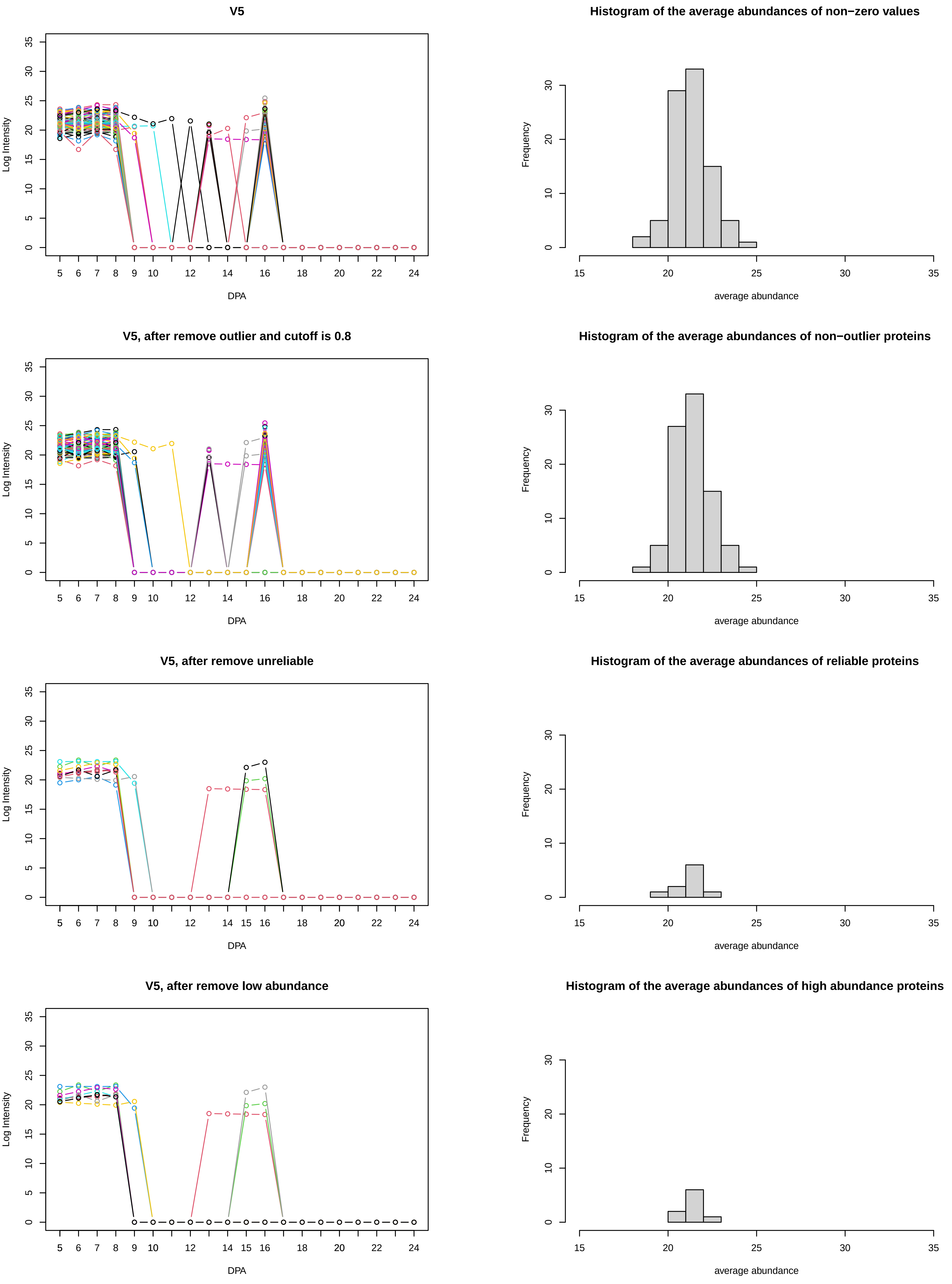
V5
Log Intensity
DPA
Histogram of the average abundances of non−zero values
Frequency
average abundance
V5, after remove outlier and cutoff is 0.8
Log Intensity
DPA
Histogram of the average abundances of non−outlier proteins
Frequency
average abundance
V5, after remove unreliable
Log Intensity
DPA
Histogram of the average abundances of reliable proteins
Frequency
average abundance
V5, after remove low abundance
Log Intensity
DPA
Histogram of the average abundances of high abundance proteins
Frequency
average abundance

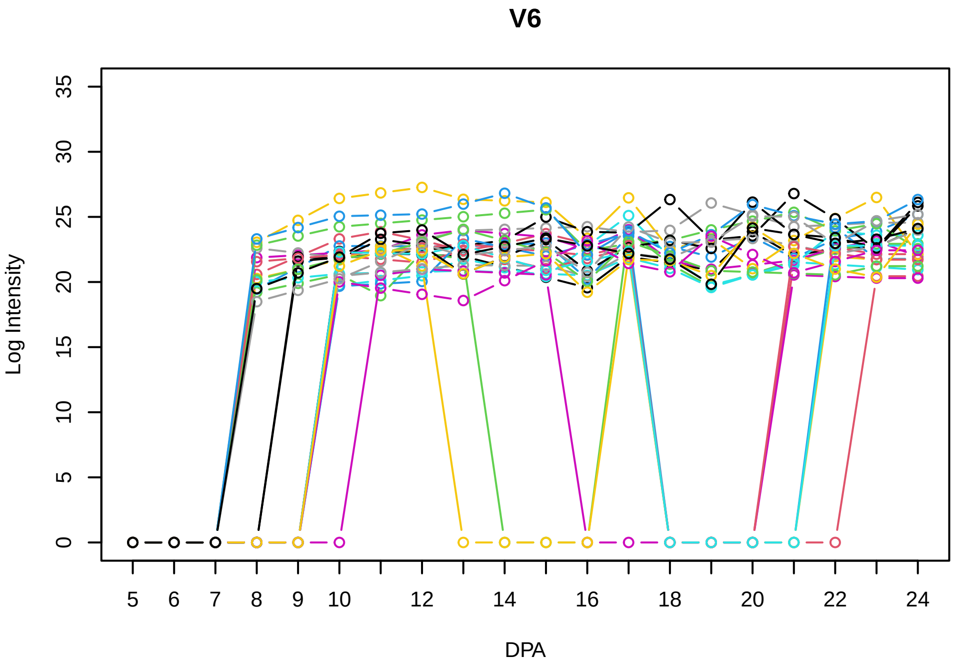
V6
Log Intensity
DPA

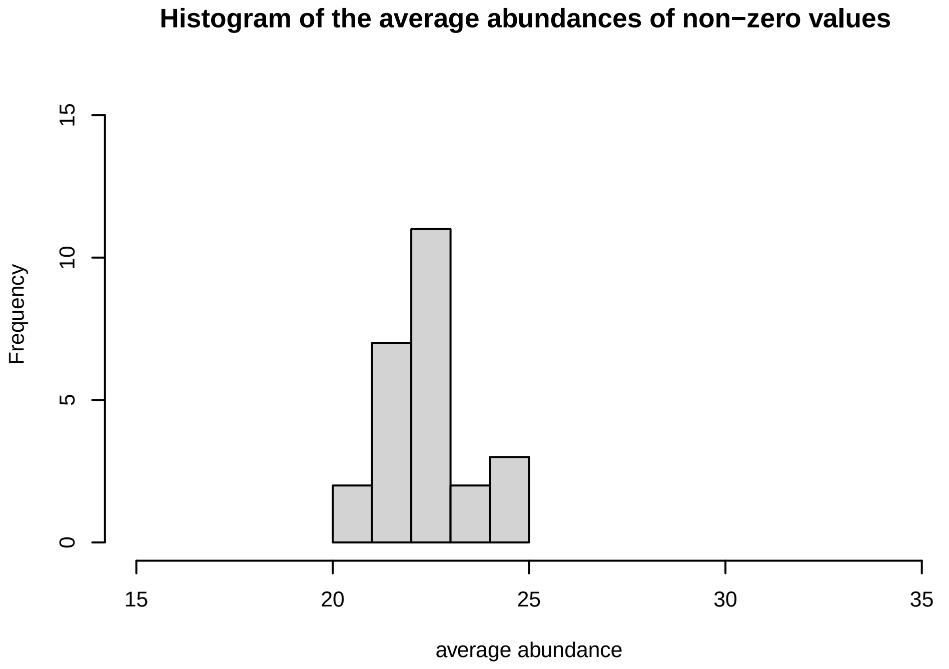
Histogram of the average abundances of non−zero values
Frequency
average abundance

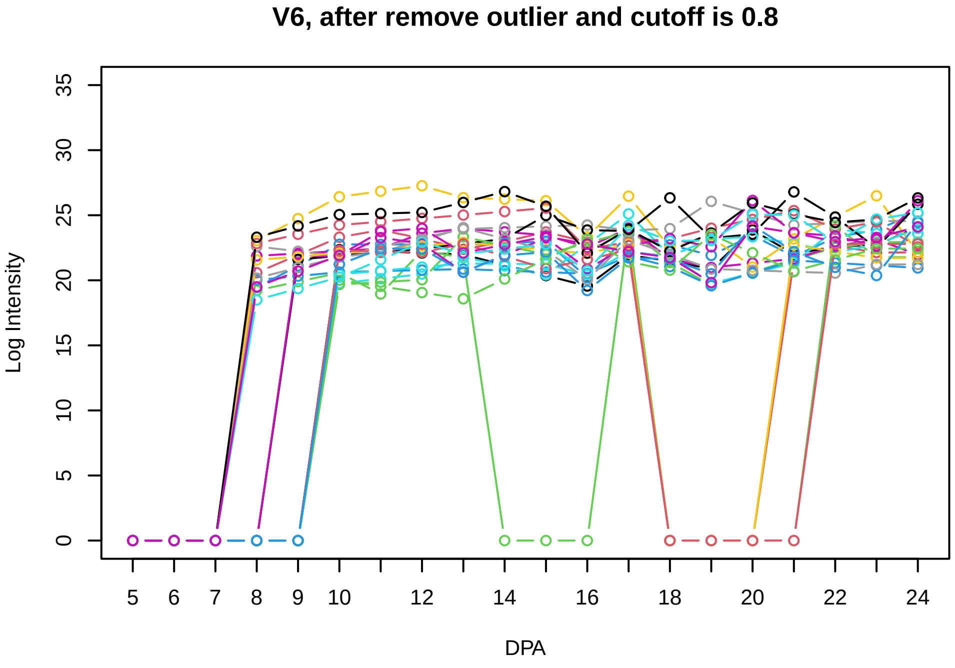
V6, after remove outlier and cutoff is 0.8
Log Intensity
DPA

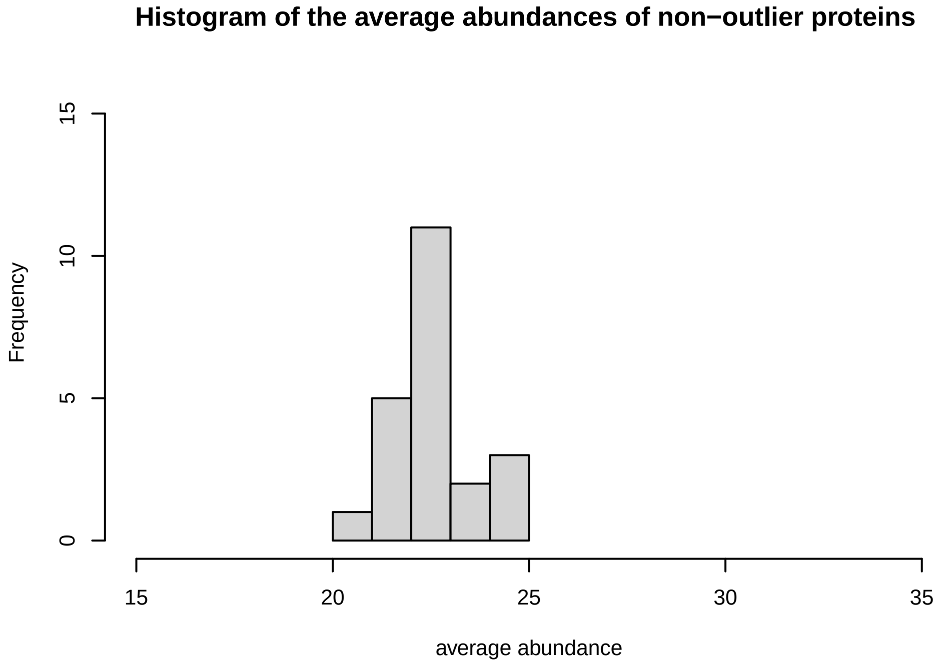
Histogram of the average abundances of non−outlier proteins
Frequency
average abundance

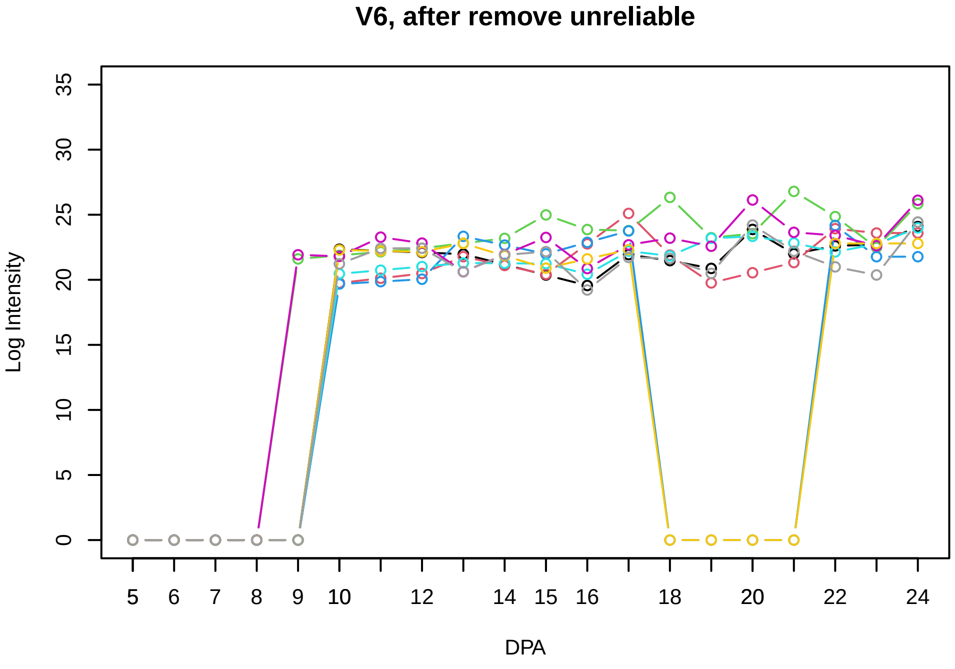
V6, after remove unreliable
Log Intensity
DPA

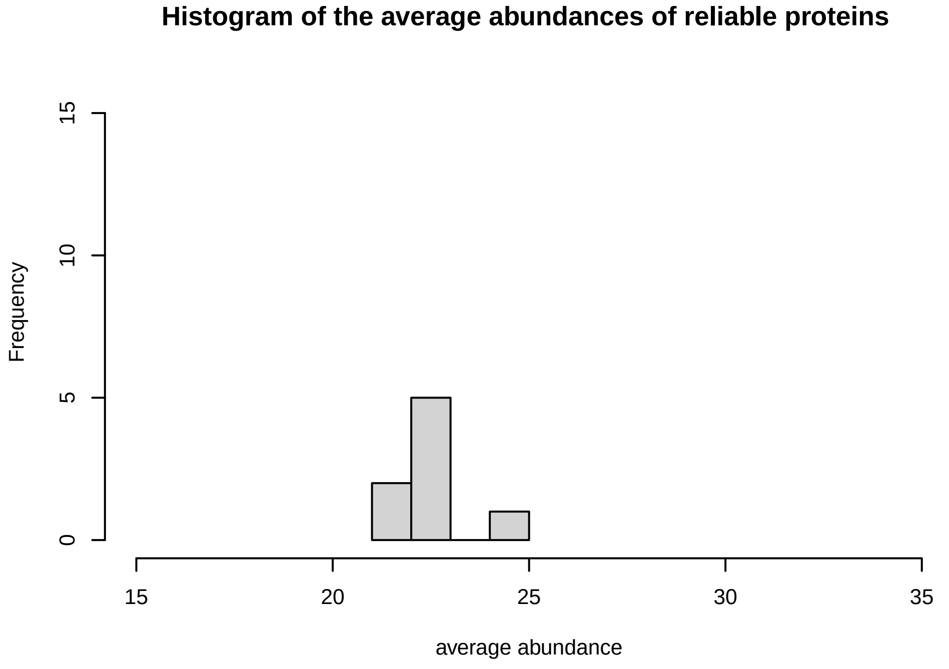
Histogram of the average abundances of reliable proteins
Frequency
average abundance

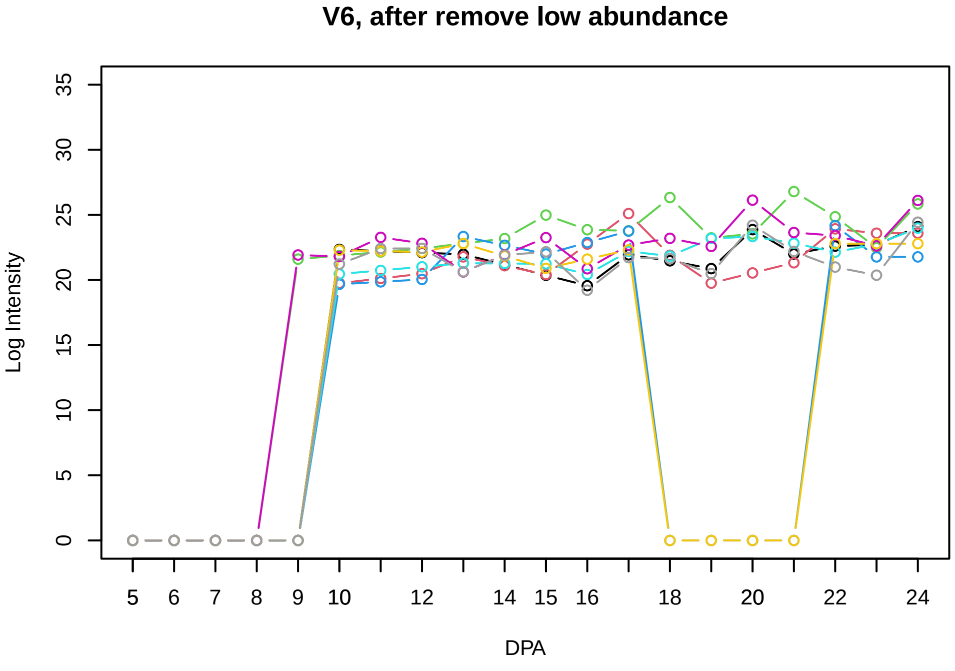
V6, after remove low abundance
Log Intensity
DPA

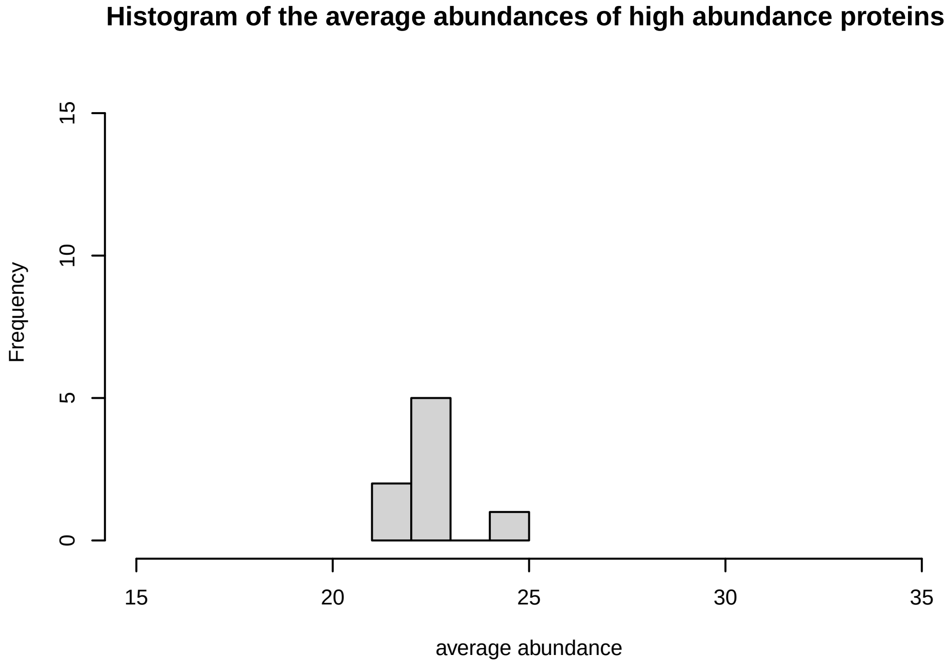
Histogram of the average abundances of high abundance proteins
Frequency
average abundance

**V7**

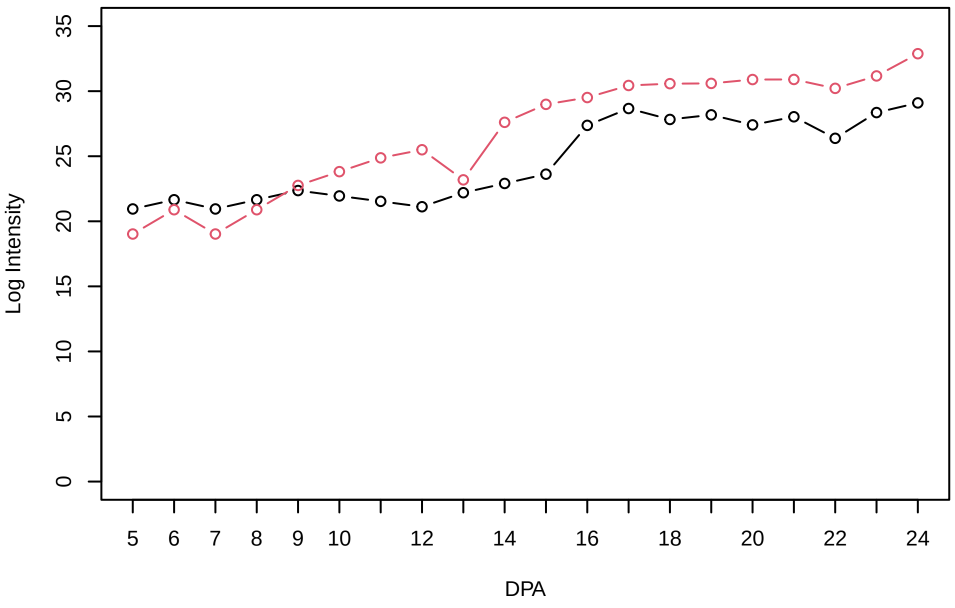


**Histogram of the average abundances of non−zero values**

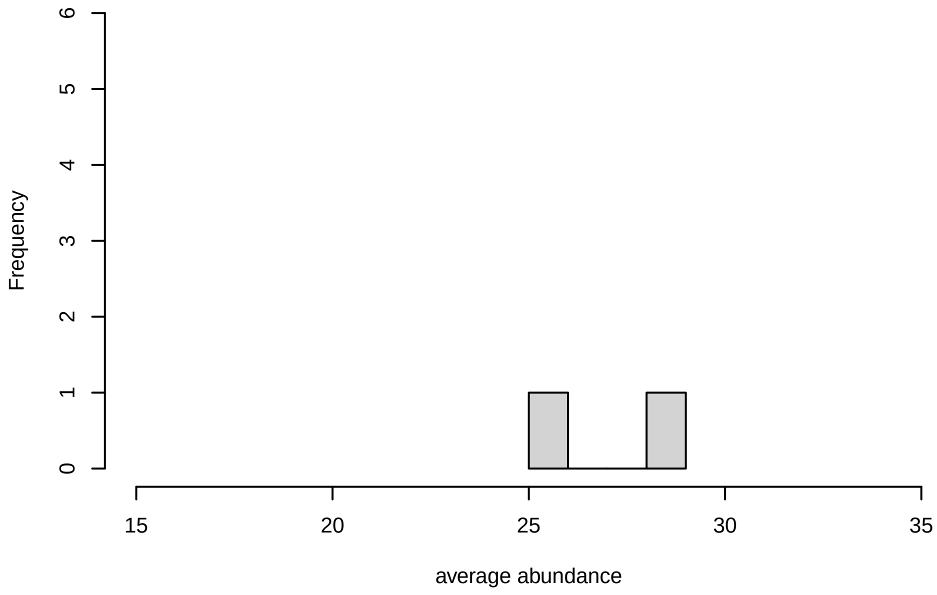


**V7, after remove outlier and cutoff is 0.8**

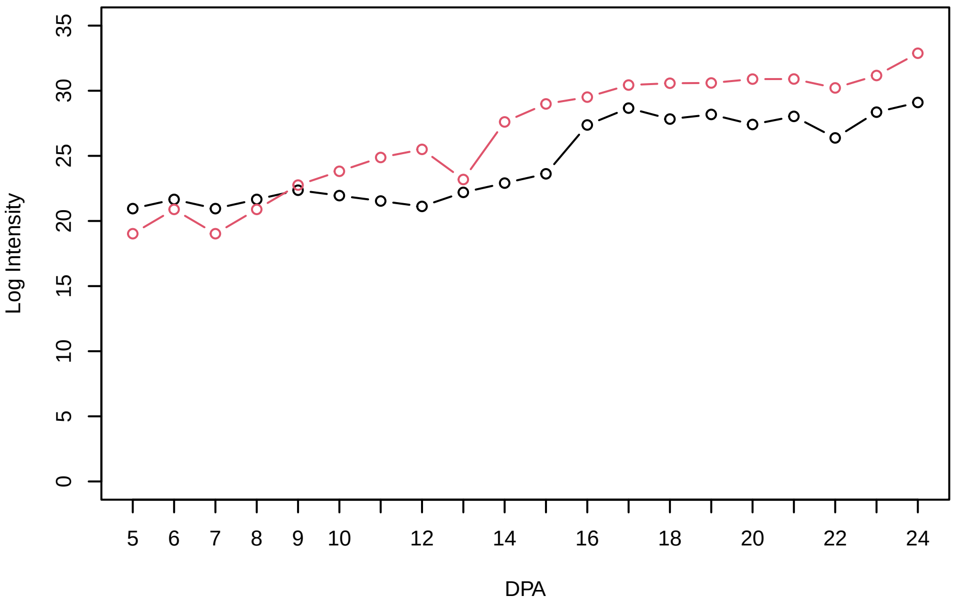


**Histogram of the average abundances of non−outlier proteins**

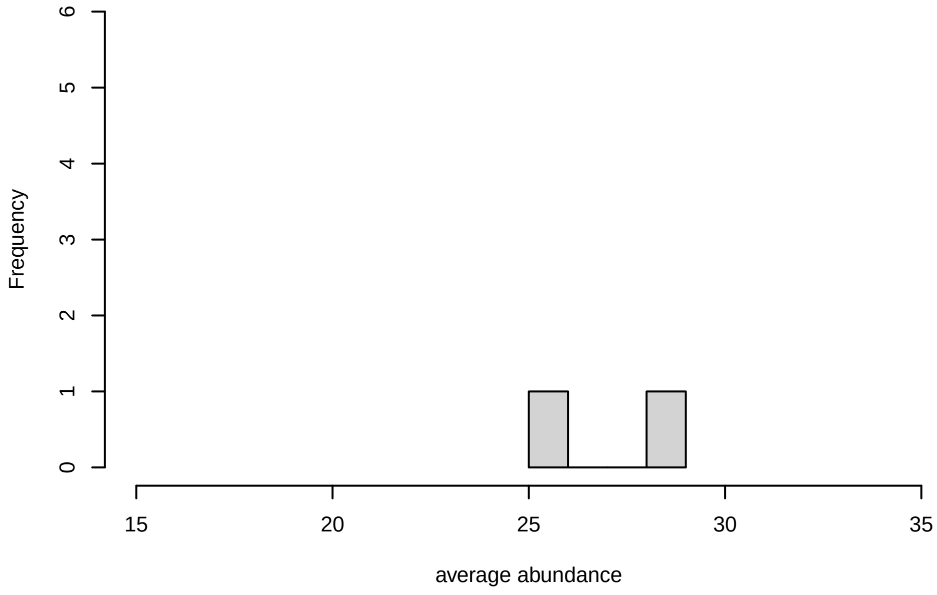


**V7, after remove unreliable**

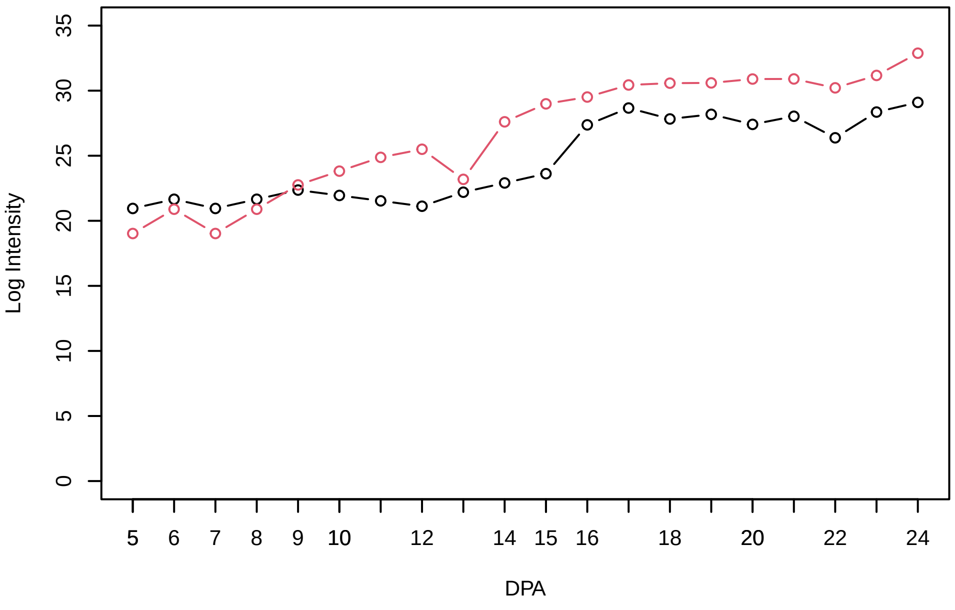


**Histogram of the average abundances of reliable proteins**

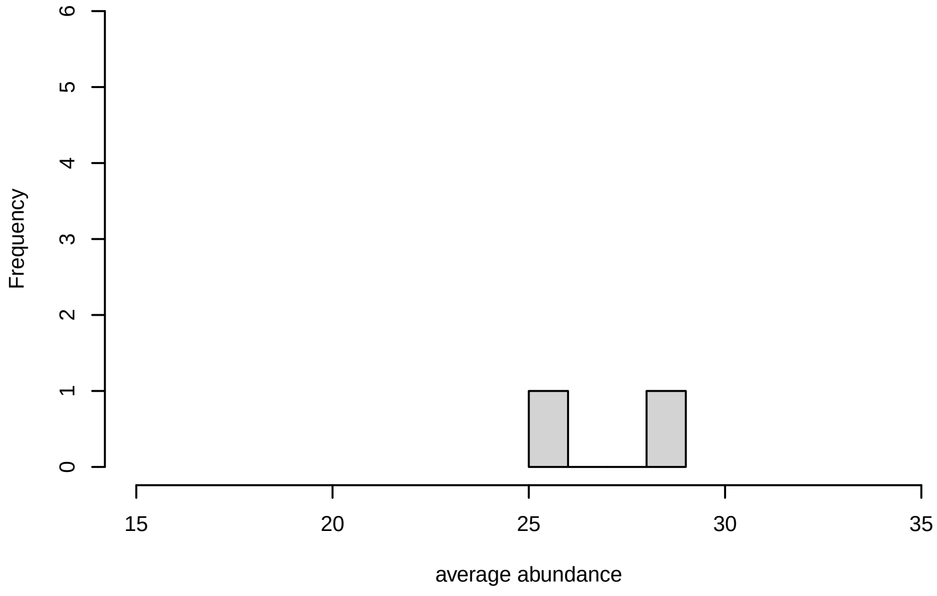


**V7, after remove low abundance**

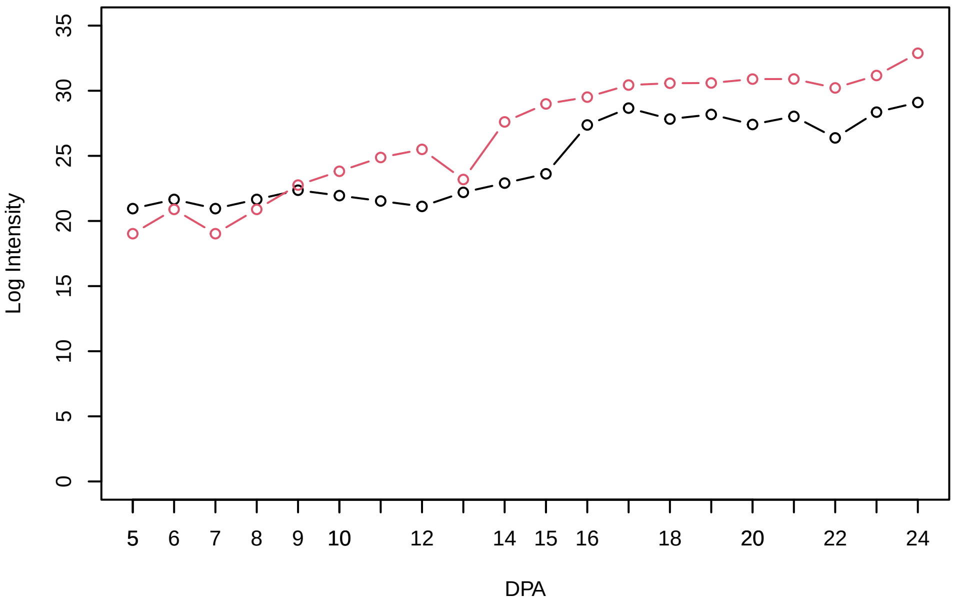


**Histogram of the average abundances of high abundance proteins**

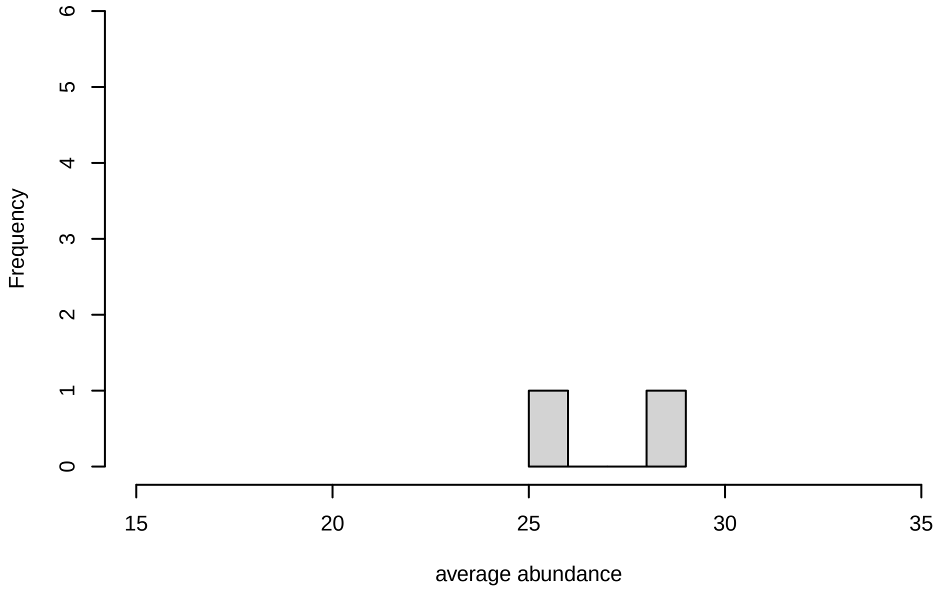

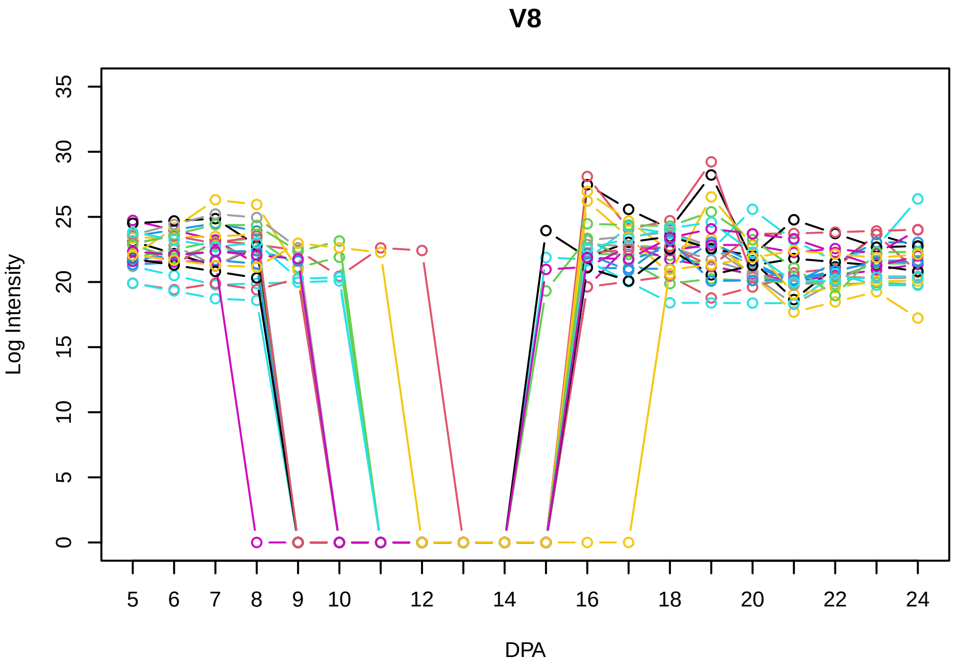
V8
Log Intensity
DPA

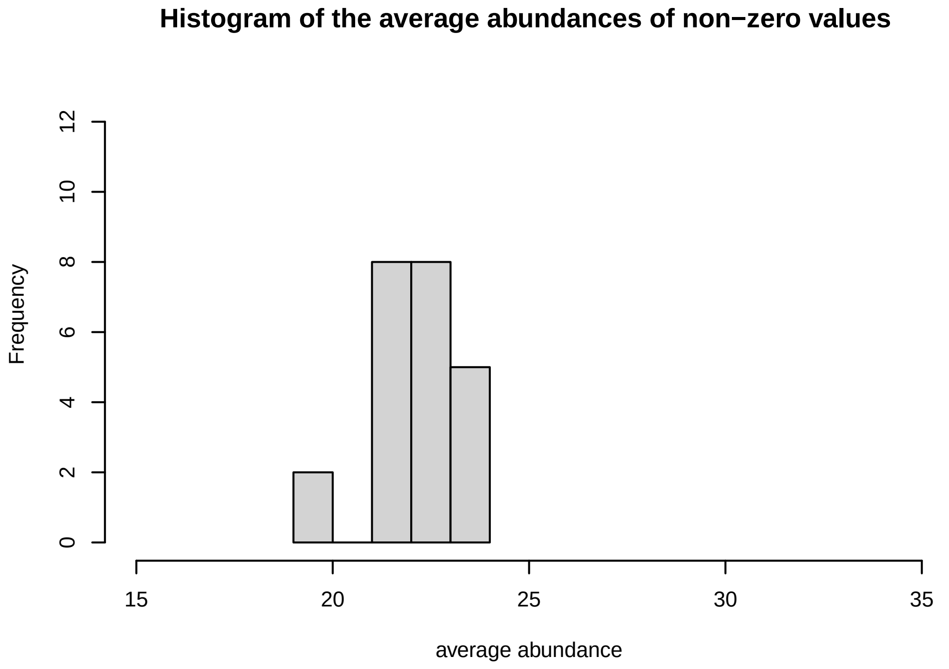
Histogram of the average abundances of non−zero values
Frequency
average abundance

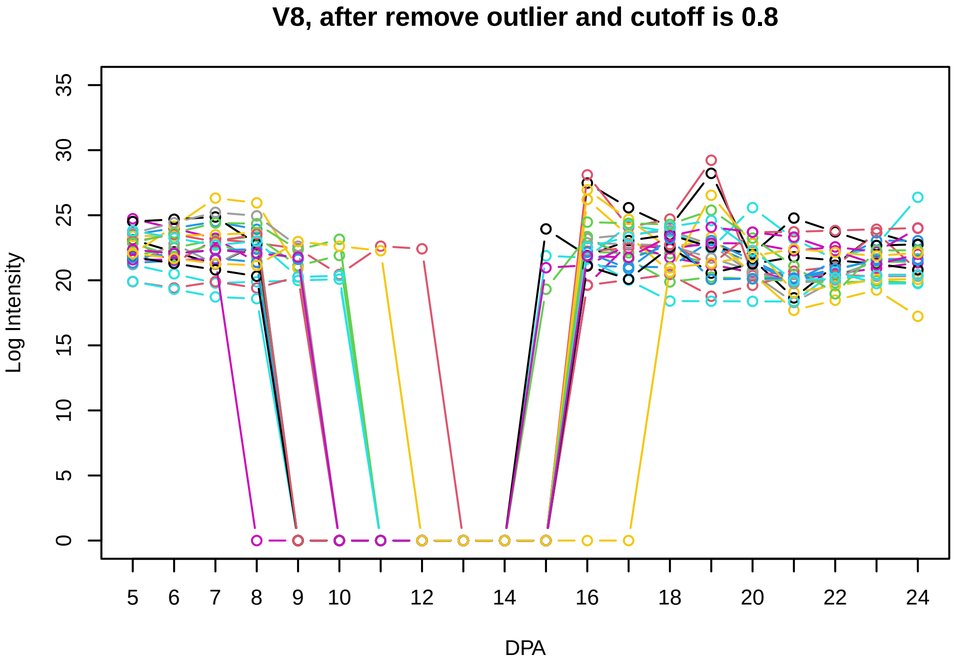
V8, after remove outlier and cutoff is 0.8
Log Intensity
DPA

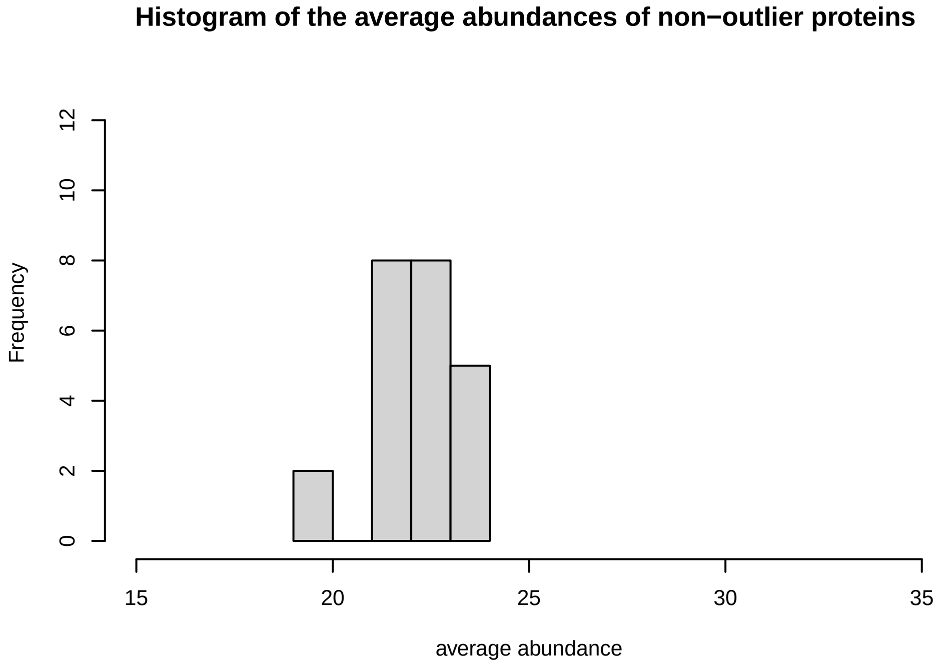
Histogram of the average abundances of non−outlier proteins
Frequency
average abundance

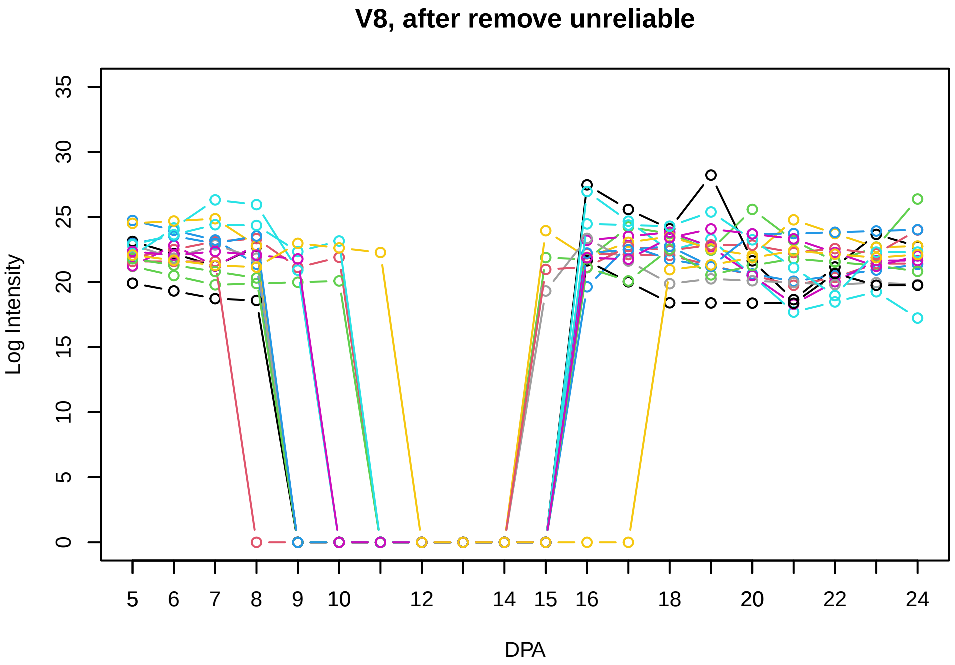
V8, after remove unreliable
Log Intensity
DPA

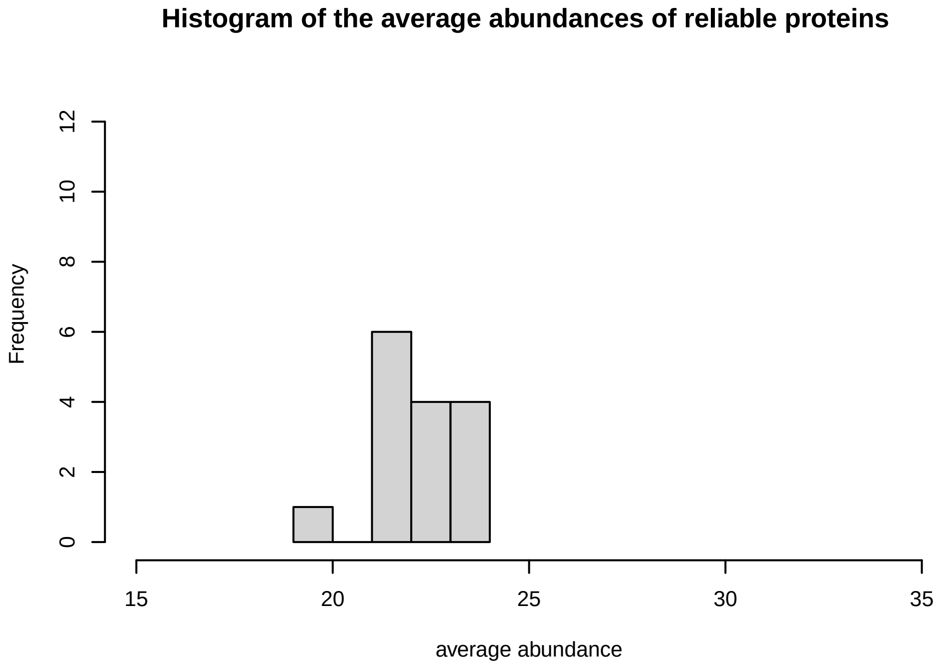
Histogram of the average abundances of reliable proteins
Frequency
average abundance

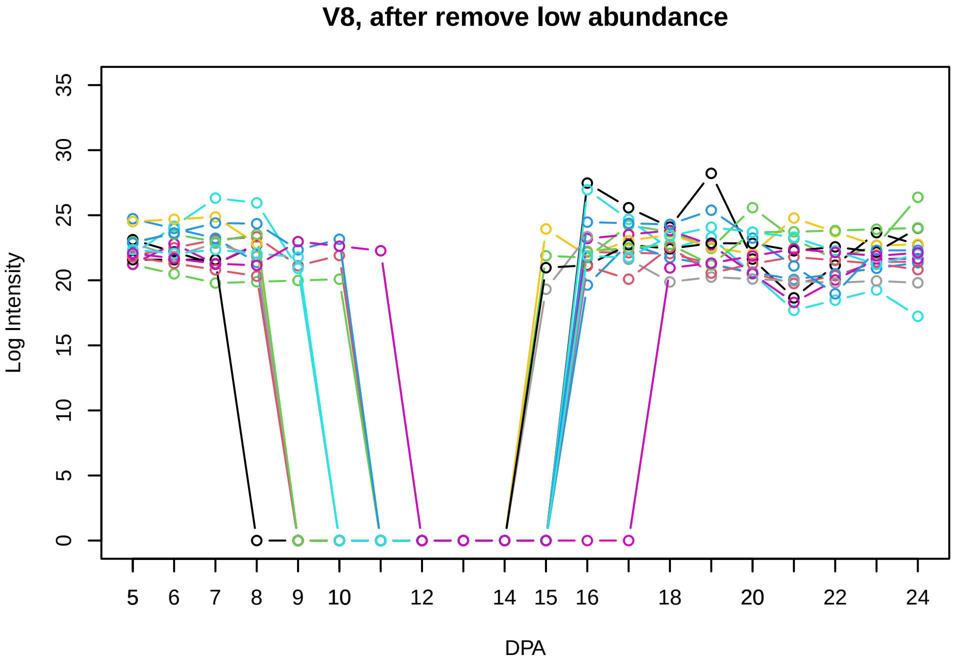
V8, after remove low abundance
Log Intensity
DPA

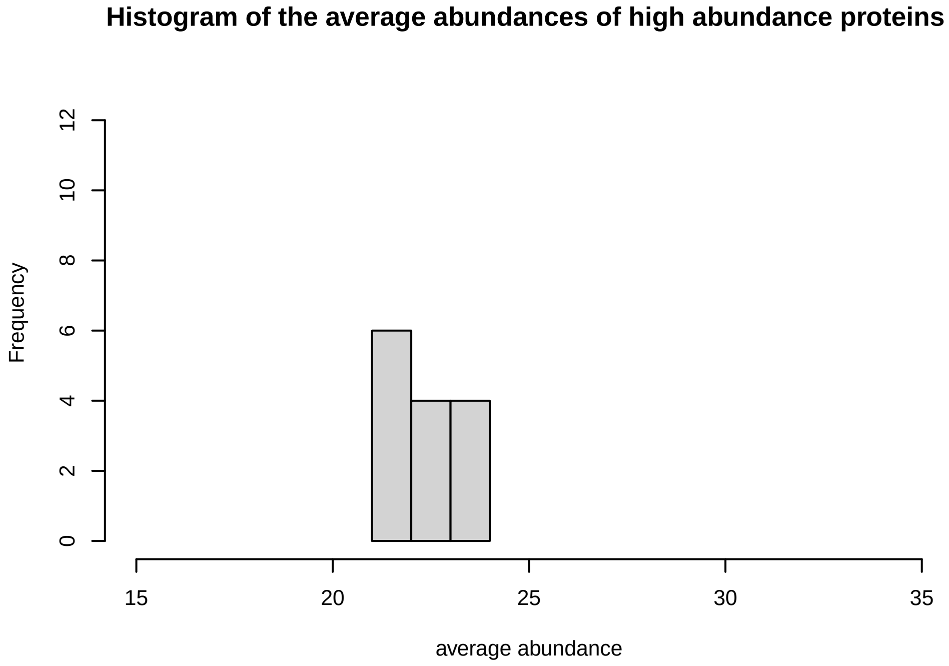
Histogram of the average abundances of high abundance proteins
Frequency
average abundance

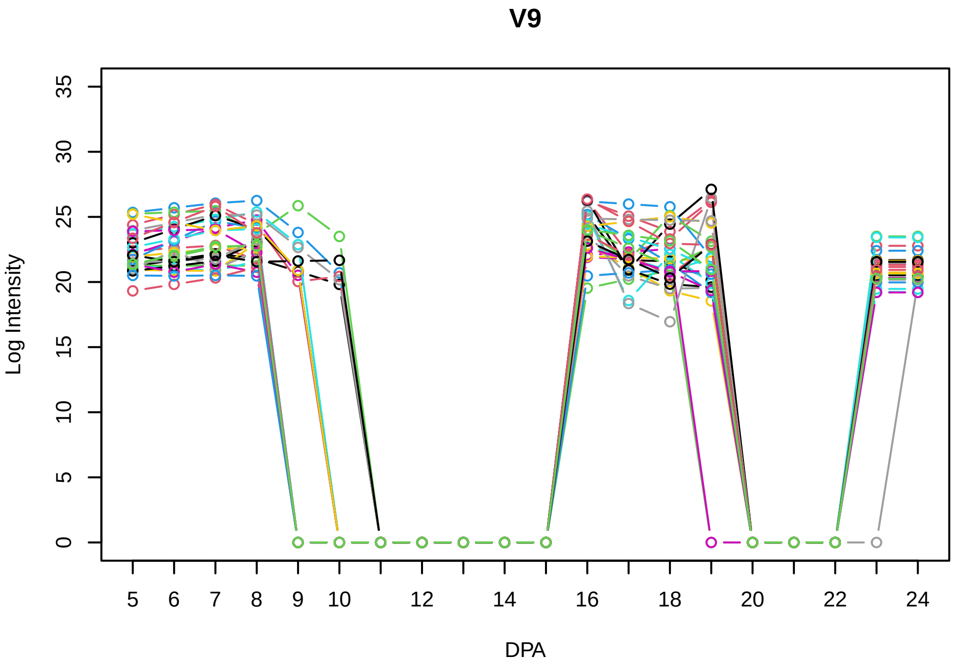
V9
Log Intensity
DPA

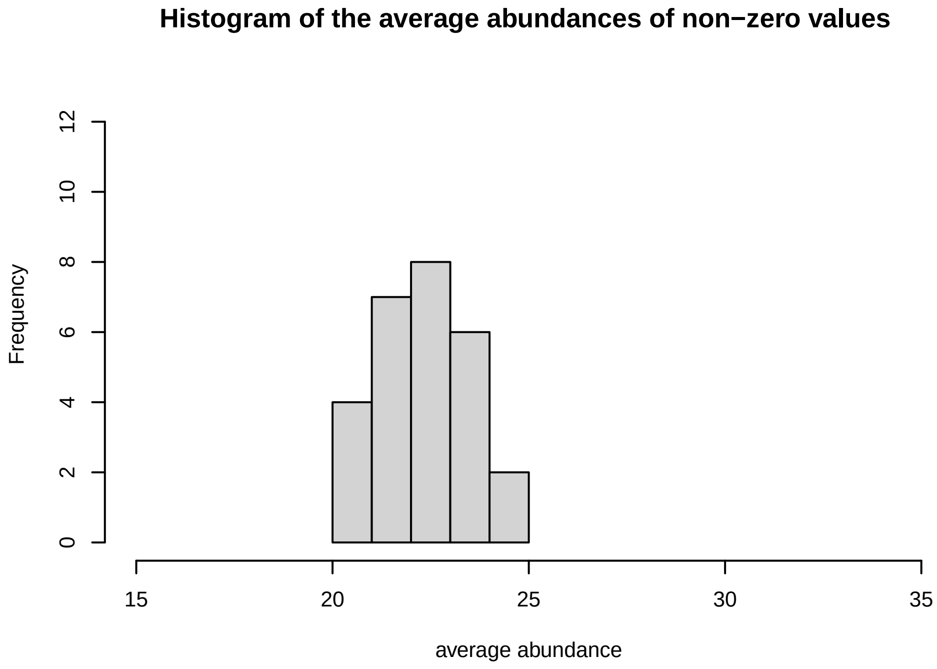
Histogram of the average abundances of non−zero values
Frequency
average abundance

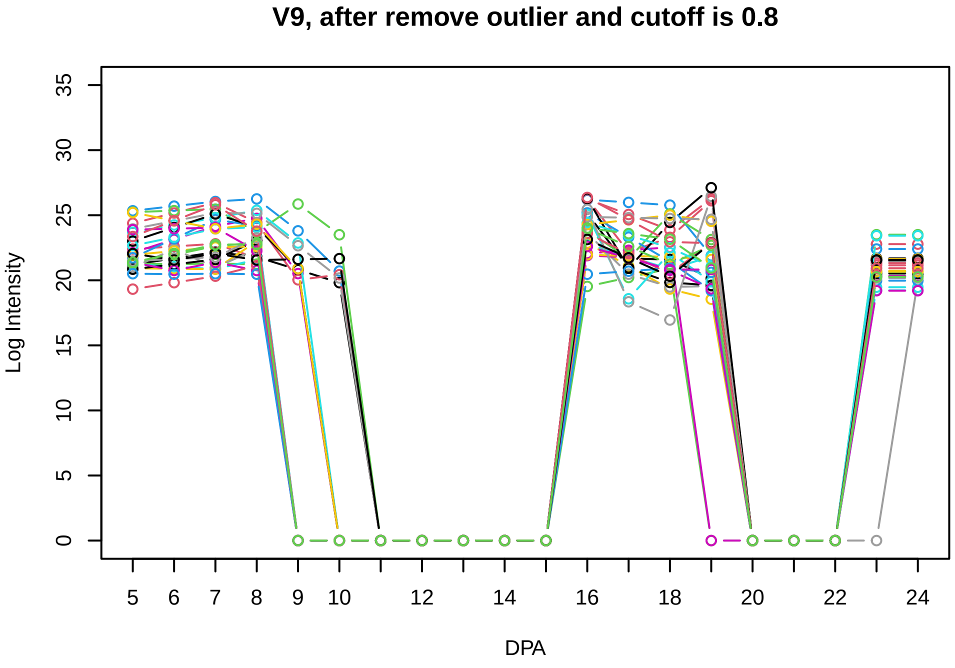
V9, after remove outlier and cutoff is 0.8
Log Intensity
DPA

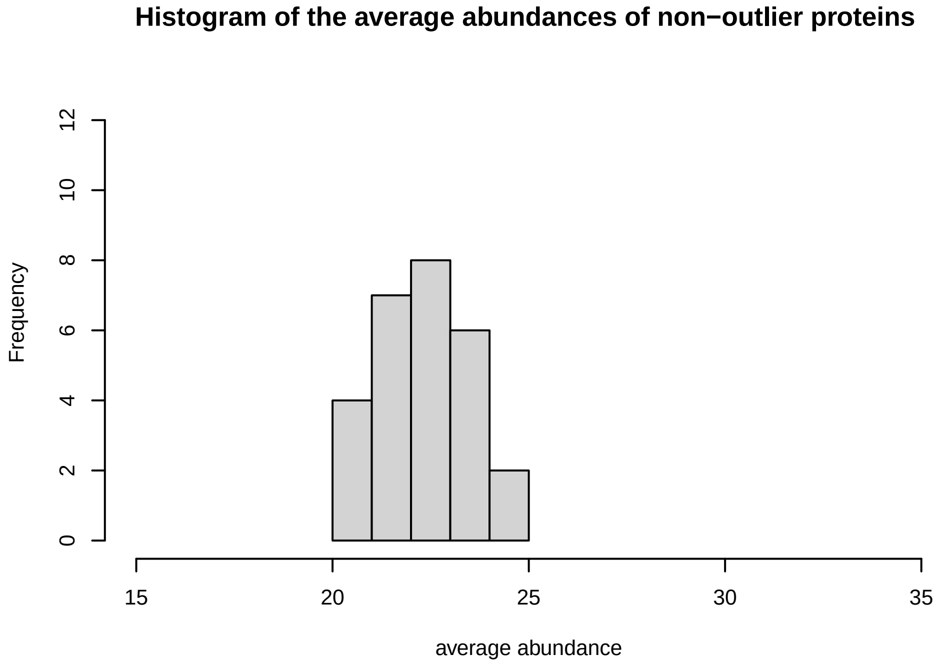
Histogram of the average abundances of non−outlier proteins
Frequency
average abundance

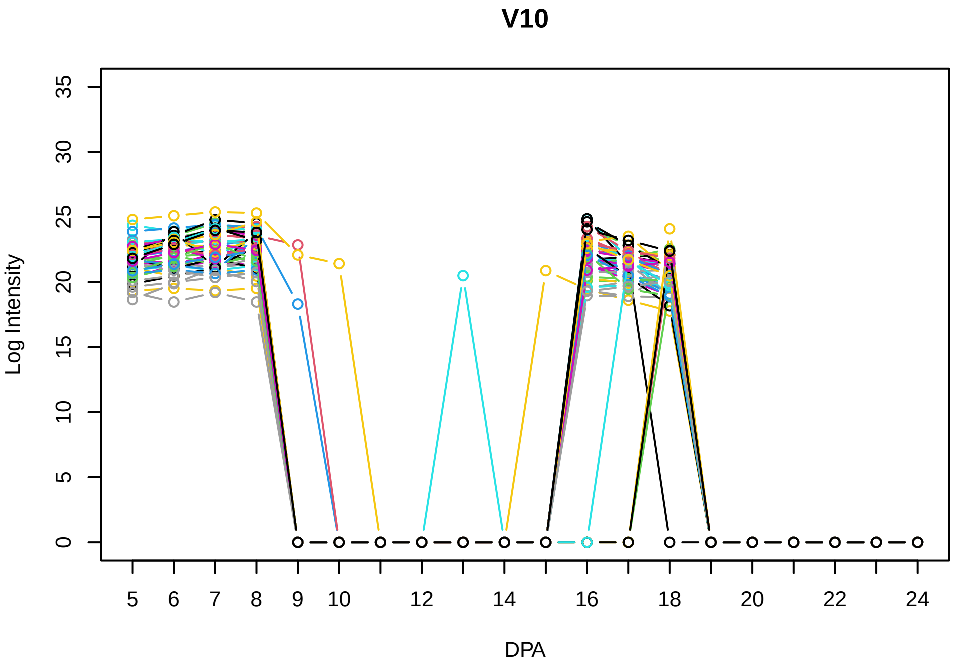
V10
Log Intensity
0 5 10 15 20 25 30 35
5 6 7 8 9 10 12 14 16 18 20 22 24
DPA

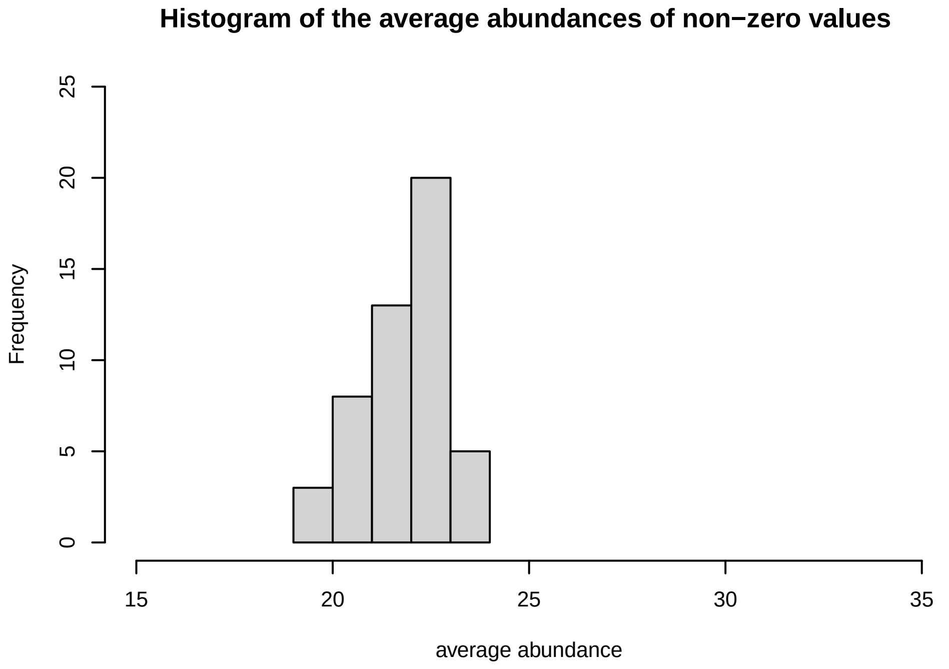
Histogram of the average abundances of non−zero values
Frequency
0 5 10 15 20 25
15 20 25 30 35
average abundance

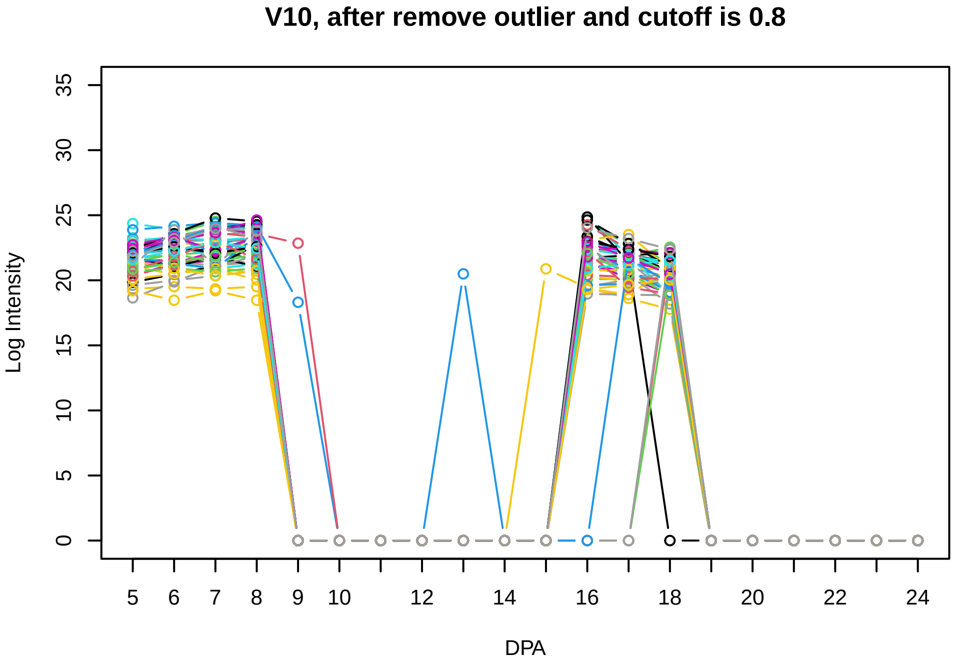
V10, after remove outlier and cutoff is 0.8
Log Intensity
0 5 10 15 20 25 30 35
5 6 7 8 9 10 12 14 16 18 20 22 24
DPA

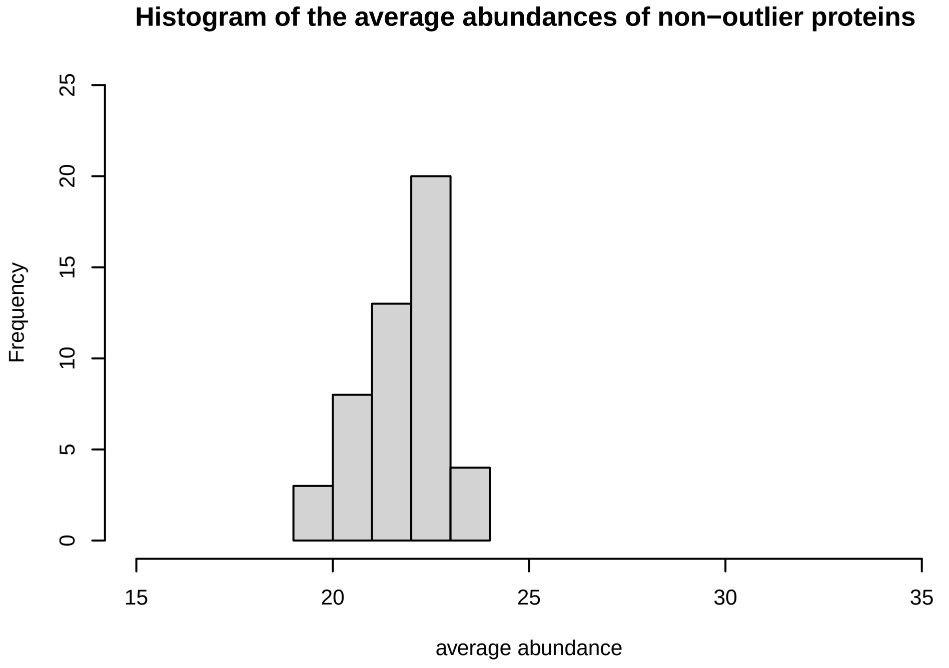
Histogram of the average abundances of non−outlier proteins
Frequency
0 5 10 15 20 25
15 20 25 30 35
average abundance

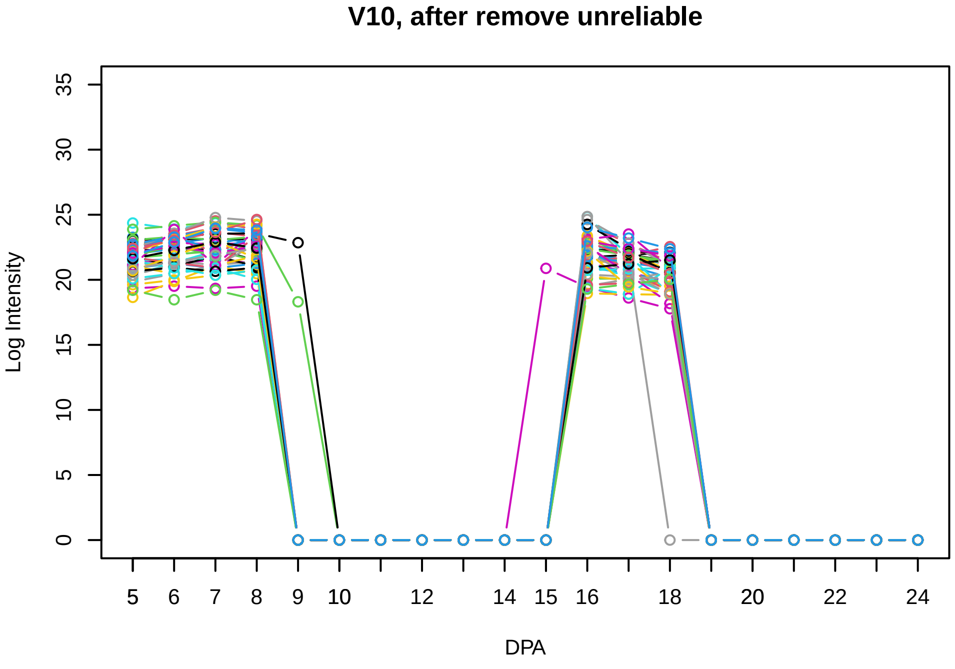
V10, after remove unreliable
Log Intensity
0 5 10 15 20 25 30 35
5 6 7 8 9 10 12 14 15 16 18 20 22 24
DPA

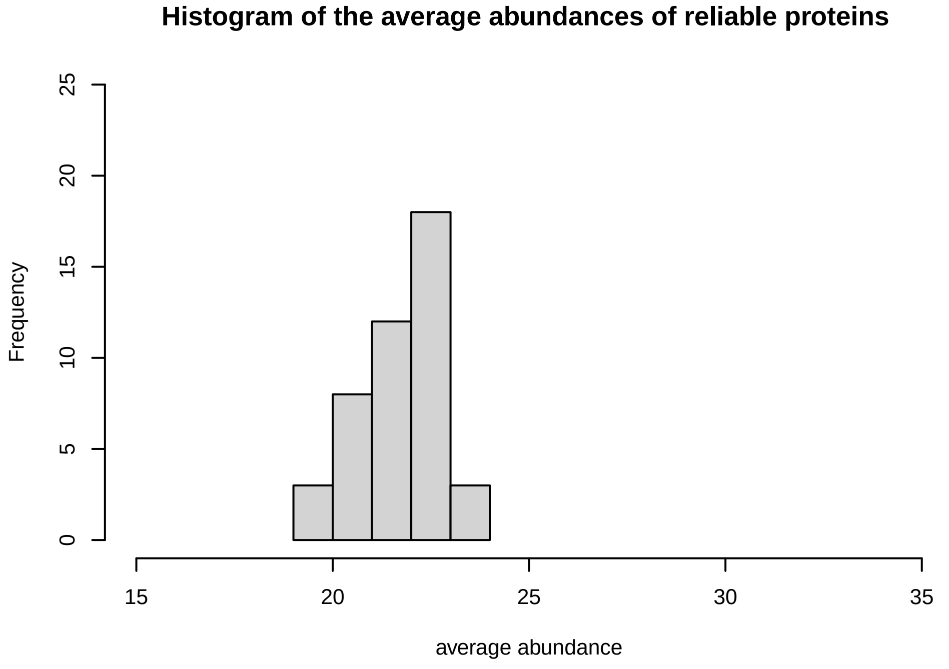
Histogram of the average abundances of reliable proteins
Frequency
0 5 10 15 20 25
15 20 25 30 35
average abundance

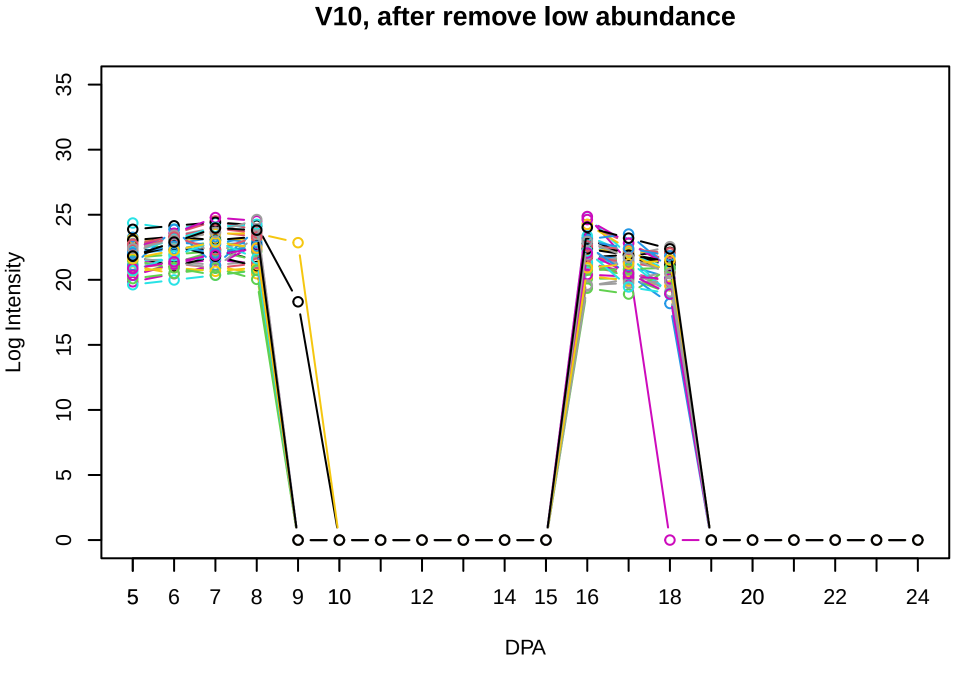
V10, after remove low abundance
Log Intensity
0 5 10 15 20 25 30 35
5 6 7 8 9 10 12 14 15 16 18 20 22 24
DPA

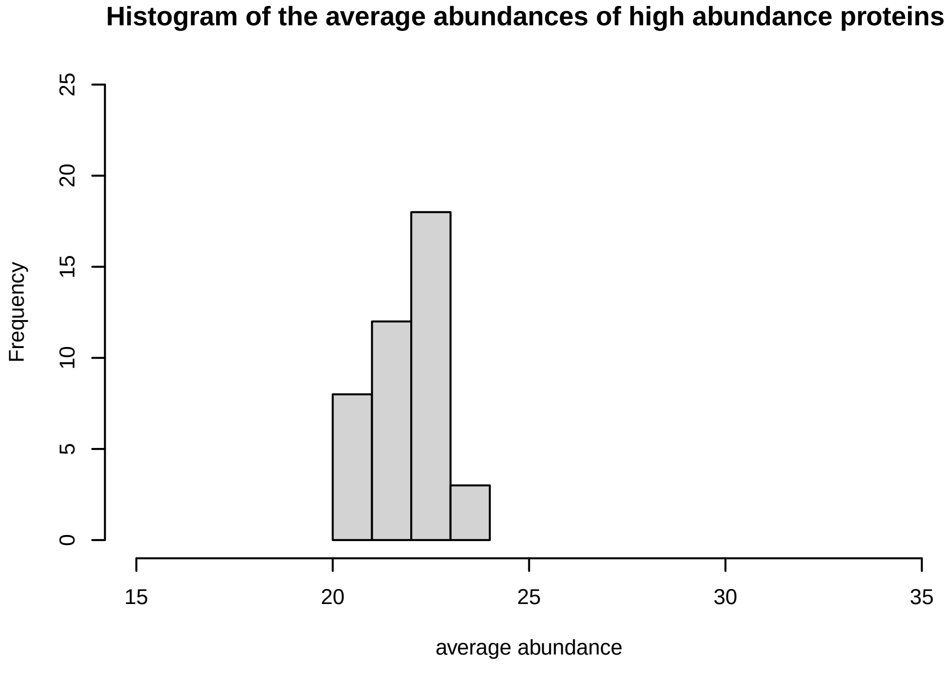
Histogram of the average abundances of high abundance proteins
Frequency
0 5 10 15 20 25
15 20 25 30 35
average abundance

**V11**

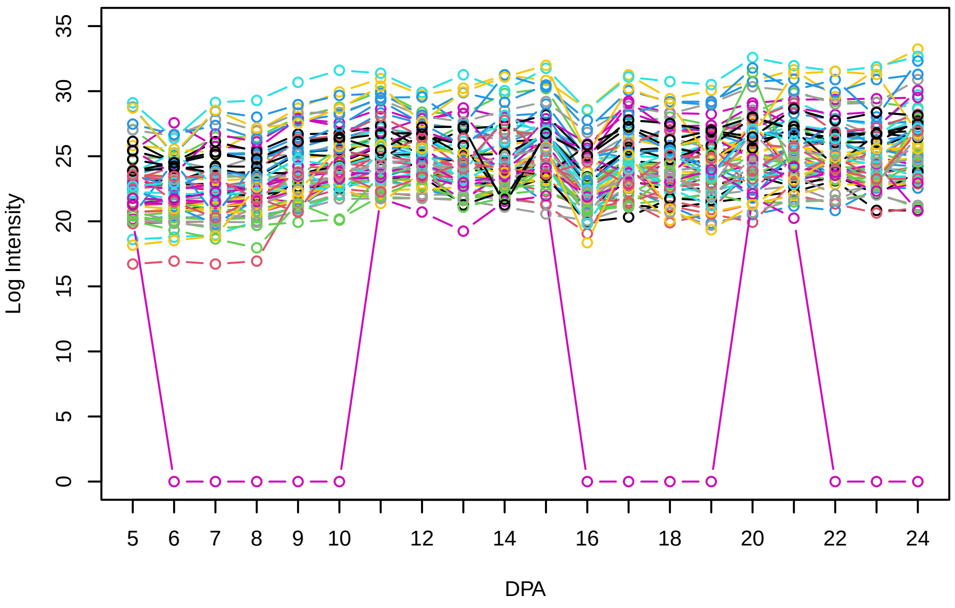


**Histogram of the average abundances of non−zero values**

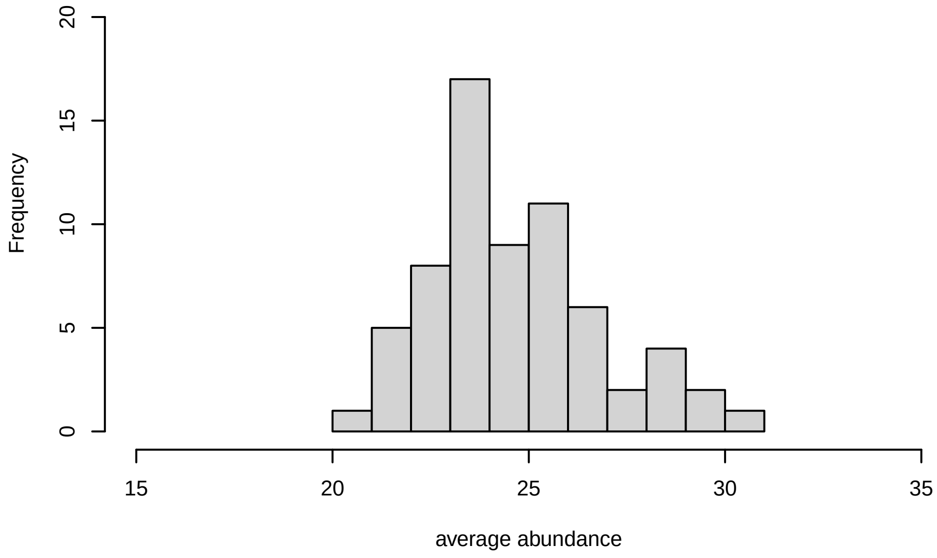


**V11, after remove outlier and cutoff is 0.8**

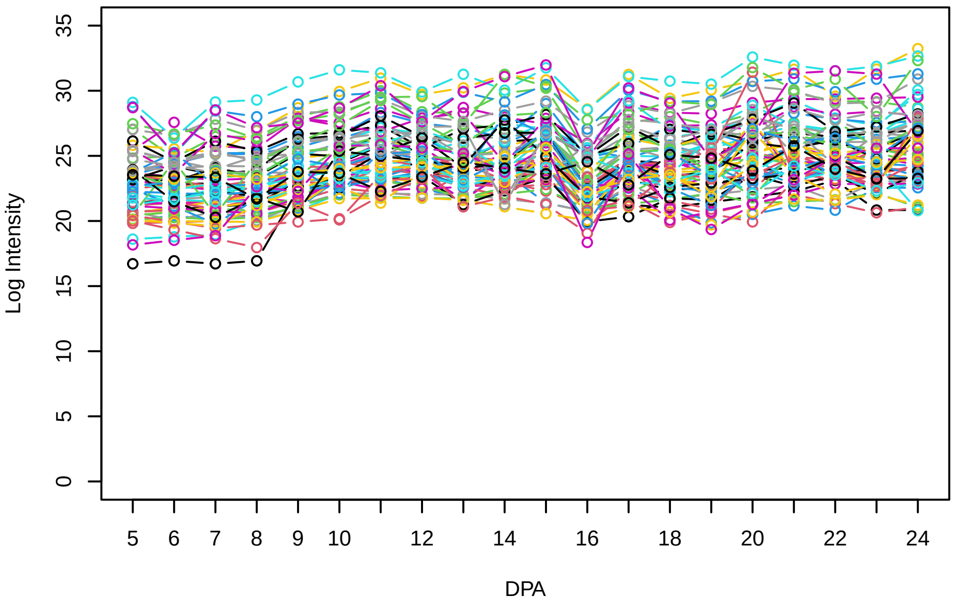


**Histogram of the average abundances of non−outlier proteins**

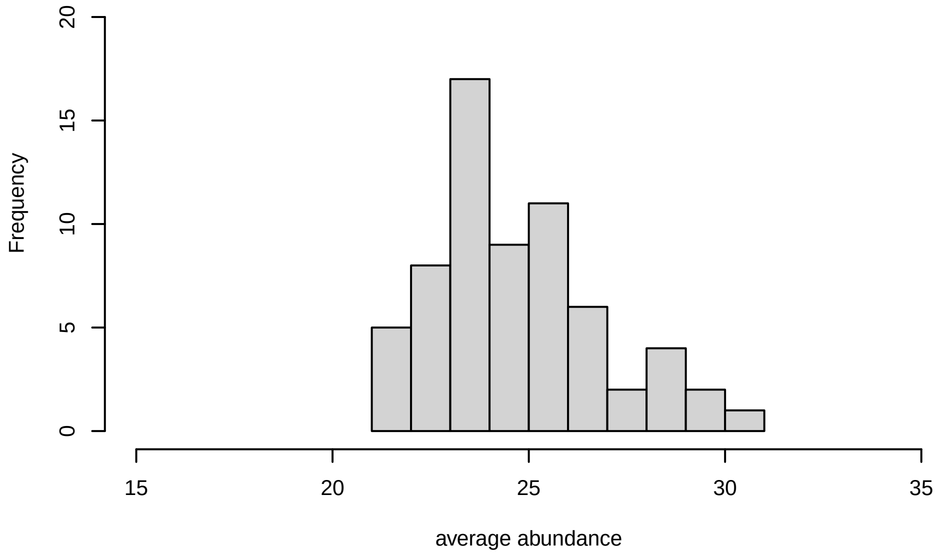


**V11, after remove unreliable**

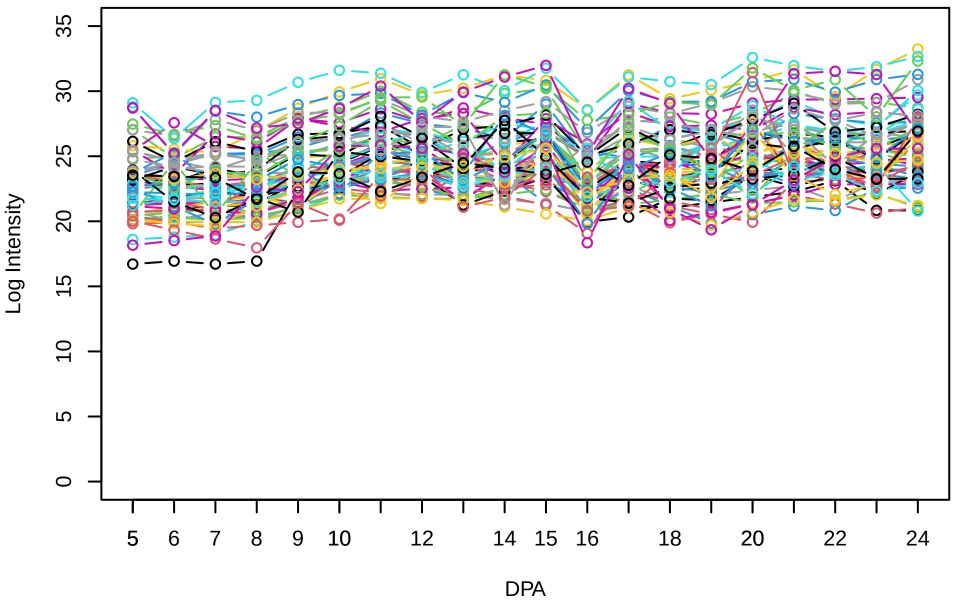


**Histogram of the average abundances of reliable proteins**

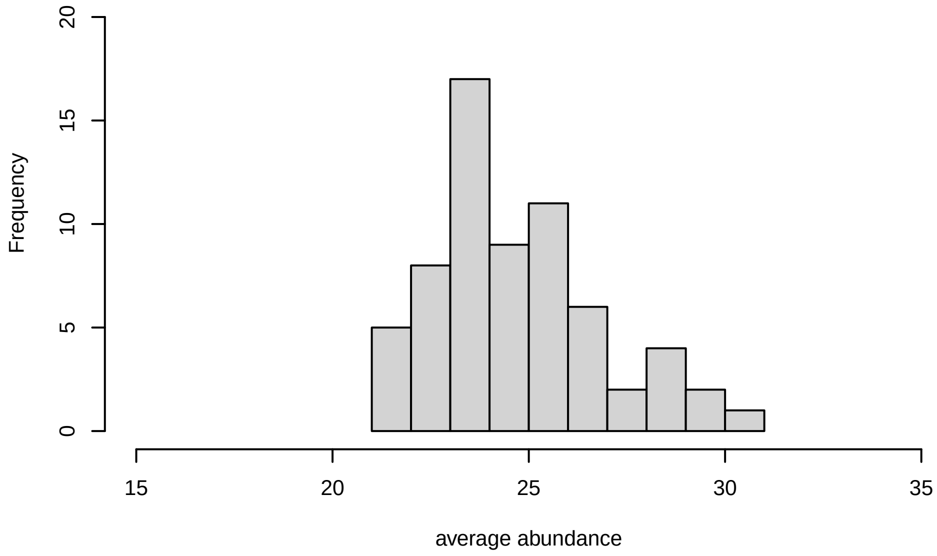


**V11, after remove low abundance**

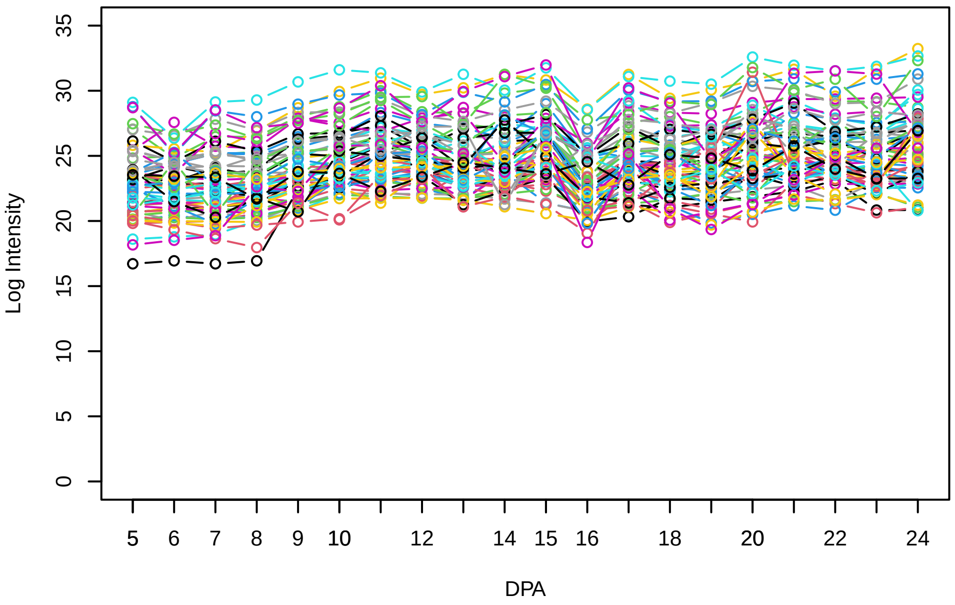


**Histogram of the average abundances of high abundance proteins**

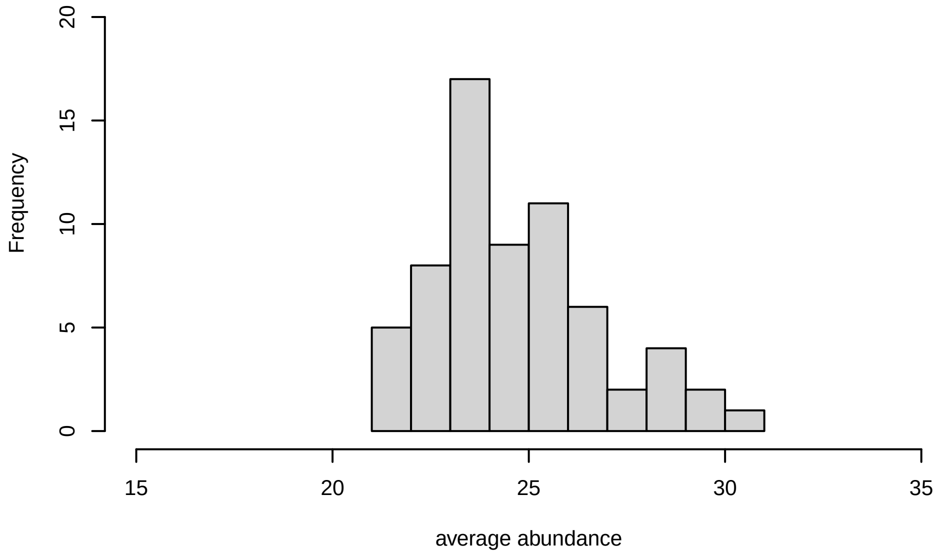

V12

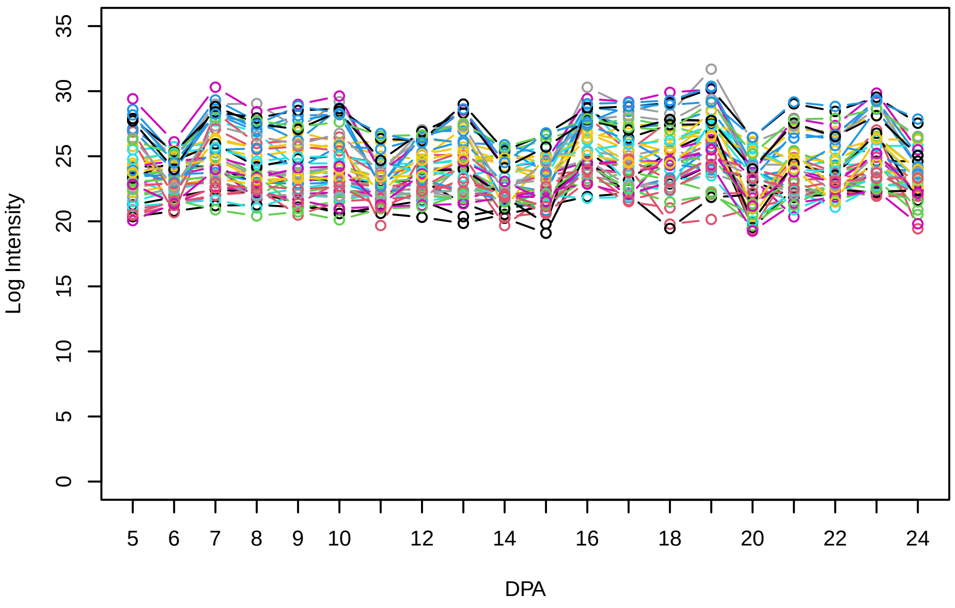


Histogram of the average abundances of non−zero values

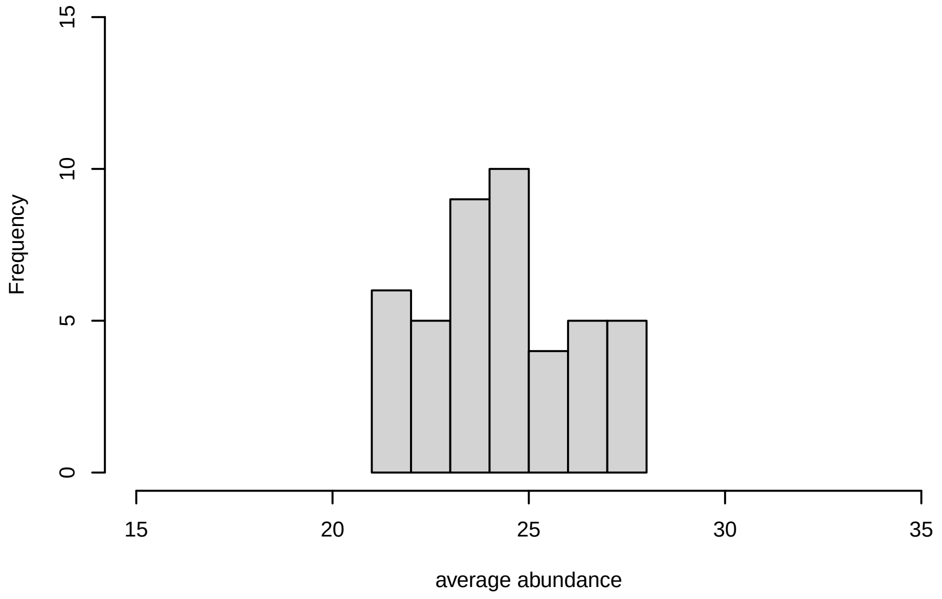


V12, after remove outlier and cutoff is 0.8

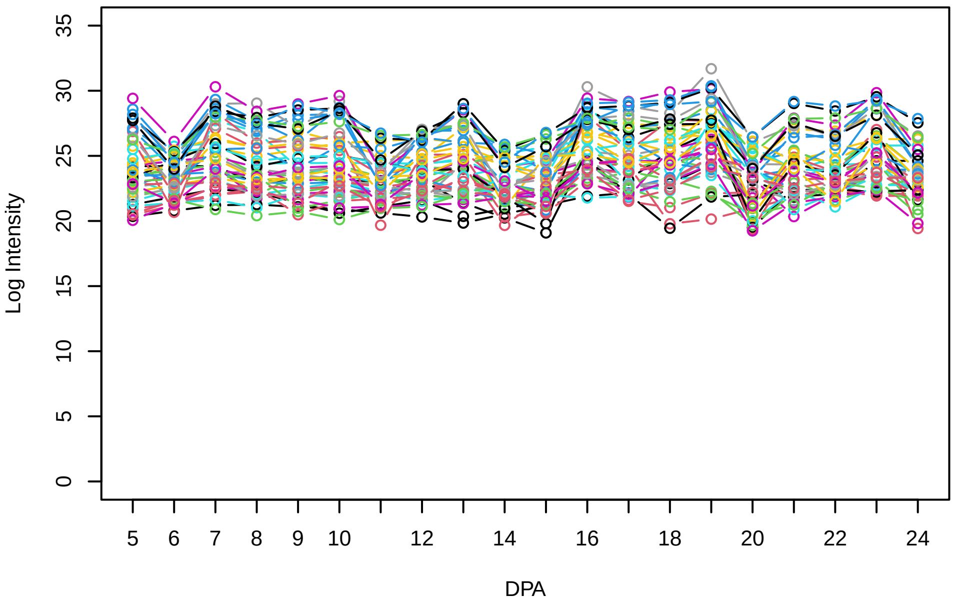


Histogram of the average abundances of non−outlier proteins

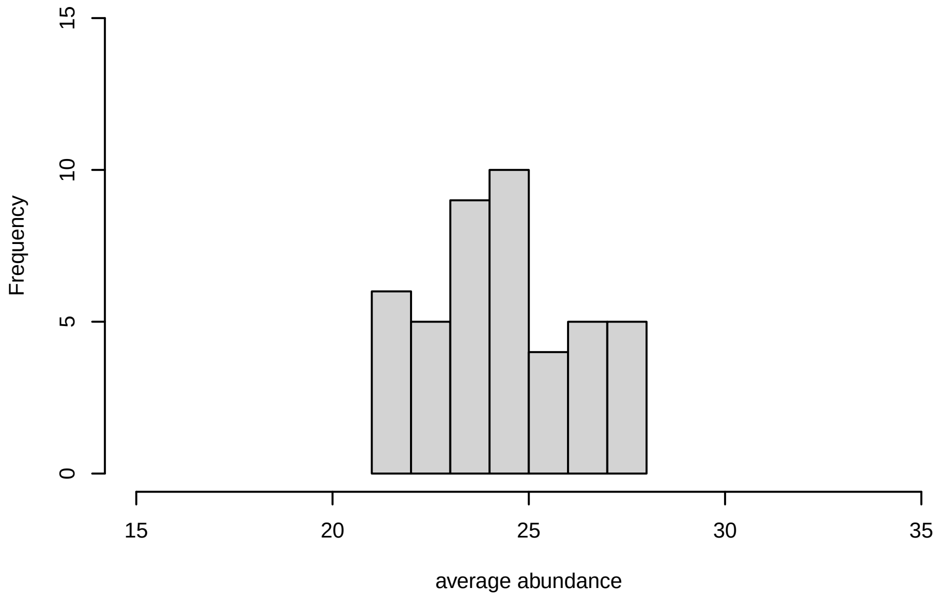


V12, after remove unreliable

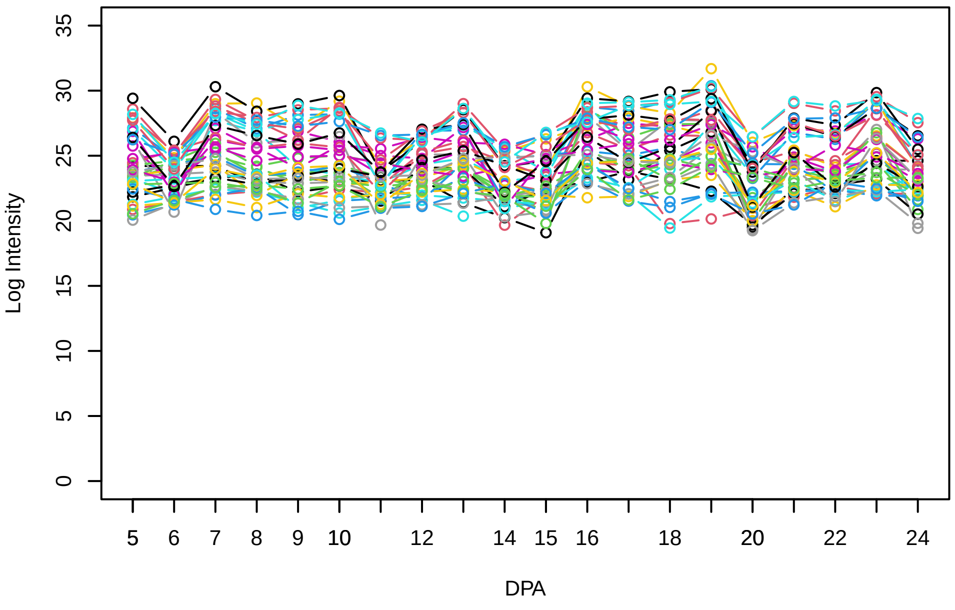


Histogram of the average abundances of reliable proteins

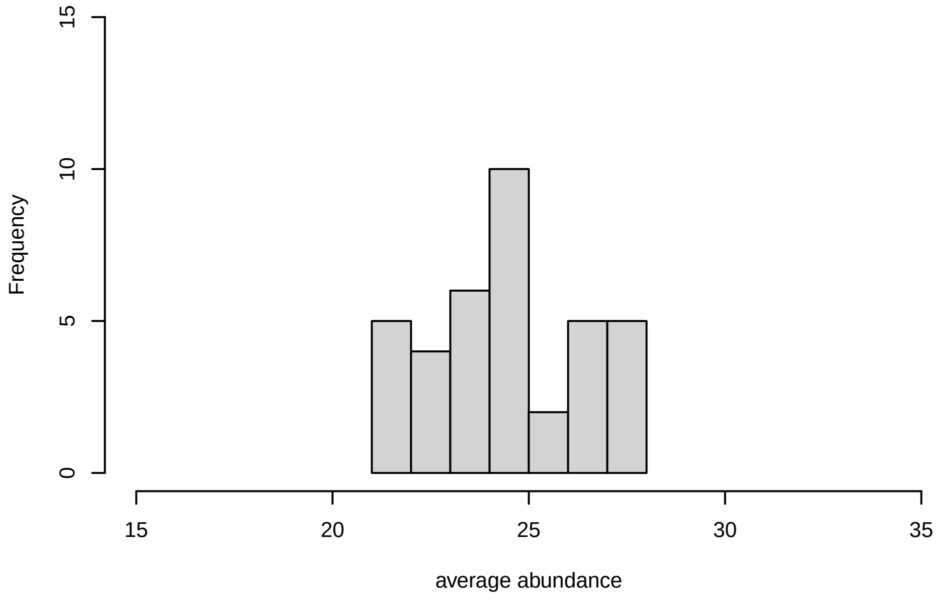


V12, after remove low abundance

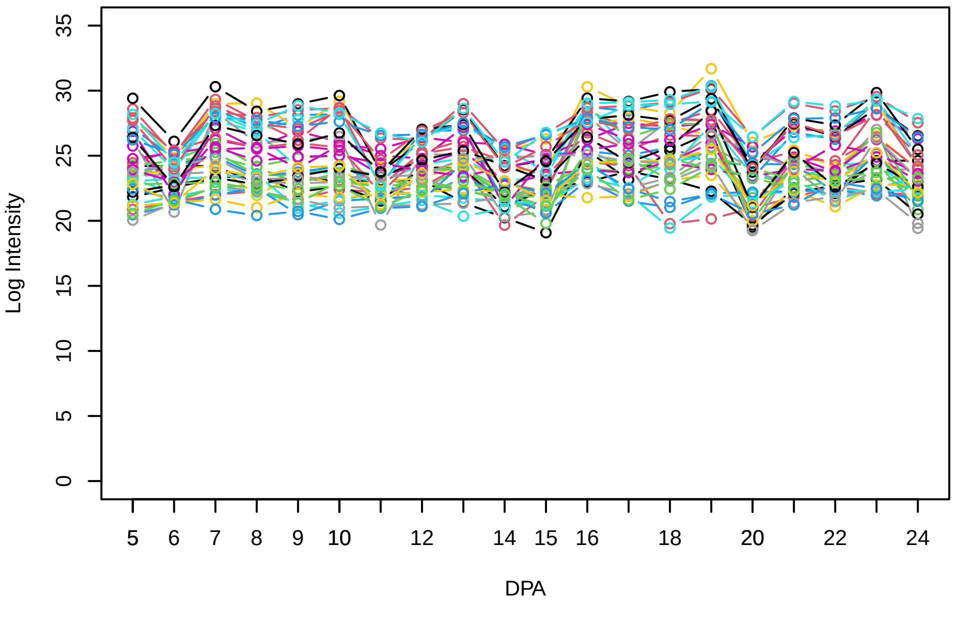


Histogram of the average abundances of high abundance proteins

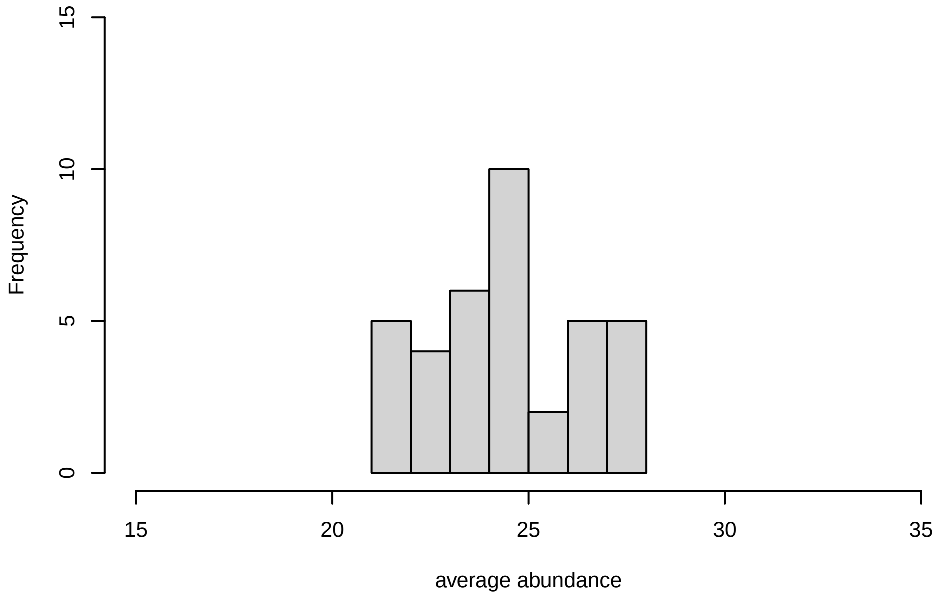

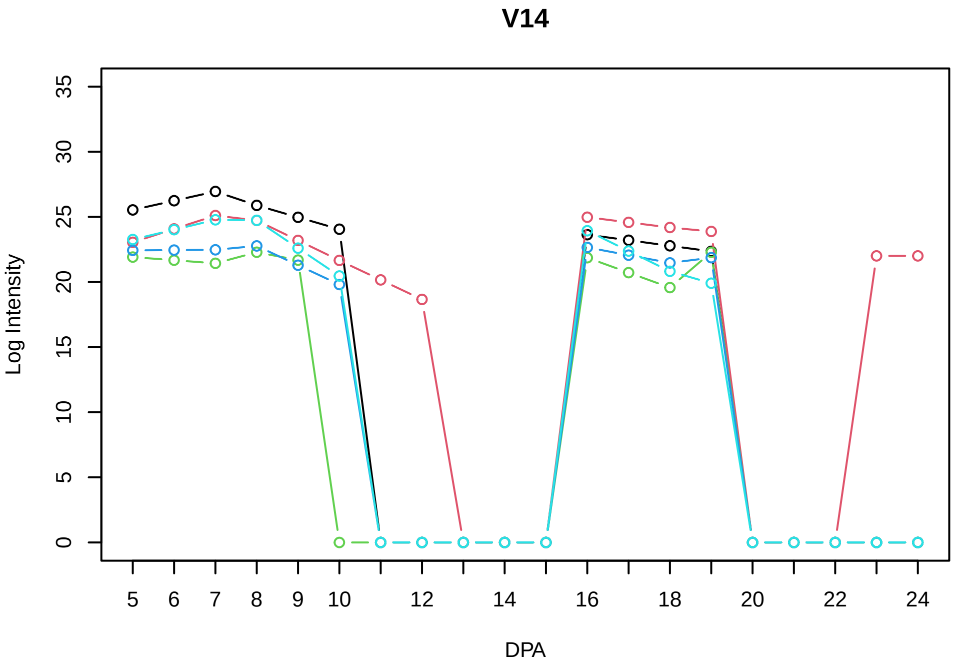
V14
Log Intensity
DPA

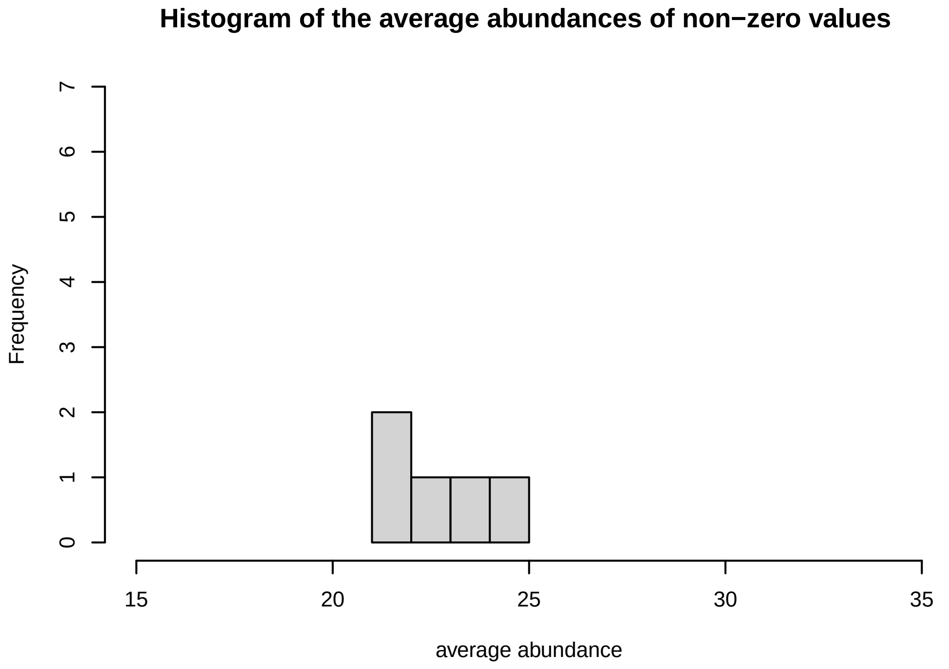
Histogram of the average abundances of non−zero values
Frequency
average abundance

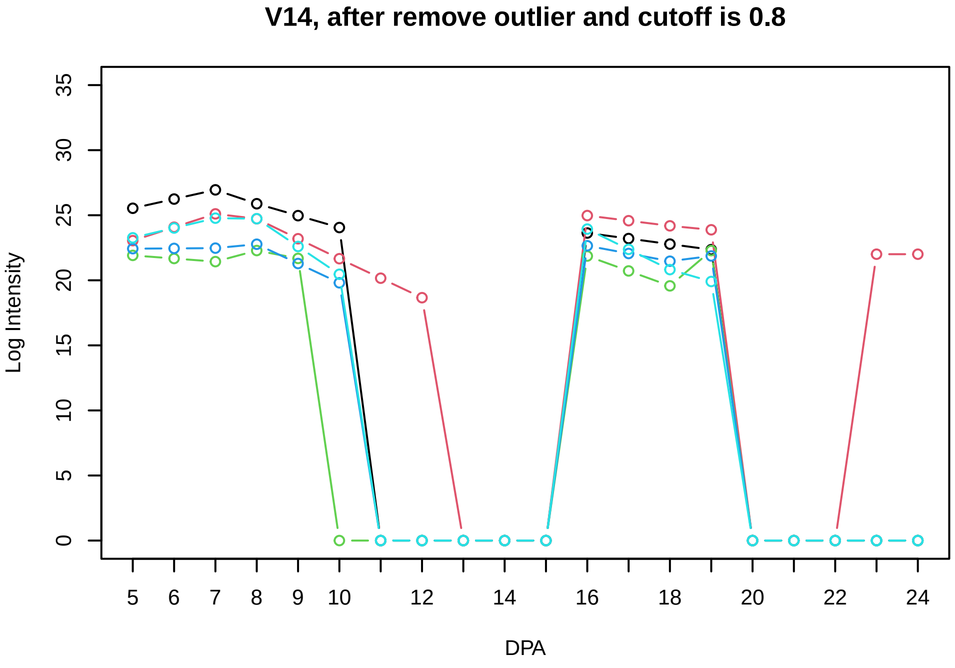
V14, after remove outlier and cutoff is 0.8
Log Intensity
DPA

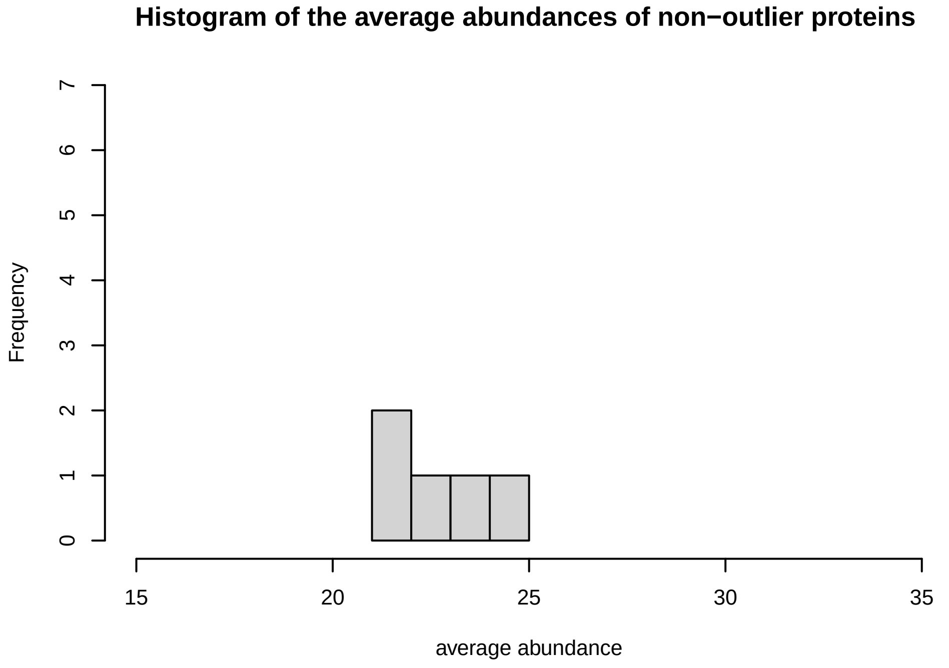
Histogram of the average abundances of non−outlier proteins
Frequency
average abundance

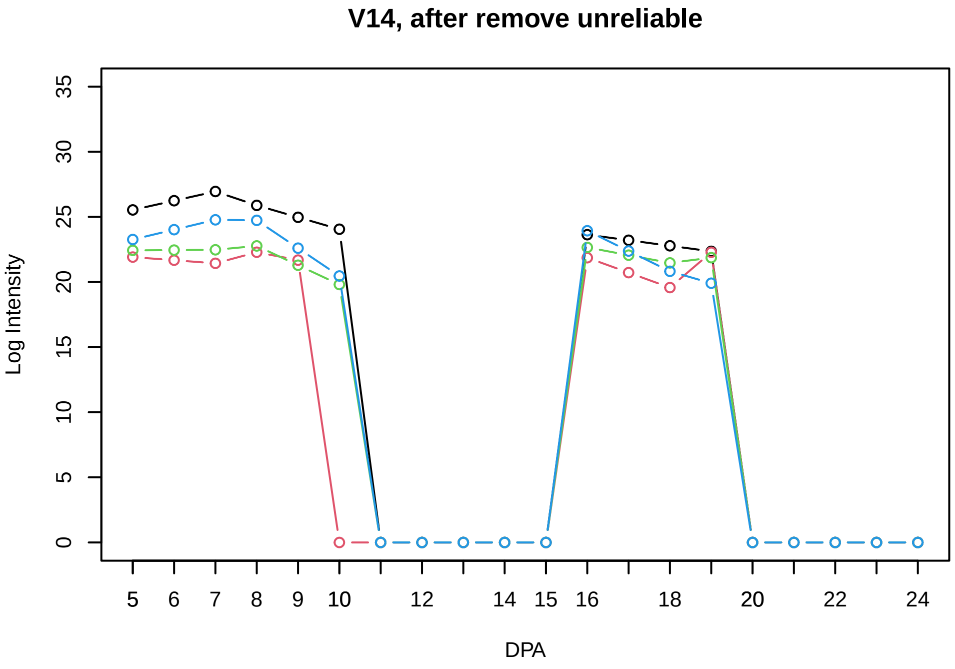
V14, after remove unreliable
Log Intensity
DPA

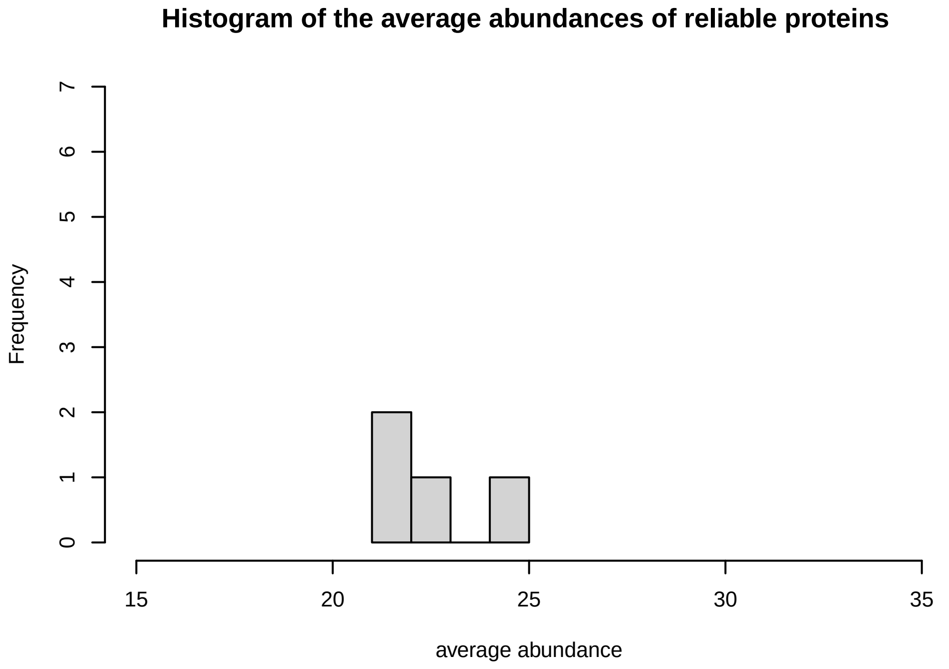
Histogram of the average abundances of reliable proteins
Frequency
average abundance

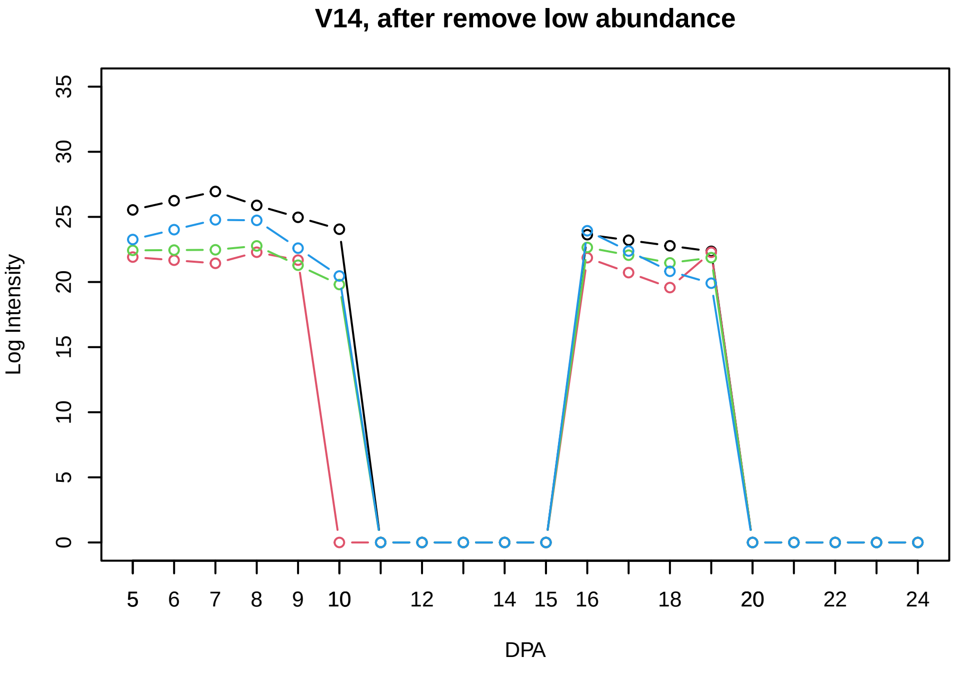
V14, after remove low abundance
Log Intensity
DPA

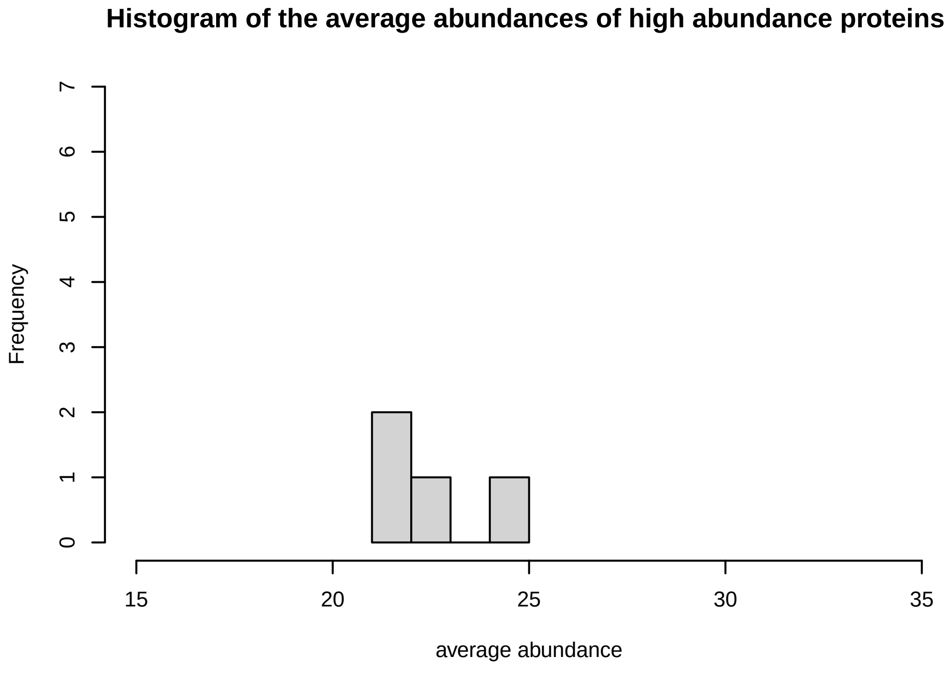
Histogram of the average abundances of high abundance proteins
Frequency
average abundance

V15

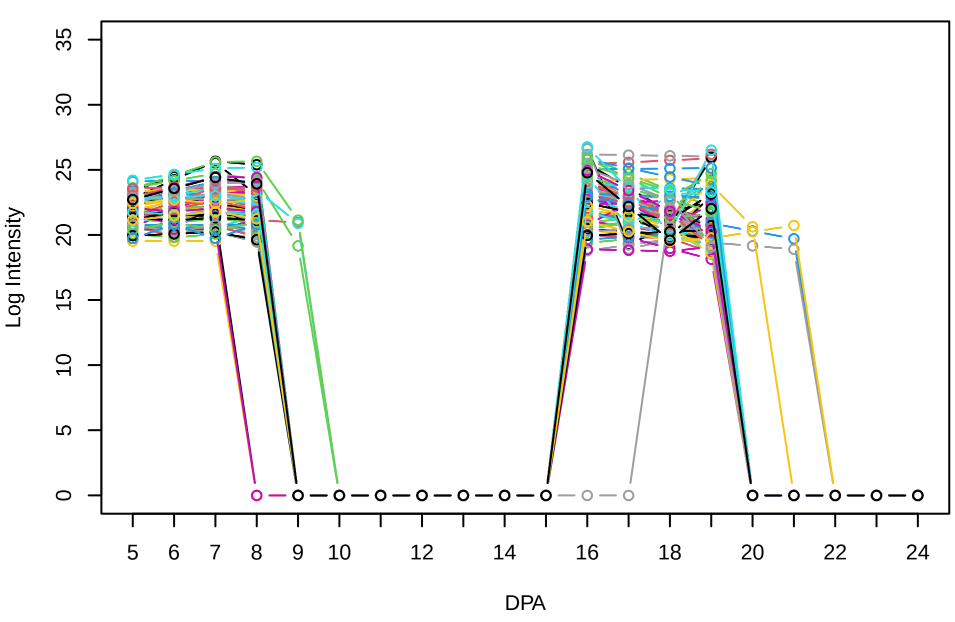


Histogram of the average abundances of non−zero values

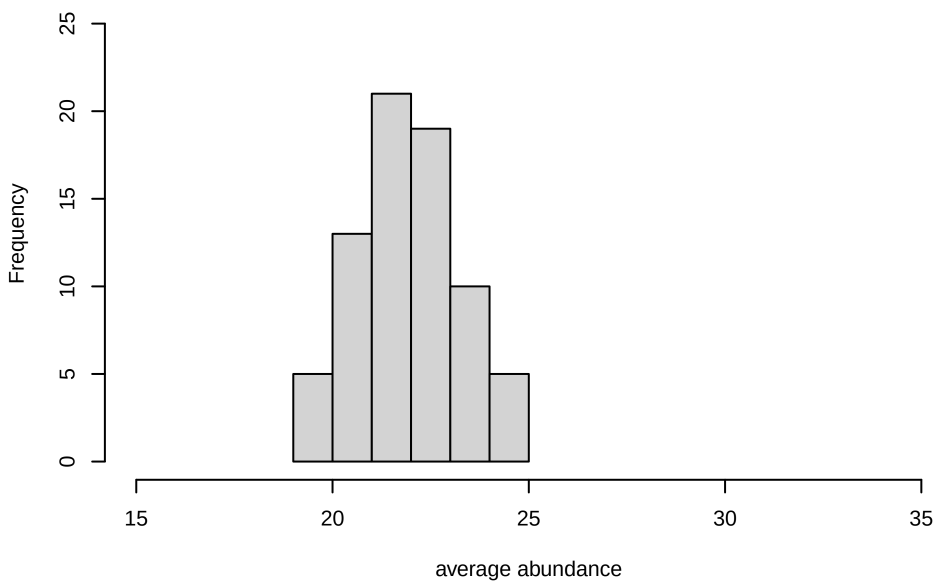


V15, after remove outlier and cutoff is 0.8

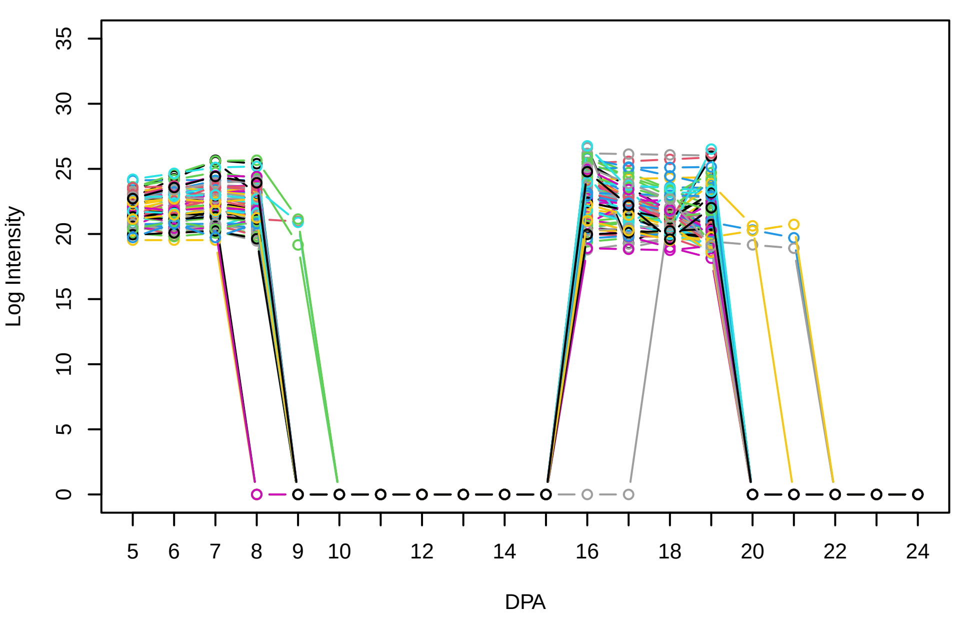


Histogram of the average abundances of non−outlier proteins

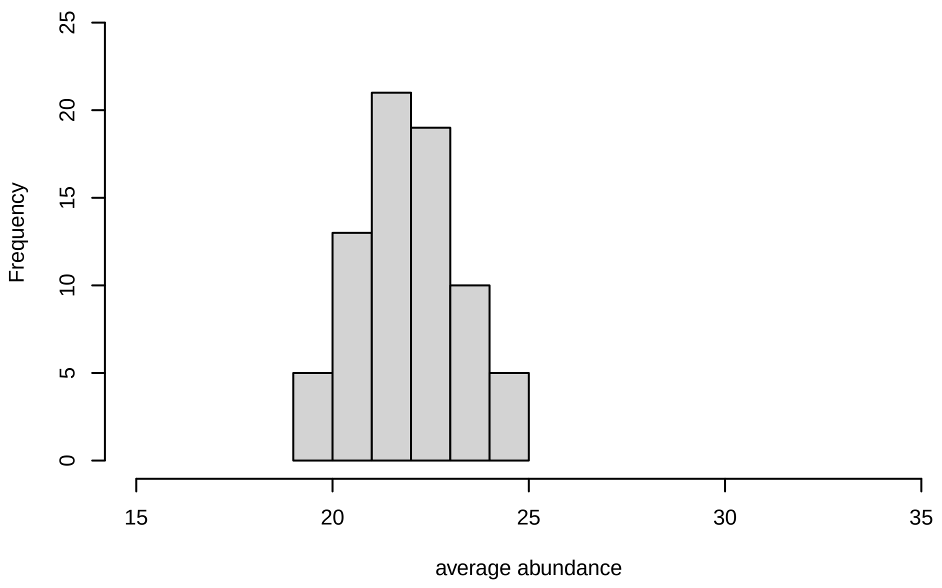


V15, after remove unreliable

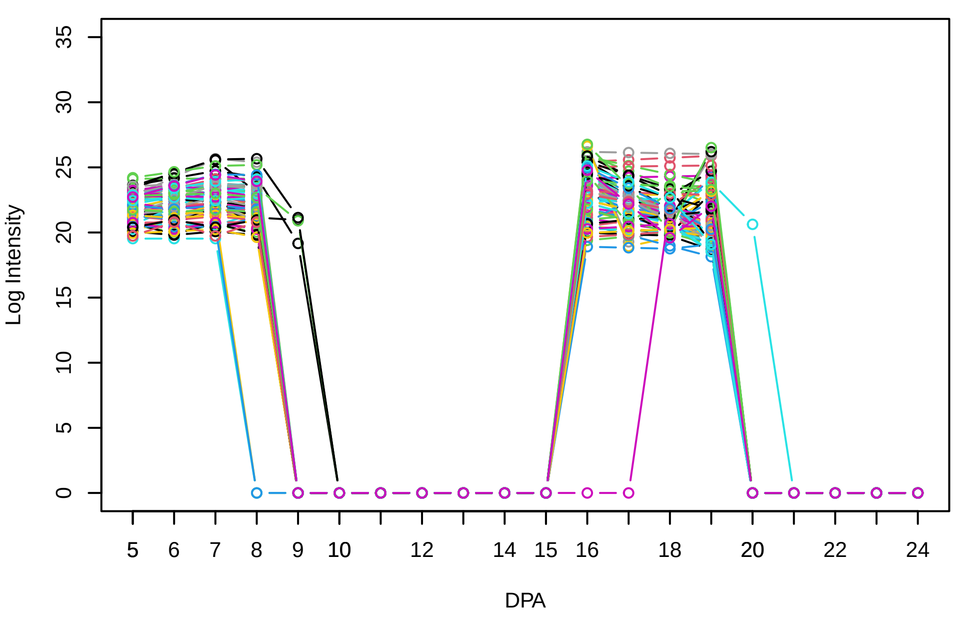


Histogram of the average abundances of reliable proteins

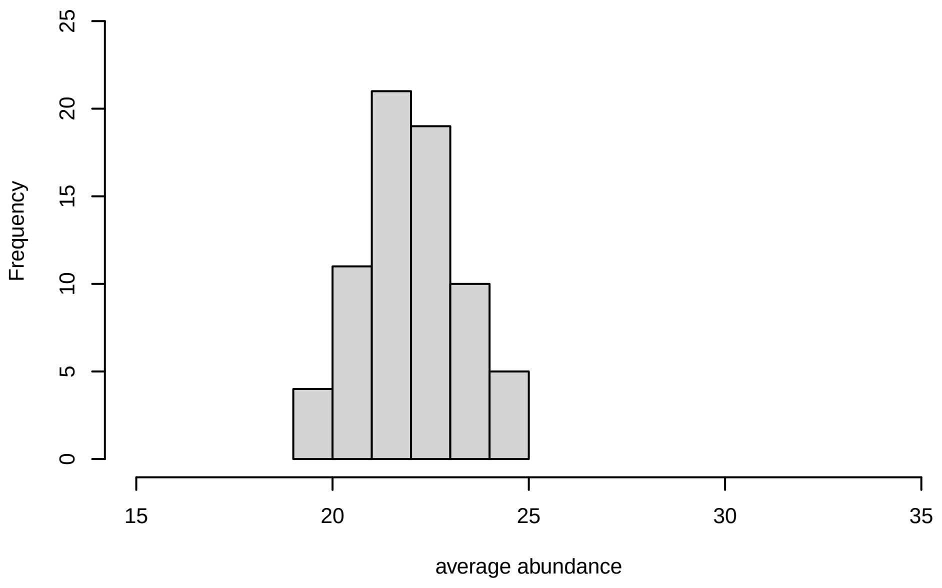


V15, after remove low abundance

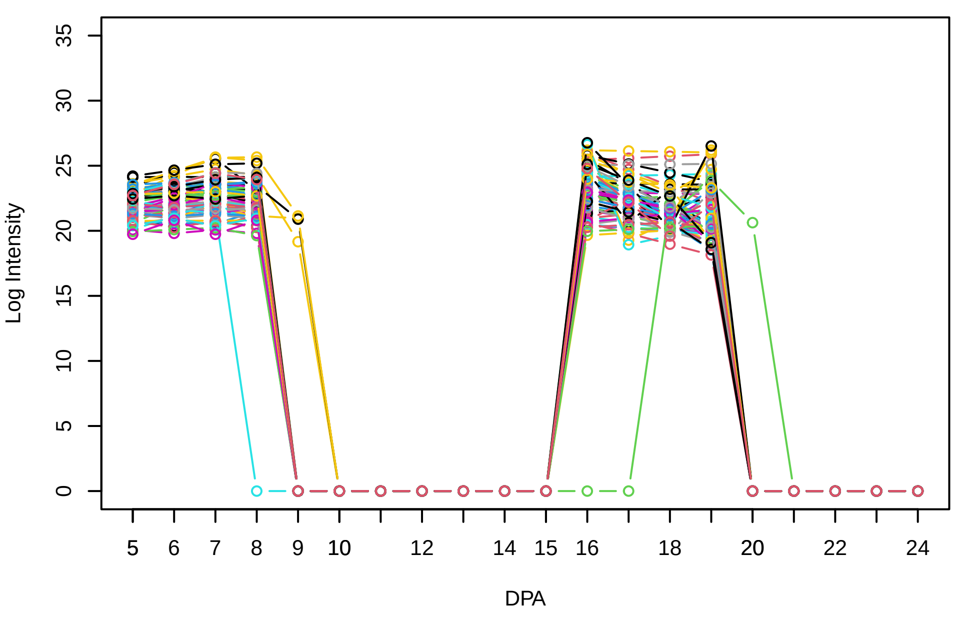


Histogram of the average abundances of high abundance proteins

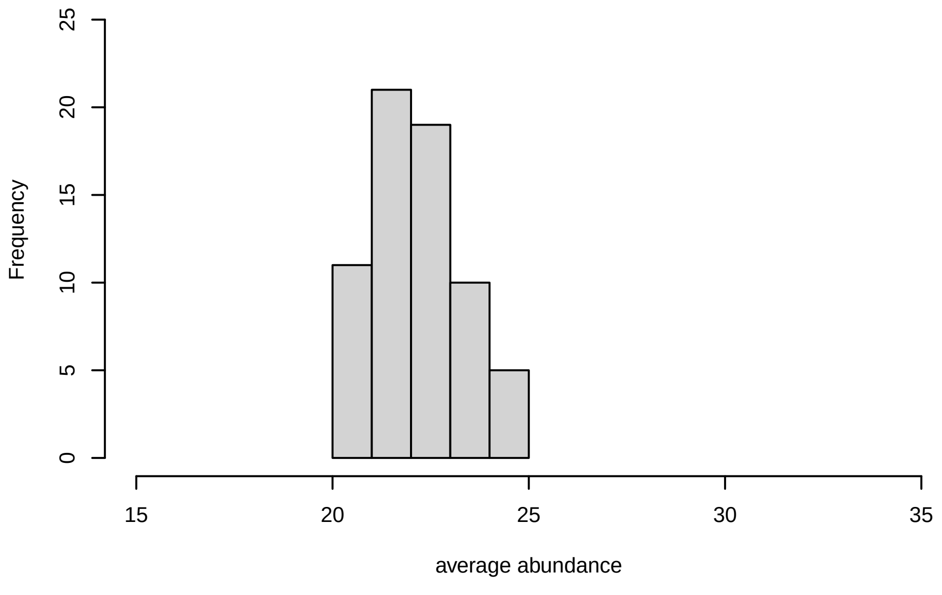

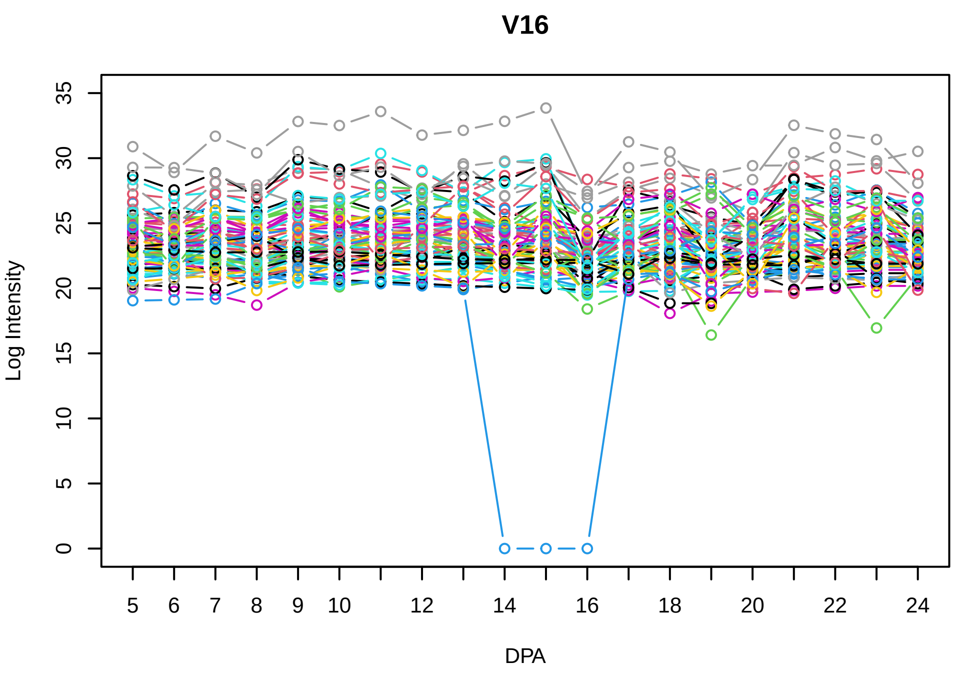
V16
Log Intensity
DPA

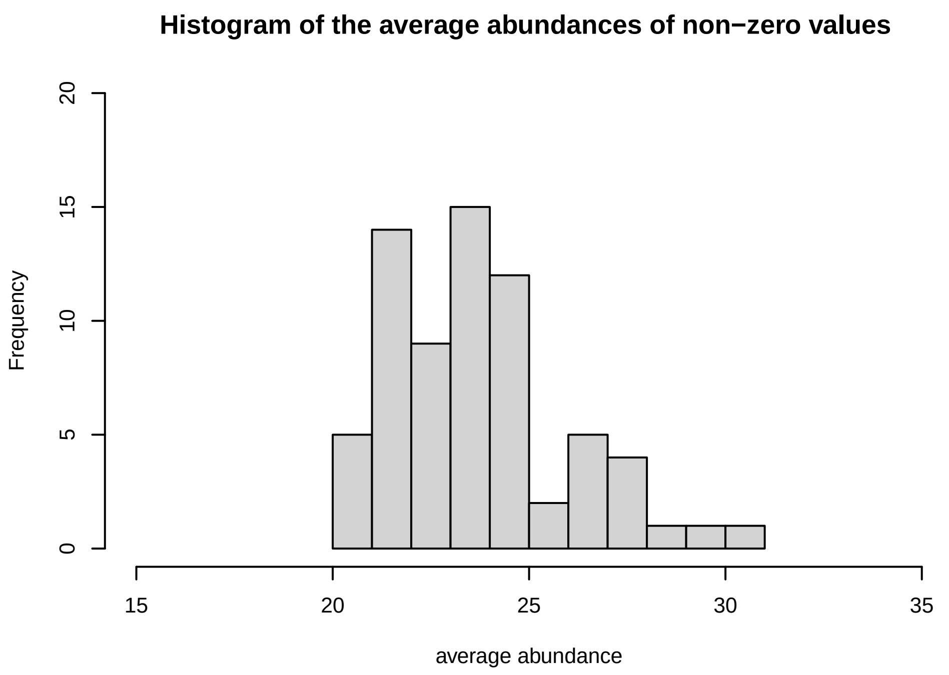
Histogram of the average abundances of non−zero values
Frequency
average abundance

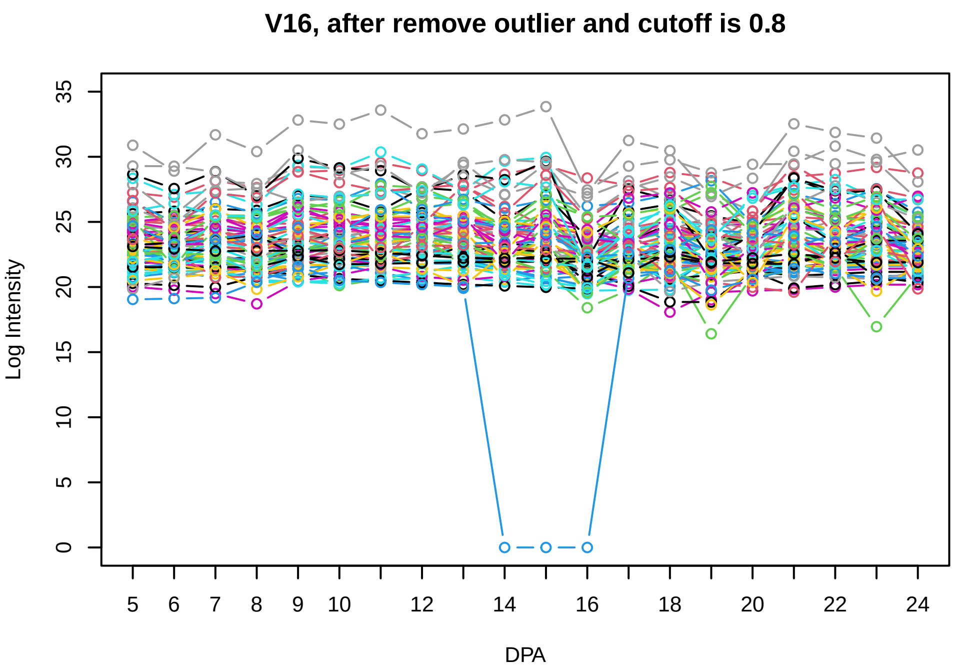
V16, after remove outlier and cutoff is 0.8
Log Intensity
DPA

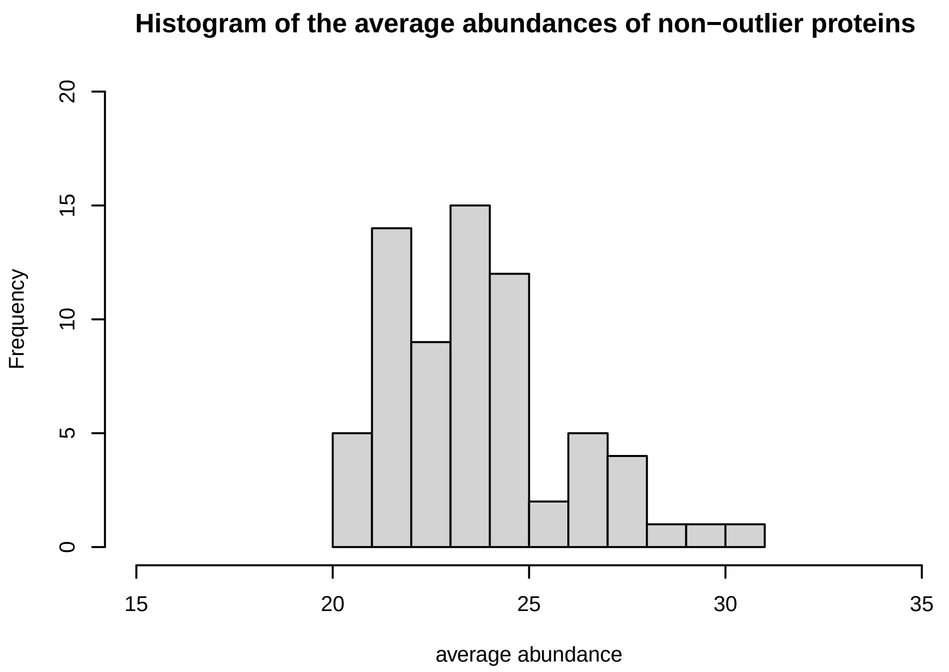
Histogram of the average abundances of non−outlier proteins
Frequency
average abundance

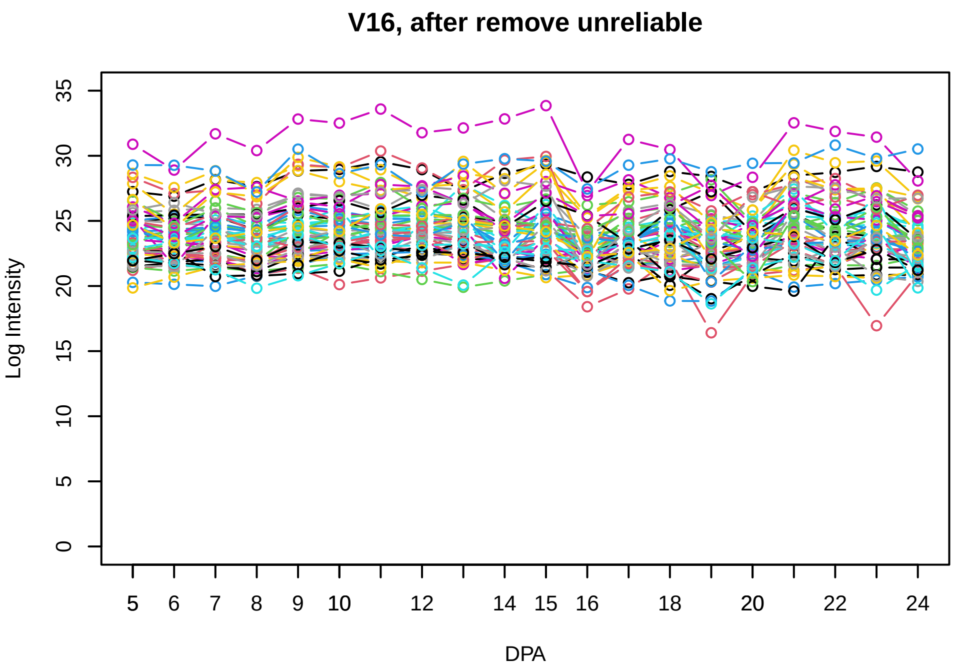
V16, after remove unreliable
Log Intensity
DPA

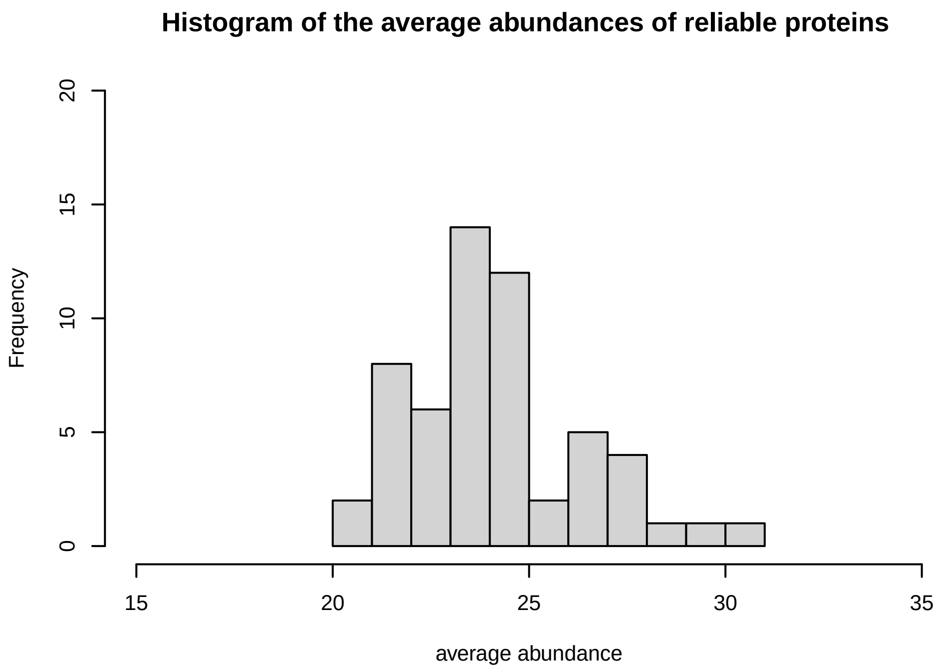
Histogram of the average abundances of reliable proteins
Frequency
average abundance

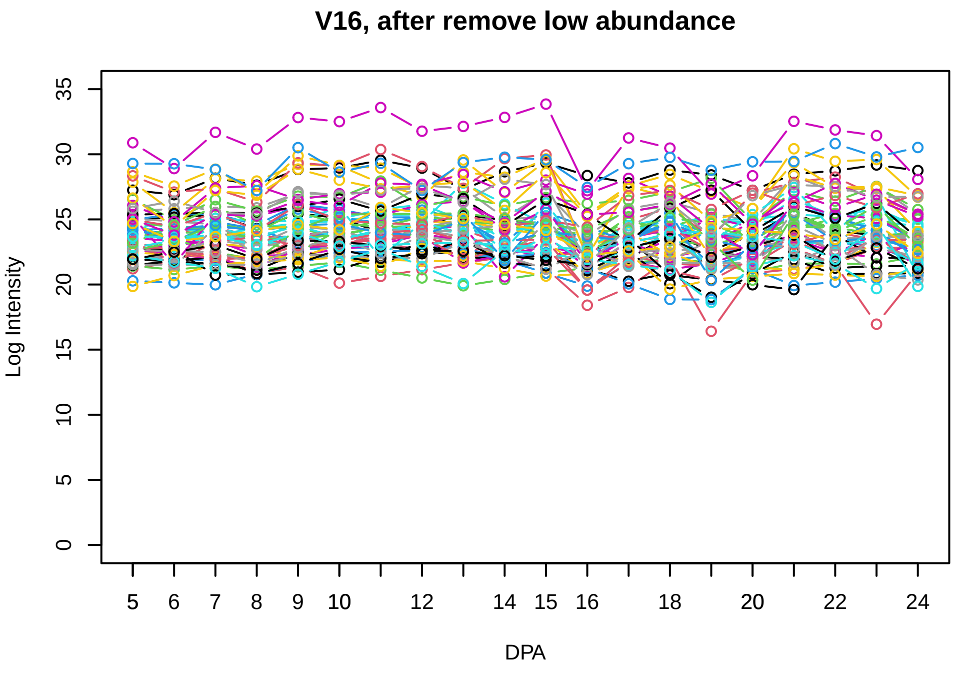
V16, after remove low abundance
Log Intensity
DPA

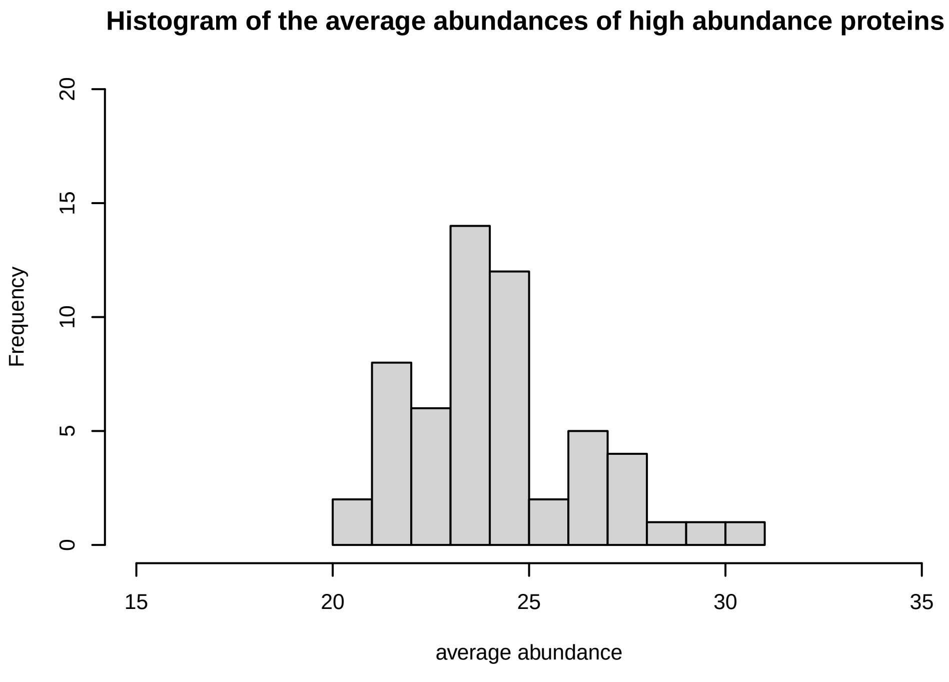
Histogram of the average abundances of high abundance proteins
Frequency
average abundance

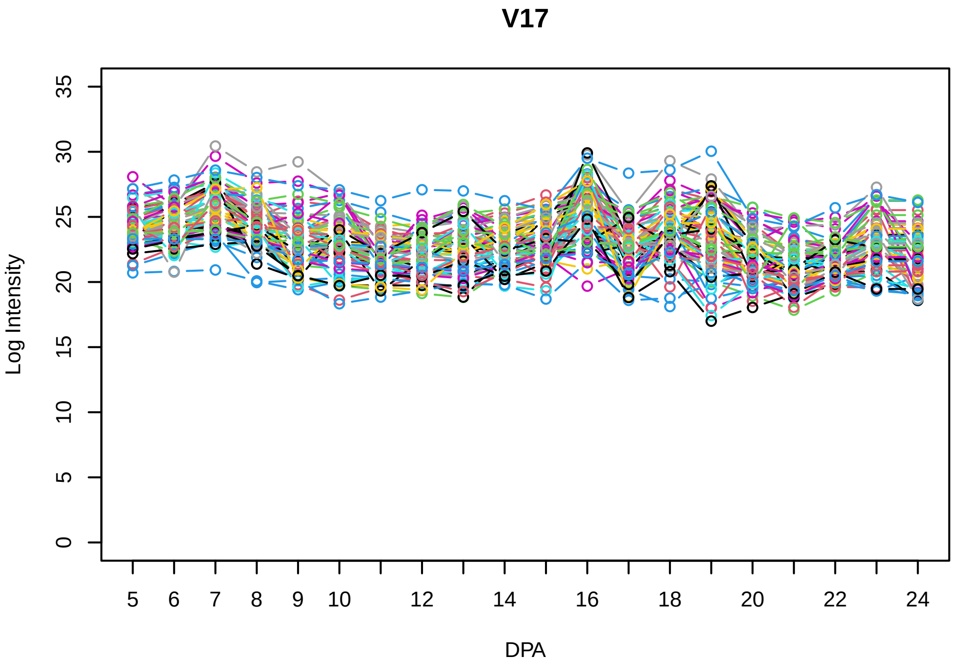
V17
Log Intensity
DPA

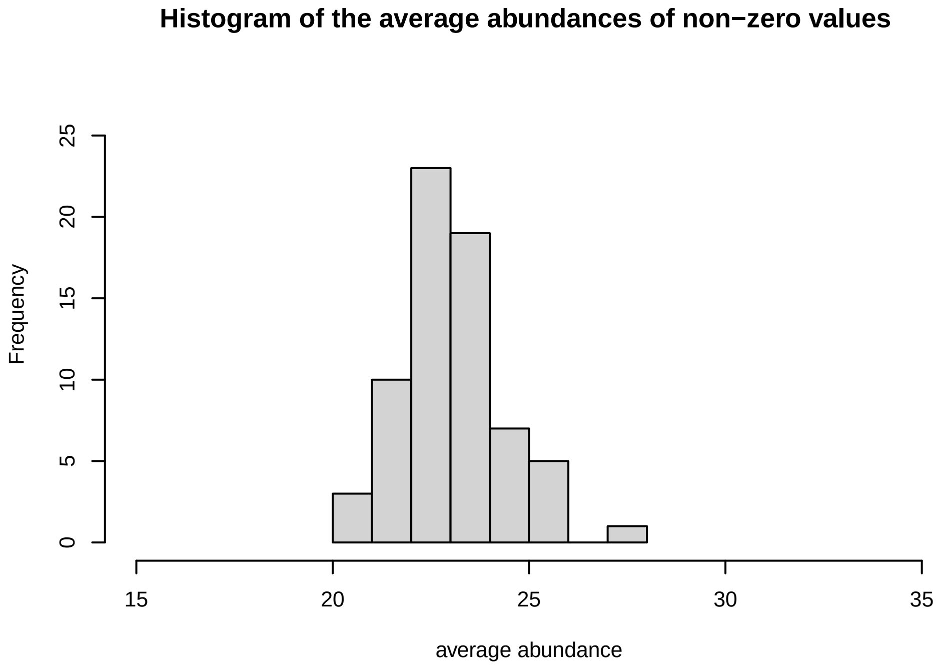
Histogram of the average abundances of non−zero values
Frequency
average abundance

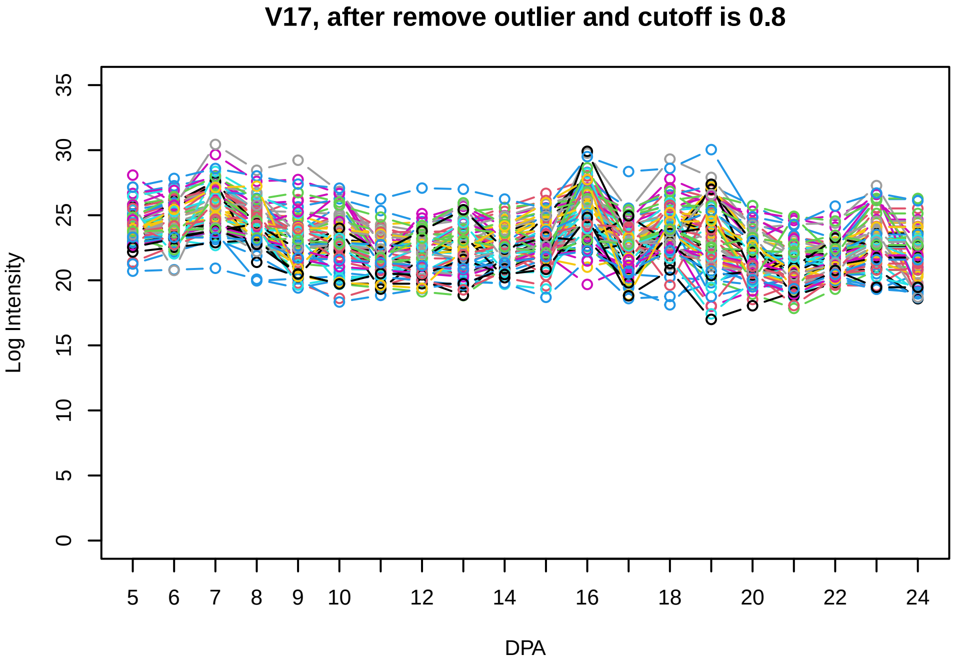
V17, after remove outlier and cutoff is 0.8
Log Intensity
DPA

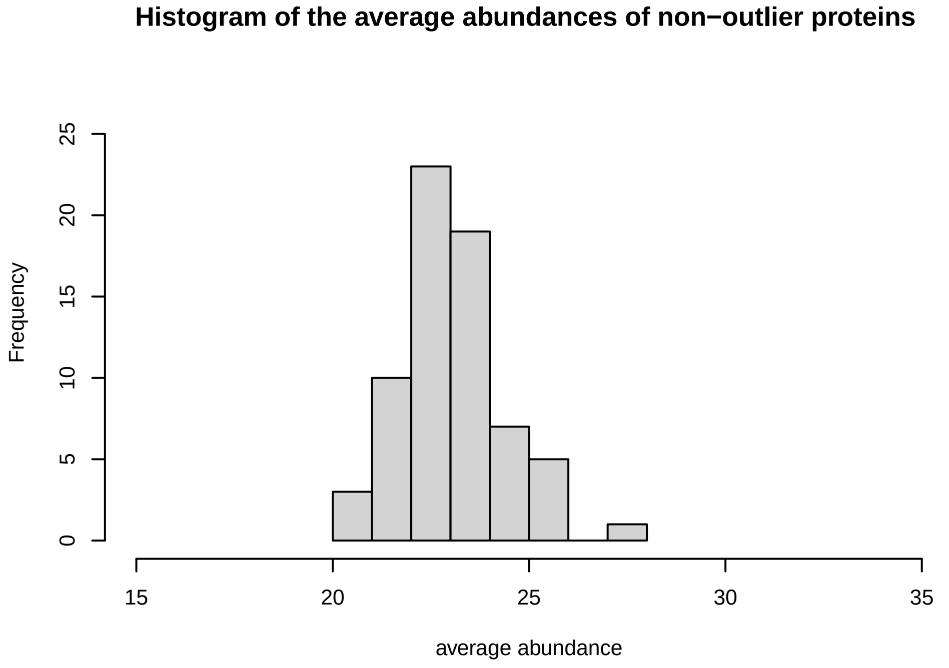
Histogram of the average abundances of non−outlier proteins
Frequency
average abundance

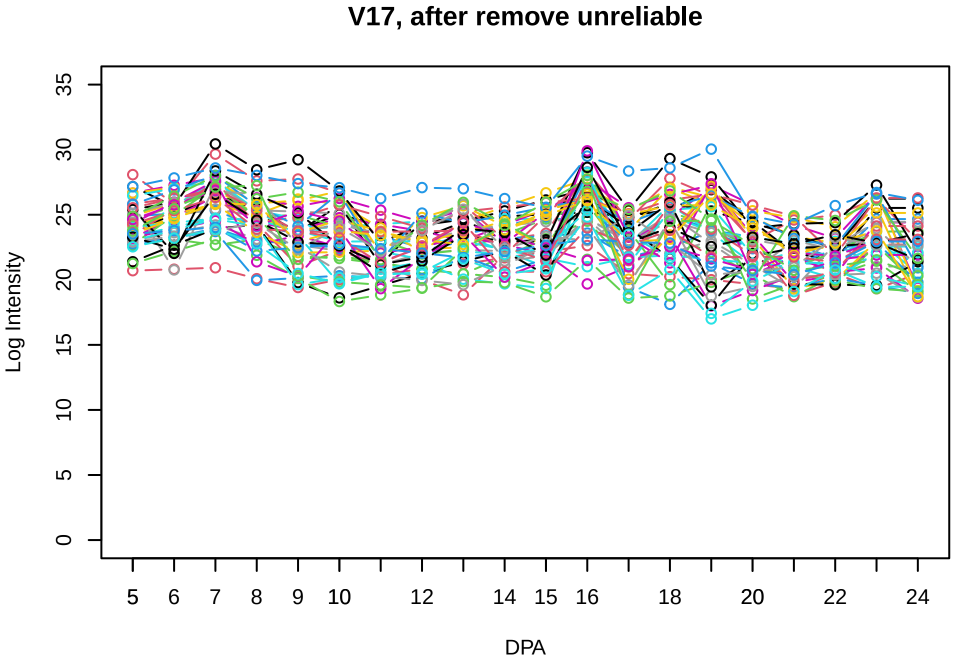
V17, after remove unreliable
Log Intensity
DPA

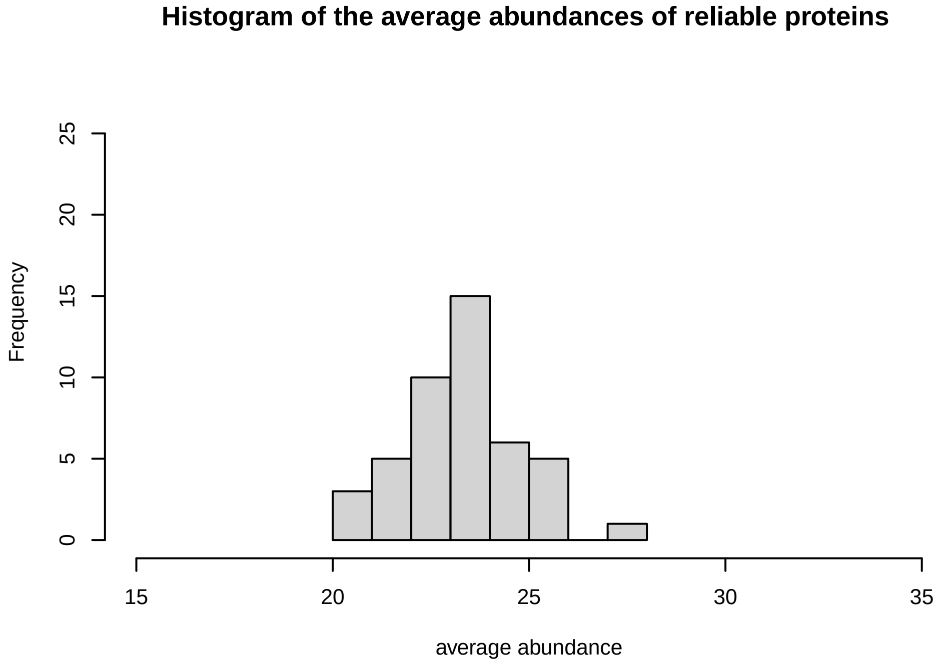
Histogram of the average abundances of reliable proteins
Frequency
average abundance

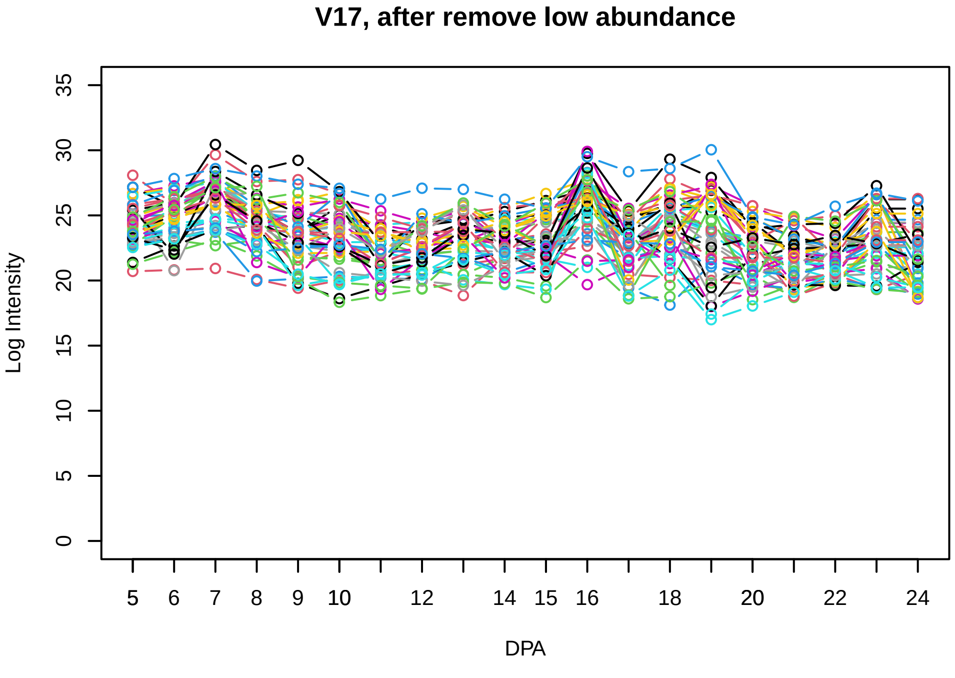
V17, after remove low abundance
Log Intensity
DPA

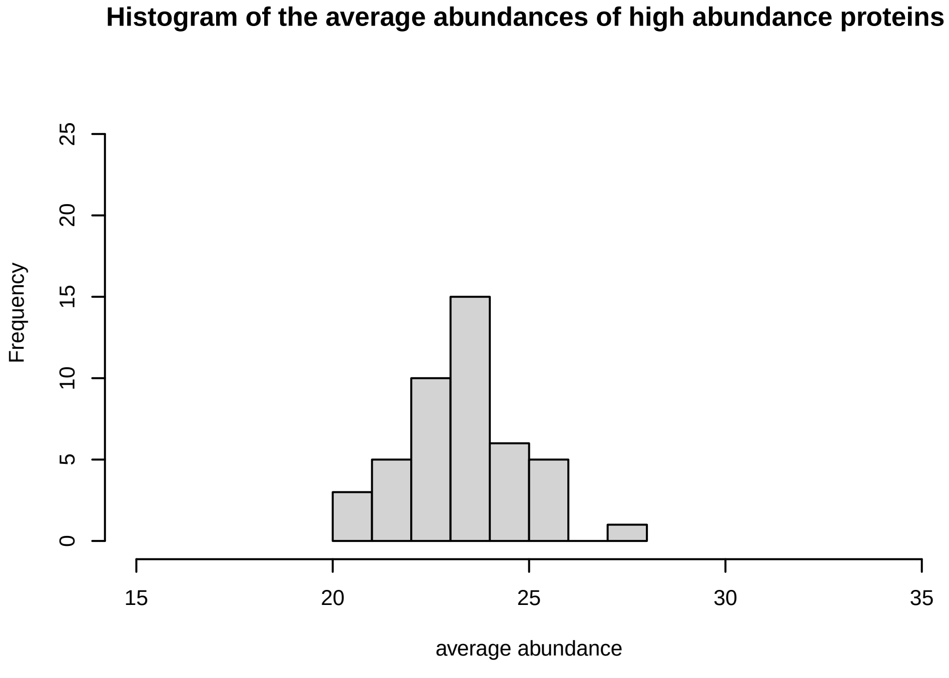
Histogram of the average abundances of high abundance proteins
Frequency
average abundance

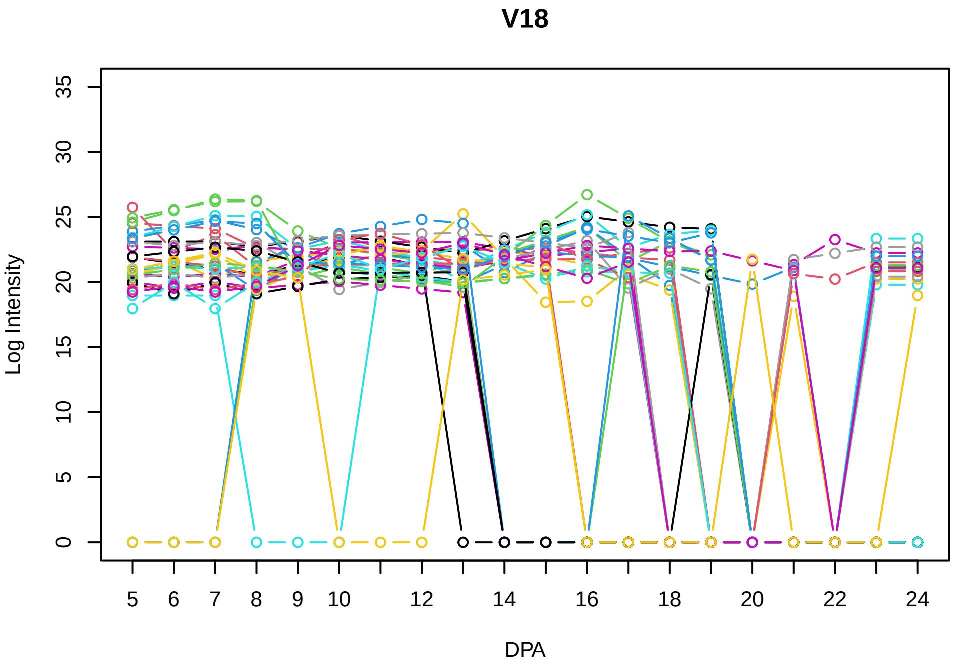
V18
Log Intensity
DPA

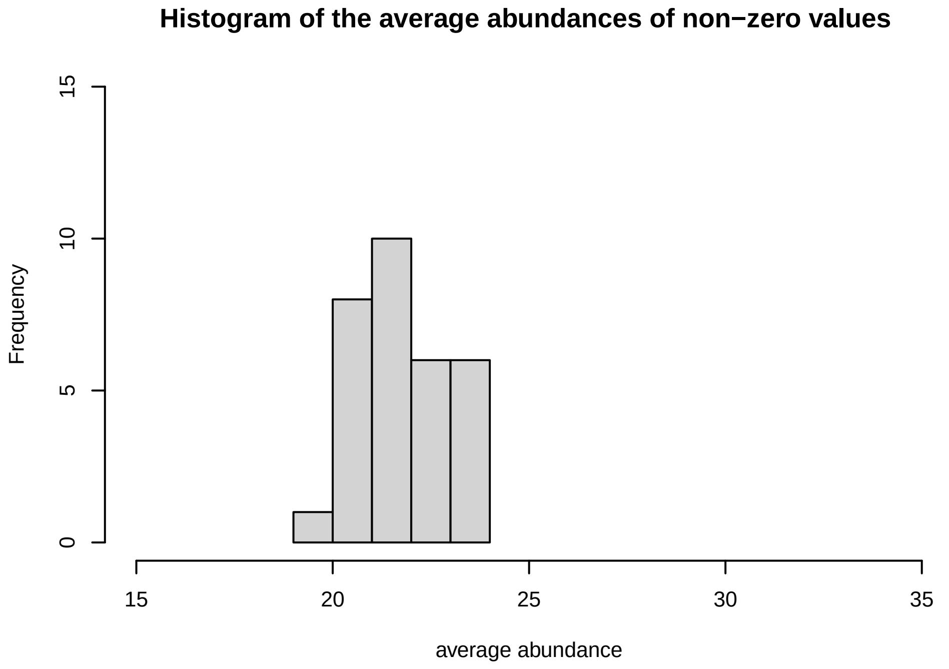
Histogram of the average abundances of non−zero values
Frequency
average abundance

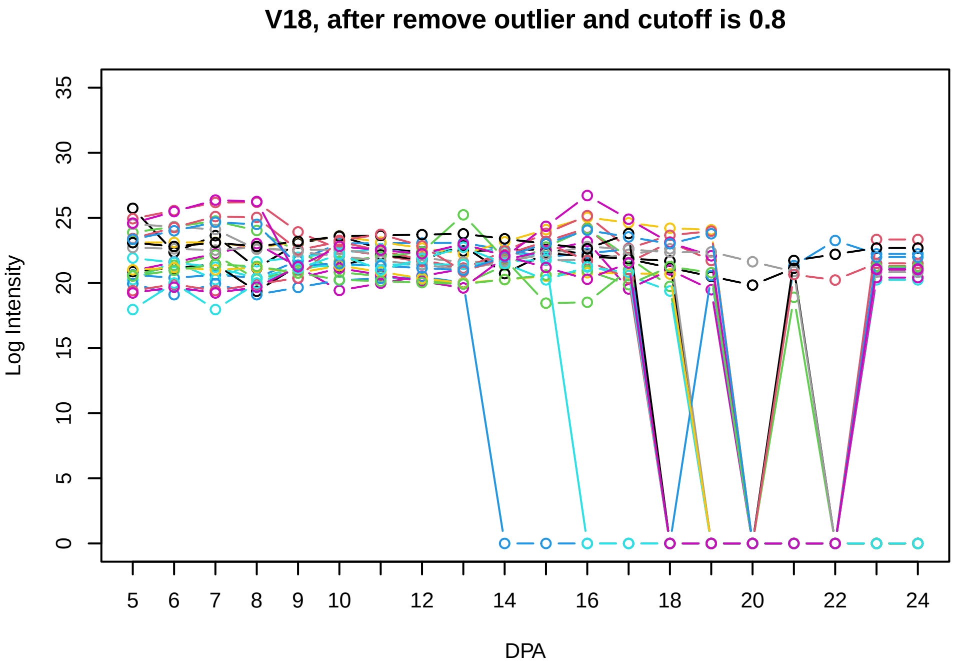
V18, after remove outlier and cutoff is 0.8
Log Intensity
DPA

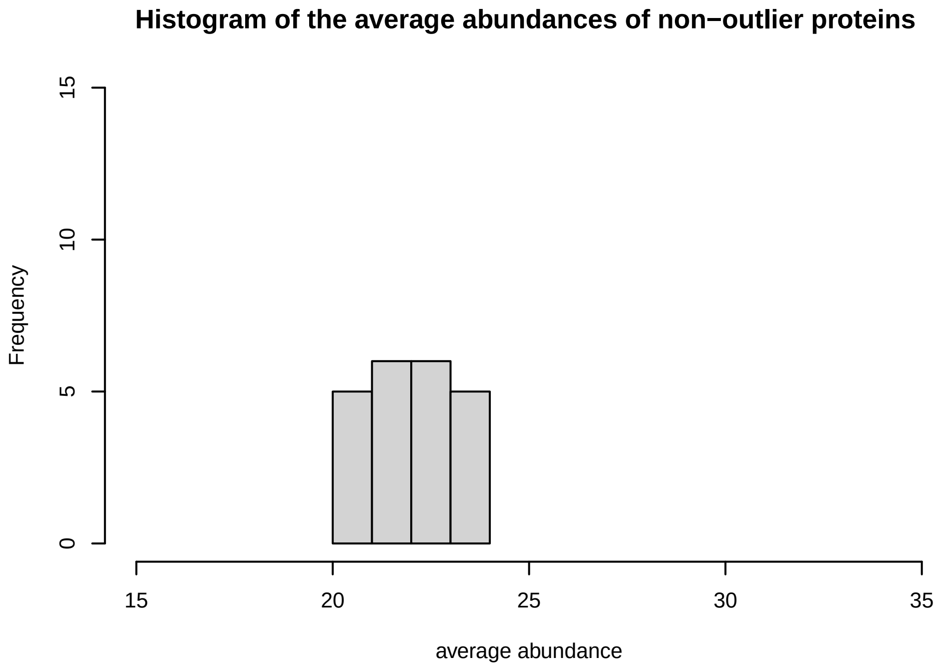
Histogram of the average abundances of non−outlier proteins
Frequency
average abundance

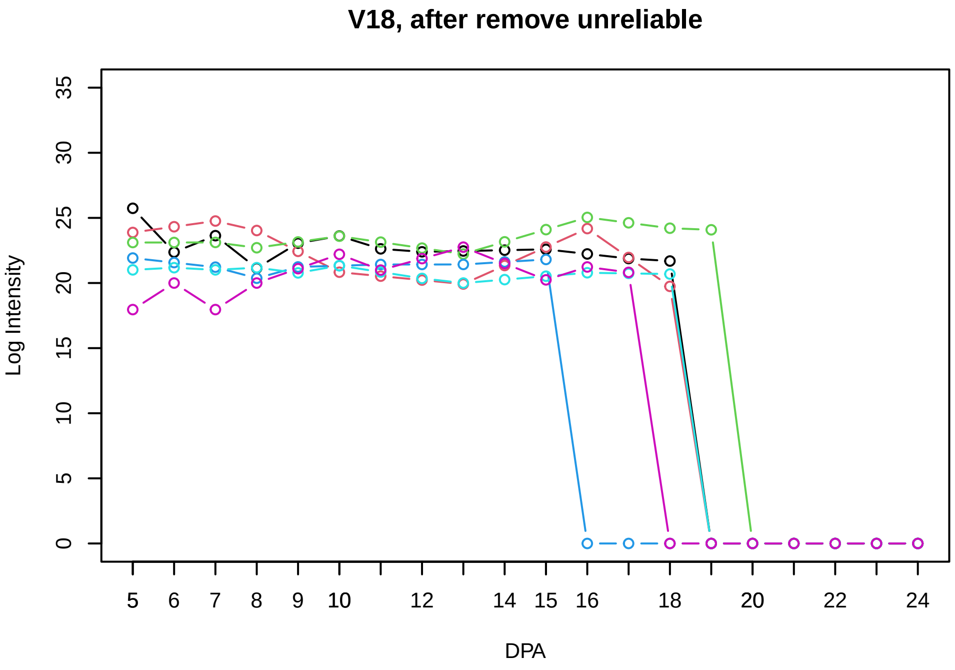
V18, after remove unreliable
Log Intensity
DPA

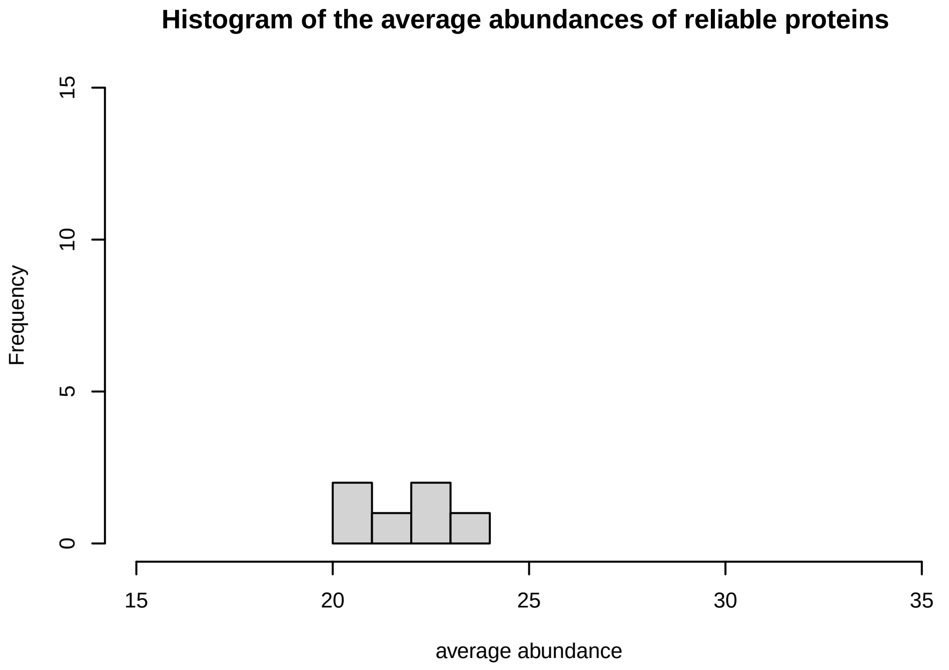
Histogram of the average abundances of reliable proteins
Frequency
average abundance

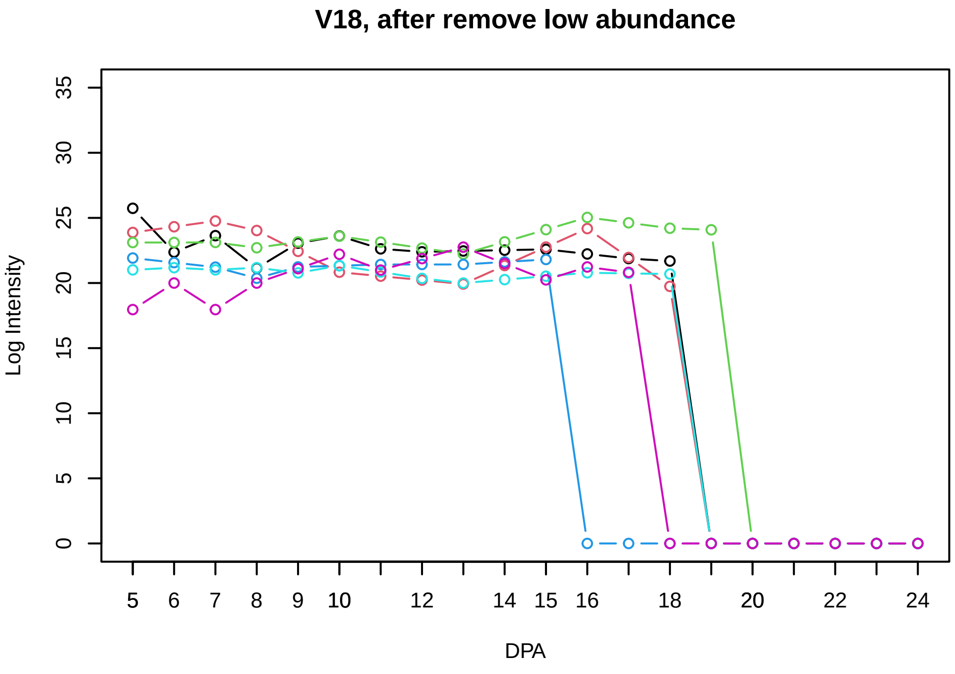
V18, after remove low abundance
Log Intensity
DPA

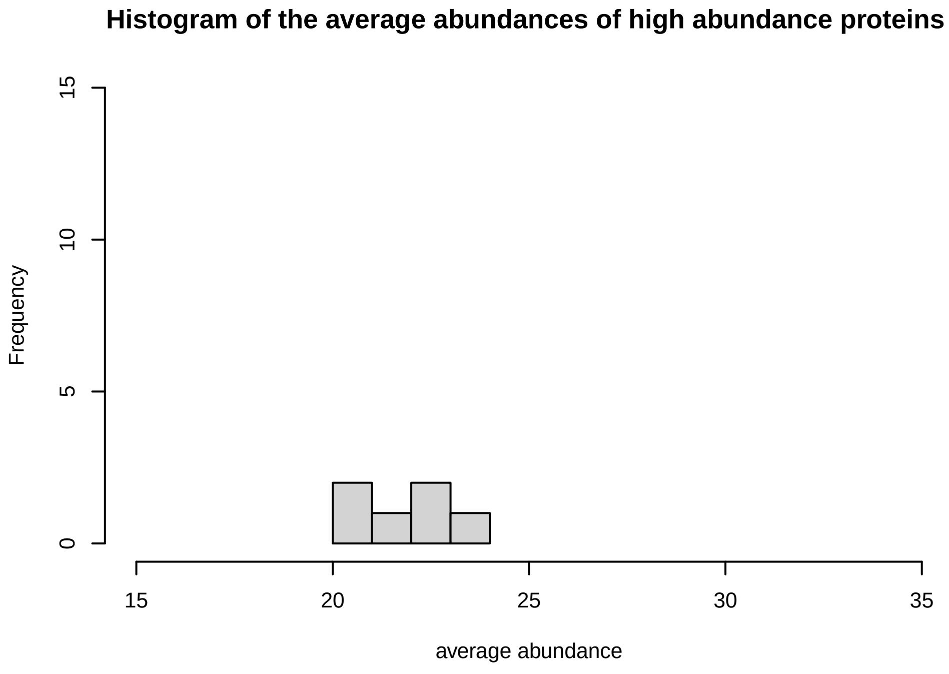
Histogram of the average abundances of high abundance proteins
Frequency
average abundance

### V19

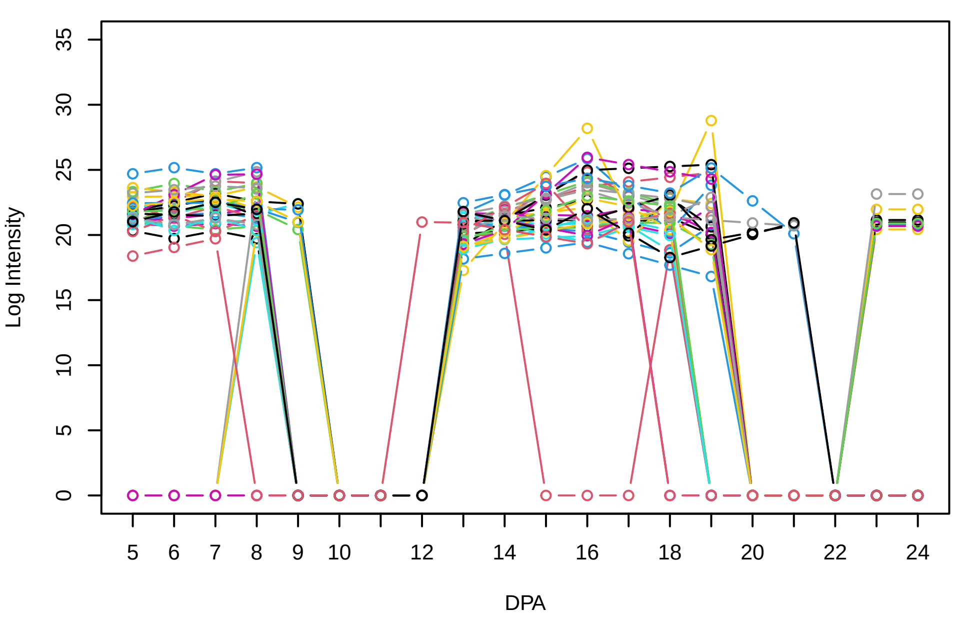


### Histogram of the average abundances of non−zero values

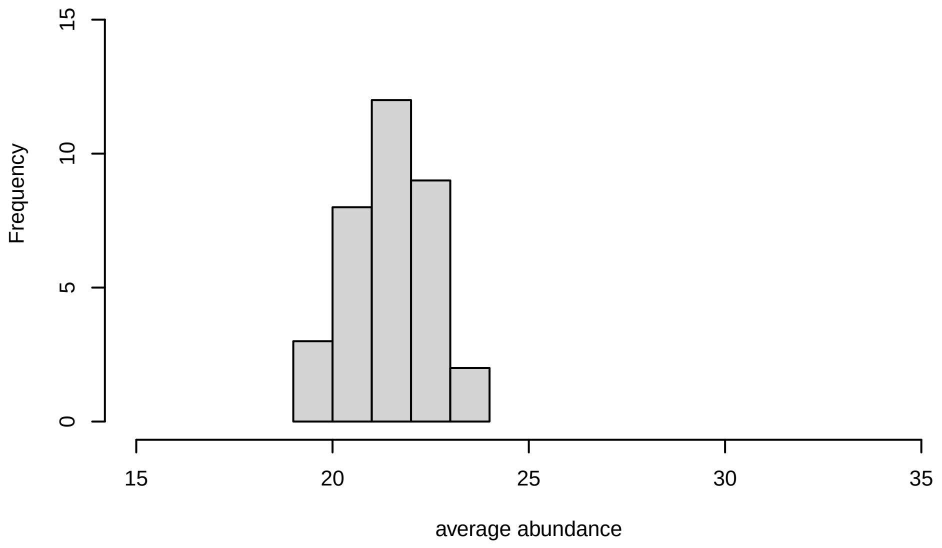


### V19, after remove outlier and cutoff is 0.8

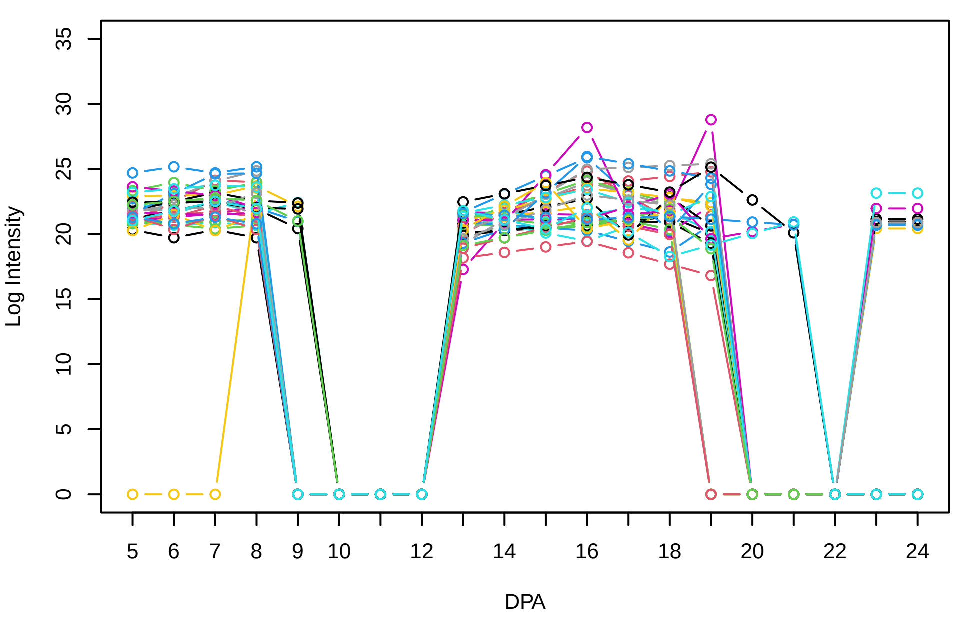


### Histogram of the average abundances of non−outlier proteins

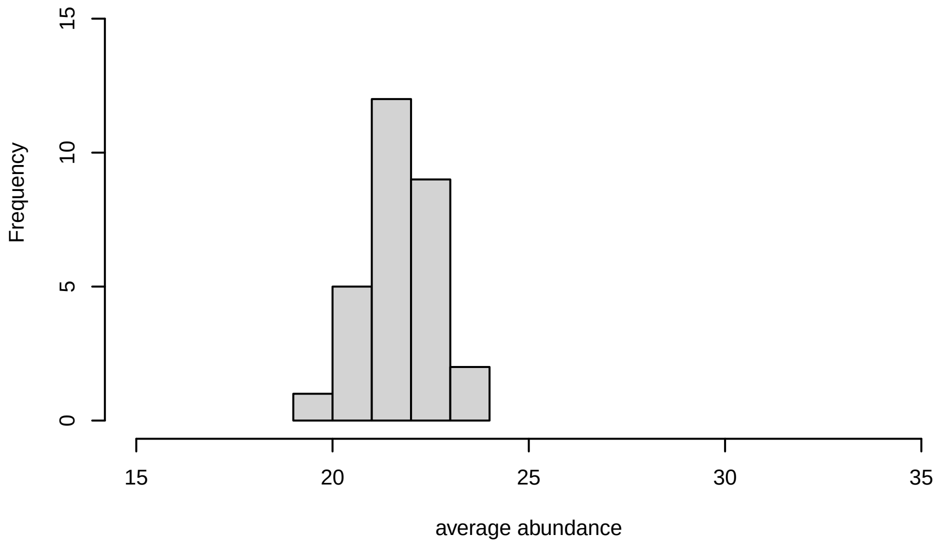


### V19, after remove unreliable

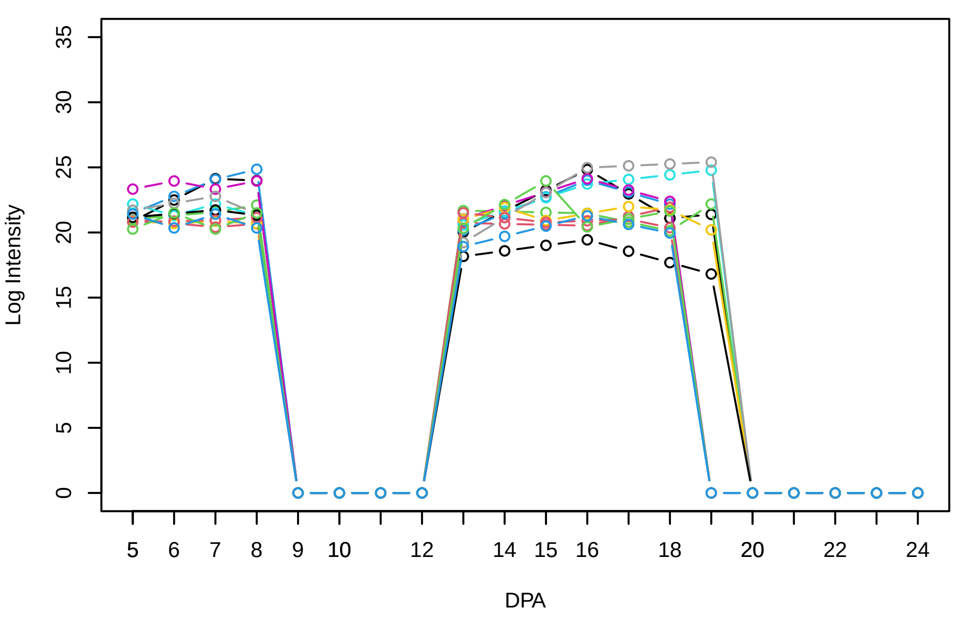


### Histogram of the average abundances of reliable proteins

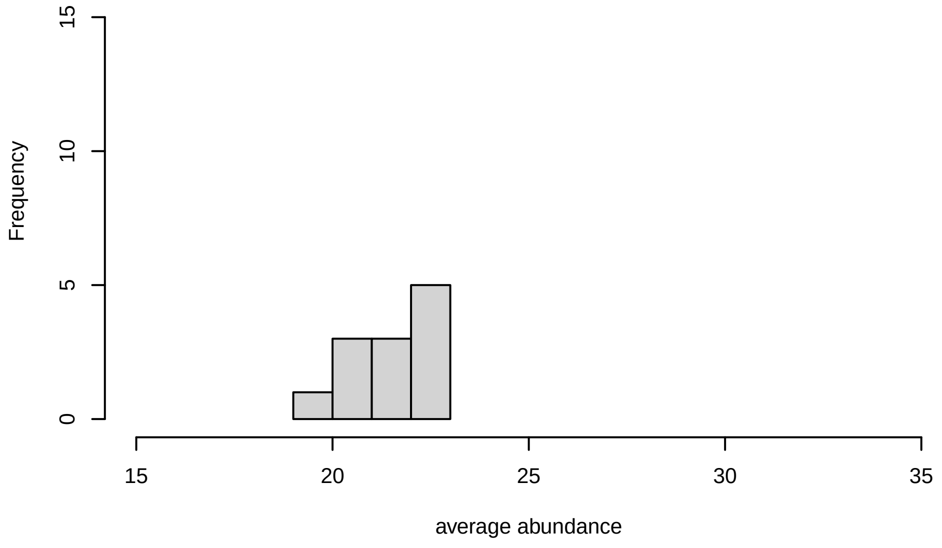


### V19, after remove low abundance

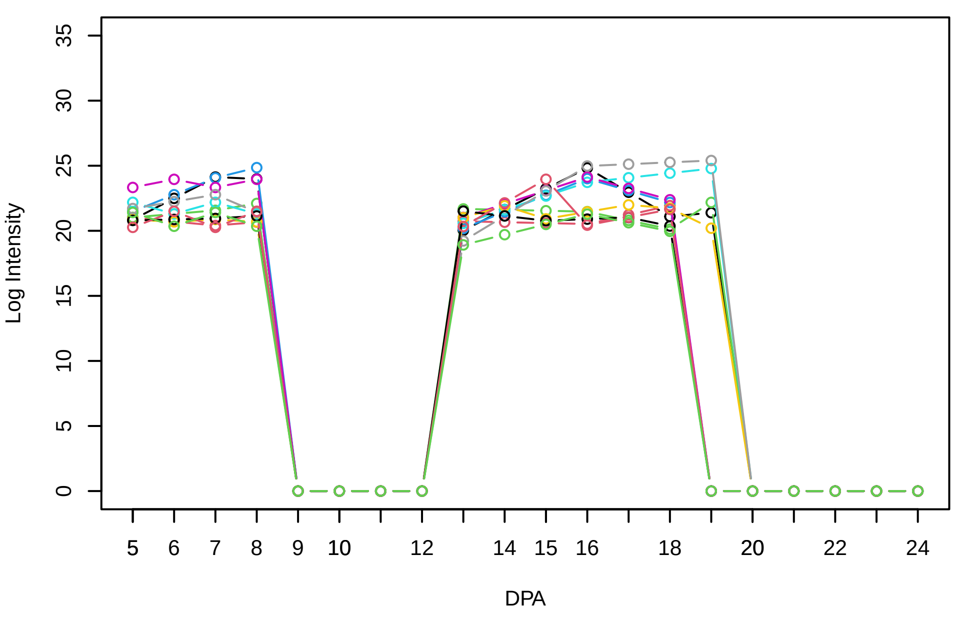


### Histogram of the average abundances of high abundance proteins

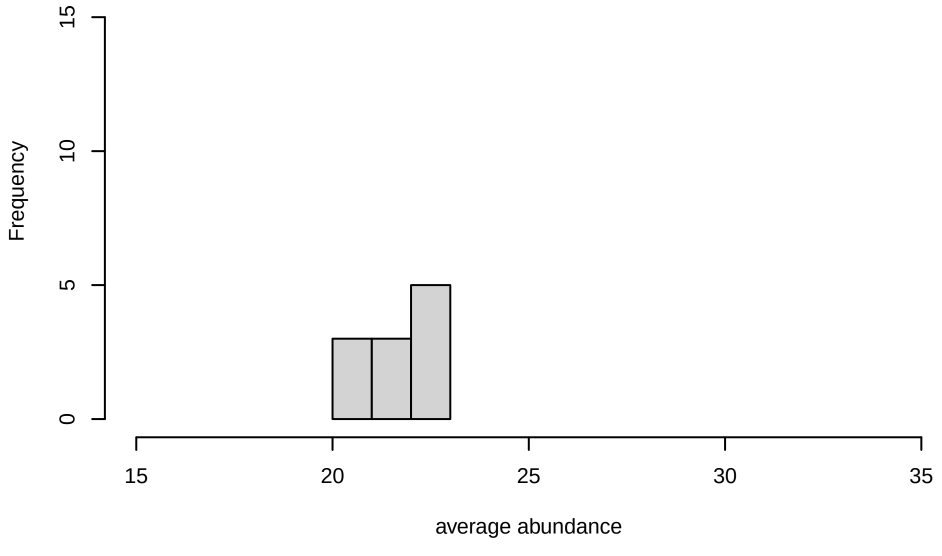

**V20**

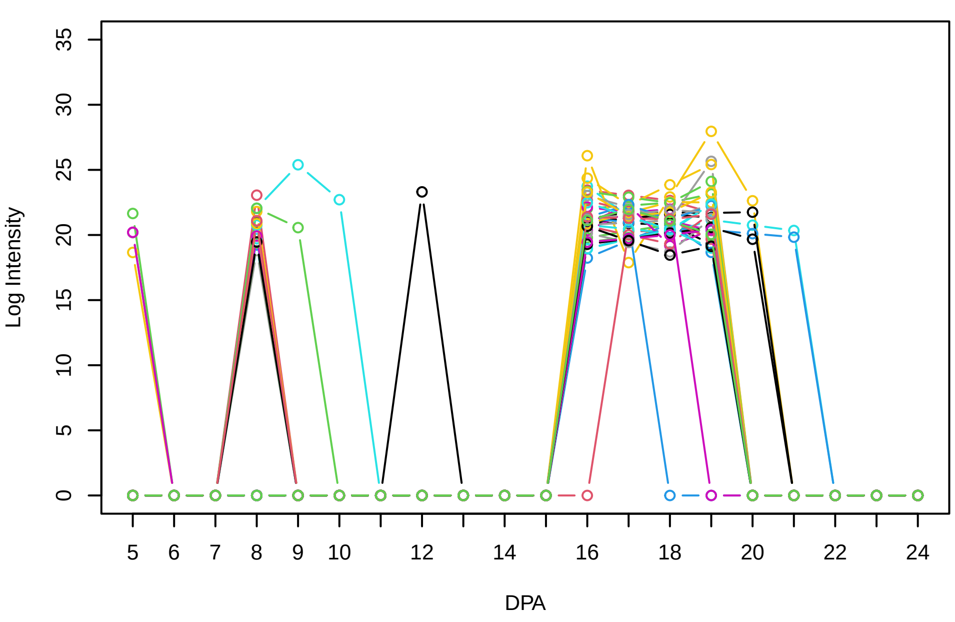


**Histogram of the average abundances of non−zero values**

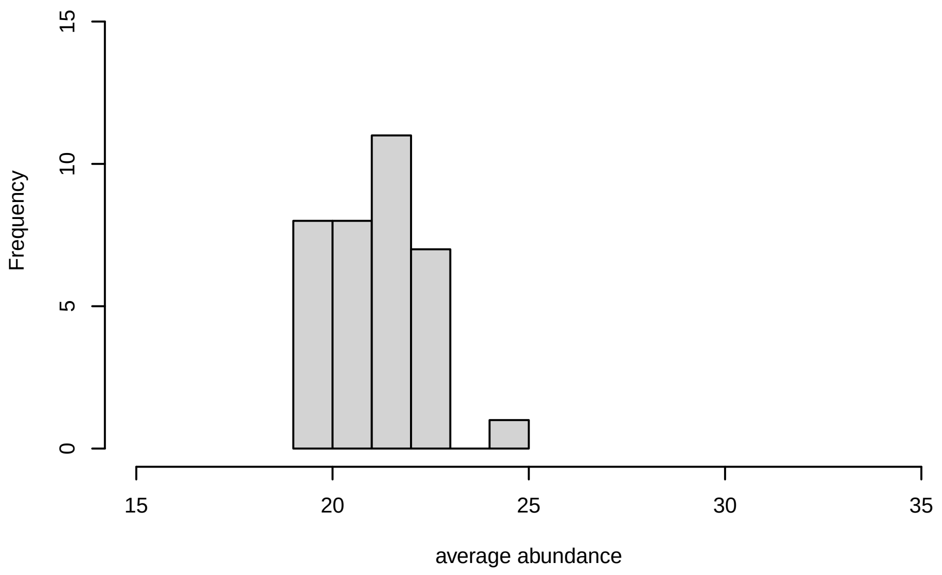


**V20, after remove outlier and cutoff is 0.8**

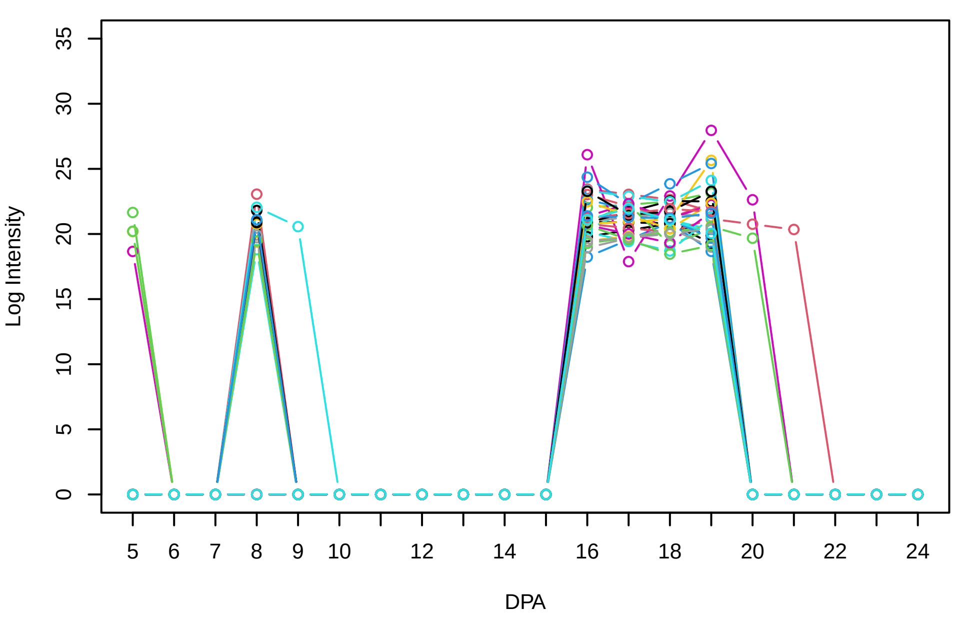


**Histogram of the average abundances of non−outlier proteins**

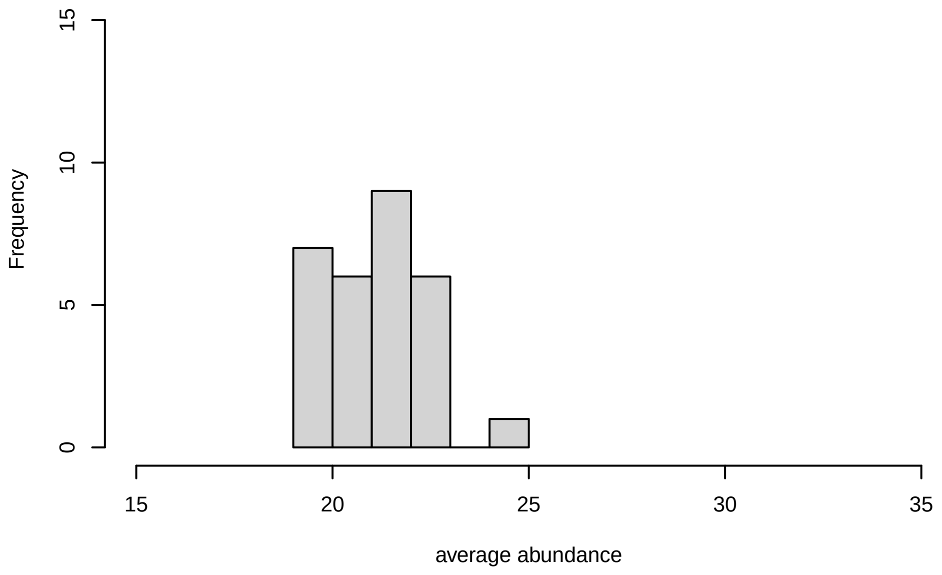


**V20, after remove unreliable**

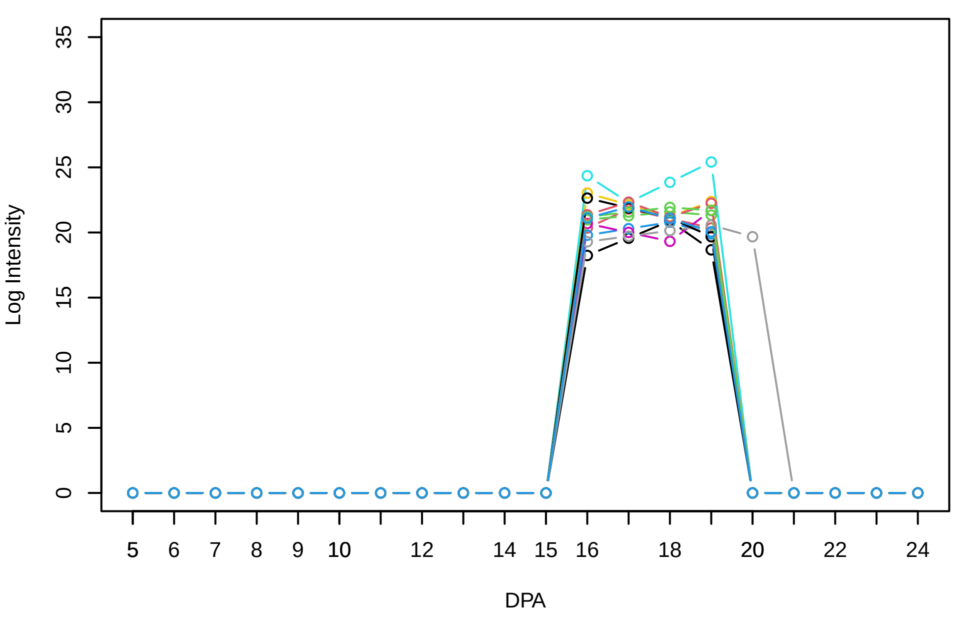


**Histogram of the average abundances of reliable proteins**

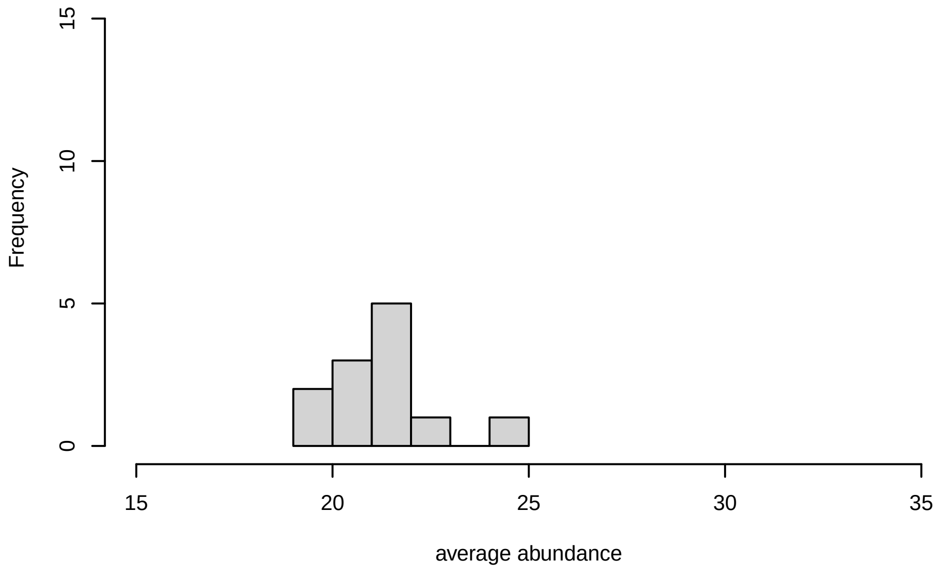


**V20, after remove low abundance**

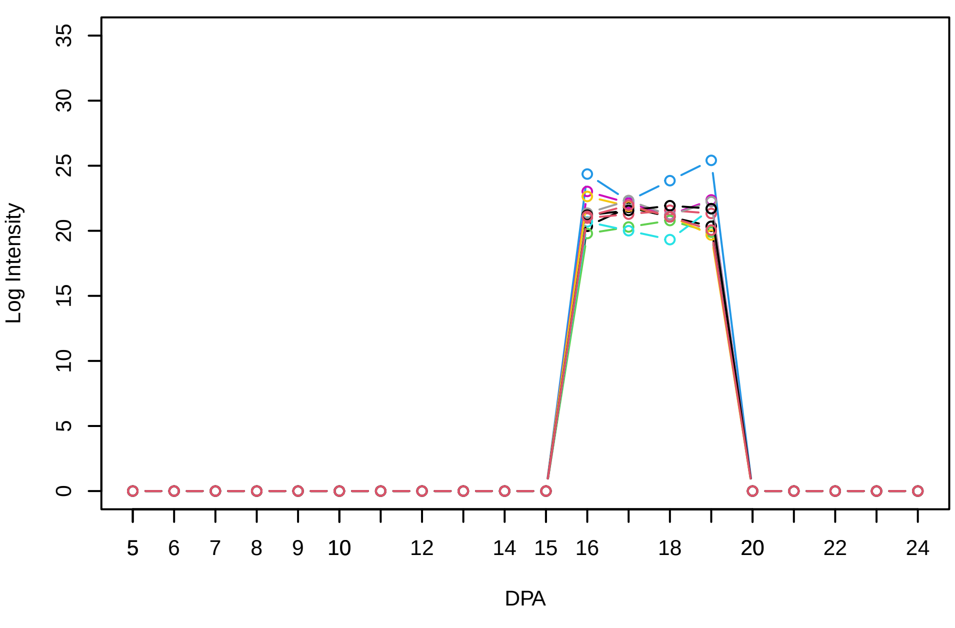


**Histogram of the average abundances of high abundance proteins**

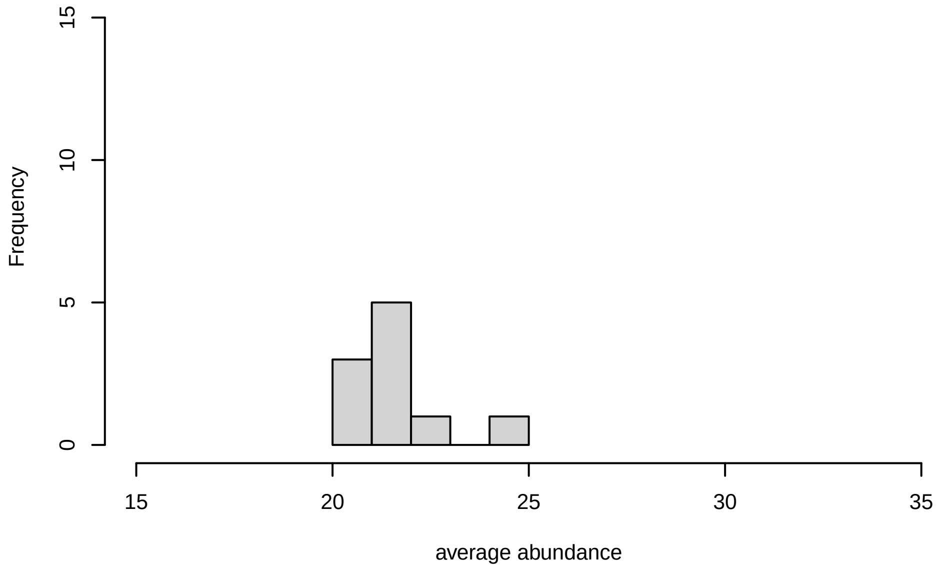
