## Supplementary Figure S5 for "Systems-level proteomic models of cotton fiber development: A high-resolution resource to analyze cell dynamics and trait engineering"

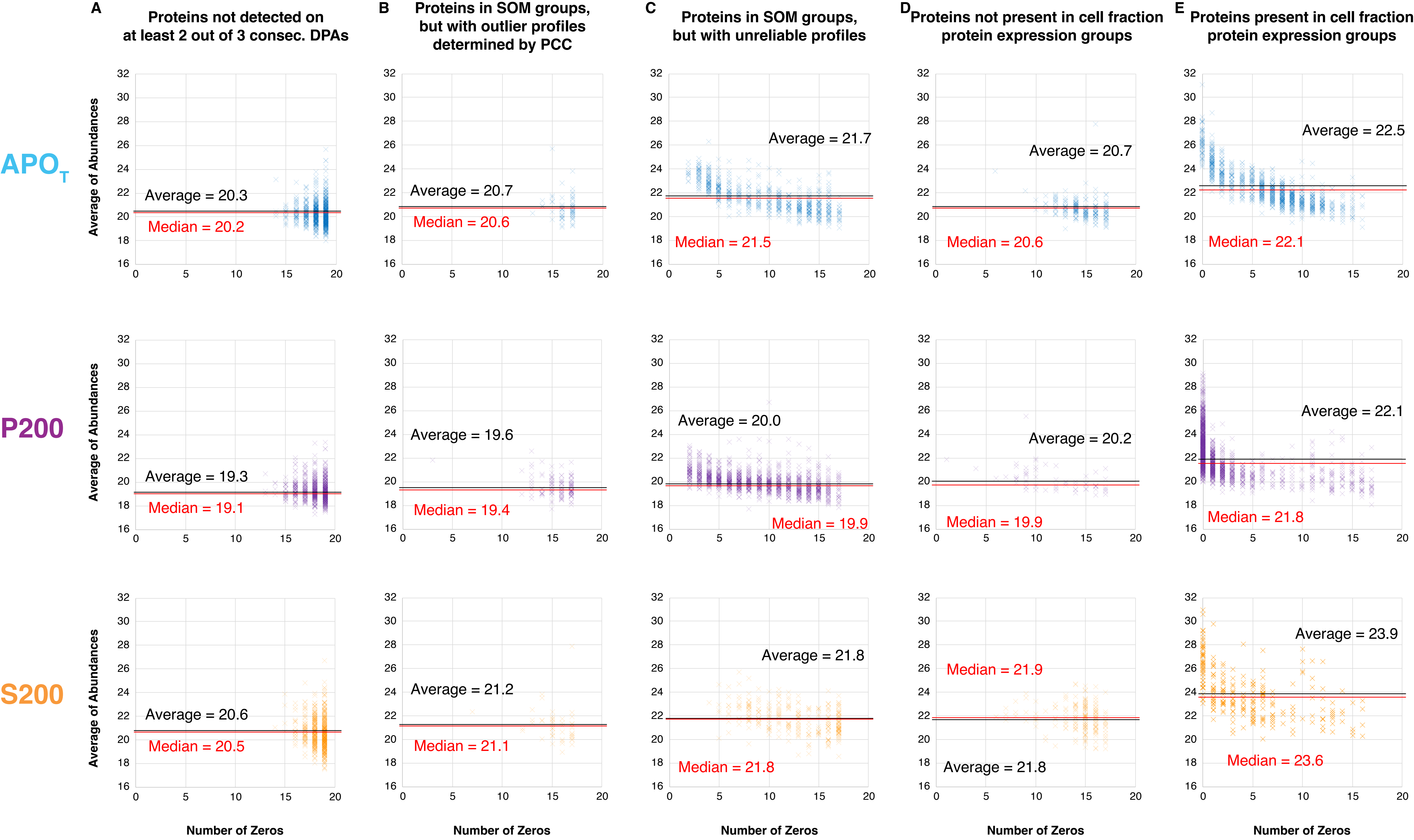


**Supplementary Figure S5.** Analysis of proteins removed at each data filtering step during the identification of reliable members.
**A,** Proteins that were not used in the SOM because they did not satisfy the criteria that 2 out 3 consecutive DPAs need to be adjacent. **B,** After SOM clustering, proteins that were identified as outliers by PCC of < 0.8 in each SOM group. **C,** Proteins that were defined as unreliable profiles. We scored a group as unreliable if the pattern reflected the overall pattern of protein coverage or if the pattern is dominated by patterns that are explained by the interpolation (e.g. 3 consecutive zeros defines the cluster). **D,** Proteins that were not present in the cell fraction protein expression groups. **E,** Proteins that are in the cell fraction protein expression groups.
