## Supplementary figures and images for "Systems-level proteomic models of cotton fiber development: A high-resolution resource to analyze cell dynamics and trait engineering"

### Supplementary Figure S6

#### Codes

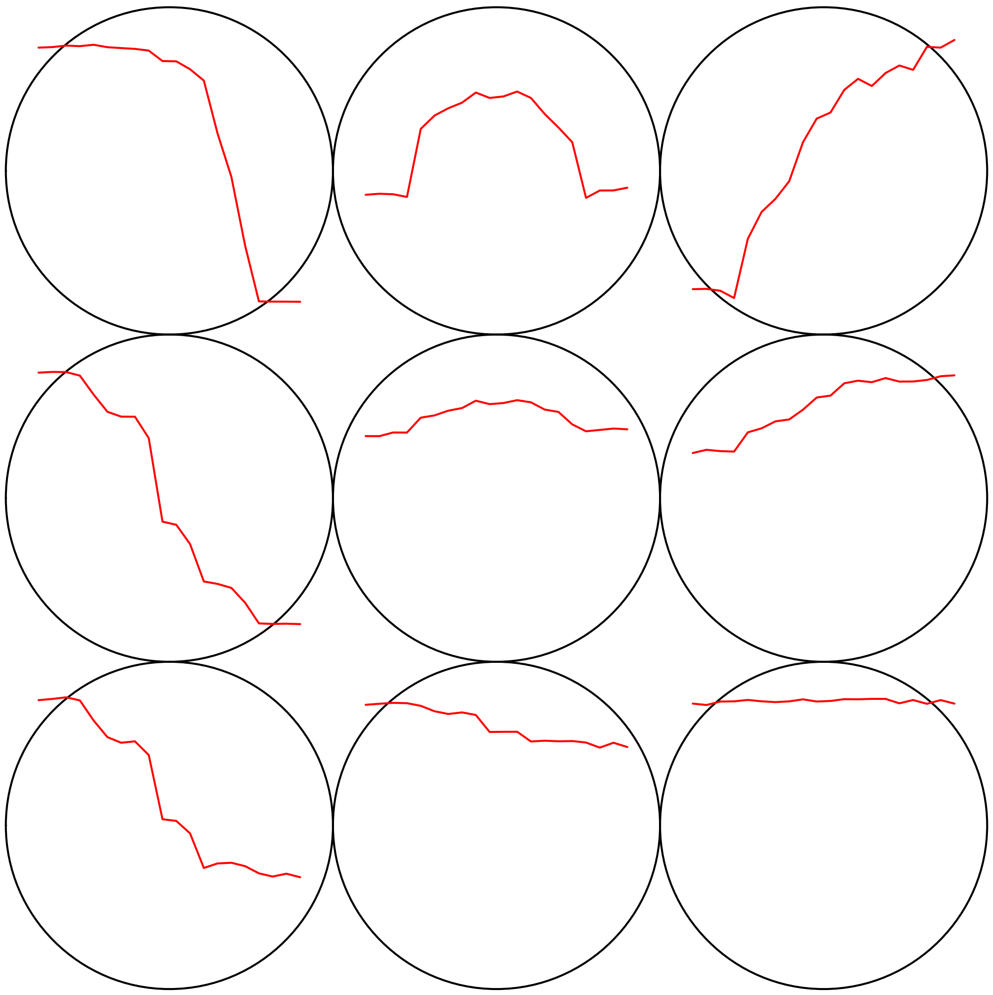

#### Count plot

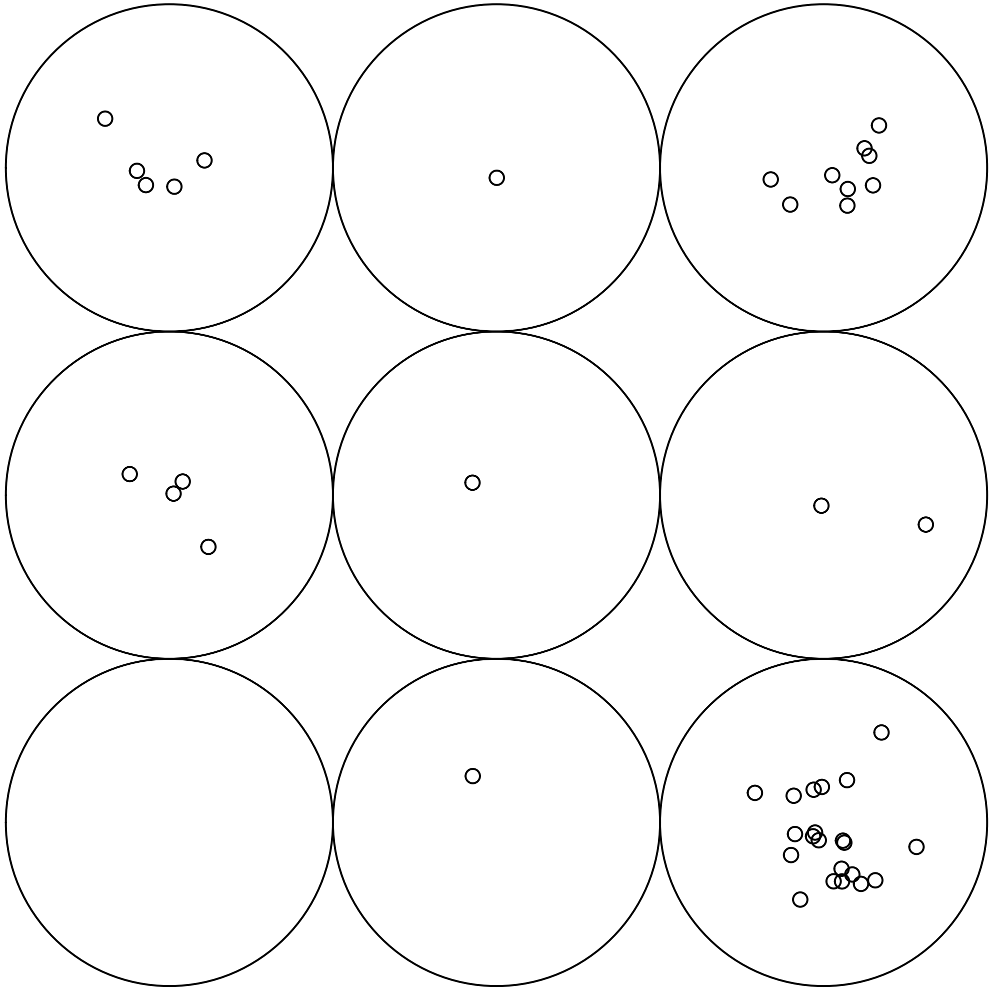

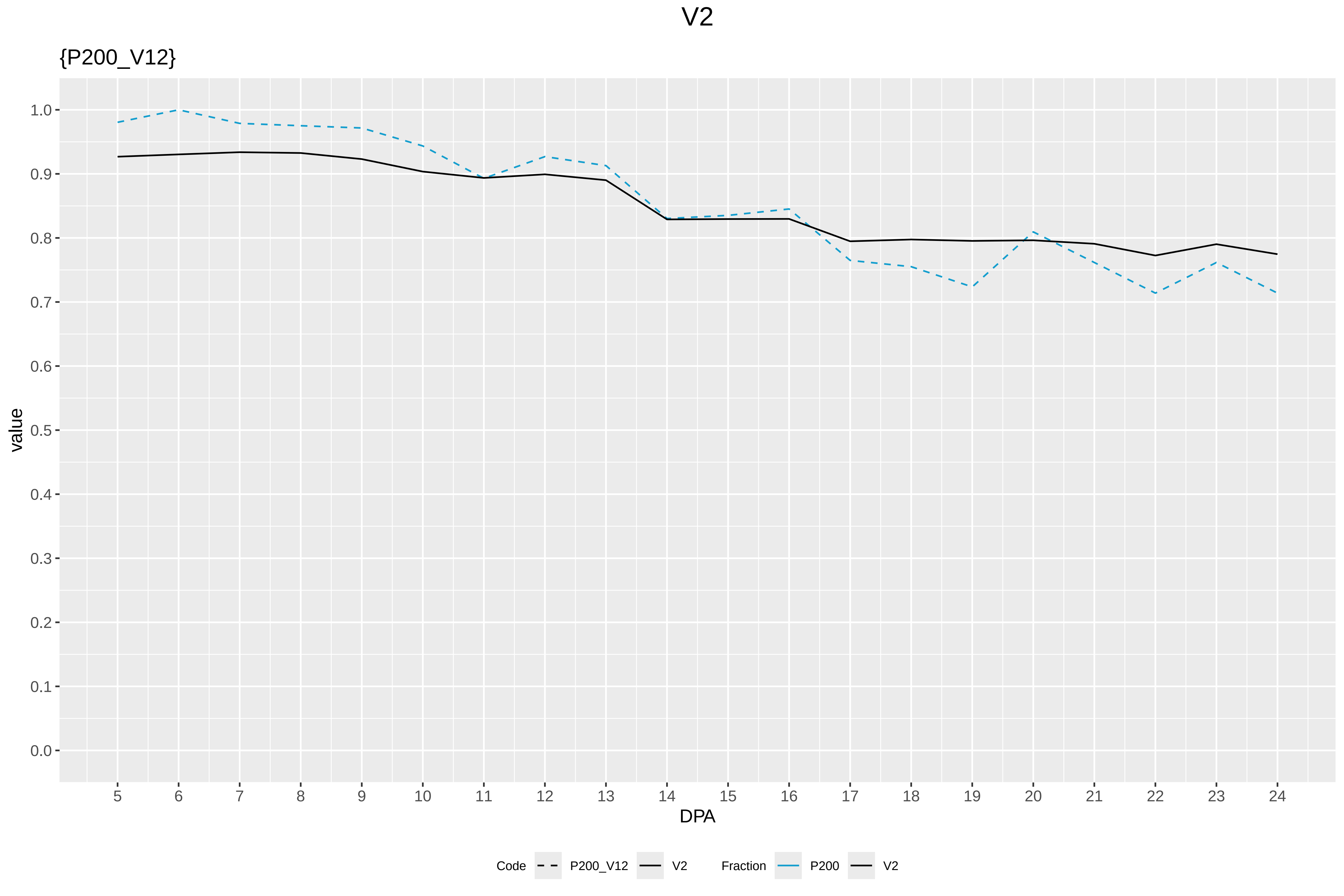

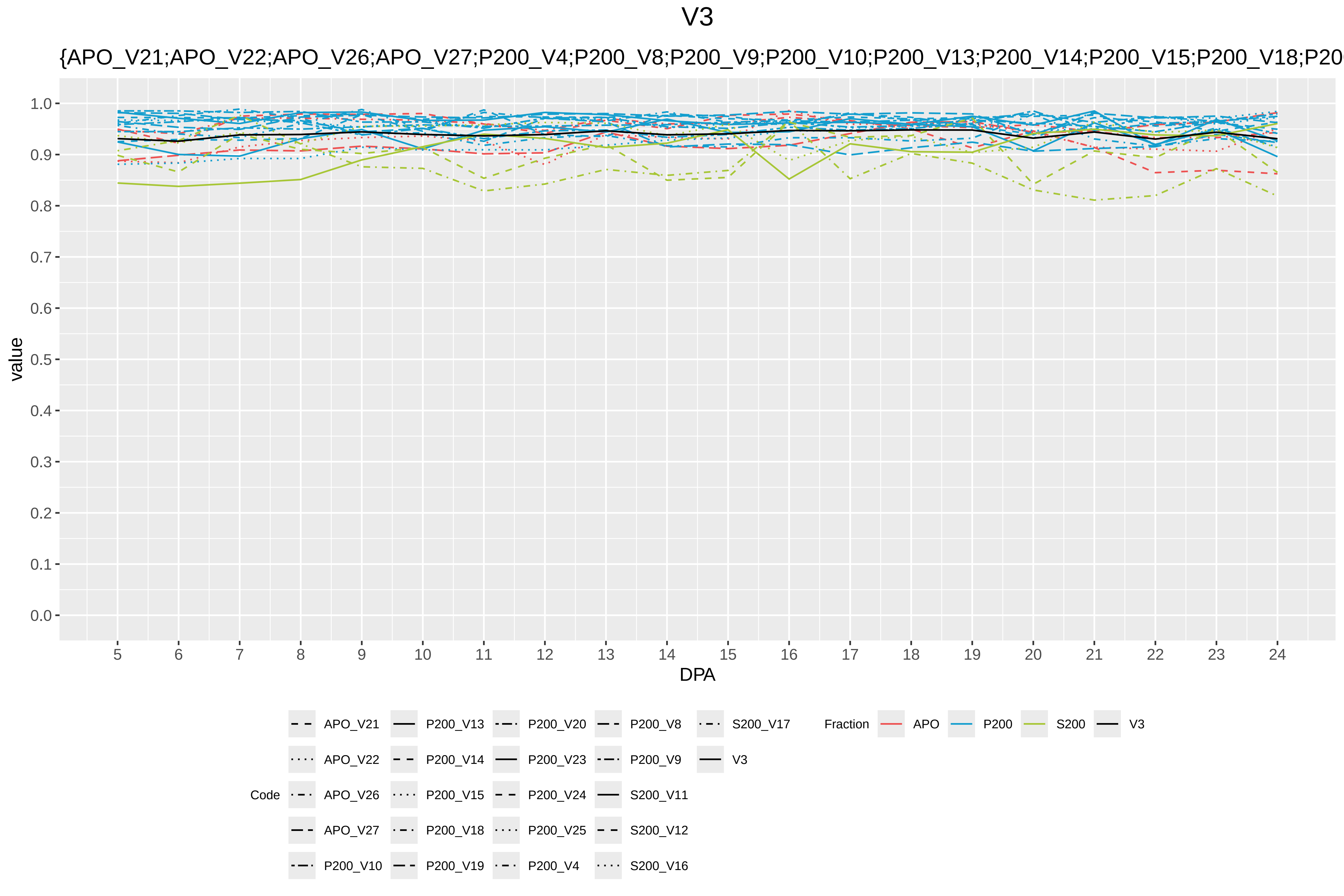

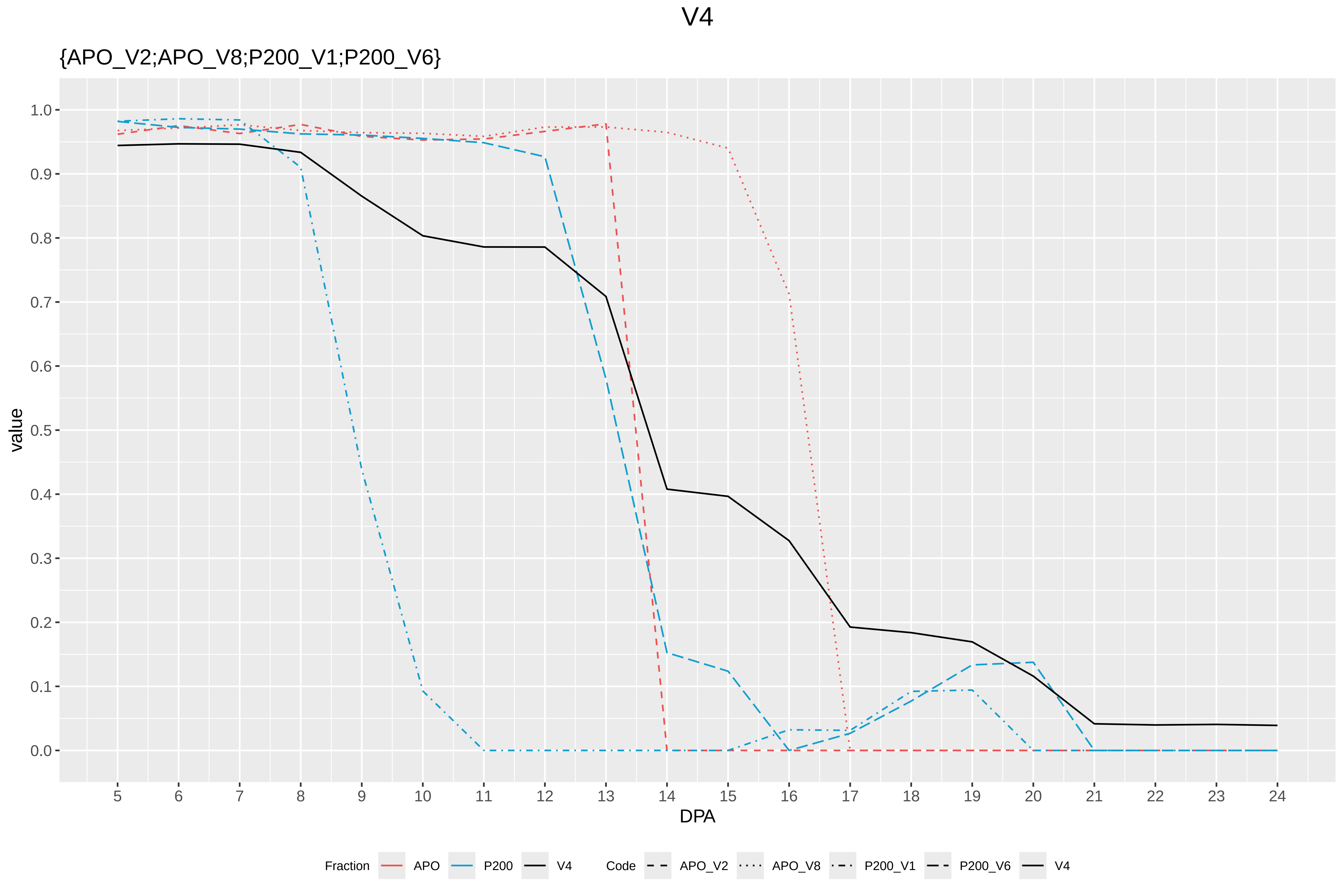

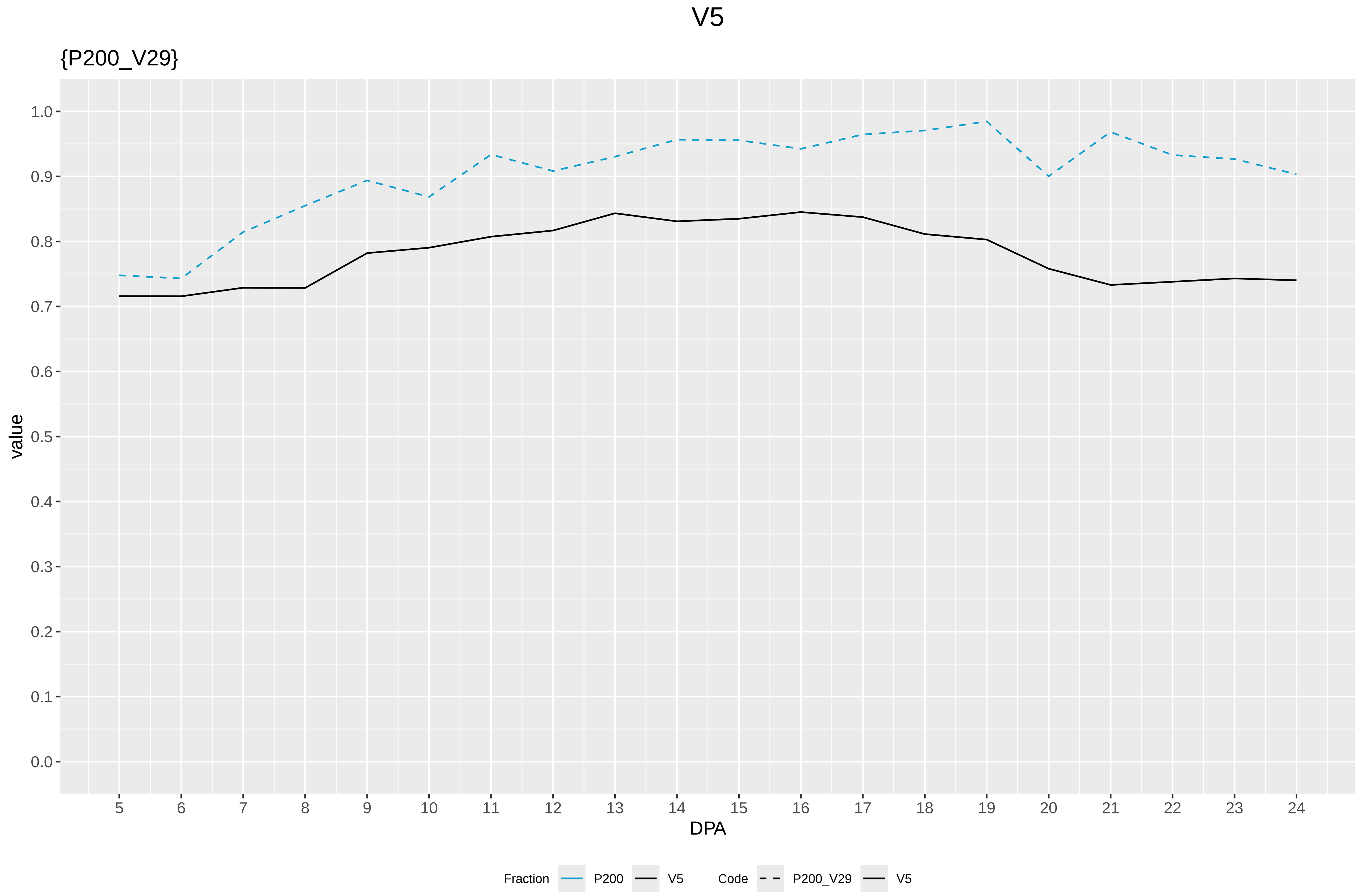

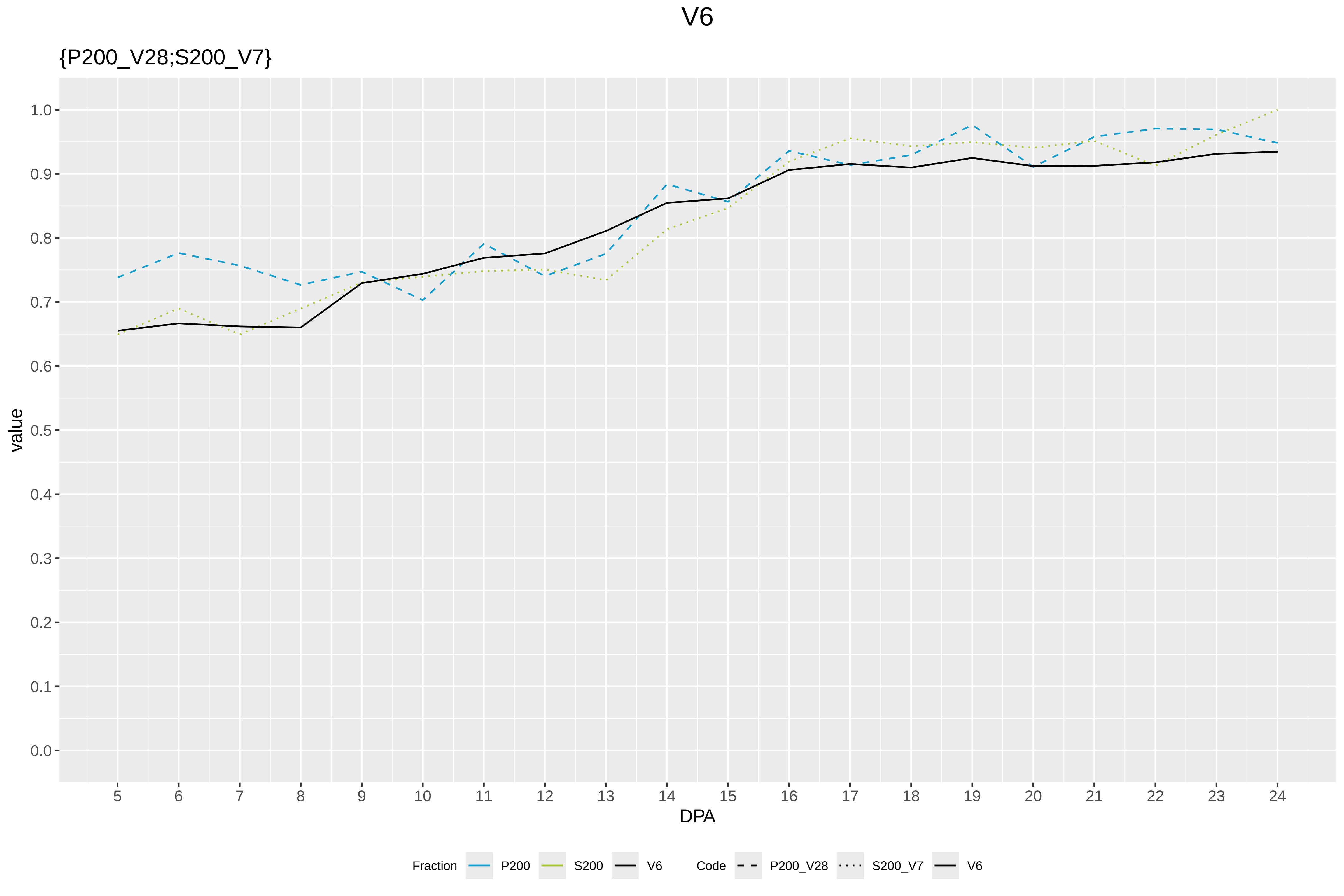

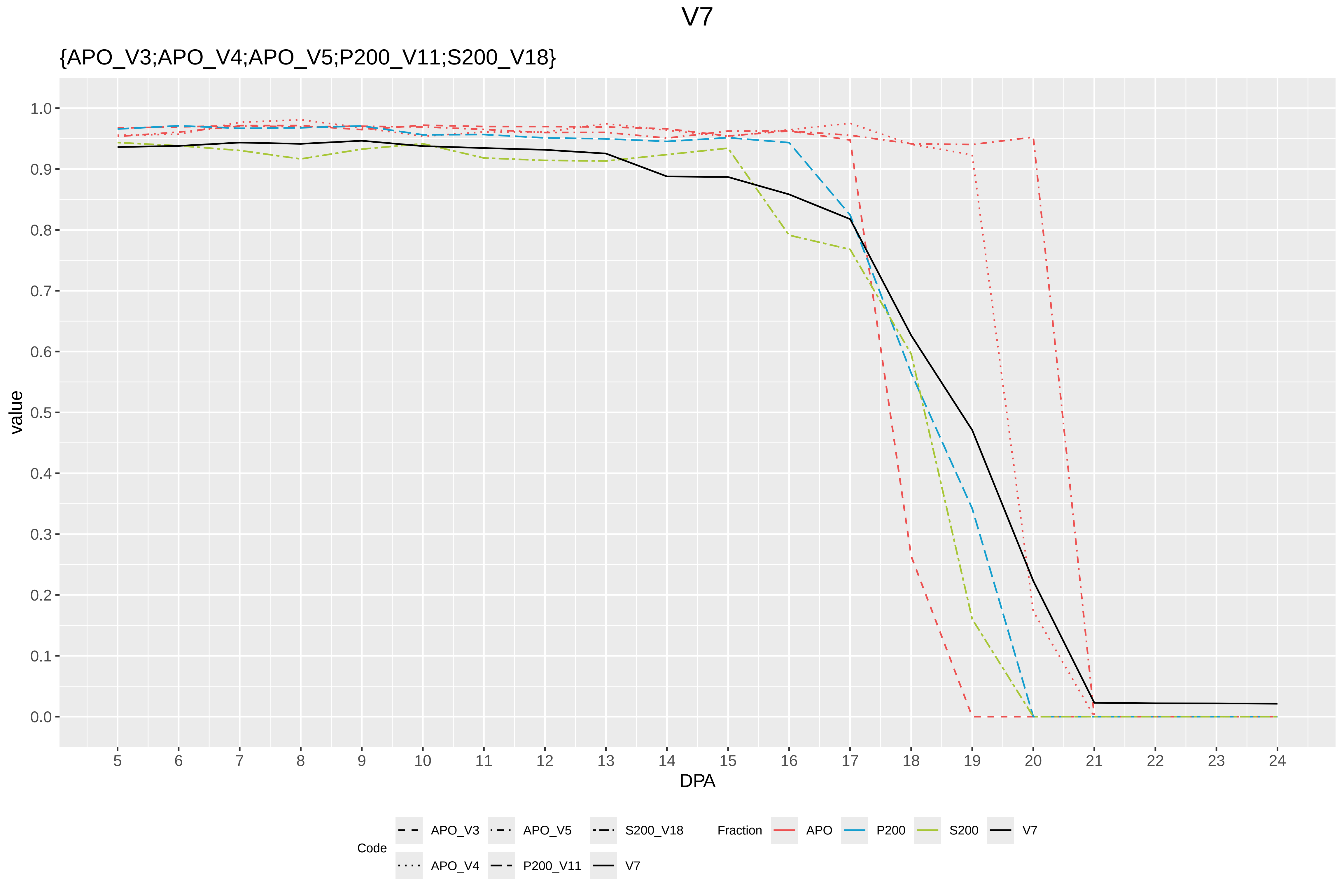

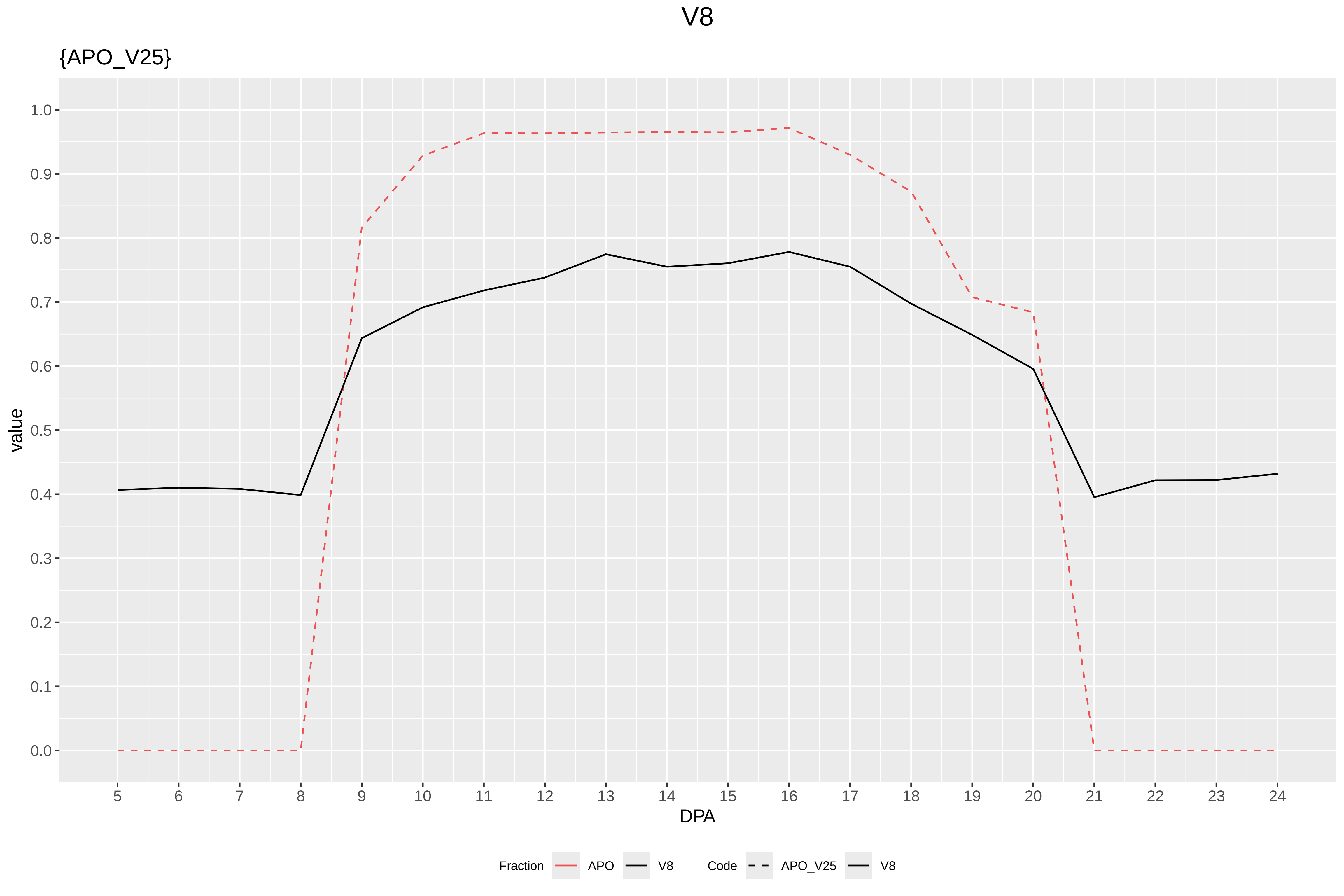

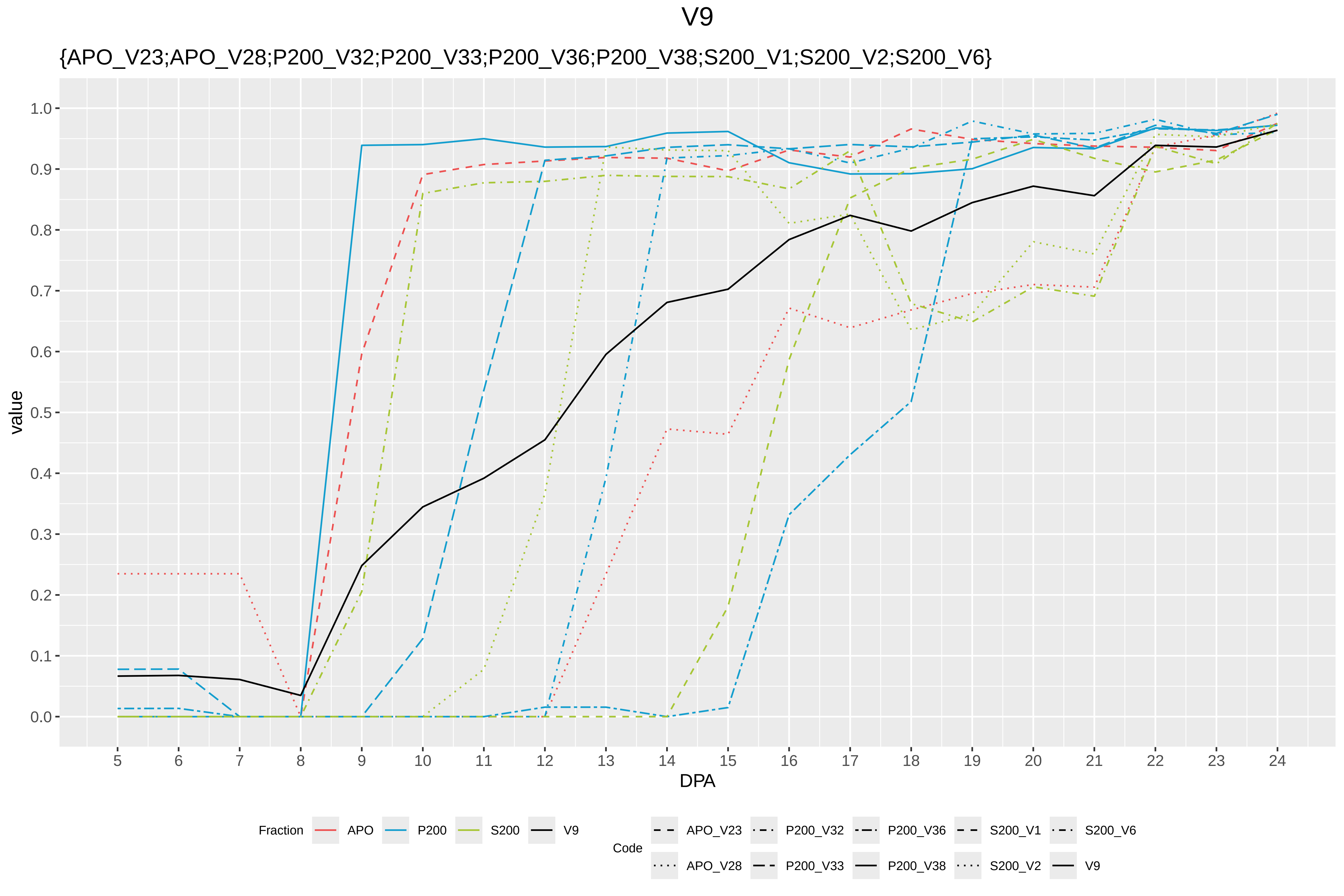
