## Supplementary Figure S7 for "Systems-level proteomic models of cotton fiber development: A high-resolution resource to analyze cell dynamics and trait engineering"

### Codes

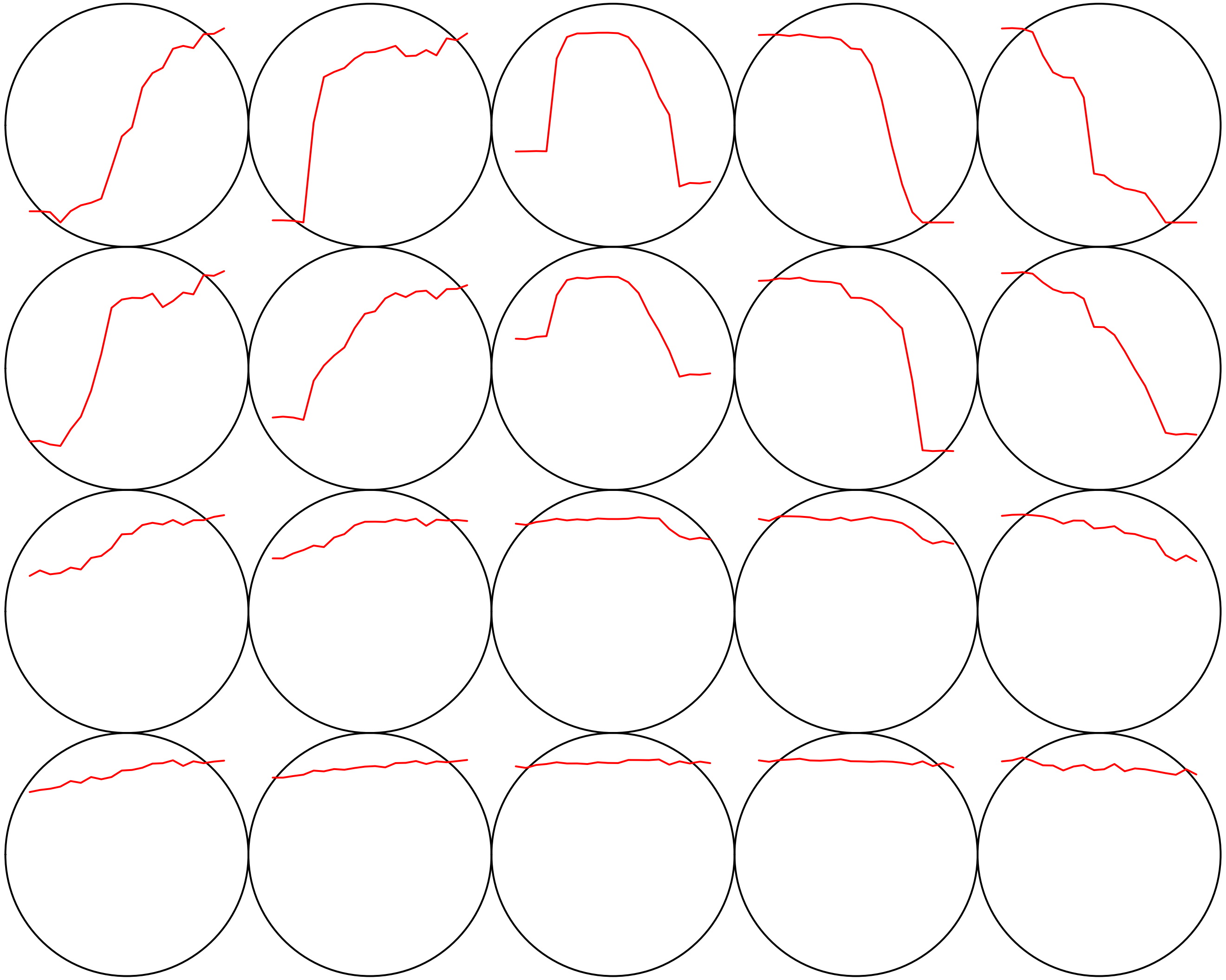

#### Count plot

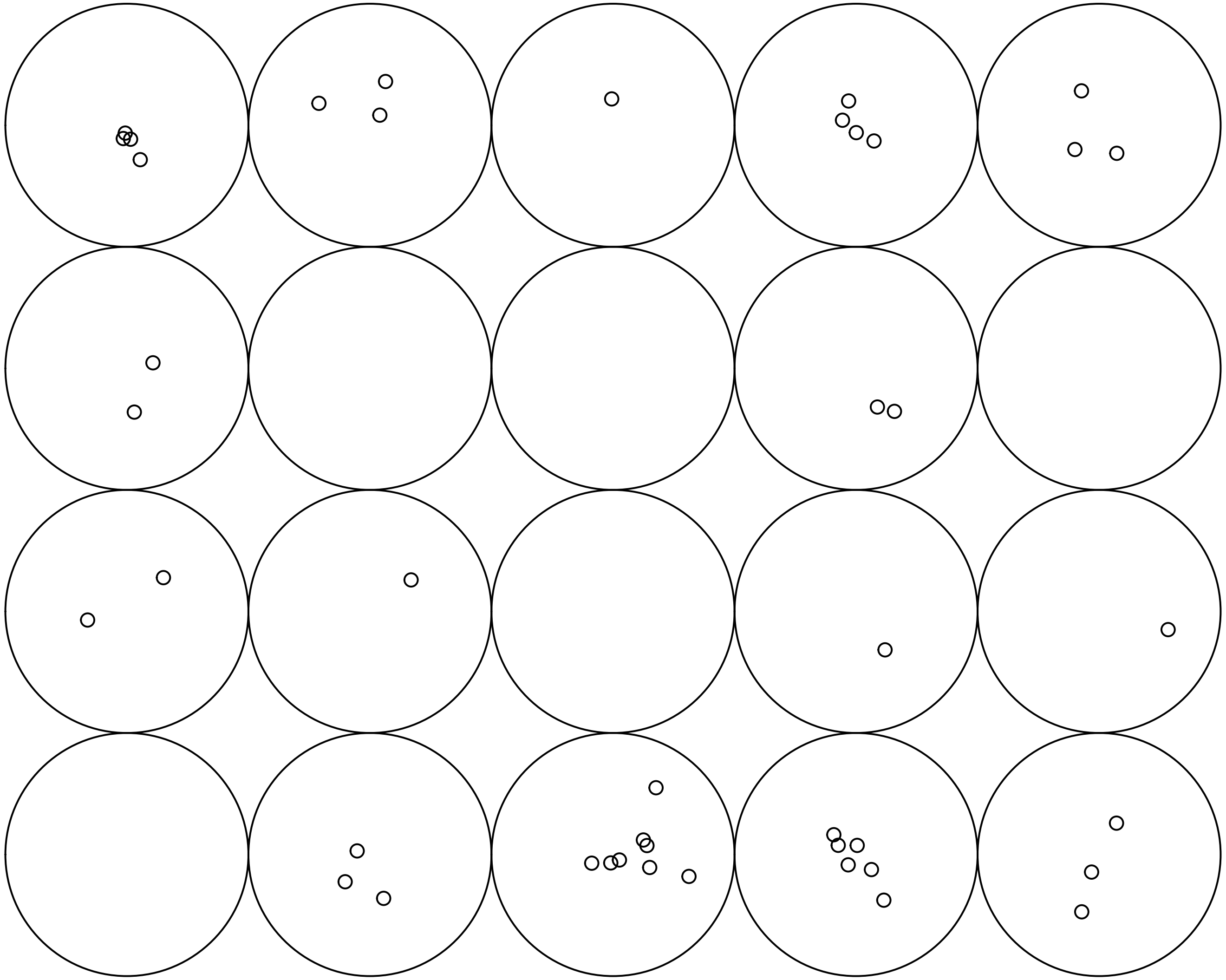

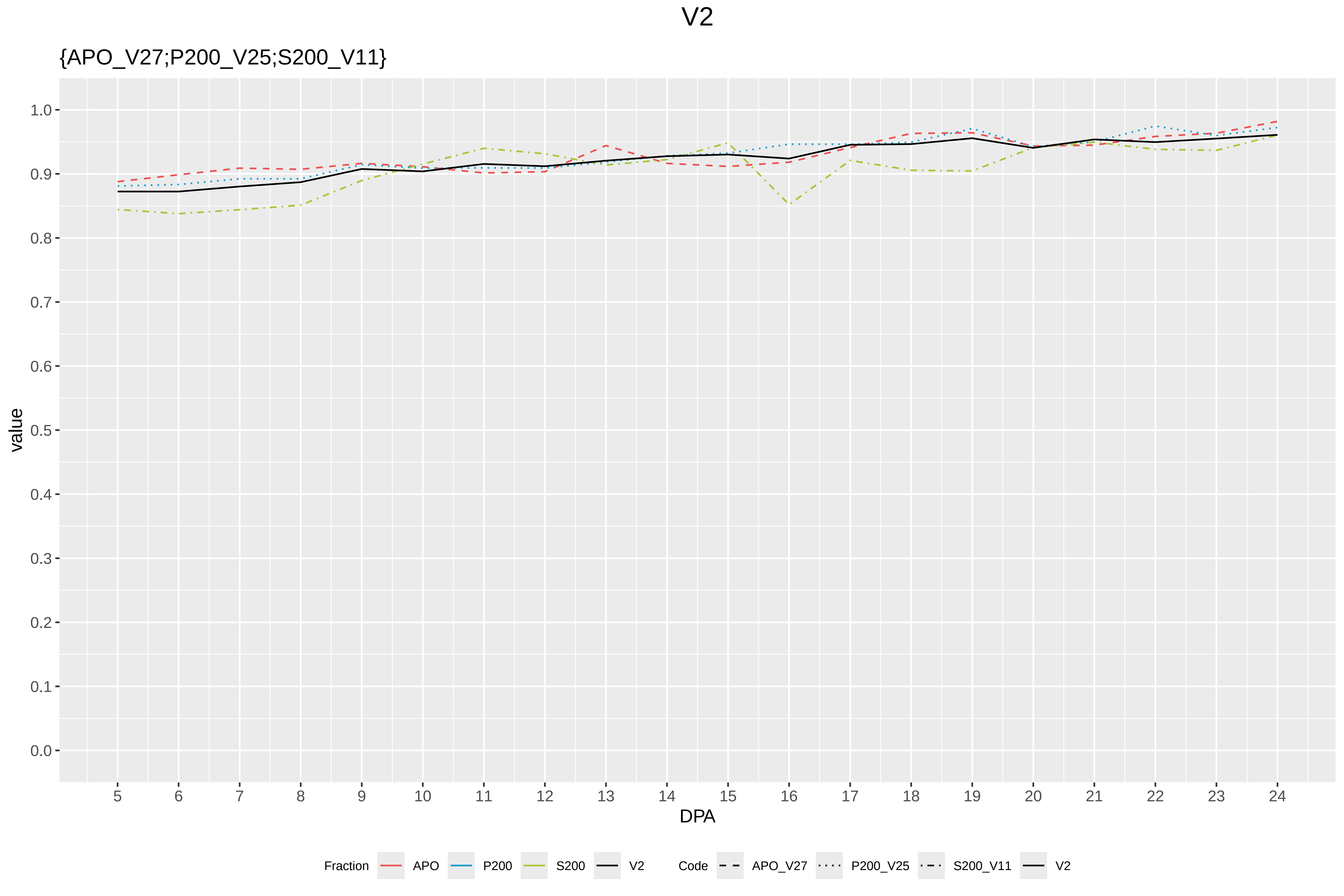

V2
{APO_V27;P200_V25;S200_V11}
value
1.0
0.9
0.8
0.7
0.6
0.5
0.4
0.3
0.2
0.1
0.0
DPA
Fraction
APO
P200
S200
V2
Code
APO_V27
P200_V25
S200_V11
V2

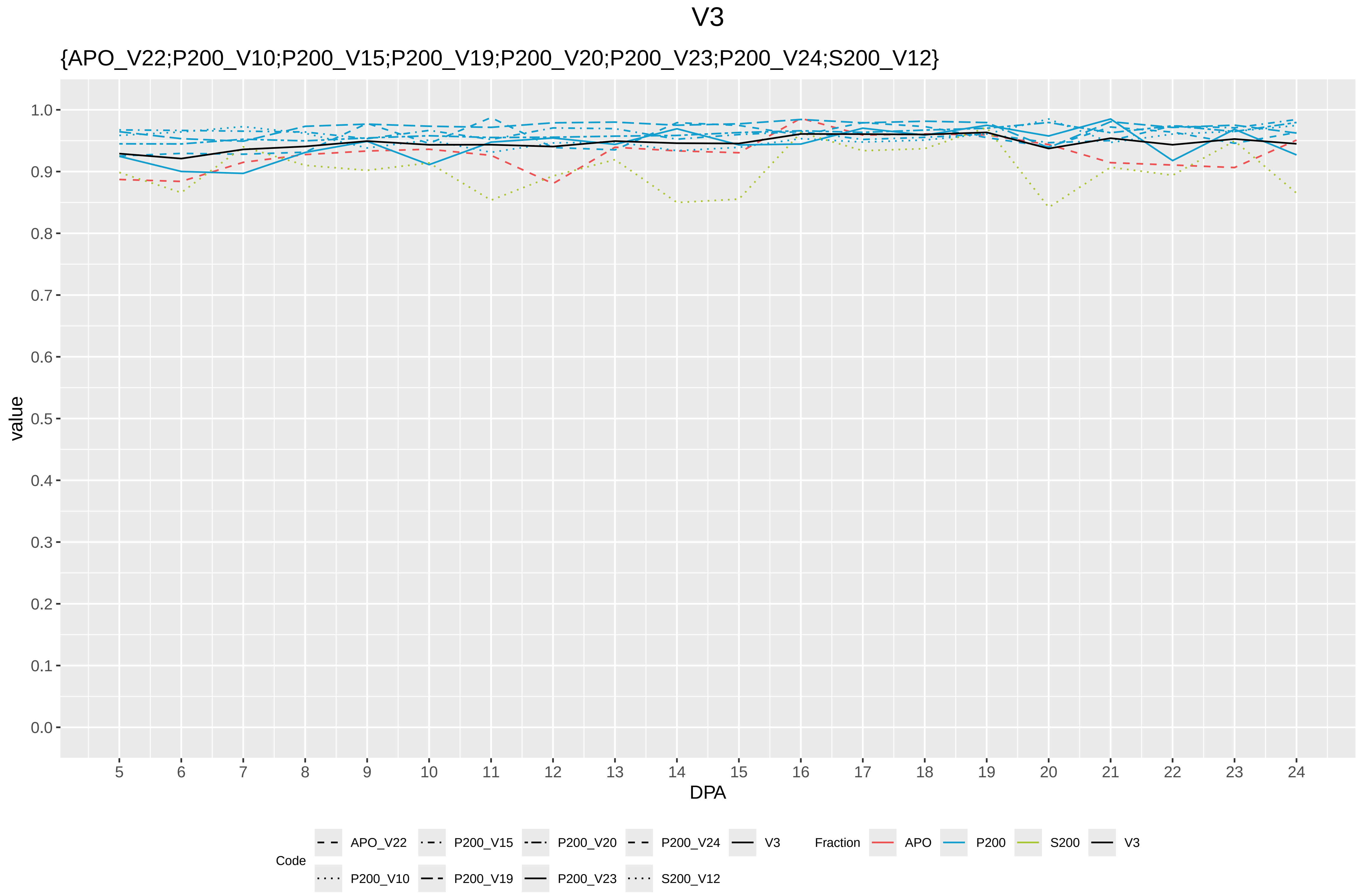
V3
{APO_V22;P200_V10;P200_V15;P200_V19;P200_V20;P200_V23;P200_V24;S200_V12}
value
1.0
0.9
0.8
0.7
0.6
0.5
0.4
0.3
0.2
0.1
0.0
DPA
Code
APO_V22
P200_V10
P200_V15
P200_V19
P200_V20
P200_V23
P200_V24
S200_V12
V3
Fraction
APO
P200
S200
V3

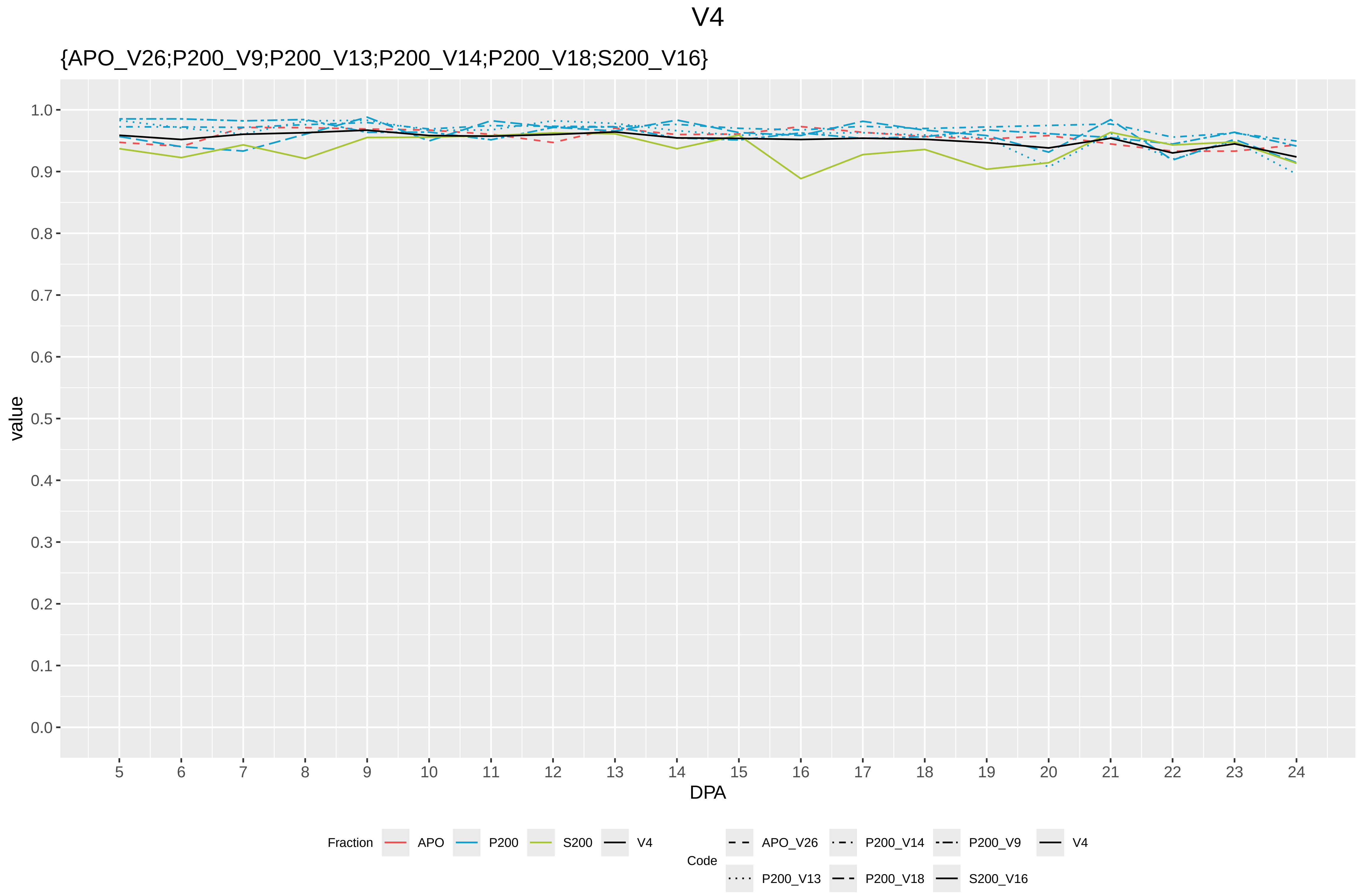
V4
{APO_V26;P200_V9;P200_V13;P200_V14;P200_V18;S200_V16}
value
1.0
0.9
0.8
0.7
0.6
0.5
0.4
0.3
0.2
0.1
0.0
DPA
Fraction
APO
P200
S200
V4
Code
APO_V26
P200_V14
P200_V9
V4
P200_V13
P200_V18
S200_V16

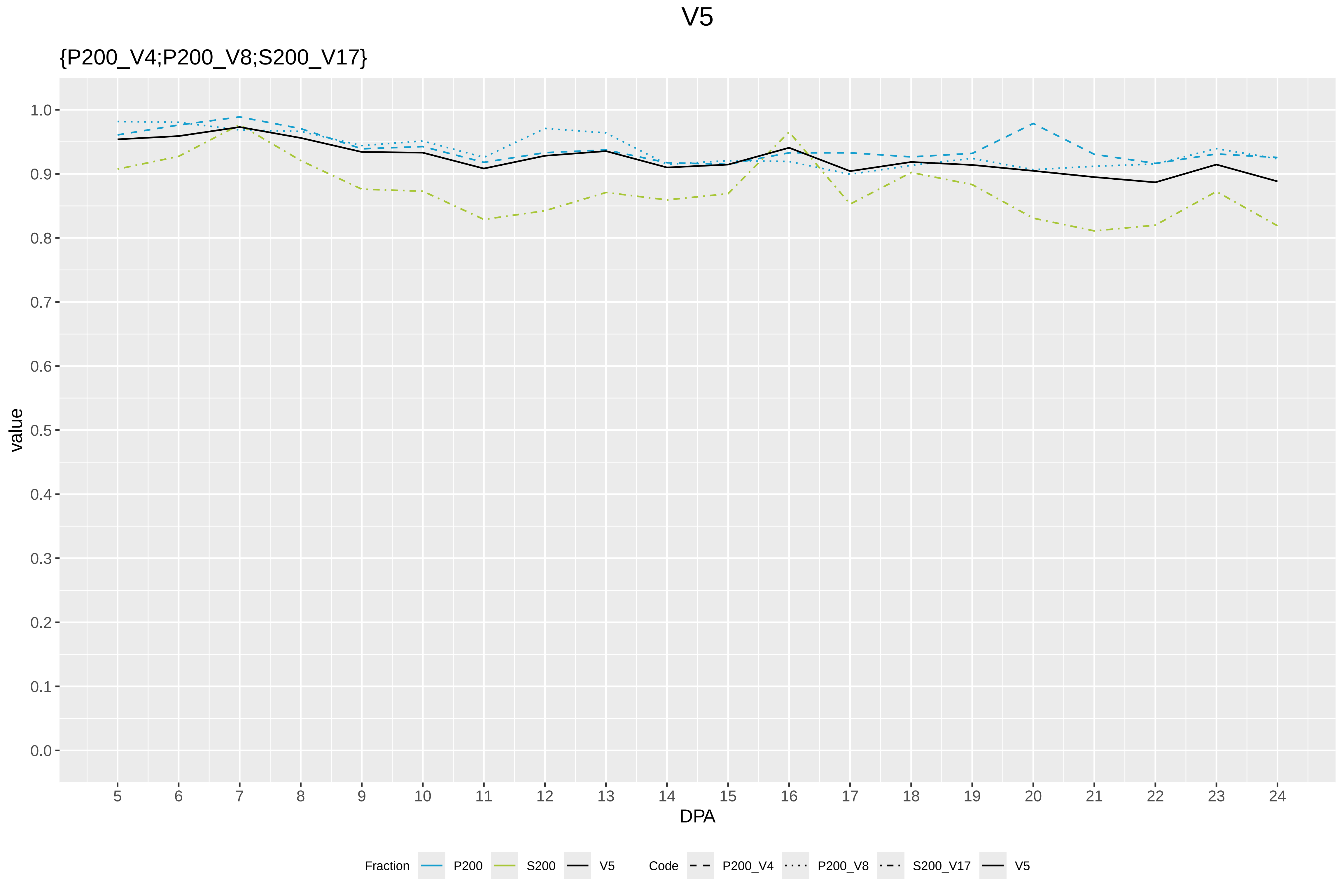

V5
{P200_V4;P200_V8;S200_V17}
value
DPA
1.0
0.9
0.8
0.7
0.6
0.5
0.4
0.3
0.2
0.1
0.0
5 6 7 8 9 10 11 12 13 14 15 16 17 18 19 20 21 22 23 24
Fraction P200 S200 V5
Code P200_V4 P200_V8 S200_V17 V5

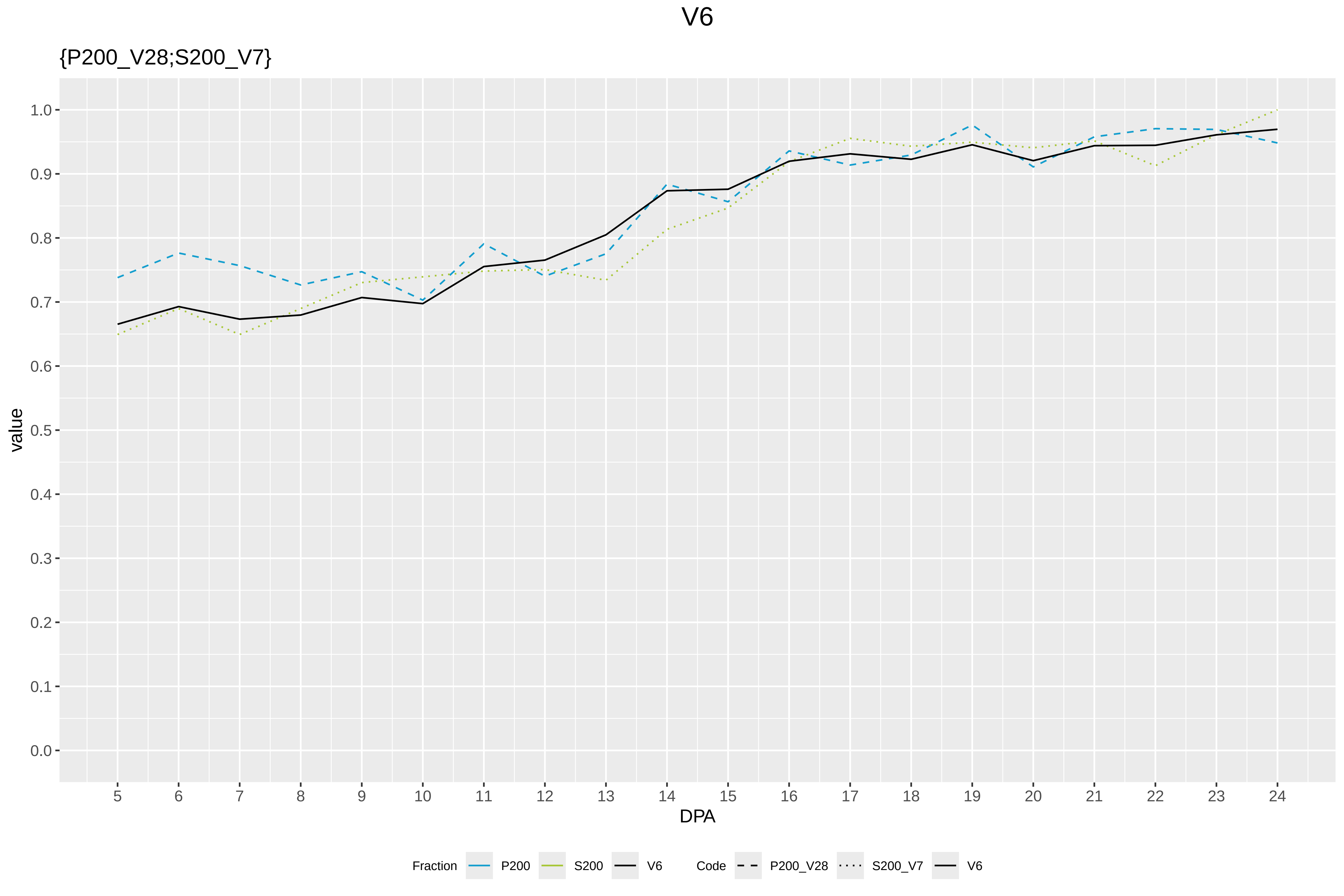
V6
{P200_V28;S200_V7}
value
1.0
0.9
0.8
0.7
0.6
0.5
0.4
0.3
0.2
0.1
0.0
DPA
Fraction
P200
S200
V6
Code
P200_V28
S200_V7
V6

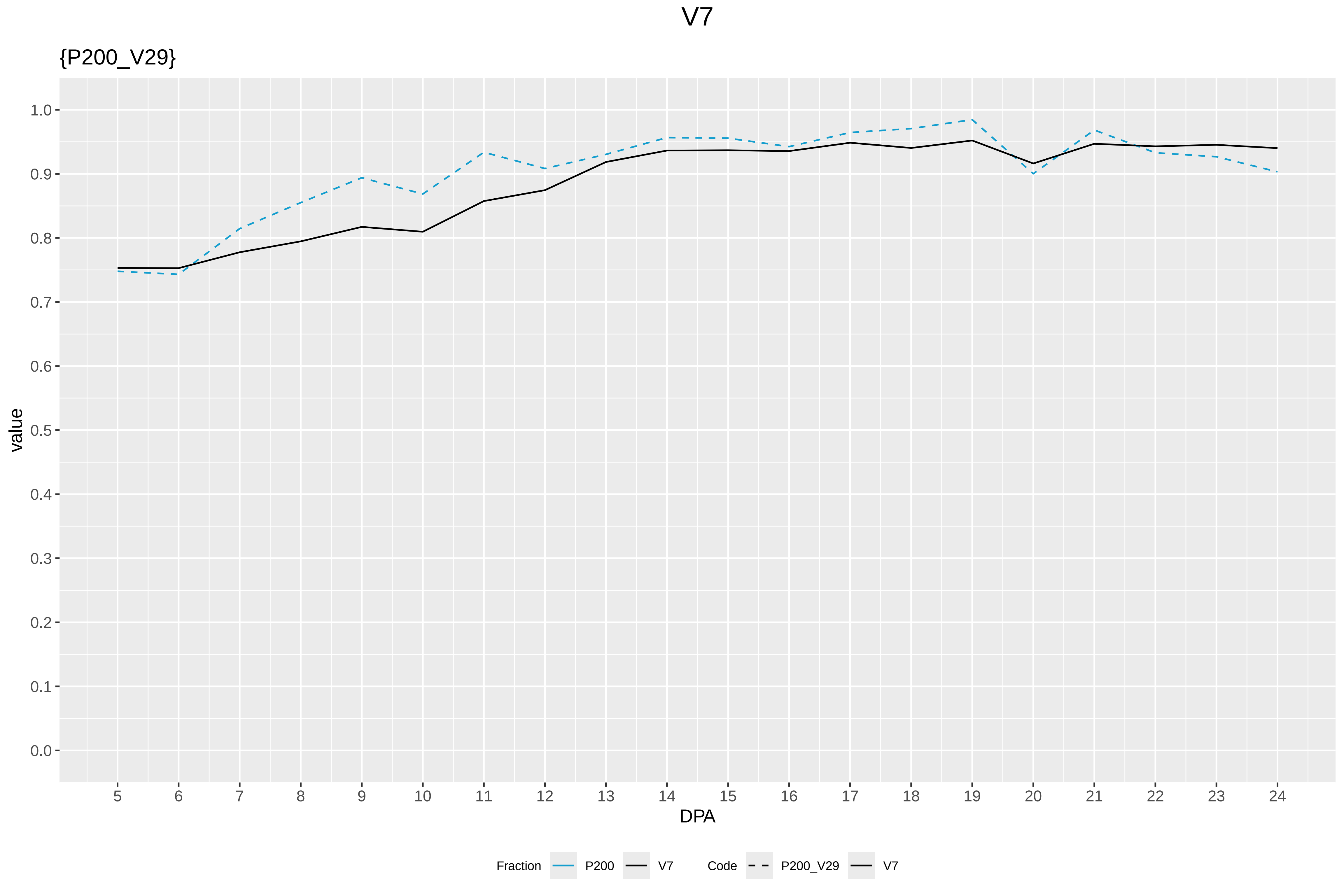

V7
{P200_V29}
value
1.0
0.9
0.8
0.7
0.6
0.5
0.4
0.3
0.2
0.1
0.0
DPA
Fraction
P200
V7
Code
P200_V29
V7

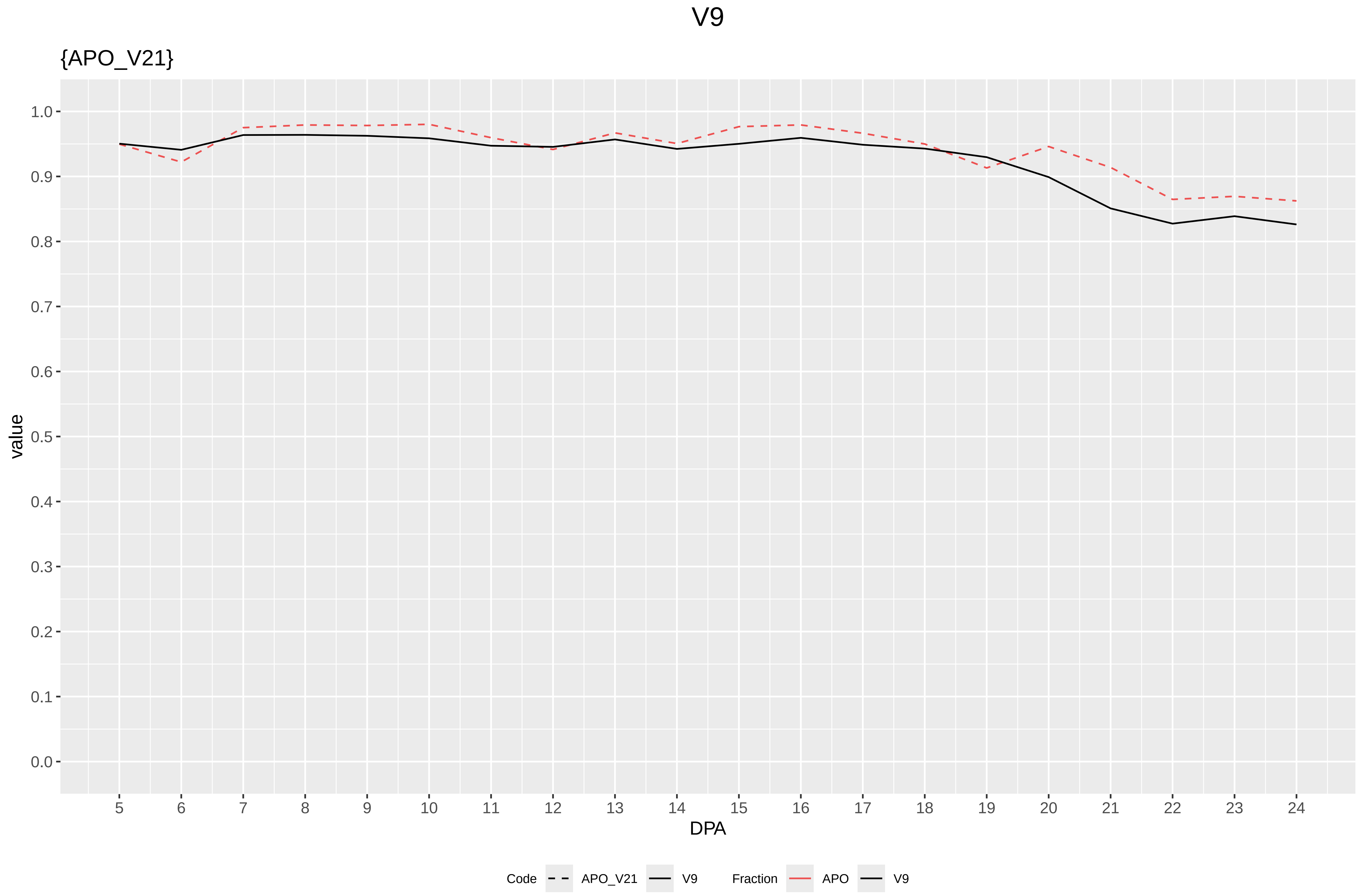

V9
{APO_V21}
value
1.0
0.9
0.8
0.7
0.6
0.5
0.4
0.3
0.2
0.1
0.0
DPA
Code
APO_V21
V9
Fraction
APO
V9

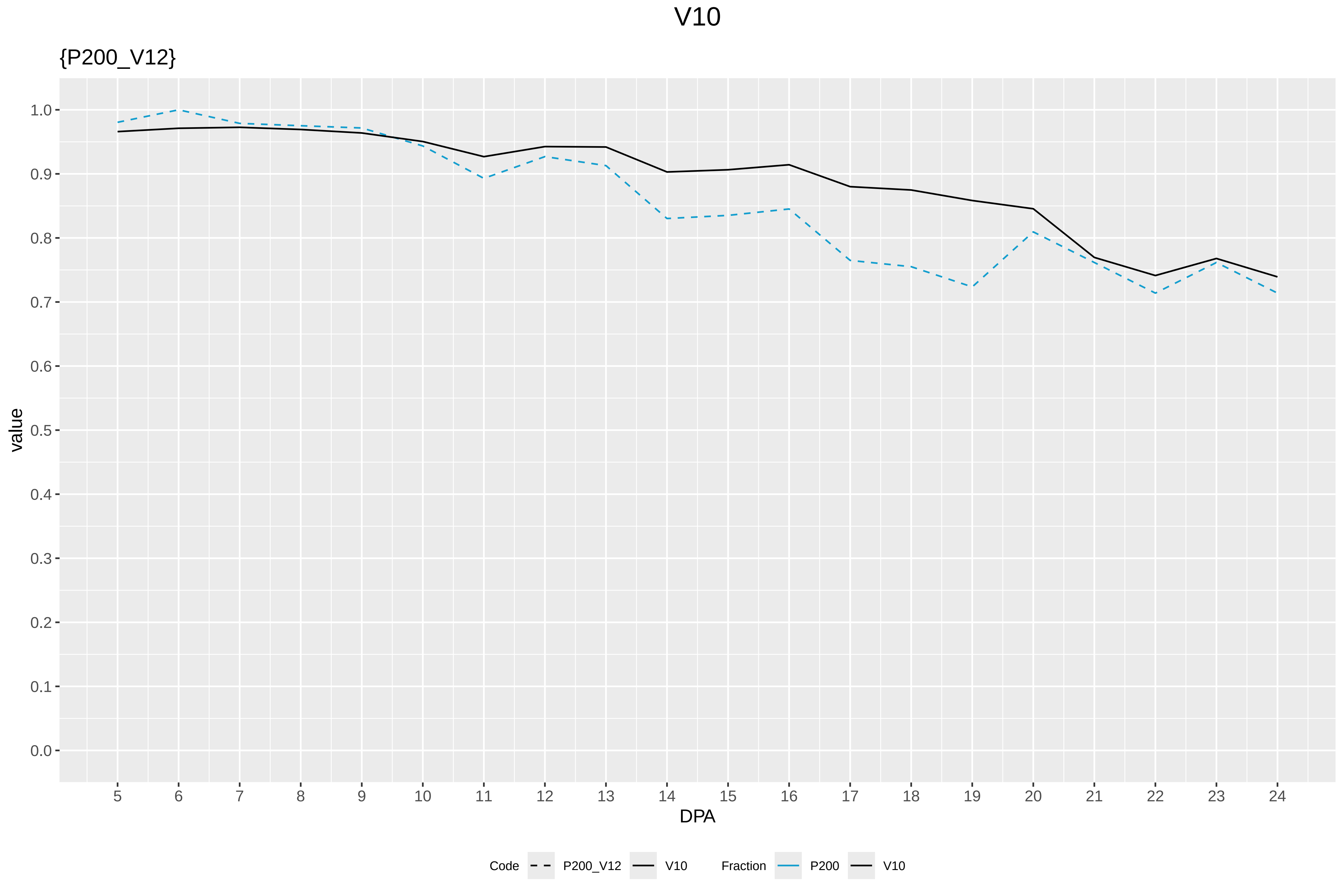

V10
{P200_V12}
value
1.0
0.9
0.8
0.7
0.6
0.5
0.4
0.3
0.2
0.1
0.0
DPA
Code
P200_V12
V10
Fraction
P200
V10

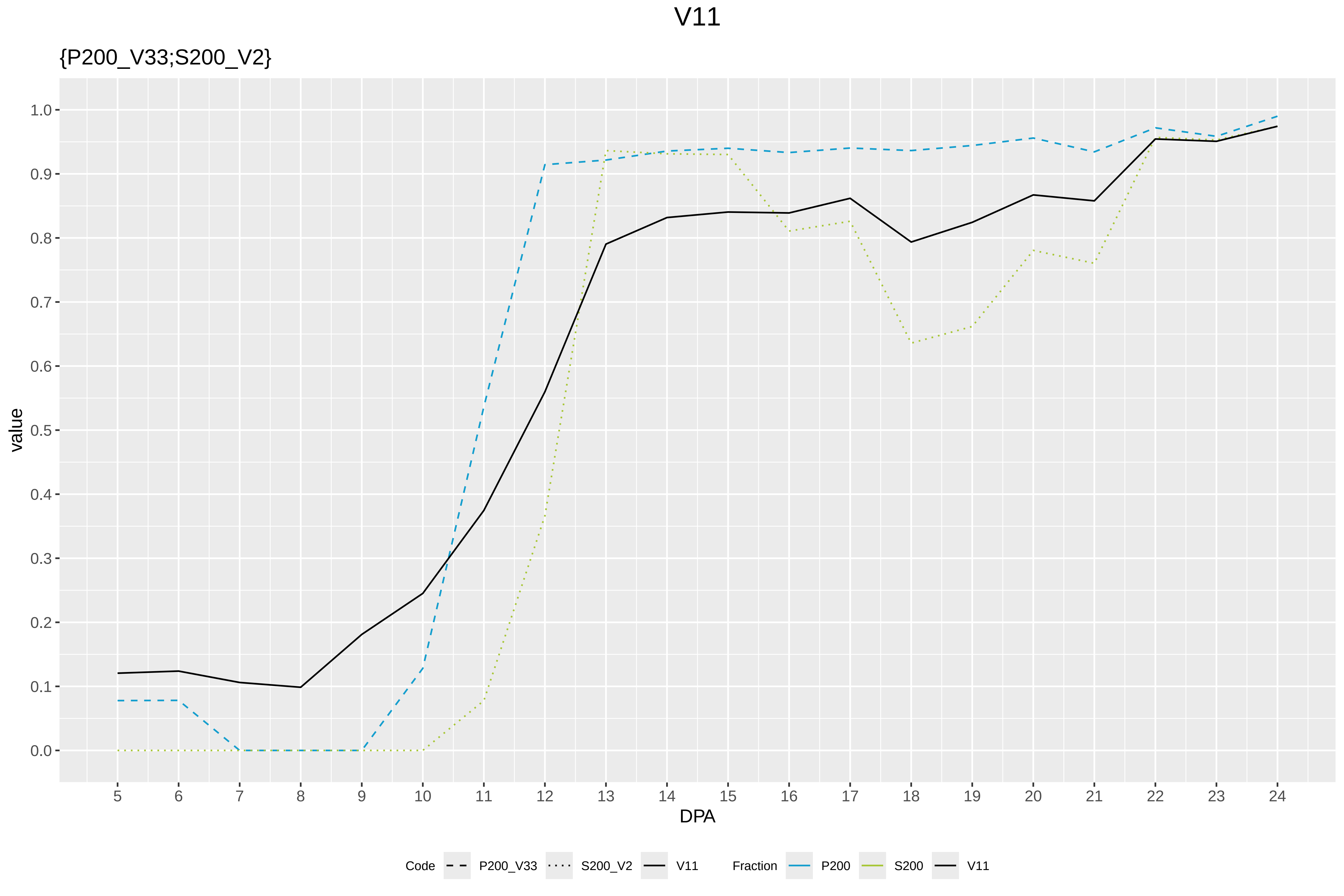

V11
{P200_V33;S200_V2}
value
1.0
0.9
0.8
0.7
0.6
0.5
0.4
0.3
0.2
0.1
0.0
DPA
Code
P200_V33
S200_V2
V11
Fraction
P200
S200
V11

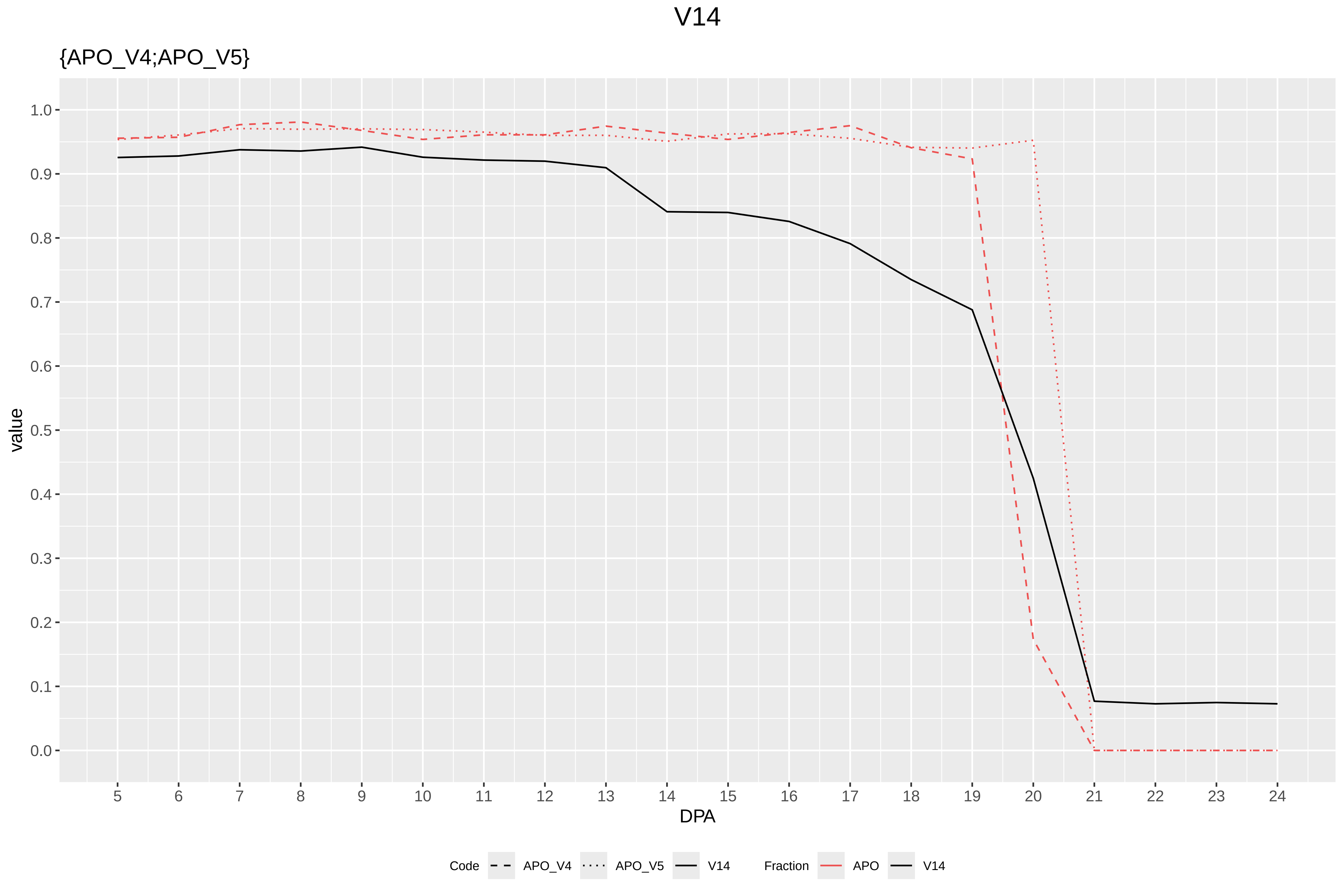

V14
{APO_V4;APO_V5}
value
1.0
0.9
0.8
0.7
0.6
0.5
0.4
0.3
0.2
0.1
0.0
DPA
Code
APO_V4
APO_V5
V14
Fraction
APO
V14

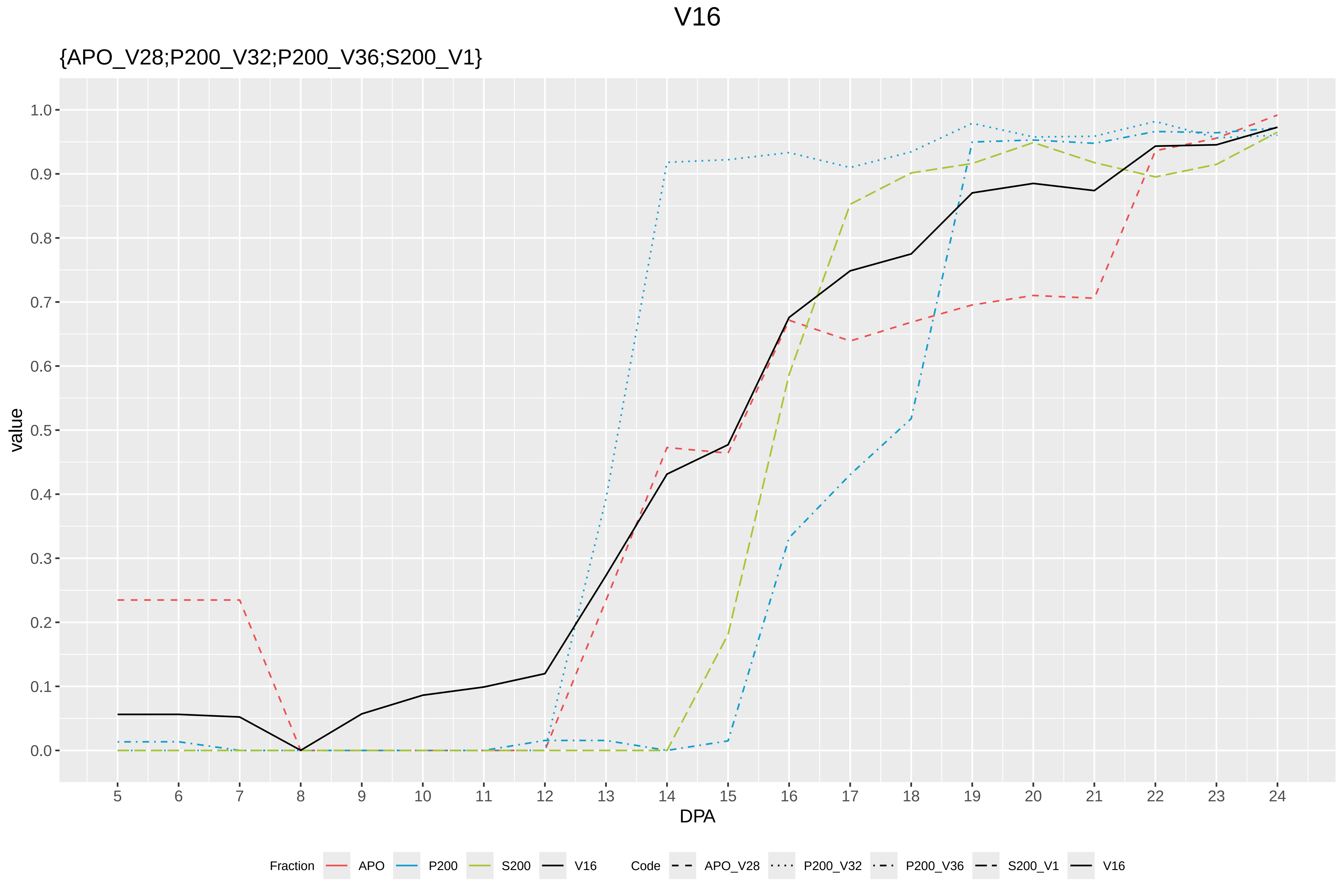

V16
{APO_V28;P200_V32;P200_V36;S200_V1}
value
1.0
0.9
0.8
0.7
0.6
0.5
0.4
0.3
0.2
0.1
0.0
DPA
Fraction
APO
P200
S200
V16
Code
APO_V28
P200_V32
P200_V36
S200_V1
V16

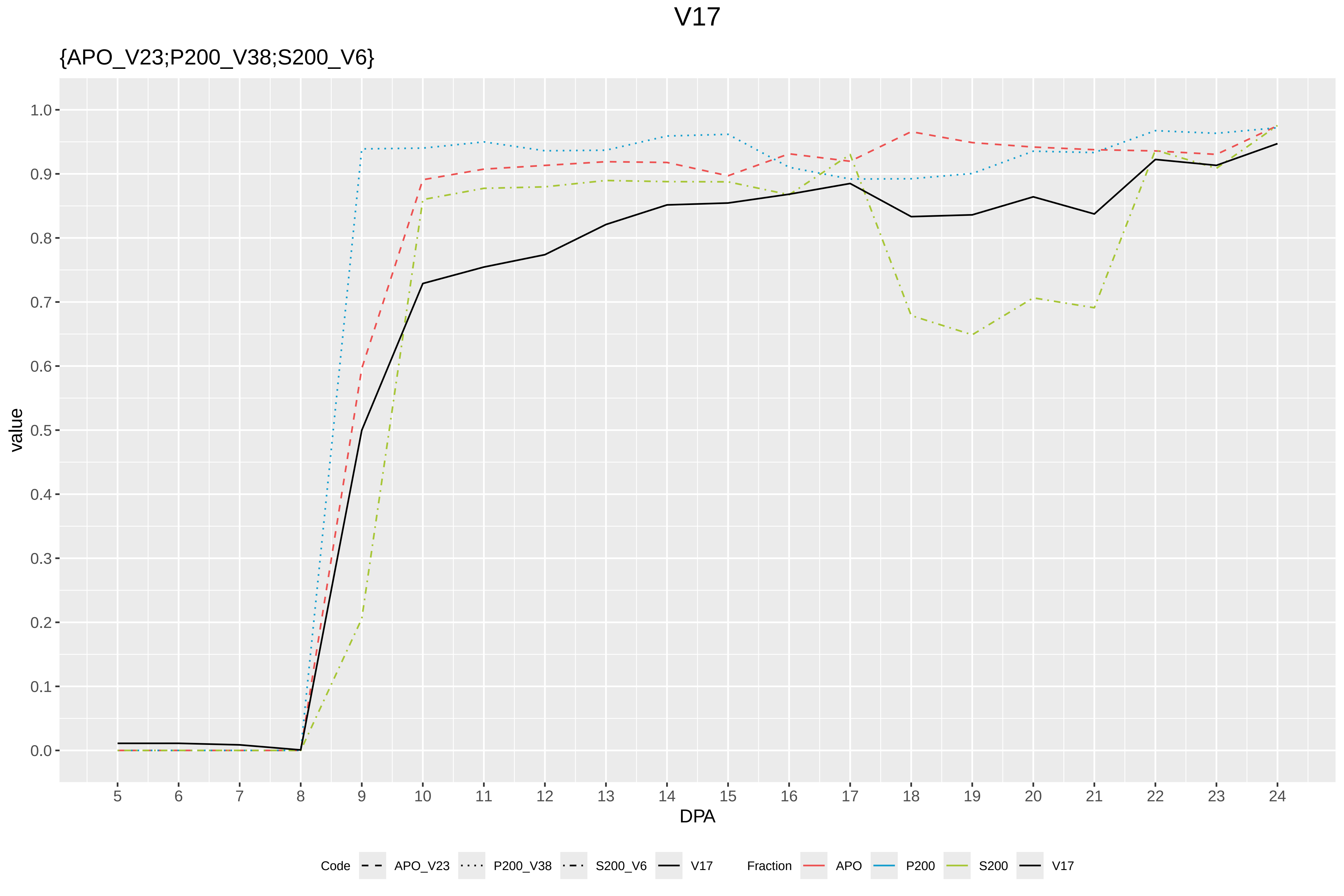

V17
{APO_V23;P200_V38;S200_V6}
value
1.0
0.9
0.8
0.7
0.6
0.5
0.4
0.3
0.2
0.1
0.0
5 6 7 8 9 10 11 12 13 14 15 16 17 18 19 20 21 22 23 24
DPA
Code APO_V23 P200_V38 S200_V6 V17
Fraction APO P200 S200 V17

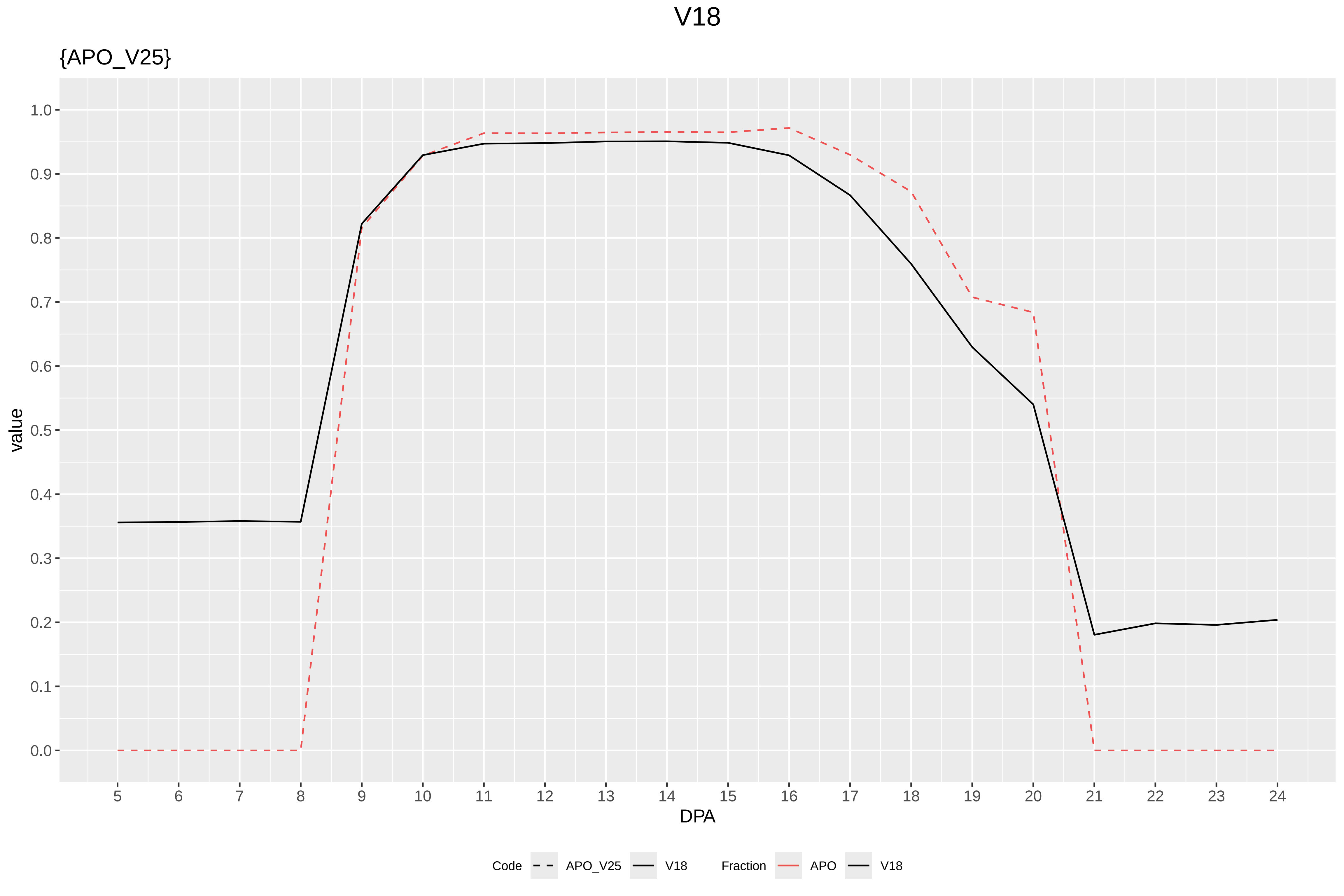
V18
{APO_V25}
value
1.0
0.9
0.8
0.7
0.6
0.5
0.4
0.3
0.2
0.1
0.0
DPA
Code
APO_V25
V18
Fraction
APO
V18

# V19

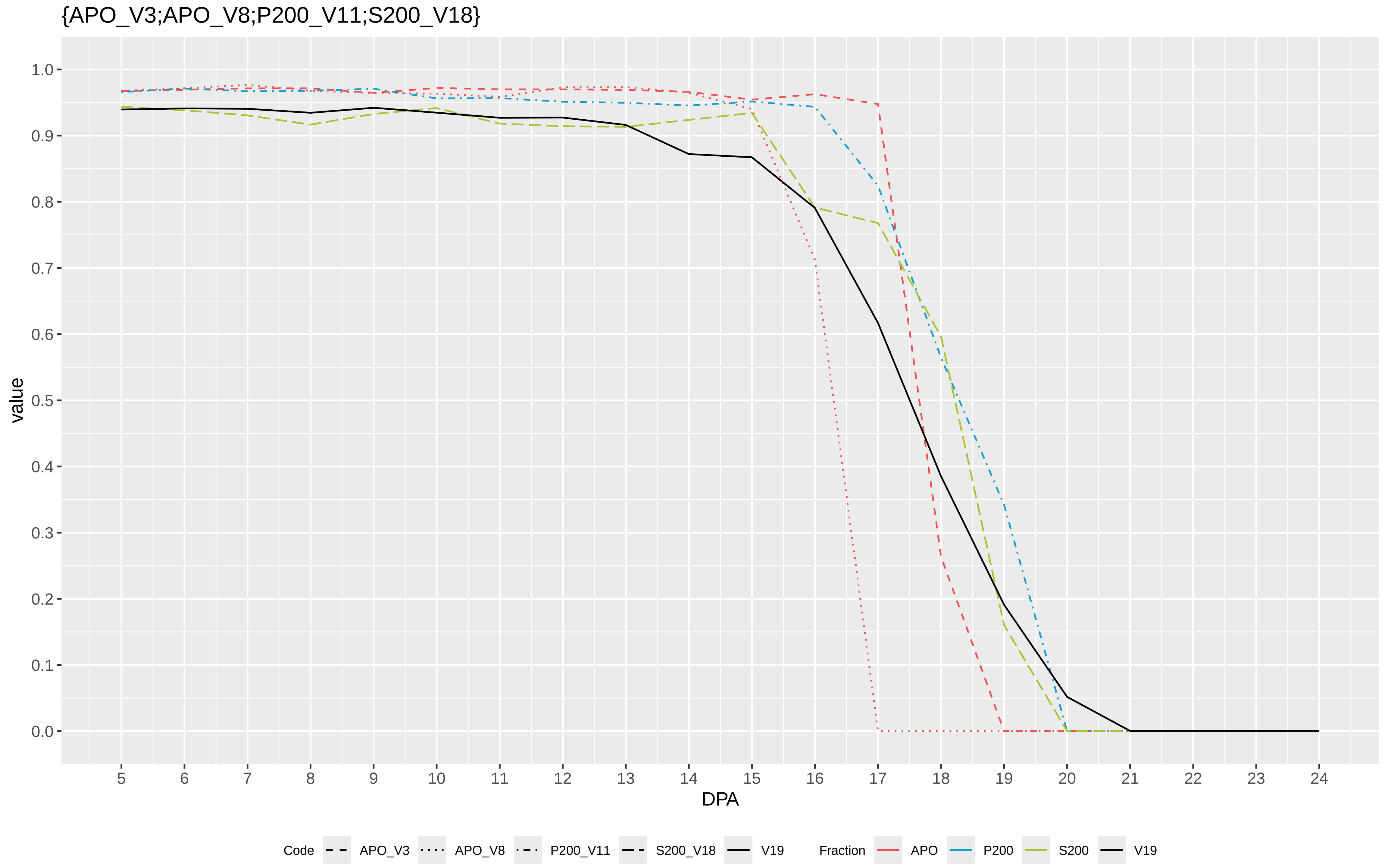

{APO_V3;APO_V8;P200_V11;S200_V18}
value
1.0
0.9
0.8
0.7
0.6
0.5
0.4
0.3
0.2
0.1
0.0
5 6 7 8 9 10 11 12 13 14 15 16 17 18 19 20 21 22 23 24
DPA
Code APO_V3 APO_V8 P200_V11 S200_V18 V19
Fraction APO P200 S200 V19

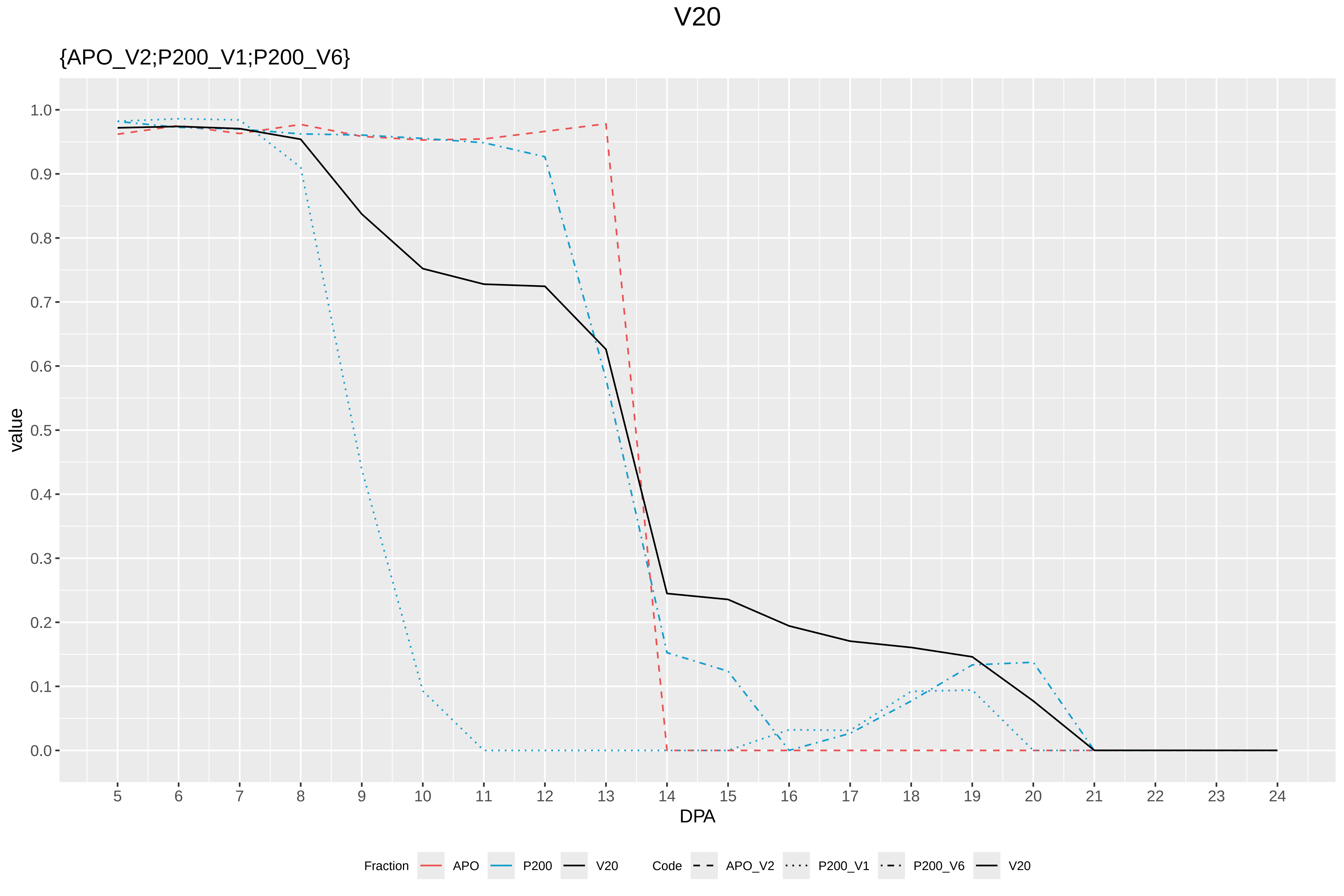
V20
{APO_V2;P200_V1;P200_V6}
value
1.0
0.9
0.8
0.7
0.6
0.5
0.4
0.3
0.2
0.1
0.0
5 6 7 8 9 10 11 12 13 14 15 16 17 18 19 20 21 22 23 24
DPA
Fraction APO P200 V20
Code APO_V2 P200_V1 P200_V6 V20
