## Supplementary Figure S8 for "Systems-level proteomic models of cotton fiber development: A high-resolution resource to analyze cell dynamics and trait engineering"

### Codes

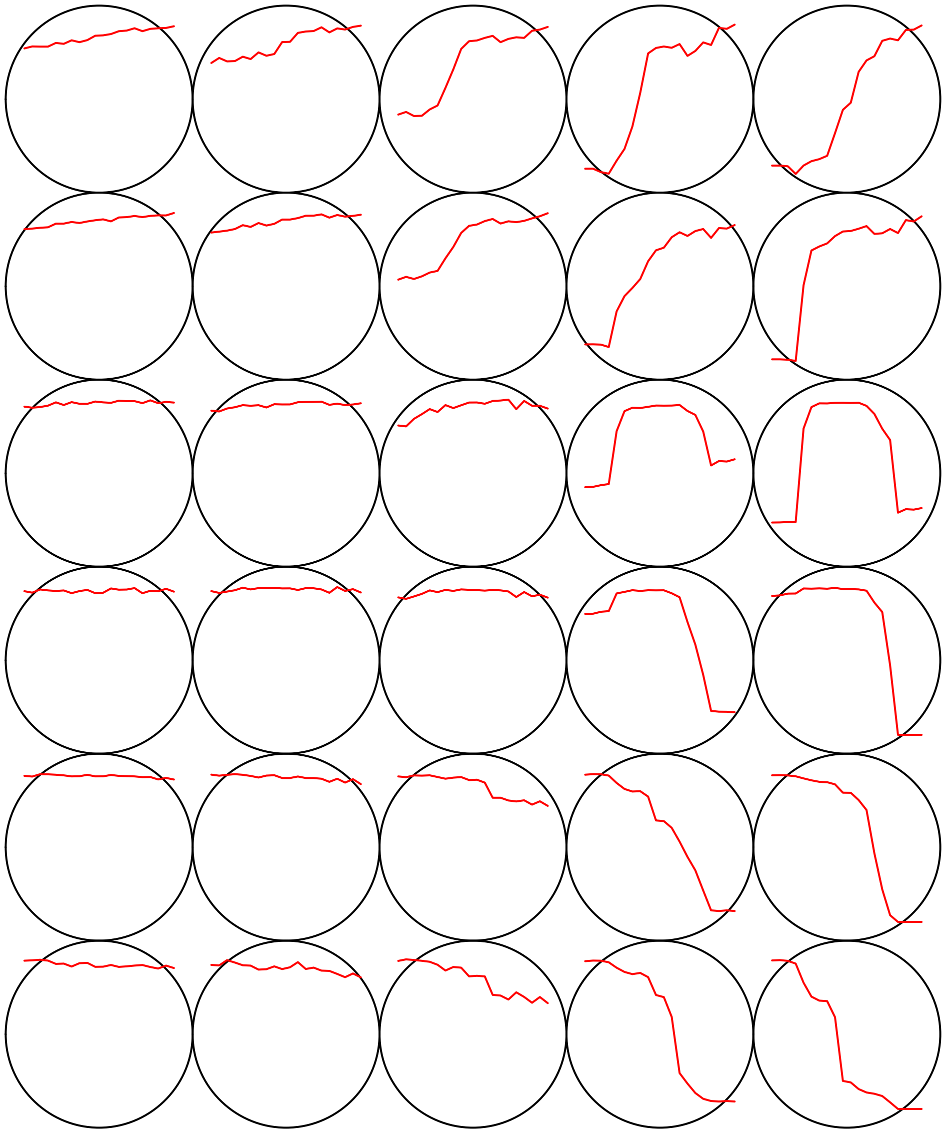

#### Count plot

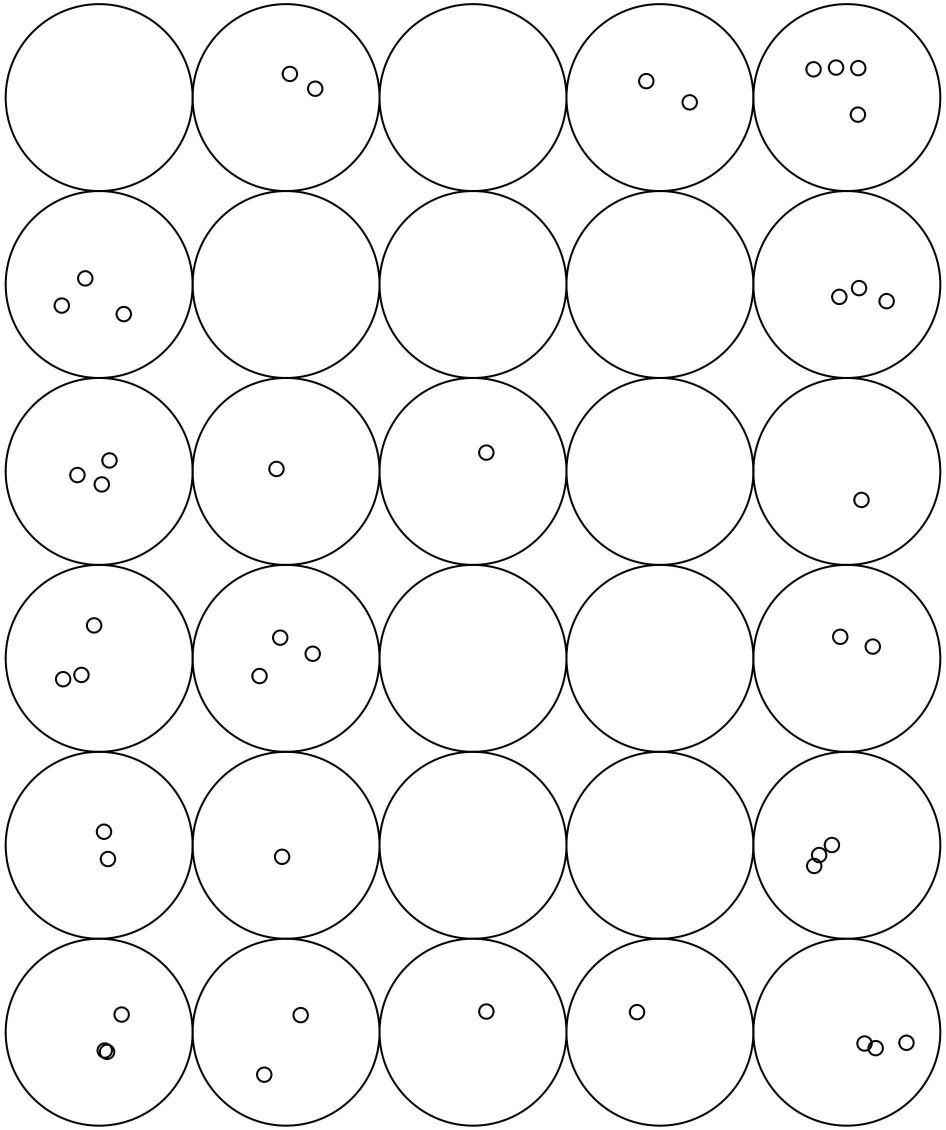

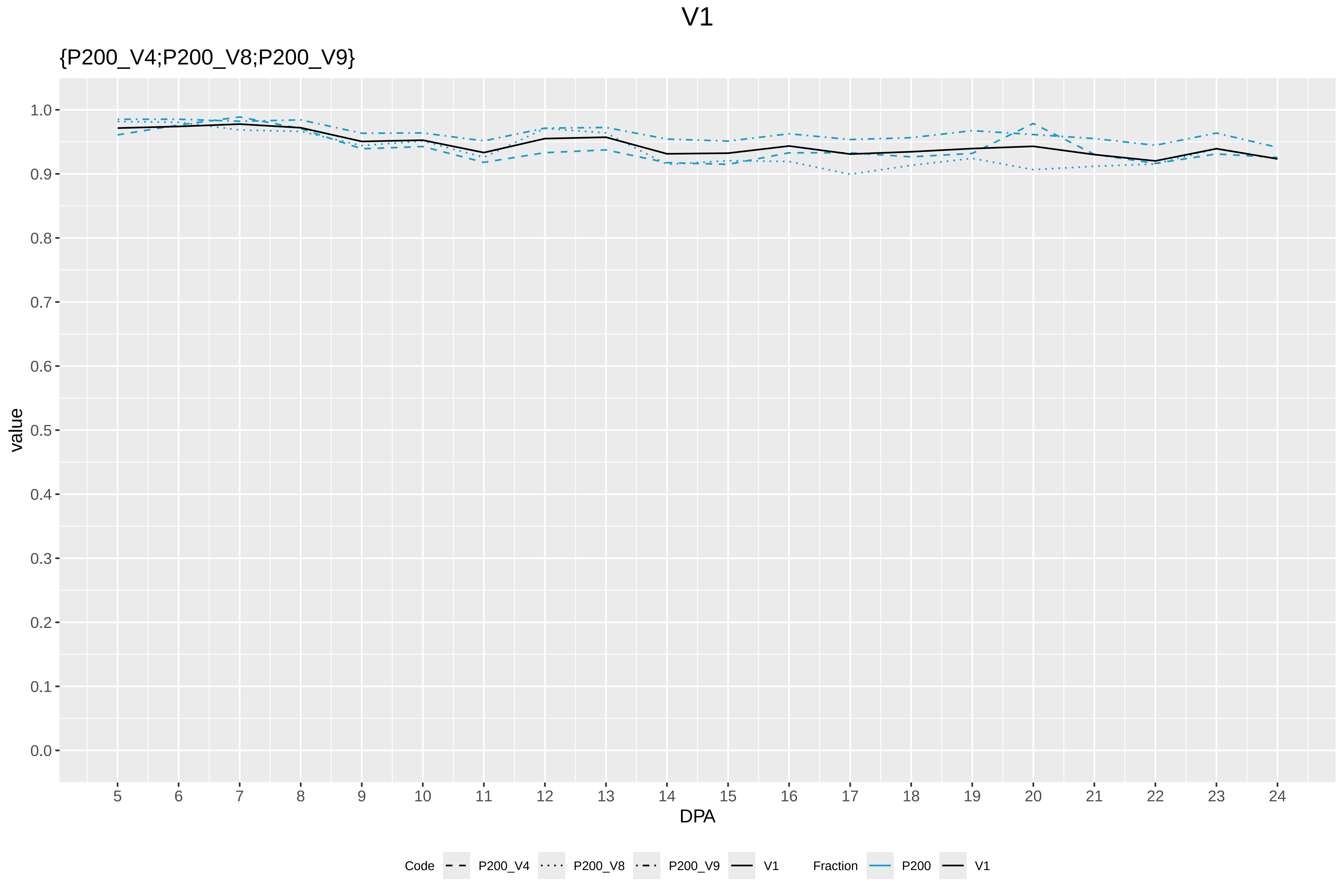

V1
{P200_V4;P200_V8;P200_V9}
value
1.0
0.9
0.8
0.7
0.6
0.5
0.4
0.3
0.2
0.1
0.0
5 6 7 8 9 10 11 12 13 14 15 16 17 18 19 20 21 22 23 24
DPA
Code P200_V4 P200_V8 P200_V9 V1
Fraction P200 V1

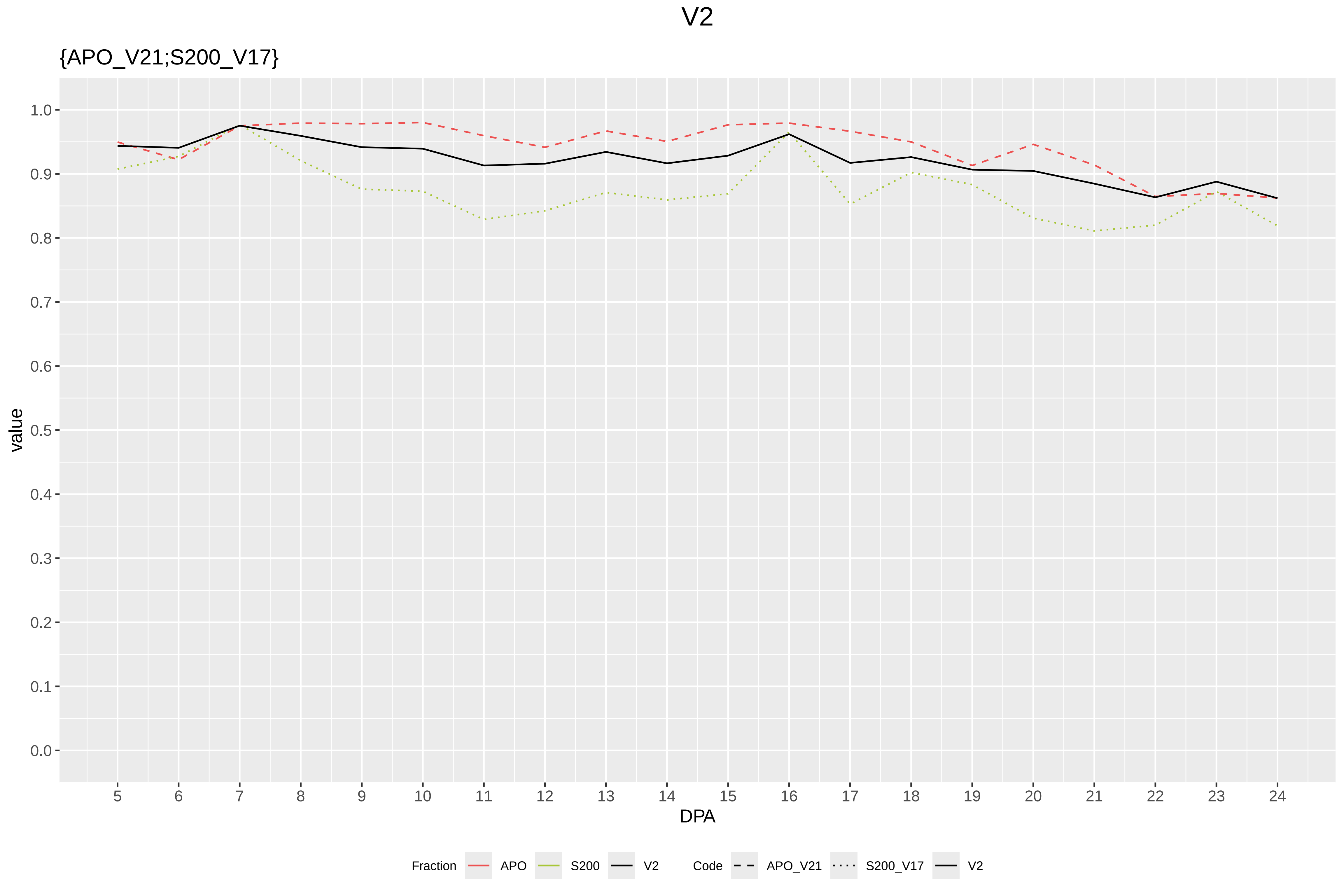

V2
{APO_V21;S200_V17}
value
1.0
0.9
0.8
0.7
0.6
0.5
0.4
0.3
0.2
0.1
0.0
DPA
Fraction
APO
S200
V2
Code
APO_V21
S200_V17
V2

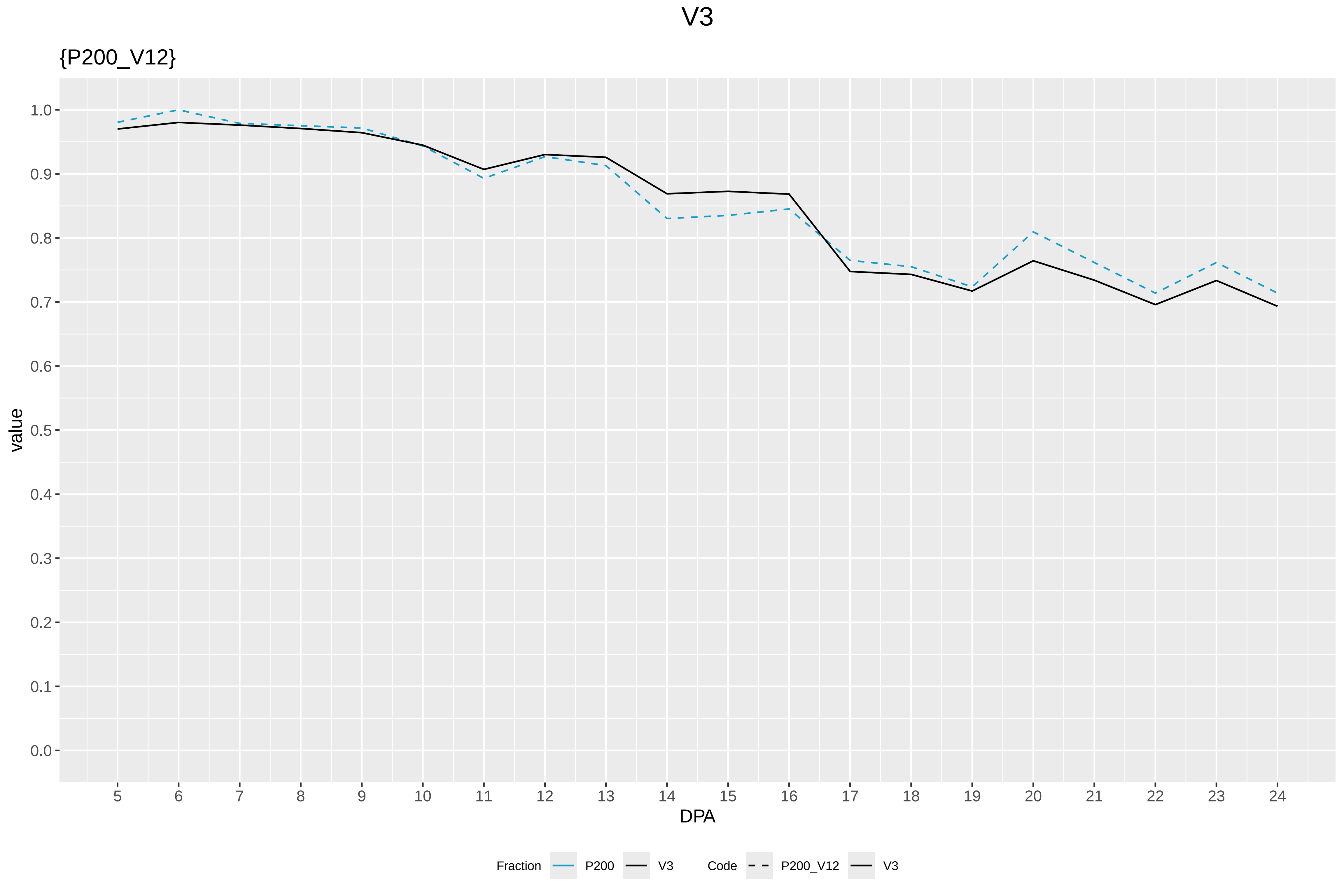

V3
{P200_V12}
value
1.0
0.9
0.8
0.7
0.6
0.5
0.4
0.3
0.2
0.1
0.0
DPA
Fraction
P200
V3
Code
P200_V12
V3

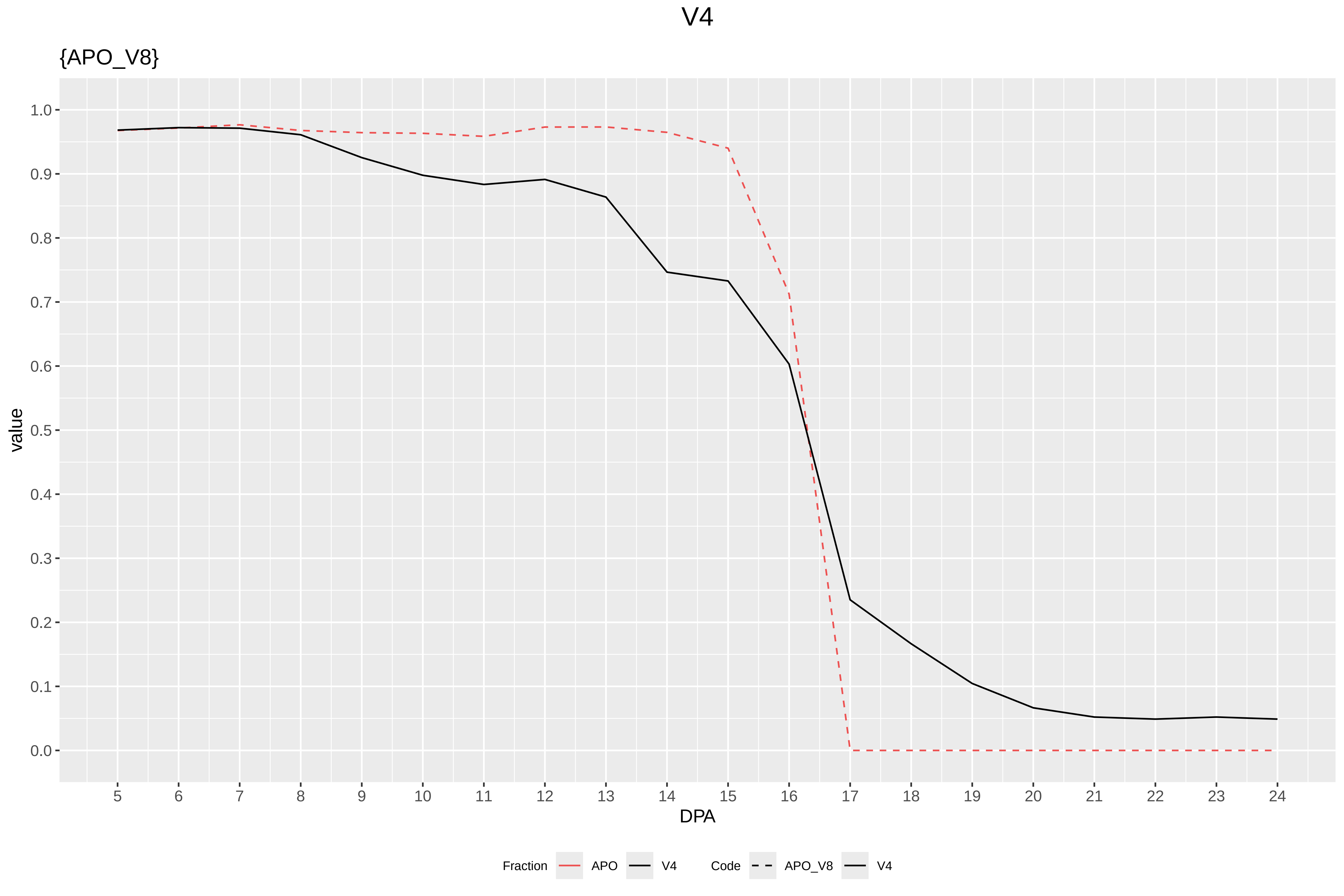

V4
{APO_V8}
value
DPA
1.0
0.9
0.8
0.7
0.6
0.5
0.4
0.3
0.2
0.1
0.0
Fraction
APO
V4
Code
APO_V8
V4

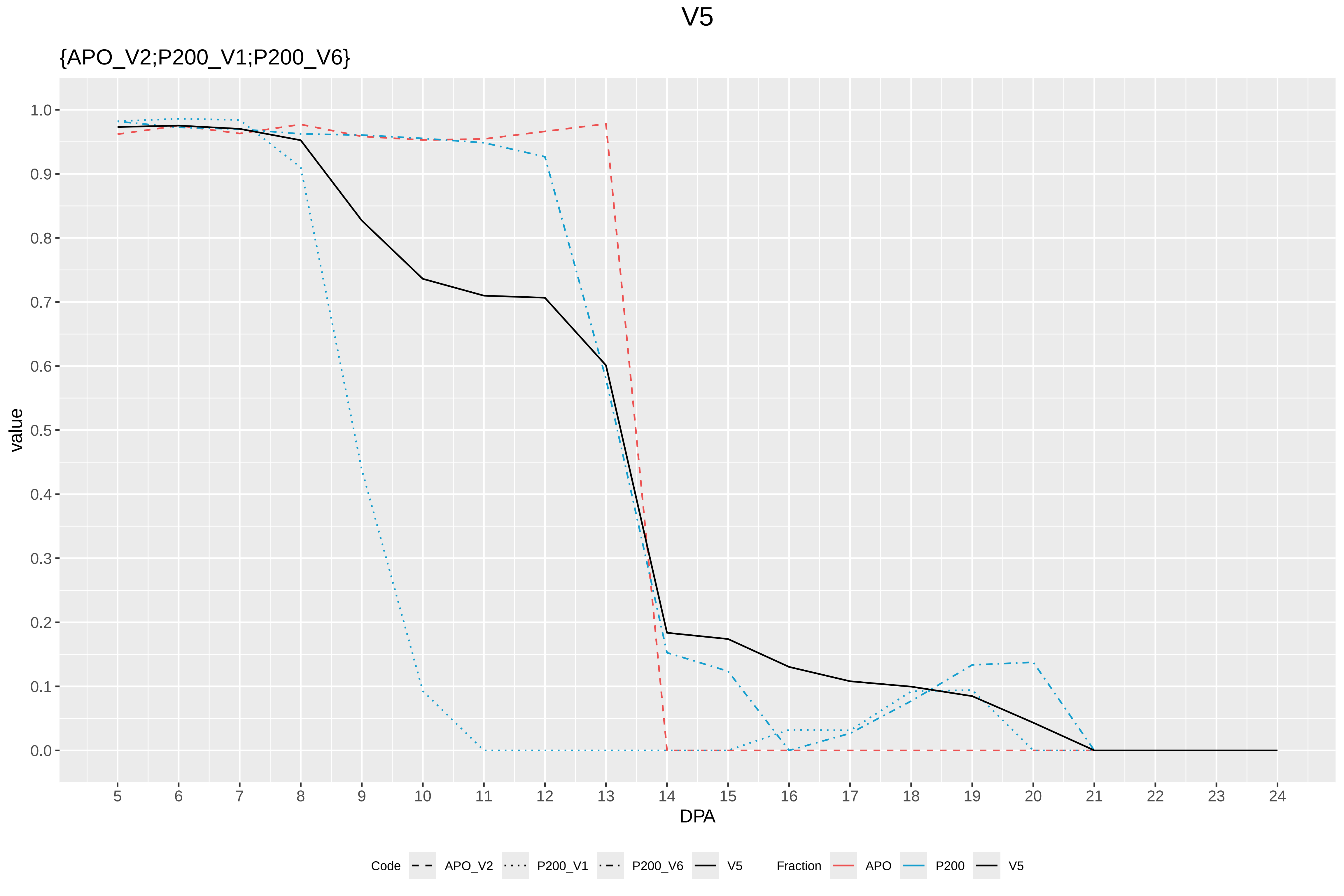
V5
{APO_V2;P200_V1;P200_V6}
value
1.0
0.9
0.8
0.7
0.6
0.5
0.4
0.3
0.2
0.1
0.0
DPA
Code
APO_V2
P200_V1
P200_V6
V5
Fraction
APO
P200
V5

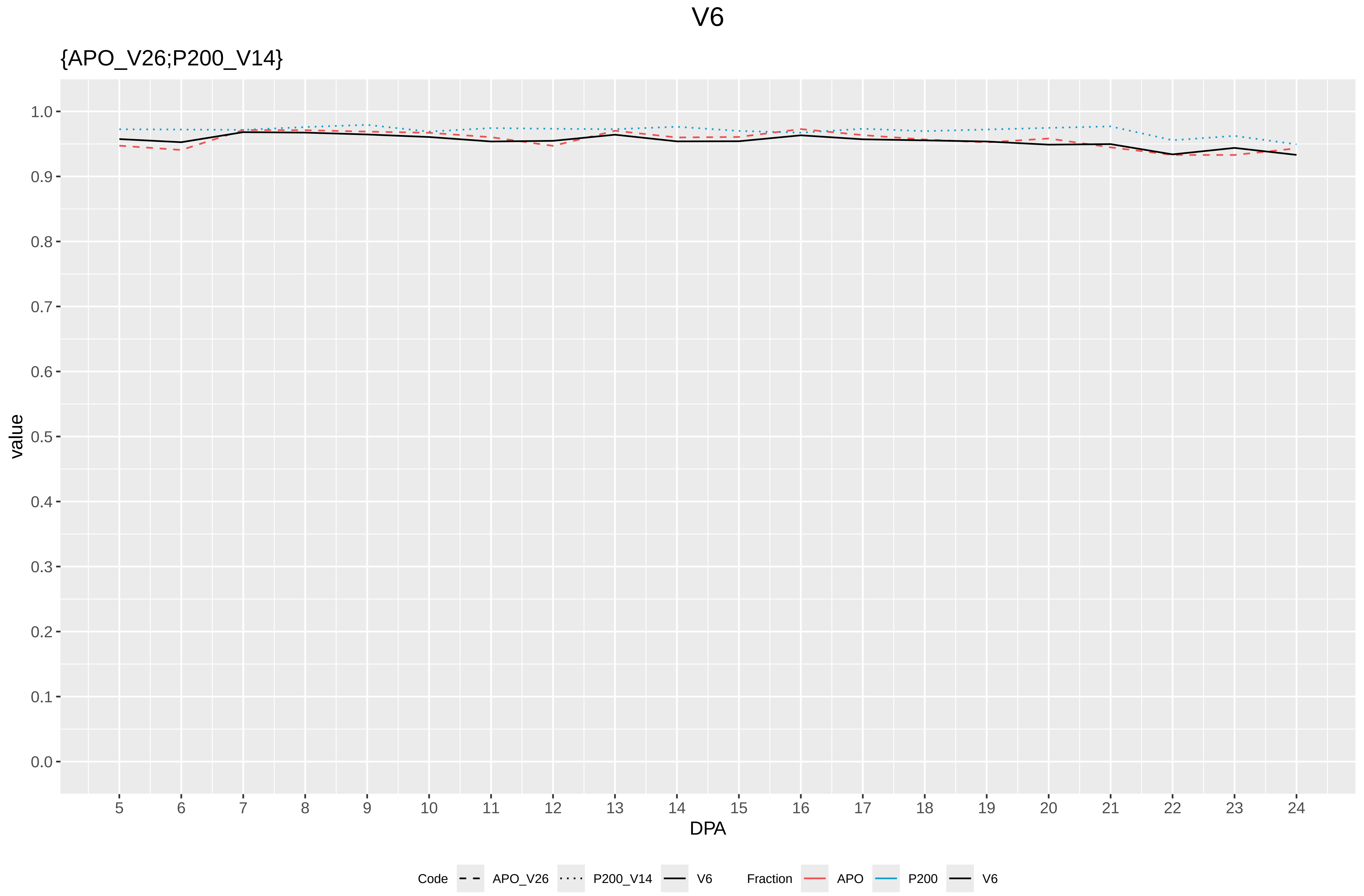

V6
{APO_V26;P200_V14}
value
1.0
0.9
0.8
0.7
0.6
0.5
0.4
0.3
0.2
0.1
0.0
DPA
Code
APO_V26
P200_V14
V6
Fraction
APO
P200
V6

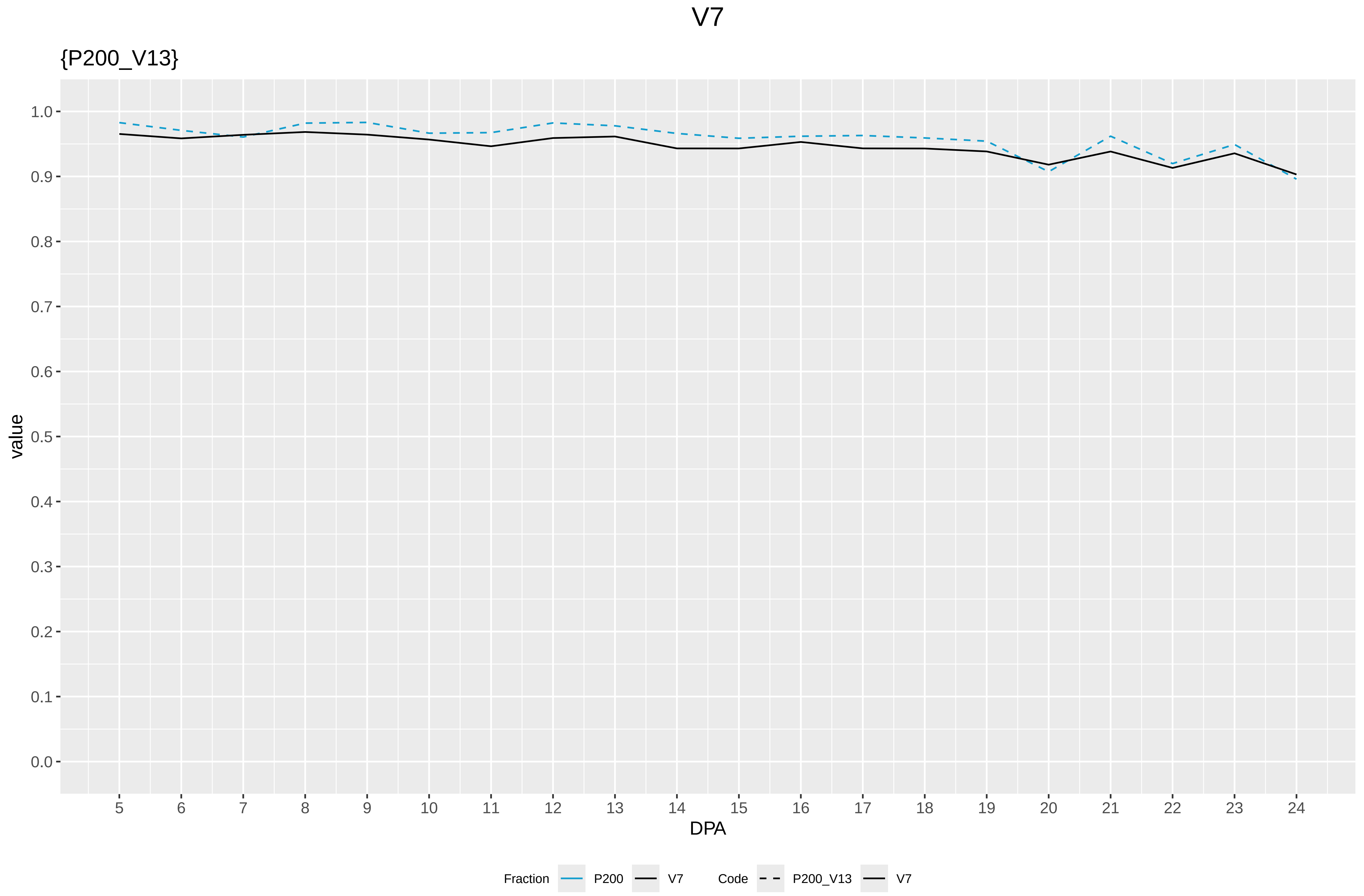

V7
{P200_V13}
value
DPA
1.0
0.9
0.8
0.7
0.6
0.5
0.4
0.3
0.2
0.1
0.0
5 6 7 8 9 10 11 12 13 14 15 16 17 18 19 20 21 22 23 24
Fraction
P200
V7
Code
P200_V13
V7

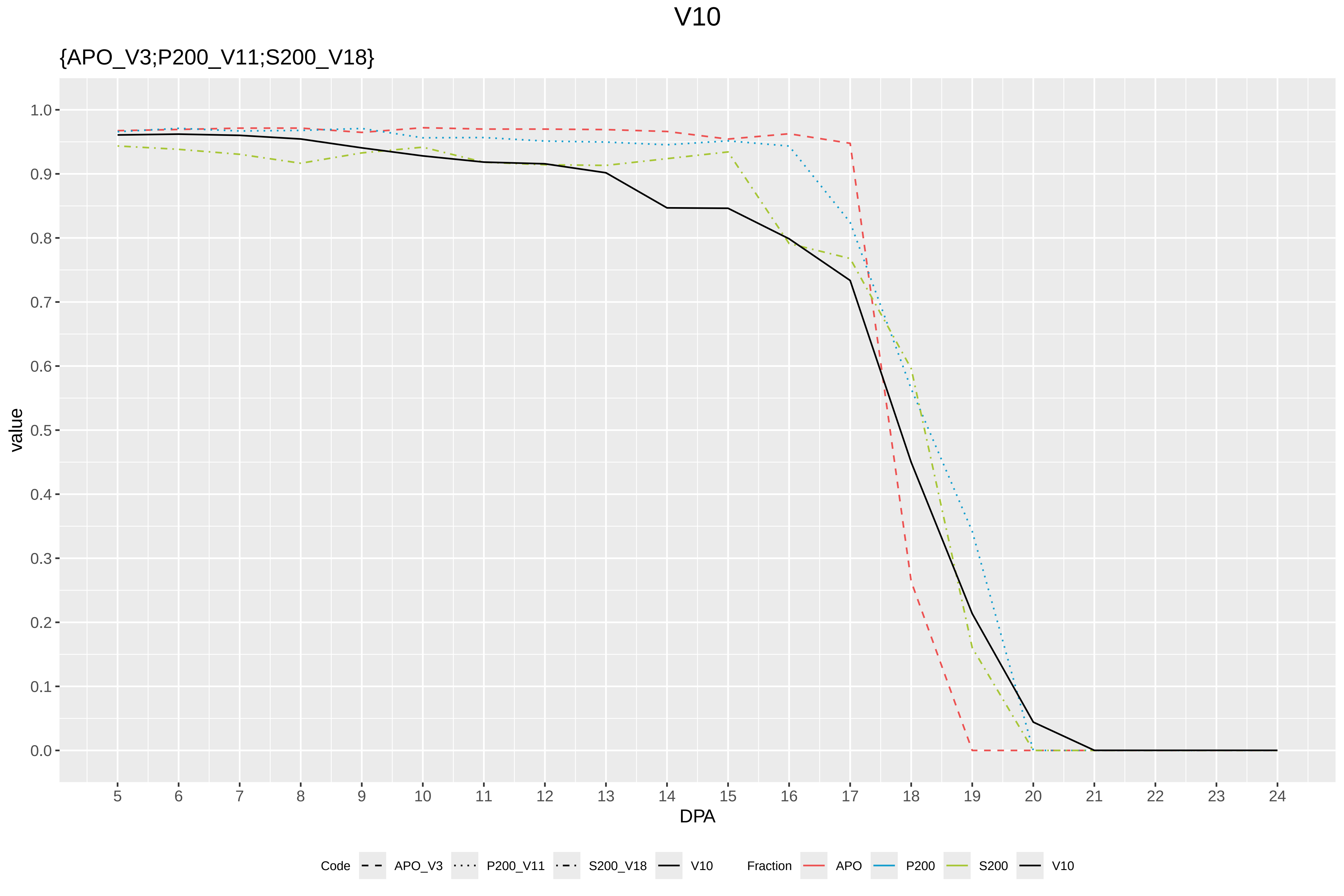

V10
{APO_V3;P200_V11;S200_V18}
value
1.0
0.9
0.8
0.7
0.6
0.5
0.4
0.3
0.2
0.1
0.0
DPA
Code
APO_V3
P200_V11
S200_V18
V10
Fraction
APO
P200
S200
V10

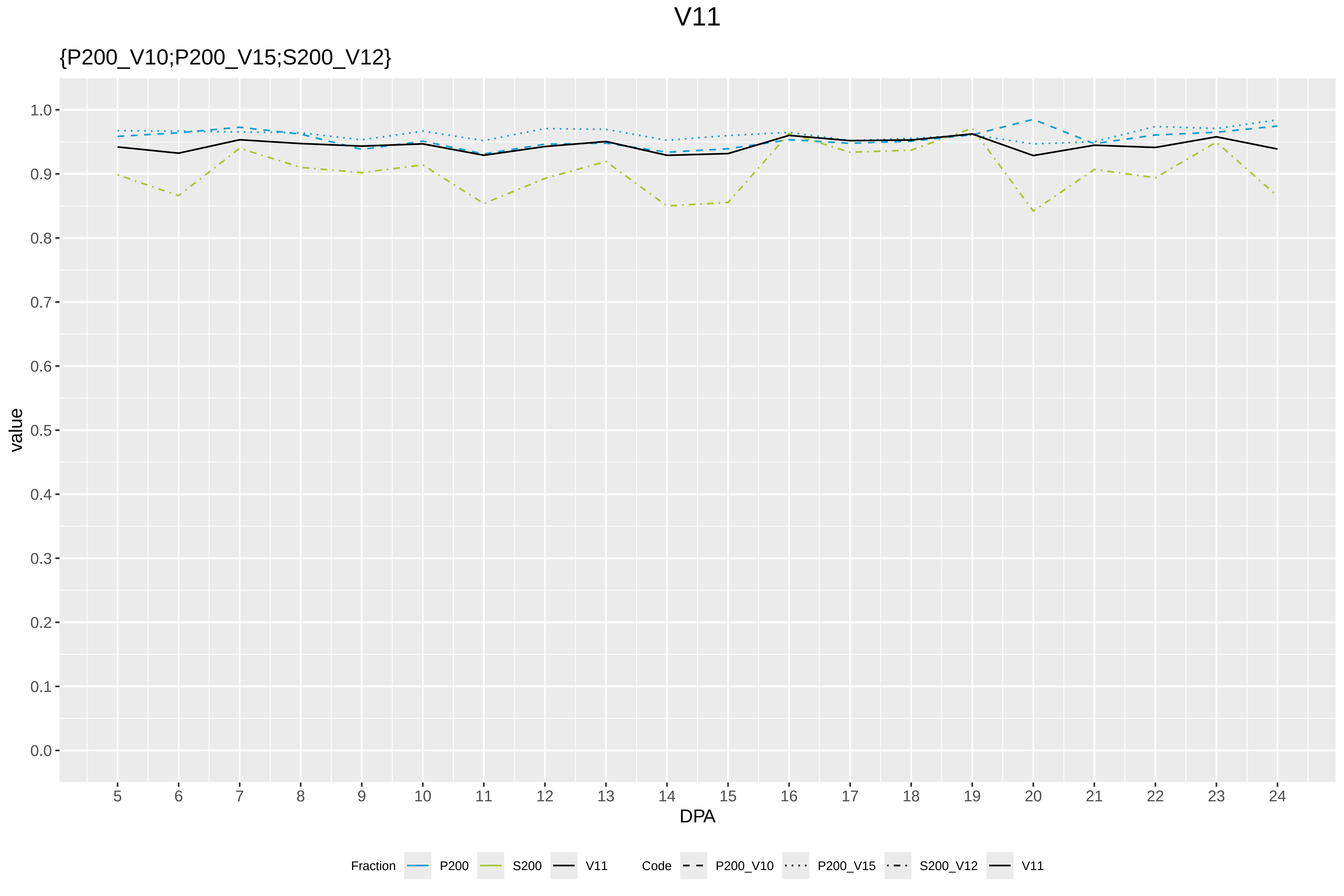
V11
{P200_V10;P200_V15;S200_V12}
value
1.0
0.9
0.8
0.7
0.6
0.5
0.4
0.3
0.2
0.1
0.0
DPA
Fraction
P200
S200
V11
Code
P200_V10
P200_V15
S200_V12
V11

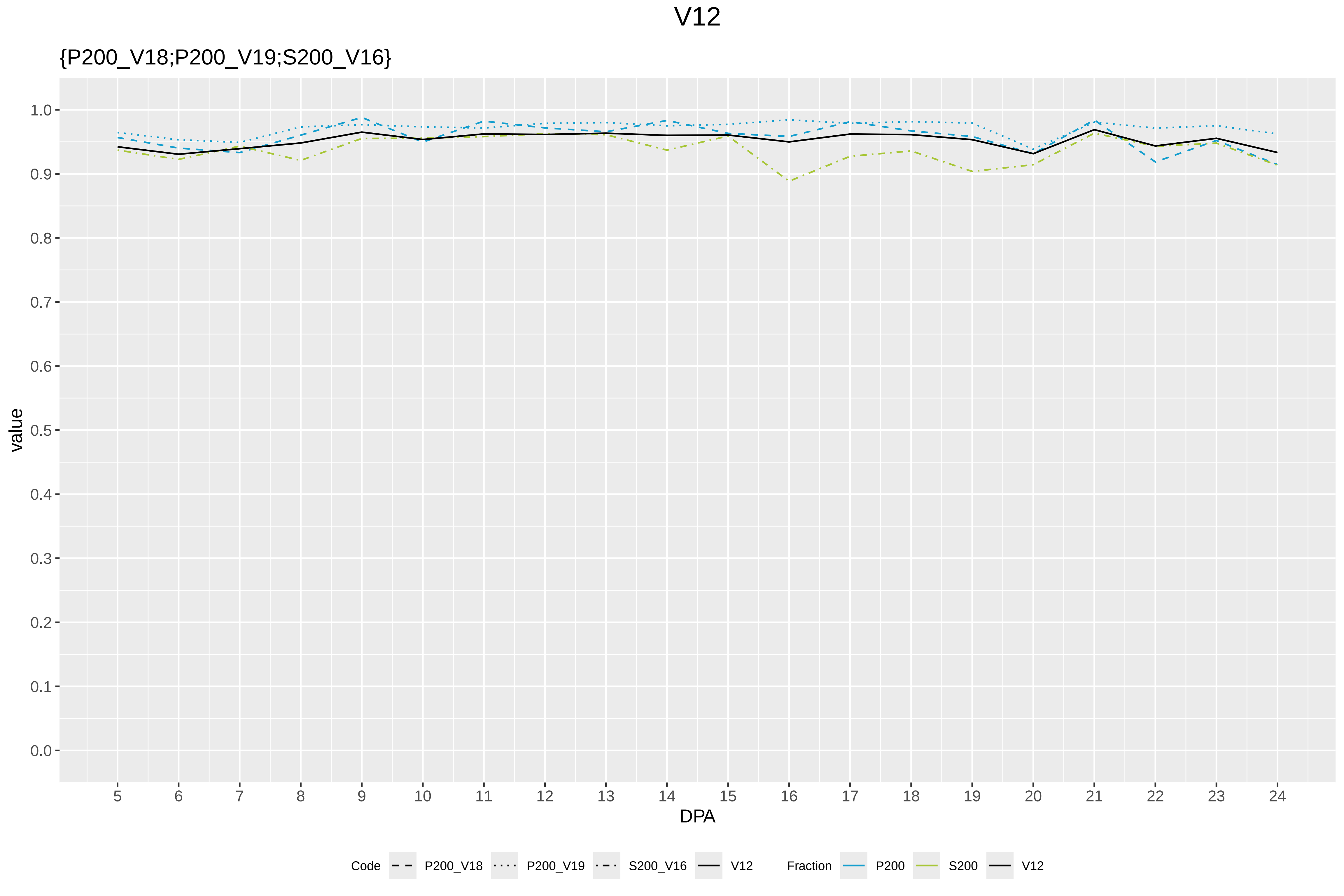

V12
{P200_V18;P200_V19;S200_V16}
1.0
0.9
0.8
0.7
0.6
0.5
0.4
0.3
0.2
0.1
0.0
value
DPA
Code
P200_V18
P200_V19
S200_V16
V12
Fraction
P200
S200
V12

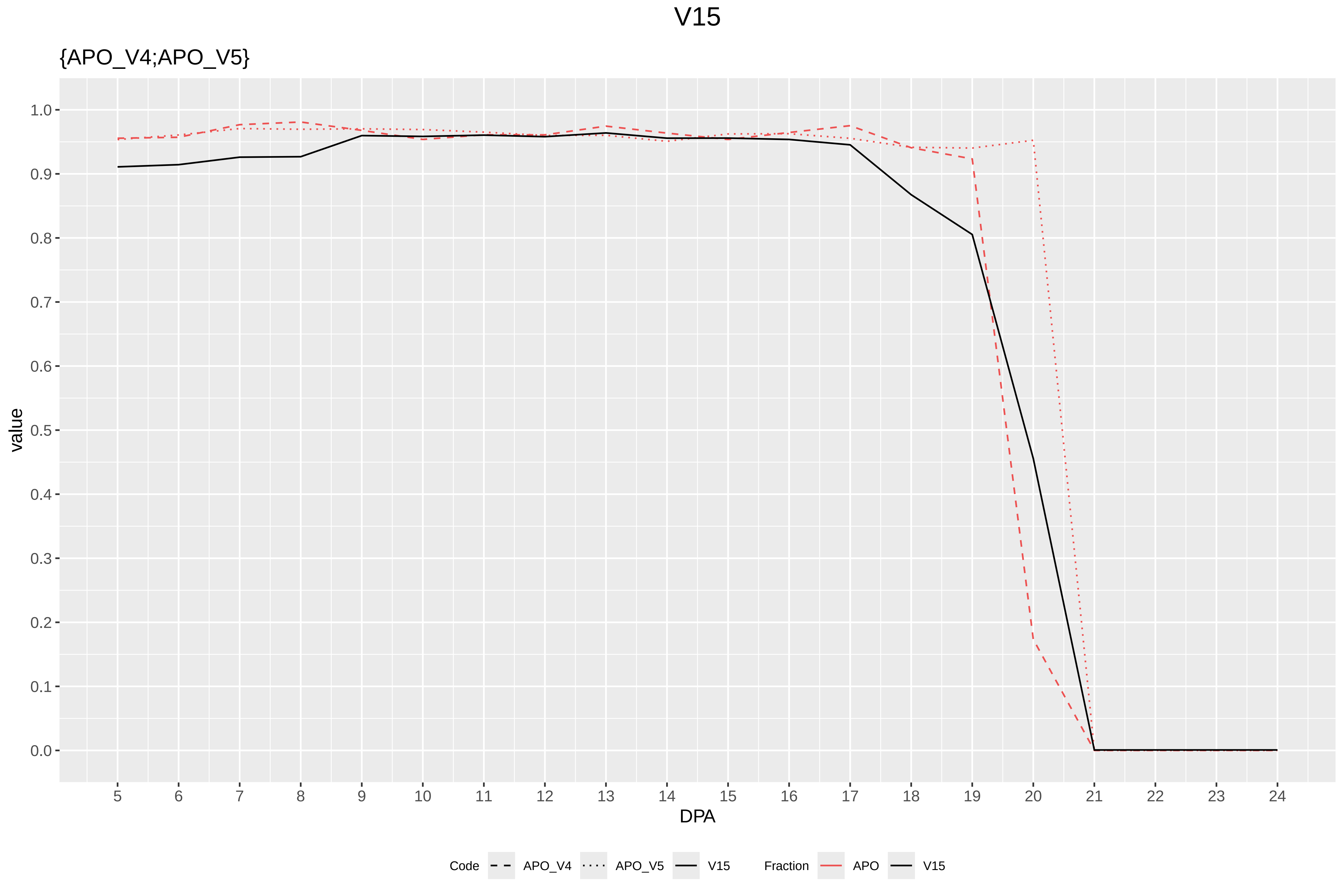

V15
{APO_V4;APO_V5}
value
1.0
0.9
0.8
0.7
0.6
0.5
0.4
0.3
0.2
0.1
0.0
DPA
Code
APO_V4
APO_V5
V15
Fraction
APO
V15

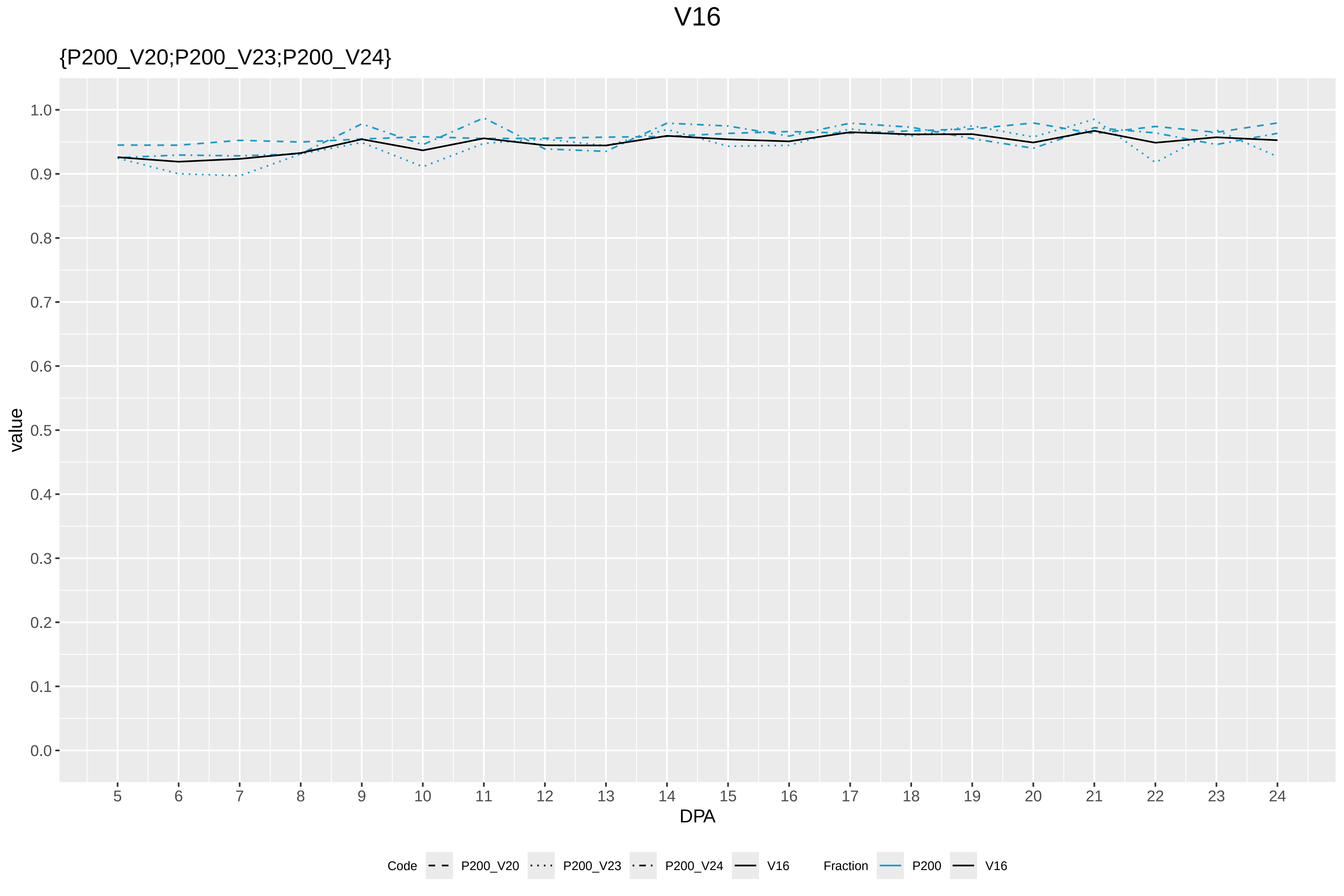
V16
{P200_V20;P200_V23;P200_V24}
value
DPA
1.0
0.9
0.8
0.7
0.6
0.5
0.4
0.3
0.2
0.1
0.0
Code
P200_V20
P200_V23
P200_V24
V16
Fraction
P200
V16

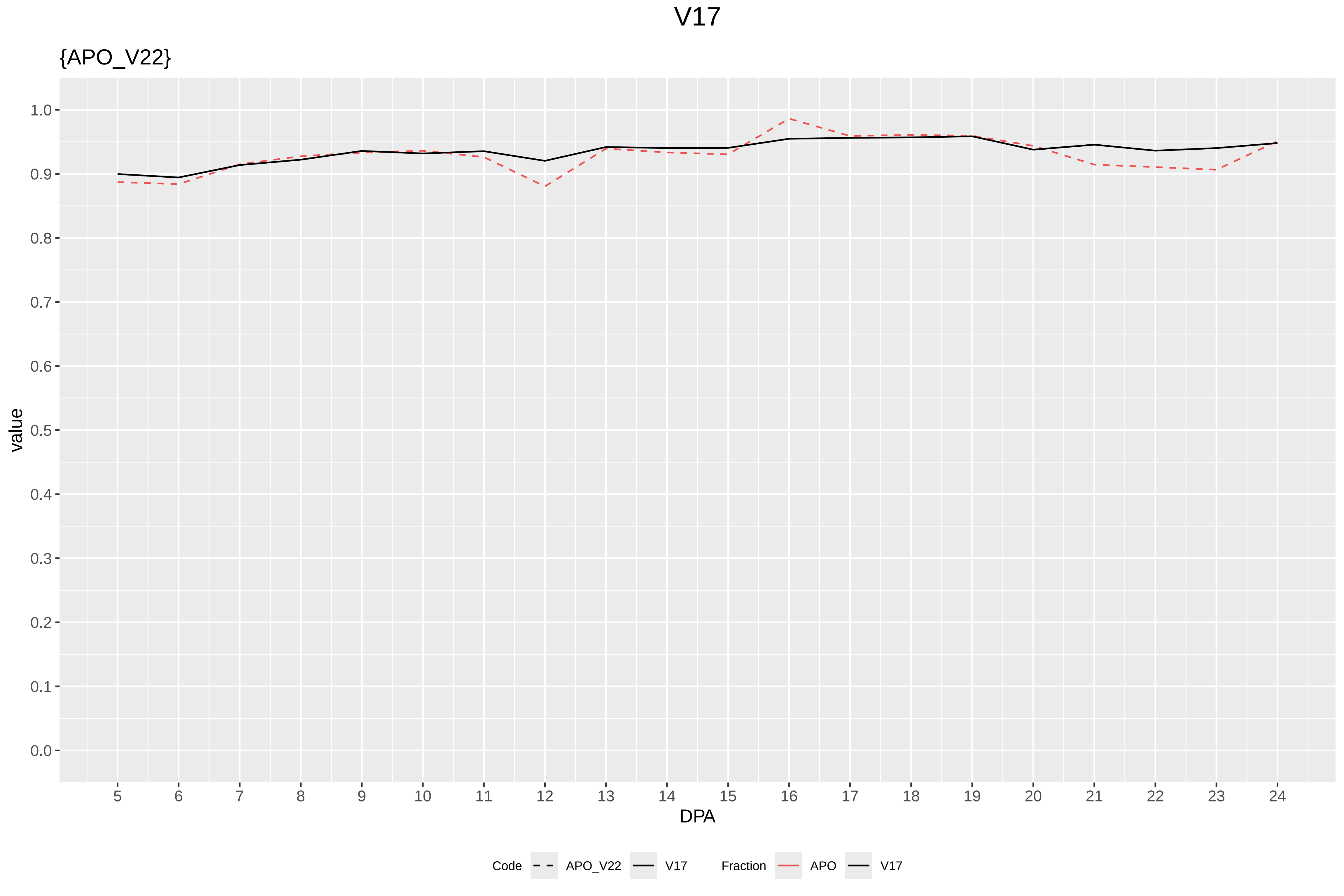
V17
{APO_V22}
value
DPA
Code
APO_V22
V17
Fraction
APO
V17

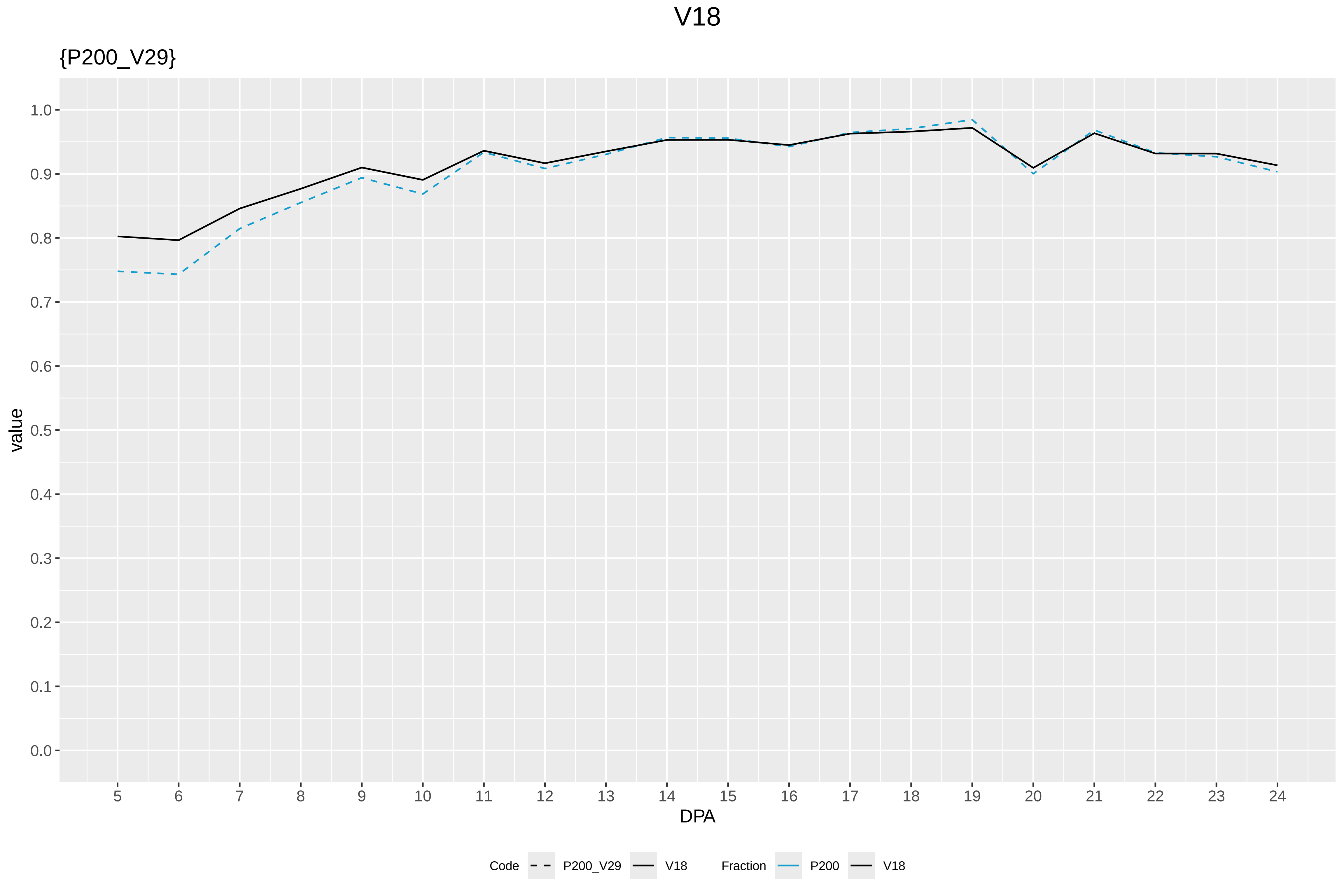
V18
{P200_V29}
value
1.0
0.9
0.8
0.7
0.6
0.5
0.4
0.3
0.2
0.1
0.0
DPA
Code
P200_V29
V18
Fraction
P200
V18

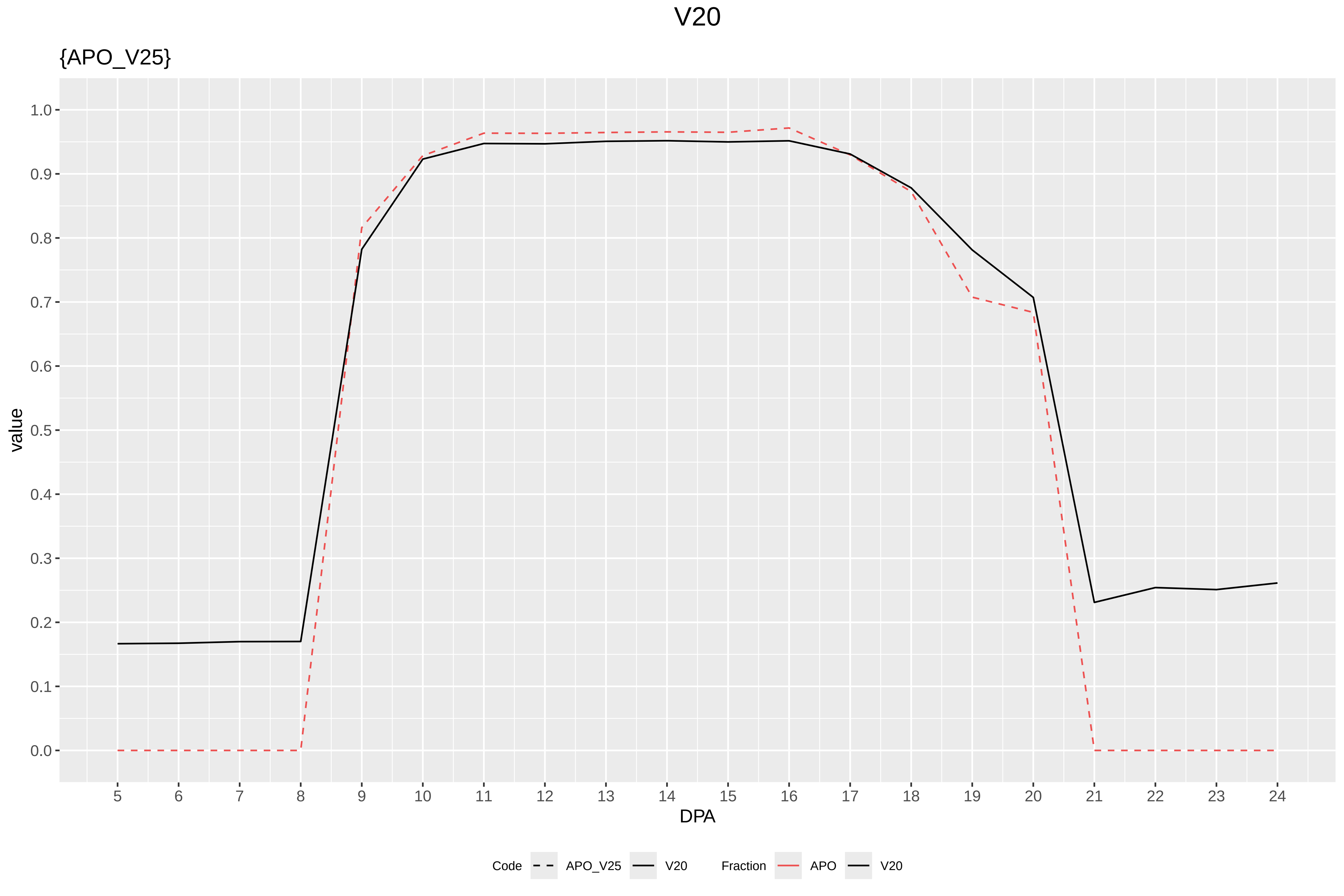
V20
{APO_V25}
value
1.0
0.9
0.8
0.7
0.6
0.5
0.4
0.3
0.2
0.1
0.0
DPA
Code
APO_V25
V20
Fraction
APO
V20

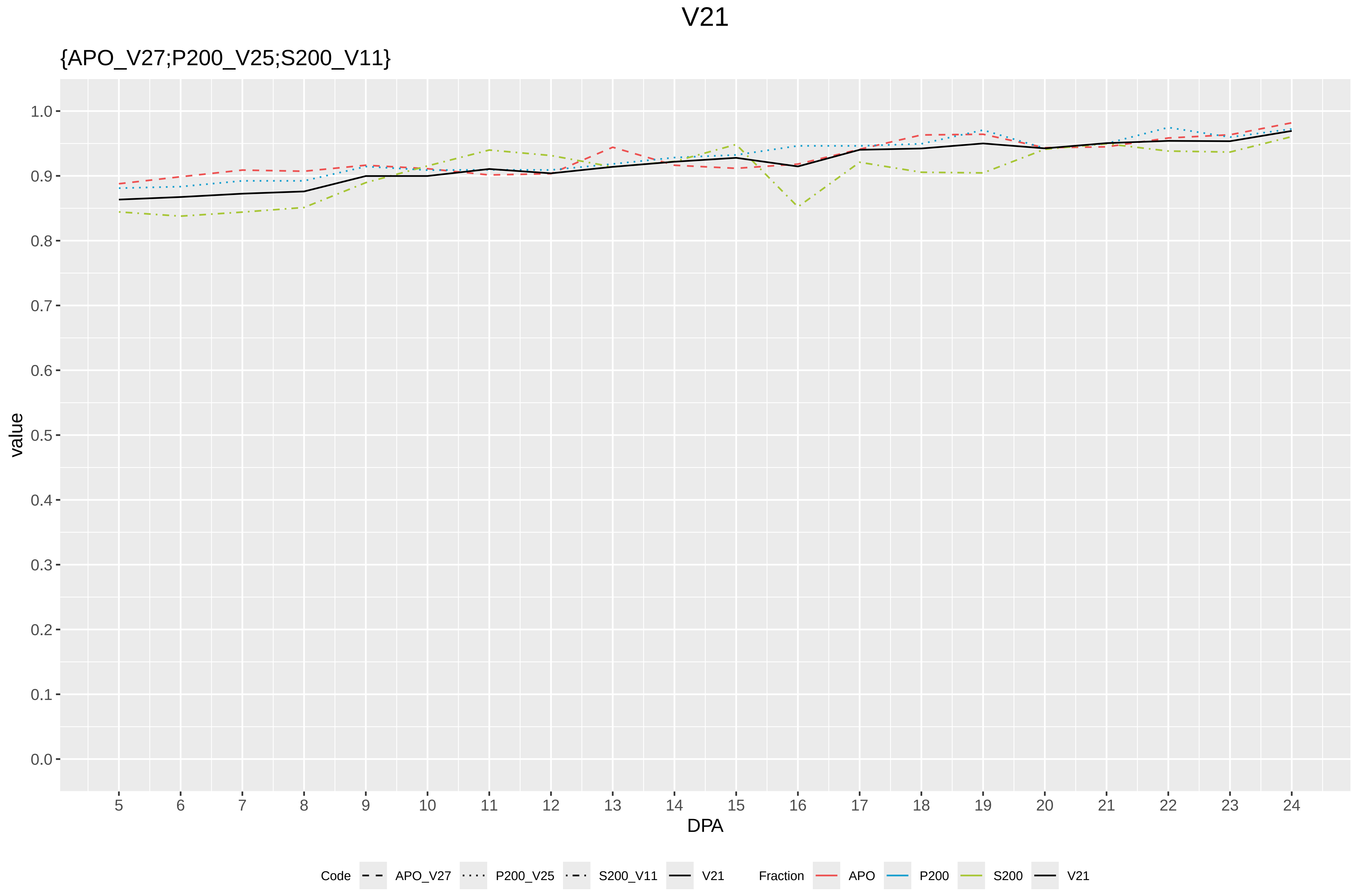
V21
{APO_V27;P200_V25;S200_V11}
value
1.0
0.9
0.8
0.7
0.6
0.5
0.4
0.3
0.2
0.1
0.0
5 6 7 8 9 10 11 12 13 14 15 16 17 18 19 20 21 22 23 24
DPA
Code
APO_V27
P200_V25
S200_V11
V21
Fraction
APO
P200
S200
V21

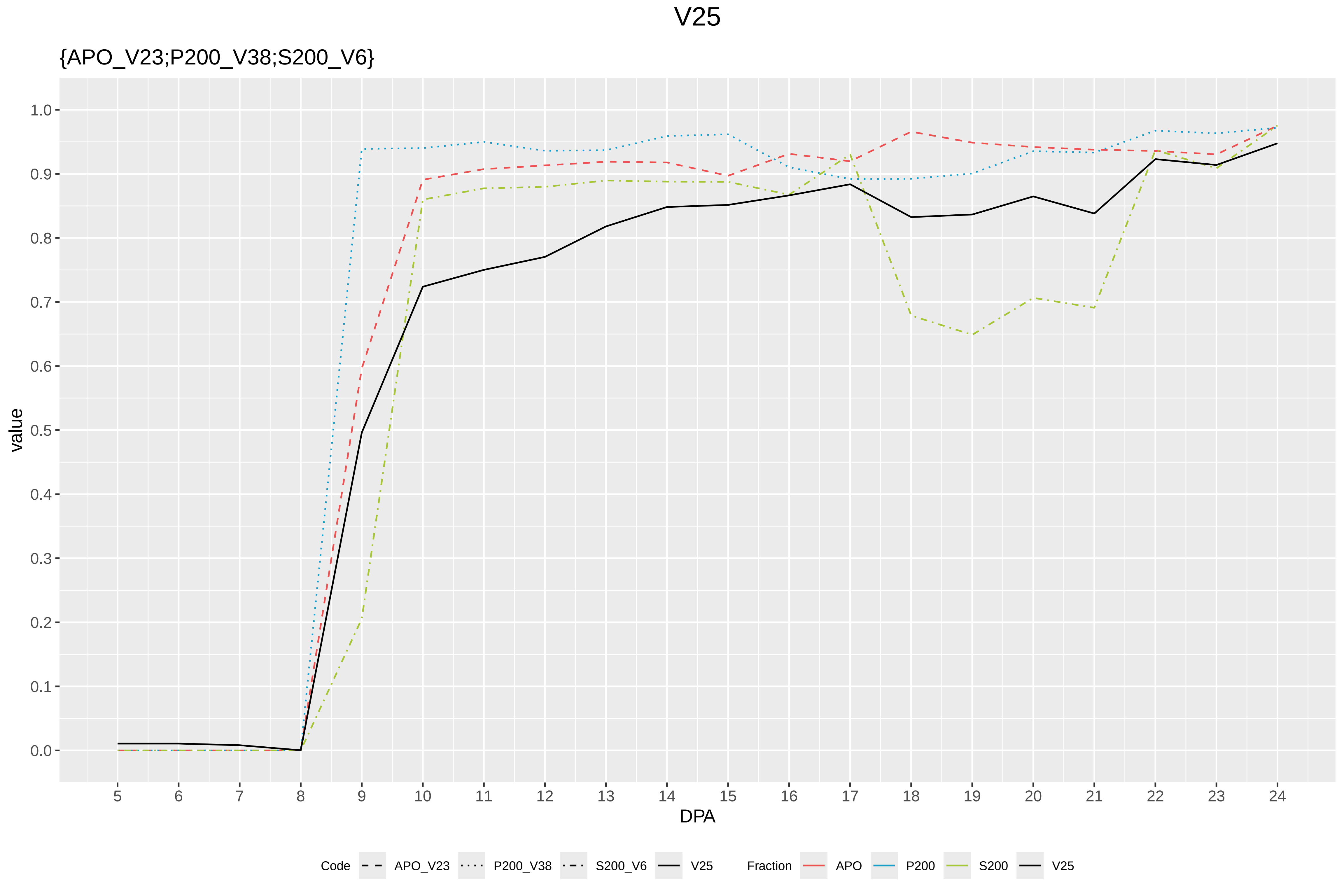
V25
{APO_V23;P200_V38;S200_V6}
value
1.0
0.9
0.8
0.7
0.6
0.5
0.4
0.3
0.2
0.1
0.0
DPA
Code
APO_V23
P200_V38
S200_V6
V25
Fraction
APO
P200
S200
V25

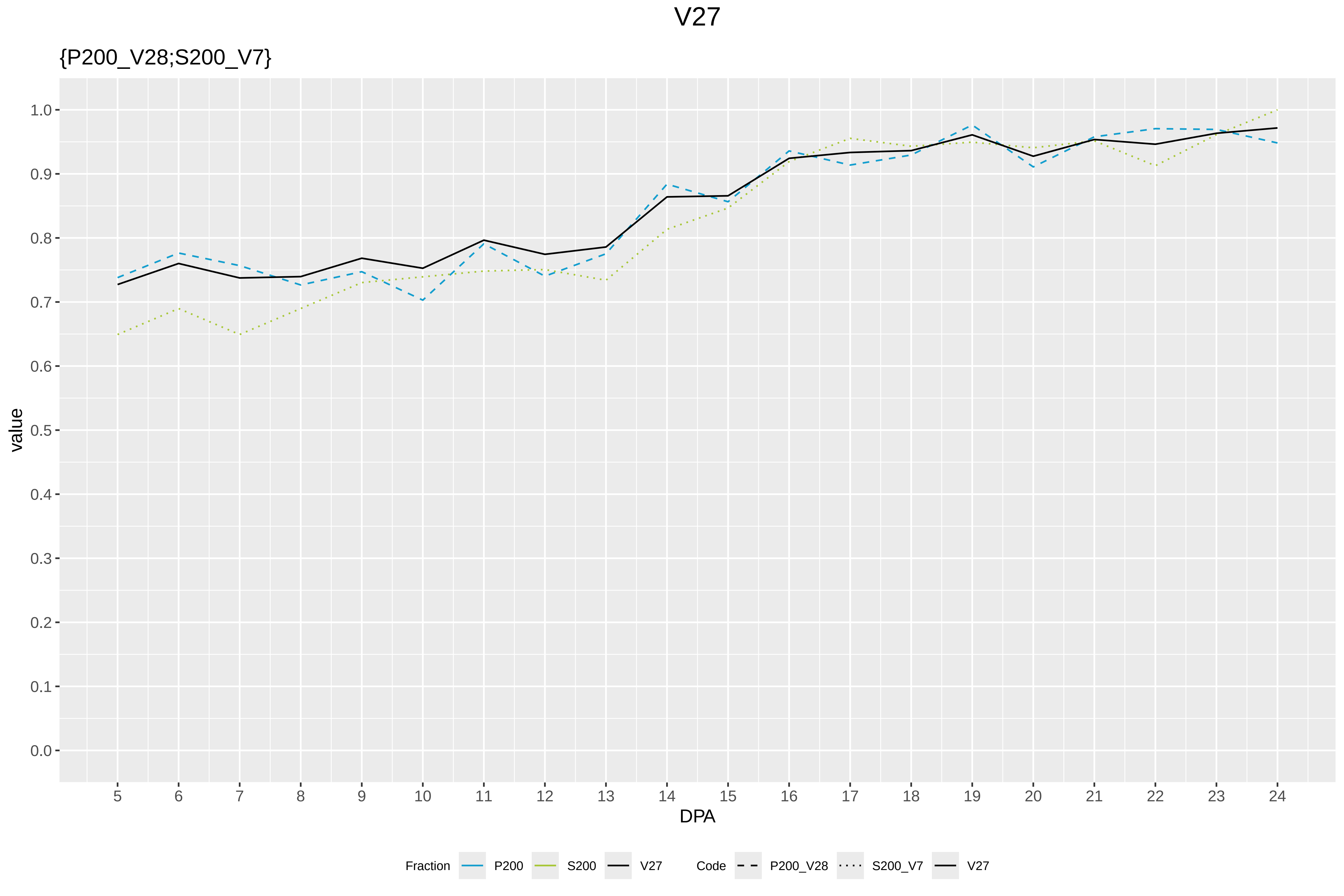
V27
{P200_V28;S200_V7}
value
1.0
0.9
0.8
0.7
0.6
0.5
0.4
0.3
0.2
0.1
0.0
DPA
Fraction
P200
S200
V27
Code
P200_V28
S200_V7
V27

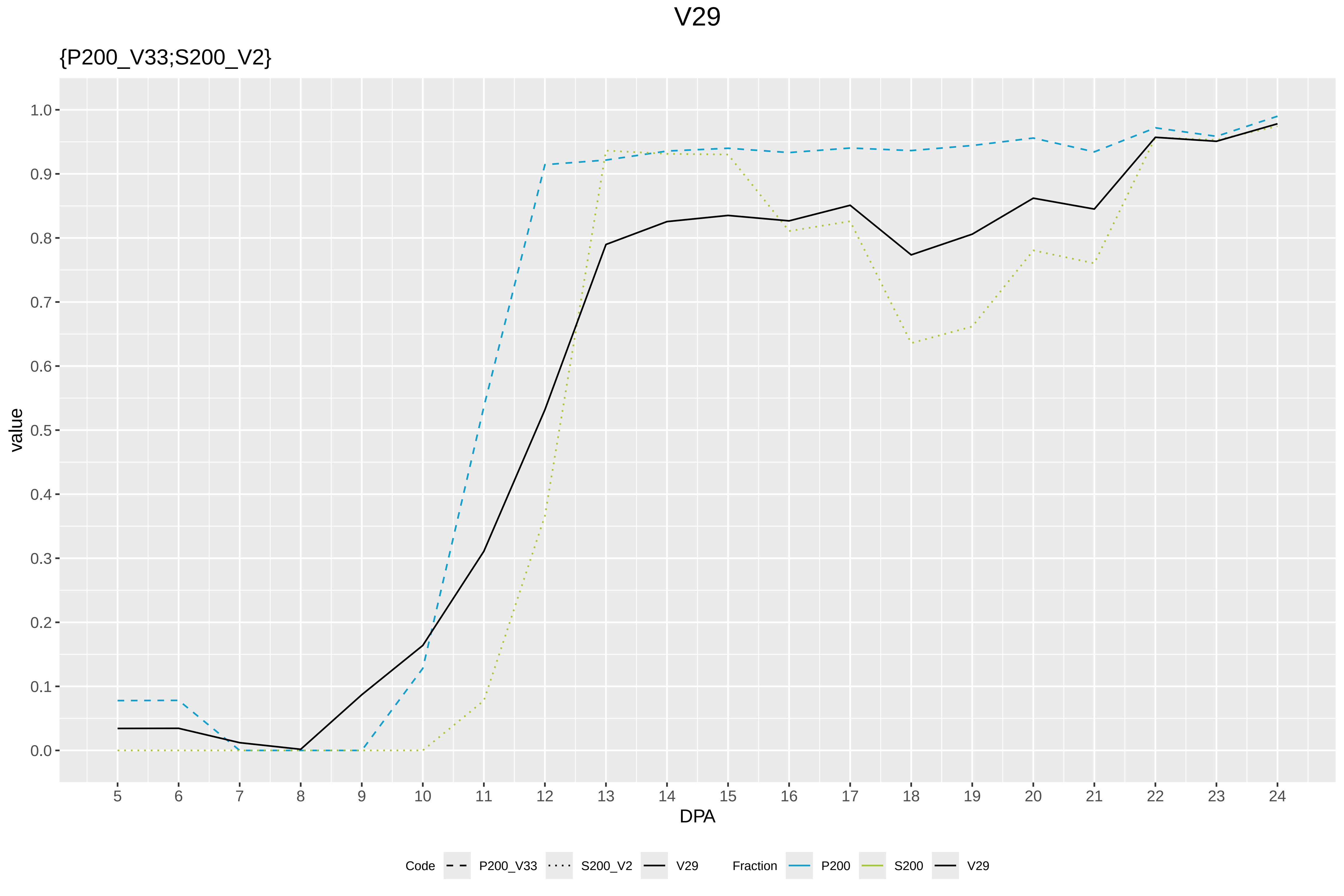

V29
{P200_V33;S200_V2}
value
1.0
0.9
0.8
0.7
0.6
0.5
0.4
0.3
0.2
0.1
0.0
DPA
Code
P200_V33
S200_V2
V29
Fraction
P200
S200
V29

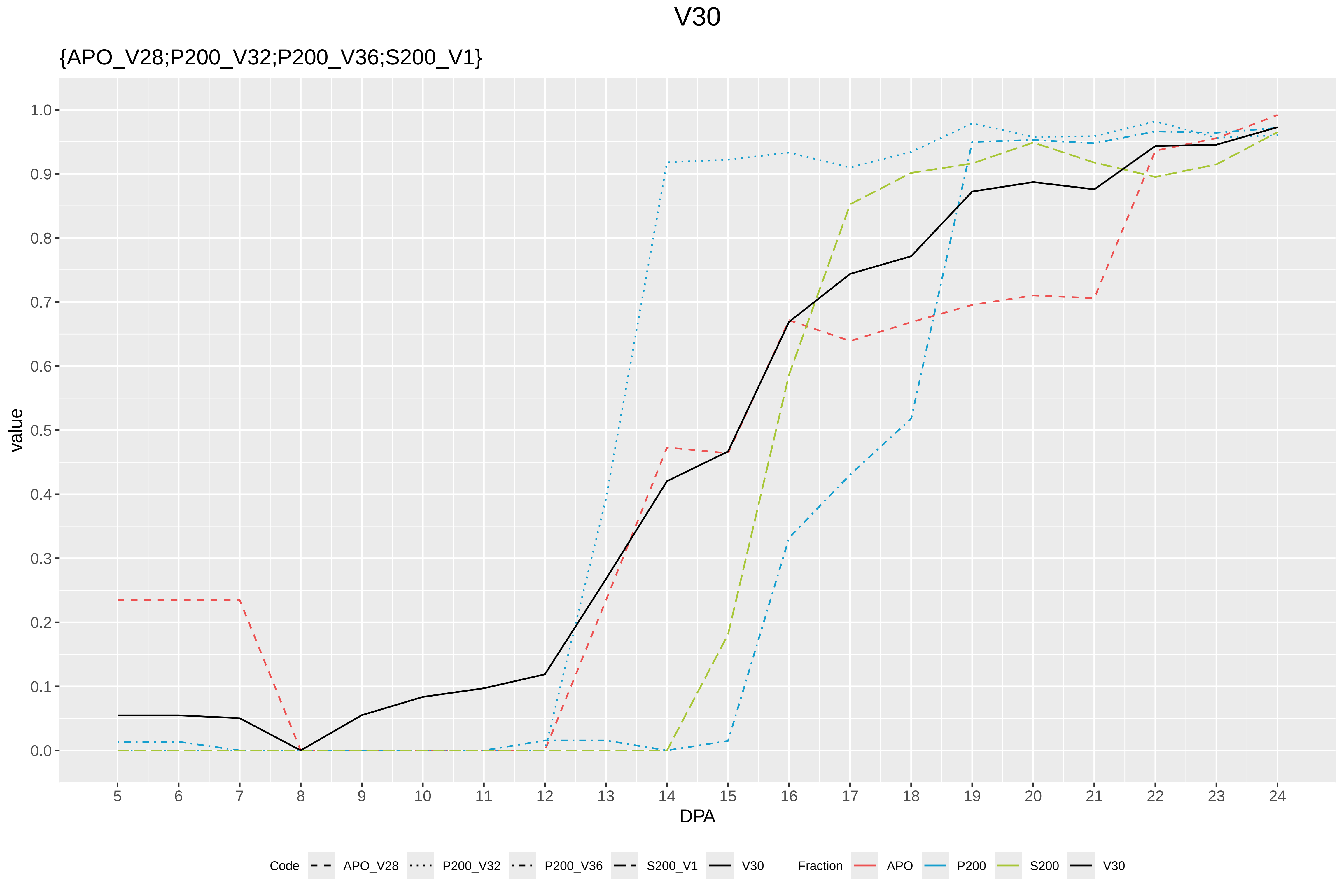

V30
{APO_V28;P200_V32;P200_V36;S200_V1}
value
1.0
0.9
0.8
0.7
0.6
0.5
0.4
0.3
0.2
0.1
0.0
DPA
Code
APO_V28
P200_V32
P200_V36
S200_V1
V30
Fraction
APO
P200
S200
V30
