## Supplementary Figure S9 for "Systems-level proteomic models of cotton fiber development: A high-resolution resource to analyze cell dynamics and trait engineering"

### Codes

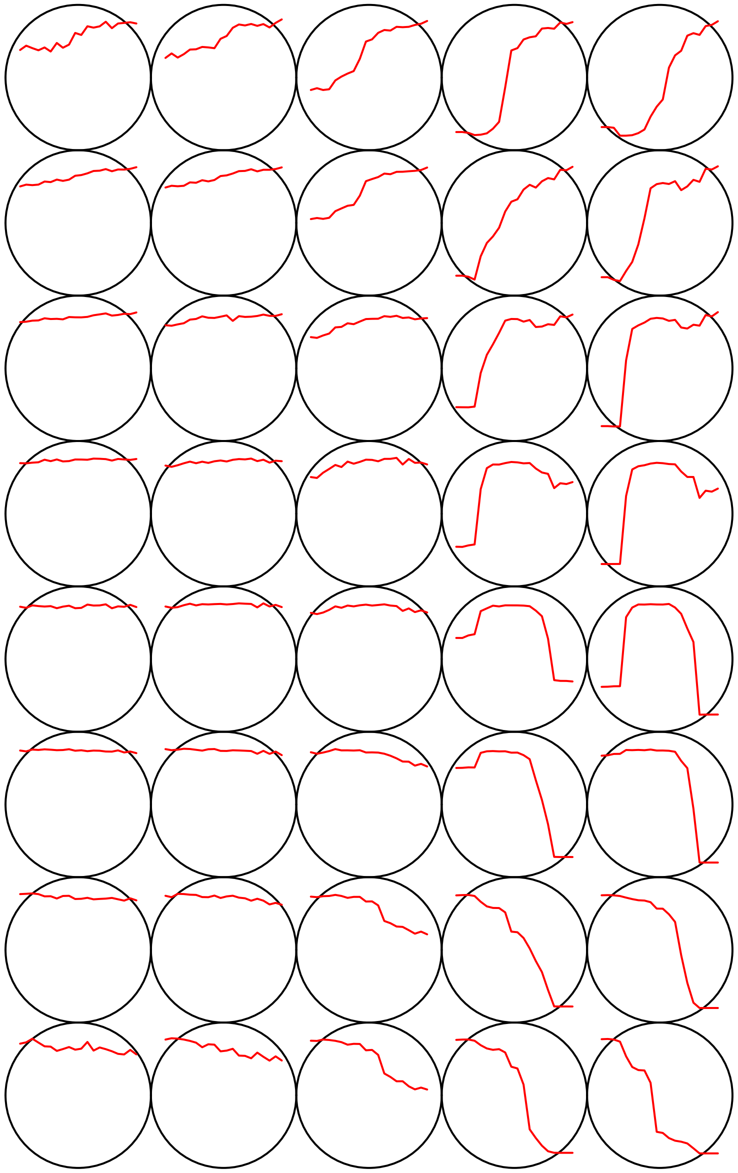

**Count plot**

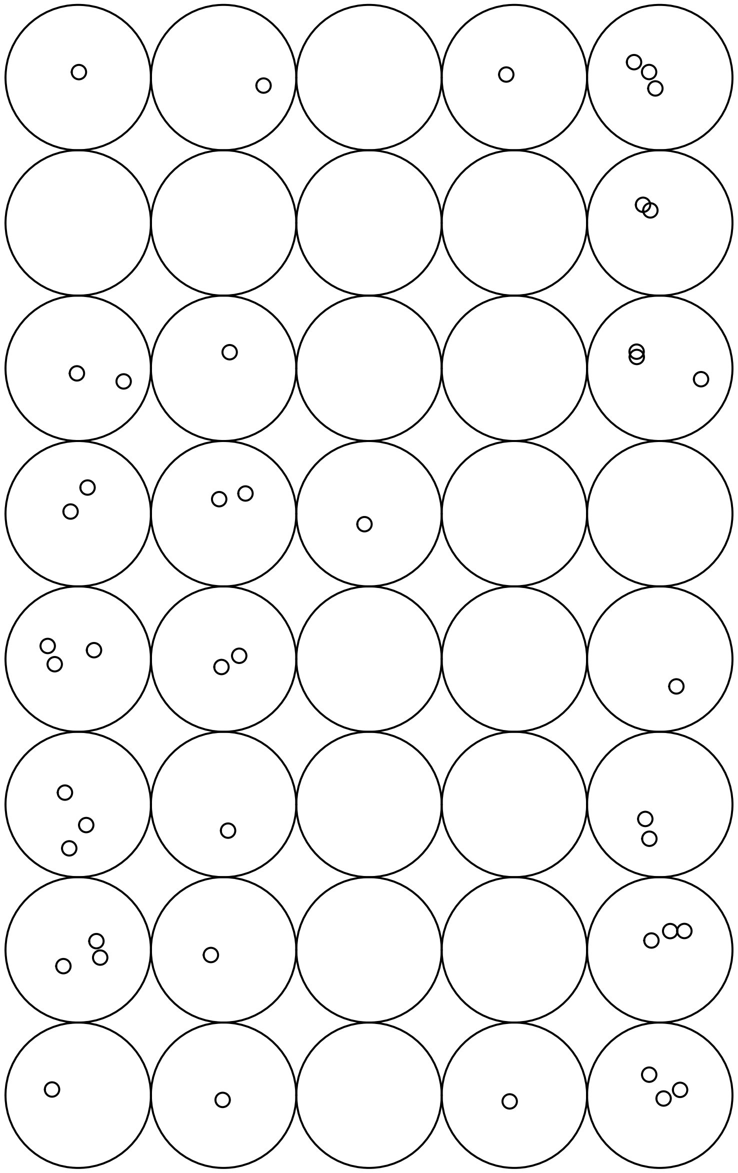

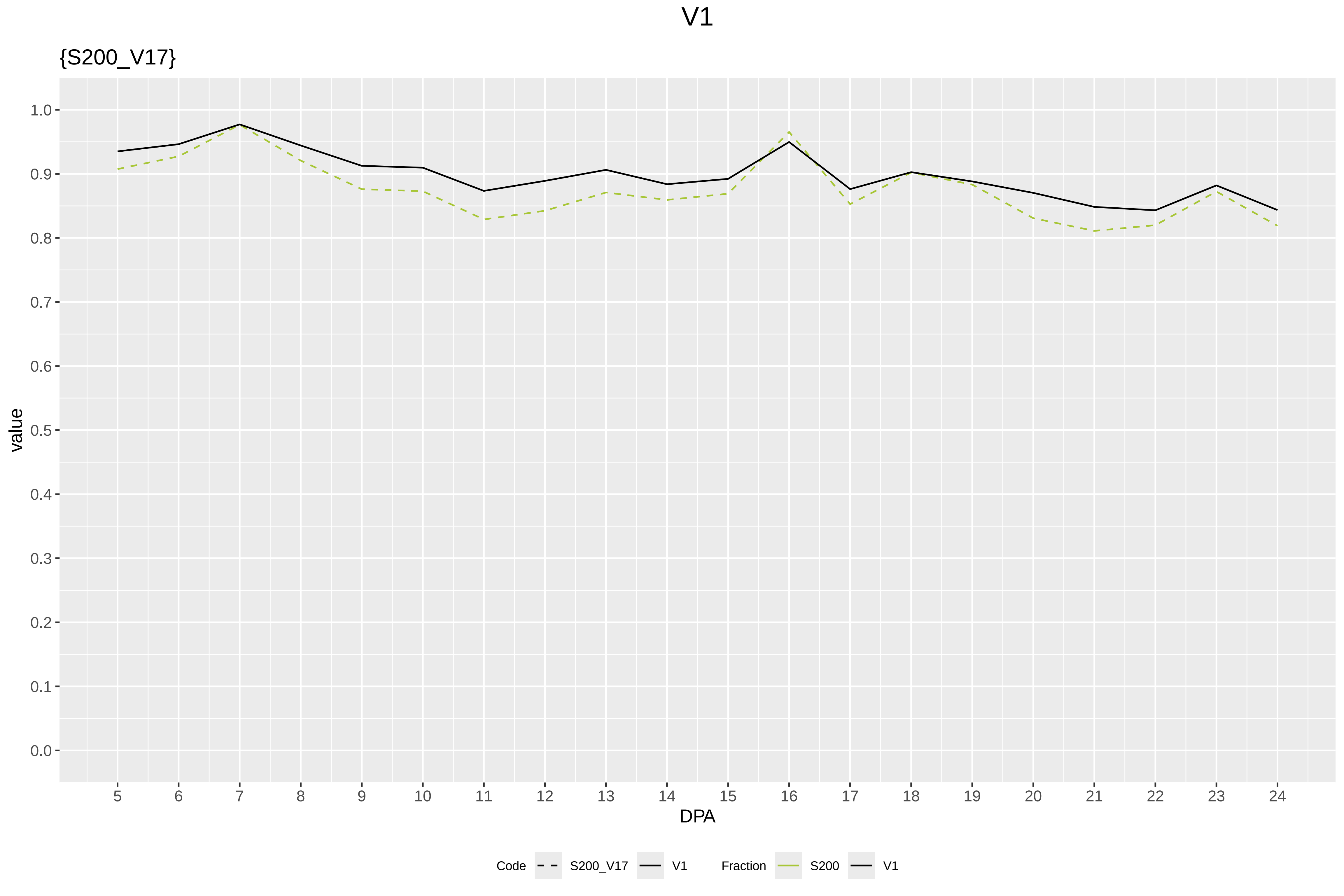
V1
{S200_V17}
value
DPA
1.0
0.9
0.8
0.7
0.6
0.5
0.4
0.3
0.2
0.1
0.0
Code
S200_V17
V1
Fraction
S200
V1

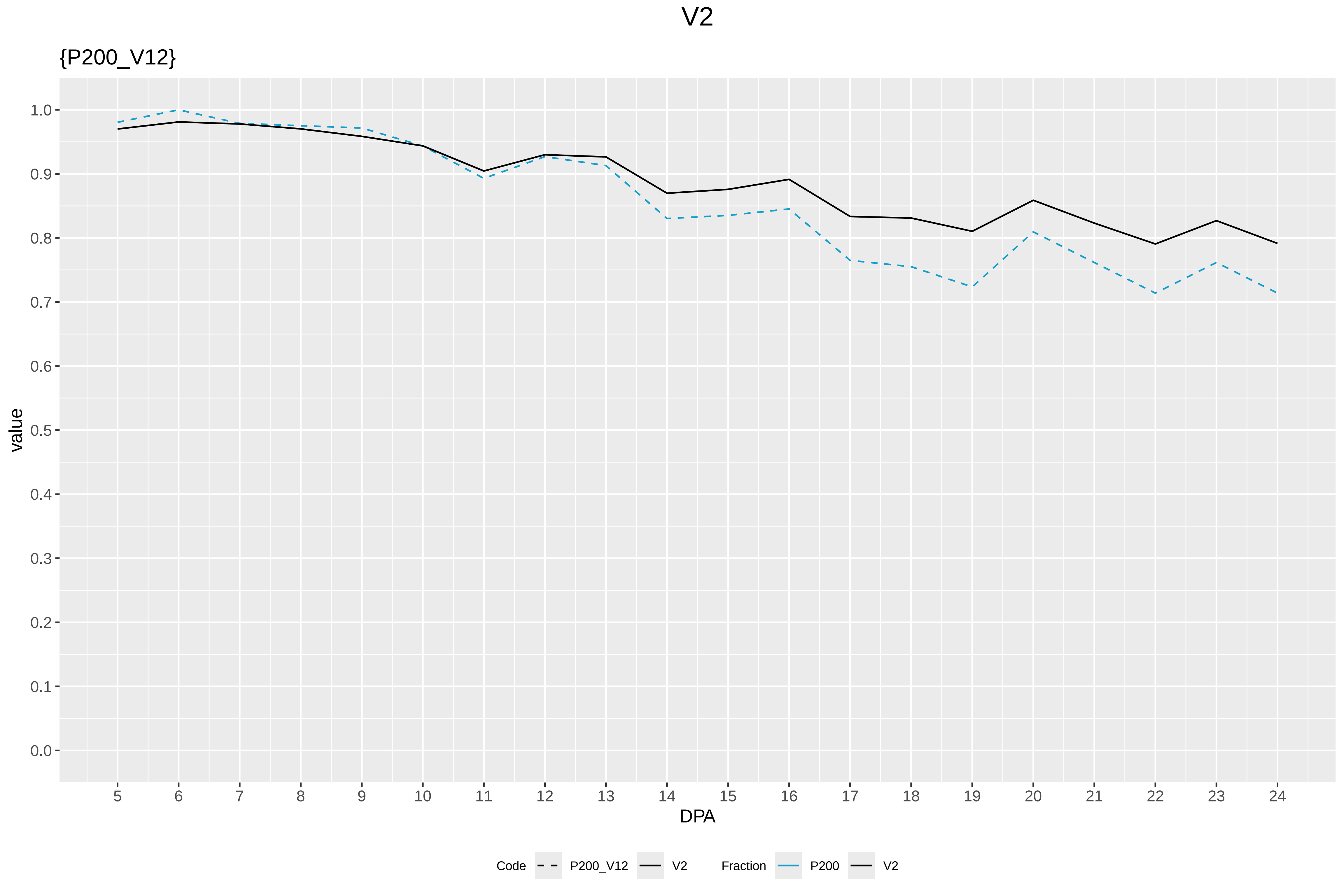
V2
{P200_V12}
value
1.0
0.9
0.8
0.7
0.6
0.5
0.4
0.3
0.2
0.1
0.0
DPA
Code
P200_V12
V2
Fraction
P200
V2

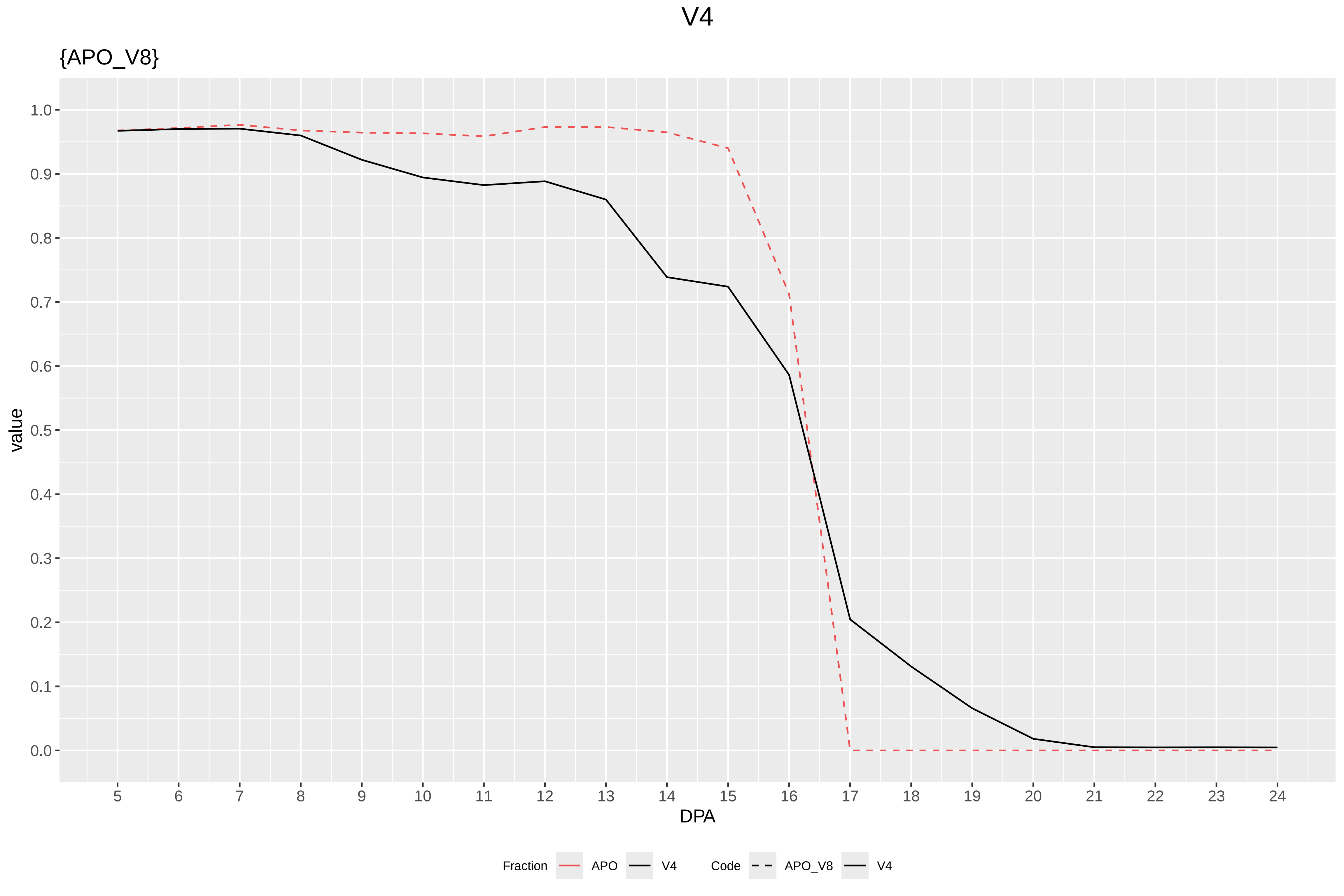
V4
{APO_V8}
value
1.0
0.9
0.8
0.7
0.6
0.5
0.4
0.3
0.2
0.1
0.0
5 6 7 8 9 10 11 12 13 14 15 16 17 18 19 20 21 22 23 24
DPA
Fraction
APO
V4
Code
APO_V8
V4

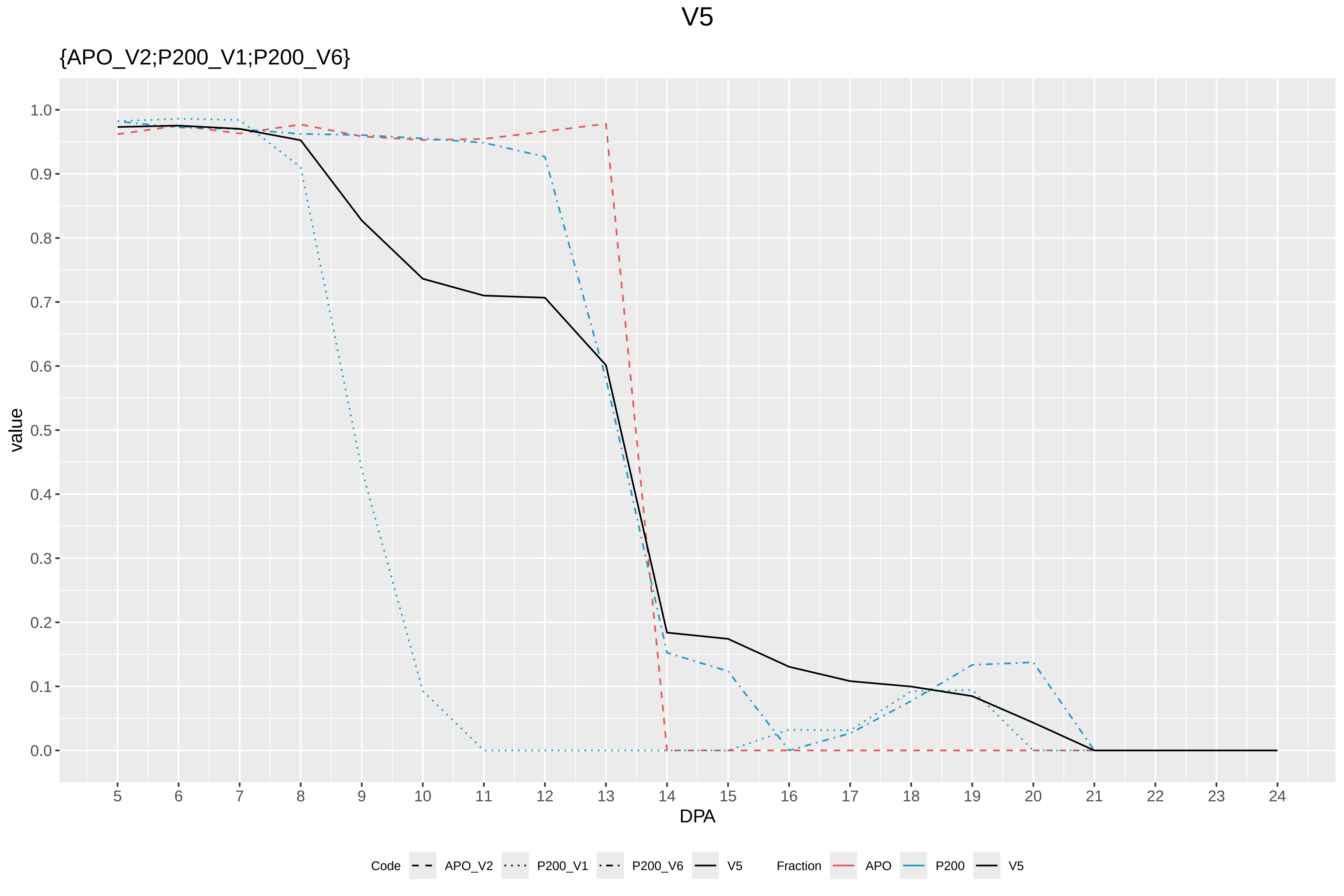

V5
{APO_V2;P200_V1;P200_V6}
value
1.0
0.9
0.8
0.7
0.6
0.5
0.4
0.3
0.2
0.1
0.0
DPA
Code
APO_V2
P200_V1
P200_V6
V5
Fraction
APO
P200
V5

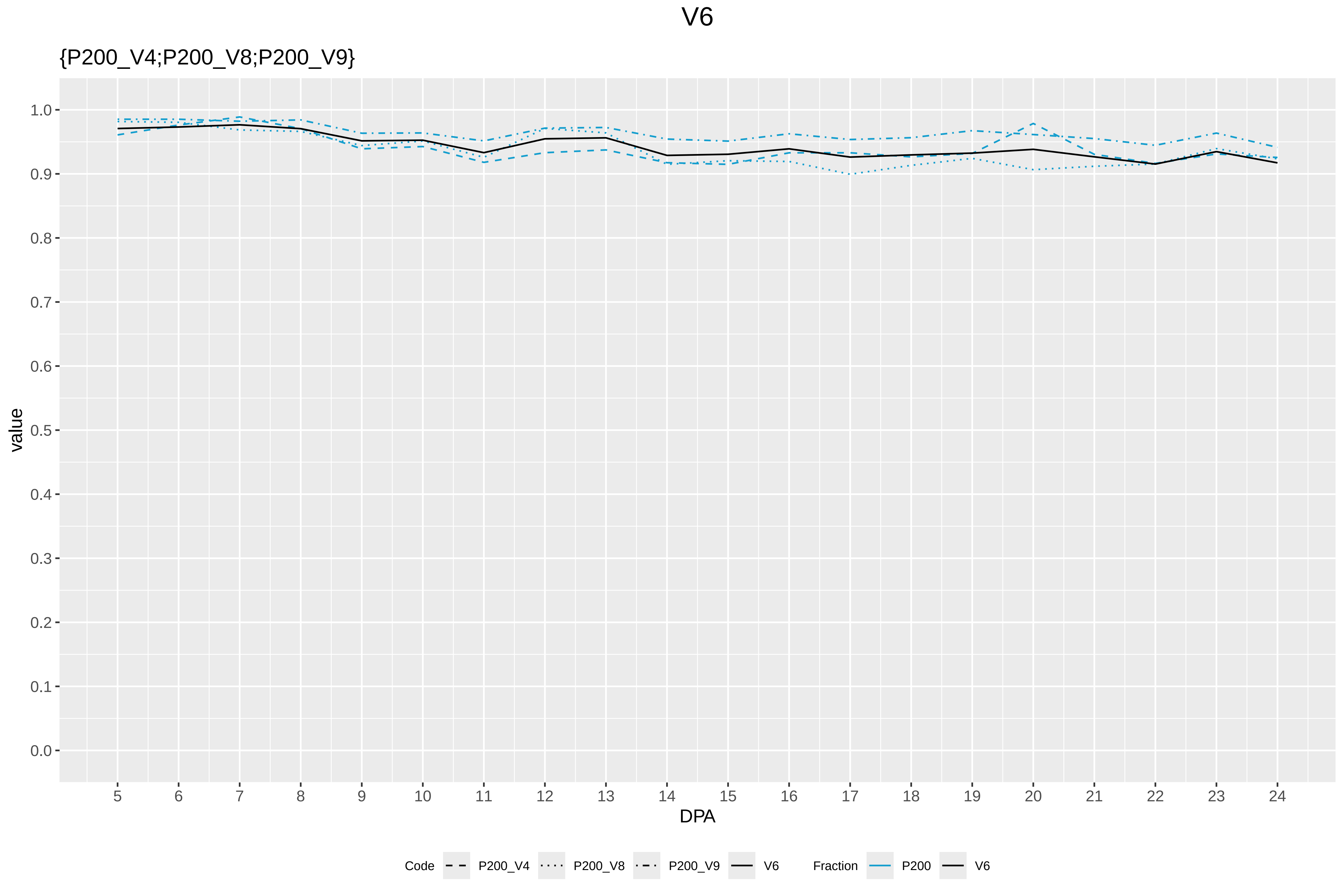
V6
{P200_V4;P200_V8;P200_V9}
value
DPA
1.0
0.9
0.8
0.7
0.6
0.5
0.4
0.3
0.2
0.1
0.0
Code
P200_V4
P200_V8
P200_V9
V6
Fraction
P200
V6

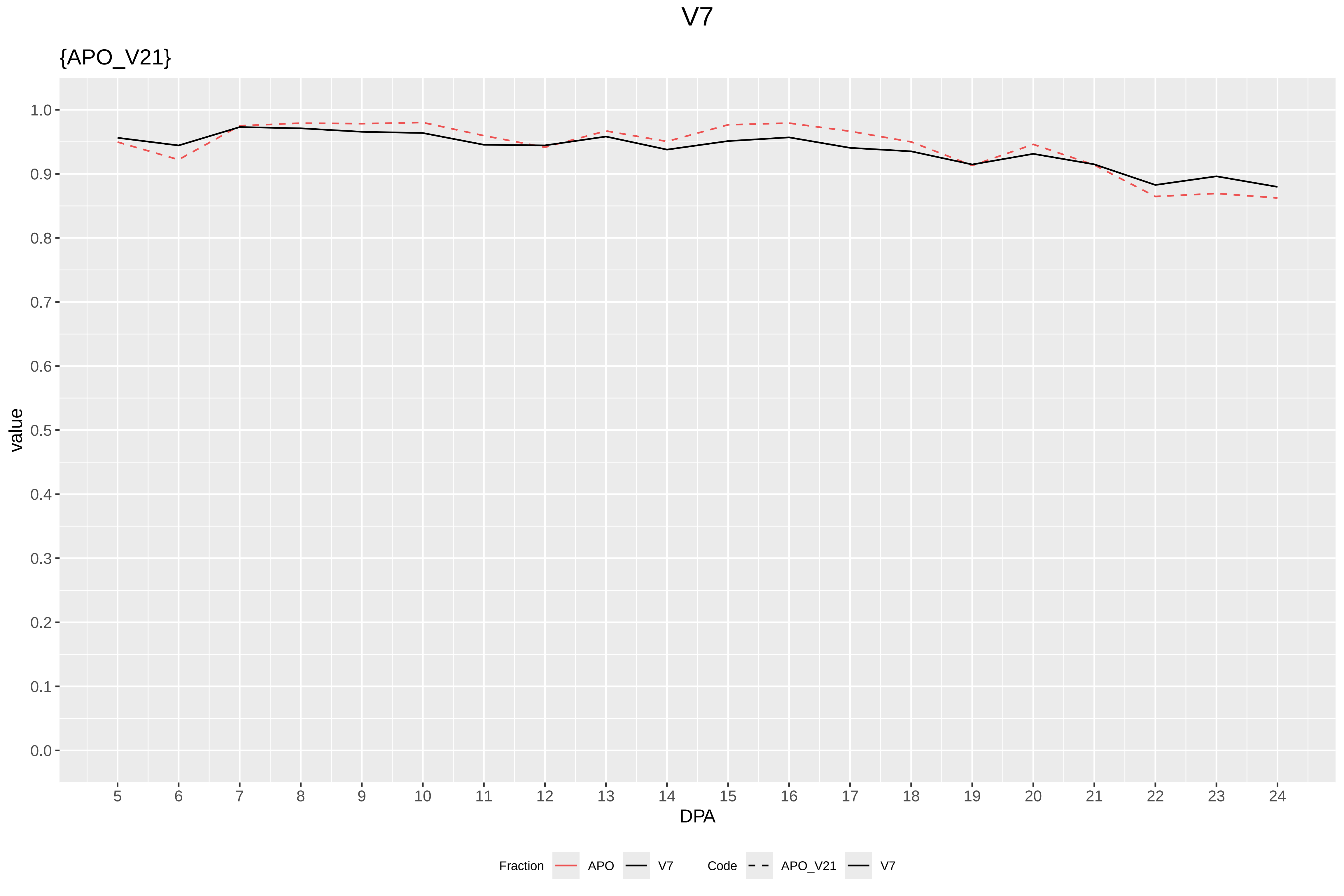

V7
{APO_V21}
value
1.0
0.9
0.8
0.7
0.6
0.5
0.4
0.3
0.2
0.1
0.0
DPA
Fraction
APO
V7
Code
APO_V21
V7

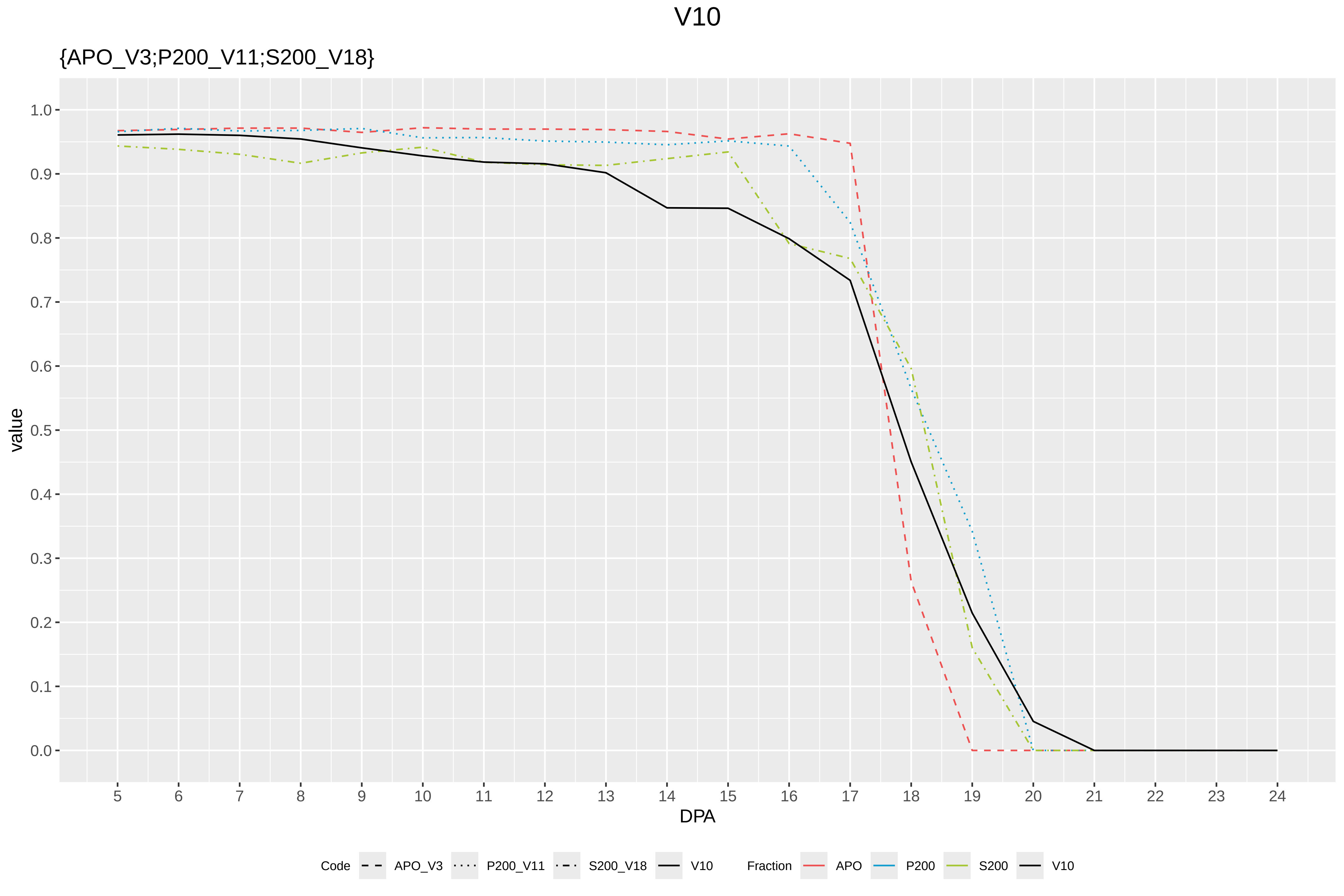

V10
{APO_V3;P200_V11;S200_V18}
value
1.0
0.9
0.8
0.7
0.6
0.5
0.4
0.3
0.2
0.1
0.0
5 6 7 8 9 10 11 12 13 14 15 16 17 18 19 20 21 22 23 24
DPA
Code APO_V3 P200_V11 S200_V18 V10
Fraction APO P200 S200 V10

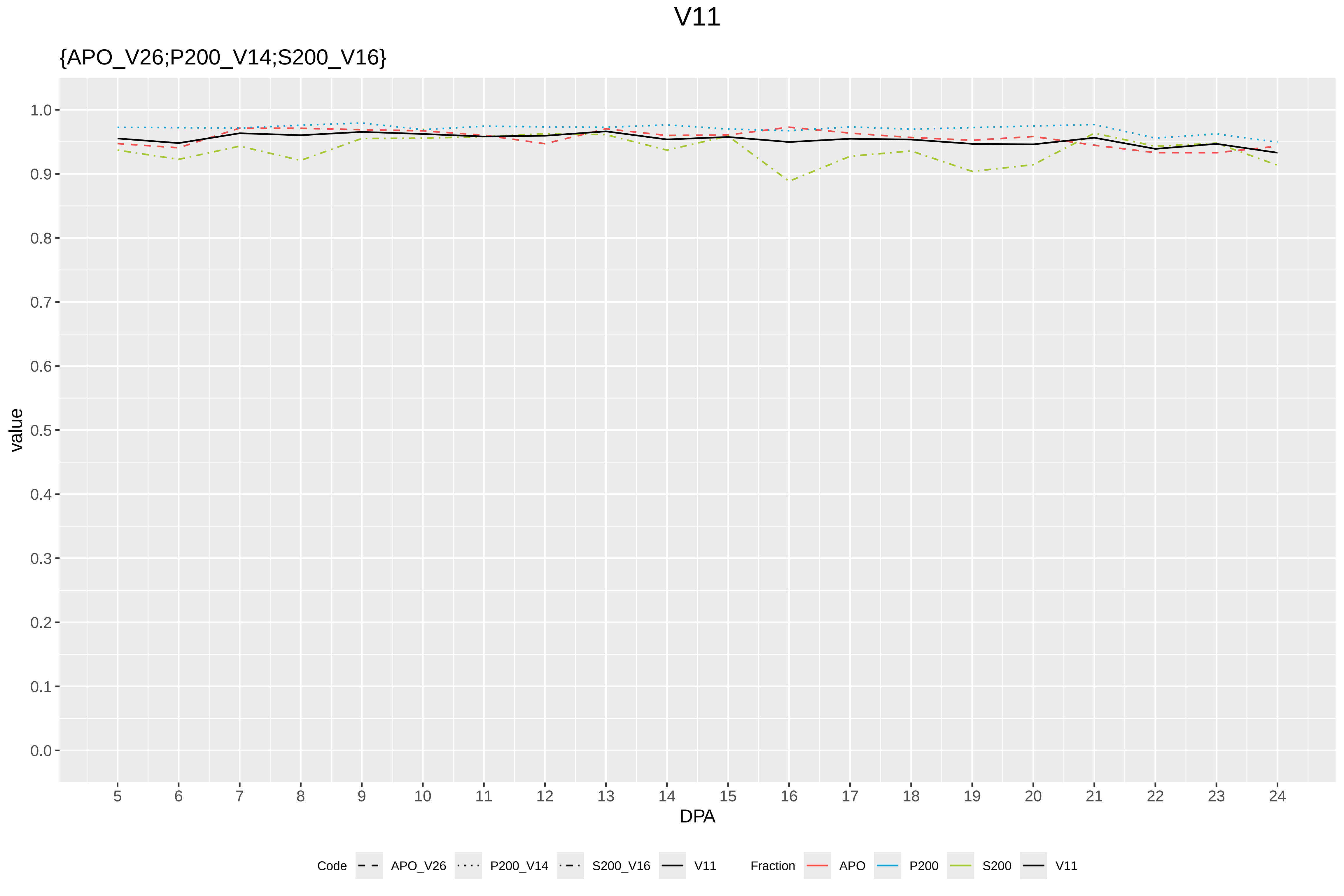
V11
{APO_V26;P200_V14;S200_V16}
value
1.0
0.9
0.8
0.7
0.6
0.5
0.4
0.3
0.2
0.1
0.0
5 6 7 8 9 10 11 12 13 14 15 16 17 18 19 20 21 22 23 24
DPA
Code
APO_V26
P200_V14
S200_V16
V11
Fraction
APO
P200
S200
V11

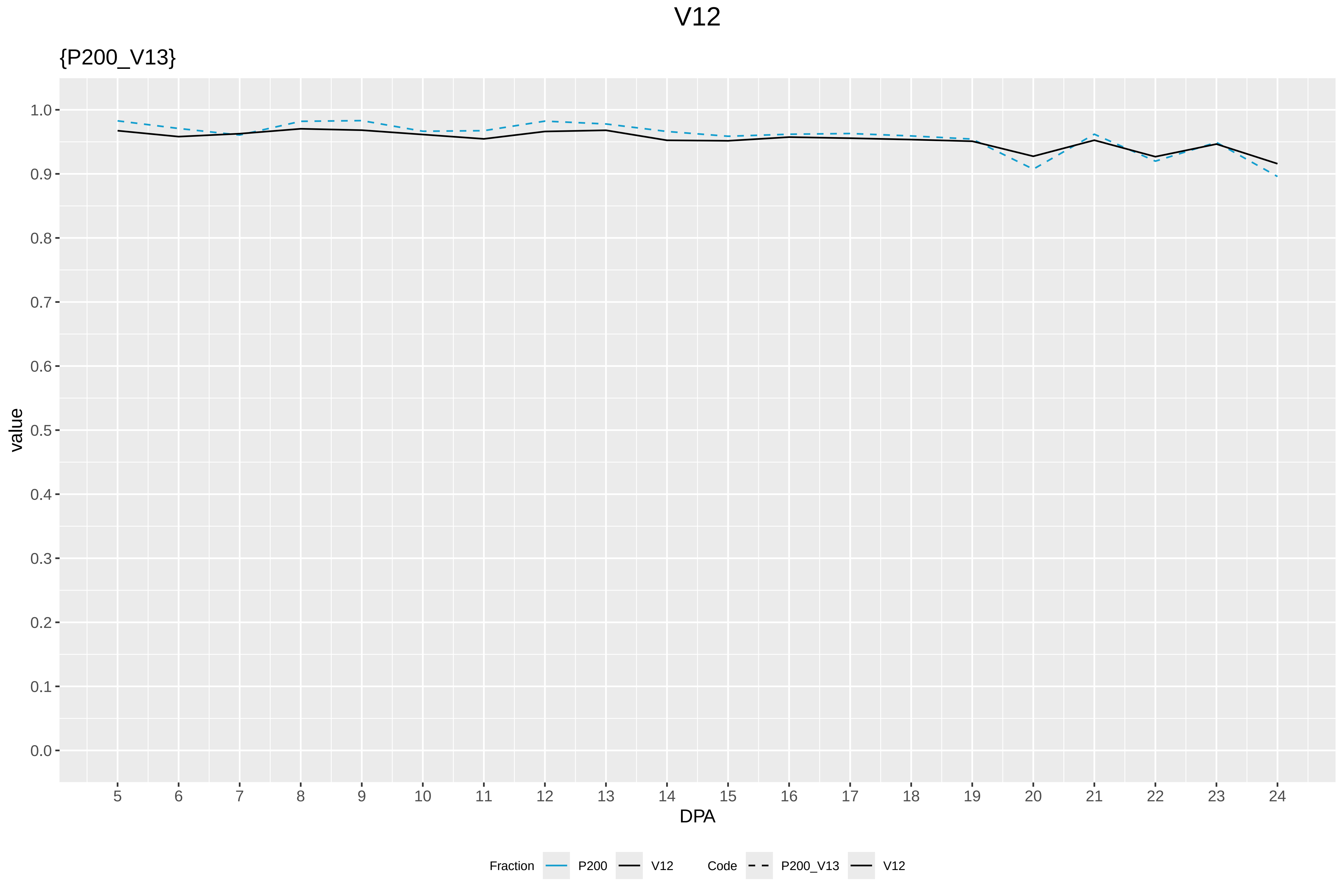

V12
{P200_V13}
value
DPA
1.0
0.9
0.8
0.7
0.6
0.5
0.4
0.3
0.2
0.1
0.0
5 6 7 8 9 10 11 12 13 14 15 16 17 18 19 20 21 22 23 24
Fraction P200 V12
Code P200_V13 V12

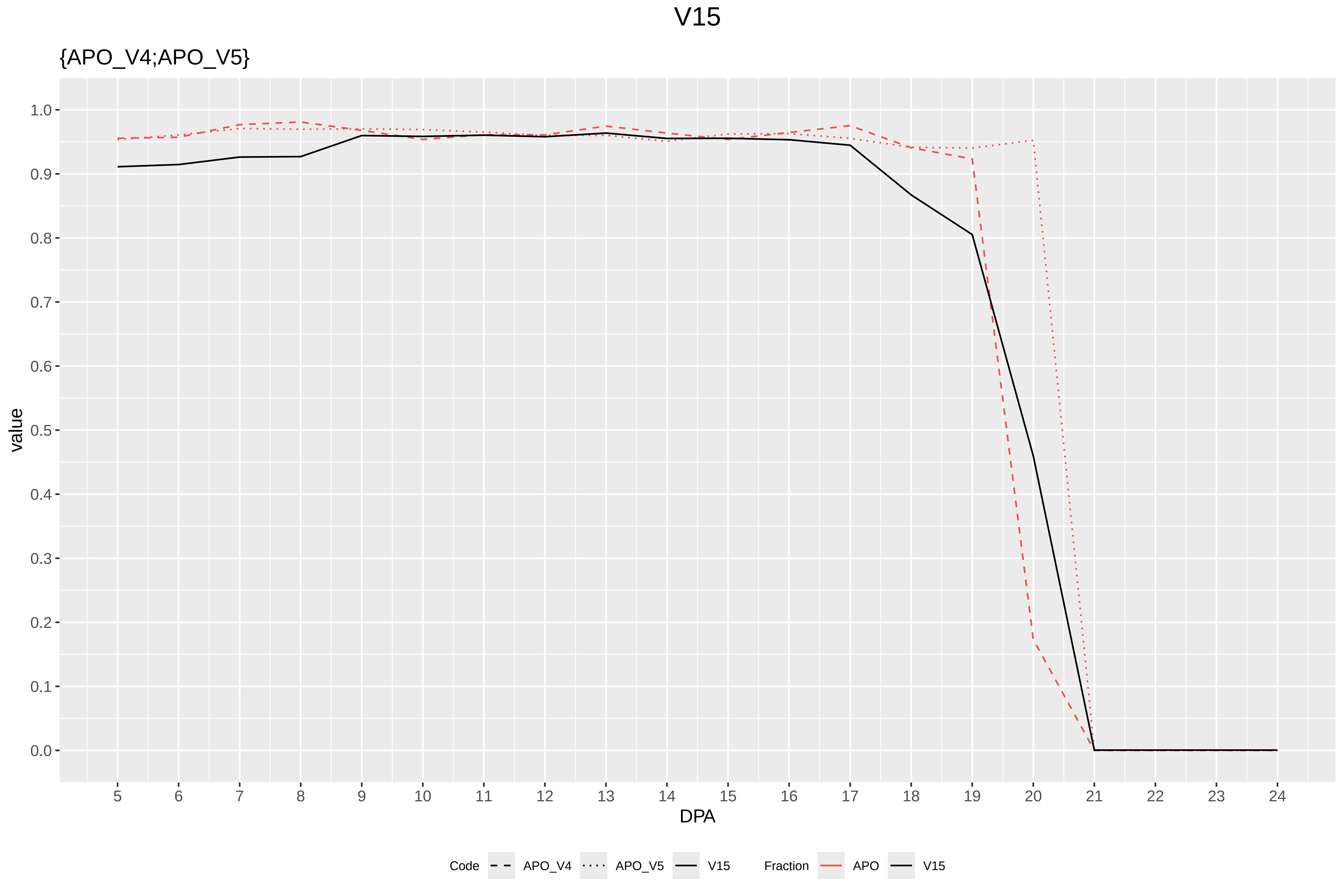
V15
{APO_V4;APO_V5}
value
DPA
1.0
0.9
0.8
0.7
0.6
0.5
0.4
0.3
0.2
0.1
0.0
Code
APO_V4
APO_V5
V15
Fraction
APO
V15

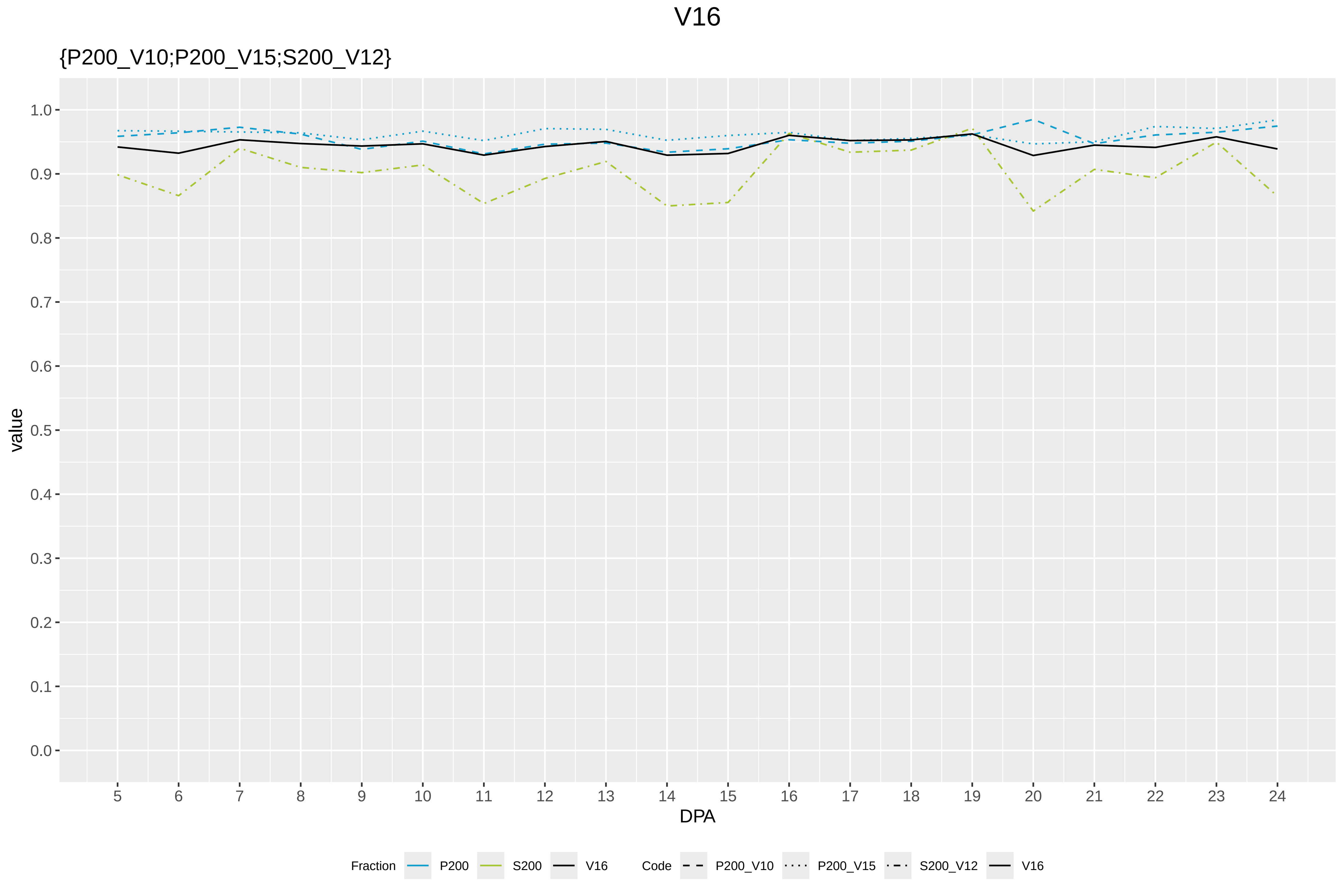

V16
{P200_V10;P200_V15;S200_V12}
value
DPA
1.0
0.9
0.8
0.7
0.6
0.5
0.4
0.3
0.2
0.1
0.0
5 6 7 8 9 10 11 12 13 14 15 16 17 18 19 20 21 22 23 24
Fraction P200 S200 V16
Code P200_V10 P200_V15 S200_V12 V16

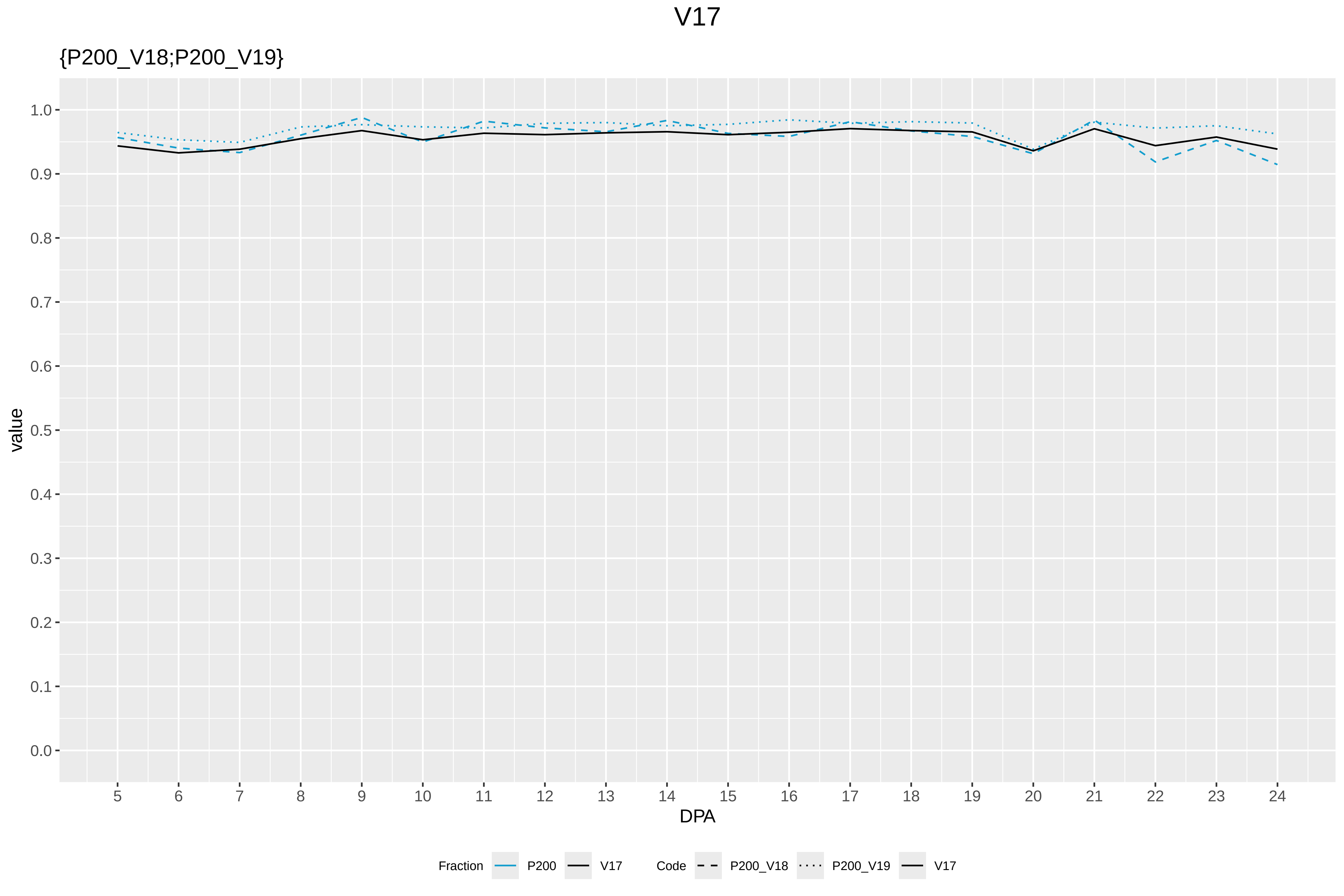

V17
{P200_V18;P200_V19}
value
1.0
0.9
0.8
0.7
0.6
0.5
0.4
0.3
0.2
0.1
0.0
DPA
Fraction
P200
V17
Code
P200_V18
P200_V19
V17

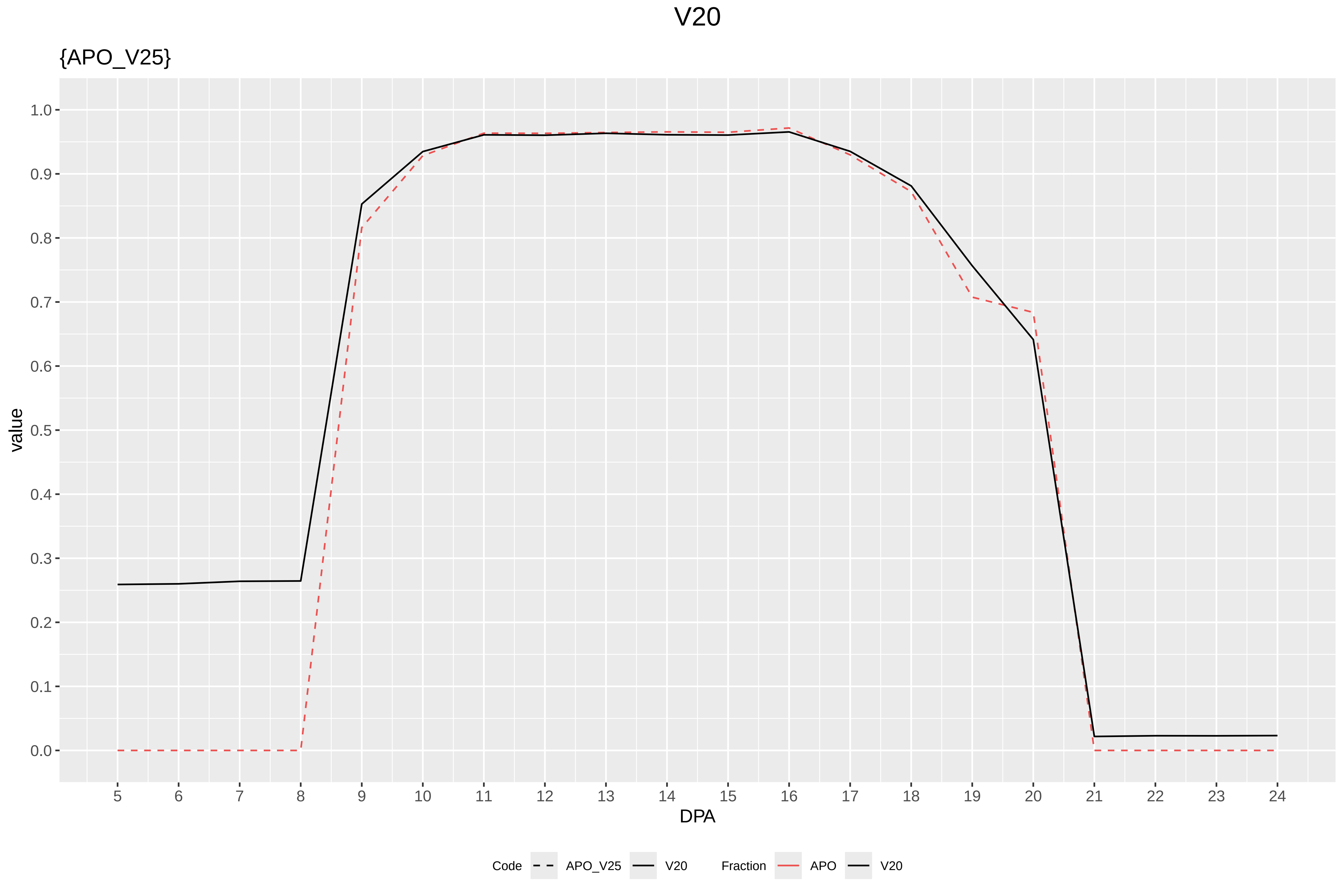

V20
{APO_V25}
value
1.0
0.9
0.8
0.7
0.6
0.5
0.4
0.3
0.2
0.1
0.0
DPA
Code
APO_V25
V20
Fraction
APO
V20

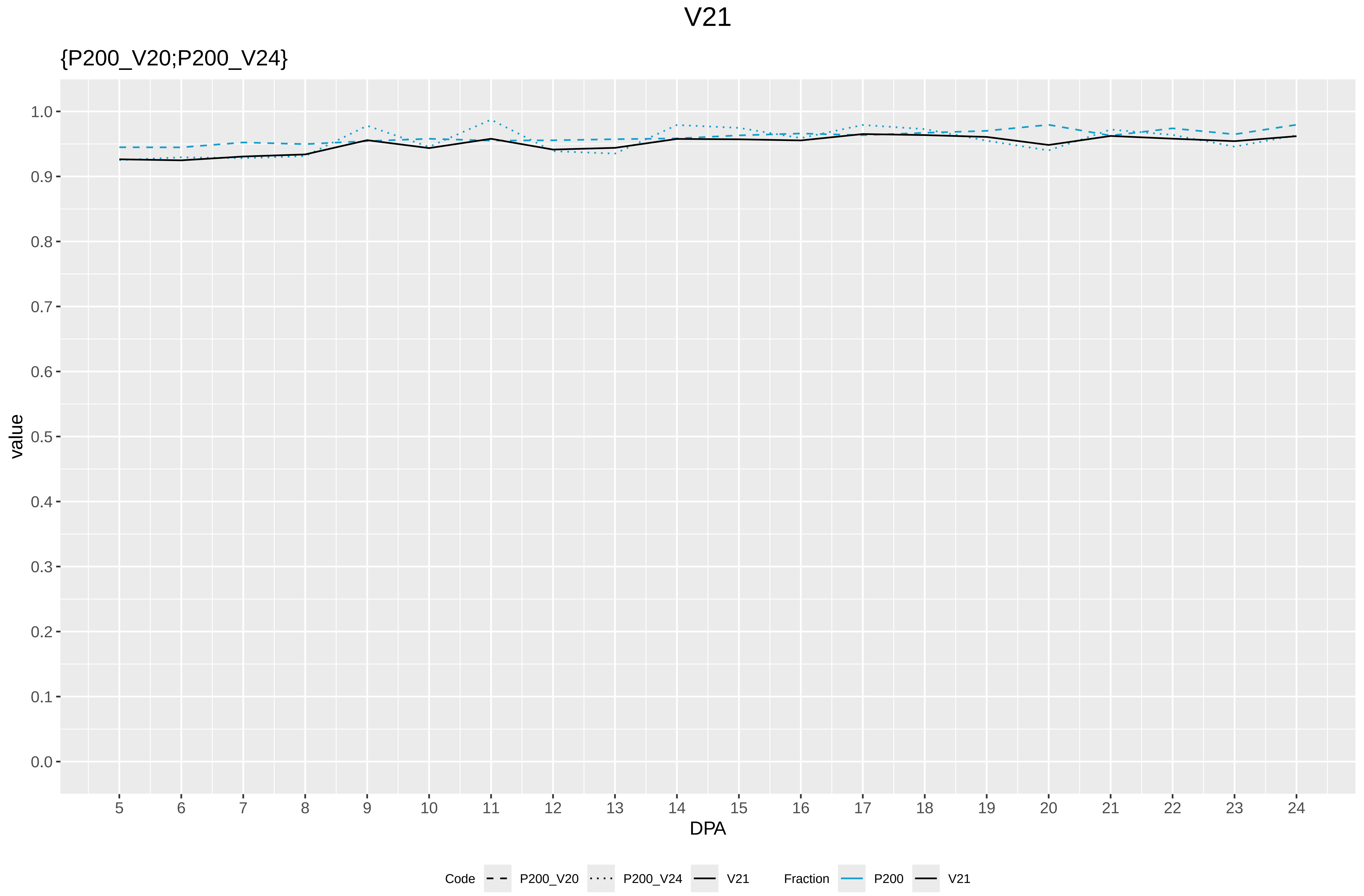

V21
{P200_V20;P200_V24}
value
1.0
0.9
0.8
0.7
0.6
0.5
0.4
0.3
0.2
0.1
0.0
DPA
Code
P200_V20
P200_V24
V21
Fraction
P200
V21

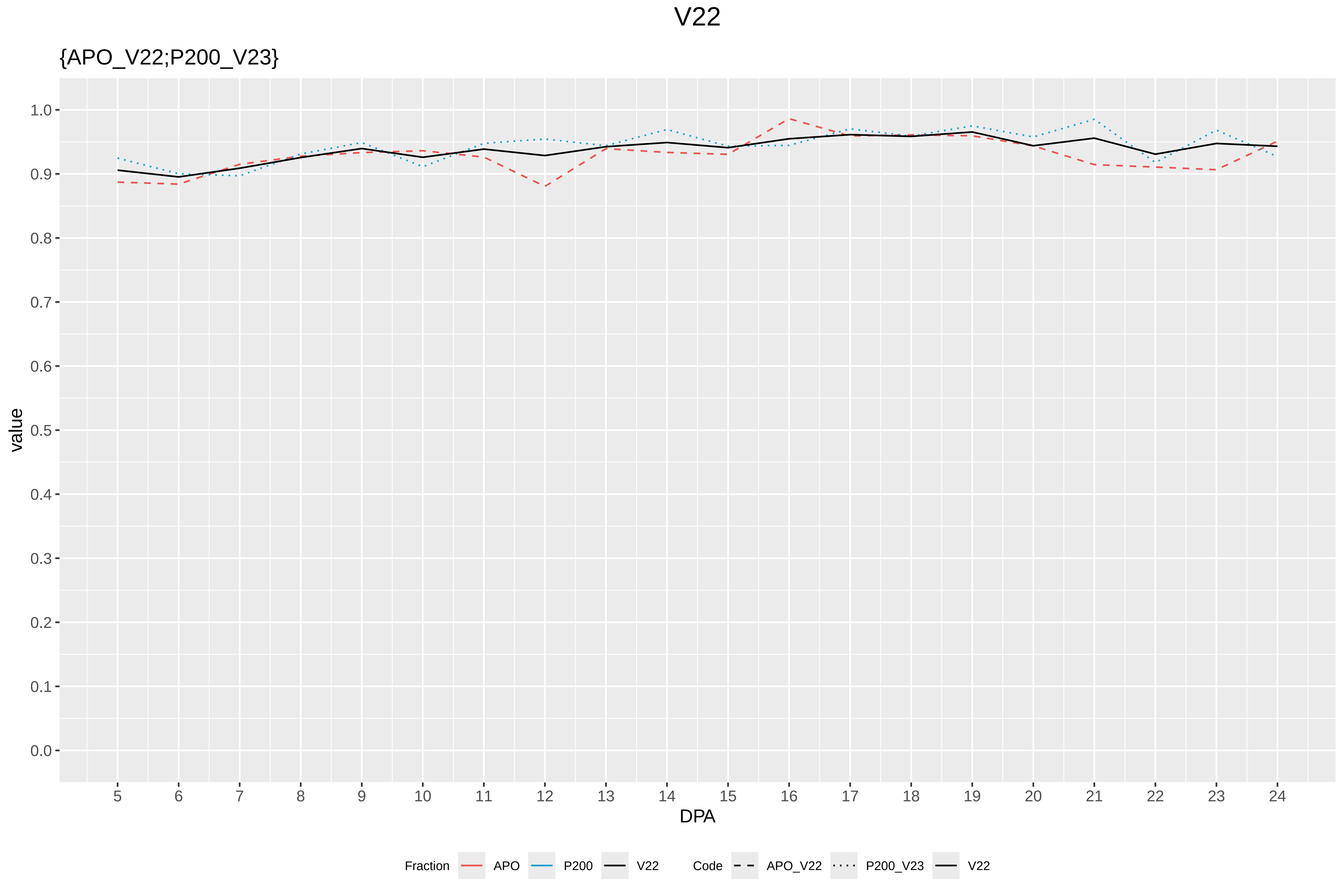

V22
{APO_V22;P200_V23}
value
DPA
1.0
0.9
0.8
0.7
0.6
0.5
0.4
0.3
0.2
0.1
0.0
5 6 7 8 9 10 11 12 13 14 15 16 17 18 19 20 21 22 23 24
Fraction APO P200 V22
Code APO_V22 P200_V23 V22

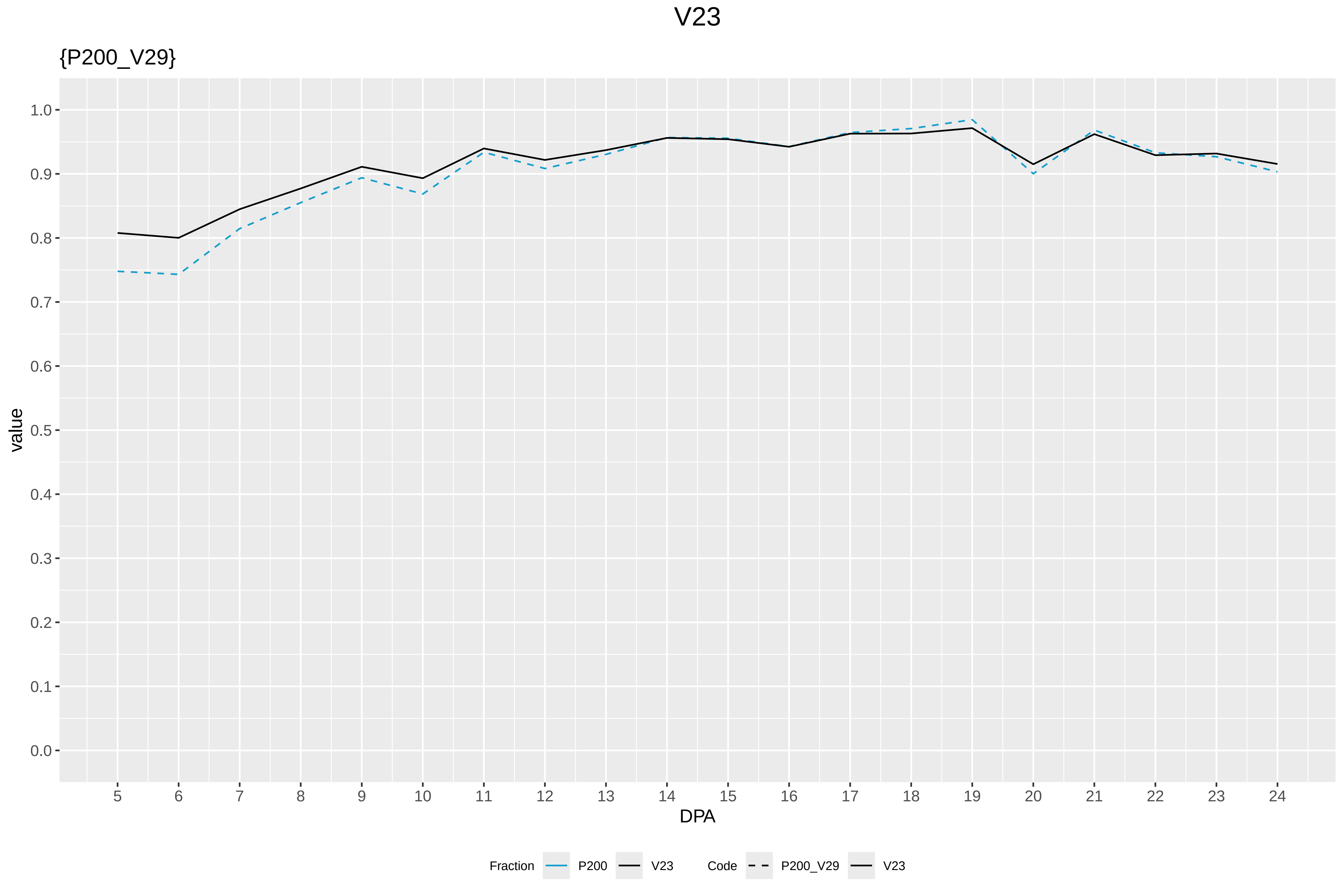

V23
{P200_V29}
value
1.0
0.9
0.8
0.7
0.6
0.5
0.4
0.3
0.2
0.1
0.0
5 6 7 8 9 10 11 12 13 14 15 16 17 18 19 20 21 22 23 24
DPA
Fraction
P200
V23
Code
P200_V29
V23

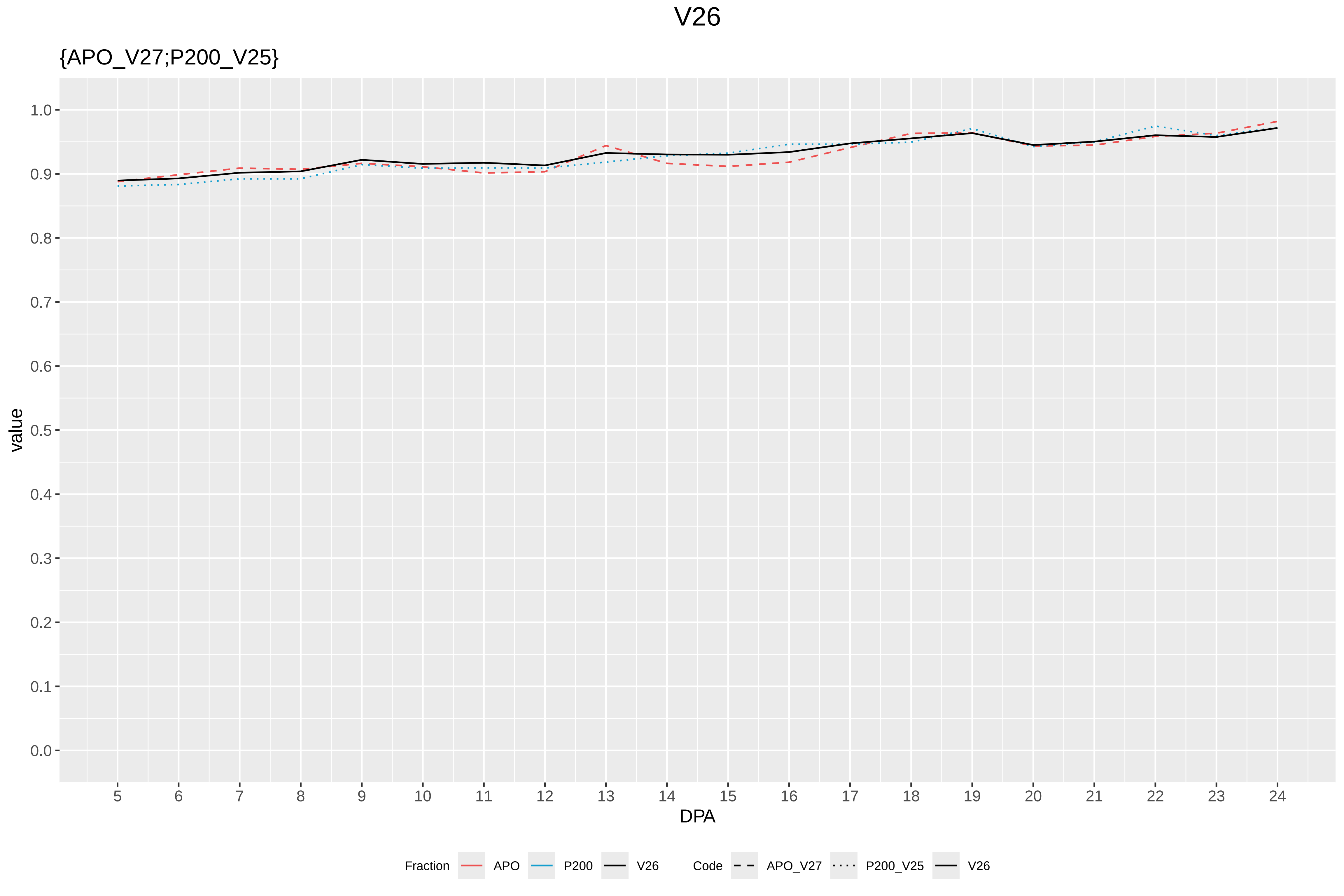
V26
{APO_V27;P200_V25}
value
1.0
0.9
0.8
0.7
0.6
0.5
0.4
0.3
0.2
0.1
0.0
DPA
Fraction
APO
P200
V26
Code
APO_V27
P200_V25
V26

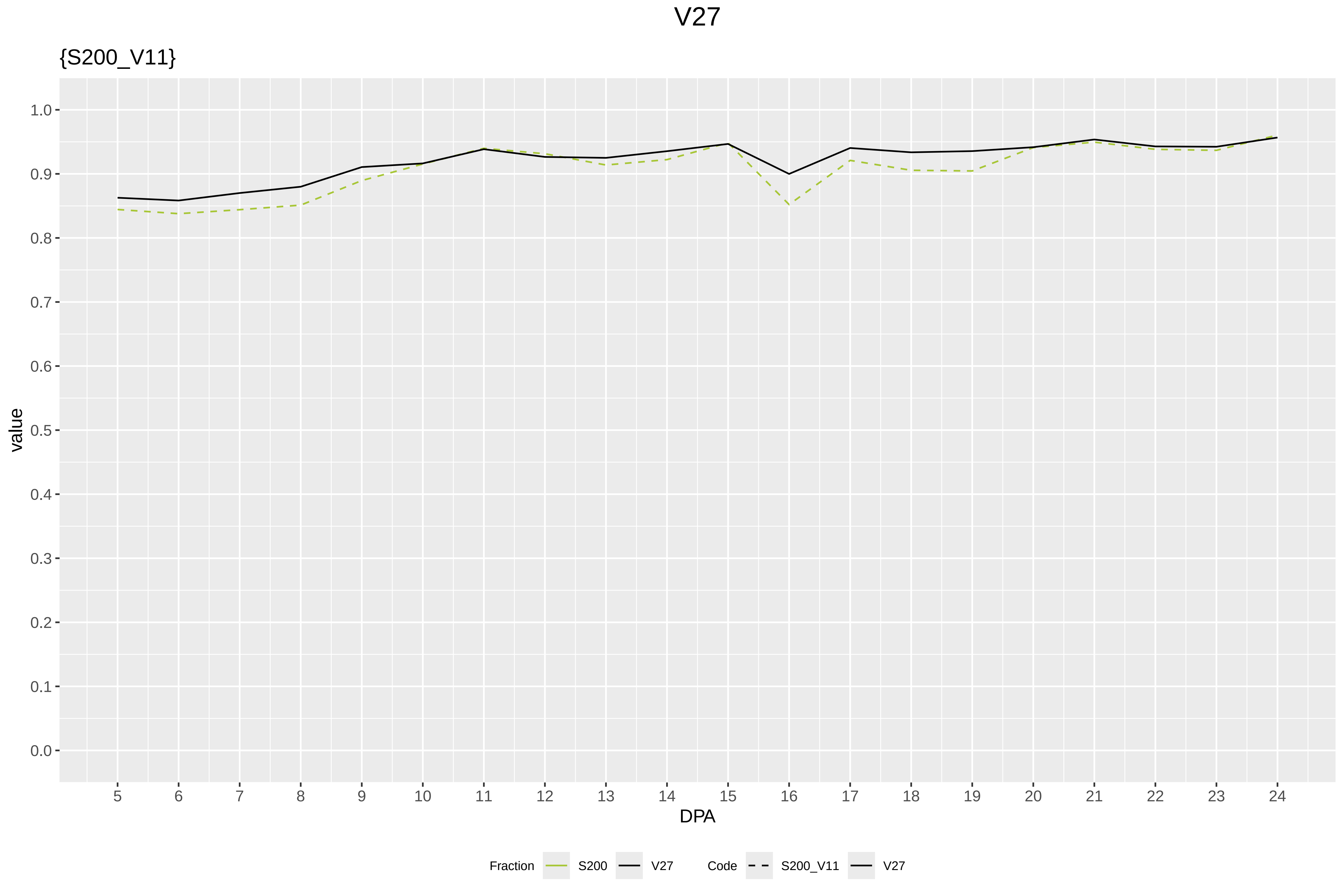

V27
{S200_V11}
value
1.0
0.9
0.8
0.7
0.6
0.5
0.4
0.3
0.2
0.1
0.0
5 6 7 8 9 10 11 12 13 14 15 16 17 18 19 20 21 22 23 24
DPA
Fraction S200 V27
Code S200_V11 V27

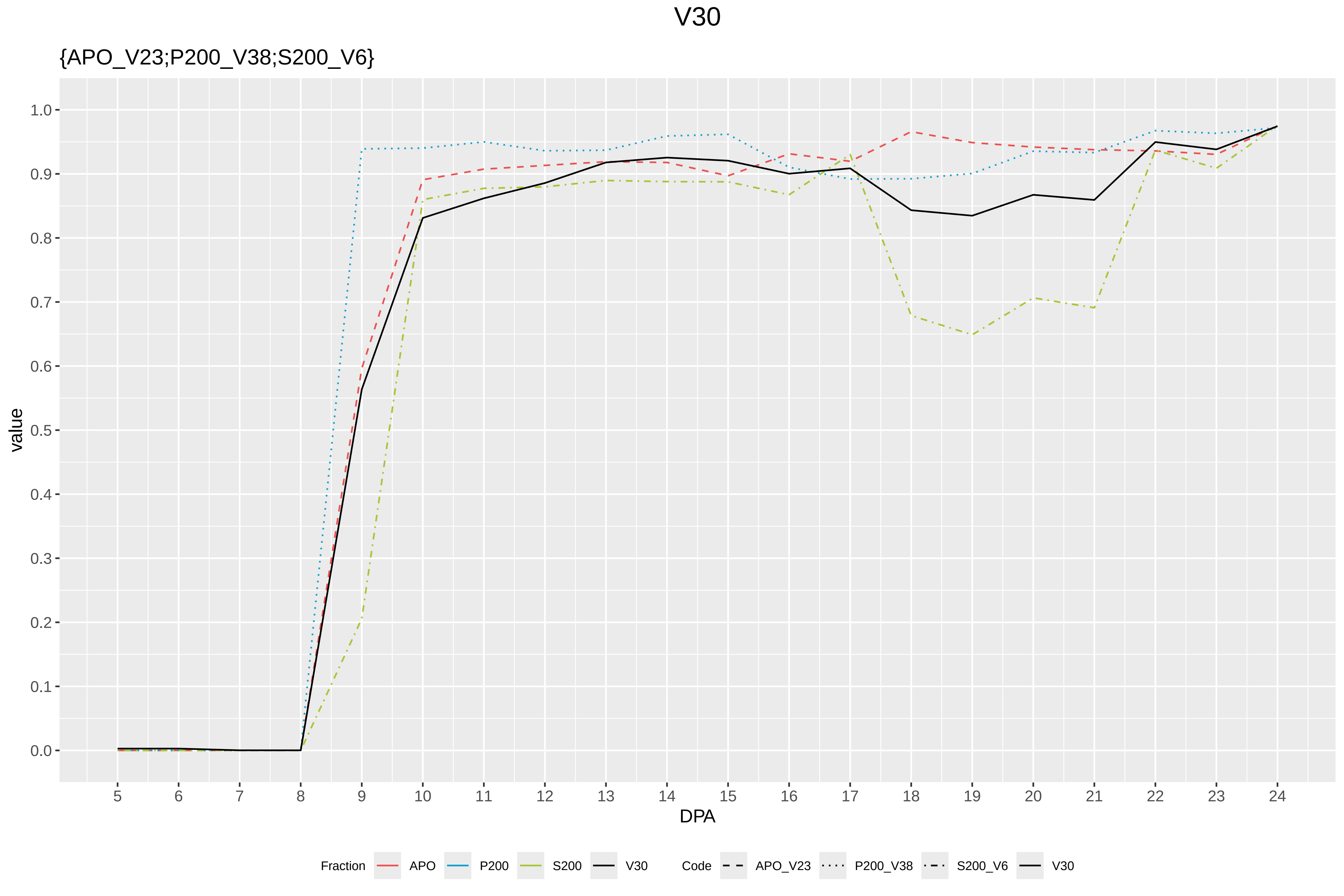

V30
{APO_V23;P200_V38;S200_V6}
value
1.0
0.9
0.8
0.7
0.6
0.5
0.4
0.3
0.2
0.1
0.0
DPA
Fraction
APO
P200
S200
V30
Code
APO_V23
P200_V38
S200_V6
V30

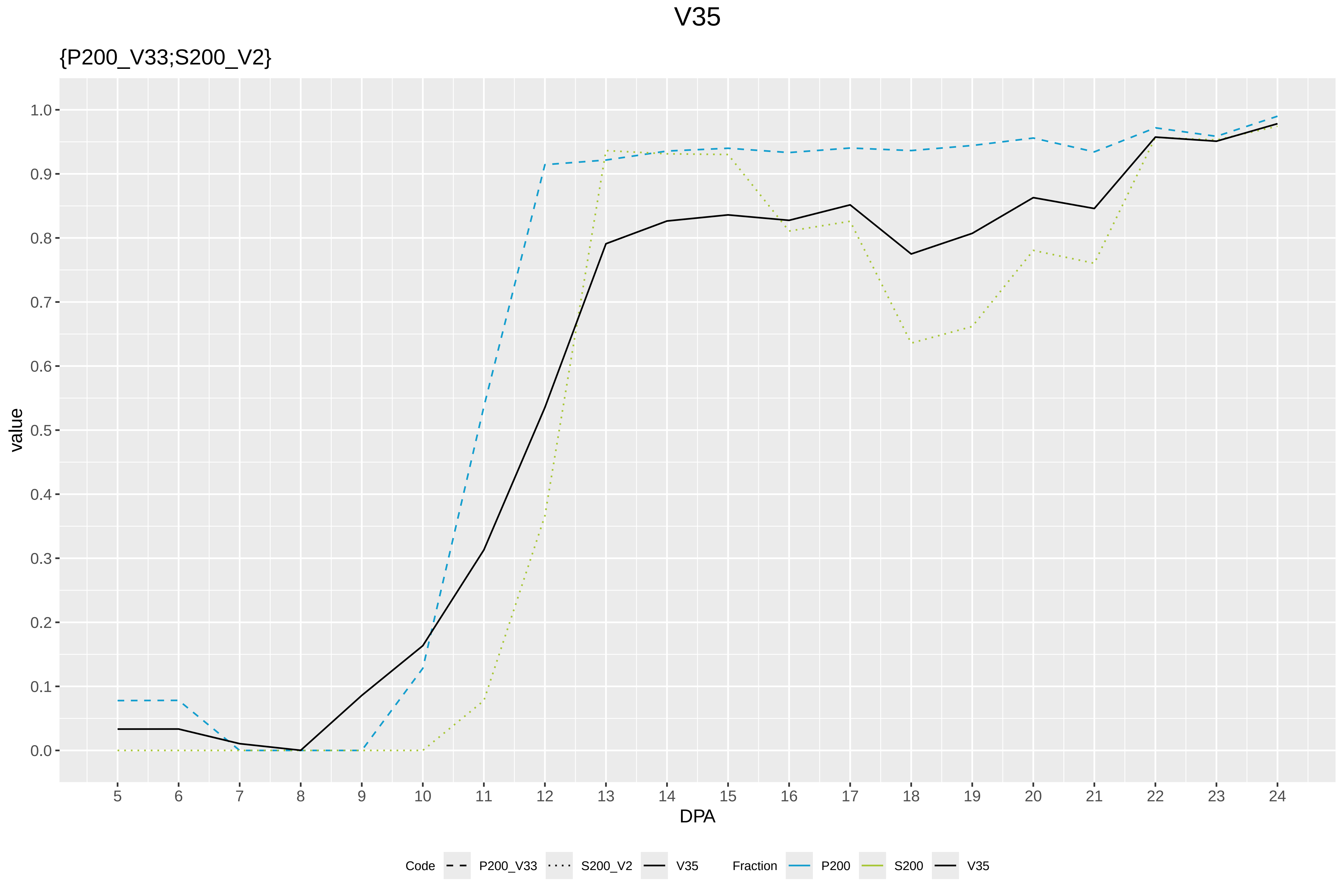
V35
{P200_V33;S200_V2}
value
1.0
0.9
0.8
0.7
0.6
0.5
0.4
0.3
0.2
0.1
0.0
5 6 7 8 9 10 11 12 13 14 15 16 17 18 19 20 21 22 23 24
DPA
Code
P200_V33
S200_V2
V35
Fraction
P200
S200
V35

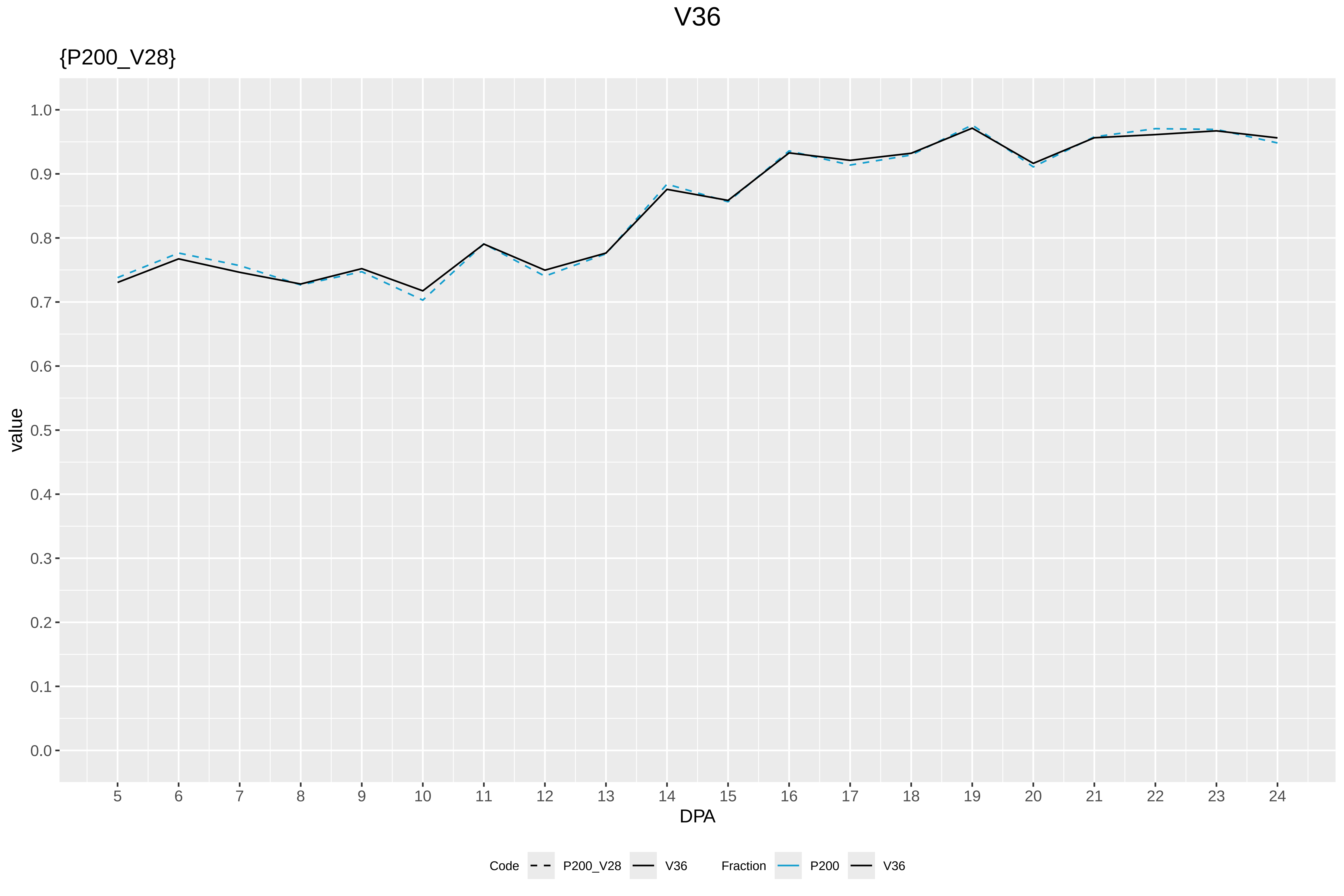

V36
{P200_V28}
value
DPA
1.0
0.9
0.8
0.7
0.6
0.5
0.4
0.3
0.2
0.1
0.0
5 6 7 8 9 10 11 12 13 14 15 16 17 18 19 20 21 22 23 24
Code
P200_V28
V36
Fraction
P200
V36

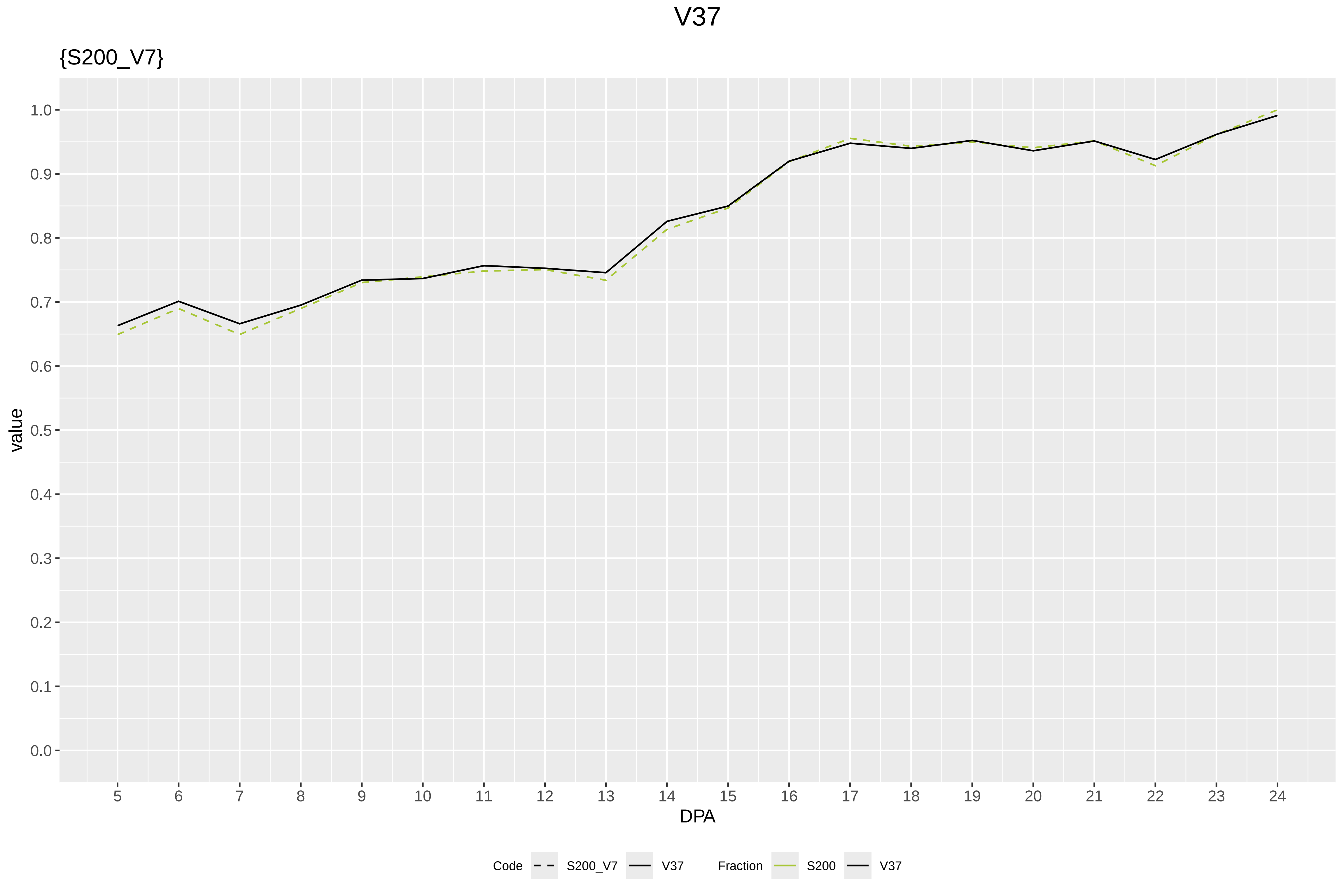

V37
{S200_V7}
value
1.0
0.9
0.8
0.7
0.6
0.5
0.4
0.3
0.2
0.1
0.0
DPA
Code
S200_V7
V37
Fraction
S200
V37

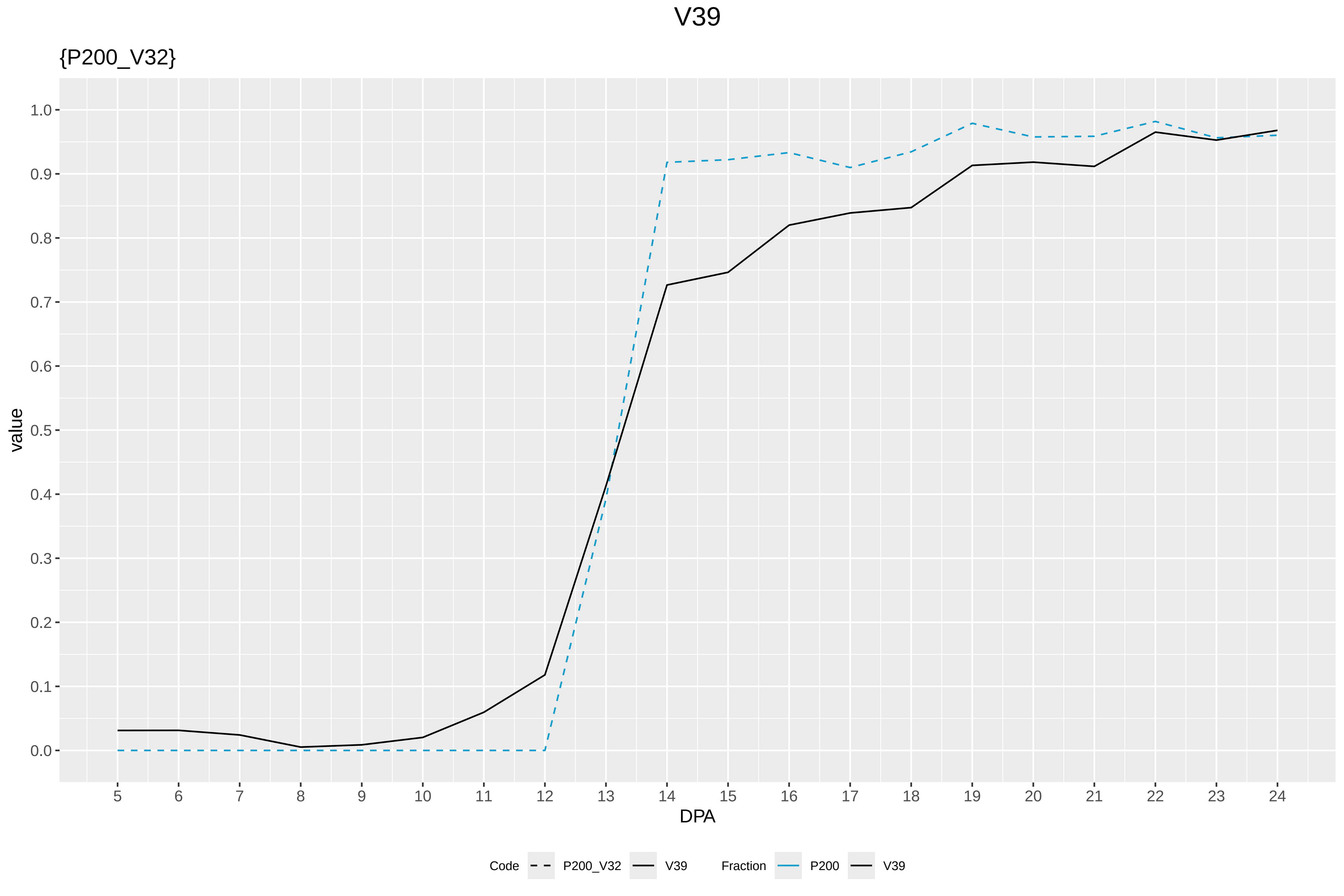

V39
{P200_V32}
value
1.0
0.9
0.8
0.7
0.6
0.5
0.4
0.3
0.2
0.1
0.0
5 6 7 8 9 10 11 12 13 14 15 16 17 18 19 20 21 22 23 24
DPA
Code
P200_V32
V39
Fraction
P200
V39

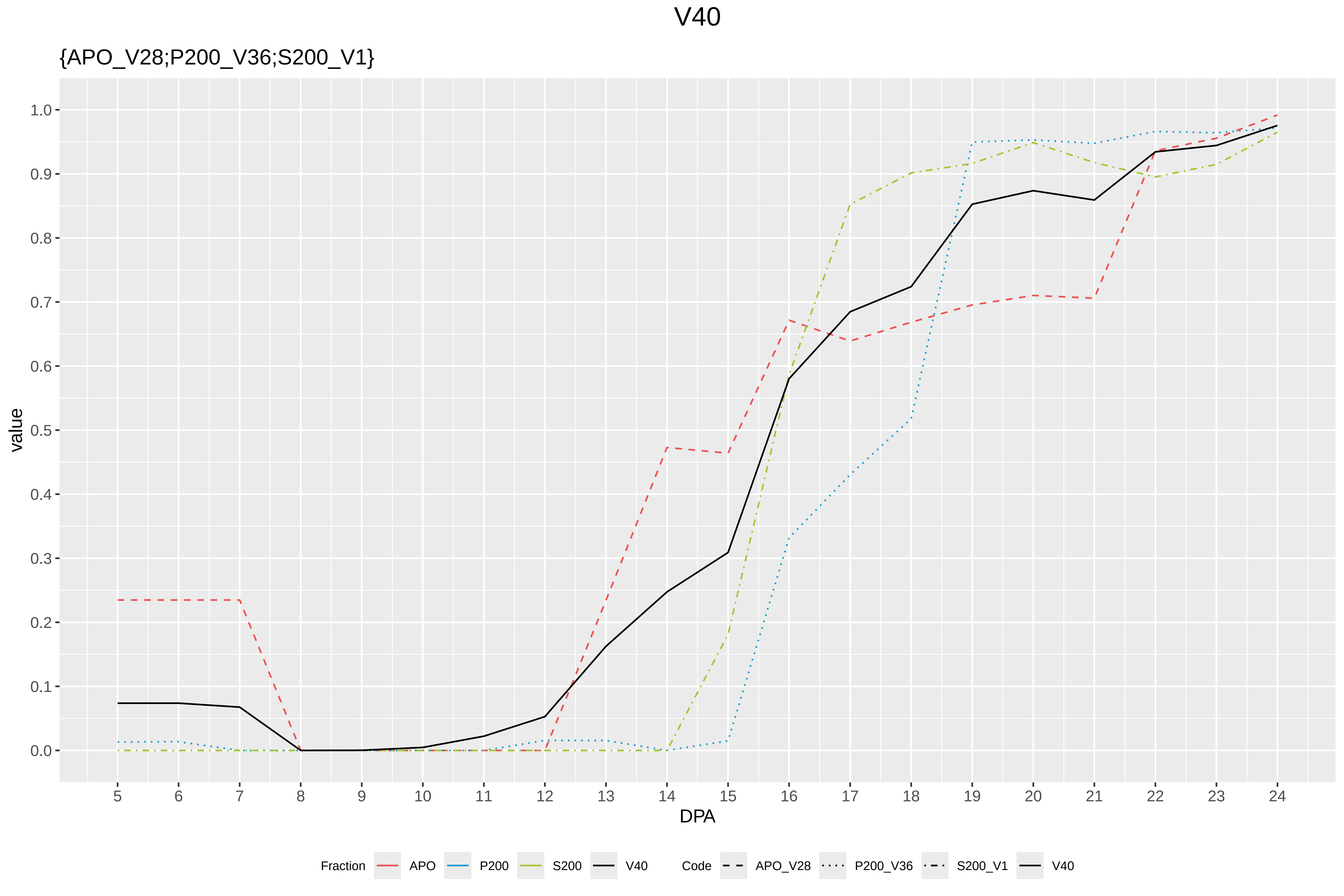

V40
{APO_V28;P200_V36;S200_V1}
value
1.0
0.9
0.8
0.7
0.6
0.5
0.4
0.3
0.2
0.1
0.0
DPA
Fraction
APO
P200
S200
V40
Code
APO_V28
P200_V36
S200_V1
V40
