## Supplementary Figure S10 for "Systems-level proteomic models of cotton fiber development: A high-resolution resource to analyze cell dynamics and trait engineering"

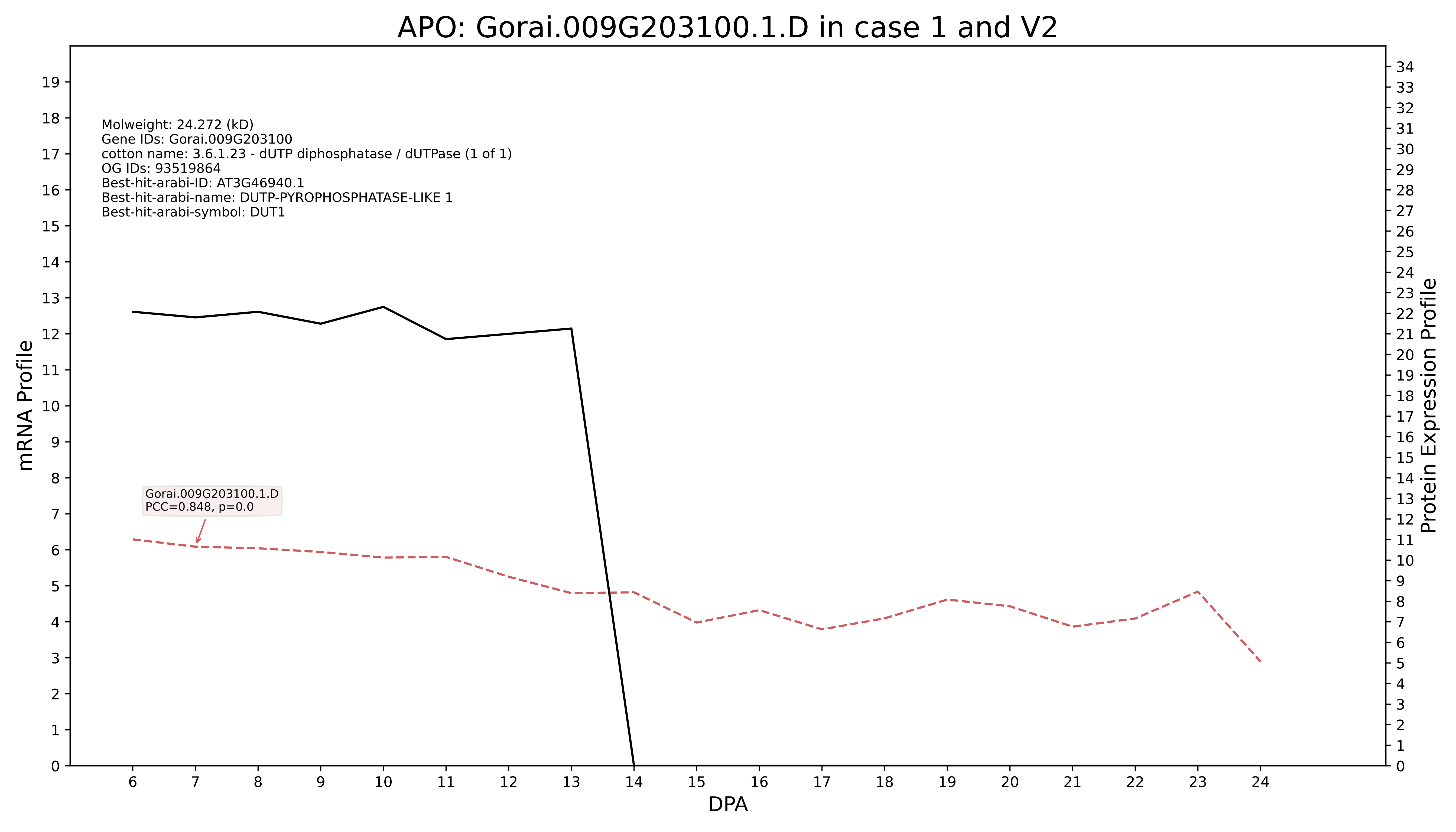
APO: Gorai.009G203100.1.D in case 1 and V2
Molweight: 24.272 (kD)
Gene IDs: Gorai.009G203100
cotton name: 3.6.1.23 - dUTP diphosphatase / dUTPase (1 of 1)
OG IDs: 93519864
Best-hit-arabi-ID: AT3G46940.1
Best-hit-arabi-name: DUTP-PYROPHOSPHATASE-LIKE 1
Best-hit-arabi-symbol: DUT1
Gorai.009G203100.1.D
PCC=0.848, p=0.0
mRNA Profile
Protein Expression Profile
DPA

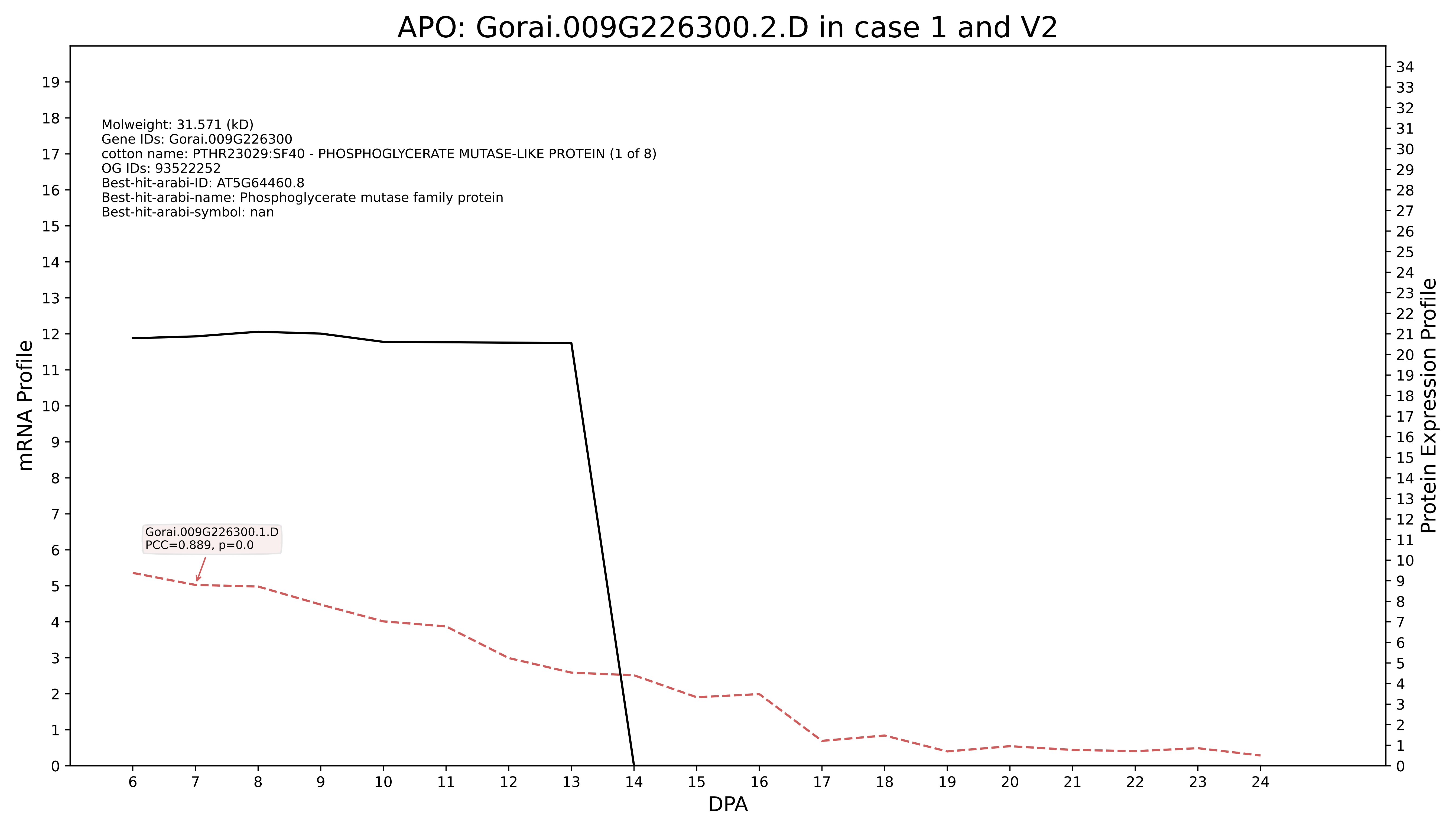
APO: Gorai.009G226300.2.D in case 1 and V2
Molweight: 31.571 (kD)
Gene IDs: Gorai.009G226300
cotton name: PTHR23029:SF40 - PHOSPHOGLYCERATE MUTASE-LIKE PROTEIN (1 of 8)
OG IDs: 93522252
Best-hit-arabi-ID: AT5G64460.8
Best-hit-arabi-name: Phosphoglycerate mutase family protein
Best-hit-arabi-symbol: nan
Gorai.009G226300.1.D
PCC=0.889, p=0.0
mRNA Profile
Protein Expression Profile
DPA

### APO: Gorai.012G139000.3.D in case 1 and V2

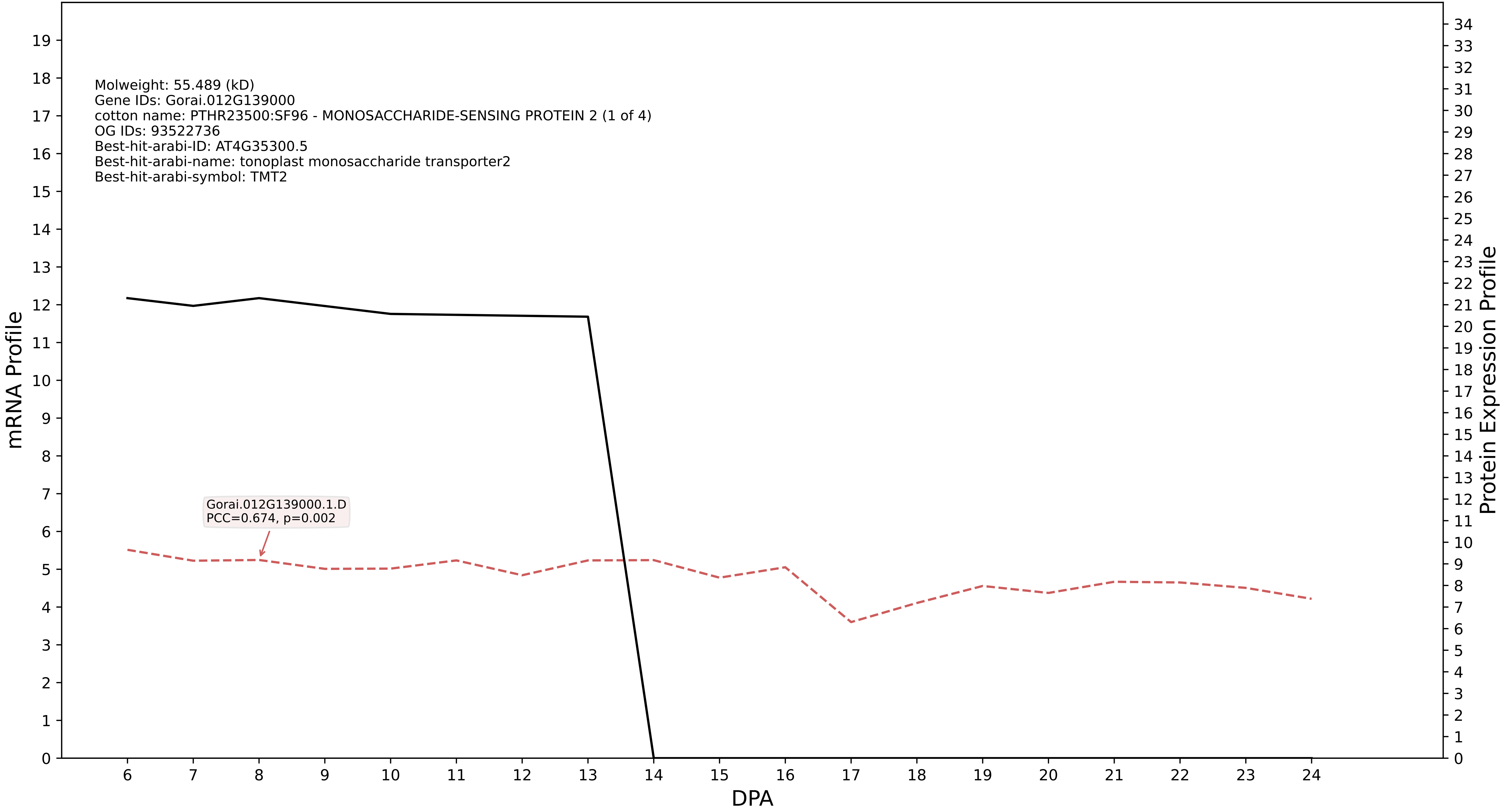

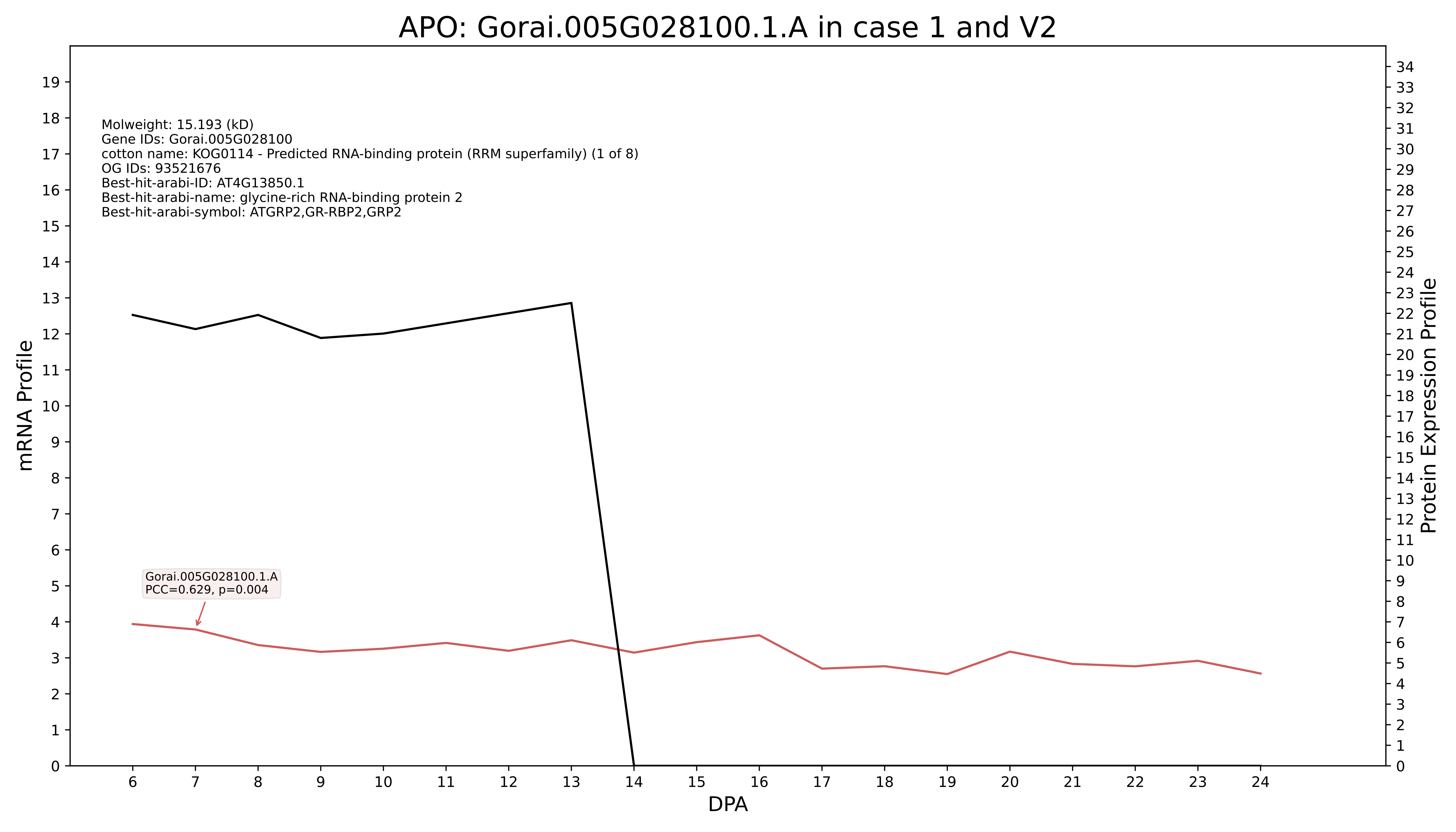

APO: Gorai.005G028100.1.A in case 1 and V2
Molweight: 15.193 (kD)
Gene IDs: Gorai.005G028100
cotton name: KOG0114 - Predicted RNA-binding protein (RRM superfamily) (1 of 8)
OG IDs: 93521676
Best-hit-arabi-ID: AT4G13850.1
Best-hit-arabi-name: glycine-rich RNA-binding protein 2
Best-hit-arabi-symbol: ATGRP2,GR-RBP2,GRP2
Gorai.005G028100.1.A
PCC=0.629, p=0.004
mRNA Profile
Protein Expression Profile
DPA

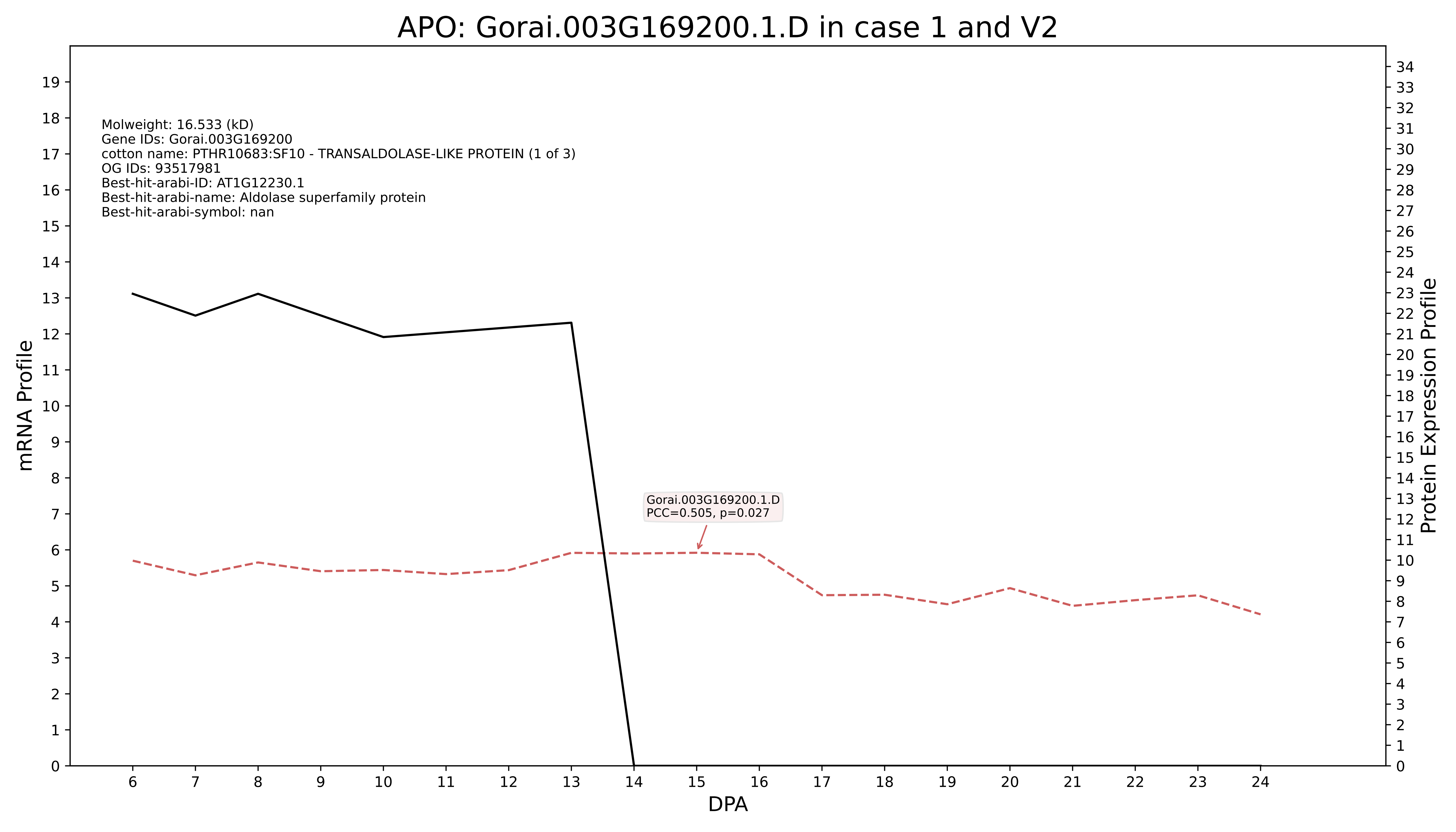
APO: Gorai.003G169200.1.D in case 1 and V2
Molweight: 16.533 (kD)
Gene IDs: Gorai.003G169200
cotton name: PTHR10683:SF10 - TRANSALDOLASE-LIKE PROTEIN (1 of 3)
OG IDs: 93517981
Best-hit-arabi-ID: AT1G12230.1
Best-hit-arabi-name: Aldolase superfamily protein
Best-hit-arabi-symbol: nan
Gorai.003G169200.1.D
PCC=0.505, p=0.027
mRNA Profile
Protein Expression Profile
DPA

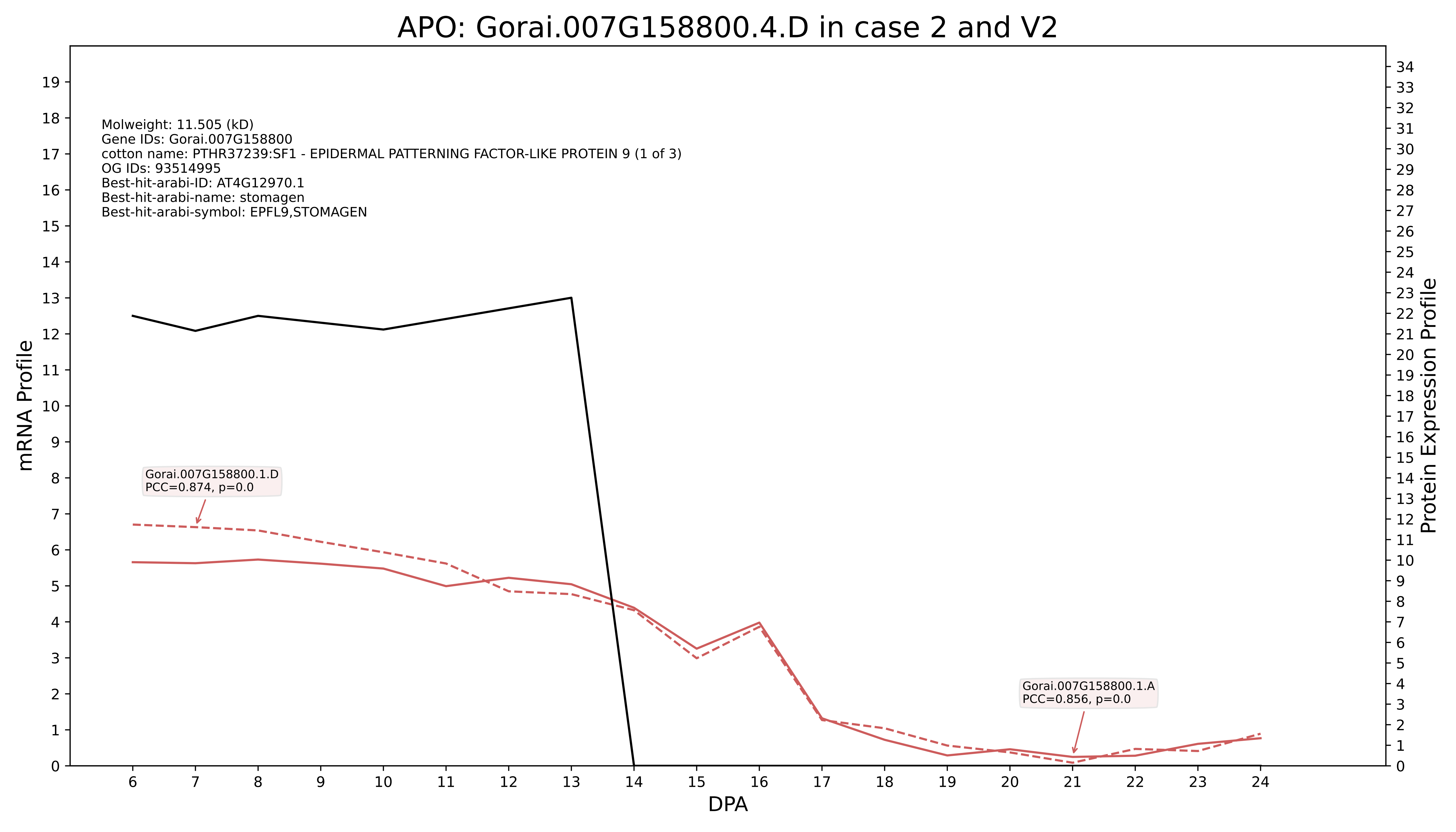
APO: Gorai.007G158800.4.D in case 2 and V2
Molweight: 11.505 (kD)
Gene IDs: Gorai.007G158800
cotton name: PTHR37239:SF1 - EPIDERMAL PATTERNING FACTOR-LIKE PROTEIN 9 (1 of 3)
OG IDs: 93514995
Best-hit-arabi-ID: AT4G12970.1
Best-hit-arabi-name: stomagen
Best-hit-arabi-symbol: EPFL9,STOMAGEN
Gorai.007G158800.1.D
PCC=0.874, p=0.0
Gorai.007G158800.1.A
PCC=0.856, p=0.0
mRNA Profile
Protein Expression Profile
DPA

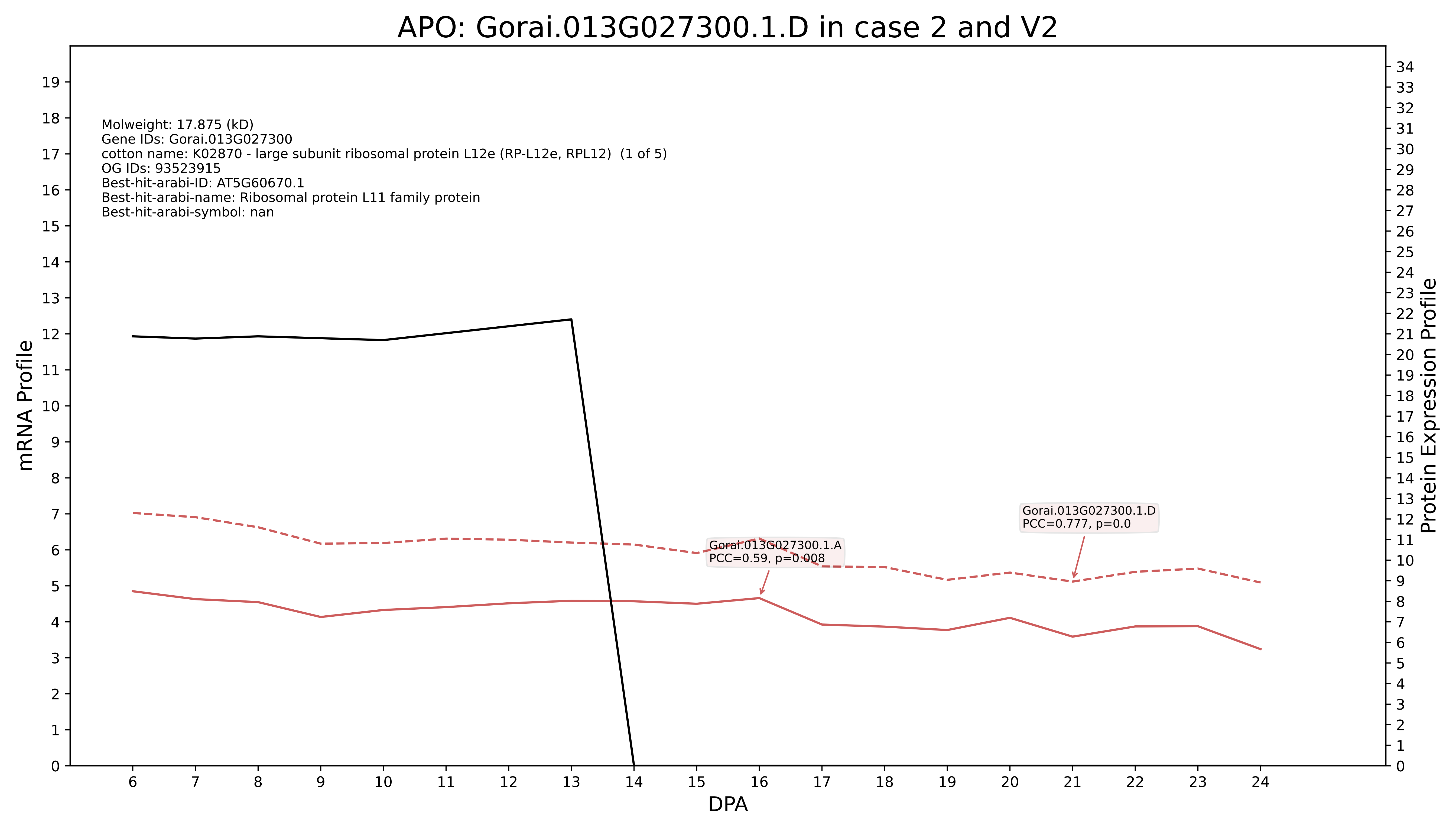
APO: Gorai.013G027300.1.D in case 2 and V2
Molweight: 17.875 (kD)
Gene IDs: Gorai.013G027300
cotton name: K02870 - large subunit ribosomal protein L12e (RP-L12e, RPL12) (1 of 5)
OG IDs: 93523915
Best-hit-arabi-ID: AT5G60670.1
Best-hit-arabi-name: Ribosomal protein L11 family protein
Best-hit-arabi-symbol: nan
Gorai.013G027300.1.A
PCC=0.59, p=0.008
Gorai.013G027300.1.D
PCC=0.777, p=0.0
mRNA Profile
Protein Expression Profile
DPA

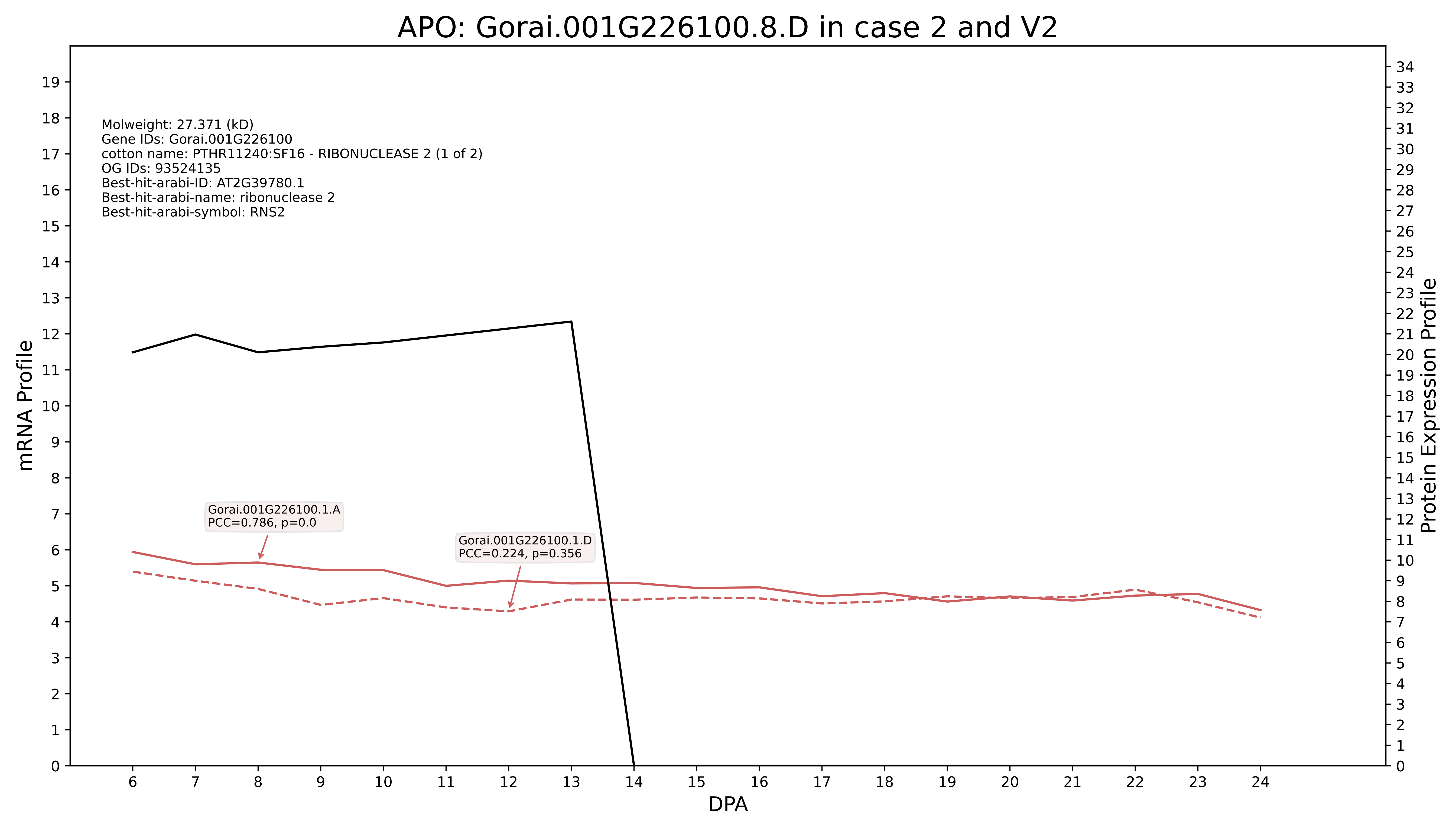

APO: Gorai.001G226100.8.D in case 2 and V2
Molweight: 27.371 (kD)
Gene IDs: Gorai.001G226100
cotton name: PTHR11240:SF16 - RIBONUCLEASE 2 (1 of 2)
OG IDs: 93524135
Best-hit-arabi-ID: AT2G39780.1
Best-hit-arabi-name: ribonuclease 2
Best-hit-arabi-symbol: RNS2
Gorai.001G226100.1.A
PCC=0.786, p=0.0
Gorai.001G226100.1.D
PCC=0.224, p=0.356
mRNA Profile
Protein Expression Profile
DPA

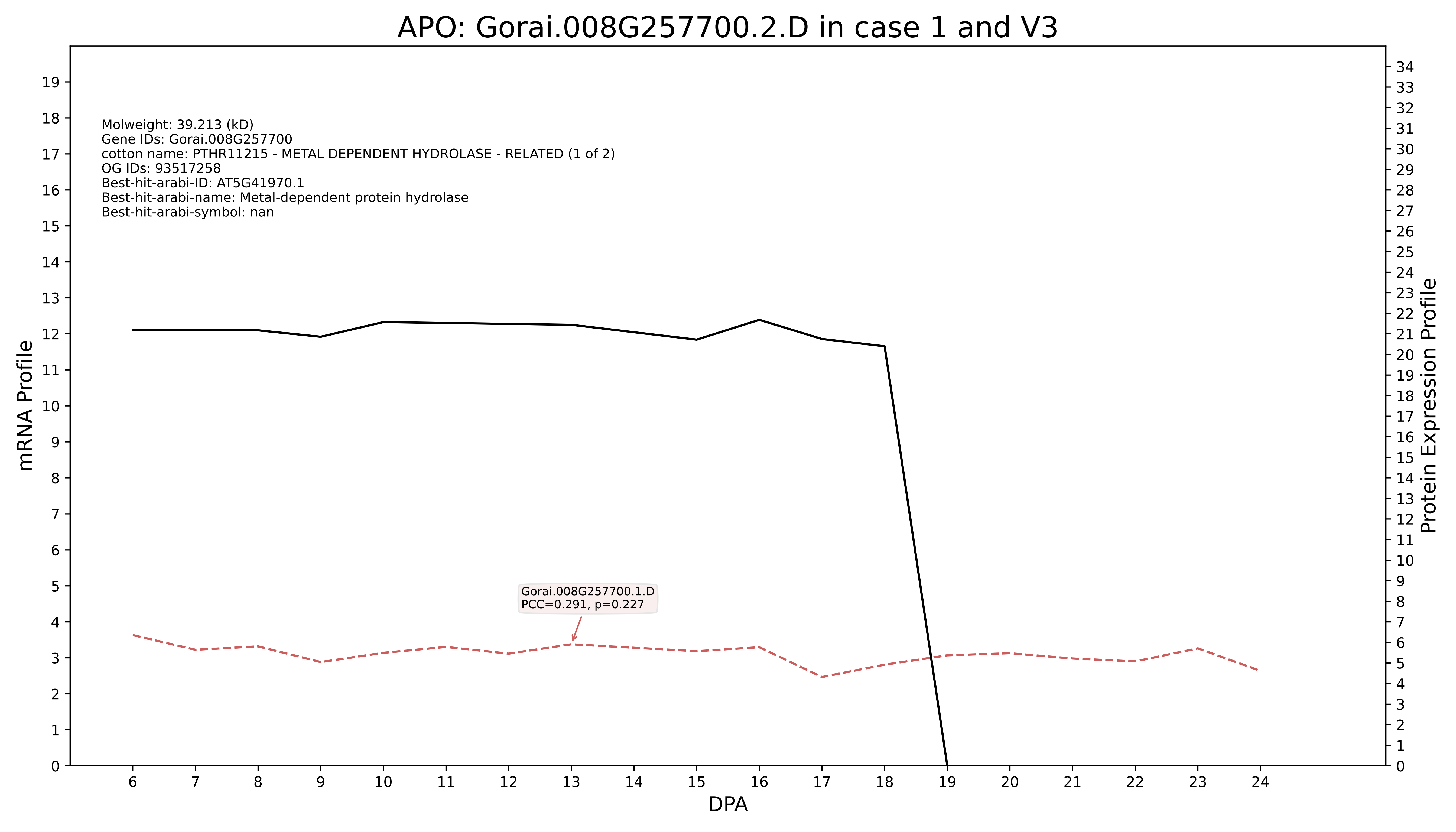
APO: Gorai.008G257700.2.D in case 1 and V3
Molweight: 39.213 (kD)
Gene IDs: Gorai.008G257700
cotton name: PTHR11215 - METAL DEPENDENT HYDROLASE - RELATED (1 of 2)
OG IDs: 93517258
Best-hit-arabi-ID: AT5G41970.1
Best-hit-arabi-name: Metal-dependent protein hydrolase
Best-hit-arabi-symbol: nan
Gorai.008G257700.1.D
PCC=0.291, p=0.227
mRNA Profile
Protein Expression Profile
DPA

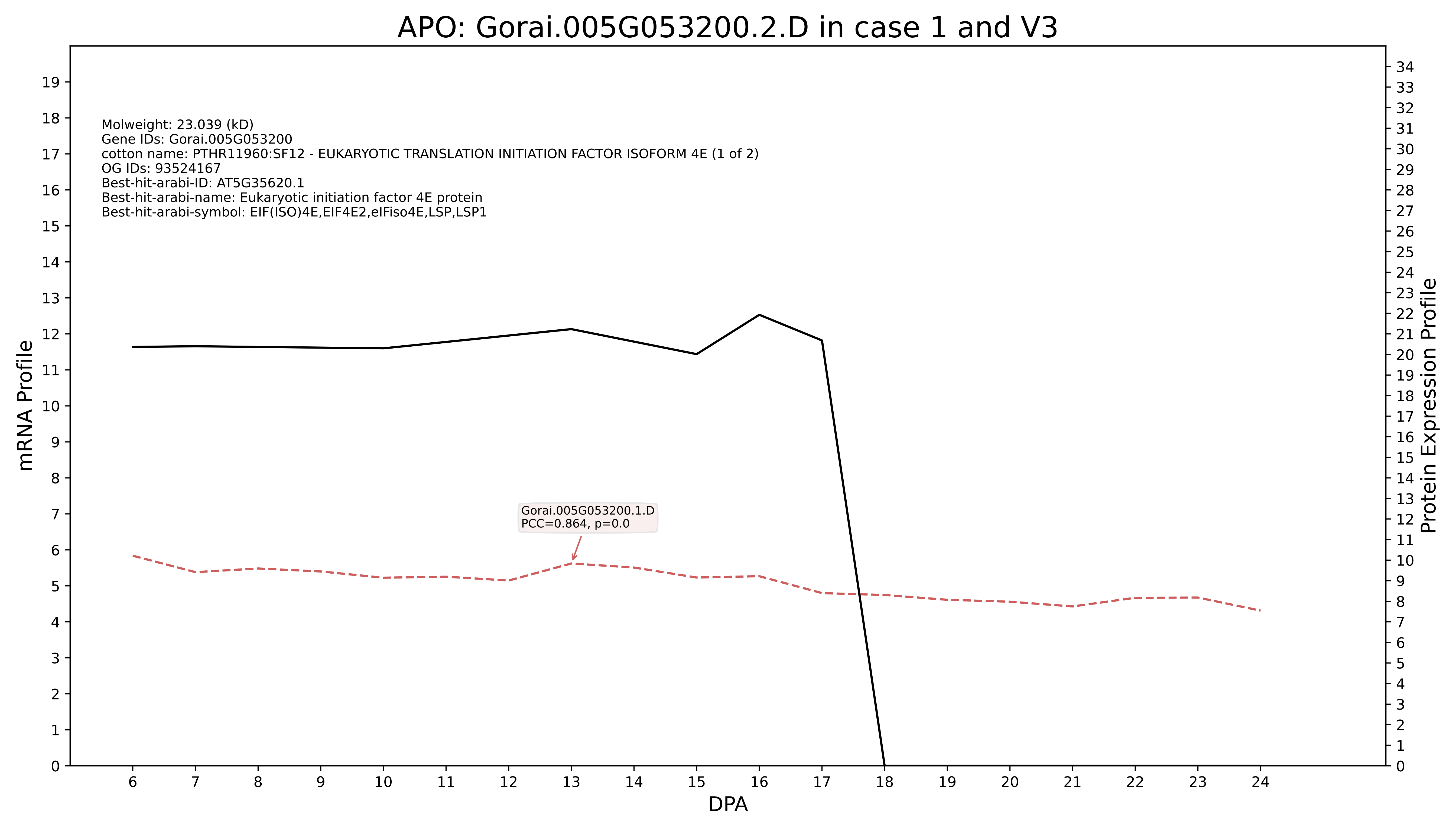
APO: Gorai.005G053200.2.D in case 1 and V3
Molweight: 23.039 (kD)
Gene IDs: Gorai.005G053200
cotton name: PTHR11960:SF12 - EUKARYOTIC TRANSLATION INITIATION FACTOR ISOFORM 4E (1 of 2)
OG IDs: 93524167
Best-hit-arabi-ID: AT5G35620.1
Best-hit-arabi-name: Eukaryotic initiation factor 4E protein
Best-hit-arabi-symbol: EIF(ISO)4E,EIF4E2,eIFiso4E,LSP,LSP1
Gorai.005G053200.1.D
PCC=0.864, p=0.0
mRNA Profile
Protein Expression Profile
DPA

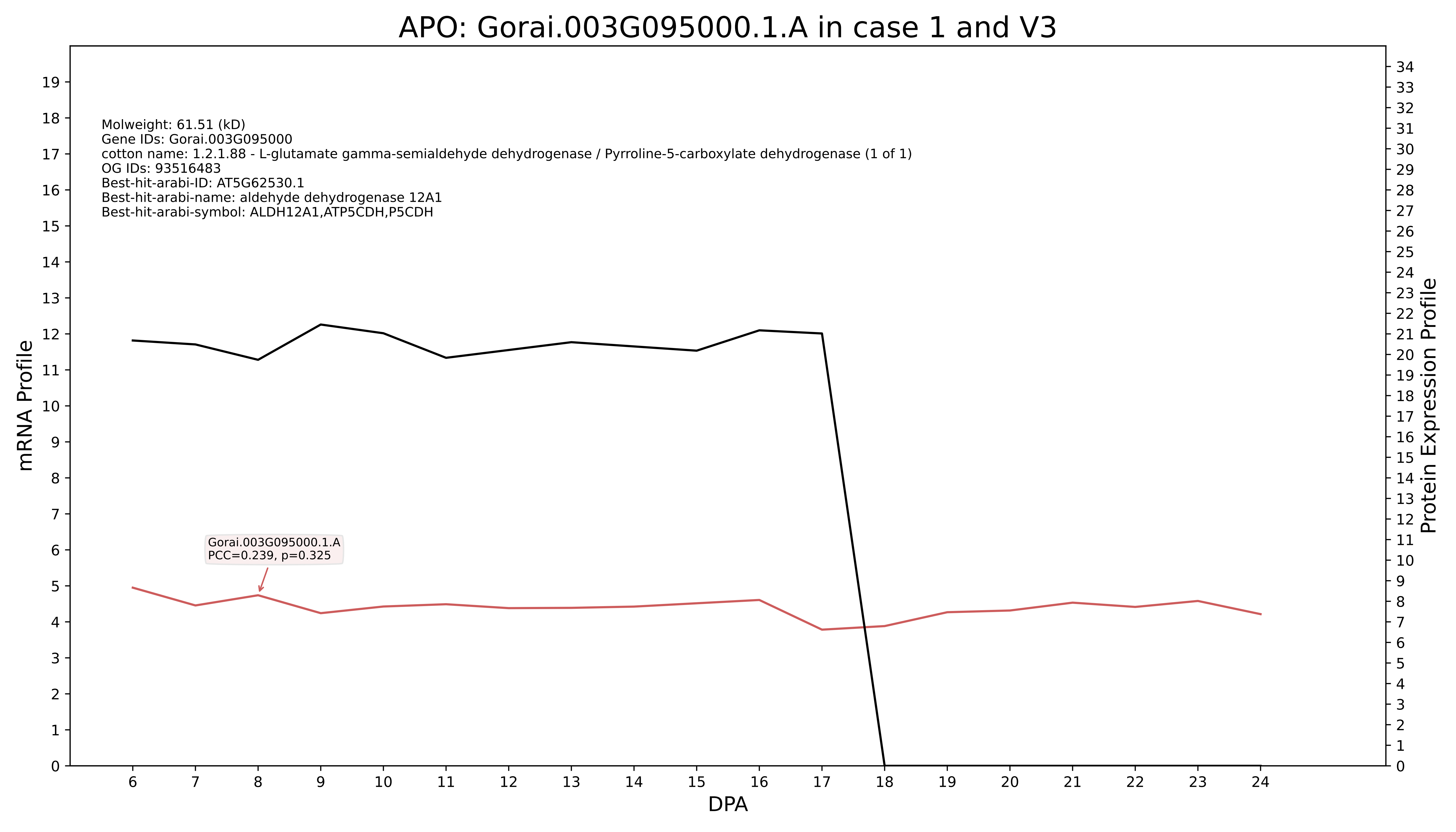
APO: Gorai.003G095000.1.A in case 1 and V3
Molweight: 61.51 (kD)
Gene IDs: Gorai.003G095000
cotton name: 1.2.1.88 - L-glutamate gamma-semialdehyde dehydrogenase / Pyrroline-5-carboxylate dehydrogenase (1 of 1)
OG IDs: 93516483
Best-hit-arabi-ID: AT5G62530.1
Best-hit-arabi-name: aldehyde dehydrogenase 12A1
Best-hit-arabi-symbol: ALDH12A1,ATP5CDH,P5CDH
Gorai.003G095000.1.A
PCC=0.239, p=0.325
mRNA Profile
Protein Expression Profile
DPA

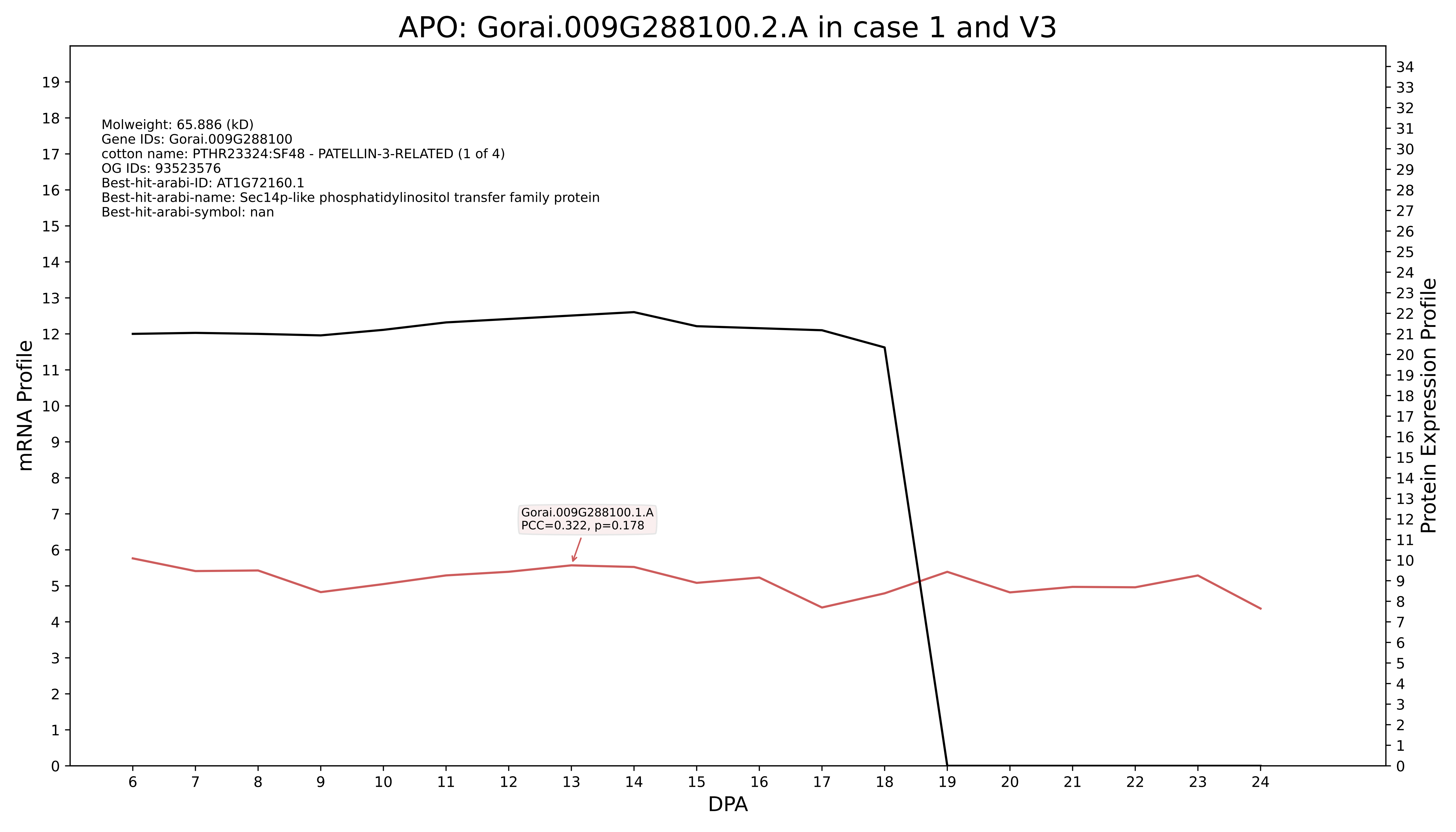
APO: Gorai.009G288100.2.A in case 1 and V3
Molweight: 65.886 (kD)
Gene IDs: Gorai.009G288100
cotton name: PTHR23324:SF48 - PATELLIN-3-RELATED (1 of 4)
OG IDs: 93523576
Best-hit-arabi-ID: AT1G72160.1
Best-hit-arabi-name: Sec14p-like phosphatidylinositol transfer family protein
Best-hit-arabi-symbol: nan
Gorai.009G288100.1.A
PCC=0.322, p=0.178
mRNA Profile
Protein Expression Profile
DPA

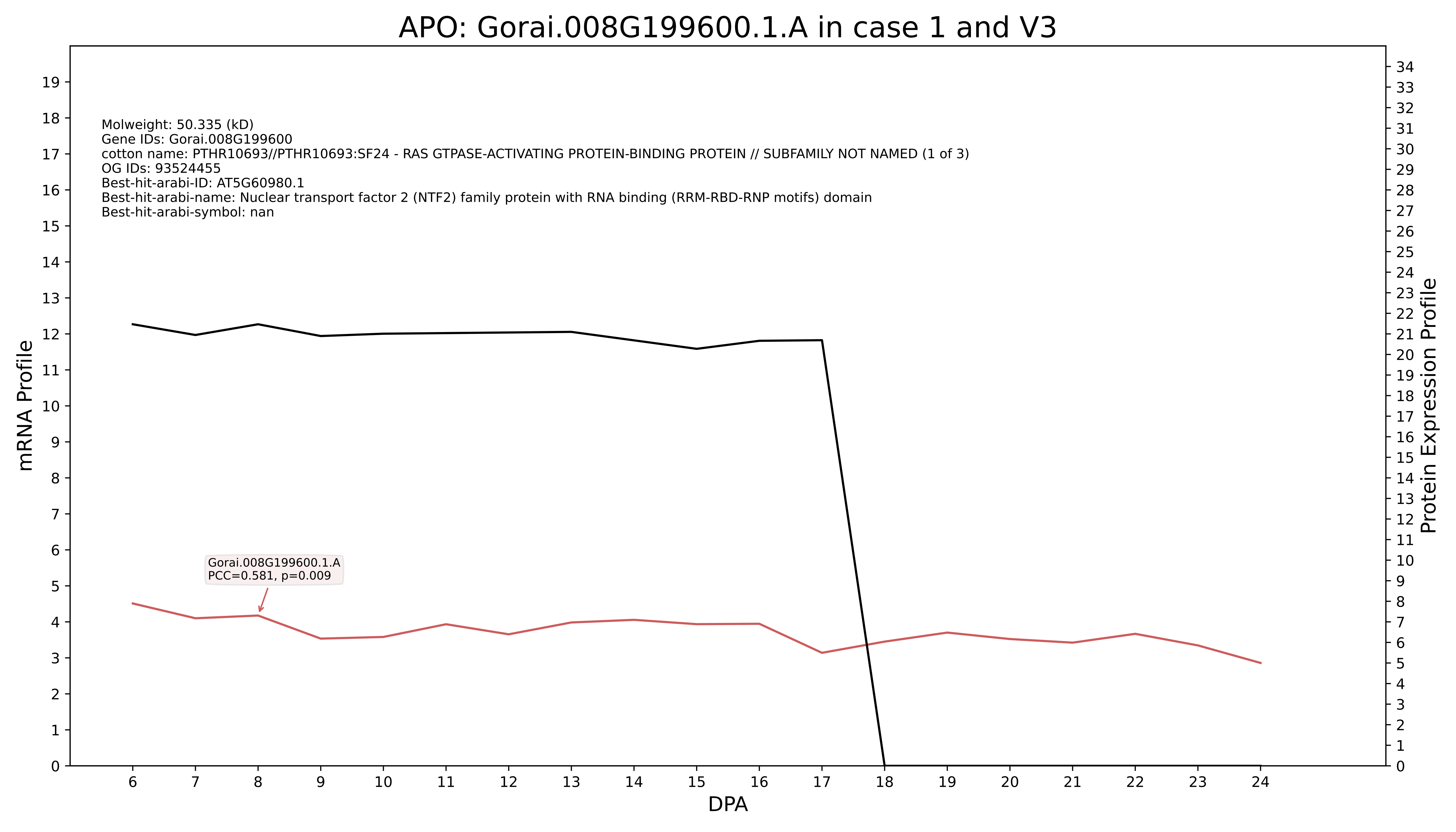

APO: Gorai.008G199600.1.A in case 1 and V3
Molweight: 50.335 (kD)
Gene IDs: Gorai.008G199600
cotton name: PTHR10693//PTHR10693:SF24 - RAS GTPASE-ACTIVATING PROTEIN-BINDING PROTEIN // SUBFAMILY NOT NAMED (1 of 3)
OG IDs: 93524455
Best-hit-arabi-ID: AT5G60980.1
Best-hit-arabi-name: Nuclear transport factor 2 (NTF2) family protein with RNA binding (RRM-RBD-RNP motifs) domain
Best-hit-arabi-symbol: nan
Gorai.008G199600.1.A
PCC=0.581, p=0.009
mRNA Profile
Protein Expression Profile
DPA

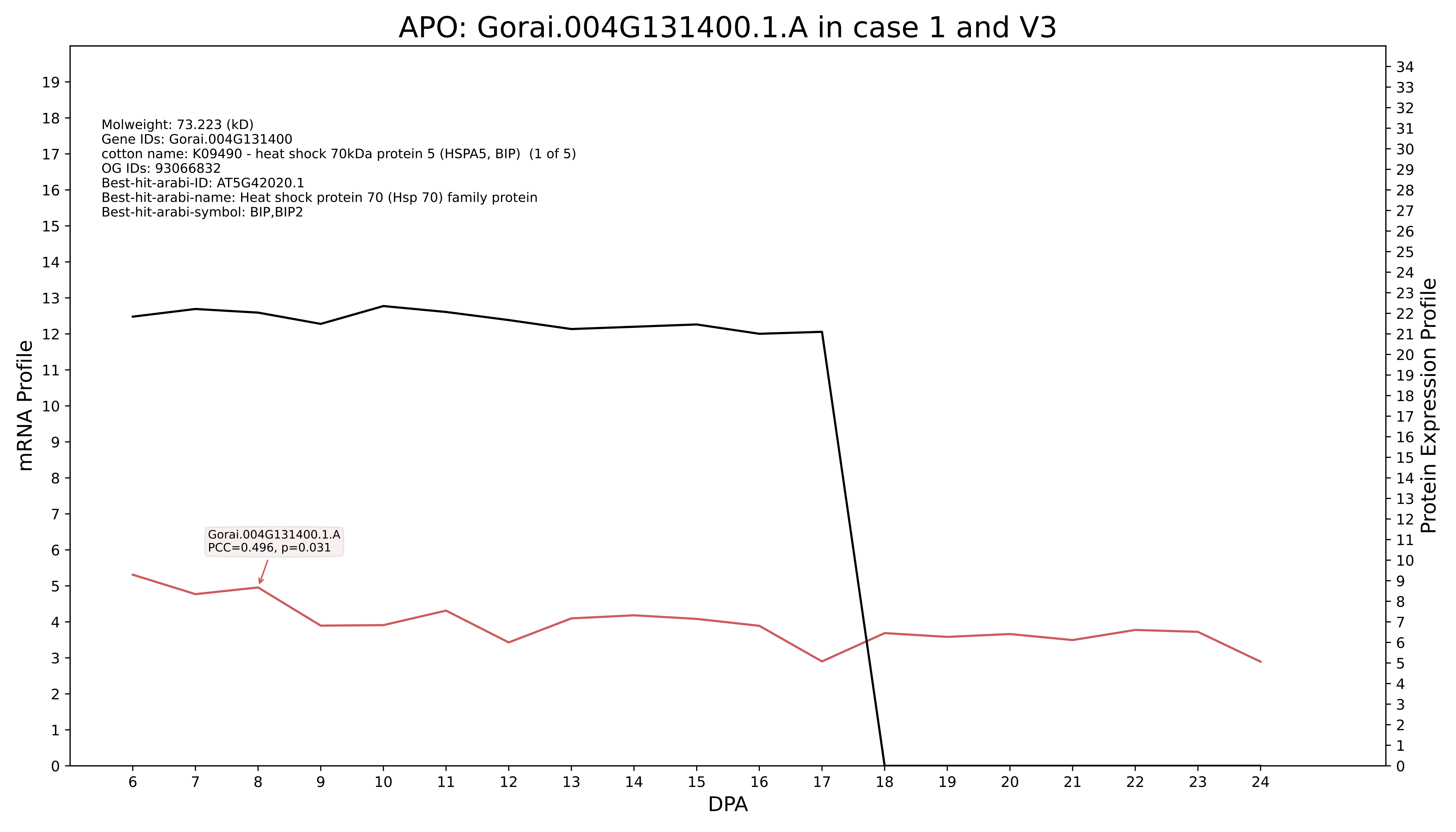
APO: Gorai.004G131400.1.A in case 1 and V3
Molweight: 73.223 (kD)
Gene IDs: Gorai.004G131400
cotton name: K09490 - heat shock 70kDa protein 5 (HSPA5, BIP) (1 of 5)
OG IDs: 93066832
Best-hit-arabi-ID: AT5G42020.1
Best-hit-arabi-name: Heat shock protein 70 (Hsp 70) family protein
Best-hit-arabi-symbol: BIP,BIP2
Gorai.004G131400.1.A
PCC=0.496, p=0.031
mRNA Profile
Protein Expression Profile
DPA

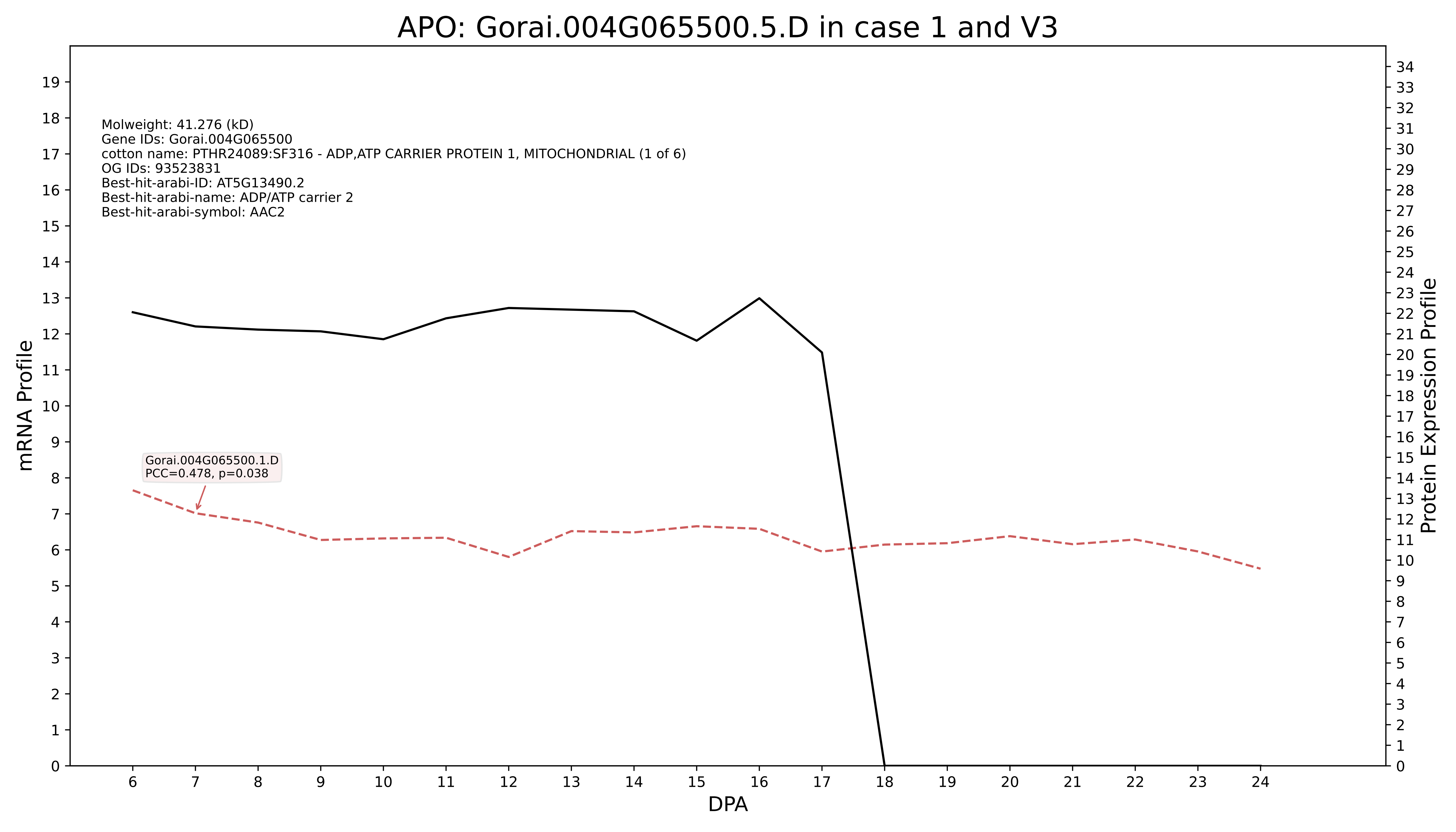

APO: Gorai.004G065500.5.D in case 1 and V3
Molweight: 41.276 (kD)
Gene IDs: Gorai.004G065500
cotton name: PTHR24089:SF316 - ADP,ATP CARRIER PROTEIN 1, MITOCHONDRIAL (1 of 6)
OG IDs: 93523831
Best-hit-arabi-ID: AT5G13490.2
Best-hit-arabi-name: ADP/ATP carrier 2
Best-hit-arabi-symbol: AAC2
Gorai.004G065500.1.D
PCC=0.478, p=0.038
mRNA Profile
Protein Expression Profile
DPA

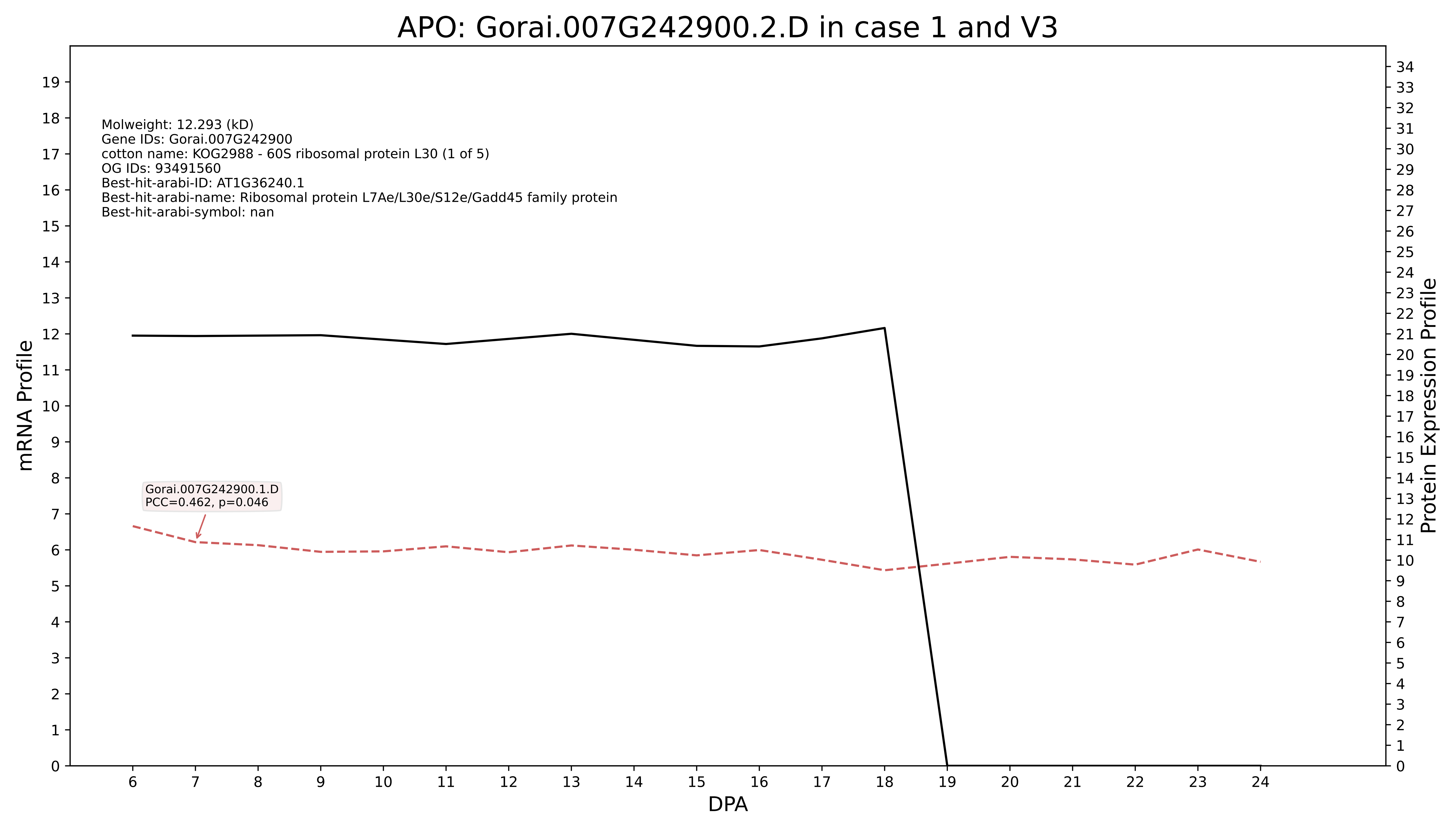
APO: Gorai.007G242900.2.D in case 1 and V3
Molweight: 12.293 (kD)
Gene IDs: Gorai.007G242900
cotton name: KOG2988 - 60S ribosomal protein L30 (1 of 5)
OG IDs: 93491560
Best-hit-arabi-ID: AT1G36240.1
Best-hit-arabi-name: Ribosomal protein L7Ae/L30e/S12e/Gadd45 family protein
Best-hit-arabi-symbol: nan
Gorai.007G242900.1.D
PCC=0.462, p=0.046
mRNA Profile
Protein Expression Profile
DPA

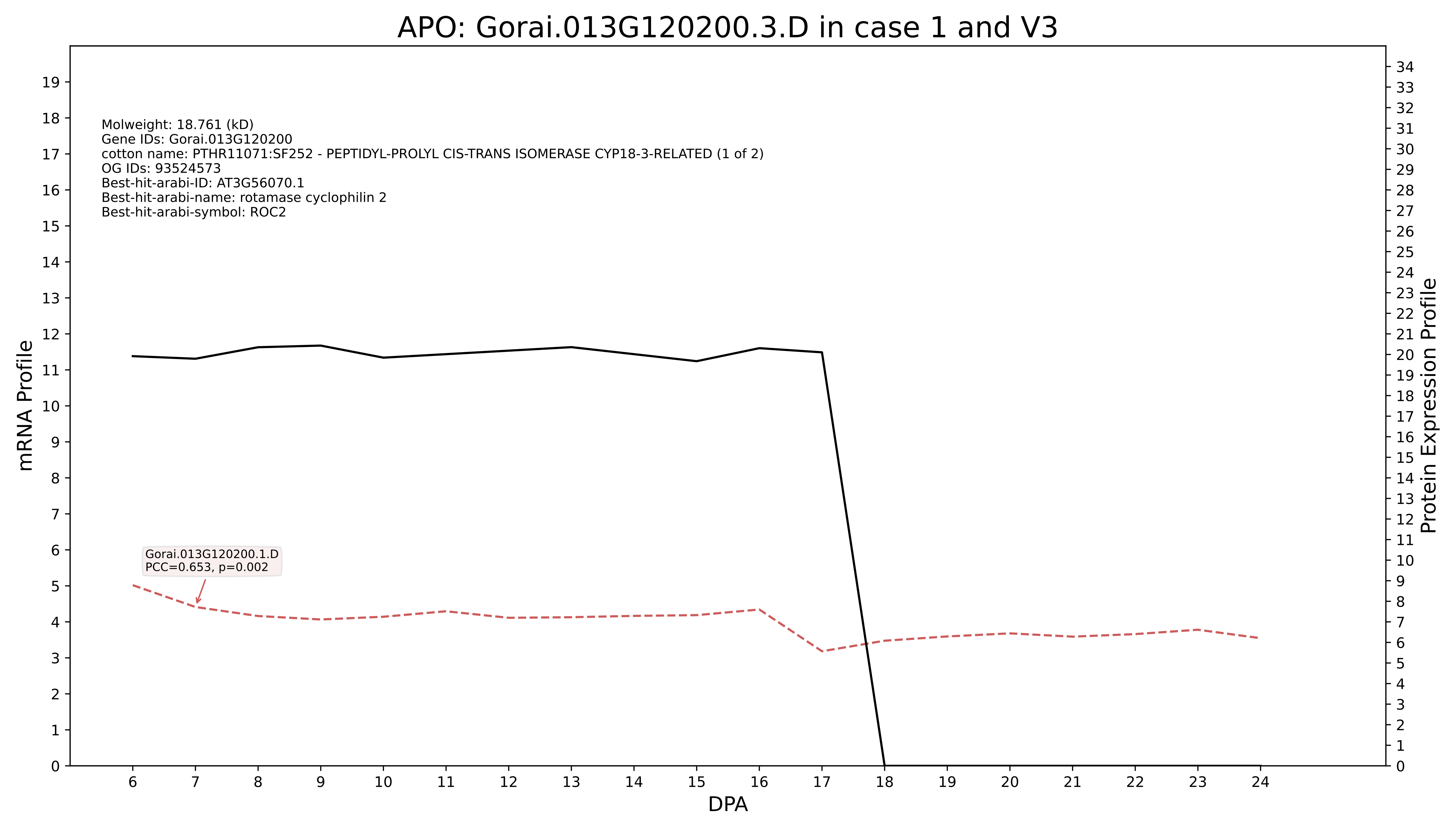
APO: Gorai.013G120200.3.D in case 1 and V3
Molweight: 18.761 (kD)
Gene IDs: Gorai.013G120200
cotton name: PTHR11071:SF252 - PEPTIDYL-PROLYL CIS-TRANS ISOMERASE CYP18-3-RELATED (1 of 2)
OG IDs: 93524573
Best-hit-arabi-ID: AT3G56070.1
Best-hit-arabi-name: rotamase cyclophilin 2
Best-hit-arabi-symbol: ROC2
Gorai.013G120200.1.D
PCC=0.653, p=0.002
mRNA Profile
Protein Expression Profile
DPA

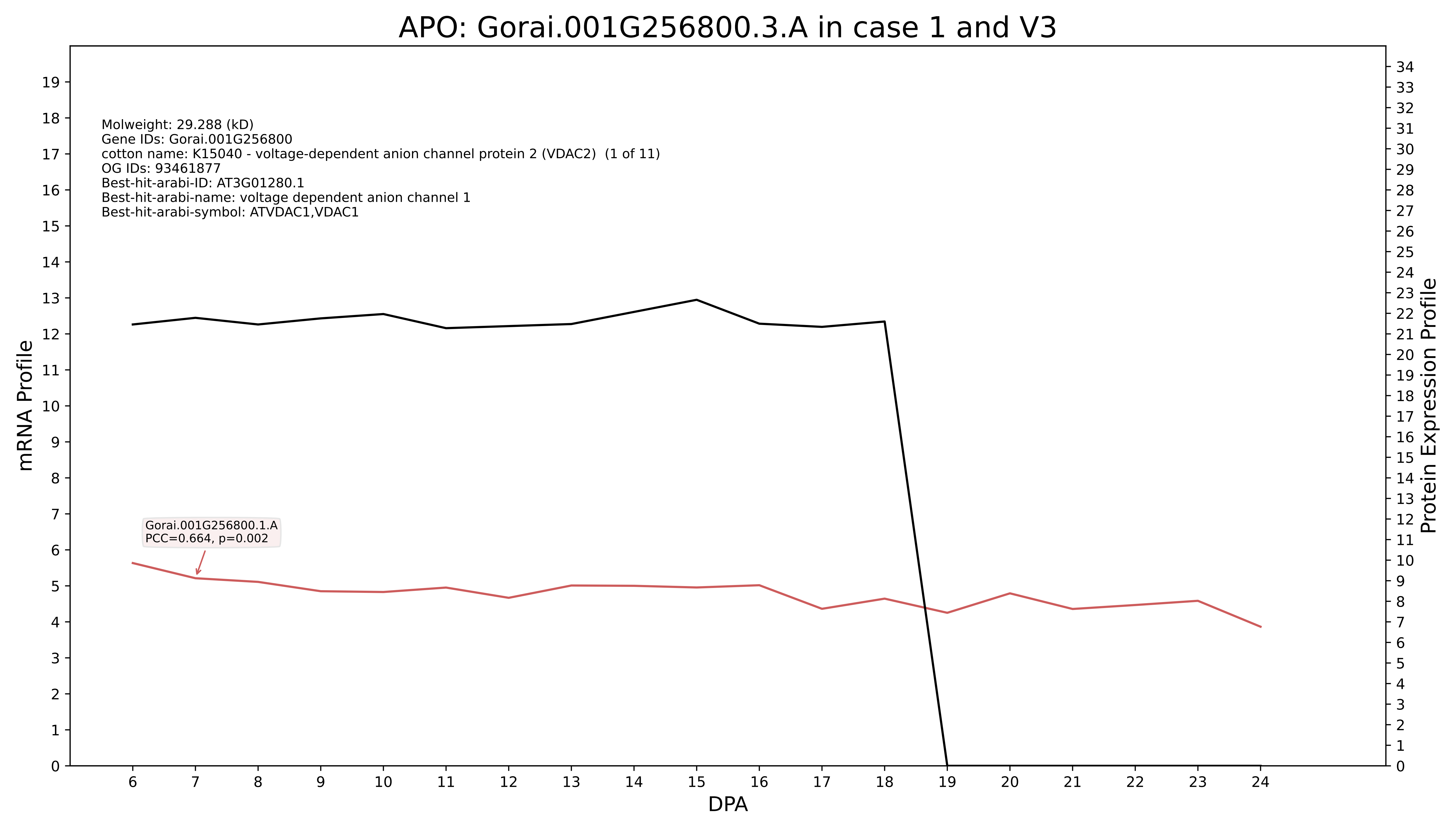
APO: Gorai.001G256800.3.A in case 1 and V3
Molweight: 29.288 (kD)
Gene IDs: Gorai.001G256800
cotton name: K15040 - voltage-dependent anion channel protein 2 (VDAC2) (1 of 11)
OG IDs: 93461877
Best-hit-arabi-ID: AT3G01280.1
Best-hit-arabi-name: voltage dependent anion channel 1
Best-hit-arabi-symbol: ATVDAC1,VDAC1
Gorai.001G256800.1.A
PCC=0.664, p=0.002
mRNA Profile
Protein Expression Profile
DPA

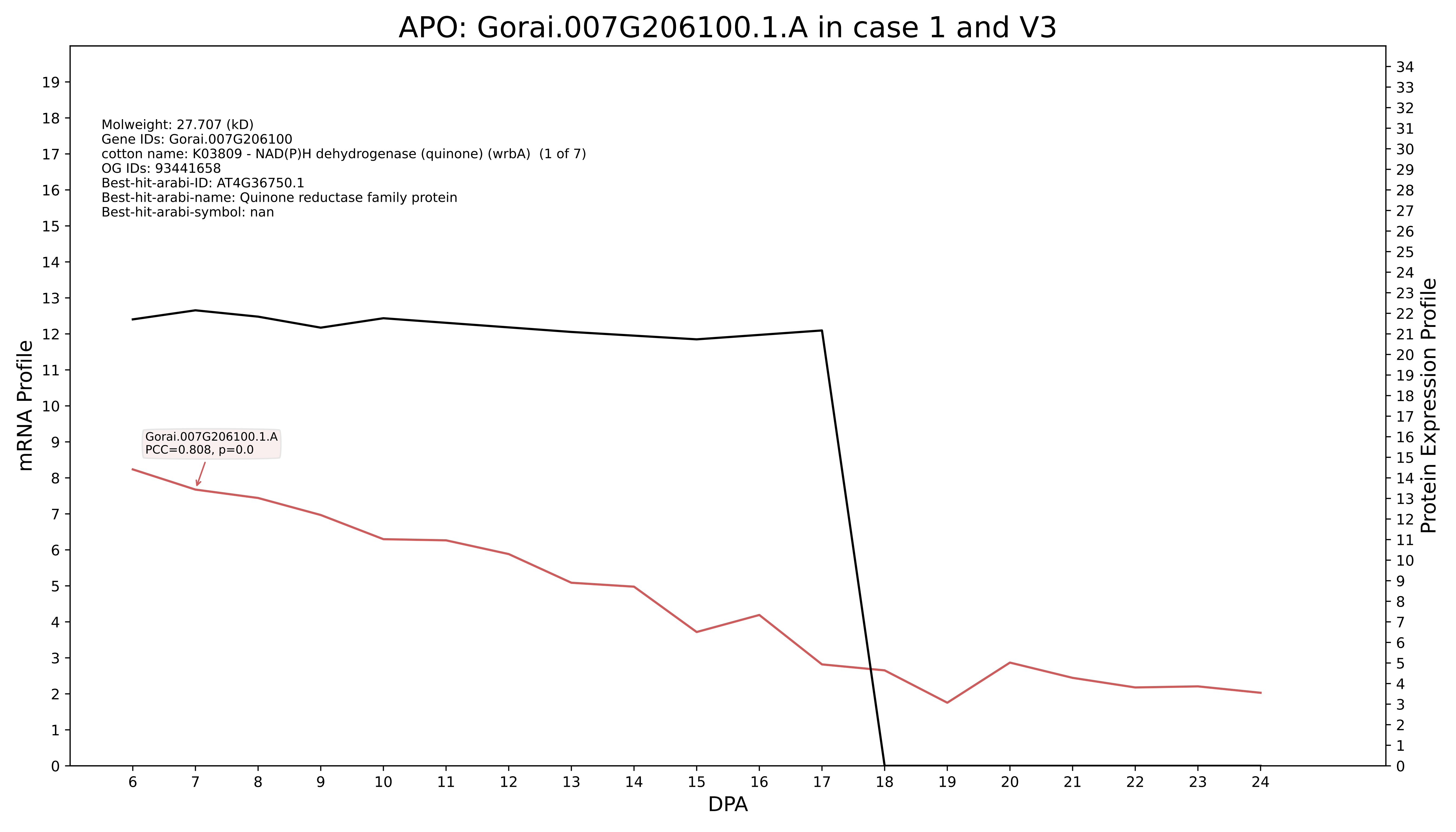
APO: Gorai.007G206100.1.A in case 1 and V3
Molweight: 27.707 (kD)
Gene IDs: Gorai.007G206100
cotton name: K03809 - NAD(P)H dehydrogenase (quinone) (wrbA) (1 of 7)
OG IDs: 93441658
Best-hit-arabi-ID: AT4G36750.1
Best-hit-arabi-name: Quinone reductase family protein
Best-hit-arabi-symbol: nan
Gorai.007G206100.1.A
PCC=0.808, p=0.0
mRNA Profile
Protein Expression Profile
DPA

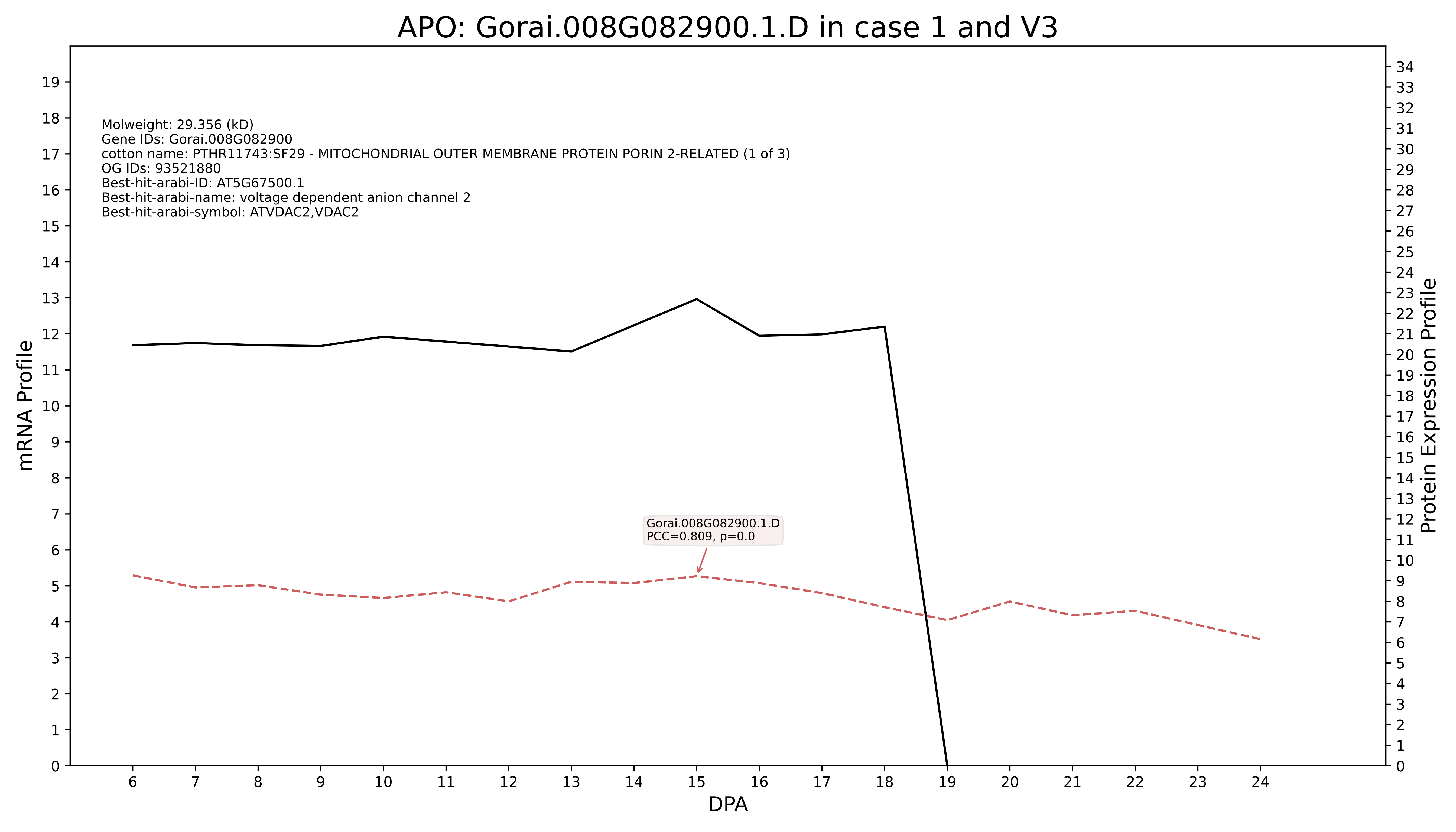
APO: Gorai.008G082900.1.D in case 1 and V3
Molweight: 29.356 (kD)
Gene IDs: Gorai.008G082900
cotton name: PTHR11743:SF29 - MITOCHONDRIAL OUTER MEMBRANE PROTEIN PORIN 2-RELATED (1 of 3)
OG IDs: 93521880
Best-hit-arabi-ID: AT5G67500.1
Best-hit-arabi-name: voltage dependent anion channel 2
Best-hit-arabi-symbol: ATVDAC2,VDAC2
Gorai.008G082900.1.D
PCC=0.809, p=0.0
mRNA Profile
Protein Expression Profile
DPA

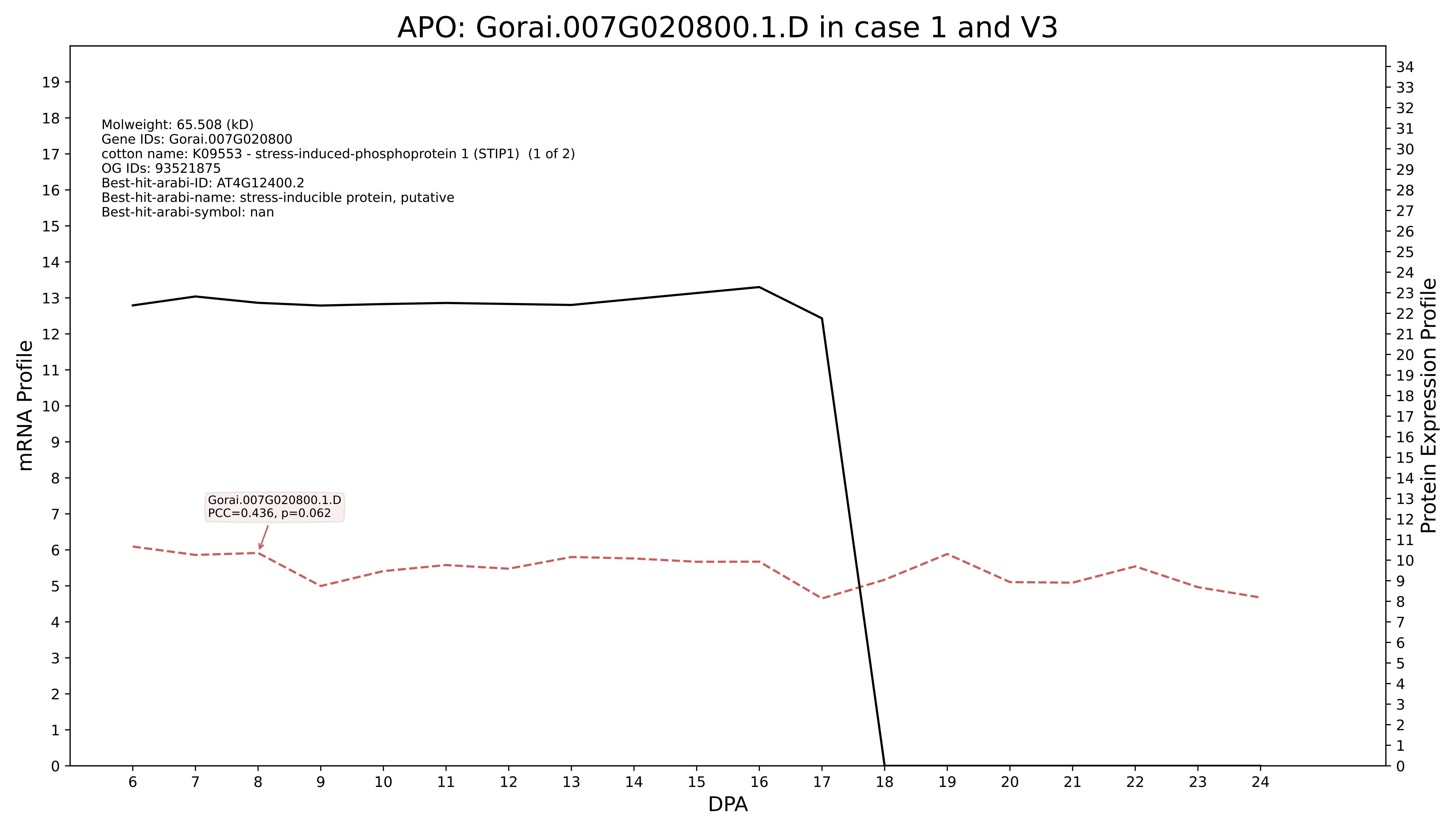
APO: Gorai.007G020800.1.D in case 1 and V3
Molweight: 65.508 (kD)
Gene IDs: Gorai.007G020800
cotton name: K09553 - stress-induced-phosphoprotein 1 (STIP1) (1 of 2)
OG IDs: 93521875
Best-hit-arabi-ID: AT4G12400.2
Best-hit-arabi-name: stress-inducible protein, putative
Best-hit-arabi-symbol: nan
Gorai.007G020800.1.D
PCC=0.436, p=0.062
mRNA Profile
Protein Expression Profile
DPA

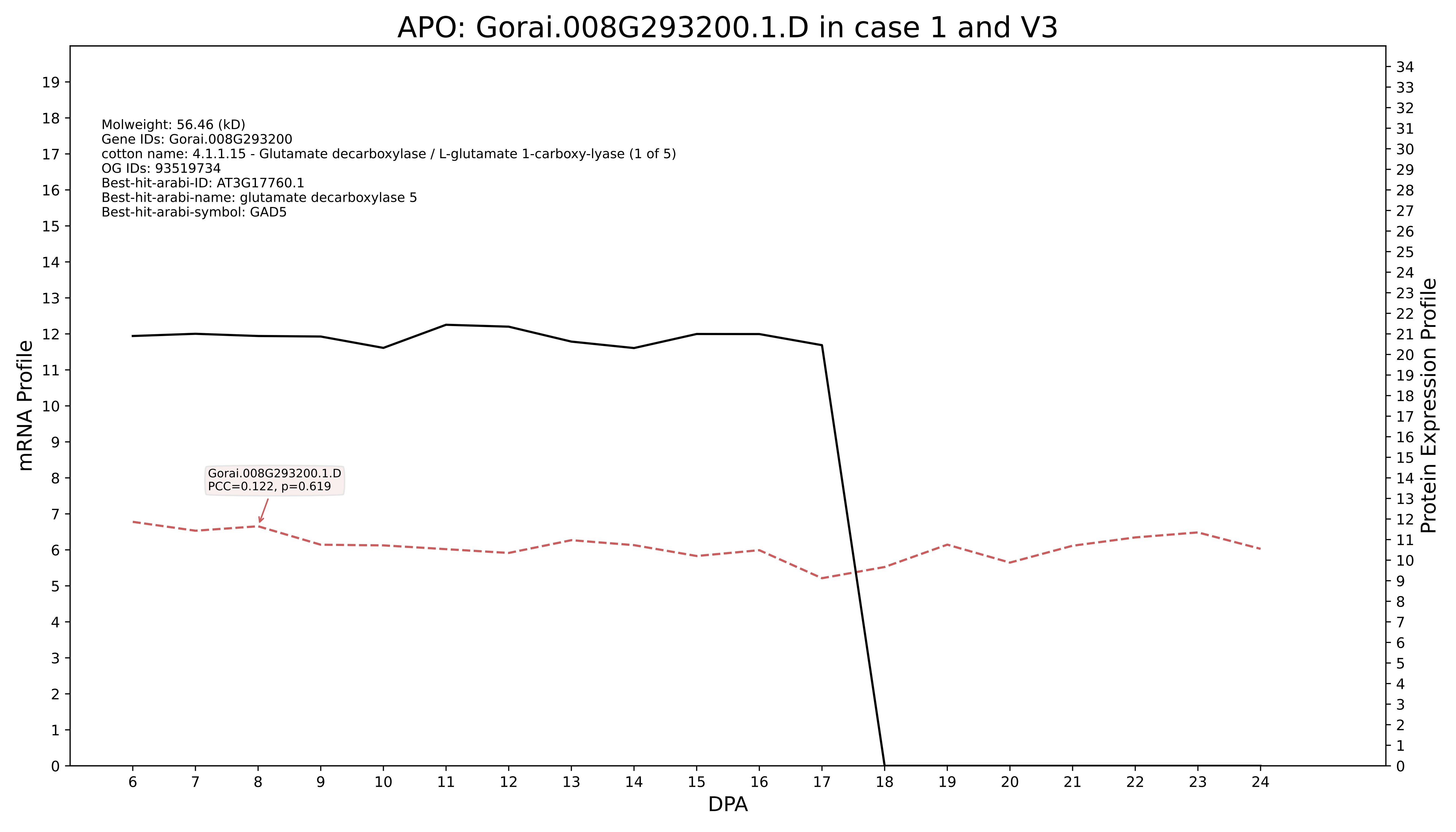
APO: Gorai.008G293200.1.D in case 1 and V3
Molweight: 56.46 (kD)
Gene IDs: Gorai.008G293200
cotton name: 4.1.1.15 - Glutamate decarboxylase / L-glutamate 1-carboxy-lyase (1 of 5)
OG IDs: 93519734
Best-hit-arabi-ID: AT3G17760.1
Best-hit-arabi-name: glutamate decarboxylase 5
Best-hit-arabi-symbol: GAD5
Gorai.008G293200.1.D
PCC=0.122, p=0.619
mRNA Profile
Protein Expression Profile
DPA

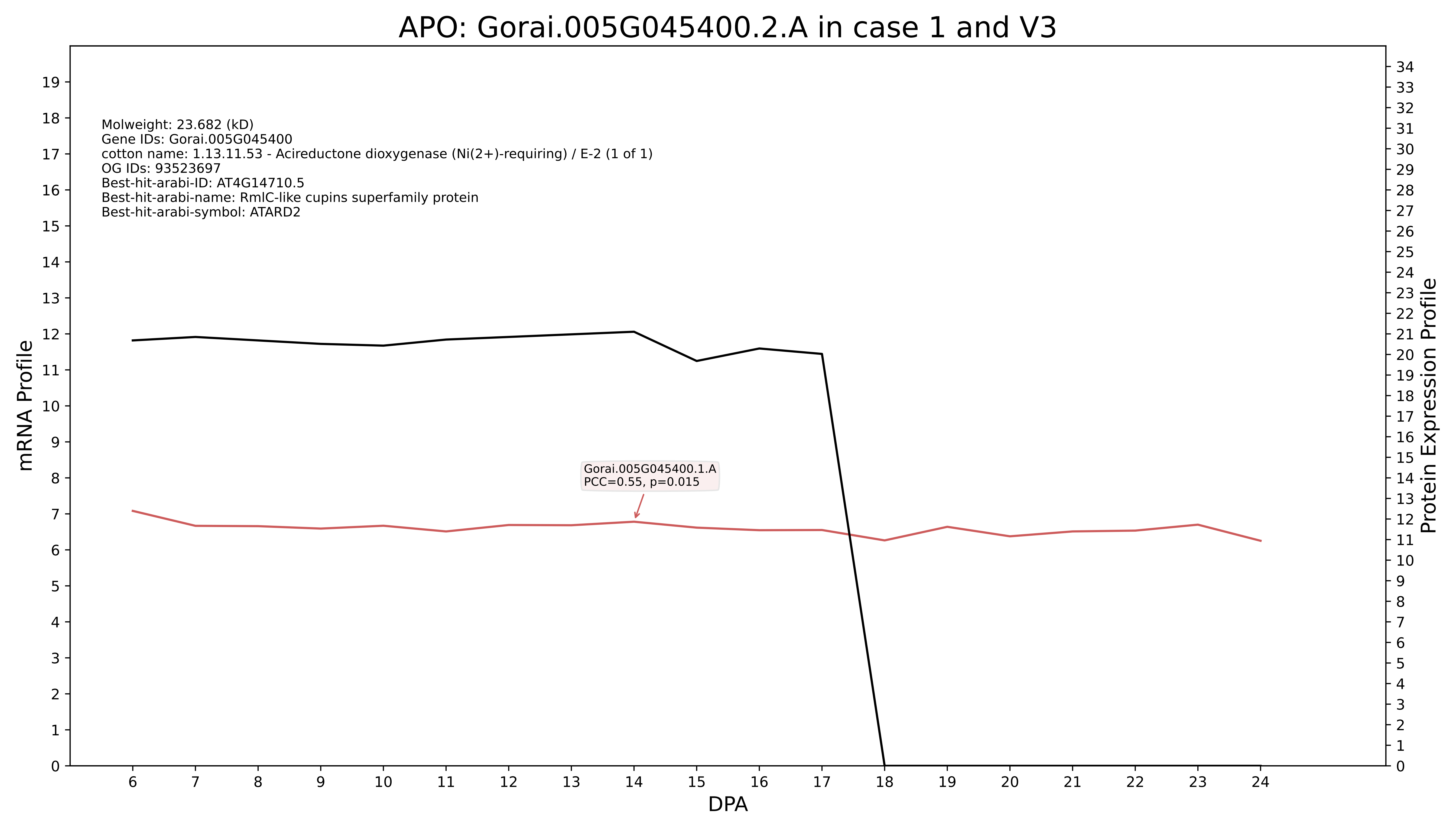
APO: Gorai.005G045400.2.A in case 1 and V3
Molweight: 23.682 (kD)
Gene IDs: Gorai.005G045400
cotton name: 1.13.11.53 - Acireductone dioxygenase (Ni(2+)-requiring) / E-2 (1 of 1)
OG IDs: 93523697
Best-hit-arabi-ID: AT4G14710.5
Best-hit-arabi-name: RmlC-like cupins superfamily protein
Best-hit-arabi-symbol: ATARD2
Gorai.005G045400.1.A
PCC=0.55, p=0.015
mRNA Profile
Protein Expression Profile
DPA

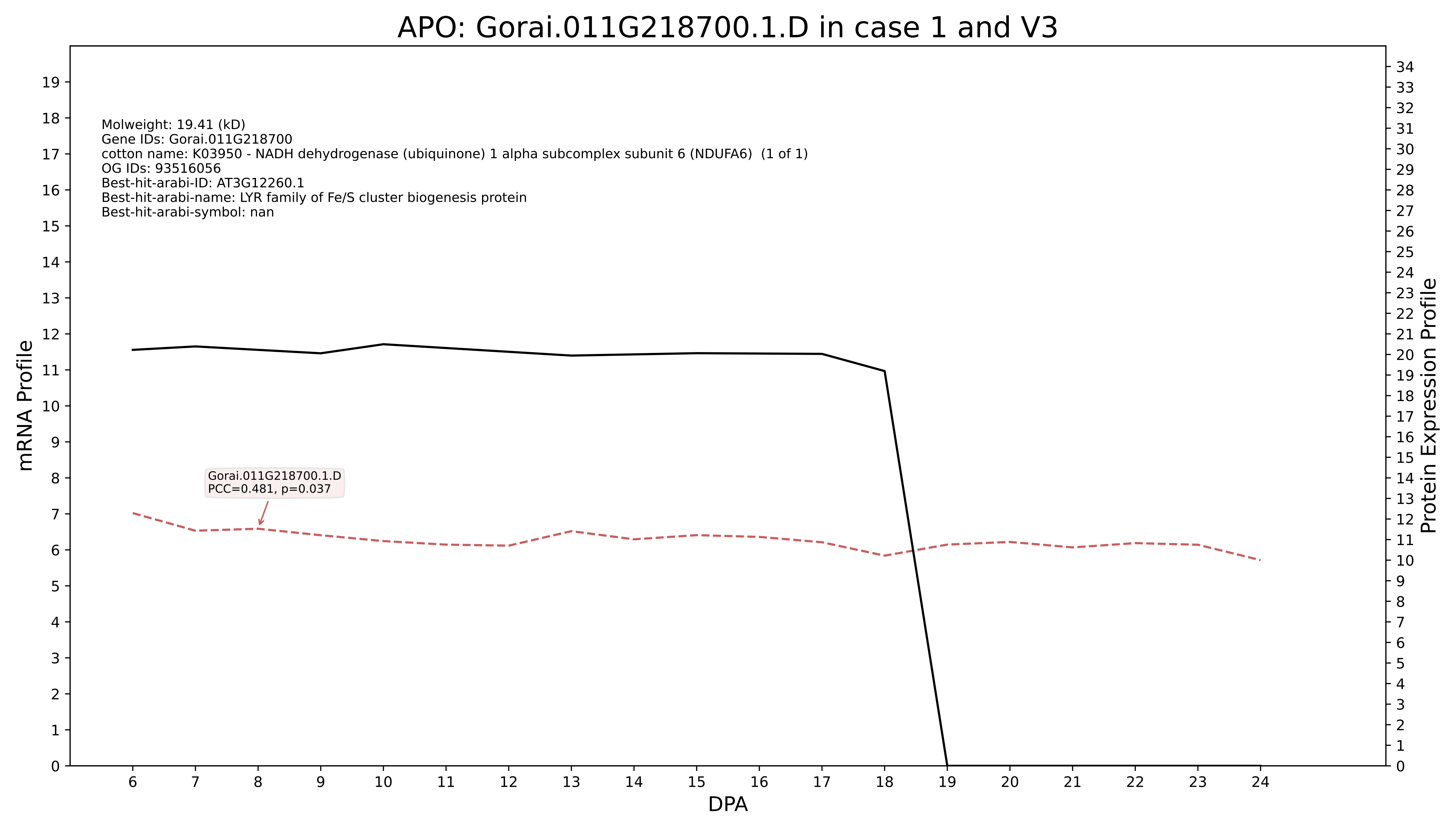
APO: Gorai.011G218700.1.D in case 1 and V3
Molweight: 19.41 (kD)
Gene IDs: Gorai.011G218700
cotton name: K03950 - NADH dehydrogenase (ubiquinone) 1 alpha subcomplex subunit 6 (NDUFA6) (1 of 1)
OG IDs: 93516056
Best-hit-arabi-ID: AT3G12260.1
Best-hit-arabi-name: LYR family of Fe/S cluster biogenesis protein
Best-hit-arabi-symbol: nan
Gorai.011G218700.1.D
PCC=0.481, p=0.037
mRNA Profile
Protein Expression Profile
DPA

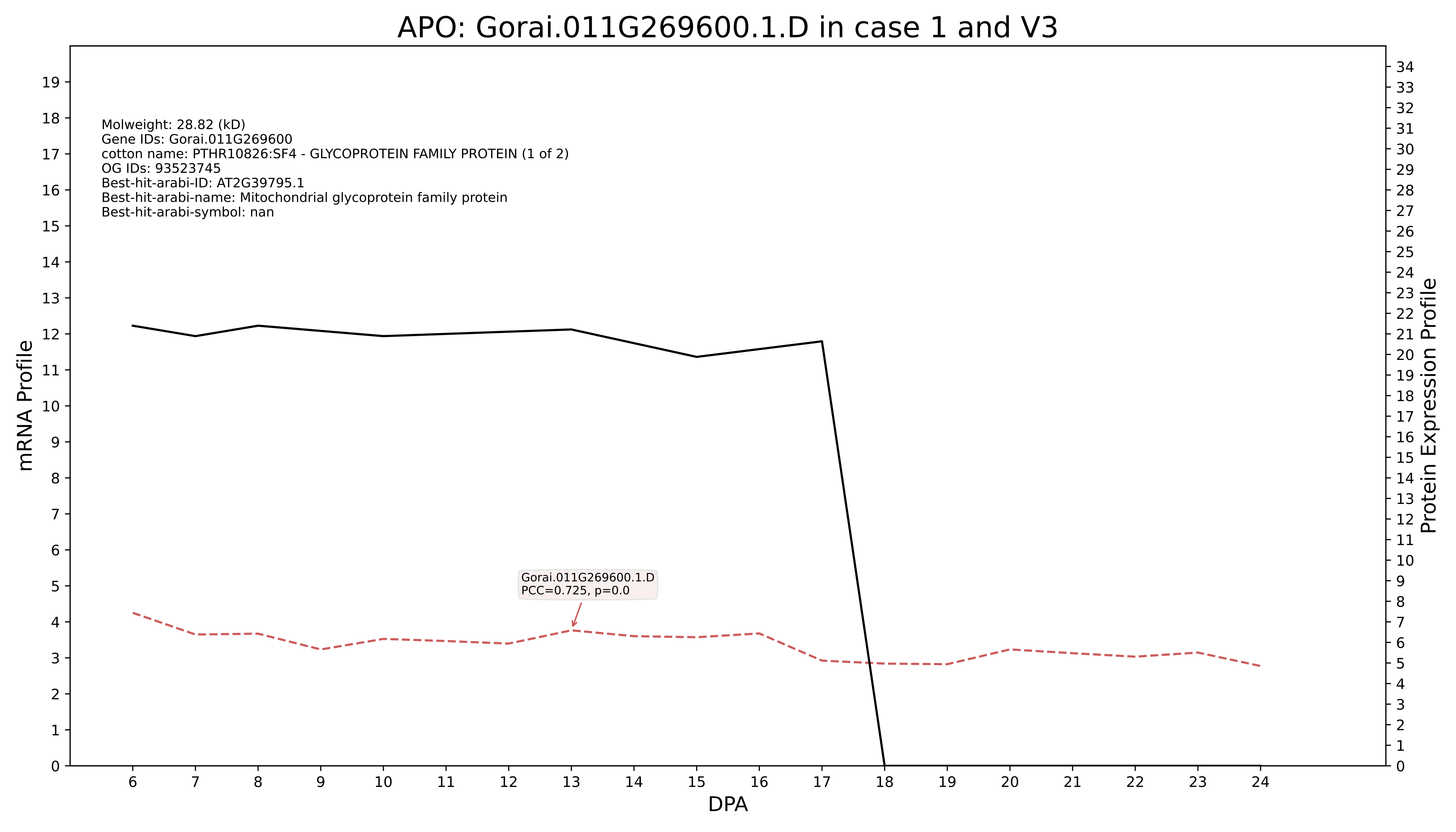
APO: Gorai.011G269600.1.D in case 1 and V3
Molweight: 28.82 (kD)
Gene IDs: Gorai.011G269600
cotton name: PTHR10826:SF4 - GLYCOPROTEIN FAMILY PROTEIN (1 of 2)
OG IDs: 93523745
Best-hit-arabi-ID: AT2G39795.1
Best-hit-arabi-name: Mitochondrial glycoprotein family protein
Best-hit-arabi-symbol: nan
Gorai.011G269600.1.D
PCC=0.725, p=0.0
mRNA Profile
Protein Expression Profile
DPA

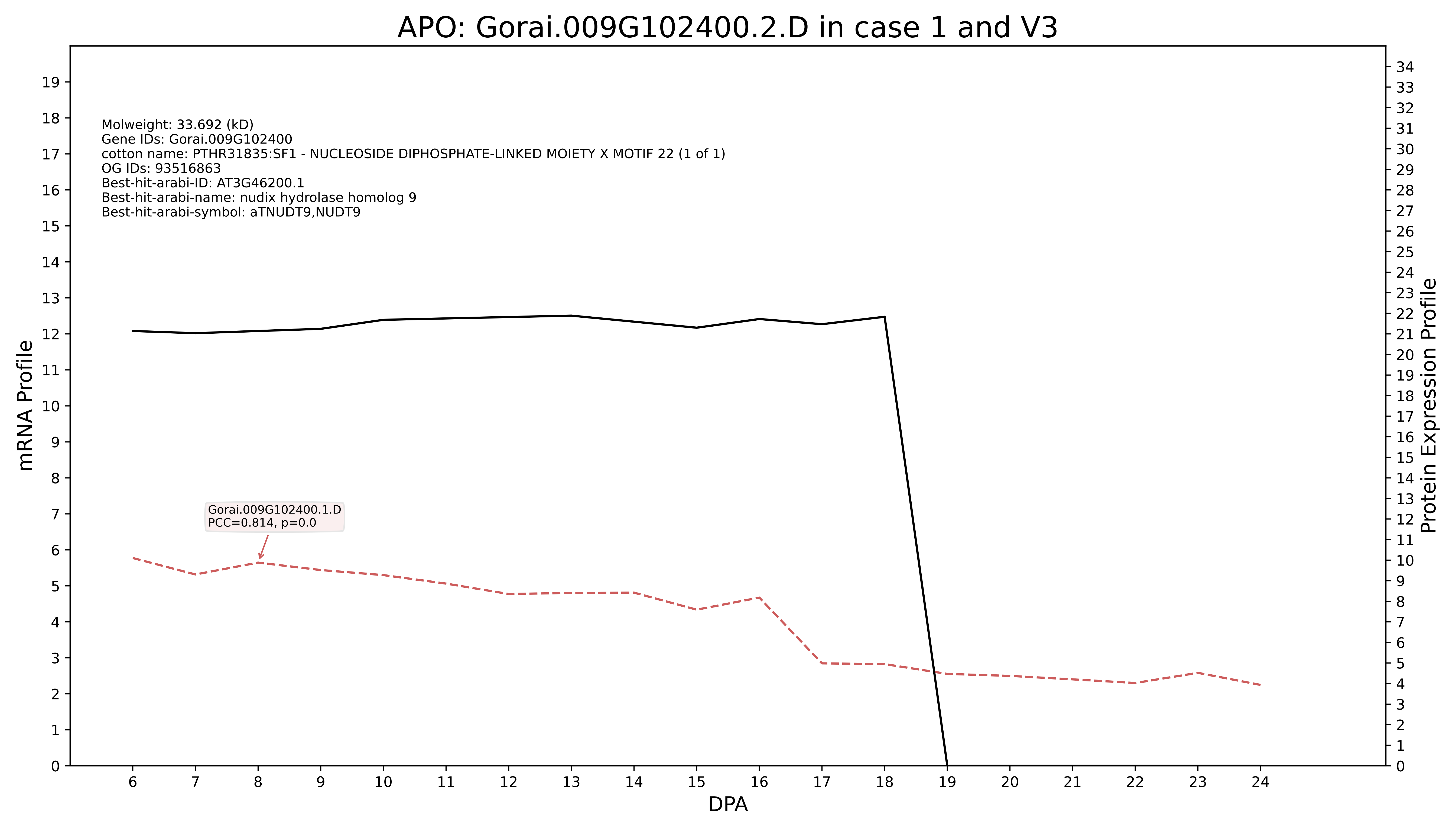

APO: Gorai.009G102400.2.D in case 1 and V3
Molweight: 33.692 (kD)
Gene IDs: Gorai.009G102400
cotton name: PTHR31835:SF1 - NUCLEOSIDE DIPHOSPHATE-LINKED MOIETY X MOTIF 22 (1 of 1)
OG IDs: 93516863
Best-hit-arabi-ID: AT3G46200.1
Best-hit-arabi-name: nudix hydrolase homolog 9
Best-hit-arabi-symbol: aTNUDT9,NUDT9
Gorai.009G102400.1.D
PCC=0.814, p=0.0
mRNA Profile
Protein Expression Profile
DPA

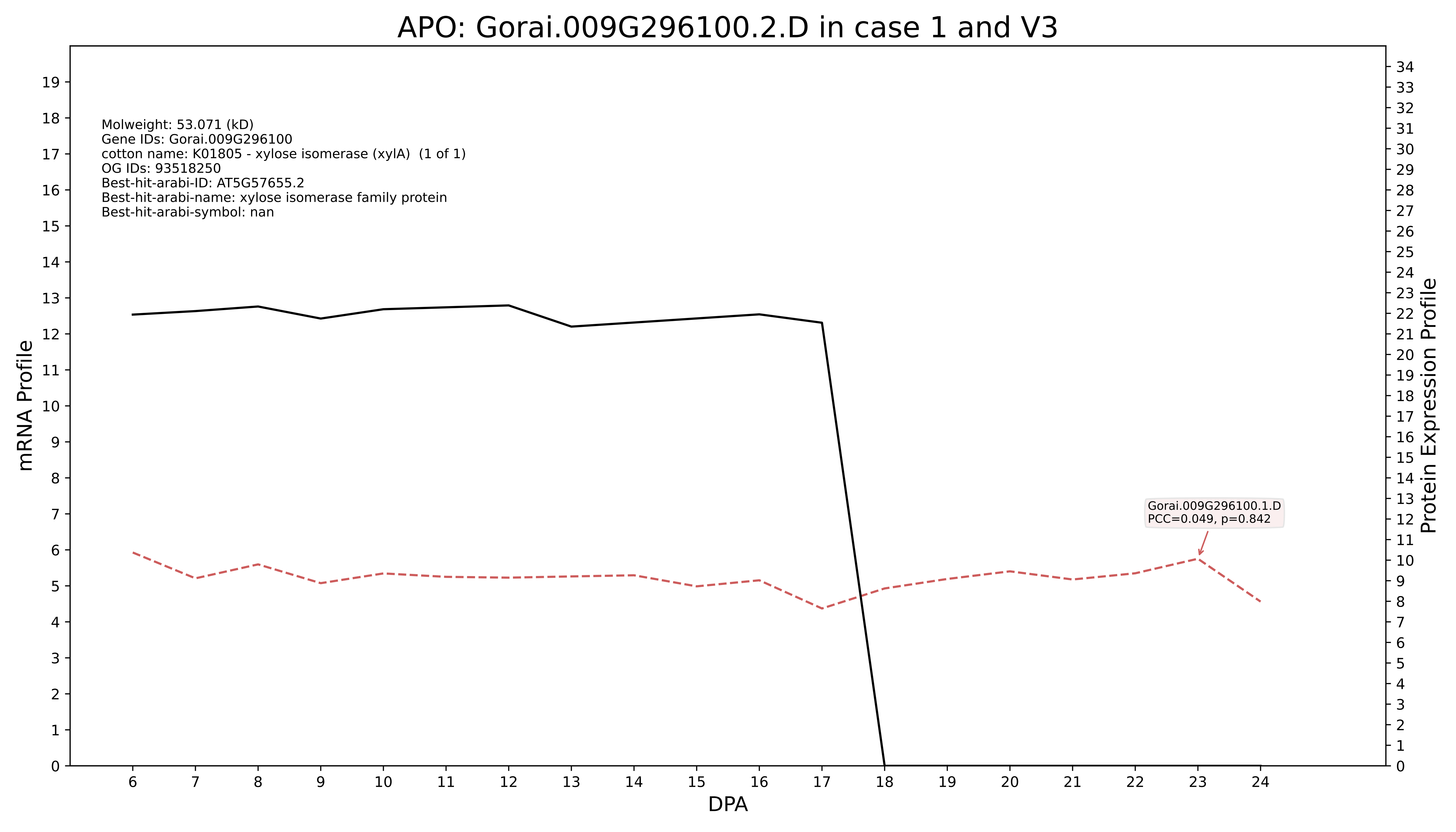

APO: Gorai.009G296100.2.D in case 1 and V3
Molweight: 53.071 (kD)
Gene IDs: Gorai.009G296100
cotton name: K01805 - xylose isomerase (xylA) (1 of 1)
OG IDs: 93518250
Best-hit-arabi-ID: AT5G57655.2
Best-hit-arabi-name: xylose isomerase family protein
Best-hit-arabi-symbol: nan
Gorai.009G296100.1.D
PCC=0.049, p=0.842
mRNA Profile
Protein Expression Profile
DPA

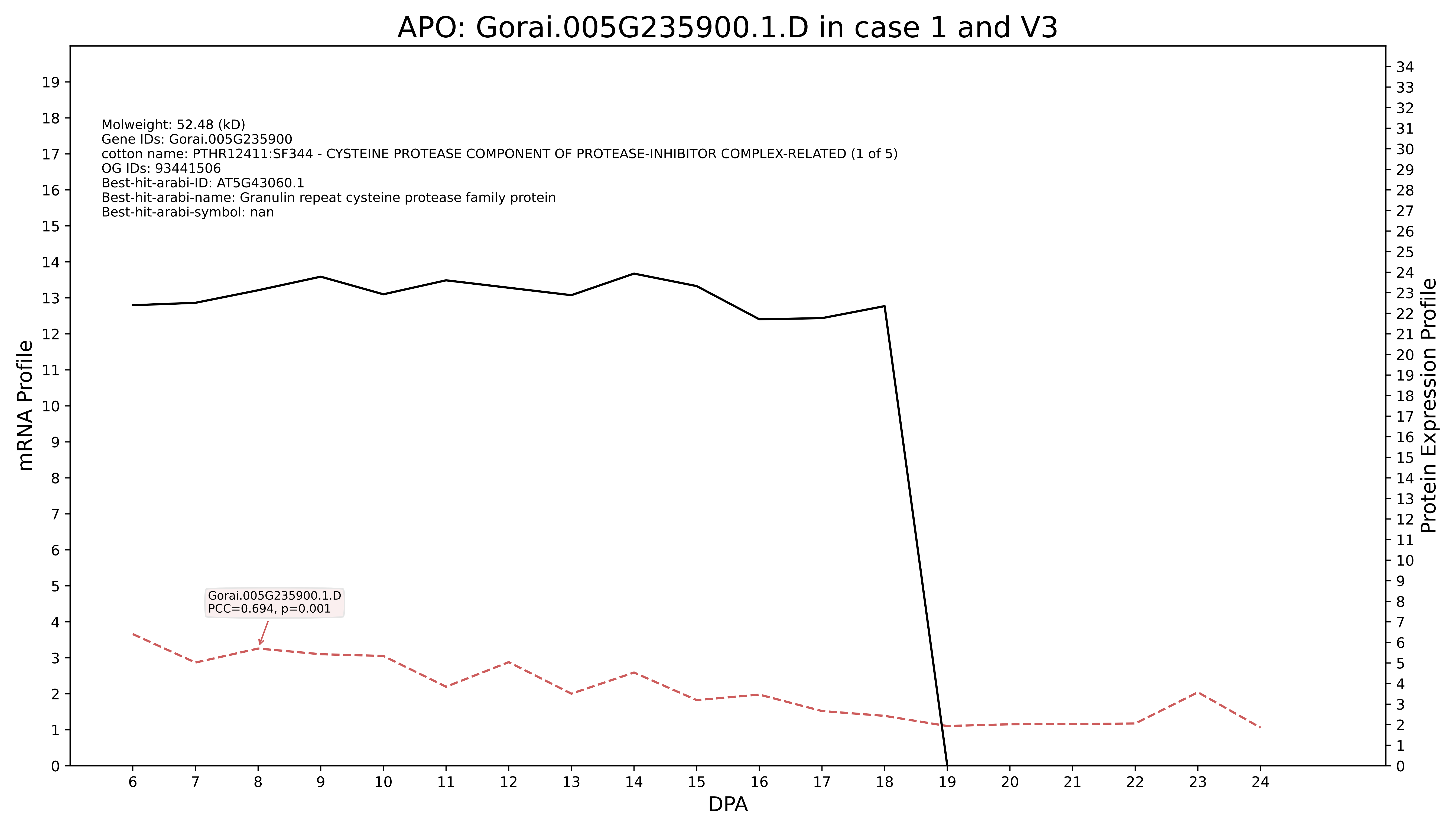
APO: Gorai.005G235900.1.D in case 1 and V3
Molweight: 52.48 (kD)
Gene IDs: Gorai.005G235900
cotton name: PTHR12411:SF344 - CYSTEINE PROTEASE COMPONENT OF PROTEASE-INHIBITOR COMPLEX-RELATED (1 of 5)
OG IDs: 93441506
Best-hit-arabi-ID: AT5G43060.1
Best-hit-arabi-name: Granulin repeat cysteine protease family protein
Best-hit-arabi-symbol: nan
Gorai.005G235900.1.D
PCC=0.694, p=0.001
mRNA Profile
Protein Expression Profile
DPA

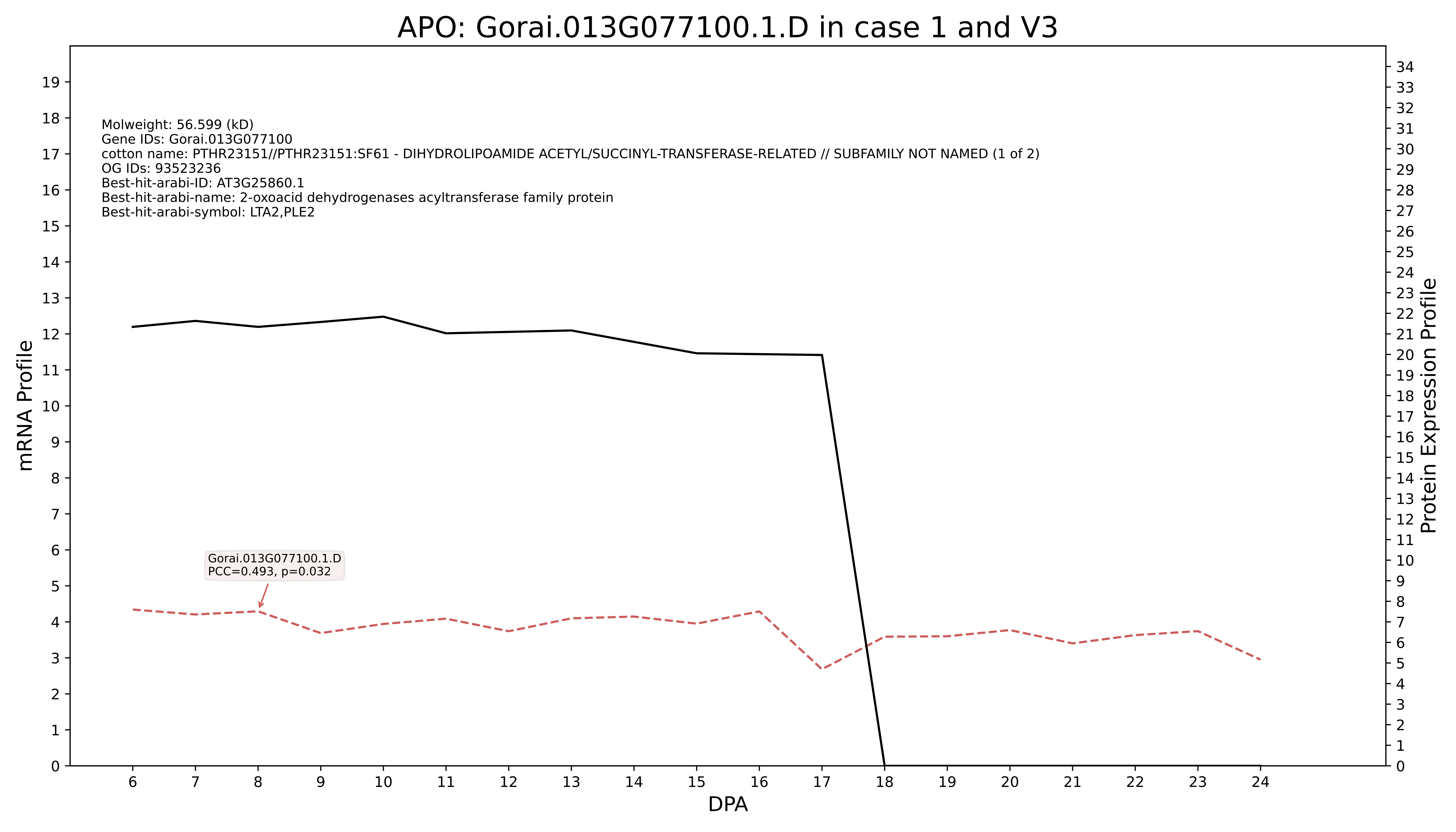
APO: Gorai.013G077100.1.D in case 1 and V3
Molweight: 56.599 (kD)
Gene IDs: Gorai.013G077100
cotton name: PTHR23151//PTHR23151:SF61 - DIHYDROLIPOAMIDE ACETYL/SUCCINYL-TRANSFERASE-RELATED // SUBFAMILY NOT NAMED (1 of 2)
OG IDs: 93523236
Best-hit-arabi-ID: AT3G25860.1
Best-hit-arabi-name: 2-oxoacid dehydrogenases acyltransferase family protein
Best-hit-arabi-symbol: LTA2,PLE2
Gorai.013G077100.1.D
PCC=0.493, p=0.032
mRNA Profile
Protein Expression Profile
DPA

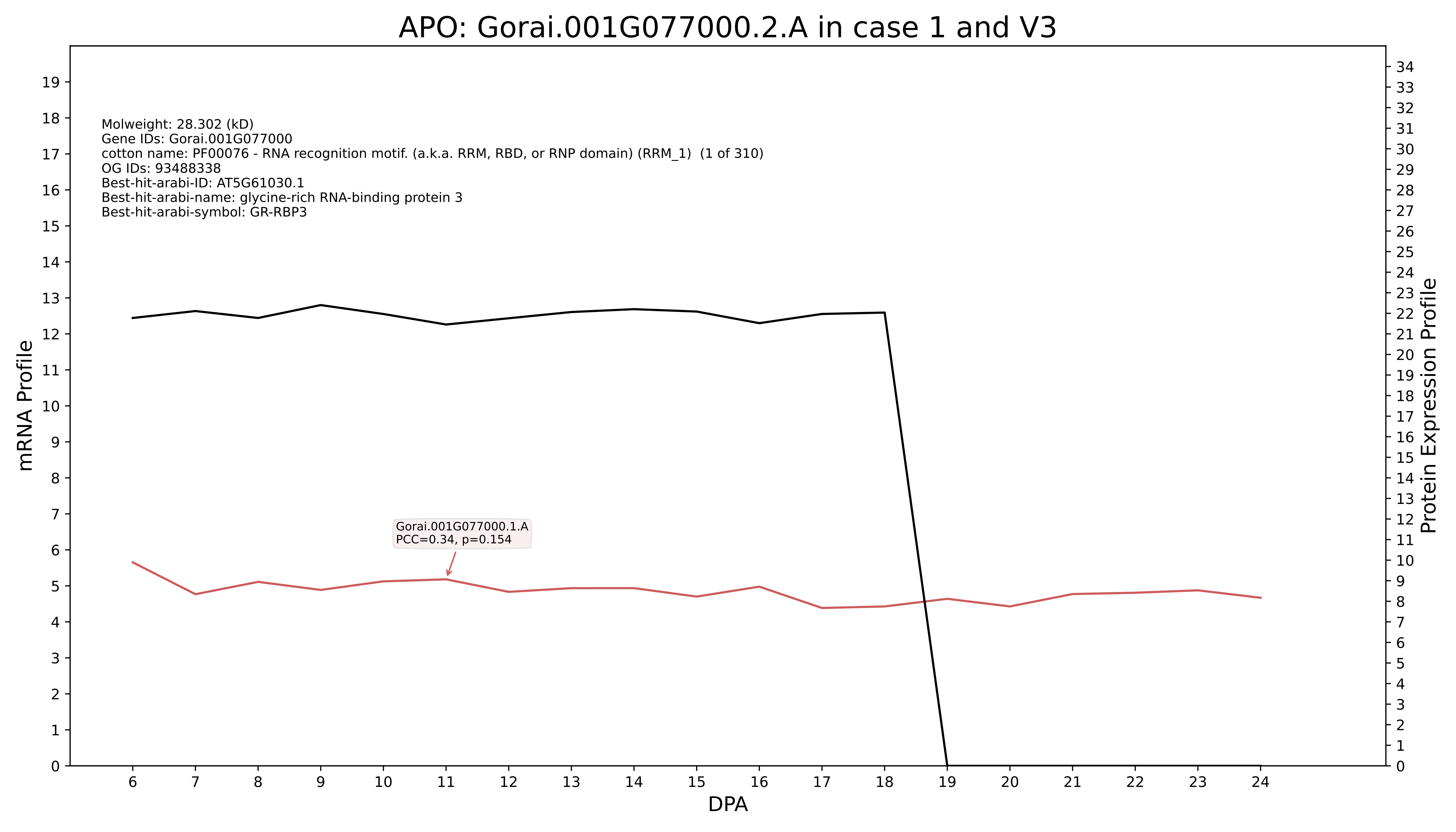
APO: Gorai.001G077000.2.A in case 1 and V3
Molweight: 28.302 (kD)
Gene IDs: Gorai.001G077000
cotton name: PF00076 - RNA recognition motif. (a.k.a. RRM, RBD, or RNP domain) (RRM_1) (1 of 310)
OG IDs: 93488338
Best-hit-arabi-ID: AT5G61030.1
Best-hit-arabi-name: glycine-rich RNA-binding protein 3
Best-hit-arabi-symbol: GR-RBP3
Gorai.001G077000.1.A
PCC=0.34, p=0.154
mRNA Profile
Protein Expression Profile
DPA

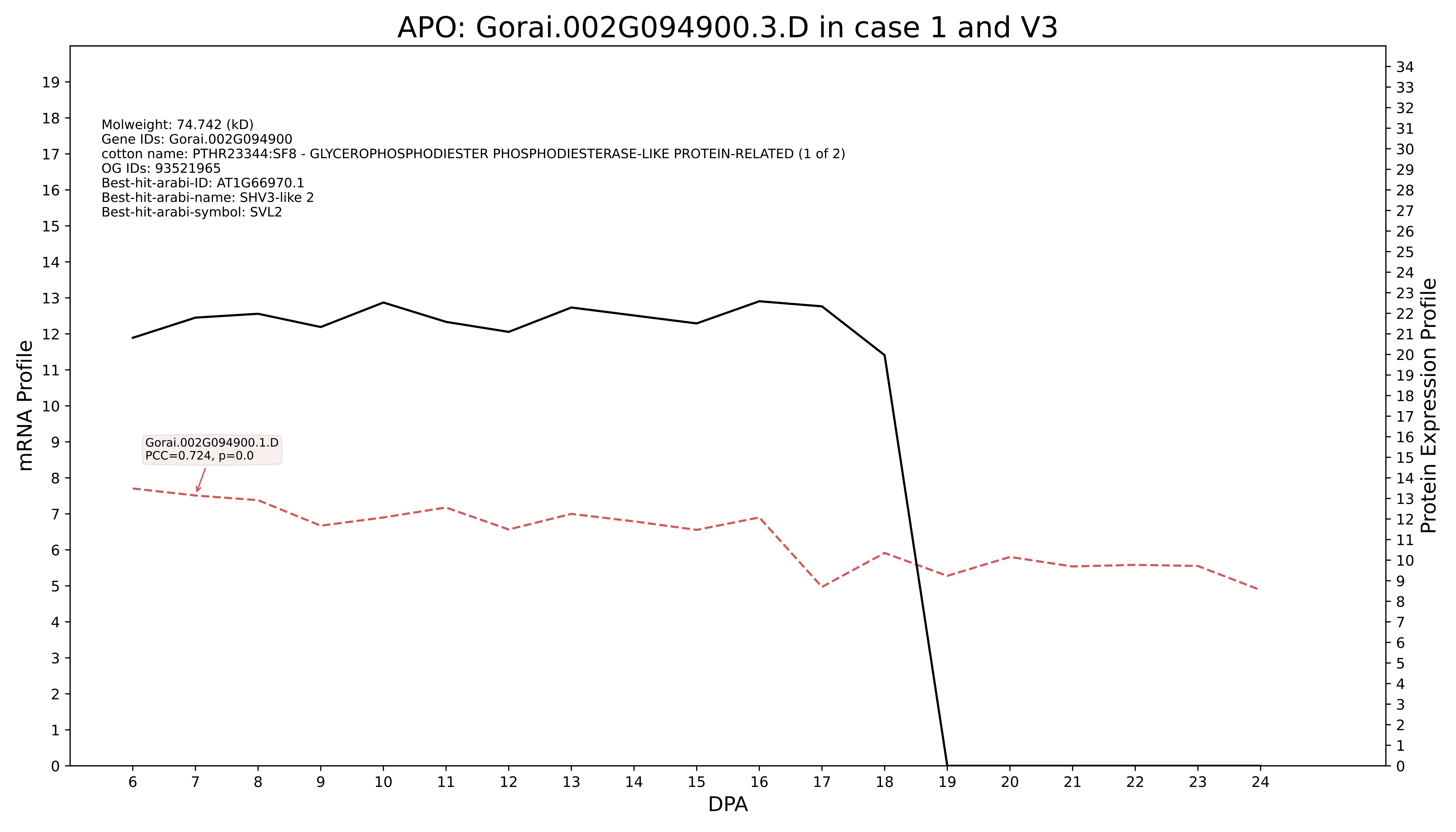

APO: Gorai.002G094900.3.D in case 1 and V3
Molweight: 74.742 (kD)
Gene IDs: Gorai.002G094900
cotton name: PTHR23344:SF8 - GLYCEROPHOSPHODIESTER PHOSPHODIESTERASE-LIKE PROTEIN-RELATED (1 of 2)
OG IDs: 93521965
Best-hit-arabi-ID: AT1G66970.1
Best-hit-arabi-name: SHV3-like 2
Best-hit-arabi-symbol: SVL2
Gorai.002G094900.1.D
PCC=0.724, p=0.0
mRNA Profile
Protein Expression Profile
DPA

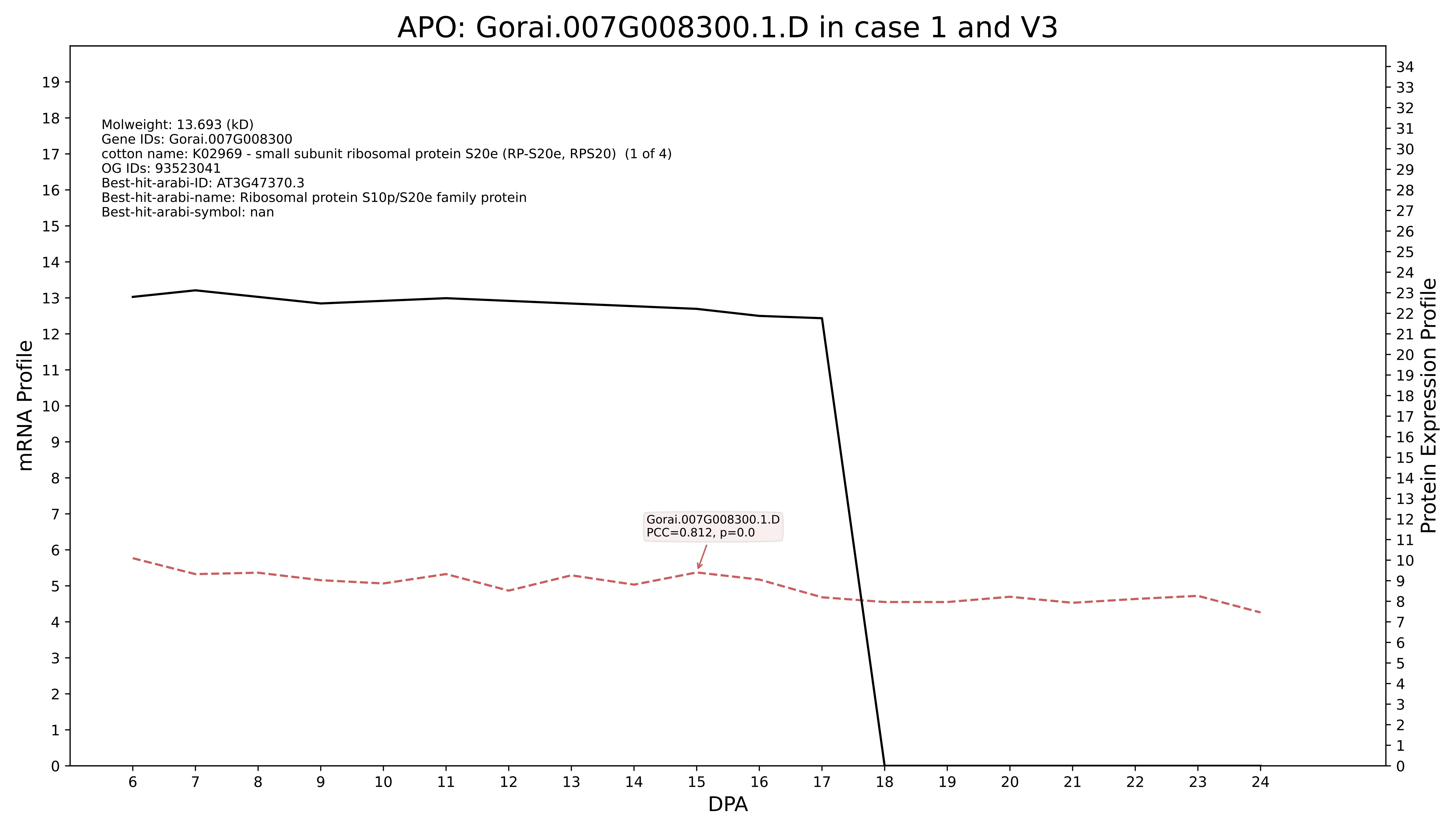
APO: Gorai.007G008300.1.D in case 1 and V3
Molweight: 13.693 (kD)
Gene IDs: Gorai.007G008300
cotton name: K02969 - small subunit ribosomal protein S20e (RP-S20e, RPS20) (1 of 4)
OG IDs: 93523041
Best-hit-arabi-ID: AT3G47370.3
Best-hit-arabi-name: Ribosomal protein S10p/S20e family protein
Best-hit-arabi-symbol: nan
Gorai.007G008300.1.D
PCC=0.812, p=0.0
mRNA Profile
Protein Expression Profile
DPA

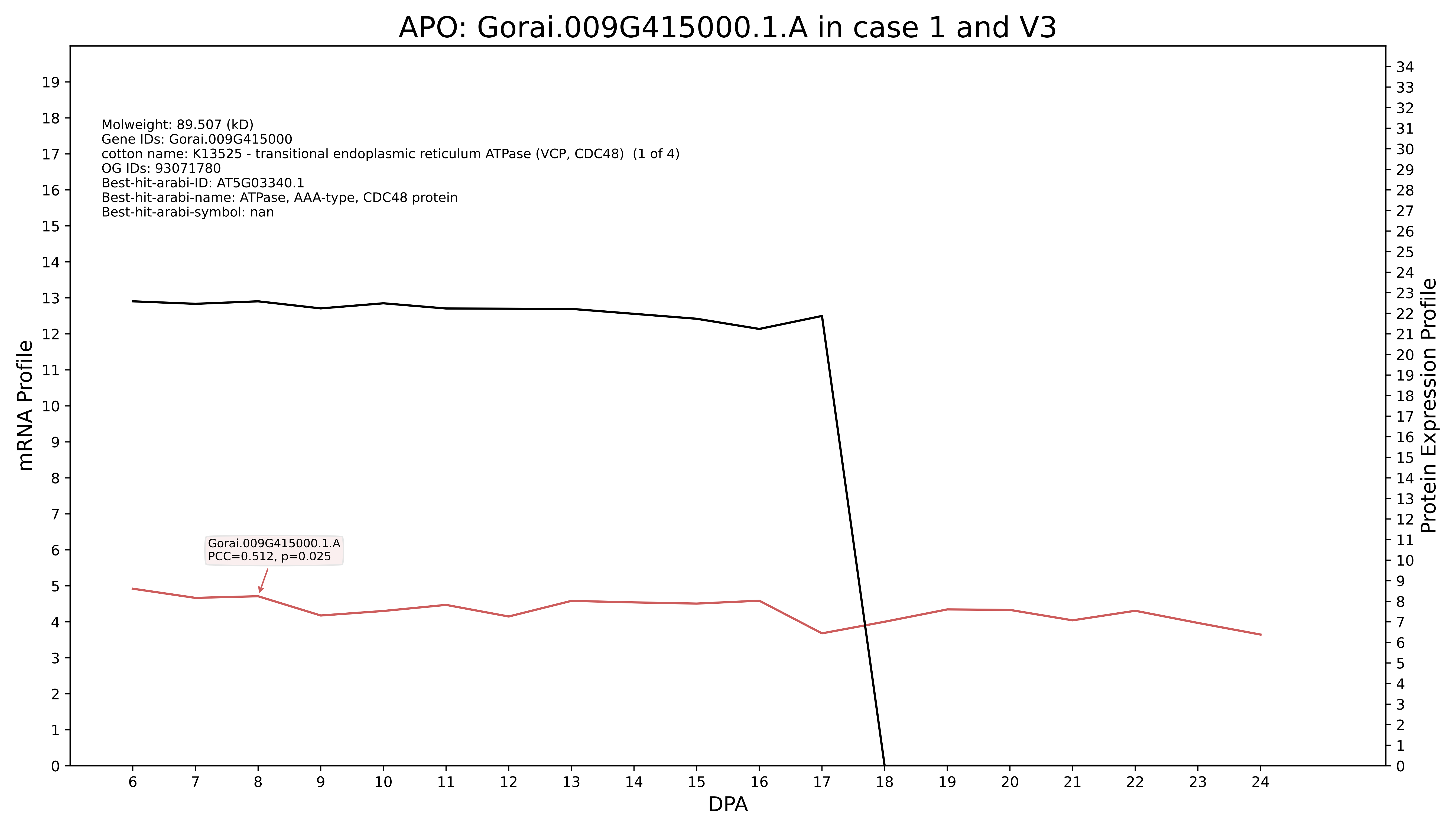

APO: Gorai.009G415000.1.A in case 1 and V3
Molweight: 89.507 (kD)
Gene IDs: Gorai.009G415000
cotton name: K13525 - transitional endoplasmic reticulum ATPase (VCP, CDC48) (1 of 4)
OG IDs: 93071780
Best-hit-arabi-ID: AT5G03340.1
Best-hit-arabi-name: ATPase, AAA-type, CDC48 protein
Best-hit-arabi-symbol: nan
Gorai.009G415000.1.A
PCC=0.512, p=0.025
mRNA Profile
Protein Expression Profile
DPA

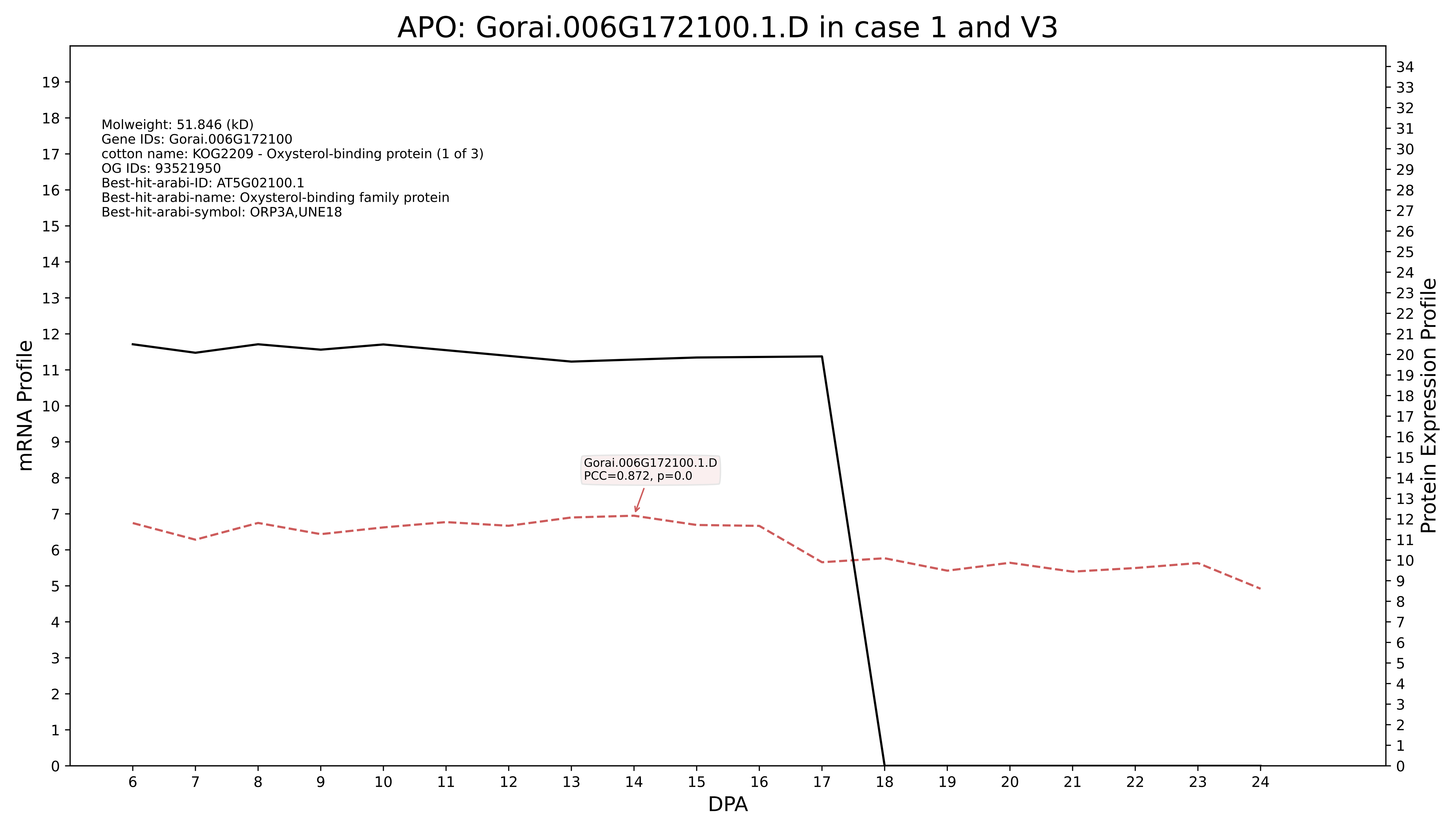
APO: Gorai.006G172100.1.D in case 1 and V3
Molweight: 51.846 (kD)
Gene IDs: Gorai.006G172100
cotton name: KOG2209 - Oxysterol-binding protein (1 of 3)
OG IDs: 93521950
Best-hit-arabi-ID: AT5G02100.1
Best-hit-arabi-name: Oxysterol-binding family protein
Best-hit-arabi-symbol: ORP3A,UNE18
Gorai.006G172100.1.D
PCC=0.872, p=0.0
mRNA Profile
Protein Expression Profile
DPA

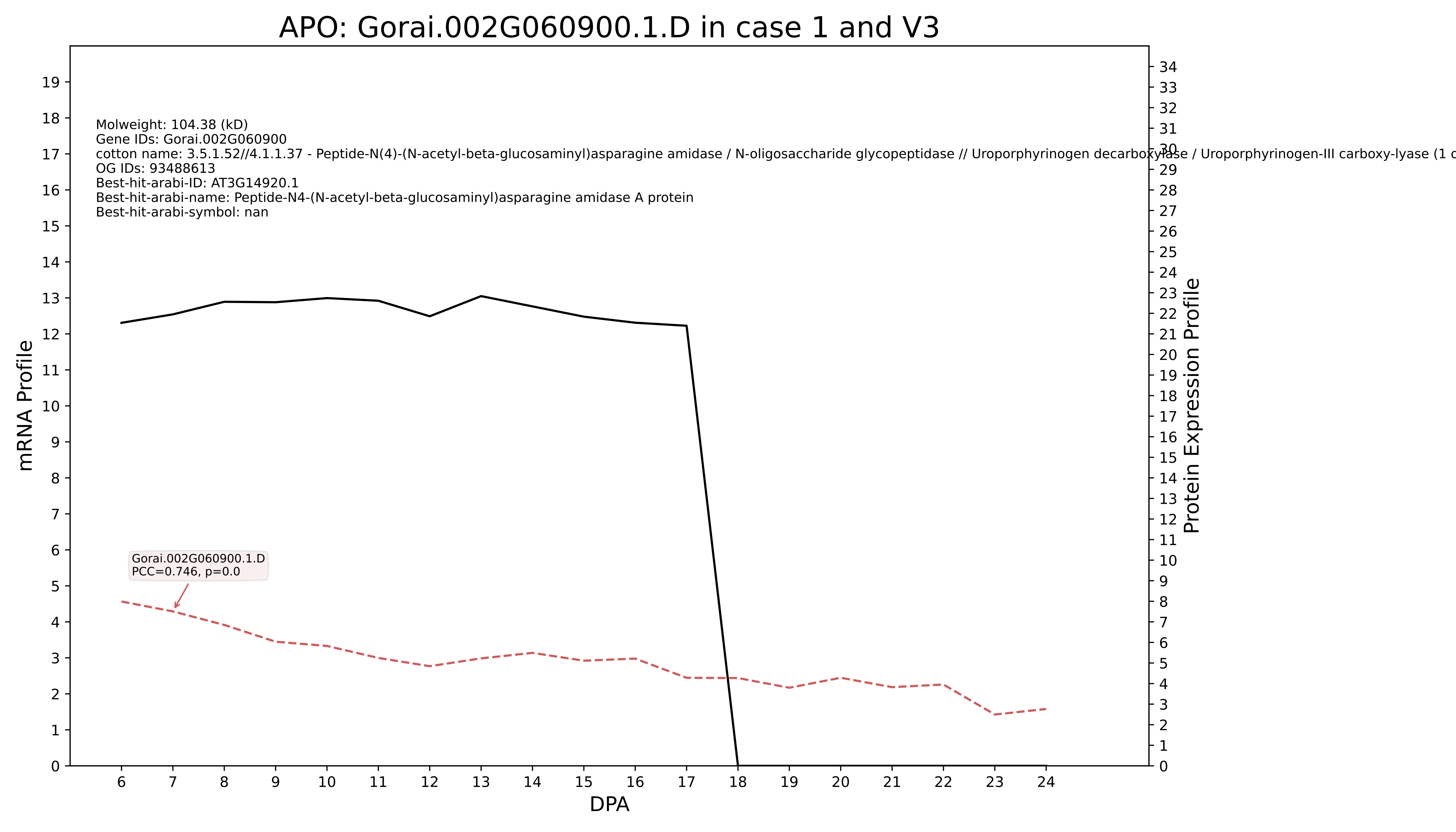
APO: Gorai.002G060900.1.D in case 1 and V3
Molweight: 104.38 (kD)
Gene IDs: Gorai.002G060900
cotton name: 3.5.1.52//4.1.1.37 - Peptide-N(4)-(N-acetyl-beta-glucosaminyl)asparagine amidase / N-oligosaccharide glycopeptidase // Uroporphyrinogen decarboxylase / Uroporphyrinogen-III carboxy-lyase (1 c
OG IDs: 93488613
Best-hit-arabi-ID: AT3G14920.1
Best-hit-arabi-name: Peptide-N4-(N-acetyl-beta-glucosaminyl)asparagine amidase A protein
Best-hit-arabi-symbol: nan
Gorai.002G060900.1.D
PCC=0.746, p=0.0
mRNA Profile
Protein Expression Profile
DPA

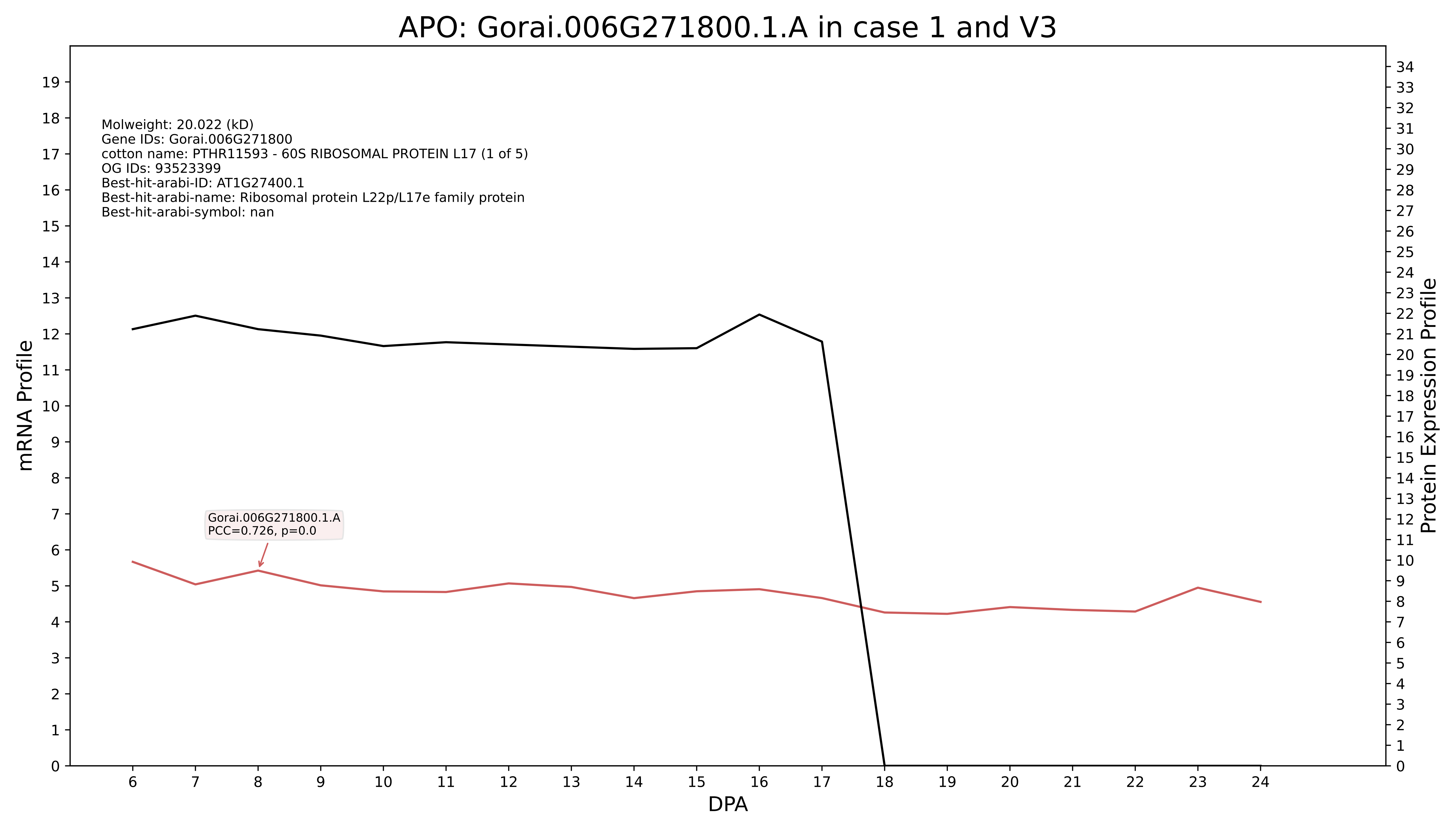
APO: Gorai.006G271800.1.A in case 1 and V3
Molweight: 20.022 (kD)
Gene IDs: Gorai.006G271800
cotton name: PTHR11593 - 60S RIBOSOMAL PROTEIN L17 (1 of 5)
OG IDs: 93523399
Best-hit-arabi-ID: AT1G27400.1
Best-hit-arabi-name: Ribosomal protein L22p/L17e family protein
Best-hit-arabi-symbol: nan
Gorai.006G271800.1.A
PCC=0.726, p=0.0
mRNA Profile
Protein Expression Profile
DPA

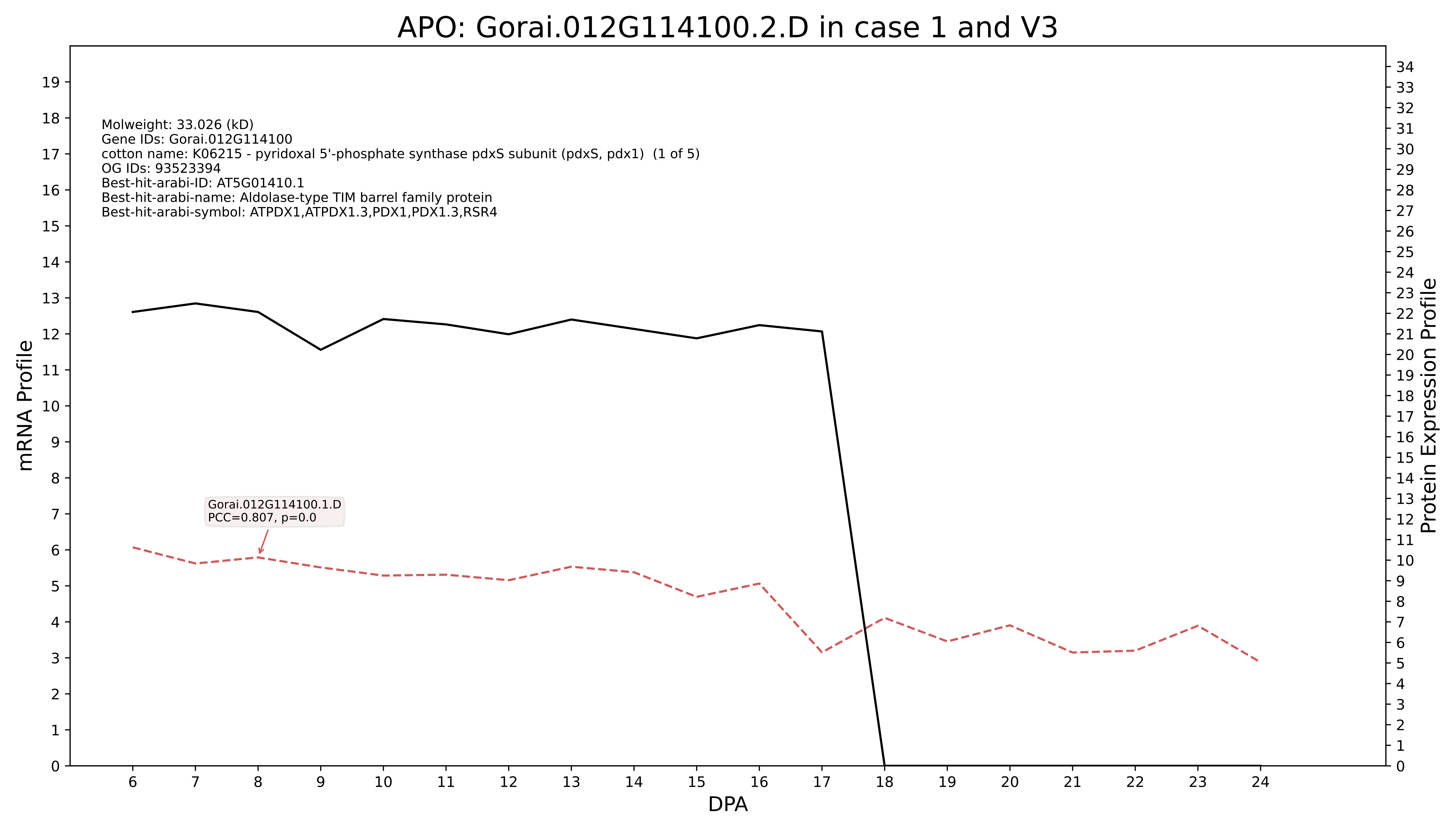
APO: Gorai.012G114100.2.D in case 1 and V3
Molweight: 33.026 (kD)
Gene IDs: Gorai.012G114100
cotton name: K06215 - pyridoxal 5'-phosphate synthase pdxS subunit (pdxS, pdx1) (1 of 5)
OG IDs: 93523394
Best-hit-arabi-ID: AT5G01410.1
Best-hit-arabi-name: Aldolase-type TIM barrel family protein
Best-hit-arabi-symbol: ATPDX1,ATPDX1.3,PDX1,PDX1.3,RSR4
Gorai.012G114100.1.D
PCC=0.807, p=0.0
mRNA Profile
Protein Expression Profile
DPA

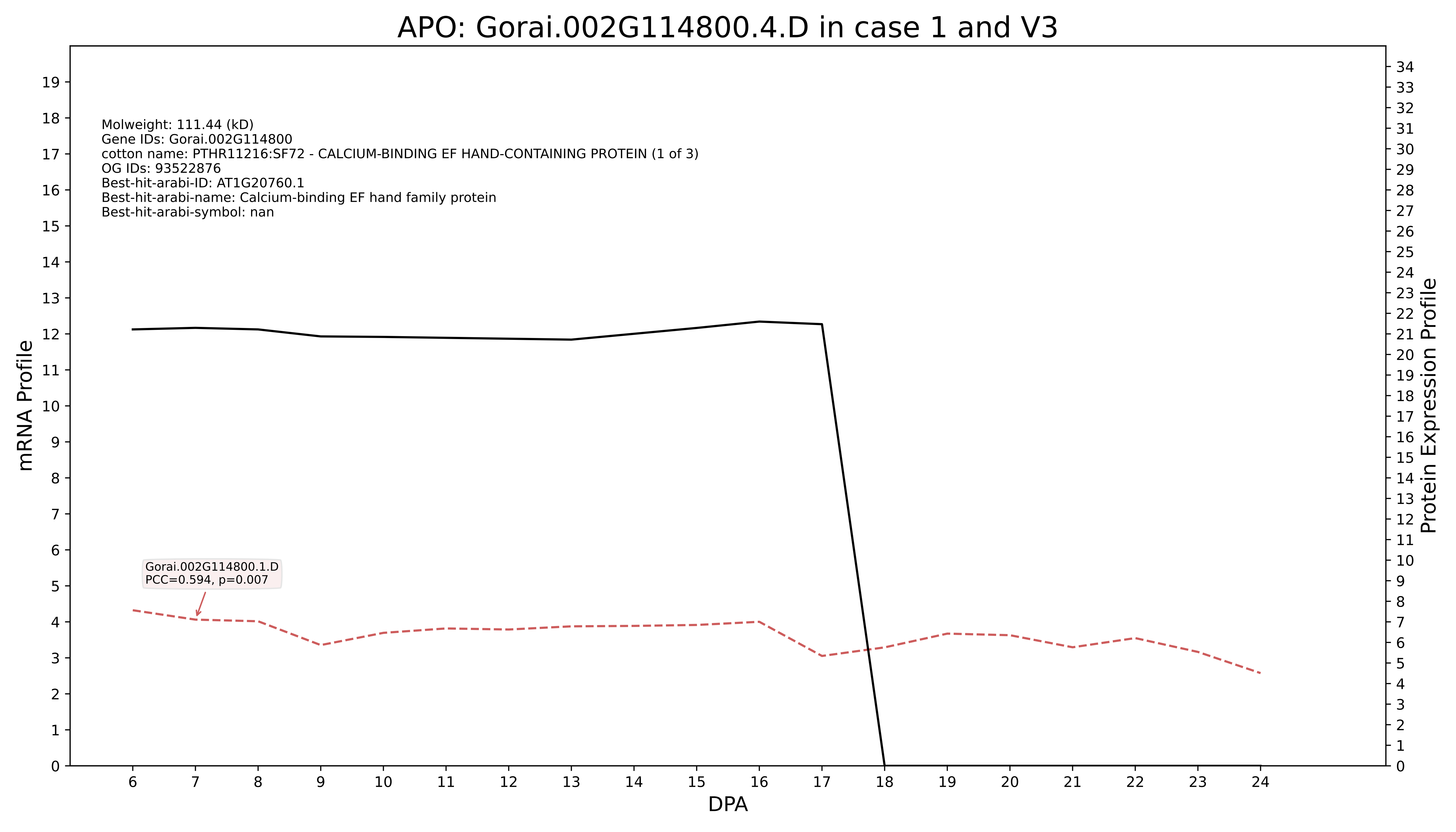

APO: Gorai.002G114800.4.D in case 1 and V3
Molweight: 111.44 (kD)
Gene IDs: Gorai.002G114800
cotton name: PTHR11216:SF72 - CALCIUM-BINDING EF HAND-CONTAINING PROTEIN (1 of 3)
OG IDs: 93522876
Best-hit-arabi-ID: AT1G20760.1
Best-hit-arabi-name: Calcium-binding EF hand family protein
Best-hit-arabi-symbol: nan
Gorai.002G114800.1.D
PCC=0.594, p=0.007
mRNA Profile
Protein Expression Profile
DPA

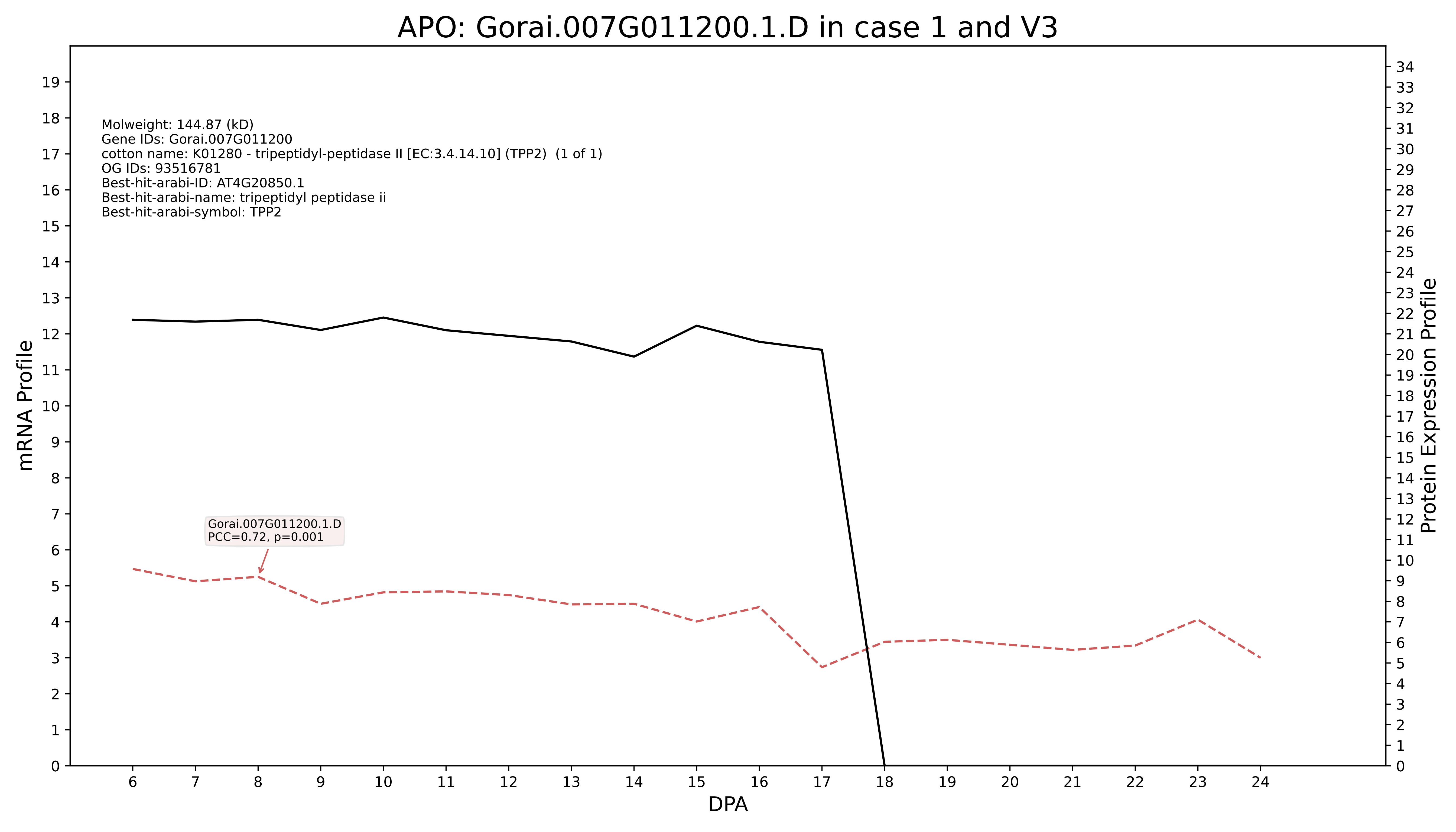
APO: Gorai.007G011200.1.D in case 1 and V3
Molweight: 144.87 (kD)
Gene IDs: Gorai.007G011200
cotton name: K01280 - tripeptidyl-peptidase II [EC:3.4.14.10] (TPP2) (1 of 1)
OG IDs: 93516781
Best-hit-arabi-ID: AT4G20850.1
Best-hit-arabi-name: tripeptidyl peptidase ii
Best-hit-arabi-symbol: TPP2
Gorai.007G011200.1.D
PCC=0.72, p=0.001
mRNA Profile
Protein Expression Profile
DPA

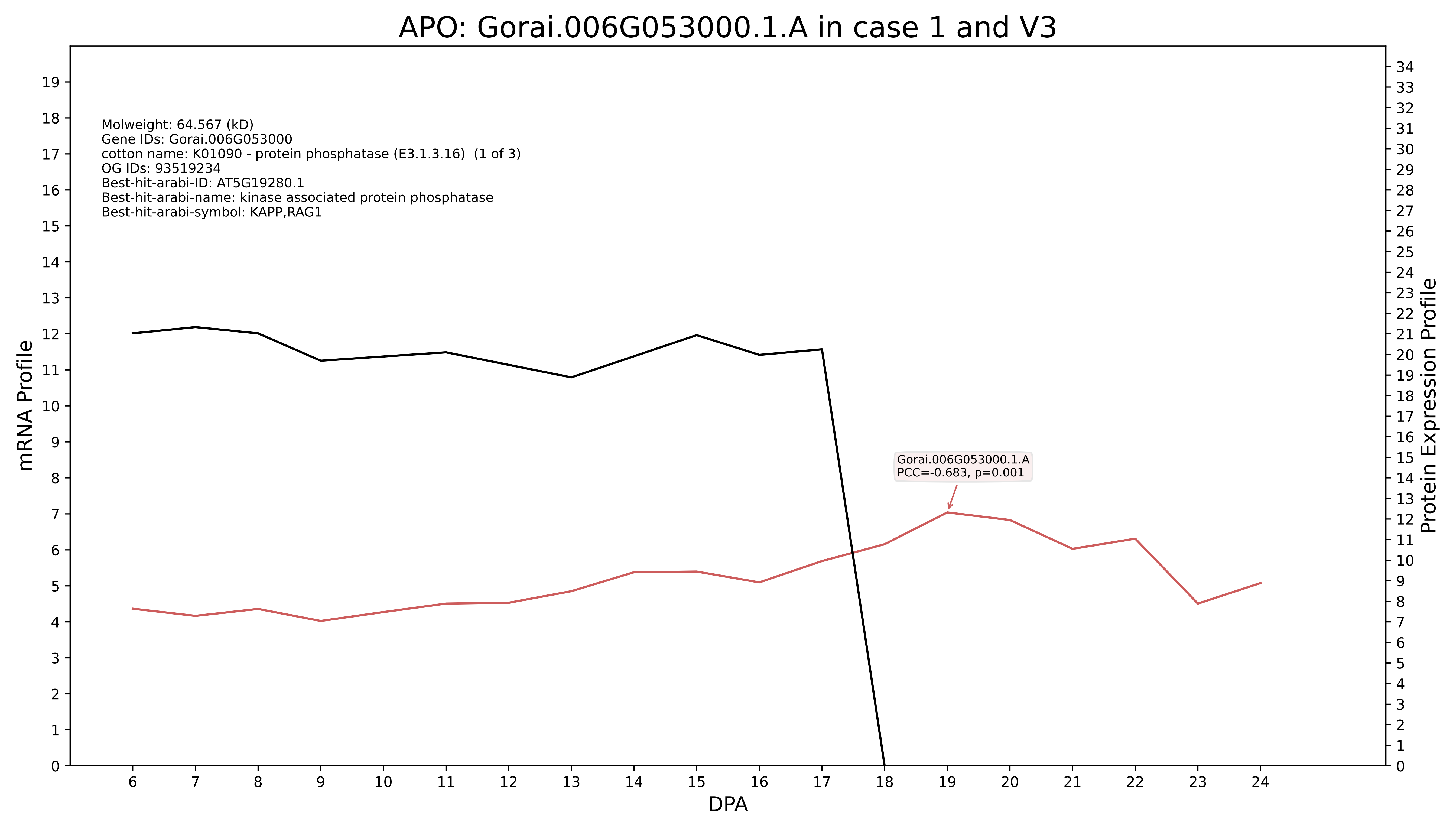

APO: Gorai.006G053000.1.A in case 1 and V3
Molweight: 64.567 (kD)
Gene IDs: Gorai.006G053000
cotton name: K01090 - protein phosphatase (E3.1.3.16) (1 of 3)
OG IDs: 93519234
Best-hit-arabi-ID: AT5G19280.1
Best-hit-arabi-name: kinase associated protein phosphatase
Best-hit-arabi-symbol: KAPP,RAG1
Gorai.006G053000.1.A
PCC=-0.683, p=0.001
mRNA Profile
Protein Expression Profile
DPA

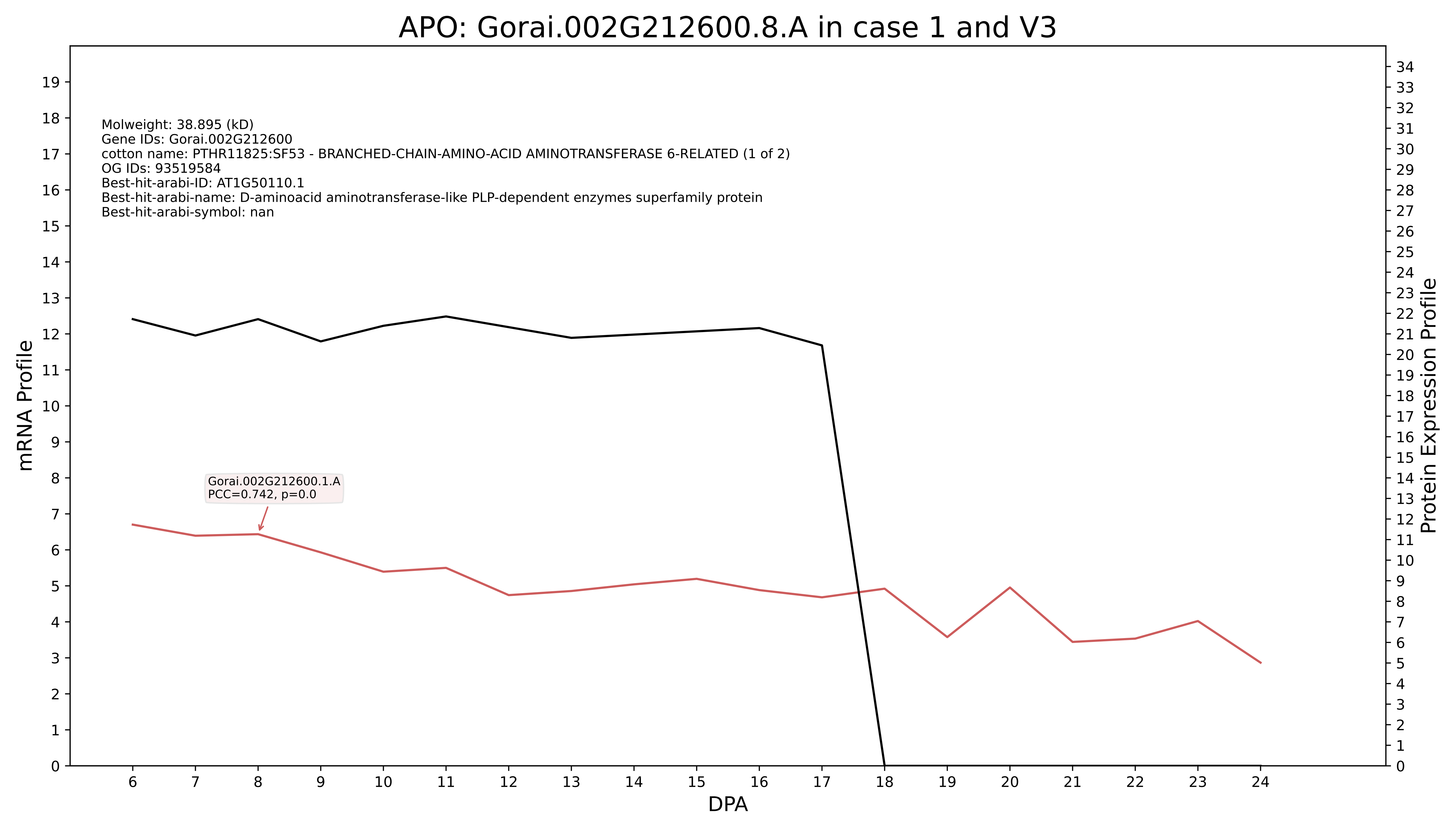

APO: Gorai.002G212600.8.A in case 1 and V3
Molweight: 38.895 (kD)
Gene IDs: Gorai.002G212600
cotton name: PTHR11825:SF53 - BRANCHED-CHAIN-AMINO-ACID AMINOTRANSFERASE 6-RELATED (1 of 2)
OG IDs: 93519584
Best-hit-arabi-ID: AT1G50110.1
Best-hit-arabi-name: D-aminoacid aminotransferase-like PLP-dependent enzymes superfamily protein
Best-hit-arabi-symbol: nan
Gorai.002G212600.1.A
PCC=0.742, p=0.0
mRNA Profile
Protein Expression Profile
DPA

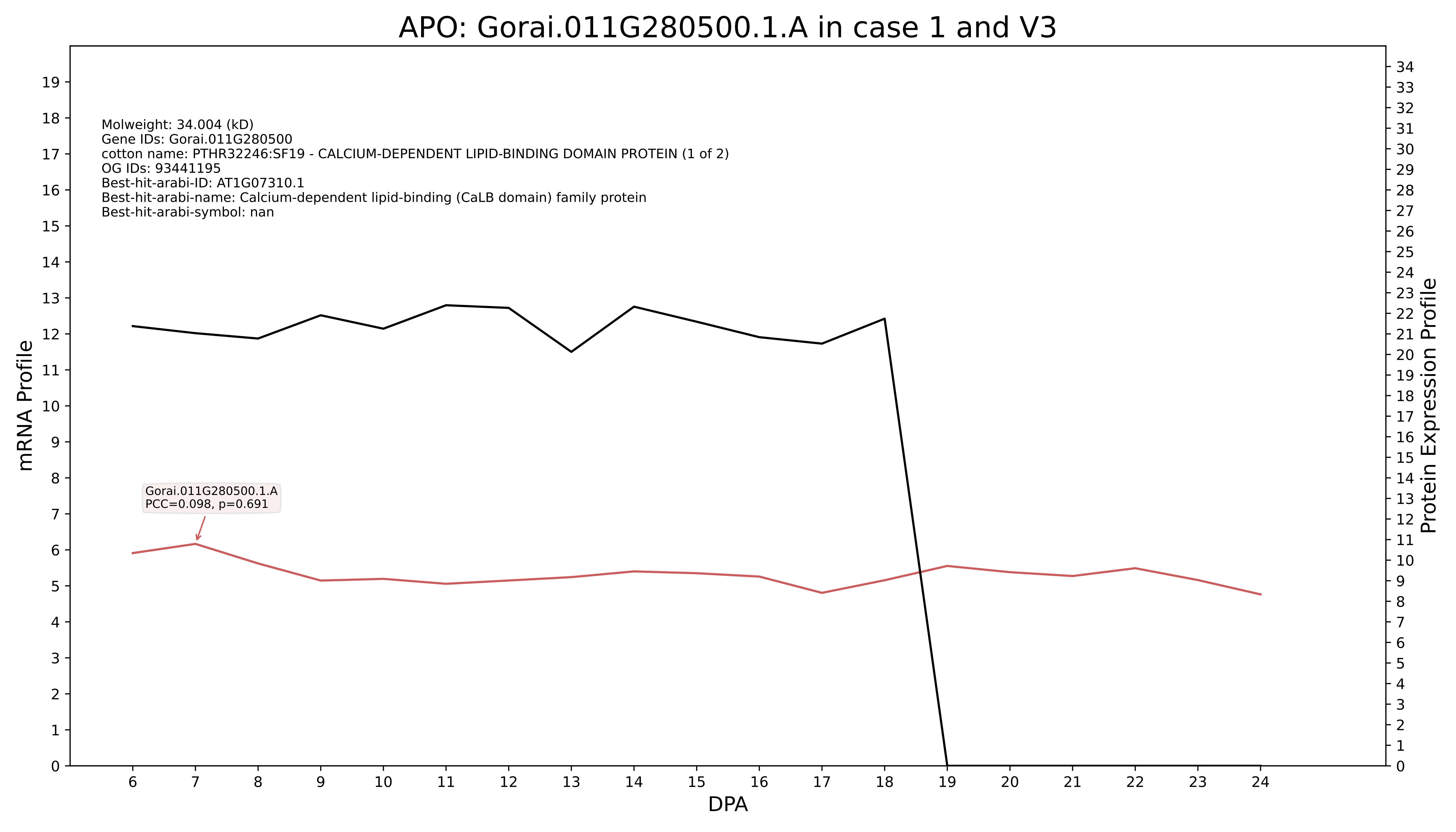

APO: Gorai.011G280500.1.A in case 1 and V3
Molweight: 34.004 (kD)
Gene IDs: Gorai.011G280500
cotton name: PTHR32246:SF19 - CALCIUM-DEPENDENT LIPID-BINDING DOMAIN PROTEIN (1 of 2)
OG IDs: 93441195
Best-hit-arabi-ID: AT1G07310.1
Best-hit-arabi-name: Calcium-dependent lipid-binding (CaLB domain) family protein
Best-hit-arabi-symbol: nan
Gorai.011G280500.1.A
PCC=0.098, p=0.691
mRNA Profile
Protein Expression Profile
DPA

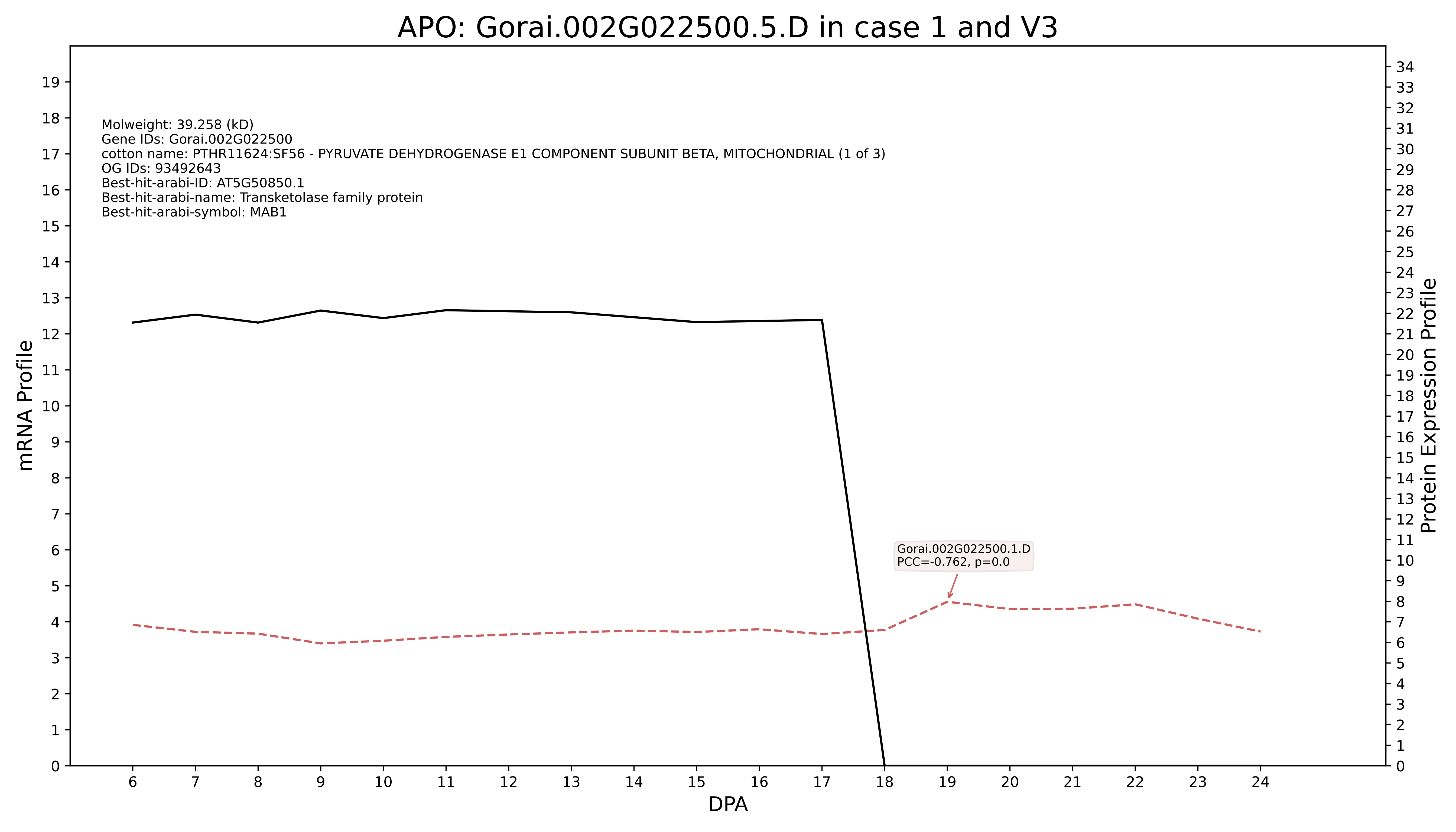
APO: Gorai.002G022500.5.D in case 1 and V3
Molweight: 39.258 (kD)
Gene IDs: Gorai.002G022500
cotton name: PTHR11624:SF56 - PYRUVATE DEHYDROGENASE E1 COMPONENT SUBUNIT BETA, MITOCHONDRIAL (1 of 3)
OG IDs: 93492643
Best-hit-arabi-ID: AT5G50850.1
Best-hit-arabi-name: Transketolase family protein
Best-hit-arabi-symbol: MAB1
Gorai.002G022500.1.D
PCC=-0.762, p=0.0
mRNA Profile
Protein Expression Profile
DPA

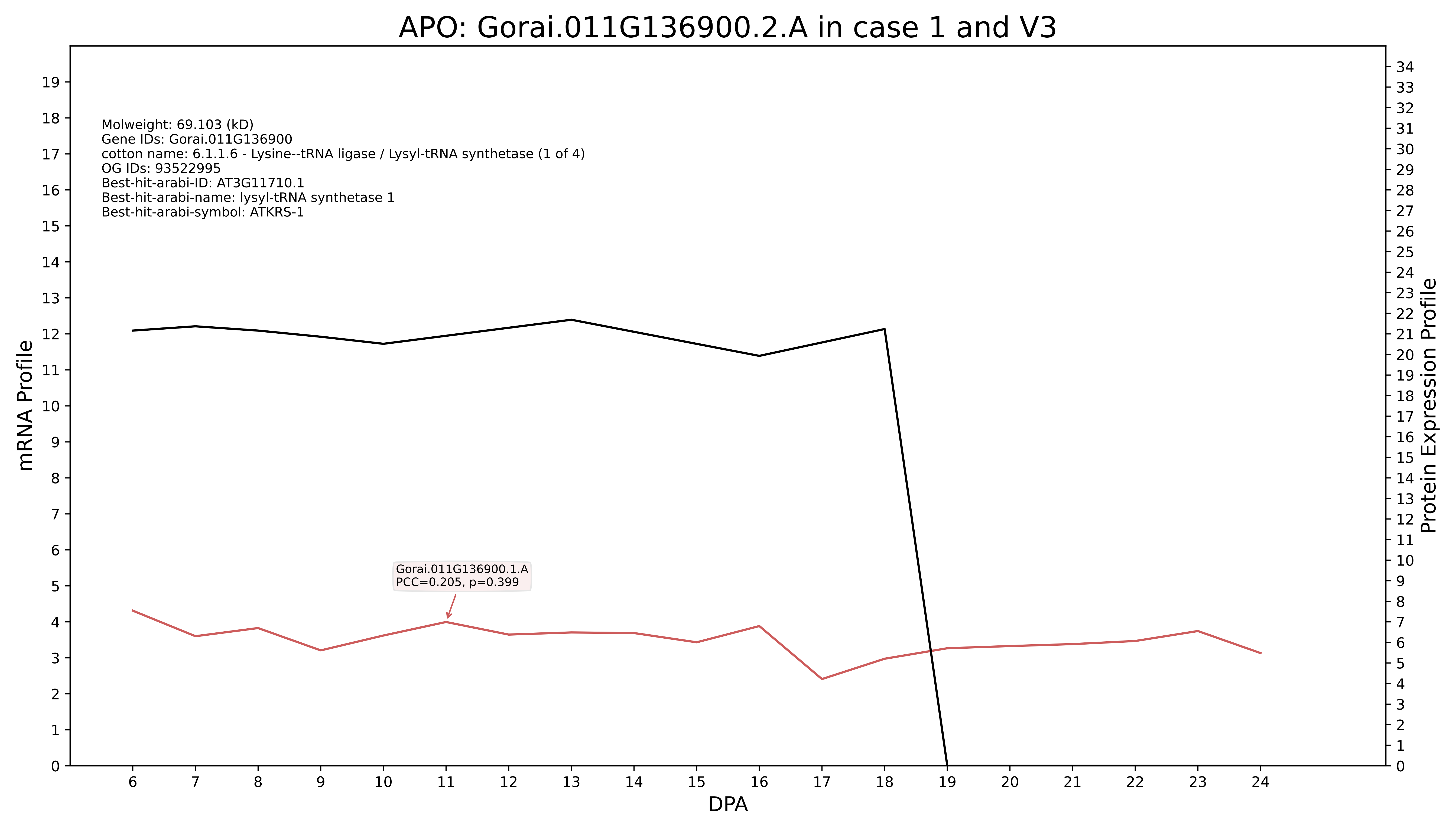
APO: Gorai.011G136900.2.A in case 1 and V3
Molweight: 69.103 (kD)
Gene IDs: Gorai.011G136900
cotton name: 6.1.1.6 - Lysine--tRNA ligase / Lysyl-tRNA synthetase (1 of 4)
OG IDs: 93522995
Best-hit-arabi-ID: AT3G11710.1
Best-hit-arabi-name: lysyl-tRNA synthetase 1
Best-hit-arabi-symbol: ATKRS-1
Gorai.011G136900.1.A
PCC=0.205, p=0.399
mRNA Profile
Protein Expression Profile
DPA

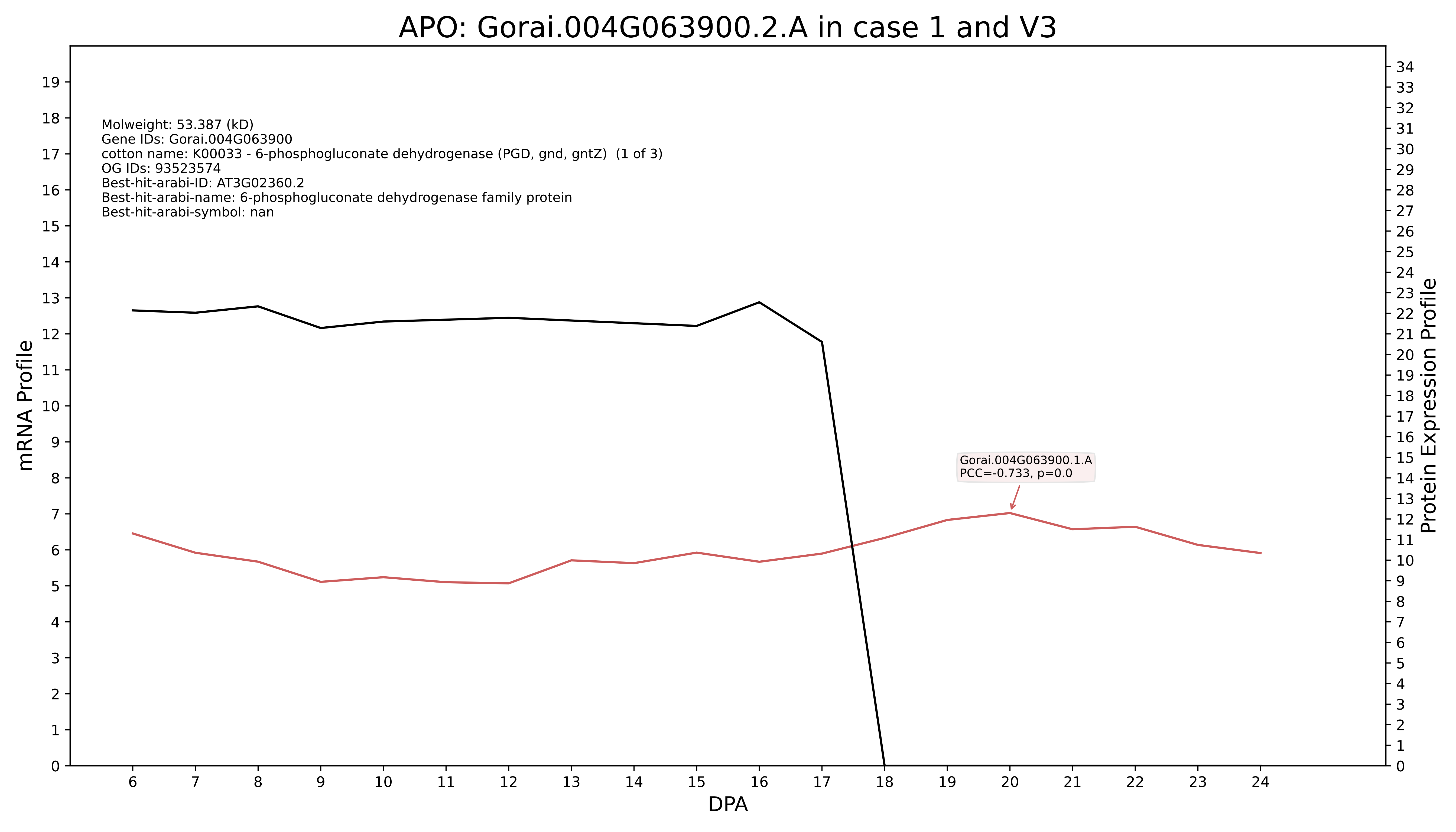
APO: Gorai.004G063900.2.A in case 1 and V3
Molweight: 53.387 (kD)
Gene IDs: Gorai.004G063900
cotton name: K00033 - 6-phosphogluconate dehydrogenase (PGD, gnd, gntZ) (1 of 3)
OG IDs: 93523574
Best-hit-arabi-ID: AT3G02360.2
Best-hit-arabi-name: 6-phosphogluconate dehydrogenase family protein
Best-hit-arabi-symbol: nan
Gorai.004G063900.1.A
PCC=-0.733, p=0.0
mRNA Profile
Protein Expression Profile
DPA

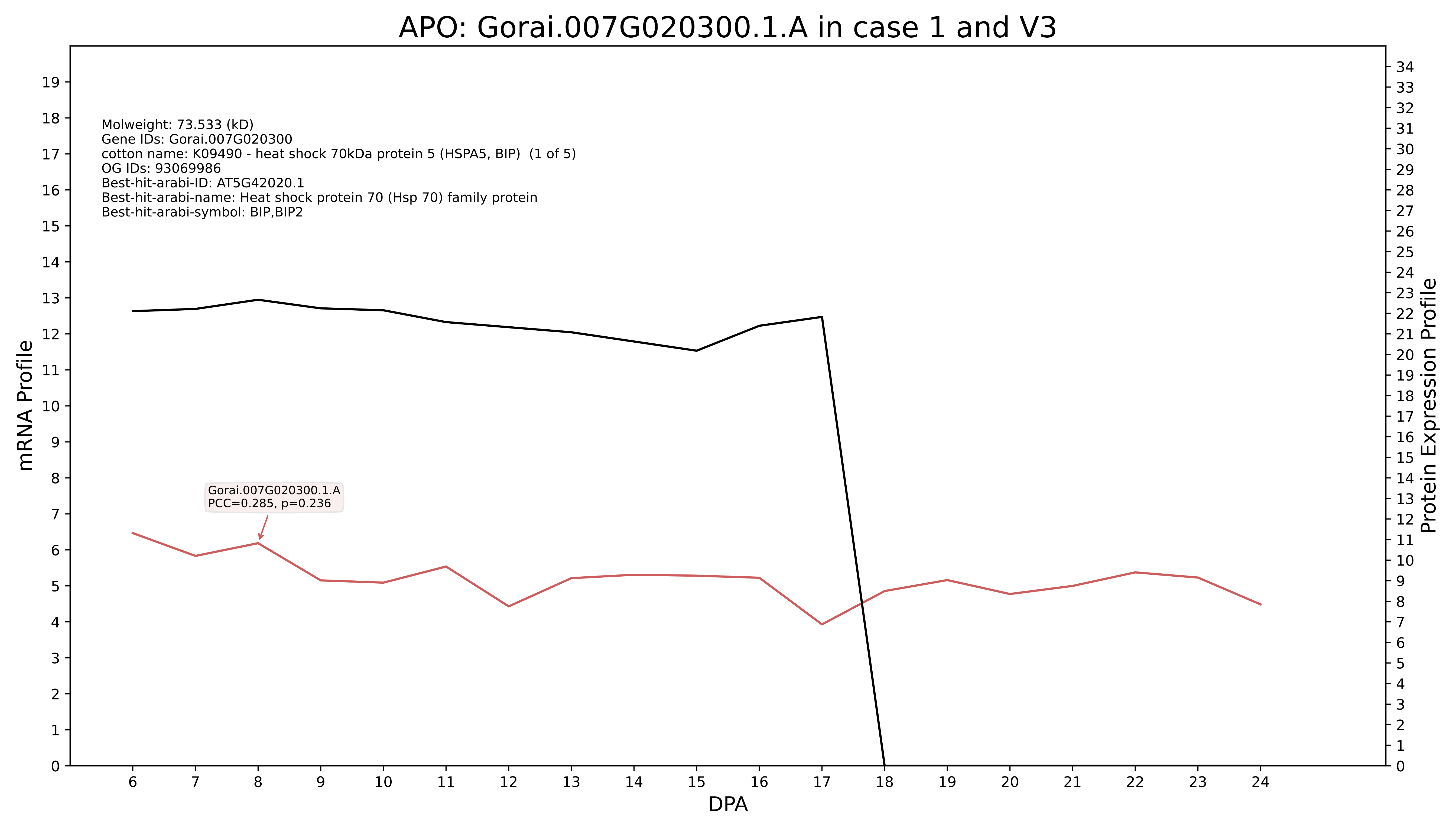
APO: Gorai.007G020300.1.A in case 1 and V3
Molweight: 73.533 (kD)
Gene IDs: Gorai.007G020300
cotton name: K09490 - heat shock 70kDa protein 5 (HSPA5, BIP) (1 of 5)
OG IDs: 93069986
Best-hit-arabi-ID: AT5G42020.1
Best-hit-arabi-name: Heat shock protein 70 (Hsp 70) family protein
Best-hit-arabi-symbol: BIP,BIP2
Gorai.007G020300.1.A
PCC=0.285, p=0.236
mRNA Profile
Protein Expression Profile
DPA

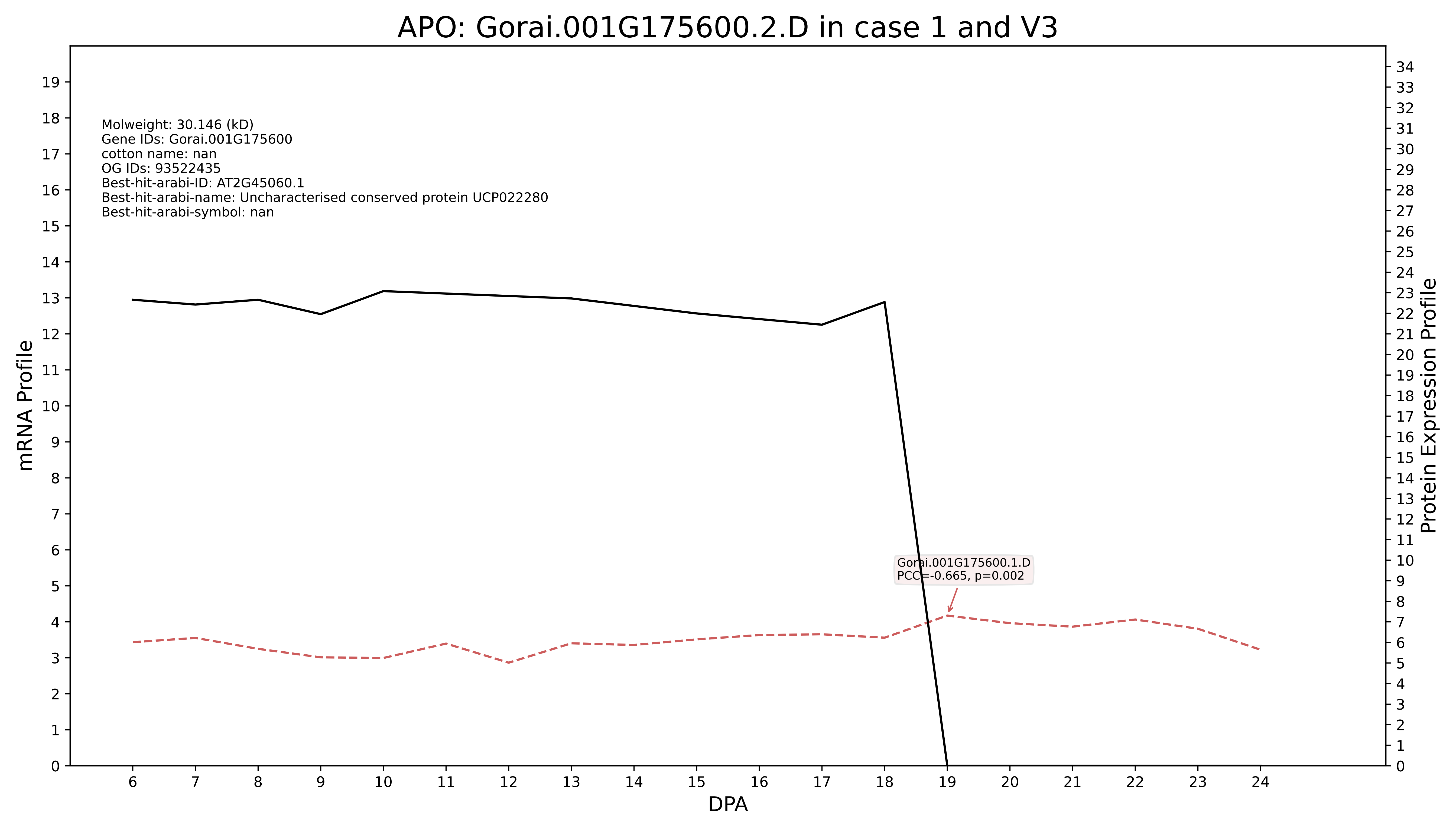
APO: Gorai.001G175600.2.D in case 1 and V3
Molweight: 30.146 (kD)
Gene IDs: Gorai.001G175600
cotton name: nan
OG IDs: 93522435
Best-hit-arabi-ID: AT2G45060.1
Best-hit-arabi-name: Uncharacterised conserved protein UCP022280
Best-hit-arabi-symbol: nan
Gorai.001G175600.1.D
PCC=-0.665, p=0.002
mRNA Profile
Protein Expression Profile
DPA

### APO: Gorai.002G221000.2.D in case 1 and V3

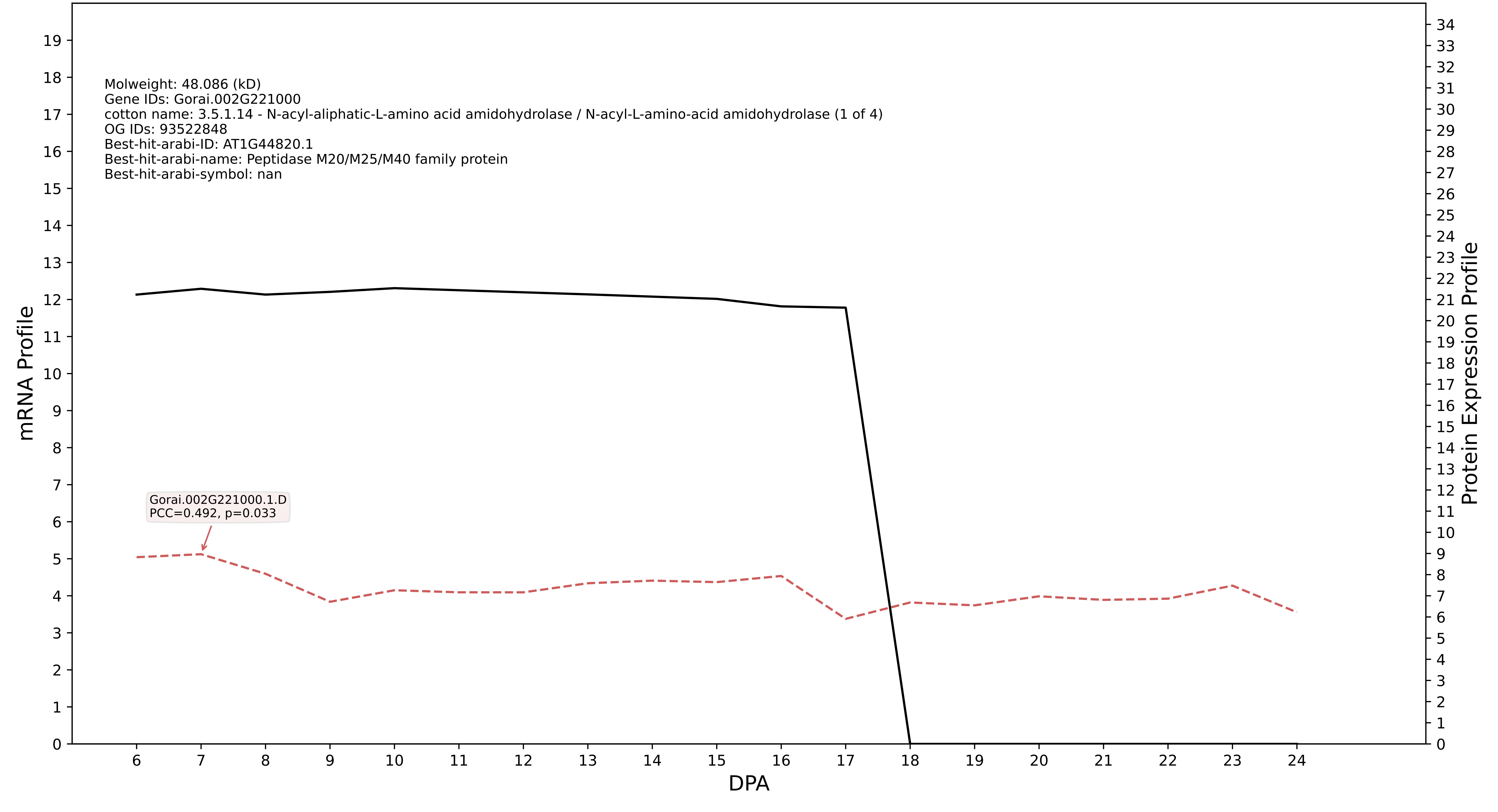

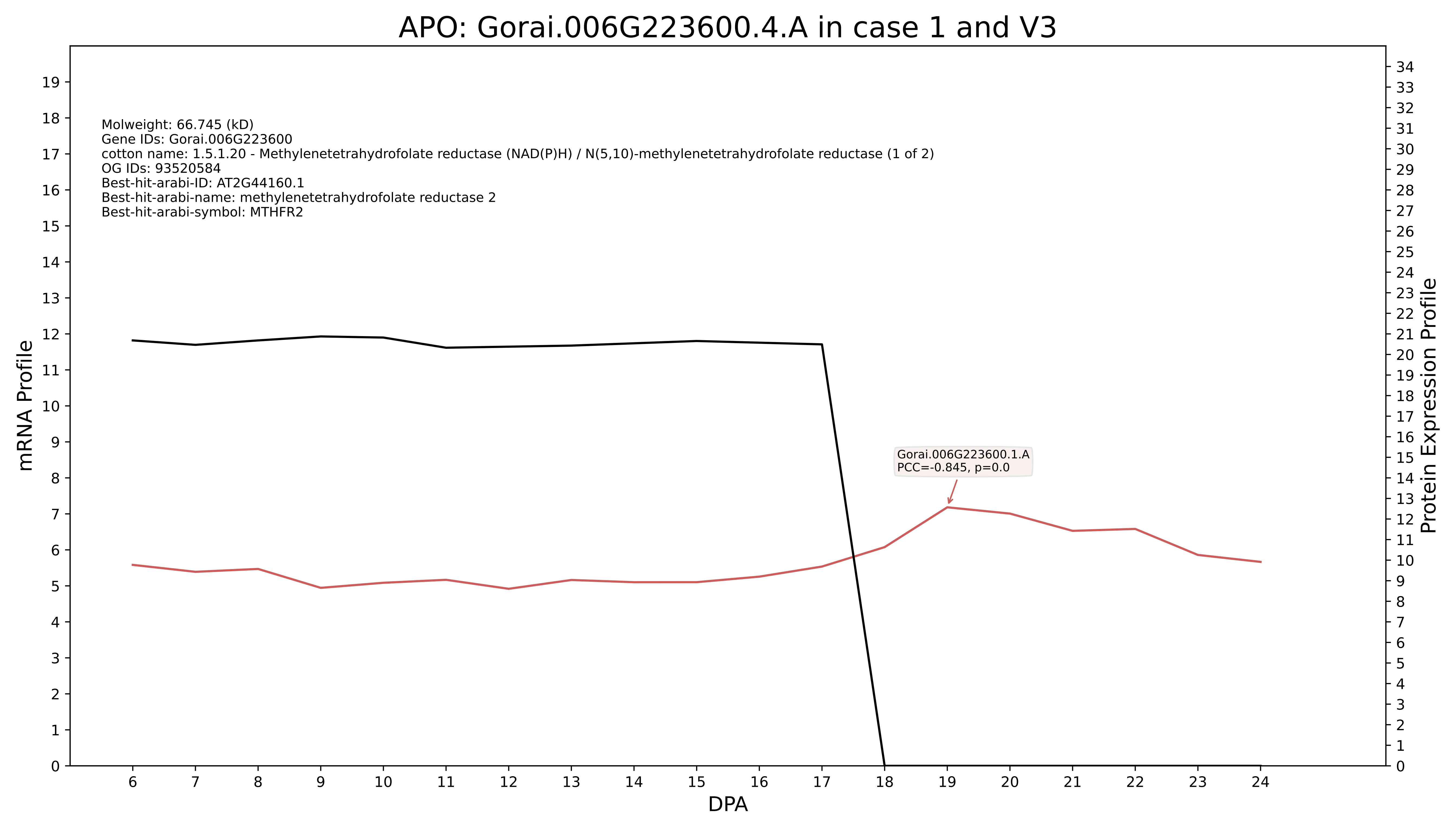
APO: Gorai.006G223600.4.A in case 1 and V3
Molweight: 66.745 (kD)
Gene IDs: Gorai.006G223600
cotton name: 1.5.1.20 - Methylenetetrahydrofolate reductase (NAD(P)H) / N(5,10)-methylenetetrahydrofolate reductase (1 of 2)
OG IDs: 93520584
Best-hit-arabi-ID: AT2G44160.1
Best-hit-arabi-name: methylenetetrahydrofolate reductase 2
Best-hit-arabi-symbol: MTHFR2
Gorai.006G223600.1.A
PCC=-0.845, p=0.0
mRNA Profile
Protein Expression Profile
DPA

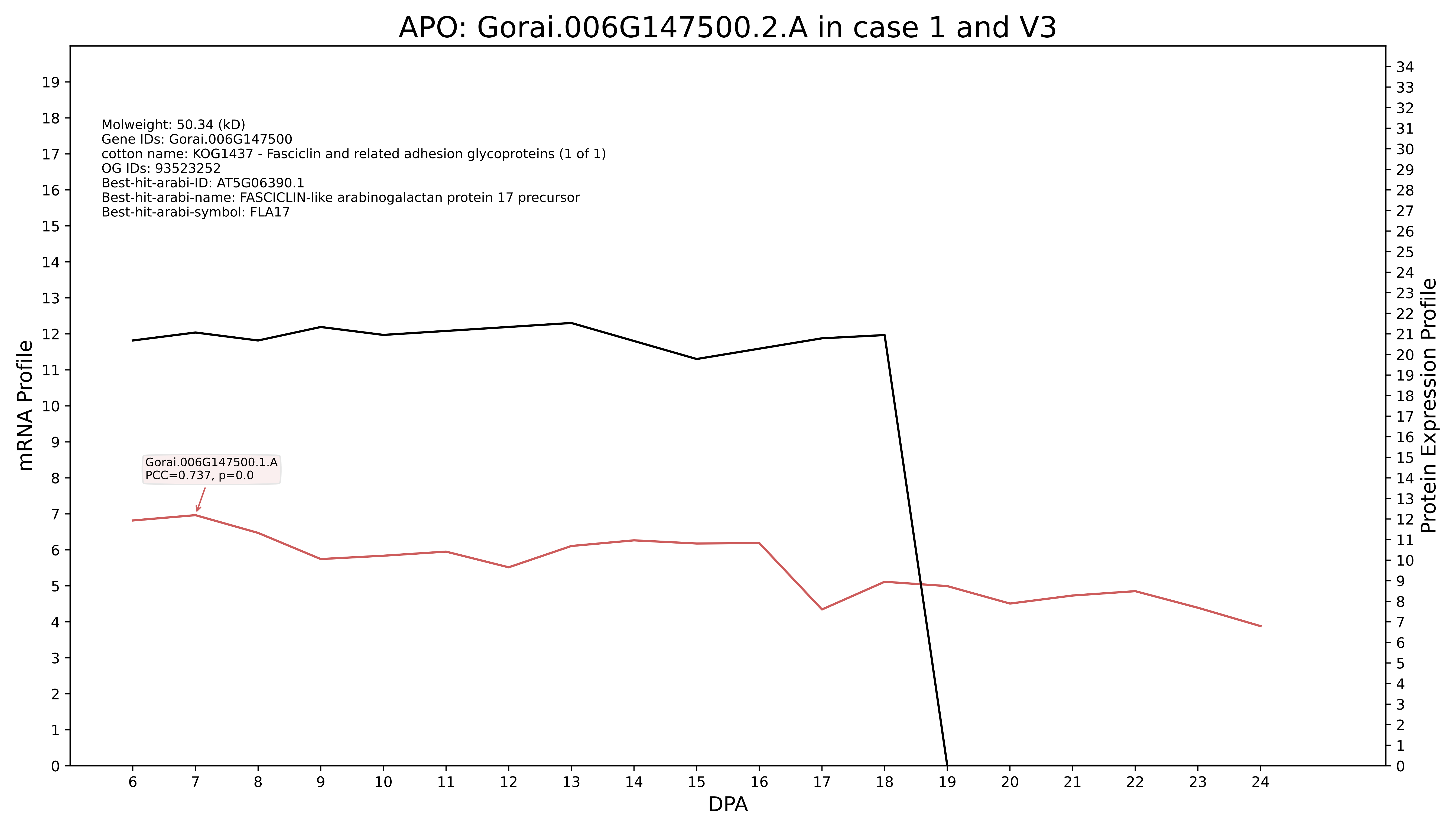
APO: Gorai.006G147500.2.A in case 1 and V3
Molweight: 50.34 (kD)
Gene IDs: Gorai.006G147500
cotton name: KOG1437 - Fasciclin and related adhesion glycoproteins (1 of 1)
OG IDs: 93523252
Best-hit-arabi-ID: AT5G06390.1
Best-hit-arabi-name: FASCICLIN-like arabinogalactan protein 17 precursor
Best-hit-arabi-symbol: FLA17
Gorai.006G147500.1.A
PCC=0.737, p=0.0
mRNA Profile
Protein Expression Profile
DPA

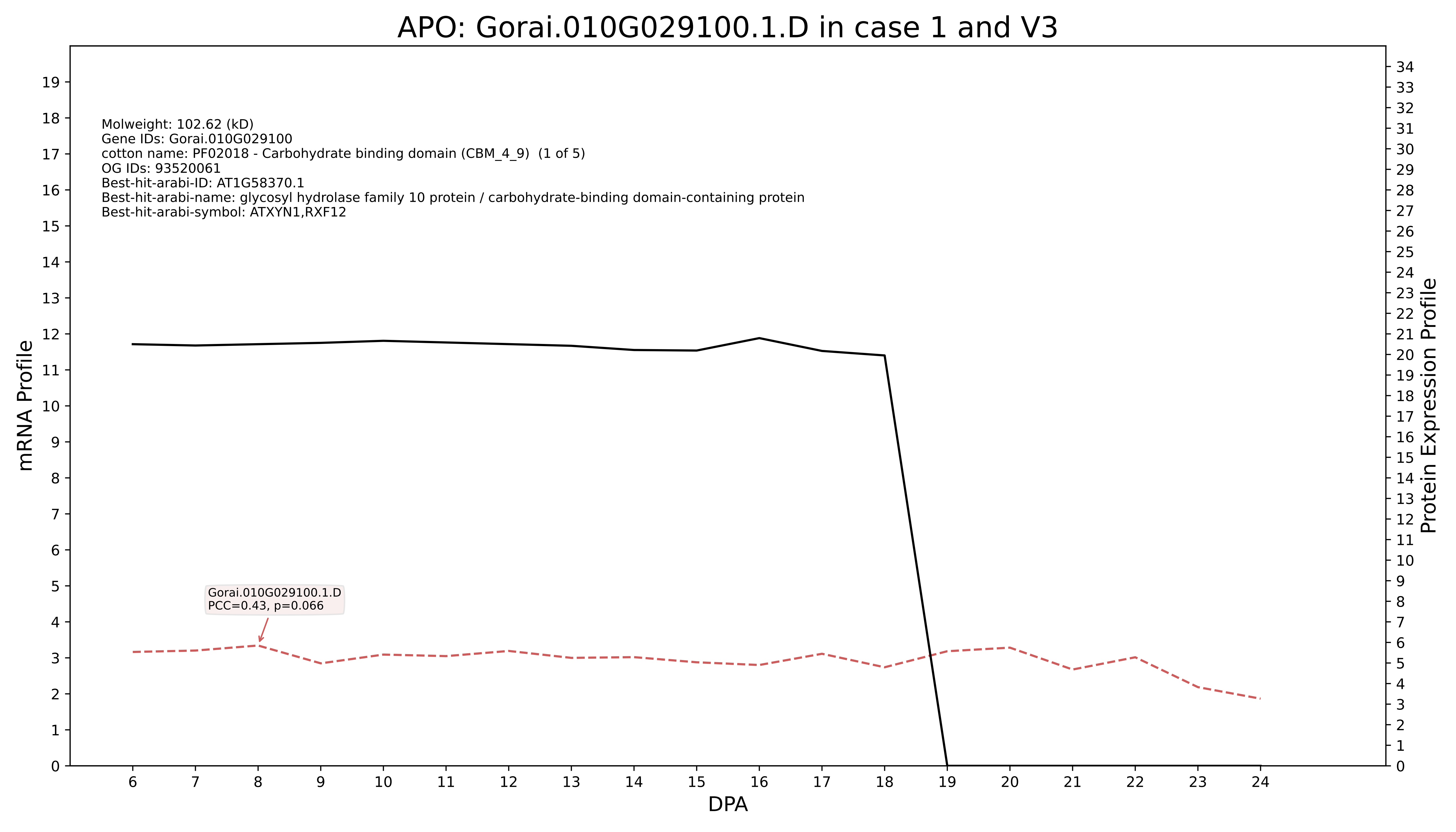
APO: Gorai.010G029100.1.D in case 1 and V3
Molweight: 102.62 (kD)
Gene IDs: Gorai.010G029100
cotton name: PF02018 - Carbohydrate binding domain (CBM_4_9) (1 of 5)
OG IDs: 93520061
Best-hit-arabi-ID: AT1G58370.1
Best-hit-arabi-name: glycosyl hydrolase family 10 protein / carbohydrate-binding domain-containing protein
Best-hit-arabi-symbol: ATXYN1,RXF12
Gorai.010G029100.1.D
PCC=0.43, p=0.066
mRNA Profile
Protein Expression Profile
DPA

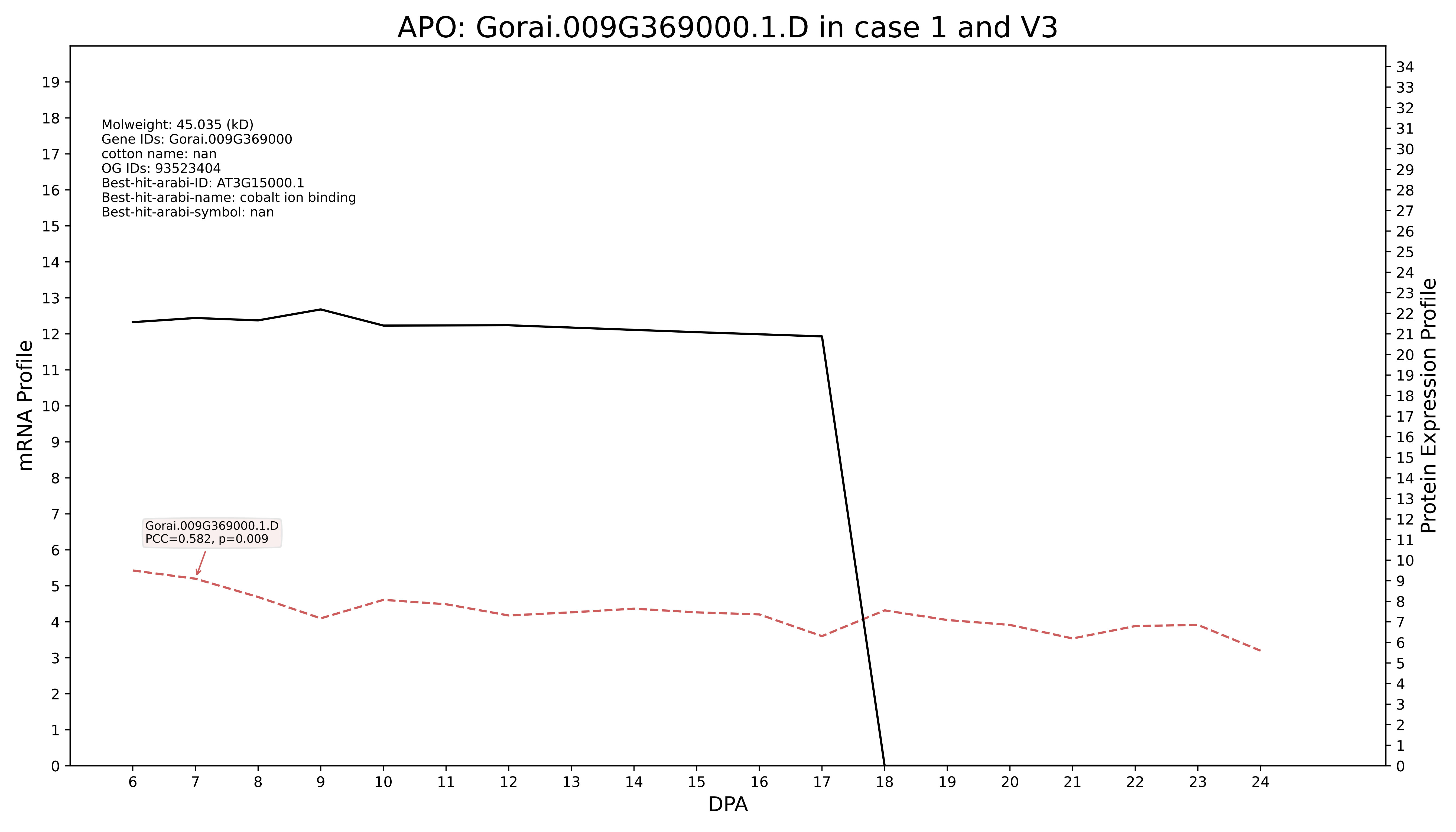
APO: Gorai.009G369000.1.D in case 1 and V3
Molweight: 45.035 (kD)
Gene IDs: Gorai.009G369000
cotton name: nan
OG IDs: 93523404
Best-hit-arabi-ID: AT3G15000.1
Best-hit-arabi-name: cobalt ion binding
Best-hit-arabi-symbol: nan
Gorai.009G369000.1.D
PCC=0.582, p=0.009
mRNA Profile
Protein Expression Profile
DPA

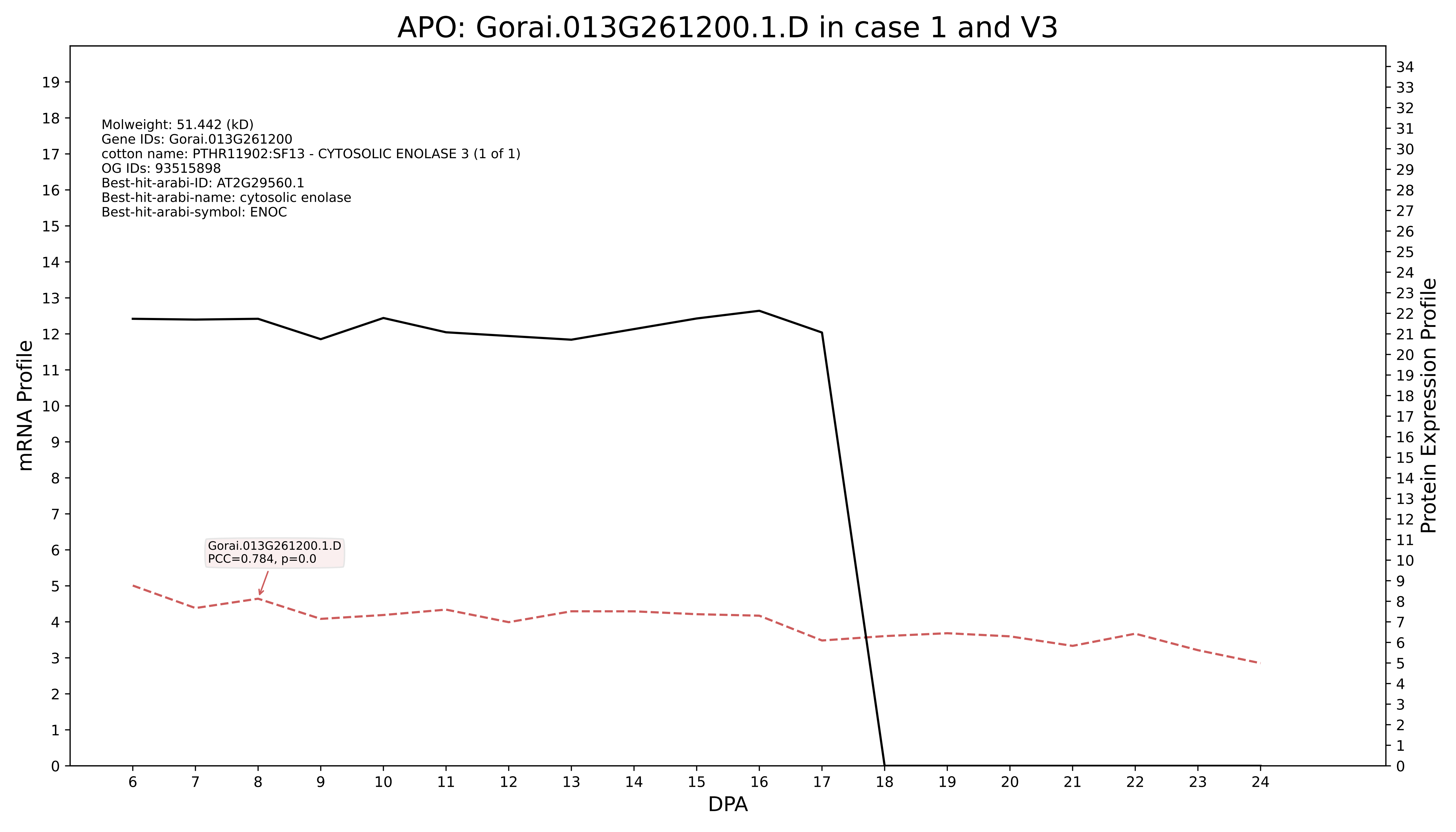
APO: Gorai.013G261200.1.D in case 1 and V3
Molweight: 51.442 (kD)
Gene IDs: Gorai.013G261200
cotton name: PTHR11902:SF13 - CYTOSOLIC ENOLASE 3 (1 of 1)
OG IDs: 93515898
Best-hit-arabi-ID: AT2G29560.1
Best-hit-arabi-name: cytosolic enolase
Best-hit-arabi-symbol: ENOC
Gorai.013G261200.1.D
PCC=0.784, p=0.0
mRNA Profile
Protein Expression Profile
DPA

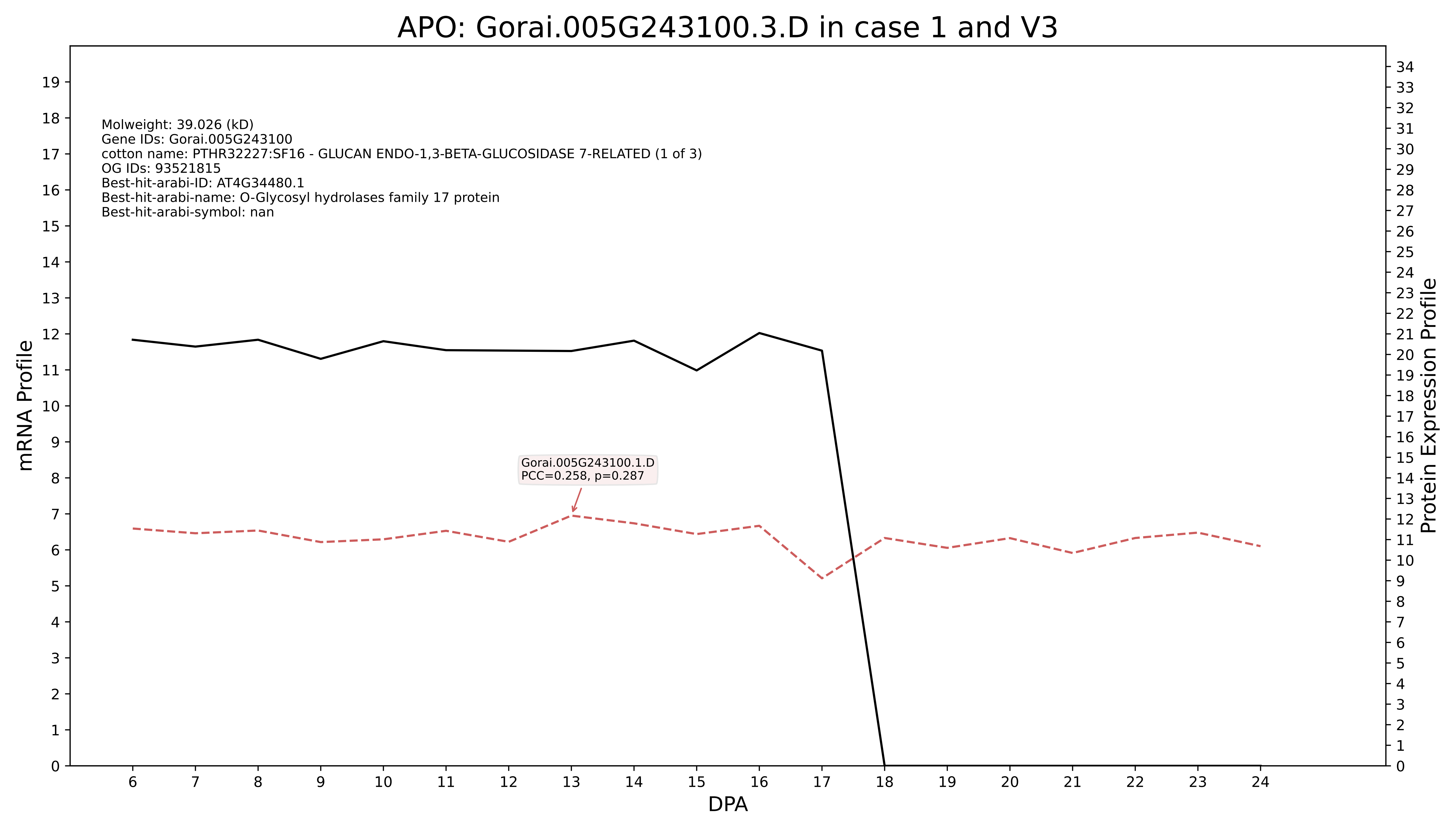
APO: Gorai.005G243100.3.D in case 1 and V3
Molweight: 39.026 (kD)
Gene IDs: Gorai.005G243100
cotton name: PTHR32227:SF16 - GLUCAN ENDO-1,3-BETA-GLUCOSIDASE 7-RELATED (1 of 3)
OG IDs: 93521815
Best-hit-arabi-ID: AT4G34480.1
Best-hit-arabi-name: O-Glycosyl hydrolases family 17 protein
Best-hit-arabi-symbol: nan
Gorai.005G243100.1.D
PCC=0.258, p=0.287
mRNA Profile
Protein Expression Profile
DPA

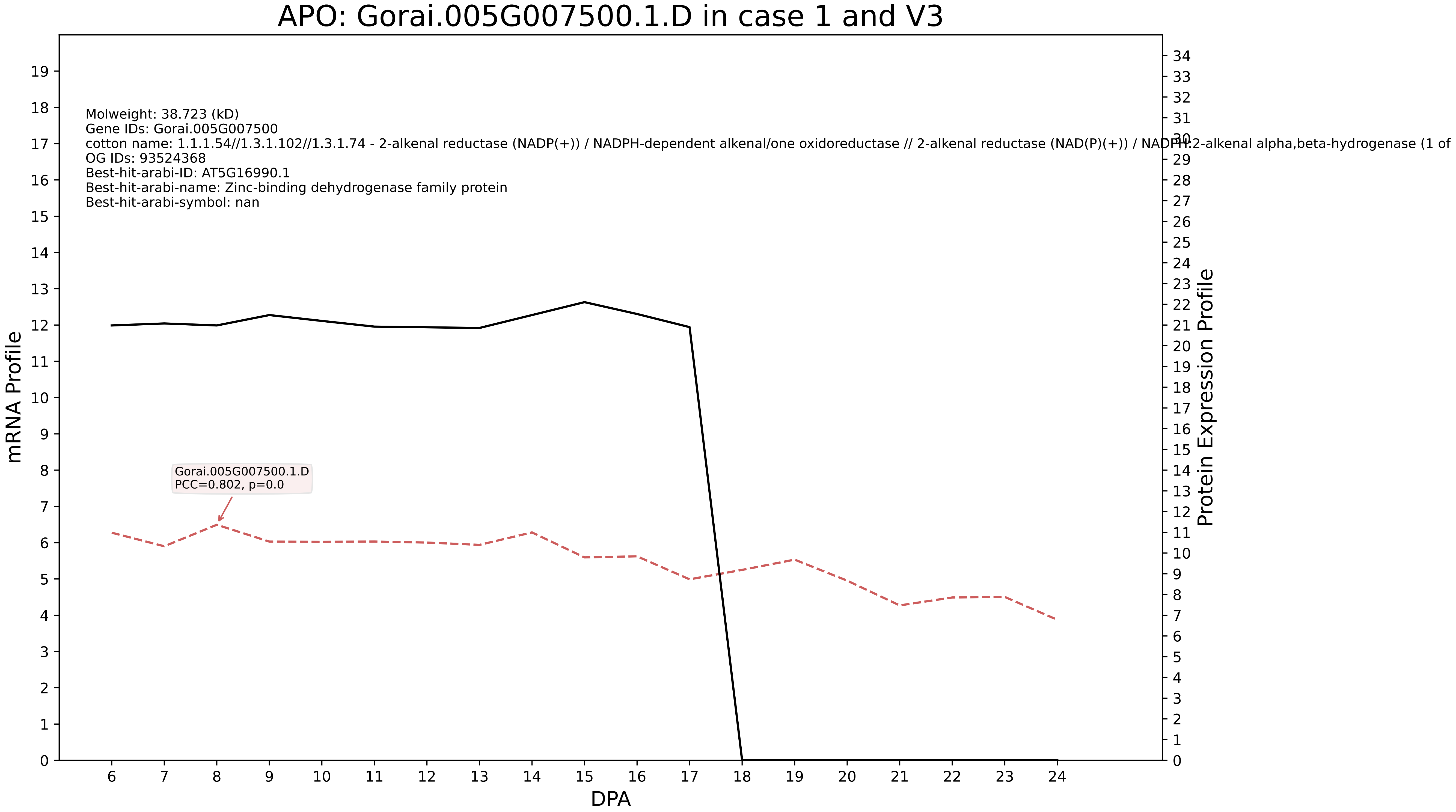
APO: Gorai.005G007500.1.D in case 1 and V3
Molweight: 38.723 (kD)
Gene IDs: Gorai.005G007500
cotton name: 1.1.1.54//1.3.1.102//1.3.1.74 - 2-alkenal reductase (NADP(+)) / NADPH-dependent alkenal/one oxidoreductase // 2-alkenal reductase (NAD(P)(+)) / NADPH:2-alkenal alpha,beta-hydrogenase (1 of
OG IDs: 93524368
Best-hit-arabi-ID: AT5G16990.1
Best-hit-arabi-name: Zinc-binding dehydrogenase family protein
Best-hit-arabi-symbol: nan
Gorai.005G007500.1.D
PCC=0.802, p=0.0
mRNA Profile
Protein Expression Profile
DPA

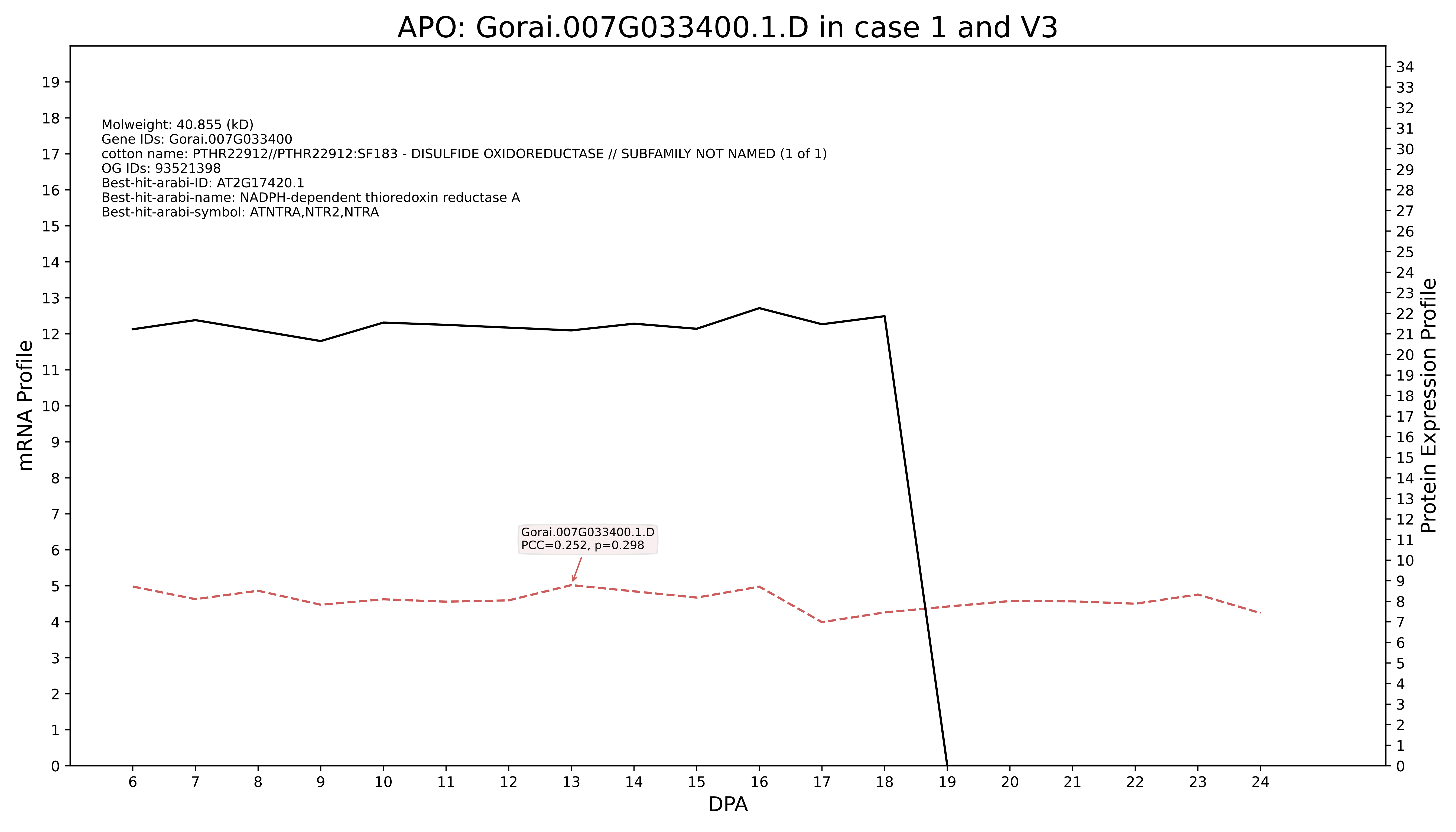

APO: Gorai.007G033400.1.D in case 1 and V3
Molweight: 40.855 (kD)
Gene IDs: Gorai.007G033400
cotton name: PTHR22912//PTHR22912:SF183 - DISULFIDE OXIDOREDUCTASE // SUBFAMILY NOT NAMED (1 of 1)
OG IDs: 93521398
Best-hit-arabi-ID: AT2G17420.1
Best-hit-arabi-name: NADPH-dependent thioredoxin reductase A
Best-hit-arabi-symbol: ATNTRA,NTR2,NTRA
Gorai.007G033400.1.D
PCC=0.252, p=0.298
mRNA Profile
Protein Expression Profile
DPA

### APO: Gorai.008G022400.3.D in case 1 and V3

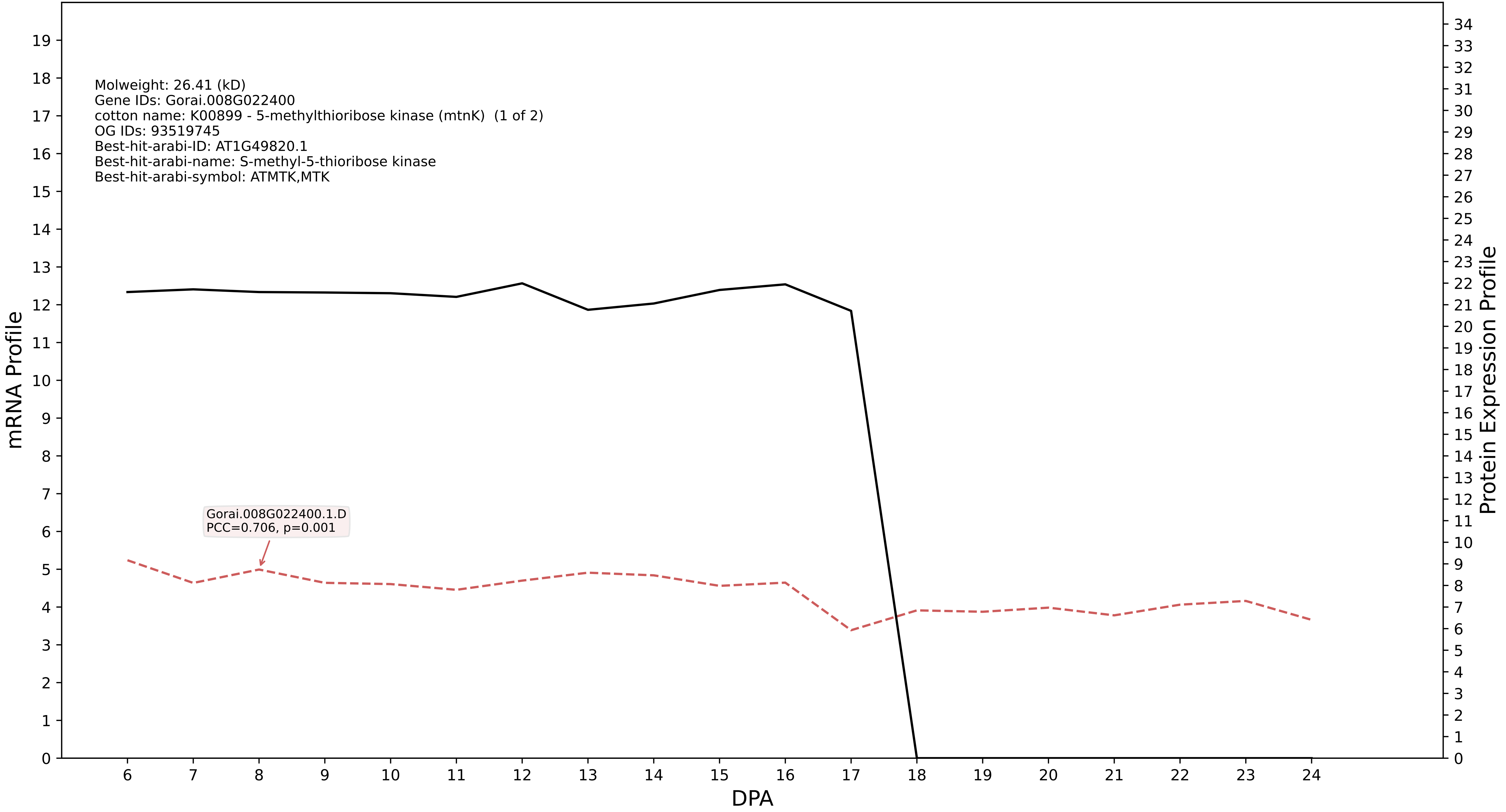

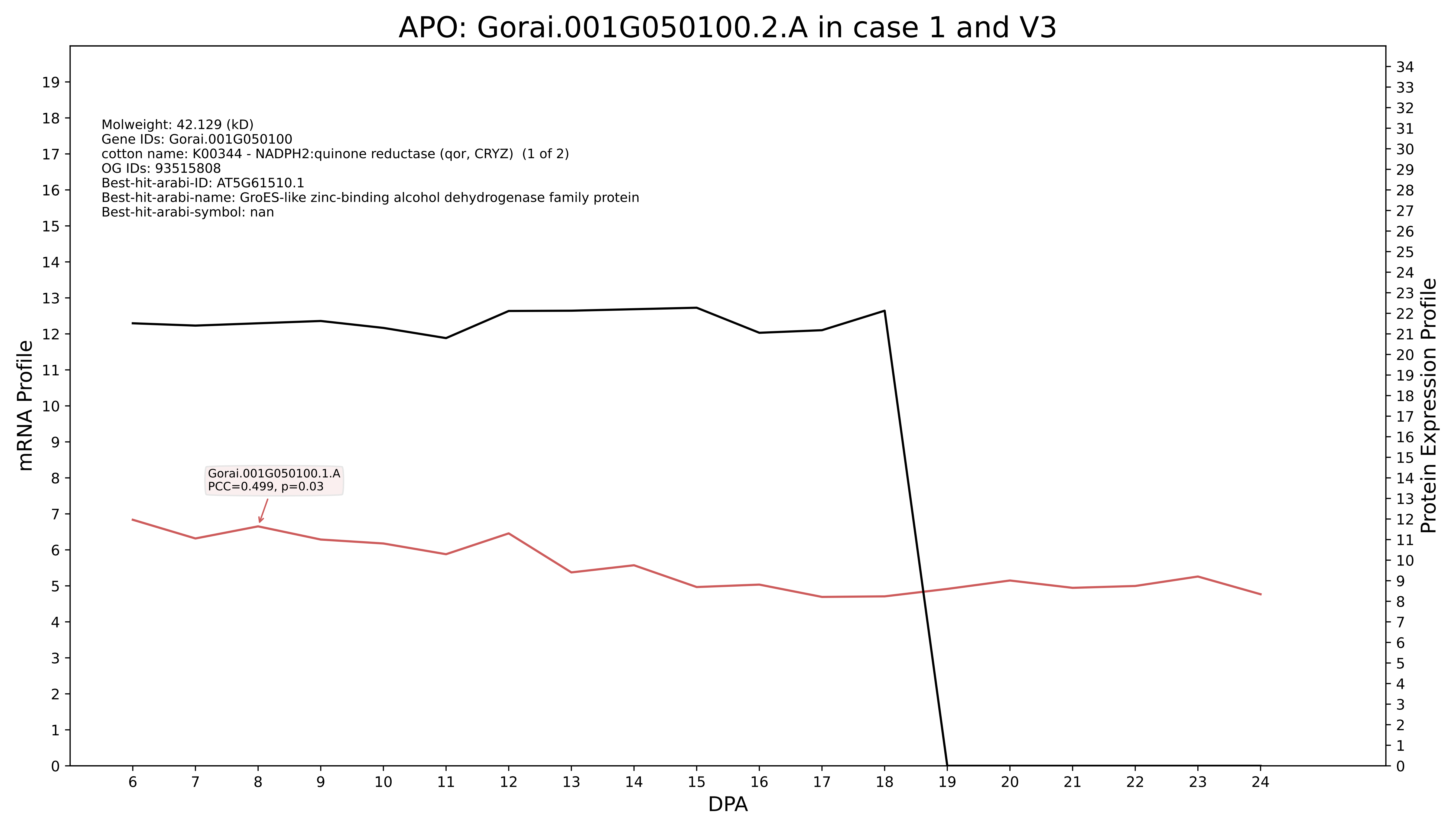
APO: Gorai.001G050100.2.A in case 1 and V3
Molweight: 42.129 (kD)
Gene IDs: Gorai.001G050100
cotton name: K00344 - NADPH2:quinone reductase (qor, CRYZ) (1 of 2)
OG IDs: 93515808
Best-hit-arabi-ID: AT5G61510.1
Best-hit-arabi-name: GroES-like zinc-binding alcohol dehydrogenase family protein
Best-hit-arabi-symbol: nan
Gorai.001G050100.1.A
PCC=0.499, p=0.03
mRNA Profile
Protein Expression Profile
DPA

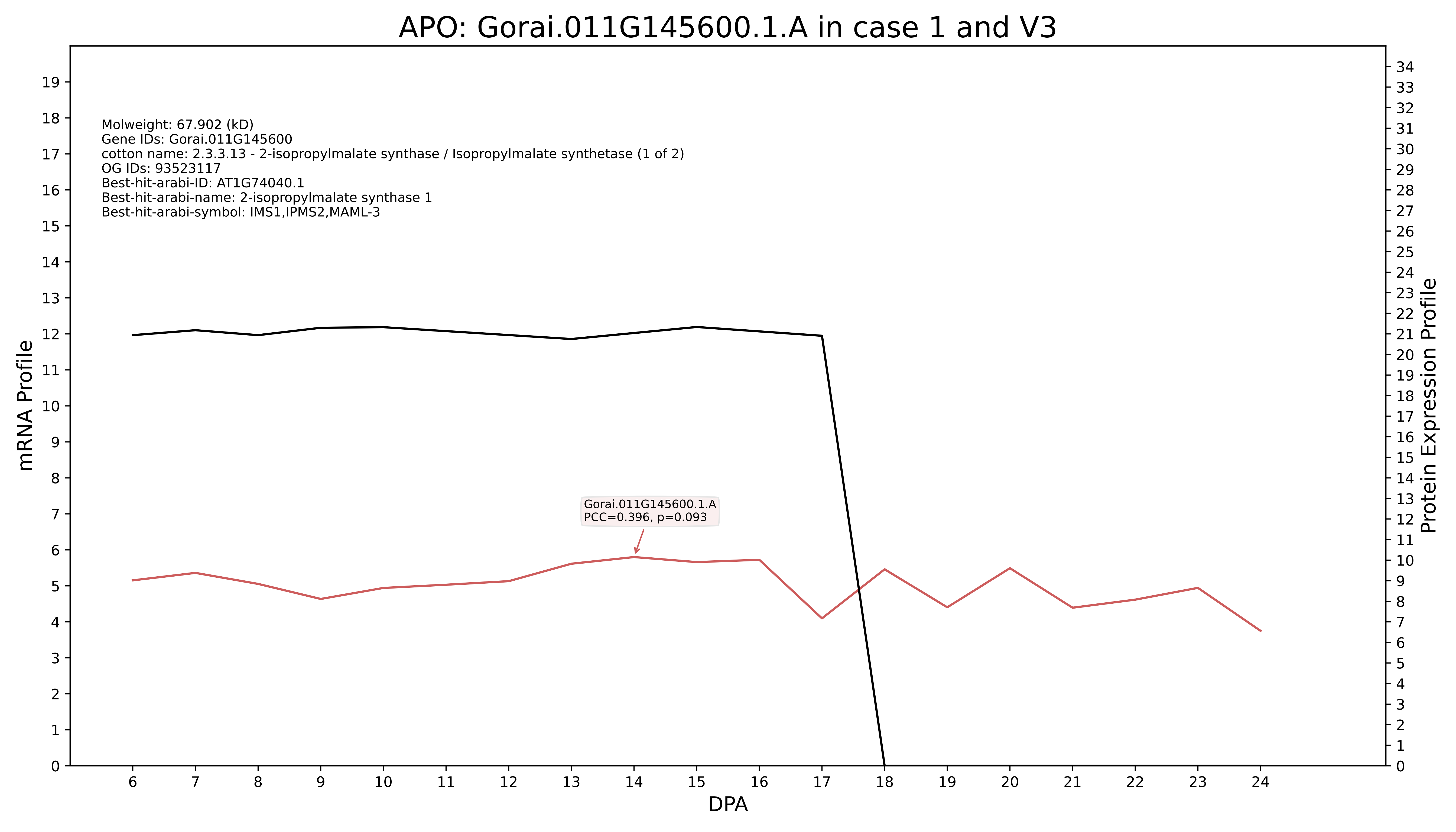

APO: Gorai.011G145600.1.A in case 1 and V3
Molweight: 67.902 (kD)
Gene IDs: Gorai.011G145600
cotton name: 2.3.3.13 - 2-isopropylmalate synthase / Isopropylmalate synthetase (1 of 2)
OG IDs: 93523117
Best-hit-arabi-ID: AT1G74040.1
Best-hit-arabi-name: 2-isopropylmalate synthase 1
Best-hit-arabi-symbol: IMS1,IPMS2,MAML-3
Gorai.011G145600.1.A
PCC=0.396, p=0.093
mRNA Profile
Protein Expression Profile
DPA

### APO: Gorai.002G257900.1.D in case 1 and V3

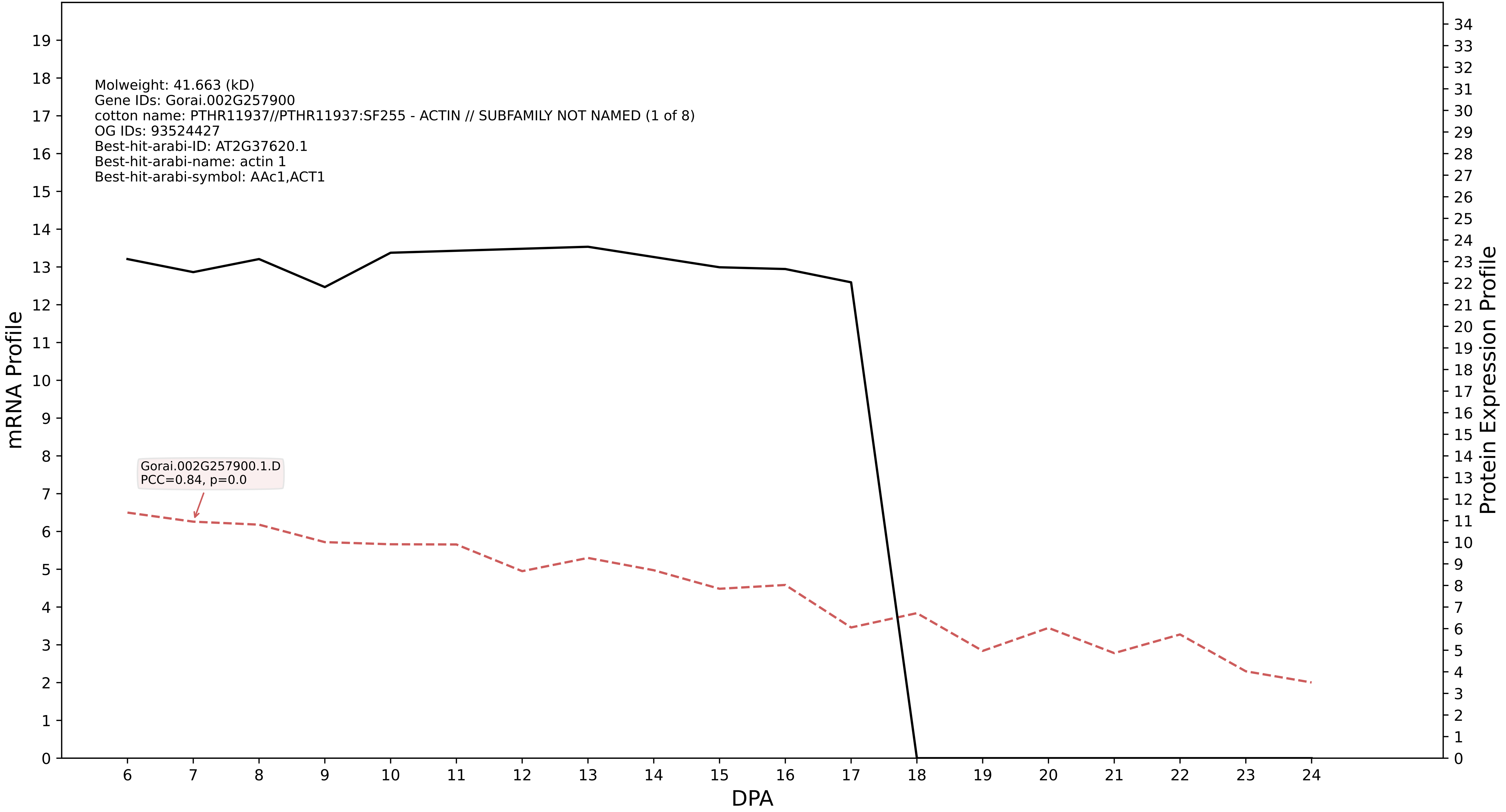

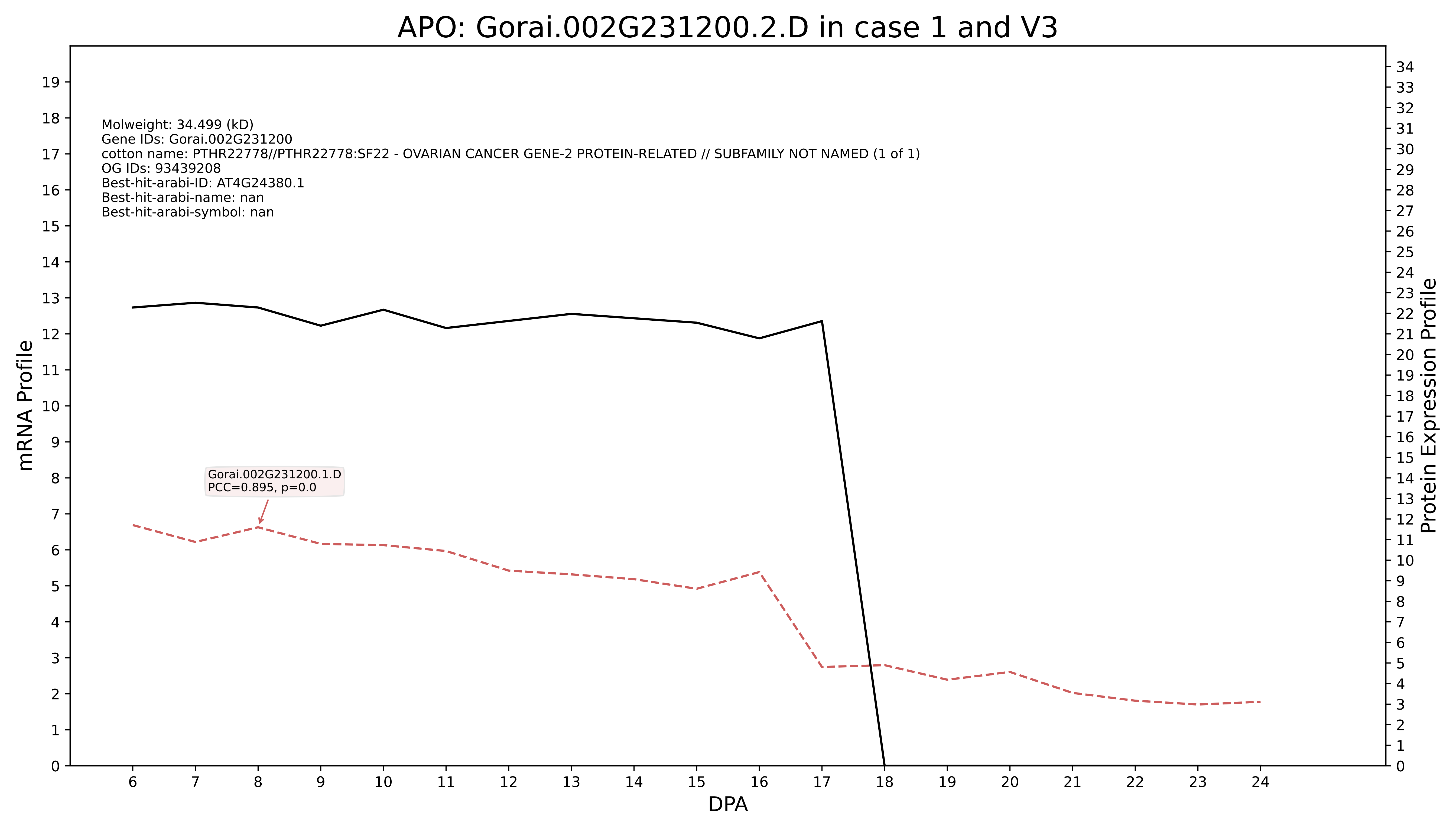
APO: Gorai.002G231200.2.D in case 1 and V3
Molweight: 34.499 (kD)
Gene IDs: Gorai.002G231200
cotton name: PTHR22778//PTHR22778:SF22 - OVARIAN CANCER GENE-2 PROTEIN-RELATED // SUBFAMILY NOT NAMED (1 of 1)
OG IDs: 93439208
Best-hit-arabi-ID: AT4G24380.1
Best-hit-arabi-name: nan
Best-hit-arabi-symbol: nan
Gorai.002G231200.1.D
PCC=0.895, p=0.0
mRNA Profile
Protein Expression Profile
DPA

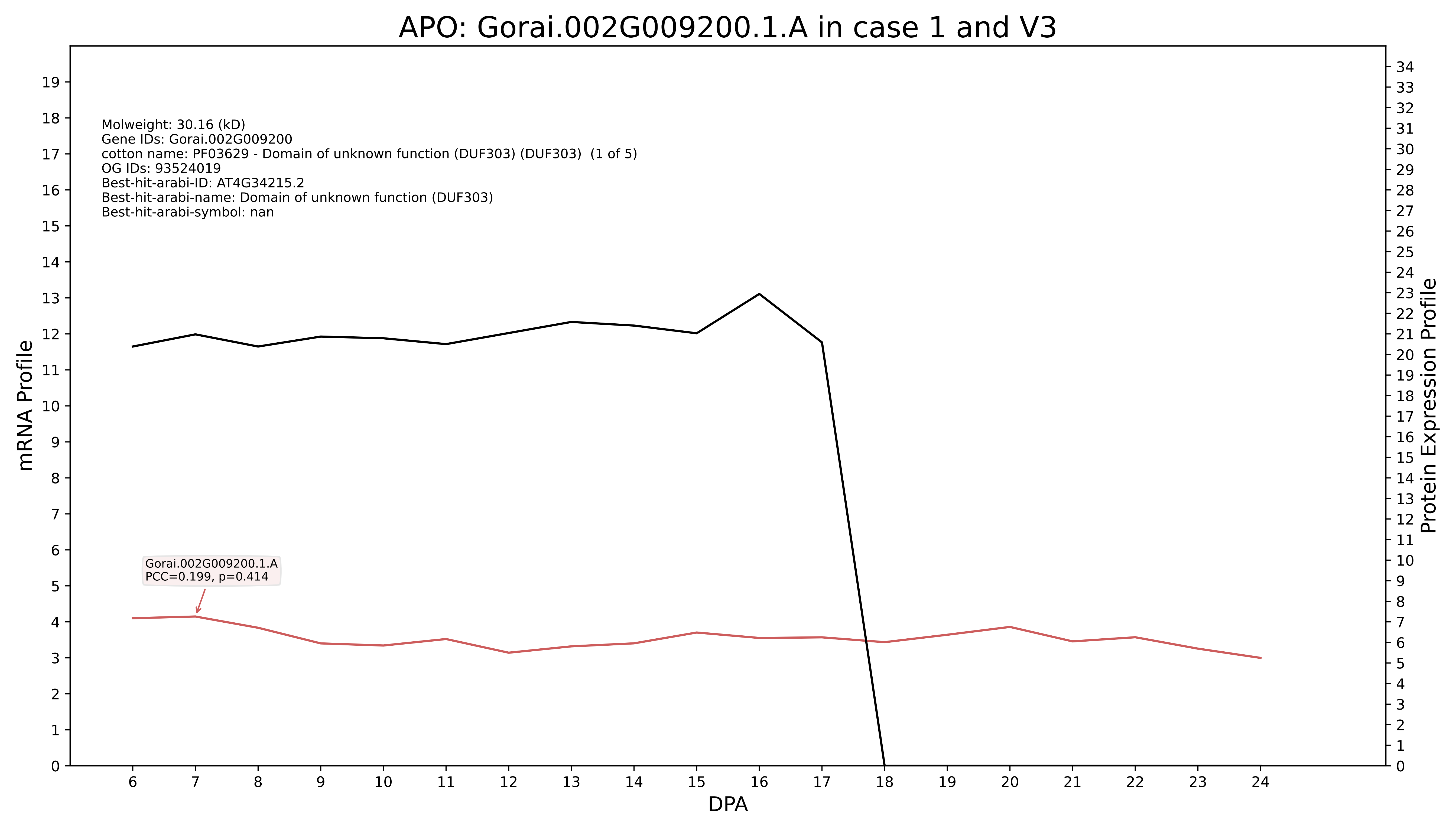
APO: Gorai.002G009200.1.A in case 1 and V3
Molweight: 30.16 (kD)
Gene IDs: Gorai.002G009200
cotton name: PF03629 - Domain of unknown function (DUF303) (DUF303) (1 of 5)
OG IDs: 93524019
Best-hit-arabi-ID: AT4G34215.2
Best-hit-arabi-name: Domain of unknown function (DUF303)
Best-hit-arabi-symbol: nan
Gorai.002G009200.1.A
PCC=0.199, p=0.414
mRNA Profile
Protein Expression Profile
DPA

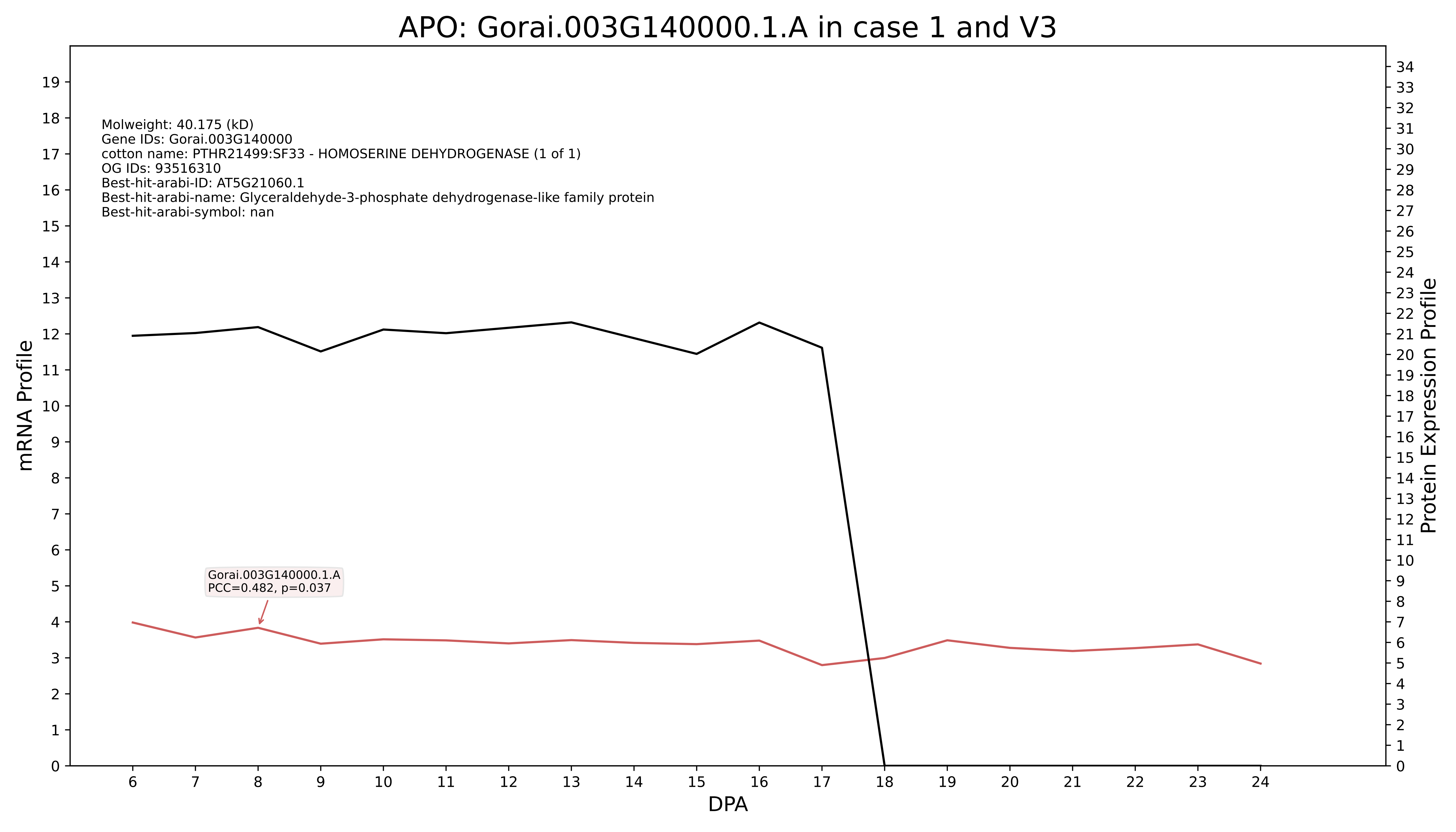
APO: Gorai.003G140000.1.A in case 1 and V3
Molweight: 40.175 (kD)
Gene IDs: Gorai.003G140000
cotton name: PTHR21499:SF33 - HOMOSERINE DEHYDROGENASE (1 of 1)
OG IDs: 93516310
Best-hit-arabi-ID: AT5G21060.1
Best-hit-arabi-name: Glyceraldehyde-3-phosphate dehydrogenase-like family protein
Best-hit-arabi-symbol: nan
Gorai.003G140000.1.A
PCC=0.482, p=0.037
mRNA Profile
Protein Expression Profile
DPA

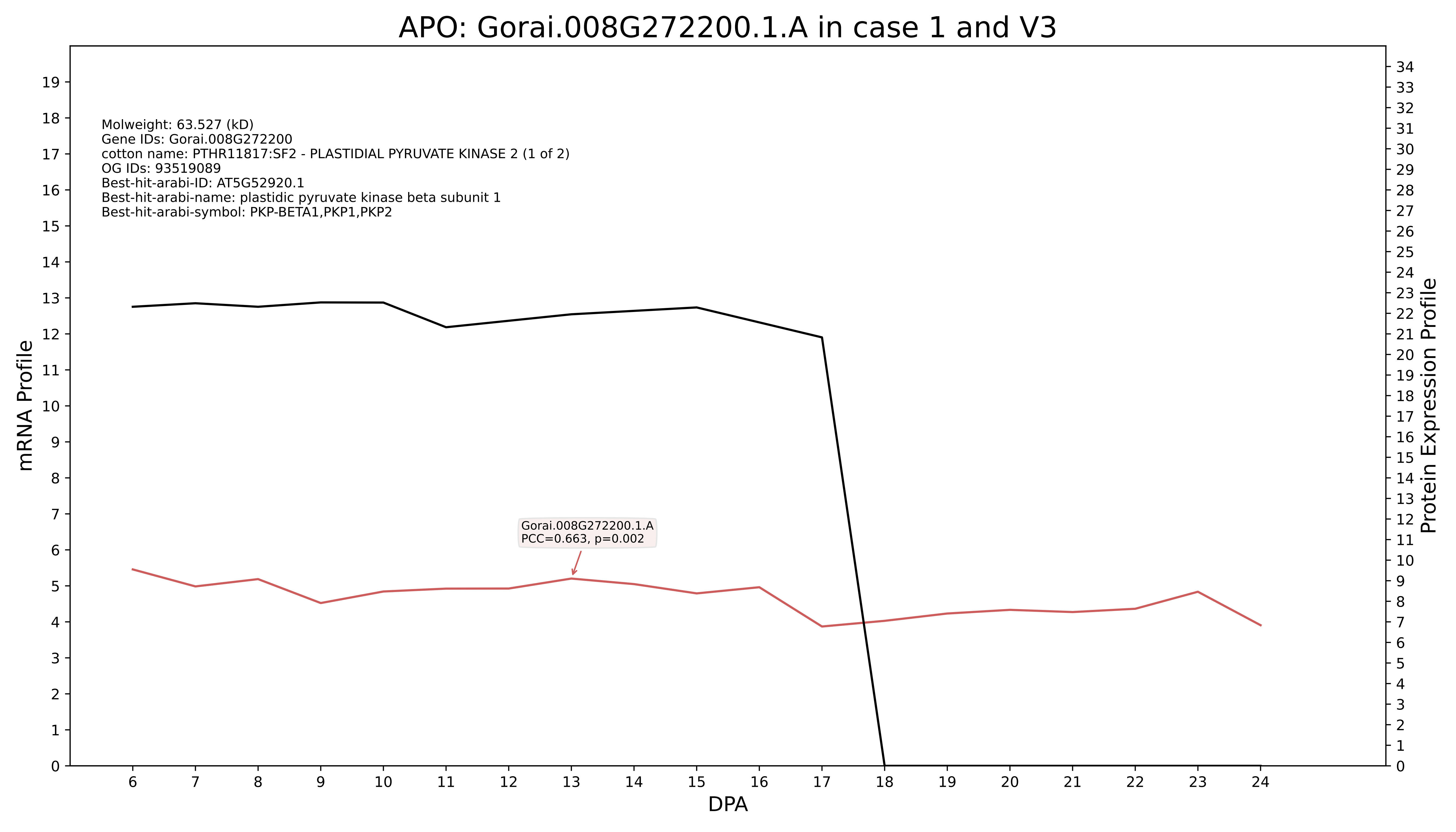

APO: Gorai.008G272200.1.A in case 1 and V3
Molweight: 63.527 (kD)
Gene IDs: Gorai.008G272200
cotton name: PTHR11817:SF2 - PLASTIDIAL PYRUVATE KINASE 2 (1 of 2)
OG IDs: 93519089
Best-hit-arabi-ID: AT5G52920.1
Best-hit-arabi-name: plastidic pyruvate kinase beta subunit 1
Best-hit-arabi-symbol: PKP-BETA1,PKP1,PKP2
Gorai.008G272200.1.A
PCC=0.663, p=0.002
mRNA Profile
Protein Expression Profile
DPA

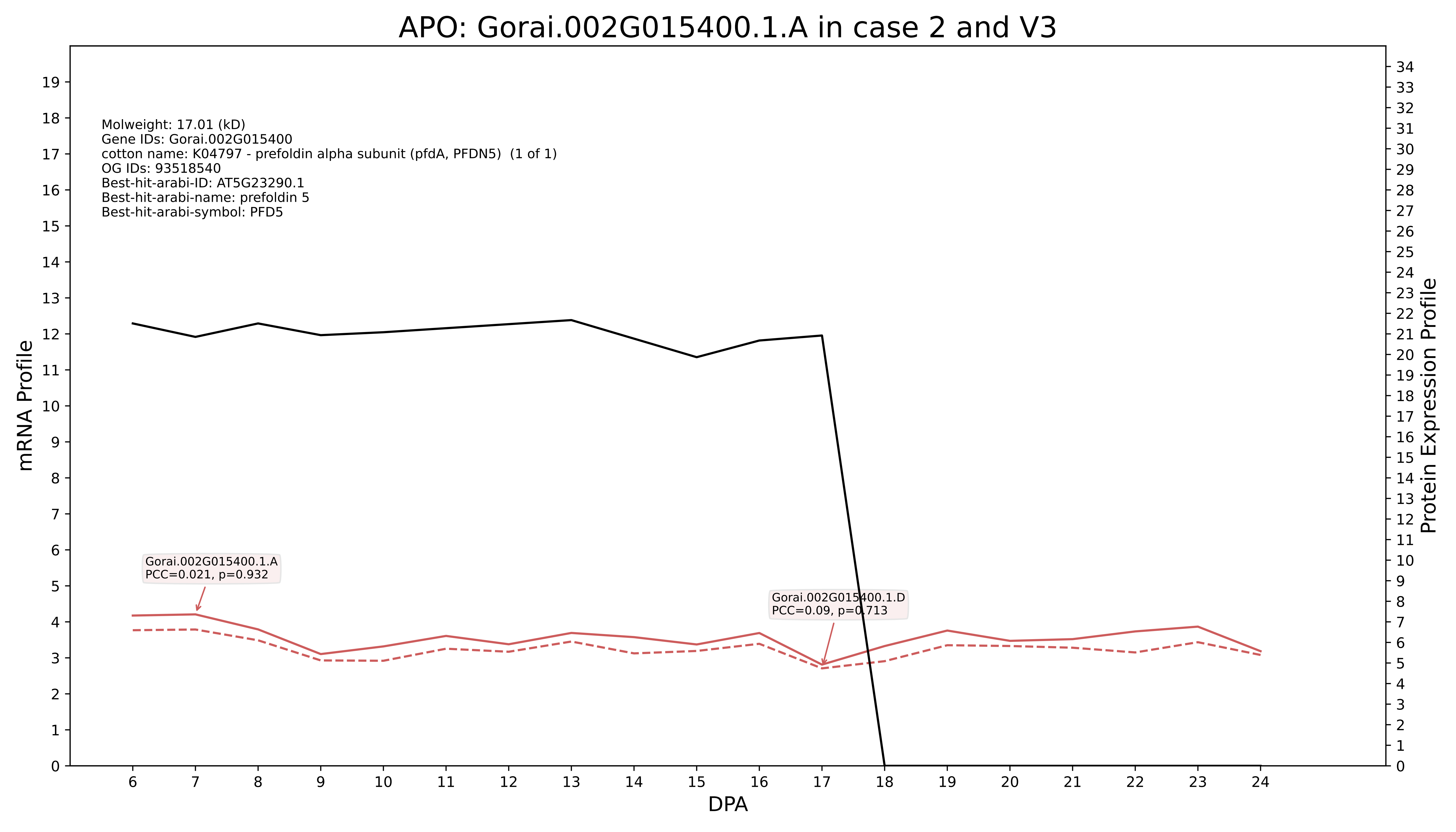
APO: Gorai.002G015400.1.A in case 2 and V3
Molweight: 17.01 (kD)
Gene IDs: Gorai.002G015400
cotton name: K04797 - prefoldin alpha subunit (pfdA, PFDN5) (1 of 1)
OG IDs: 93518540
Best-hit-arabi-ID: AT5G23290.1
Best-hit-arabi-name: prefoldin 5
Best-hit-arabi-symbol: PFD5
Gorai.002G015400.1.A
PCC=0.021, p=0.932
Gorai.002G015400.1.D
PCC=0.09, p=0.713
mRNA Profile
Protein Expression Profile
DPA

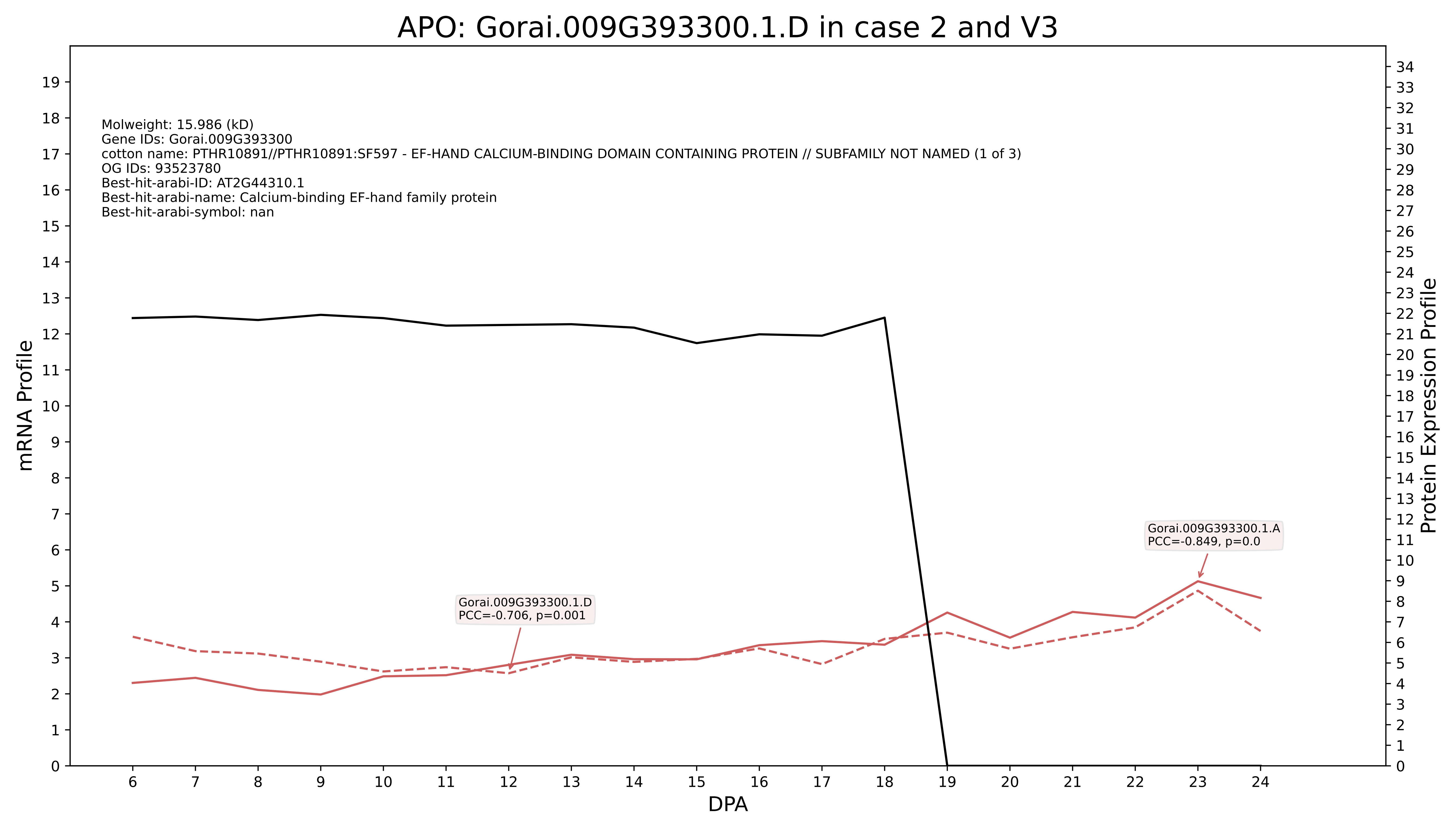
APO: Gorai.009G393300.1.D in case 2 and V3
Molweight: 15.986 (kD)
Gene IDs: Gorai.009G393300
cotton name: PTHR10891//PTHR10891:SF597 - EF-HAND CALCIUM-BINDING DOMAIN CONTAINING PROTEIN // SUBFAMILY NOT NAMED (1 of 3)
OG IDs: 93523780
Best-hit-arabi-ID: AT2G44310.1
Best-hit-arabi-name: Calcium-binding EF-hand family protein
Best-hit-arabi-symbol: nan
Gorai.009G393300.1.A
PCC=-0.849, p=0.0
Gorai.009G393300.1.D
PCC=-0.706, p=0.001
mRNA Profile
Protein Expression Profile
DPA

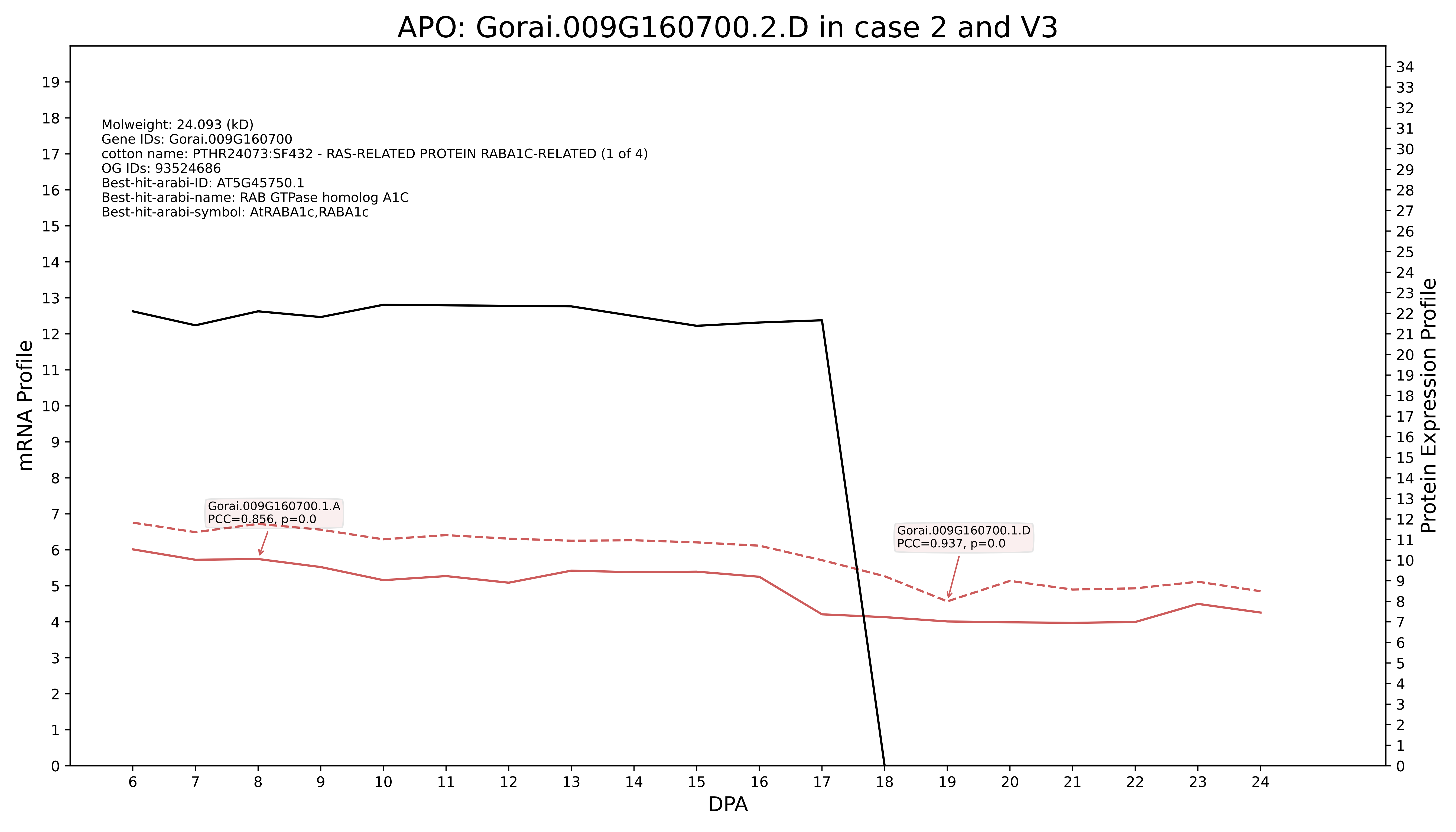
APO: Gorai.009G160700.2.D in case 2 and V3
Molweight: 24.093 (kD)
Gene IDs: Gorai.009G160700
cotton name: PTHR24073:SF432 - RAS-RELATED PROTEIN RABA1C-RELATED (1 of 4)
OG IDs: 93524686
Best-hit-arabi-ID: AT5G45750.1
Best-hit-arabi-name: RAB GTPase homolog A1C
Best-hit-arabi-symbol: AtRABA1c,RABA1c
Gorai.009G160700.1.A
PCC=0.856, p=0.0
Gorai.009G160700.1.D
PCC=0.937, p=0.0
mRNA Profile
Protein Expression Profile
DPA

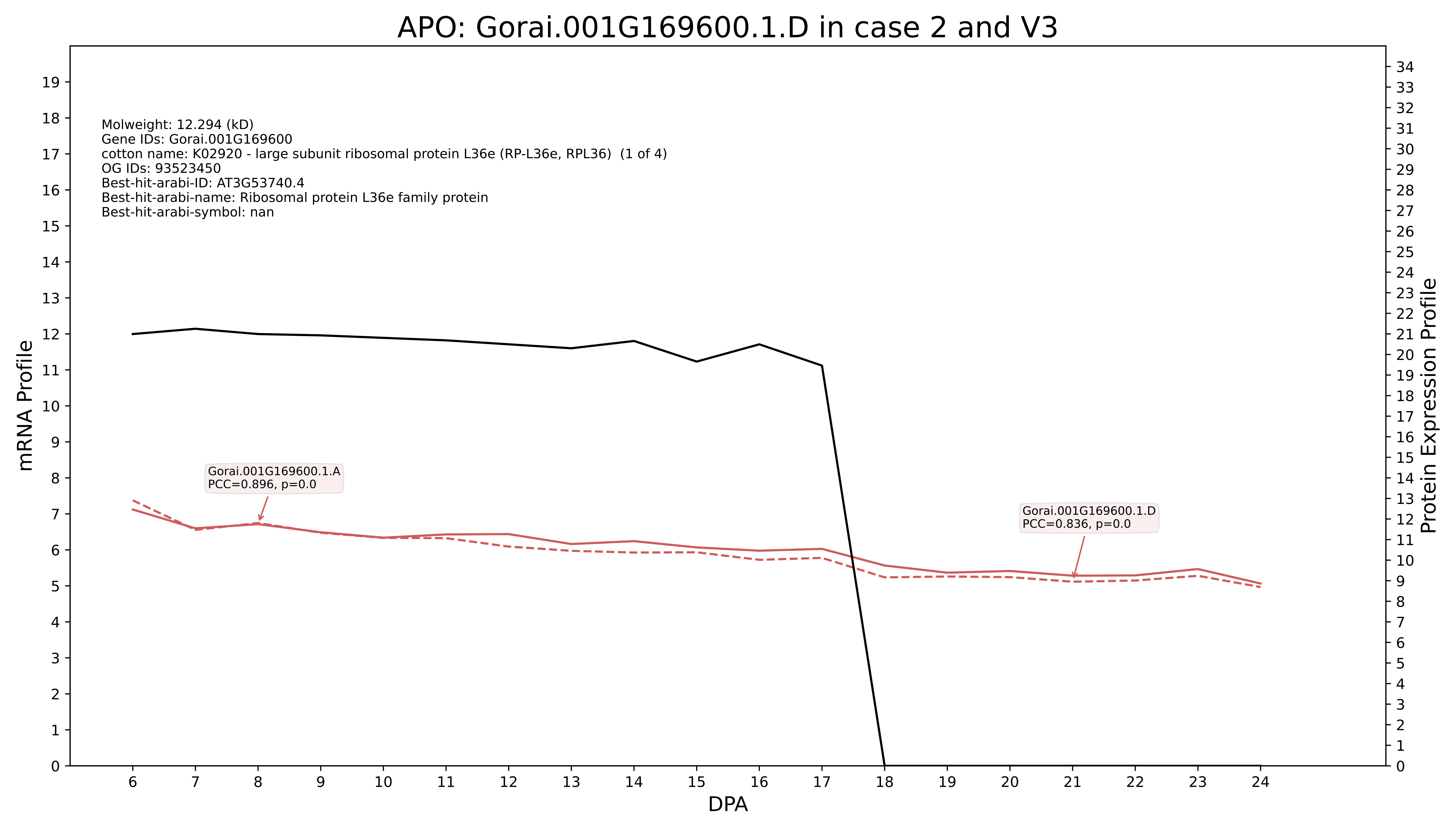

APO: Gorai.001G169600.1.D in case 2 and V3
Molweight: 12.294 (kD)
Gene IDs: Gorai.001G169600
cotton name: K02920 - large subunit ribosomal protein L36e (RP-L36e, RPL36) (1 of 4)
OG IDs: 93523450
Best-hit-arabi-ID: AT3G53740.4
Best-hit-arabi-name: Ribosomal protein L36e family protein
Best-hit-arabi-symbol: nan
Gorai.001G169600.1.A
PCC=0.896, p=0.0
Gorai.001G169600.1.D
PCC=0.836, p=0.0
mRNA Profile
Protein Expression Profile
DPA

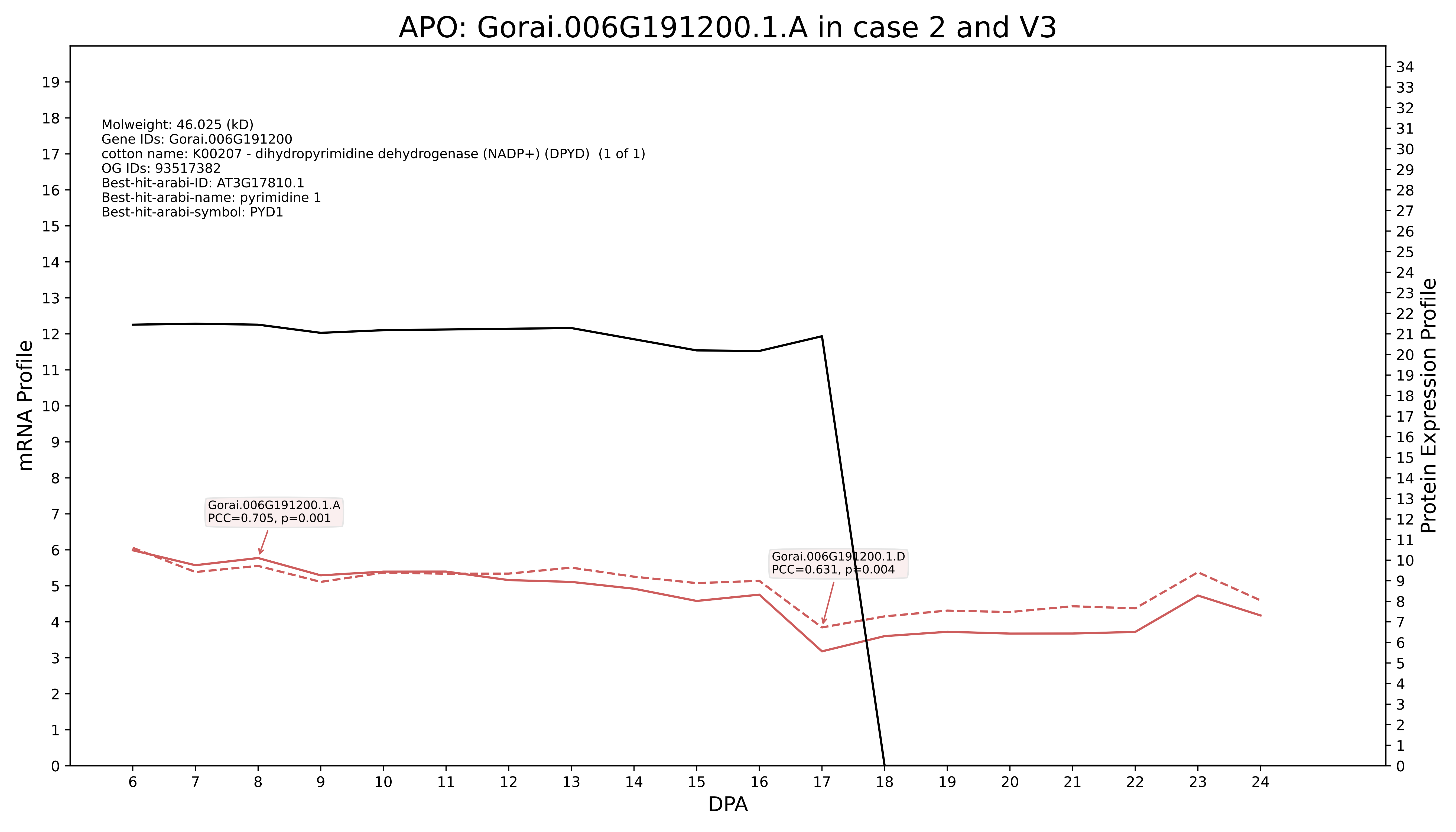
APO: Gorai.006G191200.1.A in case 2 and V3
Molweight: 46.025 (kD)
Gene IDs: Gorai.006G191200
cotton name: K00207 - dihydropyrimidine dehydrogenase (NADP+) (DPYD) (1 of 1)
OG IDs: 93517382
Best-hit-arabi-ID: AT3G17810.1
Best-hit-arabi-name: pyrimidine 1
Best-hit-arabi-symbol: PYD1
Gorai.006G191200.1.A
PCC=0.705, p=0.001
Gorai.006G191200.1.D
PCC=0.631, p=0.004
mRNA Profile
Protein Expression Profile
DPA

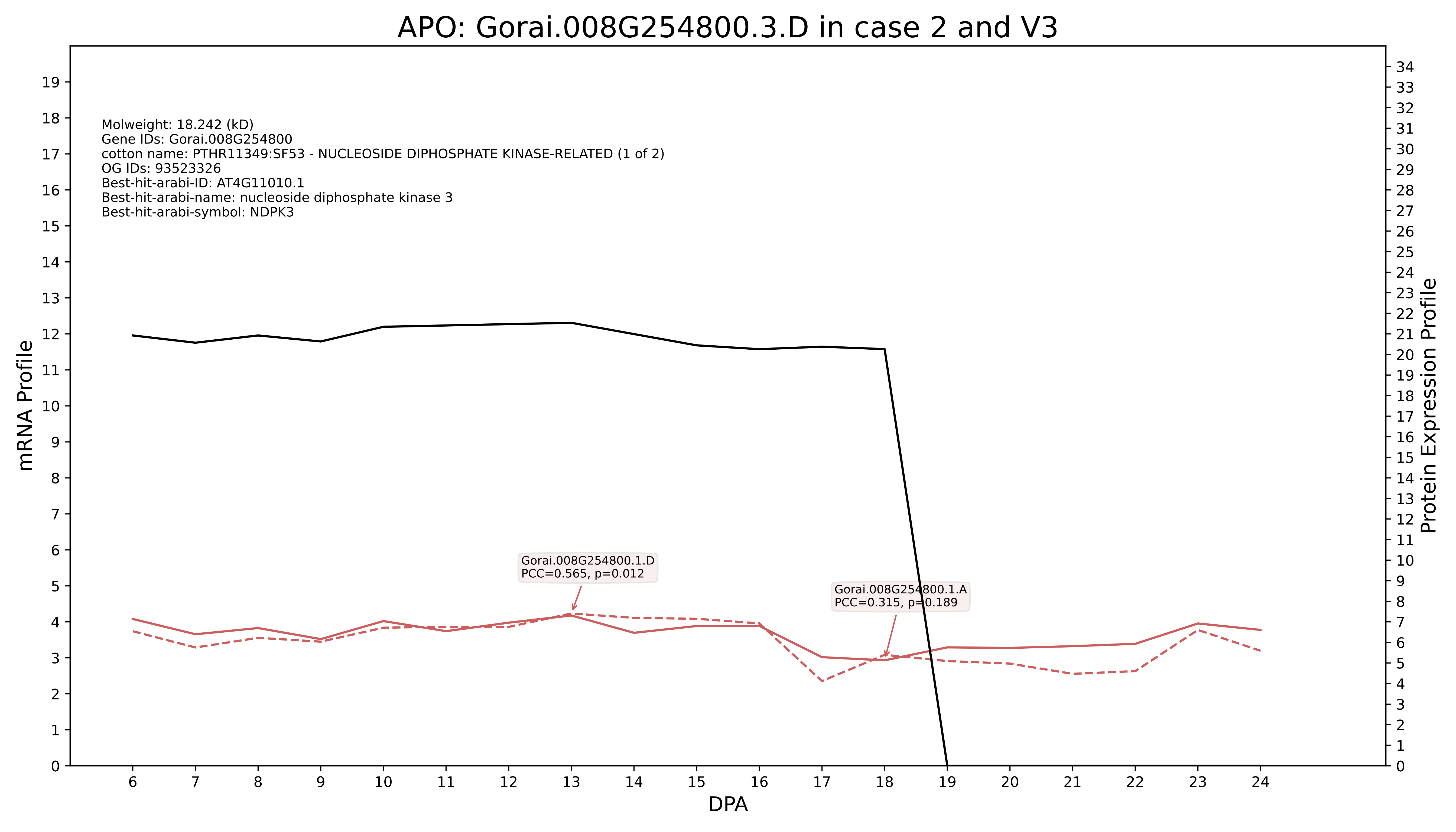

APO: Gorai.008G254800.3.D in case 2 and V3
Molweight: 18.242 (kD)
Gene IDs: Gorai.008G254800
cotton name: PTHR11349:SF53 - NUCLEOSIDE DIPHOSPHATE KINASE-RELATED (1 of 2)
OG IDs: 93523326
Best-hit-arabi-ID: AT4G11010.1
Best-hit-arabi-name: nucleoside diphosphate kinase 3
Best-hit-arabi-symbol: NDPK3
Gorai.008G254800.1.D
PCC=0.565, p=0.012
Gorai.008G254800.1.A
PCC=0.315, p=0.189
mRNA Profile
Protein Expression Profile
DPA

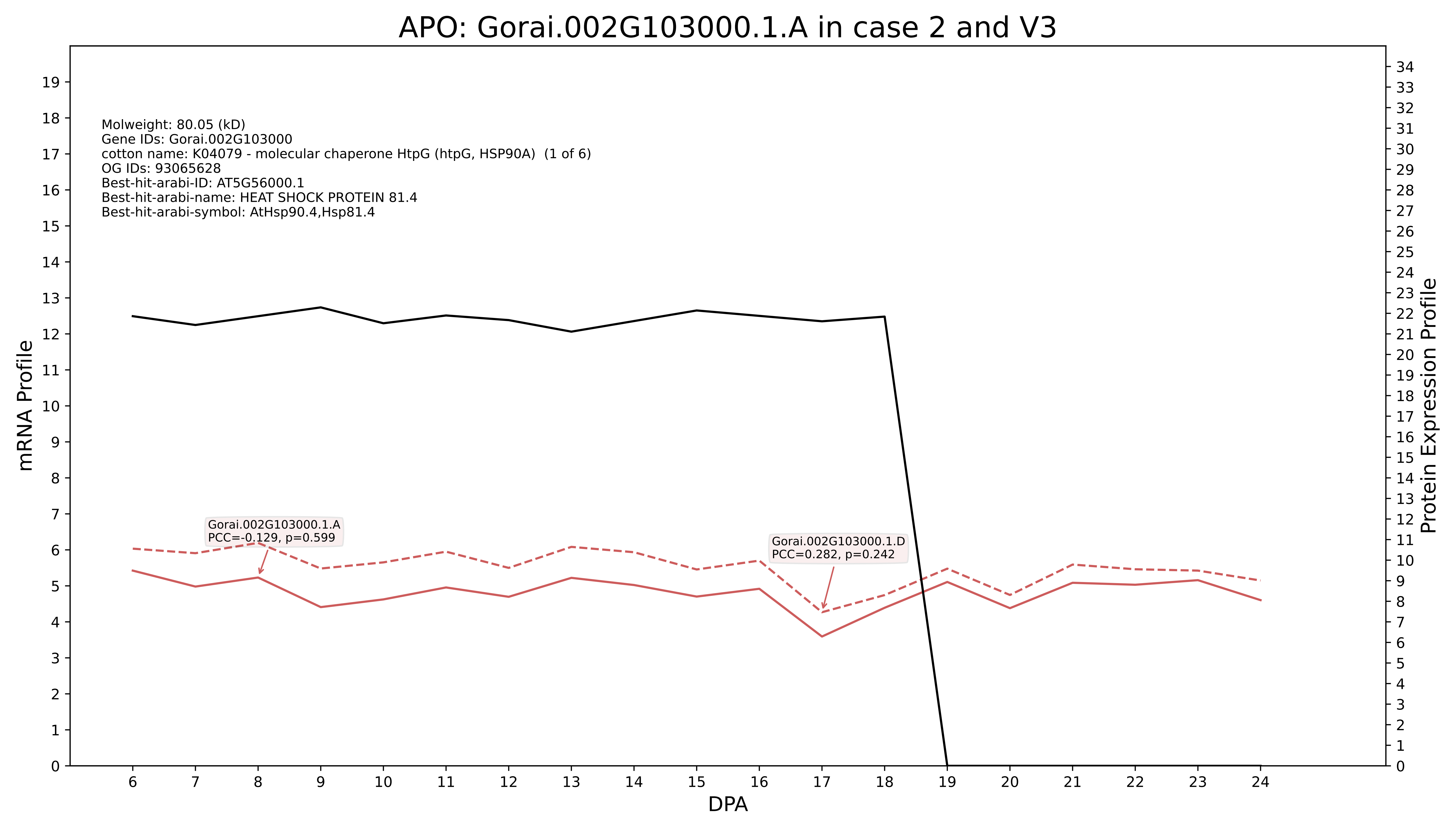

APO: Gorai.002G103000.1.A in case 2 and V3
Molweight: 80.05 (kD)
Gene IDs: Gorai.002G103000
cotton name: K04079 - molecular chaperone HtpG (htpG, HSP90A) (1 of 6)
OG IDs: 93065628
Best-hit-arabi-ID: AT5G56000.1
Best-hit-arabi-name: HEAT SHOCK PROTEIN 81.4
Best-hit-arabi-symbol: AtHsp90.4,Hsp81.4
Gorai.002G103000.1.A
PCC=-0.129, p=0.599
Gorai.002G103000.1.D
PCC=0.282, p=0.242
mRNA Profile
Protein Expression Profile
DPA

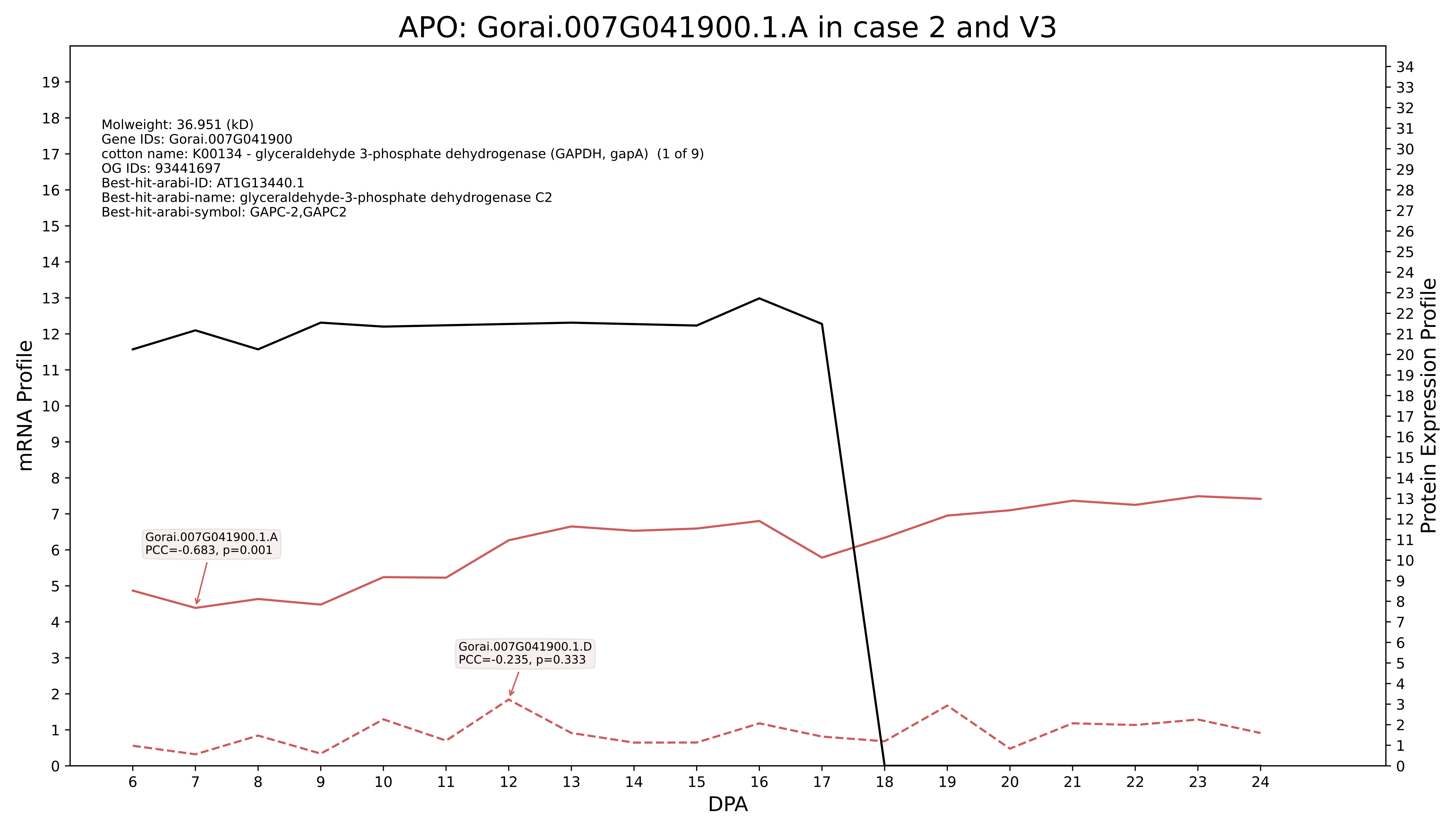
APO: Gorai.007G041900.1.A in case 2 and V3
Molweight: 36.951 (kD)
Gene IDs: Gorai.007G041900
cotton name: K00134 - glyceraldehyde 3-phosphate dehydrogenase (GAPDH, gapA) (1 of 9)
OG IDs: 93441697
Best-hit-arabi-ID: AT1G13440.1
Best-hit-arabi-name: glyceraldehyde-3-phosphate dehydrogenase C2
Best-hit-arabi-symbol: GAPC-2,GAPC2
Gorai.007G041900.1.A
PCC=-0.683, p=0.001
Gorai.007G041900.1.D
PCC=-0.235, p=0.333
mRNA Profile
Protein Expression Profile
DPA

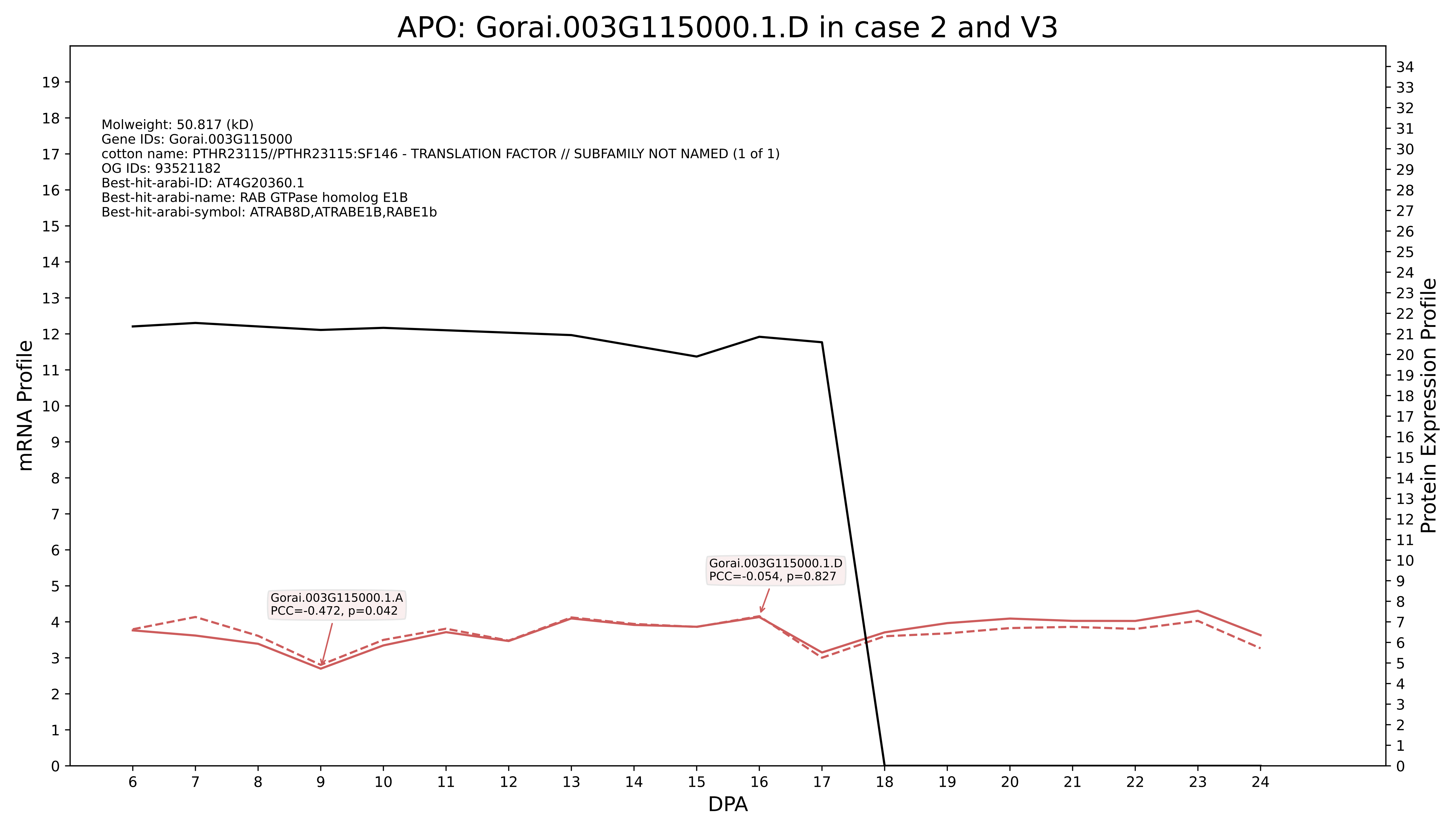
APO: Gorai.003G115000.1.D in case 2 and V3
Molweight: 50.817 (kD)
Gene IDs: Gorai.003G115000
cotton name: PTHR23115//PTHR23115:SF146 - TRANSLATION FACTOR // SUBFAMILY NOT NAMED (1 of 1)
OG IDs: 93521182
Best-hit-arabi-ID: AT4G20360.1
Best-hit-arabi-name: RAB GTPase homolog E1B
Best-hit-arabi-symbol: ATRAB8D,ATRABE1B,RABE1b
Gorai.003G115000.1.A
PCC=-0.472, p=0.042
Gorai.003G115000.1.D
PCC=-0.054, p=0.827
mRNA Profile
Protein Expression Profile
DPA

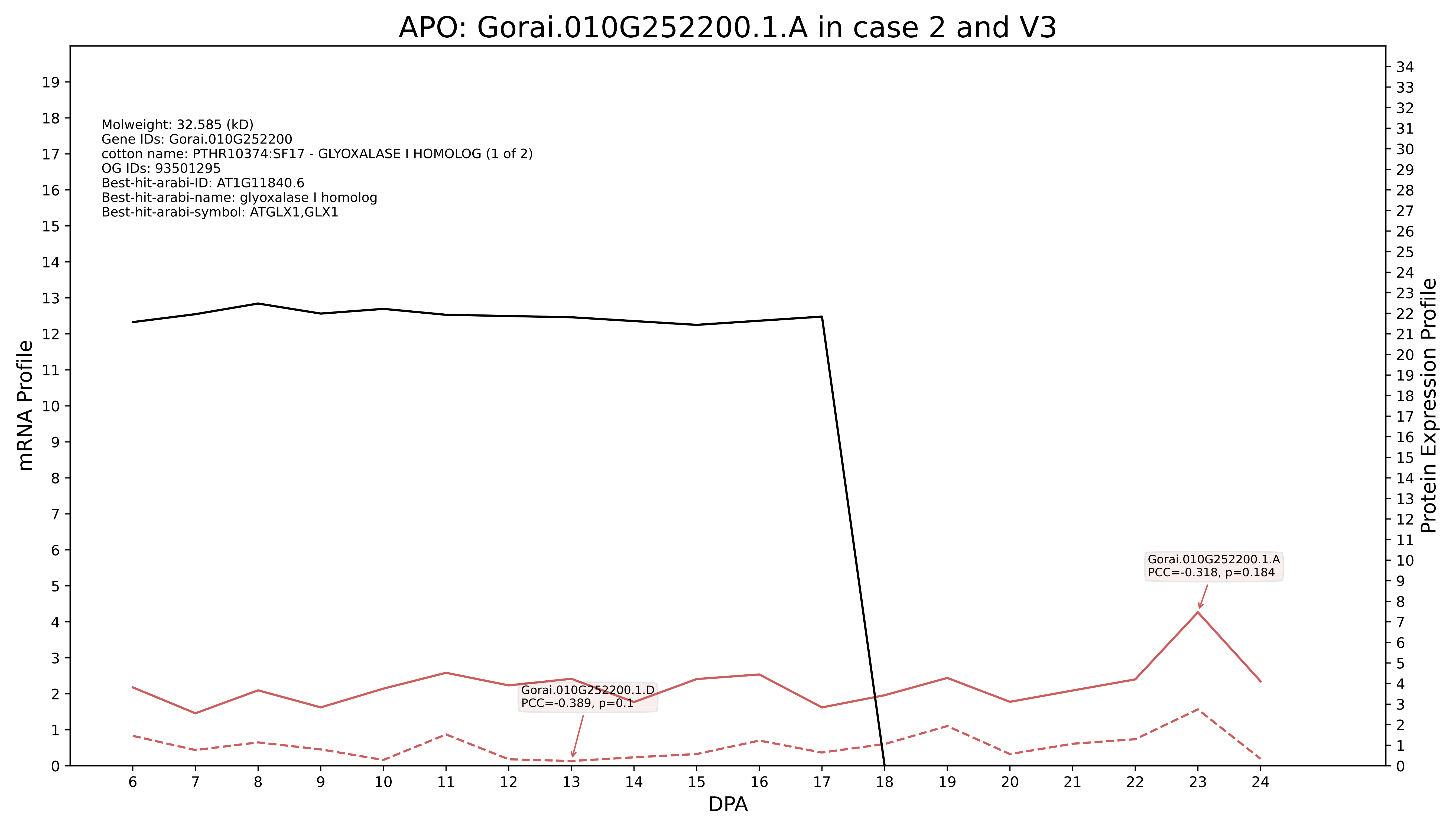

APO: Gorai.010G252200.1.A in case 2 and V3
Molweight: 32.585 (kD)
Gene IDs: Gorai.010G252200
cotton name: PTHR10374:SF17 - GLYOXALASE I HOMOLOG (1 of 2)
OG IDs: 93501295
Best-hit-arabi-ID: AT1G11840.6
Best-hit-arabi-name: glyoxalase I homolog
Best-hit-arabi-symbol: ATGLX1,GLX1
Gorai.010G252200.1.A
PCC=-0.318, p=0.184
Gorai.010G252200.1.D
PCC=-0.389, p=0.1
mRNA Profile
Protein Expression Profile
DPA

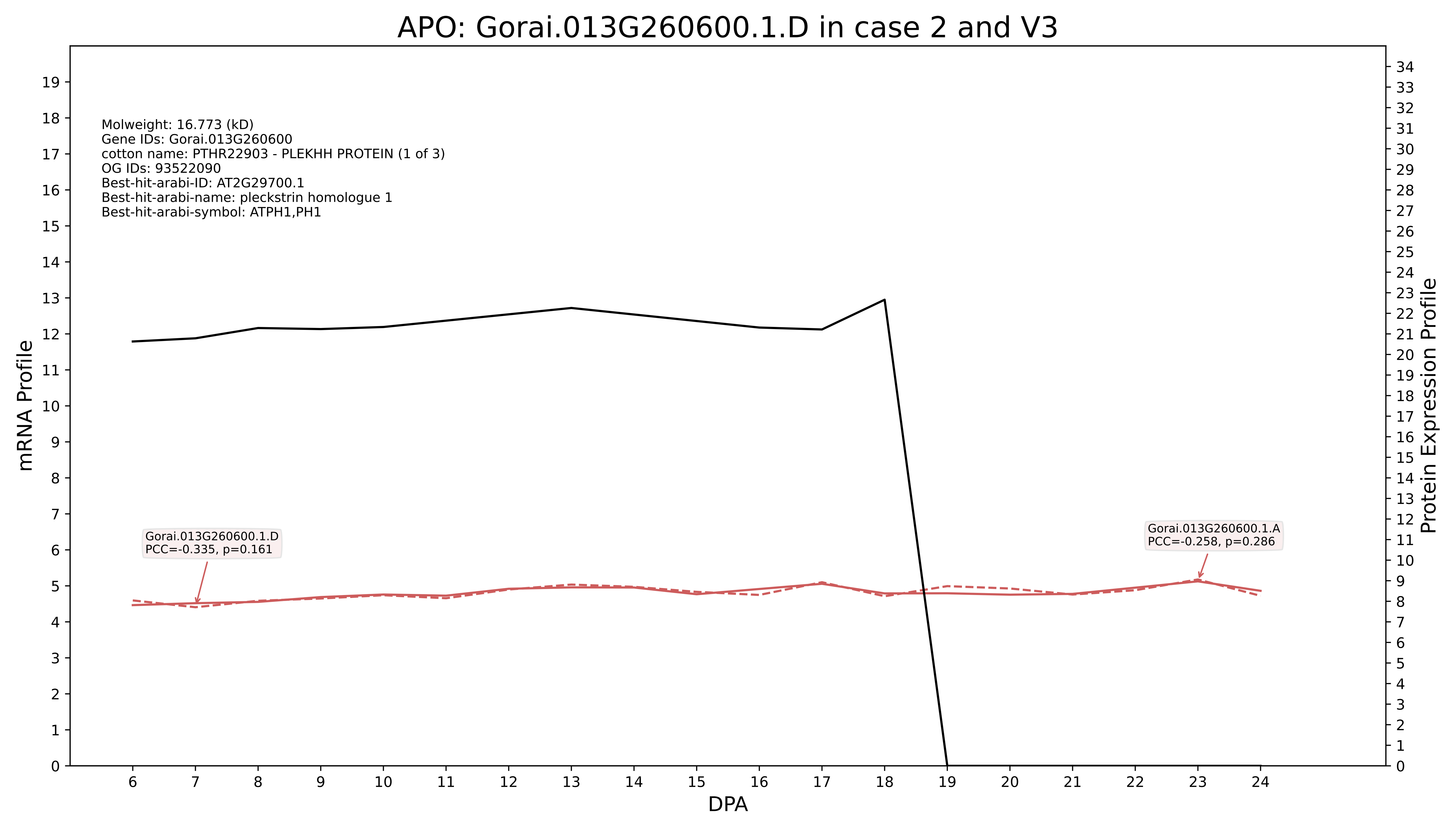

APO: Gorai.013G260600.1.D in case 2 and V3
Molweight: 16.773 (kD)
Gene IDs: Gorai.013G260600
cotton name: PTHR22903 - PLEKHH PROTEIN (1 of 3)
OG IDs: 93522090
Best-hit-arabi-ID: AT2G29700.1
Best-hit-arabi-name: pleckstrin homologue 1
Best-hit-arabi-symbol: ATPH1,PH1
Gorai.013G260600.1.D
PCC=-0.335, p=0.161
Gorai.013G260600.1.A
PCC=-0.258, p=0.286
mRNA Profile
Protein Expression Profile
DPA

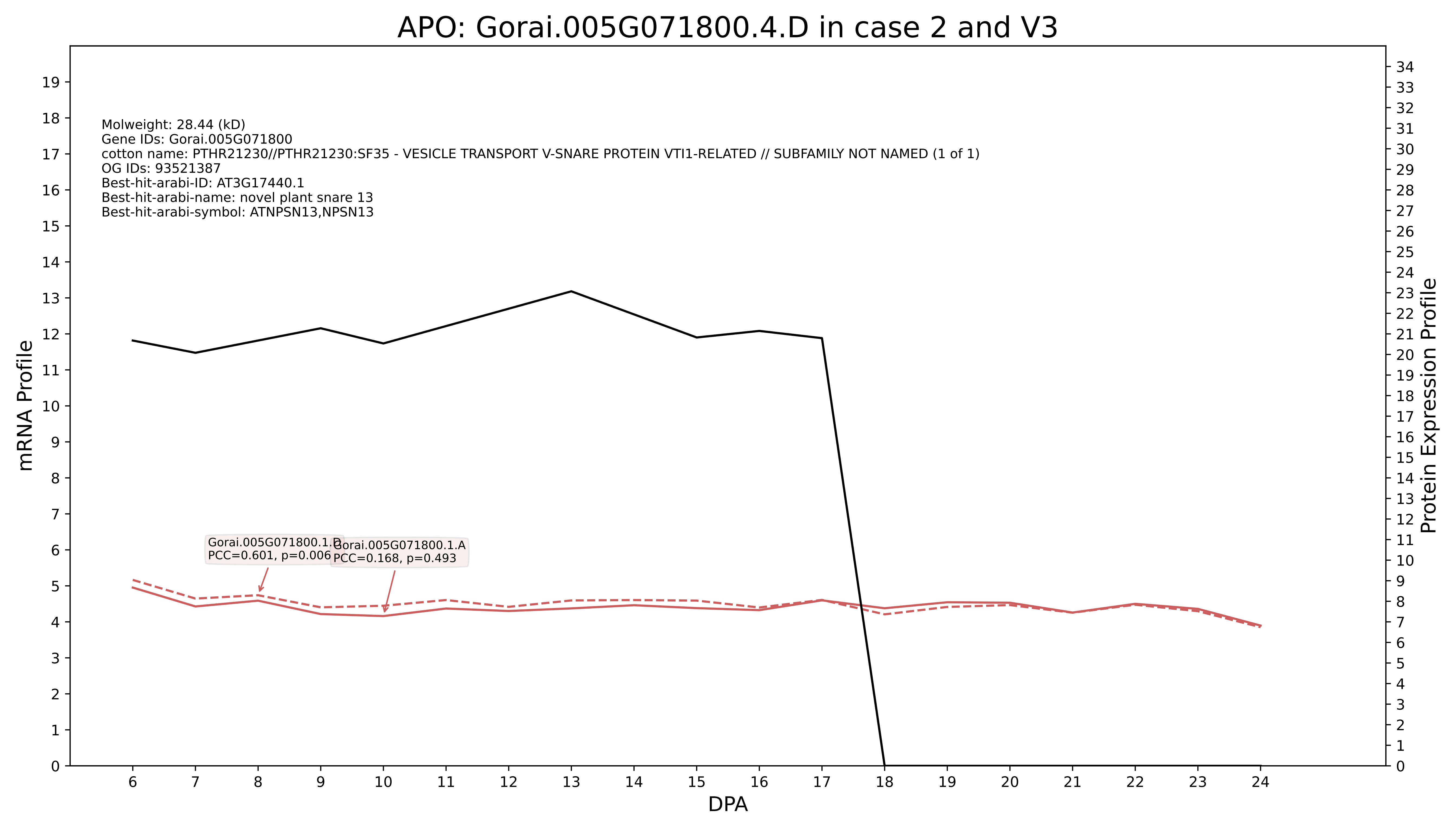
APO: Gorai.005G071800.4.D in case 2 and V3
Molweight: 28.44 (kD)
Gene IDs: Gorai.005G071800
cotton name: PTHR21230//PTHR21230:SF35 - VESICLE TRANSPORT V-SNARE PROTEIN VTI1-RELATED // SUBFAMILY NOT NAMED (1 of 1)
OG IDs: 93521387
Best-hit-arabi-ID: AT3G17440.1
Best-hit-arabi-name: novel plant snare 13
Best-hit-arabi-symbol: ATNPSN13,NPSN13
Gorai.005G071800.1.D
PCC=0.601, p=0.006
Gorai.005G071800.1.A
PCC=0.168, p=0.493
mRNA Profile
Protein Expression Profile
DPA

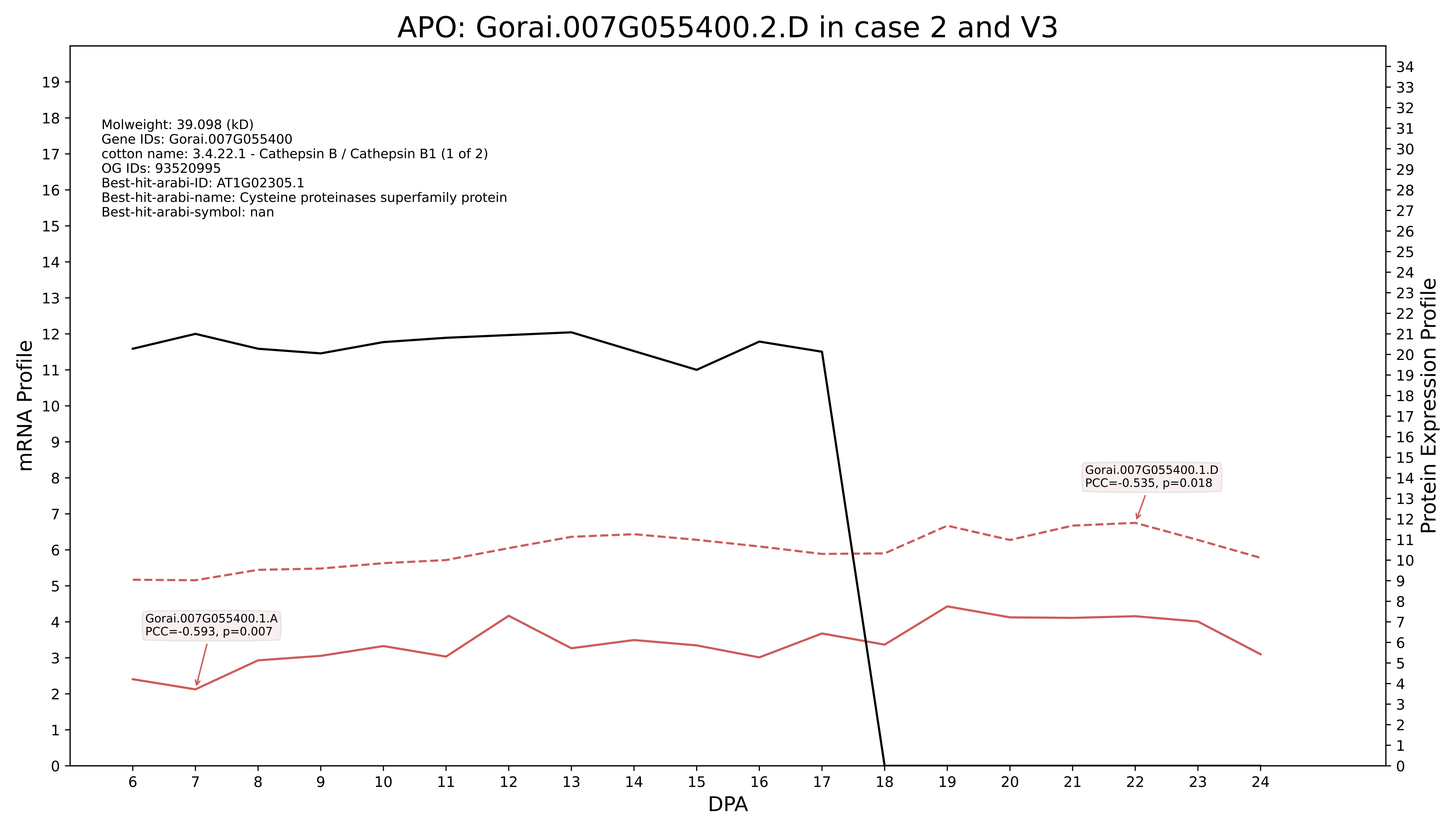

APO: Gorai.007G055400.2.D in case 2 and V3
Molweight: 39.098 (kD)
Gene IDs: Gorai.007G055400
cotton name: 3.4.22.1 - Cathepsin B / Cathepsin B1 (1 of 2)
OG IDs: 93520995
Best-hit-arabi-ID: AT1G02305.1
Best-hit-arabi-name: Cysteine proteinases superfamily protein
Best-hit-arabi-symbol: nan
Gorai.007G055400.1.D
PCC=-0.535, p=0.018
Gorai.007G055400.1.A
PCC=-0.593, p=0.007
mRNA Profile
Protein Expression Profile
DPA

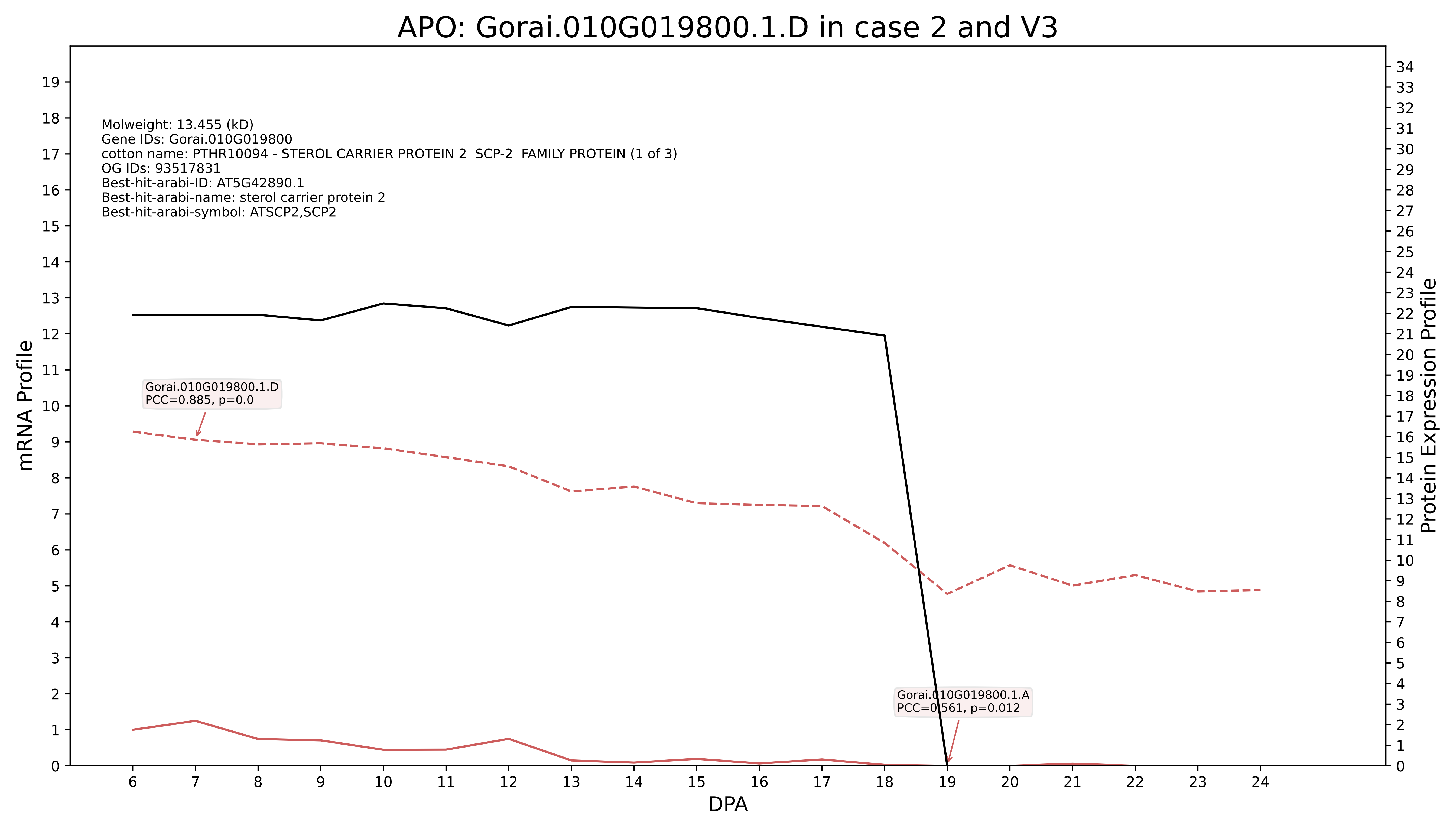
APO: Gorai.010G019800.1.D in case 2 and V3
Molweight: 13.455 (kD)
Gene IDs: Gorai.010G019800
cotton name: PTHR10094 - STEROL CARRIER PROTEIN 2 SCP-2 FAMILY PROTEIN (1 of 3)
OG IDs: 93517831
Best-hit-arabi-ID: AT5G42890.1
Best-hit-arabi-name: sterol carrier protein 2
Best-hit-arabi-symbol: ATSCP2,SCP2
Gorai.010G019800.1.D
PCC=0.885, p=0.0
Gorai.010G019800.1.A
PCC=0.561, p=0.012
mRNA Profile
Protein Expression Profile
DPA

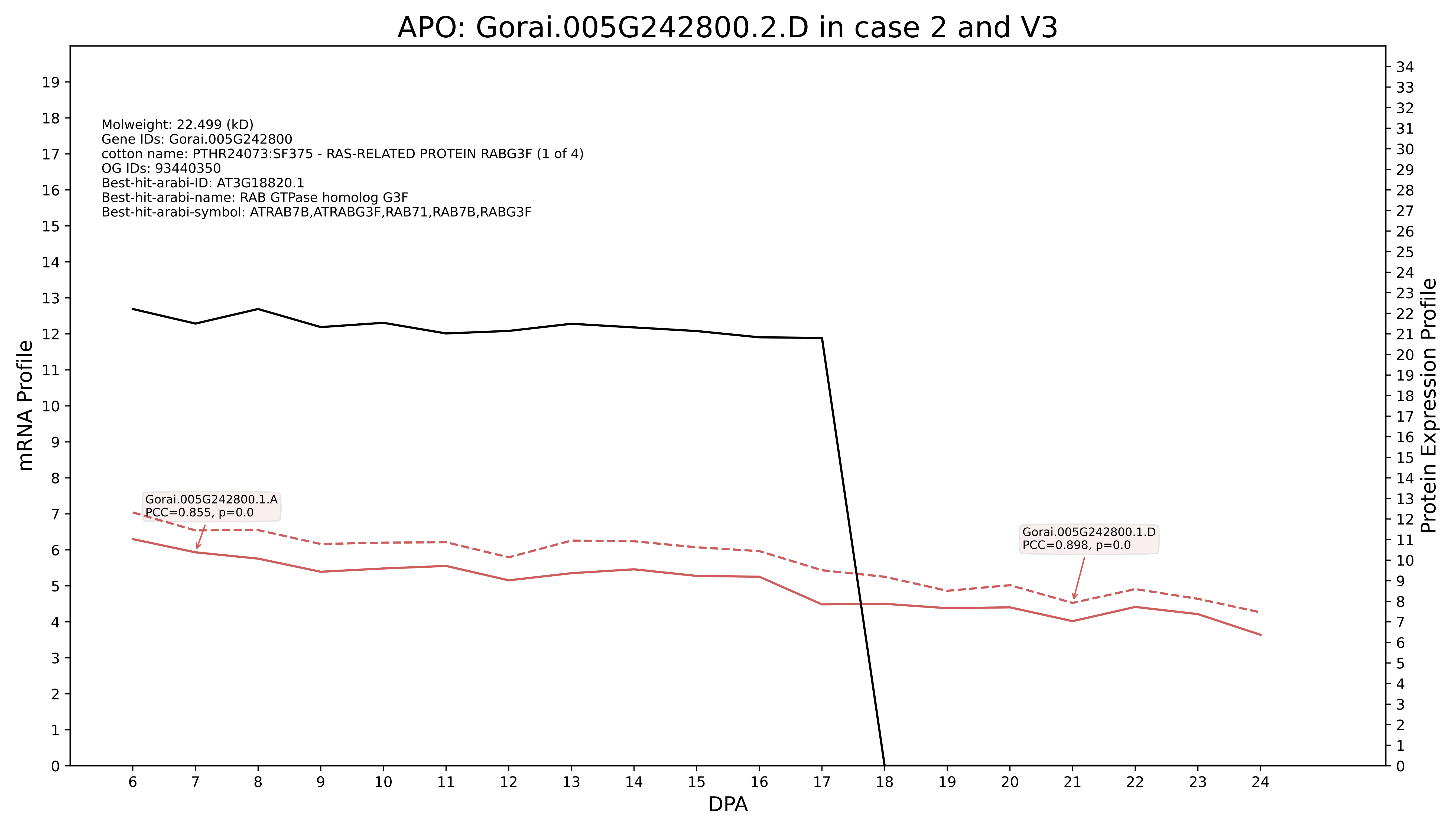

APO: Gorai.005G242800.2.D in case 2 and V3
Molweight: 22.499 (kD)
Gene IDs: Gorai.005G242800
cotton name: PTHR24073:SF375 - RAS-RELATED PROTEIN RABG3F (1 of 4)
OG IDs: 93440350
Best-hit-arabi-ID: AT3G18820.1
Best-hit-arabi-name: RAB GTPase homolog G3F
Best-hit-arabi-symbol: ATRAB7B,ATRABG3F,RAB71,RAB7B,RABG3F
Gorai.005G242800.1.A
PCC=0.855, p=0.0
Gorai.005G242800.1.D
PCC=0.898, p=0.0
mRNA Profile
Protein Expression Profile
DPA

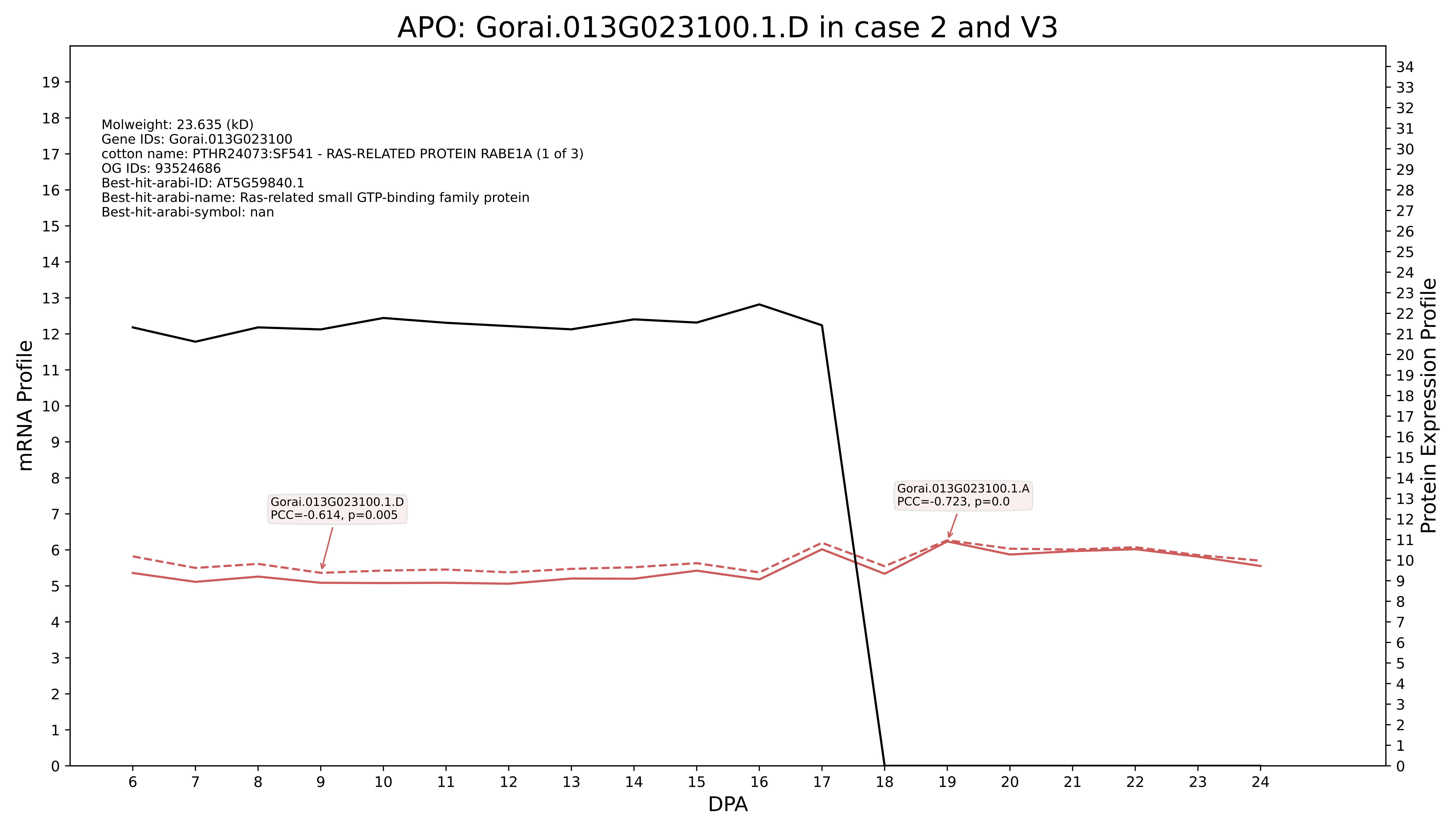

APO: Gorai.013G023100.1.D in case 2 and V3
Molweight: 23.635 (kD)
Gene IDs: Gorai.013G023100
cotton name: PTHR24073:SF541 - RAS-RELATED PROTEIN RABE1A (1 of 3)
OG IDs: 93524686
Best-hit-arabi-ID: AT5G59840.1
Best-hit-arabi-name: Ras-related small GTP-binding family protein
Best-hit-arabi-symbol: nan
Gorai.013G023100.1.D
PCC=-0.614, p=0.005
Gorai.013G023100.1.A
PCC=-0.723, p=0.0
mRNA Profile
Protein Expression Profile
DPA

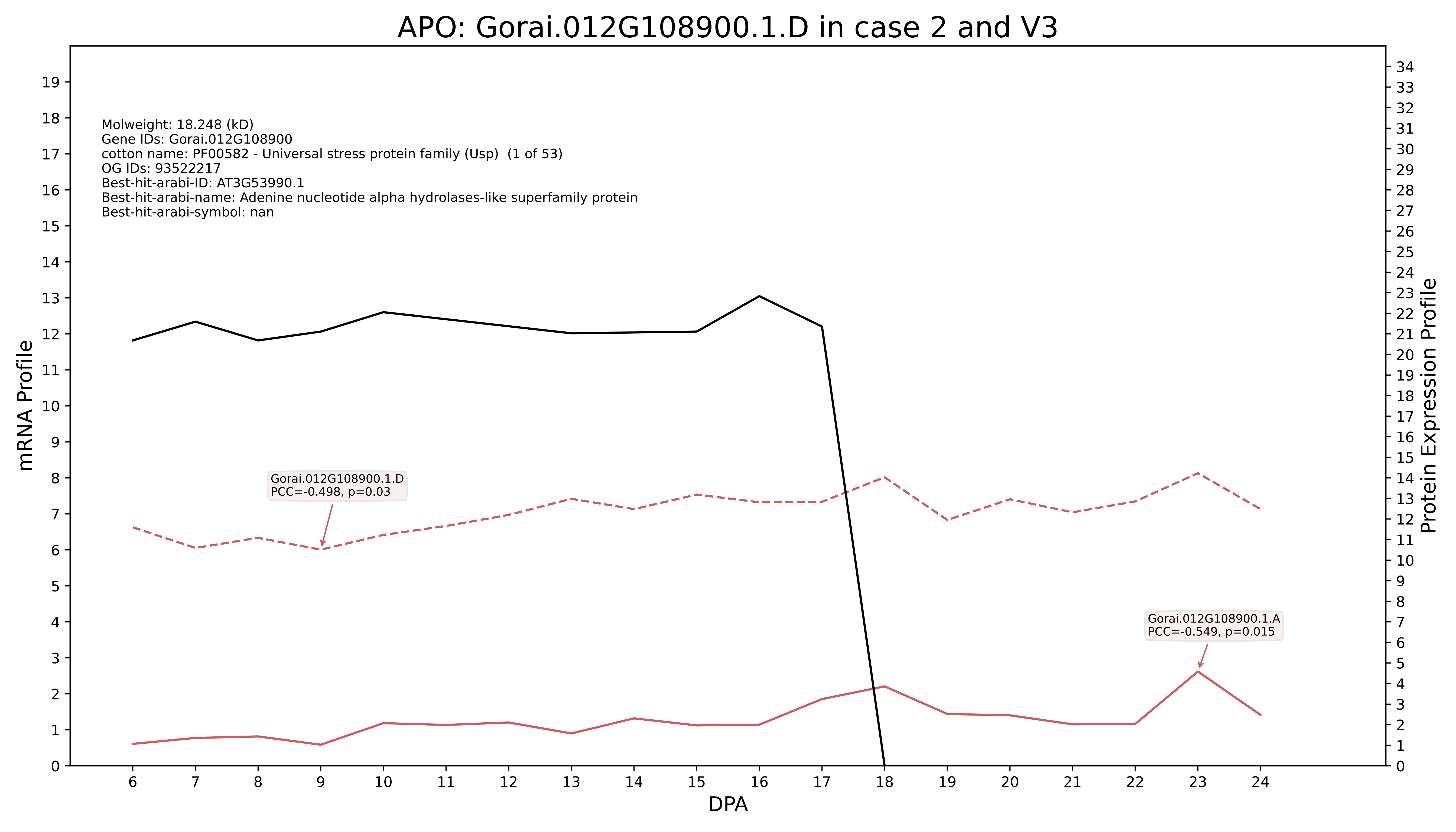

APO: Gorai.012G108900.1.D in case 2 and V3
Molweight: 18.248 (kD)
Gene IDs: Gorai.012G108900
cotton name: PF00582 - Universal stress protein family (Usp) (1 of 53)
OG IDs: 93522217
Best-hit-arabi-ID: AT3G53990.1
Best-hit-arabi-name: Adenine nucleotide alpha hydrolases-like superfamily protein
Best-hit-arabi-symbol: nan
Gorai.012G108900.1.D
PCC=-0.498, p=0.03
Gorai.012G108900.1.A
PCC=-0.549, p=0.015
mRNA Profile
Protein Expression Profile
DPA

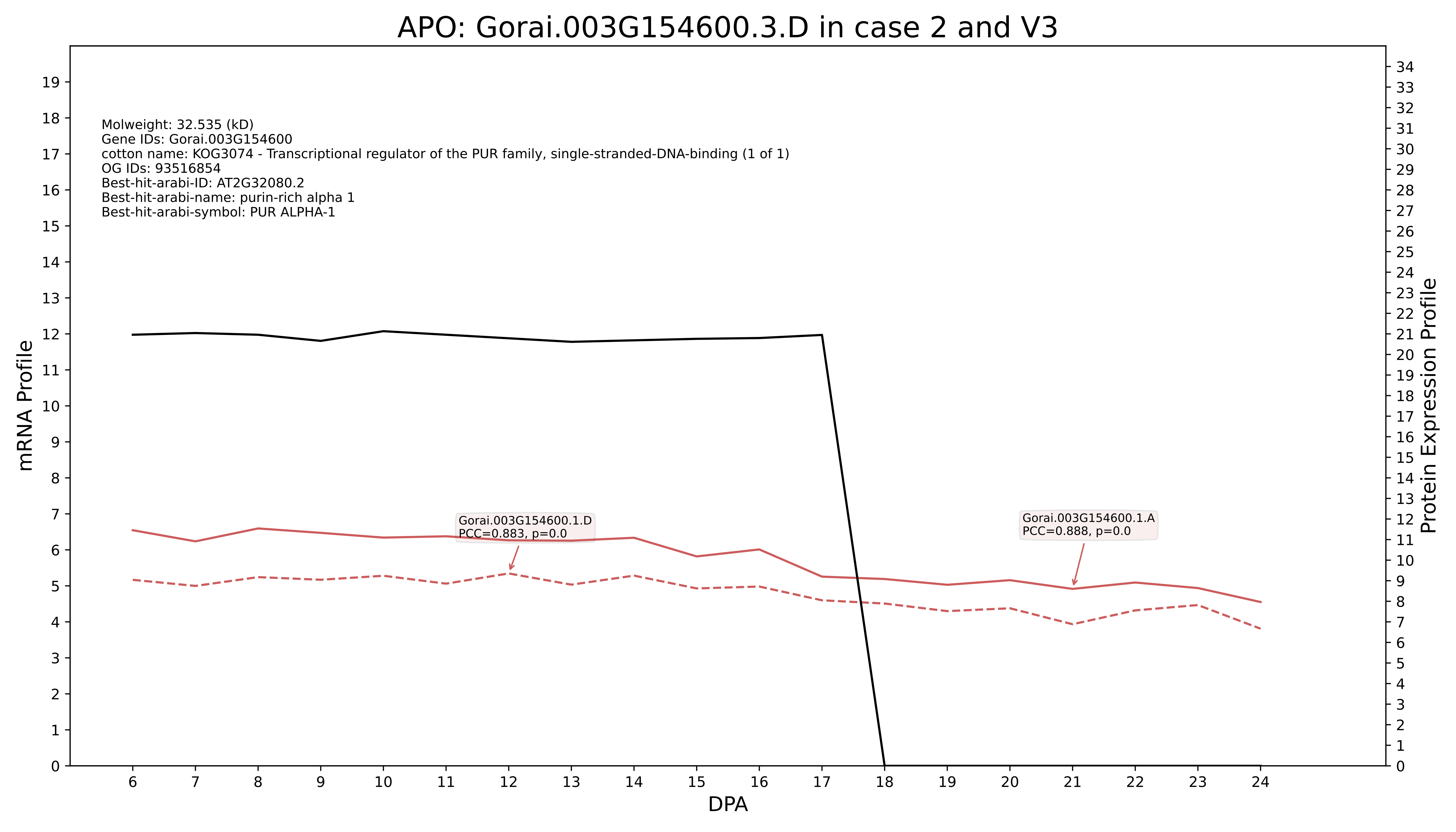

APO: Gorai.003G154600.3.D in case 2 and V3
Molweight: 32.535 (kD)
Gene IDs: Gorai.003G154600
cotton name: KOG3074 - Transcriptional regulator of the PUR family, single-stranded-DNA-binding (1 of 1)
OG IDs: 93516854
Best-hit-arabi-ID: AT2G32080.2
Best-hit-arabi-name: purin-rich alpha 1
Best-hit-arabi-symbol: PUR ALPHA-1
Gorai.003G154600.1.D
PCC=0.883, p=0.0
Gorai.003G154600.1.A
PCC=0.888, p=0.0
mRNA Profile
Protein Expression Profile
DPA

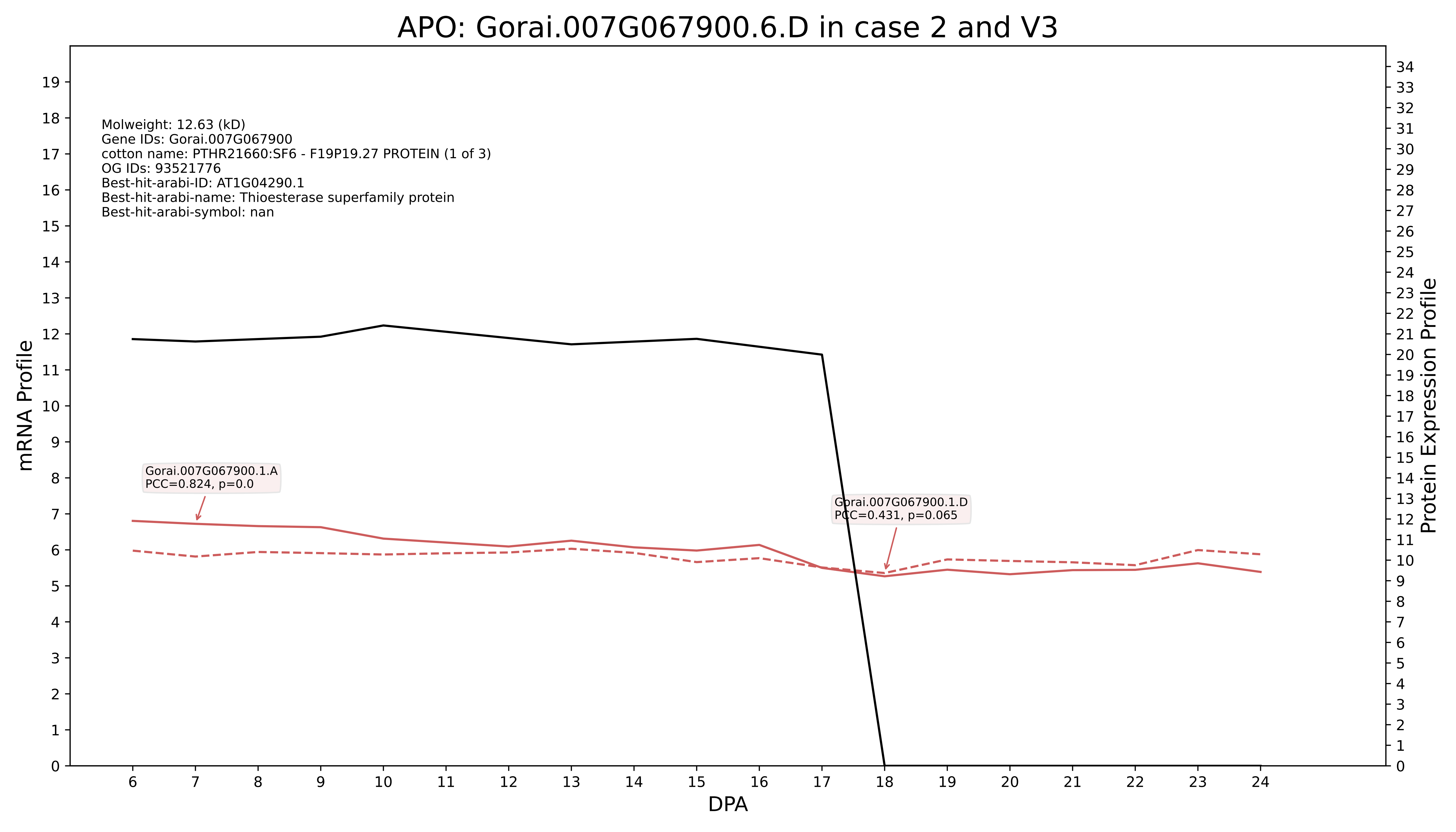
APO: Gorai.007G067900.6.D in case 2 and V3
Molweight: 12.63 (kD)
Gene IDs: Gorai.007G067900
cotton name: PTHR21660:SF6 - F19P19.27 PROTEIN (1 of 3)
OG IDs: 93521776
Best-hit-arabi-ID: AT1G04290.1
Best-hit-arabi-name: Thioesterase superfamily protein
Best-hit-arabi-symbol: nan
Gorai.007G067900.1.A
PCC=0.824, p=0.0
Gorai.007G067900.1.D
PCC=0.431, p=0.065
mRNA Profile
Protein Expression Profile
DPA

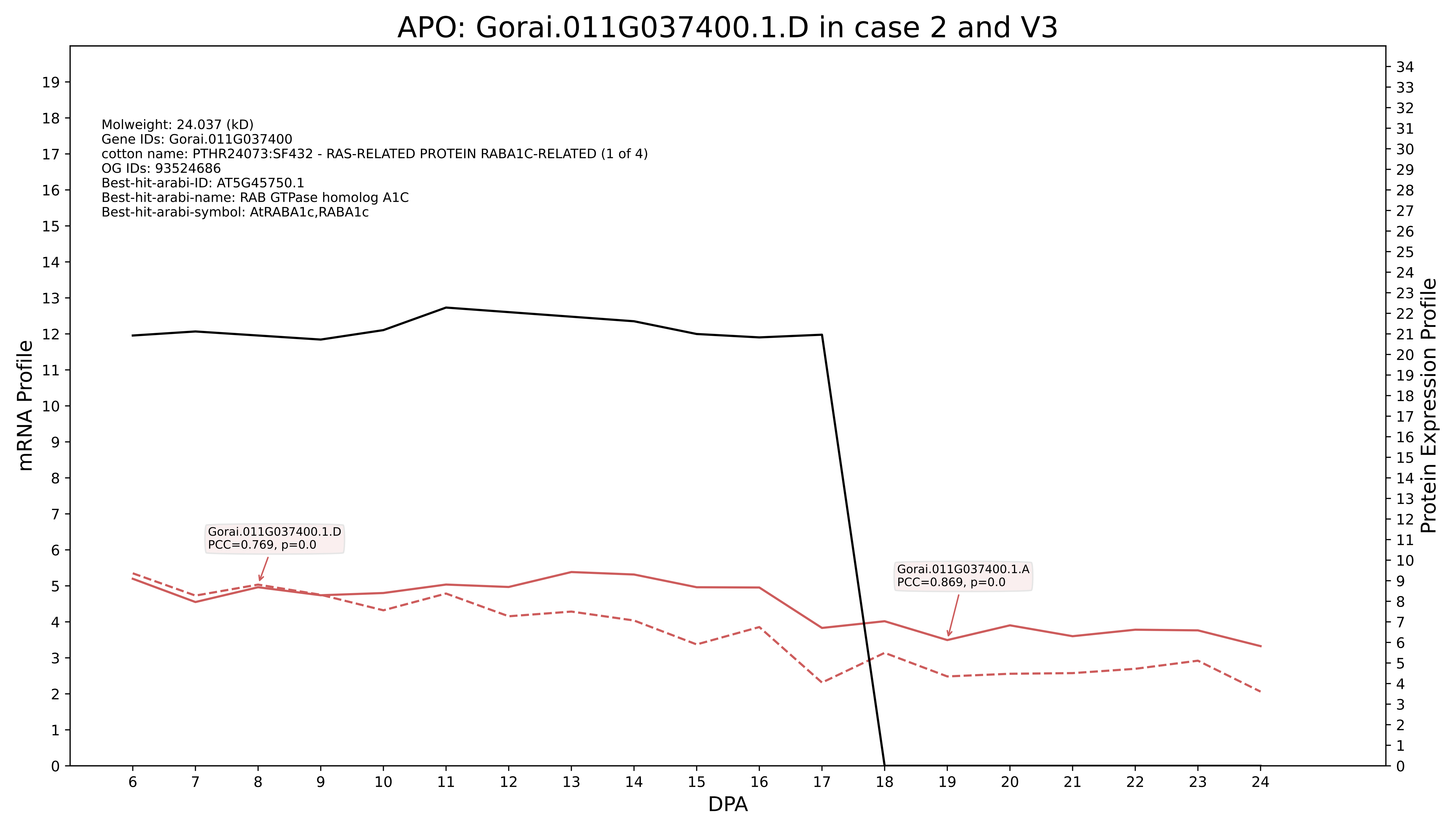
APO: Gorai.011G037400.1.D in case 2 and V3
Molweight: 24.037 (kD)
Gene IDs: Gorai.011G037400
cotton name: PTHR24073:SF432 - RAS-RELATED PROTEIN RABA1C-RELATED (1 of 4)
OG IDs: 93524686
Best-hit-arabi-ID: AT5G45750.1
Best-hit-arabi-name: RAB GTPase homolog A1C
Best-hit-arabi-symbol: AtRABA1c,RABA1c
Gorai.011G037400.1.D
PCC=0.769, p=0.0
Gorai.011G037400.1.A
PCC=0.869, p=0.0
mRNA Profile
Protein Expression Profile
DPA

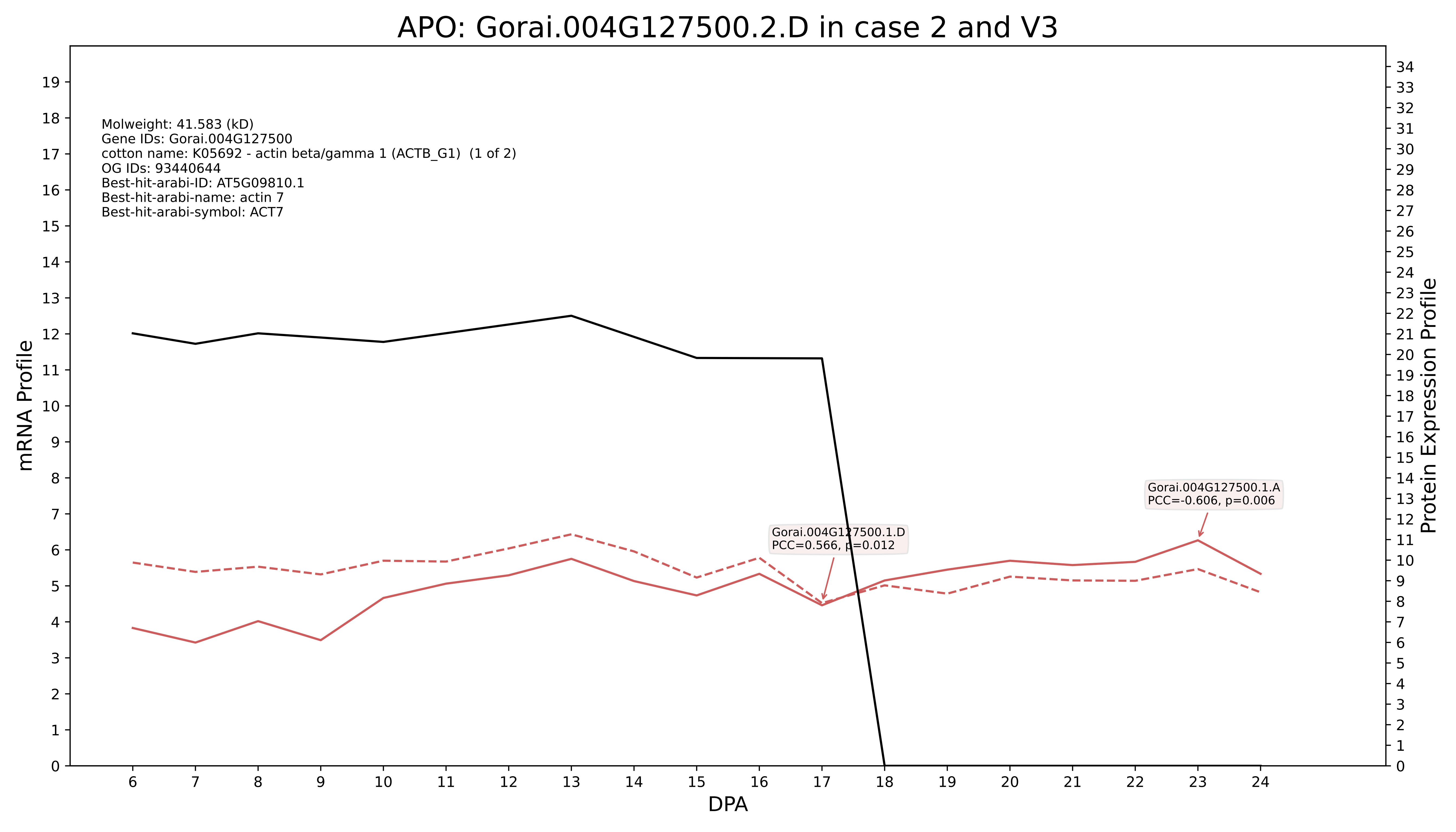
APO: Gorai.004G127500.2.D in case 2 and V3
Molweight: 41.583 (kD)
Gene IDs: Gorai.004G127500
cotton name: K05692 - actin beta/gamma 1 (ACTB_G1) (1 of 2)
OG IDs: 93440644
Best-hit-arabi-ID: AT5G09810.1
Best-hit-arabi-name: actin 7
Best-hit-arabi-symbol: ACT7
Gorai.004G127500.1.D
PCC=0.566, p=0.012
Gorai.004G127500.1.A
PCC=-0.606, p=0.006
mRNA Profile
Protein Expression Profile
DPA

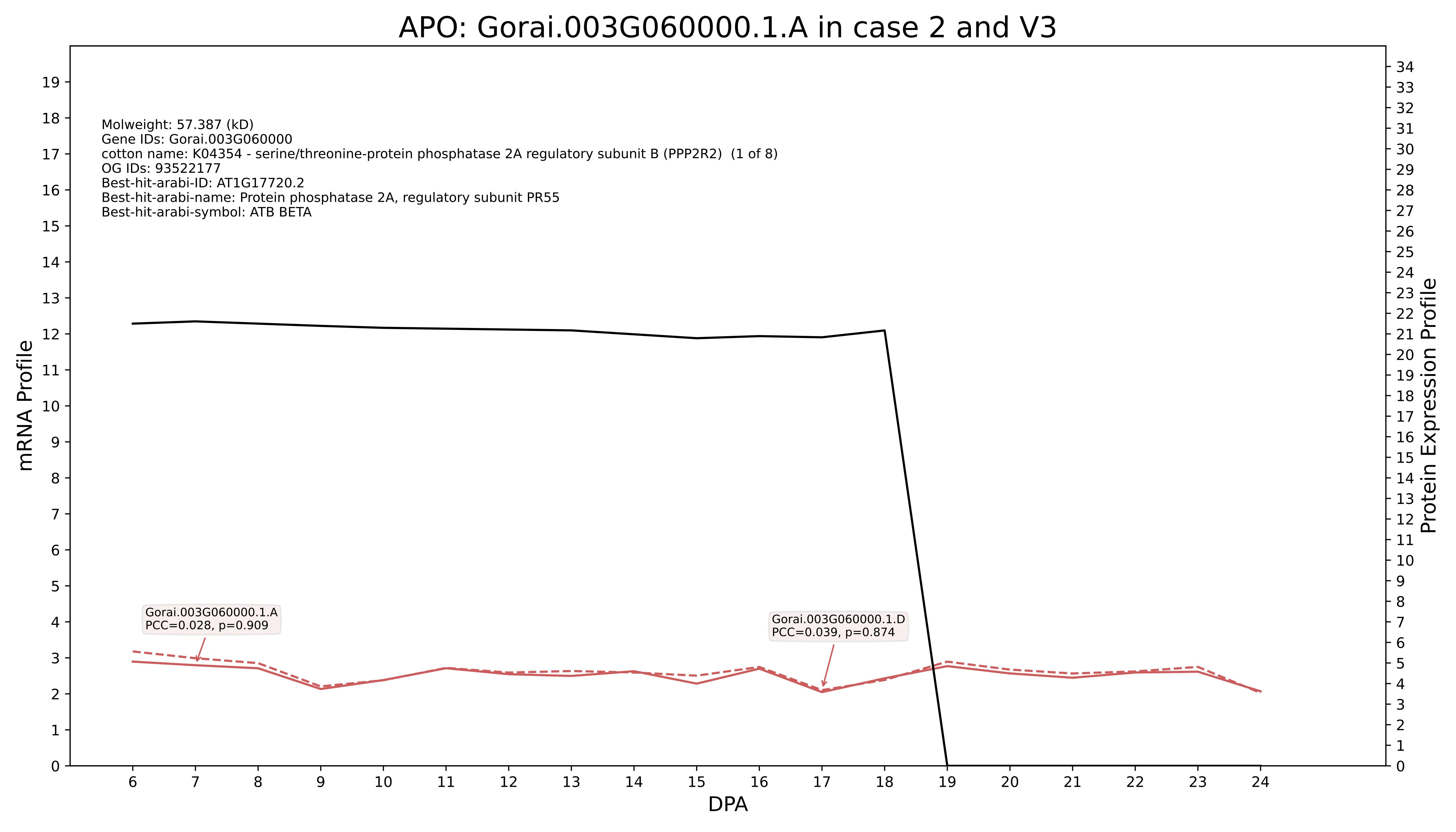
APO: Gorai.003G060000.1.A in case 2 and V3
Molweight: 57.387 (kD)
Gene IDs: Gorai.003G060000
cotton name: K04354 - serine/threonine-protein phosphatase 2A regulatory subunit B (PPP2R2) (1 of 8)
OG IDs: 93522177
Best-hit-arabi-ID: AT1G17720.2
Best-hit-arabi-name: Protein phosphatase 2A, regulatory subunit PR55
Best-hit-arabi-symbol: ATB BETA
Gorai.003G060000.1.A
PCC=0.028, p=0.909
Gorai.003G060000.1.D
PCC=0.039, p=0.874
mRNA Profile
Protein Expression Profile
DPA

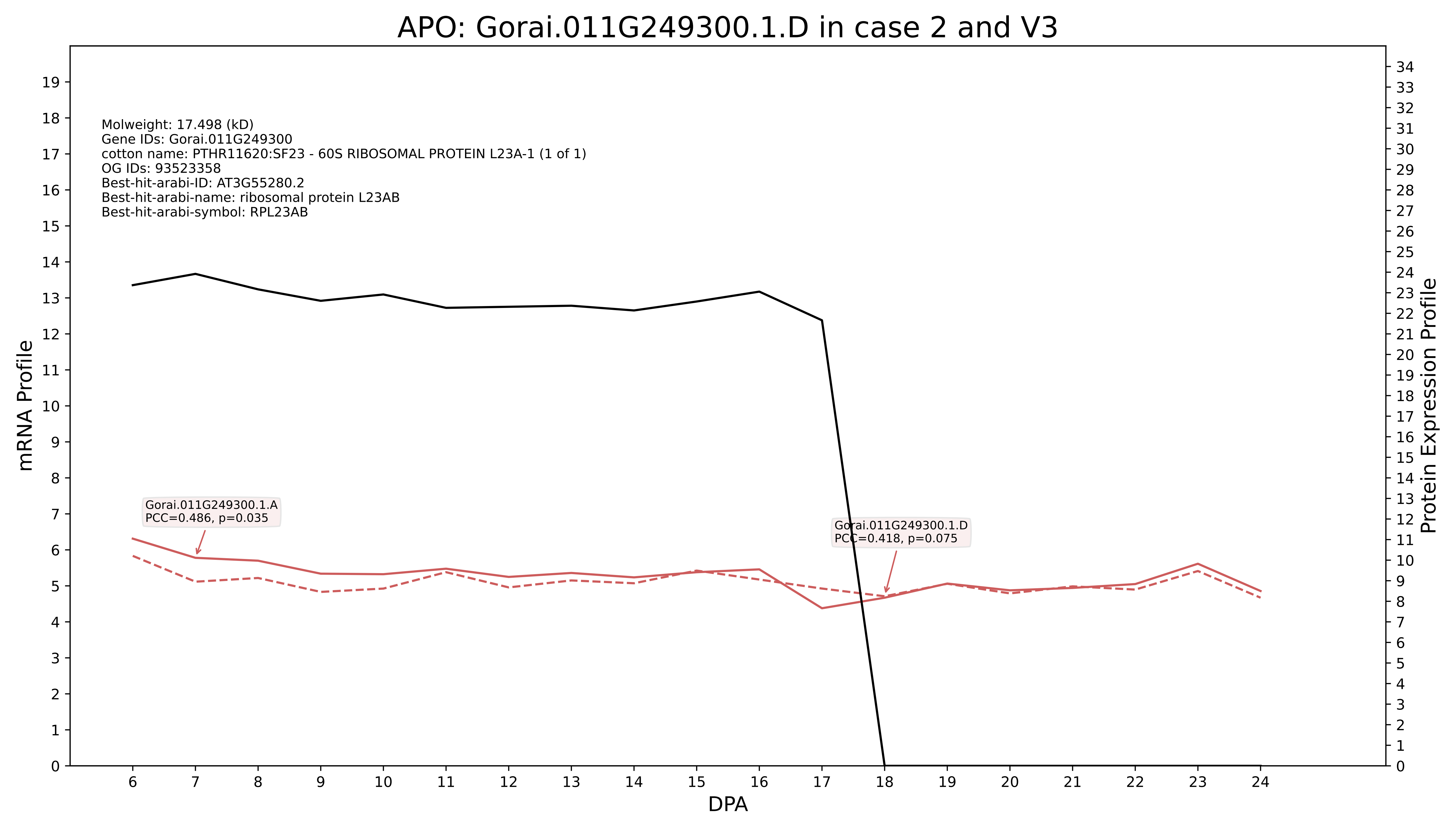

APO: Gorai.011G249300.1.D in case 2 and V3
Molweight: 17.498 (kD)
Gene IDs: Gorai.011G249300
cotton name: PTHR11620:SF23 - 60S RIBOSOMAL PROTEIN L23A-1 (1 of 1)
OG IDs: 93523358
Best-hit-arabi-ID: AT3G55280.2
Best-hit-arabi-name: ribosomal protein L23AB
Best-hit-arabi-symbol: RPL23AB
Gorai.011G249300.1.A
PCC=0.486, p=0.035
Gorai.011G249300.1.D
PCC=0.418, p=0.075
mRNA Profile
Protein Expression Profile
DPA

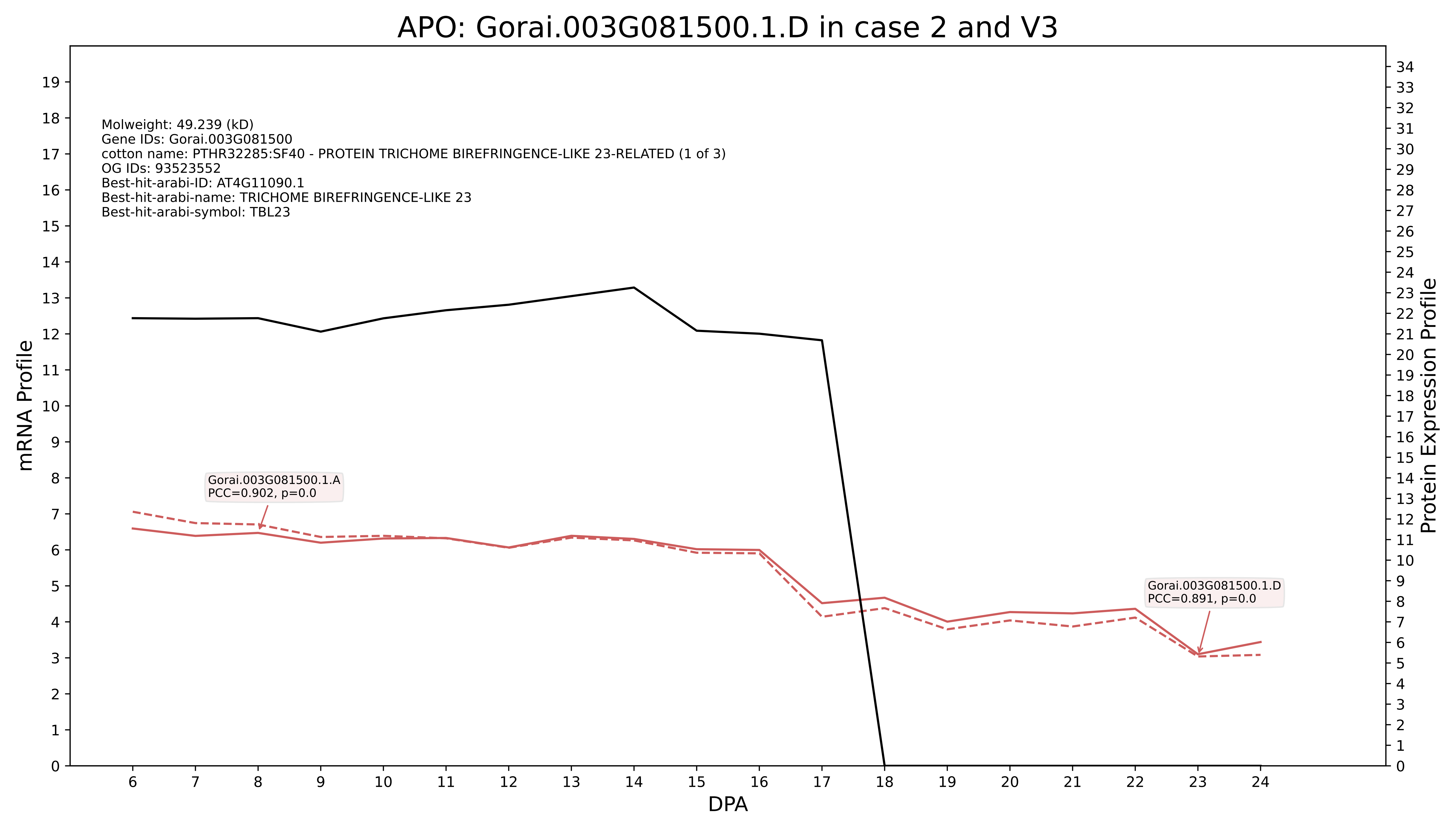
APO: Gorai.003G081500.1.D in case 2 and V3
Molweight: 49.239 (kD)
Gene IDs: Gorai.003G081500
cotton name: PTHR32285:SF40 - PROTEIN TRICHOME BIREFRINGENCE-LIKE 23-RELATED (1 of 3)
OG IDs: 93523552
Best-hit-arabi-ID: AT4G11090.1
Best-hit-arabi-name: TRICHOME BIREFRINGENCE-LIKE 23
Best-hit-arabi-symbol: TBL23
Gorai.003G081500.1.A
PCC=0.902, p=0.0
Gorai.003G081500.1.D
PCC=0.891, p=0.0
mRNA Profile
Protein Expression Profile
DPA

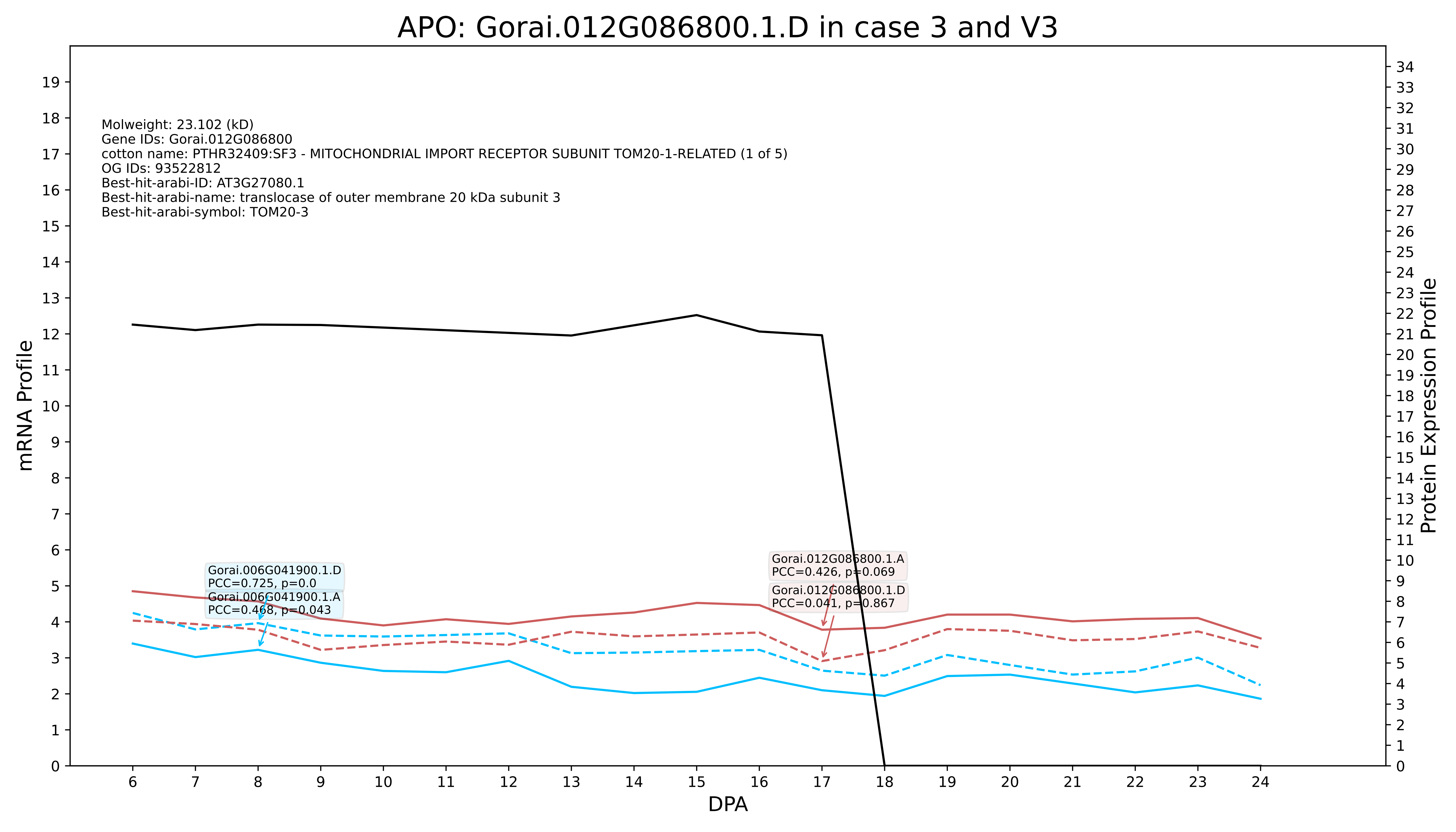
APO: Gorai.012G086800.1.D in case 3 and V3
Molweight: 23.102 (kD)
Gene IDs: Gorai.012G086800
cotton name: PTHR32409:SF3 - MITOCHONDRIAL IMPORT RECEPTOR SUBUNIT TOM20-1-RELATED (1 of 5)
OG IDs: 93522812
Best-hit-arabi-ID: AT3G27080.1
Best-hit-arabi-name: translocase of outer membrane 20 kDa subunit 3
Best-hit-arabi-symbol: TOM20-3
Gorai.006G041900.1.D
PCC=0.725, p=0.0
Gorai.006G041900.1.A
PCC=0.468, p=0.043
Gorai.012G086800.1.A
PCC=0.426, p=0.069
Gorai.012G086800.1.D
PCC=0.041, p=0.867
mRNA Profile
Protein Expression Profile
DPA

### APO: Gorai.013G154100.1.D in case 3 and V3

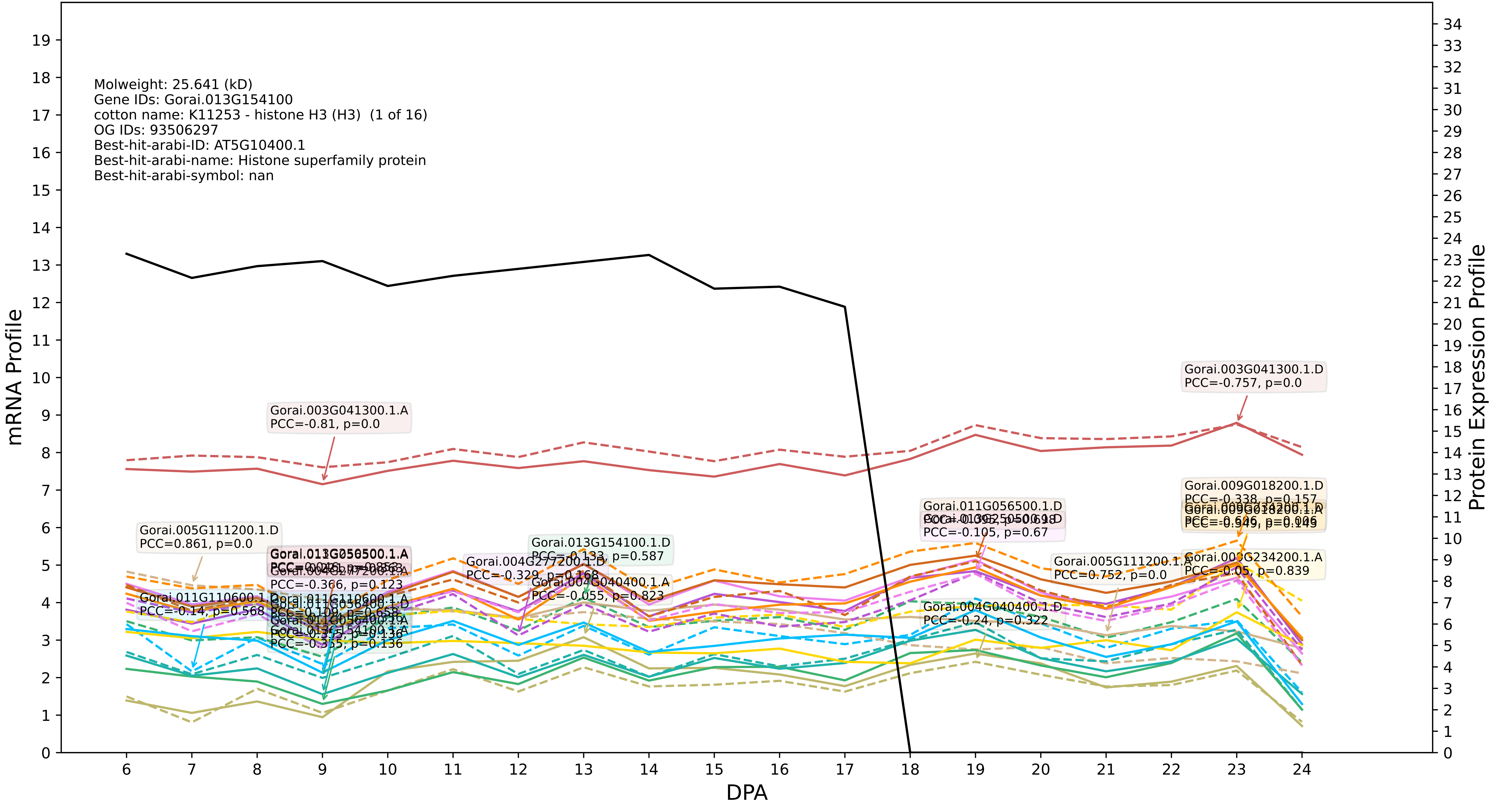

### APO: Gorai.001G103000.1.A in case 3 and V3

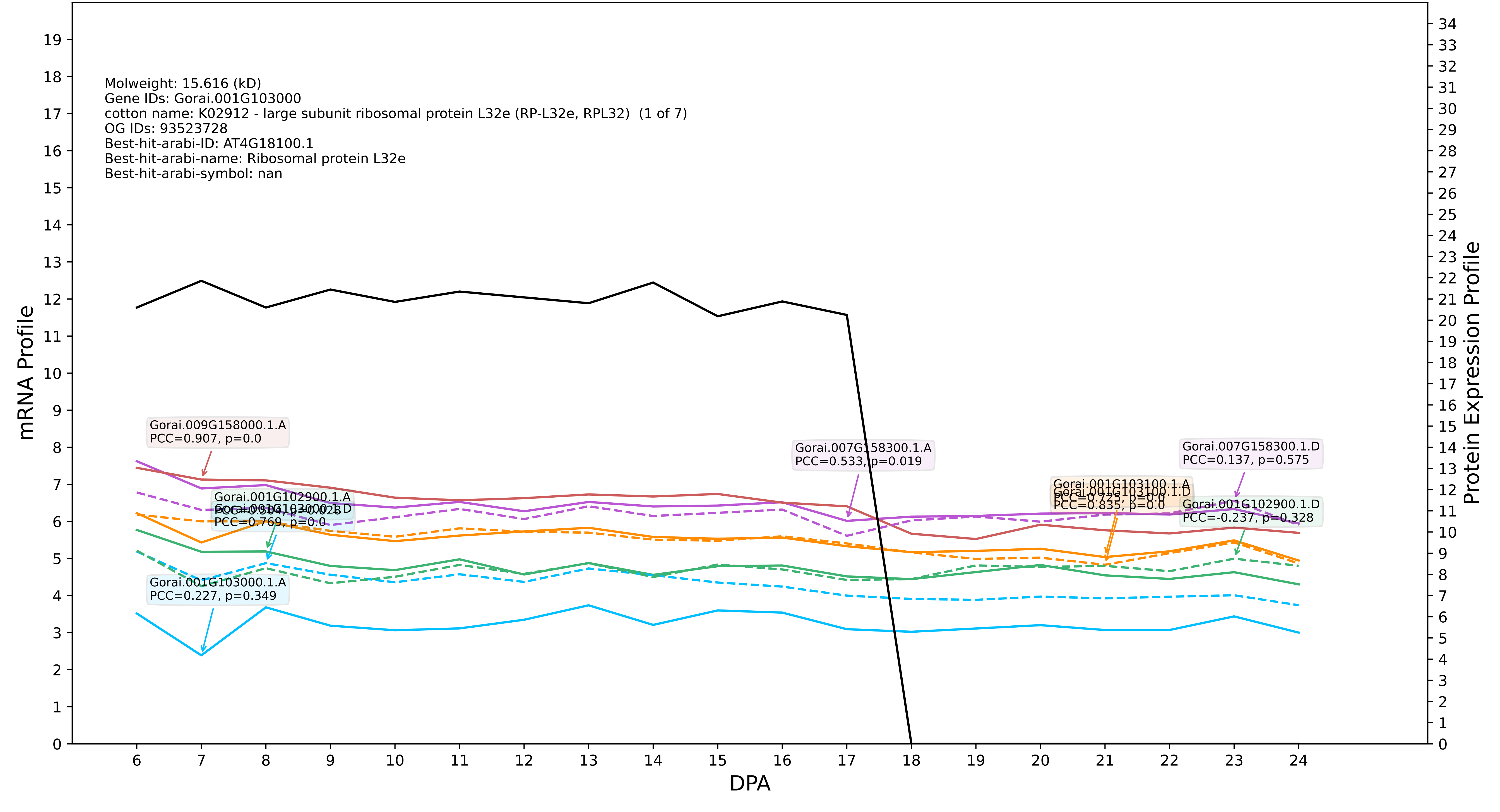

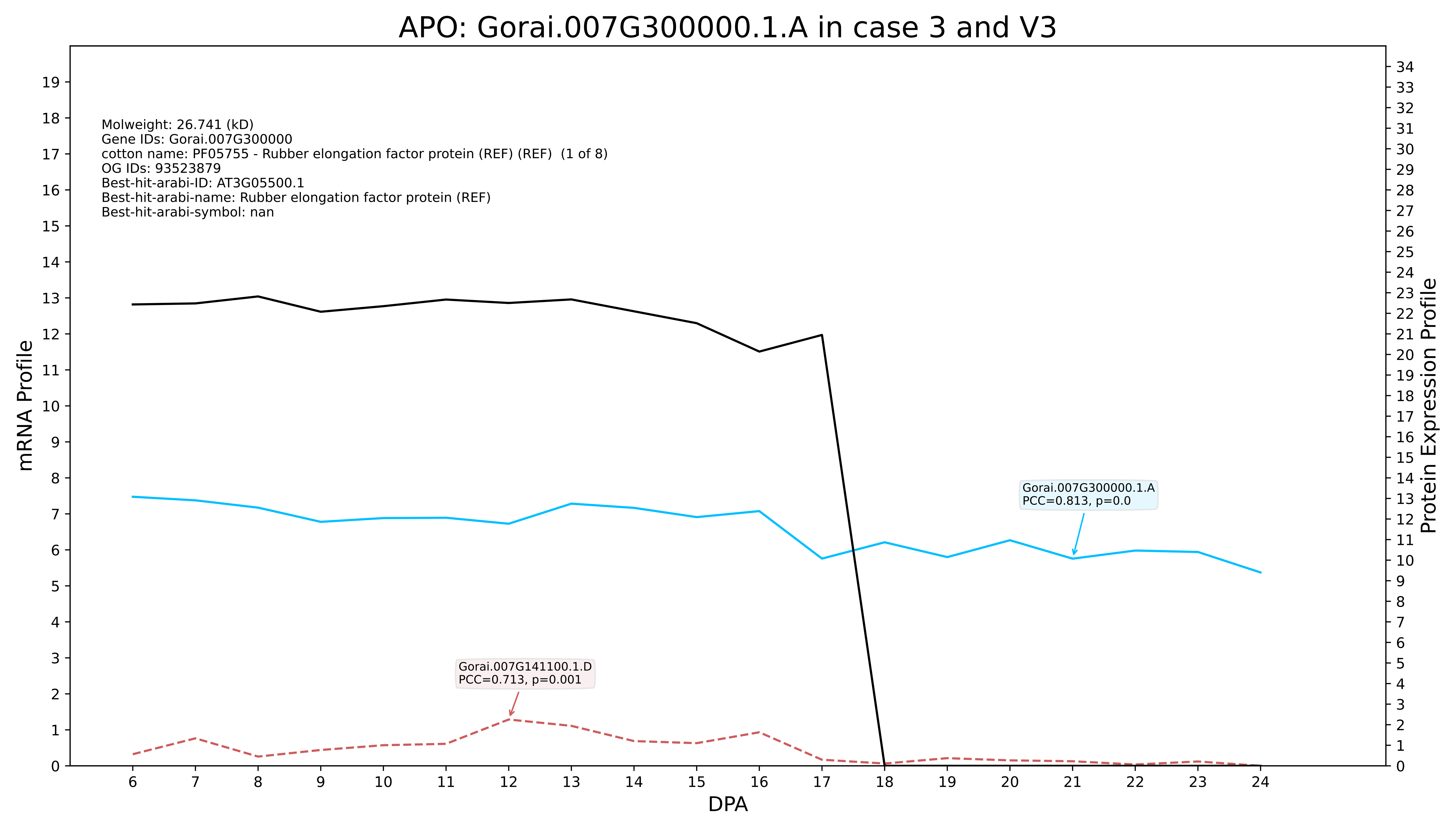

APO: Gorai.007G300000.1.A in case 3 and V3
Molweight: 26.741 (kD)
Gene IDs: Gorai.007G300000
cotton name: PF05755 - Rubber elongation factor protein (REF) (REF) (1 of 8)
OG IDs: 93523879
Best-hit-arabi-ID: AT3G05500.1
Best-hit-arabi-name: Rubber elongation factor protein (REF)
Best-hit-arabi-symbol: nan
Gorai.007G300000.1.A
PCC=0.813, p=0.0
Gorai.007G141100.1.D
PCC=0.713, p=0.001
mRNA Profile
Protein Expression Profile
DPA

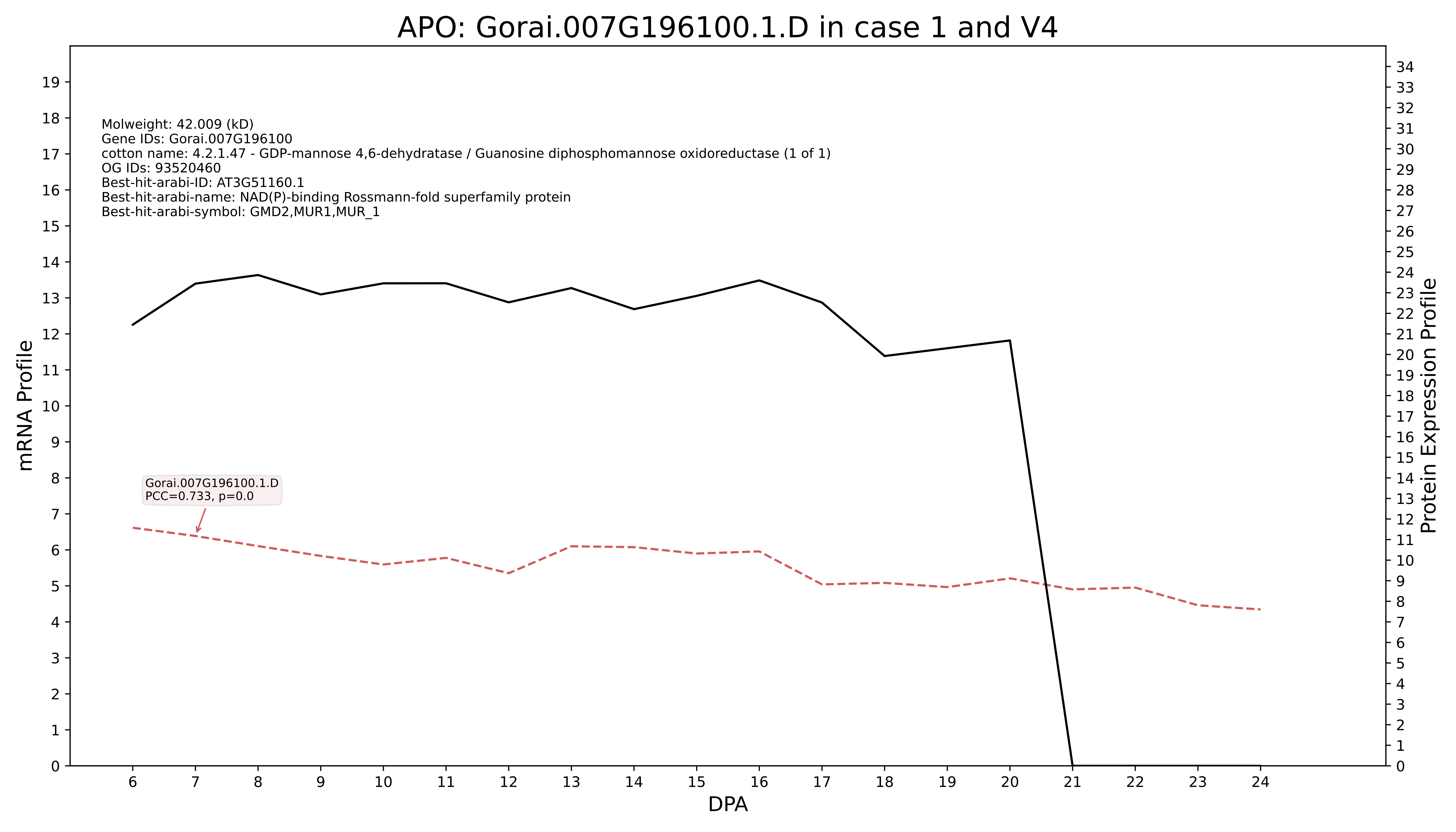
APO: Gorai.007G196100.1.D in case 1 and V4
Molweight: 42.009 (kD)
Gene IDs: Gorai.007G196100
cotton name: 4.2.1.47 - GDP-mannose 4,6-dehydratase / Guanosine diphosphomannose oxidoreductase (1 of 1)
OG IDs: 93520460
Best-hit-arabi-ID: AT3G51160.1
Best-hit-arabi-name: NAD(P)-binding Rossmann-fold superfamily protein
Best-hit-arabi-symbol: GMD2,MUR1,MUR_1
Gorai.007G196100.1.D
PCC=0.733, p=0.0
mRNA Profile
Protein Expression Profile
DPA

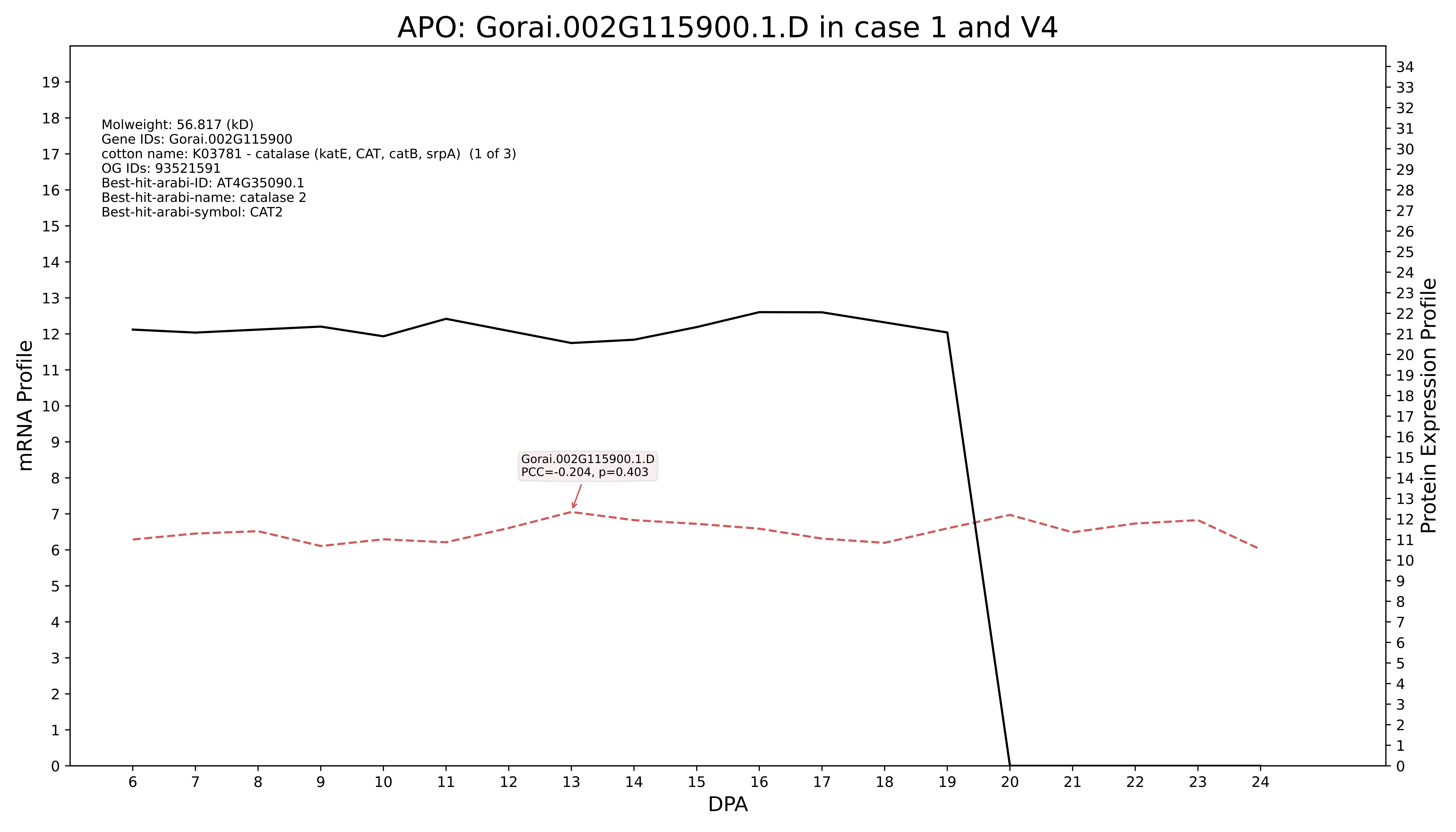

APO: Gorai.002G115900.1.D in case 1 and V4
Molweight: 56.817 (kD)
Gene IDs: Gorai.002G115900
cotton name: K03781 - catalase (katE, CAT, catB, srpA) (1 of 3)
OG IDs: 93521591
Best-hit-arabi-ID: AT4G35090.1
Best-hit-arabi-name: catalase 2
Best-hit-arabi-symbol: CAT2
Gorai.002G115900.1.D
PCC=-0.204, p=0.403
mRNA Profile
Protein Expression Profile
DPA

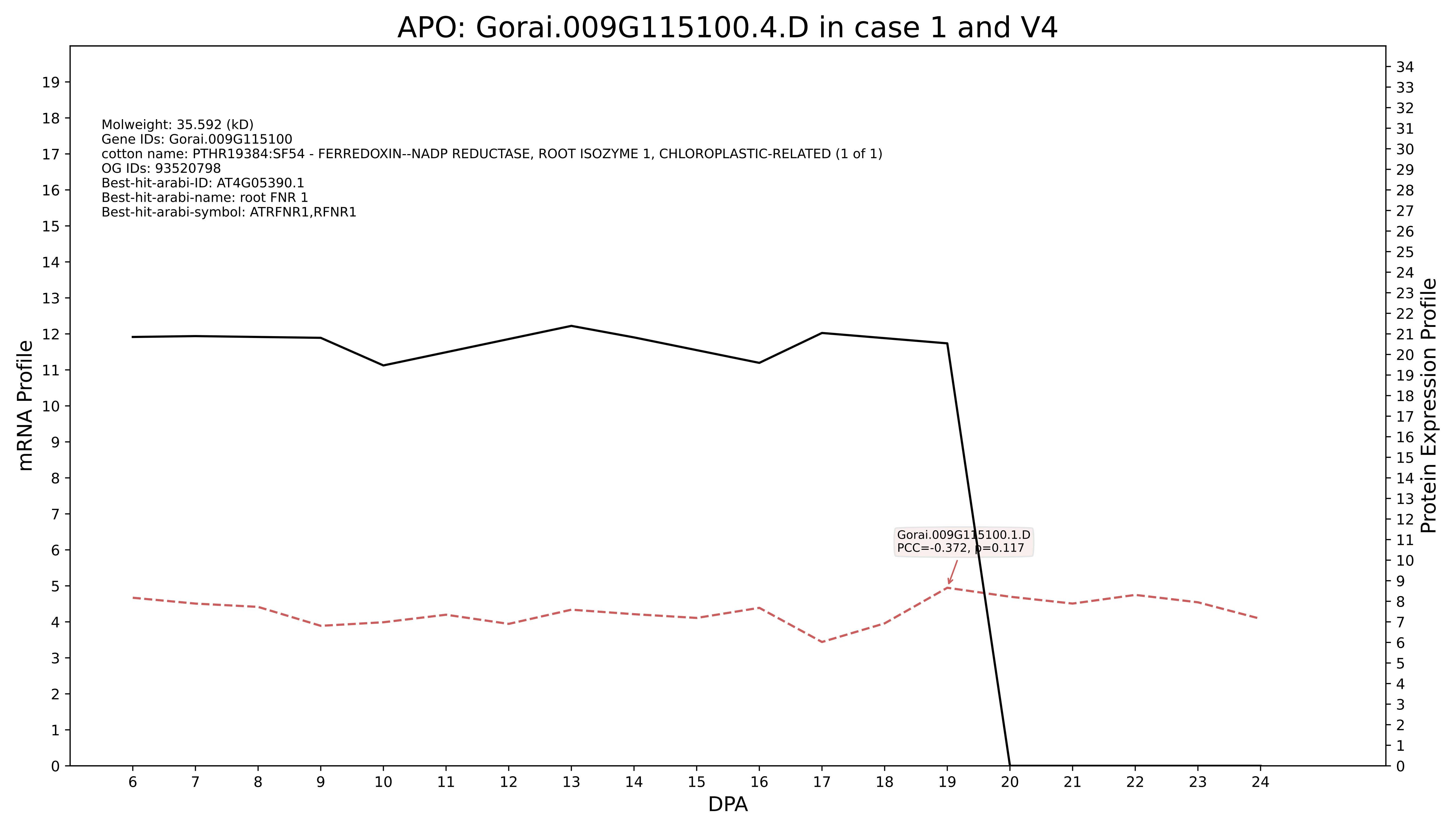

APO: Gorai.009G115100.4.D in case 1 and V4
Molweight: 35.592 (kD)
Gene IDs: Gorai.009G115100
cotton name: PTHR19384:SF54 - FERREDOXIN--NADP REDUCTASE, ROOT ISOZYME 1, CHLOROPLASTIC-RELATED (1 of 1)
OG IDs: 93520798
Best-hit-arabi-ID: AT4G05390.1
Best-hit-arabi-name: root FNR 1
Best-hit-arabi-symbol: ATRFNR1,RFNR1
Gorai.009G115100.1.D
PCC=-0.372, p=0.117
mRNA Profile
Protein Expression Profile
DPA

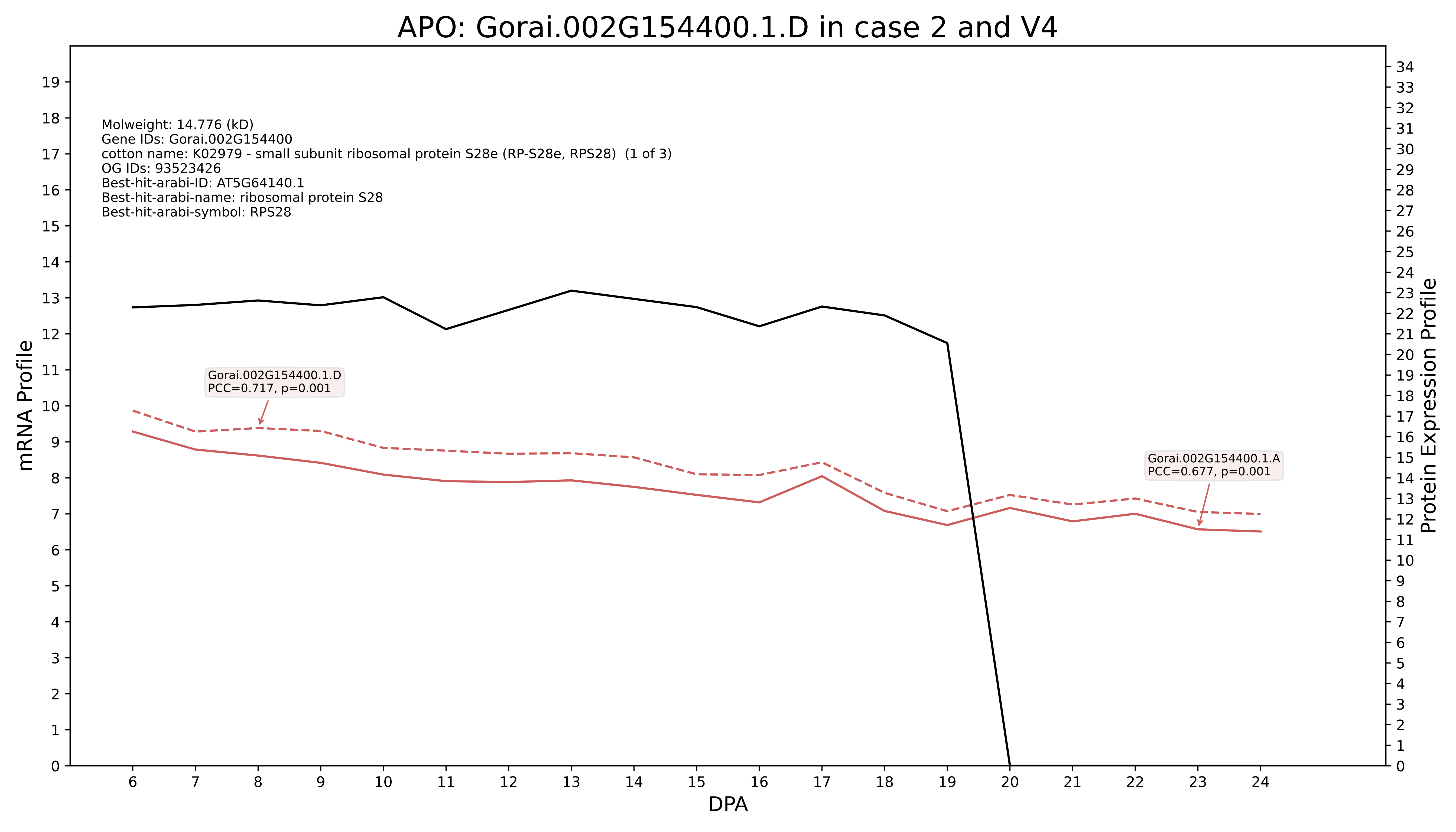

APO: Gorai.002G154400.1.D in case 2 and V4
Molweight: 14.776 (kD)
Gene IDs: Gorai.002G154400
cotton name: K02979 - small subunit ribosomal protein S28e (RP-S28e, RPS28) (1 of 3)
OG IDs: 93523426
Best-hit-arabi-ID: AT5G64140.1
Best-hit-arabi-name: ribosomal protein S28
Best-hit-arabi-symbol: RPS28
Gorai.002G154400.1.D
PCC=0.717, p=0.001
Gorai.002G154400.1.A
PCC=0.677, p=0.001
mRNA Profile
Protein Expression Profile
DPA

### APO: Gorai.001G180900.1.A in case 3 and V4

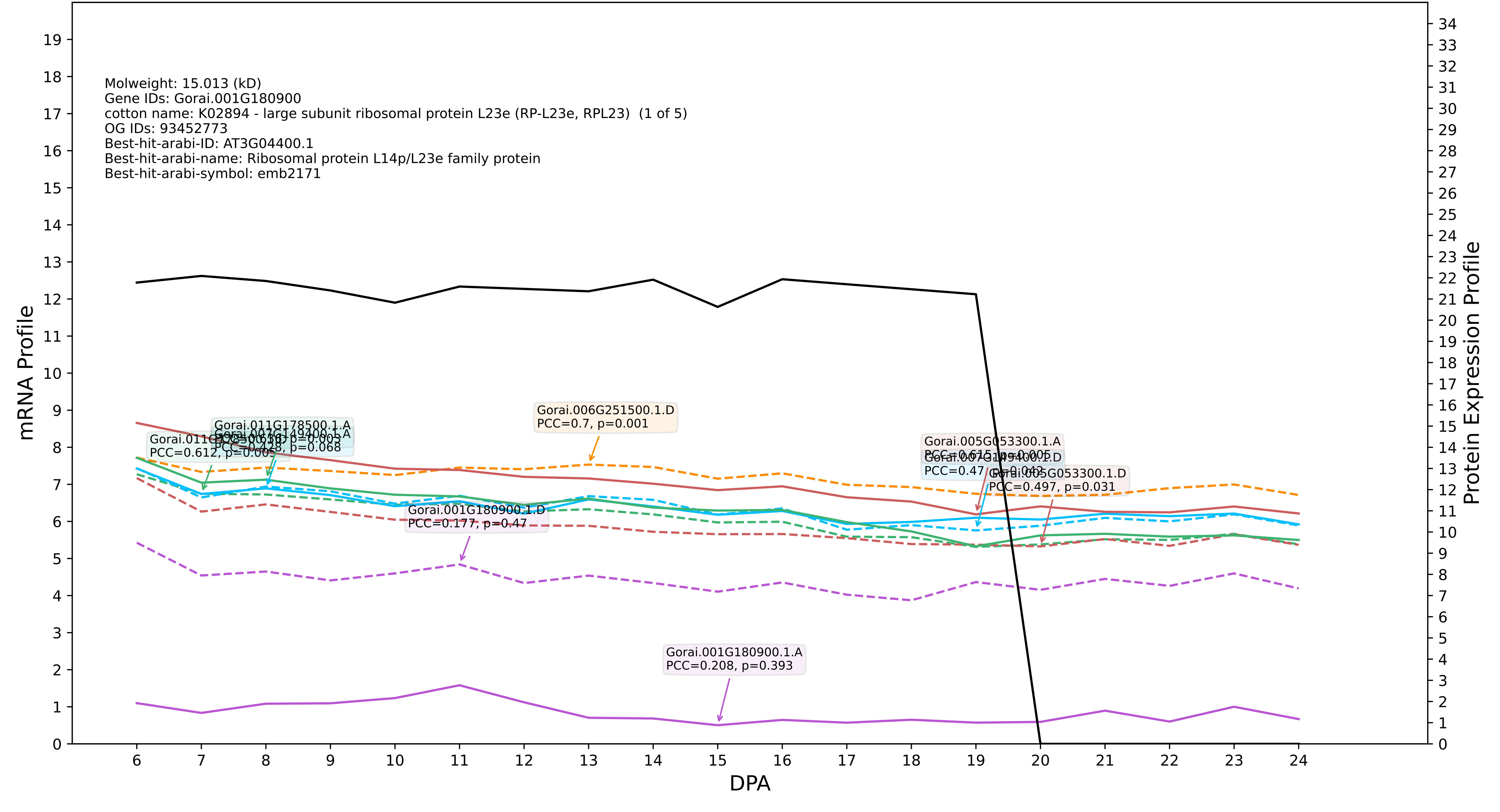

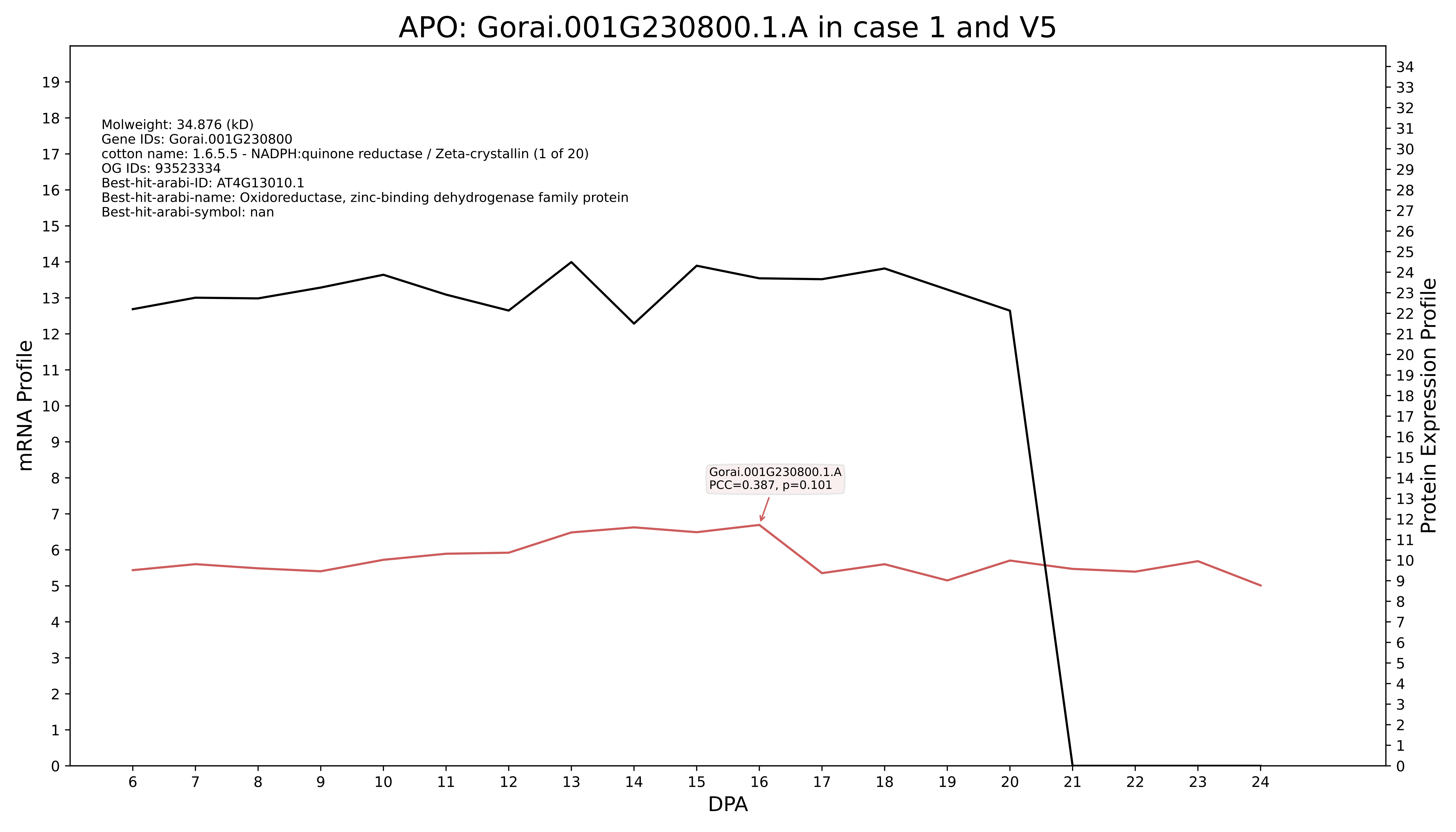

APO: Gorai.001G230800.1.A in case 1 and V5
Molweight: 34.876 (kD)
Gene IDs: Gorai.001G230800
cotton name: 1.6.5.5 - NADPH:quinone reductase / Zeta-crystallin (1 of 20)
OG IDs: 93523334
Best-hit-arabi-ID: AT4G13010.1
Best-hit-arabi-name: Oxidoreductase, zinc-binding dehydrogenase family protein
Best-hit-arabi-symbol: nan
Gorai.001G230800.1.A
PCC=0.387, p=0.101
mRNA Profile
Protein Expression Profile
DPA

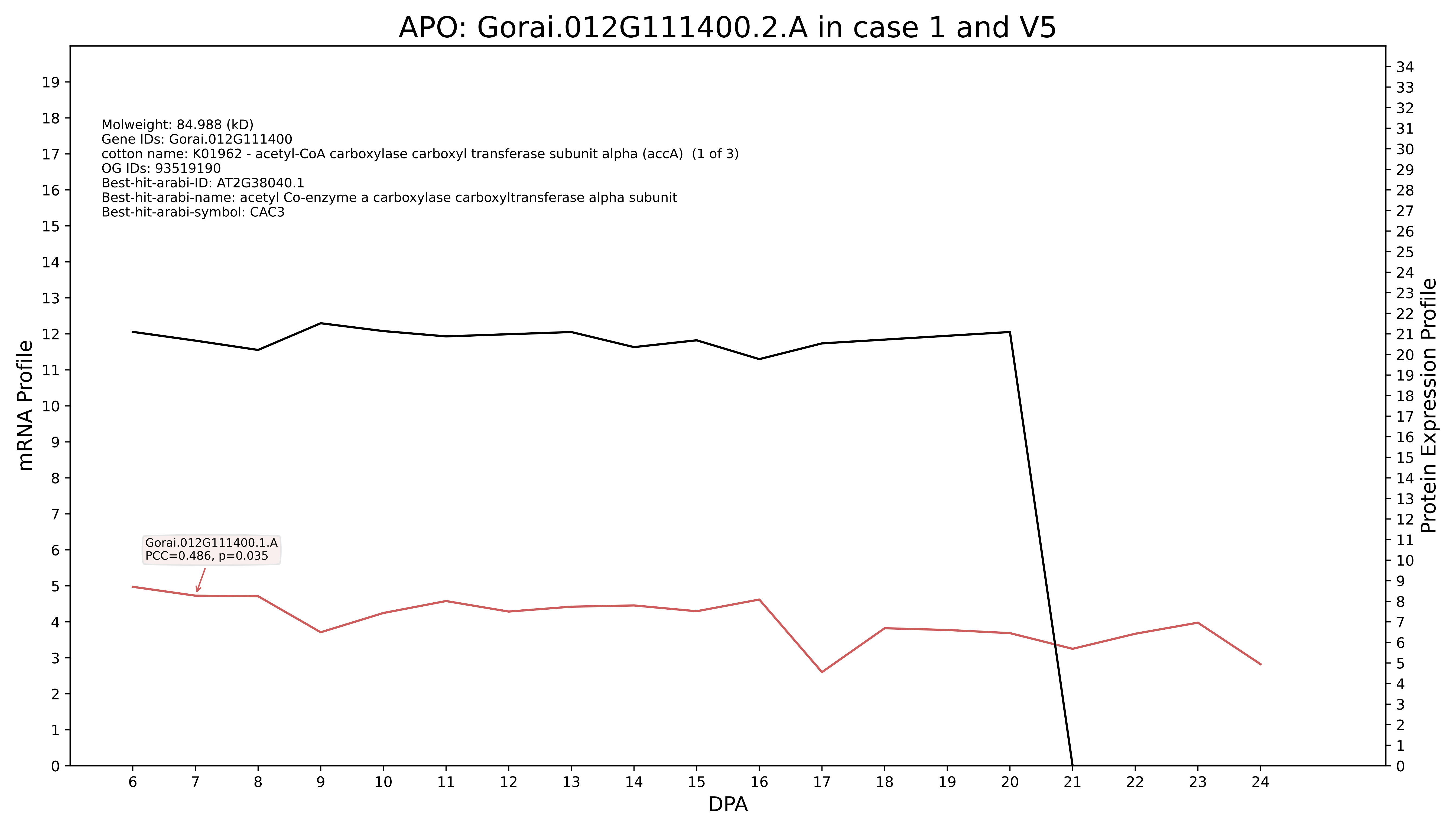

APO: Gorai.012G111400.2.A in case 1 and V5
Molweight: 84.988 (kD)
Gene IDs: Gorai.012G111400
cotton name: K01962 - acetyl-CoA carboxylase carboxyl transferase subunit alpha (accA) (1 of 3)
OG IDs: 93519190
Best-hit-arabi-ID: AT2G38040.1
Best-hit-arabi-name: acetyl Co-enzyme a carboxylase carboxyltransferase alpha subunit
Best-hit-arabi-symbol: CAC3
Gorai.012G111400.1.A
PCC=0.486, p=0.035
mRNA Profile
Protein Expression Profile
DPA

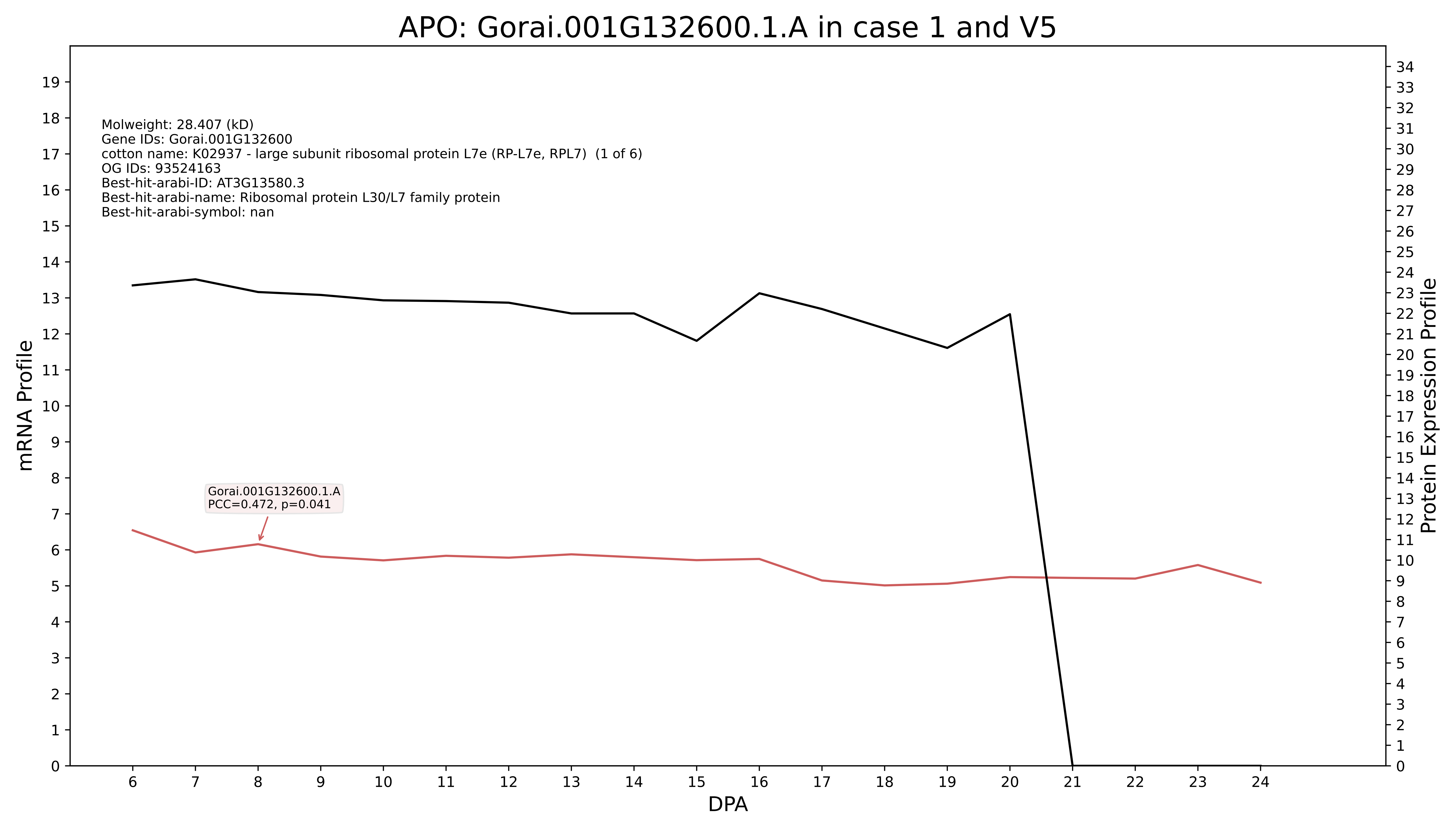
APO: Gorai.001G132600.1.A in case 1 and V5
Molweight: 28.407 (kD)
Gene IDs: Gorai.001G132600
cotton name: K02937 - large subunit ribosomal protein L7e (RP-L7e, RPL7) (1 of 6)
OG IDs: 93524163
Best-hit-arabi-ID: AT3G13580.3
Best-hit-arabi-name: Ribosomal protein L30/L7 family protein
Best-hit-arabi-symbol: nan
Gorai.001G132600.1.A
PCC=0.472, p=0.041
mRNA Profile
Protein Expression Profile
DPA

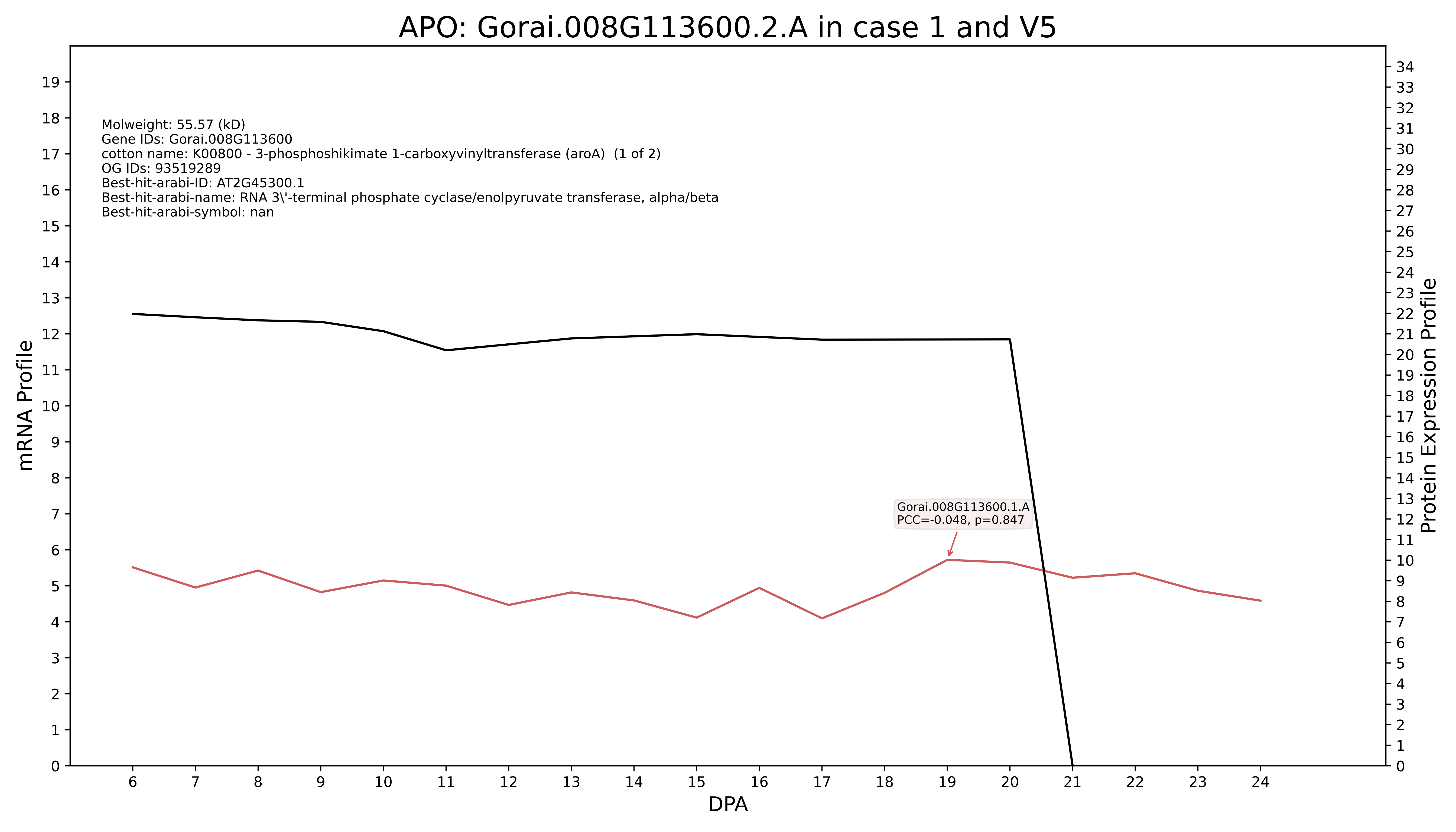

APO: Gorai.008G113600.2.A in case 1 and V5
Molweight: 55.57 (kD)
Gene IDs: Gorai.008G113600
cotton name: K00800 - 3-phosphoshikimate 1-carboxyvinyltransferase (aroA) (1 of 2)
OG IDs: 93519289
Best-hit-arabi-ID: AT2G45300.1
Best-hit-arabi-name: RNA 3\'-terminal phosphate cyclase/enolpyruvate transferase, alpha/beta
Best-hit-arabi-symbol: nan
Gorai.008G113600.1.A
PCC=-0.048, p=0.847
mRNA Profile
Protein Expression Profile
DPA

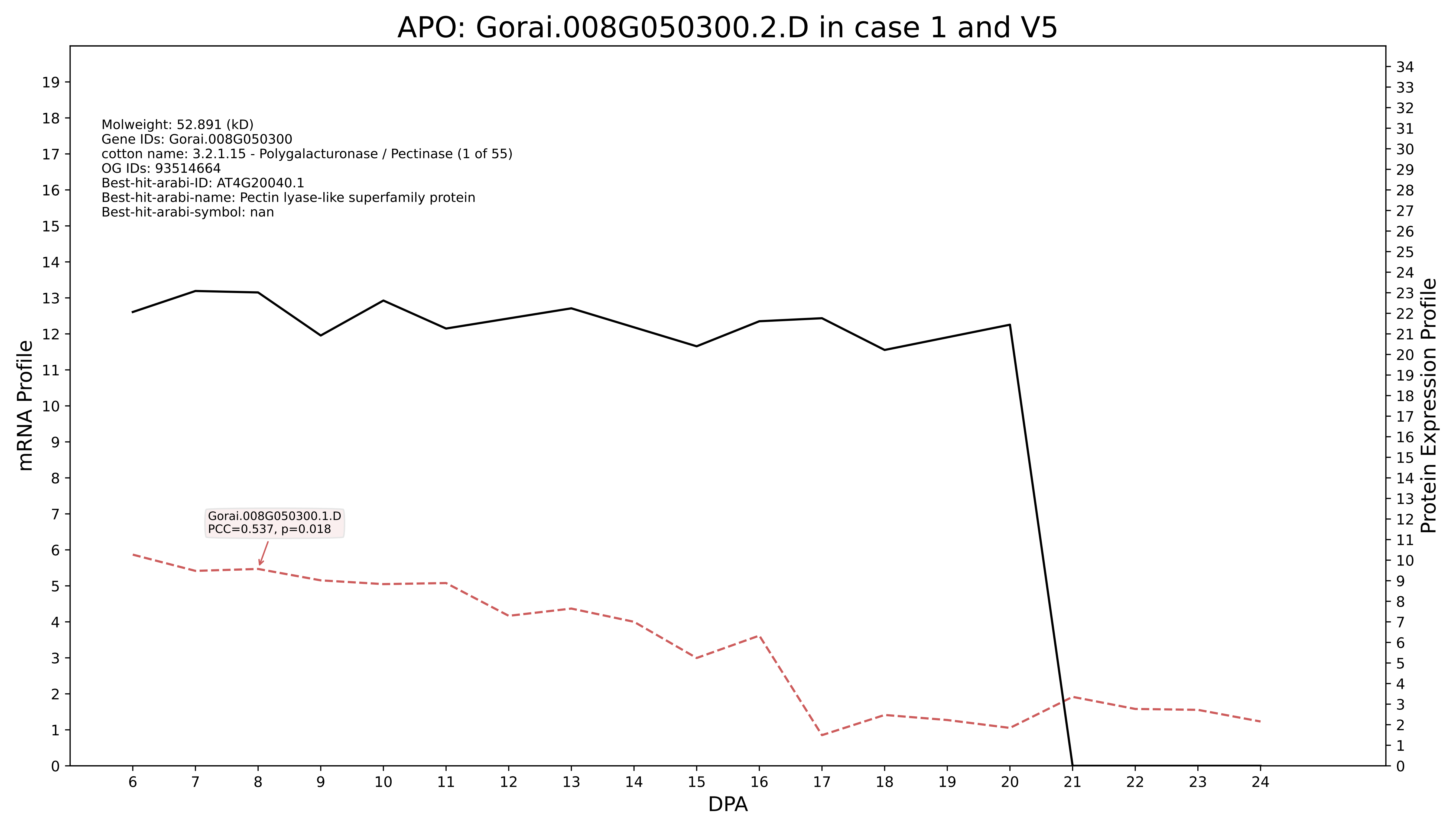
APO: Gorai.008G050300.2.D in case 1 and V5
Molweight: 52.891 (kD)
Gene IDs: Gorai.008G050300
cotton name: 3.2.1.15 - Polygalacturonase / Pectinase (1 of 55)
OG IDs: 93514664
Best-hit-arabi-ID: AT4G20040.1
Best-hit-arabi-name: Pectin lyase-like superfamily protein
Best-hit-arabi-symbol: nan
Gorai.008G050300.1.D
PCC=0.537, p=0.018
mRNA Profile
Protein Expression Profile
DPA

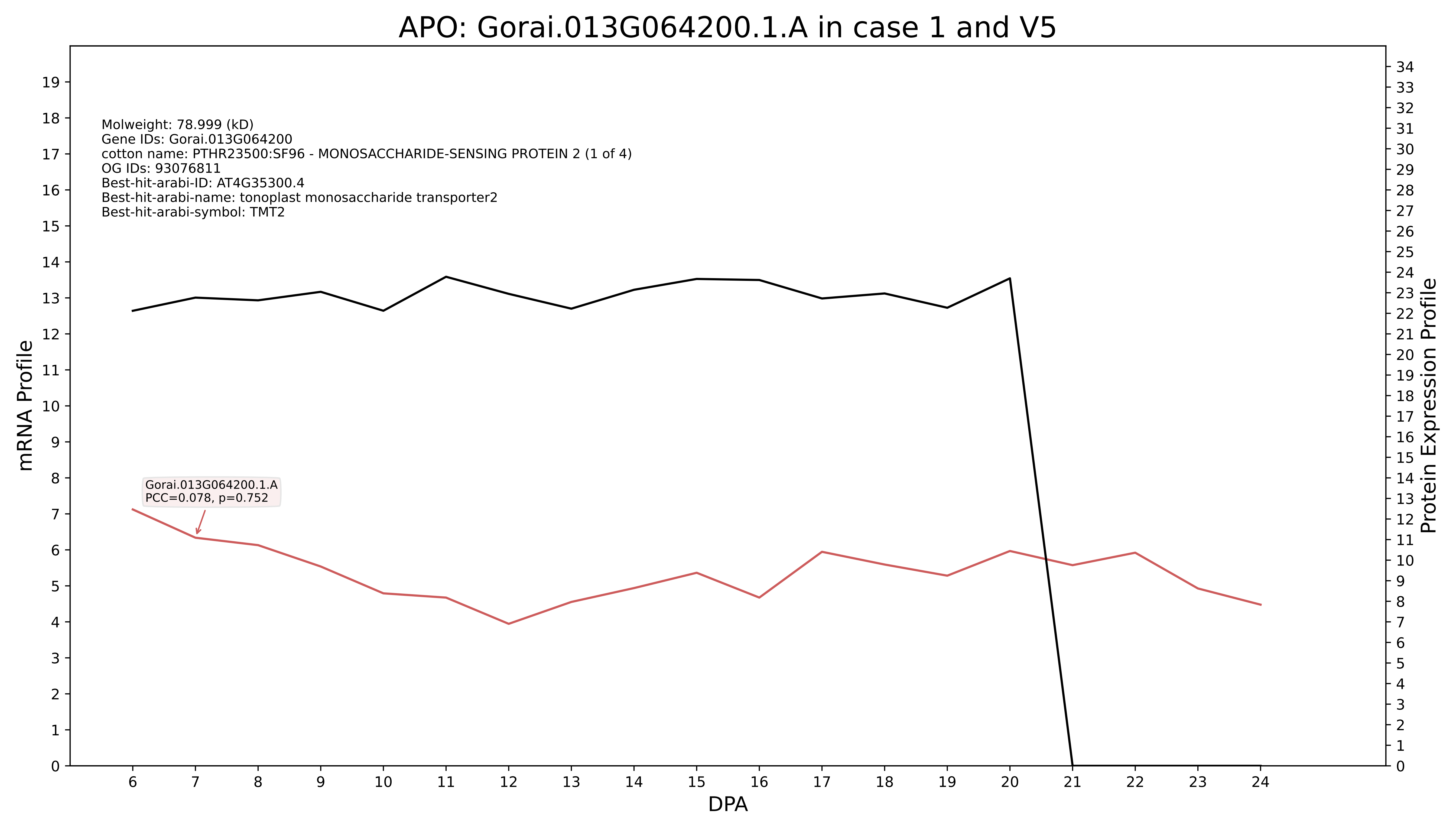

APO: Gorai.013G064200.1.A in case 1 and V5
Molweight: 78.999 (kD)
Gene IDs: Gorai.013G064200
cotton name: PTHR23500:SF96 - MONOSACCHARIDE-SENSING PROTEIN 2 (1 of 4)
OG IDs: 93076811
Best-hit-arabi-ID: AT4G35300.4
Best-hit-arabi-name: tonoplast monosaccharide transporter2
Best-hit-arabi-symbol: TMT2
Gorai.013G064200.1.A
PCC=0.078, p=0.752
mRNA Profile
Protein Expression Profile
DPA

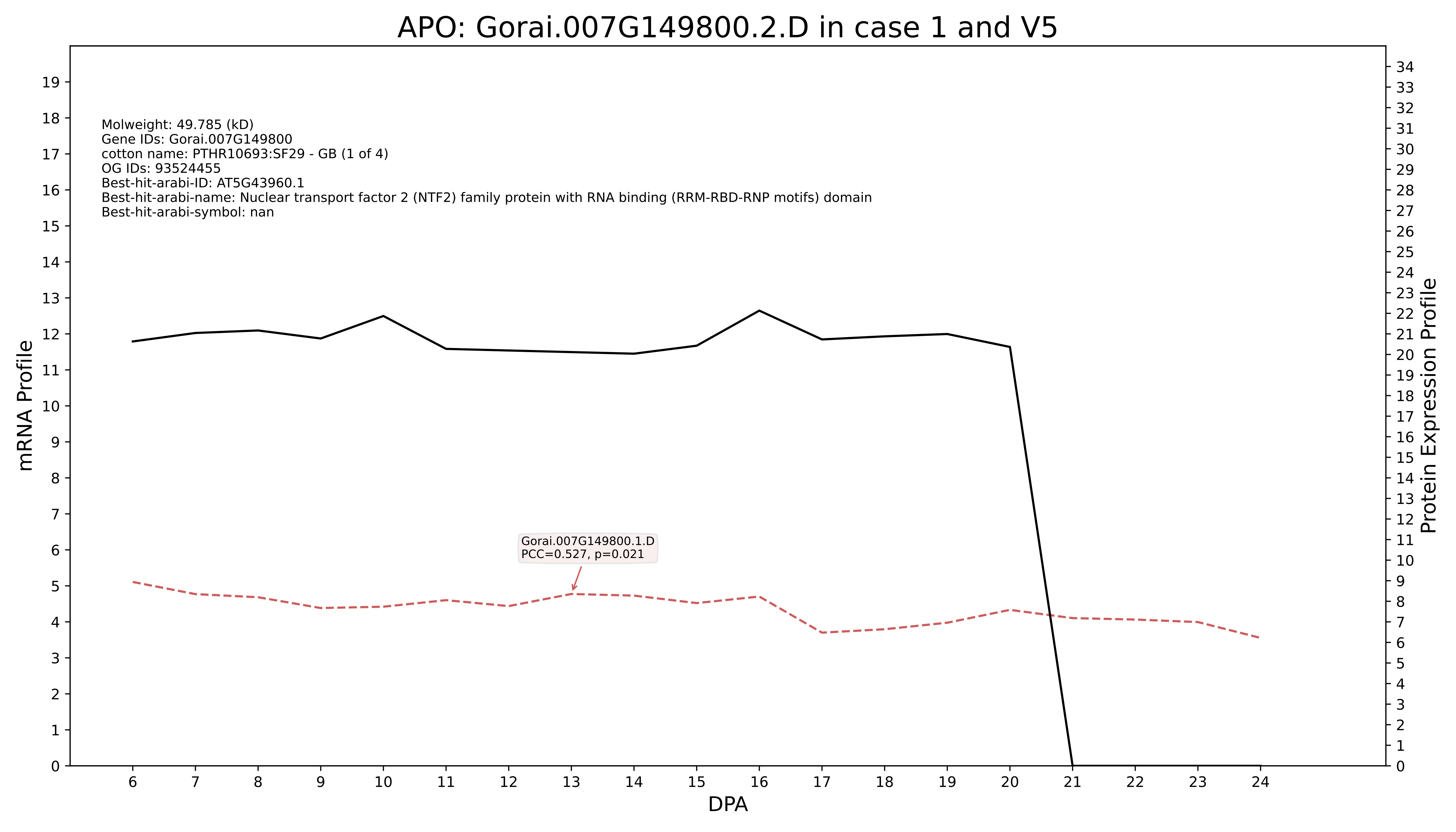
APO: Gorai.007G149800.2.D in case 1 and V5
Molweight: 49.785 (kD)
Gene IDs: Gorai.007G149800
cotton name: PTHR10693:SF29 - GB (1 of 4)
OG IDs: 93524455
Best-hit-arabi-ID: AT5G43960.1
Best-hit-arabi-name: Nuclear transport factor 2 (NTF2) family protein with RNA binding (RRM-RBD-RNP motifs) domain
Best-hit-arabi-symbol: nan
Gorai.007G149800.1.D
PCC=0.527, p=0.021
mRNA Profile
Protein Expression Profile
DPA

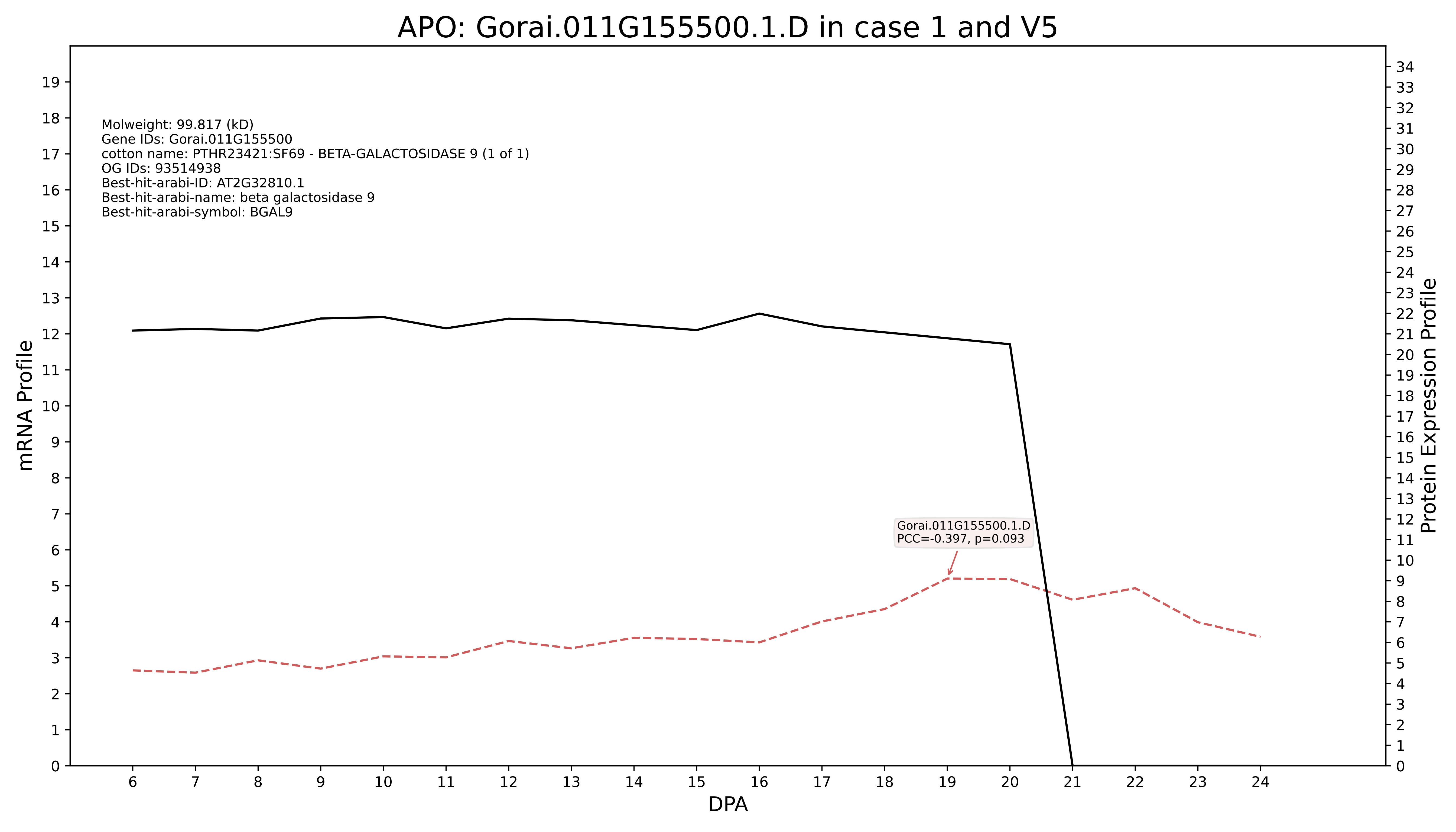

APO: Gorai.011G155500.1.D in case 1 and V5
Molweight: 99.817 (kD)
Gene IDs: Gorai.011G155500
cotton name: PTHR23421:SF69 - BETA-GALACTOSIDASE 9 (1 of 1)
OG IDs: 93514938
Best-hit-arabi-ID: AT2G32810.1
Best-hit-arabi-name: beta galactosidase 9
Best-hit-arabi-symbol: BGAL9
Gorai.011G155500.1.D
PCC=-0.397, p=0.093
mRNA Profile
Protein Expression Profile
DPA

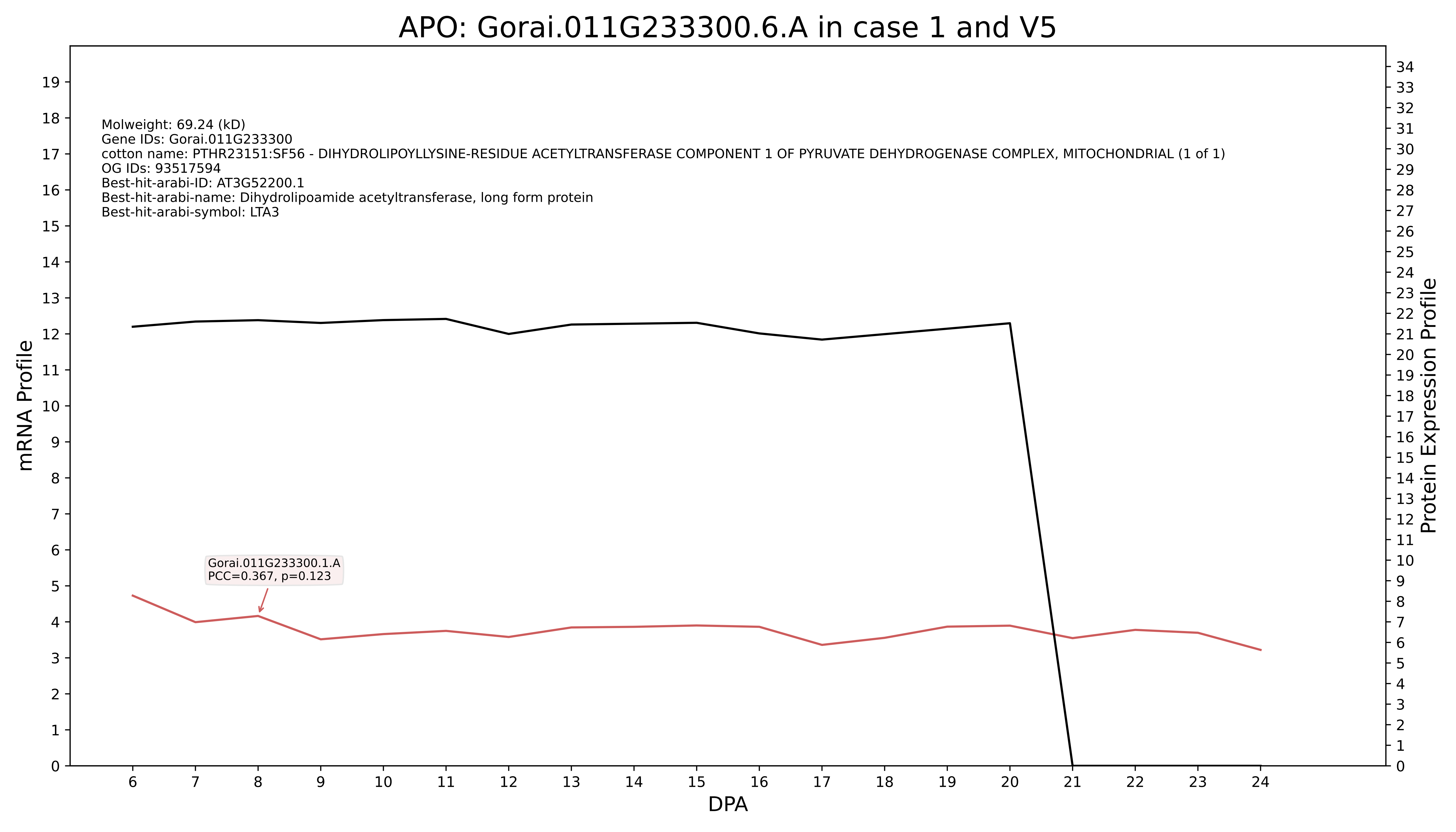
APO: Gorai.011G233300.6.A in case 1 and V5
Molweight: 69.24 (kD)
Gene IDs: Gorai.011G233300
cotton name: PTHR23151:SF56 - DIHYDROLIPOYLLYSINE-RESIDUE ACETYLTRANSFERASE COMPONENT 1 OF PYRUVATE DEHYDROGENASE COMPLEX, MITOCHONDRIAL (1 of 1)
OG IDs: 93517594
Best-hit-arabi-ID: AT3G52200.1
Best-hit-arabi-name: Dihydrolipoamide acetyltransferase, long form protein
Best-hit-arabi-symbol: LTA3
Gorai.011G233300.1.A
PCC=0.367, p=0.123
mRNA Profile
Protein Expression Profile
DPA

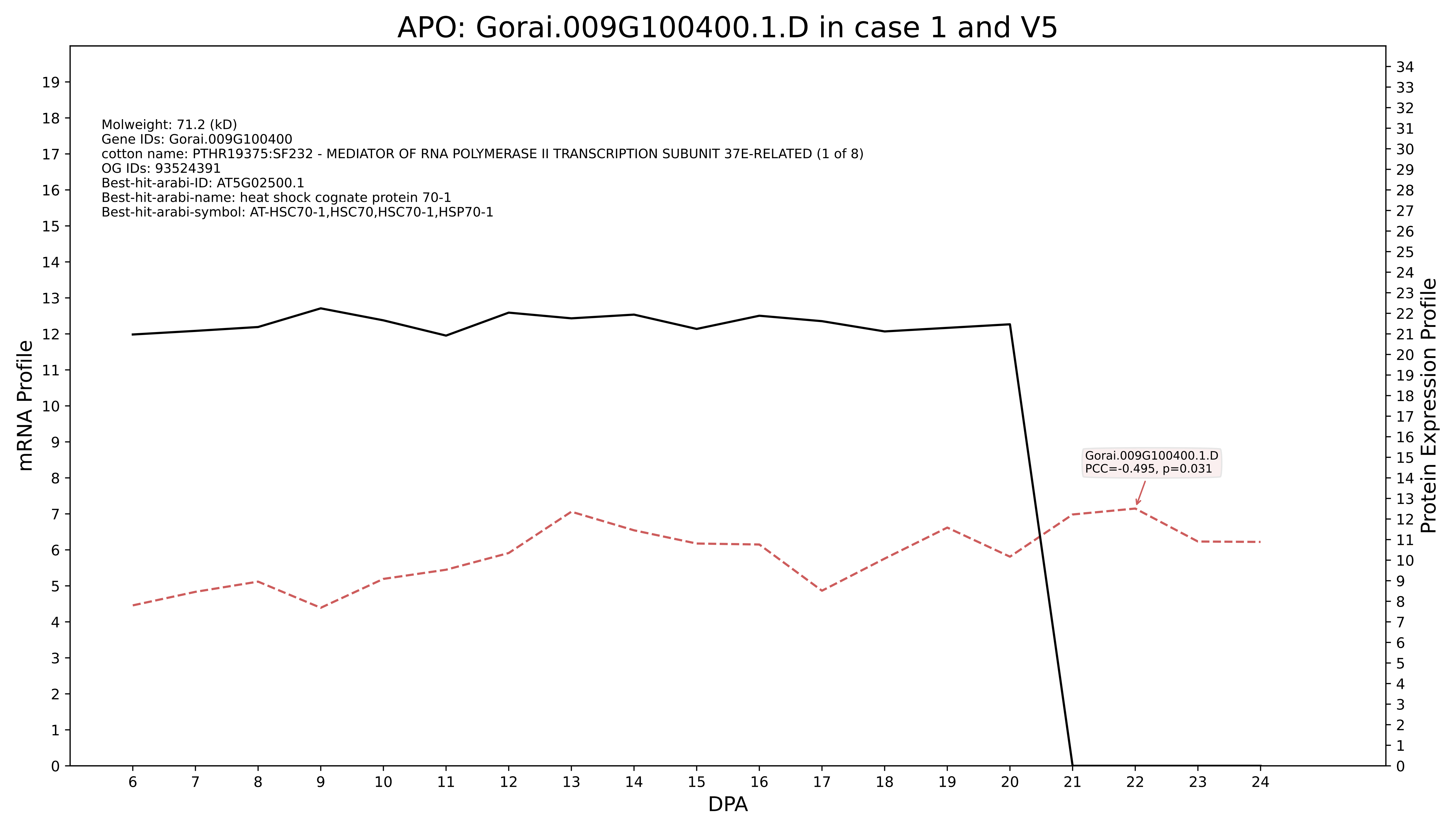
APO: Gorai.009G100400.1.D in case 1 and V5
Molweight: 71.2 (kD)
Gene IDs: Gorai.009G100400
cotton name: PTHR19375:SF232 - MEDIATOR OF RNA POLYMERASE II TRANSCRIPTION SUBUNIT 37E-RELATED (1 of 8)
OG IDs: 93524391
Best-hit-arabi-ID: AT5G02500.1
Best-hit-arabi-name: heat shock cognate protein 70-1
Best-hit-arabi-symbol: AT-HSC70-1,HSC70,HSC70-1,HSP70-1
Gorai.009G100400.1.D
PCC=-0.495, p=0.031
mRNA Profile
Protein Expression Profile
DPA

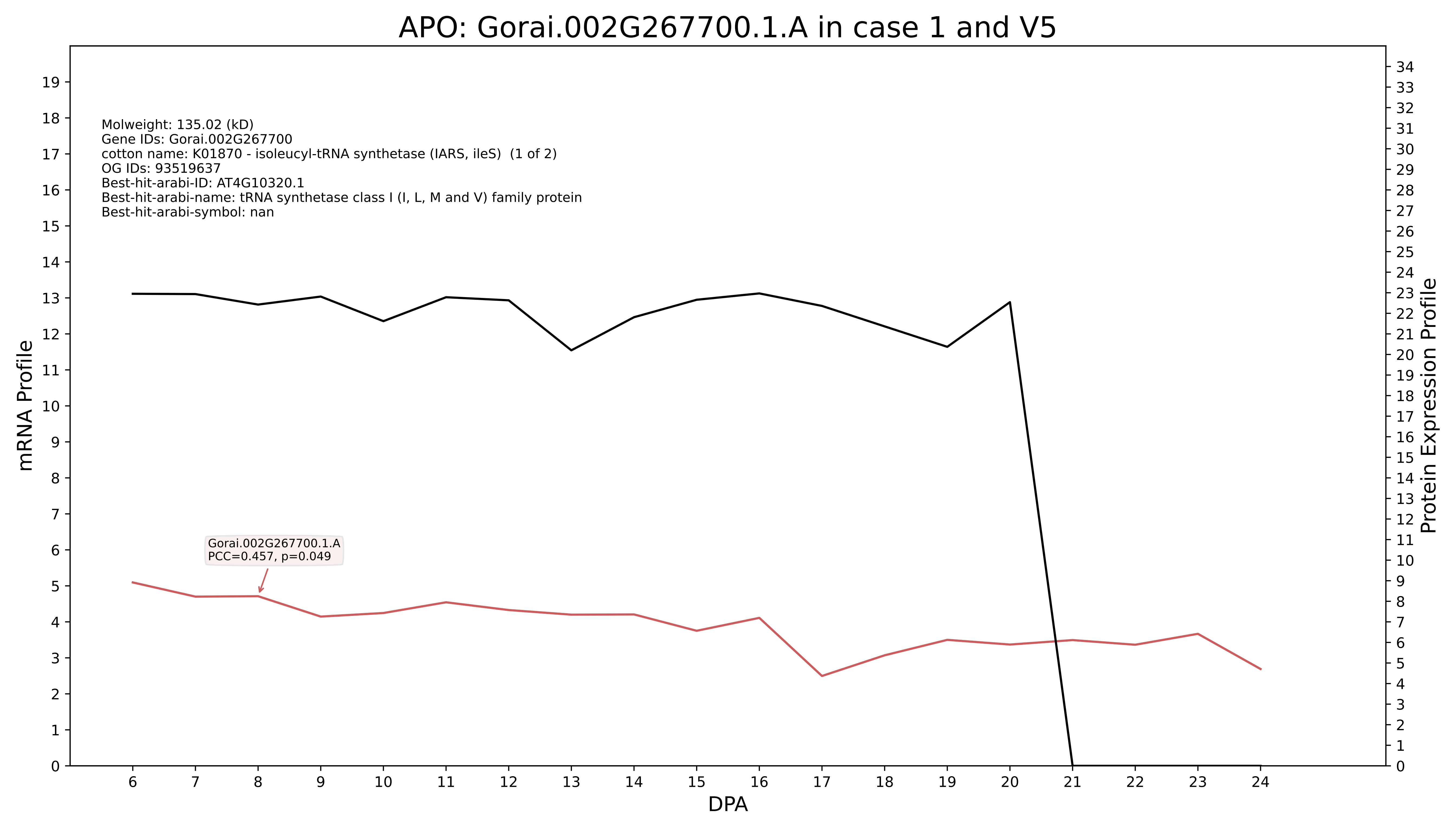
APO: Gorai.002G267700.1.A in case 1 and V5
Molweight: 135.02 (kD)
Gene IDs: Gorai.002G267700
cotton name: K01870 - isoleucyl-tRNA synthetase (IARS, ileS) (1 of 2)
OG IDs: 93519637
Best-hit-arabi-ID: AT4G10320.1
Best-hit-arabi-name: tRNA synthetase class I (I, L, M and V) family protein
Best-hit-arabi-symbol: nan
Gorai.002G267700.1.A
PCC=0.457, p=0.049
mRNA Profile
Protein Expression Profile
DPA

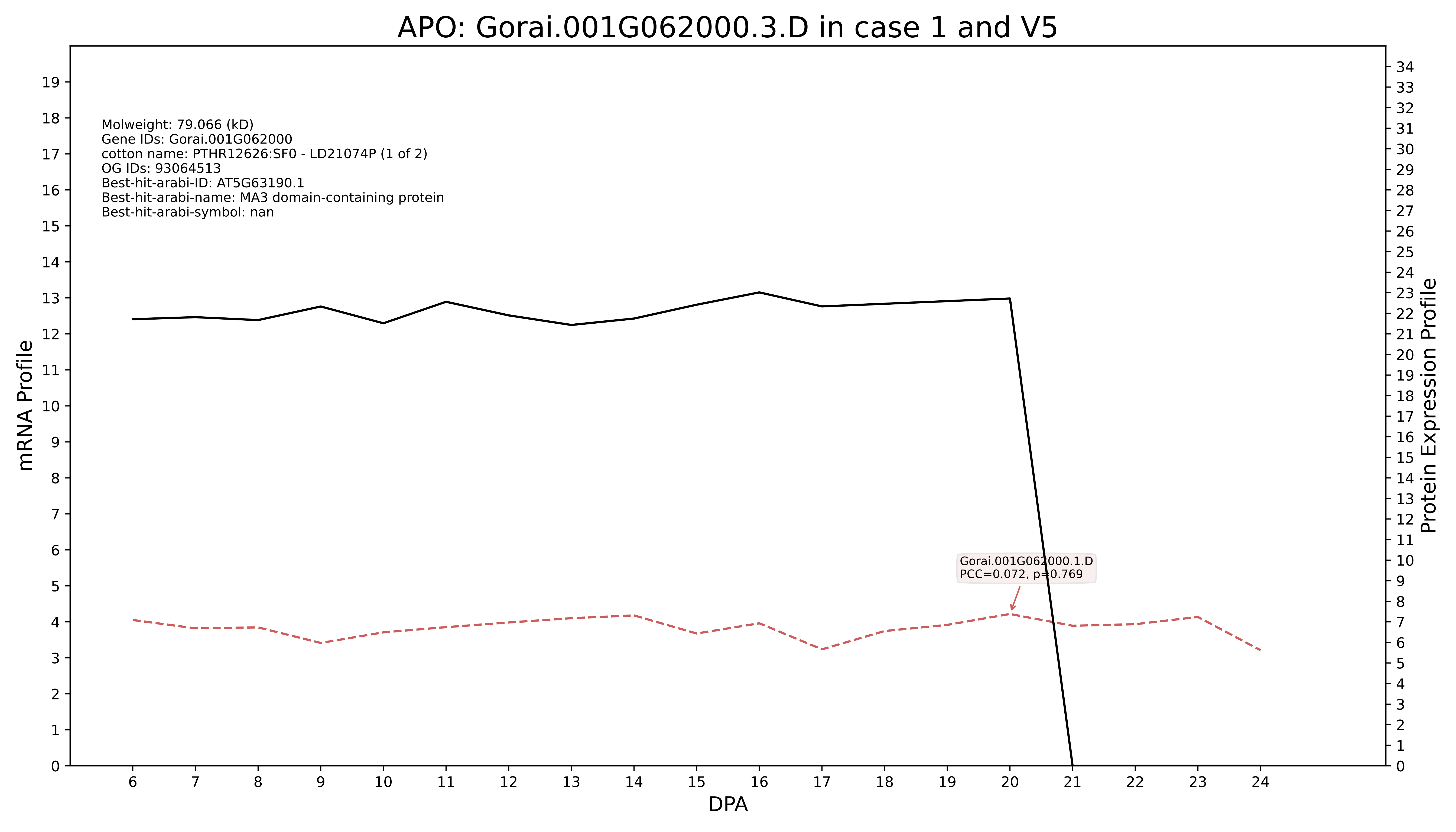
APO: Gorai.001G062000.3.D in case 1 and V5
Molweight: 79.066 (kD)
Gene IDs: Gorai.001G062000
cotton name: PTHR12626:SF0 - LD21074P (1 of 2)
OG IDs: 93064513
Best-hit-arabi-ID: AT5G63190.1
Best-hit-arabi-name: MA3 domain-containing protein
Best-hit-arabi-symbol: nan
Gorai.001G062000.1.D
PCC=0.072, p=0.769
mRNA Profile
Protein Expression Profile
DPA

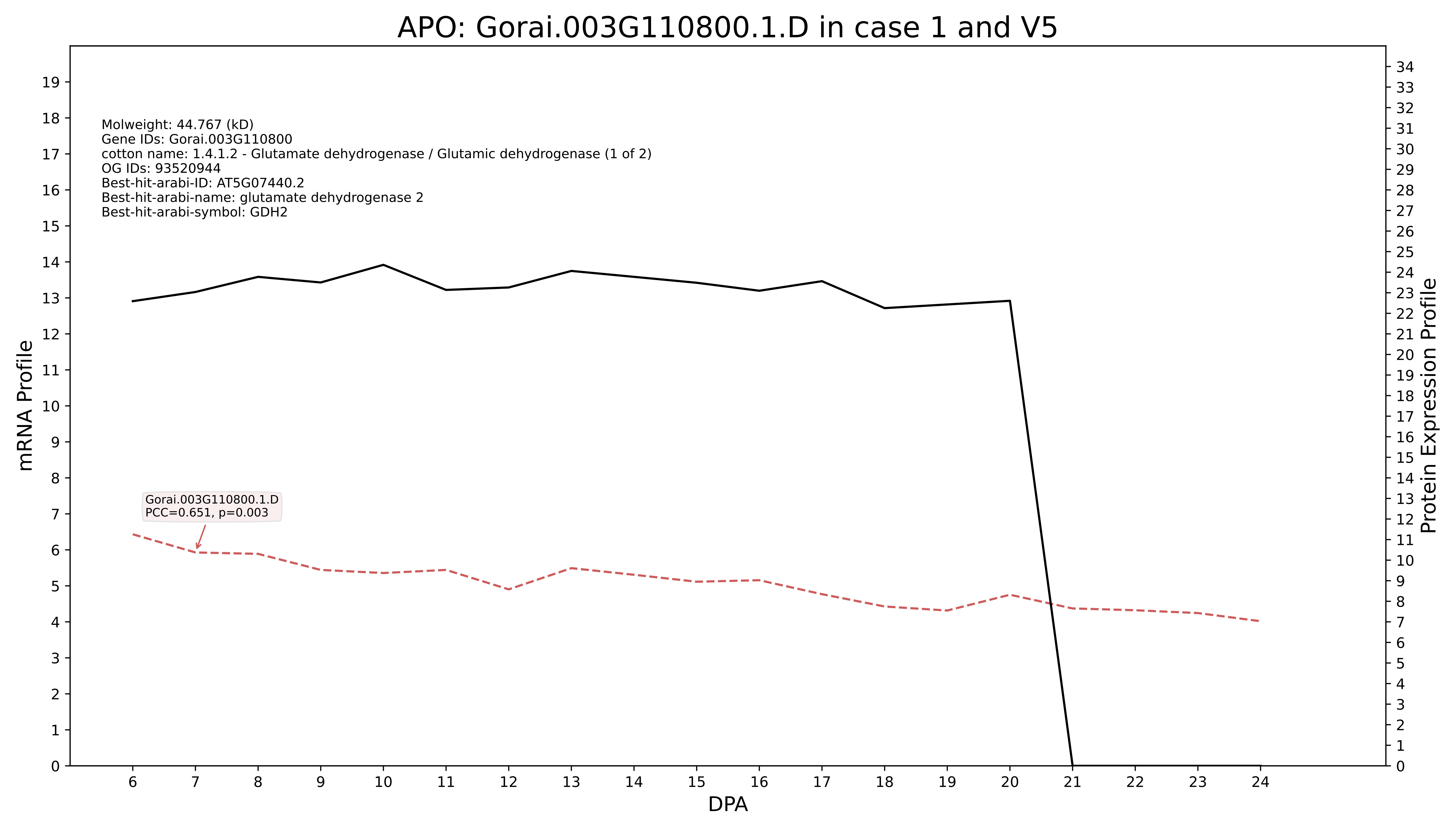

APO: Gorai.003G110800.1.D in case 1 and V5
Molweight: 44.767 (kD)
Gene IDs: Gorai.003G110800
cotton name: 1.4.1.2 - Glutamate dehydrogenase / Glutamic dehydrogenase (1 of 2)
OG IDs: 93520944
Best-hit-arabi-ID: AT5G07440.2
Best-hit-arabi-name: glutamate dehydrogenase 2
Best-hit-arabi-symbol: GDH2
Gorai.003G110800.1.D
PCC=0.651, p=0.003
mRNA Profile
Protein Expression Profile
DPA

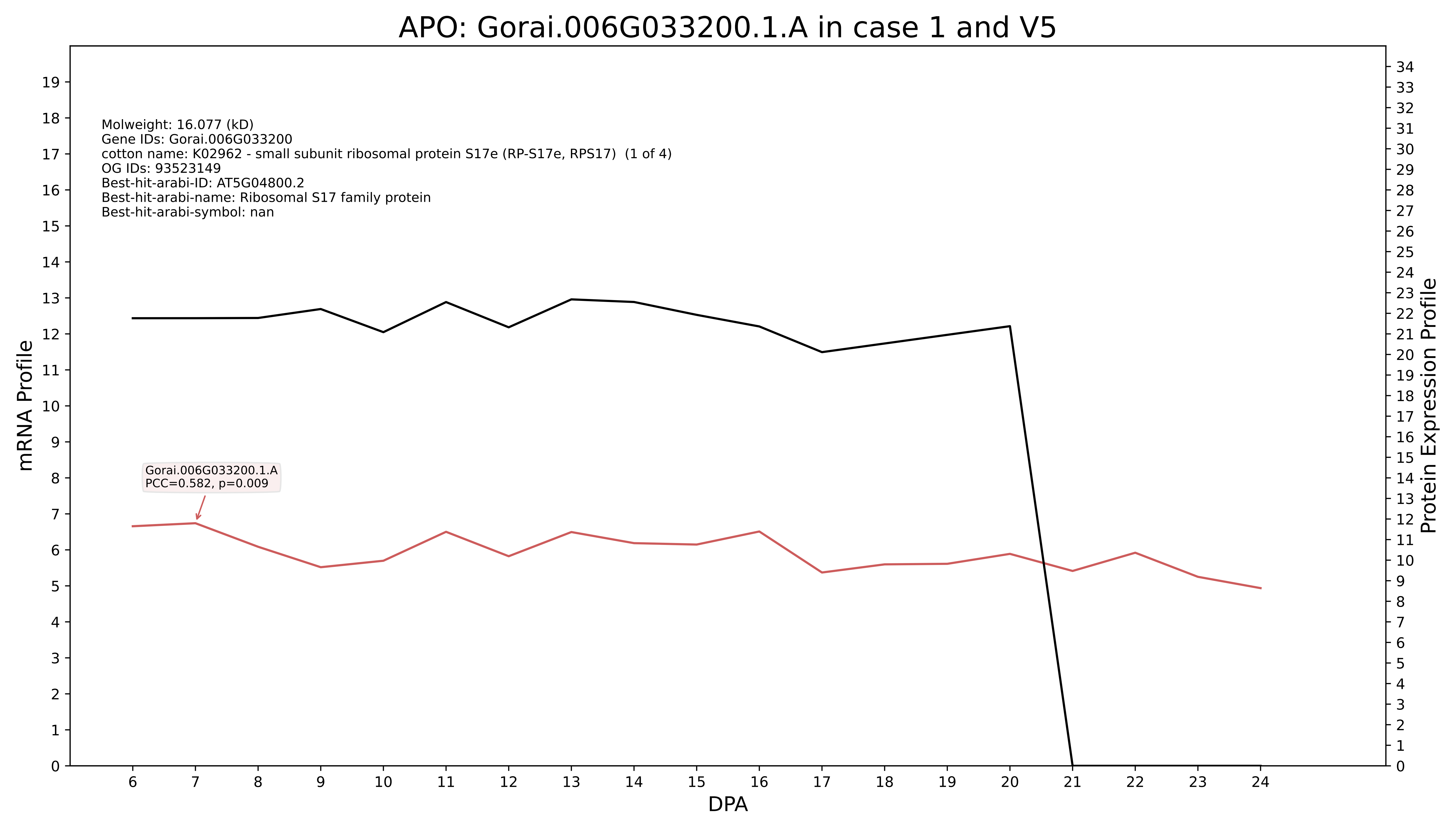

APO: Gorai.006G033200.1.A in case 1 and V5
Molweight: 16.077 (kD)
Gene IDs: Gorai.006G033200
cotton name: K02962 - small subunit ribosomal protein S17e (RP-S17e, RPS17) (1 of 4)
OG IDs: 93523149
Best-hit-arabi-ID: AT5G04800.2
Best-hit-arabi-name: Ribosomal S17 family protein
Best-hit-arabi-symbol: nan
Gorai.006G033200.1.A
PCC=0.582, p=0.009
mRNA Profile
Protein Expression Profile
DPA

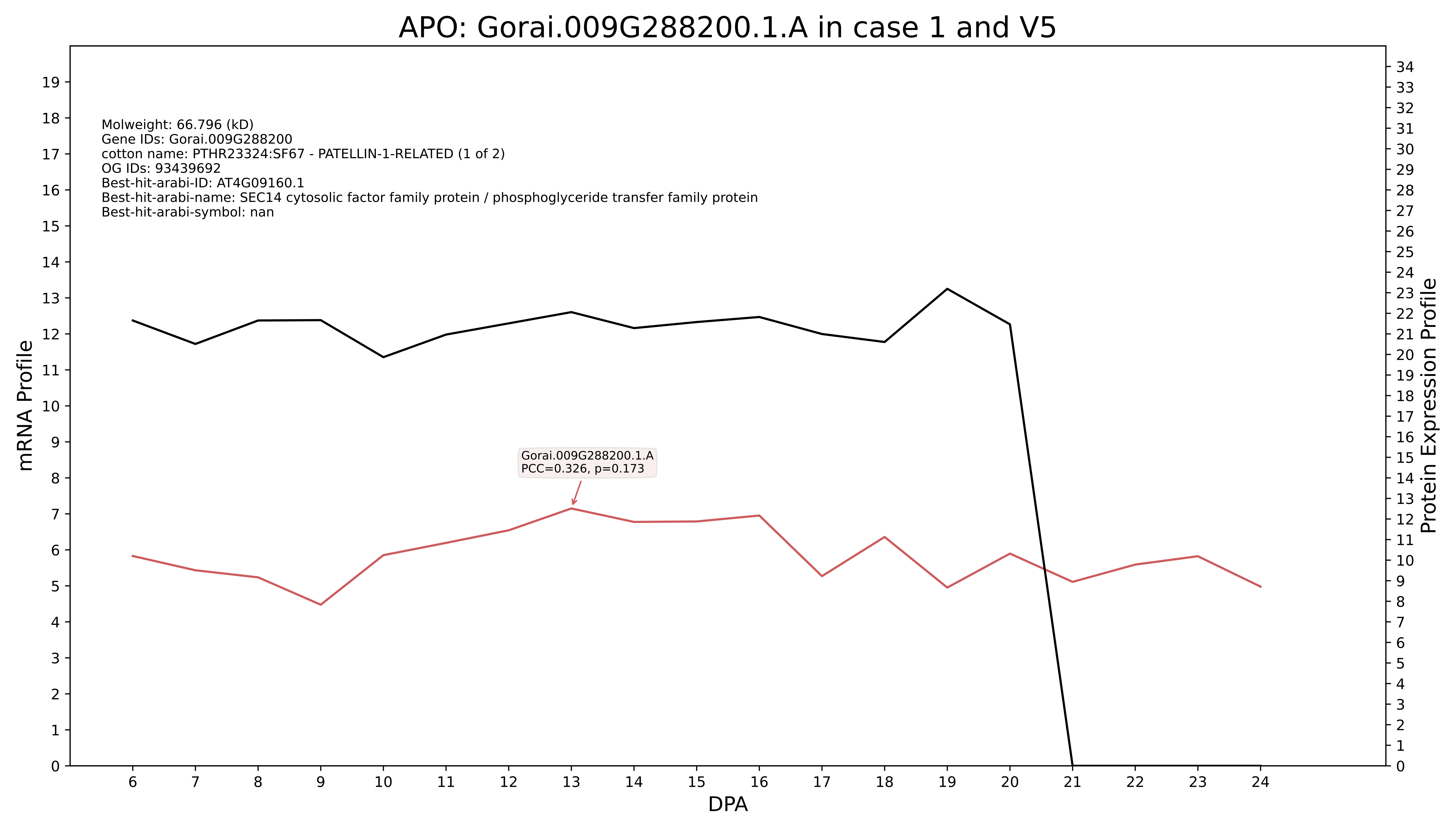
APO: Gorai.009G288200.1.A in case 1 and V5
Molweight: 66.796 (kD)
Gene IDs: Gorai.009G288200
cotton name: PTHR23324:SF67 - PATELLIN-1-RELATED (1 of 2)
OG IDs: 93439692
Best-hit-arabi-ID: AT4G09160.1
Best-hit-arabi-name: SEC14 cytosolic factor family protein / phosphoglyceride transfer family protein
Best-hit-arabi-symbol: nan
Gorai.009G288200.1.A
PCC=0.326, p=0.173
mRNA Profile
Protein Expression Profile
DPA

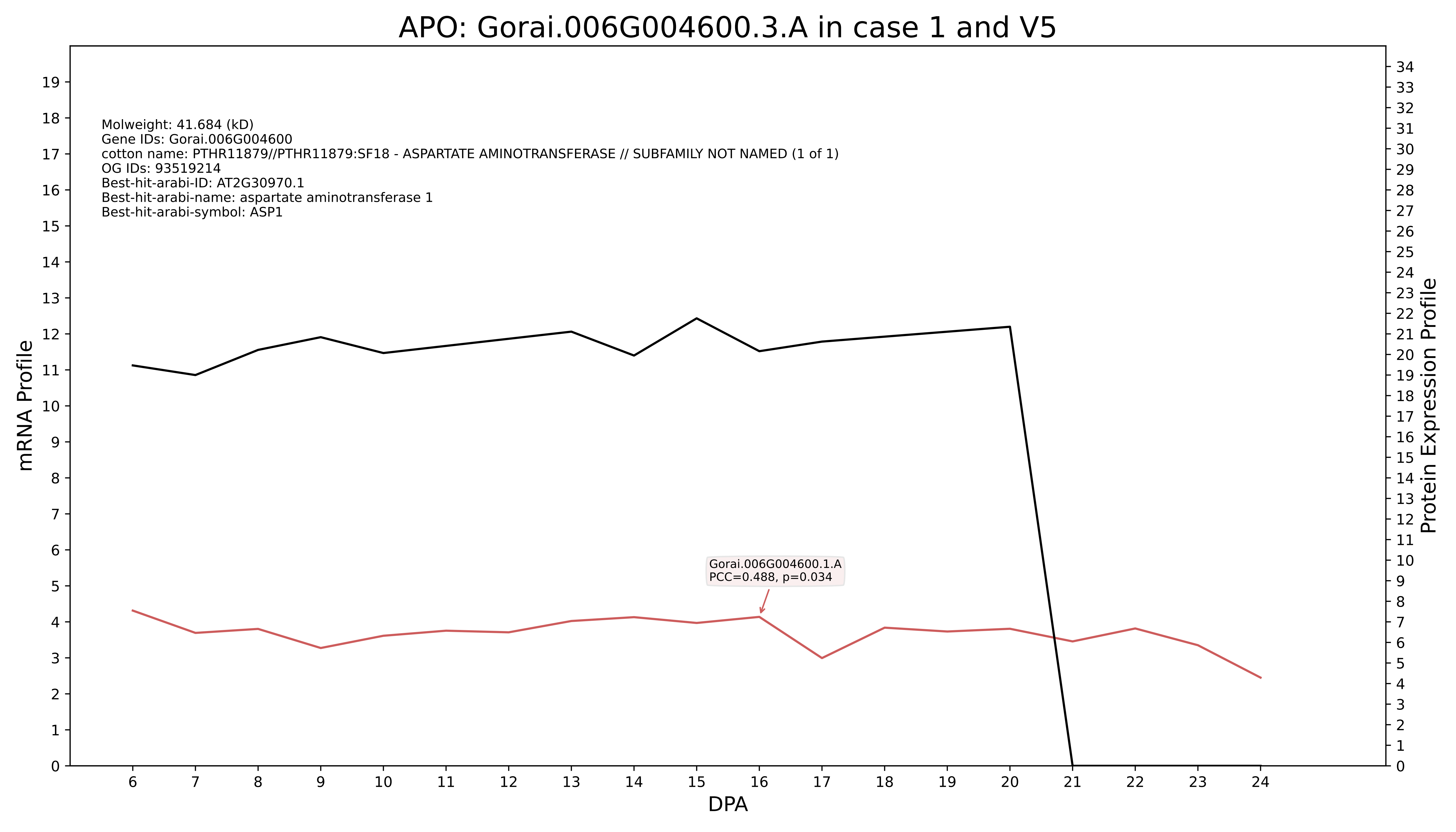
APO: Gorai.006G004600.3.A in case 1 and V5
Molweight: 41.684 (kD)
Gene IDs: Gorai.006G004600
cotton name: PTHR11879//PTHR11879:SF18 - ASPARTATE AMINOTRANSFERASE // SUBFAMILY NOT NAMED (1 of 1)
OG IDs: 93519214
Best-hit-arabi-ID: AT2G30970.1
Best-hit-arabi-name: aspartate aminotransferase 1
Best-hit-arabi-symbol: ASP1
Gorai.006G004600.1.A
PCC=0.488, p=0.034
mRNA Profile
Protein Expression Profile
DPA

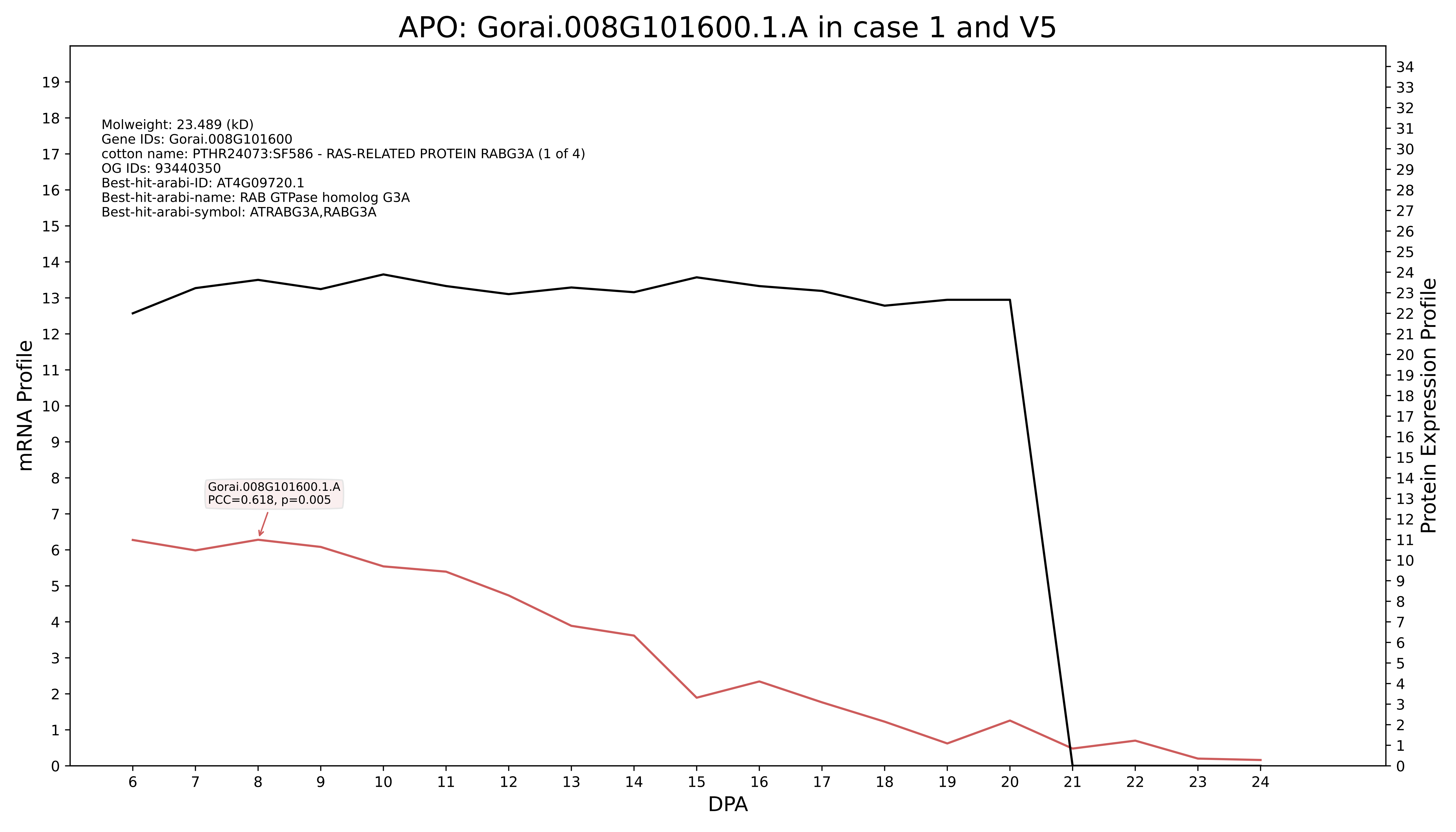
APO: Gorai.008G101600.1.A in case 1 and V5
Molweight: 23.489 (kD)
Gene IDs: Gorai.008G101600
cotton name: PTHR24073:SF586 - RAS-RELATED PROTEIN RABG3A (1 of 4)
OG IDs: 93440350
Best-hit-arabi-ID: AT4G09720.1
Best-hit-arabi-name: RAB GTPase homolog G3A
Best-hit-arabi-symbol: ATRABG3A,RABG3A
Gorai.008G101600.1.A
PCC=0.618, p=0.005
mRNA Profile
Protein Expression Profile
DPA

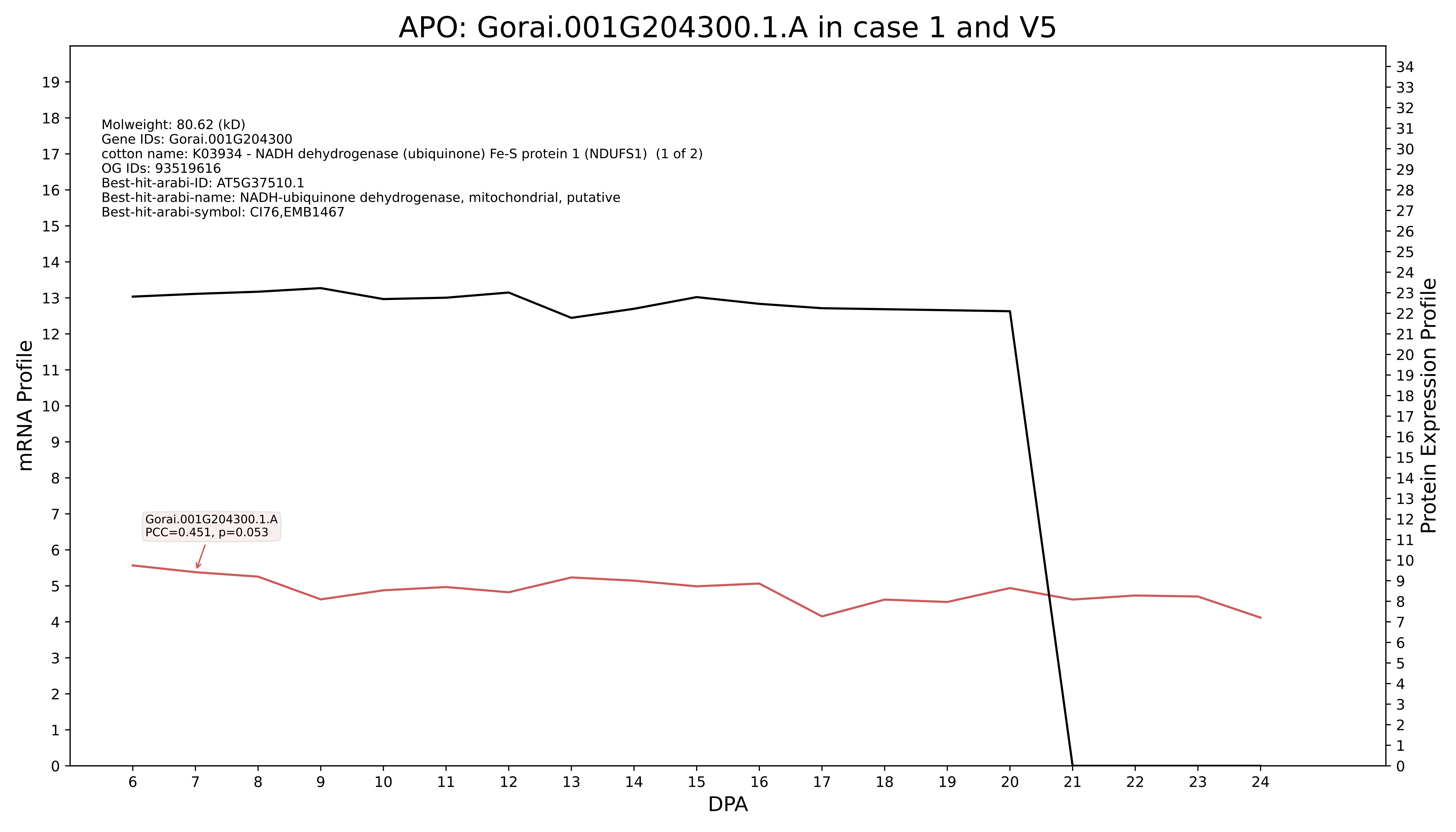

APO: Gorai.001G204300.1.A in case 1 and V5
Molweight: 80.62 (kD)
Gene IDs: Gorai.001G204300
cotton name: K03934 - NADH dehydrogenase (ubiquinone) Fe-S protein 1 (NDUFS1) (1 of 2)
OG IDs: 93519616
Best-hit-arabi-ID: AT5G37510.1
Best-hit-arabi-name: NADH-ubiquinone dehydrogenase, mitochondrial, putative
Best-hit-arabi-symbol: CI76,EMB1467
Gorai.001G204300.1.A
PCC=0.451, p=0.053
mRNA Profile
Protein Expression Profile
DPA

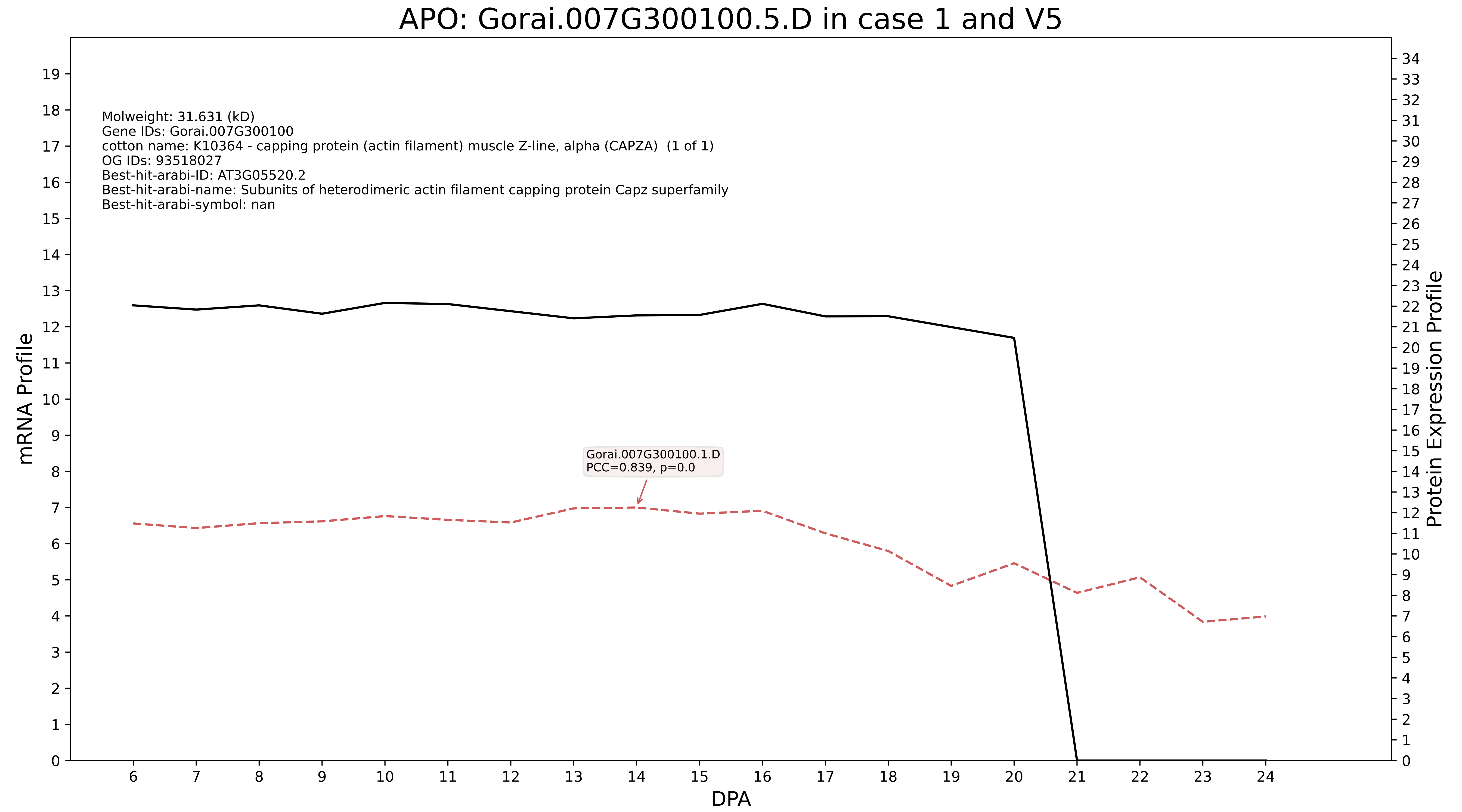

APO: Gorai.007G300100.5.D in case 1 and V5
Molweight: 31.631 (kD)
Gene IDs: Gorai.007G300100
cotton name: K10364 - capping protein (actin filament) muscle Z-line, alpha (CAPZA) (1 of 1)
OG IDs: 93518027
Best-hit-arabi-ID: AT3G05520.2
Best-hit-arabi-name: Subunits of heterodimeric actin filament capping protein Capz superfamily
Best-hit-arabi-symbol: nan
Gorai.007G300100.1.D
PCC=0.839, p=0.0
mRNA Profile
Protein Expression Profile
DPA

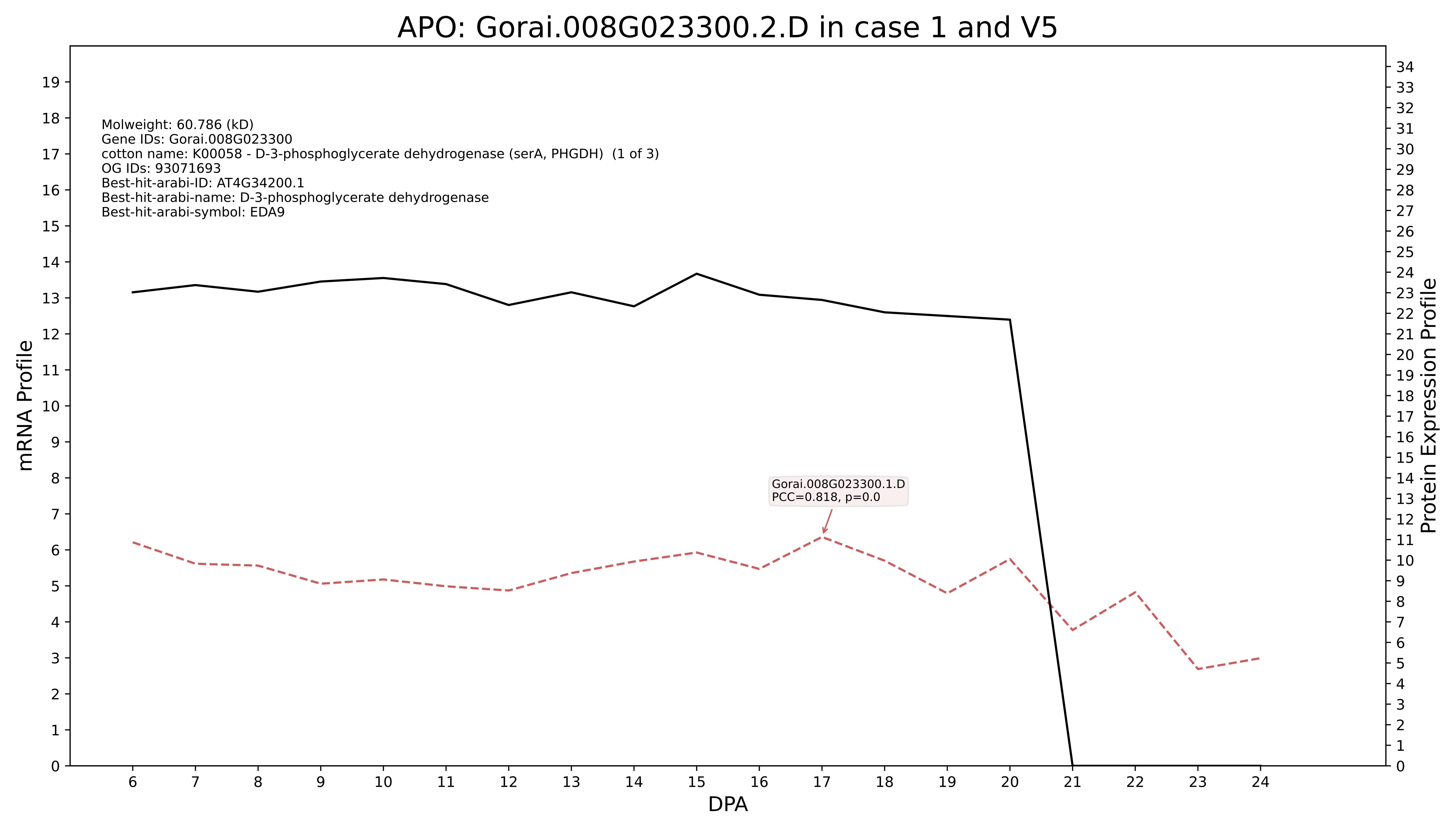

APO: Gorai.008G023300.2.D in case 1 and V5
Molweight: 60.786 (kD)
Gene IDs: Gorai.008G023300
cotton name: K00058 - D-3-phosphoglycerate dehydrogenase (serA, PHGDH) (1 of 3)
OG IDs: 93071693
Best-hit-arabi-ID: AT4G34200.1
Best-hit-arabi-name: D-3-phosphoglycerate dehydrogenase
Best-hit-arabi-symbol: EDA9
Gorai.008G023300.1.D
PCC=0.818, p=0.0
mRNA Profile
Protein Expression Profile
DPA

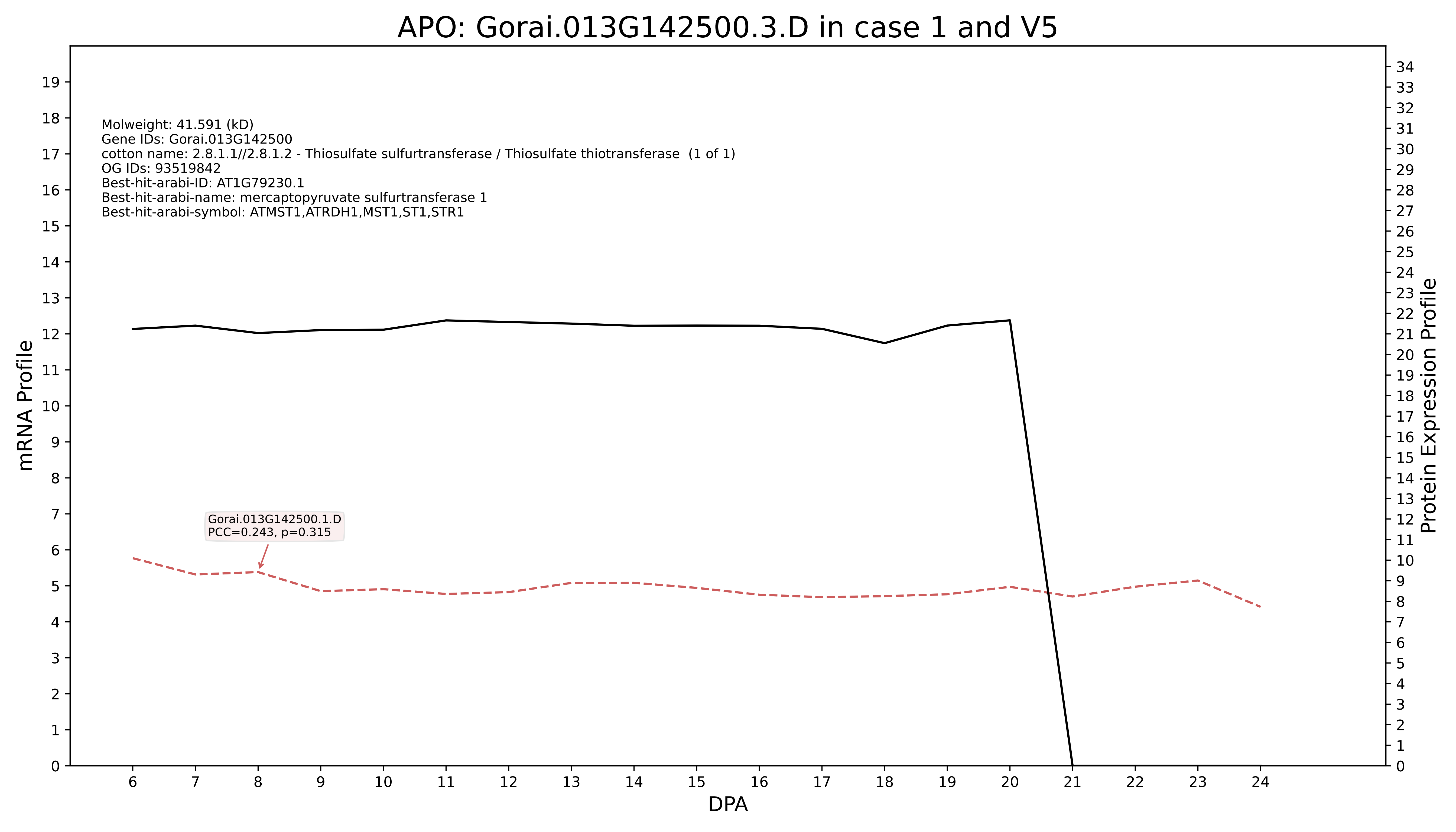
APO: Gorai.013G142500.3.D in case 1 and V5
Molweight: 41.591 (kD)
Gene IDs: Gorai.013G142500
cotton name: 2.8.1.1//2.8.1.2 - Thiosulfate sulfurtransferase / Thiosulfate thiotransferase (1 of 1)
OG IDs: 93519842
Best-hit-arabi-ID: AT1G79230.1
Best-hit-arabi-name: mercaptopyruvate sulfurtransferase 1
Best-hit-arabi-symbol: ATMST1,ATRDH1,MST1,ST1,STR1
Gorai.013G142500.1.D
PCC=0.243, p=0.315
mRNA Profile
Protein Expression Profile
DPA

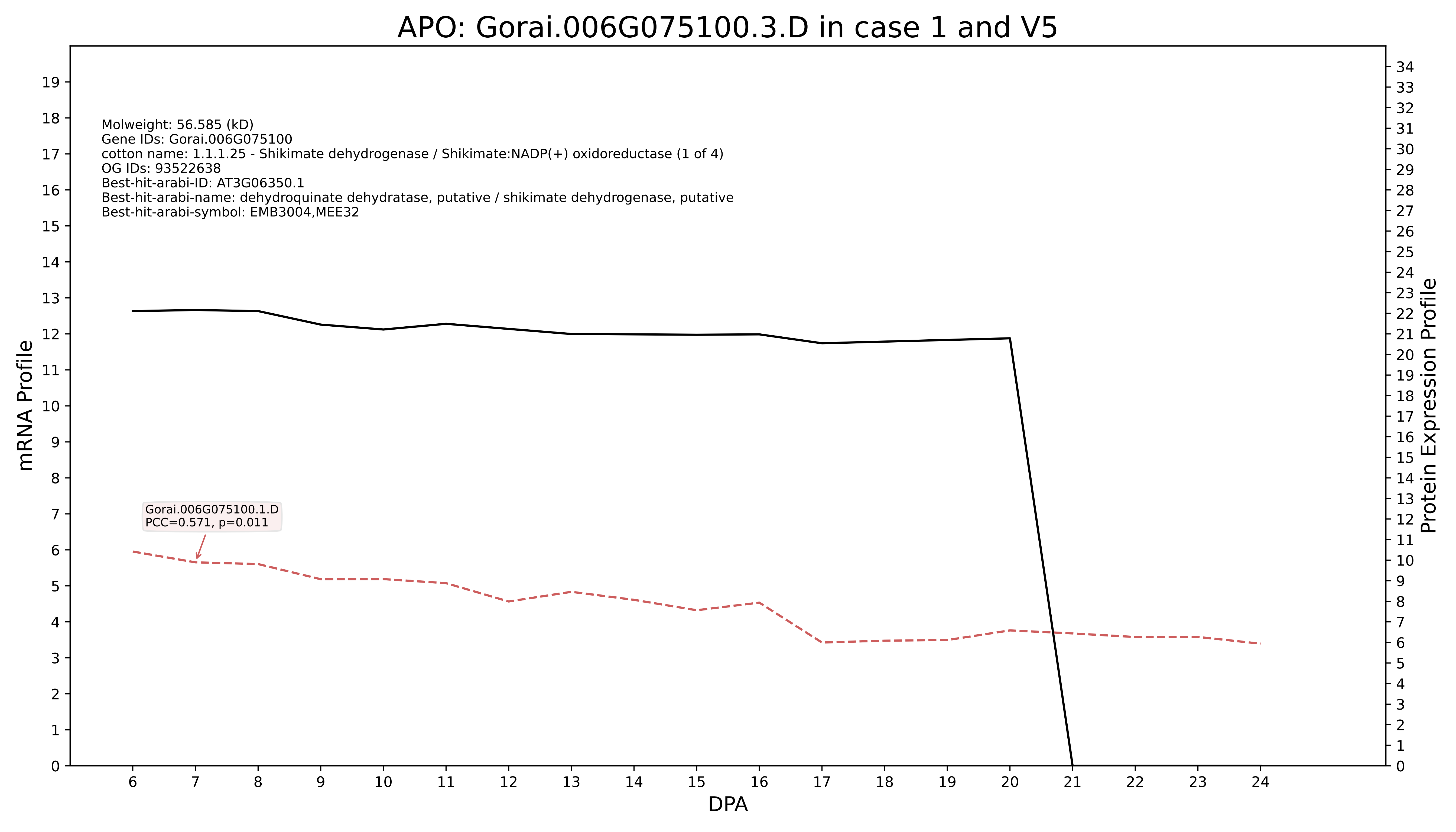

APO: Gorai.006G075100.3.D in case 1 and V5
Molweight: 56.585 (kD)
Gene IDs: Gorai.006G075100
cotton name: 1.1.1.25 - Shikimate dehydrogenase / Shikimate:NADP(+) oxidoreductase (1 of 4)
OG IDs: 93522638
Best-hit-arabi-ID: AT3G06350.1
Best-hit-arabi-name: dehydroquinate dehydratase, putative / shikimate dehydrogenase, putative
Best-hit-arabi-symbol: EMB3004,MEE32
Gorai.006G075100.1.D
PCC=0.571, p=0.011
mRNA Profile
Protein Expression Profile
DPA

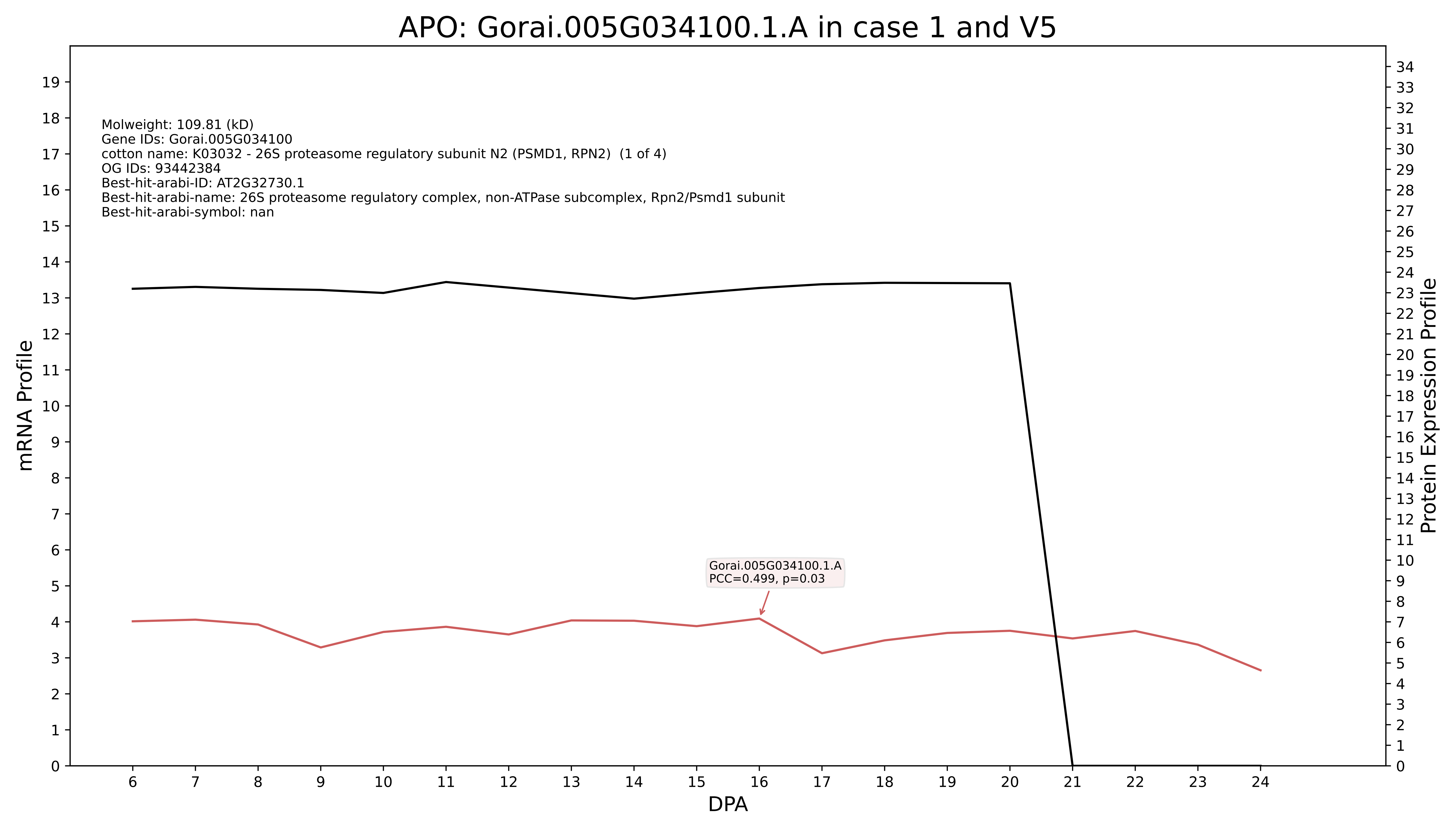

APO: Gorai.005G034100.1.A in case 1 and V5
Molweight: 109.81 (kD)
Gene IDs: Gorai.005G034100
cotton name: K03032 - 26S proteasome regulatory subunit N2 (PSMD1, RPN2) (1 of 4)
OG IDs: 93442384
Best-hit-arabi-ID: AT2G32730.1
Best-hit-arabi-name: 26S proteasome regulatory complex, non-ATPase subcomplex, Rpn2/Psmd1 subunit
Best-hit-arabi-symbol: nan
Gorai.005G034100.1.A
PCC=0.499, p=0.03
mRNA Profile
Protein Expression Profile
DPA

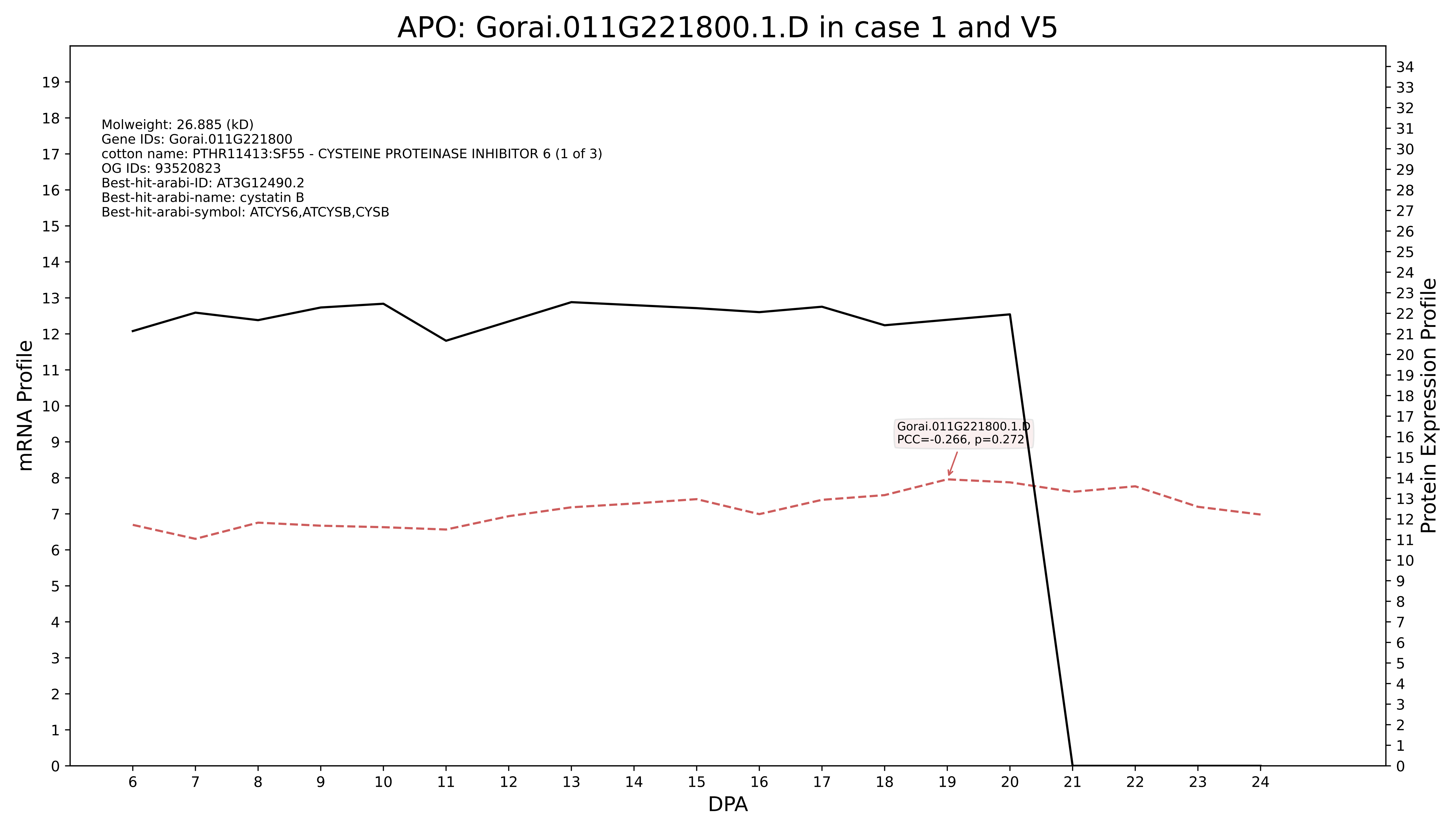

APO: Gorai.011G221800.1.D in case 1 and V5
Molweight: 26.885 (kD)
Gene IDs: Gorai.011G221800
cotton name: PTHR11413:SF55 - CYSTEINE PROTEINASE INHIBITOR 6 (1 of 3)
OG IDs: 93520823
Best-hit-arabi-ID: AT3G12490.2
Best-hit-arabi-name: cystatin B
Best-hit-arabi-symbol: ATCYS6,ATCYSB,CYSB
Gorai.011G221800.1.D
PCC=-0.266, p=0.272
mRNA Profile
Protein Expression Profile
DPA

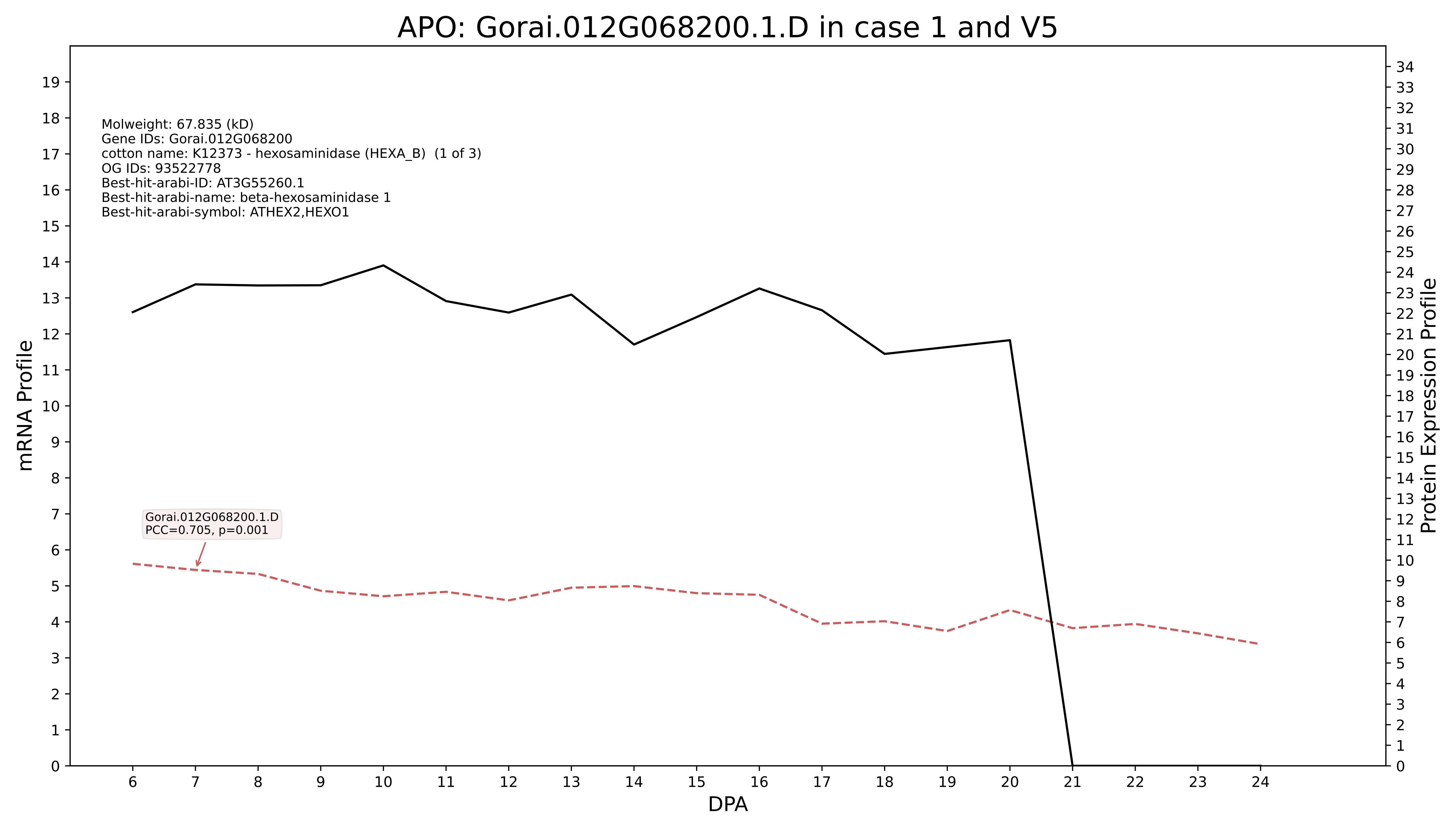

APO: Gorai.012G068200.1.D in case 1 and V5
Molweight: 67.835 (kD)
Gene IDs: Gorai.012G068200
cotton name: K12373 - hexosaminidase (HEXA_B) (1 of 3)
OG IDs: 93522778
Best-hit-arabi-ID: AT3G55260.1
Best-hit-arabi-name: beta-hexosaminidase 1
Best-hit-arabi-symbol: ATHEX2,HEXO1
Gorai.012G068200.1.D
PCC=0.705, p=0.001
mRNA Profile
Protein Expression Profile
DPA

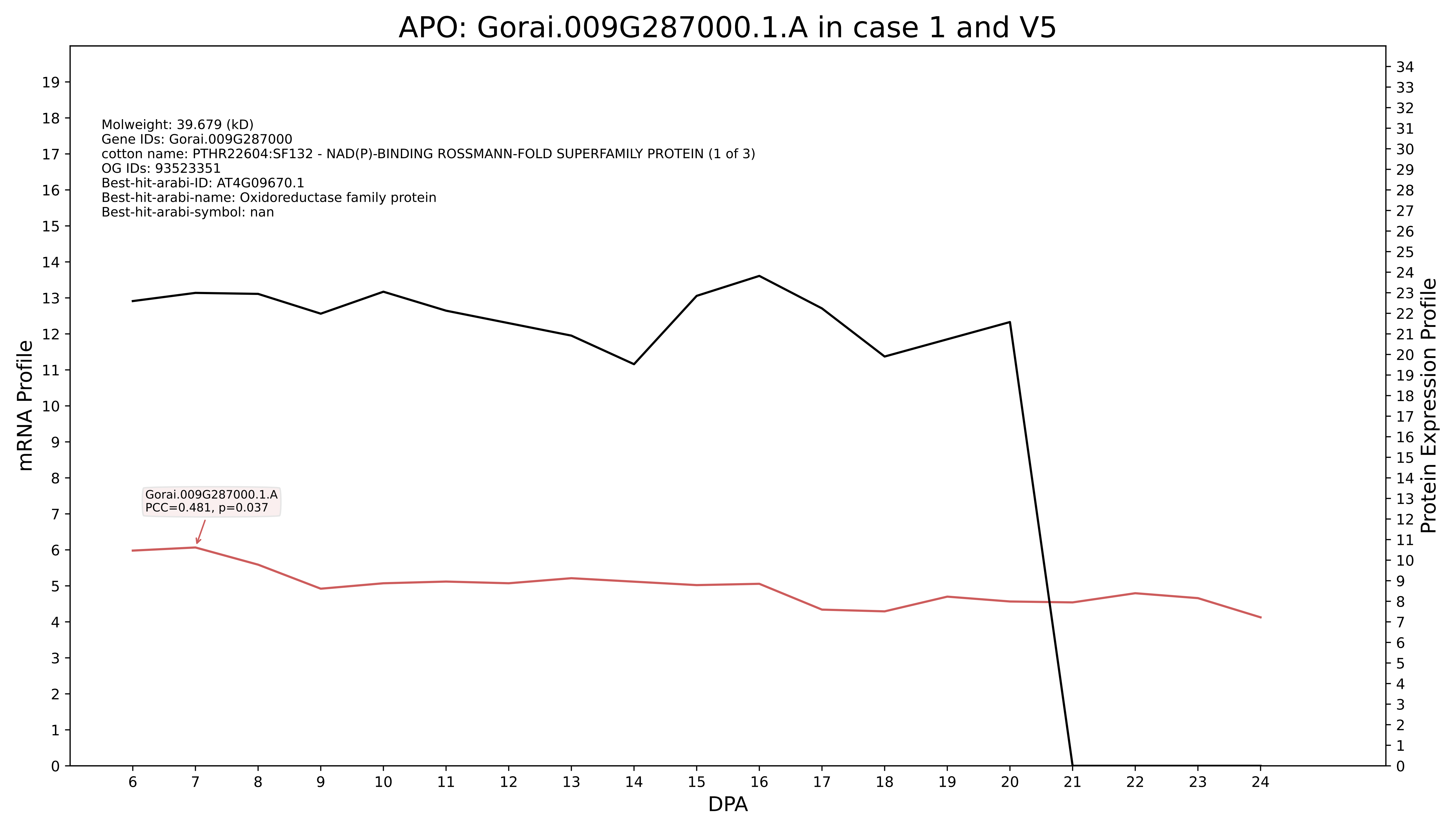

APO: Gorai.009G287000.1.A in case 1 and V5
Molweight: 39.679 (kD)
Gene IDs: Gorai.009G287000
cotton name: PTHR22604:SF132 - NAD(P)-BINDING ROSSMANN-FOLD SUPERFAMILY PROTEIN (1 of 3)
OG IDs: 93523351
Best-hit-arabi-ID: AT4G09670.1
Best-hit-arabi-name: Oxidoreductase family protein
Best-hit-arabi-symbol: nan
Gorai.009G287000.1.A
PCC=0.481, p=0.037
mRNA Profile
Protein Expression Profile
DPA

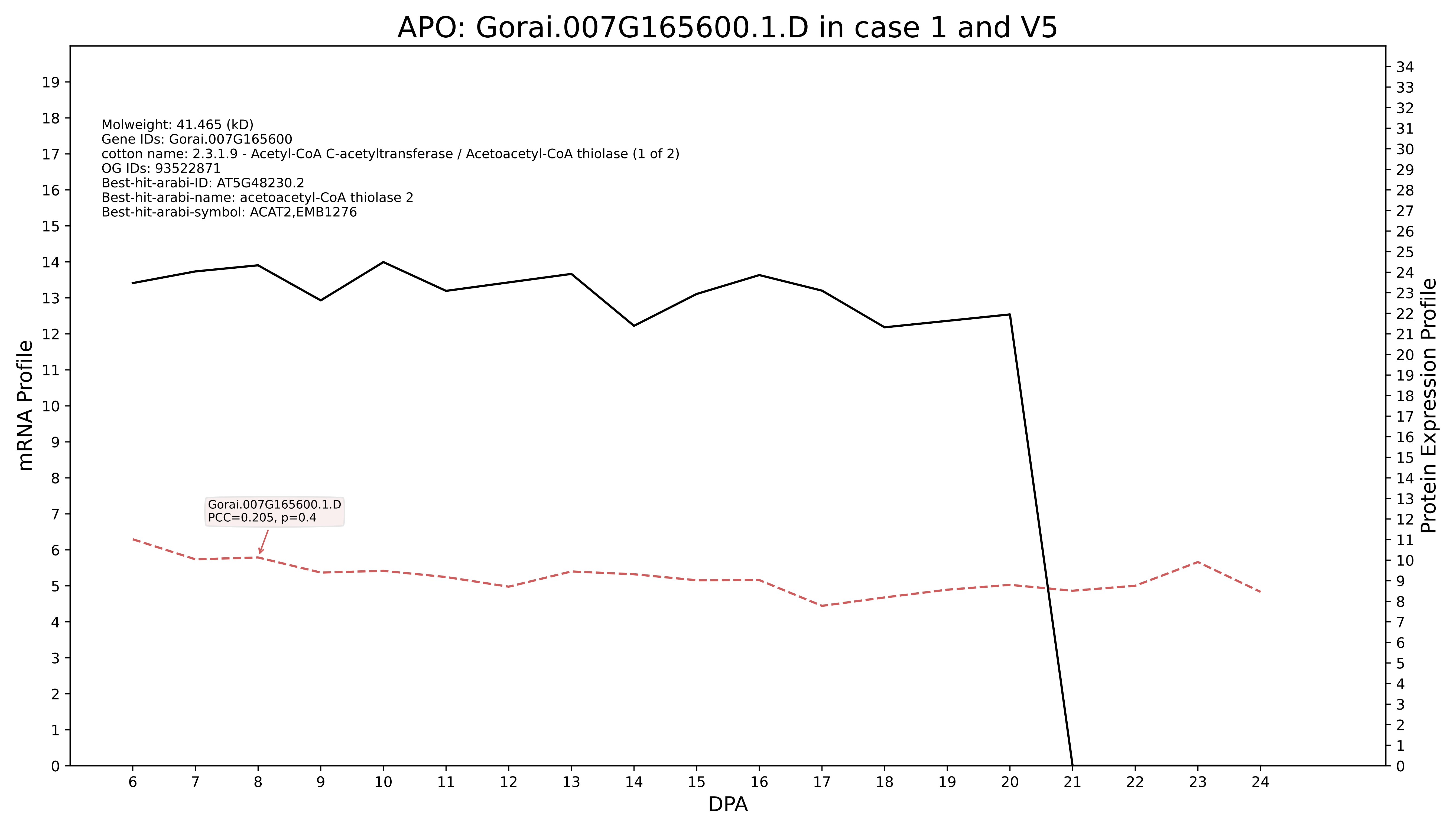
APO: Gorai.007G165600.1.D in case 1 and V5
Molweight: 41.465 (kD)
Gene IDs: Gorai.007G165600
cotton name: 2.3.1.9 - Acetyl-CoA C-acetyltransferase / Acetoacetyl-CoA thiolase (1 of 2)
OG IDs: 93522871
Best-hit-arabi-ID: AT5G48230.2
Best-hit-arabi-name: acetoacetyl-CoA thiolase 2
Best-hit-arabi-symbol: ACAT2,EMB1276
Gorai.007G165600.1.D
PCC=0.205, p=0.4
mRNA Profile
Protein Expression Profile
DPA

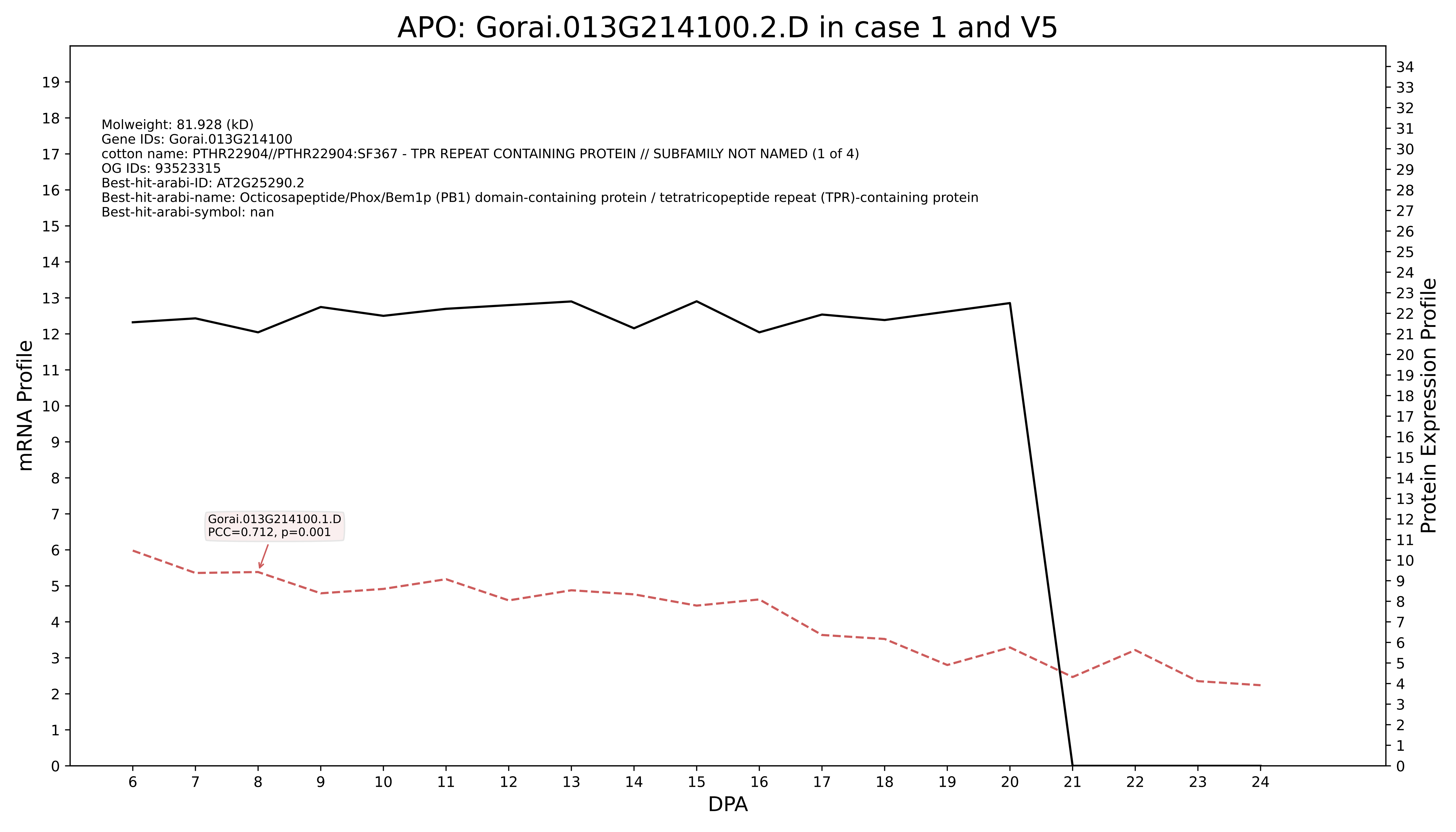
APO: Gorai.013G214100.2.D in case 1 and V5
Molweight: 81.928 (kD)
Gene IDs: Gorai.013G214100
cotton name: PTHR22904//PTHR22904:SF367 - TPR REPEAT CONTAINING PROTEIN // SUBFAMILY NOT NAMED (1 of 4)
OG IDs: 93523315
Best-hit-arabi-ID: AT2G25290.2
Best-hit-arabi-name: Octicosapeptide/Phox/Bem1p (PB1) domain-containing protein / tetratricopeptide repeat (TPR)-containing protein
Best-hit-arabi-symbol: nan
Gorai.013G214100.1.D
PCC=0.712, p=0.001
mRNA Profile
Protein Expression Profile
DPA

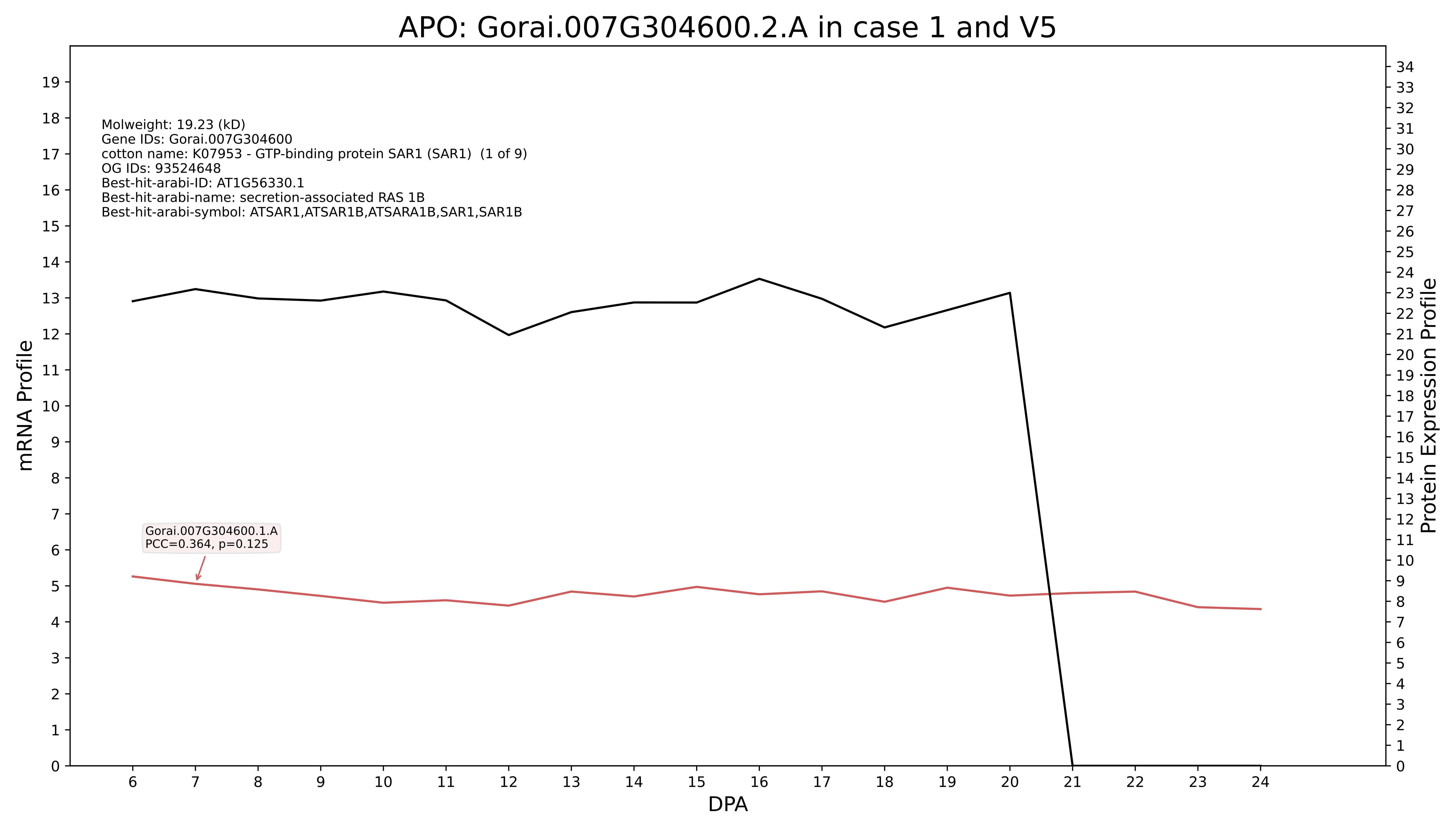
APO: Gorai.007G304600.2.A in case 1 and V5
Molweight: 19.23 (kD)
Gene IDs: Gorai.007G304600
cotton name: K07953 - GTP-binding protein SAR1 (SAR1) (1 of 9)
OG IDs: 93524648
Best-hit-arabi-ID: AT1G56330.1
Best-hit-arabi-name: secretion-associated RAS 1B
Best-hit-arabi-symbol: ATSAR1,ATSAR1B,ATSARA1B,SAR1,SAR1B
Gorai.007G304600.1.A
PCC=0.364, p=0.125
mRNA Profile
Protein Expression Profile
DPA

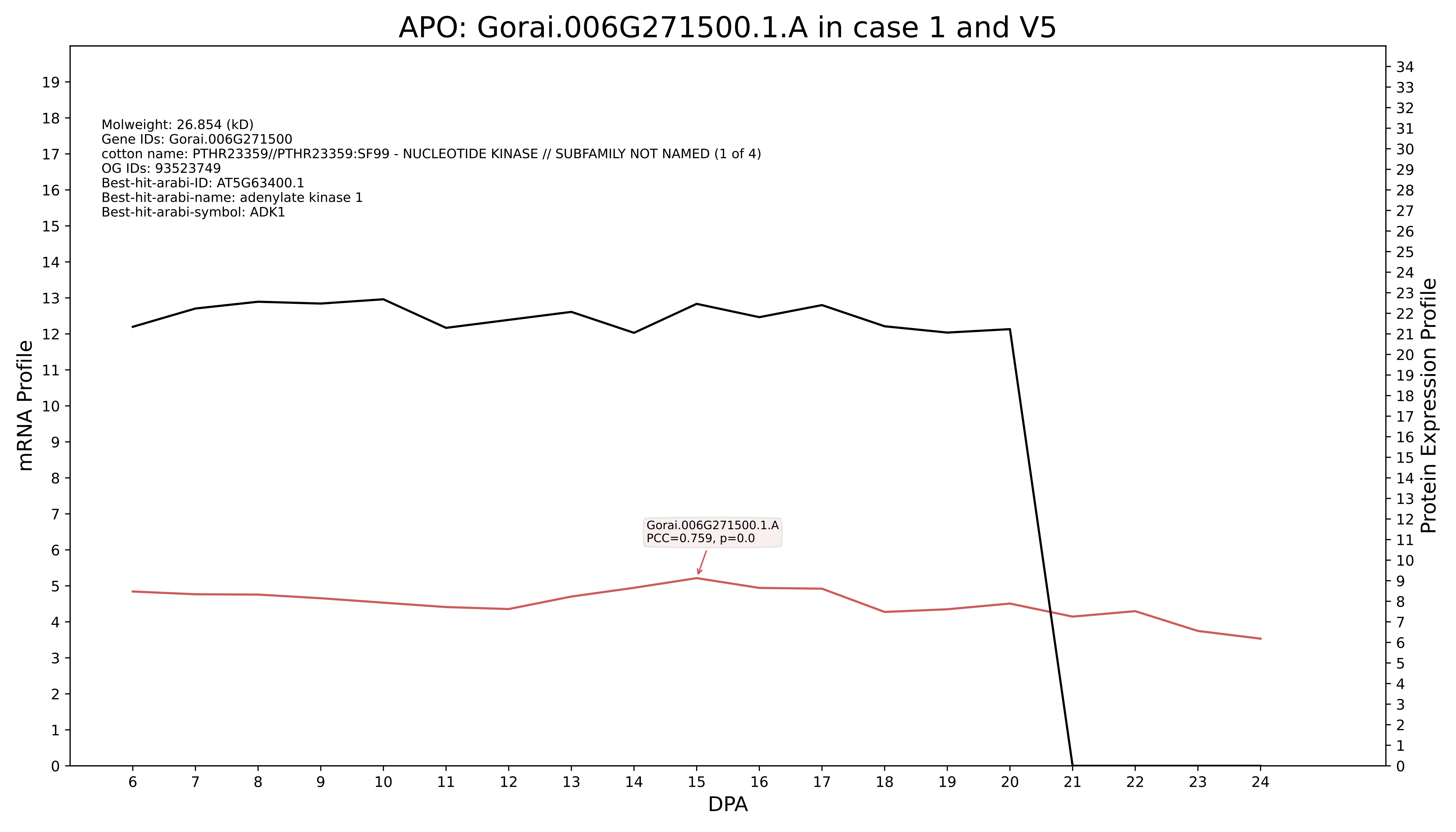

APO: Gorai.006G271500.1.A in case 1 and V5
Molweight: 26.854 (kD)
Gene IDs: Gorai.006G271500
cotton name: PTHR23359//PTHR23359:SF99 - NUCLEOTIDE KINASE // SUBFAMILY NOT NAMED (1 of 4)
OG IDs: 93523749
Best-hit-arabi-ID: AT5G63400.1
Best-hit-arabi-name: adenylate kinase 1
Best-hit-arabi-symbol: ADK1
Gorai.006G271500.1.A
PCC=0.759, p=0.0
mRNA Profile
Protein Expression Profile
DPA

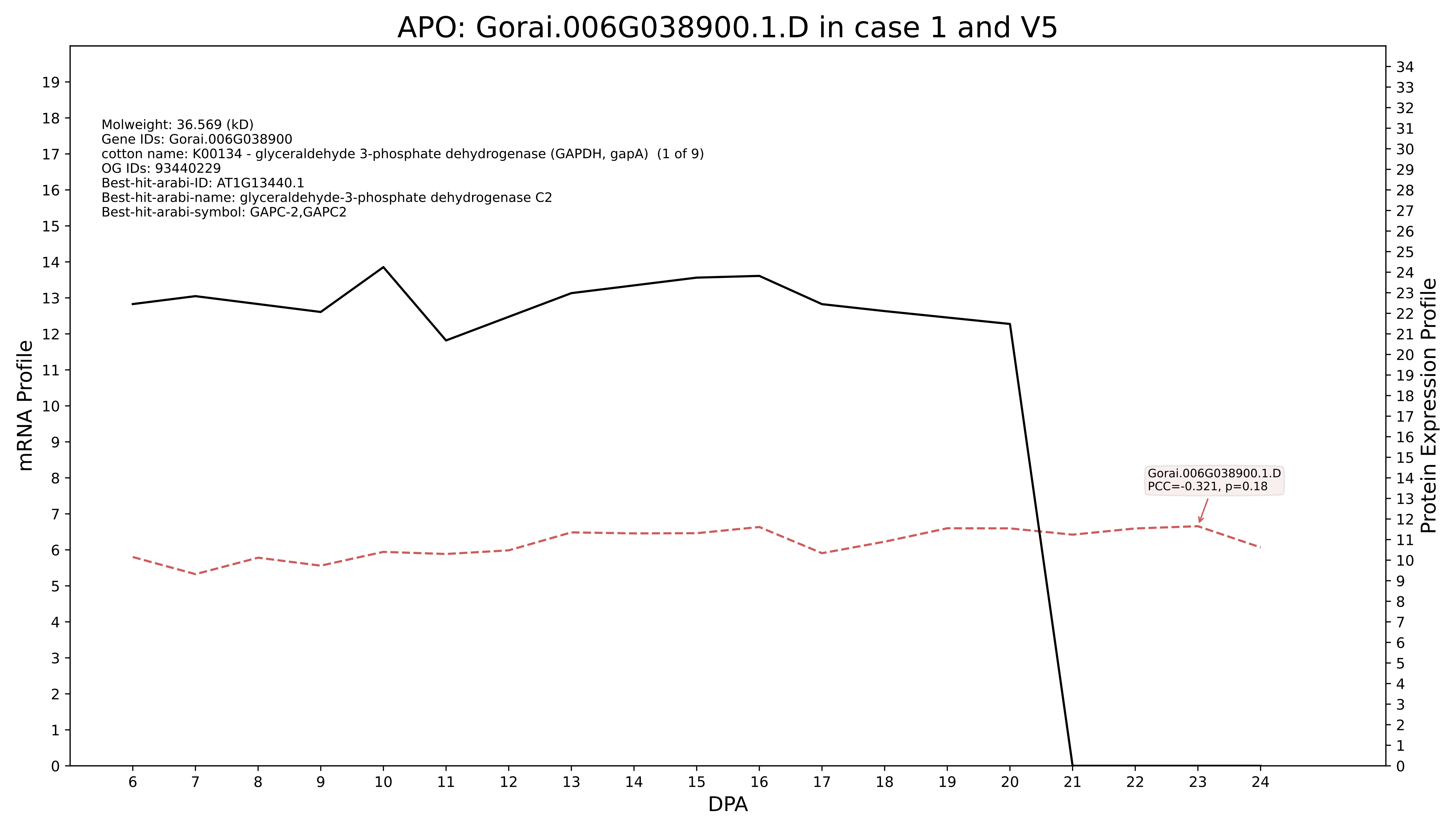
APO: Gorai.006G038900.1.D in case 1 and V5
Molweight: 36.569 (kD)
Gene IDs: Gorai.006G038900
cotton name: K00134 - glyceraldehyde 3-phosphate dehydrogenase (GAPDH, gapA) (1 of 9)
OG IDs: 93440229
Best-hit-arabi-ID: AT1G13440.1
Best-hit-arabi-name: glyceraldehyde-3-phosphate dehydrogenase C2
Best-hit-arabi-symbol: GAPC-2,GAPC2
Gorai.006G038900.1.D
PCC=-0.321, p=0.18
mRNA Profile
Protein Expression Profile
DPA

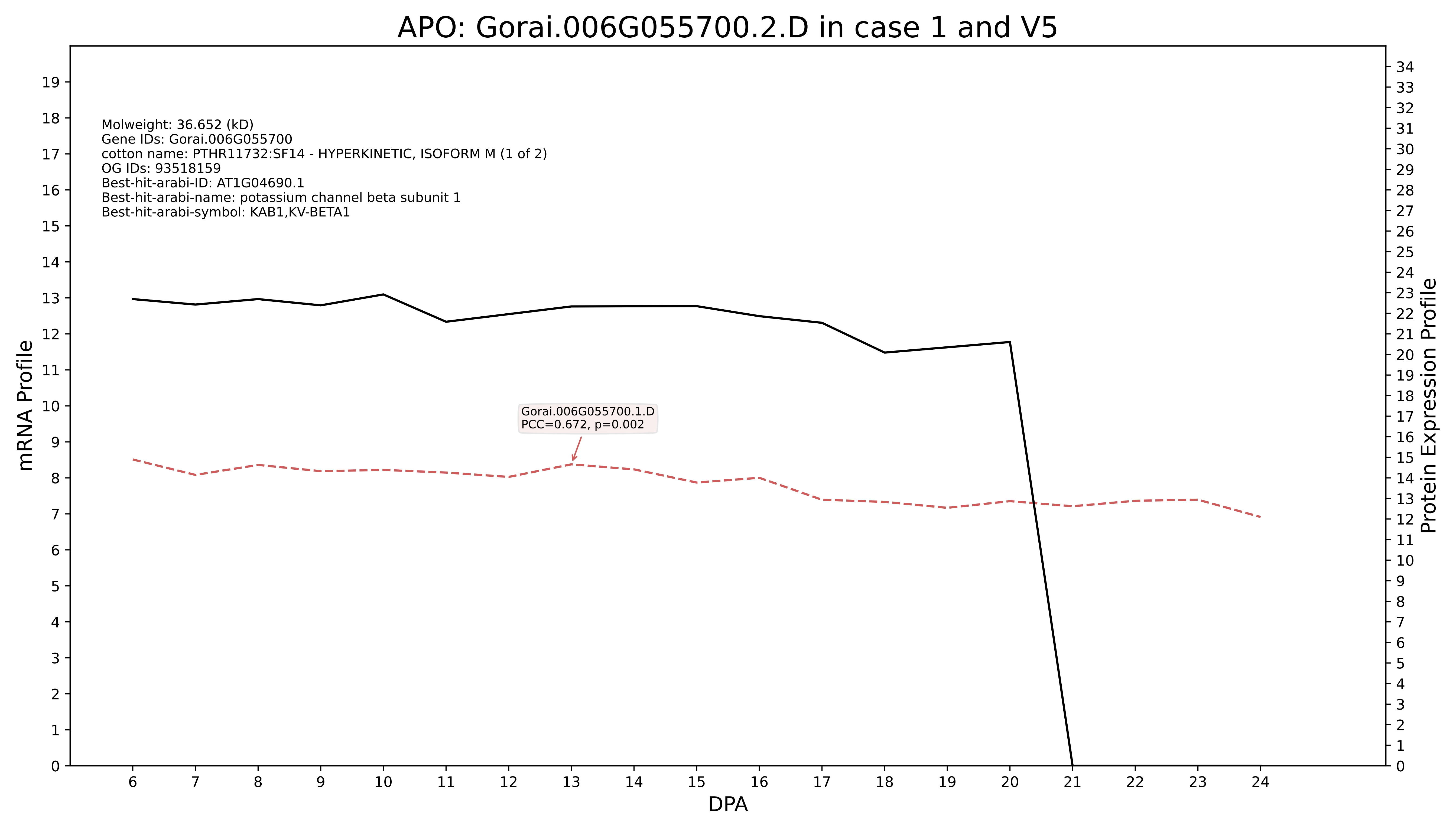
APO: Gorai.006G055700.2.D in case 1 and V5
Molweight: 36.652 (kD)
Gene IDs: Gorai.006G055700
cotton name: PTHR11732:SF14 - HYPERKINETIC, ISOFORM M (1 of 2)
OG IDs: 93518159
Best-hit-arabi-ID: AT1G04690.1
Best-hit-arabi-name: potassium channel beta subunit 1
Best-hit-arabi-symbol: KAB1,KV-BETA1
Gorai.006G055700.1.D
PCC=0.672, p=0.002
mRNA Profile
Protein Expression Profile
DPA

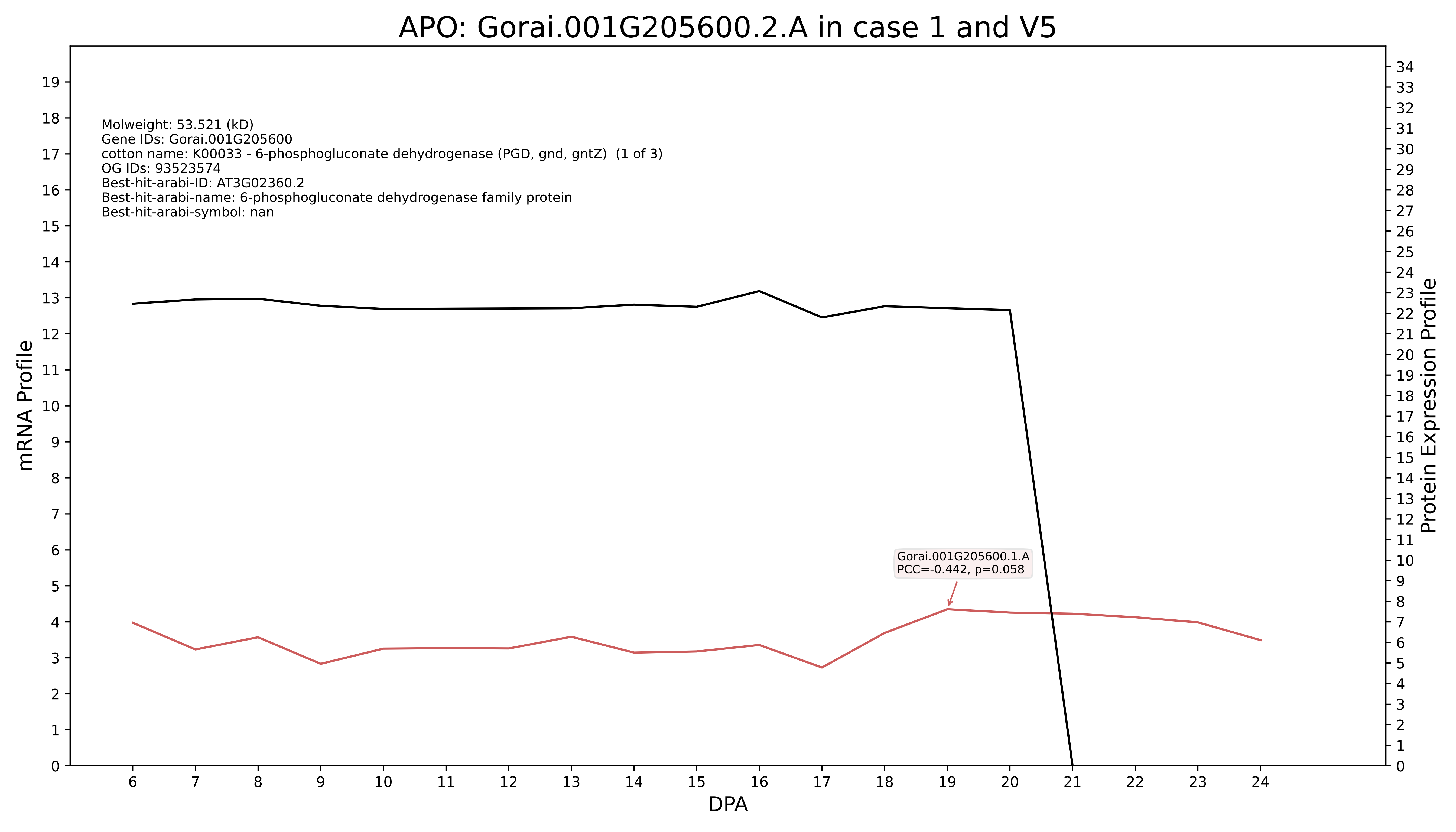

APO: Gorai.001G205600.2.A in case 1 and V5
Molweight: 53.521 (kD)
Gene IDs: Gorai.001G205600
cotton name: K00033 - 6-phosphogluconate dehydrogenase (PGD, gnd, gntZ) (1 of 3)
OG IDs: 93523574
Best-hit-arabi-ID: AT3G02360.2
Best-hit-arabi-name: 6-phosphogluconate dehydrogenase family protein
Best-hit-arabi-symbol: nan
Gorai.001G205600.1.A
PCC=-0.442, p=0.058
mRNA Profile
Protein Expression Profile
DPA

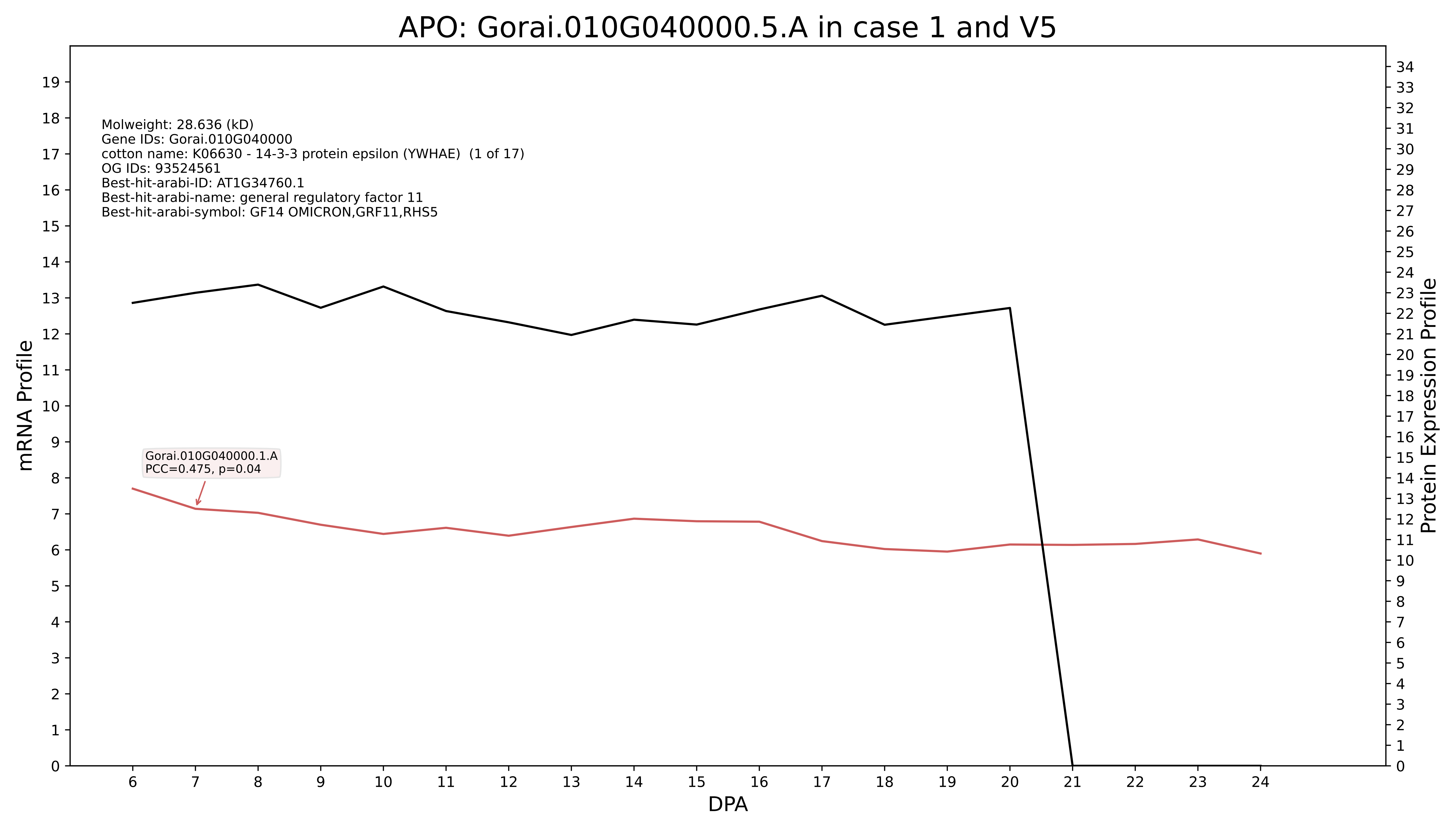
APO: Gorai.010G040000.5.A in case 1 and V5
Molweight: 28.636 (kD)
Gene IDs: Gorai.010G040000
cotton name: K06630 - 14-3-3 protein epsilon (YWHAE) (1 of 17)
OG IDs: 93524561
Best-hit-arabi-ID: AT1G34760.1
Best-hit-arabi-name: general regulatory factor 11
Best-hit-arabi-symbol: GF14 OMICRON,GRF11,RHS5
Gorai.010G040000.1.A
PCC=0.475, p=0.04
mRNA Profile
Protein Expression Profile
DPA

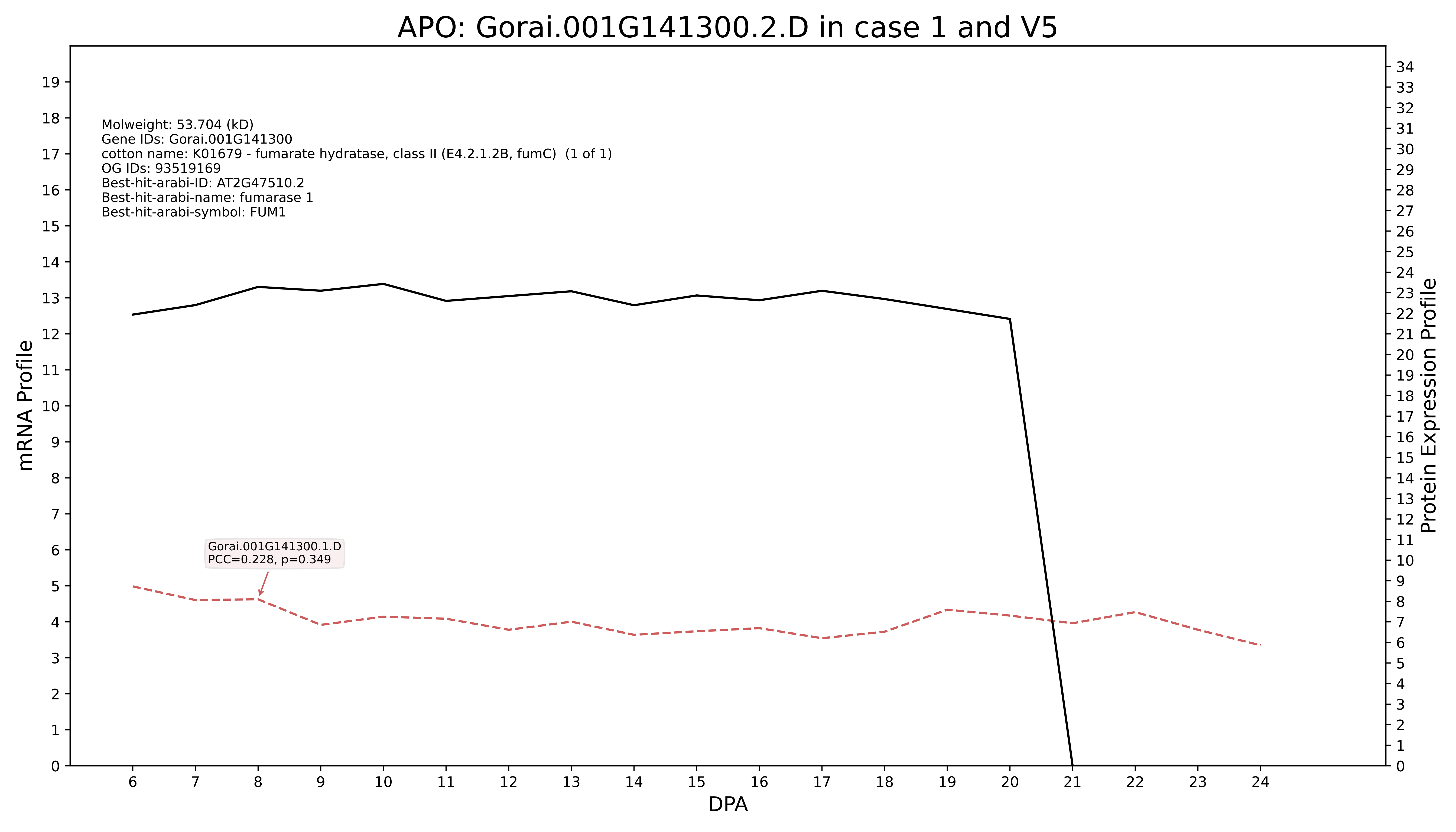
APO: Gorai.001G141300.2.D in case 1 and V5
Molweight: 53.704 (kD)
Gene IDs: Gorai.001G141300
cotton name: K01679 - fumarate hydratase, class II (E4.2.1.2B, fumC) (1 of 1)
OG IDs: 93519169
Best-hit-arabi-ID: AT2G47510.2
Best-hit-arabi-name: fumarase 1
Best-hit-arabi-symbol: FUM1
Gorai.001G141300.1.D
PCC=0.228, p=0.349
mRNA Profile
Protein Expression Profile
DPA

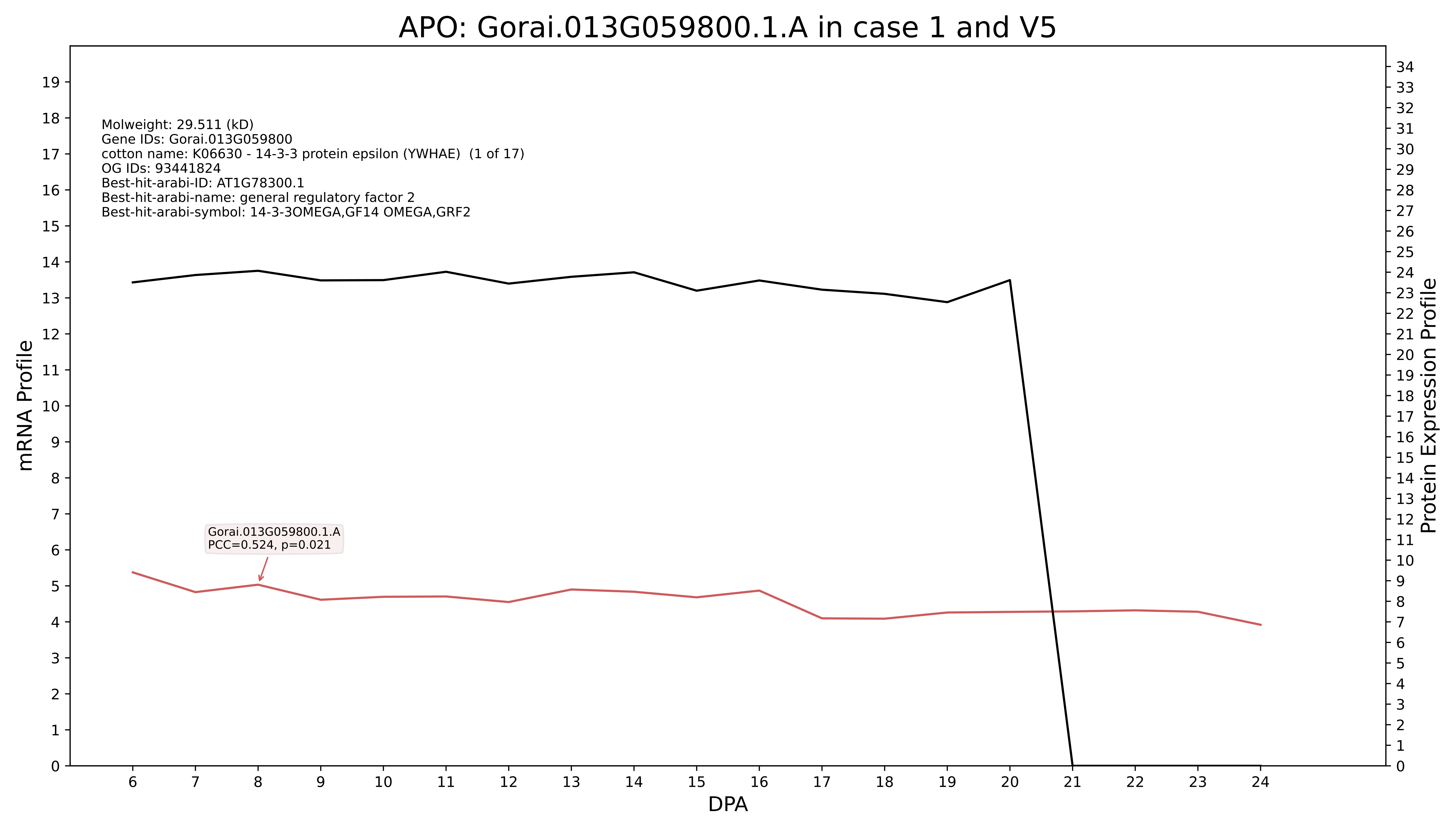

APO: Gorai.013G059800.1.A in case 1 and V5
Molweight: 29.511 (kD)
Gene IDs: Gorai.013G059800
cotton name: K06630 - 14-3-3 protein epsilon (YWHAE) (1 of 17)
OG IDs: 93441824
Best-hit-arabi-ID: AT1G78300.1
Best-hit-arabi-name: general regulatory factor 2
Best-hit-arabi-symbol: 14-3-3OMEGA,GF14 OMEGA,GRF2
Gorai.013G059800.1.A
PCC=0.524, p=0.021
mRNA Profile
Protein Expression Profile
DPA

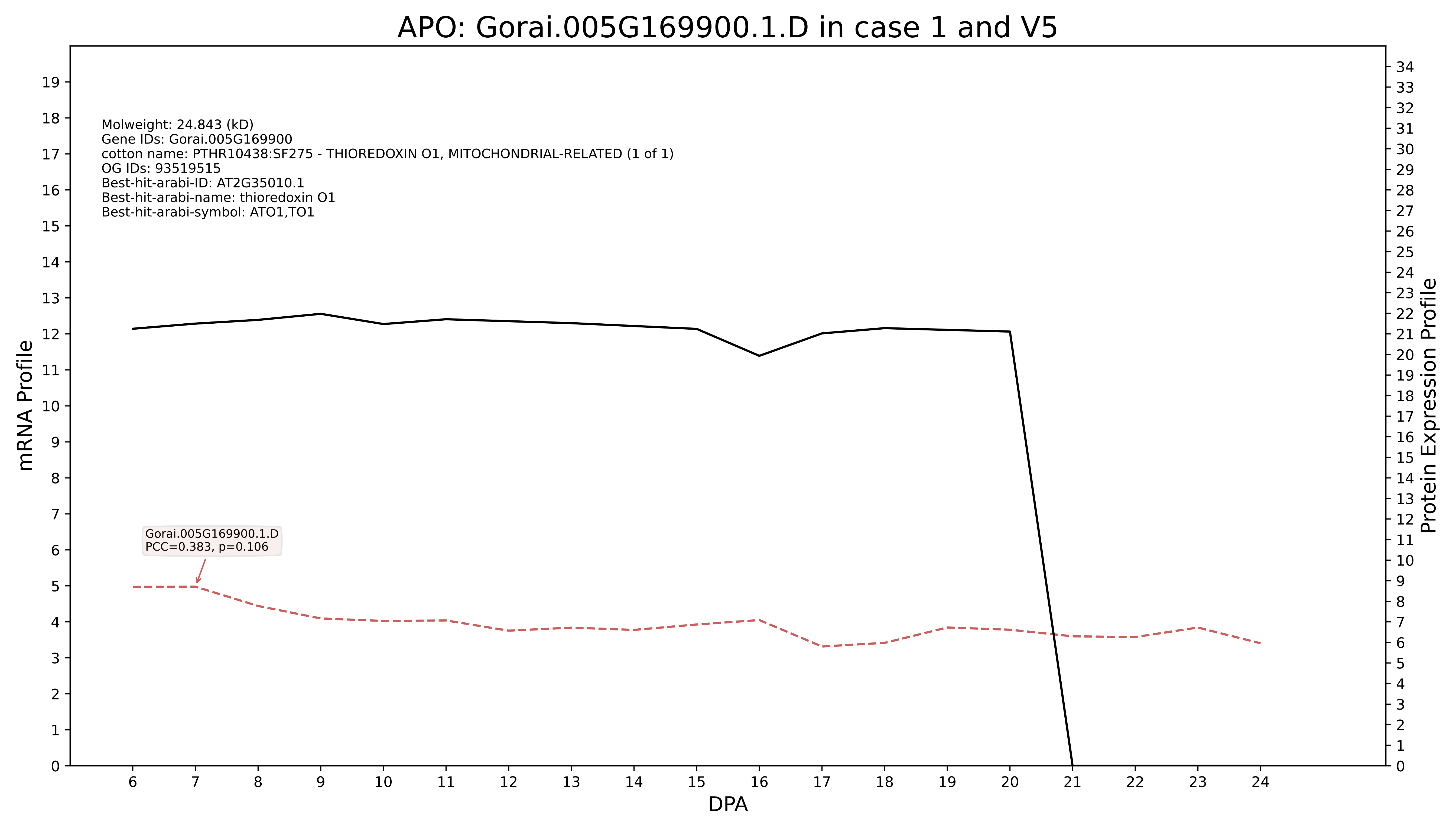

APO: Gorai.005G169900.1.D in case 1 and V5
Molweight: 24.843 (kD)
Gene IDs: Gorai.005G169900
cotton name: PTHR10438:SF275 - THIOREDOXIN O1, MITOCHONDRIAL-RELATED (1 of 1)
OG IDs: 93519515
Best-hit-arabi-ID: AT2G35010.1
Best-hit-arabi-name: thioredoxin O1
Best-hit-arabi-symbol: ATO1,TO1
Gorai.005G169900.1.D
PCC=0.383, p=0.106
mRNA Profile
Protein Expression Profile
DPA

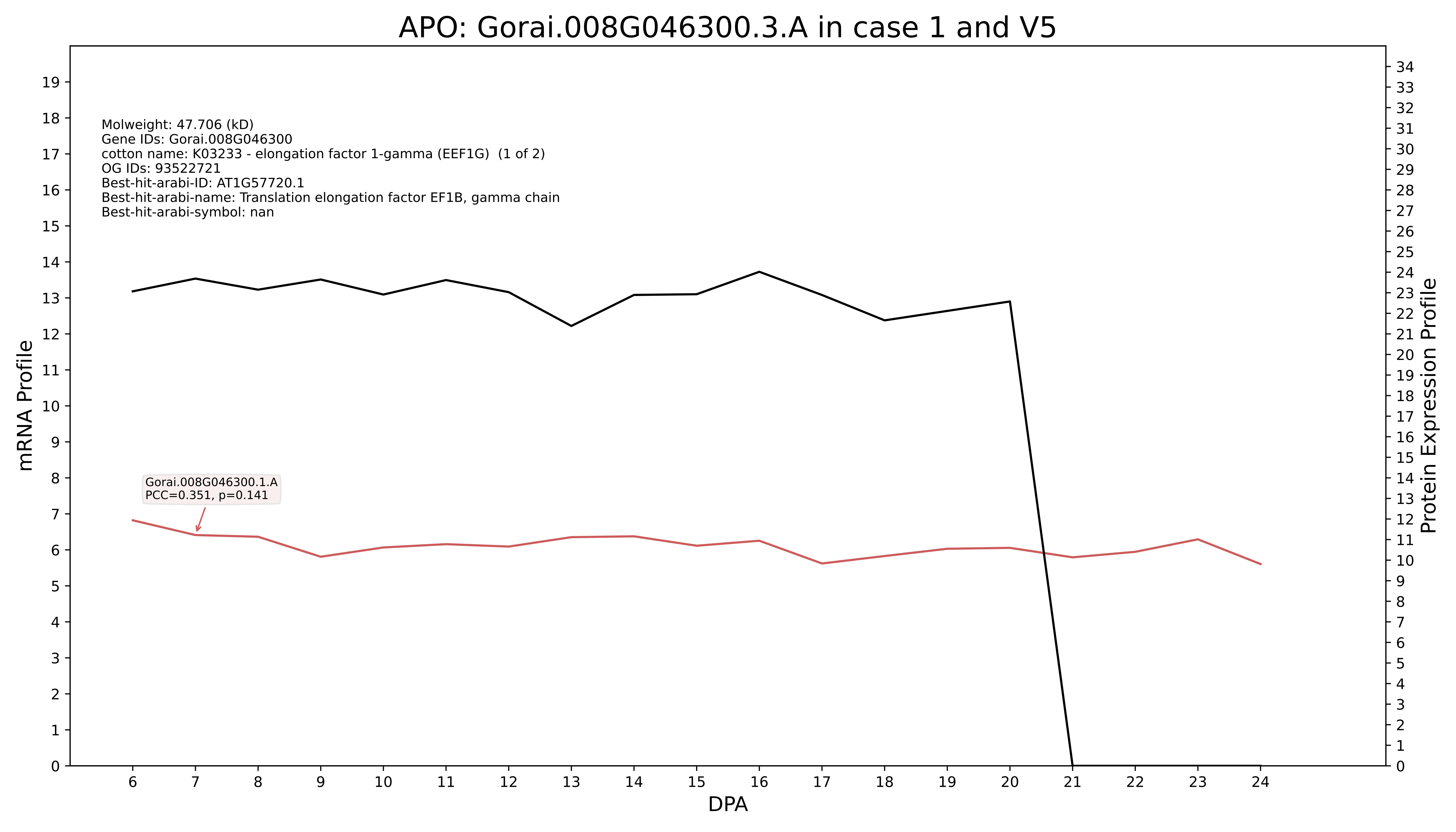
APO: Gorai.008G046300.3.A in case 1 and V5
Molweight: 47.706 (kD)
Gene IDs: Gorai.008G046300
cotton name: K03233 - elongation factor 1-gamma (EEF1G) (1 of 2)
OG IDs: 93522721
Best-hit-arabi-ID: AT1G57720.1
Best-hit-arabi-name: Translation elongation factor EF1B, gamma chain
Best-hit-arabi-symbol: nan
Gorai.008G046300.1.A
PCC=0.351, p=0.141
mRNA Profile
Protein Expression Profile
DPA

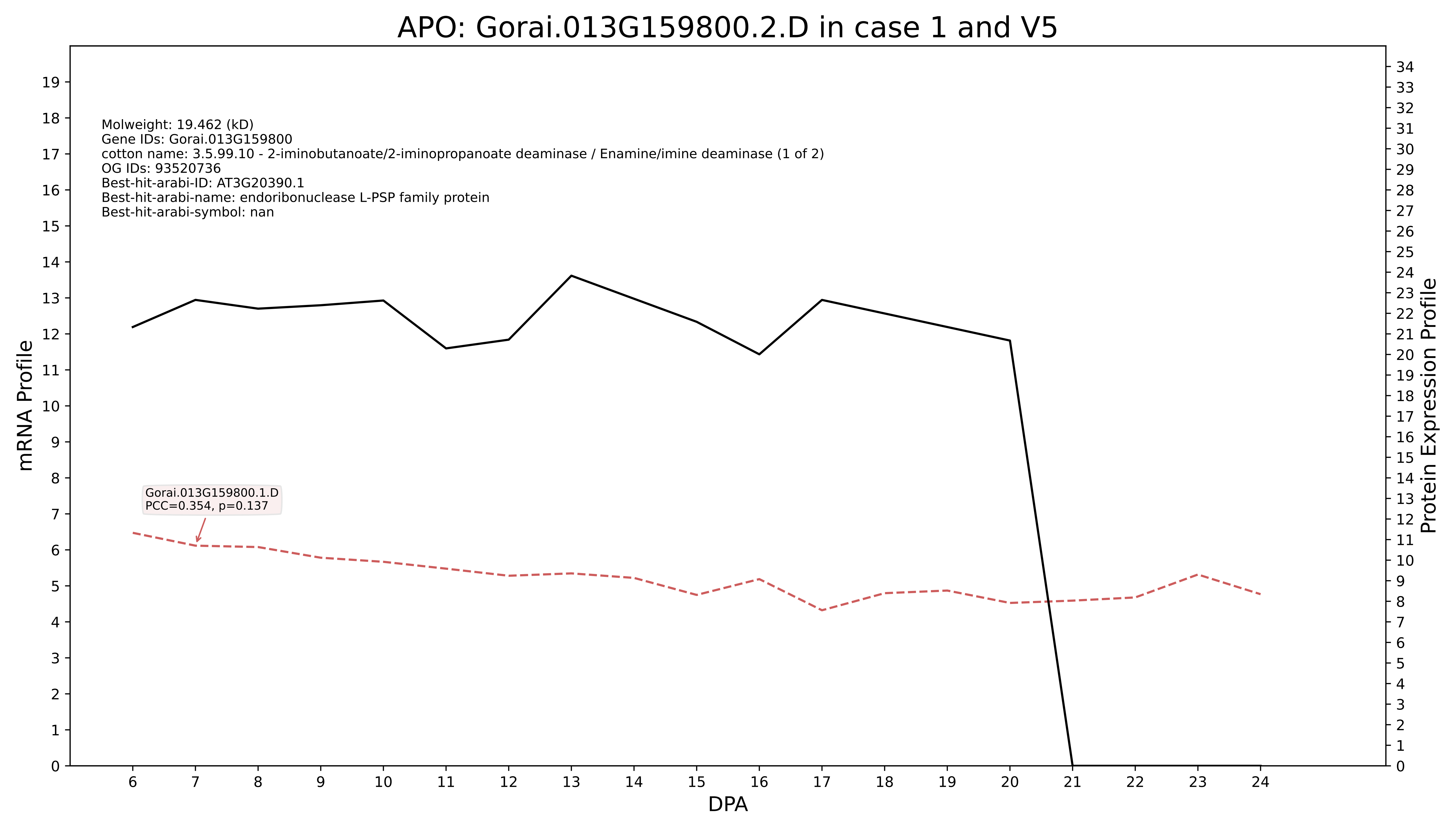
APO: Gorai.013G159800.2.D in case 1 and V5
Molweight: 19.462 (kD)
Gene IDs: Gorai.013G159800
cotton name: 3.5.99.10 - 2-iminobutanoate/2-iminopropanoate deaminase / Enamine/imine deaminase (1 of 2)
OG IDs: 93520736
Best-hit-arabi-ID: AT3G20390.1
Best-hit-arabi-name: endoribonuclease L-PSP family protein
Best-hit-arabi-symbol: nan
Gorai.013G159800.1.D
PCC=0.354, p=0.137
mRNA Profile
Protein Expression Profile
DPA

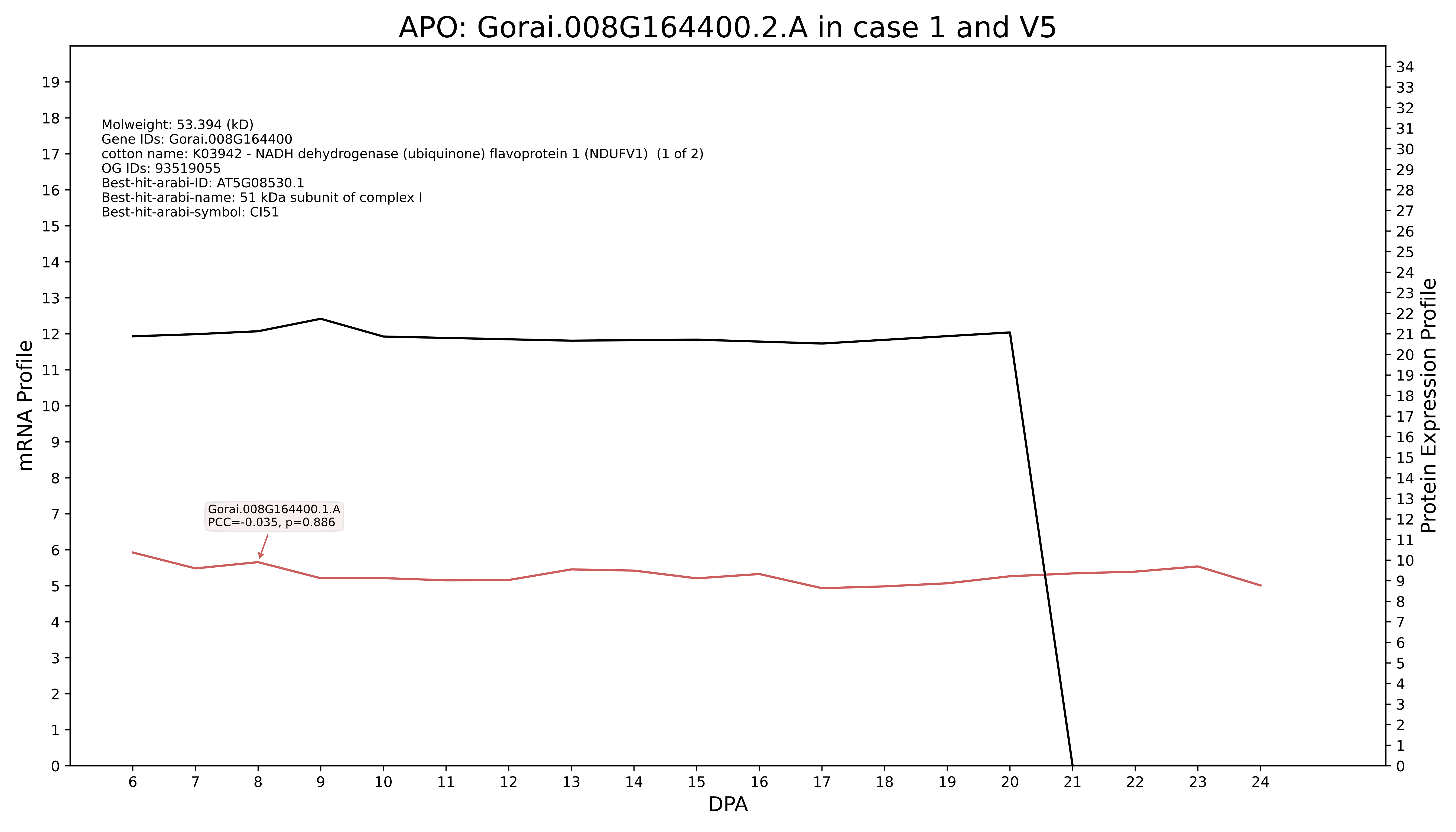
APO: Gorai.008G164400.2.A in case 1 and V5
Molweight: 53.394 (kD)
Gene IDs: Gorai.008G164400
cotton name: K03942 - NADH dehydrogenase (ubiquinone) flavoprotein 1 (NDUFV1) (1 of 2)
OG IDs: 93519055
Best-hit-arabi-ID: AT5G08530.1
Best-hit-arabi-name: 51 kDa subunit of complex I
Best-hit-arabi-symbol: CI51
Gorai.008G164400.1.A
PCC=-0.035, p=0.886
mRNA Profile
Protein Expression Profile
DPA

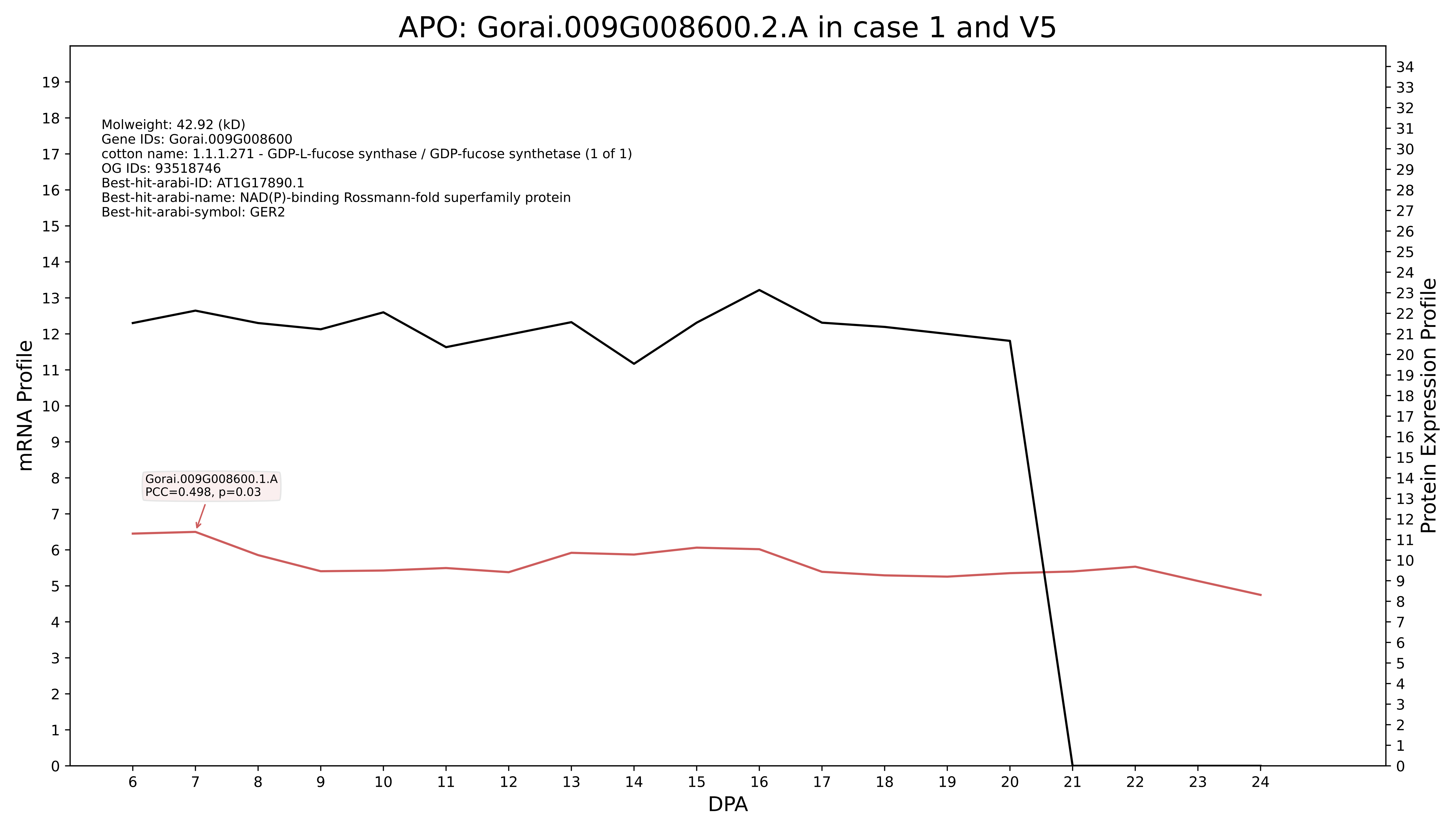
APO: Gorai.009G008600.2.A in case 1 and V5
Molweight: 42.92 (kD)
Gene IDs: Gorai.009G008600
cotton name: 1.1.1.271 - GDP-L-fucose synthase / GDP-fucose synthetase (1 of 1)
OG IDs: 93518746
Best-hit-arabi-ID: AT1G17890.1
Best-hit-arabi-name: NAD(P)-binding Rossmann-fold superfamily protein
Best-hit-arabi-symbol: GER2
Gorai.009G008600.1.A
PCC=0.498, p=0.03
mRNA Profile
Protein Expression Profile
DPA

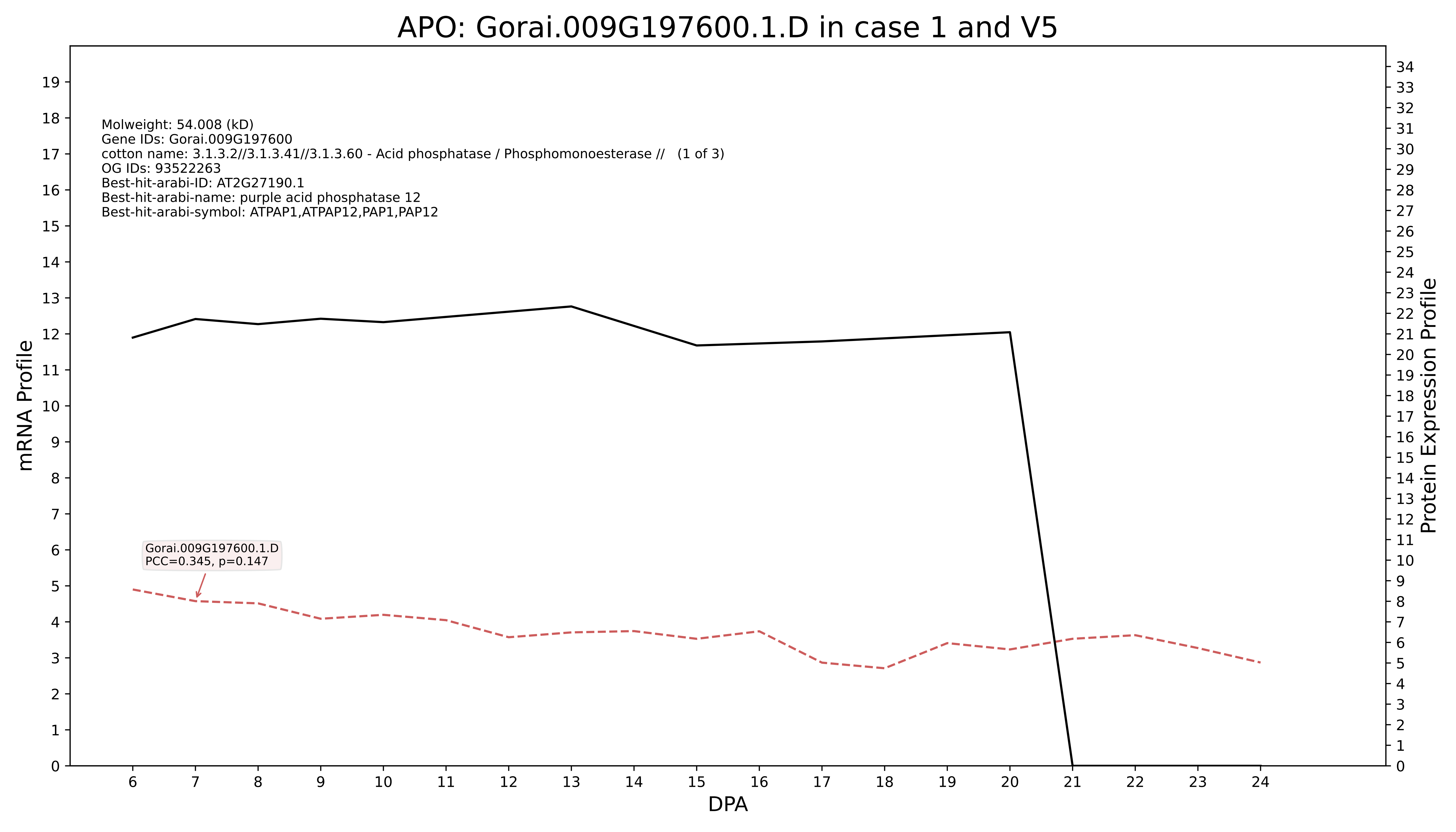
APO: Gorai.009G197600.1.D in case 1 and V5
Molweight: 54.008 (kD)
Gene IDs: Gorai.009G197600
cotton name: 3.1.3.2//3.1.3.41//3.1.3.60 - Acid phosphatase / Phosphomonoesterase // (1 of 3)
OG IDs: 93522263
Best-hit-arabi-ID: AT2G27190.1
Best-hit-arabi-name: purple acid phosphatase 12
Best-hit-arabi-symbol: ATPAP1,ATPAP12,PAP1,PAP12
Gorai.009G197600.1.D
PCC=0.345, p=0.147
mRNA Profile
Protein Expression Profile
DPA

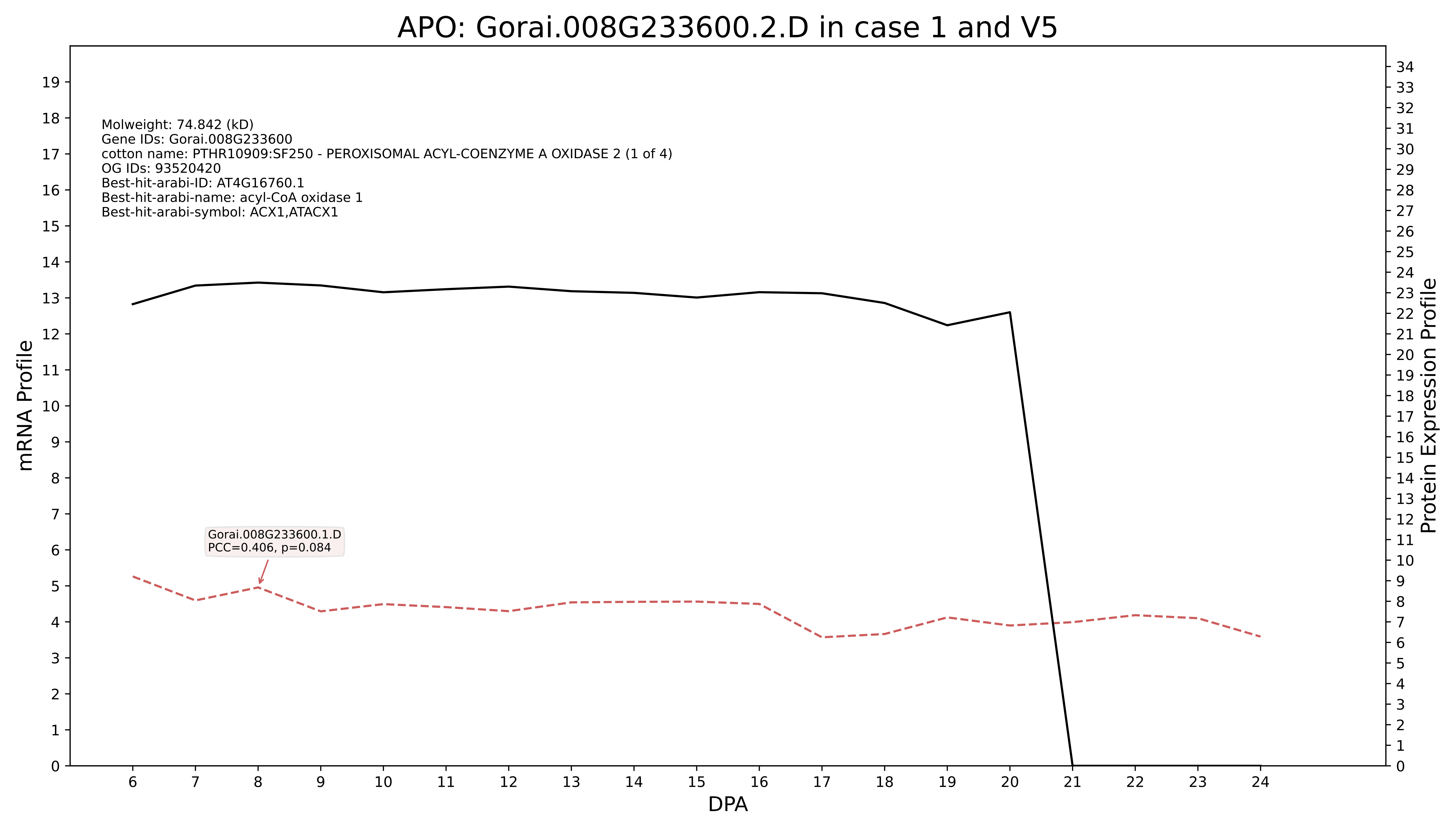
APO: Gorai.008G233600.2.D in case 1 and V5
Molweight: 74.842 (kD)
Gene IDs: Gorai.008G233600
cotton name: PTHR10909:SF250 - PEROXISOMAL ACYL-COENZYME A OXIDASE 2 (1 of 4)
OG IDs: 93520420
Best-hit-arabi-ID: AT4G16760.1
Best-hit-arabi-name: acyl-CoA oxidase 1
Best-hit-arabi-symbol: ACX1,ATACX1
Gorai.008G233600.1.D
PCC=0.406, p=0.084
mRNA Profile
Protein Expression Profile
DPA

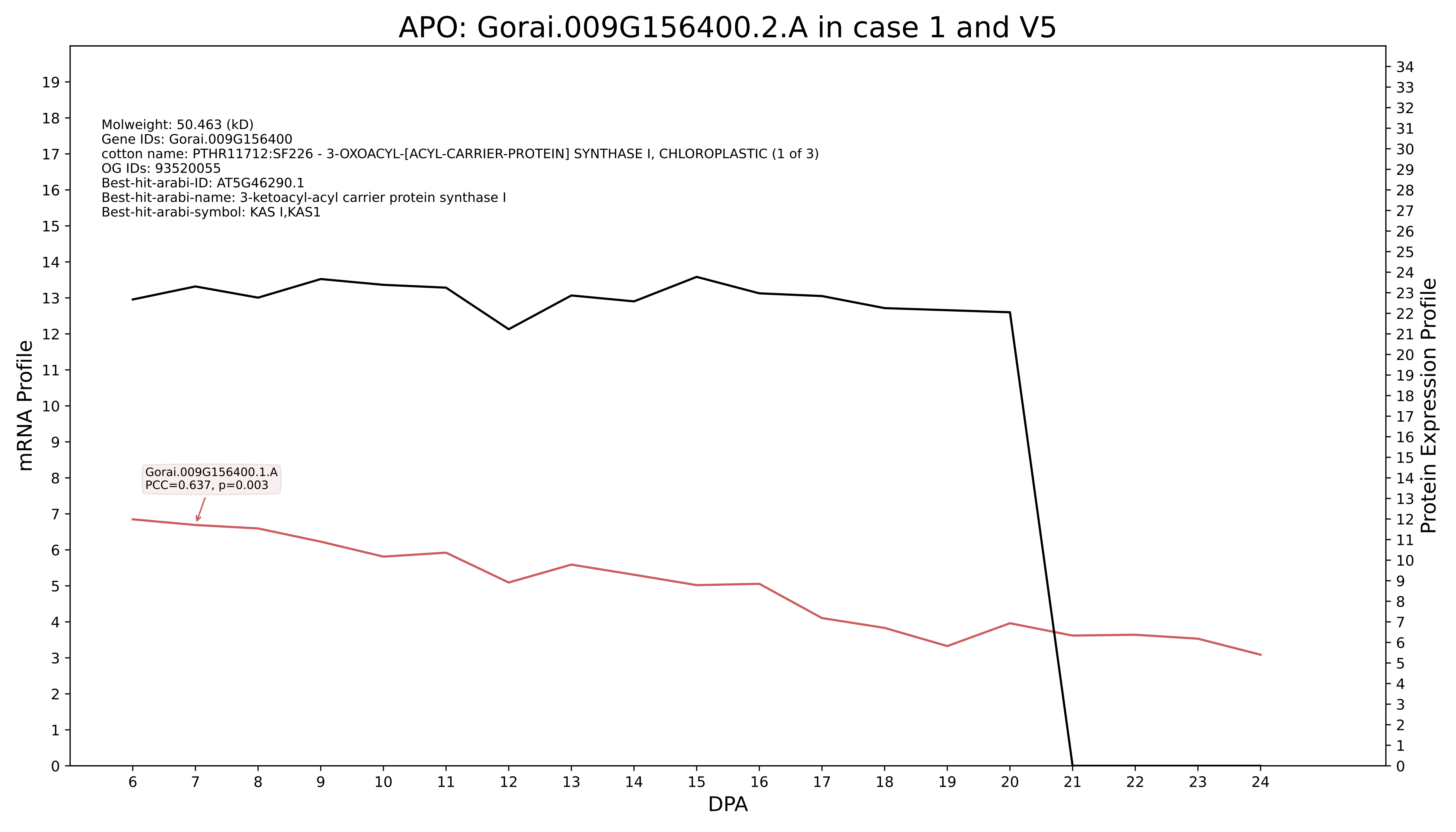

APO: Gorai.009G156400.2.A in case 1 and V5
Molweight: 50.463 (kD)
Gene IDs: Gorai.009G156400
cotton name: PTHR11712:SF226 - 3-OXOACYL-[ACYL-CARRIER-PROTEIN] SYNTHASE I, CHLOROPLASTIC (1 of 3)
OG IDs: 93520055
Best-hit-arabi-ID: AT5G46290.1
Best-hit-arabi-name: 3-ketoacyl-acyl carrier protein synthase I
Best-hit-arabi-symbol: KAS I,KAS1
Gorai.009G156400.1.A
PCC=0.637, p=0.003
mRNA Profile
Protein Expression Profile
DPA

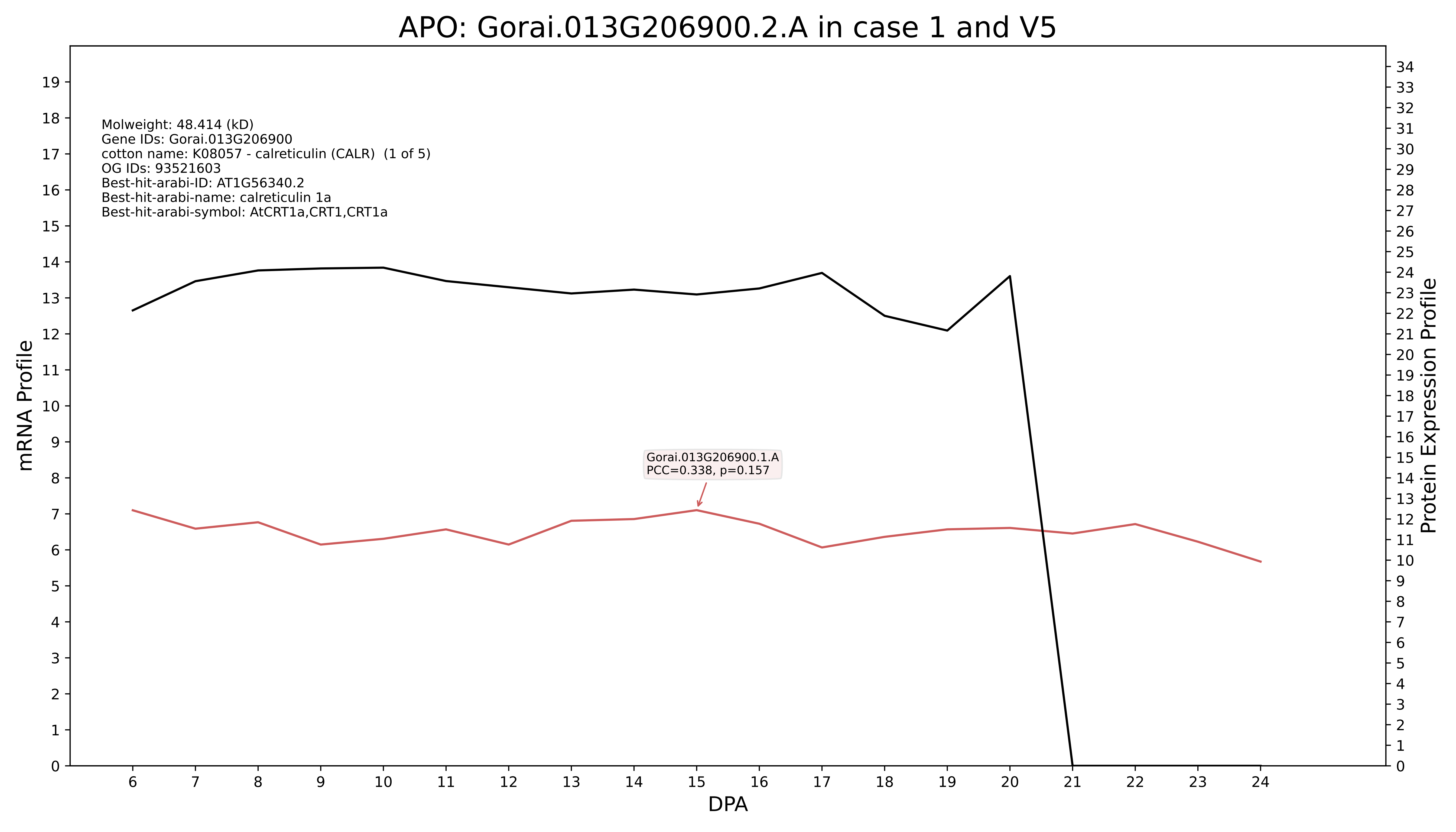
APO: Gorai.013G206900.2.A in case 1 and V5
Molweight: 48.414 (kD)
Gene IDs: Gorai.013G206900
cotton name: K08057 - calreticulin (CALR) (1 of 5)
OG IDs: 93521603
Best-hit-arabi-ID: AT1G56340.2
Best-hit-arabi-name: calreticulin 1a
Best-hit-arabi-symbol: AtCRT1a,CRT1,CRT1a
Gorai.013G206900.1.A
PCC=0.338, p=0.157
mRNA Profile
Protein Expression Profile
DPA

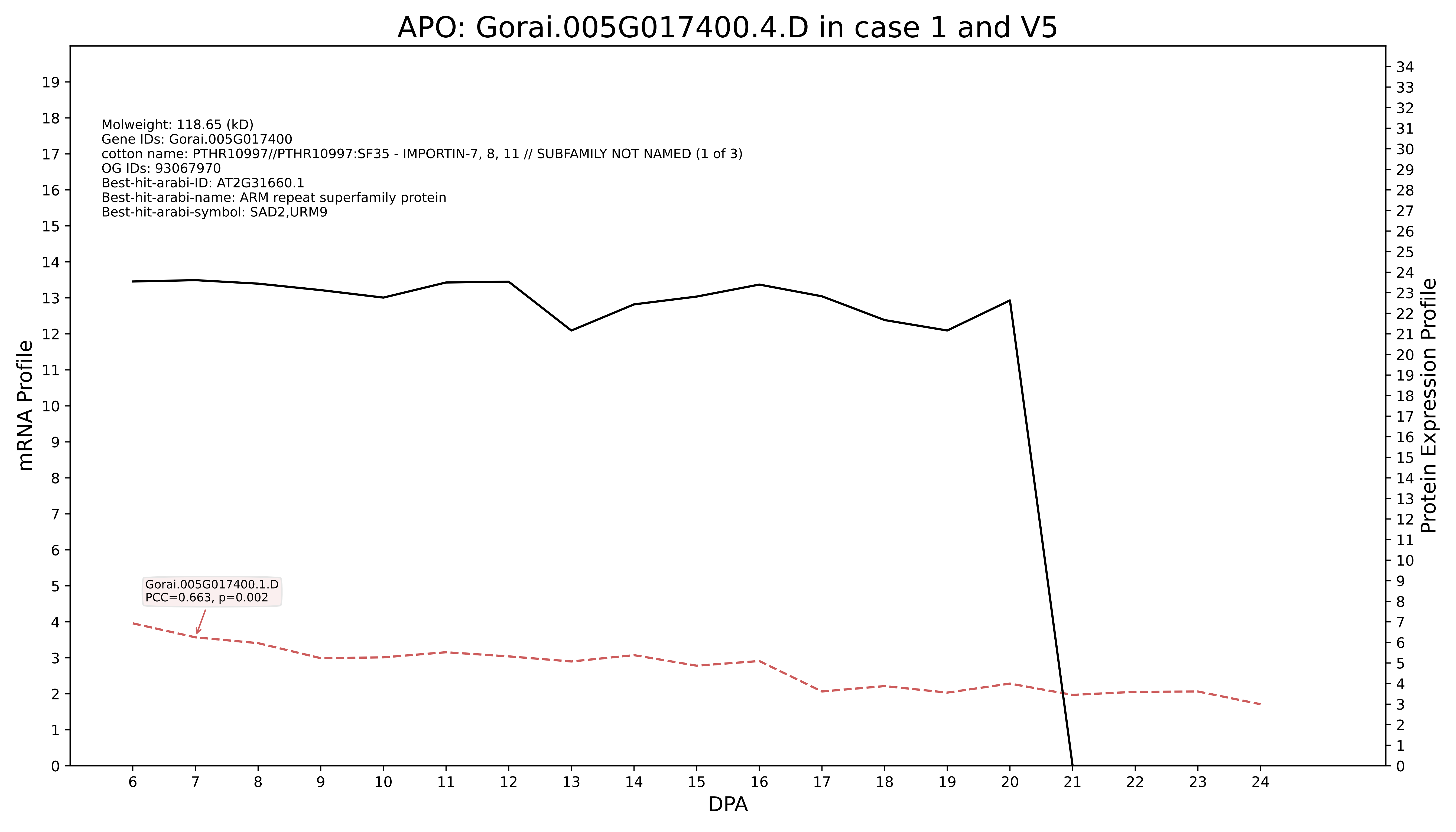
APO: Gorai.005G017400.4.D in case 1 and V5
Molweight: 118.65 (kD)
Gene IDs: Gorai.005G017400
cotton name: PTHR10997//PTHR10997:SF35 - IMPORTIN-7, 8, 11 // SUBFAMILY NOT NAMED (1 of 3)
OG IDs: 93067970
Best-hit-arabi-ID: AT2G31660.1
Best-hit-arabi-name: ARM repeat superfamily protein
Best-hit-arabi-symbol: SAD2,URM9
Gorai.005G017400.1.D
PCC=0.663, p=0.002
mRNA Profile
Protein Expression Profile
DPA

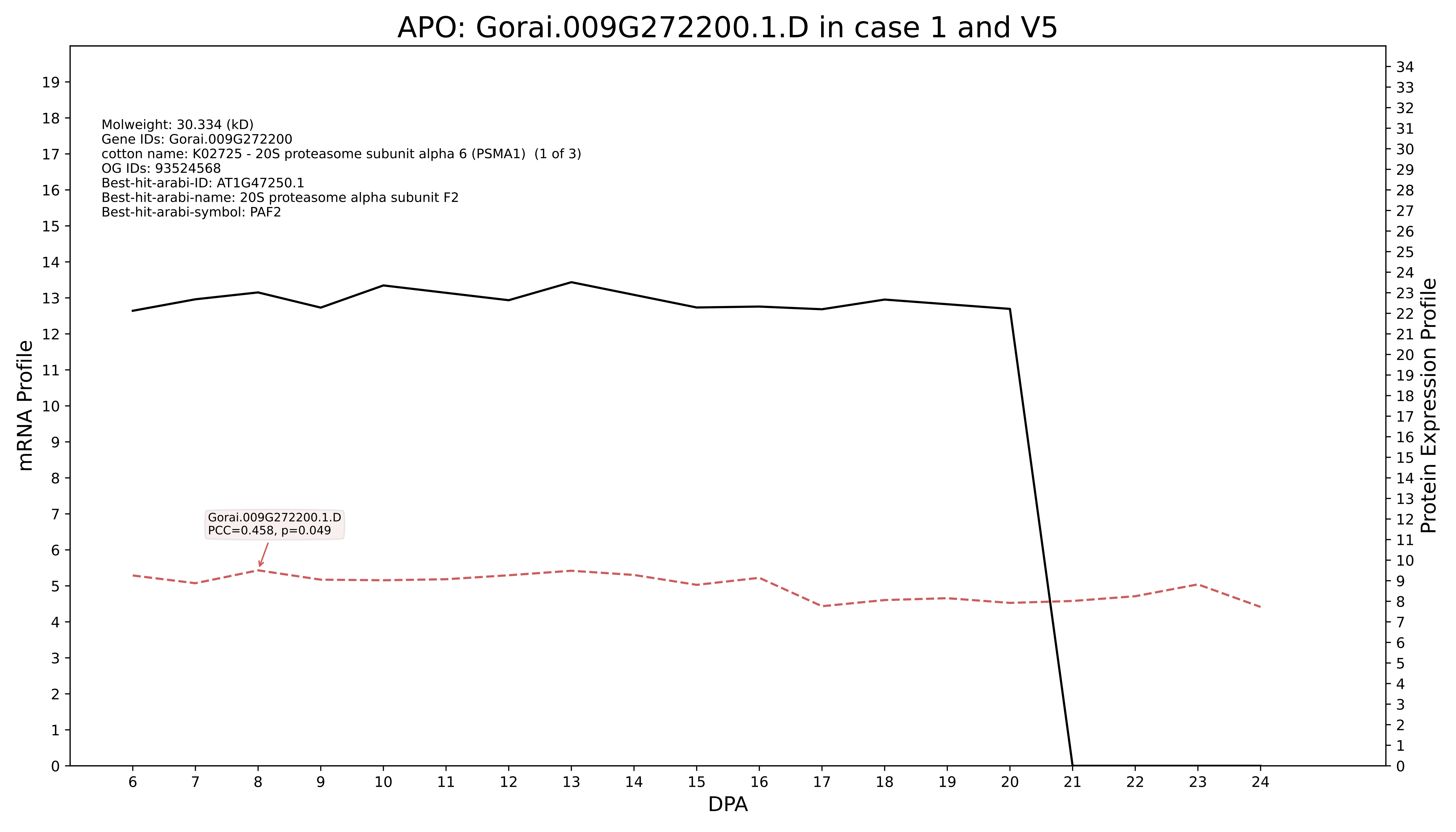

APO: Gorai.009G272200.1.D in case 1 and V5
Molweight: 30.334 (kD)
Gene IDs: Gorai.009G272200
cotton name: K02725 - 20S proteasome subunit alpha 6 (PSMA1) (1 of 3)
OG IDs: 93524568
Best-hit-arabi-ID: AT1G47250.1
Best-hit-arabi-name: 20S proteasome alpha subunit F2
Best-hit-arabi-symbol: PAF2
Gorai.009G272200.1.D
PCC=0.458, p=0.049
mRNA Profile
Protein Expression Profile
DPA

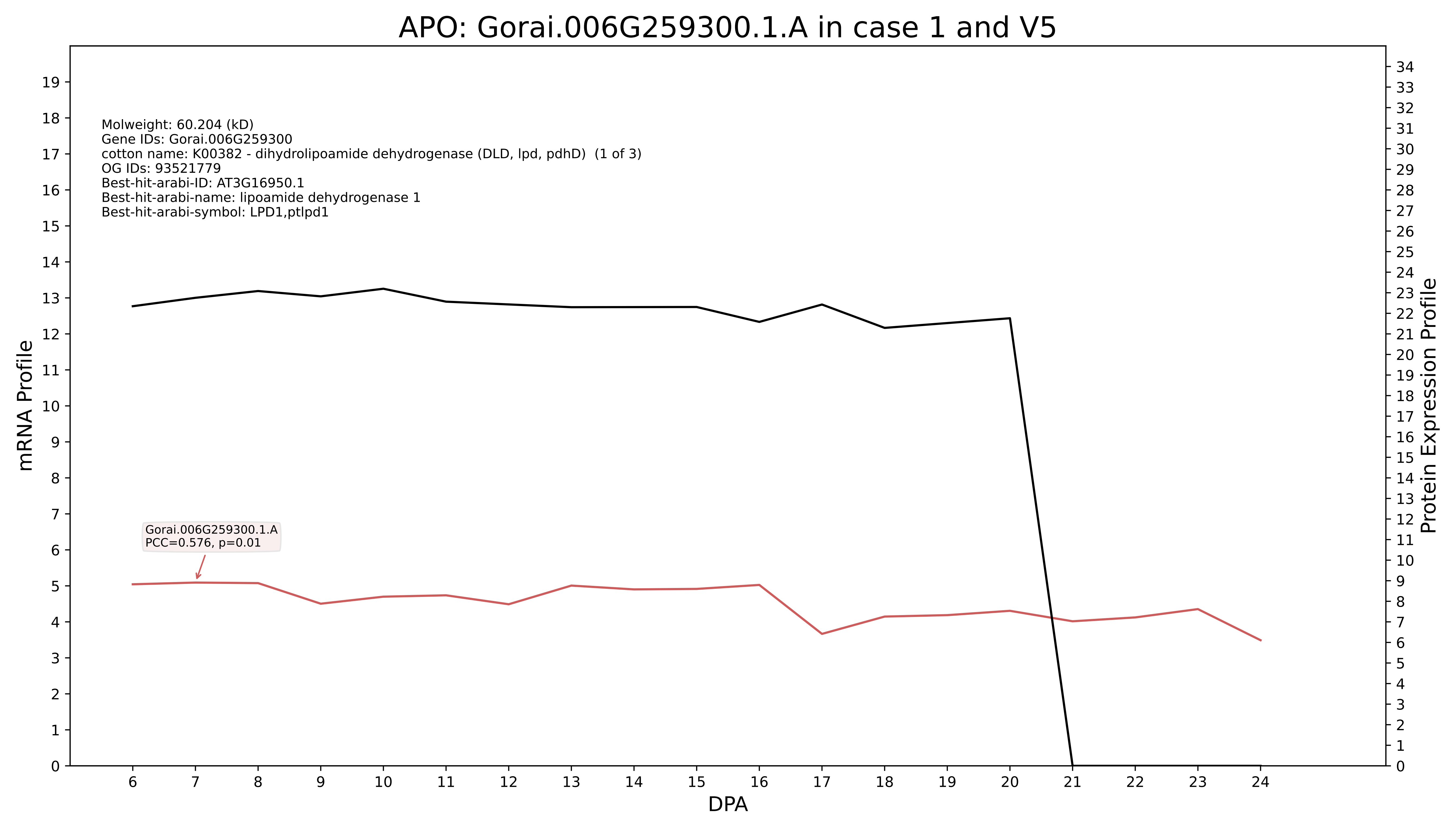
APO: Gorai.006G259300.1.A in case 1 and V5
Molweight: 60.204 (kD)
Gene IDs: Gorai.006G259300
cotton name: K00382 - dihydrolipoamide dehydrogenase (DLD, lpd, pdhD) (1 of 3)
OG IDs: 93521779
Best-hit-arabi-ID: AT3G16950.1
Best-hit-arabi-name: lipoamide dehydrogenase 1
Best-hit-arabi-symbol: LPD1,ptlpd1
Gorai.006G259300.1.A
PCC=0.576, p=0.01
mRNA Profile
Protein Expression Profile
DPA

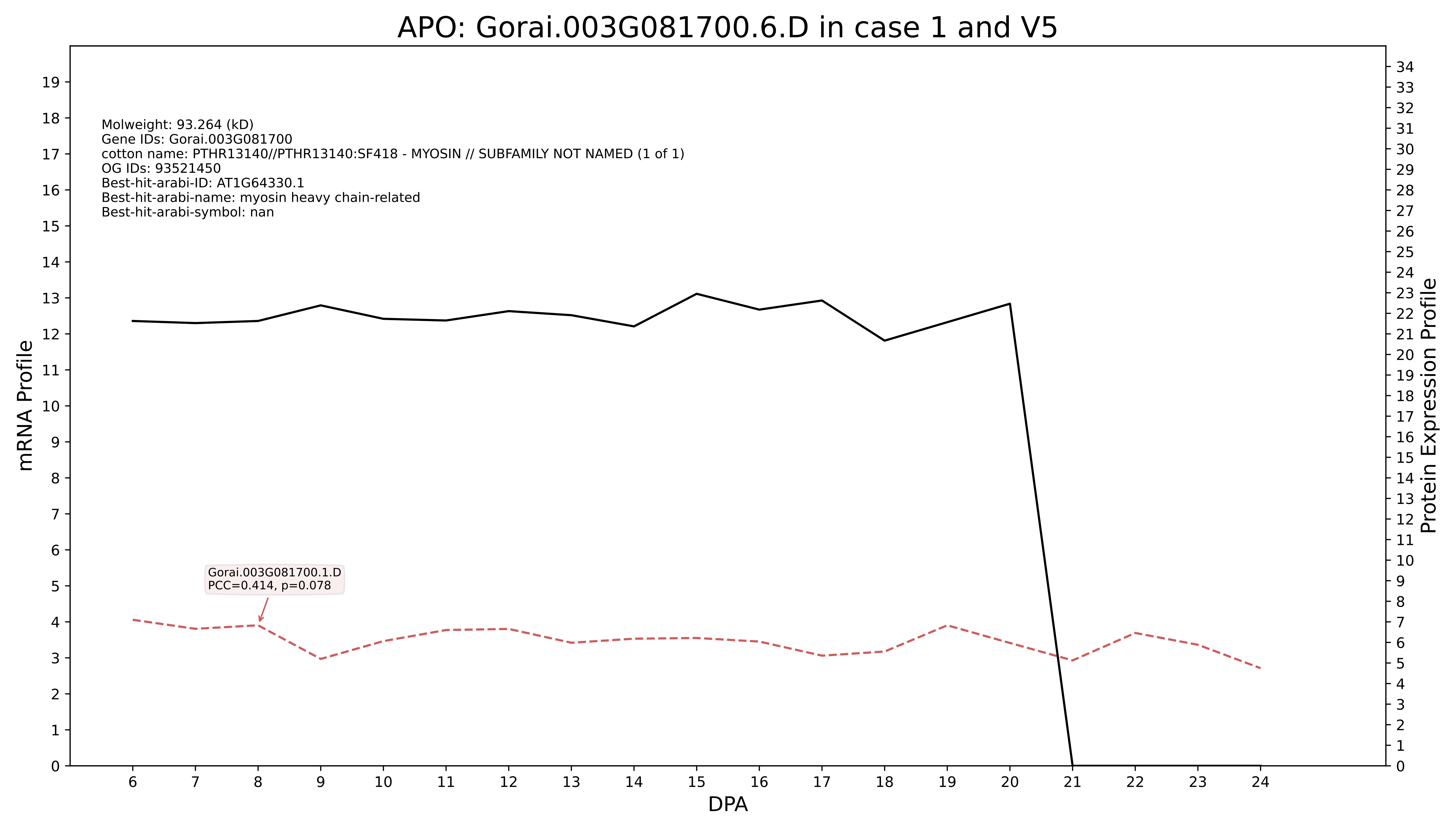
APO: Gorai.003G081700.6.D in case 1 and V5
Molweight: 93.264 (kD)
Gene IDs: Gorai.003G081700
cotton name: PTHR13140//PTHR13140:SF418 - MYOSIN // SUBFAMILY NOT NAMED (1 of 1)
OG IDs: 93521450
Best-hit-arabi-ID: AT1G64330.1
Best-hit-arabi-name: myosin heavy chain-related
Best-hit-arabi-symbol: nan
Gorai.003G081700.1.D
PCC=0.414, p=0.078
mRNA Profile
Protein Expression Profile
DPA

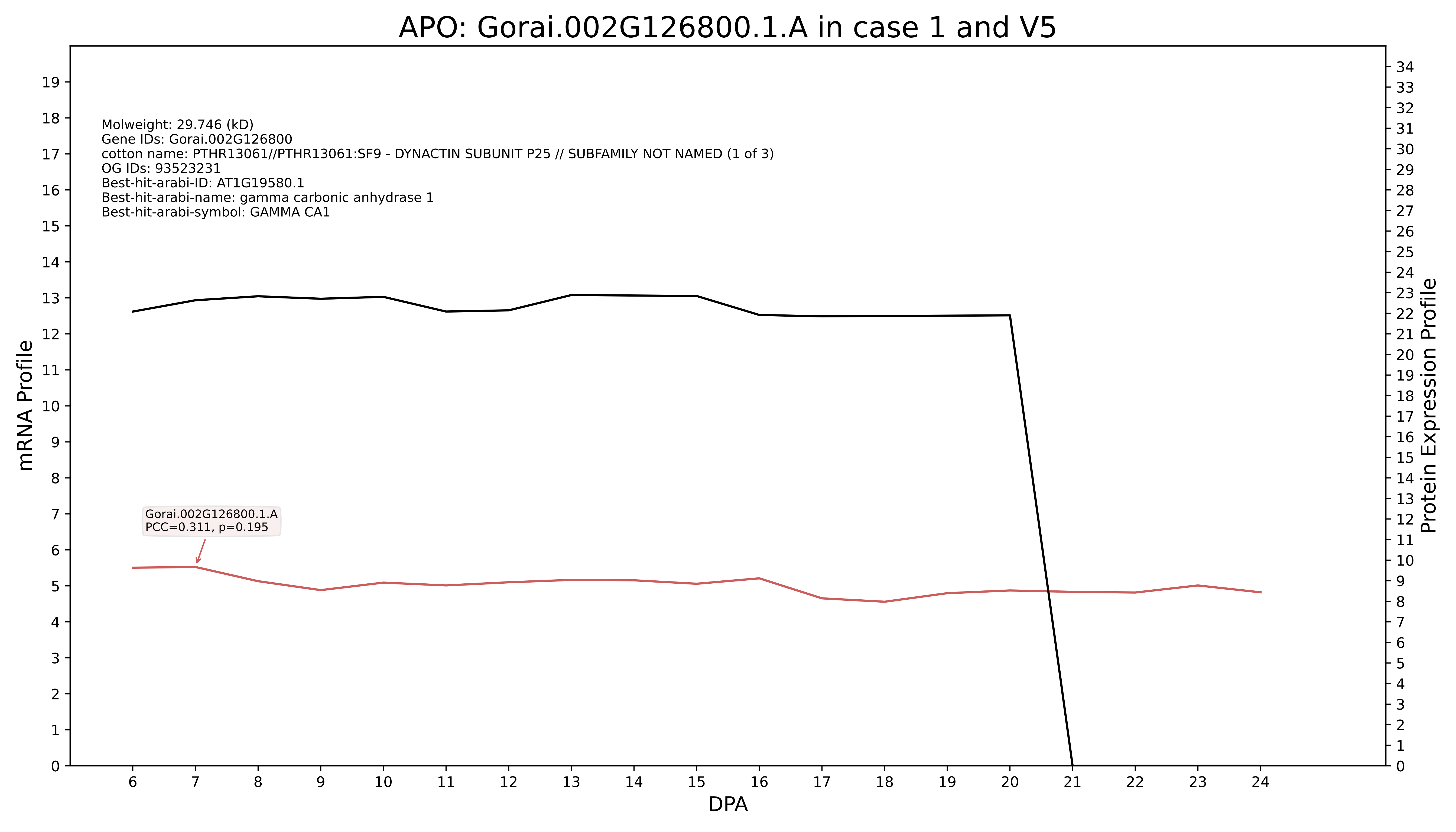
APO: Gorai.002G126800.1.A in case 1 and V5
Molweight: 29.746 (kD)
Gene IDs: Gorai.002G126800
cotton name: PTHR13061//PTHR13061:SF9 - DYNACTIN SUBUNIT P25 // SUBFAMILY NOT NAMED (1 of 3)
OG IDs: 93523231
Best-hit-arabi-ID: AT1G19580.1
Best-hit-arabi-name: gamma carbonic anhydrase 1
Best-hit-arabi-symbol: GAMMA CA1
Gorai.002G126800.1.A
PCC=0.311, p=0.195
mRNA Profile
Protein Expression Profile
DPA
0 1 2 3 4 5 6 7 8 9 10 11 12 13 14 15 16 17 18 19
0 1 2 3 4 5 6 7 8 9 10 11 12 13 14 15 16 17 18 19 20 21 22 23 24 25 26 27 28 29 30 31 32 33 34
6 7 8 9 10 11 12 13 14 15 16 17 18 19 20 21 22 23 24

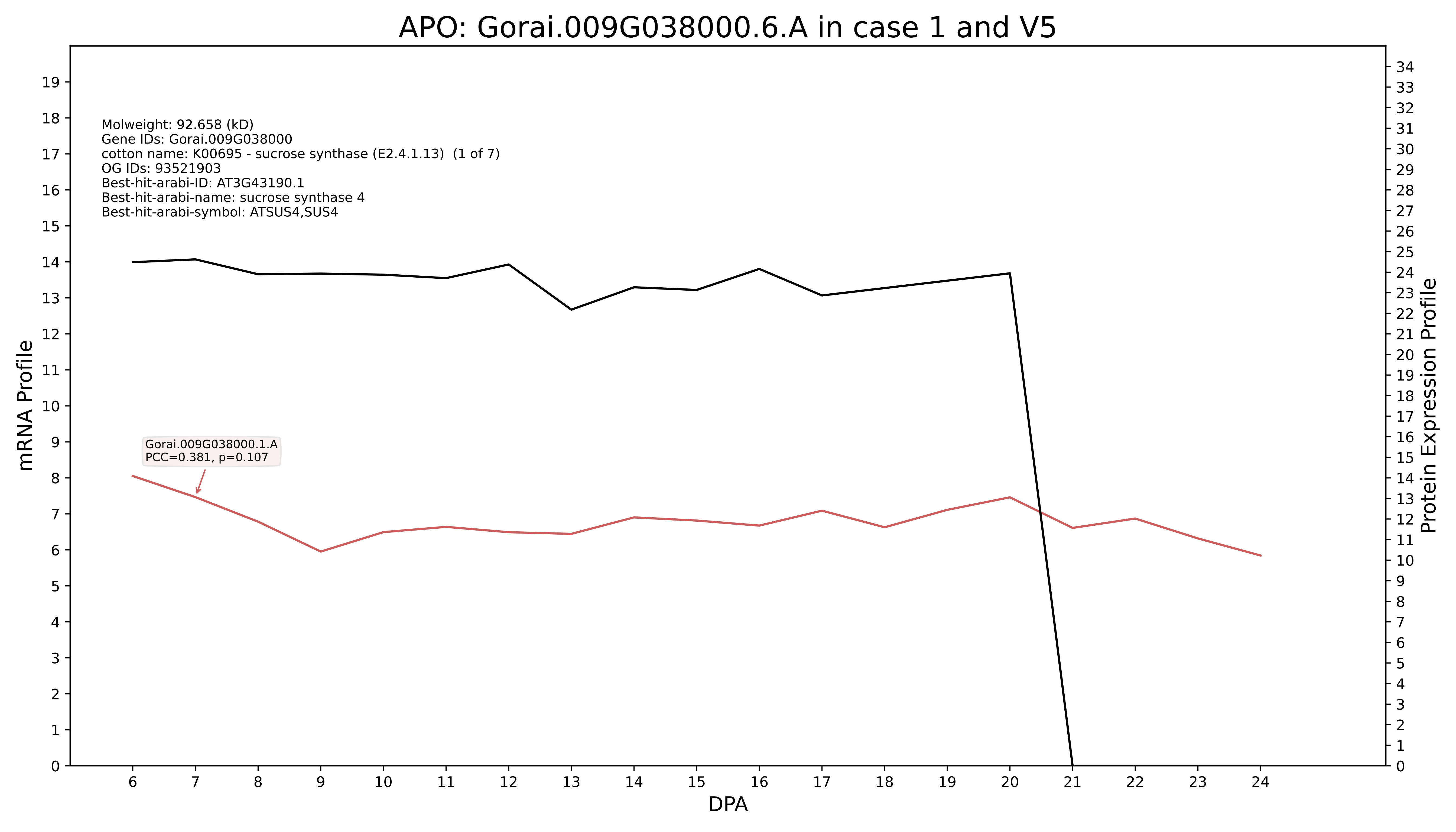

APO: Gorai.009G038000.6.A in case 1 and V5
Molweight: 92.658 (kD)
Gene IDs: Gorai.009G038000
cotton name: K00695 - sucrose synthase (E2.4.1.13) (1 of 7)
OG IDs: 93521903
Best-hit-arabi-ID: AT3G43190.1
Best-hit-arabi-name: sucrose synthase 4
Best-hit-arabi-symbol: ATSUS4,SUS4
Gorai.009G038000.1.A
PCC=0.381, p=0.107
mRNA Profile
Protein Expression Profile
DPA

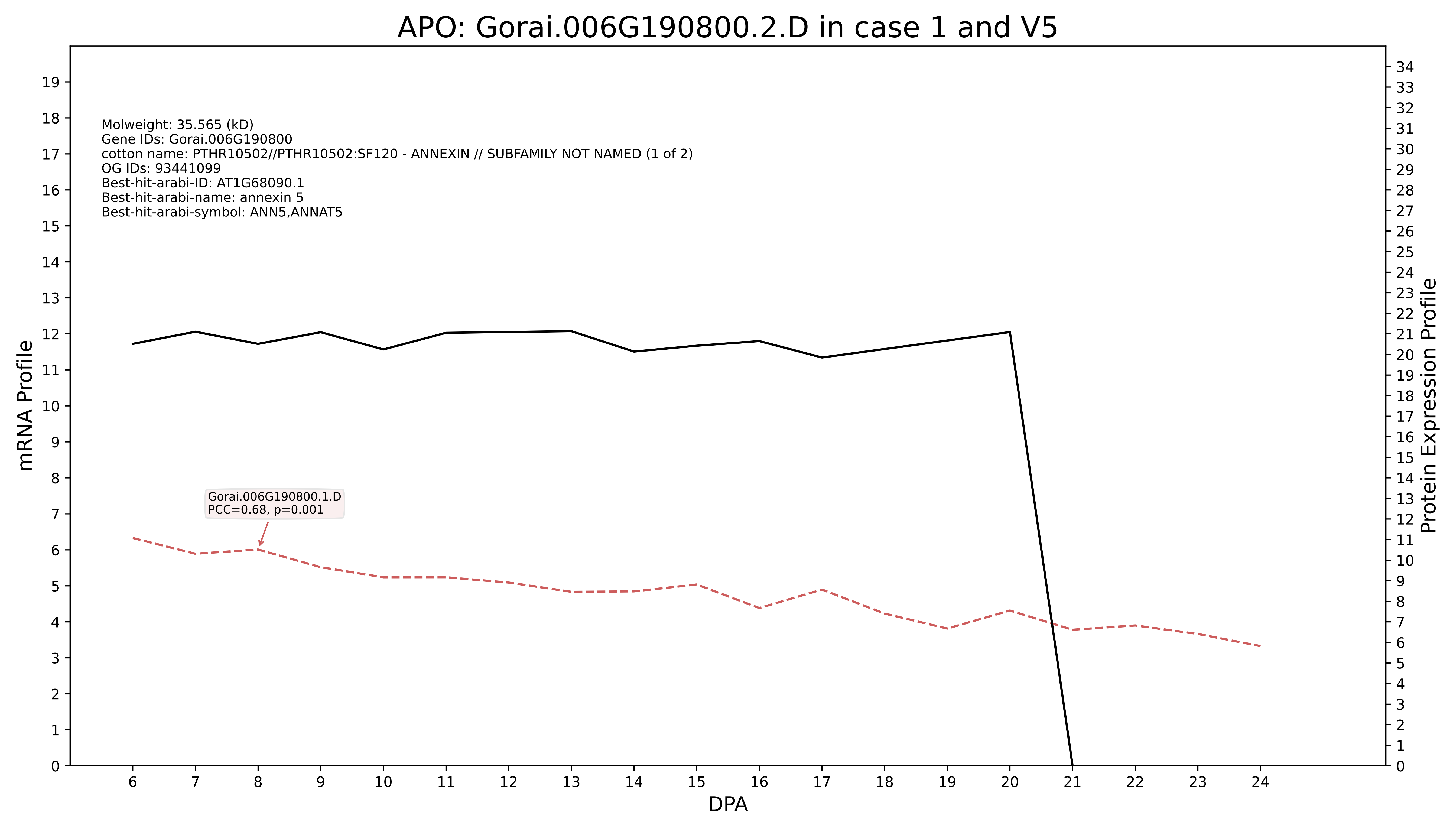
APO: Gorai.006G190800.2.D in case 1 and V5
Molweight: 35.565 (kD)
Gene IDs: Gorai.006G190800
cotton name: PTHR10502//PTHR10502:SF120 - ANNEXIN // SUBFAMILY NOT NAMED (1 of 2)
OG IDs: 93441099
Best-hit-arabi-ID: AT1G68090.1
Best-hit-arabi-name: annexin 5
Best-hit-arabi-symbol: ANN5,ANNAT5
Gorai.006G190800.1.D
PCC=0.68, p=0.001
mRNA Profile
Protein Expression Profile
DPA

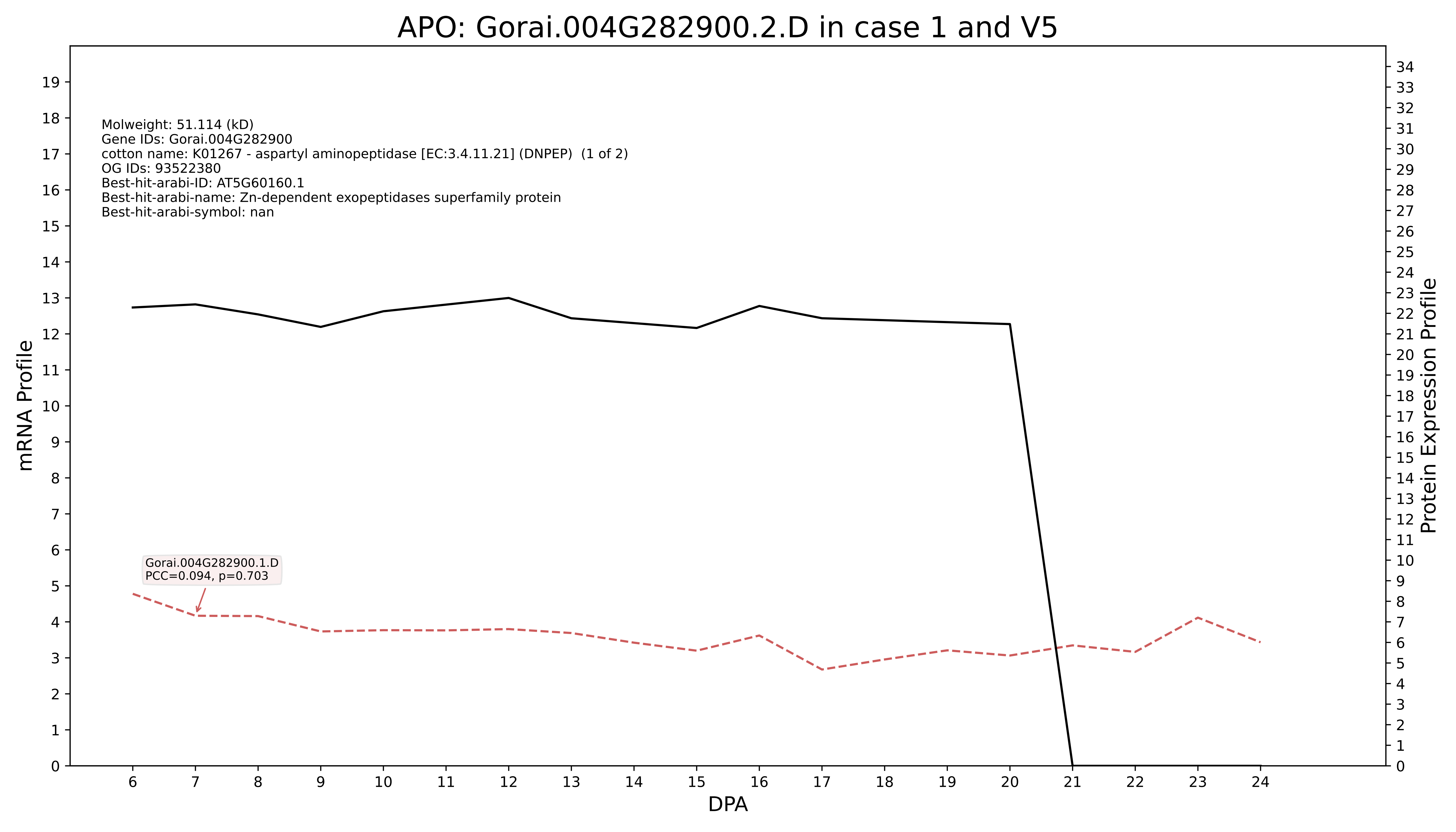
APO: Gorai.004G282900.2.D in case 1 and V5
Molweight: 51.114 (kD)
Gene IDs: Gorai.004G282900
cotton name: K01267 - aspartyl aminopeptidase [EC:3.4.11.21] (DNPEP) (1 of 2)
OG IDs: 93522380
Best-hit-arabi-ID: AT5G60160.1
Best-hit-arabi-name: Zn-dependent exopeptidases superfamily protein
Best-hit-arabi-symbol: nan
Gorai.004G282900.1.D
PCC=0.094, p=0.703
mRNA Profile
Protein Expression Profile
DPA

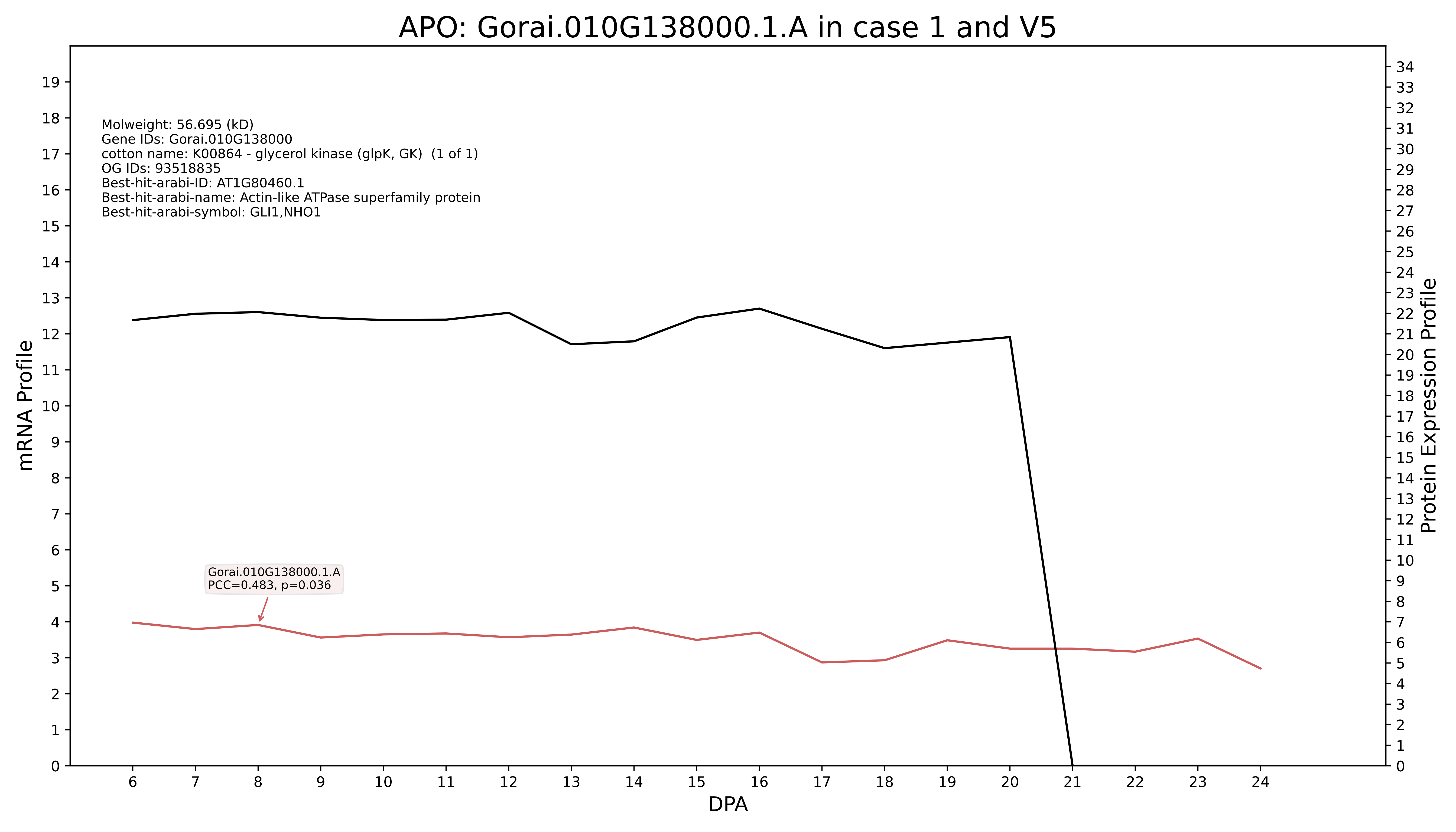
APO: Gorai.010G138000.1.A in case 1 and V5
Molweight: 56.695 (kD)
Gene IDs: Gorai.010G138000
cotton name: K00864 - glycerol kinase (glpK, GK) (1 of 1)
OG IDs: 93518835
Best-hit-arabi-ID: AT1G80460.1
Best-hit-arabi-name: Actin-like ATPase superfamily protein
Best-hit-arabi-symbol: GLI1,NHO1
Gorai.010G138000.1.A
PCC=0.483, p=0.036
mRNA Profile
Protein Expression Profile
DPA

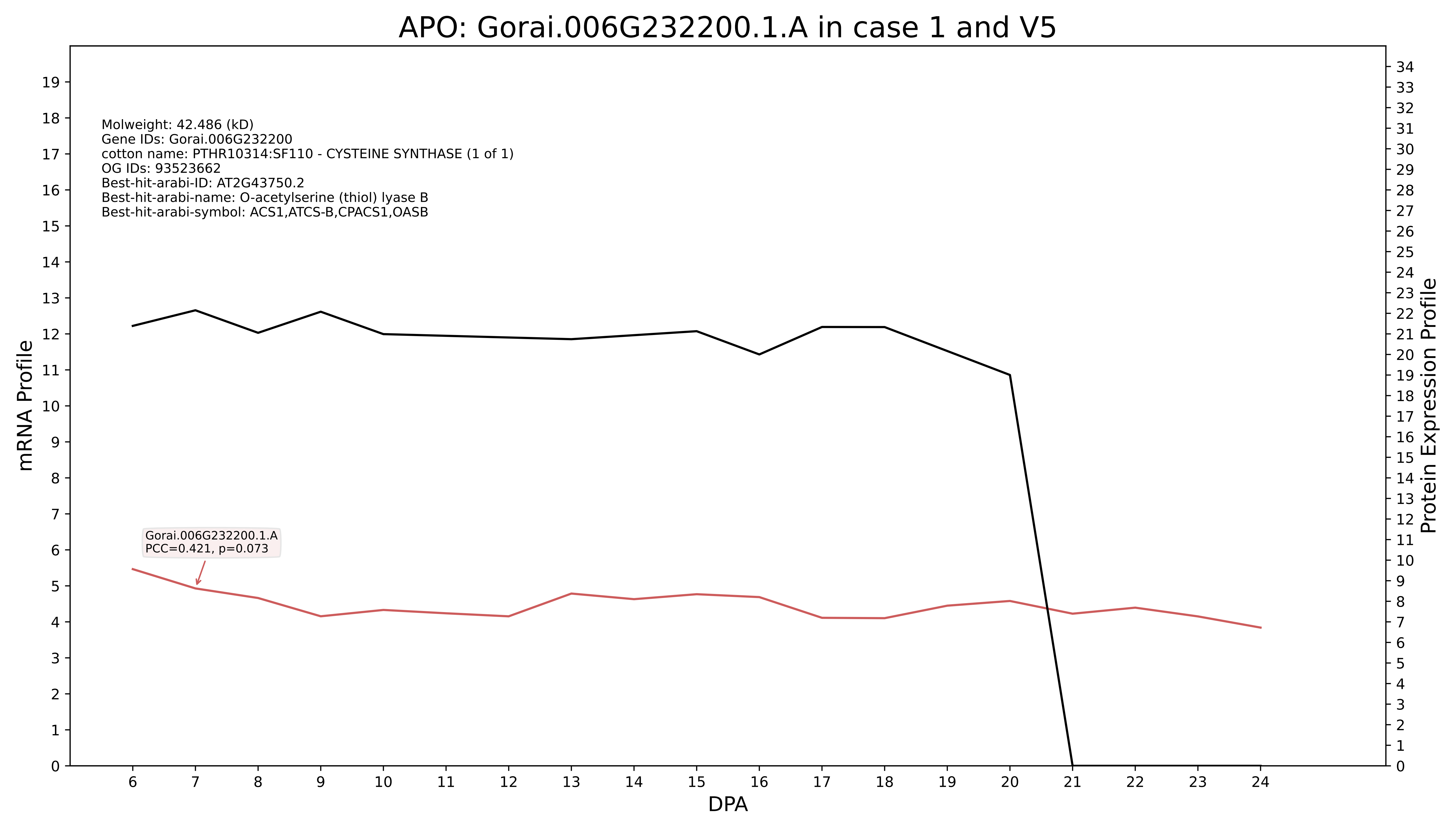

APO: Gorai.006G232200.1.A in case 1 and V5
Molweight: 42.486 (kD)
Gene IDs: Gorai.006G232200
cotton name: PTHR10314:SF110 - CYSTEINE SYNTHASE (1 of 1)
OG IDs: 93523662
Best-hit-arabi-ID: AT2G43750.2
Best-hit-arabi-name: O-acetylserine (thiol) lyase B
Best-hit-arabi-symbol: ACS1,ATCS-B,CPACS1,OASB
Gorai.006G232200.1.A
PCC=0.421, p=0.073
mRNA Profile
Protein Expression Profile
DPA

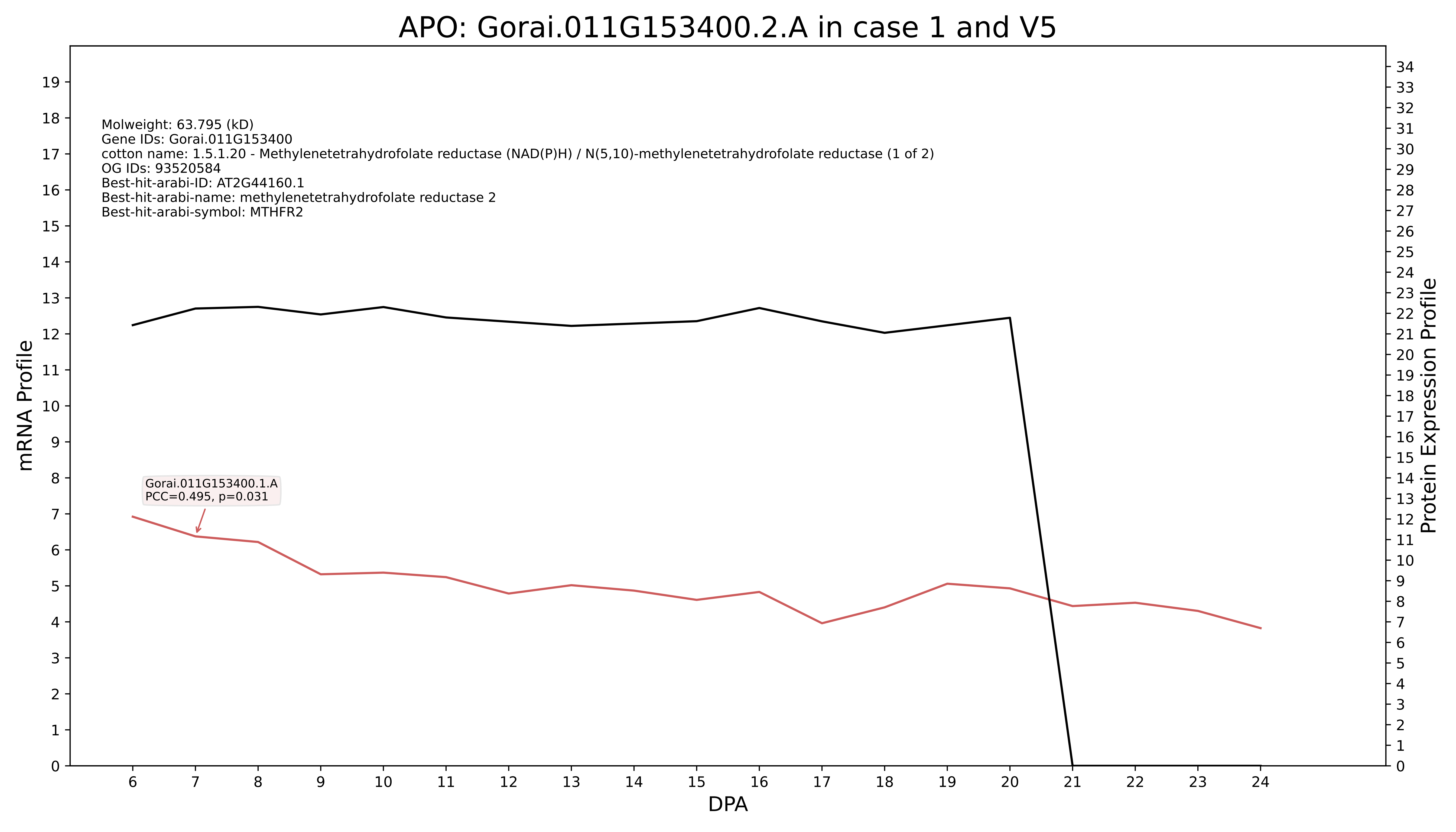
APO: Gorai.011G153400.2.A in case 1 and V5
Molweight: 63.795 (kD)
Gene IDs: Gorai.011G153400
cotton name: 1.5.1.20 - Methylenetetrahydrofolate reductase (NAD(P)H) / N(5,10)-methylenetetrahydrofolate reductase (1 of 2)
OG IDs: 93520584
Best-hit-arabi-ID: AT2G44160.1
Best-hit-arabi-name: methylenetetrahydrofolate reductase 2
Best-hit-arabi-symbol: MTHFR2
Gorai.011G153400.1.A
PCC=0.495, p=0.031
mRNA Profile
Protein Expression Profile
DPA

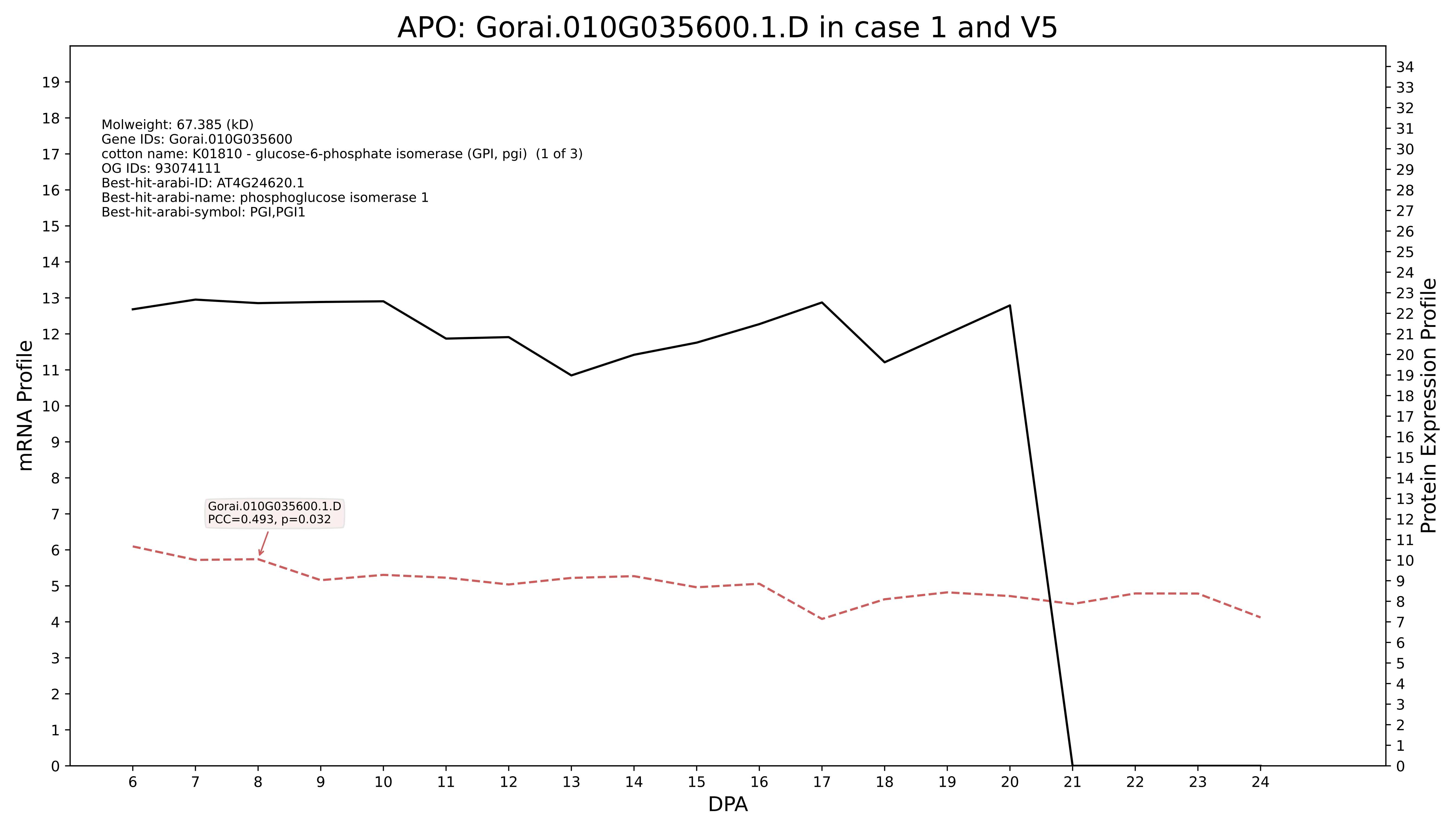
APO: Gorai.010G035600.1.D in case 1 and V5
Molweight: 67.385 (kD)
Gene IDs: Gorai.010G035600
cotton name: K01810 - glucose-6-phosphate isomerase (GPI, pgi) (1 of 3)
OG IDs: 93074111
Best-hit-arabi-ID: AT4G24620.1
Best-hit-arabi-name: phosphoglucose isomerase 1
Best-hit-arabi-symbol: PGI,PGI1
Gorai.010G035600.1.D
PCC=0.493, p=0.032
mRNA Profile
Protein Expression Profile
DPA

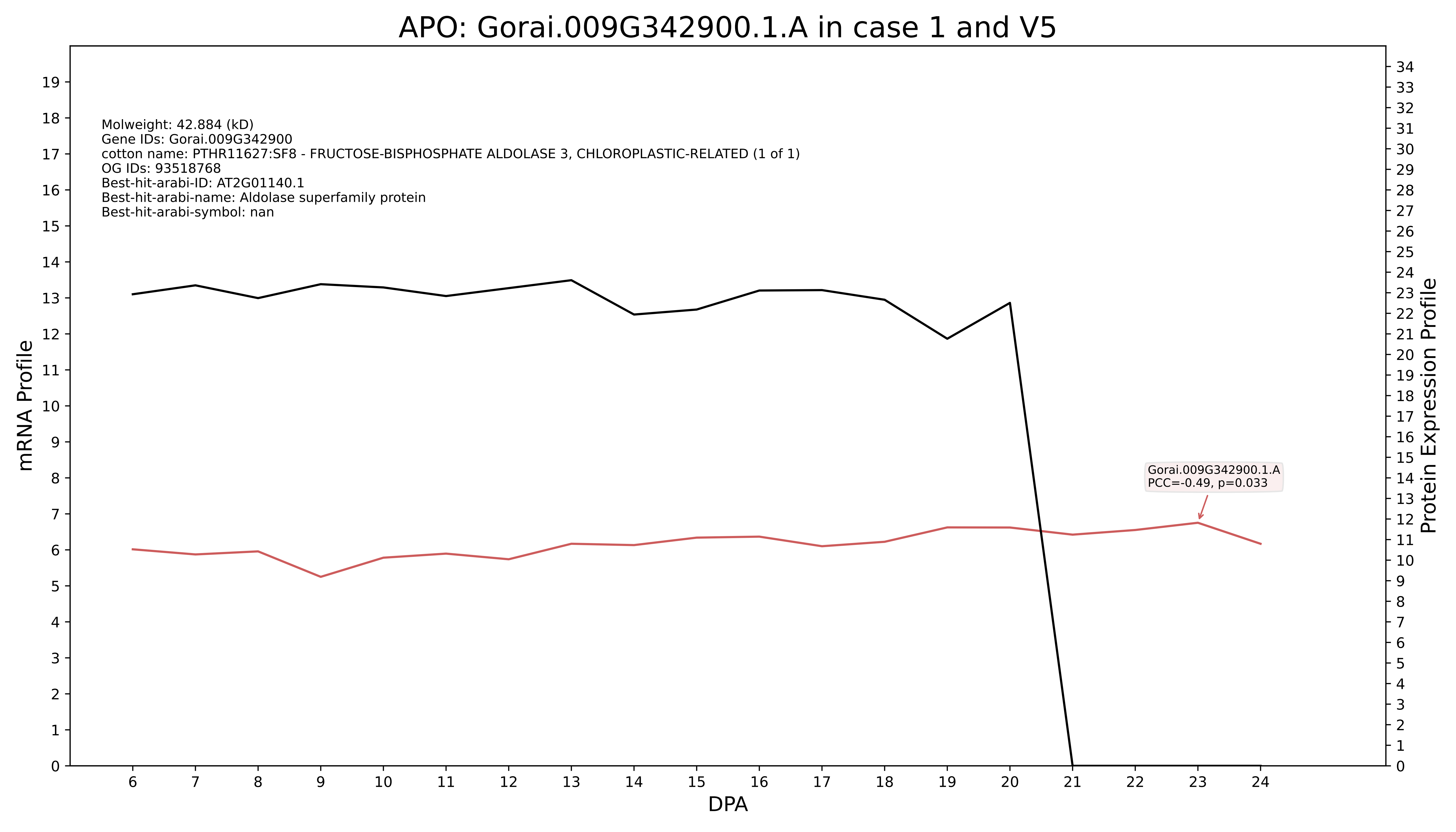
APO: Gorai.009G342900.1.A in case 1 and V5
Molweight: 42.884 (kD)
Gene IDs: Gorai.009G342900
cotton name: PTHR11627:SF8 - FRUCTOSE-BISPHOSPHATE ALDOLASE 3, CHLOROPLASTIC-RELATED (1 of 1)
OG IDs: 93518768
Best-hit-arabi-ID: AT2G01140.1
Best-hit-arabi-name: Aldolase superfamily protein
Best-hit-arabi-symbol: nan
Gorai.009G342900.1.A
PCC=-0.49, p=0.033
mRNA Profile
Protein Expression Profile
DPA

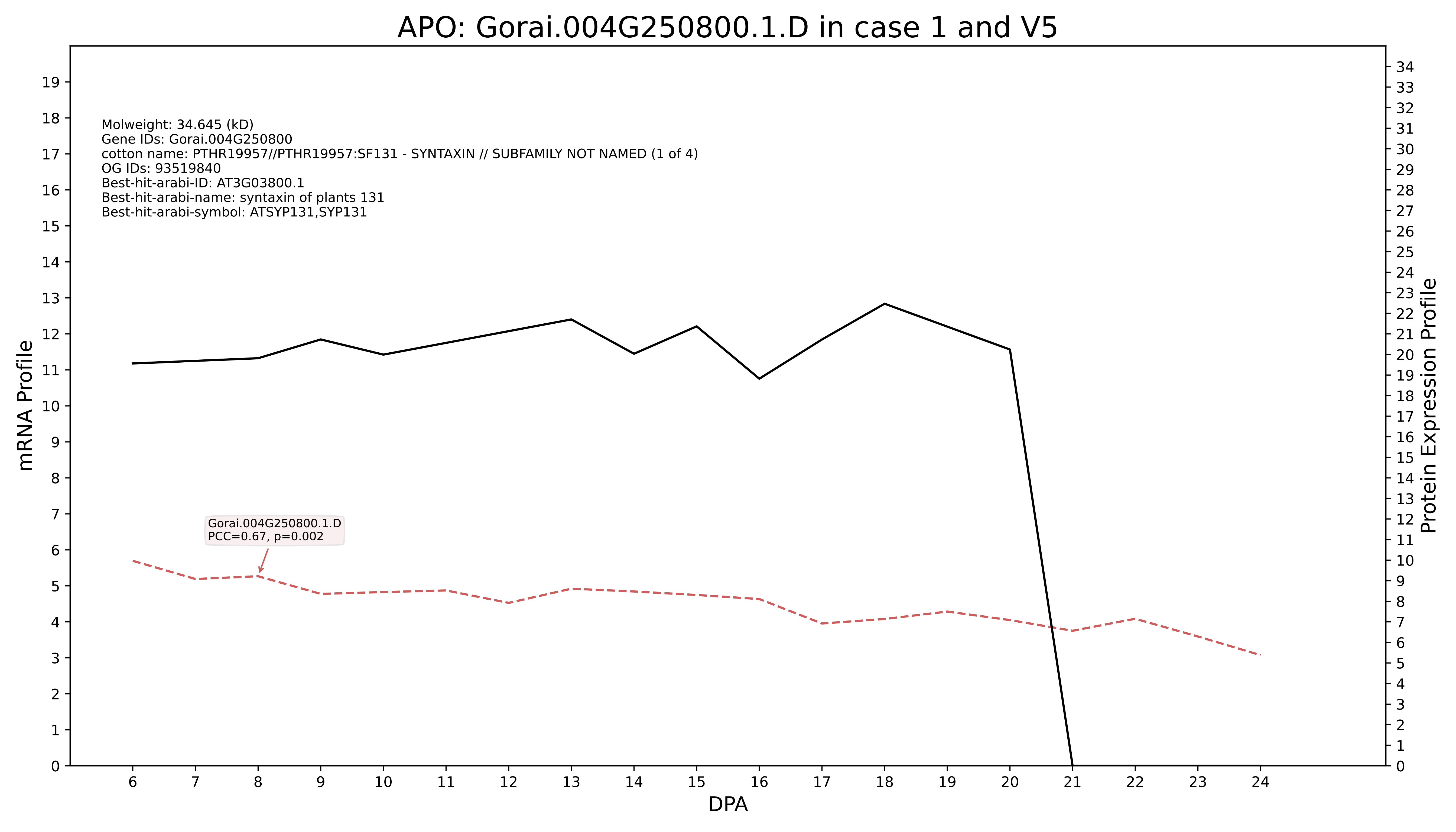
APO: Gorai.004G250800.1.D in case 1 and V5
Molweight: 34.645 (kD)
Gene IDs: Gorai.004G250800
cotton name: PTHR19957//PTHR19957:SF131 - SYNTAXIN // SUBFAMILY NOT NAMED (1 of 4)
OG IDs: 93519840
Best-hit-arabi-ID: AT3G03800.1
Best-hit-arabi-name: syntaxin of plants 131
Best-hit-arabi-symbol: ATSYP131,SYP131
Gorai.004G250800.1.D
PCC=0.67, p=0.002
mRNA Profile
Protein Expression Profile
DPA

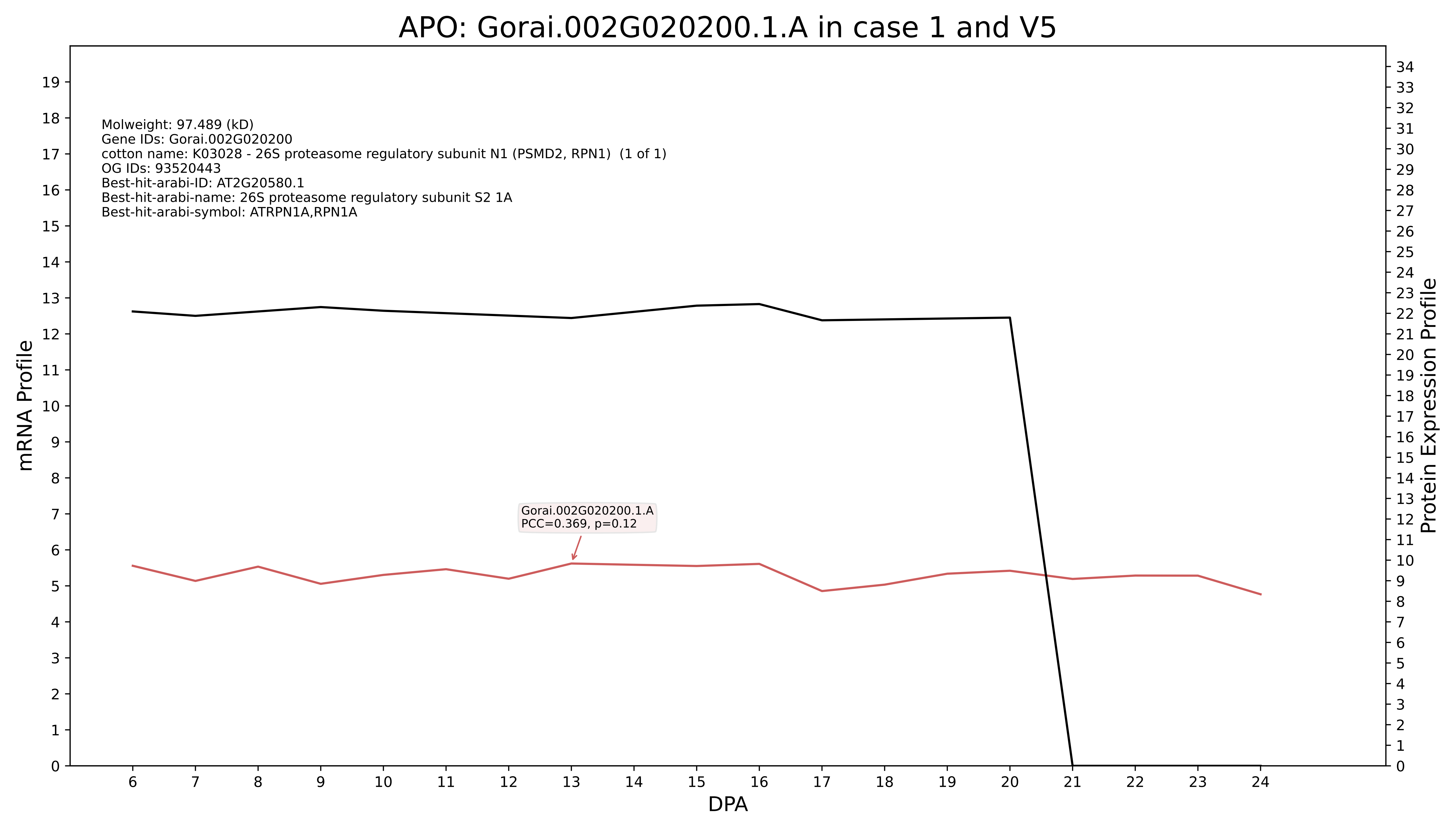

APO: Gorai.002G020200.1.A in case 1 and V5
Molweight: 97.489 (kD)
Gene IDs: Gorai.002G020200
cotton name: K03028 - 26S proteasome regulatory subunit N1 (PSMD2, RPN1) (1 of 1)
OG IDs: 93520443
Best-hit-arabi-ID: AT2G20580.1
Best-hit-arabi-name: 26S proteasome regulatory subunit S2 1A
Best-hit-arabi-symbol: ATRPN1A,RPN1A
Gorai.002G020200.1.A
PCC=0.369, p=0.12
mRNA Profile
Protein Expression Profile
DPA

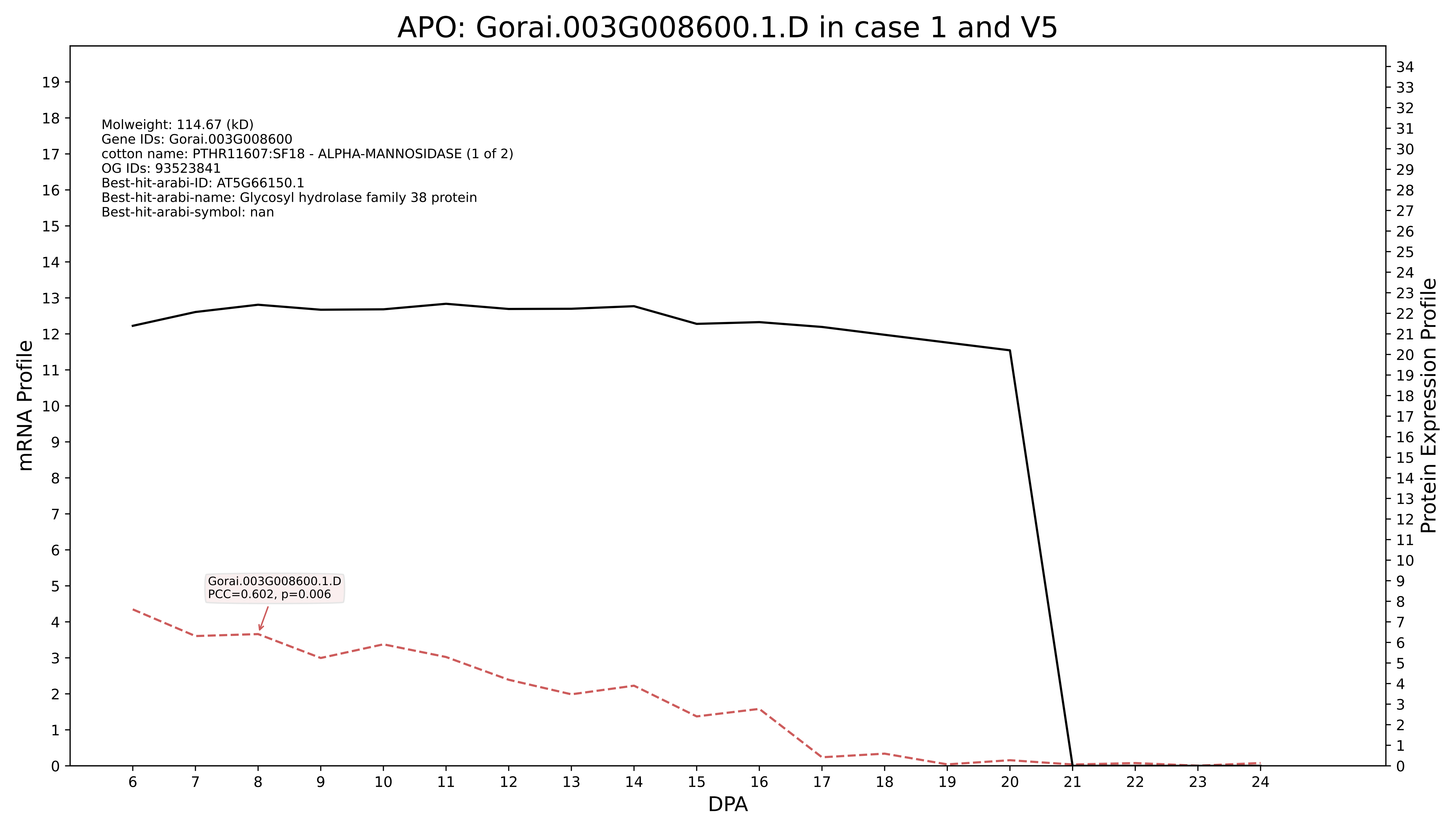
APO: Gorai.003G008600.1.D in case 1 and V5
Molweight: 114.67 (kD)
Gene IDs: Gorai.003G008600
cotton name: PTHR11607:SF18 - ALPHA-MANNOSIDASE (1 of 2)
OG IDs: 93523841
Best-hit-arabi-ID: AT5G66150.1
Best-hit-arabi-name: Glycosyl hydrolase family 38 protein
Best-hit-arabi-symbol: nan
Gorai.003G008600.1.D
PCC=0.602, p=0.006
mRNA Profile
Protein Expression Profile
DPA

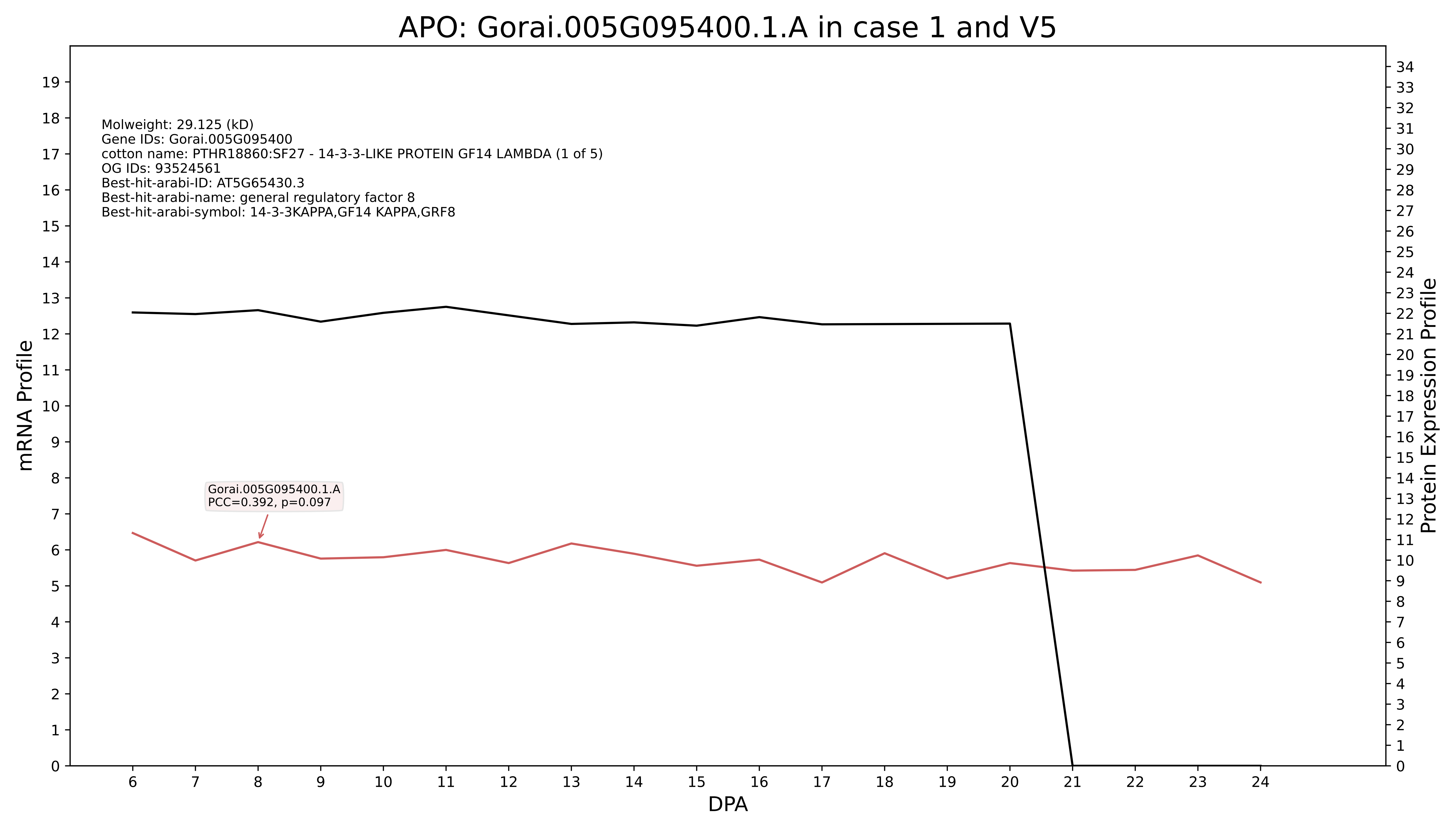

APO: Gorai.005G095400.1.A in case 1 and V5
Molweight: 29.125 (kD)
Gene IDs: Gorai.005G095400
cotton name: PTHR18860:SF27 - 14-3-3-LIKE PROTEIN GF14 LAMBDA (1 of 5)
OG IDs: 93524561
Best-hit-arabi-ID: AT5G65430.3
Best-hit-arabi-name: general regulatory factor 8
Best-hit-arabi-symbol: 14-3-3KAPPA,GF14 KAPPA,GRF8
Gorai.005G095400.1.A
PCC=0.392, p=0.097
mRNA Profile
Protein Expression Profile
DPA

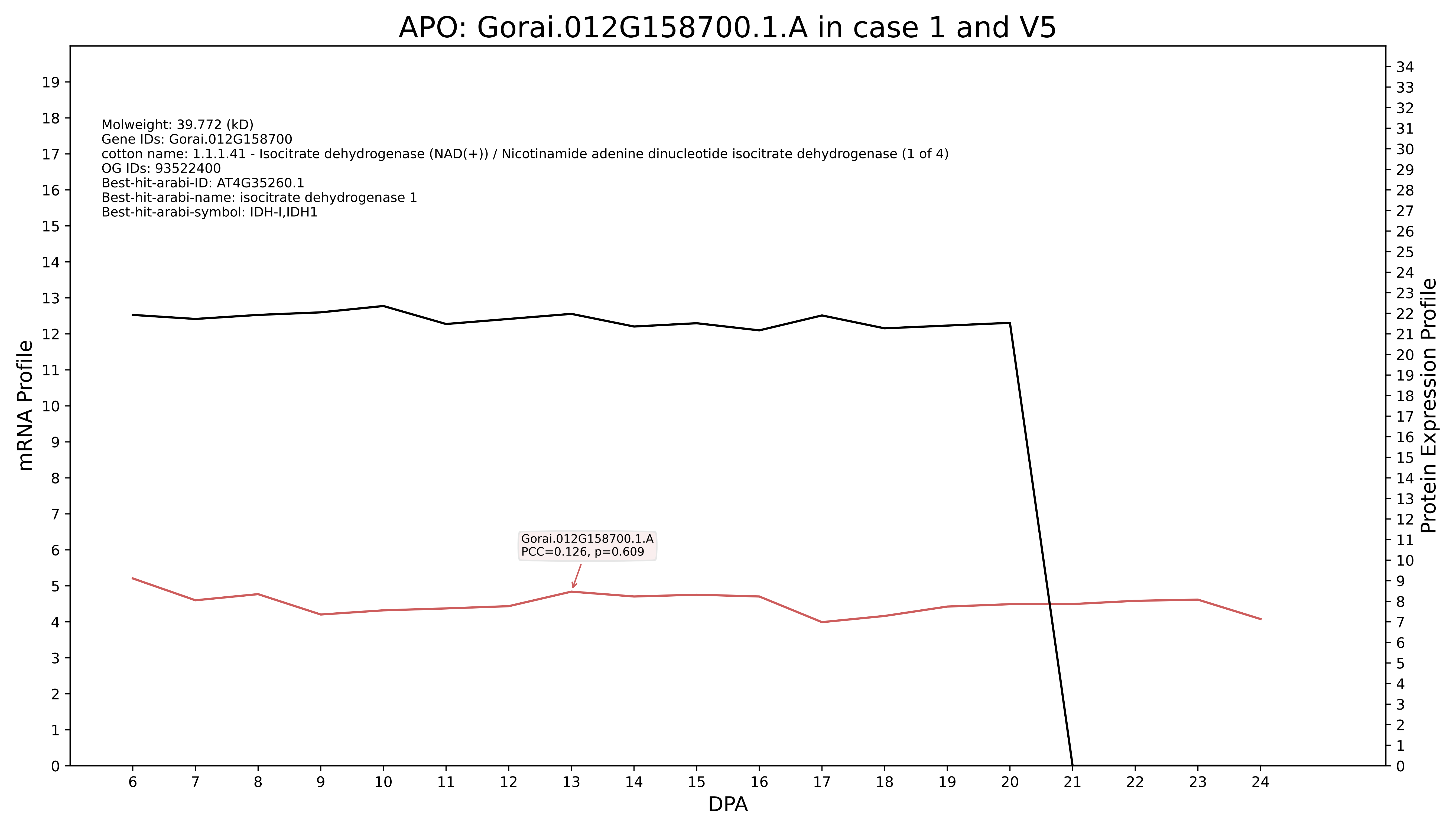

APO: Gorai.012G158700.1.A in case 1 and V5
Molweight: 39.772 (kD)
Gene IDs: Gorai.012G158700
cotton name: 1.1.1.41 - Isocitrate dehydrogenase (NAD(+)) / Nicotinamide adenine dinucleotide isocitrate dehydrogenase (1 of 4)
OG IDs: 93522400
Best-hit-arabi-ID: AT4G35260.1
Best-hit-arabi-name: isocitrate dehydrogenase 1
Best-hit-arabi-symbol: IDH-I,IDH1
Gorai.012G158700.1.A
PCC=0.126, p=0.609
mRNA Profile
Protein Expression Profile
DPA

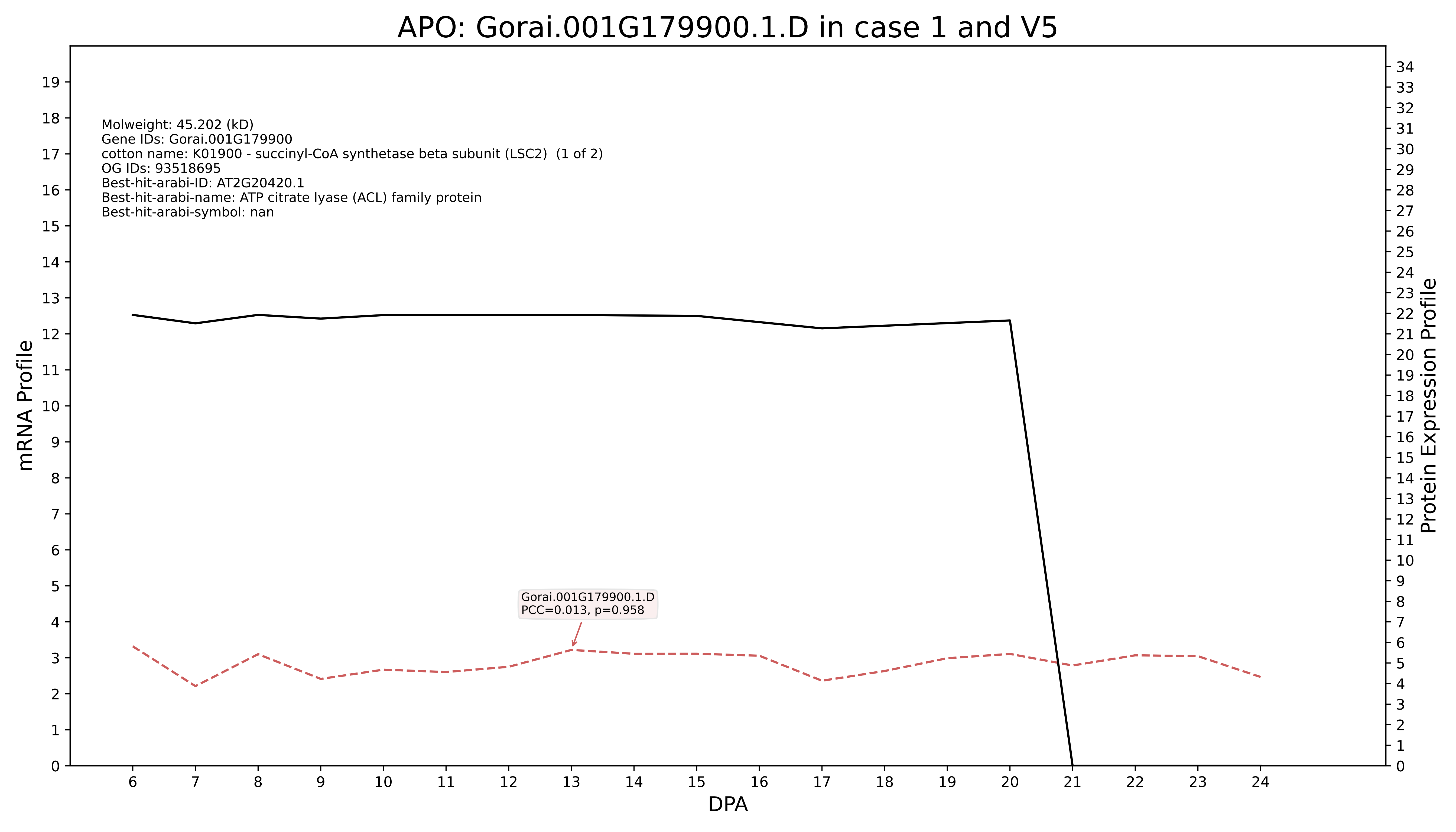
APO: Gorai.001G179900.1.D in case 1 and V5
Molweight: 45.202 (kD)
Gene IDs: Gorai.001G179900
cotton name: K01900 - succinyl-CoA synthetase beta subunit (LSC2) (1 of 2)
OG IDs: 93518695
Best-hit-arabi-ID: AT2G20420.1
Best-hit-arabi-name: ATP citrate lyase (ACL) family protein
Best-hit-arabi-symbol: nan
Gorai.001G179900.1.D
PCC=0.013, p=0.958
mRNA Profile
Protein Expression Profile
DPA

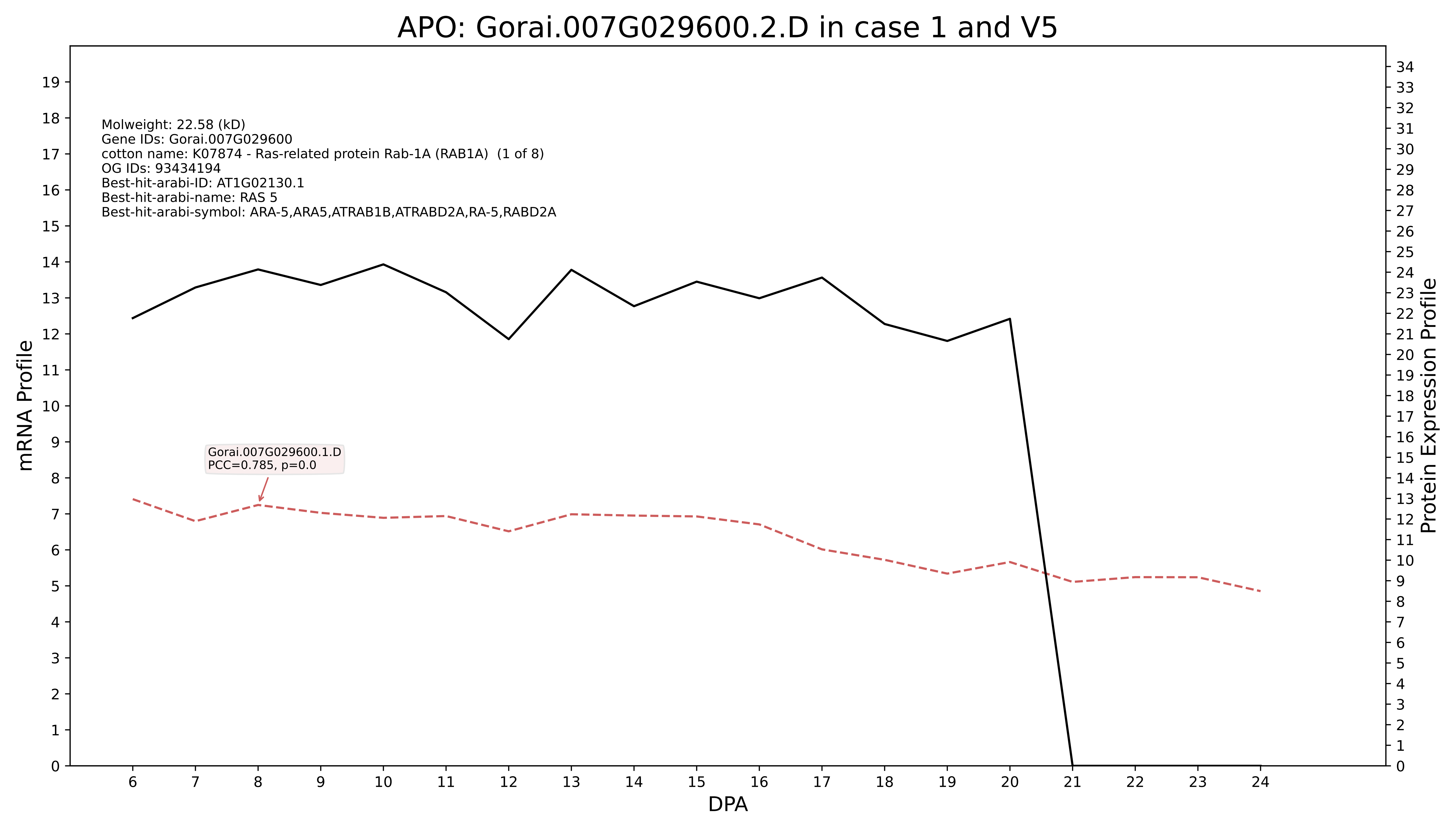
APO: Gorai.007G029600.2.D in case 1 and V5
Molweight: 22.58 (kD)
Gene IDs: Gorai.007G029600
cotton name: K07874 - Ras-related protein Rab-1A (RAB1A)  (1 of 8)
OG IDs: 93434194
Best-hit-arabi-ID: AT1G02130.1
Best-hit-arabi-name: RAS 5
Best-hit-arabi-symbol: ARA-5,ARA5,ATRAB1B,ATRABD2A,RA-5,RABD2A
Gorai.007G029600.1.D
PCC=0.785, p=0.0
mRNA Profile
Protein Expression Profile
DPA

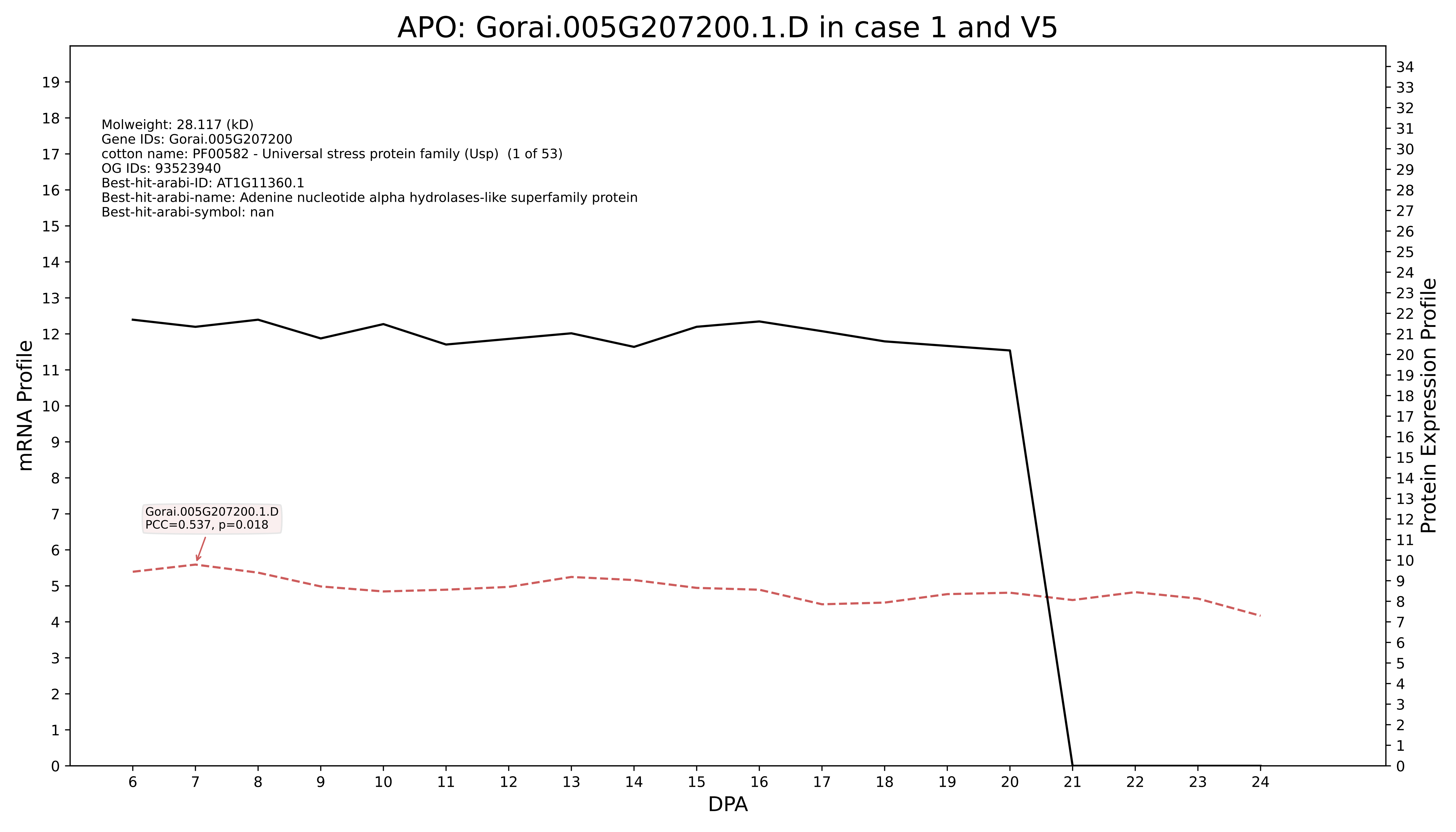

APO: Gorai.005G207200.1.D in case 1 and V5
Molweight: 28.117 (kD)
Gene IDs: Gorai.005G207200
cotton name: PF00582 - Universal stress protein family (Usp) (1 of 53)
OG IDs: 93523940
Best-hit-arabi-ID: AT1G11360.1
Best-hit-arabi-name: Adenine nucleotide alpha hydrolases-like superfamily protein
Best-hit-arabi-symbol: nan
Gorai.005G207200.1.D
PCC=0.537, p=0.018
mRNA Profile
Protein Expression Profile
DPA

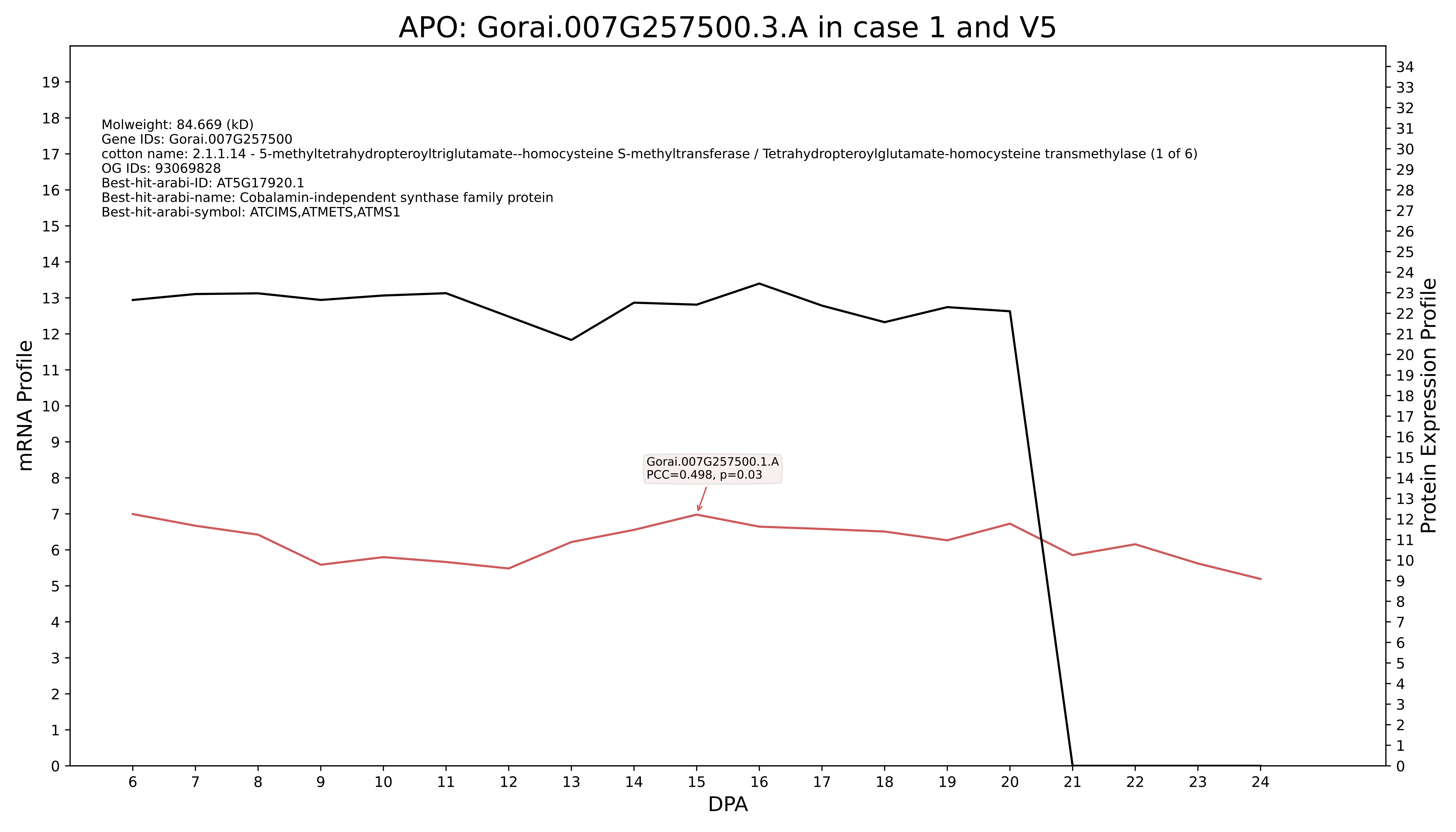
APO: Gorai.007G257500.3.A in case 1 and V5
Molweight: 84.669 (kD)
Gene IDs: Gorai.007G257500
cotton name: 2.1.1.14 - 5-methyltetrahydropteroyltriglutamate--homocysteine S-methyltransferase / Tetrahydropteroylglutamate-homocysteine transmethylase (1 of 6)
OG IDs: 93069828
Best-hit-arabi-ID: AT5G17920.1
Best-hit-arabi-name: Cobalamin-independent synthase family protein
Best-hit-arabi-symbol: ATCIMS,ATMETS,ATMS1
Gorai.007G257500.1.A
PCC=0.498, p=0.03
mRNA Profile
Protein Expression Profile
DPA

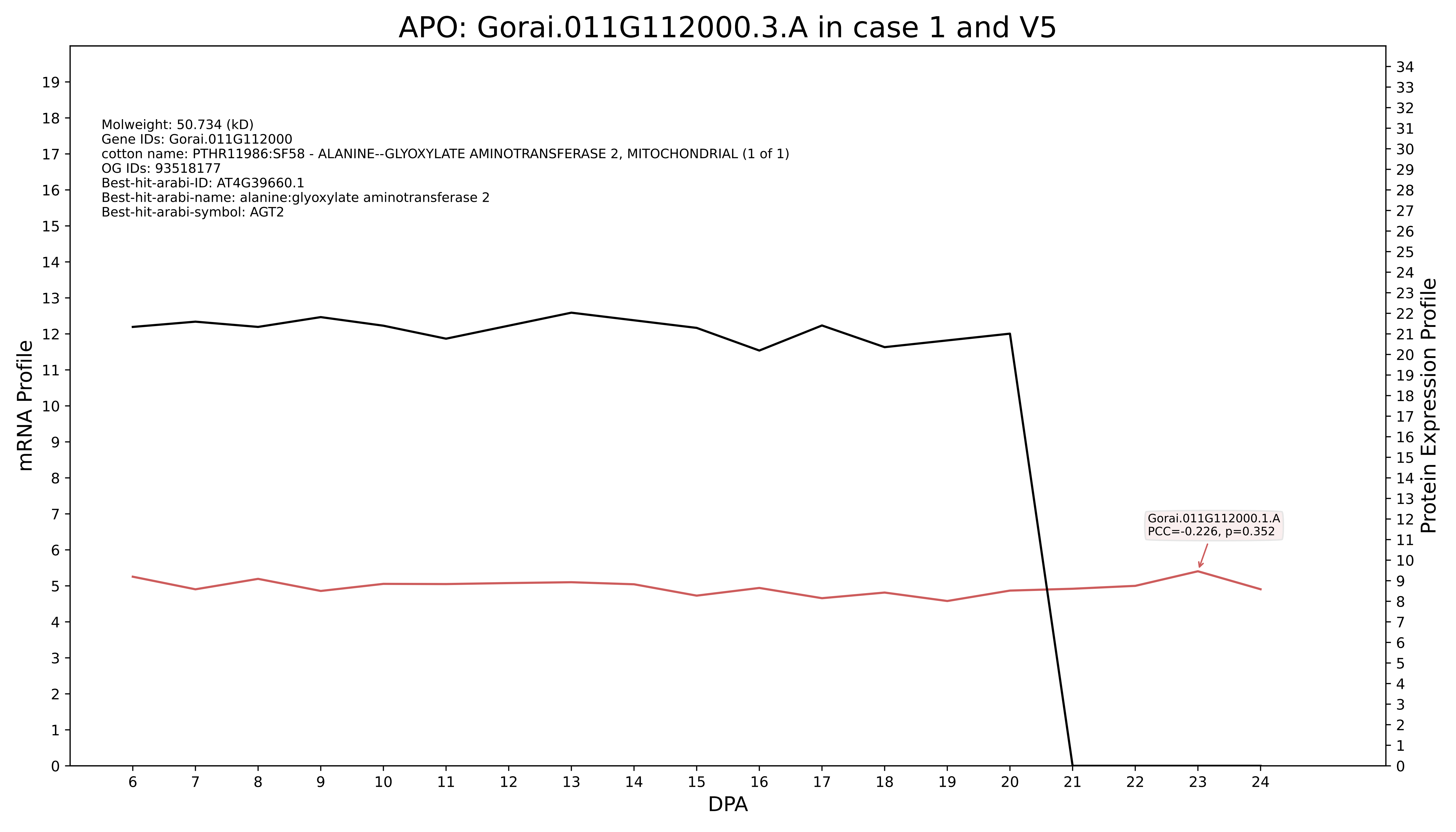
APO: Gorai.011G112000.3.A in case 1 and V5
Molweight: 50.734 (kD)
Gene IDs: Gorai.011G112000
cotton name: PTHR11986:SF58 - ALANINE--GLYOXYLATE AMINOTRANSFERASE 2, MITOCHONDRIAL (1 of 1)
OG IDs: 93518177
Best-hit-arabi-ID: AT4G39660.1
Best-hit-arabi-name: alanine:glyoxylate aminotransferase 2
Best-hit-arabi-symbol: AGT2
Gorai.011G112000.1.A
PCC=-0.226, p=0.352
mRNA Profile
Protein Expression Profile
DPA

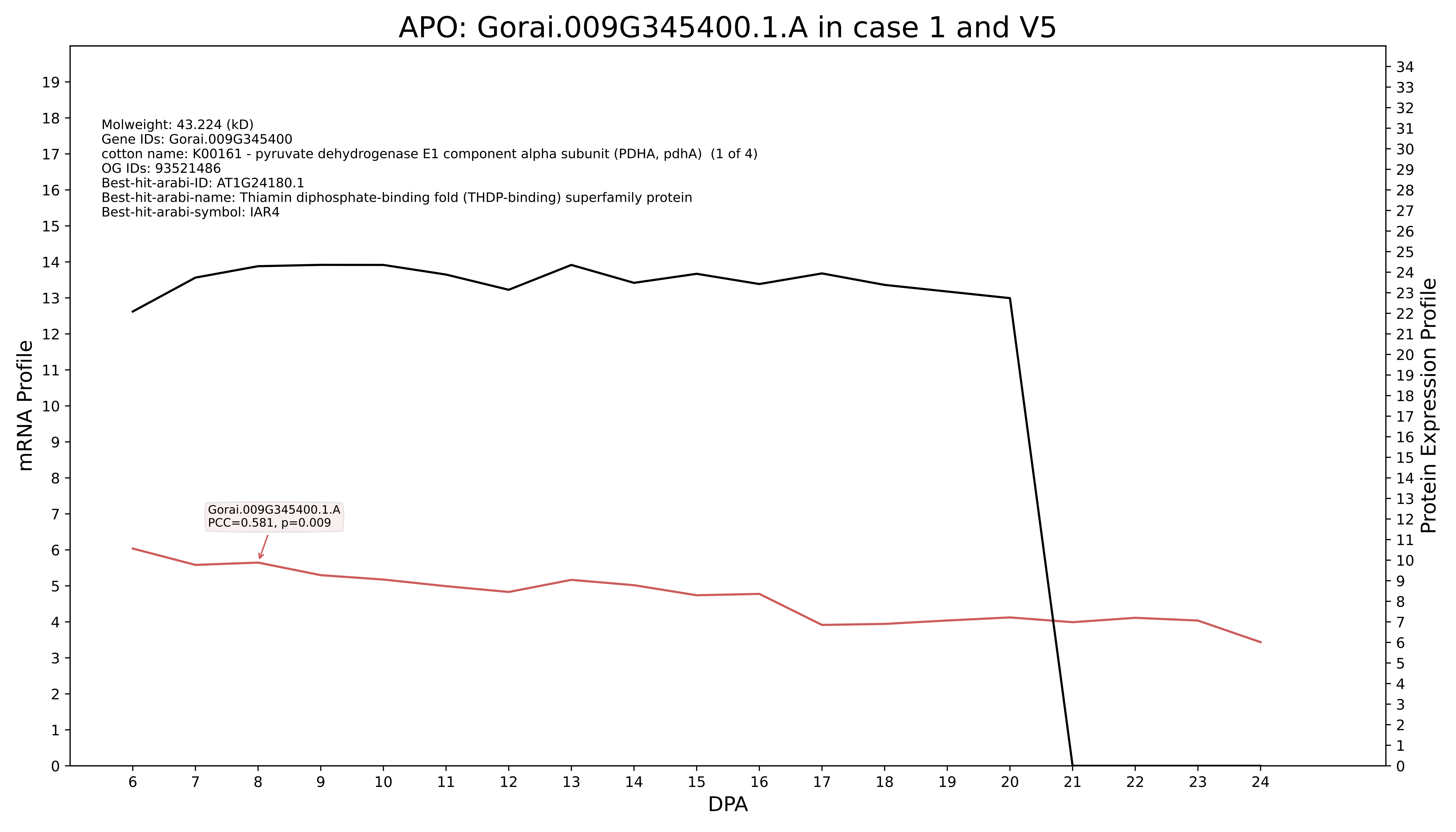
APO: Gorai.009G345400.1.A in case 1 and V5
Molweight: 43.224 (kD)
Gene IDs: Gorai.009G345400
cotton name: K00161 - pyruvate dehydrogenase E1 component alpha subunit (PDHA, pdhA) (1 of 4)
OG IDs: 93521486
Best-hit-arabi-ID: AT1G24180.1
Best-hit-arabi-name: Thiamin diphosphate-binding fold (THDP-binding) superfamily protein
Best-hit-arabi-symbol: IAR4
Gorai.009G345400.1.A
PCC=0.581, p=0.009
mRNA Profile
Protein Expression Profile
DPA

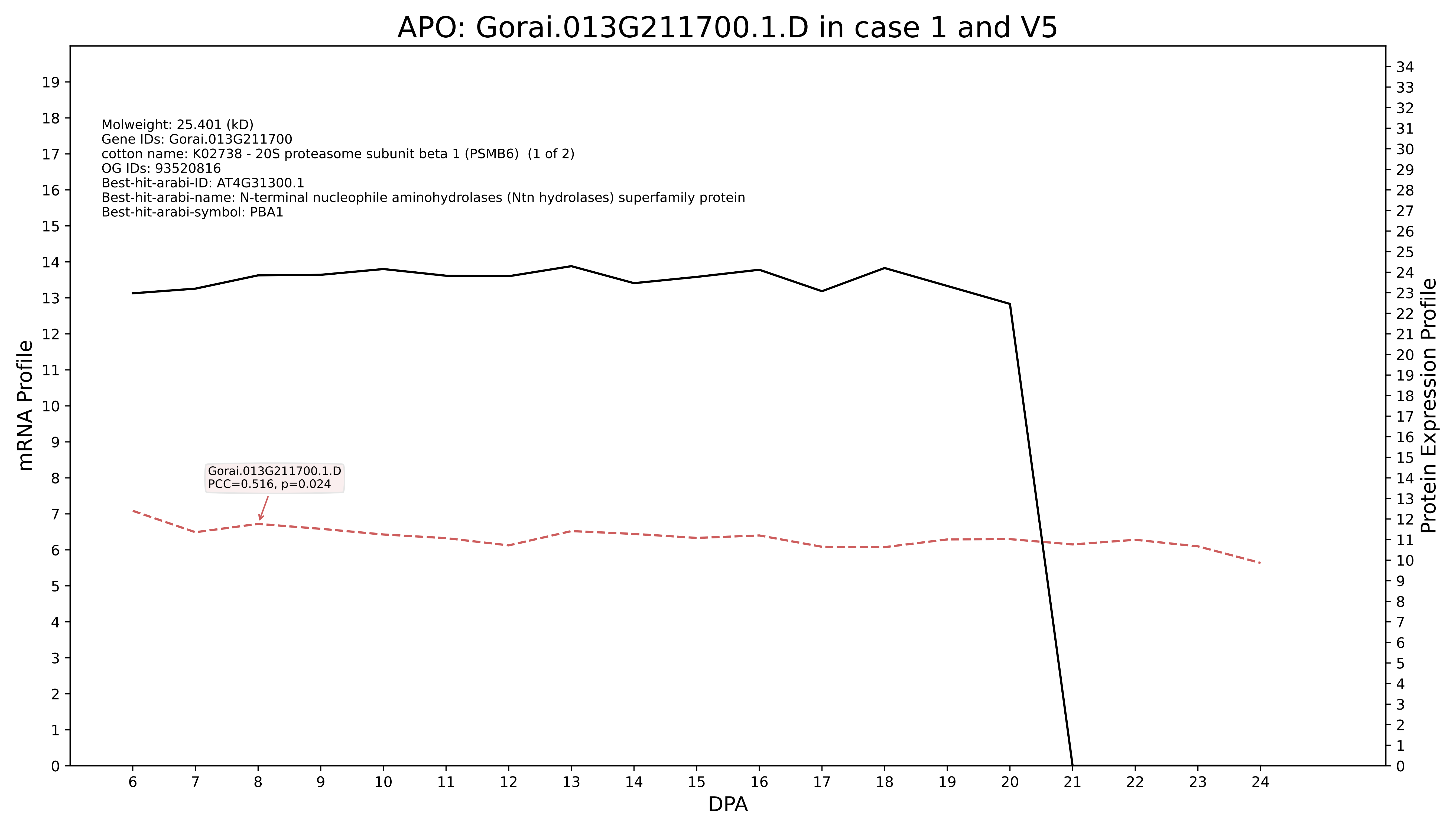
APO: Gorai.013G211700.1.D in case 1 and V5
Molweight: 25.401 (kD)
Gene IDs: Gorai.013G211700
cotton name: K02738 - 20S proteasome subunit beta 1 (PSMB6) (1 of 2)
OG IDs: 93520816
Best-hit-arabi-ID: AT4G31300.1
Best-hit-arabi-name: N-terminal nucleophile aminohydrolases (Ntn hydrolases) superfamily protein
Best-hit-arabi-symbol: PBA1
Gorai.013G211700.1.D
PCC=0.516, p=0.024
mRNA Profile
Protein Expression Profile
DPA

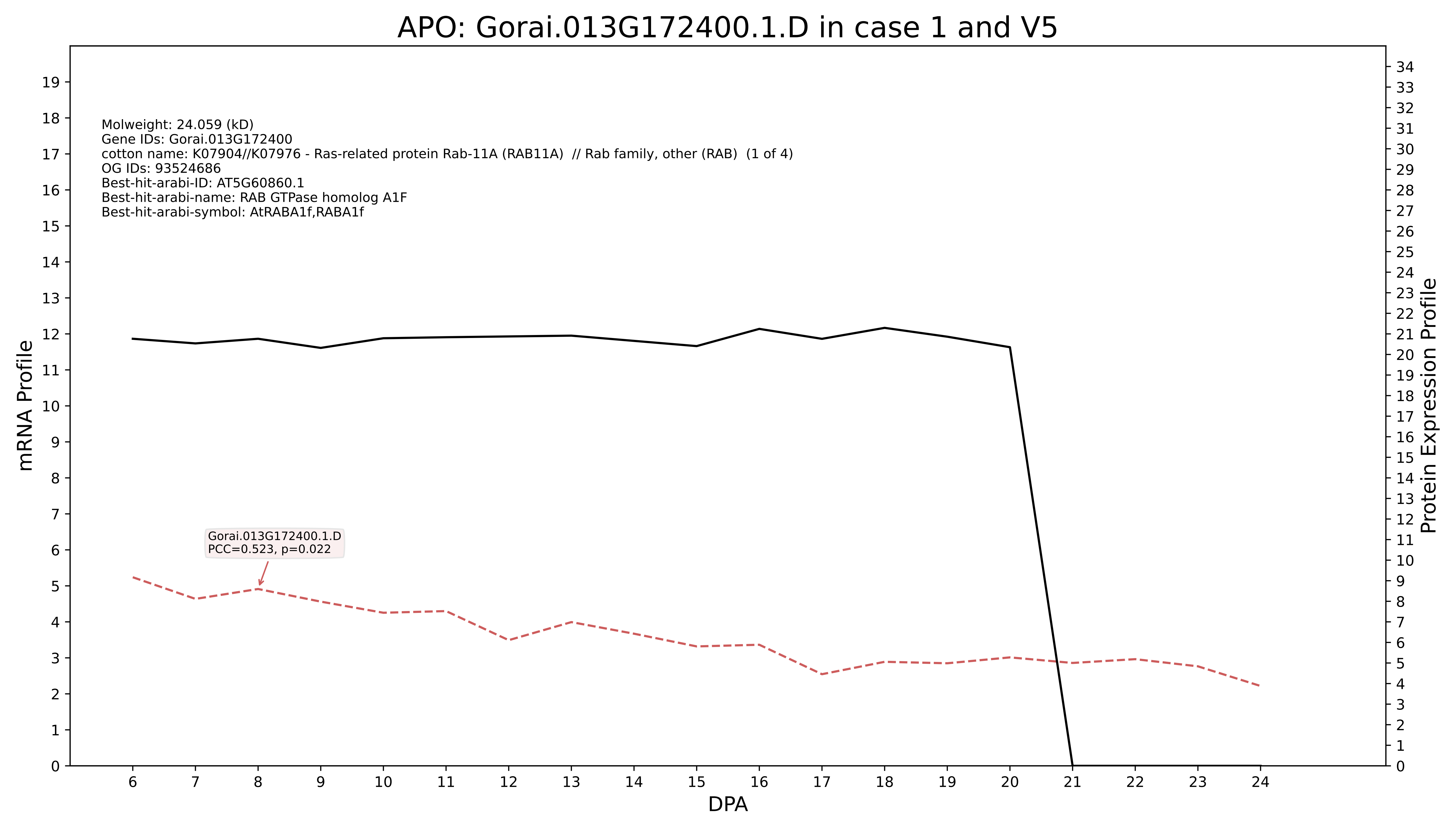
APO: Gorai.013G172400.1.D in case 1 and V5
Molweight: 24.059 (kD)
Gene IDs: Gorai.013G172400
cotton name: K07904//K07976 - Ras-related protein Rab-11A (RAB11A) // Rab family, other (RAB) (1 of 4)
OG IDs: 93524686
Best-hit-arabi-ID: AT5G60860.1
Best-hit-arabi-name: RAB GTPase homolog A1F
Best-hit-arabi-symbol: AtRABA1f,RABA1f
Gorai.013G172400.1.D
PCC=0.523, p=0.022
mRNA Profile
Protein Expression Profile
DPA

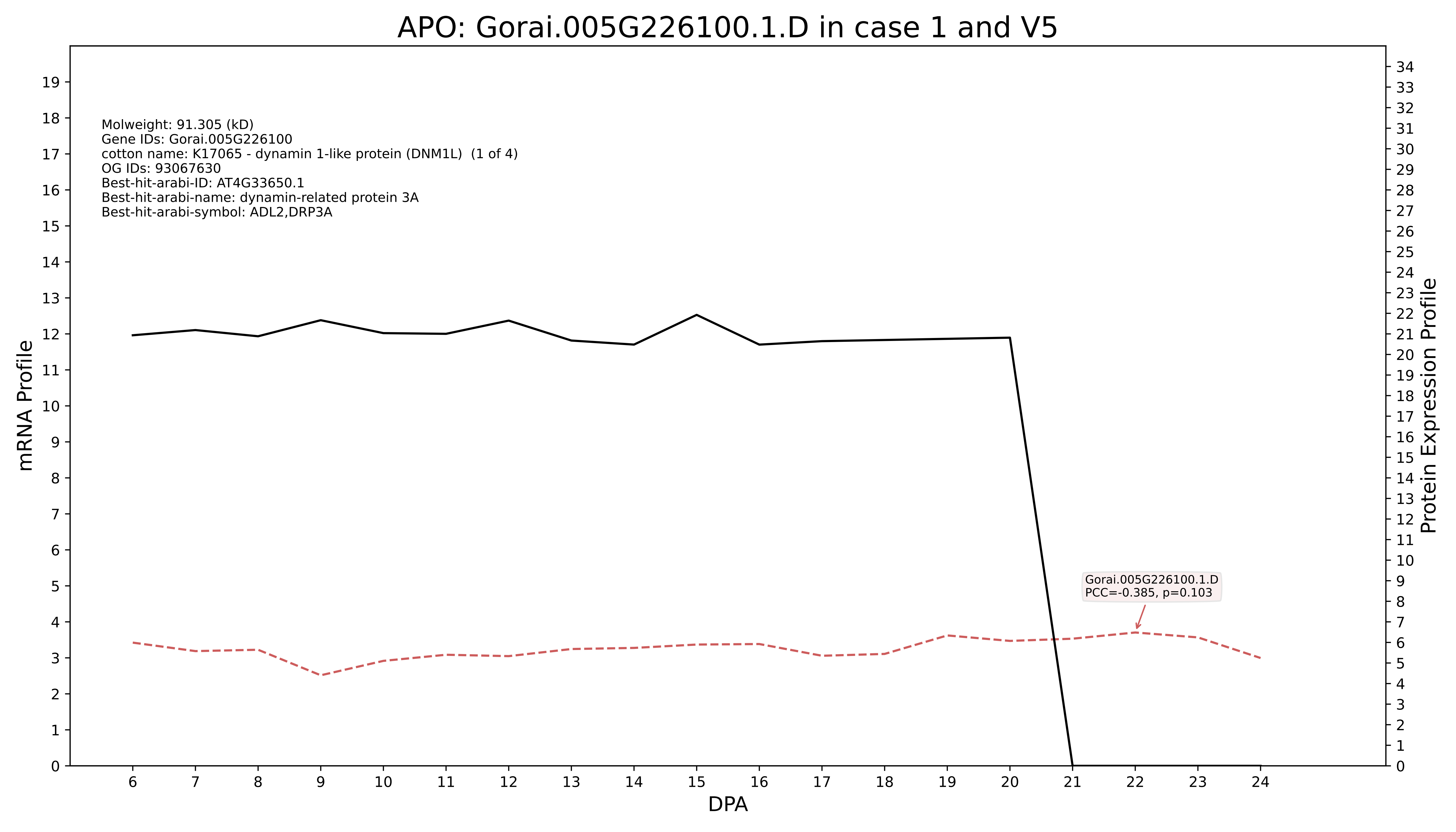

APO: Gorai.005G226100.1.D in case 1 and V5
Molweight: 91.305 (kD)
Gene IDs: Gorai.005G226100
cotton name: K17065 - dynamin 1-like protein (DNM1L) (1 of 4)
OG IDs: 93067630
Best-hit-arabi-ID: AT4G33650.1
Best-hit-arabi-name: dynamin-related protein 3A
Best-hit-arabi-symbol: ADL2,DRP3A
Gorai.005G226100.1.D
PCC=-0.385, p=0.103
mRNA Profile
Protein Expression Profile
DPA

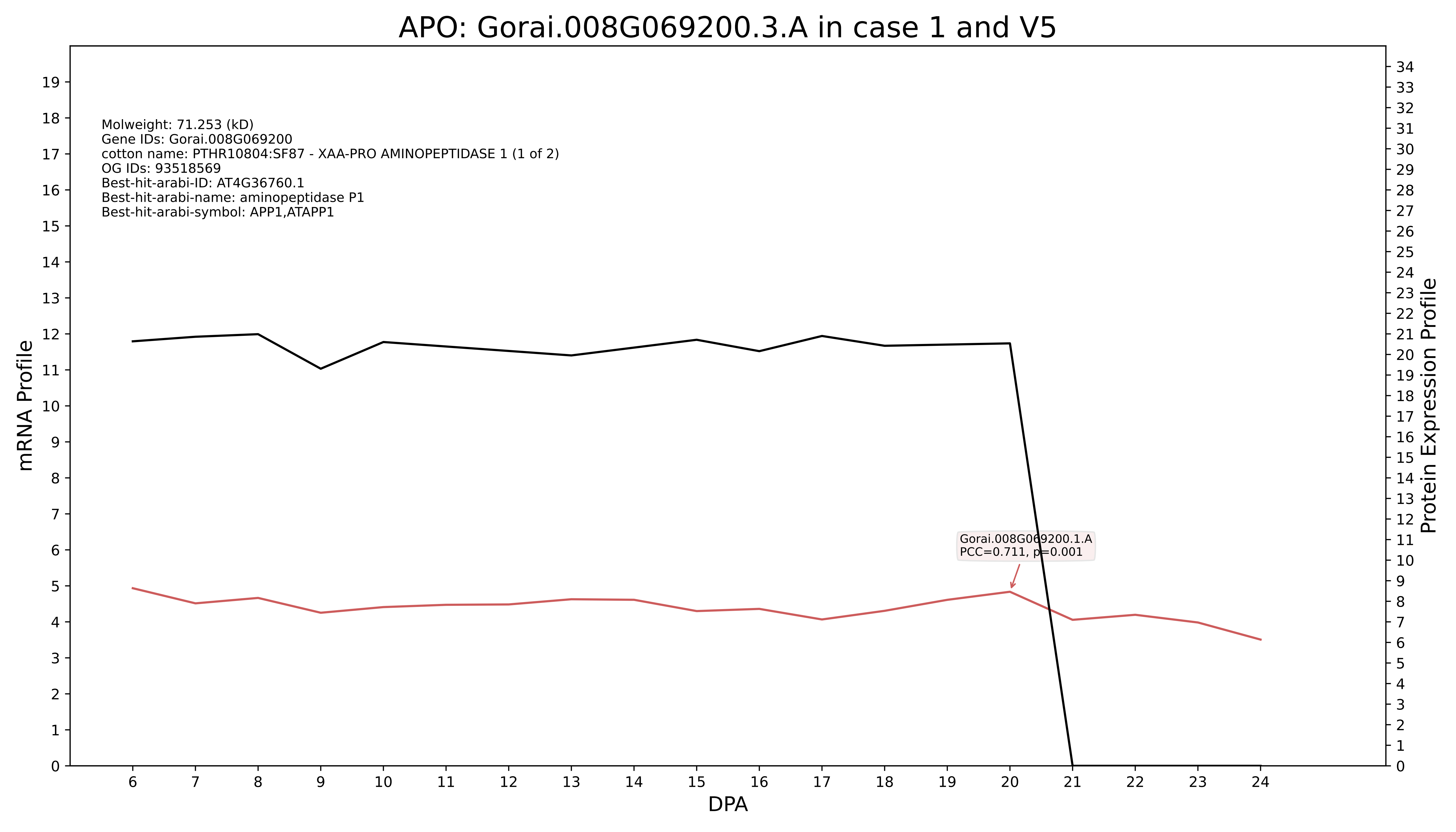
APO: Gorai.008G069200.3.A in case 1 and V5
Molweight: 71.253 (kD)
Gene IDs: Gorai.008G069200
cotton name: PTHR10804:SF87 - XAA-PRO AMINOPEPTIDASE 1 (1 of 2)
OG IDs: 93518569
Best-hit-arabi-ID: AT4G36760.1
Best-hit-arabi-name: aminopeptidase P1
Best-hit-arabi-symbol: APP1,ATAPP1
Gorai.008G069200.1.A
PCC=0.711, p=0.001
mRNA Profile
Protein Expression Profile
DPA

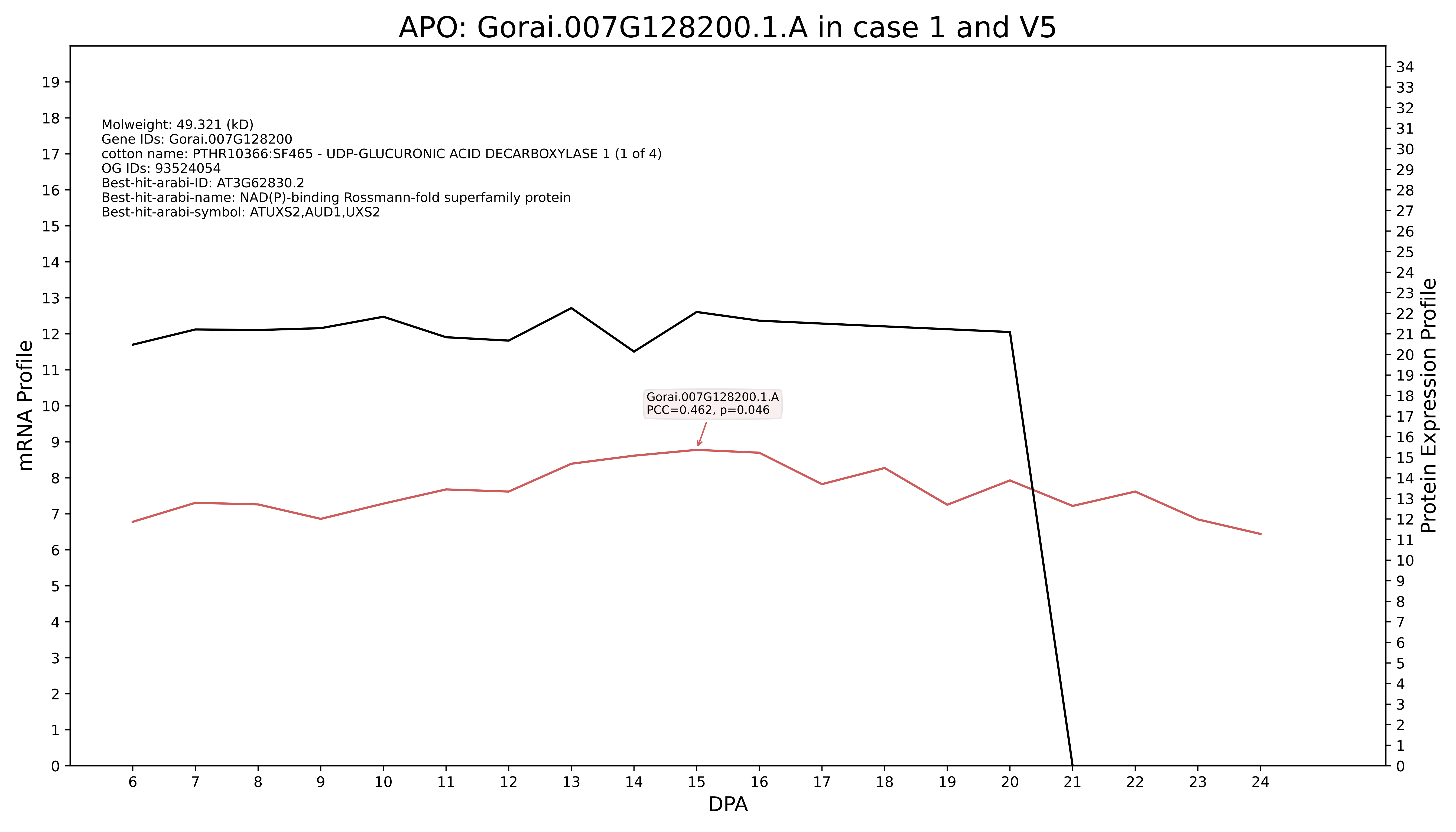

APO: Gorai.007G128200.1.A in case 1 and V5
Molweight: 49.321 (kD)
Gene IDs: Gorai.007G128200
cotton name: PTHR10366:SF465 - UDP-GLUCURONIC ACID DECARBOXYLASE 1 (1 of 4)
OG IDs: 93524054
Best-hit-arabi-ID: AT3G62830.2
Best-hit-arabi-name: NAD(P)-binding Rossmann-fold superfamily protein
Best-hit-arabi-symbol: ATUXS2,AUD1,UXS2
Gorai.007G128200.1.A
PCC=0.462, p=0.046
mRNA Profile
Protein Expression Profile
DPA

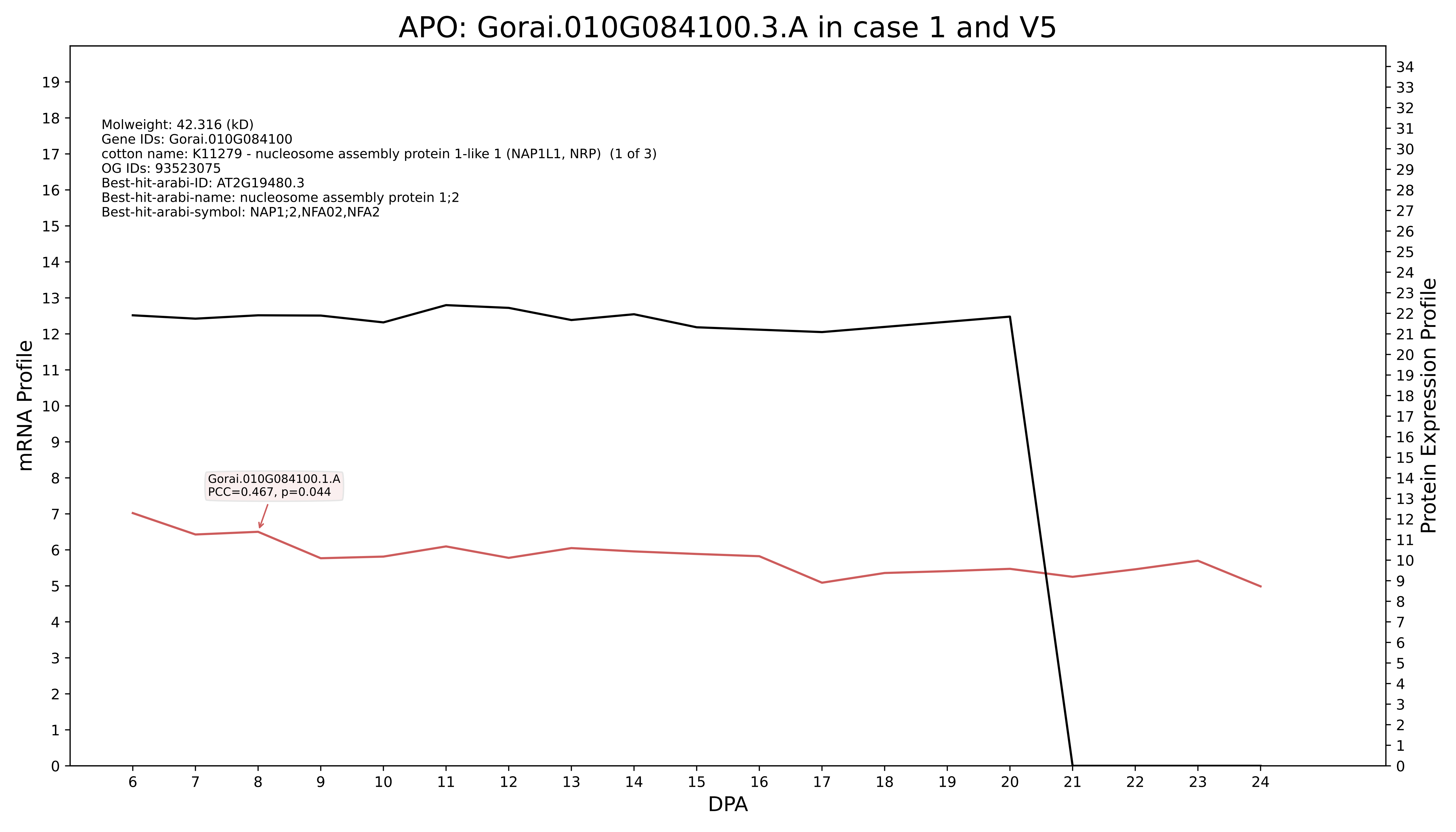

APO: Gorai.010G084100.3.A in case 1 and V5
Molweight: 42.316 (kD)
Gene IDs: Gorai.010G084100
cotton name: K11279 - nucleosome assembly protein 1-like 1 (NAP1L1, NRP) (1 of 3)
OG IDs: 93523075
Best-hit-arabi-ID: AT2G19480.3
Best-hit-arabi-name: nucleosome assembly protein 1;2
Best-hit-arabi-symbol: NAP1;2,NFA02,NFA2
Gorai.010G084100.1.A
PCC=0.467, p=0.044
mRNA Profile
Protein Expression Profile
DPA

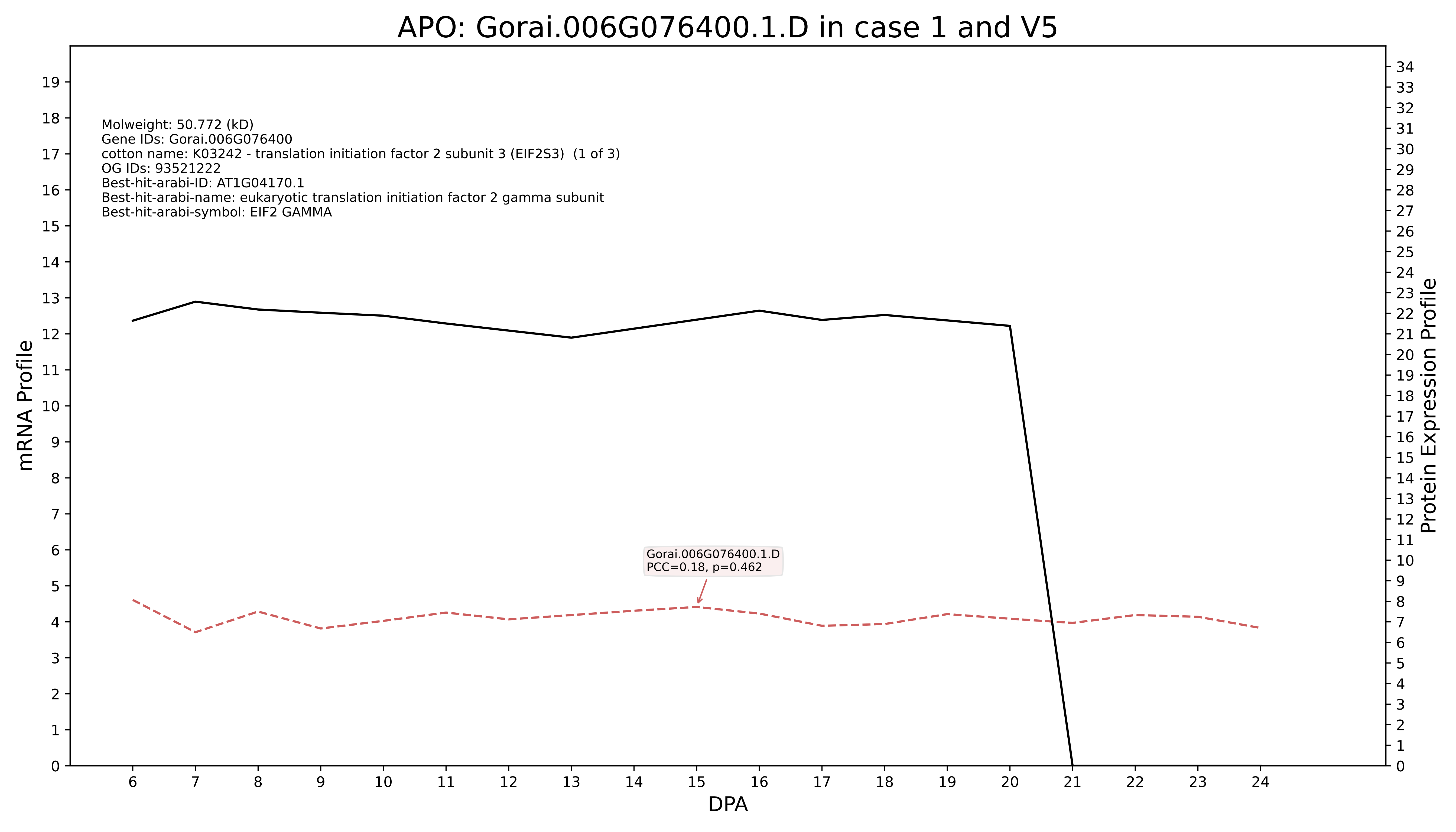
APO: Gorai.006G076400.1.D in case 1 and V5
Molweight: 50.772 (kD)
Gene IDs: Gorai.006G076400
cotton name: K03242 - translation initiation factor 2 subunit 3 (EIF2S3) (1 of 3)
OG IDs: 93521222
Best-hit-arabi-ID: AT1G04170.1
Best-hit-arabi-name: eukaryotic translation initiation factor 2 gamma subunit
Best-hit-arabi-symbol: EIF2 GAMMA
Gorai.006G076400.1.D
PCC=0.18, p=0.462
mRNA Profile
Protein Expression Profile
DPA

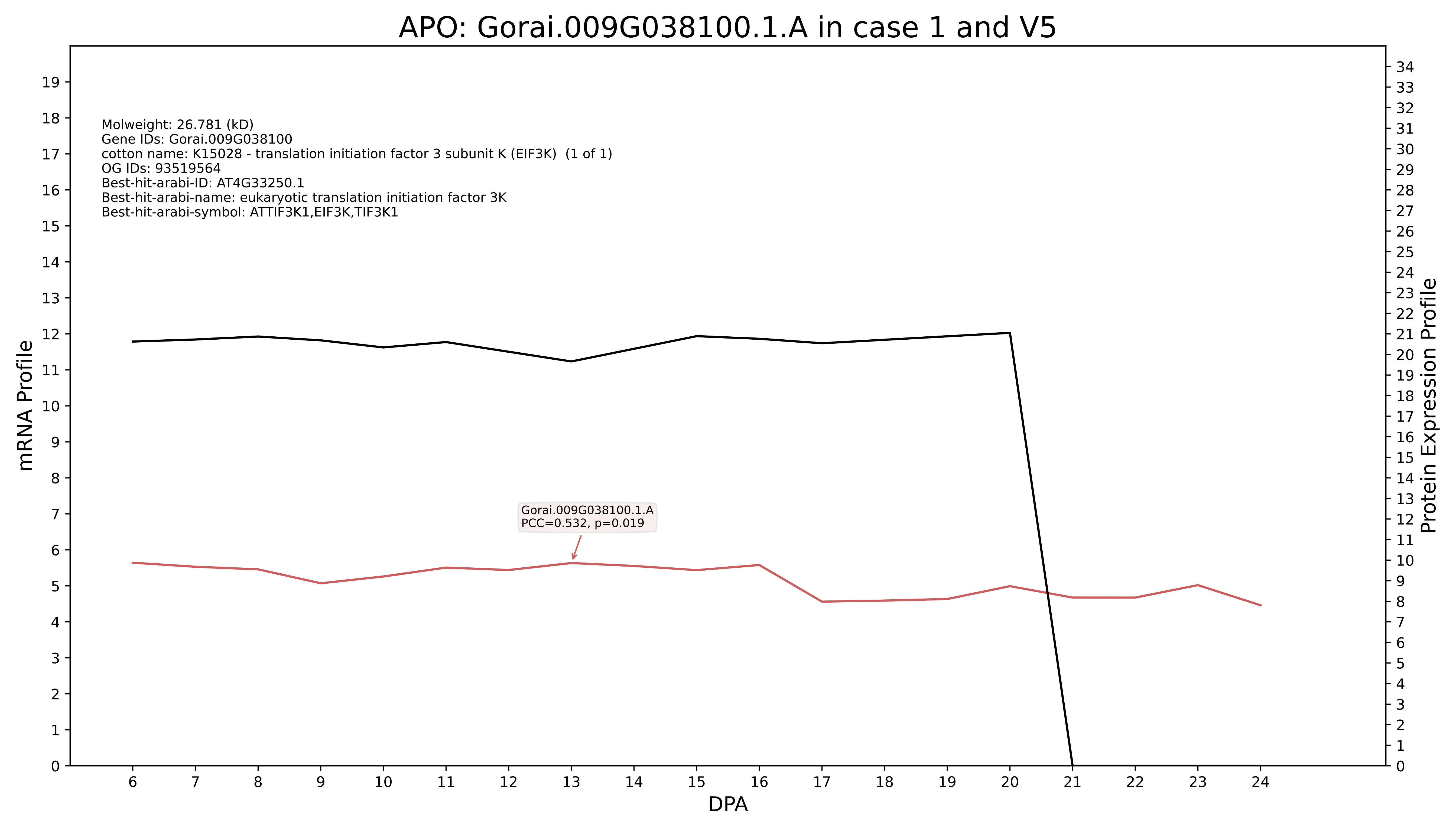
APO: Gorai.009G038100.1.A in case 1 and V5
Molweight: 26.781 (kD)
Gene IDs: Gorai.009G038100
cotton name: K15028 - translation initiation factor 3 subunit K (EIF3K) (1 of 1)
OG IDs: 93519564
Best-hit-arabi-ID: AT4G33250.1
Best-hit-arabi-name: eukaryotic translation initiation factor 3K
Best-hit-arabi-symbol: ATTIF3K1,EIF3K,TIF3K1
Gorai.009G038100.1.A
PCC=0.532, p=0.019
mRNA Profile
Protein Expression Profile
DPA

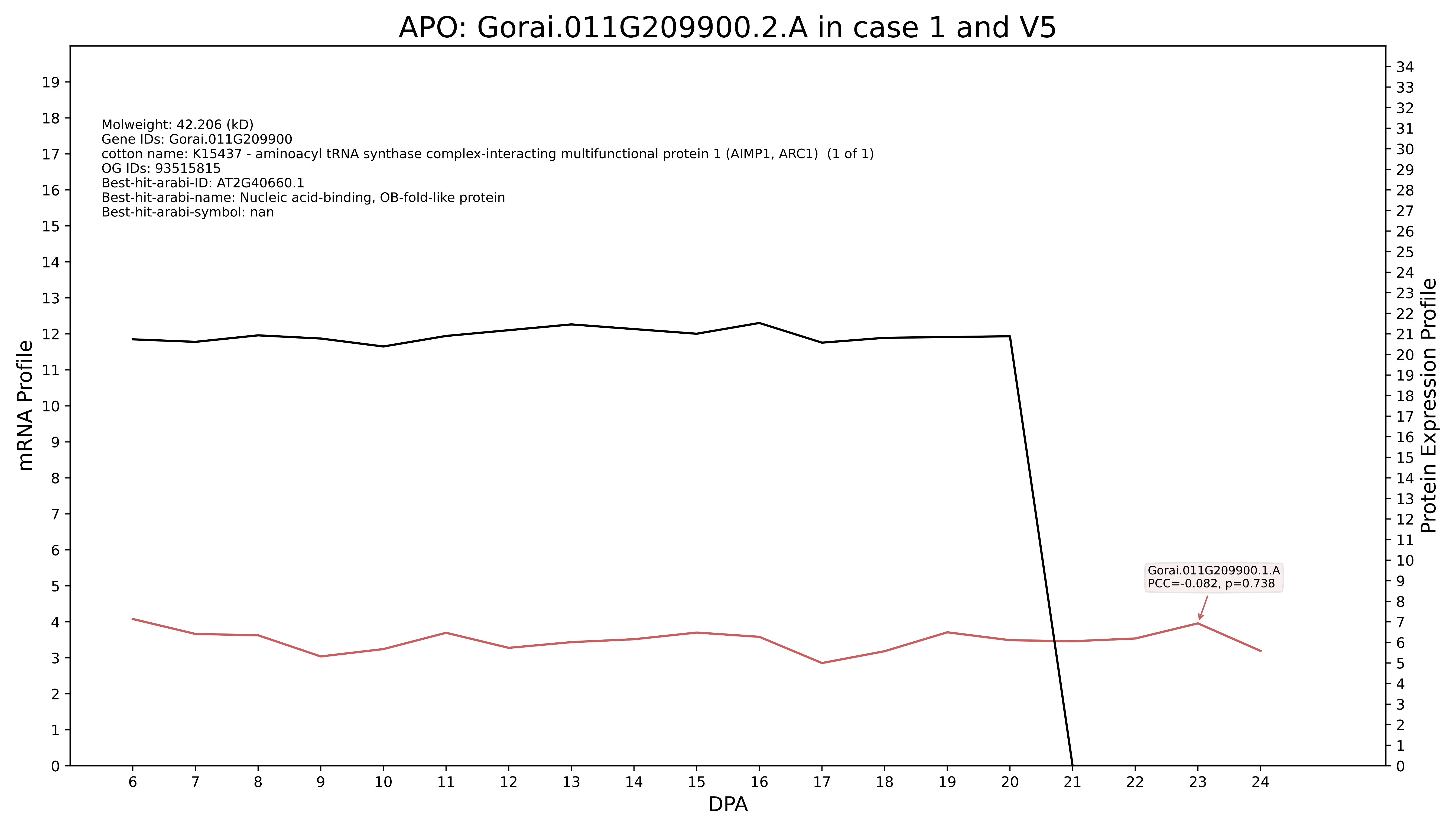

APO: Gorai.011G209900.2.A in case 1 and V5
Molweight: 42.206 (kD)
Gene IDs: Gorai.011G209900
cotton name: K15437 - aminoacyl tRNA synthase complex-interacting multifunctional protein 1 (AIMP1, ARC1) (1 of 1)
OG IDs: 93515815
Best-hit-arabi-ID: AT2G40660.1
Best-hit-arabi-name: Nucleic acid-binding, OB-fold-like protein
Best-hit-arabi-symbol: nan
Gorai.011G209900.1.A
PCC=-0.082, p=0.738
mRNA Profile
Protein Expression Profile
DPA

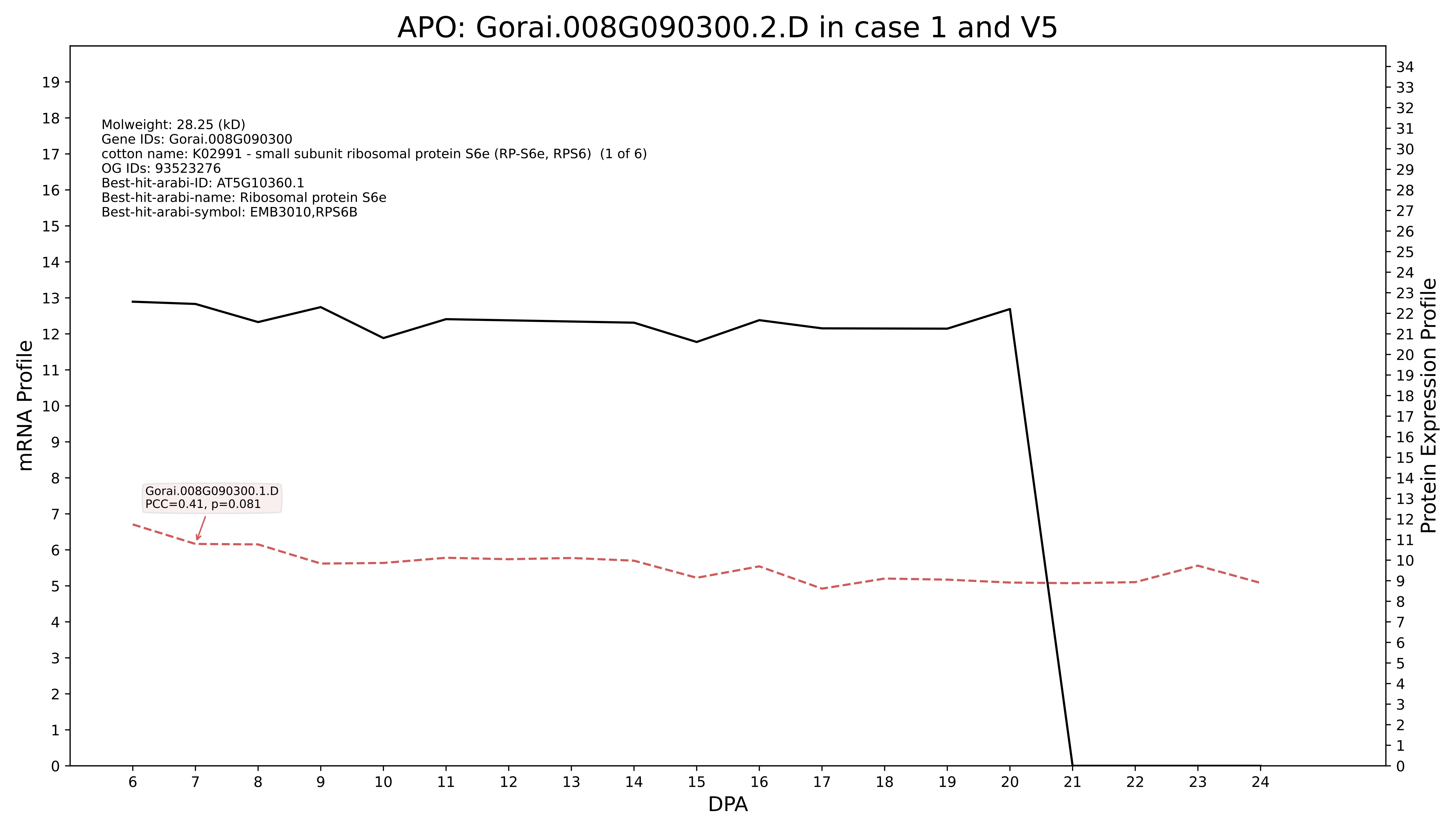
APO: Gorai.008G090300.2.D in case 1 and V5
Molweight: 28.25 (kD)
Gene IDs: Gorai.008G090300
cotton name: K02991 - small subunit ribosomal protein S6e (RP-S6e, RPS6) (1 of 6)
OG IDs: 93523276
Best-hit-arabi-ID: AT5G10360.1
Best-hit-arabi-name: Ribosomal protein S6e
Best-hit-arabi-symbol: EMB3010,RPS6B
Gorai.008G090300.1.D
PCC=0.41, p=0.081
mRNA Profile
Protein Expression Profile
DPA

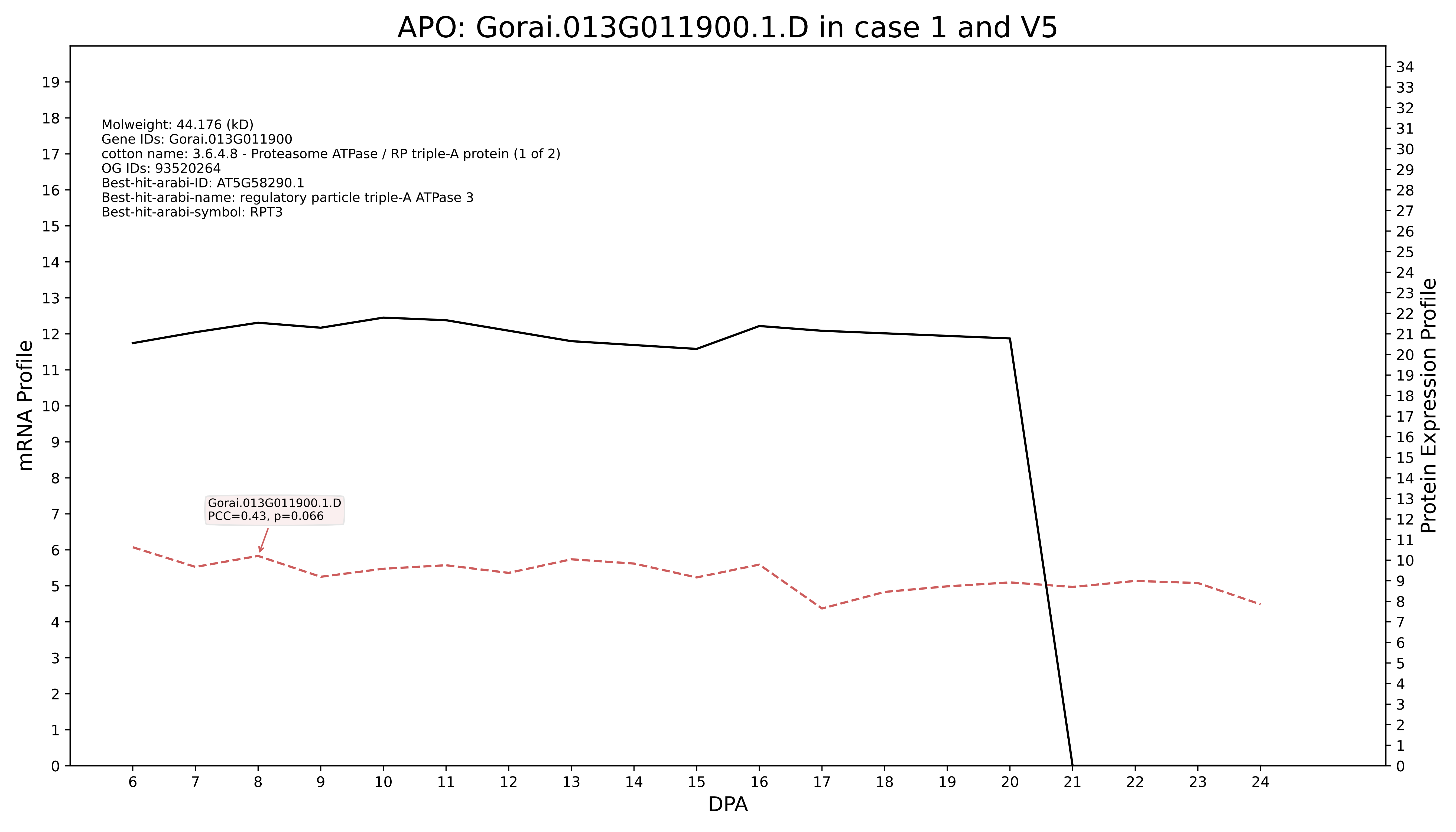

APO: Gorai.013G011900.1.D in case 1 and V5
Molweight: 44.176 (kD)
Gene IDs: Gorai.013G011900
cotton name: 3.6.4.8 - Proteasome ATPase / RP triple-A protein (1 of 2)
OG IDs: 93520264
Best-hit-arabi-ID: AT5G58290.1
Best-hit-arabi-name: regulatory particle triple-A ATPase 3
Best-hit-arabi-symbol: RPT3
Gorai.013G011900.1.D
PCC=0.43, p=0.066
mRNA Profile
Protein Expression Profile
DPA

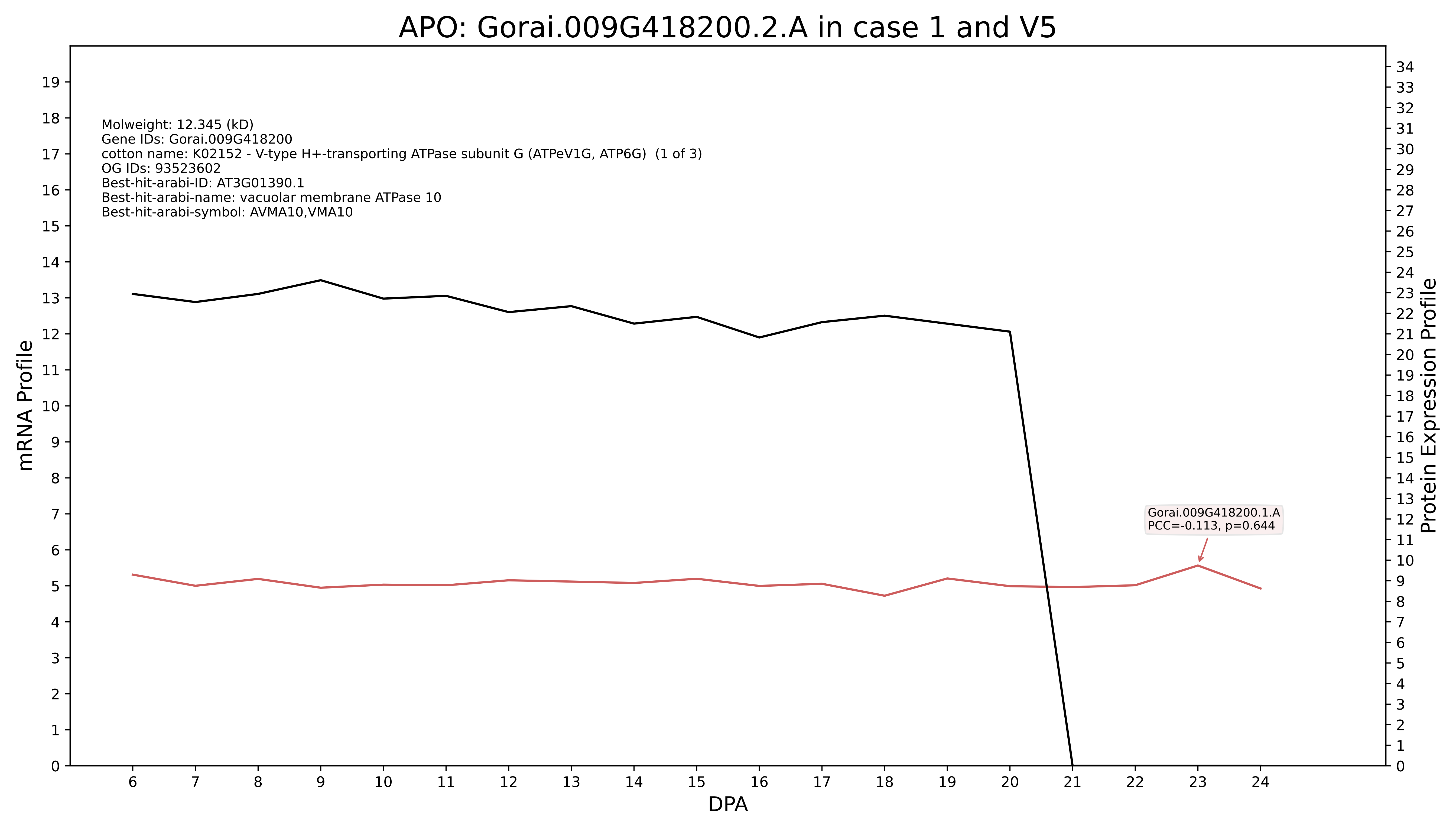

APO: Gorai.009G418200.2.A in case 1 and V5
Molweight: 12.345 (kD)
Gene IDs: Gorai.009G418200
cotton name: K02152 - V-type H+-transporting ATPase subunit G (ATPeV1G, ATP6G) (1 of 3)
OG IDs: 93523602
Best-hit-arabi-ID: AT3G01390.1
Best-hit-arabi-name: vacuolar membrane ATPase 10
Best-hit-arabi-symbol: AVMA10,VMA10
Gorai.009G418200.1.A
PCC=-0.113, p=0.644
mRNA Profile
Protein Expression Profile
DPA

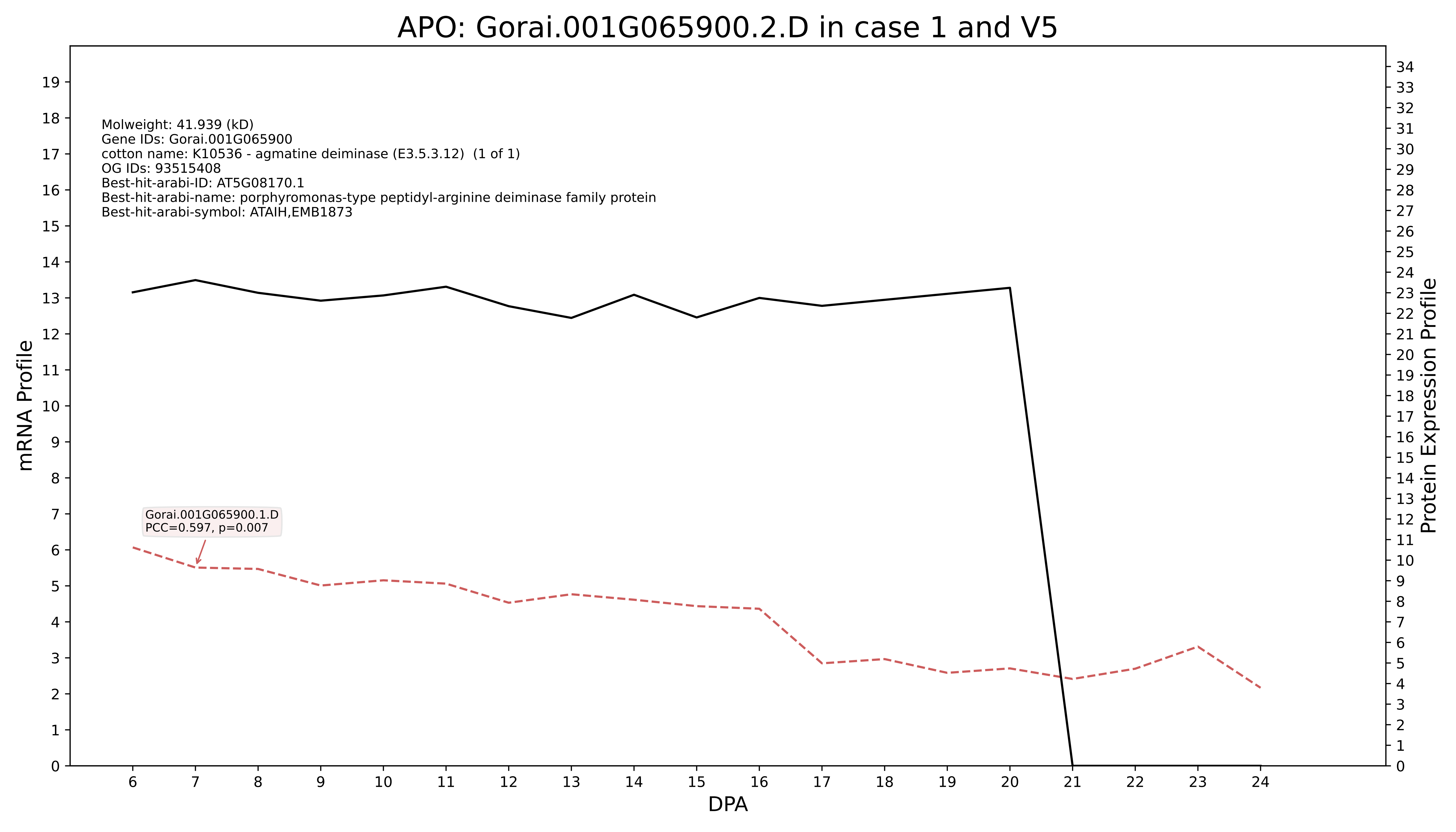

APO: Gorai.001G065900.2.D in case 1 and V5
Molweight: 41.939 (kD)
Gene IDs: Gorai.001G065900
cotton name: K10536 - agmatine deiminase (E3.5.3.12) (1 of 1)
OG IDs: 93515408
Best-hit-arabi-ID: AT5G08170.1
Best-hit-arabi-name: porphyromonas-type peptidyl-arginine deiminase family protein
Best-hit-arabi-symbol: ATAIH,EMB1873
Gorai.001G065900.1.D
PCC=0.597, p=0.007
mRNA Profile
Protein Expression Profile
DPA

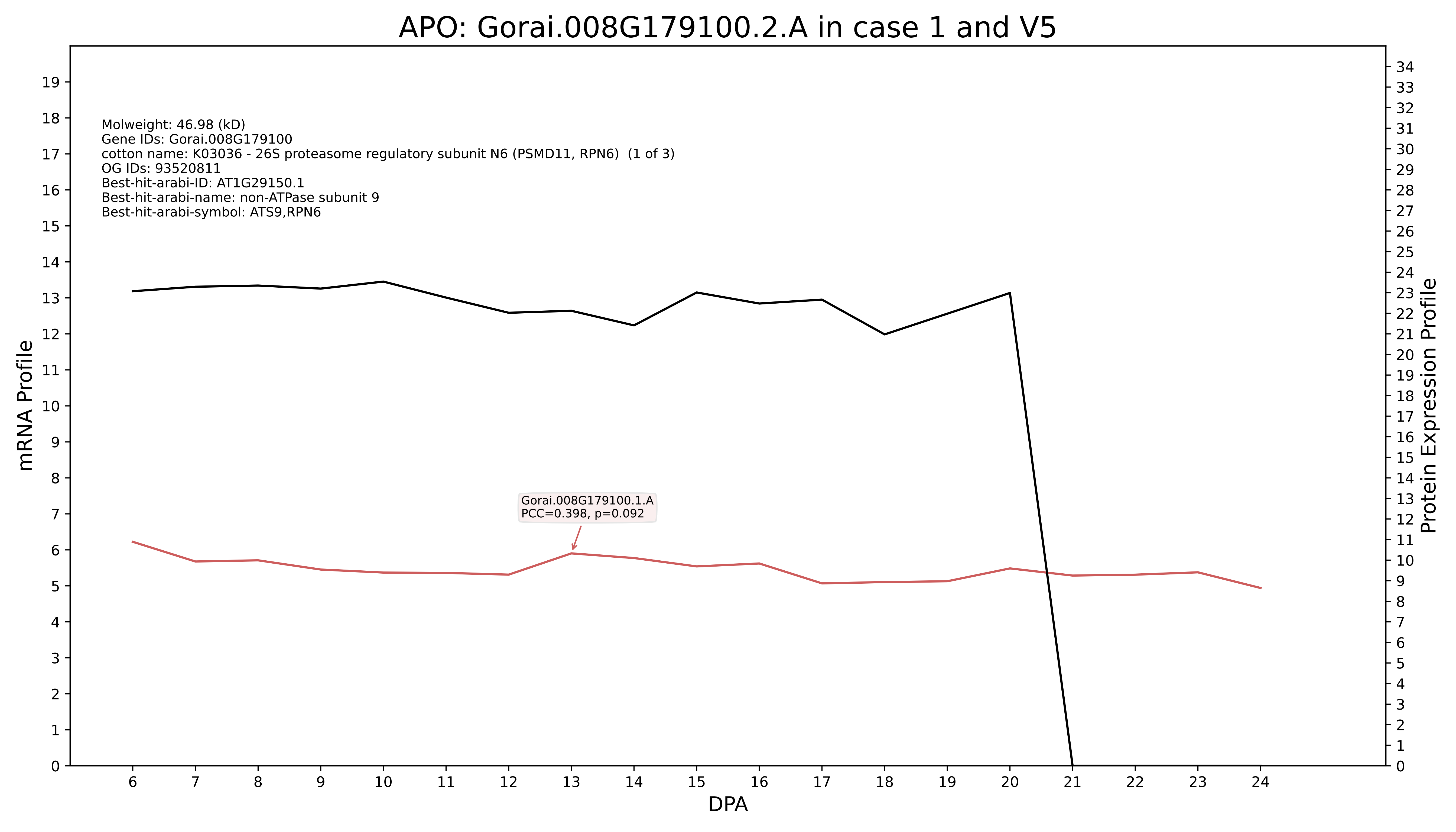

APO: Gorai.008G179100.2.A in case 1 and V5
Molweight: 46.98 (kD)
Gene IDs: Gorai.008G179100
cotton name: K03036 - 26S proteasome regulatory subunit N6 (PSMD11, RPN6) (1 of 3)
OG IDs: 93520811
Best-hit-arabi-ID: AT1G29150.1
Best-hit-arabi-name: non-ATPase subunit 9
Best-hit-arabi-symbol: ATS9,RPN6
Gorai.008G179100.1.A
PCC=0.398, p=0.092
mRNA Profile
Protein Expression Profile
DPA

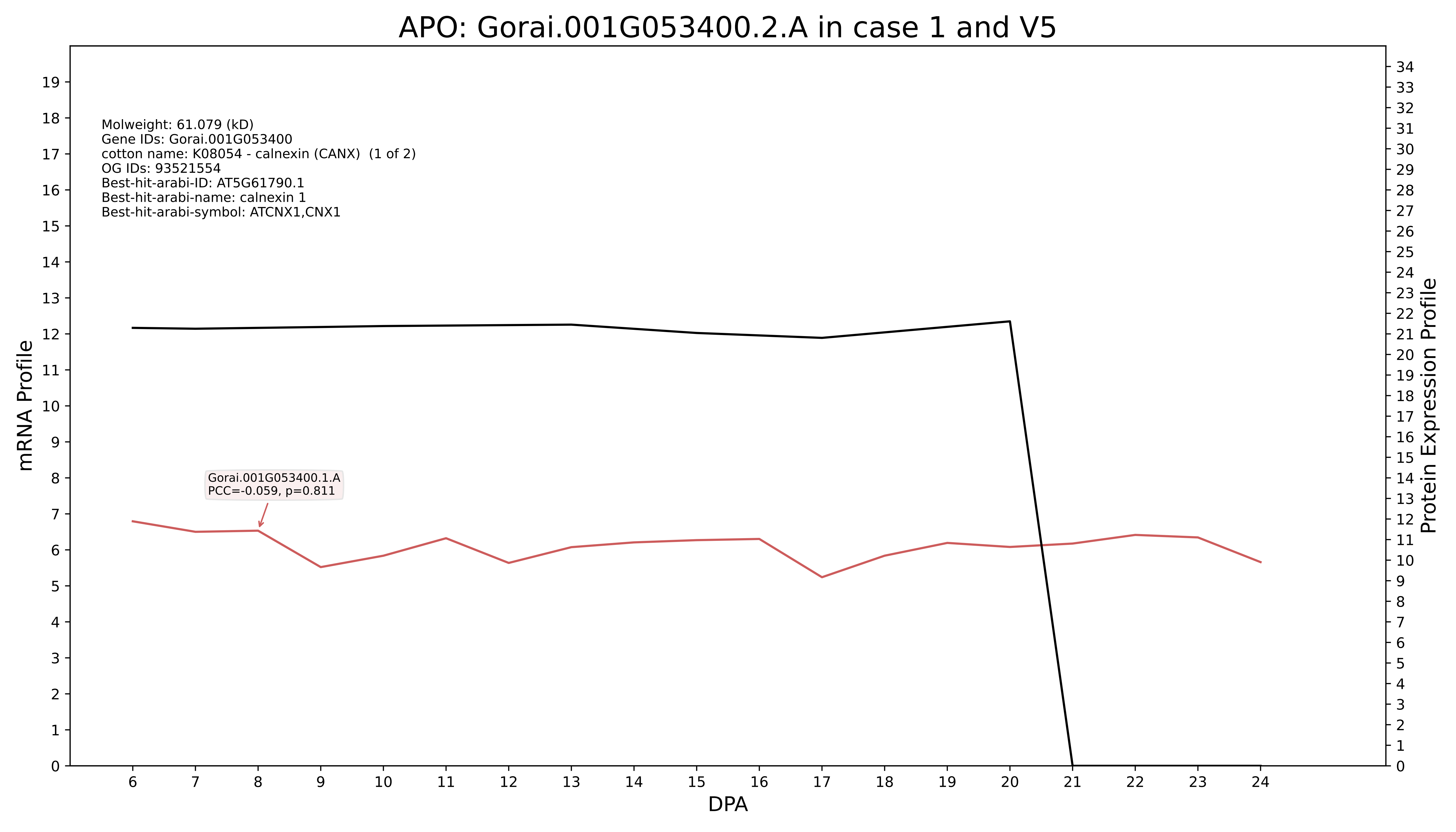

APO: Gorai.001G053400.2.A in case 1 and V5
Molweight: 61.079 (kD)
Gene IDs: Gorai.001G053400
cotton name: K08054 - calnexin (CANX) (1 of 2)
OG IDs: 93521554
Best-hit-arabi-ID: AT5G61790.1
Best-hit-arabi-name: calnexin 1
Best-hit-arabi-symbol: ATCNX1,CNX1
Gorai.001G053400.1.A
PCC=-0.059, p=0.811
mRNA Profile
Protein Expression Profile
DPA

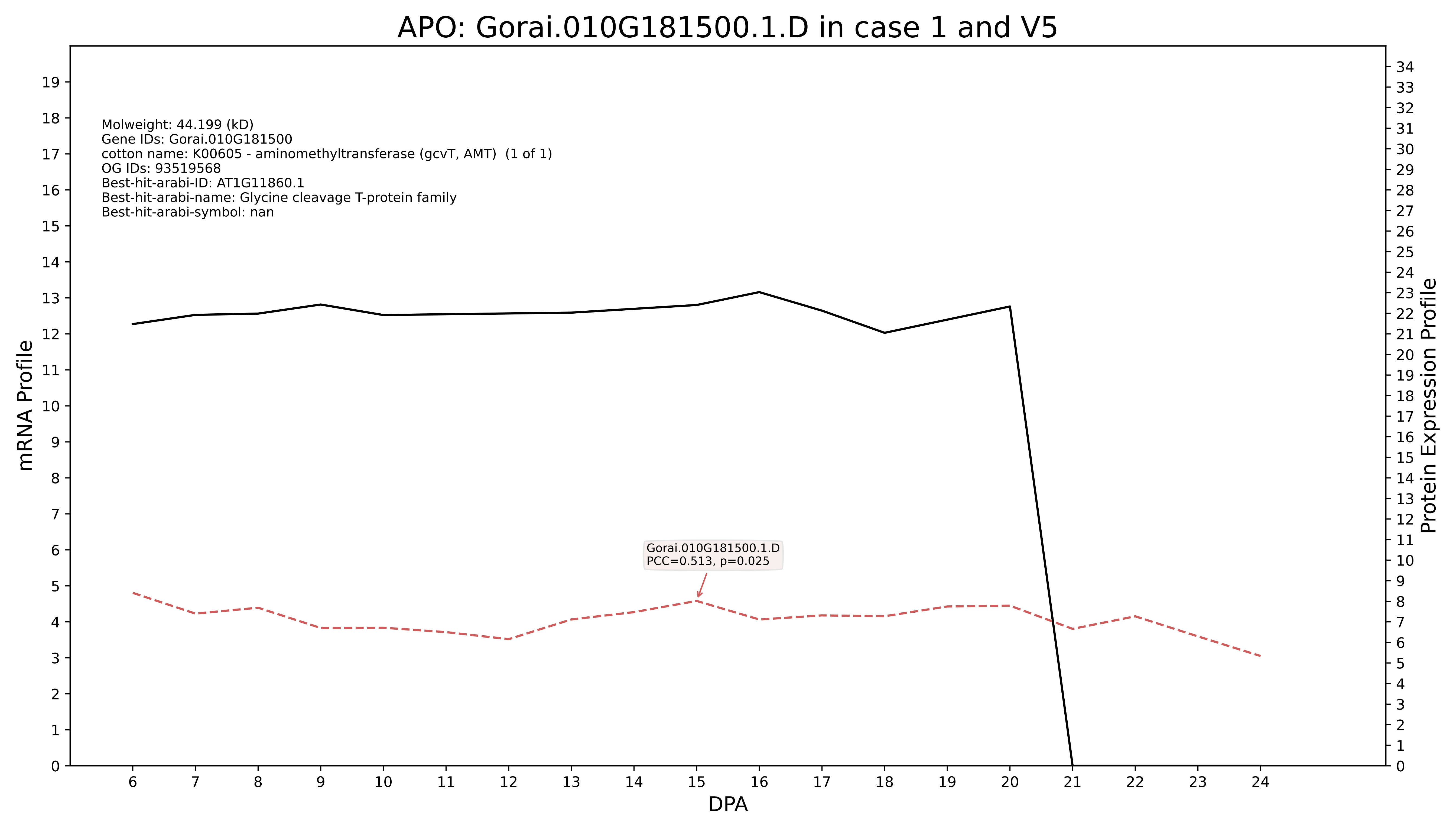
APO: Gorai.010G181500.1.D in case 1 and V5
Molweight: 44.199 (kD)
Gene IDs: Gorai.010G181500
cotton name: K00605 - aminomethyltransferase (gcvT, AMT) (1 of 1)
OG IDs: 93519568
Best-hit-arabi-ID: AT1G11860.1
Best-hit-arabi-name: Glycine cleavage T-protein family
Best-hit-arabi-symbol: nan
Gorai.010G181500.1.D
PCC=0.513, p=0.025
mRNA Profile
Protein Expression Profile
DPA

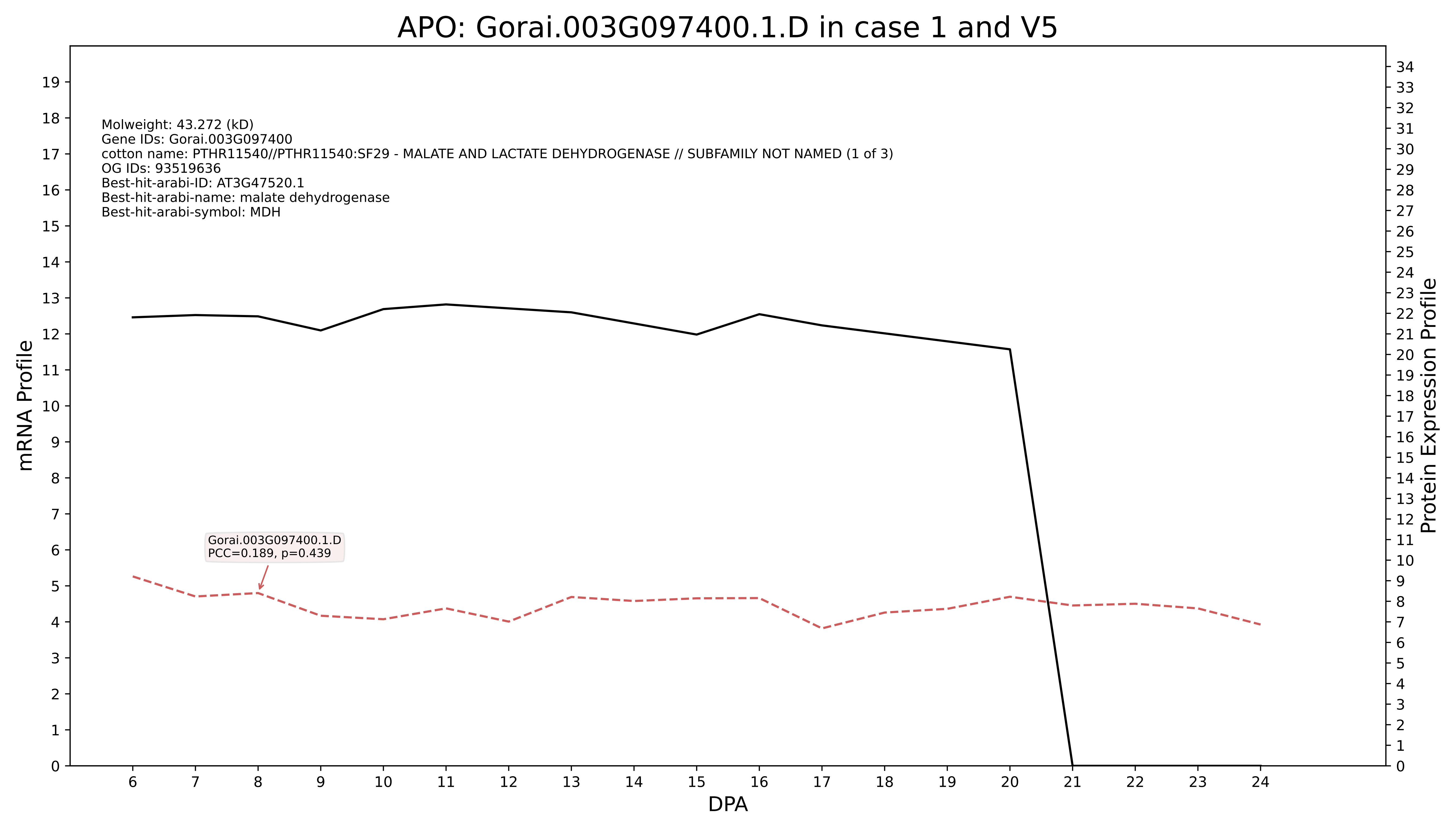

APO: Gorai.003G097400.1.D in case 1 and V5
Molweight: 43.272 (kD)
Gene IDs: Gorai.003G097400
cotton name: PTHR11540//PTHR11540:SF29 - MALATE AND LACTATE DEHYDROGENASE // SUBFAMILY NOT NAMED (1 of 3)
OG IDs: 93519636
Best-hit-arabi-ID: AT3G47520.1
Best-hit-arabi-name: malate dehydrogenase
Best-hit-arabi-symbol: MDH
Gorai.003G097400.1.D
PCC=0.189, p=0.439
mRNA Profile
Protein Expression Profile
DPA

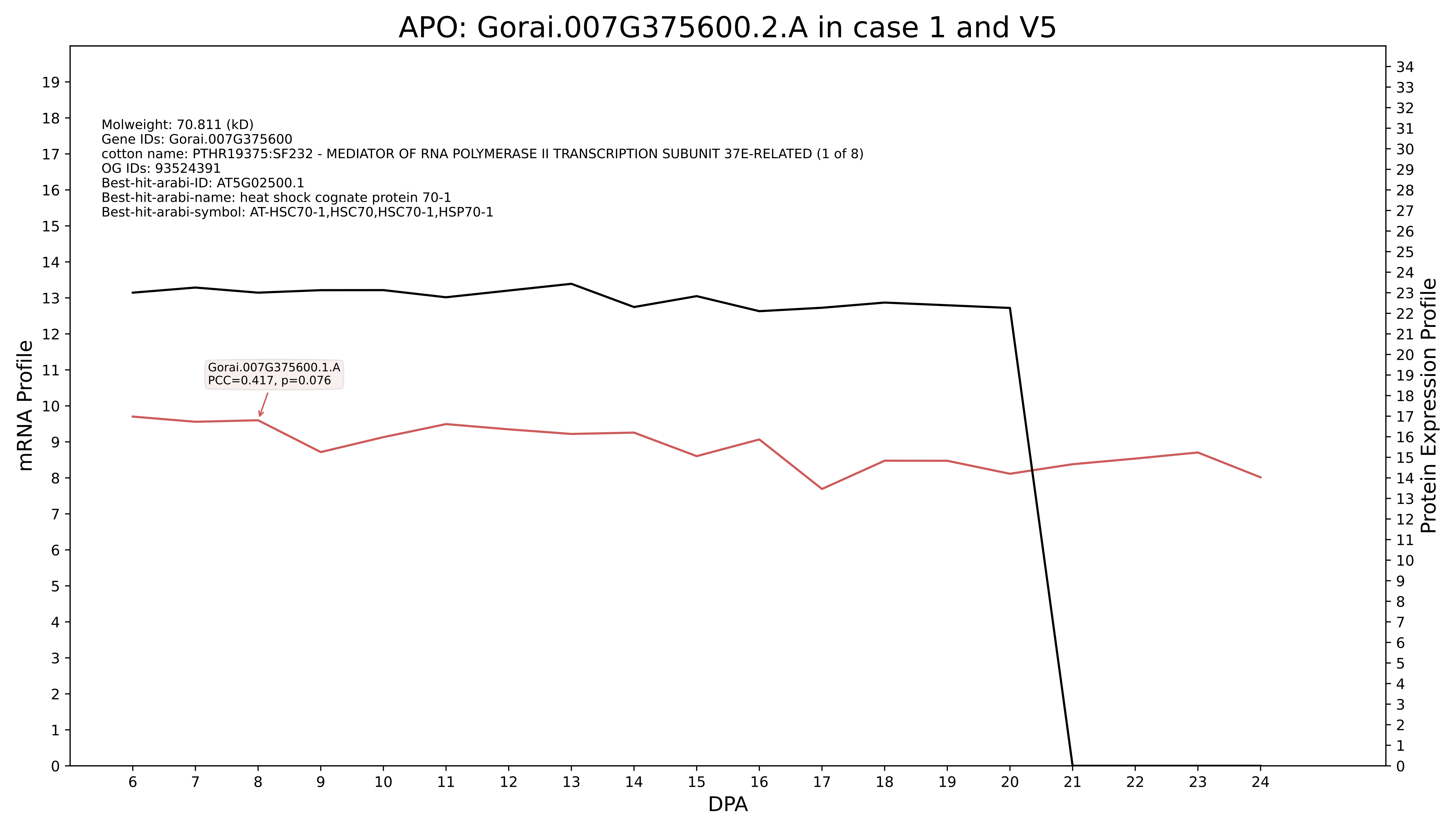
APO: Gorai.007G375600.2.A in case 1 and V5
Molweight: 70.811 (kD)
Gene IDs: Gorai.007G375600
cotton name: PTHR19375:SF232 - MEDIATOR OF RNA POLYMERASE II TRANSCRIPTION SUBUNIT 37E-RELATED (1 of 8)
OG IDs: 93524391
Best-hit-arabi-ID: AT5G02500.1
Best-hit-arabi-name: heat shock cognate protein 70-1
Best-hit-arabi-symbol: AT-HSC70-1,HSC70,HSC70-1,HSP70-1
Gorai.007G375600.1.A
PCC=0.417, p=0.076
mRNA Profile
Protein Expression Profile
DPA

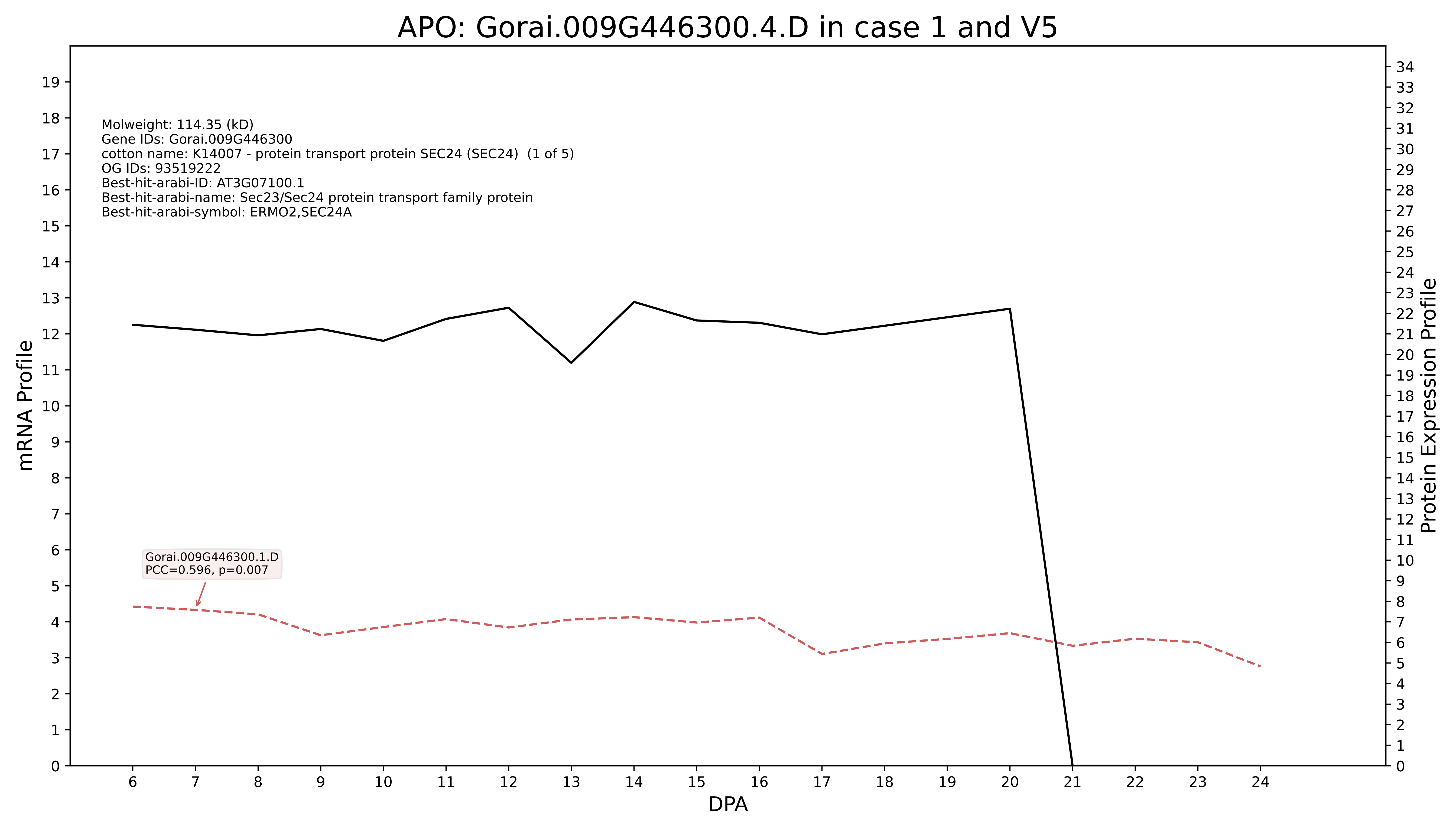
APO: Gorai.009G446300.4.D in case 1 and V5
Molweight: 114.35 (kD)
Gene IDs: Gorai.009G446300
cotton name: K14007 - protein transport protein SEC24 (SEC24) (1 of 5)
OG IDs: 93519222
Best-hit-arabi-ID: AT3G07100.1
Best-hit-arabi-name: Sec23/Sec24 protein transport family protein
Best-hit-arabi-symbol: ERMO2,SEC24A
Gorai.009G446300.1.D
PCC=0.596, p=0.007
mRNA Profile
Protein Expression Profile
DPA

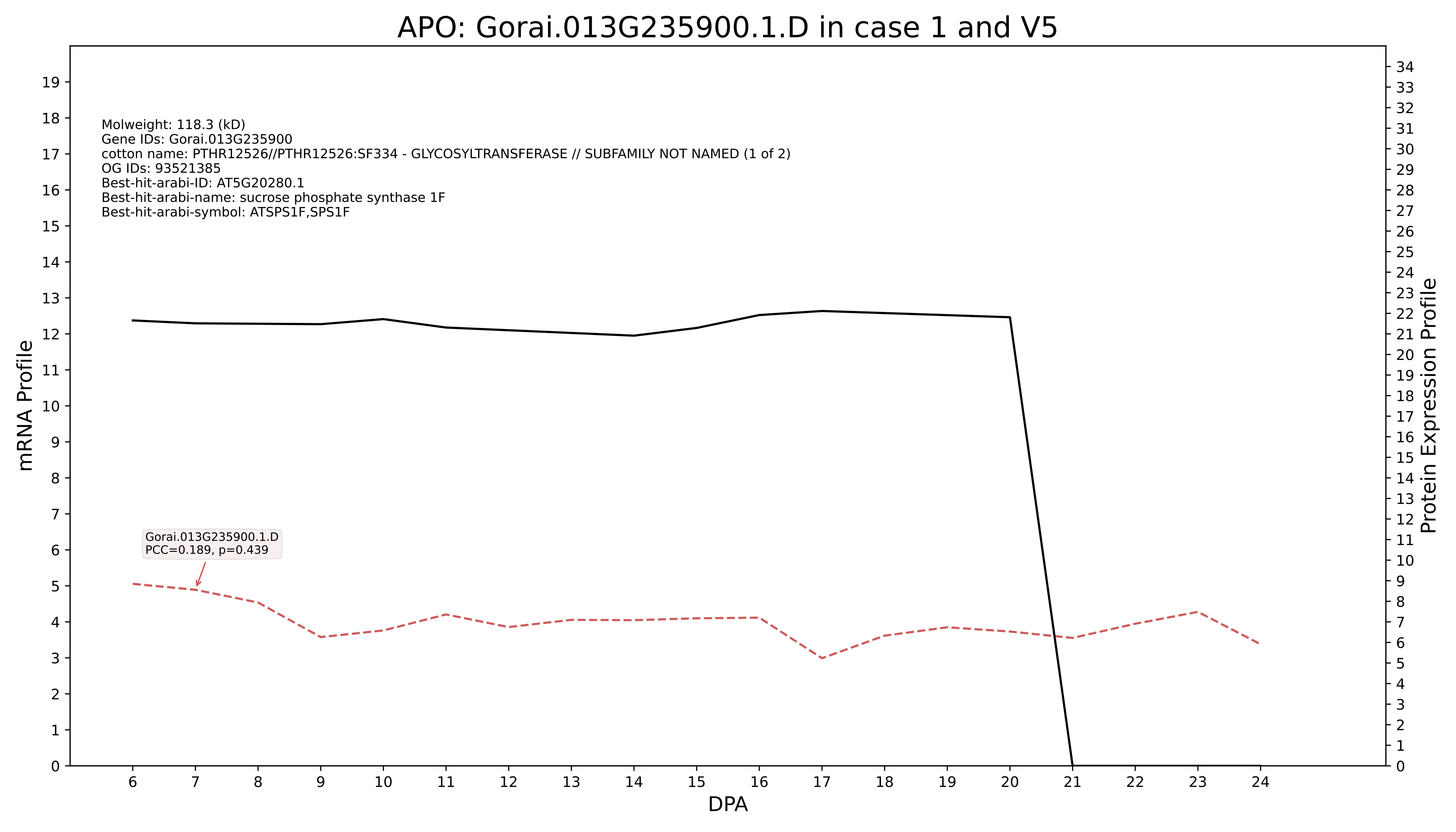
APO: Gorai.013G235900.1.D in case 1 and V5
Molweight: 118.3 (kD)
Gene IDs: Gorai.013G235900
cotton name: PTHR12526//PTHR12526:SF334 - GLYCOSYLTRANSFERASE // SUBFAMILY NOT NAMED (1 of 2)
OG IDs: 93521385
Best-hit-arabi-ID: AT5G20280.1
Best-hit-arabi-name: sucrose phosphate synthase 1F
Best-hit-arabi-symbol: ATSPS1F,SPS1F
Gorai.013G235900.1.D
PCC=0.189, p=0.439
mRNA Profile
Protein Expression Profile
DPA

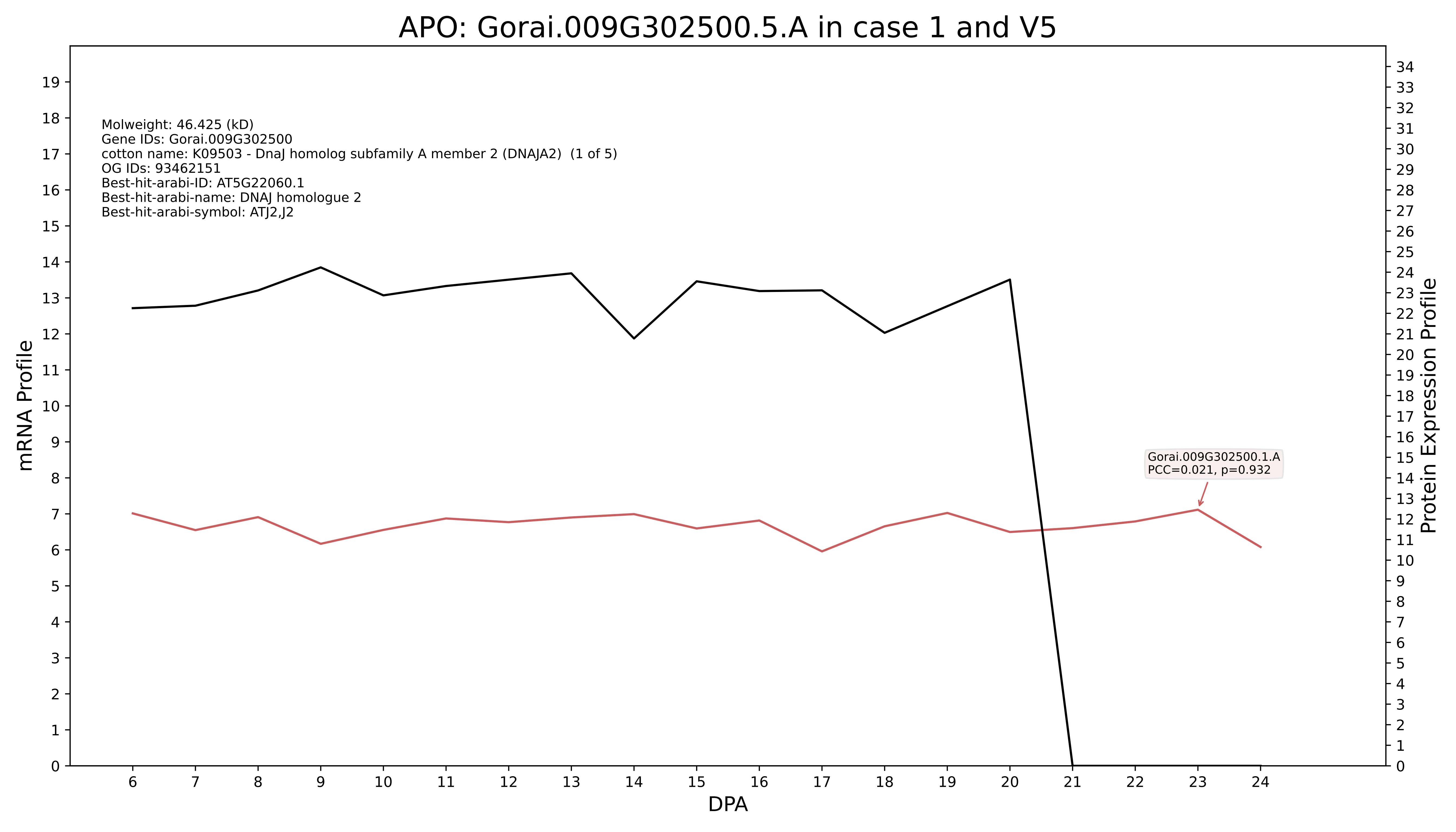

APO: Gorai.009G302500.5.A in case 1 and V5
Molweight: 46.425 (kD)
Gene IDs: Gorai.009G302500
cotton name: K09503 - DnaJ homolog subfamily A member 2 (DNAJA2) (1 of 5)
OG IDs: 93462151
Best-hit-arabi-ID: AT5G22060.1
Best-hit-arabi-name: DNAJ homologue 2
Best-hit-arabi-symbol: ATJ2,J2
Gorai.009G302500.1.A
PCC=0.021, p=0.932
mRNA Profile
Protein Expression Profile
DPA

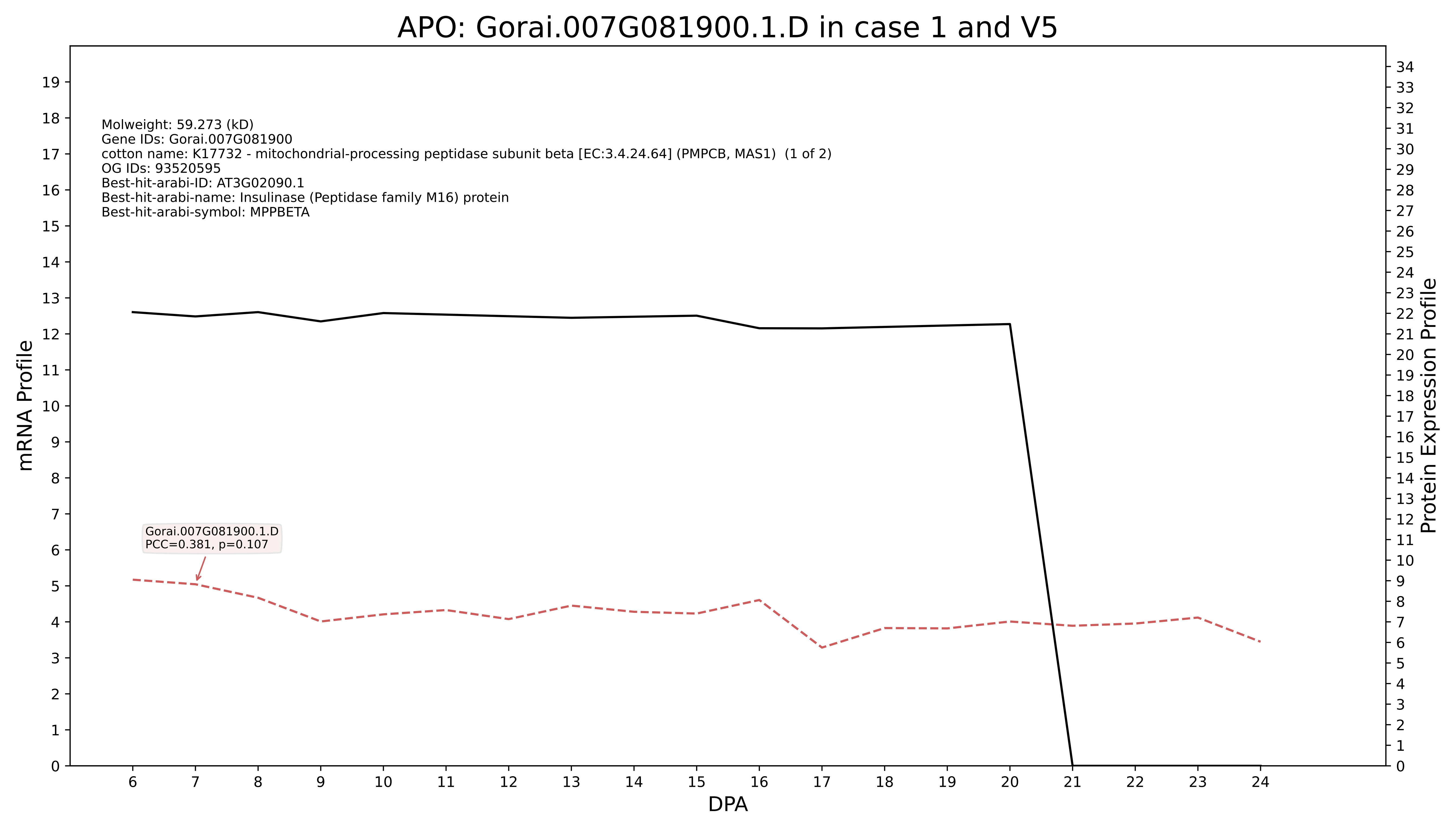
APO: Gorai.007G081900.1.D in case 1 and V5
Molweight: 59.273 (kD)
Gene IDs: Gorai.007G081900
cotton name: K17732 - mitochondrial-processing peptidase subunit beta [EC:3.4.24.64] (PMPCB, MAS1) (1 of 2)
OG IDs: 93520595
Best-hit-arabi-ID: AT3G02090.1
Best-hit-arabi-name: Insulinase (Peptidase family M16) protein
Best-hit-arabi-symbol: MPPBETA
Gorai.007G081900.1.D
PCC=0.381, p=0.107
mRNA Profile
Protein Expression Profile
DPA

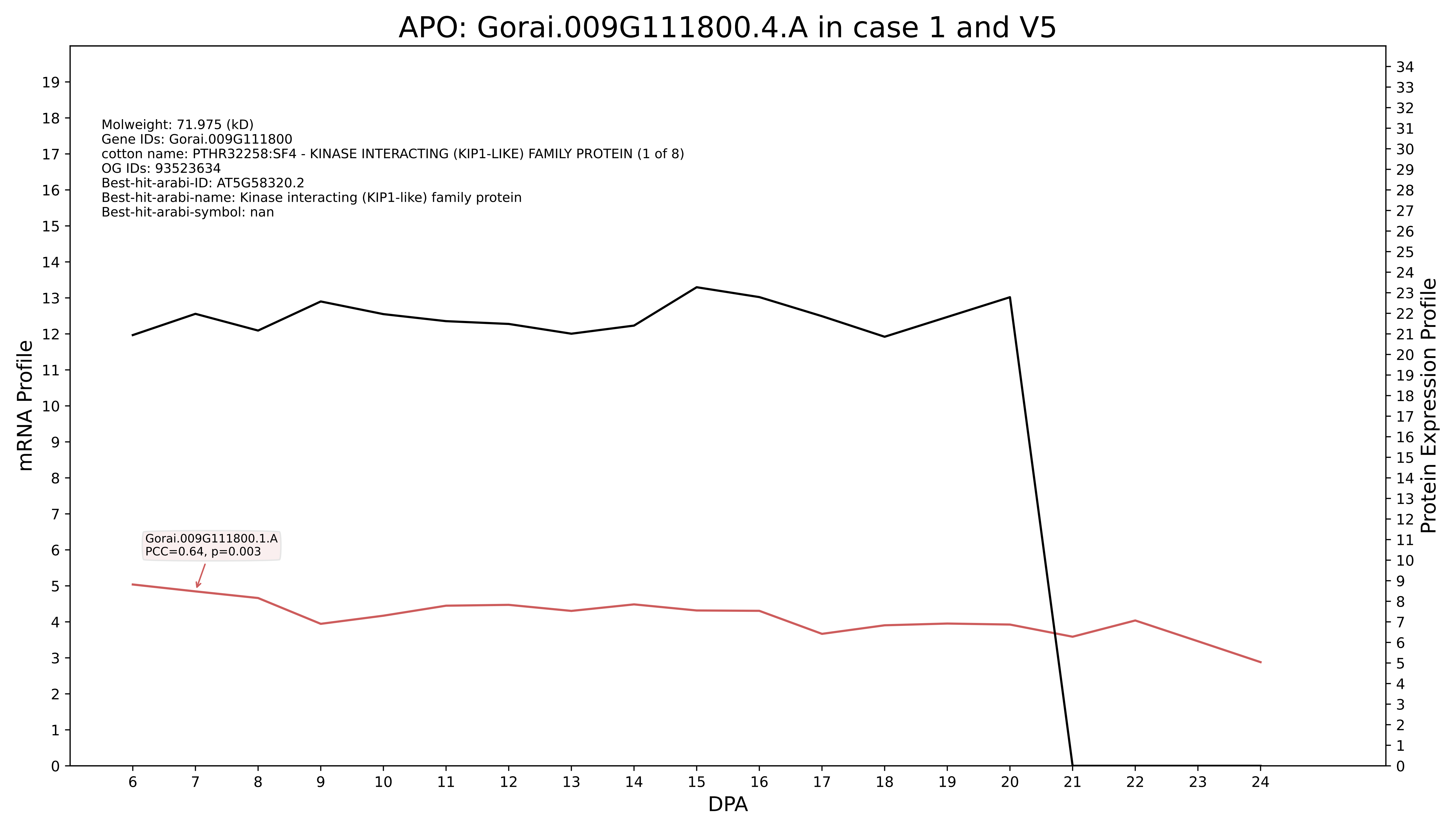

APO: Gorai.009G111800.4.A in case 1 and V5
Molweight: 71.975 (kD)
Gene IDs: Gorai.009G111800
cotton name: PTHR32258:SF4 - KINASE INTERACTING (KIP1-LIKE) FAMILY PROTEIN (1 of 8)
OG IDs: 93523634
Best-hit-arabi-ID: AT5G58320.2
Best-hit-arabi-name: Kinase interacting (KIP1-like) family protein
Best-hit-arabi-symbol: nan
Gorai.009G111800.1.A
PCC=0.64, p=0.003
mRNA Profile
Protein Expression Profile
DPA

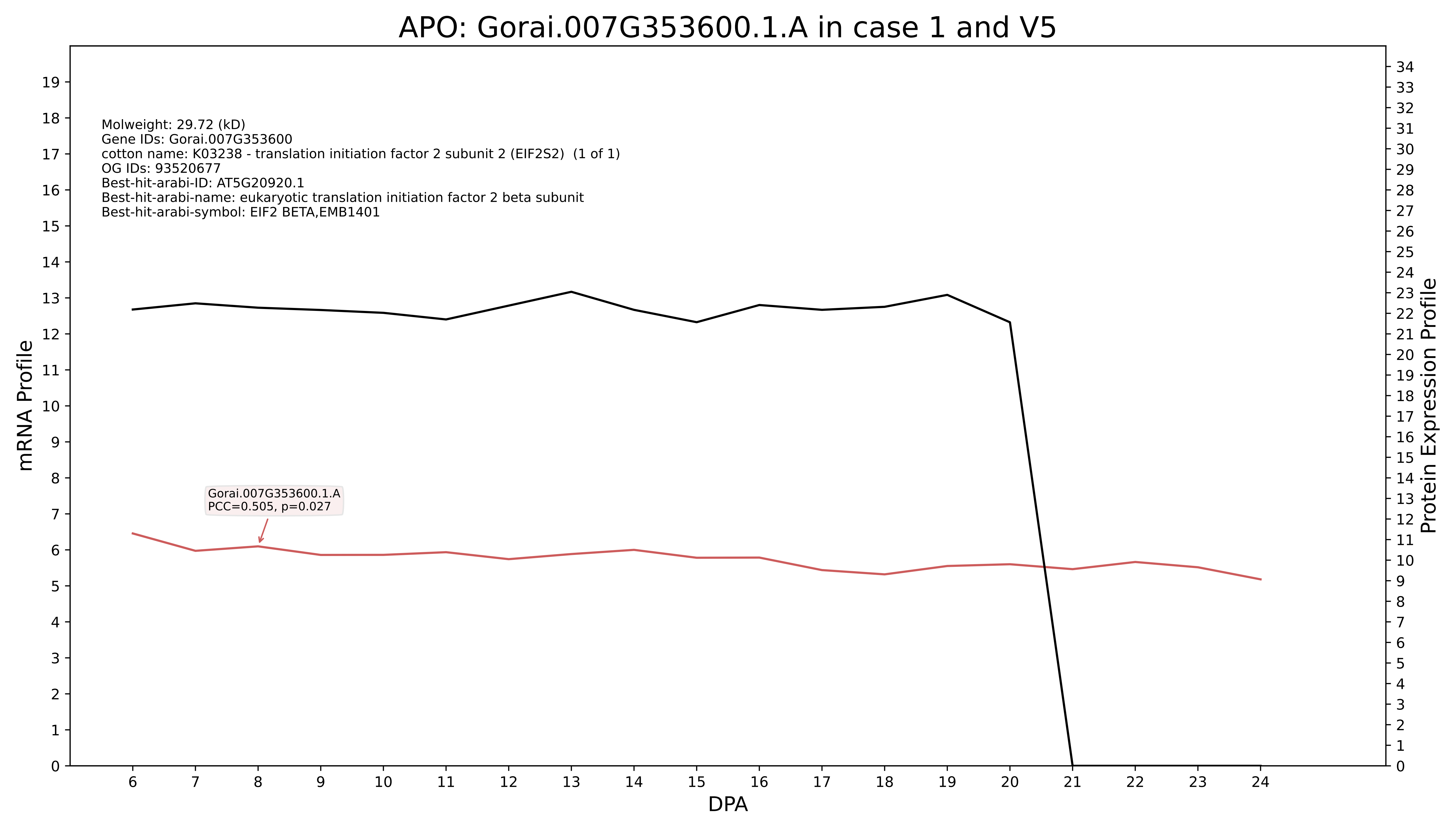
APO: Gorai.007G353600.1.A in case 1 and V5
Molweight: 29.72 (kD)
Gene IDs: Gorai.007G353600
cotton name: K03238 - translation initiation factor 2 subunit 2 (EIF2S2) (1 of 1)
OG IDs: 93520677
Best-hit-arabi-ID: AT5G20920.1
Best-hit-arabi-name: eukaryotic translation initiation factor 2 beta subunit
Best-hit-arabi-symbol: EIF2 BETA,EMB1401
Gorai.007G353600.1.A
PCC=0.505, p=0.027
mRNA Profile
Protein Expression Profile
DPA

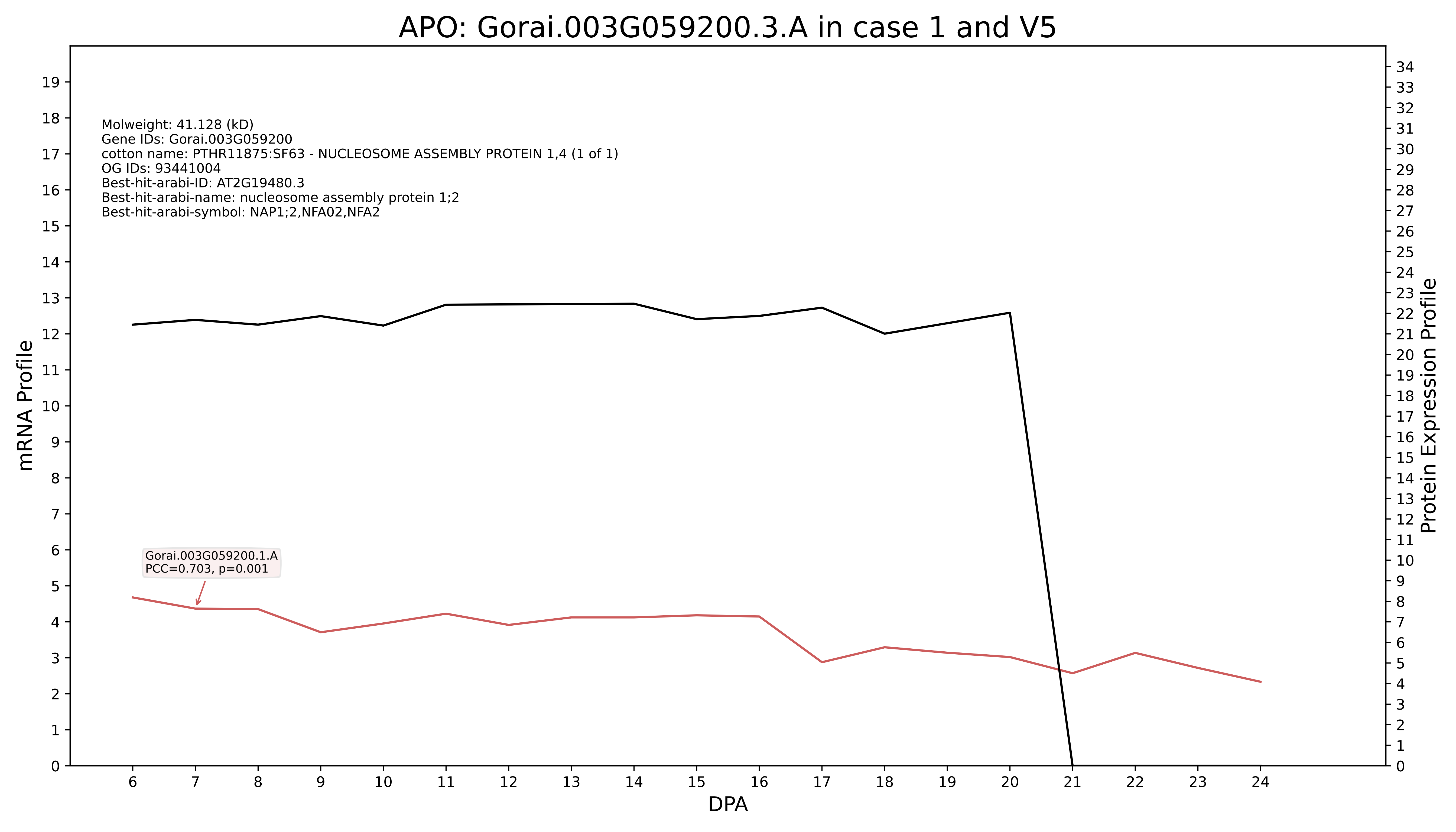

APO: Gorai.003G059200.3.A in case 1 and V5
Molweight: 41.128 (kD)
Gene IDs: Gorai.003G059200
cotton name: PTHR11875:SF63 - NUCLEOSOME ASSEMBLY PROTEIN 1,4 (1 of 1)
OG IDs: 93441004
Best-hit-arabi-ID: AT2G19480.3
Best-hit-arabi-name: nucleosome assembly protein 1;2
Best-hit-arabi-symbol: NAP1;2,NFA02,NFA2
Gorai.003G059200.1.A
PCC=0.703, p=0.001
mRNA Profile
Protein Expression Profile
DPA

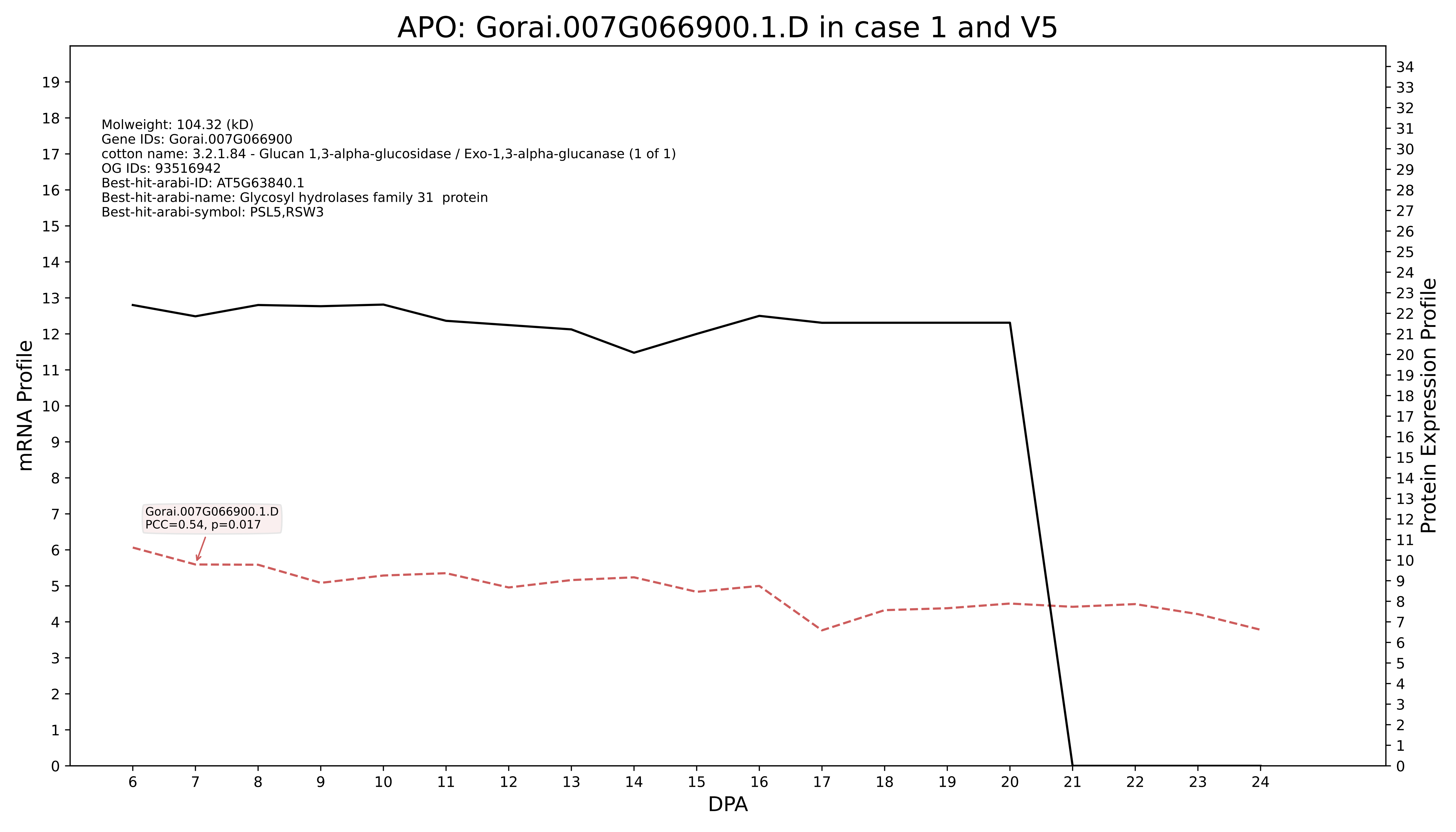
APO: Gorai.007G066900.1.D in case 1 and V5
Molweight: 104.32 (kD)
Gene IDs: Gorai.007G066900
cotton name: 3.2.1.84 - Glucan 1,3-alpha-glucosidase / Exo-1,3-alpha-glucanase (1 of 1)
OG IDs: 93516942
Best-hit-arabi-ID: AT5G63840.1
Best-hit-arabi-name: Glycosyl hydrolases family 31 protein
Best-hit-arabi-symbol: PSL5,RSW3
Gorai.007G066900.1.D
PCC=0.54, p=0.017
mRNA Profile
Protein Expression Profile
DPA

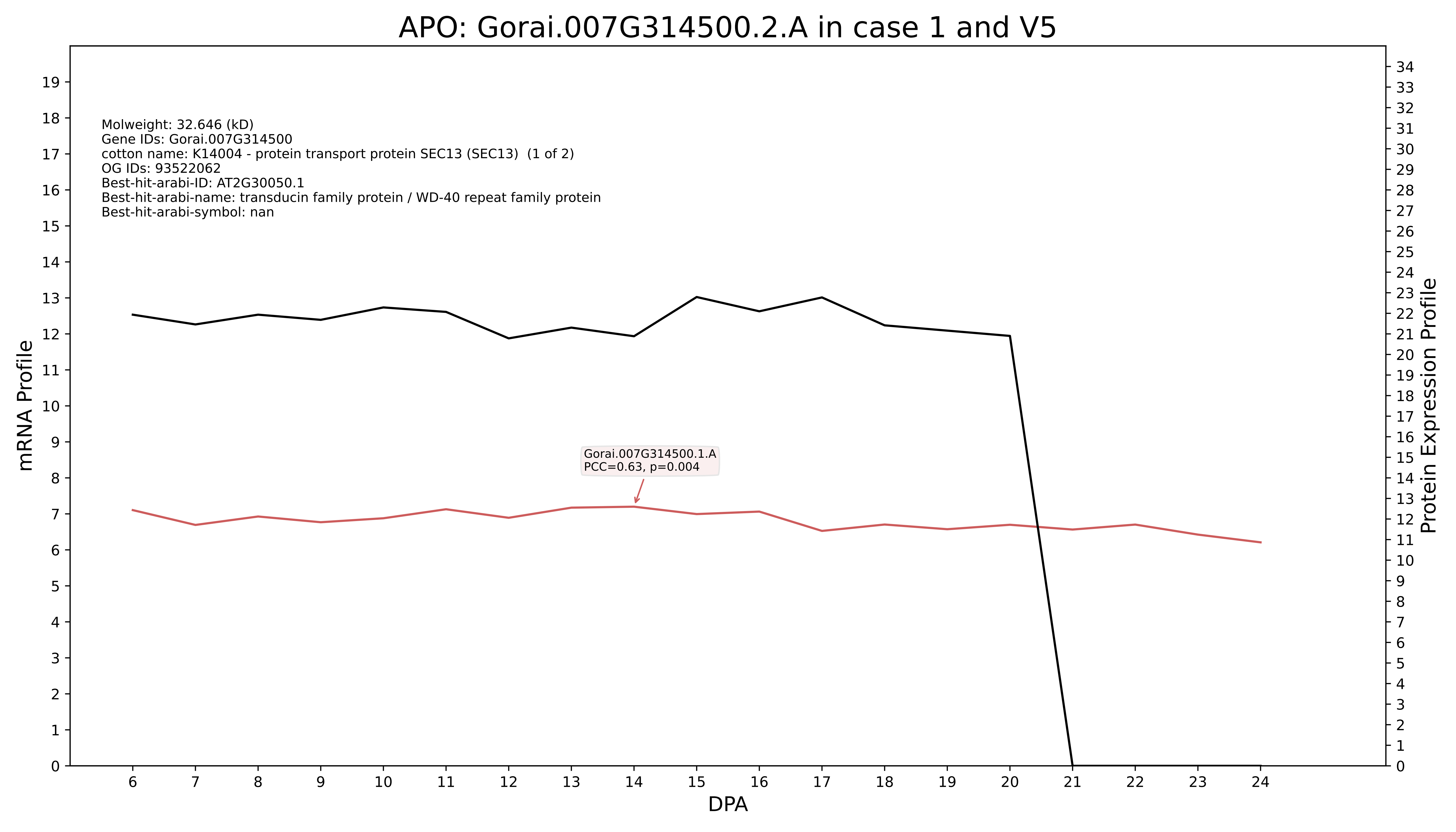

APO: Gorai.007G314500.2.A in case 1 and V5
Molweight: 32.646 (kD)
Gene IDs: Gorai.007G314500
cotton name: K14004 - protein transport protein SEC13 (SEC13) (1 of 2)
OG IDs: 93522062
Best-hit-arabi-ID: AT2G30050.1
Best-hit-arabi-name: transducin family protein / WD-40 repeat family protein
Best-hit-arabi-symbol: nan
Gorai.007G314500.1.A
PCC=0.63, p=0.004
mRNA Profile
Protein Expression Profile
DPA

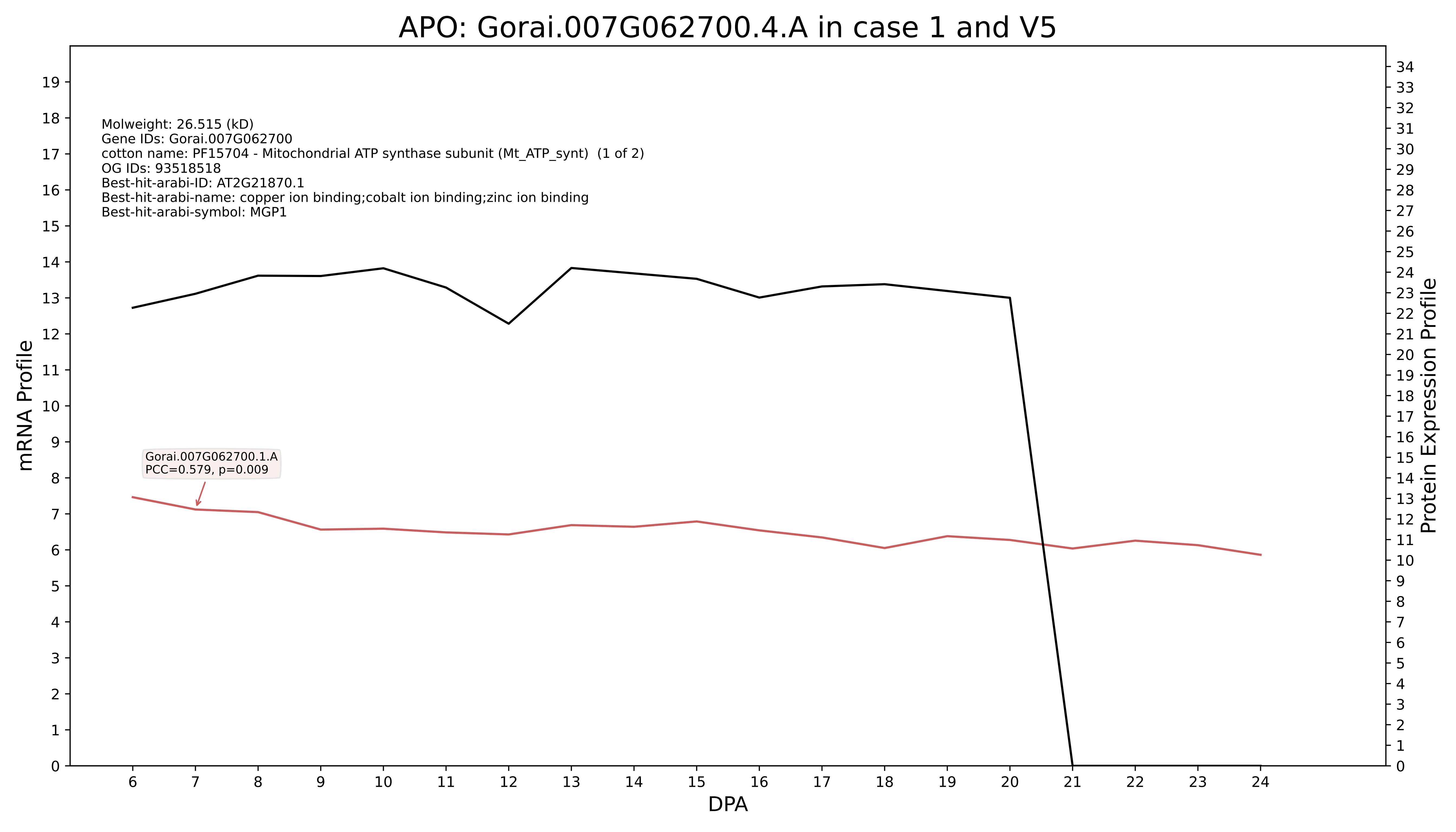
APO: Gorai.007G062700.4.A in case 1 and V5
Molweight: 26.515 (kD)
Gene IDs: Gorai.007G062700
cotton name: PF15704 - Mitochondrial ATP synthase subunit (Mt_ATP_synt) (1 of 2)
OG IDs: 93518518
Best-hit-arabi-ID: AT2G21870.1
Best-hit-arabi-name: copper ion binding;cobalt ion binding;zinc ion binding
Best-hit-arabi-symbol: MGP1
Gorai.007G062700.1.A
PCC=0.579, p=0.009
mRNA Profile
Protein Expression Profile
DPA

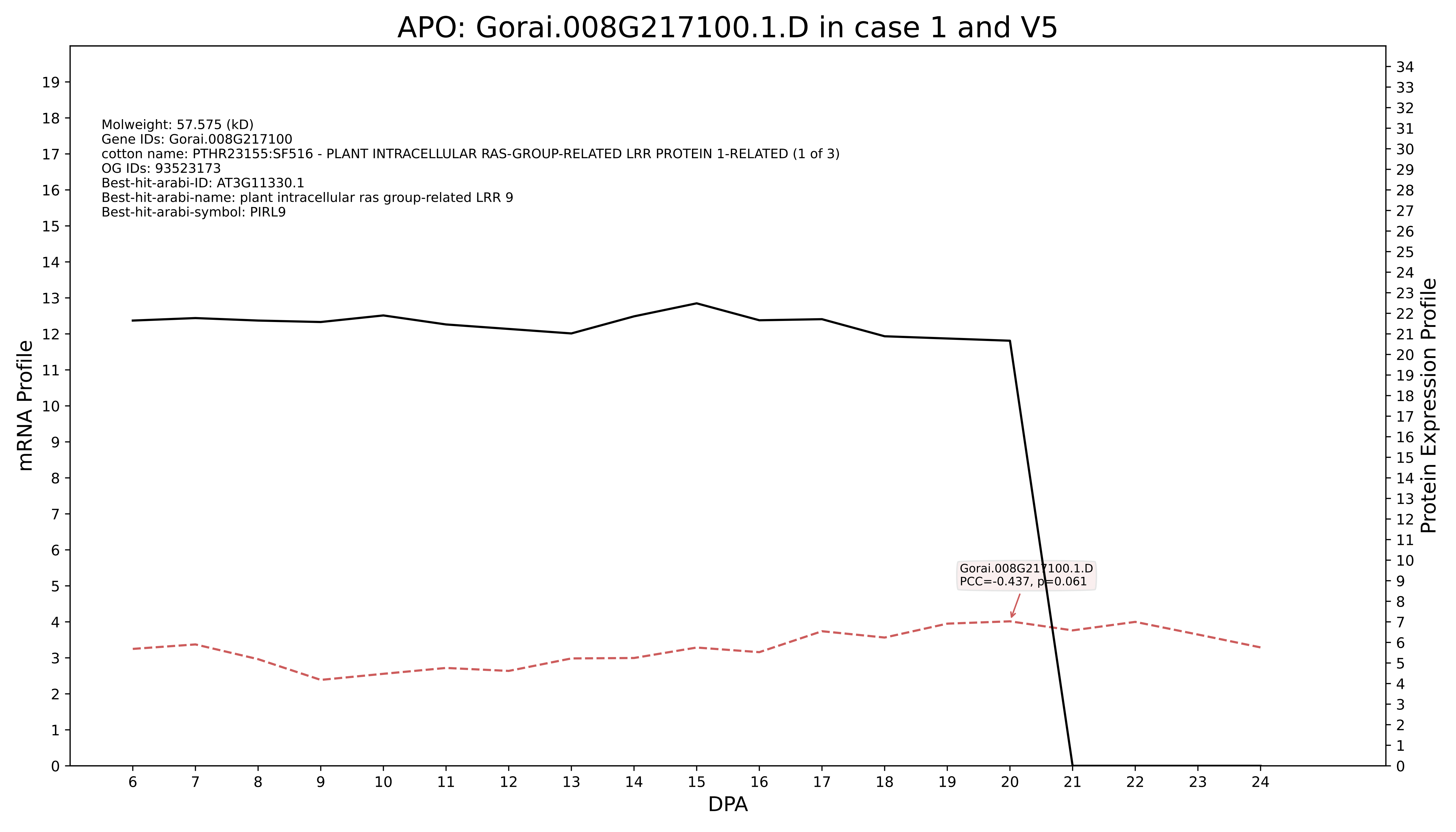
APO: Gorai.008G217100.1.D in case 1 and V5
Molweight: 57.575 (kD)
Gene IDs: Gorai.008G217100
cotton name: PTHR23155:SF516 - PLANT INTRACELLULAR RAS-GROUP-RELATED LRR PROTEIN 1-RELATED (1 of 3)
OG IDs: 93523173
Best-hit-arabi-ID: AT3G11330.1
Best-hit-arabi-name: plant intracellular ras group-related LRR 9
Best-hit-arabi-symbol: PIRL9
Gorai.008G217100.1.D
PCC=-0.437, p=0.061
mRNA Profile
Protein Expression Profile
DPA

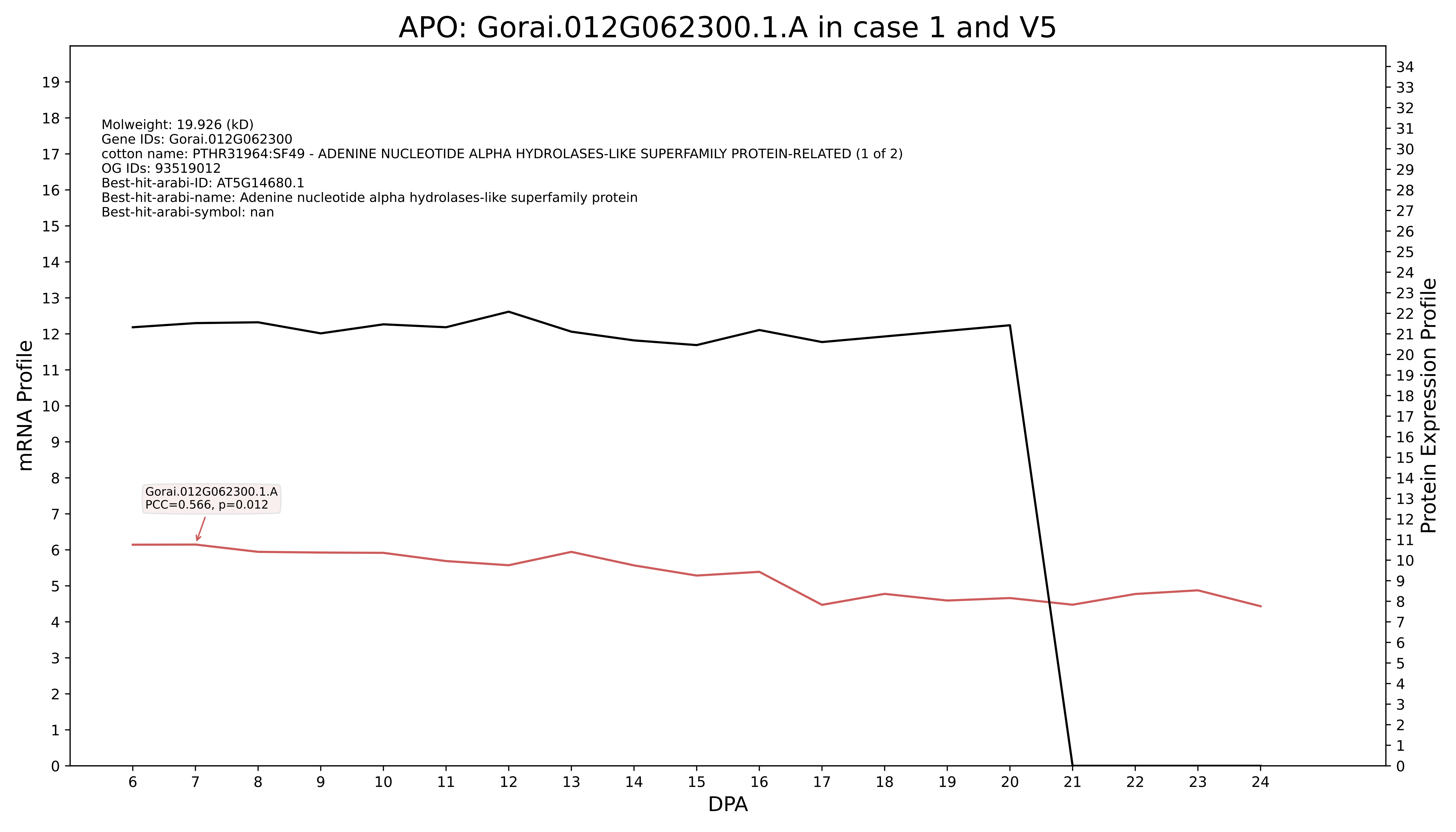

APO: Gorai.012G062300.1.A in case 1 and V5
Molweight: 19.926 (kD)
Gene IDs: Gorai.012G062300
cotton name: PTHR31964:SF49 - ADENINE NUCLEOTIDE ALPHA HYDROLASES-LIKE SUPERFAMILY PROTEIN-RELATED (1 of 2)
OG IDs: 93519012
Best-hit-arabi-ID: AT5G14680.1
Best-hit-arabi-name: Adenine nucleotide alpha hydrolases-like superfamily protein
Best-hit-arabi-symbol: nan
Gorai.012G062300.1.A
PCC=0.566, p=0.012
mRNA Profile
Protein Expression Profile
DPA

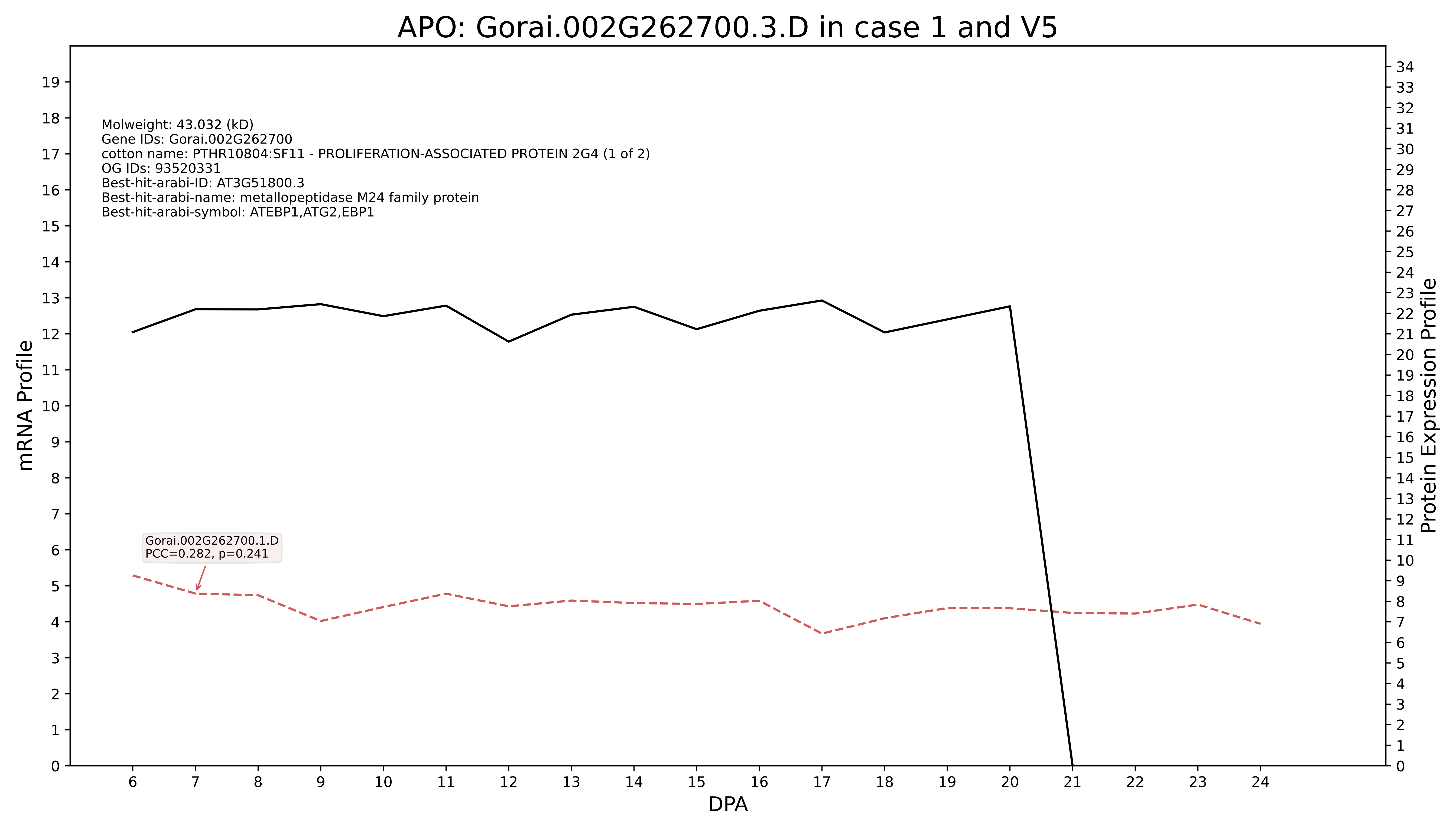

APO: Gorai.002G262700.3.D in case 1 and V5
Molweight: 43.032 (kD)
Gene IDs: Gorai.002G262700
cotton name: PTHR10804:SF11 - PROLIFERATION-ASSOCIATED PROTEIN 2G4 (1 of 2)
OG IDs: 93520331
Best-hit-arabi-ID: AT3G51800.3
Best-hit-arabi-name: metallopeptidase M24 family protein
Best-hit-arabi-symbol: ATEBP1,ATG2,EBP1
Gorai.002G262700.1.D
PCC=0.282, p=0.241
mRNA Profile
Protein Expression Profile
DPA

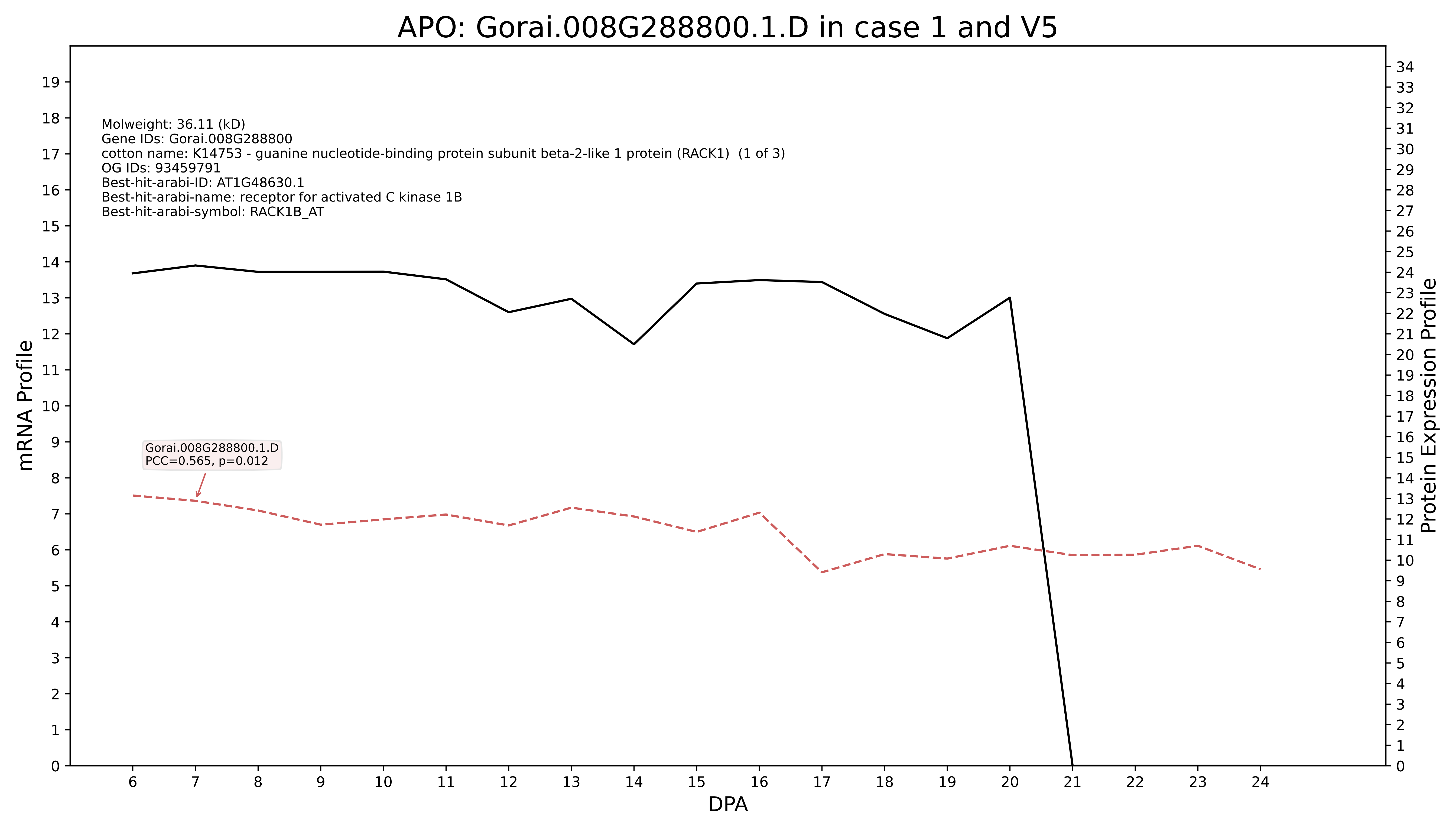

APO: Gorai.008G288800.1.D in case 1 and V5
Molweight: 36.11 (kD)
Gene IDs: Gorai.008G288800
cotton name: K14753 - guanine nucleotide-binding protein subunit beta-2-like 1 protein (RACK1) (1 of 3)
OG IDs: 93459791
Best-hit-arabi-ID: AT1G48630.1
Best-hit-arabi-name: receptor for activated C kinase 1B
Best-hit-arabi-symbol: RACK1B_AT
Gorai.008G288800.1.D
PCC=0.565, p=0.012
mRNA Profile
Protein Expression Profile
DPA

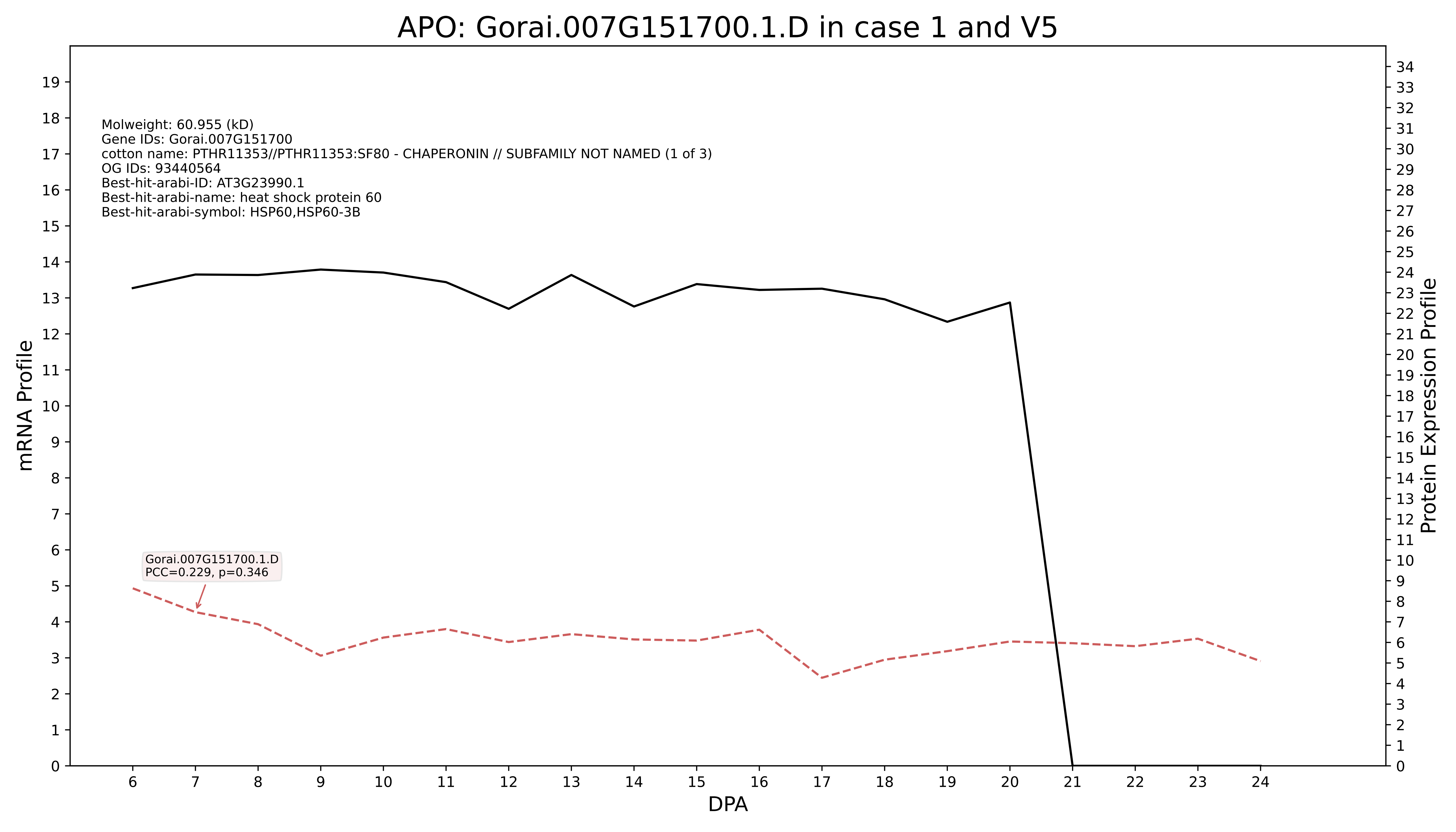

APO: Gorai.007G151700.1.D in case 1 and V5
Molweight: 60.955 (kD)
Gene IDs: Gorai.007G151700
cotton name: PTHR11353//PTHR11353:SF80 - CHAPERONIN // SUBFAMILY NOT NAMED (1 of 3)
OG IDs: 93440564
Best-hit-arabi-ID: AT3G23990.1
Best-hit-arabi-name: heat shock protein 60
Best-hit-arabi-symbol: HSP60,HSP60-3B
Gorai.007G151700.1.D
PCC=0.229, p=0.346
mRNA Profile
Protein Expression Profile
DPA

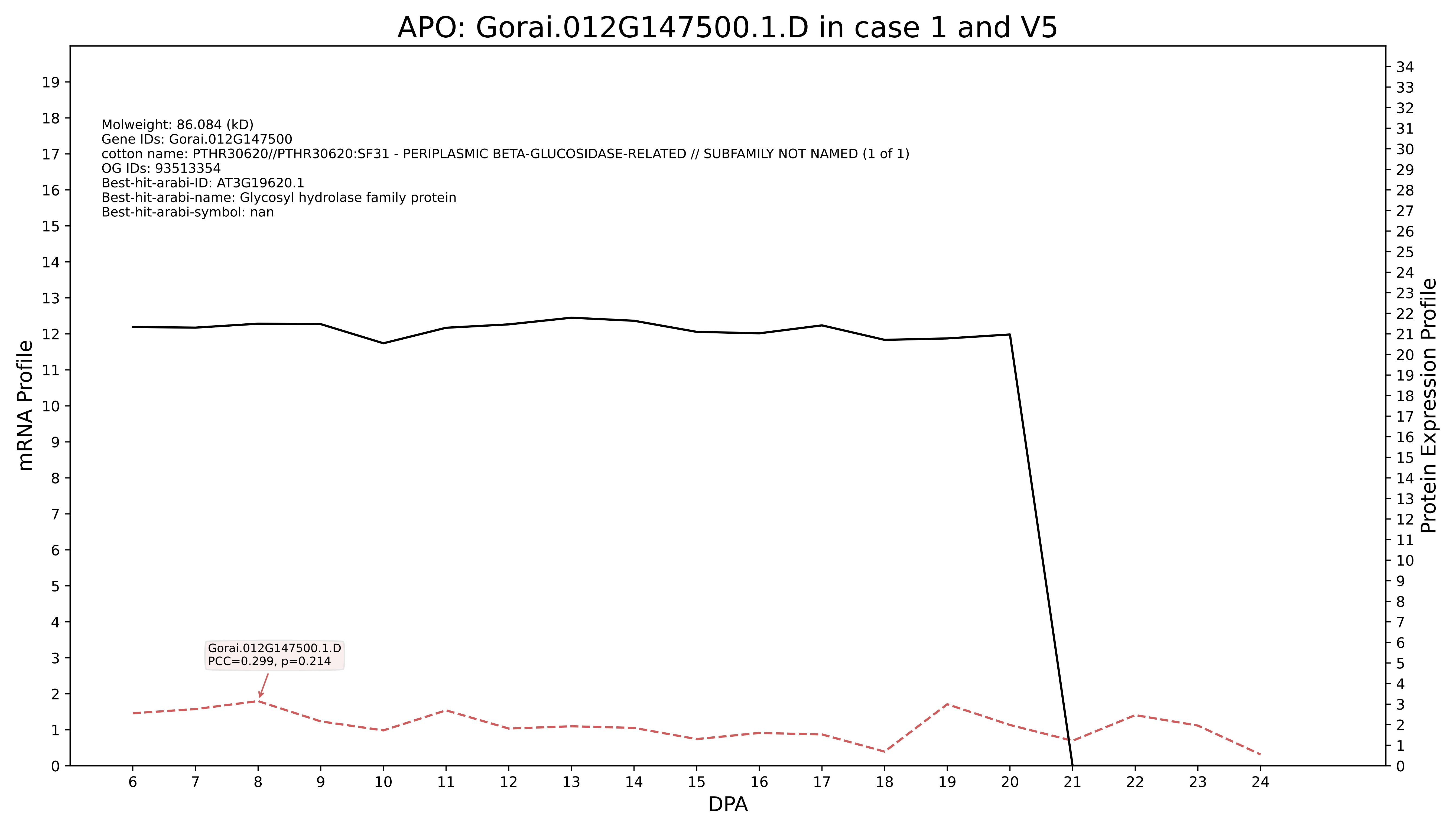
APO: Gorai.012G147500.1.D in case 1 and V5
Molweight: 86.084 (kD)
Gene IDs: Gorai.012G147500
cotton name: PTHR30620//PTHR30620:SF31 - PERIPLASMIC BETA-GLUCOSIDASE-RELATED // SUBFAMILY NOT NAMED (1 of 1)
OG IDs: 93513354
Best-hit-arabi-ID: AT3G19620.1
Best-hit-arabi-name: Glycosyl hydrolase family protein
Best-hit-arabi-symbol: nan
Gorai.012G147500.1.D
PCC=0.299, p=0.214
mRNA Profile
Protein Expression Profile
DPA

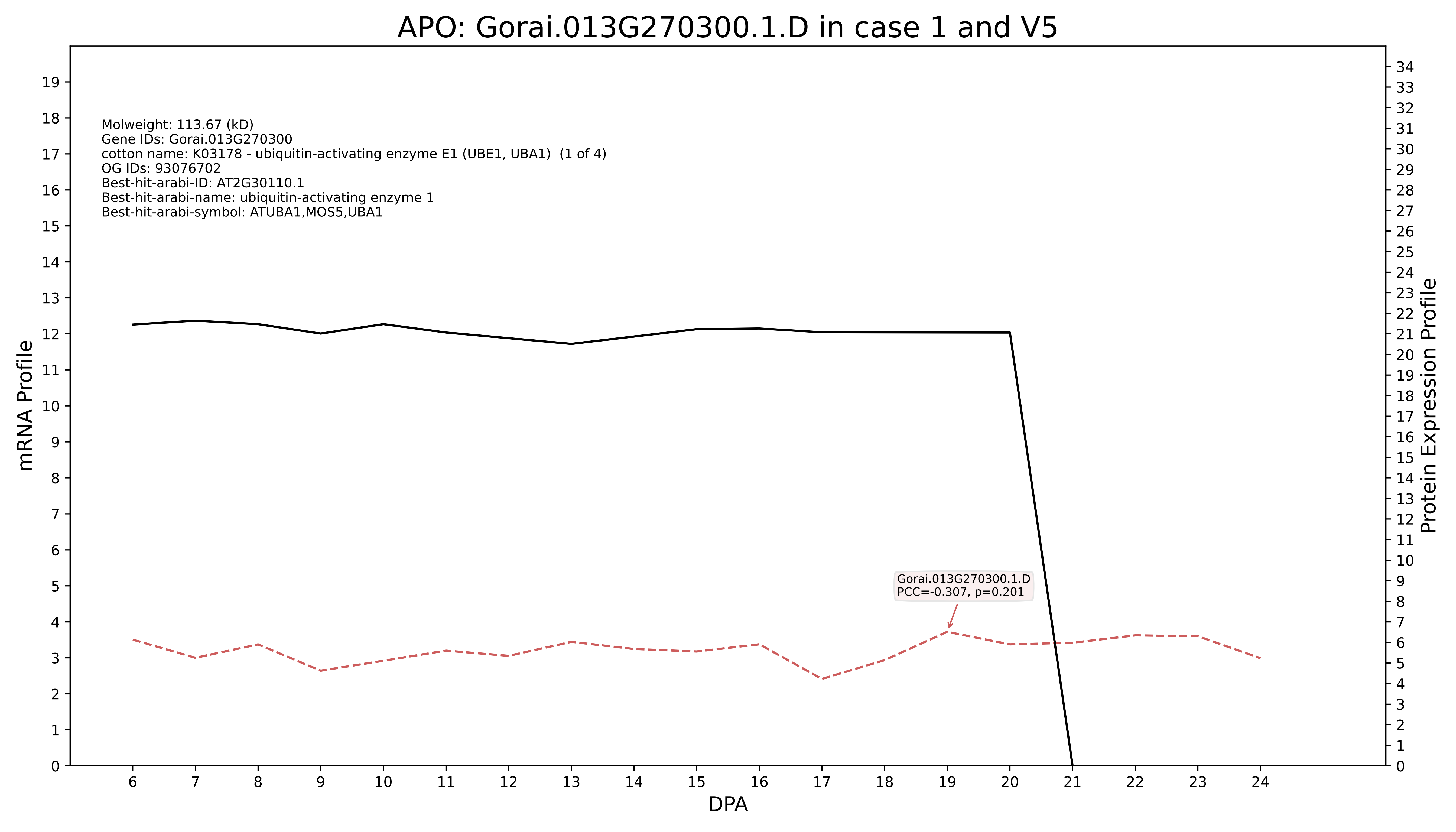
APO: Gorai.013G270300.1.D in case 1 and V5
Molweight: 113.67 (kD)
Gene IDs: Gorai.013G270300
cotton name: K03178 - ubiquitin-activating enzyme E1 (UBE1, UBA1) (1 of 4)
OG IDs: 93076702
Best-hit-arabi-ID: AT2G30110.1
Best-hit-arabi-name: ubiquitin-activating enzyme 1
Best-hit-arabi-symbol: ATUBA1,MOS5,UBA1
Gorai.013G270300.1.D
PCC=-0.307, p=0.201
mRNA Profile
Protein Expression Profile
DPA

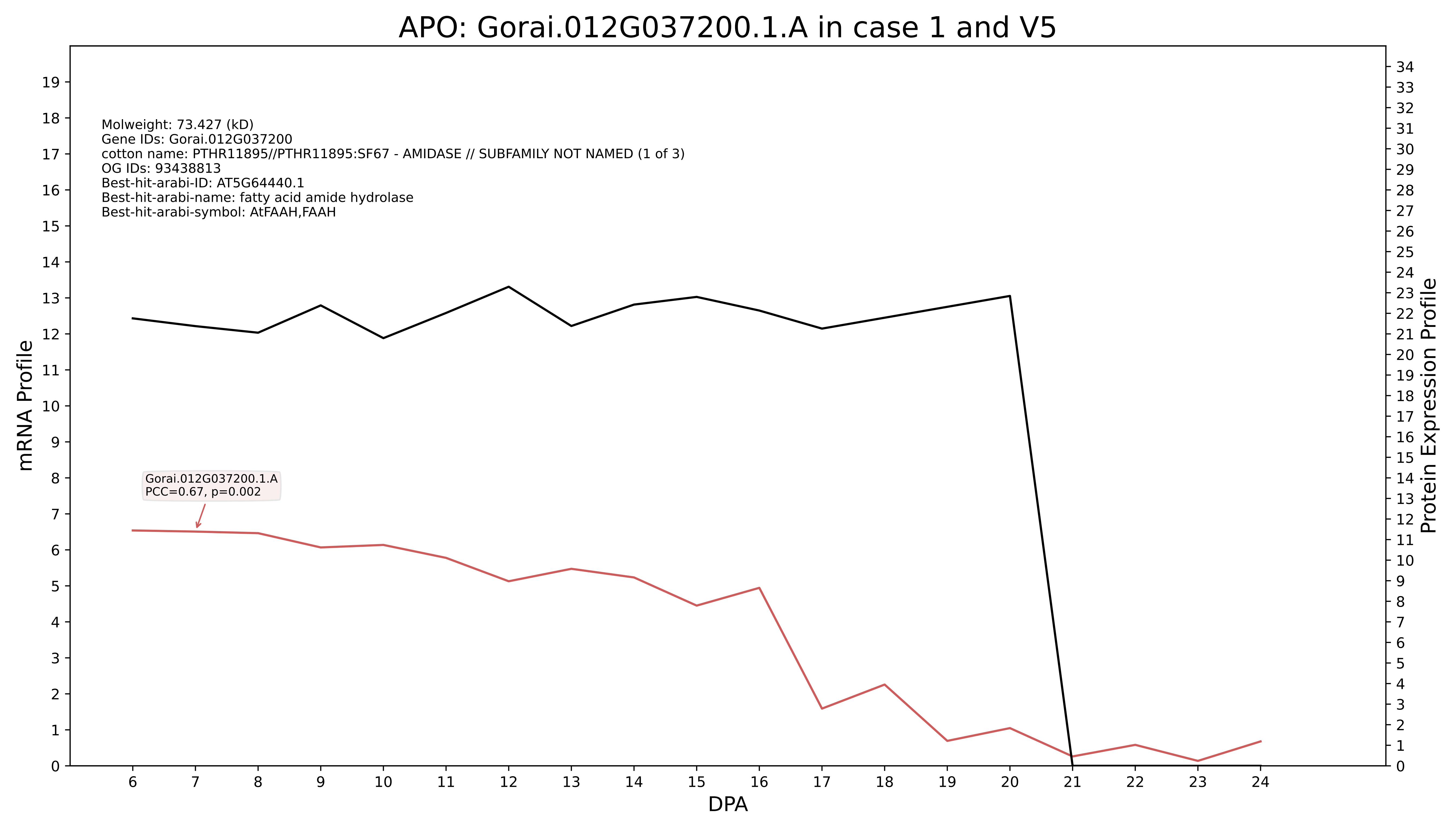

APO: Gorai.012G037200.1.A in case 1 and V5
Molweight: 73.427 (kD)
Gene IDs: Gorai.012G037200
cotton name: PTHR11895//PTHR11895:SF67 - AMIDASE // SUBFAMILY NOT NAMED (1 of 3)
OG IDs: 93438813
Best-hit-arabi-ID: AT5G64440.1
Best-hit-arabi-name: fatty acid amide hydrolase
Best-hit-arabi-symbol: AtFAAH,FAAH
Gorai.012G037200.1.A
PCC=0.67, p=0.002
mRNA Profile
Protein Expression Profile
DPA

### APO: Gorai.001G079900.2.D in case 1 and V5

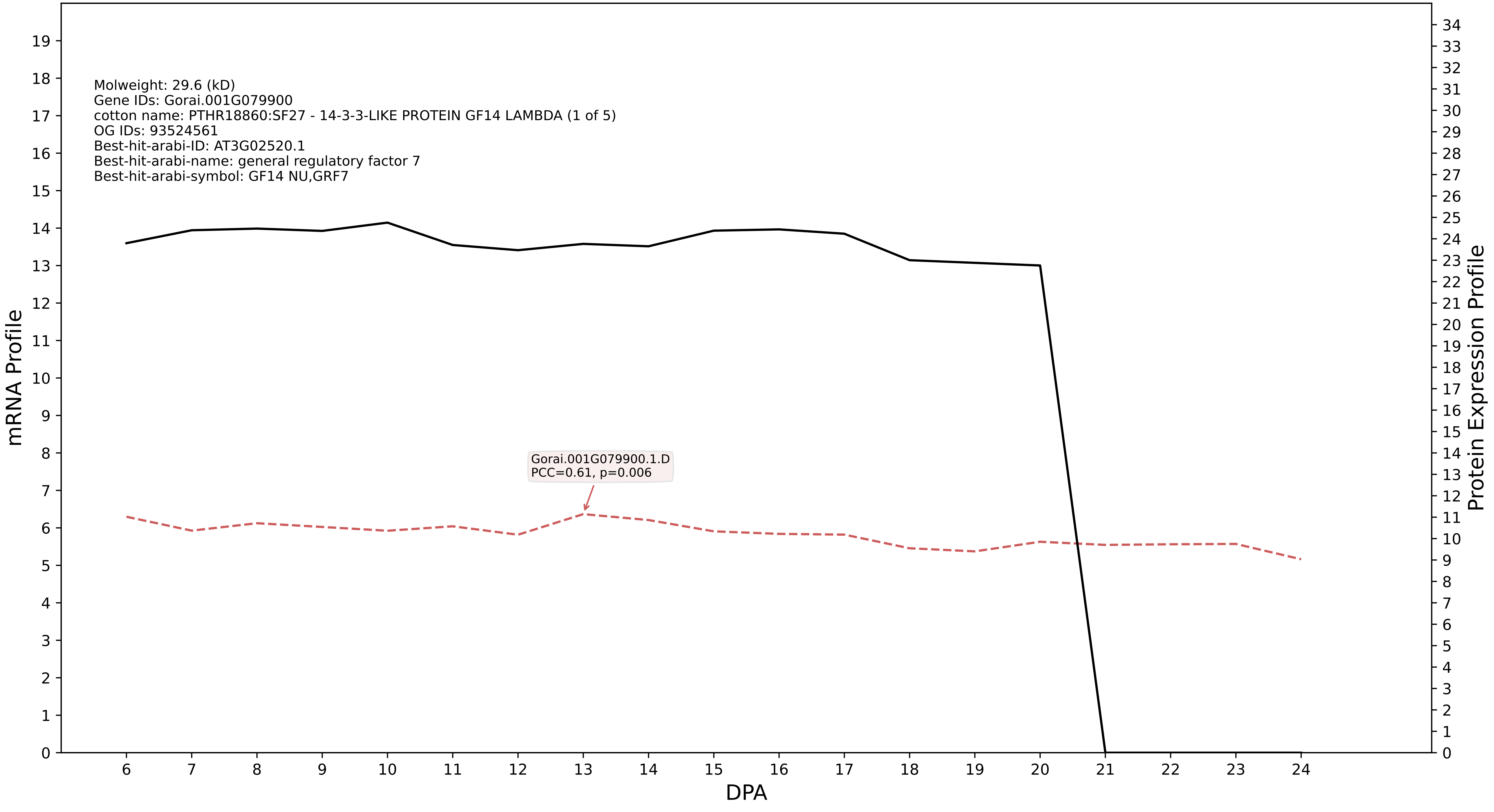

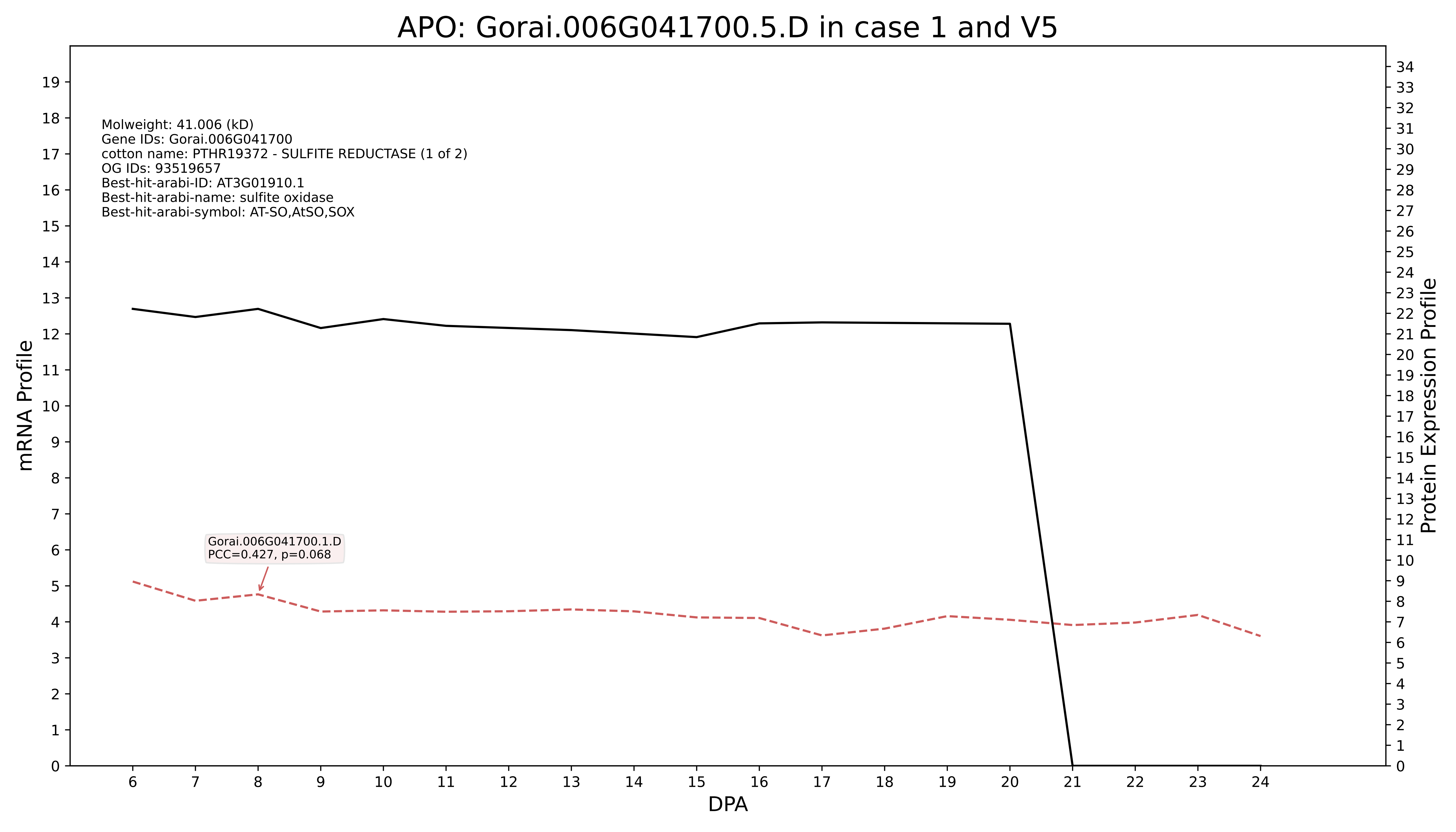

APO: Gorai.006G041700.5.D in case 1 and V5
Molweight: 41.006 (kD)
Gene IDs: Gorai.006G041700
cotton name: PTHR19372 - SULFITE REDUCTASE (1 of 2)
OG IDs: 93519657
Best-hit-arabi-ID: AT3G01910.1
Best-hit-arabi-name: sulfite oxidase
Best-hit-arabi-symbol: AT-SO,AtSO,SOX
Gorai.006G041700.1.D
PCC=0.427, p=0.068
mRNA Profile
Protein Expression Profile
DPA

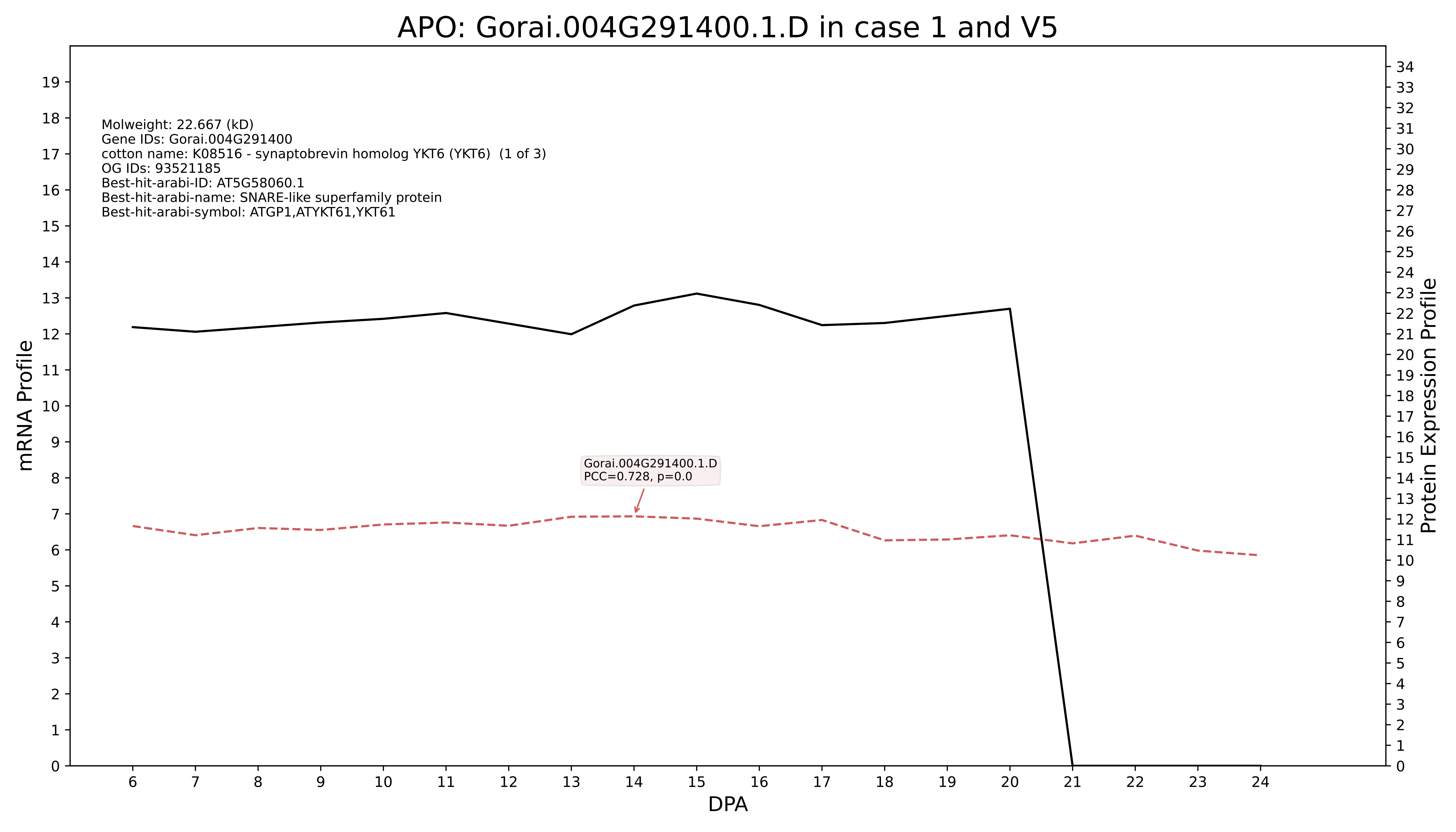

APO: Gorai.004G291400.1.D in case 1 and V5
Molweight: 22.667 (kD)
Gene IDs: Gorai.004G291400
cotton name: K08516 - synaptobrevin homolog YKT6 (YKT6) (1 of 3)
OG IDs: 93521185
Best-hit-arabi-ID: AT5G58060.1
Best-hit-arabi-name: SNARE-like superfamily protein
Best-hit-arabi-symbol: ATGP1,ATYKT61,YKT61
Gorai.004G291400.1.D
PCC=0.728, p=0.0
mRNA Profile
Protein Expression Profile
DPA

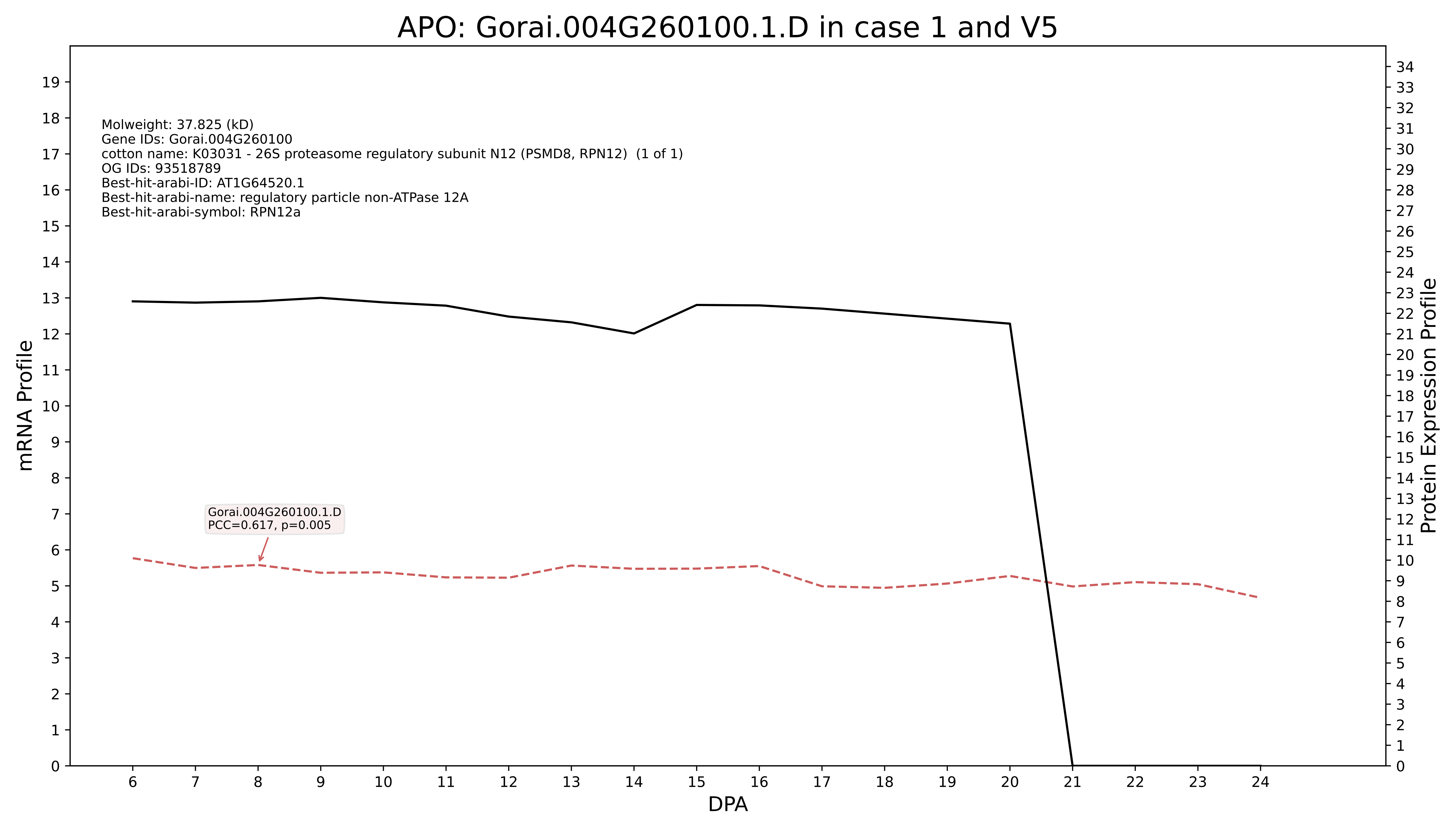

APO: Gorai.004G260100.1.D in case 1 and V5
Molweight: 37.825 (kD)
Gene IDs: Gorai.004G260100
cotton name: K03031 - 26S proteasome regulatory subunit N12 (PSMD8, RPN12) (1 of 1)
OG IDs: 93518789
Best-hit-arabi-ID: AT1G64520.1
Best-hit-arabi-name: regulatory particle non-ATPase 12A
Best-hit-arabi-symbol: RPN12a
Gorai.004G260100.1.D
PCC=0.617, p=0.005
mRNA Profile
Protein Expression Profile
DPA

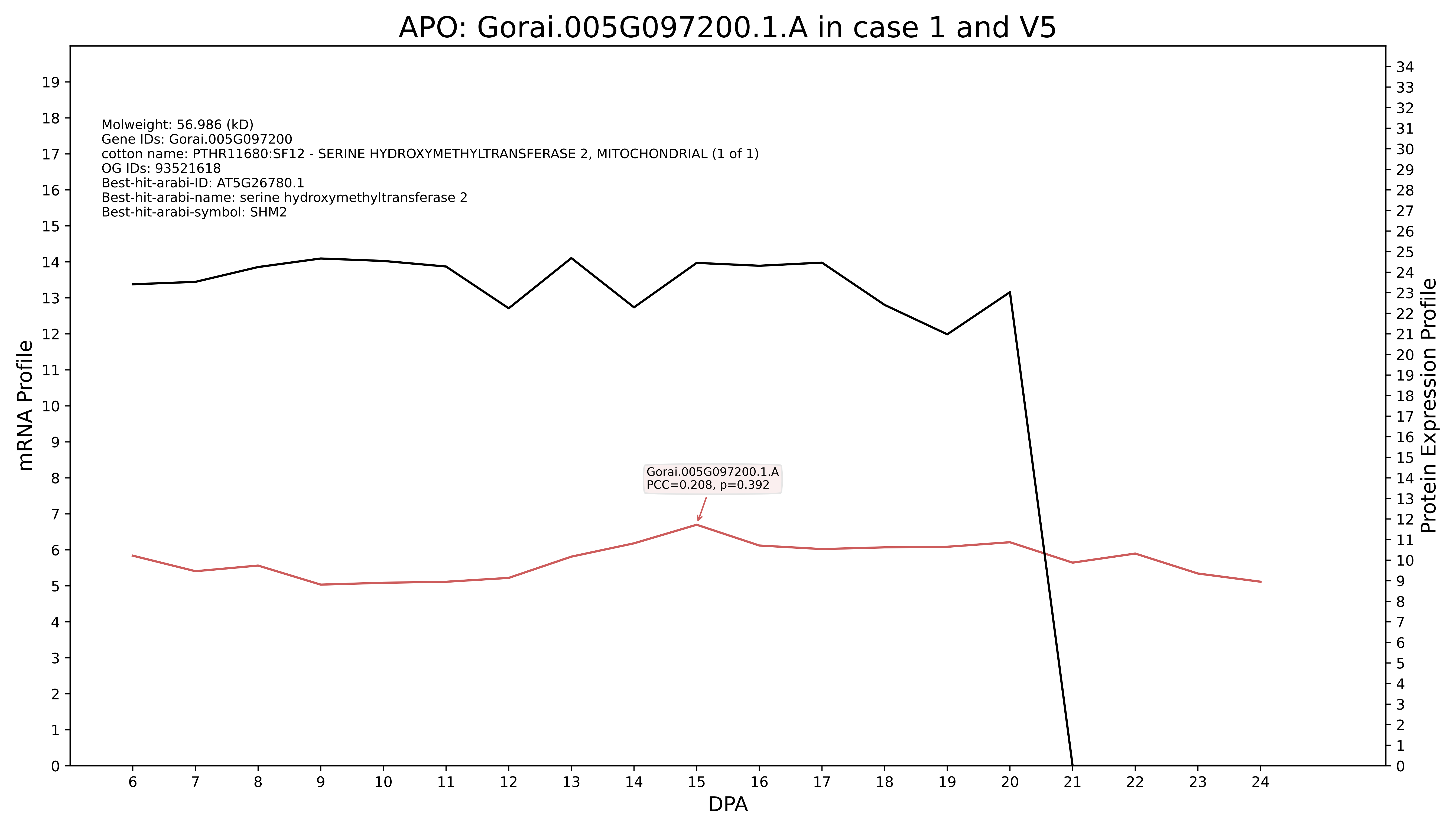
APO: Gorai.005G097200.1.A in case 1 and V5
Molweight: 56.986 (kD)
Gene IDs: Gorai.005G097200
cotton name: PTHR11680:SF12 - SERINE HYDROXYMETHYLTRANSFERASE 2, MITOCHONDRIAL (1 of 1)
OG IDs: 93521618
Best-hit-arabi-ID: AT5G26780.1
Best-hit-arabi-name: serine hydroxymethyltransferase 2
Best-hit-arabi-symbol: SHM2
Gorai.005G097200.1.A
PCC=0.208, p=0.392
mRNA Profile
Protein Expression Profile
DPA

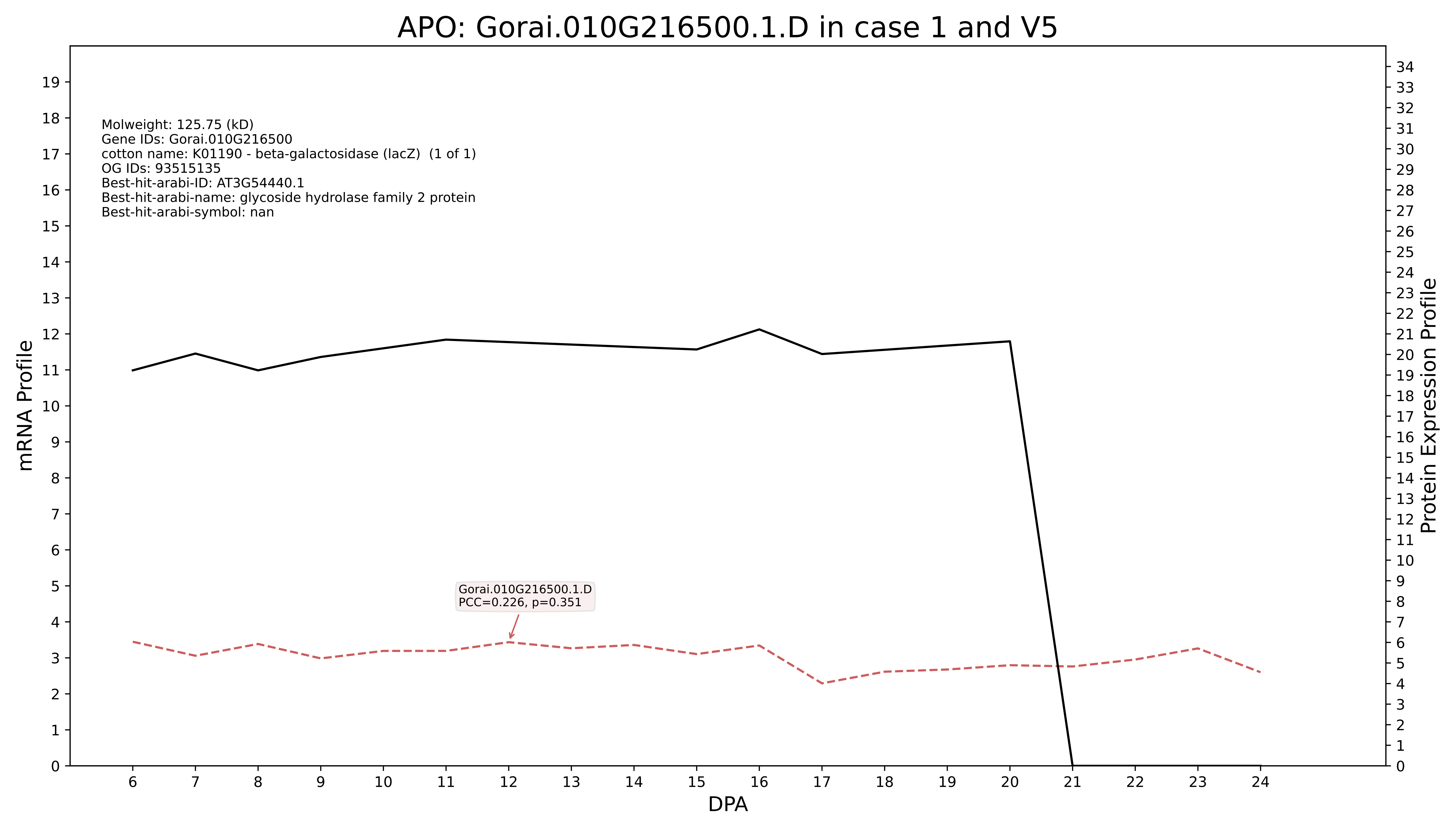

APO: Gorai.010G216500.1.D in case 1 and V5
Molweight: 125.75 (kD)
Gene IDs: Gorai.010G216500
cotton name: K01190 - beta-galactosidase (lacZ) (1 of 1)
OG IDs: 93515135
Best-hit-arabi-ID: AT3G54440.1
Best-hit-arabi-name: glycoside hydrolase family 2 protein
Best-hit-arabi-symbol: nan
Gorai.010G216500.1.D
PCC=0.226, p=0.351
mRNA Profile
Protein Expression Profile
DPA

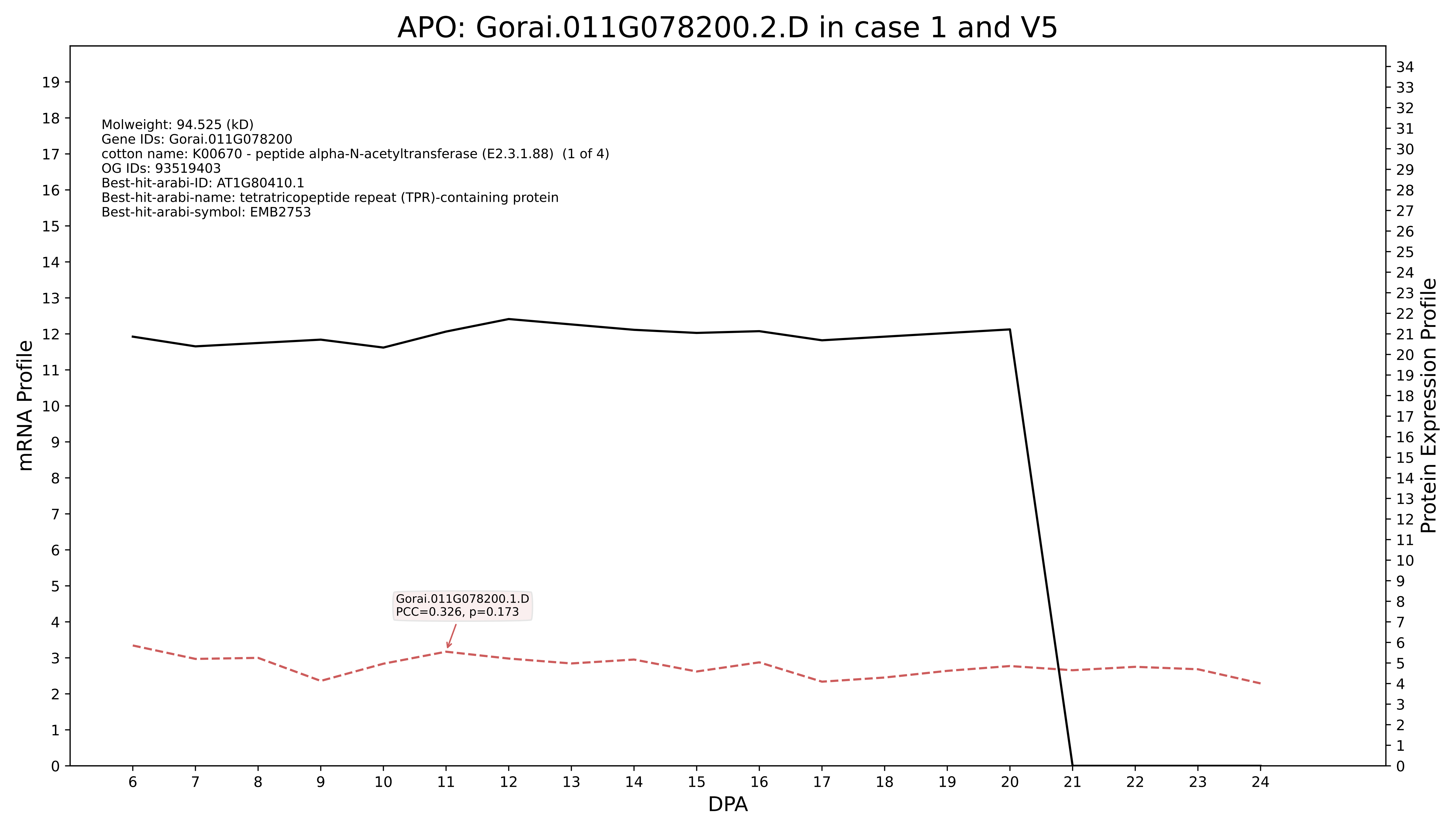
APO: Gorai.011G078200.2.D in case 1 and V5
Molweight: 94.525 (kD)
Gene IDs: Gorai.011G078200
cotton name: K00670 - peptide alpha-N-acetyltransferase (E2.3.1.88) (1 of 4)
OG IDs: 93519403
Best-hit-arabi-ID: AT1G80410.1
Best-hit-arabi-name: tetratricopeptide repeat (TPR)-containing protein
Best-hit-arabi-symbol: EMB2753
Gorai.011G078200.1.D
PCC=0.326, p=0.173
mRNA Profile
Protein Expression Profile
DPA

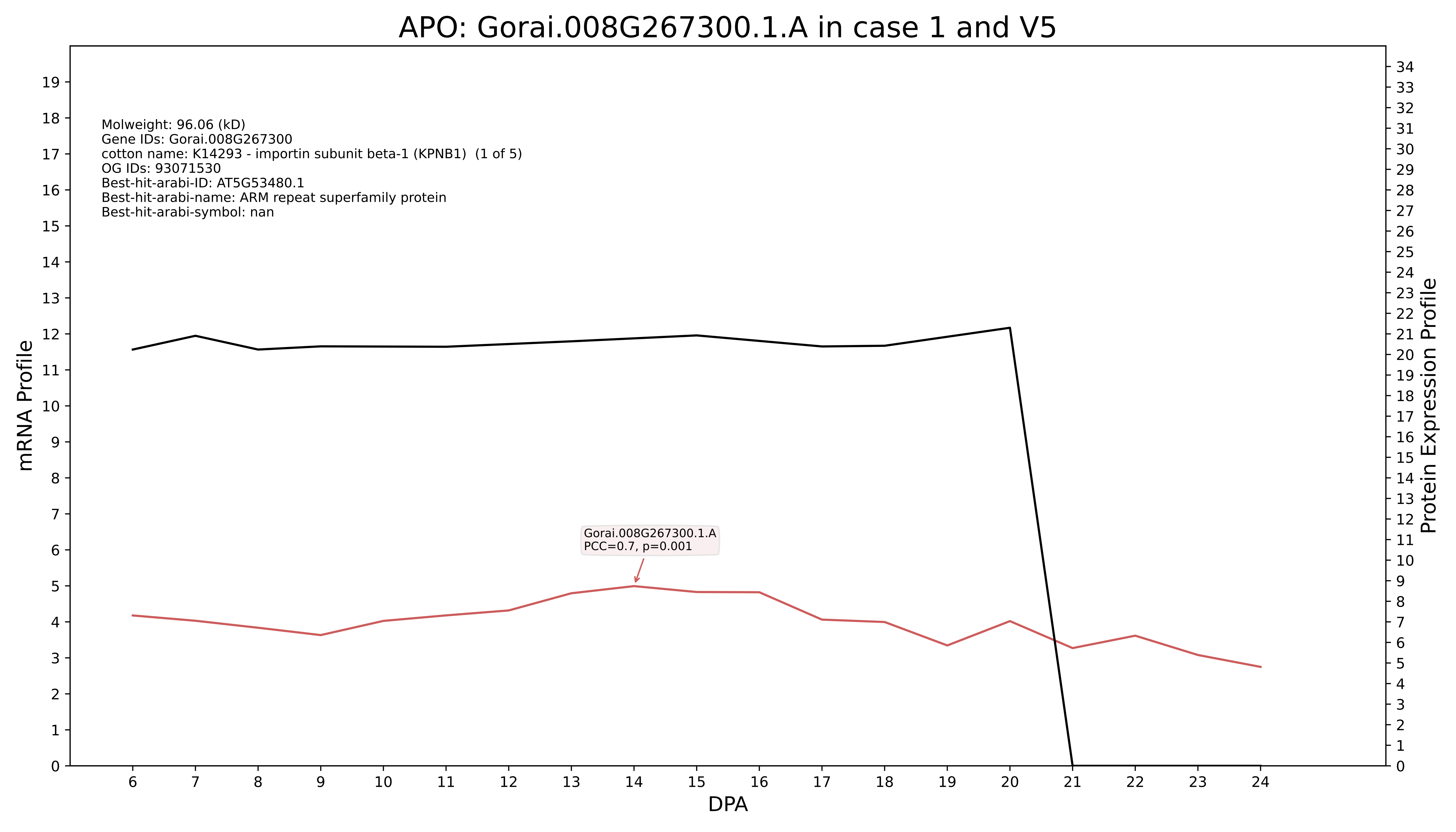
APO: Gorai.008G267300.1.A in case 1 and V5
Molweight: 96.06 (kD)
Gene IDs: Gorai.008G267300
cotton name: K14293 - importin subunit beta-1 (KPNB1) (1 of 5)
OG IDs: 93071530
Best-hit-arabi-ID: AT5G53480.1
Best-hit-arabi-name: ARM repeat superfamily protein
Best-hit-arabi-symbol: nan
Gorai.008G267300.1.A
PCC=0.7, p=0.001
mRNA Profile
Protein Expression Profile
DPA

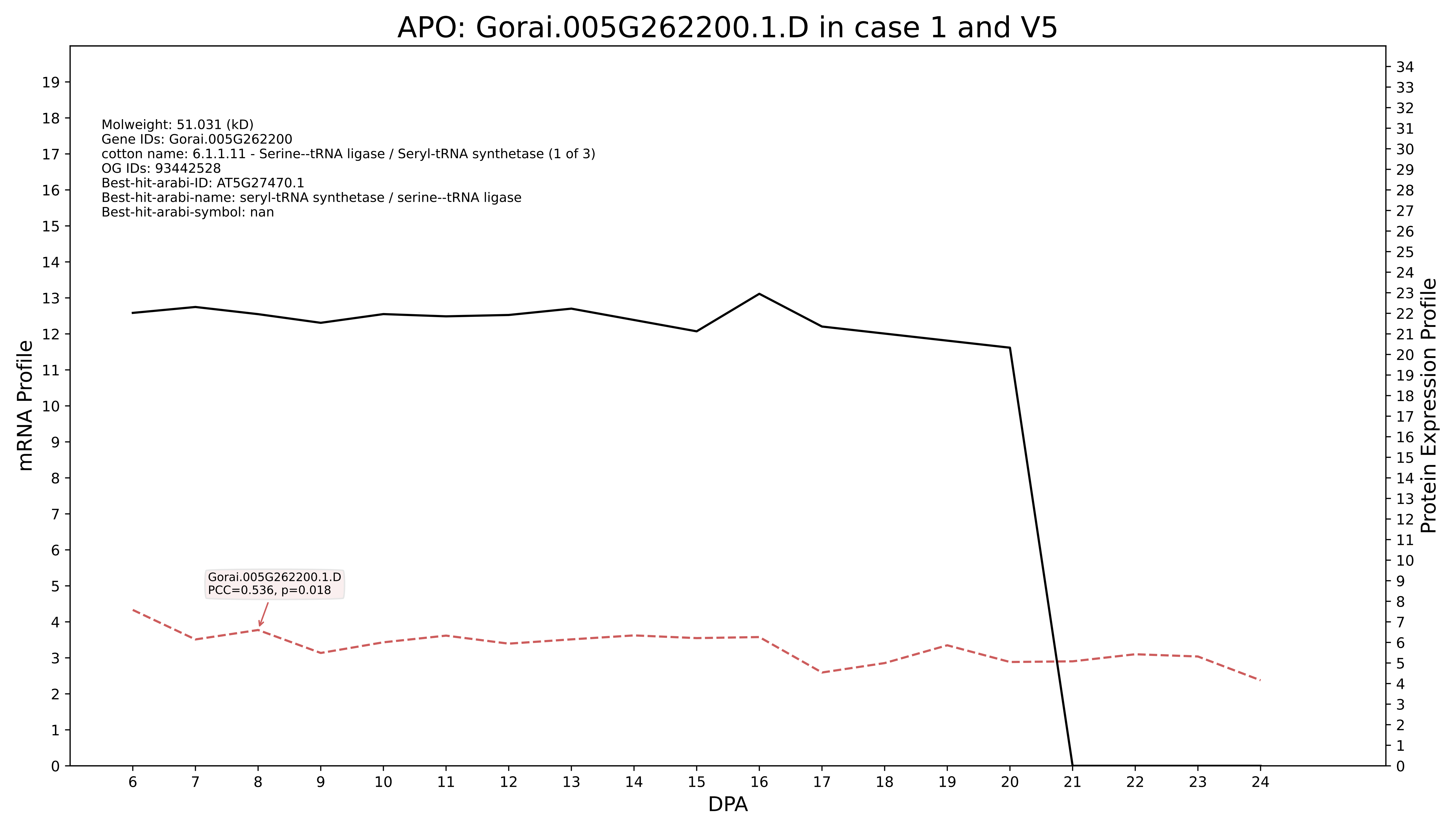
APO: Gorai.005G262200.1.D in case 1 and V5
Molweight: 51.031 (kD)
Gene IDs: Gorai.005G262200
cotton name: 6.1.1.11 - Serine--tRNA ligase / Seryl-tRNA synthetase (1 of 3)
OG IDs: 93442528
Best-hit-arabi-ID: AT5G27470.1
Best-hit-arabi-name: seryl-tRNA synthetase / serine--tRNA ligase
Best-hit-arabi-symbol: nan
Gorai.005G262200.1.D
PCC=0.536, p=0.018
mRNA Profile
Protein Expression Profile
DPA

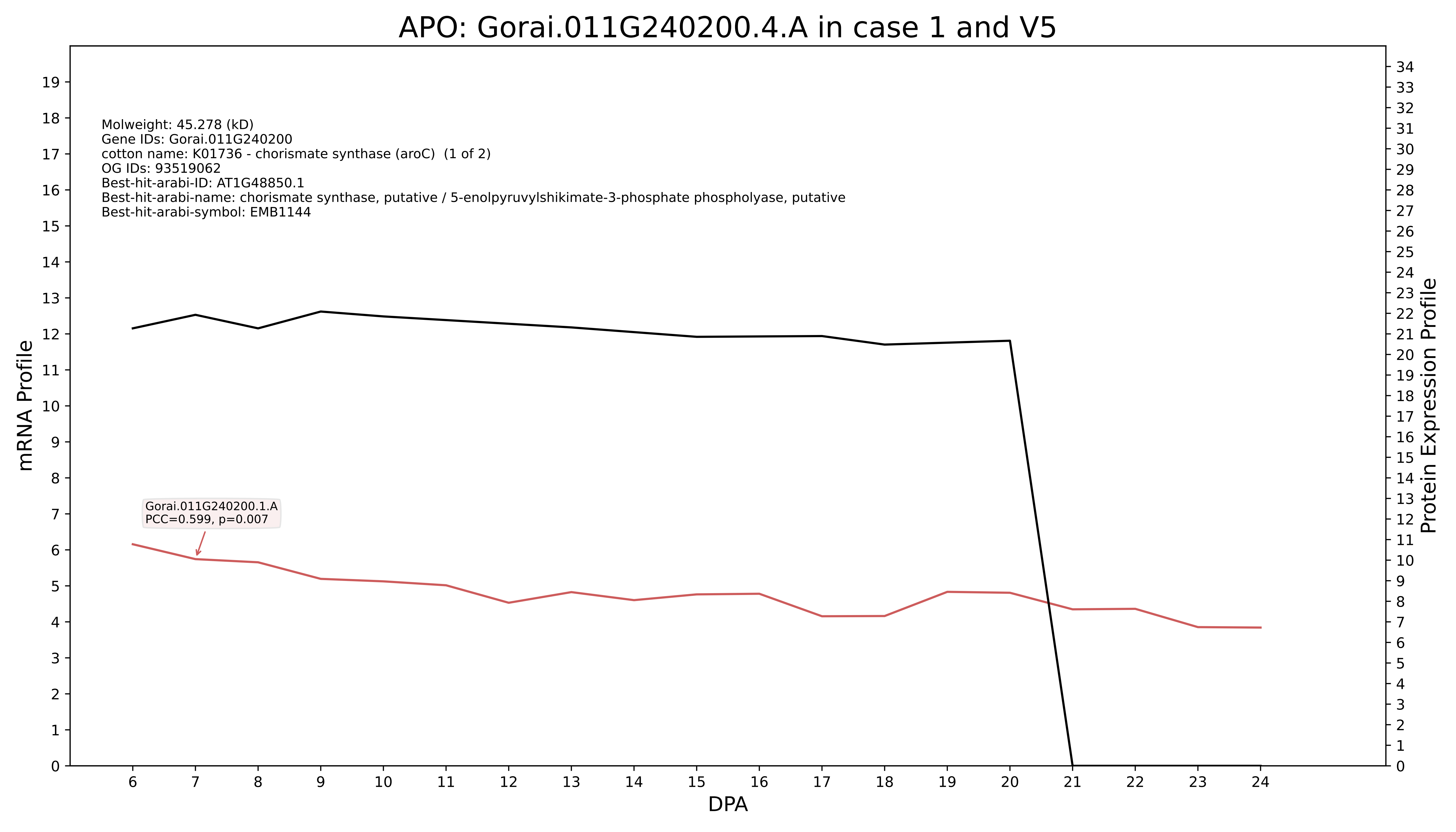
APO: Gorai.011G240200.4.A in case 1 and V5
Molweight: 45.278 (kD)
Gene IDs: Gorai.011G240200
cotton name: K01736 - chorismate synthase (aroC) (1 of 2)
OG IDs: 93519062
Best-hit-arabi-ID: AT1G48850.1
Best-hit-arabi-name: chorismate synthase, putative / 5-enolpyruvylshikimate-3-phosphate phospholyase, putative
Best-hit-arabi-symbol: EMB1144
Gorai.011G240200.1.A
PCC=0.599, p=0.007
mRNA Profile
Protein Expression Profile
DPA

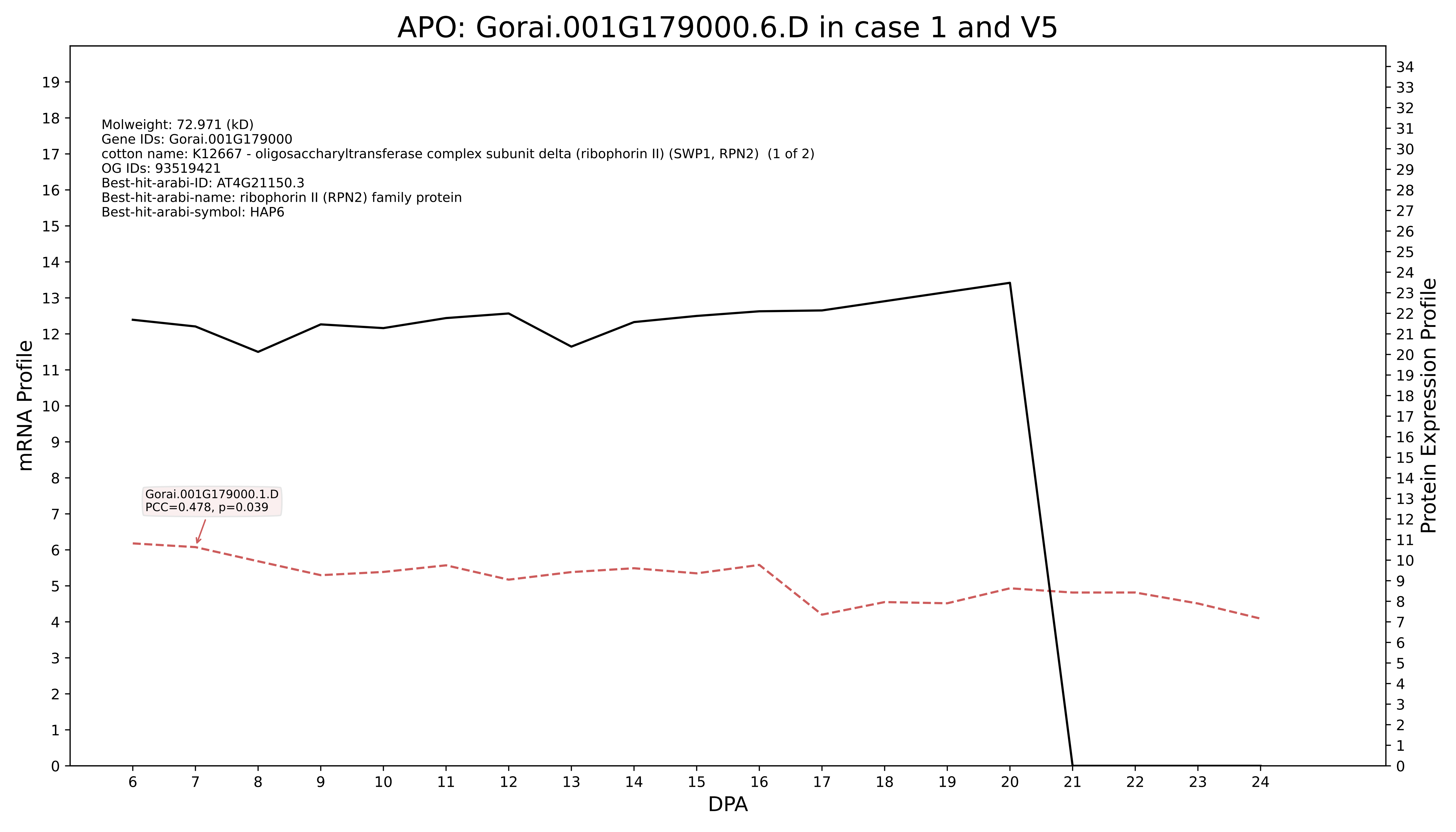
APO: Gorai.001G179000.6.D in case 1 and V5
Molweight: 72.971 (kD)
Gene IDs: Gorai.001G179000
cotton name: K12667 - oligosaccharyltransferase complex subunit delta (ribophorin II) (SWP1, RPN2) (1 of 2)
OG IDs: 93519421
Best-hit-arabi-ID: AT4G21150.3
Best-hit-arabi-name: ribophorin II (RPN2) family protein
Best-hit-arabi-symbol: HAP6
Gorai.001G179000.1.D
PCC=0.478, p=0.039
mRNA Profile
Protein Expression Profile
DPA

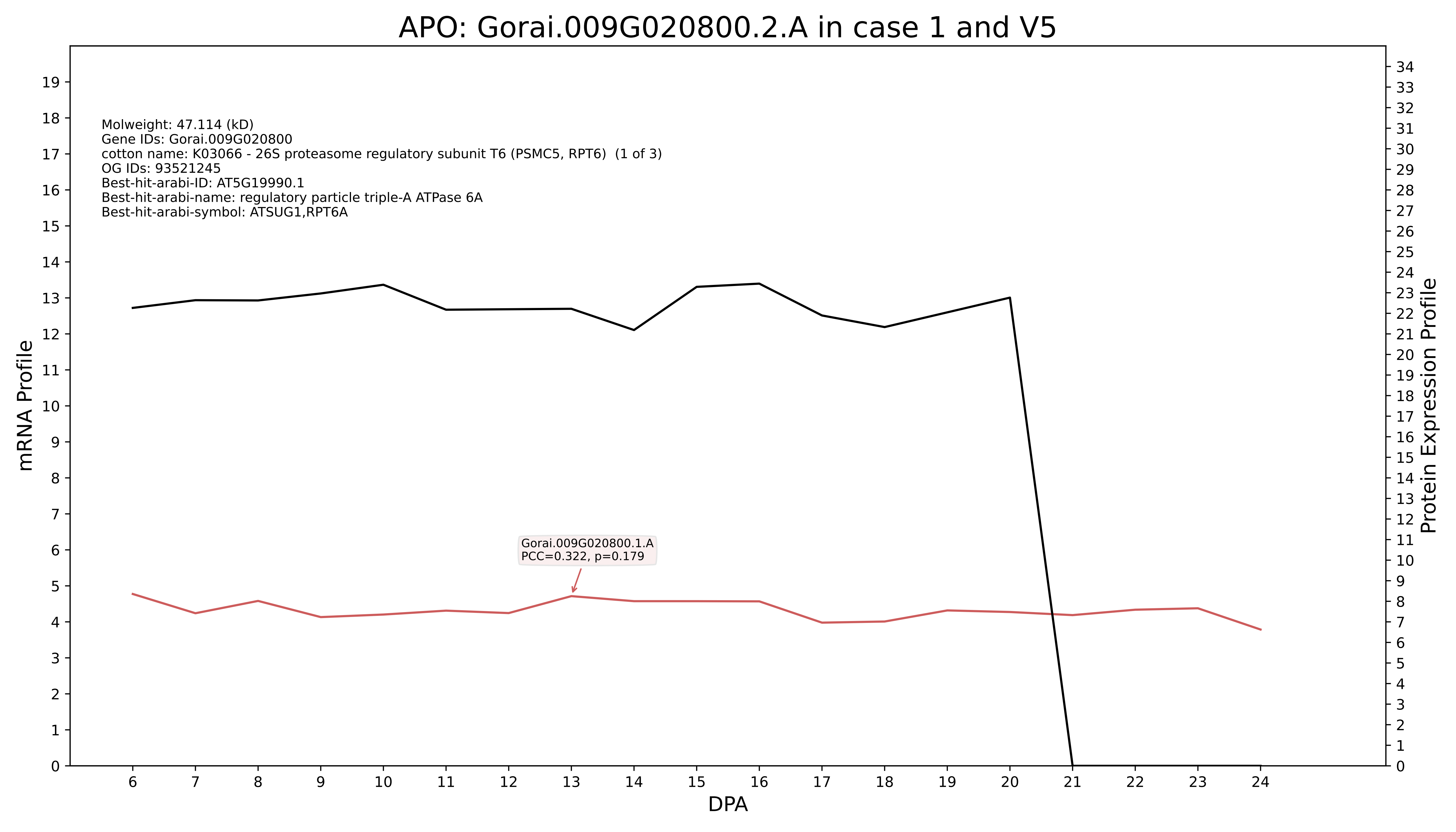

APO: Gorai.009G020800.2.A in case 1 and V5
Molweight: 47.114 (kD)
Gene IDs: Gorai.009G020800
cotton name: K03066 - 26S proteasome regulatory subunit T6 (PSMC5, RPT6) (1 of 3)
OG IDs: 93521245
Best-hit-arabi-ID: AT5G19990.1
Best-hit-arabi-name: regulatory particle triple-A ATPase 6A
Best-hit-arabi-symbol: ATSUG1,RPT6A
Gorai.009G020800.1.A
PCC=0.322, p=0.179
mRNA Profile
Protein Expression Profile
DPA

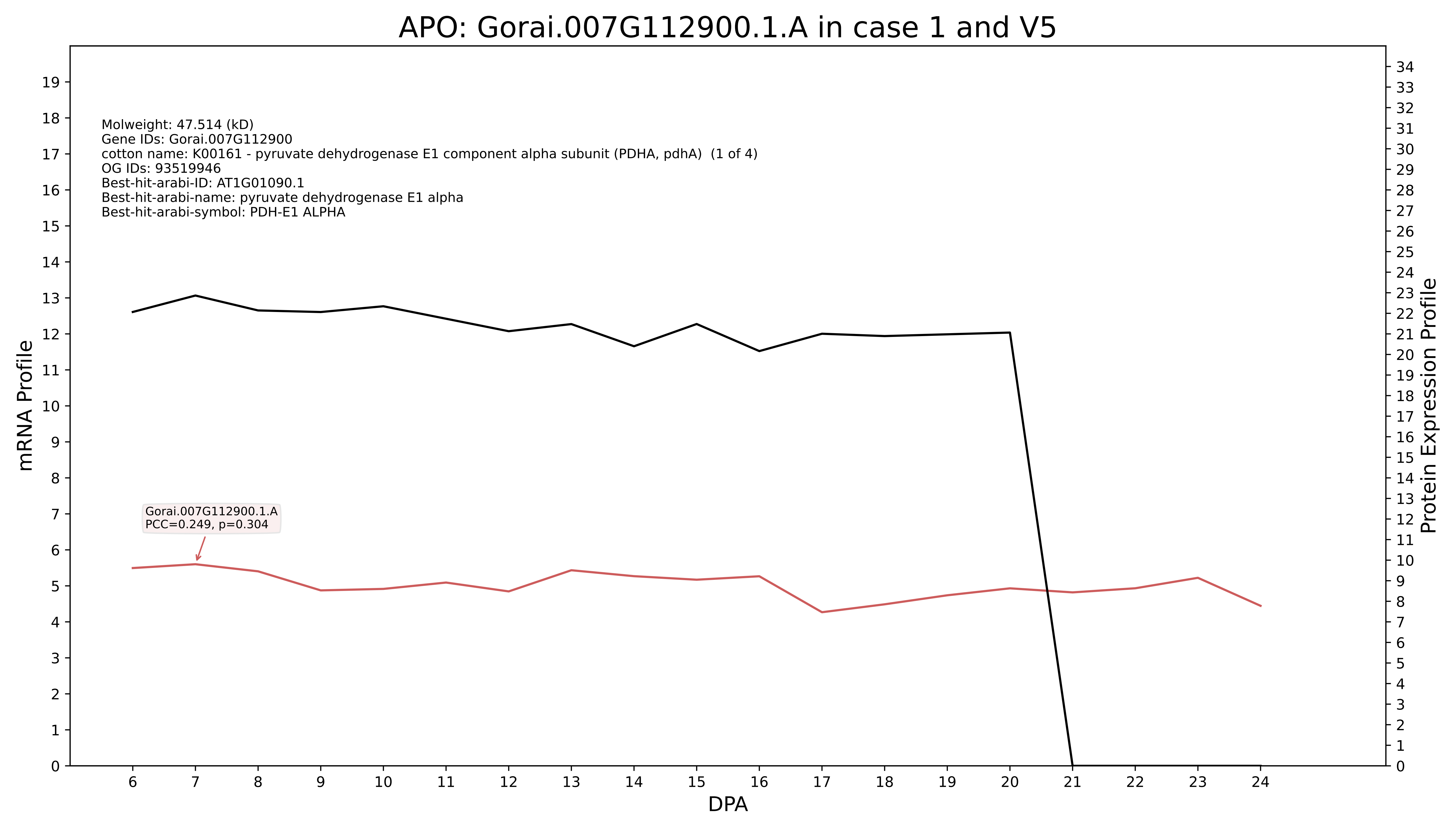
APO: Gorai.007G112900.1.A in case 1 and V5
Molweight: 47.514 (kD)
Gene IDs: Gorai.007G112900
cotton name: K00161 - pyruvate dehydrogenase E1 component alpha subunit (PDHA, pdhA) (1 of 4)
OG IDs: 93519946
Best-hit-arabi-ID: AT1G01090.1
Best-hit-arabi-name: pyruvate dehydrogenase E1 alpha
Best-hit-arabi-symbol: PDH-E1 ALPHA
Gorai.007G112900.1.A
PCC=0.249, p=0.304
mRNA Profile
Protein Expression Profile
DPA

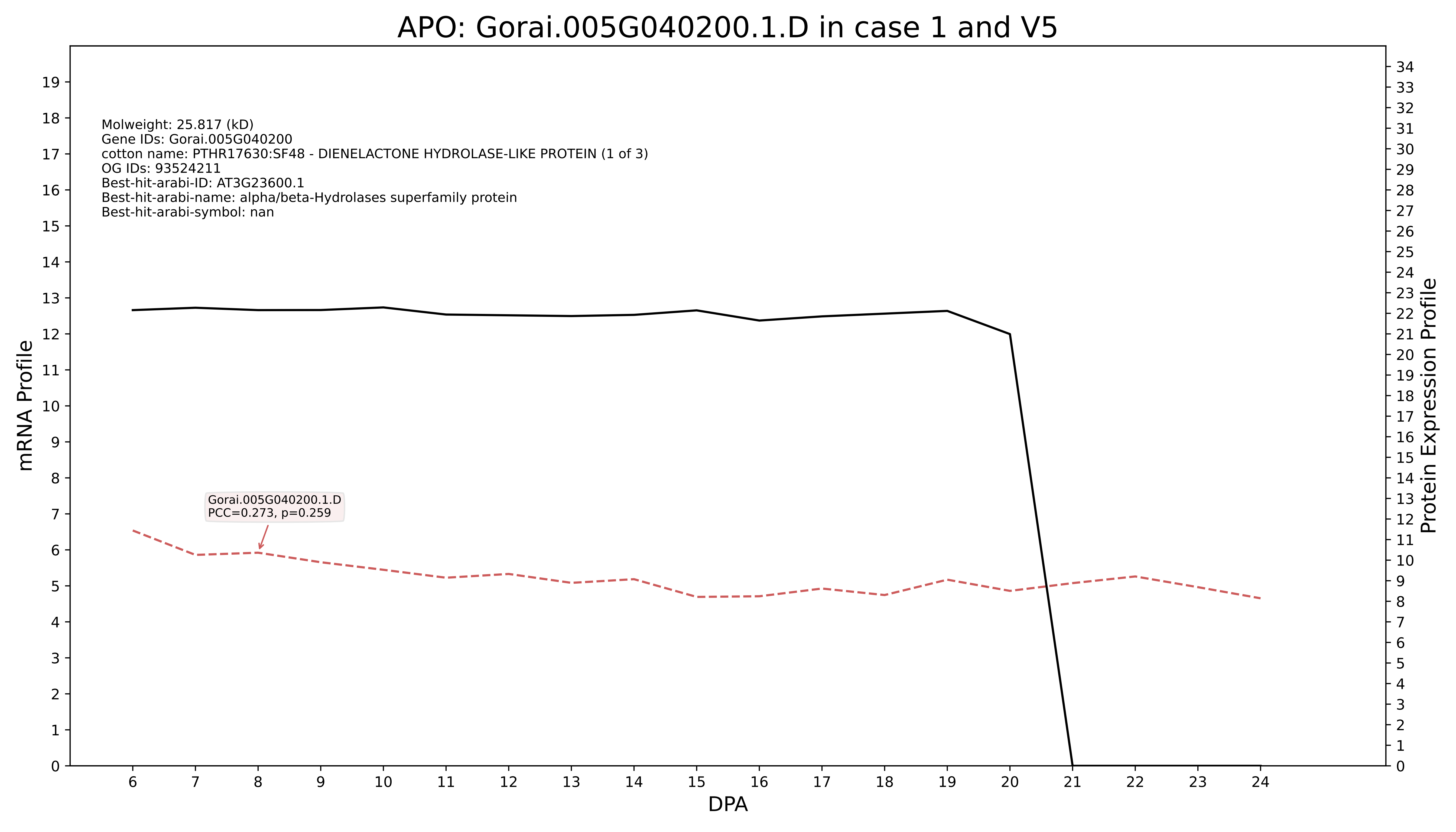
APO: Gorai.005G040200.1.D in case 1 and V5
Molweight: 25.817 (kD)
Gene IDs: Gorai.005G040200
cotton name: PTHR17630:SF48 - DIENELACTONE HYDROLASE-LIKE PROTEIN (1 of 3)
OG IDs: 93524211
Best-hit-arabi-ID: AT3G23600.1
Best-hit-arabi-name: alpha/beta-Hydrolases superfamily protein
Best-hit-arabi-symbol: nan
Gorai.005G040200.1.D
PCC=0.273, p=0.259
mRNA Profile
Protein Expression Profile
DPA

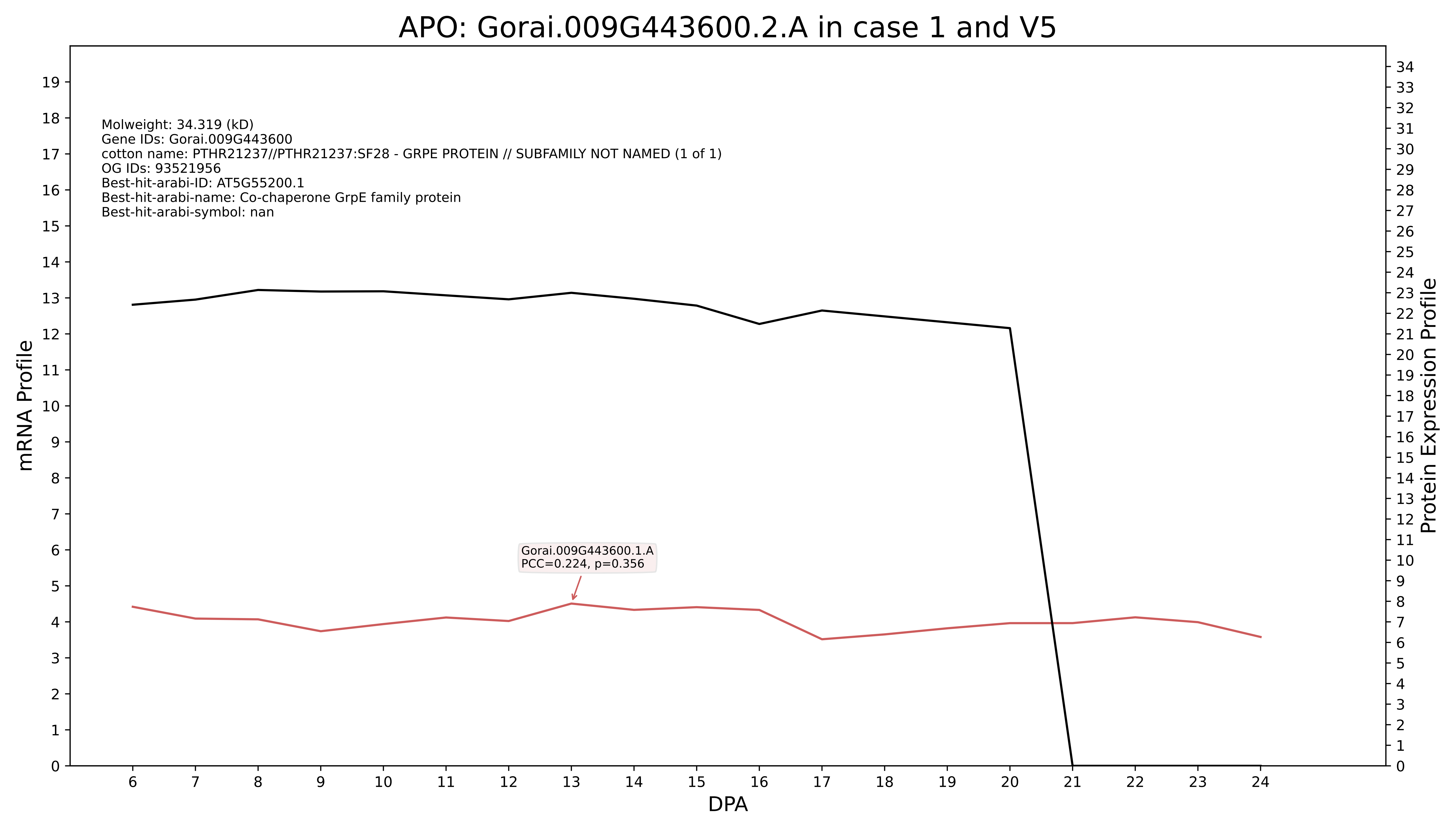
APO: Gorai.009G443600.2.A in case 1 and V5
Molweight: 34.319 (kD)
Gene IDs: Gorai.009G443600
cotton name: PTHR21237//PTHR21237:SF28 - GRPE PROTEIN // SUBFAMILY NOT NAMED (1 of 1)
OG IDs: 93521956
Best-hit-arabi-ID: AT5G55200.1
Best-hit-arabi-name: Co-chaperone GrpE family protein
Best-hit-arabi-symbol: nan
Gorai.009G443600.1.A
PCC=0.224, p=0.356
mRNA Profile
Protein Expression Profile
DPA

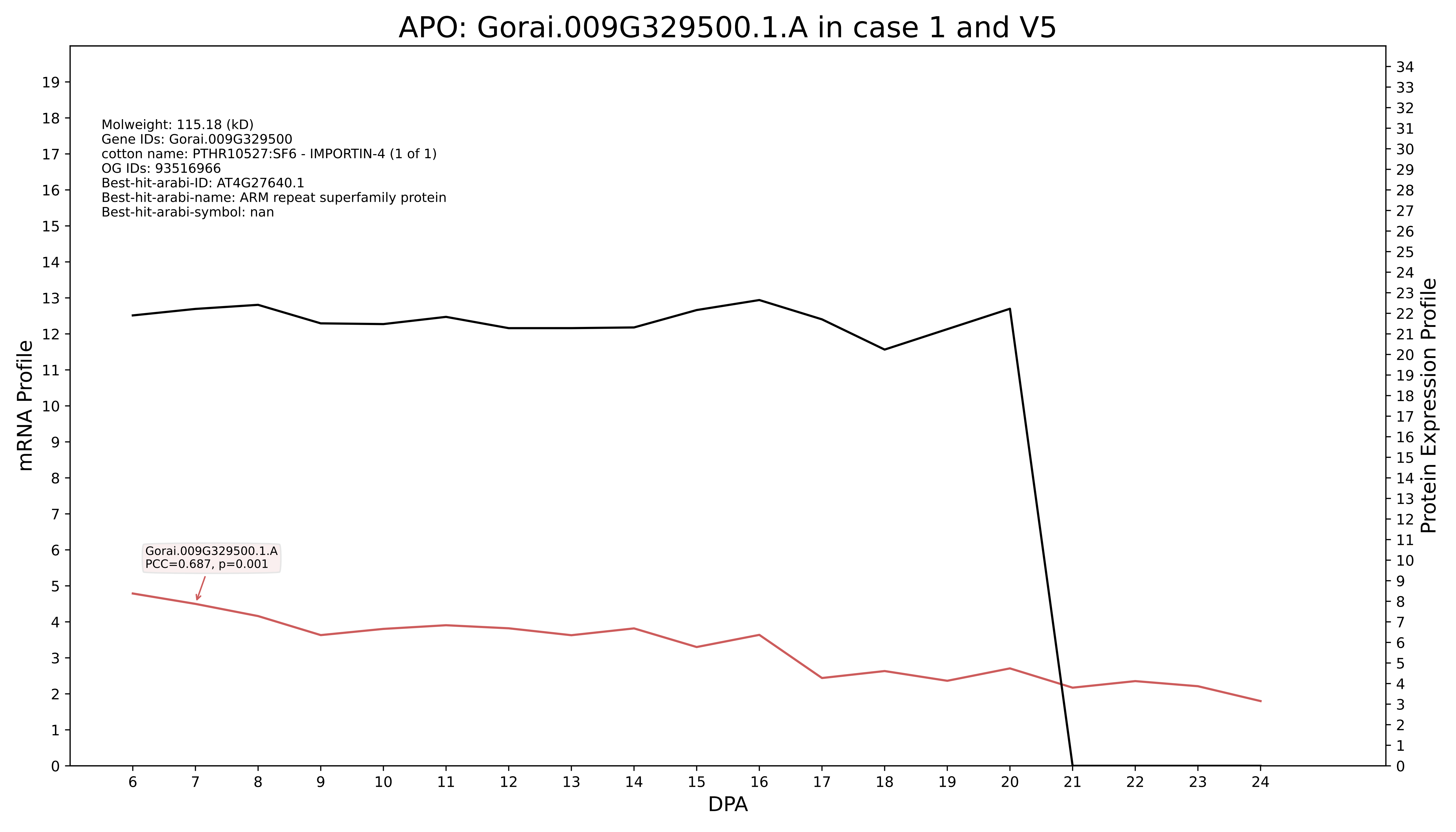
APO: Gorai.009G329500.1.A in case 1 and V5
Molweight: 115.18 (kD)
Gene IDs: Gorai.009G329500
cotton name: PTHR10527:SF6 - IMPORTIN-4 (1 of 1)
OG IDs: 93516966
Best-hit-arabi-ID: AT4G27640.1
Best-hit-arabi-name: ARM repeat superfamily protein
Best-hit-arabi-symbol: nan
Gorai.009G329500.1.A
PCC=0.687, p=0.001
mRNA Profile
Protein Expression Profile
DPA

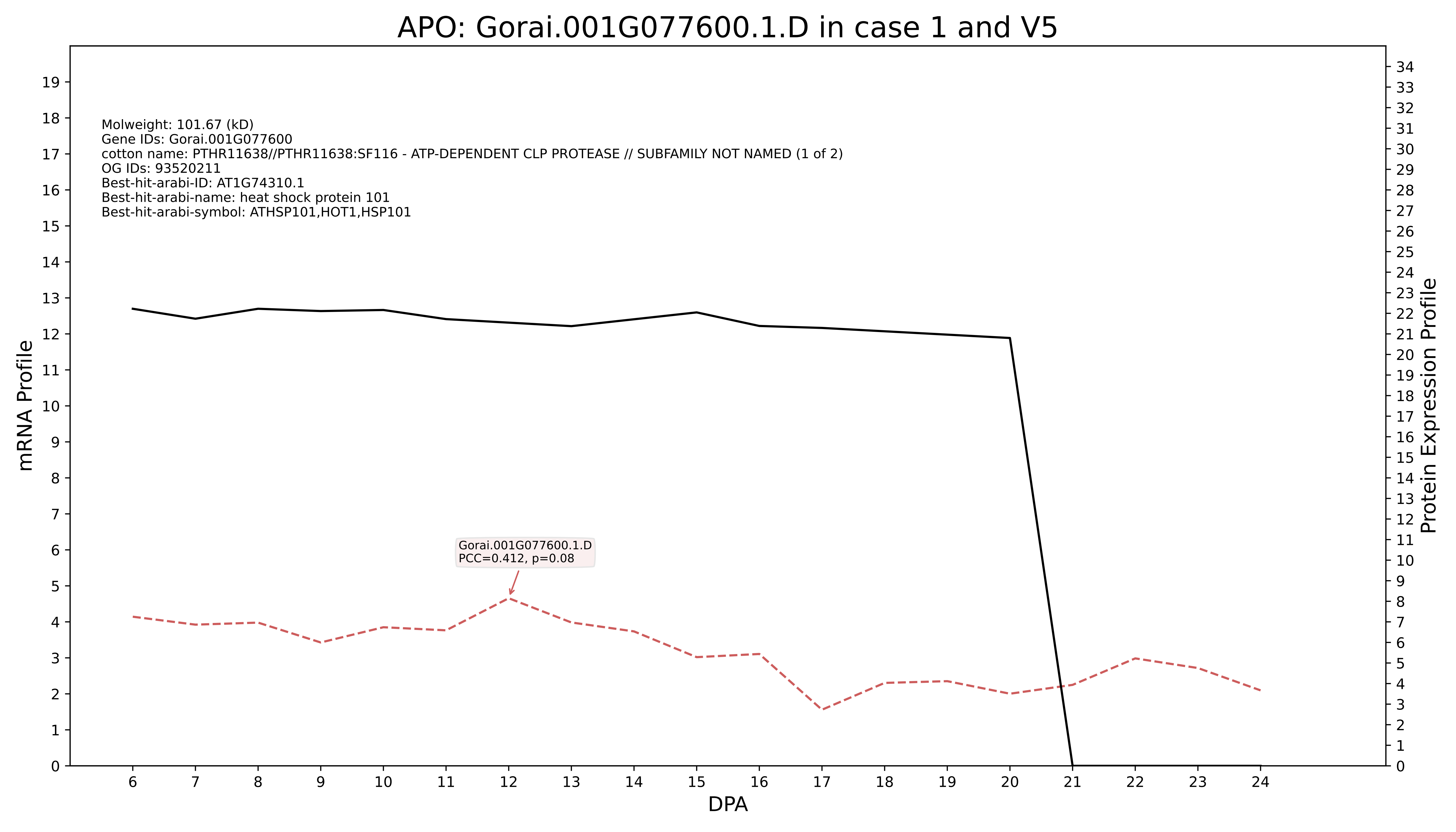
APO: Gorai.001G077600.1.D in case 1 and V5
Molweight: 101.67 (kD)
Gene IDs: Gorai.001G077600
cotton name: PTHR11638//PTHR11638:SF116 - ATP-DEPENDENT CLP PROTEASE // SUBFAMILY NOT NAMED (1 of 2)
OG IDs: 93520211
Best-hit-arabi-ID: AT1G74310.1
Best-hit-arabi-name: heat shock protein 101
Best-hit-arabi-symbol: ATHSP101,HOT1,HSP101
Gorai.001G077600.1.D
PCC=0.412, p=0.08
mRNA Profile
Protein Expression Profile
DPA

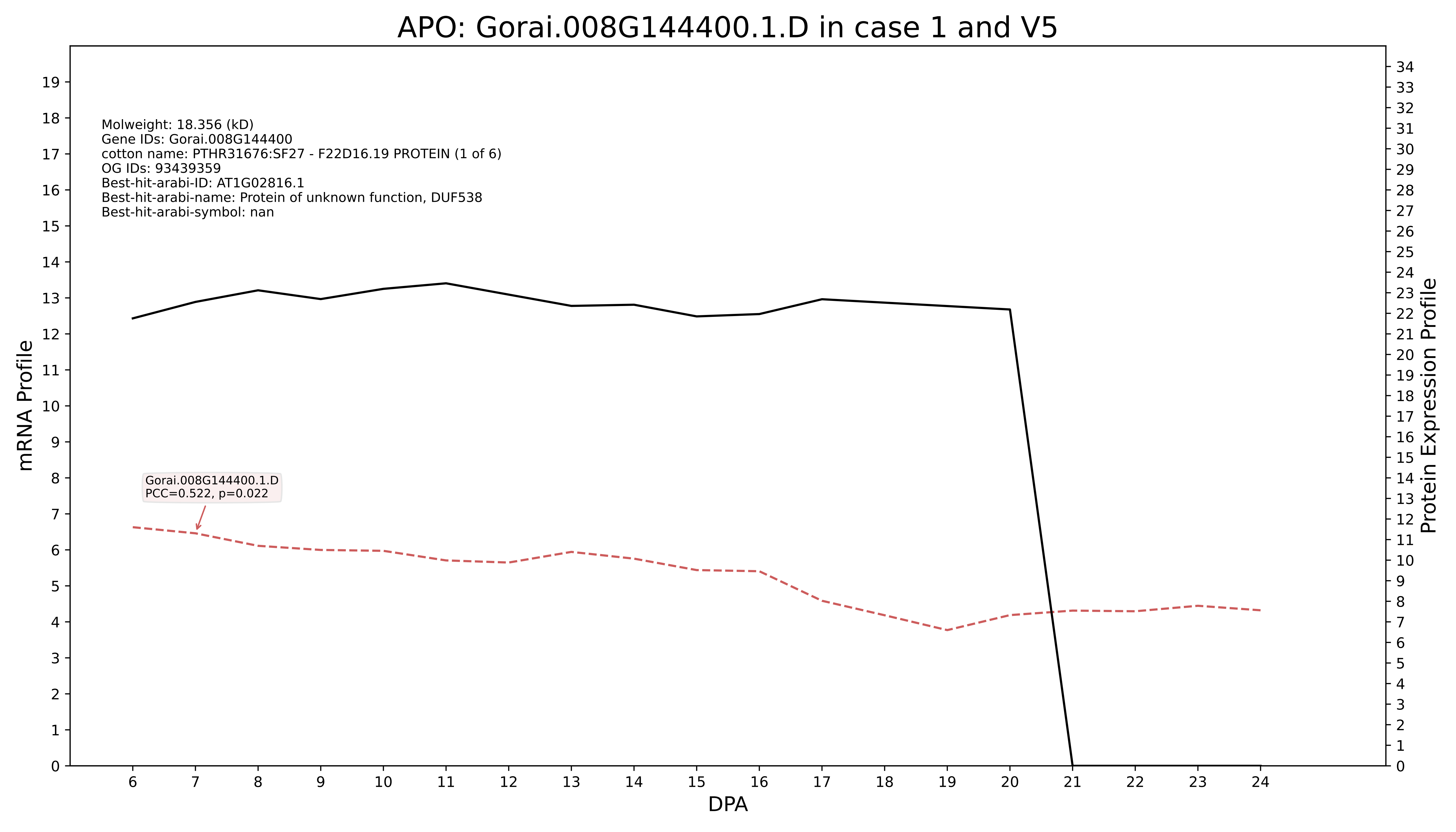
APO: Gorai.008G144400.1.D in case 1 and V5
Molweight: 18.356 (kD)
Gene IDs: Gorai.008G144400
cotton name: PTHR31676:SF27 - F22D16.19 PROTEIN (1 of 6)
OG IDs: 93439359
Best-hit-arabi-ID: AT1G02816.1
Best-hit-arabi-name: Protein of unknown function, DUF538
Best-hit-arabi-symbol: nan
Gorai.008G144400.1.D
PCC=0.522, p=0.022
mRNA Profile
Protein Expression Profile
DPA

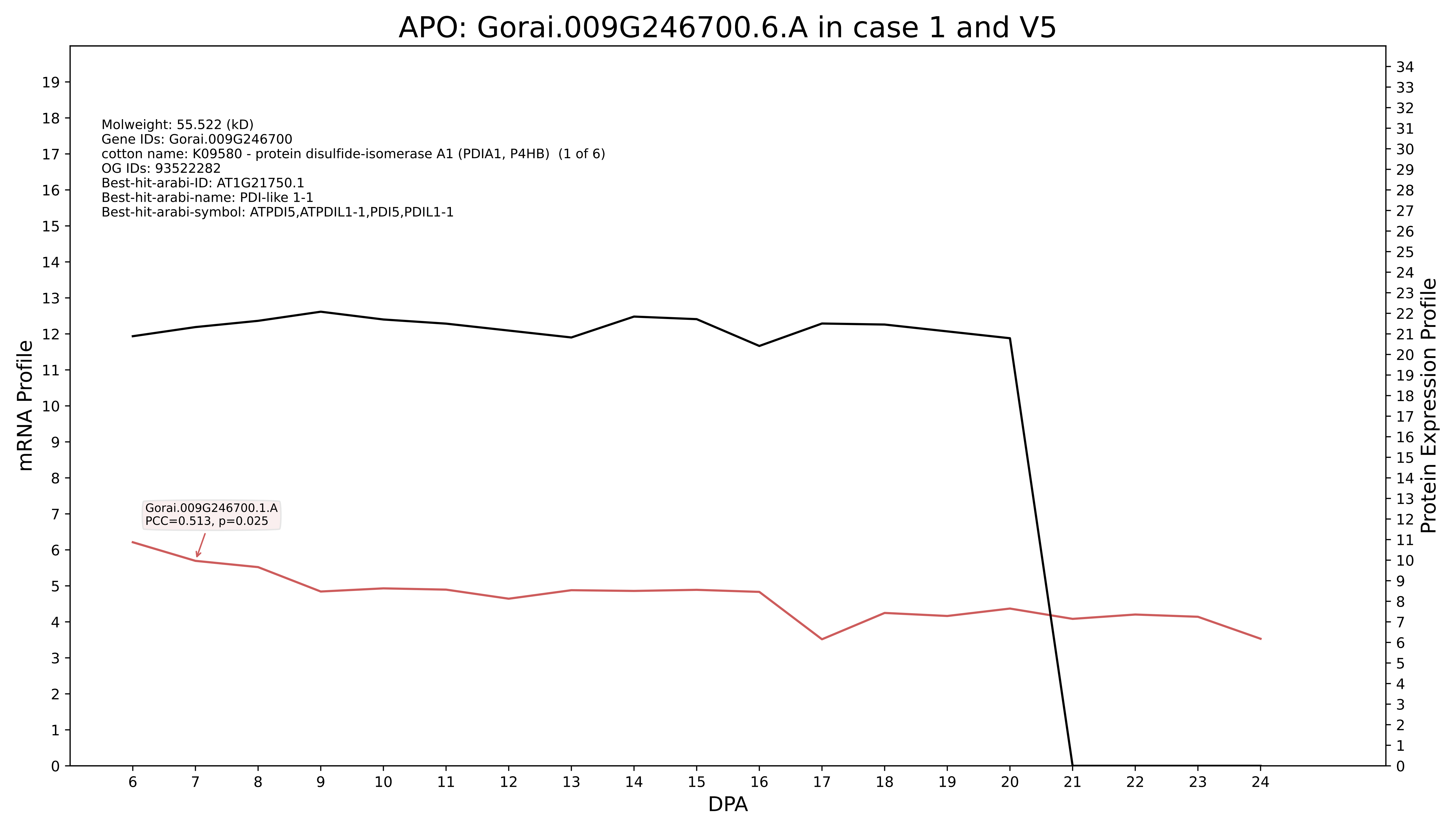
APO: Gorai.009G246700.6.A in case 1 and V5
Molweight: 55.522 (kD)
Gene IDs: Gorai.009G246700
cotton name: K09580 - protein disulfide-isomerase A1 (PDIA1, P4HB) (1 of 6)
OG IDs: 93522282
Best-hit-arabi-ID: AT1G21750.1
Best-hit-arabi-name: PDI-like 1-1
Best-hit-arabi-symbol: ATPDI5,ATPDIL1-1,PDI5,PDIL1-1
Gorai.009G246700.1.A
PCC=0.513, p=0.025
mRNA Profile
Protein Expression Profile
DPA

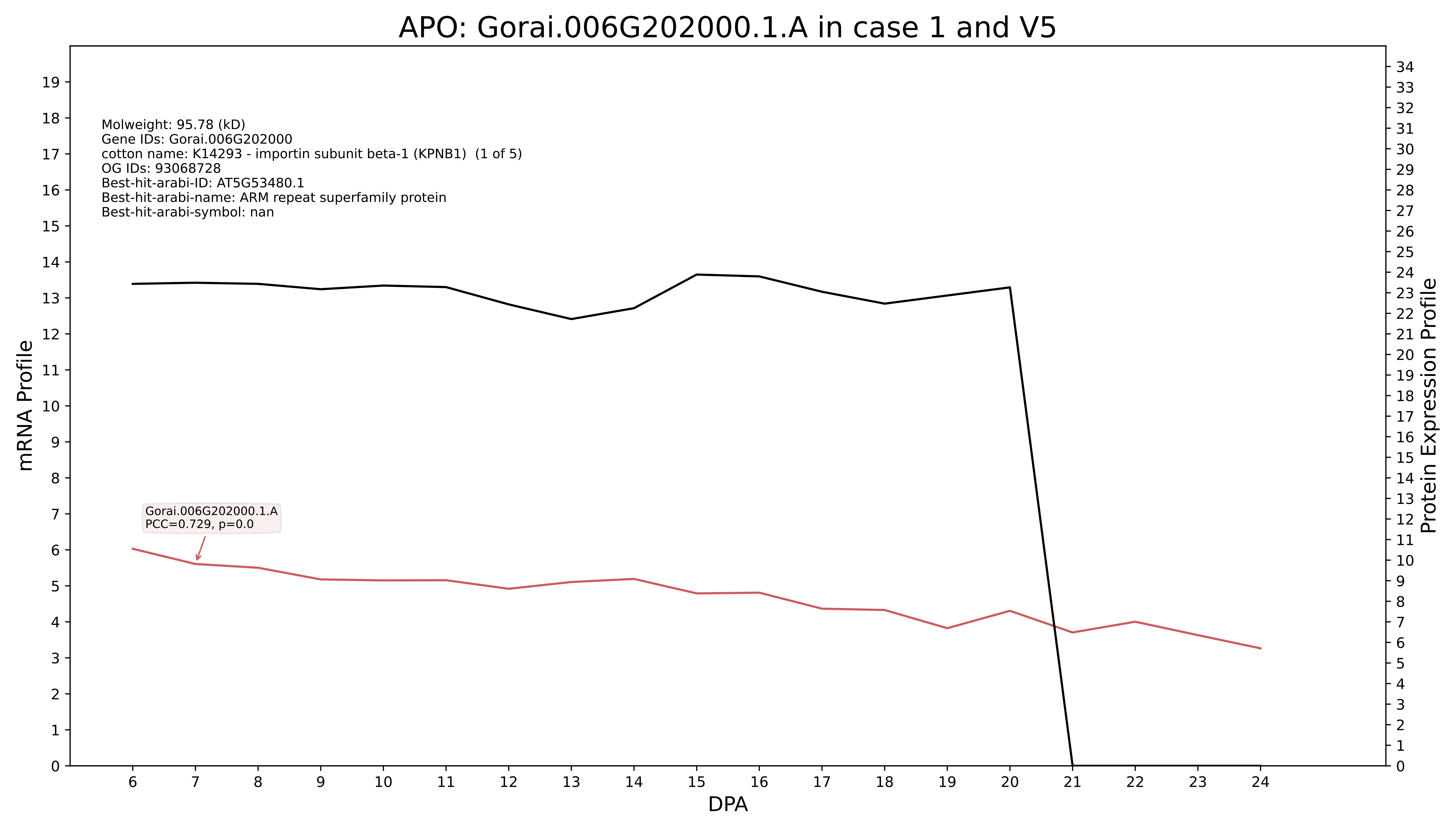
APO: Gorai.006G202000.1.A in case 1 and V5
Molweight: 95.78 (kD)
Gene IDs: Gorai.006G202000
cotton name: K14293 - importin subunit beta-1 (KPNB1) (1 of 5)
OG IDs: 93068728
Best-hit-arabi-ID: AT5G53480.1
Best-hit-arabi-name: ARM repeat superfamily protein
Best-hit-arabi-symbol: nan
Gorai.006G202000.1.A
PCC=0.729, p=0.0
mRNA Profile
Protein Expression Profile
DPA

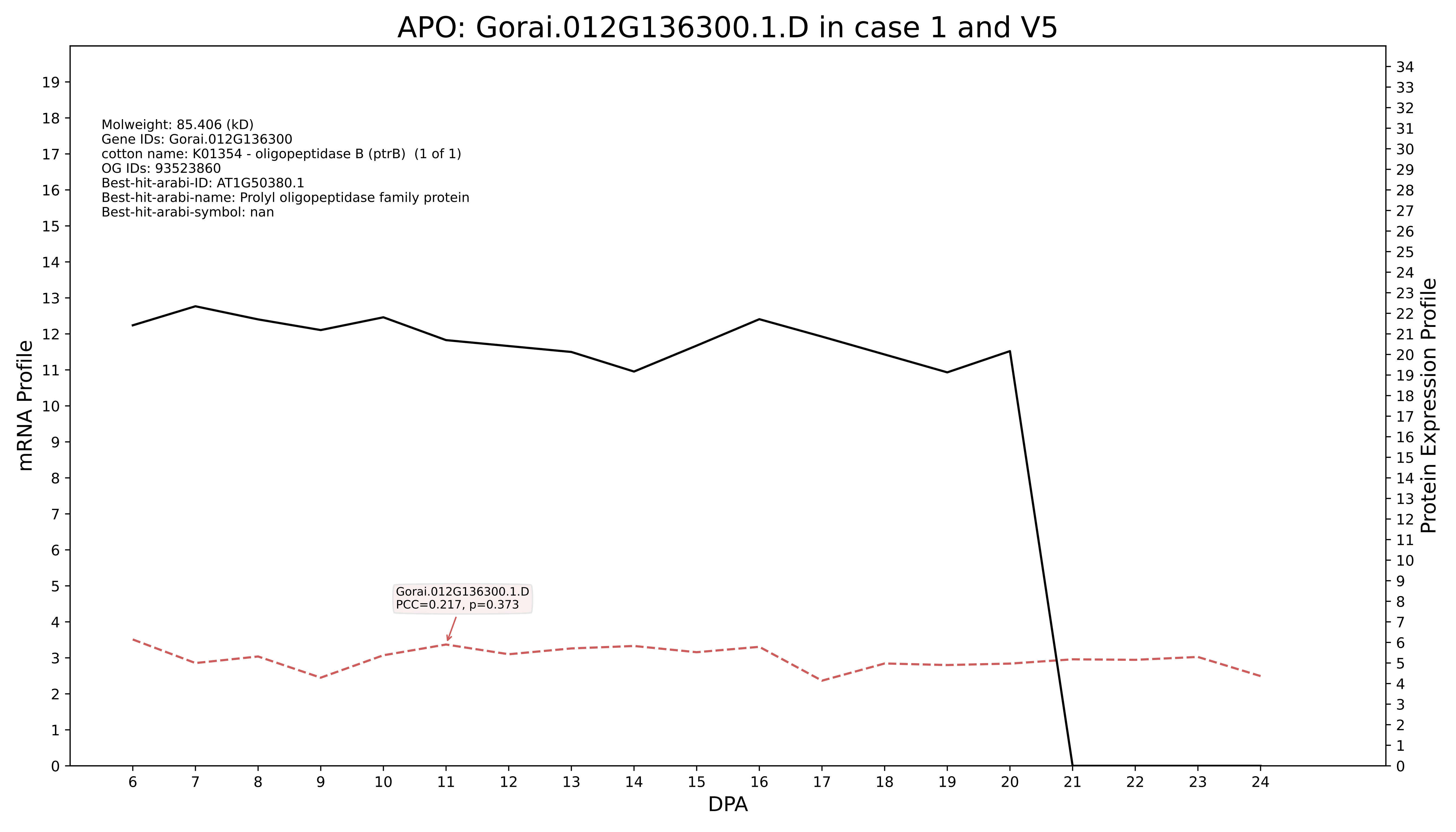

APO: Gorai.012G136300.1.D in case 1 and V5
Molweight: 85.406 (kD)
Gene IDs: Gorai.012G136300
cotton name: K01354 - oligopeptidase B (ptrB) (1 of 1)
OG IDs: 93523860
Best-hit-arabi-ID: AT1G50380.1
Best-hit-arabi-name: Prolyl oligopeptidase family protein
Best-hit-arabi-symbol: nan
Gorai.012G136300.1.D
PCC=0.217, p=0.373
mRNA Profile
Protein Expression Profile
DPA

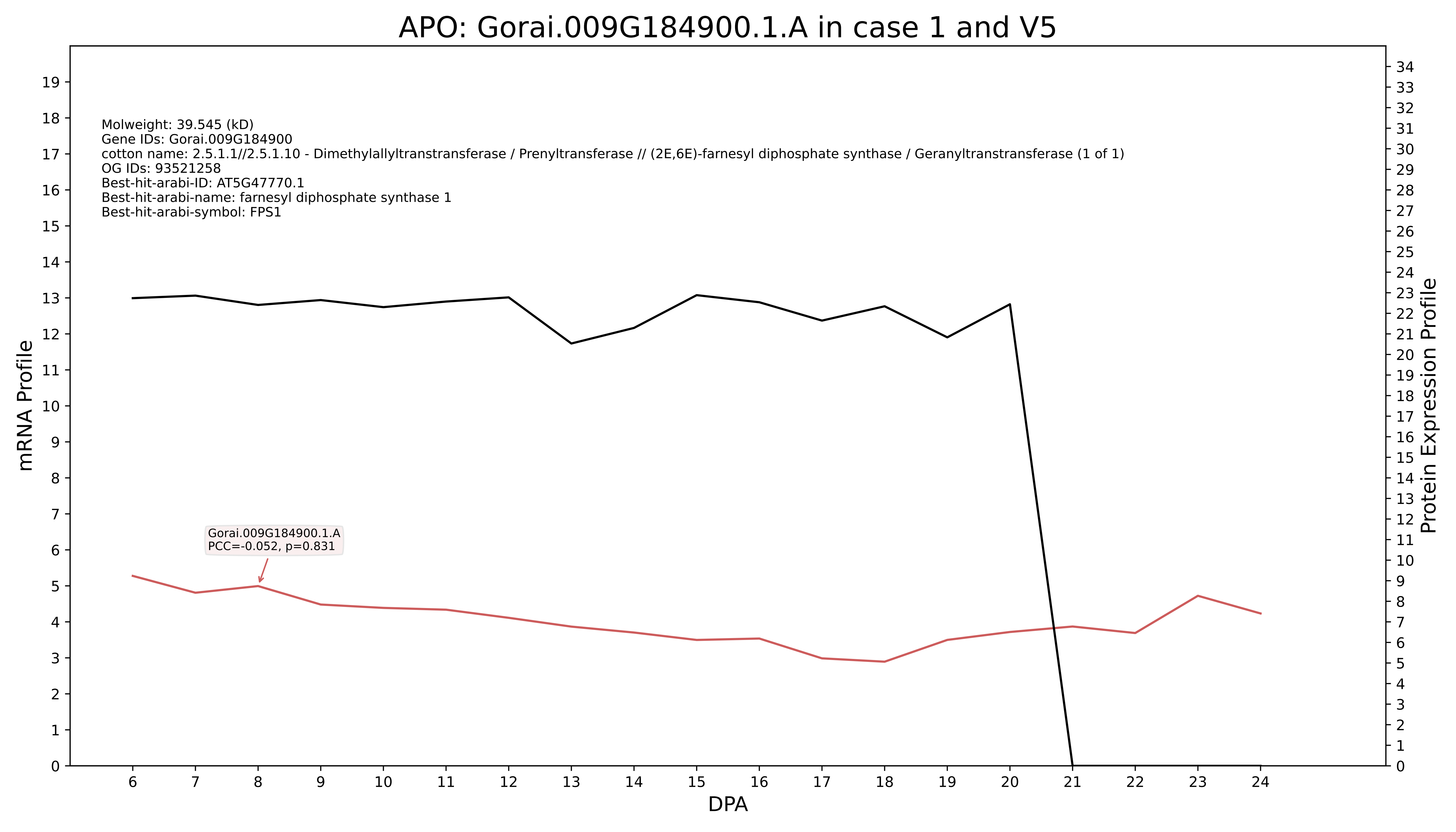
APO: Gorai.009G184900.1.A in case 1 and V5
Molweight: 39.545 (kD)
Gene IDs: Gorai.009G184900
cotton name: 2.5.1.1//2.5.1.10 - Dimethylallyltranstransferase / Prenyltransferase // (2E,6E)-farnesyl diphosphate synthase / Geranyltranstransferase (1 of 1)
OG IDs: 93521258
Best-hit-arabi-ID: AT5G47770.1
Best-hit-arabi-name: farnesyl diphosphate synthase 1
Best-hit-arabi-symbol: FPS1
Gorai.009G184900.1.A
PCC=-0.052, p=0.831
mRNA Profile
Protein Expression Profile
DPA

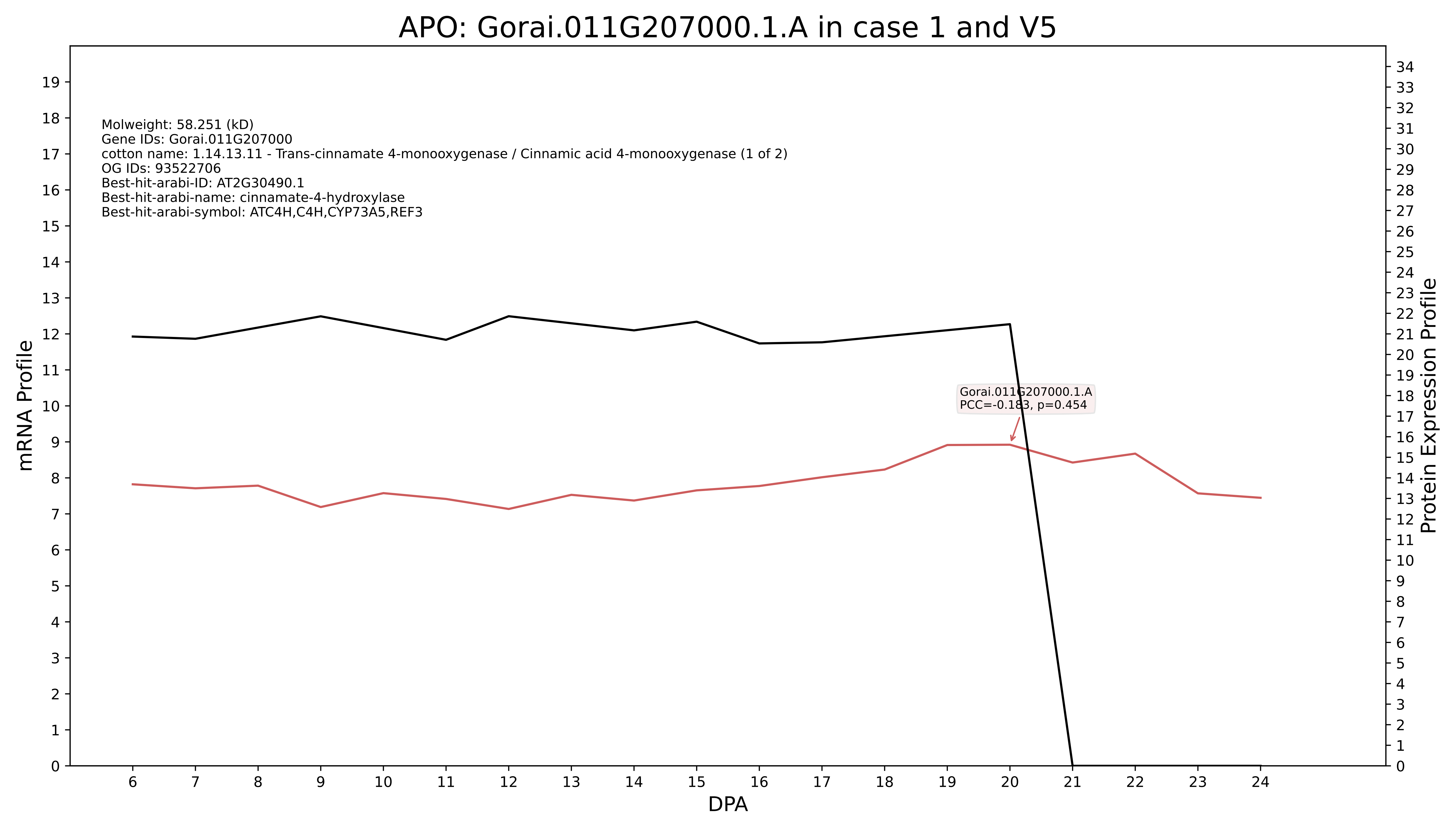

APO: Gorai.011G207000.1.A in case 1 and V5
Molweight: 58.251 (kD)
Gene IDs: Gorai.011G207000
cotton name: 1.14.13.11 - Trans-cinnamate 4-monooxygenase / Cinnamic acid 4-monooxygenase (1 of 2)
OG IDs: 93522706
Best-hit-arabi-ID: AT2G30490.1
Best-hit-arabi-name: cinnamate-4-hydroxylase
Best-hit-arabi-symbol: ATC4H,C4H,CYP73A5,REF3
Gorai.011G207000.1.A
PCC=-0.183, p=0.454
mRNA Profile
Protein Expression Profile
DPA

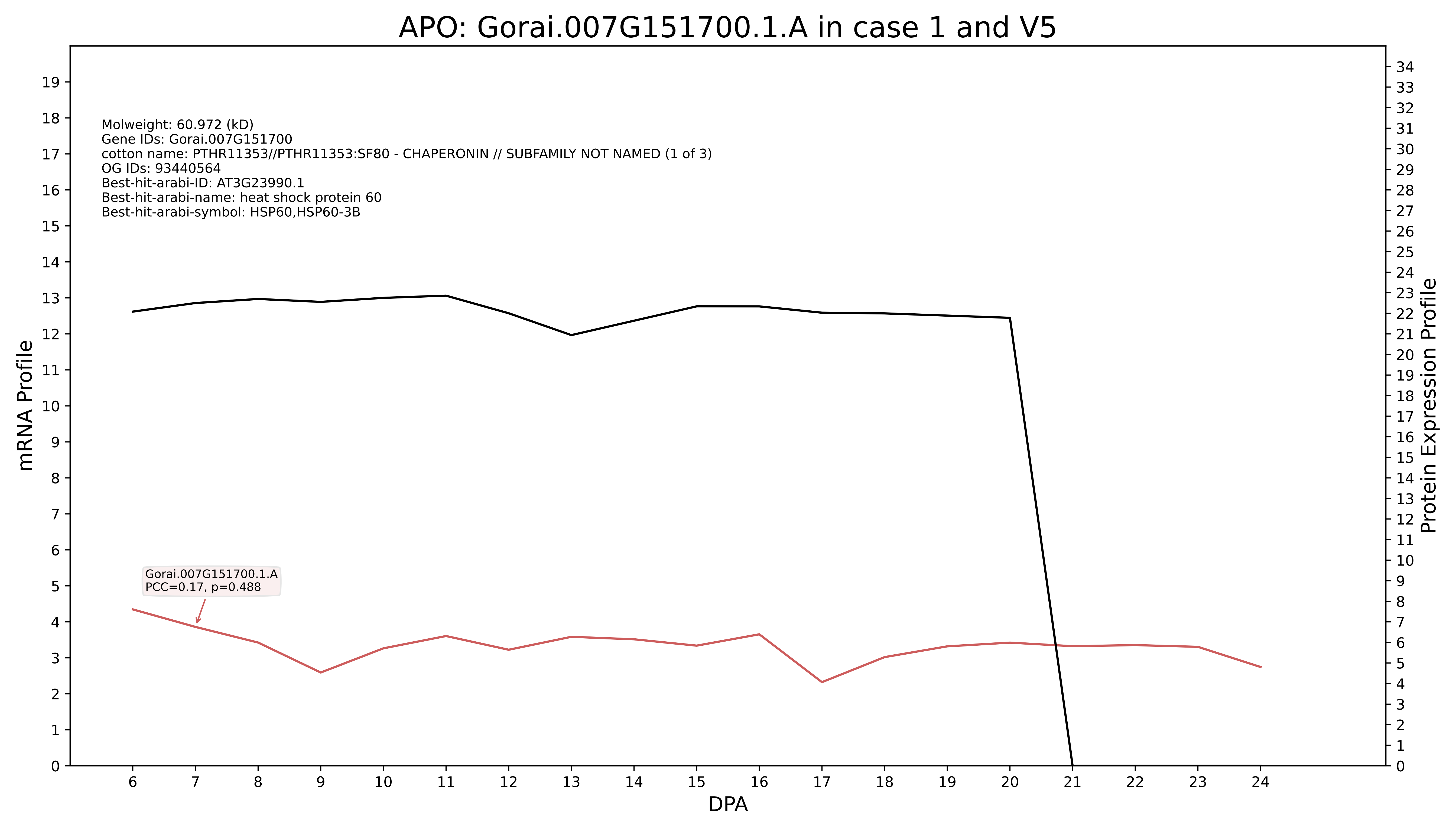

APO: Gorai.007G151700.1.A in case 1 and V5
Molweight: 60.972 (kD)
Gene IDs: Gorai.007G151700
cotton name: PTHR11353//PTHR11353:SF80 - CHAPERONIN // SUBFAMILY NOT NAMED (1 of 3)
OG IDs: 93440564
Best-hit-arabi-ID: AT3G23990.1
Best-hit-arabi-name: heat shock protein 60
Best-hit-arabi-symbol: HSP60,HSP60-3B
Gorai.007G151700.1.A
PCC=0.17, p=0.488
mRNA Profile
Protein Expression Profile
DPA

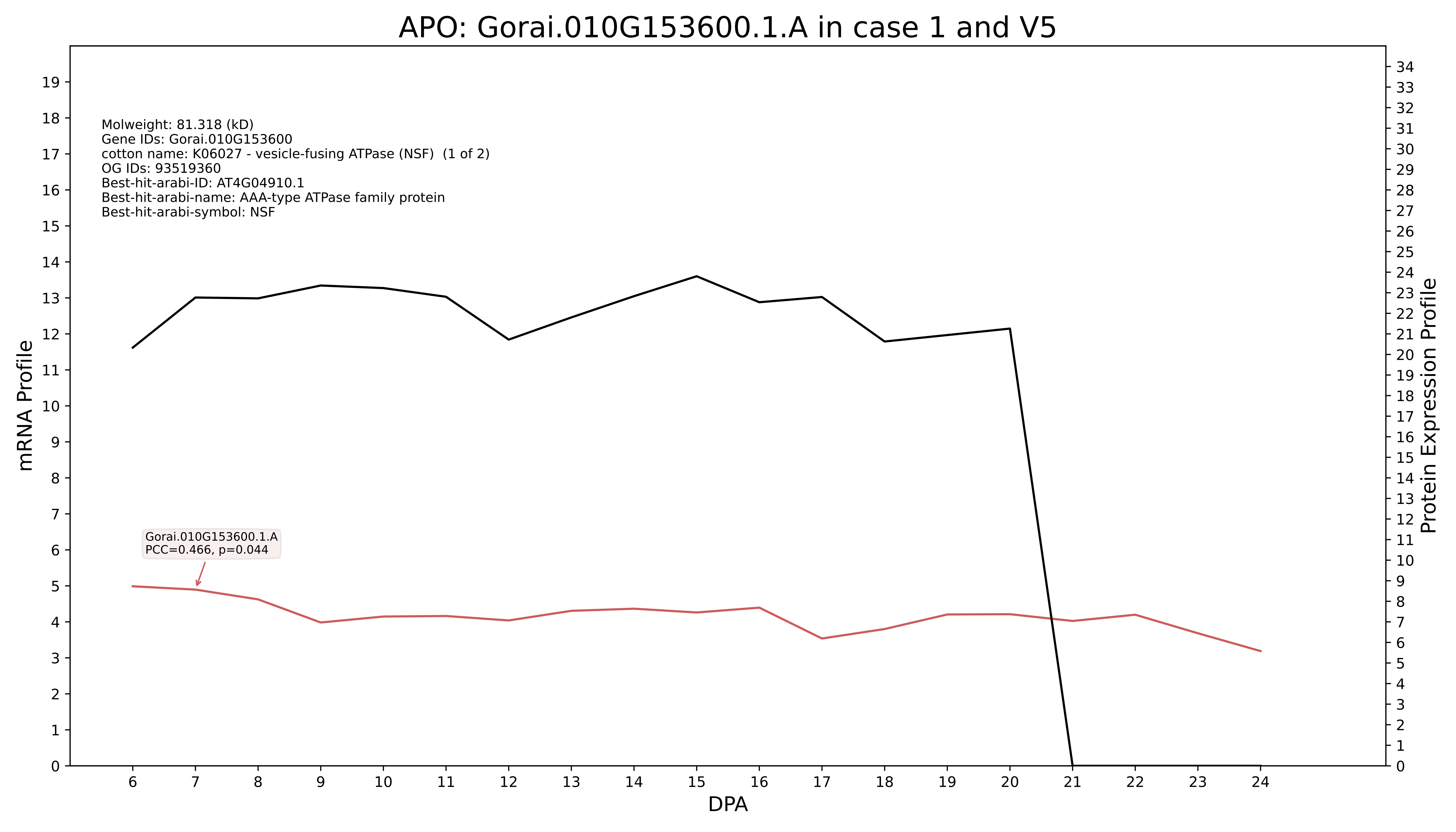

APO: Gorai.010G153600.1.A in case 1 and V5
Molweight: 81.318 (kD)
Gene IDs: Gorai.010G153600
cotton name: K06027 - vesicle-fusing ATPase (NSF) (1 of 2)
OG IDs: 93519360
Best-hit-arabi-ID: AT4G04910.1
Best-hit-arabi-name: AAA-type ATPase family protein
Best-hit-arabi-symbol: NSF
Gorai.010G153600.1.A
PCC=0.466, p=0.044
mRNA Profile
Protein Expression Profile
DPA

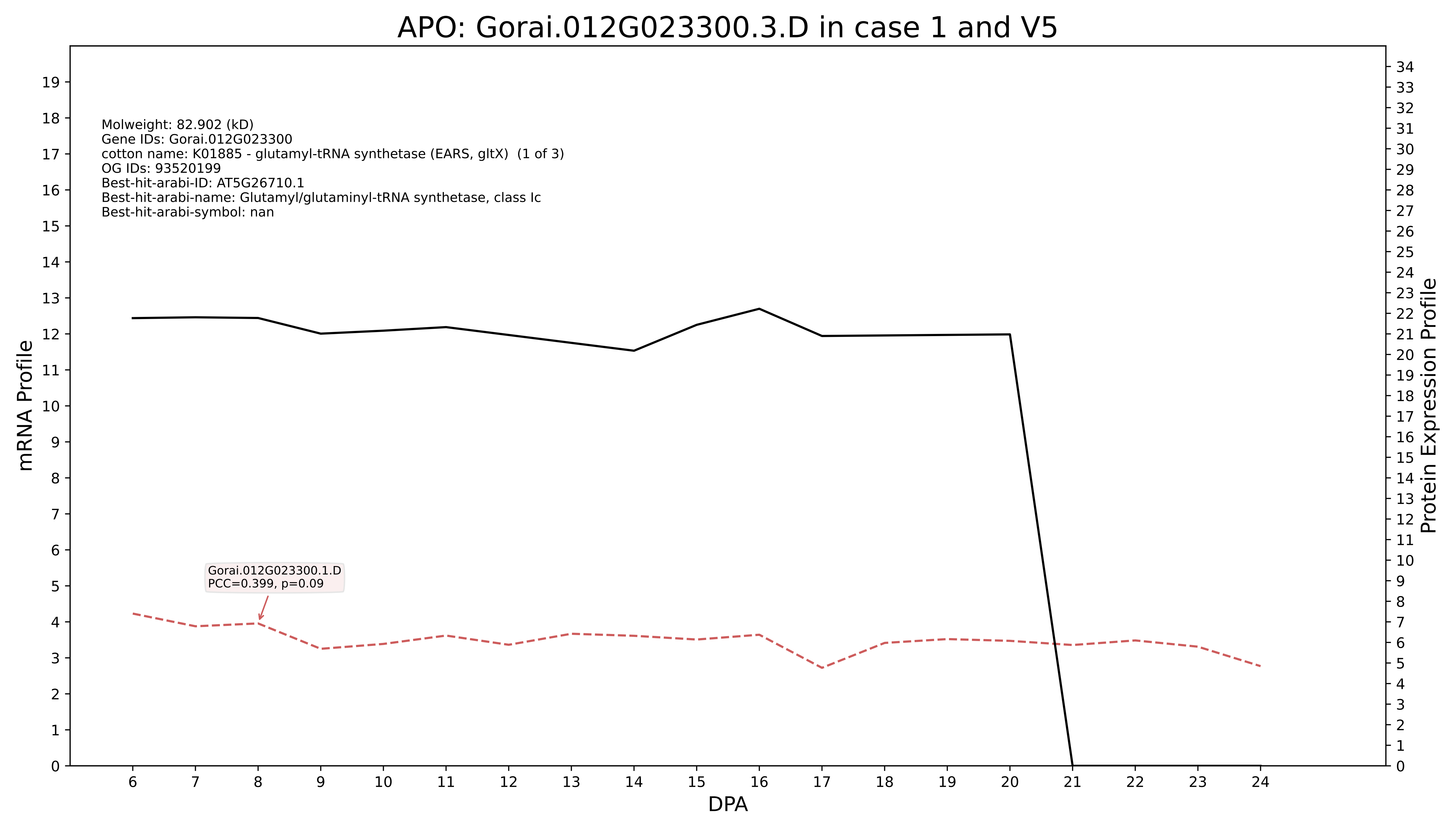

APO: Gorai.012G023300.3.D in case 1 and V5
Molweight: 82.902 (kD)
Gene IDs: Gorai.012G023300
cotton name: K01885 - glutamyl-tRNA synthetase (EARS, gltX) (1 of 3)
OG IDs: 93520199
Best-hit-arabi-ID: AT5G26710.1
Best-hit-arabi-name: Glutamyl/glutaminyl-tRNA synthetase, class Ic
Best-hit-arabi-symbol: nan
Gorai.012G023300.1.D
PCC=0.399, p=0.09
mRNA Profile
Protein Expression Profile
DPA

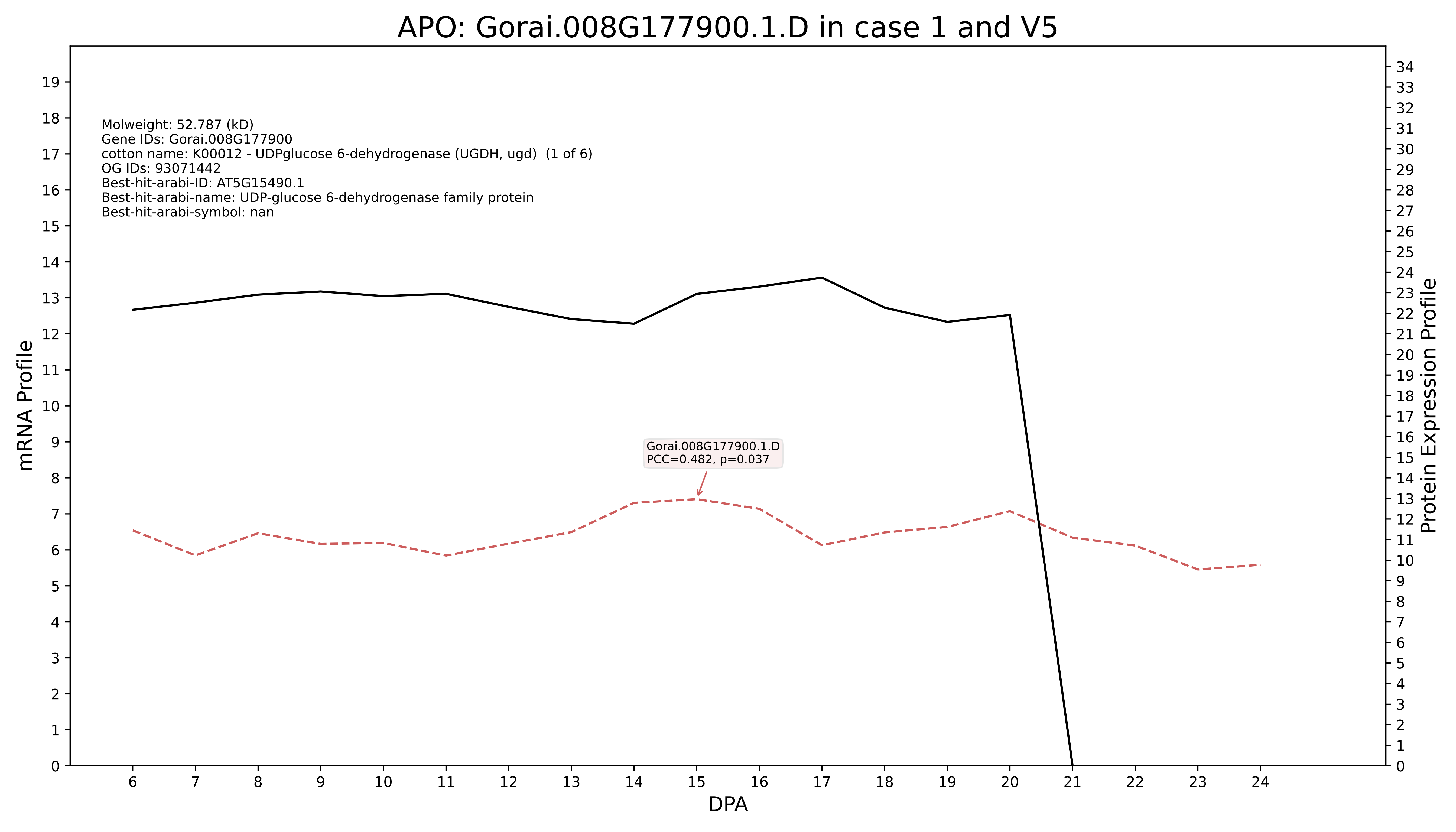

APO: Gorai.008G177900.1.D in case 1 and V5
Molweight: 52.787 (kD)
Gene IDs: Gorai.008G177900
cotton name: K00012 - UDPglucose 6-dehydrogenase (UGDH, ugd) (1 of 6)
OG IDs: 93071442
Best-hit-arabi-ID: AT5G15490.1
Best-hit-arabi-name: UDP-glucose 6-dehydrogenase family protein
Best-hit-arabi-symbol: nan
Gorai.008G177900.1.D
PCC=0.482, p=0.037
mRNA Profile
Protein Expression Profile
DPA

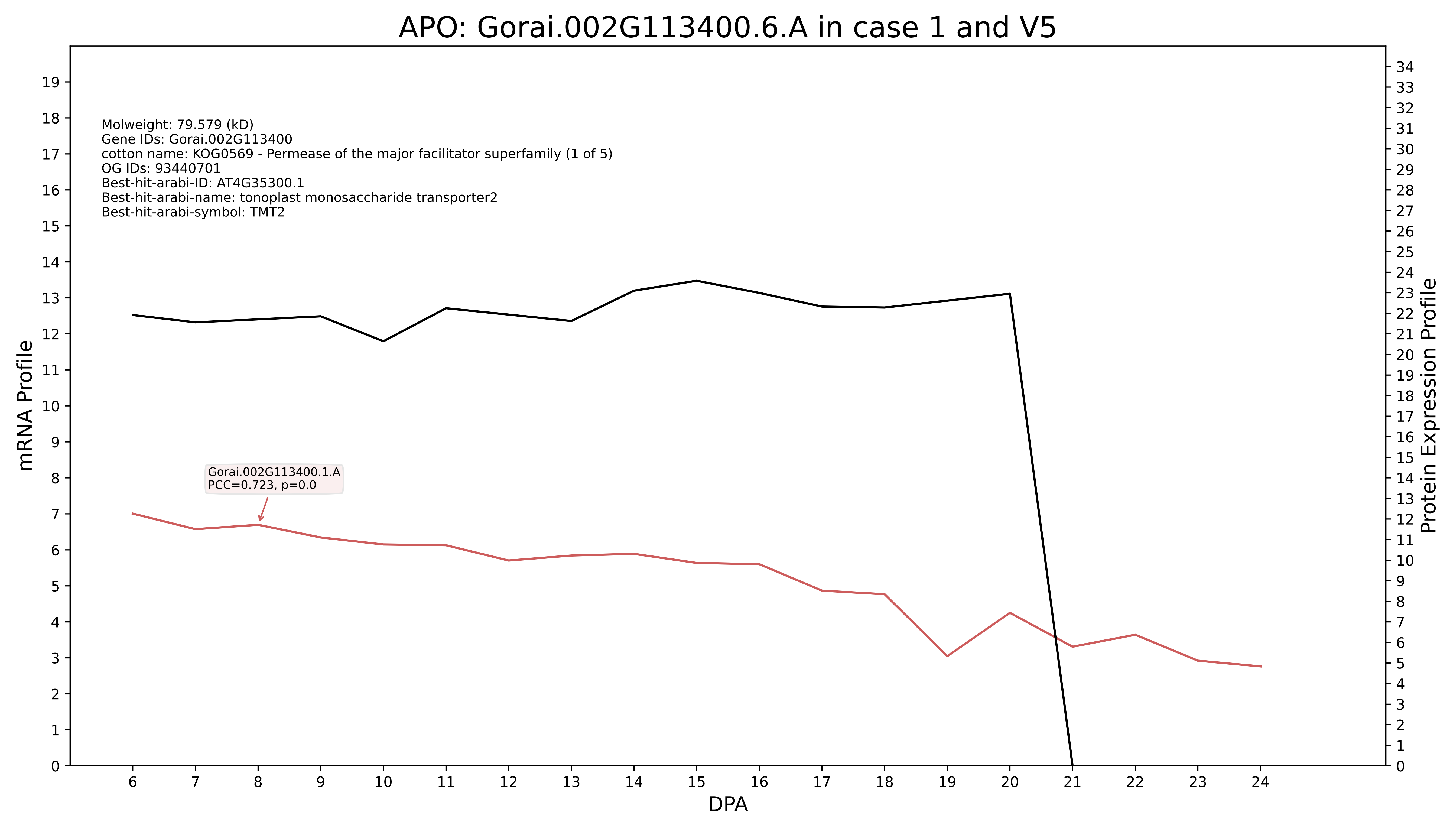
APO: Gorai.002G113400.6.A in case 1 and V5
Molweight: 79.579 (kD)
Gene IDs: Gorai.002G113400
cotton name: KOG0569 - Permease of the major facilitator superfamily (1 of 5)
OG IDs: 93440701
Best-hit-arabi-ID: AT4G35300.1
Best-hit-arabi-name: tonoplast monosaccharide transporter2
Best-hit-arabi-symbol: TMT2
Gorai.002G113400.1.A
PCC=0.723, p=0.0
mRNA Profile
Protein Expression Profile
DPA

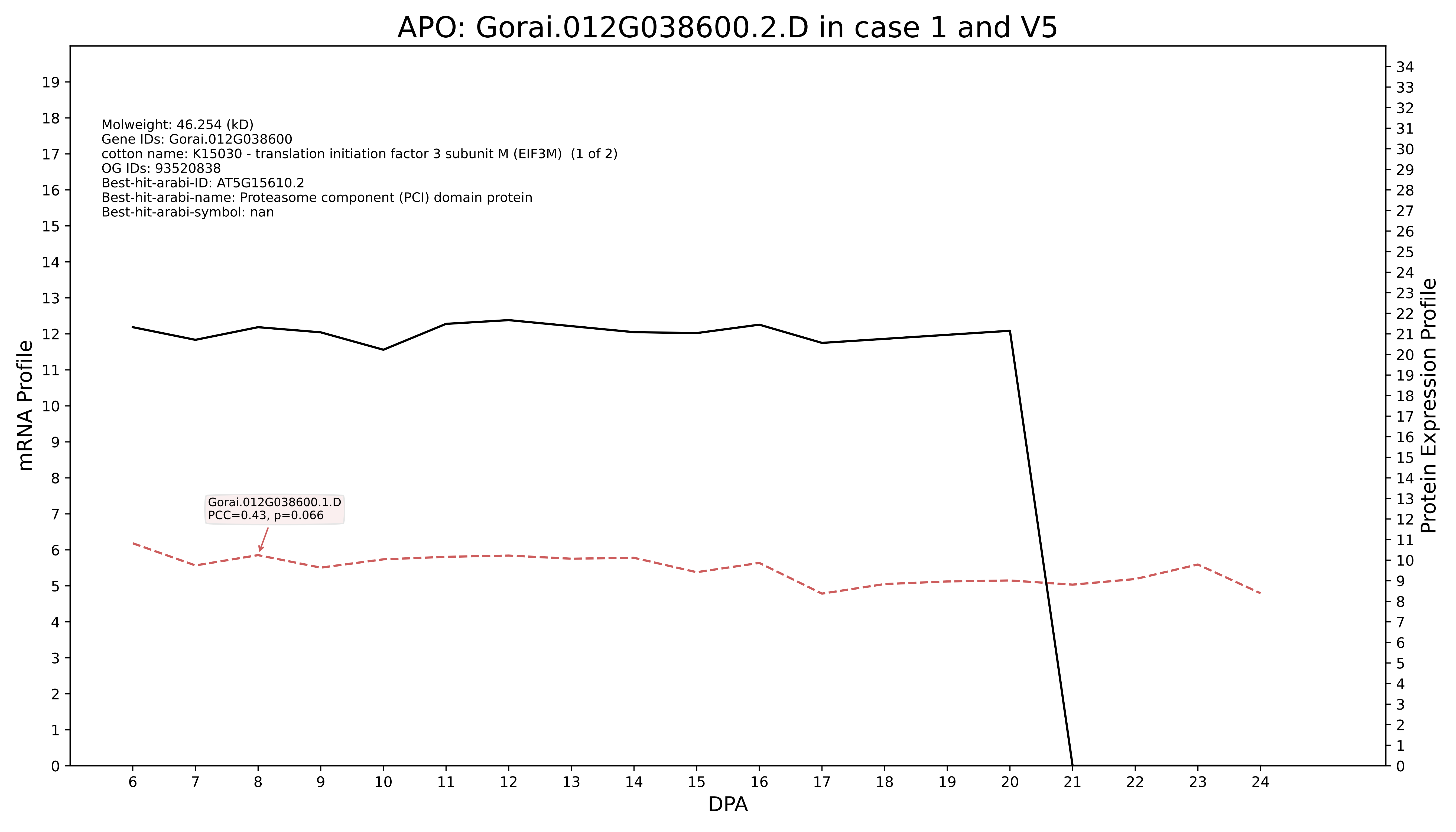

APO: Gorai.012G038600.2.D in case 1 and V5
Molweight: 46.254 (kD)
Gene IDs: Gorai.012G038600
cotton name: K15030 - translation initiation factor 3 subunit M (EIF3M) (1 of 2)
OG IDs: 93520838
Best-hit-arabi-ID: AT5G15610.2
Best-hit-arabi-name: Proteasome component (PCI) domain protein
Best-hit-arabi-symbol: nan
Gorai.012G038600.1.D
PCC=0.43, p=0.066
mRNA Profile
Protein Expression Profile
DPA

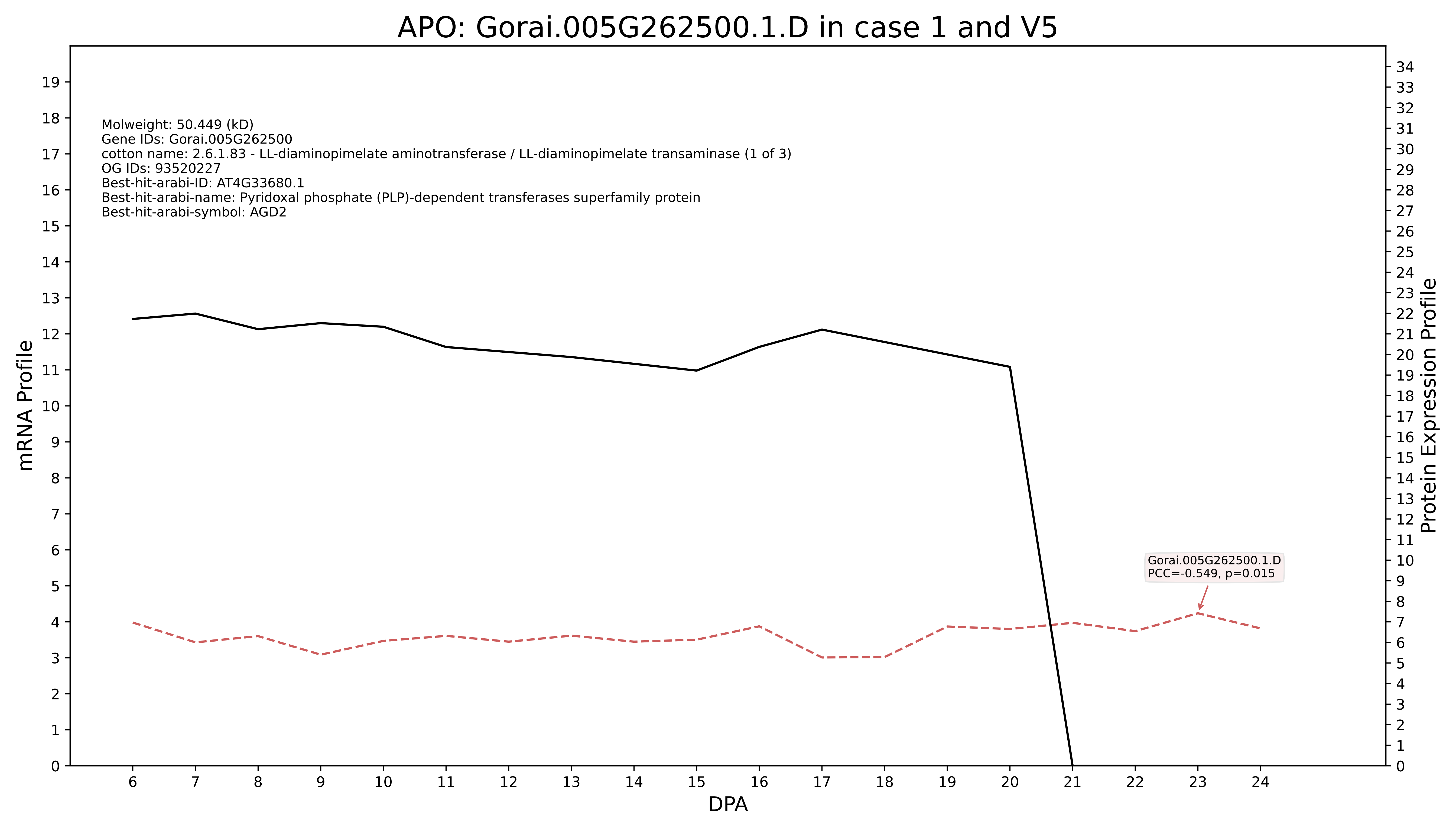
APO: Gorai.005G262500.1.D in case 1 and V5
Molweight: 50.449 (kD)
Gene IDs: Gorai.005G262500
cotton name: 2.6.1.83 - LL-diaminopimelate aminotransferase / LL-diaminopimelate transaminase (1 of 3)
OG IDs: 93520227
Best-hit-arabi-ID: AT4G33680.1
Best-hit-arabi-name: Pyridoxal phosphate (PLP)-dependent transferases superfamily protein
Best-hit-arabi-symbol: AGD2
Gorai.005G262500.1.D
PCC=-0.549, p=0.015
mRNA Profile
Protein Expression Profile
DPA

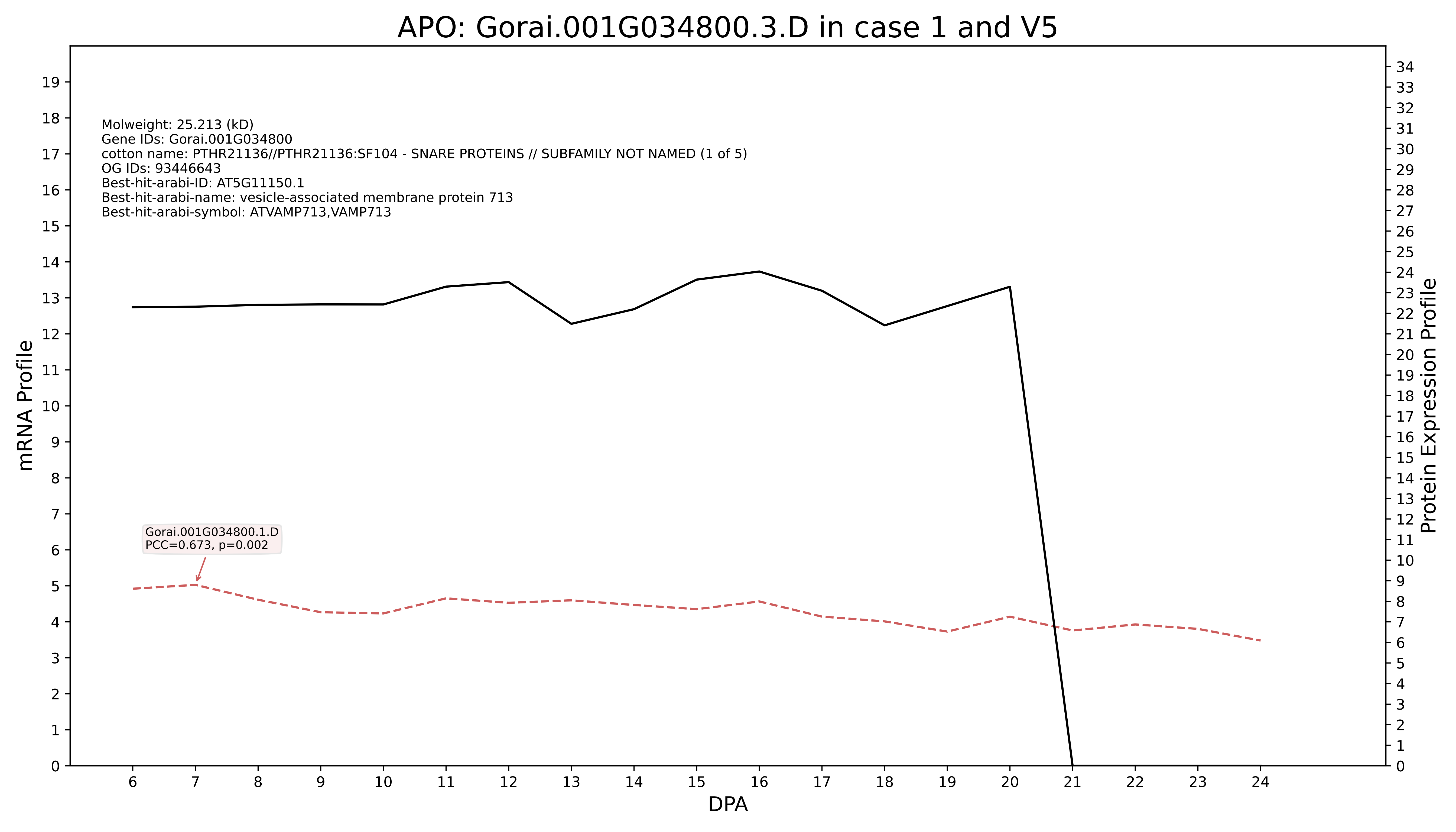
APO: Gorai.001G034800.3.D in case 1 and V5
Molweight: 25.213 (kD)
Gene IDs: Gorai.001G034800
cotton name: PTHR21136//PTHR21136:SF104 - SNARE PROTEINS // SUBFAMILY NOT NAMED (1 of 5)
OG IDs: 93446643
Best-hit-arabi-ID: AT5G11150.1
Best-hit-arabi-name: vesicle-associated membrane protein 713
Best-hit-arabi-symbol: ATVAMP713,VAMP713
Gorai.001G034800.1.D
PCC=0.673, p=0.002
mRNA Profile
Protein Expression Profile
DPA

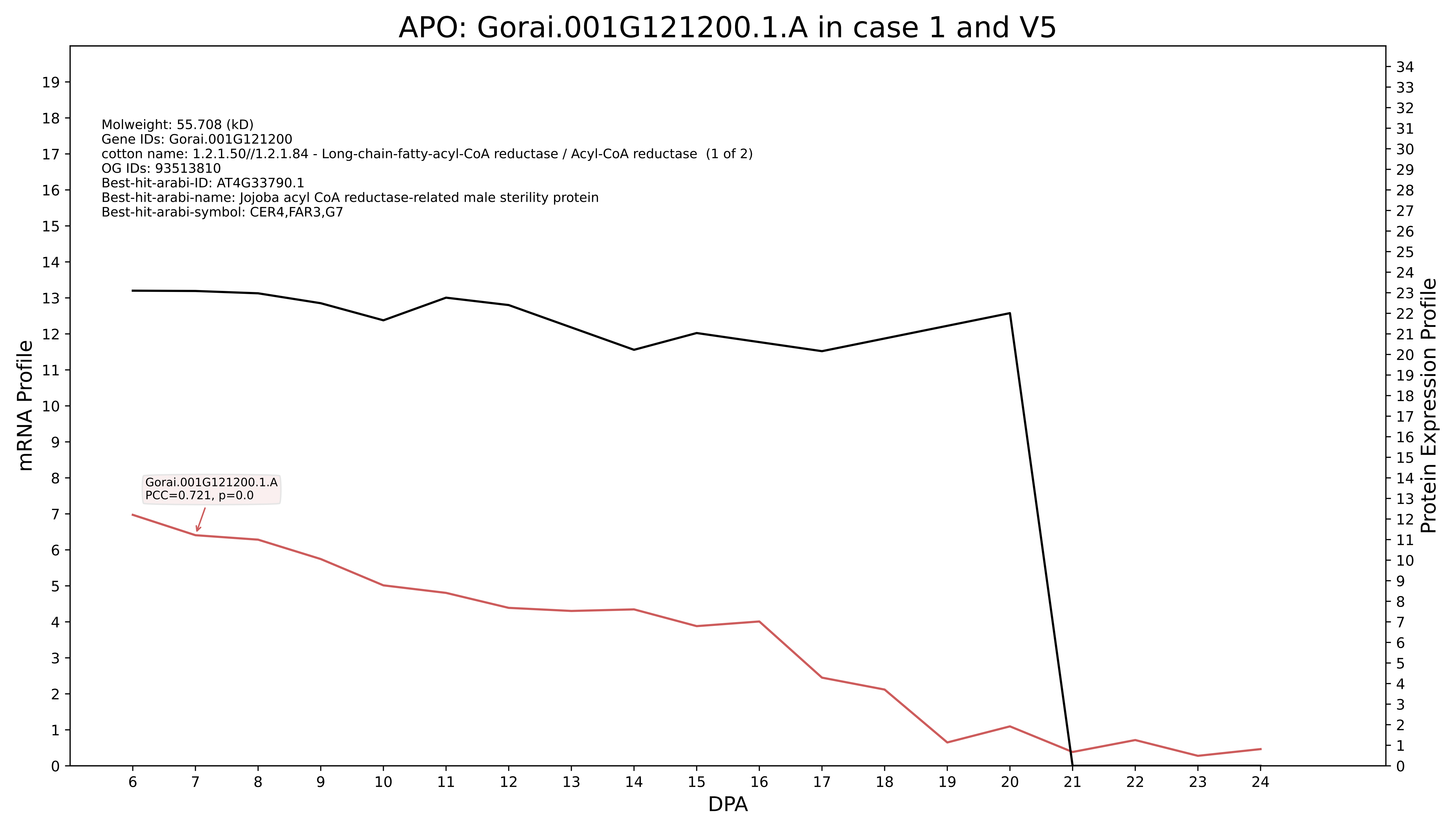
APO: Gorai.001G121200.1.A in case 1 and V5
Molweight: 55.708 (kD)
Gene IDs: Gorai.001G121200
cotton name: 1.2.1.50//1.2.1.84 - Long-chain-fatty-acyl-CoA reductase / Acyl-CoA reductase (1 of 2)
OG IDs: 93513810
Best-hit-arabi-ID: AT4G33790.1
Best-hit-arabi-name: Jojoba acyl CoA reductase-related male sterility protein
Best-hit-arabi-symbol: CER4,FAR3,G7
Gorai.001G121200.1.A
PCC=0.721, p=0.0
mRNA Profile
Protein Expression Profile
DPA

### APO: Gorai.002G110500.5.D in case 1 and V5

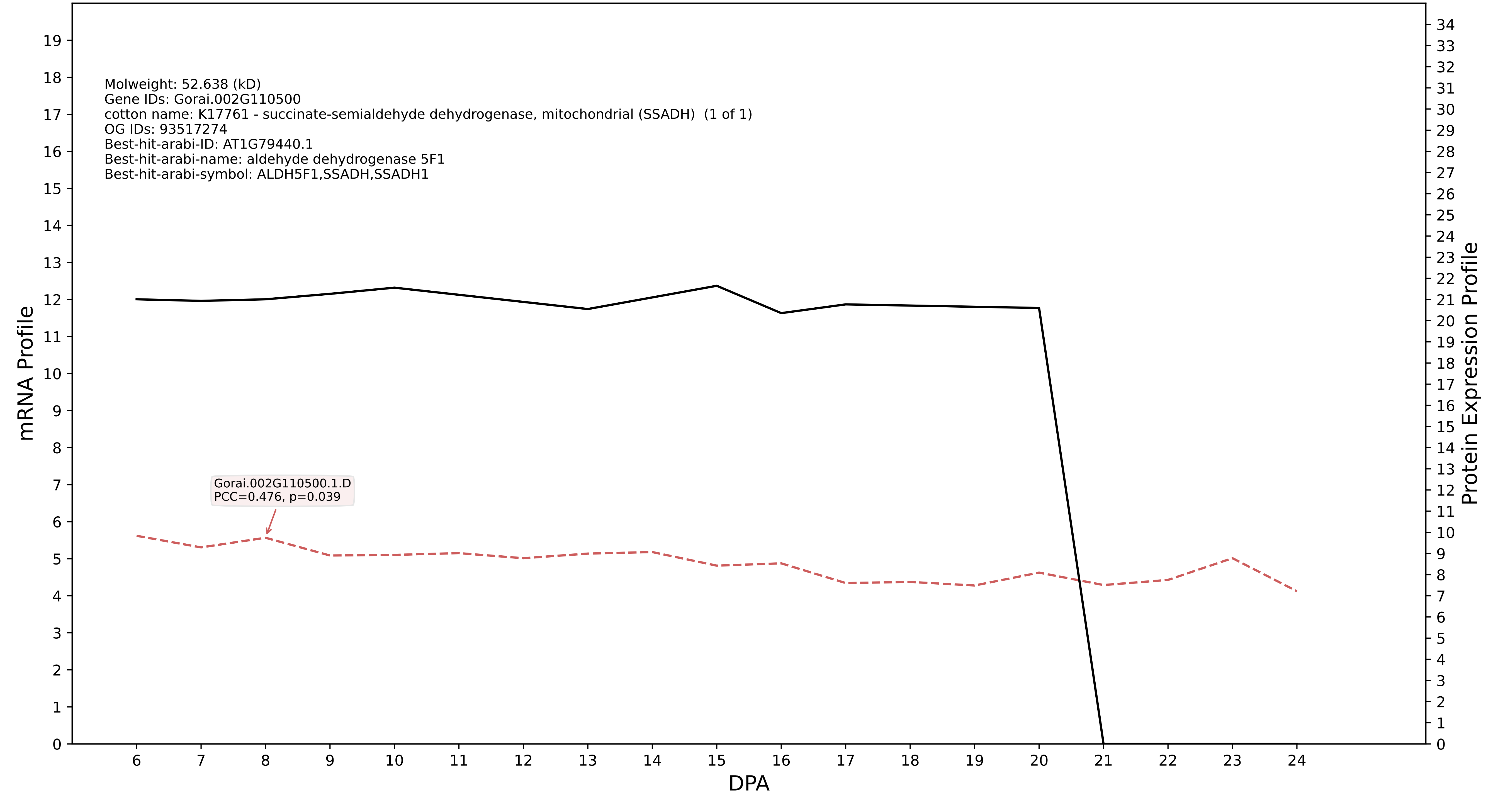

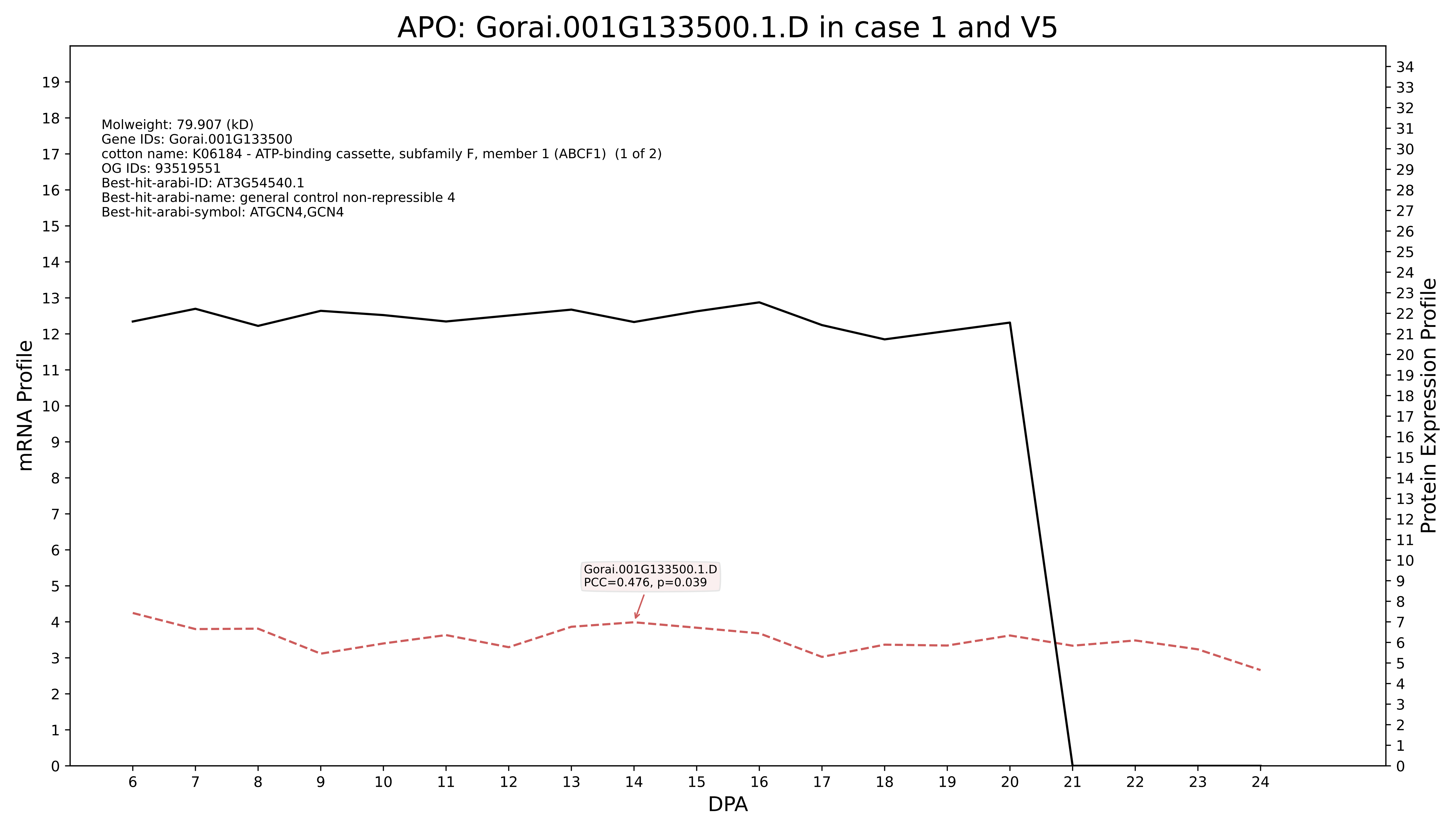
APO: Gorai.001G133500.1.D in case 1 and V5
Molweight: 79.907 (kD)
Gene IDs: Gorai.001G133500
cotton name: K06184 - ATP-binding cassette, subfamily F, member 1 (ABCF1) (1 of 2)
OG IDs: 93519551
Best-hit-arabi-ID: AT3G54540.1
Best-hit-arabi-name: general control non-repressible 4
Best-hit-arabi-symbol: ATGCN4,GCN4
Gorai.001G133500.1.D
PCC=0.476, p=0.039
mRNA Profile
Protein Expression Profile
DPA

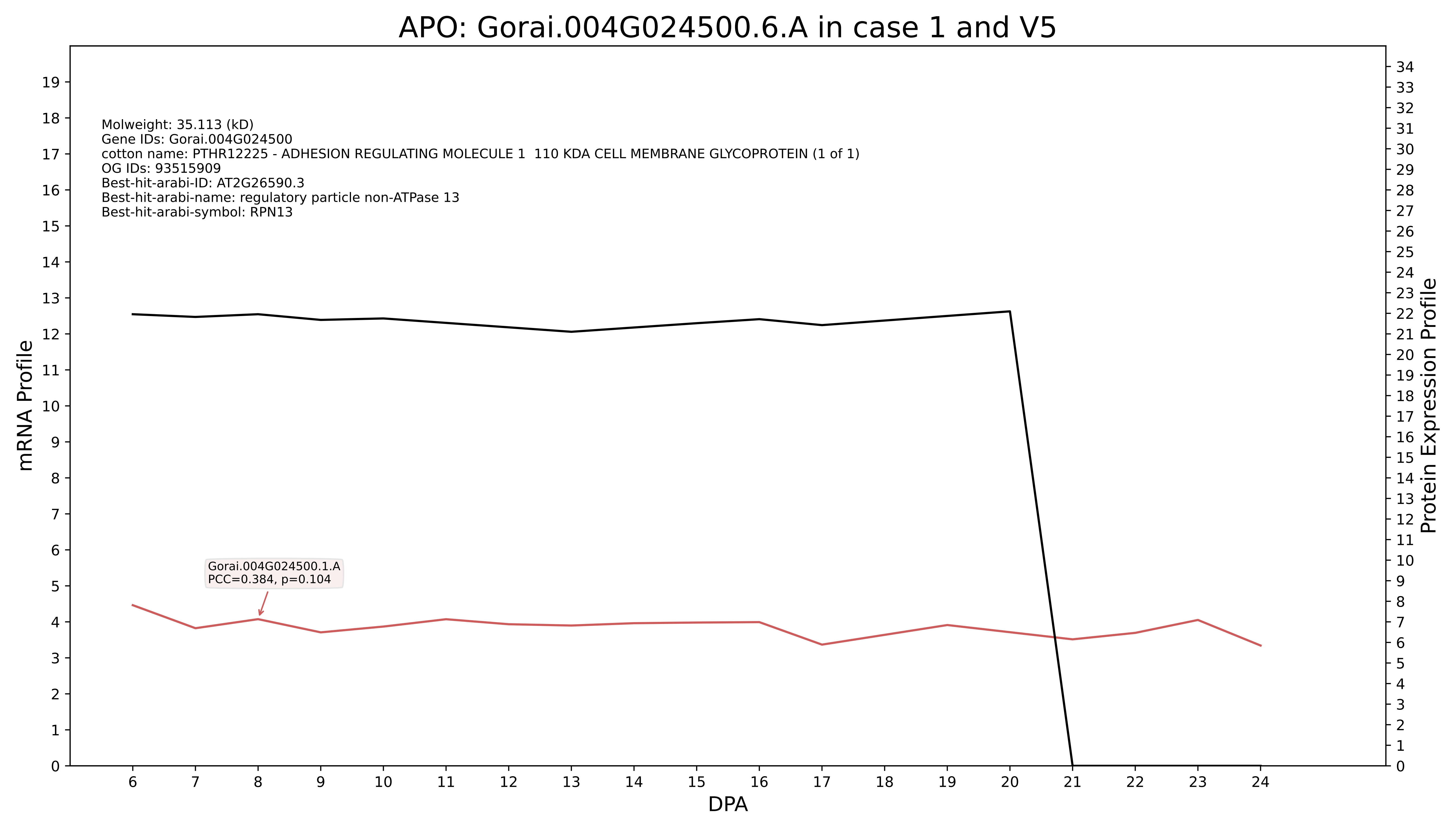
APO: Gorai.004G024500.6.A in case 1 and V5
Molweight: 35.113 (kD)
Gene IDs: Gorai.004G024500
cotton name: PTHR12225 - ADHESION REGULATING MOLECULE 1 110 KDA CELL MEMBRANE GLYCOPROTEIN (1 of 1)
OG IDs: 93515909
Best-hit-arabi-ID: AT2G26590.3
Best-hit-arabi-name: regulatory particle non-ATPase 13
Best-hit-arabi-symbol: RPN13
Gorai.004G024500.1.A
PCC=0.384, p=0.104
mRNA Profile
Protein Expression Profile
DPA

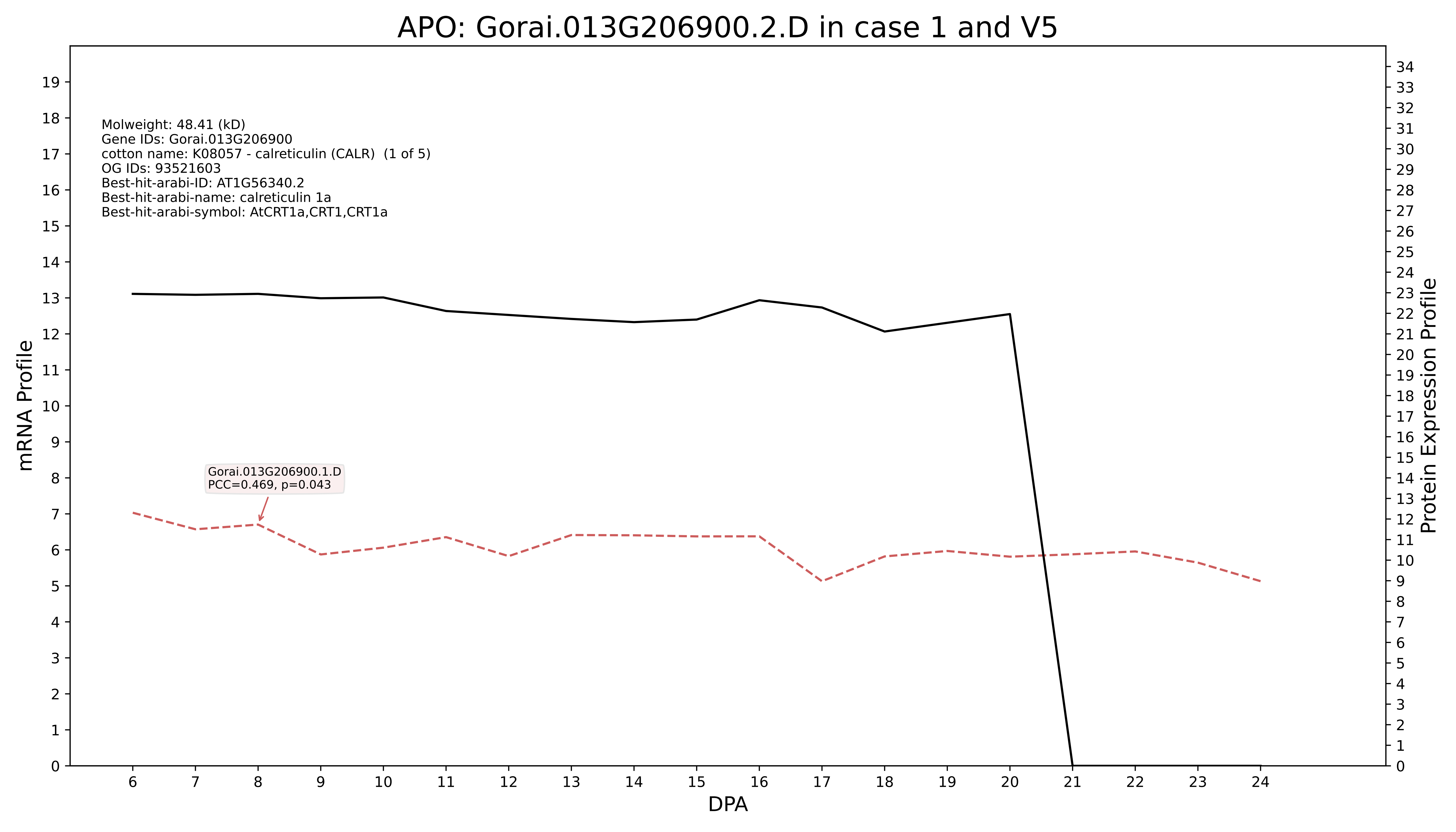

APO: Gorai.013G206900.2.D in case 1 and V5
Molweight: 48.41 (kD)
Gene IDs: Gorai.013G206900
cotton name: K08057 - calreticulin (CALR) (1 of 5)
OG IDs: 93521603
Best-hit-arabi-ID: AT1G56340.2
Best-hit-arabi-name: calreticulin 1a
Best-hit-arabi-symbol: AtCRT1a,CRT1,CRT1a
Gorai.013G206900.1.D
PCC=0.469, p=0.043
mRNA Profile
Protein Expression Profile
DPA

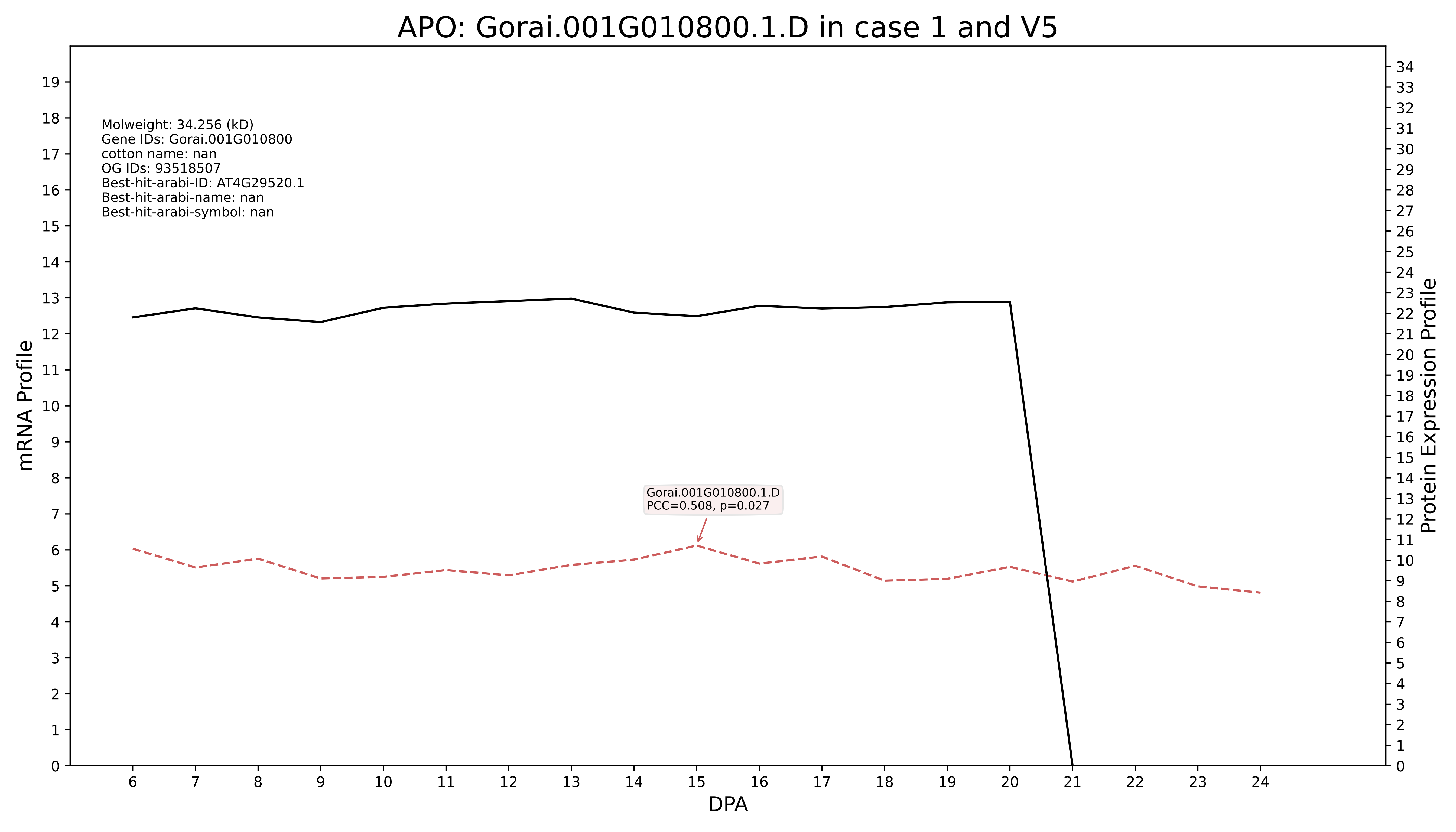

APO: Gorai.001G010800.1.D in case 1 and V5
Molweight: 34.256 (kD)
Gene IDs: Gorai.001G010800
cotton name: nan
OG IDs: 93518507
Best-hit-arabi-ID: AT4G29520.1
Best-hit-arabi-name: nan
Best-hit-arabi-symbol: nan
Gorai.001G010800.1.D
PCC=0.508, p=0.027
mRNA Profile
Protein Expression Profile
DPA

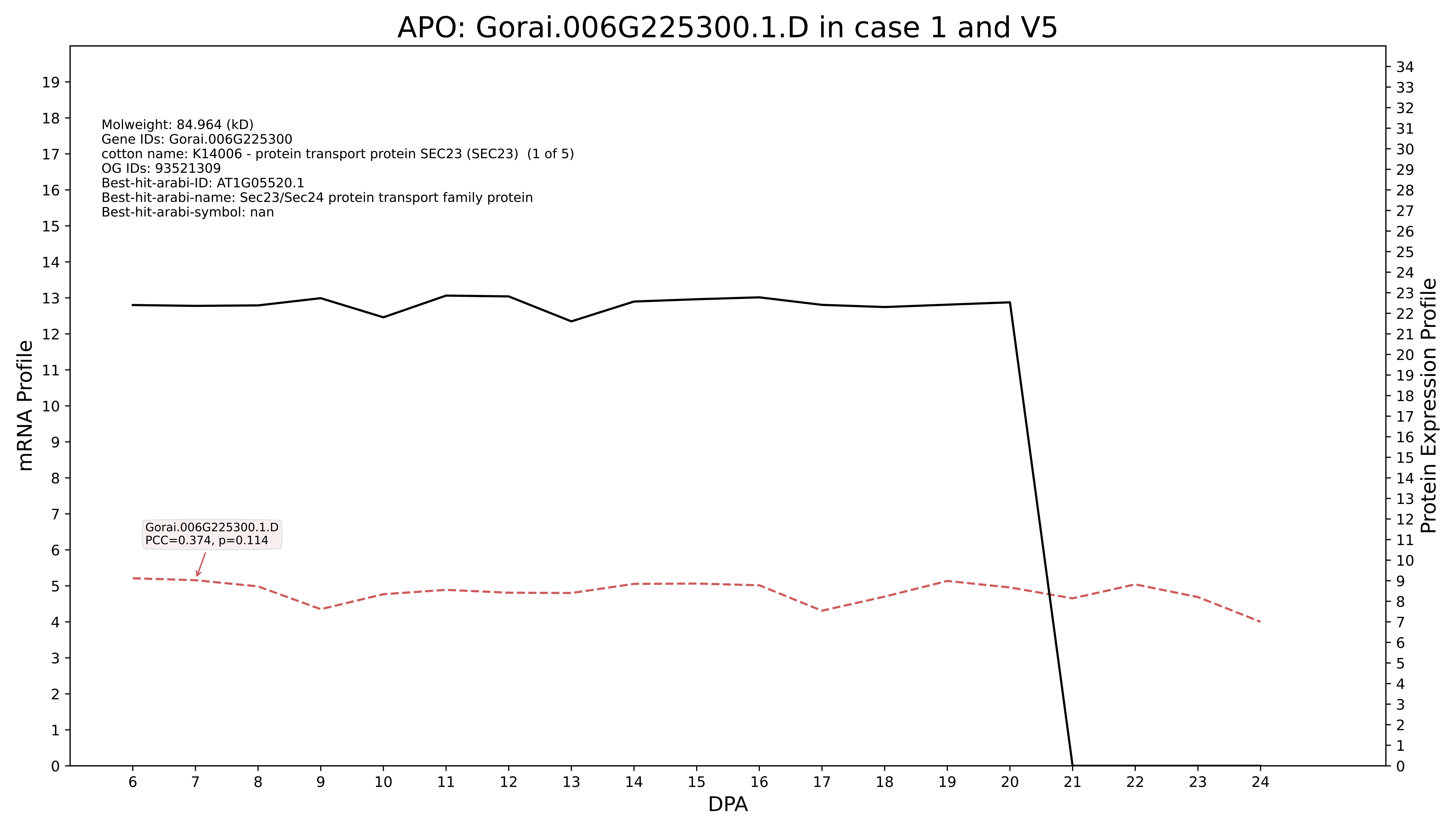

APO: Gorai.006G225300.1.D in case 1 and V5
Molweight: 84.964 (kD)
Gene IDs: Gorai.006G225300
cotton name: K14006 - protein transport protein SEC23 (SEC23) (1 of 5)
OG IDs: 93521309
Best-hit-arabi-ID: AT1G05520.1
Best-hit-arabi-name: Sec23/Sec24 protein transport family protein
Best-hit-arabi-symbol: nan
Gorai.006G225300.1.D
PCC=0.374, p=0.114
mRNA Profile
Protein Expression Profile
DPA

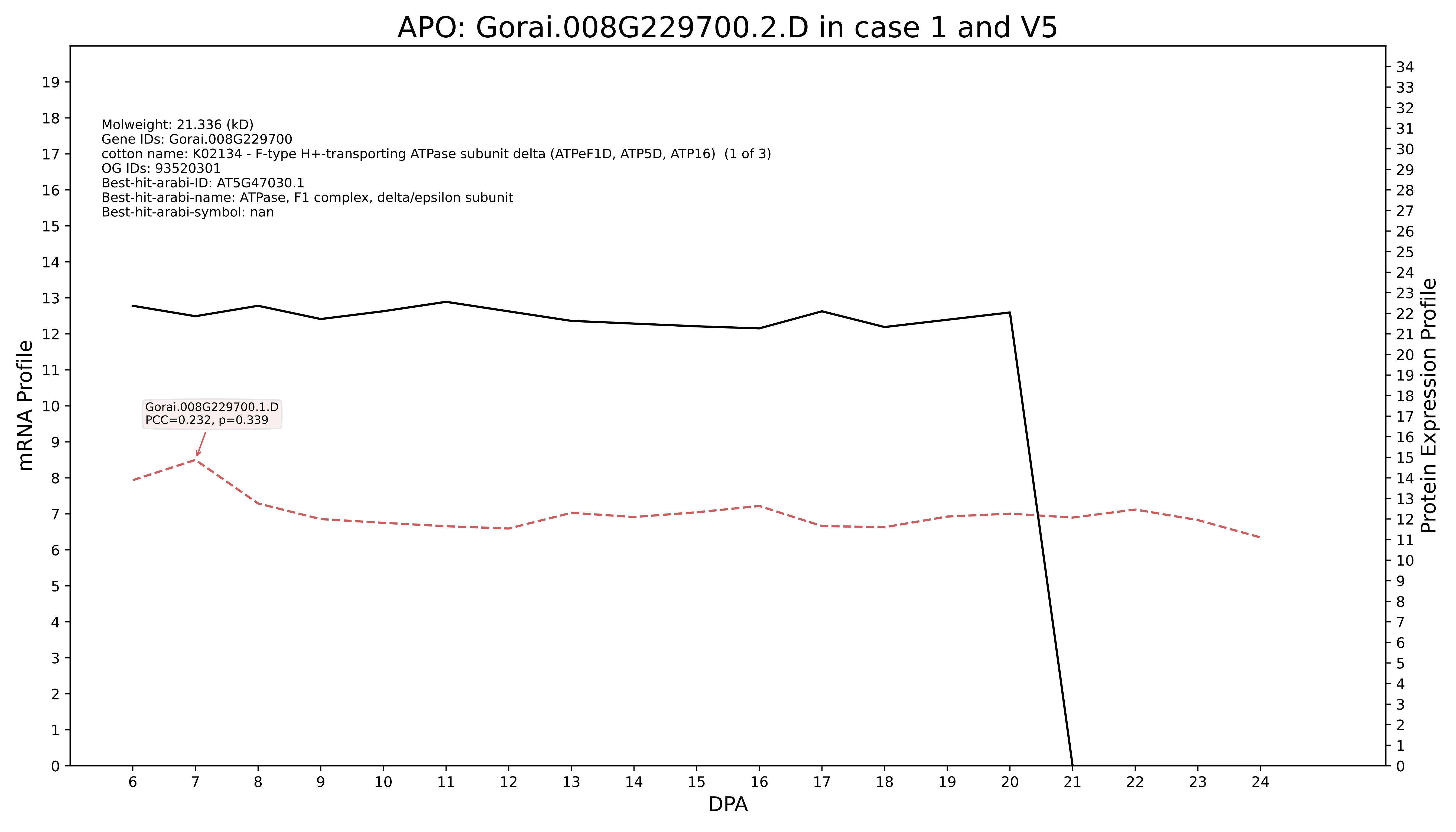

APO: Gorai.008G229700.2.D in case 1 and V5
Molweight: 21.336 (kD)
Gene IDs: Gorai.008G229700
cotton name: K02134 - F-type H+-transporting ATPase subunit delta (ATPeF1D, ATP5D, ATP16) (1 of 3)
OG IDs: 93520301
Best-hit-arabi-ID: AT5G47030.1
Best-hit-arabi-name: ATPase, F1 complex, delta/epsilon subunit
Best-hit-arabi-symbol: nan
Gorai.008G229700.1.D
PCC=0.232, p=0.339
mRNA Profile
Protein Expression Profile
DPA

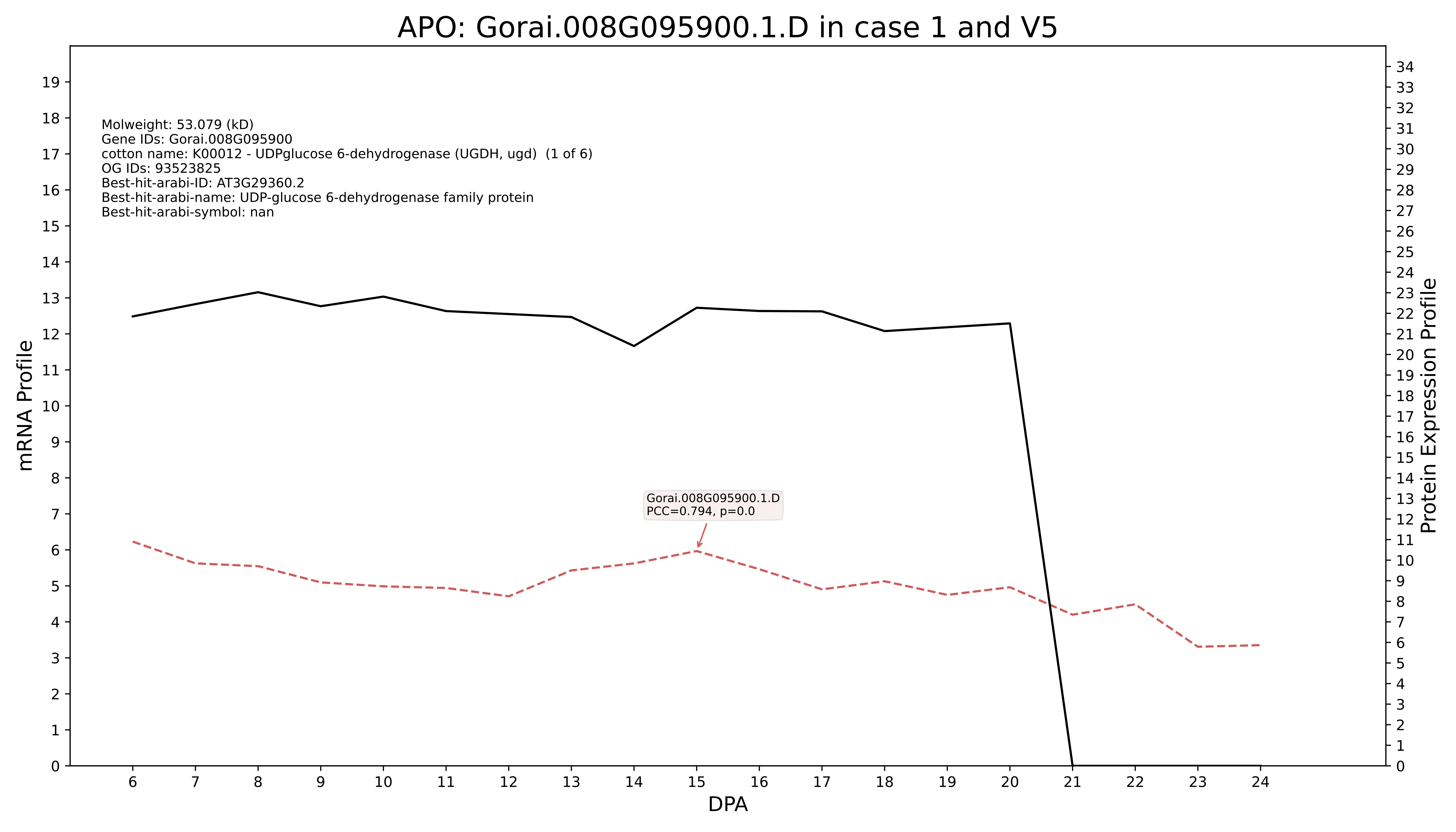

APO: Gorai.008G095900.1.D in case 1 and V5
Molweight: 53.079 (kD)
Gene IDs: Gorai.008G095900
cotton name: K00012 - UDPglucose 6-dehydrogenase (UGDH, ugd) (1 of 6)
OG IDs: 93523825
Best-hit-arabi-ID: AT3G29360.2
Best-hit-arabi-name: UDP-glucose 6-dehydrogenase family protein
Best-hit-arabi-symbol: nan
Gorai.008G095900.1.D
PCC=0.794, p=0.0
mRNA Profile
Protein Expression Profile
DPA

### APO: Gorai.006G183900.2.D in case 1 and V5

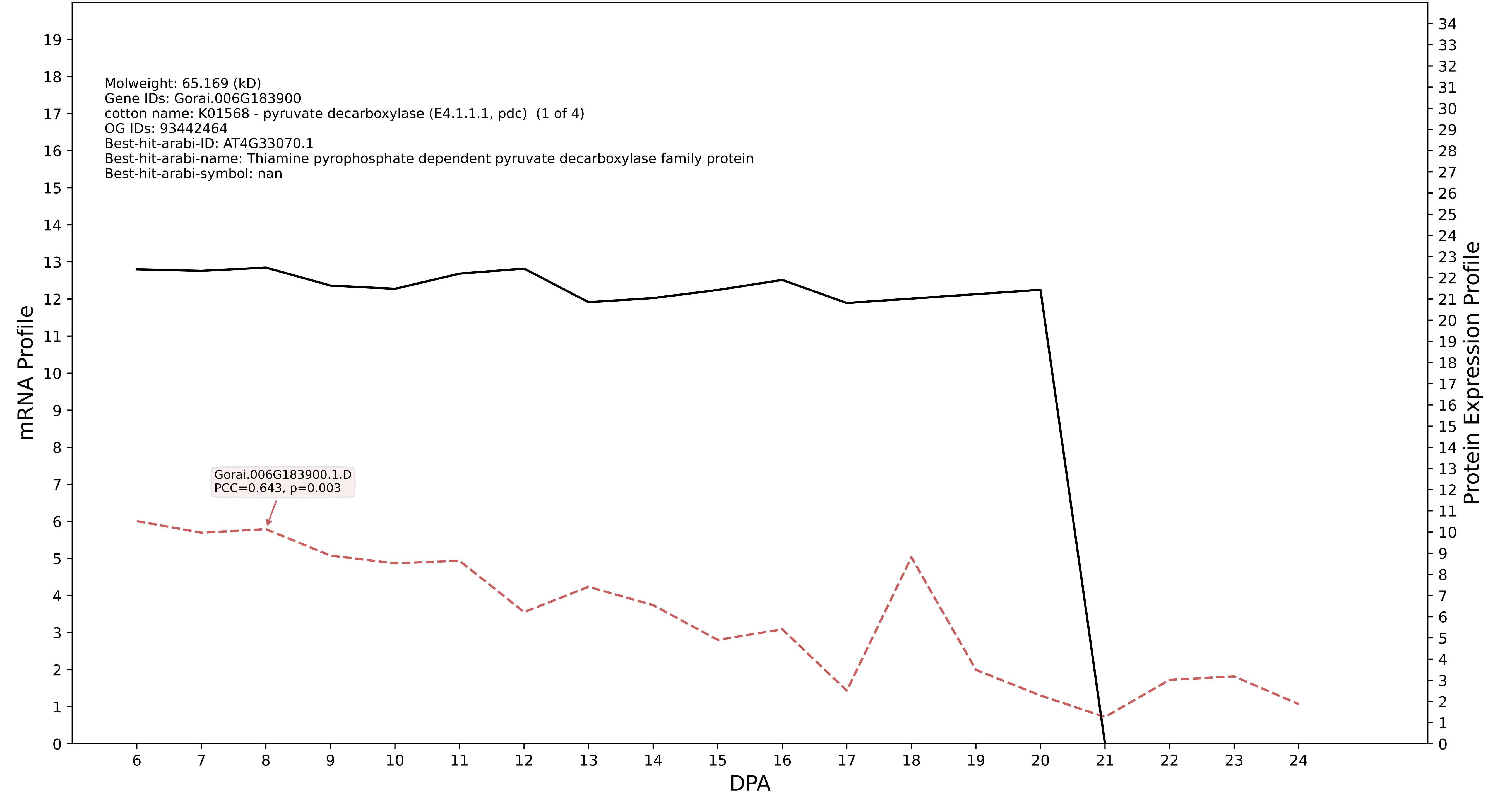

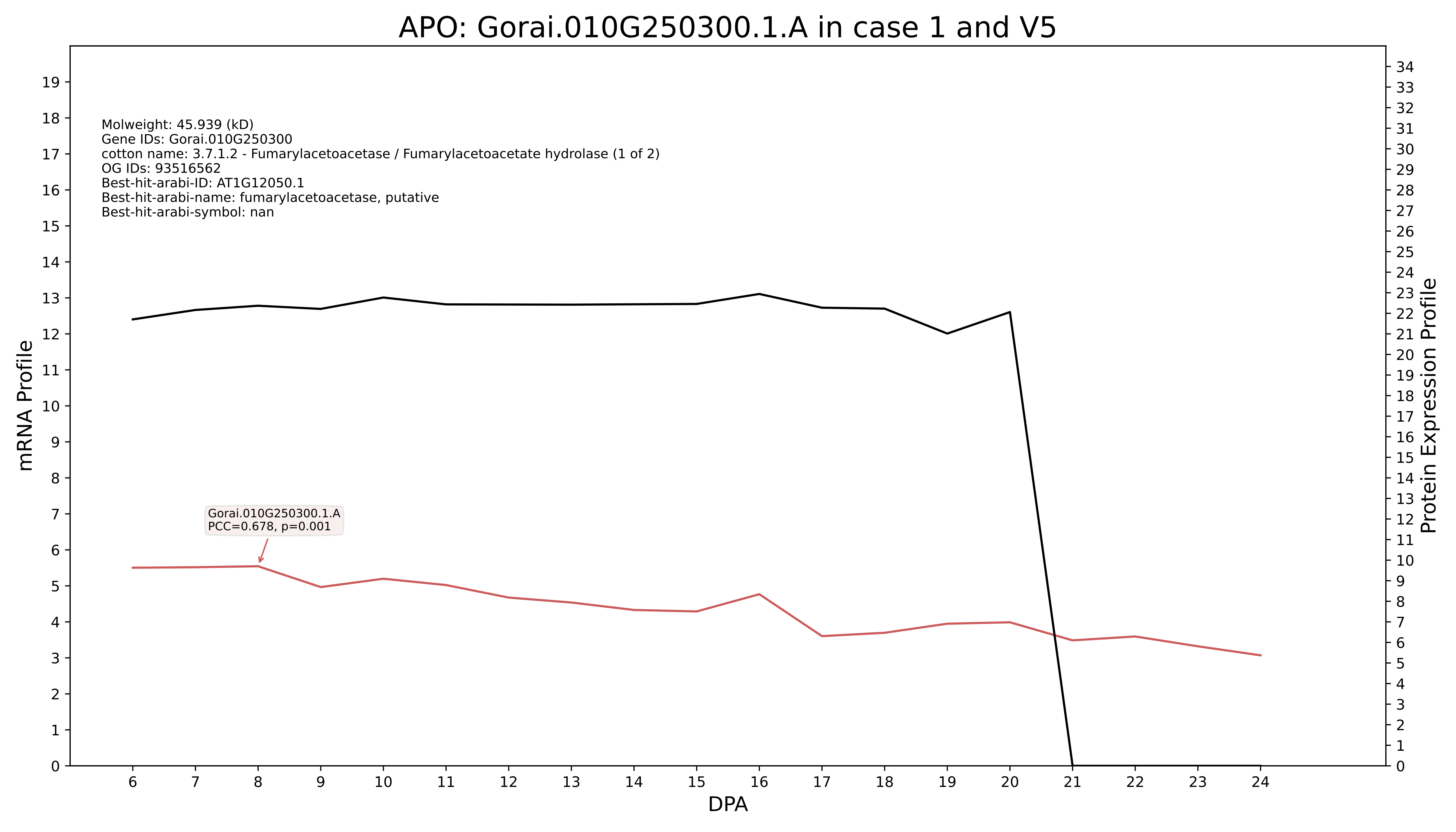

APO: Gorai.010G250300.1.A in case 1 and V5
Molweight: 45.939 (kD)
Gene IDs: Gorai.010G250300
cotton name: 3.7.1.2 - Fumarylacetoacetase / Fumarylacetoacetate hydrolase (1 of 2)
OG IDs: 93516562
Best-hit-arabi-ID: AT1G12050.1
Best-hit-arabi-name: fumarylacetoacetase, putative
Best-hit-arabi-symbol: nan
Gorai.010G250300.1.A
PCC=0.678, p=0.001
mRNA Profile
Protein Expression Profile
DPA

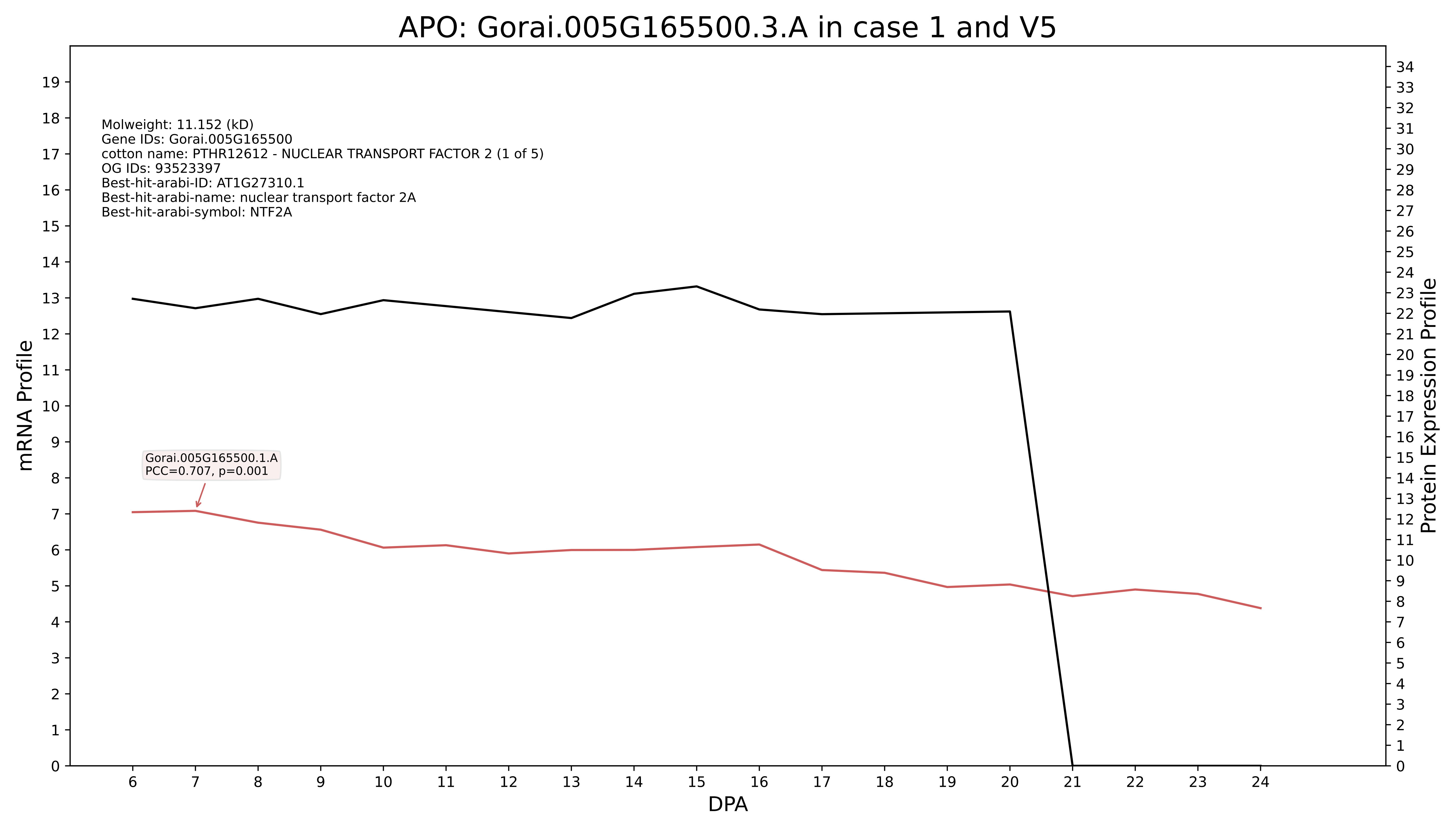

APO: Gorai.005G165500.3.A in case 1 and V5
Molweight: 11.152 (kD)
Gene IDs: Gorai.005G165500
cotton name: PTHR12612 - NUCLEAR TRANSPORT FACTOR 2 (1 of 5)
OG IDs: 93523397
Best-hit-arabi-ID: AT1G27310.1
Best-hit-arabi-name: nuclear transport factor 2A
Best-hit-arabi-symbol: NTF2A
Gorai.005G165500.1.A
PCC=0.707, p=0.001
mRNA Profile
Protein Expression Profile
DPA

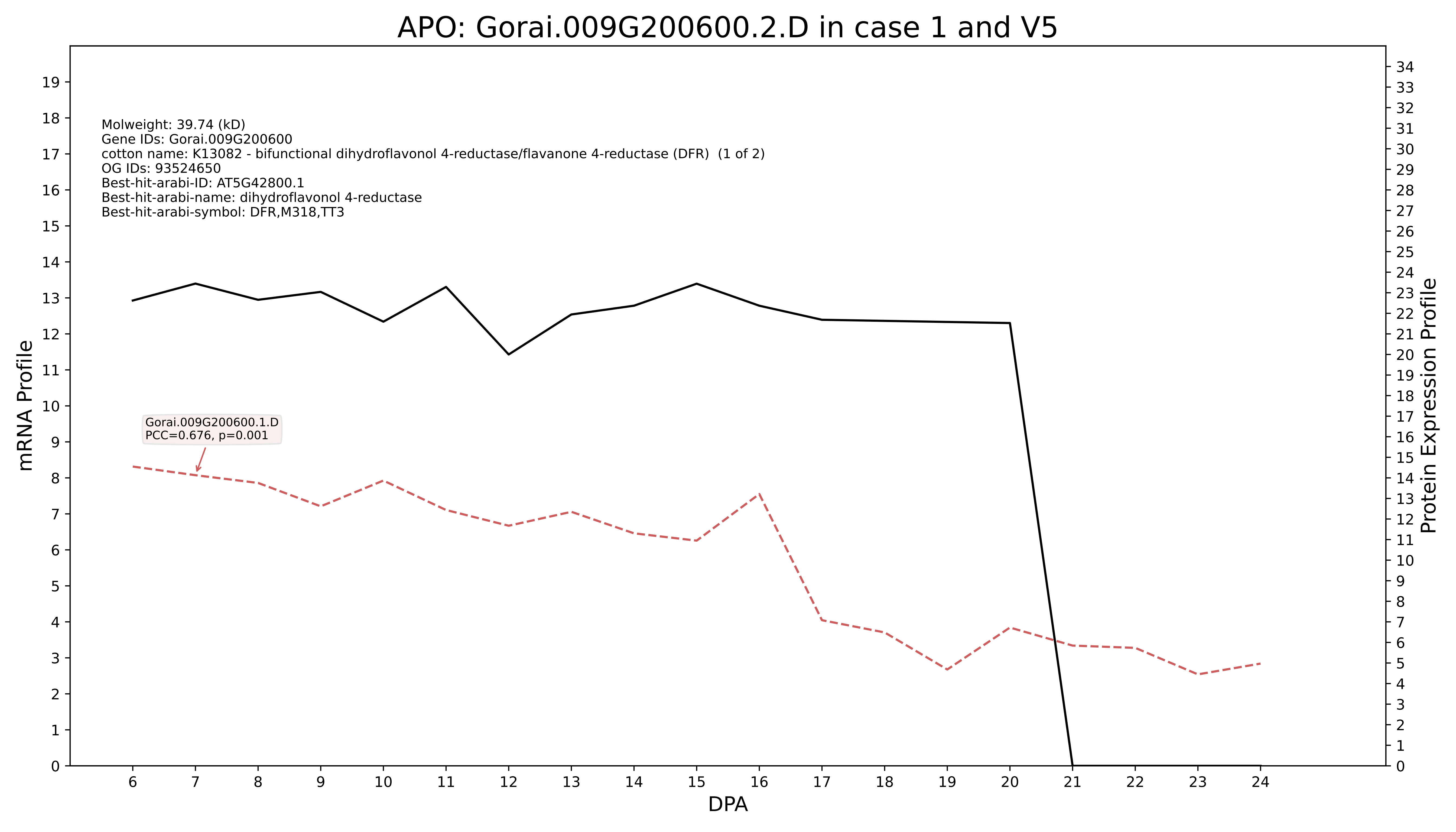
APO: Gorai.009G200600.2.D in case 1 and V5
Molweight: 39.74 (kD)
Gene IDs: Gorai.009G200600
cotton name: K13082 - bifunctional dihydroflavonol 4-reductase/flavanone 4-reductase (DFR) (1 of 2)
OG IDs: 93524650
Best-hit-arabi-ID: AT5G42800.1
Best-hit-arabi-name: dihydroflavonol 4-reductase
Best-hit-arabi-symbol: DFR,M318,TT3
Gorai.009G200600.1.D
PCC=0.676, p=0.001
mRNA Profile
Protein Expression Profile
DPA

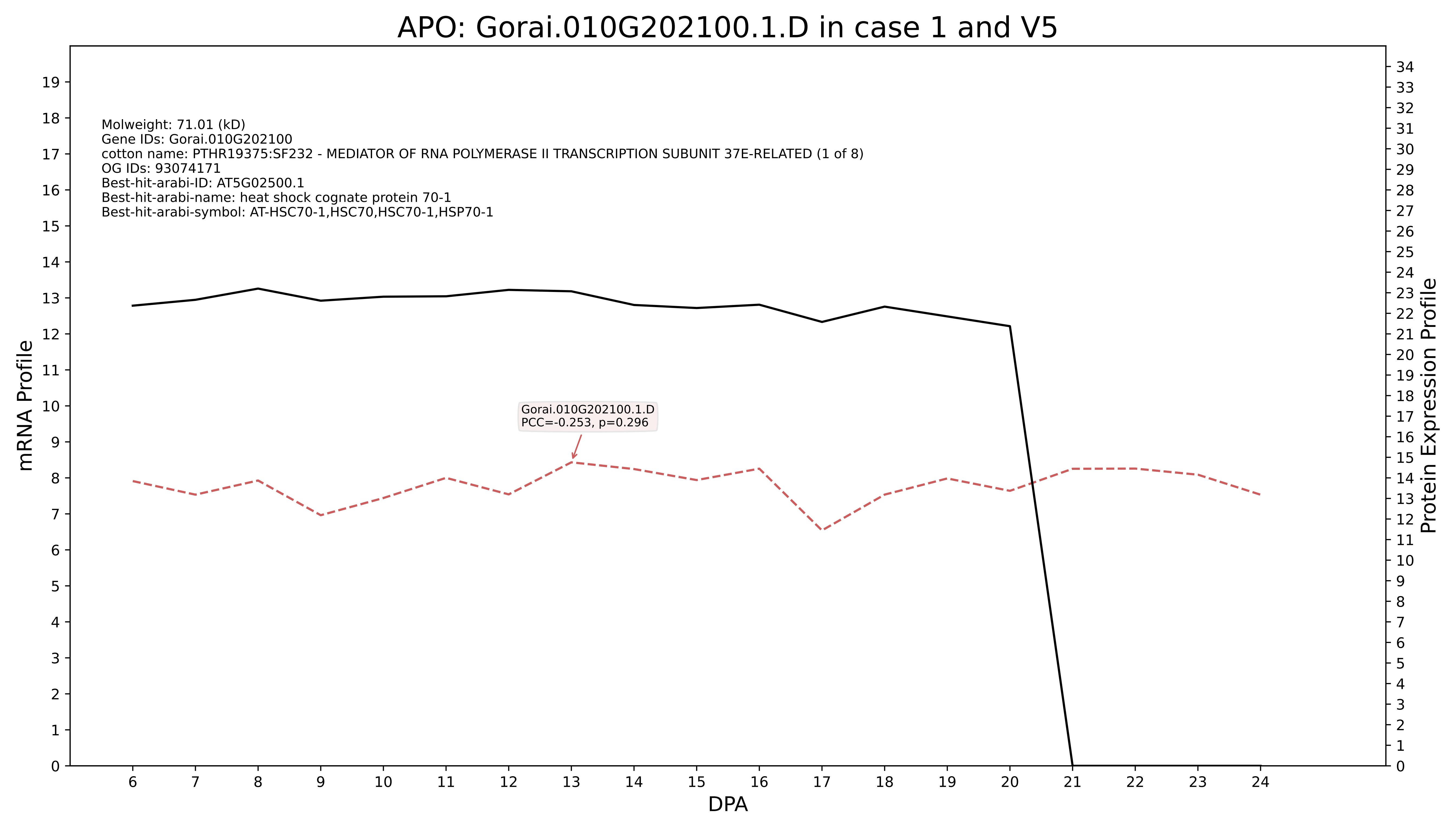

APO: Gorai.010G202100.1.D in case 1 and V5
Molweight: 71.01 (kD)
Gene IDs: Gorai.010G202100
cotton name: PTHR19375:SF232 - MEDIATOR OF RNA POLYMERASE II TRANSCRIPTION SUBUNIT 37E-RELATED (1 of 8)
OG IDs: 93074171
Best-hit-arabi-ID: AT5G02500.1
Best-hit-arabi-name: heat shock cognate protein 70-1
Best-hit-arabi-symbol: AT-HSC70-1,HSC70,HSC70-1,HSP70-1
Gorai.010G202100.1.D
PCC=-0.253, p=0.296
mRNA Profile
Protein Expression Profile
DPA

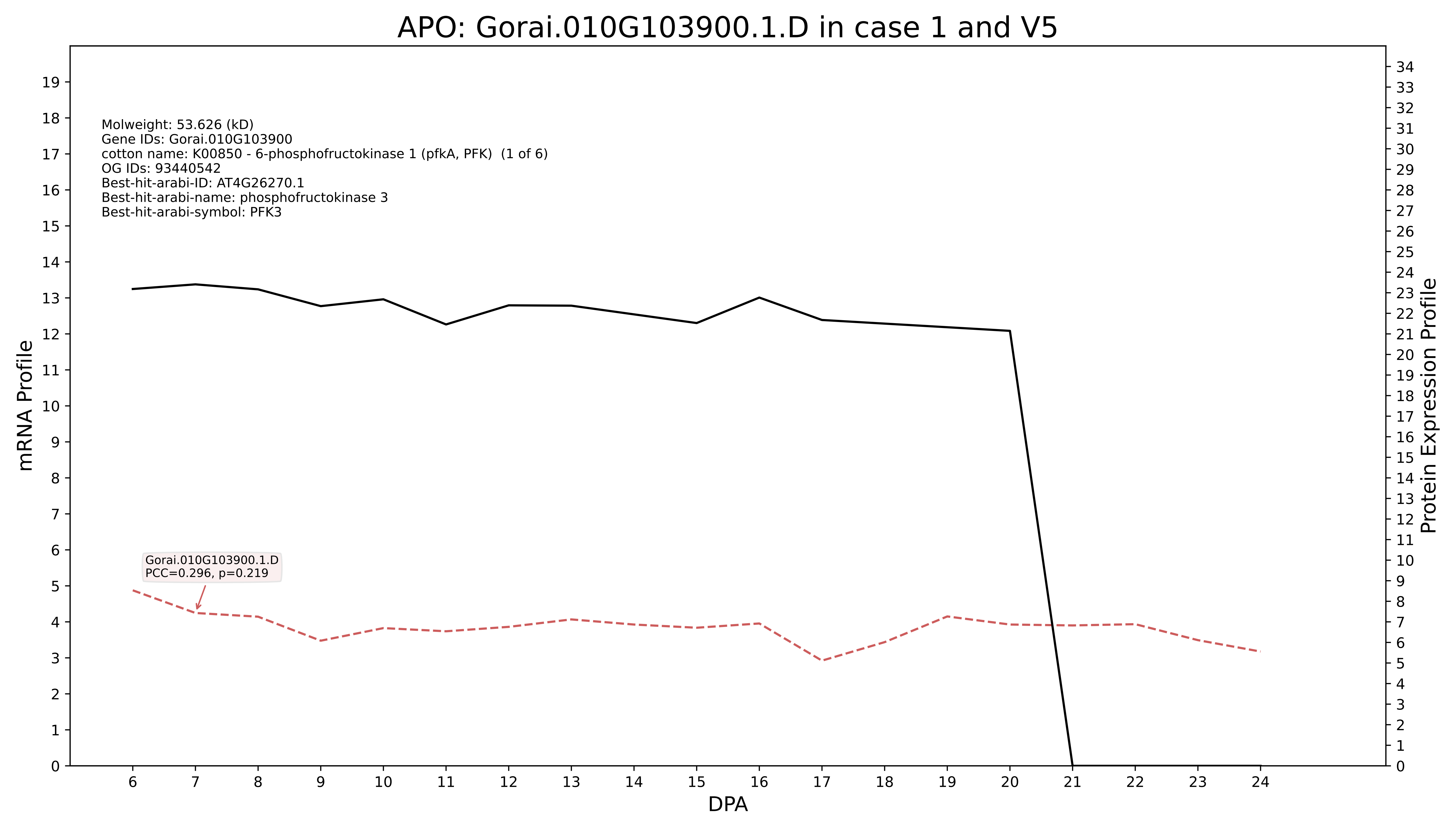
APO: Gorai.010G103900.1.D in case 1 and V5
Molweight: 53.626 (kD)
Gene IDs: Gorai.010G103900
cotton name: K00850 - 6-phosphofructokinase 1 (pfkA, PFK) (1 of 6)
OG IDs: 93440542
Best-hit-arabi-ID: AT4G26270.1
Best-hit-arabi-name: phosphofructokinase 3
Best-hit-arabi-symbol: PFK3
Gorai.010G103900.1.D
PCC=0.296, p=0.219
mRNA Profile
Protein Expression Profile
DPA

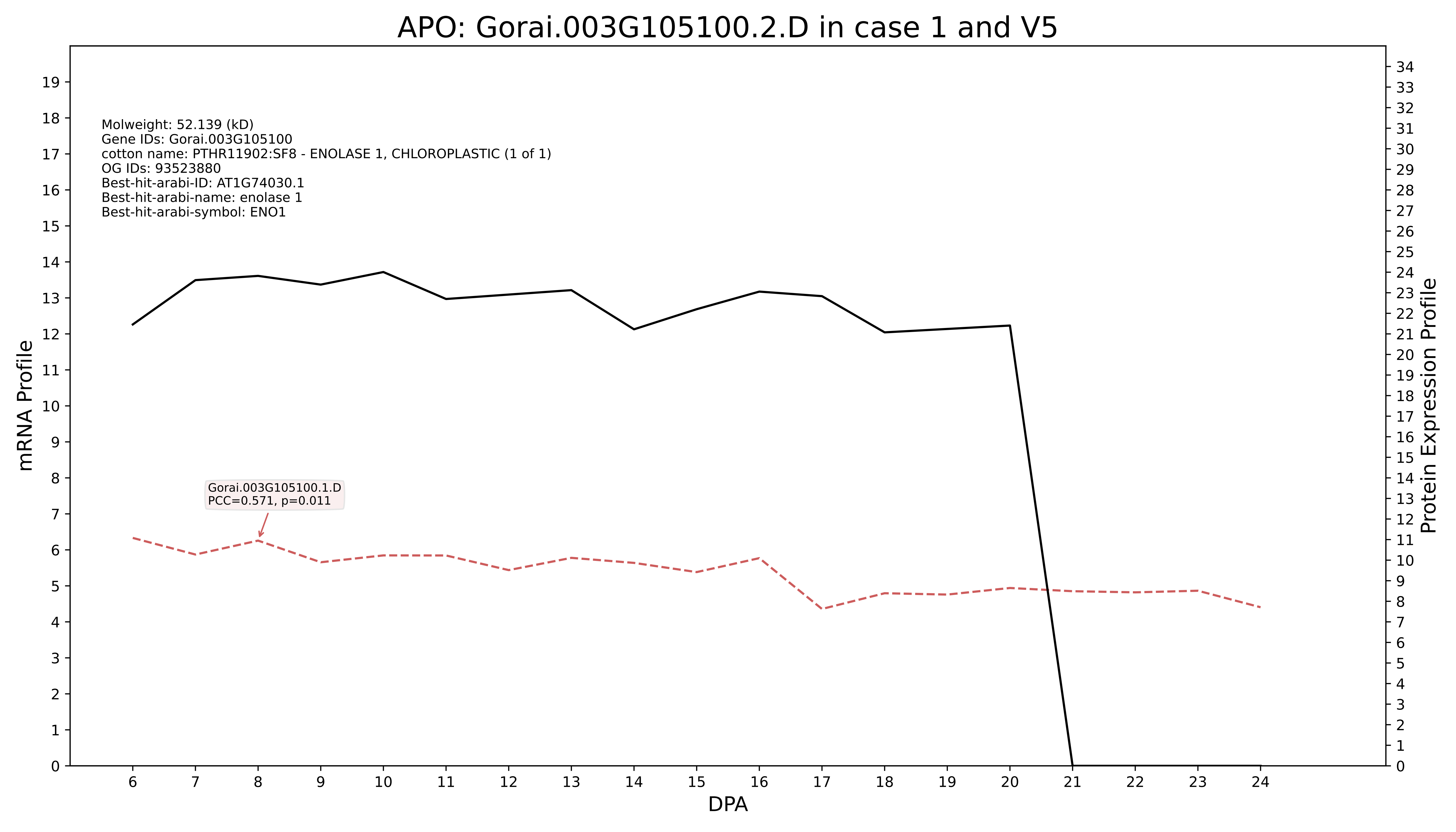

APO: Gorai.003G105100.2.D in case 1 and V5
Molweight: 52.139 (kD)
Gene IDs: Gorai.003G105100
cotton name: PTHR11902:SF8 - ENOLASE 1, CHLOROPLASTIC (1 of 1)
OG IDs: 93523880
Best-hit-arabi-ID: AT1G74030.1
Best-hit-arabi-name: enolase 1
Best-hit-arabi-symbol: ENO1
Gorai.003G105100.1.D
PCC=0.571, p=0.011
mRNA Profile
Protein Expression Profile
DPA

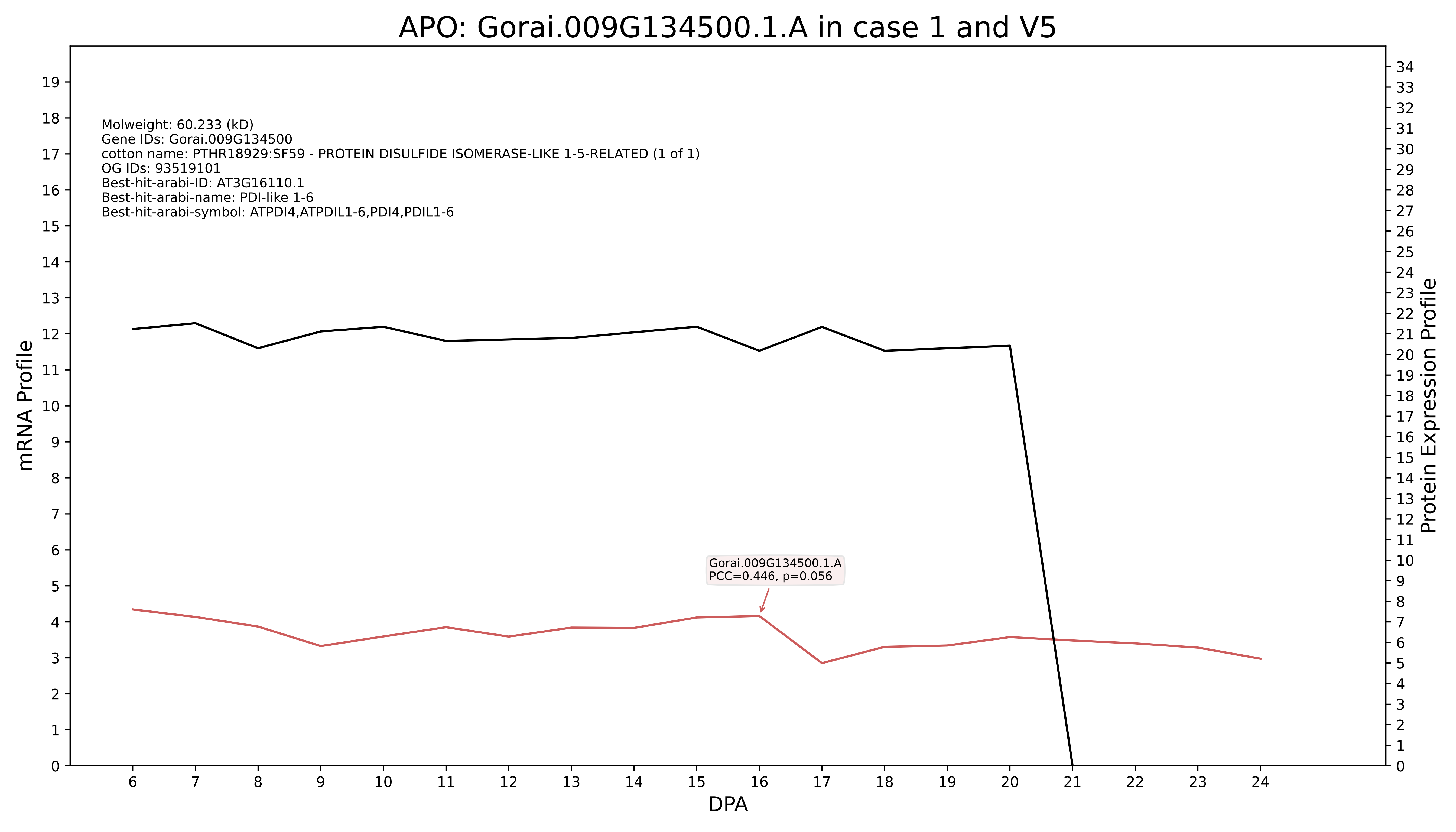
APO: Gorai.009G134500.1.A in case 1 and V5
Molweight: 60.233 (kD)
Gene IDs: Gorai.009G134500
cotton name: PTHR18929:SF59 - PROTEIN DISULFIDE ISOMERASE-LIKE 1-5-RELATED (1 of 1)
OG IDs: 93519101
Best-hit-arabi-ID: AT3G16110.1
Best-hit-arabi-name: PDI-like 1-6
Best-hit-arabi-symbol: ATPDI4,ATPDIL1-6,PDI4,PDIL1-6
Gorai.009G134500.1.A
PCC=0.446, p=0.056
mRNA Profile
Protein Expression Profile
DPA

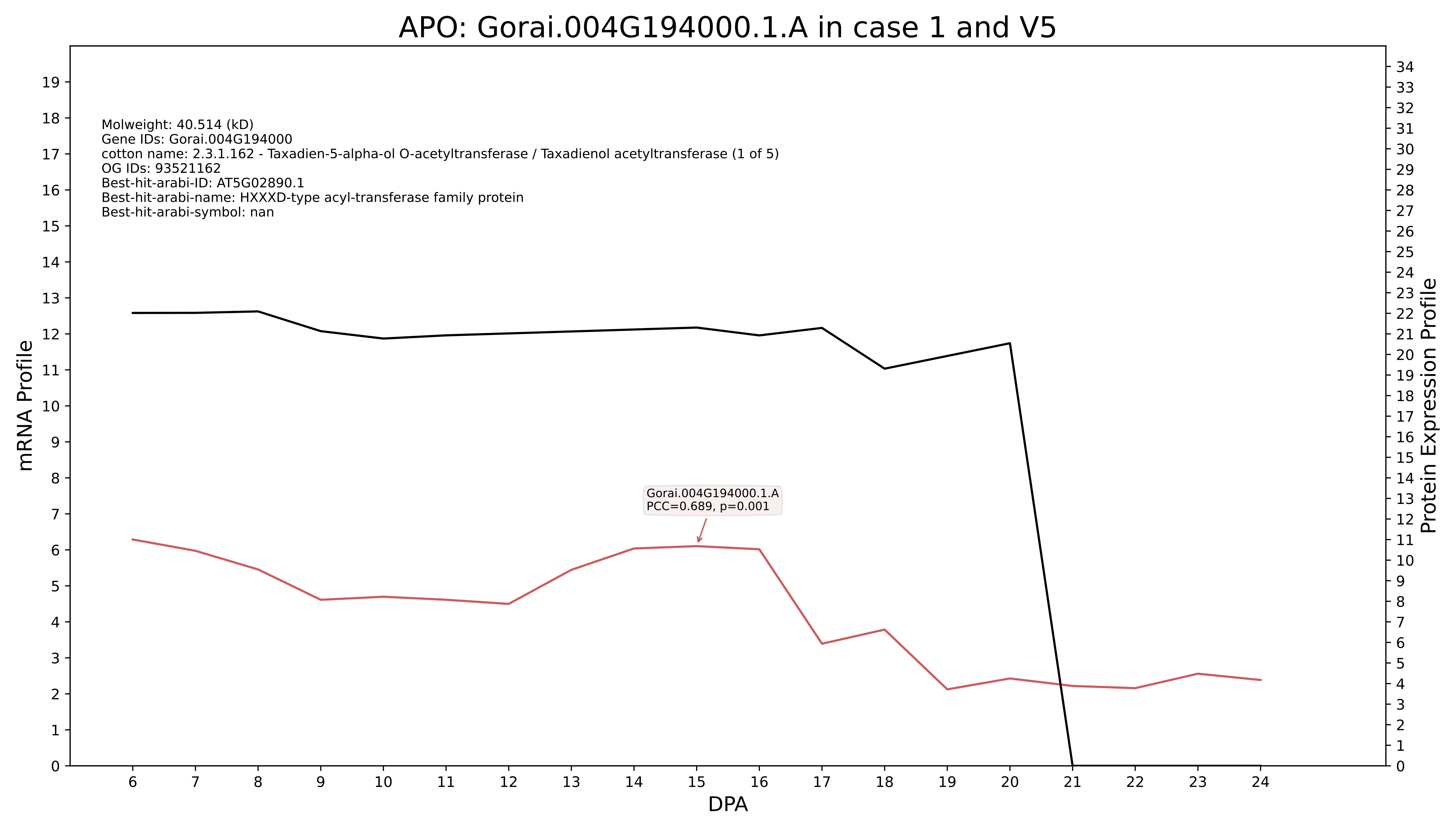

APO: Gorai.004G194000.1.A in case 1 and V5
Molweight: 40.514 (kD)
Gene IDs: Gorai.004G194000
cotton name: 2.3.1.162 - Taxadien-5-alpha-ol O-acetyltransferase / Taxadienol acetyltransferase (1 of 5)
OG IDs: 93521162
Best-hit-arabi-ID: AT5G02890.1
Best-hit-arabi-name: HXXXD-type acyl-transferase family protein
Best-hit-arabi-symbol: nan
Gorai.004G194000.1.A
PCC=0.689, p=0.001
mRNA Profile
Protein Expression Profile
DPA

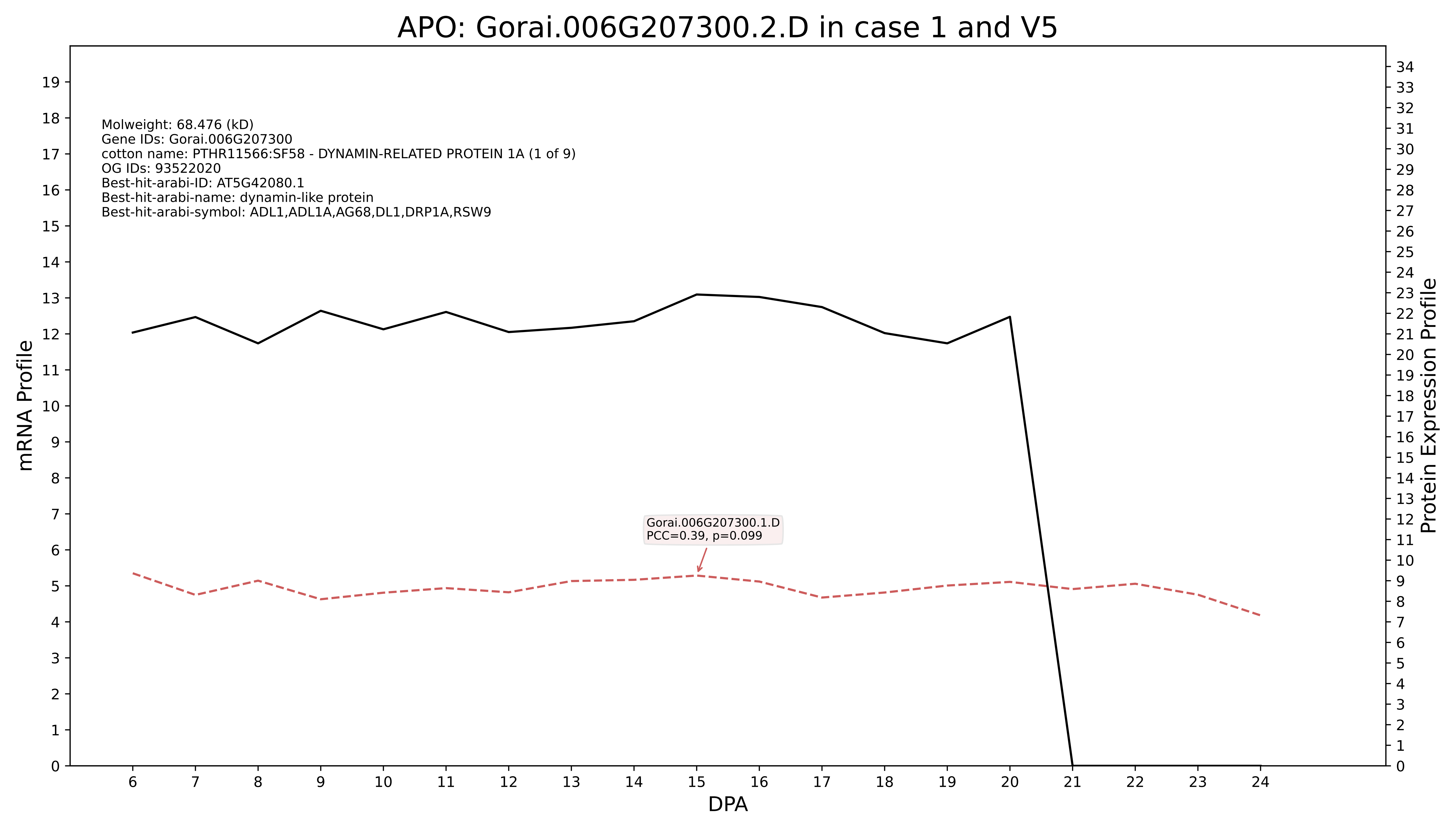
APO: Gorai.006G207300.2.D in case 1 and V5
Molweight: 68.476 (kD)
Gene IDs: Gorai.006G207300
cotton name: PTHR11566:SF58 - DYNAMIN-RELATED PROTEIN 1A (1 of 9)
OG IDs: 93522020
Best-hit-arabi-ID: AT5G42080.1
Best-hit-arabi-name: dynamin-like protein
Best-hit-arabi-symbol: ADL1,ADL1A,AG68,DL1,DRP1A,RSW9
Gorai.006G207300.1.D
PCC=0.39, p=0.099
mRNA Profile
Protein Expression Profile
DPA

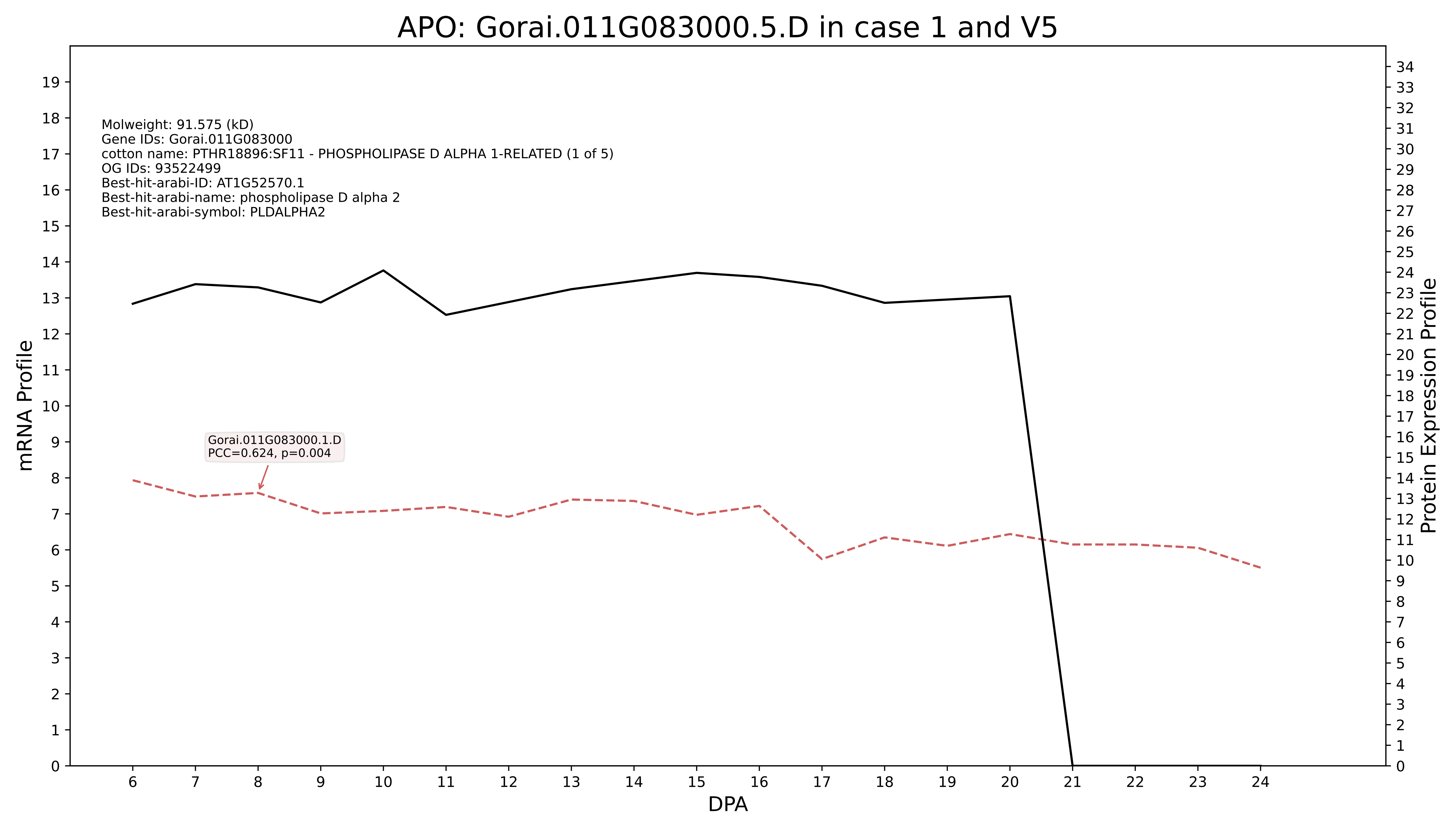
APO: Gorai.011G083000.5.D in case 1 and V5
Molweight: 91.575 (kD)
Gene IDs: Gorai.011G083000
cotton name: PTHR18896:SF11 - PHOSPHOLIPASE D ALPHA 1-RELATED (1 of 5)
OG IDs: 93522499
Best-hit-arabi-ID: AT1G52570.1
Best-hit-arabi-name: phospholipase D alpha 2
Best-hit-arabi-symbol: PLDALPHA2
Gorai.011G083000.1.D
PCC=0.624, p=0.004
mRNA Profile
Protein Expression Profile
DPA

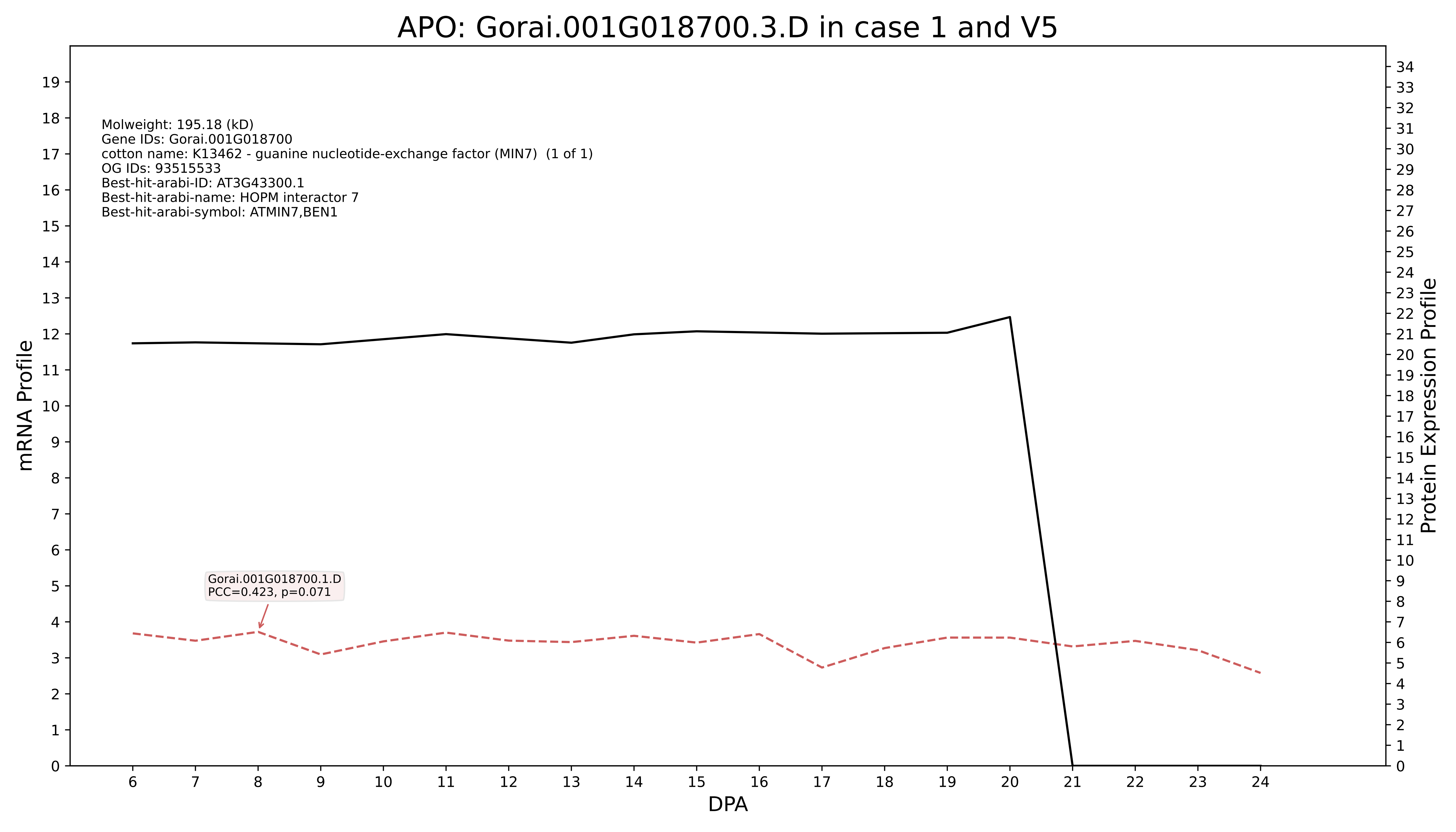

APO: Gorai.001G018700.3.D in case 1 and V5
Molweight: 195.18 (kD)
Gene IDs: Gorai.001G018700
cotton name: K13462 - guanine nucleotide-exchange factor (MIN7) (1 of 1)
OG IDs: 93515533
Best-hit-arabi-ID: AT3G43300.1
Best-hit-arabi-name: HOPM interactor 7
Best-hit-arabi-symbol: ATMIN7,BEN1
Gorai.001G018700.1.D
PCC=0.423, p=0.071
mRNA Profile
Protein Expression Profile
DPA

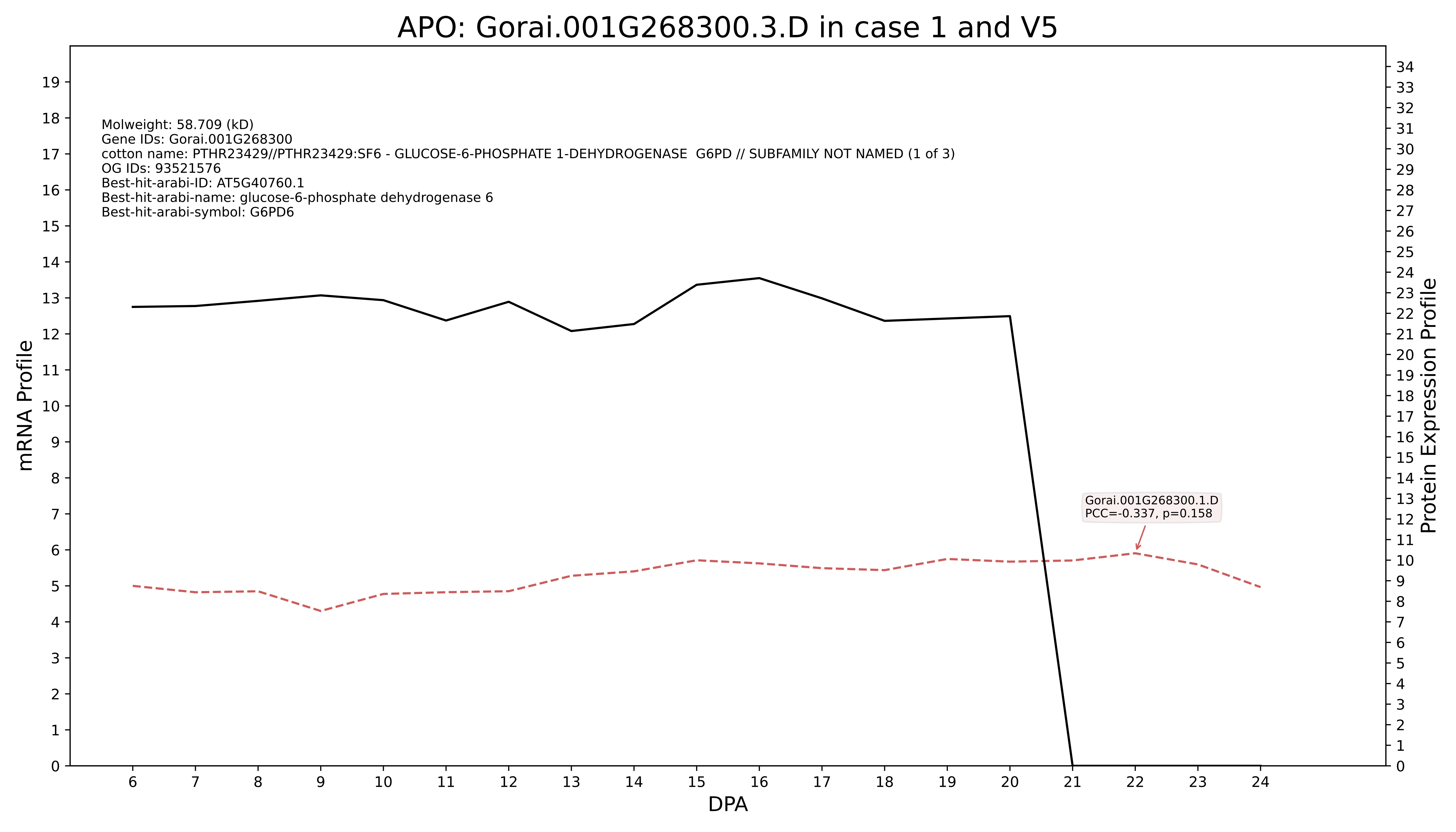
APO: Gorai.001G268300.3.D in case 1 and V5
Molweight: 58.709 (kD)
Gene IDs: Gorai.001G268300
cotton name: PTHR23429//PTHR23429:SF6 - GLUCOSE-6-PHOSPHATE 1-DEHYDROGENASE G6PD // SUBFAMILY NOT NAMED (1 of 3)
OG IDs: 93521576
Best-hit-arabi-ID: AT5G40760.1
Best-hit-arabi-name: glucose-6-phosphate dehydrogenase 6
Best-hit-arabi-symbol: G6PD6
Gorai.001G268300.1.D
PCC=-0.337, p=0.158
mRNA Profile
Protein Expression Profile
DPA

### APO: Gorai.002G255500.5.D in case 1 and V5

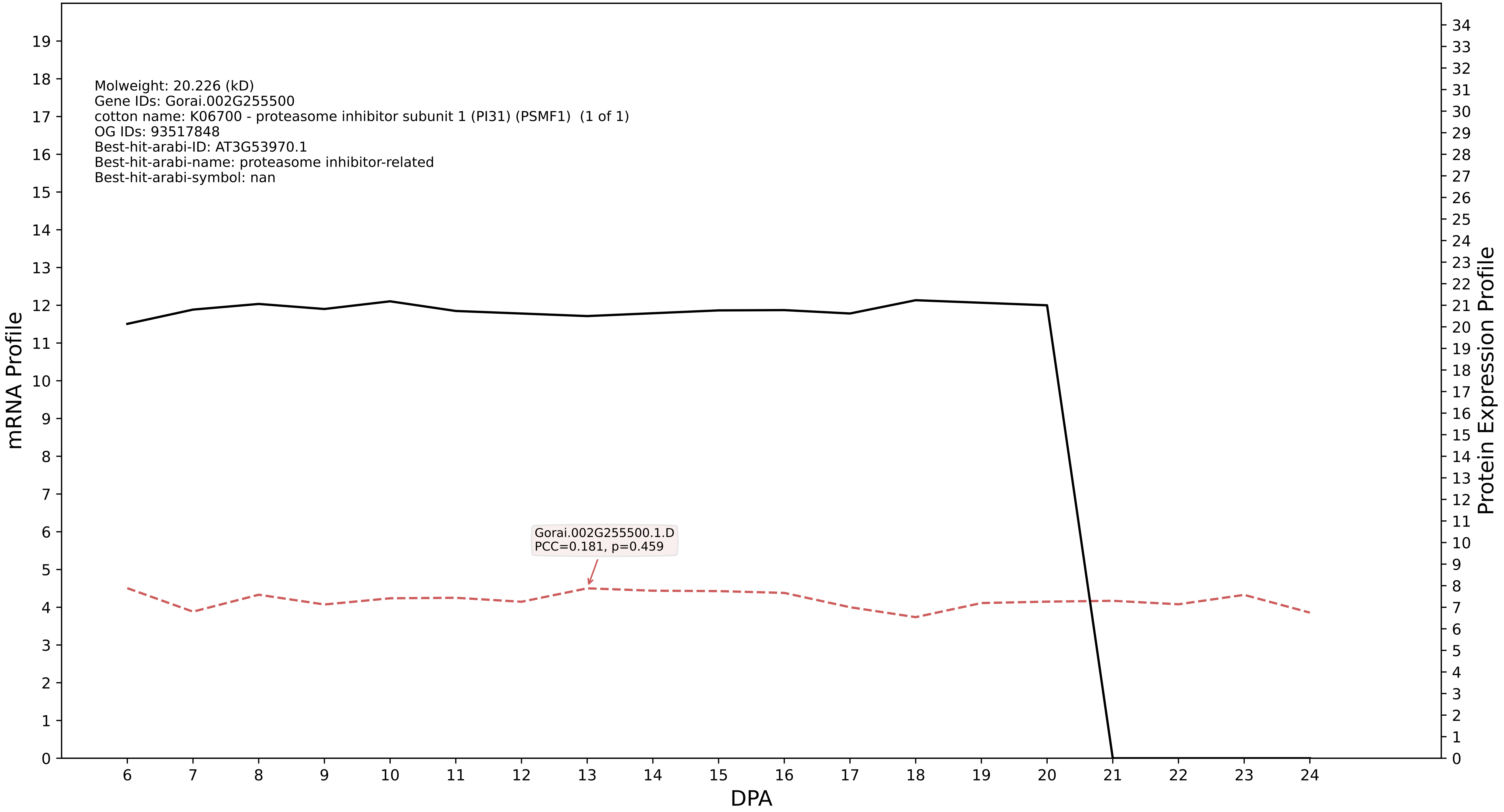

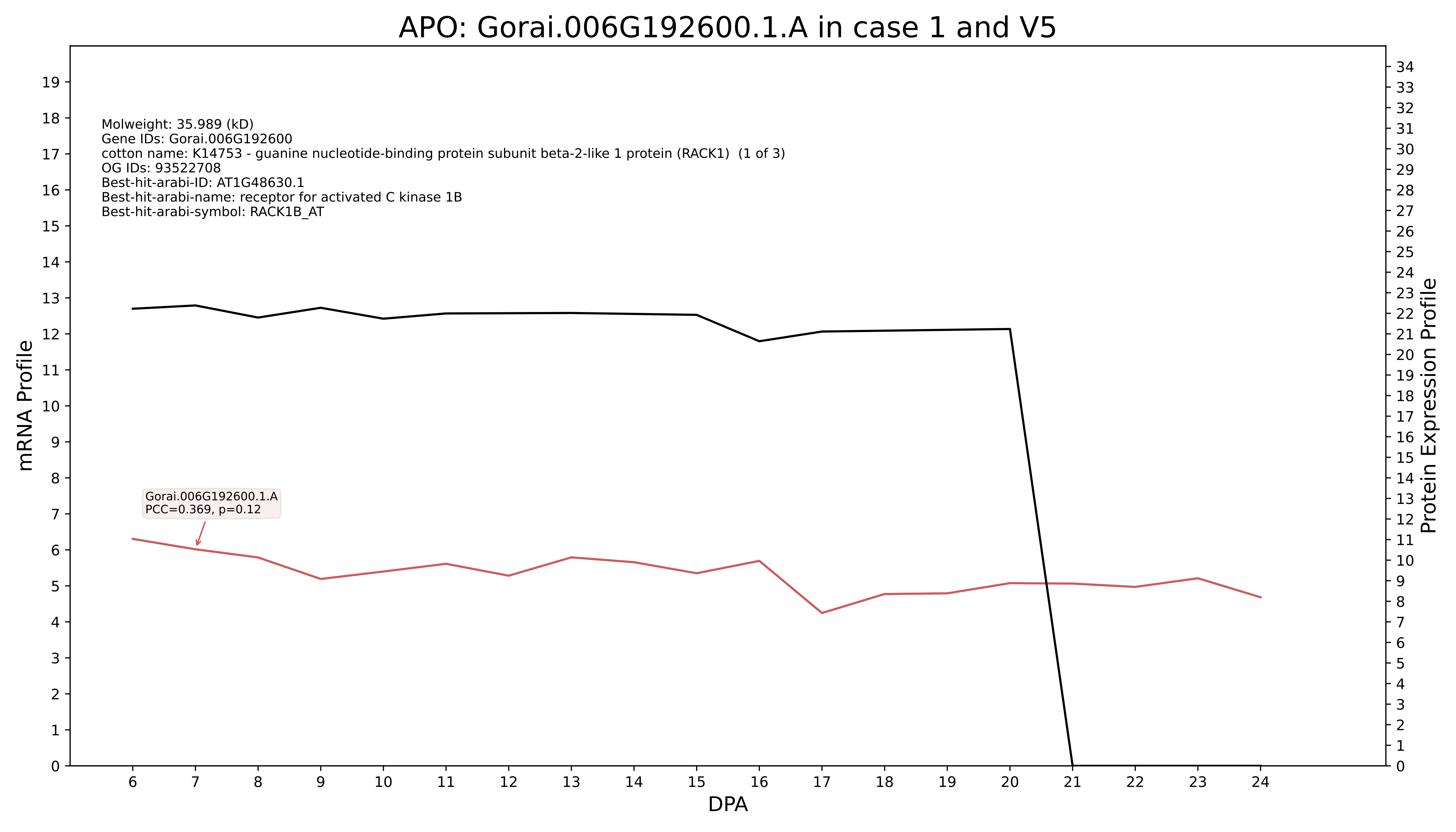
APO: Gorai.006G192600.1.A in case 1 and V5
Molweight: 35.989 (kD)
Gene IDs: Gorai.006G192600
cotton name: K14753 - guanine nucleotide-binding protein subunit beta-2-like 1 protein (RACK1) (1 of 3)
OG IDs: 93522708
Best-hit-arabi-ID: AT1G48630.1
Best-hit-arabi-name: receptor for activated C kinase 1B
Best-hit-arabi-symbol: RACK1B_AT
Gorai.006G192600.1.A
PCC=0.369, p=0.12
mRNA Profile
Protein Expression Profile
DPA

### APO: Gorai.013G129200.2.D in case 1 and V5

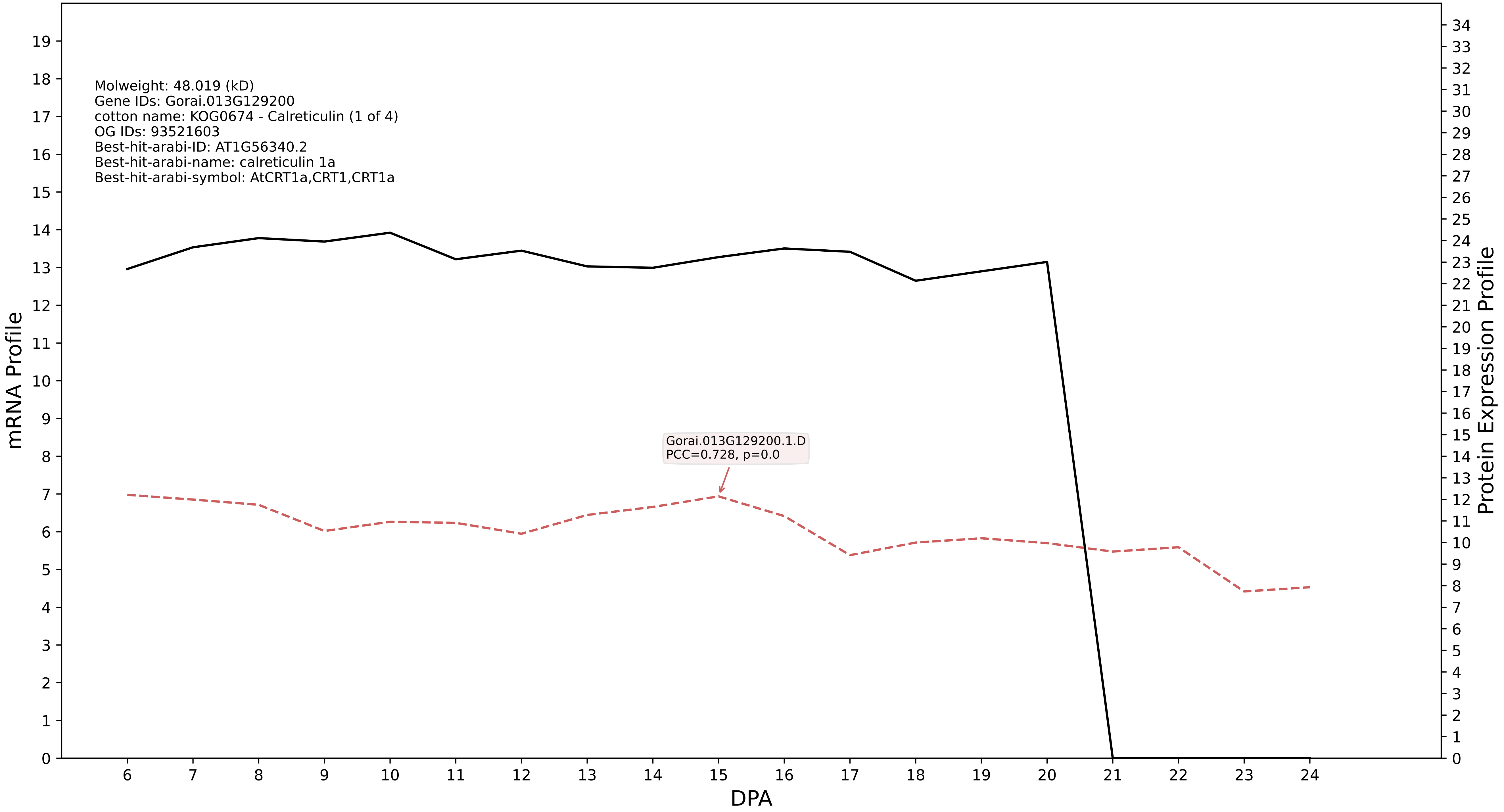

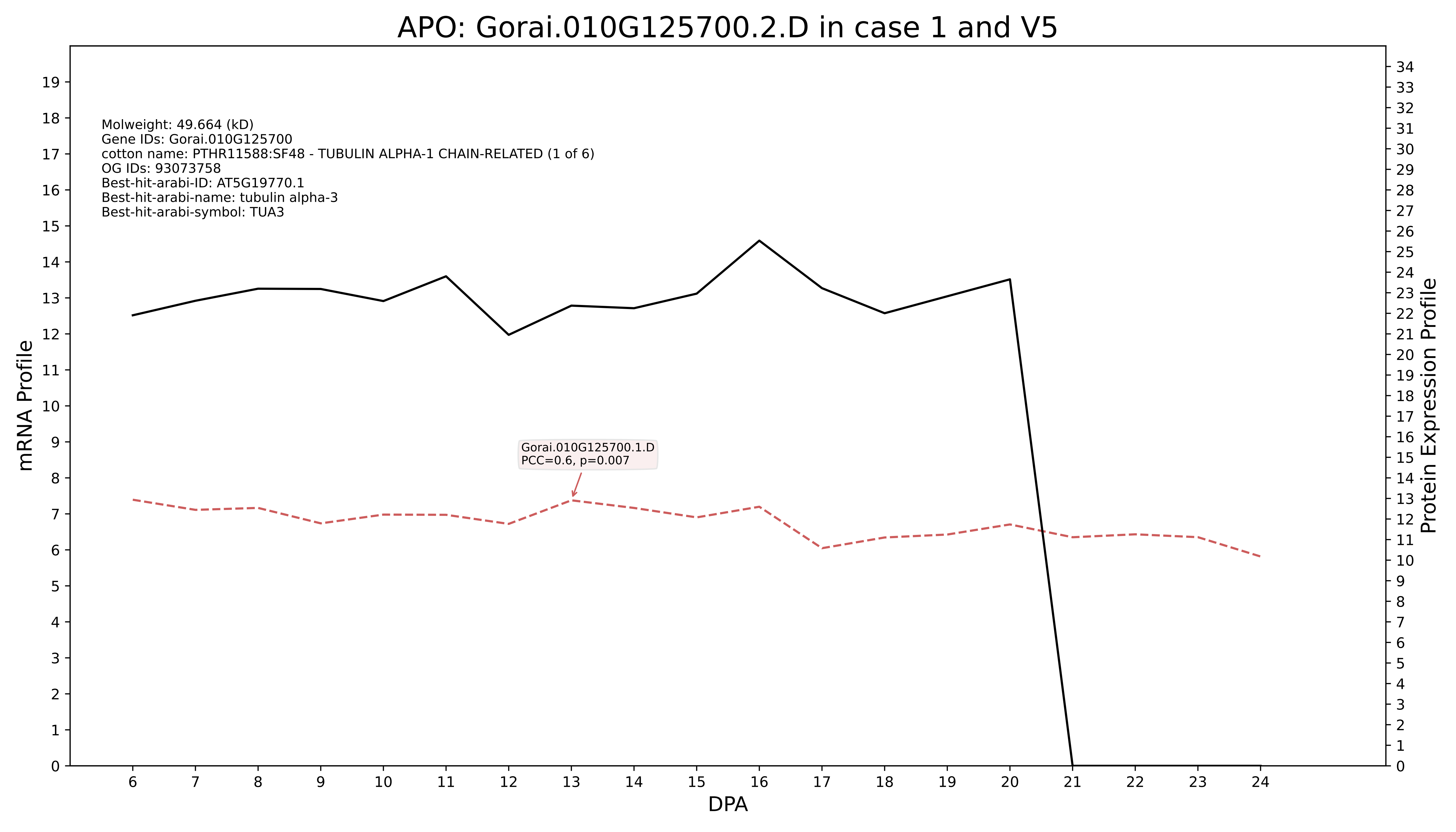

APO: Gorai.010G125700.2.D in case 1 and V5
Molweight: 49.664 (kD)
Gene IDs: Gorai.010G125700
cotton name: PTHR11588:SF48 - TUBULIN ALPHA-1 CHAIN-RELATED (1 of 6)
OG IDs: 93073758
Best-hit-arabi-ID: AT5G19770.1
Best-hit-arabi-name: tubulin alpha-3
Best-hit-arabi-symbol: TUA3
Gorai.010G125700.1.D
PCC=0.6, p=0.007
mRNA Profile
Protein Expression Profile
DPA

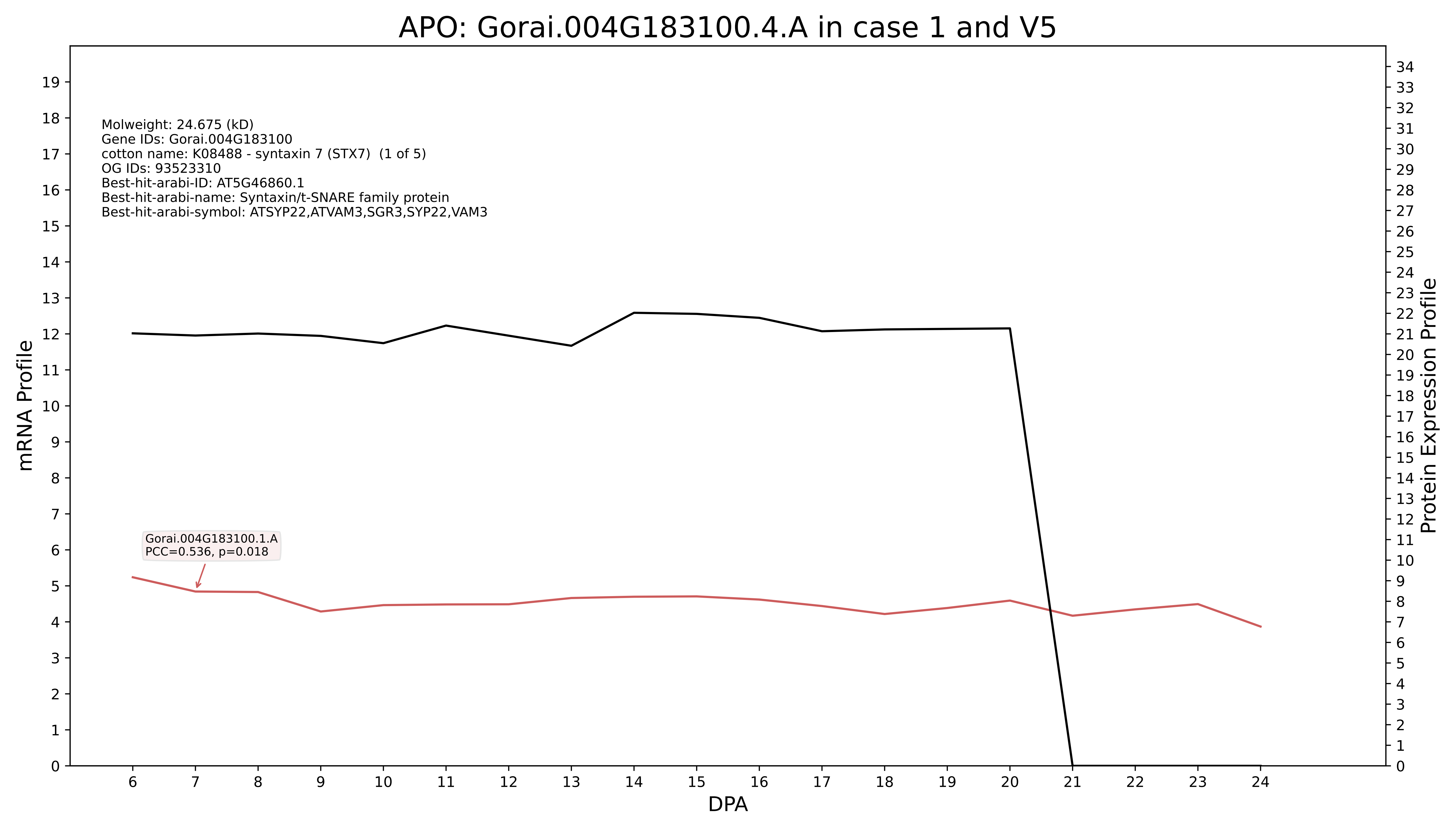
APO: Gorai.004G183100.4.A in case 1 and V5
Molweight: 24.675 (kD)
Gene IDs: Gorai.004G183100
cotton name: K08488 - syntaxin 7 (STX7) (1 of 5)
OG IDs: 93523310
Best-hit-arabi-ID: AT5G46860.1
Best-hit-arabi-name: Syntaxin/t-SNARE family protein
Best-hit-arabi-symbol: ATSYP22,ATVAM3,SGR3,SYP22,VAM3
Gorai.004G183100.1.A
PCC=0.536, p=0.018
mRNA Profile
Protein Expression Profile
DPA

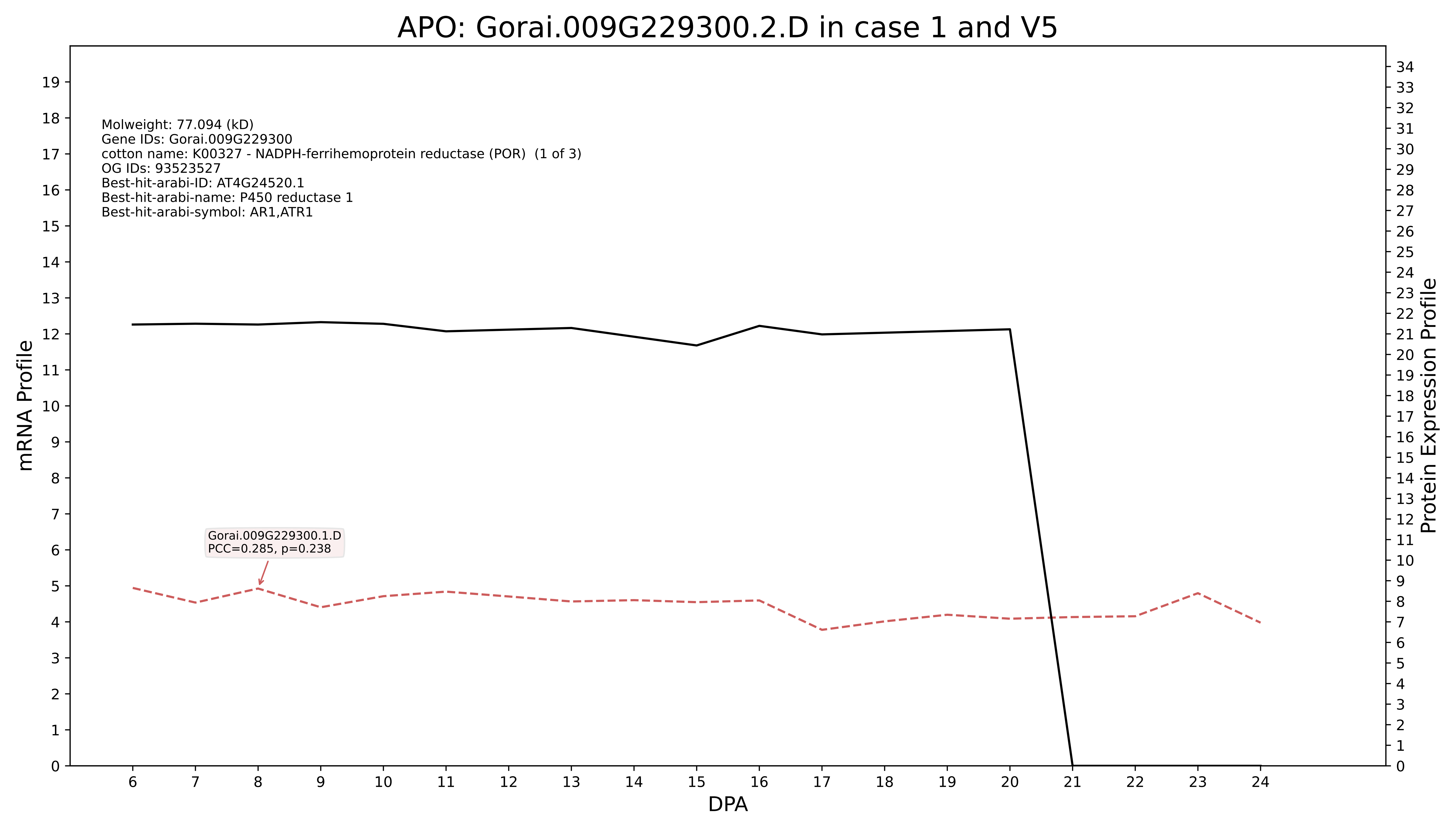
APO: Gorai.009G229300.2.D in case 1 and V5
Molweight: 77.094 (kD)
Gene IDs: Gorai.009G229300
cotton name: K00327 - NADPH-ferrihemoprotein reductase (POR) (1 of 3)
OG IDs: 93523527
Best-hit-arabi-ID: AT4G24520.1
Best-hit-arabi-name: P450 reductase 1
Best-hit-arabi-symbol: AR1,ATR1
Gorai.009G229300.1.D
PCC=0.285, p=0.238
mRNA Profile
Protein Expression Profile
DPA

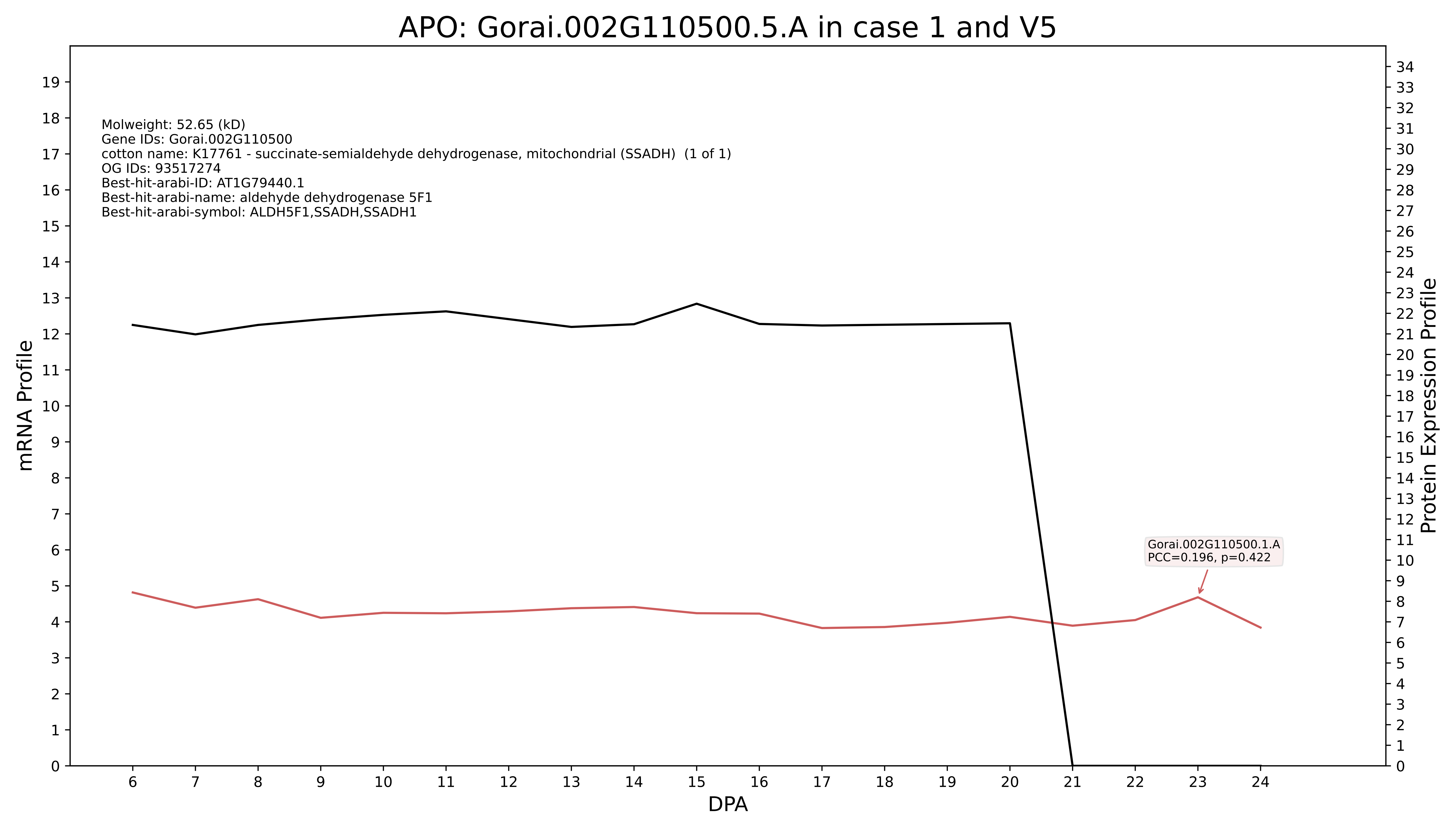

APO: Gorai.002G110500.5.A in case 1 and V5
Molweight: 52.65 (kD)
Gene IDs: Gorai.002G110500
cotton name: K17761 - succinate-semialdehyde dehydrogenase, mitochondrial (SSADH) (1 of 1)
OG IDs: 93517274
Best-hit-arabi-ID: AT1G79440.1
Best-hit-arabi-name: aldehyde dehydrogenase 5F1
Best-hit-arabi-symbol: ALDH5F1,SSADH,SSADH1
Gorai.002G110500.1.A
PCC=0.196, p=0.422
mRNA Profile
Protein Expression Profile
DPA

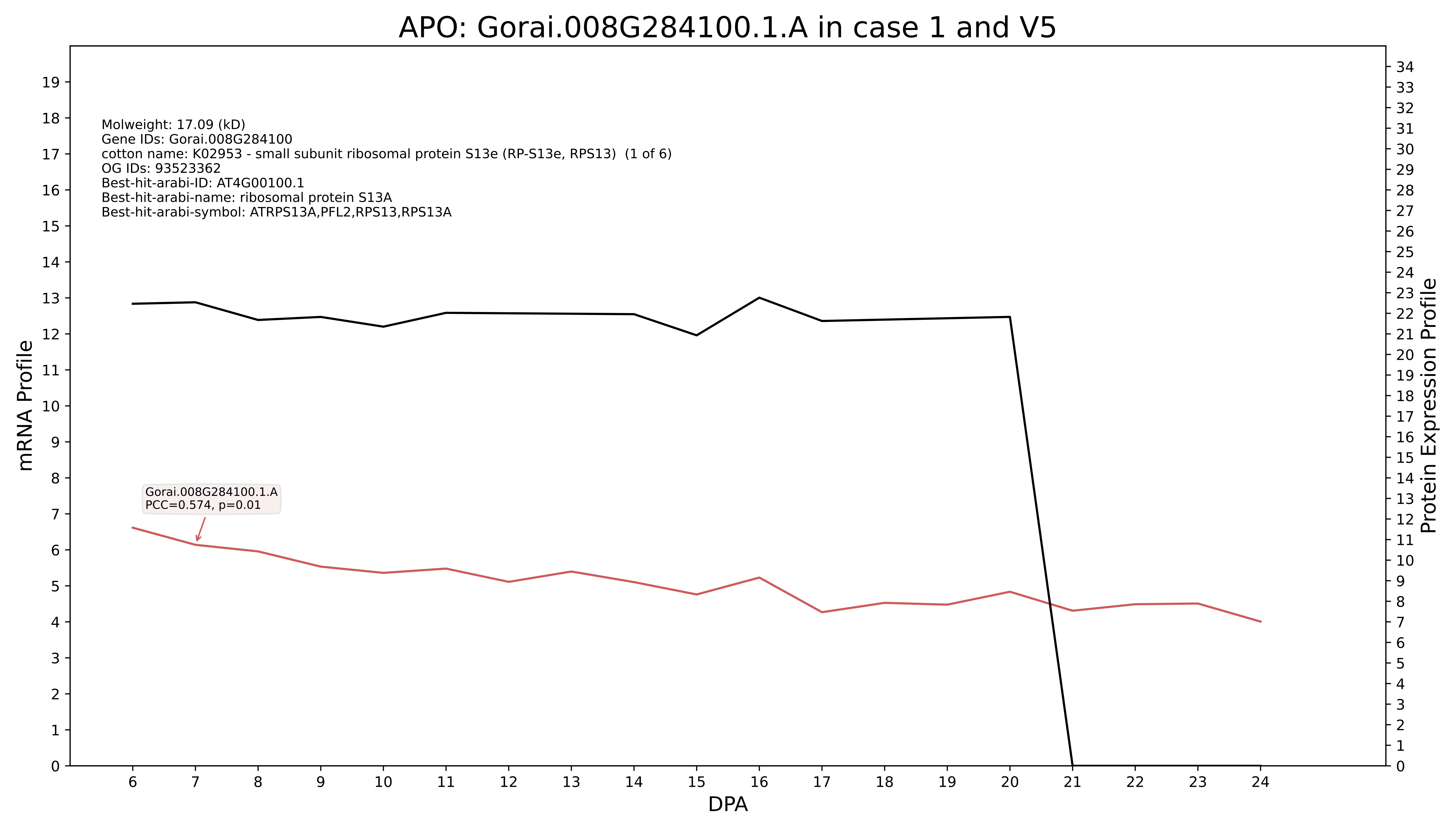
APO: Gorai.008G284100.1.A in case 1 and V5
Molweight: 17.09 (kD)
Gene IDs: Gorai.008G284100
cotton name: K02953 - small subunit ribosomal protein S13e (RP-S13e, RPS13) (1 of 6)
OG IDs: 93523362
Best-hit-arabi-ID: AT4G00100.1
Best-hit-arabi-name: ribosomal protein S13A
Best-hit-arabi-symbol: ATRPS13A,PFL2,RPS13,RPS13A
Gorai.008G284100.1.A
PCC=0.574, p=0.01
mRNA Profile
Protein Expression Profile
DPA

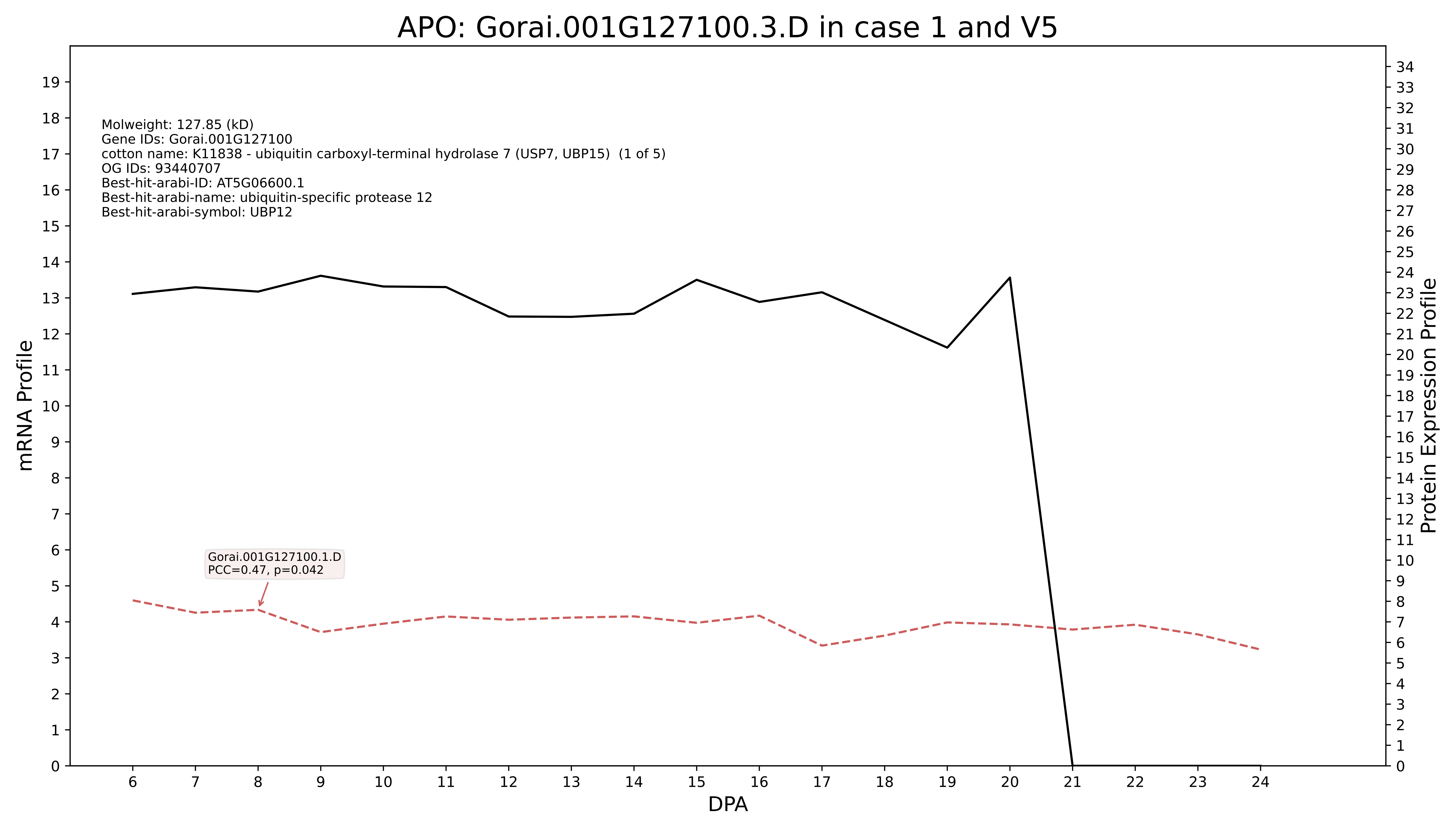
APO: Gorai.001G127100.3.D in case 1 and V5
Molweight: 127.85 (kD)
Gene IDs: Gorai.001G127100
cotton name: K11838 - ubiquitin carboxyl-terminal hydrolase 7 (USP7, UBP15) (1 of 5)
OG IDs: 93440707
Best-hit-arabi-ID: AT5G06600.1
Best-hit-arabi-name: ubiquitin-specific protease 12
Best-hit-arabi-symbol: UBP12
Gorai.001G127100.1.D
PCC=0.47, p=0.042
mRNA Profile
Protein Expression Profile
DPA

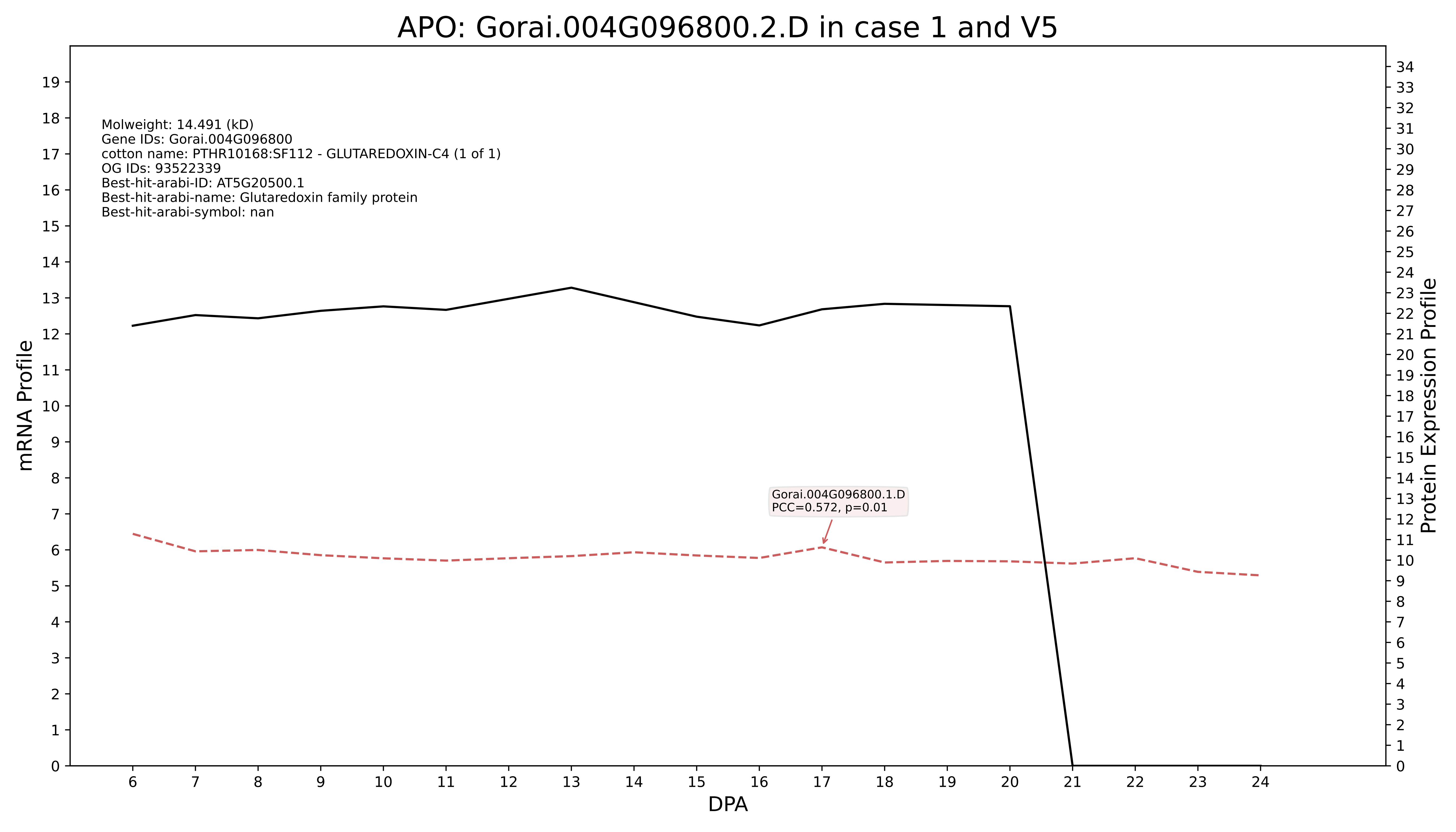

APO: Gorai.004G096800.2.D in case 1 and V5
Molweight: 14.491 (kD)
Gene IDs: Gorai.004G096800
cotton name: PTHR10168:SF112 - GLUTAREDOXIN-C4 (1 of 1)
OG IDs: 93522339
Best-hit-arabi-ID: AT5G20500.1
Best-hit-arabi-name: Glutaredoxin family protein
Best-hit-arabi-symbol: nan
Gorai.004G096800.1.D
PCC=0.572, p=0.01
mRNA Profile
Protein Expression Profile
DPA

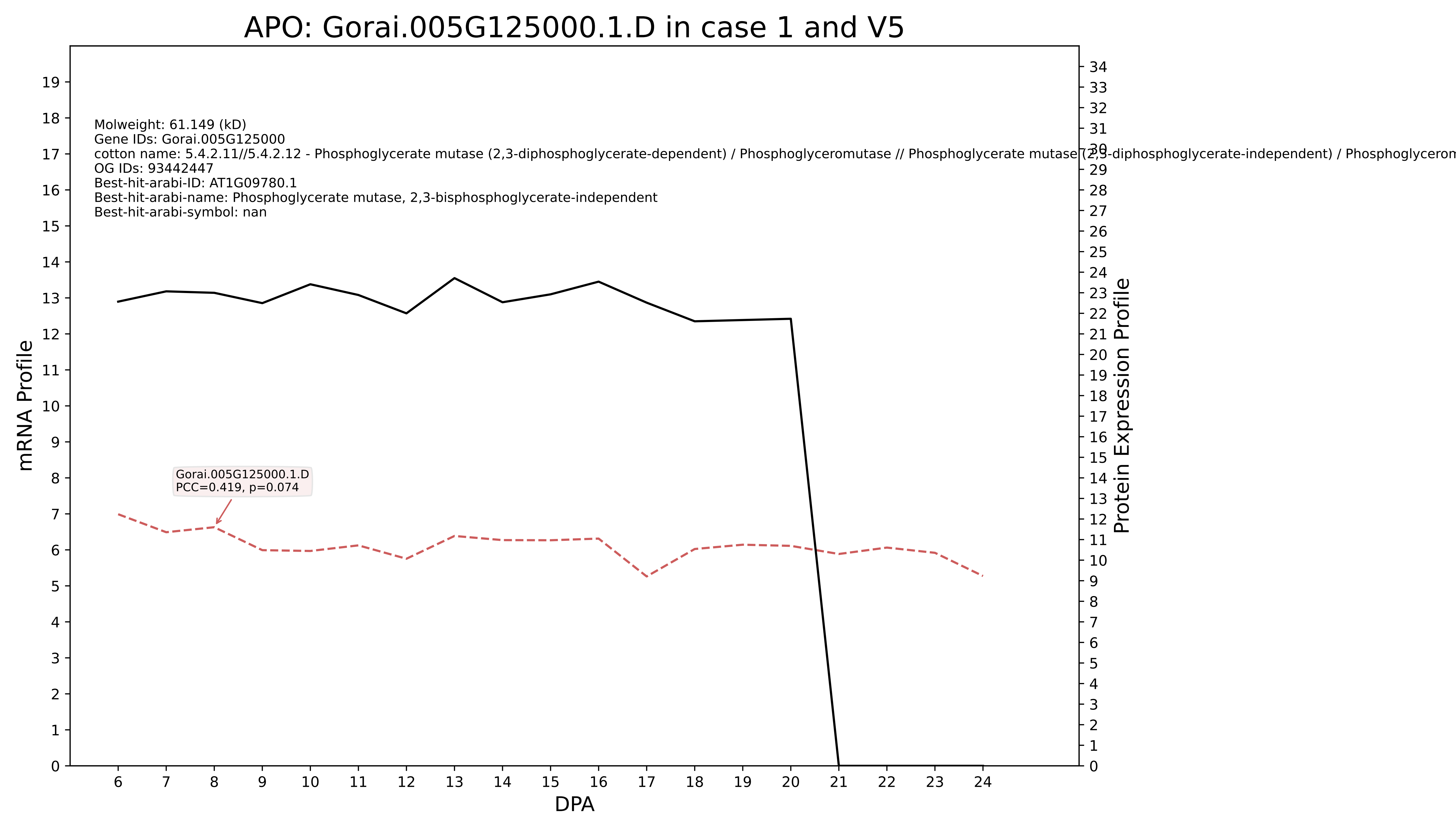
APO: Gorai.005G125000.1.D in case 1 and V5
Molweight: 61.149 (kD)
Gene IDs: Gorai.005G125000
cotton name: 5.4.2.11//5.4.2.12 - Phosphoglycerate mutase (2,3-diphosphoglycerate-dependent) / Phosphoglyceromutase // Phosphoglycerate mutase (2,3-diphosphoglycerate-independent) / Phosphoglyceron
OG IDs: 93442447
Best-hit-arabi-ID: AT1G09780.1
Best-hit-arabi-name: Phosphoglycerate mutase, 2,3-bisphosphoglycerate-independent
Best-hit-arabi-symbol: nan
Gorai.005G125000.1.D
PCC=0.419, p=0.074
mRNA Profile
Protein Expression Profile
DPA

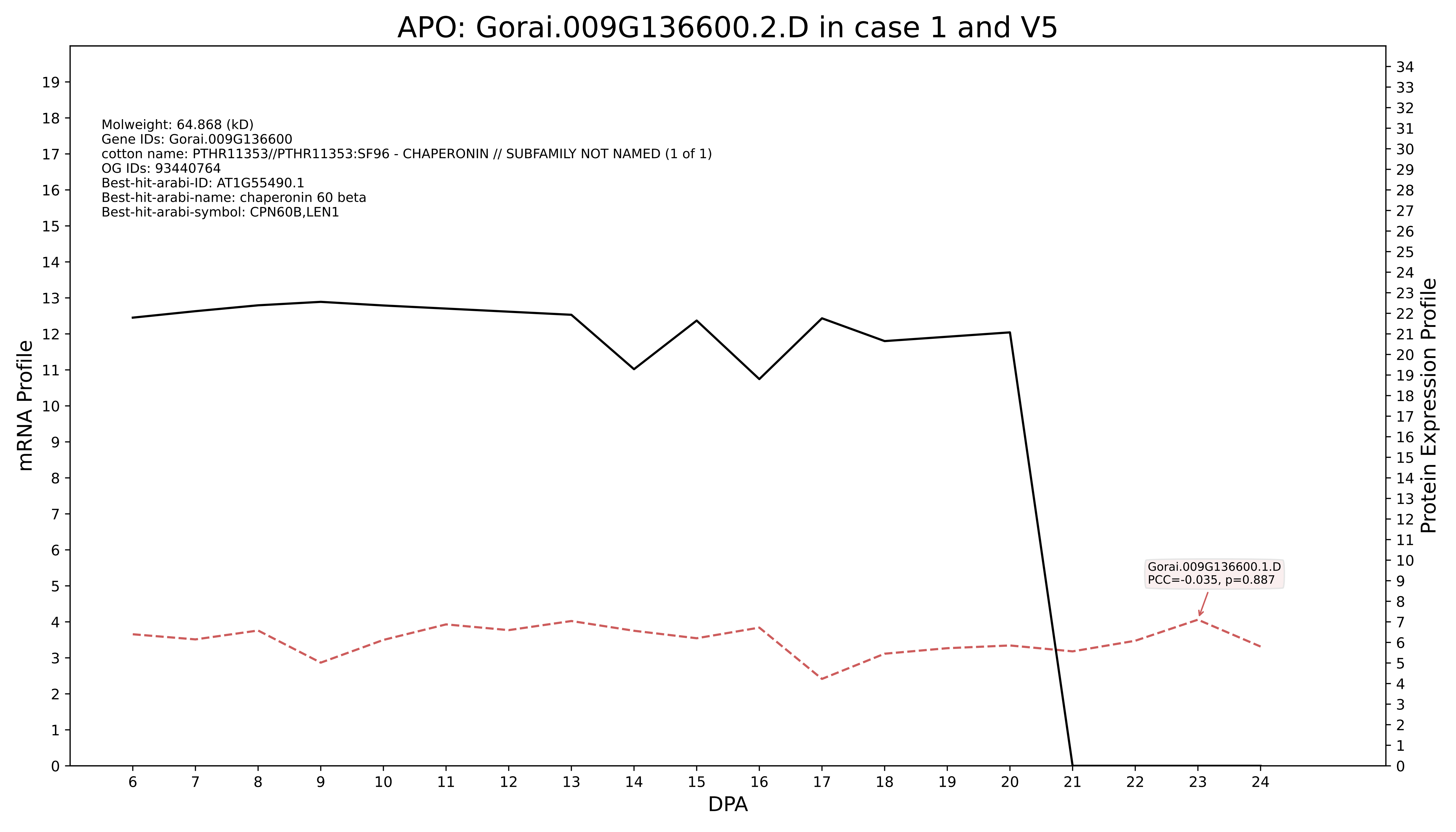
APO: Gorai.009G136600.2.D in case 1 and V5
Molweight: 64.868 (kD)
Gene IDs: Gorai.009G136600
cotton name: PTHR11353//PTHR11353:SF96 - CHAPERONIN // SUBFAMILY NOT NAMED (1 of 1)
OG IDs: 93440764
Best-hit-arabi-ID: AT1G55490.1
Best-hit-arabi-name: chaperonin 60 beta
Best-hit-arabi-symbol: CPN60B,LEN1
Gorai.009G136600.1.D
PCC=-0.035, p=0.887
mRNA Profile
Protein Expression Profile
DPA

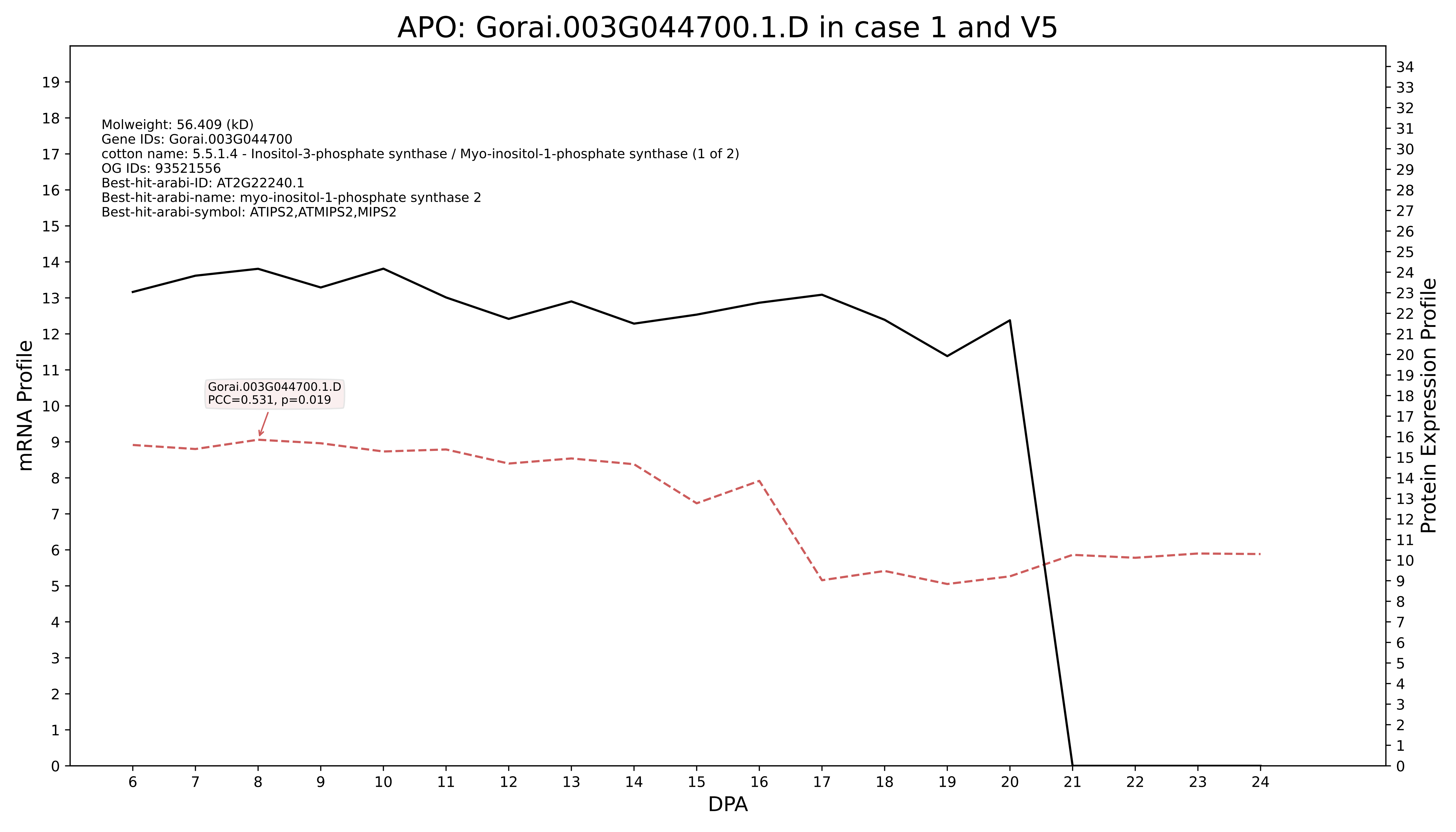
APO: Gorai.003G044700.1.D in case 1 and V5
Molweight: 56.409 (kD)
Gene IDs: Gorai.003G044700
cotton name: 5.5.1.4 - Inositol-3-phosphate synthase / Myo-inositol-1-phosphate synthase (1 of 2)
OG IDs: 93521556
Best-hit-arabi-ID: AT2G22240.1
Best-hit-arabi-name: myo-inositol-1-phosphate synthase 2
Best-hit-arabi-symbol: ATIPS2,ATMIPS2,MIPS2
Gorai.003G044700.1.D
PCC=0.531, p=0.019
mRNA Profile
Protein Expression Profile
DPA

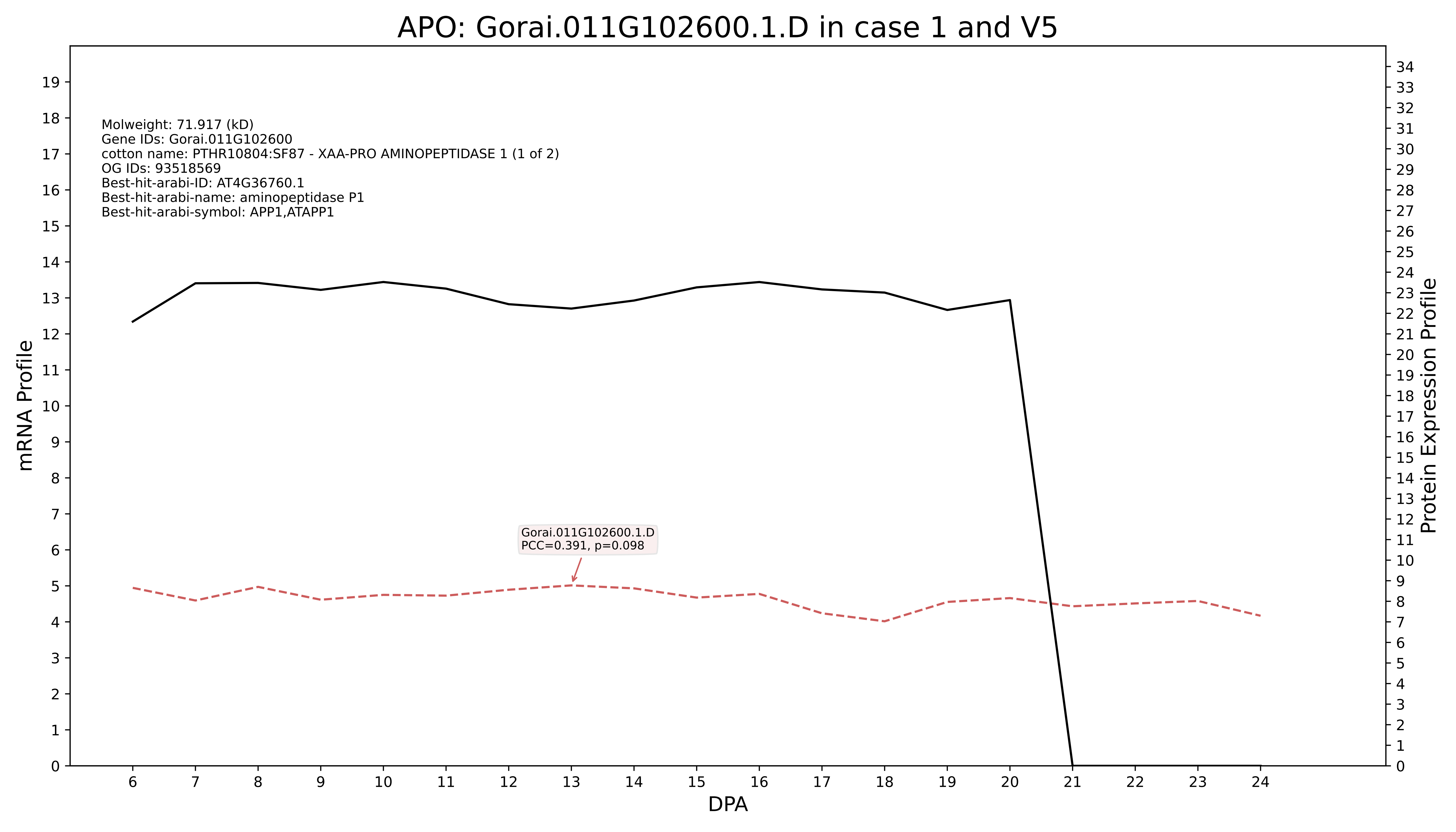

APO: Gorai.011G102600.1.D in case 1 and V5
Molweight: 71.917 (kD)
Gene IDs: Gorai.011G102600
cotton name: PTHR10804:SF87 - XAA-PRO AMINOPEPTIDASE 1 (1 of 2)
OG IDs: 93518569
Best-hit-arabi-ID: AT4G36760.1
Best-hit-arabi-name: aminopeptidase P1
Best-hit-arabi-symbol: APP1,ATAPP1
Gorai.011G102600.1.D
PCC=0.391, p=0.098
mRNA Profile
Protein Expression Profile
DPA

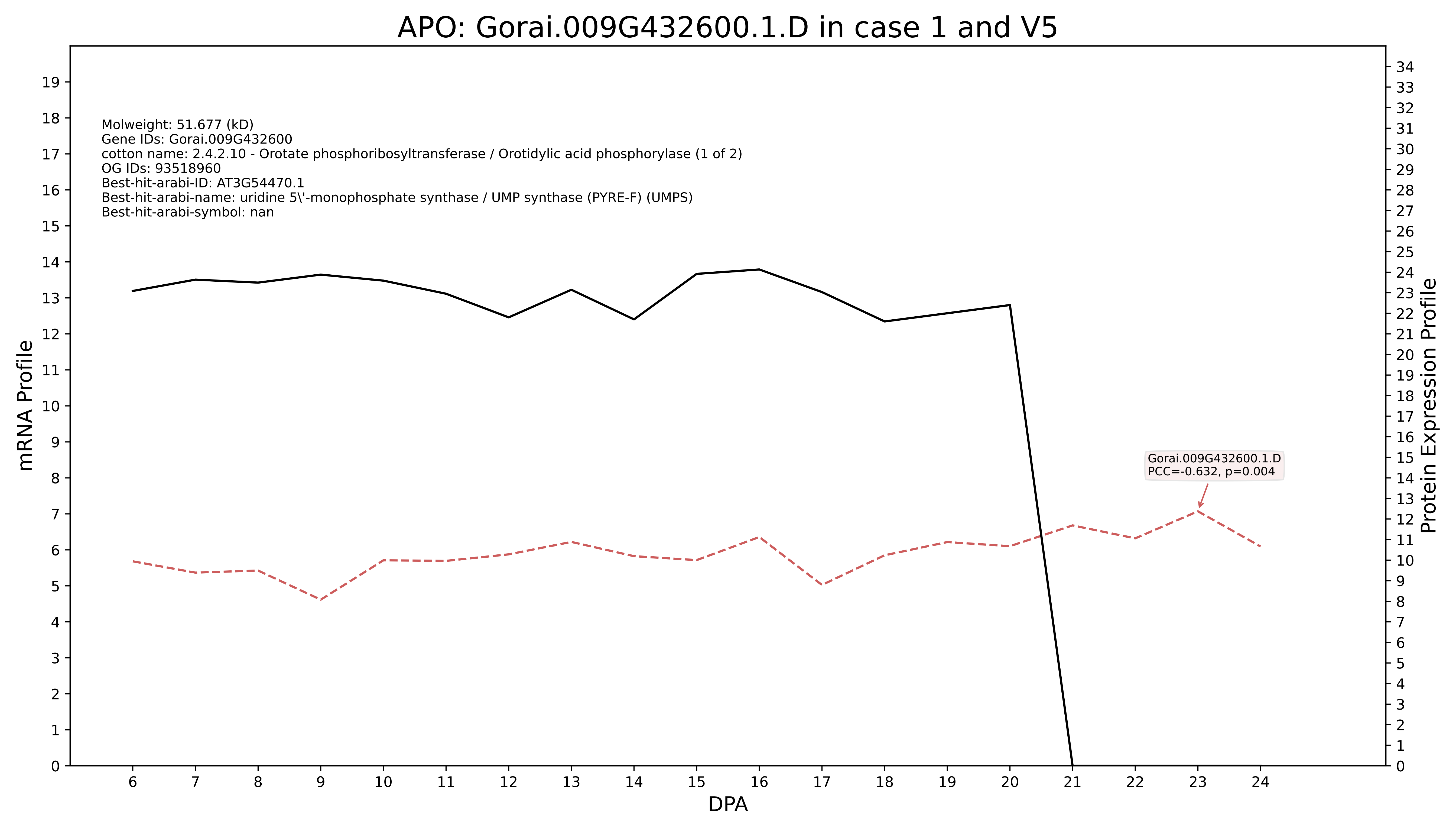

APO: Gorai.009G432600.1.D in case 1 and V5
Molweight: 51.677 (kD)
Gene IDs: Gorai.009G432600
cotton name: 2.4.2.10 - Orotate phosphoribosyltransferase / Orotidylic acid phosphorylase (1 of 2)
OG IDs: 93518960
Best-hit-arabi-ID: AT3G54470.1
Best-hit-arabi-name: uridine 5\'-monophosphate synthase / UMP synthase (PYRE-F) (UMPS)
Best-hit-arabi-symbol: nan
Gorai.009G432600.1.D
PCC=-0.632, p=0.004
mRNA Profile
Protein Expression Profile
DPA

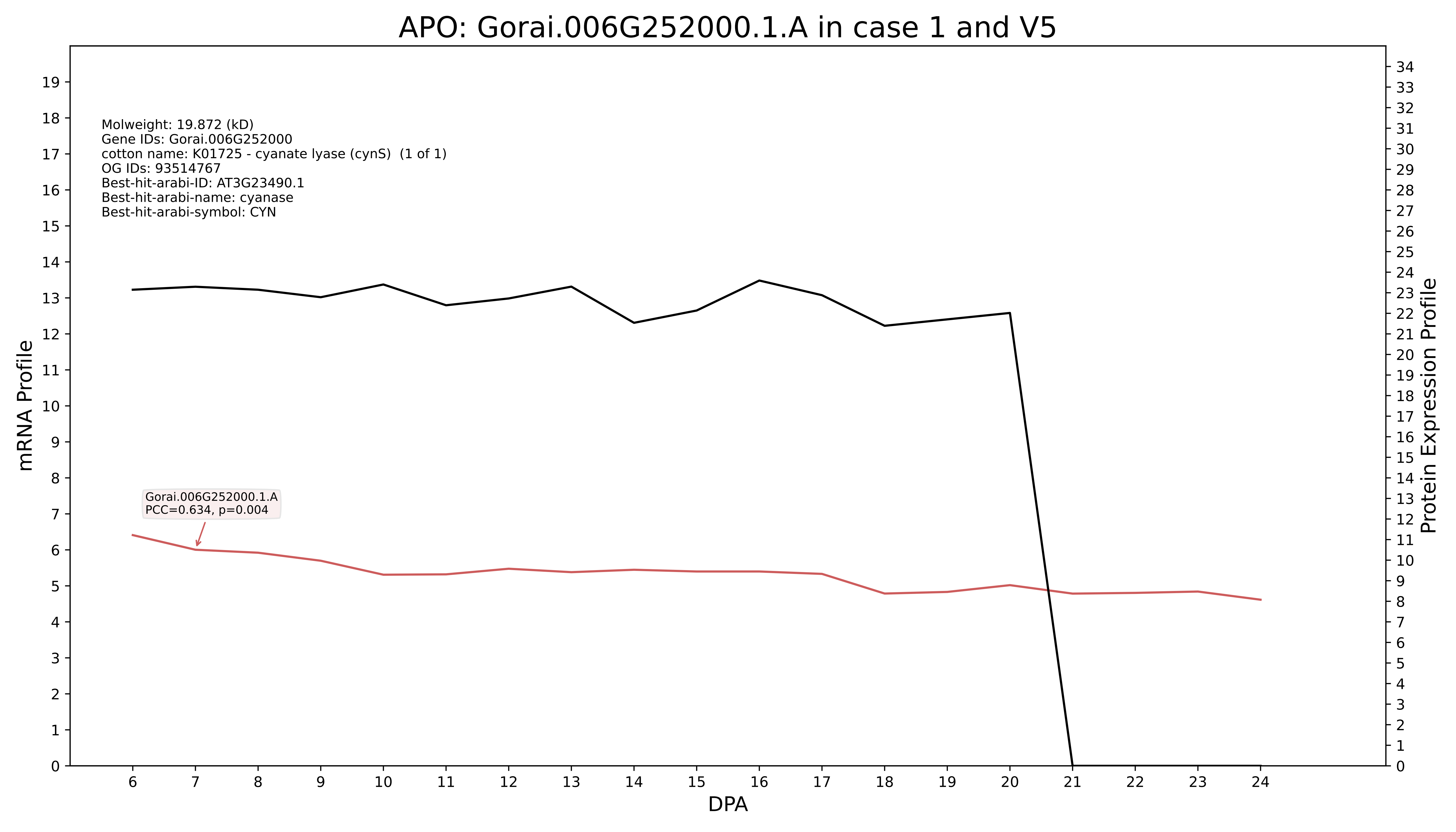

APO: Gorai.006G252000.1.A in case 1 and V5
Molweight: 19.872 (kD)
Gene IDs: Gorai.006G252000
cotton name: K01725 - cyanate lyase (cynS) (1 of 1)
OG IDs: 93514767
Best-hit-arabi-ID: AT3G23490.1
Best-hit-arabi-name: cyanase
Best-hit-arabi-symbol: CYN
Gorai.006G252000.1.A
PCC=0.634, p=0.004
mRNA Profile
Protein Expression Profile
DPA

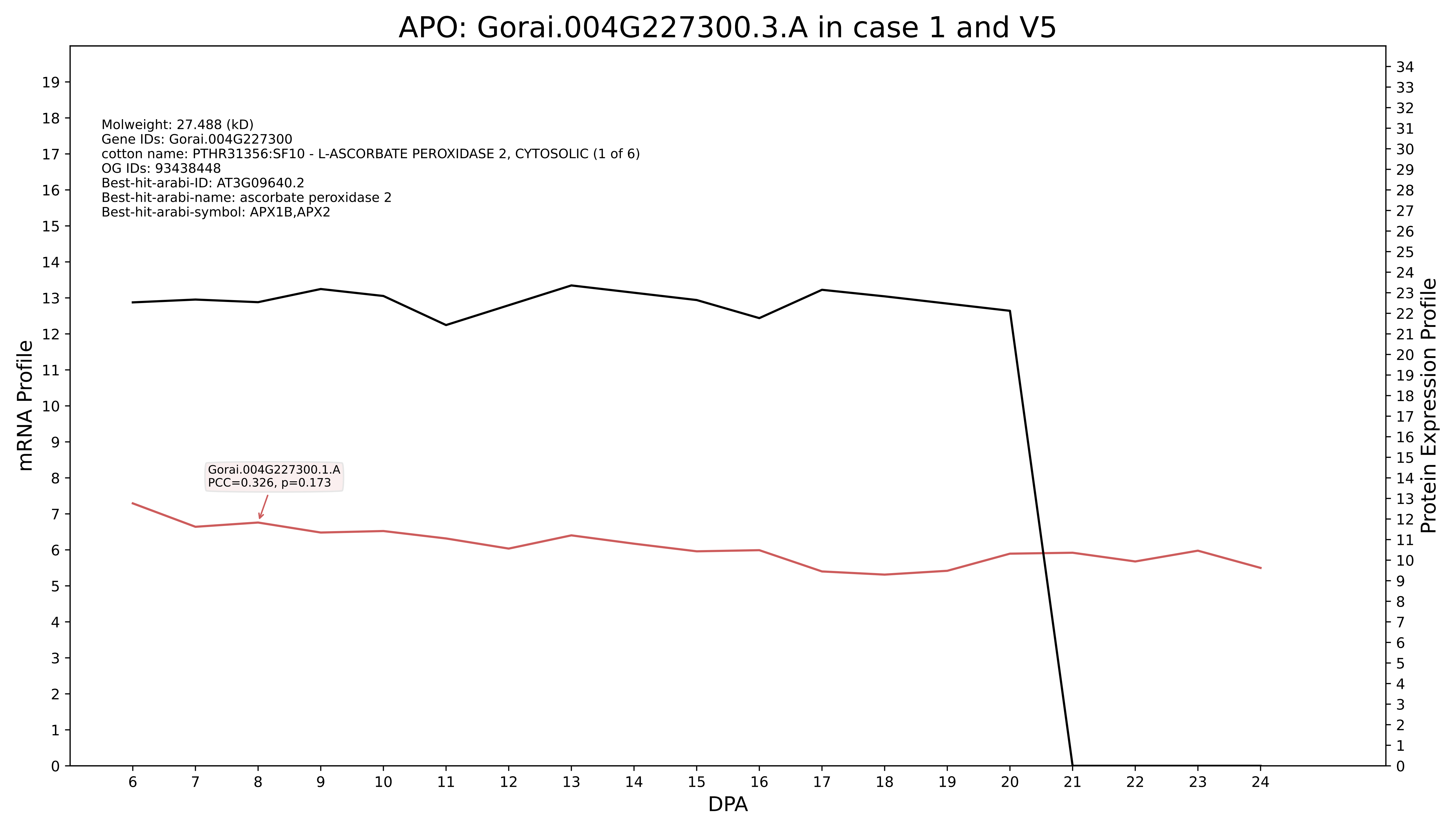
APO: Gorai.004G227300.3.A in case 1 and V5
Molweight: 27.488 (kD)
Gene IDs: Gorai.004G227300
cotton name: PTHR31356:SF10 - L-ASCORBATE PEROXIDASE 2, CYTOSOLIC (1 of 6)
OG IDs: 93438448
Best-hit-arabi-ID: AT3G09640.2
Best-hit-arabi-name: ascorbate peroxidase 2
Best-hit-arabi-symbol: APX1B,APX2
Gorai.004G227300.1.A
PCC=0.326, p=0.173
mRNA Profile
Protein Expression Profile
DPA

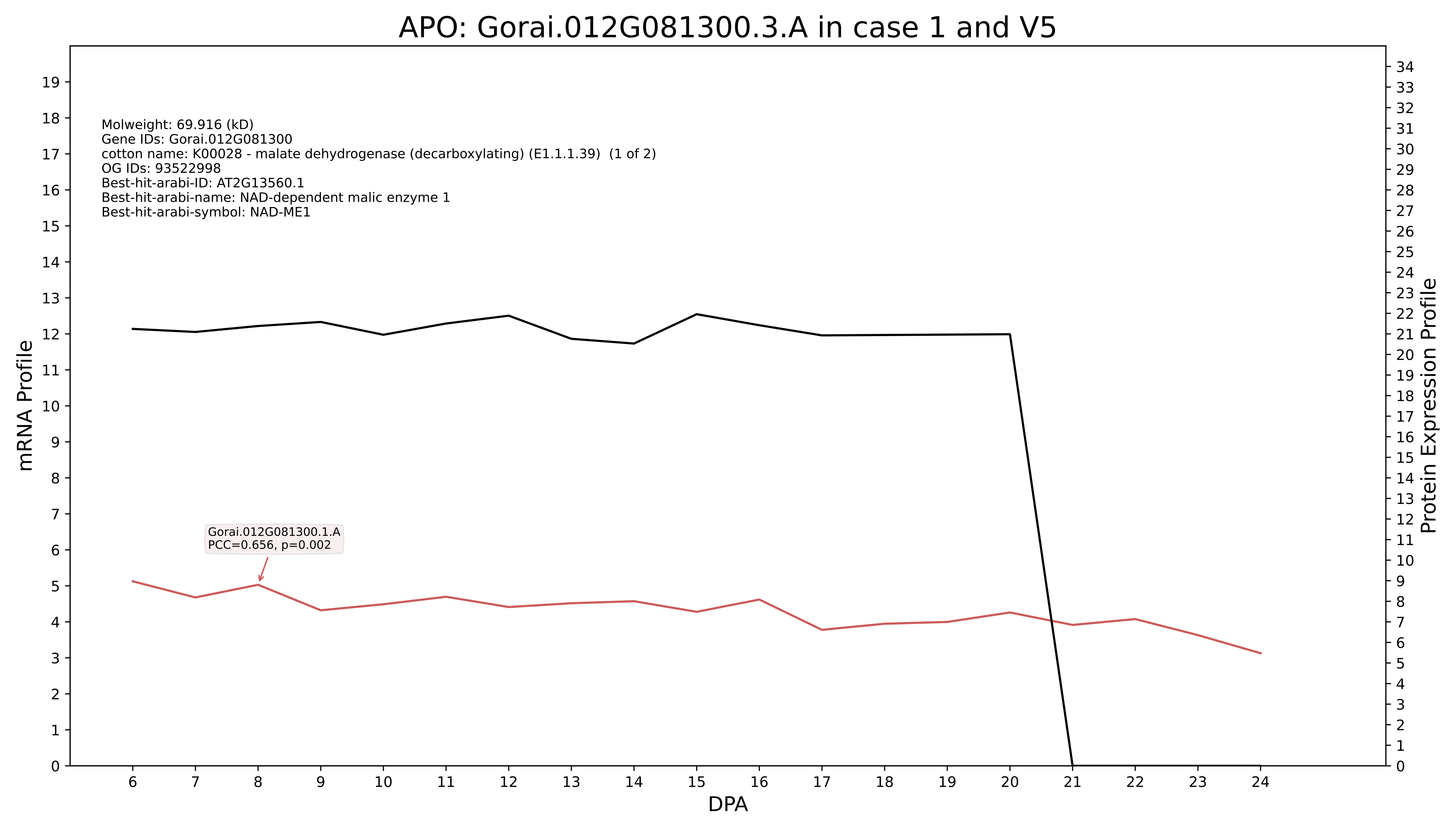
APO: Gorai.012G081300.3.A in case 1 and V5
Molweight: 69.916 (kD)
Gene IDs: Gorai.012G081300
cotton name: K00028 - malate dehydrogenase (decarboxylating) (E1.1.1.39) (1 of 2)
OG IDs: 93522998
Best-hit-arabi-ID: AT2G13560.1
Best-hit-arabi-name: NAD-dependent malic enzyme 1
Best-hit-arabi-symbol: NAD-ME1
Gorai.012G081300.1.A
PCC=0.656, p=0.002
mRNA Profile
Protein Expression Profile
DPA

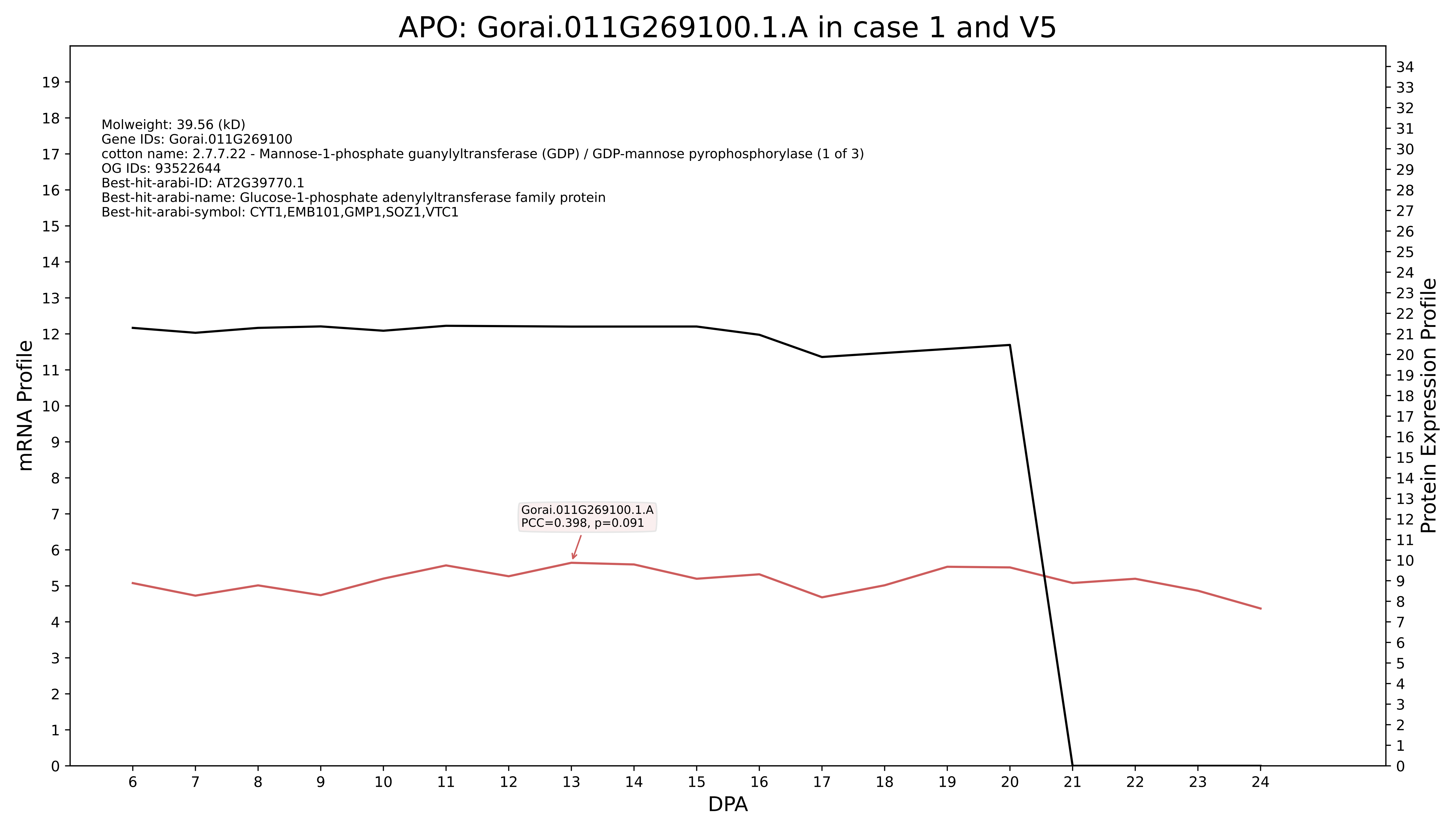
APO: Gorai.011G269100.1.A in case 1 and V5
Molweight: 39.56 (kD)
Gene IDs: Gorai.011G269100
cotton name: 2.7.7.22 - Mannose-1-phosphate guanylyltransferase (GDP) / GDP-mannose pyrophosphorylase (1 of 3)
OG IDs: 93522644
Best-hit-arabi-ID: AT2G39770.1
Best-hit-arabi-name: Glucose-1-phosphate adenylyltransferase family protein
Best-hit-arabi-symbol: CYT1,EMB101,GMP1,SOZ1,VTC1
Gorai.011G269100.1.A
PCC=0.398, p=0.091
mRNA Profile
Protein Expression Profile
DPA

### APO: Gorai.001G240900.4.A in case 1 and V5

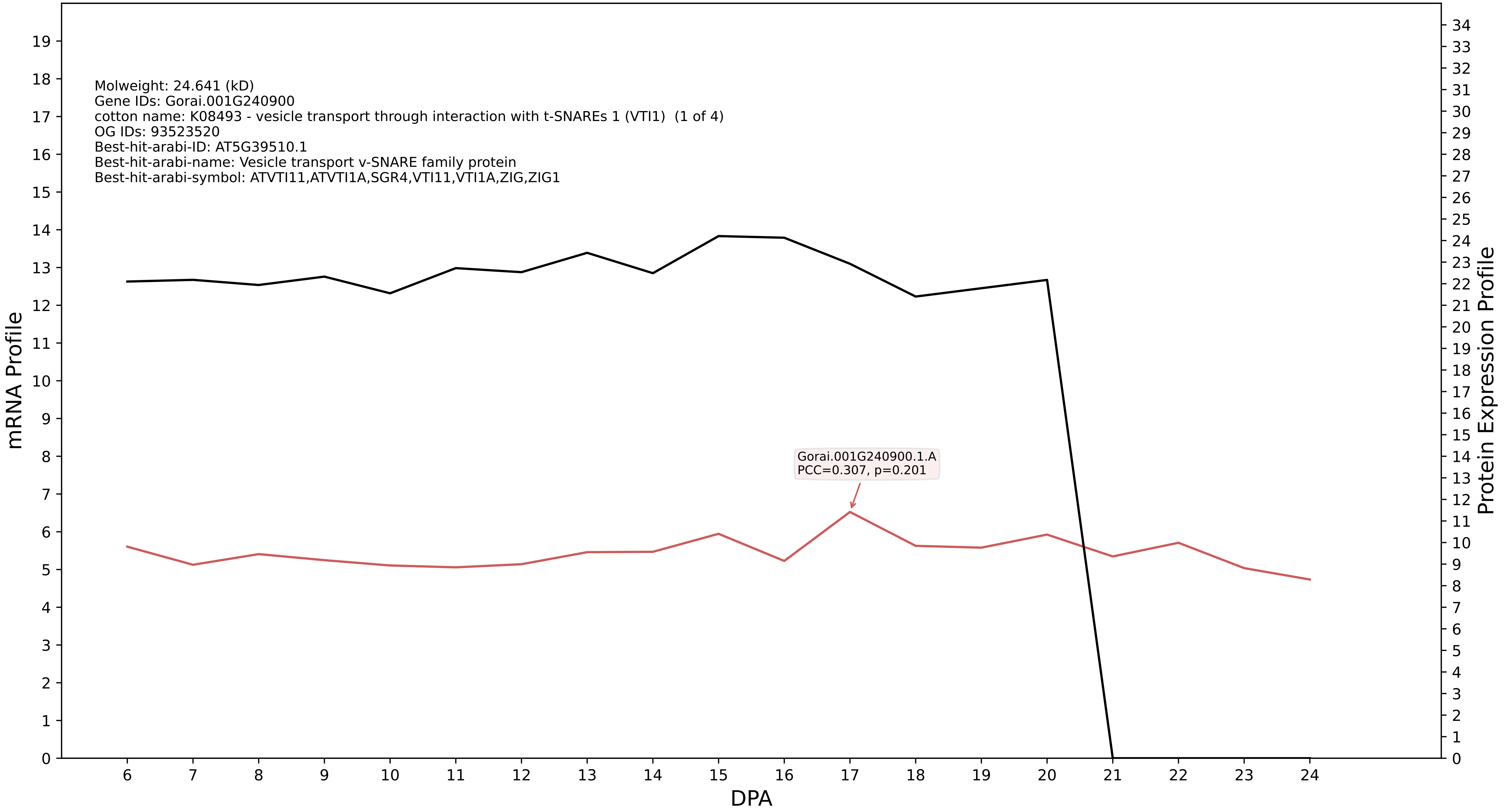

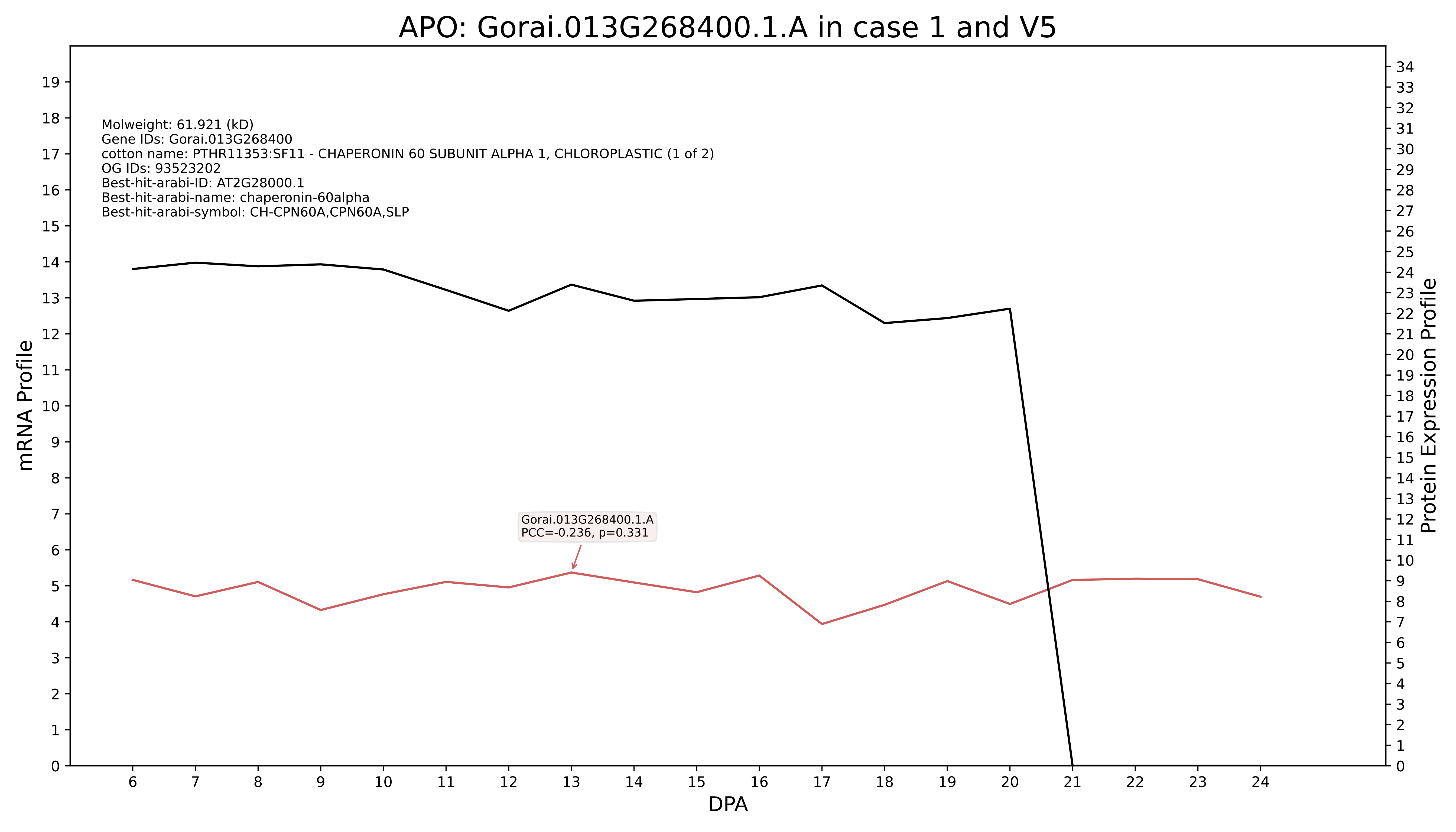
APO: Gorai.013G268400.1.A in case 1 and V5
Molweight: 61.921 (kD)
Gene IDs: Gorai.013G268400
cotton name: PTHR11353:SF11 - CHAPERONIN 60 SUBUNIT ALPHA 1, CHLOROPLASTIC (1 of 2)
OG IDs: 93523202
Best-hit-arabi-ID: AT2G28000.1
Best-hit-arabi-name: chaperonin-60alpha
Best-hit-arabi-symbol: CH-CPN60A,CPN60A,SLP
Gorai.013G268400.1.A
PCC=-0.236, p=0.331
mRNA Profile
Protein Expression Profile
DPA

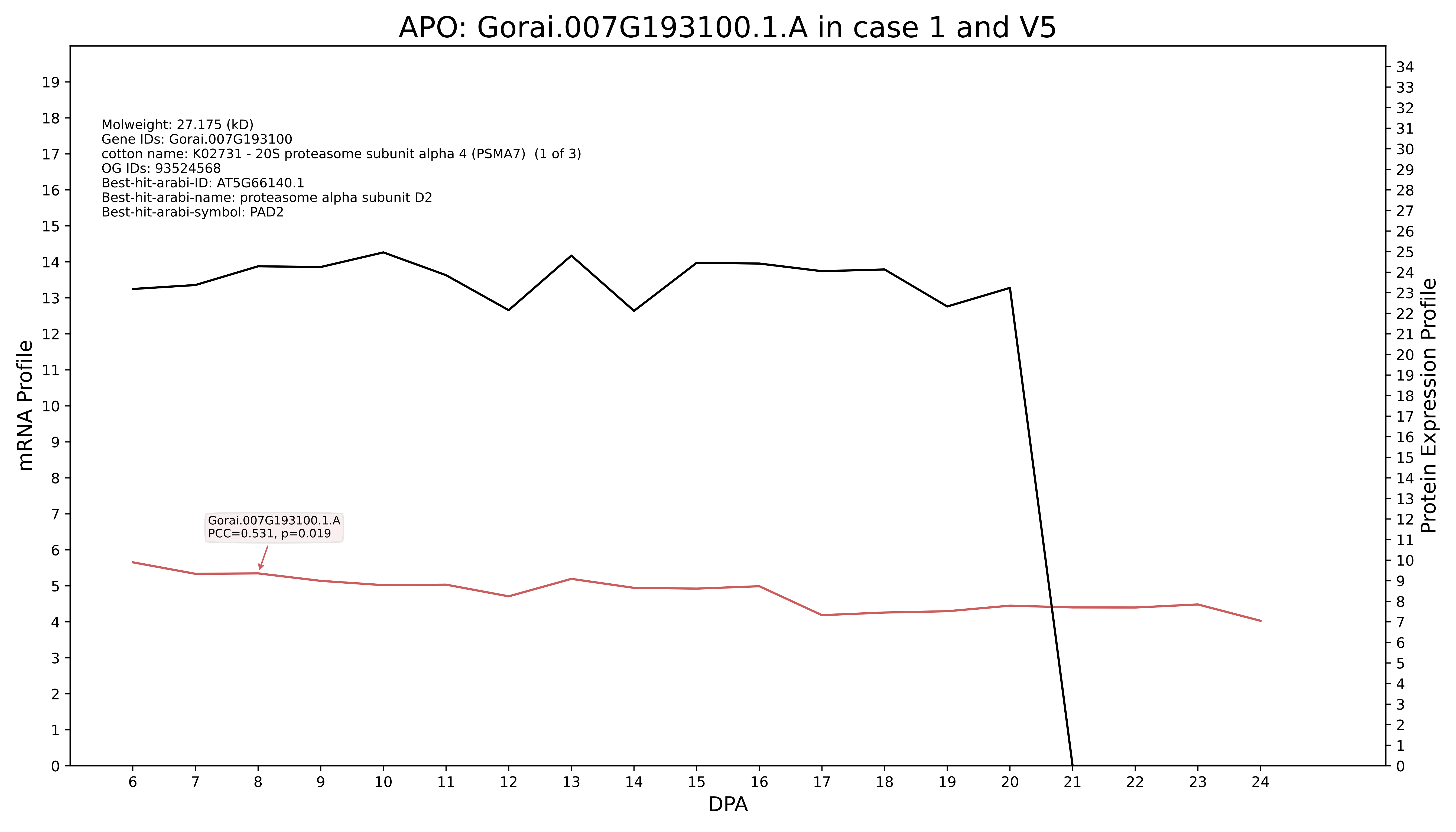
APO: Gorai.007G193100.1.A in case 1 and V5
Molweight: 27.175 (kD)
Gene IDs: Gorai.007G193100
cotton name: K02731 - 20S proteasome subunit alpha 4 (PSMA7) (1 of 3)
OG IDs: 93524568
Best-hit-arabi-ID: AT5G66140.1
Best-hit-arabi-name: proteasome alpha subunit D2
Best-hit-arabi-symbol: PAD2
Gorai.007G193100.1.A
PCC=0.531, p=0.019
mRNA Profile
Protein Expression Profile
DPA

### APO: Gorai.013G168900.2.A in case 1 and V5

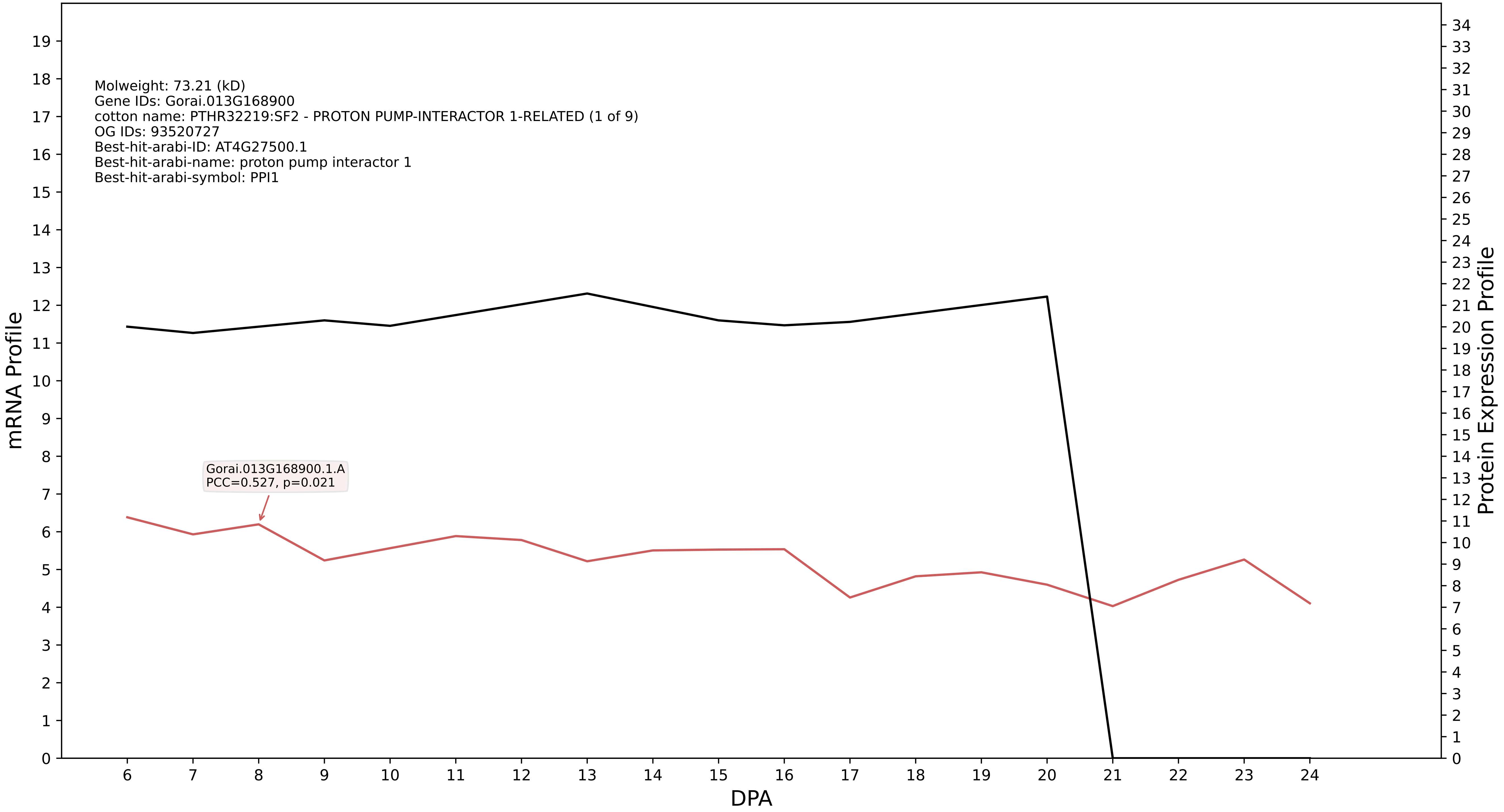

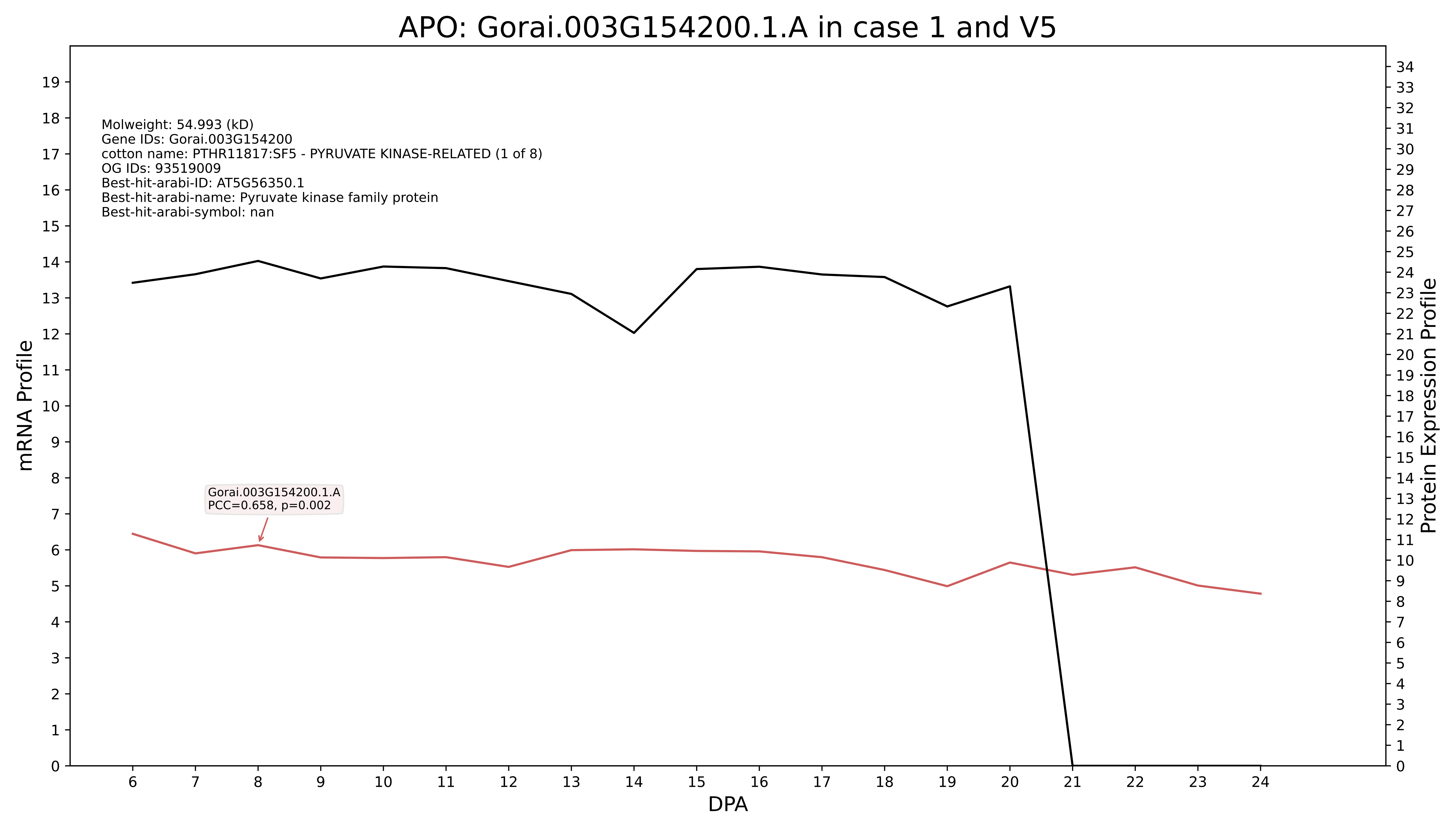
APO: Gorai.003G154200.1.A in case 1 and V5
Molweight: 54.993 (kD)
Gene IDs: Gorai.003G154200
cotton name: PTHR11817:SF5 - PYRUVATE KINASE-RELATED (1 of 8)
OG IDs: 93519009
Best-hit-arabi-ID: AT5G56350.1
Best-hit-arabi-name: Pyruvate kinase family protein
Best-hit-arabi-symbol: nan
Gorai.003G154200.1.A
PCC=0.658, p=0.002
mRNA Profile
Protein Expression Profile
DPA

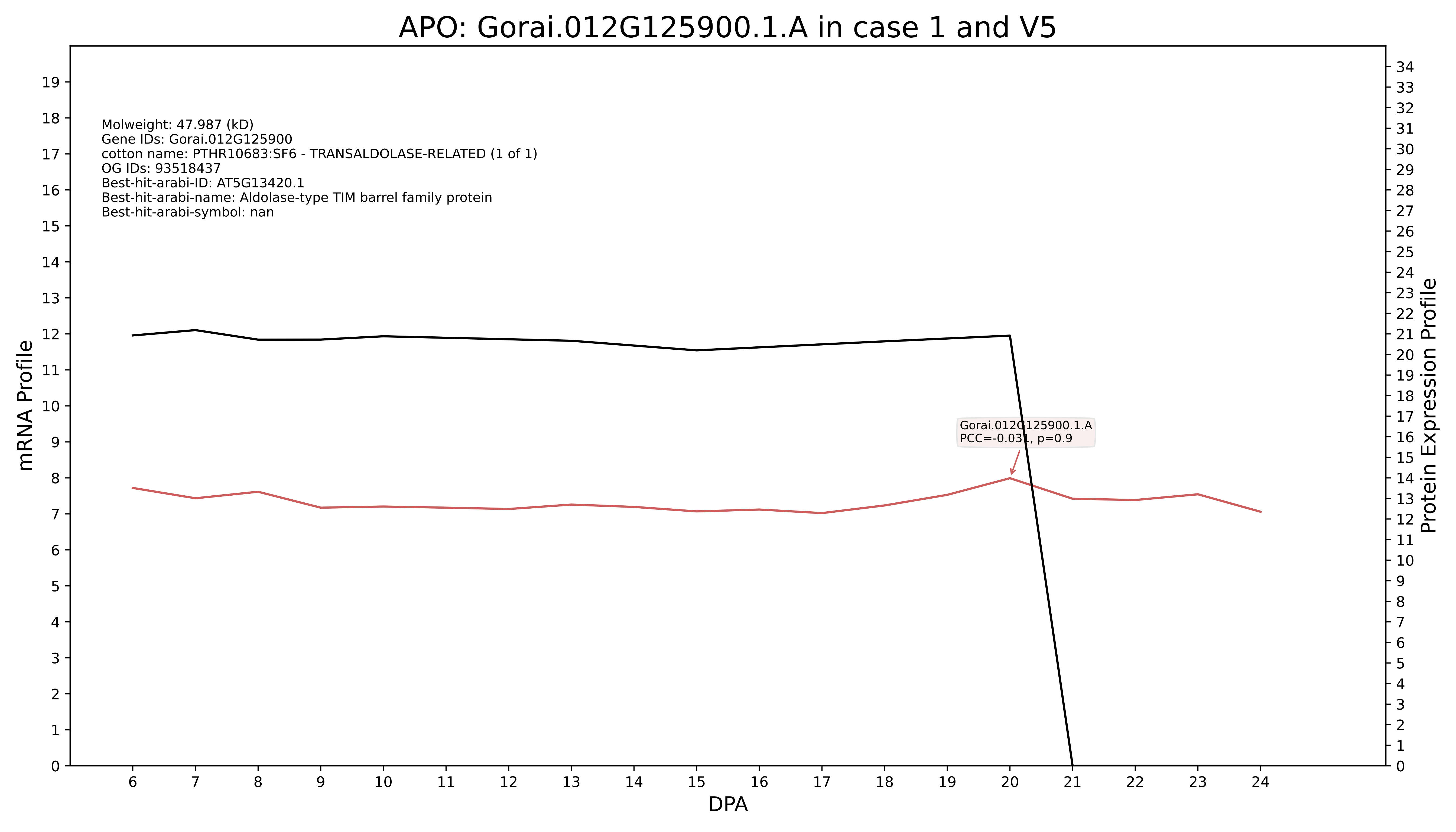

APO: Gorai.012G125900.1.A in case 1 and V5
Molweight: 47.987 (kD)
Gene IDs: Gorai.012G125900
cotton name: PTHR10683:SF6 - TRANSALDOLASE-RELATED (1 of 1)
OG IDs: 93518437
Best-hit-arabi-ID: AT5G13420.1
Best-hit-arabi-name: Aldolase-type TIM barrel family protein
Best-hit-arabi-symbol: nan
Gorai.012G125900.1.A
PCC=-0.031, p=0.9
mRNA Profile
Protein Expression Profile
DPA

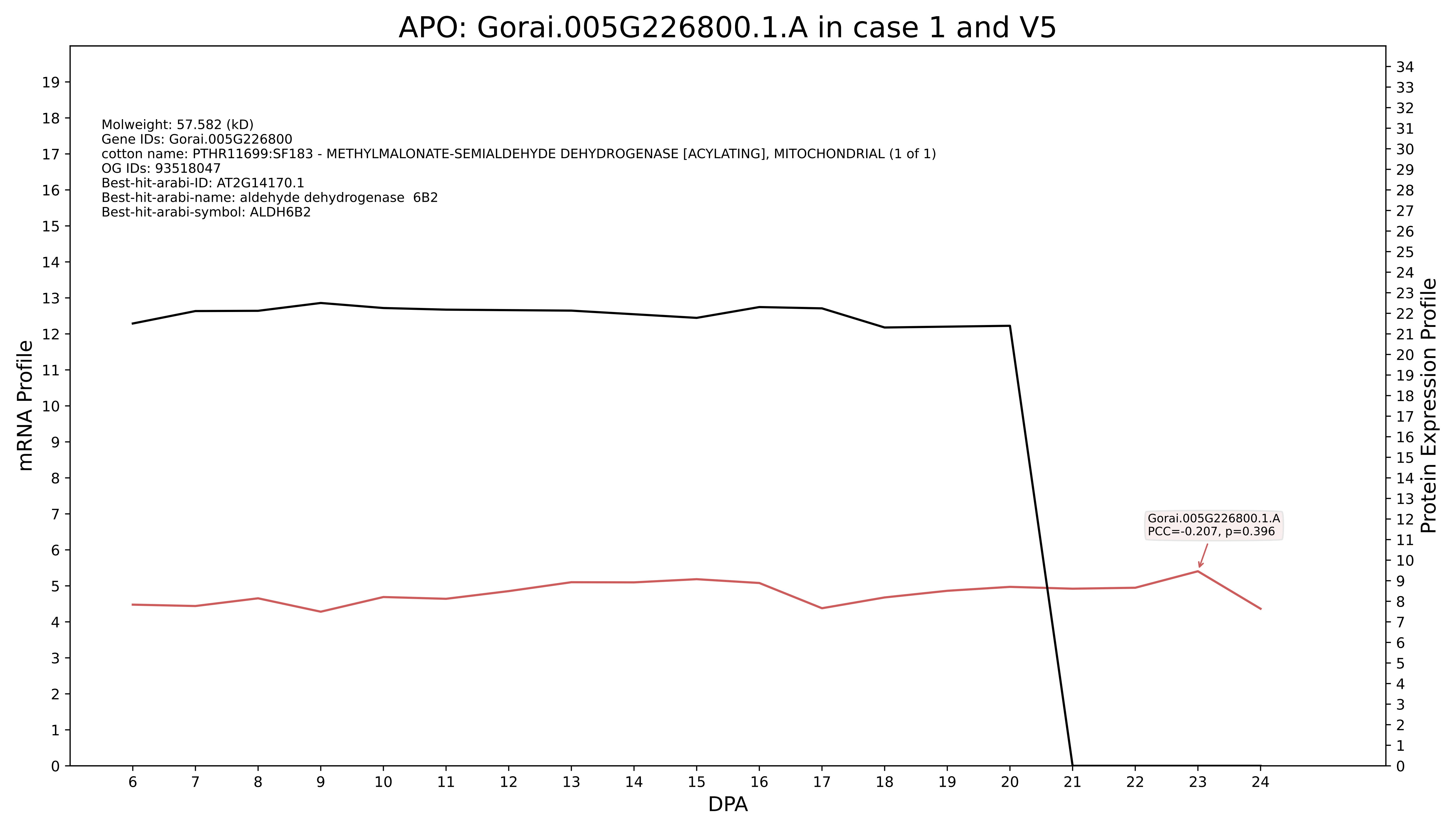
APO: Gorai.005G226800.1.A in case 1 and V5
Molweight: 57.582 (kD)
Gene IDs: Gorai.005G226800
cotton name: PTHR11699:SF183 - METHYLMALONATE-SEMIALDEHYDE DEHYDROGENASE [ACYLATING], MITOCHONDRIAL (1 of 1)
OG IDs: 93518047
Best-hit-arabi-ID: AT2G14170.1
Best-hit-arabi-name: aldehyde dehydrogenase 6B2
Best-hit-arabi-symbol: ALDH6B2
Gorai.005G226800.1.A
PCC=-0.207, p=0.396
mRNA Profile
Protein Expression Profile
DPA

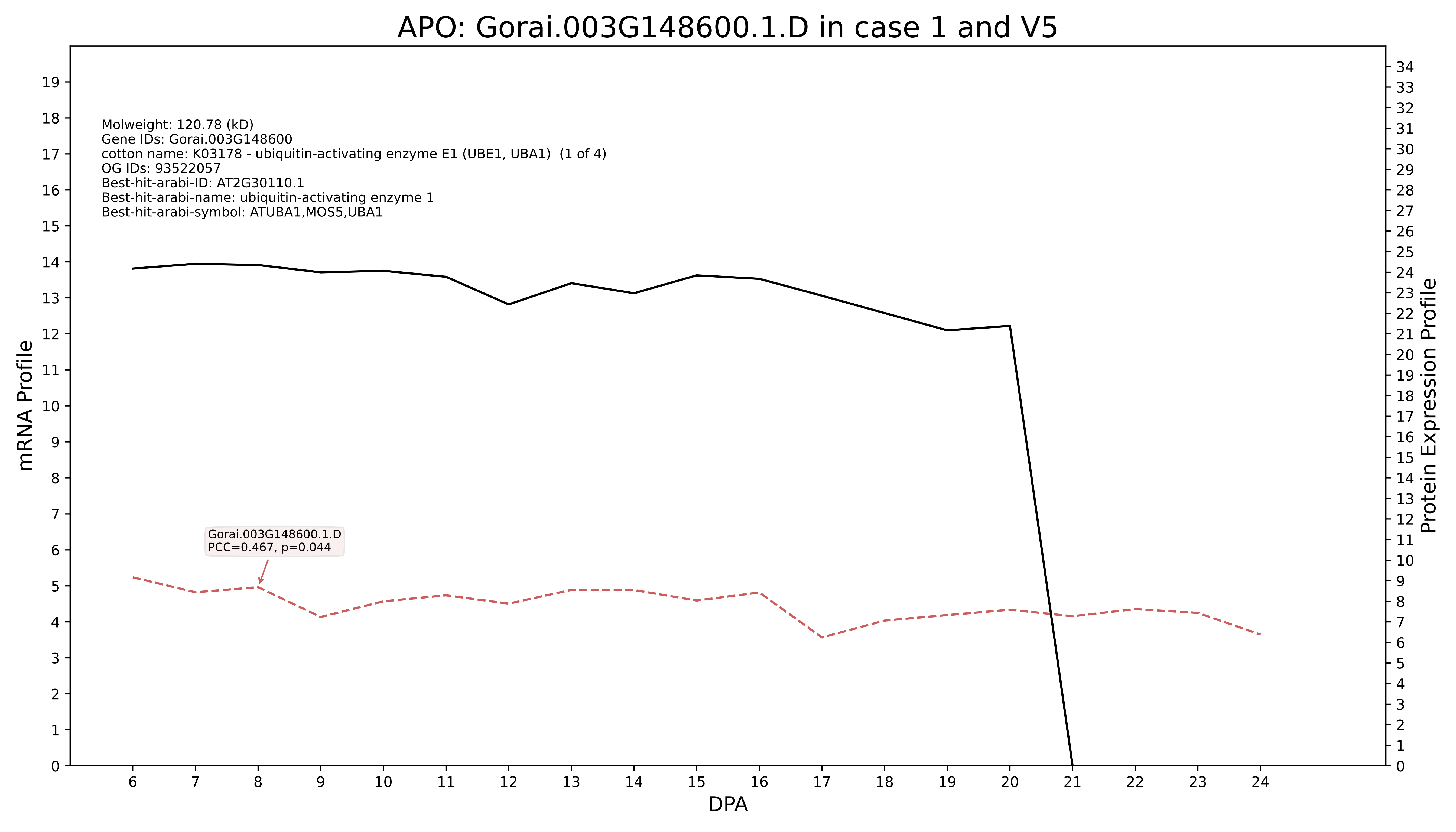

APO: Gorai.003G148600.1.D in case 1 and V5
Molweight: 120.78 (kD)
Gene IDs: Gorai.003G148600
cotton name: K03178 - ubiquitin-activating enzyme E1 (UBE1, UBA1) (1 of 4)
OG IDs: 93522057
Best-hit-arabi-ID: AT2G30110.1
Best-hit-arabi-name: ubiquitin-activating enzyme 1
Best-hit-arabi-symbol: ATUBA1,MOS5,UBA1
Gorai.003G148600.1.D
PCC=0.467, p=0.044
mRNA Profile
Protein Expression Profile
DPA

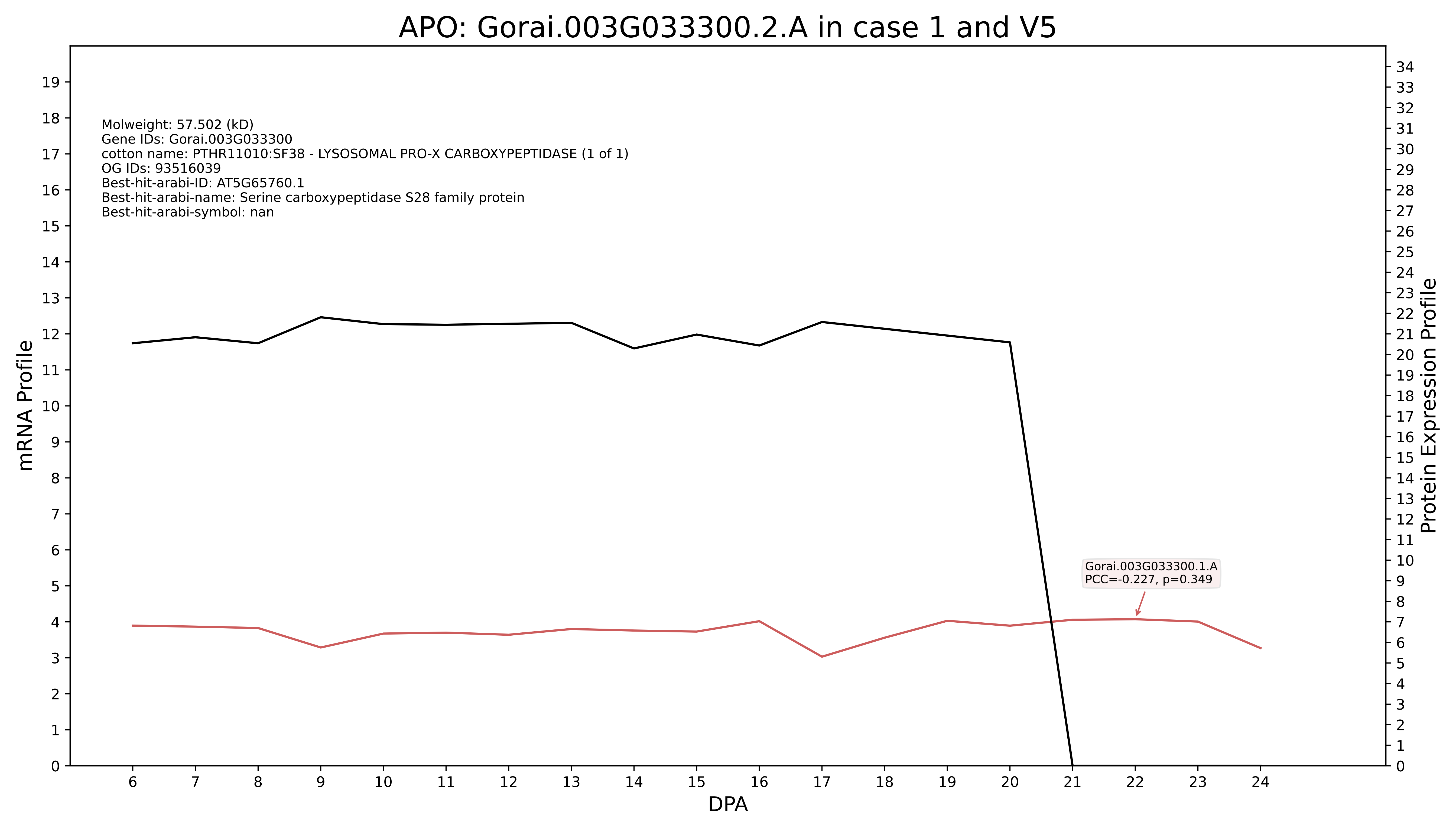

APO: Gorai.003G033300.2.A in case 1 and V5
Molweight: 57.502 (kD)
Gene IDs: Gorai.003G033300
cotton name: PTHR11010:SF38 - LYSOSOMAL PRO-X CARBOXYPEPTIDASE (1 of 1)
OG IDs: 93516039
Best-hit-arabi-ID: AT5G65760.1
Best-hit-arabi-name: Serine carboxypeptidase S28 family protein
Best-hit-arabi-symbol: nan
Gorai.003G033300.1.A
PCC=-0.227, p=0.349
mRNA Profile
Protein Expression Profile
DPA

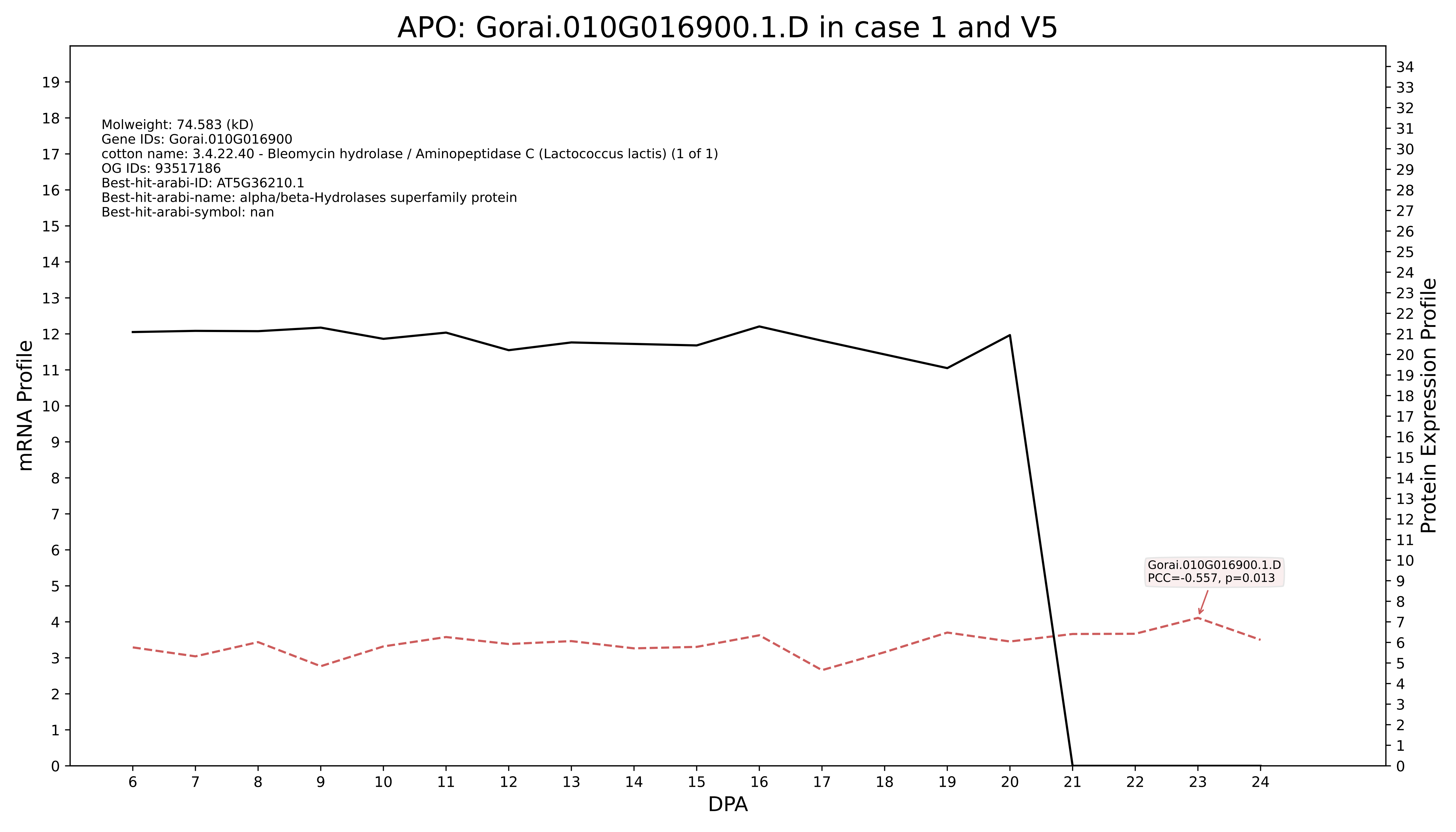
APO: Gorai.010G016900.1.D in case 1 and V5
Molweight: 74.583 (kD)
Gene IDs: Gorai.010G016900
cotton name: 3.4.22.40 - Bleomycin hydrolase / Aminopeptidase C (Lactococcus lactis) (1 of 1)
OG IDs: 93517186
Best-hit-arabi-ID: AT5G36210.1
Best-hit-arabi-name: alpha/beta-Hydrolases superfamily protein
Best-hit-arabi-symbol: nan
Gorai.010G016900.1.D
PCC=-0.557, p=0.013
mRNA Profile
Protein Expression Profile
DPA

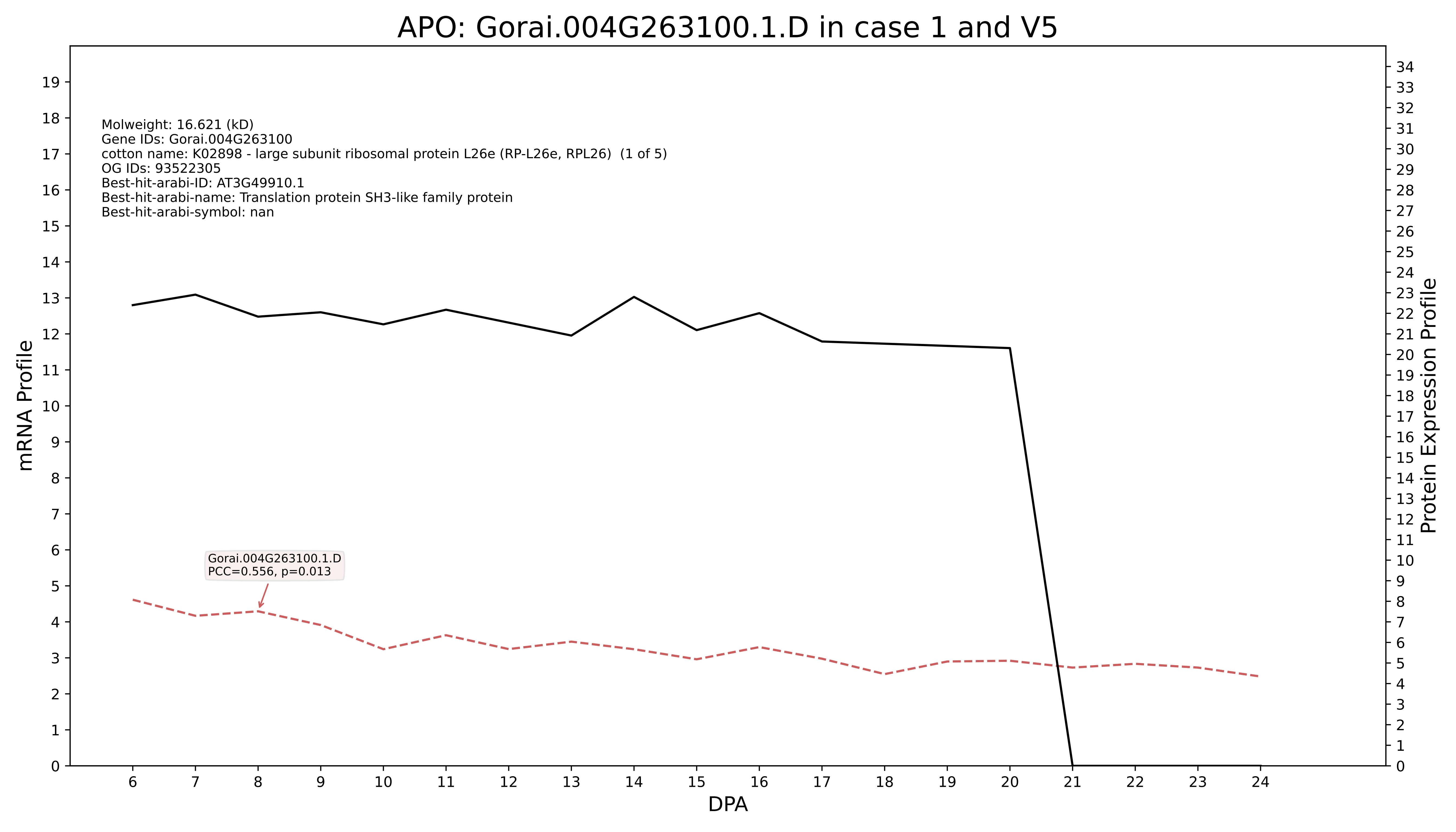
APO: Gorai.004G263100.1.D in case 1 and V5
Molweight: 16.621 (kD)
Gene IDs: Gorai.004G263100
cotton name: K02898 - large subunit ribosomal protein L26e (RP-L26e, RPL26) (1 of 5)
OG IDs: 93522305
Best-hit-arabi-ID: AT3G49910.1
Best-hit-arabi-name: Translation protein SH3-like family protein
Best-hit-arabi-symbol: nan
Gorai.004G263100.1.D
PCC=0.556, p=0.013
mRNA Profile
Protein Expression Profile
DPA

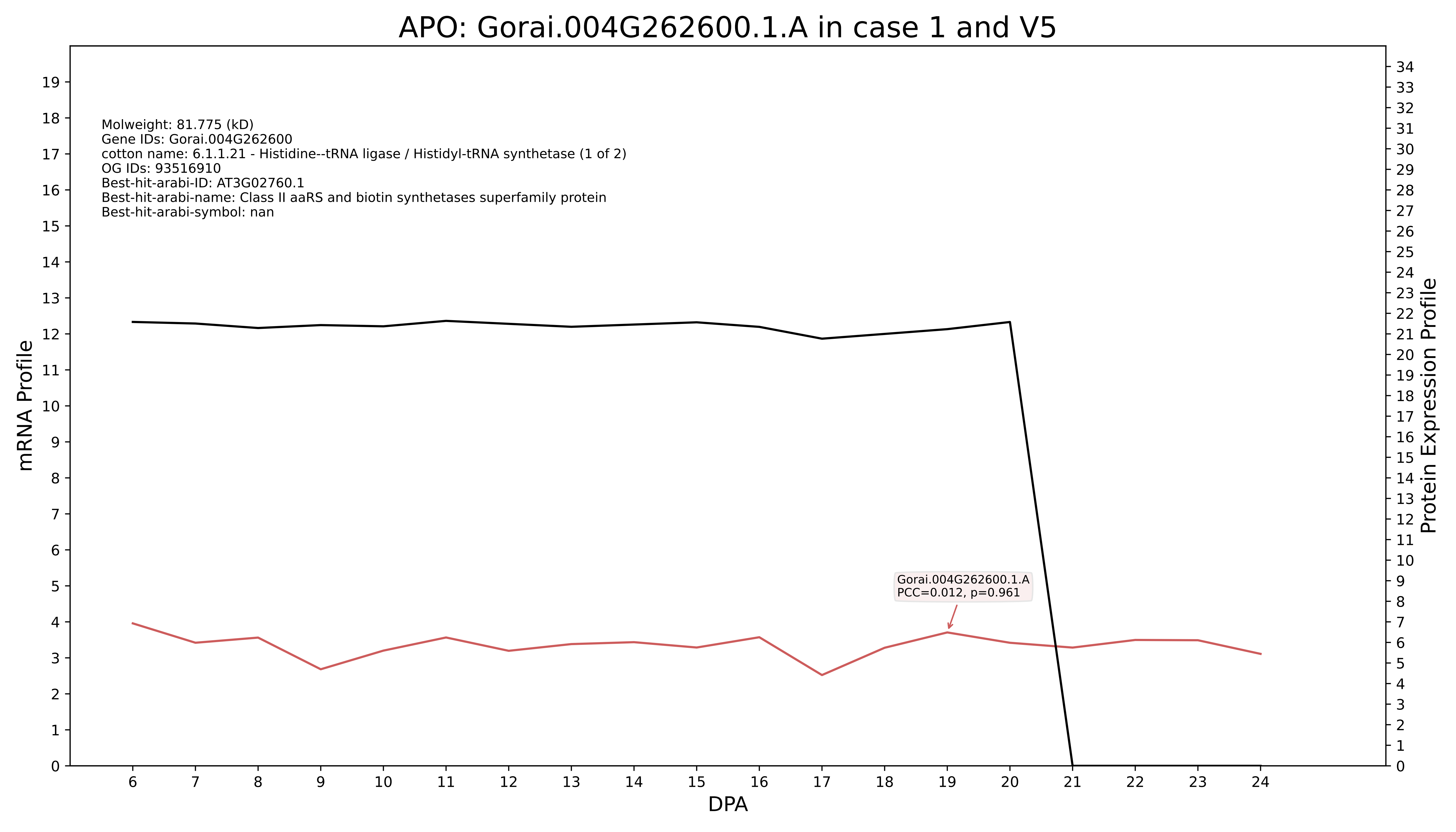

APO: Gorai.004G262600.1.A in case 1 and V5
Molweight: 81.775 (kD)
Gene IDs: Gorai.004G262600
cotton name: 6.1.1.21 - Histidine--tRNA ligase / Histidyl-tRNA synthetase (1 of 2)
OG IDs: 93516910
Best-hit-arabi-ID: AT3G02760.1
Best-hit-arabi-name: Class II aaRS and biotin synthetases superfamily protein
Best-hit-arabi-symbol: nan
Gorai.004G262600.1.A
PCC=0.012, p=0.961
mRNA Profile
Protein Expression Profile
DPA

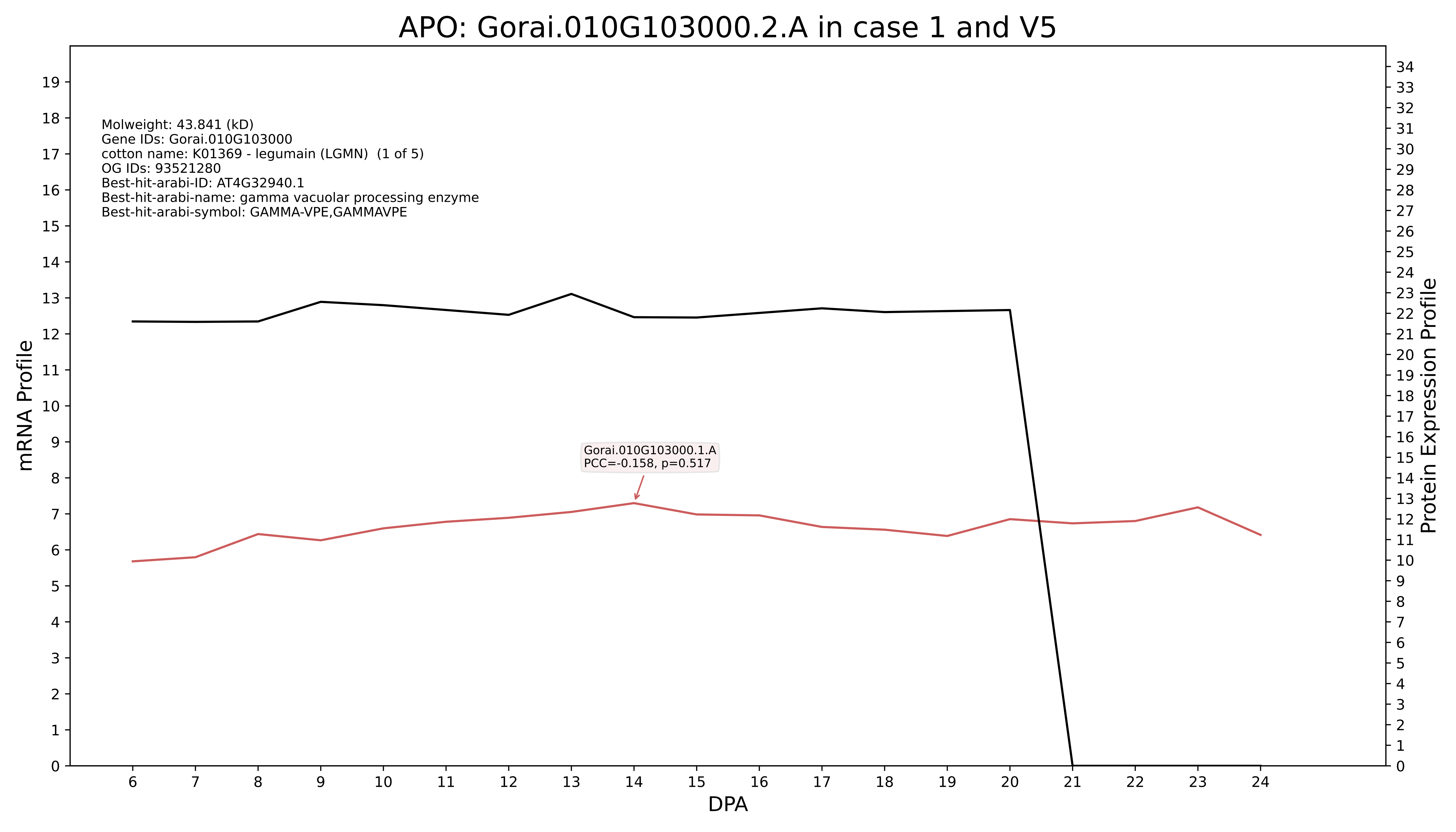

APO: Gorai.010G103000.2.A in case 1 and V5
Molweight: 43.841 (kD)
Gene IDs: Gorai.010G103000
cotton name: K01369 - legumain (LGMN) (1 of 5)
OG IDs: 93521280
Best-hit-arabi-ID: AT4G32940.1
Best-hit-arabi-name: gamma vacuolar processing enzyme
Best-hit-arabi-symbol: GAMMA-VPE,GAMMAVPE
Gorai.010G103000.1.A
PCC=-0.158, p=0.517
mRNA Profile
Protein Expression Profile
DPA
6 7 8 9 10 11 12 13 14 15 16 17 18 19 20 21 22 23 24

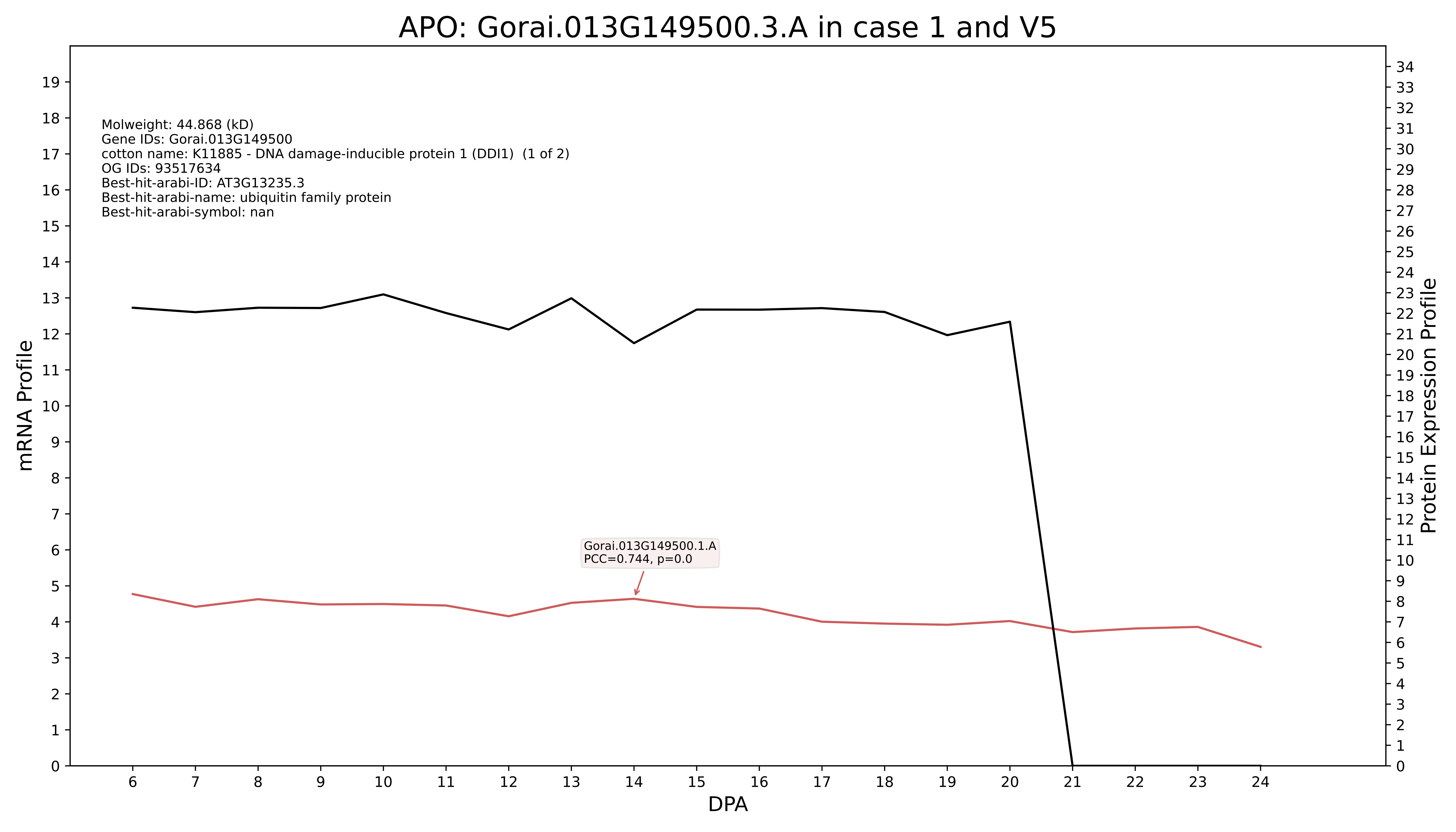

APO: Gorai.013G149500.3.A in case 1 and V5
Molweight: 44.868 (kD)
Gene IDs: Gorai.013G149500
cotton name: K11885 - DNA damage-inducible protein 1 (DDI1) (1 of 2)
OG IDs: 93517634
Best-hit-arabi-ID: AT3G13235.3
Best-hit-arabi-name: ubiquitin family protein
Best-hit-arabi-symbol: nan
Gorai.013G149500.1.A
PCC=0.744, p=0.0
mRNA Profile
Protein Expression Profile
DPA

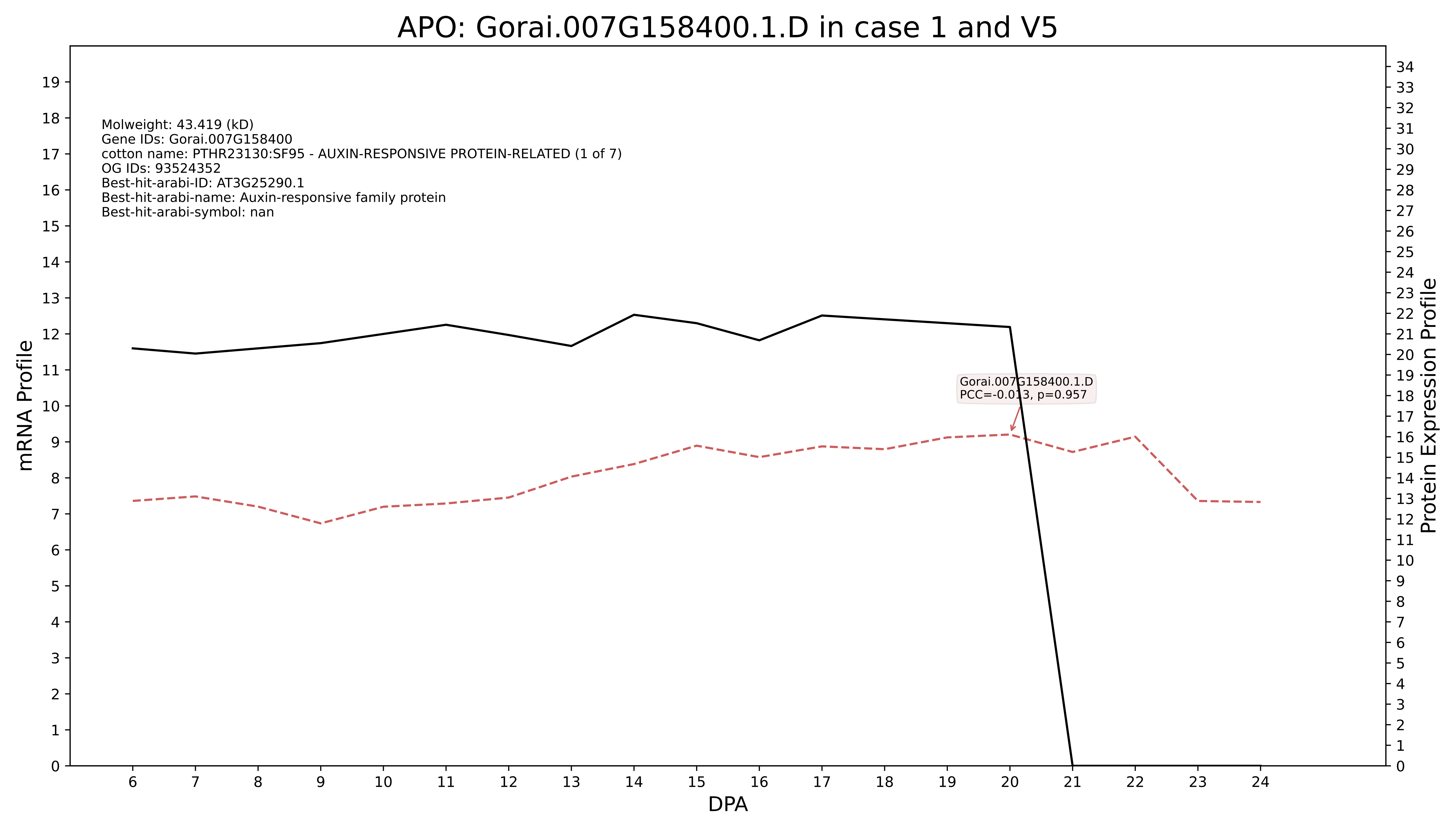
APO: Gorai.007G158400.1.D in case 1 and V5
Molweight: 43.419 (kD)
Gene IDs: Gorai.007G158400
cotton name: PTHR23130:SF95 - AUXIN-RESPONSIVE PROTEIN-RELATED (1 of 7)
OG IDs: 93524352
Best-hit-arabi-ID: AT3G25290.1
Best-hit-arabi-name: Auxin-responsive family protein
Best-hit-arabi-symbol: nan
Gorai.007G158400.1.D
PCC=-0.013, p=0.957
mRNA Profile
Protein Expression Profile
DPA

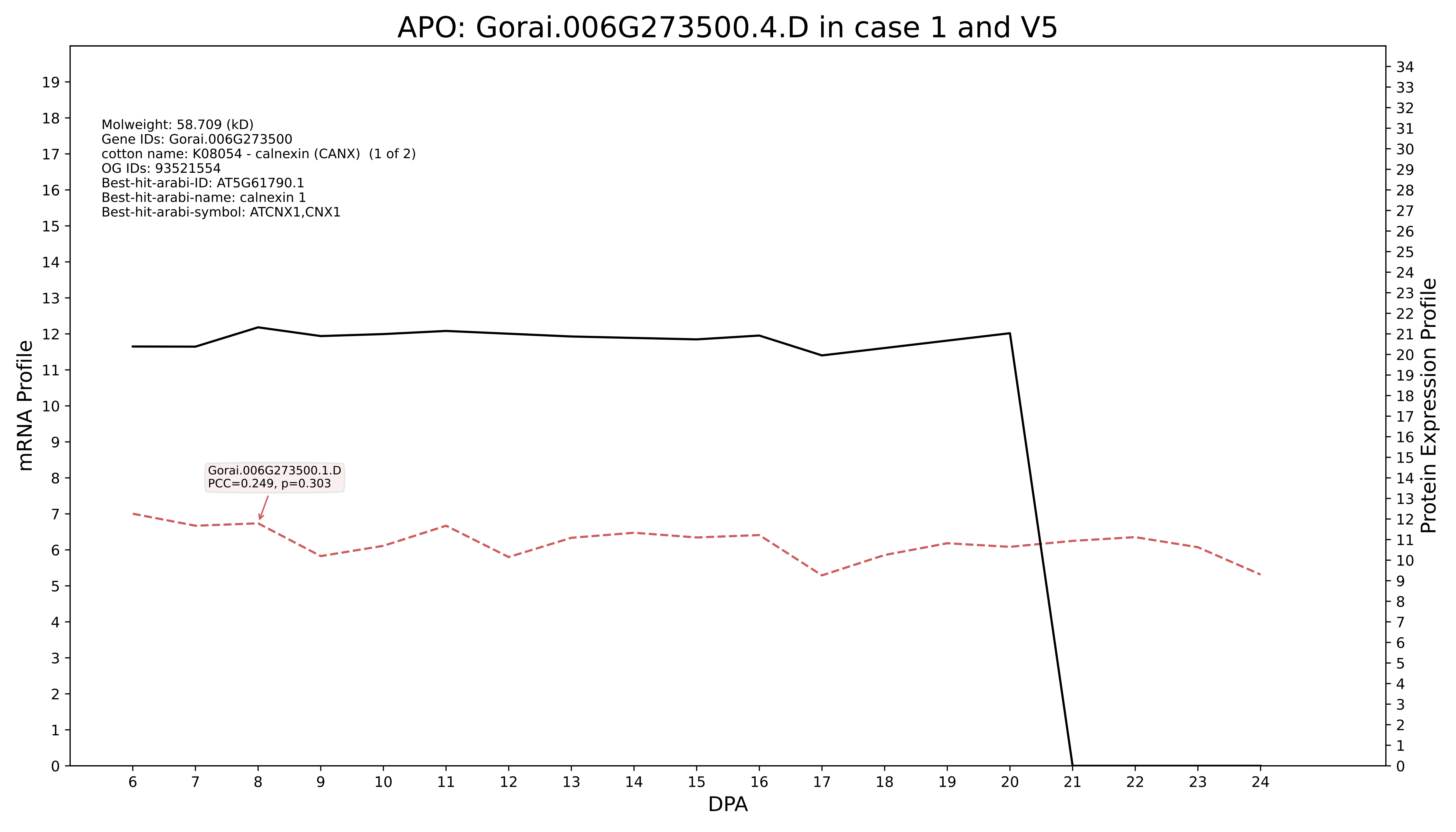

APO: Gorai.006G273500.4.D in case 1 and V5
Molweight: 58.709 (kD)
Gene IDs: Gorai.006G273500
cotton name: K08054 - calnexin (CANX) (1 of 2)
OG IDs: 93521554
Best-hit-arabi-ID: AT5G61790.1
Best-hit-arabi-name: calnexin 1
Best-hit-arabi-symbol: ATCNX1,CNX1
Gorai.006G273500.1.D
PCC=0.249, p=0.303
mRNA Profile
Protein Expression Profile
DPA

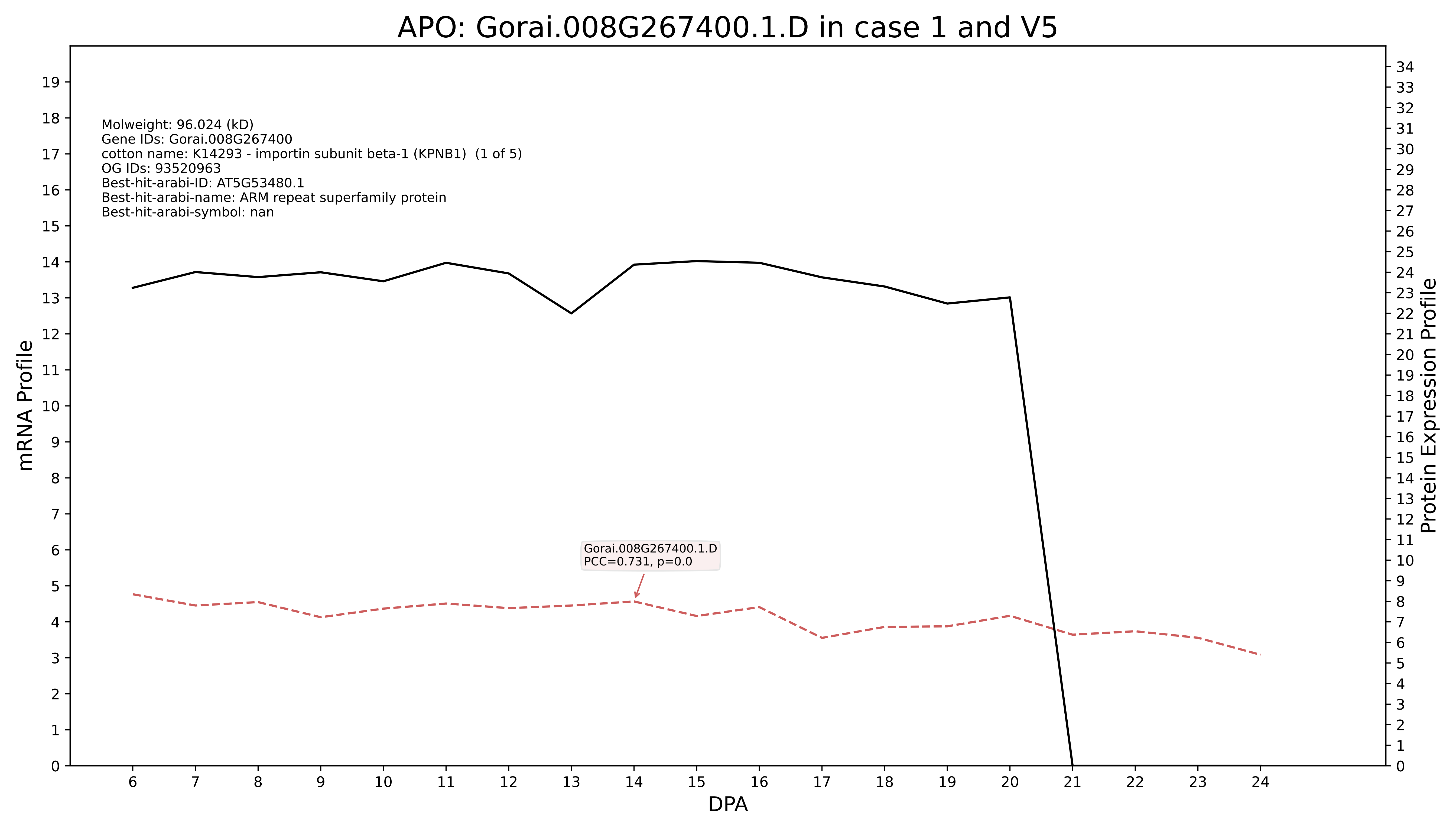
APO: Gorai.008G267400.1.D in case 1 and V5
Molweight: 96.024 (kD)
Gene IDs: Gorai.008G267400
cotton name: K14293 - importin subunit beta-1 (KPNB1) (1 of 5)
OG IDs: 93520963
Best-hit-arabi-ID: AT5G53480.1
Best-hit-arabi-name: ARM repeat superfamily protein
Best-hit-arabi-symbol: nan
Gorai.008G267400.1.D
PCC=0.731, p=0.0
mRNA Profile
Protein Expression Profile
DPA

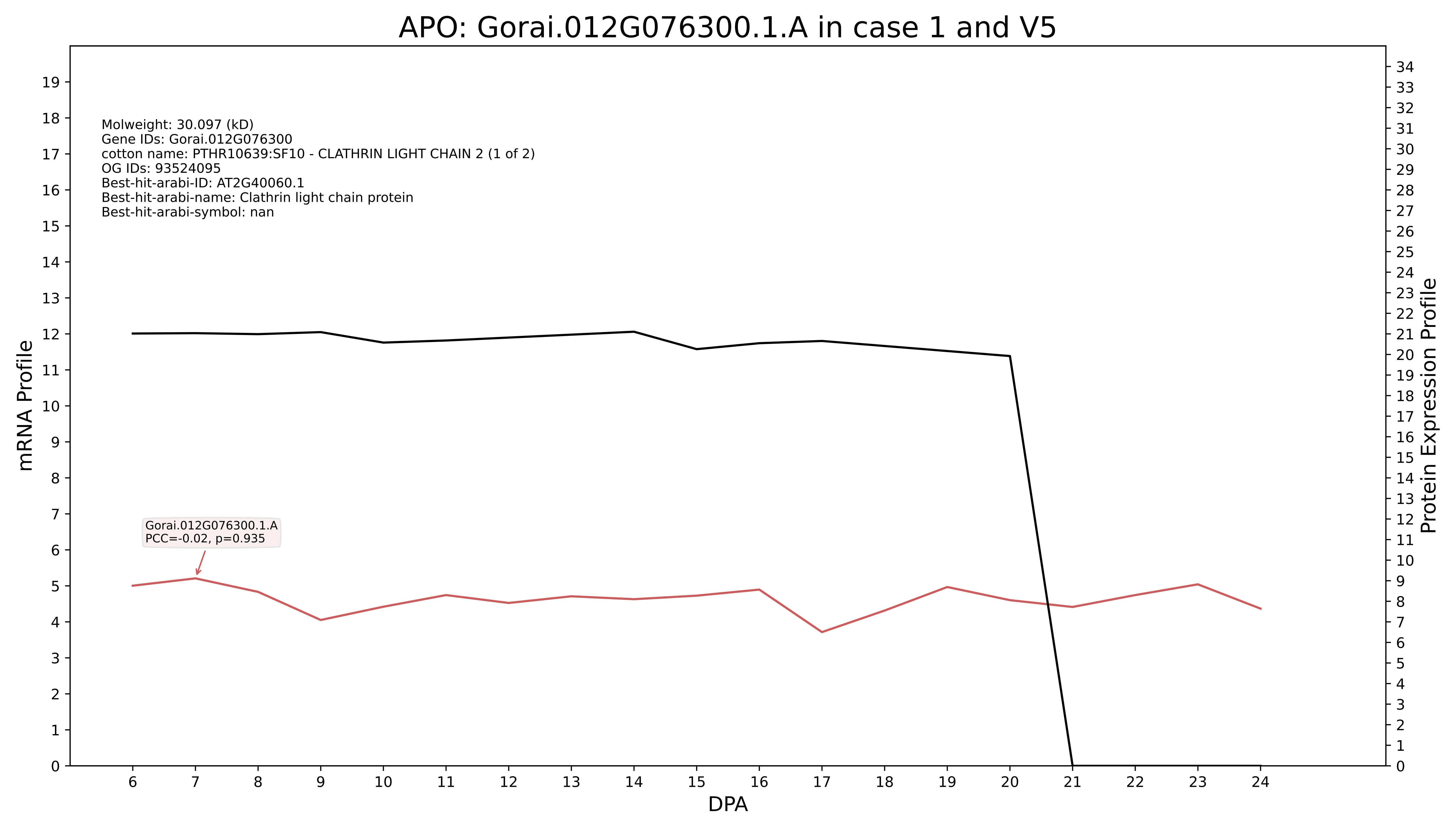

APO: Gorai.012G076300.1.A in case 1 and V5
Molweight: 30.097 (kD)
Gene IDs: Gorai.012G076300
cotton name: PTHR10639:SF10 - CLATHRIN LIGHT CHAIN 2 (1 of 2)
OG IDs: 93524095
Best-hit-arabi-ID: AT2G40060.1
Best-hit-arabi-name: Clathrin light chain protein
Best-hit-arabi-symbol: nan
Gorai.012G076300.1.A
PCC=-0.02, p=0.935
mRNA Profile
Protein Expression Profile
DPA

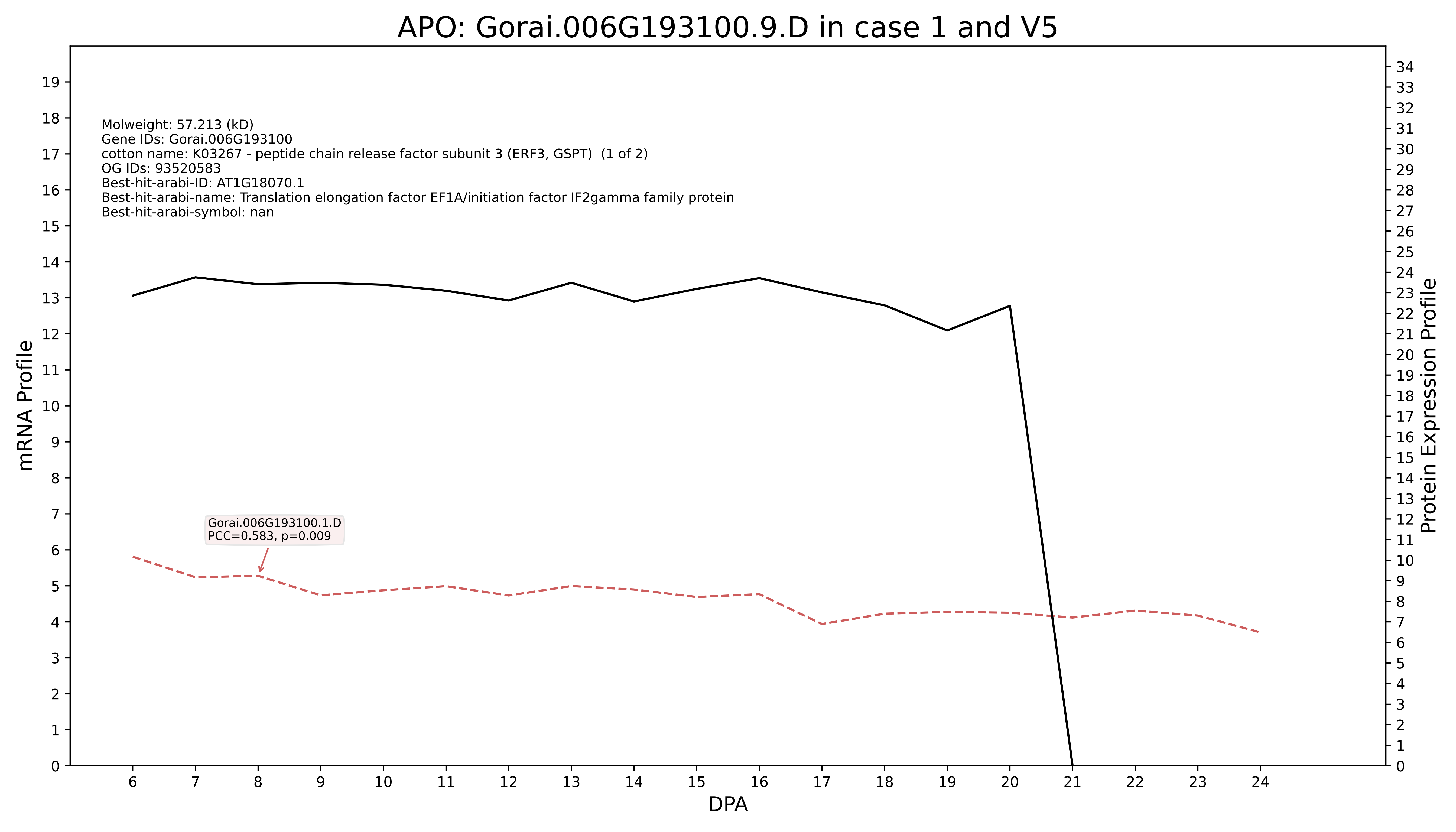

APO: Gorai.006G193100.9.D in case 1 and V5
Molweight: 57.213 (kD)
Gene IDs: Gorai.006G193100
cotton name: K03267 - peptide chain release factor subunit 3 (ERF3, GSPT) (1 of 2)
OG IDs: 93520583
Best-hit-arabi-ID: AT1G18070.1
Best-hit-arabi-name: Translation elongation factor EF1A/initiation factor IF2gamma family protein
Best-hit-arabi-symbol: nan
Gorai.006G193100.1.D
PCC=0.583, p=0.009
mRNA Profile
Protein Expression Profile
DPA

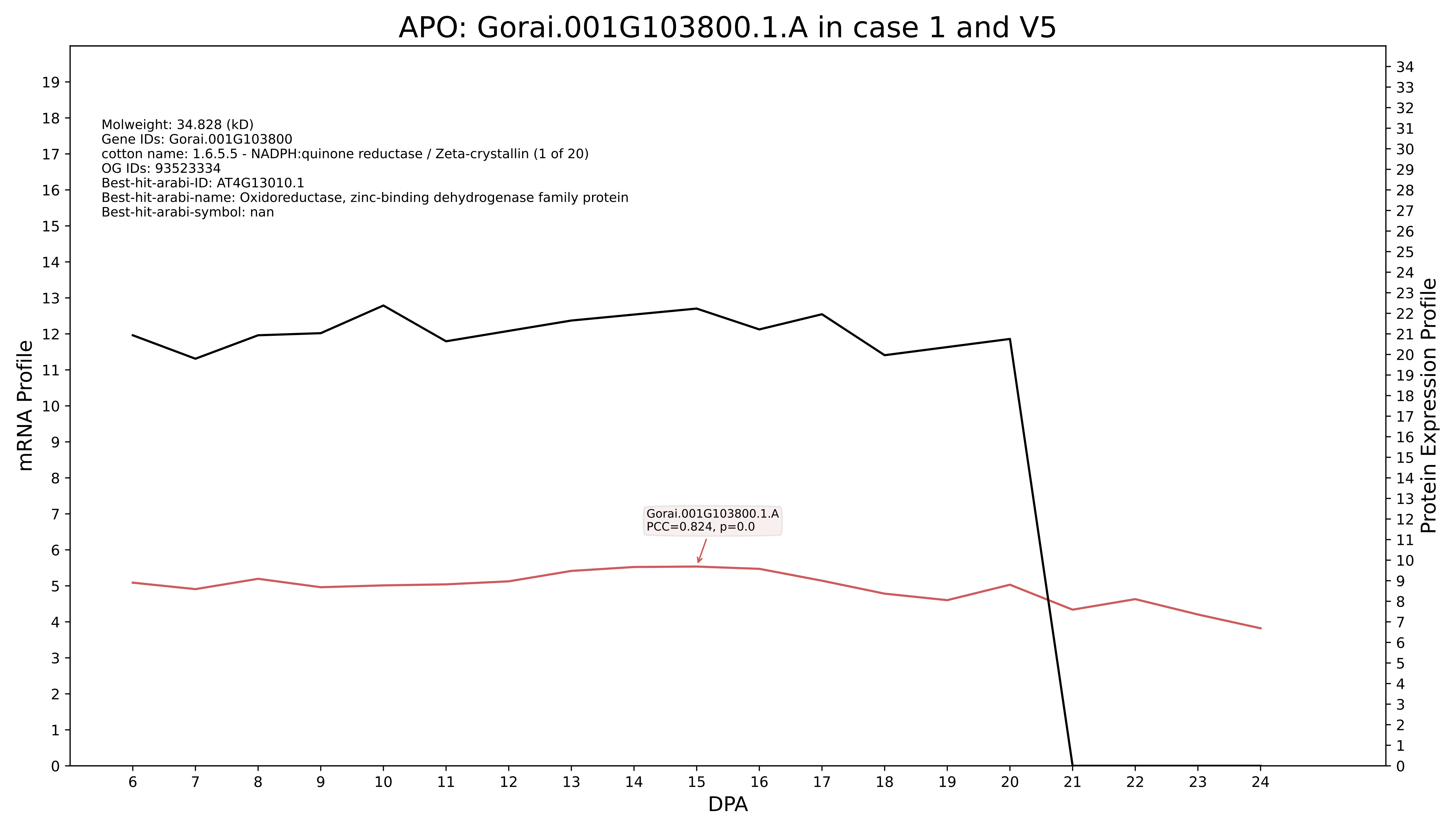
APO: Gorai.001G103800.1.A in case 1 and V5
Molweight: 34.828 (kD)
Gene IDs: Gorai.001G103800
cotton name: 1.6.5.5 - NADPH:quinone reductase / Zeta-crystallin (1 of 20)
OG IDs: 93523334
Best-hit-arabi-ID: AT4G13010.1
Best-hit-arabi-name: Oxidoreductase, zinc-binding dehydrogenase family protein
Best-hit-arabi-symbol: nan
Gorai.001G103800.1.A
PCC=0.824, p=0.0
mRNA Profile
Protein Expression Profile
DPA

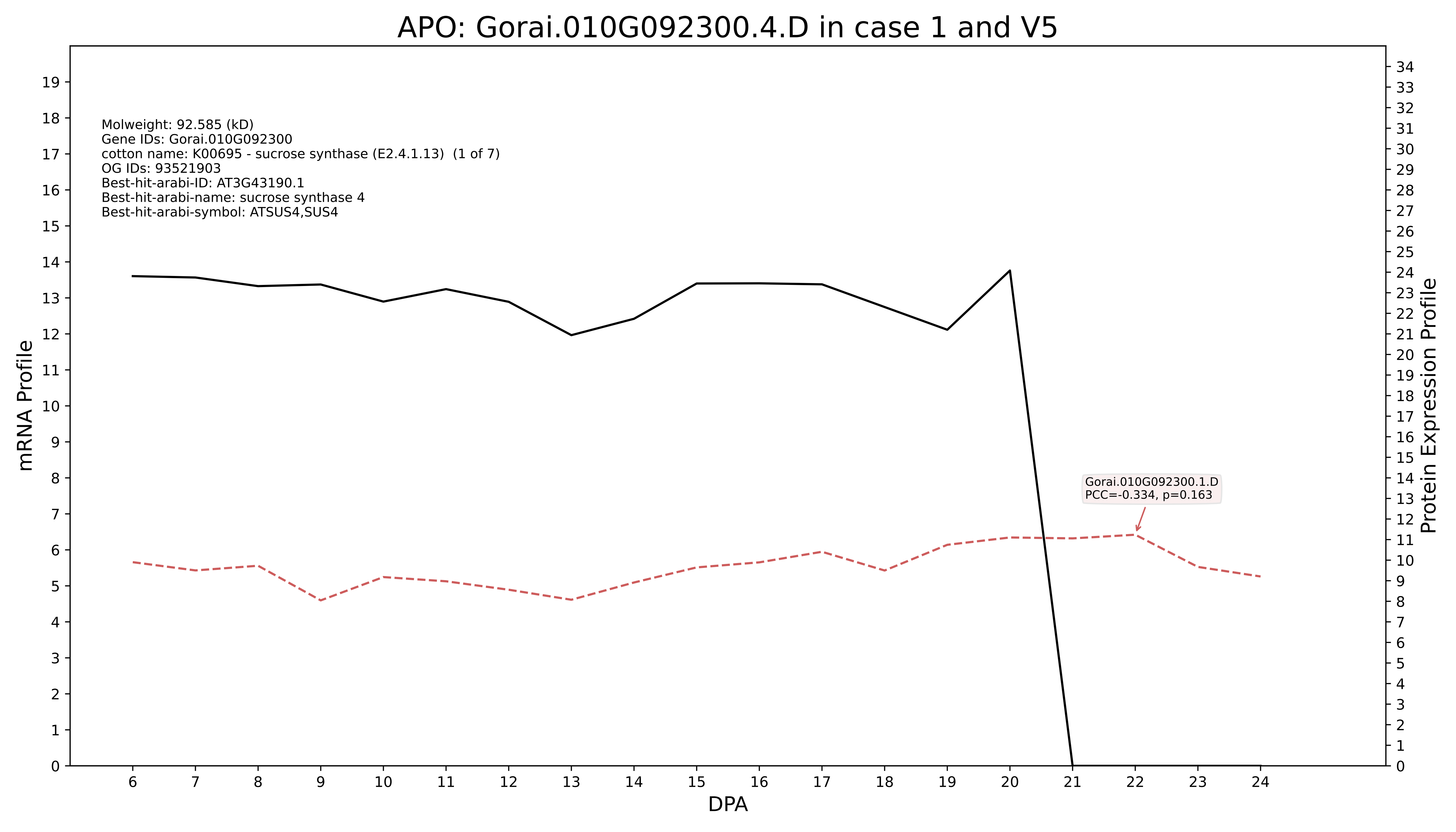
APO: Gorai.010G092300.4.D in case 1 and V5
Molweight: 92.585 (kD)
Gene IDs: Gorai.010G092300
cotton name: K00695 - sucrose synthase (E2.4.1.13) (1 of 7)
OG IDs: 93521903
Best-hit-arabi-ID: AT3G43190.1
Best-hit-arabi-name: sucrose synthase 4
Best-hit-arabi-symbol: ATSUS4,SUS4
Gorai.010G092300.1.D
PCC=-0.334, p=0.163
mRNA Profile
Protein Expression Profile
DPA

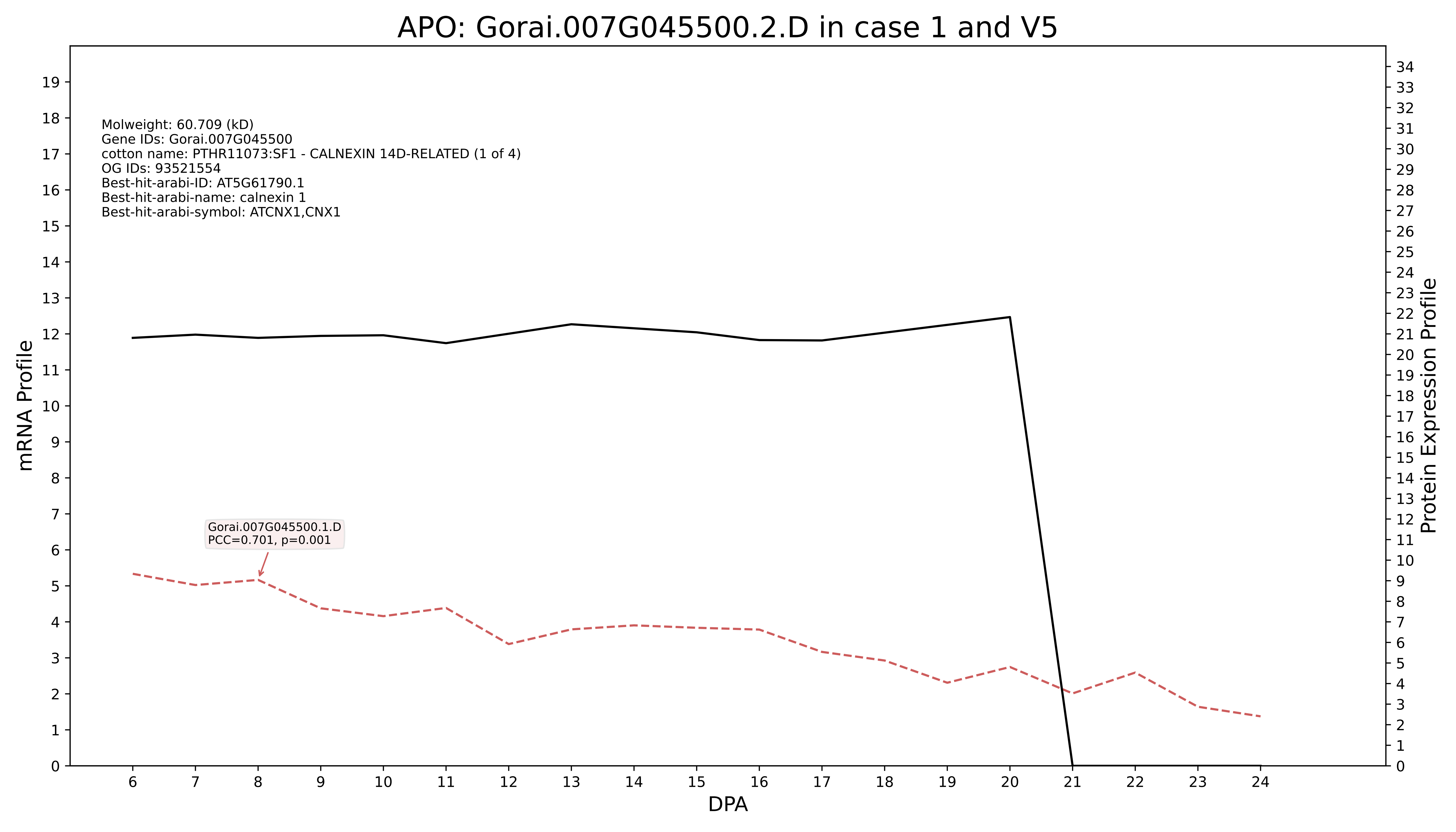
APO: Gorai.007G045500.2.D in case 1 and V5
Molweight: 60.709 (kD)
Gene IDs: Gorai.007G045500
cotton name: PTHR11073:SF1 - CALNEXIN 14D-RELATED (1 of 4)
OG IDs: 93521554
Best-hit-arabi-ID: AT5G61790.1
Best-hit-arabi-name: calnexin 1
Best-hit-arabi-symbol: ATCNX1,CNX1
Gorai.007G045500.1.D
PCC=0.701, p=0.001
mRNA Profile
Protein Expression Profile
DPA

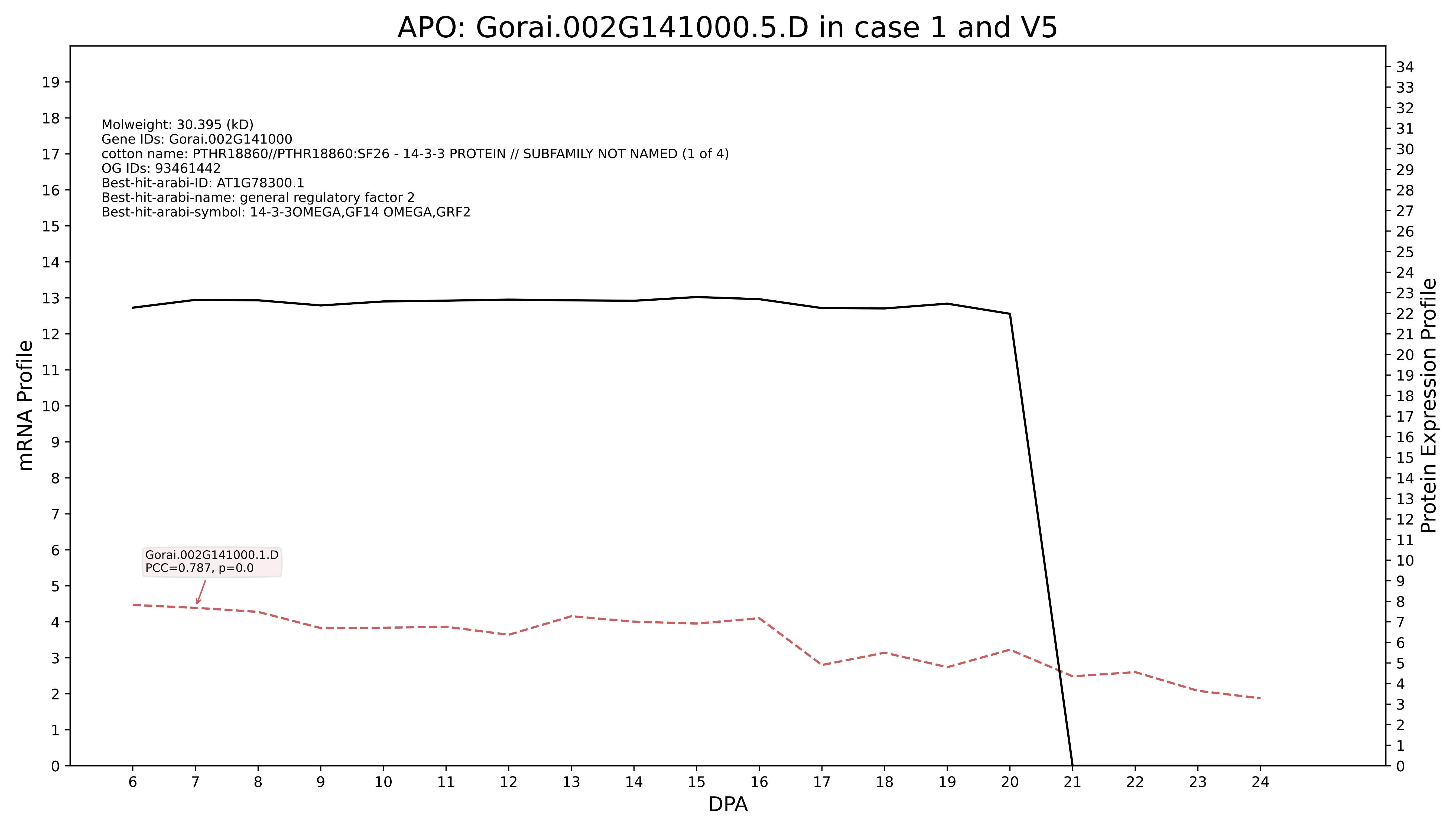
APO: Gorai.002G141000.5.D in case 1 and V5
Molweight: 30.395 (kD)
Gene IDs: Gorai.002G141000
cotton name: PTHR18860//PTHR18860:SF26 - 14-3-3 PROTEIN // SUBFAMILY NOT NAMED (1 of 4)
OG IDs: 93461442
Best-hit-arabi-ID: AT1G78300.1
Best-hit-arabi-name: general regulatory factor 2
Best-hit-arabi-symbol: 14-3-3OMEGA,GF14 OMEGA,GRF2
Gorai.002G141000.1.D
PCC=0.787, p=0.0
mRNA Profile
Protein Expression Profile
DPA

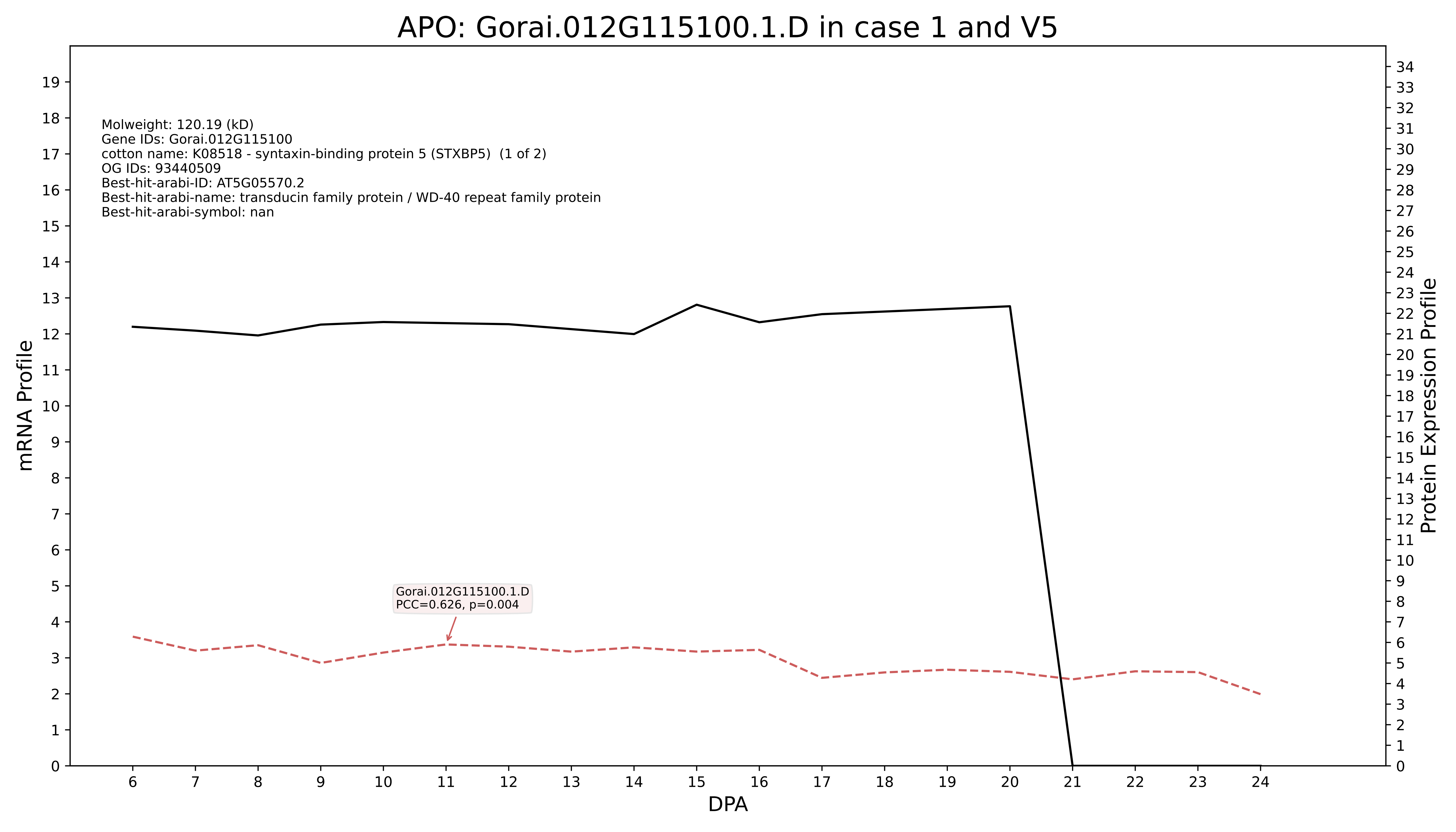

APO: Gorai.012G115100.1.D in case 1 and V5
Molweight: 120.19 (kD)
Gene IDs: Gorai.012G115100
cotton name: K08518 - syntaxin-binding protein 5 (STXBP5) (1 of 2)
OG IDs: 93440509
Best-hit-arabi-ID: AT5G05570.2
Best-hit-arabi-name: transducin family protein / WD-40 repeat family protein
Best-hit-arabi-symbol: nan
Gorai.012G115100.1.D
PCC=0.626, p=0.004
mRNA Profile
Protein Expression Profile
DPA

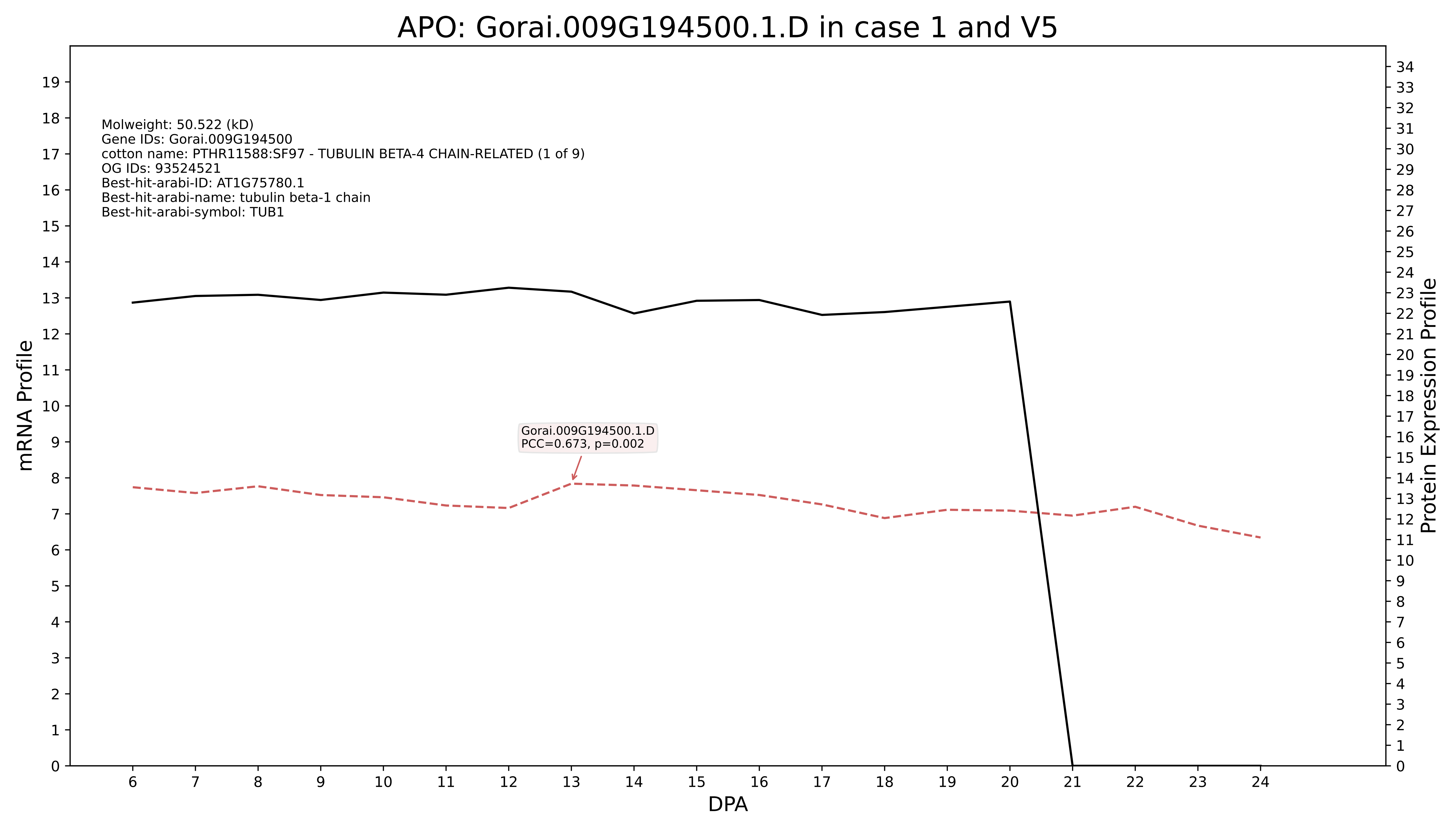

APO: Gorai.009G194500.1.D in case 1 and V5
Molweight: 50.522 (kD)
Gene IDs: Gorai.009G194500
cotton name: PTHR11588:SF97 - TUBULIN BETA-4 CHAIN-RELATED (1 of 9)
OG IDs: 93524521
Best-hit-arabi-ID: AT1G75780.1
Best-hit-arabi-name: tubulin beta-1 chain
Best-hit-arabi-symbol: TUB1
Gorai.009G194500.1.D
PCC=0.673, p=0.002
mRNA Profile
Protein Expression Profile
DPA

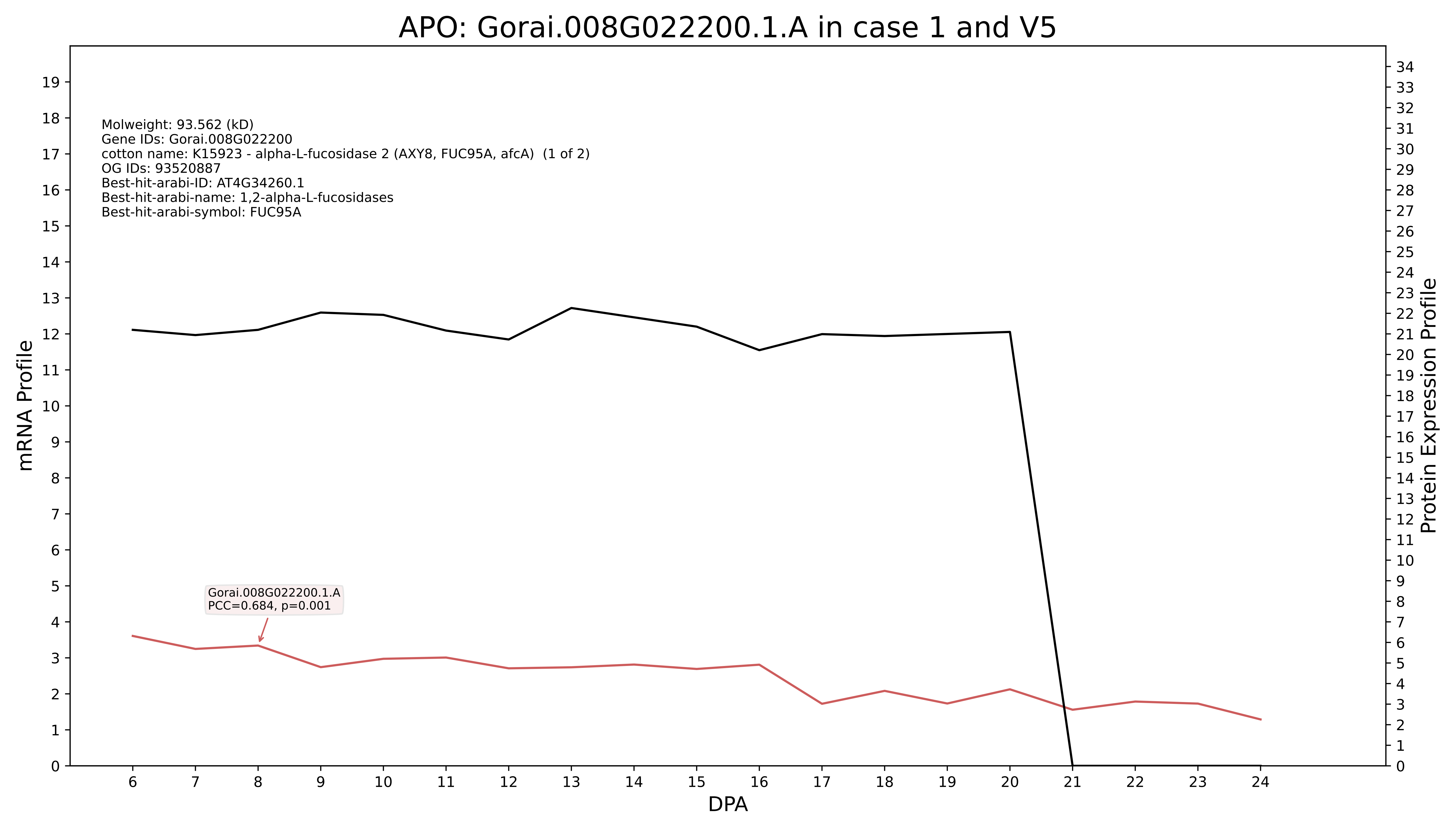
APO: Gorai.008G022200.1.A in case 1 and V5
Molweight: 93.562 (kD)
Gene IDs: Gorai.008G022200
cotton name: K15923 - alpha-L-fucosidase 2 (AXY8, FUC95A, afcA) (1 of 2)
OG IDs: 93520887
Best-hit-arabi-ID: AT4G34260.1
Best-hit-arabi-name: 1,2-alpha-L-fucosidases
Best-hit-arabi-symbol: FUC95A
Gorai.008G022200.1.A
PCC=0.684, p=0.001
mRNA Profile
Protein Expression Profile
DPA

### APO: Gorai.002G262000.1.D in case 1 and V5

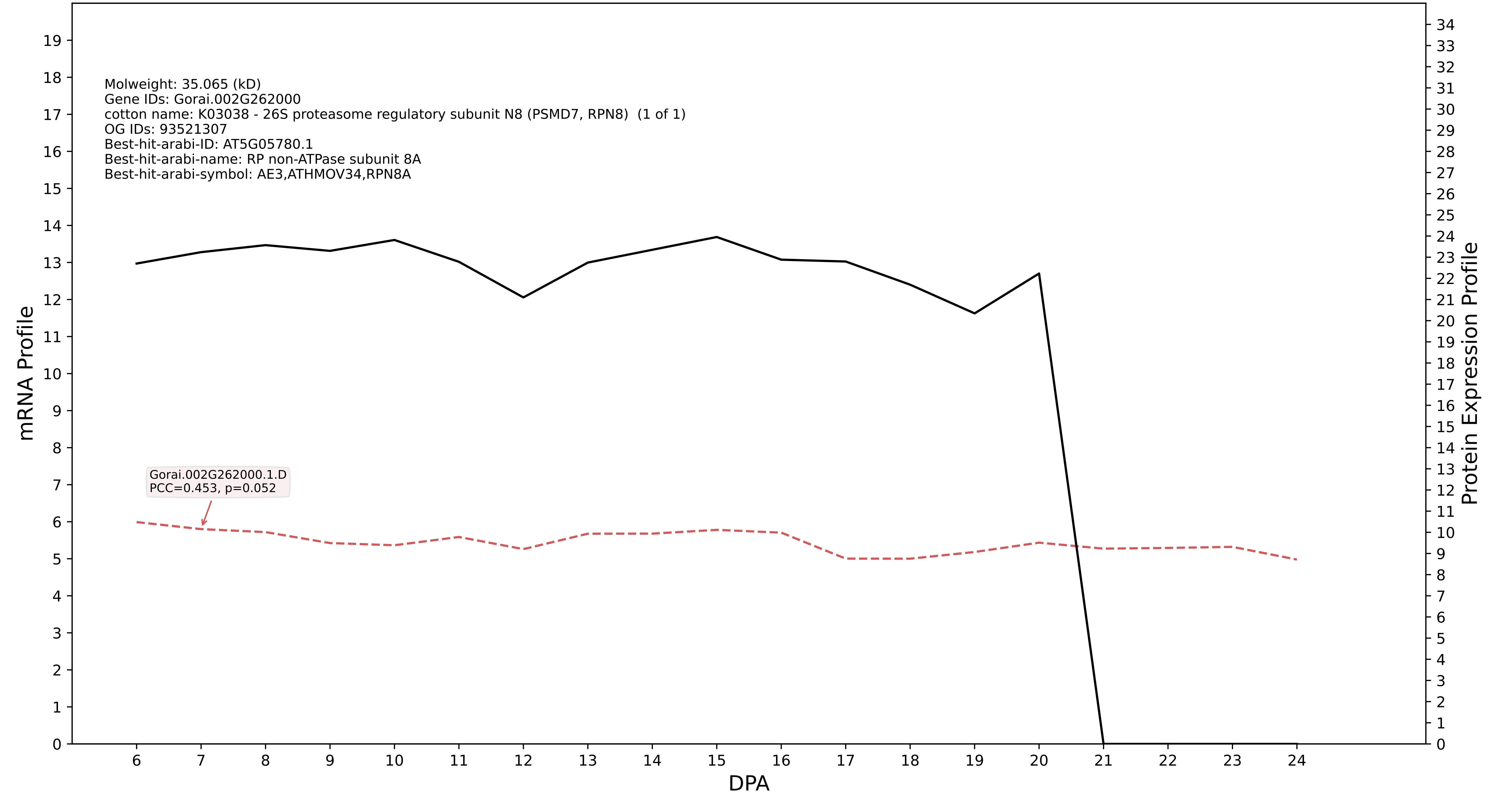

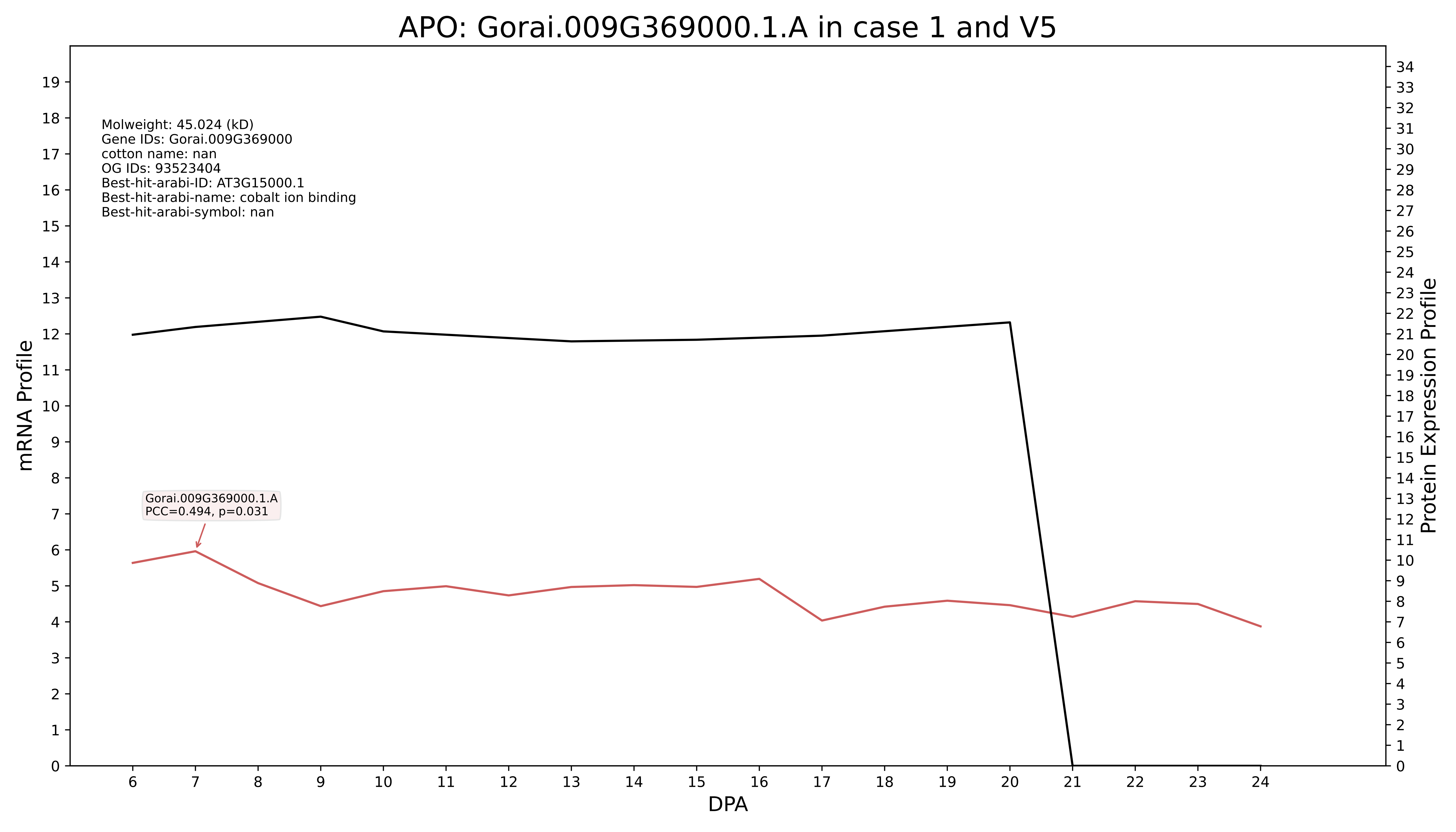
APO: Gorai.009G369000.1.A in case 1 and V5
Molweight: 45.024 (kD)
Gene IDs: Gorai.009G369000
cotton name: nan
OG IDs: 93523404
Best-hit-arabi-ID: AT3G15000.1
Best-hit-arabi-name: cobalt ion binding
Best-hit-arabi-symbol: nan
Gorai.009G369000.1.A
PCC=0.494, p=0.031
mRNA Profile
Protein Expression Profile
DPA

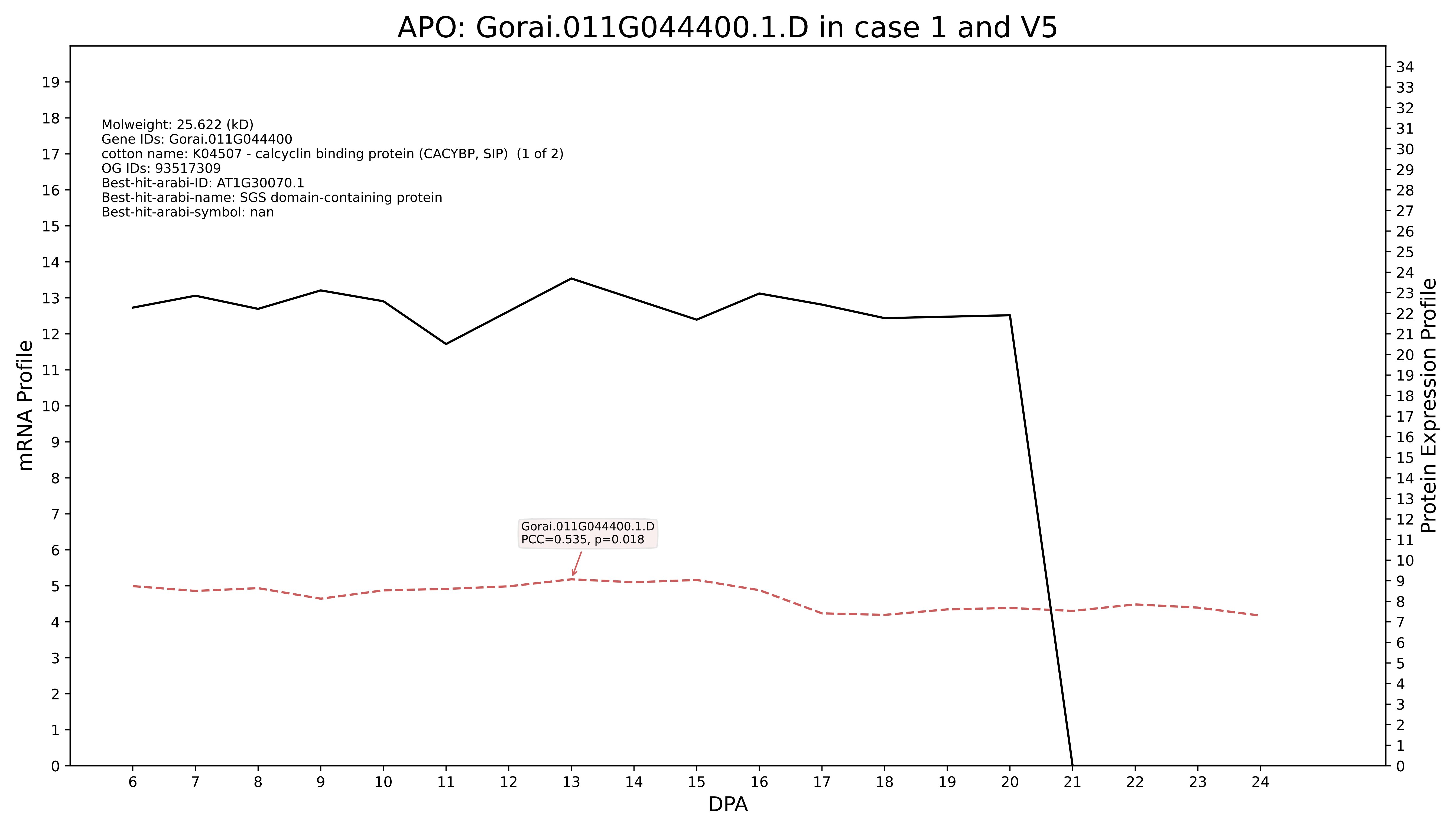
APO: Gorai.011G044400.1.D in case 1 and V5
Molweight: 25.622 (kD)
Gene IDs: Gorai.011G044400
cotton name: K04507 - calcyclin binding protein (CACYBP, SIP) (1 of 2)
OG IDs: 93517309
Best-hit-arabi-ID: AT1G30070.1
Best-hit-arabi-name: SGS domain-containing protein
Best-hit-arabi-symbol: nan
Gorai.011G044400.1.D
PCC=0.535, p=0.018
mRNA Profile
Protein Expression Profile
DPA

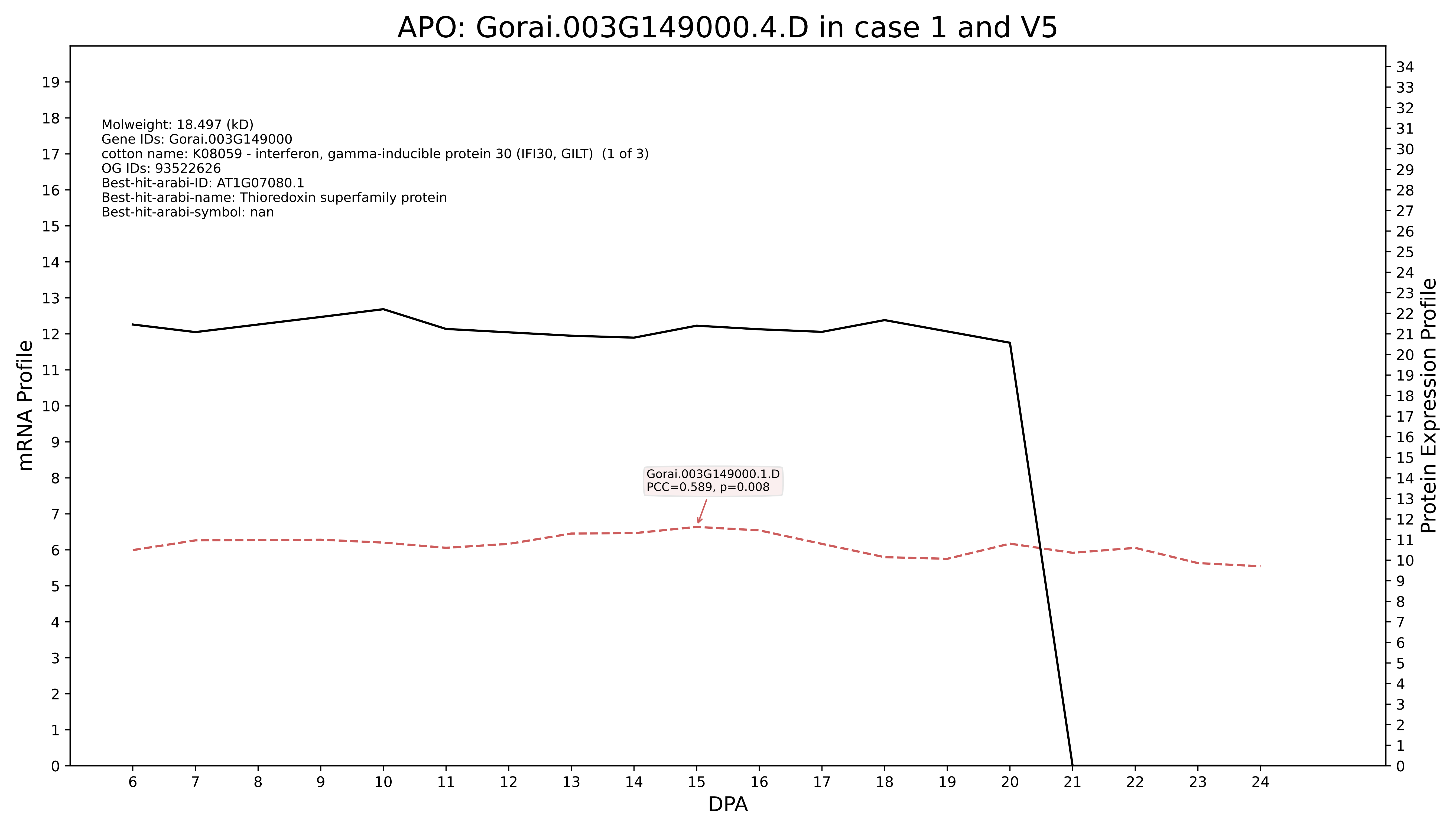

APO: Gorai.003G149000.4.D in case 1 and V5
Molweight: 18.497 (kD)
Gene IDs: Gorai.003G149000
cotton name: K08059 - interferon, gamma-inducible protein 30 (IFI30, GILT) (1 of 3)
OG IDs: 93522626
Best-hit-arabi-ID: AT1G07080.1
Best-hit-arabi-name: Thioredoxin superfamily protein
Best-hit-arabi-symbol: nan
Gorai.003G149000.1.D
PCC=0.589, p=0.008
mRNA Profile
Protein Expression Profile
DPA

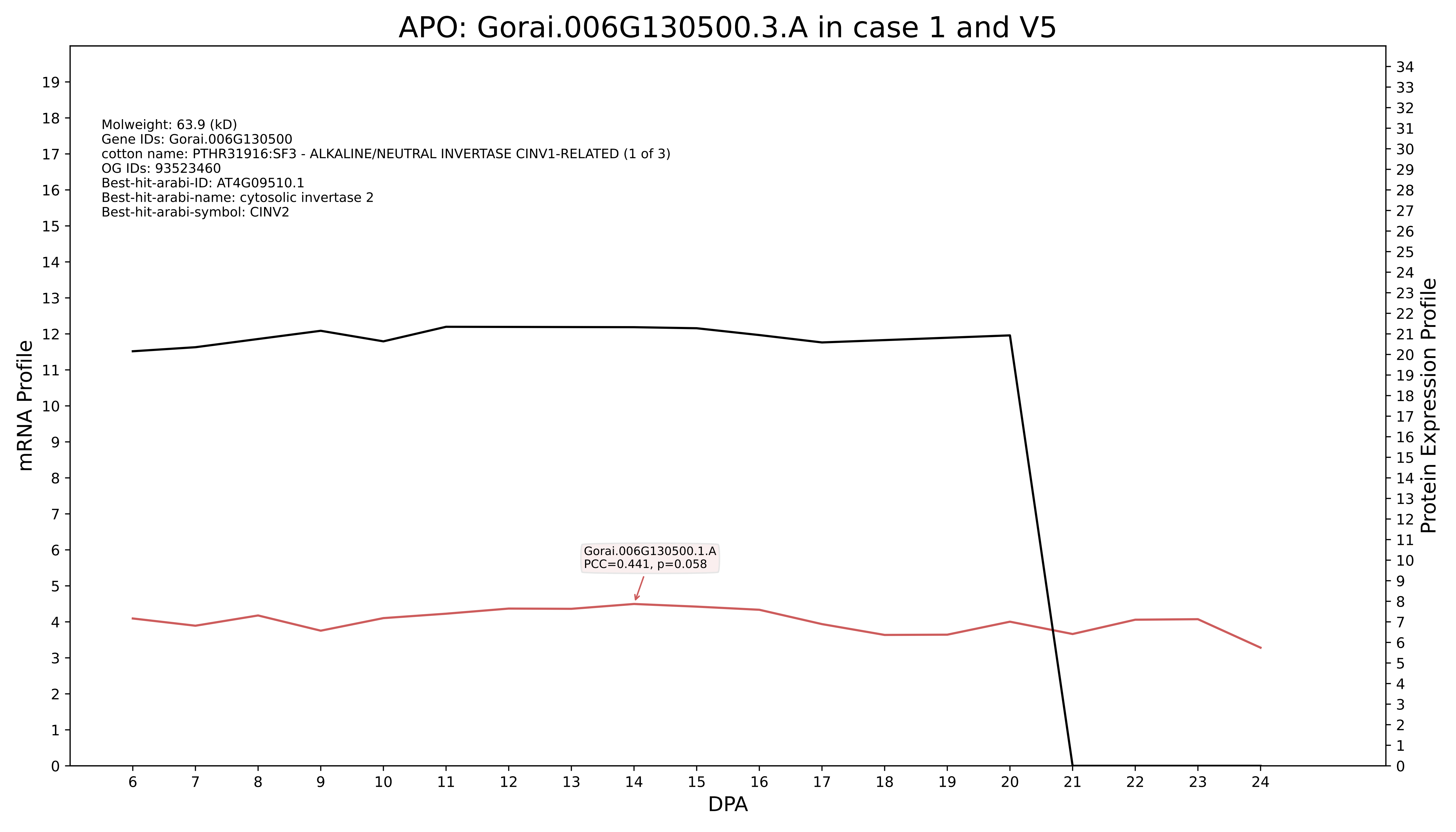

APO: Gorai.006G130500.3.A in case 1 and V5
Molweight: 63.9 (kD)
Gene IDs: Gorai.006G130500
cotton name: PTHR31916:SF3 - ALKALINE/NEUTRAL INVERTASE CINV1-RELATED (1 of 3)
OG IDs: 93523460
Best-hit-arabi-ID: AT4G09510.1
Best-hit-arabi-name: cytosolic invertase 2
Best-hit-arabi-symbol: CINV2
Gorai.006G130500.1.A
PCC=0.441, p=0.058
mRNA Profile
Protein Expression Profile
DPA

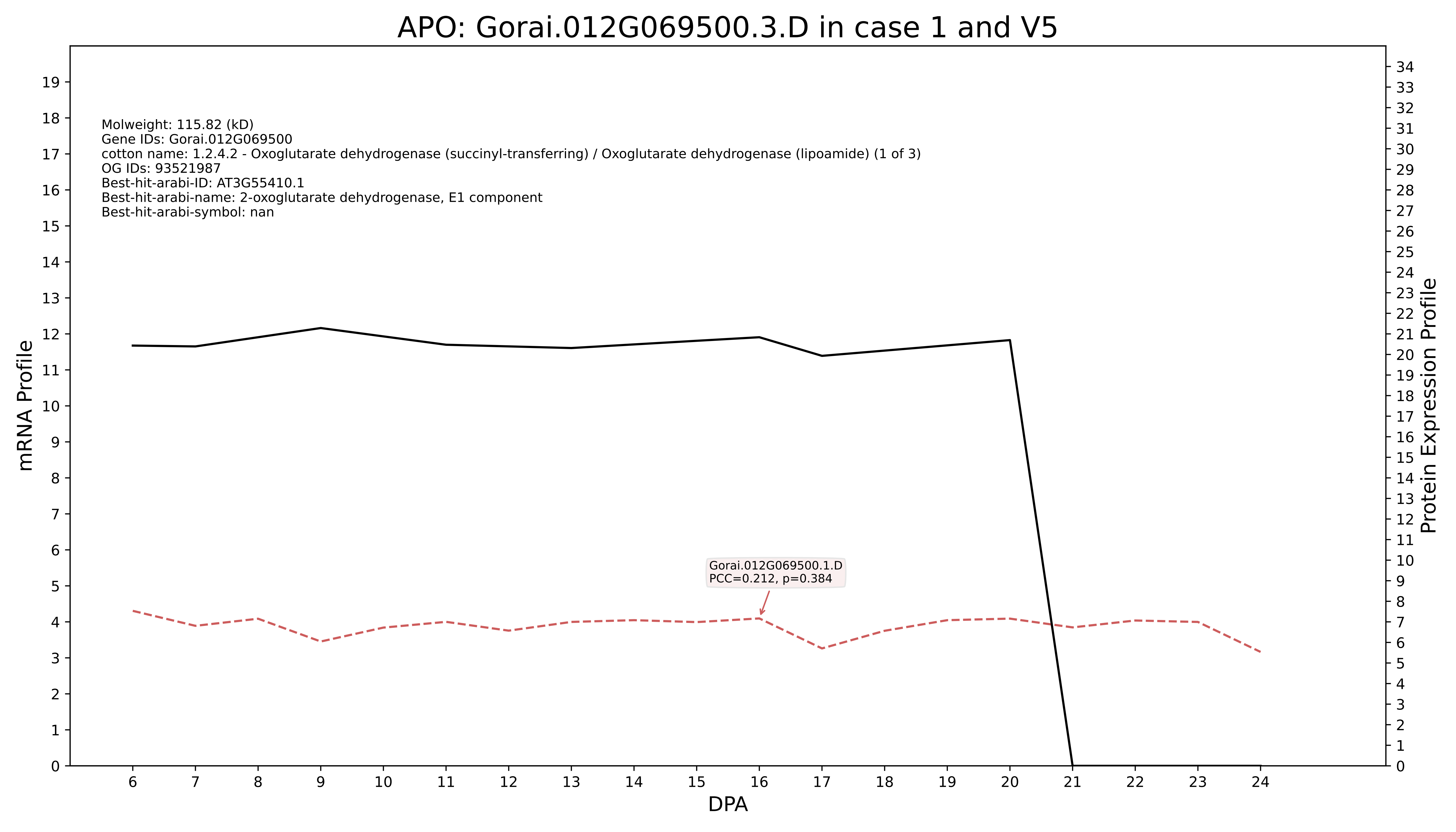
APO: Gorai.012G069500.3.D in case 1 and V5
Molweight: 115.82 (kD)
Gene IDs: Gorai.012G069500
cotton name: 1.2.4.2 - Oxoglutarate dehydrogenase (succinyl-transferring) / Oxoglutarate dehydrogenase (lipoamide) (1 of 3)
OG IDs: 93521987
Best-hit-arabi-ID: AT3G55410.1
Best-hit-arabi-name: 2-oxoglutarate dehydrogenase, E1 component
Best-hit-arabi-symbol: nan
Gorai.012G069500.1.D
PCC=0.212, p=0.384
mRNA Profile
Protein Expression Profile
DPA

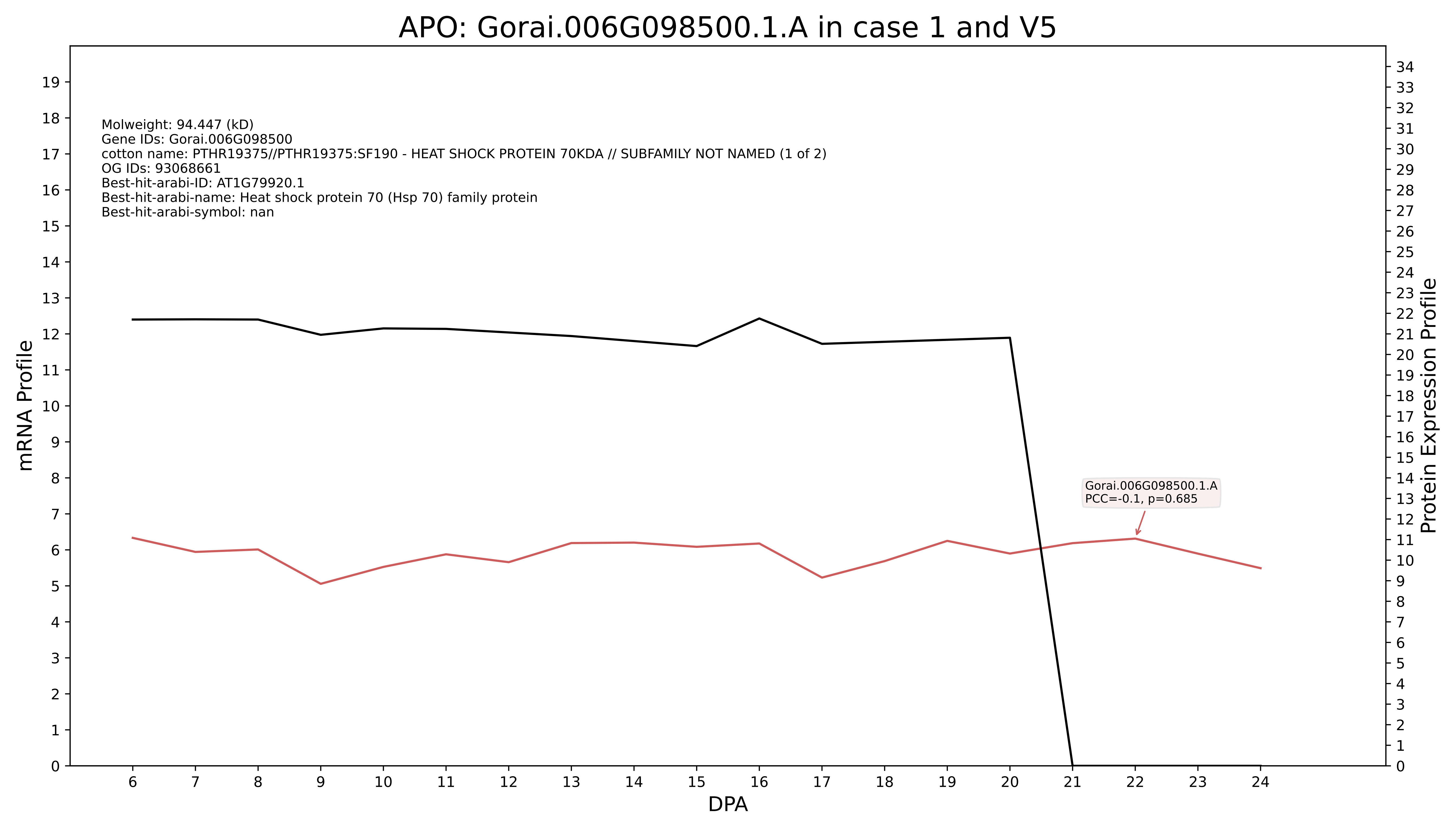

APO: Gorai.006G098500.1.A in case 1 and V5
Molweight: 94.447 (kD)
Gene IDs: Gorai.006G098500
cotton name: PTHR19375//PTHR19375:SF190 - HEAT SHOCK PROTEIN 70KDA // SUBFAMILY NOT NAMED (1 of 2)
OG IDs: 93068661
Best-hit-arabi-ID: AT1G79920.1
Best-hit-arabi-name: Heat shock protein 70 (Hsp 70) family protein
Best-hit-arabi-symbol: nan
Gorai.006G098500.1.A
PCC=-0.1, p=0.685
mRNA Profile
Protein Expression Profile
DPA

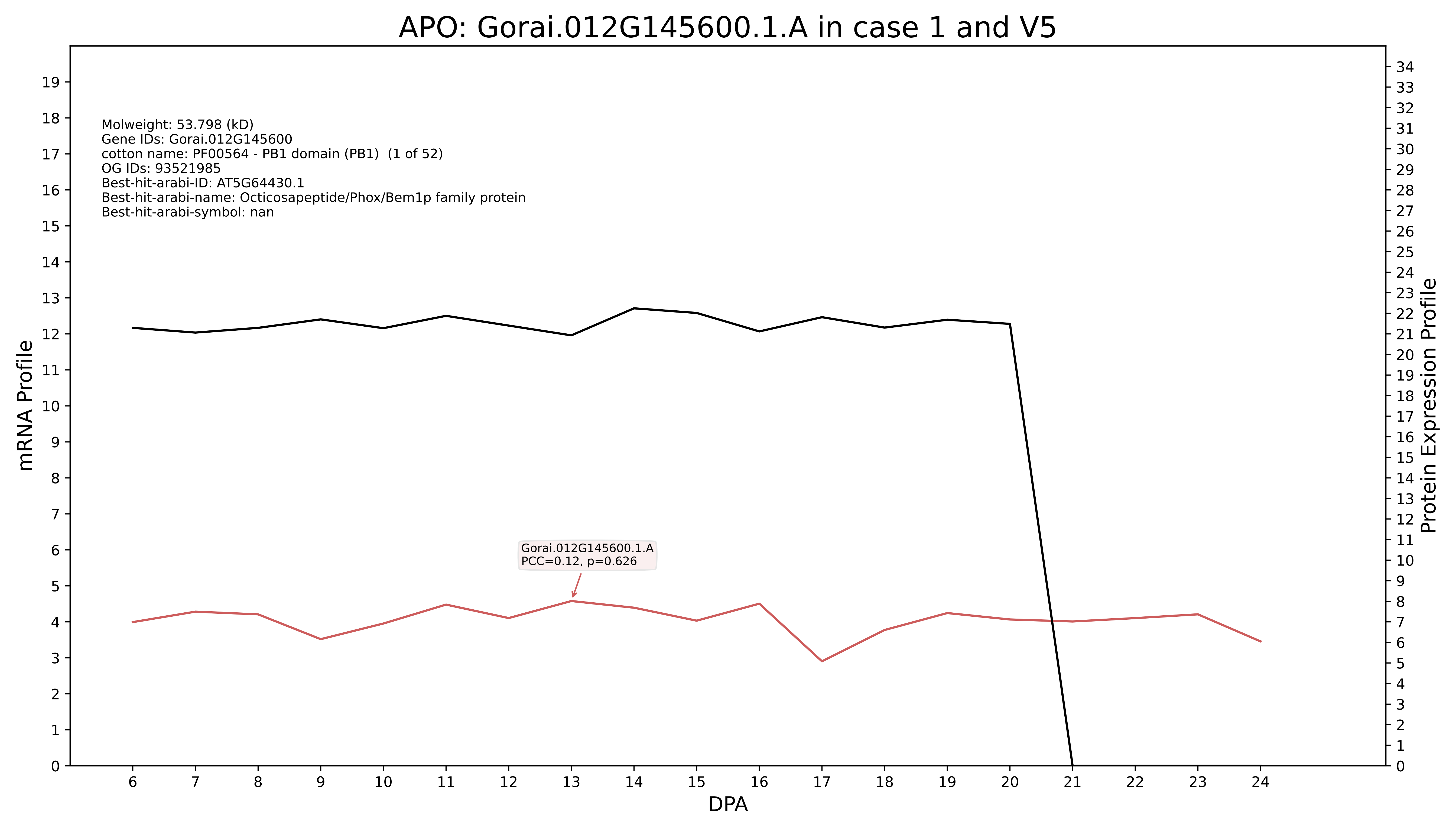
APO: Gorai.012G145600.1.A in case 1 and V5
Molweight: 53.798 (kD)
Gene IDs: Gorai.012G145600
cotton name: PF00564 - PB1 domain (PB1) (1 of 52)
OG IDs: 93521985
Best-hit-arabi-ID: AT5G64430.1
Best-hit-arabi-name: Octicosapeptide/Phox/Bem1p family protein
Best-hit-arabi-symbol: nan
Gorai.012G145600.1.A
PCC=0.12, p=0.626
mRNA Profile
Protein Expression Profile
DPA

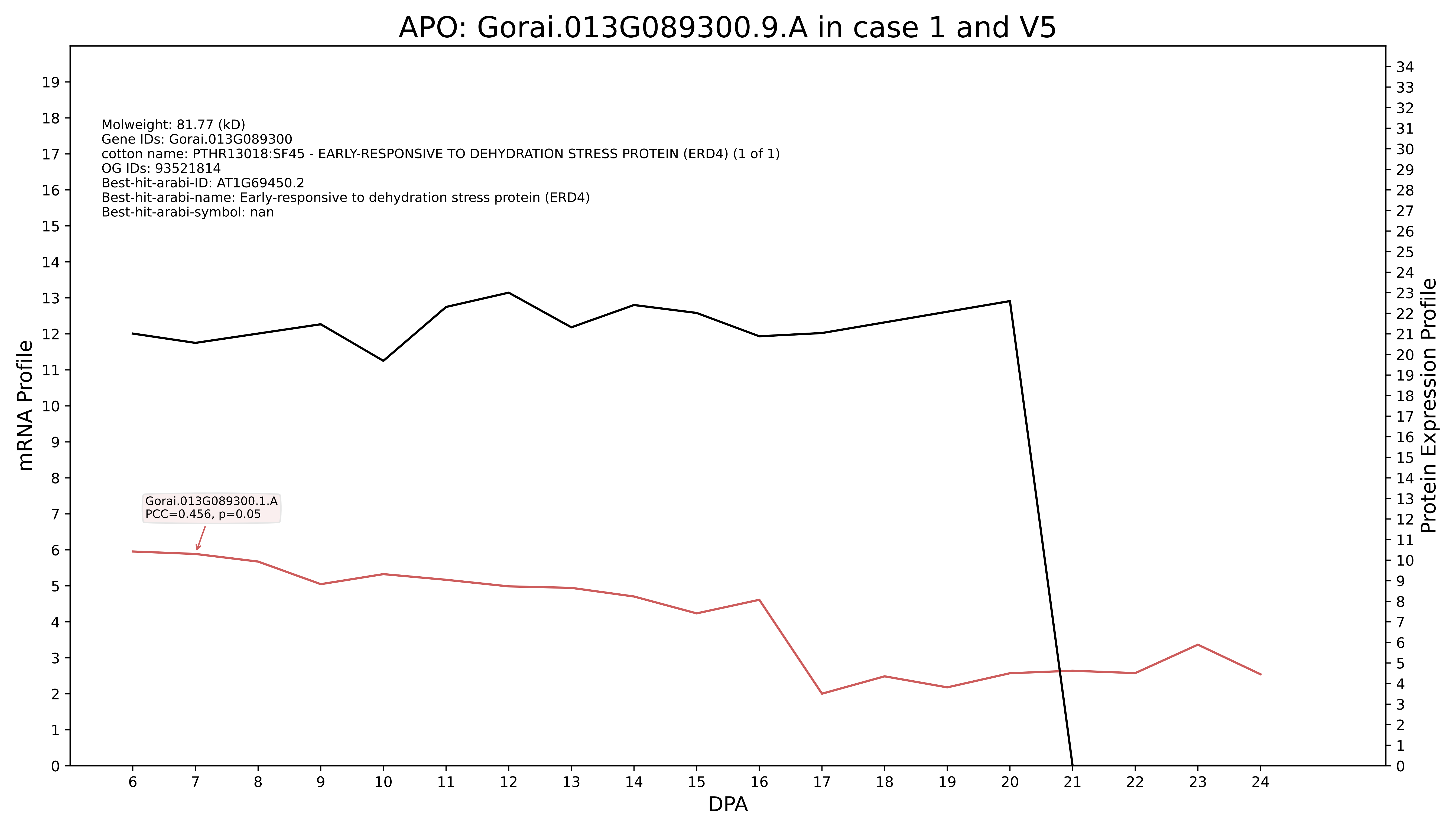
APO: Gorai.013G089300.9.A in case 1 and V5
Molweight: 81.77 (kD)
Gene IDs: Gorai.013G089300
cotton name: PTHR13018:SF45 - EARLY-RESPONSIVE TO DEHYDRATION STRESS PROTEIN (ERD4) (1 of 1)
OG IDs: 93521814
Best-hit-arabi-ID: AT1G69450.2
Best-hit-arabi-name: Early-responsive to dehydration stress protein (ERD4)
Best-hit-arabi-symbol: nan
Gorai.013G089300.1.A
PCC=0.456, p=0.05
mRNA Profile
Protein Expression Profile
DPA

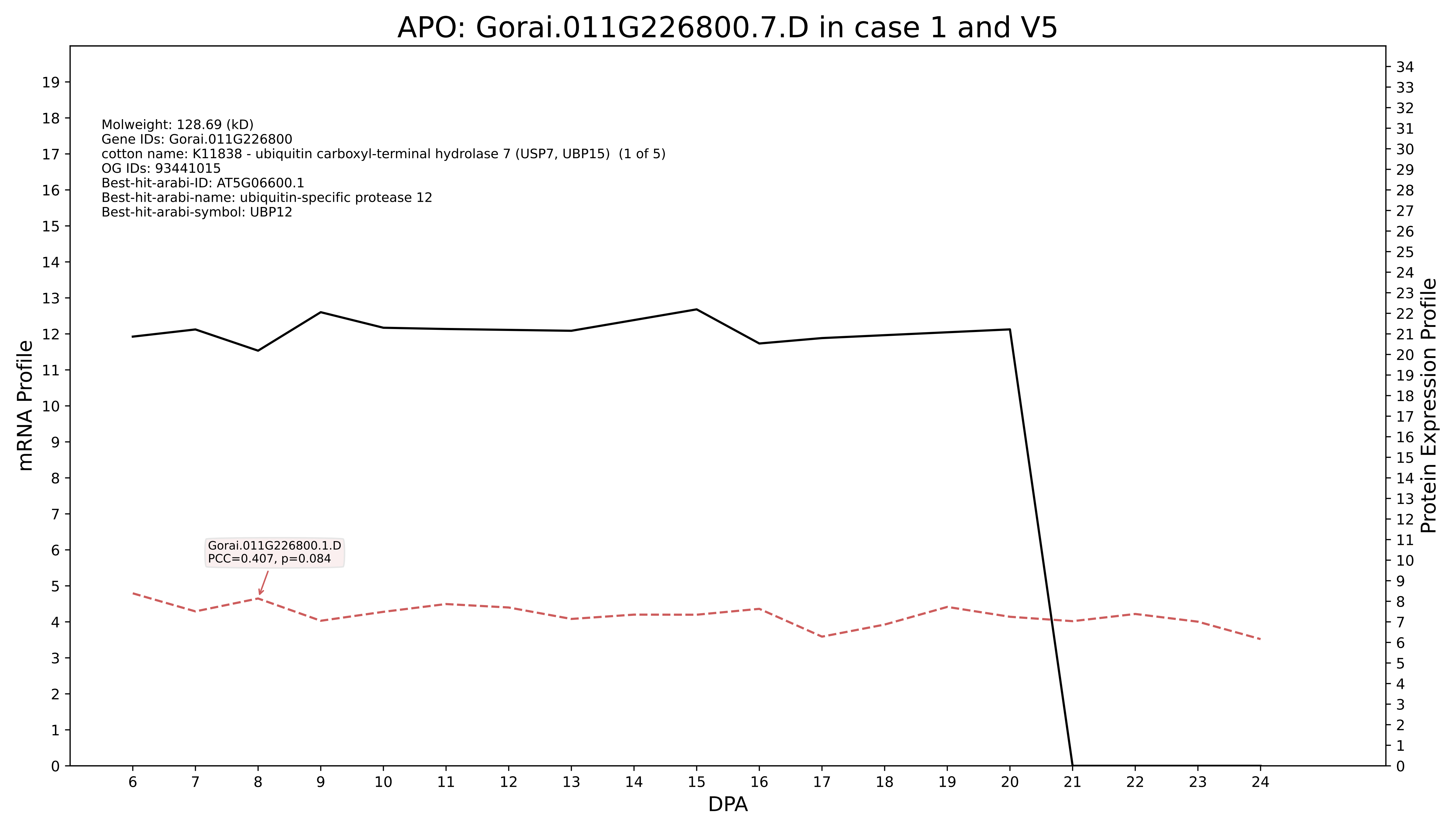

APO: Gorai.011G226800.7.D in case 1 and V5
Molweight: 128.69 (kD)
Gene IDs: Gorai.011G226800
cotton name: K11838 - ubiquitin carboxyl-terminal hydrolase 7 (USP7, UBP15) (1 of 5)
OG IDs: 93441015
Best-hit-arabi-ID: AT5G06600.1
Best-hit-arabi-name: ubiquitin-specific protease 12
Best-hit-arabi-symbol: UBP12
Gorai.011G226800.1.D
PCC=0.407, p=0.084
mRNA Profile
Protein Expression Profile
DPA

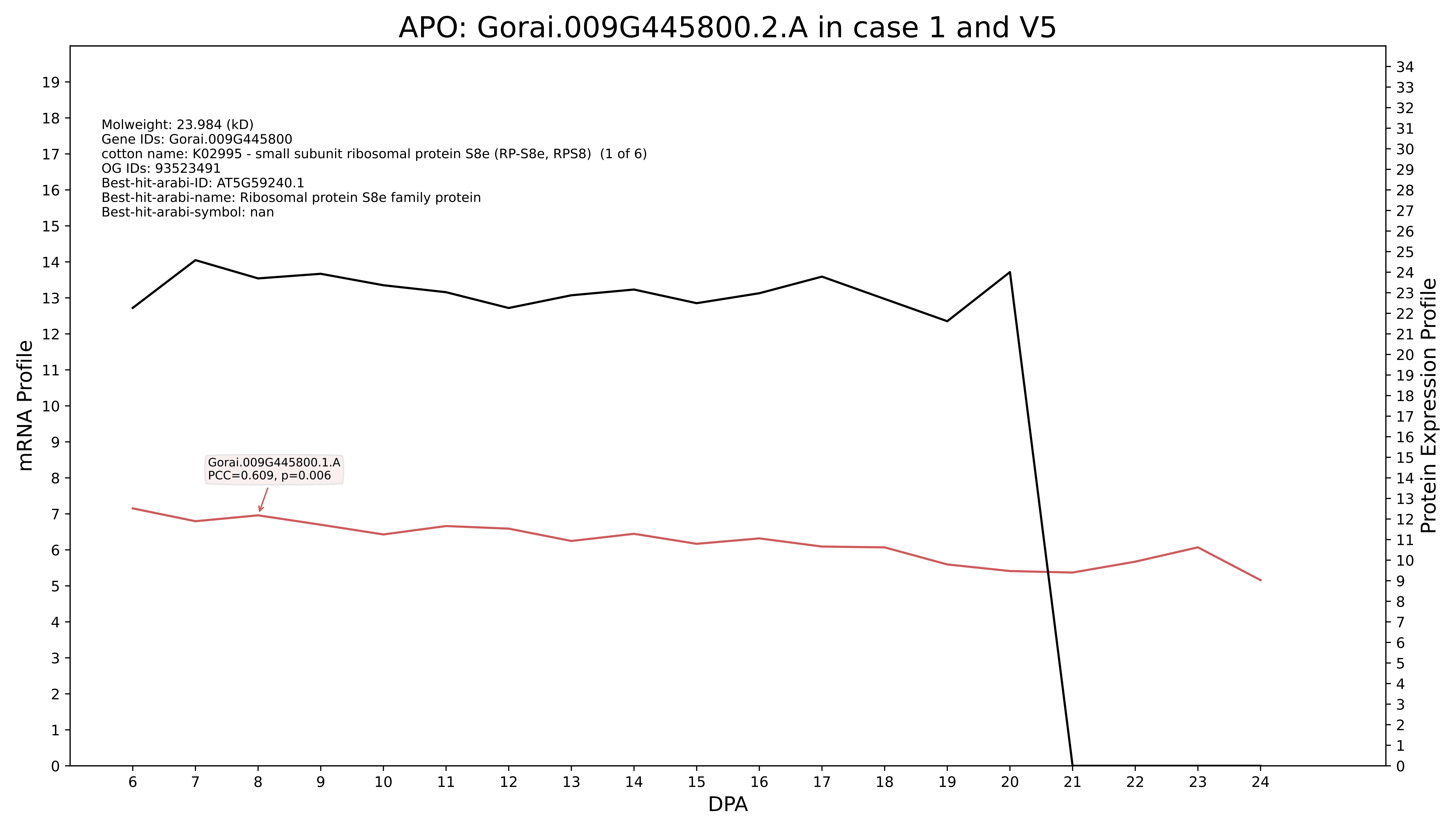
APO: Gorai.009G445800.2.A in case 1 and V5
Molweight: 23.984 (kD)
Gene IDs: Gorai.009G445800
cotton name: K02995 - small subunit ribosomal protein S8e (RP-S8e, RPS8) (1 of 6)
OG IDs: 93523491
Best-hit-arabi-ID: AT5G59240.1
Best-hit-arabi-name: Ribosomal protein S8e family protein
Best-hit-arabi-symbol: nan
Gorai.009G445800.1.A
PCC=0.609, p=0.006
mRNA Profile
Protein Expression Profile
DPA

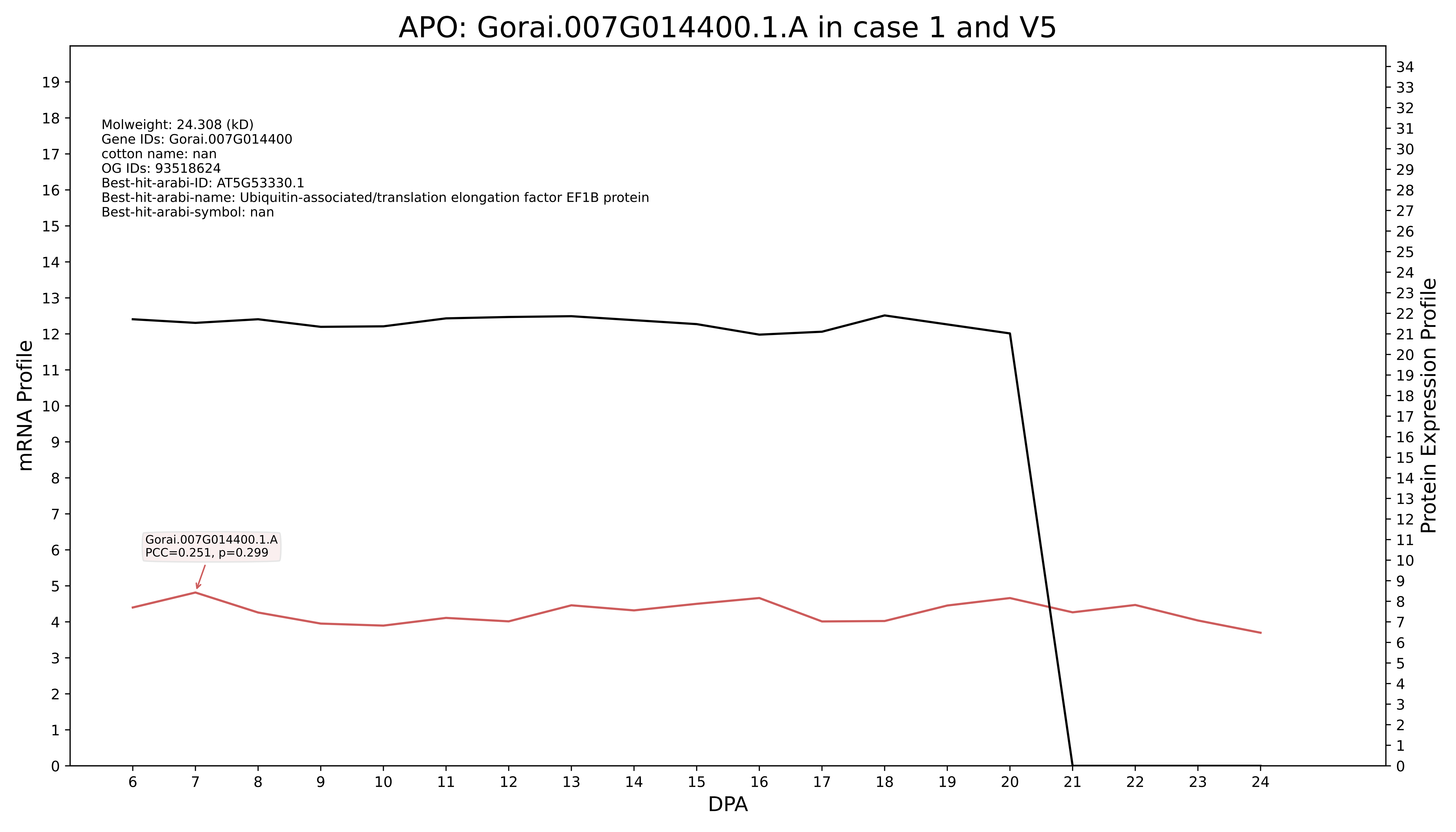
APO: Gorai.007G014400.1.A in case 1 and V5
Molweight: 24.308 (kD)
Gene IDs: Gorai.007G014400
cotton name: nan
OG IDs: 93518624
Best-hit-arabi-ID: AT5G53330.1
Best-hit-arabi-name: Ubiquitin-associated/translation elongation factor EF1B protein
Best-hit-arabi-symbol: nan
Gorai.007G014400.1.A
PCC=0.251, p=0.299
mRNA Profile
Protein Expression Profile
DPA

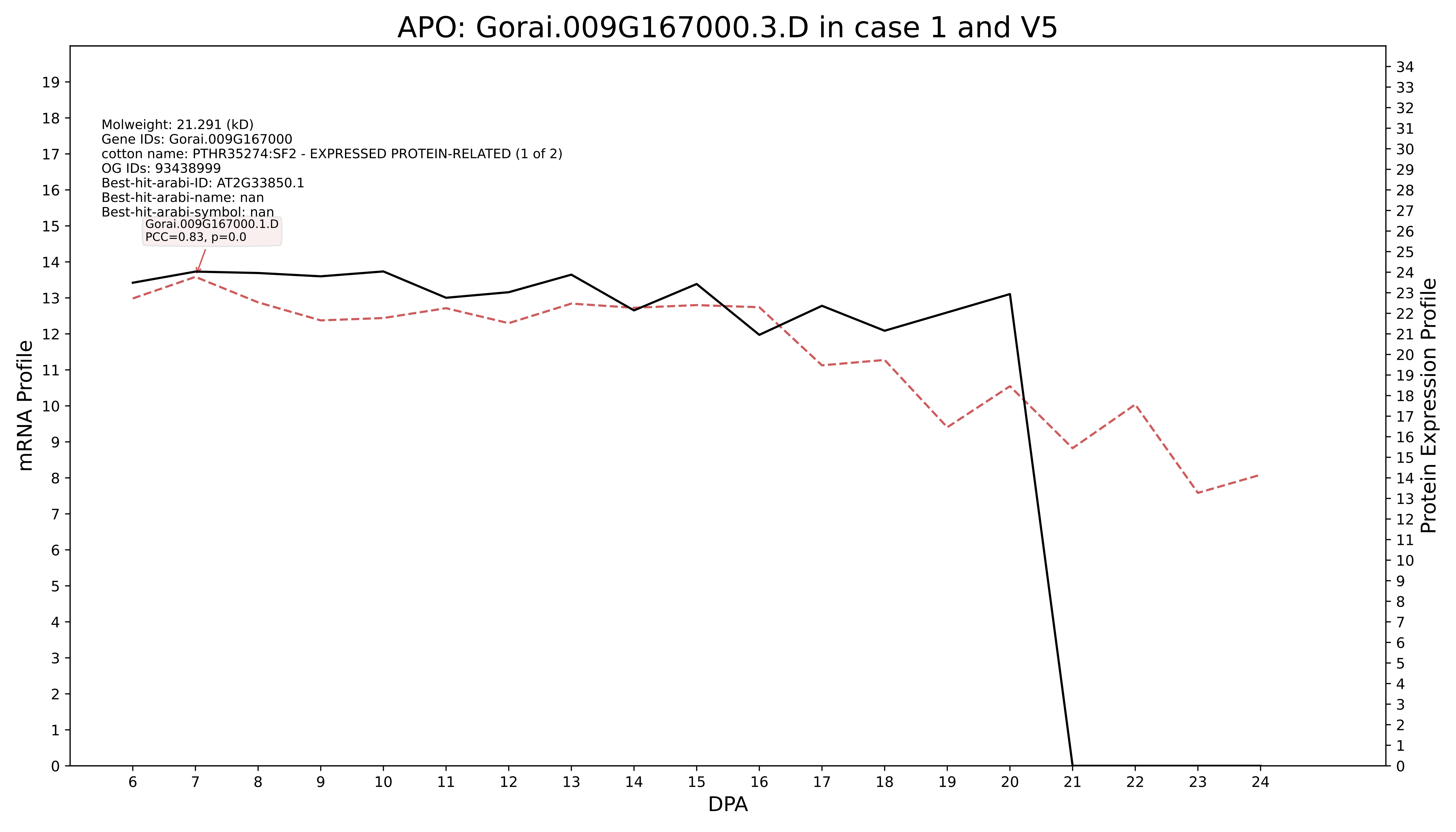

APO: Gorai.009G167000.3.D in case 1 and V5
Molweight: 21.291 (kD)
Gene IDs: Gorai.009G167000
cotton name: PTHR35274:SF2 - EXPRESSED PROTEIN-RELATED (1 of 2)
OG IDs: 93438999
Best-hit-arabi-ID: AT2G33850.1
Best-hit-arabi-name: nan
Best-hit-arabi-symbol: nan
Gorai.009G167000.1.D
PCC=0.83, p=0.0
mRNA Profile
Protein Expression Profile
DPA

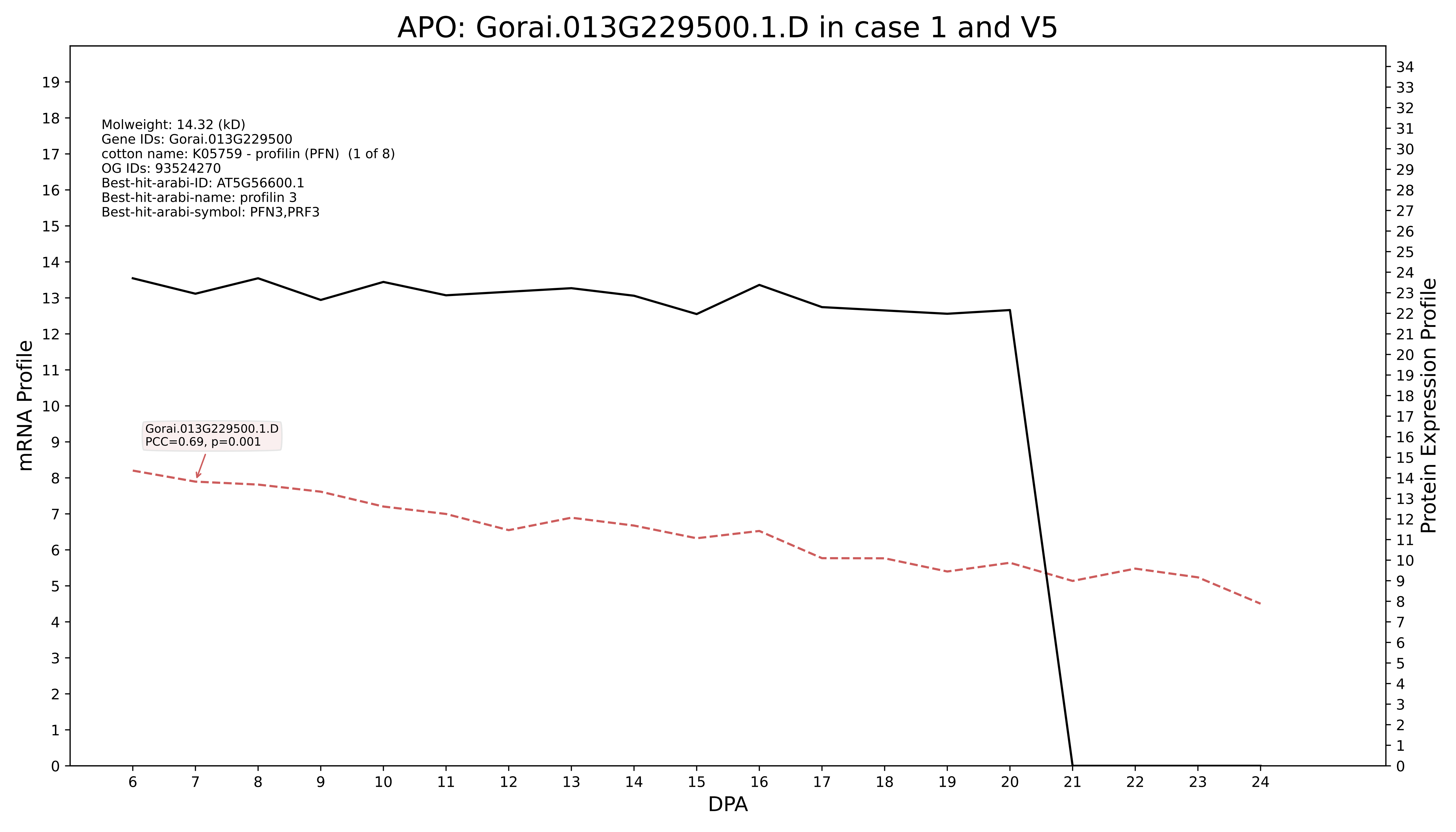
APO: Gorai.013G229500.1.D in case 1 and V5
Molweight: 14.32 (kD)
Gene IDs: Gorai.013G229500
cotton name: K05759 - profilin (PFN) (1 of 8)
OG IDs: 93524270
Best-hit-arabi-ID: AT5G56600.1
Best-hit-arabi-name: profilin 3
Best-hit-arabi-symbol: PFN3,PRF3
Gorai.013G229500.1.D
PCC=0.69, p=0.001
mRNA Profile
Protein Expression Profile
DPA

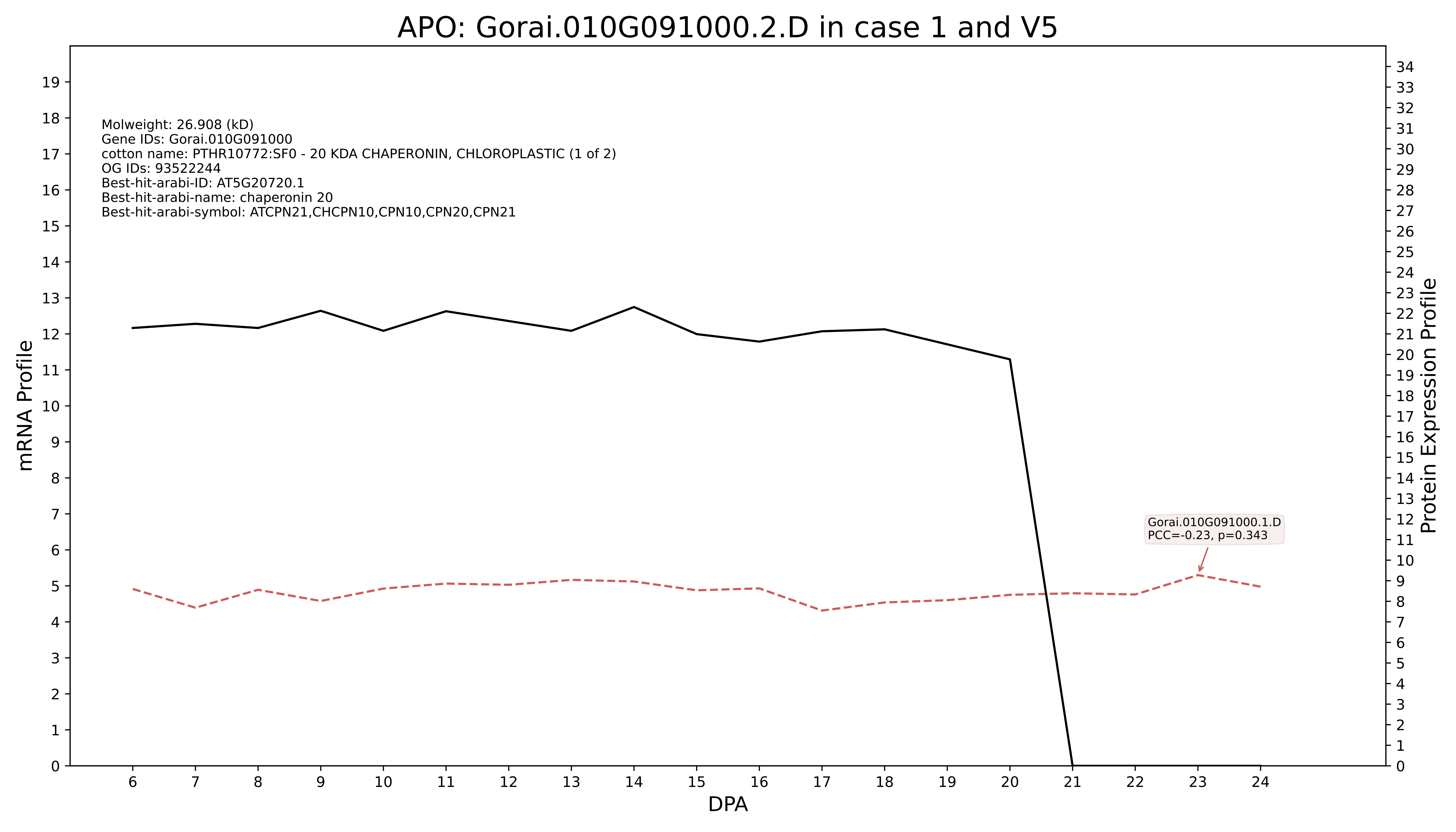

APO: Gorai.010G091000.2.D in case 1 and V5
Molweight: 26.908 (kD)
Gene IDs: Gorai.010G091000
cotton name: PTHR10772:SF0 - 20 KDA CHAPERONIN, CHLOROPLASTIC (1 of 2)
OG IDs: 93522244
Best-hit-arabi-ID: AT5G20720.1
Best-hit-arabi-name: chaperonin 20
Best-hit-arabi-symbol: ATCPN21,CHCPN10,CPN10,CPN20,CPN21
Gorai.010G091000.1.D
PCC=-0.23, p=0.343
mRNA Profile
Protein Expression Profile
DPA

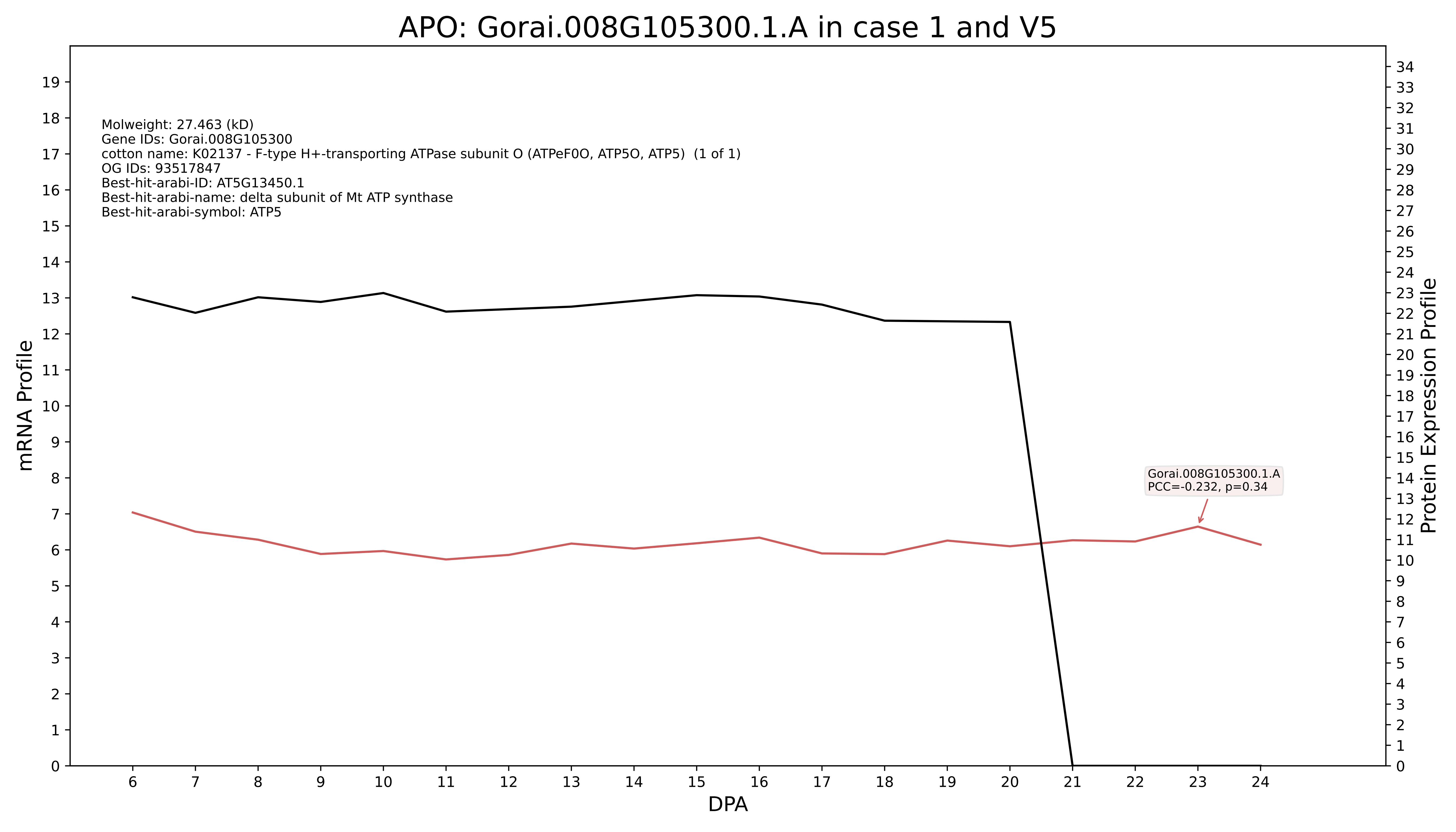
APO: Gorai.008G105300.1.A in case 1 and V5
Molweight: 27.463 (kD)
Gene IDs: Gorai.008G105300
cotton name: K02137 - F-type H+-transporting ATPase subunit O (ATPeF0O, ATP5O, ATP5) (1 of 1)
OG IDs: 93517847
Best-hit-arabi-ID: AT5G13450.1
Best-hit-arabi-name: delta subunit of Mt ATP synthase
Best-hit-arabi-symbol: ATP5
Gorai.008G105300.1.A
PCC=-0.232, p=0.34
mRNA Profile
Protein Expression Profile
DPA

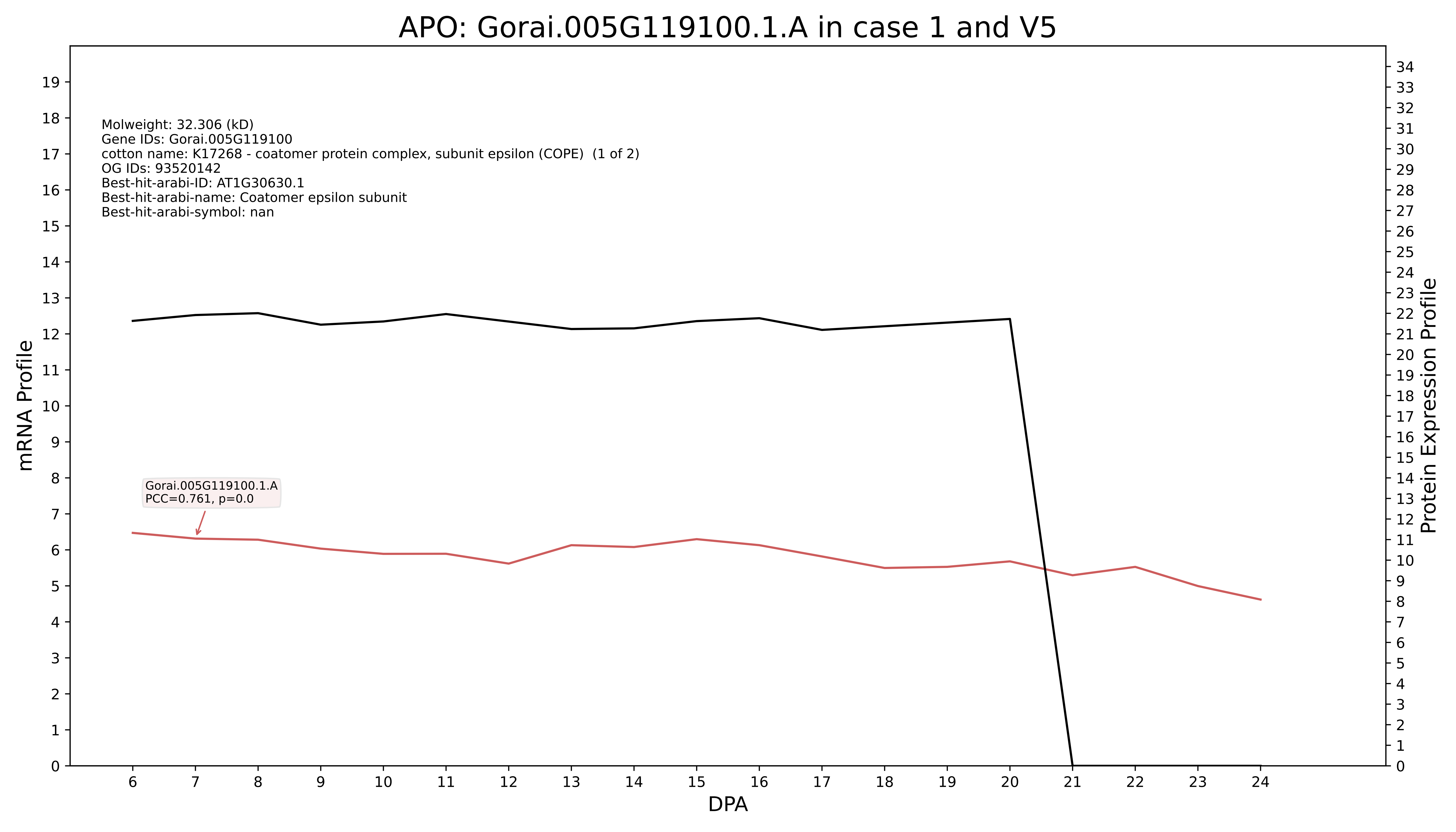

APO: Gorai.005G119100.1.A in case 1 and V5
Molweight: 32.306 (kD)
Gene IDs: Gorai.005G119100
cotton name: K17268 - coatomer protein complex, subunit epsilon (COPE) (1 of 2)
OG IDs: 93520142
Best-hit-arabi-ID: AT1G30630.1
Best-hit-arabi-name: Coatomer epsilon subunit
Best-hit-arabi-symbol: nan
Gorai.005G119100.1.A
PCC=0.761, p=0.0
mRNA Profile
Protein Expression Profile
DPA

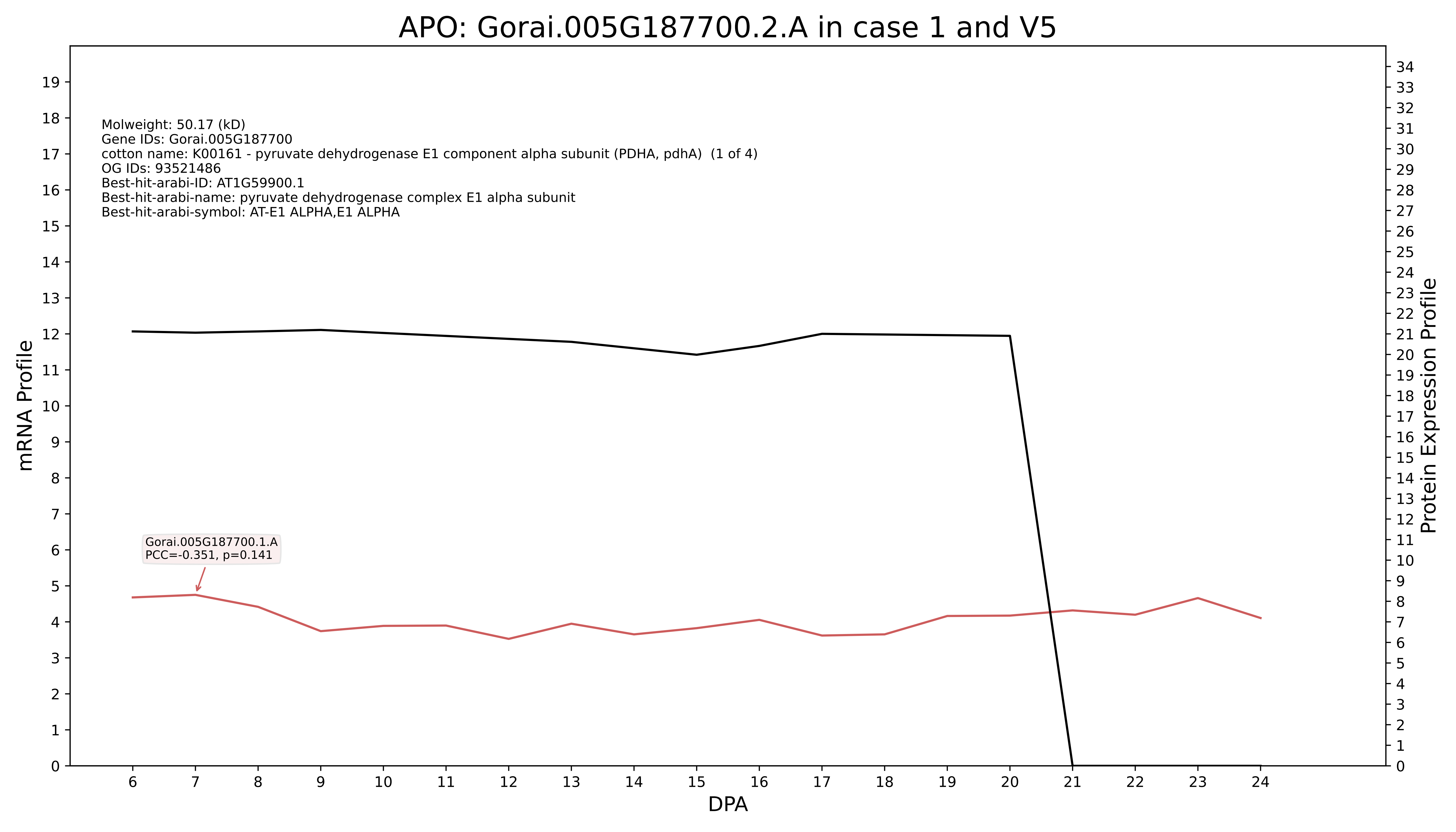
APO: Gorai.005G187700.2.A in case 1 and V5
Molweight: 50.17 (kD)
Gene IDs: Gorai.005G187700
cotton name: K00161 - pyruvate dehydrogenase E1 component alpha subunit (PDHA, pdhA) (1 of 4)
OG IDs: 93521486
Best-hit-arabi-ID: AT1G59900.1
Best-hit-arabi-name: pyruvate dehydrogenase complex E1 alpha subunit
Best-hit-arabi-symbol: AT-E1 ALPHA,E1 ALPHA
Gorai.005G187700.1.A
PCC=-0.351, p=0.141
mRNA Profile
Protein Expression Profile
DPA

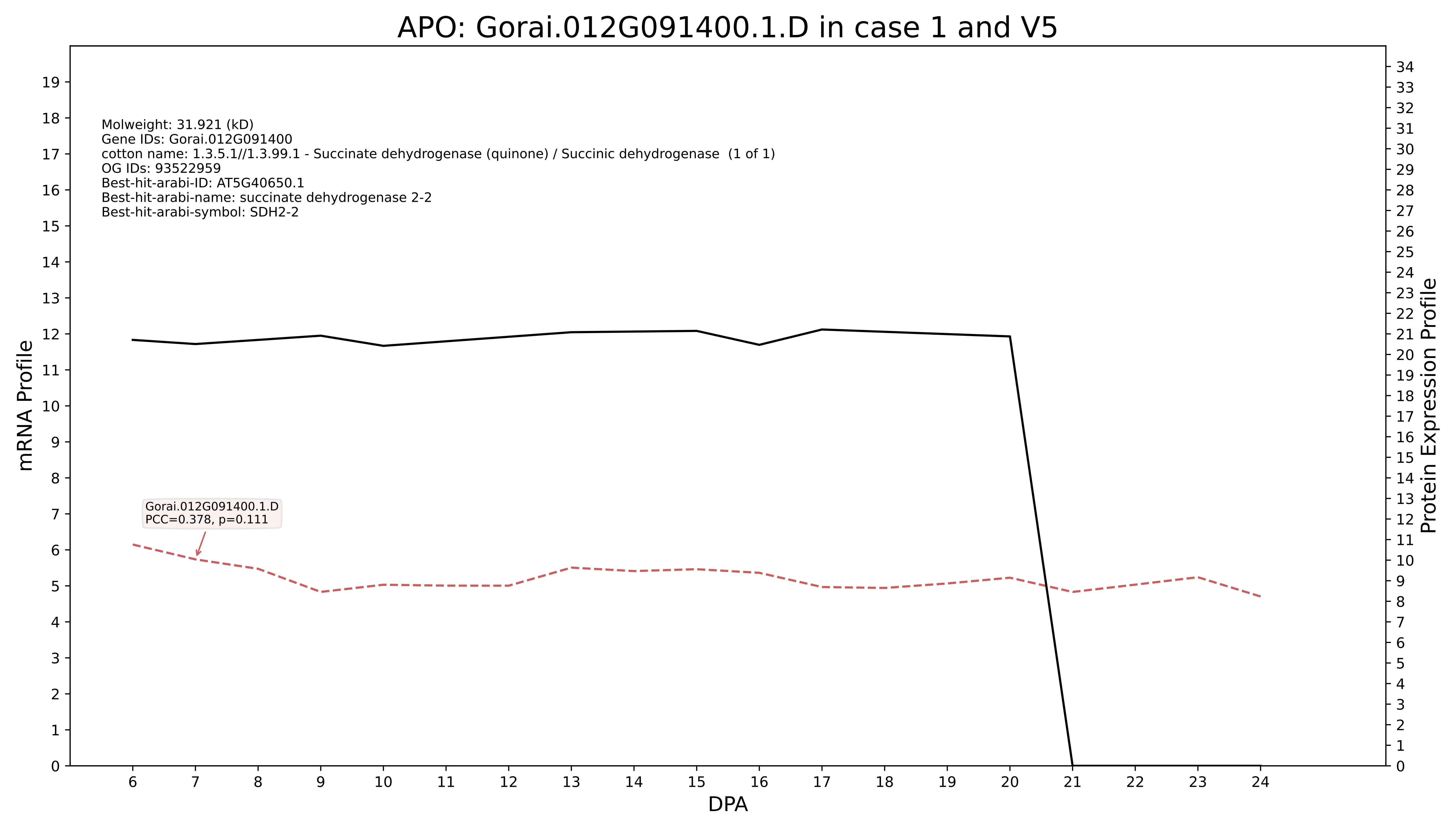
APO: Gorai.012G091400.1.D in case 1 and V5
Molweight: 31.921 (kD)
Gene IDs: Gorai.012G091400
cotton name: 1.3.5.1//1.3.99.1 - Succinate dehydrogenase (quinone) / Succinic dehydrogenase (1 of 1)
OG IDs: 93522959
Best-hit-arabi-ID: AT5G40650.1
Best-hit-arabi-name: succinate dehydrogenase 2-2
Best-hit-arabi-symbol: SDH2-2
Gorai.012G091400.1.D
PCC=0.378, p=0.111
mRNA Profile
Protein Expression Profile
DPA

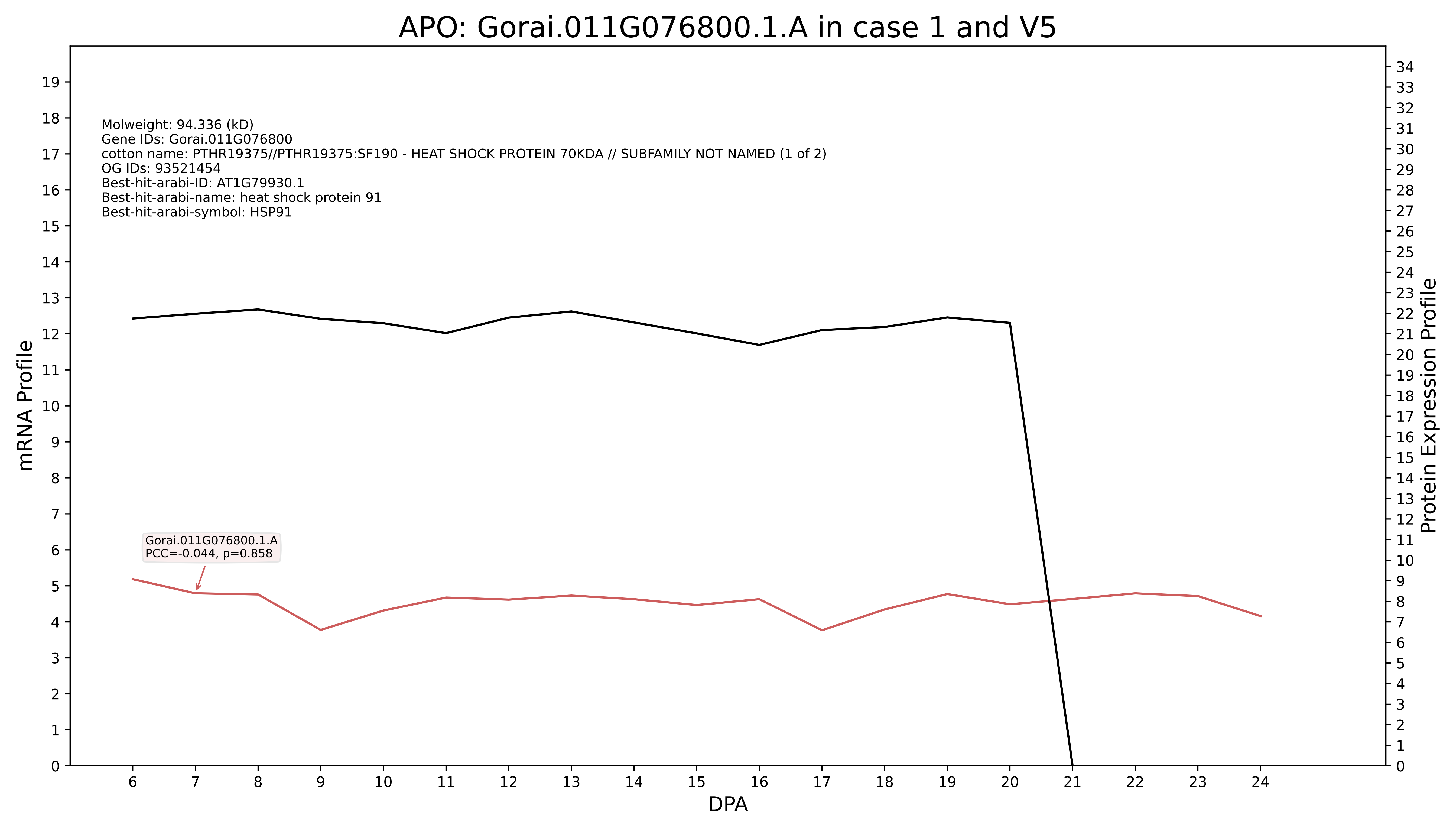
APO: Gorai.011G076800.1.A in case 1 and V5
Molweight: 94.336 (kD)
Gene IDs: Gorai.011G076800
cotton name: PTHR19375//PTHR19375:SF190 - HEAT SHOCK PROTEIN 70KDA // SUBFAMILY NOT NAMED (1 of 2)
OG IDs: 93521454
Best-hit-arabi-ID: AT1G79930.1
Best-hit-arabi-name: heat shock protein 91
Best-hit-arabi-symbol: HSP91
Gorai.011G076800.1.A
PCC=-0.044, p=0.858
mRNA Profile
Protein Expression Profile
DPA

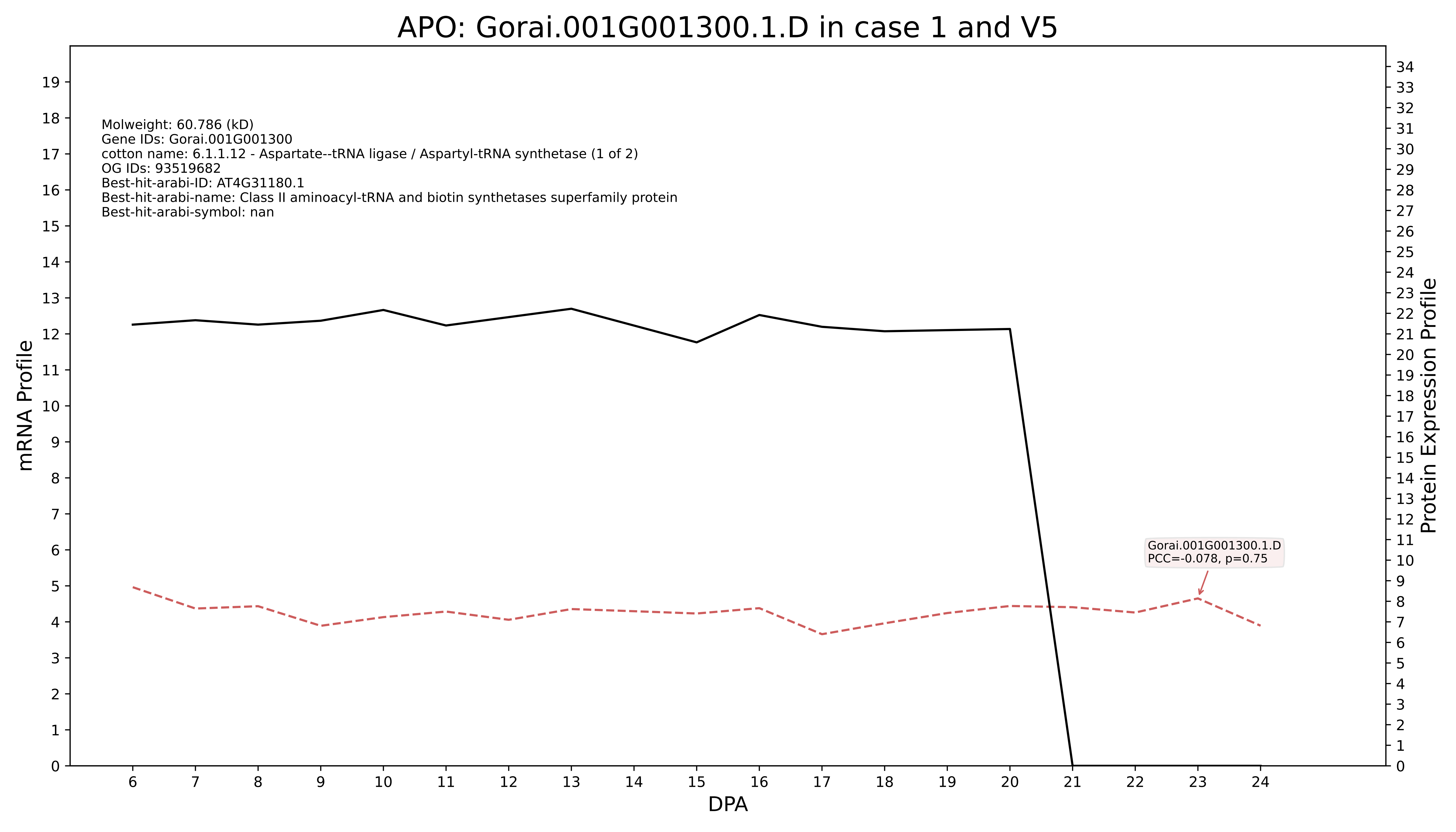
APO: Gorai.001G001300.1.D in case 1 and V5
Molweight: 60.786 (kD)
Gene IDs: Gorai.001G001300
cotton name: 6.1.1.12 - Aspartate--tRNA ligase / Aspartyl-tRNA synthetase (1 of 2)
OG IDs: 93519682
Best-hit-arabi-ID: AT4G31180.1
Best-hit-arabi-name: Class II aminoacyl-tRNA and biotin synthetases superfamily protein
Best-hit-arabi-symbol: nan
Gorai.001G001300.1.D
PCC=-0.078, p=0.75
mRNA Profile
Protein Expression Profile
DPA

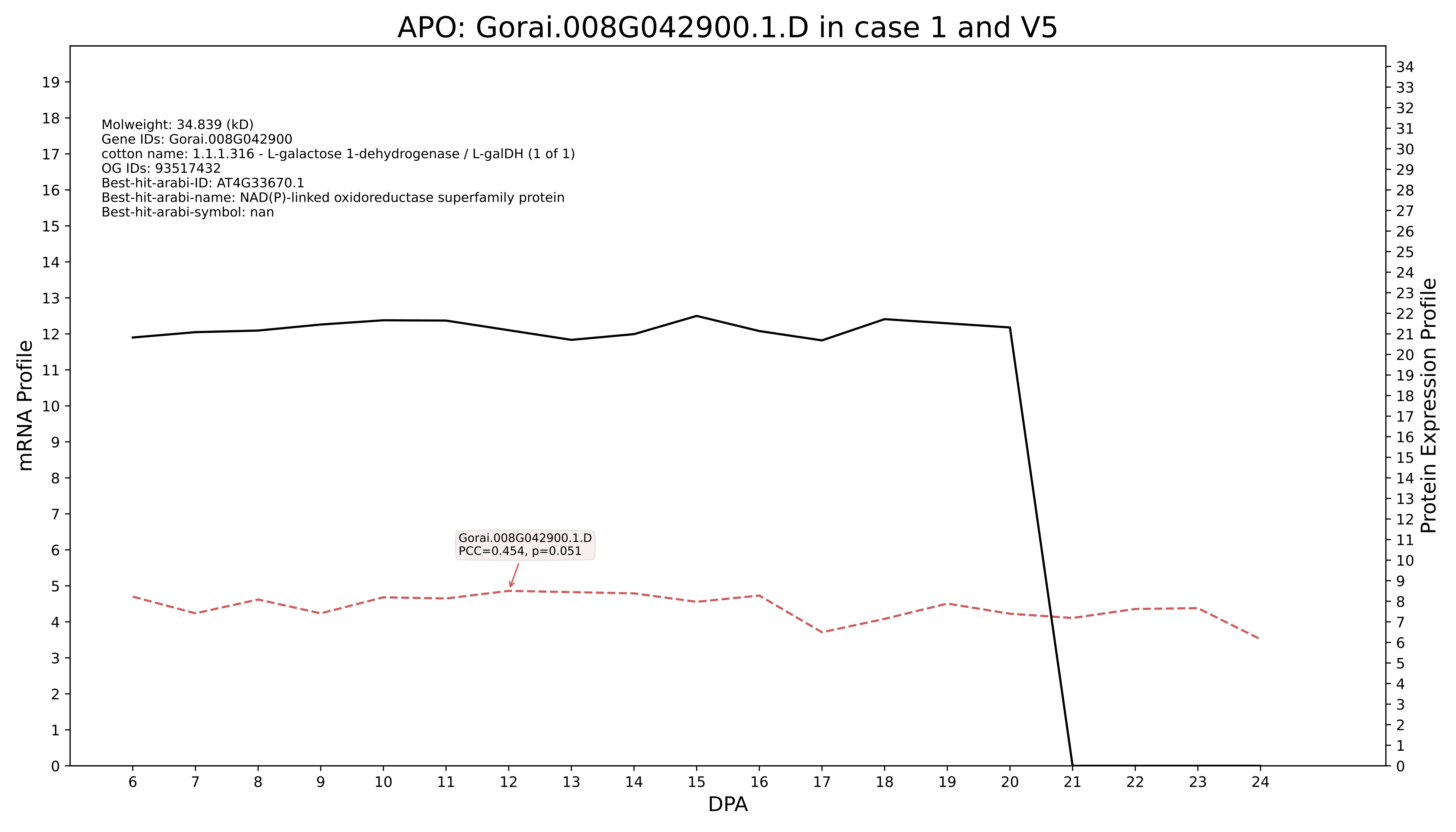

APO: Gorai.008G042900.1.D in case 1 and V5
Molweight: 34.839 (kD)
Gene IDs: Gorai.008G042900
cotton name: 1.1.1.316 - L-galactose 1-dehydrogenase / L-galDH (1 of 1)
OG IDs: 93517432
Best-hit-arabi-ID: AT4G33670.1
Best-hit-arabi-name: NAD(P)-linked oxidoreductase superfamily protein
Best-hit-arabi-symbol: nan
Gorai.008G042900.1.D
PCC=0.454, p=0.051
mRNA Profile
Protein Expression Profile
DPA

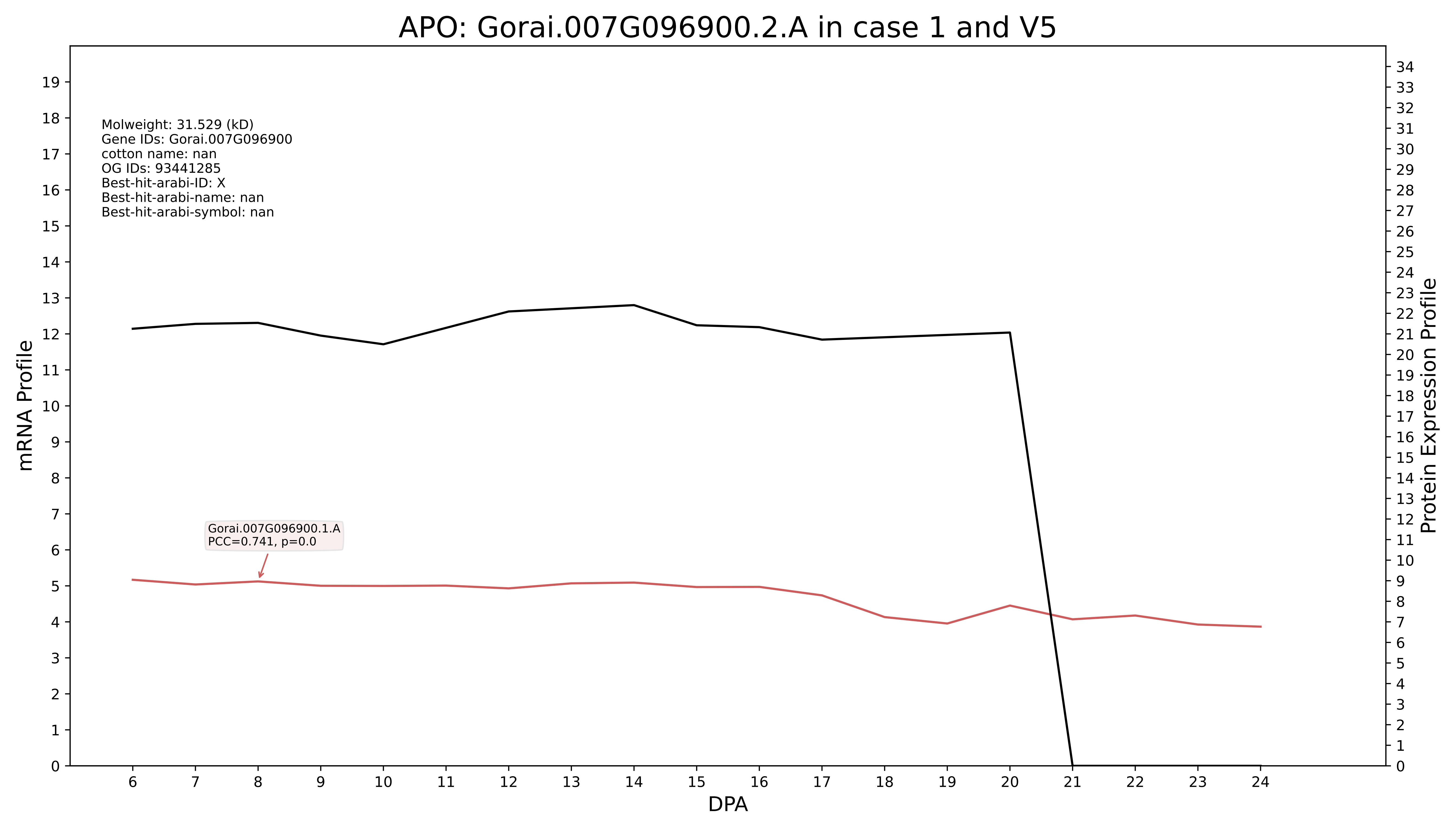

APO: Gorai.007G096900.2.A in case 1 and V5
Molweight: 31.529 (kD)
Gene IDs: Gorai.007G096900
cotton name: nan
OG IDs: 93441285
Best-hit-arabi-ID: X
Best-hit-arabi-name: nan
Best-hit-arabi-symbol: nan
Gorai.007G096900.1.A
PCC=0.741, p=0.0
mRNA Profile
Protein Expression Profile
DPA

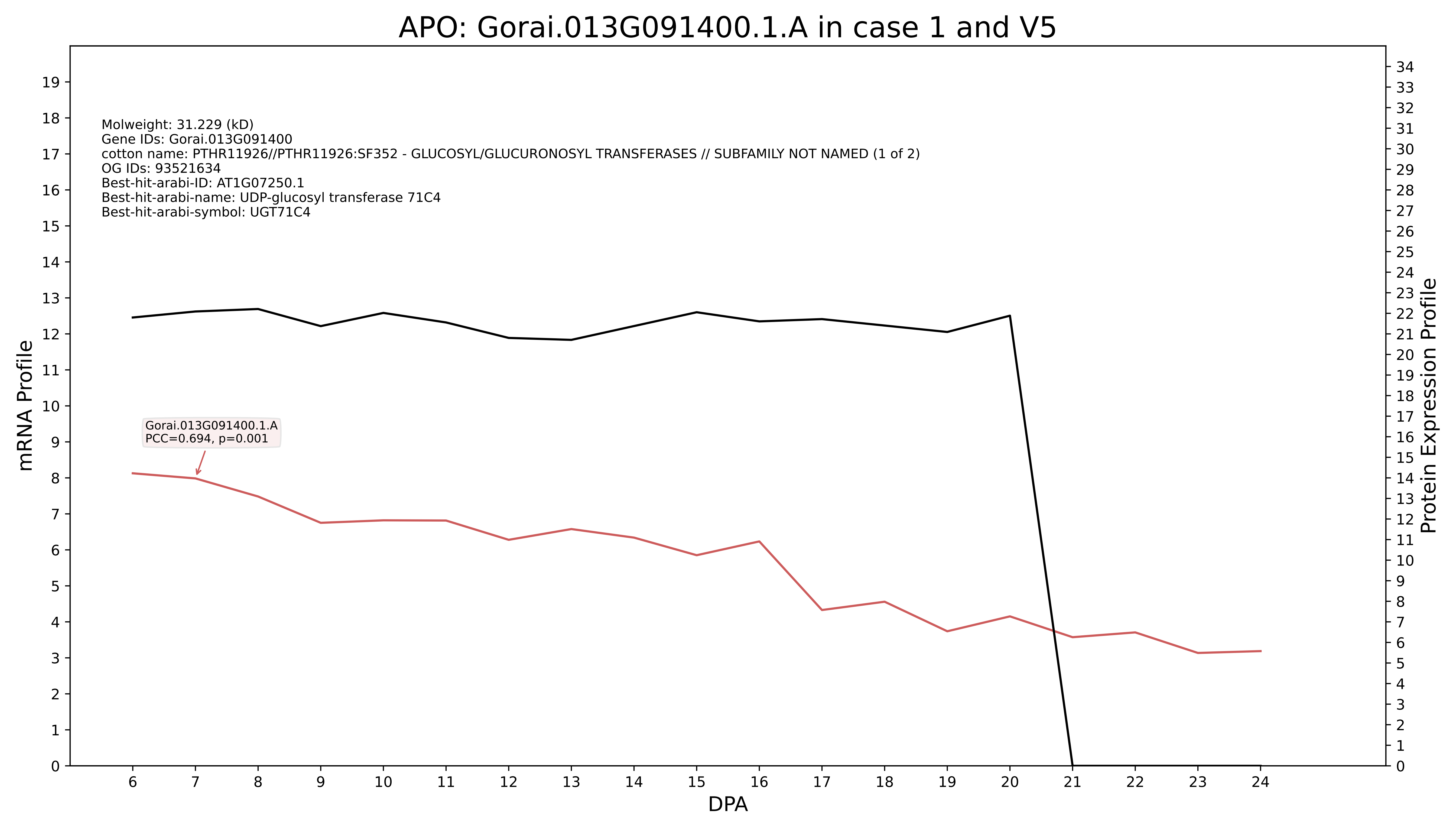
APO: Gorai.013G091400.1.A in case 1 and V5
Molweight: 31.229 (kD)
Gene IDs: Gorai.013G091400
cotton name: PTHR11926//PTHR11926:SF352 - GLUCOSYL/GLUCURONOSYL TRANSFERASES // SUBFAMILY NOT NAMED (1 of 2)
OG IDs: 93521634
Best-hit-arabi-ID: AT1G07250.1
Best-hit-arabi-name: UDP-glucosyl transferase 71C4
Best-hit-arabi-symbol: UGT71C4
Gorai.013G091400.1.A
PCC=0.694, p=0.001
mRNA Profile
Protein Expression Profile
DPA

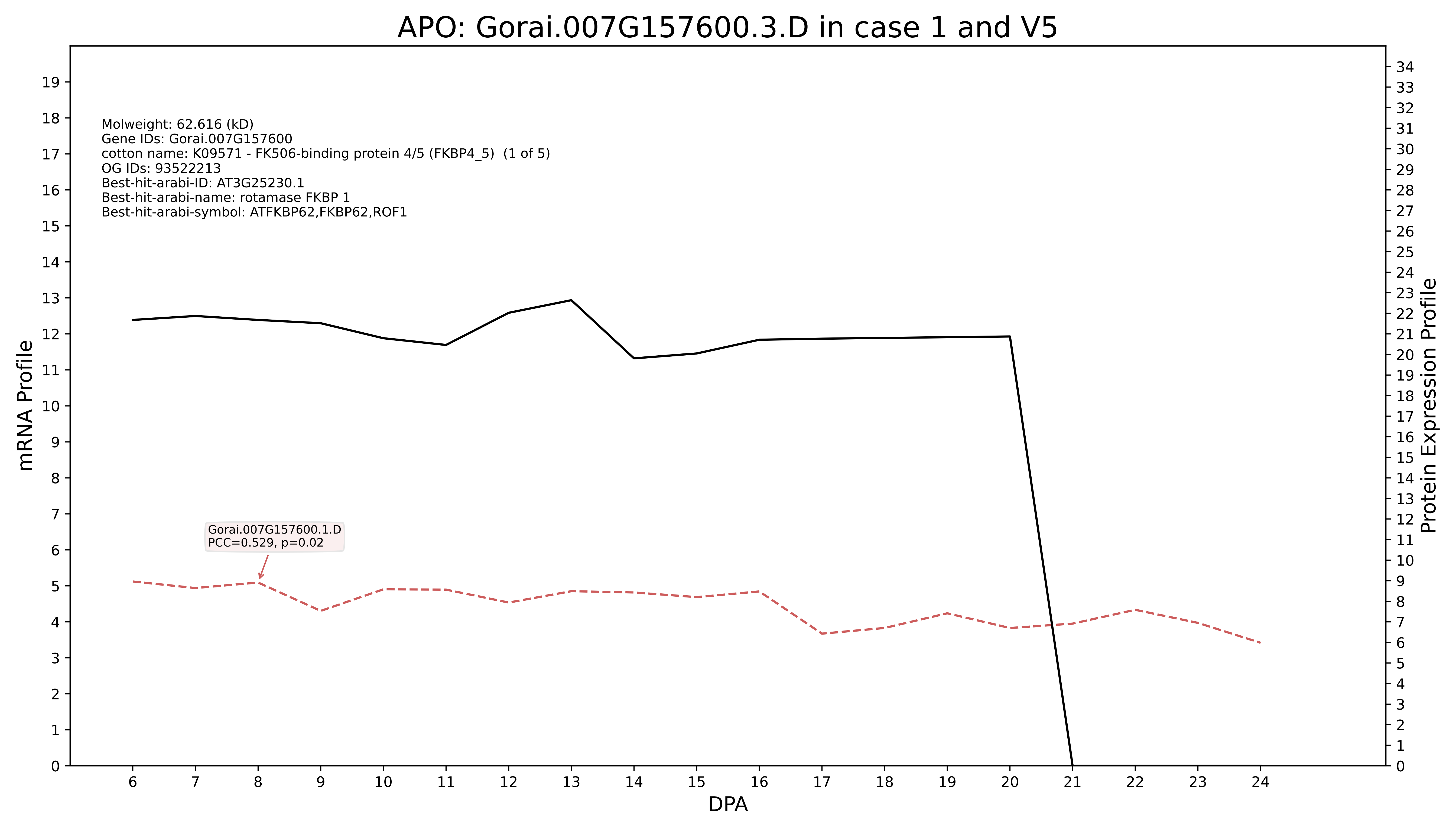

APO: Gorai.007G157600.3.D in case 1 and V5
Molweight: 62.616 (kD)
Gene IDs: Gorai.007G157600
cotton name: K09571 - FK506-binding protein 4/5 (FKBP4_5) (1 of 5)
OG IDs: 93522213
Best-hit-arabi-ID: AT3G25230.1
Best-hit-arabi-name: rotamase FKBP 1
Best-hit-arabi-symbol: ATFKBP62,FKBP62,ROF1
Gorai.007G157600.1.D
PCC=0.529, p=0.02
mRNA Profile
Protein Expression Profile
DPA

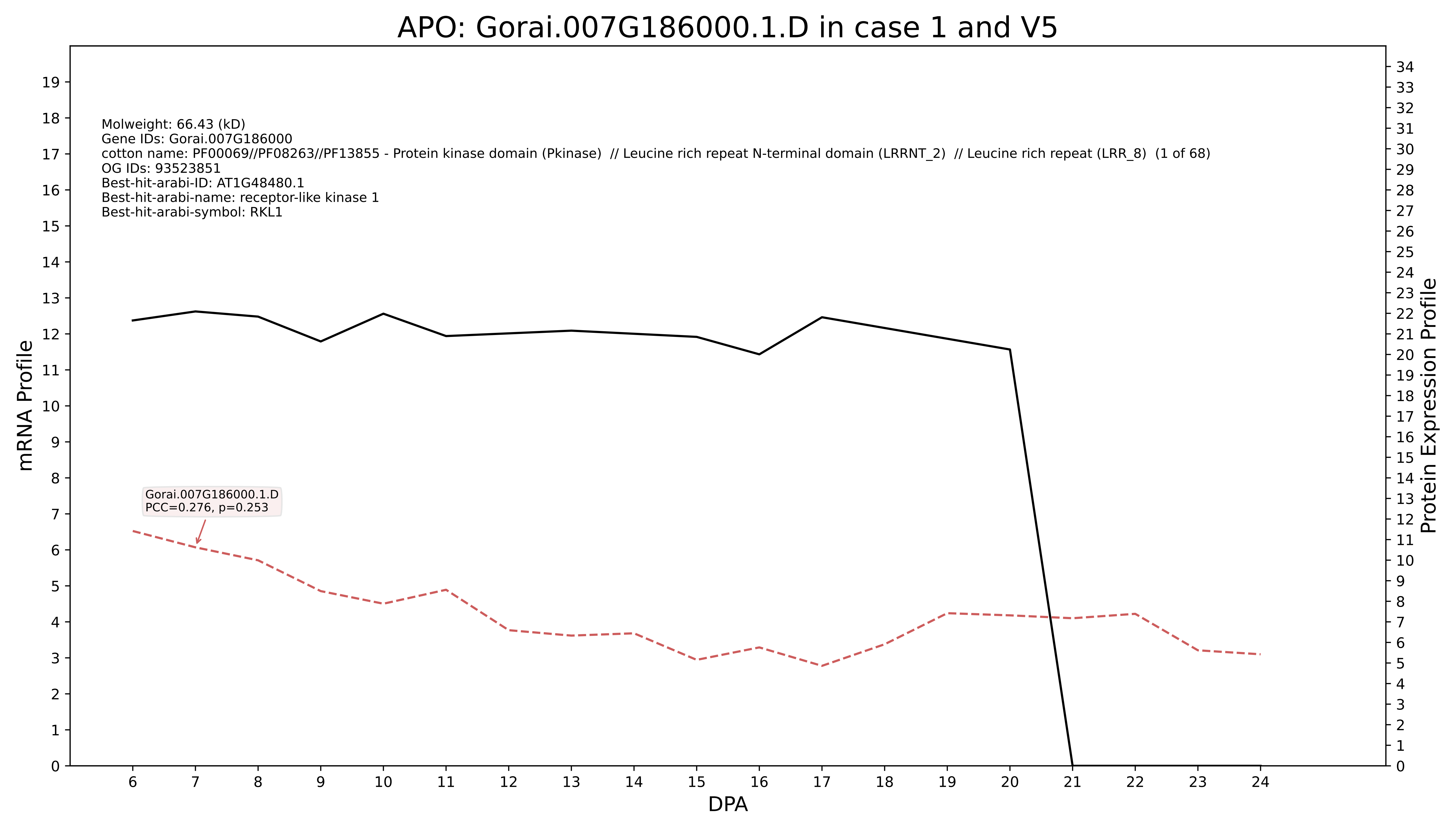

APO: Gorai.007G186000.1.D in case 1 and V5
Molweight: 66.43 (kD)
Gene IDs: Gorai.007G186000
cotton name: PF00069//PF08263//PF13855 - Protein kinase domain (Pkinase) // Leucine rich repeat N-terminal domain (LRRNT_2) // Leucine rich repeat (LRR_8) (1 of 68)
OG IDs: 93523851
Best-hit-arabi-ID: AT1G48480.1
Best-hit-arabi-name: receptor-like kinase 1
Best-hit-arabi-symbol: RKL1
Gorai.007G186000.1.D
PCC=0.276, p=0.253
mRNA Profile
Protein Expression Profile
DPA

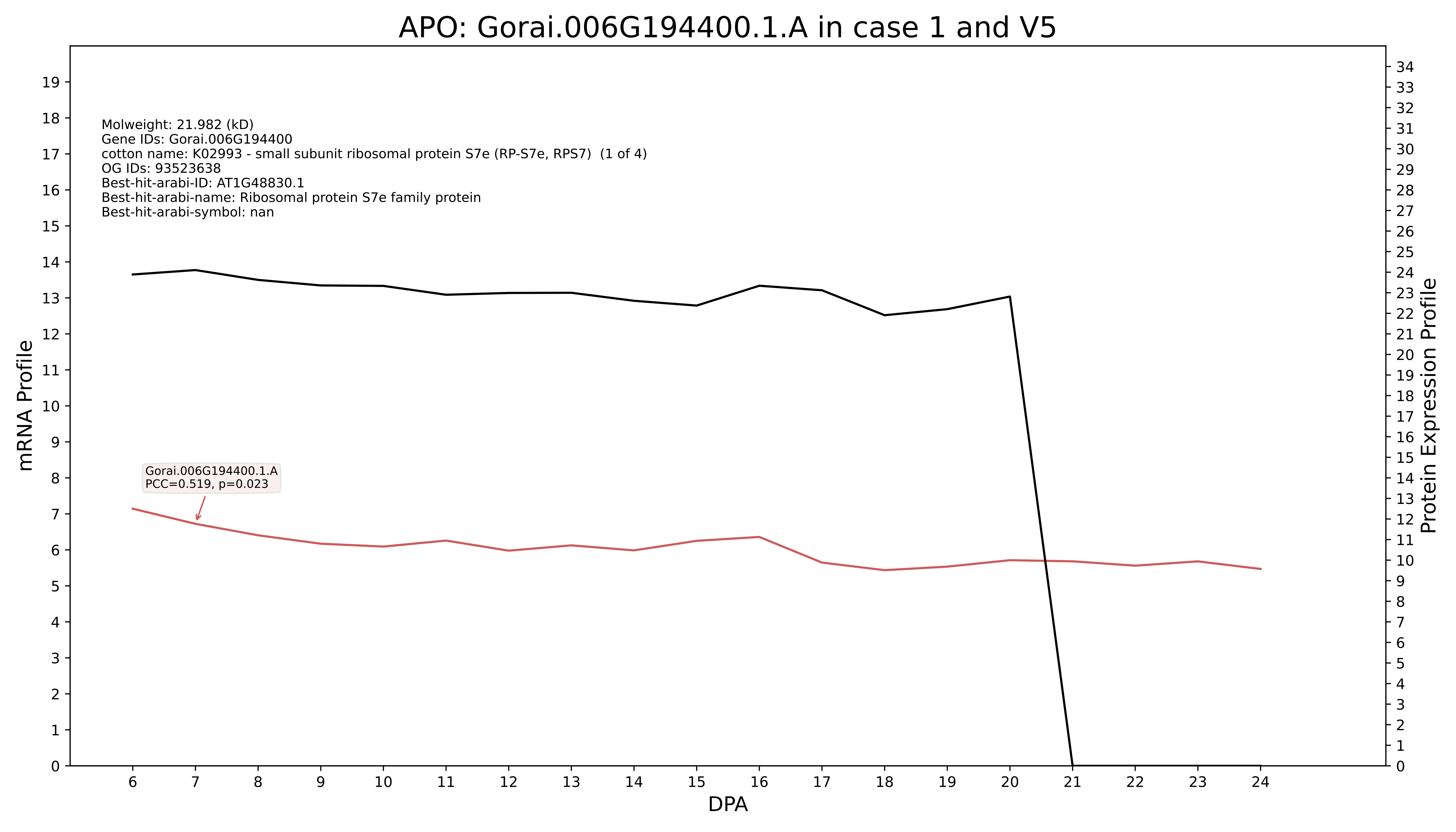
APO: Gorai.006G194400.1.A in case 1 and V5
Molweight: 21.982 (kD)
Gene IDs: Gorai.006G194400
cotton name: K02993 - small subunit ribosomal protein S7e (RP-S7e, RPS7) (1 of 4)
OG IDs: 93523638
Best-hit-arabi-ID: AT1G48830.1
Best-hit-arabi-name: Ribosomal protein S7e family protein
Best-hit-arabi-symbol: nan
Gorai.006G194400.1.A
PCC=0.519, p=0.023
mRNA Profile
Protein Expression Profile
DPA

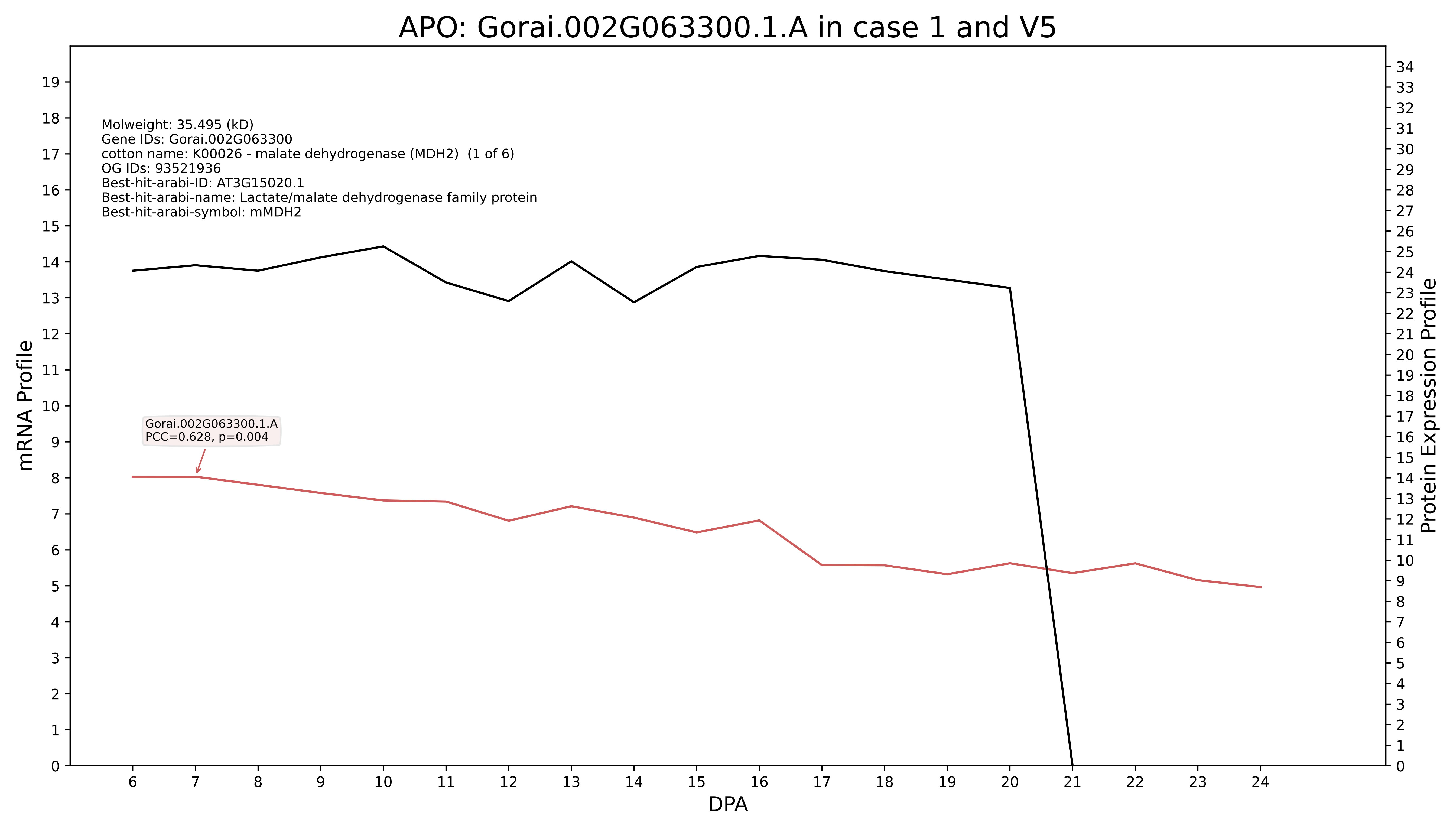

APO: Gorai.002G063300.1.A in case 1 and V5
Molweight: 35.495 (kD)
Gene IDs: Gorai.002G063300
cotton name: K00026 - malate dehydrogenase (MDH2) (1 of 6)
OG IDs: 93521936
Best-hit-arabi-ID: AT3G15020.1
Best-hit-arabi-name: Lactate/malate dehydrogenase family protein
Best-hit-arabi-symbol: mMDH2
Gorai.002G063300.1.A
PCC=0.628, p=0.004
mRNA Profile
Protein Expression Profile
DPA

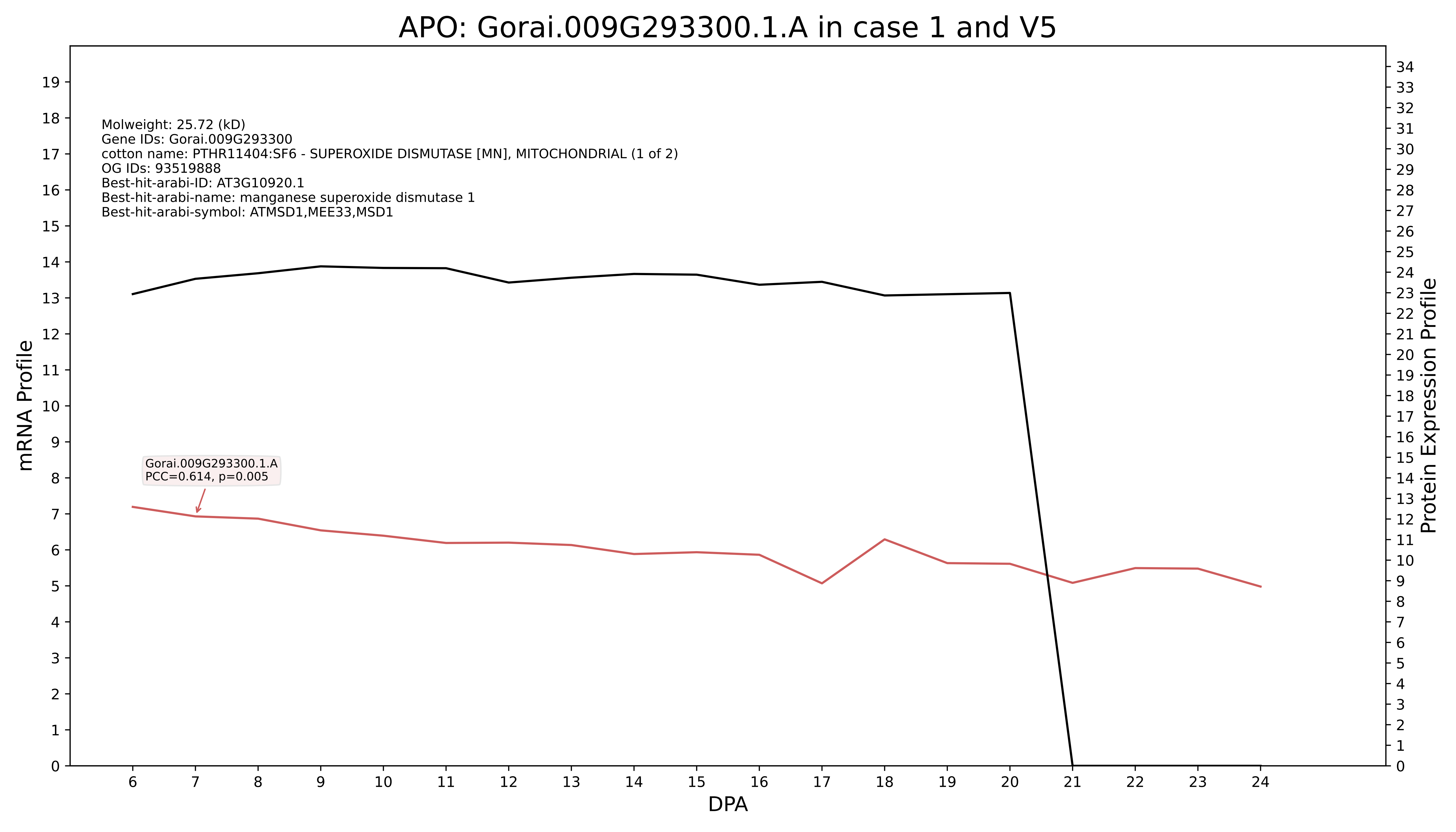

APO: Gorai.009G293300.1.A in case 1 and V5
Molweight: 25.72 (kD)
Gene IDs: Gorai.009G293300
cotton name: PTHR11404:SF6 - SUPEROXIDE DISMUTASE [MN], MITOCHONDRIAL (1 of 2)
OG IDs: 93519888
Best-hit-arabi-ID: AT3G10920.1
Best-hit-arabi-name: manganese superoxide dismutase 1
Best-hit-arabi-symbol: ATMSD1,MEE33,MSD1
Gorai.009G293300.1.A
PCC=0.614, p=0.005
mRNA Profile
Protein Expression Profile
DPA

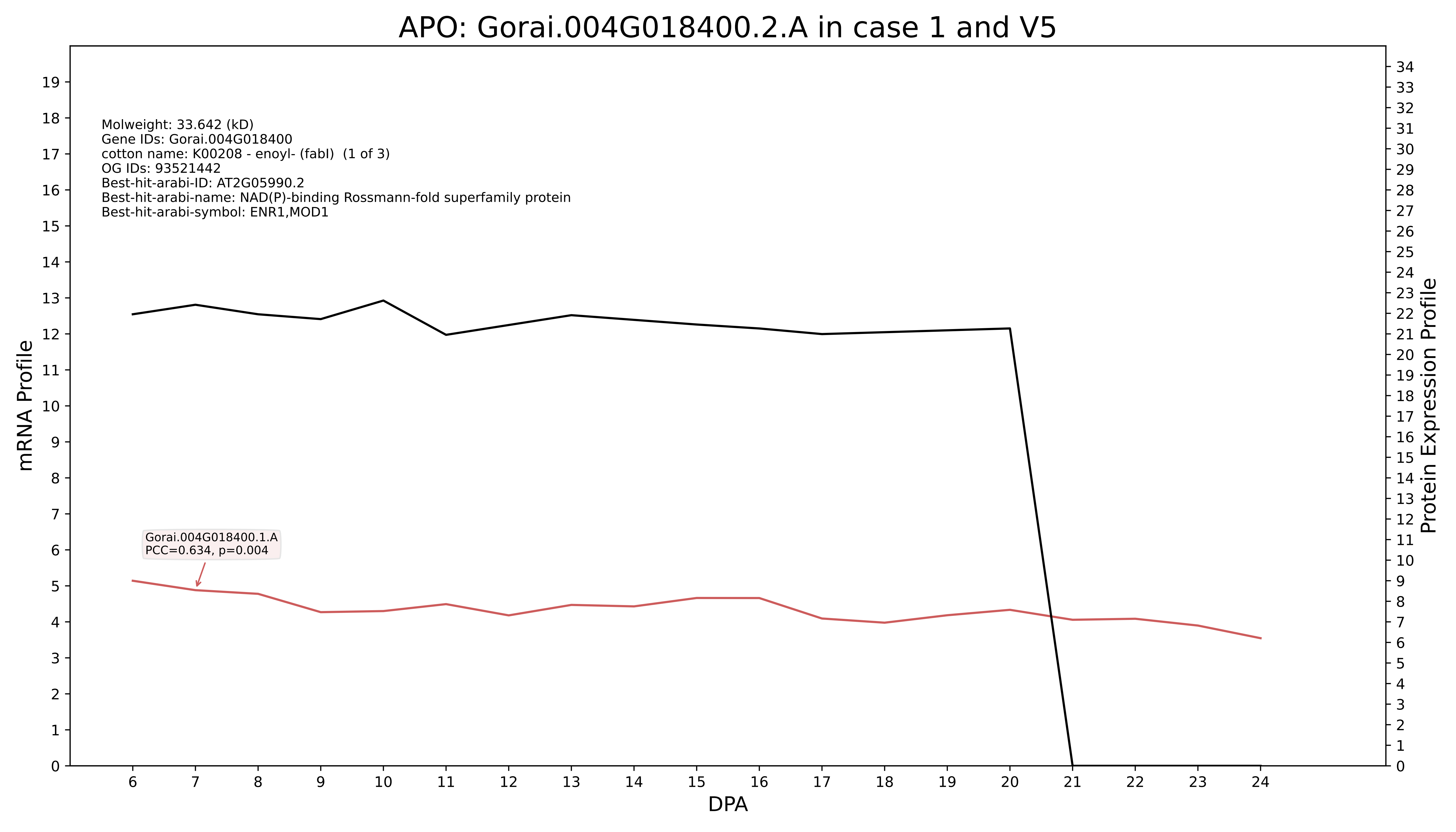
APO: Gorai.004G018400.2.A in case 1 and V5
Molweight: 33.642 (kD)
Gene IDs: Gorai.004G018400
cotton name: K00208 - enoyl- (fabI) (1 of 3)
OG IDs: 93521442
Best-hit-arabi-ID: AT2G05990.2
Best-hit-arabi-name: NAD(P)-binding Rossmann-fold superfamily protein
Best-hit-arabi-symbol: ENR1,MOD1
Gorai.004G018400.1.A
PCC=0.634, p=0.004
mRNA Profile
Protein Expression Profile
DPA

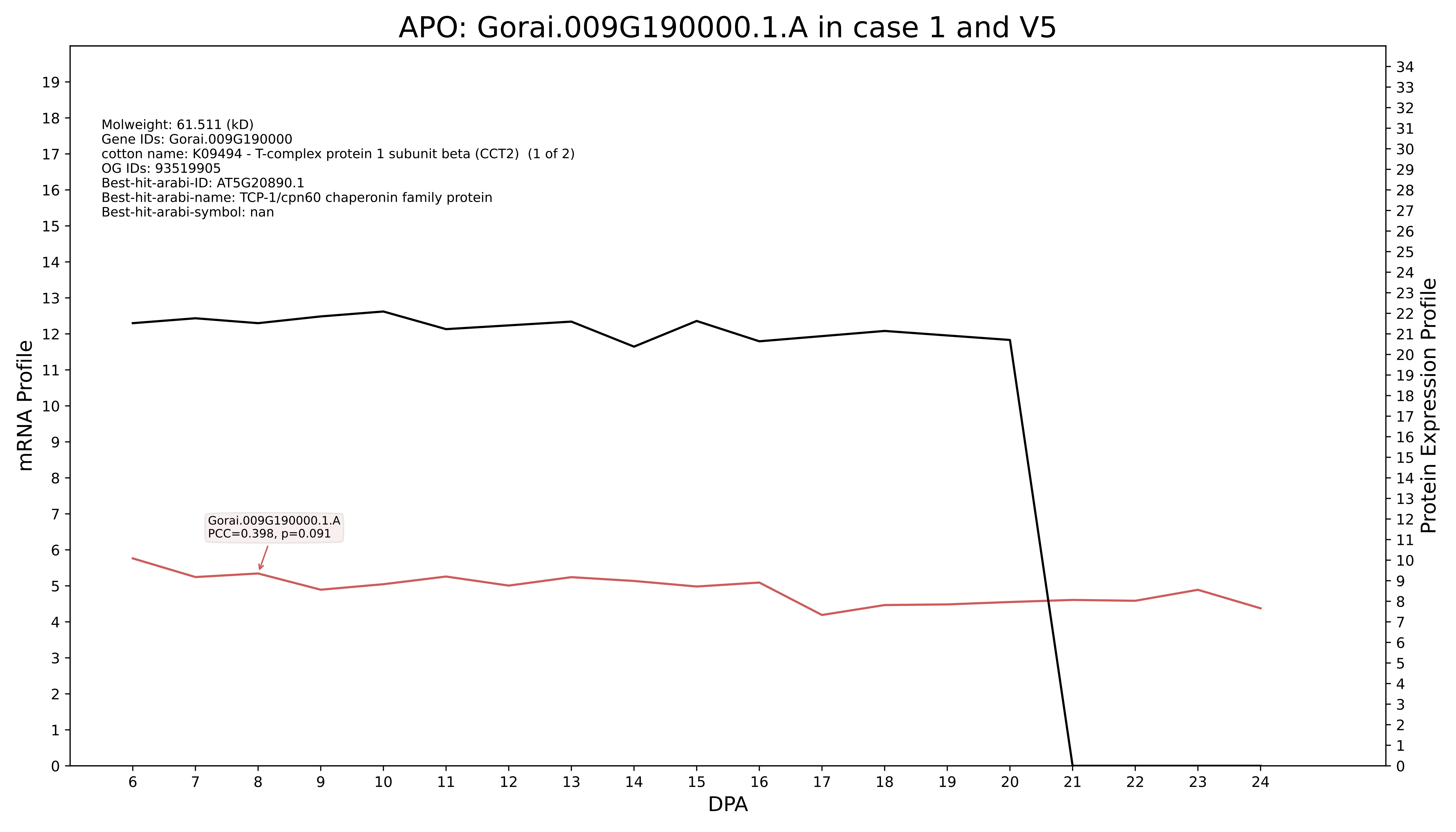
APO: Gorai.009G190000.1.A in case 1 and V5
Molweight: 61.511 (kD)
Gene IDs: Gorai.009G190000
cotton name: K09494 - T-complex protein 1 subunit beta (CCT2) (1 of 2)
OG IDs: 93519905
Best-hit-arabi-ID: AT5G20890.1
Best-hit-arabi-name: TCP-1/cpn60 chaperonin family protein
Best-hit-arabi-symbol: nan
Gorai.009G190000.1.A
PCC=0.398, p=0.091
mRNA Profile
Protein Expression Profile
DPA

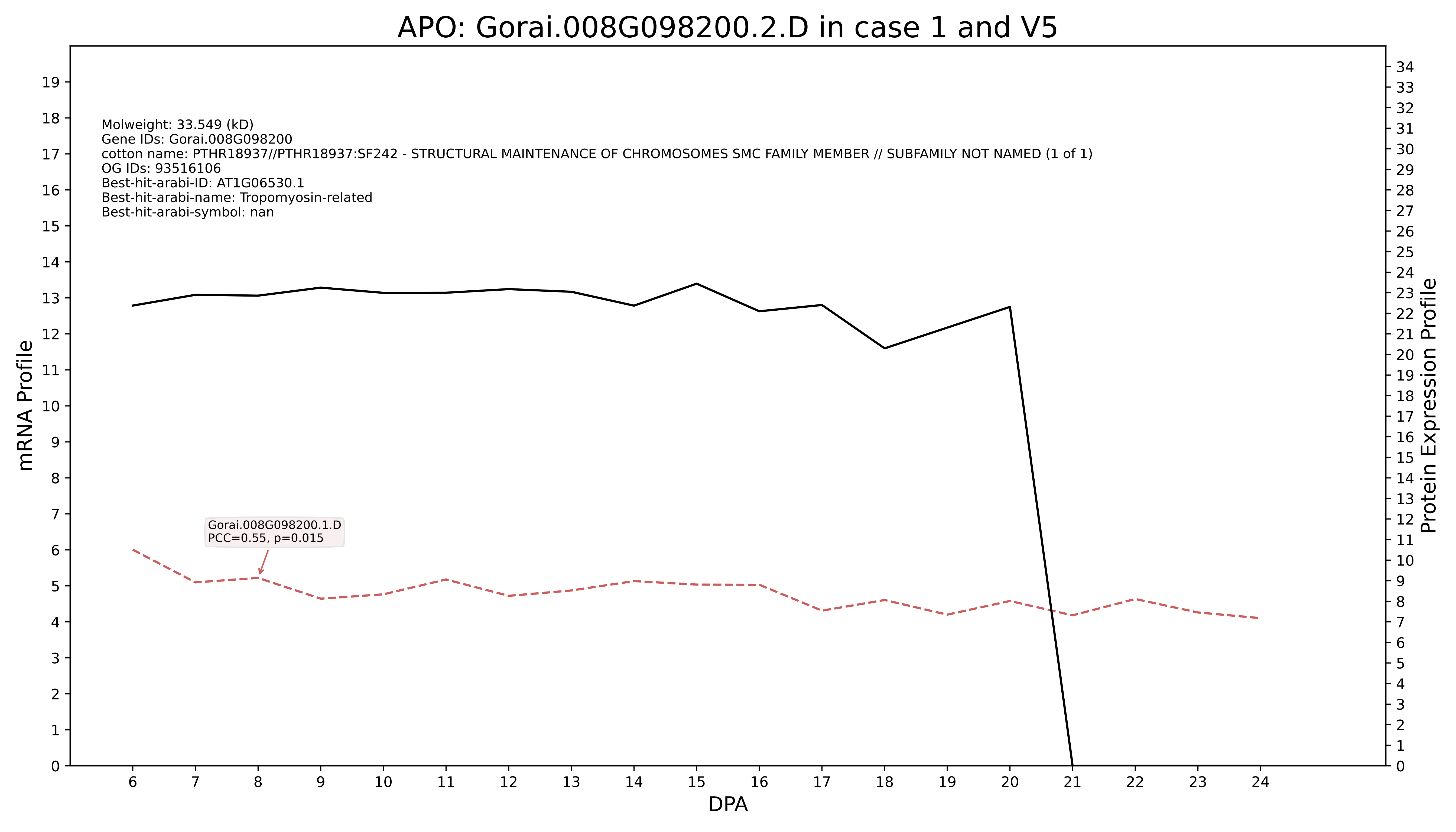
APO: Gorai.008G098200.2.D in case 1 and V5
Molweight: 33.549 (kD)
Gene IDs: Gorai.008G098200
cotton name: PTHR18937//PTHR18937:SF242 - STRUCTURAL MAINTENANCE OF CHROMOSOMES SMC FAMILY MEMBER // SUBFAMILY NOT NAMED (1 of 1)
OG IDs: 93516106
Best-hit-arabi-ID: AT1G06530.1
Best-hit-arabi-name: Tropomyosin-related
Best-hit-arabi-symbol: nan
Gorai.008G098200.1.D
PCC=0.55, p=0.015
mRNA Profile
Protein Expression Profile
DPA

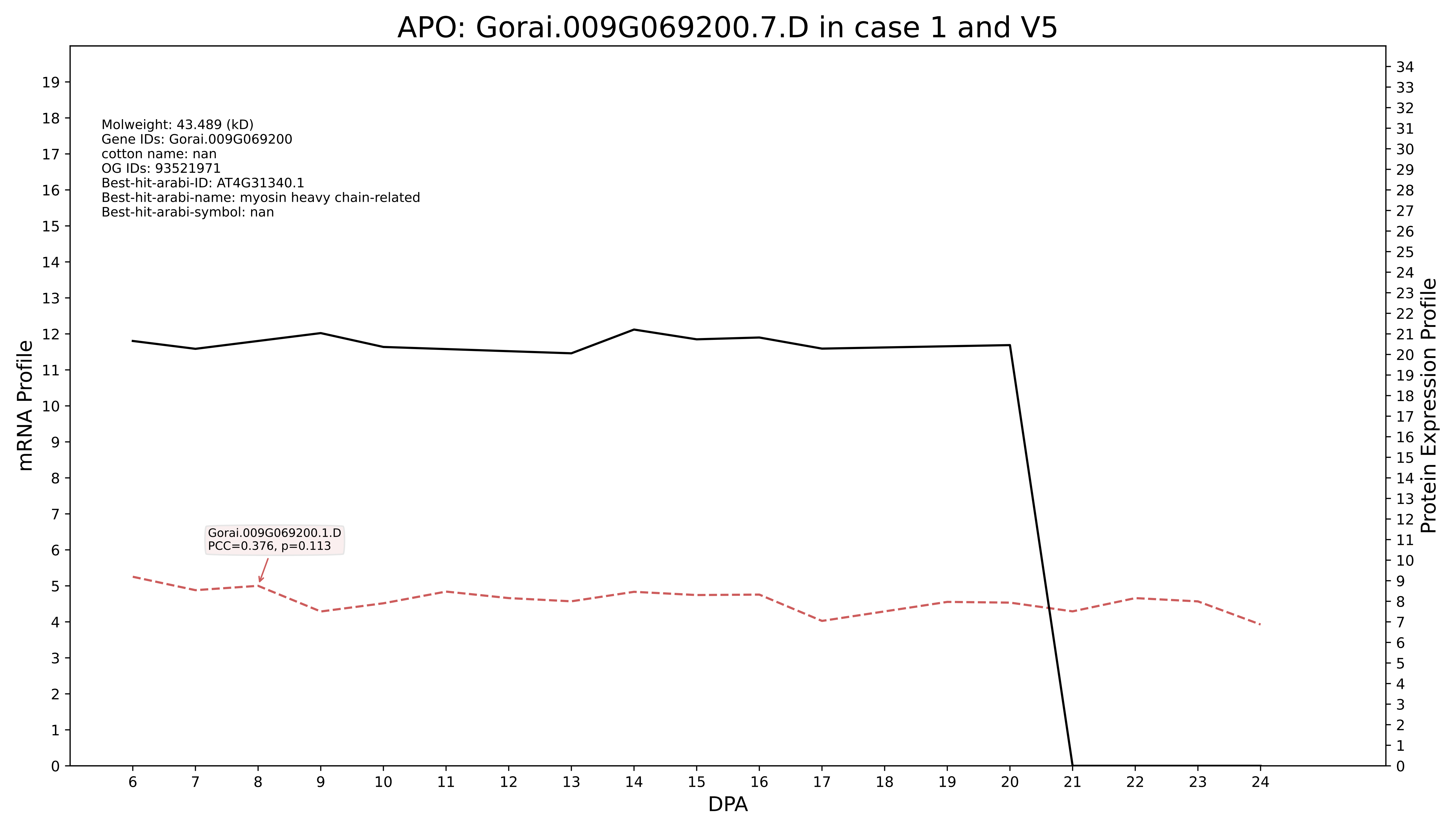

APO: Gorai.009G069200.7.D in case 1 and V5
Molweight: 43.489 (kD)
Gene IDs: Gorai.009G069200
cotton name: nan
OG IDs: 93521971
Best-hit-arabi-ID: AT4G31340.1
Best-hit-arabi-name: myosin heavy chain-related
Best-hit-arabi-symbol: nan
Gorai.009G069200.1.D
PCC=0.376, p=0.113
mRNA Profile
Protein Expression Profile
DPA

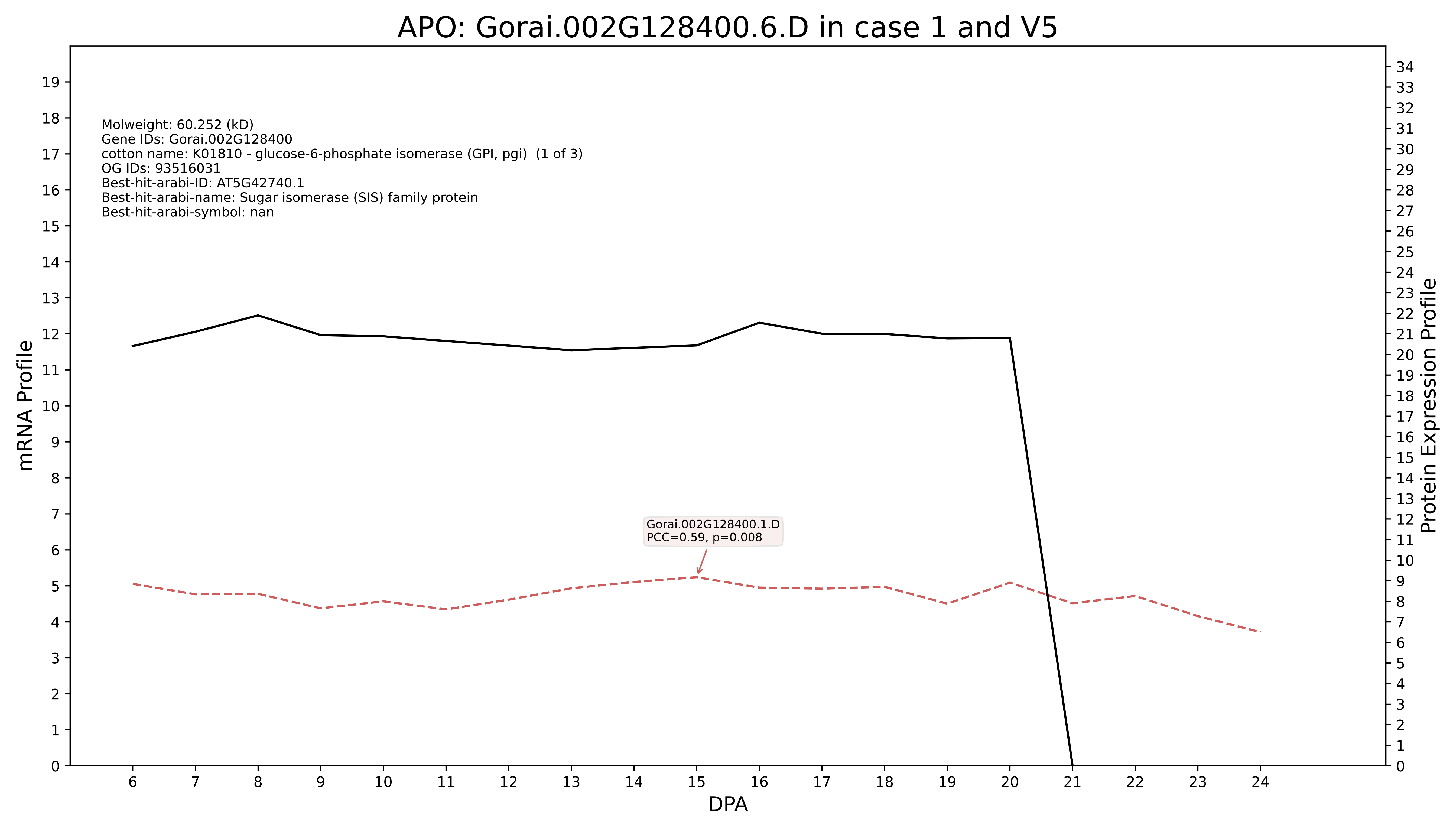
APO: Gorai.002G128400.6.D in case 1 and V5
Molweight: 60.252 (kD)
Gene IDs: Gorai.002G128400
cotton name: K01810 - glucose-6-phosphate isomerase (GPI, pgi) (1 of 3)
OG IDs: 93516031
Best-hit-arabi-ID: AT5G42740.1
Best-hit-arabi-name: Sugar isomerase (SIS) family protein
Best-hit-arabi-symbol: nan
Gorai.002G128400.1.D
PCC=0.59, p=0.008
mRNA Profile
Protein Expression Profile
DPA

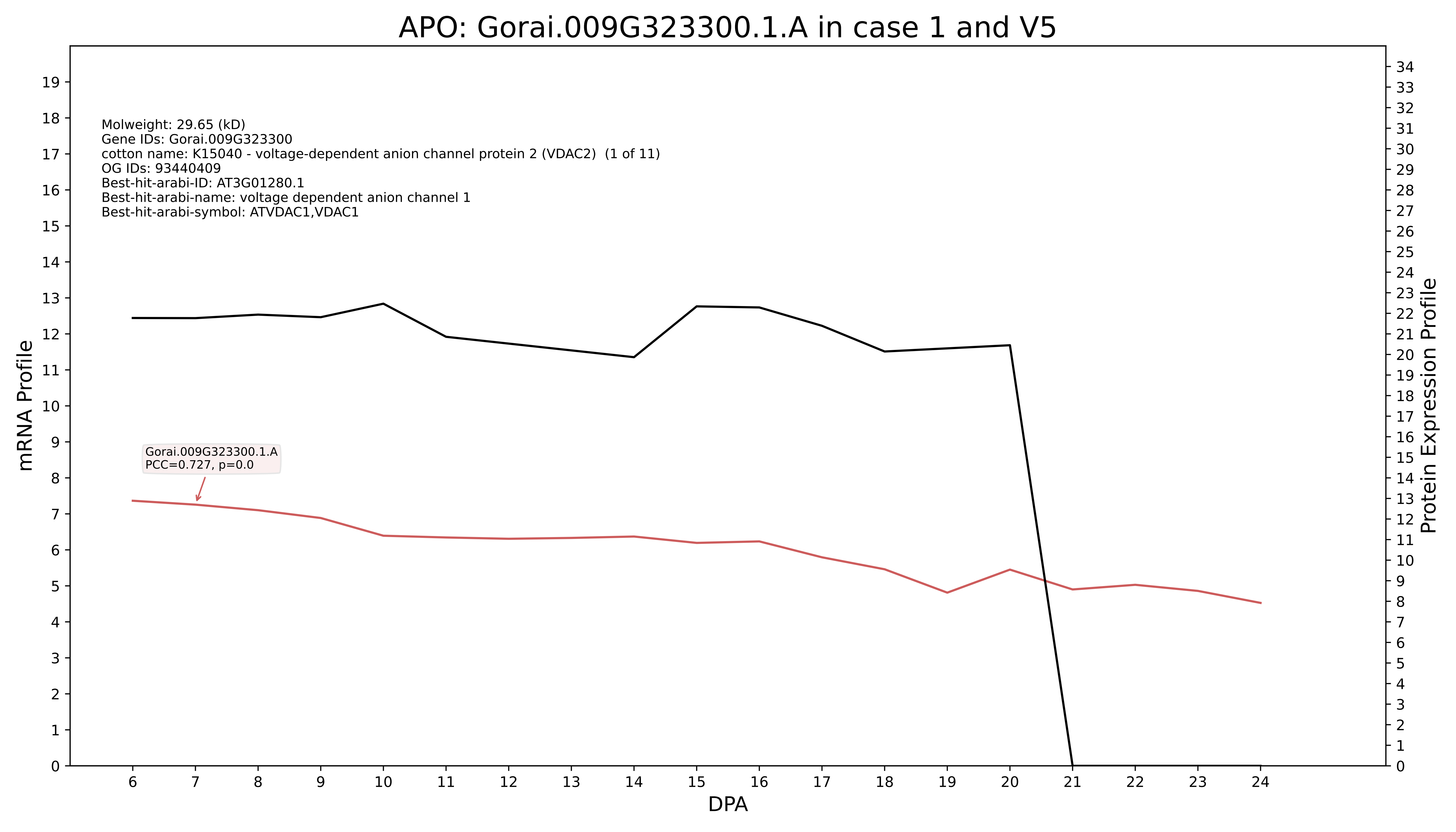
APO: Gorai.009G323300.1.A in case 1 and V5
Molweight: 29.65 (kD)
Gene IDs: Gorai.009G323300
cotton name: K15040 - voltage-dependent anion channel protein 2 (VDAC2) (1 of 11)
OG IDs: 93440409
Best-hit-arabi-ID: AT3G01280.1
Best-hit-arabi-name: voltage dependent anion channel 1
Best-hit-arabi-symbol: ATVDAC1,VDAC1
Gorai.009G323300.1.A
PCC=0.727, p=0.0
mRNA Profile
Protein Expression Profile
DPA

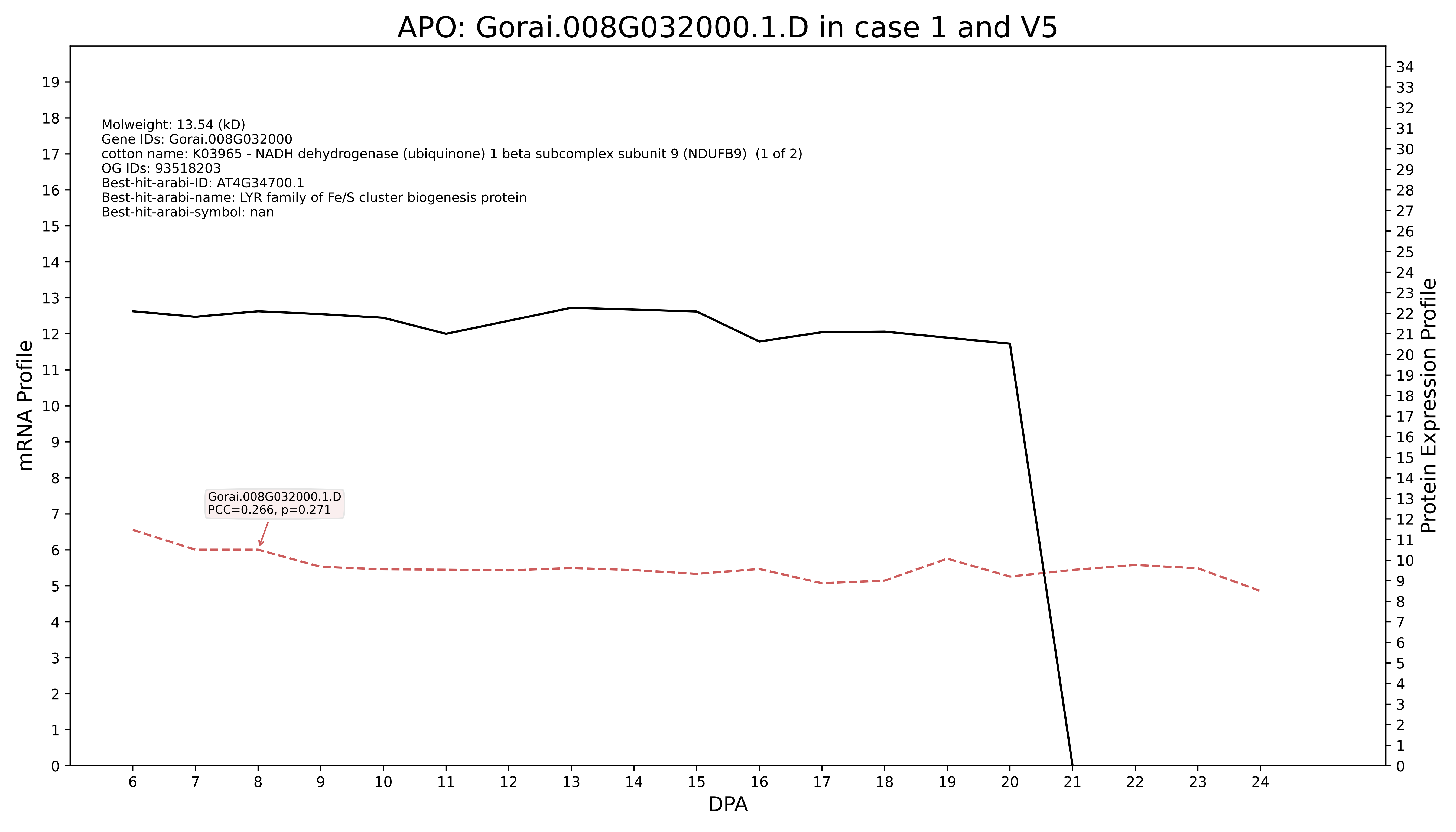

APO: Gorai.008G032000.1.D in case 1 and V5
Molweight: 13.54 (kD)
Gene IDs: Gorai.008G032000
cotton name: K03965 - NADH dehydrogenase (ubiquinone) 1 beta subcomplex subunit 9 (NDUFB9) (1 of 2)
OG IDs: 93518203
Best-hit-arabi-ID: AT4G34700.1
Best-hit-arabi-name: LYR family of Fe/S cluster biogenesis protein
Best-hit-arabi-symbol: nan
Gorai.008G032000.1.D
PCC=0.266, p=0.271
mRNA Profile
Protein Expression Profile
DPA

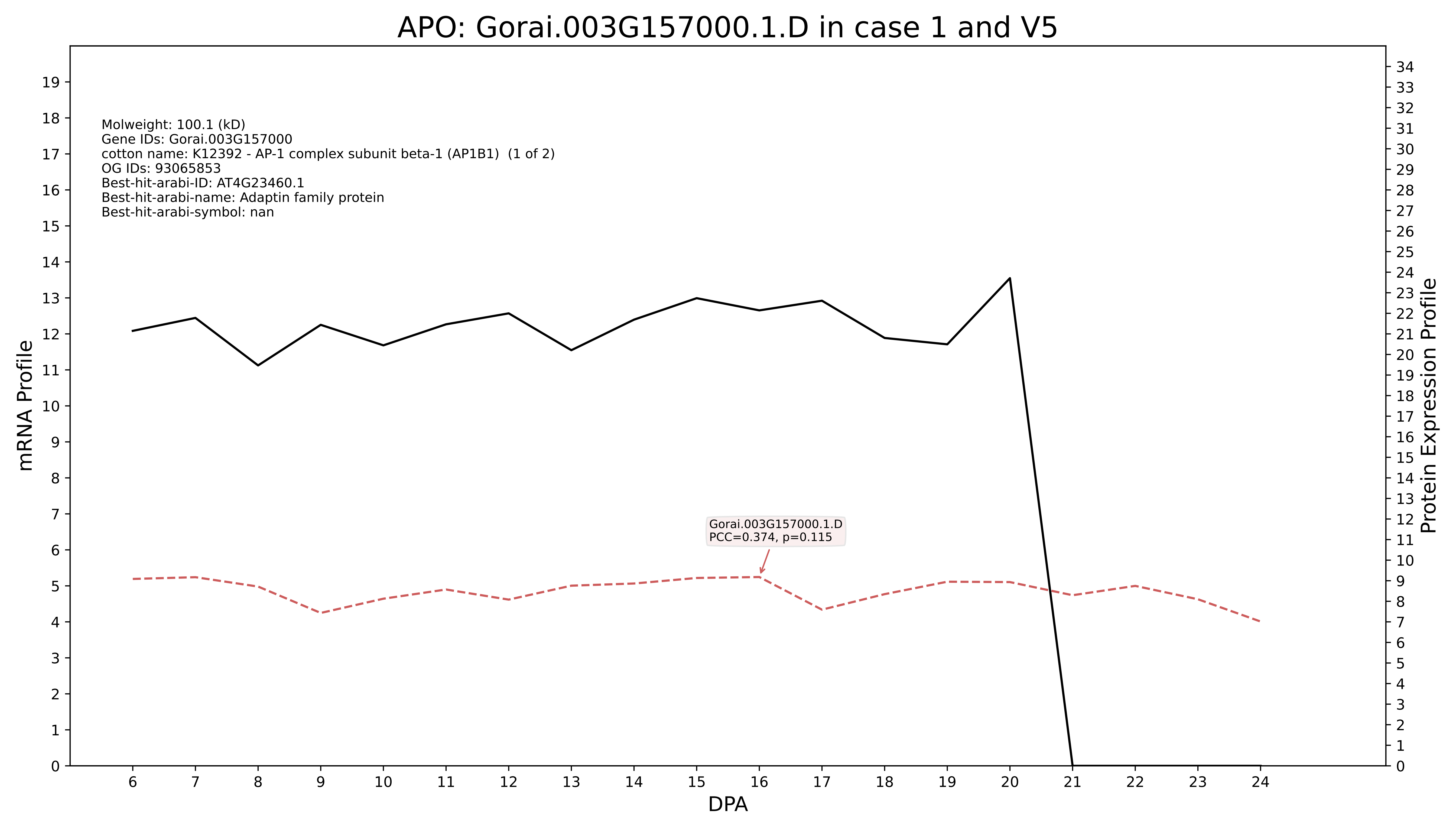
APO: Gorai.003G157000.1.D in case 1 and V5
Molweight: 100.1 (kD)
Gene IDs: Gorai.003G157000
cotton name: K12392 - AP-1 complex subunit beta-1 (AP1B1) (1 of 2)
OG IDs: 93065853
Best-hit-arabi-ID: AT4G23460.1
Best-hit-arabi-name: Adaptin family protein
Best-hit-arabi-symbol: nan
Gorai.003G157000.1.D
PCC=0.374, p=0.115
mRNA Profile
Protein Expression Profile
DPA

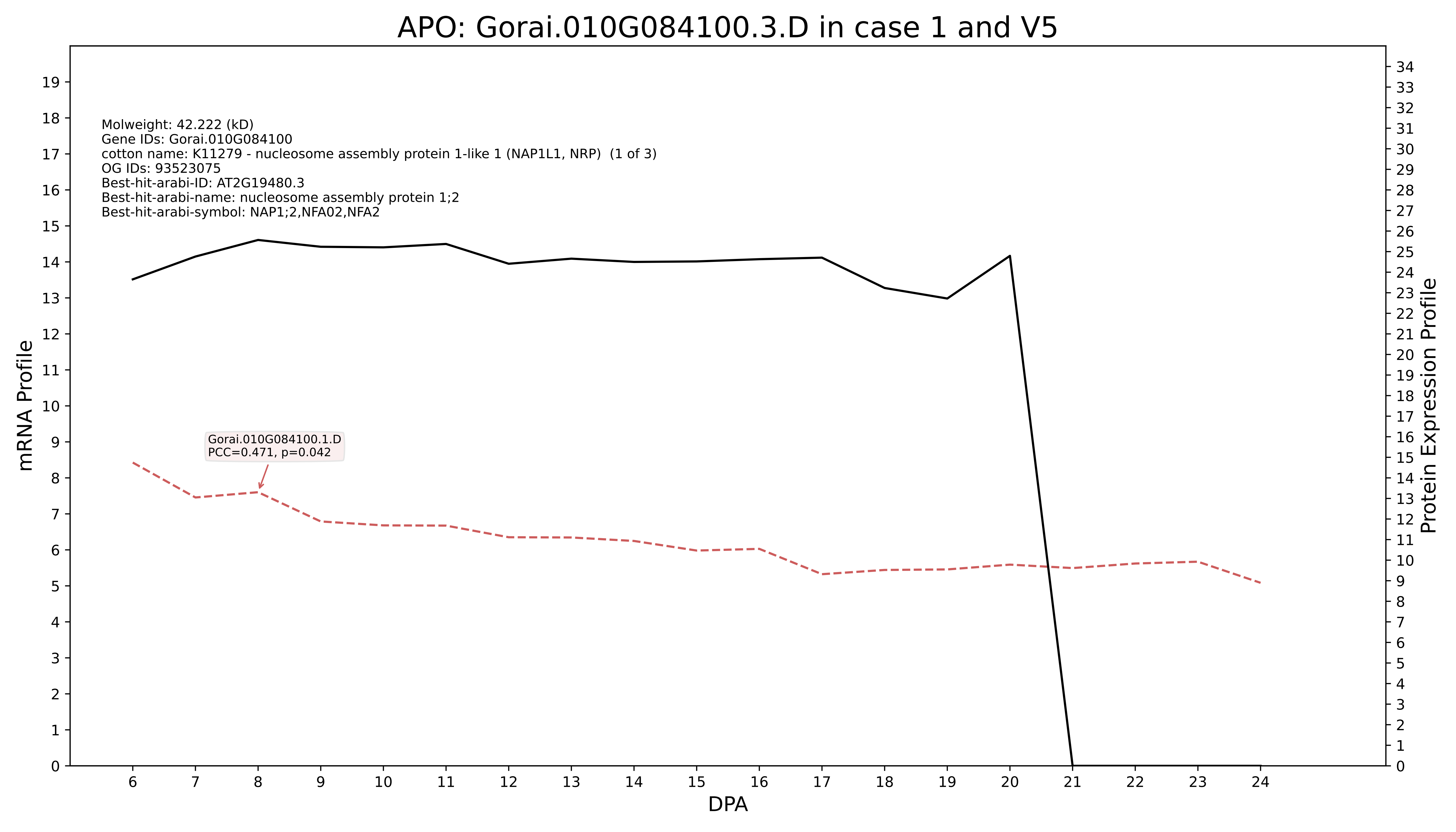
APO: Gorai.010G084100.3.D in case 1 and V5
Molweight: 42.222 (kD)
Gene IDs: Gorai.010G084100
cotton name: K11279 - nucleosome assembly protein 1-like 1 (NAP1L1, NRP) (1 of 3)
OG IDs: 93523075
Best-hit-arabi-ID: AT2G19480.3
Best-hit-arabi-name: nucleosome assembly protein 1;2
Best-hit-arabi-symbol: NAP1;2,NFA02,NFA2
Gorai.010G084100.1.D
PCC=0.471, p=0.042
mRNA Profile
Protein Expression Profile
DPA

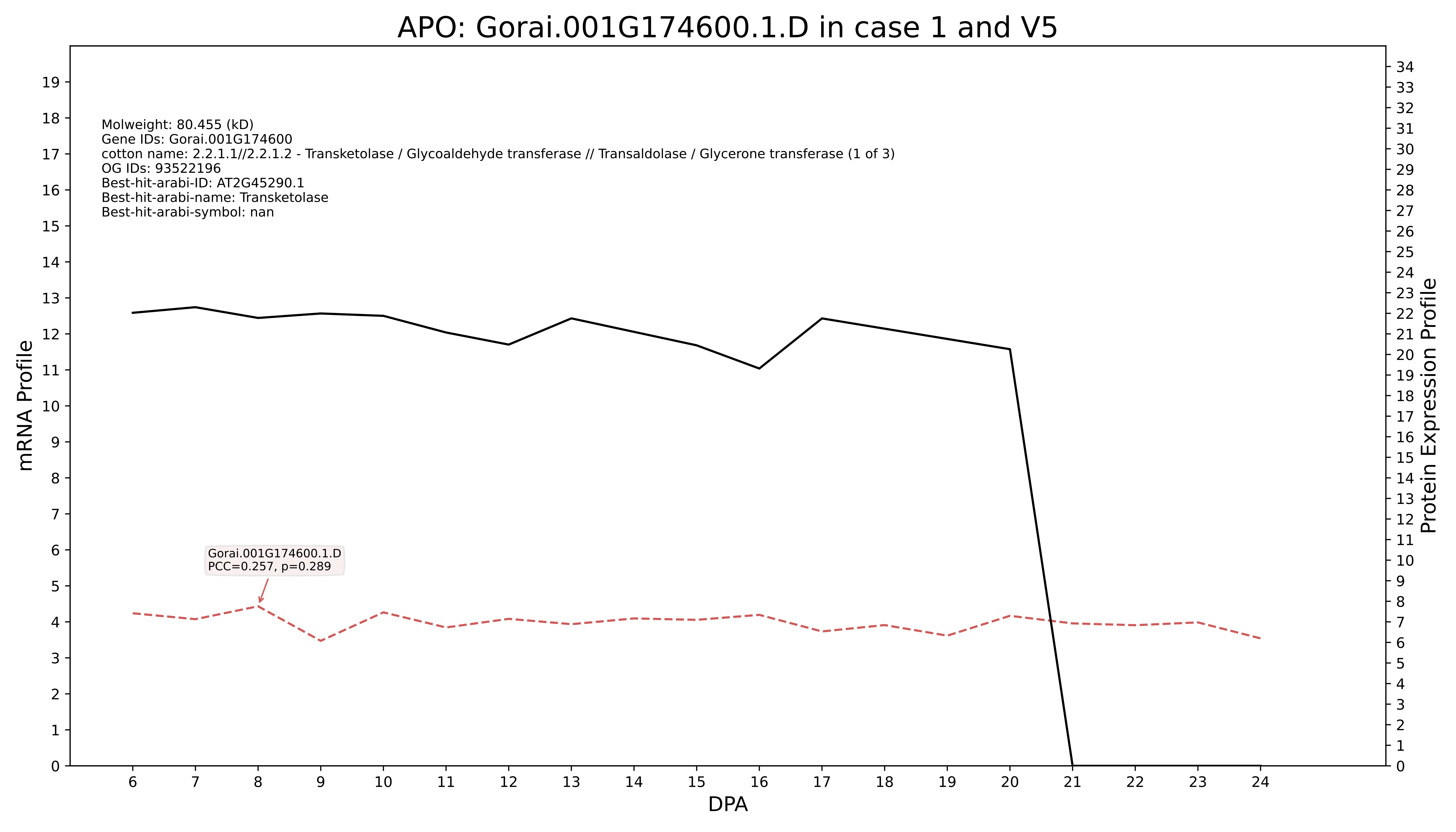
APO: Gorai.001G174600.1.D in case 1 and V5
Molweight: 80.455 (kD)
Gene IDs: Gorai.001G174600
cotton name: 2.2.1.1//2.2.1.2 - Transketolase / Glycoaldehyde transferase // Transaldolase / Glycerone transferase (1 of 3)
OG IDs: 93522196
Best-hit-arabi-ID: AT2G45290.1
Best-hit-arabi-name: Transketolase
Best-hit-arabi-symbol: nan
Gorai.001G174600.1.D
PCC=0.257, p=0.289
mRNA Profile
Protein Expression Profile
DPA

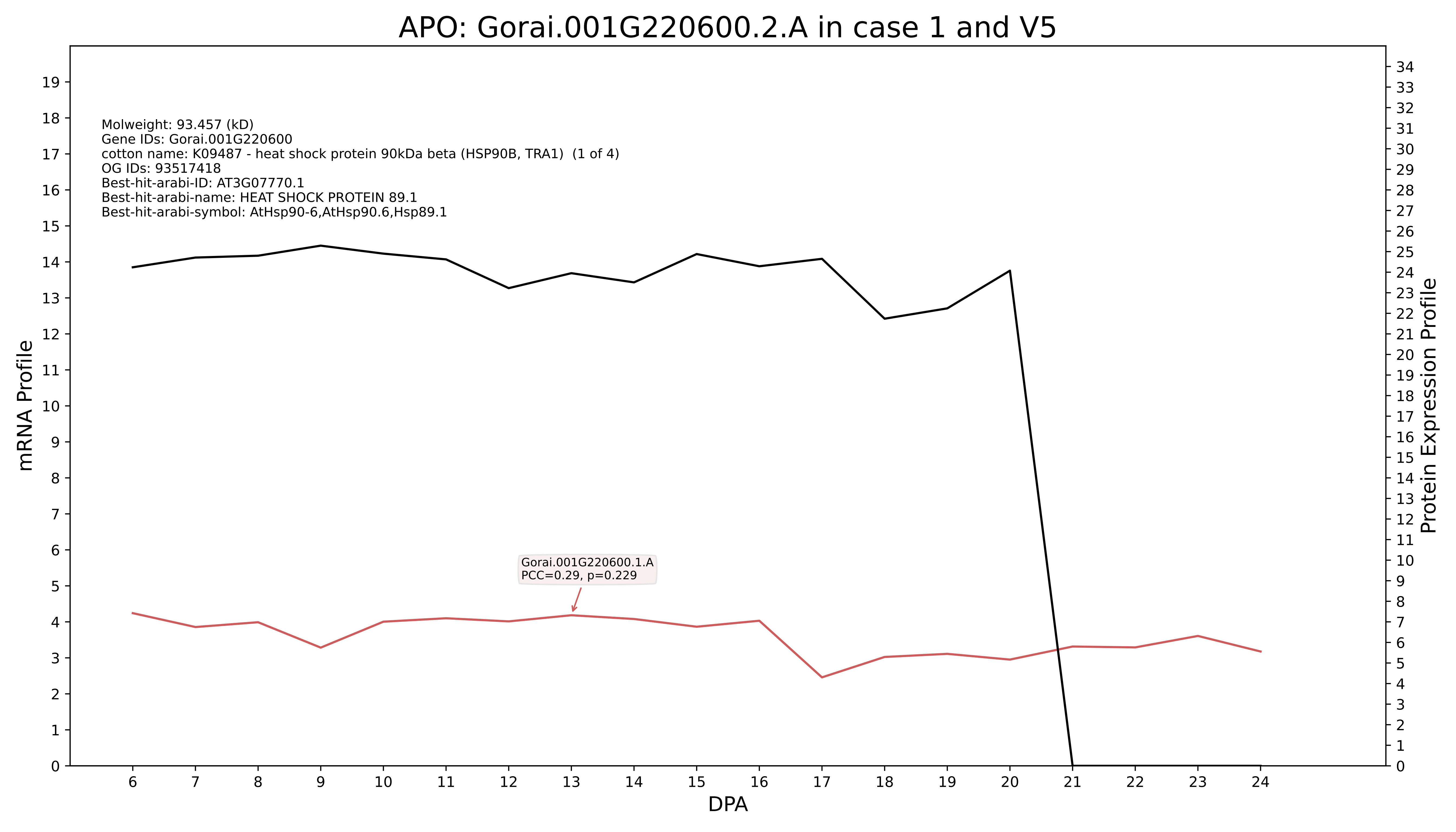

APO: Gorai.001G220600.2.A in case 1 and V5
Molweight: 93.457 (kD)
Gene IDs: Gorai.001G220600
cotton name: K09487 - heat shock protein 90kDa beta (HSP90B, TRA1) (1 of 4)
OG IDs: 93517418
Best-hit-arabi-ID: AT3G07770.1
Best-hit-arabi-name: HEAT SHOCK PROTEIN 89.1
Best-hit-arabi-symbol: AtHsp90-6,AtHsp90.6,Hsp89.1
Gorai.001G220600.1.A
PCC=0.29, p=0.229
mRNA Profile
Protein Expression Profile
DPA

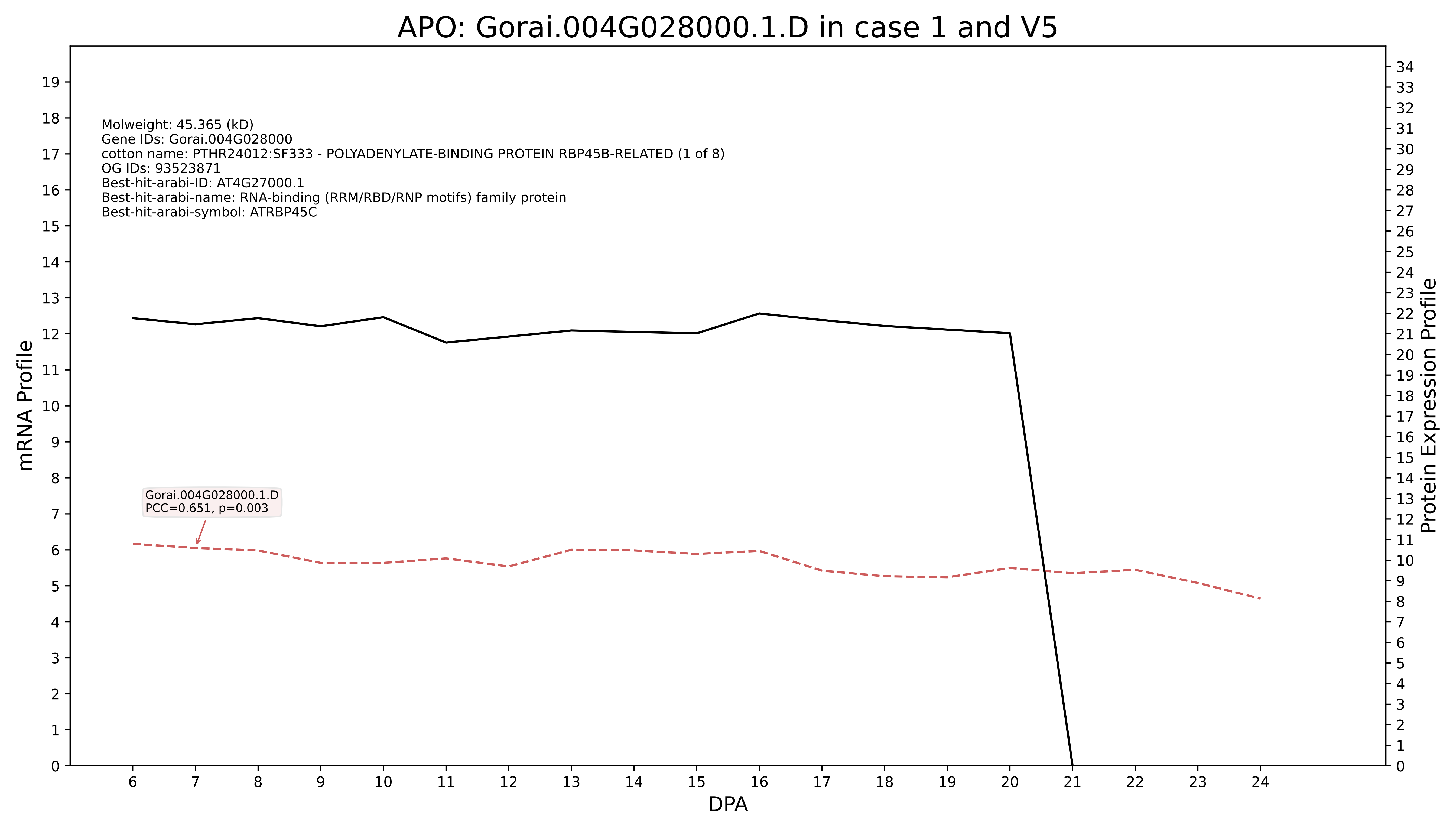

APO: Gorai.004G028000.1.D in case 1 and V5
Molweight: 45.365 (kD)
Gene IDs: Gorai.004G028000
cotton name: PTHR24012:SF333 - POLYADENYLATE-BINDING PROTEIN RBP45B-RELATED (1 of 8)
OG IDs: 93523871
Best-hit-arabi-ID: AT4G27000.1
Best-hit-arabi-name: RNA-binding (RRM/RBD/RNP motifs) family protein
Best-hit-arabi-symbol: ATRBP45C
Gorai.004G028000.1.D
PCC=0.651, p=0.003
mRNA Profile
Protein Expression Profile
DPA

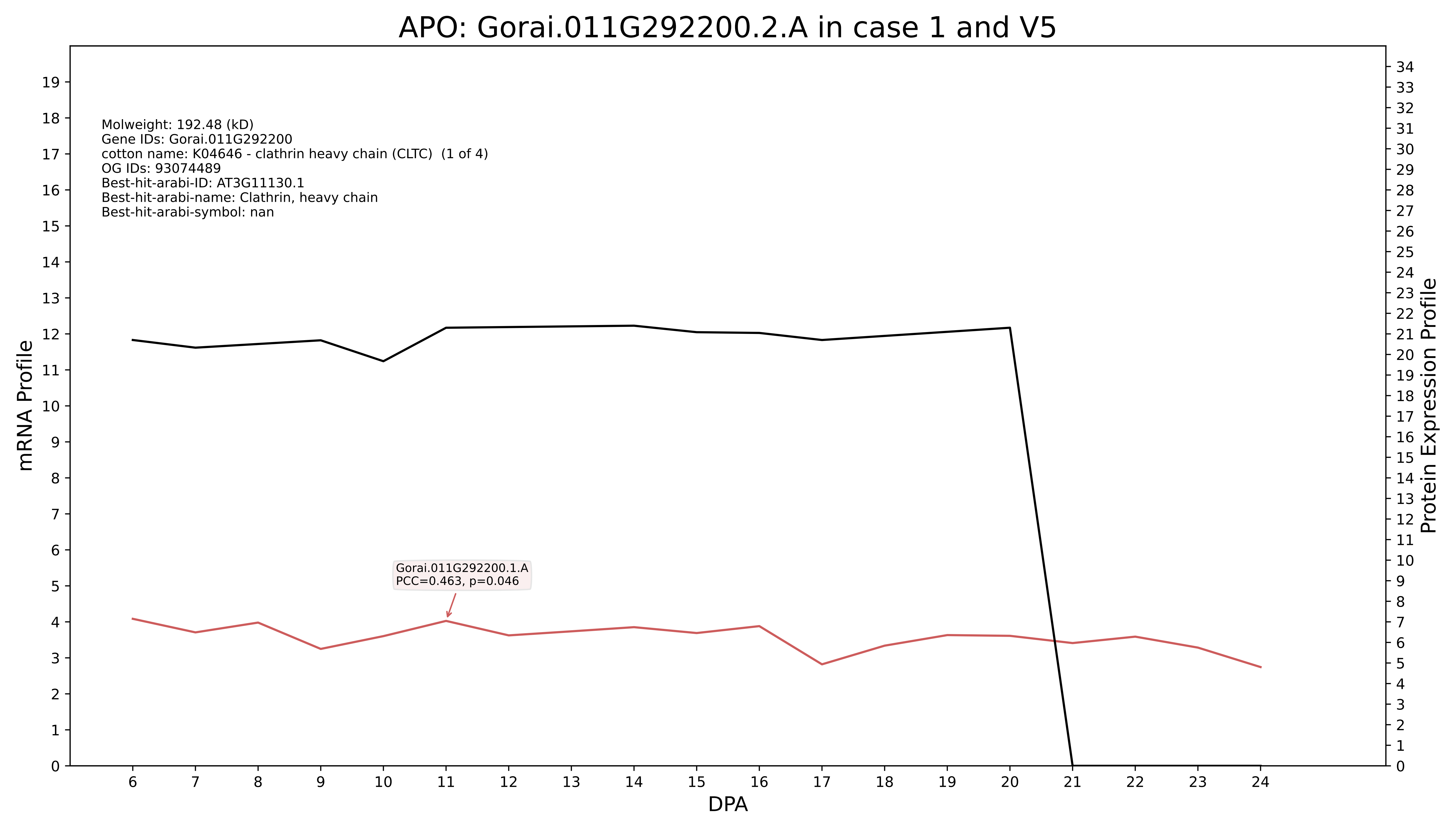

APO: Gorai.011G292200.2.A in case 1 and V5
Molweight: 192.48 (kD)
Gene IDs: Gorai.011G292200
cotton name: K04646 - clathrin heavy chain (CLTC) (1 of 4)
OG IDs: 93074489
Best-hit-arabi-ID: AT3G11130.1
Best-hit-arabi-name: Clathrin, heavy chain
Best-hit-arabi-symbol: nan
Gorai.011G292200.1.A
PCC=0.463, p=0.046
mRNA Profile
Protein Expression Profile
DPA

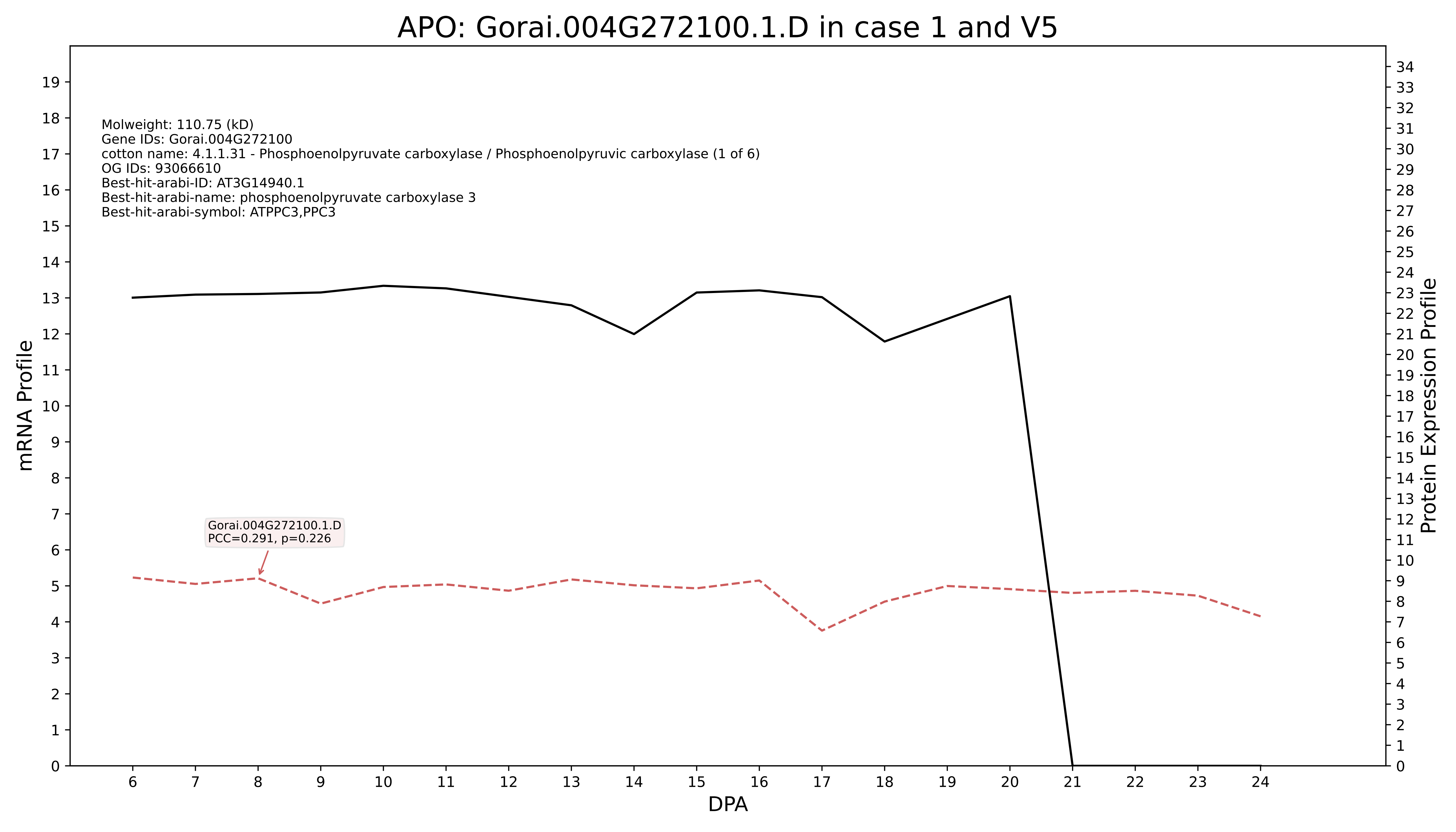
APO: Gorai.004G272100.1.D in case 1 and V5
Molweight: 110.75 (kD)
Gene IDs: Gorai.004G272100
cotton name: 4.1.1.31 - Phosphoenolpyruvate carboxylase / Phosphoenolpyruvic carboxylase (1 of 6)
OG IDs: 93066610
Best-hit-arabi-ID: AT3G14940.1
Best-hit-arabi-name: phosphoenolpyruvate carboxylase 3
Best-hit-arabi-symbol: ATPPC3,PPC3
Gorai.004G272100.1.D
PCC=0.291, p=0.226
mRNA Profile
Protein Expression Profile
DPA

#### APO: Gorai.013G255200.1.D in case 1 and V5

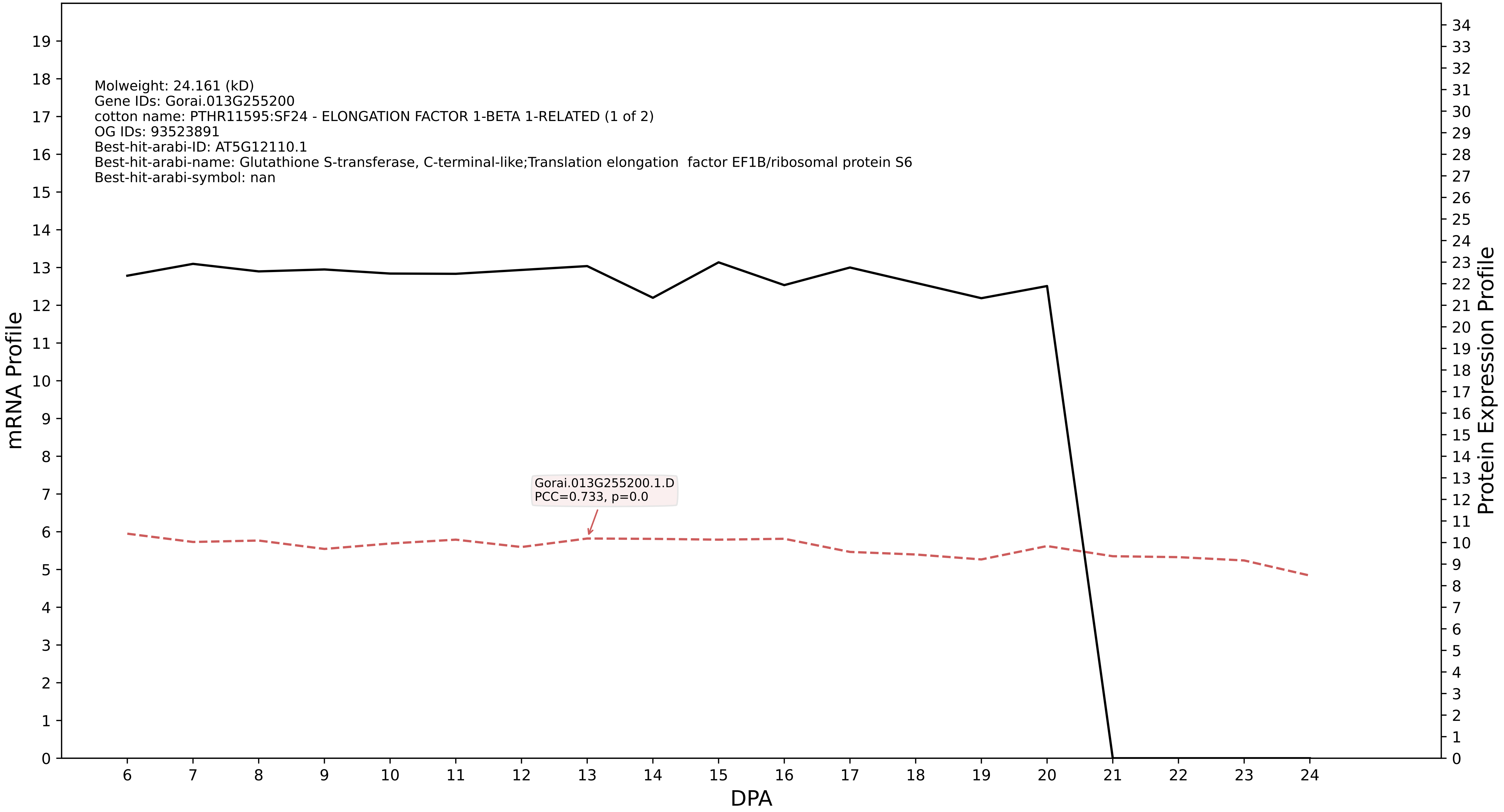

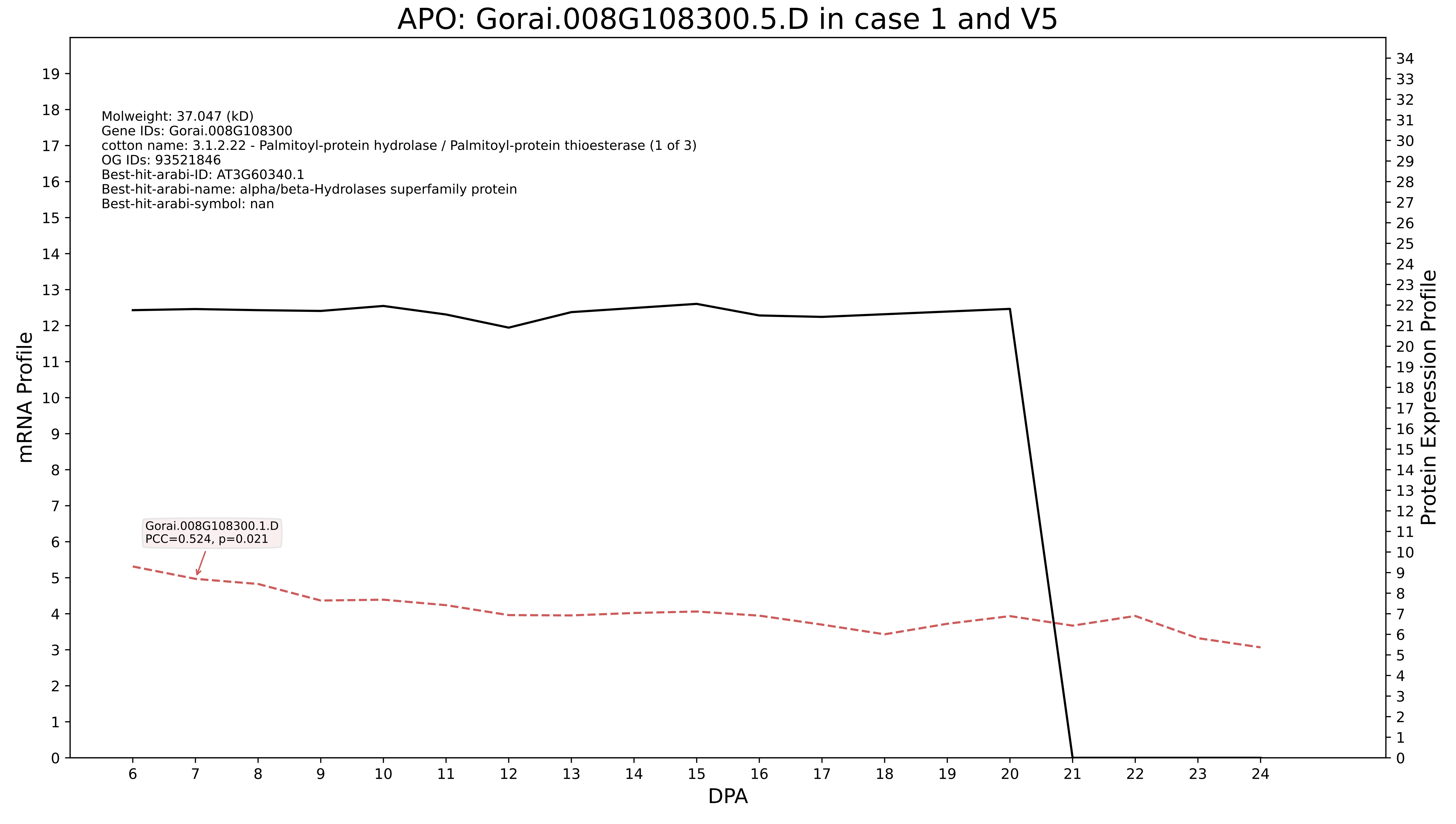
APO: Gorai.008G108300.5.D in case 1 and V5
Molweight: 37.047 (kD)
Gene IDs: Gorai.008G108300
cotton name: 3.1.2.22 - Palmitoyl-protein hydrolase / Palmitoyl-protein thioesterase (1 of 3)
OG IDs: 93521846
Best-hit-arabi-ID: AT3G60340.1
Best-hit-arabi-name: alpha/beta-Hydrolases superfamily protein
Best-hit-arabi-symbol: nan
Gorai.008G108300.1.D
PCC=0.524, p=0.021
mRNA Profile
Protein Expression Profile
DPA

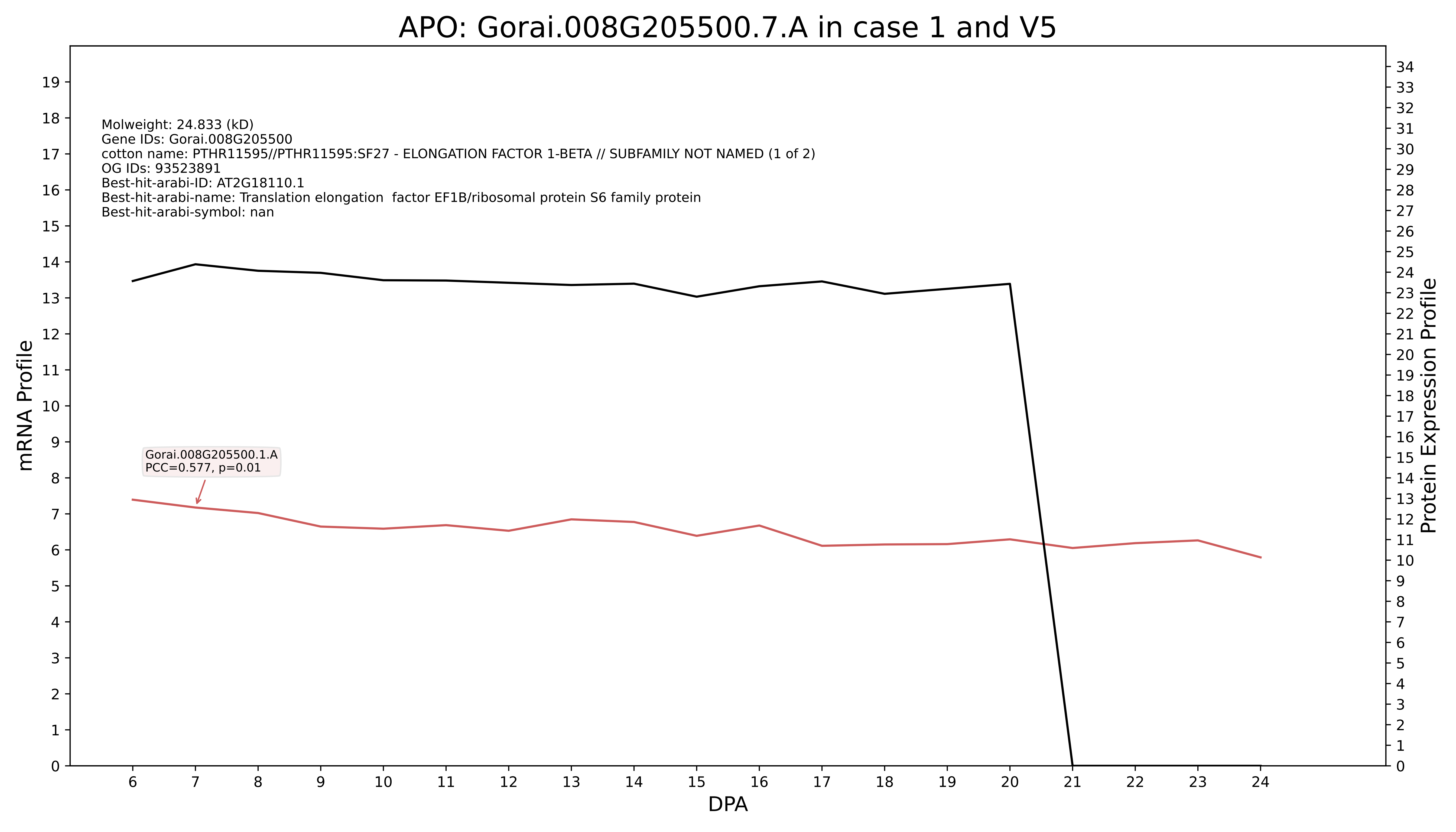
APO: Gorai.008G205500.7.A in case 1 and V5
Molweight: 24.833 (kD)
Gene IDs: Gorai.008G205500
cotton name: PTHR11595//PTHR11595:SF27 - ELONGATION FACTOR 1-BETA // SUBFAMILY NOT NAMED (1 of 2)
OG IDs: 93523891
Best-hit-arabi-ID: AT2G18110.1
Best-hit-arabi-name: Translation elongation factor EF1B/ribosomal protein S6 family protein
Best-hit-arabi-symbol: nan
Gorai.008G205500.1.A
PCC=0.577, p=0.01
mRNA Profile
Protein Expression Profile
DPA

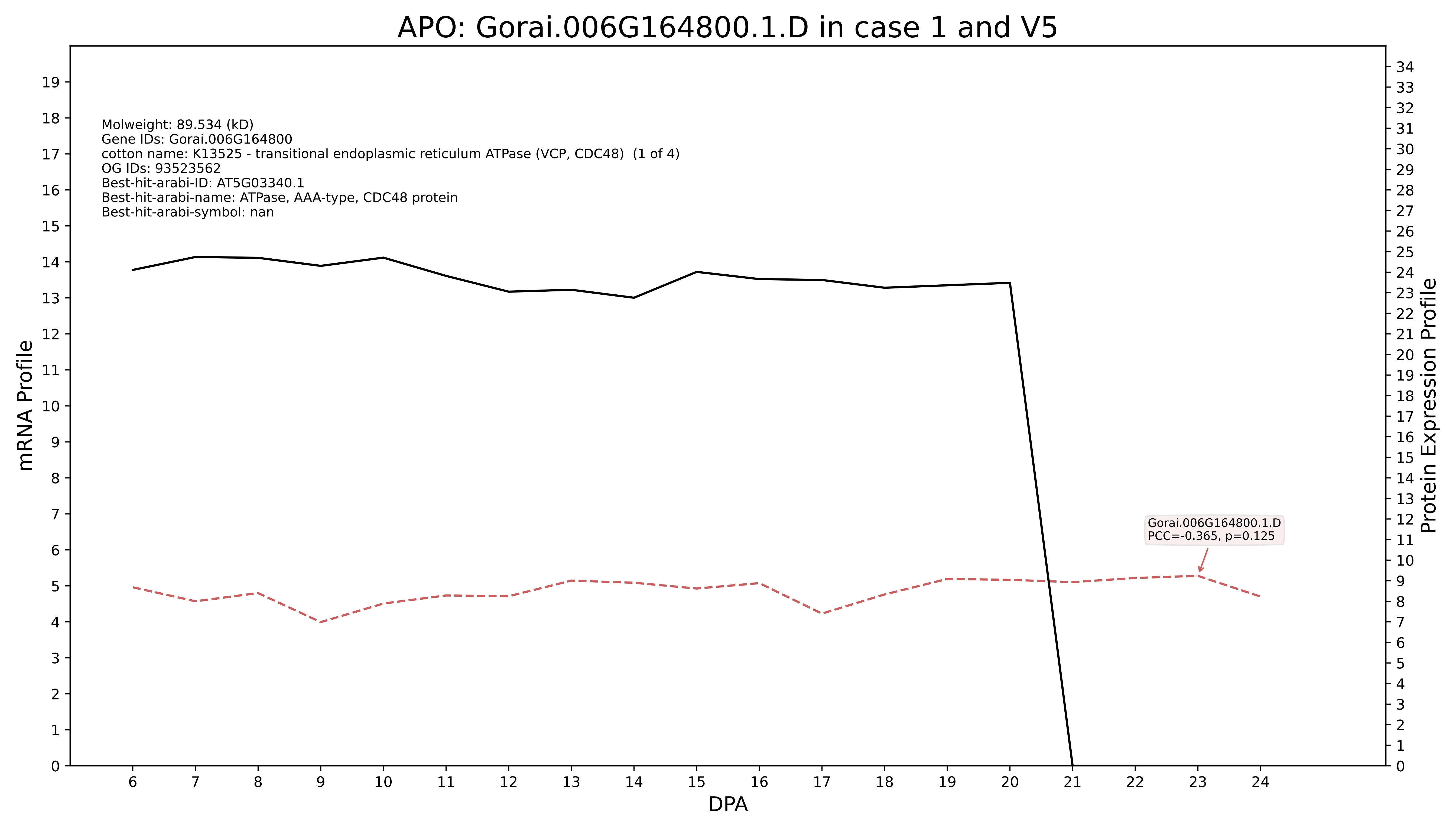

APO: Gorai.006G164800.1.D in case 1 and V5
Molweight: 89.534 (kD)
Gene IDs: Gorai.006G164800
cotton name: K13525 - transitional endoplasmic reticulum ATPase (VCP, CDC48) (1 of 4)
OG IDs: 93523562
Best-hit-arabi-ID: AT5G03340.1
Best-hit-arabi-name: ATPase, AAA-type, CDC48 protein
Best-hit-arabi-symbol: nan
Gorai.006G164800.1.D
PCC=-0.365, p=0.125
mRNA Profile
Protein Expression Profile
DPA

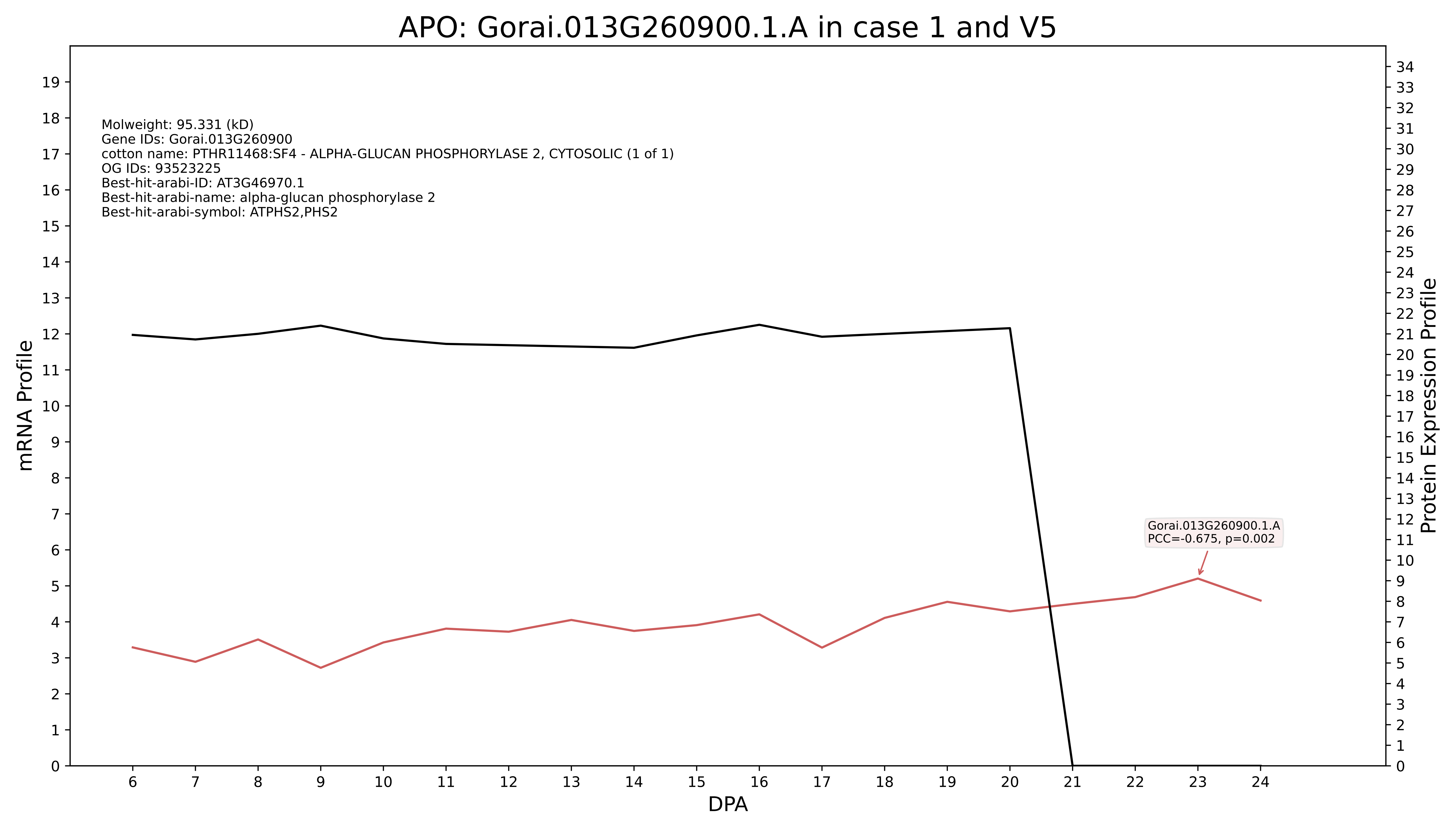
APO: Gorai.013G260900.1.A in case 1 and V5
Molweight: 95.331 (kD)
Gene IDs: Gorai.013G260900
cotton name: PTHR11468:SF4 - ALPHA-GLUCAN PHOSPHORYLASE 2, CYTOSOLIC (1 of 1)
OG IDs: 93523225
Best-hit-arabi-ID: AT3G46970.1
Best-hit-arabi-name: alpha-glucan phosphorylase 2
Best-hit-arabi-symbol: ATPHS2,PHS2
Gorai.013G260900.1.A
PCC=-0.675, p=0.002
mRNA Profile
Protein Expression Profile
DPA

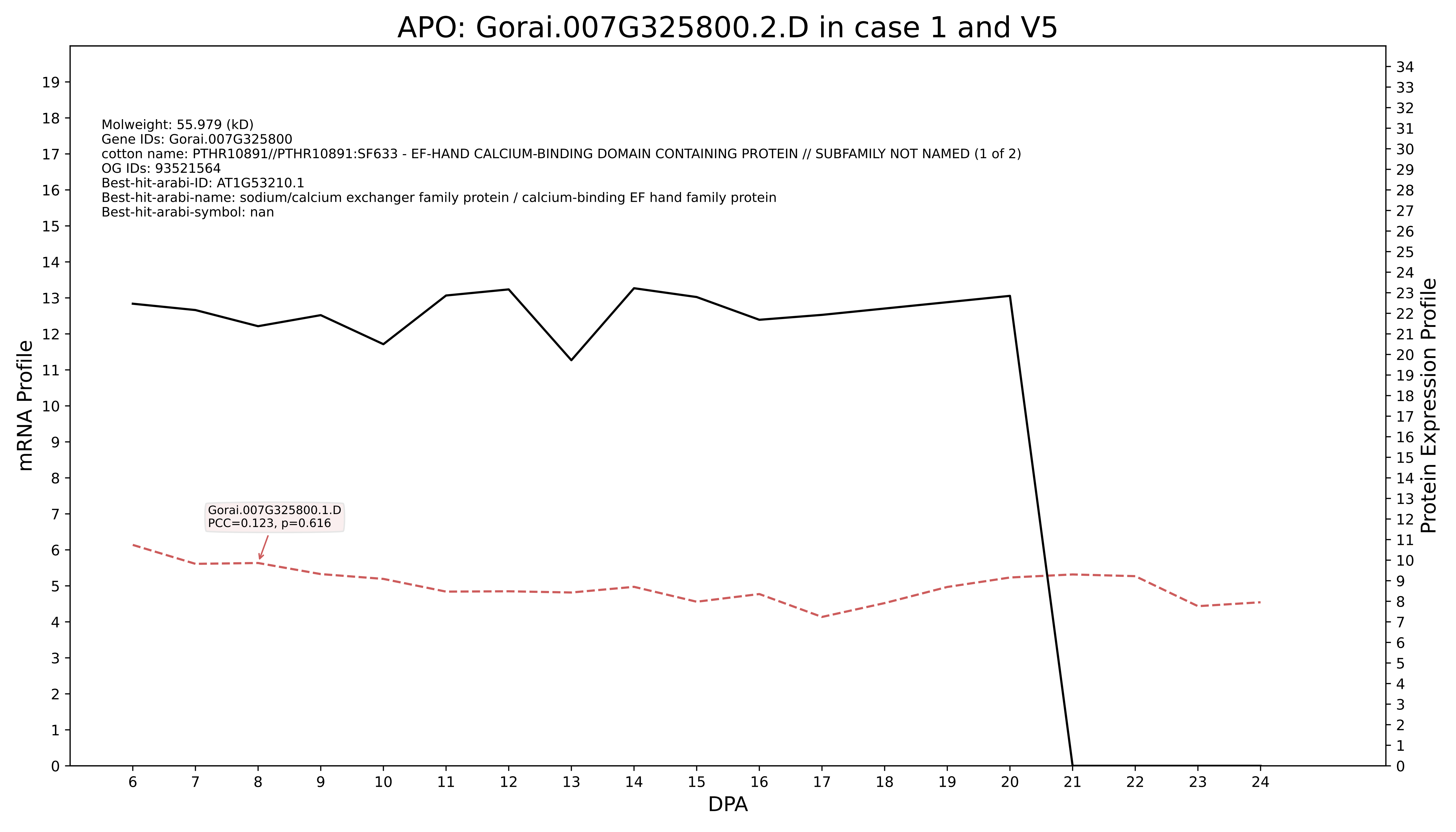
APO: Gorai.007G325800.2.D in case 1 and V5
Molweight: 55.979 (kD)
Gene IDs: Gorai.007G325800
cotton name: PTHR10891//PTHR10891:SF633 - EF-HAND CALCIUM-BINDING DOMAIN CONTAINING PROTEIN // SUBFAMILY NOT NAMED (1 of 2)
OG IDs: 93521564
Best-hit-arabi-ID: AT1G53210.1
Best-hit-arabi-name: sodium/calcium exchanger family protein / calcium-binding EF hand family protein
Best-hit-arabi-symbol: nan
Gorai.007G325800.1.D
PCC=0.123, p=0.616
mRNA Profile
Protein Expression Profile
DPA

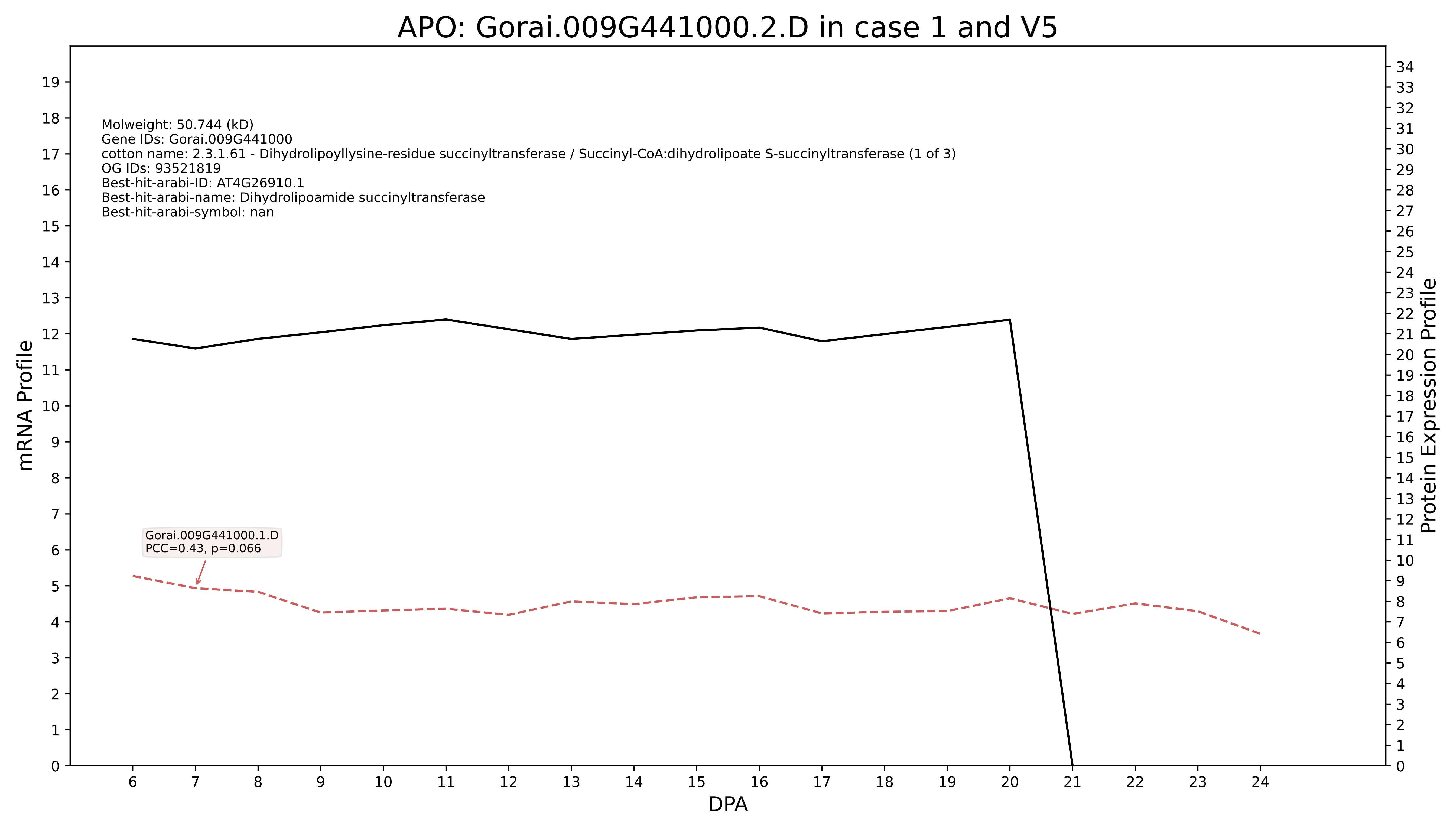

APO: Gorai.009G441000.2.D in case 1 and V5
Molweight: 50.744 (kD)
Gene IDs: Gorai.009G441000
cotton name: 2.3.1.61 - Dihydrolipoyllysine-residue succinyltransferase / Succinyl-CoA:dihydrolipoate S-succinyltransferase (1 of 3)
OG IDs: 93521819
Best-hit-arabi-ID: AT4G26910.1
Best-hit-arabi-name: Dihydrolipoamide succinyltransferase
Best-hit-arabi-symbol: nan
Gorai.009G441000.1.D
PCC=0.43, p=0.066
mRNA Profile
Protein Expression Profile
DPA

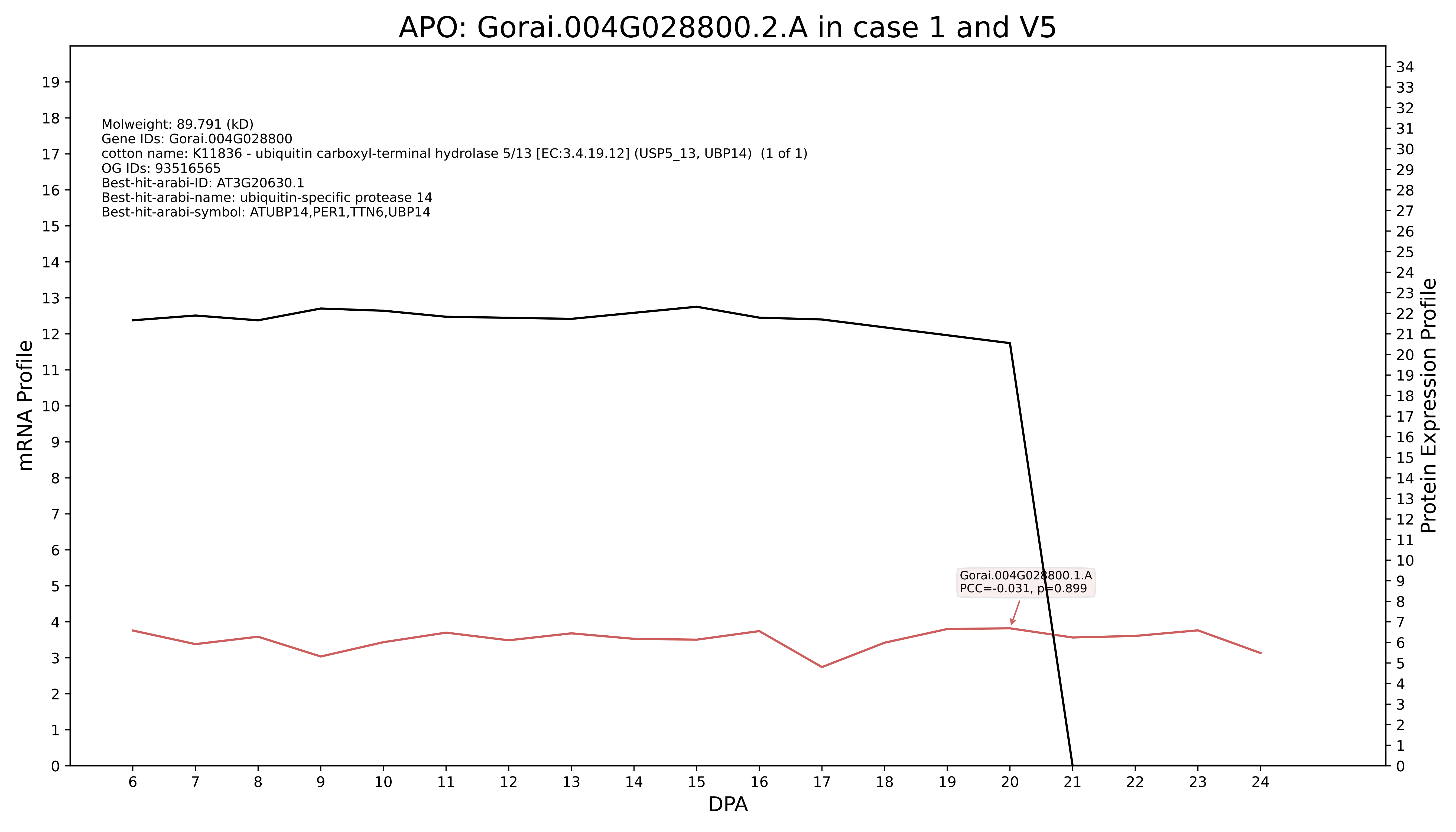
APO: Gorai.004G028800.2.A in case 1 and V5
Molweight: 89.791 (kD)
Gene IDs: Gorai.004G028800
cotton name: K11836 - ubiquitin carboxyl-terminal hydrolase 5/13 [EC:3.4.19.12] (USP5_13, UBP14) (1 of 1)
OG IDs: 93516565
Best-hit-arabi-ID: AT3G20630.1
Best-hit-arabi-name: ubiquitin-specific protease 14
Best-hit-arabi-symbol: ATUBP14,PER1,TTN6,UBP14
Gorai.004G028800.1.A
PCC=-0.031, p=0.899
mRNA Profile
Protein Expression Profile
DPA

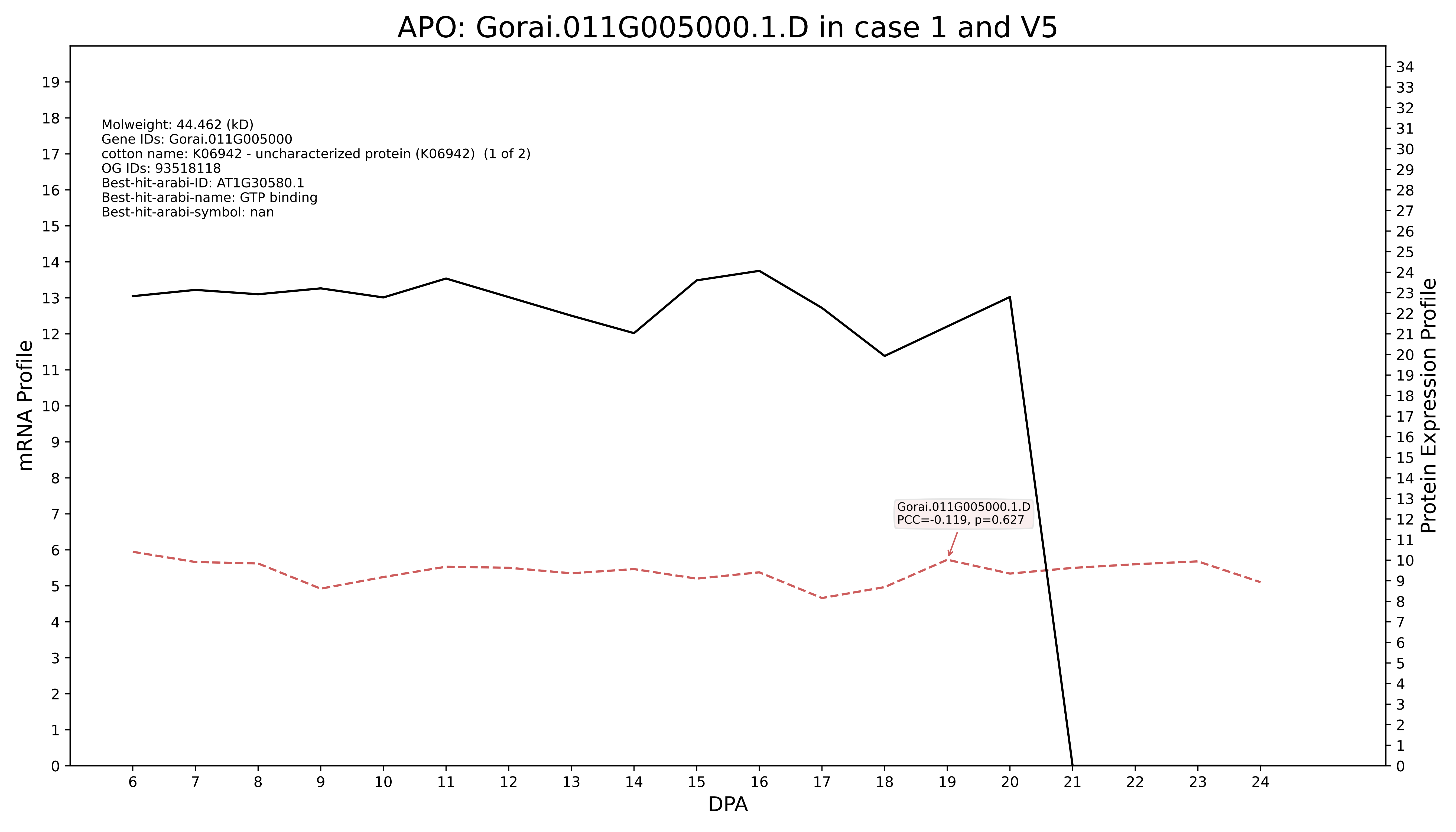

APO: Gorai.011G005000.1.D in case 1 and V5
Molweight: 44.462 (kD)
Gene IDs: Gorai.011G005000
cotton name: K06942 - uncharacterized protein (K06942) (1 of 2)
OG IDs: 93518118
Best-hit-arabi-ID: AT1G30580.1
Best-hit-arabi-name: GTP binding
Best-hit-arabi-symbol: nan
Gorai.011G005000.1.D
PCC=-0.119, p=0.627
mRNA Profile
Protein Expression Profile
DPA

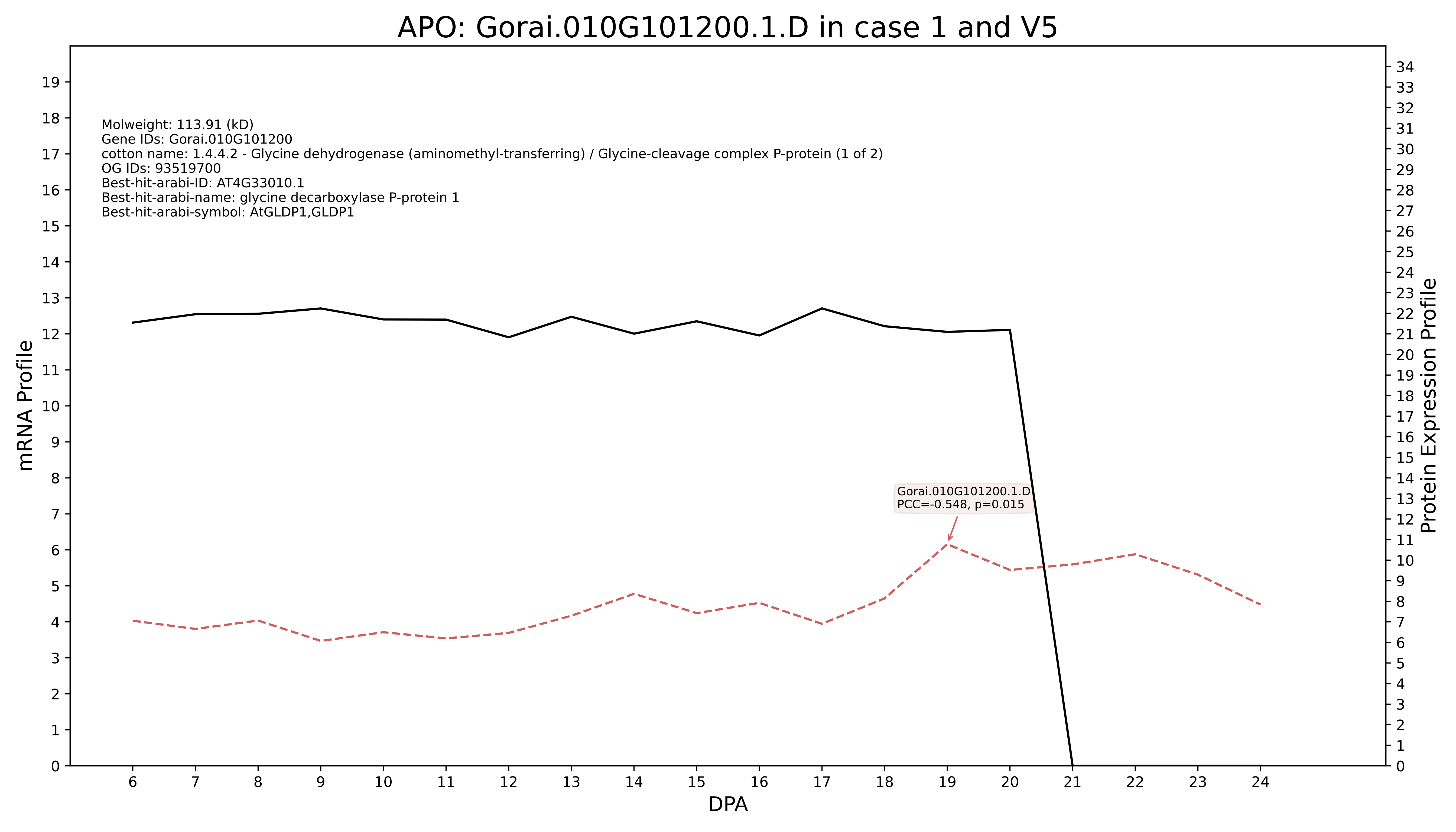

APO: Gorai.010G101200.1.D in case 1 and V5
Molweight: 113.91 (kD)
Gene IDs: Gorai.010G101200
cotton name: 1.4.4.2 - Glycine dehydrogenase (aminomethyl-transferring) / Glycine-cleavage complex P-protein (1 of 2)
OG IDs: 93519700
Best-hit-arabi-ID: AT4G33010.1
Best-hit-arabi-name: glycine decarboxylase P-protein 1
Best-hit-arabi-symbol: AtGLDP1,GLDP1
Gorai.010G101200.1.D
PCC=-0.548, p=0.015
mRNA Profile
Protein Expression Profile
DPA

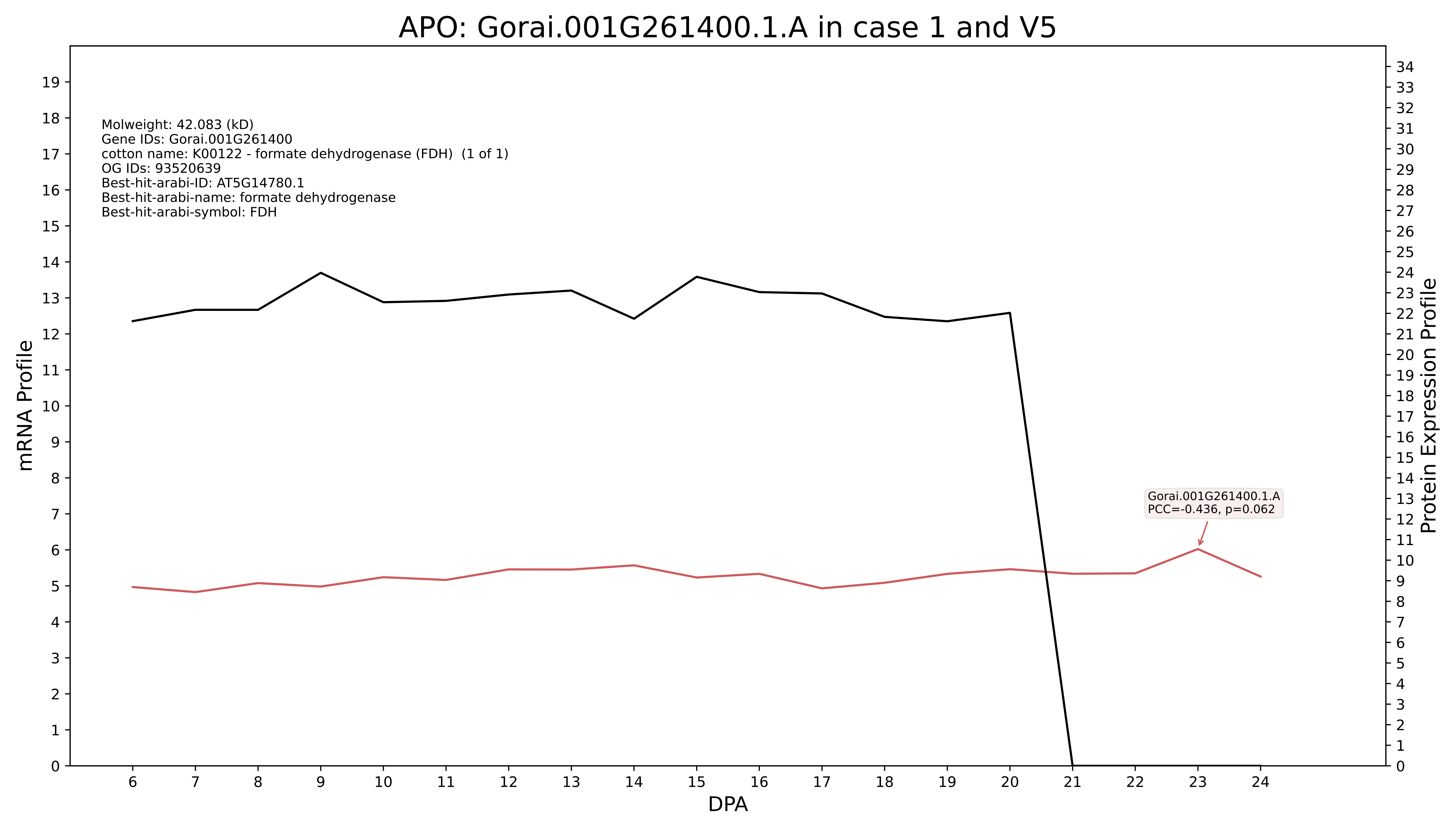
APO: Gorai.001G261400.1.A in case 1 and V5
Molweight: 42.083 (kD)
Gene IDs: Gorai.001G261400
cotton name: K00122 - formate dehydrogenase (FDH) (1 of 1)
OG IDs: 93520639
Best-hit-arabi-ID: AT5G14780.1
Best-hit-arabi-name: formate dehydrogenase
Best-hit-arabi-symbol: FDH
Gorai.001G261400.1.A
PCC=-0.436, p=0.062
mRNA Profile
Protein Expression Profile
DPA

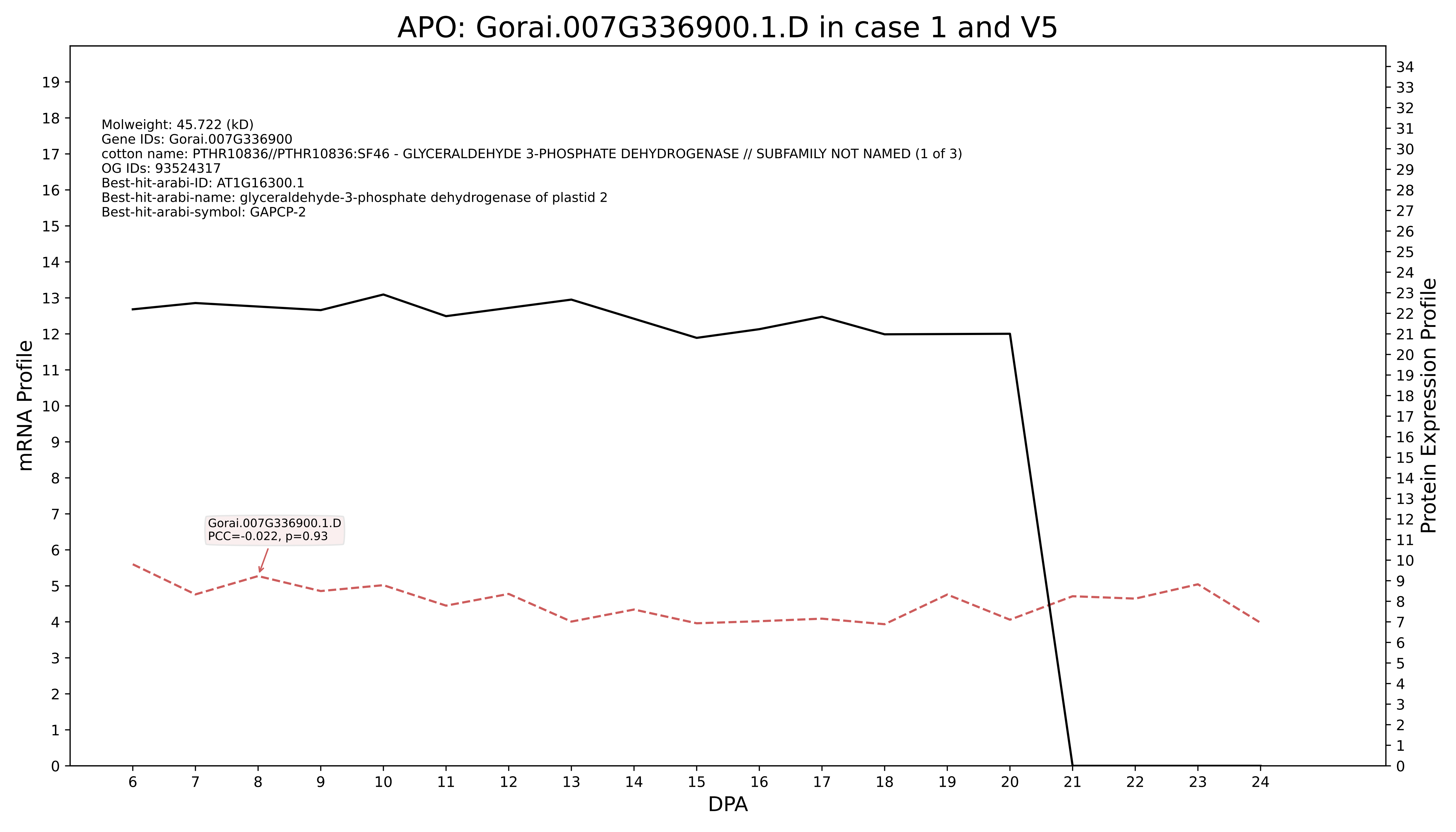
APO: Gorai.007G336900.1.D in case 1 and V5
Molweight: 45.722 (kD)
Gene IDs: Gorai.007G336900
cotton name: PTHR10836//PTHR10836:SF46 - GLYCERALDEHYDE 3-PHOSPHATE DEHYDROGENASE // SUBFAMILY NOT NAMED (1 of 3)
OG IDs: 93524317
Best-hit-arabi-ID: AT1G16300.1
Best-hit-arabi-name: glyceraldehyde-3-phosphate dehydrogenase of plastid 2
Best-hit-arabi-symbol: GAPCP-2
Gorai.007G336900.1.D
PCC=-0.022, p=0.93
mRNA Profile
Protein Expression Profile
DPA

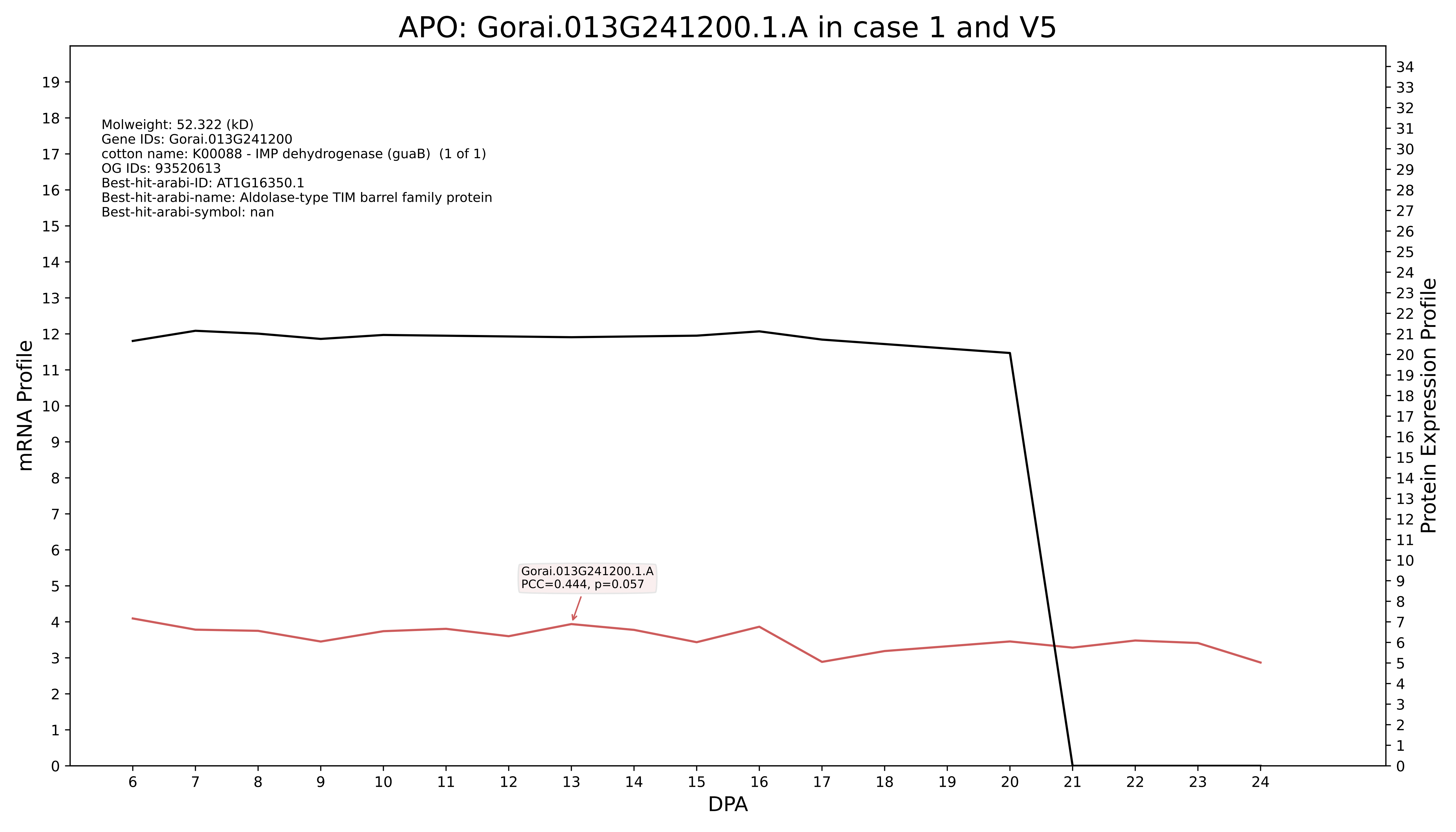

APO: Gorai.013G241200.1.A in case 1 and V5
Molweight: 52.322 (kD)
Gene IDs: Gorai.013G241200
cotton name: K00088 - IMP dehydrogenase (guaB) (1 of 1)
OG IDs: 93520613
Best-hit-arabi-ID: AT1G16350.1
Best-hit-arabi-name: Aldolase-type TIM barrel family protein
Best-hit-arabi-symbol: nan
Gorai.013G241200.1.A
PCC=0.444, p=0.057
mRNA Profile
Protein Expression Profile
DPA

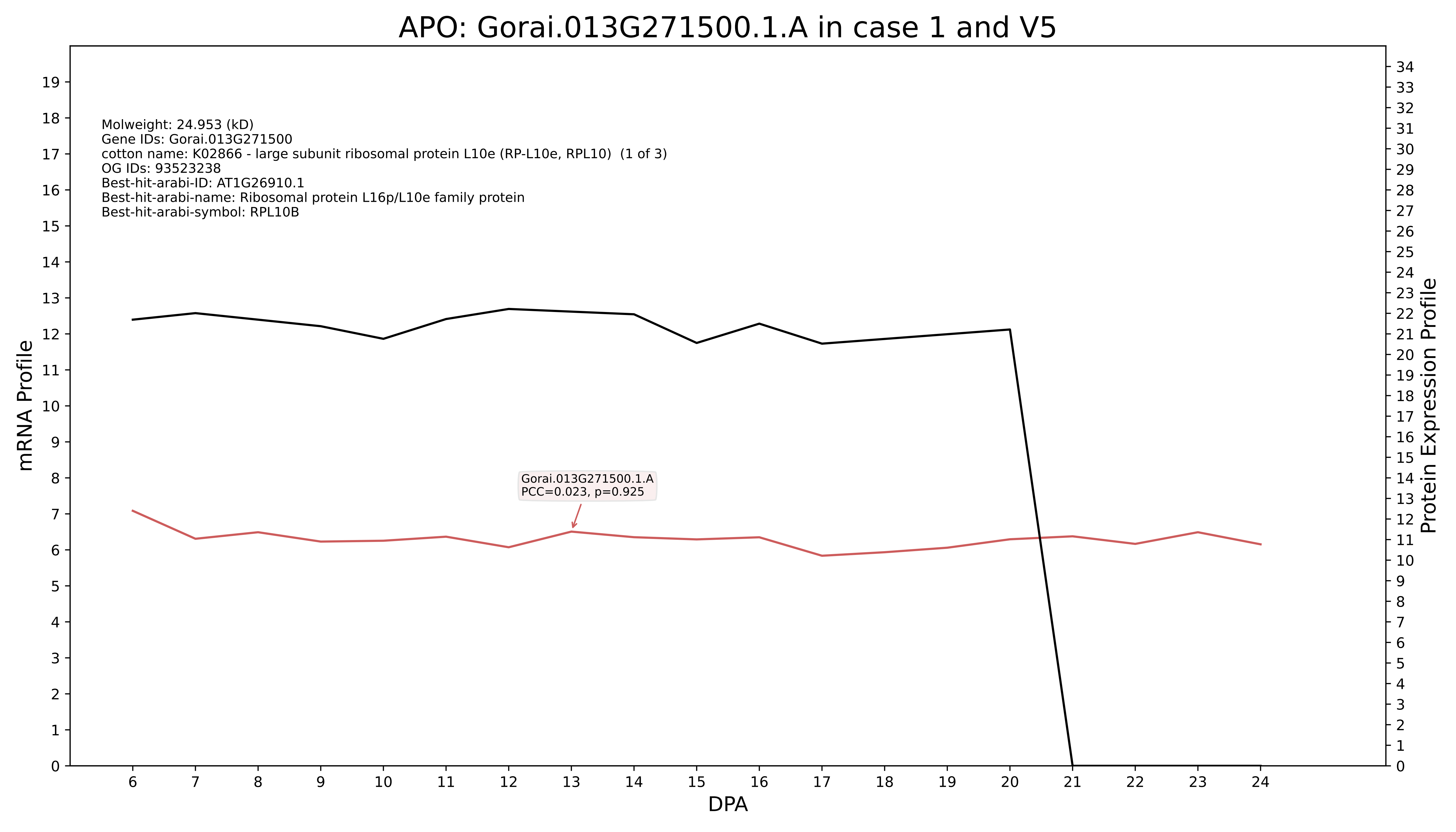
APO: Gorai.013G271500.1.A in case 1 and V5
Molweight: 24.953 (kD)
Gene IDs: Gorai.013G271500
cotton name: K02866 - large subunit ribosomal protein L10e (RP-L10e, RPL10) (1 of 3)
OG IDs: 93523238
Best-hit-arabi-ID: AT1G26910.1
Best-hit-arabi-name: Ribosomal protein L16p/L10e family protein
Best-hit-arabi-symbol: RPL10B
Gorai.013G271500.1.A
PCC=0.023, p=0.925
mRNA Profile
Protein Expression Profile
DPA

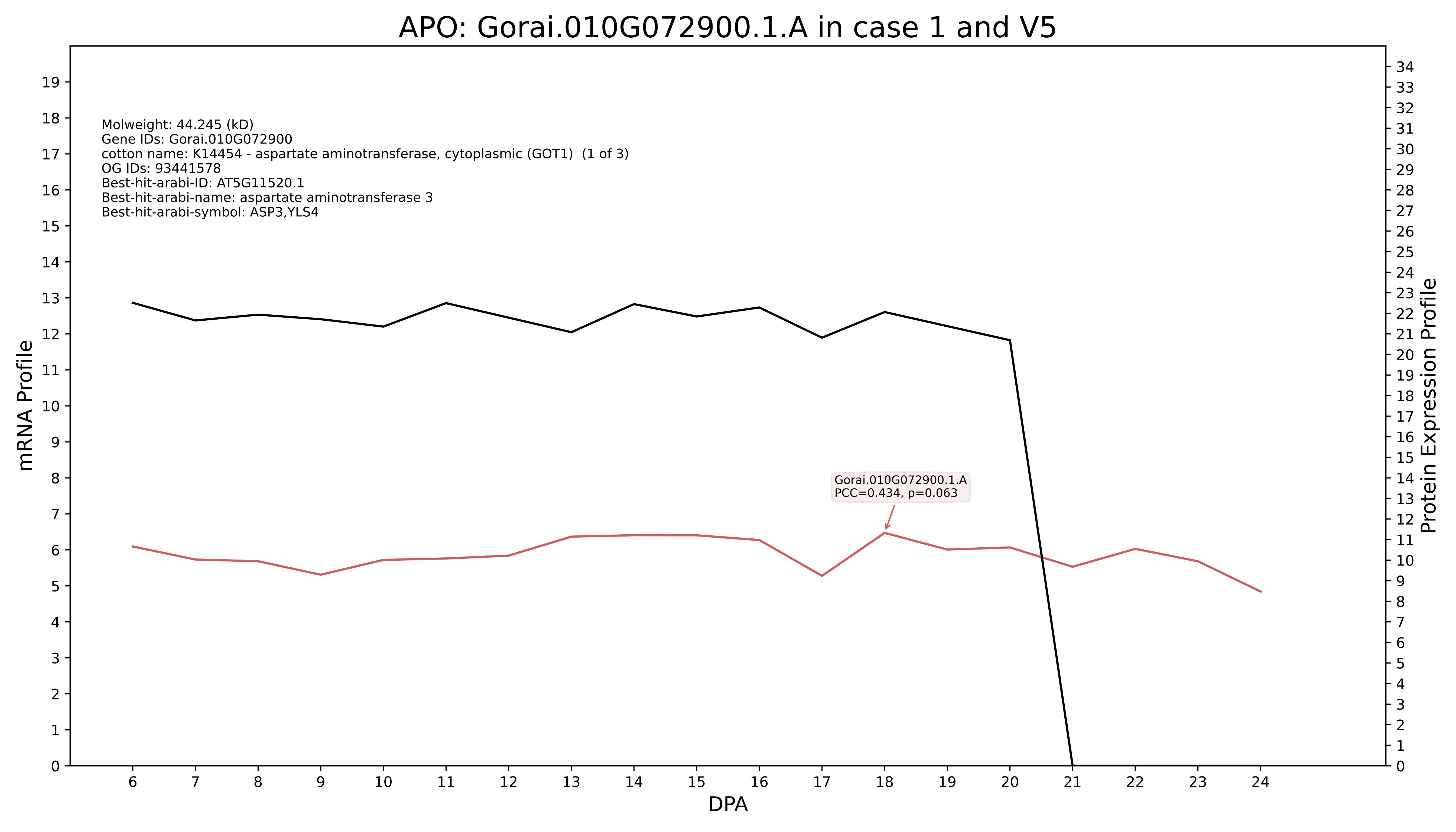
APO: Gorai.010G072900.1.A in case 1 and V5
Molweight: 44.245 (kD)
Gene IDs: Gorai.010G072900
cotton name: K14454 - aspartate aminotransferase, cytoplasmic (GOT1) (1 of 3)
OG IDs: 93441578
Best-hit-arabi-ID: AT5G11520.1
Best-hit-arabi-name: aspartate aminotransferase 3
Best-hit-arabi-symbol: ASP3,YLS4
Gorai.010G072900.1.A
PCC=0.434, p=0.063
mRNA Profile
Protein Expression Profile
DPA

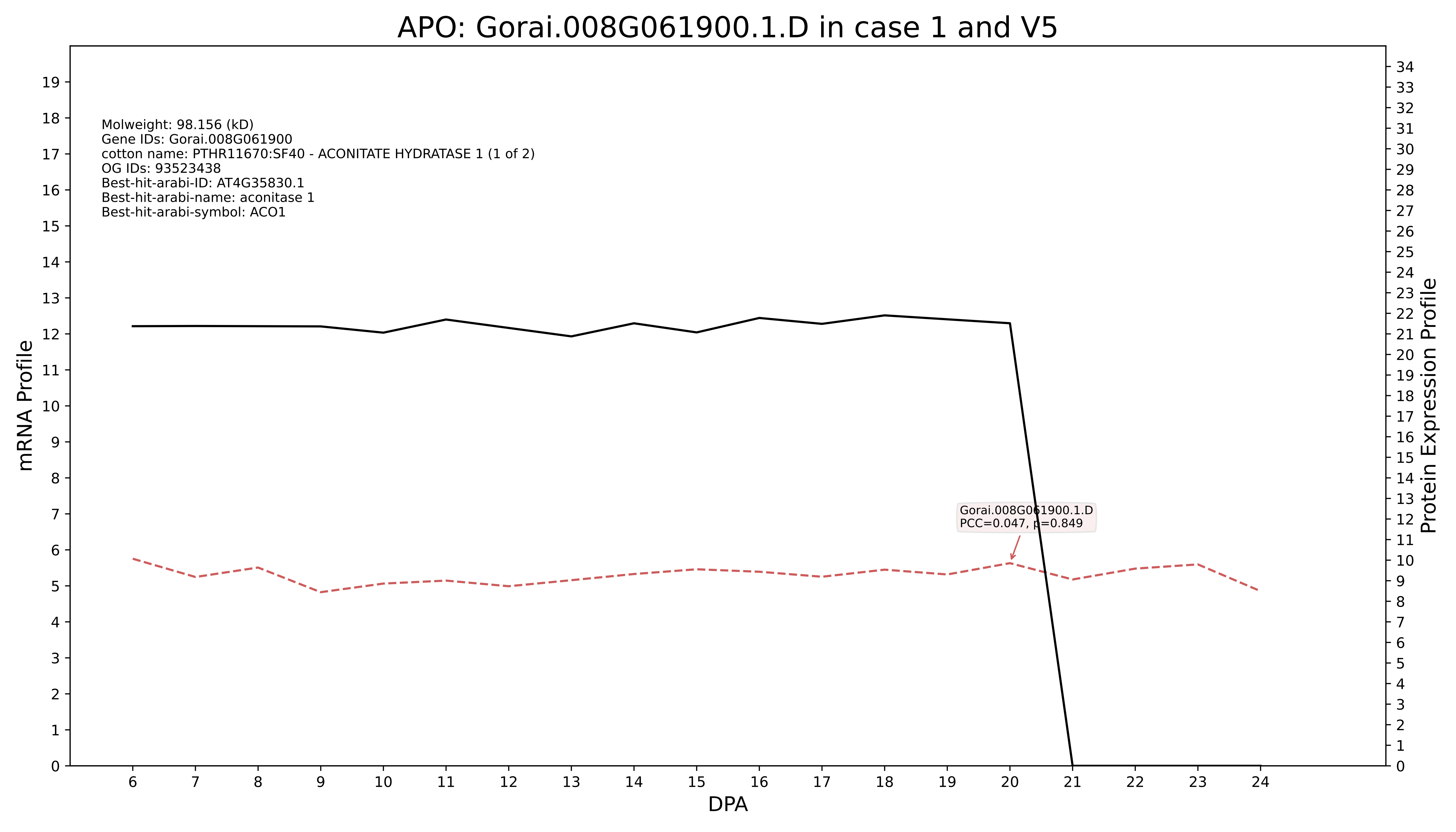

APO: Gorai.008G061900.1.D in case 1 and V5
Molweight: 98.156 (kD)
Gene IDs: Gorai.008G061900
cotton name: PTHR11670:SF40 - ACONITATE HYDRATASE 1 (1 of 2)
OG IDs: 93523438
Best-hit-arabi-ID: AT4G35830.1
Best-hit-arabi-name: aconitase 1
Best-hit-arabi-symbol: ACO1
Gorai.008G061900.1.D
PCC=0.047, p=0.849
mRNA Profile
Protein Expression Profile
DPA

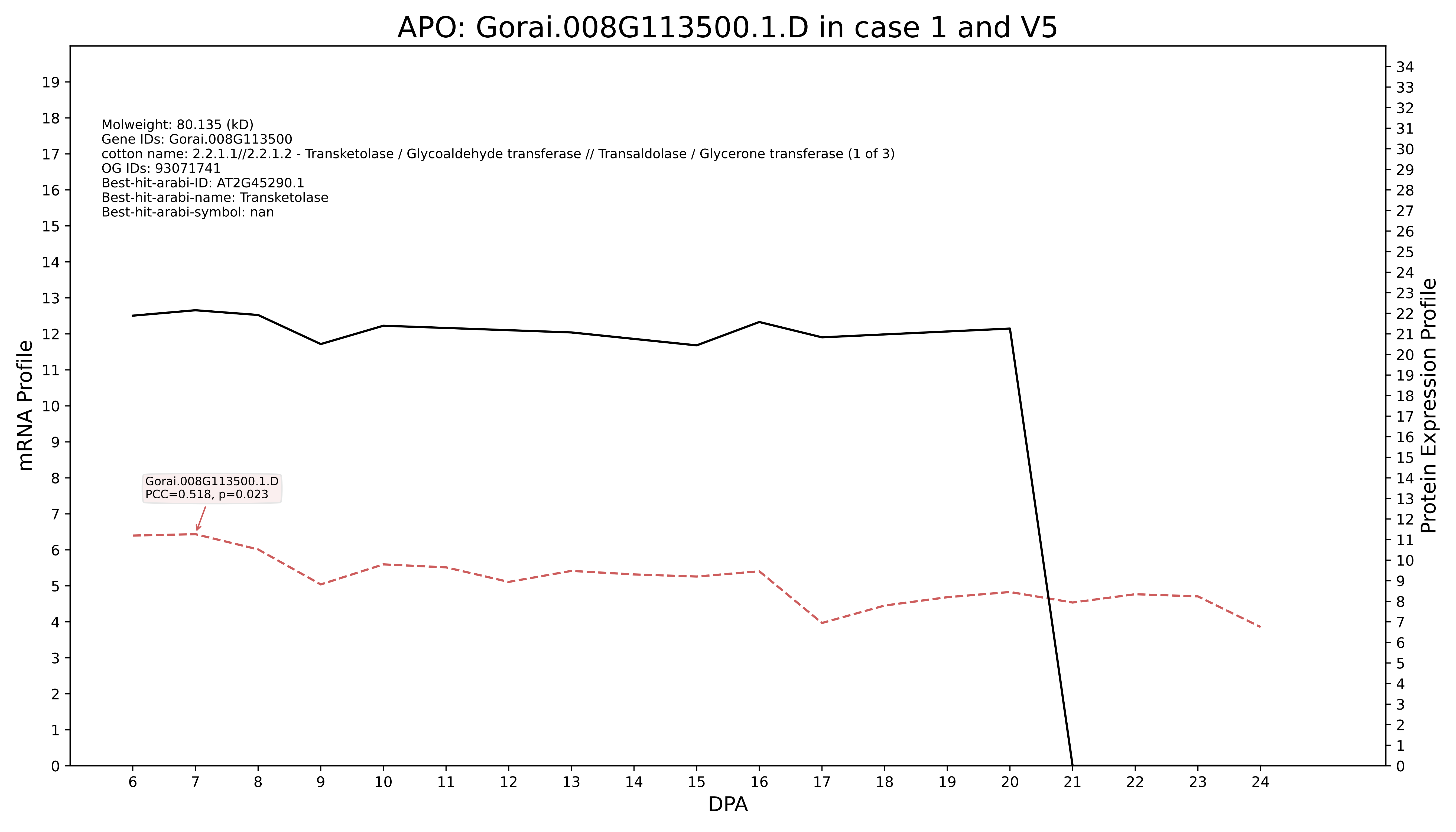
APO: Gorai.008G113500.1.D in case 1 and V5
Molweight: 80.135 (kD)
Gene IDs: Gorai.008G113500
cotton name: 2.2.1.1//2.2.1.2 - Transketolase / Glycoaldehyde transferase // Transaldolase / Glycerone transferase (1 of 3)
OG IDs: 93071741
Best-hit-arabi-ID: AT2G45290.1
Best-hit-arabi-name: Transketolase
Best-hit-arabi-symbol: nan
Gorai.008G113500.1.D
PCC=0.518, p=0.023
mRNA Profile
Protein Expression Profile
DPA

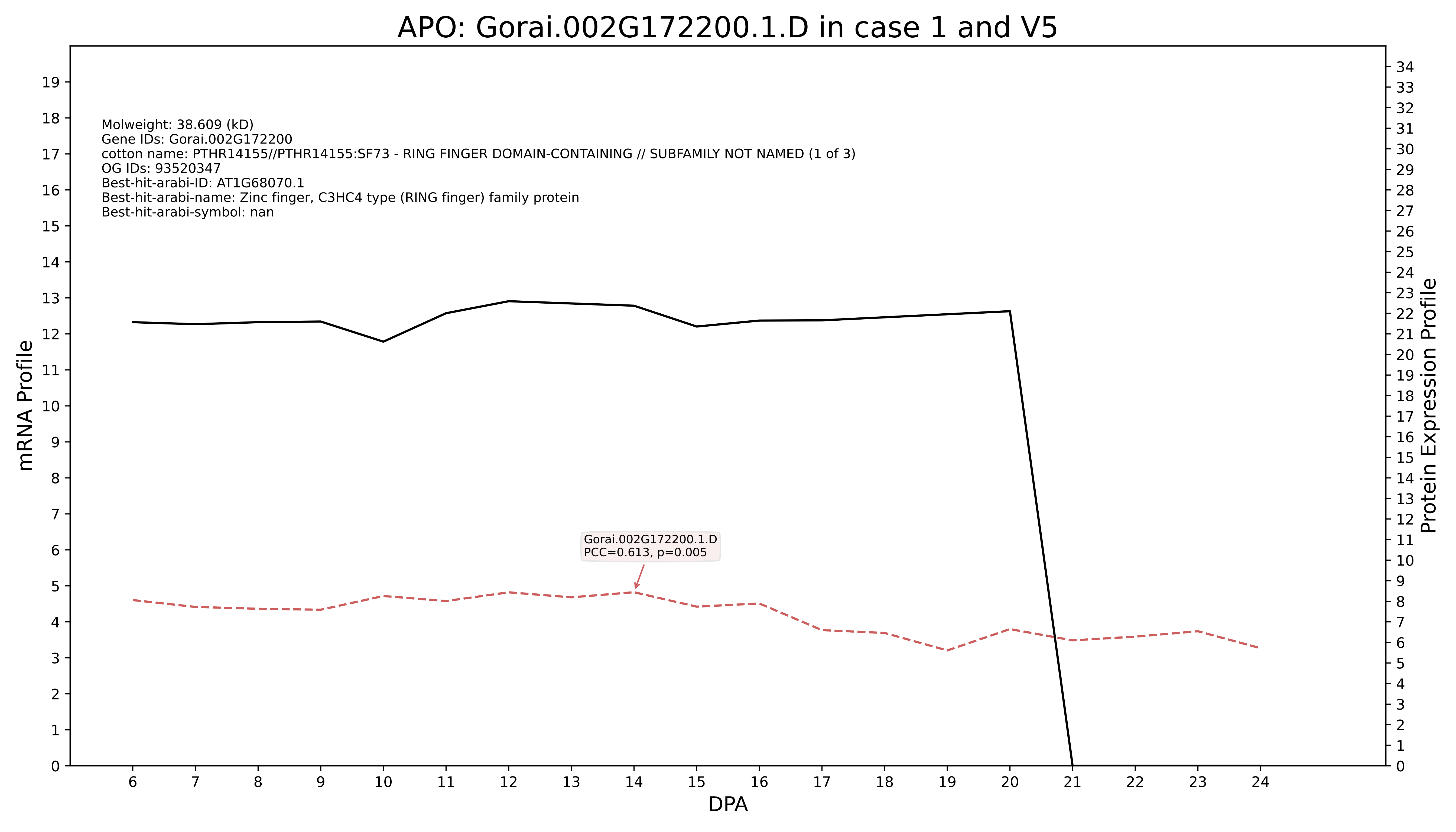
APO: Gorai.002G172200.1.D in case 1 and V5
Molweight: 38.609 (kD)
Gene IDs: Gorai.002G172200
cotton name: PTHR14155//PTHR14155:SF73 - RING FINGER DOMAIN-CONTAINING // SUBFAMILY NOT NAMED (1 of 3)
OG IDs: 93520347
Best-hit-arabi-ID: AT1G68070.1
Best-hit-arabi-name: Zinc finger, C3HC4 type (RING finger) family protein
Best-hit-arabi-symbol: nan
Gorai.002G172200.1.D
PCC=0.613, p=0.005
mRNA Profile
Protein Expression Profile
DPA

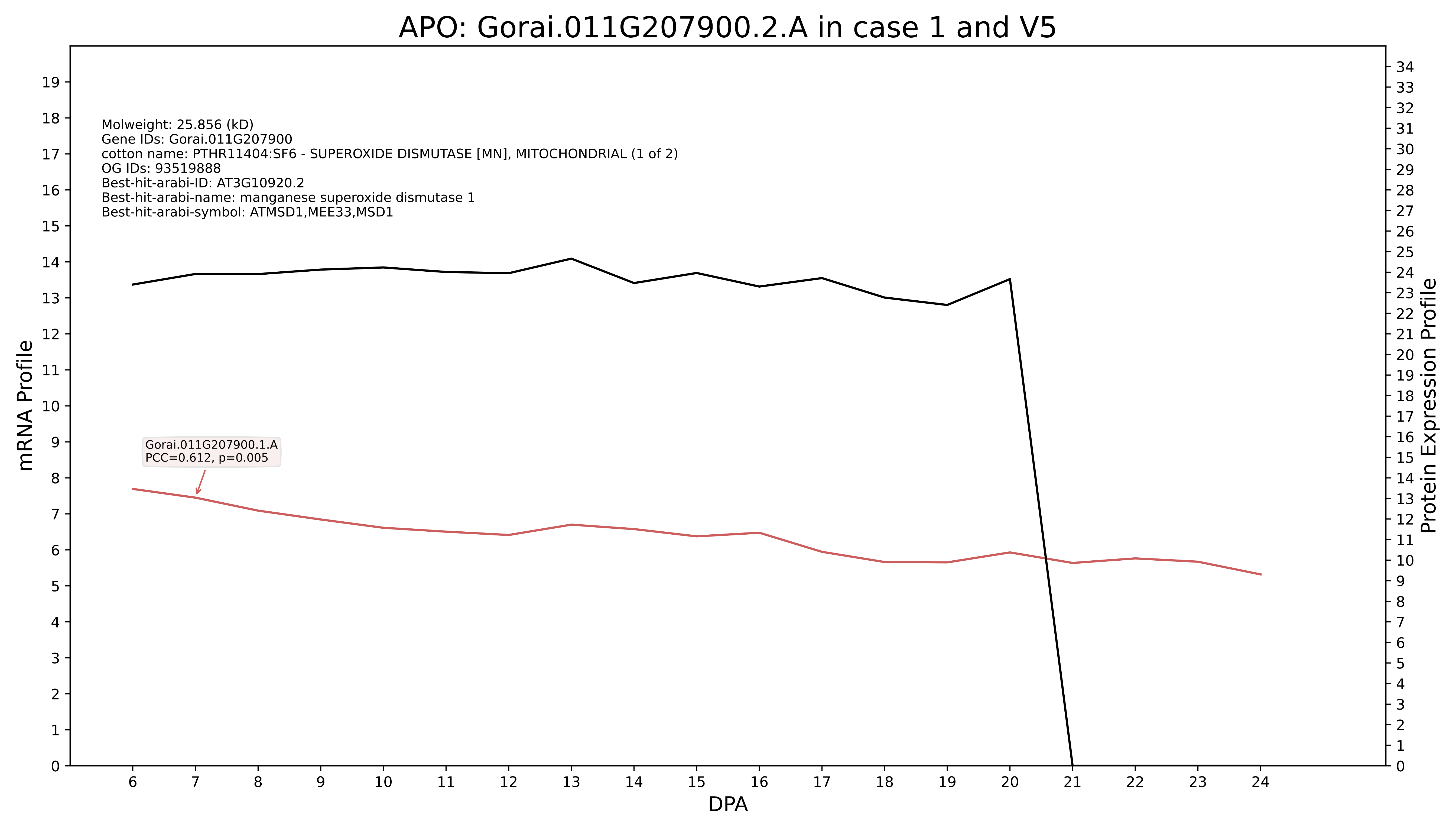
APO: Gorai.011G207900.2.A in case 1 and V5
Molweight: 25.856 (kD)
Gene IDs: Gorai.011G207900
cotton name: PTHR11404:SF6 - SUPEROXIDE DISMUTASE [MN], MITOCHONDRIAL (1 of 2)
OG IDs: 93519888
Best-hit-arabi-ID: AT3G10920.2
Best-hit-arabi-name: manganese superoxide dismutase 1
Best-hit-arabi-symbol: ATMSD1,MEE33,MSD1
Gorai.011G207900.1.A
PCC=0.612, p=0.005
mRNA Profile
Protein Expression Profile
DPA

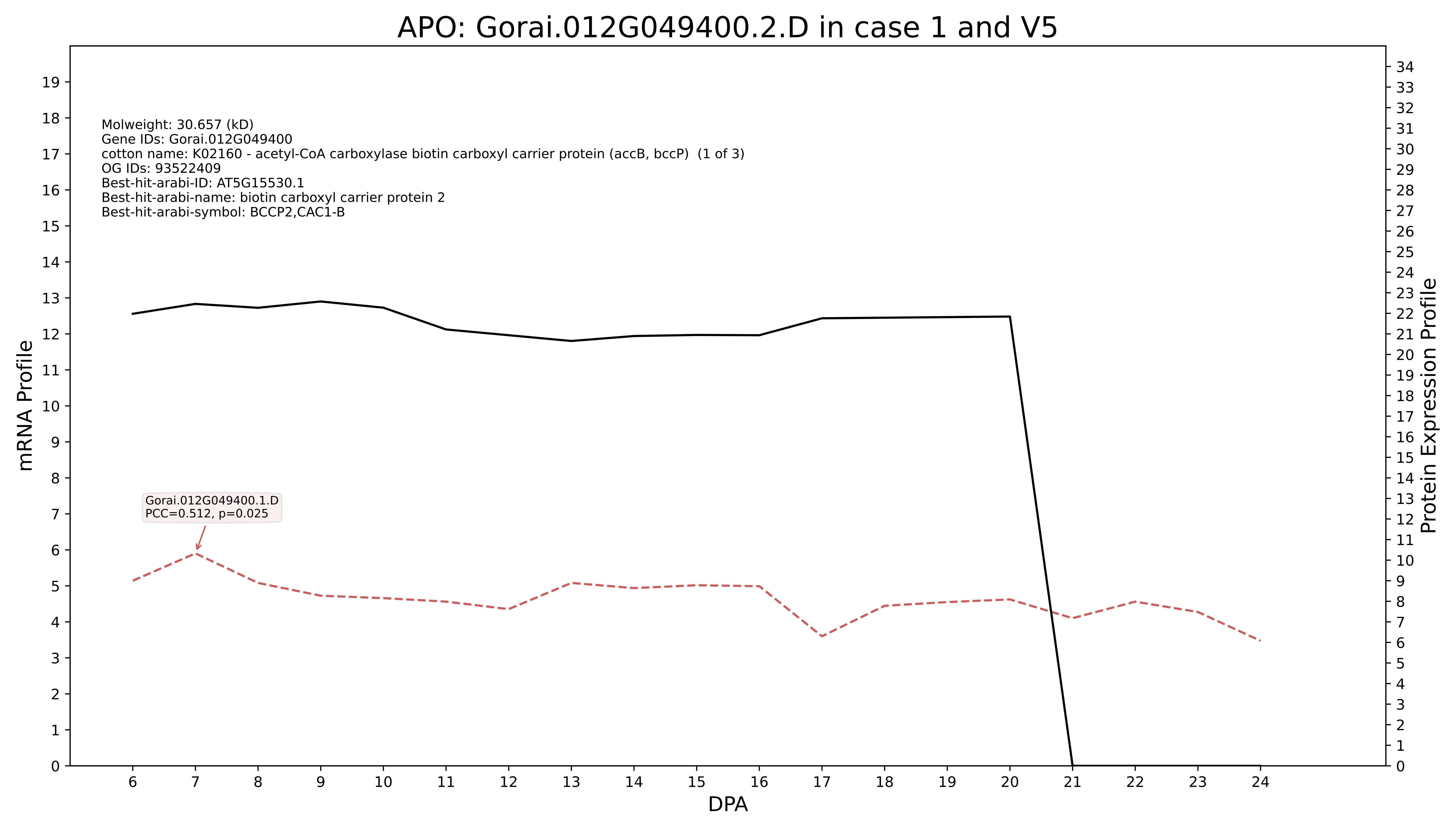

APO: Gorai.012G049400.2.D in case 1 and V5
Molweight: 30.657 (kD)
Gene IDs: Gorai.012G049400
cotton name: K02160 - acetyl-CoA carboxylase biotin carboxyl carrier protein (accB, bccP) (1 of 3)
OG IDs: 93522409
Best-hit-arabi-ID: AT5G15530.1
Best-hit-arabi-name: biotin carboxyl carrier protein 2
Best-hit-arabi-symbol: BCCP2,CAC1-B
Gorai.012G049400.1.D
PCC=0.512, p=0.025
mRNA Profile
Protein Expression Profile
DPA

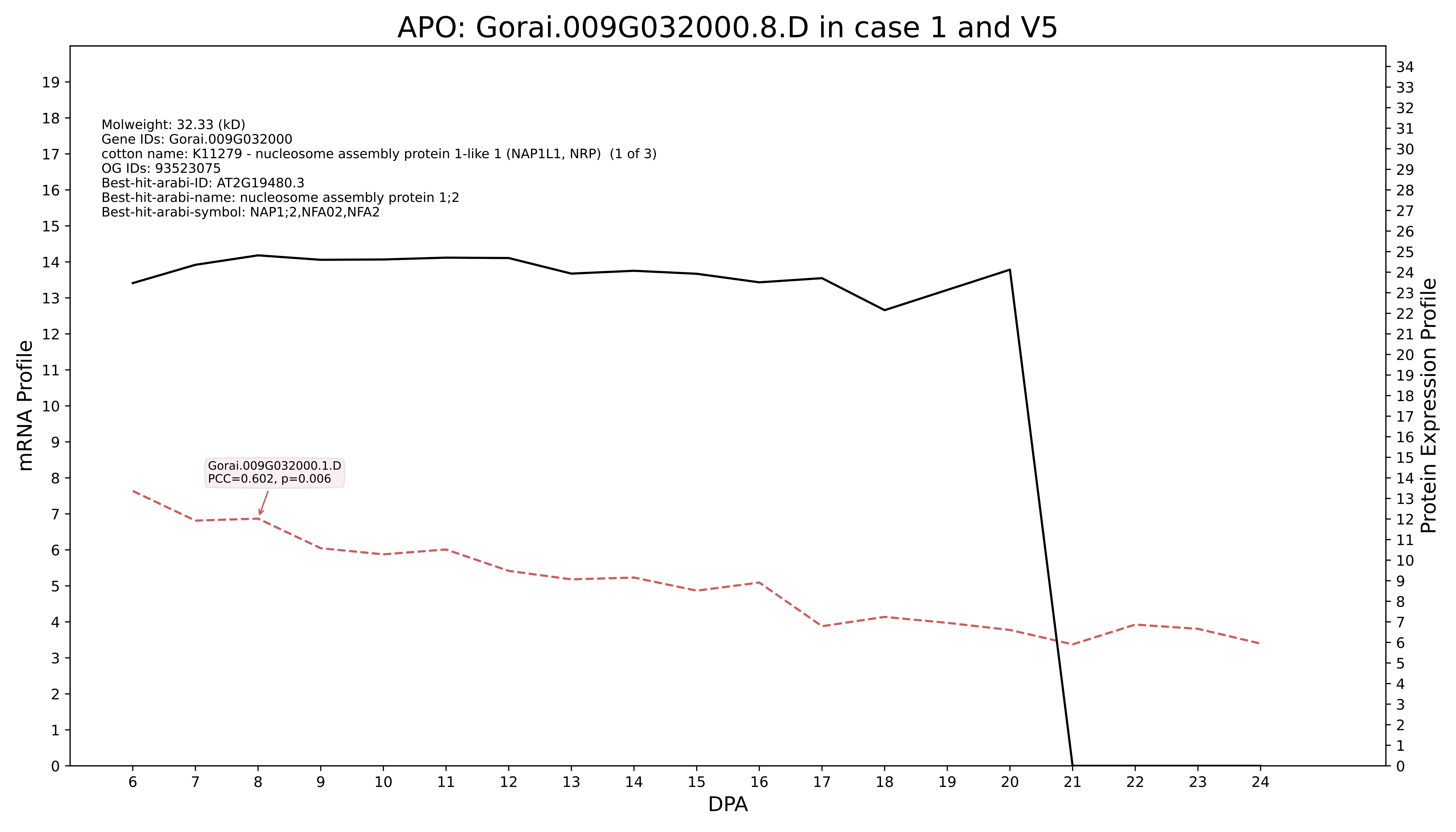
APO: Gorai.009G032000.8.D in case 1 and V5
Molweight: 32.33 (kD)
Gene IDs: Gorai.009G032000
cotton name: K11279 - nucleosome assembly protein 1-like 1 (NAP1L1, NRP) (1 of 3)
OG IDs: 93523075
Best-hit-arabi-ID: AT2G19480.3
Best-hit-arabi-name: nucleosome assembly protein 1;2
Best-hit-arabi-symbol: NAP1;2,NFA02,NFA2
Gorai.009G032000.1.D
PCC=0.602, p=0.006
mRNA Profile
Protein Expression Profile
DPA

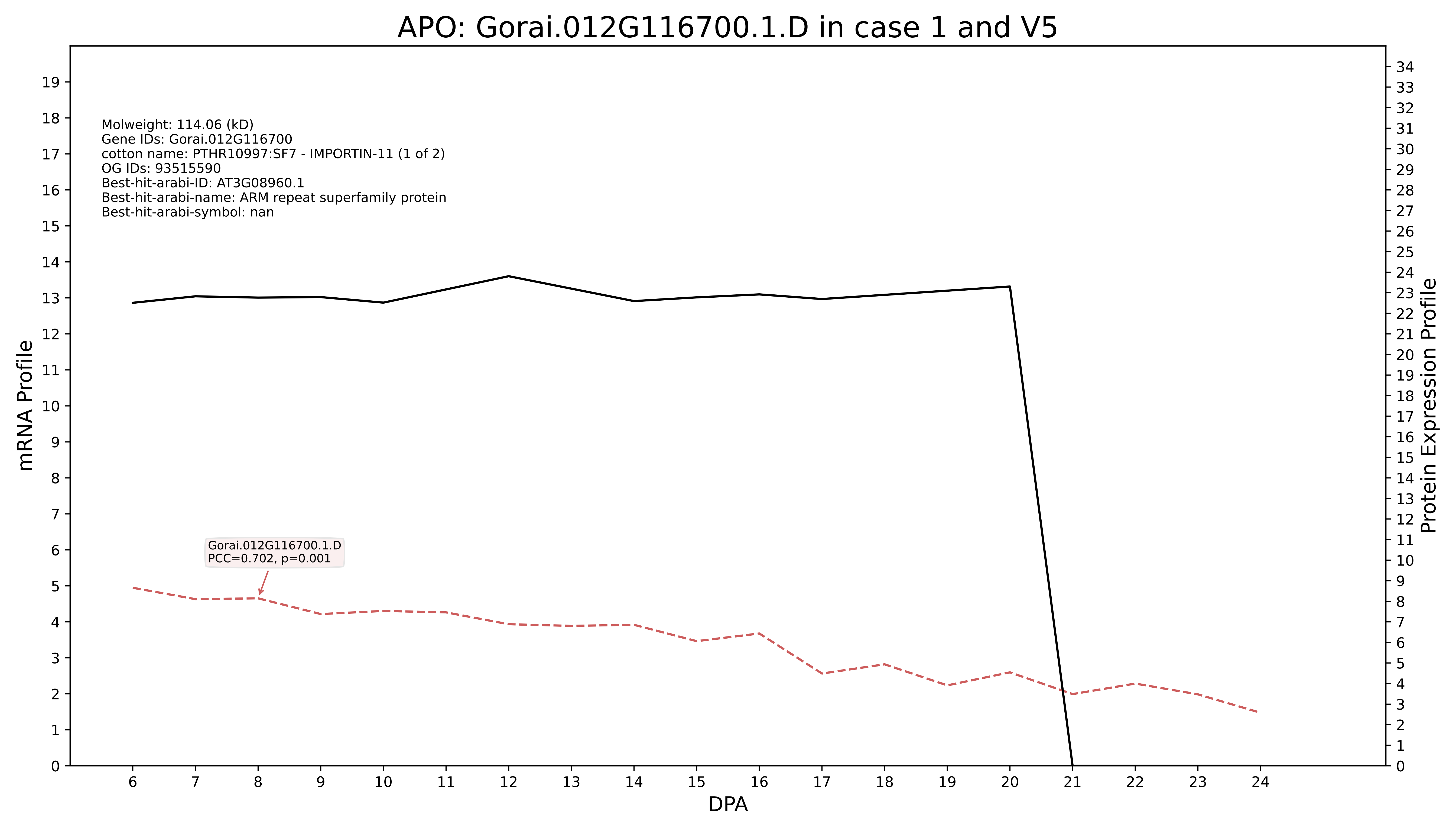
APO: Gorai.012G116700.1.D in case 1 and V5
Molweight: 114.06 (kD)
Gene IDs: Gorai.012G116700
cotton name: PTHR10997:SF7 - IMPORTIN-11 (1 of 2)
OG IDs: 93515590
Best-hit-arabi-ID: AT3G08960.1
Best-hit-arabi-name: ARM repeat superfamily protein
Best-hit-arabi-symbol: nan
Gorai.012G116700.1.D
PCC=0.702, p=0.001
mRNA Profile
Protein Expression Profile
DPA

APO: Gorai.011G076800.1.D in case 1 and V5

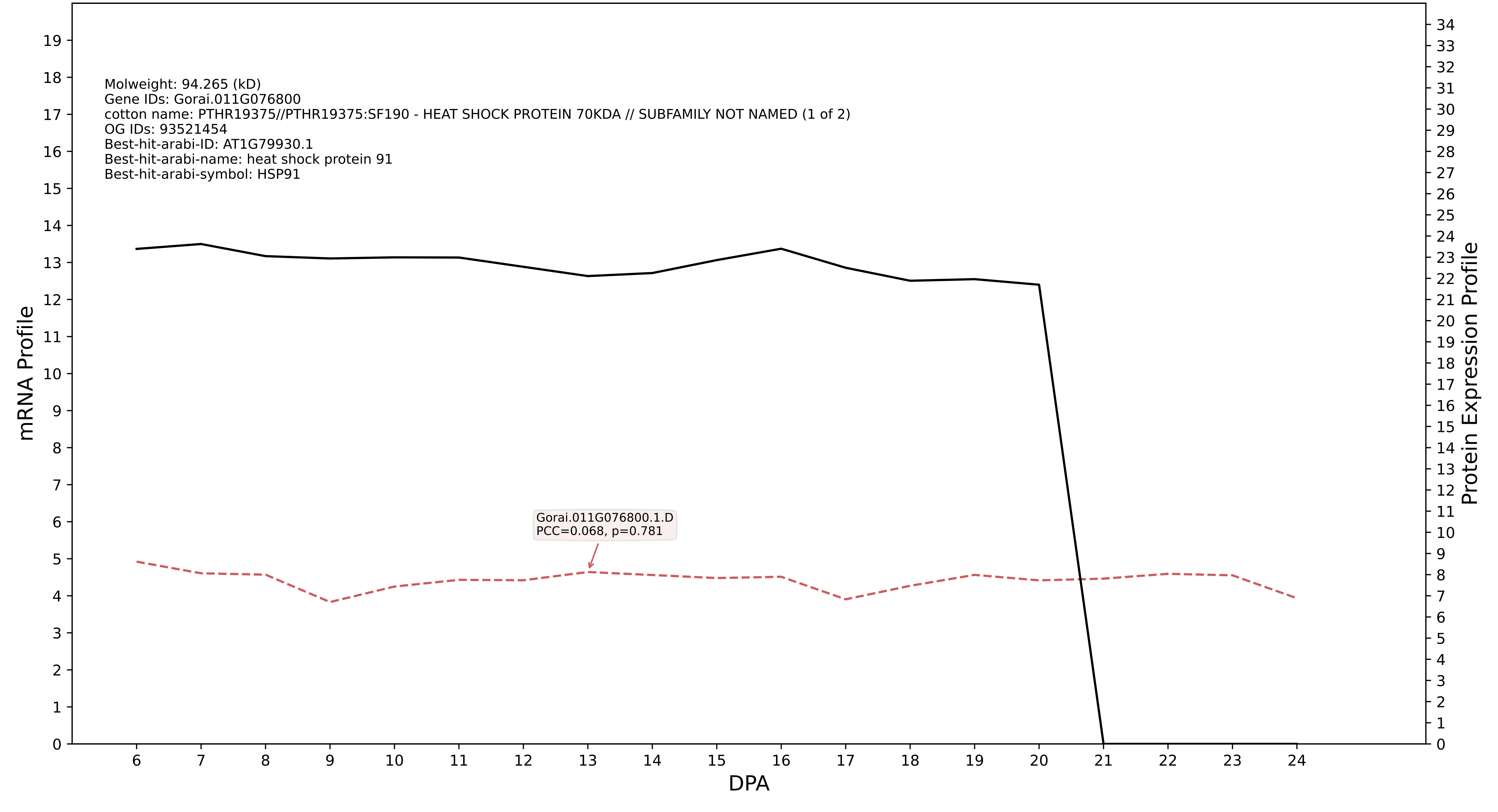

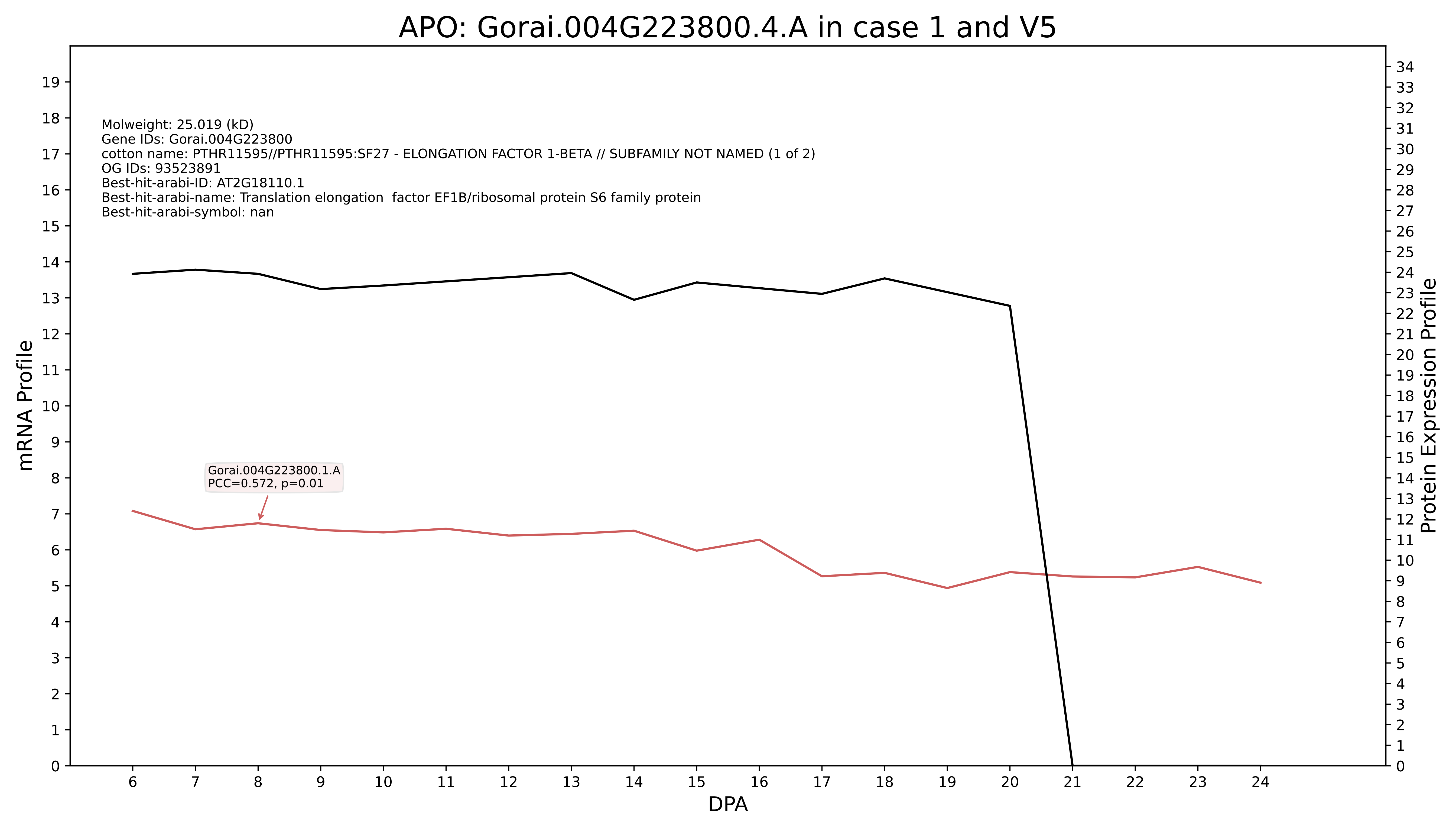

APO: Gorai.004G223800.4.A in case 1 and V5
Molweight: 25.019 (kD)
Gene IDs: Gorai.004G223800
cotton name: PTHR11595//PTHR11595:SF27 - ELONGATION FACTOR 1-BETA // SUBFAMILY NOT NAMED (1 of 2)
OG IDs: 93523891
Best-hit-arabi-ID: AT2G18110.1
Best-hit-arabi-name: Translation elongation factor EF1B/ribosomal protein S6 family protein
Best-hit-arabi-symbol: nan
Gorai.004G223800.1.A
PCC=0.572, p=0.01
mRNA Profile
Protein Expression Profile
DPA

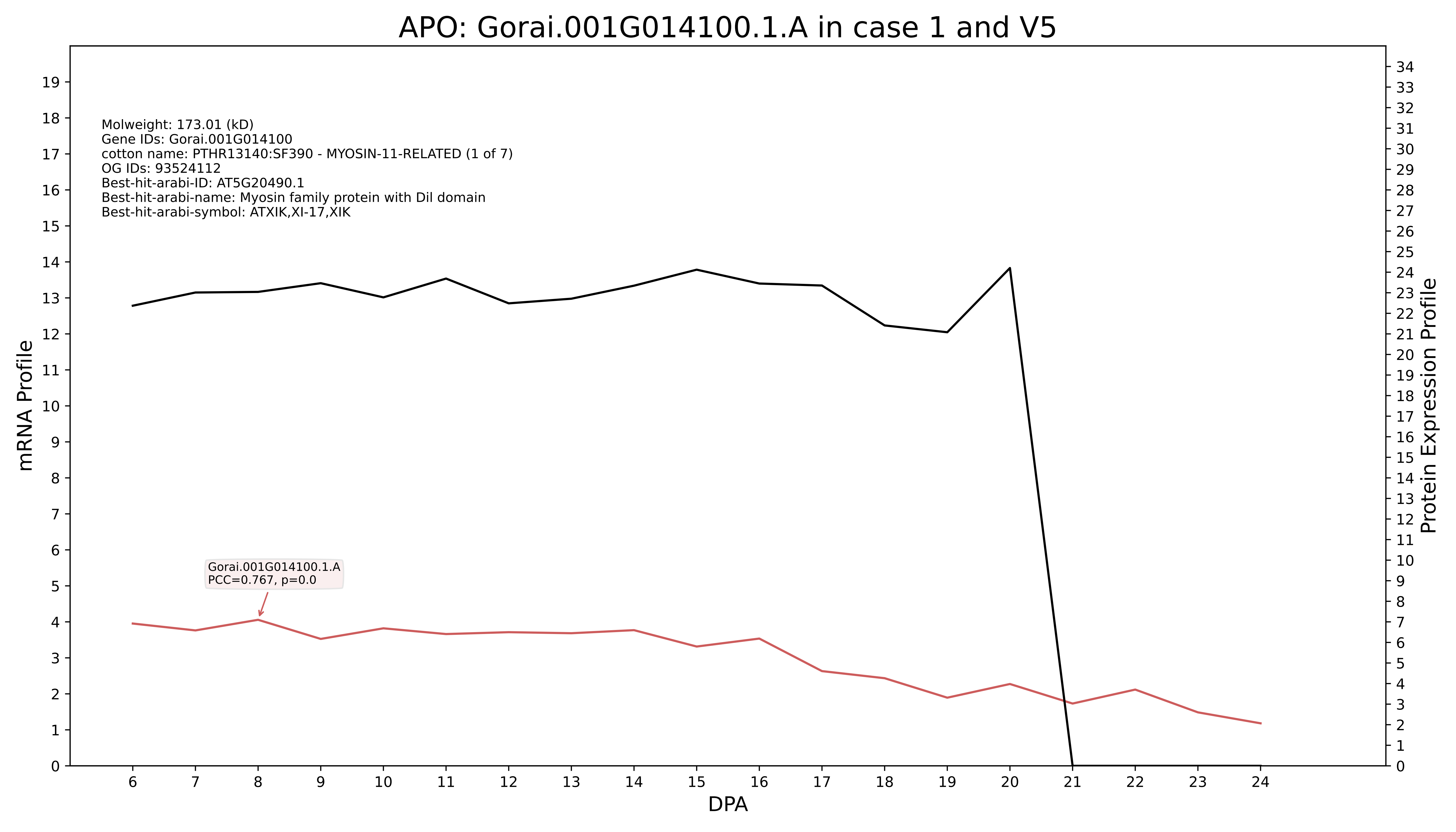
APO: Gorai.001G014100.1.A in case 1 and V5
Molweight: 173.01 (kD)
Gene IDs: Gorai.001G014100
cotton name: PTHR13140:SF390 - MYOSIN-11-RELATED (1 of 7)
OG IDs: 93524112
Best-hit-arabi-ID: AT5G20490.1
Best-hit-arabi-name: Myosin family protein with Dil domain
Best-hit-arabi-symbol: ATXIK,XI-17,XIK
Gorai.001G014100.1.A
PCC=0.767, p=0.0
mRNA Profile
Protein Expression Profile
DPA

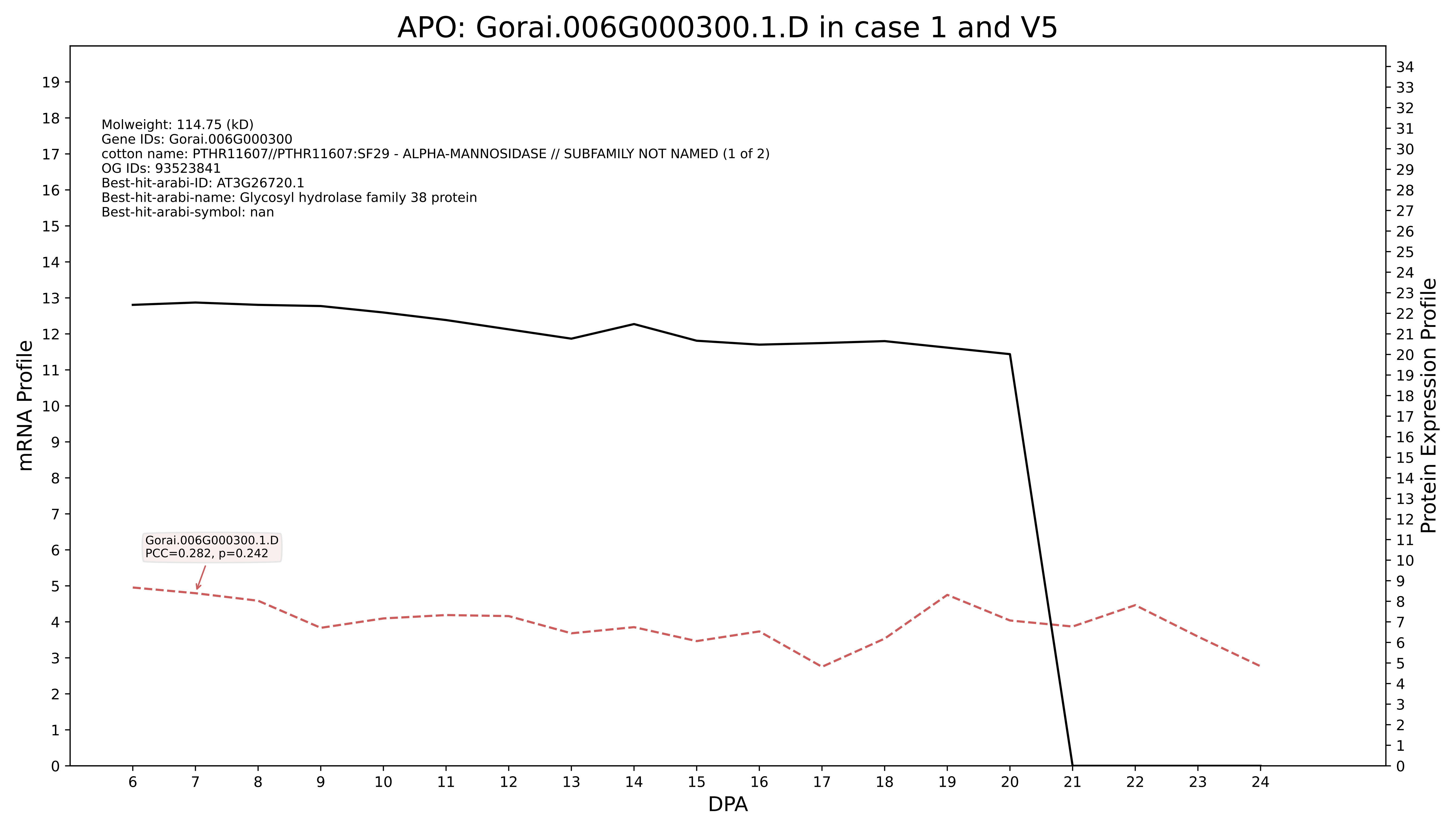

APO: Gorai.006G000300.1.D in case 1 and V5
Molweight: 114.75 (kD)
Gene IDs: Gorai.006G000300
cotton name: PTHR11607//PTHR11607:SF29 - ALPHA-MANNOSIDASE // SUBFAMILY NOT NAMED (1 of 2)
OG IDs: 93523841
Best-hit-arabi-ID: AT3G26720.1
Best-hit-arabi-name: Glycosyl hydrolase family 38 protein
Best-hit-arabi-symbol: nan
Gorai.006G000300.1.D
PCC=0.282, p=0.242
mRNA Profile
Protein Expression Profile
DPA

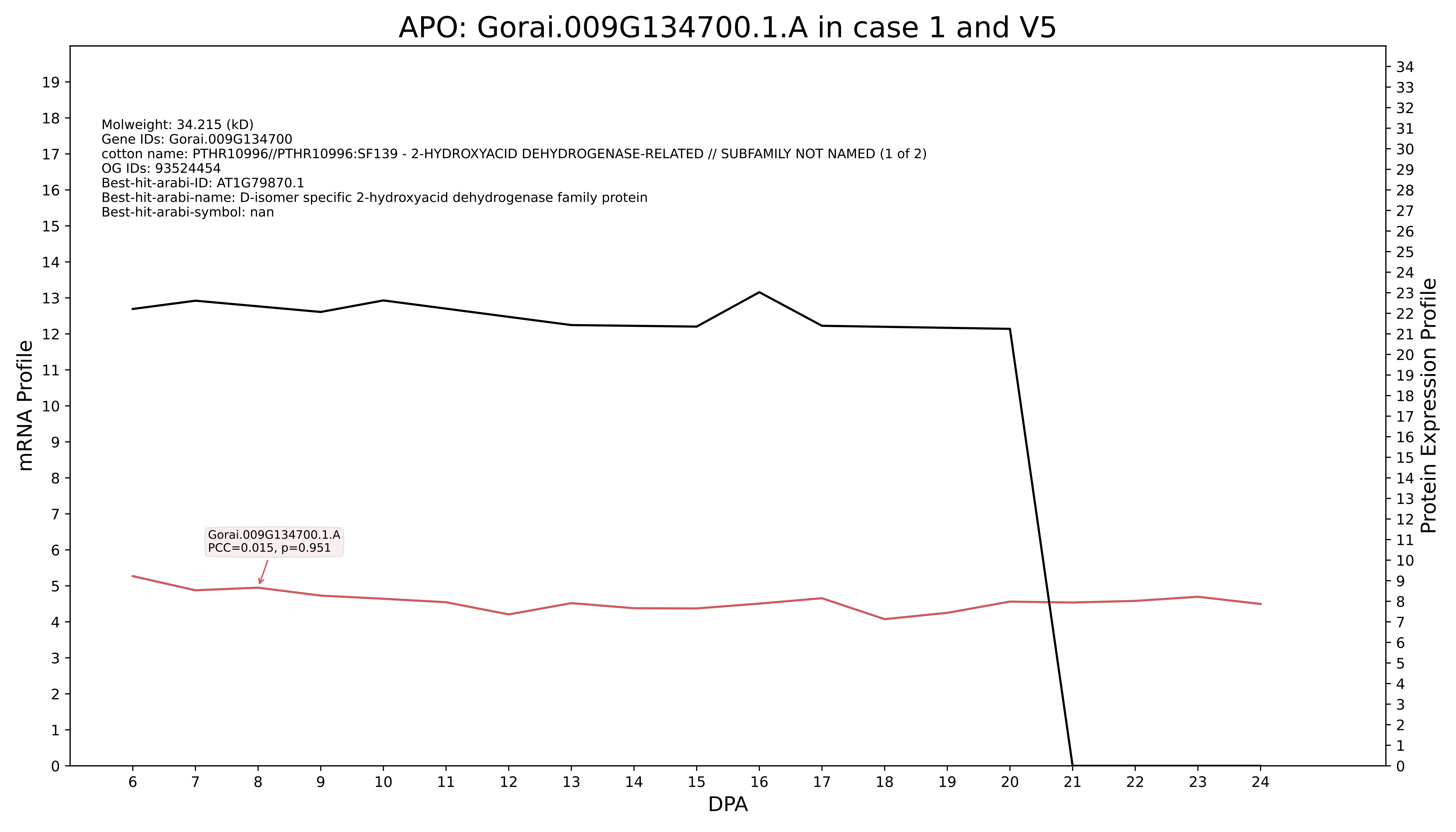

APO: Gorai.009G134700.1.A in case 1 and V5
Molweight: 34.215 (kD)
Gene IDs: Gorai.009G134700
cotton name: PTHR10996//PTHR10996:SF139 - 2-HYDROXYACID DEHYDROGENASE-RELATED // SUBFAMILY NOT NAMED (1 of 2)
OG IDs: 93524454
Best-hit-arabi-ID: AT1G79870.1
Best-hit-arabi-name: D-isomer specific 2-hydroxyacid dehydrogenase family protein
Best-hit-arabi-symbol: nan
Gorai.009G134700.1.A
PCC=0.015, p=0.951
mRNA Profile
Protein Expression Profile
DPA

### APO: Gorai.011G011600.1.D in case 1 and V5

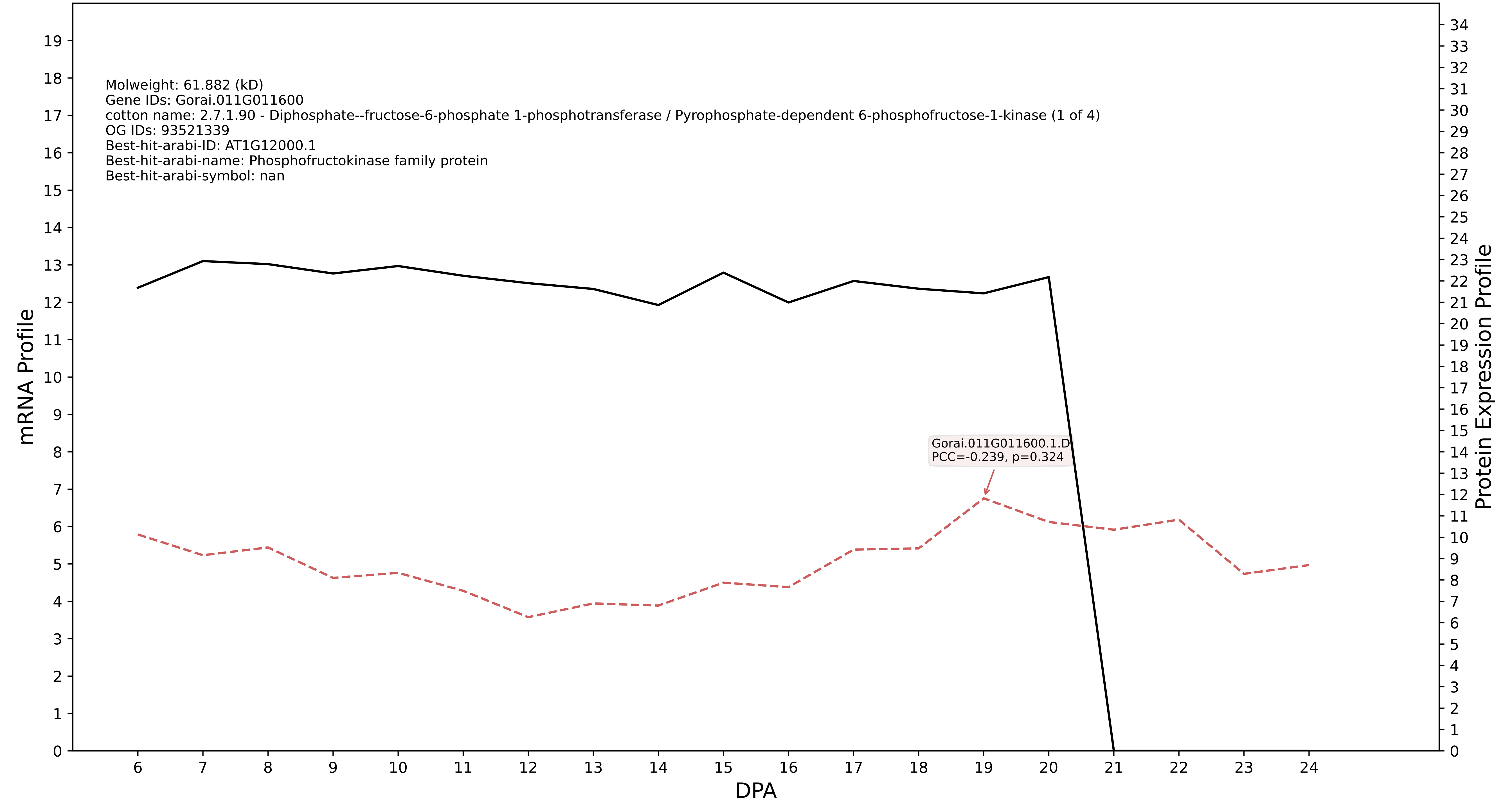

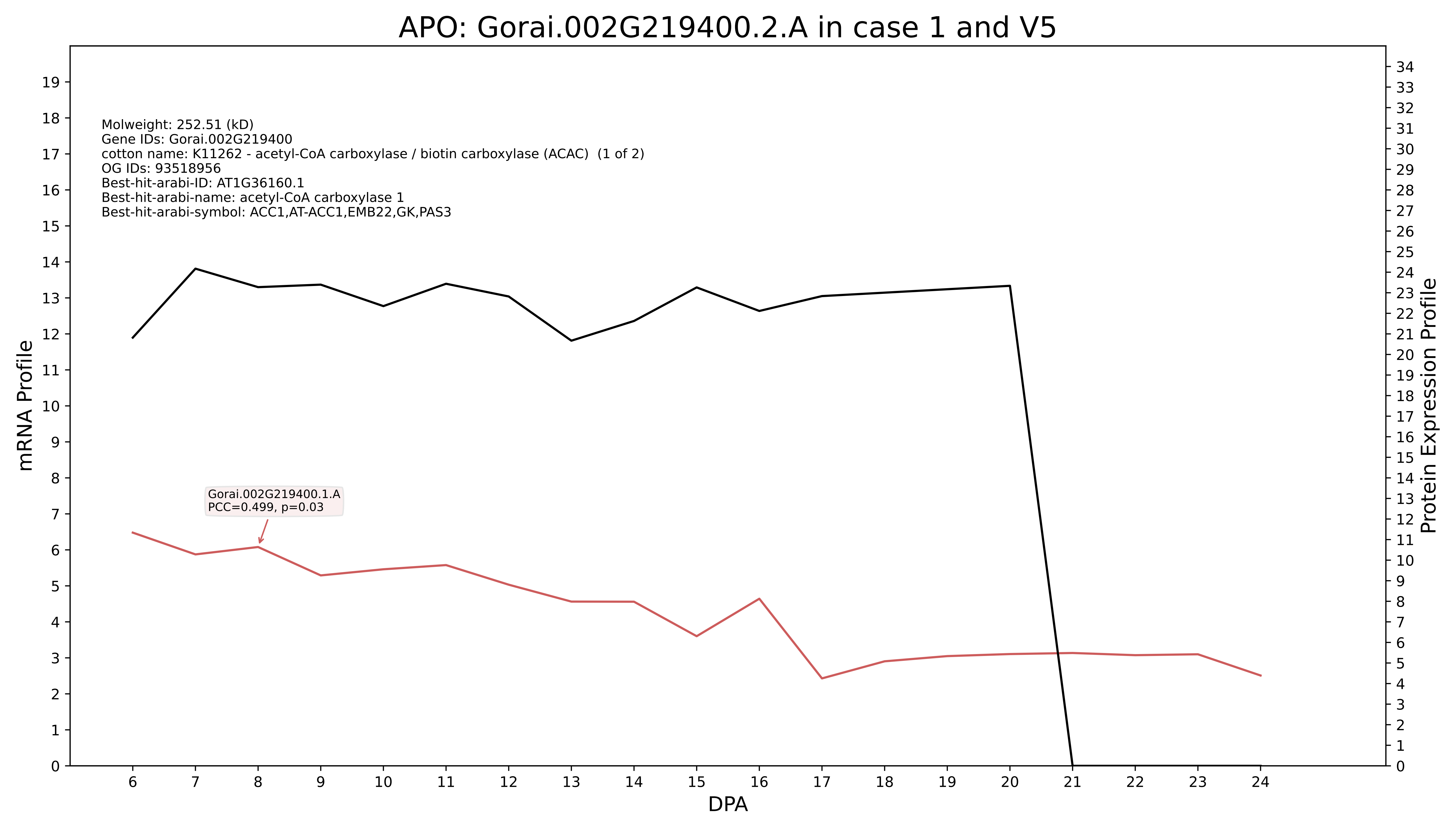
APO: Gorai.002G219400.2.A in case 1 and V5
Molweight: 252.51 (kD)
Gene IDs: Gorai.002G219400
cotton name: K11262 - acetyl-CoA carboxylase / biotin carboxylase (ACAC) (1 of 2)
OG IDs: 93518956
Best-hit-arabi-ID: AT1G36160.1
Best-hit-arabi-name: acetyl-CoA carboxylase 1
Best-hit-arabi-symbol: ACC1,AT-ACC1,EMB22,GK,PAS3
Gorai.002G219400.1.A
PCC=0.499, p=0.03
mRNA Profile
Protein Expression Profile
DPA

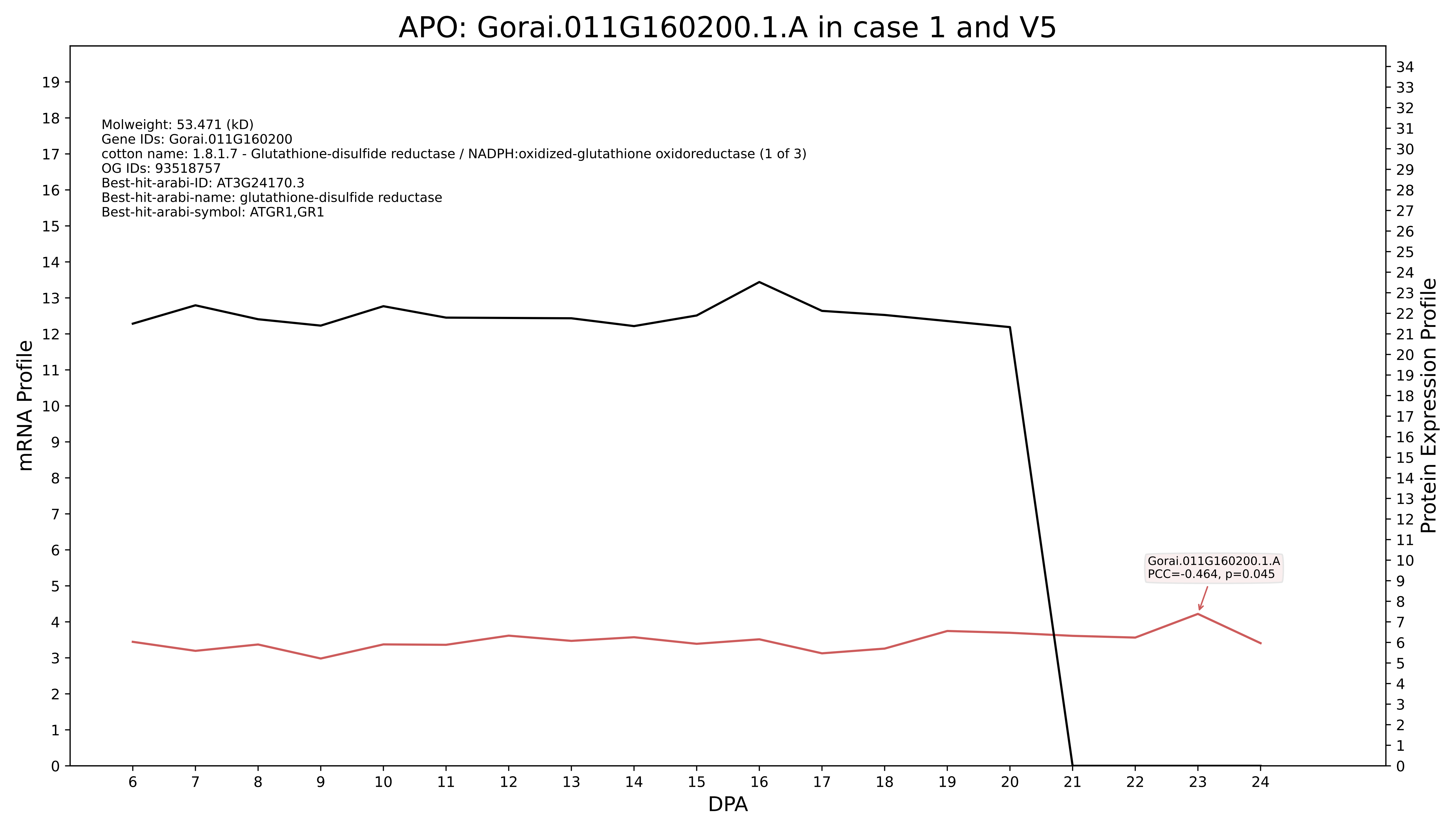
APO: Gorai.011G160200.1.A in case 1 and V5
Molweight: 53.471 (kD)
Gene IDs: Gorai.011G160200
cotton name: 1.8.1.7 - Glutathione-disulfide reductase / NADPH:oxidized-glutathione oxidoreductase (1 of 3)
OG IDs: 93518757
Best-hit-arabi-ID: AT3G24170.3
Best-hit-arabi-name: glutathione-disulfide reductase
Best-hit-arabi-symbol: ATGR1,GR1
Gorai.011G160200.1.A
PCC=-0.464, p=0.045
mRNA Profile
Protein Expression Profile
DPA

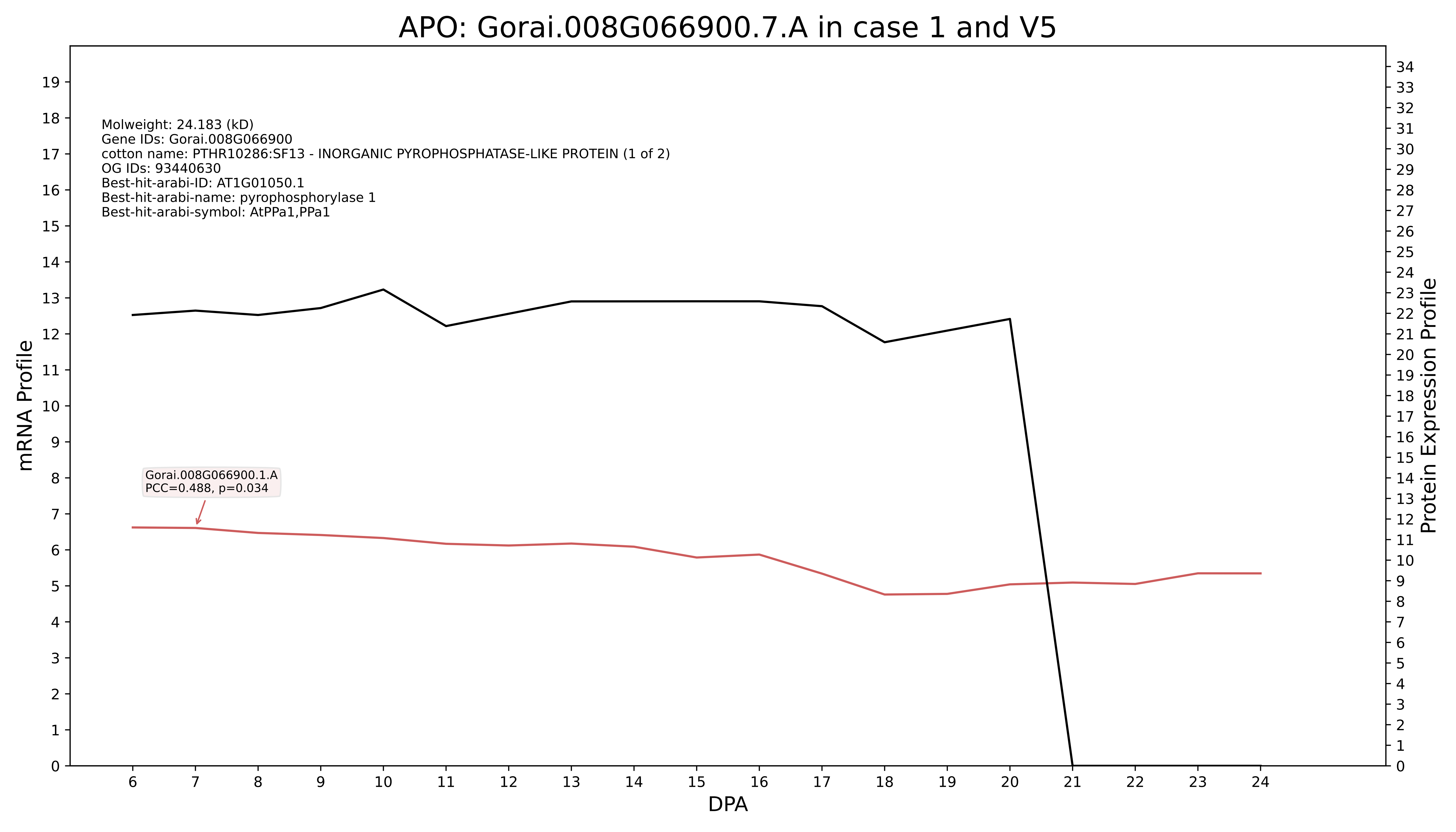

APO: Gorai.008G066900.7.A in case 1 and V5
Molweight: 24.183 (kD)
Gene IDs: Gorai.008G066900
cotton name: PTHR10286:SF13 - INORGANIC PYROPHOSPHATASE-LIKE PROTEIN (1 of 2)
OG IDs: 93440630
Best-hit-arabi-ID: AT1G01050.1
Best-hit-arabi-name: pyrophosphorylase 1
Best-hit-arabi-symbol: AtPPa1,PPa1
Gorai.008G066900.1.A
PCC=0.488, p=0.034
mRNA Profile
Protein Expression Profile
DPA

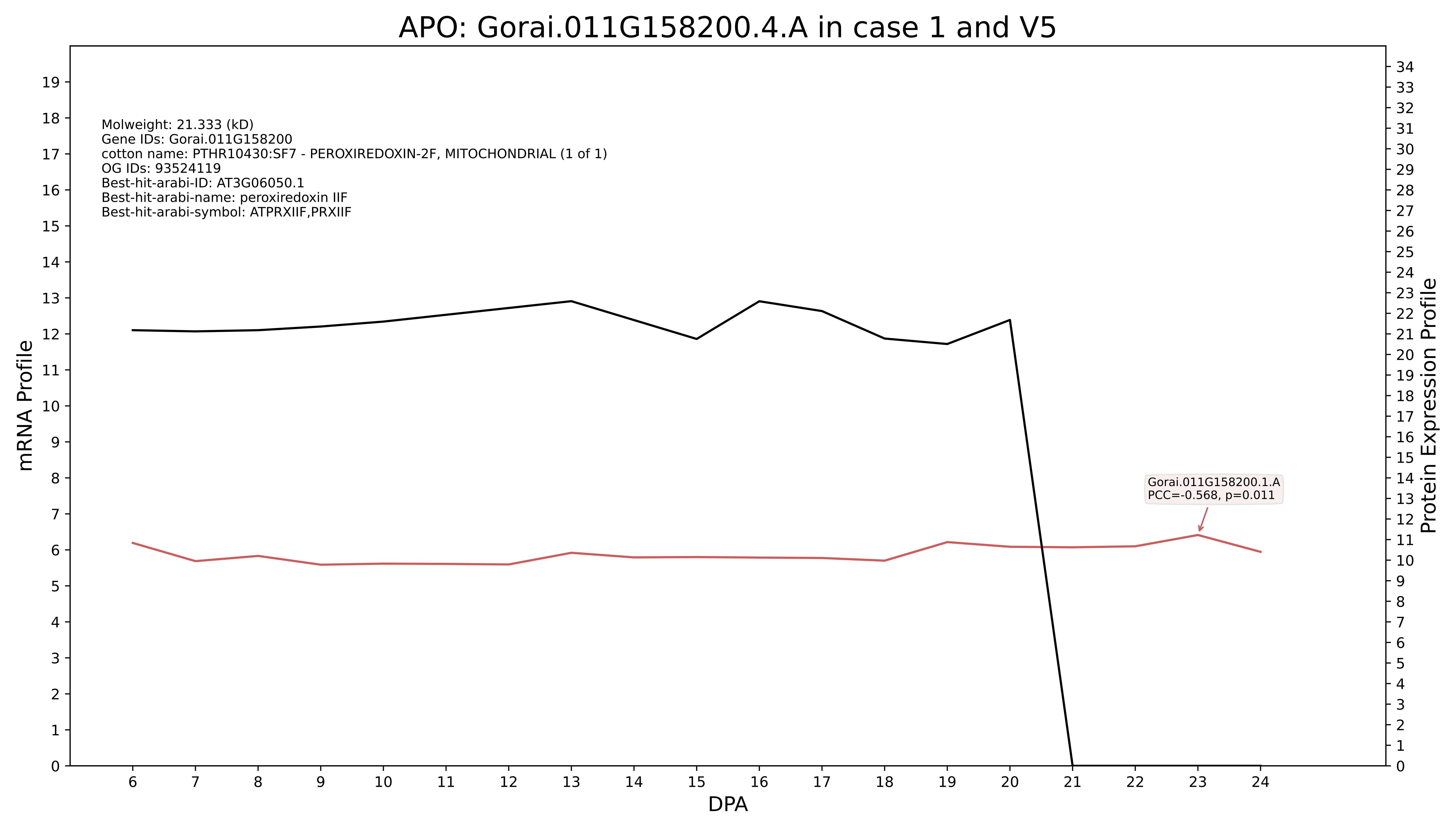

APO: Gorai.011G158200.4.A in case 1 and V5
Molweight: 21.333 (kD)
Gene IDs: Gorai.011G158200
cotton name: PTHR10430:SF7 - PEROXIREDOXIN-2F, MITOCHONDRIAL (1 of 1)
OG IDs: 93524119
Best-hit-arabi-ID: AT3G06050.1
Best-hit-arabi-name: peroxiredoxin IIF
Best-hit-arabi-symbol: ATPRXIIF,PRXIIF
Gorai.011G158200.1.A
PCC=-0.568, p=0.011
mRNA Profile
Protein Expression Profile
DPA

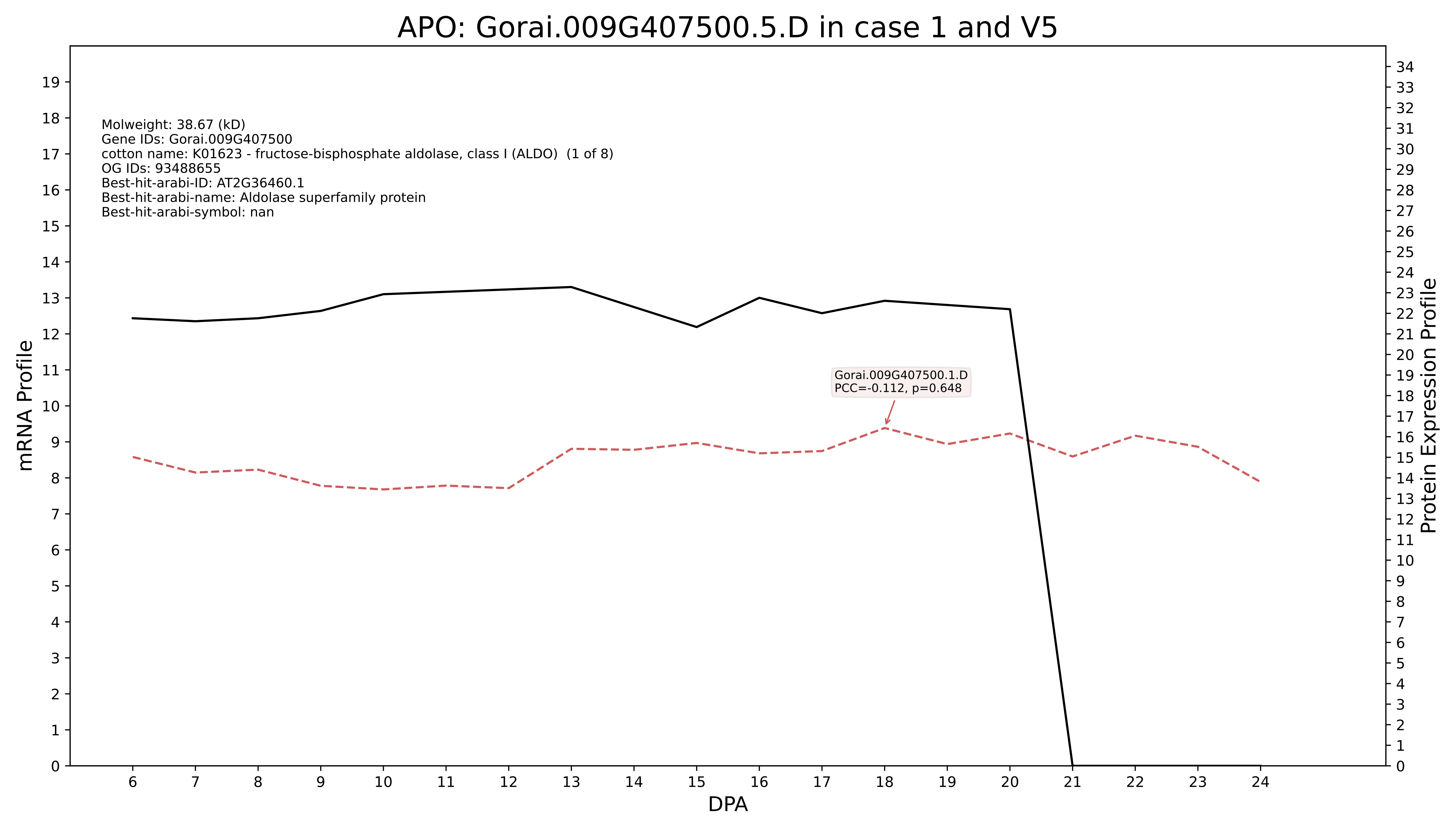
APO: Gorai.009G407500.5.D in case 1 and V5
Molweight: 38.67 (kD)
Gene IDs: Gorai.009G407500
cotton name: K01623 - fructose-bisphosphate aldolase, class I (ALDO) (1 of 8)
OG IDs: 93488655
Best-hit-arabi-ID: AT2G36460.1
Best-hit-arabi-name: Aldolase superfamily protein
Best-hit-arabi-symbol: nan
Gorai.009G407500.1.D
PCC=-0.112, p=0.648
mRNA Profile
Protein Expression Profile
DPA

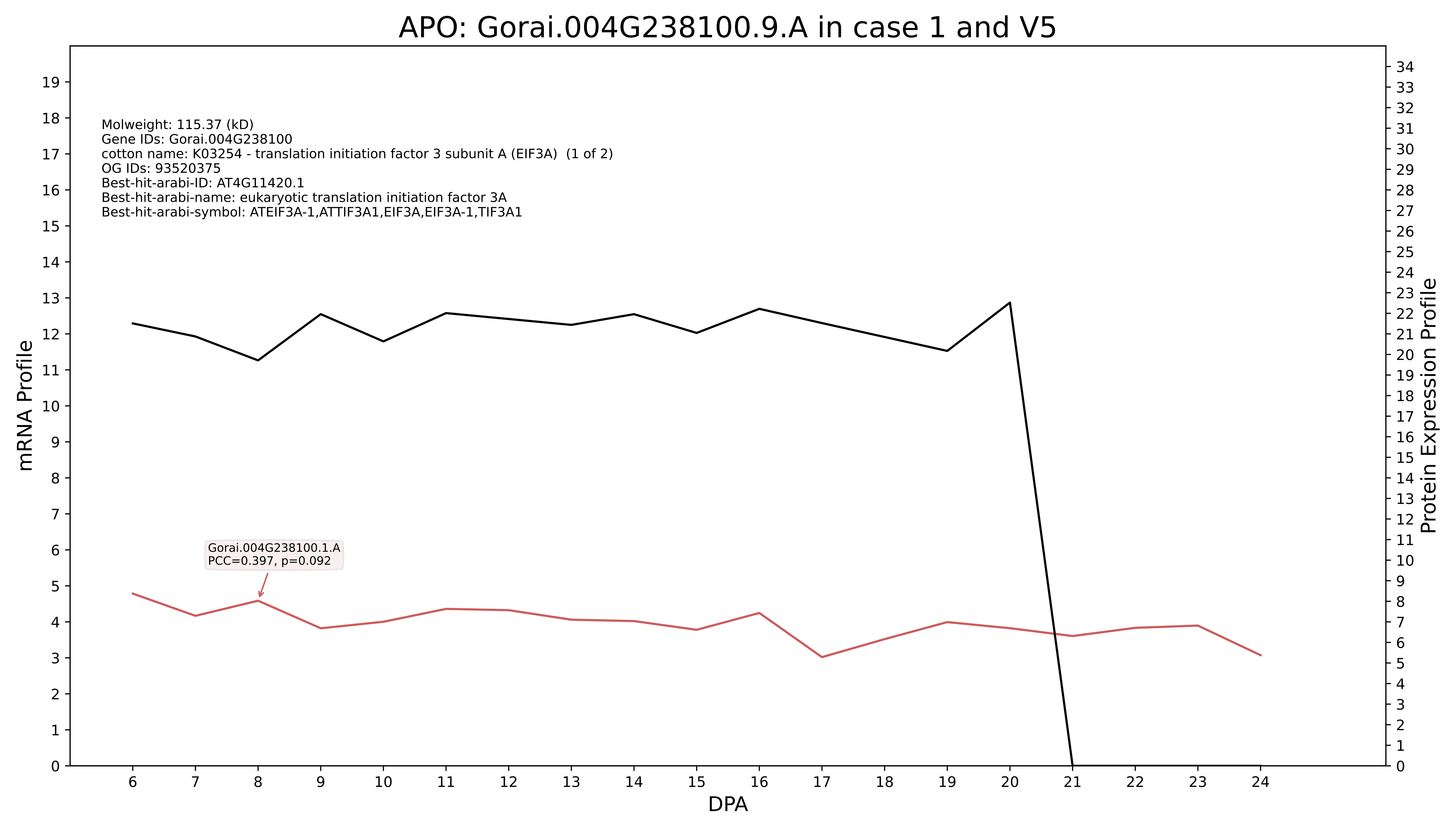

APO: Gorai.004G238100.9.A in case 1 and V5
Molweight: 115.37 (kD)
Gene IDs: Gorai.004G238100
cotton name: K03254 - translation initiation factor 3 subunit A (EIF3A) (1 of 2)
OG IDs: 93520375
Best-hit-arabi-ID: AT4G11420.1
Best-hit-arabi-name: eukaryotic translation initiation factor 3A
Best-hit-arabi-symbol: ATEIF3A-1,ATTIF3A1,EIF3A,EIF3A-1,TIF3A1
Gorai.004G238100.1.A
PCC=0.397, p=0.092
mRNA Profile
Protein Expression Profile
DPA

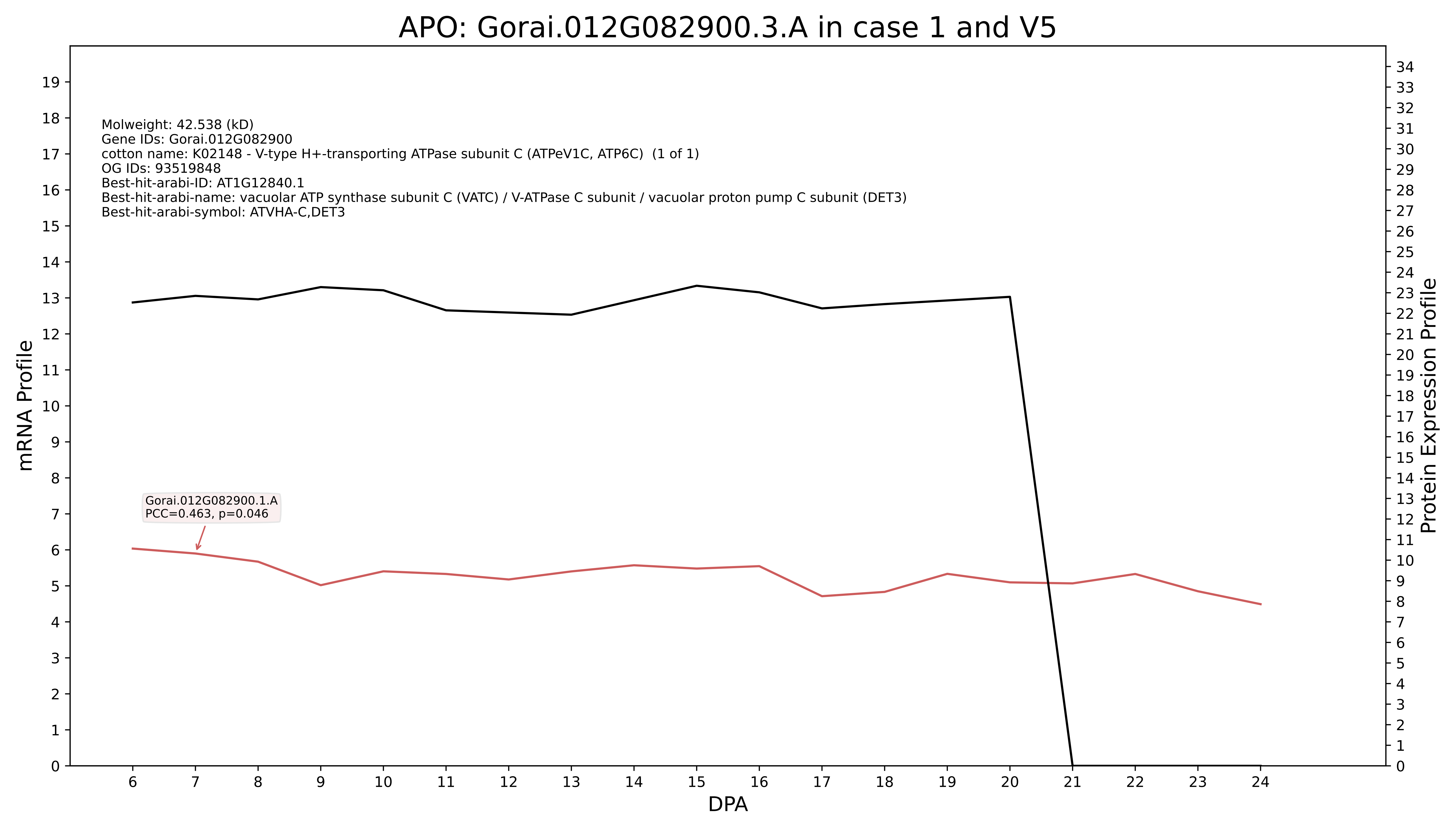
APO: Gorai.012G082900.3.A in case 1 and V5
Molweight: 42.538 (kD)
Gene IDs: Gorai.012G082900
cotton name: K02148 - V-type H+-transporting ATPase subunit C (ATPeV1C, ATP6C) (1 of 1)
OG IDs: 93519848
Best-hit-arabi-ID: AT1G12840.1
Best-hit-arabi-name: vacuolar ATP synthase subunit C (VATC) / V-ATPase C subunit / vacuolar proton pump C subunit (DET3)
Best-hit-arabi-symbol: ATVHA-C,DET3
Gorai.012G082900.1.A
PCC=0.463, p=0.046
mRNA Profile
Protein Expression Profile
DPA

### APO: Gorai.009G347600.1.A in case 1 and V5

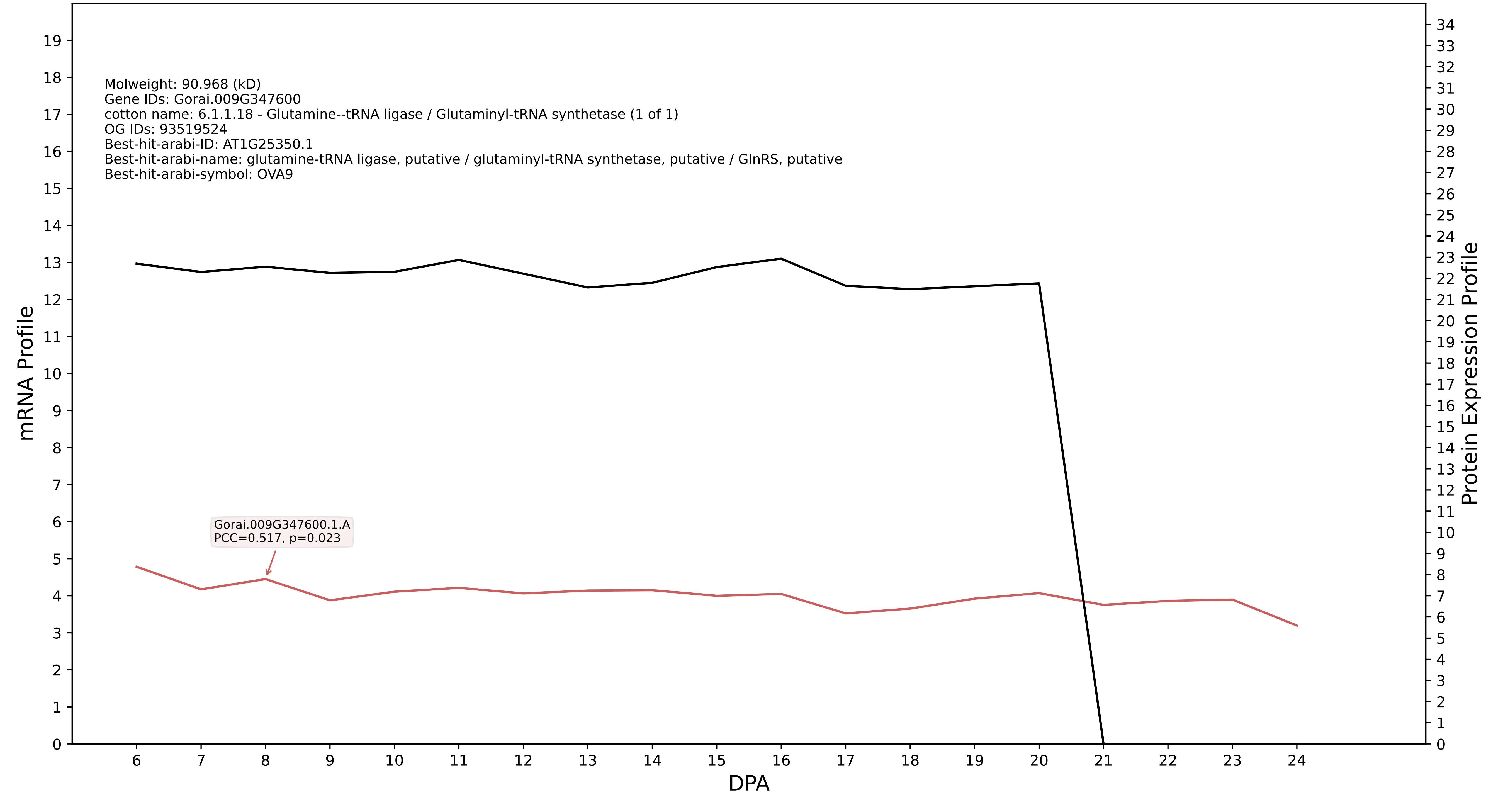

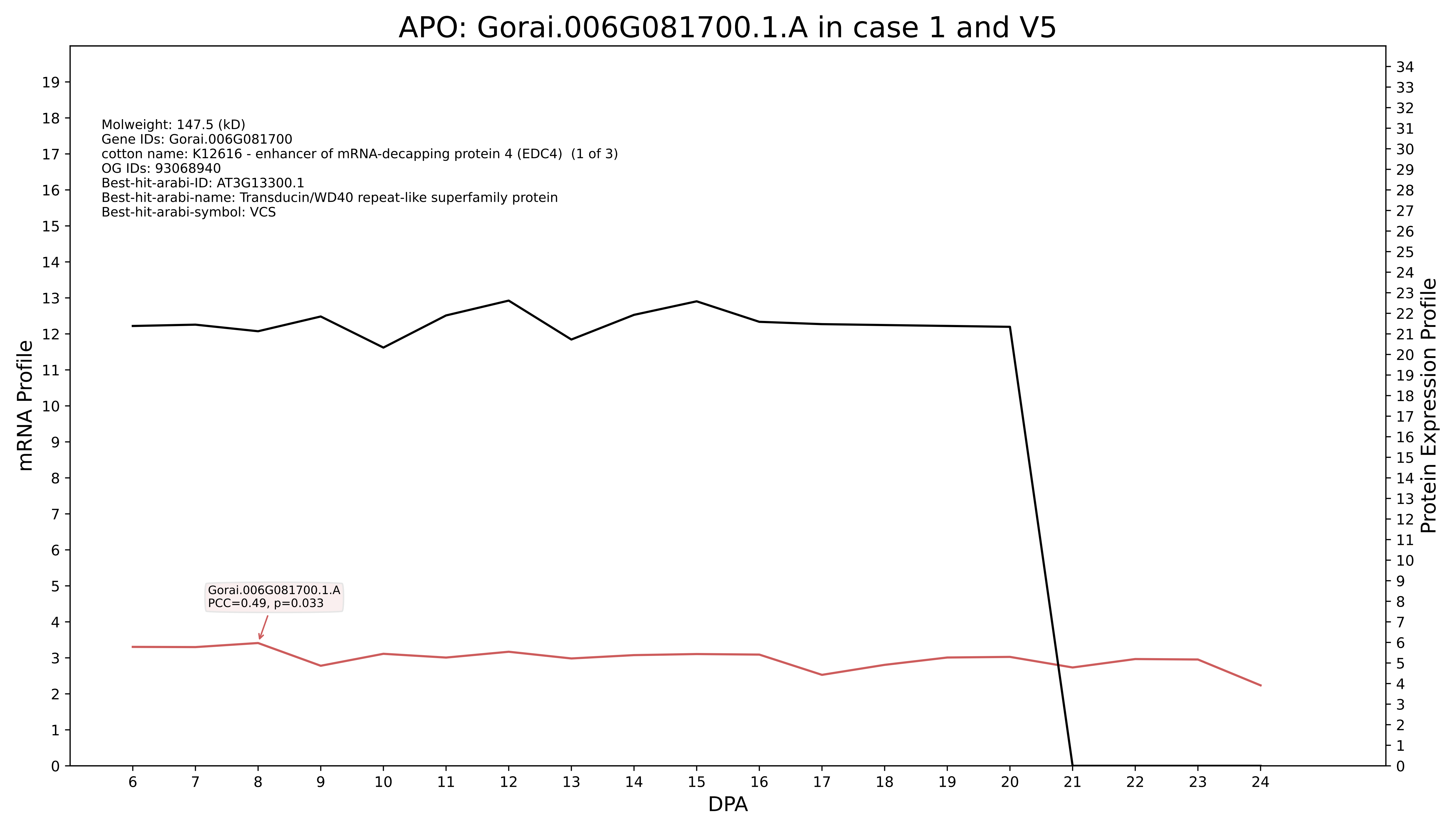

APO: Gorai.006G081700.1.A in case 1 and V5
Molweight: 147.5 (kD)
Gene IDs: Gorai.006G081700
cotton name: K12616 - enhancer of mRNA-decapping protein 4 (EDC4) (1 of 3)
OG IDs: 93068940
Best-hit-arabi-ID: AT3G13300.1
Best-hit-arabi-name: Transducin/WD40 repeat-like superfamily protein
Best-hit-arabi-symbol: VCS
Gorai.006G081700.1.A
PCC=0.49, p=0.033
mRNA Profile
Protein Expression Profile
DPA
6 7 8 9 10 11 12 13 14 15 16 17 18 19 20 21 22 23 24

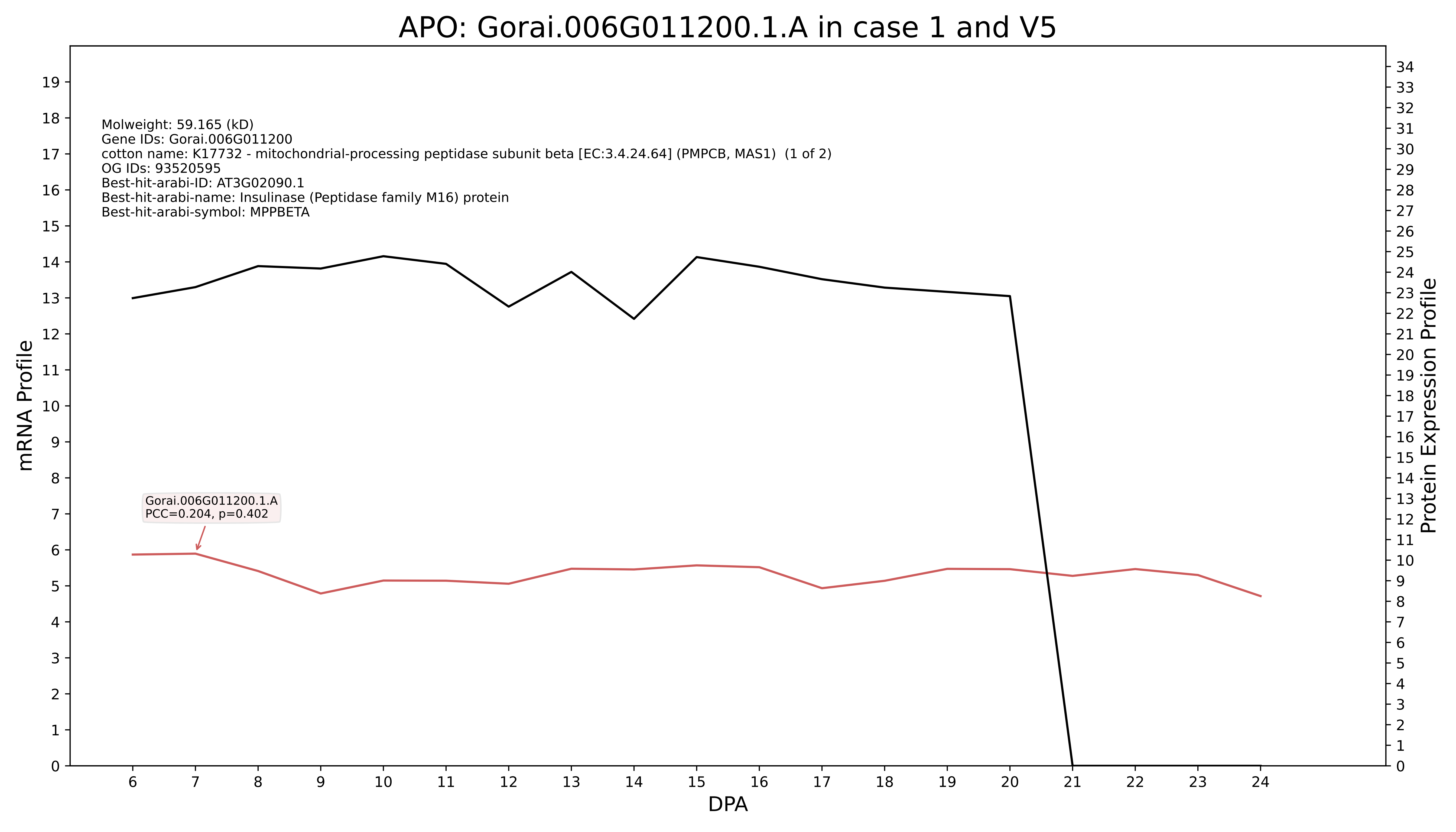
APO: Gorai.006G011200.1.A in case 1 and V5
Molweight: 59.165 (kD)
Gene IDs: Gorai.006G011200
cotton name: K17732 - mitochondrial-processing peptidase subunit beta [EC:3.4.24.64] (PMPCB, MAS1) (1 of 2)
OG IDs: 93520595
Best-hit-arabi-ID: AT3G02090.1
Best-hit-arabi-name: Insulinase (Peptidase family M16) protein
Best-hit-arabi-symbol: MPPBETA
Gorai.006G011200.1.A
PCC=0.204, p=0.402
mRNA Profile
Protein Expression Profile
DPA

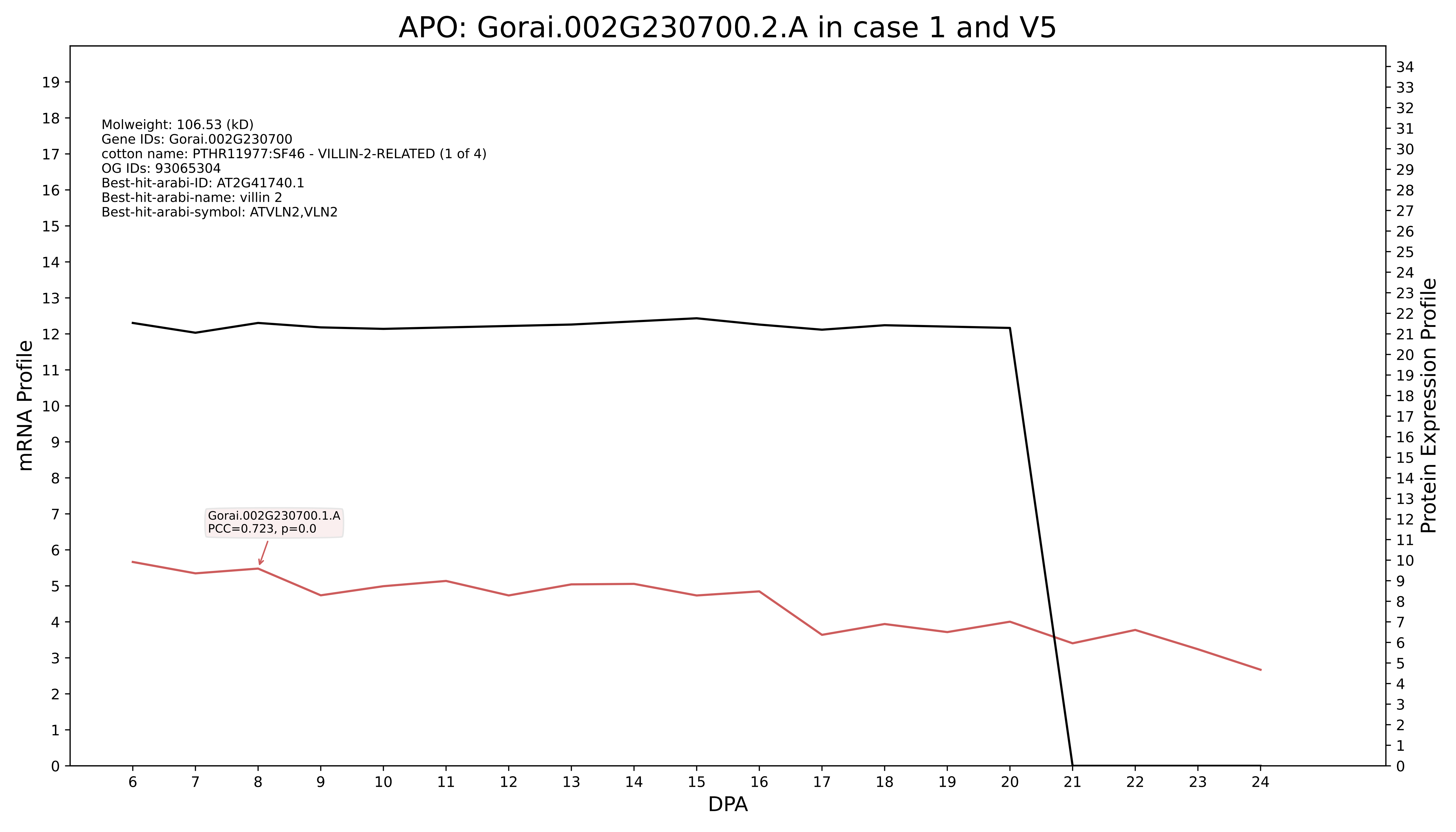

APO: Gorai.002G230700.2.A in case 1 and V5
Molweight: 106.53 (kD)
Gene IDs: Gorai.002G230700
cotton name: PTHR11977:SF46 - VILLIN-2-RELATED (1 of 4)
OG IDs: 93065304
Best-hit-arabi-ID: AT2G41740.1
Best-hit-arabi-name: villin 2
Best-hit-arabi-symbol: ATVLN2,VLN2
Gorai.002G230700.1.A
PCC=0.723, p=0.0
mRNA Profile
Protein Expression Profile
DPA

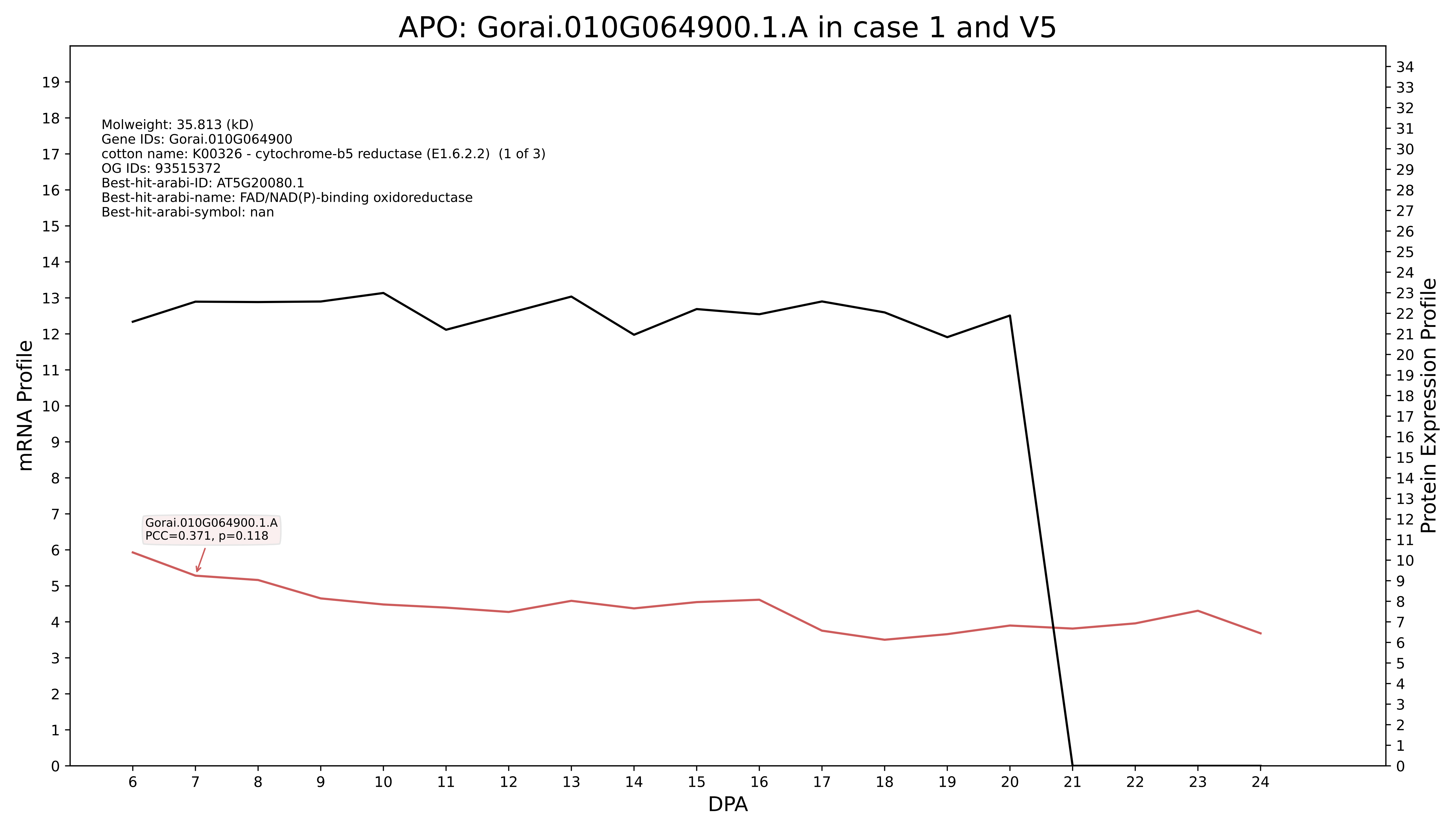

APO: Gorai.010G064900.1.A in case 1 and V5
Molweight: 35.813 (kD)
Gene IDs: Gorai.010G064900
cotton name: K00326 - cytochrome-b5 reductase (E1.6.2.2) (1 of 3)
OG IDs: 93515372
Best-hit-arabi-ID: AT5G20080.1
Best-hit-arabi-name: FAD/NAD(P)-binding oxidoreductase
Best-hit-arabi-symbol: nan
Gorai.010G064900.1.A
PCC=0.371, p=0.118
mRNA Profile
Protein Expression Profile
DPA

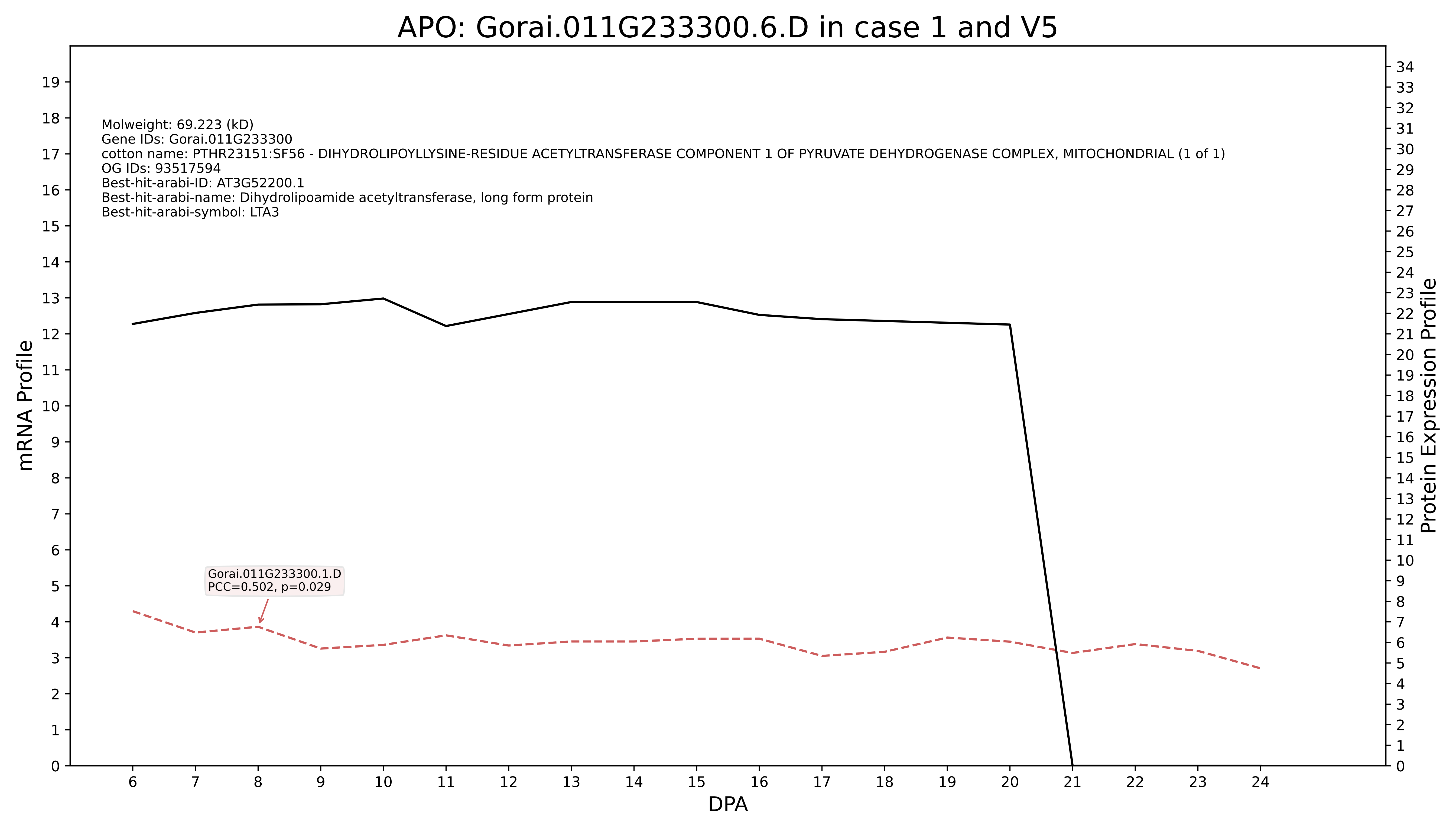

APO: Gorai.011G233300.6.D in case 1 and V5
Molweight: 69.223 (kD)
Gene IDs: Gorai.011G233300
cotton name: PTHR23151:SF56 - DIHYDROLIPOYLLYSINE-RESIDUE ACETYLTRANSFERASE COMPONENT 1 OF PYRUVATE DEHYDROGENASE COMPLEX, MITOCHONDRIAL (1 of 1)
OG IDs: 93517594
Best-hit-arabi-ID: AT3G52200.1
Best-hit-arabi-name: Dihydrolipoamide acetyltransferase, long form protein
Best-hit-arabi-symbol: LTA3
Gorai.011G233300.1.D
PCC=0.502, p=0.029
mRNA Profile
Protein Expression Profile
DPA

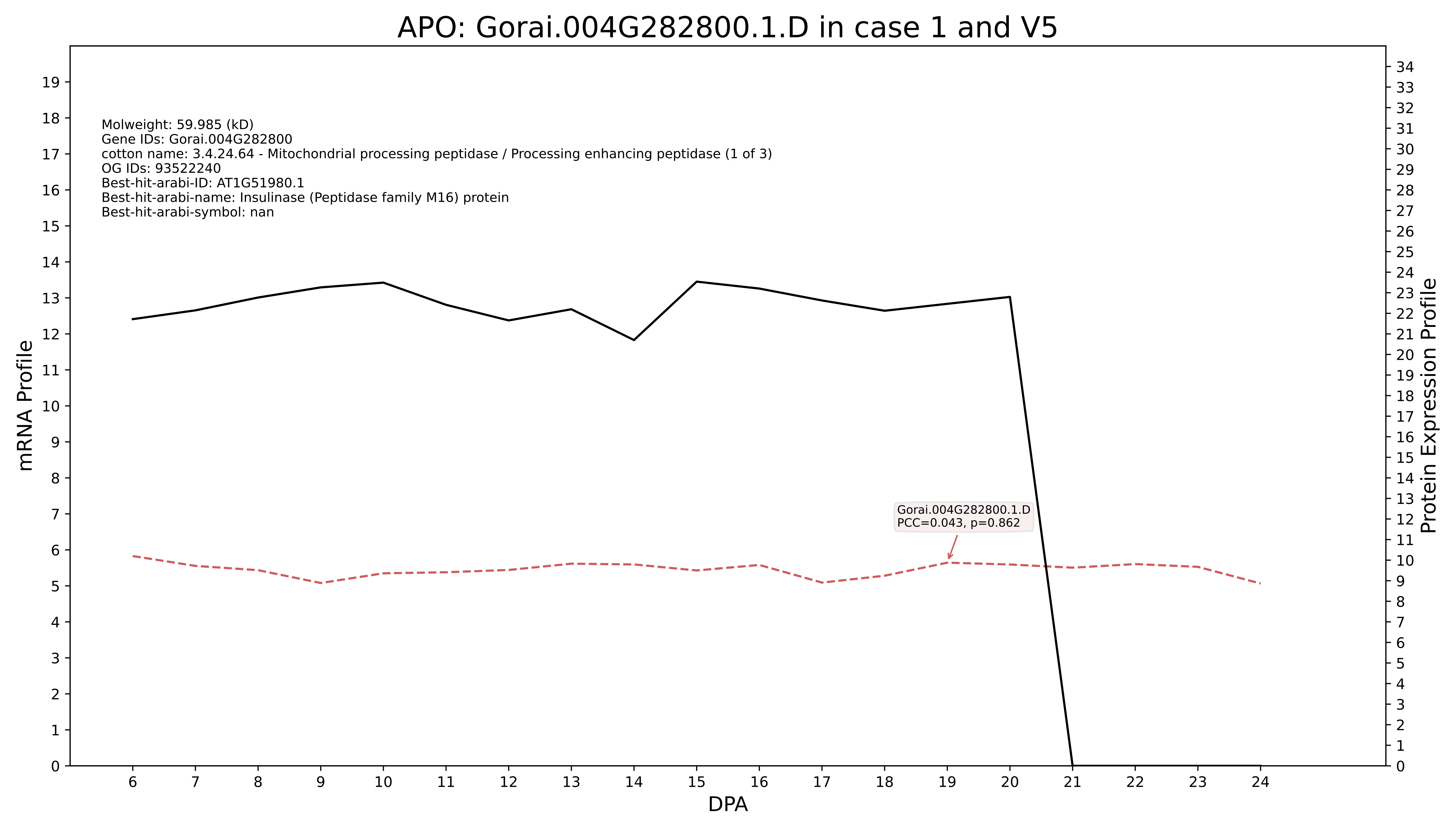
APO: Gorai.004G282800.1.D in case 1 and V5
Molweight: 59.985 (kD)
Gene IDs: Gorai.004G282800
cotton name: 3.4.24.64 - Mitochondrial processing peptidase / Processing enhancing peptidase (1 of 3)
OG IDs: 93522240
Best-hit-arabi-ID: AT1G51980.1
Best-hit-arabi-name: Insulinase (Peptidase family M16) protein
Best-hit-arabi-symbol: nan
Gorai.004G282800.1.D
PCC=0.043, p=0.862
mRNA Profile
Protein Expression Profile
DPA

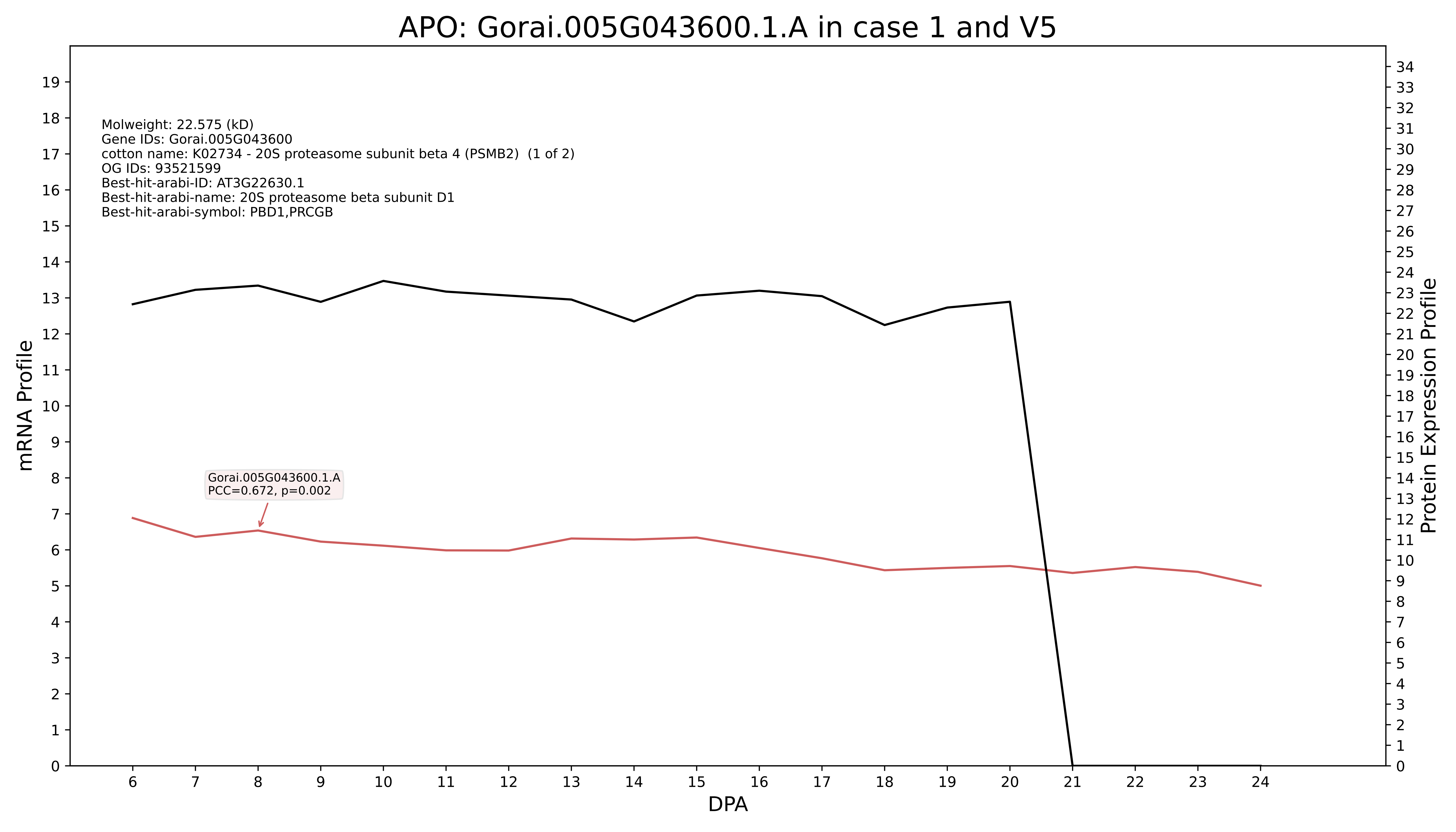
APO: Gorai.005G043600.1.A in case 1 and V5
Molweight: 22.575 (kD)
Gene IDs: Gorai.005G043600
cotton name: K02734 - 20S proteasome subunit beta 4 (PSMB2) (1 of 2)
OG IDs: 93521599
Best-hit-arabi-ID: AT3G22630.1
Best-hit-arabi-name: 20S proteasome beta subunit D1
Best-hit-arabi-symbol: PBD1,PRCGB
Gorai.005G043600.1.A
PCC=0.672, p=0.002
mRNA Profile
Protein Expression Profile
DPA

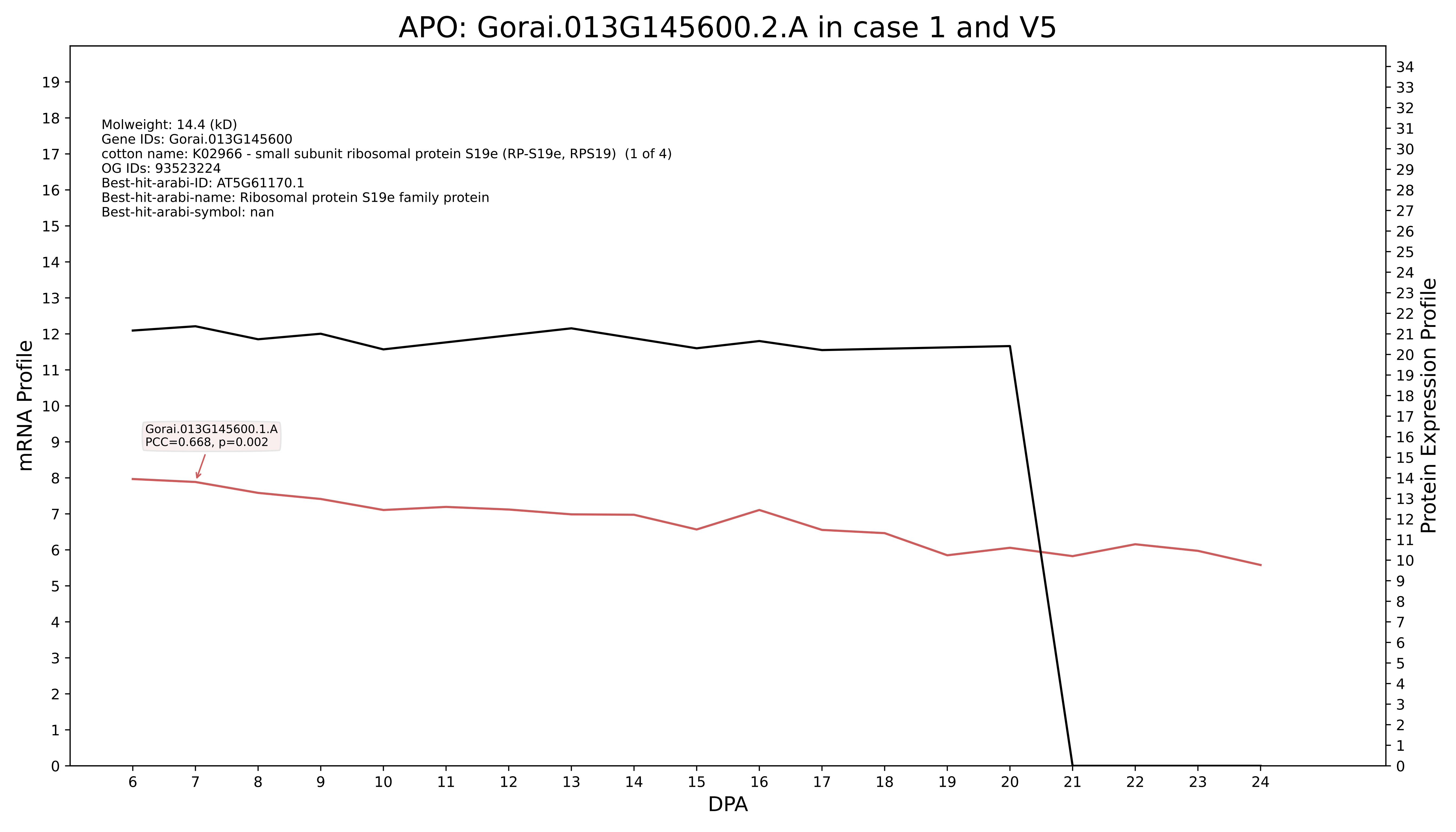
APO: Gorai.013G145600.2.A in case 1 and V5
Molweight: 14.4 (kD)
Gene IDs: Gorai.013G145600
cotton name: K02966 - small subunit ribosomal protein S19e (RP-S19e, RPS19) (1 of 4)
OG IDs: 93523224
Best-hit-arabi-ID: AT5G61170.1
Best-hit-arabi-name: Ribosomal protein S19e family protein
Best-hit-arabi-symbol: nan
Gorai.013G145600.1.A
PCC=0.668, p=0.002
mRNA Profile
Protein Expression Profile
DPA

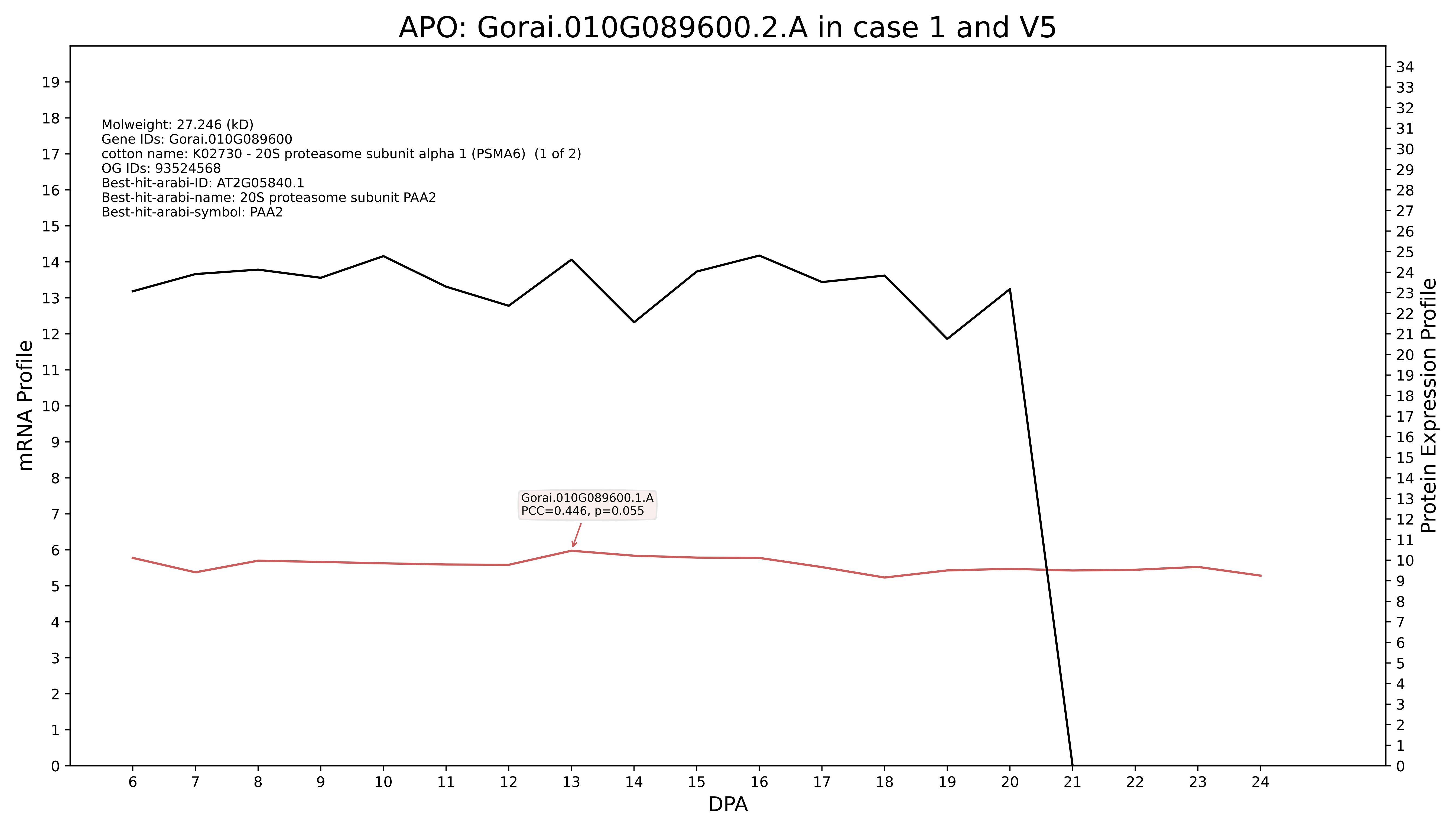

APO: Gorai.010G089600.2.A in case 1 and V5
Molweight: 27.246 (kD)
Gene IDs: Gorai.010G089600
cotton name: K02730 - 20S proteasome subunit alpha 1 (PSMA6) (1 of 2)
OG IDs: 93524568
Best-hit-arabi-ID: AT2G05840.1
Best-hit-arabi-name: 20S proteasome subunit PAA2
Best-hit-arabi-symbol: PAA2
Gorai.010G089600.1.A
PCC=0.446, p=0.055
mRNA Profile
Protein Expression Profile
DPA

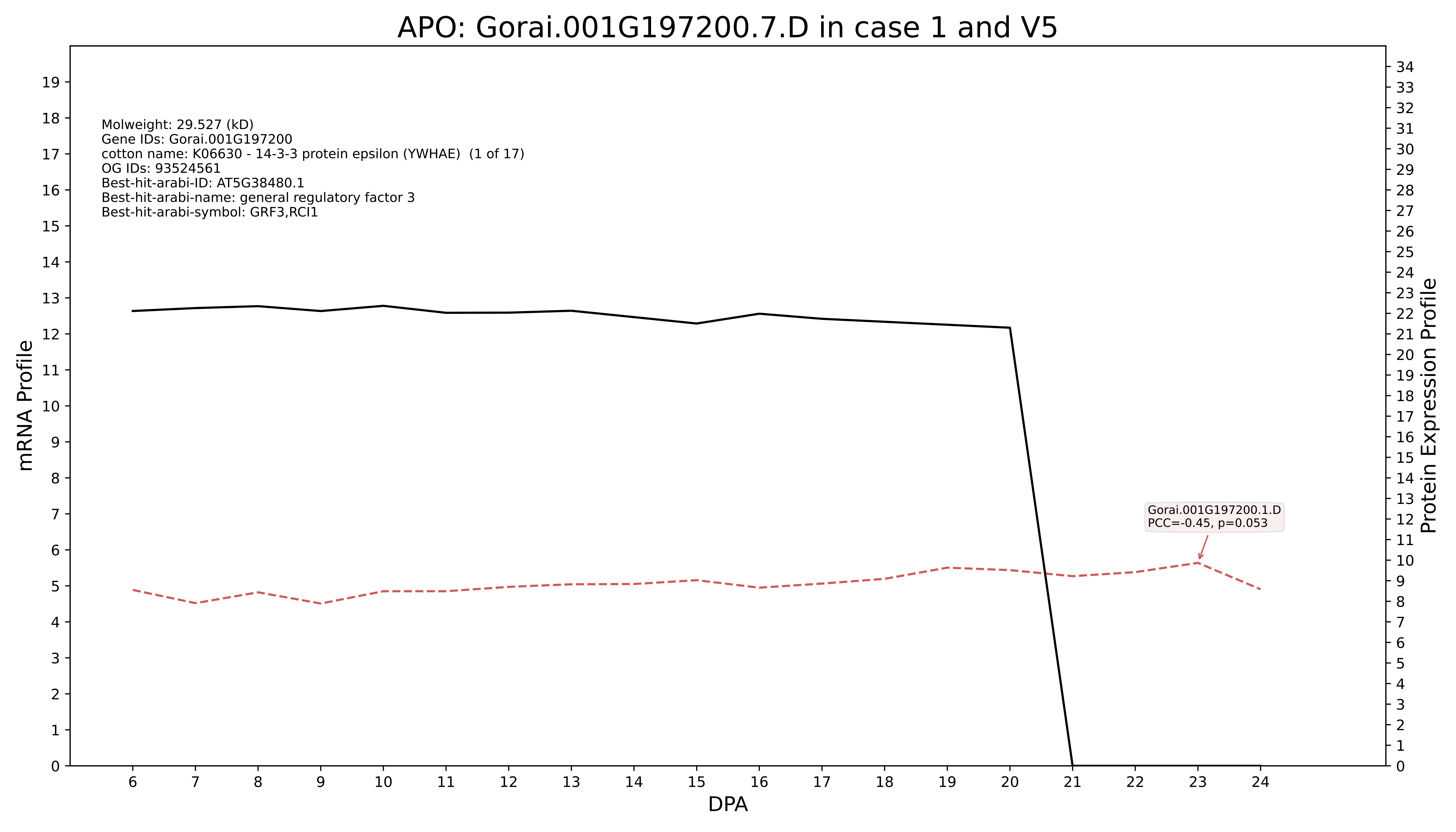

APO: Gorai.001G197200.7.D in case 1 and V5
Molweight: 29.527 (kD)
Gene IDs: Gorai.001G197200
cotton name: K06630 - 14-3-3 protein epsilon (YWHAE) (1 of 17)
OG IDs: 93524561
Best-hit-arabi-ID: AT5G38480.1
Best-hit-arabi-name: general regulatory factor 3
Best-hit-arabi-symbol: GRF3,RCI1
Gorai.001G197200.1.D
PCC=-0.45, p=0.053
mRNA Profile
Protein Expression Profile
DPA

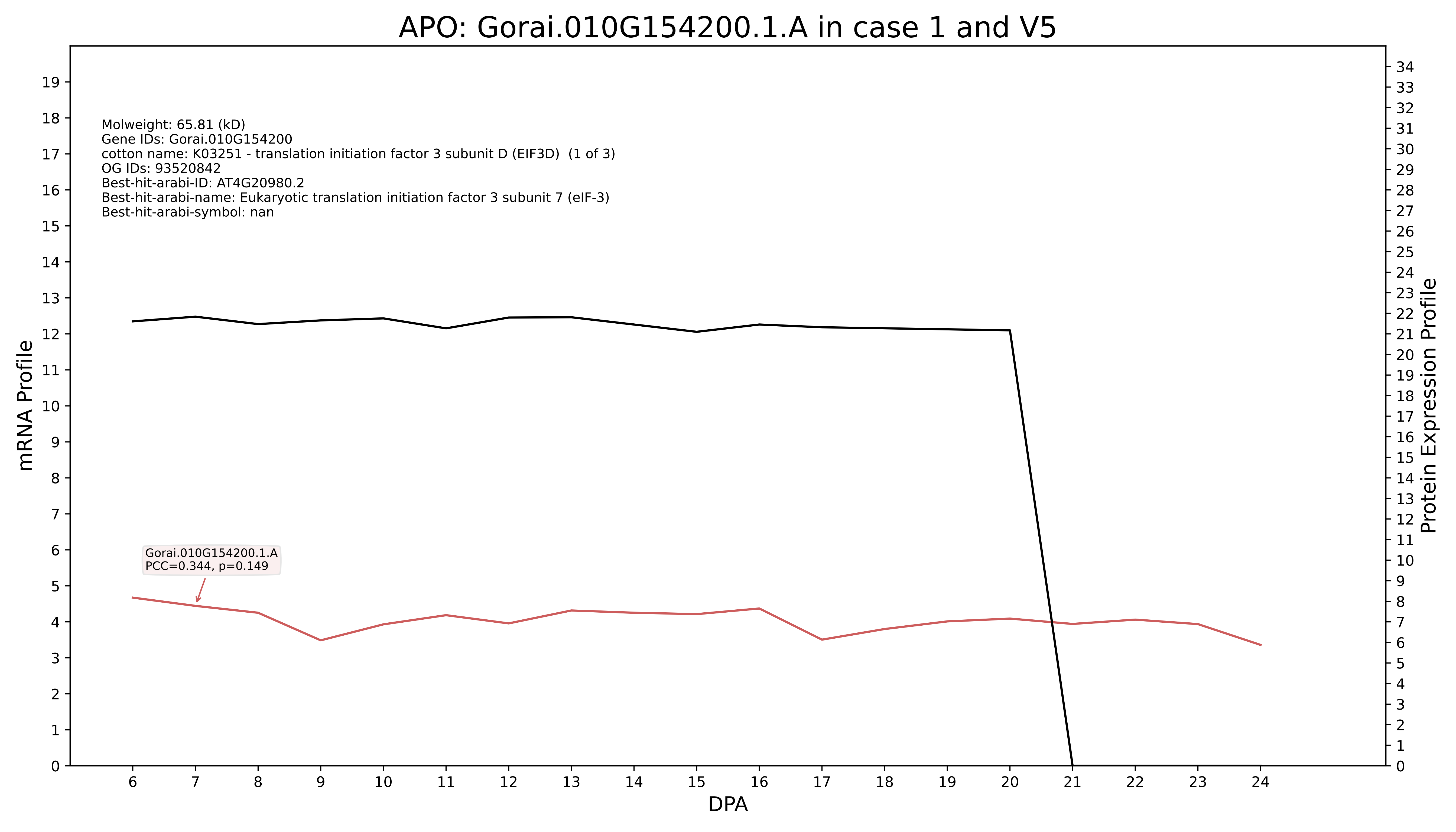
APO: Gorai.010G154200.1.A in case 1 and V5
Molweight: 65.81 (kD)
Gene IDs: Gorai.010G154200
cotton name: K03251 - translation initiation factor 3 subunit D (EIF3D) (1 of 3)
OG IDs: 93520842
Best-hit-arabi-ID: AT4G20980.2
Best-hit-arabi-name: Eukaryotic translation initiation factor 3 subunit 7 (eIF-3)
Best-hit-arabi-symbol: nan
Gorai.010G154200.1.A
PCC=0.344, p=0.149
mRNA Profile
Protein Expression Profile
DPA

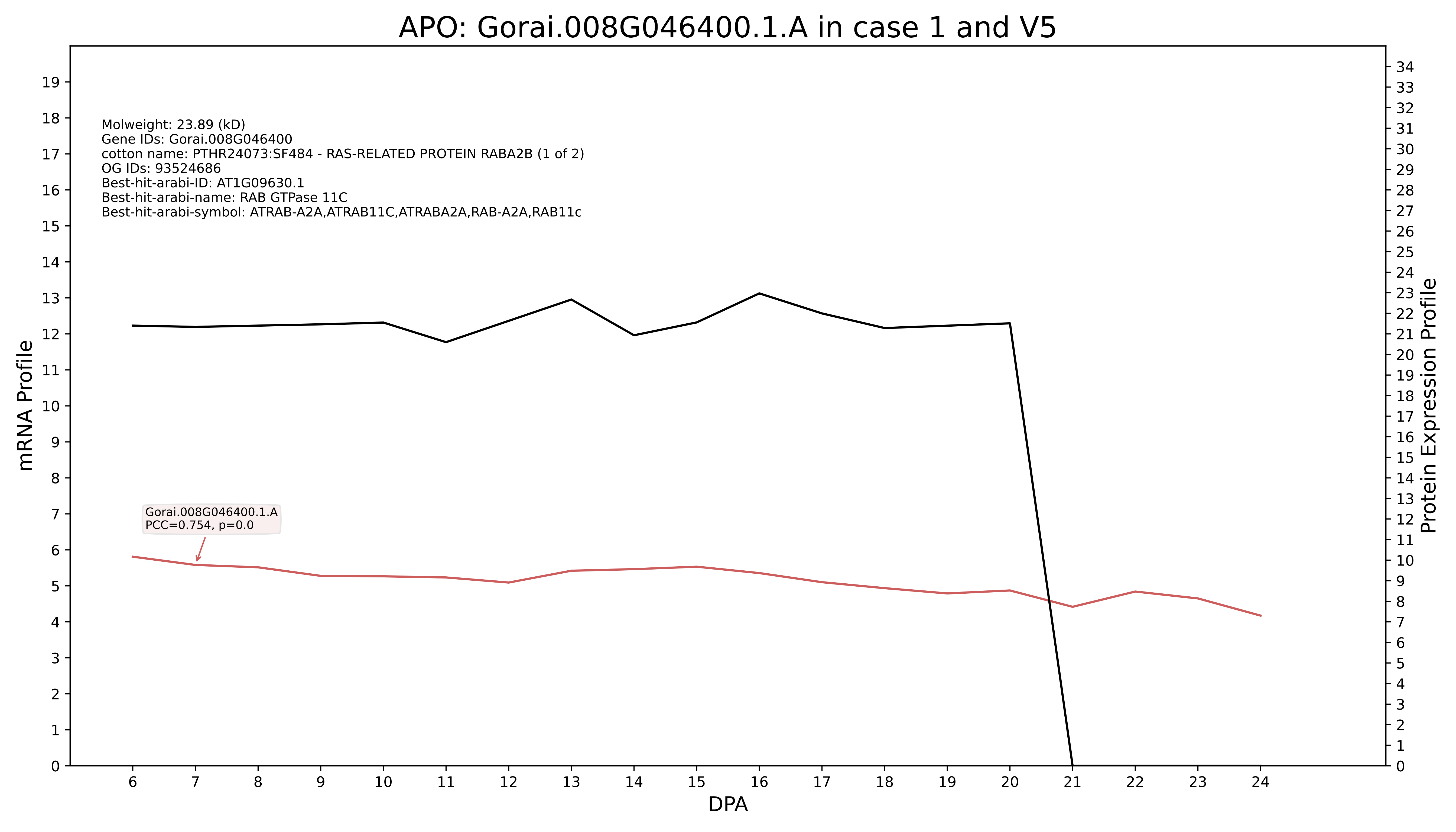
APO: Gorai.008G046400.1.A in case 1 and V5
Molweight: 23.89 (kD)
Gene IDs: Gorai.008G046400
cotton name: PTHR24073:SF484 - RAS-RELATED PROTEIN RABA2B (1 of 2)
OG IDs: 93524686
Best-hit-arabi-ID: AT1G09630.1
Best-hit-arabi-name: RAB GTPase 11C
Best-hit-arabi-symbol: ATRAB-A2A,ATRAB11C,ATRABA2A,RAB-A2A,RAB11c
Gorai.008G046400.1.A
PCC=0.754, p=0.0
mRNA Profile
Protein Expression Profile
DPA

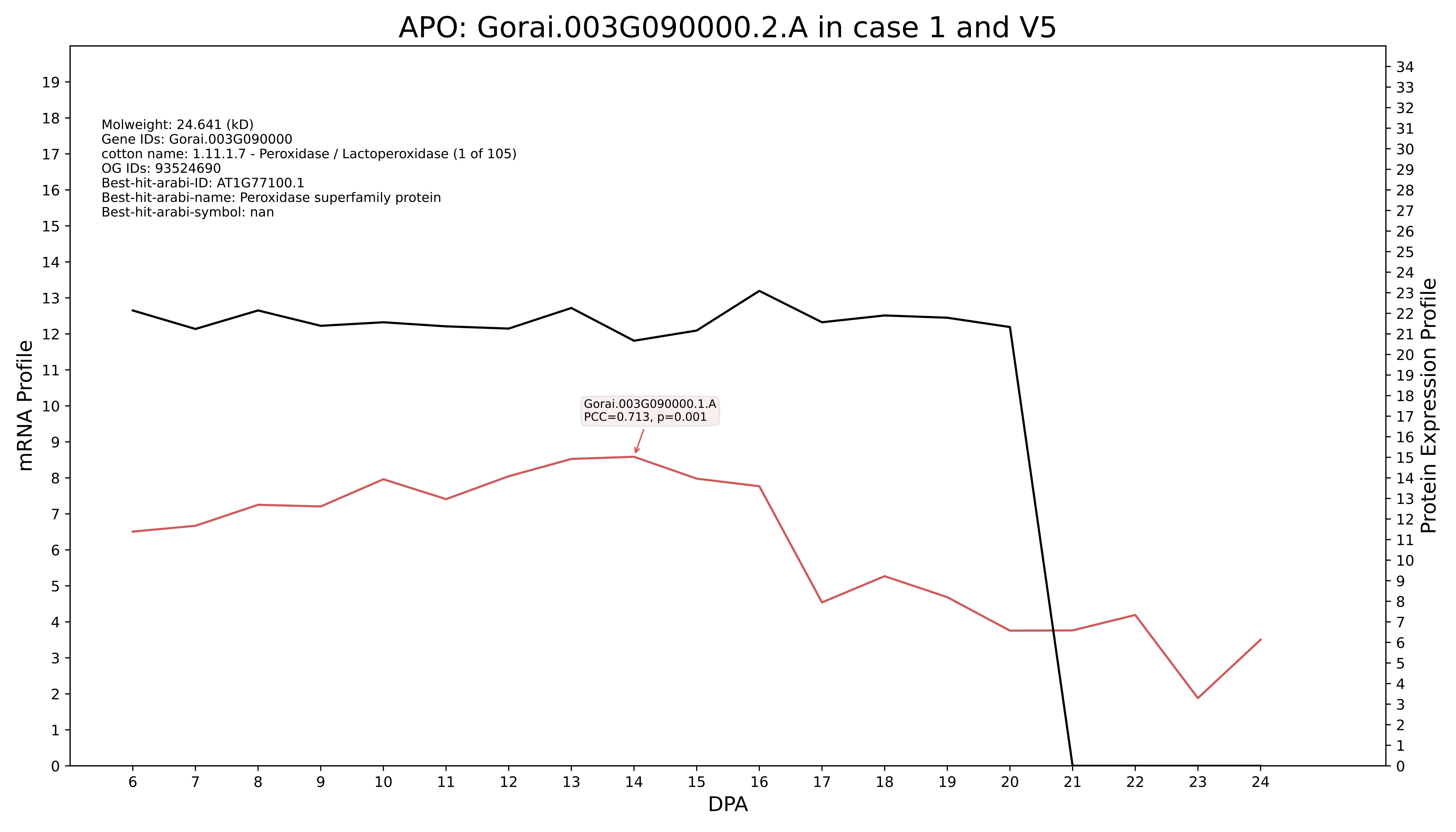
APO: Gorai.003G090000.2.A in case 1 and V5
Molweight: 24.641 (kD)
Gene IDs: Gorai.003G090000
cotton name: 1.11.1.7 - Peroxidase / Lactoperoxidase (1 of 105)
OG IDs: 93524690
Best-hit-arabi-ID: AT1G77100.1
Best-hit-arabi-name: Peroxidase superfamily protein
Best-hit-arabi-symbol: nan
Gorai.003G090000.1.A
PCC=0.713, p=0.001
mRNA Profile
Protein Expression Profile
DPA

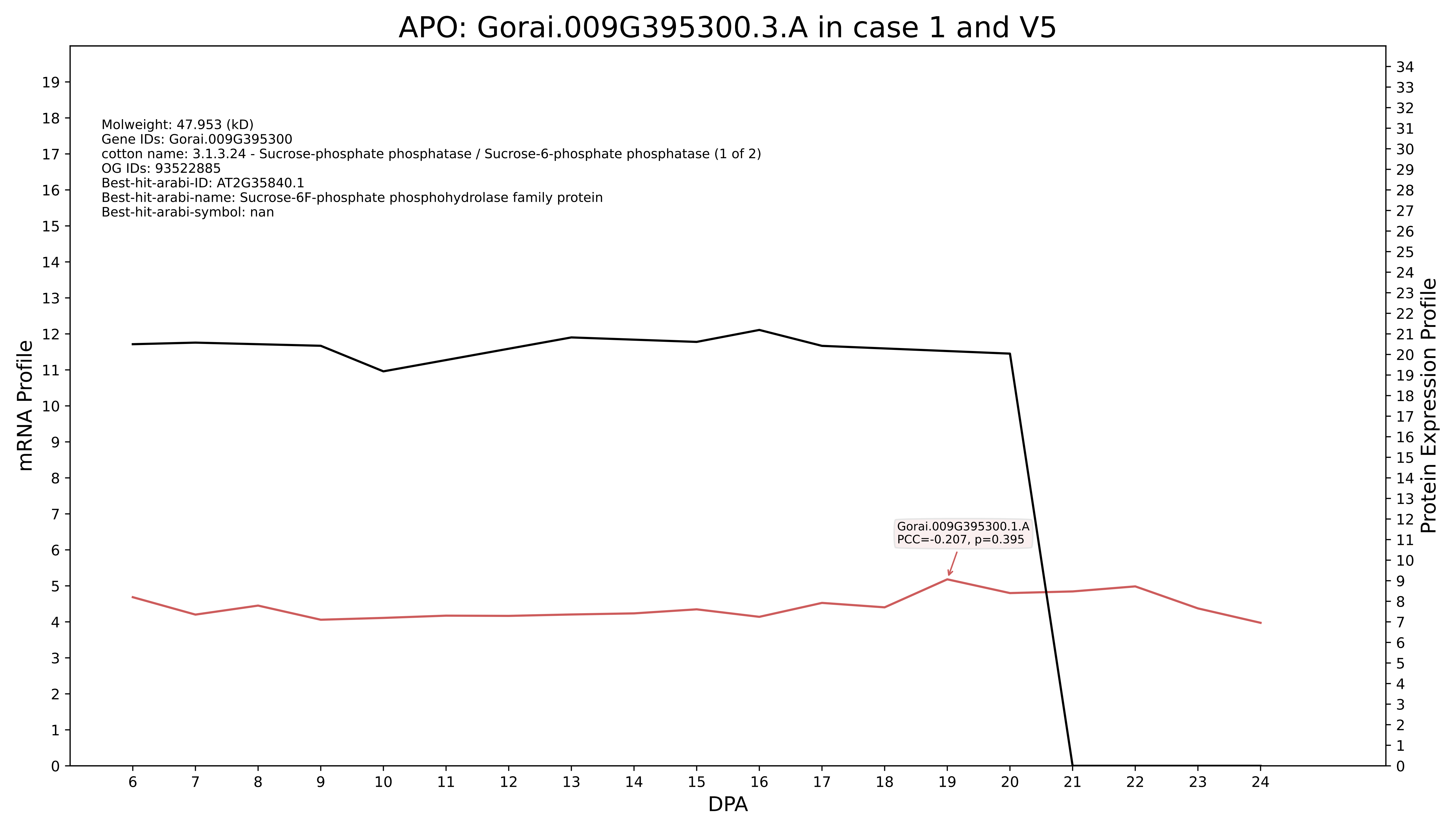
APO: Gorai.009G395300.3.A in case 1 and V5
Molweight: 47.953 (kD)
Gene IDs: Gorai.009G395300
cotton name: 3.1.3.24 - Sucrose-phosphate phosphatase / Sucrose-6-phosphate phosphatase (1 of 2)
OG IDs: 93522885
Best-hit-arabi-ID: AT2G35840.1
Best-hit-arabi-name: Sucrose-6F-phosphate phosphohydrolase family protein
Best-hit-arabi-symbol: nan
Gorai.009G395300.1.A
PCC=-0.207, p=0.395
mRNA Profile
Protein Expression Profile
DPA

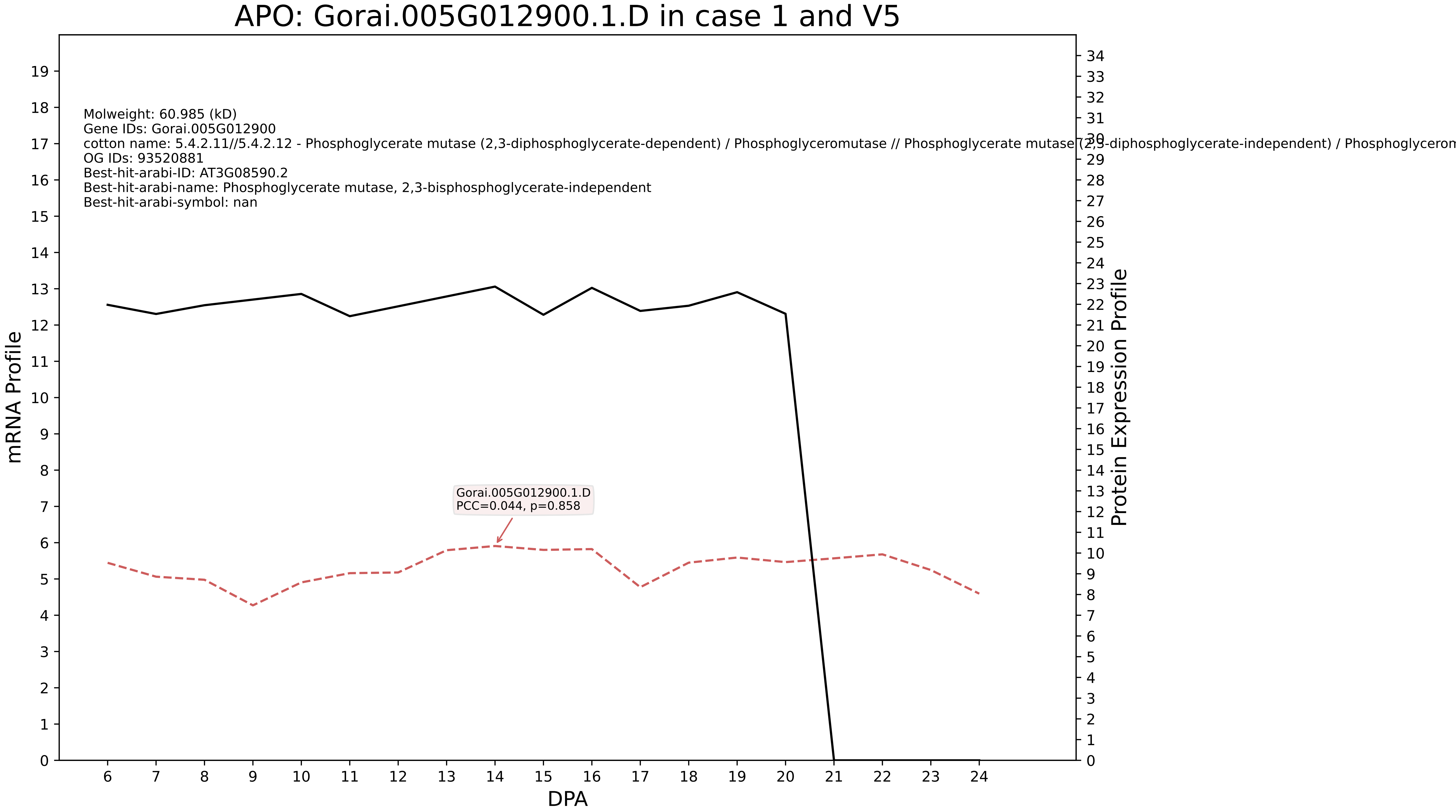
APO: Gorai.005G012900.1.D in case 1 and V5
Molweight: 60.985 (kD)
Gene IDs: Gorai.005G012900
cotton name: 5.4.2.11//5.4.2.12 - Phosphoglycerate mutase (2,3-diphosphoglycerate-dependent) / Phosphoglyceromutase // Phosphoglycerate mutase (2,3-diphosphoglycerate-independent) / Phosphoglyceron
OG IDs: 93520881
Best-hit-arabi-ID: AT3G08590.2
Best-hit-arabi-name: Phosphoglycerate mutase, 2,3-bisphosphoglycerate-independent
Best-hit-arabi-symbol: nan
Gorai.005G012900.1.D
PCC=0.044, p=0.858
mRNA Profile
Protein Expression Profile
DPA

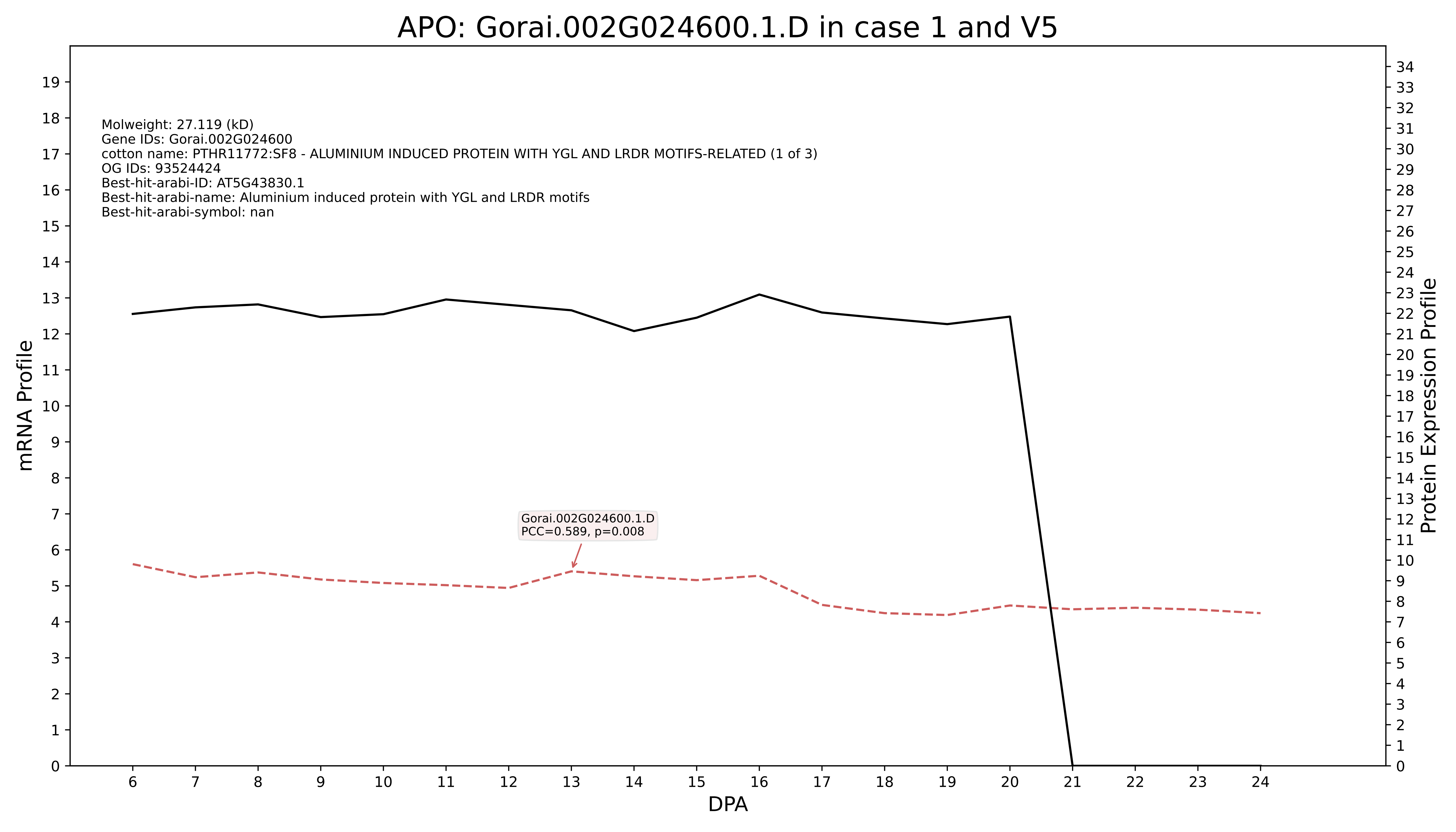
APO: Gorai.002G024600.1.D in case 1 and V5
Molweight: 27.119 (kD)
Gene IDs: Gorai.002G024600
cotton name: PTHR11772:SF8 - ALUMINIUM INDUCED PROTEIN WITH YGL AND LRDR MOTIFS-RELATED (1 of 3)
OG IDs: 93524424
Best-hit-arabi-ID: AT5G43830.1
Best-hit-arabi-name: Aluminium induced protein with YGL and LRDR motifs
Best-hit-arabi-symbol: nan
Gorai.002G024600.1.D
PCC=0.589, p=0.008
mRNA Profile
Protein Expression Profile
DPA

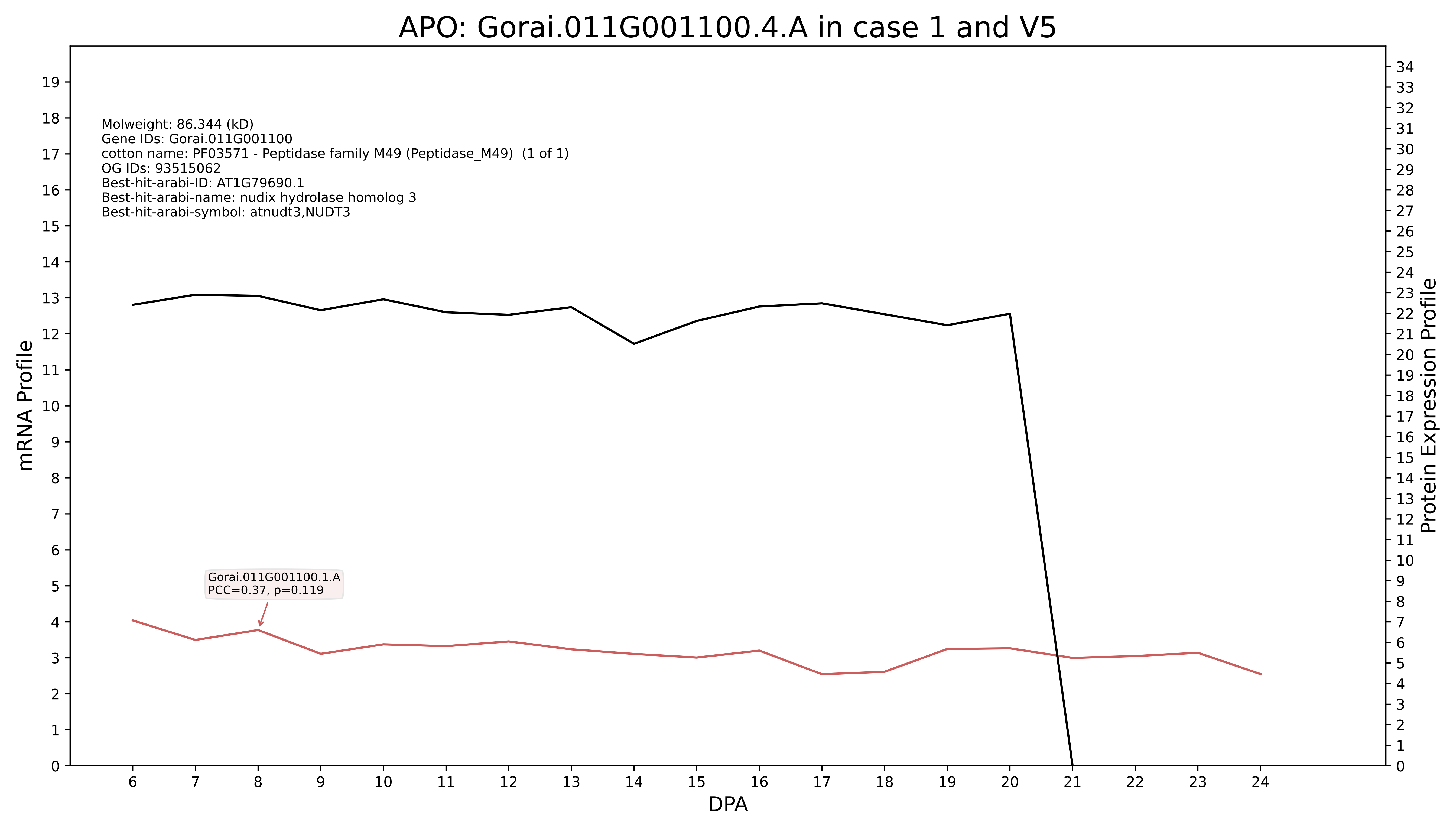
APO: Gorai.011G001100.4.A in case 1 and V5
Molweight: 86.344 (kD)
Gene IDs: Gorai.011G001100
cotton name: PF03571 - Peptidase family M49 (Peptidase_M49) (1 of 1)
OG IDs: 93515062
Best-hit-arabi-ID: AT1G79690.1
Best-hit-arabi-name: nudix hydrolase homolog 3
Best-hit-arabi-symbol: atnudt3,NUDT3
Gorai.011G001100.1.A
PCC=0.37, p=0.119
mRNA Profile
Protein Expression Profile
DPA

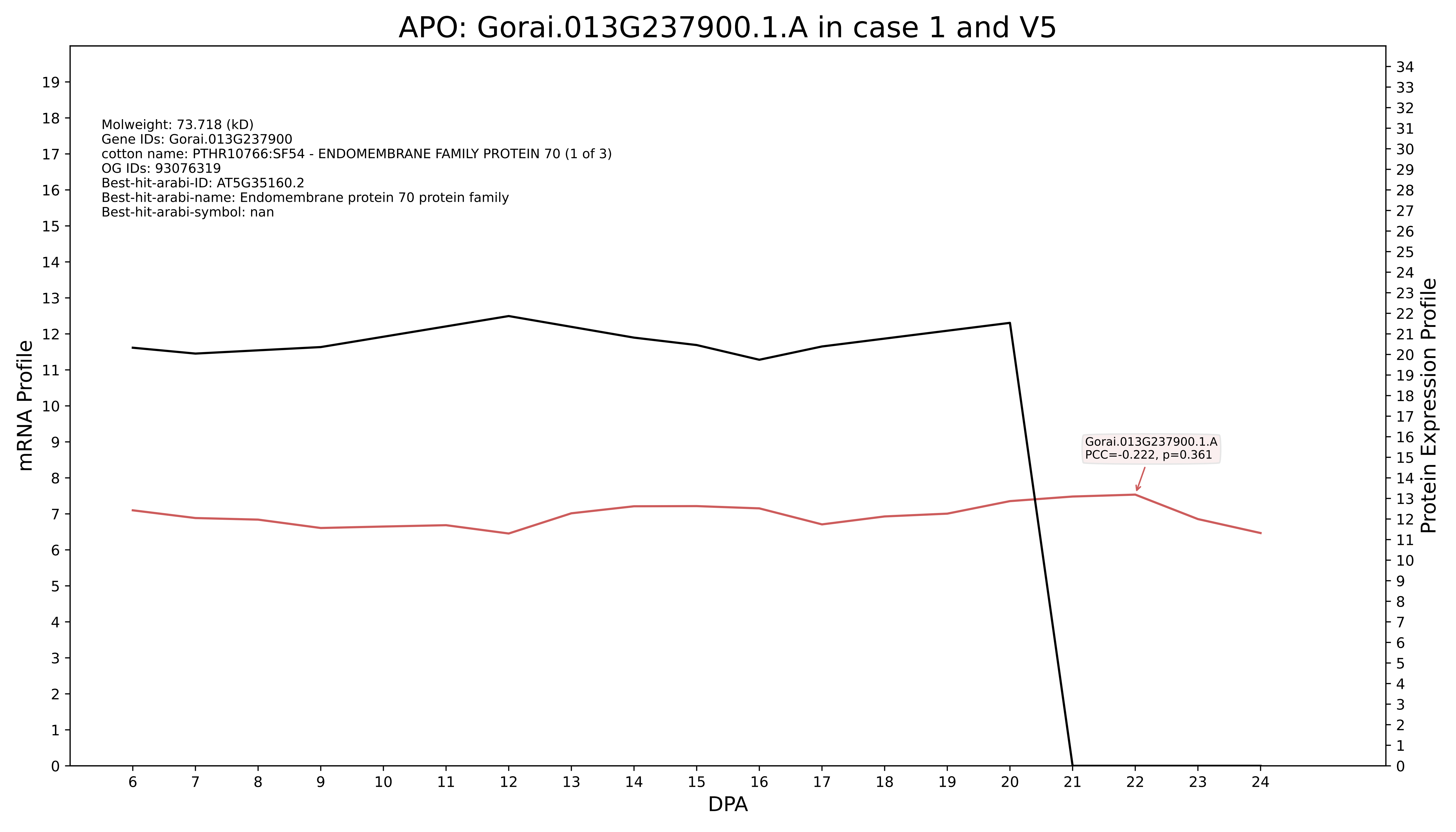

APO: Gorai.013G237900.1.A in case 1 and V5
Molweight: 73.718 (kD)
Gene IDs: Gorai.013G237900
cotton name: PTHR10766:SF54 - ENDOMEMBRANE FAMILY PROTEIN 70 (1 of 3)
OG IDs: 93076319
Best-hit-arabi-ID: AT5G35160.2
Best-hit-arabi-name: Endomembrane protein 70 protein family
Best-hit-arabi-symbol: nan
Gorai.013G237900.1.A
PCC=-0.222, p=0.361
mRNA Profile
Protein Expression Profile
DPA

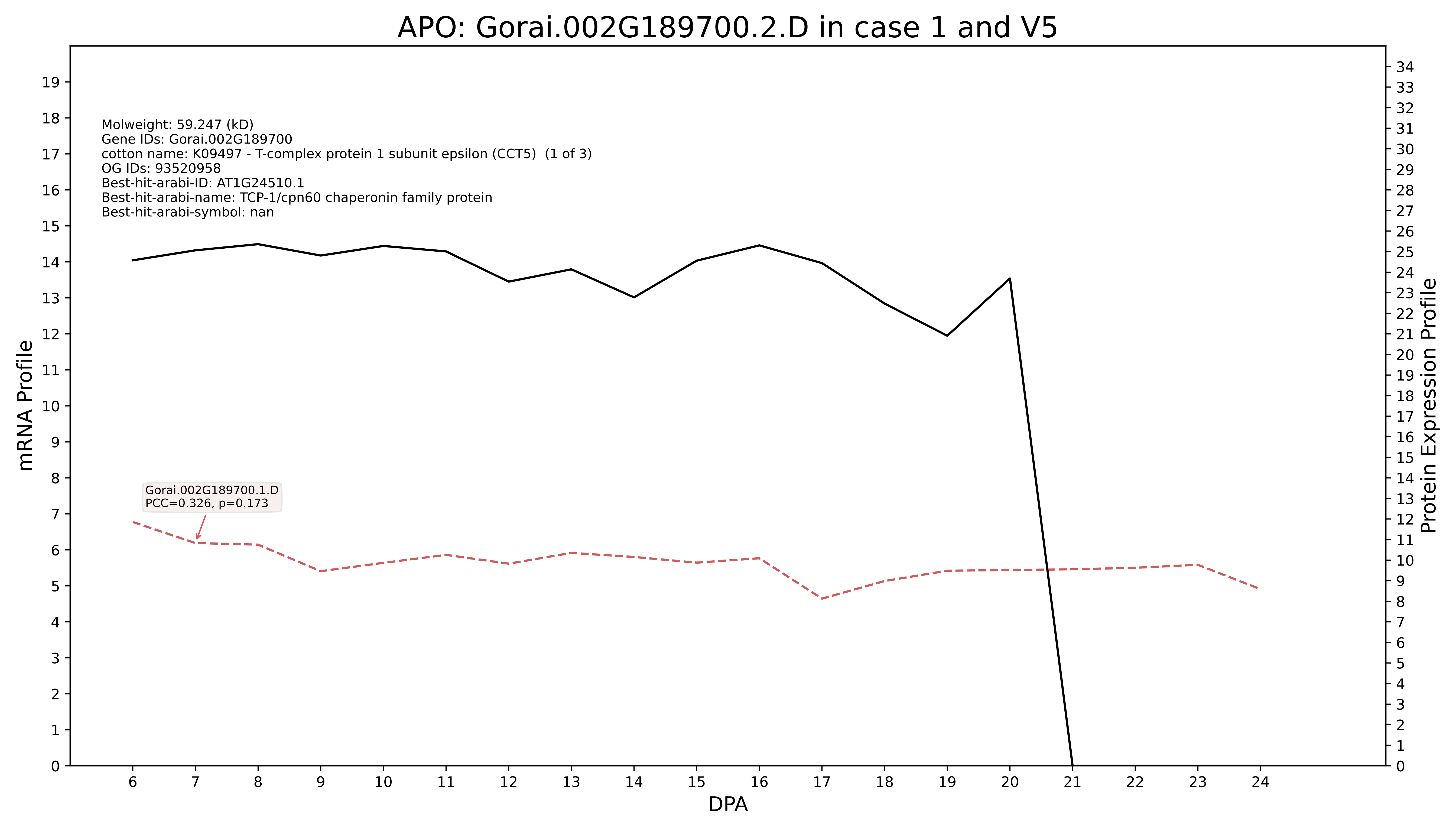

APO: Gorai.002G189700.2.D in case 1 and V5
Molweight: 59.247 (kD)
Gene IDs: Gorai.002G189700
cotton name: K09497 - T-complex protein 1 subunit epsilon (CCT5) (1 of 3)
OG IDs: 93520958
Best-hit-arabi-ID: AT1G24510.1
Best-hit-arabi-name: TCP-1/cpn60 chaperonin family protein
Best-hit-arabi-symbol: nan
Gorai.002G189700.1.D
PCC=0.326, p=0.173
mRNA Profile
Protein Expression Profile
DPA

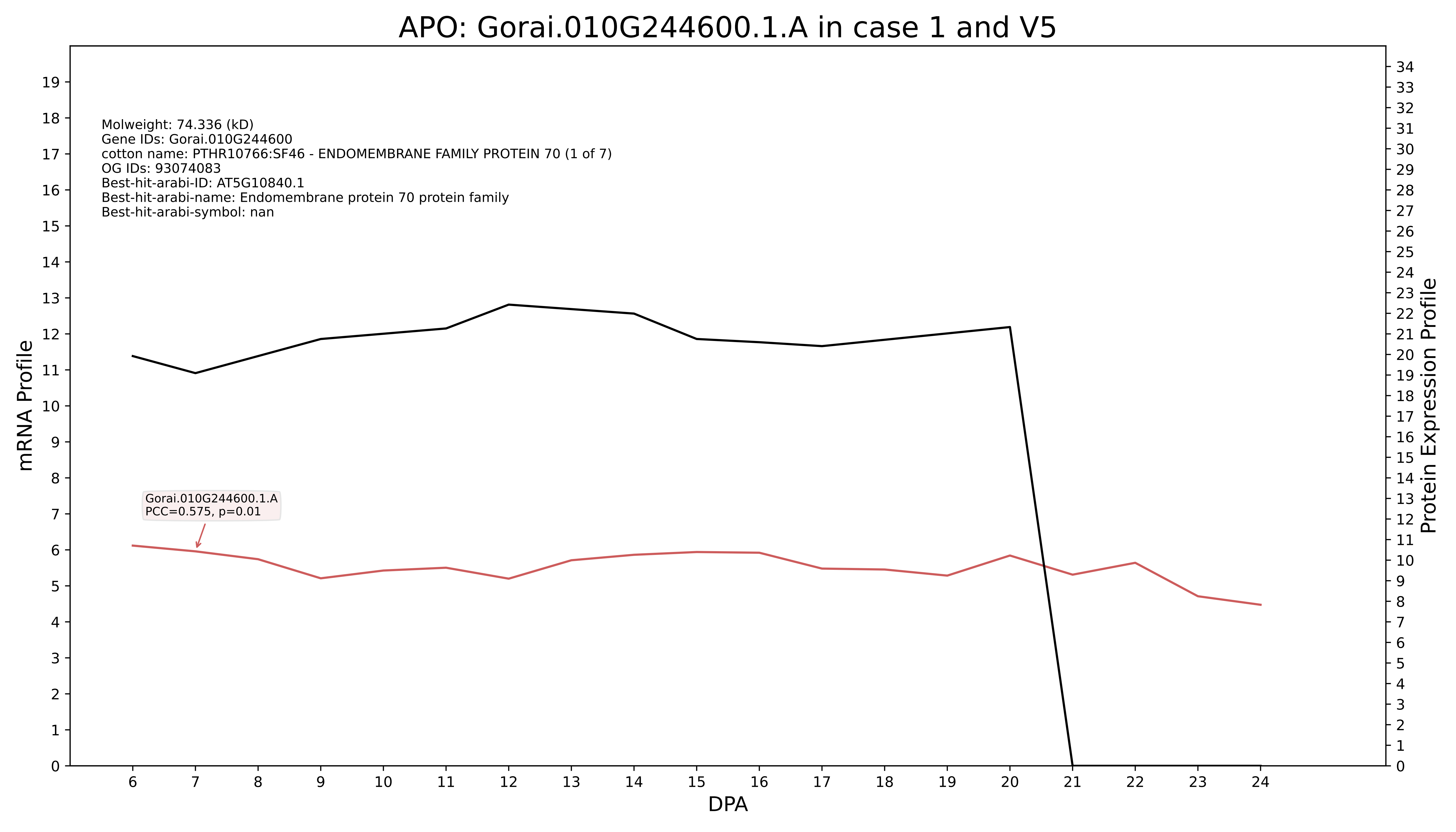
APO: Gorai.010G244600.1.A in case 1 and V5
Molweight: 74.336 (kD)
Gene IDs: Gorai.010G244600
cotton name: PTHR10766:SF46 - ENDOMEMBRANE FAMILY PROTEIN 70 (1 of 7)
OG IDs: 93074083
Best-hit-arabi-ID: AT5G10840.1
Best-hit-arabi-name: Endomembrane protein 70 protein family
Best-hit-arabi-symbol: nan
Gorai.010G244600.1.A
PCC=0.575, p=0.01
mRNA Profile
Protein Expression Profile
DPA

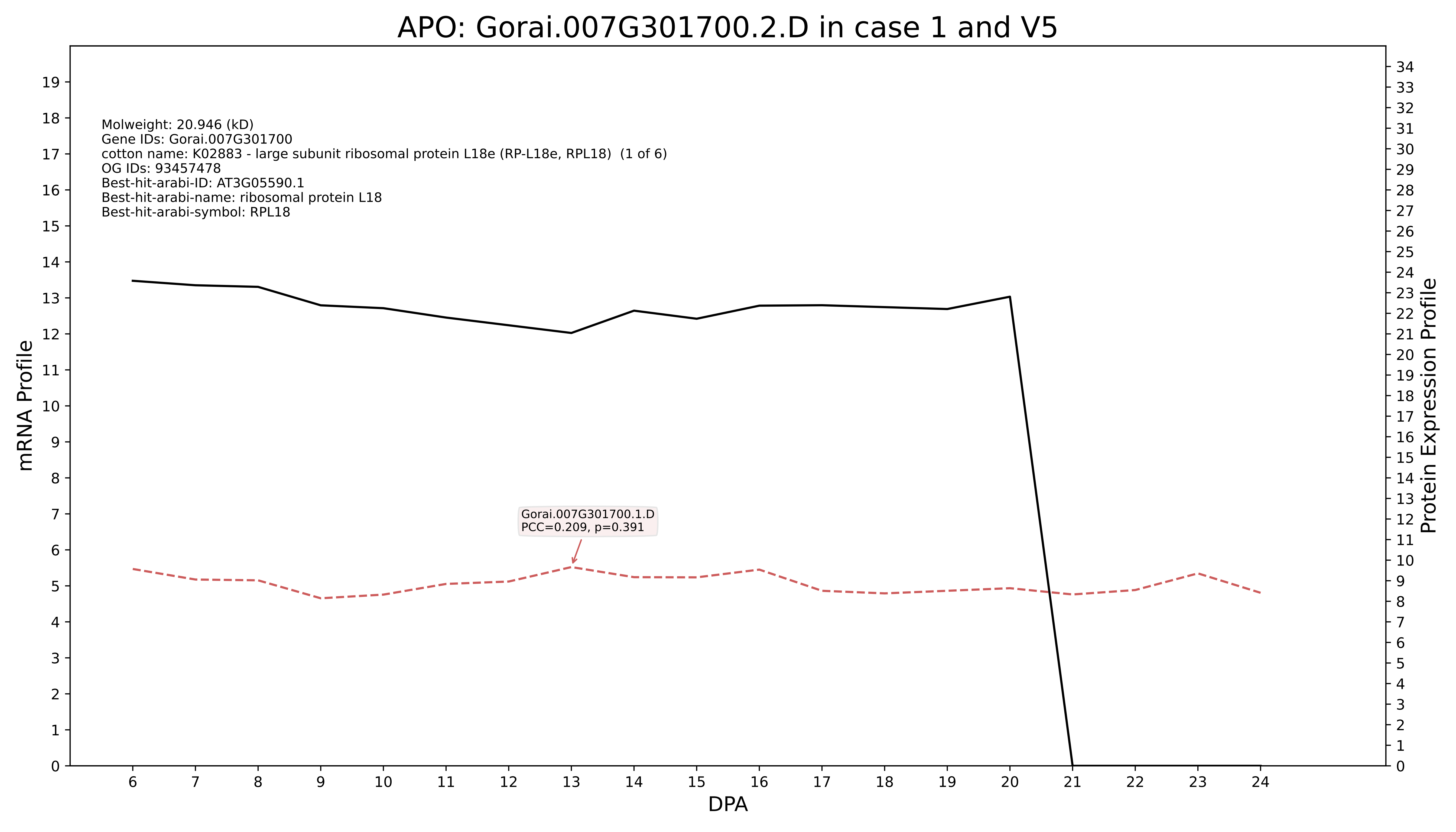
APO: Gorai.007G301700.2.D in case 1 and V5
Molweight: 20.946 (kD)
Gene IDs: Gorai.007G301700
cotton name: K02883 - large subunit ribosomal protein L18e (RP-L18e, RPL18) (1 of 6)
OG IDs: 93457478
Best-hit-arabi-ID: AT3G05590.1
Best-hit-arabi-name: ribosomal protein L18
Best-hit-arabi-symbol: RPL18
Gorai.007G301700.1.D
PCC=0.209, p=0.391
mRNA Profile
Protein Expression Profile
DPA

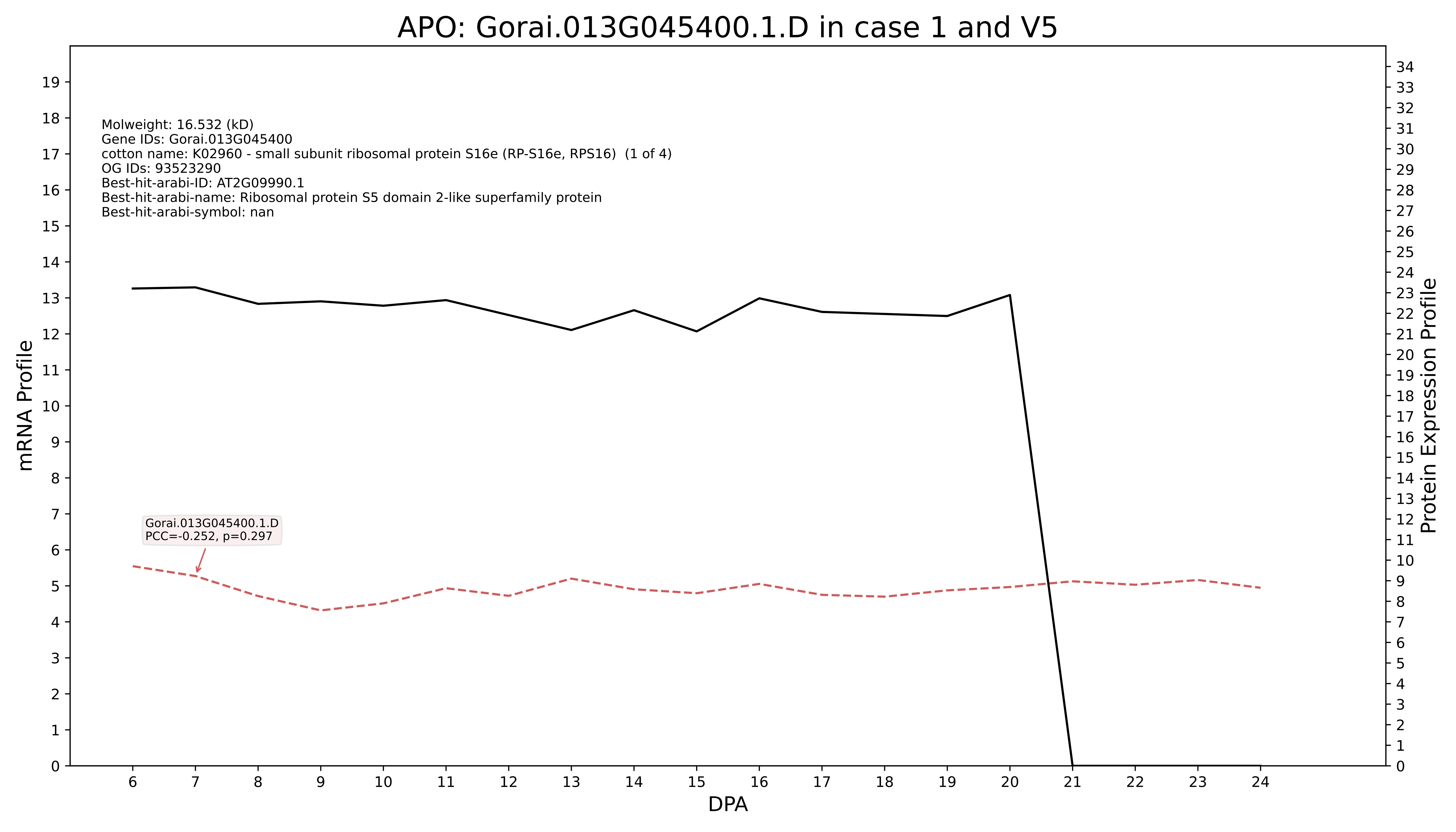
APO: Gorai.013G045400.1.D in case 1 and V5
Molweight: 16.532 (kD)
Gene IDs: Gorai.013G045400
cotton name: K02960 - small subunit ribosomal protein S16e (RP-S16e, RPS16) (1 of 4)
OG IDs: 93523290
Best-hit-arabi-ID: AT2G09990.1
Best-hit-arabi-name: Ribosomal protein S5 domain 2-like superfamily protein
Best-hit-arabi-symbol: nan
Gorai.013G045400.1.D
PCC=-0.252, p=0.297
mRNA Profile
Protein Expression Profile
DPA

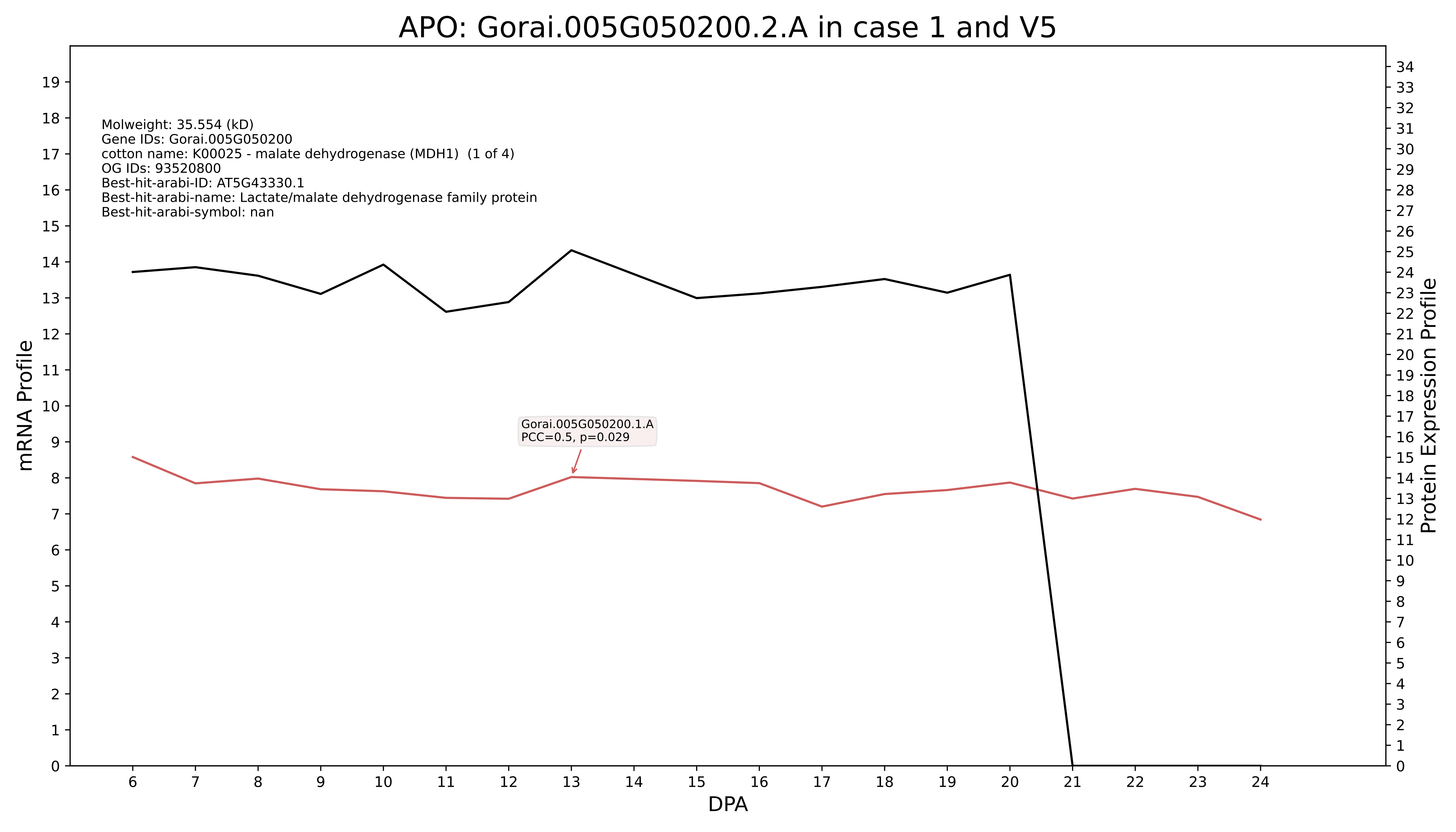

APO: Gorai.005G050200.2.A in case 1 and V5
Molweight: 35.554 (kD)
Gene IDs: Gorai.005G050200
cotton name: K00025 - malate dehydrogenase (MDH1) (1 of 4)
OG IDs: 93520800
Best-hit-arabi-ID: AT5G43330.1
Best-hit-arabi-name: Lactate/malate dehydrogenase family protein
Best-hit-arabi-symbol: nan
Gorai.005G050200.1.A
PCC=0.5, p=0.029
mRNA Profile
Protein Expression Profile
DPA

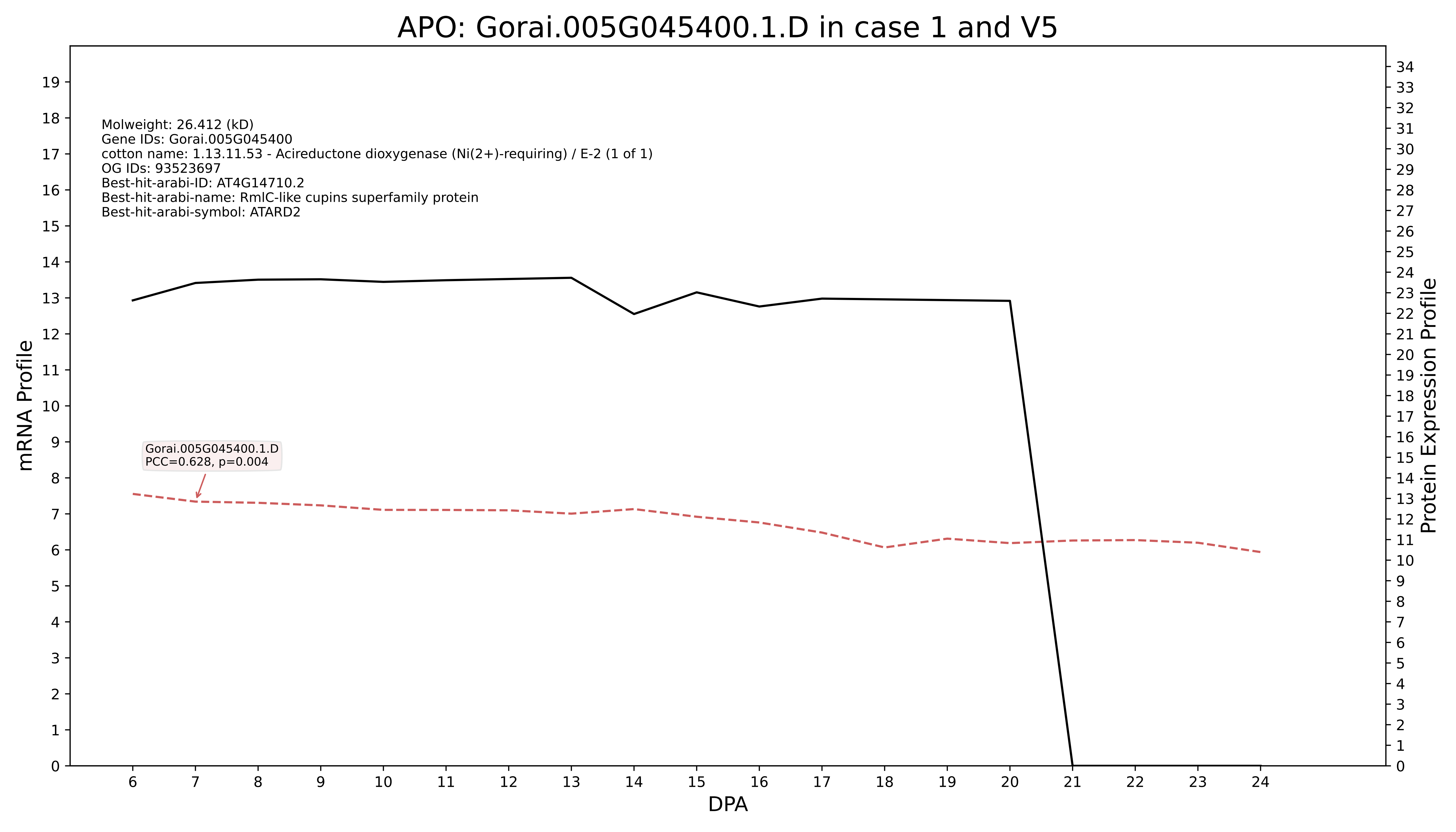
APO: Gorai.005G045400.1.D in case 1 and V5
Molweight: 26.412 (kD)
Gene IDs: Gorai.005G045400
cotton name: 1.13.11.53 - Acireductone dioxygenase (Ni(2+)-requiring) / E-2 (1 of 1)
OG IDs: 93523697
Best-hit-arabi-ID: AT4G14710.2
Best-hit-arabi-name: RmlC-like cupins superfamily protein
Best-hit-arabi-symbol: ATARD2
Gorai.005G045400.1.D
PCC=0.628, p=0.004
mRNA Profile
Protein Expression Profile
DPA

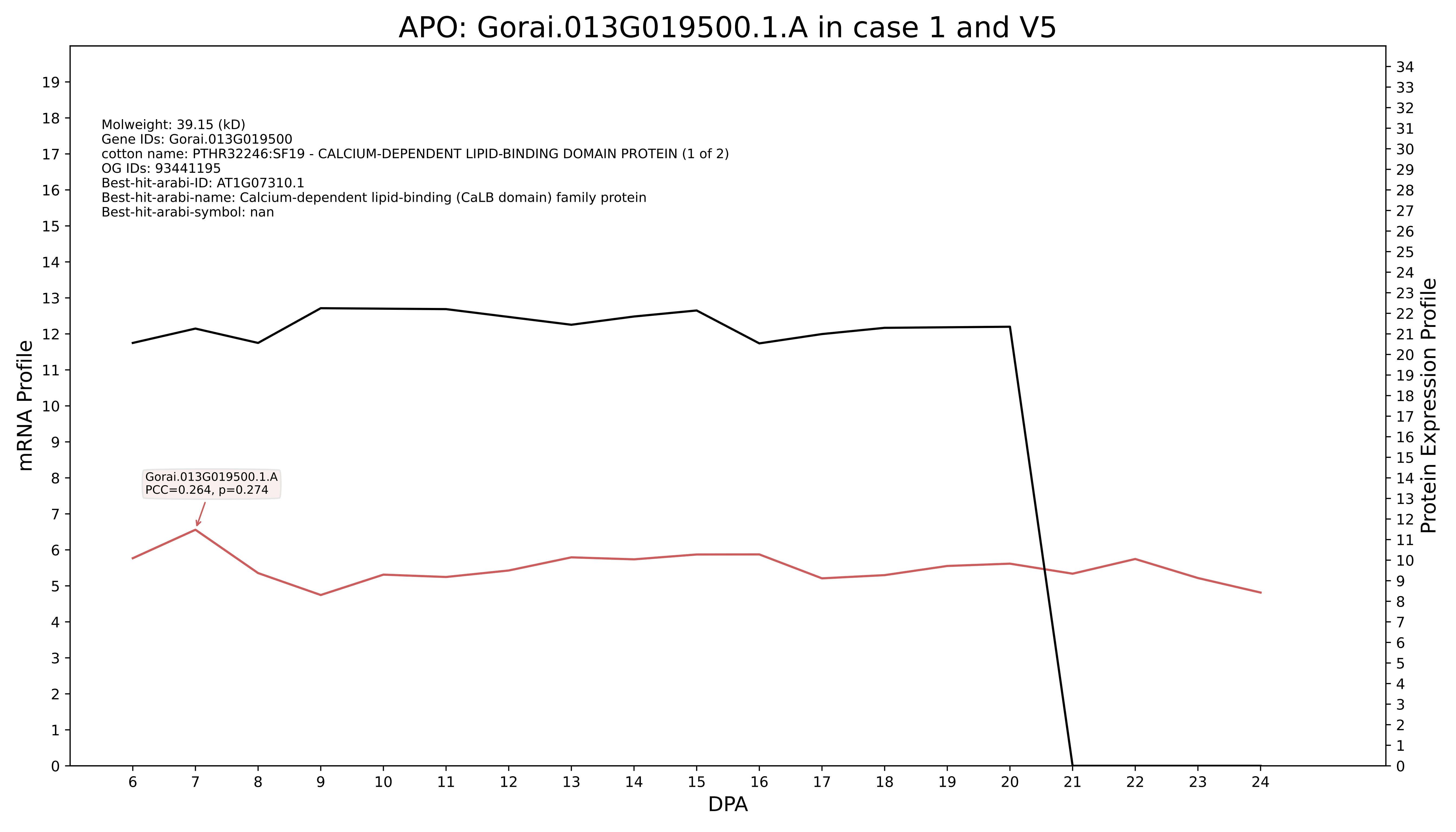
APO: Gorai.013G019500.1.A in case 1 and V5
Molweight: 39.15 (kD)
Gene IDs: Gorai.013G019500
cotton name: PTHR32246:SF19 - CALCIUM-DEPENDENT LIPID-BINDING DOMAIN PROTEIN (1 of 2)
OG IDs: 93441195
Best-hit-arabi-ID: AT1G07310.1
Best-hit-arabi-name: Calcium-dependent lipid-binding (CaLB domain) family protein
Best-hit-arabi-symbol: nan
Gorai.013G019500.1.A
PCC=0.264, p=0.274
mRNA Profile
Protein Expression Profile
DPA

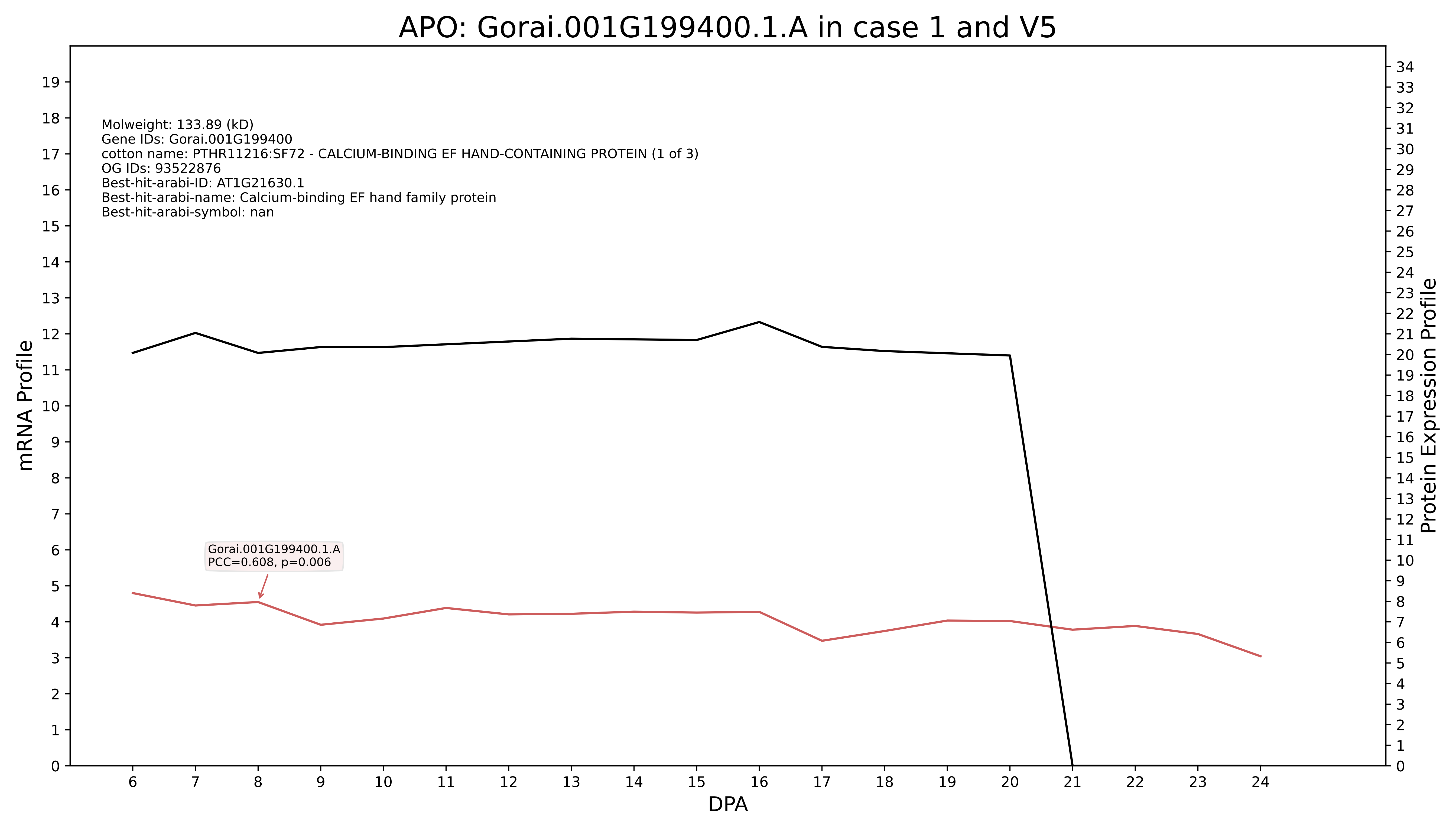
APO: Gorai.001G199400.1.A in case 1 and V5
Molweight: 133.89 (kD)
Gene IDs: Gorai.001G199400
cotton name: PTHR11216:SF72 - CALCIUM-BINDING EF HAND-CONTAINING PROTEIN (1 of 3)
OG IDs: 93522876
Best-hit-arabi-ID: AT1G21630.1
Best-hit-arabi-name: Calcium-binding EF hand family protein
Best-hit-arabi-symbol: nan
Gorai.001G199400.1.A
PCC=0.608, p=0.006
mRNA Profile
Protein Expression Profile
DPA

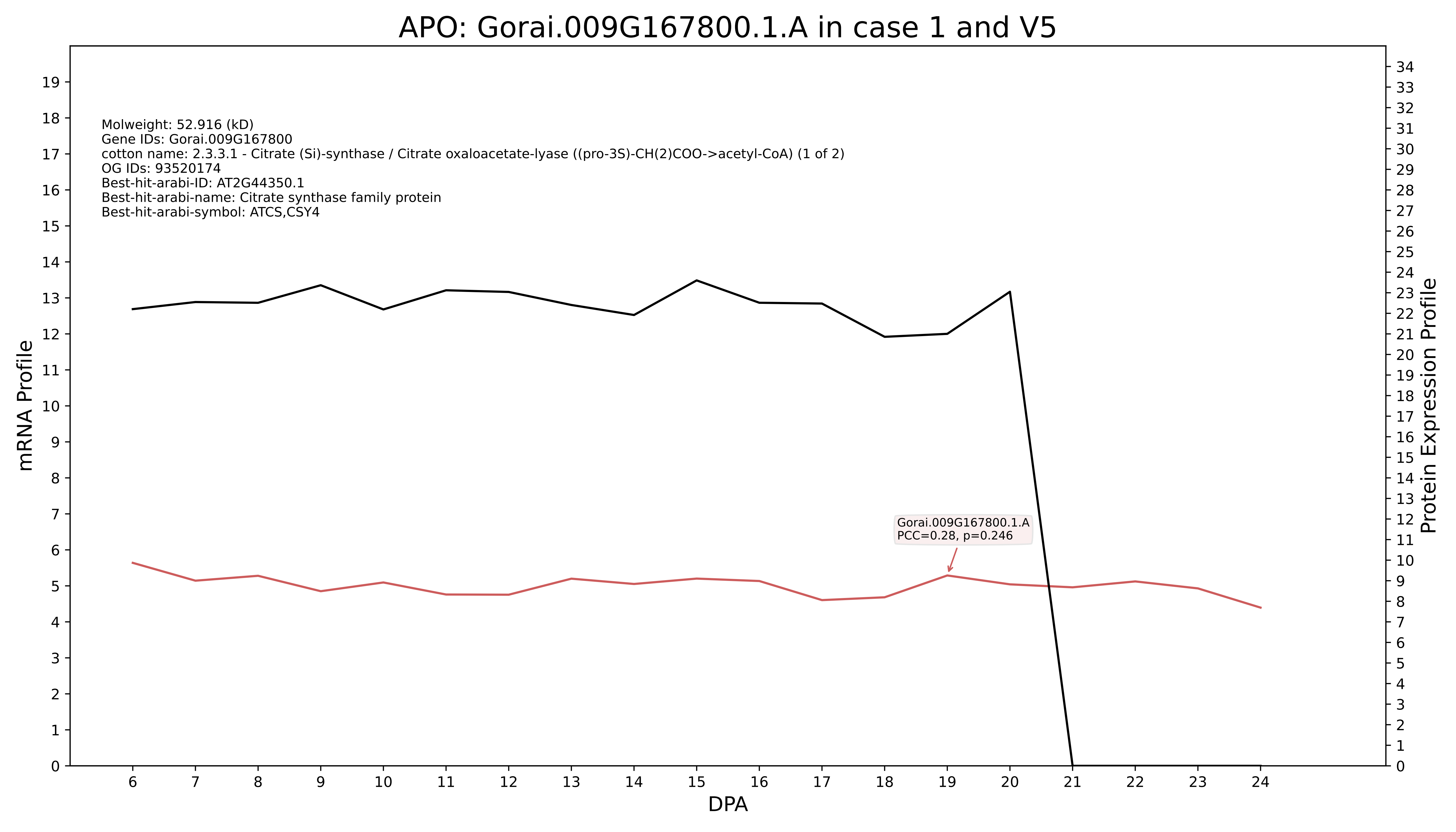

APO: Gorai.009G167800.1.A in case 1 and V5
Molweight: 52.916 (kD)
Gene IDs: Gorai.009G167800
cotton name: 2.3.3.1 - Citrate (Si)-synthase / Citrate oxaloacetate-lyase ((pro-3S)-CH(2)COO->acetyl-CoA) (1 of 2)
OG IDs: 93520174
Best-hit-arabi-ID: AT2G44350.1
Best-hit-arabi-name: Citrate synthase family protein
Best-hit-arabi-symbol: ATCS,CSY4
Gorai.009G167800.1.A
PCC=0.28, p=0.246
mRNA Profile
Protein Expression Profile
DPA

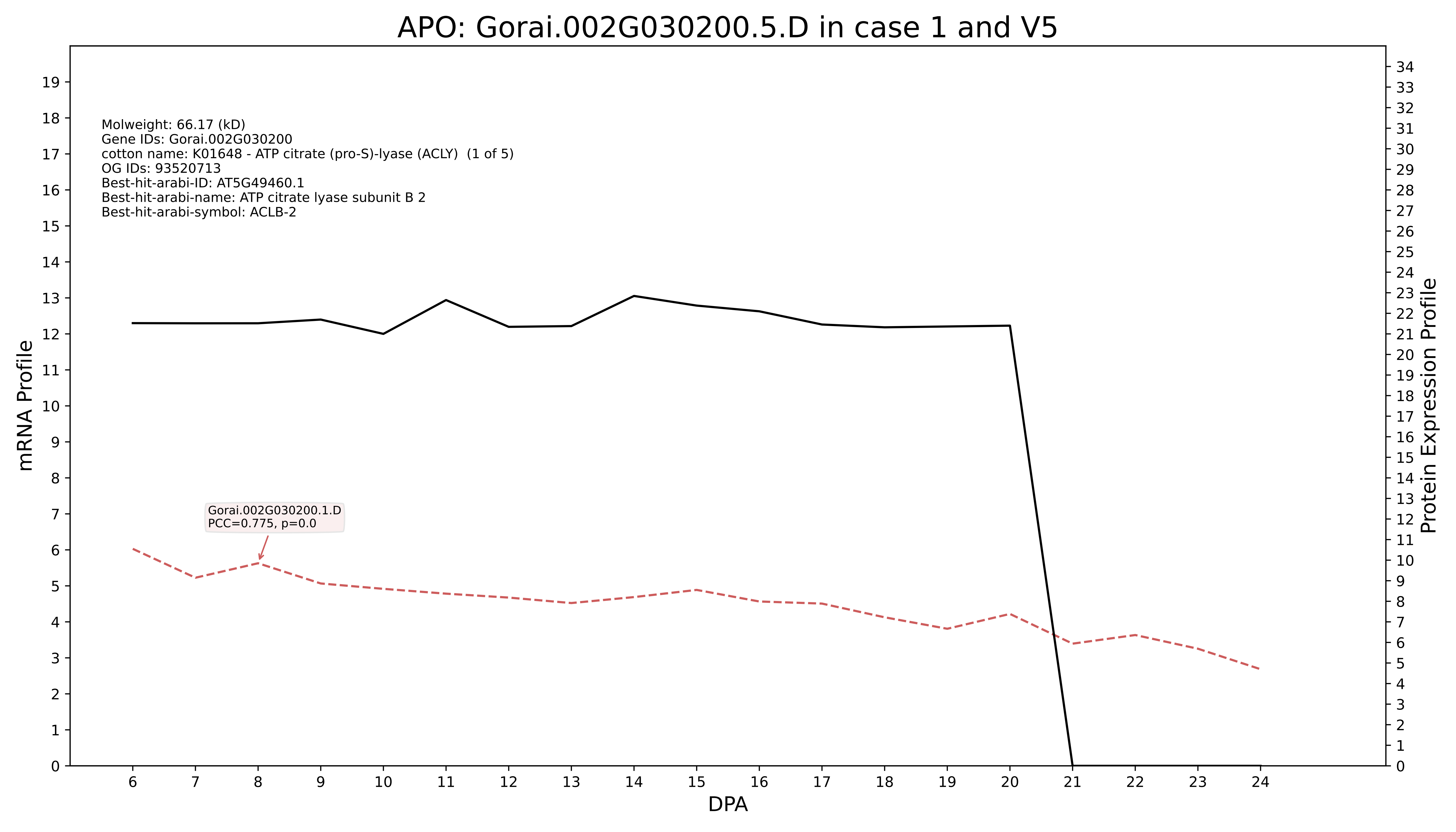

APO: Gorai.002G030200.5.D in case 1 and V5
Molweight: 66.17 (kD)
Gene IDs: Gorai.002G030200
cotton name: K01648 - ATP citrate (pro-S)-lyase (ACLY) (1 of 5)
OG IDs: 93520713
Best-hit-arabi-ID: AT5G49460.1
Best-hit-arabi-name: ATP citrate lyase subunit B 2
Best-hit-arabi-symbol: ACLB-2
Gorai.002G030200.1.D
PCC=0.775, p=0.0
mRNA Profile
Protein Expression Profile
DPA

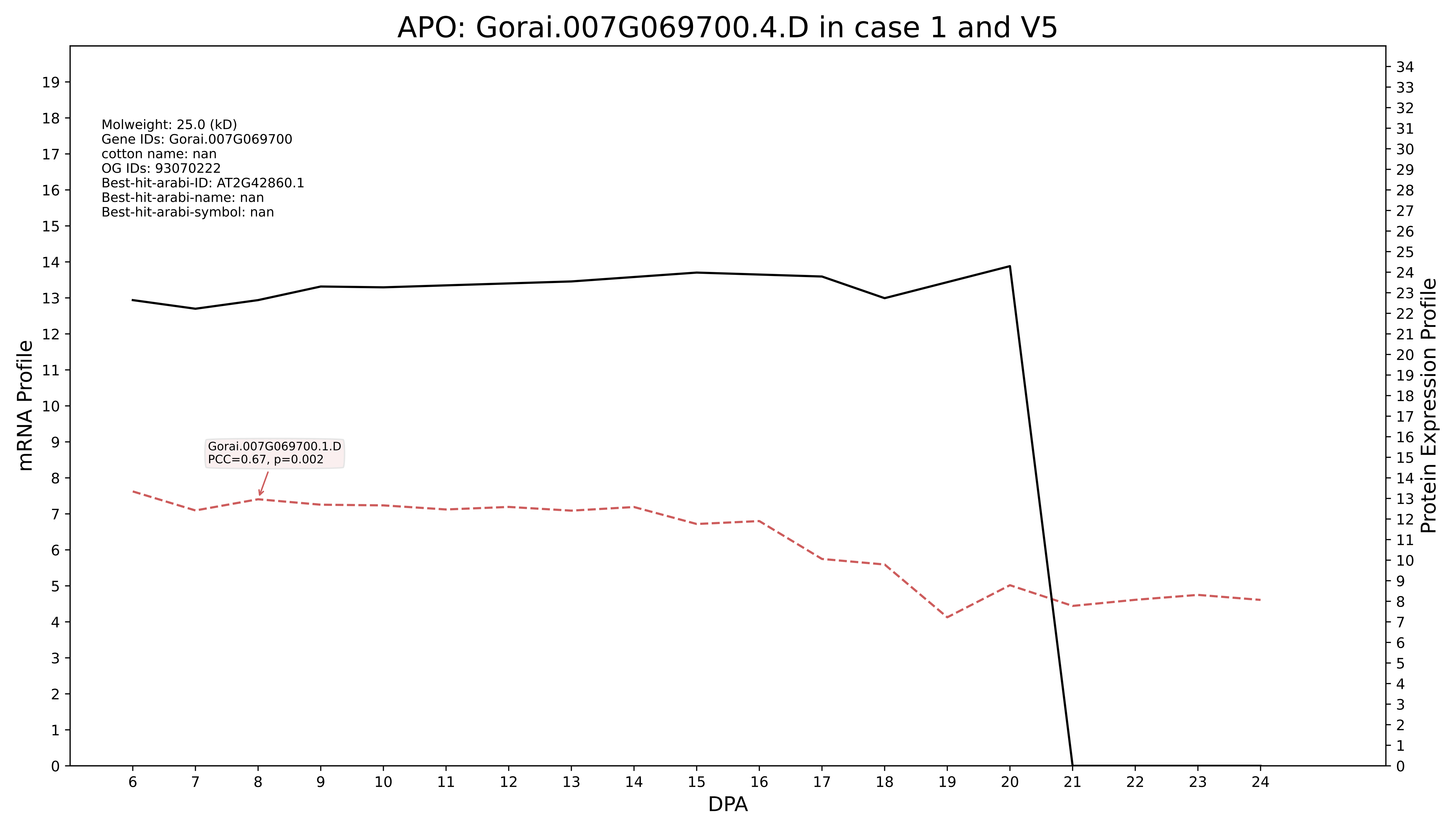
APO: Gorai.007G069700.4.D in case 1 and V5
Molweight: 25.0 (kD)
Gene IDs: Gorai.007G069700
cotton name: nan
OG IDs: 93070222
Best-hit-arabi-ID: AT2G42860.1
Best-hit-arabi-name: nan
Best-hit-arabi-symbol: nan
Gorai.007G069700.1.D
PCC=0.67, p=0.002
mRNA Profile
Protein Expression Profile
DPA

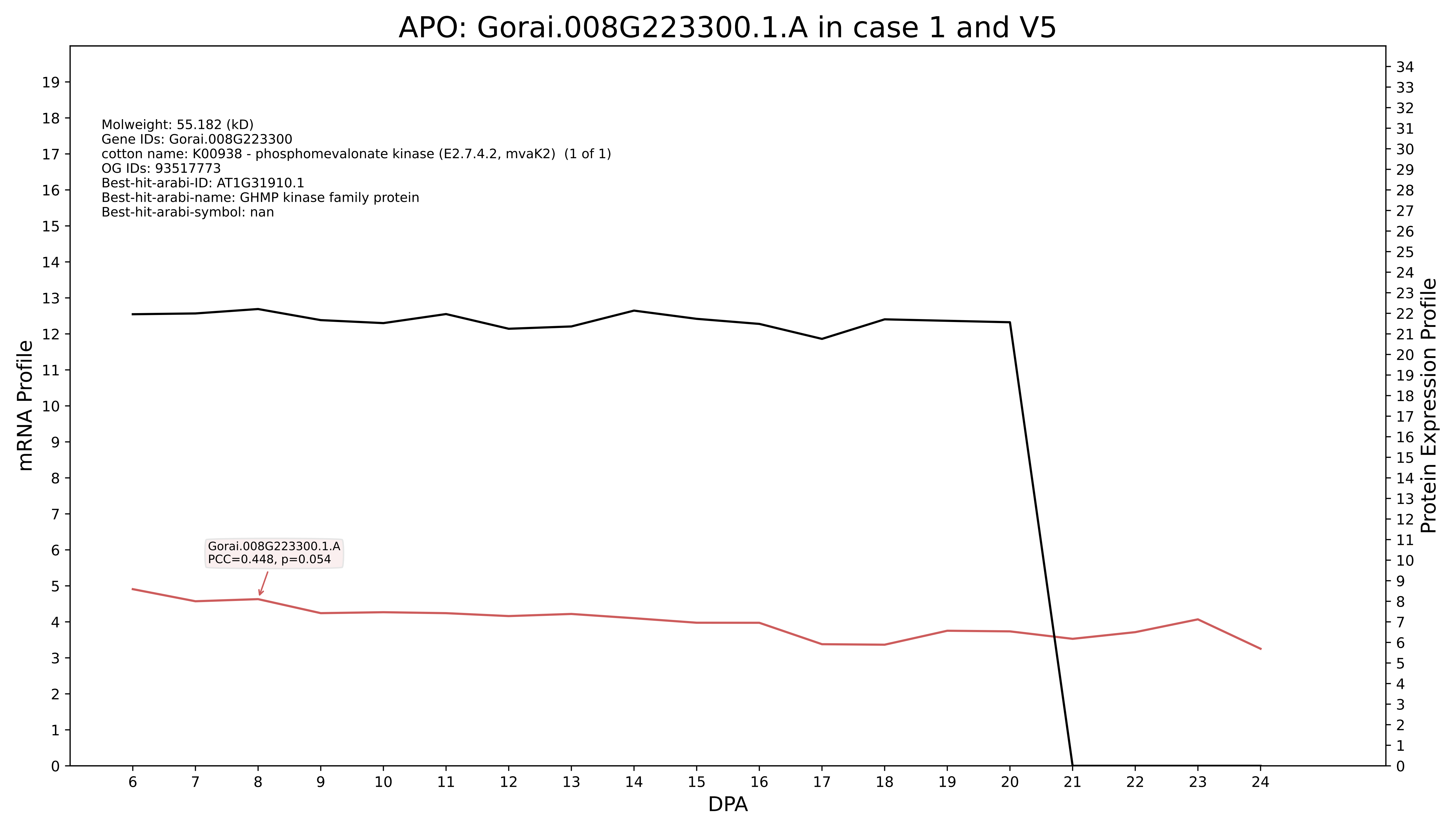
APO: Gorai.008G223300.1.A in case 1 and V5
Molweight: 55.182 (kD)
Gene IDs: Gorai.008G223300
cotton name: K00938 - phosphomevalonate kinase (E2.7.4.2, mvaK2) (1 of 1)
OG IDs: 93517773
Best-hit-arabi-ID: AT1G31910.1
Best-hit-arabi-name: GHMP kinase family protein
Best-hit-arabi-symbol: nan
Gorai.008G223300.1.A
PCC=0.448, p=0.054
mRNA Profile
Protein Expression Profile
DPA

### APO: Gorai.006G233600.1.D in case 1 and V5

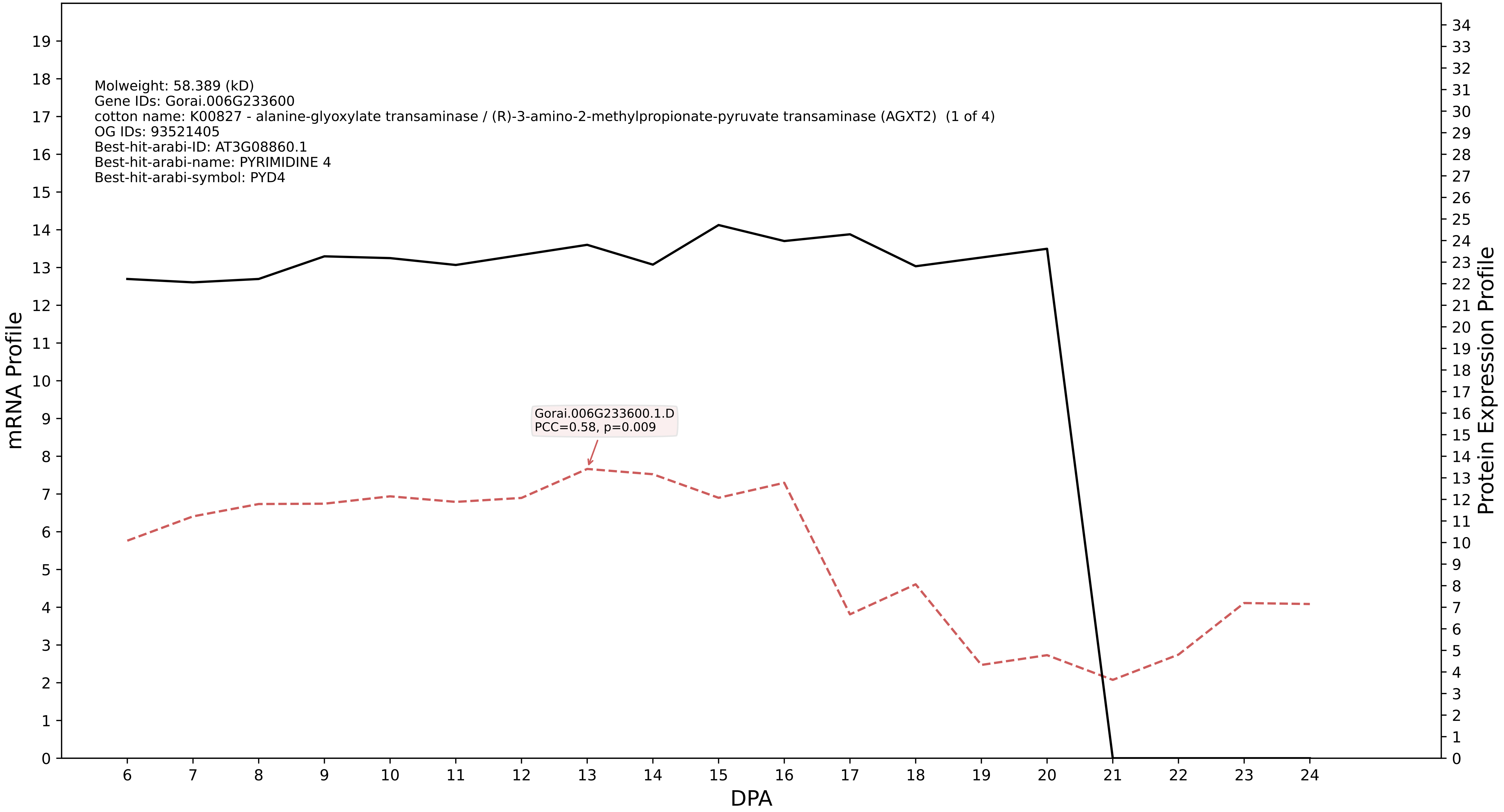

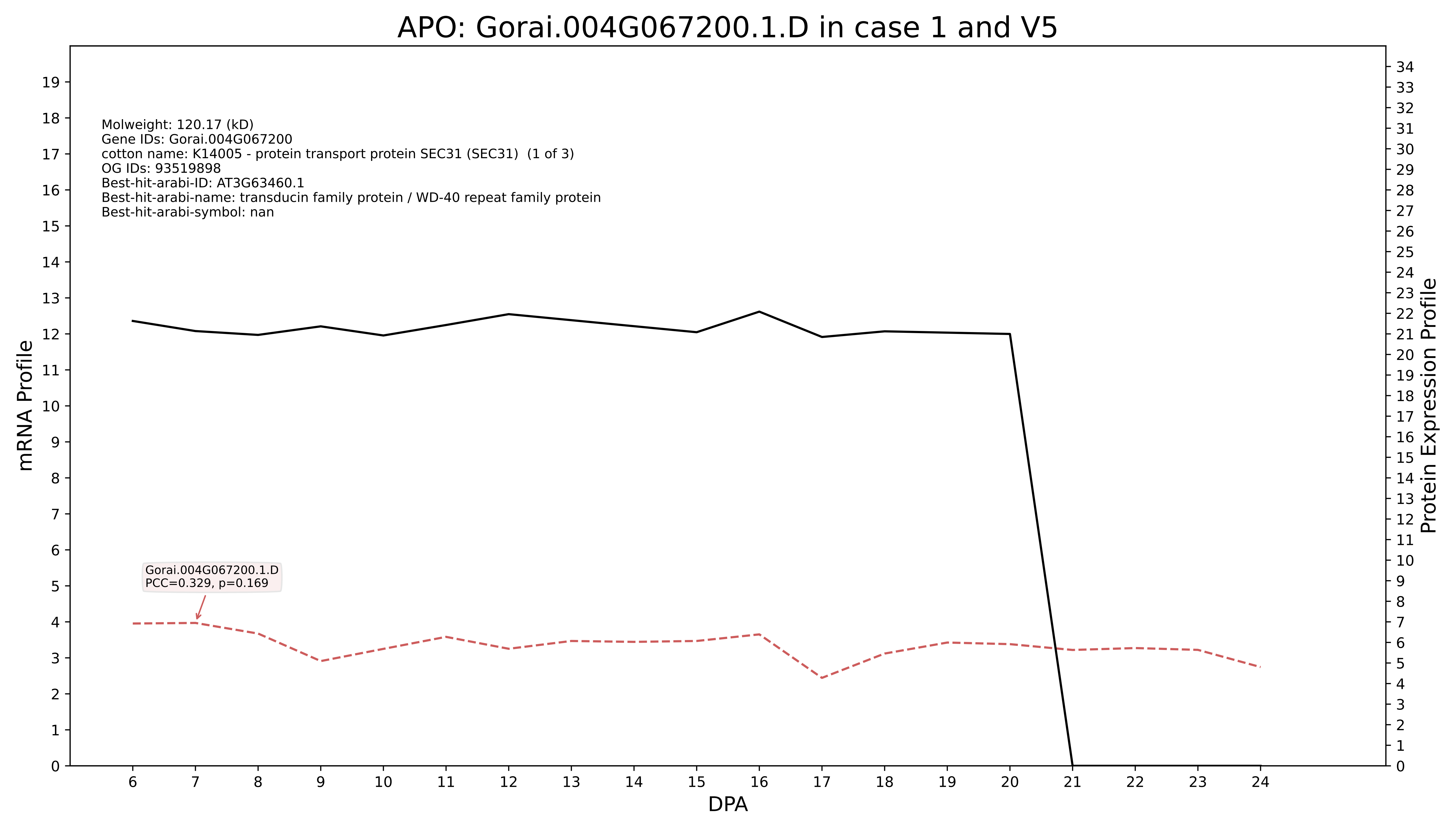
APO: Gorai.004G067200.1.D in case 1 and V5
Molweight: 120.17 (kD)
Gene IDs: Gorai.004G067200
cotton name: K14005 - protein transport protein SEC31 (SEC31) (1 of 3)
OG IDs: 93519898
Best-hit-arabi-ID: AT3G63460.1
Best-hit-arabi-name: transducin family protein / WD-40 repeat family protein
Best-hit-arabi-symbol: nan
Gorai.004G067200.1.D
PCC=0.329, p=0.169
mRNA Profile
Protein Expression Profile
DPA

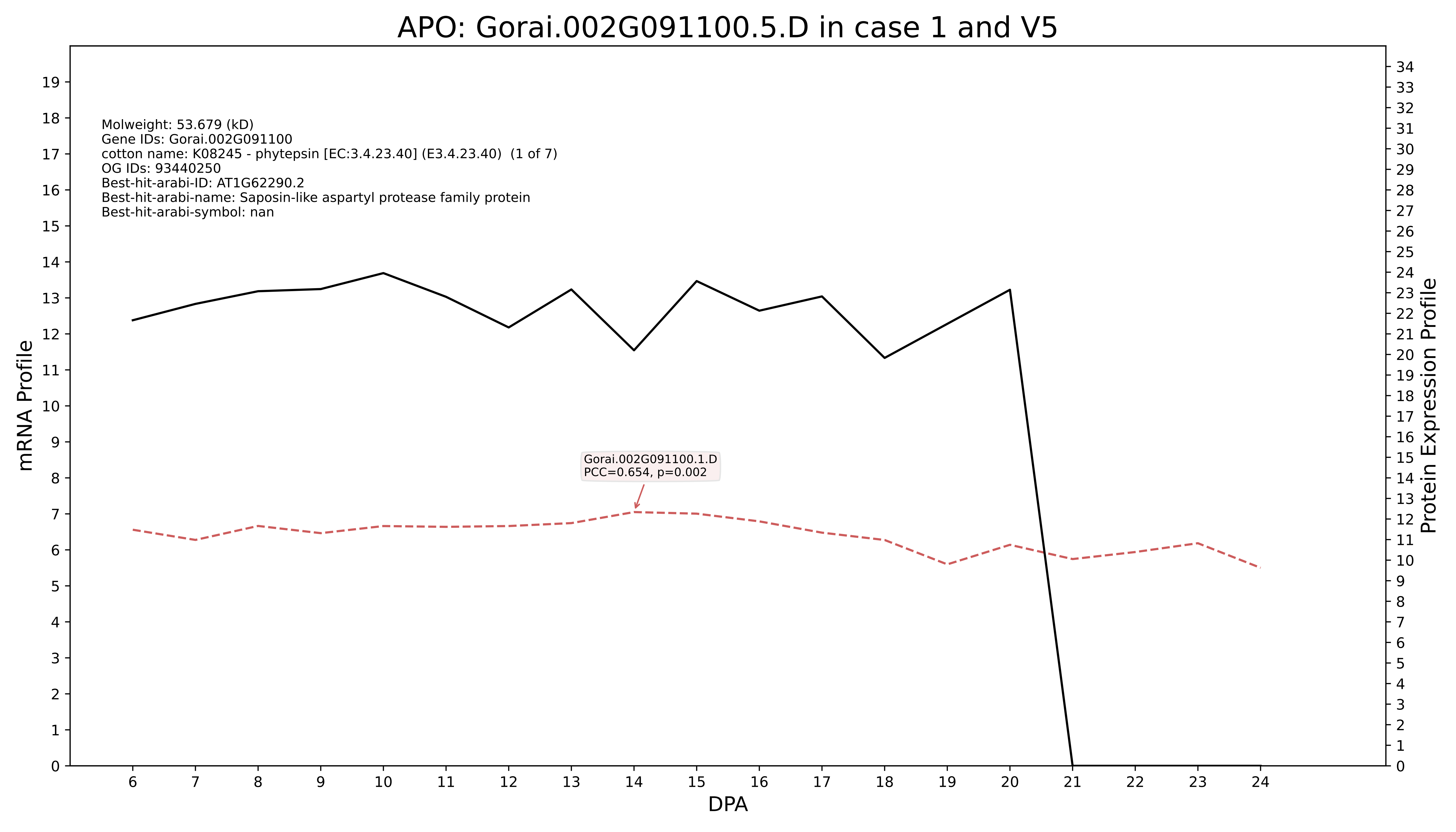
APO: Gorai.002G091100.5.D in case 1 and V5
Molweight: 53.679 (kD)
Gene IDs: Gorai.002G091100
cotton name: K08245 - phytepsin [EC:3.4.23.40] (E3.4.23.40) (1 of 7)
OG IDs: 93440250
Best-hit-arabi-ID: AT1G62290.2
Best-hit-arabi-name: Saposin-like aspartyl protease family protein
Best-hit-arabi-symbol: nan
Gorai.002G091100.1.D
PCC=0.654, p=0.002
mRNA Profile
Protein Expression Profile
DPA

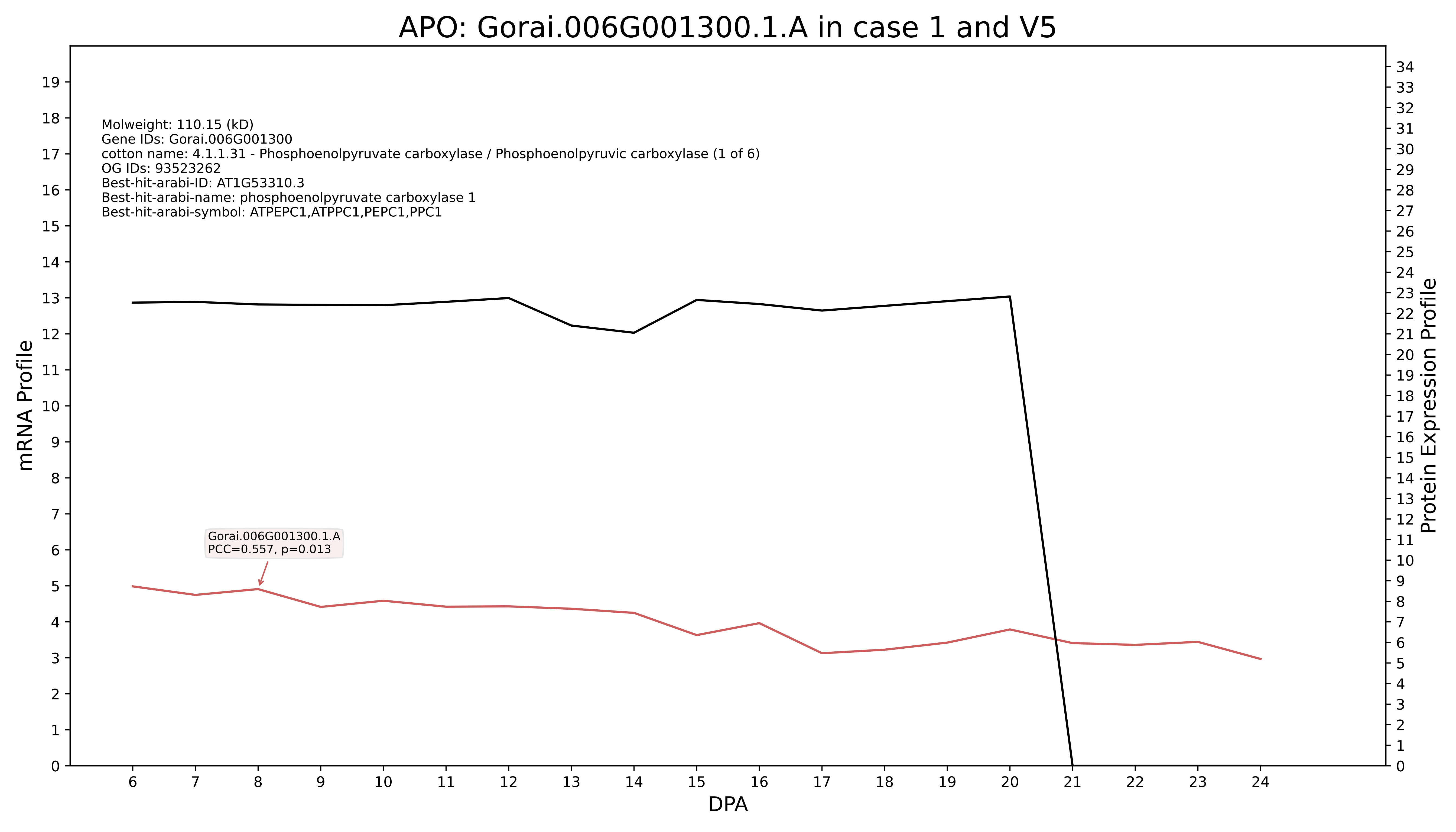

APO: Gorai.006G001300.1.A in case 1 and V5
Molweight: 110.15 (kD)
Gene IDs: Gorai.006G001300
cotton name: 4.1.1.31 - Phosphoenolpyruvate carboxylase / Phosphoenolpyruvic carboxylase (1 of 6)
OG IDs: 93523262
Best-hit-arabi-ID: AT1G53310.3
Best-hit-arabi-name: phosphoenolpyruvate carboxylase 1
Best-hit-arabi-symbol: ATPEPC1,ATPPC1,PEPC1,PPC1
Gorai.006G001300.1.A
PCC=0.557, p=0.013
mRNA Profile
Protein Expression Profile
DPA

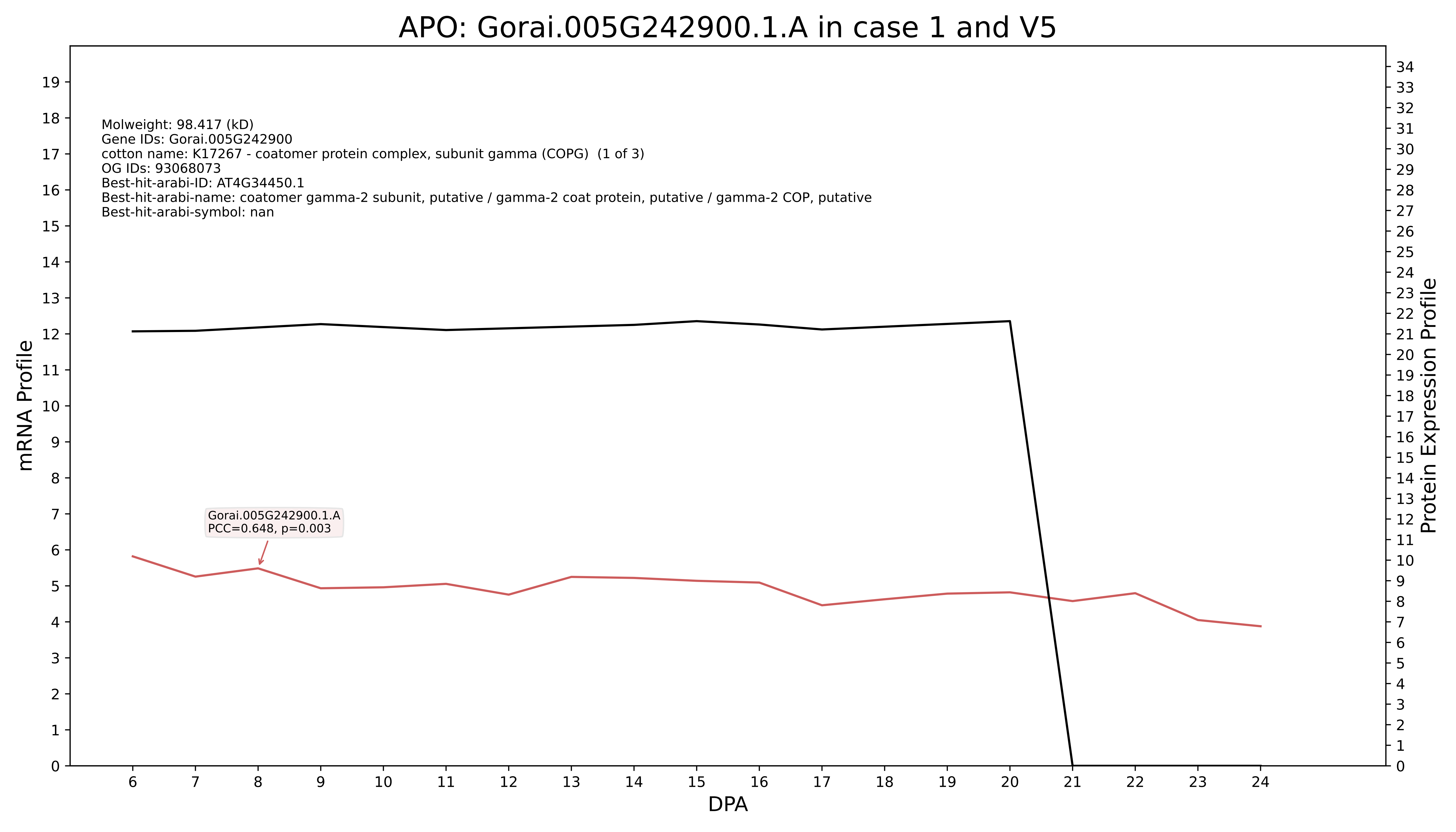
APO: Gorai.005G242900.1.A in case 1 and V5
Molweight: 98.417 (kD)
Gene IDs: Gorai.005G242900
cotton name: K17267 - coatomer protein complex, subunit gamma (COPG) (1 of 3)
OG IDs: 93068073
Best-hit-arabi-ID: AT4G34450.1
Best-hit-arabi-name: coatomer gamma-2 subunit, putative / gamma-2 coat protein, putative / gamma-2 COP, putative
Best-hit-arabi-symbol: nan
Gorai.005G242900.1.A
PCC=0.648, p=0.003
mRNA Profile
Protein Expression Profile
DPA

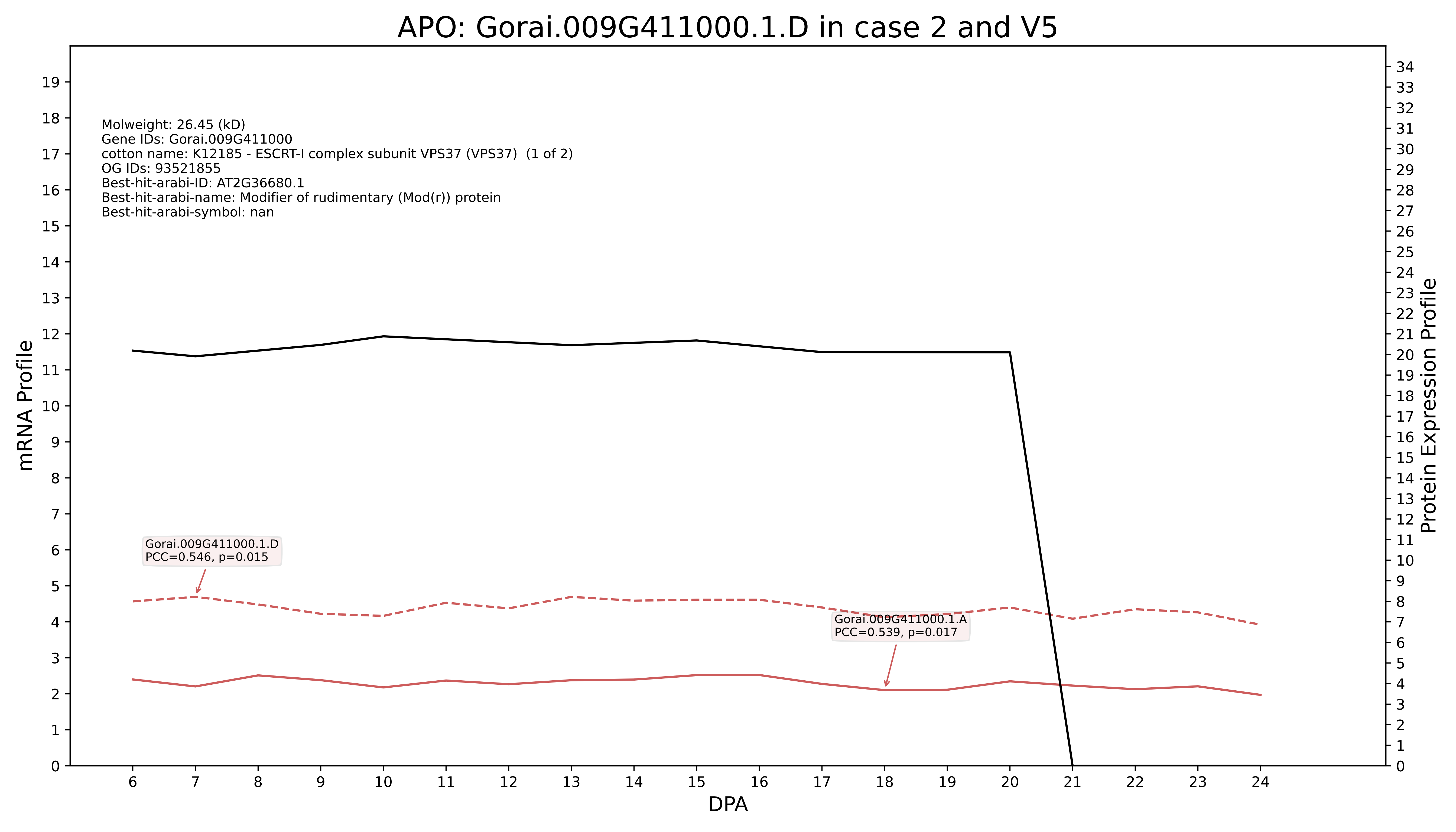

APO: Gorai.009G411000.1.D in case 2 and V5
Molweight: 26.45 (kD)
Gene IDs: Gorai.009G411000
cotton name: K12185 - ESCRT-I complex subunit VPS37 (VPS37) (1 of 2)
OG IDs: 93521855
Best-hit-arabi-ID: AT2G36680.1
Best-hit-arabi-name: Modifier of rudimentary (Mod(r)) protein
Best-hit-arabi-symbol: nan
Gorai.009G411000.1.D
PCC=0.546, p=0.015
Gorai.009G411000.1.A
PCC=0.539, p=0.017
mRNA Profile
Protein Expression Profile
DPA

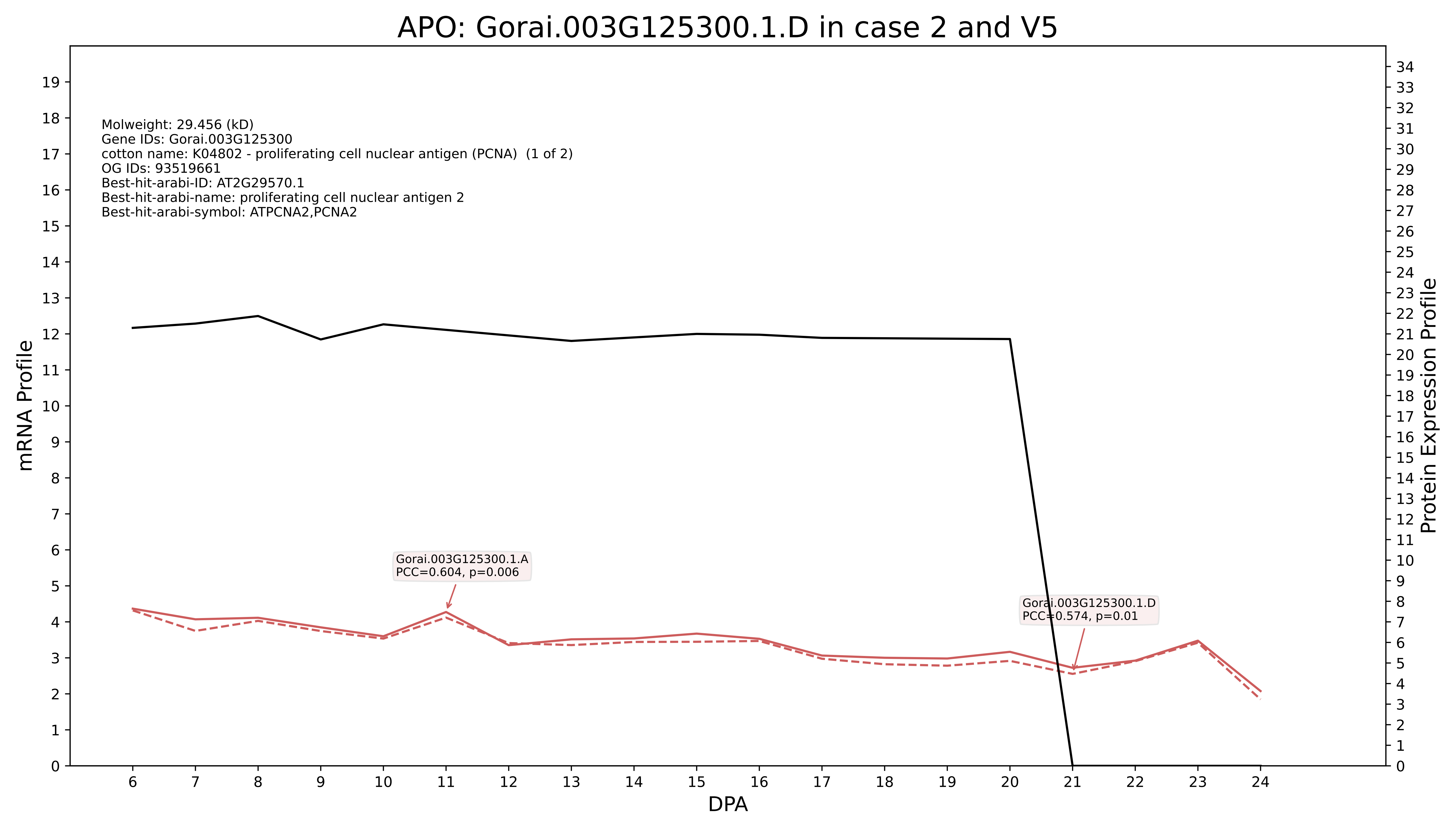
APO: Gorai.003G125300.1.D in case 2 and V5
Molweight: 29.456 (kD)
Gene IDs: Gorai.003G125300
cotton name: K04802 - proliferating cell nuclear antigen (PCNA) (1 of 2)
OG IDs: 93519661
Best-hit-arabi-ID: AT2G29570.1
Best-hit-arabi-name: proliferating cell nuclear antigen 2
Best-hit-arabi-symbol: ATPCNA2,PCNA2
Gorai.003G125300.1.A
PCC=0.604, p=0.006
Gorai.003G125300.1.D
PCC=0.574, p=0.01
mRNA Profile
Protein Expression Profile
DPA

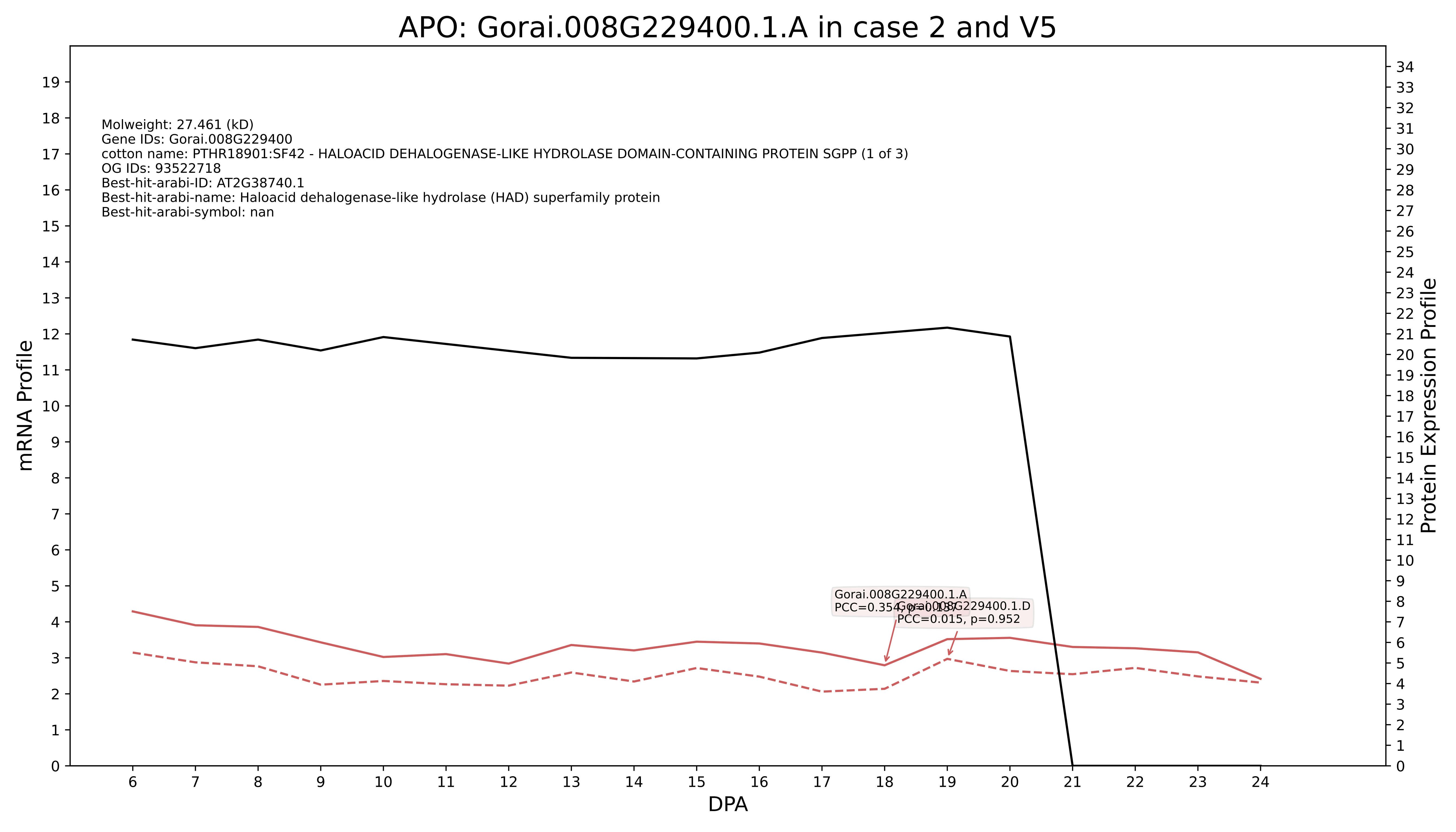

APO: Gorai.008G229400.1.A in case 2 and V5
Molweight: 27.461 (kD)
Gene IDs: Gorai.008G229400
cotton name: PTHR18901:SF42 - HALOACID DEHALOGENASE-LIKE HYDROLASE DOMAIN-CONTAINING PROTEIN SGPP (1 of 3)
OG IDs: 93522718
Best-hit-arabi-ID: AT2G38740.1
Best-hit-arabi-name: Haloacid dehalogenase-like hydrolase (HAD) superfamily protein
Best-hit-arabi-symbol: nan
Gorai.008G229400.1.A
PCC=0.354, p=0.137
Gorai.008G229400.1.D
PCC=0.015, p=0.952
mRNA Profile
Protein Expression Profile
DPA

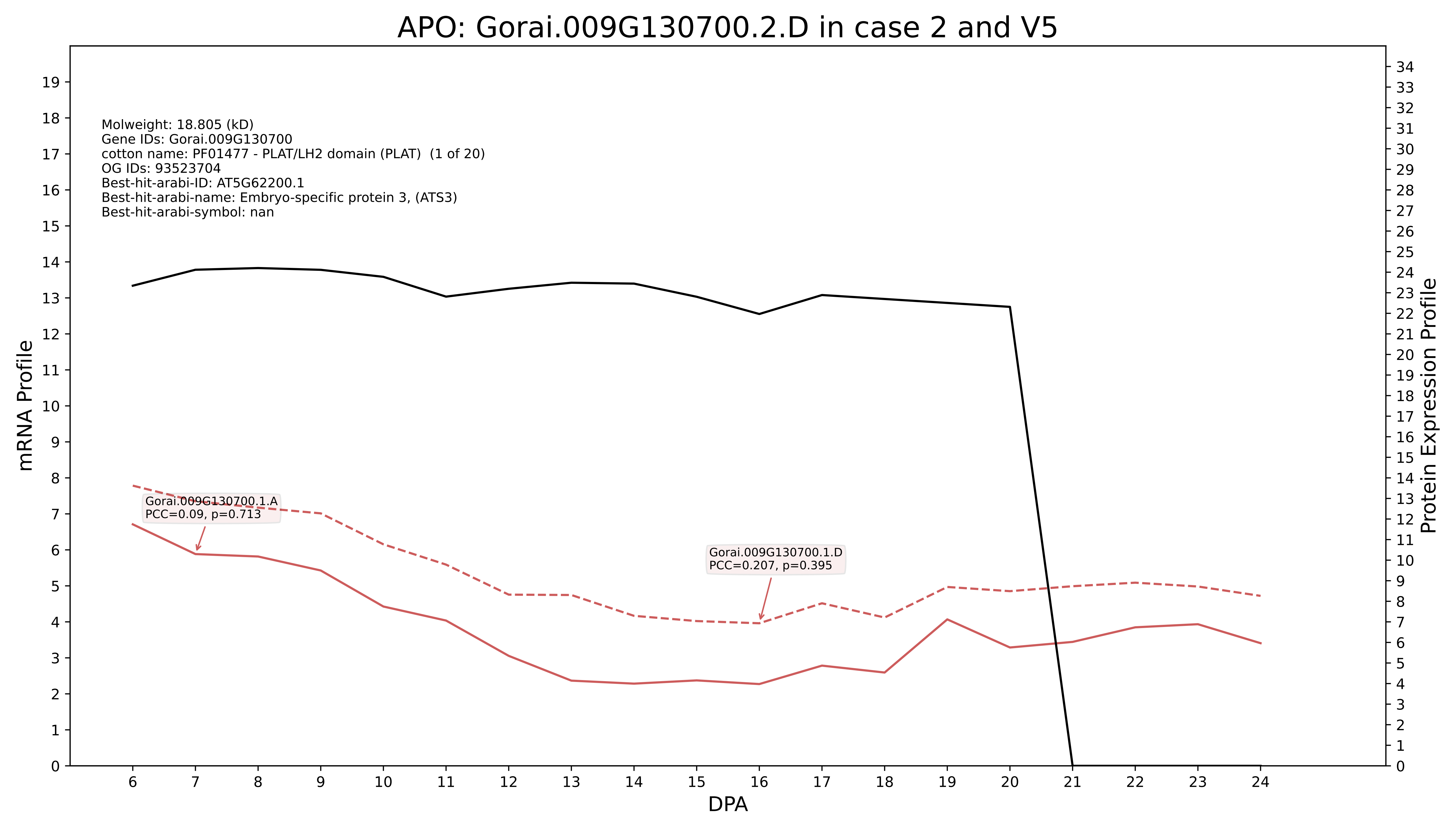
APO: Gorai.009G130700.2.D in case 2 and V5
Molweight: 18.805 (kD)
Gene IDs: Gorai.009G130700
cotton name: PF01477 - PLAT/LH2 domain (PLAT) (1 of 20)
OG IDs: 93523704
Best-hit-arabi-ID: AT5G62200.1
Best-hit-arabi-name: Embryo-specific protein 3, (ATS3)
Best-hit-arabi-symbol: nan
Gorai.009G130700.1.A
PCC=0.09, p=0.713
Gorai.009G130700.1.D
PCC=0.207, p=0.395
mRNA Profile
Protein Expression Profile
DPA

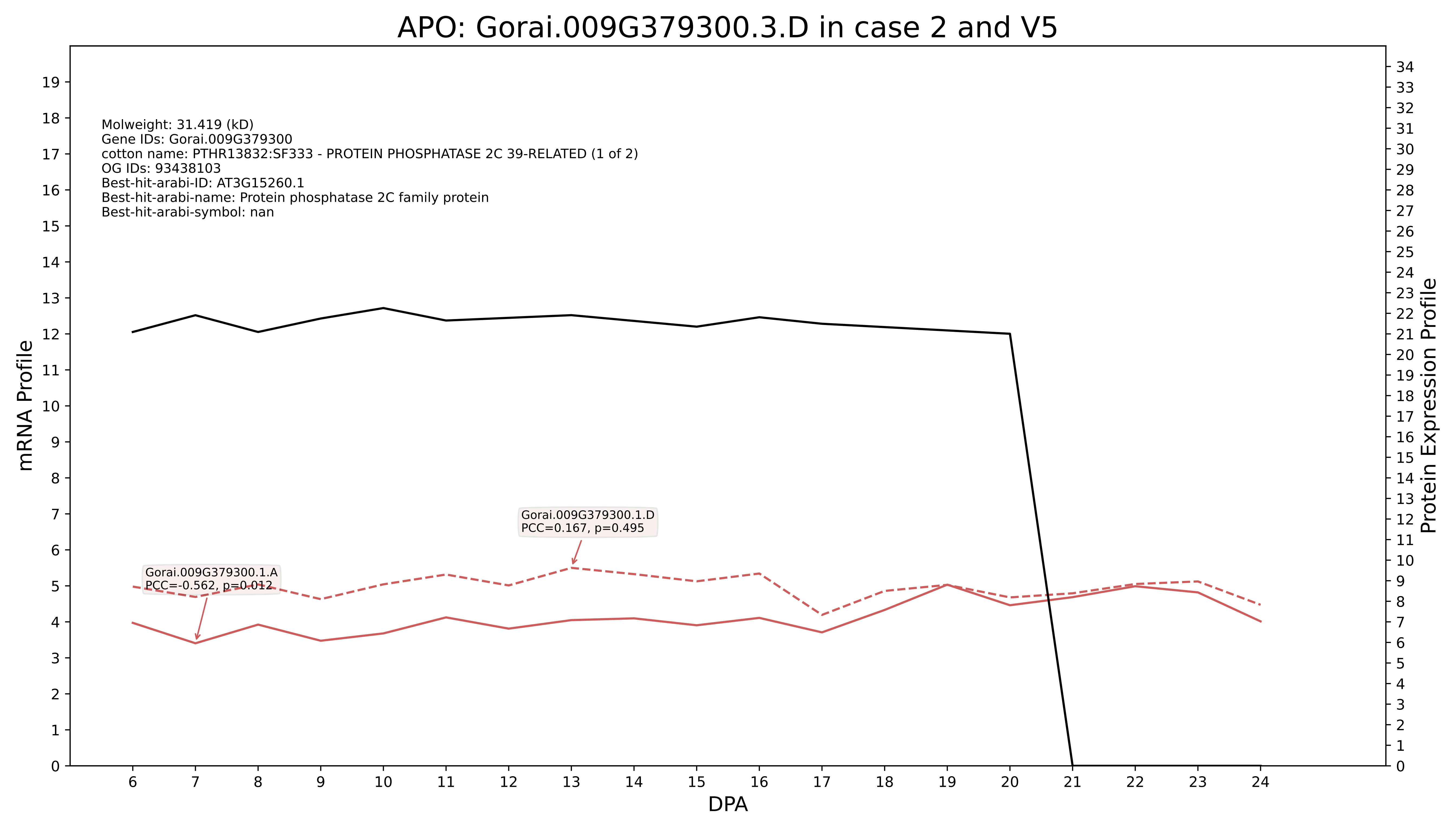
APO: Gorai.009G379300.3.D in case 2 and V5
Molweight: 31.419 (kD)
Gene IDs: Gorai.009G379300
cotton name: PTHR13832:SF333 - PROTEIN PHOSPHATASE 2C 39-RELATED (1 of 2)
OG IDs: 93438103
Best-hit-arabi-ID: AT3G15260.1
Best-hit-arabi-name: Protein phosphatase 2C family protein
Best-hit-arabi-symbol: nan
Gorai.009G379300.1.D
PCC=0.167, p=0.495
Gorai.009G379300.1.A
PCC=-0.562, p=0.012
mRNA Profile
Protein Expression Profile
DPA

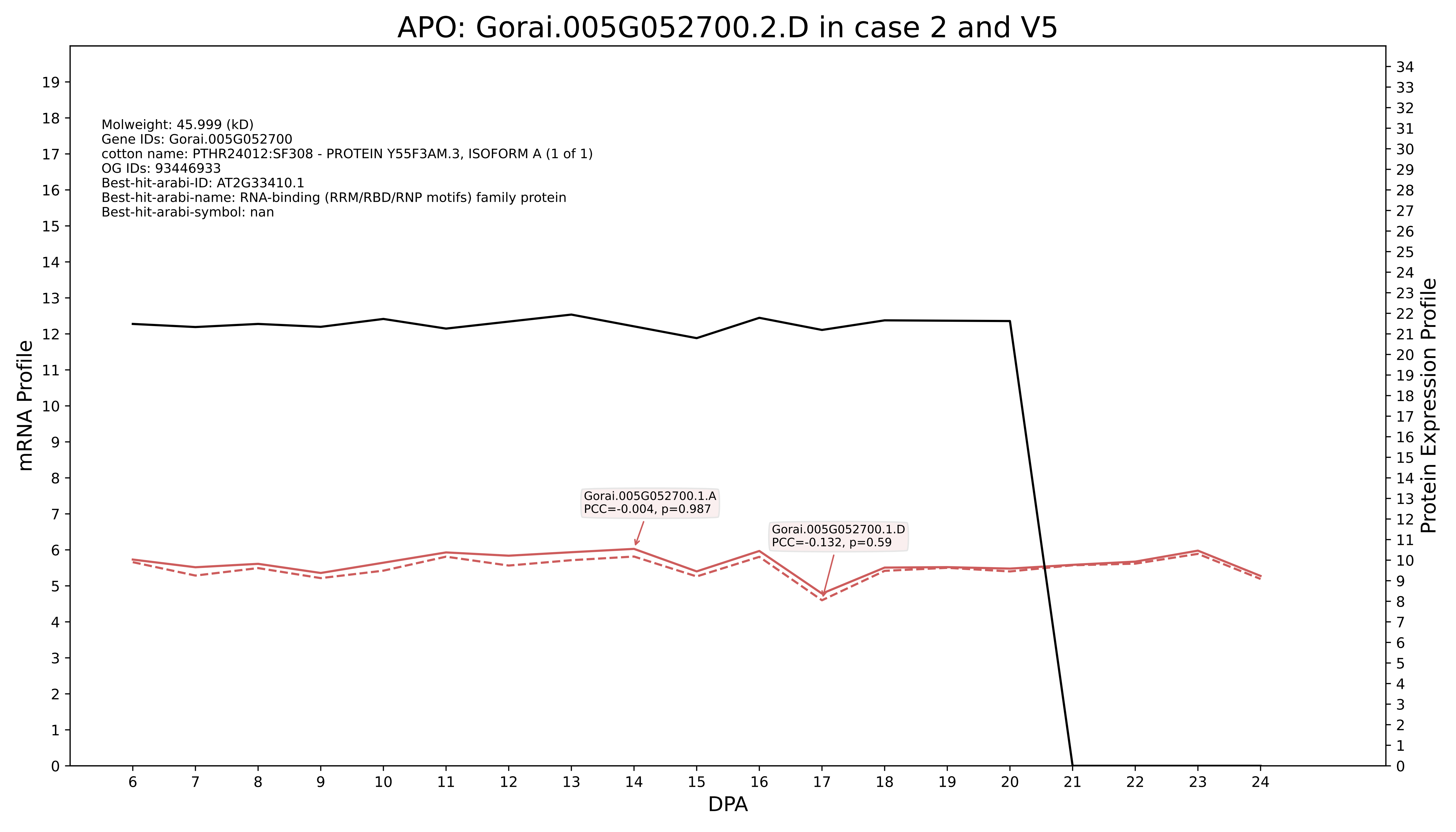
APO: Gorai.005G052700.2.D in case 2 and V5
Molweight: 45.999 (kD)
Gene IDs: Gorai.005G052700
cotton name: PTHR24012:SF308 - PROTEIN Y55F3AM.3, ISOFORM A (1 of 1)
OG IDs: 93446933
Best-hit-arabi-ID: AT2G33410.1
Best-hit-arabi-name: RNA-binding (RRM/RBD/RNP motifs) family protein
Best-hit-arabi-symbol: nan
Gorai.005G052700.1.A
PCC=-0.004, p=0.987
Gorai.005G052700.1.D
PCC=-0.132, p=0.59
mRNA Profile
Protein Expression Profile
DPA

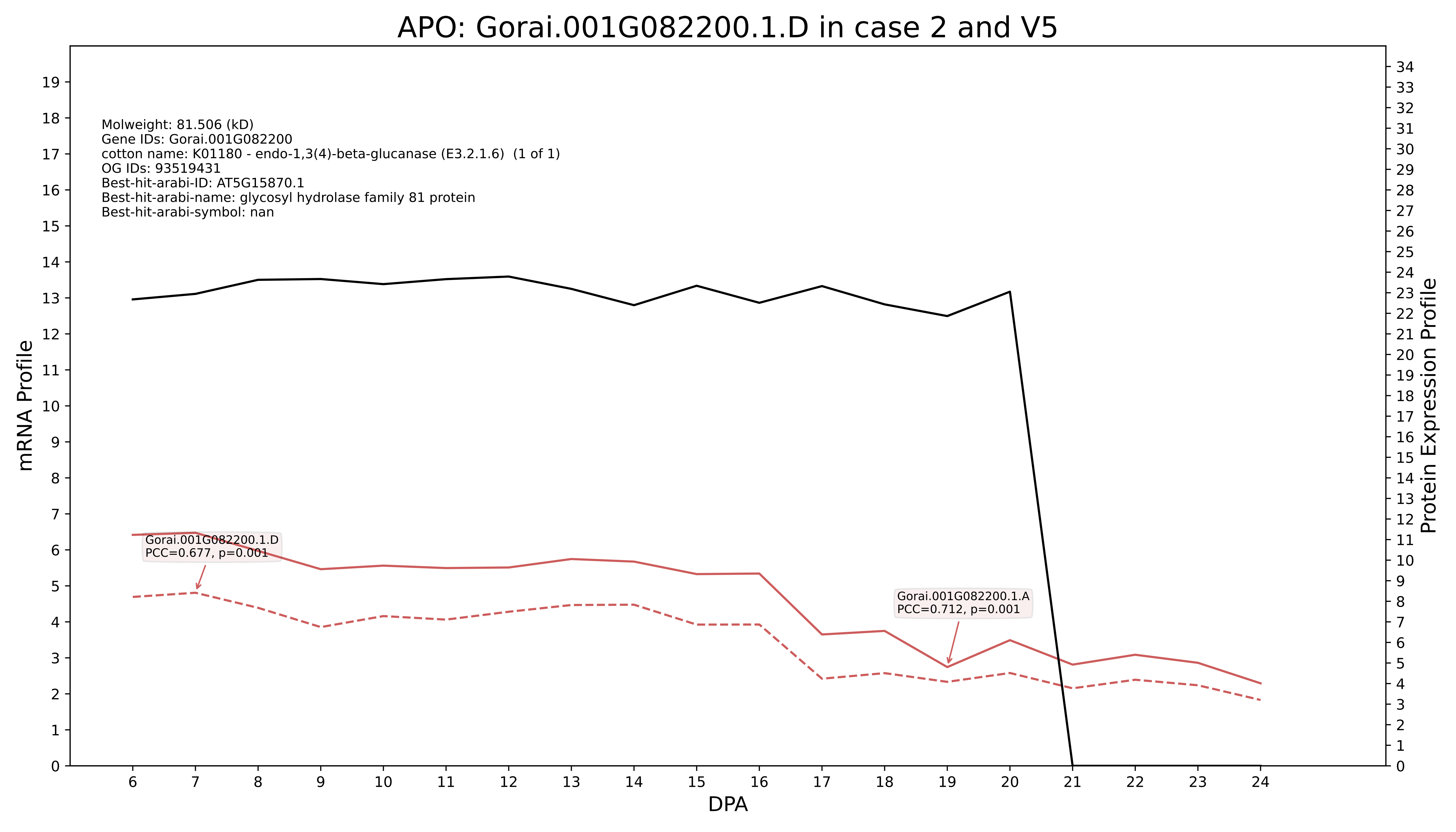
APO: Gorai.001G082200.1.D in case 2 and V5
Molweight: 81.506 (kD)
Gene IDs: Gorai.001G082200
cotton name: K01180 - endo-1,3(4)-beta-glucanase (E3.2.1.6) (1 of 1)
OG IDs: 93519431
Best-hit-arabi-ID: AT5G15870.1
Best-hit-arabi-name: glycosyl hydrolase family 81 protein
Best-hit-arabi-symbol: nan
Gorai.001G082200.1.D
PCC=0.677, p=0.001
Gorai.001G082200.1.A
PCC=0.712, p=0.001
mRNA Profile
Protein Expression Profile
DPA

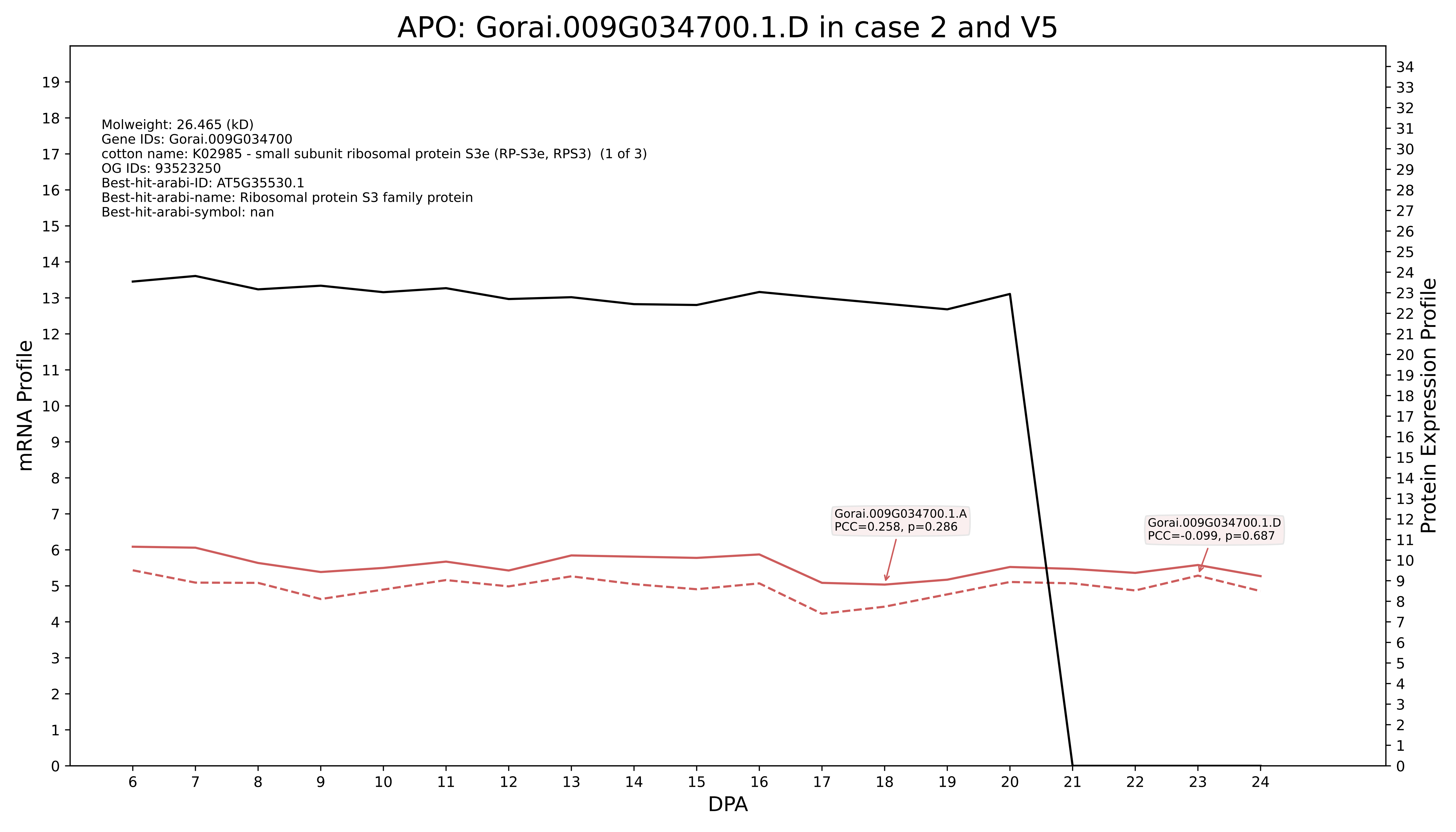

APO: Gorai.009G034700.1.D in case 2 and V5
Molweight: 26.465 (kD)
Gene IDs: Gorai.009G034700
cotton name: K02985 - small subunit ribosomal protein S3e (RP-S3e, RPS3) (1 of 3)
OG IDs: 93523250
Best-hit-arabi-ID: AT5G35530.1
Best-hit-arabi-name: Ribosomal protein S3 family protein
Best-hit-arabi-symbol: nan
Gorai.009G034700.1.A
PCC=0.258, p=0.286
Gorai.009G034700.1.D
PCC=-0.099, p=0.687
mRNA Profile
Protein Expression Profile
DPA

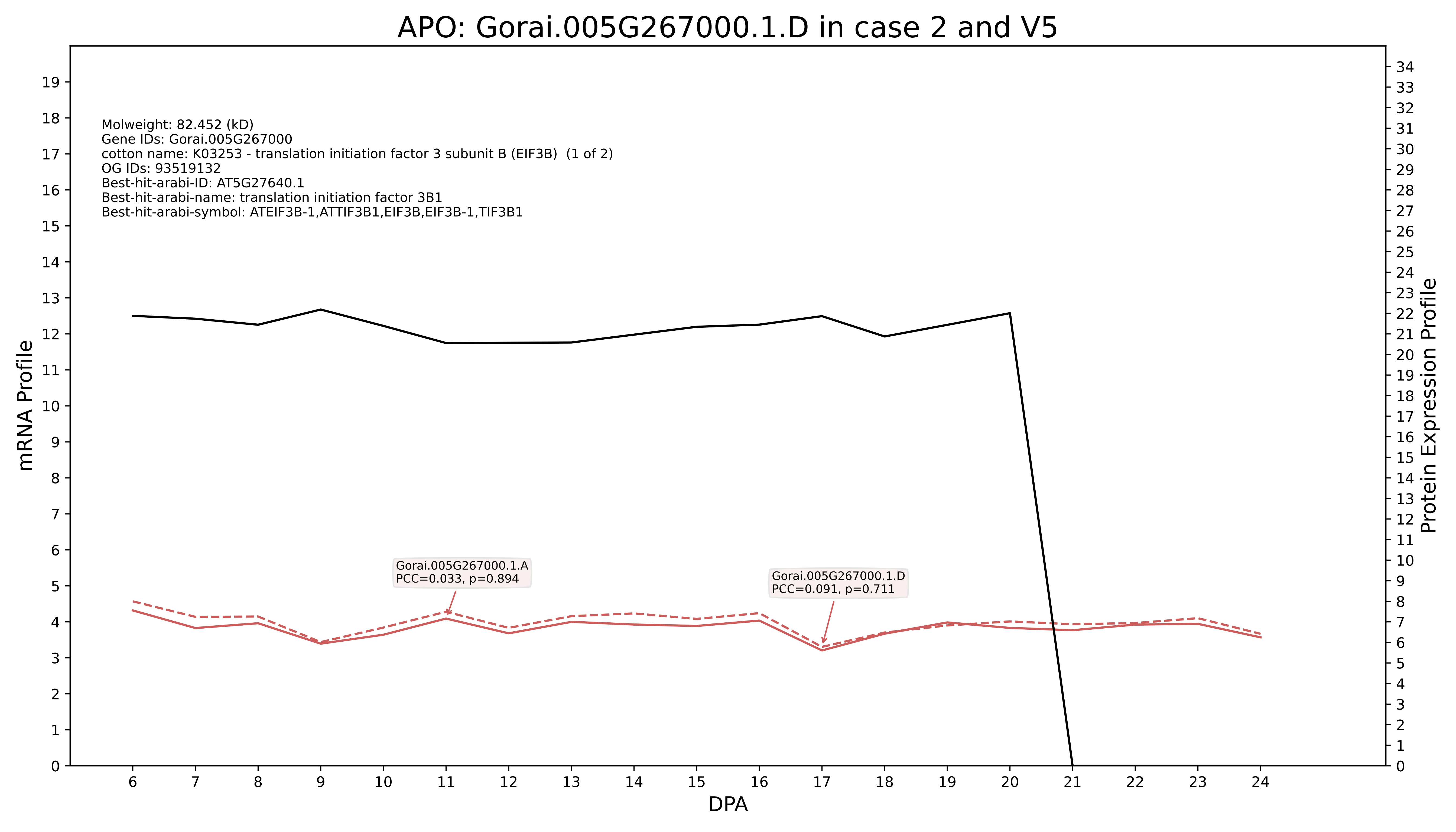

APO: Gorai.005G267000.1.D in case 2 and V5
Molweight: 82.452 (kD)
Gene IDs: Gorai.005G267000
cotton name: K03253 - translation initiation factor 3 subunit B (EIF3B) (1 of 2)
OG IDs: 93519132
Best-hit-arabi-ID: AT5G27640.1
Best-hit-arabi-name: translation initiation factor 3B1
Best-hit-arabi-symbol: ATEIF3B-1,ATTIF3B1,EIF3B,EIF3B-1,TIF3B1
Gorai.005G267000.1.A
PCC=0.033, p=0.894
Gorai.005G267000.1.D
PCC=0.091, p=0.711
mRNA Profile
Protein Expression Profile
DPA

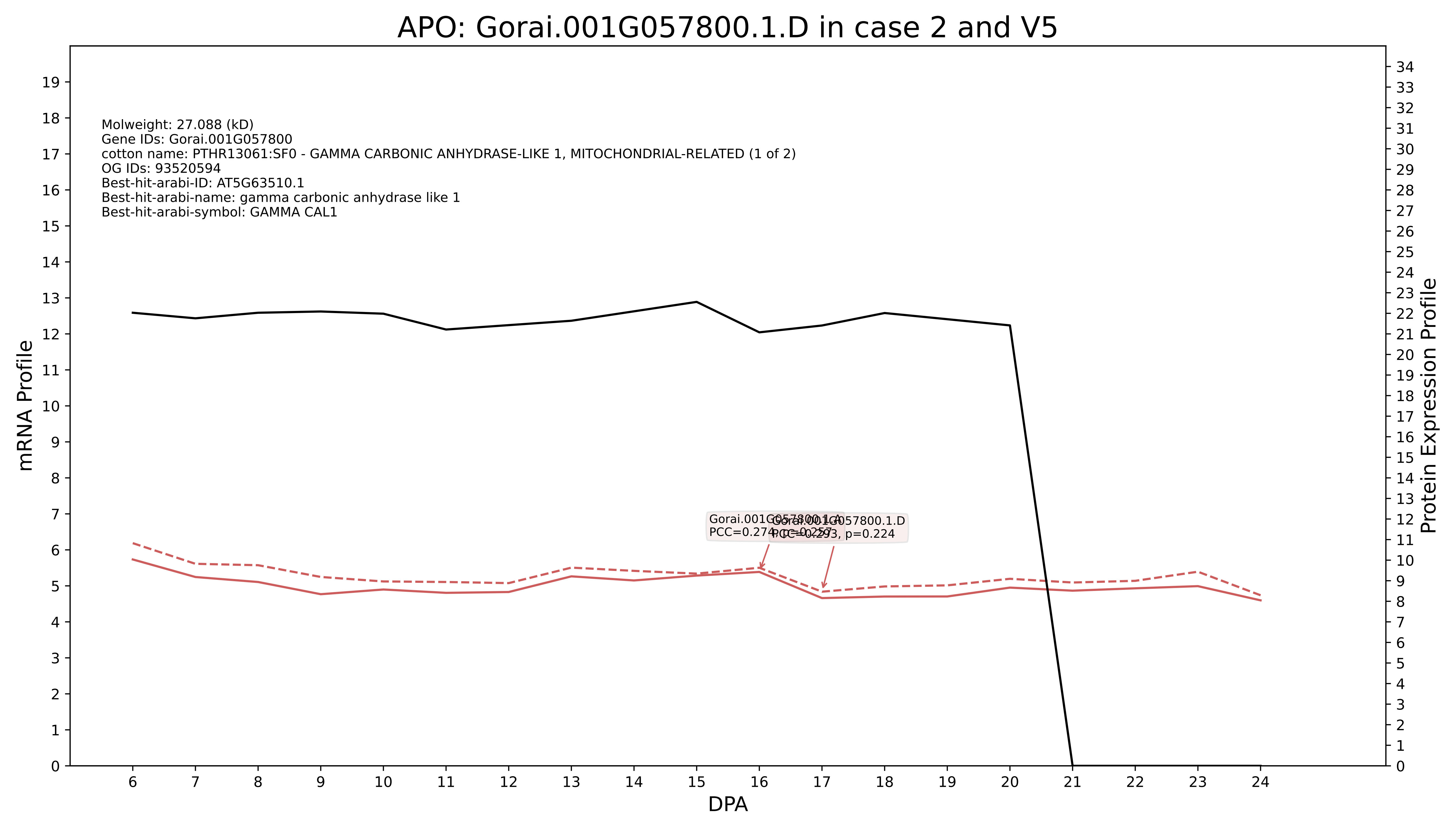

APO: Gorai.001G057800.1.D in case 2 and V5
Molweight: 27.088 (kD)
Gene IDs: Gorai.001G057800
cotton name: PTHR13061:SF0 - GAMMA CARBONIC ANHYDRASE-LIKE 1, MITOCHONDRIAL-RELATED (1 of 2)
OG IDs: 93520594
Best-hit-arabi-ID: AT5G63510.1
Best-hit-arabi-name: gamma carbonic anhydrase like 1
Best-hit-arabi-symbol: GAMMA CAL1
Gorai.001G057800.1.D
PCC=0.293, p=0.224
mRNA Profile
Protein Expression Profile
DPA

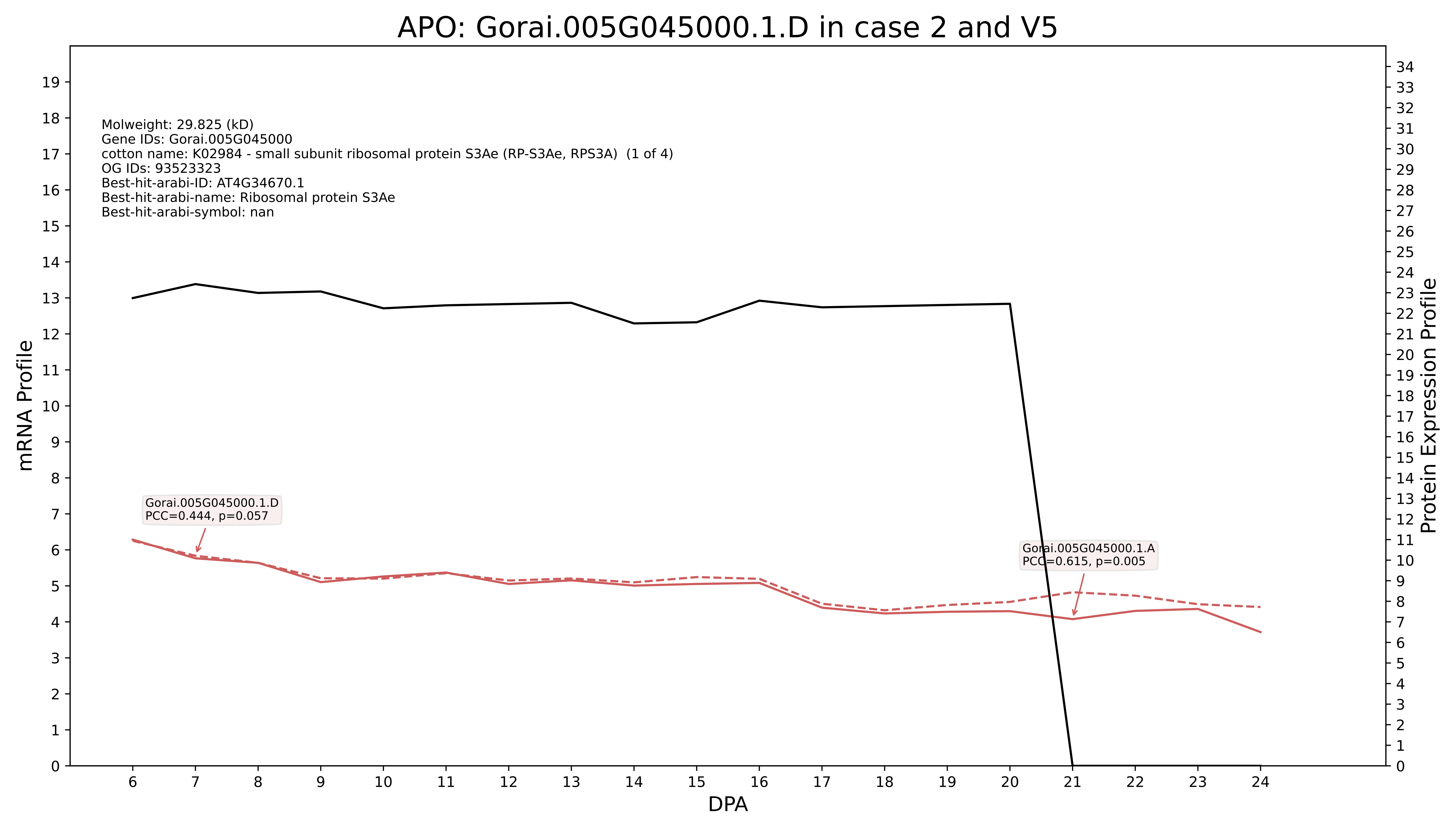

APO: Gorai.005G045000.1.D in case 2 and V5
Molweight: 29.825 (kD)
Gene IDs: Gorai.005G045000
cotton name: K02984 - small subunit ribosomal protein S3Ae (RP-S3Ae, RPS3A) (1 of 4)
OG IDs: 93523323
Best-hit-arabi-ID: AT4G34670.1
Best-hit-arabi-name: Ribosomal protein S3Ae
Best-hit-arabi-symbol: nan
Gorai.005G045000.1.D
PCC=0.444, p=0.057
Gorai.005G045000.1.A
PCC=0.615, p=0.005
mRNA Profile
Protein Expression Profile
DPA

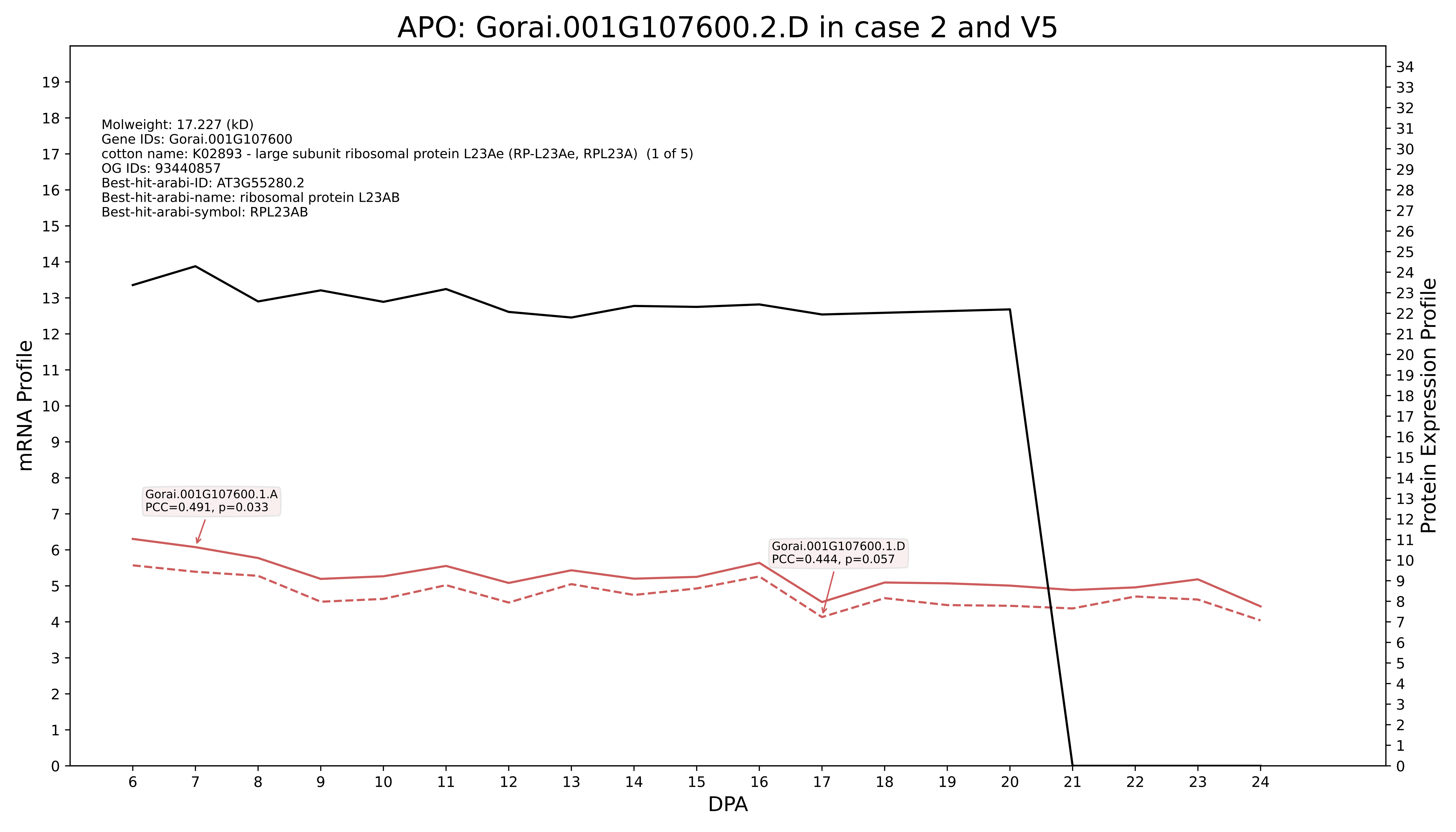

APO: Gorai.001G107600.2.D in case 2 and V5
Molweight: 17.227 (kD)
Gene IDs: Gorai.001G107600
cotton name: K02893 - large subunit ribosomal protein L23Ae (RP-L23Ae, RPL23A) (1 of 5)
OG IDs: 93440857
Best-hit-arabi-ID: AT3G55280.2
Best-hit-arabi-name: ribosomal protein L23AB
Best-hit-arabi-symbol: RPL23AB
Gorai.001G107600.1.A
PCC=0.491, p=0.033
Gorai.001G107600.1.D
PCC=0.444, p=0.057
mRNA Profile
Protein Expression Profile
DPA

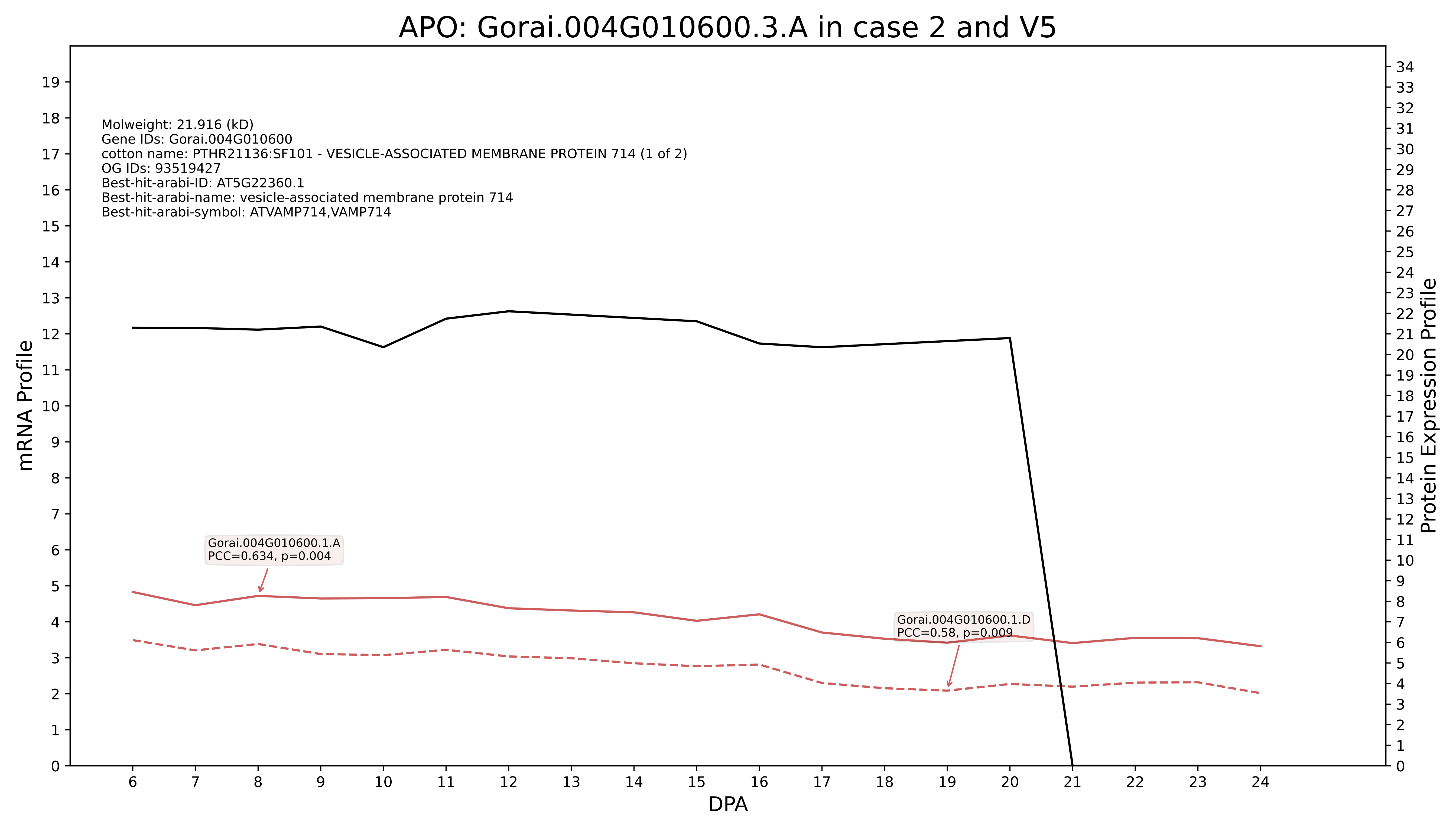
APO: Gorai.004G010600.3.A in case 2 and V5
Molweight: 21.916 (kD)
Gene IDs: Gorai.004G010600
cotton name: PTHR21136:SF101 - VESICLE-ASSOCIATED MEMBRANE PROTEIN 714 (1 of 2)
OG IDs: 93519427
Best-hit-arabi-ID: AT5G22360.1
Best-hit-arabi-name: vesicle-associated membrane protein 714
Best-hit-arabi-symbol: ATVAMP714,VAMP714
Gorai.004G010600.1.A
PCC=0.634, p=0.004
Gorai.004G010600.1.D
PCC=0.58, p=0.009
mRNA Profile
Protein Expression Profile
DPA

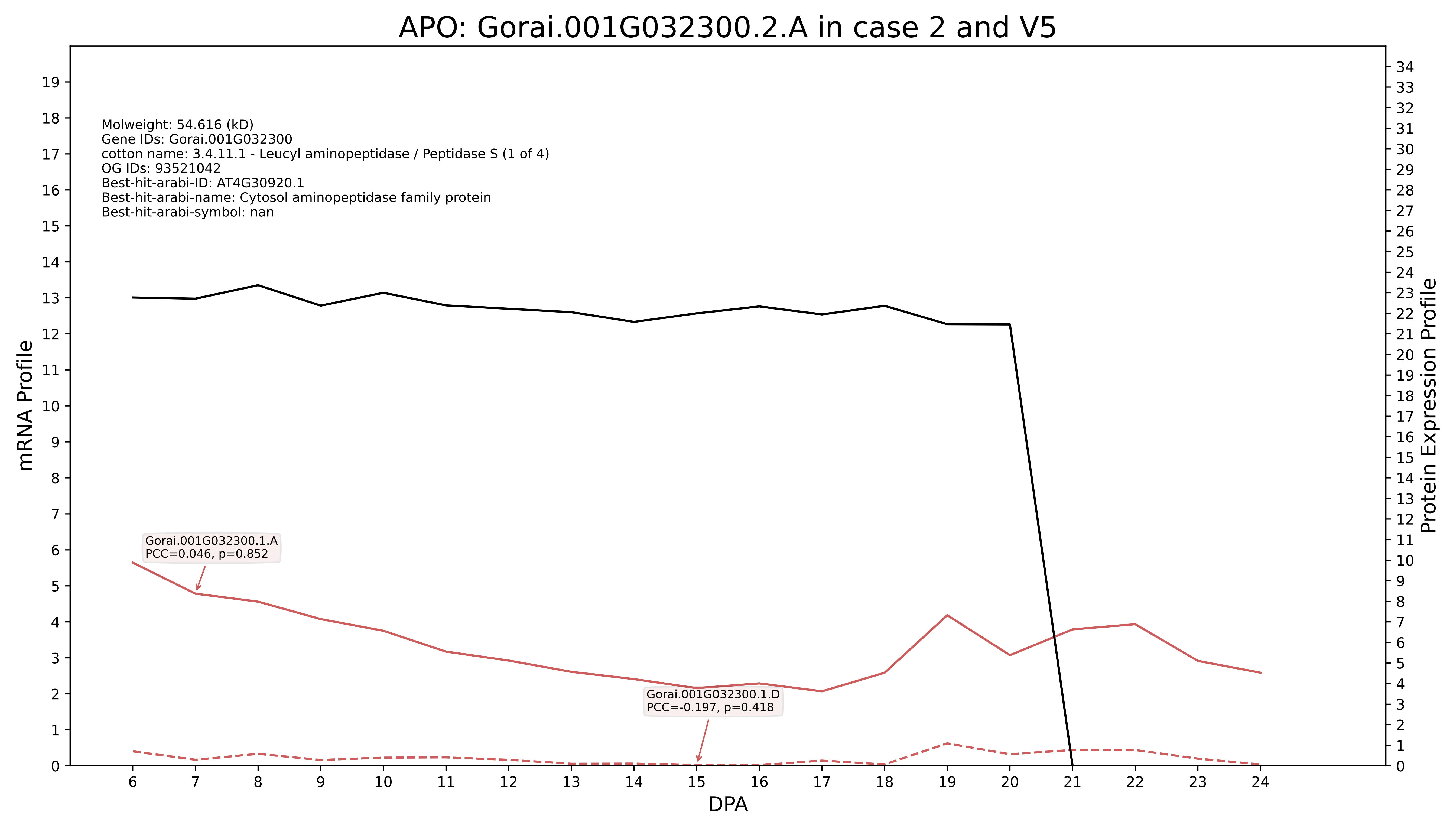
APO: Gorai.001G032300.2.A in case 2 and V5
Molweight: 54.616 (kD)
Gene IDs: Gorai.001G032300
cotton name: 3.4.11.1 - Leucyl aminopeptidase / Peptidase S (1 of 4)
OG IDs: 93521042
Best-hit-arabi-ID: AT4G30920.1
Best-hit-arabi-name: Cytosol aminopeptidase family protein
Best-hit-arabi-symbol: nan
Gorai.001G032300.1.A
PCC=0.046, p=0.852
Gorai.001G032300.1.D
PCC=-0.197, p=0.418
mRNA Profile
Protein Expression Profile
DPA

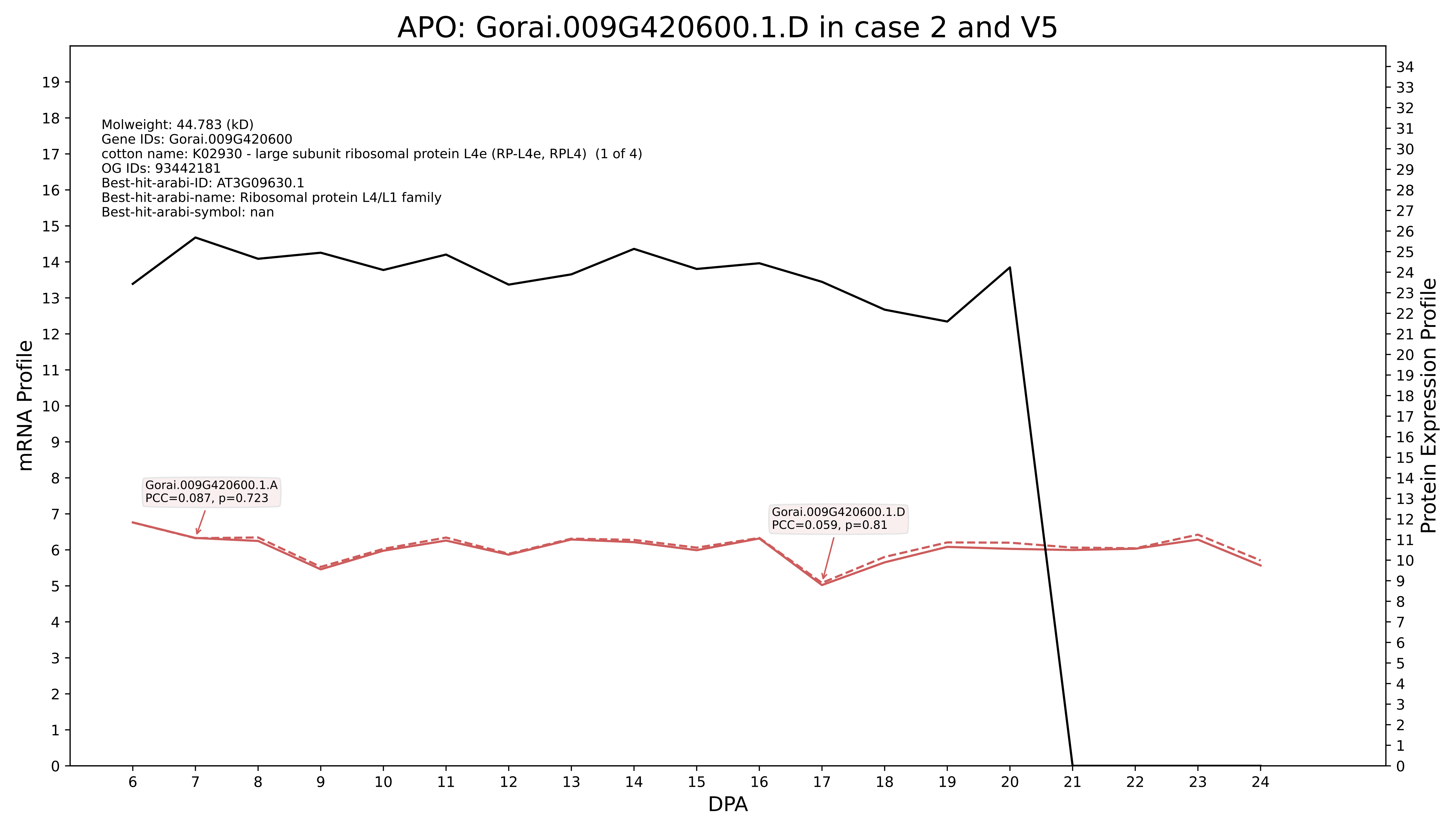

APO: Gorai.009G420600.1.D in case 2 and V5
Molweight: 44.783 (kD)
Gene IDs: Gorai.009G420600
cotton name: K02930 - large subunit ribosomal protein L4e (RP-L4e, RPL4) (1 of 4)
OG IDs: 93442181
Best-hit-arabi-ID: AT3G09630.1
Best-hit-arabi-name: Ribosomal protein L4/L1 family
Best-hit-arabi-symbol: nan
Gorai.009G420600.1.A
PCC=0.087, p=0.723
Gorai.009G420600.1.D
PCC=0.059, p=0.81
mRNA Profile
Protein Expression Profile
DPA

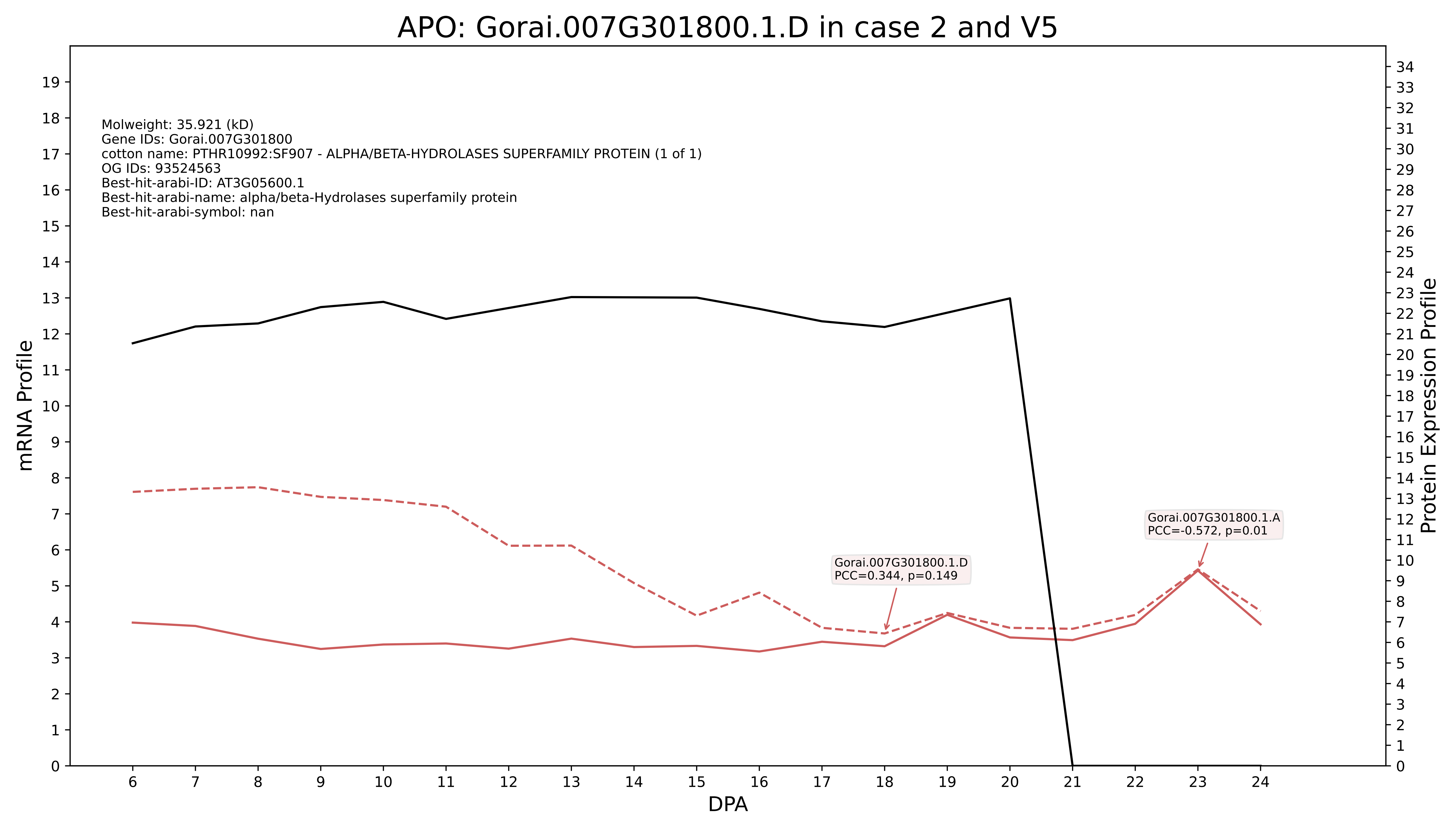

APO: Gorai.007G301800.1.D in case 2 and V5
Molweight: 35.921 (kD)
Gene IDs: Gorai.007G301800
cotton name: PTHR10992:SF907 - ALPHA/BETA-HYDROLASES SUPERFAMILY PROTEIN (1 of 1)
OG IDs: 93524563
Best-hit-arabi-ID: AT3G05600.1
Best-hit-arabi-name: alpha/beta-Hydrolases superfamily protein
Best-hit-arabi-symbol: nan
Gorai.007G301800.1.D
PCC=0.344, p=0.149
Gorai.007G301800.1.A
PCC=-0.572, p=0.01
mRNA Profile
Protein Expression Profile
DPA

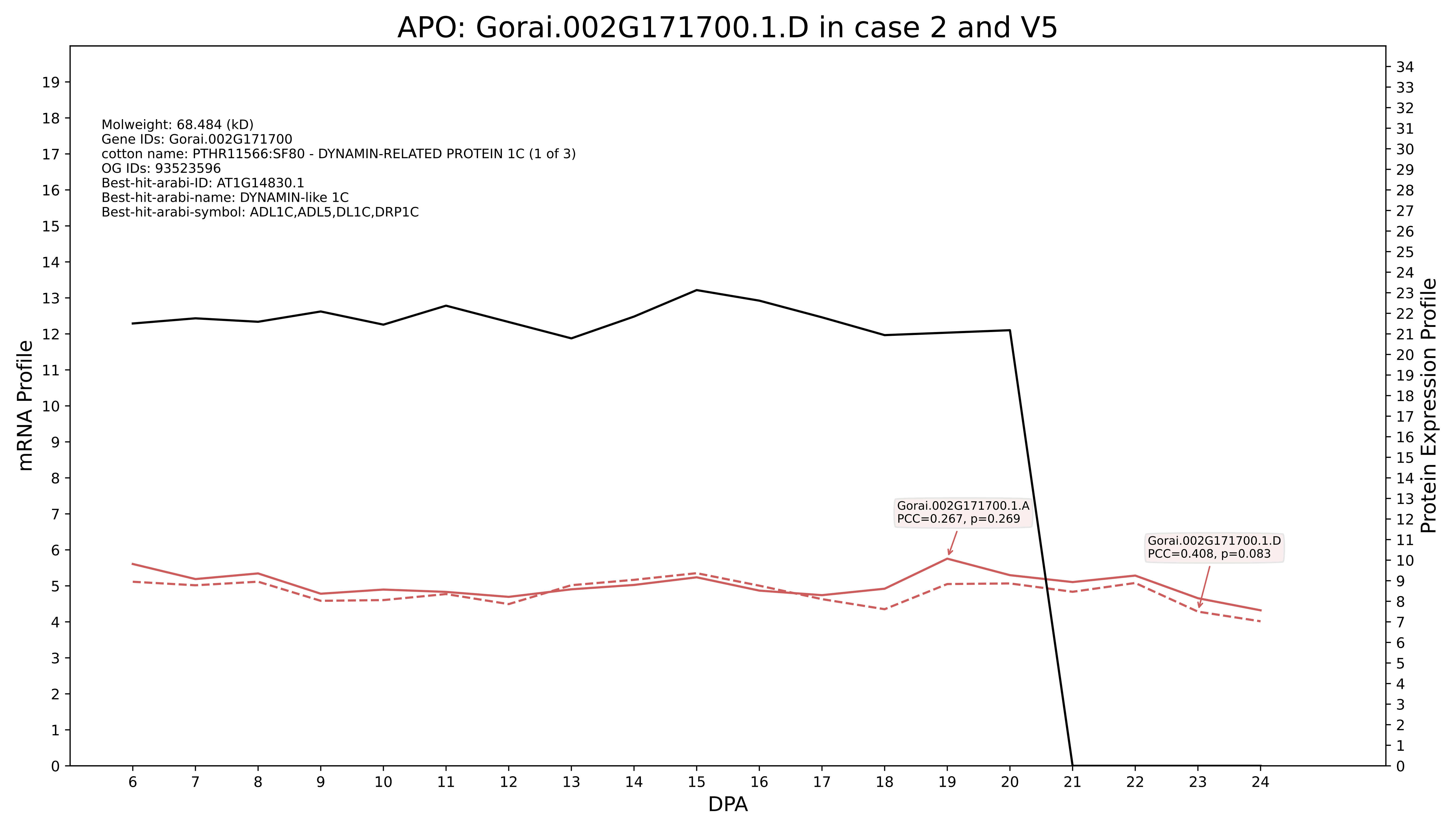
APO: Gorai.002G171700.1.D in case 2 and V5
Molweight: 68.484 (kD)
Gene IDs: Gorai.002G171700
cotton name: PTHR11566:SF80 - DYNAMIN-RELATED PROTEIN 1C (1 of 3)
OG IDs: 93523596
Best-hit-arabi-ID: AT1G14830.1
Best-hit-arabi-name: DYNAMIN-like 1C
Best-hit-arabi-symbol: ADL1C,ADL5,DL1C,DRP1C
Gorai.002G171700.1.A
PCC=0.267, p=0.269
Gorai.002G171700.1.D
PCC=0.408, p=0.083
mRNA Profile
Protein Expression Profile
DPA

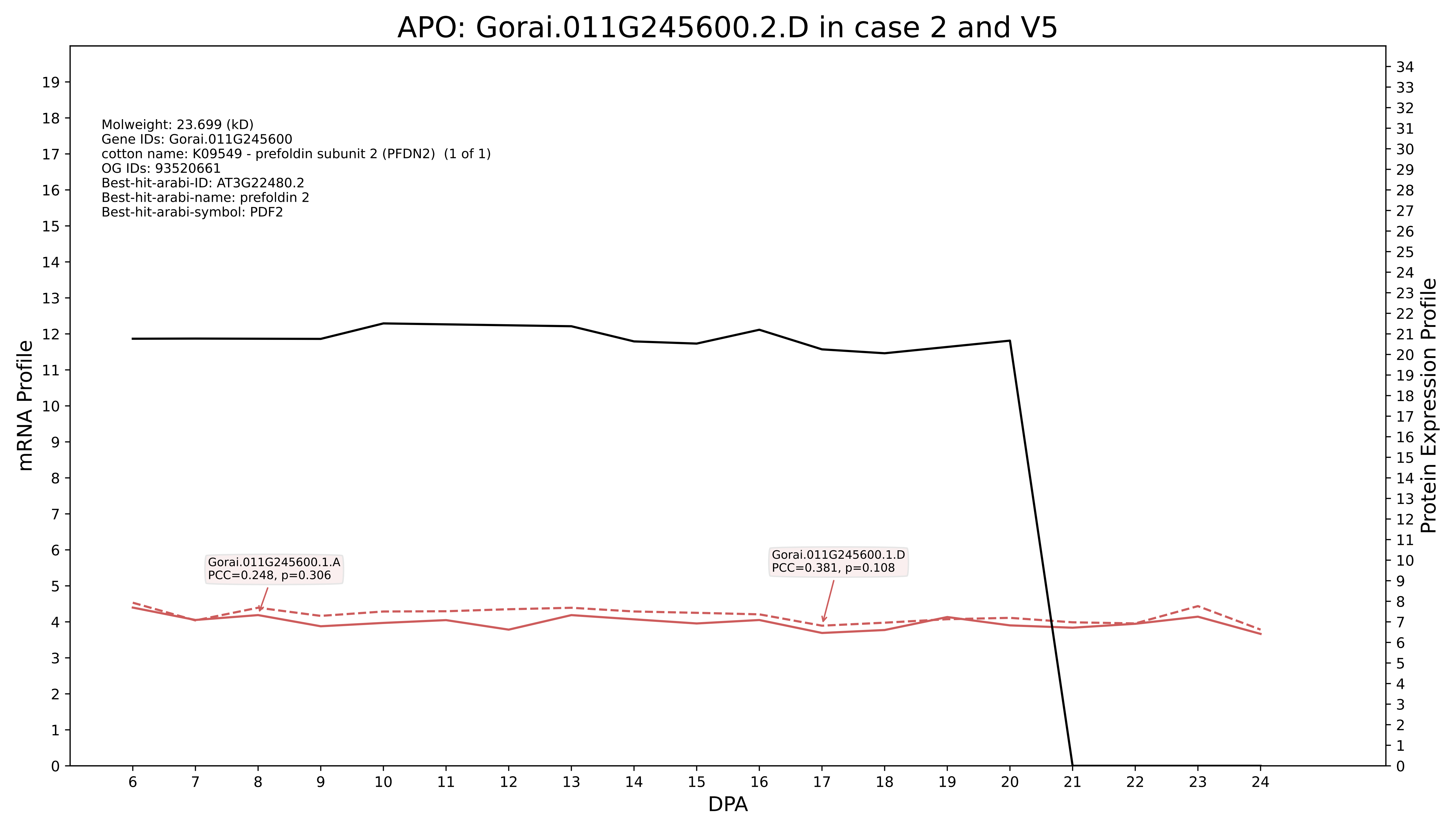

APO: Gorai.011G245600.2.D in case 2 and V5
Molweight: 23.699 (kD)
Gene IDs: Gorai.011G245600
cotton name: K09549 - prefoldin subunit 2 (PFDN2) (1 of 1)
OG IDs: 93520661
Best-hit-arabi-ID: AT3G22480.2
Best-hit-arabi-name: prefoldin 2
Best-hit-arabi-symbol: PDF2
Gorai.011G245600.1.A
PCC=0.248, p=0.306
Gorai.011G245600.1.D
PCC=0.381, p=0.108
mRNA Profile
Protein Expression Profile
DPA

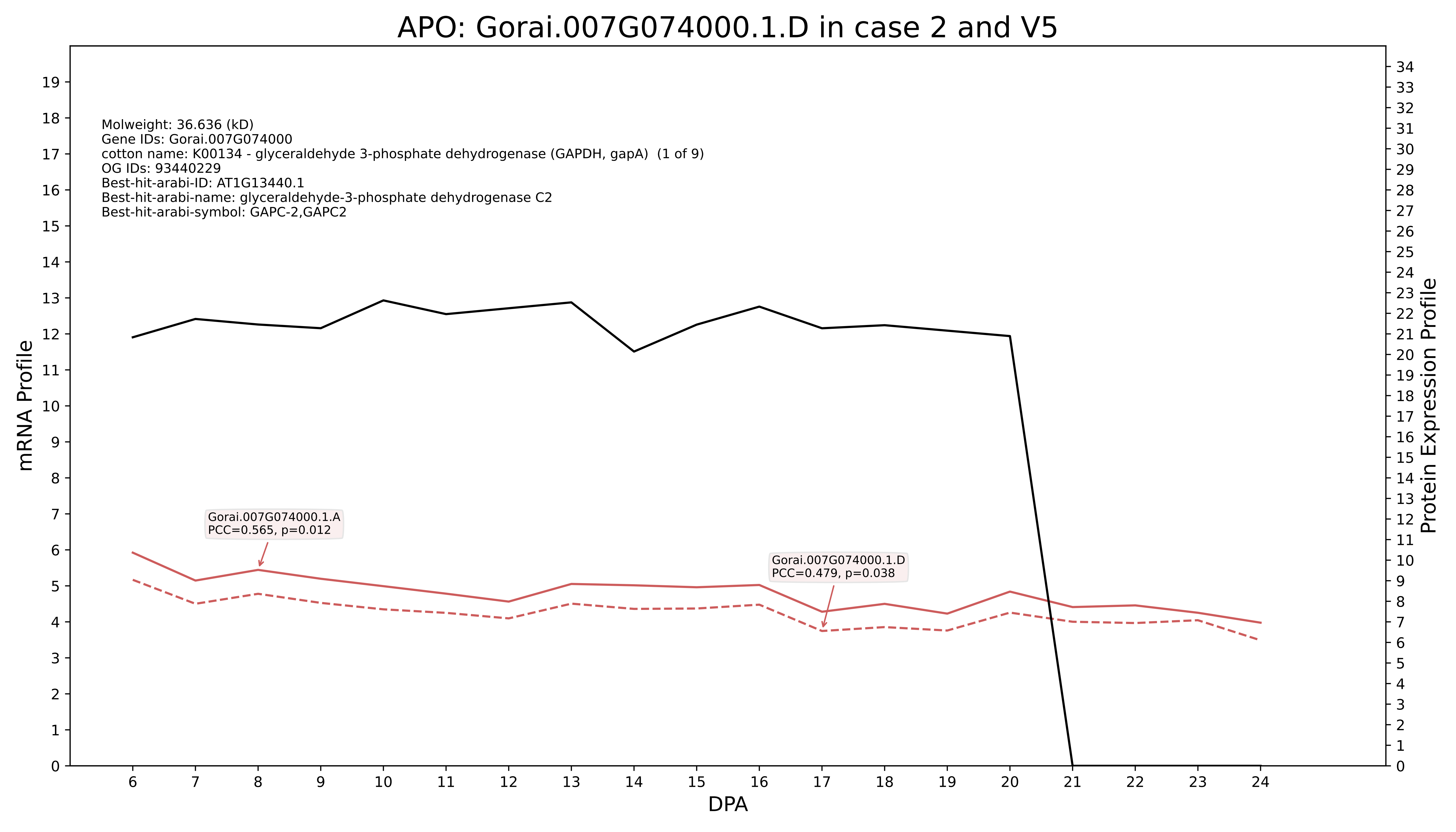
APO: Gorai.007G074000.1.D in case 2 and V5
Molweight: 36.636 (kD)
Gene IDs: Gorai.007G074000
cotton name: K00134 - glyceraldehyde 3-phosphate dehydrogenase (GAPDH, gapA) (1 of 9)
OG IDs: 93440229
Best-hit-arabi-ID: AT1G13440.1
Best-hit-arabi-name: glyceraldehyde-3-phosphate dehydrogenase C2
Best-hit-arabi-symbol: GAPC-2,GAPC2
Gorai.007G074000.1.A
PCC=0.565, p=0.012
Gorai.007G074000.1.D
PCC=0.479, p=0.038
mRNA Profile
Protein Expression Profile
DPA

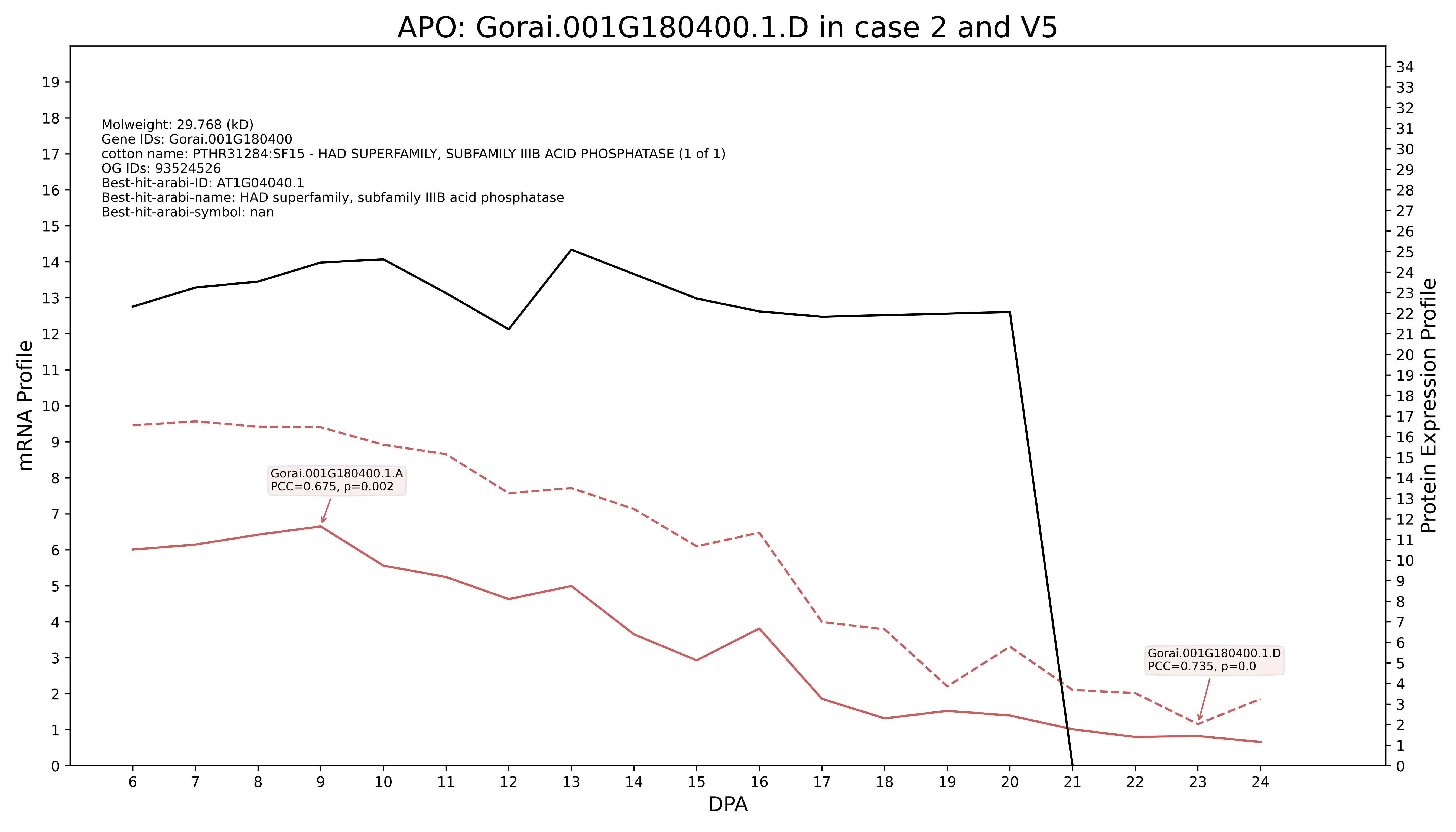
APO: Gorai.001G180400.1.D in case 2 and V5
Molweight: 29.768 (kD)
Gene IDs: Gorai.001G180400
cotton name: PTHR31284:SF15 - HAD SUPERFAMILY, SUBFAMILY IIIB ACID PHOSPHATASE (1 of 1)
OG IDs: 93524526
Best-hit-arabi-ID: AT1G04040.1
Best-hit-arabi-name: HAD superfamily, subfamily IIIB acid phosphatase
Best-hit-arabi-symbol: nan
Gorai.001G180400.1.A
PCC=0.675, p=0.002
Gorai.001G180400.1.D
PCC=0.735, p=0.0
mRNA Profile
Protein Expression Profile
DPA

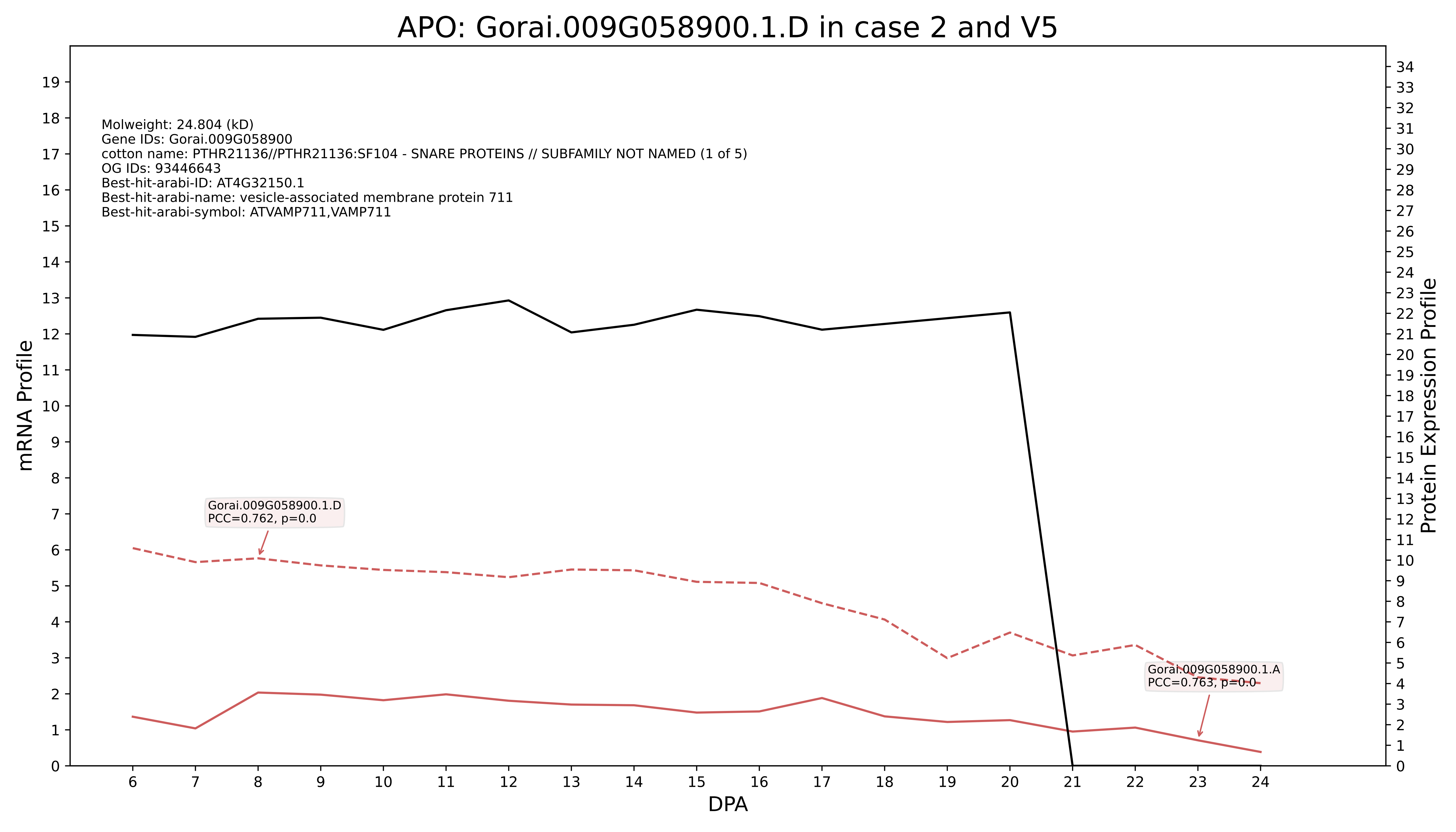

APO: Gorai.009G058900.1.D in case 2 and V5
Molweight: 24.804 (kD)
Gene IDs: Gorai.009G058900
cotton name: PTHR21136//PTHR21136:SF104 - SNARE PROTEINS // SUBFAMILY NOT NAMED (1 of 5)
OG IDs: 93446643
Best-hit-arabi-ID: AT4G32150.1
Best-hit-arabi-name: vesicle-associated membrane protein 711
Best-hit-arabi-symbol: ATVAMP711,VAMP711
Gorai.009G058900.1.D
PCC=0.762, p=0.0
Gorai.009G058900.1.A
PCC=0.763, p=0.0
mRNA Profile
Protein Expression Profile
DPA

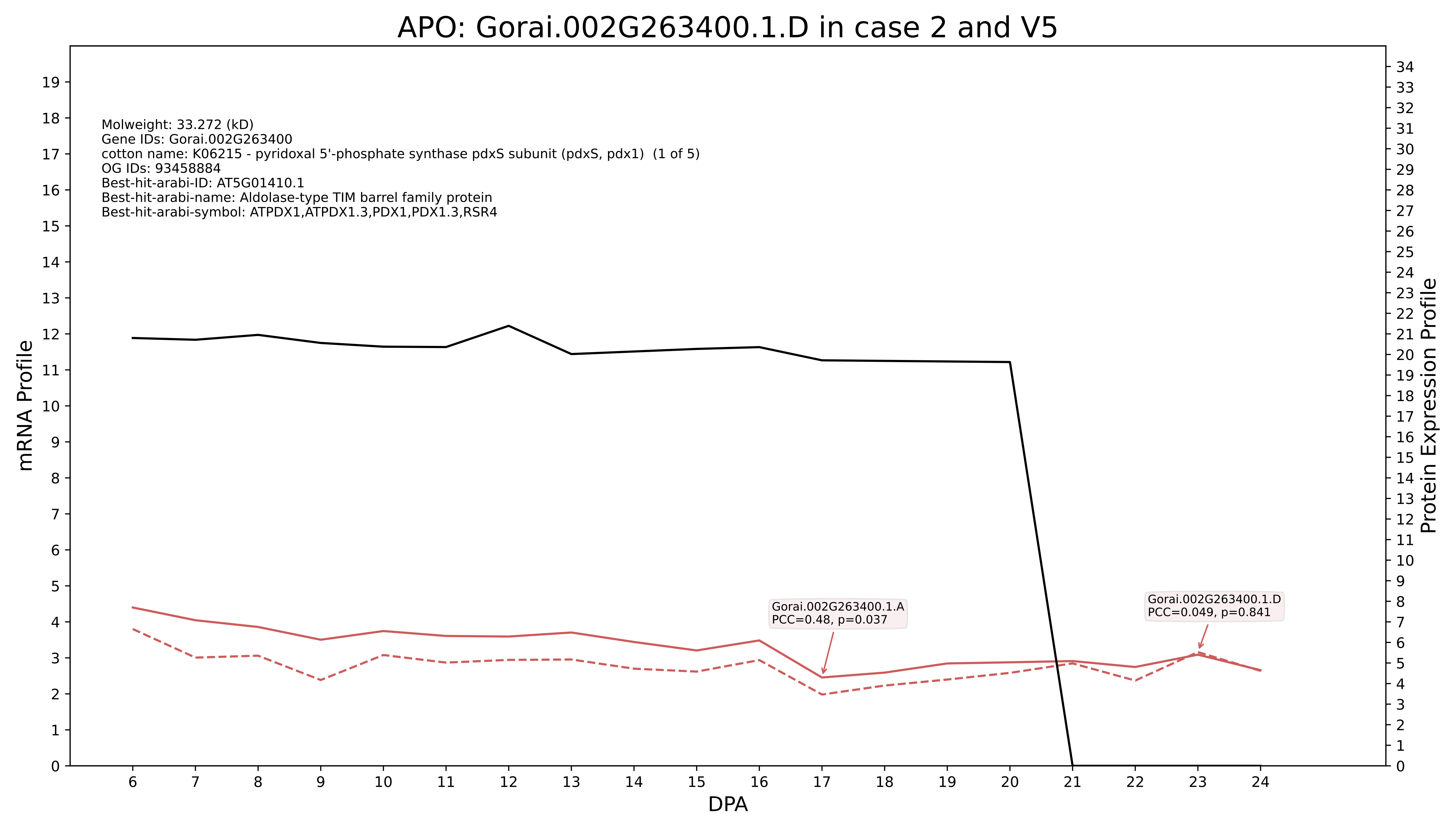

APO: Gorai.002G263400.1.D in case 2 and V5
Molweight: 33.272 (kD)
Gene IDs: Gorai.002G263400
cotton name: K06215 - pyridoxal 5'-phosphate synthase pdxS subunit (pdxS, pdx1) (1 of 5)
OG IDs: 93458884
Best-hit-arabi-ID: AT5G01410.1
Best-hit-arabi-name: Aldolase-type TIM barrel family protein
Best-hit-arabi-symbol: ATPDX1,ATPDX1.3,PDX1,PDX1.3,RSR4
Gorai.002G263400.1.A
PCC=0.48, p=0.037
Gorai.002G263400.1.D
PCC=0.049, p=0.841
mRNA Profile
Protein Expression Profile
DPA

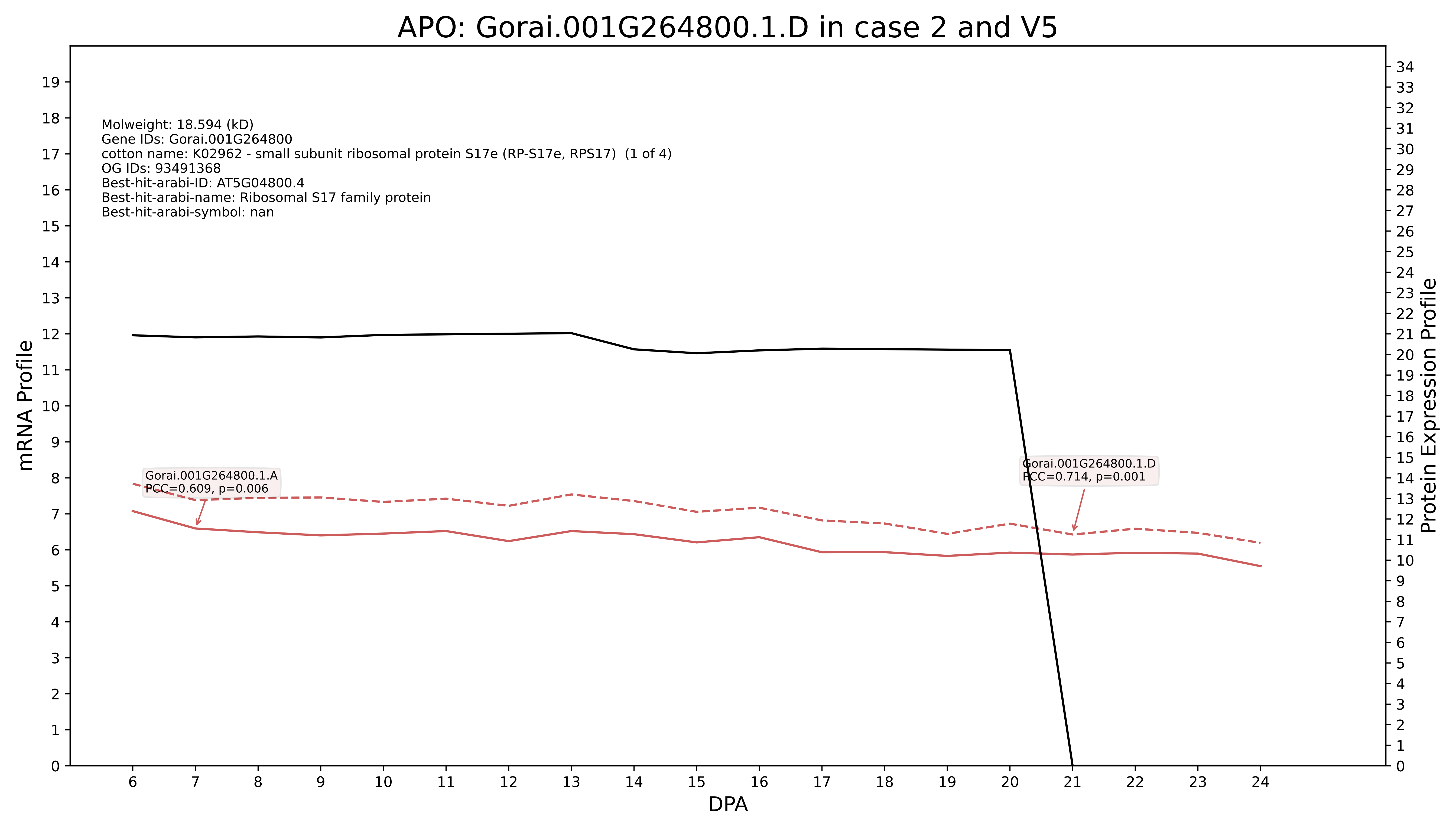

APO: Gorai.001G264800.1.D in case 2 and V5
Molweight: 18.594 (kD)
Gene IDs: Gorai.001G264800
cotton name: K02962 - small subunit ribosomal protein S17e (RP-S17e, RPS17) (1 of 4)
OG IDs: 93491368
Best-hit-arabi-ID: AT5G04800.4
Best-hit-arabi-name: Ribosomal S17 family protein
Best-hit-arabi-symbol: nan
Gorai.001G264800.1.A
PCC=0.609, p=0.006
Gorai.001G264800.1.D
PCC=0.714, p=0.001
mRNA Profile
Protein Expression Profile
DPA

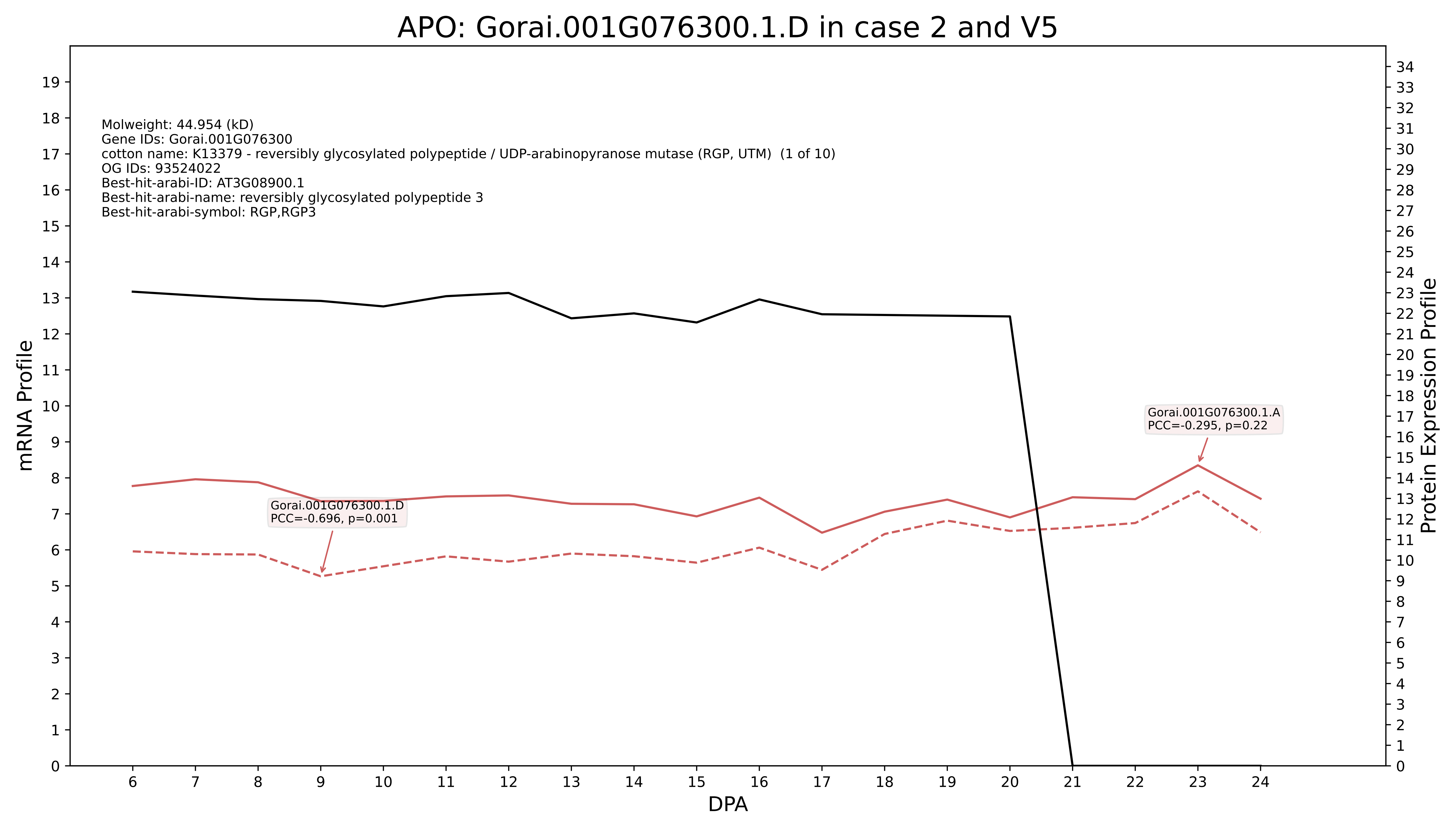
APO: Gorai.001G076300.1.D in case 2 and V5
Molweight: 44.954 (kD)
Gene IDs: Gorai.001G076300
cotton name: K13379 - reversibly glycosylated polypeptide / UDP-arabinopyranose mutase (RGP, UTM) (1 of 10)
OG IDs: 93524022
Best-hit-arabi-ID: AT3G08900.1
Best-hit-arabi-name: reversibly glycosylated polypeptide 3
Best-hit-arabi-symbol: RGP,RGP3
Gorai.001G076300.1.A
PCC=-0.295, p=0.22
Gorai.001G076300.1.D
PCC=-0.696, p=0.001
mRNA Profile
Protein Expression Profile
DPA

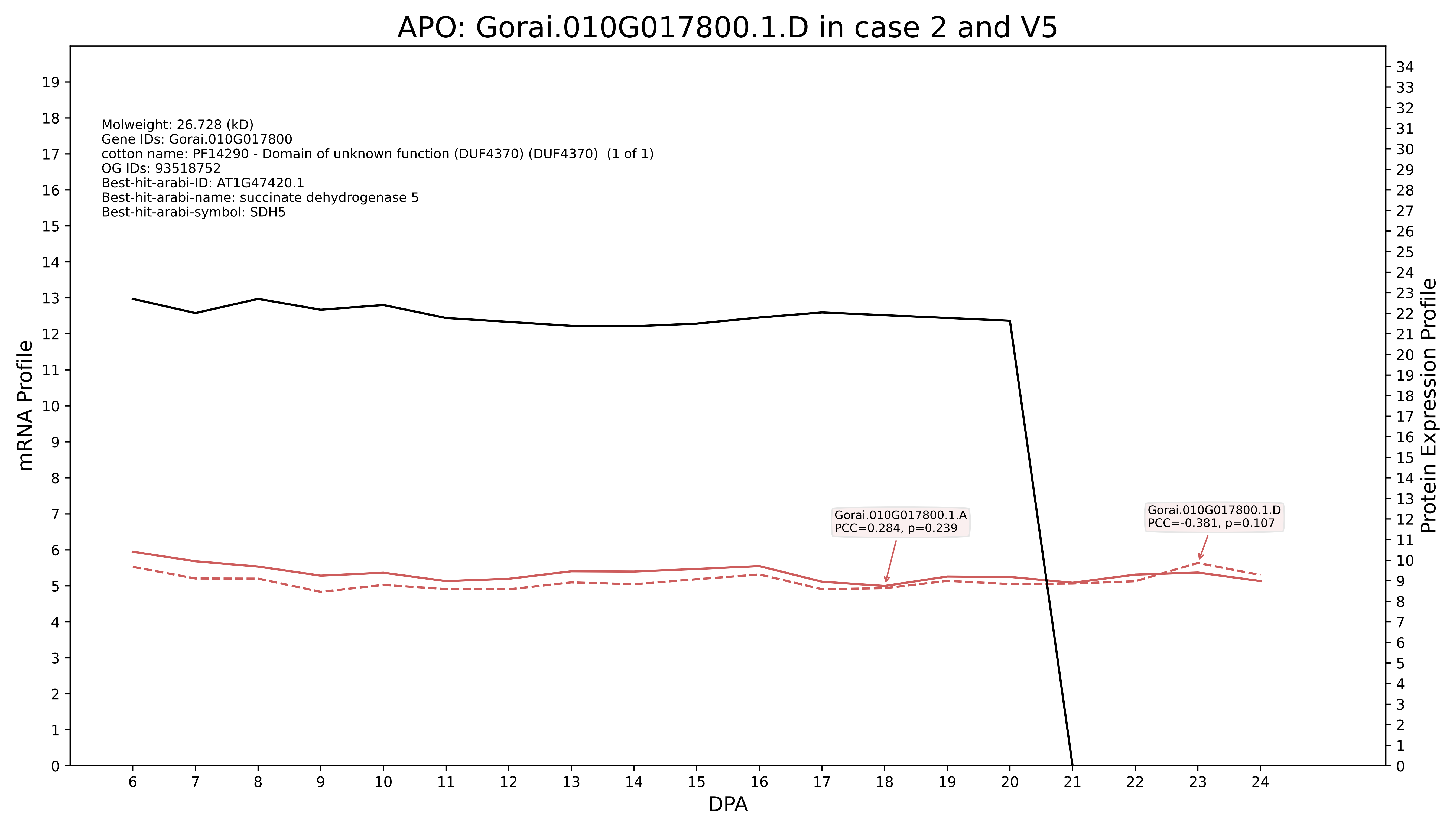
APO: Gorai.010G017800.1.D in case 2 and V5
Molweight: 26.728 (kD)
Gene IDs: Gorai.010G017800
cotton name: PF14290 - Domain of unknown function (DUF4370) (DUF4370) (1 of 1)
OG IDs: 93518752
Best-hit-arabi-ID: AT1G47420.1
Best-hit-arabi-name: succinate dehydrogenase 5
Best-hit-arabi-symbol: SDH5
Gorai.010G017800.1.A
PCC=0.284, p=0.239
Gorai.010G017800.1.D
PCC=-0.381, p=0.107
mRNA Profile
Protein Expression Profile
DPA

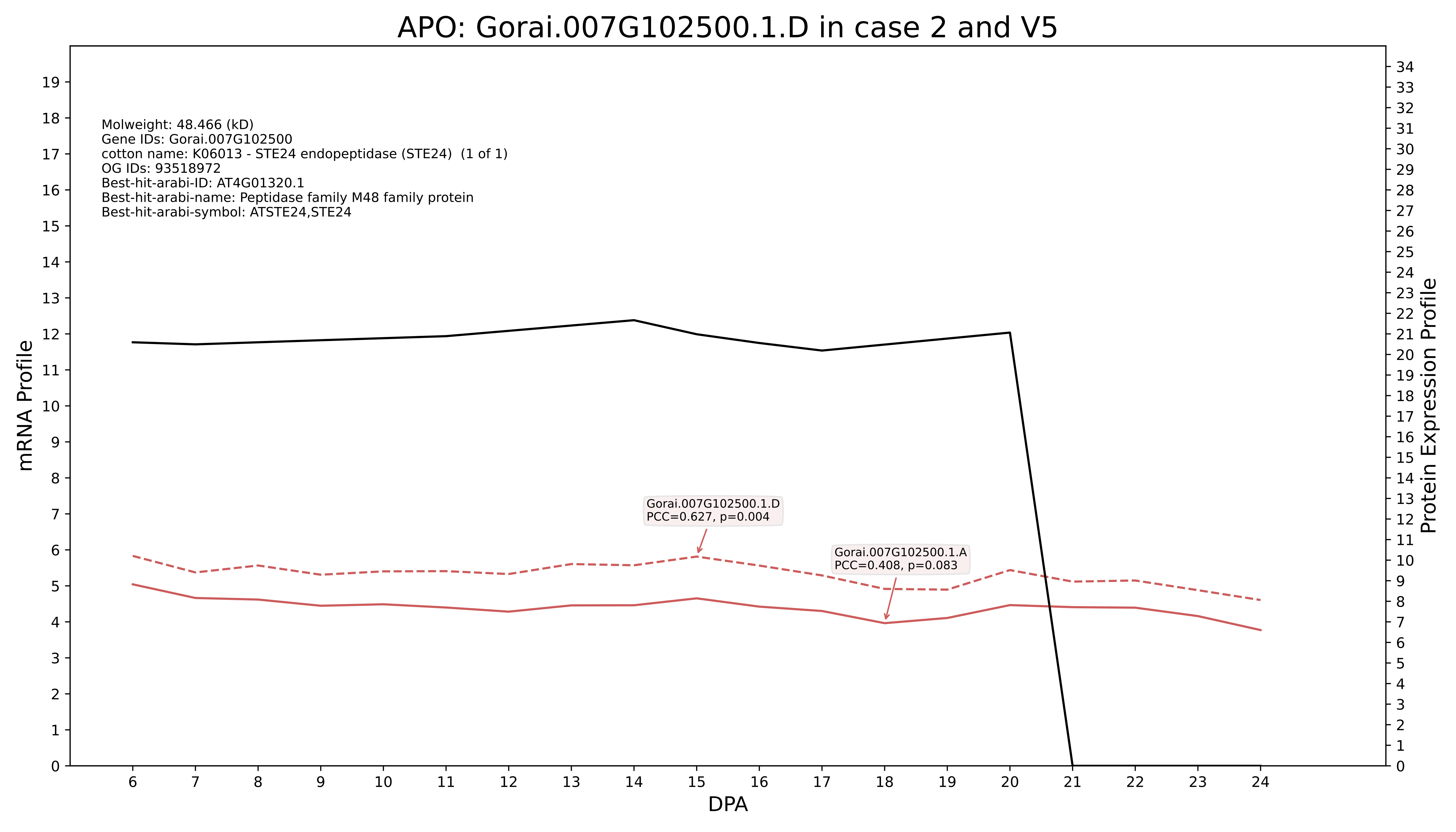

APO: Gorai.007G102500.1.D in case 2 and V5
Molweight: 48.466 (kD)
Gene IDs: Gorai.007G102500
cotton name: K06013 - STE24 endopeptidase (STE24) (1 of 1)
OG IDs: 93518972
Best-hit-arabi-ID: AT4G01320.1
Best-hit-arabi-name: Peptidase family M48 family protein
Best-hit-arabi-symbol: ATSTE24,STE24
Gorai.007G102500.1.D
PCC=0.627, p=0.004
Gorai.007G102500.1.A
PCC=0.408, p=0.083
mRNA Profile
Protein Expression Profile
DPA

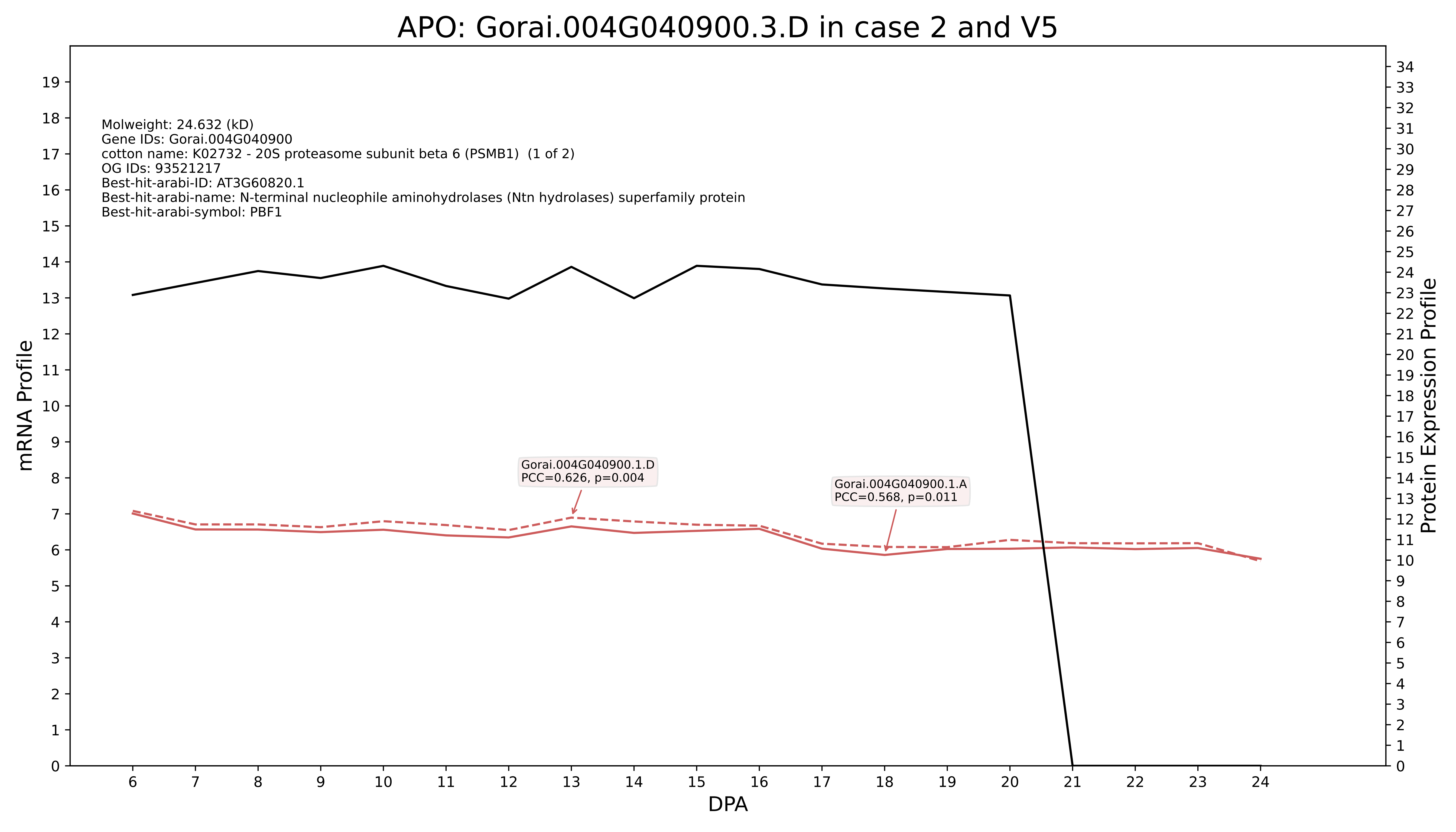

APO: Gorai.004G040900.3.D in case 2 and V5
Molweight: 24.632 (kD)
Gene IDs: Gorai.004G040900
cotton name: K02732 - 20S proteasome subunit beta 6 (PSMB1) (1 of 2)
OG IDs: 93521217
Best-hit-arabi-ID: AT3G60820.1
Best-hit-arabi-name: N-terminal nucleophile aminohydrolases (Ntn hydrolases) superfamily protein
Best-hit-arabi-symbol: PBF1
Gorai.004G040900.1.D
PCC=0.626, p=0.004
Gorai.004G040900.1.A
PCC=0.568, p=0.011
mRNA Profile
Protein Expression Profile
DPA

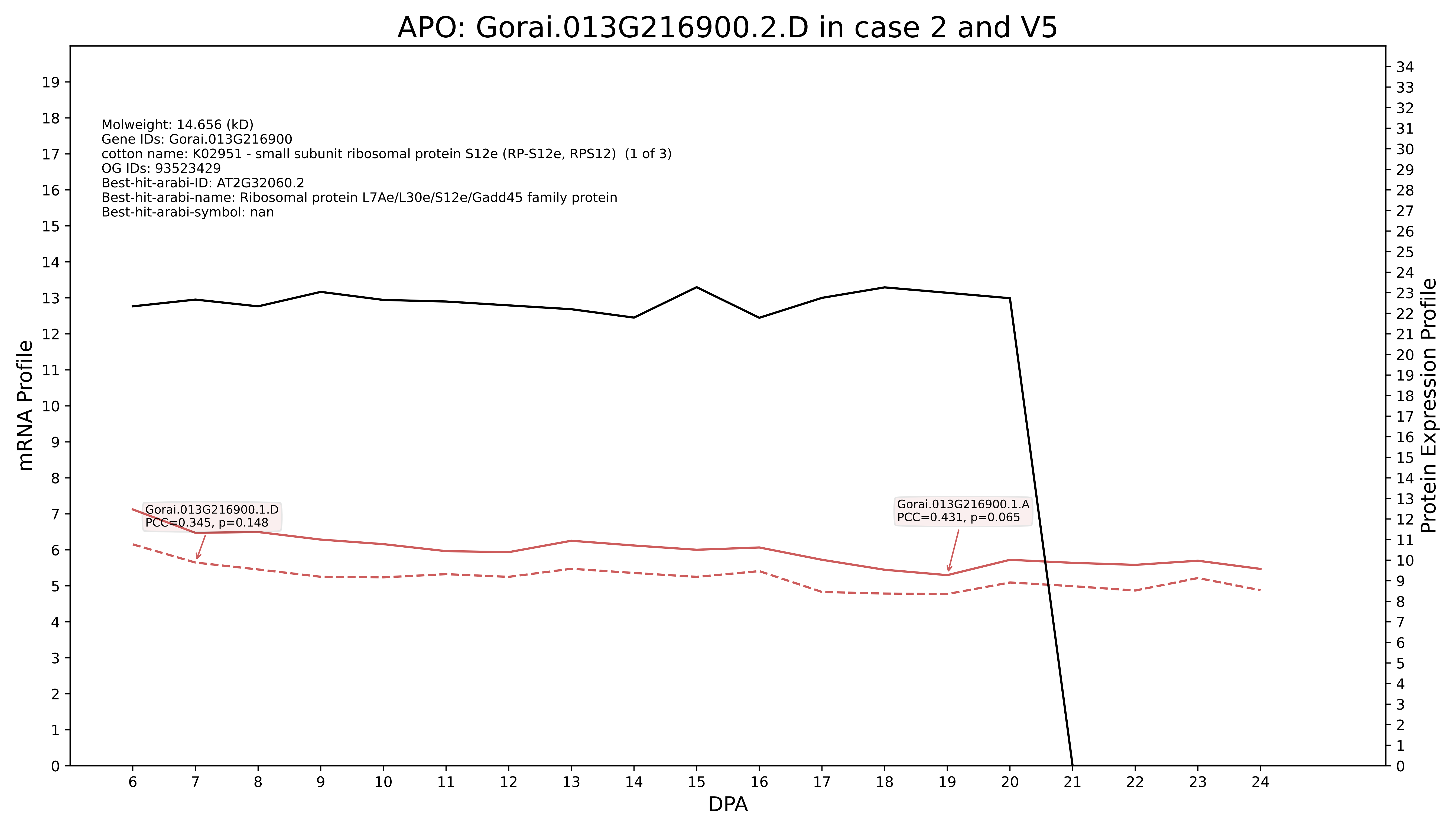
APO: Gorai.013G216900.2.D in case 2 and V5
Molweight: 14.656 (kD)
Gene IDs: Gorai.013G216900
cotton name: K02951 - small subunit ribosomal protein S12e (RP-S12e, RPS12) (1 of 3)
OG IDs: 93523429
Best-hit-arabi-ID: AT2G32060.2
Best-hit-arabi-name: Ribosomal protein L7Ae/L30e/S12e/Gadd45 family protein
Best-hit-arabi-symbol: nan
Gorai.013G216900.1.D
PCC=0.345, p=0.148
Gorai.013G216900.1.A
PCC=0.431, p=0.065
mRNA Profile
Protein Expression Profile
DPA

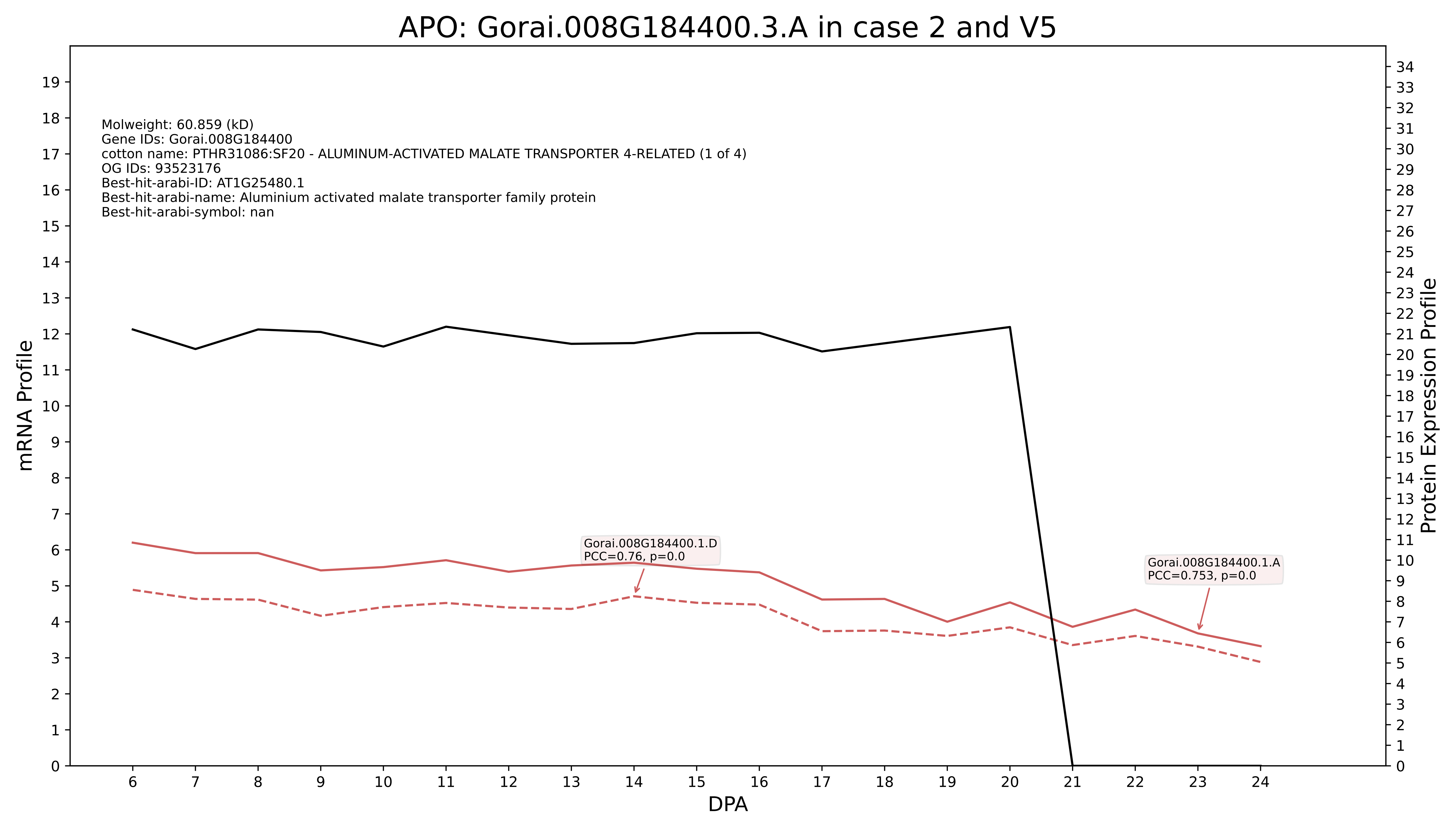
APO: Gorai.008G184400.3.A in case 2 and V5
Molweight: 60.859 (kD)
Gene IDs: Gorai.008G184400
cotton name: PTHR31086:SF20 - ALUMINUM-ACTIVATED MALATE TRANSPORTER 4-RELATED (1 of 4)
OG IDs: 93523176
Best-hit-arabi-ID: AT1G25480.1
Best-hit-arabi-name: Aluminium activated malate transporter family protein
Best-hit-arabi-symbol: nan
Gorai.008G184400.1.D
PCC=0.76, p=0.0
Gorai.008G184400.1.A
PCC=0.753, p=0.0
mRNA Profile
Protein Expression Profile
DPA

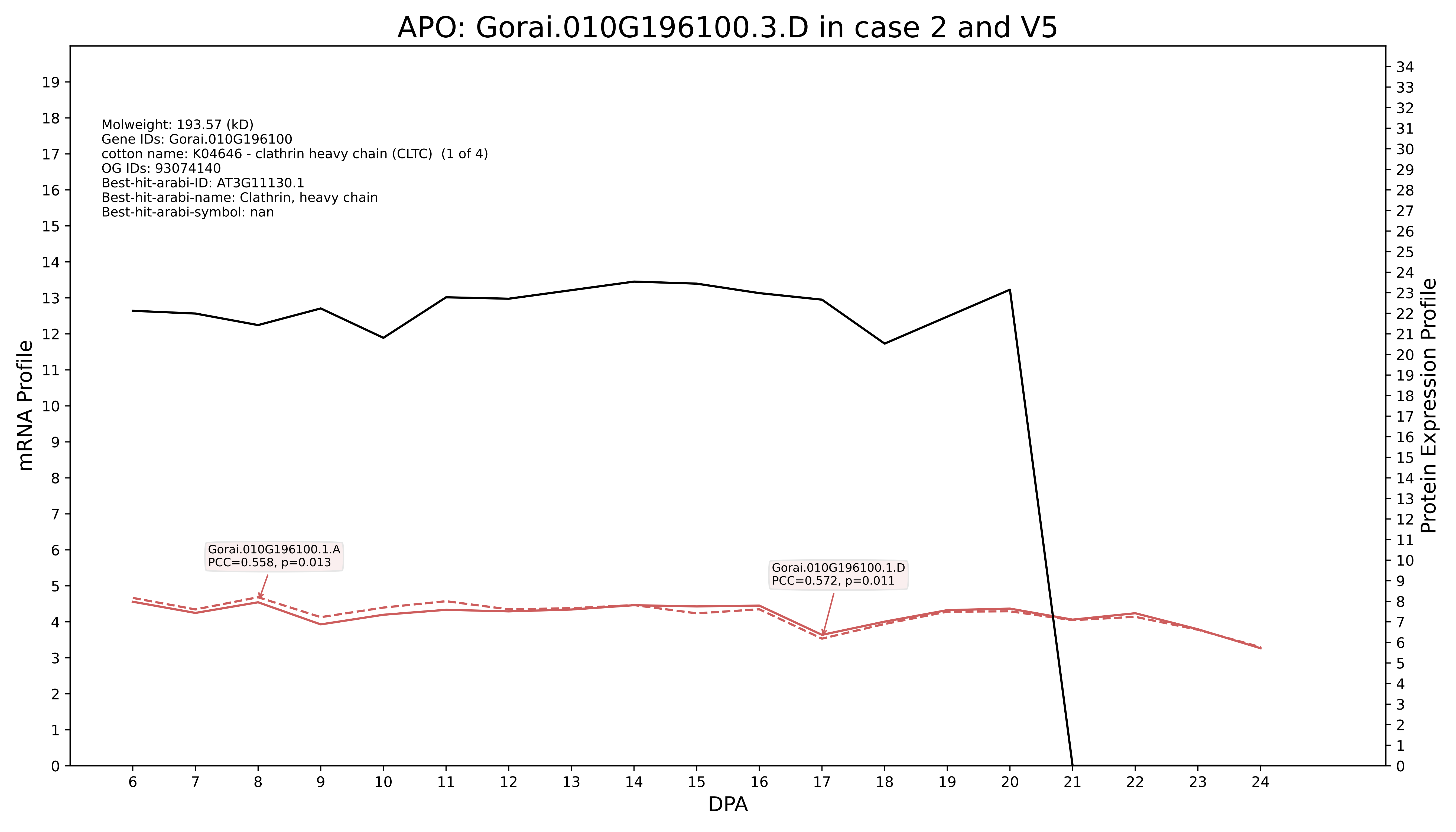
APO: Gorai.010G196100.3.D in case 2 and V5
Molweight: 193.57 (kD)
Gene IDs: Gorai.010G196100
cotton name: K04646 - clathrin heavy chain (CLTC) (1 of 4)
OG IDs: 93074140
Best-hit-arabi-ID: AT3G11130.1
Best-hit-arabi-name: Clathrin, heavy chain
Best-hit-arabi-symbol: nan
Gorai.010G196100.1.A
PCC=0.558, p=0.013
Gorai.010G196100.1.D
PCC=0.572, p=0.011
mRNA Profile
Protein Expression Profile
DPA

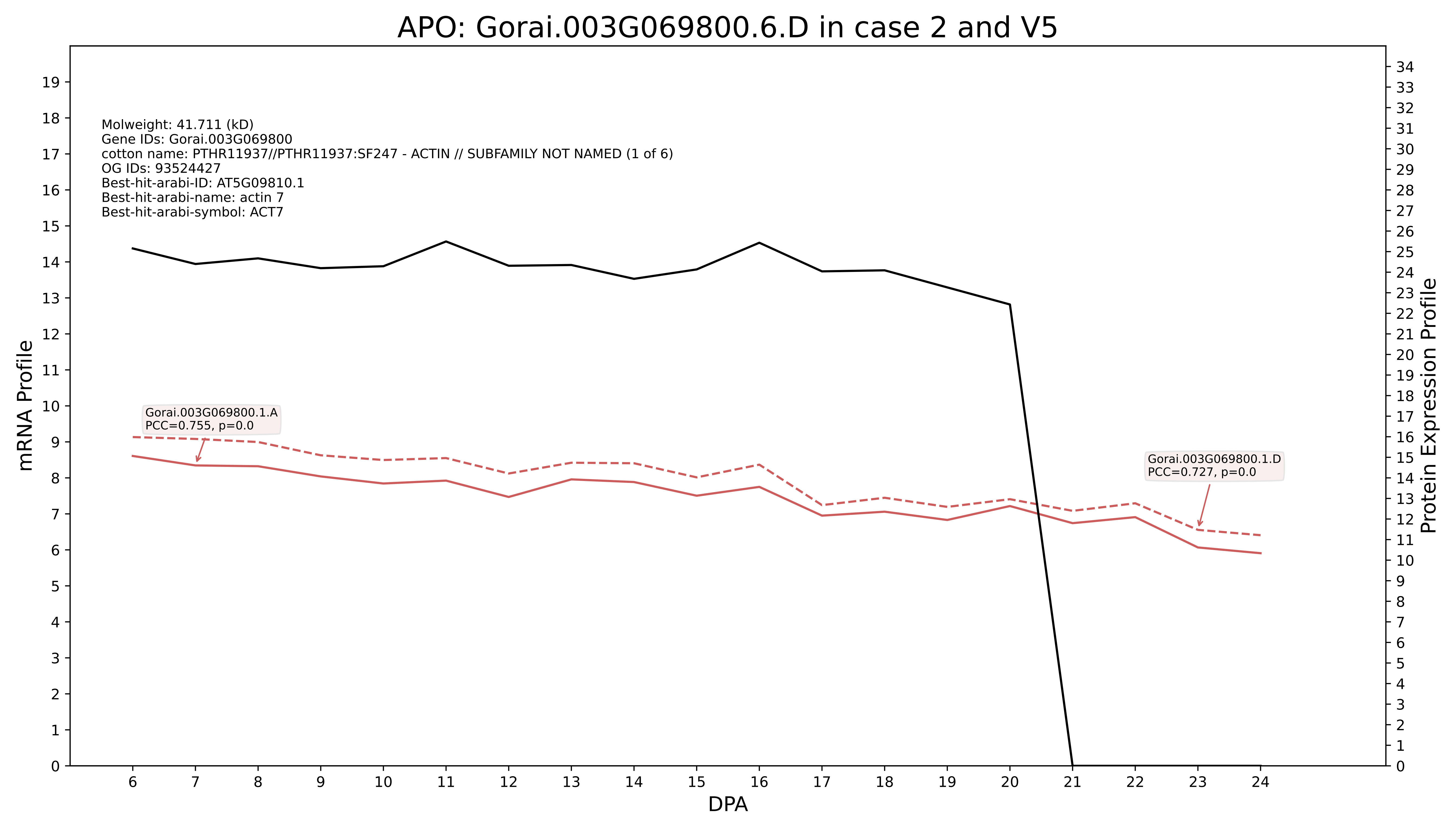
APO: Gorai.003G069800.6.D in case 2 and V5
Molweight: 41.711 (kD)
Gene IDs: Gorai.003G069800
cotton name: PTHR11937//PTHR11937:SF247 - ACTIN // SUBFAMILY NOT NAMED (1 of 6)
OG IDs: 93524427
Best-hit-arabi-ID: AT5G09810.1
Best-hit-arabi-name: actin 7
Best-hit-arabi-symbol: ACT7
Gorai.003G069800.1.A
PCC=0.755, p=0.0
Gorai.003G069800.1.D
PCC=0.727, p=0.0
mRNA Profile
Protein Expression Profile
DPA

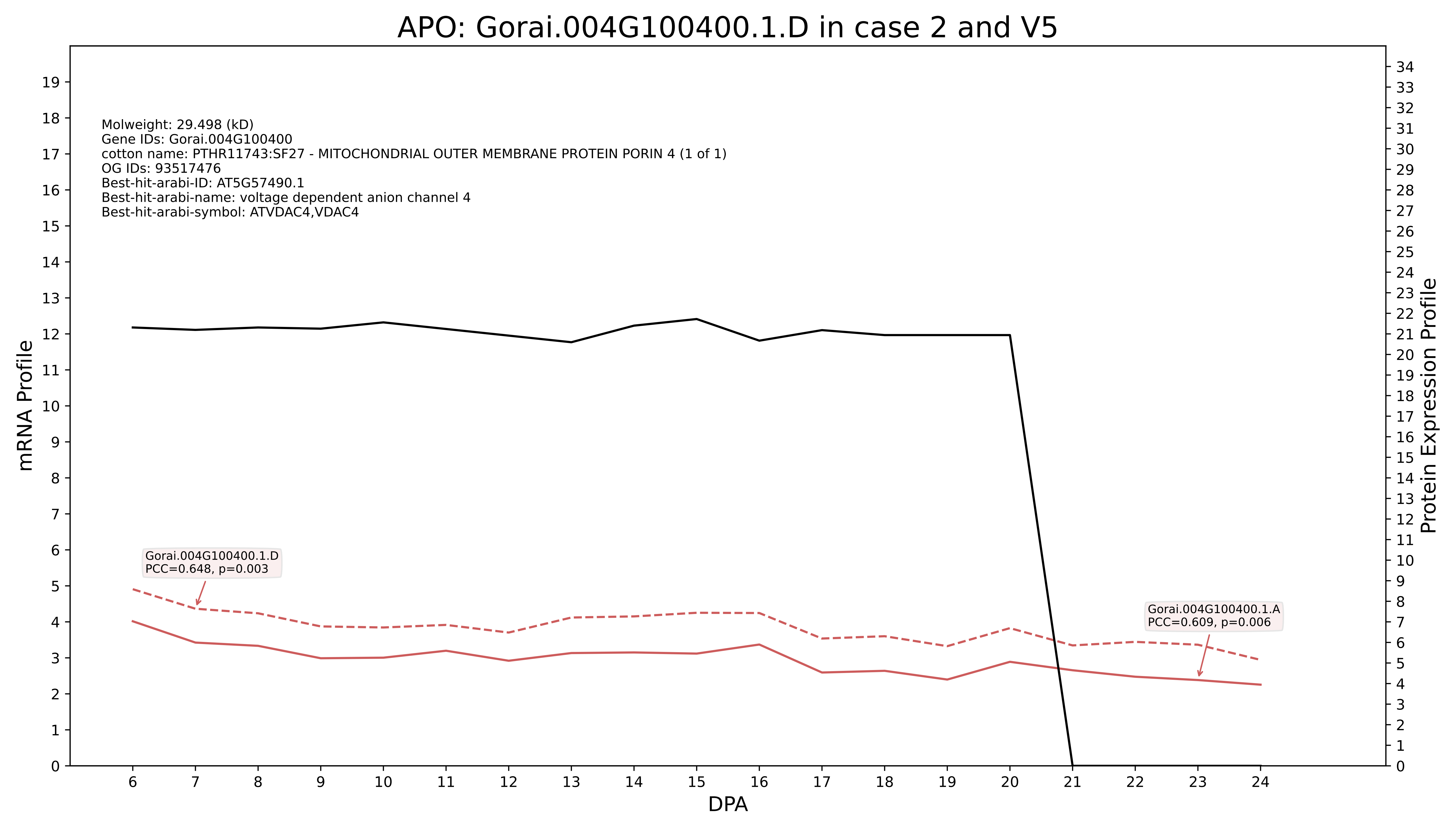
APO: Gorai.004G100400.1.D in case 2 and V5
Molweight: 29.498 (kD)
Gene IDs: Gorai.004G100400
cotton name: PTHR11743:SF27 - MITOCHONDRIAL OUTER MEMBRANE PROTEIN PORIN 4 (1 of 1)
OG IDs: 93517476
Best-hit-arabi-ID: AT5G57490.1
Best-hit-arabi-name: voltage dependent anion channel 4
Best-hit-arabi-symbol: ATVDAC4,VDAC4
Gorai.004G100400.1.D
PCC=0.648, p=0.003
Gorai.004G100400.1.A
PCC=0.609, p=0.006
mRNA Profile
Protein Expression Profile
DPA

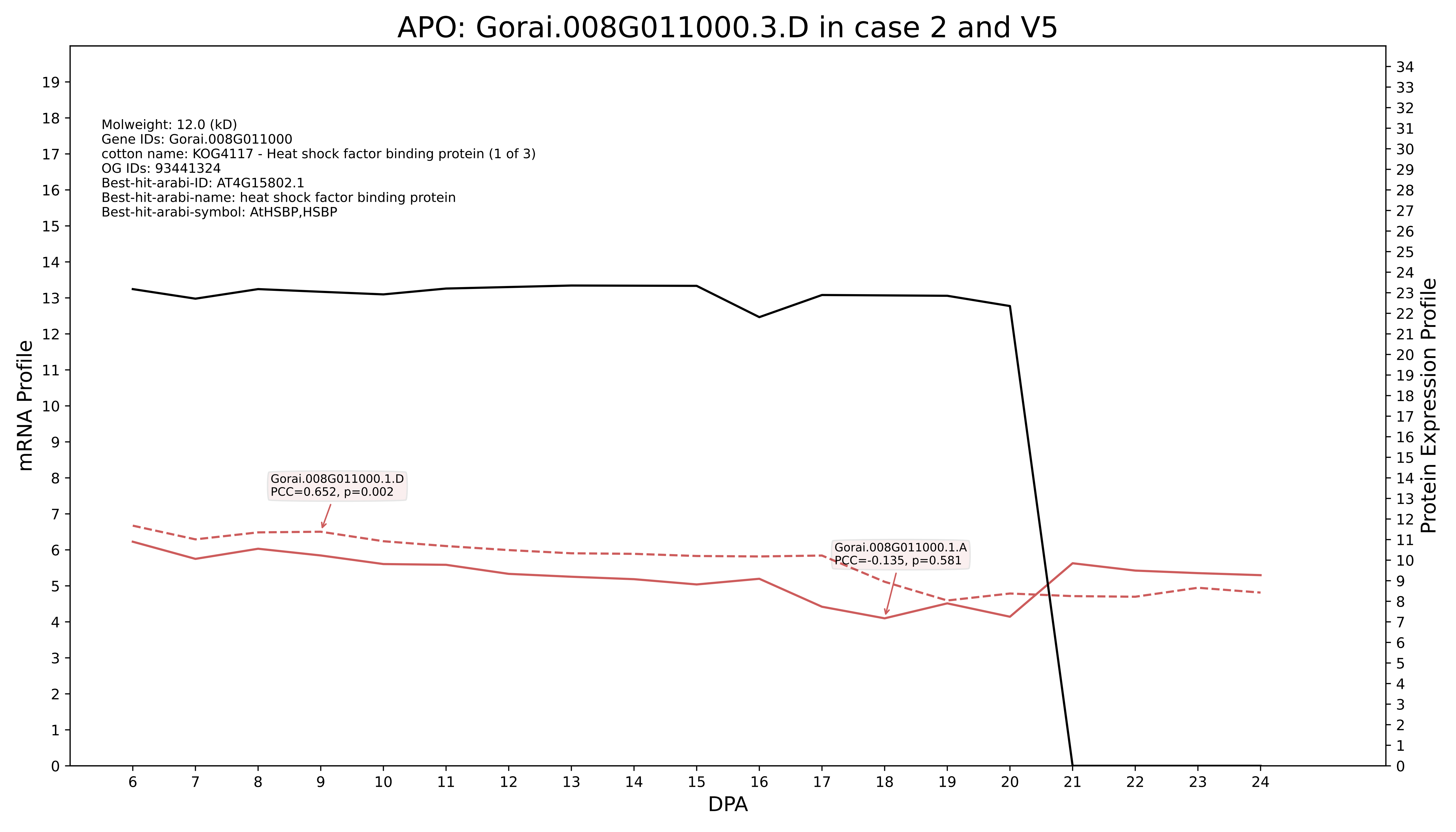

APO: Gorai.008G011000.3.D in case 2 and V5
Molweight: 12.0 (kD)
Gene IDs: Gorai.008G011000
cotton name: KOG4117 - Heat shock factor binding protein (1 of 3)
OG IDs: 93441324
Best-hit-arabi-ID: AT4G15802.1
Best-hit-arabi-name: heat shock factor binding protein
Best-hit-arabi-symbol: AtHSBP,HSBP
Gorai.008G011000.1.D
PCC=0.652, p=0.002
Gorai.008G011000.1.A
PCC=-0.135, p=0.581
mRNA Profile
Protein Expression Profile
DPA

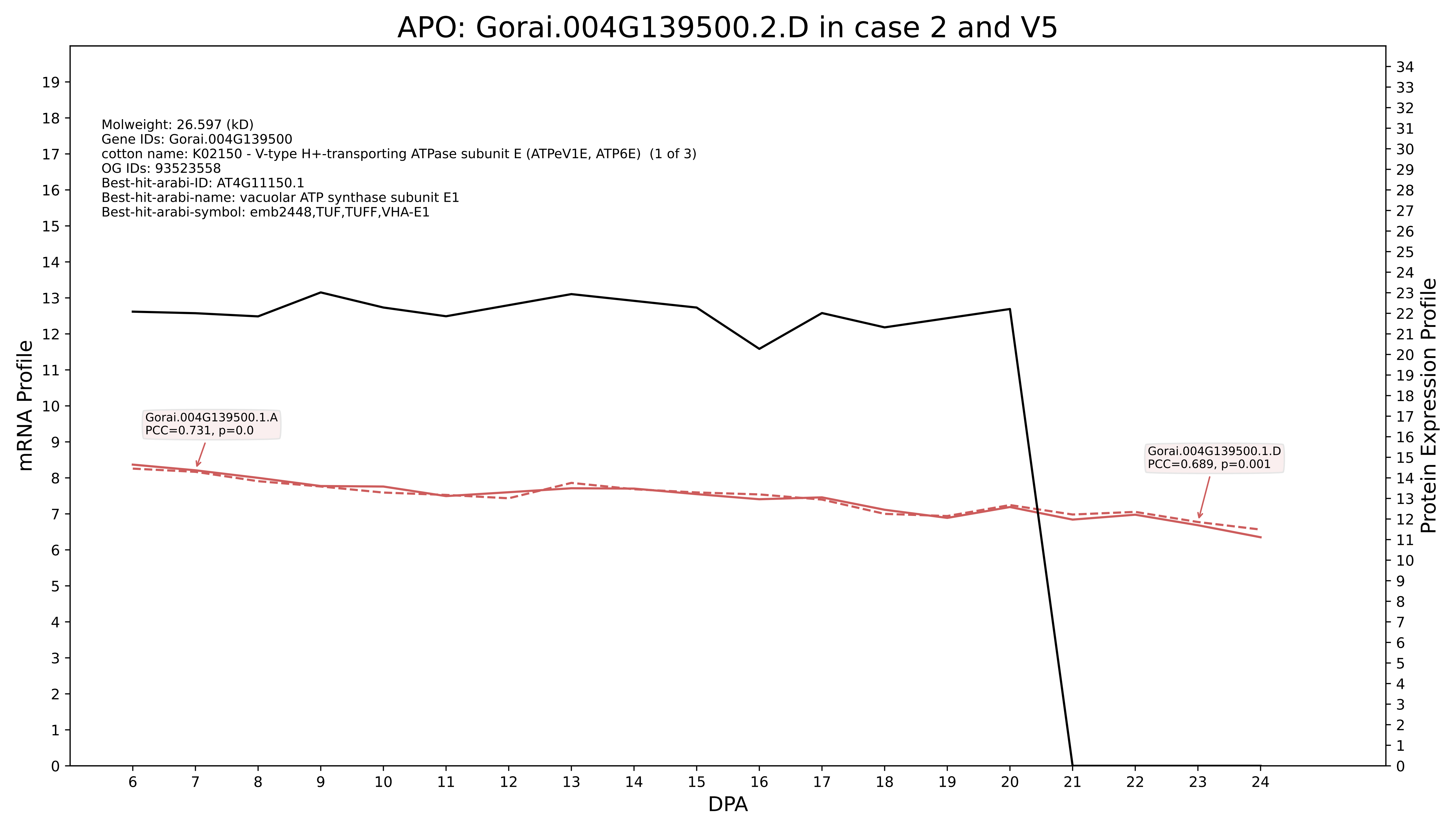

APO: Gorai.004G139500.2.D in case 2 and V5
Molweight: 26.597 (kD)
Gene IDs: Gorai.004G139500
cotton name: K02150 - V-type H+-transporting ATPase subunit E (ATPeV1E, ATP6E) (1 of 3)
OG IDs: 93523558
Best-hit-arabi-ID: AT4G11150.1
Best-hit-arabi-name: vacuolar ATP synthase subunit E1
Best-hit-arabi-symbol: emb2448,TUF,TUFF,VHA-E1
Gorai.004G139500.1.A
PCC=0.731, p=0.0
Gorai.004G139500.1.D
PCC=0.689, p=0.001
mRNA Profile
Protein Expression Profile
DPA

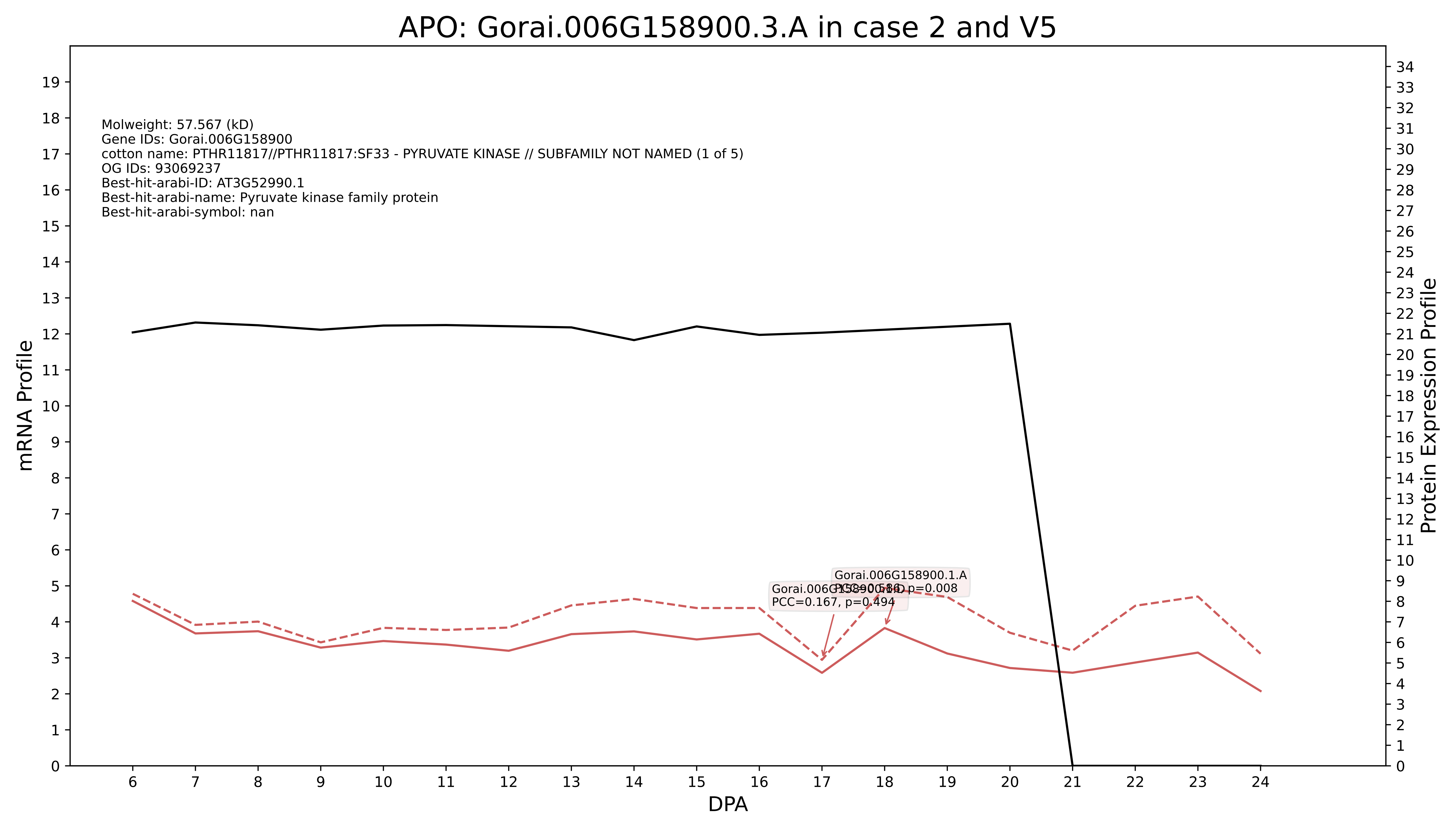
APO: Gorai.006G158900.3.A in case 2 and V5
Molweight: 57.567 (kD)
Gene IDs: Gorai.006G158900
cotton name: PTHR11817//PTHR11817:SF33 - PYRUVATE KINASE // SUBFAMILY NOT NAMED (1 of 5)
OG IDs: 93069237
Best-hit-arabi-ID: AT3G52990.1
Best-hit-arabi-name: Pyruvate kinase family protein
Best-hit-arabi-symbol: nan
Gorai.006G158900.1.A
PCC=0.167, p=0.494
mRNA Profile
Protein Expression Profile
DPA

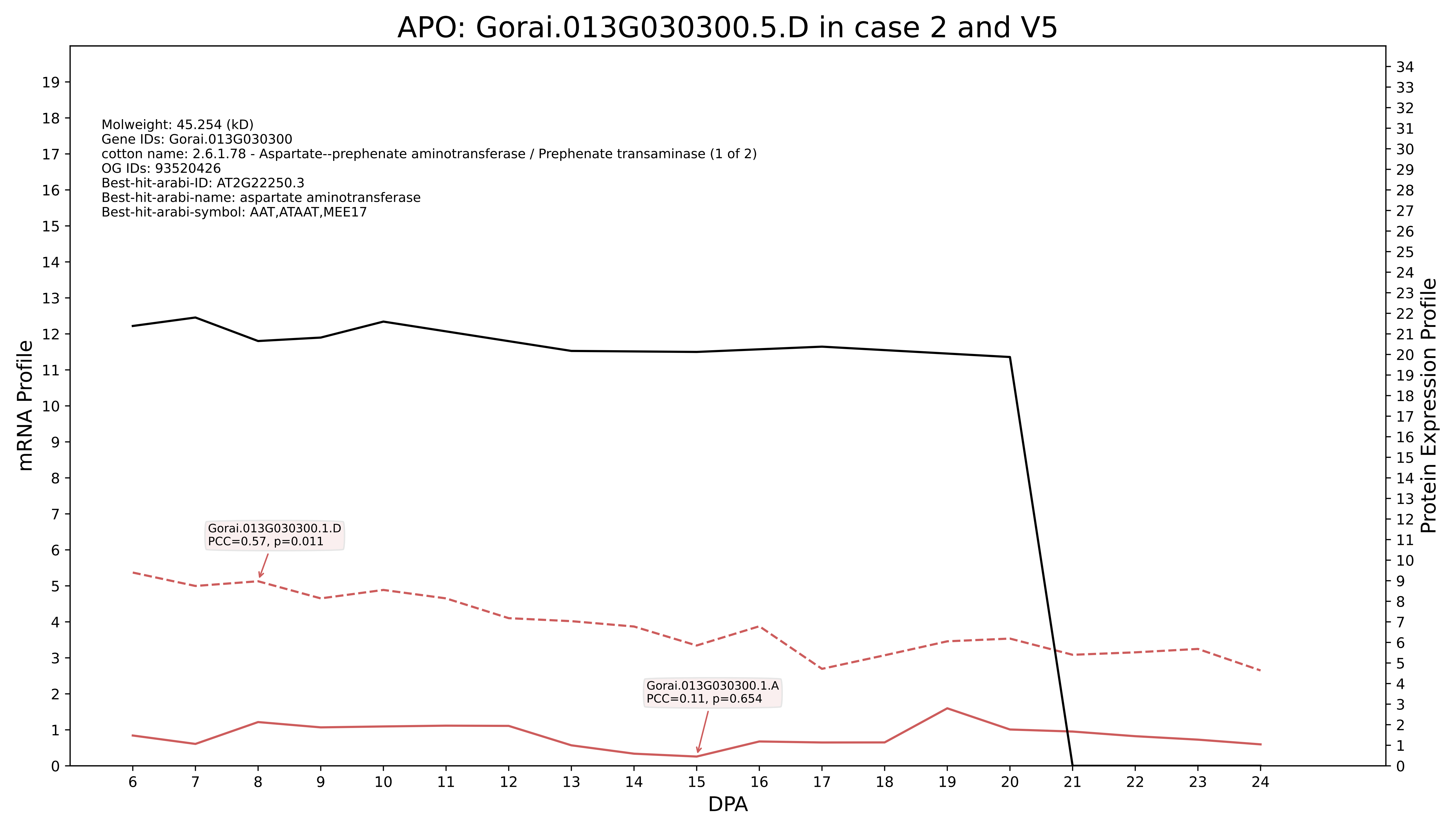
APO: Gorai.013G030300.5.D in case 2 and V5
Molweight: 45.254 (kD)
Gene IDs: Gorai.013G030300
cotton name: 2.6.1.78 - Aspartate--prephenate aminotransferase / Prephenate transaminase (1 of 2)
OG IDs: 93520426
Best-hit-arabi-ID: AT2G22250.3
Best-hit-arabi-name: aspartate aminotransferase
Best-hit-arabi-symbol: AAT,ATAAT,MEE17
Gorai.013G030300.1.D
PCC=0.57, p=0.011
Gorai.013G030300.1.A
PCC=0.11, p=0.654
mRNA Profile
Protein Expression Profile
DPA

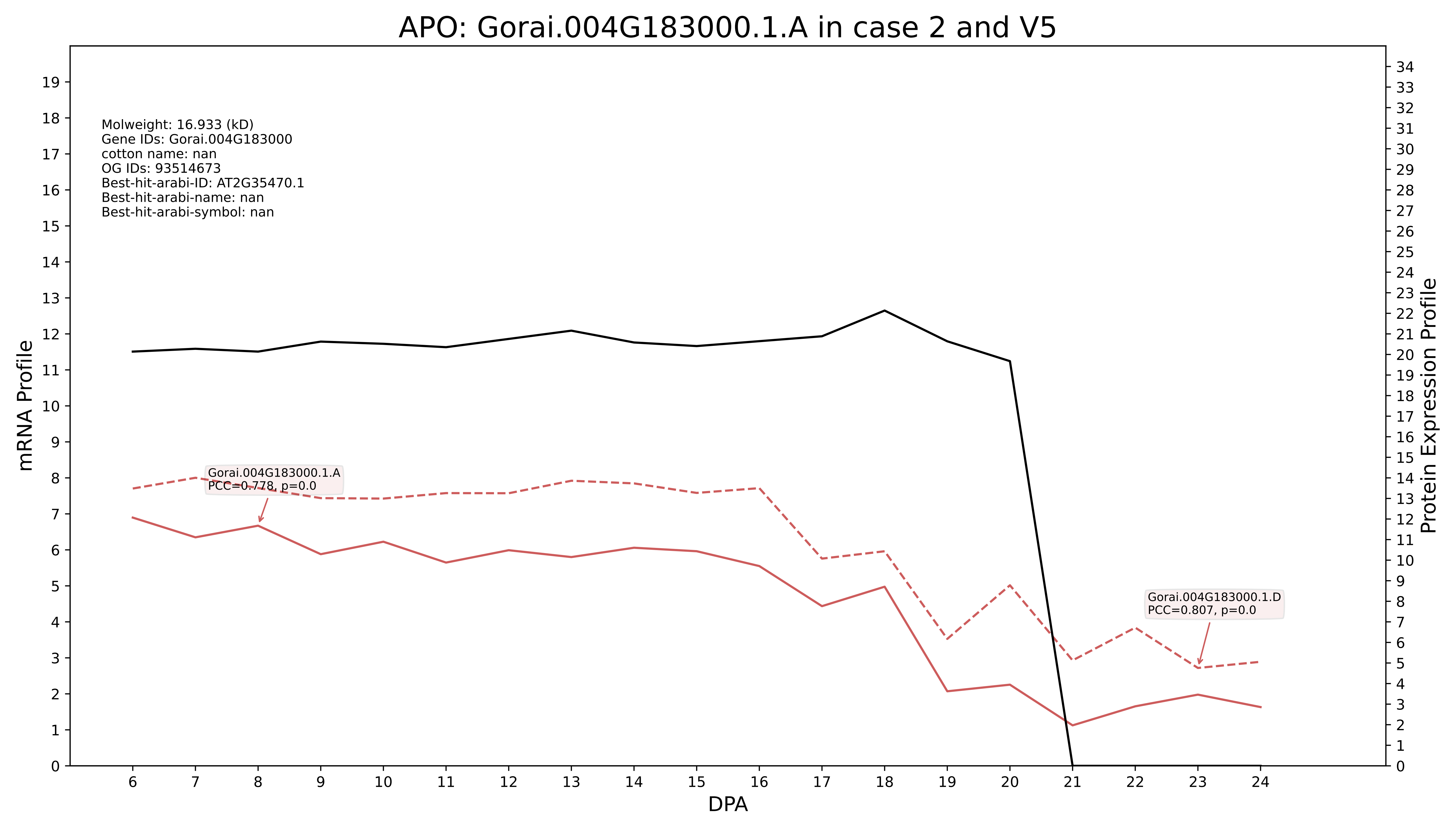
APO: Gorai.004G183000.1.A in case 2 and V5
Molweight: 16.933 (kD)
Gene IDs: Gorai.004G183000
cotton name: nan
OG IDs: 93514673
Best-hit-arabi-ID: AT2G35470.1
Best-hit-arabi-name: nan
Best-hit-arabi-symbol: nan
Gorai.004G183000.1.A
PCC=0.778, p=0.0
Gorai.004G183000.1.D
PCC=0.807, p=0.0
mRNA Profile
Protein Expression Profile
DPA

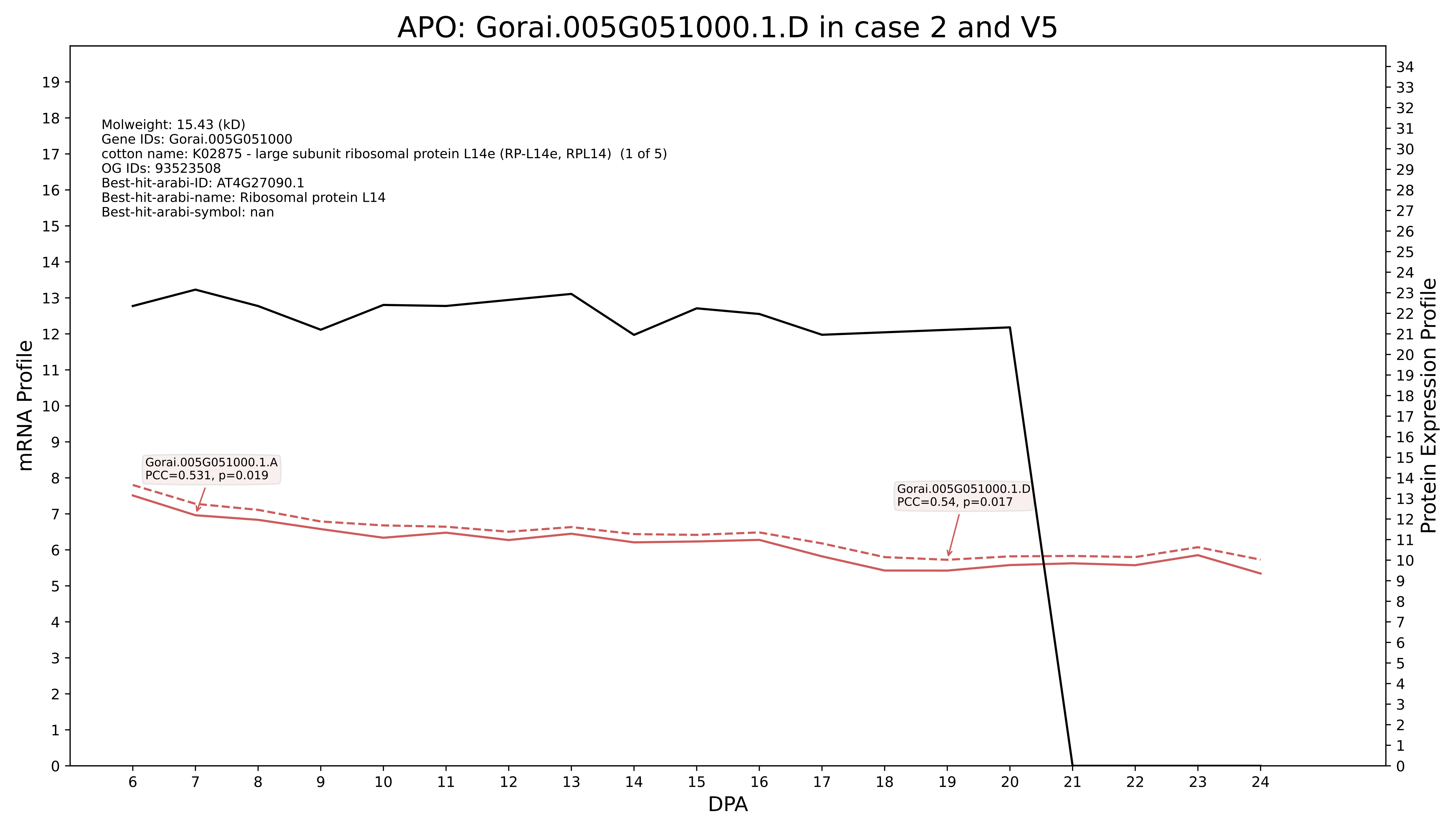
APO: Gorai.005G051000.1.D in case 2 and V5
Molweight: 15.43 (kD)
Gene IDs: Gorai.005G051000
cotton name: K02875 - large subunit ribosomal protein L14e (RP-L14e, RPL14) (1 of 5)
OG IDs: 93523508
Best-hit-arabi-ID: AT4G27090.1
Best-hit-arabi-name: Ribosomal protein L14
Best-hit-arabi-symbol: nan
Gorai.005G051000.1.A
PCC=0.531, p=0.019
Gorai.005G051000.1.D
PCC=0.54, p=0.017
mRNA Profile
Protein Expression Profile
DPA

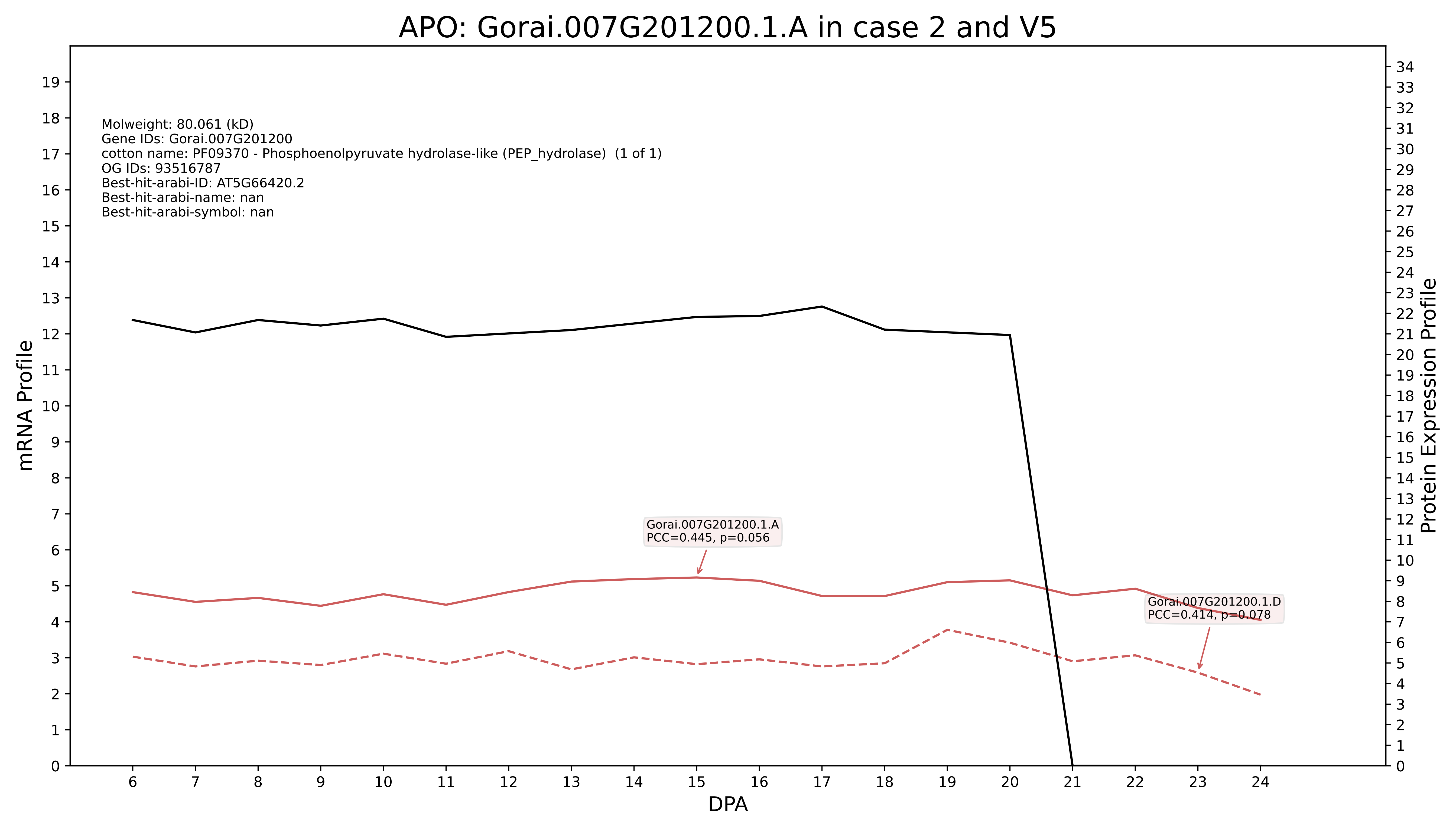
APO: Gorai.007G201200.1.A in case 2 and V5
Molweight: 80.061 (kD)
Gene IDs: Gorai.007G201200
cotton name: PF09370 - Phosphoenolpyruvate hydrolase-like (PEP_hydrolase) (1 of 1)
OG IDs: 93516787
Best-hit-arabi-ID: AT5G66420.2
Best-hit-arabi-name: nan
Best-hit-arabi-symbol: nan
Gorai.007G201200.1.A
PCC=0.445, p=0.056
Gorai.007G201200.1.D
PCC=0.414, p=0.078
mRNA Profile
Protein Expression Profile
DPA

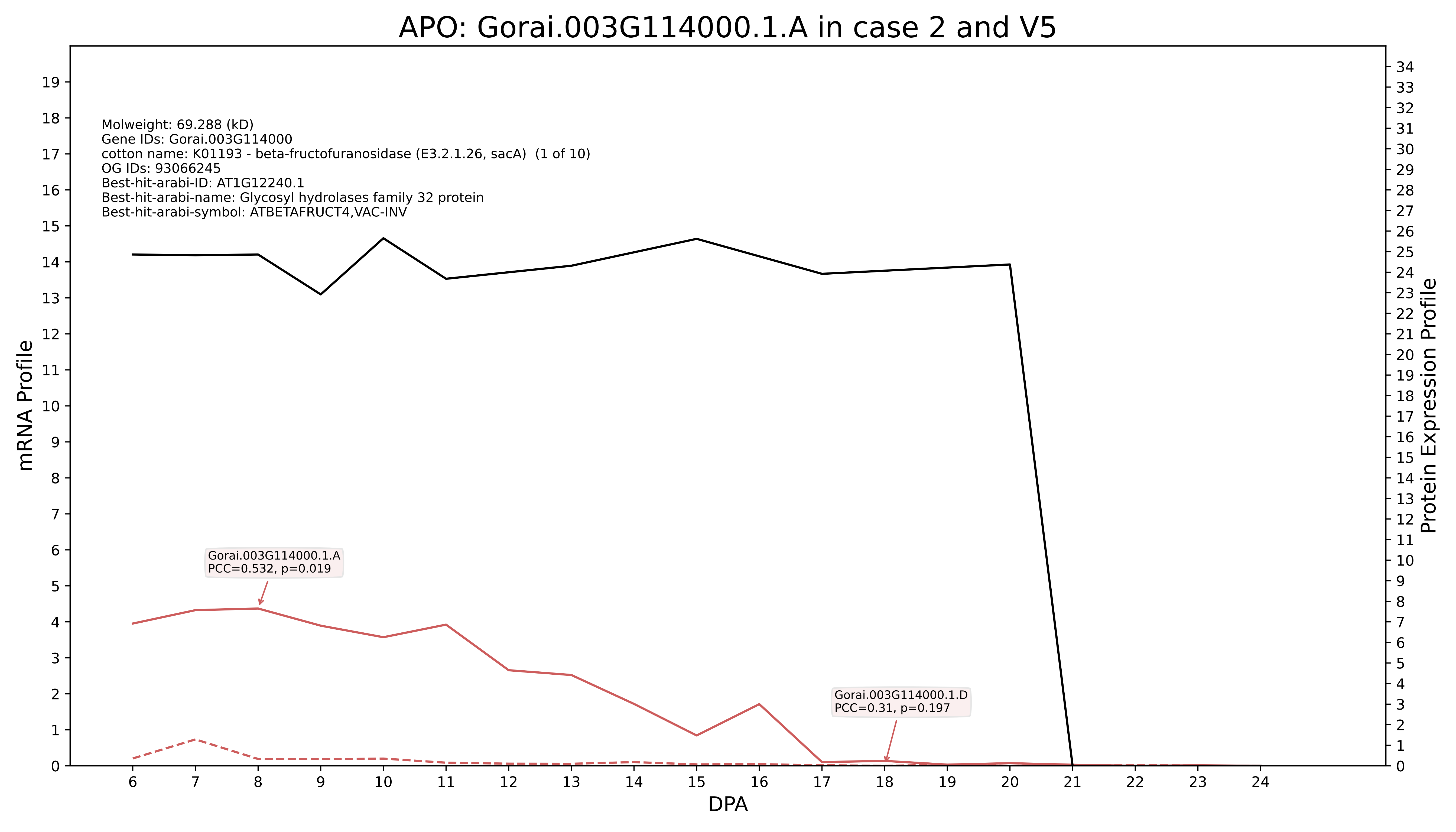

APO: Gorai.003G114000.1.A in case 2 and V5
Molweight: 69.288 (kD)
Gene IDs: Gorai.003G114000
cotton name: K01193 - beta-fructofuranosidase (E3.2.1.26, sacA) (1 of 10)
OG IDs: 93066245
Best-hit-arabi-ID: AT1G12240.1
Best-hit-arabi-name: Glycosyl hydrolases family 32 protein
Best-hit-arabi-symbol: ATBETAFRUCT4,VAC-INV
Gorai.003G114000.1.A
PCC=0.532, p=0.019
Gorai.003G114000.1.D
PCC=0.31, p=0.197
mRNA Profile
Protein Expression Profile
DPA

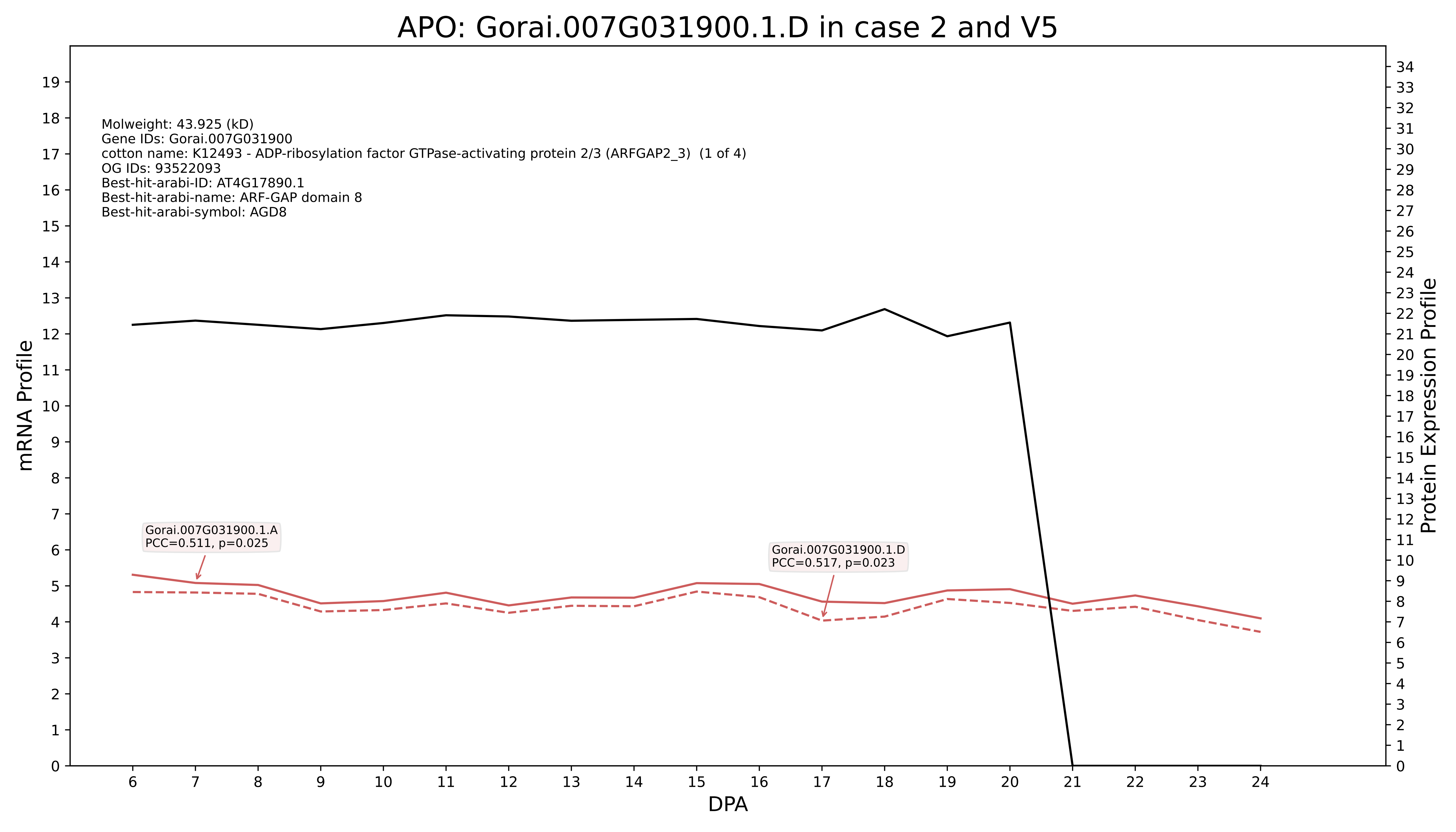
APO: Gorai.007G031900.1.D in case 2 and V5
Molweight: 43.925 (kD)
Gene IDs: Gorai.007G031900
cotton name: K12493 - ADP-ribosylation factor GTPase-activating protein 2/3 (ARFGAP2_3) (1 of 4)
OG IDs: 93522093
Best-hit-arabi-ID: AT4G17890.1
Best-hit-arabi-name: ARF-GAP domain 8
Best-hit-arabi-symbol: AGD8
Gorai.007G031900.1.A
PCC=0.511, p=0.025
Gorai.007G031900.1.D
PCC=0.517, p=0.023
mRNA Profile
Protein Expression Profile
DPA

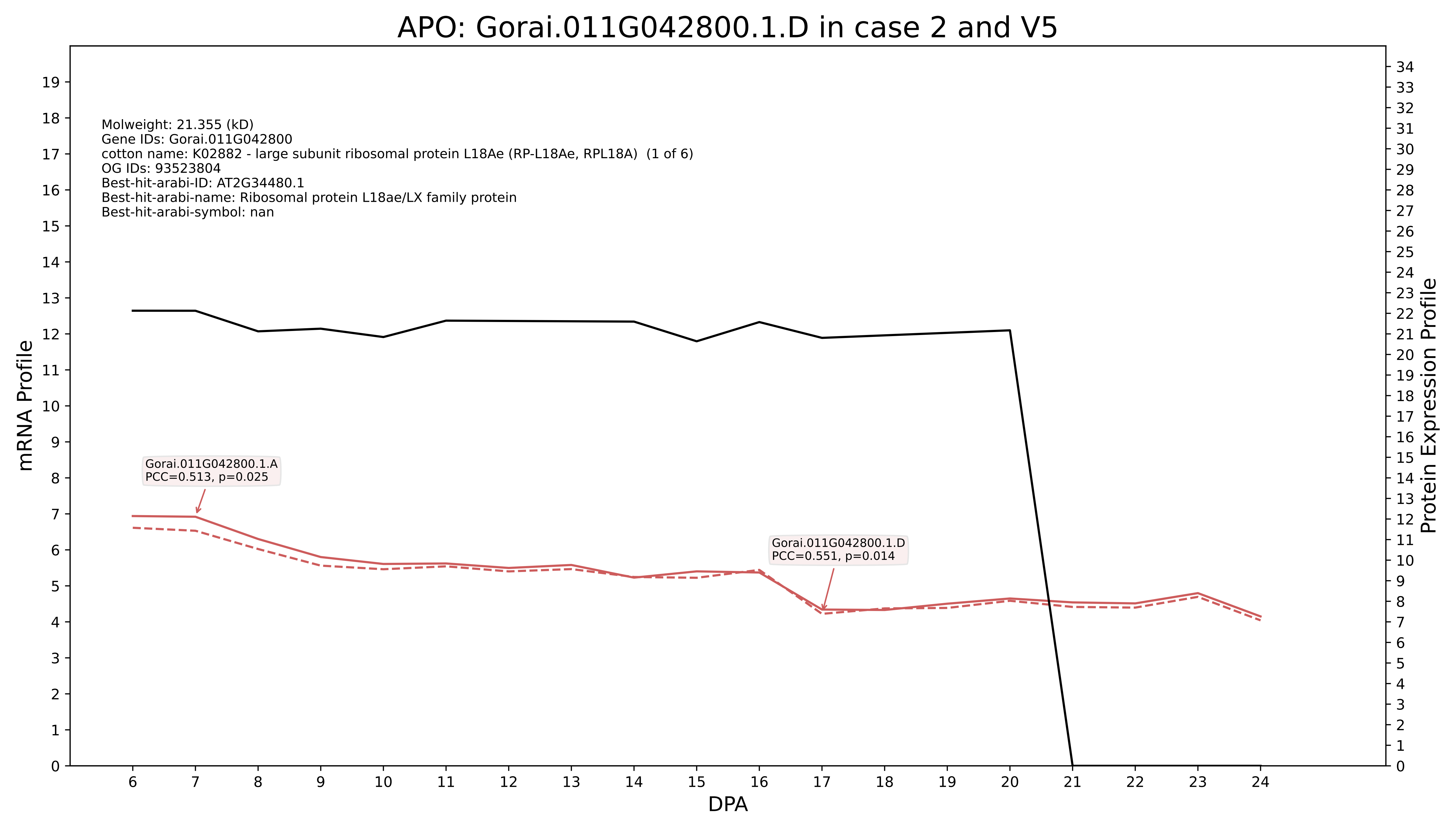

APO: Gorai.011G042800.1.D in case 2 and V5
Molweight: 21.355 (kD)
Gene IDs: Gorai.011G042800
cotton name: K02882 - large subunit ribosomal protein L18Ae (RP-L18Ae, RPL18A) (1 of 6)
OG IDs: 93523804
Best-hit-arabi-ID: AT2G34480.1
Best-hit-arabi-name: Ribosomal protein L18ae/LX family protein
Best-hit-arabi-symbol: nan
Gorai.011G042800.1.A
PCC=0.513, p=0.025
Gorai.011G042800.1.D
PCC=0.551, p=0.014
mRNA Profile
Protein Expression Profile
DPA

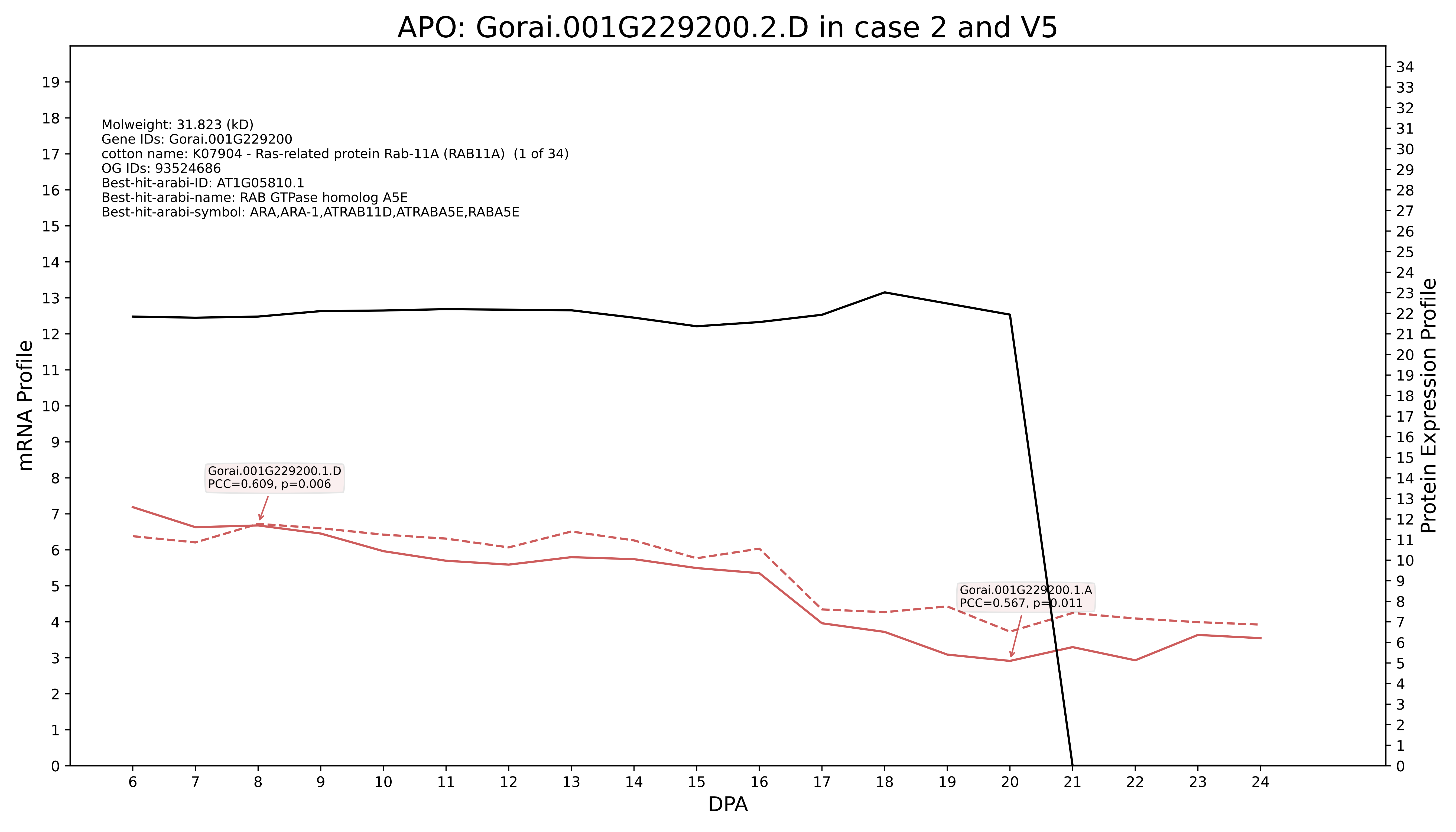
APO: Gorai.001G229200.2.D in case 2 and V5
Molweight: 31.823 (kD)
Gene IDs: Gorai.001G229200
cotton name: K07904 - Ras-related protein Rab-11A (RAB11A) (1 of 34)
OG IDs: 93524686
Best-hit-arabi-ID: AT1G05810.1
Best-hit-arabi-name: RAB GTPase homolog A5E
Best-hit-arabi-symbol: ARA,ARA-1,ATRAB11D,ATRABA5E,RABA5E
Gorai.001G229200.1.D
PCC=0.609, p=0.006
Gorai.001G229200.1.A
PCC=0.567, p=0.011
mRNA Profile
Protein Expression Profile
DPA

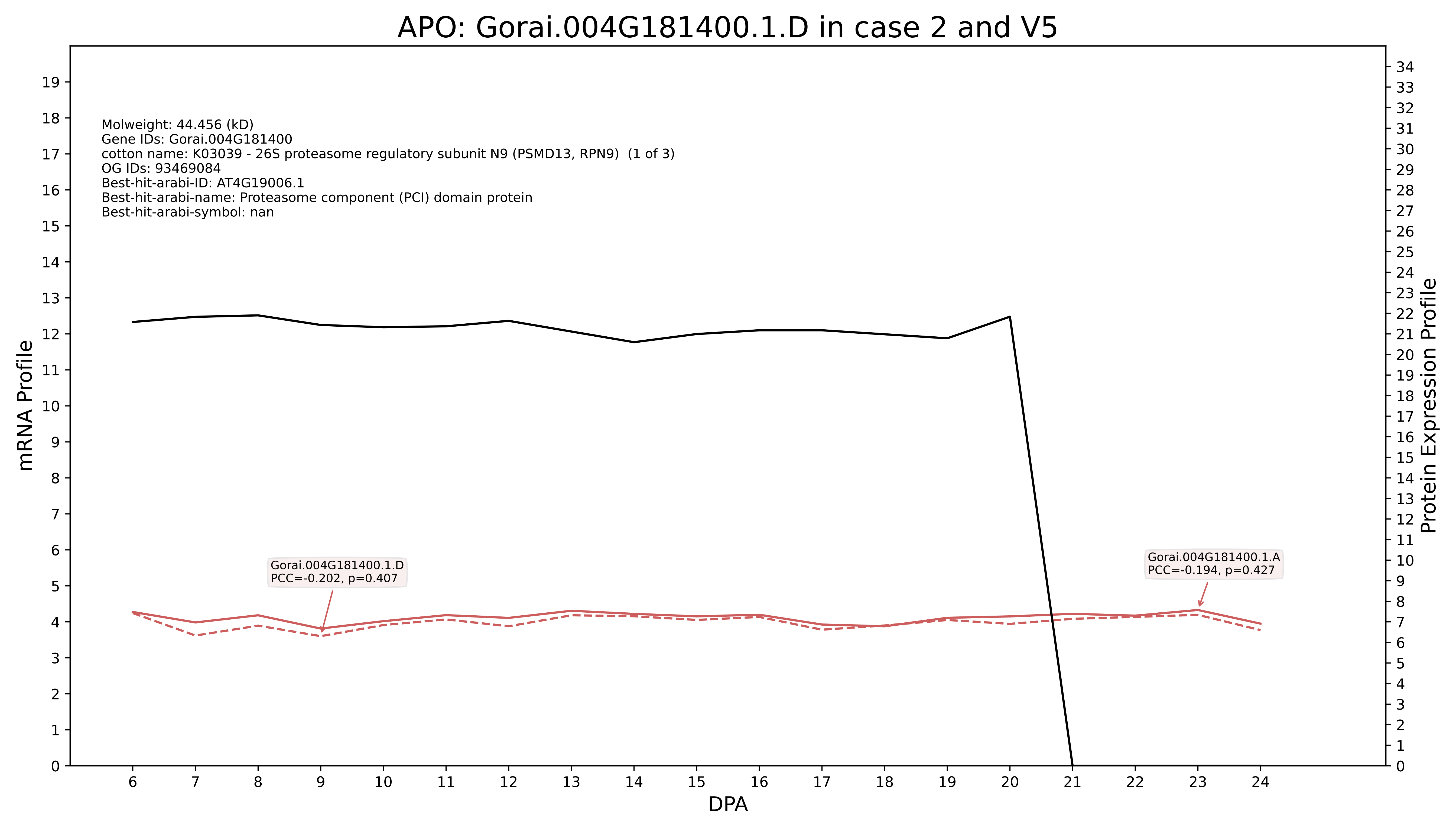

APO: Gorai.004G181400.1.D in case 2 and V5
Molweight: 44.456 (kD)
Gene IDs: Gorai.004G181400
cotton name: K03039 - 26S proteasome regulatory subunit N9 (PSMD13, RPN9) (1 of 3)
OG IDs: 93469084
Best-hit-arabi-ID: AT4G19006.1
Best-hit-arabi-name: Proteasome component (PCI) domain protein
Best-hit-arabi-symbol: nan
Gorai.004G181400.1.D
PCC=-0.202, p=0.407
Gorai.004G181400.1.A
PCC=-0.194, p=0.427
mRNA Profile
Protein Expression Profile
DPA

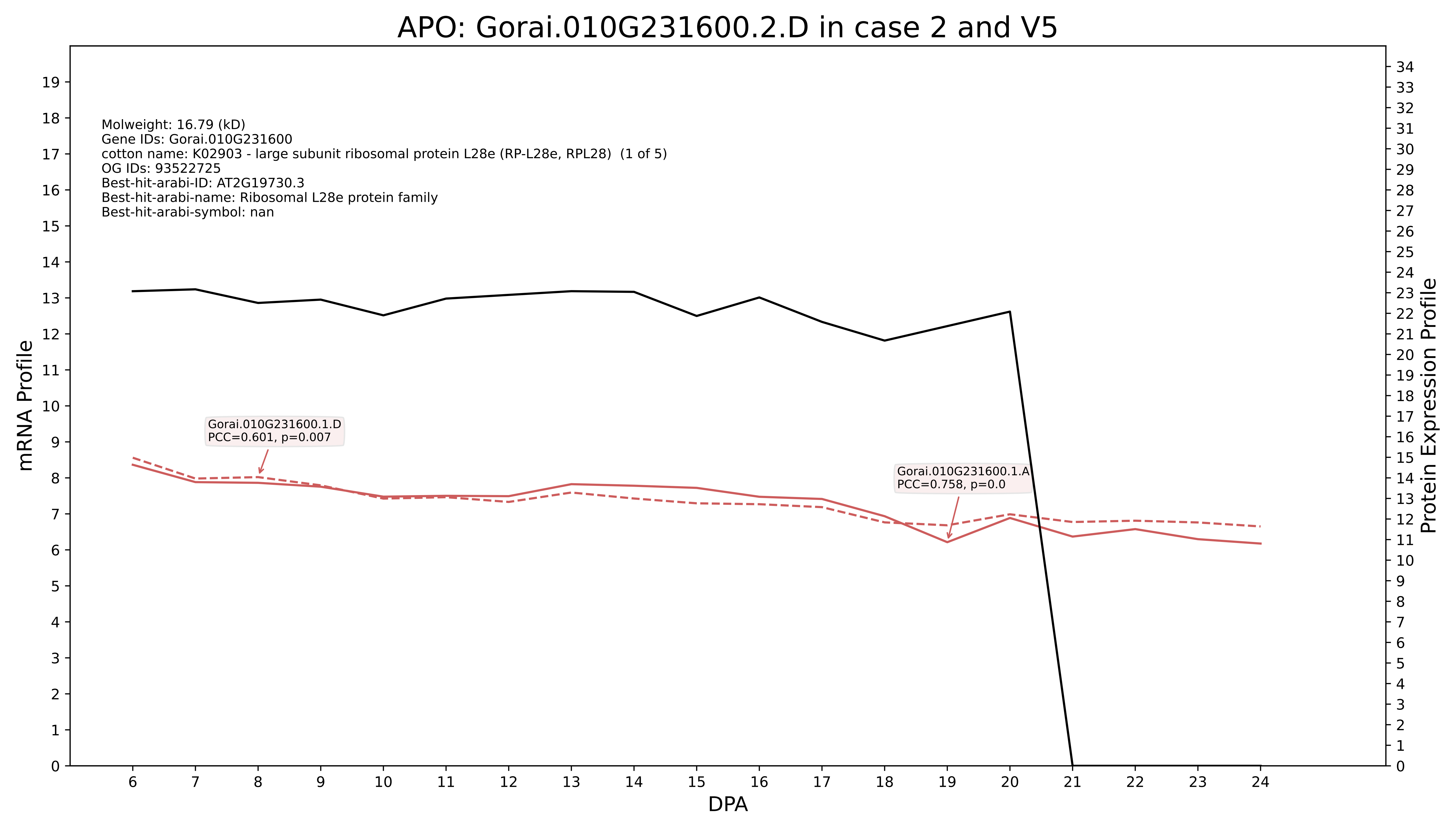
APO: Gorai.010G231600.2.D in case 2 and V5
Molweight: 16.79 (kD)
Gene IDs: Gorai.010G231600
cotton name: K02903 - large subunit ribosomal protein L28e (RP-L28e, RPL28) (1 of 5)
OG IDs: 93522725
Best-hit-arabi-ID: AT2G19730.3
Best-hit-arabi-name: Ribosomal L28e protein family
Best-hit-arabi-symbol: nan
Gorai.010G231600.1.D
PCC=0.601, p=0.007
Gorai.010G231600.1.A
PCC=0.758, p=0.0
mRNA Profile
Protein Expression Profile
DPA

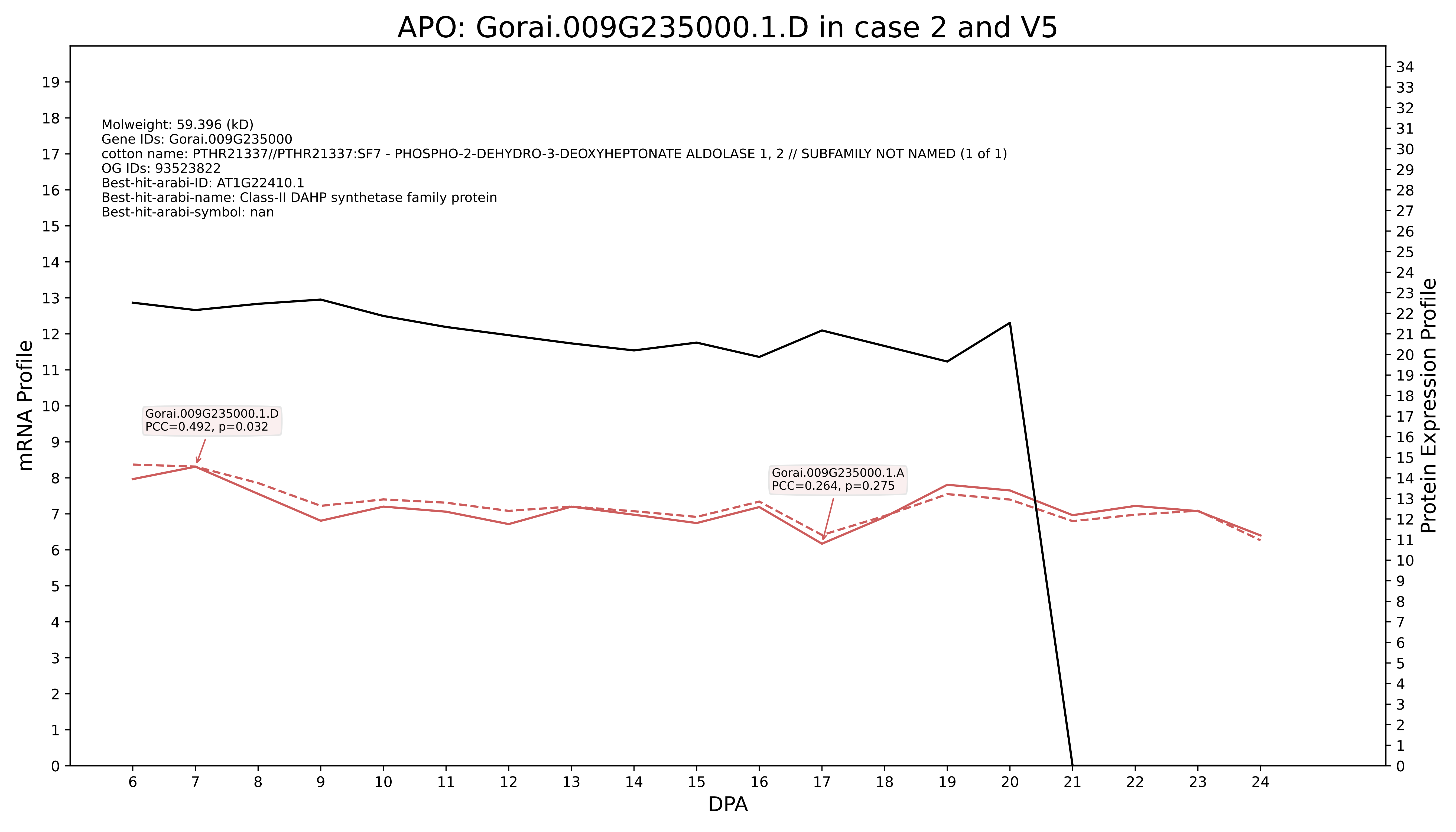

APO: Gorai.009G235000.1.D in case 2 and V5
Molweight: 59.396 (kD)
Gene IDs: Gorai.009G235000
cotton name: PTHR21337//PTHR21337:SF7 - PHOSPHO-2-DEHYDRO-3-DEOXYHEPTONATE ALDOLASE 1, 2 // SUBFAMILY NOT NAMED (1 of 1)
OG IDs: 93523822
Best-hit-arabi-ID: AT1G22410.1
Best-hit-arabi-name: Class-II DAHP synthetase family protein
Best-hit-arabi-symbol: nan
Gorai.009G235000.1.D
PCC=0.492, p=0.032
Gorai.009G235000.1.A
PCC=0.264, p=0.275
mRNA Profile
Protein Expression Profile
DPA

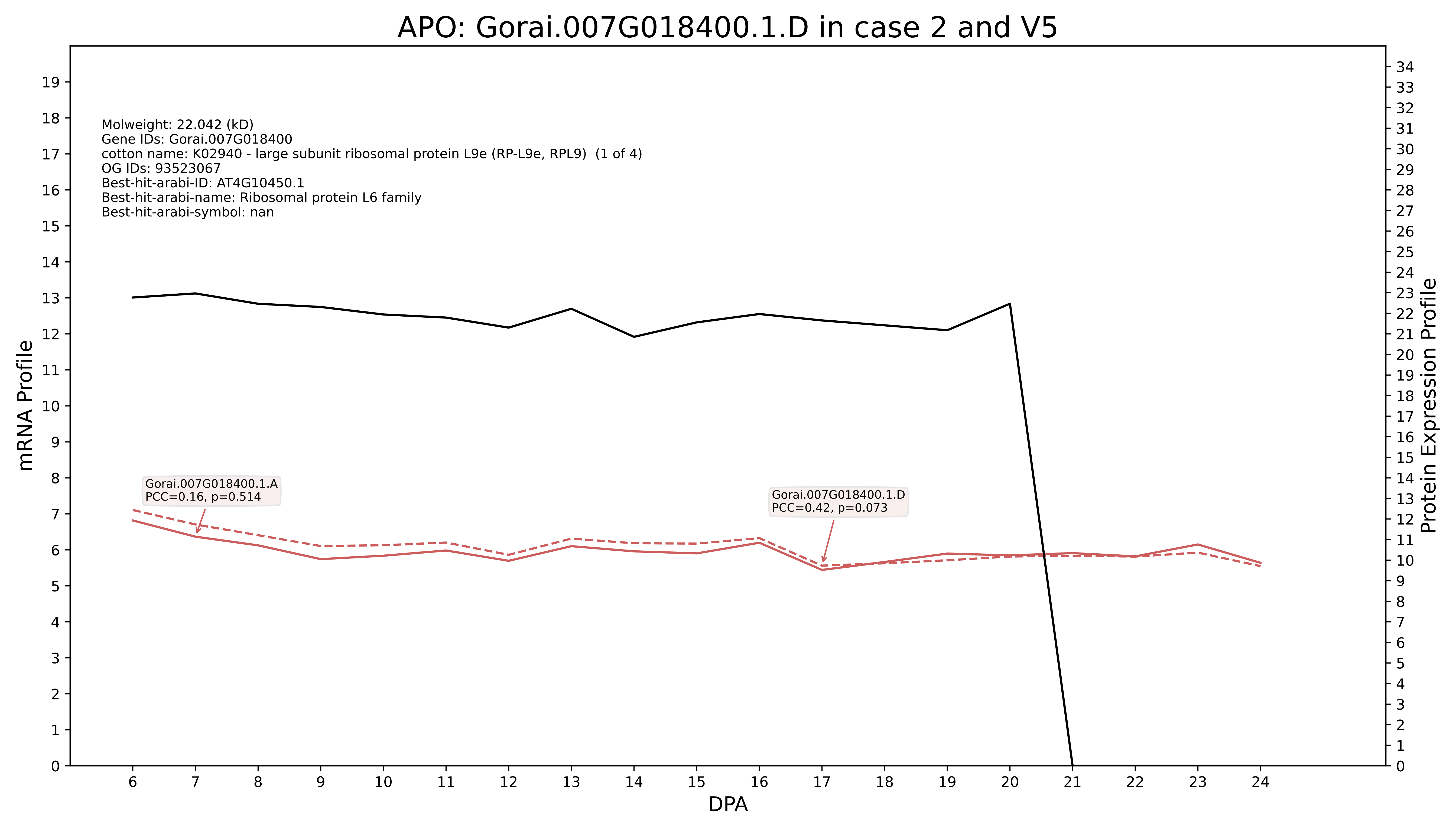

APO: Gorai.007G018400.1.D in case 2 and V5
Molweight: 22.042 (kD)
Gene IDs: Gorai.007G018400
cotton name: K02940 - large subunit ribosomal protein L9e (RP-L9e, RPL9) (1 of 4)
OG IDs: 93523067
Best-hit-arabi-ID: AT4G10450.1
Best-hit-arabi-name: Ribosomal protein L6 family
Best-hit-arabi-symbol: nan
Gorai.007G018400.1.A
PCC=0.16, p=0.514
Gorai.007G018400.1.D
PCC=0.42, p=0.073
mRNA Profile
Protein Expression Profile
DPA

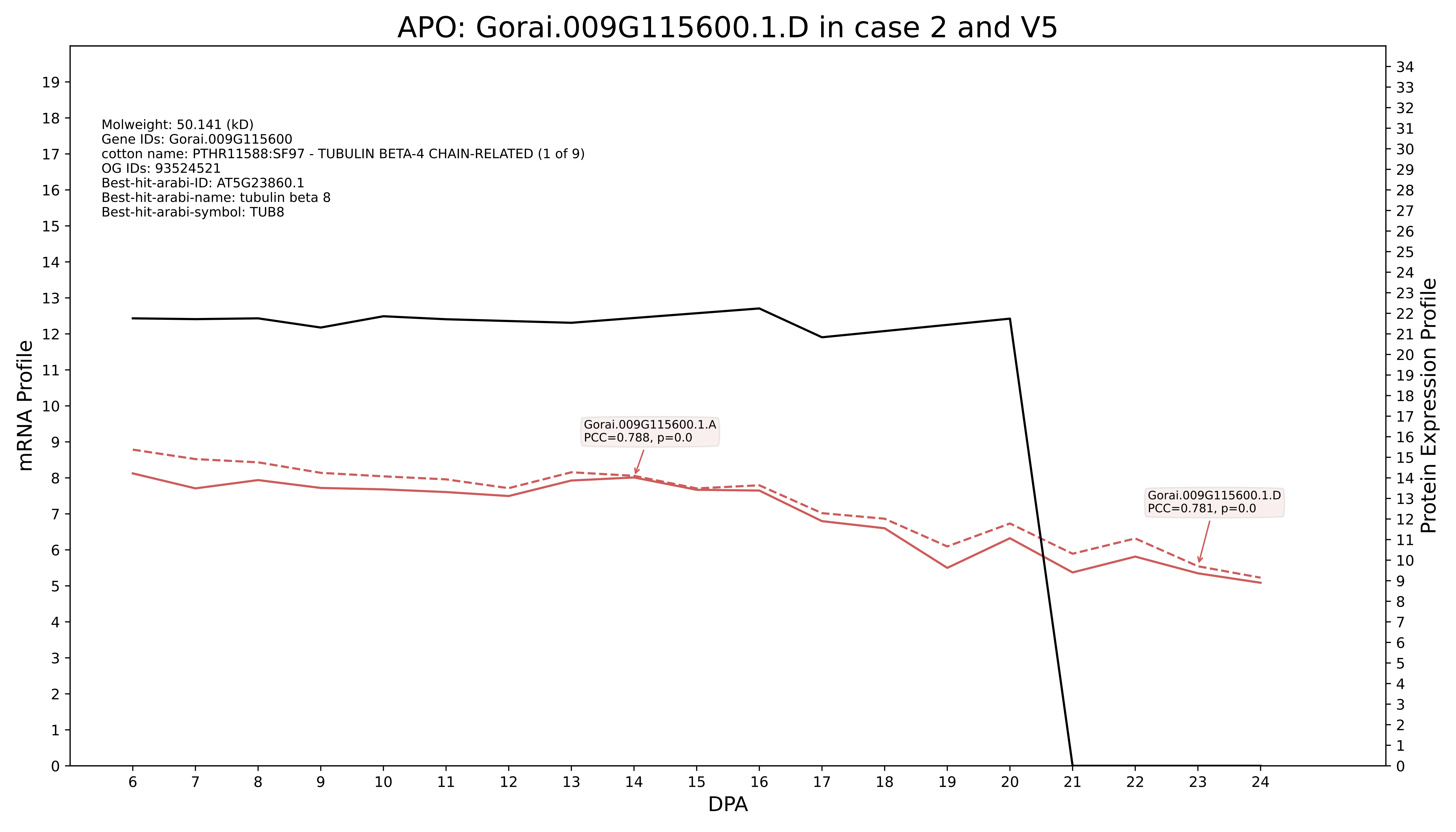

APO: Gorai.009G115600.1.D in case 2 and V5
Molweight: 50.141 (kD)
Gene IDs: Gorai.009G115600
cotton name: PTHR11588:SF97 - TUBULIN BETA-4 CHAIN-RELATED (1 of 9)
OG IDs: 93524521
Best-hit-arabi-ID: AT5G23860.1
Best-hit-arabi-name: tubulin beta 8
Best-hit-arabi-symbol: TUB8
Gorai.009G115600.1.A
PCC=0.788, p=0.0
Gorai.009G115600.1.D
PCC=0.781, p=0.0
mRNA Profile
Protein Expression Profile
DPA

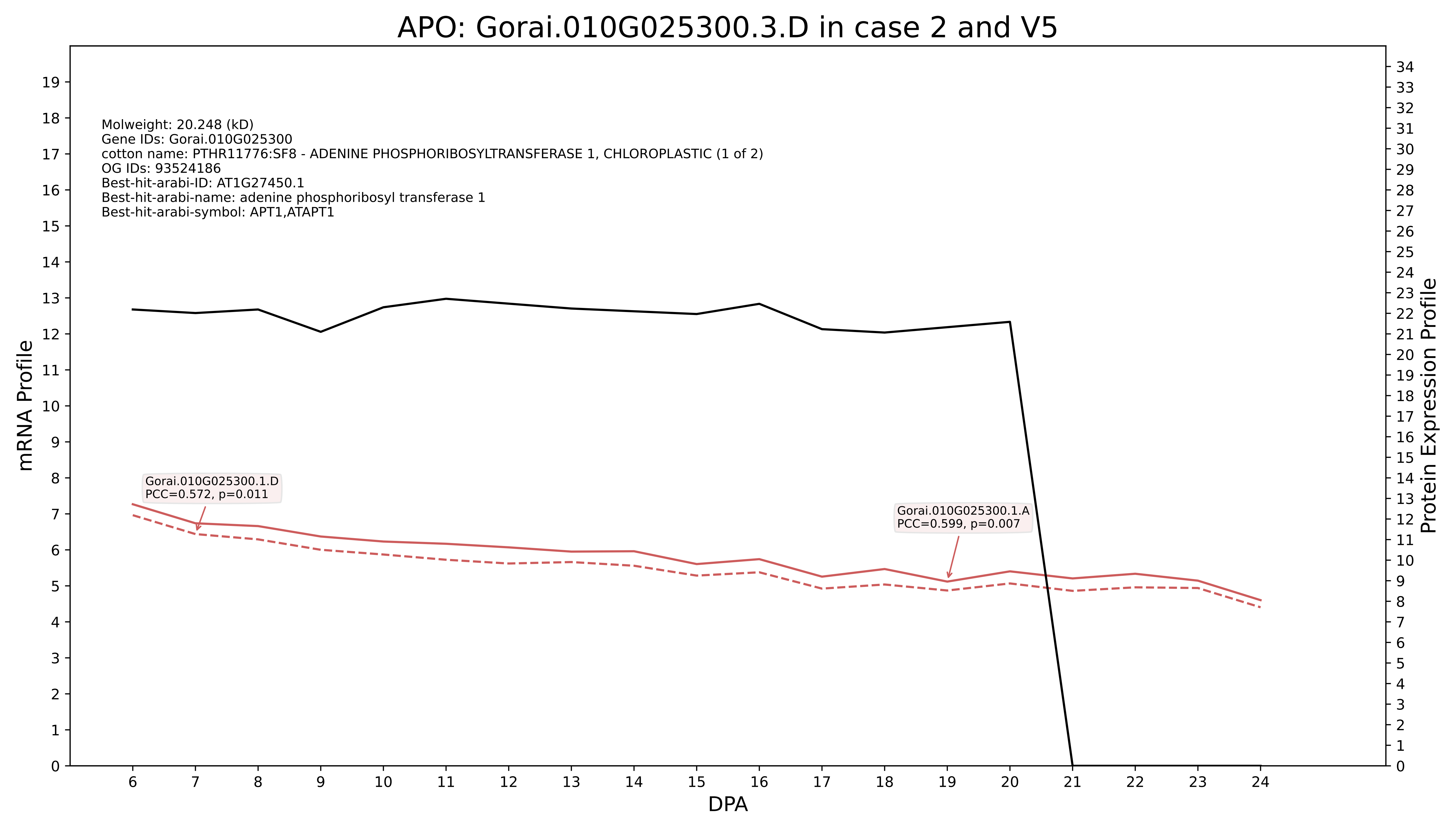

APO: Gorai.010G025300.3.D in case 2 and V5
Molweight: 20.248 (kD)
Gene IDs: Gorai.010G025300
cotton name: PTHR11776:SF8 - ADENINE PHOSPHORIBOSYLTRANSFERASE 1, CHLOROPLASTIC (1 of 2)
OG IDs: 93524186
Best-hit-arabi-ID: AT1G27450.1
Best-hit-arabi-name: adenine phosphoribosyl transferase 1
Best-hit-arabi-symbol: APT1,ATAPT1
Gorai.010G025300.1.D
PCC=0.572, p=0.011
Gorai.010G025300.1.A
PCC=0.599, p=0.007
mRNA Profile
Protein Expression Profile
DPA

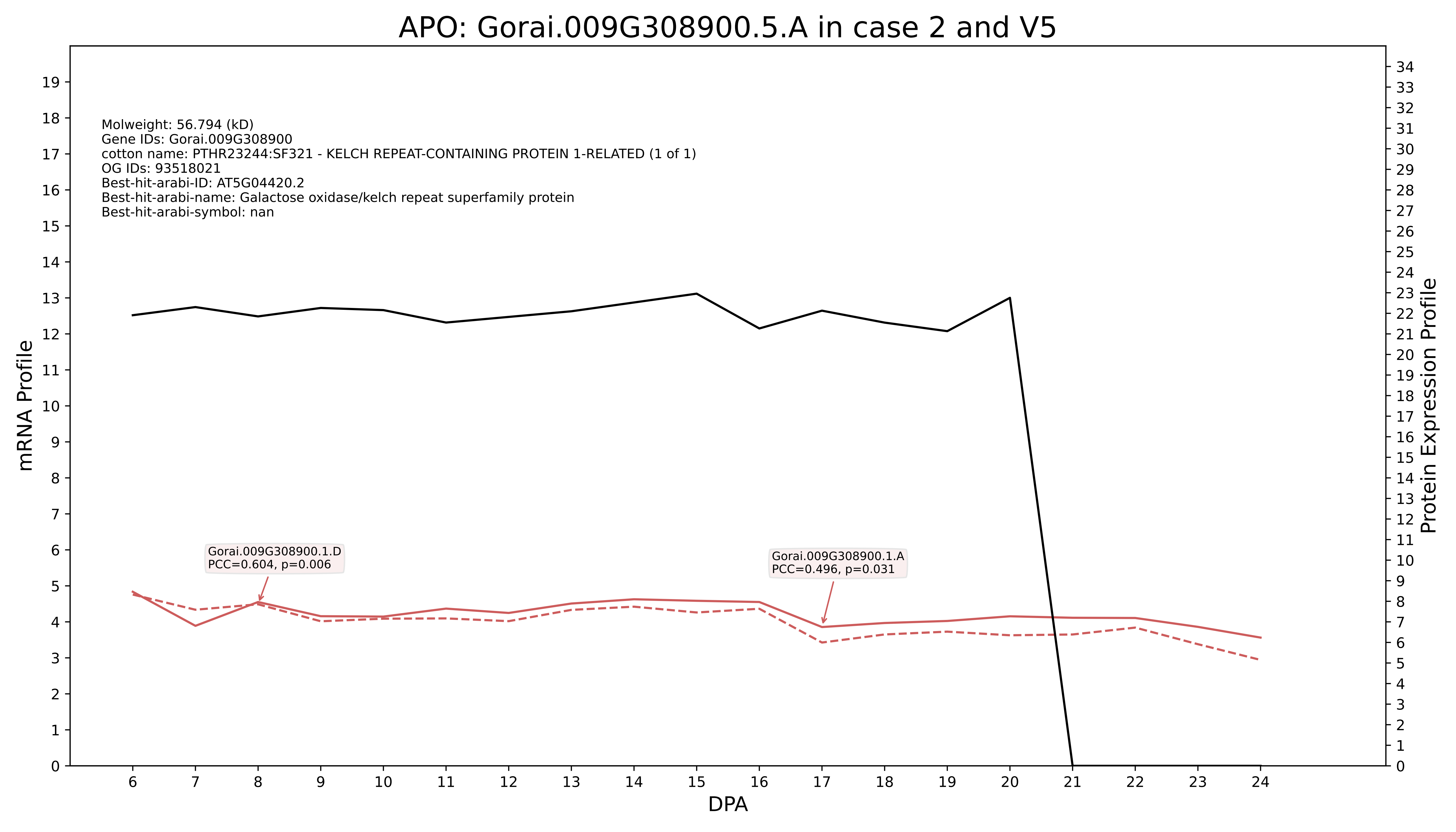
APO: Gorai.009G308900.5.A in case 2 and V5
Molweight: 56.794 (kD)
Gene IDs: Gorai.009G308900
cotton name: PTHR23244:SF321 - KELCH REPEAT-CONTAINING PROTEIN 1-RELATED (1 of 1)
OG IDs: 93518021
Best-hit-arabi-ID: AT5G04420.2
Best-hit-arabi-name: Galactose oxidase/kelch repeat superfamily protein
Best-hit-arabi-symbol: nan
Gorai.009G308900.1.D
PCC=0.604, p=0.006
Gorai.009G308900.1.A
PCC=0.496, p=0.031
mRNA Profile
Protein Expression Profile
DPA

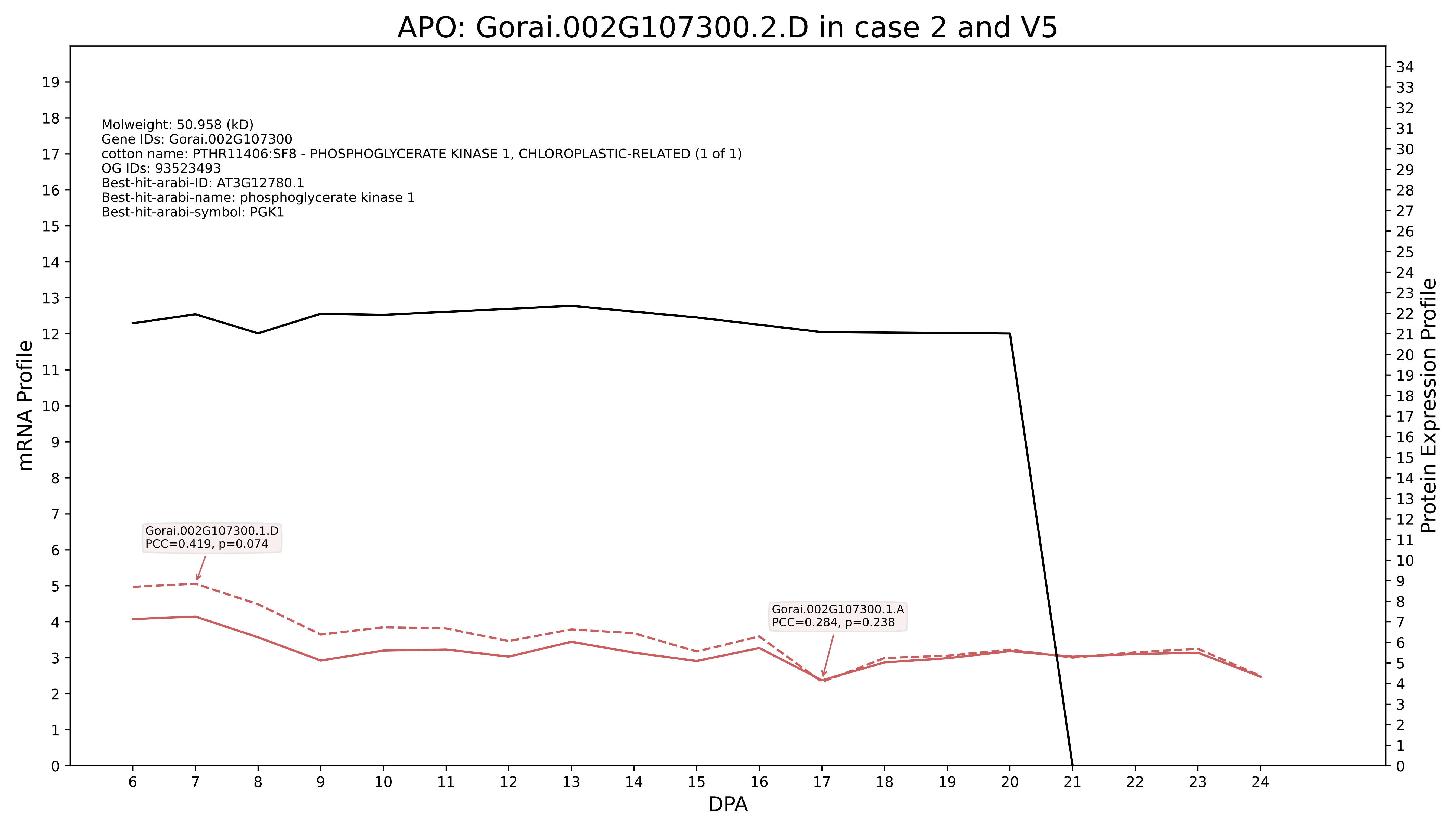

APO: Gorai.002G107300.2.D in case 2 and V5
Molweight: 50.958 (kD)
Gene IDs: Gorai.002G107300
cotton name: PTHR11406:SF8 - PHOSPHOGLYCERATE KINASE 1, CHLOROPLASTIC-RELATED (1 of 1)
OG IDs: 93523493
Best-hit-arabi-ID: AT3G12780.1
Best-hit-arabi-name: phosphoglycerate kinase 1
Best-hit-arabi-symbol: PGK1
Gorai.002G107300.1.D
PCC=0.419, p=0.074
Gorai.002G107300.1.A
PCC=0.284, p=0.238
mRNA Profile
Protein Expression Profile
DPA

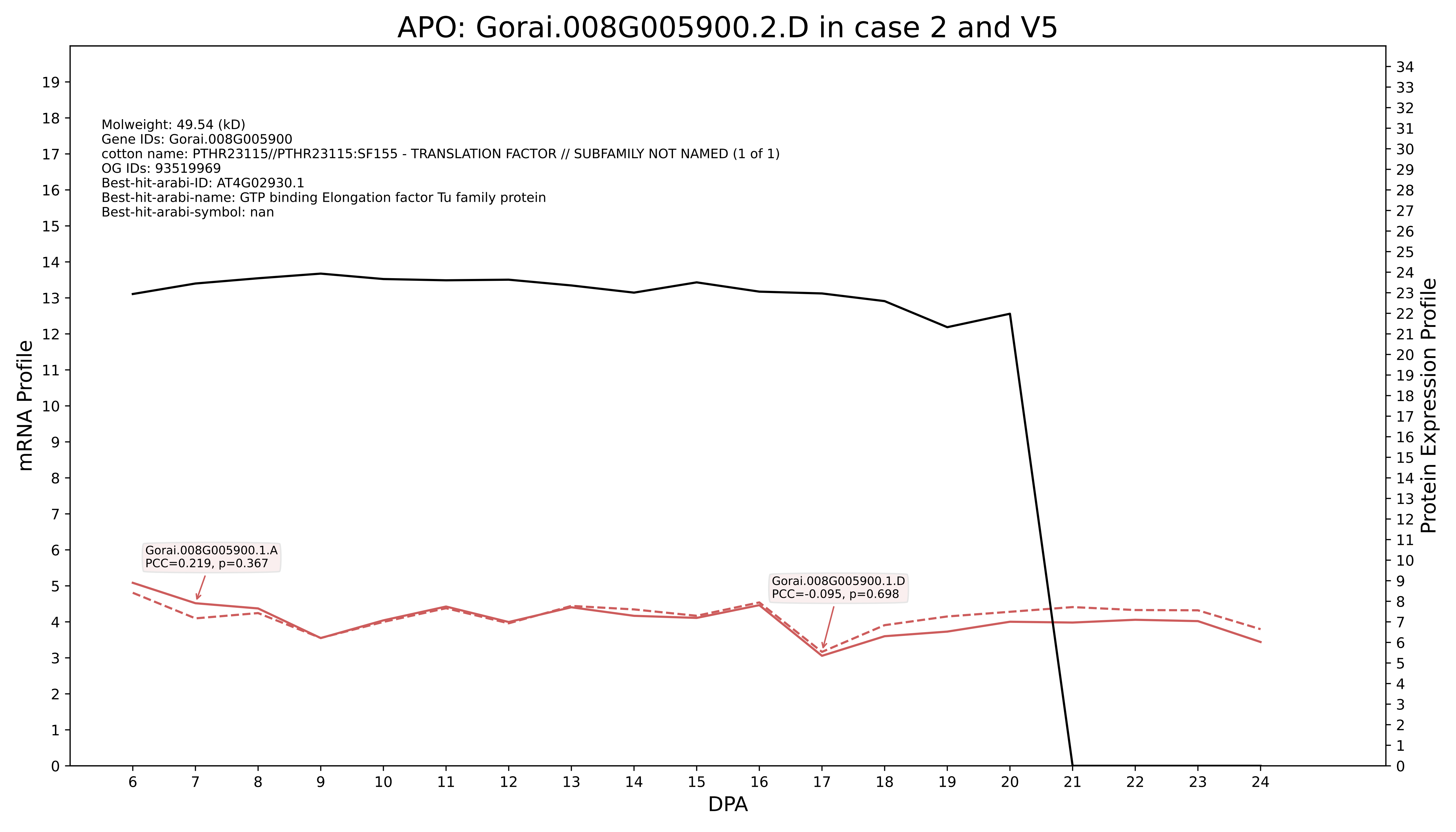
APO: Gorai.008G005900.2.D in case 2 and V5
Molweight: 49.54 (kD)
Gene IDs: Gorai.008G005900
cotton name: PTHR23115//PTHR23115:SF155 - TRANSLATION FACTOR // SUBFAMILY NOT NAMED (1 of 1)
OG IDs: 93519969
Best-hit-arabi-ID: AT4G02930.1
Best-hit-arabi-name: GTP binding Elongation factor Tu family protein
Best-hit-arabi-symbol: nan
Gorai.008G005900.1.A
PCC=0.219, p=0.367
Gorai.008G005900.1.D
PCC=-0.095, p=0.698
mRNA Profile
Protein Expression Profile
DPA

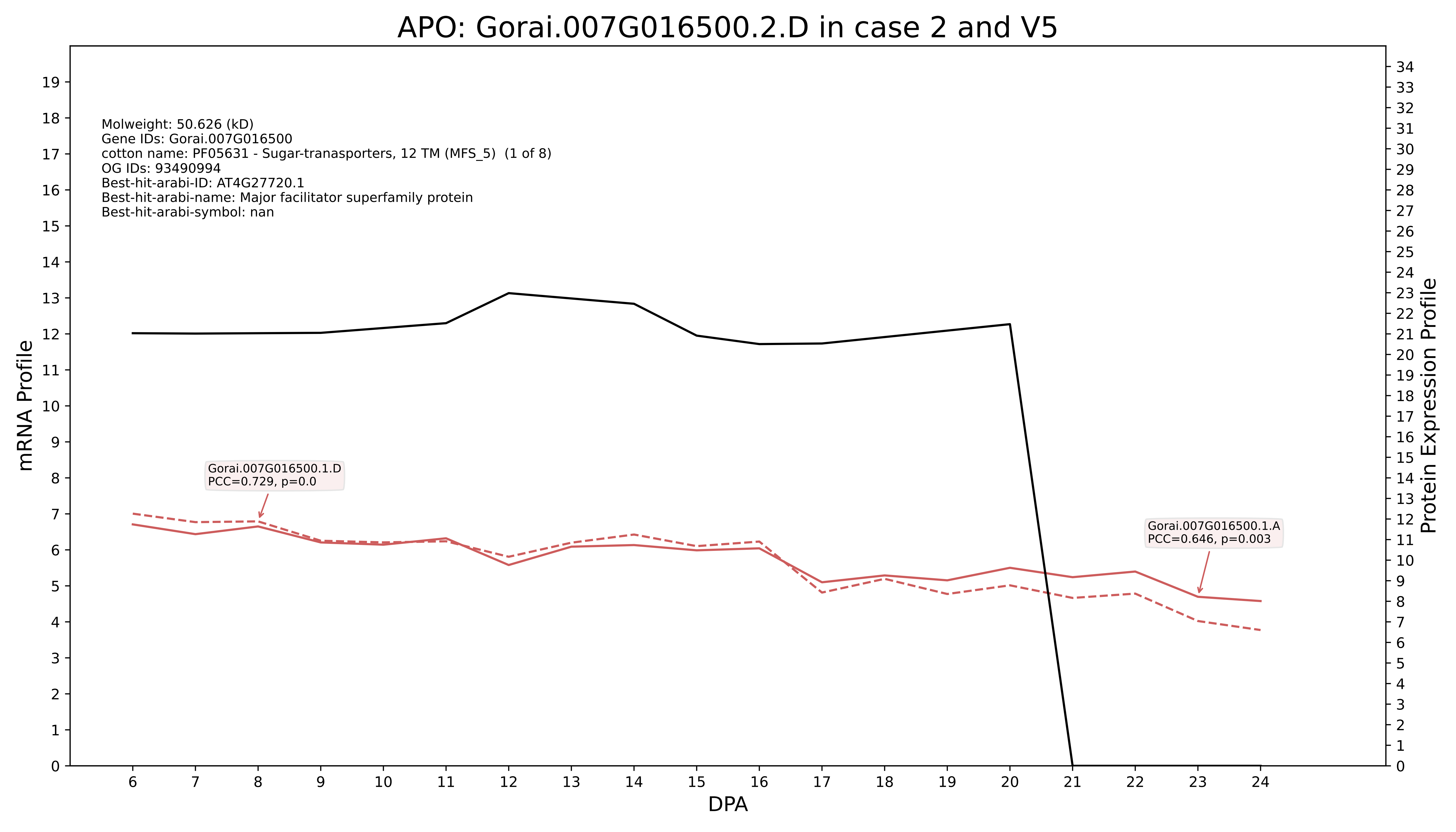

APO: Gorai.007G016500.2.D in case 2 and V5
Molweight: 50.626 (kD)
Gene IDs: Gorai.007G016500
cotton name: PF05631 - Sugar-tranasporters, 12 TM (MFS_5) (1 of 8)
OG IDs: 93490994
Best-hit-arabi-ID: AT4G27720.1
Best-hit-arabi-name: Major facilitator superfamily protein
Best-hit-arabi-symbol: nan
Gorai.007G016500.1.D
PCC=0.729, p=0.0
Gorai.007G016500.1.A
PCC=0.646, p=0.003
mRNA Profile
Protein Expression Profile
DPA

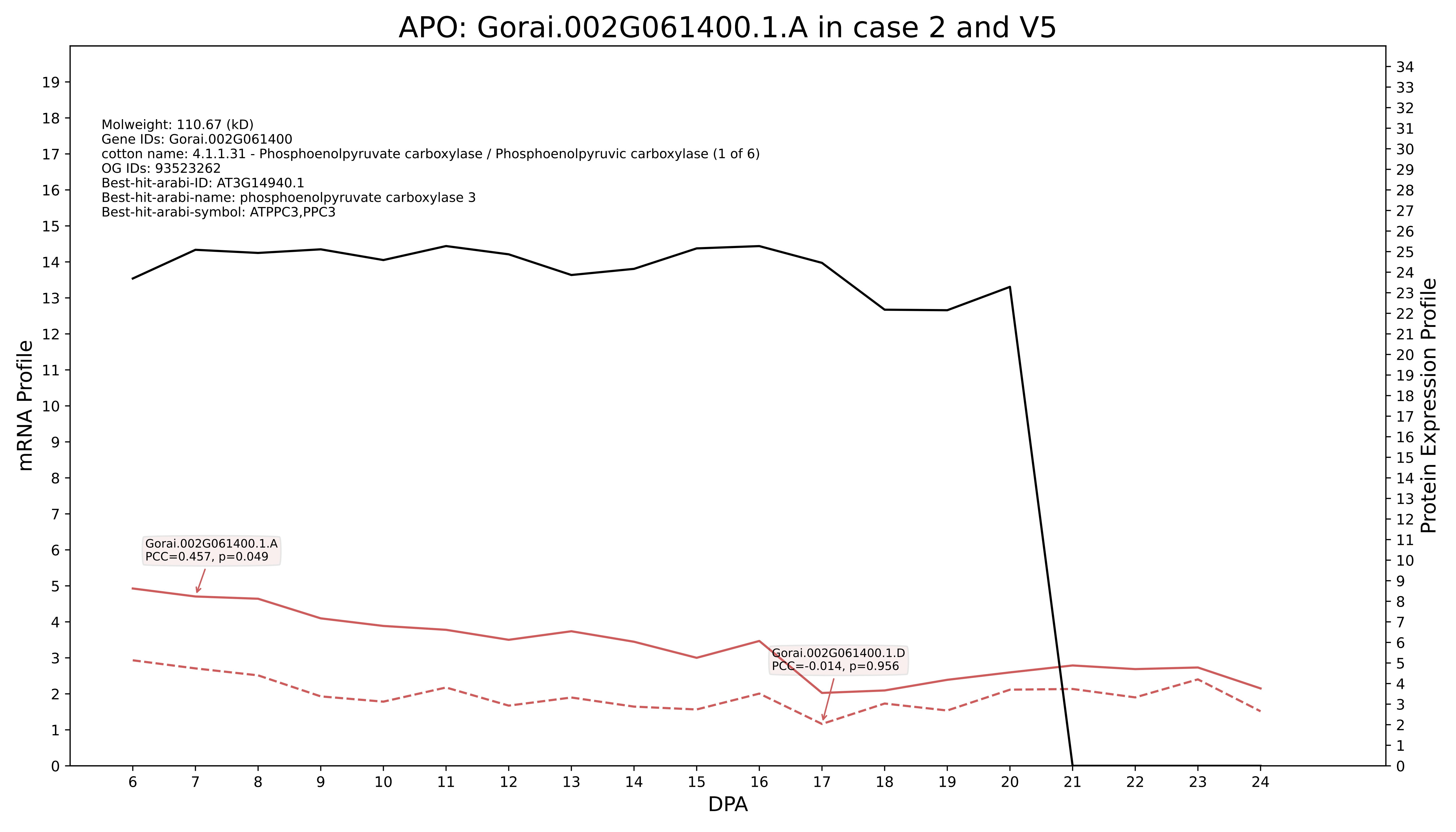
APO: Gorai.002G061400.1.A in case 2 and V5
Molweight: 110.67 (kD)
Gene IDs: Gorai.002G061400
cotton name: 4.1.1.31 - Phosphoenolpyruvate carboxylase / Phosphoenolpyruvic carboxylase (1 of 6)
OG IDs: 93523262
Best-hit-arabi-ID: AT3G14940.1
Best-hit-arabi-name: phosphoenolpyruvate carboxylase 3
Best-hit-arabi-symbol: ATPPC3,PPC3
Gorai.002G061400.1.A
PCC=0.457, p=0.049
Gorai.002G061400.1.D
PCC=-0.014, p=0.956
mRNA Profile
Protein Expression Profile
DPA

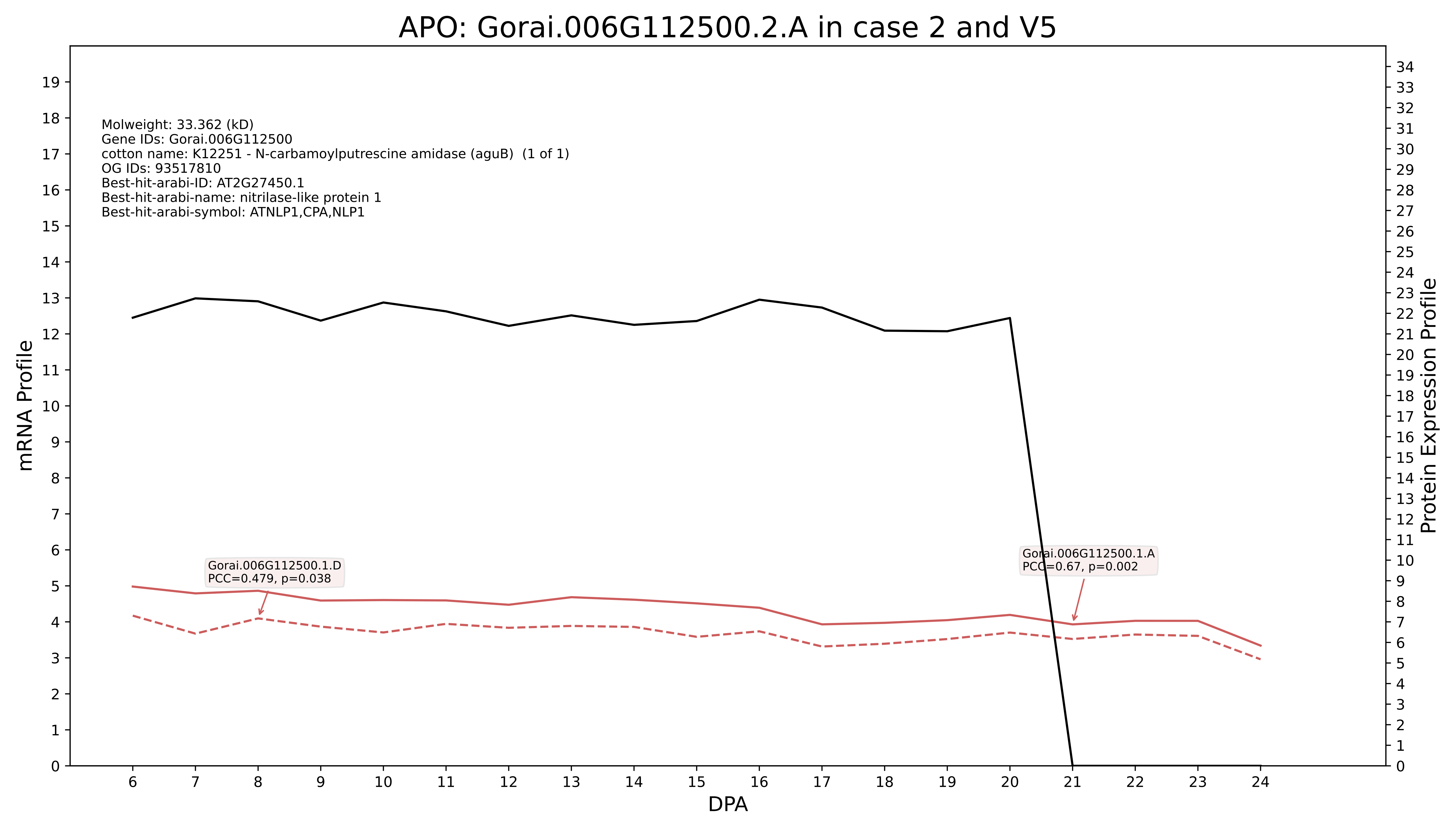
APO: Gorai.006G112500.2.A in case 2 and V5
Molweight: 33.362 (kD)
Gene IDs: Gorai.006G112500
cotton name: K12251 - N-carbamoylputrescine amidase (aguB) (1 of 1)
OG IDs: 93517810
Best-hit-arabi-ID: AT2G27450.1
Best-hit-arabi-name: nitrilase-like protein 1
Best-hit-arabi-symbol: ATNLP1,CPA,NLP1
Gorai.006G112500.1.D
PCC=0.479, p=0.038
Gorai.006G112500.1.A
PCC=0.67, p=0.002
mRNA Profile
Protein Expression Profile
DPA

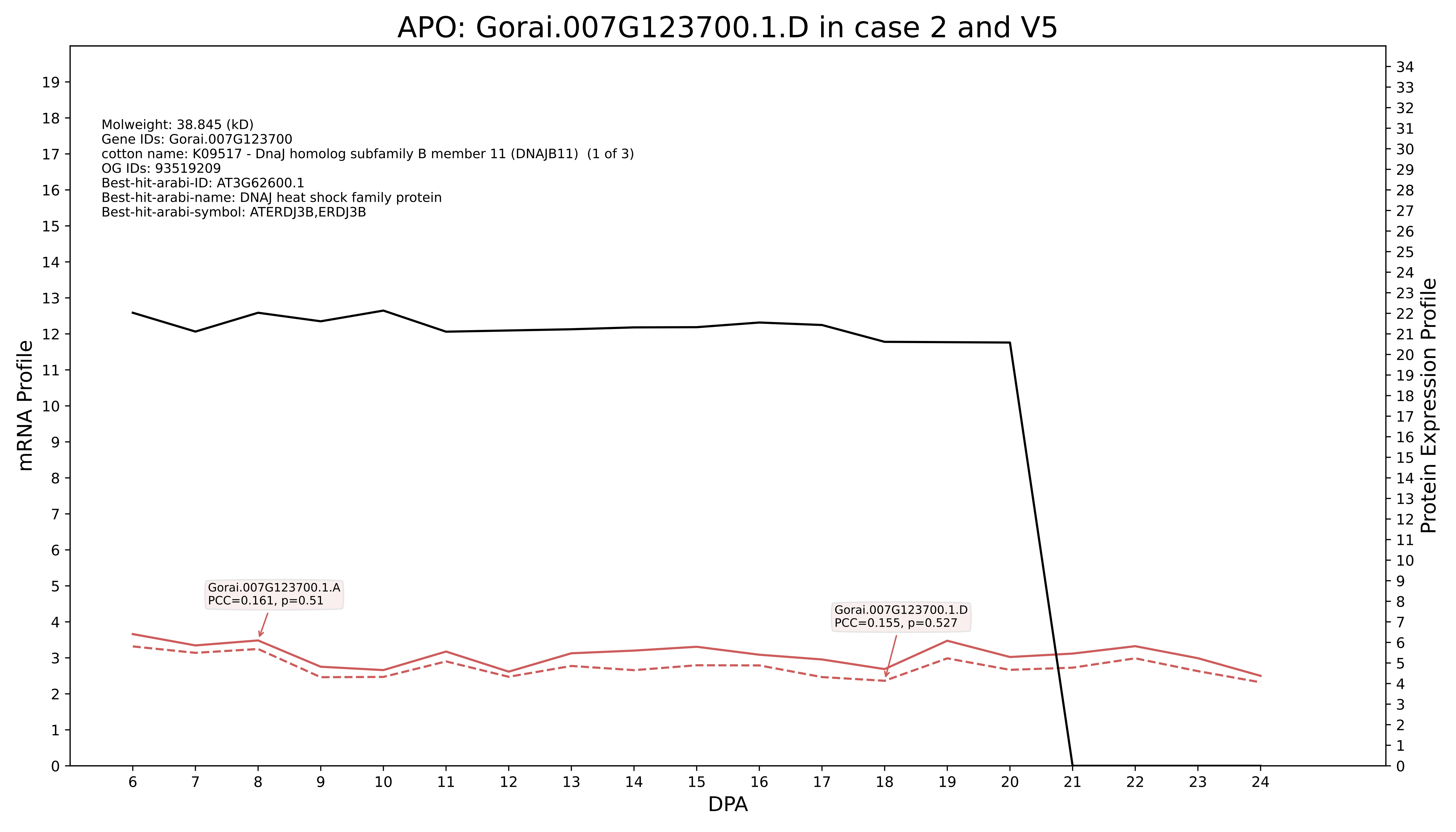

APO: Gorai.007G123700.1.D in case 2 and V5
Molweight: 38.845 (kD)
Gene IDs: Gorai.007G123700
cotton name: K09517 - DnaJ homolog subfamily B member 11 (DNAJB11) (1 of 3)
OG IDs: 93519209
Best-hit-arabi-ID: AT3G62600.1
Best-hit-arabi-name: DNAJ heat shock family protein
Best-hit-arabi-symbol: ATERDJ3B,ERDJ3B
Gorai.007G123700.1.A
PCC=0.161, p=0.51
Gorai.007G123700.1.D
PCC=0.155, p=0.527
mRNA Profile
Protein Expression Profile
DPA

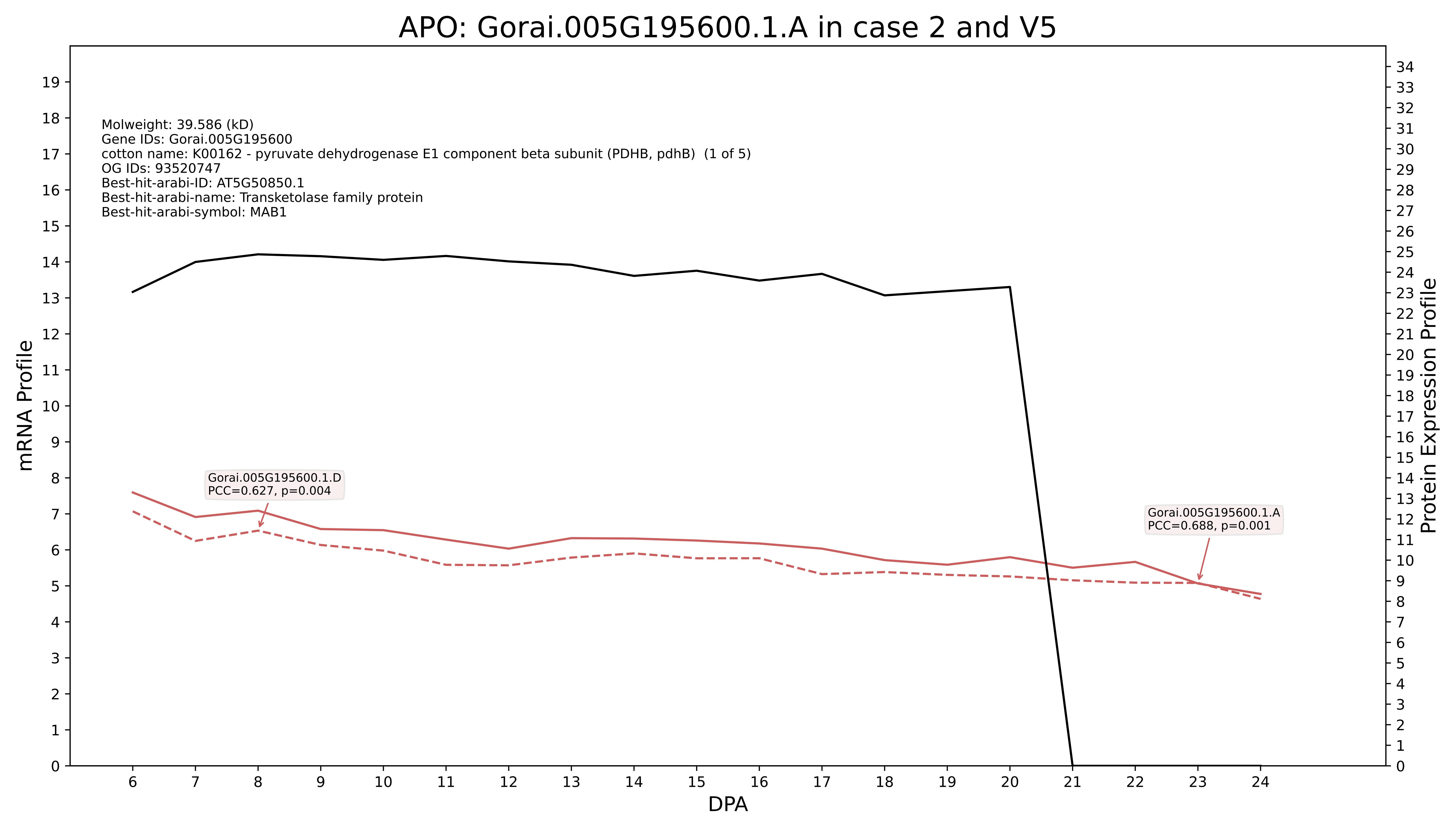

APO: Gorai.005G195600.1.A in case 2 and V5
Molweight: 39.586 (kD)
Gene IDs: Gorai.005G195600
cotton name: K00162 - pyruvate dehydrogenase E1 component beta subunit (PDHB, pdhB) (1 of 5)
OG IDs: 93520747
Best-hit-arabi-ID: AT5G50850.1
Best-hit-arabi-name: Transketolase family protein
Best-hit-arabi-symbol: MAB1
Gorai.005G195600.1.D
PCC=0.627, p=0.004
Gorai.005G195600.1.A
PCC=0.688, p=0.001
mRNA Profile
Protein Expression Profile
DPA

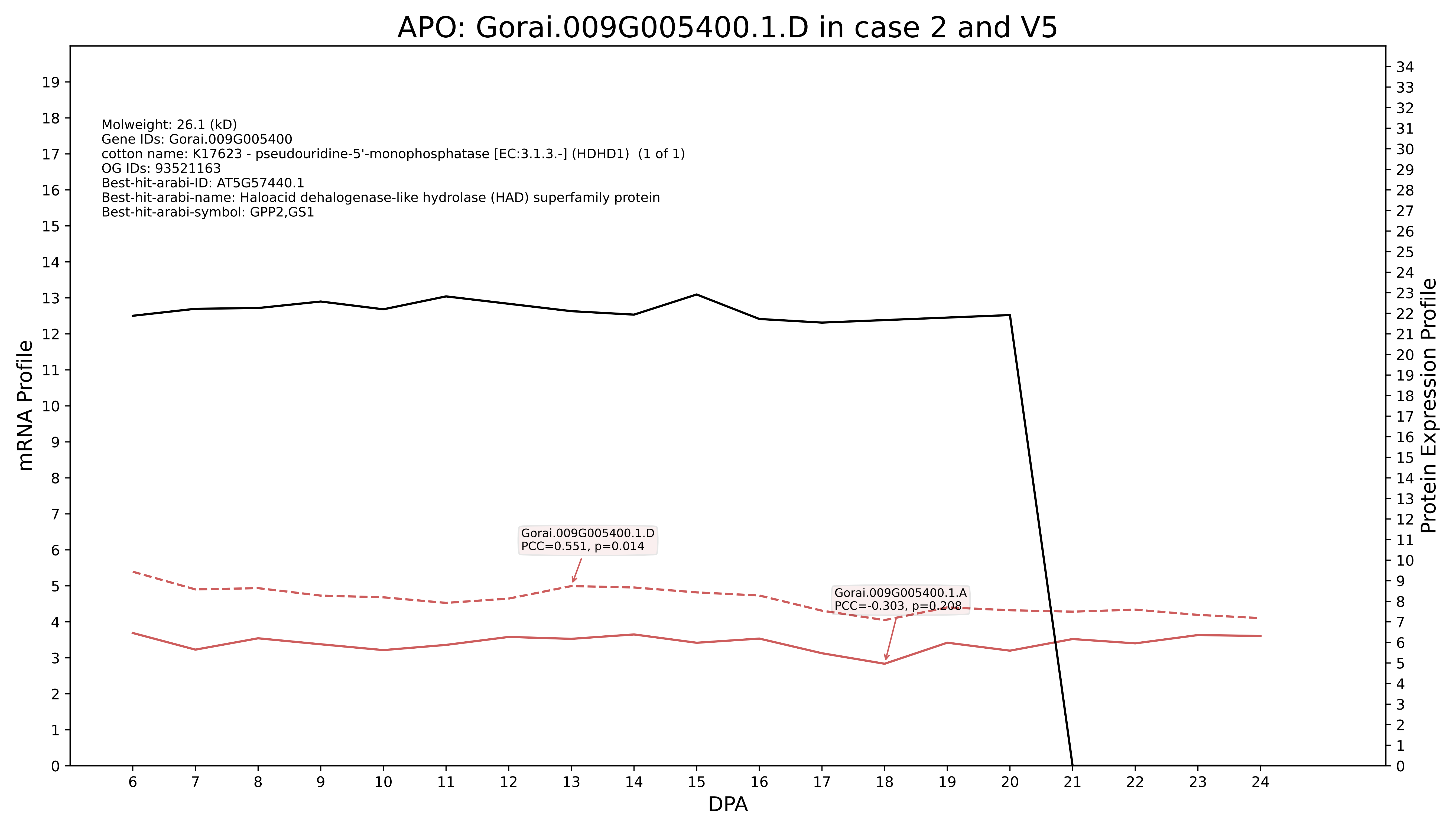
APO: Gorai.009G005400.1.D in case 2 and V5
Molweight: 26.1 (kD)
Gene IDs: Gorai.009G005400
cotton name: K17623 - pseudouridine-5'-monophosphatase [EC:3.1.3.-] (HDHD1) (1 of 1)
OG IDs: 93521163
Best-hit-arabi-ID: AT5G57440.1
Best-hit-arabi-name: Haloacid dehalogenase-like hydrolase (HAD) superfamily protein
Best-hit-arabi-symbol: GPP2,GS1
Gorai.009G005400.1.D
PCC=0.551, p=0.014
Gorai.009G005400.1.A
PCC=-0.303, p=0.208
mRNA Profile
Protein Expression Profile
DPA

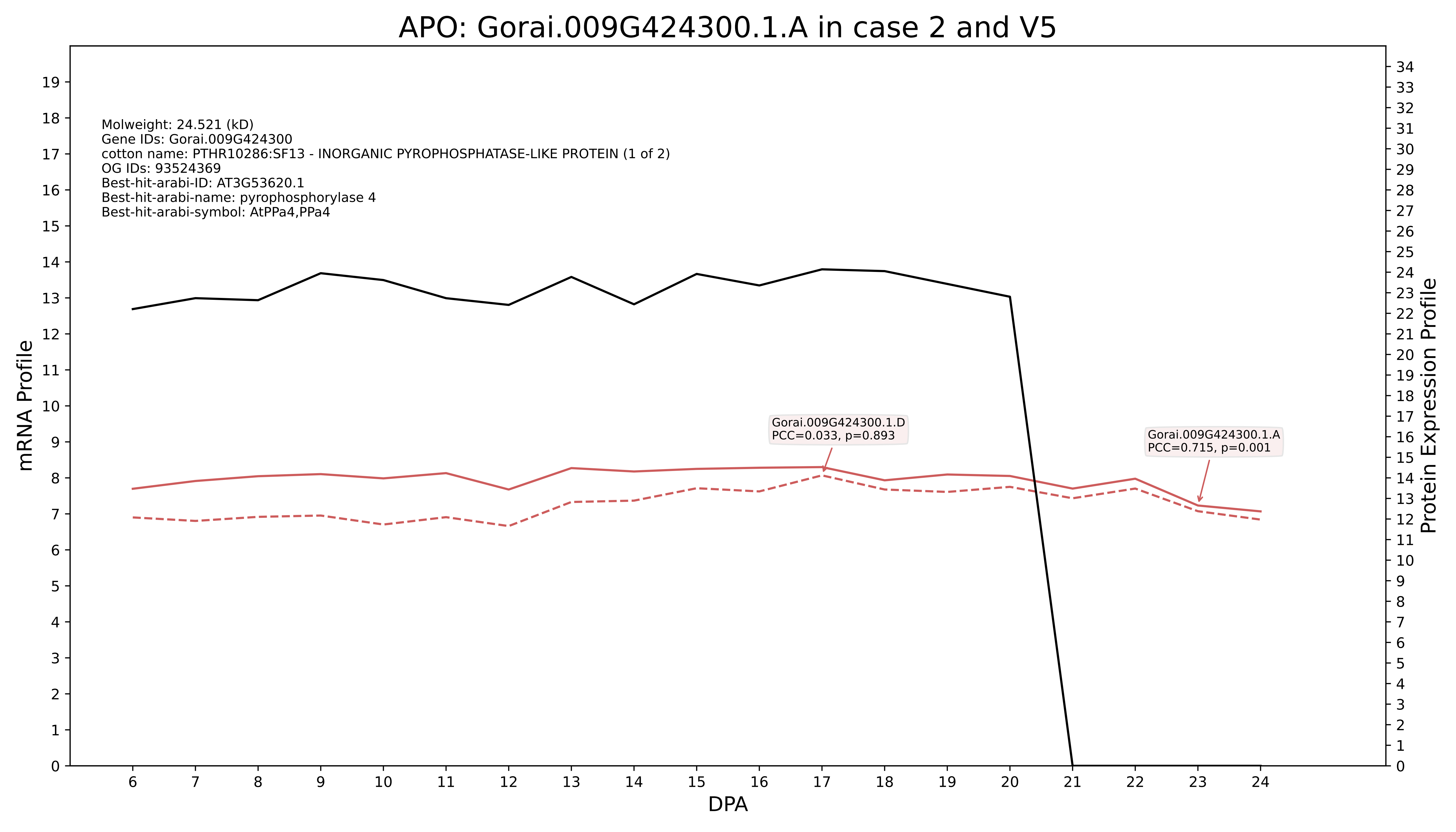

APO: Gorai.009G424300.1.A in case 2 and V5
Molweight: 24.521 (kD)
Gene IDs: Gorai.009G424300
cotton name: PTHR10286:SF13 - INORGANIC PYROPHOSPHATASE-LIKE PROTEIN (1 of 2)
OG IDs: 93524369
Best-hit-arabi-ID: AT3G53620.1
Best-hit-arabi-name: pyrophosphorylase 4
Best-hit-arabi-symbol: AtPPa4,PPa4
Gorai.009G424300.1.D
PCC=0.033, p=0.893
Gorai.009G424300.1.A
PCC=0.715, p=0.001
mRNA Profile
Protein Expression Profile
DPA

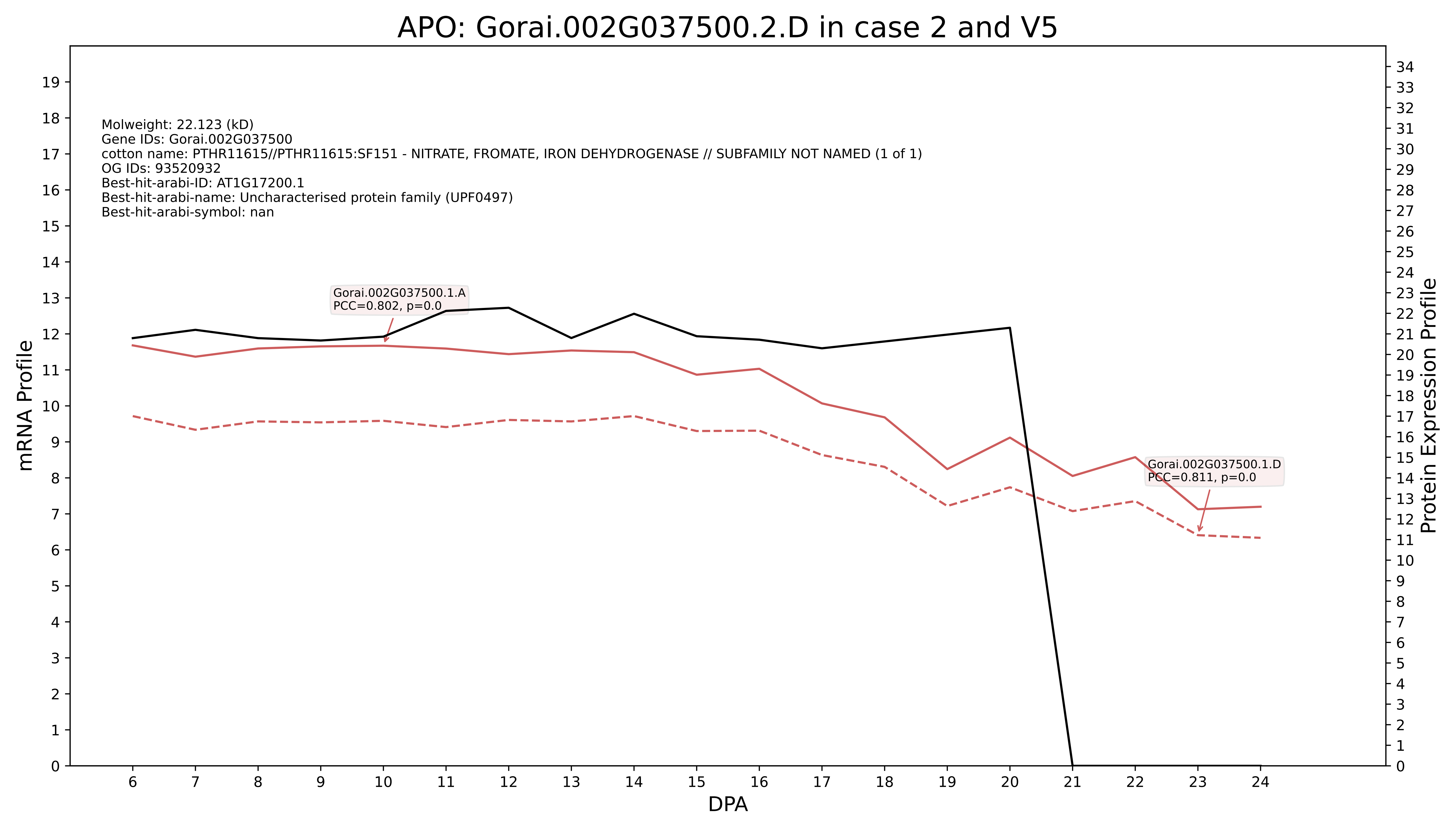

APO: Gorai.002G037500.2.D in case 2 and V5
Molweight: 22.123 (kD)
Gene IDs: Gorai.002G037500
cotton name: PTHR11615//PTHR11615:SF151 - NITRATE, FROMATE, IRON DEHYDROGENASE // SUBFAMILY NOT NAMED (1 of 1)
OG IDs: 93520932
Best-hit-arabi-ID: AT1G17200.1
Best-hit-arabi-name: Uncharacterised protein family (UPF0497)
Best-hit-arabi-symbol: nan
Gorai.002G037500.1.A
PCC=0.802, p=0.0
Gorai.002G037500.1.D
PCC=0.811, p=0.0
mRNA Profile
Protein Expression Profile
DPA

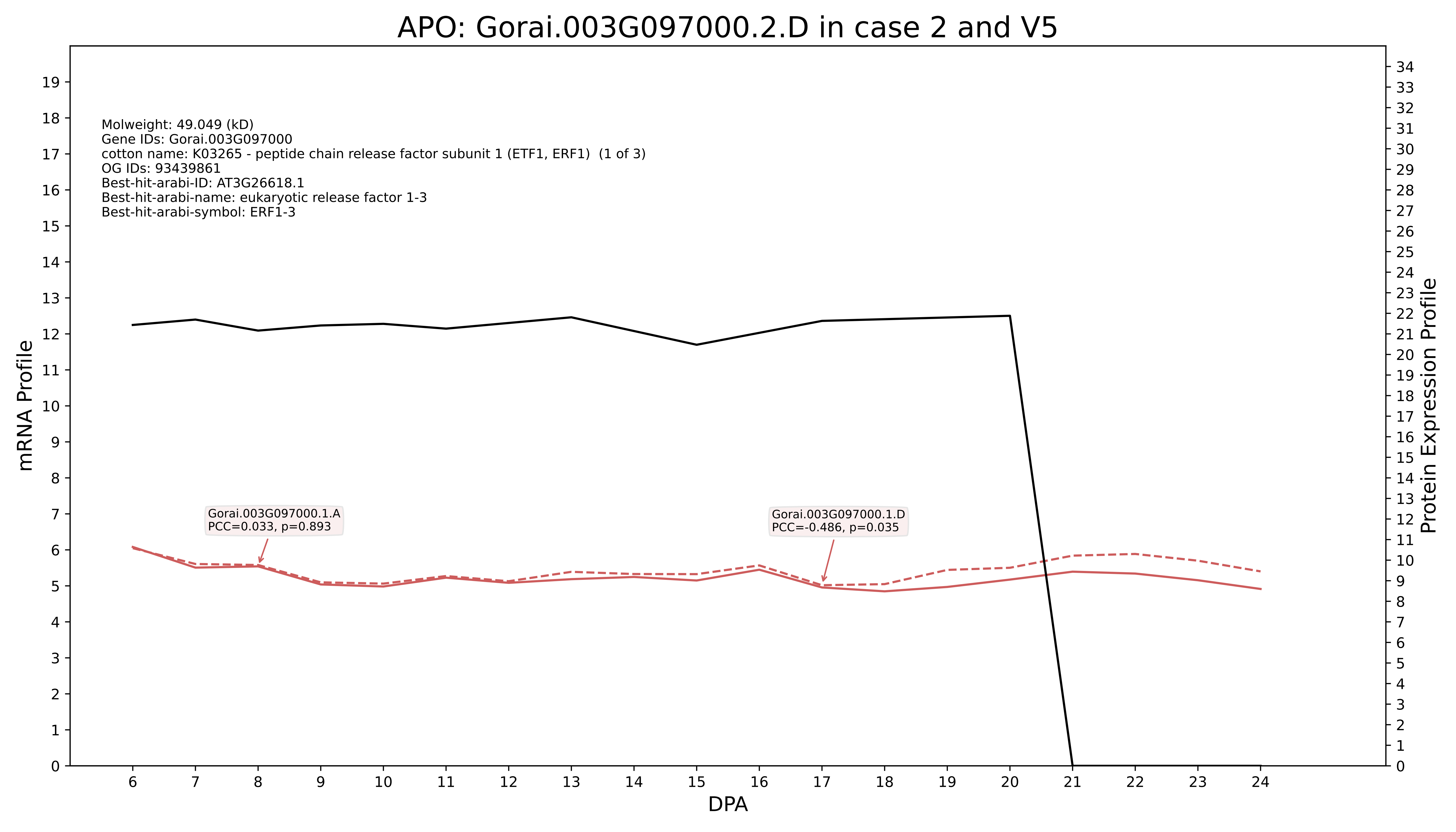
APO: Gorai.003G097000.2.D in case 2 and V5
Molweight: 49.049 (kD)
Gene IDs: Gorai.003G097000
cotton name: K03265 - peptide chain release factor subunit 1 (ETF1, ERF1) (1 of 3)
OG IDs: 93439861
Best-hit-arabi-ID: AT3G26618.1
Best-hit-arabi-name: eukaryotic release factor 1-3
Best-hit-arabi-symbol: ERF1-3
Gorai.003G097000.1.A
PCC=0.033, p=0.893
Gorai.003G097000.1.D
PCC=-0.486, p=0.035
mRNA Profile
Protein Expression Profile
DPA

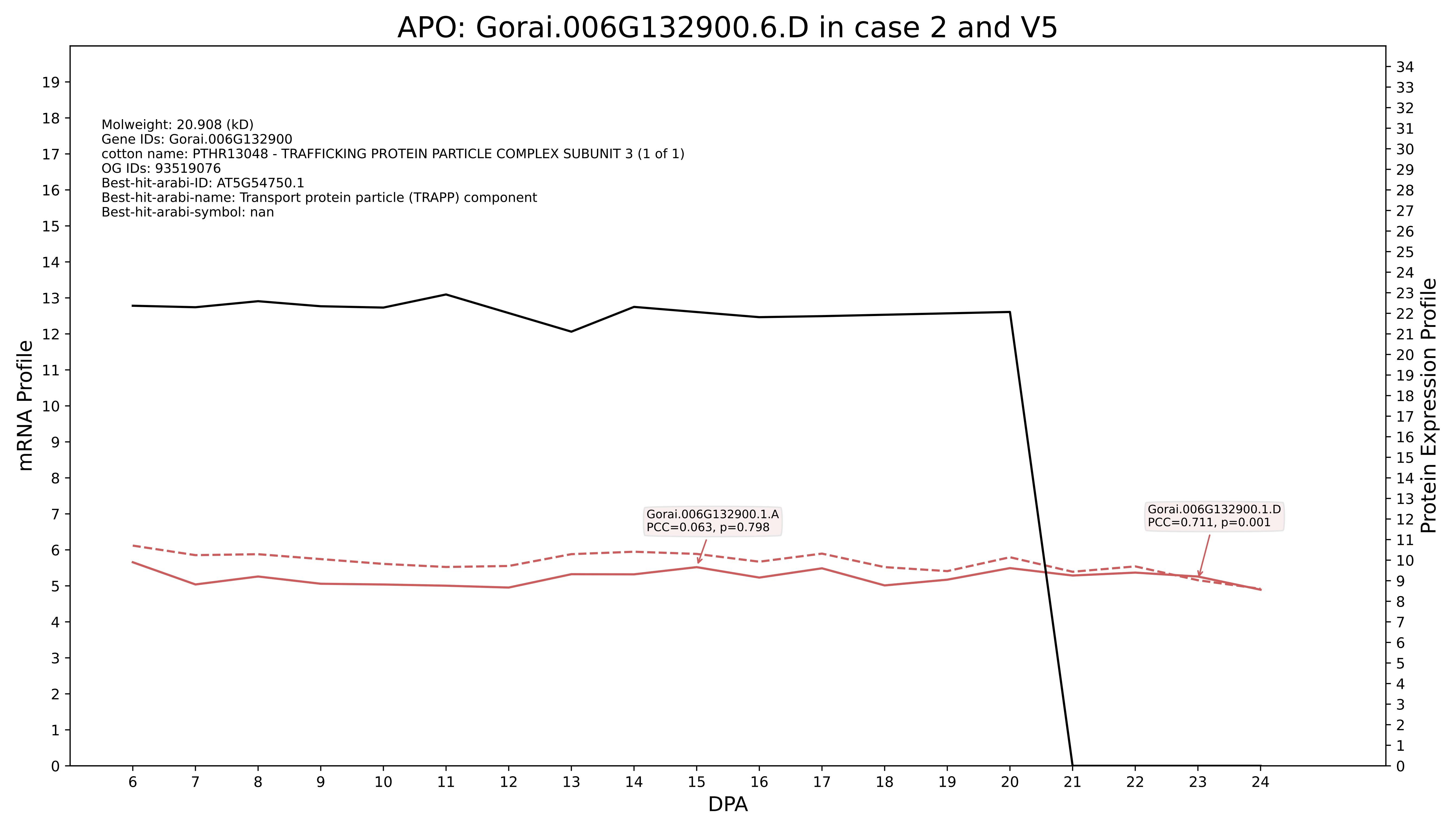
APO: Gorai.006G132900.6.D in case 2 and V5
Molweight: 20.908 (kD)
Gene IDs: Gorai.006G132900
cotton name: PTHR13048 - TRAFFICKING PROTEIN PARTICLE COMPLEX SUBUNIT 3 (1 of 1)
OG IDs: 93519076
Best-hit-arabi-ID: AT5G54750.1
Best-hit-arabi-name: Transport protein particle (TRAPP) component
Best-hit-arabi-symbol: nan
Gorai.006G132900.1.A
PCC=0.063, p=0.798
Gorai.006G132900.1.D
PCC=0.711, p=0.001
mRNA Profile
Protein Expression Profile
DPA

### APO: Gorai.011G133700.1.D in case 2 and V5

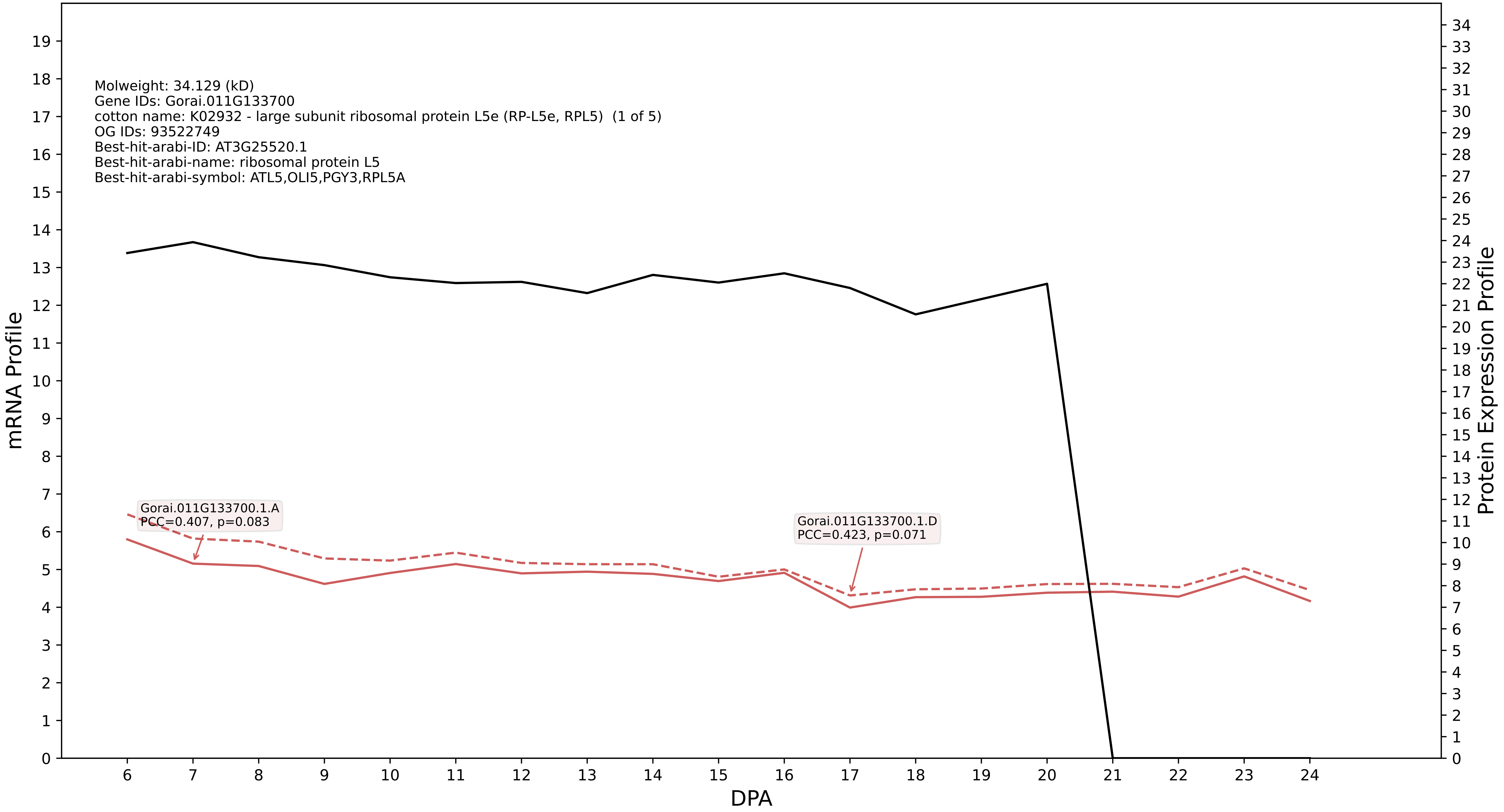

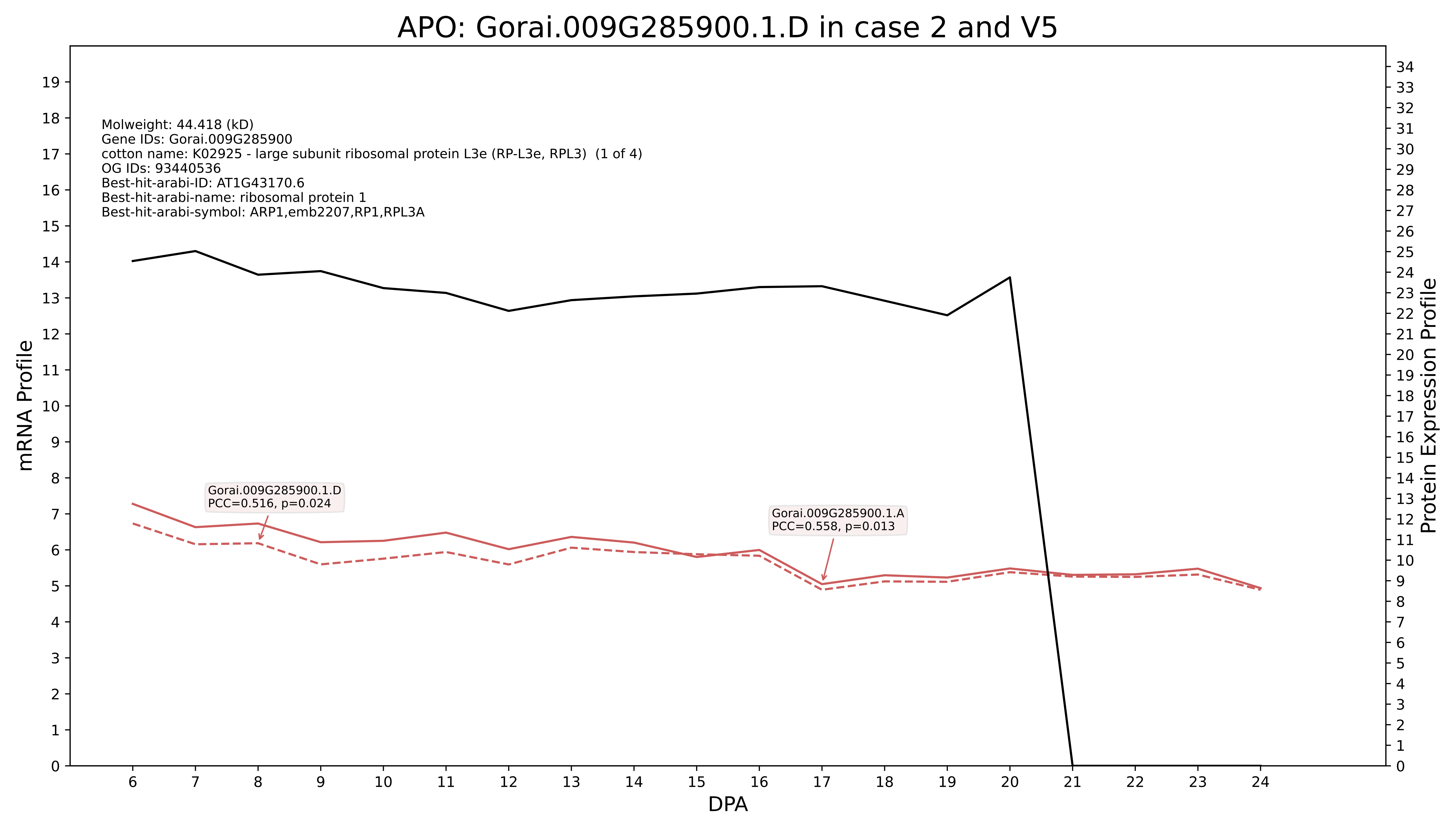
APO: Gorai.009G285900.1.D in case 2 and V5
Molweight: 44.418 (kD)
Gene IDs: Gorai.009G285900
cotton name: K02925 - large subunit ribosomal protein L3e (RP-L3e, RPL3) (1 of 4)
OG IDs: 93440536
Best-hit-arabi-ID: AT1G43170.6
Best-hit-arabi-name: ribosomal protein 1
Best-hit-arabi-symbol: ARP1,emb2207,RP1,RPL3A
Gorai.009G285900.1.D
PCC=0.516, p=0.024
Gorai.009G285900.1.A
PCC=0.558, p=0.013
mRNA Profile
Protein Expression Profile
DPA

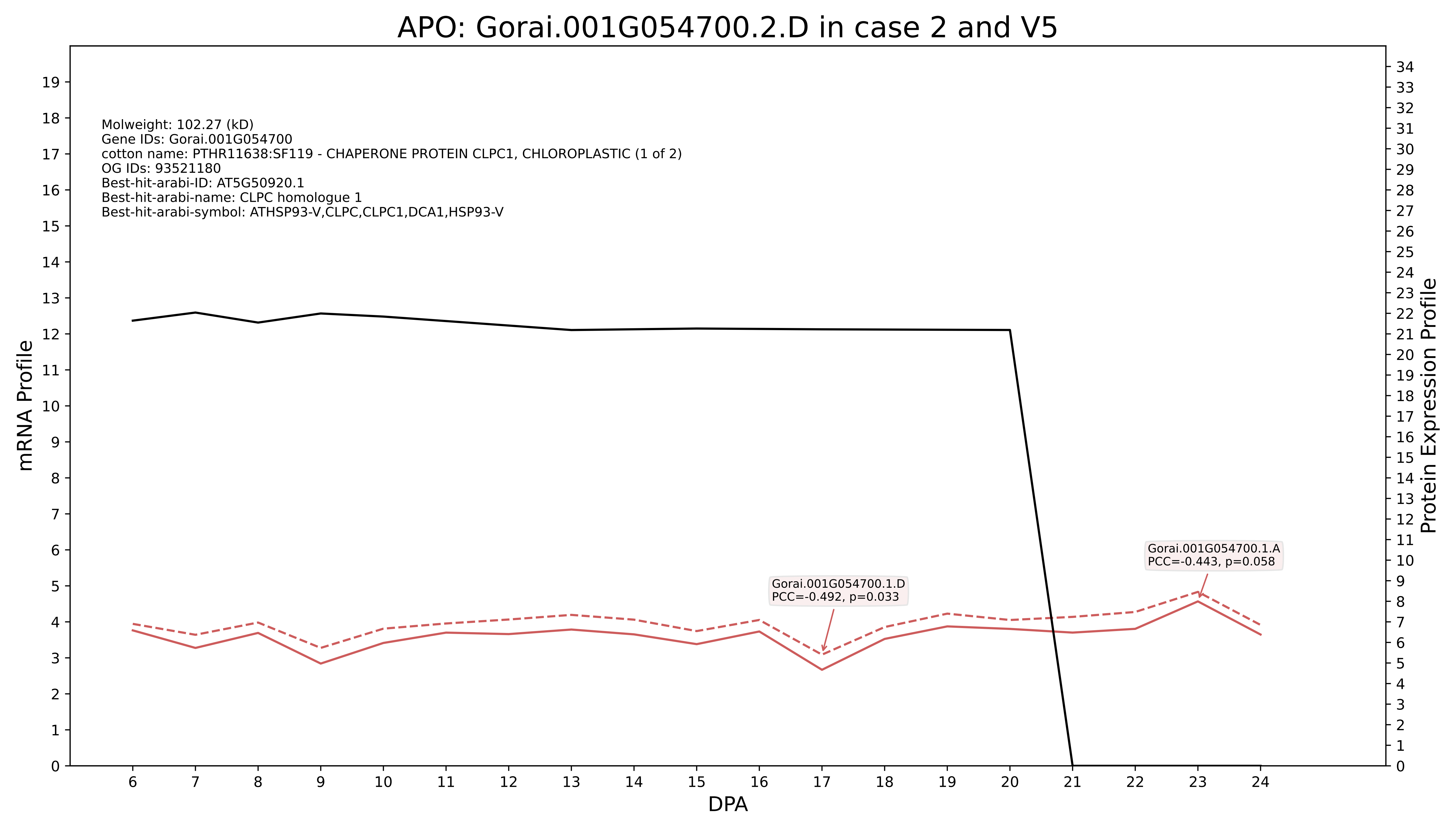

APO: Gorai.001G054700.2.D in case 2 and V5
Molweight: 102.27 (kD)
Gene IDs: Gorai.001G054700
cotton name: PTHR11638:SF119 - CHAPERONE PROTEIN CLPC1, CHLOROPLASTIC (1 of 2)
OG IDs: 93521180
Best-hit-arabi-ID: AT5G50920.1
Best-hit-arabi-name: CLPC homologue 1
Best-hit-arabi-symbol: ATHSP93-V,CLPC,CLPC1,DCA1,HSP93-V
Gorai.001G054700.1.D
PCC=-0.492, p=0.033
Gorai.001G054700.1.A
PCC=-0.443, p=0.058
mRNA Profile
Protein Expression Profile
DPA

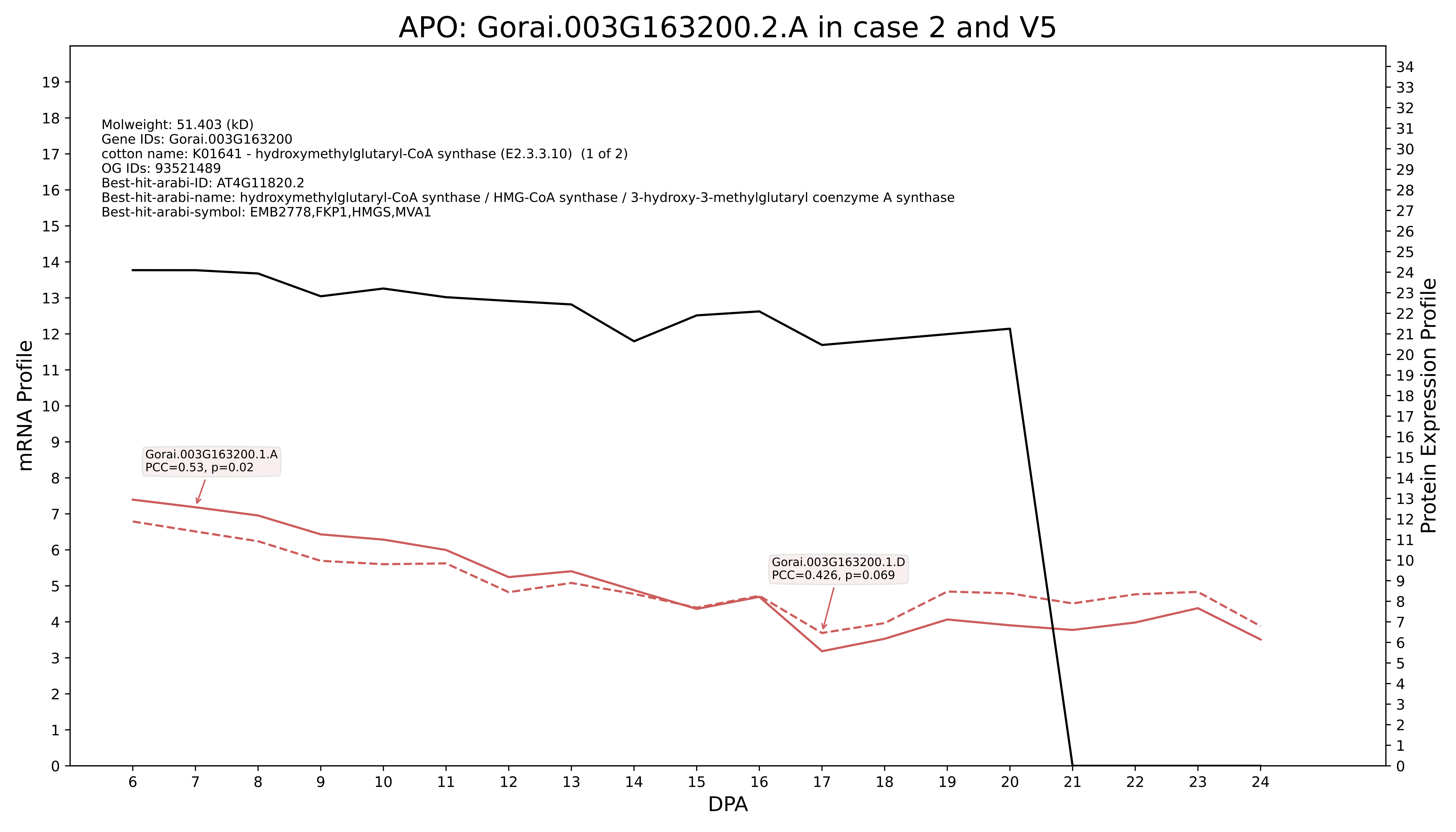
APO: Gorai.003G163200.2.A in case 2 and V5
Molweight: 51.403 (kD)
Gene IDs: Gorai.003G163200
cotton name: K01641 - hydroxymethylglutaryl-CoA synthase (E2.3.3.10) (1 of 2)
OG IDs: 93521489
Best-hit-arabi-ID: AT4G11820.2
Best-hit-arabi-name: hydroxymethylglutaryl-CoA synthase / HMG-CoA synthase / 3-hydroxy-3-methylglutaryl coenzyme A synthase
Best-hit-arabi-symbol: EMB2778,FKP1,HMGS,MVA1
Gorai.003G163200.1.A
PCC=0.53, p=0.02
Gorai.003G163200.1.D
PCC=0.426, p=0.069
mRNA Profile
Protein Expression Profile
DPA

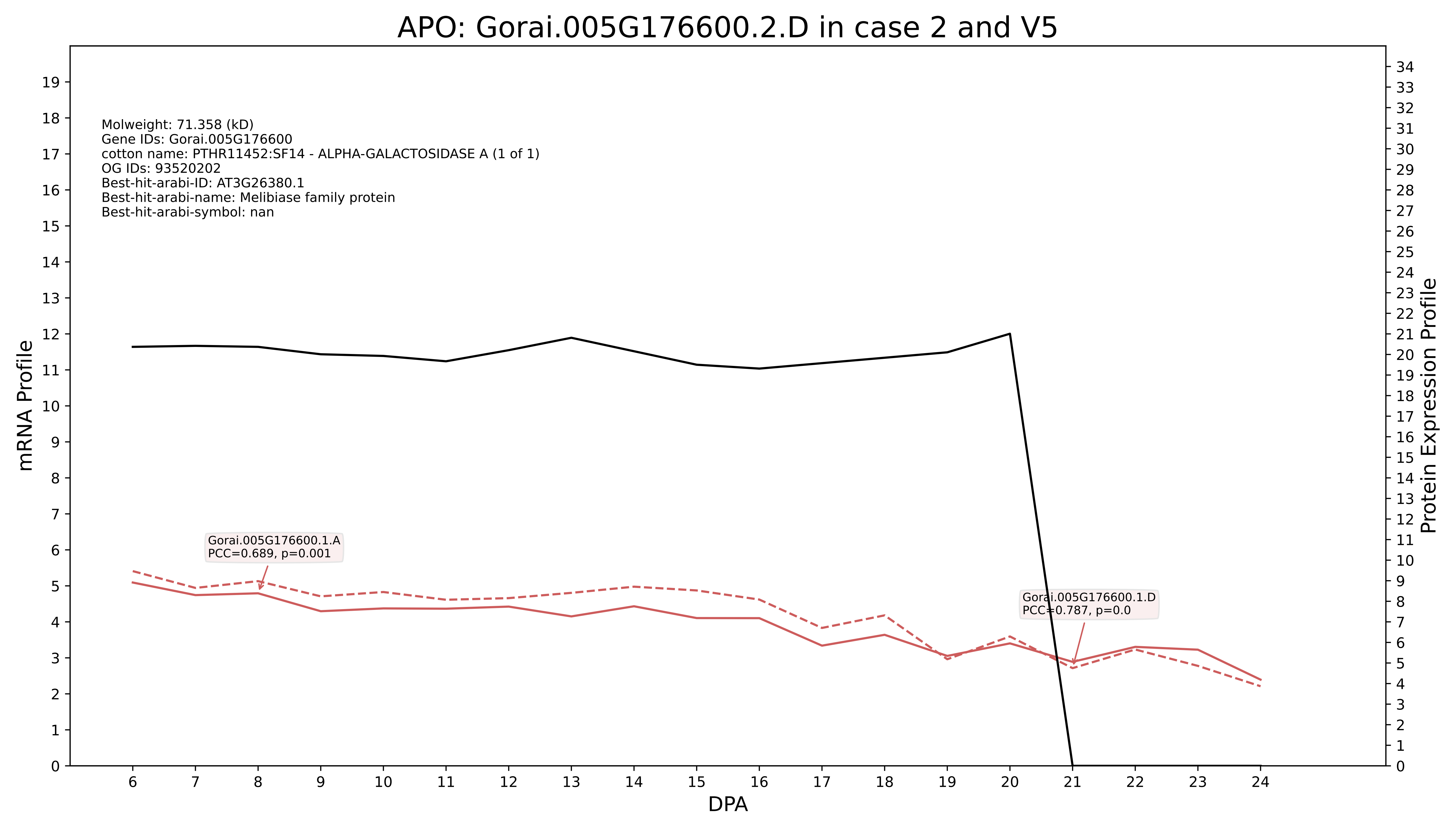
APO: Gorai.005G176600.2.D in case 2 and V5
Molweight: 71.358 (kD)
Gene IDs: Gorai.005G176600
cotton name: PTHR11452:SF14 - ALPHA-GALACTOSIDASE A (1 of 1)
OG IDs: 93520202
Best-hit-arabi-ID: AT3G26380.1
Best-hit-arabi-name: Melibiase family protein
Best-hit-arabi-symbol: nan
Gorai.005G176600.1.A
PCC=0.689, p=0.001
Gorai.005G176600.1.D
PCC=0.787, p=0.0
mRNA Profile
Protein Expression Profile
DPA

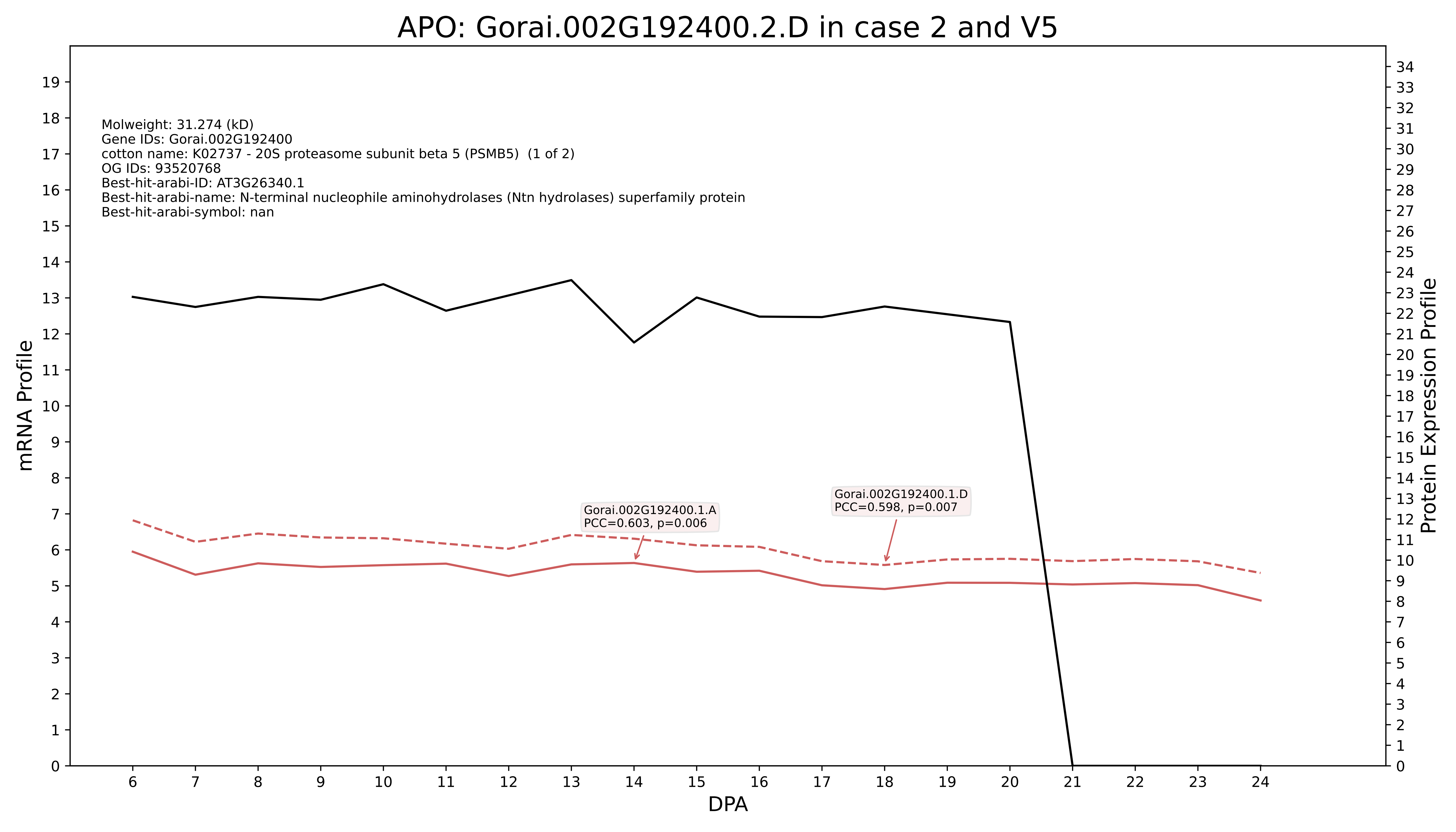
APO: Gorai.002G192400.2.D in case 2 and V5
Molweight: 31.274 (kD)
Gene IDs: Gorai.002G192400
cotton name: K02737 - 20S proteasome subunit beta 5 (PSMB5) (1 of 2)
OG IDs: 93520768
Best-hit-arabi-ID: AT3G26340.1
Best-hit-arabi-name: N-terminal nucleophile aminohydrolases (Ntn hydrolases) superfamily protein
Best-hit-arabi-symbol: nan
Gorai.002G192400.1.A
PCC=0.603, p=0.006
Gorai.002G192400.1.D
PCC=0.598, p=0.007
mRNA Profile
Protein Expression Profile
DPA

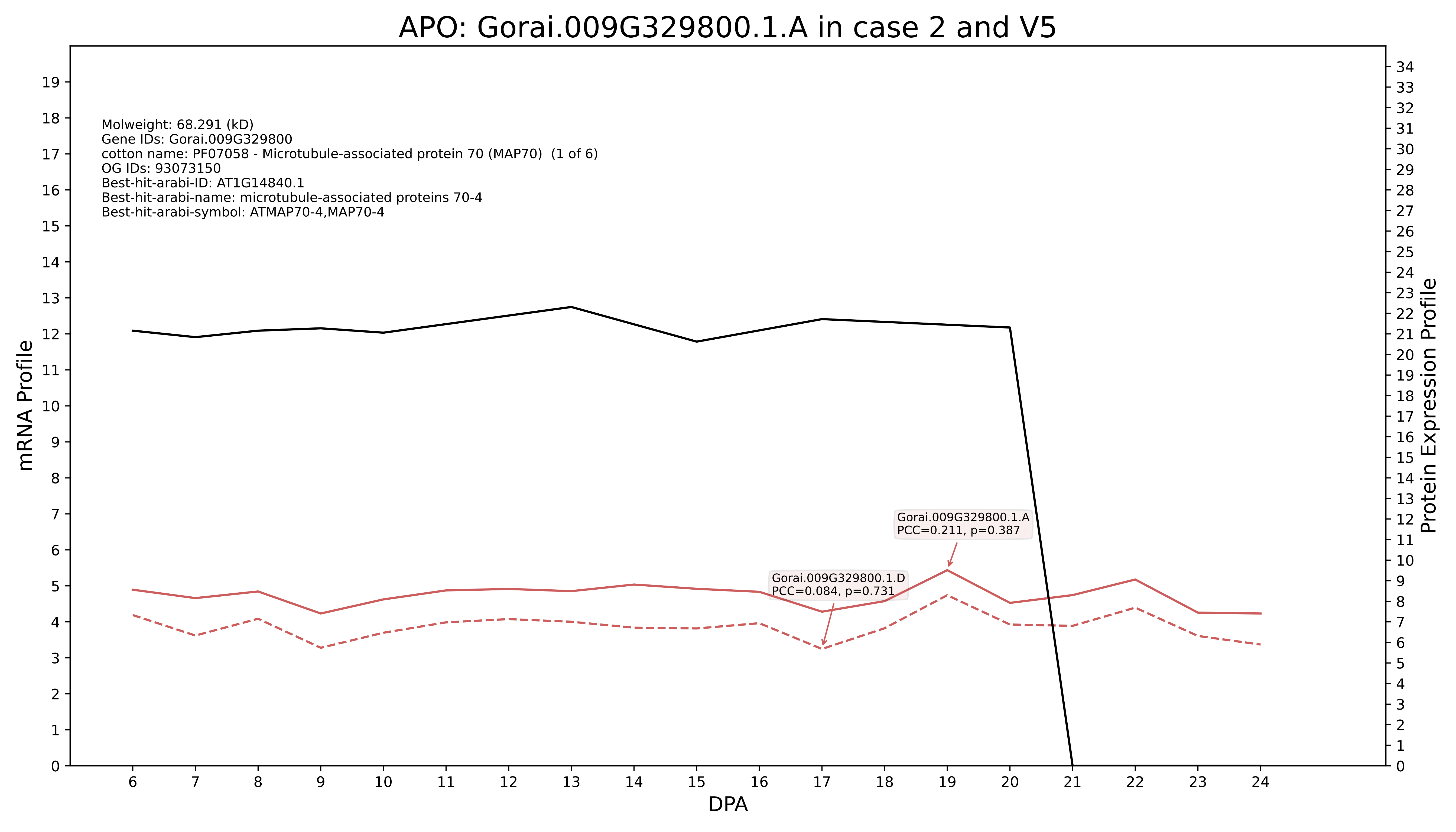

APO: Gorai.009G329800.1.A in case 2 and V5
Molweight: 68.291 (kD)
Gene IDs: Gorai.009G329800
cotton name: PF07058 - Microtubule-associated protein 70 (MAP70) (1 of 6)
OG IDs: 93073150
Best-hit-arabi-ID: AT1G14840.1
Best-hit-arabi-name: microtubule-associated proteins 70-4
Best-hit-arabi-symbol: ATMAP70-4,MAP70-4
Gorai.009G329800.1.A
PCC=0.211, p=0.387
Gorai.009G329800.1.D
PCC=0.084, p=0.731
mRNA Profile
Protein Expression Profile
DPA

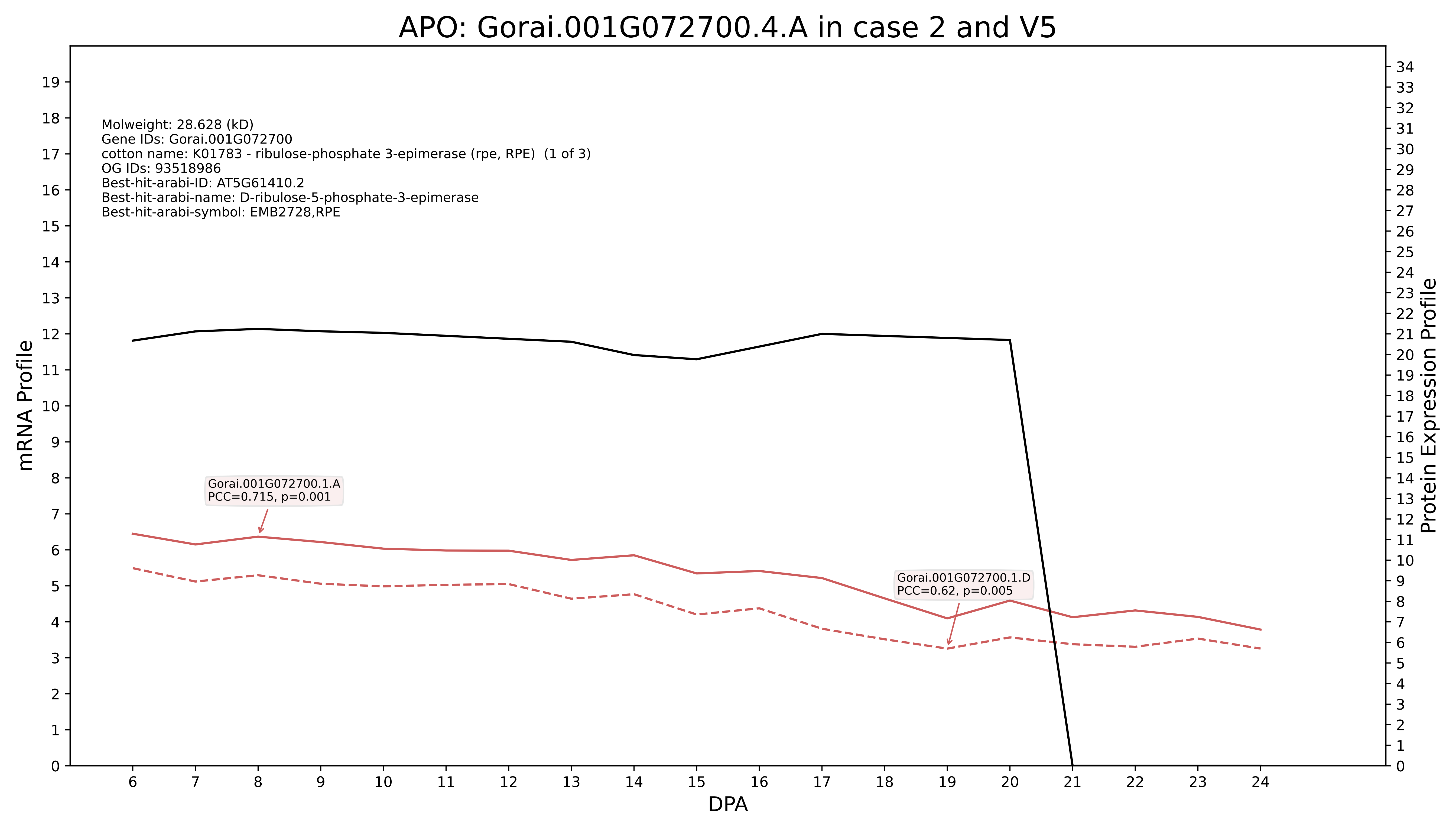
APO: Gorai.001G072700.4.A in case 2 and V5
Molweight: 28.628 (kD)
Gene IDs: Gorai.001G072700
cotton name: K01783 - ribulose-phosphate 3-epimerase (rpe, RPE) (1 of 3)
OG IDs: 93518986
Best-hit-arabi-ID: AT5G61410.2
Best-hit-arabi-name: D-ribulose-5-phosphate-3-epimerase
Best-hit-arabi-symbol: EMB2728,RPE
Gorai.001G072700.1.A
PCC=0.715, p=0.001
Gorai.001G072700.1.D
PCC=0.62, p=0.005
mRNA Profile
Protein Expression Profile
DPA

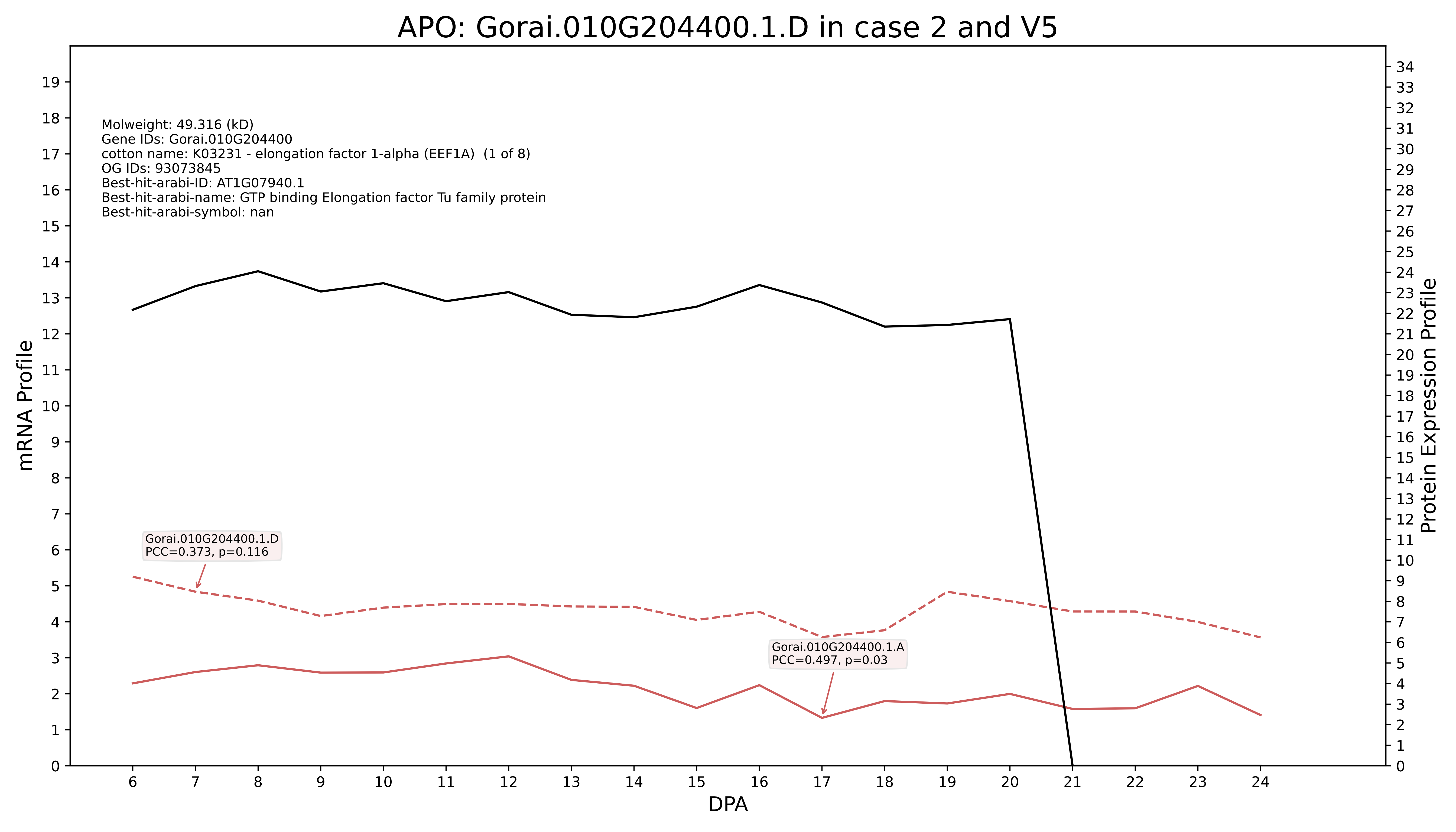
APO: Gorai.010G204400.1.D in case 2 and V5
Molweight: 49.316 (kD)
Gene IDs: Gorai.010G204400
cotton name: K03231 - elongation factor 1-alpha (EEF1A) (1 of 8)
OG IDs: 93073845
Best-hit-arabi-ID: AT1G07940.1
Best-hit-arabi-name: GTP binding Elongation factor Tu family protein
Best-hit-arabi-symbol: nan
Gorai.010G204400.1.D
PCC=0.373, p=0.116
Gorai.010G204400.1.A
PCC=0.497, p=0.03
mRNA Profile
Protein Expression Profile
DPA

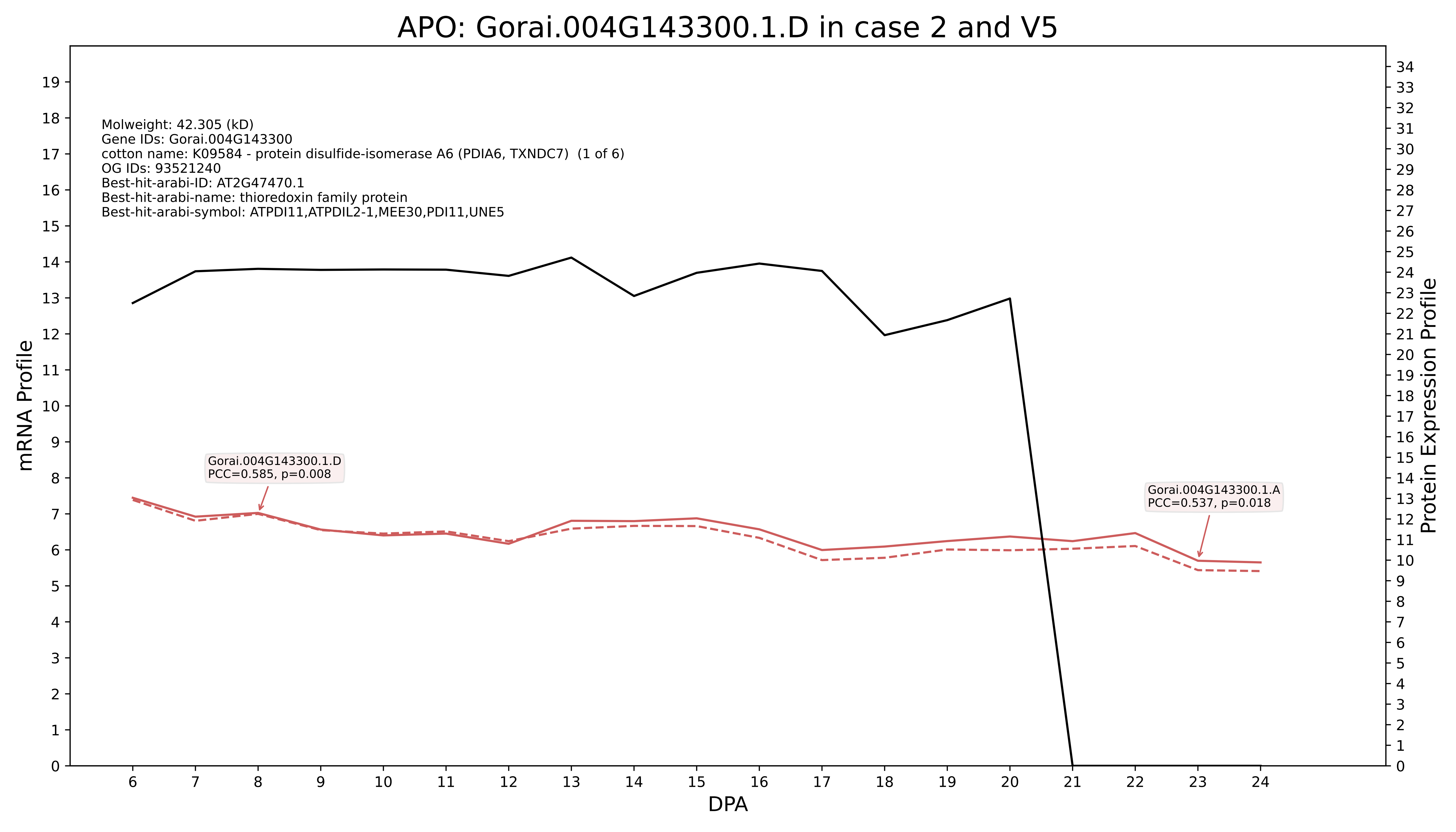

APO: Gorai.004G143300.1.D in case 2 and V5
Molweight: 42.305 (kD)
Gene IDs: Gorai.004G143300
cotton name: K09584 - protein disulfide-isomerase A6 (PDIA6, TXNDC7) (1 of 6)
OG IDs: 93521240
Best-hit-arabi-ID: AT2G47470.1
Best-hit-arabi-name: thioredoxin family protein
Best-hit-arabi-symbol: ATPDI11,ATPDIL2-1,MEE30,PDI11,UNE5
Gorai.004G143300.1.D
PCC=0.585, p=0.008
Gorai.004G143300.1.A
PCC=0.537, p=0.018
mRNA Profile
Protein Expression Profile
DPA

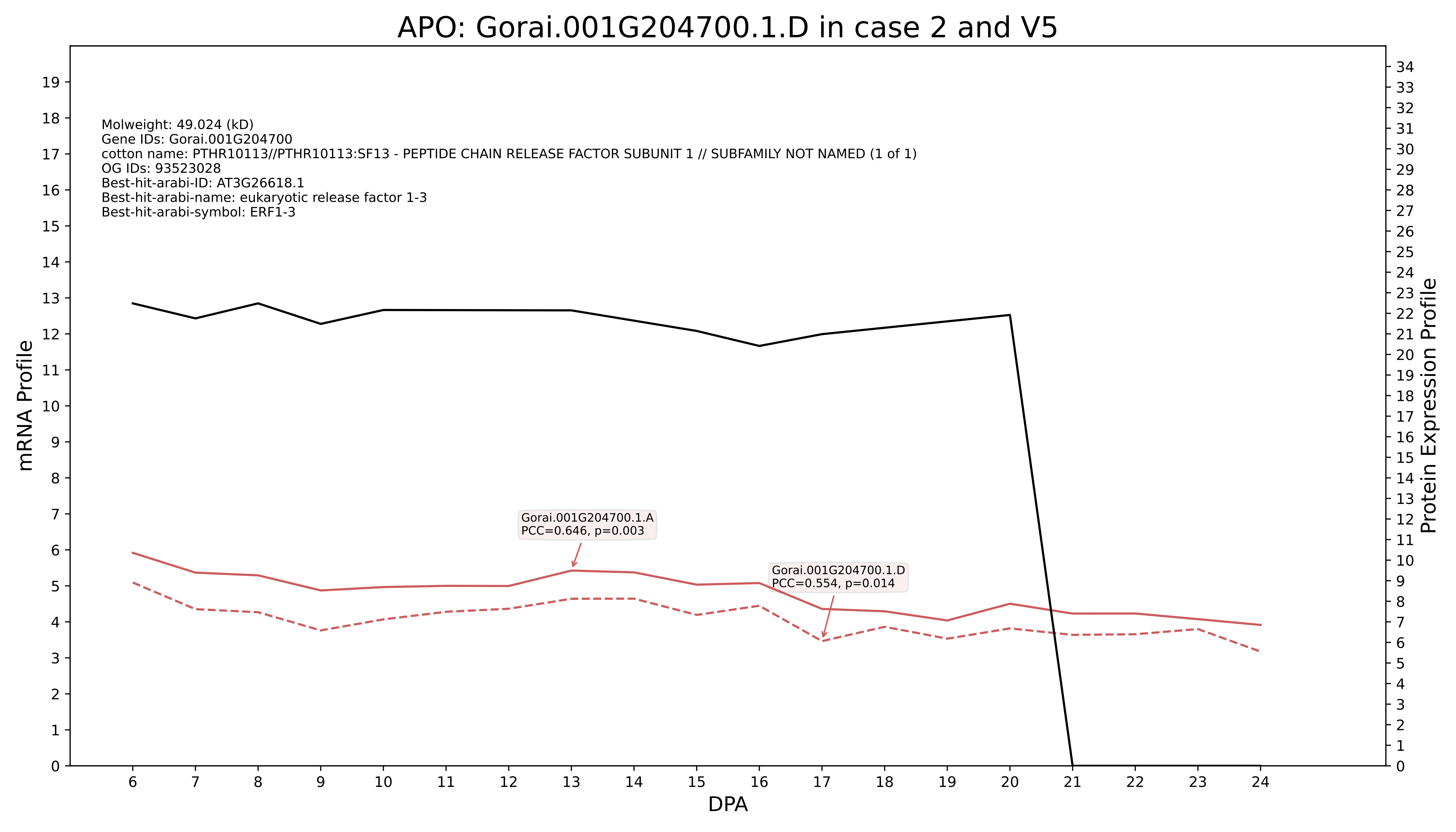
APO: Gorai.001G204700.1.D in case 2 and V5
Molweight: 49.024 (kD)
Gene IDs: Gorai.001G204700
cotton name: PTHR10113//PTHR10113:SF13 - PEPTIDE CHAIN RELEASE FACTOR SUBUNIT 1 // SUBFAMILY NOT NAMED (1 of 1)
OG IDs: 93523028
Best-hit-arabi-ID: AT3G26618.1
Best-hit-arabi-name: eukaryotic release factor 1-3
Best-hit-arabi-symbol: ERF1-3
Gorai.001G204700.1.A
PCC=0.646, p=0.003
Gorai.001G204700.1.D
PCC=0.554, p=0.014
mRNA Profile
Protein Expression Profile
DPA

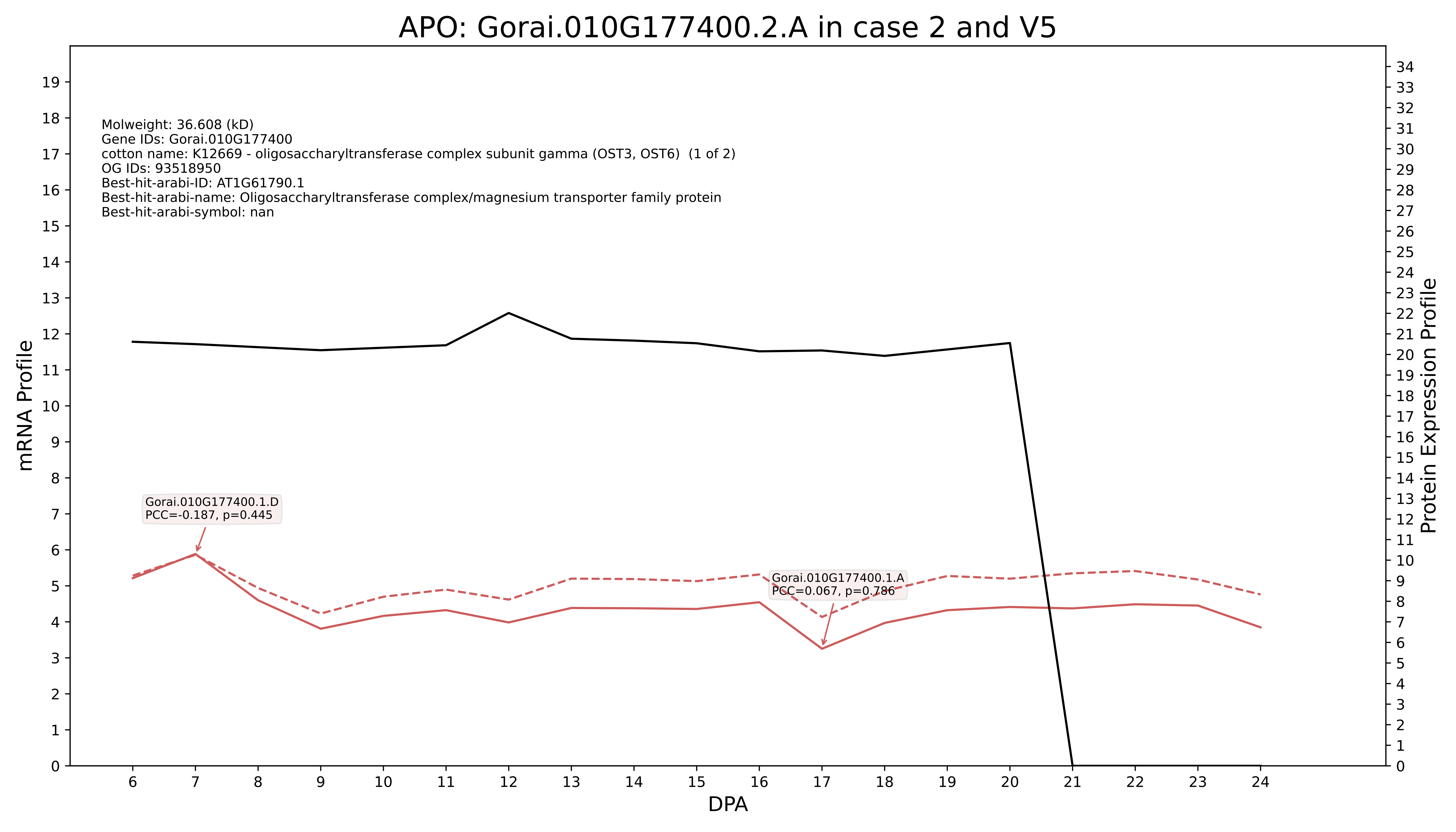
APO: Gorai.010G177400.2.A in case 2 and V5
Molweight: 36.608 (kD)
Gene IDs: Gorai.010G177400
cotton name: K12669 - oligosaccharyltransferase complex subunit gamma (OST3, OST6) (1 of 2)
OG IDs: 93518950
Best-hit-arabi-ID: AT1G61790.1
Best-hit-arabi-name: Oligosaccharyltransferase complex/magnesium transporter family protein
Best-hit-arabi-symbol: nan
Gorai.010G177400.1.D
PCC=-0.187, p=0.445
Gorai.010G177400.1.A
PCC=0.067, p=0.786
mRNA Profile
Protein Expression Profile
DPA

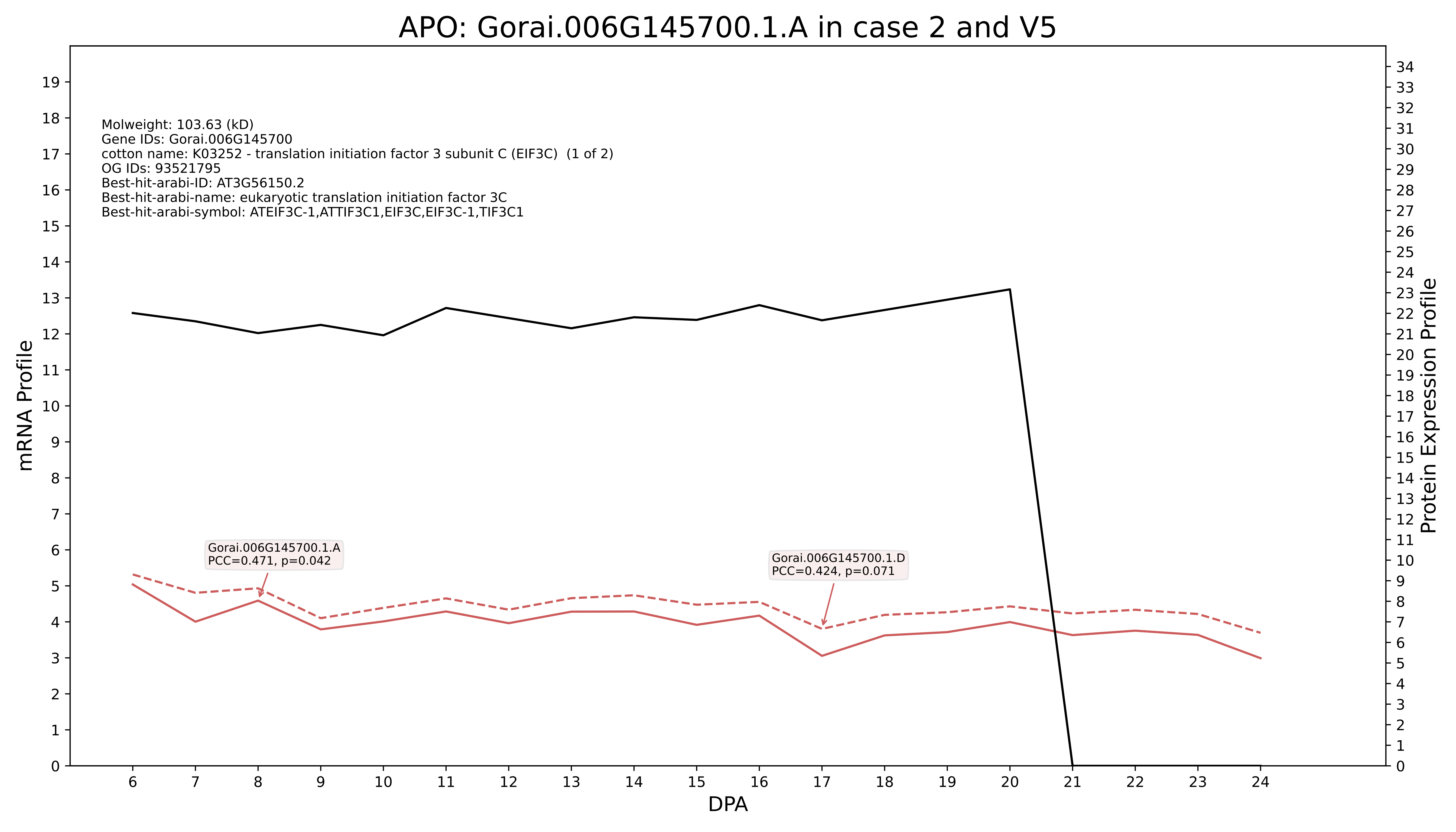
APO: Gorai.006G145700.1.A in case 2 and V5
Molweight: 103.63 (kD)
Gene IDs: Gorai.006G145700
cotton name: K03252 - translation initiation factor 3 subunit C (EIF3C) (1 of 2)
OG IDs: 93521795
Best-hit-arabi-ID: AT3G56150.2
Best-hit-arabi-name: eukaryotic translation initiation factor 3C
Best-hit-arabi-symbol: ATEIF3C-1,ATTIF3C1,EIF3C,EIF3C-1,TIF3C1
Gorai.006G145700.1.A
PCC=0.471, p=0.042
Gorai.006G145700.1.D
PCC=0.424, p=0.071
mRNA Profile
Protein Expression Profile
DPA

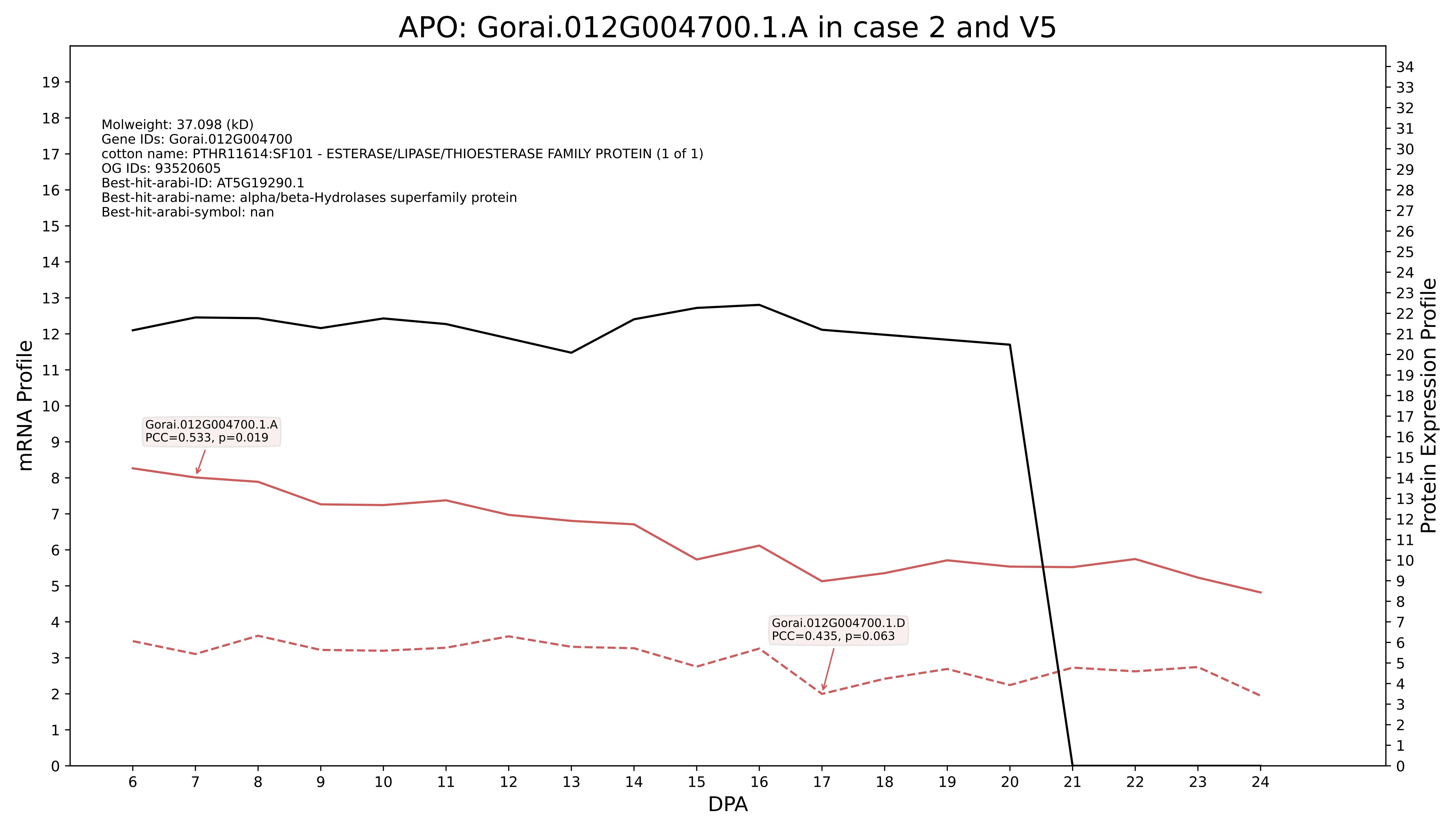
APO: Gorai.012G004700.1.A in case 2 and V5
Molweight: 37.098 (kD)
Gene IDs: Gorai.012G004700
cotton name: PTHR11614:SF101 - ESTERASE/LIPASE/THIOESTERASE FAMILY PROTEIN (1 of 1)
OG IDs: 93520605
Best-hit-arabi-ID: AT5G19290.1
Best-hit-arabi-name: alpha/beta-Hydrolases superfamily protein
Best-hit-arabi-symbol: nan
Gorai.012G004700.1.A
PCC=0.533, p=0.019
Gorai.012G004700.1.D
PCC=0.435, p=0.063
mRNA Profile
Protein Expression Profile
DPA

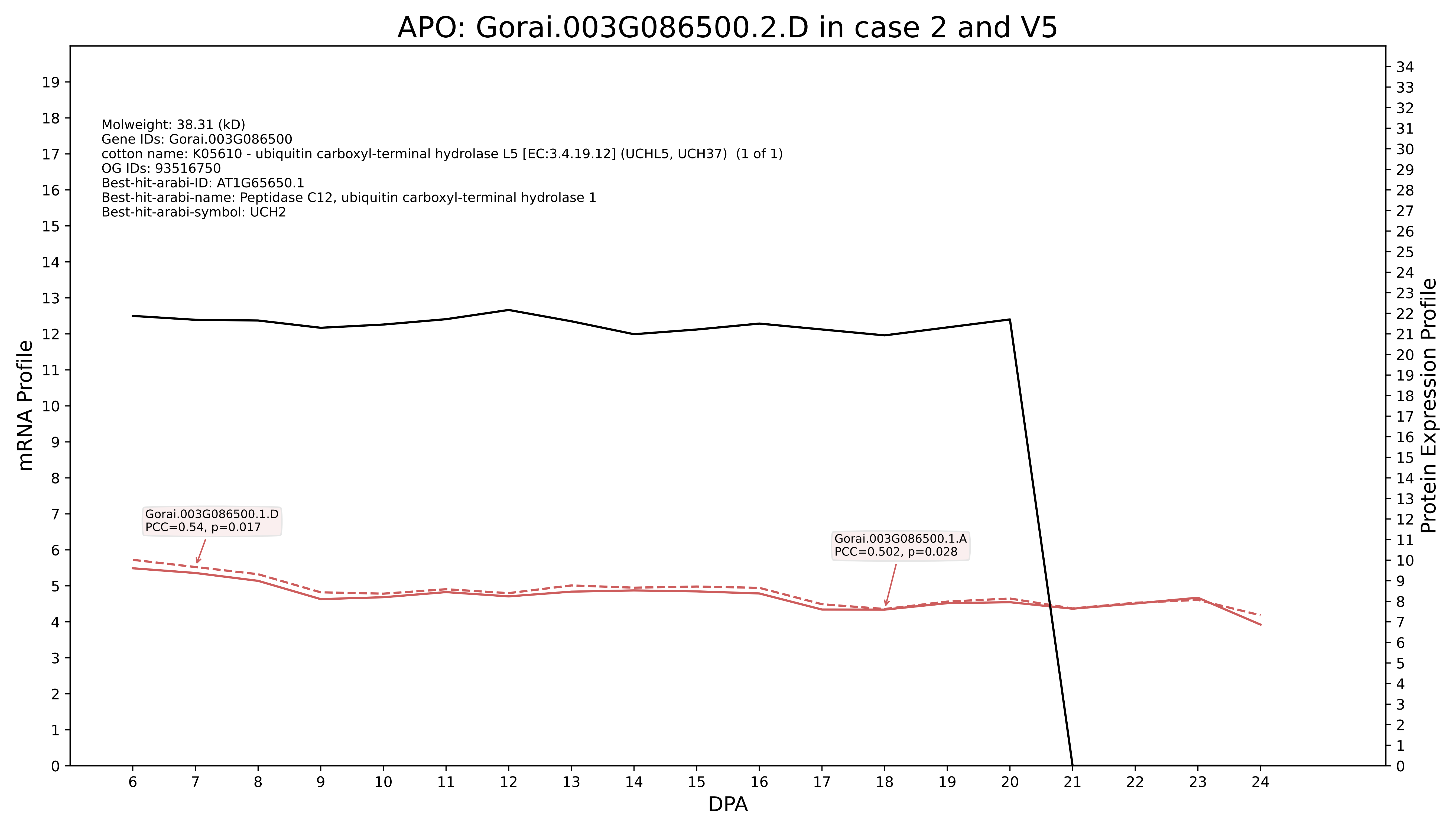
APO: Gorai.003G086500.2.D in case 2 and V5
Molweight: 38.31 (kD)
Gene IDs: Gorai.003G086500
cotton name: K05610 - ubiquitin carboxyl-terminal hydrolase L5 [EC:3.4.19.12] (UCHL5, UCH37) (1 of 1)
OG IDs: 93516750
Best-hit-arabi-ID: AT1G65650.1
Best-hit-arabi-name: Peptidase C12, ubiquitin carboxyl-terminal hydrolase 1
Best-hit-arabi-symbol: UCH2
Gorai.003G086500.1.D
PCC=0.54, p=0.017
Gorai.003G086500.1.A
PCC=0.502, p=0.028
mRNA Profile
Protein Expression Profile
DPA

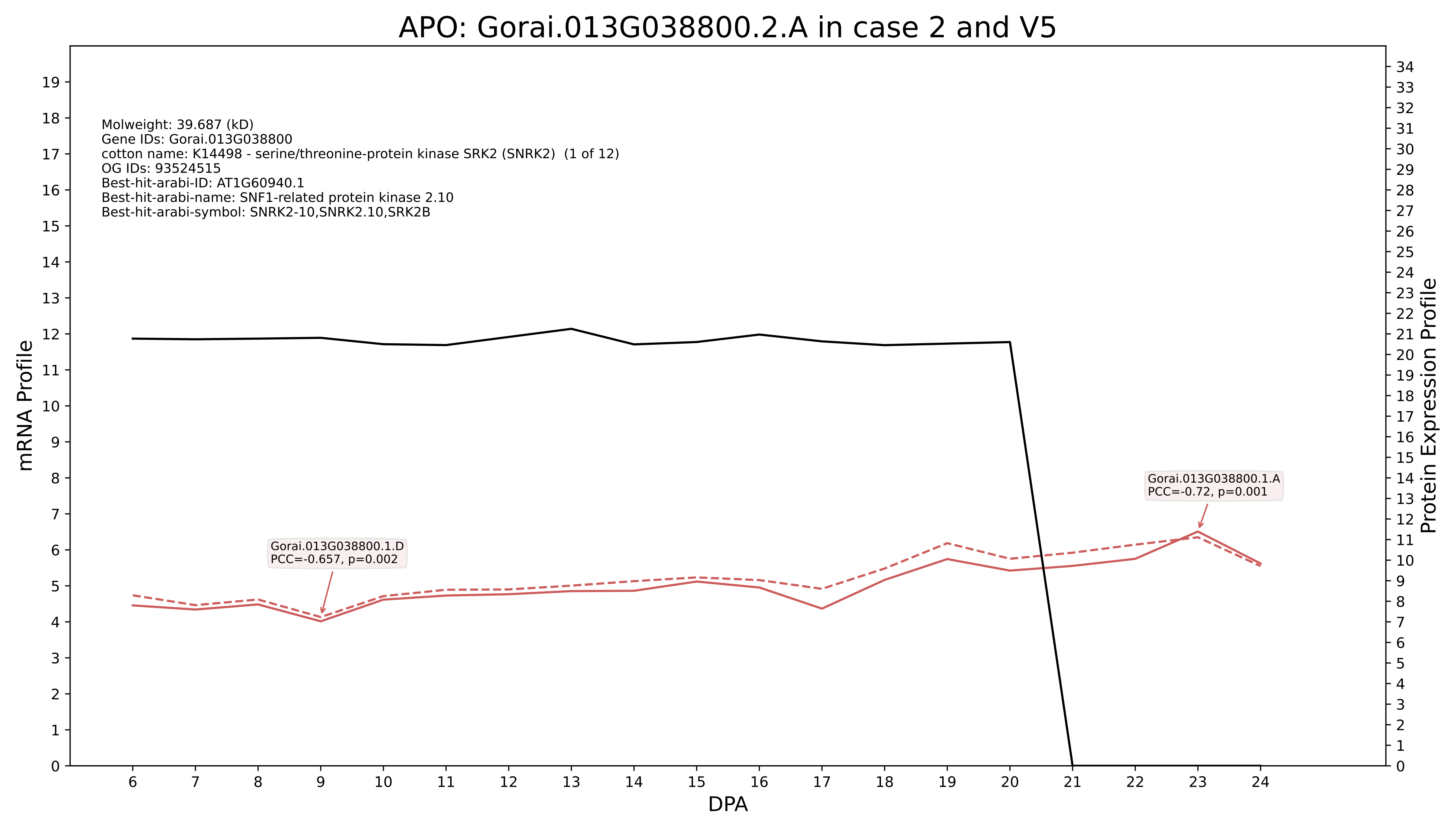
APO: Gorai.013G038800.2.A in case 2 and V5
Molweight: 39.687 (kD)
Gene IDs: Gorai.013G038800
cotton name: K14498 - serine/threonine-protein kinase SRK2 (SNRK2) (1 of 12)
OG IDs: 93524515
Best-hit-arabi-ID: AT1G60940.1
Best-hit-arabi-name: SNF1-related protein kinase 2.10
Best-hit-arabi-symbol: SNRK2-10,SNRK2.10,SRK2B
Gorai.013G038800.1.D
PCC=-0.657, p=0.002
Gorai.013G038800.1.A
PCC=-0.72, p=0.001
mRNA Profile
Protein Expression Profile
DPA

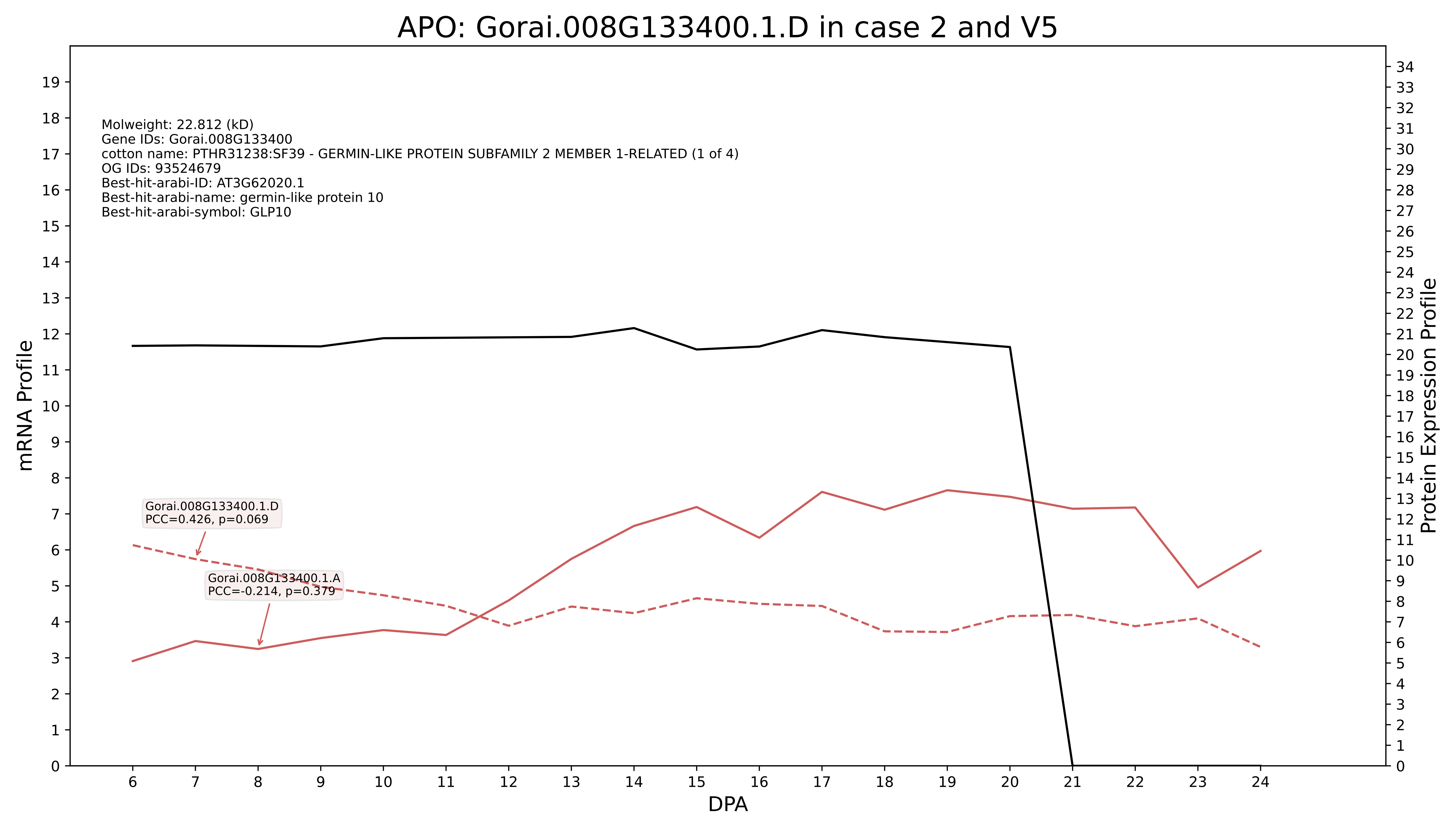
APO: Gorai.008G133400.1.D in case 2 and V5
Molweight: 22.812 (kD)
Gene IDs: Gorai.008G133400
cotton name: PTHR31238:SF39 - GERMIN-LIKE PROTEIN SUBFAMILY 2 MEMBER 1-RELATED (1 of 4)
OG IDs: 93524679
Best-hit-arabi-ID: AT3G62020.1
Best-hit-arabi-name: germin-like protein 10
Best-hit-arabi-symbol: GLP10
Gorai.008G133400.1.D
PCC=0.426, p=0.069
Gorai.008G133400.1.A
PCC=-0.214, p=0.379
mRNA Profile
Protein Expression Profile
DPA

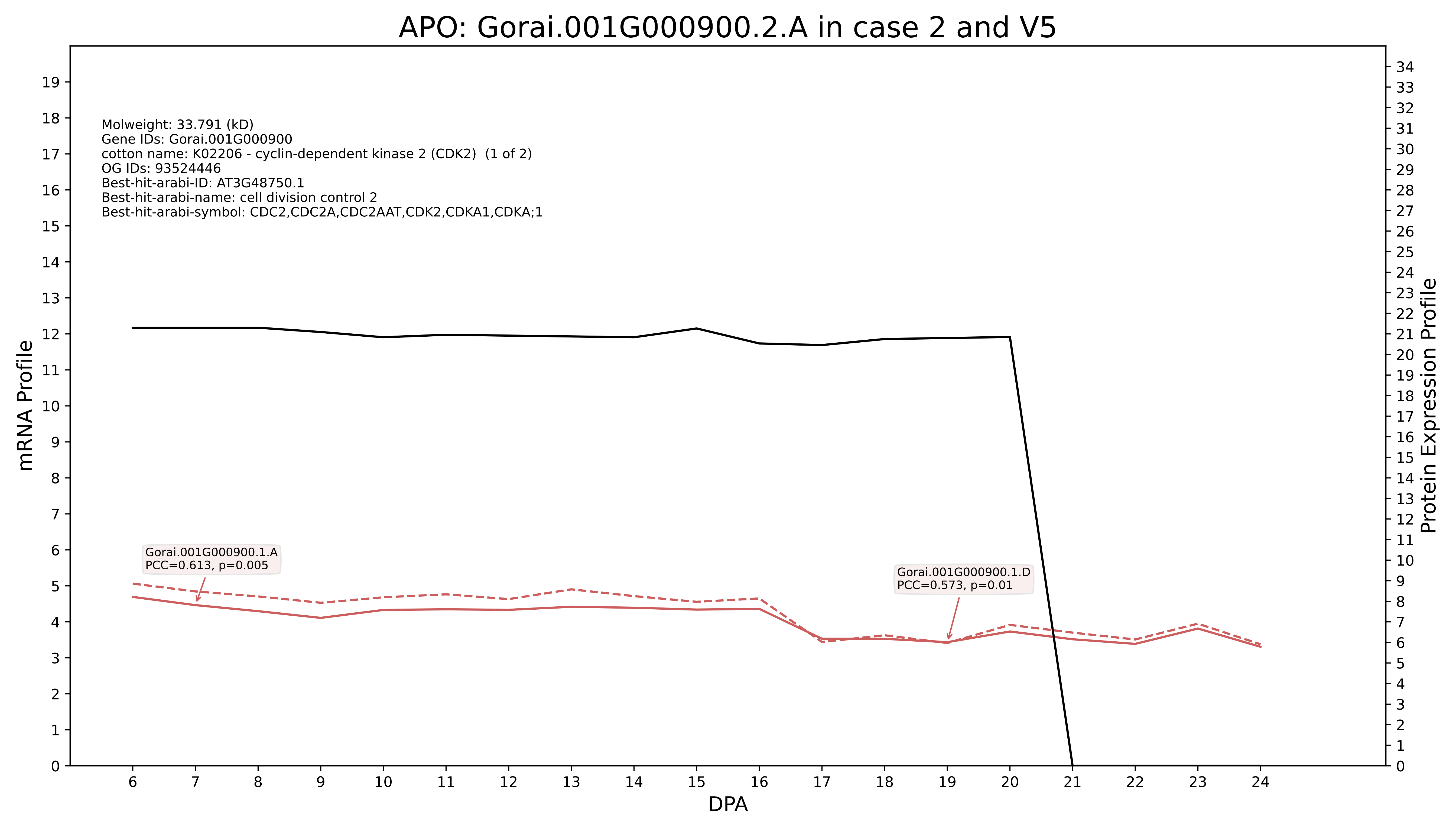
APO: Gorai.001G000900.2.A in case 2 and V5
Molweight: 33.791 (kD)
Gene IDs: Gorai.001G000900
cotton name: K02206 - cyclin-dependent kinase 2 (CDK2) (1 of 2)
OG IDs: 93524446
Best-hit-arabi-ID: AT3G48750.1
Best-hit-arabi-name: cell division control 2
Best-hit-arabi-symbol: CDC2,CDC2A,CDC2AAT,CDK2,CDKA1,CDKA;1
Gorai.001G000900.1.A
PCC=0.613, p=0.005
Gorai.001G000900.1.D
PCC=0.573, p=0.01
mRNA Profile
Protein Expression Profile
DPA

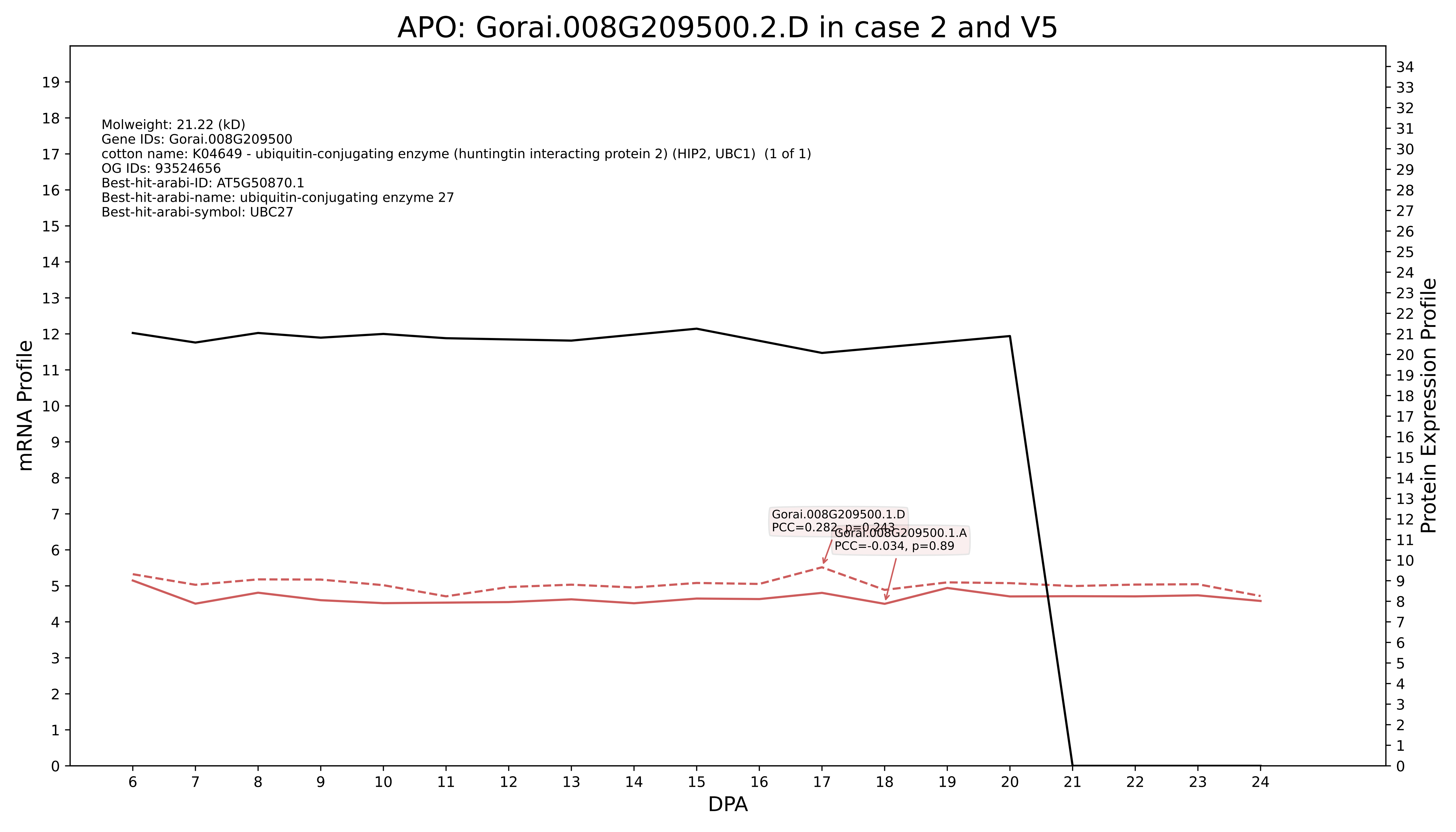

APO: Gorai.008G209500.2.D in case 2 and V5
Molweight: 21.22 (kD)
Gene IDs: Gorai.008G209500
cotton name: K04649 - ubiquitin-conjugating enzyme (huntingtin interacting protein 2) (HIP2, UBC1) (1 of 1)
OG IDs: 93524656
Best-hit-arabi-ID: AT5G50870.1
Best-hit-arabi-name: ubiquitin-conjugating enzyme 27
Best-hit-arabi-symbol: UBC27
Gorai.008G209500.1.D
PCC=0.282, p=0.243
Gorai.008G209500.1.A
PCC=-0.034, p=0.89
mRNA Profile
Protein Expression Profile
DPA

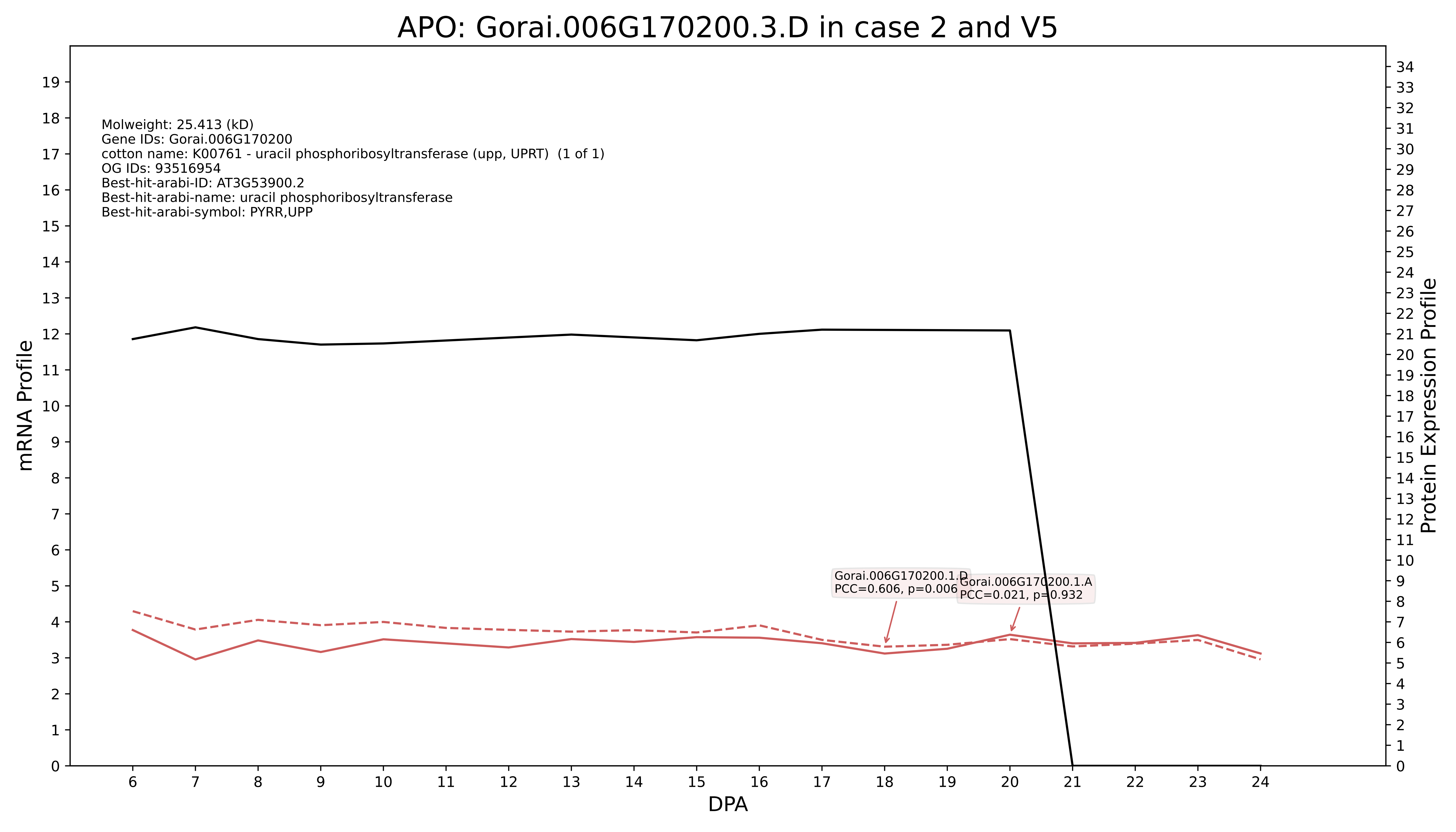

APO: Gorai.006G170200.3.D in case 2 and V5
Molweight: 25.413 (kD)
Gene IDs: Gorai.006G170200
cotton name: K00761 - uracil phosphoribosyltransferase (upp, UPRT) (1 of 1)
OG IDs: 93516954
Best-hit-arabi-ID: AT3G53900.2
Best-hit-arabi-name: uracil phosphoribosyltransferase
Best-hit-arabi-symbol: PYRR,UPP
Gorai.006G170200.1.D
PCC=0.606, p=0.006
Gorai.006G170200.1.A
PCC=0.021, p=0.932
mRNA Profile
Protein Expression Profile
DPA

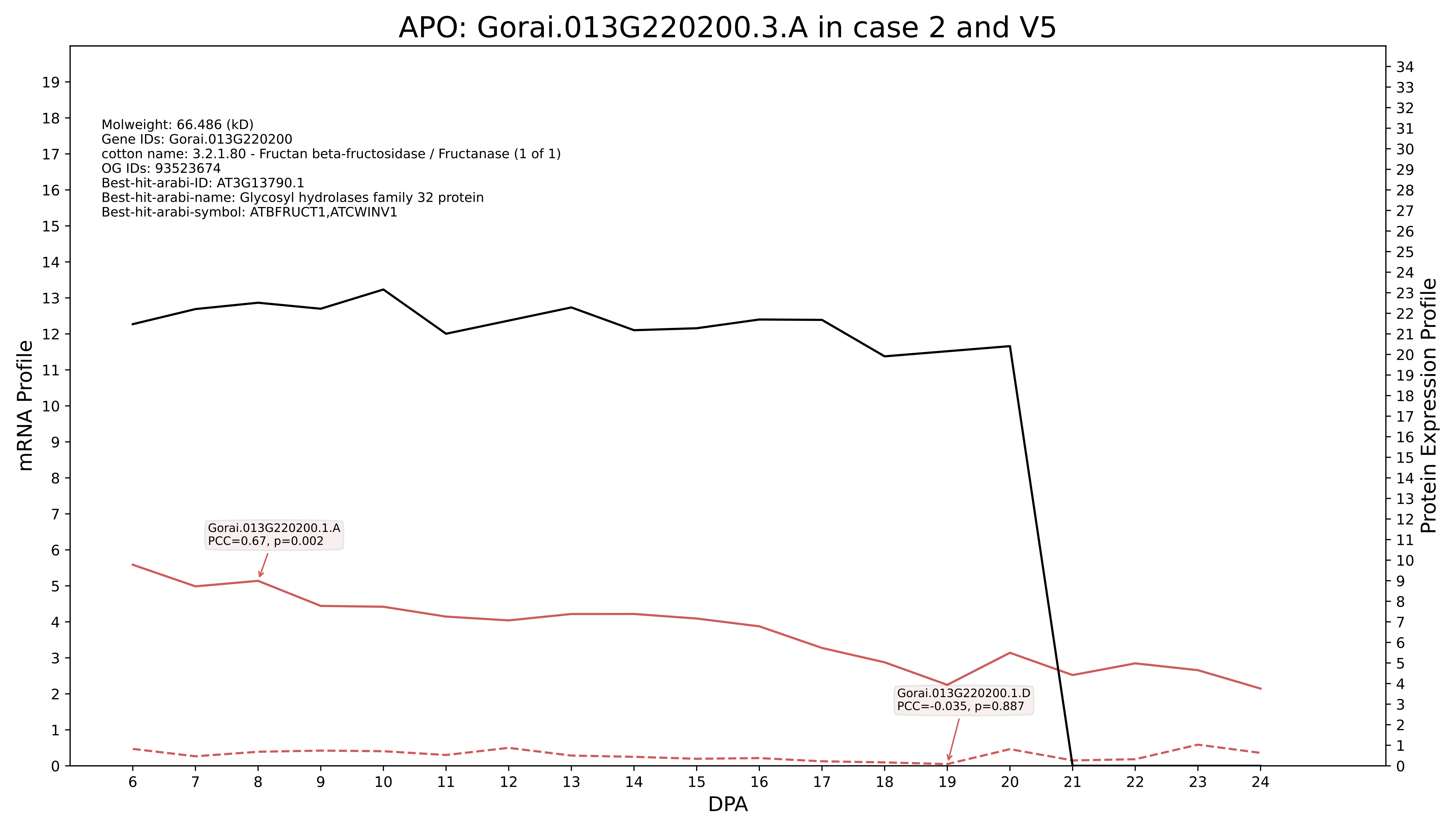
APO: Gorai.013G220200.3.A in case 2 and V5
Molweight: 66.486 (kD)
Gene IDs: Gorai.013G220200
cotton name: 3.2.1.80 - Fructan beta-fructosidase / Fructanase (1 of 1)
OG IDs: 93523674
Best-hit-arabi-ID: AT3G13790.1
Best-hit-arabi-name: Glycosyl hydrolases family 32 protein
Best-hit-arabi-symbol: ATBFRUCT1,ATCWINV1
Gorai.013G220200.1.A
PCC=0.67, p=0.002
Gorai.013G220200.1.D
PCC=-0.035, p=0.887
mRNA Profile
Protein Expression Profile
DPA

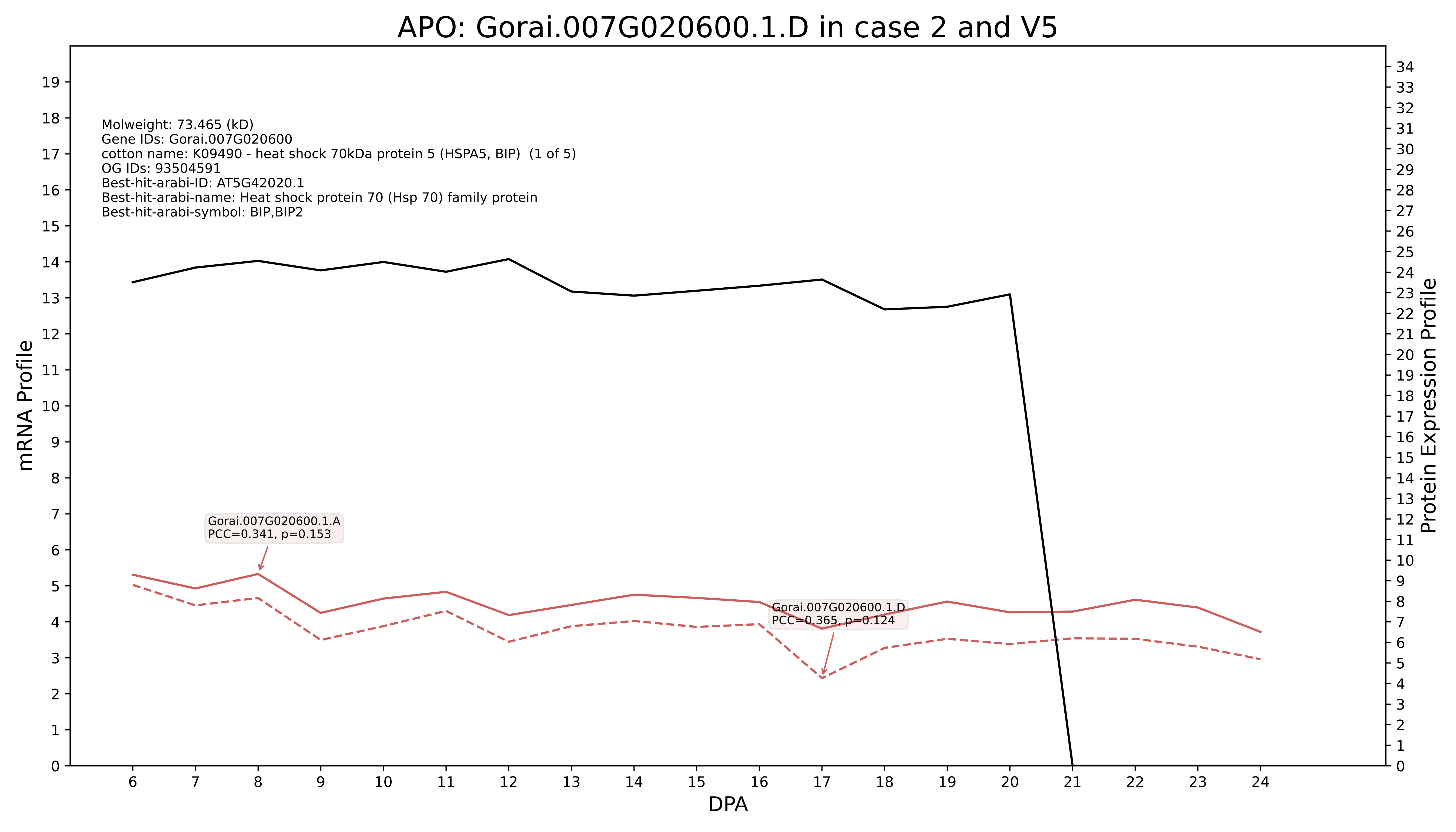

APO: Gorai.007G020600.1.D in case 2 and V5
Molweight: 73.465 (kD)
Gene IDs: Gorai.007G020600
cotton name: K09490 - heat shock 70kDa protein 5 (HSPA5, BIP) (1 of 5)
OG IDs: 93504591
Best-hit-arabi-ID: AT5G42020.1
Best-hit-arabi-name: Heat shock protein 70 (Hsp 70) family protein
Best-hit-arabi-symbol: BIP,BIP2
Gorai.007G020600.1.A
PCC=0.341, p=0.153
Gorai.007G020600.1.D
PCC=0.365, p=0.124
mRNA Profile
Protein Expression Profile
DPA

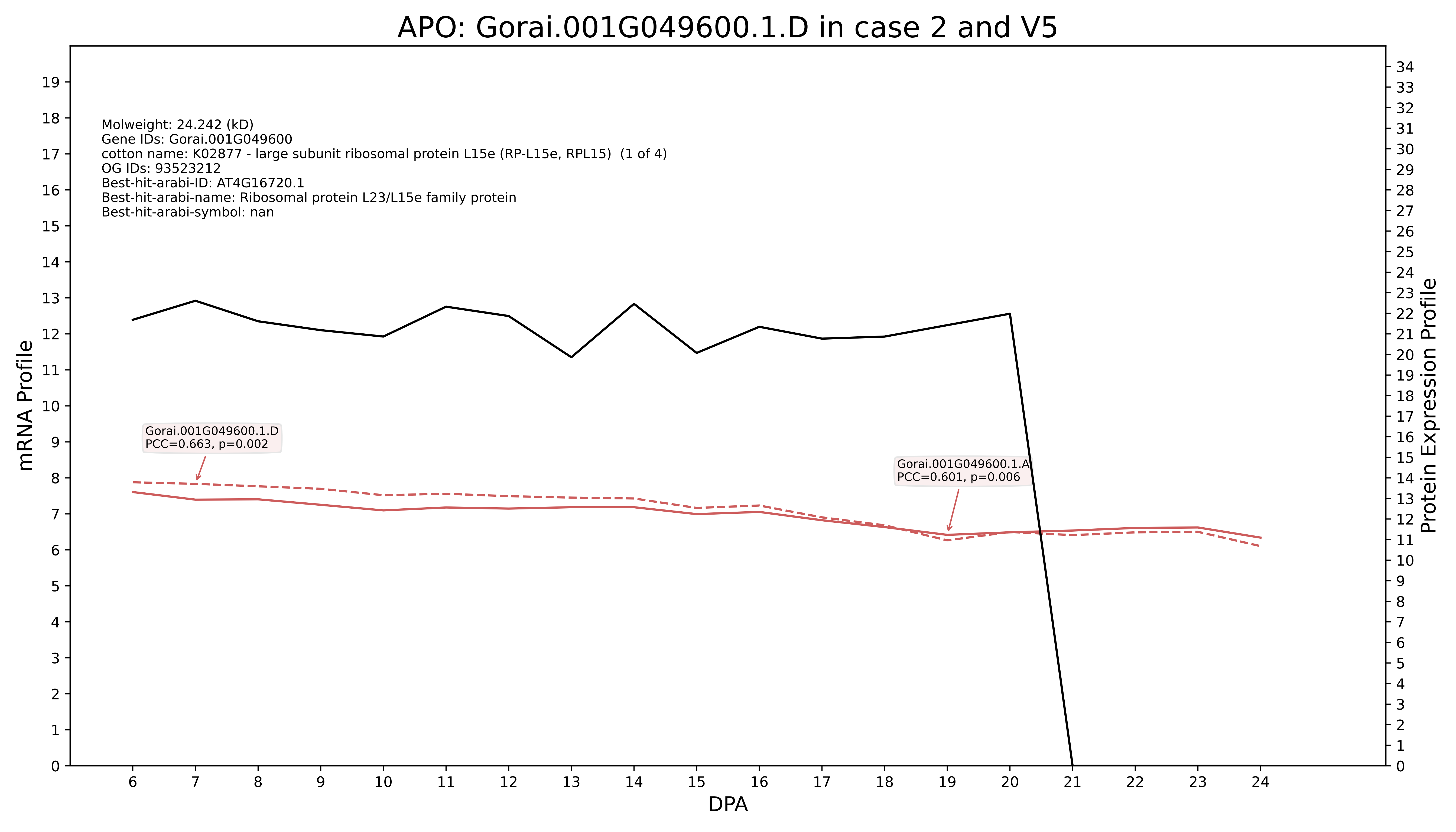
APO: Gorai.001G049600.1.D in case 2 and V5
Molweight: 24.242 (kD)
Gene IDs: Gorai.001G049600
cotton name: K02877 - large subunit ribosomal protein L15e (RP-L15e, RPL15) (1 of 4)
OG IDs: 93523212
Best-hit-arabi-ID: AT4G16720.1
Best-hit-arabi-name: Ribosomal protein L23/L15e family protein
Best-hit-arabi-symbol: nan
Gorai.001G049600.1.D
PCC=0.663, p=0.002
Gorai.001G049600.1.A
PCC=0.601, p=0.006
mRNA Profile
Protein Expression Profile
DPA

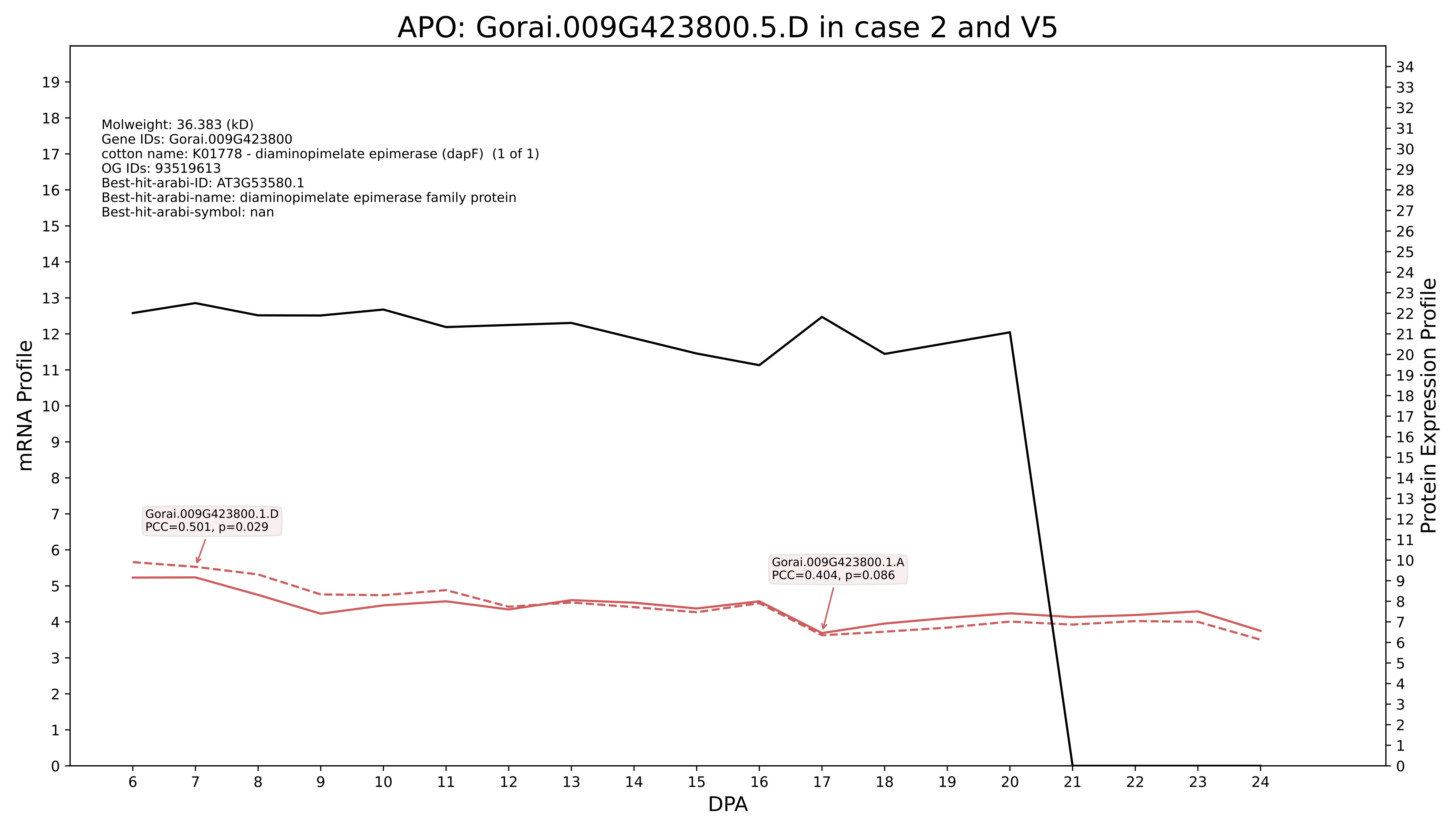
APO: Gorai.009G423800.5.D in case 2 and V5
Molweight: 36.383 (kD)
Gene IDs: Gorai.009G423800
cotton name: K01778 - diaminopimelate epimerase (dapF) (1 of 1)
OG IDs: 93519613
Best-hit-arabi-ID: AT3G53580.1
Best-hit-arabi-name: diaminopimelate epimerase family protein
Best-hit-arabi-symbol: nan
Gorai.009G423800.1.D
PCC=0.501, p=0.029
Gorai.009G423800.1.A
PCC=0.404, p=0.086
mRNA Profile
Protein Expression Profile
DPA

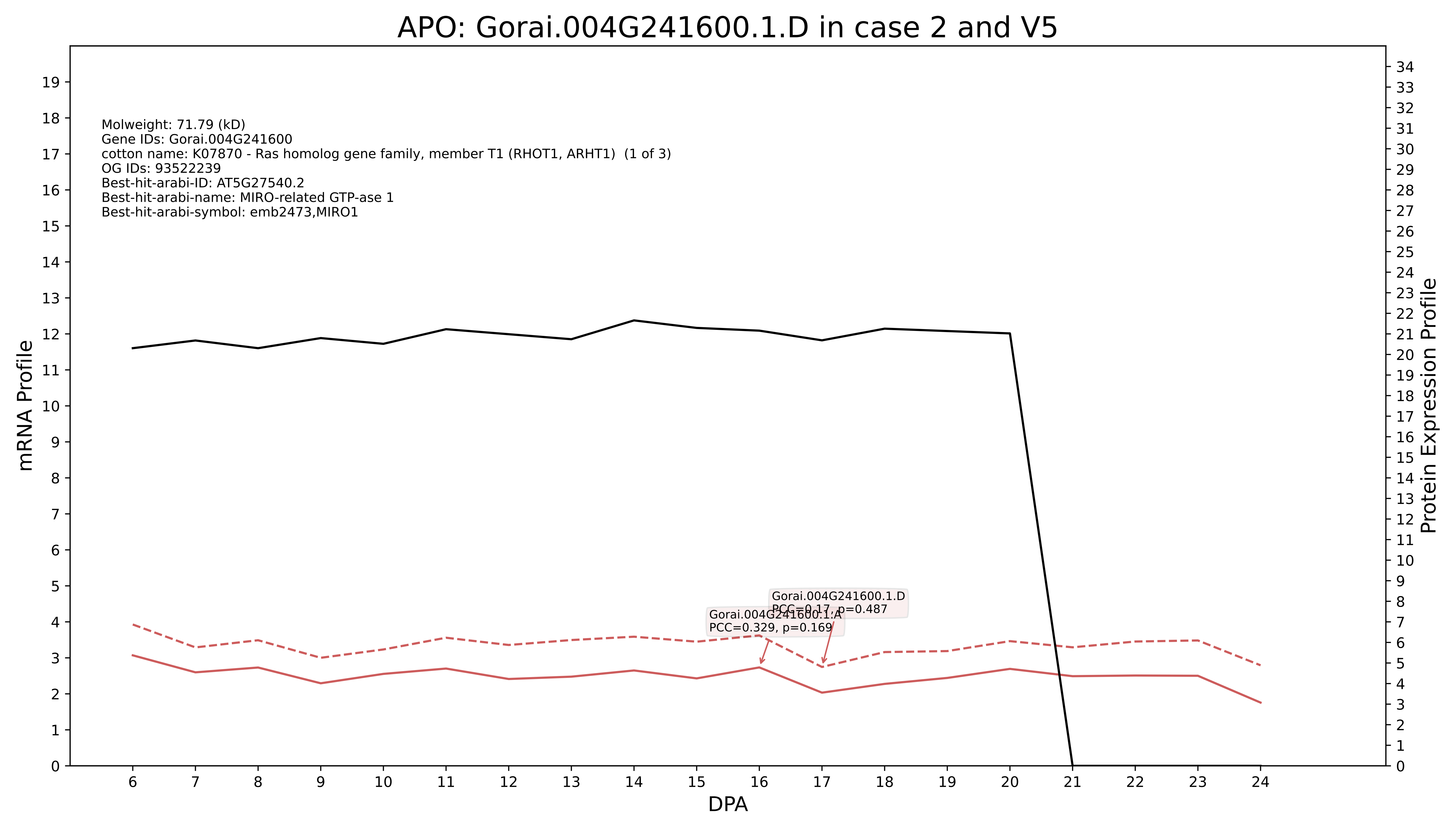
APO: Gorai.004G241600.1.D in case 2 and V5
Molweight: 71.79 (kD)
Gene IDs: Gorai.004G241600
cotton name: K07870 - Ras homolog gene family, member T1 (RHOT1, ARHT1) (1 of 3)
OG IDs: 93522239
Best-hit-arabi-ID: AT5G27540.2
Best-hit-arabi-name: MIRO-related GTP-ase 1
Best-hit-arabi-symbol: emb2473,MIRO1
Gorai.004G241600.1.D
PCC=0.17, p=0.487
Gorai.004G241600.1.A
PCC=0.329, p=0.169
mRNA Profile
Protein Expression Profile
DPA

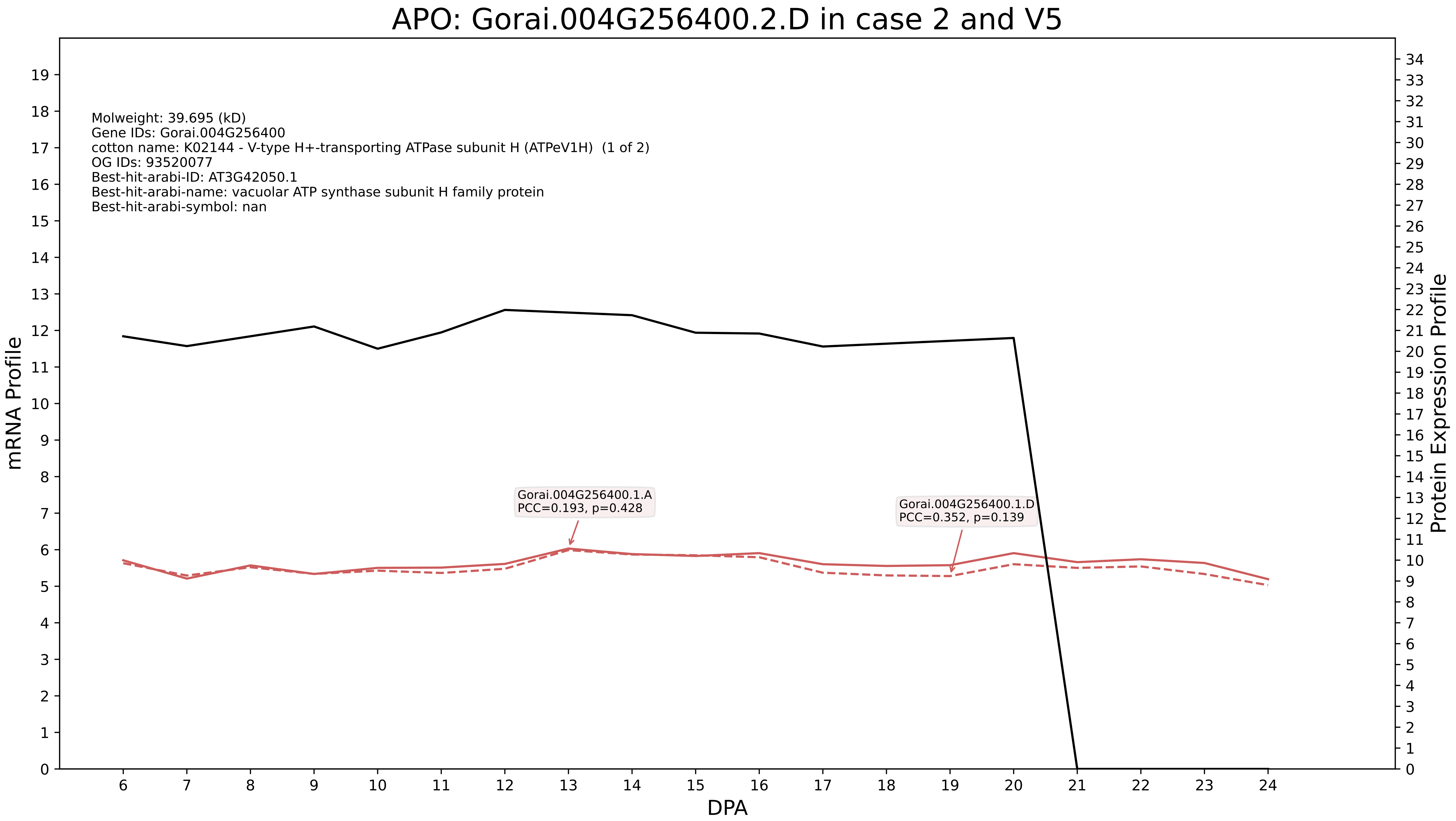
APO: Gorai.004G256400.2.D in case 2 and V5
Molweight: 39.695 (kD)
Gene IDs: Gorai.004G256400
cotton name: K02144 - V-type H+-transporting ATPase subunit H (ATPeV1H) (1 of 2)
OG IDs: 93520077
Best-hit-arabi-ID: AT3G42050.1
Best-hit-arabi-name: vacuolar ATP synthase subunit H family protein
Best-hit-arabi-symbol: nan
Gorai.004G256400.1.A
PCC=0.193, p=0.428
Gorai.004G256400.1.D
PCC=0.352, p=0.139
mRNA Profile
Protein Expression Profile
DPA

### APO: Gorai.001G056300.1.D in case 2 and V5

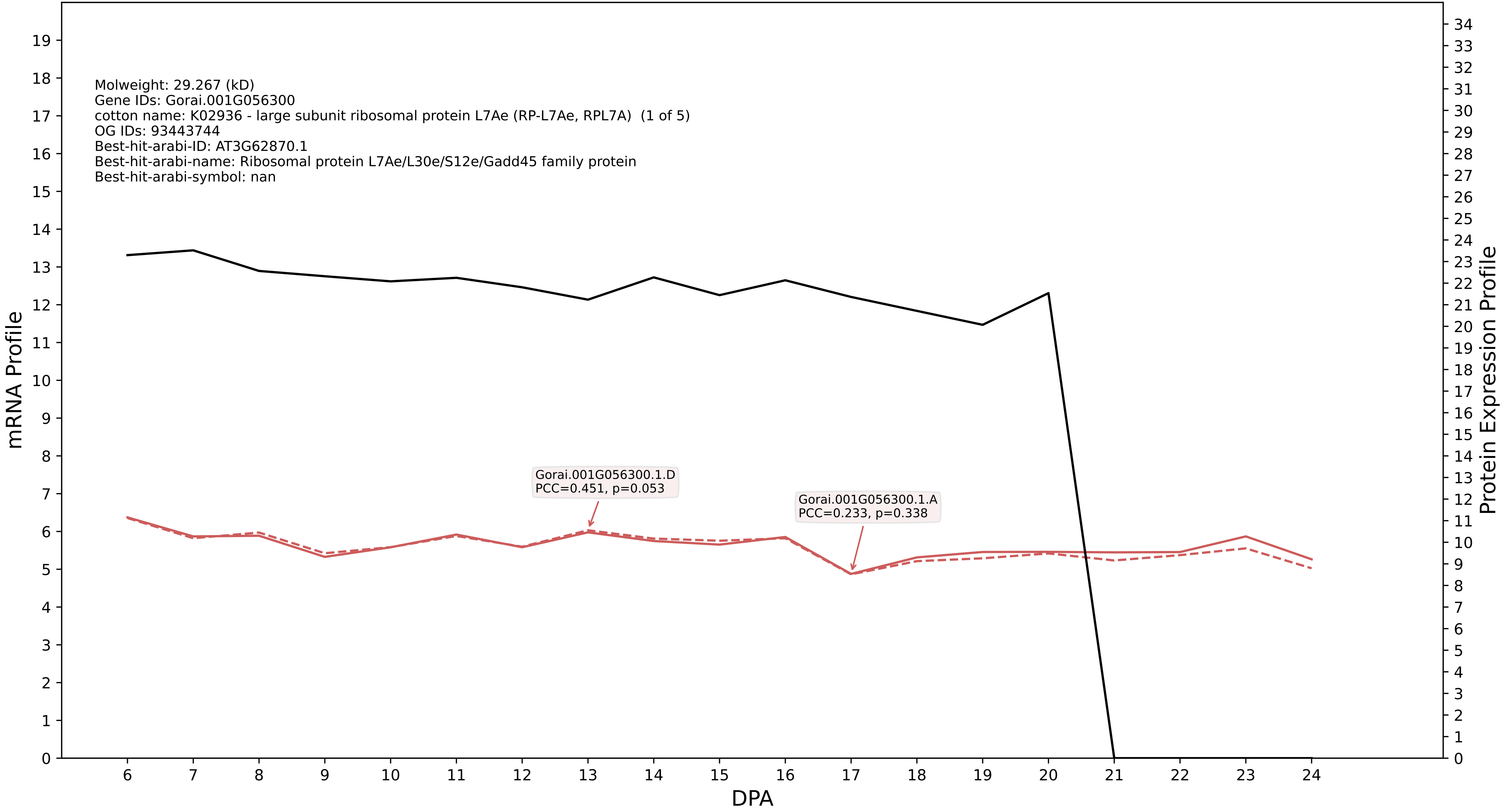

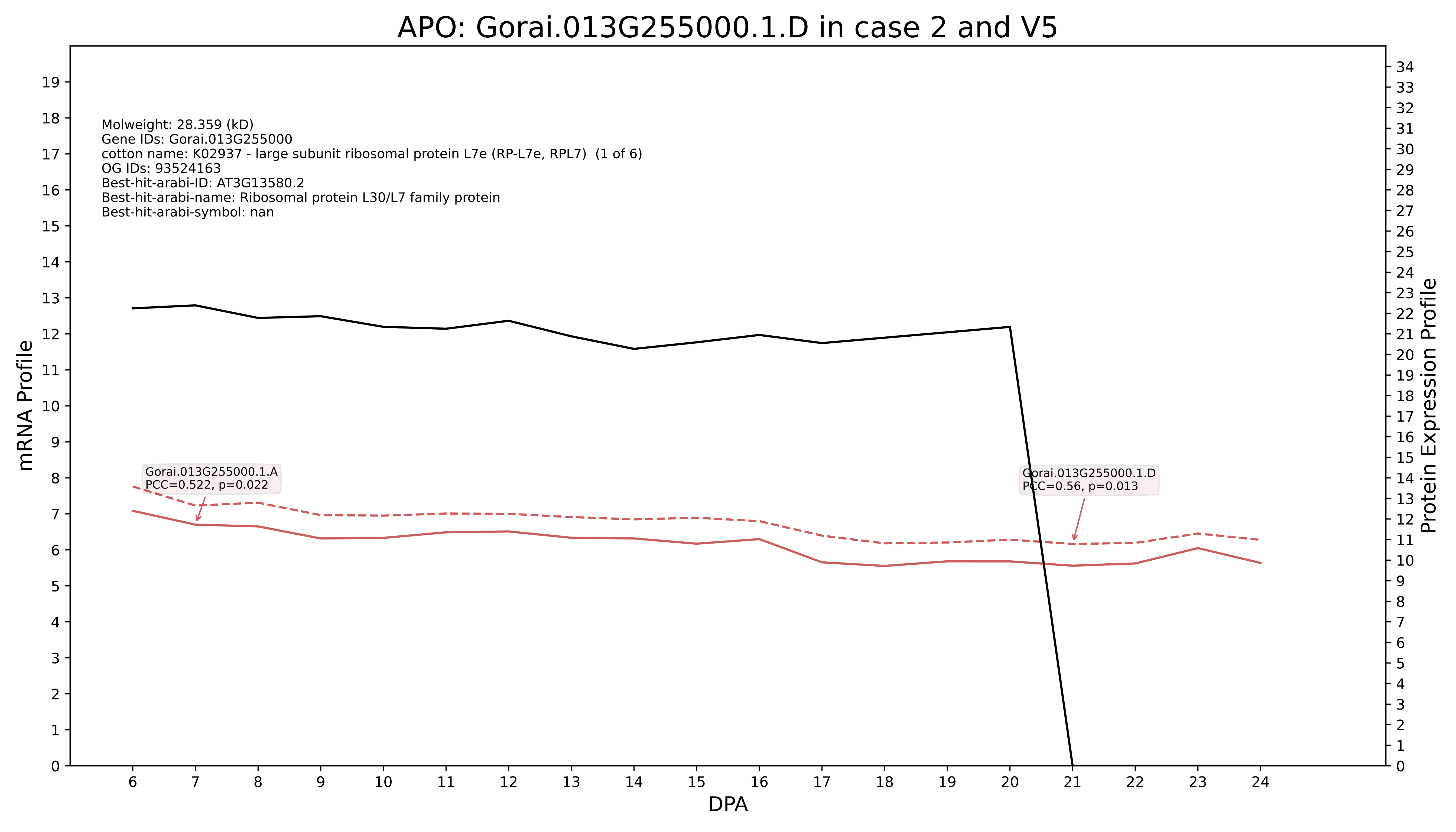

APO: Gorai.013G255000.1.D in case 2 and V5
Molweight: 28.359 (kD)
Gene IDs: Gorai.013G255000
cotton name: K02937 - large subunit ribosomal protein L7e (RP-L7e, RPL7) (1 of 6)
OG IDs: 93524163
Best-hit-arabi-ID: AT3G13580.2
Best-hit-arabi-name: Ribosomal protein L30/L7 family protein
Best-hit-arabi-symbol: nan
Gorai.013G255000.1.A
PCC=0.522, p=0.022
Gorai.013G255000.1.D
PCC=0.56, p=0.013
mRNA Profile
Protein Expression Profile
DPA

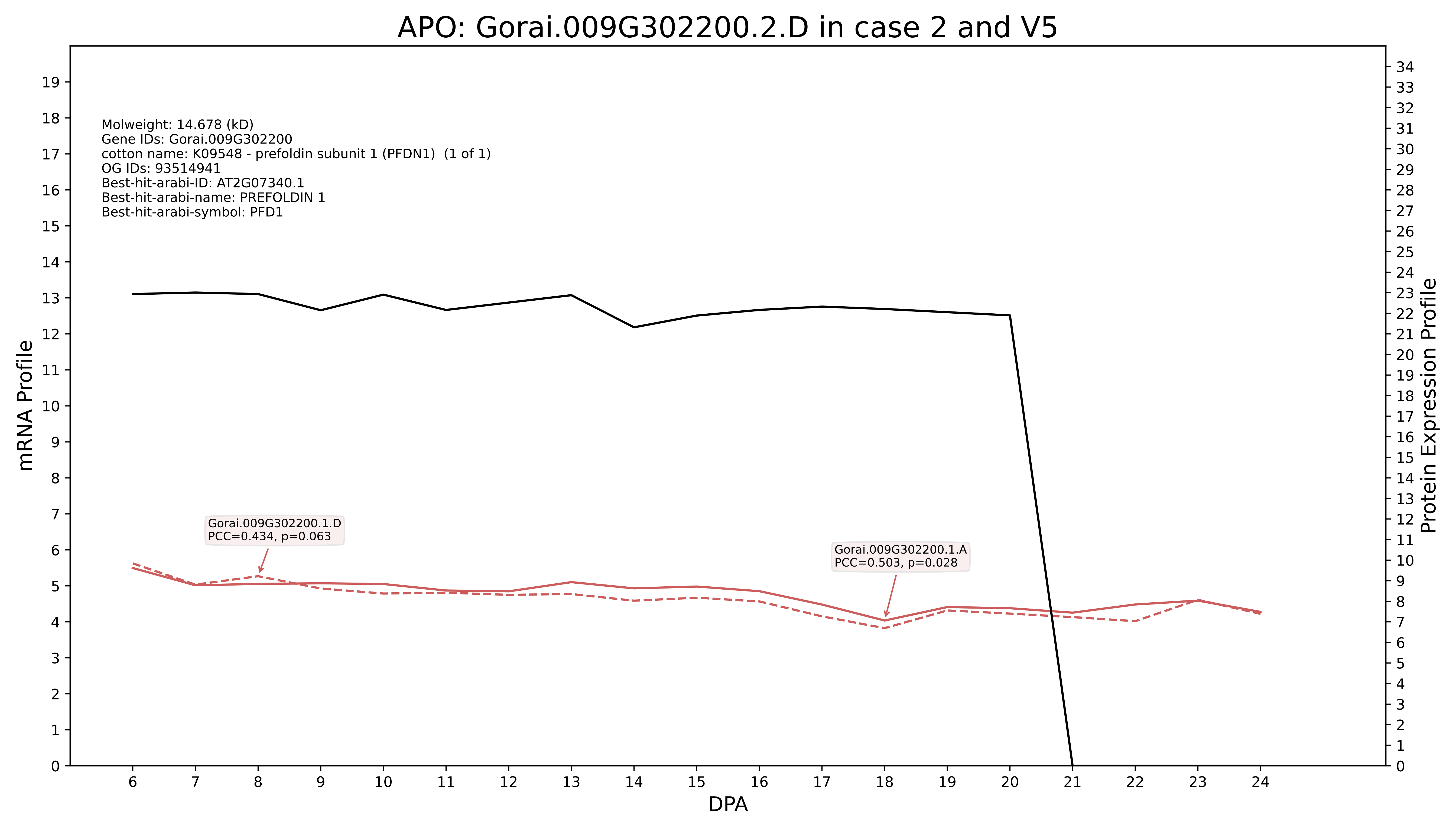
APO: Gorai.009G302200.2.D in case 2 and V5
Molweight: 14.678 (kD)
Gene IDs: Gorai.009G302200
cotton name: K09548 - prefoldin subunit 1 (PFDN1) (1 of 1)
OG IDs: 93514941
Best-hit-arabi-ID: AT2G07340.1
Best-hit-arabi-name: PREFOLDIN 1
Best-hit-arabi-symbol: PFD1
Gorai.009G302200.1.D
PCC=0.434, p=0.063
Gorai.009G302200.1.A
PCC=0.503, p=0.028
mRNA Profile
Protein Expression Profile
DPA

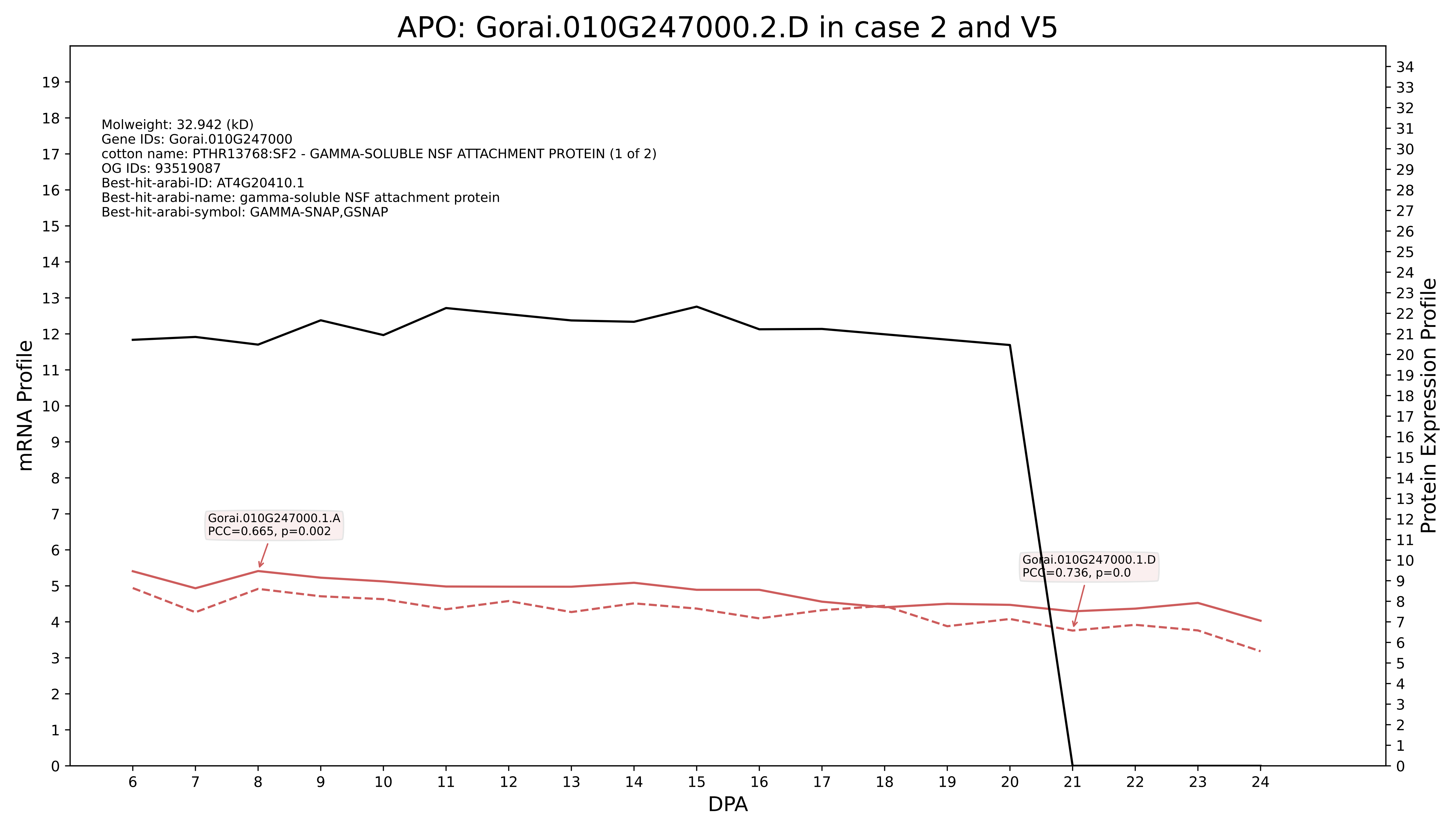

APO: Gorai.010G247000.2.D in case 2 and V5
Molweight: 32.942 (kD)
Gene IDs: Gorai.010G247000
cotton name: PTHR13768:SF2 - GAMMA-SOLUBLE NSF ATTACHMENT PROTEIN (1 of 2)
OG IDs: 93519087
Best-hit-arabi-ID: AT4G20410.1
Best-hit-arabi-name: gamma-soluble NSF attachment protein
Best-hit-arabi-symbol: GAMMA-SNAP,GSNAP
Gorai.010G247000.1.A
PCC=0.665, p=0.002
Gorai.010G247000.1.D
PCC=0.736, p=0.0
mRNA Profile
Protein Expression Profile
DPA

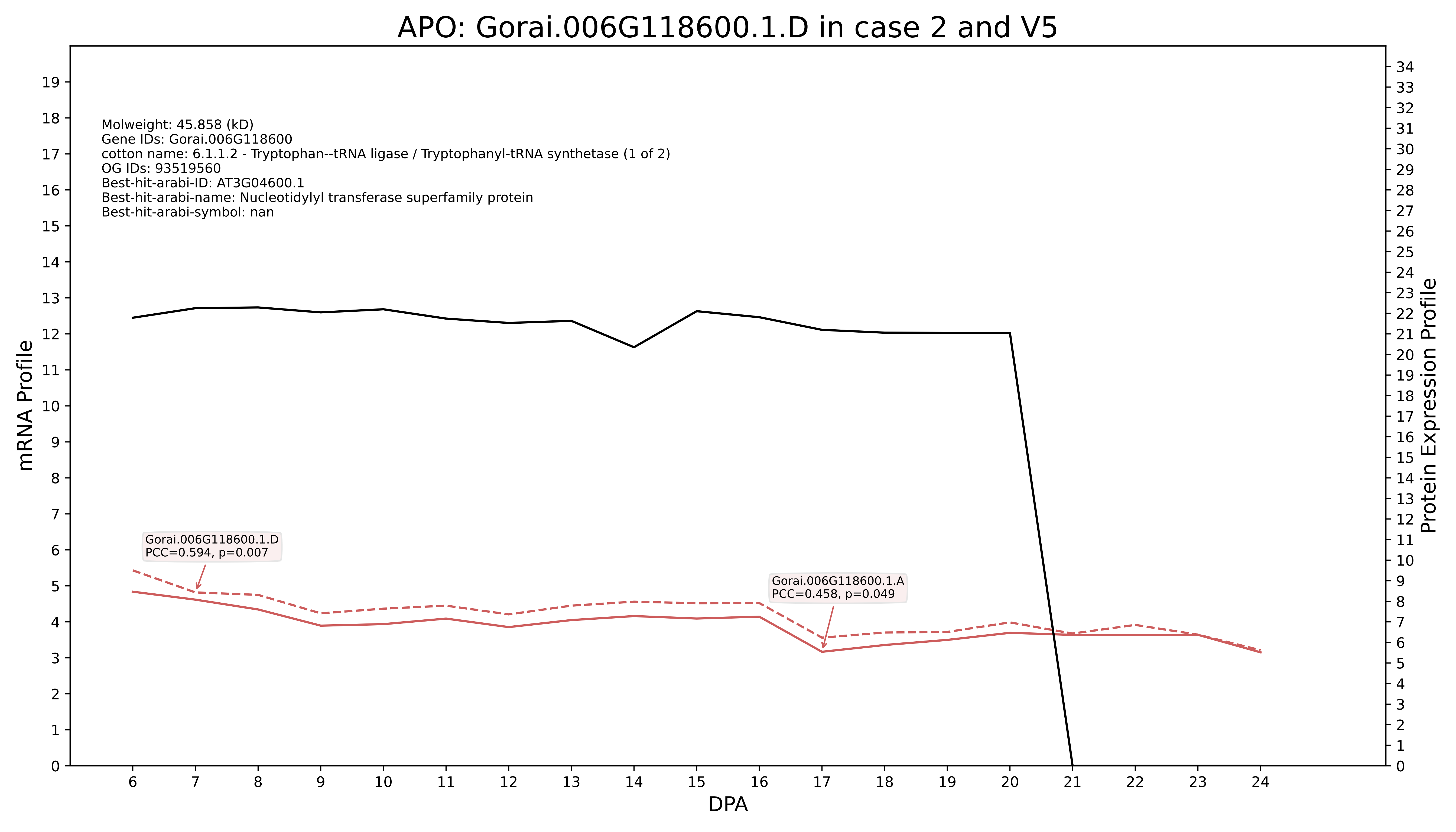
APO: Gorai.006G118600.1.D in case 2 and V5
Molweight: 45.858 (kD)
Gene IDs: Gorai.006G118600
cotton name: 6.1.1.2 - Tryptophan--tRNA ligase / Tryptophanyl-tRNA synthetase (1 of 2)
OG IDs: 93519560
Best-hit-arabi-ID: AT3G04600.1
Best-hit-arabi-name: Nucleotidylyl transferase superfamily protein
Best-hit-arabi-symbol: nan
Gorai.006G118600.1.D
PCC=0.594, p=0.007
Gorai.006G118600.1.A
PCC=0.458, p=0.049
mRNA Profile
Protein Expression Profile
DPA

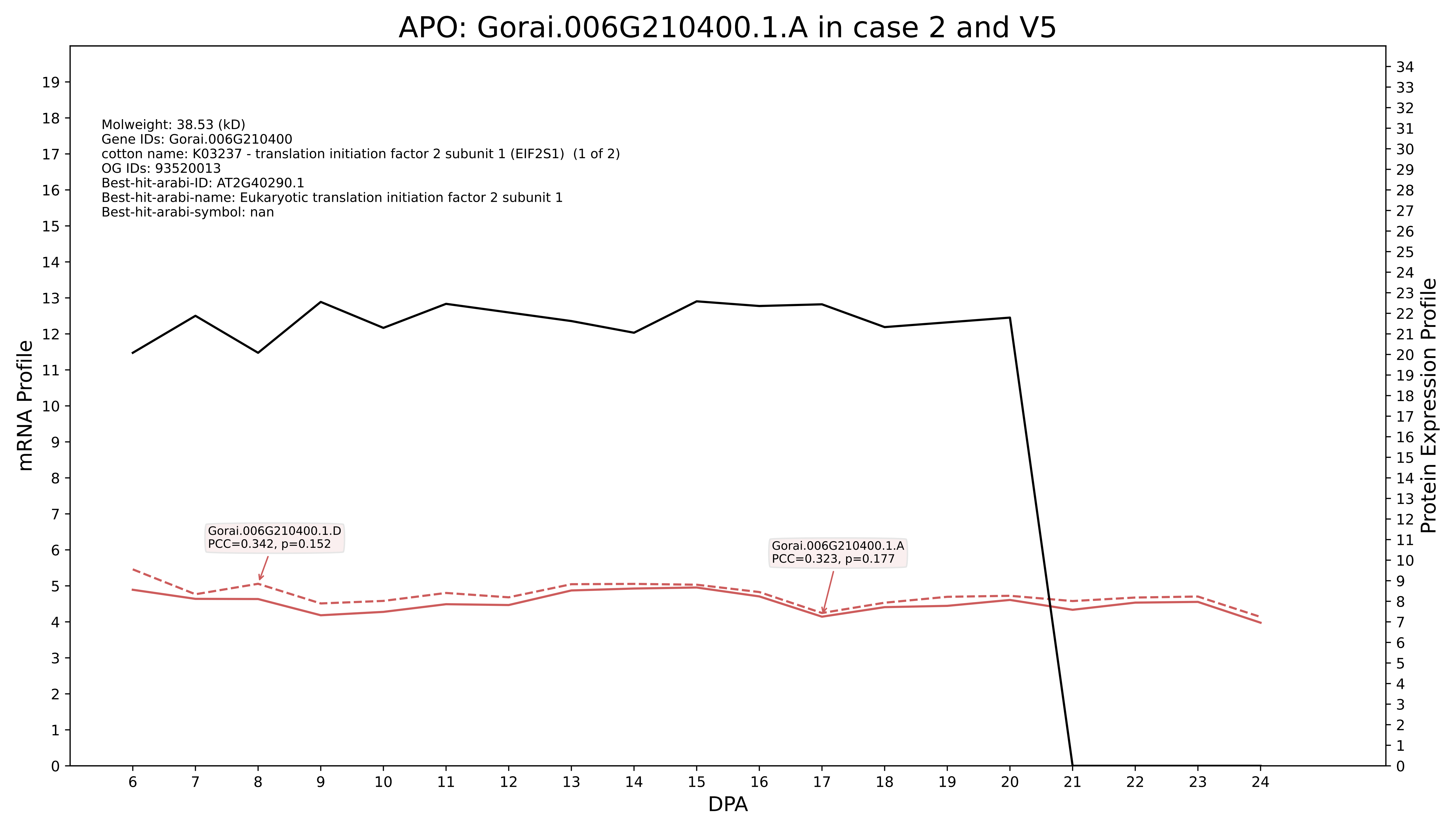

APO: Gorai.006G210400.1.A in case 2 and V5
Molweight: 38.53 (kD)
Gene IDs: Gorai.006G210400
cotton name: K03237 - translation initiation factor 2 subunit 1 (EIF2S1) (1 of 2)
OG IDs: 93520013
Best-hit-arabi-ID: AT2G40290.1
Best-hit-arabi-name: Eukaryotic translation initiation factor 2 subunit 1
Best-hit-arabi-symbol: nan
Gorai.006G210400.1.D
PCC=0.342, p=0.152
Gorai.006G210400.1.A
PCC=0.323, p=0.177
mRNA Profile
Protein Expression Profile
DPA

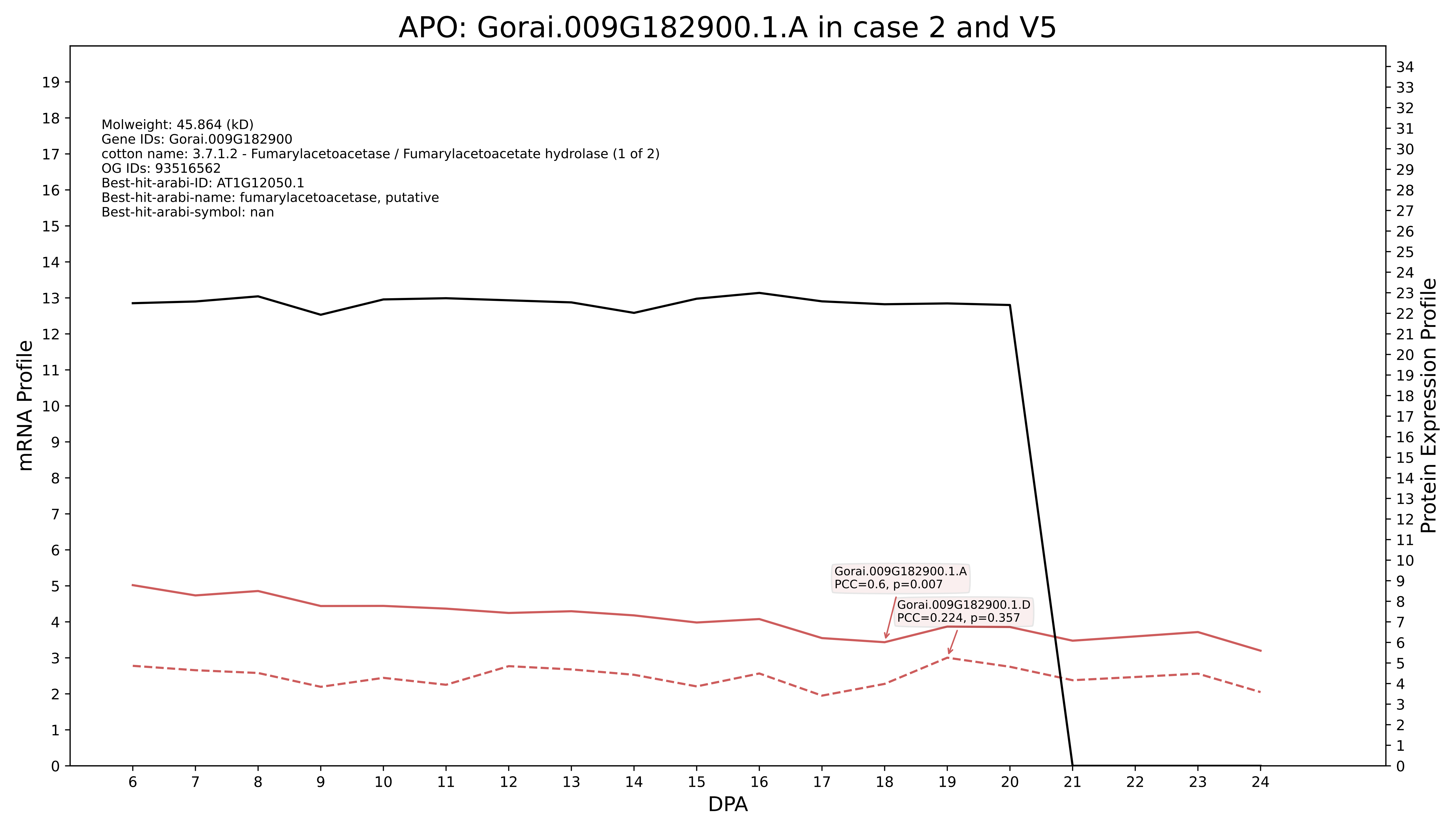
APO: Gorai.009G182900.1.A in case 2 and V5
Molweight: 45.864 (kD)
Gene IDs: Gorai.009G182900
cotton name: 3.7.1.2 - Fumarylacetoacetase / Fumarylacetoacetate hydrolase (1 of 2)
OG IDs: 93516562
Best-hit-arabi-ID: AT1G12050.1
Best-hit-arabi-name: fumarylacetoacetase, putative
Best-hit-arabi-symbol: nan
Gorai.009G182900.1.A
PCC=0.6, p=0.007
Gorai.009G182900.1.D
PCC=0.224, p=0.357
mRNA Profile
Protein Expression Profile
DPA

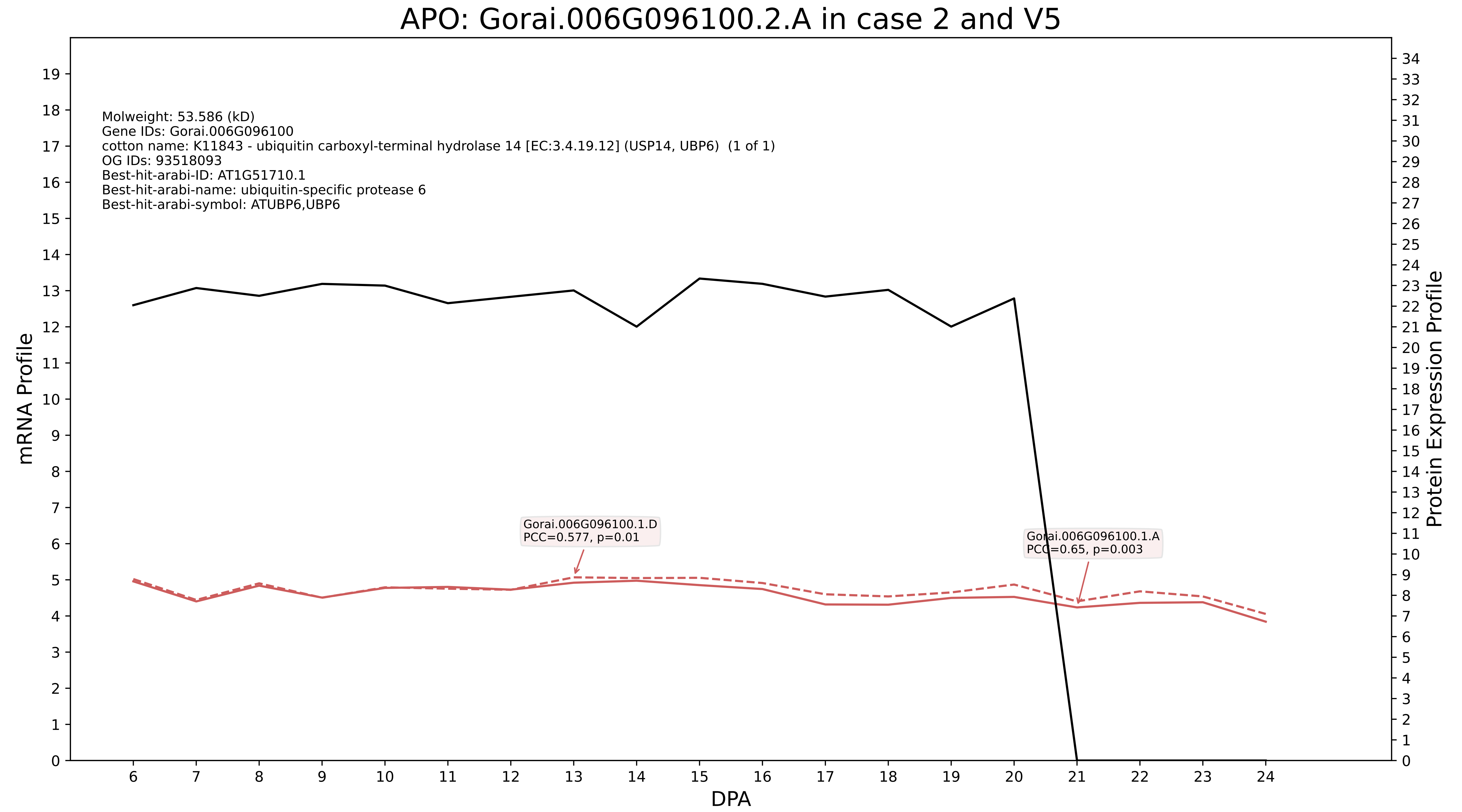

APO: Gorai.006G096100.2.A in case 2 and V5
Molweight: 53.586 (kD)
Gene IDs: Gorai.006G096100
cotton name: K11843 - ubiquitin carboxyl-terminal hydrolase 14 [EC:3.4.19.12] (USP14, UBP6) (1 of 1)
OG IDs: 93518093
Best-hit-arabi-ID: AT1G51710.1
Best-hit-arabi-name: ubiquitin-specific protease 6
Best-hit-arabi-symbol: ATUBP6,UBP6
Gorai.006G096100.1.D
PCC=0.577, p=0.01
Gorai.006G096100.1.A
PCC=0.65, p=0.003
mRNA Profile
Protein Expression Profile
DPA

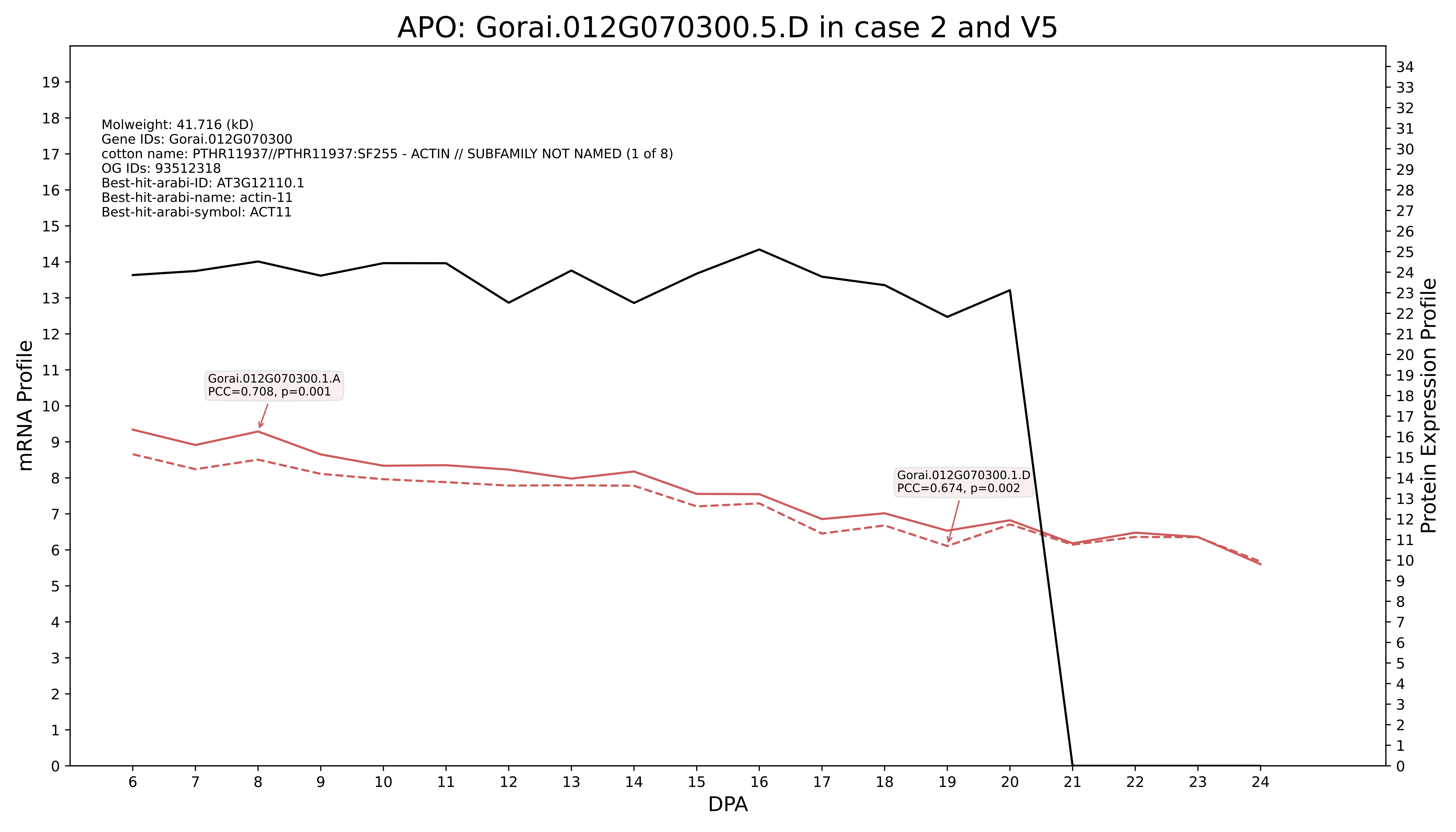

APO: Gorai.012G070300.5.D in case 2 and V5
Molweight: 41.716 (kD)
Gene IDs: Gorai.012G070300
cotton name: PTHR11937//PTHR11937:SF255 - ACTIN // SUBFAMILY NOT NAMED (1 of 8)
OG IDs: 93512318
Best-hit-arabi-ID: AT3G12110.1
Best-hit-arabi-name: actin-11
Best-hit-arabi-symbol: ACT11
Gorai.012G070300.1.A
PCC=0.708, p=0.001
Gorai.012G070300.1.D
PCC=0.674, p=0.002
mRNA Profile
Protein Expression Profile
DPA

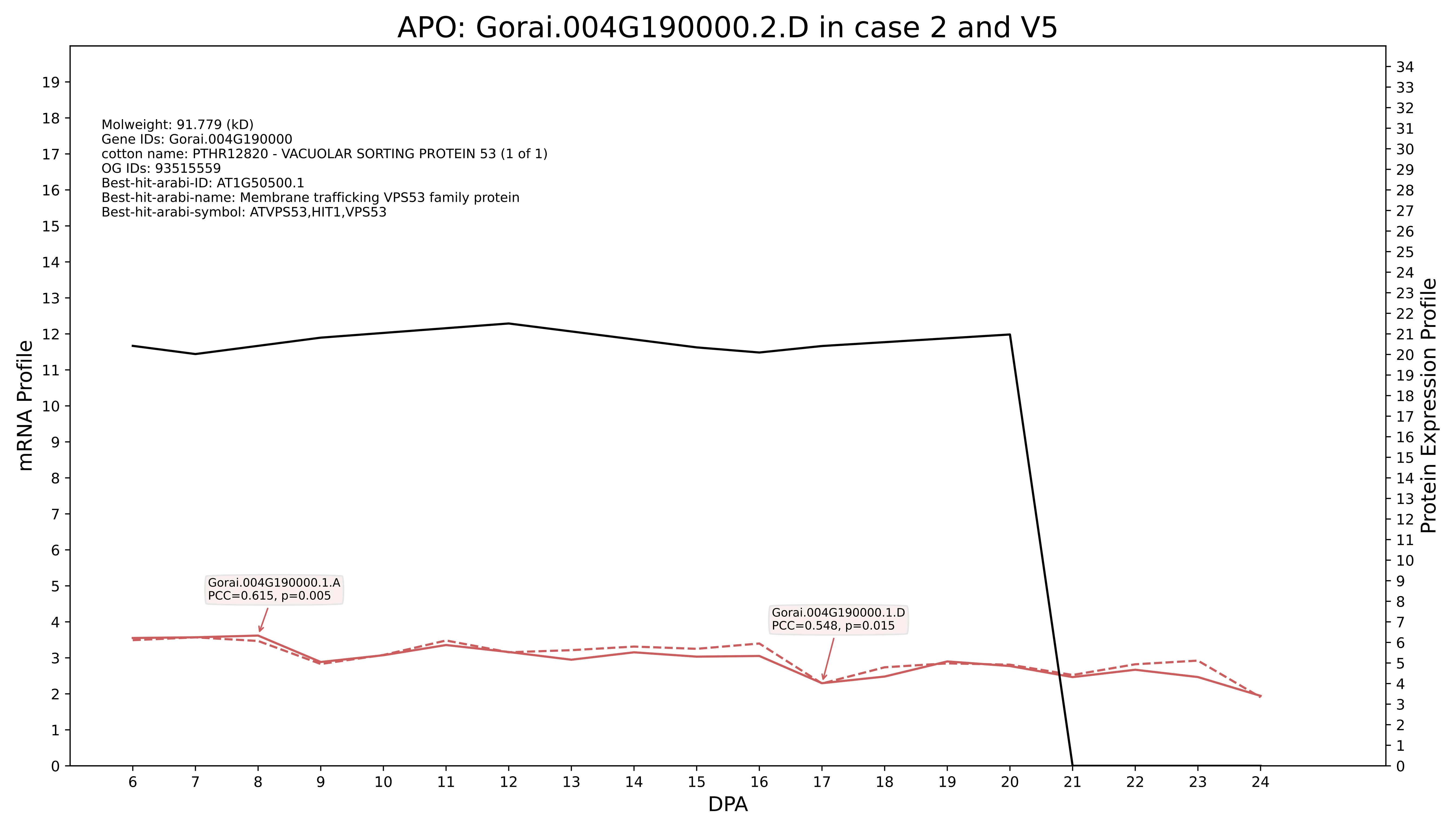

APO: Gorai.004G190000.2.D in case 2 and V5
Molweight: 91.779 (kD)
Gene IDs: Gorai.004G190000
cotton name: PTHR12820 - VACUOLAR SORTING PROTEIN 53 (1 of 1)
OG IDs: 93515559
Best-hit-arabi-ID: AT1G50500.1
Best-hit-arabi-name: Membrane trafficking VPS53 family protein
Best-hit-arabi-symbol: ATVPS53,HIT1,VPS53
Gorai.004G190000.1.A
PCC=0.615, p=0.005
Gorai.004G190000.1.D
PCC=0.548, p=0.015
mRNA Profile
Protein Expression Profile
DPA

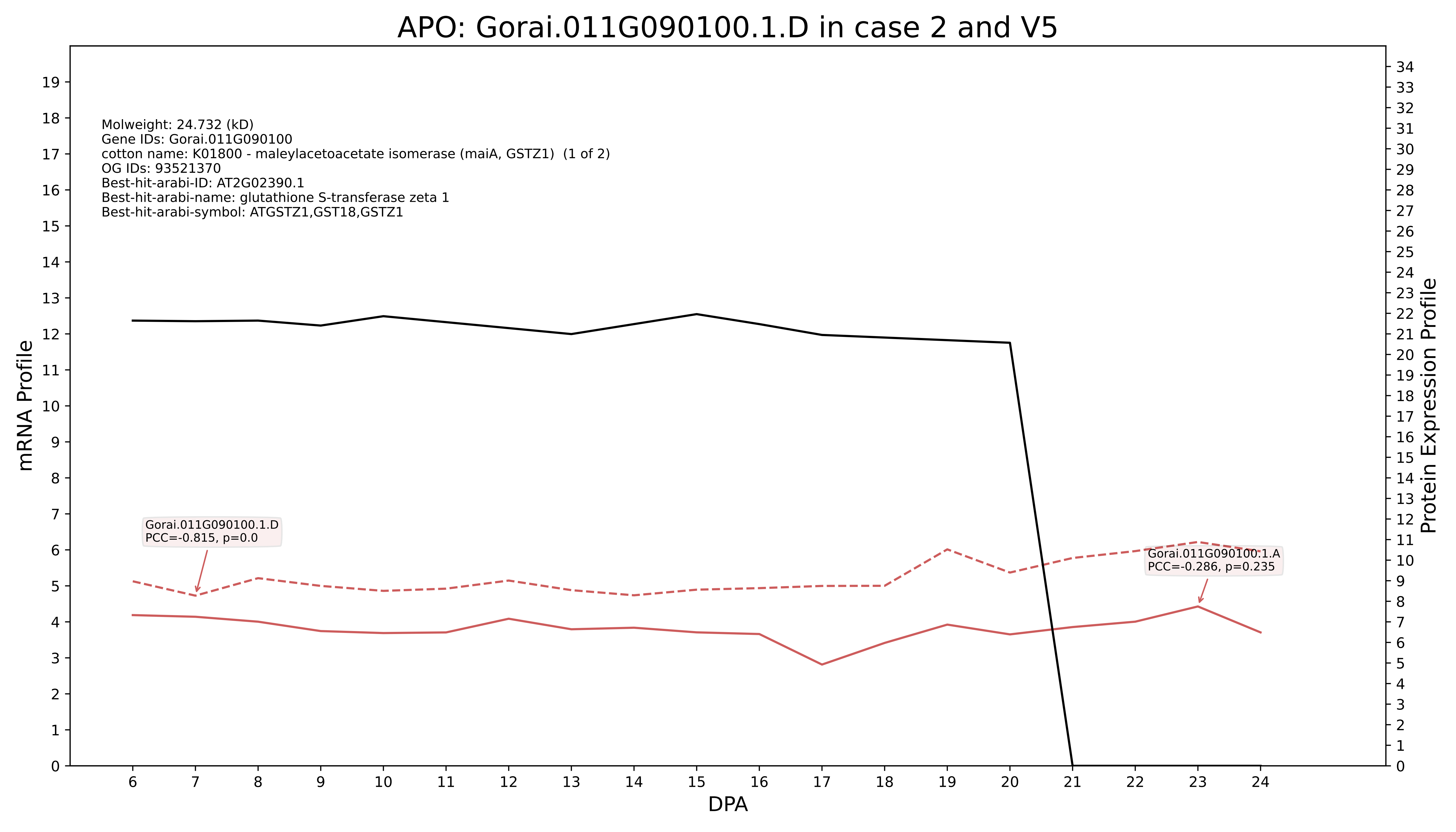

APO: Gorai.011G090100.1.D in case 2 and V5
Molweight: 24.732 (kD)
Gene IDs: Gorai.011G090100
cotton name: K01800 - maleylacetoacetate isomerase (maiA, GSTZ1) (1 of 2)
OG IDs: 93521370
Best-hit-arabi-ID: AT2G02390.1
Best-hit-arabi-name: glutathione S-transferase zeta 1
Best-hit-arabi-symbol: ATGSTZ1,GST18,GSTZ1
Gorai.011G090100.1.D
PCC=-0.815, p=0.0
Gorai.011G090100.1.A
PCC=-0.286, p=0.235
mRNA Profile
Protein Expression Profile
DPA

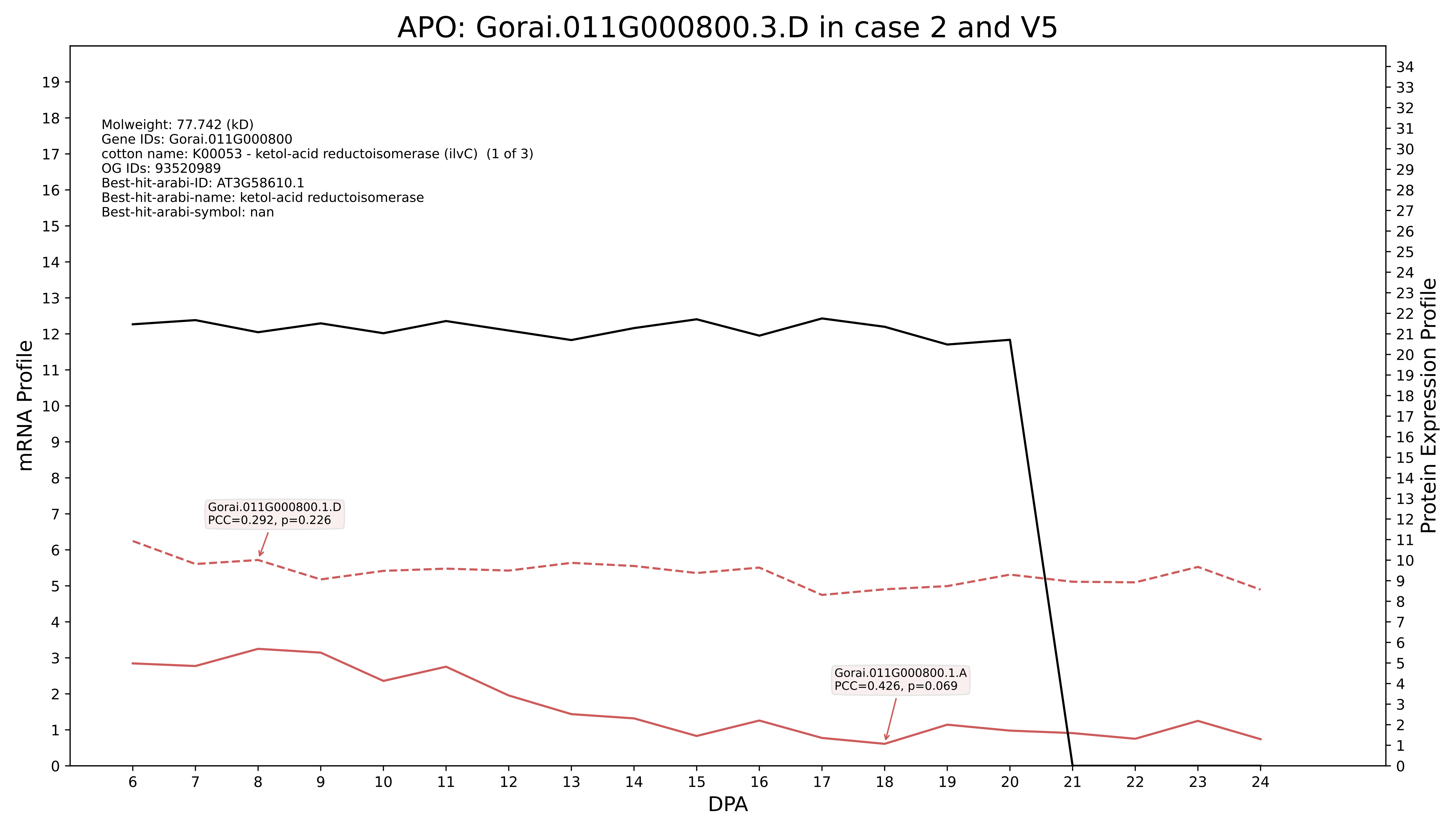

APO: Gorai.011G000800.3.D in case 2 and V5
Molweight: 77.742 (kD)
Gene IDs: Gorai.011G000800
cotton name: K00053 - ketol-acid reductoisomerase (ilvC) (1 of 3)
OG IDs: 93520989
Best-hit-arabi-ID: AT3G58610.1
Best-hit-arabi-name: ketol-acid reductoisomerase
Best-hit-arabi-symbol: nan
Gorai.011G000800.1.D
PCC=0.292, p=0.226
Gorai.011G000800.1.A
PCC=0.426, p=0.069
mRNA Profile
Protein Expression Profile
DPA

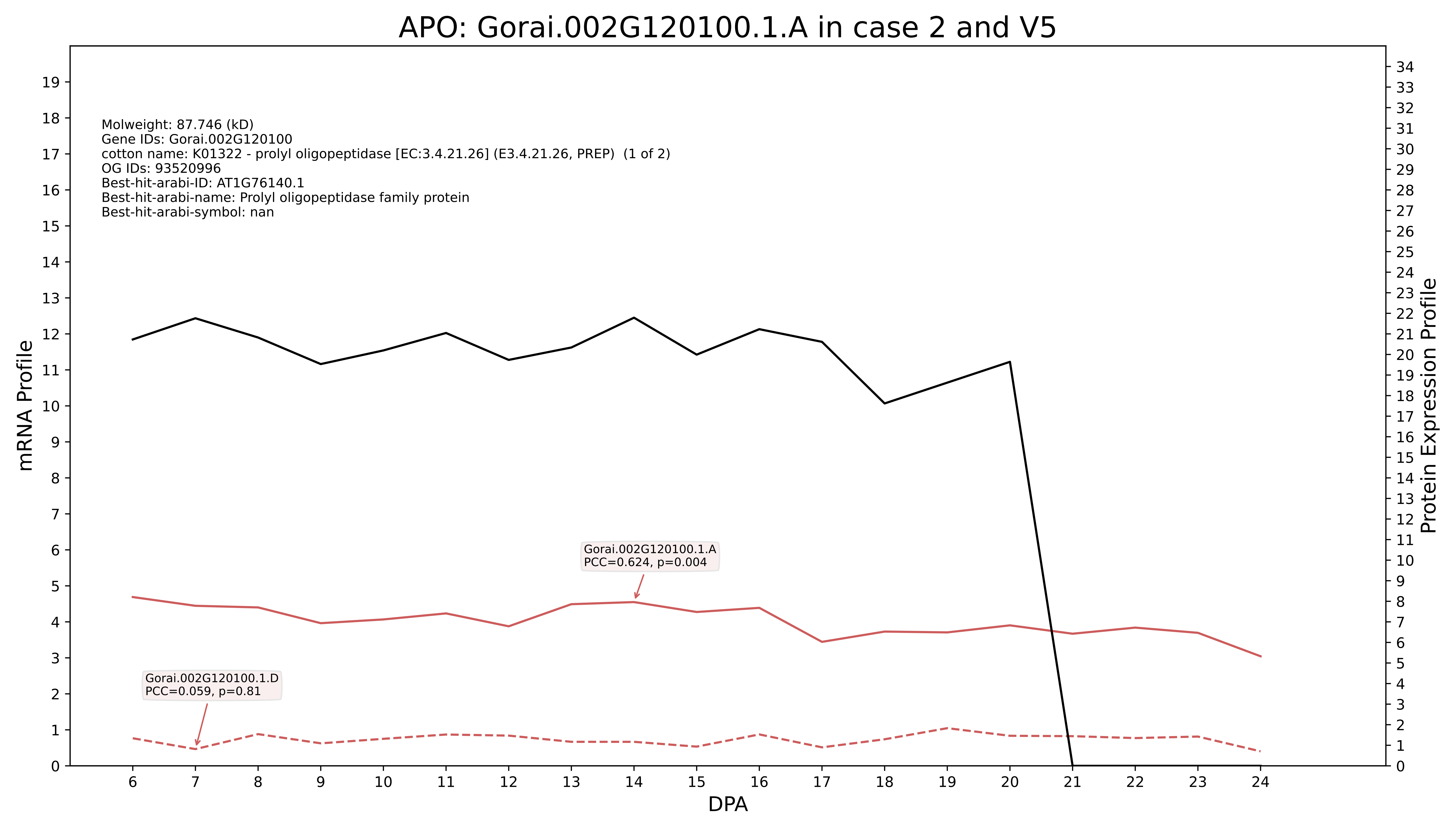

APO: Gorai.002G120100.1.A in case 2 and V5
Molweight: 87.746 (kD)
Gene IDs: Gorai.002G120100
cotton name: K01322 - prolyl oligopeptidase [EC:3.4.21.26] (E3.4.21.26, PREP) (1 of 2)
OG IDs: 93520996
Best-hit-arabi-ID: AT1G76140.1
Best-hit-arabi-name: Prolyl oligopeptidase family protein
Best-hit-arabi-symbol: nan
Gorai.002G120100.1.A
PCC=0.624, p=0.004
Gorai.002G120100.1.D
PCC=0.059, p=0.81
mRNA Profile
Protein Expression Profile
DPA

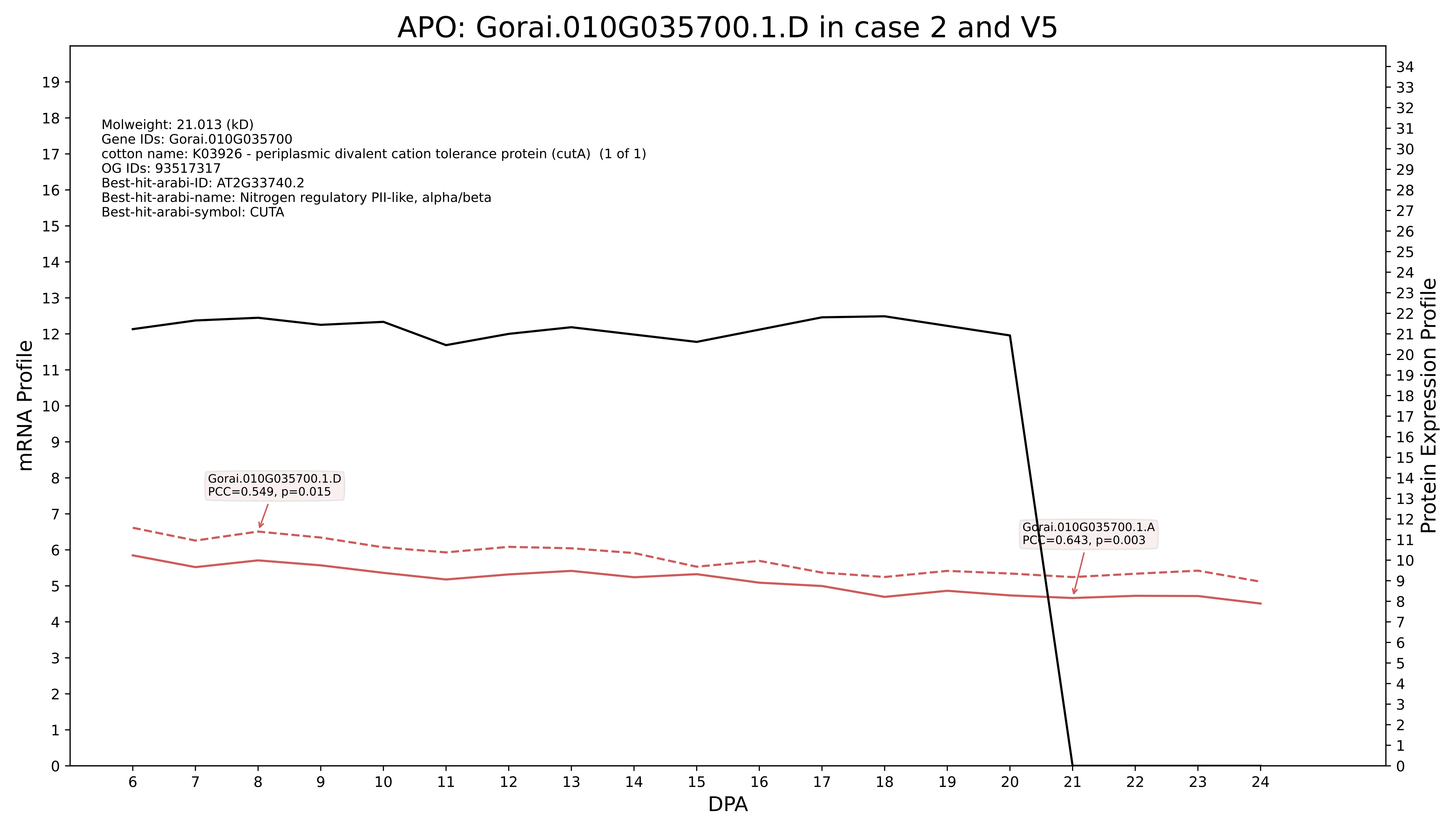
APO: Gorai.010G035700.1.D in case 2 and V5
Molweight: 21.013 (kD)
Gene IDs: Gorai.010G035700
cotton name: K03926 - periplasmic divalent cation tolerance protein (cutA) (1 of 1)
OG IDs: 93517317
Best-hit-arabi-ID: AT2G33740.2
Best-hit-arabi-name: Nitrogen regulatory PII-like, alpha/beta
Best-hit-arabi-symbol: CUTA
Gorai.010G035700.1.D
PCC=0.549, p=0.015
Gorai.010G035700.1.A
PCC=0.643, p=0.003
mRNA Profile
Protein Expression Profile
DPA

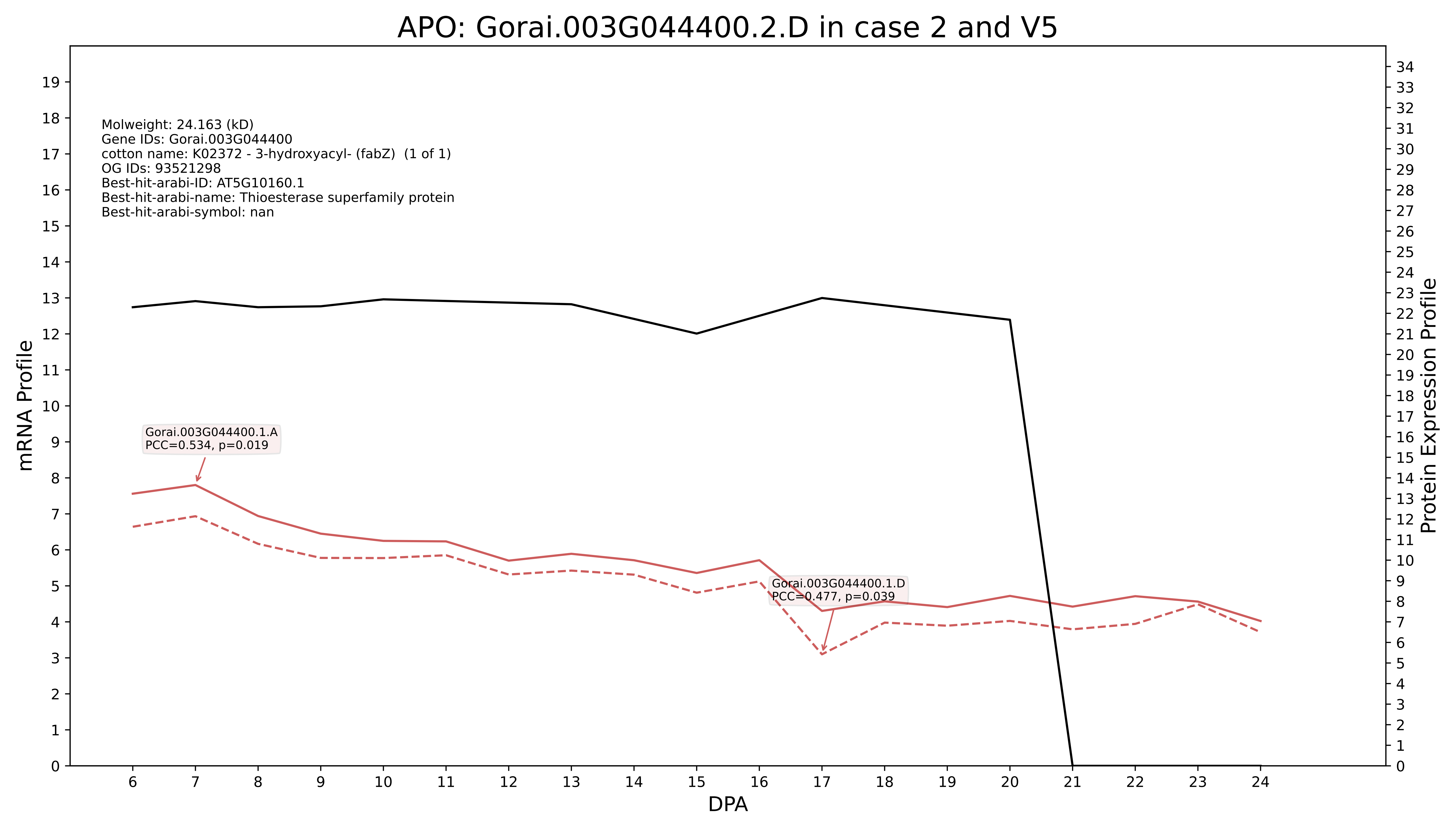
APO: Gorai.003G044400.2.D in case 2 and V5
Molweight: 24.163 (kD)
Gene IDs: Gorai.003G044400
cotton name: K02372 - 3-hydroxyacyl- (fabZ) (1 of 1)
OG IDs: 93521298
Best-hit-arabi-ID: AT5G10160.1
Best-hit-arabi-name: Thioesterase superfamily protein
Best-hit-arabi-symbol: nan
Gorai.003G044400.1.A
PCC=0.534, p=0.019
Gorai.003G044400.1.D
PCC=0.477, p=0.039
mRNA Profile
Protein Expression Profile
DPA

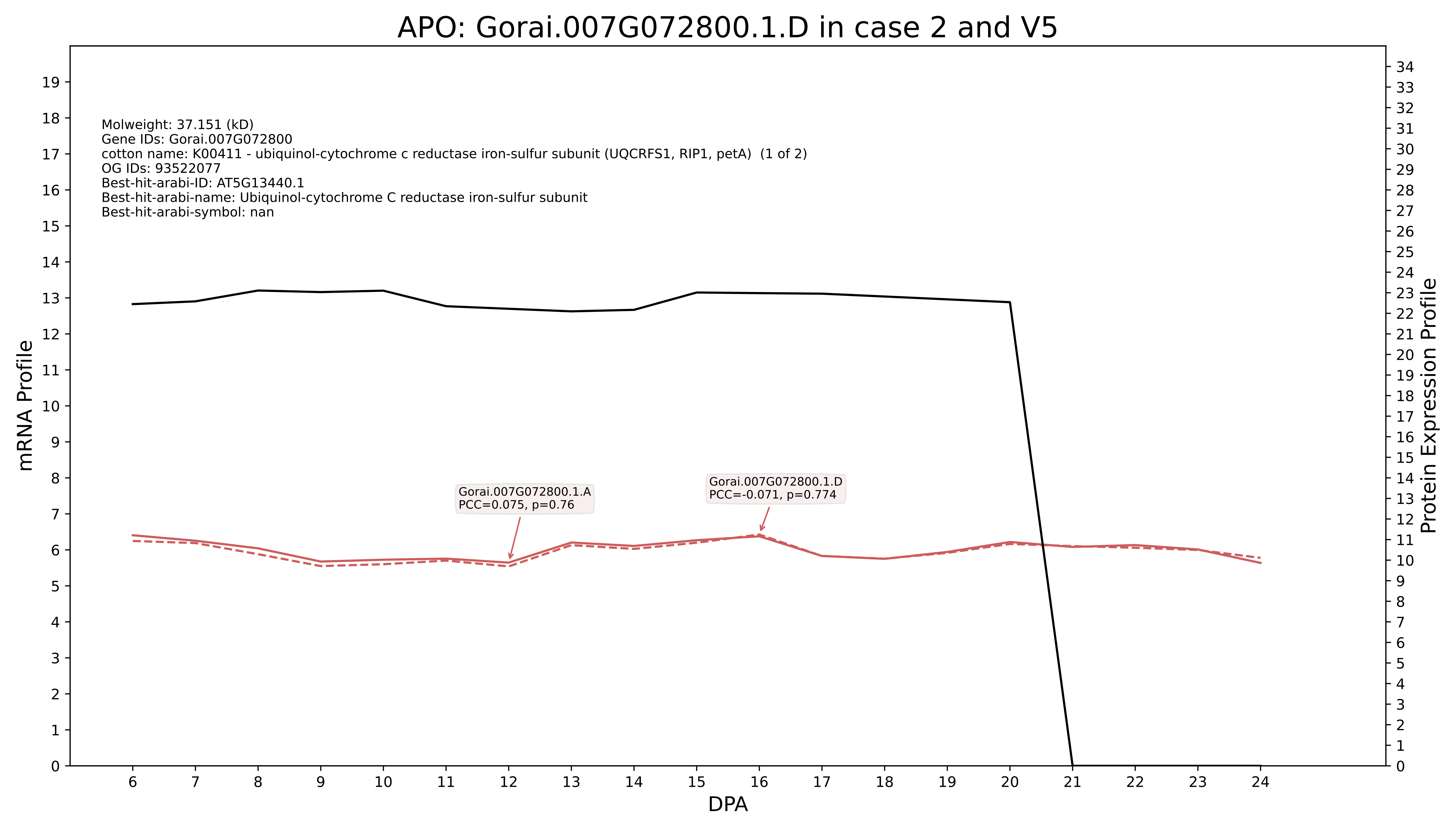

APO: Gorai.007G072800.1.D in case 2 and V5
Molweight: 37.151 (kD)
Gene IDs: Gorai.007G072800
cotton name: K00411 - ubiquinol-cytochrome c reductase iron-sulfur subunit (UQCRFS1, RIP1, petA) (1 of 2)
OG IDs: 93522077
Best-hit-arabi-ID: AT5G13440.1
Best-hit-arabi-name: Ubiquinol-cytochrome C reductase iron-sulfur subunit
Best-hit-arabi-symbol: nan
Gorai.007G072800.1.A
PCC=0.075, p=0.76
Gorai.007G072800.1.D
PCC=-0.071, p=0.774
mRNA Profile
Protein Expression Profile
DPA

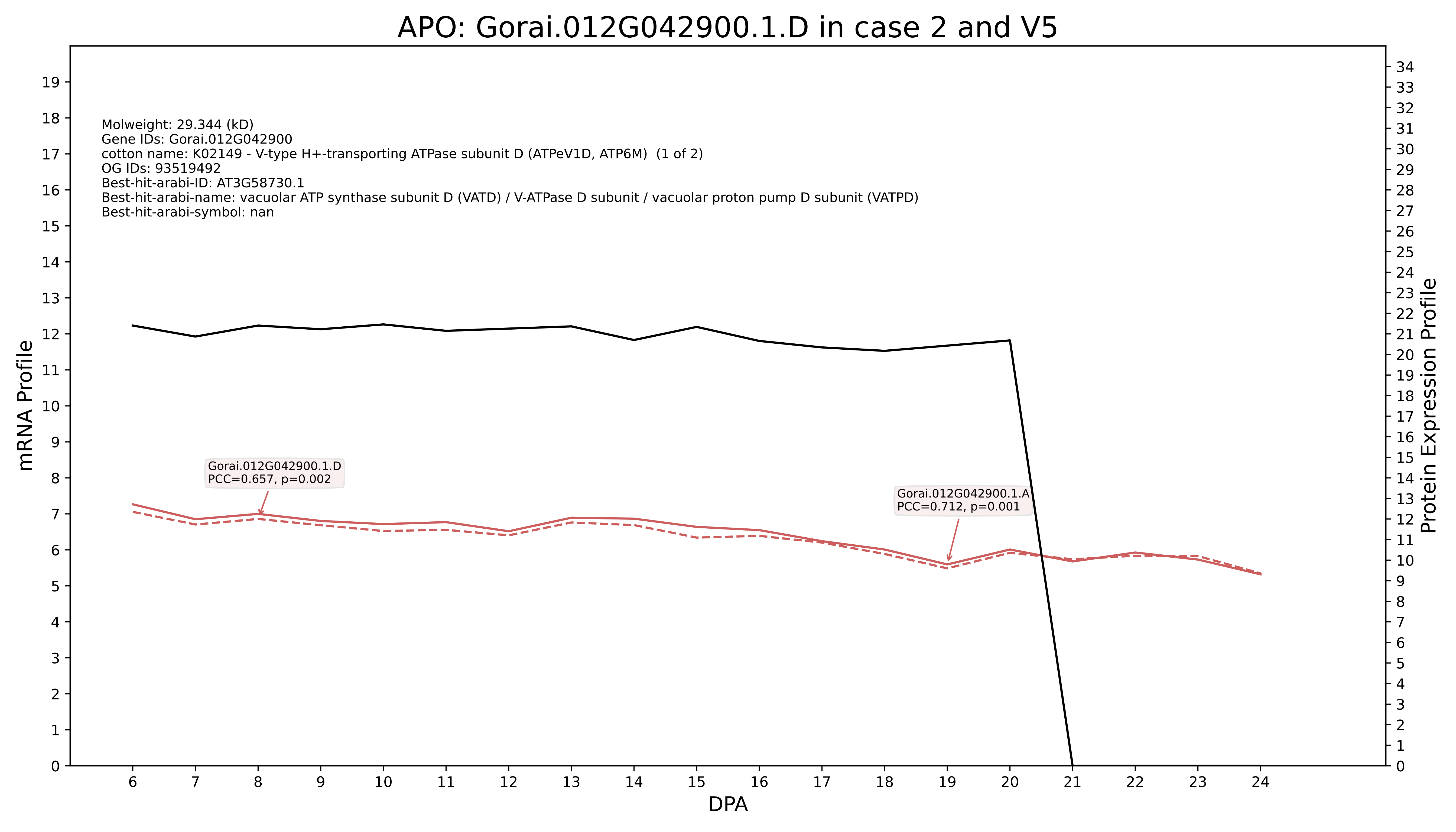

APO: Gorai.012G042900.1.D in case 2 and V5
Molweight: 29.344 (kD)
Gene IDs: Gorai.012G042900
cotton name: K02149 - V-type H+-transporting ATPase subunit D (ATPeV1D, ATP6M) (1 of 2)
OG IDs: 93519492
Best-hit-arabi-ID: AT3G58730.1
Best-hit-arabi-name: vacuolar ATP synthase subunit D (VATD) / V-ATPase D subunit / vacuolar proton pump D subunit (VATPD)
Best-hit-arabi-symbol: nan
Gorai.012G042900.1.D
PCC=0.657, p=0.002
Gorai.012G042900.1.A
PCC=0.712, p=0.001
mRNA Profile
Protein Expression Profile
DPA

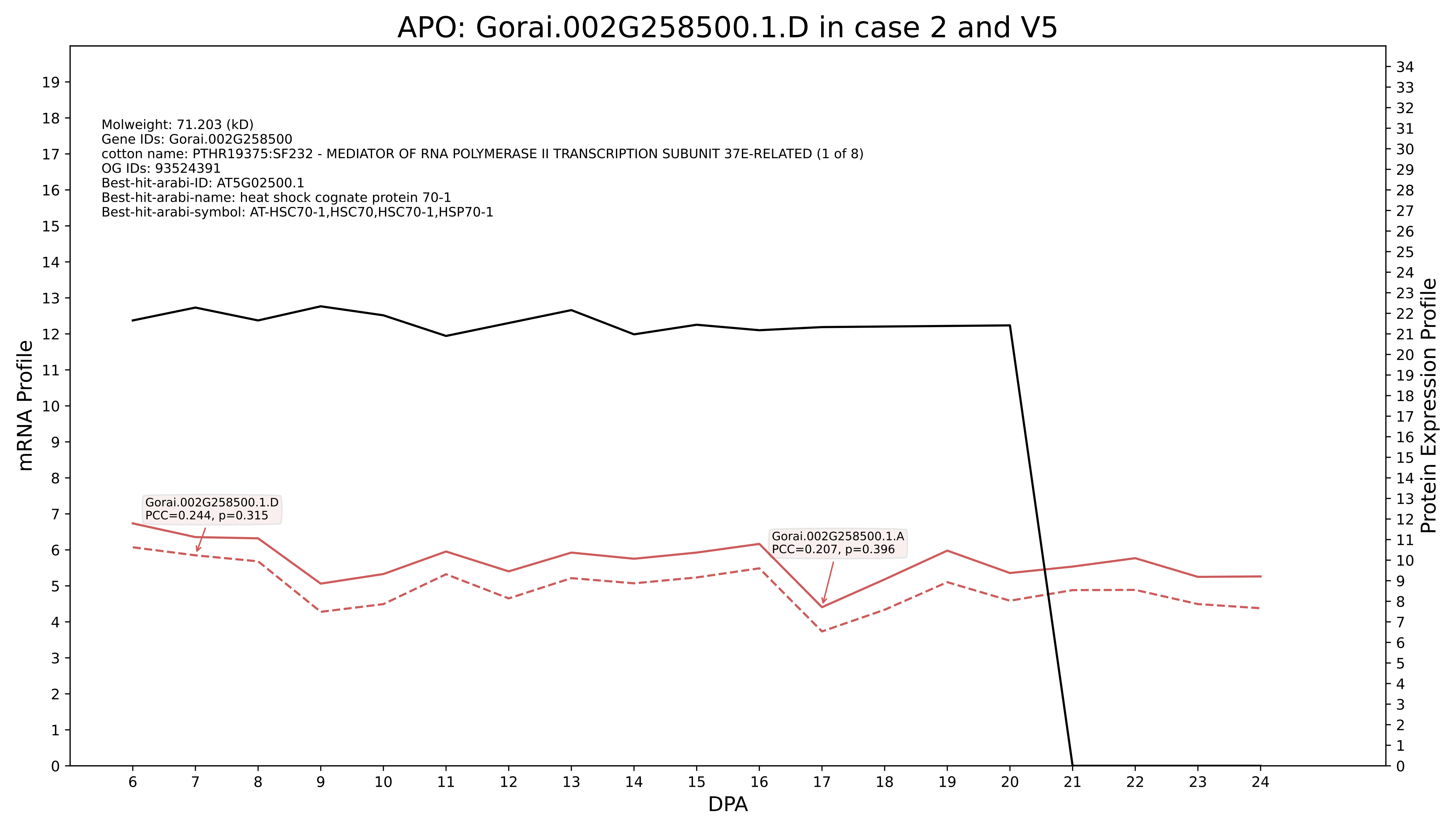
APO: Gorai.002G258500.1.D in case 2 and V5
Molweight: 71.203 (kD)
Gene IDs: Gorai.002G258500
cotton name: PTHR19375:SF232 - MEDIATOR OF RNA POLYMERASE II TRANSCRIPTION SUBUNIT 37E-RELATED (1 of 8)
OG IDs: 93524391
Best-hit-arabi-ID: AT5G02500.1
Best-hit-arabi-name: heat shock cognate protein 70-1
Best-hit-arabi-symbol: AT-HSC70-1,HSC70,HSC70-1,HSP70-1
Gorai.002G258500.1.D
PCC=0.244, p=0.315
Gorai.002G258500.1.A
PCC=0.207, p=0.396
mRNA Profile
Protein Expression Profile
DPA

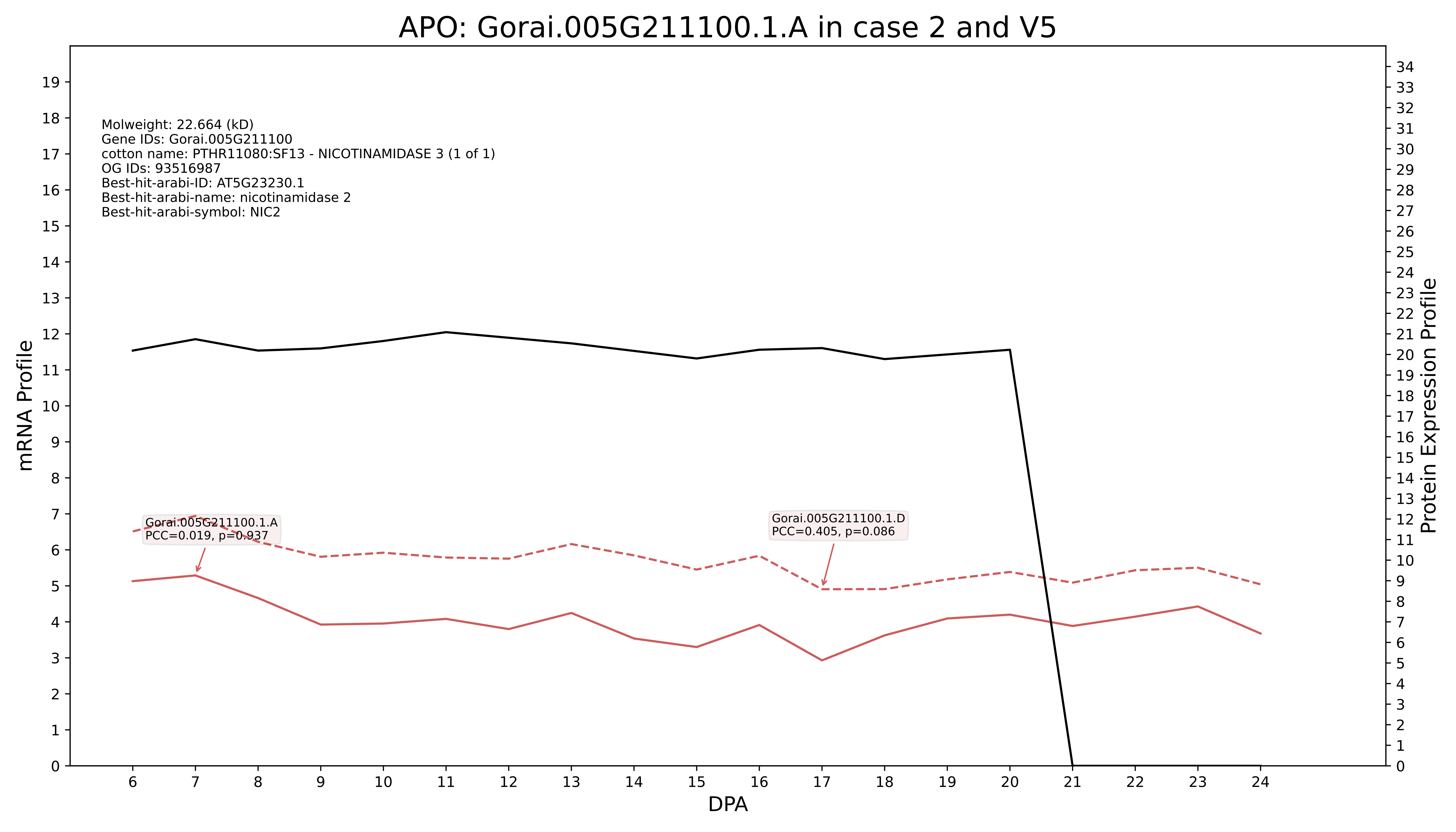
APO: Gorai.005G211100.1.A in case 2 and V5
Molweight: 22.664 (kD)
Gene IDs: Gorai.005G211100
cotton name: PTHR11080:SF13 - NICOTINAMIDASE 3 (1 of 1)
OG IDs: 93516987
Best-hit-arabi-ID: AT5G23230.1
Best-hit-arabi-name: nicotinamidase 2
Best-hit-arabi-symbol: NIC2
Gorai.005G211100.1.A
PCC=0.019, p=0.937
Gorai.005G211100.1.D
PCC=0.405, p=0.086
mRNA Profile
Protein Expression Profile
DPA

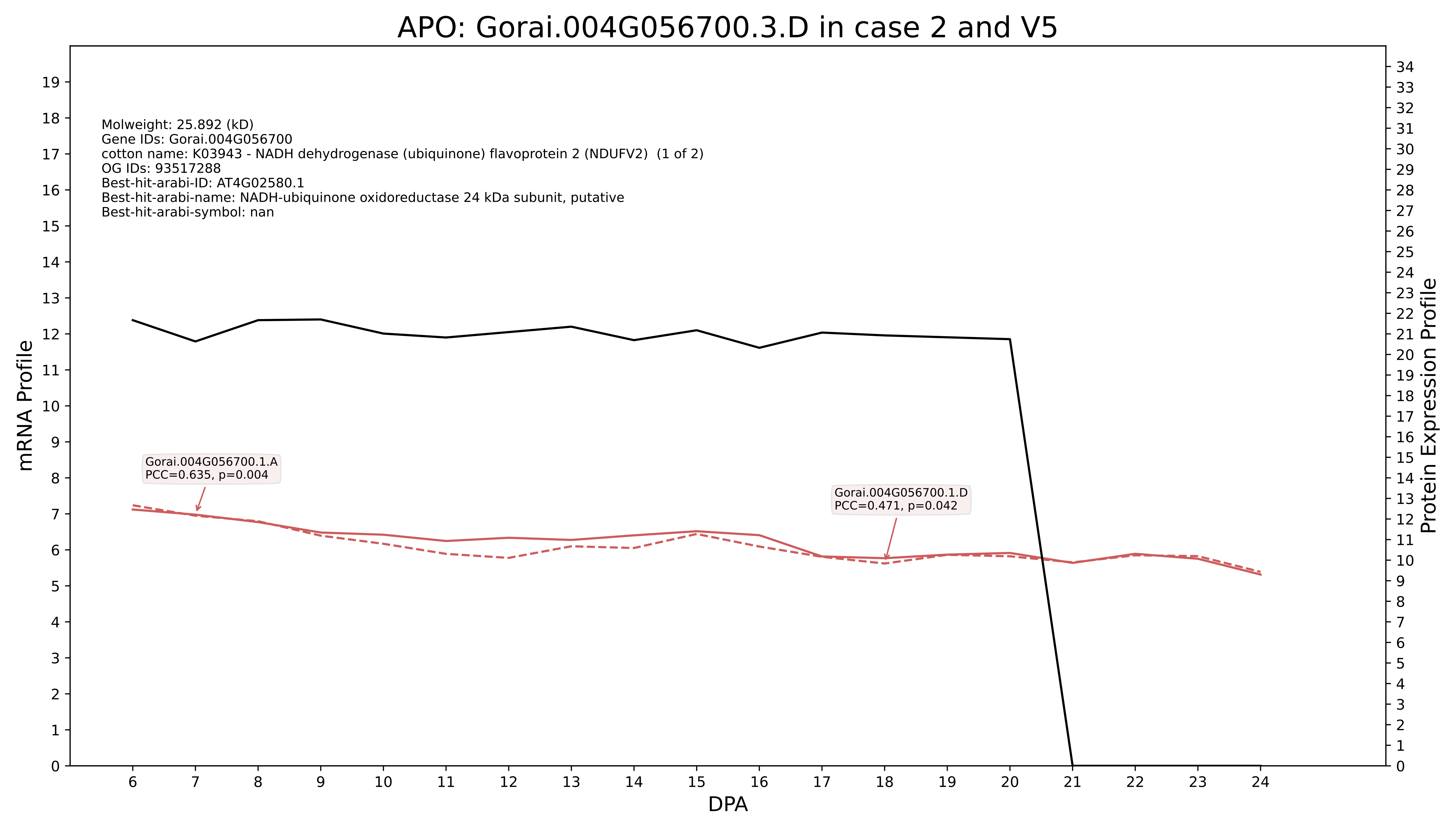

APO: Gorai.004G056700.3.D in case 2 and V5
Molweight: 25.892 (kD)
Gene IDs: Gorai.004G056700
cotton name: K03943 - NADH dehydrogenase (ubiquinone) flavoprotein 2 (NDUFV2) (1 of 2)
OG IDs: 93517288
Best-hit-arabi-ID: AT4G02580.1
Best-hit-arabi-name: NADH-ubiquinone oxidoreductase 24 kDa subunit, putative
Best-hit-arabi-symbol: nan
Gorai.004G056700.1.A
PCC=0.635, p=0.004
Gorai.004G056700.1.D
PCC=0.471, p=0.042
mRNA Profile
Protein Expression Profile
DPA

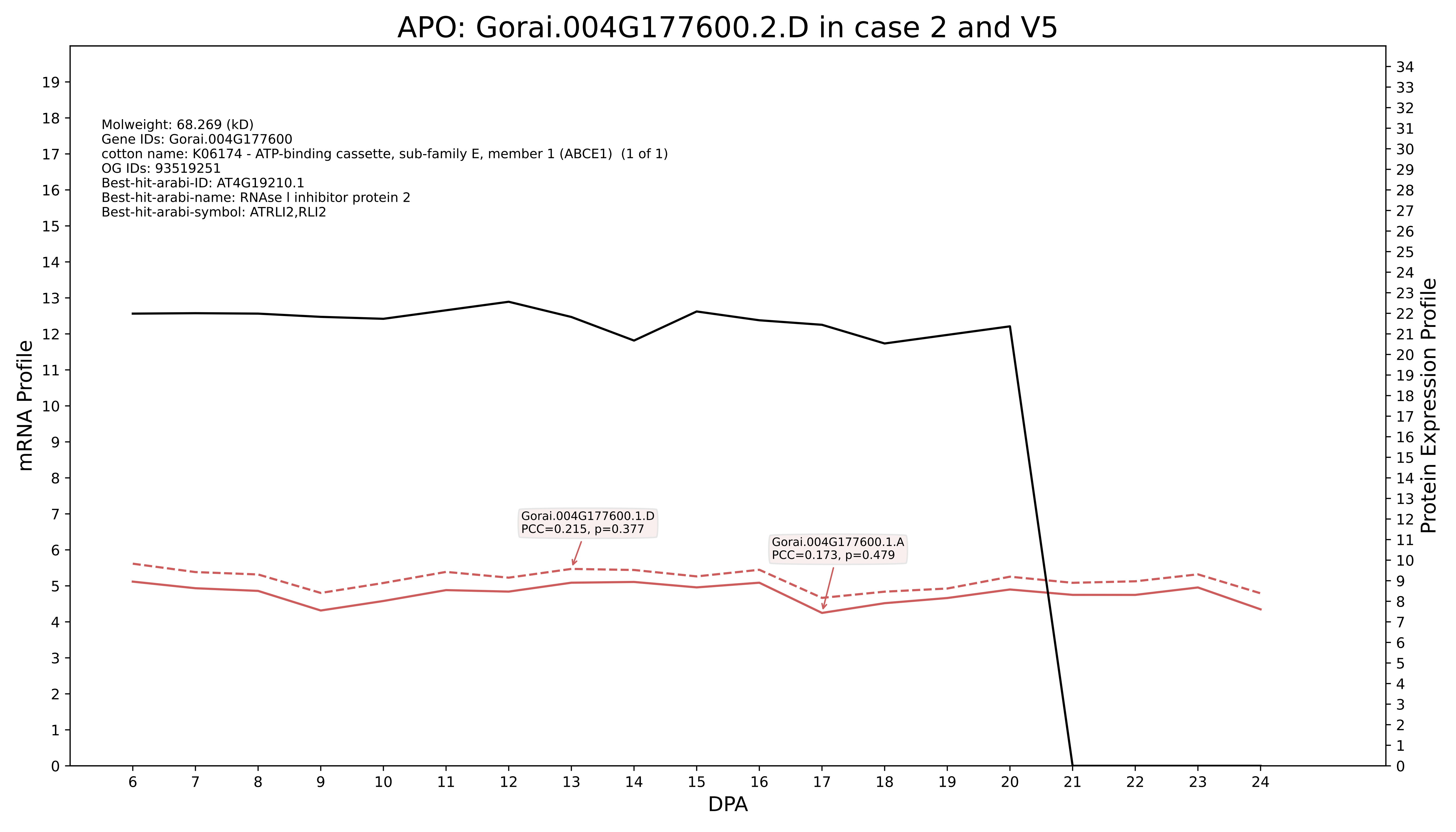
APO: Gorai.004G177600.2.D in case 2 and V5
Molweight: 68.269 (kD)
Gene IDs: Gorai.004G177600
cotton name: K06174 - ATP-binding cassette, sub-family E, member 1 (ABCE1) (1 of 1)
OG IDs: 93519251
Best-hit-arabi-ID: AT4G19210.1
Best-hit-arabi-name: RNAse l inhibitor protein 2
Best-hit-arabi-symbol: ATRLI2,RLI2
Gorai.004G177600.1.D
PCC=0.215, p=0.377
Gorai.004G177600.1.A
PCC=0.173, p=0.479
mRNA Profile
Protein Expression Profile
DPA

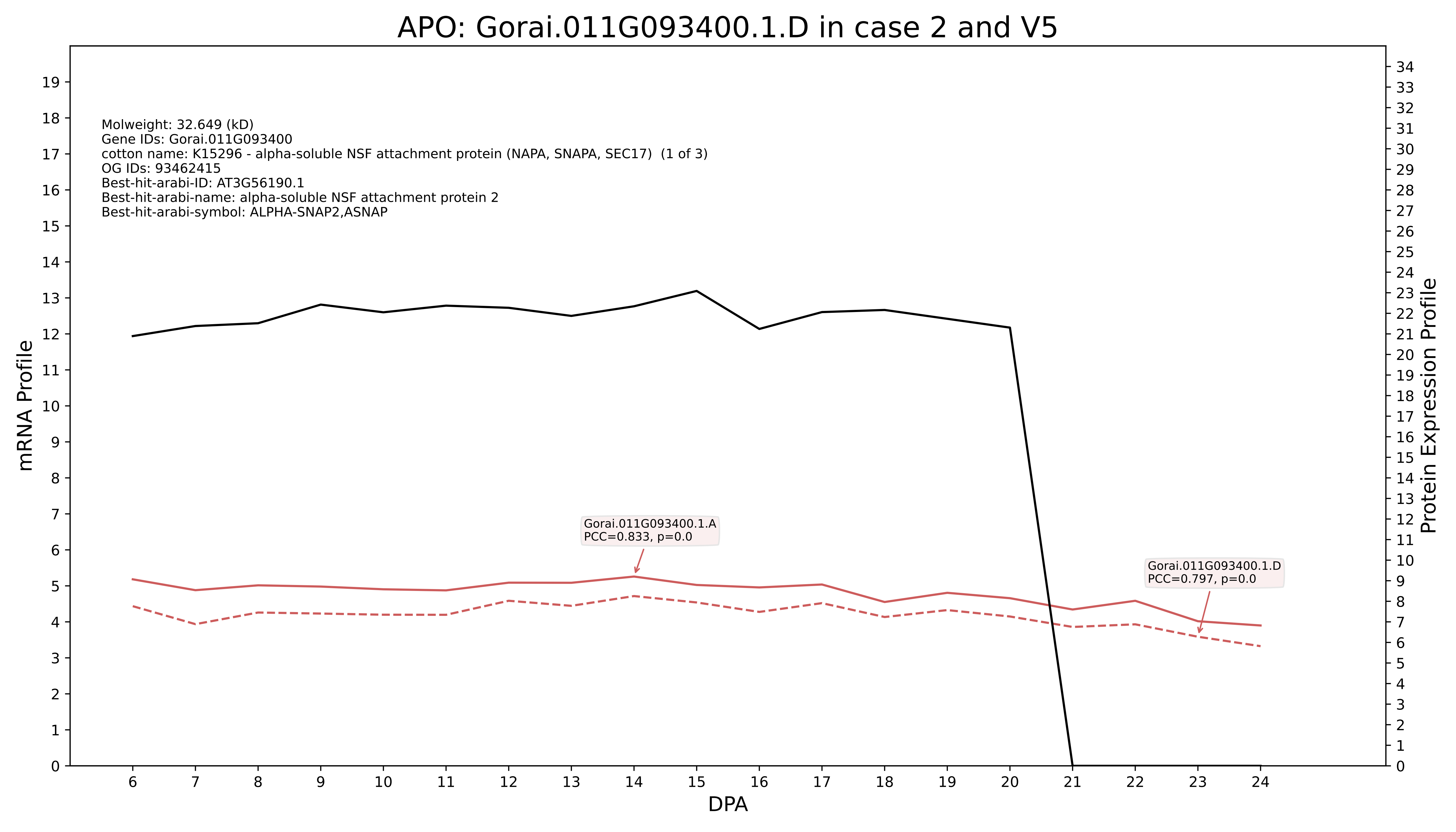

APO: Gorai.011G093400.1.D in case 2 and V5
Molweight: 32.649 (kD)
Gene IDs: Gorai.011G093400
cotton name: K15296 - alpha-soluble NSF attachment protein (NAPA, SNAPA, SEC17) (1 of 3)
OG IDs: 93462415
Best-hit-arabi-ID: AT3G56190.1
Best-hit-arabi-name: alpha-soluble NSF attachment protein 2
Best-hit-arabi-symbol: ALPHA-SNAP2,ASNAP
Gorai.011G093400.1.A
PCC=0.833, p=0.0
Gorai.011G093400.1.D
PCC=0.797, p=0.0
mRNA Profile
Protein Expression Profile
DPA

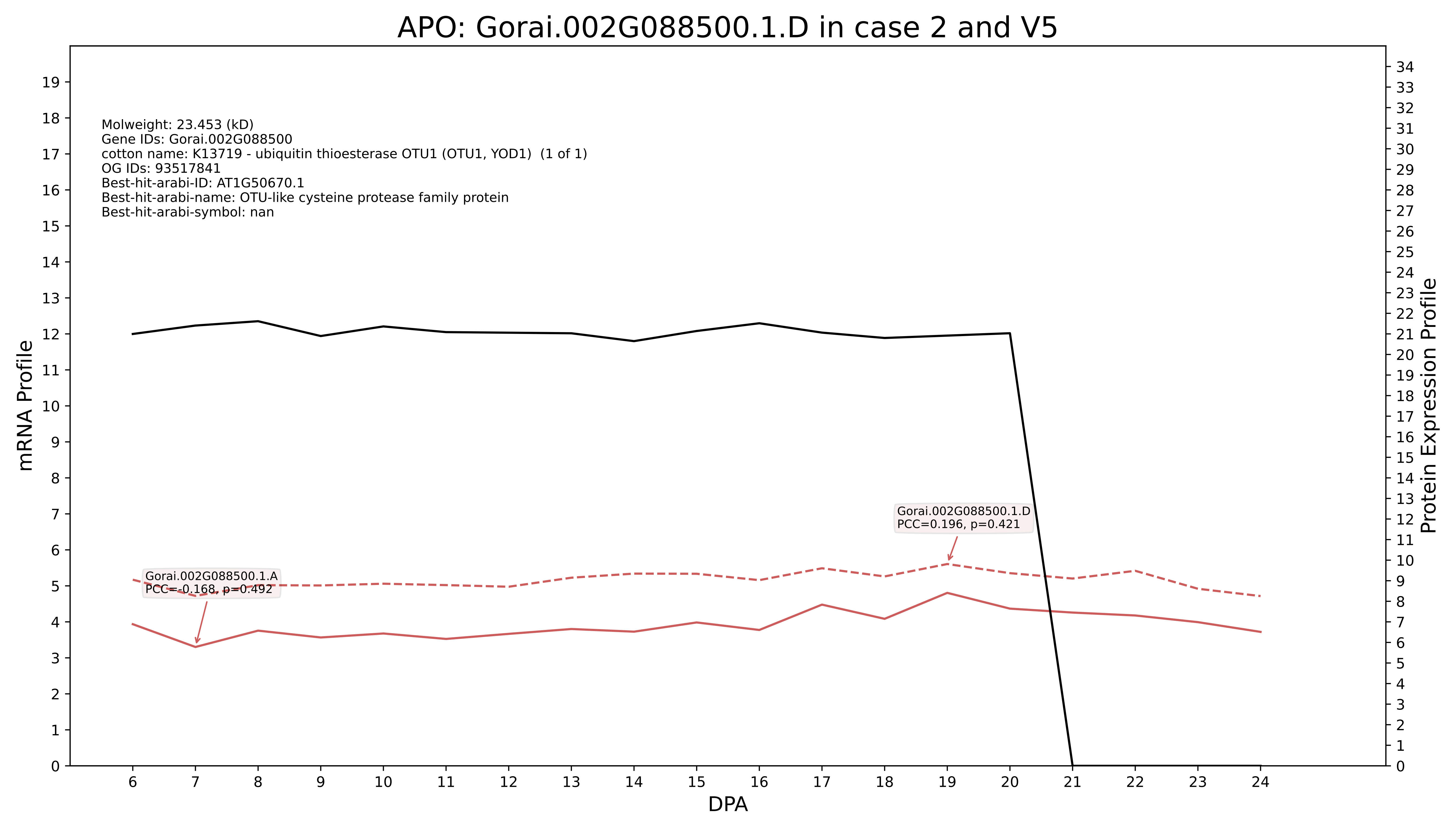
APO: Gorai.002G088500.1.D in case 2 and V5
Molweight: 23.453 (kD)
Gene IDs: Gorai.002G088500
cotton name: K13719 - ubiquitin thioesterase OTU1 (OTU1, YOD1) (1 of 1)
OG IDs: 93517841
Best-hit-arabi-ID: AT1G50670.1
Best-hit-arabi-name: OTU-like cysteine protease family protein
Best-hit-arabi-symbol: nan
Gorai.002G088500.1.D
PCC=0.196, p=0.421
Gorai.002G088500.1.A
PCC=-0.168, p=0.492
mRNA Profile
Protein Expression Profile
DPA

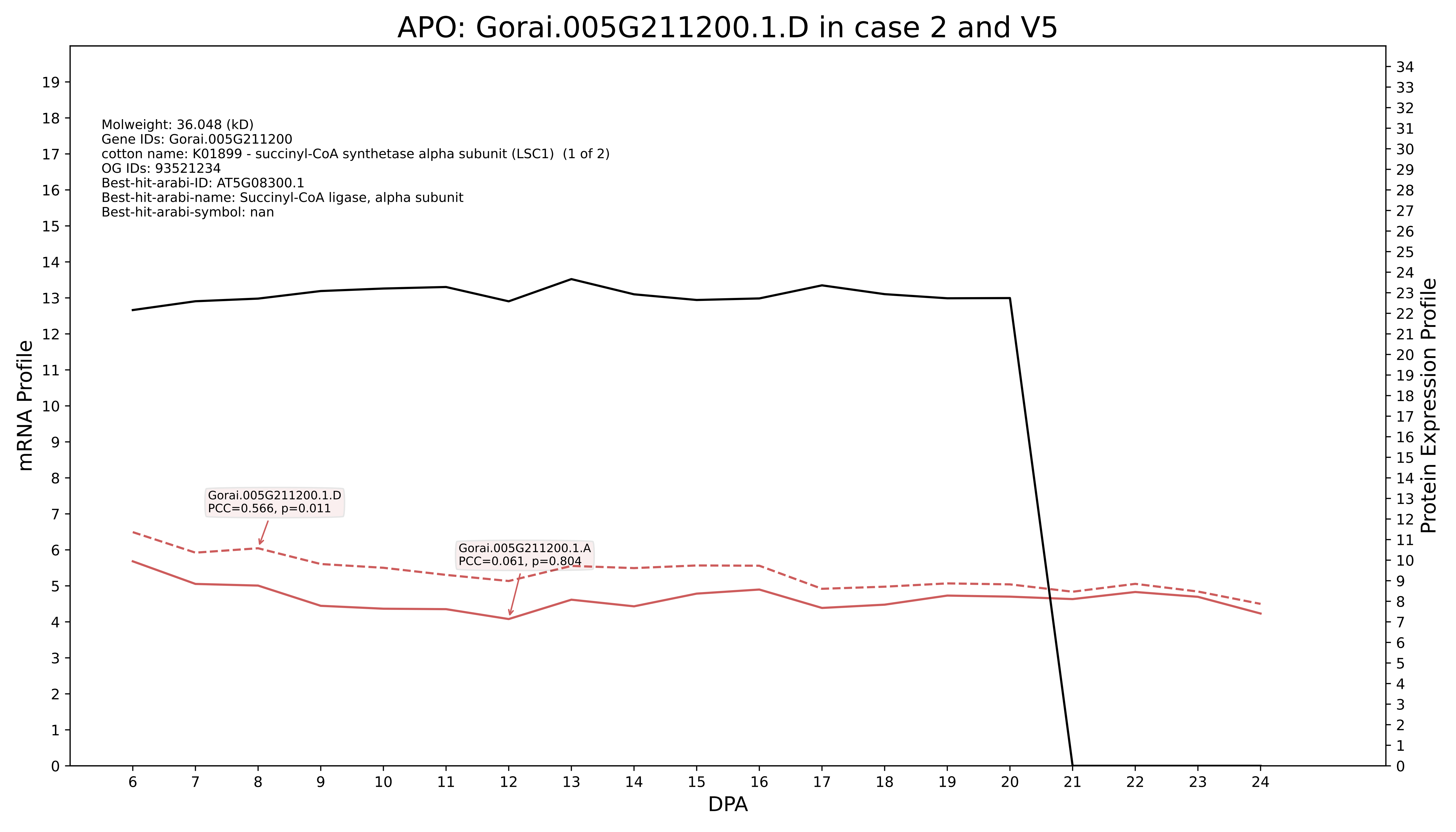
APO: Gorai.005G211200.1.D in case 2 and V5
Molweight: 36.048 (kD)
Gene IDs: Gorai.005G211200
cotton name: K01899 - succinyl-CoA synthetase alpha subunit (LSC1) (1 of 2)
OG IDs: 93521234
Best-hit-arabi-ID: AT5G08300.1
Best-hit-arabi-name: Succinyl-CoA ligase, alpha subunit
Best-hit-arabi-symbol: nan
Gorai.005G211200.1.D
PCC=0.566, p=0.011
Gorai.005G211200.1.A
PCC=0.061, p=0.804
mRNA Profile
Protein Expression Profile
DPA

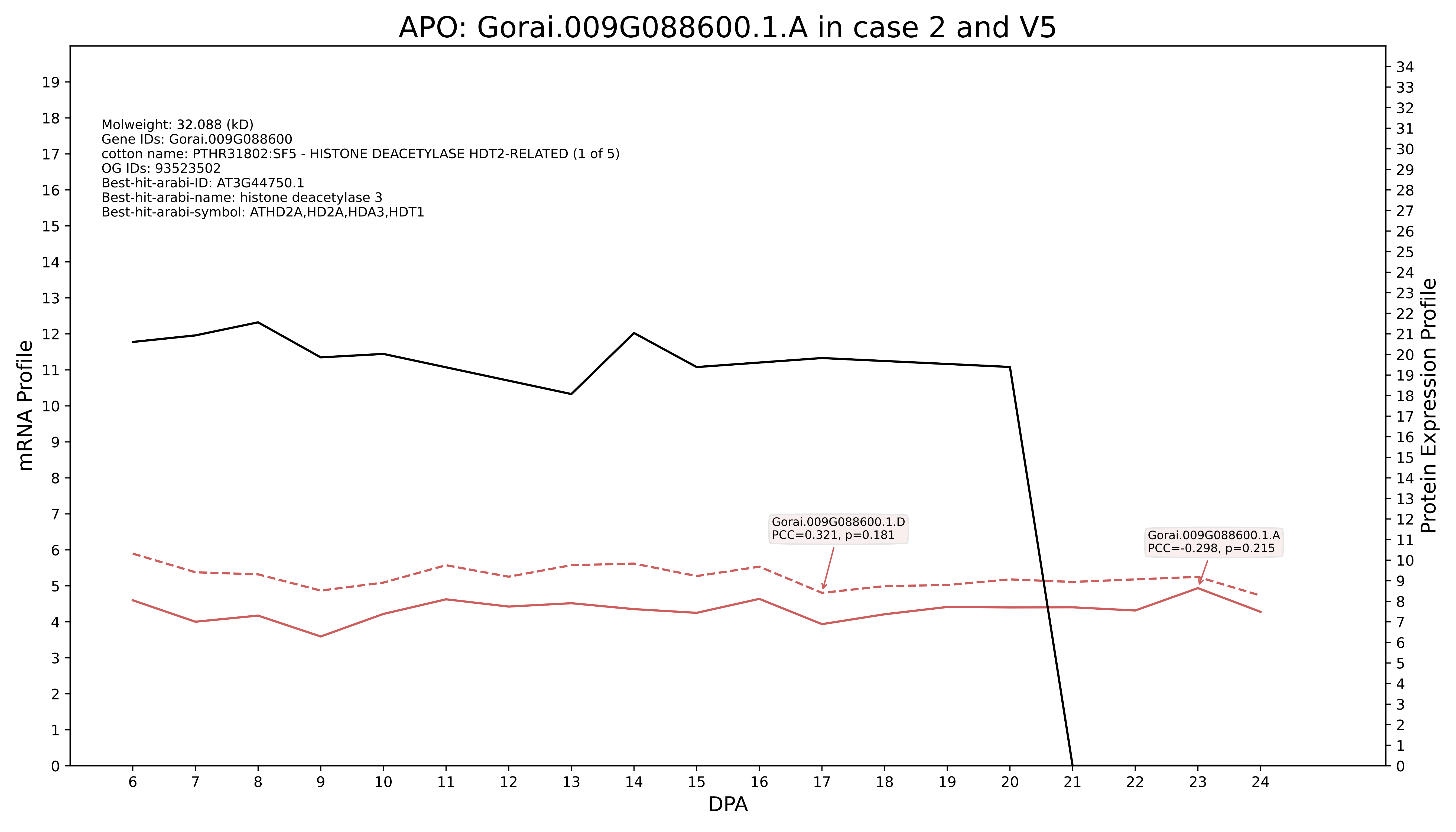
APO: Gorai.009G088600.1.A in case 2 and V5
Molweight: 32.088 (kD)
Gene IDs: Gorai.009G088600
cotton name: PTHR31802:SF5 - HISTONE DEACETYLASE HDT2-RELATED (1 of 5)
OG IDs: 93523502
Best-hit-arabi-ID: AT3G44750.1
Best-hit-arabi-name: histone deacetylase 3
Best-hit-arabi-symbol: ATHD2A,HD2A,HDA3,HDT1
Gorai.009G088600.1.D
PCC=0.321, p=0.181
Gorai.009G088600.1.A
PCC=-0.298, p=0.215
mRNA Profile
Protein Expression Profile
DPA

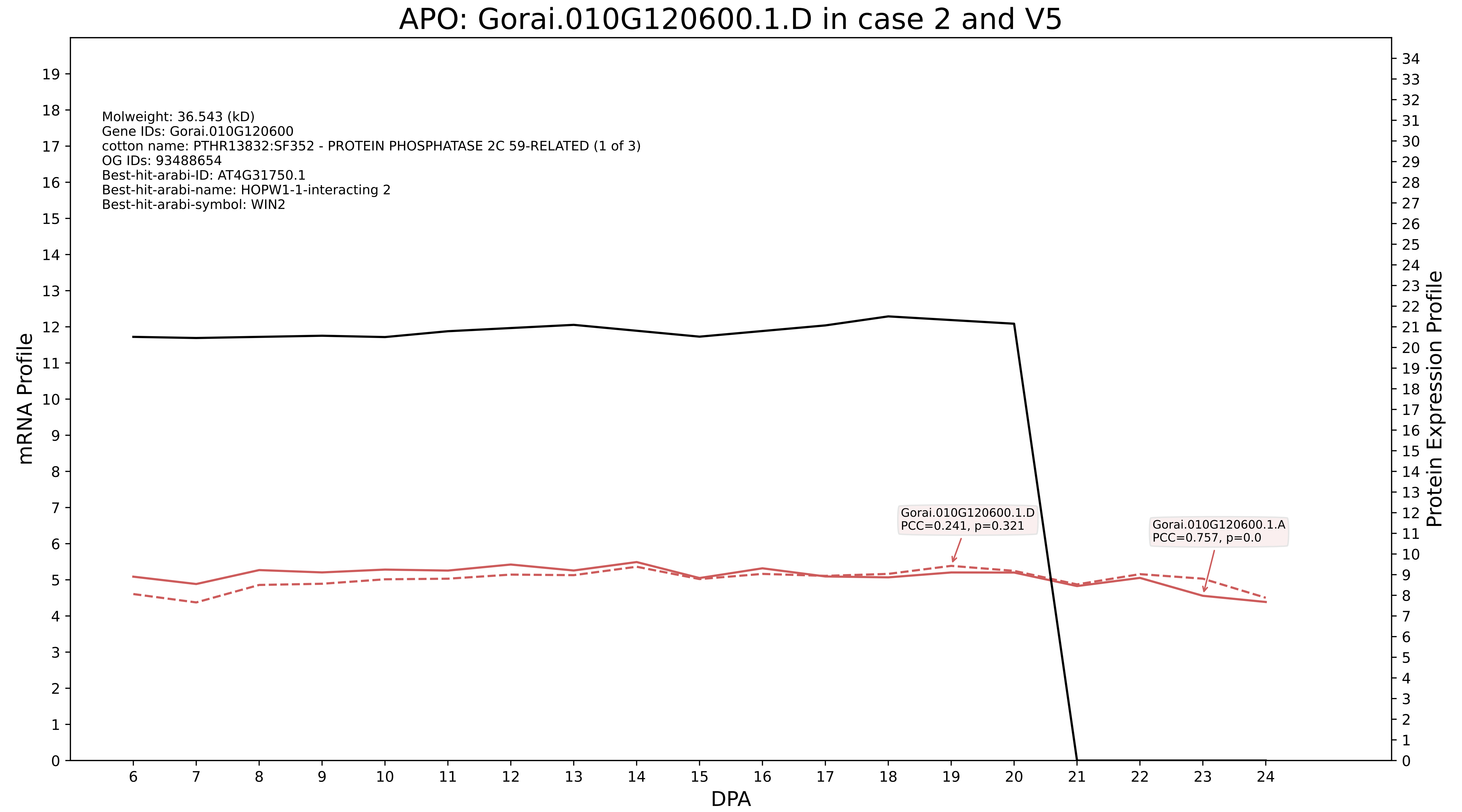
APO: Gorai.010G120600.1.D in case 2 and V5
Molweight: 36.543 (kD)
Gene IDs: Gorai.010G120600
cotton name: PTHR13832:SF352 - PROTEIN PHOSPHATASE 2C 59-RELATED (1 of 3)
OG IDs: 93488654
Best-hit-arabi-ID: AT4G31750.1
Best-hit-arabi-name: HOPW1-1-interacting 2
Best-hit-arabi-symbol: WIN2
Gorai.010G120600.1.D
PCC=0.241, p=0.321
Gorai.010G120600.1.A
PCC=0.757, p=0.0
mRNA Profile
Protein Expression Profile
DPA
6 7 8 9 10 11 12 13 14 15 16 17 18 19 20 21 22 23 24
0 1 2 3 4 5 6 7 8 9 10 11 12 13 14 15 16 17 18 19
0 1 2 3 4 5 6 7 8 9 10 11 12 13 14 15 16 17 18 19 20 21 22 23 24 25 26 27 28 29 30 31 32 33 34

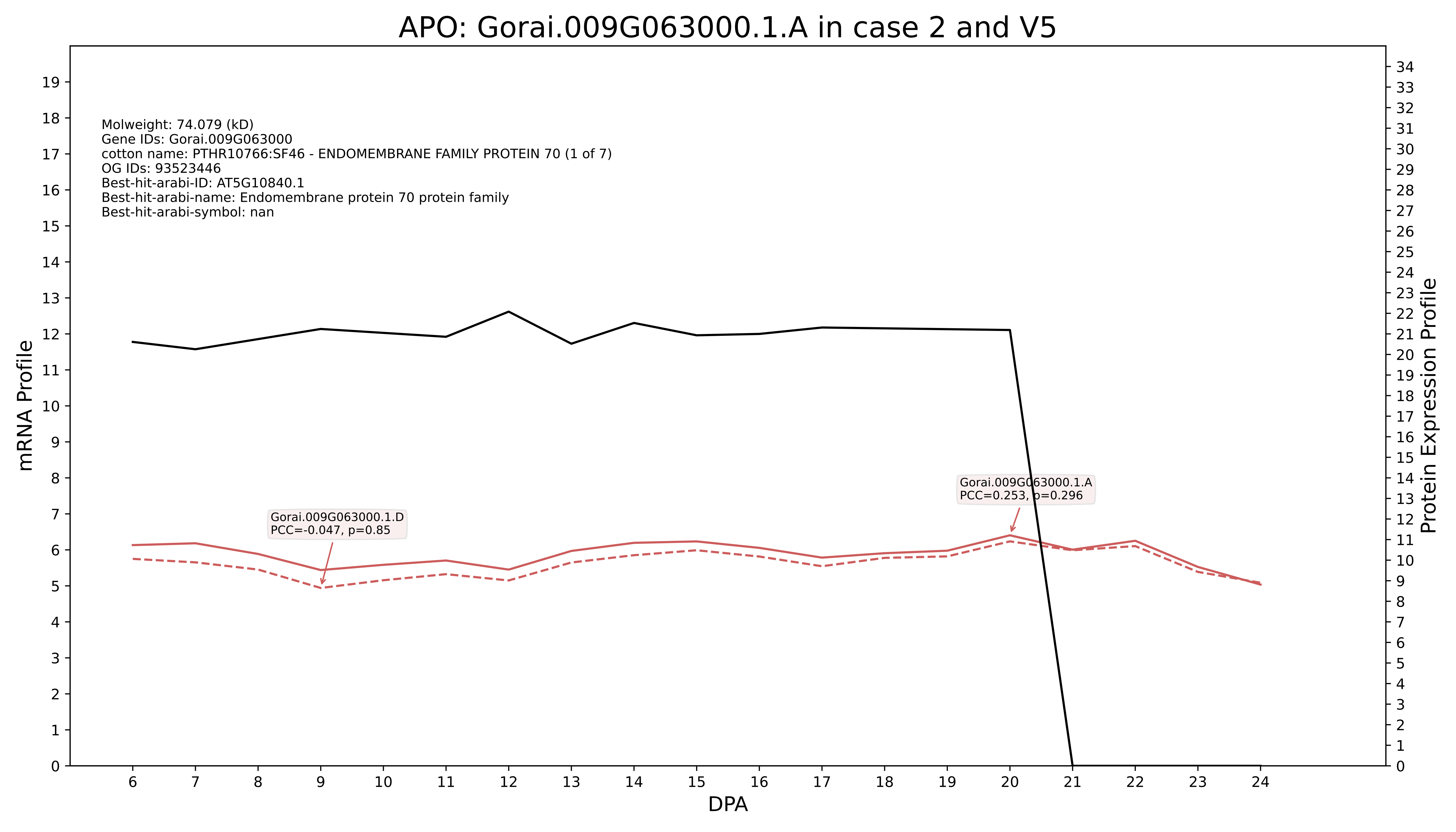
APO: Gorai.009G063000.1.A in case 2 and V5
Molweight: 74.079 (kD)
Gene IDs: Gorai.009G063000
cotton name: PTHR10766:SF46 - ENDOMEMBRANE FAMILY PROTEIN 70 (1 of 7)
OG IDs: 93523446
Best-hit-arabi-ID: AT5G10840.1
Best-hit-arabi-name: Endomembrane protein 70 protein family
Best-hit-arabi-symbol: nan
Gorai.009G063000.1.D
PCC=-0.047, p=0.85
Gorai.009G063000.1.A
PCC=0.253, p=0.296
mRNA Profile
Protein Expression Profile
DPA

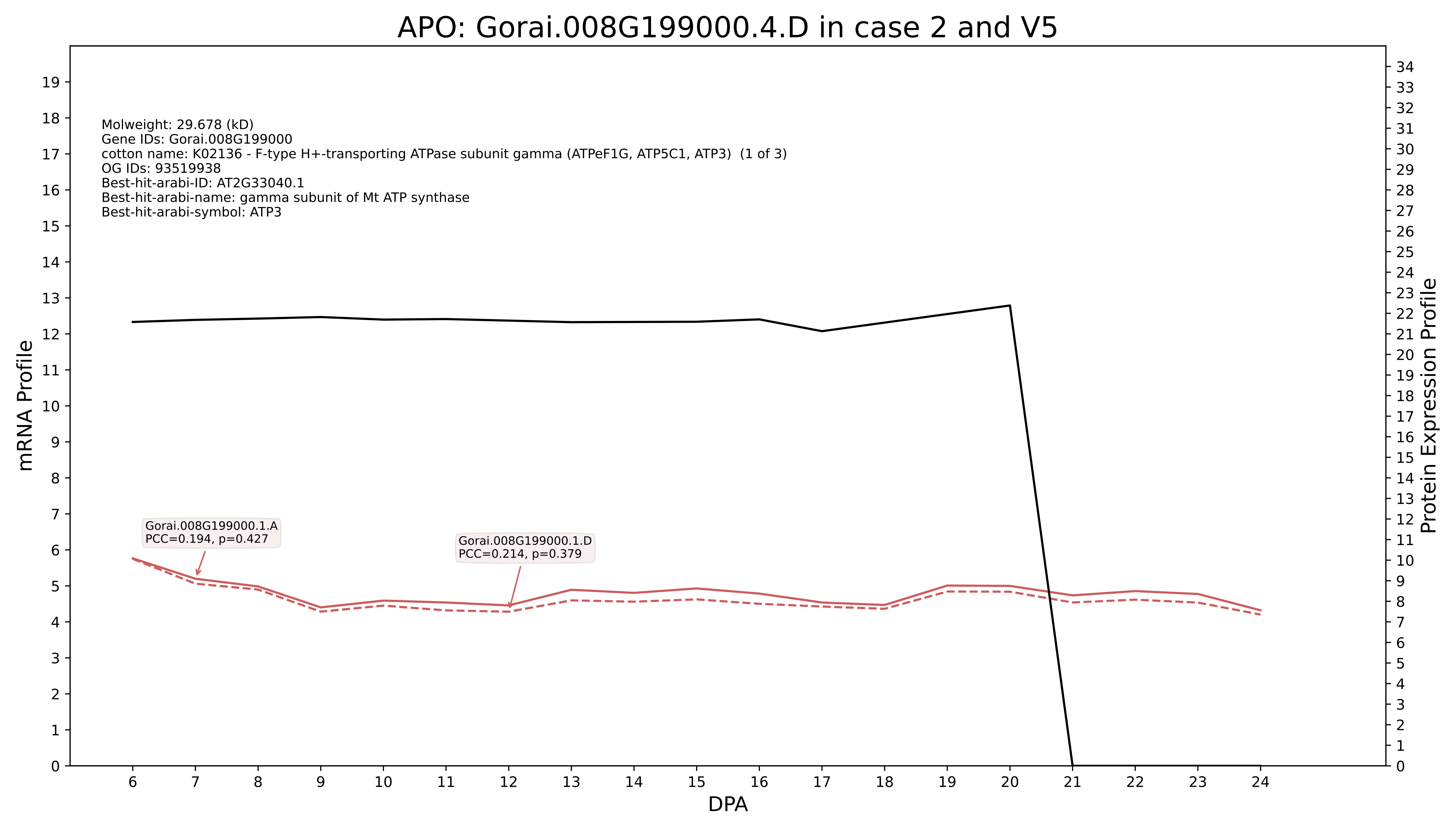
APO: Gorai.008G199000.4.D in case 2 and V5
Molweight: 29.678 (kD)
Gene IDs: Gorai.008G199000
cotton name: K02136 - F-type H+-transporting ATPase subunit gamma (ATPeF1G, ATP5C1, ATP3) (1 of 3)
OG IDs: 93519938
Best-hit-arabi-ID: AT2G33040.1
Best-hit-arabi-name: gamma subunit of Mt ATP synthase
Best-hit-arabi-symbol: ATP3
Gorai.008G199000.1.A
PCC=0.194, p=0.427
Gorai.008G199000.1.D
PCC=0.214, p=0.379
mRNA Profile
Protein Expression Profile
DPA

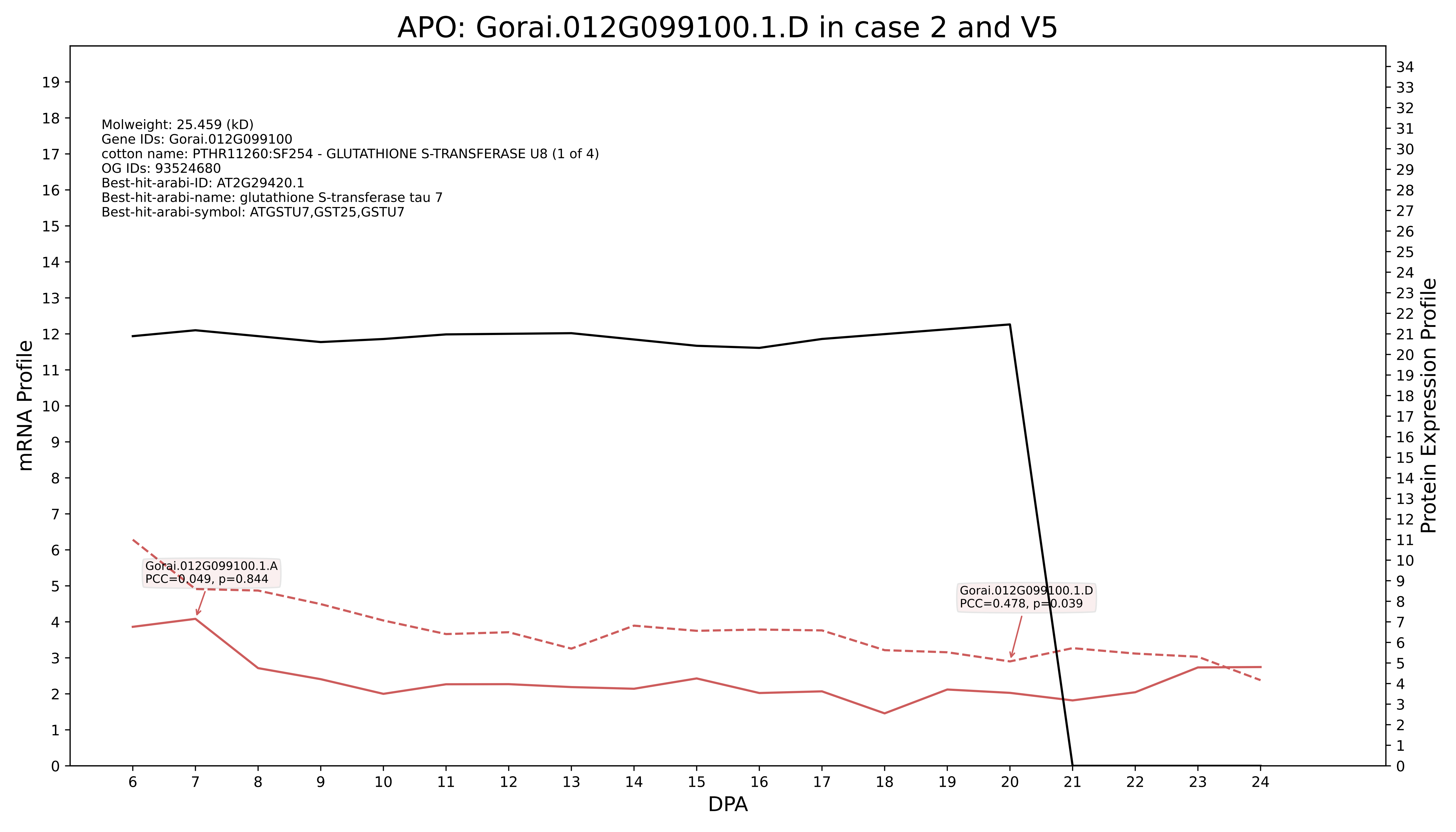

APO: Gorai.012G099100.1.D in case 2 and V5
Molweight: 25.459 (kD)
Gene IDs: Gorai.012G099100
cotton name: PTHR11260:SF254 - GLUTATHIONE S-TRANSFERASE U8 (1 of 4)
OG IDs: 93524680
Best-hit-arabi-ID: AT2G29420.1
Best-hit-arabi-name: glutathione S-transferase tau 7
Best-hit-arabi-symbol: ATGSTU7,GST25,GSTU7
Gorai.012G099100.1.A
PCC=0.049, p=0.844
Gorai.012G099100.1.D
PCC=0.478, p=0.039
mRNA Profile
Protein Expression Profile
DPA

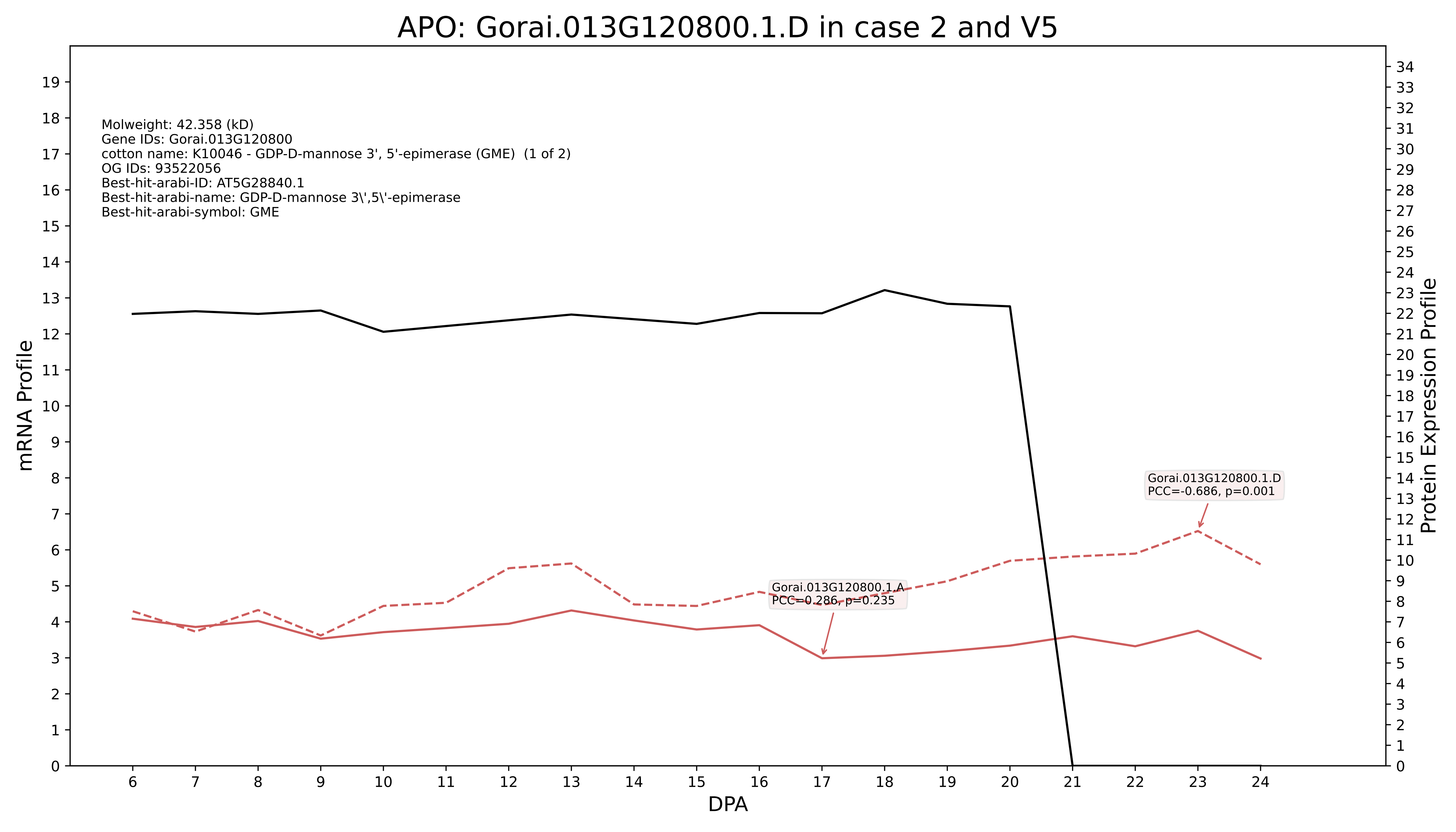

APO: Gorai.013G120800.1.D in case 2 and V5
Molweight: 42.358 (kD)
Gene IDs: Gorai.013G120800
cotton name: K10046 - GDP-D-mannose 3', 5'-epimerase (GME) (1 of 2)
OG IDs: 93522056
Best-hit-arabi-ID: AT5G28840.1
Best-hit-arabi-name: GDP-D-mannose 3\',5\'-epimerase
Best-hit-arabi-symbol: GME
Gorai.013G120800.1.D
PCC=-0.686, p=0.001
Gorai.013G120800.1.A
PCC=0.286, p=0.235
mRNA Profile
Protein Expression Profile
DPA

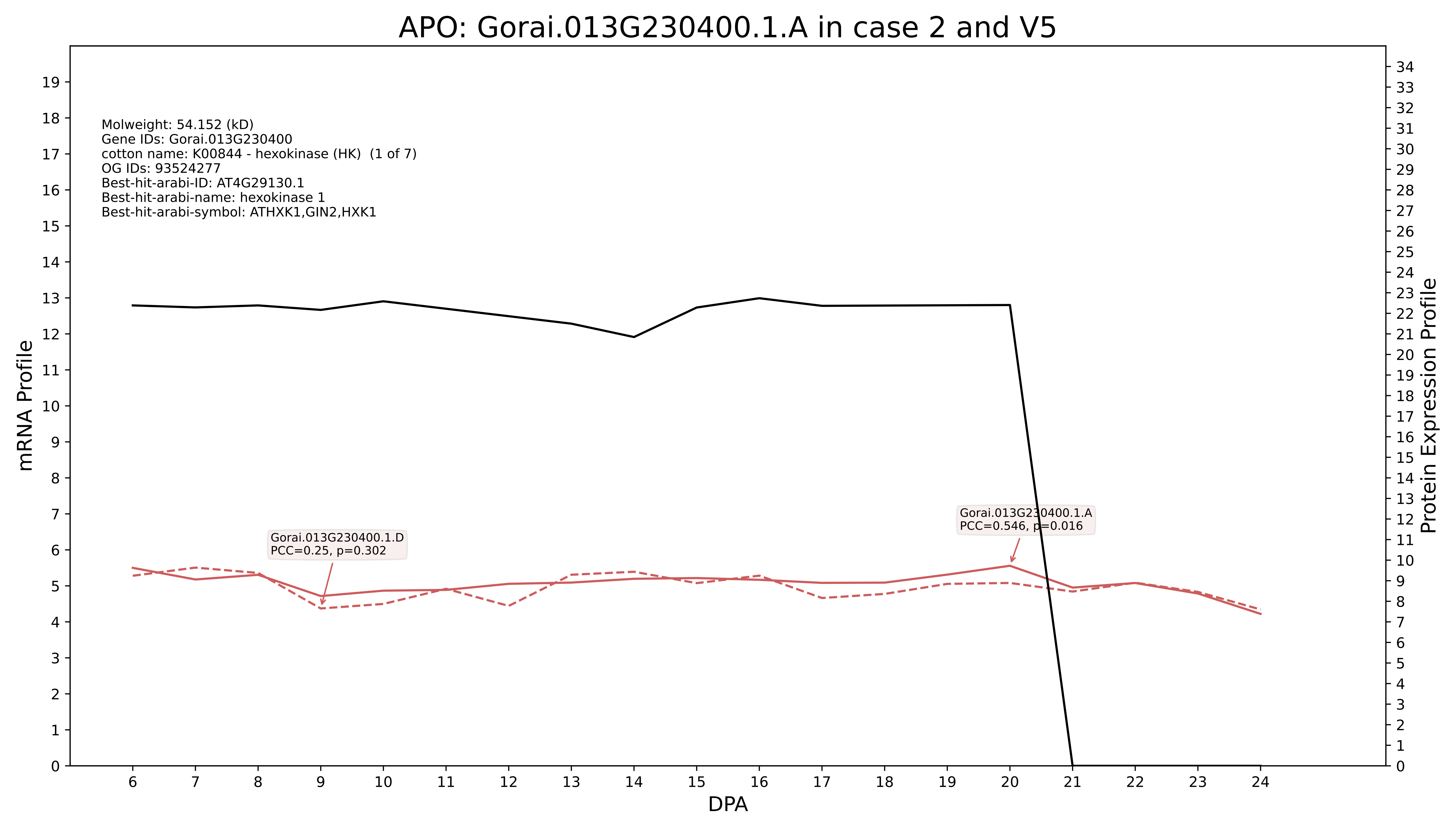
APO: Gorai.013G230400.1.A in case 2 and V5
Molweight: 54.152 (kD)
Gene IDs: Gorai.013G230400
cotton name: K00844 - hexokinase (HK) (1 of 7)
OG IDs: 93524277
Best-hit-arabi-ID: AT4G29130.1
Best-hit-arabi-name: hexokinase 1
Best-hit-arabi-symbol: ATHXK1,GIN2,HXK1
Gorai.013G230400.1.D
PCC=0.25, p=0.302
Gorai.013G230400.1.A
PCC=0.546, p=0.016
mRNA Profile
Protein Expression Profile
DPA
0 1 2 3 4 5 6 7 8 9 10 11 12 13 14 15 16 17 18 19
0 1 2 3 4 5 6 7 8 9 10 11 12 13 14 15 16 17 18 19 20 21 22 23 24 25 26 27 28 29 30 31 32 33 34
6 7 8 9 10 11 12 13 14 15 16 17 18 19 20 21 22 23 24

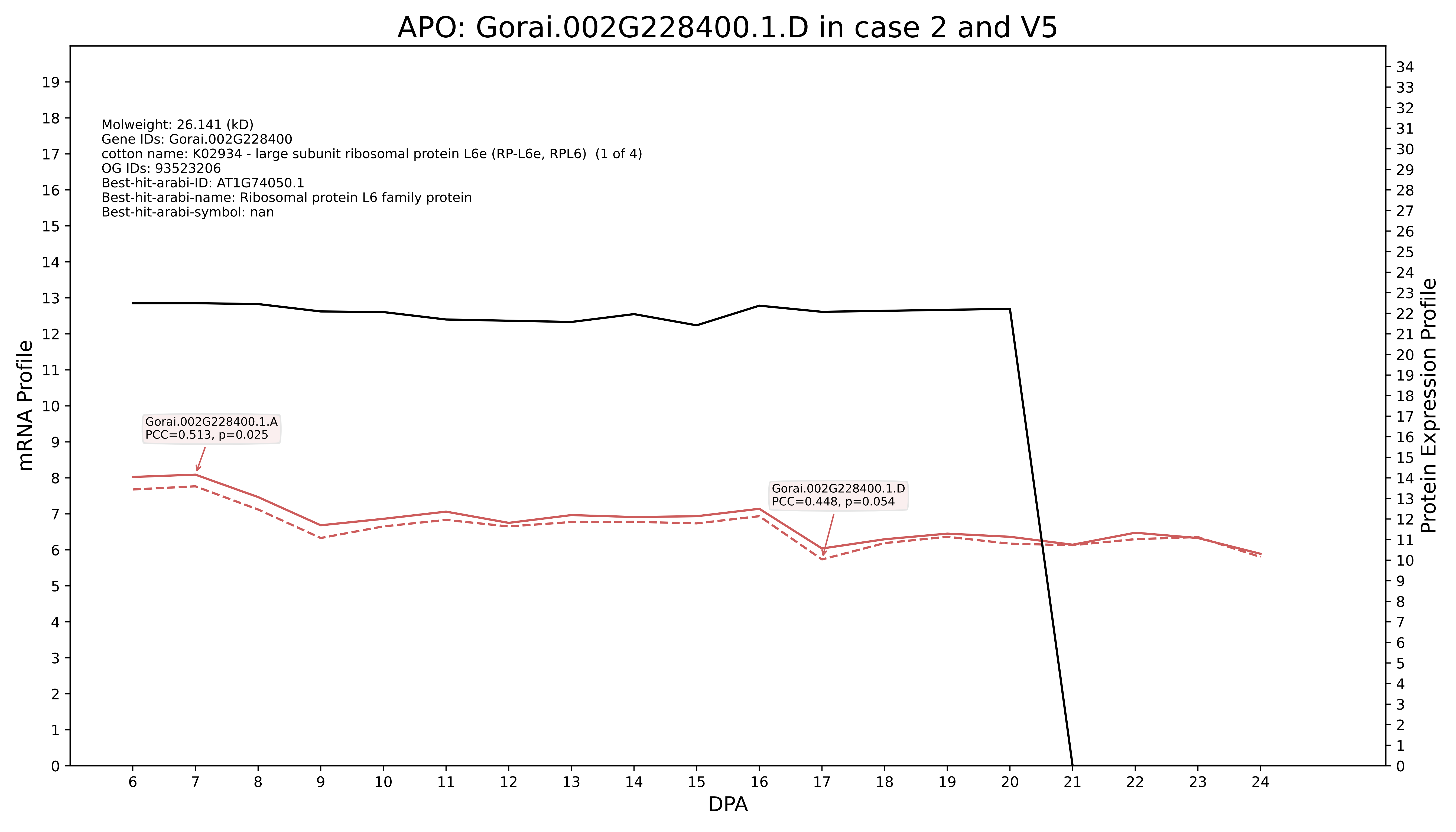

APO: Gorai.002G228400.1.D in case 2 and V5
Molweight: 26.141 (kD)
Gene IDs: Gorai.002G228400
cotton name: K02934 - large subunit ribosomal protein L6e (RP-L6e, RPL6) (1 of 4)
OG IDs: 93523206
Best-hit-arabi-ID: AT1G74050.1
Best-hit-arabi-name: Ribosomal protein L6 family protein
Best-hit-arabi-symbol: nan
Gorai.002G228400.1.A
PCC=0.513, p=0.025
Gorai.002G228400.1.D
PCC=0.448, p=0.054
mRNA Profile
Protein Expression Profile
DPA

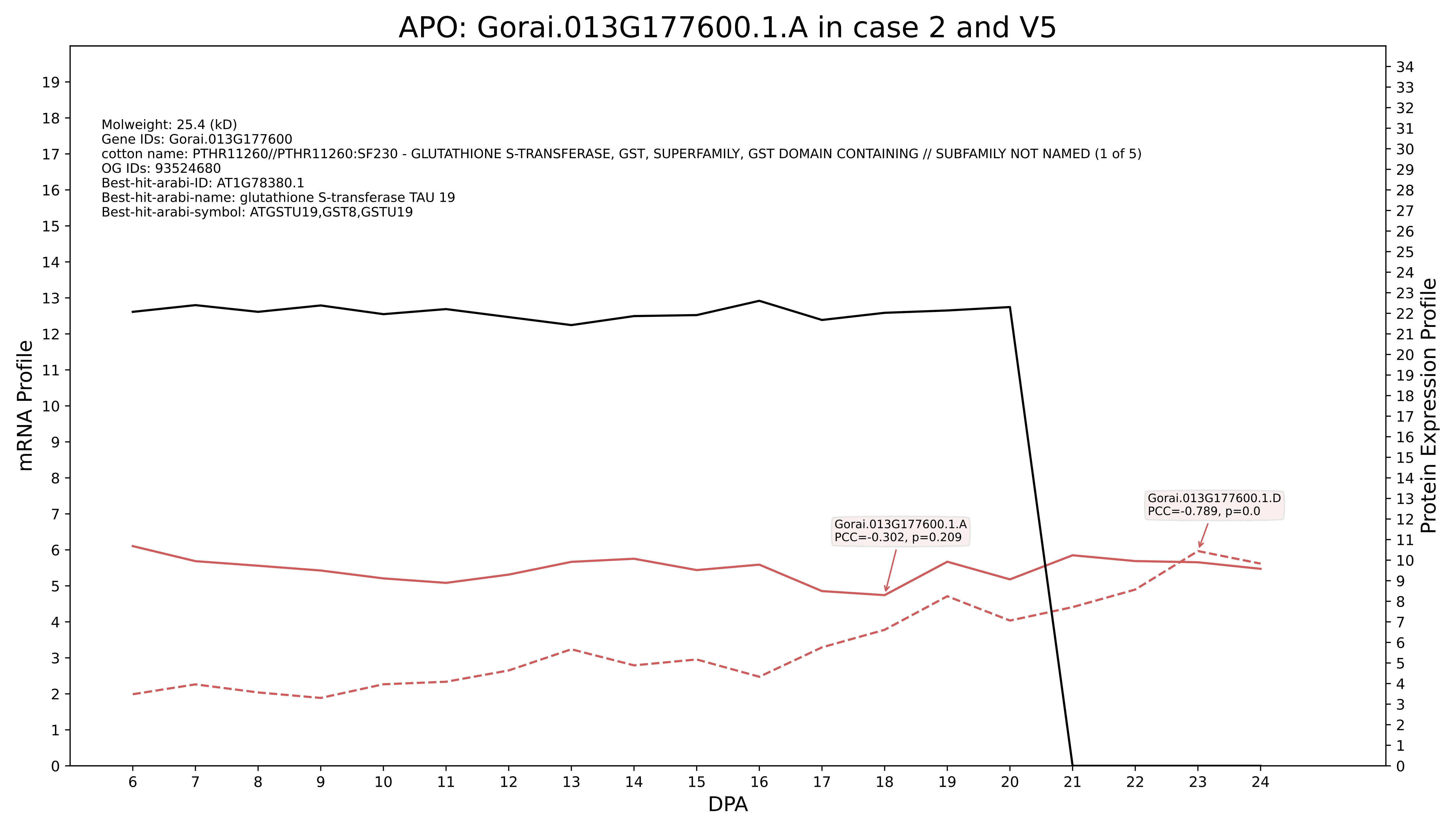

APO: Gorai.013G177600.1.A in case 2 and V5
Molweight: 25.4 (kD)
Gene IDs: Gorai.013G177600
cotton name: PTHR11260//PTHR11260:SF230 - GLUTATHIONE S-TRANSFERASE, GST, SUPERFAMILY, GST DOMAIN CONTAINING // SUBFAMILY NOT NAMED (1 of 5)
OG IDs: 93524680
Best-hit-arabi-ID: AT1G78380.1
Best-hit-arabi-name: glutathione S-transferase TAU 19
Best-hit-arabi-symbol: ATGSTU19,GST8,GSTU19
Gorai.013G177600.1.A
PCC=-0.302, p=0.209
Gorai.013G177600.1.D
PCC=-0.789, p=0.0
mRNA Profile
Protein Expression Profile
DPA

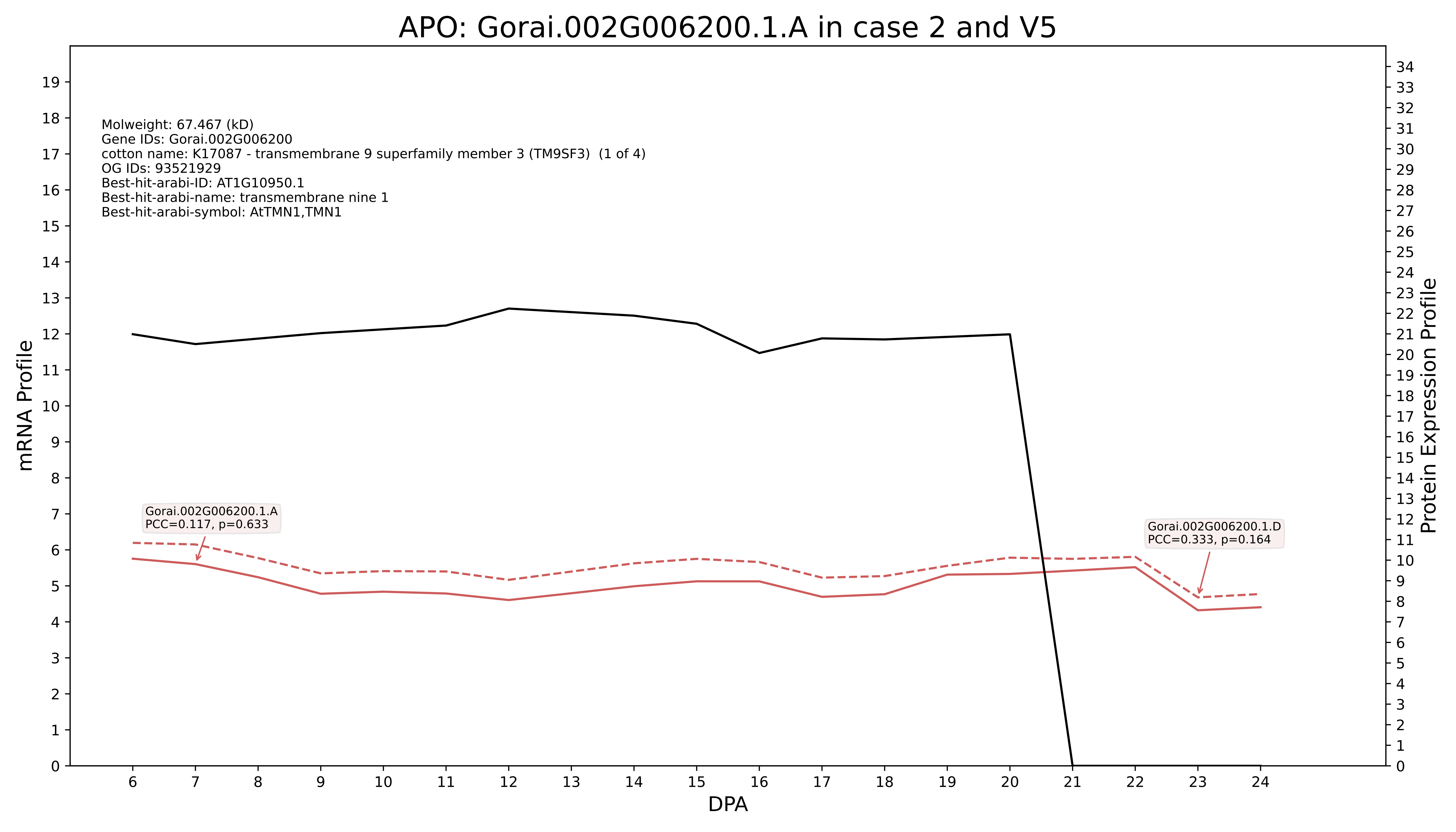

APO: Gorai.002G006200.1.A in case 2 and V5
Molweight: 67.467 (kD)
Gene IDs: Gorai.002G006200
cotton name: K17087 - transmembrane 9 superfamily member 3 (TM9SF3) (1 of 4)
OG IDs: 93521929
Best-hit-arabi-ID: AT1G10950.1
Best-hit-arabi-name: transmembrane nine 1
Best-hit-arabi-symbol: AtTMN1,TMN1
Gorai.002G006200.1.A
PCC=0.117, p=0.633
Gorai.002G006200.1.D
PCC=0.333, p=0.164
mRNA Profile
Protein Expression Profile
DPA

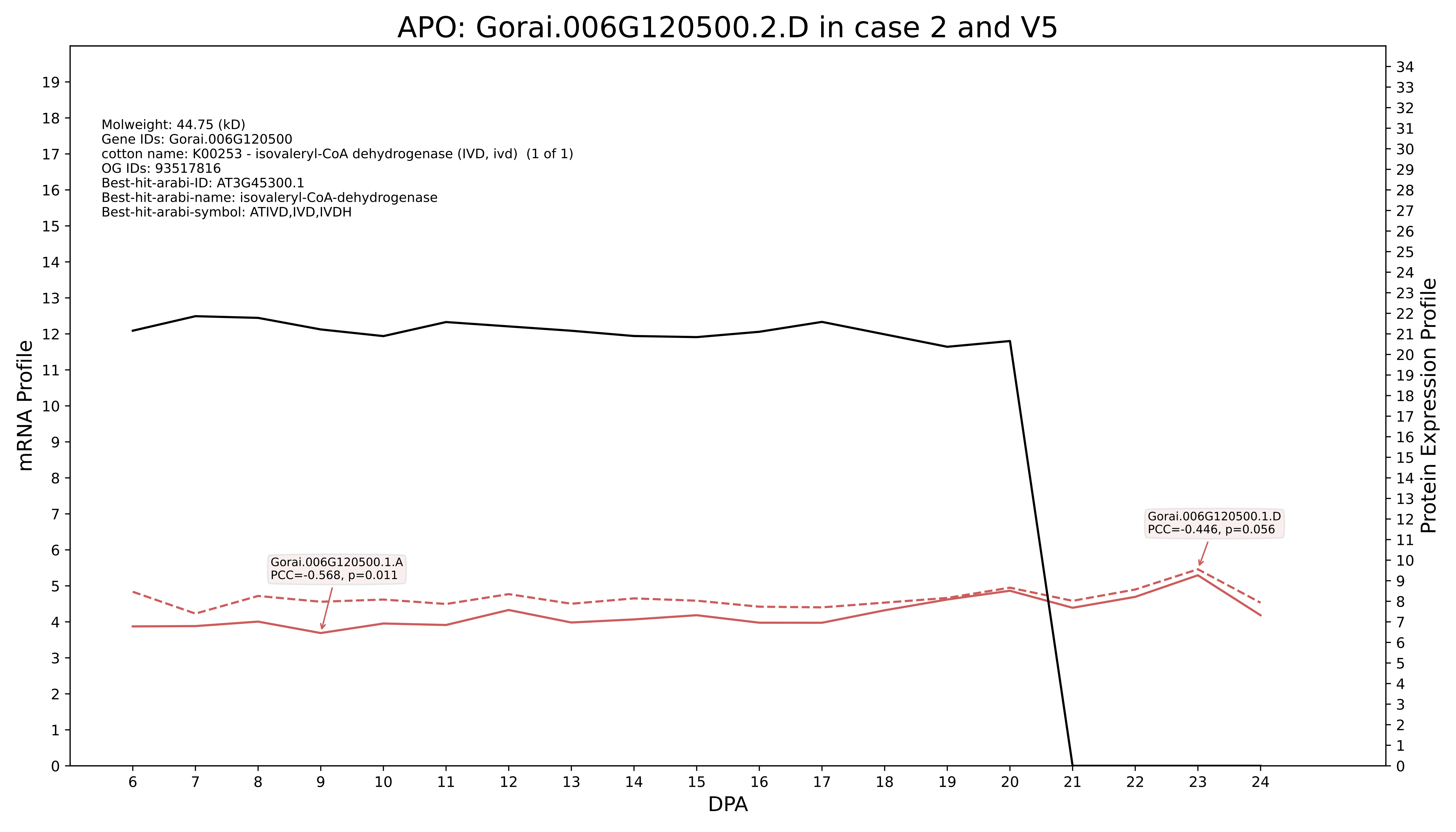

APO: Gorai.006G120500.2.D in case 2 and V5
Molweight: 44.75 (kD)
Gene IDs: Gorai.006G120500
cotton name: K00253 - isovaleryl-CoA dehydrogenase (IVD, ivd) (1 of 1)
OG IDs: 93517816
Best-hit-arabi-ID: AT3G45300.1
Best-hit-arabi-name: isovaleryl-CoA-dehydrogenase
Best-hit-arabi-symbol: ATIVD,IVD,IVDH
Gorai.006G120500.1.A
PCC=-0.568, p=0.011
Gorai.006G120500.1.D
PCC=-0.446, p=0.056
mRNA Profile
Protein Expression Profile
DPA

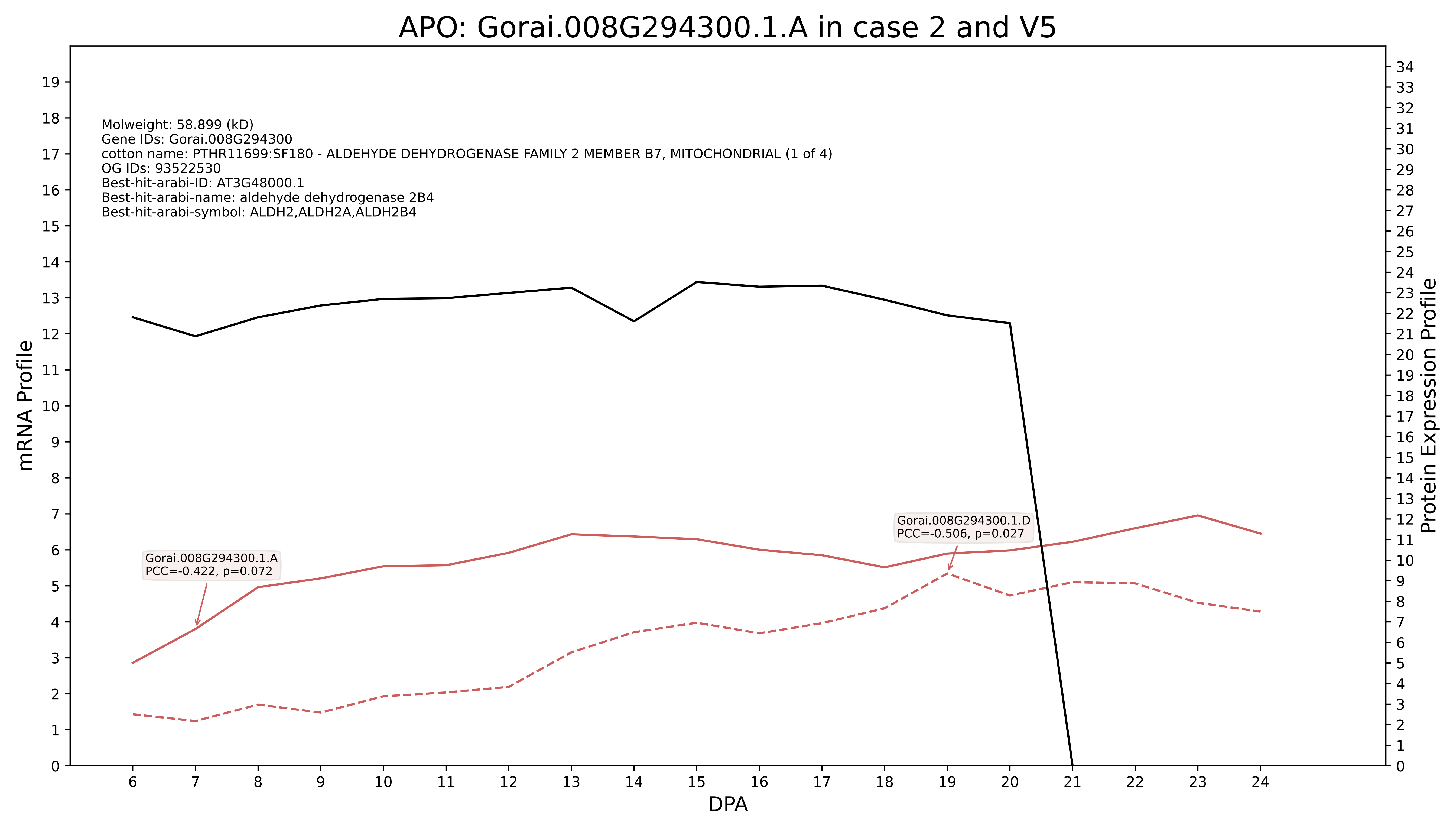
APO: Gorai.008G294300.1.A in case 2 and V5
Molweight: 58.899 (kD)
Gene IDs: Gorai.008G294300
cotton name: PTHR11699:SF180 - ALDEHYDE DEHYDROGENASE FAMILY 2 MEMBER B7, MITOCHONDRIAL (1 of 4)
OG IDs: 93522530
Best-hit-arabi-ID: AT3G48000.1
Best-hit-arabi-name: aldehyde dehydrogenase 2B4
Best-hit-arabi-symbol: ALDH2,ALDH2A,ALDH2B4
Gorai.008G294300.1.D
PCC=-0.506, p=0.027
Gorai.008G294300.1.A
PCC=-0.422, p=0.072
mRNA Profile
Protein Expression Profile
DPA

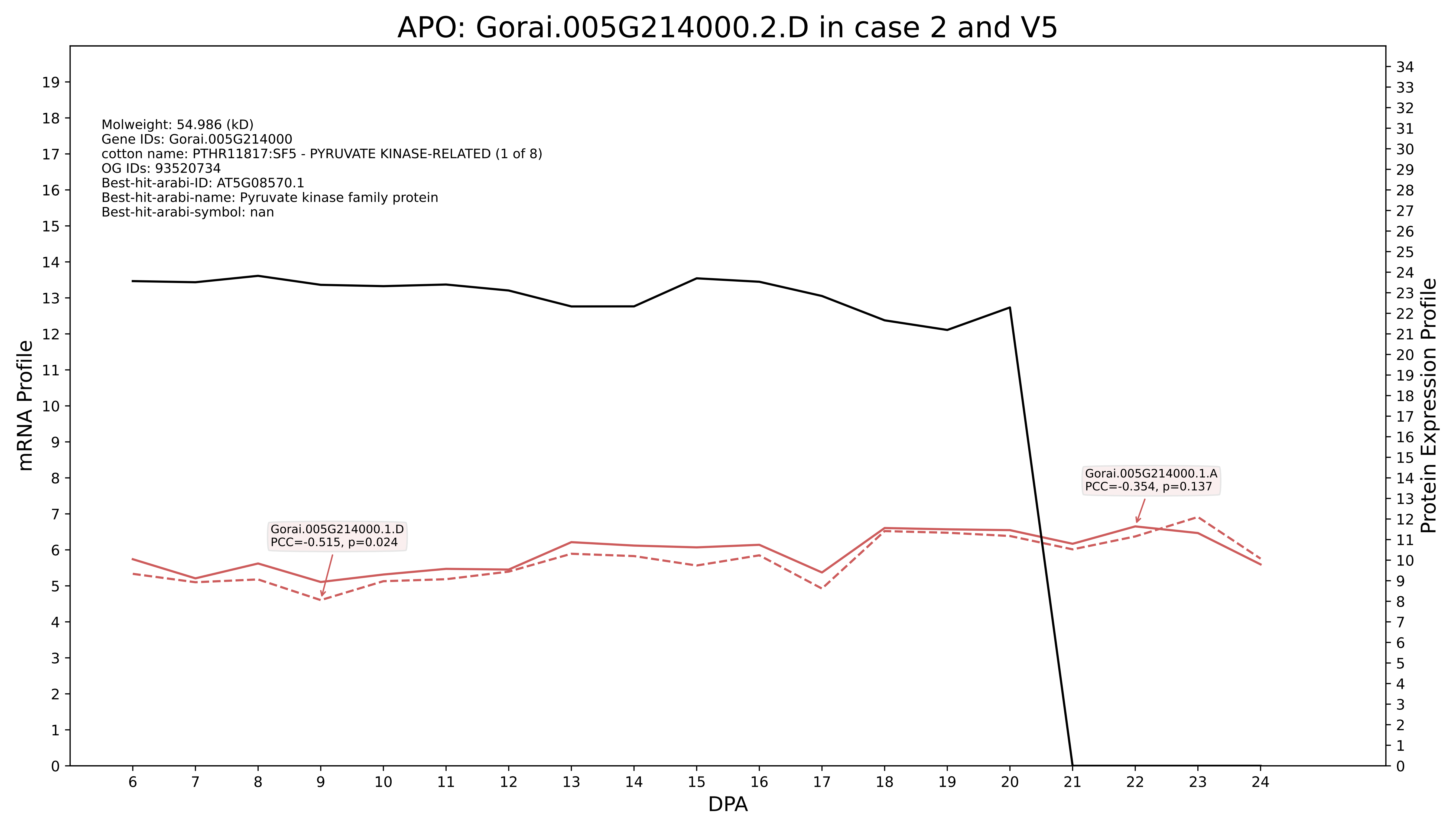

APO: Gorai.005G214000.2.D in case 2 and V5
Molweight: 54.986 (kD)
Gene IDs: Gorai.005G214000
cotton name: PTHR11817:SF5 - PYRUVATE KINASE-RELATED (1 of 8)
OG IDs: 93520734
Best-hit-arabi-ID: AT5G08570.1
Best-hit-arabi-name: Pyruvate kinase family protein
Best-hit-arabi-symbol: nan
Gorai.005G214000.1.D
PCC=-0.515, p=0.024
Gorai.005G214000.1.A
PCC=-0.354, p=0.137
mRNA Profile
Protein Expression Profile
DPA

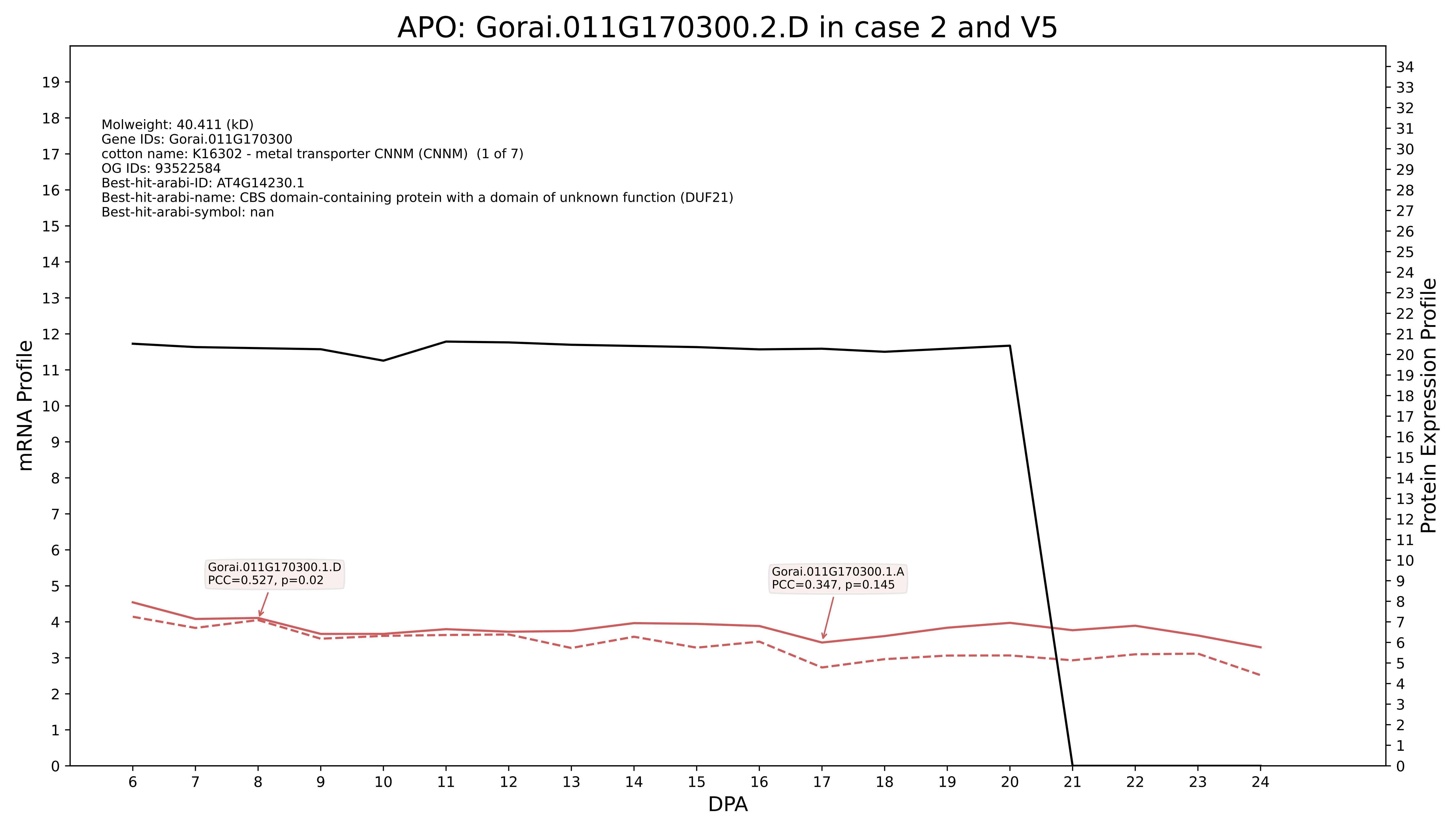

APO: Gorai.011G170300.2.D in case 2 and V5
Molweight: 40.411 (kD)
Gene IDs: Gorai.011G170300
cotton name: K16302 - metal transporter CNNM (CNNM) (1 of 7)
OG IDs: 93522584
Best-hit-arabi-ID: AT4G14230.1
Best-hit-arabi-name: CBS domain-containing protein with a domain of unknown function (DUF21)
Best-hit-arabi-symbol: nan
Gorai.011G170300.1.D
PCC=0.527, p=0.02
Gorai.011G170300.1.A
PCC=0.347, p=0.145
mRNA Profile
Protein Expression Profile
DPA

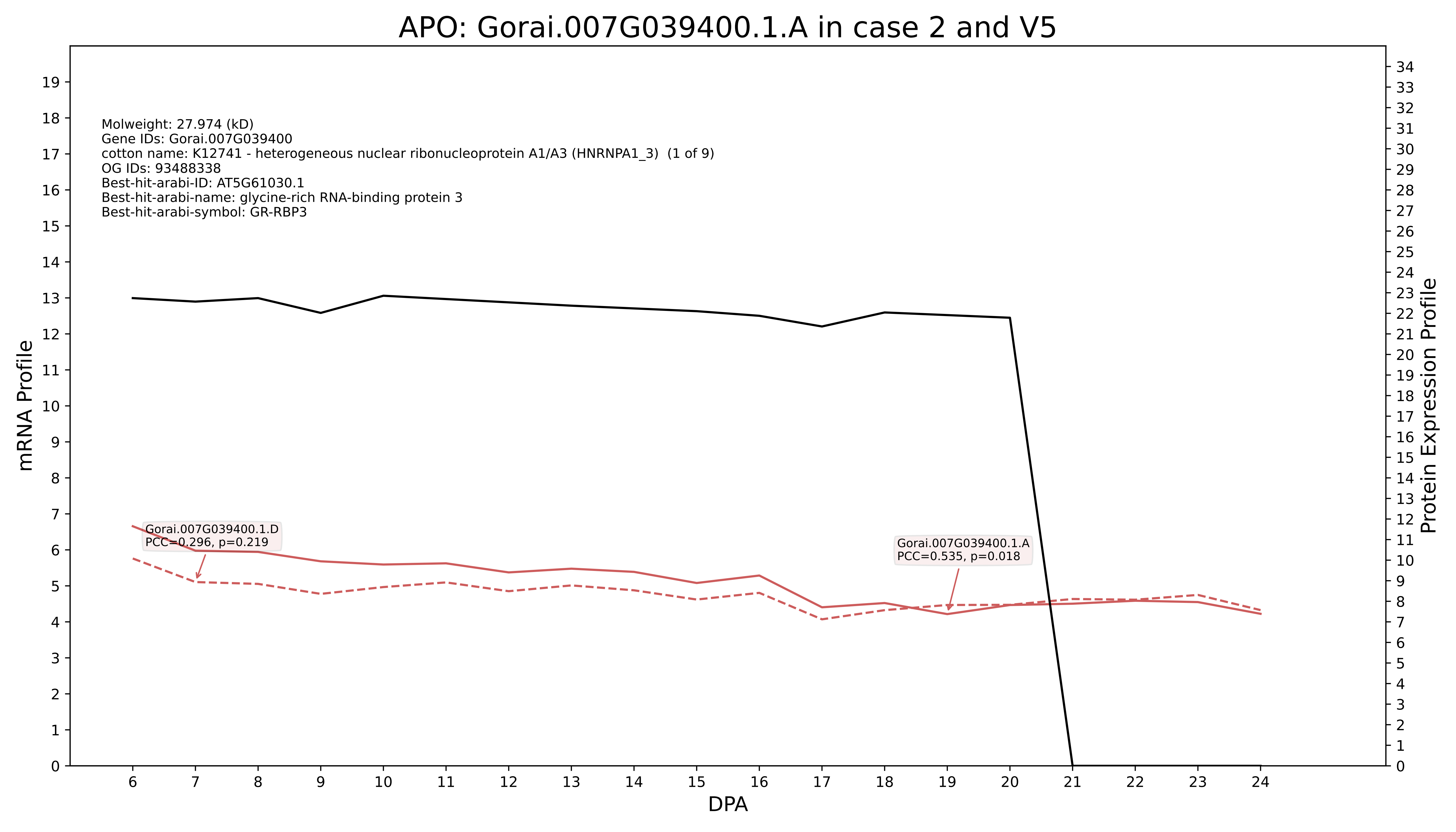
APO: Gorai.007G039400.1.A in case 2 and V5
Molweight: 27.974 (kD)
Gene IDs: Gorai.007G039400
cotton name: K12741 - heterogeneous nuclear ribonucleoprotein A1/A3 (HNRNPA1_3) (1 of 9)
OG IDs: 93488338
Best-hit-arabi-ID: AT5G61030.1
Best-hit-arabi-name: glycine-rich RNA-binding protein 3
Best-hit-arabi-symbol: GR-RBP3
Gorai.007G039400.1.D
PCC=0.296, p=0.219
Gorai.007G039400.1.A
PCC=0.535, p=0.018
mRNA Profile
Protein Expression Profile
DPA

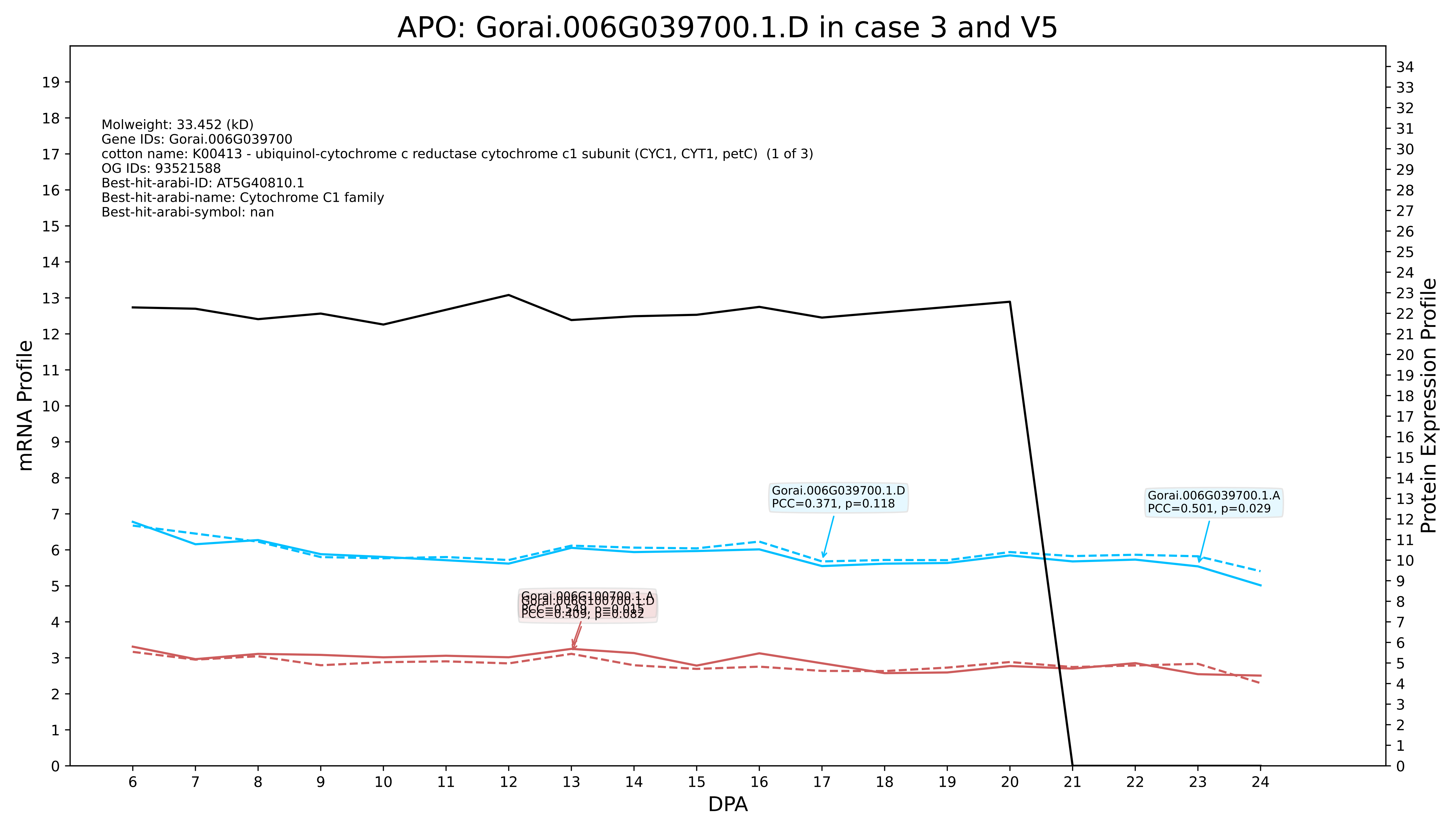

APO: Gorai.006G039700.1.D in case 3 and V5
Molweight: 33.452 (kD)
Gene IDs: Gorai.006G039700
cotton name: K00413 - ubiquinol-cytochrome c reductase cytochrome c1 subunit (CYC1, CYT1, petC) (1 of 3)
OG IDs: 93521588
Best-hit-arabi-ID: AT5G40810.1
Best-hit-arabi-name: Cytochrome C1 family
Best-hit-arabi-symbol: nan
Gorai.006G039700.1.D
PCC=0.371, p=0.118
Gorai.006G039700.1.A
PCC=0.501, p=0.029
mRNA Profile
Protein Expression Profile
DPA

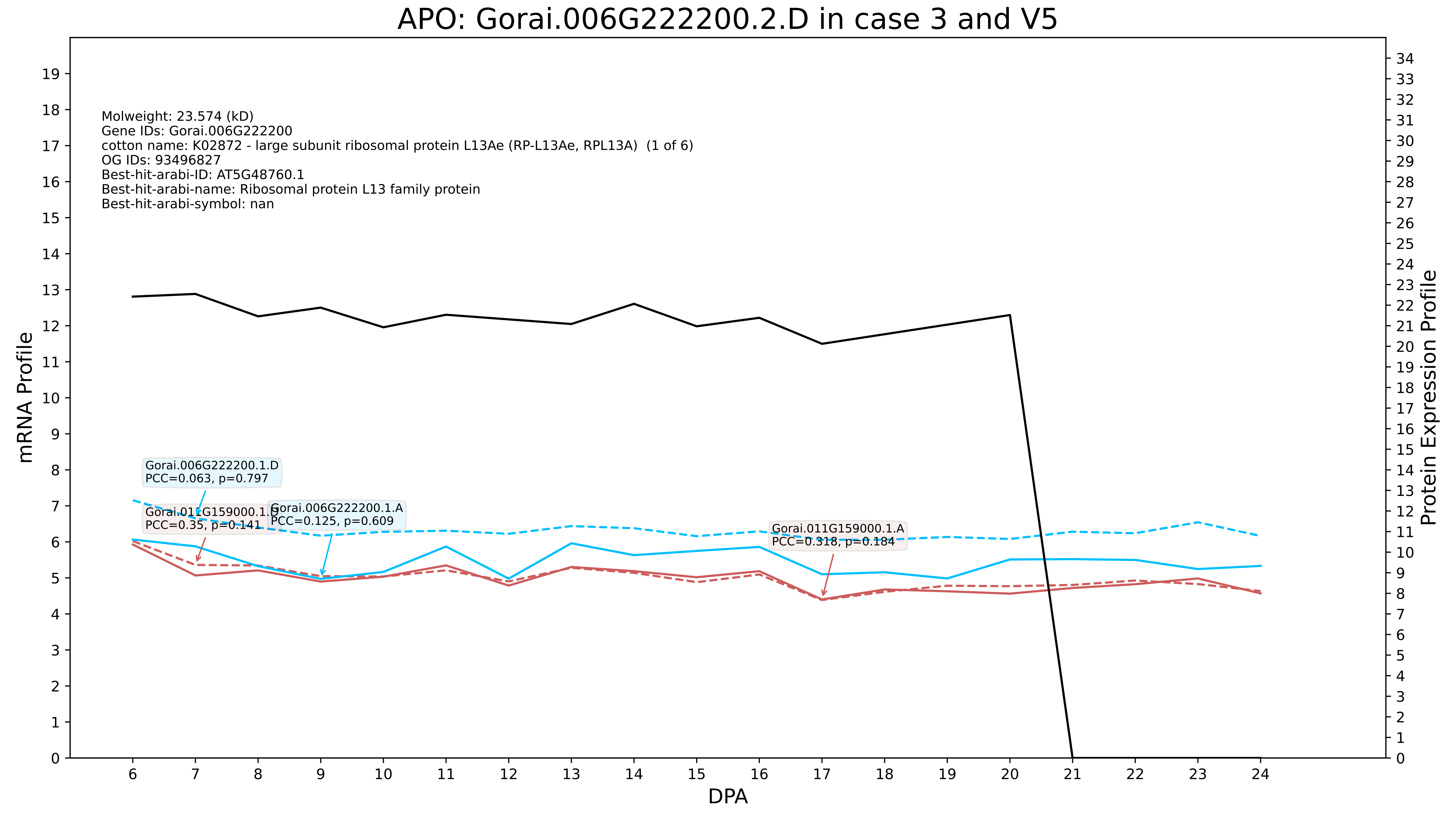
APO: Gorai.006G222200.2.D in case 3 and V5
Molweight: 23.574 (kD)
Gene IDs: Gorai.006G222200
cotton name: K02872 - large subunit ribosomal protein L13Ae (RP-L13Ae, RPL13A) (1 of 6)
OG IDs: 93496827
Best-hit-arabi-ID: AT5G48760.1
Best-hit-arabi-name: Ribosomal protein L13 family protein
Best-hit-arabi-symbol: nan
Gorai.006G222200.1.D
PCC=0.063, p=0.797
Gorai.011G159000.1.D
PCC=0.35, p=0.141
Gorai.006G222200.1.A
PCC=0.125, p=0.609
Gorai.011G159000.1.A
PCC=0.318, p=0.184
mRNA Profile
Protein Expression Profile
DPA

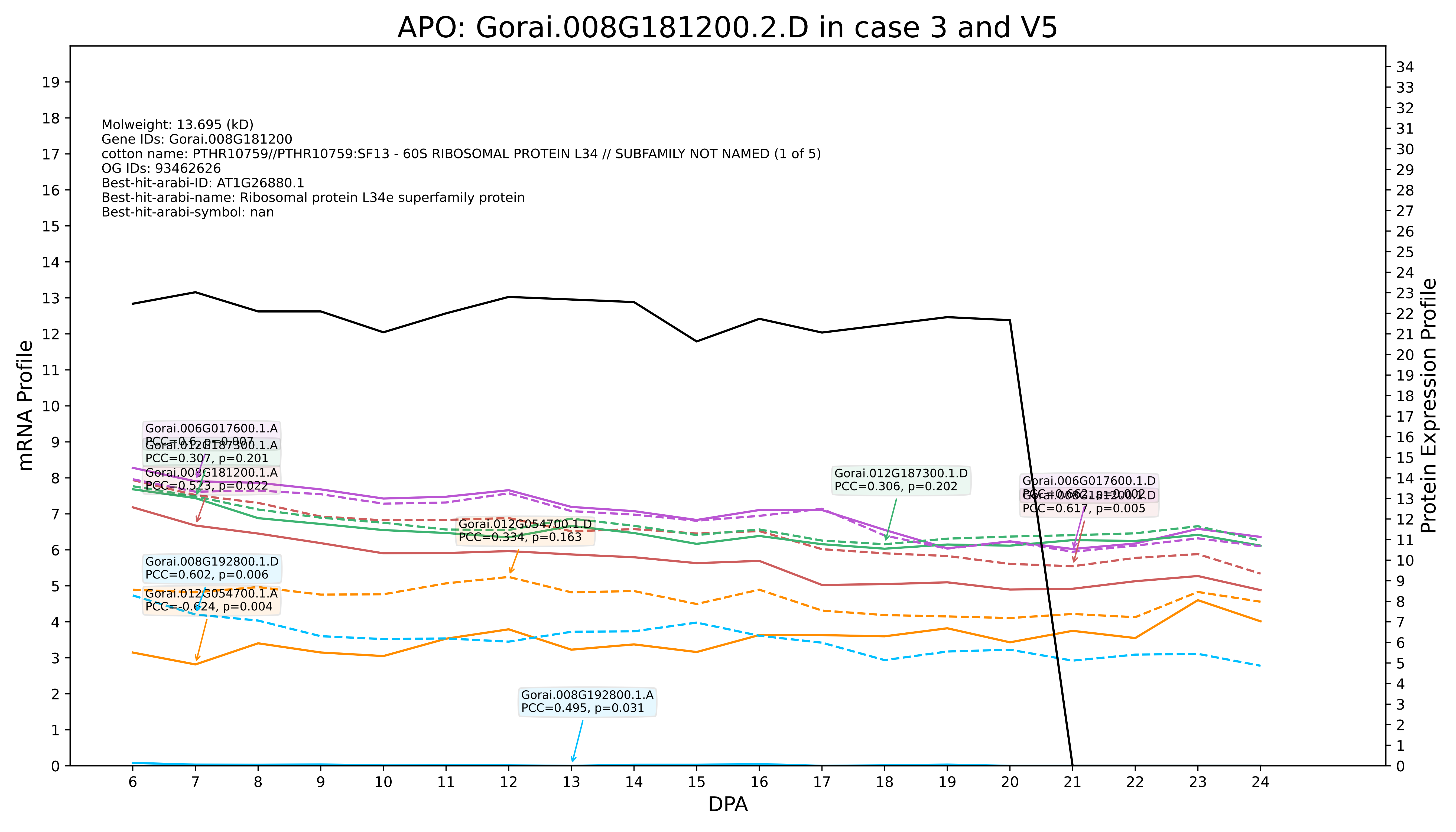

APO: Gorai.008G181200.2.D in case 3 and V5
Molweight: 13.695 (kD)
Gene IDs: Gorai.008G181200
cotton name: PTHR10759//PTHR10759:SF13 - 60S RIBOSOMAL PROTEIN L34 // SUBFAMILY NOT NAMED (1 of 5)
OG IDs: 93462626
Best-hit-arabi-ID: AT1G26880.1
Best-hit-arabi-name: Ribosomal protein L34e superfamily protein
Best-hit-arabi-symbol: nan
Gorai.006G017600.1.A
Gorai.012G187300.1.A
PCC=0.307, p=0.201
Gorai.008G181200.1.A
PCC=0.523, p=0.022
Gorai.012G187300.1.D
PCC=0.306, p=0.202
Gorai.006G017600.1.D
Gorai.008G181200.1.D
PCC=0.617, p=0.005
Gorai.012G054700.1.D
PCC=0.334, p=0.163
Gorai.008G192800.1.D
PCC=0.602, p=0.006
Gorai.012G054700.1.A
PCC=-0.624, p=0.004
Gorai.008G192800.1.A
PCC=0.495, p=0.031
mRNA Profile
Protein Expression Profile
DPA

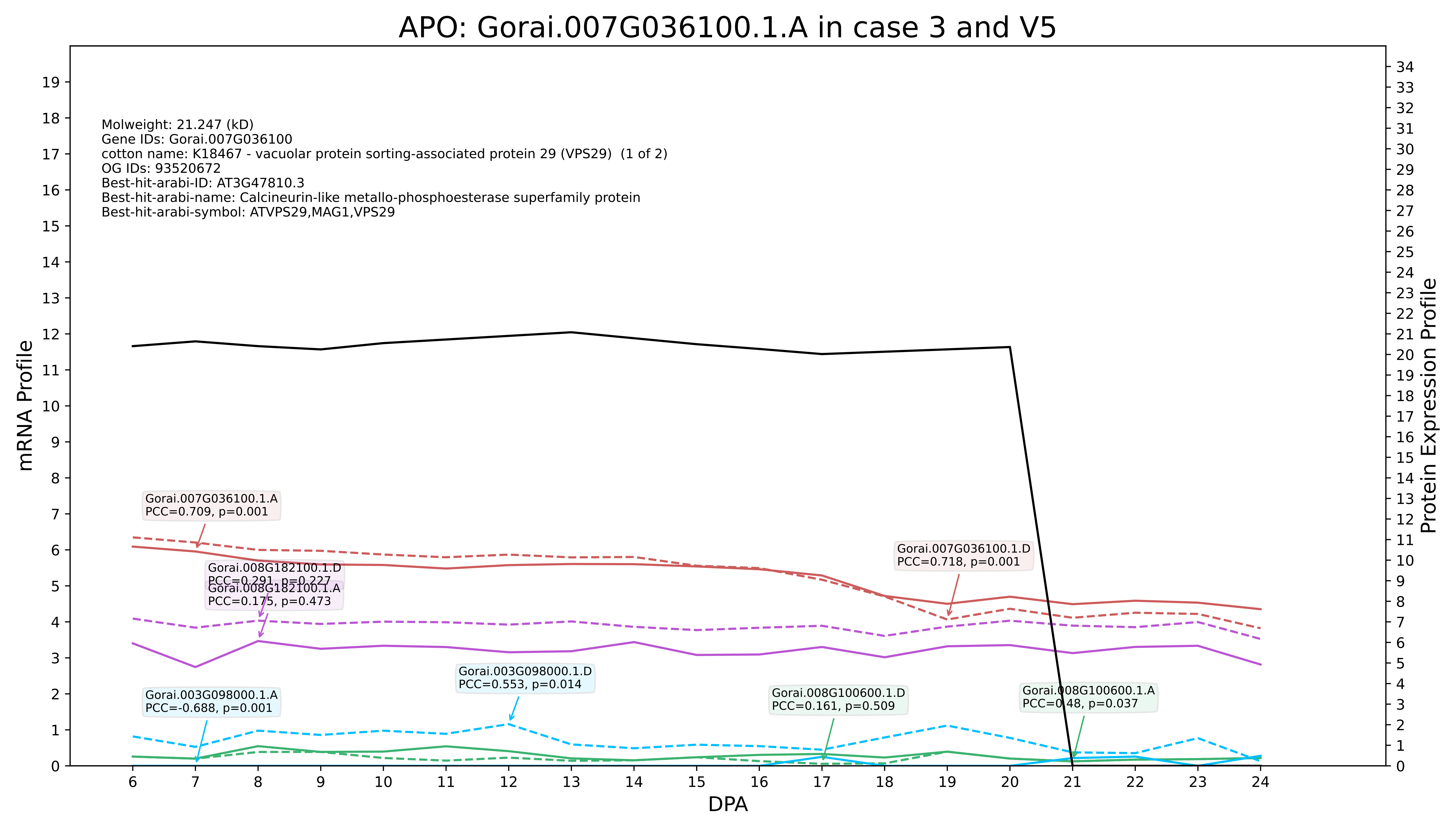

APO: Gorai.007G036100.1.A in case 3 and V5
Molweight: 21.247 (kD)
Gene IDs: Gorai.007G036100
cotton name: K18467 - vacuolar protein sorting-associated protein 29 (VPS29) (1 of 2)
OG IDs: 93520672
Best-hit-arabi-ID: AT3G47810.3
Best-hit-arabi-name: Calcineurin-like metallo-phosphoesterase superfamily protein
Best-hit-arabi-symbol: ATVPS29,MAG1,VPS29
Gorai.007G036100.1.A
PCC=0.709, p=0.001
Gorai.007G036100.1.D
PCC=0.718, p=0.001
Gorai.008G182100.1.D
PCC=0.291, p=0.227
Gorai.008G182100.1.A
PCC=0.175, p=0.473
Gorai.003G098000.1.D
PCC=0.553, p=0.014
Gorai.003G098000.1.A
PCC=-0.688, p=0.001
Gorai.008G100600.1.D
PCC=0.161, p=0.509
Gorai.008G100600.1.A
PCC=0.48, p=0.037
mRNA Profile
Protein Expression Profile
DPA

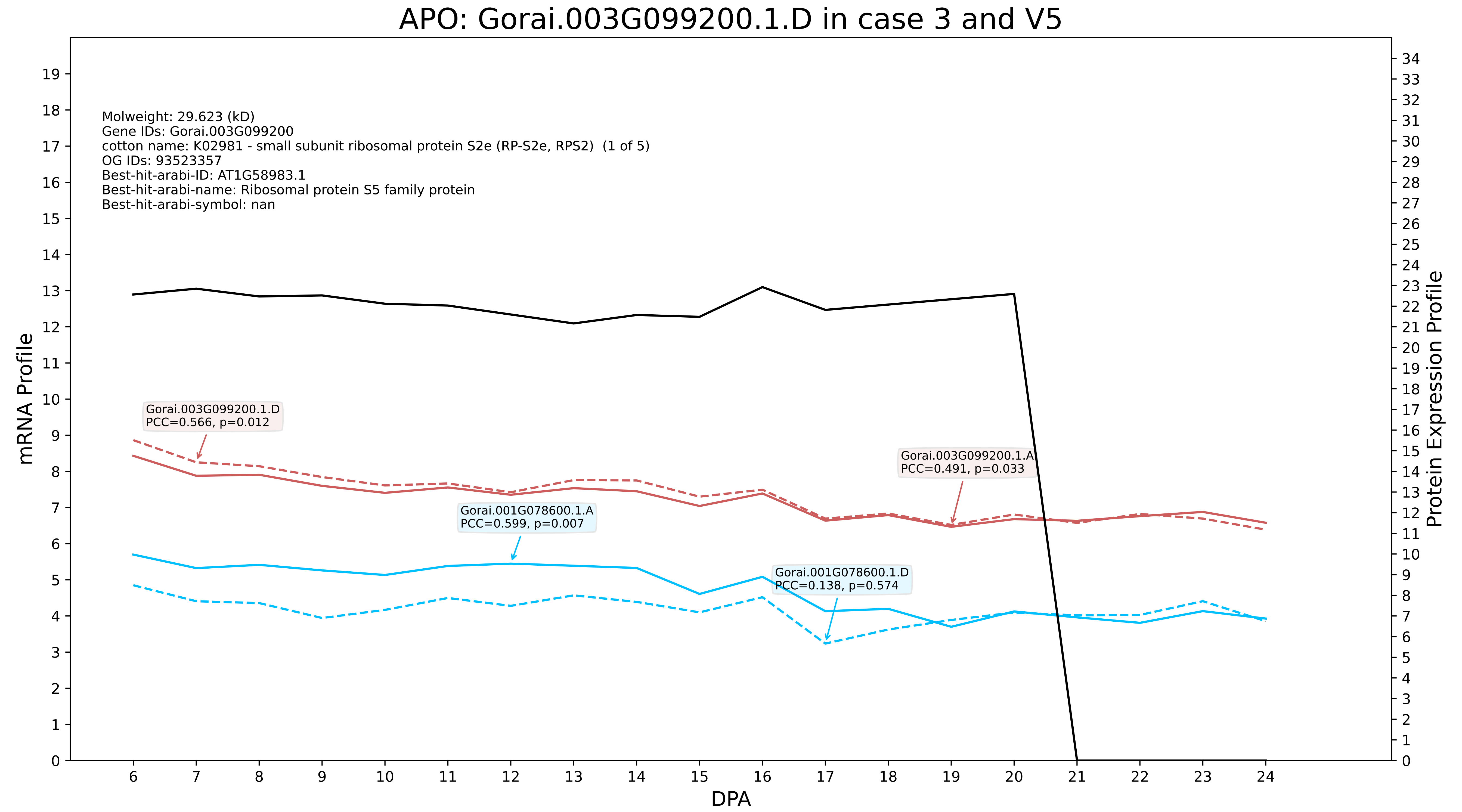

APO: Gorai.003G099200.1.D in case 3 and V5
Molweight: 29.623 (kD)
Gene IDs: Gorai.003G099200
cotton name: K02981 - small subunit ribosomal protein S2e (RP-S2e, RPS2) (1 of 5)
OG IDs: 93523357
Best-hit-arabi-ID: AT1G58983.1
Best-hit-arabi-name: Ribosomal protein S5 family protein
Best-hit-arabi-symbol: nan
Gorai.003G099200.1.D
PCC=0.566, p=0.012
Gorai.003G099200.1.A
PCC=0.491, p=0.033
Gorai.001G078600.1.A
PCC=0.599, p=0.007
Gorai.001G078600.1.D
PCC=0.138, p=0.574
mRNA Profile
Protein Expression Profile
DPA

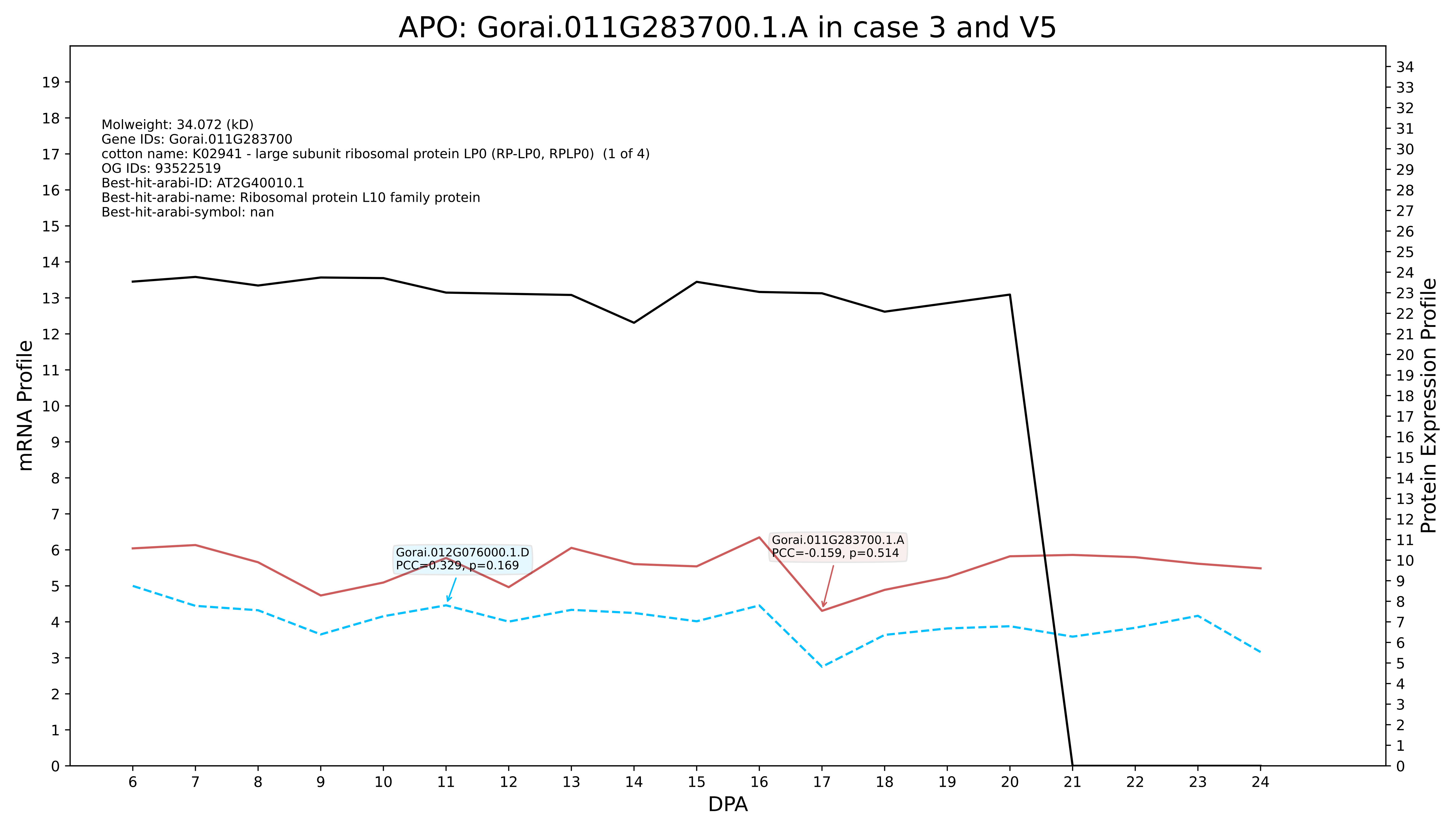

APO: Gorai.011G283700.1.A in case 3 and V5
Molweight: 34.072 (kD)
Gene IDs: Gorai.011G283700
cotton name: K02941 - large subunit ribosomal protein LP0 (RP-LP0, RPLP0) (1 of 4)
OG IDs: 93522519
Best-hit-arabi-ID: AT2G40010.1
Best-hit-arabi-name: Ribosomal protein L10 family protein
Best-hit-arabi-symbol: nan
Gorai.012G076000.1.D
PCC=0.329, p=0.169
Gorai.011G283700.1.A
PCC=-0.159, p=0.514
mRNA Profile
Protein Expression Profile
DPA

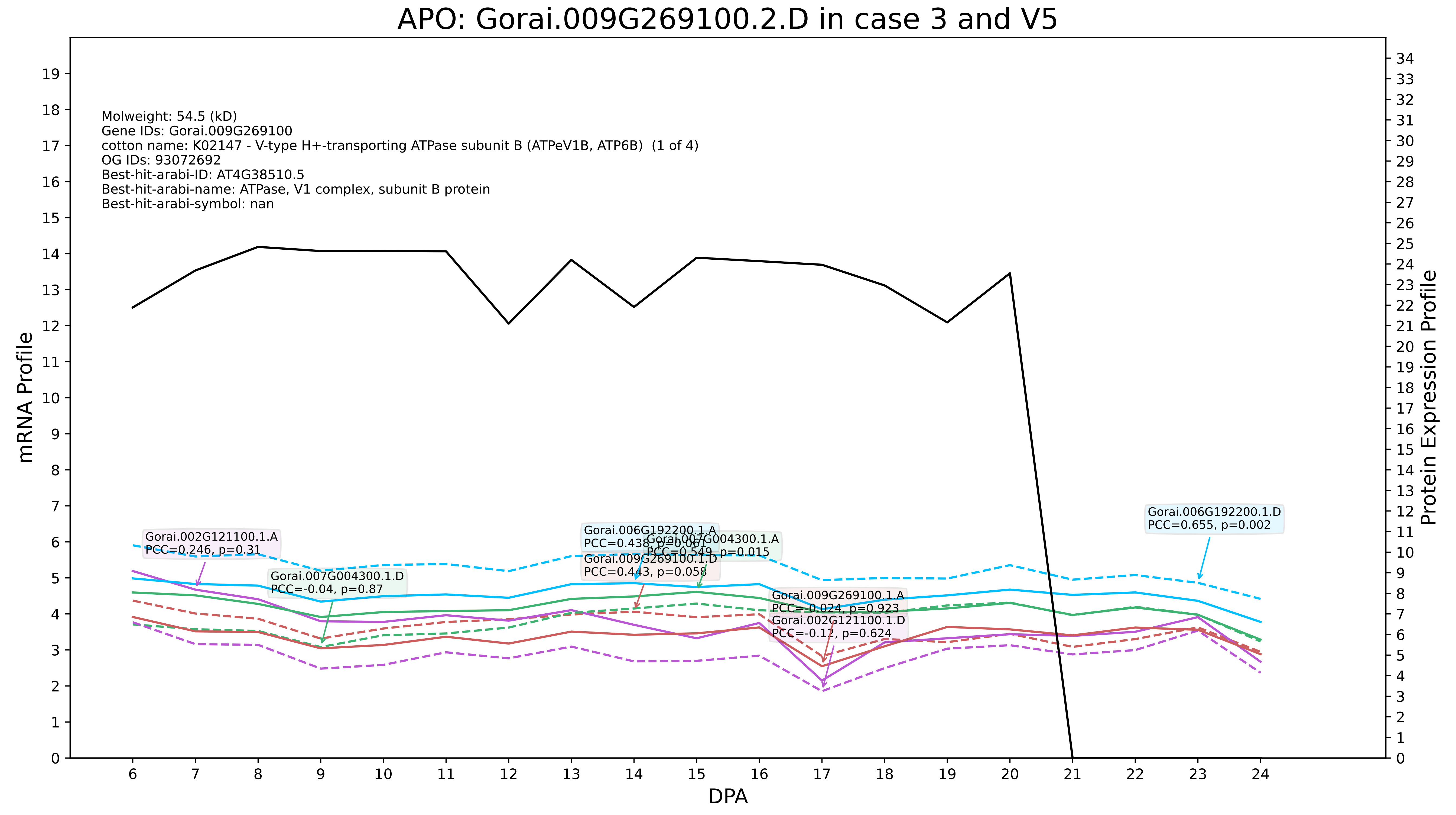
APO: Gorai.009G269100.2.D in case 3 and V5
Molweight: 54.5 (kD)
Gene IDs: Gorai.009G269100
cotton name: K02147 - V-type H+-transporting ATPase subunit B (ATPeV1B, ATP6B) (1 of 4)
OG IDs: 93072692
Best-hit-arabi-ID: AT4G38510.5
Best-hit-arabi-name: ATPase, V1 complex, subunit B protein
Best-hit-arabi-symbol: nan
Gorai.002G121100.1.A
PCC=0.246, p=0.31
Gorai.007G004300.1.D
PCC=-0.04, p=0.87
Gorai.006G192200.1.A
Gorai.007G004300.1.A
PCC=0.549, p=0.015
Gorai.009G269100.1.D
PCC=0.443, p=0.058
Gorai.009G269100.1.A
PCC=-0.024, p=0.923
Gorai.002G121100.1.D
PCC=-0.12, p=0.624
Gorai.006G192200.1.D
PCC=0.655, p=0.002
mRNA Profile
Protein Expression Profile
DPA

### APO: Gorai.008G149500.1.D in case 3 and V5

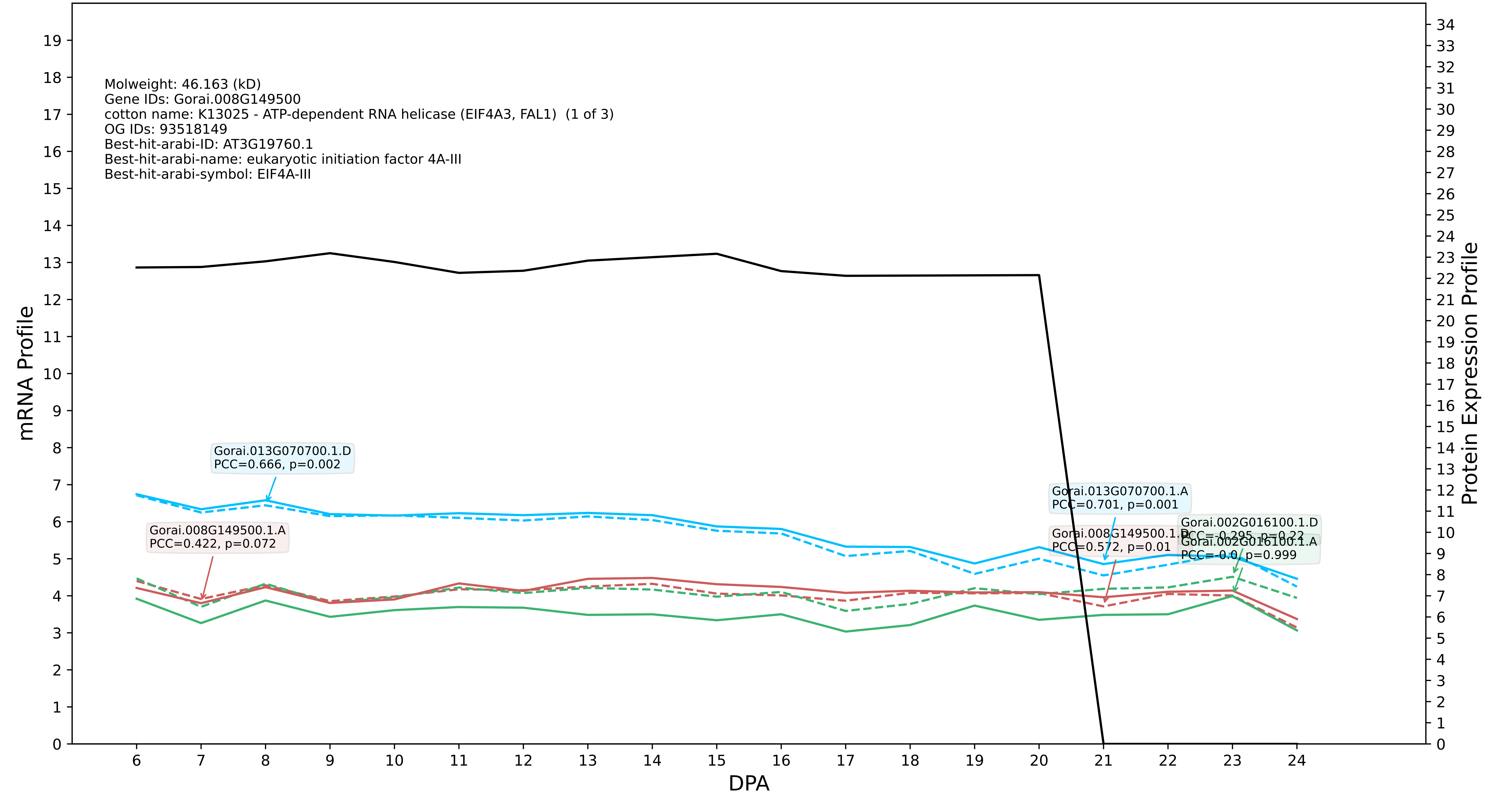

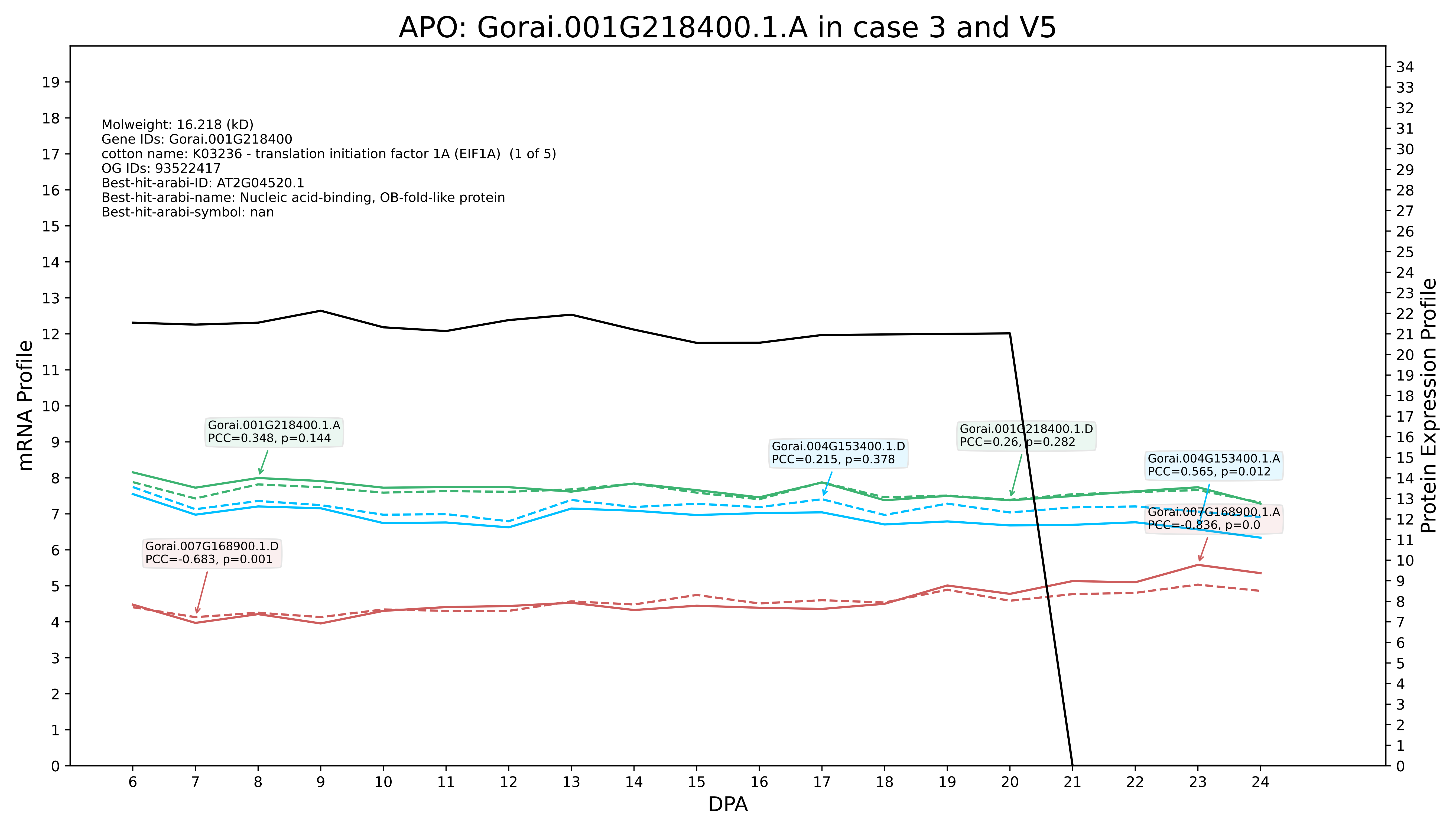

APO: Gorai.001G218400.1.A in case 3 and V5
Molweight: 16.218 (kD)
Gene IDs: Gorai.001G218400
cotton name: K03236 - translation initiation factor 1A (EIF1A) (1 of 5)
OG IDs: 93522417
Best-hit-arabi-ID: AT2G04520.1
Best-hit-arabi-name: Nucleic acid-binding, OB-fold-like protein
Best-hit-arabi-symbol: nan
Gorai.001G218400.1.A
PCC=0.348, p=0.144
Gorai.001G218400.1.D
PCC=0.26, p=0.282
Gorai.004G153400.1.D
PCC=0.215, p=0.378
Gorai.004G153400.1.A
PCC=0.565, p=0.012
Gorai.007G168900.1.A
PCC=-0.836, p=0.0
Gorai.007G168900.1.D
PCC=-0.683, p=0.001
mRNA Profile
Protein Expression Profile
DPA

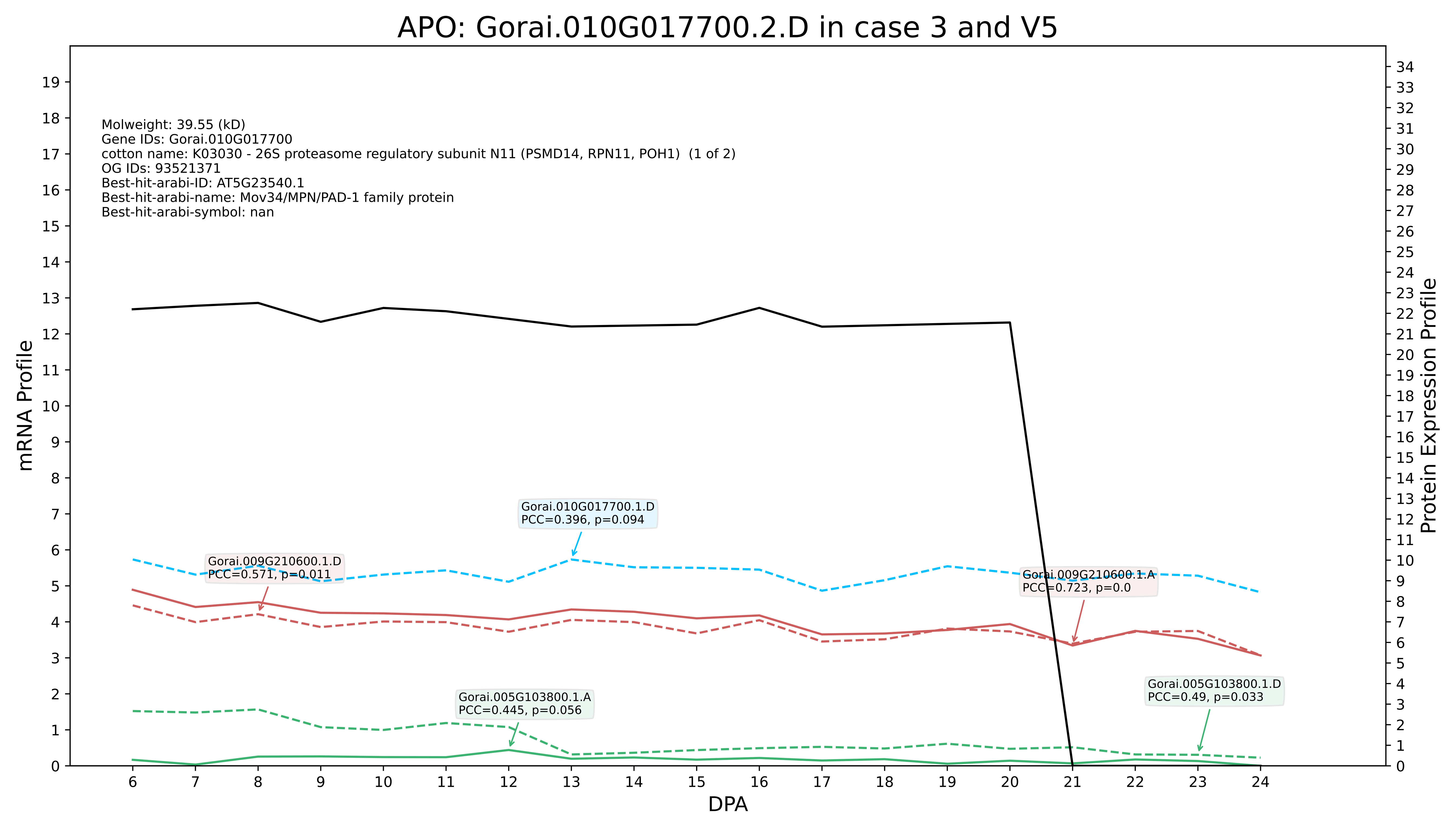
APO: Gorai.010G017700.2.D in case 3 and V5
Molweight: 39.55 (kD)
Gene IDs: Gorai.010G017700
cotton name: K03030 - 26S proteasome regulatory subunit N11 (PSMD14, RPN11, POH1) (1 of 2)
OG IDs: 93521371
Best-hit-arabi-ID: AT5G23540.1
Best-hit-arabi-name: Mov34/MPN/PAD-1 family protein
Best-hit-arabi-symbol: nan
Gorai.010G017700.1.D
PCC=0.396, p=0.094
Gorai.009G210600.1.D
PCC=0.571, p=0.011
Gorai.009G210600.1.A
PCC=0.723, p=0.0
Gorai.005G103800.1.A
PCC=0.445, p=0.056
Gorai.005G103800.1.D
PCC=0.49, p=0.033
mRNA Profile
Protein Expression Profile
DPA

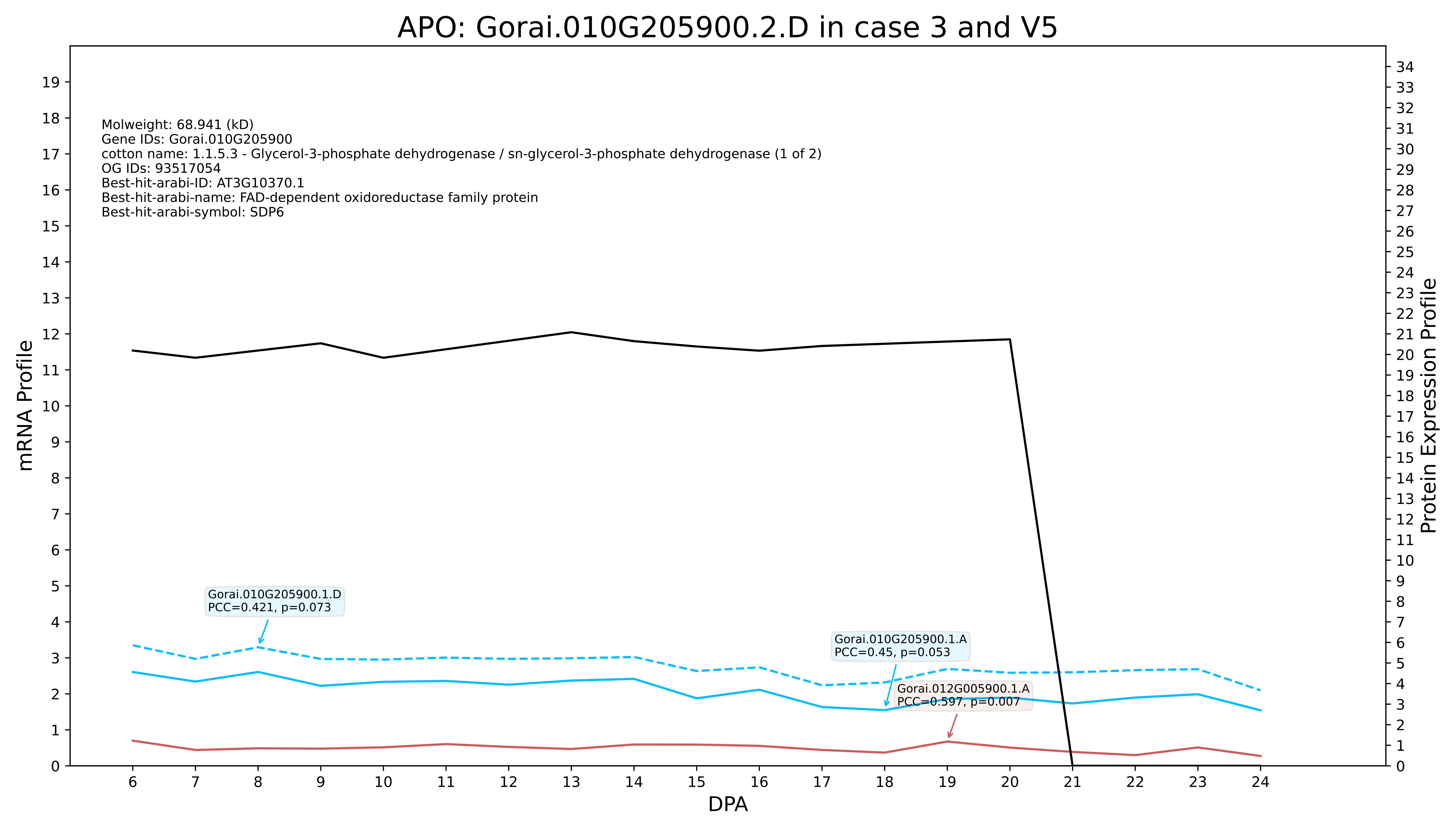
APO: Gorai.010G205900.2.D in case 3 and V5
Molweight: 68.941 (kD)
Gene IDs: Gorai.010G205900
cotton name: 1.1.5.3 - Glycerol-3-phosphate dehydrogenase / sn-glycerol-3-phosphate dehydrogenase (1 of 2)
OG IDs: 93517054
Best-hit-arabi-ID: AT3G10370.1
Best-hit-arabi-name: FAD-dependent oxidoreductase family protein
Best-hit-arabi-symbol: SDP6
Gorai.010G205900.1.D
PCC=0.421, p=0.073
Gorai.010G205900.1.A
PCC=0.45, p=0.053
Gorai.012G005900.1.A
PCC=0.597, p=0.007
mRNA Profile
Protein Expression Profile
DPA

### APO: Gorai.009G082000.1.D in case 3 and V5

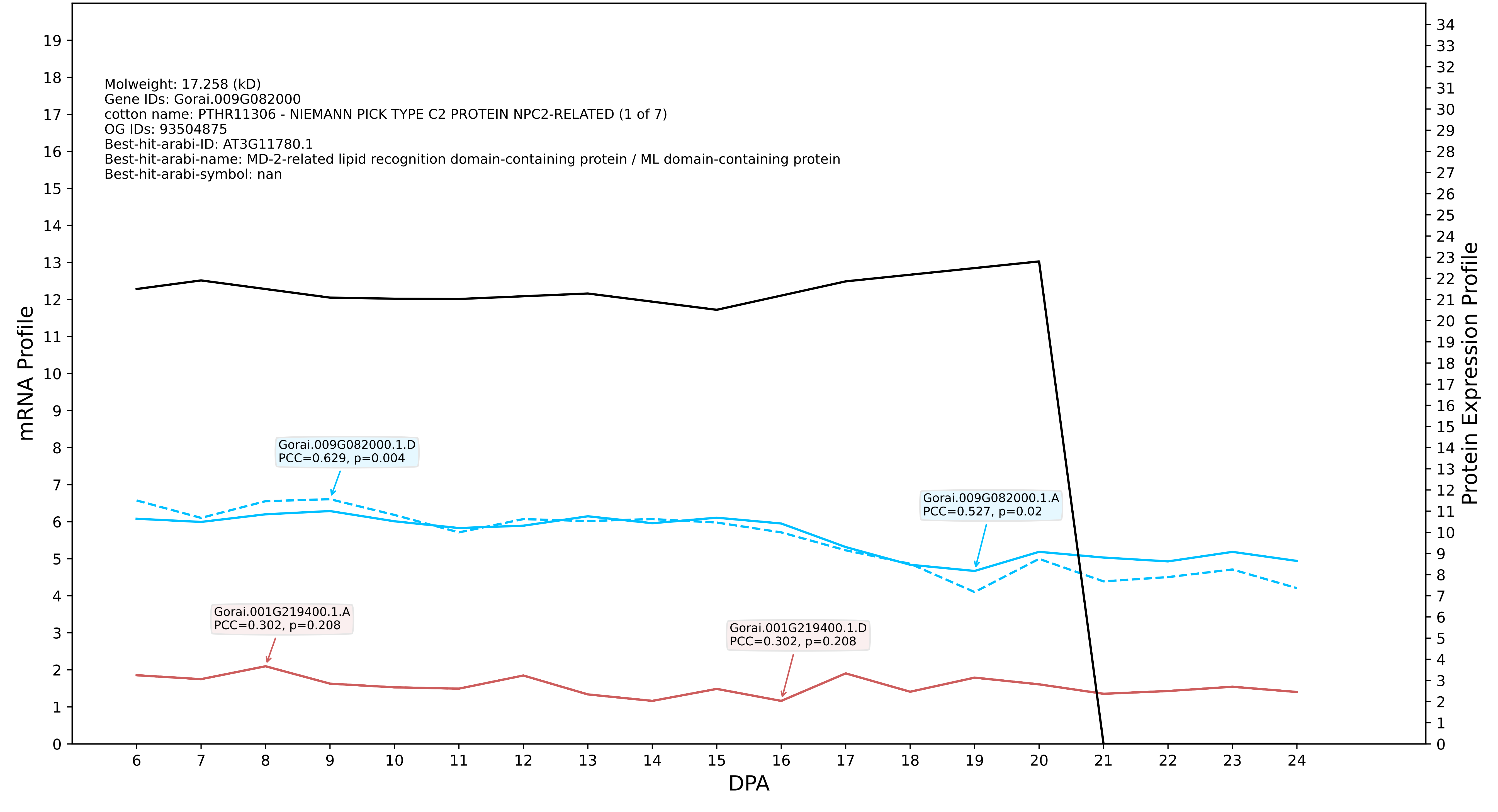

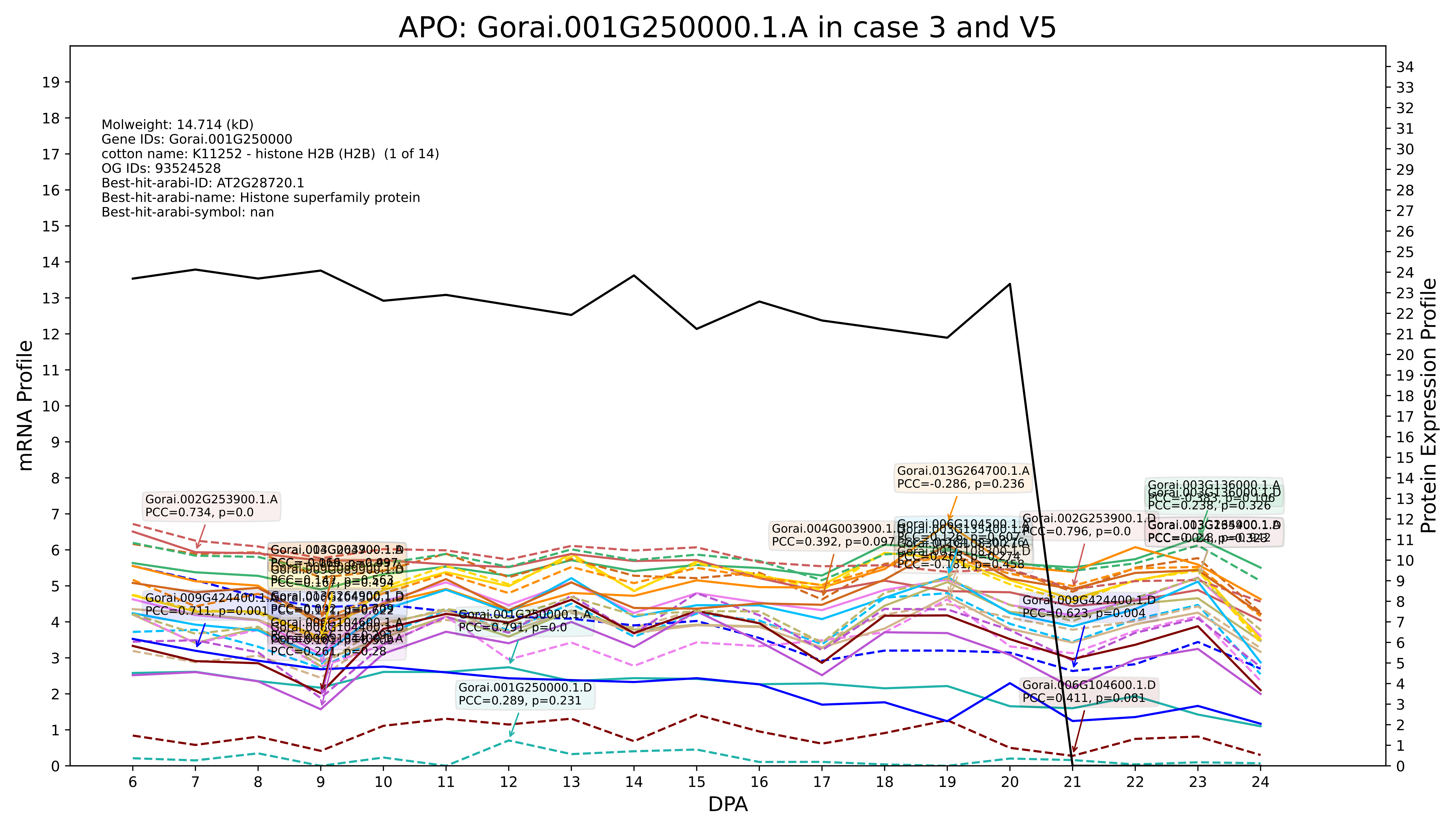

APO: Gorai.001G250000.1.A in case 3 and V5
Molweight: 14.714 (kD)
Gene IDs: Gorai.001G250000
cotton name: K11252 - histone H2B (H2B) (1 of 14)
OG IDs: 93524528
Best-hit-arabi-ID: AT2G28720.1
Best-hit-arabi-name: Histone superfamily protein
Best-hit-arabi-symbol: nan
Gorai.002G253900.1.A
PCC=0.734, p=0.0
Gorai.009G424400.1.A
PCC=0.711, p=0.001
Gorai.001G250000.1.A
PCC=0.791, p=0.0
Gorai.001G250000.1.D
PCC=0.289, p=0.231
Gorai.013G264700.1.A
PCC=-0.286, p=0.236
Gorai.004G003900.1.D
PCC=0.392, p=0.097
Gorai.002G253900.1.D
PCC=0.796, p=0.0
Gorai.009G424400.1.D
PCC=0.623, p=0.004
Gorai.006G104600.1.D
PCC=0.411, p=0.081
Gorai.003G136000.1.A
Gorai.003G136000.1.D
PCC=0.238, p=0.326
mRNA Profile
Protein Expression Profile
DPA

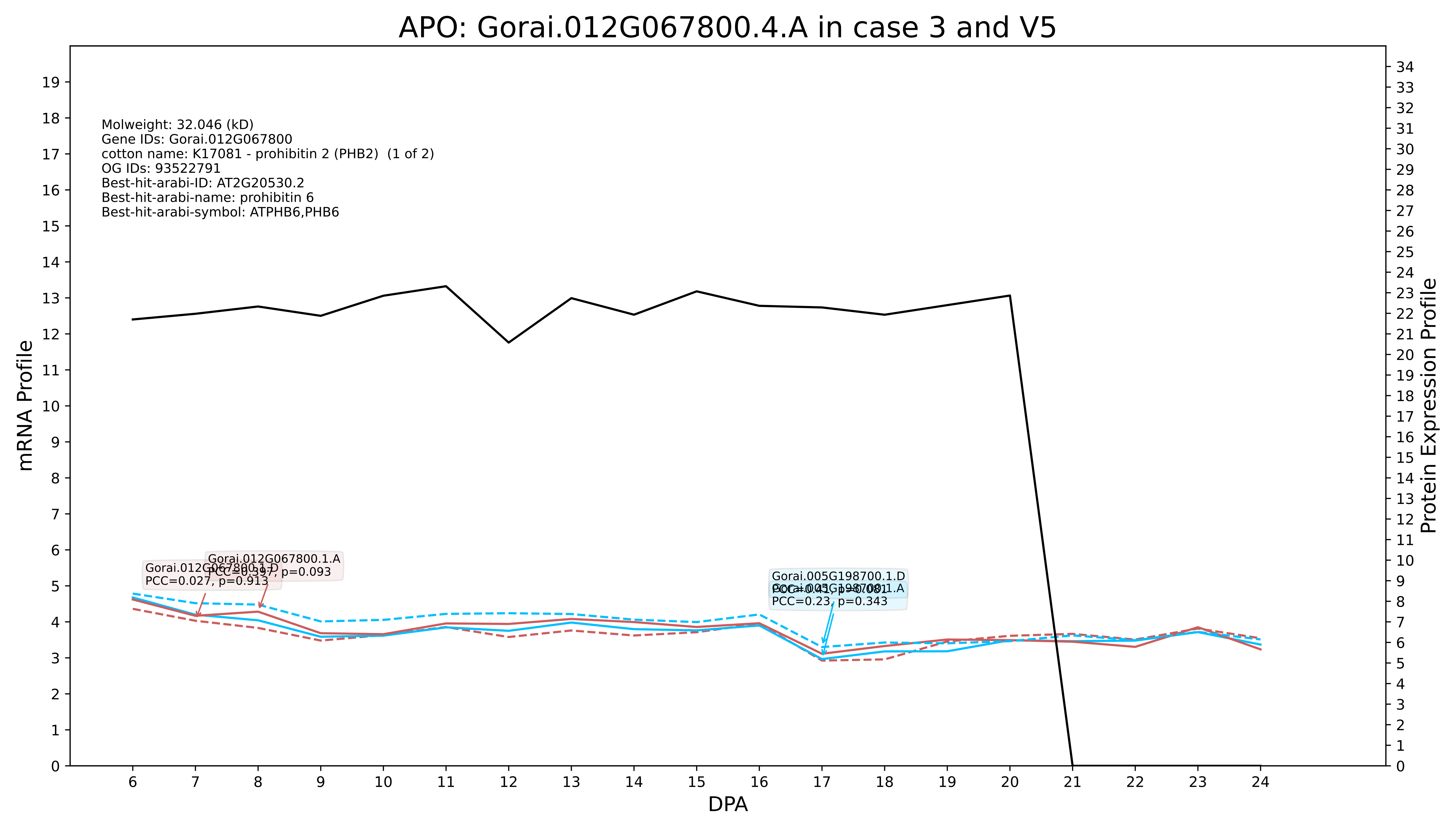
APO: Gorai.012G067800.4.A in case 3 and V5
Molweight: 32.046 (kD)
Gene IDs: Gorai.012G067800
cotton name: K17081 - prohibitin 2 (PHB2) (1 of 2)
OG IDs: 93522791
Best-hit-arabi-ID: AT2G20530.2
Best-hit-arabi-name: prohibitin 6
Best-hit-arabi-symbol: ATPHB6,PHB6
Gorai.012G067800.1.D
PCC=0.027, p=0.913
Gorai.012G067800.1.A
PCC=0.397, p=0.093
Gorai.005G198700.1.D
PCC=0.41, p=0.081
Gorai.005G198700.1.A
PCC=0.23, p=0.343
mRNA Profile
Protein Expression Profile
DPA

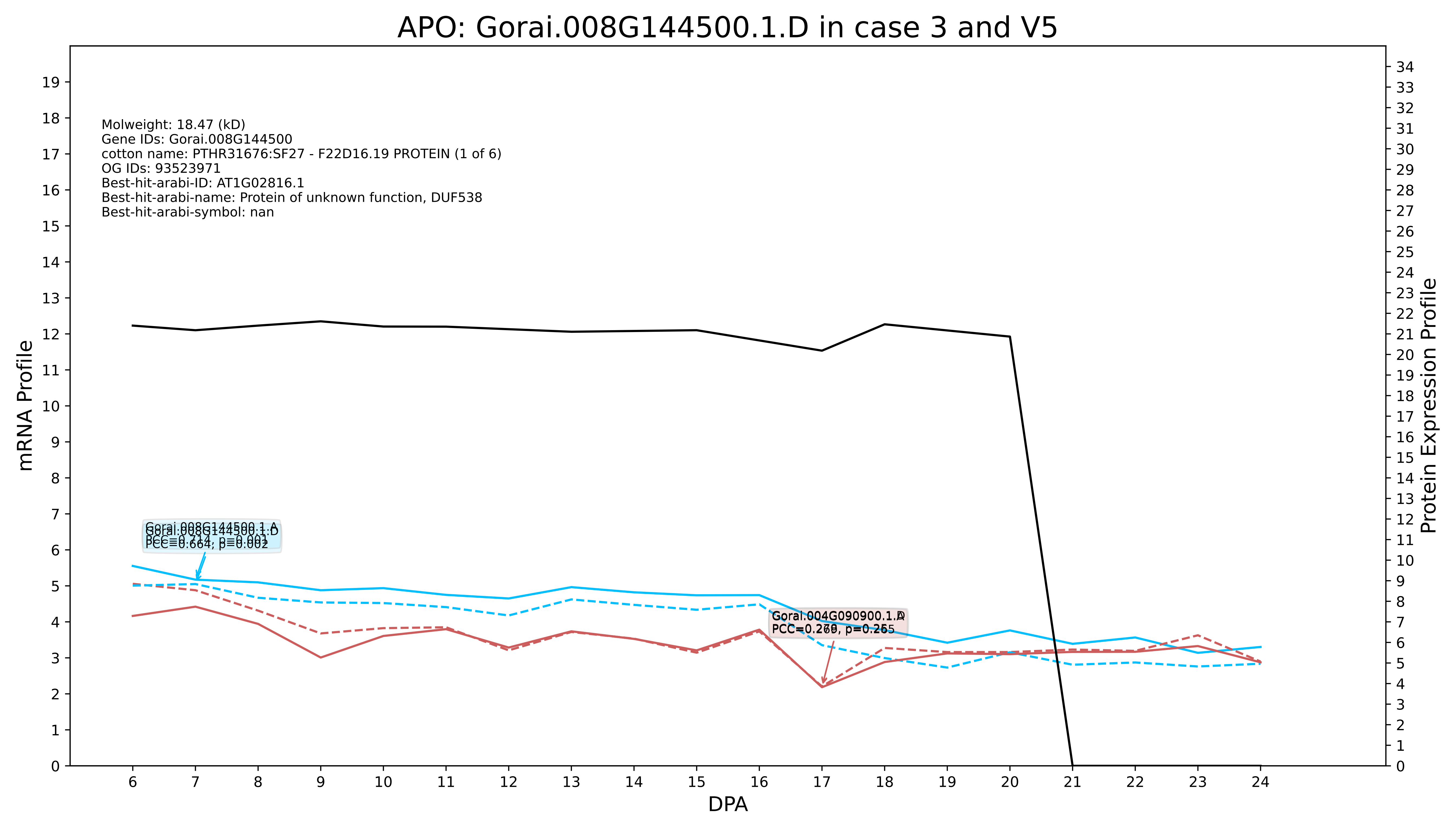

APO: Gorai.008G144500.1.D in case 3 and V5
Molweight: 18.47 (kD)
Gene IDs: Gorai.008G144500
cotton name: PTHR31676:SF27 - F22D16.19 PROTEIN (1 of 6)
OG IDs: 93523971
Best-hit-arabi-ID: AT1G02816.1
Best-hit-arabi-name: Protein of unknown function, DUF538
Best-hit-arabi-symbol: nan
Gorai.008G144500.1.A
Gorai.008G144500.1.D
PCC=0.714, p=0.001
PCC=0.664, p=0.002
Gorai.004G090900.1.A
Gorai.004G090900.1.D
PCC=0.279, p=0.26
PCC=0.265
mRNA Profile
Protein Expression Profile
DPA

### APO: Gorai.006G208700.1.D in case 3 and V5

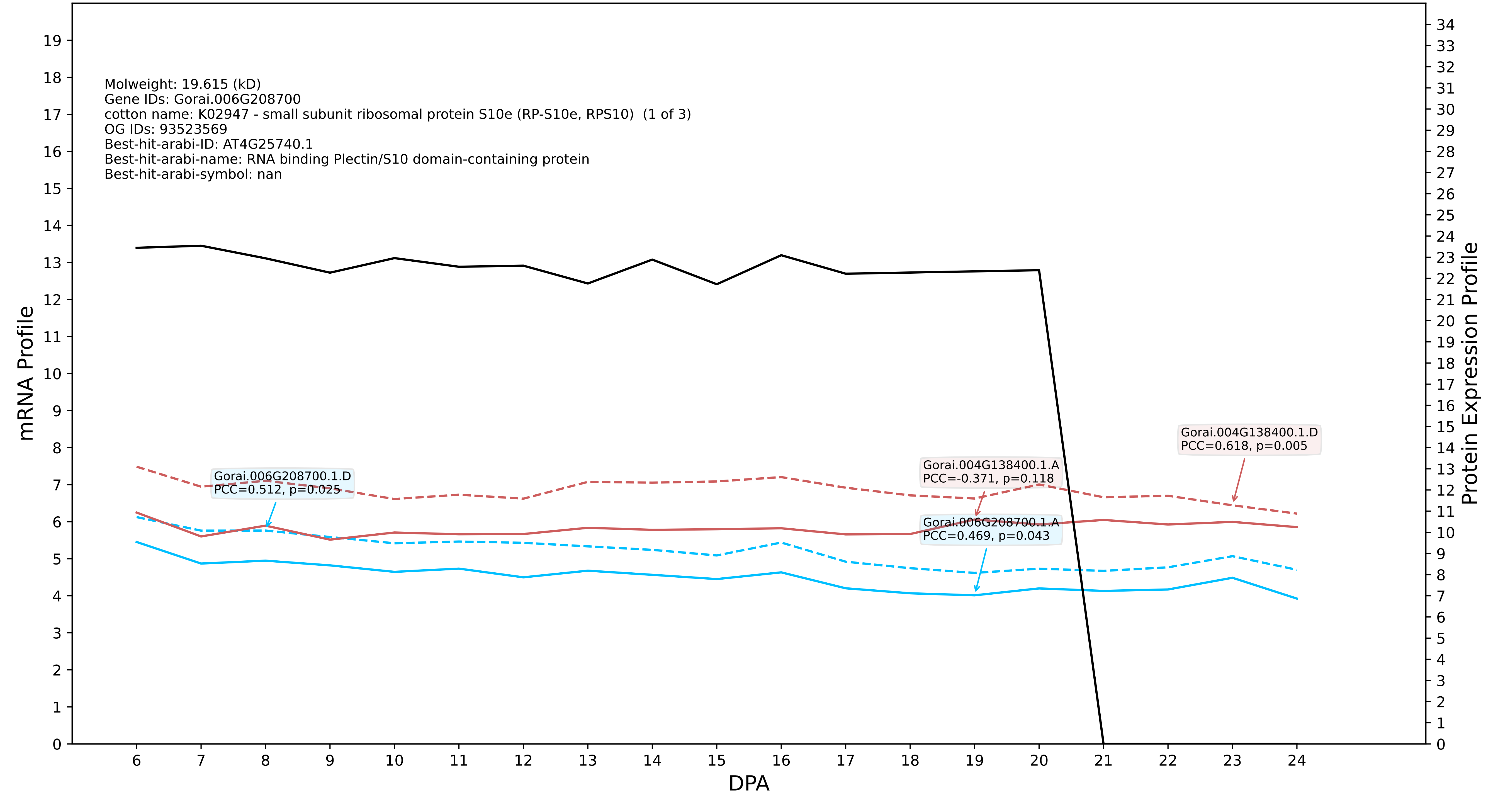

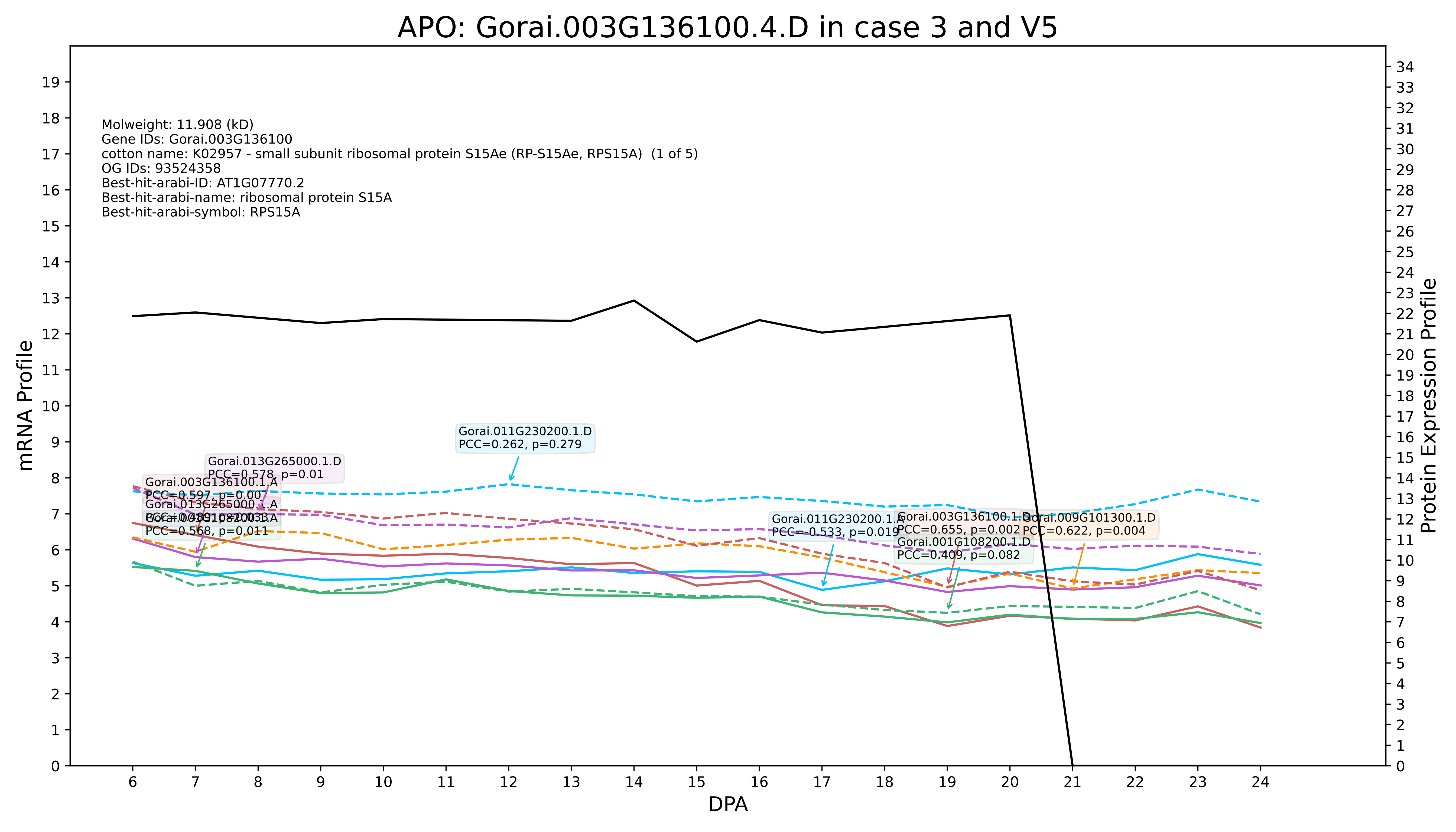
APO: Gorai.003G136100.4.D in case 3 and V5
Molweight: 11.908 (kD)
Gene IDs: Gorai.003G136100
cotton name: K02957 - small subunit ribosomal protein S15Ae (RP-S15Ae, RPS15A) (1 of 5)
OG IDs: 93524358
Best-hit-arabi-ID: AT1G07770.2
Best-hit-arabi-name: ribosomal protein S15A
Best-hit-arabi-symbol: RPS15A
Gorai.011G230200.1.D
PCC=0.262, p=0.279
Gorai.013G265000.1.D
PCC=0.578, p=0.01
Gorai.003G136100.1.A
PCC=0.597, p=0.007
Gorai.013G265000.1.A
Gorai.011G230200.1.A
PCC=-0.533, p=0.019
Gorai.003G136100.1.D
PCC=0.655, p=0.002
Gorai.009G101300.1.D
PCC=0.622, p=0.004
Gorai.001G108200.1.D
PCC=0.409, p=0.082
PCC=0.568, p=0.011
mRNA Profile
Protein Expression Profile
DPA

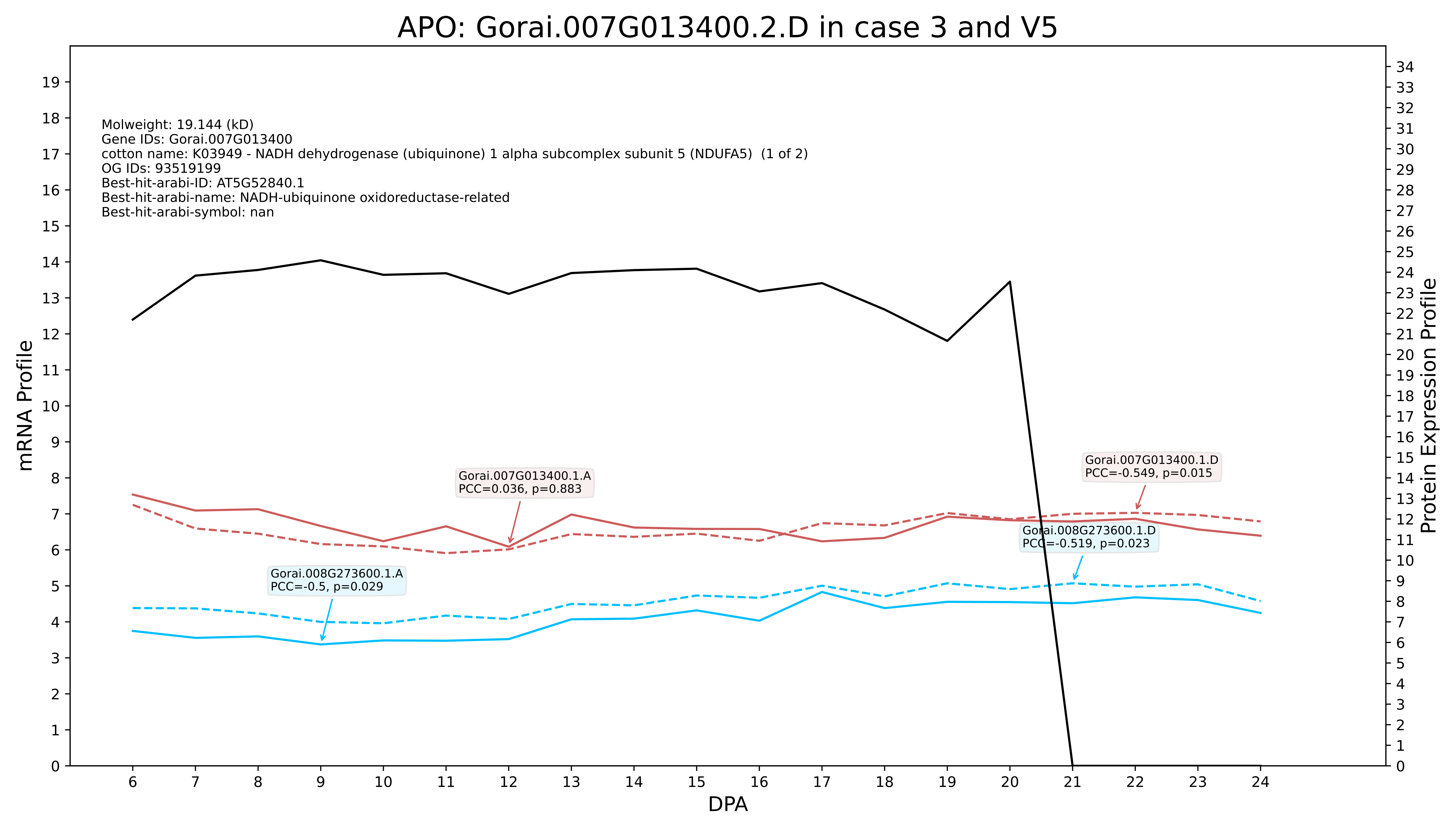
APO: Gorai.007G013400.2.D in case 3 and V5
Molweight: 19.144 (kD)
Gene IDs: Gorai.007G013400
cotton name: K03949 - NADH dehydrogenase (ubiquinone) 1 alpha subcomplex subunit 5 (NDUFA5) (1 of 2)
OG IDs: 93519199
Best-hit-arabi-ID: AT5G52840.1
Best-hit-arabi-name: NADH-ubiquinone oxidoreductase-related
Best-hit-arabi-symbol: nan
Gorai.007G013400.1.A
PCC=0.036, p=0.883
Gorai.007G013400.1.D
PCC=-0.549, p=0.015
Gorai.008G273600.1.D
PCC=-0.519, p=0.023
Gorai.008G273600.1.A
PCC=-0.5, p=0.029
mRNA Profile
Protein Expression Profile
DPA

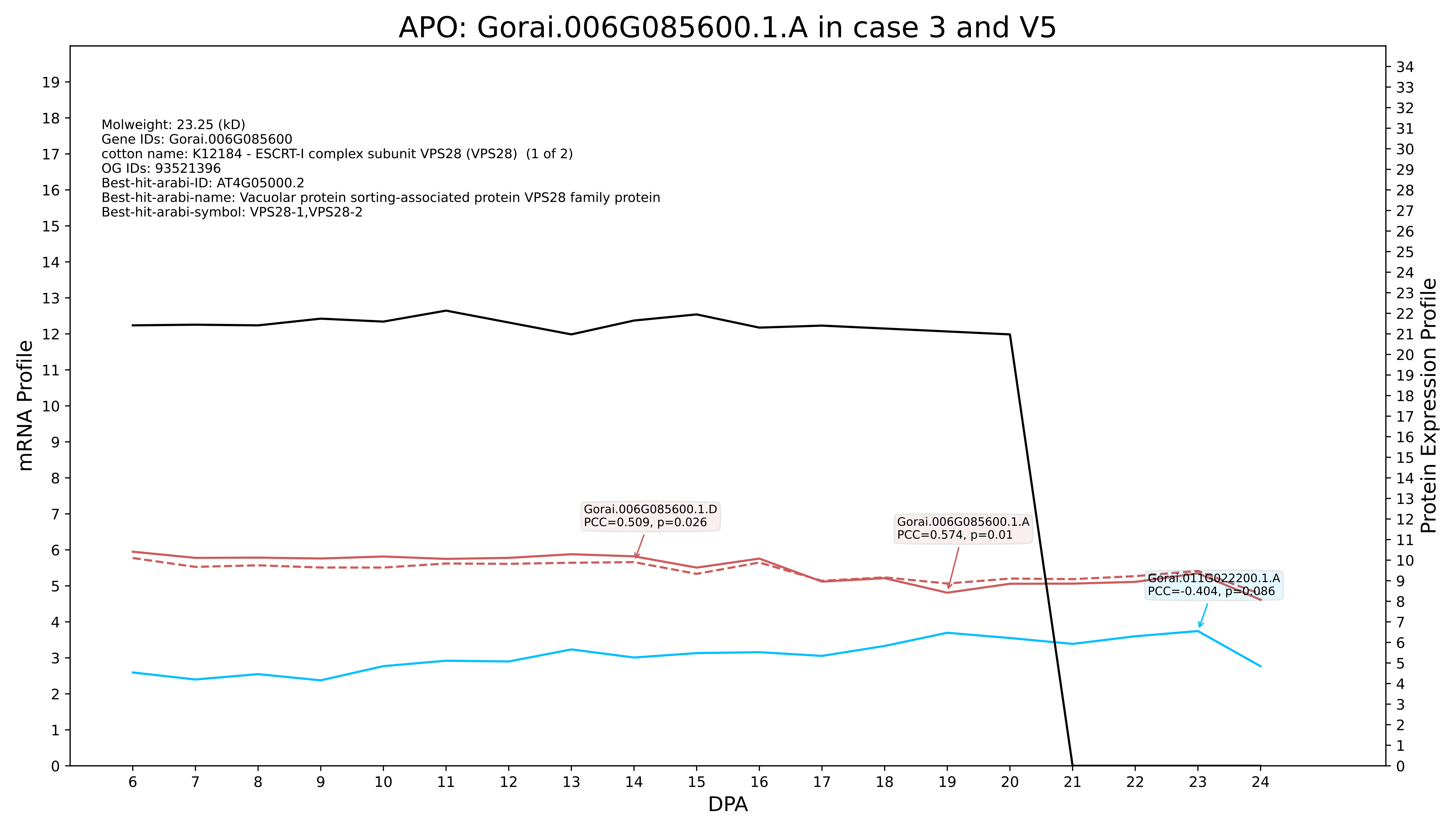
APO: Gorai.006G085600.1.A in case 3 and V5
Molweight: 23.25 (kD)
Gene IDs: Gorai.006G085600
cotton name: K12184 - ESCRT-I complex subunit VPS28 (VPS28) (1 of 2)
OG IDs: 93521396
Best-hit-arabi-ID: AT4G05000.2
Best-hit-arabi-name: Vacuolar protein sorting-associated protein VPS28 family protein
Best-hit-arabi-symbol: VPS28-1,VPS28-2
Gorai.006G085600.1.D
PCC=0.509, p=0.026
Gorai.006G085600.1.A
PCC=0.574, p=0.01
Gorai.011G022200.1.A
PCC=-0.404, p=0.086
mRNA Profile
Protein Expression Profile
DPA

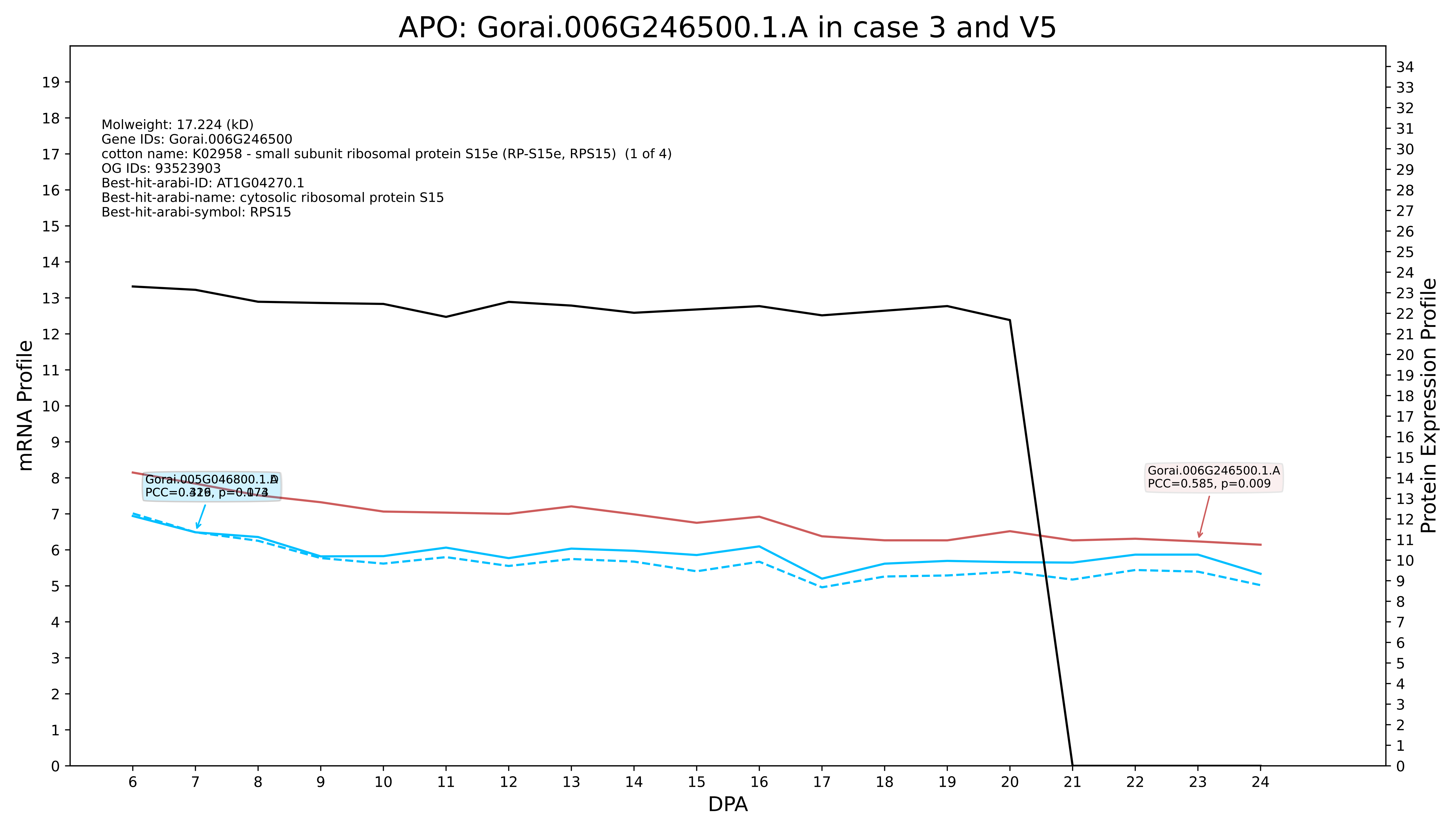

APO: Gorai.006G246500.1.A in case 3 and V5
Molweight: 17.224 (kD)
Gene IDs: Gorai.006G246500
cotton name: K02958 - small subunit ribosomal protein S15e (RP-S15e, RPS15) (1 of 4)
OG IDs: 93523903
Best-hit-arabi-ID: AT1G04270.1
Best-hit-arabi-name: cytosolic ribosomal protein S15
Best-hit-arabi-symbol: RPS15
Gorai.006G246500.1.A
PCC=0.585, p=0.009
mRNA Profile
Protein Expression Profile
DPA

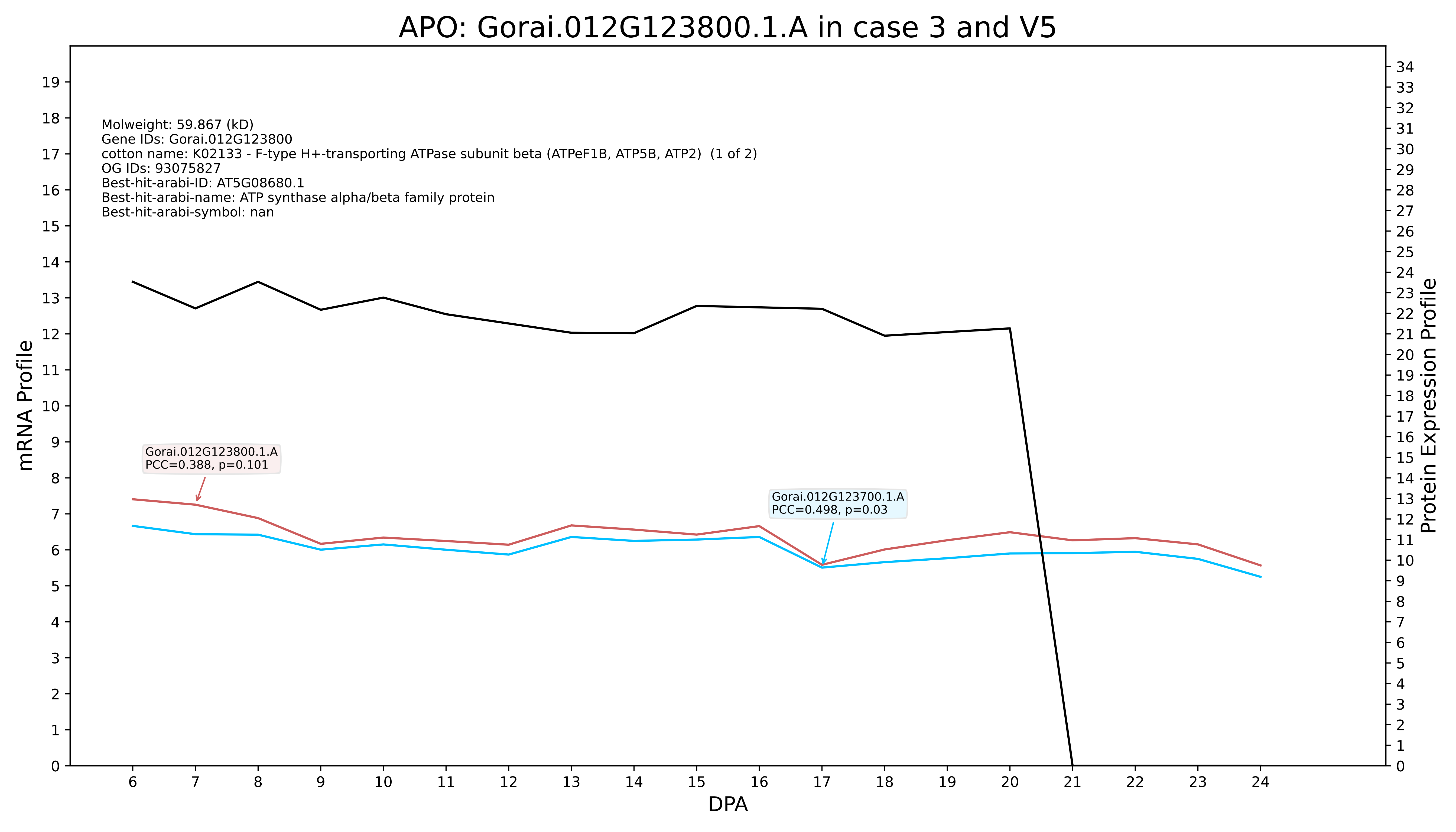
APO: Gorai.012G123800.1.A in case 3 and V5
Molweight: 59.867 (kD)
Gene IDs: Gorai.012G123800
cotton name: K02133 - F-type H+-transporting ATPase subunit beta (ATPeF1B, ATP5B, ATP2) (1 of 2)
OG IDs: 93075827
Best-hit-arabi-ID: AT5G08680.1
Best-hit-arabi-name: ATP synthase alpha/beta family protein
Best-hit-arabi-symbol: nan
Gorai.012G123800.1.A
PCC=0.388, p=0.101
Gorai.012G123700.1.A
PCC=0.498, p=0.03
mRNA Profile
Protein Expression Profile
DPA

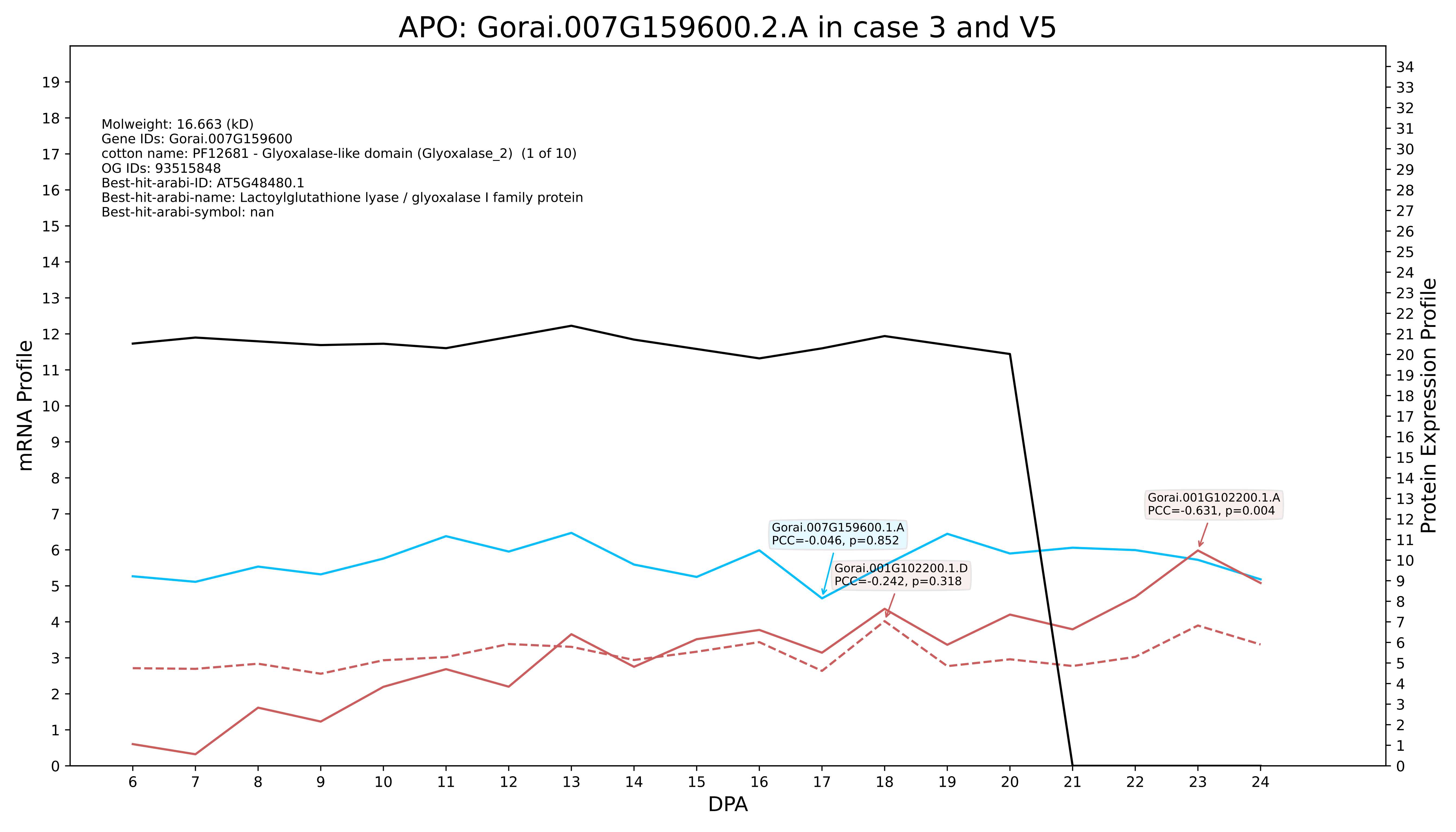

APO: Gorai.007G159600.2.A in case 3 and V5
Molweight: 16.663 (kD)
Gene IDs: Gorai.007G159600
cotton name: PF12681 - Glyoxalase-like domain (Glyoxalase_2) (1 of 10)
OG IDs: 93515848
Best-hit-arabi-ID: AT5G48480.1
Best-hit-arabi-name: Lactoylglutathione lyase / glyoxalase I family protein
Best-hit-arabi-symbol: nan
Gorai.007G159600.1.A
PCC=-0.046, p=0.852
Gorai.001G102200.1.D
PCC=-0.242, p=0.318
Gorai.001G102200.1.A
PCC=-0.631, p=0.004
mRNA Profile
Protein Expression Profile
DPA

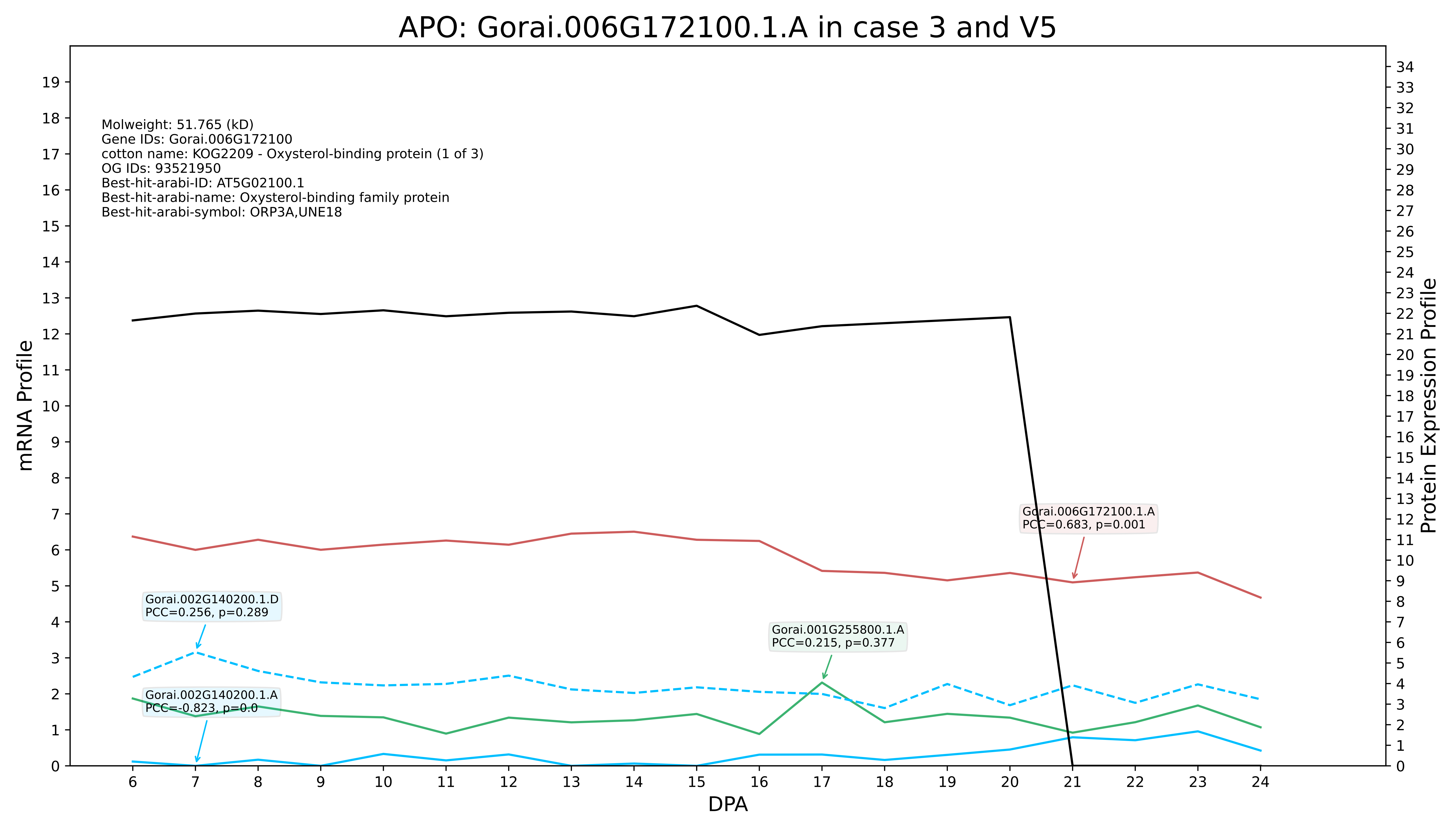
APO: Gorai.006G172100.1.A in case 3 and V5
Molweight: 51.765 (kD)
Gene IDs: Gorai.006G172100
cotton name: KOG2209 - Oxysterol-binding protein (1 of 3)
OG IDs: 93521950
Best-hit-arabi-ID: AT5G02100.1
Best-hit-arabi-name: Oxysterol-binding family protein
Best-hit-arabi-symbol: ORP3A,UNE18
Gorai.006G172100.1.A
PCC=0.683, p=0.001
Gorai.002G140200.1.D
PCC=0.256, p=0.289
Gorai.001G255800.1.A
PCC=0.215, p=0.377
Gorai.002G140200.1.A
PCC=-0.823, p=0.0
mRNA Profile
Protein Expression Profile
DPA

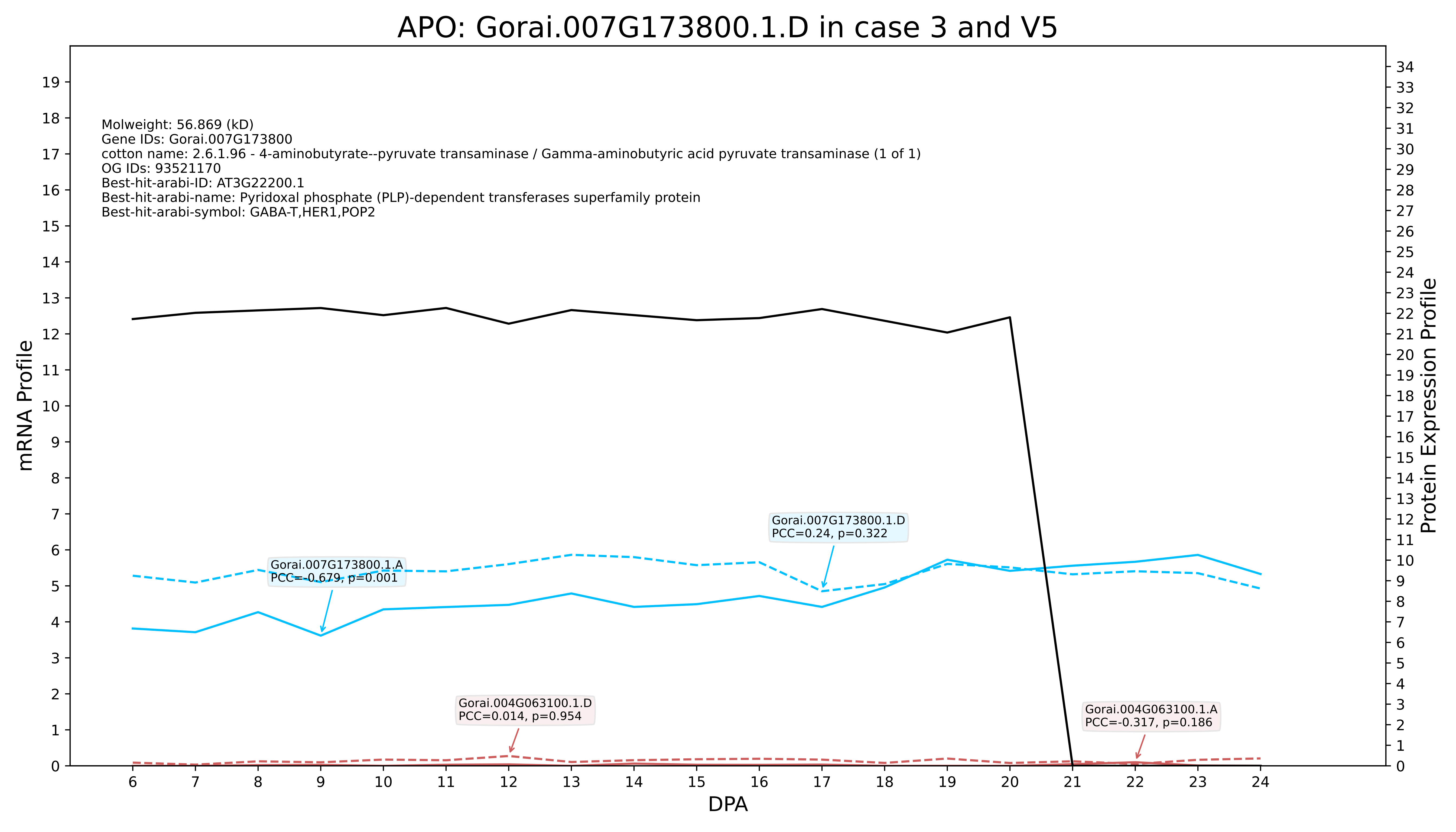
APO: Gorai.007G173800.1.D in case 3 and V5
Molweight: 56.869 (kD)
Gene IDs: Gorai.007G173800
cotton name: 2.6.1.96 - 4-aminobutyrate--pyruvate transaminase / Gamma-aminobutyric acid pyruvate transaminase (1 of 1)
OG IDs: 93521170
Best-hit-arabi-ID: AT3G22200.1
Best-hit-arabi-name: Pyridoxal phosphate (PLP)-dependent transferases superfamily protein
Best-hit-arabi-symbol: GABA-T,HER1,POP2
Gorai.007G173800.1.D
PCC=0.24, p=0.322
Gorai.007G173800.1.A
PCC=-0.679, p=0.001
Gorai.004G063100.1.D
PCC=0.014, p=0.954
Gorai.004G063100.1.A
PCC=-0.317, p=0.186
mRNA Profile
Protein Expression Profile
DPA

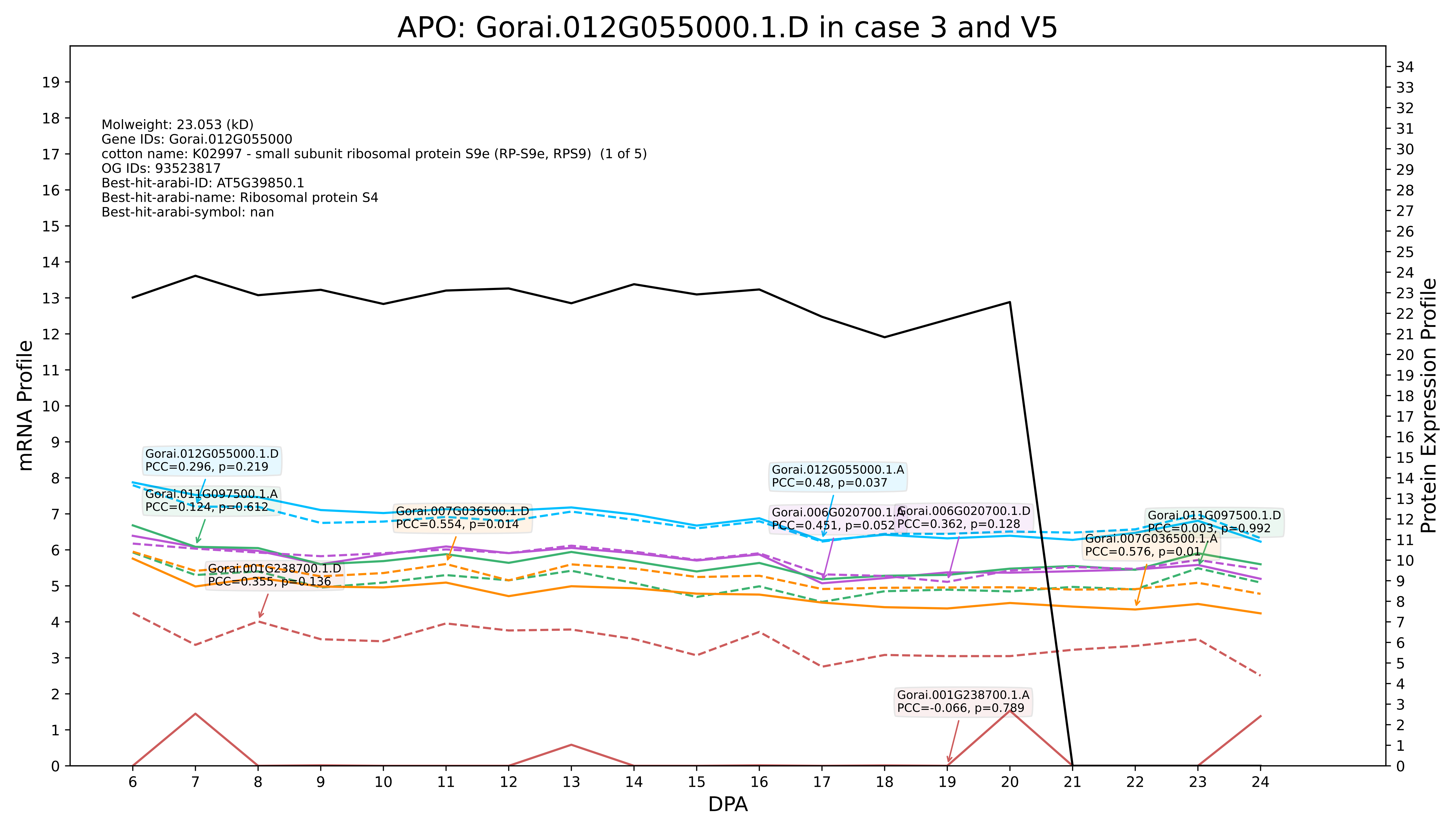

APO: Gorai.012G055000.1.D in case 3 and V5
Molweight: 23.053 (kD)
Gene IDs: Gorai.012G055000
cotton name: K02997 - small subunit ribosomal protein S9e (RP-S9e, RPS9) (1 of 5)
OG IDs: 93523817
Best-hit-arabi-ID: AT5G39850.1
Best-hit-arabi-name: Ribosomal protein S4
Best-hit-arabi-symbol: nan
Gorai.012G055000.1.D
PCC=0.296, p=0.219
Gorai.011G097500.1.A
PCC=0.124, p=0.612
Gorai.007G036500.1.D
PCC=0.554, p=0.014
Gorai.001G238700.1.D
PCC=0.355, p=0.136
Gorai.012G055000.1.A
PCC=0.48, p=0.037
Gorai.006G020700.1.A
PCC=0.451, p=0.052
Gorai.006G020700.1.D
PCC=0.362, p=0.128
Gorai.011G097500.1.D
PCC=0.003, p=0.992
Gorai.007G036500.1.A
PCC=0.576, p=0.01
Gorai.001G238700.1.A
PCC=-0.066, p=0.789
mRNA Profile
Protein Expression Profile
DPA

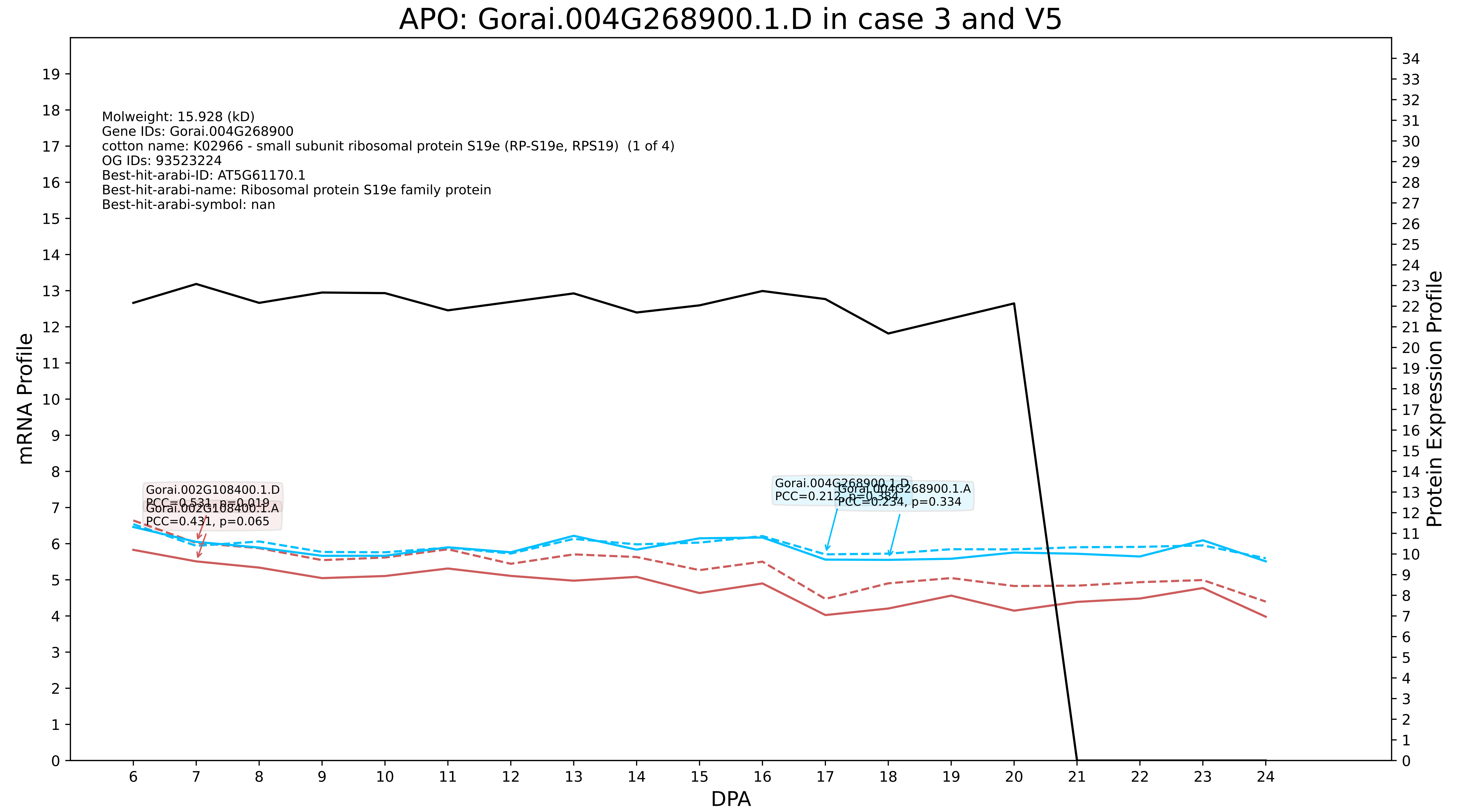

APO: Gorai.004G268900.1.D in case 3 and V5
Molweight: 15.928 (kD)
Gene IDs: Gorai.004G268900
cotton name: K02966 - small subunit ribosomal protein S19e (RP-S19e, RPS19) (1 of 4)
OG IDs: 93523224
Best-hit-arabi-ID: AT5G61170.1
Best-hit-arabi-name: Ribosomal protein S19e family protein
Best-hit-arabi-symbol: nan
Gorai.002G108400.1.D
PCC=0.531, p=0.019
Gorai.002G108400.1.A
PCC=0.431, p=0.065
Gorai.004G268900.1.D
PCC=0.212, p=0.384
Gorai.004G268900.1.A
PCC=0.234, p=0.334
mRNA Profile
Protein Expression Profile
DPA

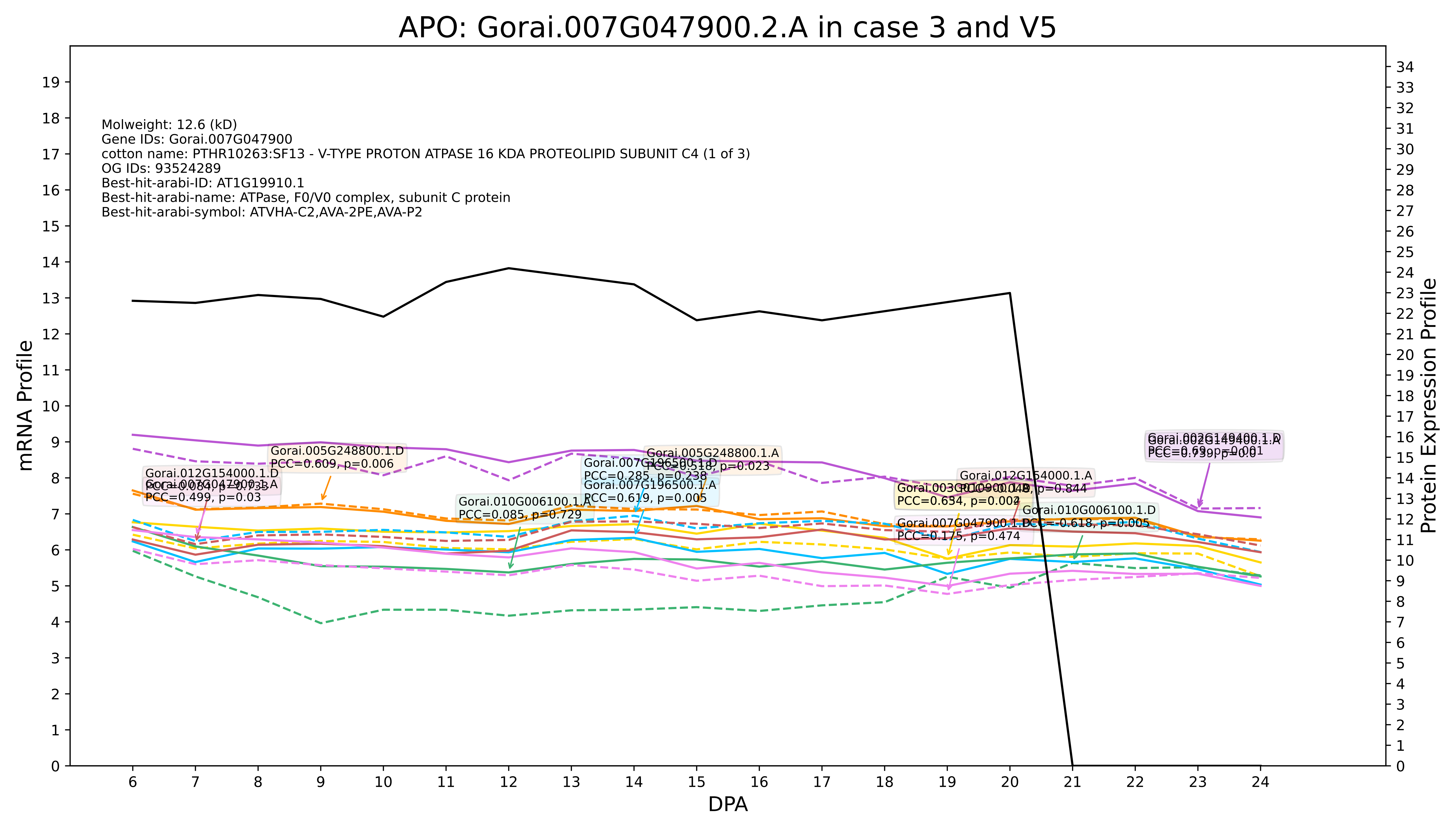

APO: Gorai.007G047900.2.A in case 3 and V5
Molweight: 12.6 (kD)
Gene IDs: Gorai.007G047900
cotton name: PTHR10263:SF13 - V-TYPE PROTON ATPASE 16 KDA PROTEOLIPID SUBUNIT C4 (1 of 3)
OG IDs: 93524289
Best-hit-arabi-ID: AT1G19910.1
Best-hit-arabi-name: ATPase, F0/V0 complex, subunit C protein
Best-hit-arabi-symbol: ATVHA-C2,AVA-2PE,AVA-P2
Gorai.005G248800.1.D
PCC=0.609, p=0.006
Gorai.012G154000.1.D
PCC=0.499, p=0.03
Gorai.005G248800.1.A
Gorai.010G006100.1.A
PCC=0.085, p=0.729
Gorai.007G196500.1.A
PCC=0.619, p=0.005
Gorai.012G154000.1.A
PCC=0.634, p=0.004
Gorai.010G006100.1.D
PCC=0.618, p=0.005
PCC=0.175, p=0.474
mRNA Profile
Protein Expression Profile
DPA

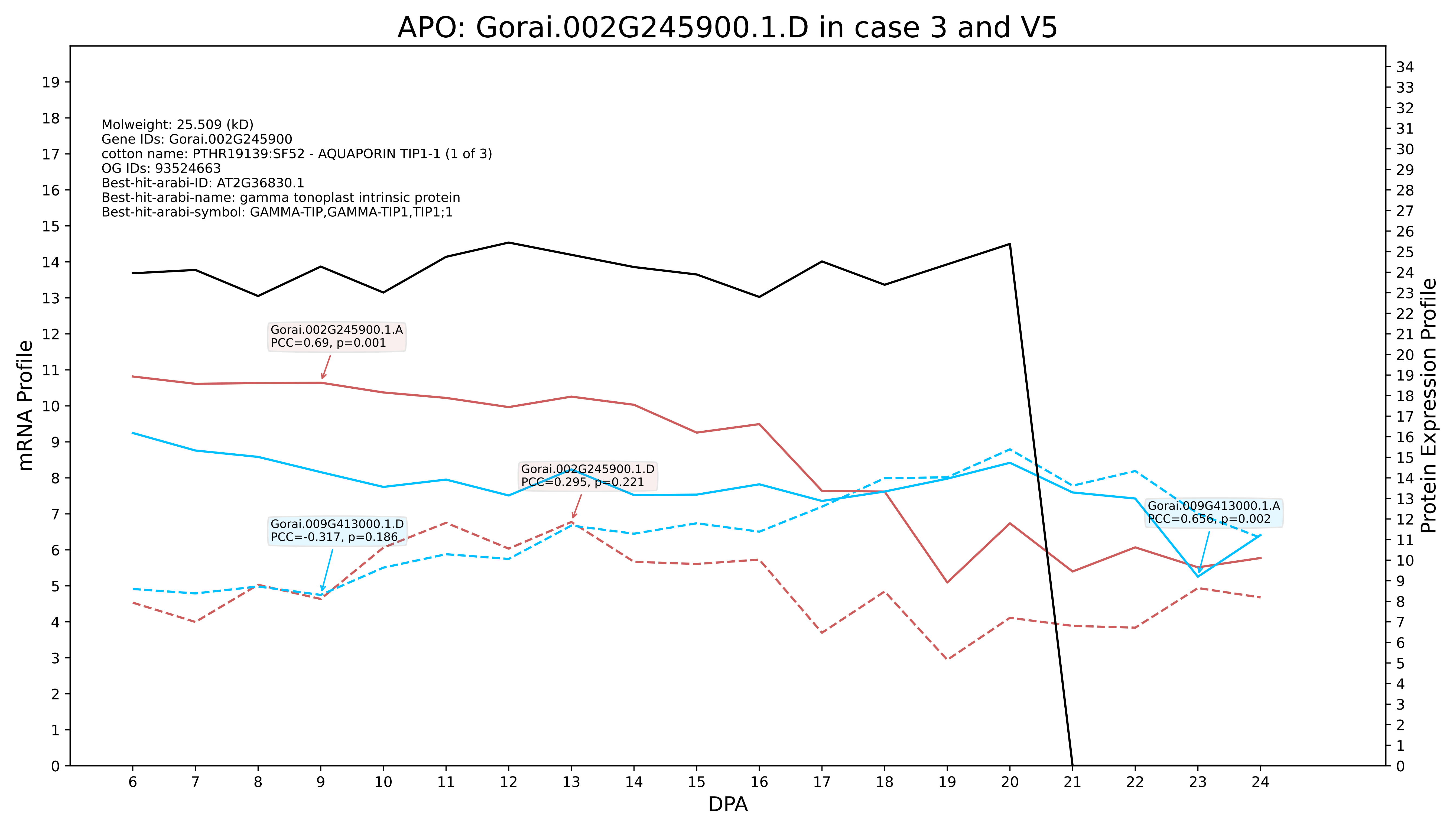
APO: Gorai.002G245900.1.D in case 3 and V5
Molweight: 25.509 (kD)
Gene IDs: Gorai.002G245900
cotton name: PTHR19139:SF52 - AQUAPORIN TIP1-1 (1 of 3)
OG IDs: 93524663
Best-hit-arabi-ID: AT2G36830.1
Best-hit-arabi-name: gamma tonoplast intrinsic protein
Best-hit-arabi-symbol: GAMMA-TIP,GAMMA-TIP1,TIP1;1
Gorai.002G245900.1.A
PCC=0.69, p=0.001
Gorai.002G245900.1.D
PCC=0.295, p=0.221
Gorai.009G413000.1.D
PCC=-0.317, p=0.186
Gorai.009G413000.1.A
PCC=0.656, p=0.002
mRNA Profile
Protein Expression Profile
DPA

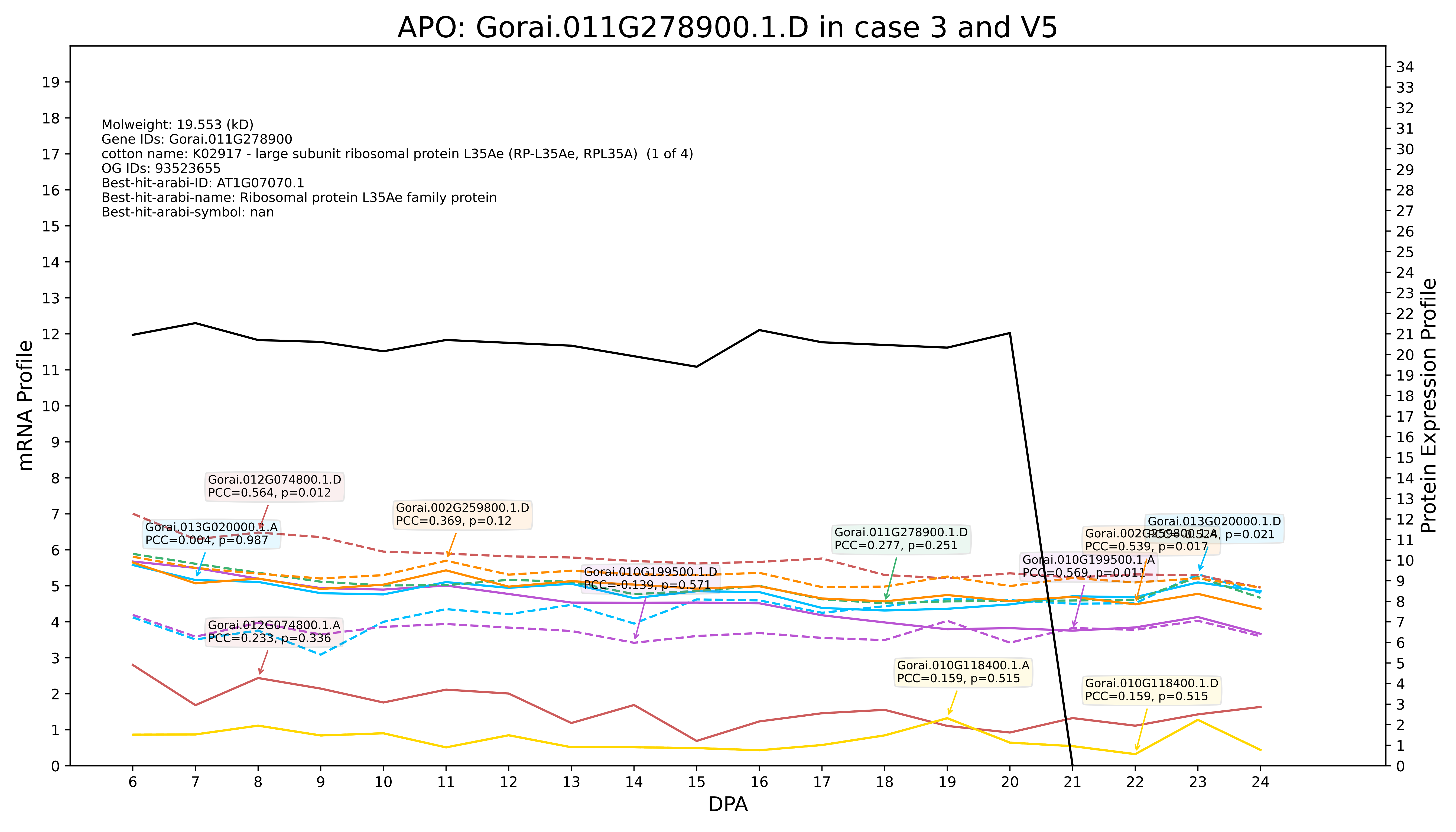
APO: Gorai.011G278900.1.D in case 3 and V5
Molweight: 19.553 (kD)
Gene IDs: Gorai.011G278900
cotton name: K02917 - large subunit ribosomal protein L35Ae (RP-L35Ae, RPL35A) (1 of 4)
OG IDs: 93523655
Best-hit-arabi-ID: AT1G07070.1
Best-hit-arabi-name: Ribosomal protein L35Ae family protein
Best-hit-arabi-symbol: nan
Gorai.012G074800.1.D
PCC=0.564, p=0.012
Gorai.002G259800.1.D
PCC=0.369, p=0.12
Gorai.013G020000.1.A
PCC=0.004, p=0.987
Gorai.011G278900.1.D
PCC=0.277, p=0.251
Gorai.013G020000.1.D
Gorai.010G199500.1.A
PCC=0.569, p=0.011
Gorai.010G199500.1.D
PCC=-0.139, p=0.571
Gorai.012G074800.1.A
PCC=0.233, p=0.336
Gorai.010G118400.1.A
PCC=0.159, p=0.515
Gorai.010G118400.1.D
PCC=0.159, p=0.515
mRNA Profile
Protein Expression Profile
DPA

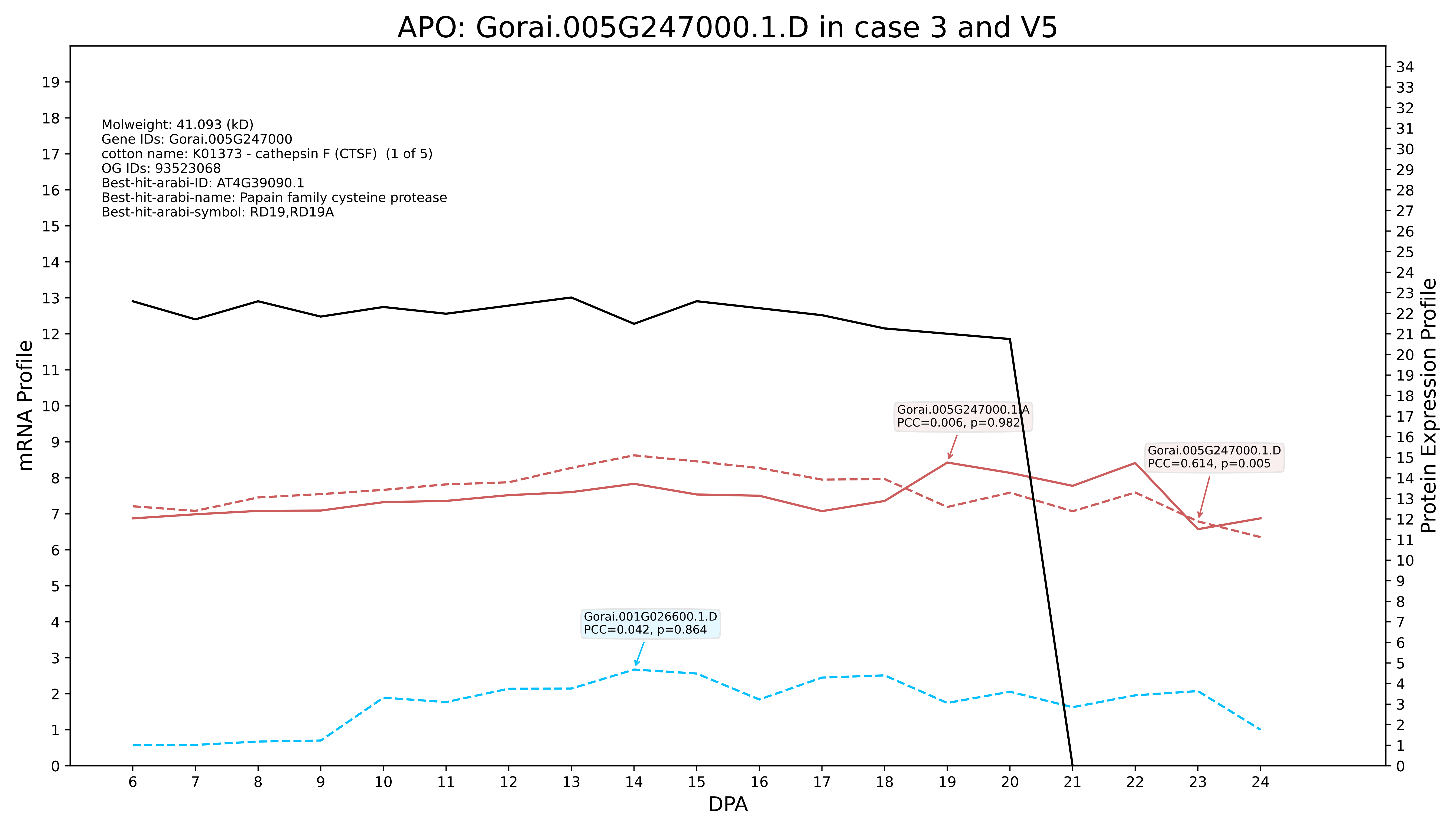

APO: Gorai.005G247000.1.D in case 3 and V5
Molweight: 41.093 (kD)
Gene IDs: Gorai.005G247000
cotton name: K01373 - cathepsin F (CTSF) (1 of 5)
OG IDs: 93523068
Best-hit-arabi-ID: AT4G39090.1
Best-hit-arabi-name: Papain family cysteine protease
Best-hit-arabi-symbol: RD19,RD19A
Gorai.005G247000.1.A
PCC=0.006, p=0.982
Gorai.005G247000.1.D
PCC=0.614, p=0.005
Gorai.001G026600.1.D
PCC=0.042, p=0.864
mRNA Profile
Protein Expression Profile
DPA

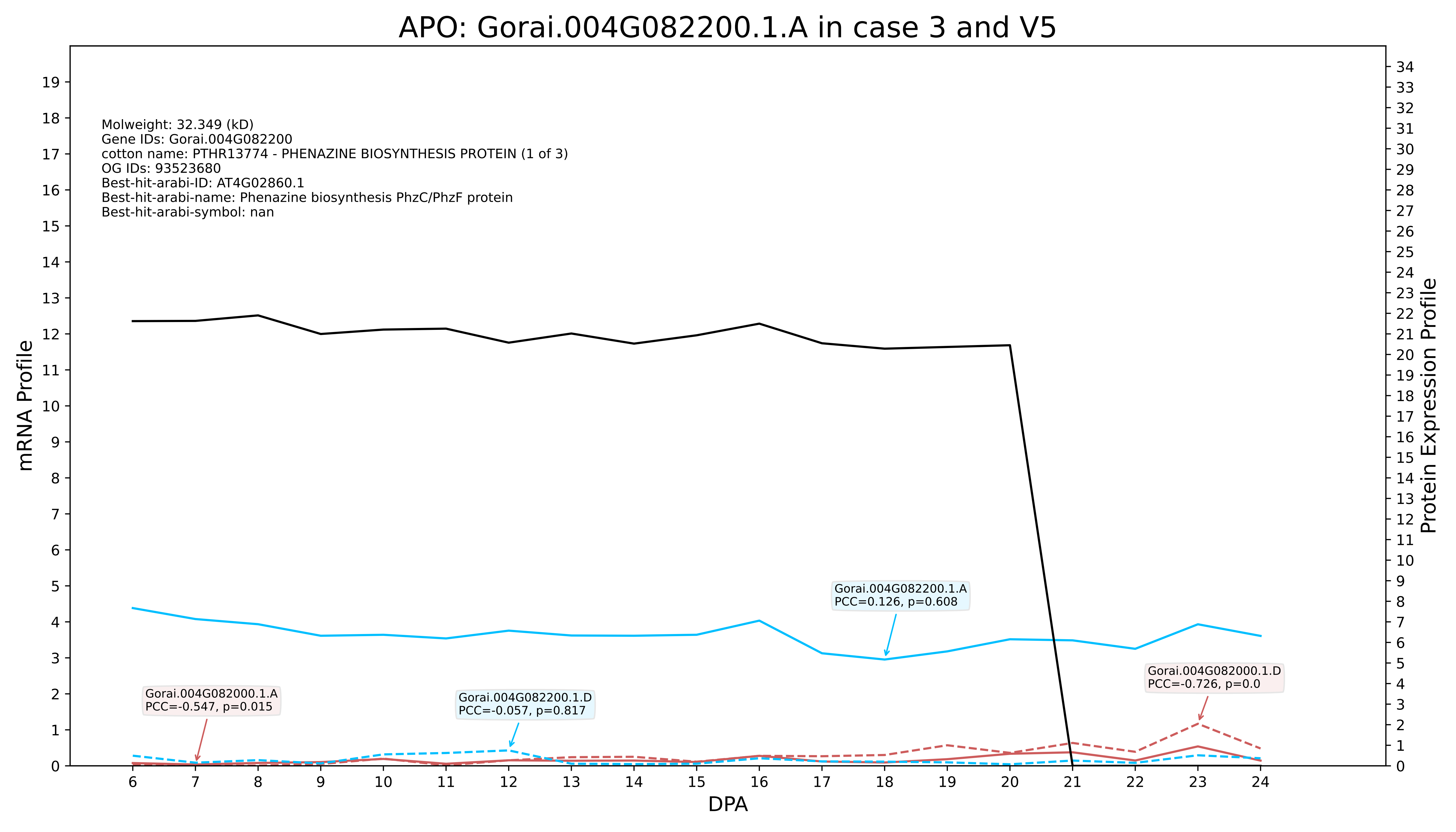
APO: Gorai.004G082200.1.A in case 3 and V5
Molweight: 32.349 (kD)
Gene IDs: Gorai.004G082200
cotton name: PTHR13774 - PHENAZINE BIOSYNTHESIS PROTEIN (1 of 3)
OG IDs: 93523680
Best-hit-arabi-ID: AT4G02860.1
Best-hit-arabi-name: Phenazine biosynthesis PhzC/PhzF protein
Best-hit-arabi-symbol: nan
Gorai.004G082200.1.A
PCC=0.126, p=0.608
Gorai.004G082000.1.D
PCC=-0.726, p=0.0
Gorai.004G082000.1.A
PCC=-0.547, p=0.015
Gorai.004G082200.1.D
PCC=-0.057, p=0.817
mRNA Profile
Protein Expression Profile
DPA

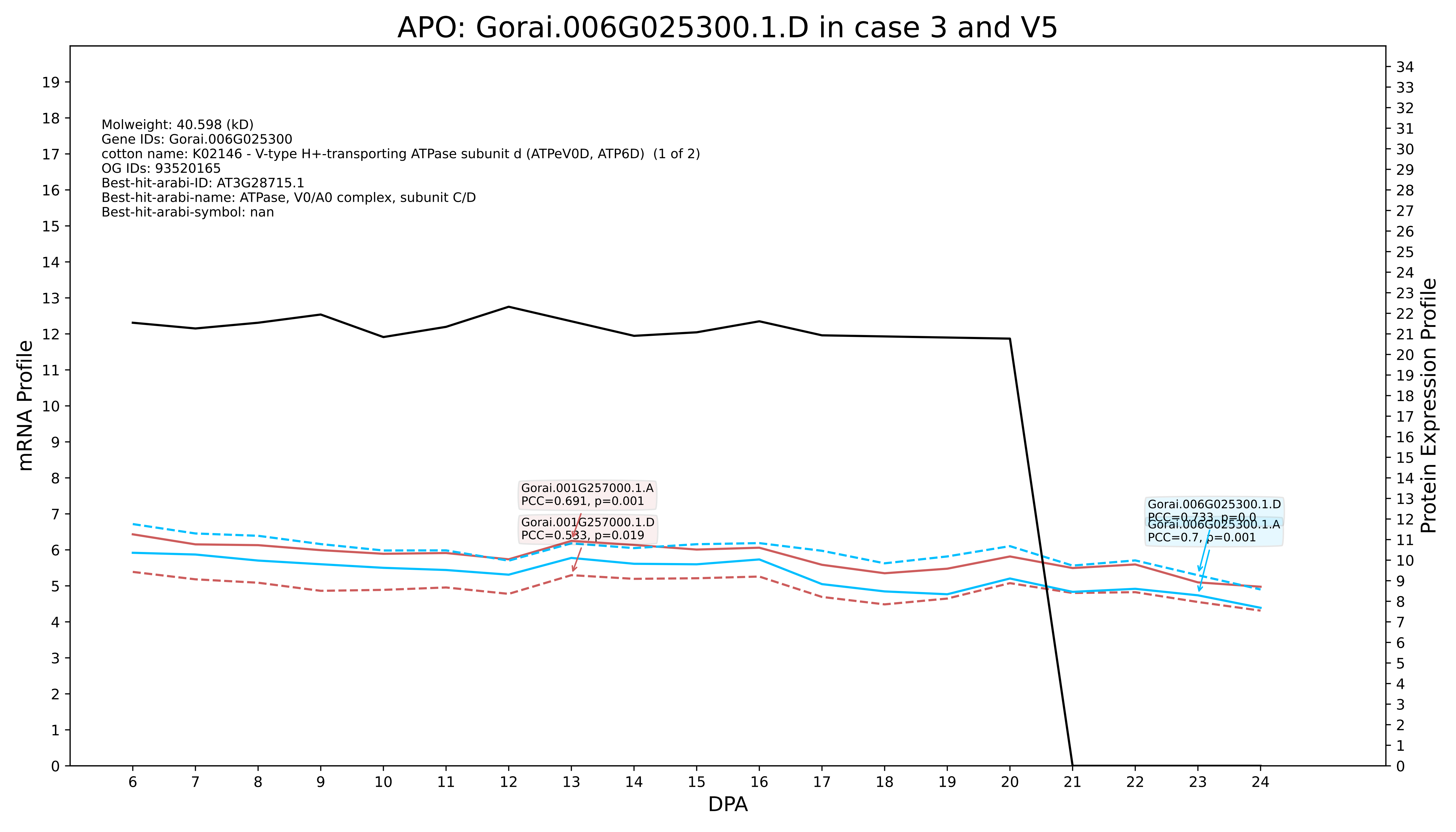

APO: Gorai.006G025300.1.D in case 3 and V5
Molweight: 40.598 (kD)
Gene IDs: Gorai.006G025300
cotton name: K02146 - V-type H+-transporting ATPase subunit d (ATPeV0D, ATP6D) (1 of 2)
OG IDs: 93520165
Best-hit-arabi-ID: AT3G28715.1
Best-hit-arabi-name: ATPase, V0/A0 complex, subunit C/D
Best-hit-arabi-symbol: nan
Gorai.001G257000.1.A
PCC=0.691, p=0.001
Gorai.001G257000.1.D
PCC=0.533, p=0.019
Gorai.006G025300.1.D
PCC=0.733, p=0.0
Gorai.006G025300.1.A
PCC=0.7, p=0.001
mRNA Profile
Protein Expression Profile
DPA

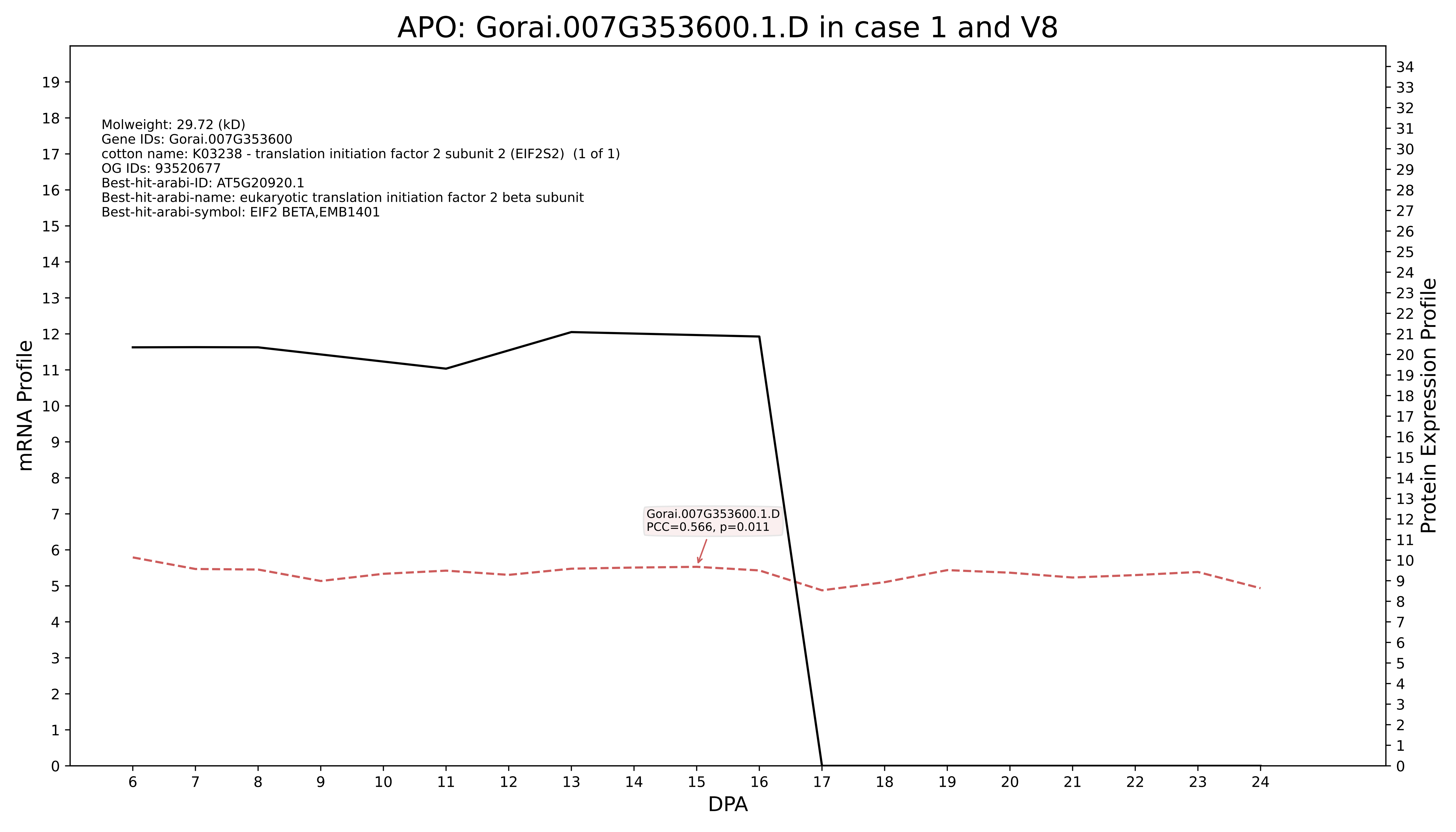
APO: Gorai.007G353600.1.D in case 1 and V8
Molweight: 29.72 (kD)
Gene IDs: Gorai.007G353600
cotton name: K03238 - translation initiation factor 2 subunit 2 (EIF2S2) (1 of 1)
OG IDs: 93520677
Best-hit-arabi-ID: AT5G20920.1
Best-hit-arabi-name: eukaryotic translation initiation factor 2 beta subunit
Best-hit-arabi-symbol: EIF2 BETA,EMB1401
Gorai.007G353600.1.D
PCC=0.566, p=0.011
mRNA Profile
Protein Expression Profile
DPA

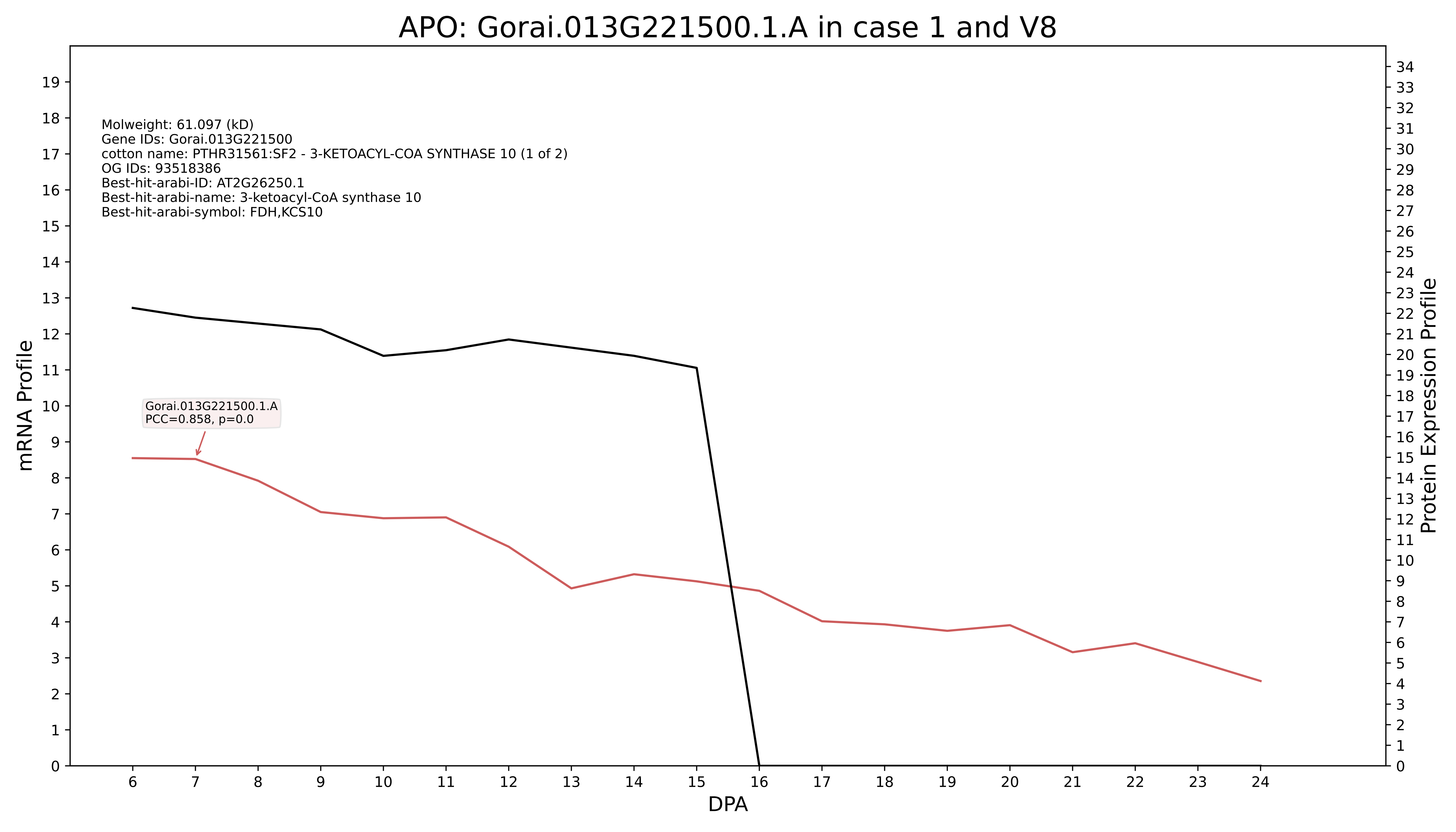
APO: Gorai.013G221500.1.A in case 1 and V8
Molweight: 61.097 (kD)
Gene IDs: Gorai.013G221500
cotton name: PTHR31561:SF2 - 3-KETOACYL-COA SYNTHASE 10 (1 of 2)
OG IDs: 93518386
Best-hit-arabi-ID: AT2G26250.1
Best-hit-arabi-name: 3-ketoacyl-CoA synthase 10
Best-hit-arabi-symbol: FDH,KCS10
Gorai.013G221500.1.A
PCC=0.858, p=0.0
mRNA Profile
Protein Expression Profile
DPA

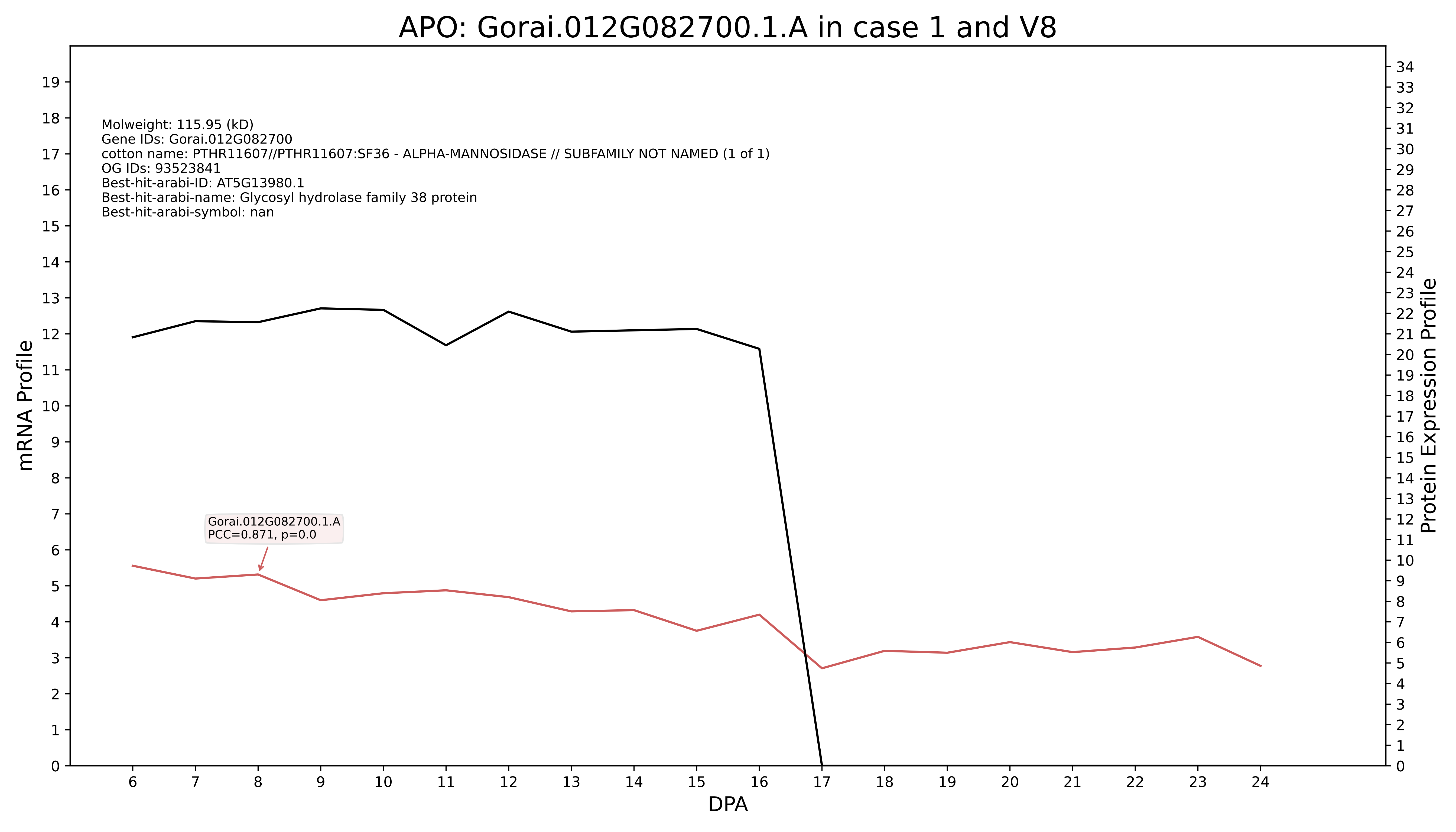

APO: Gorai.012G082700.1.A in case 1 and V8
Molweight: 115.95 (kD)
Gene IDs: Gorai.012G082700
cotton name: PTHR11607//PTHR11607:SF36 - ALPHA-MANNOSIDASE // SUBFAMILY NOT NAMED (1 of 1)
OG IDs: 93523841
Best-hit-arabi-ID: AT5G13980.1
Best-hit-arabi-name: Glycosyl hydrolase family 38 protein
Best-hit-arabi-symbol: nan
Gorai.012G082700.1.A
PCC=0.871, p=0.0
mRNA Profile
Protein Expression Profile
DPA

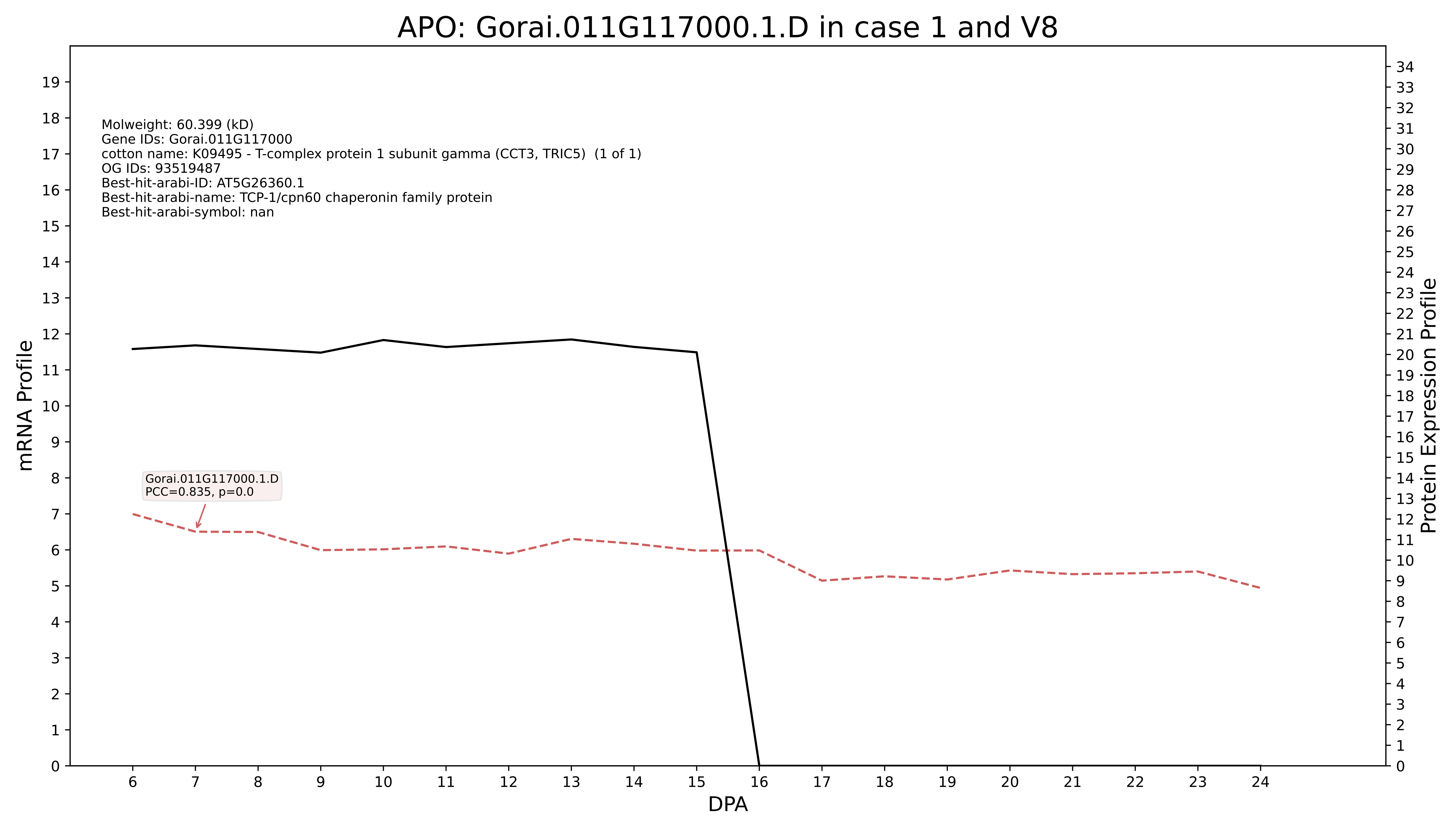

APO: Gorai.011G117000.1.D in case 1 and V8
Molweight: 60.399 (kD)
Gene IDs: Gorai.011G117000
cotton name: K09495 - T-complex protein 1 subunit gamma (CCT3, TRIC5) (1 of 1)
OG IDs: 93519487
Best-hit-arabi-ID: AT5G26360.1
Best-hit-arabi-name: TCP-1/cpn60 chaperonin family protein
Best-hit-arabi-symbol: nan
Gorai.011G117000.1.D
PCC=0.835, p=0.0
mRNA Profile
Protein Expression Profile
DPA

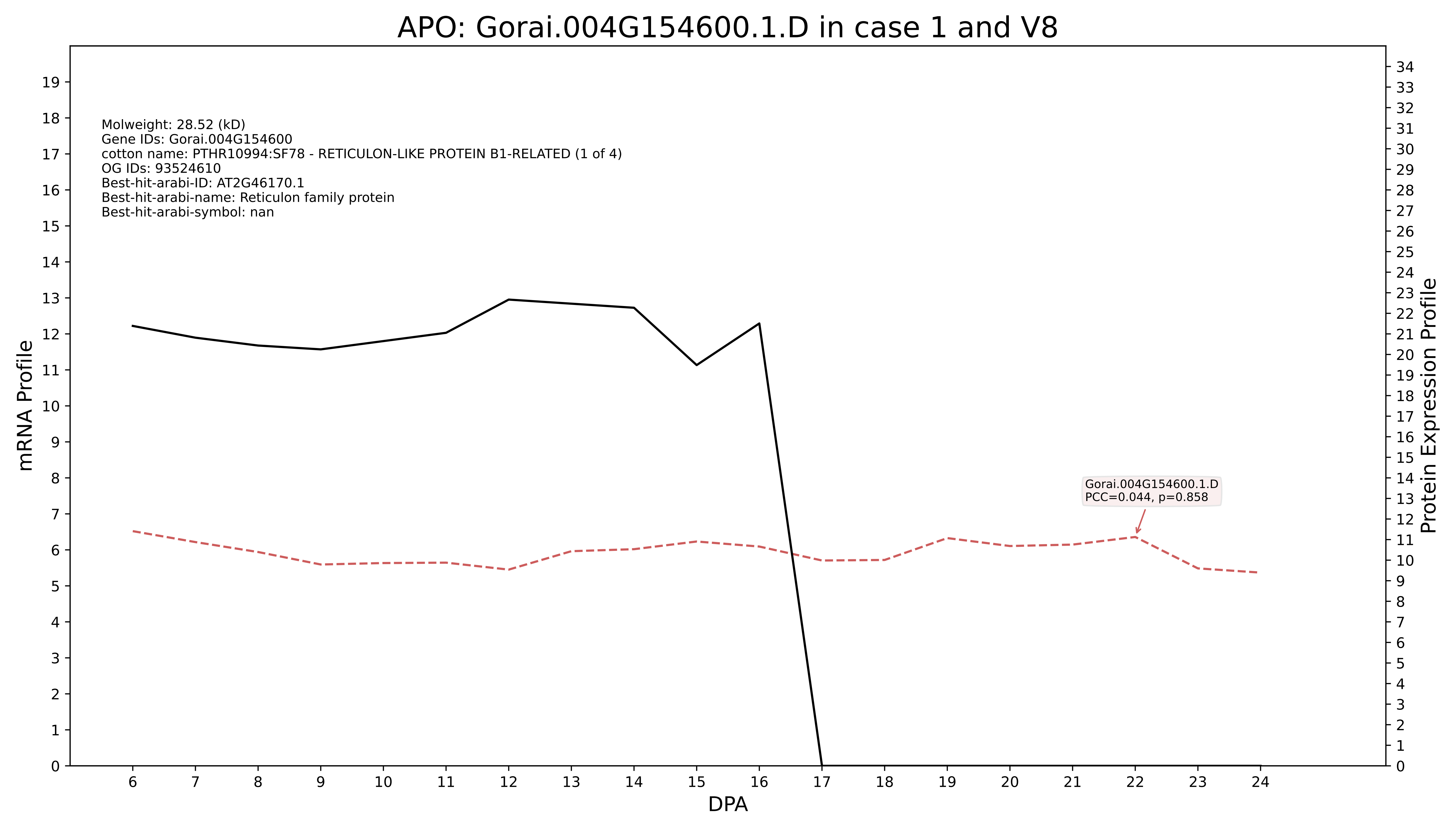
APO: Gorai.004G154600.1.D in case 1 and V8
Molweight: 28.52 (kD)
Gene IDs: Gorai.004G154600
cotton name: PTHR10994:SF78 - RETICULON-LIKE PROTEIN B1-RELATED (1 of 4)
OG IDs: 93524610
Best-hit-arabi-ID: AT2G46170.1
Best-hit-arabi-name: Reticulon family protein
Best-hit-arabi-symbol: nan
Gorai.004G154600.1.D
PCC=0.044, p=0.858
mRNA Profile
Protein Expression Profile
DPA

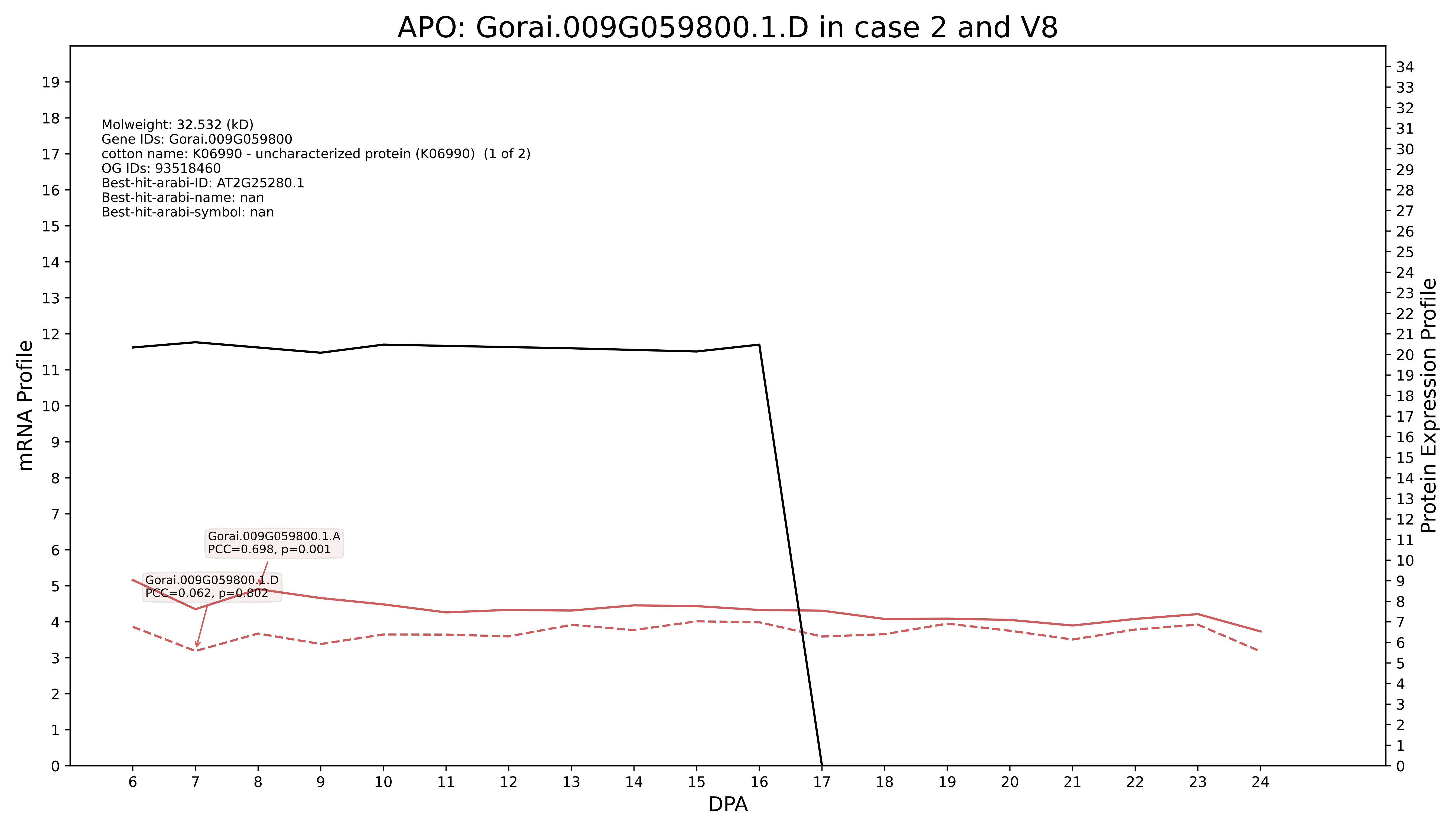
APO: Gorai.009G059800.1.D in case 2 and V8
Molweight: 32.532 (kD)
Gene IDs: Gorai.009G059800
cotton name: K06990 - uncharacterized protein (K06990) (1 of 2)
OG IDs: 93518460
Best-hit-arabi-ID: AT2G25280.1
Best-hit-arabi-name: nan
Best-hit-arabi-symbol: nan
Gorai.009G059800.1.A
PCC=0.698, p=0.001
Gorai.009G059800.1.D
PCC=0.062, p=0.802
mRNA Profile
Protein Expression Profile
DPA

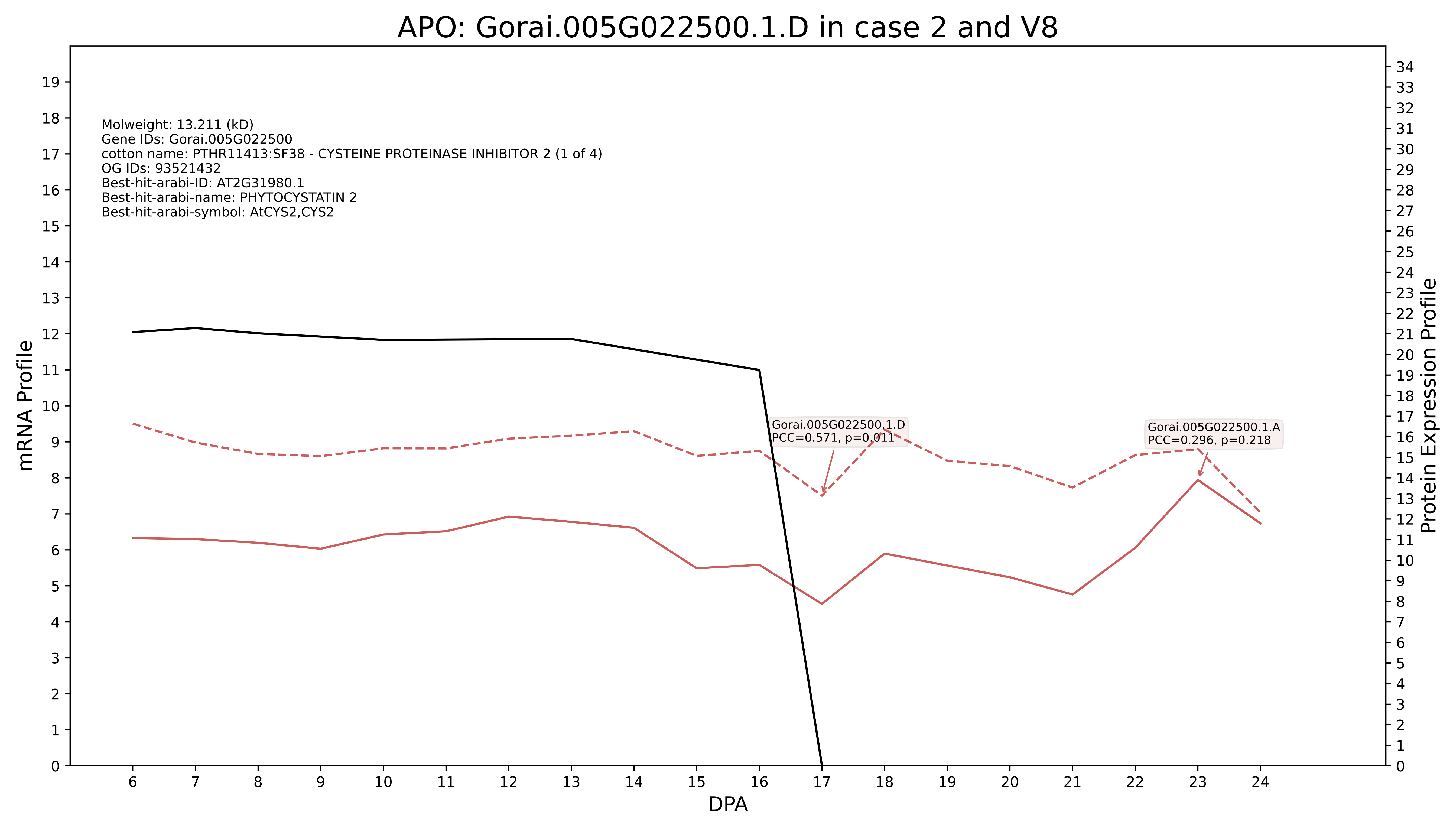
APO: Gorai.005G022500.1.D in case 2 and V8
Molweight: 13.211 (kD)
Gene IDs: Gorai.005G022500
cotton name: PTHR11413:SF38 - CYSTEINE PROTEINASE INHIBITOR 2 (1 of 4)
OG IDs: 93521432
Best-hit-arabi-ID: AT2G31980.1
Best-hit-arabi-name: PHYTOCYSTATIN 2
Best-hit-arabi-symbol: AtCYS2,CYS2
Gorai.005G022500.1.D
PCC=0.571, p=0.011
Gorai.005G022500.1.A
PCC=0.296, p=0.218
mRNA Profile
Protein Expression Profile
DPA

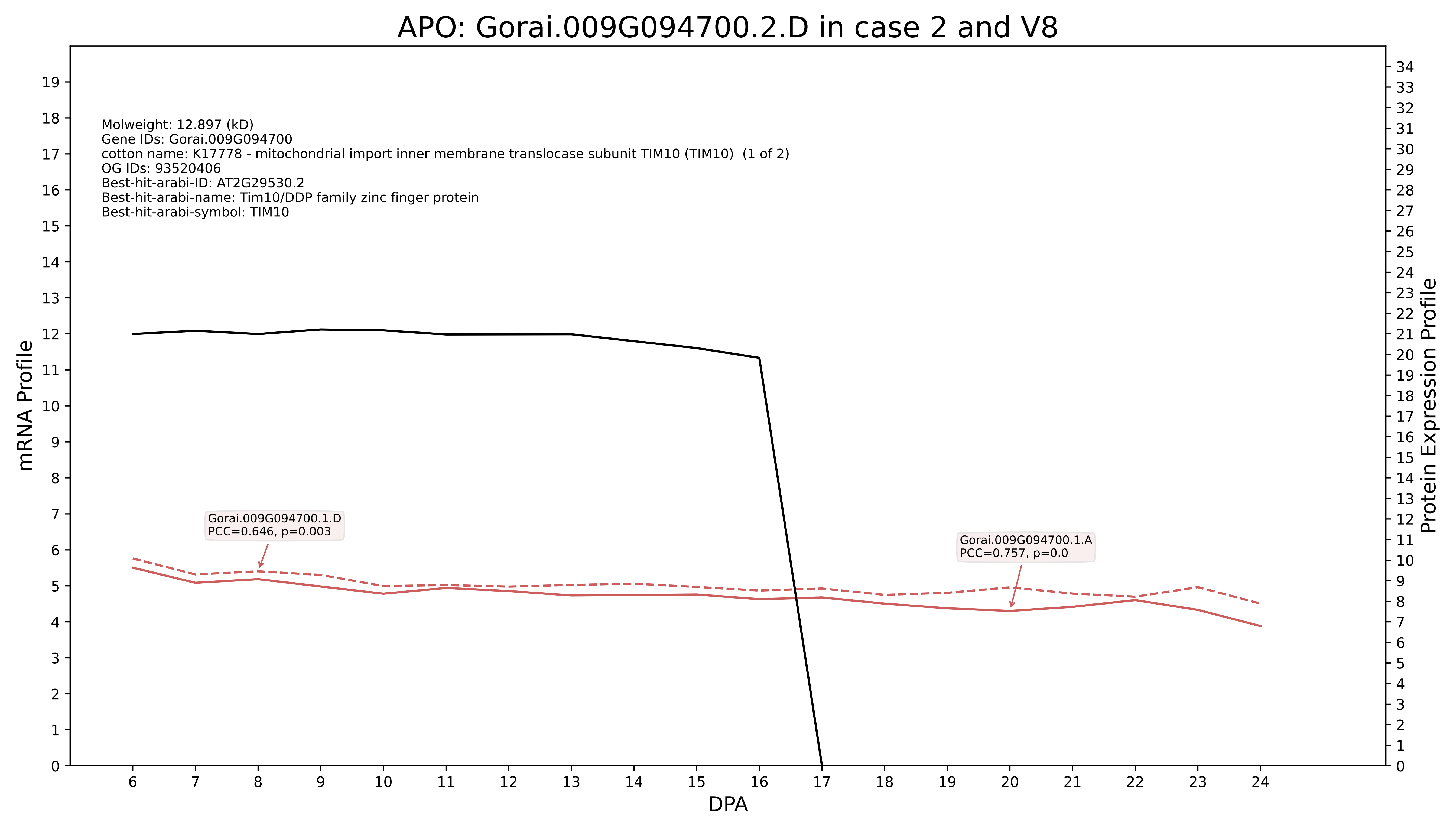
APO: Gorai.009G094700.2.D in case 2 and V8
Molweight: 12.897 (kD)
Gene IDs: Gorai.009G094700
cotton name: K17778 - mitochondrial import inner membrane translocase subunit TIM10 (TIM10) (1 of 2)
OG IDs: 93520406
Best-hit-arabi-ID: AT2G29530.2
Best-hit-arabi-name: Tim10/DDP family zinc finger protein
Best-hit-arabi-symbol: TIM10
Gorai.009G094700.1.D
PCC=0.646, p=0.003
Gorai.009G094700.1.A
PCC=0.757, p=0.0
mRNA Profile
Protein Expression Profile
DPA

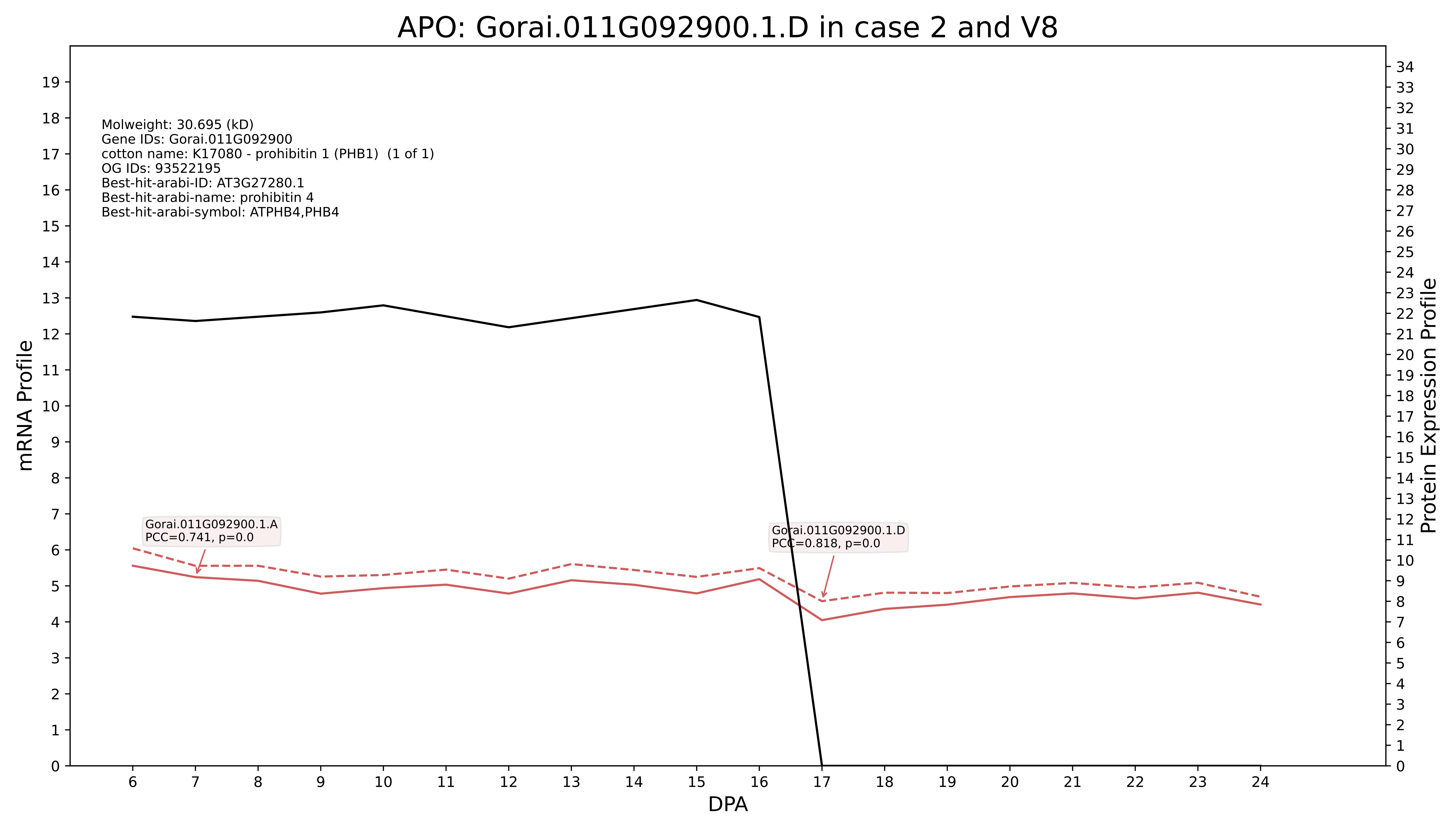
APO: Gorai.011G092900.1.D in case 2 and V8
Molweight: 30.695 (kD)
Gene IDs: Gorai.011G092900
cotton name: K17080 - prohibitin 1 (PHB1) (1 of 1)
OG IDs: 93522195
Best-hit-arabi-ID: AT3G27280.1
Best-hit-arabi-name: prohibitin 4
Best-hit-arabi-symbol: ATPHB4,PHB4
Gorai.011G092900.1.A
PCC=0.741, p=0.0
Gorai.011G092900.1.D
PCC=0.818, p=0.0
mRNA Profile
Protein Expression Profile
DPA

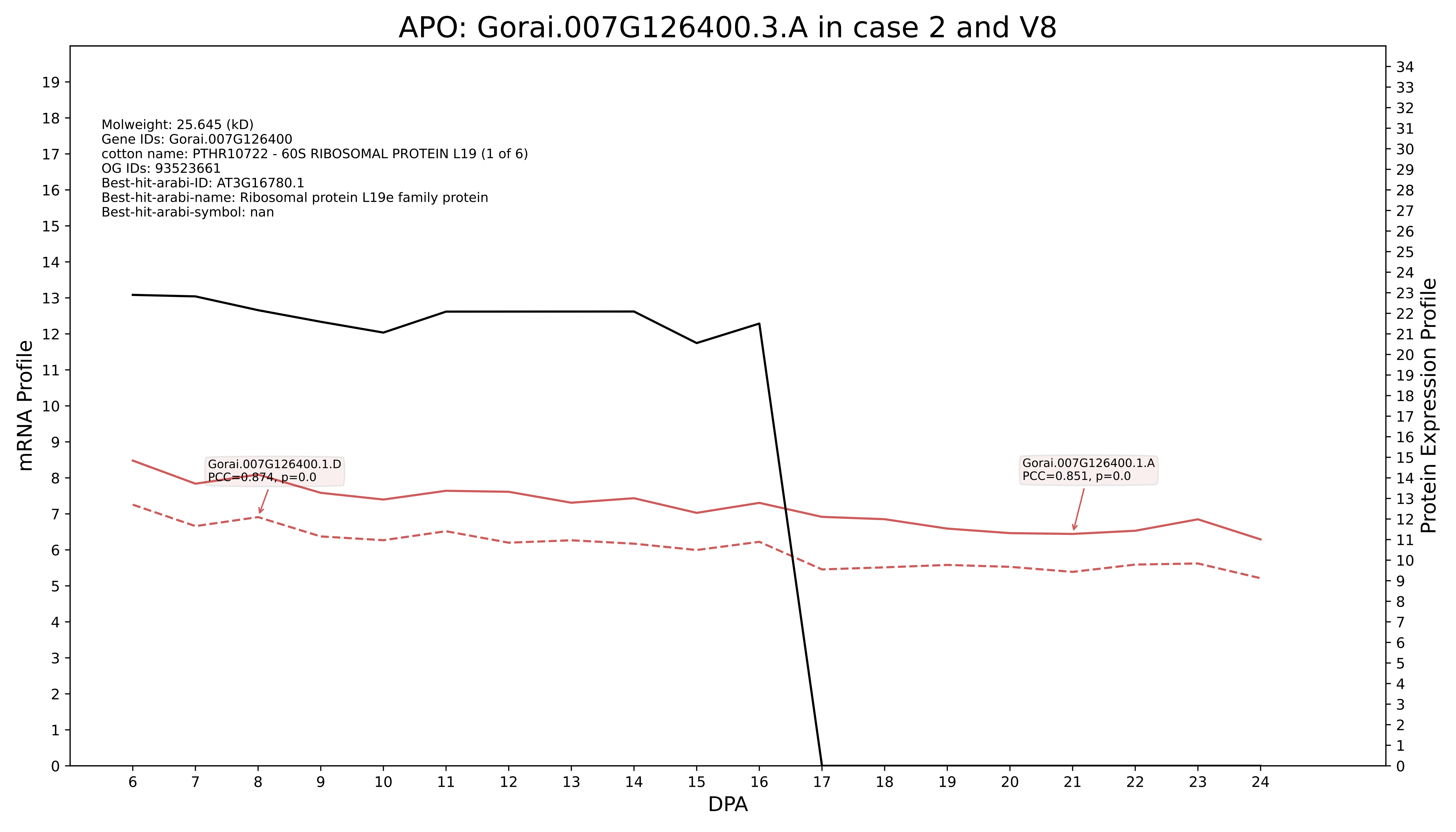
APO: Gorai.007G126400.3.A in case 2 and V8
Molweight: 25.645 (kD)
Gene IDs: Gorai.007G126400
cotton name: PTHR10722 - 60S RIBOSOMAL PROTEIN L19 (1 of 6)
OG IDs: 93523661
Best-hit-arabi-ID: AT3G16780.1
Best-hit-arabi-name: Ribosomal protein L19e family protein
Best-hit-arabi-symbol: nan
Gorai.007G126400.1.D
PCC=0.874, p=0.0
Gorai.007G126400.1.A
PCC=0.851, p=0.0
mRNA Profile
Protein Expression Profile
DPA

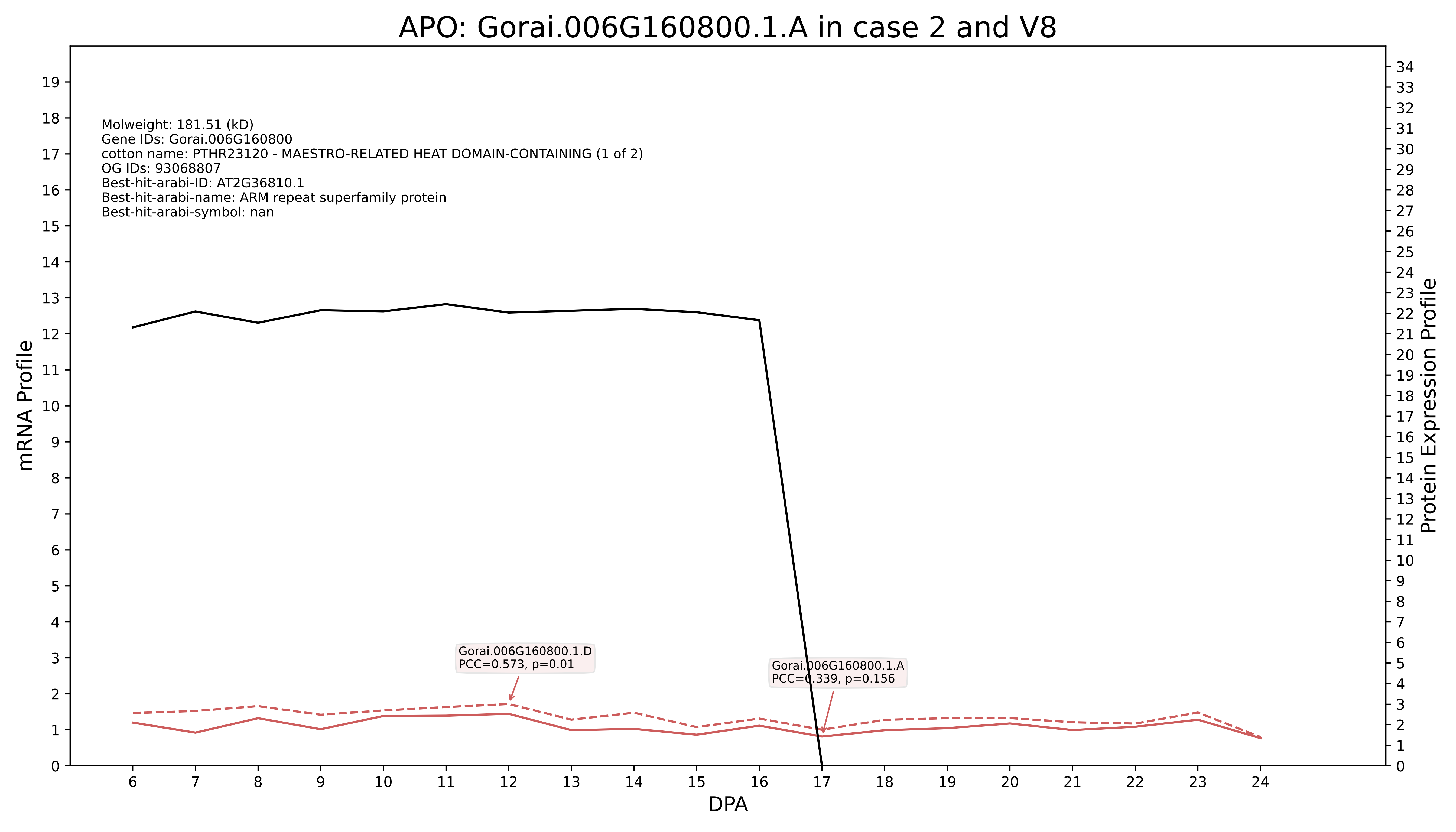
APO: Gorai.006G160800.1.A in case 2 and V8
Molweight: 181.51 (kD)
Gene IDs: Gorai.006G160800
cotton name: PTHR23120 - MAESTRO-RELATED HEAT DOMAIN-CONTAINING (1 of 2)
OG IDs: 93068807
Best-hit-arabi-ID: AT2G36810.1
Best-hit-arabi-name: ARM repeat superfamily protein
Best-hit-arabi-symbol: nan
Gorai.006G160800.1.D
PCC=0.573, p=0.01
Gorai.006G160800.1.A
PCC=0.339, p=0.156
mRNA Profile
Protein Expression Profile
DPA

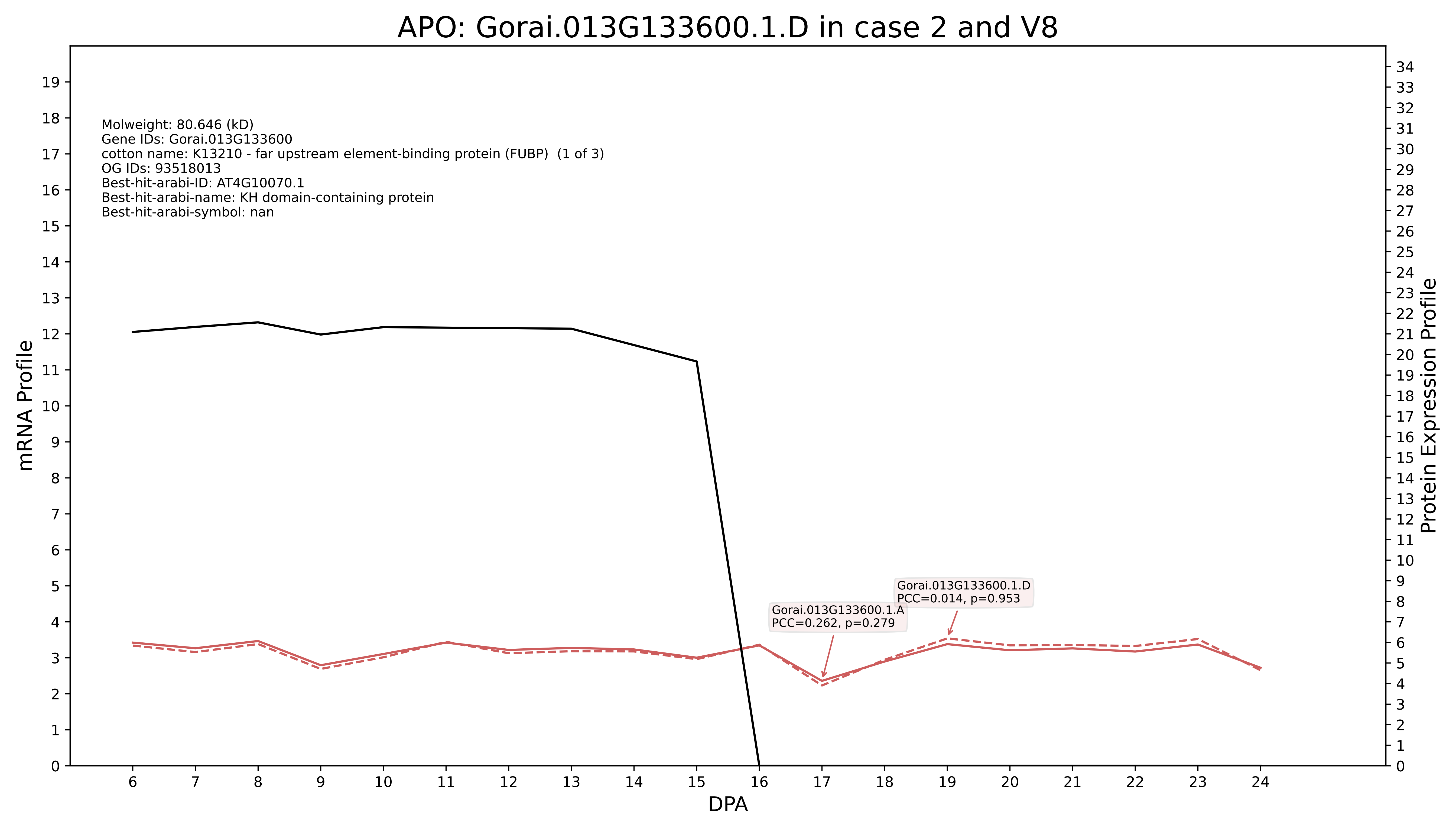
APO: Gorai.013G133600.1.D in case 2 and V8
Molweight: 80.646 (kD)
Gene IDs: Gorai.013G133600
cotton name: K13210 - far upstream element-binding protein (FUBP) (1 of 3)
OG IDs: 93518013
Best-hit-arabi-ID: AT4G10070.1
Best-hit-arabi-name: KH domain-containing protein
Best-hit-arabi-symbol: nan
Gorai.013G133600.1.D
PCC=0.014, p=0.953
Gorai.013G133600.1.A
PCC=0.262, p=0.279
mRNA Profile
Protein Expression Profile
DPA

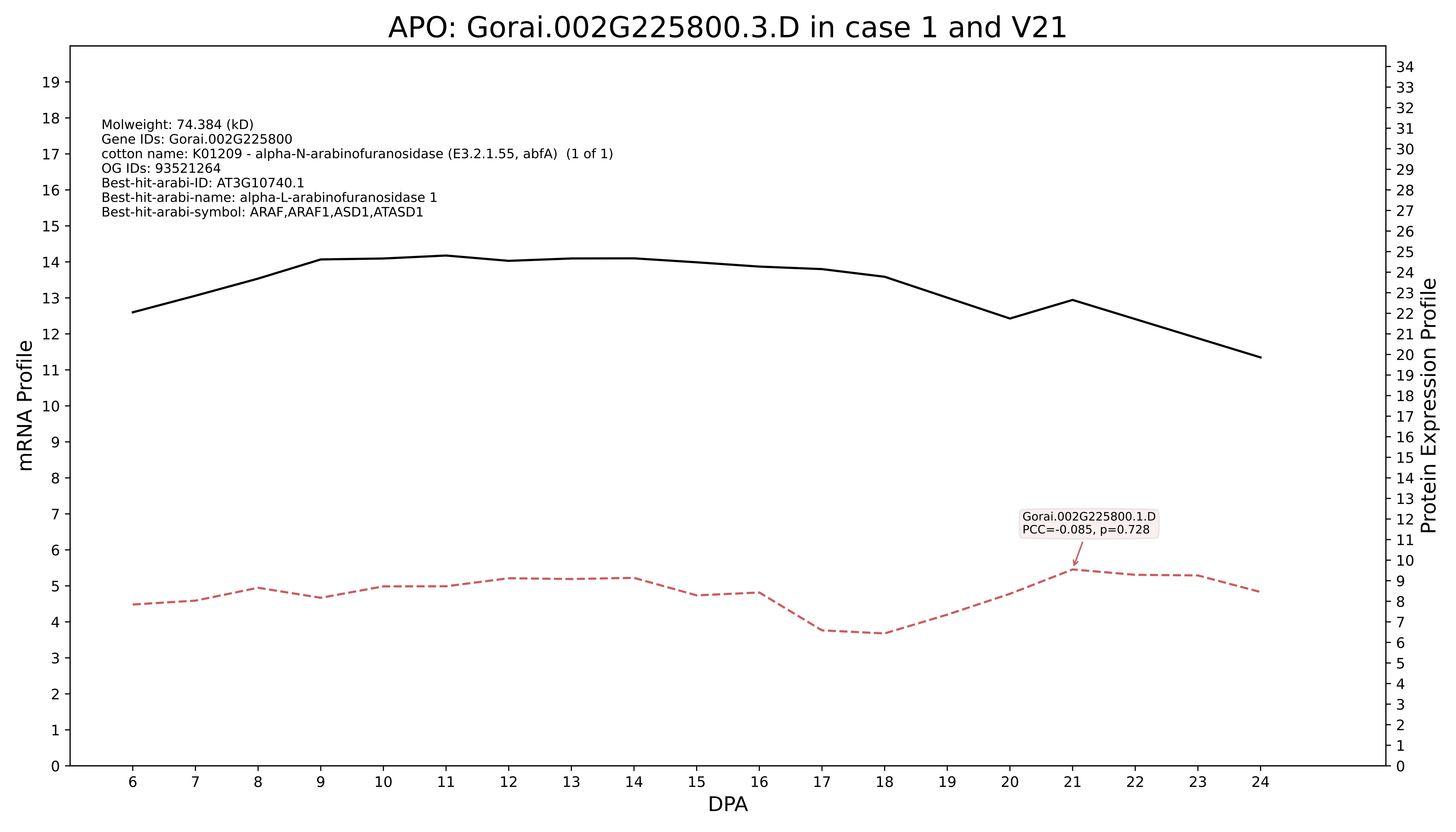
APO: Gorai.002G225800.3.D in case 1 and V21
Molweight: 74.384 (kD)
Gene IDs: Gorai.002G225800
cotton name: K01209 - alpha-N-arabinofuranosidase (E3.2.1.55, abfA) (1 of 1)
OG IDs: 93521264
Best-hit-arabi-ID: AT3G10740.1
Best-hit-arabi-name: alpha-L-arabinofuranosidase 1
Best-hit-arabi-symbol: ARAF,ARAF1,ASD1,ATASD1
Gorai.002G225800.1.D
PCC=-0.085, p=0.728
mRNA Profile
Protein Expression Profile
DPA

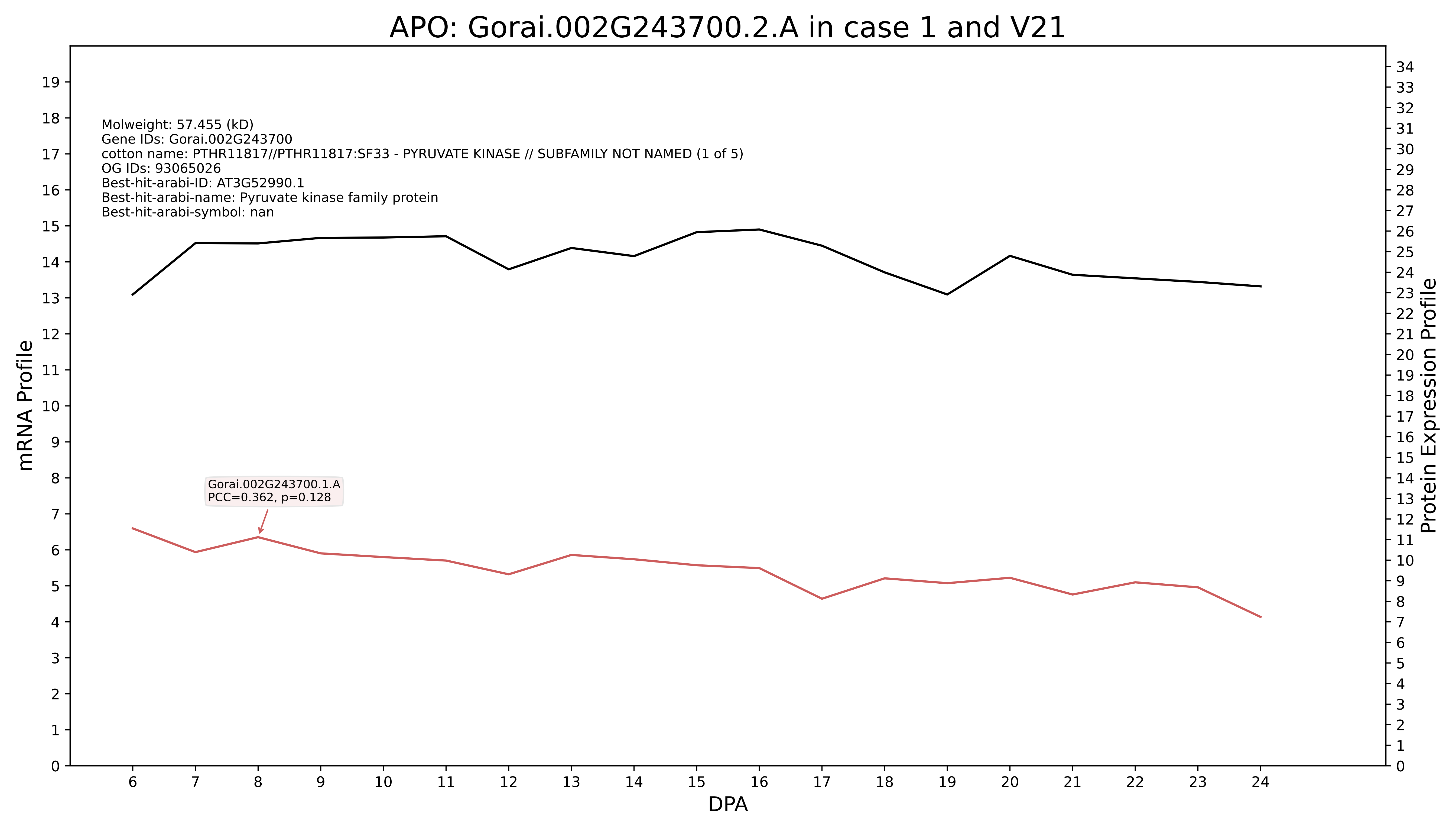

APO: Gorai.002G243700.2.A in case 1 and V21
Molweight: 57.455 (kD)
Gene IDs: Gorai.002G243700
cotton name: PTHR11817//PTHR11817:SF33 - PYRUVATE KINASE // SUBFAMILY NOT NAMED (1 of 5)
OG IDs: 93065026
Best-hit-arabi-ID: AT3G52990.1
Best-hit-arabi-name: Pyruvate kinase family protein
Best-hit-arabi-symbol: nan
Gorai.002G243700.1.A
PCC=0.362, p=0.128
mRNA Profile
Protein Expression Profile
DPA

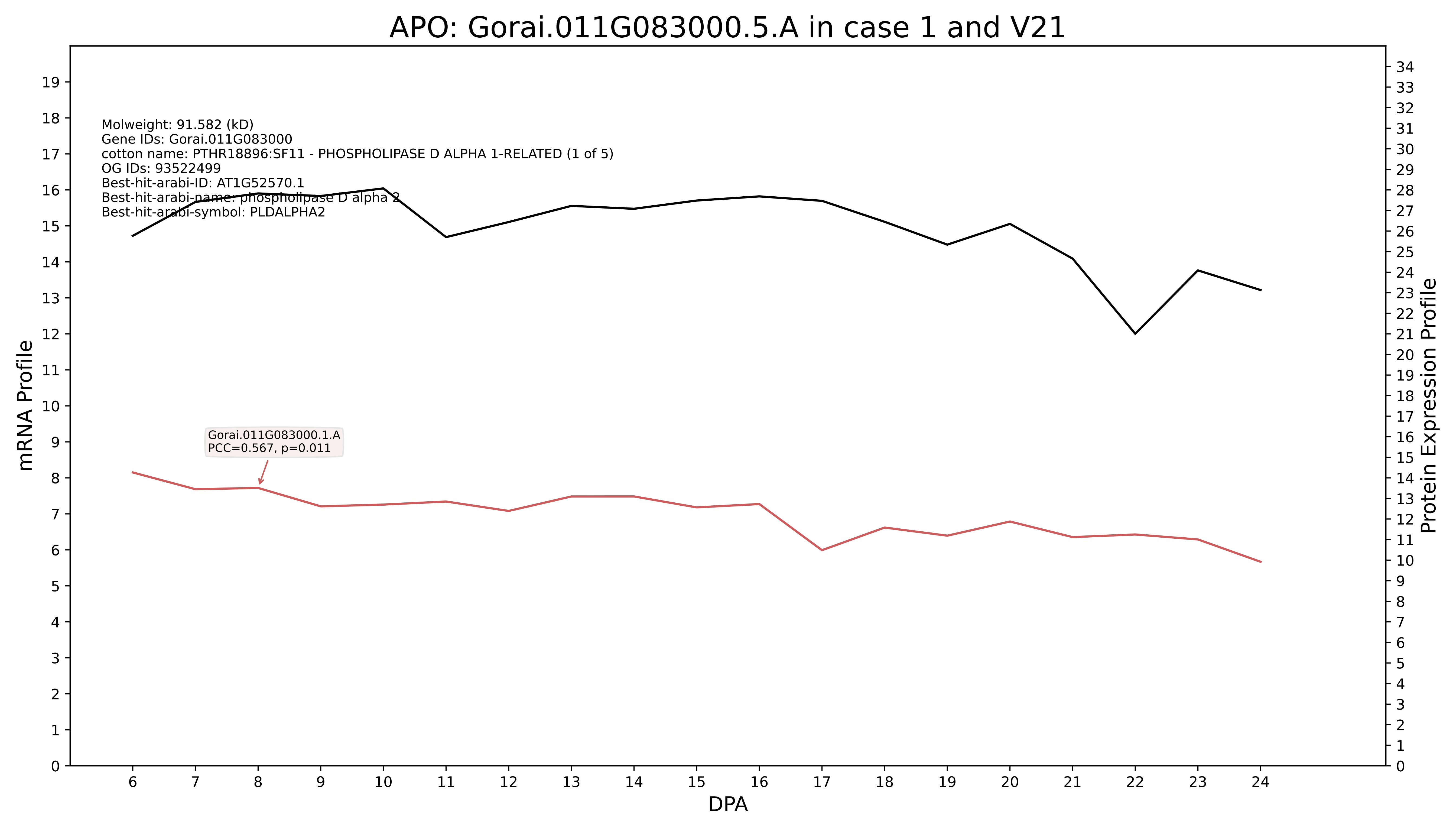
APO: Gorai.011G083000.5.A in case 1 and V21
Molweight: 91.582 (kD)
Gene IDs: Gorai.011G083000
cotton name: PTHR18896:SF11 - PHOSPHOLIPASE D ALPHA 1-RELATED (1 of 5)
OG IDs: 93522499
Best-hit-arabi-ID: AT1G52570.1
Best-hit-arabi-name: phospholipase D alpha 2
Best-hit-arabi-symbol: PLDALPHA2
Gorai.011G083000.1.A
PCC=0.567, p=0.011
mRNA Profile
Protein Expression Profile
DPA

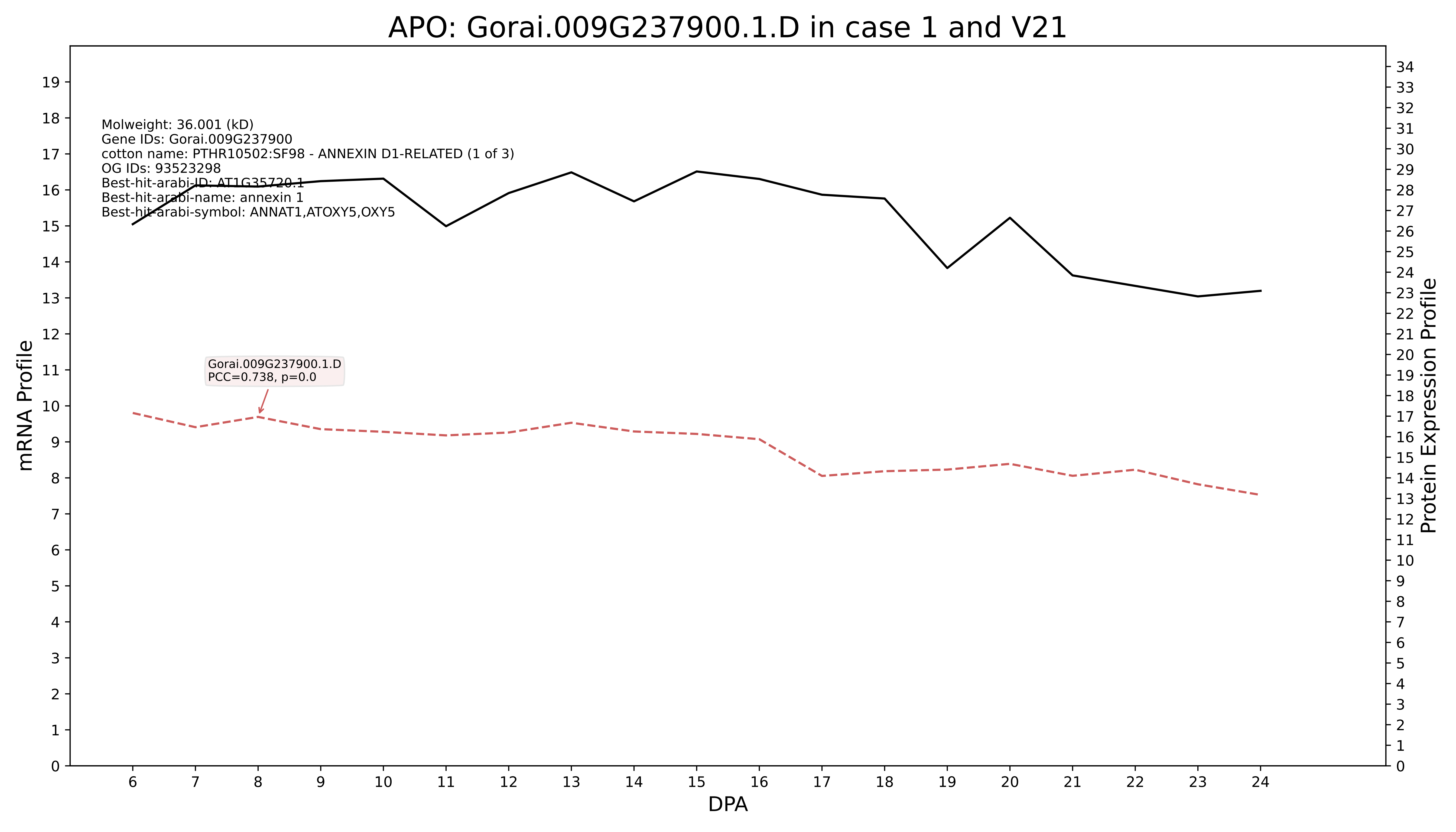
APO: Gorai.009G237900.1.D in case 1 and V21
Molweight: 36.001 (kD)
Gene IDs: Gorai.009G237900
cotton name: PTHR10502:SF98 - ANNEXIN D1-RELATED (1 of 3)
OG IDs: 93523298
Best-hit-arabi-ID: AT1G35720.1
Best-hit-arabi-name: annexin 1
Best-hit-arabi-symbol: ANNAT1,ATOXY5,OXY5
Gorai.009G237900.1.D
PCC=0.738, p=0.0
mRNA Profile
Protein Expression Profile
DPA

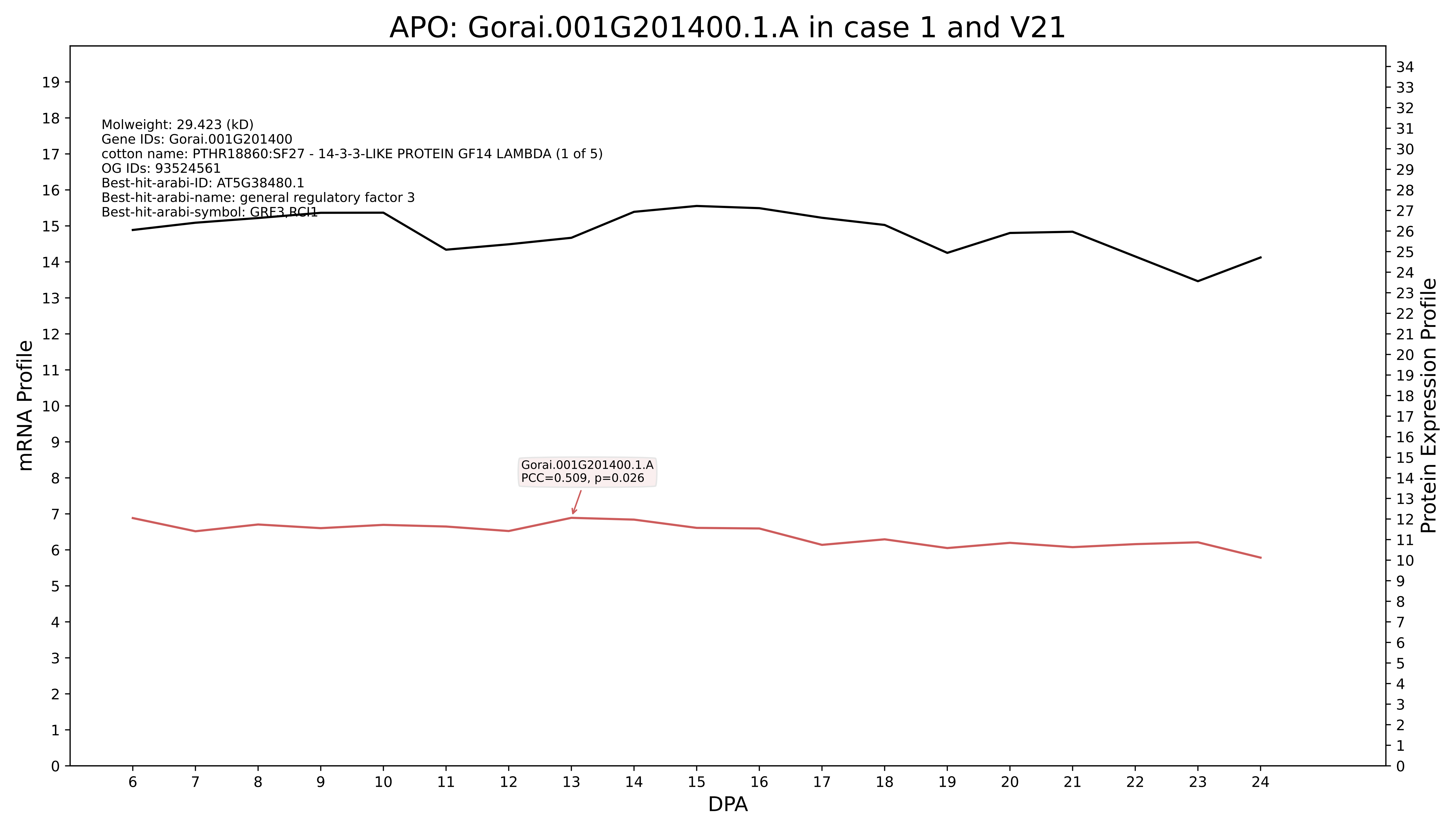
APO: Gorai.001G201400.1.A in case 1 and V21
Molweight: 29.423 (kD)
Gene IDs: Gorai.001G201400
cotton name: PTHR18860:SF27 - 14-3-3-LIKE PROTEIN GF14 LAMBDA (1 of 5)
OG IDs: 93524561
Best-hit-arabi-ID: AT5G38480.1
Best-hit-arabi-name: general regulatory factor 3
Best-hit-arabi-symbol: GRF3,RCI1
Gorai.001G201400.1.A
PCC=0.509, p=0.026
mRNA Profile
Protein Expression Profile
DPA

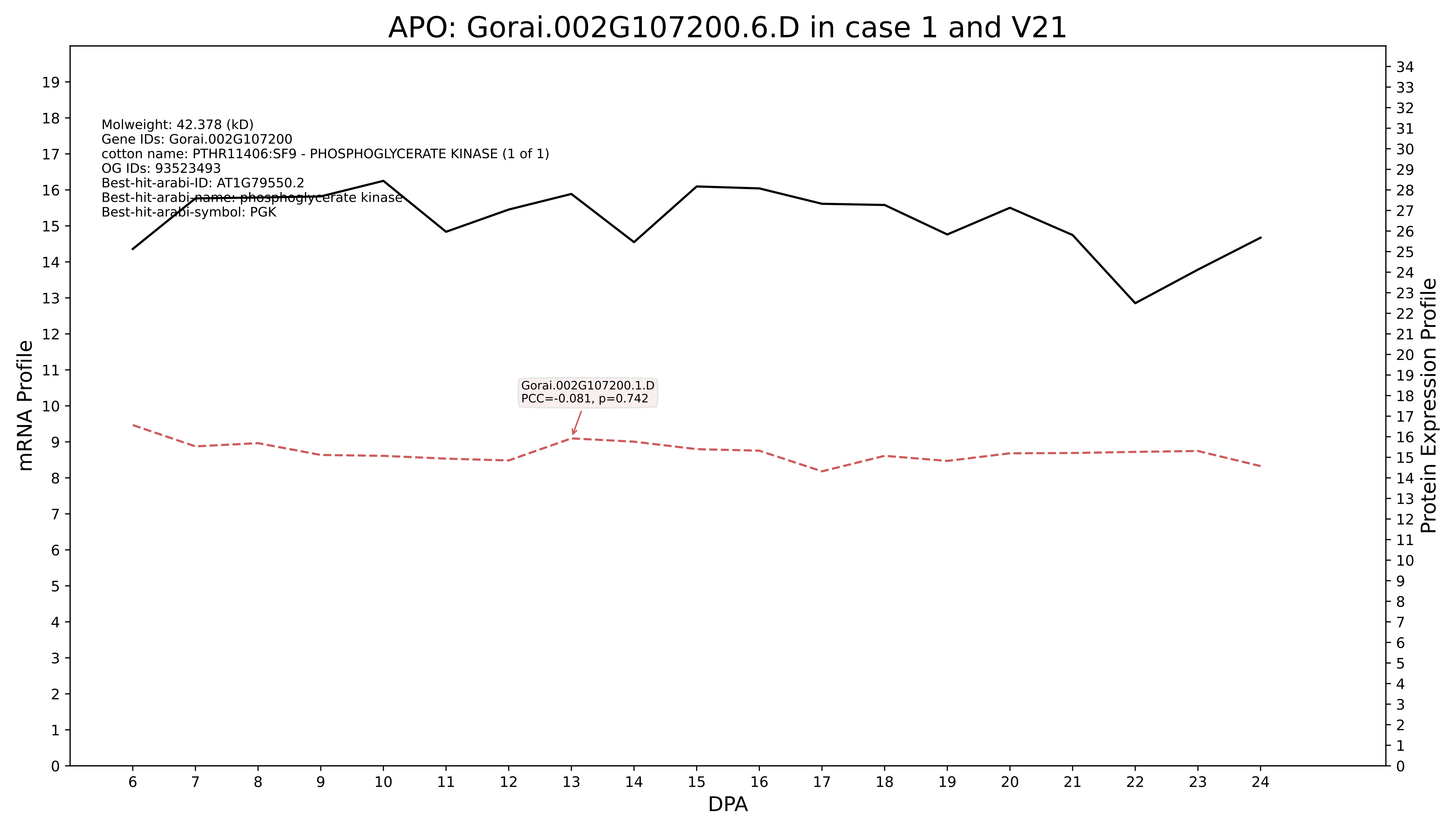

APO: Gorai.002G107200.6.D in case 1 and V21
Molweight: 42.378 (kD)
Gene IDs: Gorai.002G107200
cotton name: PTHR11406:SF9 - PHOSPHOGLYCERATE KINASE (1 of 1)
OG IDs: 93523493
Best-hit-arabi-ID: AT1G79550.2
Best-hit-arabi-name: phosphoglycerate kinase
Best-hit-arabi-symbol: PGK
Gorai.002G107200.1.D
PCC=-0.081, p=0.742
mRNA Profile
Protein Expression Profile
DPA

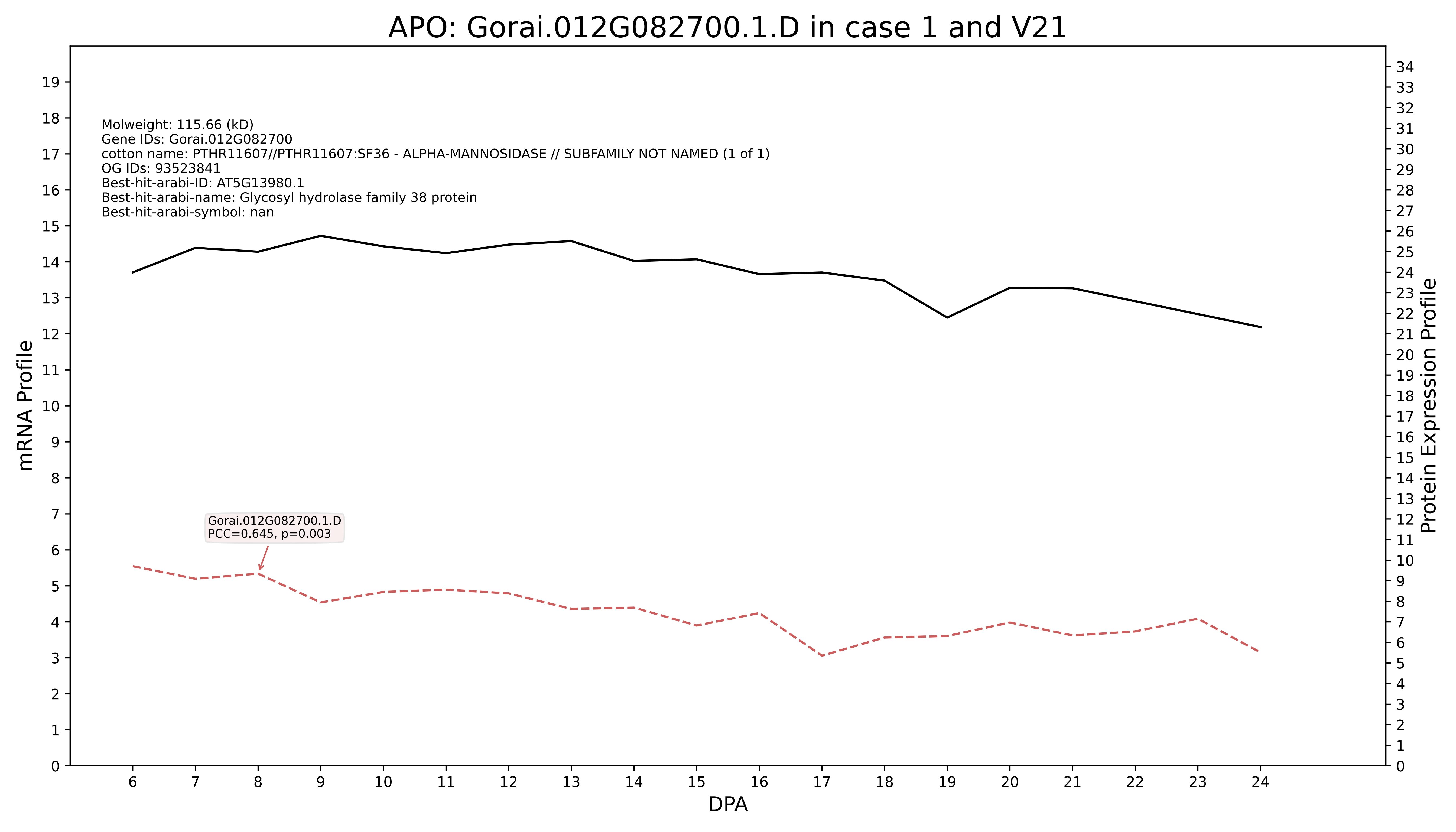
APO: Gorai.012G082700.1.D in case 1 and V21
Molweight: 115.66 (kD)
Gene IDs: Gorai.012G082700
cotton name: PTHR11607//PTHR11607:SF36 - ALPHA-MANNOSIDASE // SUBFAMILY NOT NAMED (1 of 1)
OG IDs: 93523841
Best-hit-arabi-ID: AT5G13980.1
Best-hit-arabi-name: Glycosyl hydrolase family 38 protein
Best-hit-arabi-symbol: nan
Gorai.012G082700.1.D
PCC=0.645, p=0.003
mRNA Profile
Protein Expression Profile
DPA

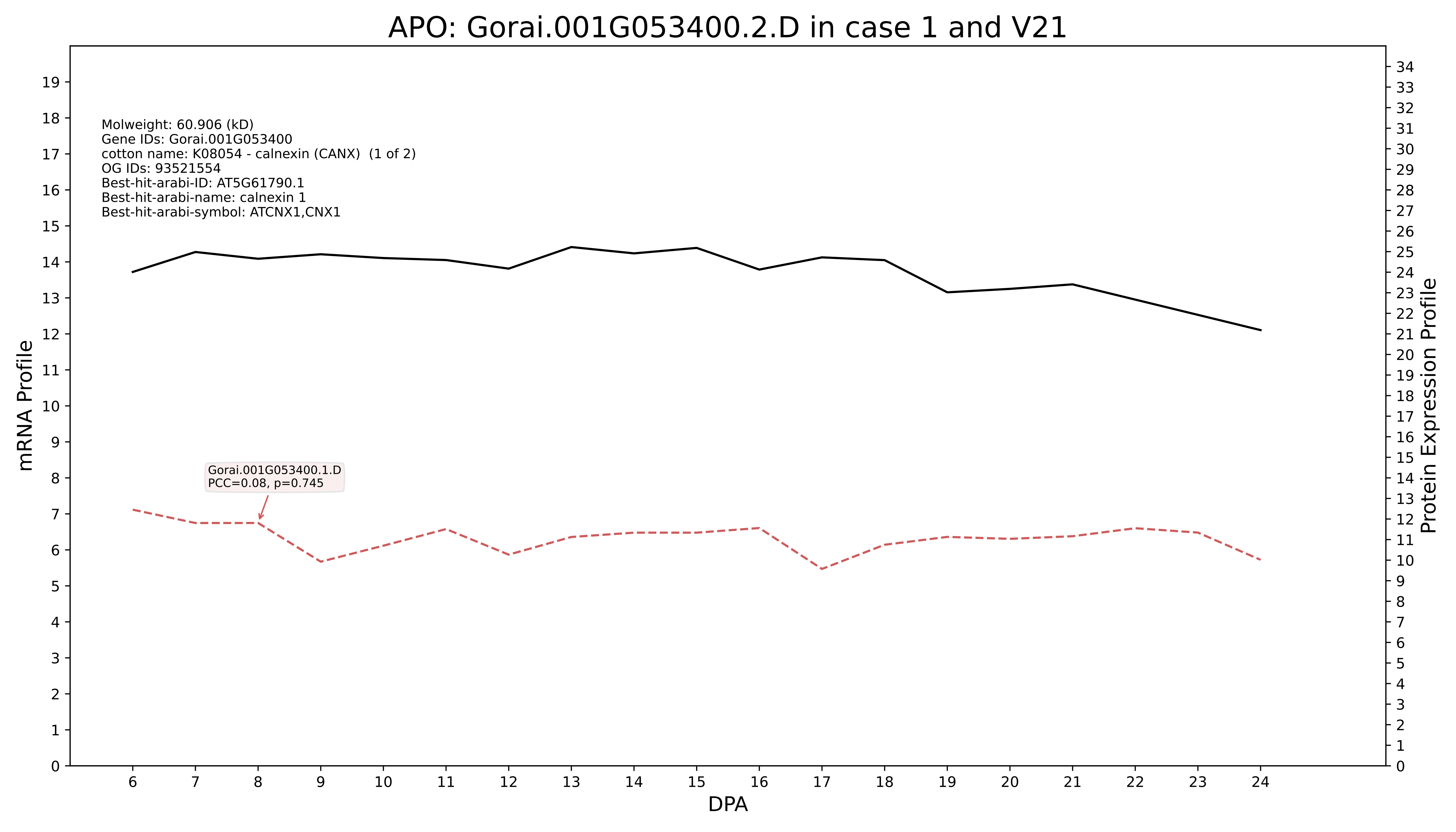

APO: Gorai.001G053400.2.D in case 1 and V21
Molweight: 60.906 (kD)
Gene IDs: Gorai.001G053400
cotton name: K08054 - calnexin (CANX) (1 of 2)
OG IDs: 93521554
Best-hit-arabi-ID: AT5G61790.1
Best-hit-arabi-name: calnexin 1
Best-hit-arabi-symbol: ATCNX1,CNX1
Gorai.001G053400.1.D
PCC=0.08, p=0.745
mRNA Profile
Protein Expression Profile
DPA

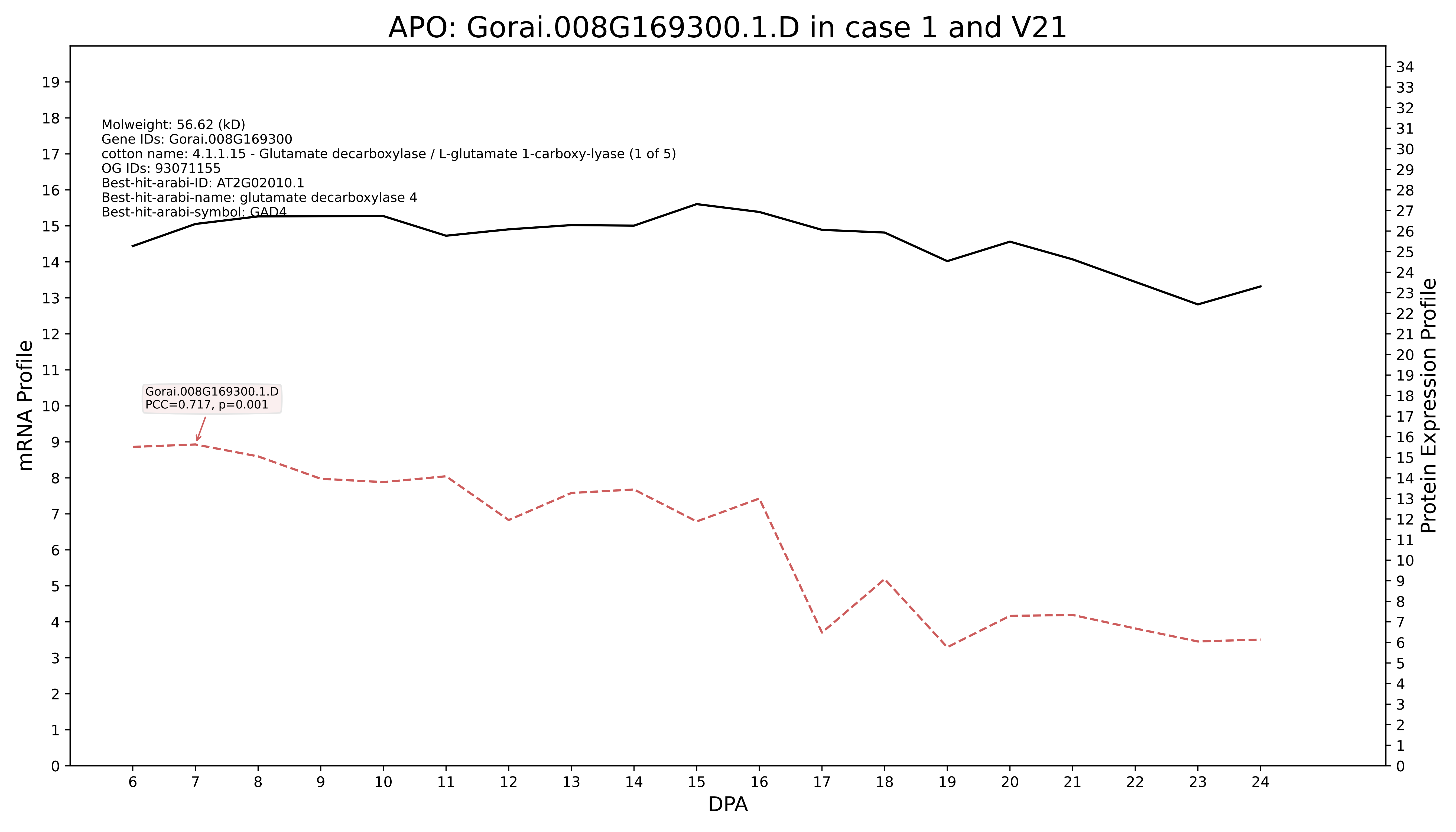

APO: Gorai.008G169300.1.D in case 1 and V21
Molweight: 56.62 (kD)
Gene IDs: Gorai.008G169300
cotton name: 4.1.1.15 - Glutamate decarboxylase / L-glutamate 1-carboxy-lyase (1 of 5)
OG IDs: 93071155
Best-hit-arabi-ID: AT2G02010.1
Best-hit-arabi-name: glutamate decarboxylase 4
Best-hit-arabi-symbol: GAD4
Gorai.008G169300.1.D
PCC=0.717, p=0.001
mRNA Profile
Protein Expression Profile
DPA

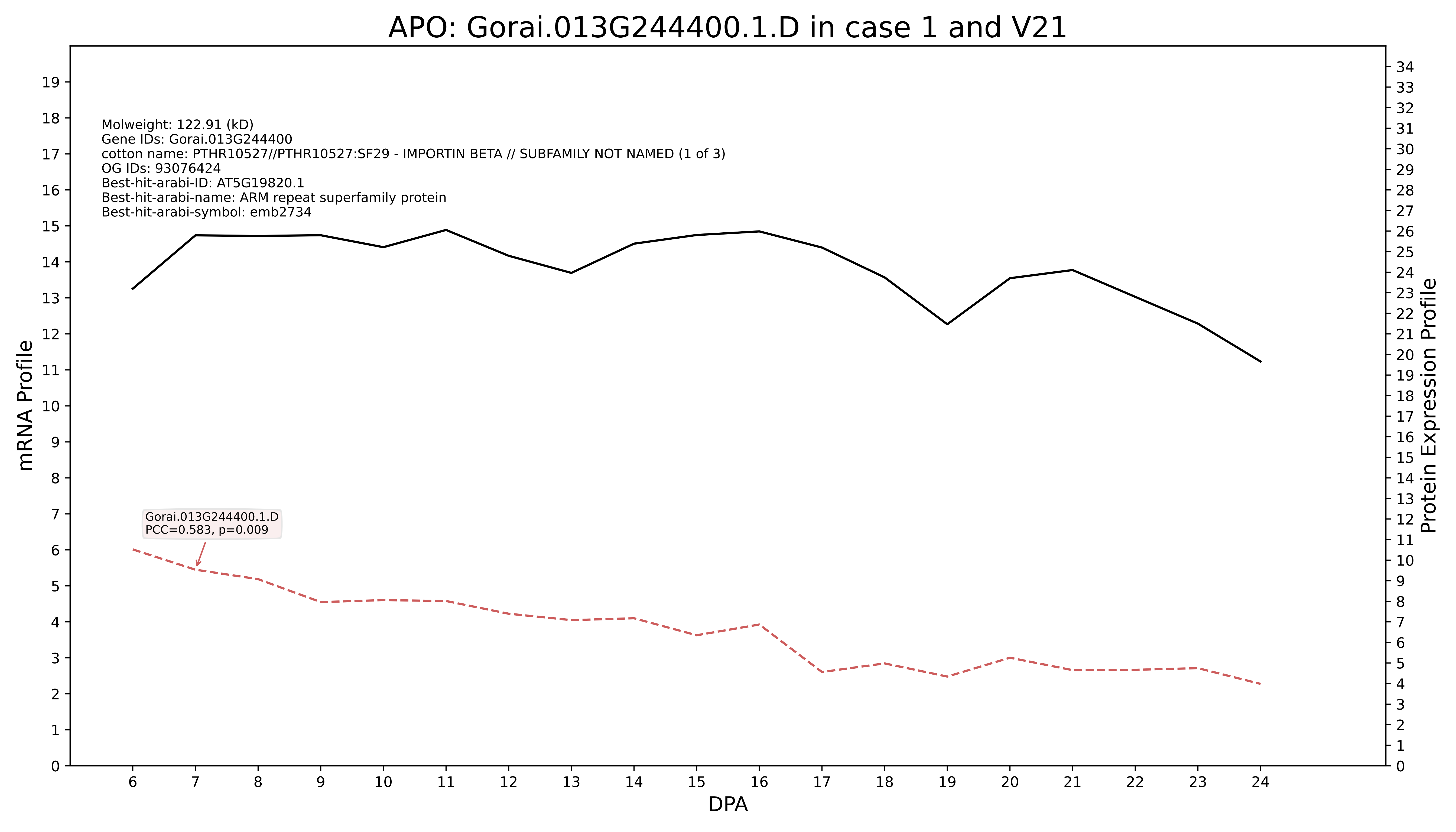

APO: Gorai.013G244400.1.D in case 1 and V21
Molweight: 122.91 (kD)
Gene IDs: Gorai.013G244400
cotton name: PTHR10527//PTHR10527:SF29 - IMPORTIN BETA // SUBFAMILY NOT NAMED (1 of 3)
OG IDs: 93076424
Best-hit-arabi-ID: AT5G19820.1
Best-hit-arabi-name: ARM repeat superfamily protein
Best-hit-arabi-symbol: emb2734
Gorai.013G244400.1.D
PCC=0.583, p=0.009
mRNA Profile
Protein Expression Profile
DPA

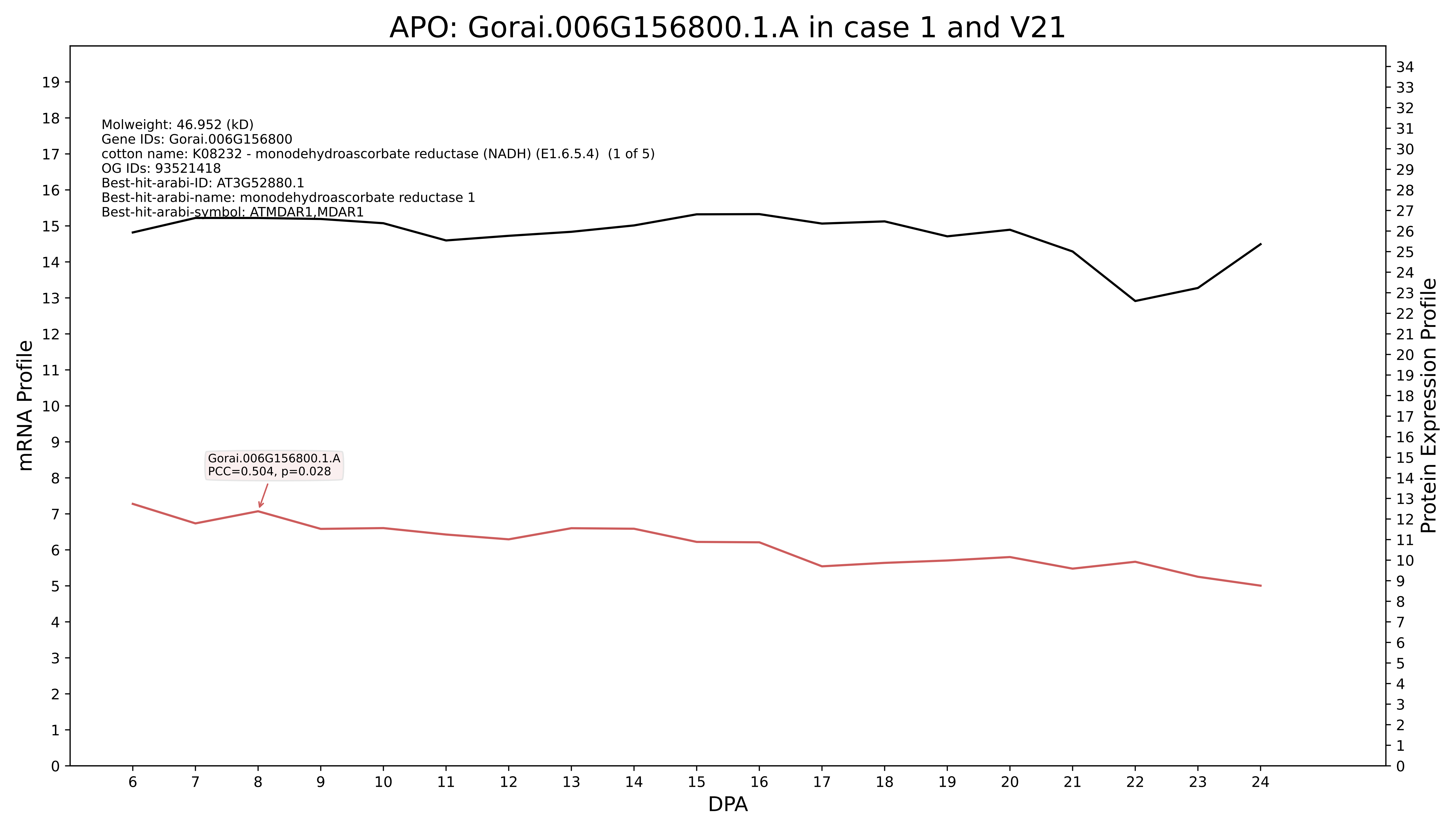
APO: Gorai.006G156800.1.A in case 1 and V21
Molweight: 46.952 (kD)
Gene IDs: Gorai.006G156800
cotton name: K08232 - monodehydroascorbate reductase (NADH) (E1.6.5.4) (1 of 5)
OG IDs: 93521418
Best-hit-arabi-ID: AT3G52880.1
Best-hit-arabi-name: monodehydroascorbate reductase 1
Best-hit-arabi-symbol: ATMDAR1,MDAR1
Gorai.006G156800.1.A
PCC=0.504, p=0.028
mRNA Profile
Protein Expression Profile
DPA

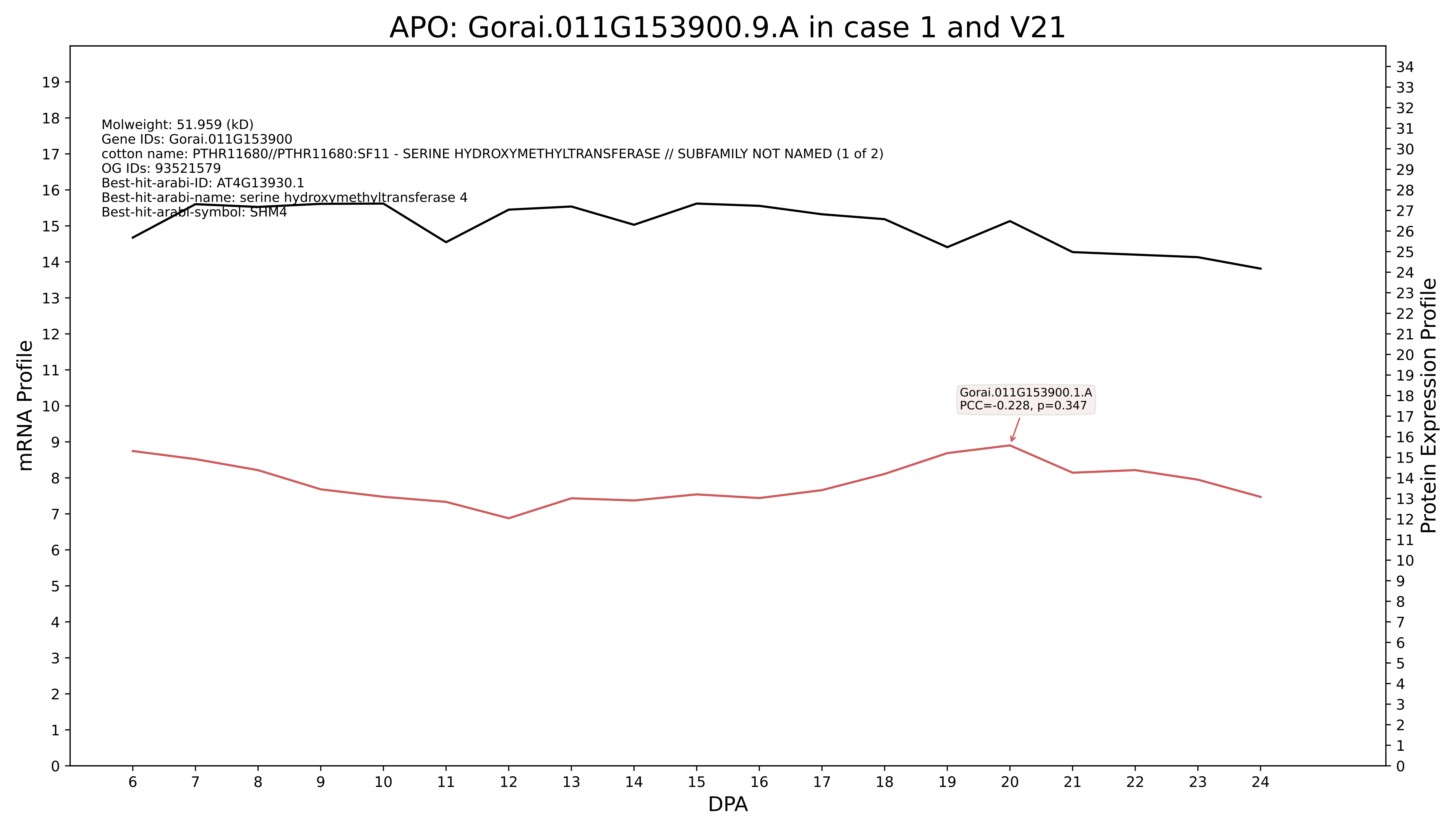
APO: Gorai.011G153900.9.A in case 1 and V21
Molweight: 51.959 (kD)
Gene IDs: Gorai.011G153900
cotton name: PTHR11680//PTHR11680:SF11 - SERINE HYDROXYMETHYLTRANSFERASE // SUBFAMILY NOT NAMED (1 of 2)
OG IDs: 93521579
Best-hit-arabi-ID: AT4G13930.1
Best-hit-arabi-name: serine hydroxymethyltransferase 4
Best-hit-arabi-symbol: SHM4
Gorai.011G153900.1.A
PCC=-0.228, p=0.347
mRNA Profile
Protein Expression Profile
DPA

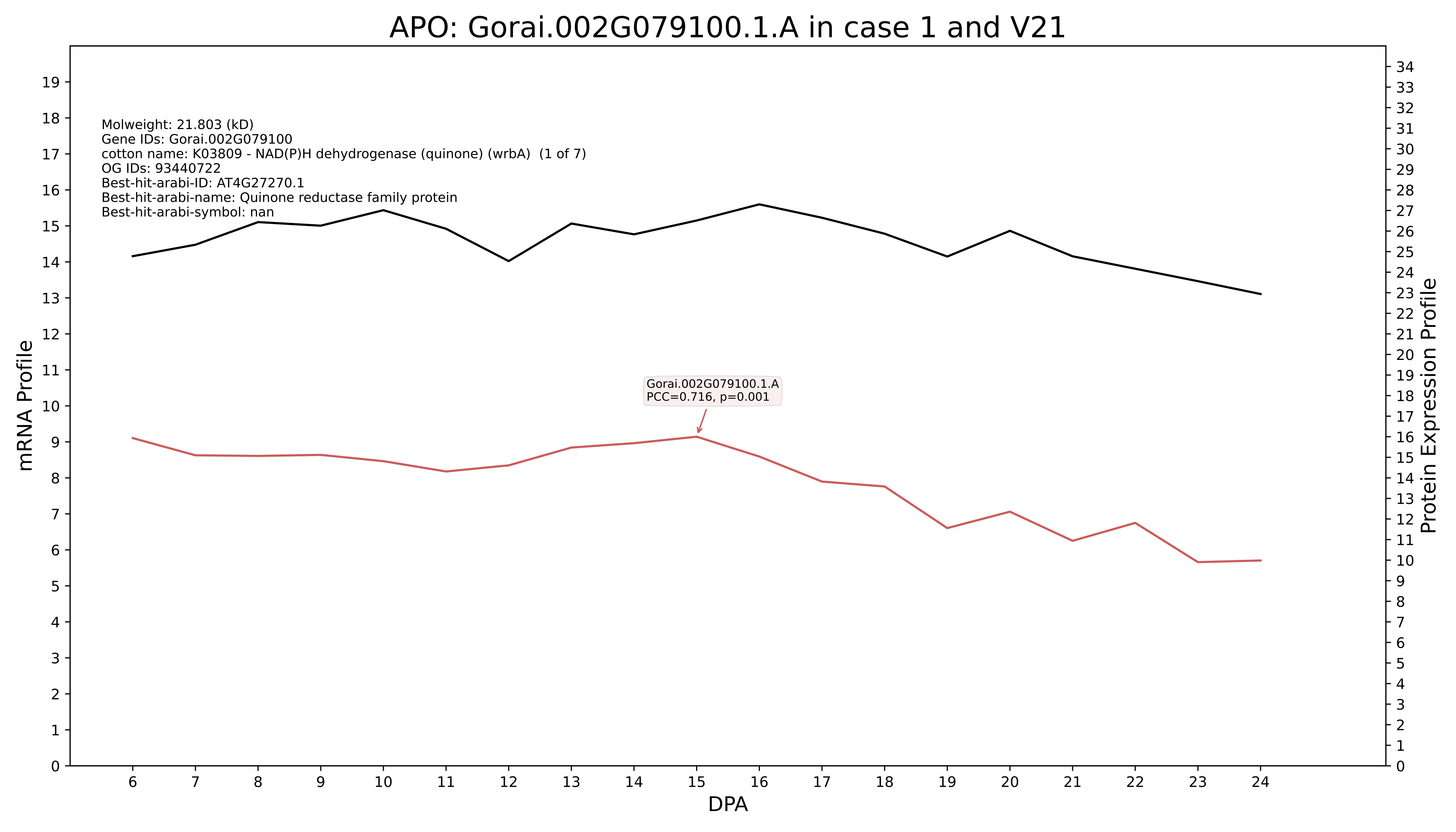
APO: Gorai.002G079100.1.A in case 1 and V21
Molweight: 21.803 (kD)
Gene IDs: Gorai.002G079100
cotton name: K03809 - NAD(P)H dehydrogenase (quinone) (wrbA) (1 of 7)
OG IDs: 93440722
Best-hit-arabi-ID: AT4G27270.1
Best-hit-arabi-name: Quinone reductase family protein
Best-hit-arabi-symbol: nan
Gorai.002G079100.1.A
PCC=0.716, p=0.001
mRNA Profile
Protein Expression Profile
DPA

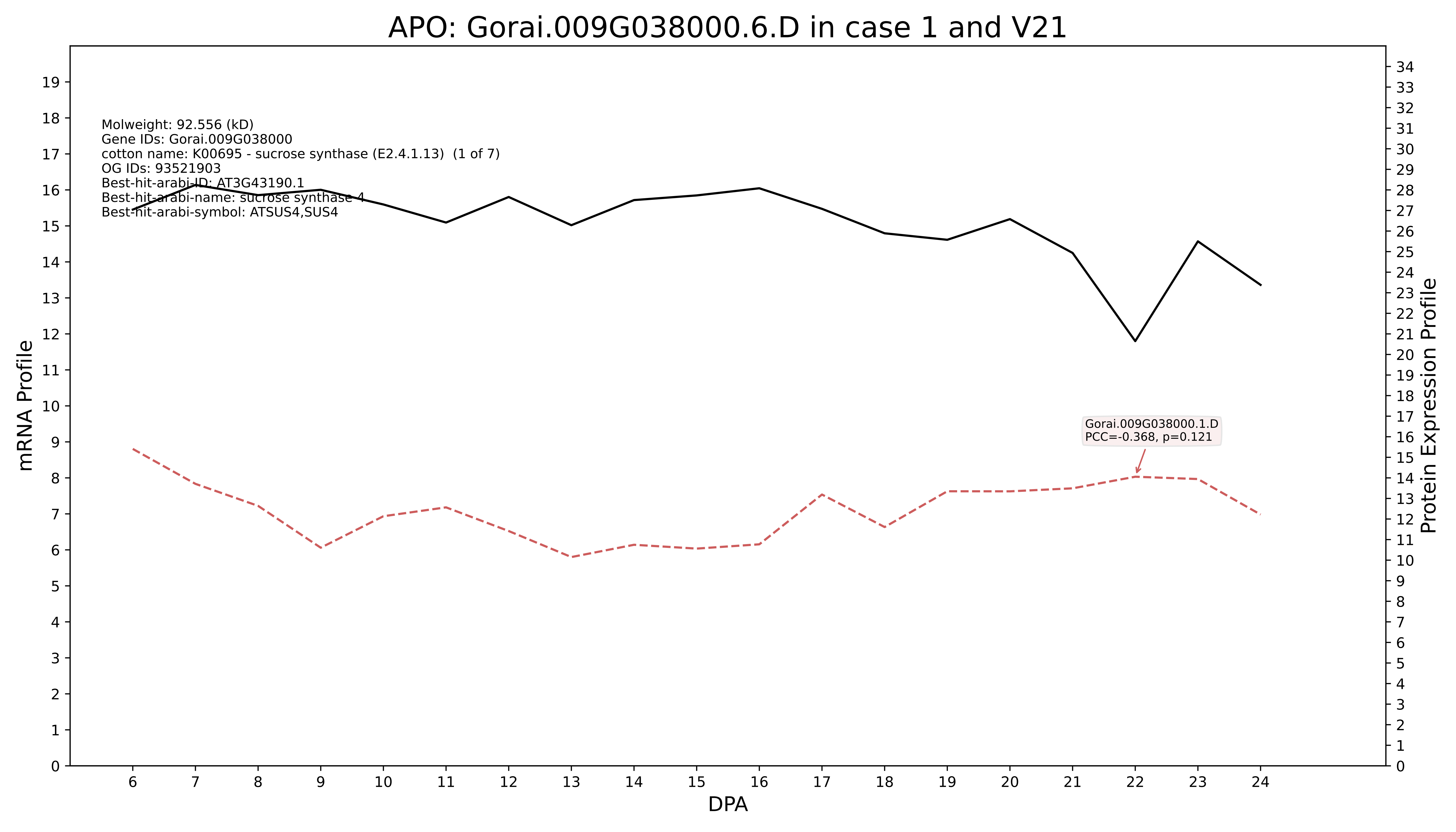

APO: Gorai.009G038000.6.D in case 1 and V21
Molweight: 92.556 (kD)
Gene IDs: Gorai.009G038000
cotton name: K00695 - sucrose synthase (E2.4.1.13) (1 of 7)
OG IDs: 93521903
Best-hit-arabi-ID: AT3G43190.1
Best-hit-arabi-name: sucrose synthase 4
Best-hit-arabi-symbol: ATSUS4,SUS4
Gorai.009G038000.1.D
PCC=-0.368, p=0.121
mRNA Profile
Protein Expression Profile
DPA

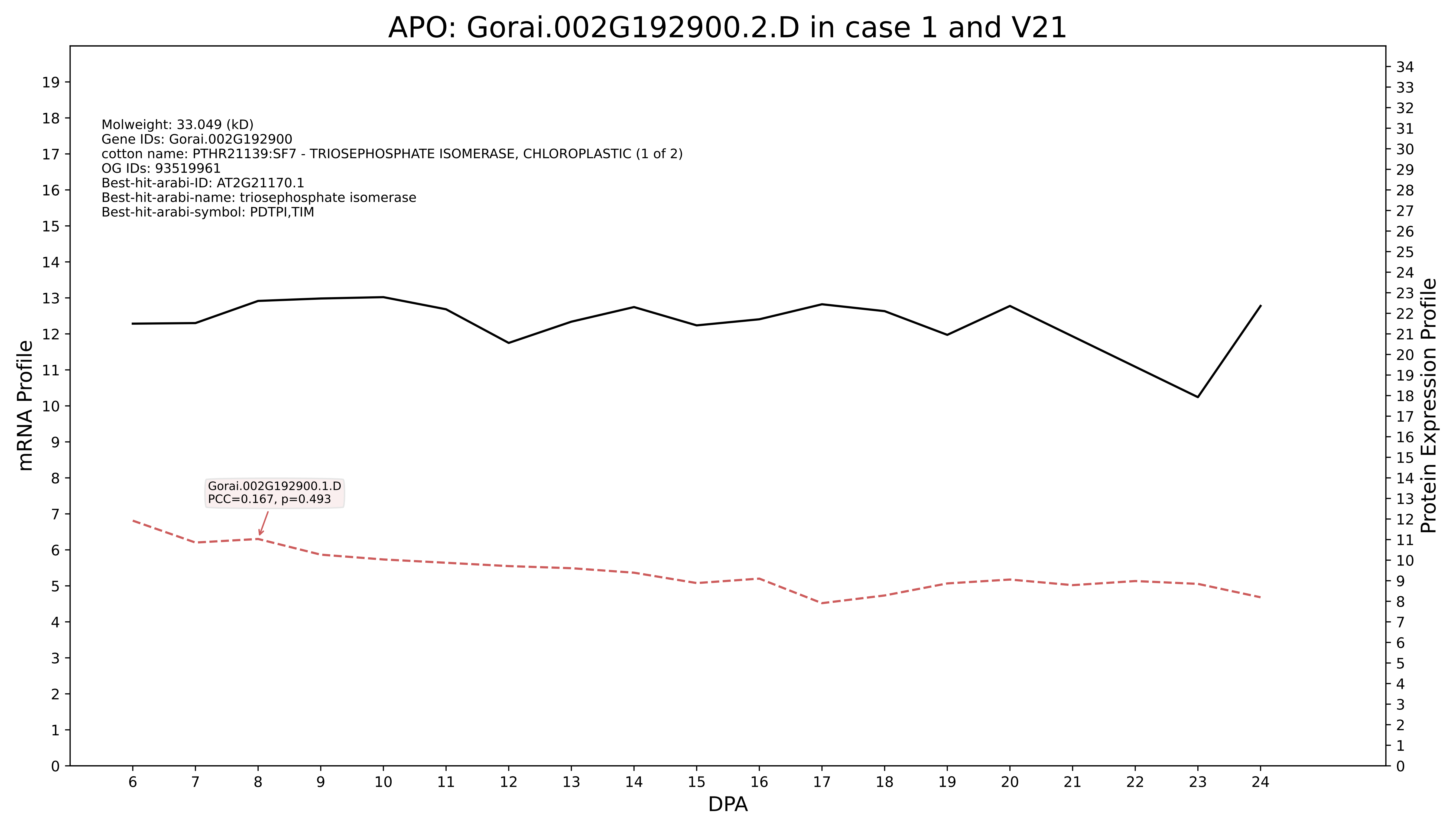

APO: Gorai.002G192900.2.D in case 1 and V21
Molweight: 33.049 (kD)
Gene IDs: Gorai.002G192900
cotton name: PTHR21139:SF7 - TRIOSEPHOSPHATE ISOMERASE, CHLOROPLASTIC (1 of 2)
OG IDs: 93519961
Best-hit-arabi-ID: AT2G21170.1
Best-hit-arabi-name: triosephosphate isomerase
Best-hit-arabi-symbol: PDTPI,TIM
Gorai.002G192900.1.D
PCC=0.167, p=0.493
mRNA Profile
Protein Expression Profile
DPA

#### APO: Gorai.009G234600.3.D in case 1 and V21

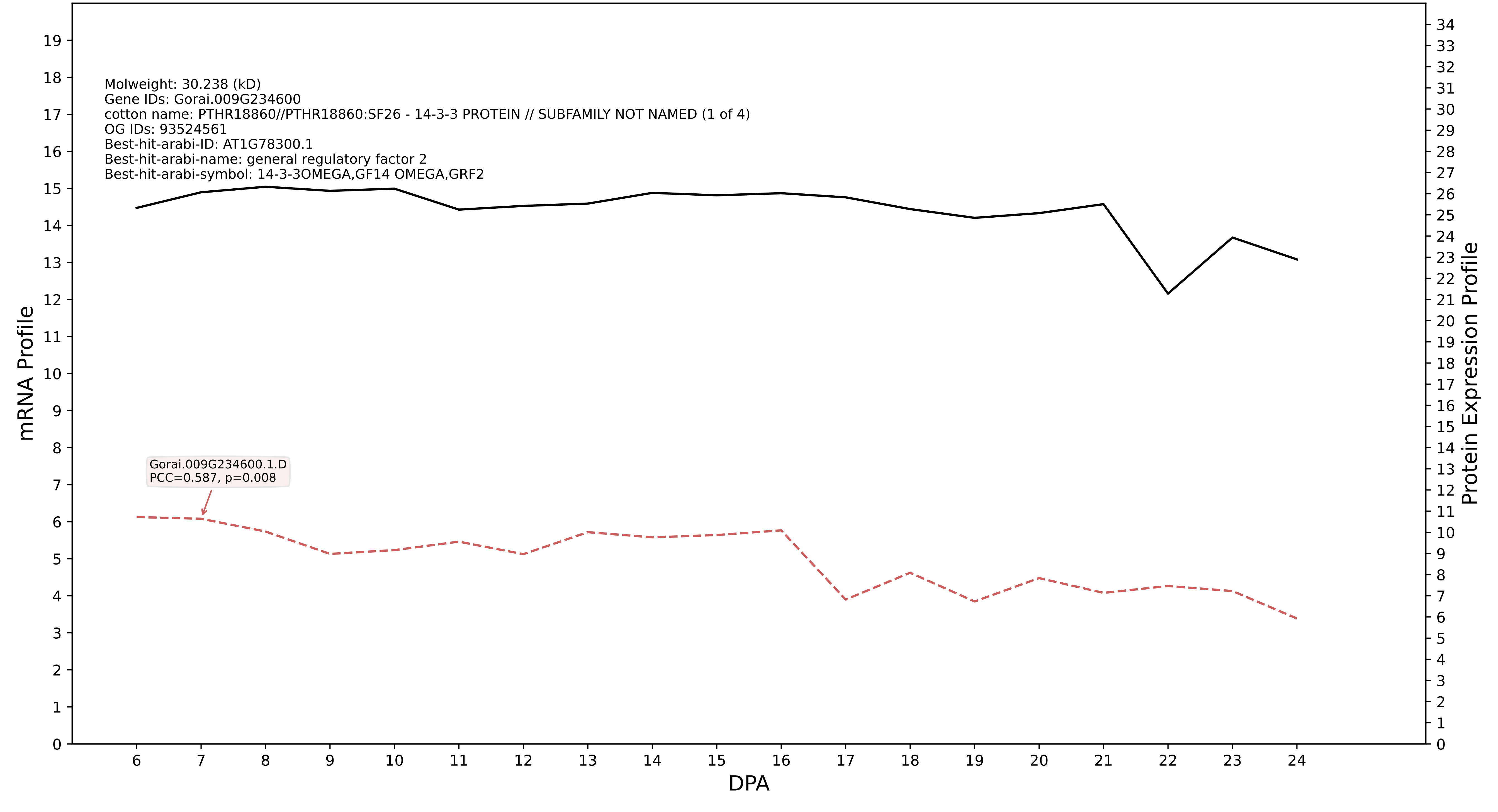

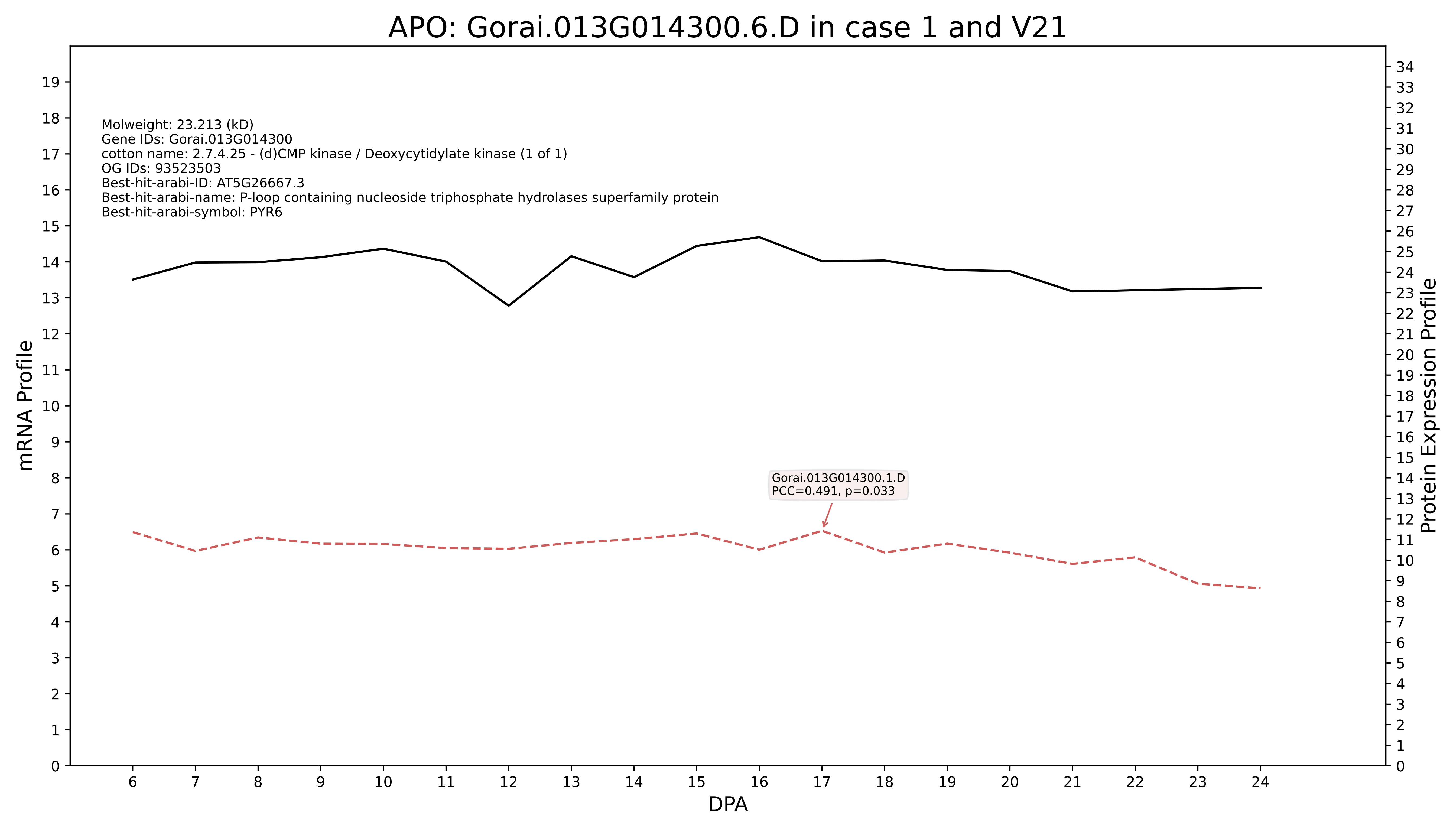
APO: Gorai.013G014300.6.D in case 1 and V21
Molweight: 23.213 (kD)
Gene IDs: Gorai.013G014300
cotton name: 2.7.4.25 - (d)CMP kinase / Deoxycytidylate kinase (1 of 1)
OG IDs: 93523503
Best-hit-arabi-ID: AT5G26667.3
Best-hit-arabi-name: P-loop containing nucleoside triphosphate hydrolases superfamily protein
Best-hit-arabi-symbol: PYR6
Gorai.013G014300.1.D
PCC=0.491, p=0.033
mRNA Profile
Protein Expression Profile
DPA

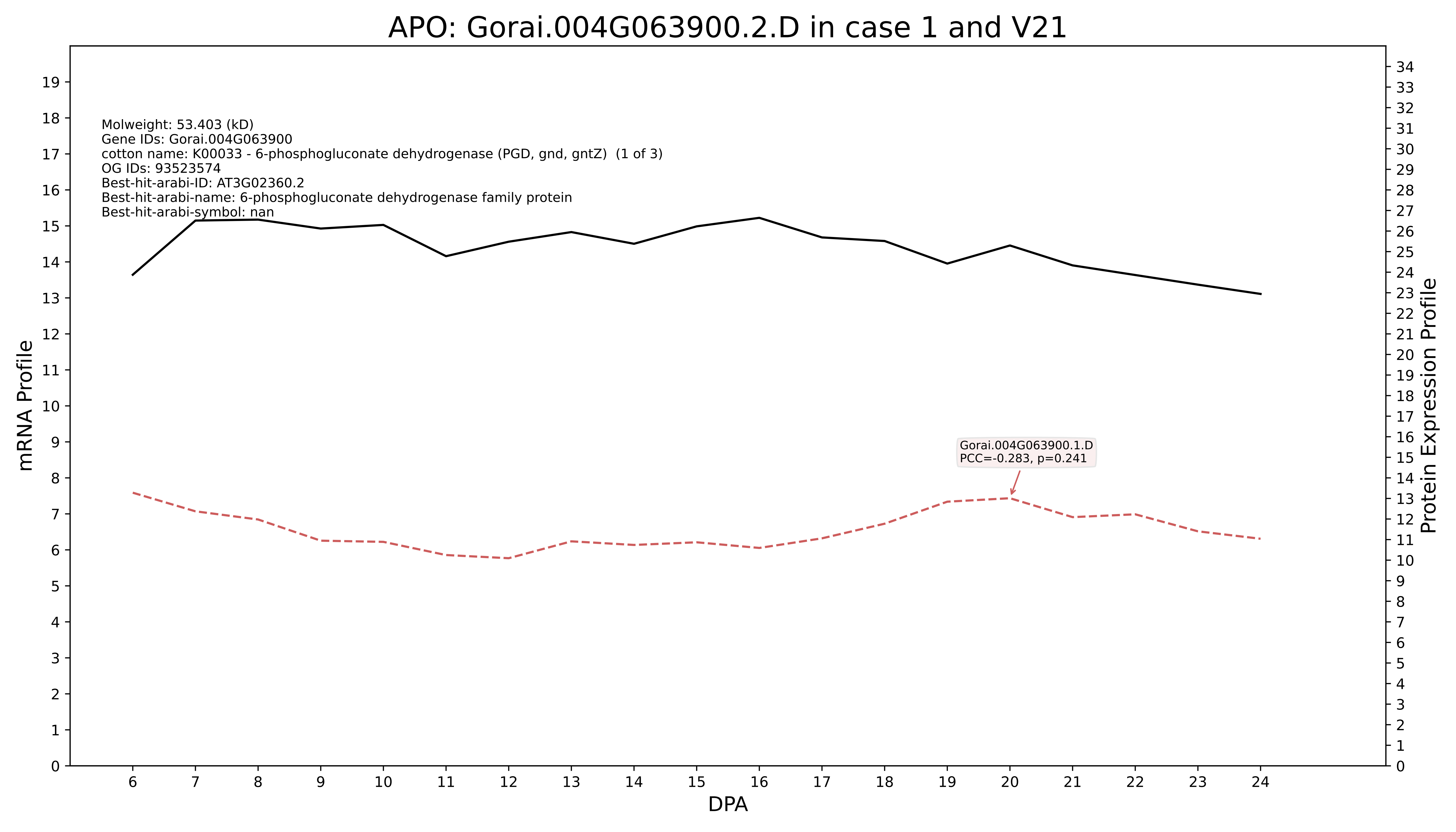
APO: Gorai.004G063900.2.D in case 1 and V21
Molweight: 53.403 (kD)
Gene IDs: Gorai.004G063900
cotton name: K00033 - 6-phosphogluconate dehydrogenase (PGD, gnd, gntZ) (1 of 3)
OG IDs: 93523574
Best-hit-arabi-ID: AT3G02360.2
Best-hit-arabi-name: 6-phosphogluconate dehydrogenase family protein
Best-hit-arabi-symbol: nan
Gorai.004G063900.1.D
PCC=-0.283, p=0.241
mRNA Profile
Protein Expression Profile
DPA

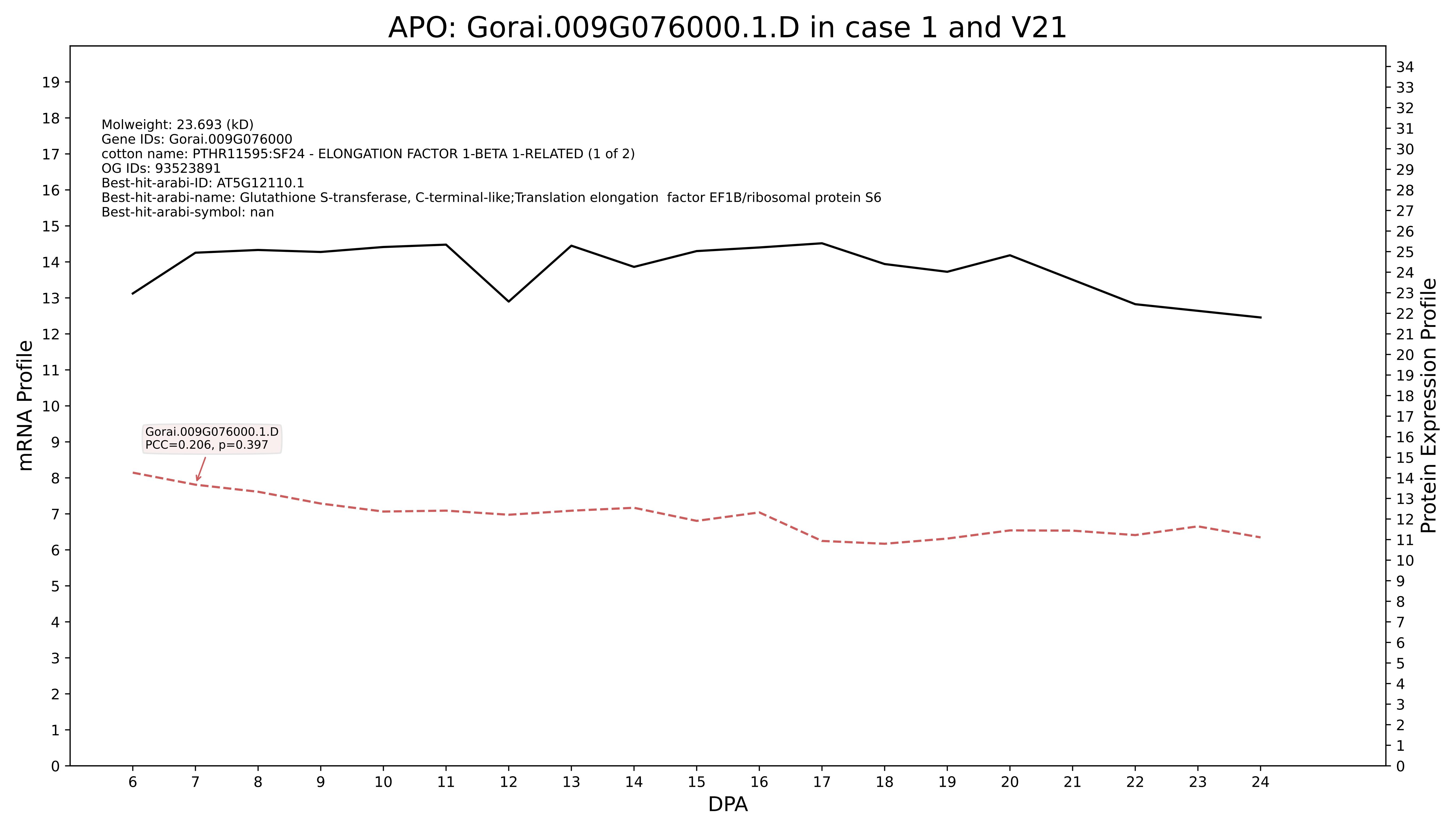

APO: Gorai.009G076000.1.D in case 1 and V21
Molweight: 23.693 (kD)
Gene IDs: Gorai.009G076000
cotton name: PTHR11595:SF24 - ELONGATION FACTOR 1-BETA 1-RELATED (1 of 2)
OG IDs: 93523891
Best-hit-arabi-ID: AT5G12110.1
Best-hit-arabi-name: Glutathione S-transferase, C-terminal-like;Translation elongation factor EF1B/ribosomal protein S6
Best-hit-arabi-symbol: nan
Gorai.009G076000.1.D
PCC=0.206, p=0.397
mRNA Profile
Protein Expression Profile
DPA

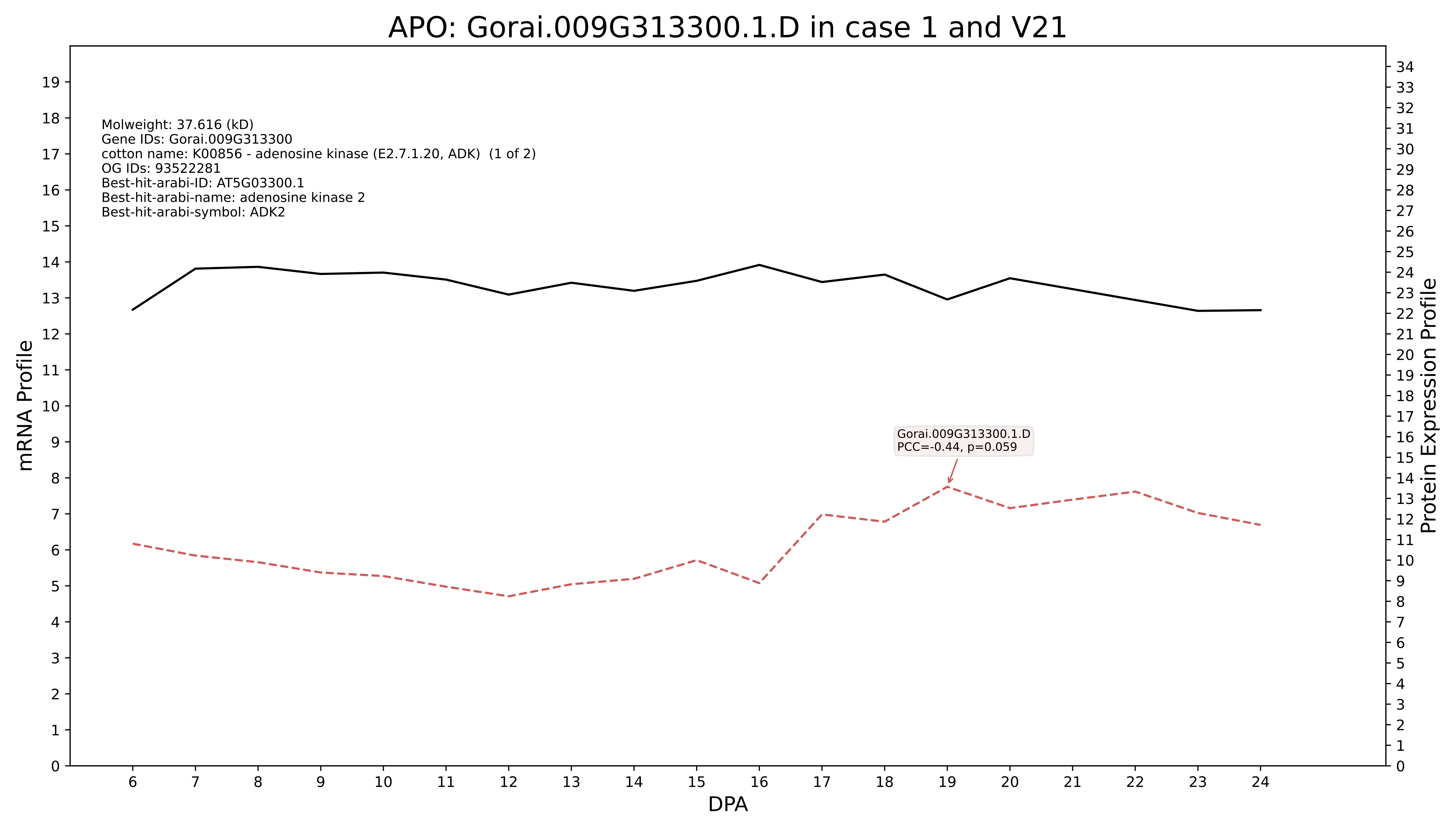

APO: Gorai.009G313300.1.D in case 1 and V21
Molweight: 37.616 (kD)
Gene IDs: Gorai.009G313300
cotton name: K00856 - adenosine kinase (E2.7.1.20, ADK) (1 of 2)
OG IDs: 93522281
Best-hit-arabi-ID: AT5G03300.1
Best-hit-arabi-name: adenosine kinase 2
Best-hit-arabi-symbol: ADK2
Gorai.009G313300.1.D
PCC=-0.44, p=0.059
mRNA Profile
Protein Expression Profile
DPA

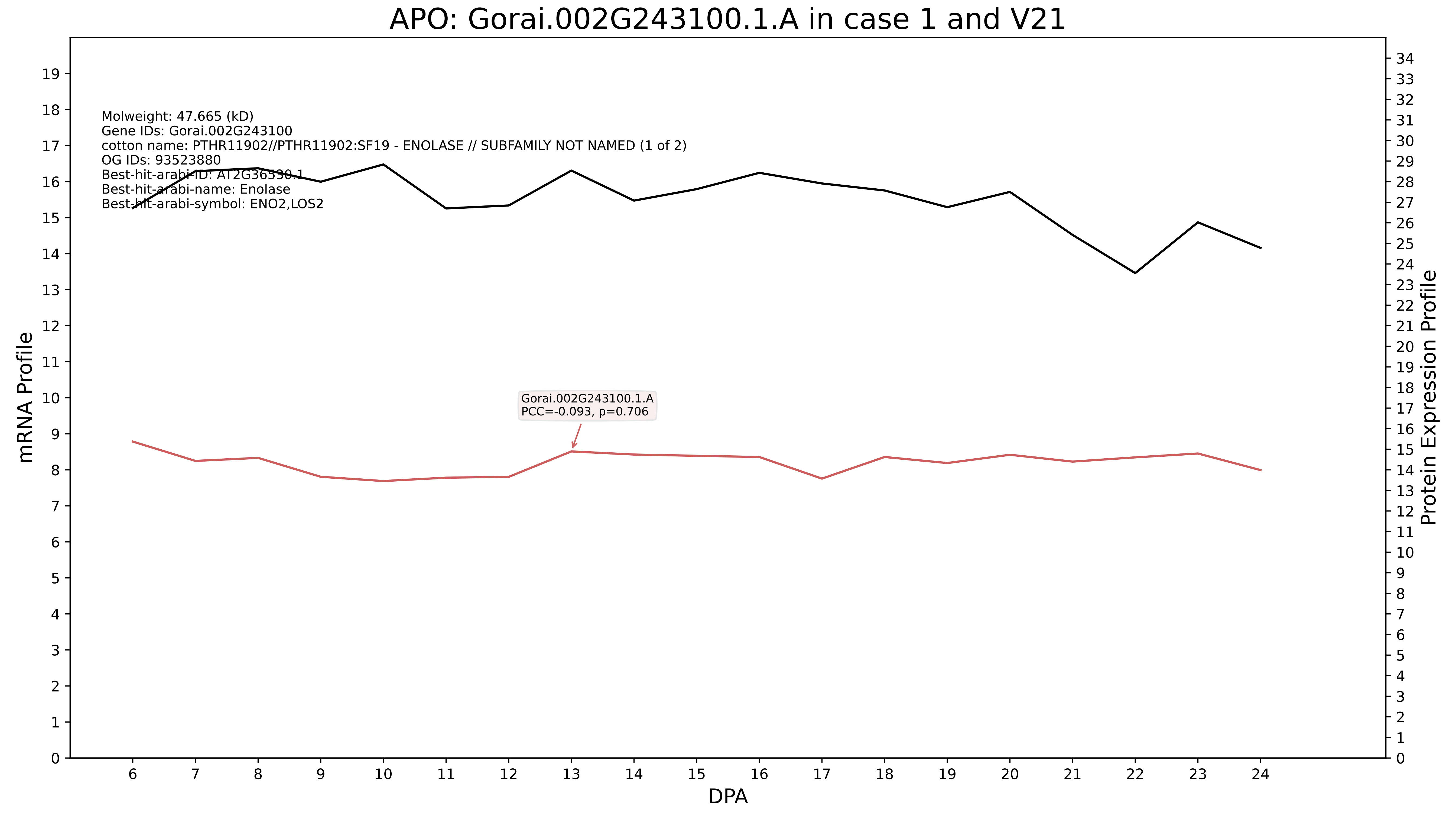
APO: Gorai.002G243100.1.A in case 1 and V21
Molweight: 47.665 (kD)
Gene IDs: Gorai.002G243100
cotton name: PTHR11902//PTHR11902:SF19 - ENOLASE // SUBFAMILY NOT NAMED (1 of 2)
OG IDs: 93523880
Best-hit-arabi-ID: AT2G36530.1
Best-hit-arabi-name: Enolase
Best-hit-arabi-symbol: ENO2,LOS2
Gorai.002G243100.1.A
PCC=-0.093, p=0.706
mRNA Profile
Protein Expression Profile
DPA

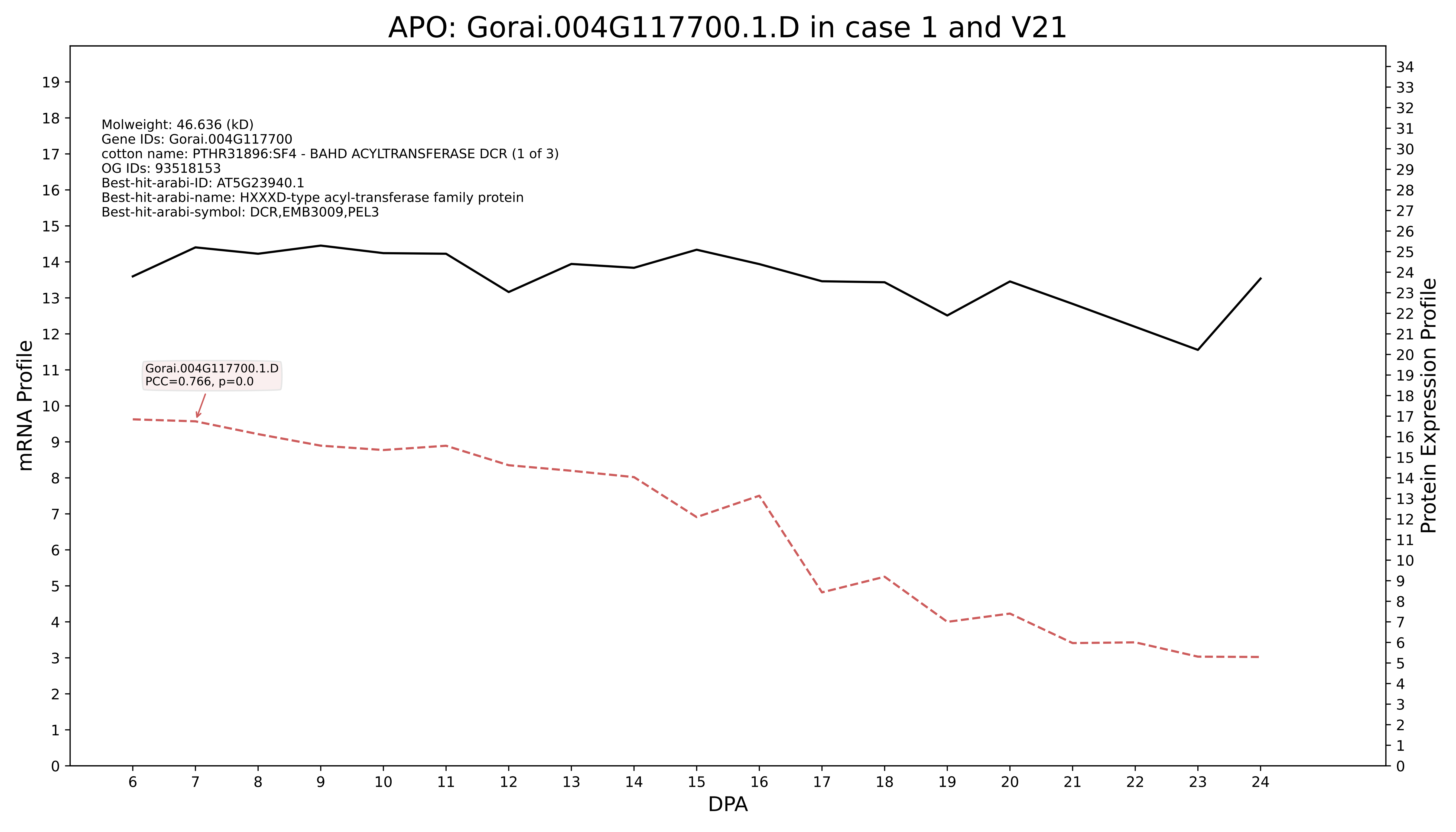
APO: Gorai.004G117700.1.D in case 1 and V21
Molweight: 46.636 (kD)
Gene IDs: Gorai.004G117700
cotton name: PTHR31896:SF4 - BAHD ACYLTRANSFERASE DCR (1 of 3)
OG IDs: 93518153
Best-hit-arabi-ID: AT5G23940.1
Best-hit-arabi-name: HXXXD-type acyl-transferase family protein
Best-hit-arabi-symbol: DCR,EMB3009,PEL3
Gorai.004G117700.1.D
PCC=0.766, p=0.0
mRNA Profile
Protein Expression Profile
DPA

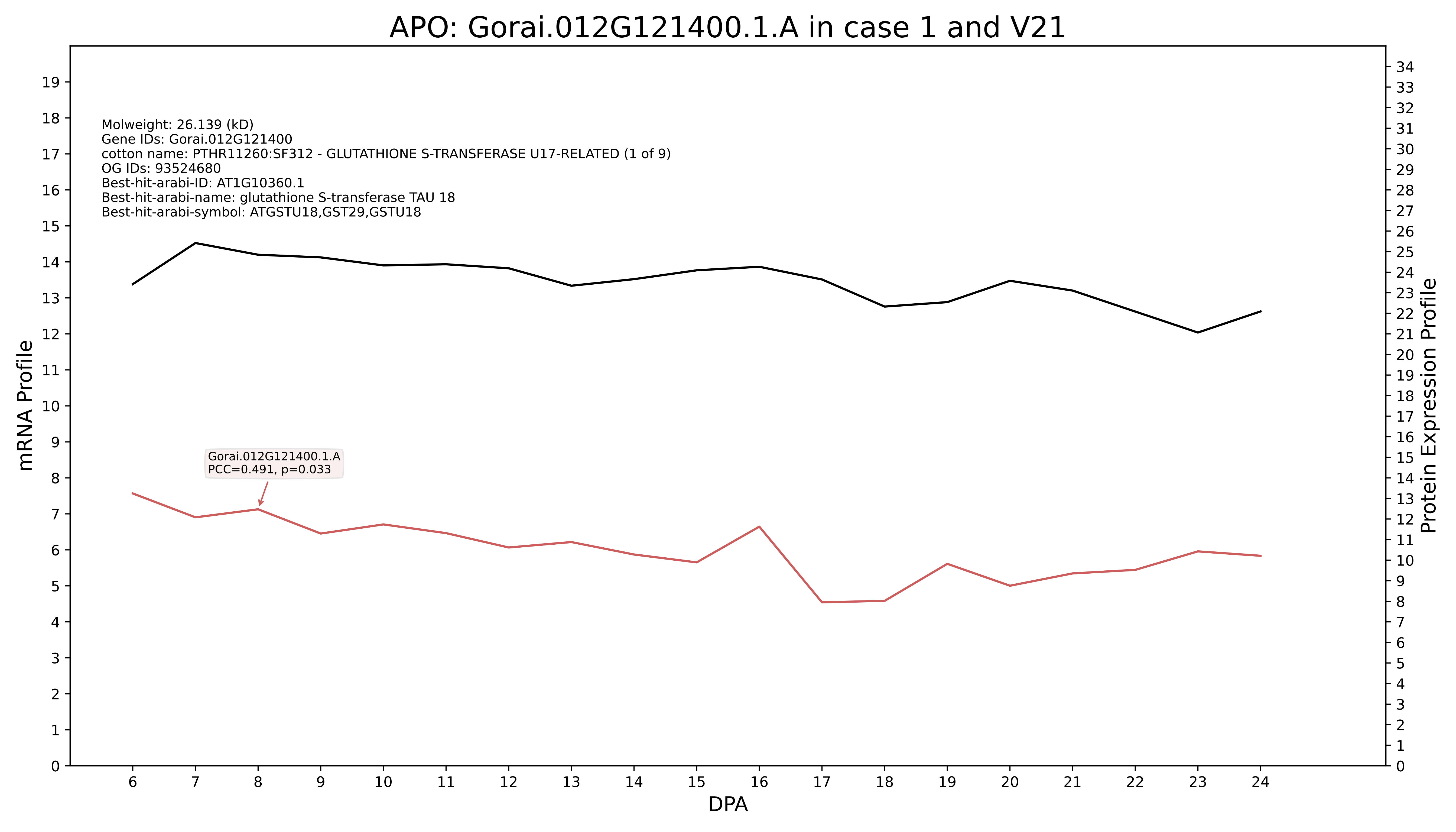
APO: Gorai.012G121400.1.A in case 1 and V21
Molweight: 26.139 (kD)
Gene IDs: Gorai.012G121400
cotton name: PTHR11260:SF312 - GLUTATHIONE S-TRANSFERASE U17-RELATED (1 of 9)
OG IDs: 93524680
Best-hit-arabi-ID: AT1G10360.1
Best-hit-arabi-name: glutathione S-transferase TAU 18
Best-hit-arabi-symbol: ATGSTU18,GST29,GSTU18
Gorai.012G121400.1.A
PCC=0.491, p=0.033
mRNA Profile
Protein Expression Profile
DPA

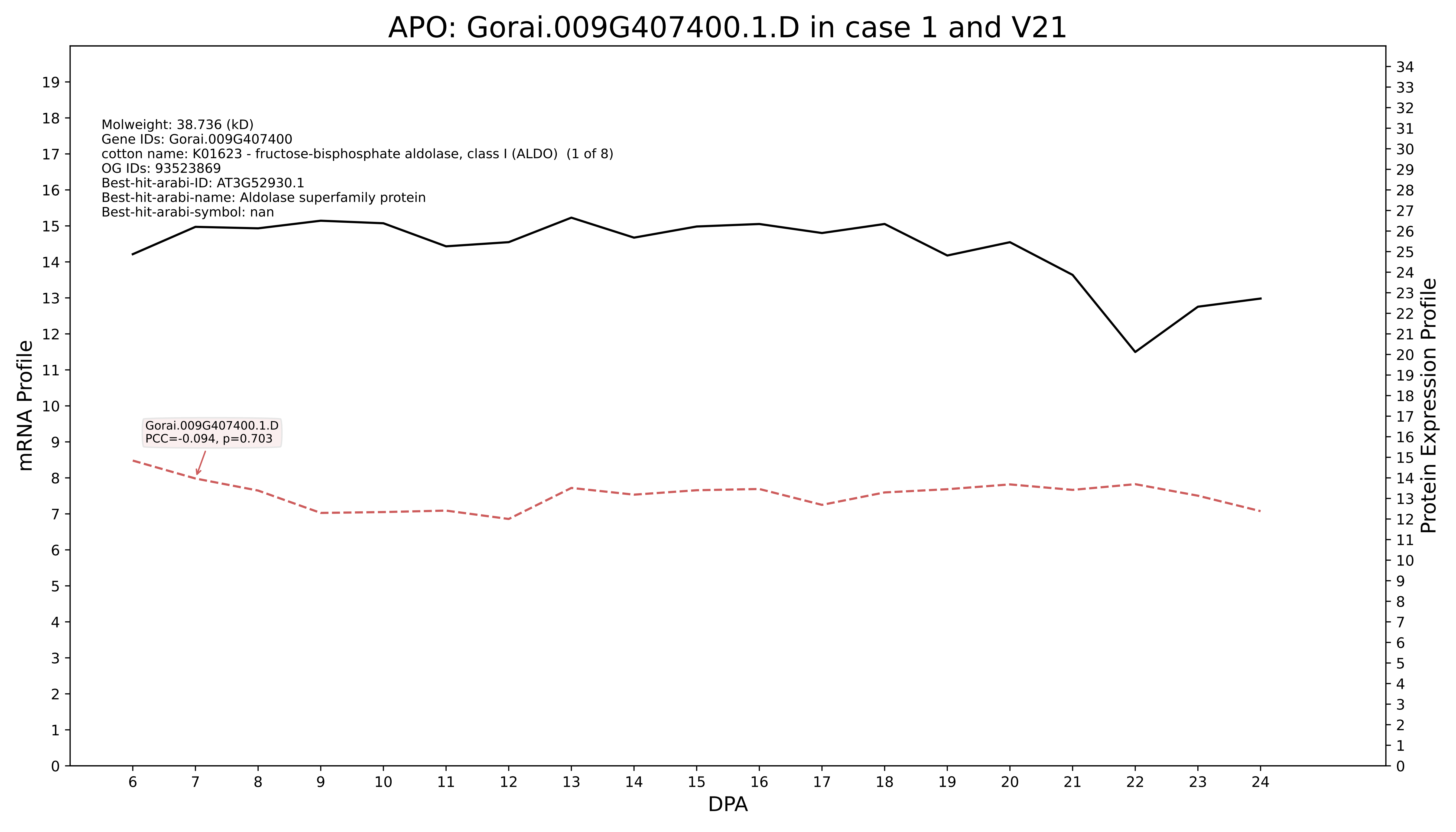

APO: Gorai.009G407400.1.D in case 1 and V21
Molweight: 38.736 (kD)
Gene IDs: Gorai.009G407400
cotton name: K01623 - fructose-bisphosphate aldolase, class I (ALDO) (1 of 8)
OG IDs: 93523869
Best-hit-arabi-ID: AT3G52930.1
Best-hit-arabi-name: Aldolase superfamily protein
Best-hit-arabi-symbol: nan
Gorai.009G407400.1.D
PCC=-0.094, p=0.703
mRNA Profile
Protein Expression Profile
DPA

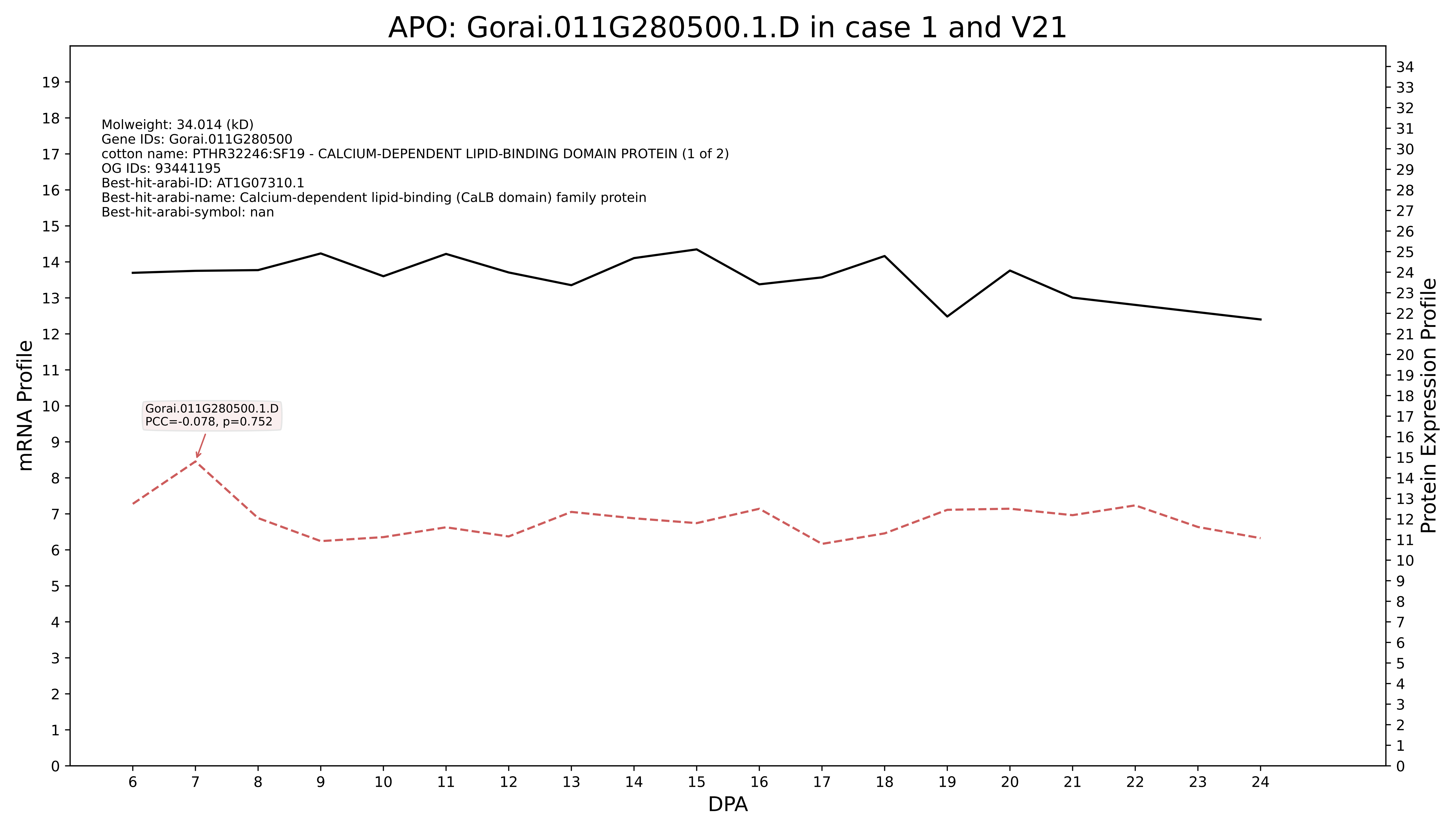

APO: Gorai.011G280500.1.D in case 1 and V21
Molweight: 34.014 (kD)
Gene IDs: Gorai.011G280500
cotton name: PTHR32246:SF19 - CALCIUM-DEPENDENT LIPID-BINDING DOMAIN PROTEIN (1 of 2)
OG IDs: 93441195
Best-hit-arabi-ID: AT1G07310.1
Best-hit-arabi-name: Calcium-dependent lipid-binding (CaLB domain) family protein
Best-hit-arabi-symbol: nan
Gorai.011G280500.1.D
PCC=-0.078, p=0.752
mRNA Profile
Protein Expression Profile
DPA

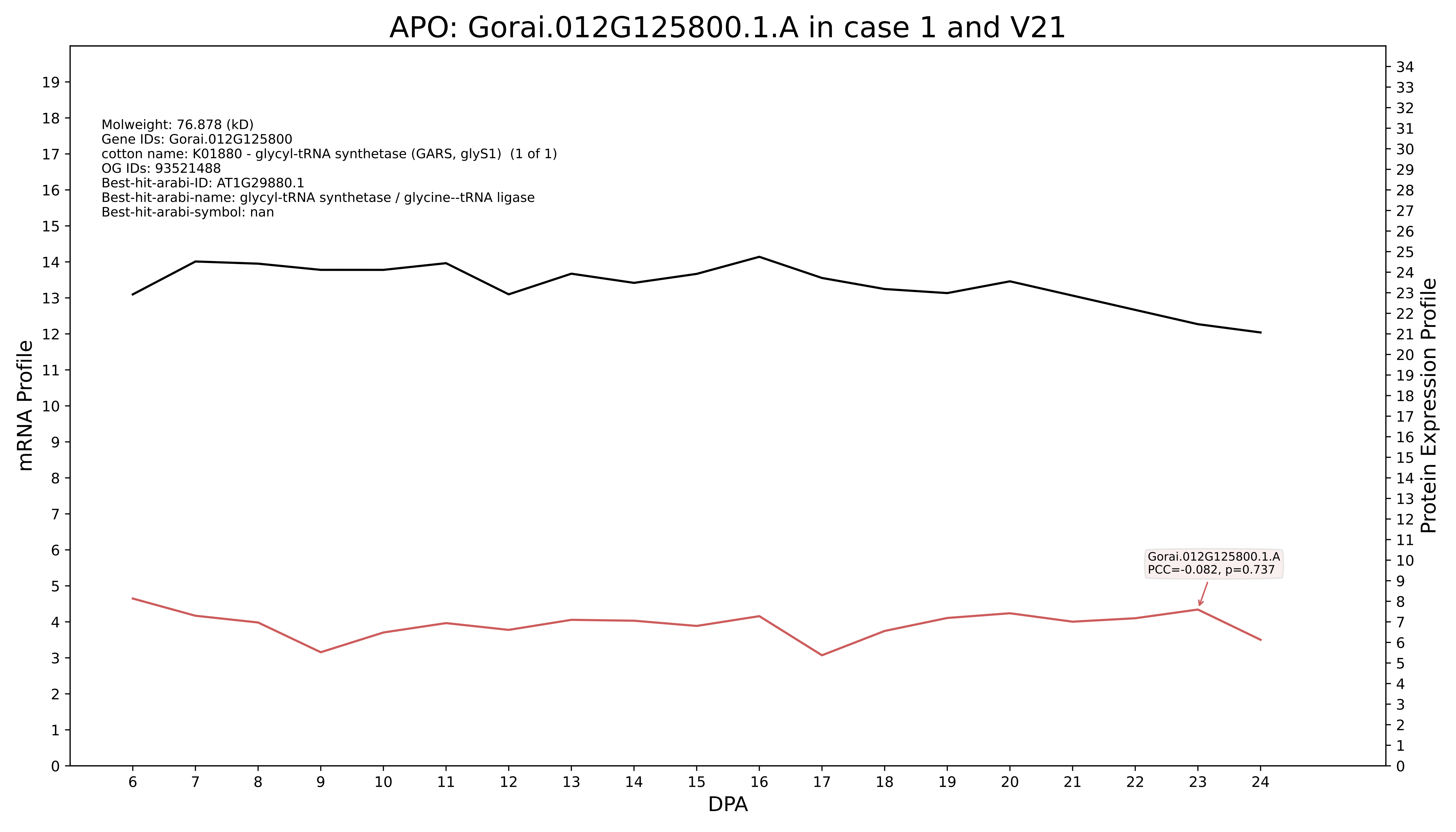
APO: Gorai.012G125800.1.A in case 1 and V21
Molweight: 76.878 (kD)
Gene IDs: Gorai.012G125800
cotton name: K01880 - glycyl-tRNA synthetase (GARS, glyS1) (1 of 1)
OG IDs: 93521488
Best-hit-arabi-ID: AT1G29880.1
Best-hit-arabi-name: glycyl-tRNA synthetase / glycine--tRNA ligase
Best-hit-arabi-symbol: nan
Gorai.012G125800.1.A
PCC=-0.082, p=0.737
mRNA Profile
Protein Expression Profile
DPA

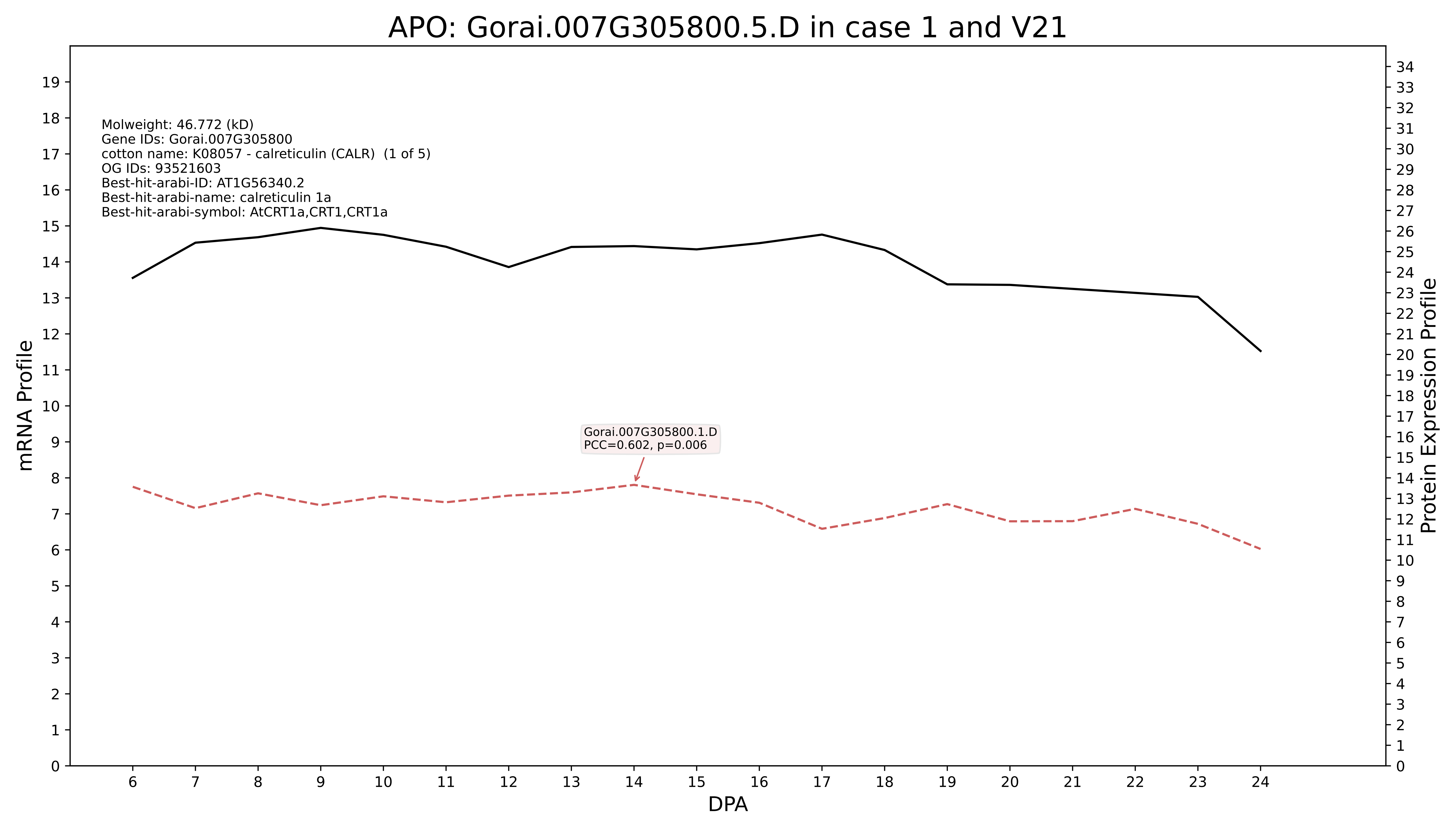

APO: Gorai.007G305800.5.D in case 1 and V21
Molweight: 46.772 (kD)
Gene IDs: Gorai.007G305800
cotton name: K08057 - calreticulin (CALR) (1 of 5)
OG IDs: 93521603
Best-hit-arabi-ID: AT1G56340.2
Best-hit-arabi-name: calreticulin 1a
Best-hit-arabi-symbol: AtCRT1a,CRT1,CRT1a
Gorai.007G305800.1.D
PCC=0.602, p=0.006
mRNA Profile
Protein Expression Profile
DPA

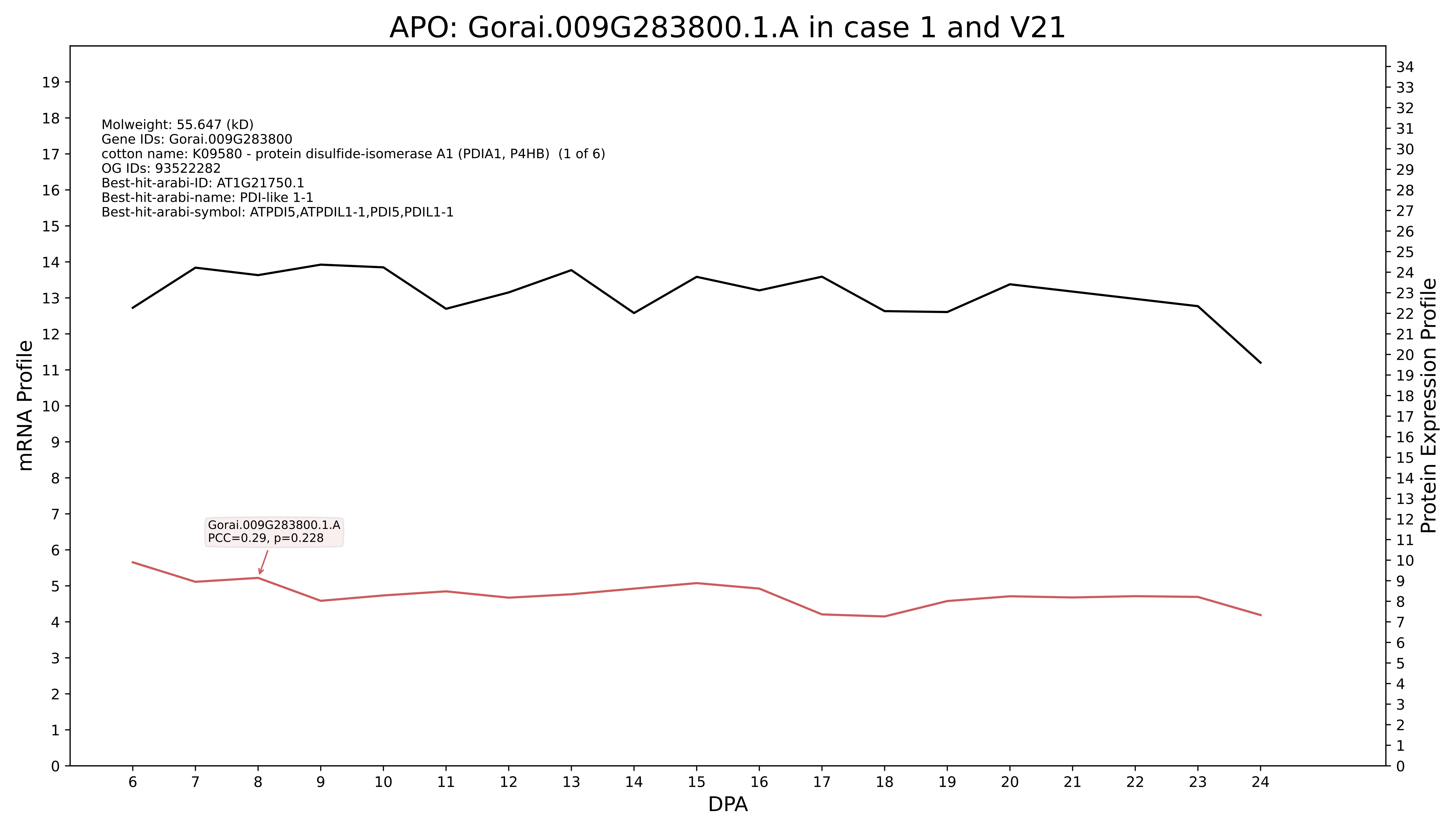
APO: Gorai.009G283800.1.A in case 1 and V21
Molweight: 55.647 (kD)
Gene IDs: Gorai.009G283800
cotton name: K09580 - protein disulfide-isomerase A1 (PDIA1, P4HB) (1 of 6)
OG IDs: 93522282
Best-hit-arabi-ID: AT1G21750.1
Best-hit-arabi-name: PDI-like 1-1
Best-hit-arabi-symbol: ATPDI5,ATPDIL1-1,PDI5,PDIL1-1
Gorai.009G283800.1.A
PCC=0.29, p=0.228
mRNA Profile
Protein Expression Profile
DPA

APO: Gorai.007G188300.2.A in case 1 and V21

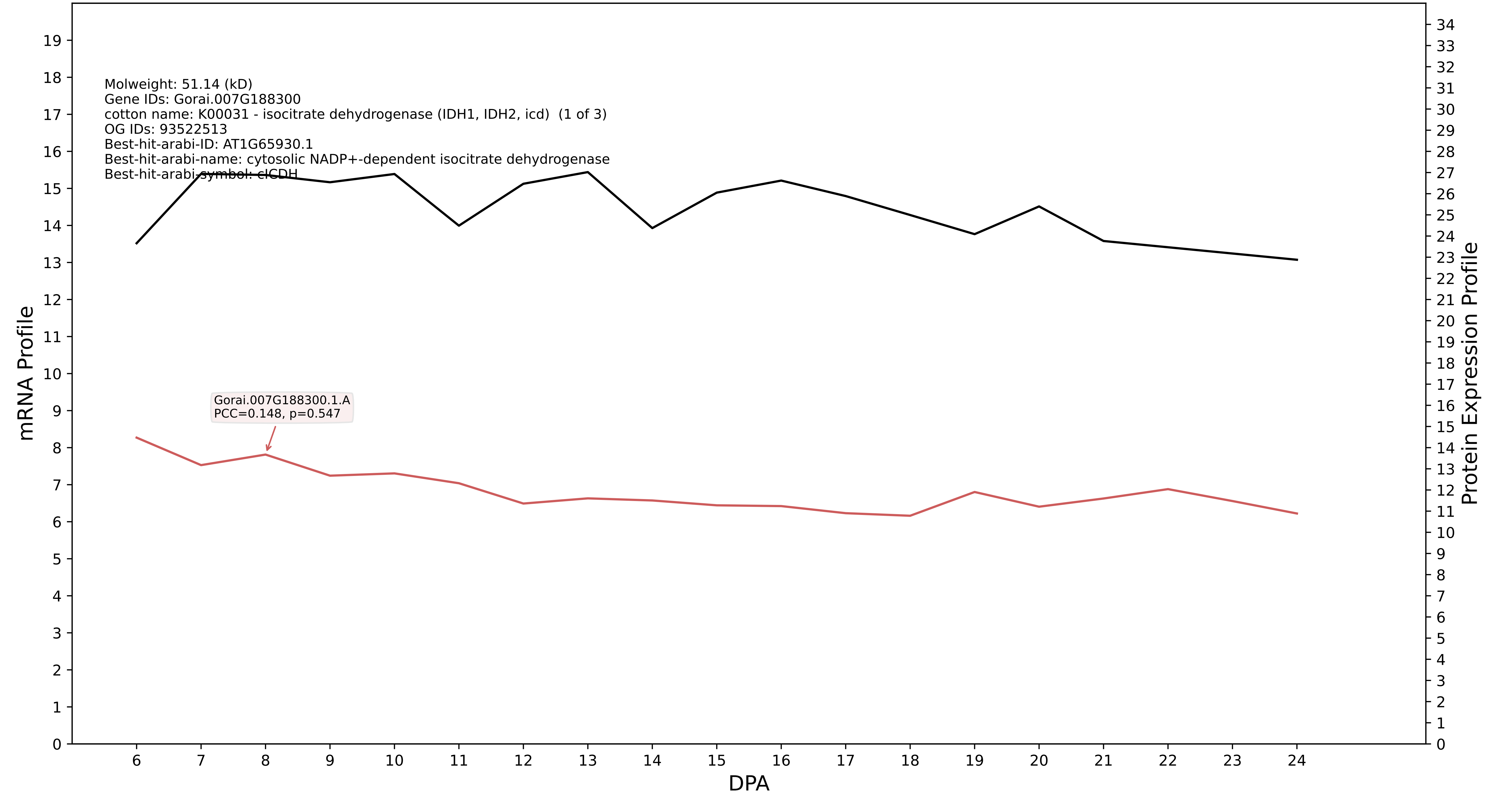

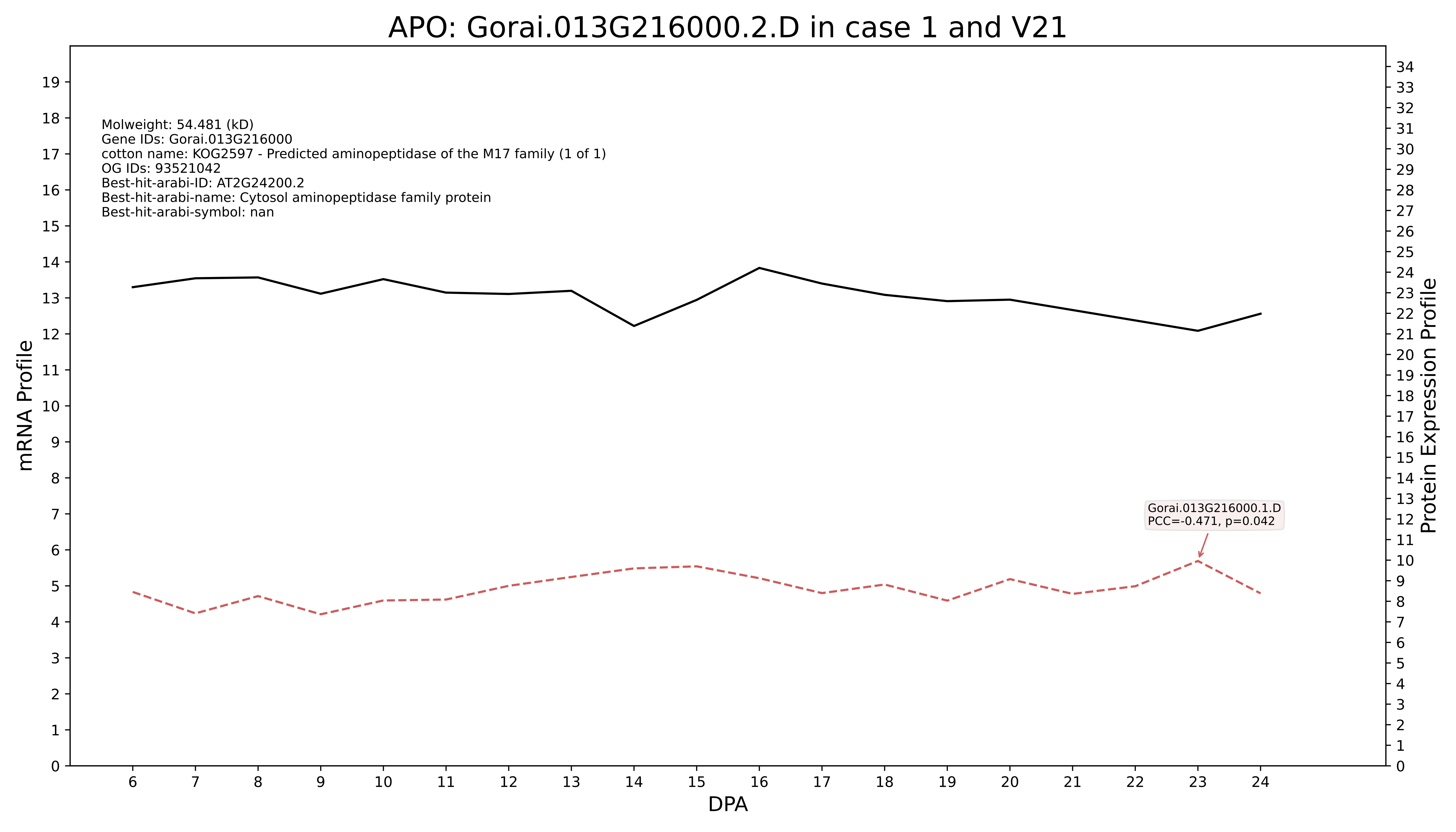
APO: Gorai.013G216000.2.D in case 1 and V21
Molweight: 54.481 (kD)
Gene IDs: Gorai.013G216000
cotton name: KOG2597 - Predicted aminopeptidase of the M17 family (1 of 1)
OG IDs: 93521042
Best-hit-arabi-ID: AT2G24200.2
Best-hit-arabi-name: Cytosol aminopeptidase family protein
Best-hit-arabi-symbol: nan
Gorai.013G216000.1.D
PCC=-0.471, p=0.042
mRNA Profile
Protein Expression Profile
DPA

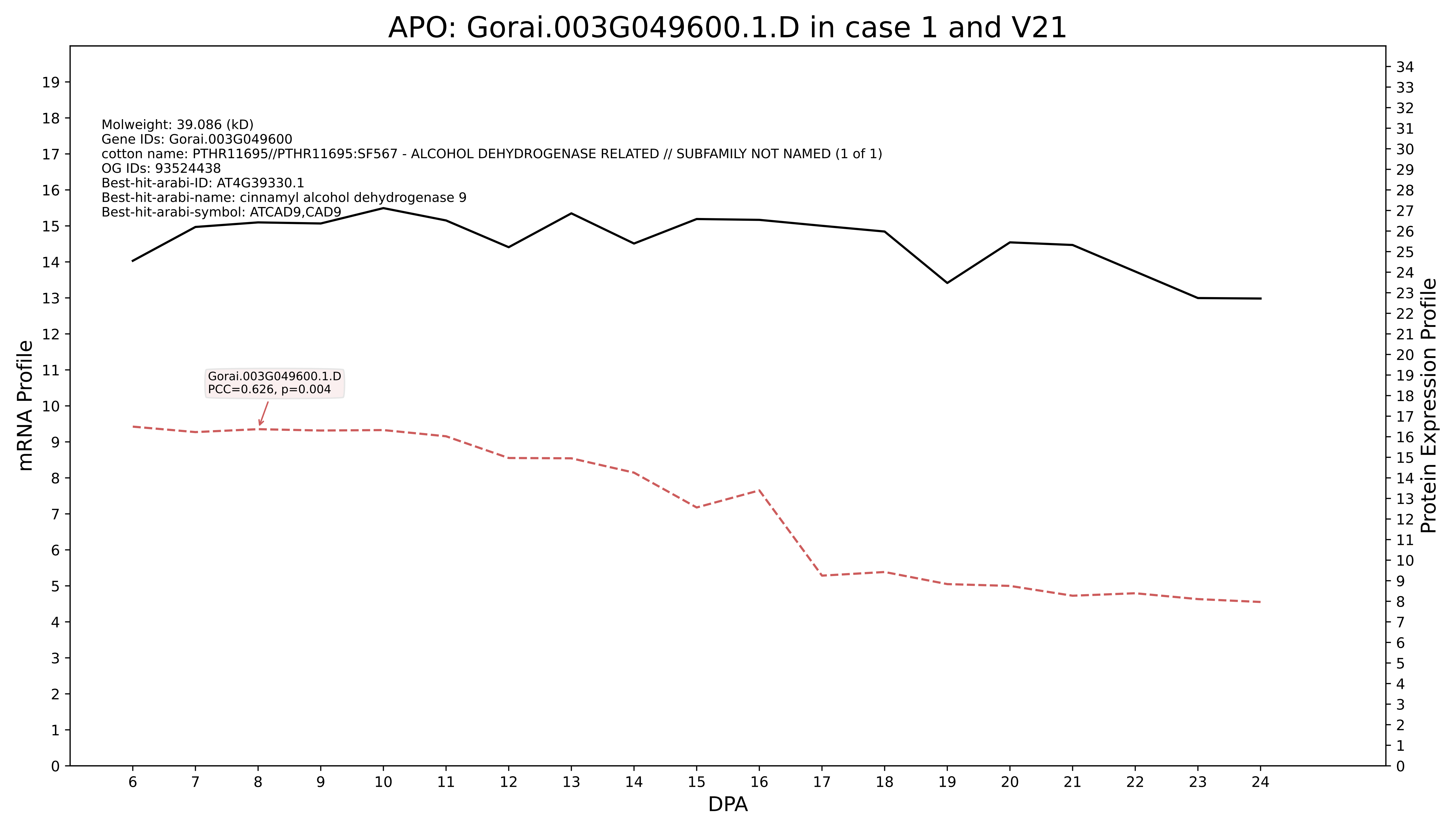

APO: Gorai.003G049600.1.D in case 1 and V21
Molweight: 39.086 (kD)
Gene IDs: Gorai.003G049600
cotton name: PTHR11695//PTHR11695:SF567 - ALCOHOL DEHYDROGENASE RELATED // SUBFAMILY NOT NAMED (1 of 1)
OG IDs: 93524438
Best-hit-arabi-ID: AT4G39330.1
Best-hit-arabi-name: cinnamyl alcohol dehydrogenase 9
Best-hit-arabi-symbol: ATCAD9,CAD9
Gorai.003G049600.1.D
PCC=0.626, p=0.004
mRNA Profile
Protein Expression Profile
DPA

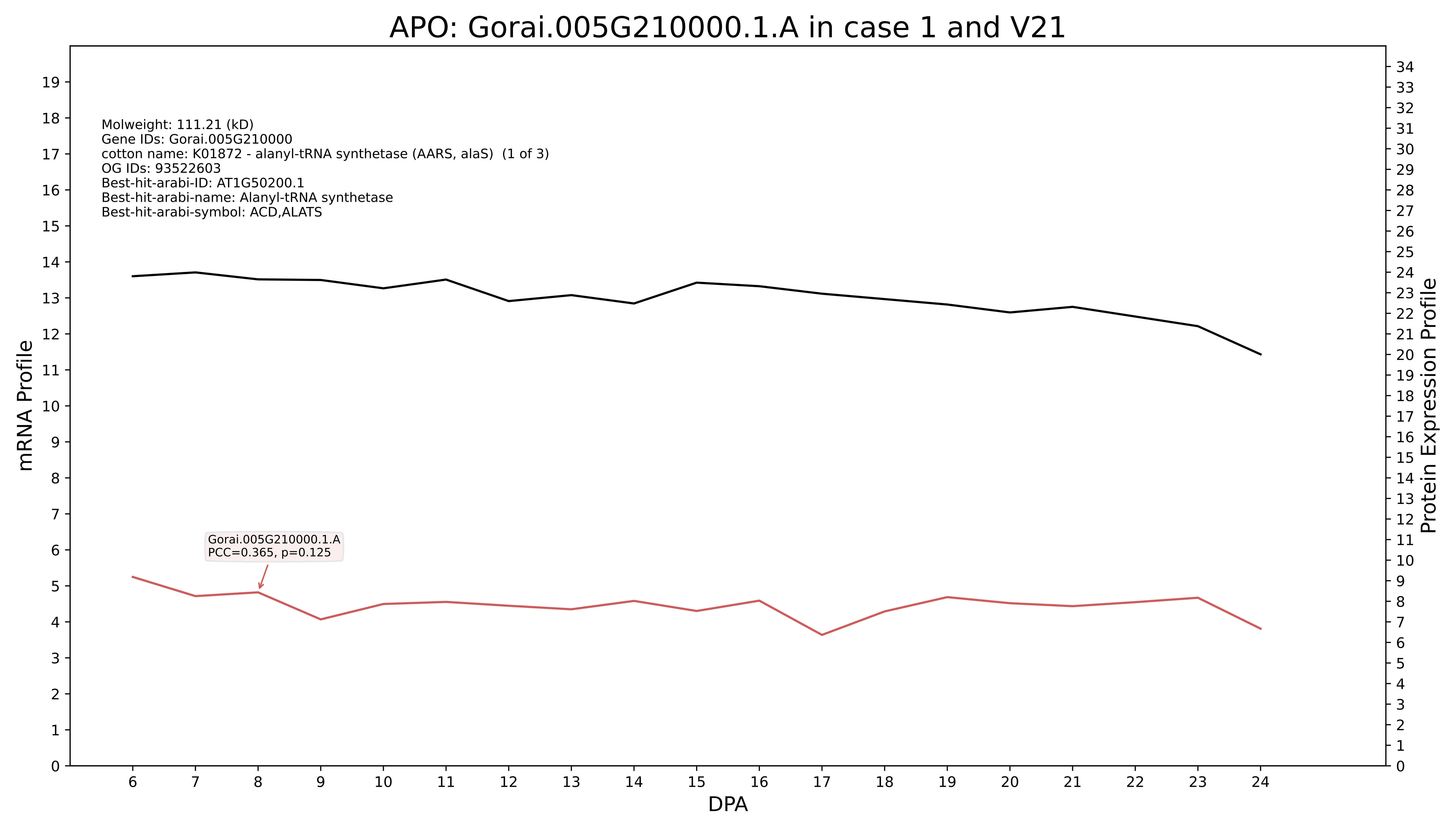
APO: Gorai.005G210000.1.A in case 1 and V21
Molweight: 111.21 (kD)
Gene IDs: Gorai.005G210000
cotton name: K01872 - alanyl-tRNA synthetase (AARS, alaS) (1 of 3)
OG IDs: 93522603
Best-hit-arabi-ID: AT1G50200.1
Best-hit-arabi-name: Alanyl-tRNA synthetase
Best-hit-arabi-symbol: ACD,ALATS
Gorai.005G210000.1.A
PCC=0.365, p=0.125
mRNA Profile
Protein Expression Profile
DPA

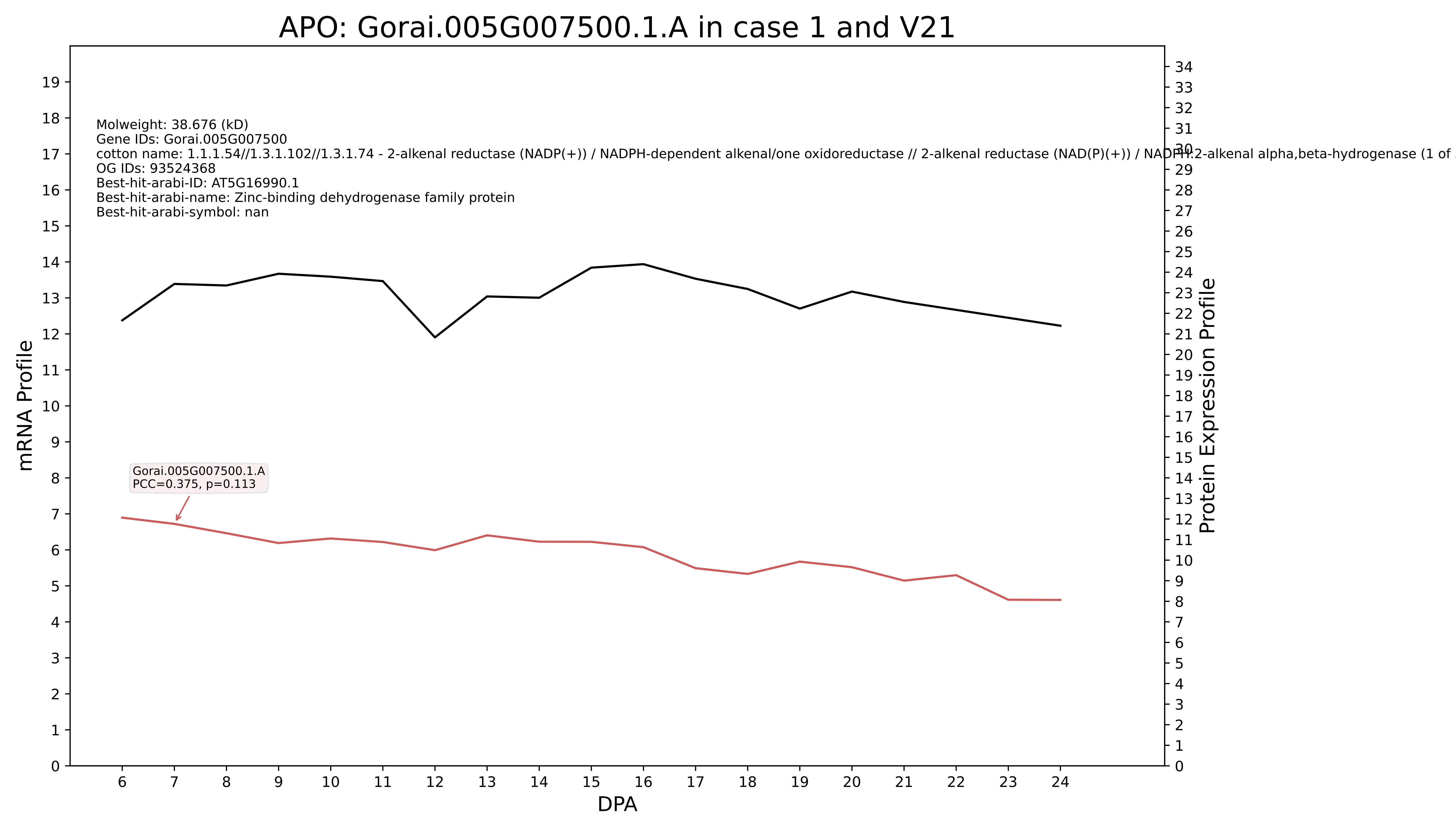

APO: Gorai.005G007500.1.A in case 1 and V21
Molweight: 38.676 (kD)
Gene IDs: Gorai.005G007500
cotton name: 1.1.1.54//1.3.1.102//1.3.1.74 - 2-alkenal reductase (NADP(+)) / NADPH-dependent alkenal/one oxidoreductase // 2-alkenal reductase (NAD(P)(+)) / NADPH:2-alkenal alpha,beta-hydrogenase (1 of
OG IDs: 93524368
Best-hit-arabi-ID: AT5G16990.1
Best-hit-arabi-name: Zinc-binding dehydrogenase family protein
Best-hit-arabi-symbol: nan
Gorai.005G007500.1.A
PCC=0.375, p=0.113
mRNA Profile
Protein Expression Profile
DPA

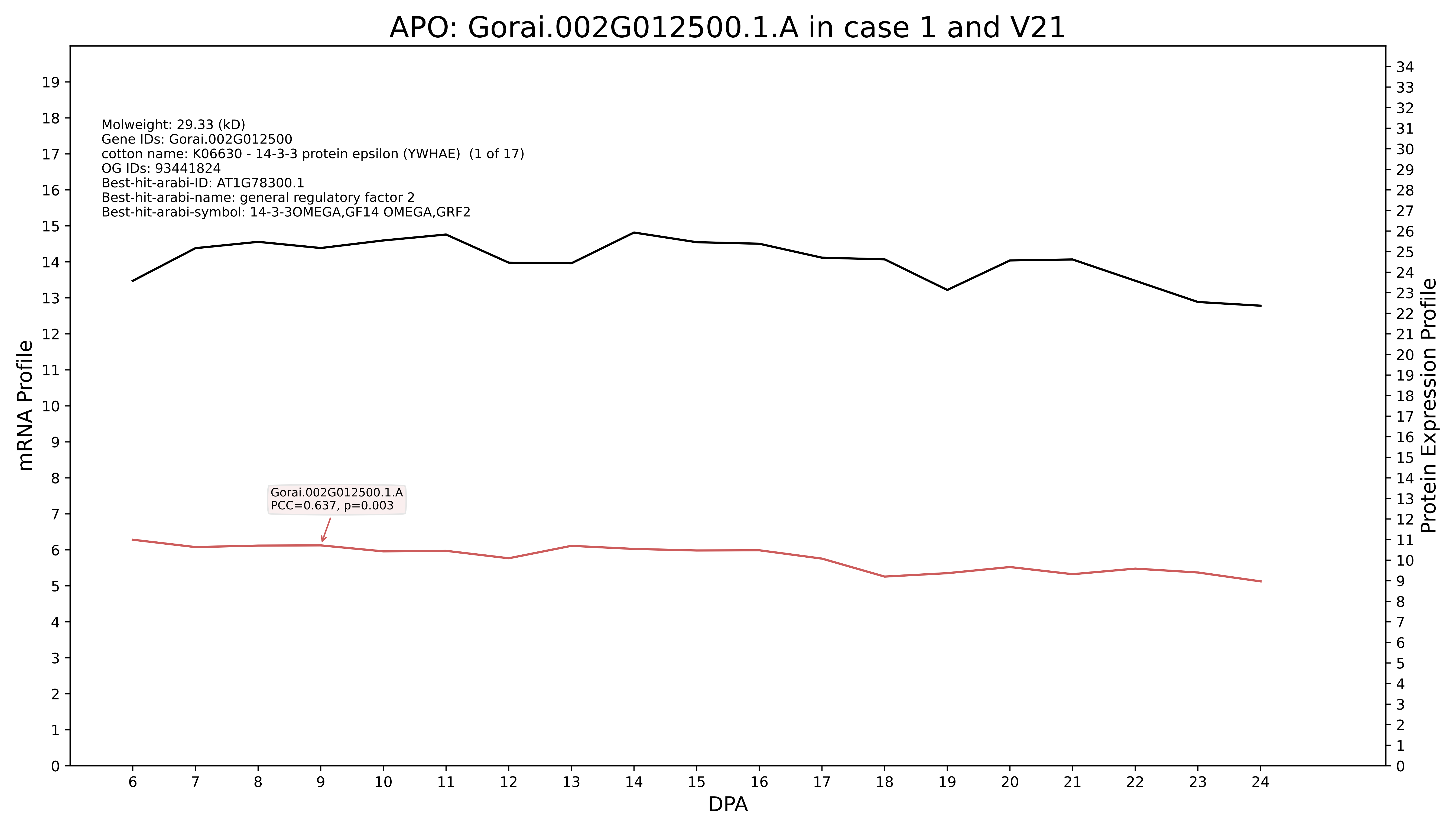
APO: Gorai.002G012500.1.A in case 1 and V21
Molweight: 29.33 (kD)
Gene IDs: Gorai.002G012500
cotton name: K06630 - 14-3-3 protein epsilon (YWHAE) (1 of 17)
OG IDs: 93441824
Best-hit-arabi-ID: AT1G78300.1
Best-hit-arabi-name: general regulatory factor 2
Best-hit-arabi-symbol: 14-3-3OMEGA,GF14 OMEGA,GRF2
Gorai.002G012500.1.A
PCC=0.637, p=0.003
mRNA Profile
Protein Expression Profile
DPA

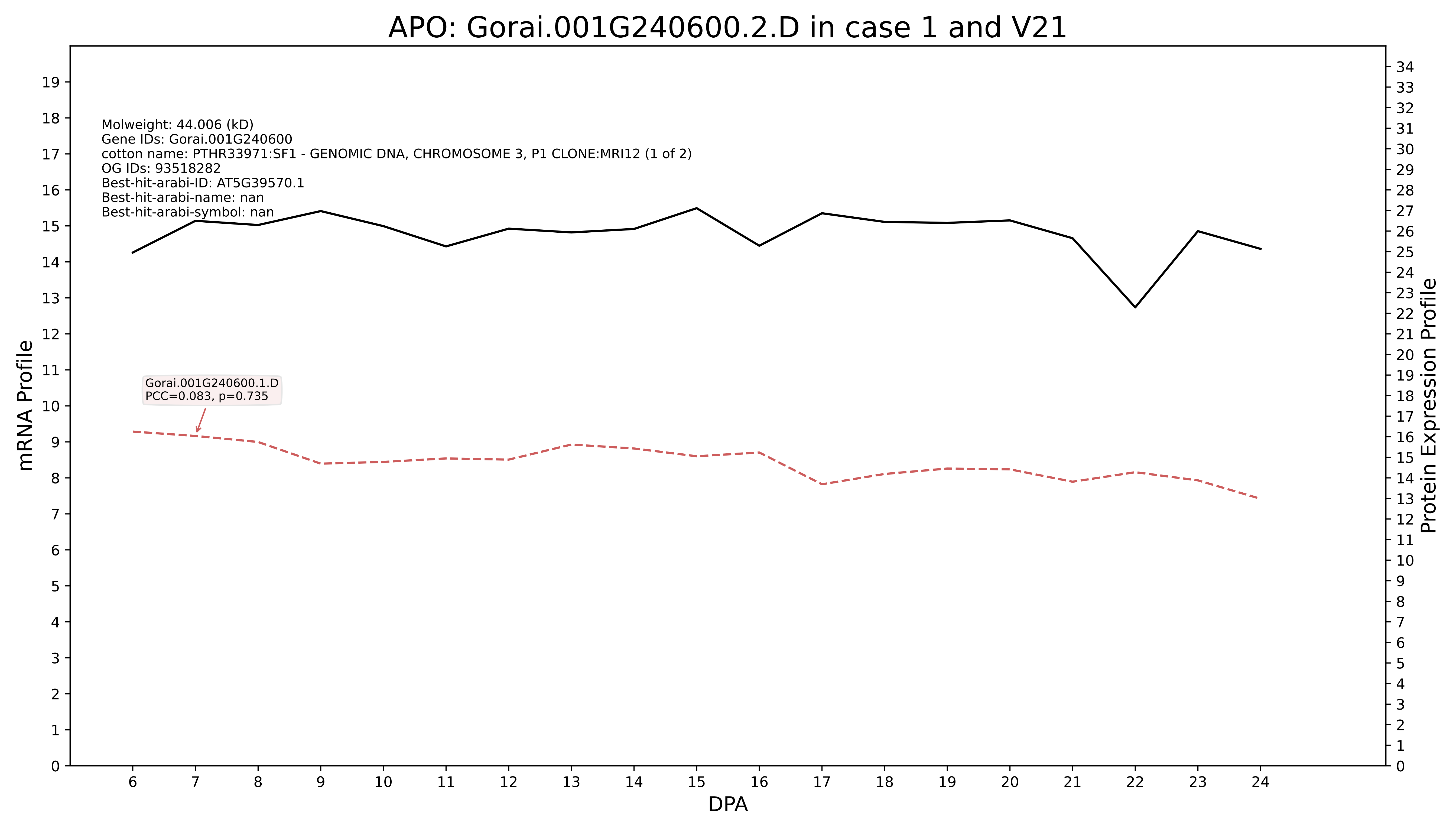
APO: Gorai.001G240600.2.D in case 1 and V21
Molweight: 44.006 (kD)
Gene IDs: Gorai.001G240600
cotton name: PTHR33971:SF1 - GENOMIC DNA, CHROMOSOME 3, P1 CLONE:MRI12 (1 of 2)
OG IDs: 93518282
Best-hit-arabi-ID: AT5G39570.1
Best-hit-arabi-name: nan
Best-hit-arabi-symbol: nan
Gorai.001G240600.1.D
PCC=0.083, p=0.735
mRNA Profile
Protein Expression Profile
DPA

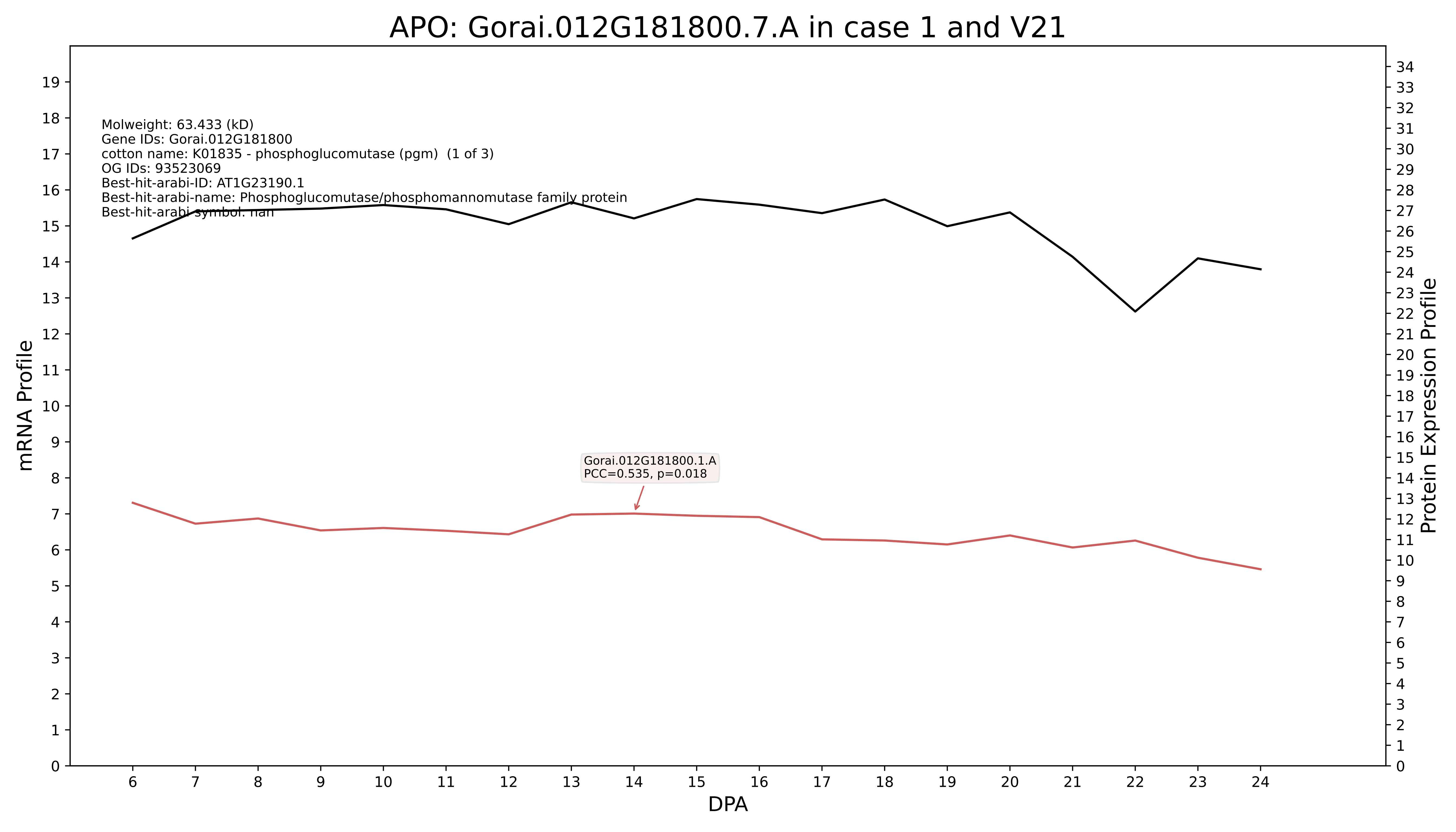

APO: Gorai.012G181800.7.A in case 1 and V21
Molweight: 63.433 (kD)
Gene IDs: Gorai.012G181800
cotton name: K01835 - phosphoglucomutase (pgm) (1 of 3)
OG IDs: 93523069
Best-hit-arabi-ID: AT1G23190.1
Best-hit-arabi-name: Phosphoglucomutase/phosphomannomutase family protein
Best-hit-arabi-symbol: nan
Gorai.012G181800.1.A
PCC=0.535, p=0.018
mRNA Profile
Protein Expression Profile
DPA

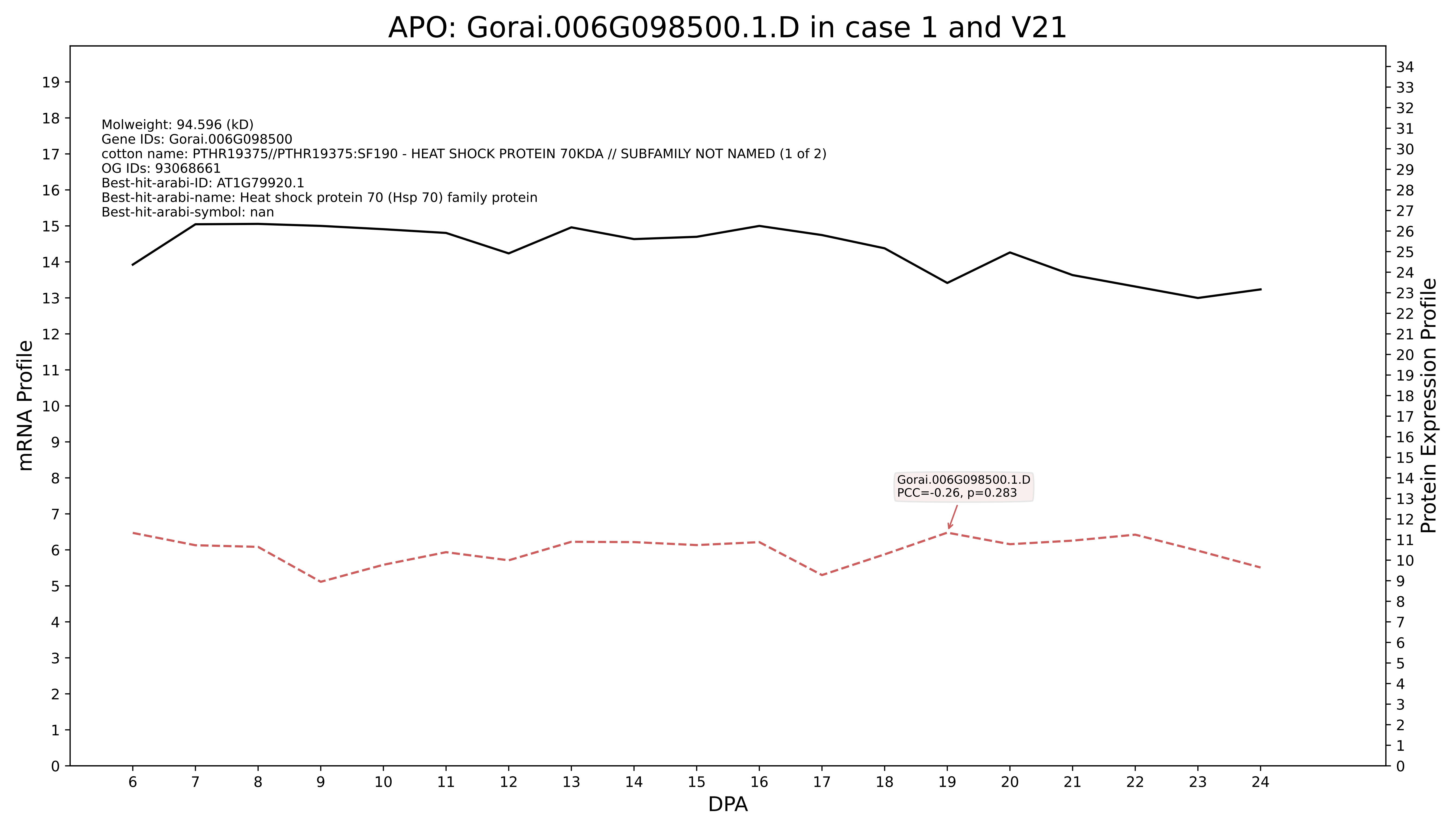
APO: Gorai.006G098500.1.D in case 1 and V21
Molweight: 94.596 (kD)
Gene IDs: Gorai.006G098500
cotton name: PTHR19375//PTHR19375:SF190 - HEAT SHOCK PROTEIN 70KDA // SUBFAMILY NOT NAMED (1 of 2)
OG IDs: 93068661
Best-hit-arabi-ID: AT1G79920.1
Best-hit-arabi-name: Heat shock protein 70 (Hsp 70) family protein
Best-hit-arabi-symbol: nan
Gorai.006G098500.1.D
PCC=-0.26, p=0.283
mRNA Profile
Protein Expression Profile
DPA

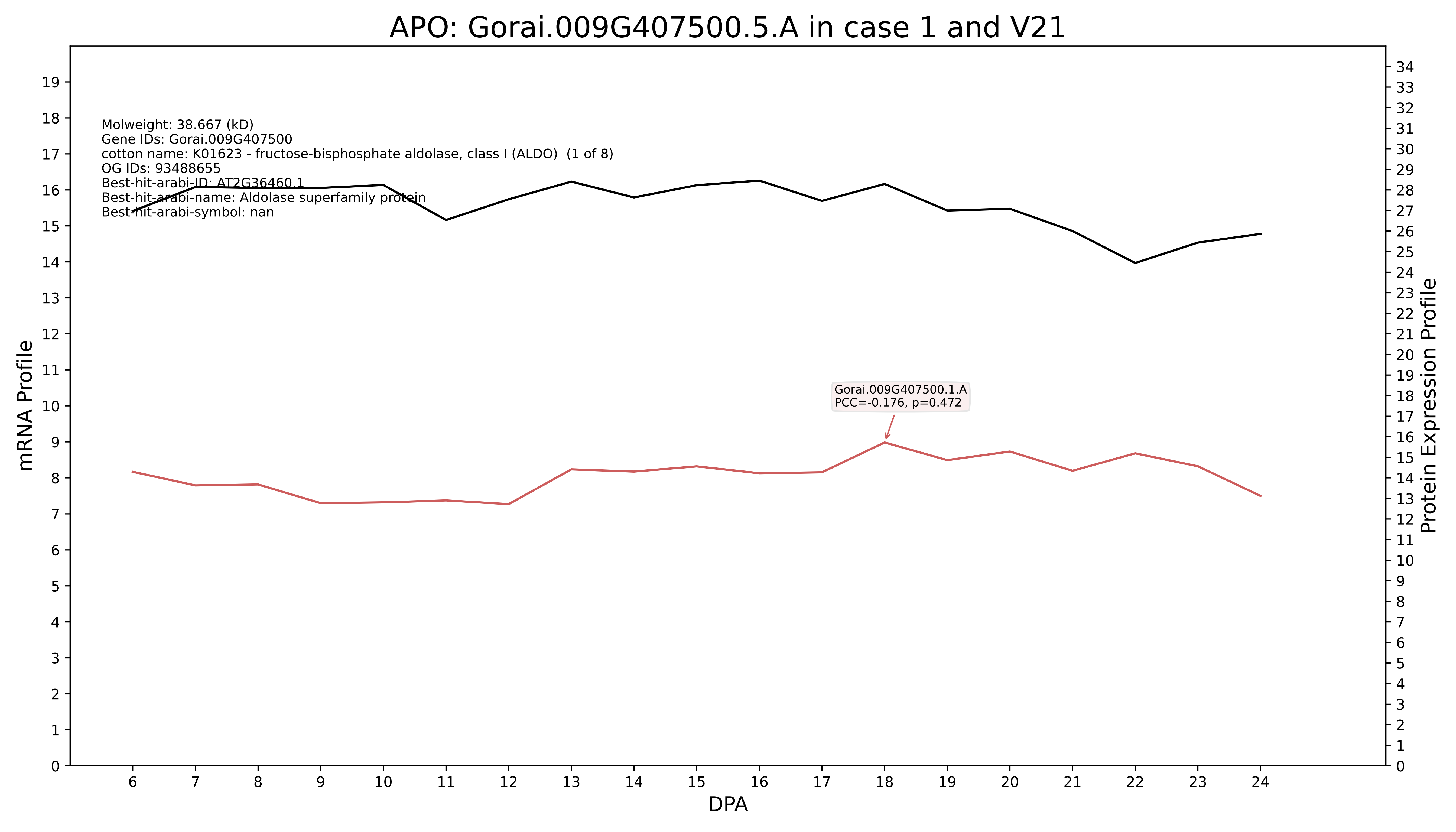
APO: Gorai.009G407500.5.A in case 1 and V21
Molweight: 38.667 (kD)
Gene IDs: Gorai.009G407500
cotton name: K01623 - fructose-bisphosphate aldolase, class I (ALDO) (1 of 8)
OG IDs: 93488655
Best-hit-arabi-ID: AT2G36460.1
Best-hit-arabi-name: Aldolase superfamily protein
Best-hit-arabi-symbol: nan
Gorai.009G407500.1.A
PCC=-0.176, p=0.472
mRNA Profile
Protein Expression Profile
DPA

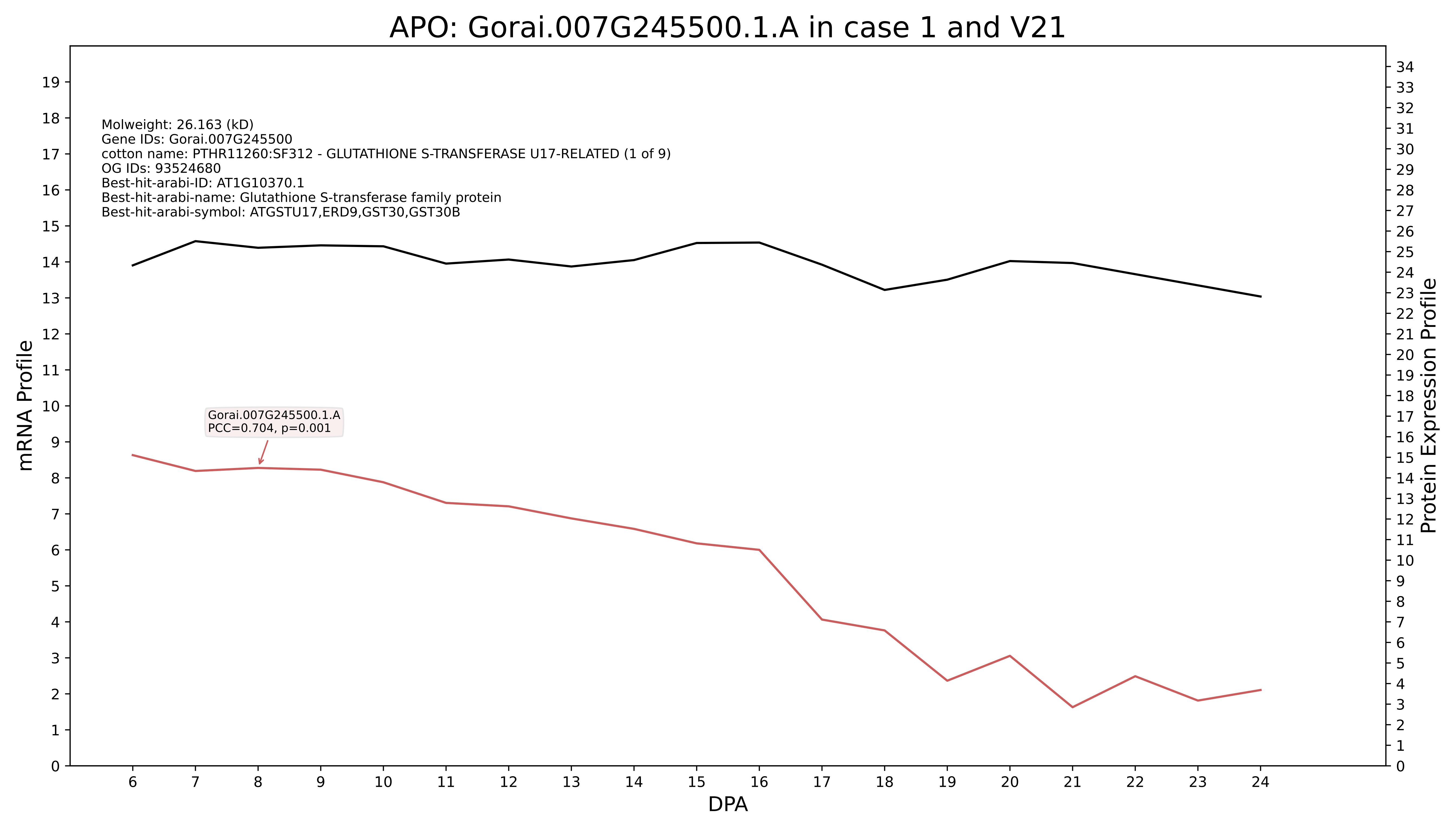
APO: Gorai.007G245500.1.A in case 1 and V21
Molweight: 26.163 (kD)
Gene IDs: Gorai.007G245500
cotton name: PTHR11260:SF312 - GLUTATHIONE S-TRANSFERASE U17-RELATED (1 of 9)
OG IDs: 93524680
Best-hit-arabi-ID: AT1G10370.1
Best-hit-arabi-name: Glutathione S-transferase family protein
Best-hit-arabi-symbol: ATGSTU17,ERD9,GST30,GST30B
Gorai.007G245500.1.A
PCC=0.704, p=0.001
mRNA Profile
Protein Expression Profile
DPA

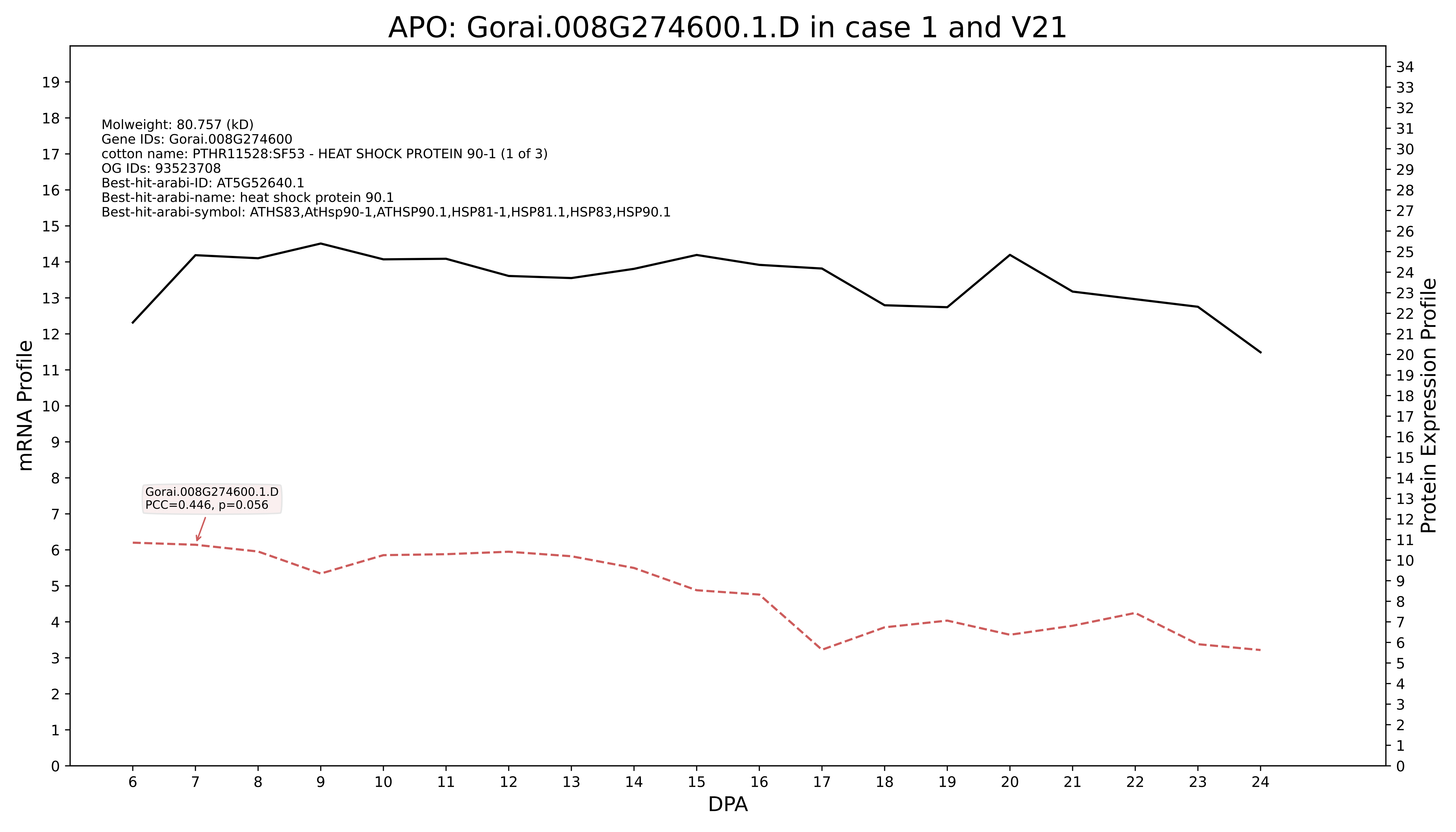

APO: Gorai.008G274600.1.D in case 1 and V21
Molweight: 80.757 (kD)
Gene IDs: Gorai.008G274600
cotton name: PTHR11528:SF53 - HEAT SHOCK PROTEIN 90-1 (1 of 3)
OG IDs: 93523708
Best-hit-arabi-ID: AT5G52640.1
Best-hit-arabi-name: heat shock protein 90.1
Best-hit-arabi-symbol: ATHS83,AtHsp90-1,ATHSP90.1,HSP81-1,HSP81.1,HSP83,HSP90.1
Gorai.008G274600.1.D
PCC=0.446, p=0.056
mRNA Profile
Protein Expression Profile
DPA

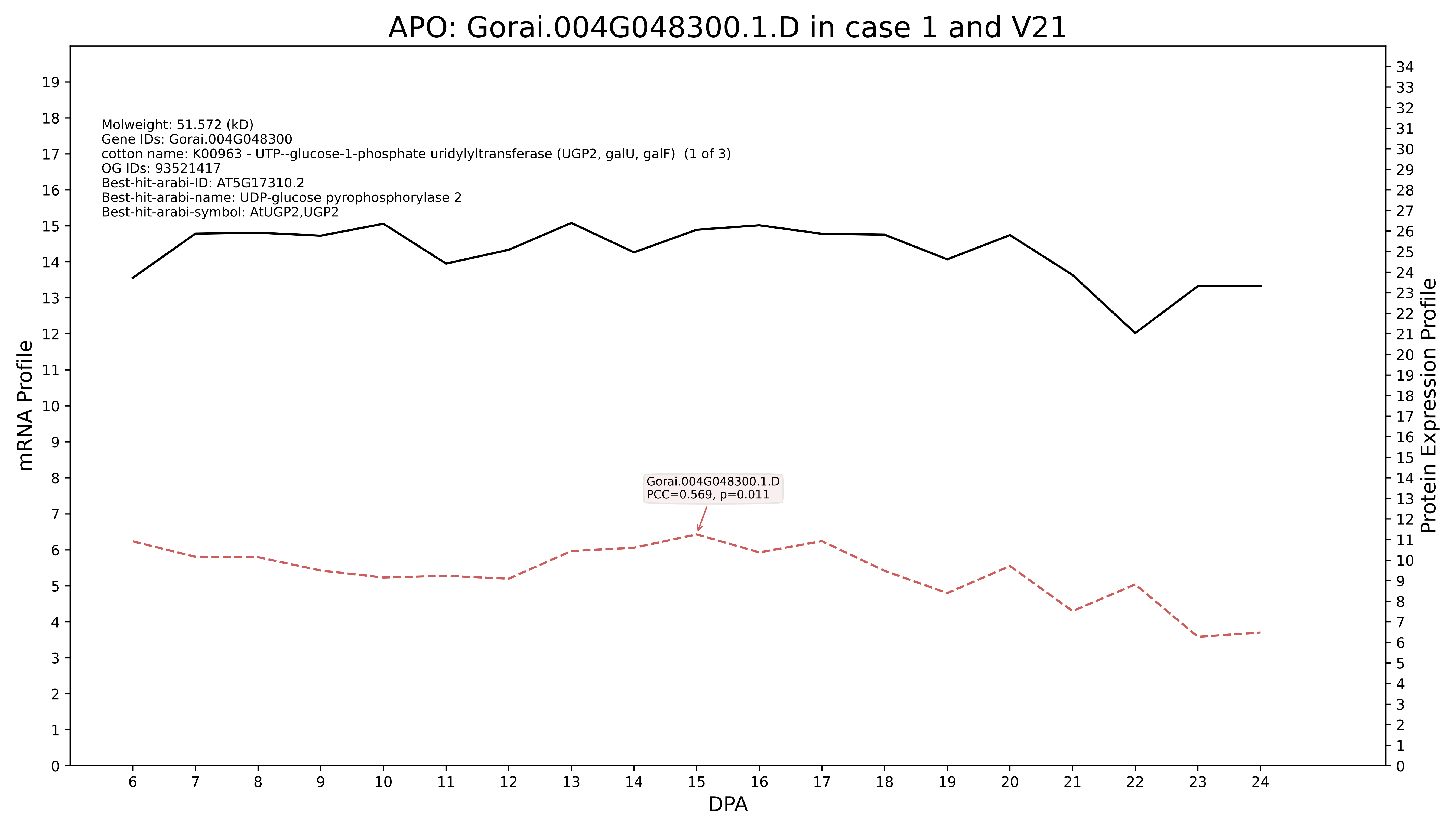
APO: Gorai.004G048300.1.D in case 1 and V21
Molweight: 51.572 (kD)
Gene IDs: Gorai.004G048300
cotton name: K00963 - UTP--glucose-1-phosphate uridylyltransferase (UGP2, galU, galF) (1 of 3)
OG IDs: 93521417
Best-hit-arabi-ID: AT5G17310.2
Best-hit-arabi-name: UDP-glucose pyrophosphorylase 2
Best-hit-arabi-symbol: AtUGP2,UGP2
Gorai.004G048300.1.D
PCC=0.569, p=0.011
mRNA Profile
Protein Expression Profile
DPA

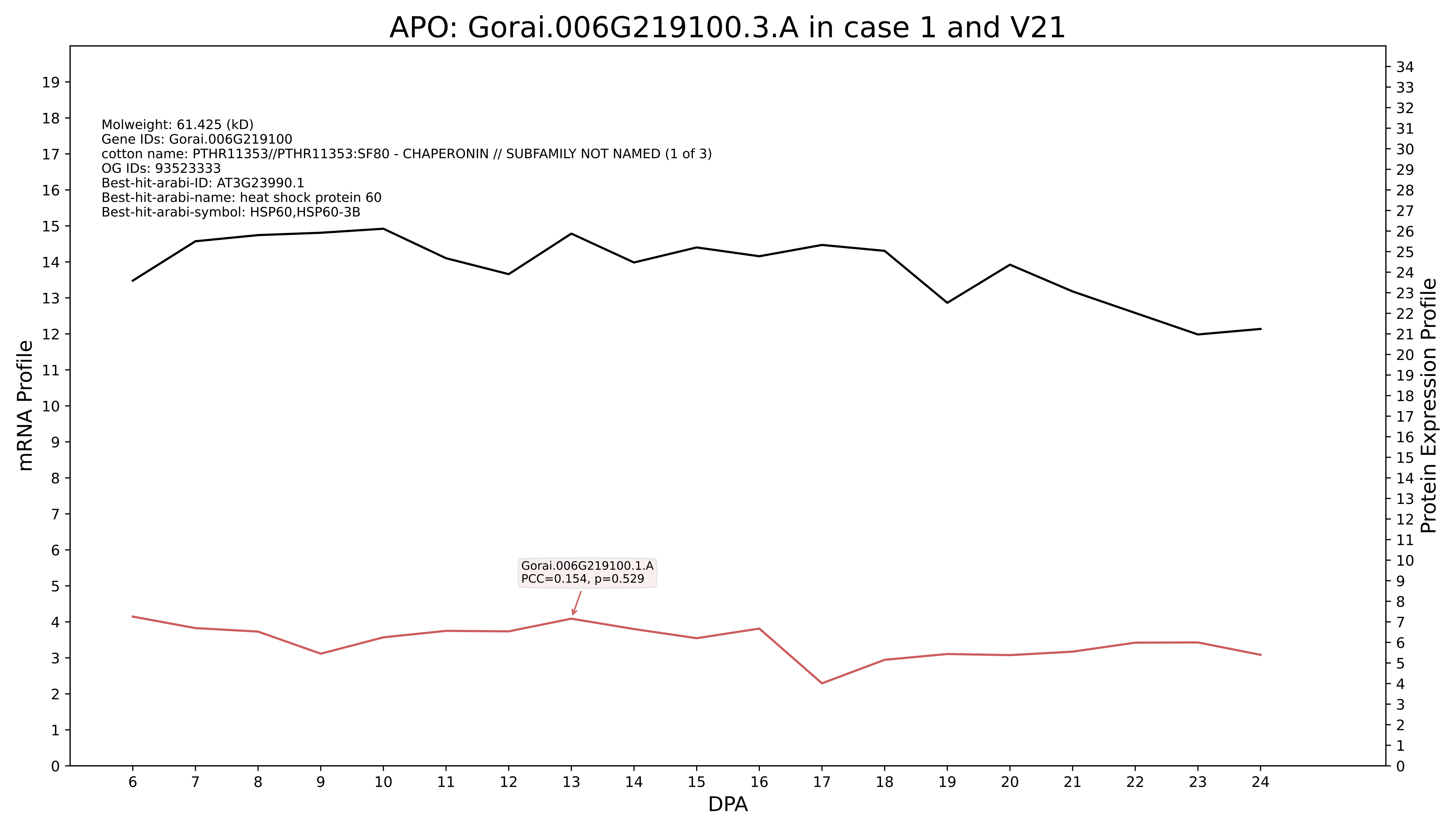
APO: Gorai.006G219100.3.A in case 1 and V21
Molweight: 61.425 (kD)
Gene IDs: Gorai.006G219100
cotton name: PTHR11353//PTHR11353:SF80 - CHAPERONIN // SUBFAMILY NOT NAMED (1 of 3)
OG IDs: 93523333
Best-hit-arabi-ID: AT3G23990.1
Best-hit-arabi-name: heat shock protein 60
Best-hit-arabi-symbol: HSP60,HSP60-3B
Gorai.006G219100.1.A
PCC=0.154, p=0.529
mRNA Profile
Protein Expression Profile
DPA

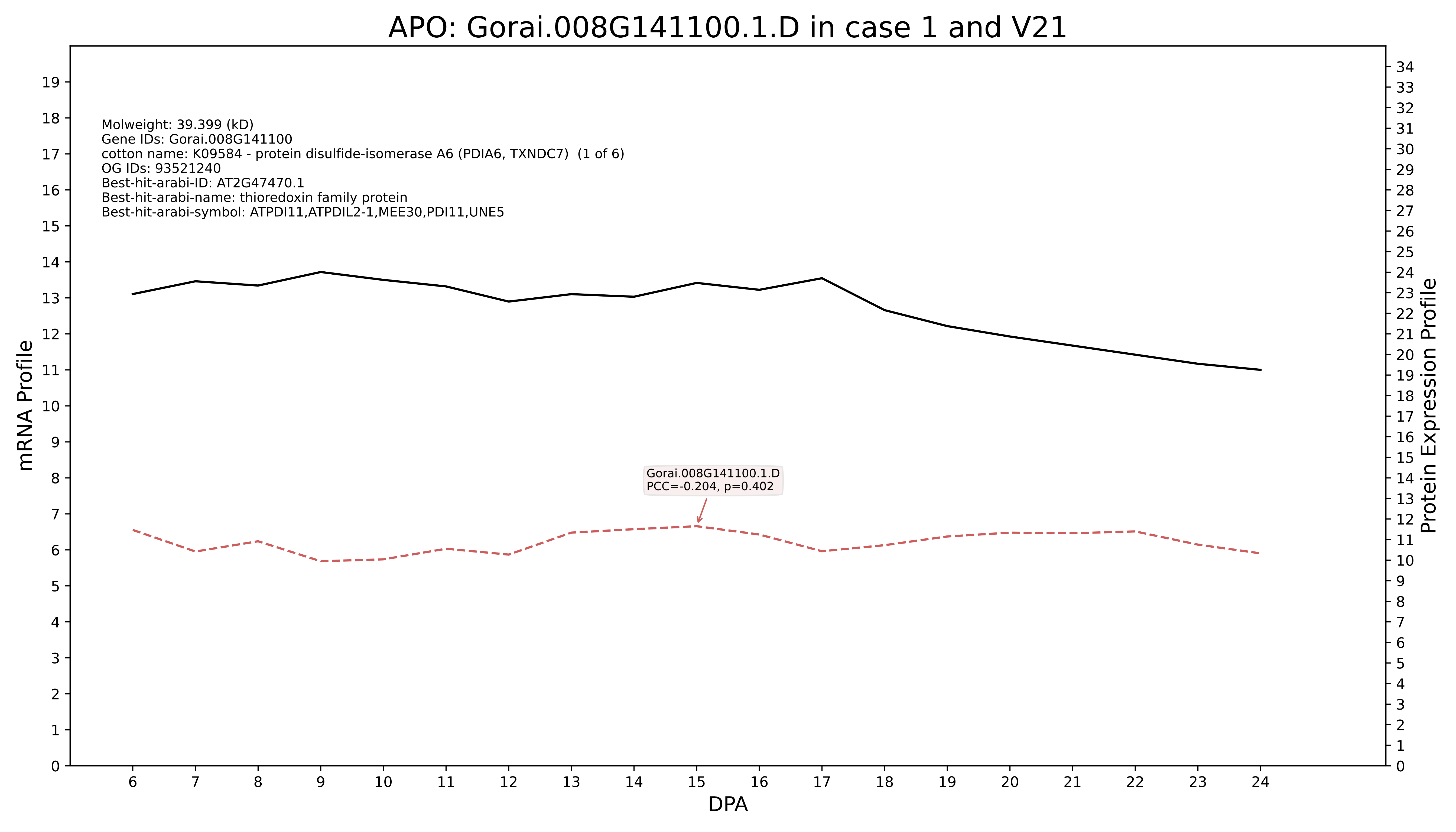
APO: Gorai.008G141100.1.D in case 1 and V21
Molweight: 39.399 (kD)
Gene IDs: Gorai.008G141100
cotton name: K09584 - protein disulfide-isomerase A6 (PDIA6, TXNDC7) (1 of 6)
OG IDs: 93521240
Best-hit-arabi-ID: AT2G47470.1
Best-hit-arabi-name: thioredoxin family protein
Best-hit-arabi-symbol: ATPDI11,ATPDIL2-1,MEE30,PDI11,UNE5
Gorai.008G141100.1.D
PCC=-0.204, p=0.402
mRNA Profile
Protein Expression Profile
DPA

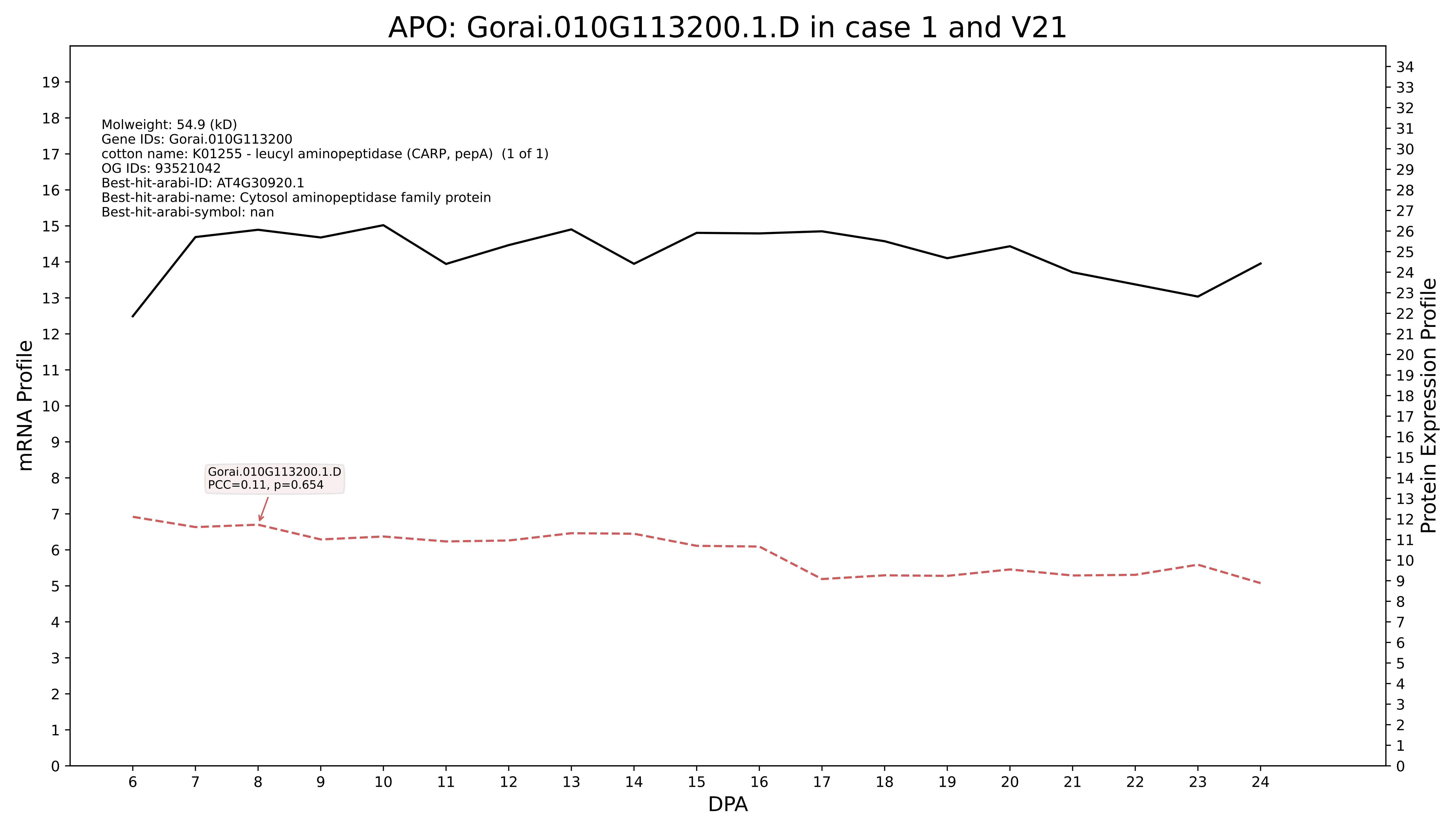
APO: Gorai.010G113200.1.D in case 1 and V21
Molweight: 54.9 (kD)
Gene IDs: Gorai.010G113200
cotton name: K01255 - leucyl aminopeptidase (CARP, pepA) (1 of 1)
OG IDs: 93521042
Best-hit-arabi-ID: AT4G30920.1
Best-hit-arabi-name: Cytosol aminopeptidase family protein
Best-hit-arabi-symbol: nan
Gorai.010G113200.1.D
PCC=0.11, p=0.654
mRNA Profile
Protein Expression Profile
DPA

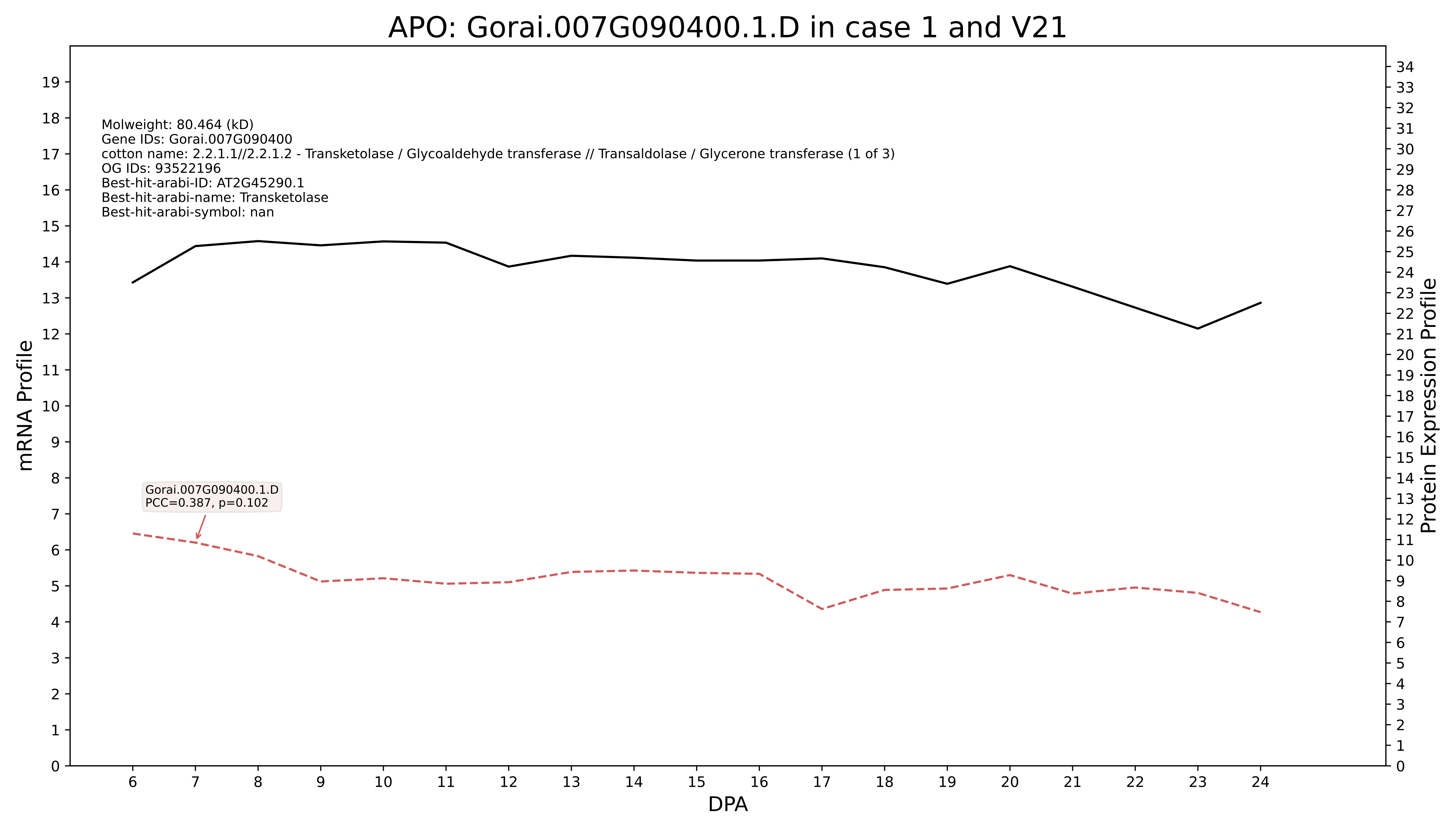

APO: Gorai.007G090400.1.D in case 1 and V21
Molweight: 80.464 (kD)
Gene IDs: Gorai.007G090400
cotton name: 2.2.1.1//2.2.1.2 - Transketolase / Glycoaldehyde transferase // Transaldolase / Glycerone transferase (1 of 3)
OG IDs: 93522196
Best-hit-arabi-ID: AT2G45290.1
Best-hit-arabi-name: Transketolase
Best-hit-arabi-symbol: nan
Gorai.007G090400.1.D
PCC=0.387, p=0.102
mRNA Profile
Protein Expression Profile
DPA

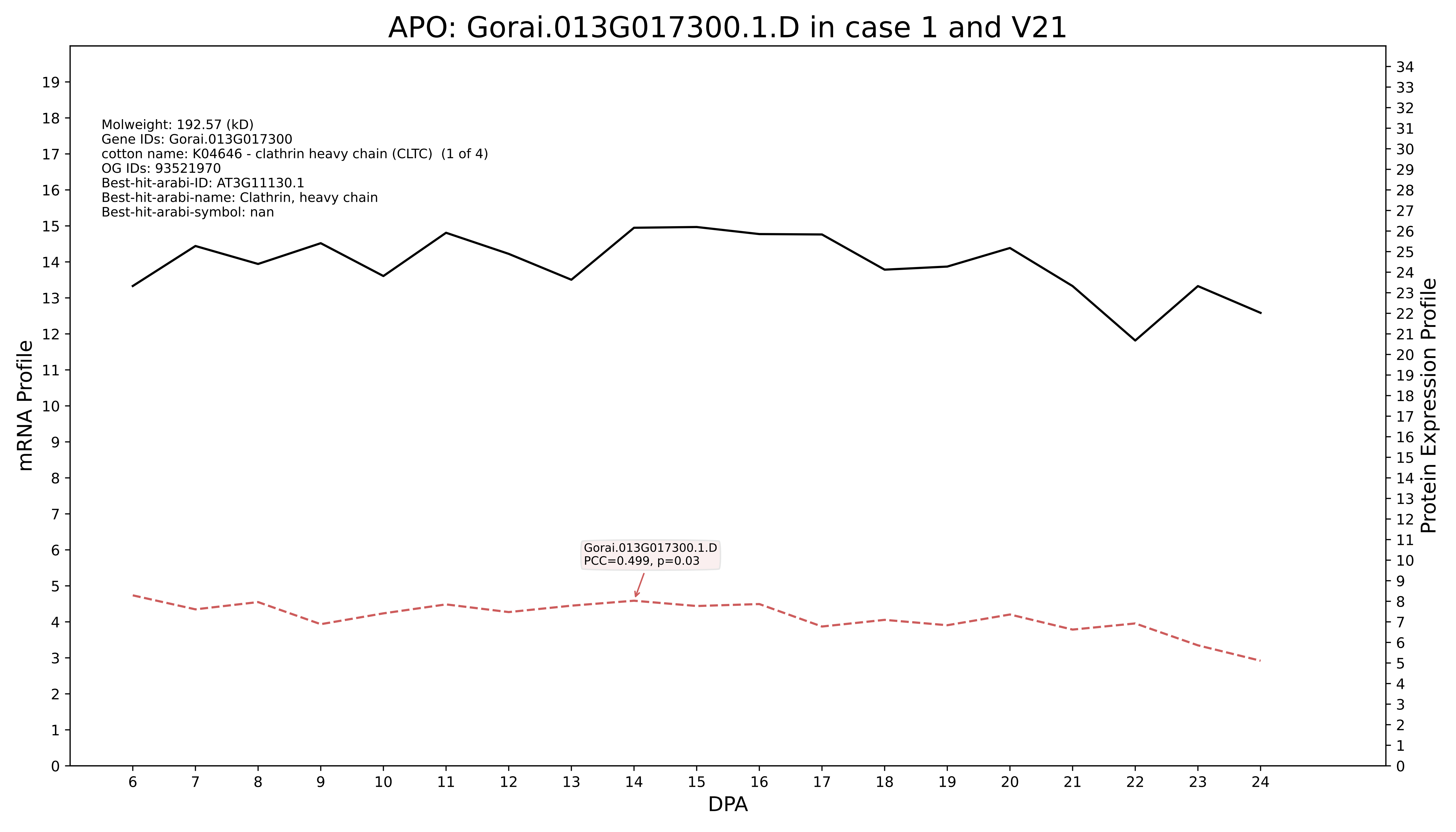

APO: Gorai.013G017300.1.D in case 1 and V21
Molweight: 192.57 (kD)
Gene IDs: Gorai.013G017300
cotton name: K04646 - clathrin heavy chain (CLTC) (1 of 4)
OG IDs: 93521970
Best-hit-arabi-ID: AT3G11130.1
Best-hit-arabi-name: Clathrin, heavy chain
Best-hit-arabi-symbol: nan
Gorai.013G017300.1.D
PCC=0.499, p=0.03
mRNA Profile
Protein Expression Profile
DPA

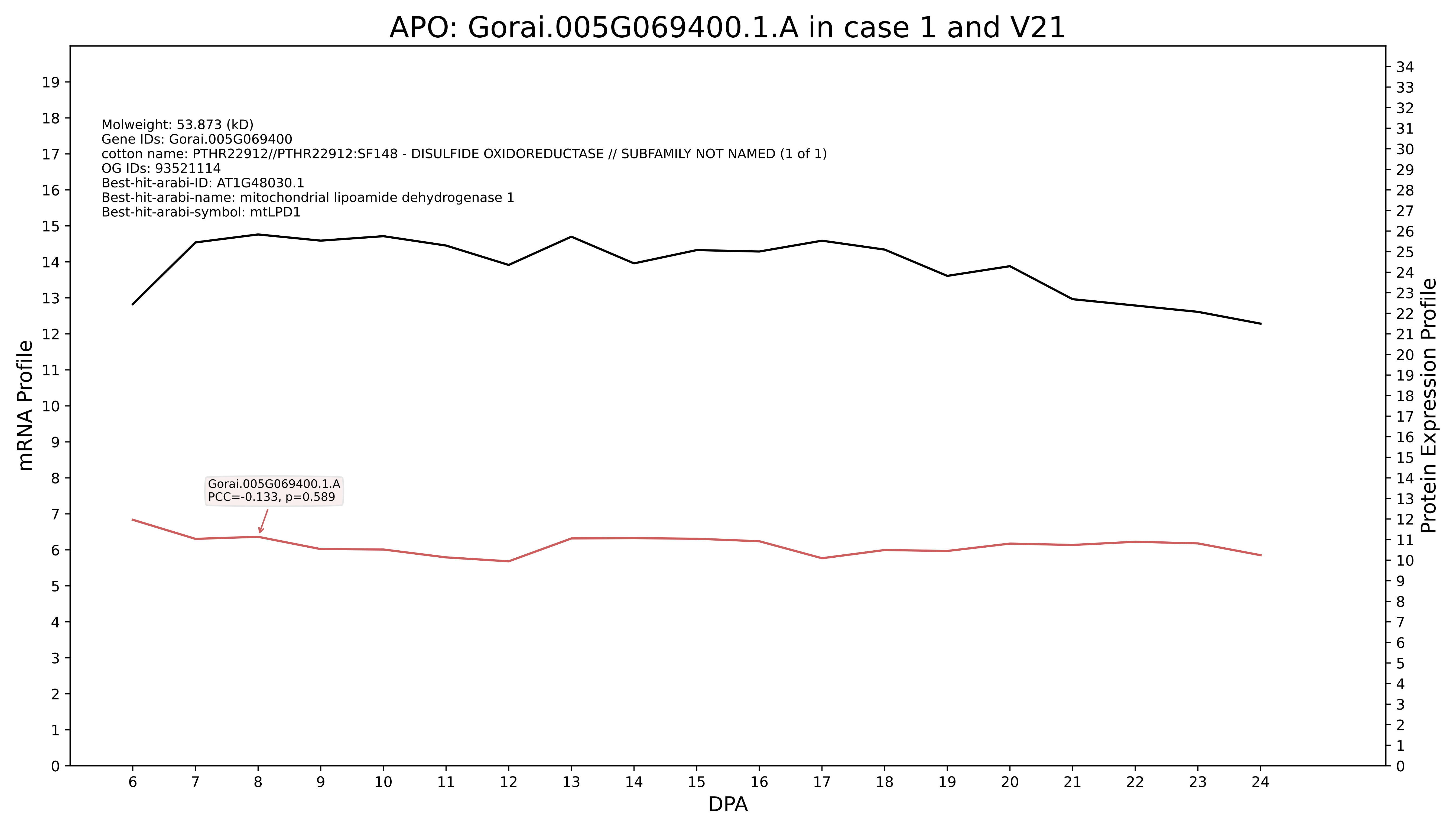

APO: Gorai.005G069400.1.A in case 1 and V21
Molweight: 53.873 (kD)
Gene IDs: Gorai.005G069400
cotton name: PTHR22912//PTHR22912:SF148 - DISULFIDE OXIDOREDUCTASE // SUBFAMILY NOT NAMED (1 of 1)
OG IDs: 93521114
Best-hit-arabi-ID: AT1G48030.1
Best-hit-arabi-name: mitochondrial lipoamide dehydrogenase 1
Best-hit-arabi-symbol: mtLPD1
Gorai.005G069400.1.A
PCC=-0.133, p=0.589
mRNA Profile
Protein Expression Profile
DPA

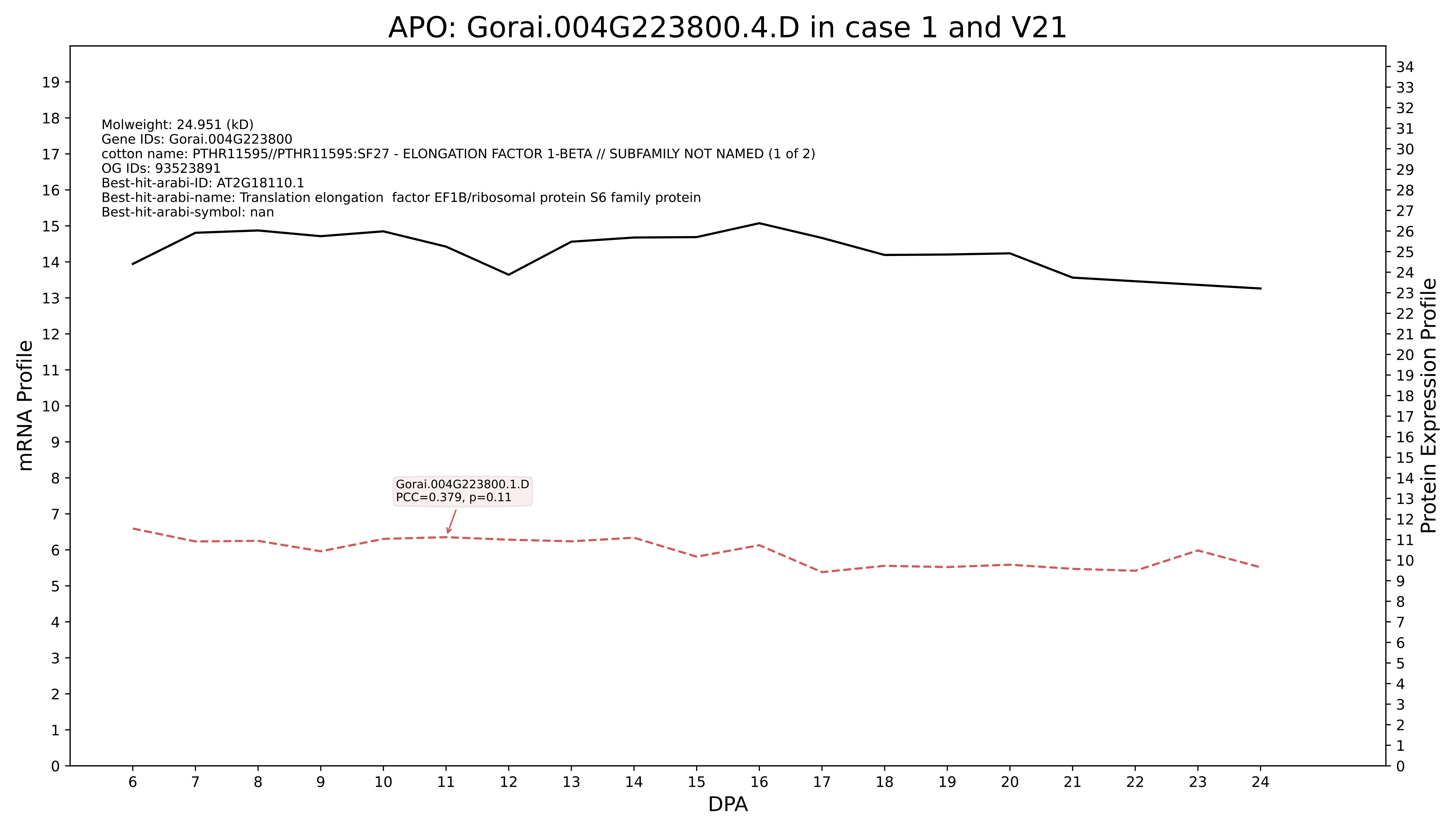
APO: Gorai.004G223800.4.D in case 1 and V21
Molweight: 24.951 (kD)
Gene IDs: Gorai.004G223800
cotton name: PTHR11595//PTHR11595:SF27 - ELONGATION FACTOR 1-BETA // SUBFAMILY NOT NAMED (1 of 2)
OG IDs: 93523891
Best-hit-arabi-ID: AT2G18110.1
Best-hit-arabi-name: Translation elongation factor EF1B/ribosomal protein S6 family protein
Best-hit-arabi-symbol: nan
Gorai.004G223800.1.D
PCC=0.379, p=0.11
mRNA Profile
Protein Expression Profile
DPA

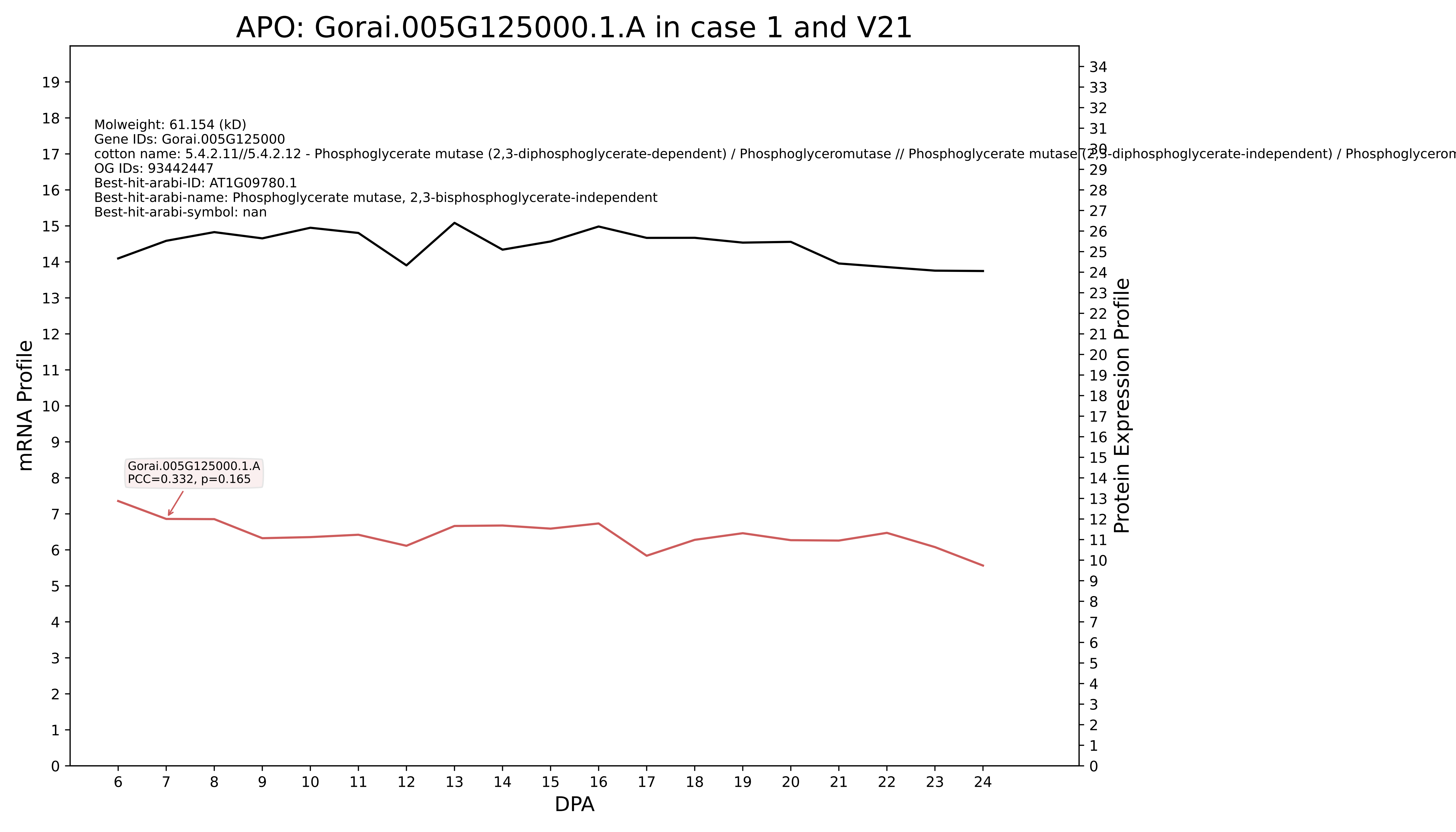

APO: Gorai.005G125000.1.A in case 1 and V21
Molweight: 61.154 (kD)
Gene IDs: Gorai.005G125000
cotton name: 5.4.2.11//5.4.2.12 - Phosphoglycerate mutase (2,3-diphosphoglycerate-dependent) / Phosphoglyceromutase // Phosphoglycerate mutase (2,3-diphosphoglycerate-independent) / Phosphoglyceron
OG IDs: 93442447
Best-hit-arabi-ID: AT1G09780.1
Best-hit-arabi-name: Phosphoglycerate mutase, 2,3-bisphosphoglycerate-independent
Best-hit-arabi-symbol: nan
Gorai.005G125000.1.A
PCC=0.332, p=0.165
mRNA Profile
Protein Expression Profile
DPA

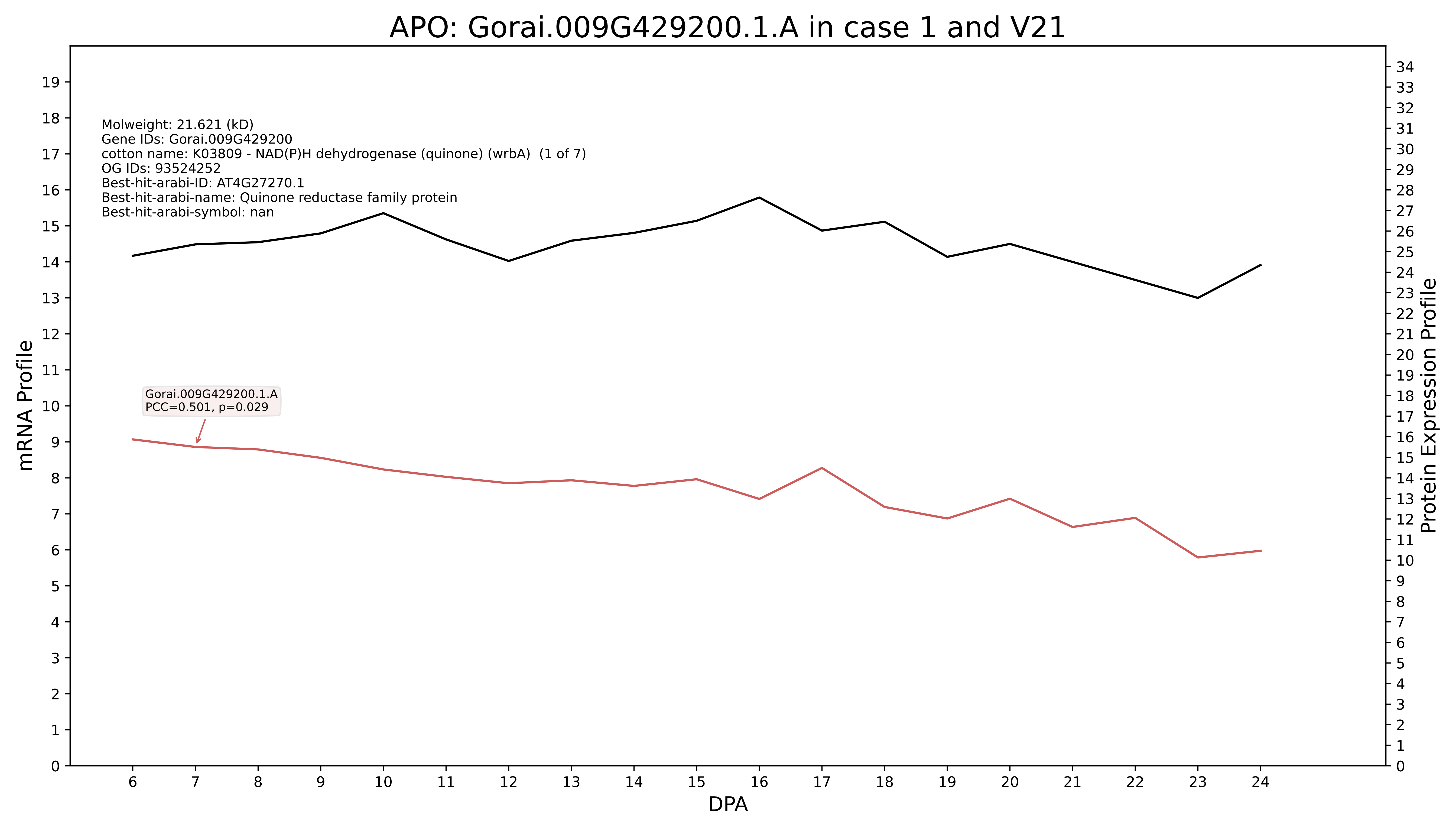
APO: Gorai.009G429200.1.A in case 1 and V21
Molweight: 21.621 (kD)
Gene IDs: Gorai.009G429200
cotton name: K03809 - NAD(P)H dehydrogenase (quinone) (wrbA) (1 of 7)
OG IDs: 93524252
Best-hit-arabi-ID: AT4G27270.1
Best-hit-arabi-name: Quinone reductase family protein
Best-hit-arabi-symbol: nan
Gorai.009G429200.1.A
PCC=0.501, p=0.029
mRNA Profile
Protein Expression Profile
DPA

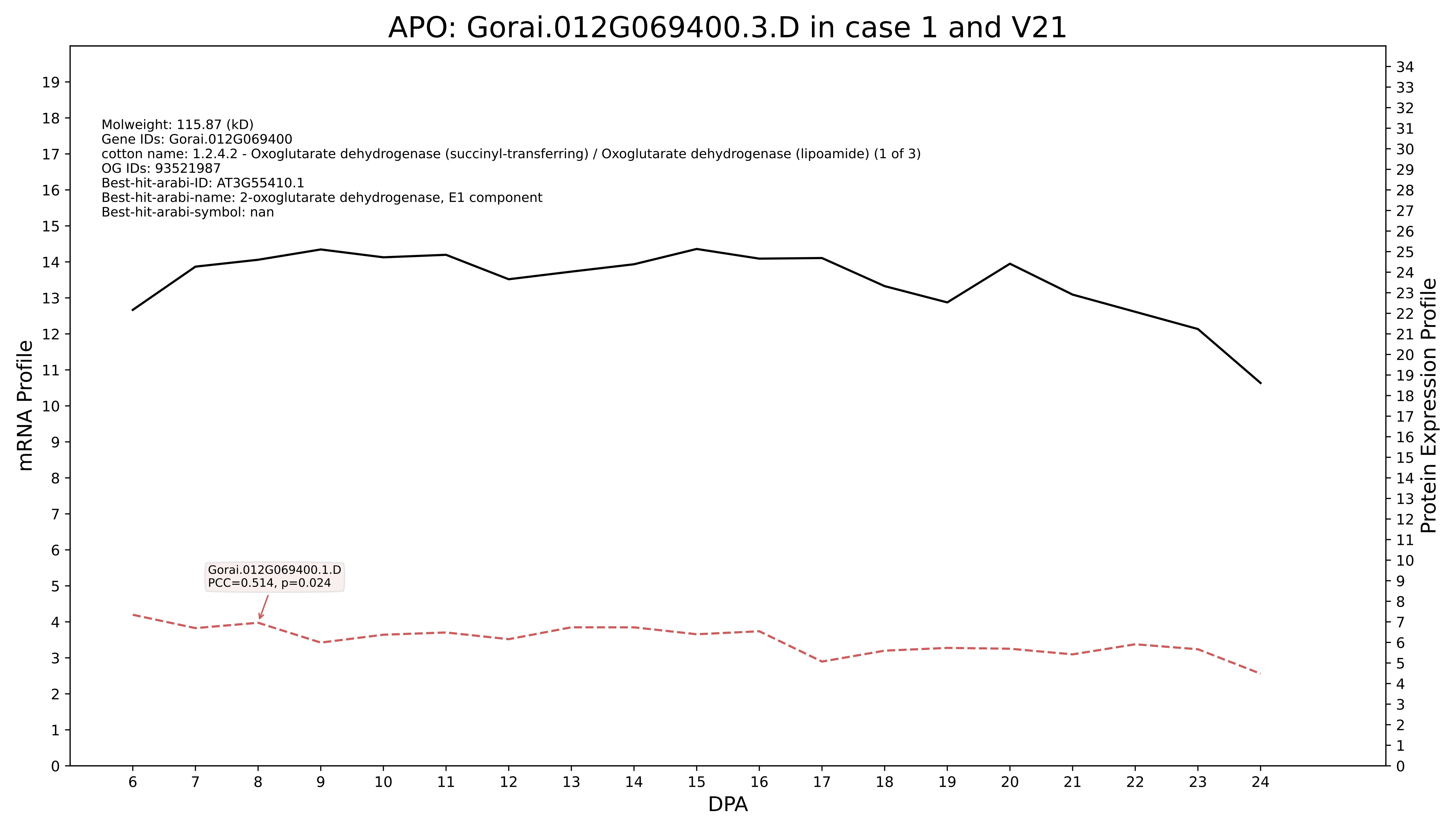

APO: Gorai.012G069400.3.D in case 1 and V21
Molweight: 115.87 (kD)
Gene IDs: Gorai.012G069400
cotton name: 1.2.4.2 - Oxoglutarate dehydrogenase (succinyl-transferring) / Oxoglutarate dehydrogenase (lipoamide) (1 of 3)
OG IDs: 93521987
Best-hit-arabi-ID: AT3G55410.1
Best-hit-arabi-name: 2-oxoglutarate dehydrogenase, E1 component
Best-hit-arabi-symbol: nan
Gorai.012G069400.1.D
PCC=0.514, p=0.024
mRNA Profile
Protein Expression Profile
DPA

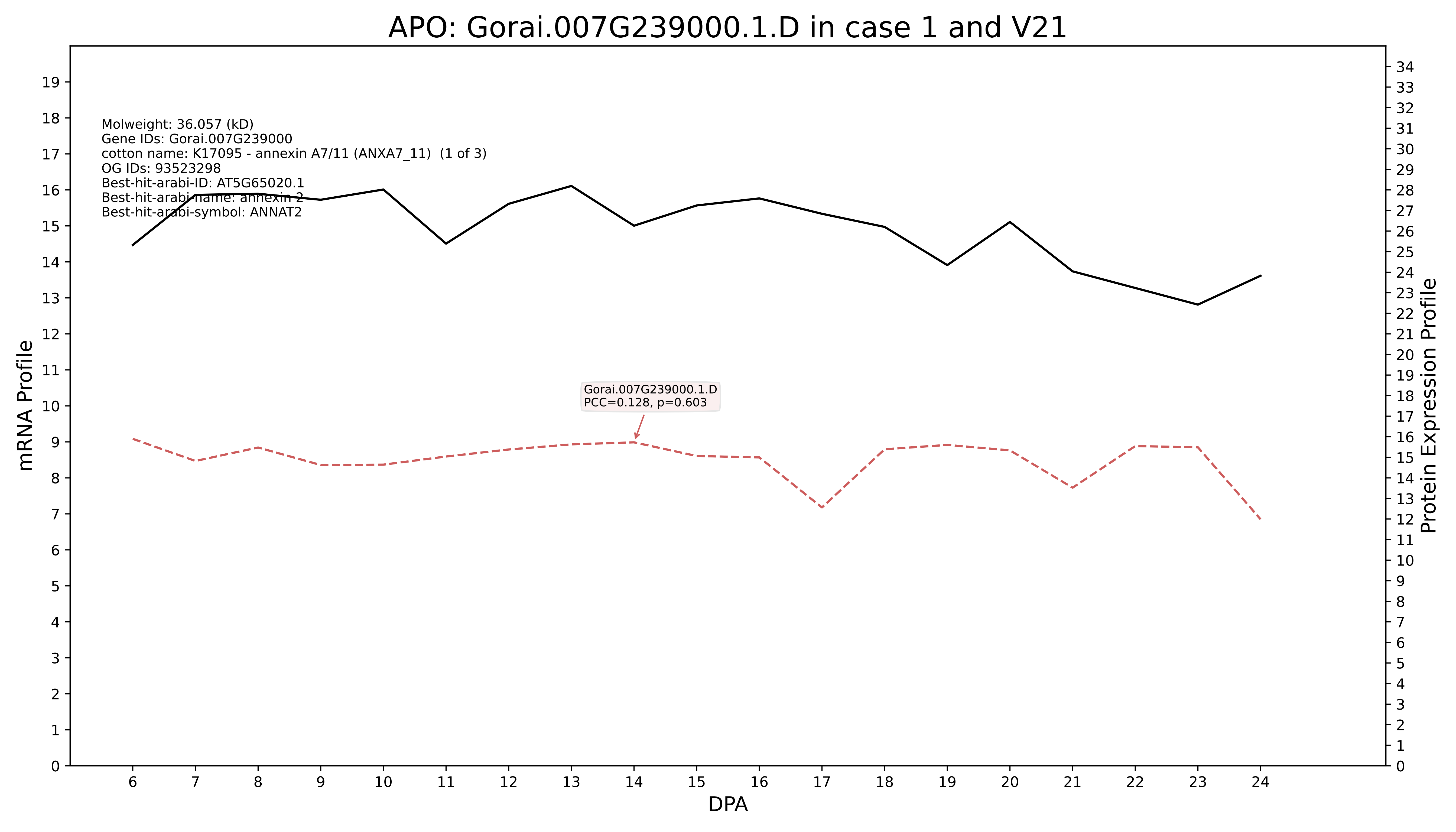
APO: Gorai.007G239000.1.D in case 1 and V21
Molweight: 36.057 (kD)
Gene IDs: Gorai.007G239000
cotton name: K17095 - annexin A7/11 (ANXA7_11) (1 of 3)
OG IDs: 93523298
Best-hit-arabi-ID: AT5G65020.1
Best-hit-arabi-name: annexin 2
Best-hit-arabi-symbol: ANNAT2
Gorai.007G239000.1.D
PCC=0.128, p=0.603
mRNA Profile
Protein Expression Profile
DPA

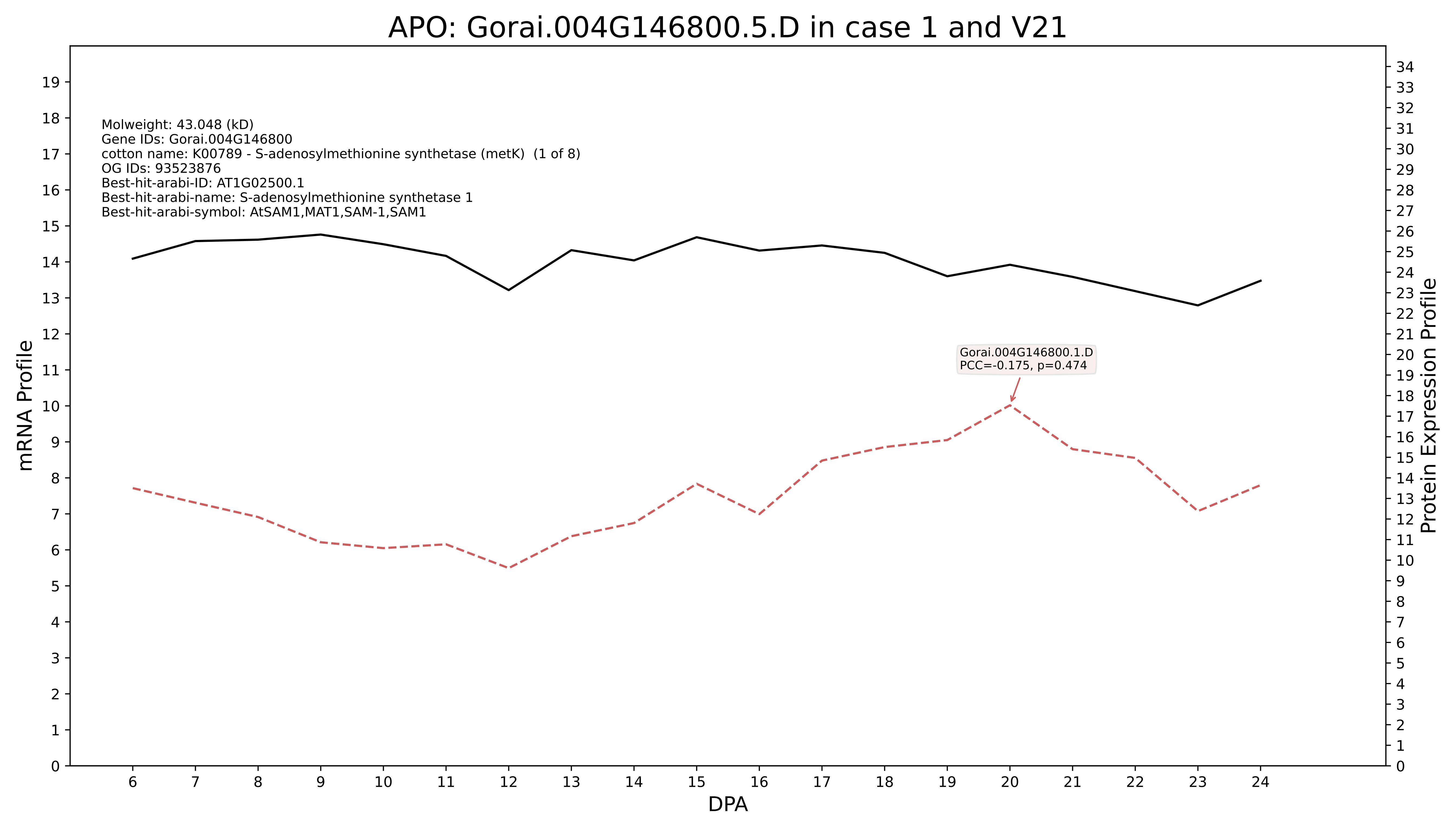

APO: Gorai.004G146800.5.D in case 1 and V21
Molweight: 43.048 (kD)
Gene IDs: Gorai.004G146800
cotton name: K00789 - S-adenosylmethionine synthetase (metK) (1 of 8)
OG IDs: 93523876
Best-hit-arabi-ID: AT1G02500.1
Best-hit-arabi-name: S-adenosylmethionine synthetase 1
Best-hit-arabi-symbol: AtSAM1,MAT1,SAM-1,SAM1
Gorai.004G146800.1.D
PCC=-0.175, p=0.474
mRNA Profile
Protein Expression Profile
DPA

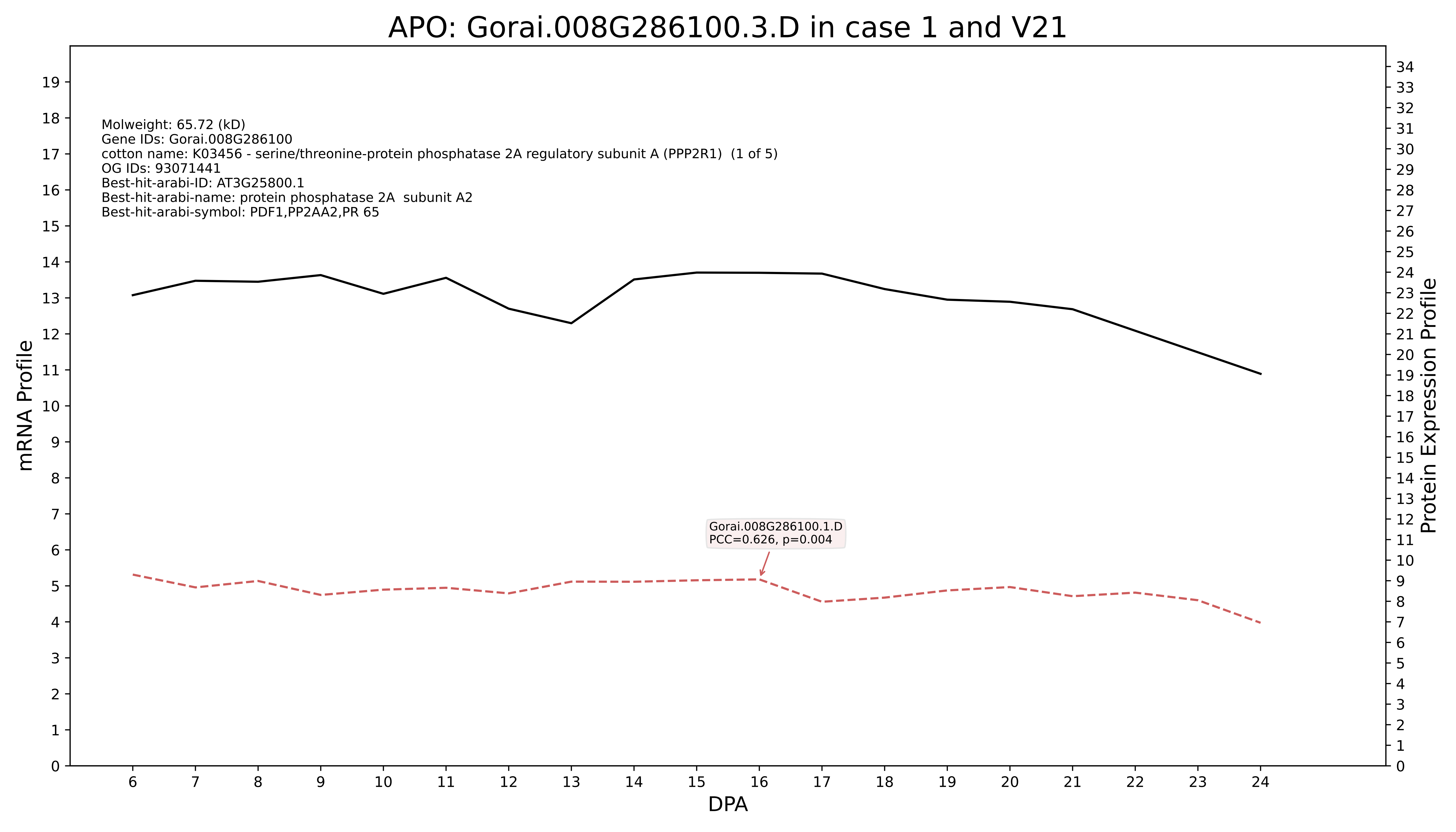
APO: Gorai.008G286100.3.D in case 1 and V21
Molweight: 65.72 (kD)
Gene IDs: Gorai.008G286100
cotton name: K03456 - serine/threonine-protein phosphatase 2A regulatory subunit A (PPP2R1) (1 of 5)
OG IDs: 93071441
Best-hit-arabi-ID: AT3G25800.1
Best-hit-arabi-name: protein phosphatase 2A subunit A2
Best-hit-arabi-symbol: PDF1,PP2AA2,PR 65
Gorai.008G286100.1.D
PCC=0.626, p=0.004
mRNA Profile
Protein Expression Profile
DPA

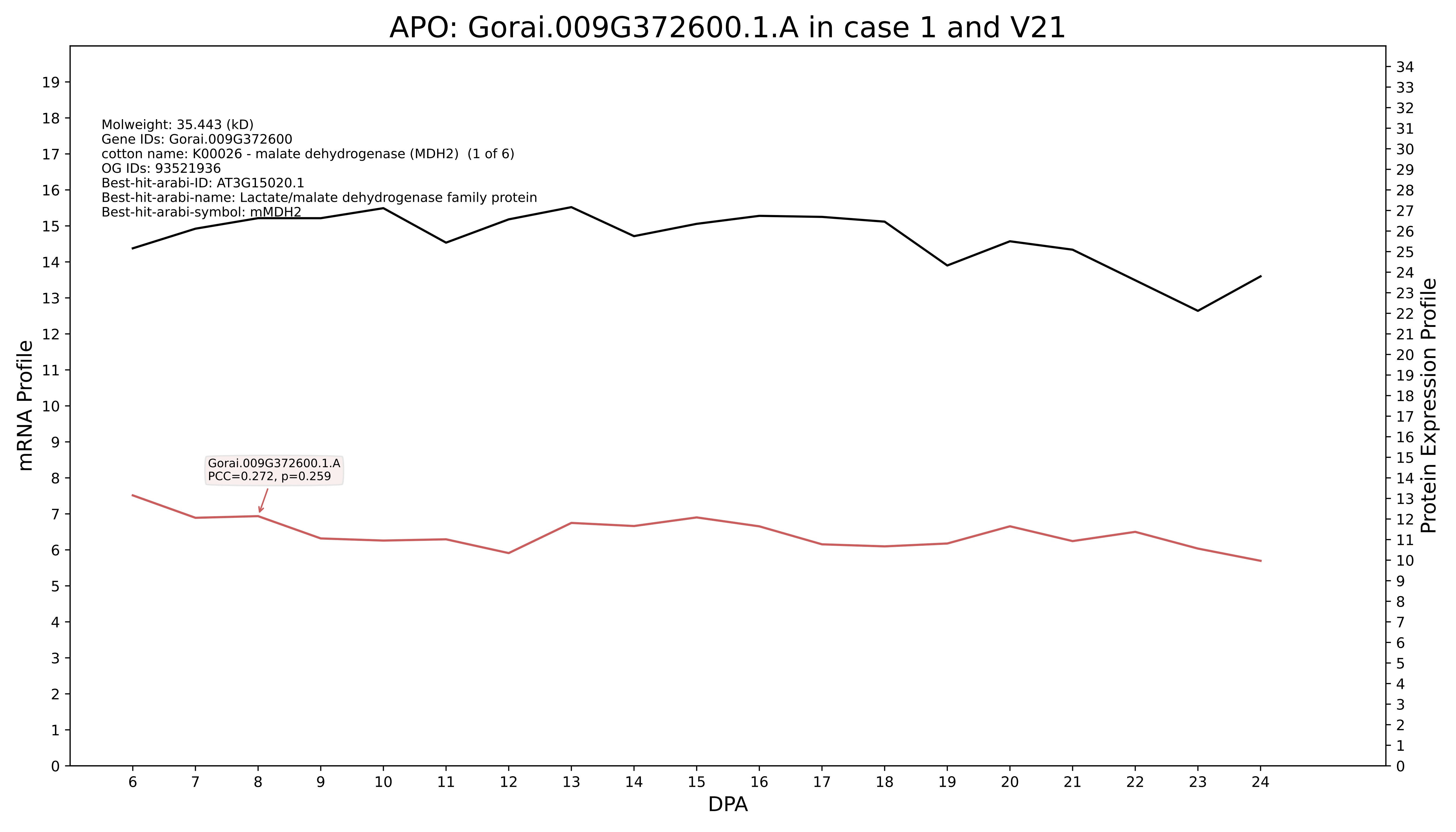
APO: Gorai.009G372600.1.A in case 1 and V21
Molweight: 35.443 (kD)
Gene IDs: Gorai.009G372600
cotton name: K00026 - malate dehydrogenase (MDH2) (1 of 6)
OG IDs: 93521936
Best-hit-arabi-ID: AT3G15020.1
Best-hit-arabi-name: Lactate/malate dehydrogenase family protein
Best-hit-arabi-symbol: mMDH2
Gorai.009G372600.1.A
PCC=0.272, p=0.259
mRNA Profile
Protein Expression Profile
DPA

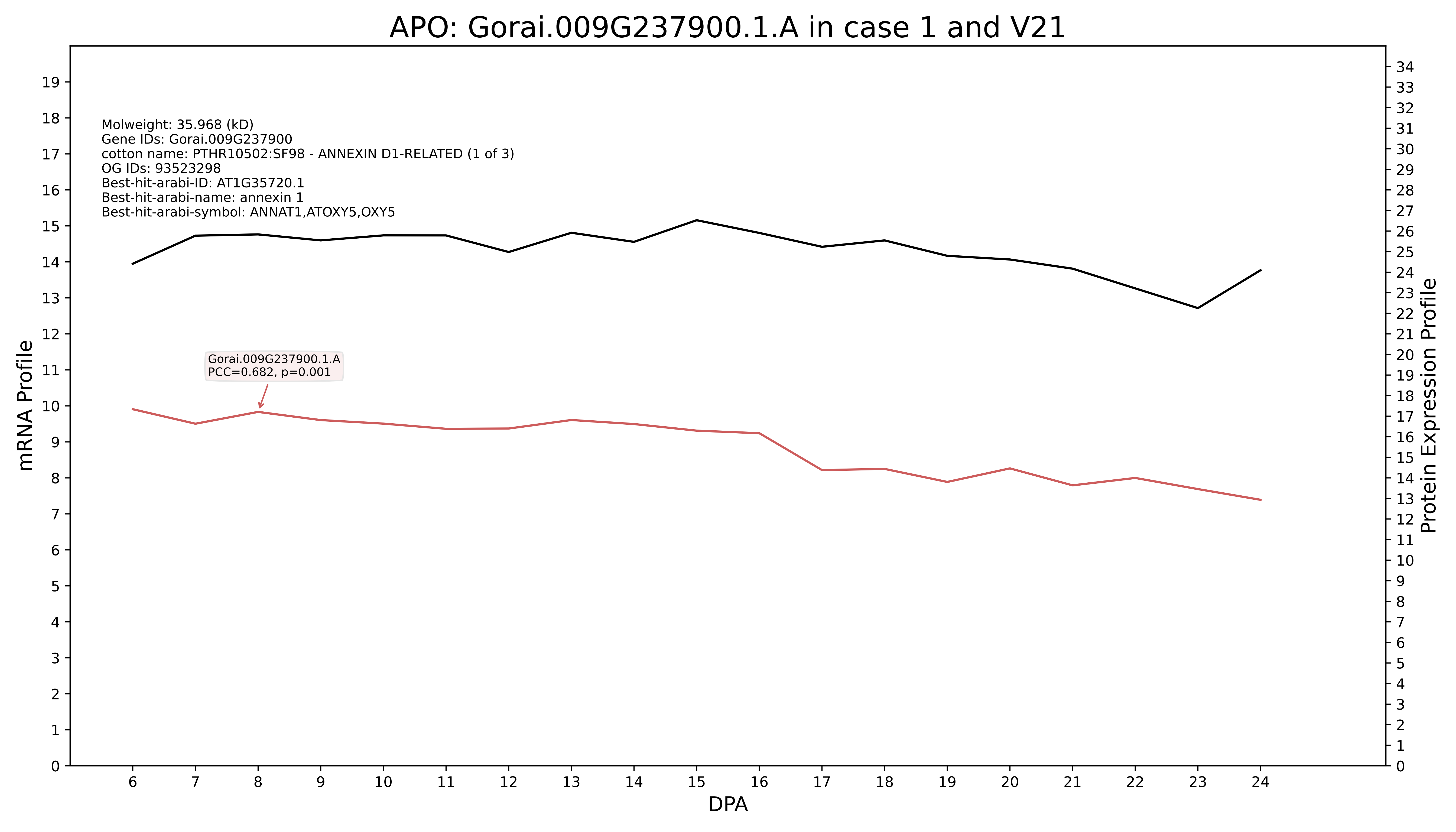

APO: Gorai.009G237900.1.A in case 1 and V21
Molweight: 35.968 (kD)
Gene IDs: Gorai.009G237900
cotton name: PTHR10502:SF98 - ANNEXIN D1-RELATED (1 of 3)
OG IDs: 93523298
Best-hit-arabi-ID: AT1G35720.1
Best-hit-arabi-name: annexin 1
Best-hit-arabi-symbol: ANNAT1,ATOXY5,OXY5
Gorai.009G237900.1.A
PCC=0.682, p=0.001
mRNA Profile
Protein Expression Profile
DPA

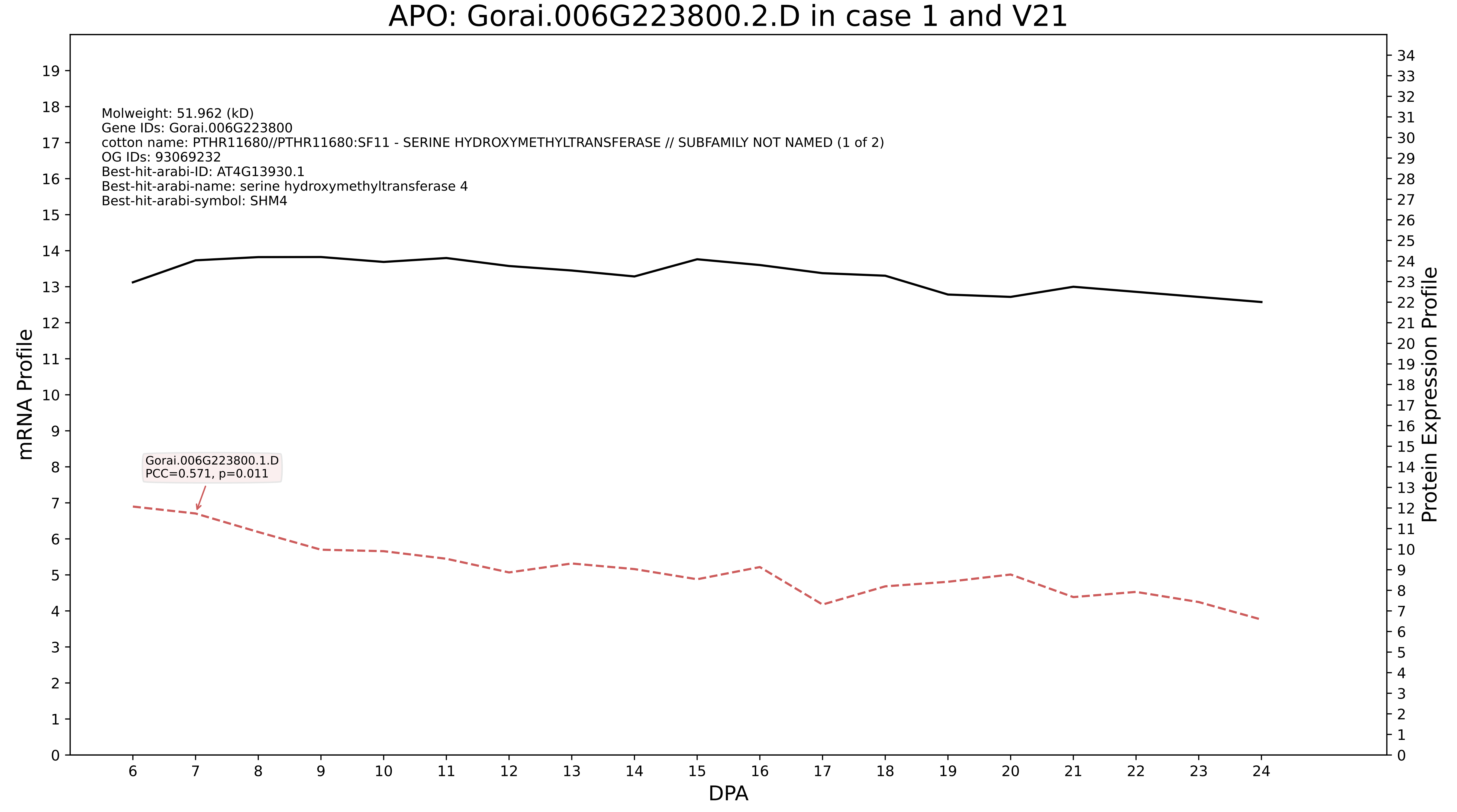

APO: Gorai.006G223800.2.D in case 1 and V21
Molweight: 51.962 (kD)
Gene IDs: Gorai.006G223800
cotton name: PTHR11680//PTHR11680:SF11 - SERINE HYDROXYMETHYLTRANSFERASE // SUBFAMILY NOT NAMED (1 of 2)
OG IDs: 93069232
Best-hit-arabi-ID: AT4G13930.1
Best-hit-arabi-name: serine hydroxymethyltransferase 4
Best-hit-arabi-symbol: SHM4
Gorai.006G223800.1.D
PCC=0.571, p=0.011
mRNA Profile
Protein Expression Profile
DPA

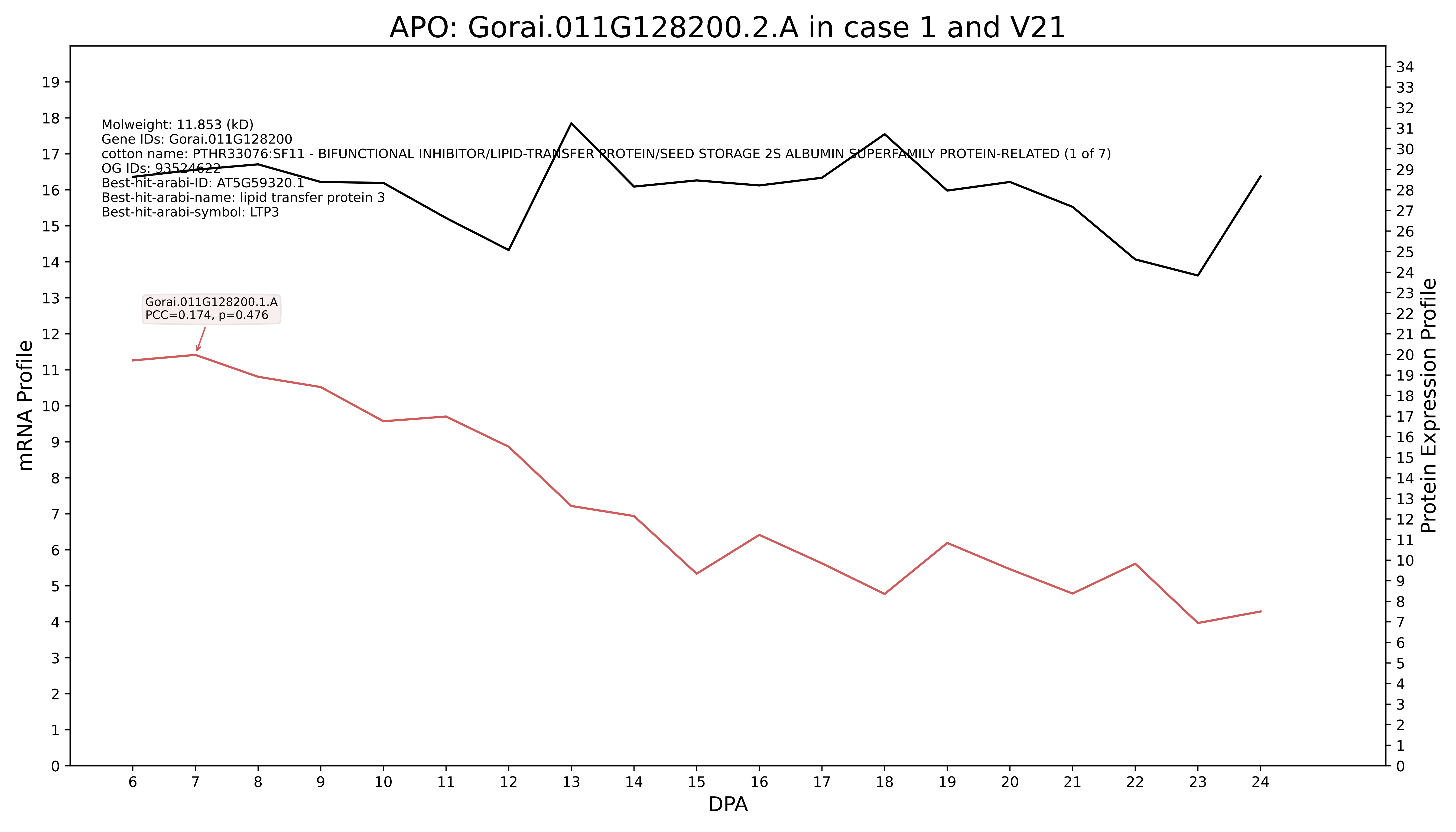
APO: Gorai.011G128200.2.A in case 1 and V21
Molweight: 11.853 (kD)
Gene IDs: Gorai.011G128200
cotton name: PTHR33076:SF11 - BIFUNCTIONAL INHIBITOR/LIPID-TRANSFER PROTEIN/SEED STORAGE 2S ALBUMIN SUPERFAMILY PROTEIN-RELATED (1 of 7)
OG IDs: 93524622
Best-hit-arabi-ID: AT5G59320.1
Best-hit-arabi-name: lipid transfer protein 3
Best-hit-arabi-symbol: LTP3
Gorai.011G128200.1.A
PCC=0.174, p=0.476
mRNA Profile
Protein Expression Profile
DPA

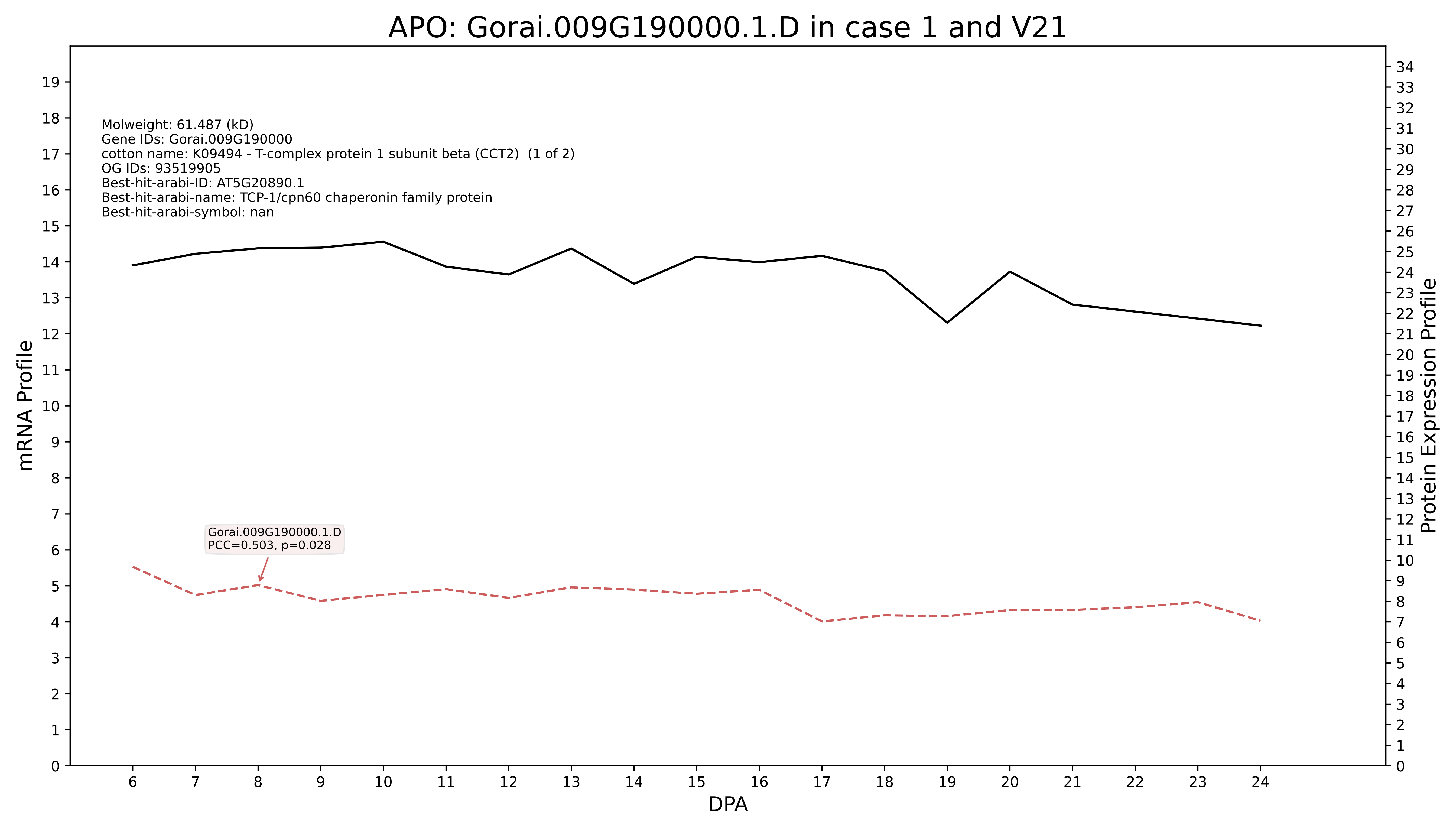

APO: Gorai.009G190000.1.D in case 1 and V21
Molweight: 61.487 (kD)
Gene IDs: Gorai.009G190000
cotton name: K09494 - T-complex protein 1 subunit beta (CCT2) (1 of 2)
OG IDs: 93519905
Best-hit-arabi-ID: AT5G20890.1
Best-hit-arabi-name: TCP-1/cpn60 chaperonin family protein
Best-hit-arabi-symbol: nan
Gorai.009G190000.1.D
PCC=0.503, p=0.028
mRNA Profile
Protein Expression Profile
DPA

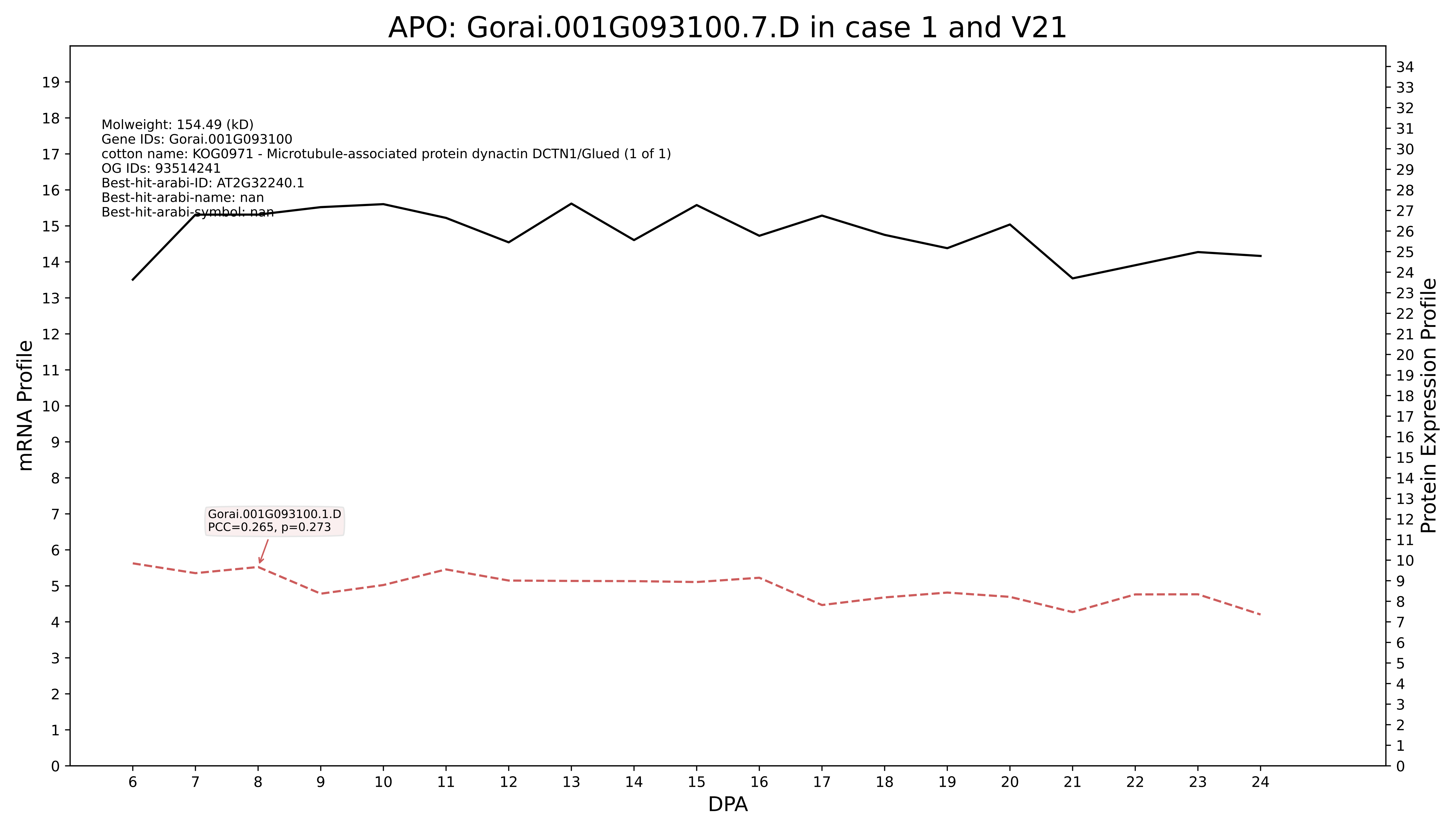
APO: Gorai.001G093100.7.D in case 1 and V21
Molweight: 154.49 (kD)
Gene IDs: Gorai.001G093100
cotton name: KOG0971 - Microtubule-associated protein dynactin DCTN1/Glued (1 of 1)
OG IDs: 93514241
Best-hit-arabi-ID: AT2G32240.1
Best-hit-arabi-name: nan
Best-hit-arabi-symbol: nan
Gorai.001G093100.1.D
PCC=0.265, p=0.273
mRNA Profile
Protein Expression Profile
DPA

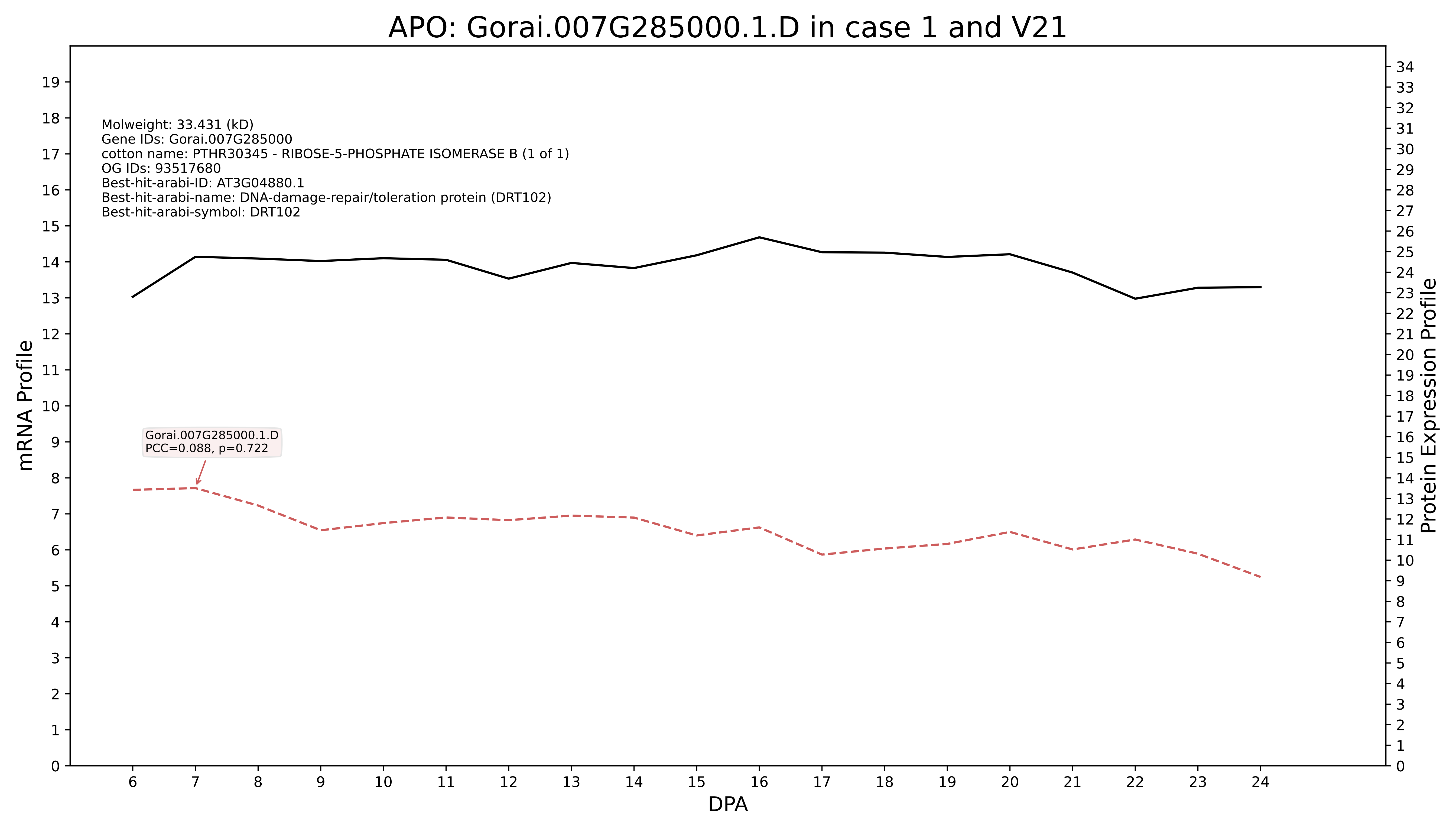

APO: Gorai.007G285000.1.D in case 1 and V21
Molweight: 33.431 (kD)
Gene IDs: Gorai.007G285000
cotton name: PTHR30345 - RIBOSE-5-PHOSPHATE ISOMERASE B (1 of 1)
OG IDs: 93517680
Best-hit-arabi-ID: AT3G04880.1
Best-hit-arabi-name: DNA-damage-repair/toleration protein (DRT102)
Best-hit-arabi-symbol: DRT102
Gorai.007G285000.1.D
PCC=0.088, p=0.722
mRNA Profile
Protein Expression Profile
DPA

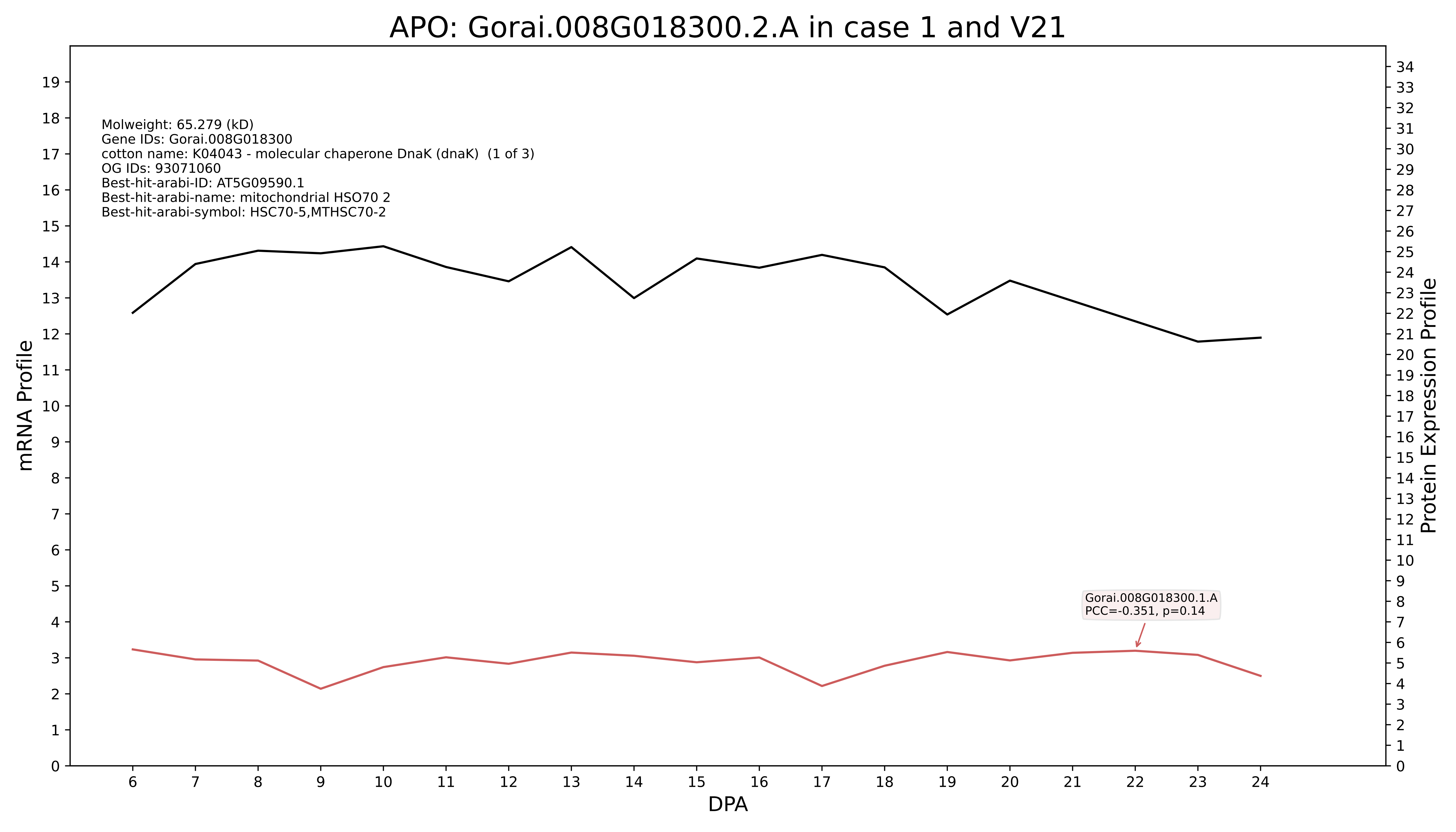
APO: Gorai.008G018300.2.A in case 1 and V21
Molweight: 65.279 (kD)
Gene IDs: Gorai.008G018300
cotton name: K04043 - molecular chaperone DnaK (dnaK) (1 of 3)
OG IDs: 93071060
Best-hit-arabi-ID: AT5G09590.1
Best-hit-arabi-name: mitochondrial HSO70 2
Best-hit-arabi-symbol: HSC70-5,MTHSC70-2
Gorai.008G018300.1.A
PCC=-0.351, p=0.14
mRNA Profile
Protein Expression Profile
DPA

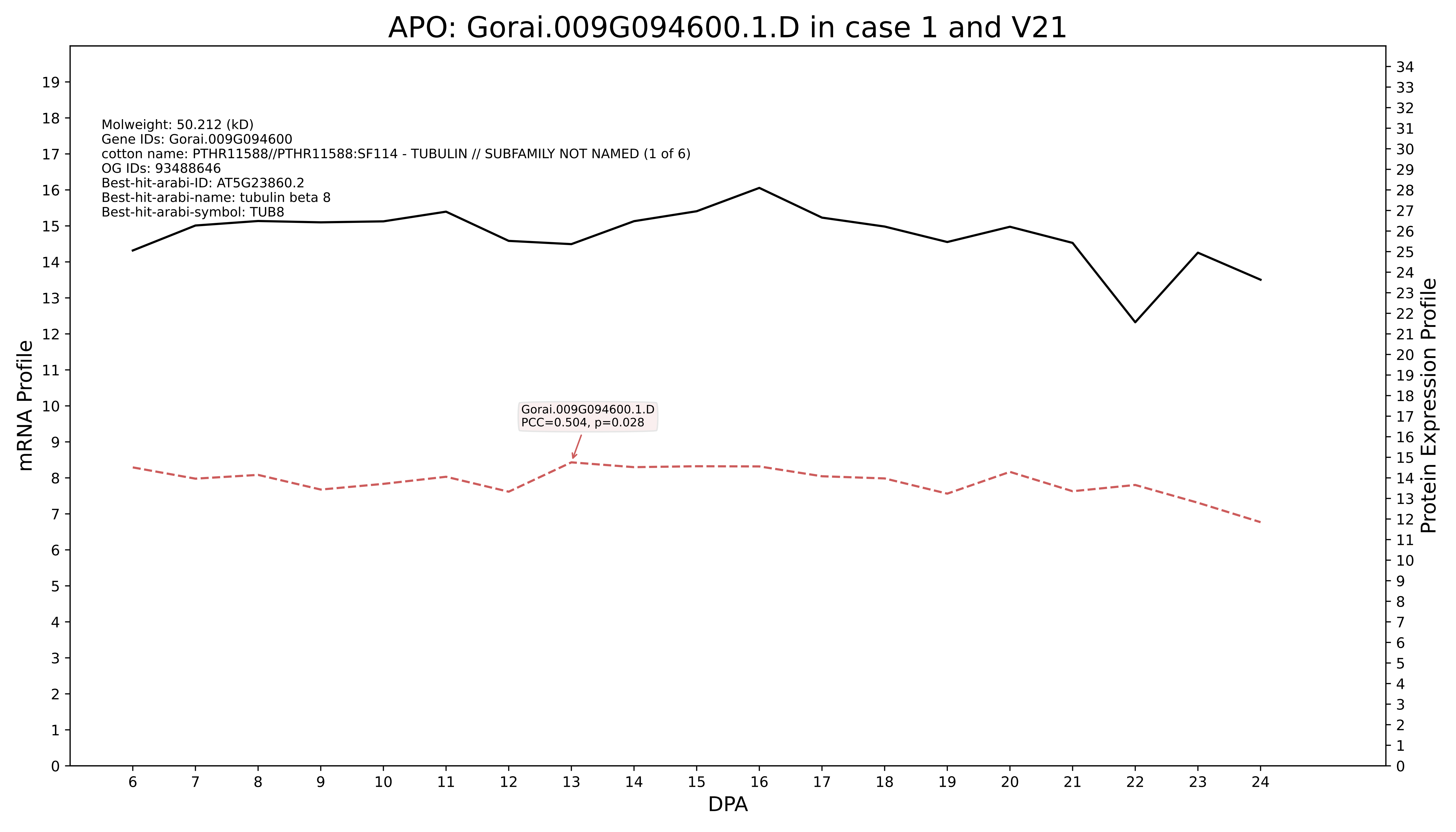

APO: Gorai.009G094600.1.D in case 1 and V21
Molweight: 50.212 (kD)
Gene IDs: Gorai.009G094600
cotton name: PTHR11588//PTHR11588:SF114 - TUBULIN // SUBFAMILY NOT NAMED (1 of 6)
OG IDs: 93488646
Best-hit-arabi-ID: AT5G23860.2
Best-hit-arabi-name: tubulin beta 8
Best-hit-arabi-symbol: TUB8
Gorai.009G094600.1.D
PCC=0.504, p=0.028
mRNA Profile
Protein Expression Profile
DPA

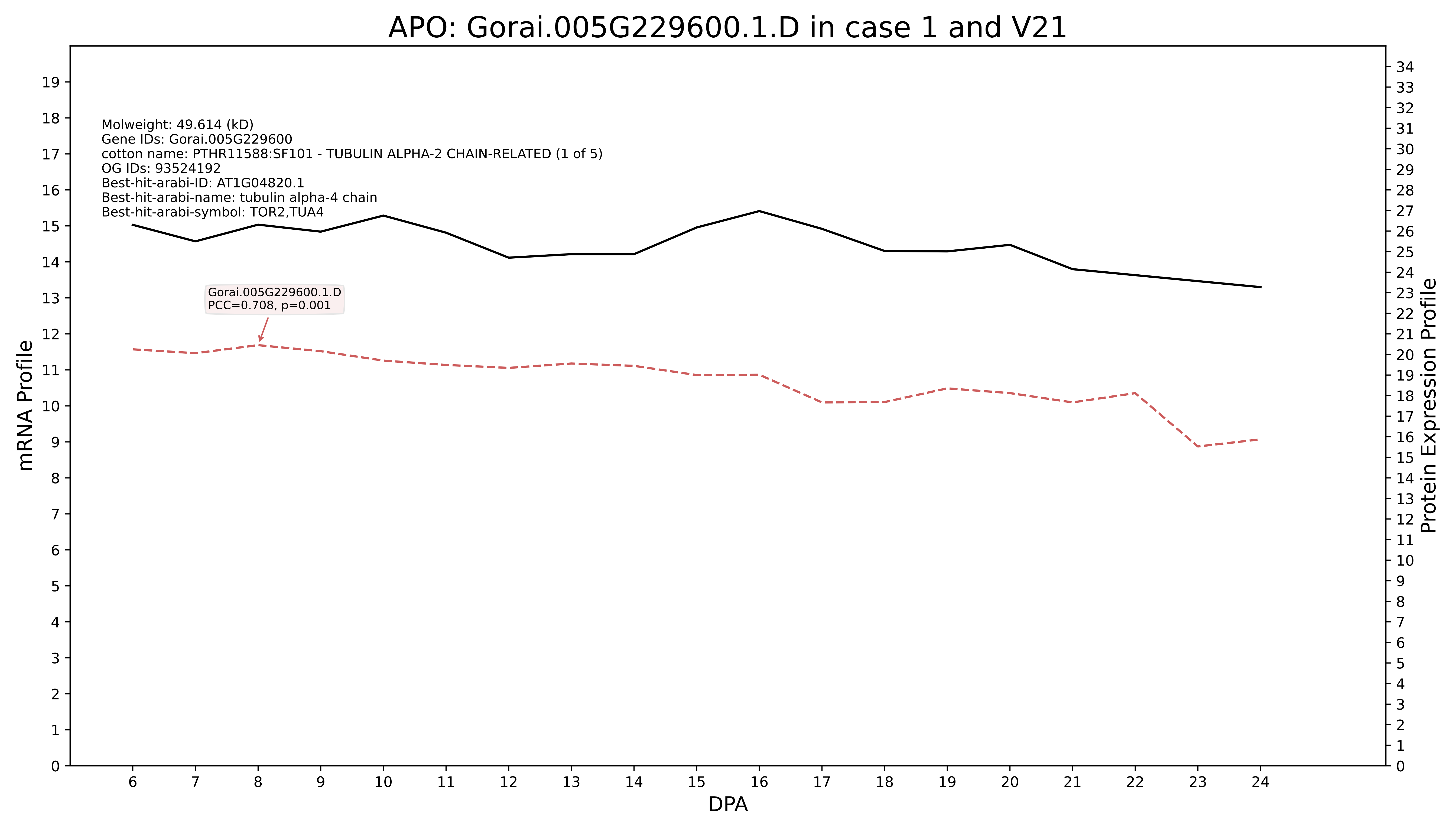
APO: Gorai.005G229600.1.D in case 1 and V21
Molweight: 49.614 (kD)
Gene IDs: Gorai.005G229600
cotton name: PTHR11588:SF101 - TUBULIN ALPHA-2 CHAIN-RELATED (1 of 5)
OG IDs: 93524192
Best-hit-arabi-ID: AT1G04820.1
Best-hit-arabi-name: tubulin alpha-4 chain
Best-hit-arabi-symbol: TOR2,TUA4
Gorai.005G229600.1.D
PCC=0.708, p=0.001
mRNA Profile
Protein Expression Profile
DPA

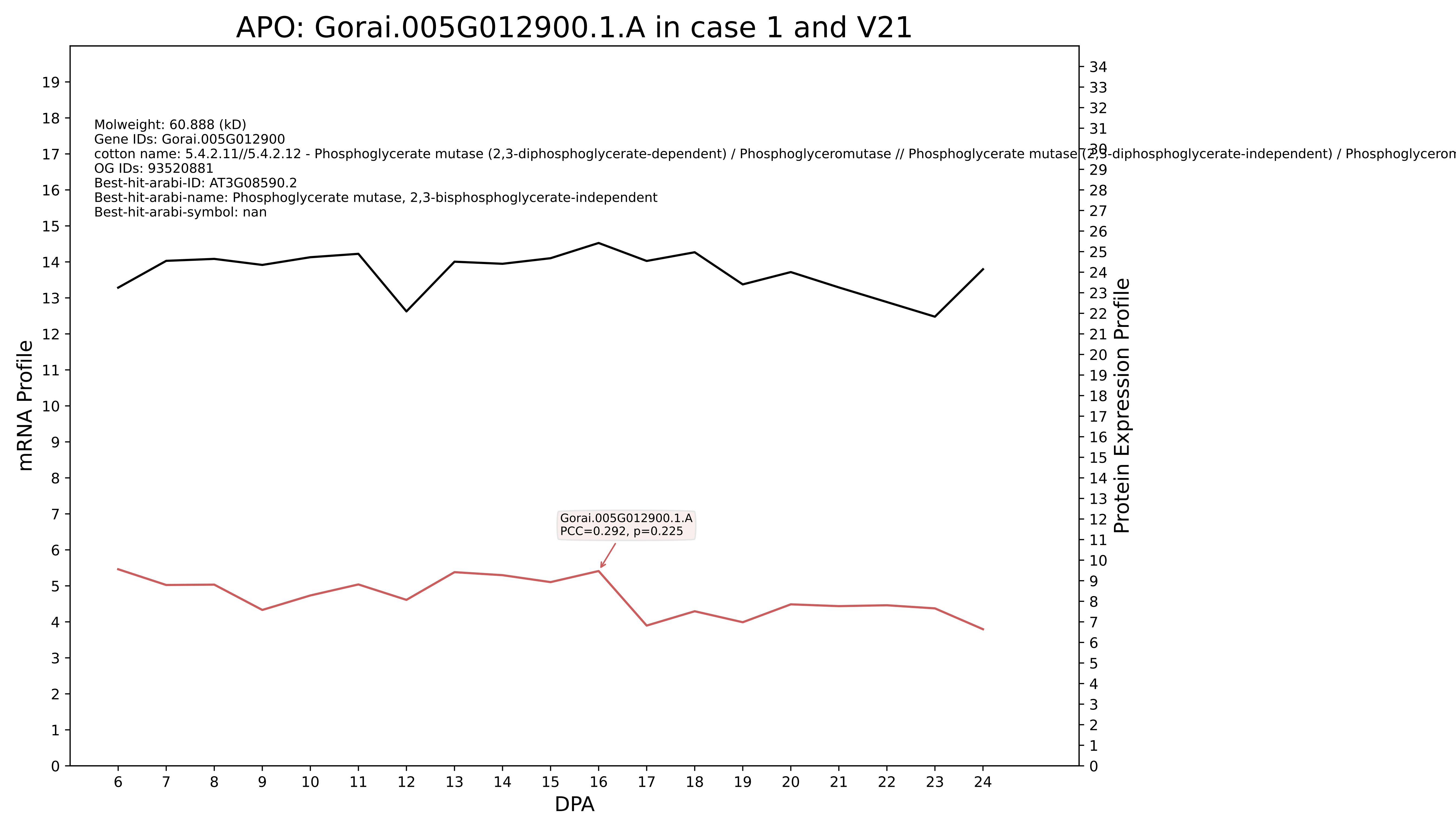
APO: Gorai.005G012900.1.A in case 1 and V21
Molweight: 60.888 (kD)
Gene IDs: Gorai.005G012900
cotton name: 5.4.2.11//5.4.2.12 - Phosphoglycerate mutase (2,3-diphosphoglycerate-dependent) / Phosphoglyceromutase // Phosphoglycerate mutase (2,3-diphosphoglycerate-independent) / Phosphoglyceron
OG IDs: 93520881
Best-hit-arabi-ID: AT3G08590.2
Best-hit-arabi-name: Phosphoglycerate mutase, 2,3-bisphosphoglycerate-independent
Best-hit-arabi-symbol: nan
Gorai.005G012900.1.A
PCC=0.292, p=0.225
mRNA Profile
Protein Expression Profile
DPA

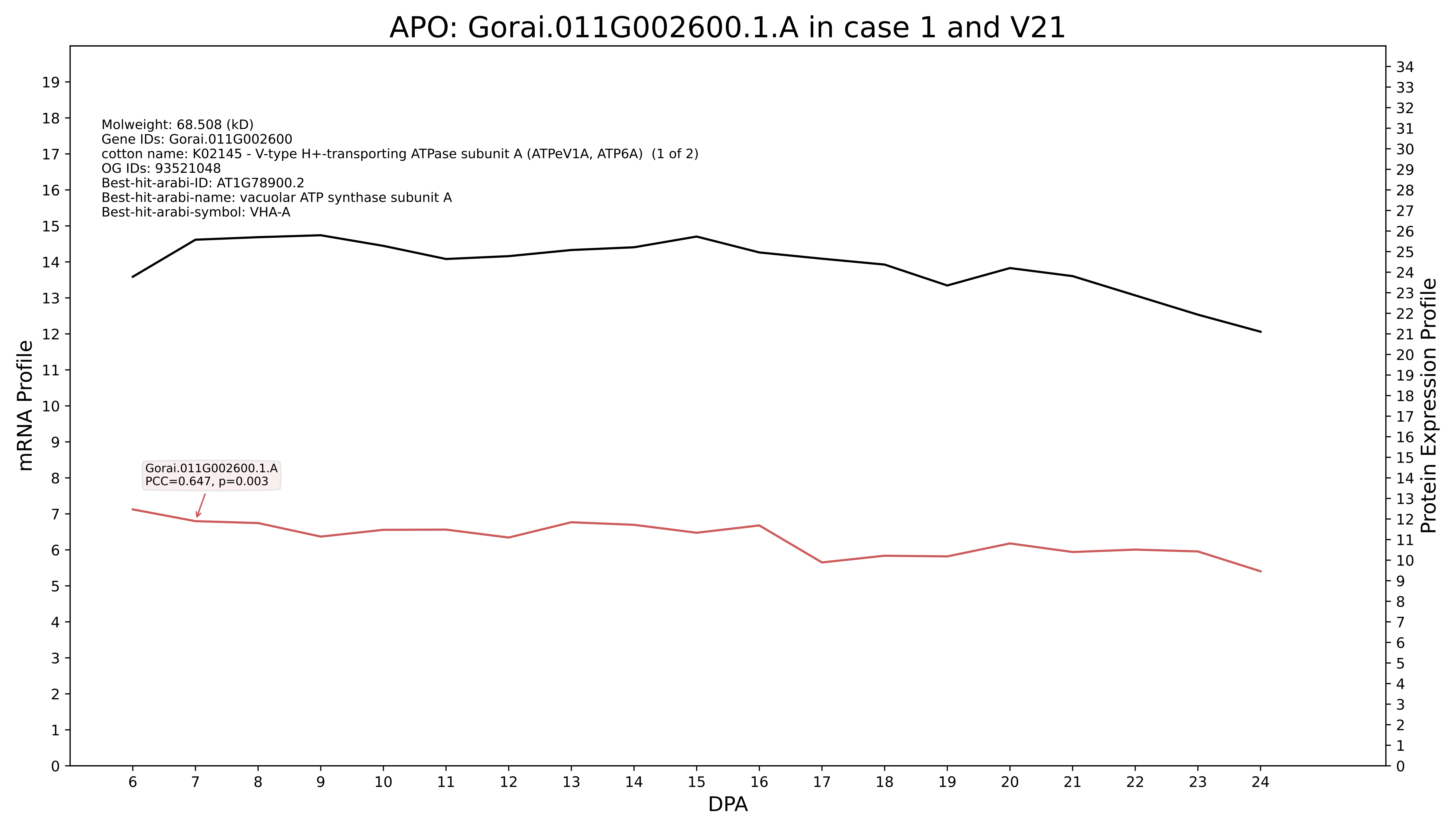
APO: Gorai.011G002600.1.A in case 1 and V21
Molweight: 68.508 (kD)
Gene IDs: Gorai.011G002600
cotton name: K02145 - V-type H+-transporting ATPase subunit A (ATPeV1A, ATP6A) (1 of 2)
OG IDs: 93521048
Best-hit-arabi-ID: AT1G78900.2
Best-hit-arabi-name: vacuolar ATP synthase subunit A
Best-hit-arabi-symbol: VHA-A
Gorai.011G002600.1.A
PCC=0.647, p=0.003
mRNA Profile
Protein Expression Profile
DPA

### APO: Gorai.011G161700.1.D in case 1 and V21

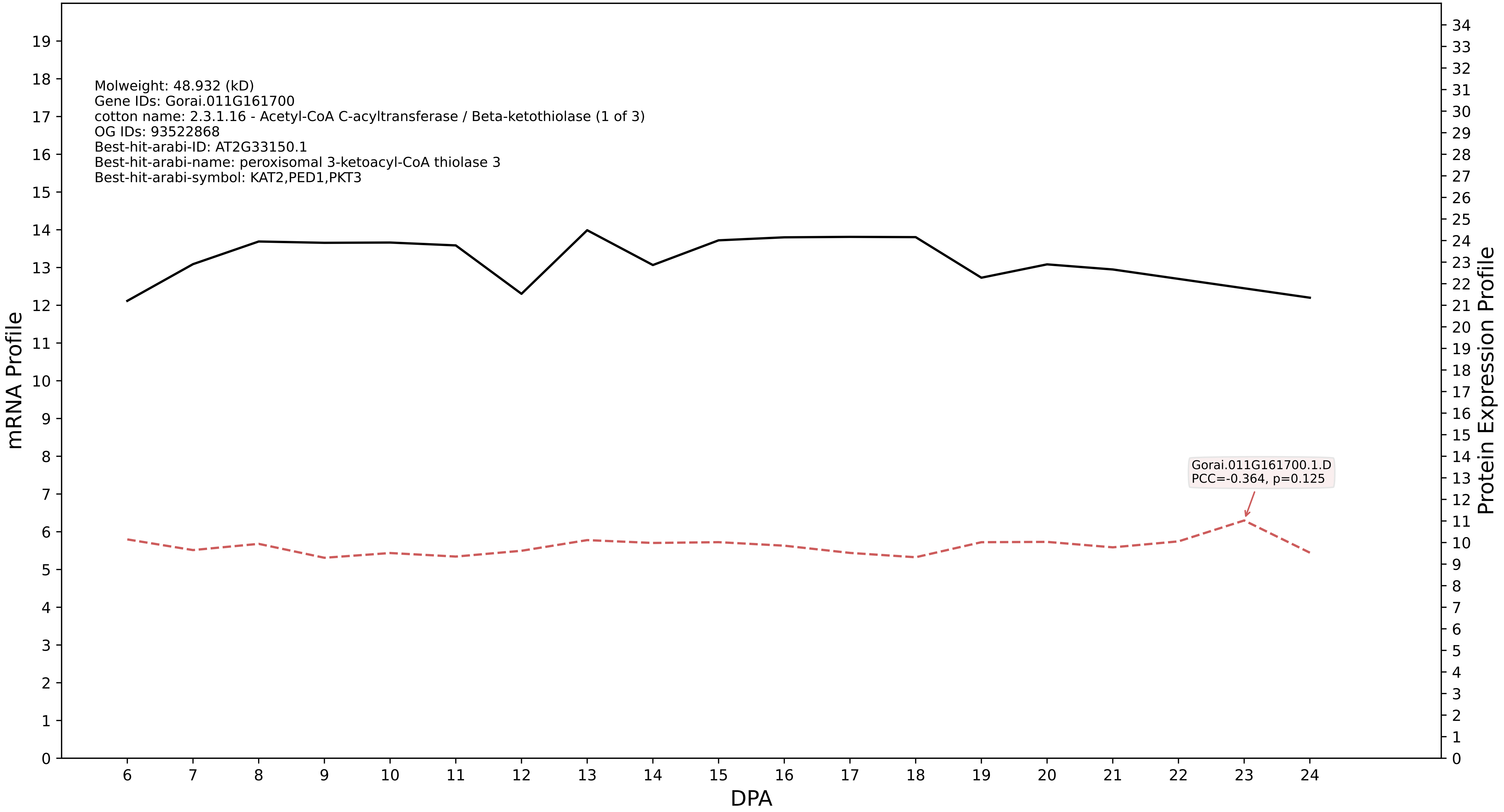

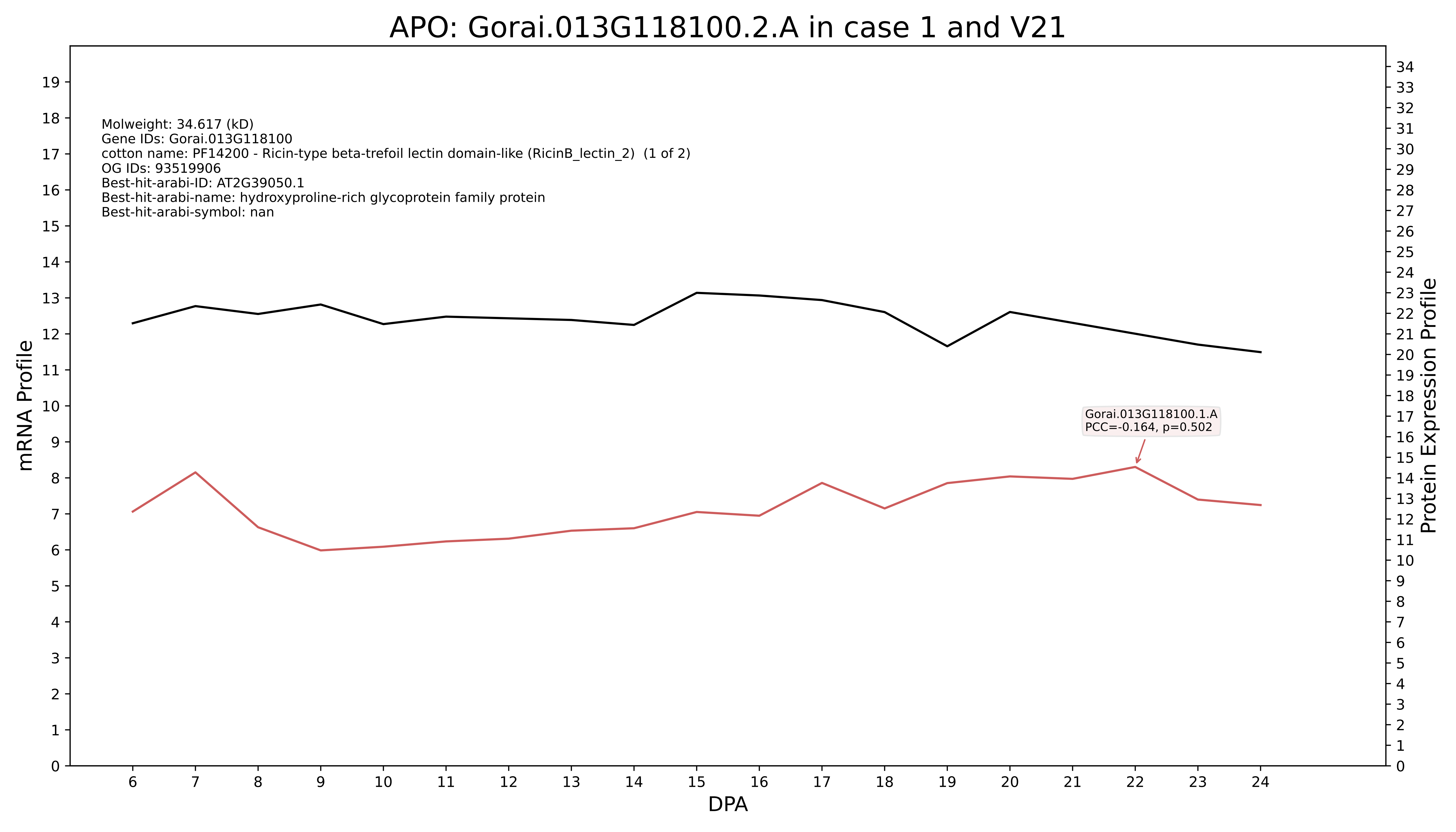
APO: Gorai.013G118100.2.A in case 1 and V21
Molweight: 34.617 (kD)
Gene IDs: Gorai.013G118100
cotton name: PF14200 - Ricin-type beta-trefoil lectin domain-like (RicinB_lectin_2) (1 of 2)
OG IDs: 93519906
Best-hit-arabi-ID: AT2G39050.1
Best-hit-arabi-name: hydroxyproline-rich glycoprotein family protein
Best-hit-arabi-symbol: nan
Gorai.013G118100.1.A
PCC=-0.164, p=0.502
mRNA Profile
Protein Expression Profile
DPA

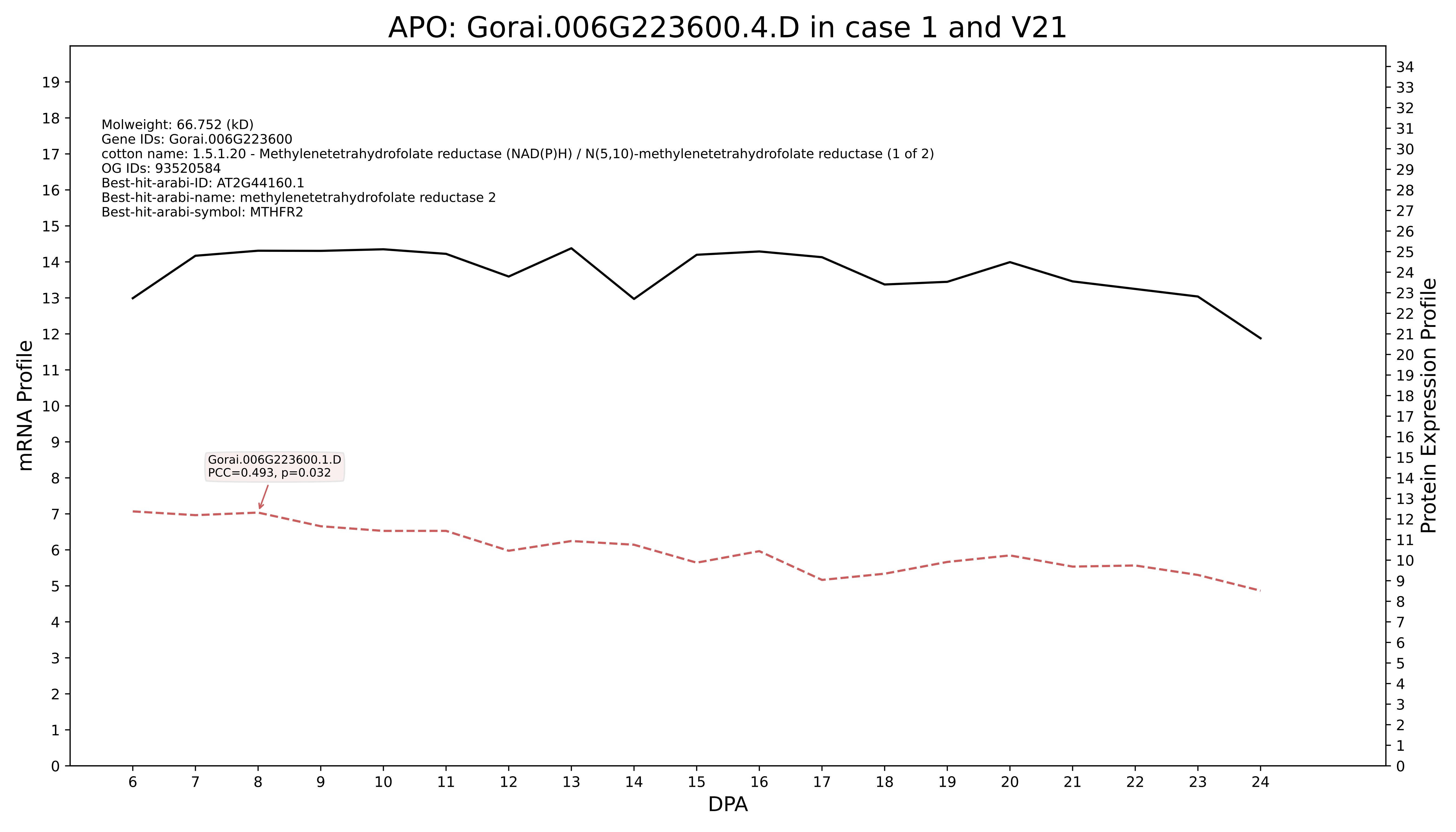

APO: Gorai.006G223600.4.D in case 1 and V21
Molweight: 66.752 (kD)
Gene IDs: Gorai.006G223600
cotton name: 1.5.1.20 - Methylenetetrahydrofolate reductase (NAD(P)H) / N(5,10)-methylenetetrahydrofolate reductase (1 of 2)
OG IDs: 93520584
Best-hit-arabi-ID: AT2G44160.1
Best-hit-arabi-name: methylenetetrahydrofolate reductase 2
Best-hit-arabi-symbol: MTHFR2
Gorai.006G223600.1.D
PCC=0.493, p=0.032
mRNA Profile
Protein Expression Profile
DPA

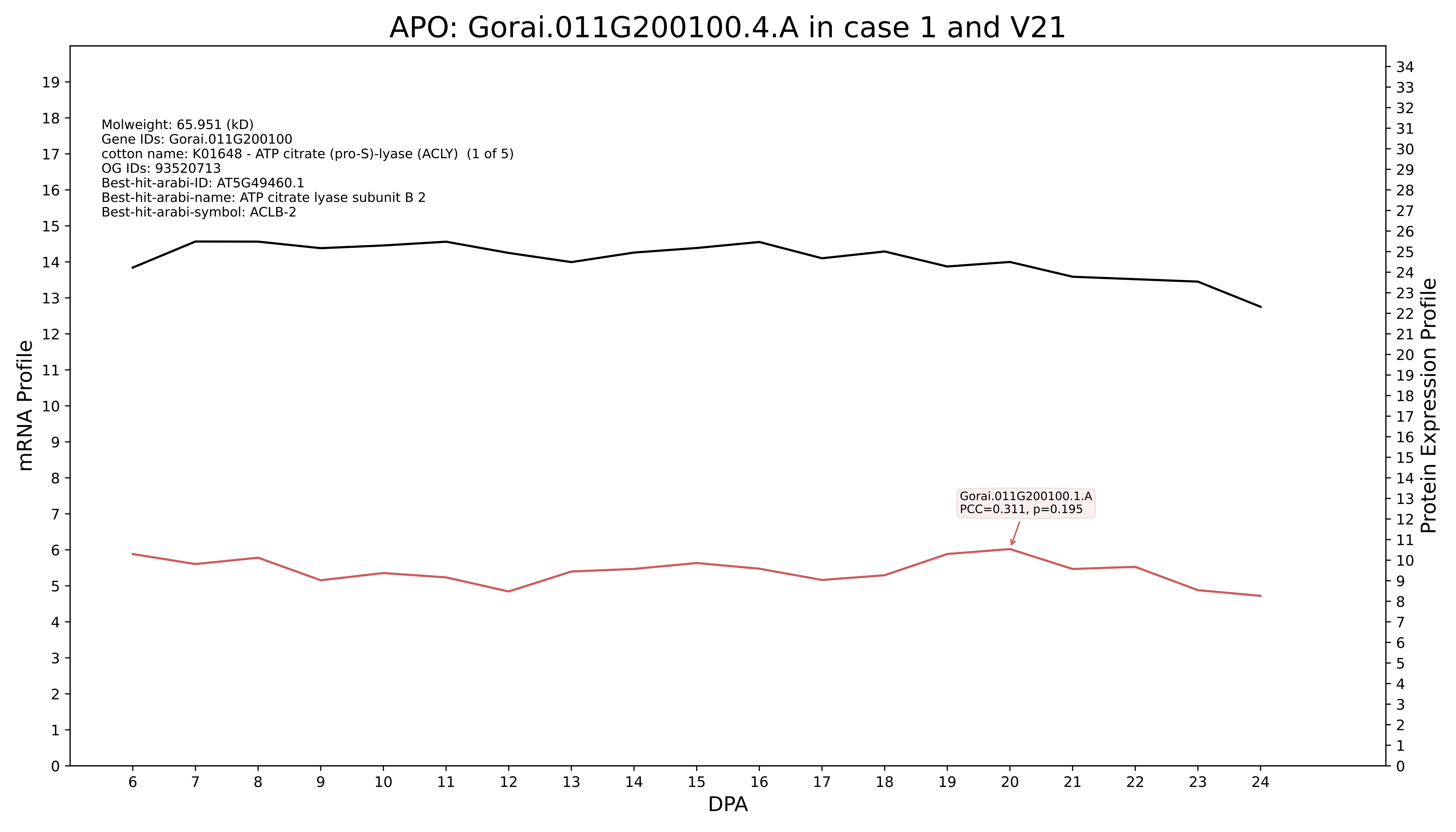
APO: Gorai.011G200100.4.A in case 1 and V21
Molweight: 65.951 (kD)
Gene IDs: Gorai.011G200100
cotton name: K01648 - ATP citrate (pro-S)-lyase (ACLY) (1 of 5)
OG IDs: 93520713
Best-hit-arabi-ID: AT5G49460.1
Best-hit-arabi-name: ATP citrate lyase subunit B 2
Best-hit-arabi-symbol: ACLB-2
Gorai.011G200100.1.A
PCC=0.311, p=0.195
mRNA Profile
Protein Expression Profile
DPA

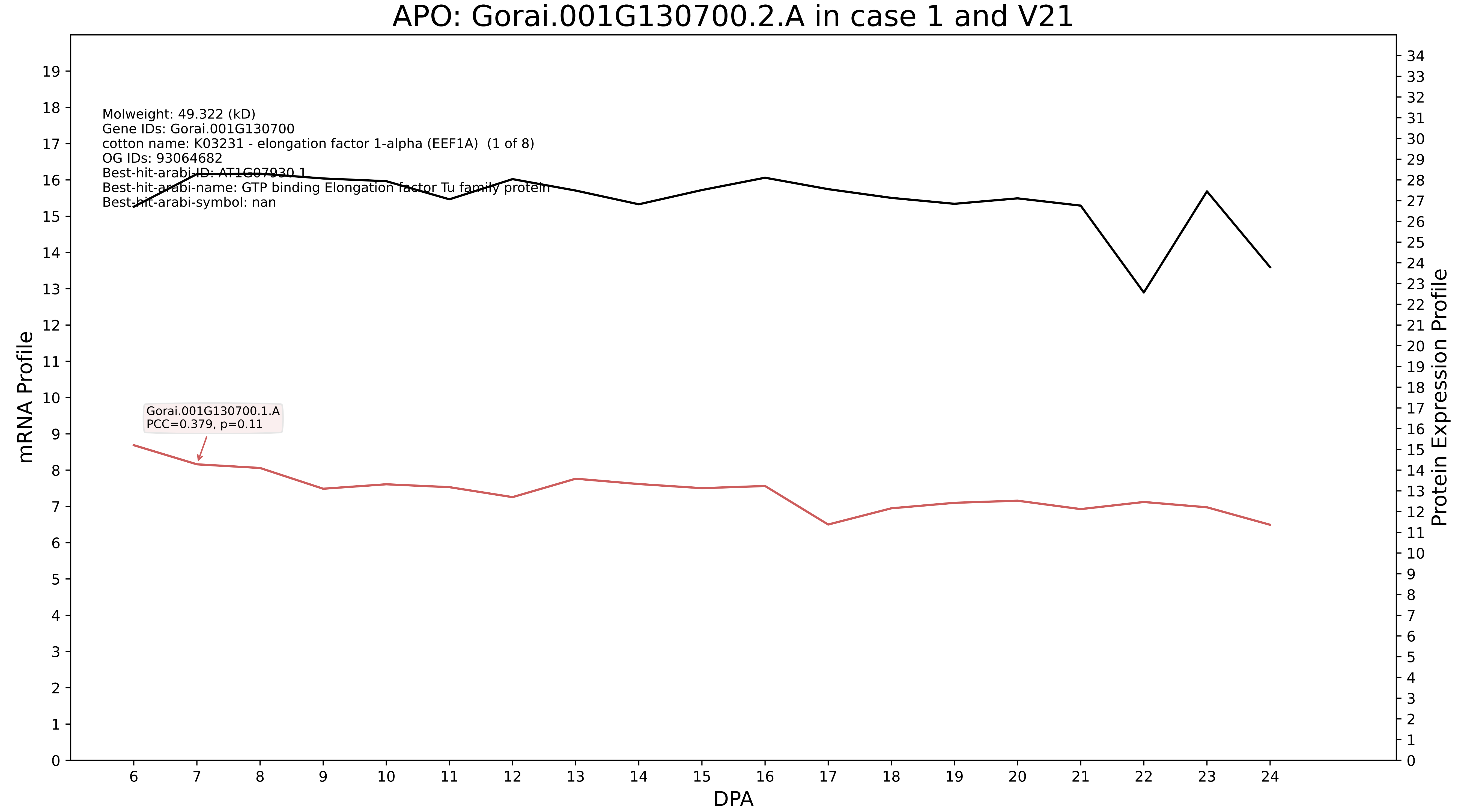

APO: Gorai.001G130700.2.A in case 1 and V21
Molweight: 49.322 (kD)
Gene IDs: Gorai.001G130700
cotton name: K03231 - elongation factor 1-alpha (EEF1A) (1 of 8)
OG IDs: 93064682
Best-hit-arabi-ID: AT1G07930.1
Best-hit-arabi-name: GTP binding Elongation factor Tu family protein
Best-hit-arabi-symbol: nan
Gorai.001G130700.1.A
PCC=0.379, p=0.11
mRNA Profile
Protein Expression Profile
DPA

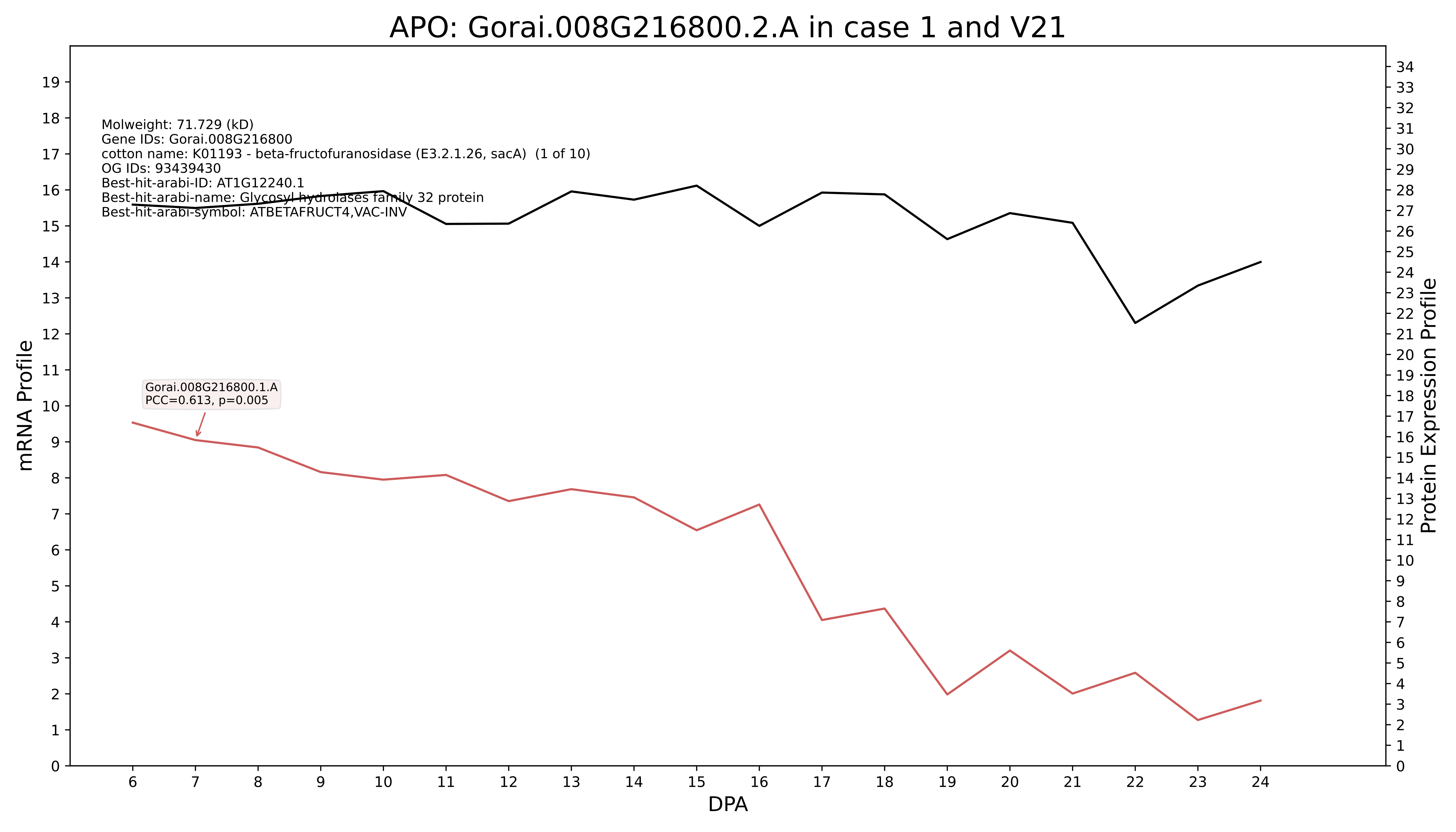
APO: Gorai.008G216800.2.A in case 1 and V21
Molweight: 71.729 (kD)
Gene IDs: Gorai.008G216800
cotton name: K01193 - beta-fructofuranosidase (E3.2.1.26, sacA) (1 of 10)
OG IDs: 93439430
Best-hit-arabi-ID: AT1G12240.1
Best-hit-arabi-name: Glycosyl hydrolases family 32 protein
Best-hit-arabi-symbol: ATBETAFRUCT4,VAC-INV
Gorai.008G216800.1.A
PCC=0.613, p=0.005
mRNA Profile
Protein Expression Profile
DPA

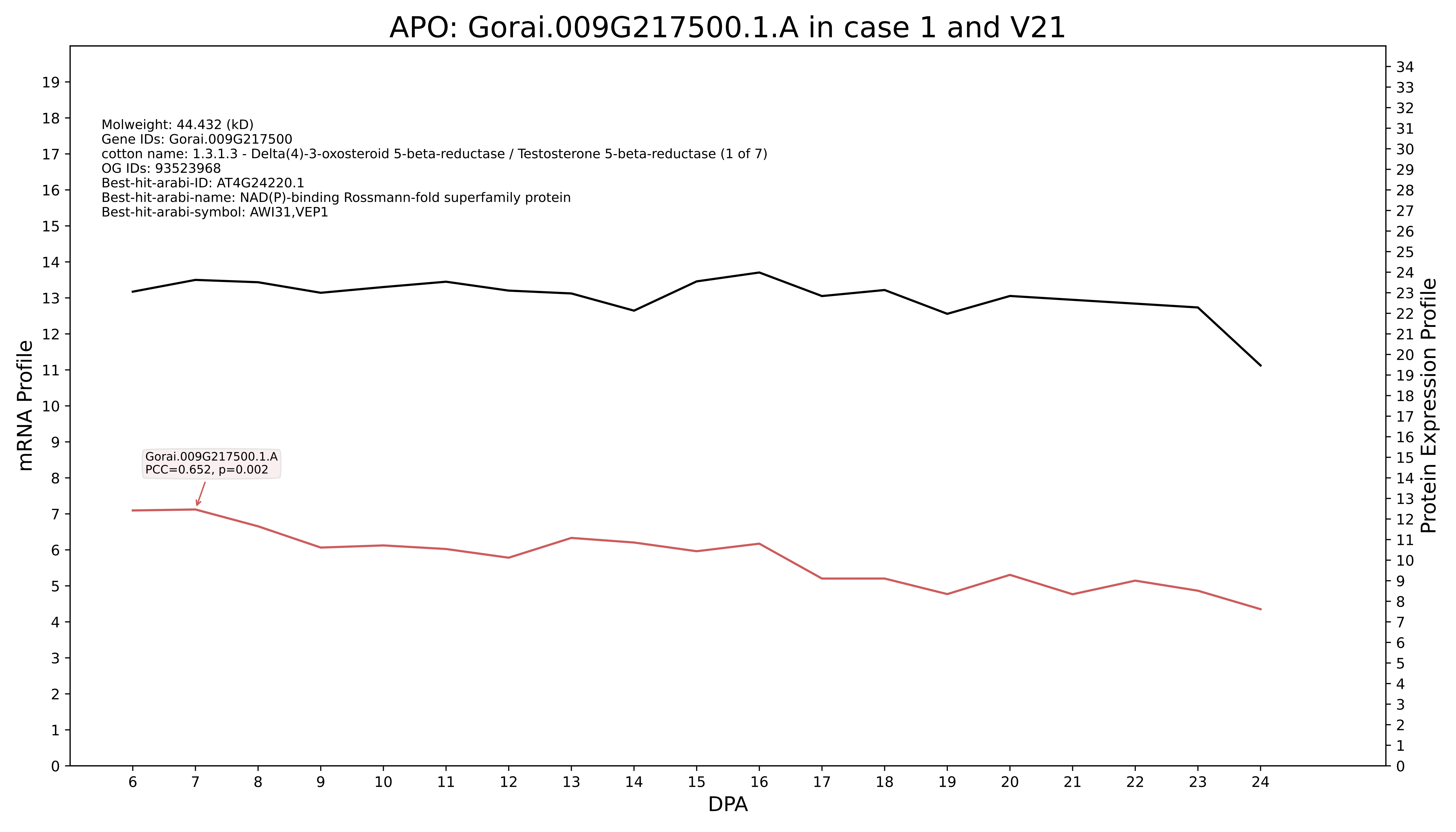

APO: Gorai.009G217500.1.A in case 1 and V21
Molweight: 44.432 (kD)
Gene IDs: Gorai.009G217500
cotton name: 1.3.1.3 - Delta(4)-3-oxosteroid 5-beta-reductase / Testosterone 5-beta-reductase (1 of 7)
OG IDs: 93523968
Best-hit-arabi-ID: AT4G24220.1
Best-hit-arabi-name: NAD(P)-binding Rossmann-fold superfamily protein
Best-hit-arabi-symbol: AWI31,VEP1
Gorai.009G217500.1.A
PCC=0.652, p=0.002
mRNA Profile
Protein Expression Profile
DPA

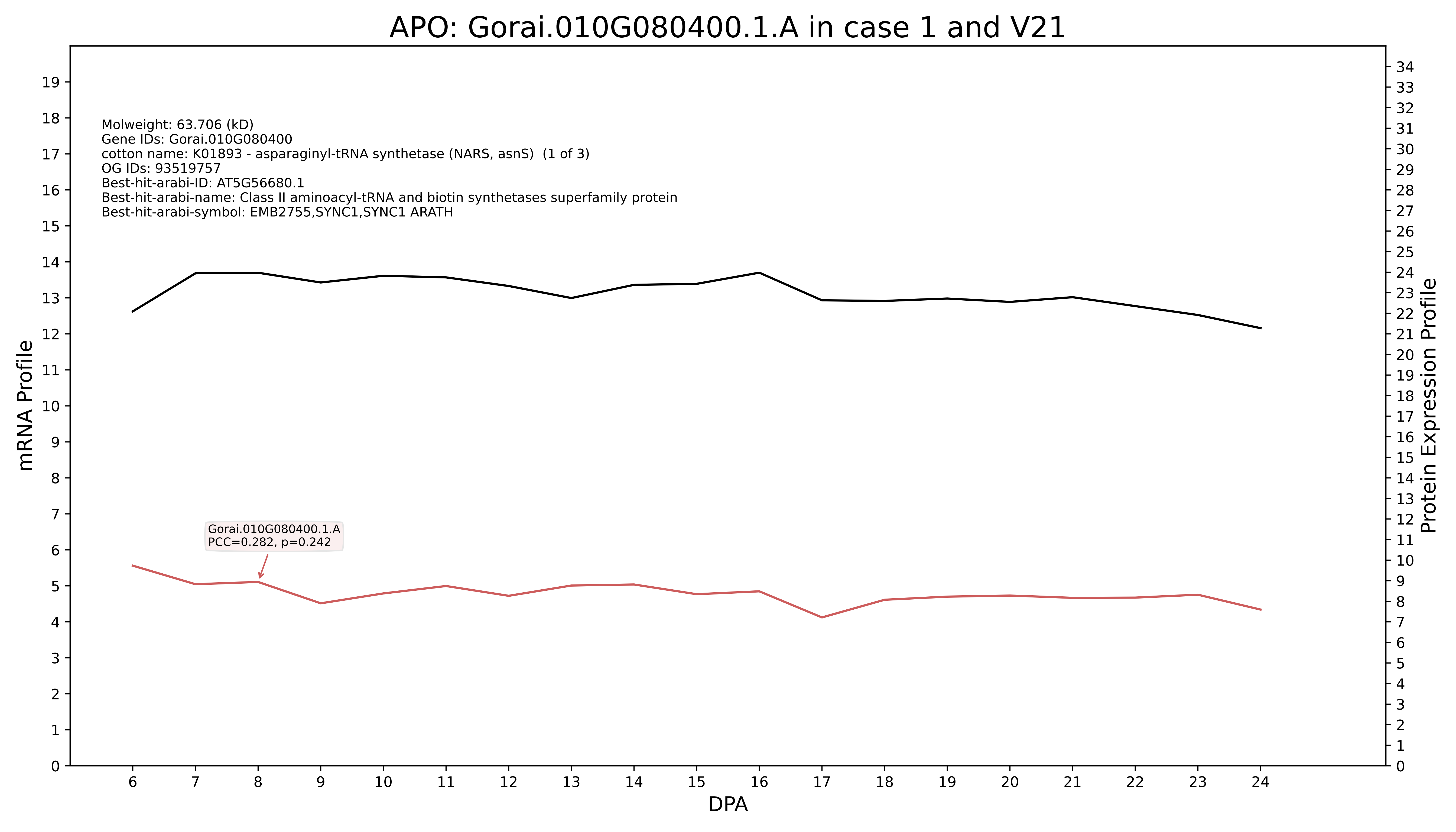
APO: Gorai.010G080400.1.A in case 1 and V21
Molweight: 63.706 (kD)
Gene IDs: Gorai.010G080400
cotton name: K01893 - asparaginyl-tRNA synthetase (NARS, asnS) (1 of 3)
OG IDs: 93519757
Best-hit-arabi-ID: AT5G56680.1
Best-hit-arabi-name: Class II aminoacyl-tRNA and biotin synthetases superfamily protein
Best-hit-arabi-symbol: EMB2755,SYNC1,SYNC1 ARATH
Gorai.010G080400.1.A
PCC=0.282, p=0.242
mRNA Profile
Protein Expression Profile
DPA

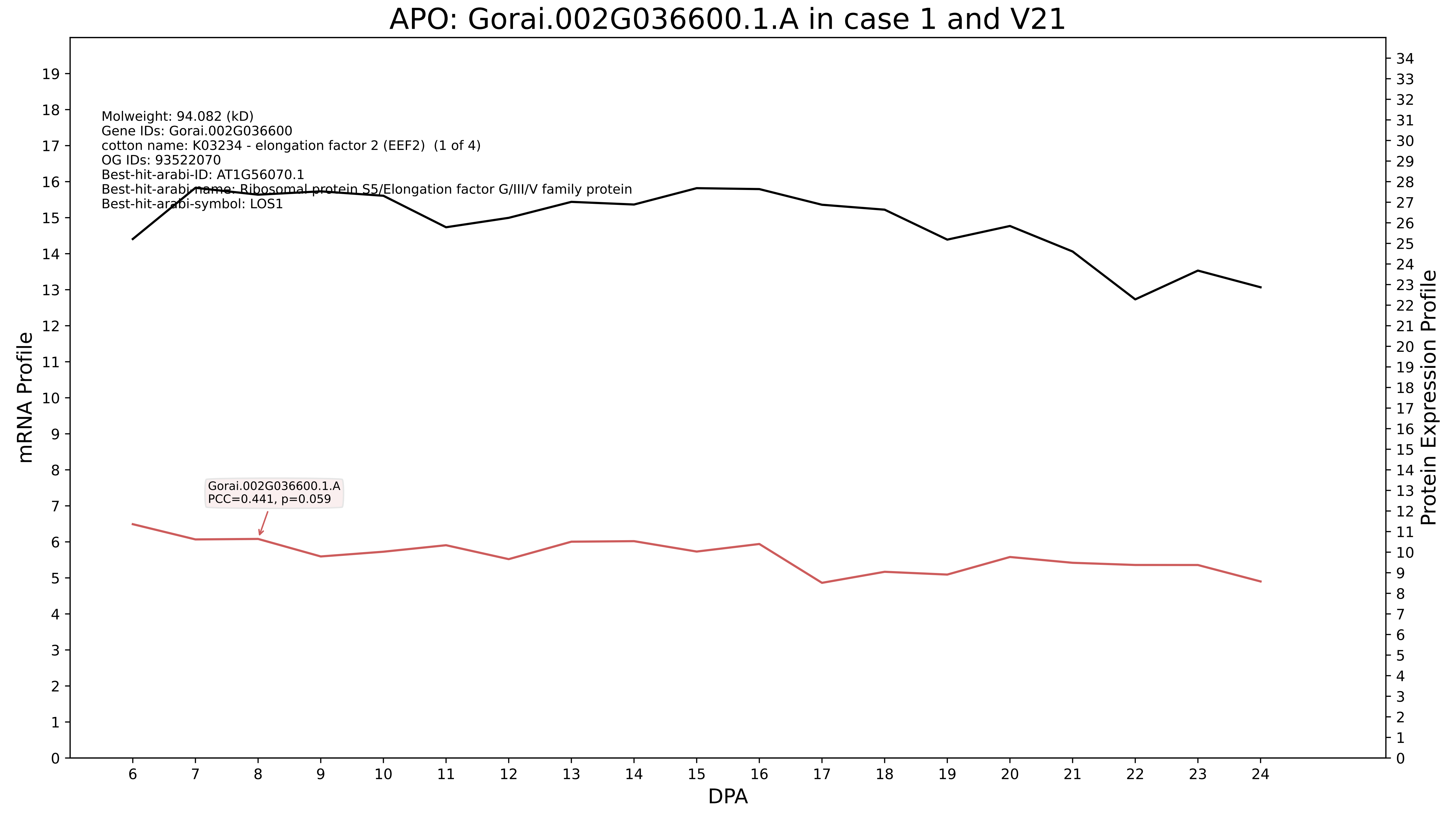

APO: Gorai.002G036600.1.A in case 1 and V21
Molweight: 94.082 (kD)
Gene IDs: Gorai.002G036600
cotton name: K03234 - elongation factor 2 (EEF2) (1 of 4)
OG IDs: 93522070
Best-hit-arabi-ID: AT1G56070.1
Best-hit-arabi-name: Ribosomal protein S5/Elongation factor G/III/V family protein
Best-hit-arabi-symbol: LOS1
Gorai.002G036600.1.A
PCC=0.441, p=0.059
mRNA Profile
Protein Expression Profile
DPA

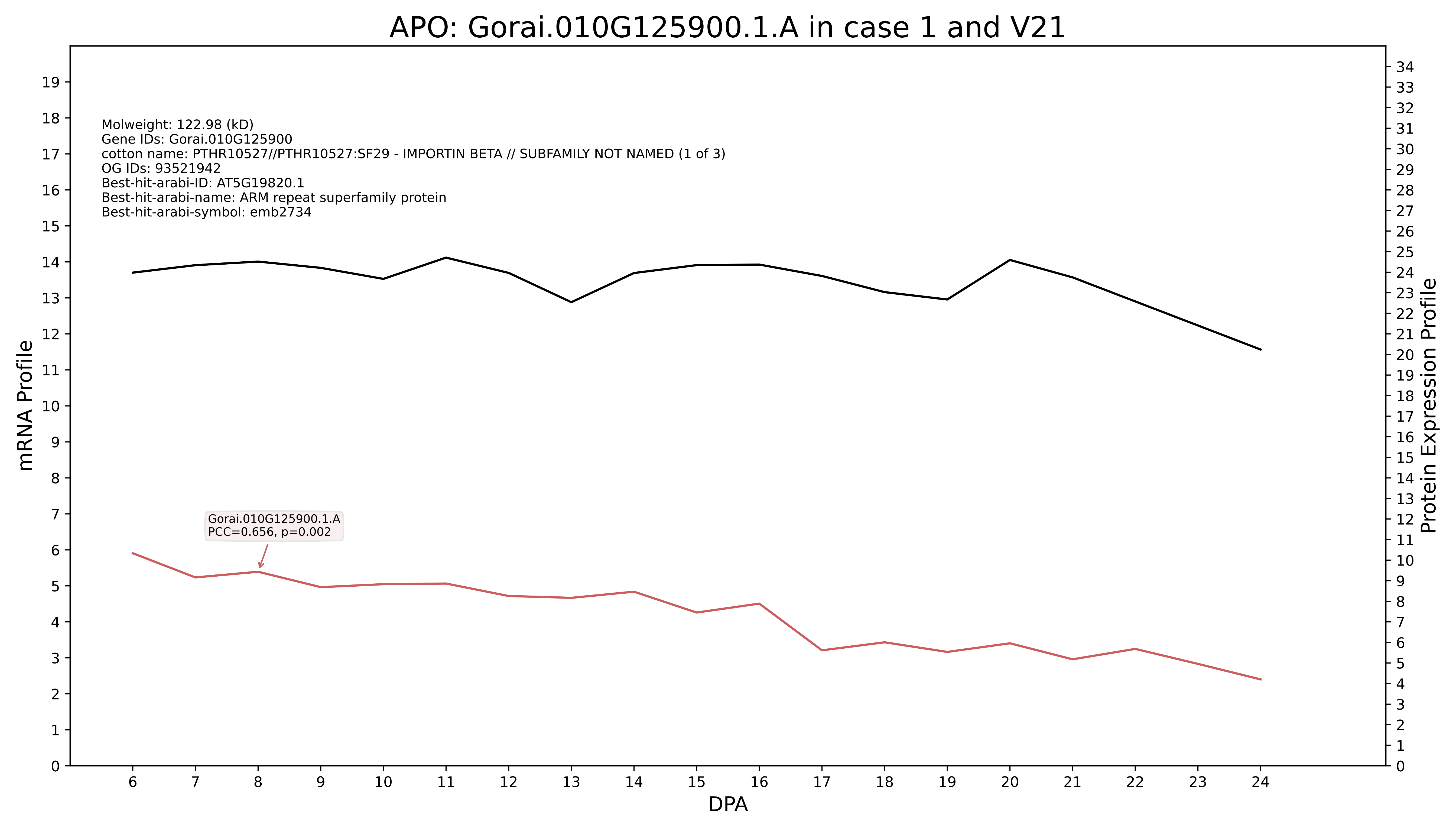
APO: Gorai.010G125900.1.A in case 1 and V21
Molweight: 122.98 (kD)
Gene IDs: Gorai.010G125900
cotton name: PTHR10527//PTHR10527:SF29 - IMPORTIN BETA // SUBFAMILY NOT NAMED (1 of 3)
OG IDs: 93521942
Best-hit-arabi-ID: AT5G19820.1
Best-hit-arabi-name: ARM repeat superfamily protein
Best-hit-arabi-symbol: emb2734
Gorai.010G125900.1.A
PCC=0.656, p=0.002
mRNA Profile
Protein Expression Profile
DPA

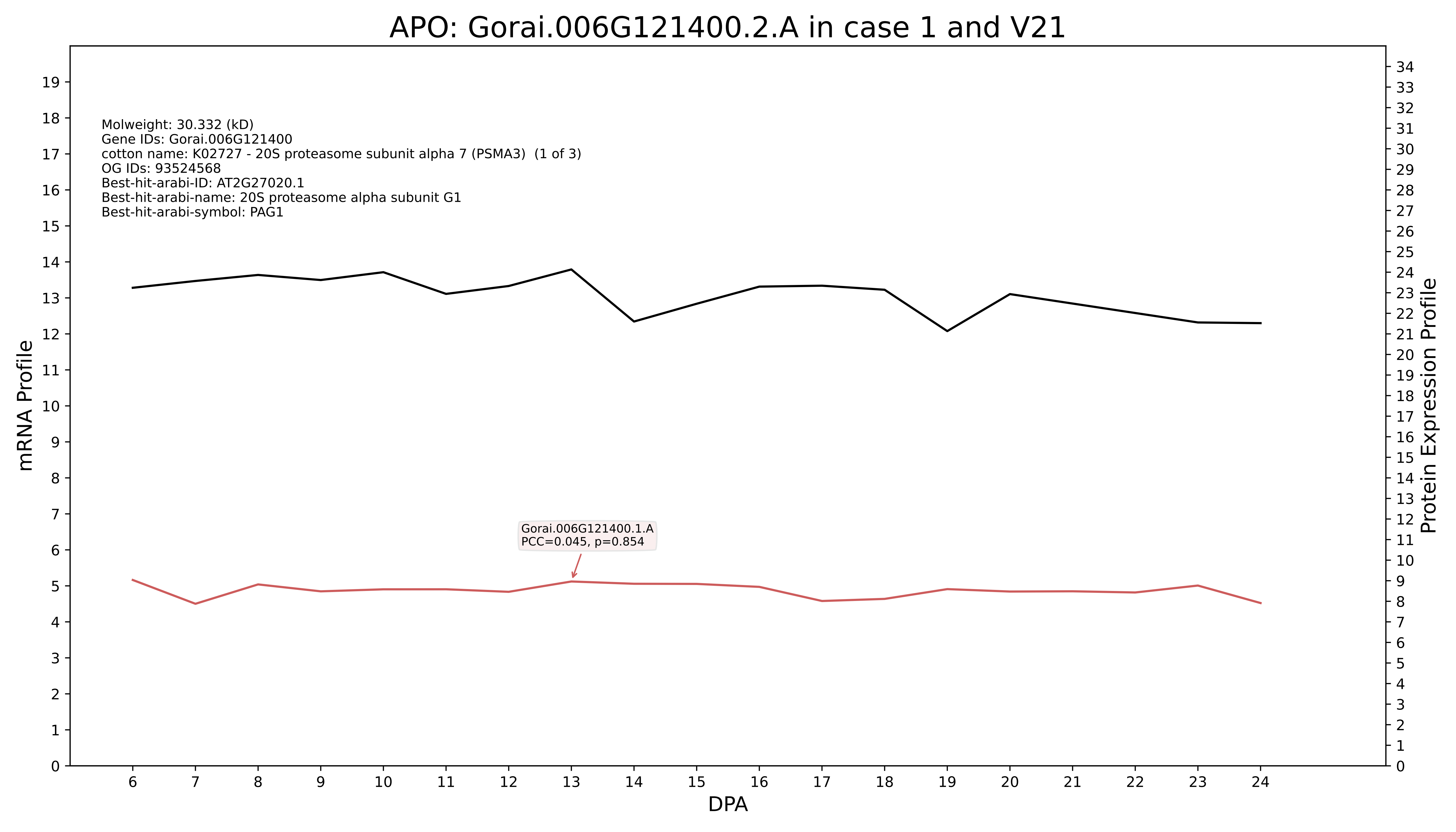

APO: Gorai.006G121400.2.A in case 1 and V21
Molweight: 30.332 (kD)
Gene IDs: Gorai.006G121400
cotton name: K02727 - 20S proteasome subunit alpha 7 (PSMA3) (1 of 3)
OG IDs: 93524568
Best-hit-arabi-ID: AT2G27020.1
Best-hit-arabi-name: 20S proteasome alpha subunit G1
Best-hit-arabi-symbol: PAG1
Gorai.006G121400.1.A
PCC=0.045, p=0.854
mRNA Profile
Protein Expression Profile
DPA

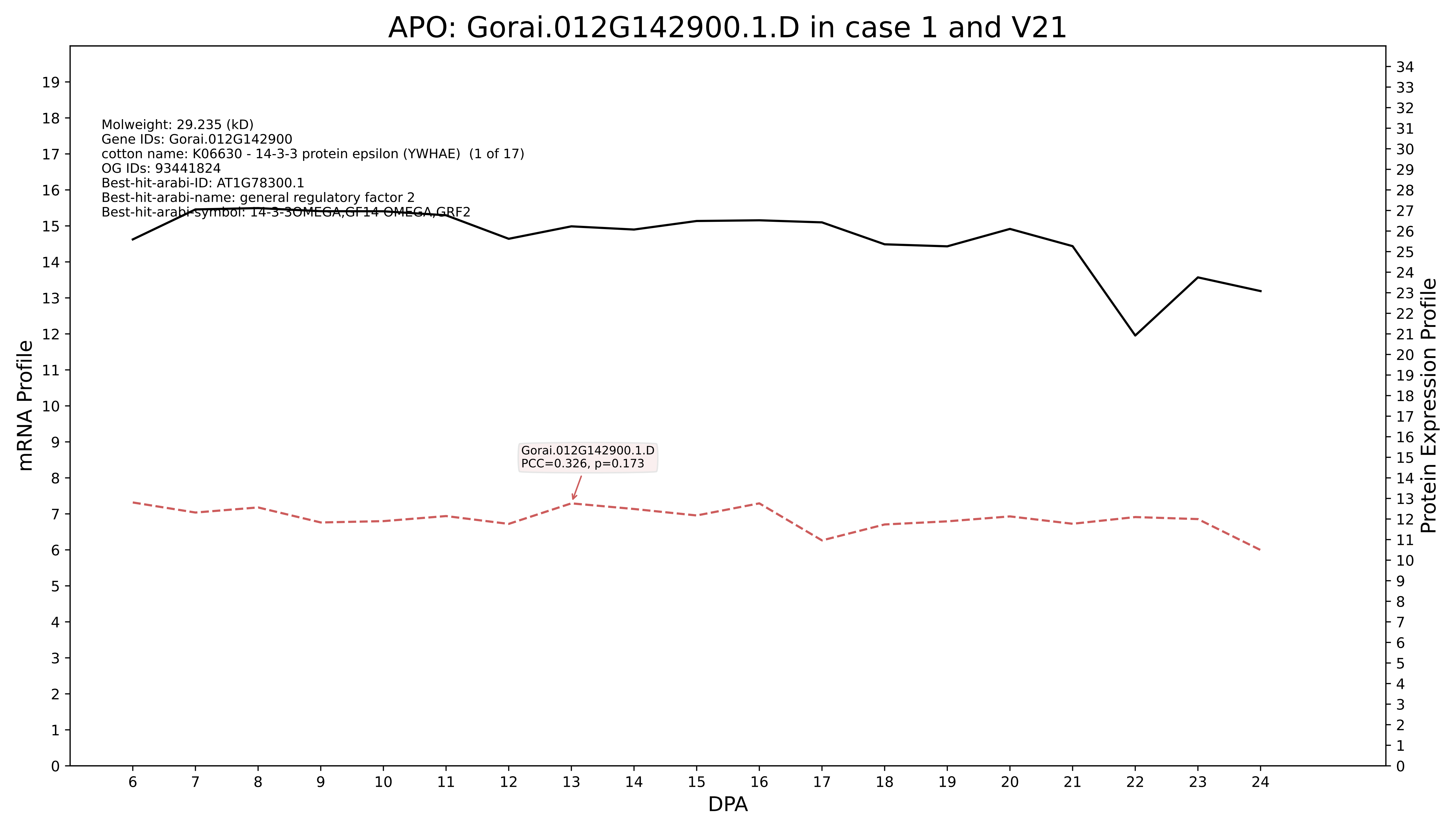
APO: Gorai.012G142900.1.D in case 1 and V21
Molweight: 29.235 (kD)
Gene IDs: Gorai.012G142900
cotton name: K06630 - 14-3-3 protein epsilon (YWHAE) (1 of 17)
OG IDs: 93441824
Best-hit-arabi-ID: AT1G78300.1
Best-hit-arabi-name: general regulatory factor 2
Best-hit-arabi-symbol: 14-3-3OMEGA,GF14 OMEGA,GRF2
Gorai.012G142900.1.D
PCC=0.326, p=0.173
mRNA Profile
Protein Expression Profile
DPA

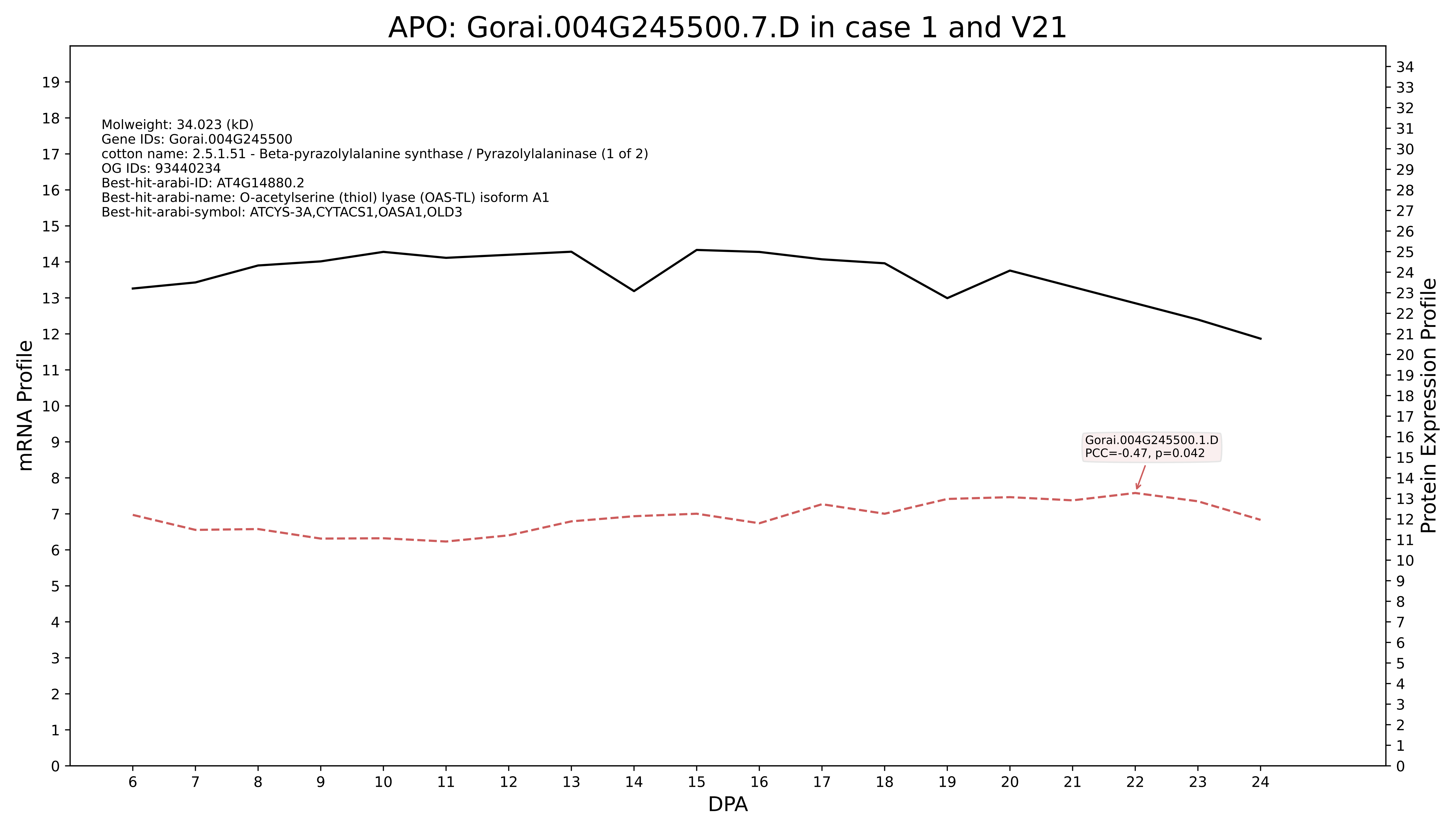

APO: Gorai.004G245500.7.D in case 1 and V21
Molweight: 34.023 (kD)
Gene IDs: Gorai.004G245500
cotton name: 2.5.1.51 - Beta-pyrazolylalanine synthase / Pyrazolylalaninase (1 of 2)
OG IDs: 93440234
Best-hit-arabi-ID: AT4G14880.2
Best-hit-arabi-name: O-acetylserine (thiol) lyase (OAS-TL) isoform A1
Best-hit-arabi-symbol: ATCYS-3A,CYTACS1,OASA1,OLD3
Gorai.004G245500.1.D
PCC=-0.47, p=0.042
mRNA Profile
Protein Expression Profile
DPA

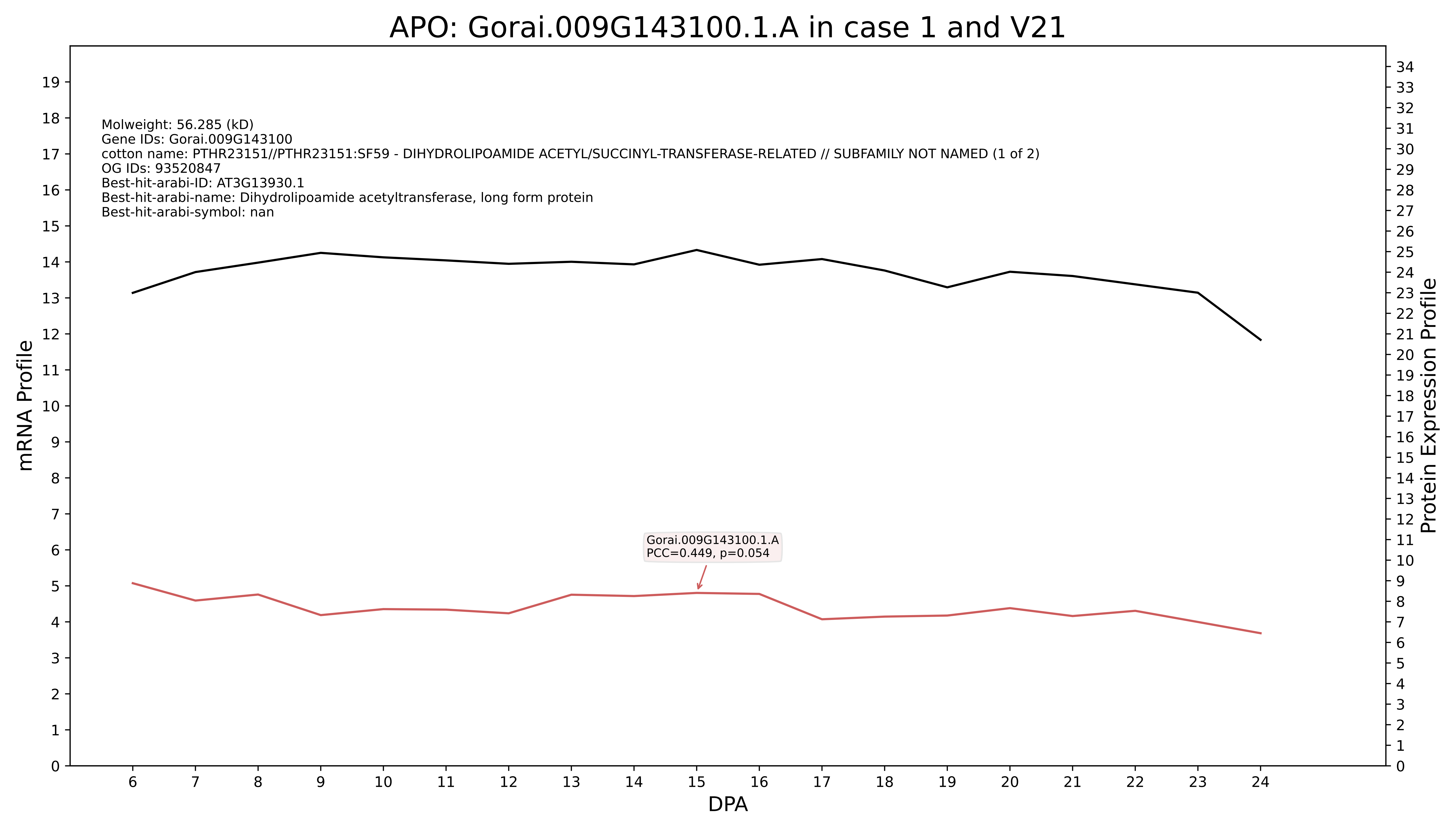
APO: Gorai.009G143100.1.A in case 1 and V21
Molweight: 56.285 (kD)
Gene IDs: Gorai.009G143100
cotton name: PTHR23151//PTHR23151:SF59 - DIHYDROLIPOAMIDE ACETYL/SUCCINYL-TRANSFERASE-RELATED // SUBFAMILY NOT NAMED (1 of 2)
OG IDs: 93520847
Best-hit-arabi-ID: AT3G13930.1
Best-hit-arabi-name: Dihydrolipoamide acetyltransferase, long form protein
Best-hit-arabi-symbol: nan
Gorai.009G143100.1.A
PCC=0.449, p=0.054
mRNA Profile
Protein Expression Profile
DPA

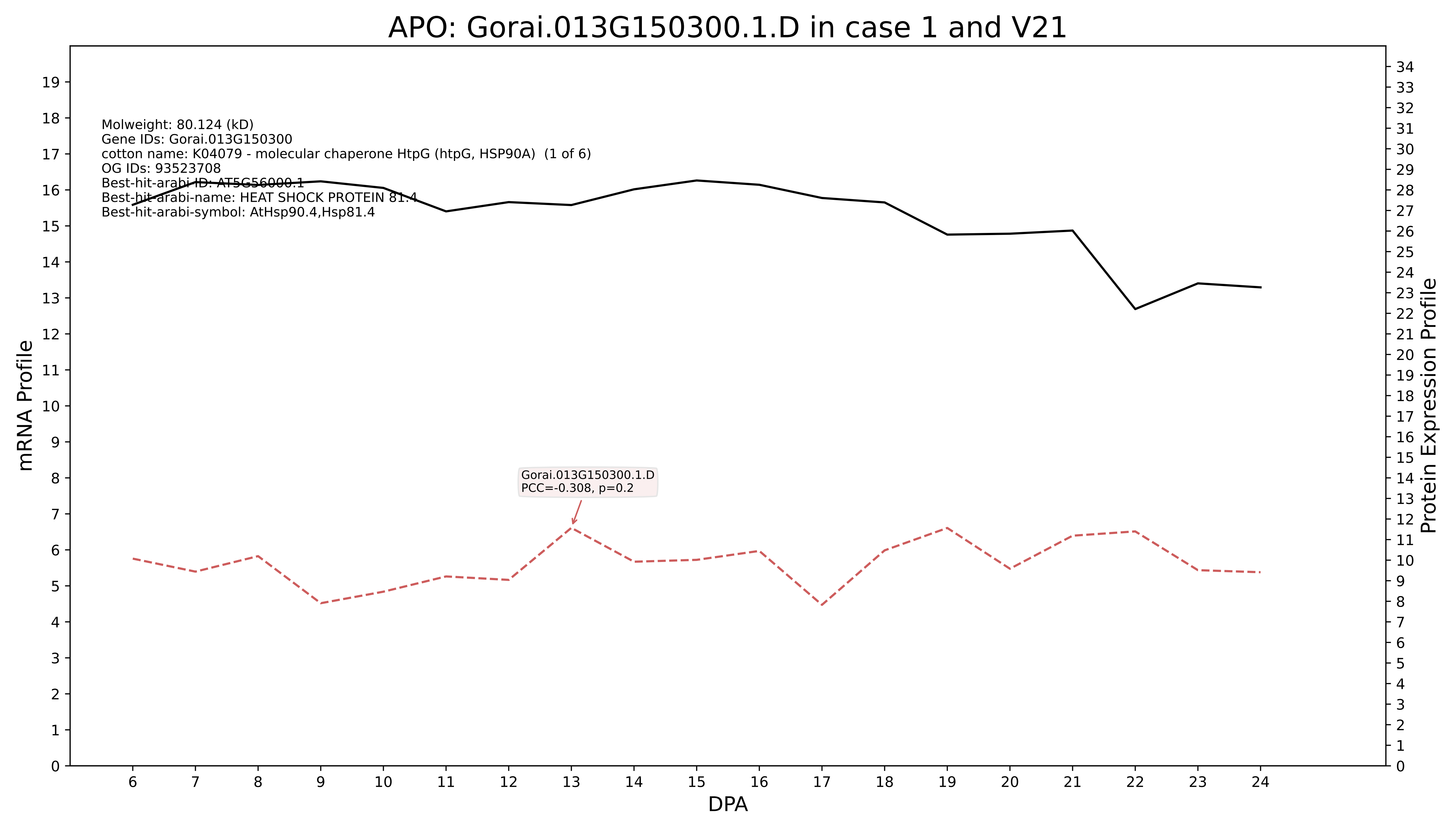
APO: Gorai.013G150300.1.D in case 1 and V21
Molweight: 80.124 (kD)
Gene IDs: Gorai.013G150300
cotton name: K04079 - molecular chaperone HtpG (htpG, HSP90A) (1 of 6)
OG IDs: 93523708
Best-hit-arabi-ID: AT5G56000.1
Best-hit-arabi-name: HEAT SHOCK PROTEIN 81.4
Best-hit-arabi-symbol: AtHsp90.4,Hsp81.4
Gorai.013G150300.1.D
PCC=-0.308, p=0.2
mRNA Profile
Protein Expression Profile
DPA

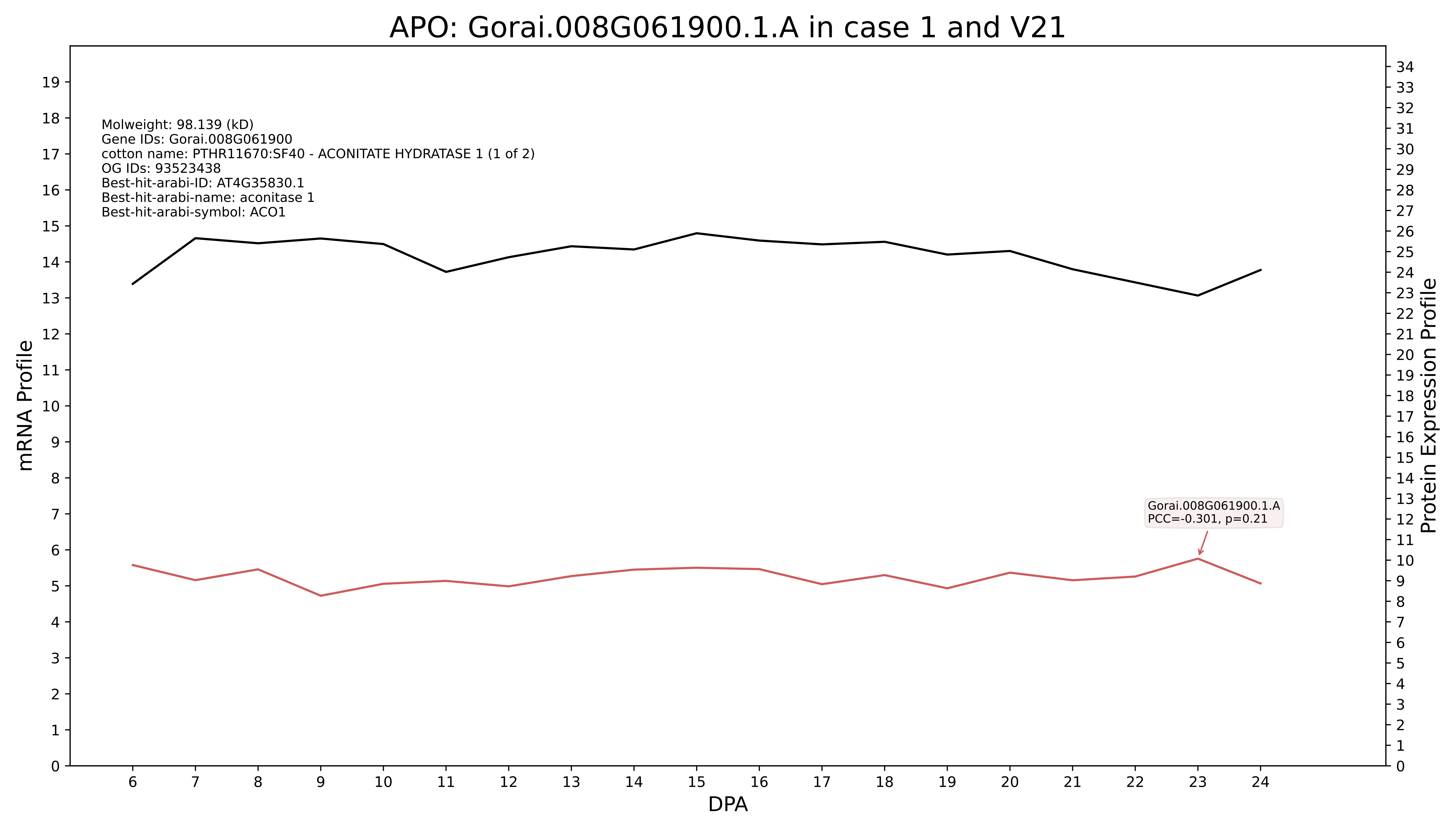

APO: Gorai.008G061900.1.A in case 1 and V21
Molweight: 98.139 (kD)
Gene IDs: Gorai.008G061900
cotton name: PTHR11670:SF40 - ACONITATE HYDRATASE 1 (1 of 2)
OG IDs: 93523438
Best-hit-arabi-ID: AT4G35830.1
Best-hit-arabi-name: aconitase 1
Best-hit-arabi-symbol: ACO1
Gorai.008G061900.1.A
PCC=-0.301, p=0.21
mRNA Profile
Protein Expression Profile
DPA

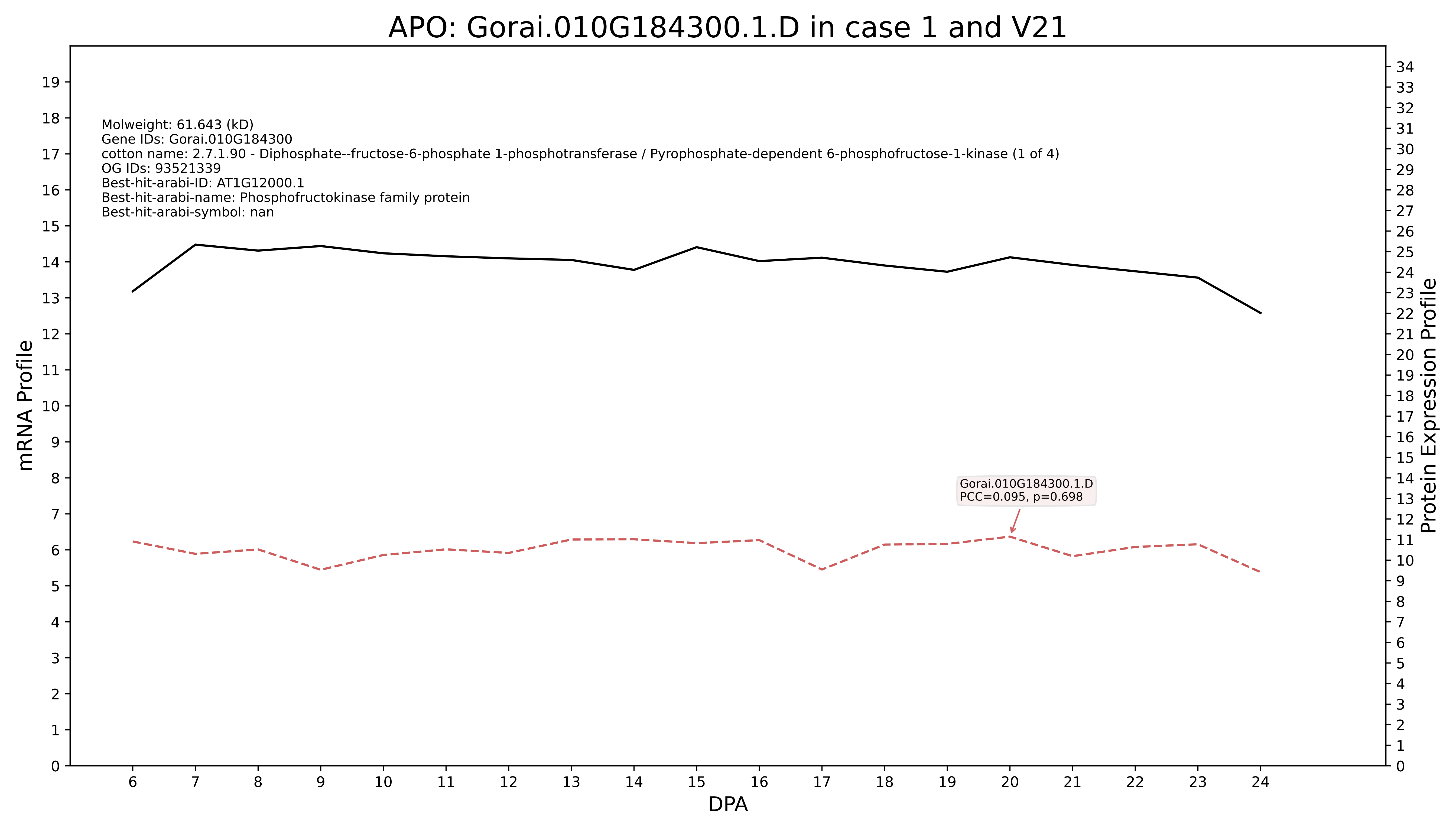

APO: Gorai.010G184300.1.D in case 1 and V21
Molweight: 61.643 (kD)
Gene IDs: Gorai.010G184300
cotton name: 2.7.1.90 - Diphosphate--fructose-6-phosphate 1-phosphotransferase / Pyrophosphate-dependent 6-phosphofructose-1-kinase (1 of 4)
OG IDs: 93521339
Best-hit-arabi-ID: AT1G12000.1
Best-hit-arabi-name: Phosphofructokinase family protein
Best-hit-arabi-symbol: nan
Gorai.010G184300.1.D
PCC=0.095, p=0.698
mRNA Profile
Protein Expression Profile
DPA

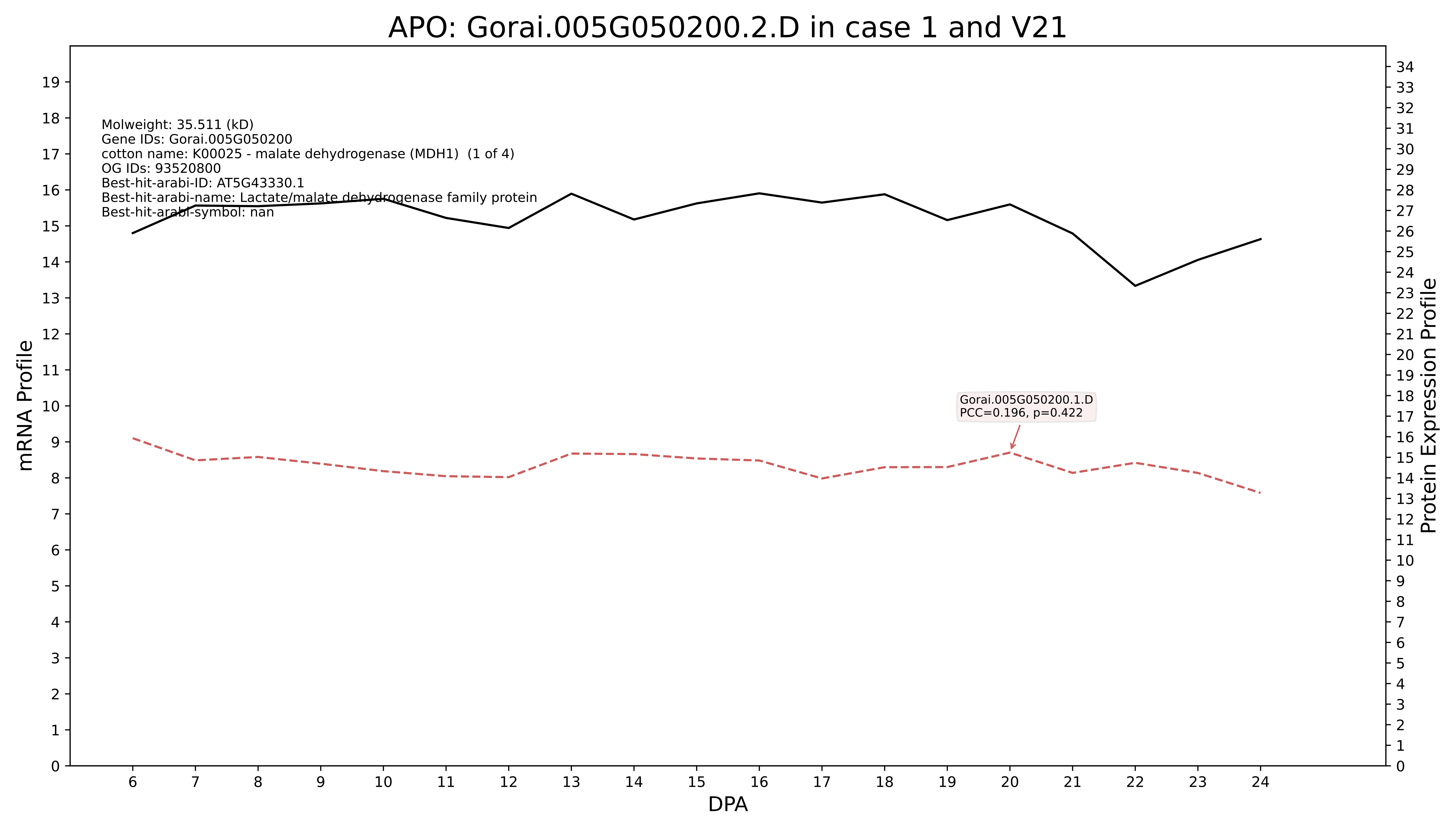
APO: Gorai.005G050200.2.D in case 1 and V21
Molweight: 35.511 (kD)
Gene IDs: Gorai.005G050200
cotton name: K00025 - malate dehydrogenase (MDH1) (1 of 4)
OG IDs: 93520800
Best-hit-arabi-ID: AT5G43330.1
Best-hit-arabi-name: Lactate/malate dehydrogenase family protein
Best-hit-arabi-symbol: nan
Gorai.005G050200.1.D
PCC=0.196, p=0.422
mRNA Profile
Protein Expression Profile
DPA

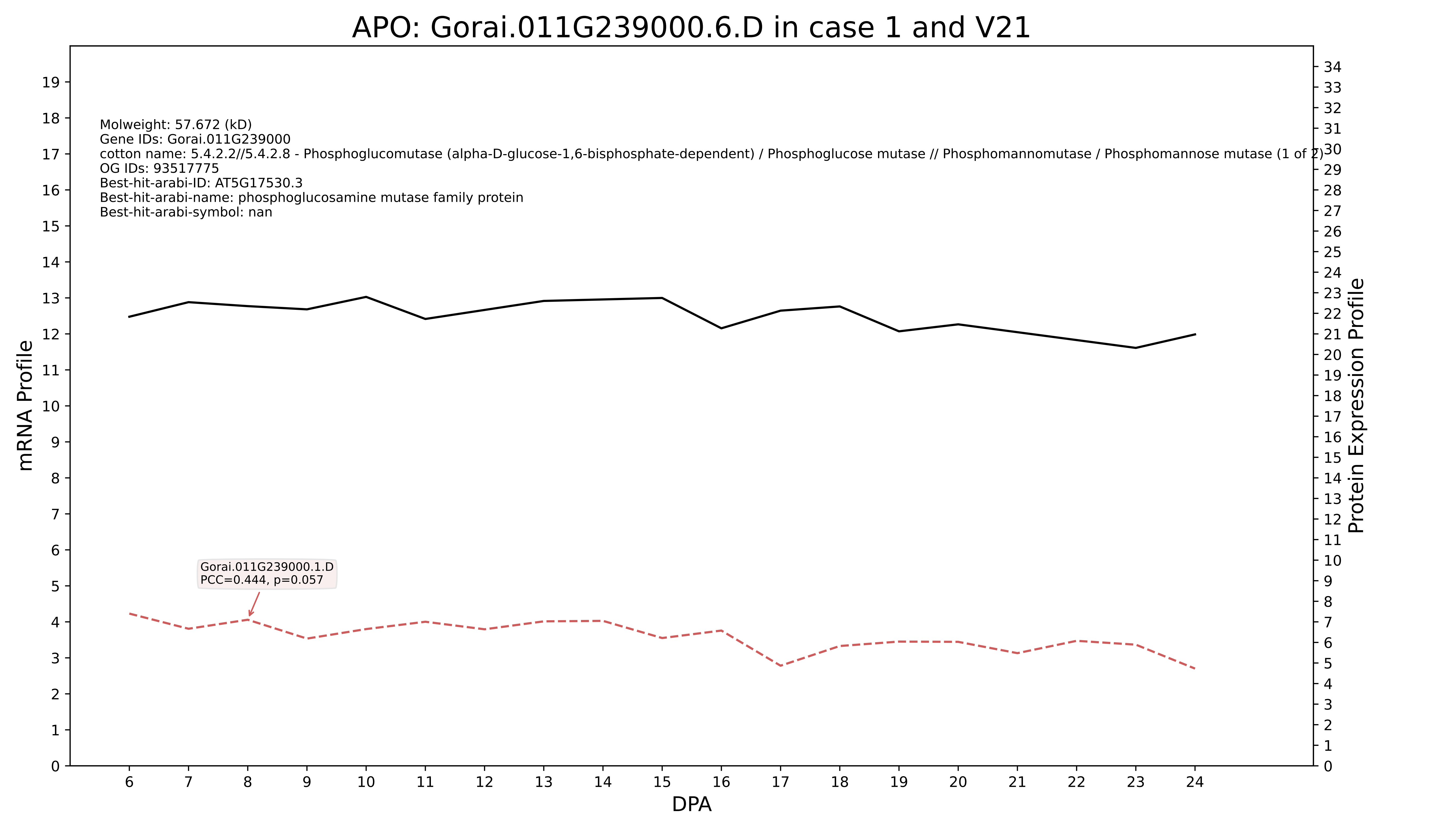

APO: Gorai.011G239000.6.D in case 1 and V21
Molweight: 57.672 (kD)
Gene IDs: Gorai.011G239000
cotton name: 5.4.2.2//5.4.2.8 - Phosphoglucomutase (alpha-D-glucose-1,6-bisphosphate-dependent) / Phosphoglucose mutase // Phosphomannomutase / Phosphomannose mutase (1 of 2)
OG IDs: 93517775
Best-hit-arabi-ID: AT5G17530.3
Best-hit-arabi-name: phosphoglucosamine mutase family protein
Best-hit-arabi-symbol: nan
Gorai.011G239000.1.D
PCC=0.444, p=0.057
mRNA Profile
Protein Expression Profile
DPA

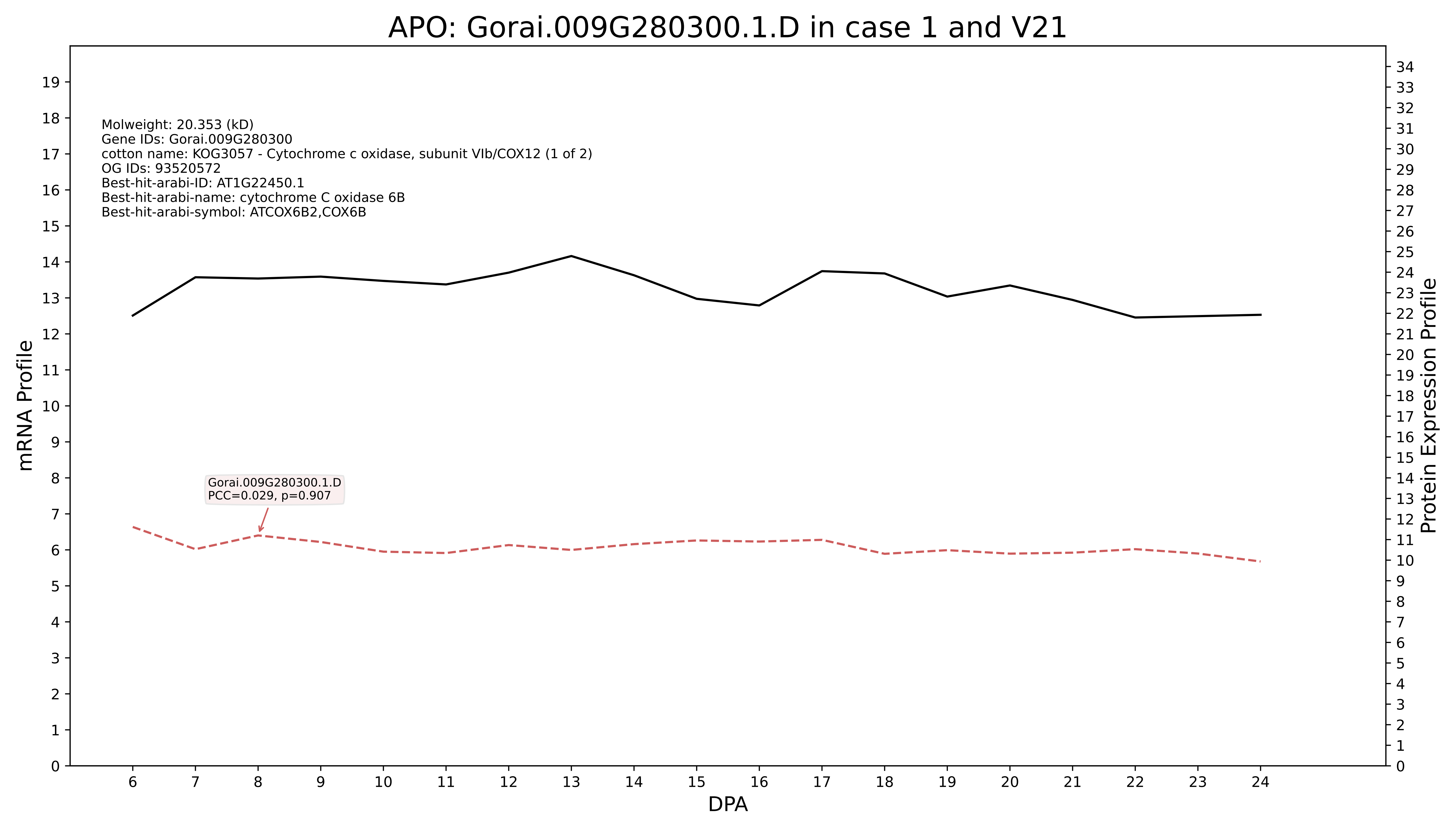

APO: Gorai.009G280300.1.D in case 1 and V21
Molweight: 20.353 (kD)
Gene IDs: Gorai.009G280300
cotton name: KOG3057 - Cytochrome c oxidase, subunit VIb/COX12 (1 of 2)
OG IDs: 93520572
Best-hit-arabi-ID: AT1G22450.1
Best-hit-arabi-name: cytochrome C oxidase 6B
Best-hit-arabi-symbol: ATCOX6B2,COX6B
Gorai.009G280300.1.D
PCC=0.029, p=0.907
mRNA Profile
Protein Expression Profile
DPA

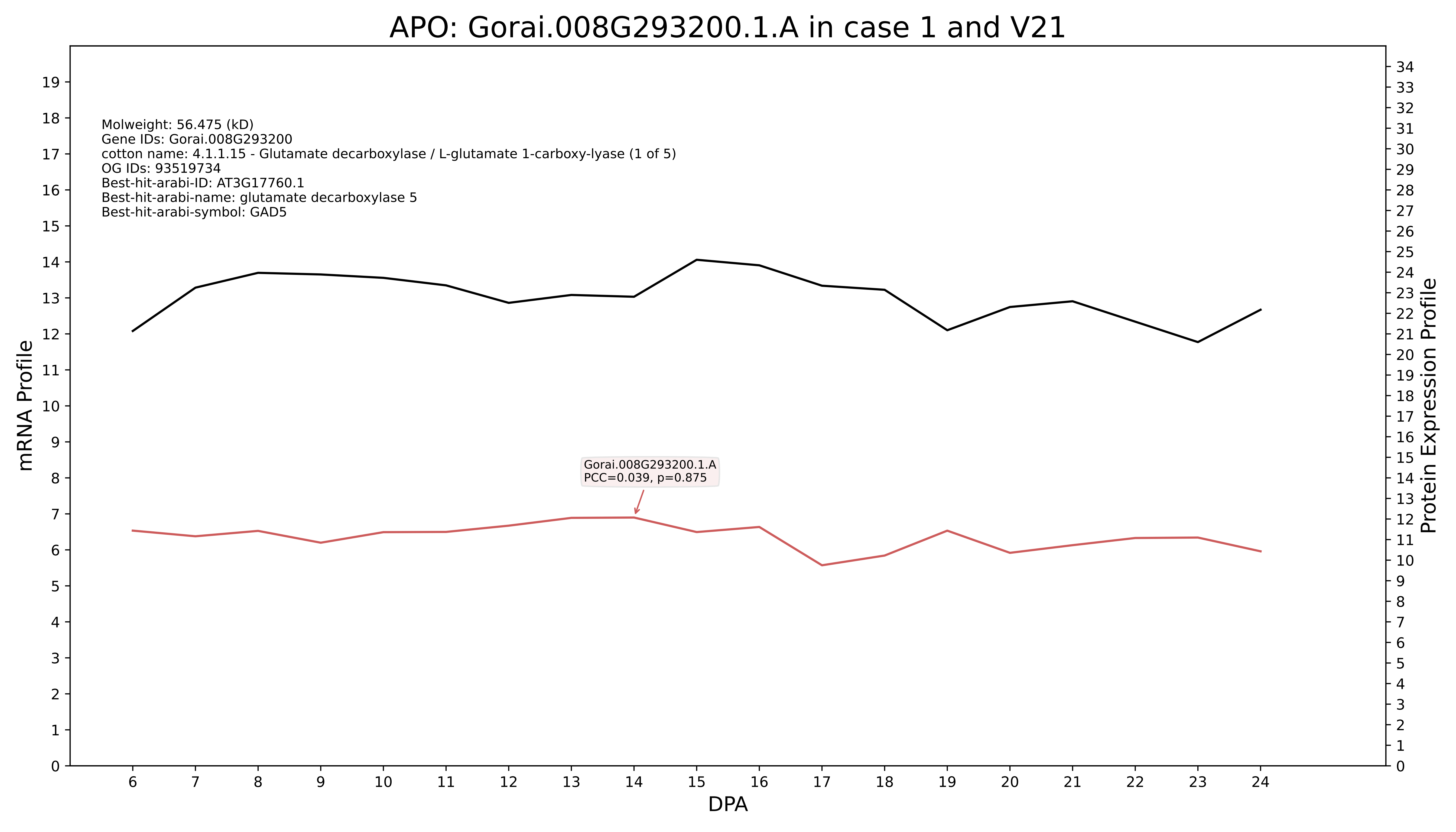

APO: Gorai.008G293200.1.A in case 1 and V21
Molweight: 56.475 (kD)
Gene IDs: Gorai.008G293200
cotton name: 4.1.1.15 - Glutamate decarboxylase / L-glutamate 1-carboxy-lyase (1 of 5)
OG IDs: 93519734
Best-hit-arabi-ID: AT3G17760.1
Best-hit-arabi-name: glutamate decarboxylase 5
Best-hit-arabi-symbol: GAD5
Gorai.008G293200.1.A
PCC=0.039, p=0.875
mRNA Profile
Protein Expression Profile
DPA

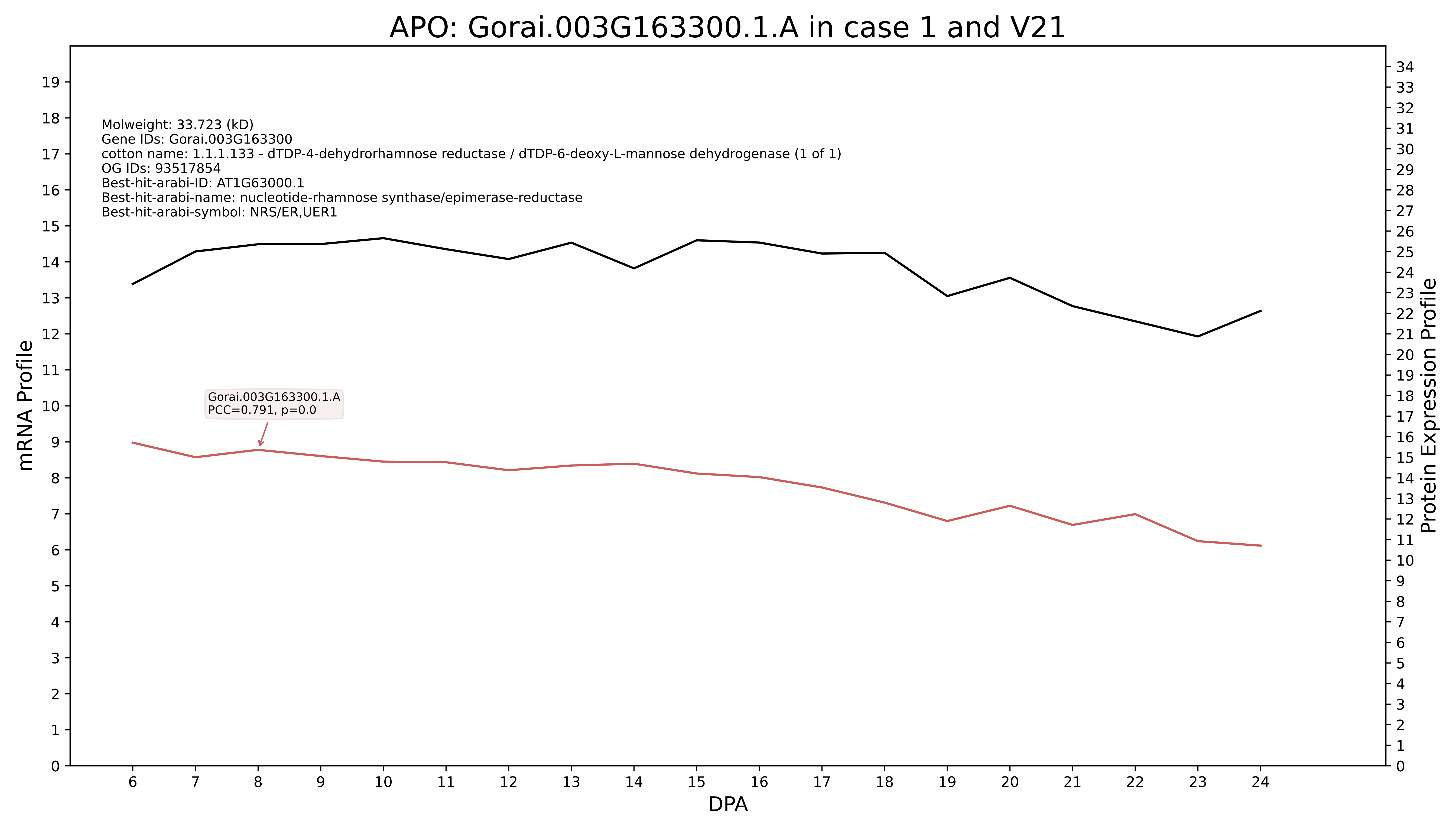
APO: Gorai.003G163300.1.A in case 1 and V21
Molweight: 33.723 (kD)
Gene IDs: Gorai.003G163300
cotton name: 1.1.1.133 - dTDP-4-dehydrorhamnose reductase / dTDP-6-deoxy-L-mannose dehydrogenase (1 of 1)
OG IDs: 93517854
Best-hit-arabi-ID: AT1G63000.1
Best-hit-arabi-name: nucleotide-rhamnose synthase/epimerase-reductase
Best-hit-arabi-symbol: NRS/ER,UER1
Gorai.003G163300.1.A
PCC=0.791, p=0.0
mRNA Profile
Protein Expression Profile
DPA

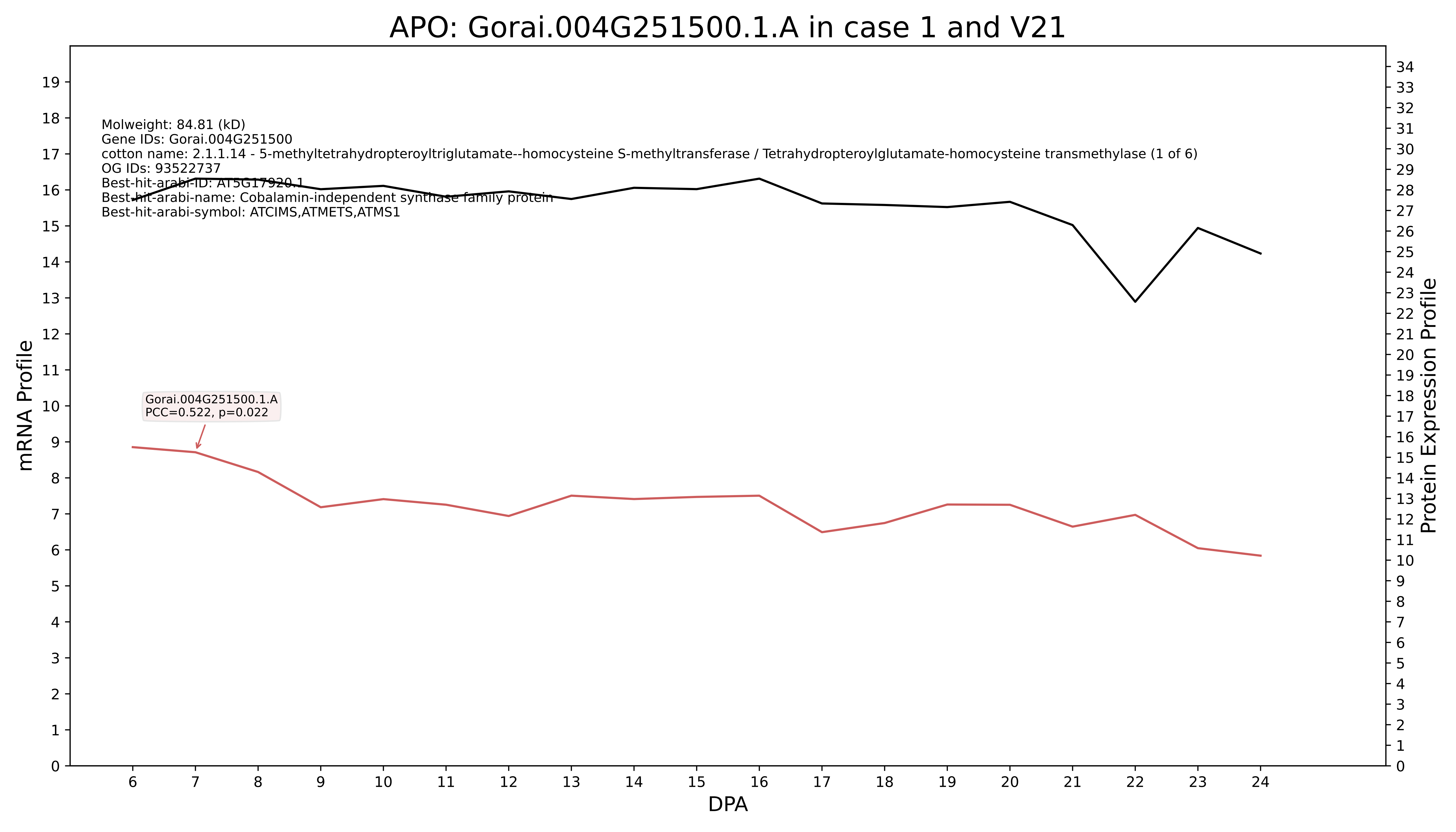
APO: Gorai.004G251500.1.A in case 1 and V21
Molweight: 84.81 (kD)
Gene IDs: Gorai.004G251500
cotton name: 2.1.1.14 - 5-methyltetrahydropteroyltriglutamate--homocysteine S-methyltransferase / Tetrahydropteroylglutamate-homocysteine transmethylase (1 of 6)
OG IDs: 93522737
Best-hit-arabi-ID: AT5G17920.1
Best-hit-arabi-name: Cobalamin-independent synthase family protein
Best-hit-arabi-symbol: ATCIMS,ATMETS,ATMS1
Gorai.004G251500.1.A
PCC=0.522, p=0.022
mRNA Profile
Protein Expression Profile
DPA

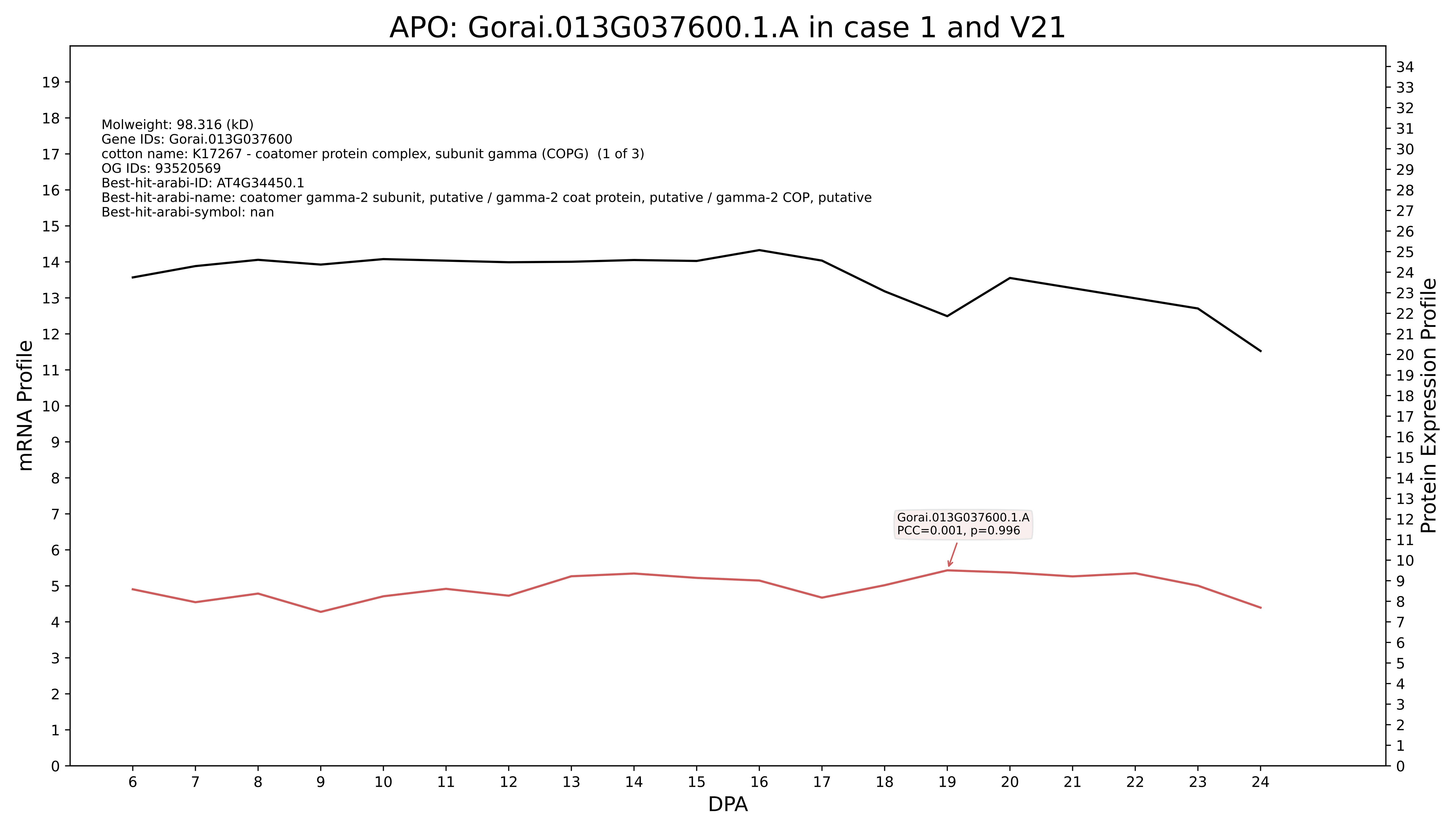
APO: Gorai.013G037600.1.A in case 1 and V21
Molweight: 98.316 (kD)
Gene IDs: Gorai.013G037600
cotton name: K17267 - coatomer protein complex, subunit gamma (COPG) (1 of 3)
OG IDs: 93520569
Best-hit-arabi-ID: AT4G34450.1
Best-hit-arabi-name: coatomer gamma-2 subunit, putative / gamma-2 coat protein, putative / gamma-2 COP, putative
Best-hit-arabi-symbol: nan
Gorai.013G037600.1.A
PCC=0.001, p=0.996
mRNA Profile
Protein Expression Profile
DPA

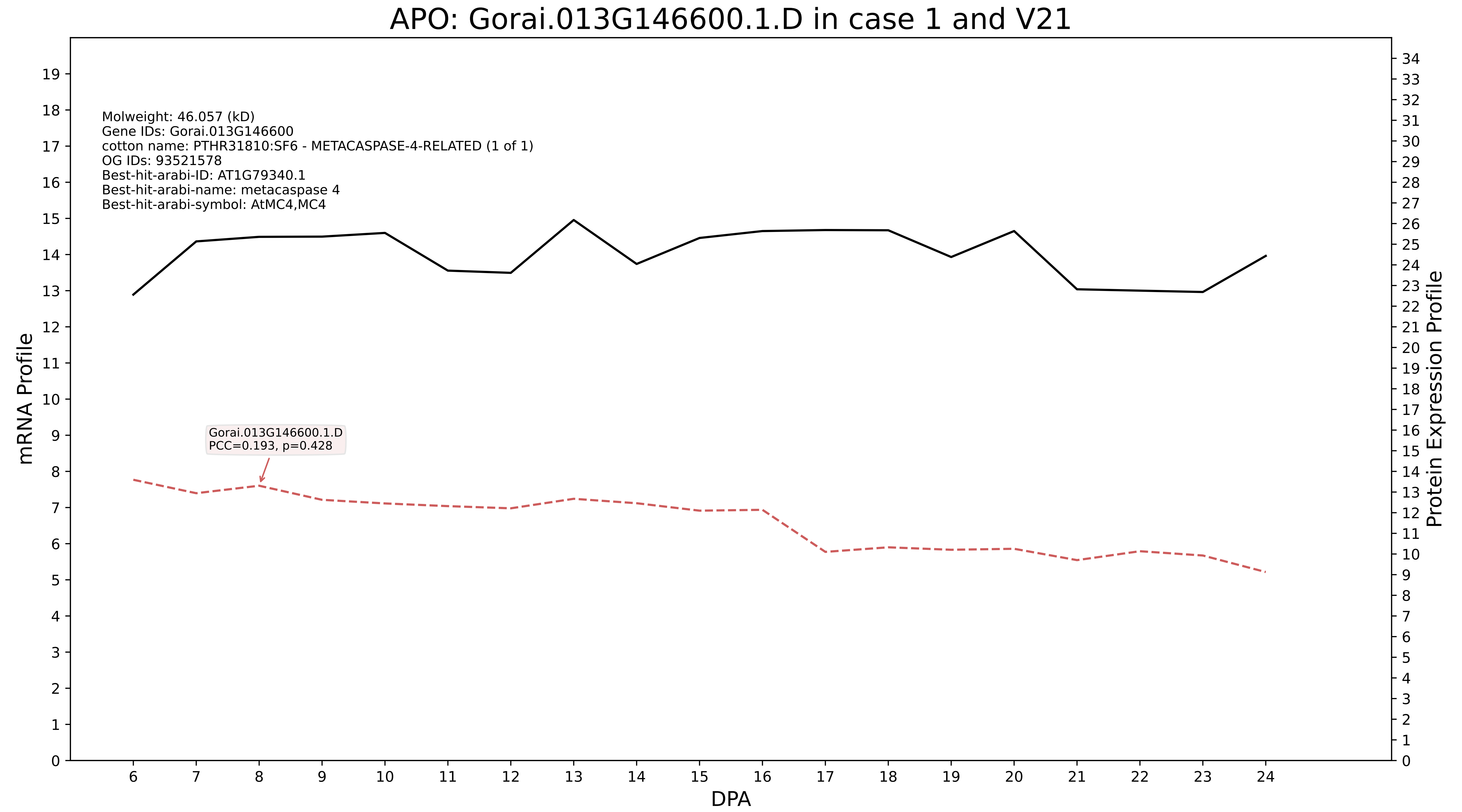

APO: Gorai.013G146600.1.D in case 1 and V21
Molweight: 46.057 (kD)
Gene IDs: Gorai.013G146600
cotton name: PTHR31810:SF6 - METACASPASE-4-RELATED (1 of 1)
OG IDs: 93521578
Best-hit-arabi-ID: AT1G79340.1
Best-hit-arabi-name: metacaspase 4
Best-hit-arabi-symbol: AtMC4,MC4
Gorai.013G146600.1.D
PCC=0.193, p=0.428
mRNA Profile
Protein Expression Profile
DPA

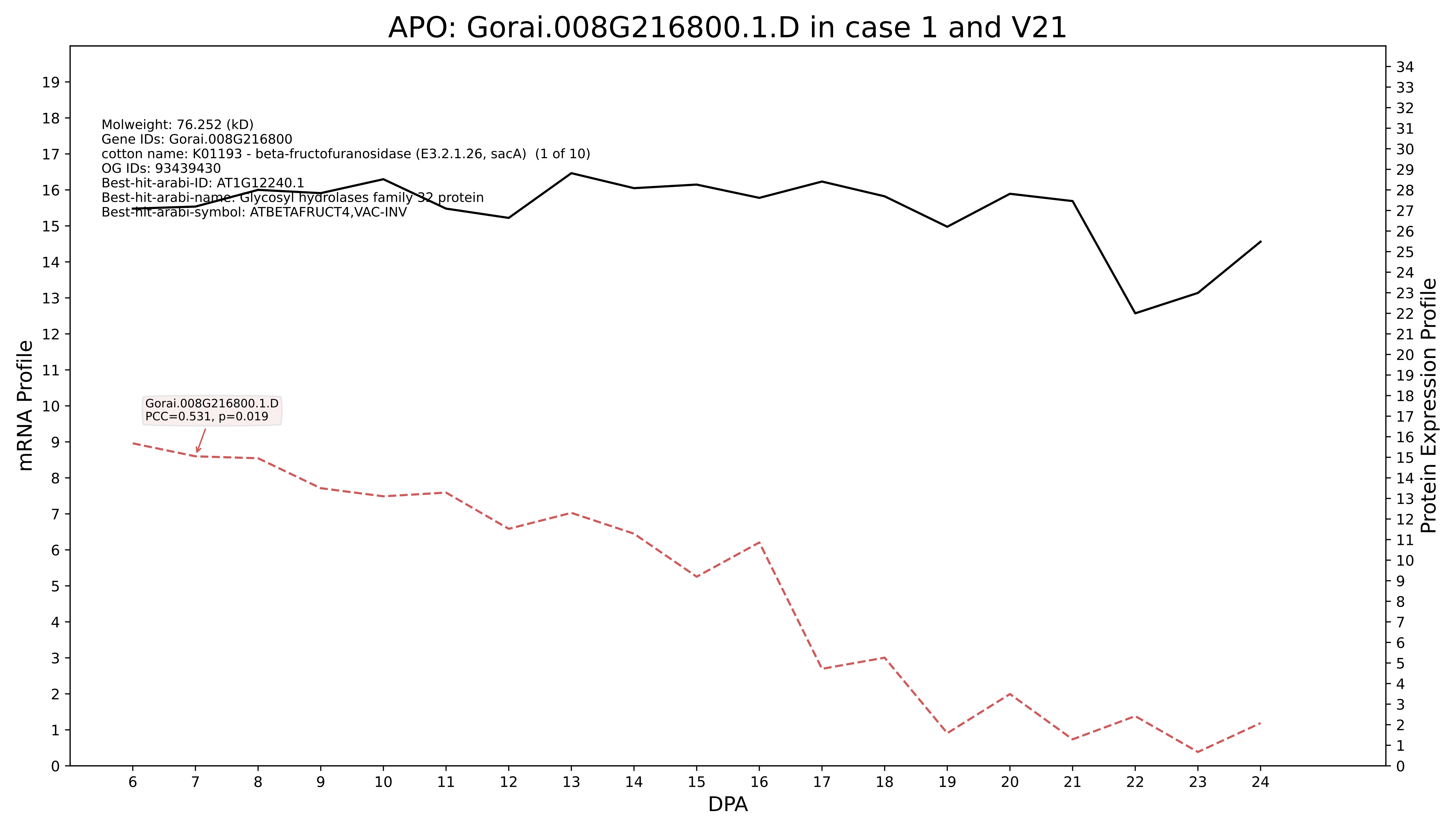
APO: Gorai.008G216800.1.D in case 1 and V21
Molweight: 76.252 (kD)
Gene IDs: Gorai.008G216800
cotton name: K01193 - beta-fructofuranosidase (E3.2.1.26, sacA) (1 of 10)
OG IDs: 93439430
Best-hit-arabi-ID: AT1G12240.1
Best-hit-arabi-name: Glycosyl hydrolases family 32 protein
Best-hit-arabi-symbol: ATBETAFRUCT4,VAC-INV
Gorai.008G216800.1.D
PCC=0.531, p=0.019
mRNA Profile
Protein Expression Profile
DPA

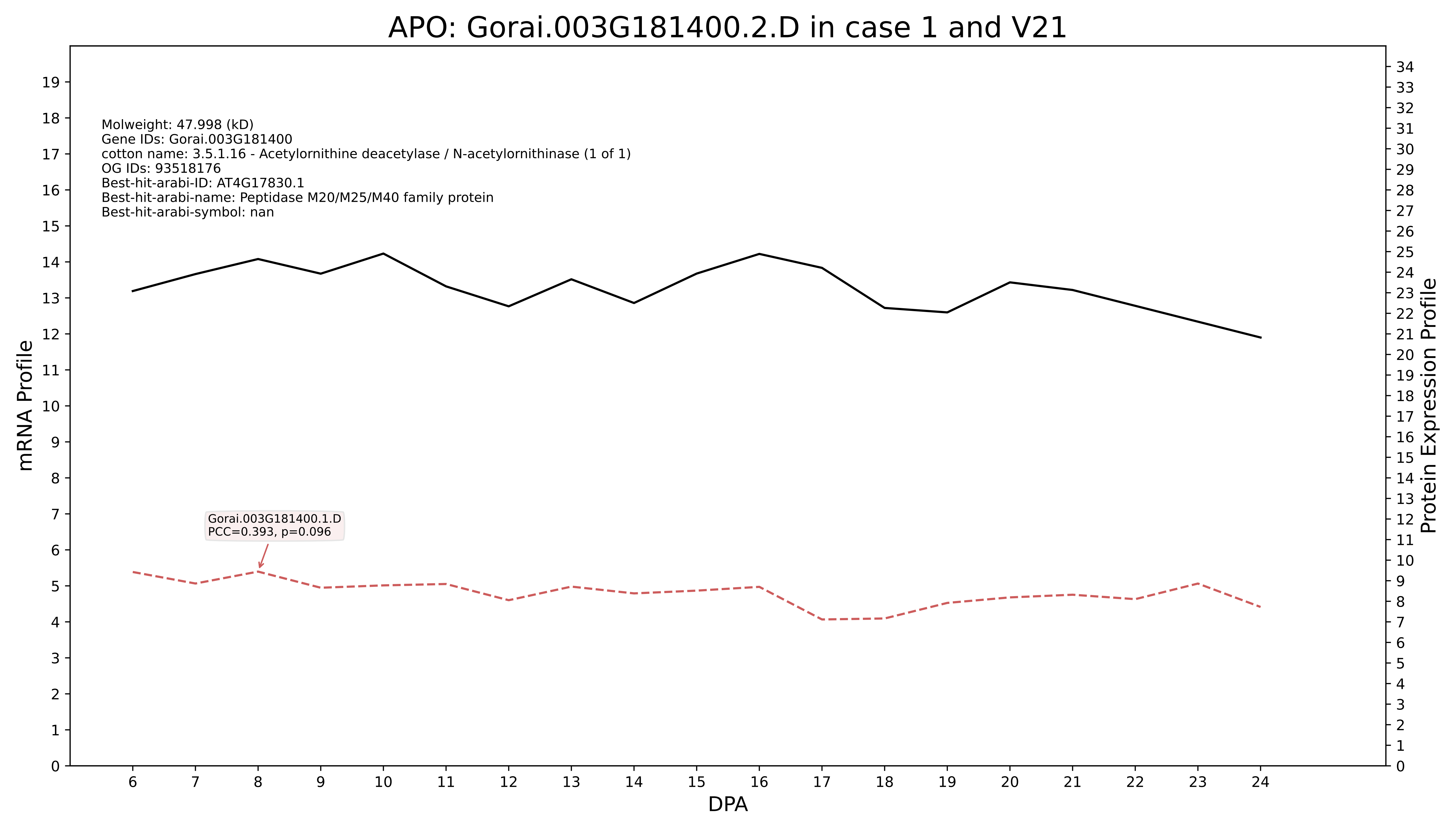

APO: Gorai.003G181400.2.D in case 1 and V21
Molweight: 47.998 (kD)
Gene IDs: Gorai.003G181400
cotton name: 3.5.1.16 - Acetylornithine deacetylase / N-acetylornithinase (1 of 1)
OG IDs: 93518176
Best-hit-arabi-ID: AT4G17830.1
Best-hit-arabi-name: Peptidase M20/M25/M40 family protein
Best-hit-arabi-symbol: nan
Gorai.003G181400.1.D
PCC=0.393, p=0.096
mRNA Profile
Protein Expression Profile
DPA

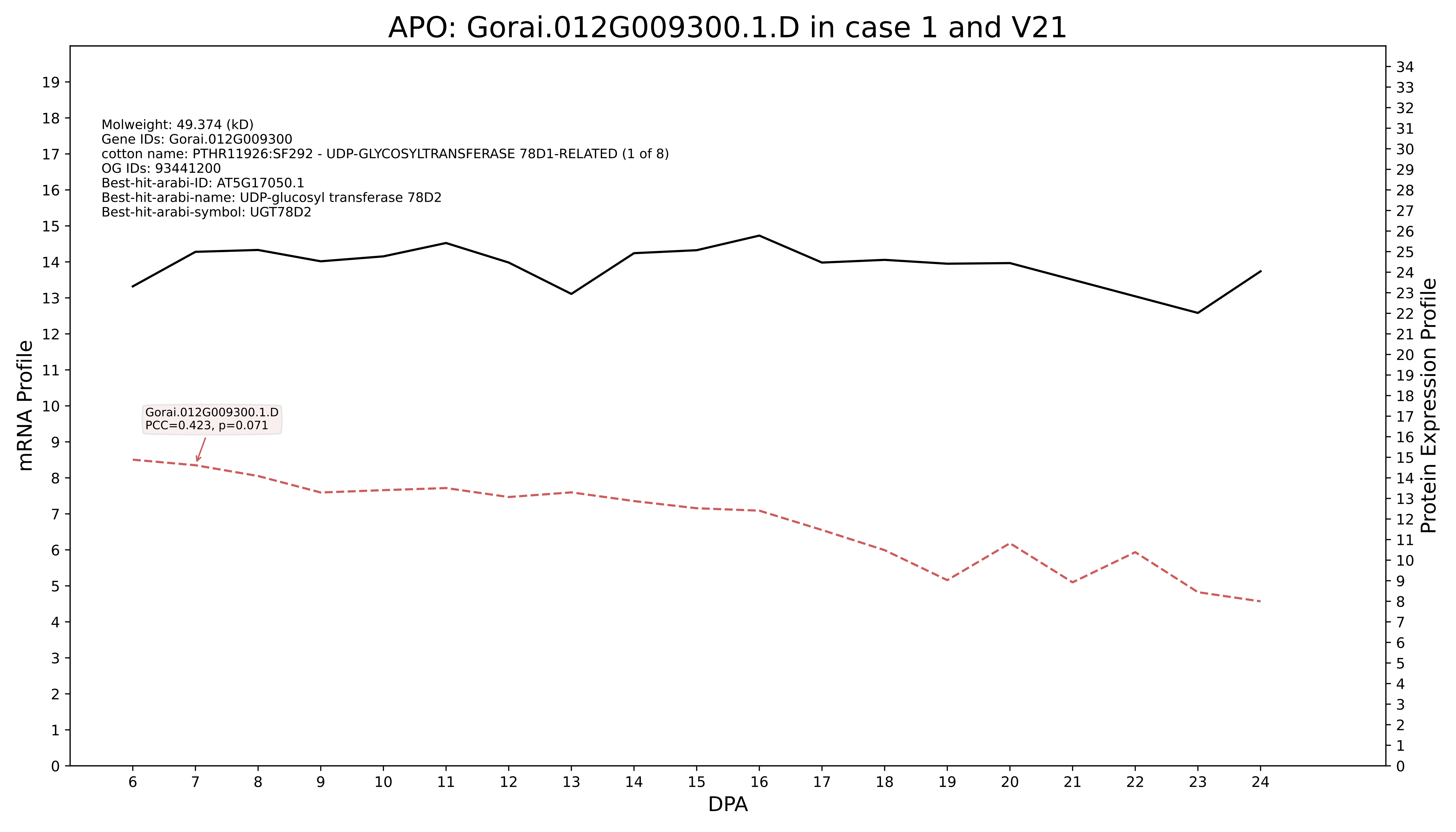
APO: Gorai.012G009300.1.D in case 1 and V21
Molweight: 49.374 (kD)
Gene IDs: Gorai.012G009300
cotton name: PTHR11926:SF292 - UDP-GLYCOSYLTRANSFERASE 78D1-RELATED (1 of 8)
OG IDs: 93441200
Best-hit-arabi-ID: AT5G17050.1
Best-hit-arabi-name: UDP-glucosyl transferase 78D2
Best-hit-arabi-symbol: UGT78D2
Gorai.012G009300.1.D
PCC=0.423, p=0.071
mRNA Profile
Protein Expression Profile
DPA

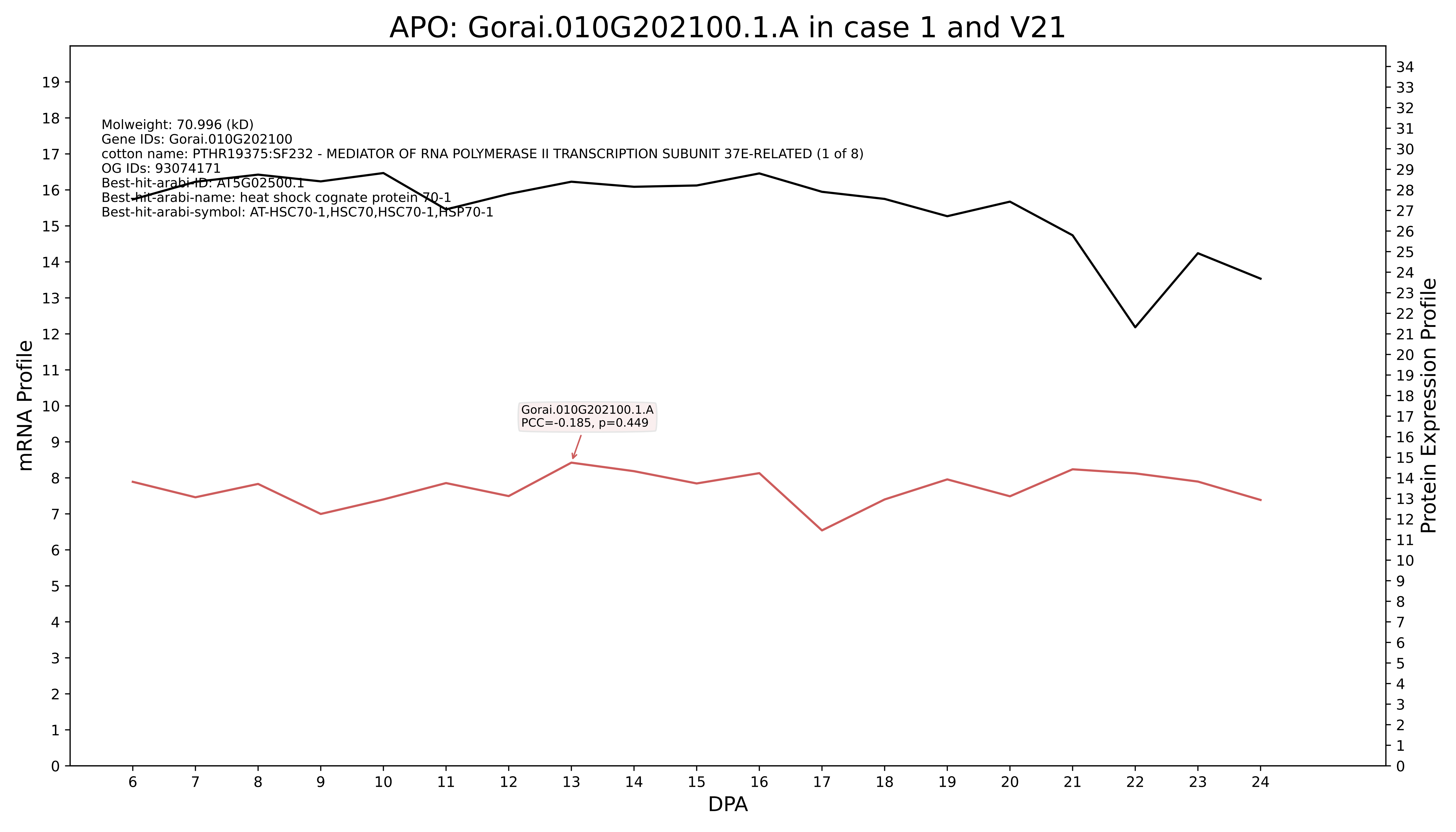
APO: Gorai.010G202100.1.A in case 1 and V21
Molweight: 70.996 (kD)
Gene IDs: Gorai.010G202100
cotton name: PTHR19375:SF232 - MEDIATOR OF RNA POLYMERASE II TRANSCRIPTION SUBUNIT 37E-RELATED (1 of 8)
OG IDs: 93074171
Best-hit-arabi-ID: AT5G02500.1
Best-hit-arabi-name: heat shock cognate protein 70-1
Best-hit-arabi-symbol: AT-HSC70-1,HSC70,HSC70-1,HSP70-1
Gorai.010G202100.1.A
PCC=-0.185, p=0.449
mRNA Profile
Protein Expression Profile
DPA

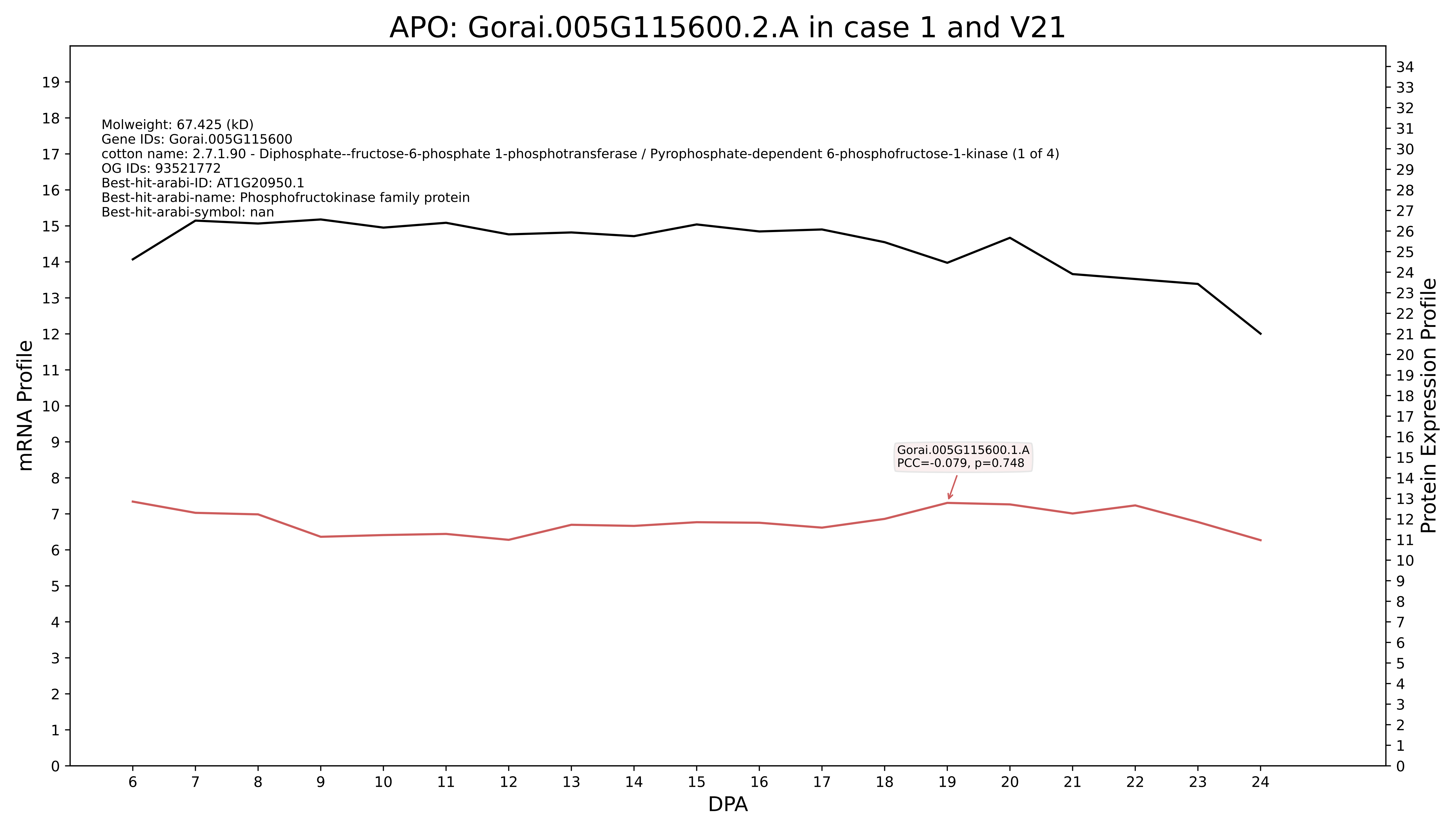
APO: Gorai.005G115600.2.A in case 1 and V21
Molweight: 67.425 (kD)
Gene IDs: Gorai.005G115600
cotton name: 2.7.1.90 - Diphosphate--fructose-6-phosphate 1-phosphotransferase / Pyrophosphate-dependent 6-phosphofructose-1-kinase (1 of 4)
OG IDs: 93521772
Best-hit-arabi-ID: AT1G20950.1
Best-hit-arabi-name: Phosphofructokinase family protein
Best-hit-arabi-symbol: nan
Gorai.005G115600.1.A
PCC=-0.079, p=0.748
mRNA Profile
Protein Expression Profile
DPA

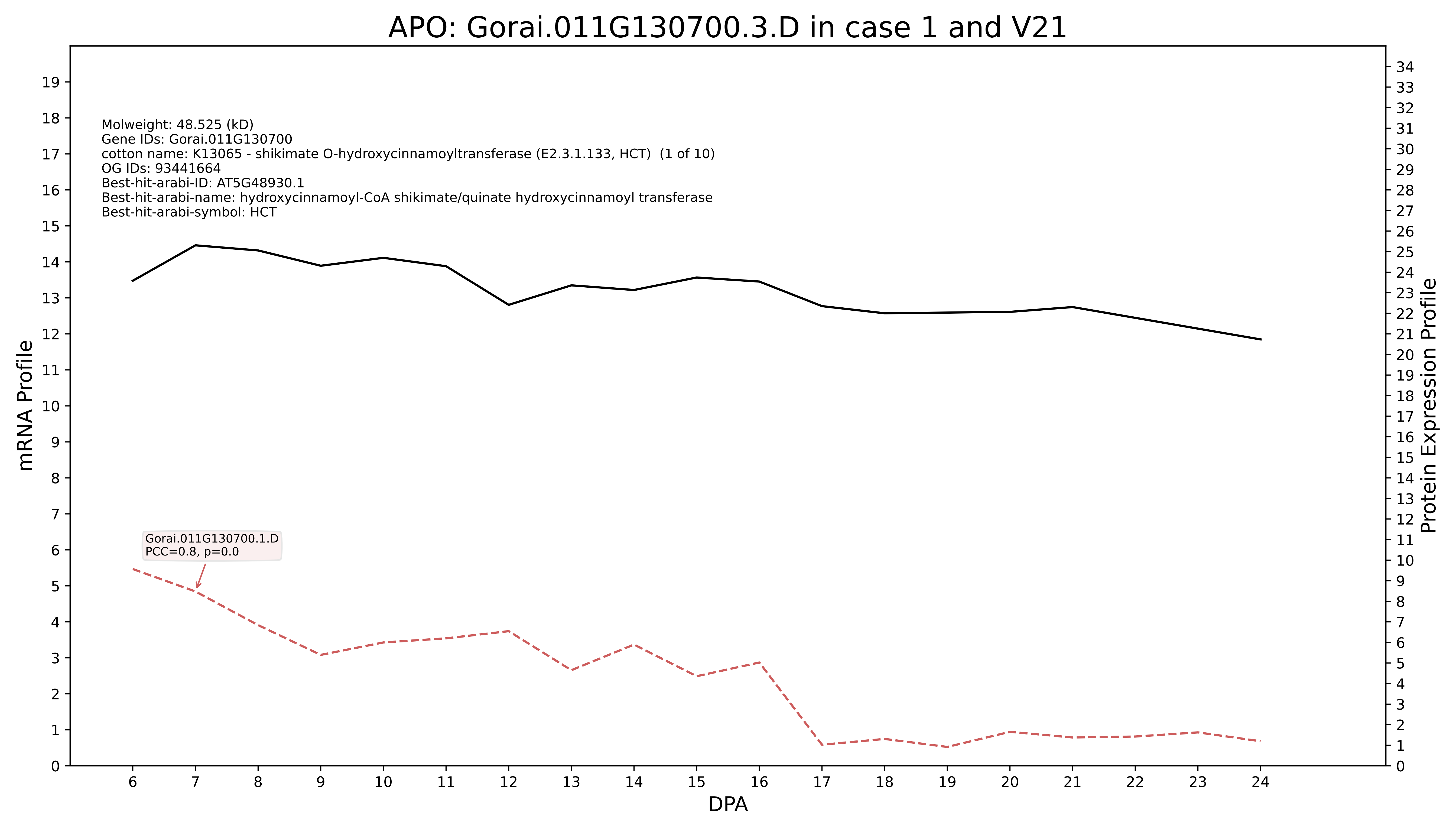
APO: Gorai.011G130700.3.D in case 1 and V21
Molweight: 48.525 (kD)
Gene IDs: Gorai.011G130700
cotton name: K13065 - shikimate O-hydroxycinnamoyltransferase (E2.3.1.133, HCT) (1 of 10)
OG IDs: 93441664
Best-hit-arabi-ID: AT5G48930.1
Best-hit-arabi-name: hydroxycinnamoyl-CoA shikimate/quinate hydroxycinnamoyl transferase
Best-hit-arabi-symbol: HCT
Gorai.011G130700.1.D
PCC=0.8, p=0.0
mRNA Profile
Protein Expression Profile
DPA
0 1 2 3 4 5 6 7 8 9 10 11 12 13 14 15 16 17 18 19
0 1 2 3 4 5 6 7 8 9 10 11 12 13 14 15 16 17 18 19 20 21 22 23 24 25 26 27 28 29 30 31 32 33 34
6 7 8 9 10 11 12 13 14 15 16 17 18 19 20 21 22 23 24

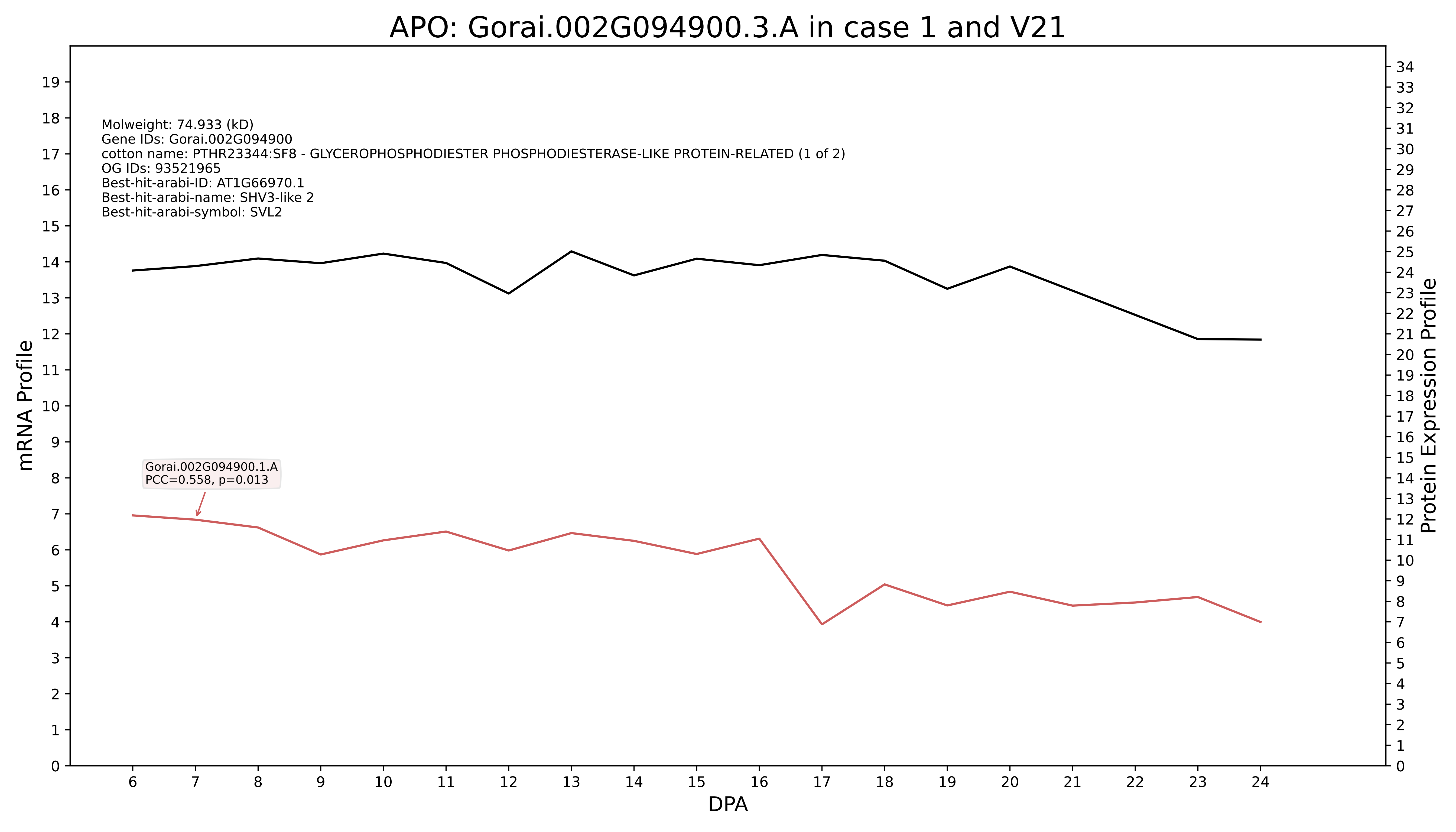

APO: Gorai.002G094900.3.A in case 1 and V21
Molweight: 74.933 (kD)
Gene IDs: Gorai.002G094900
cotton name: PTHR23344:SF8 - GLYCEROPHOSPHODIESTER PHOSPHODIESTERASE-LIKE PROTEIN-RELATED (1 of 2)
OG IDs: 93521965
Best-hit-arabi-ID: AT1G66970.1
Best-hit-arabi-name: SHV3-like 2
Best-hit-arabi-symbol: SVL2
Gorai.002G094900.1.A
PCC=0.558, p=0.013
mRNA Profile
Protein Expression Profile
DPA

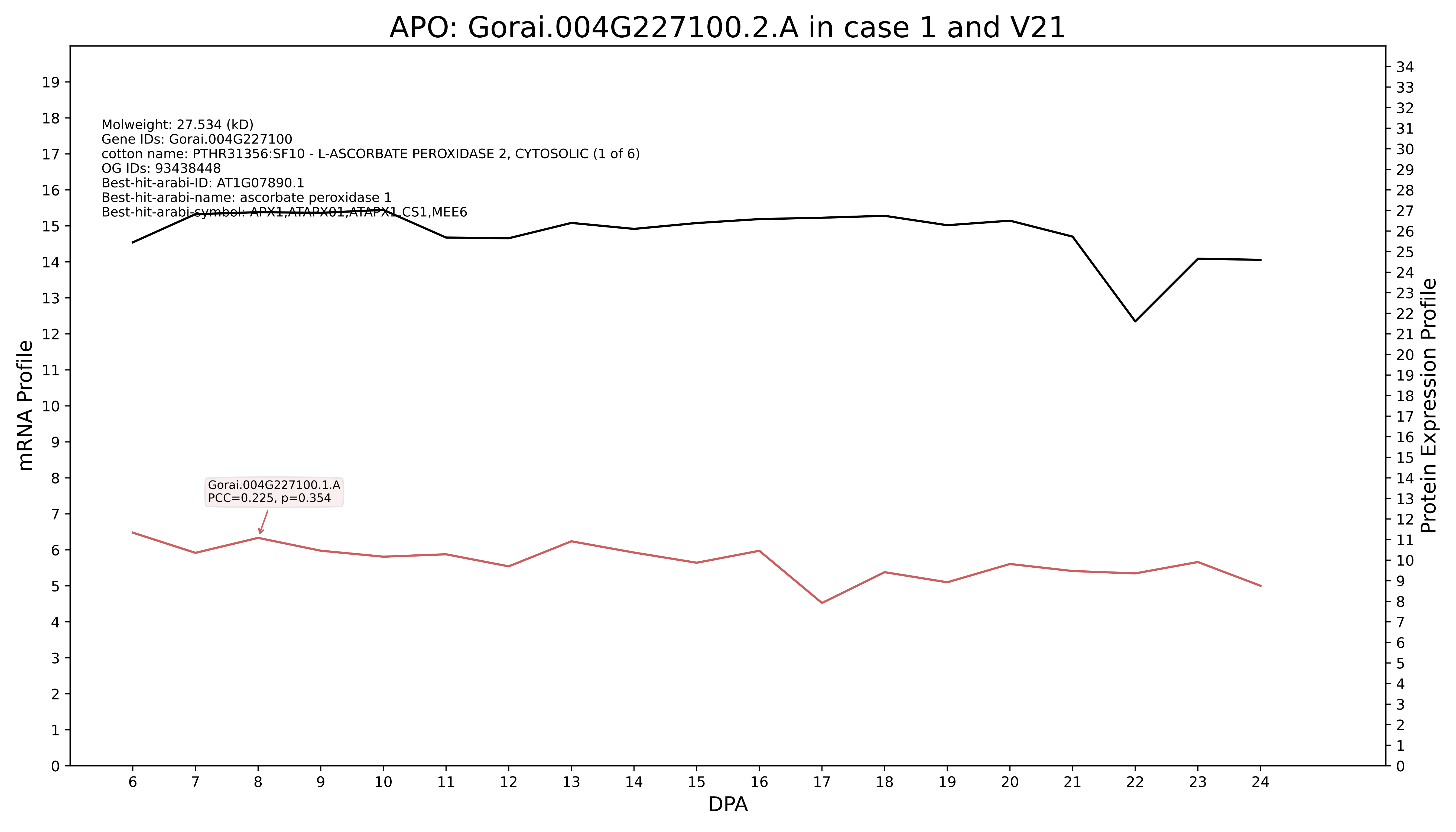
APO: Gorai.004G227100.2.A in case 1 and V21
Molweight: 27.534 (kD)
Gene IDs: Gorai.004G227100
cotton name: PTHR31356:SF10 - L-ASCORBATE PEROXIDASE 2, CYTOSOLIC (1 of 6)
OG IDs: 93438448
Best-hit-arabi-ID: AT1G07890.1
Best-hit-arabi-name: ascorbate peroxidase 1
Best-hit-arabi-symbol: APX1,ATAPX01,ATAPX1,CS1,MEE6
Gorai.004G227100.1.A
PCC=0.225, p=0.354
mRNA Profile
Protein Expression Profile
DPA

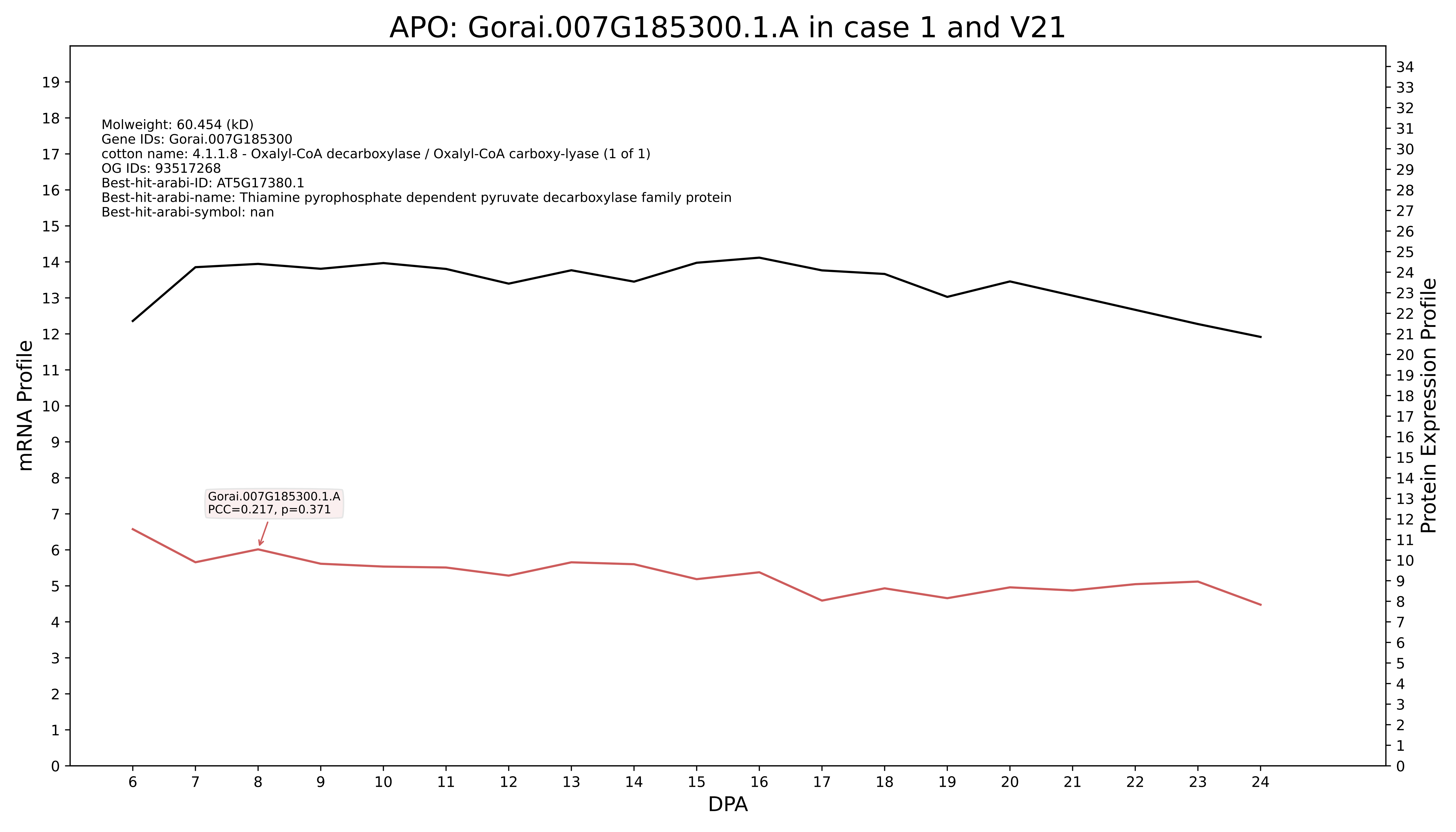
APO: Gorai.007G185300.1.A in case 1 and V21
Molweight: 60.454 (kD)
Gene IDs: Gorai.007G185300
cotton name: 4.1.1.8 - Oxalyl-CoA decarboxylase / Oxalyl-CoA carboxy-lyase (1 of 1)
OG IDs: 93517268
Best-hit-arabi-ID: AT5G17380.1
Best-hit-arabi-name: Thiamine pyrophosphate dependent pyruvate decarboxylase family protein
Best-hit-arabi-symbol: nan
Gorai.007G185300.1.A
PCC=0.217, p=0.371
mRNA Profile
Protein Expression Profile
DPA

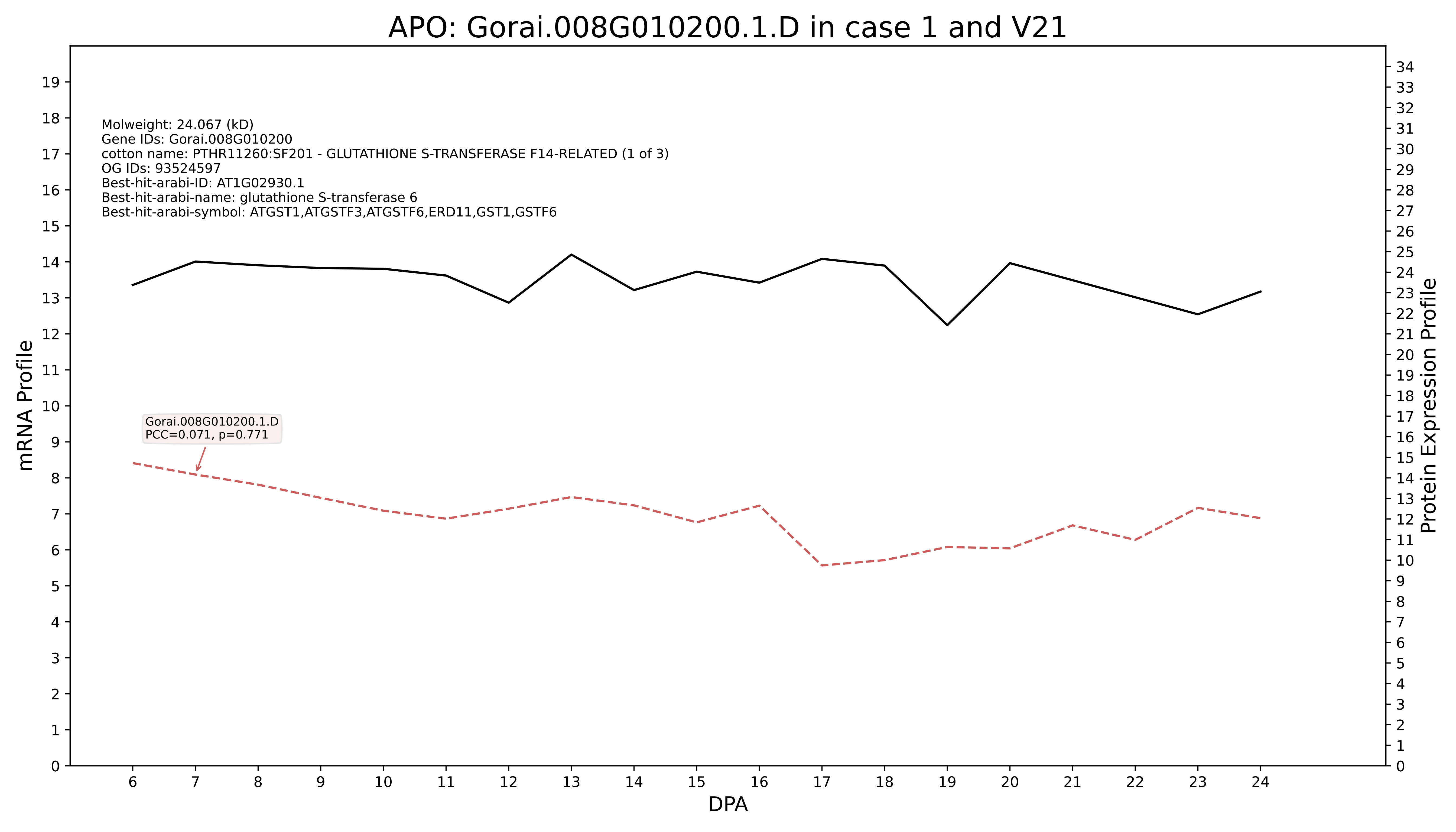
APO: Gorai.008G010200.1.D in case 1 and V21
Molweight: 24.067 (kD)
Gene IDs: Gorai.008G010200
cotton name: PTHR11260:SF201 - GLUTATHIONE S-TRANSFERASE F14-RELATED (1 of 3)
OG IDs: 93524597
Best-hit-arabi-ID: AT1G02930.1
Best-hit-arabi-name: glutathione S-transferase 6
Best-hit-arabi-symbol: ATGST1,ATGSTF3,ATGSTF6,ERD11,GST1,GSTF6
Gorai.008G010200.1.D
PCC=0.071, p=0.771
mRNA Profile
Protein Expression Profile
DPA

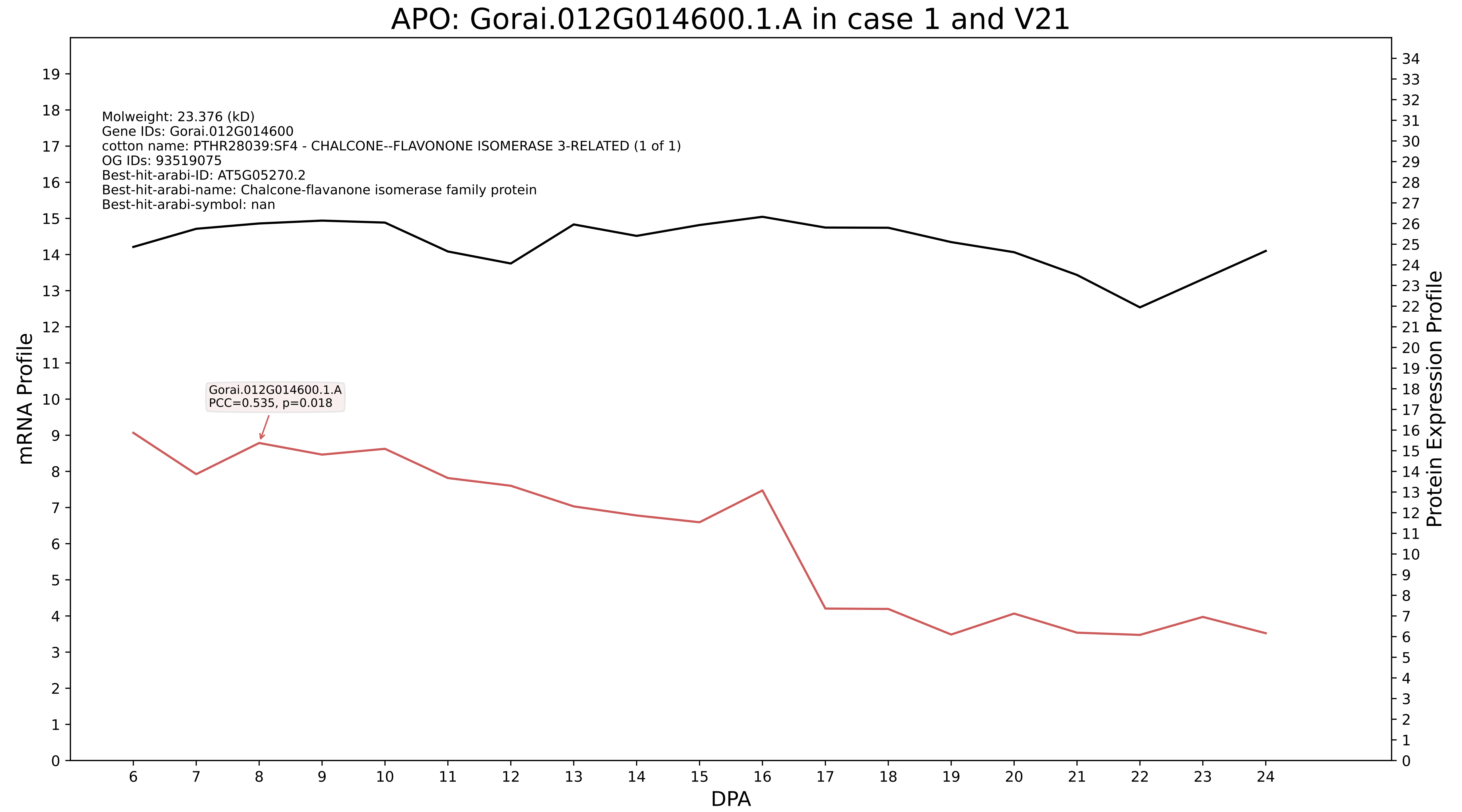
APO: Gorai.012G014600.1.A in case 1 and V21
Molweight: 23.376 (kD)
Gene IDs: Gorai.012G014600
cotton name: PTHR28039:SF4 - CHALCONE--FLAVONONE ISOMERASE 3-RELATED (1 of 1)
OG IDs: 93519075
Best-hit-arabi-ID: AT5G05270.2
Best-hit-arabi-name: Chalcone-flavanone isomerase family protein
Best-hit-arabi-symbol: nan
Gorai.012G014600.1.A
PCC=0.535, p=0.018
mRNA Profile
Protein Expression Profile
DPA

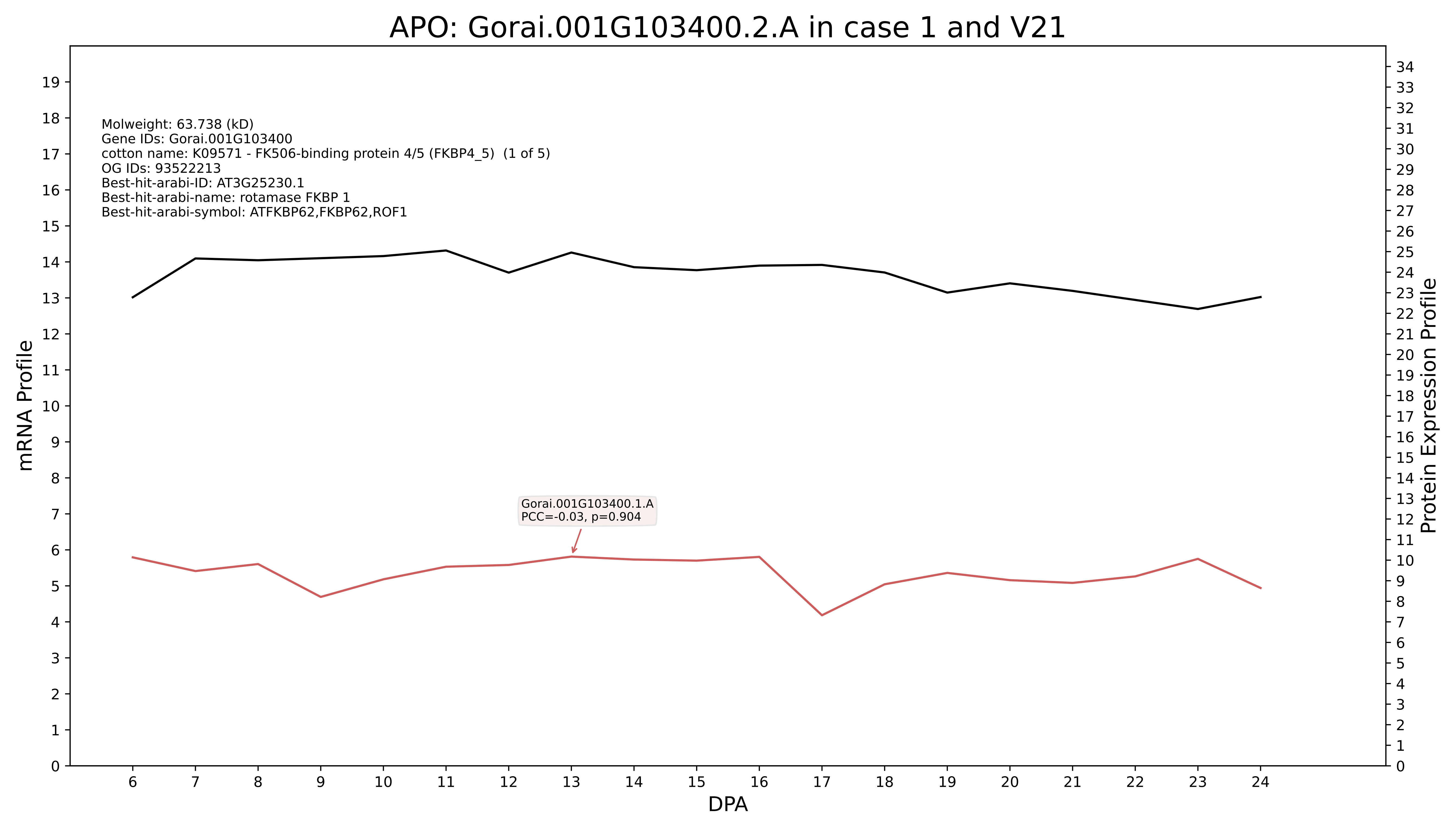

APO: Gorai.001G103400.2.A in case 1 and V21
Molweight: 63.738 (kD)
Gene IDs: Gorai.001G103400
cotton name: K09571 - FK506-binding protein 4/5 (FKBP4_5) (1 of 5)
OG IDs: 93522213
Best-hit-arabi-ID: AT3G25230.1
Best-hit-arabi-name: rotamase FKBP 1
Best-hit-arabi-symbol: ATFKBP62,FKBP62,ROF1
Gorai.001G103400.1.A
PCC=-0.03, p=0.904
mRNA Profile
Protein Expression Profile
DPA

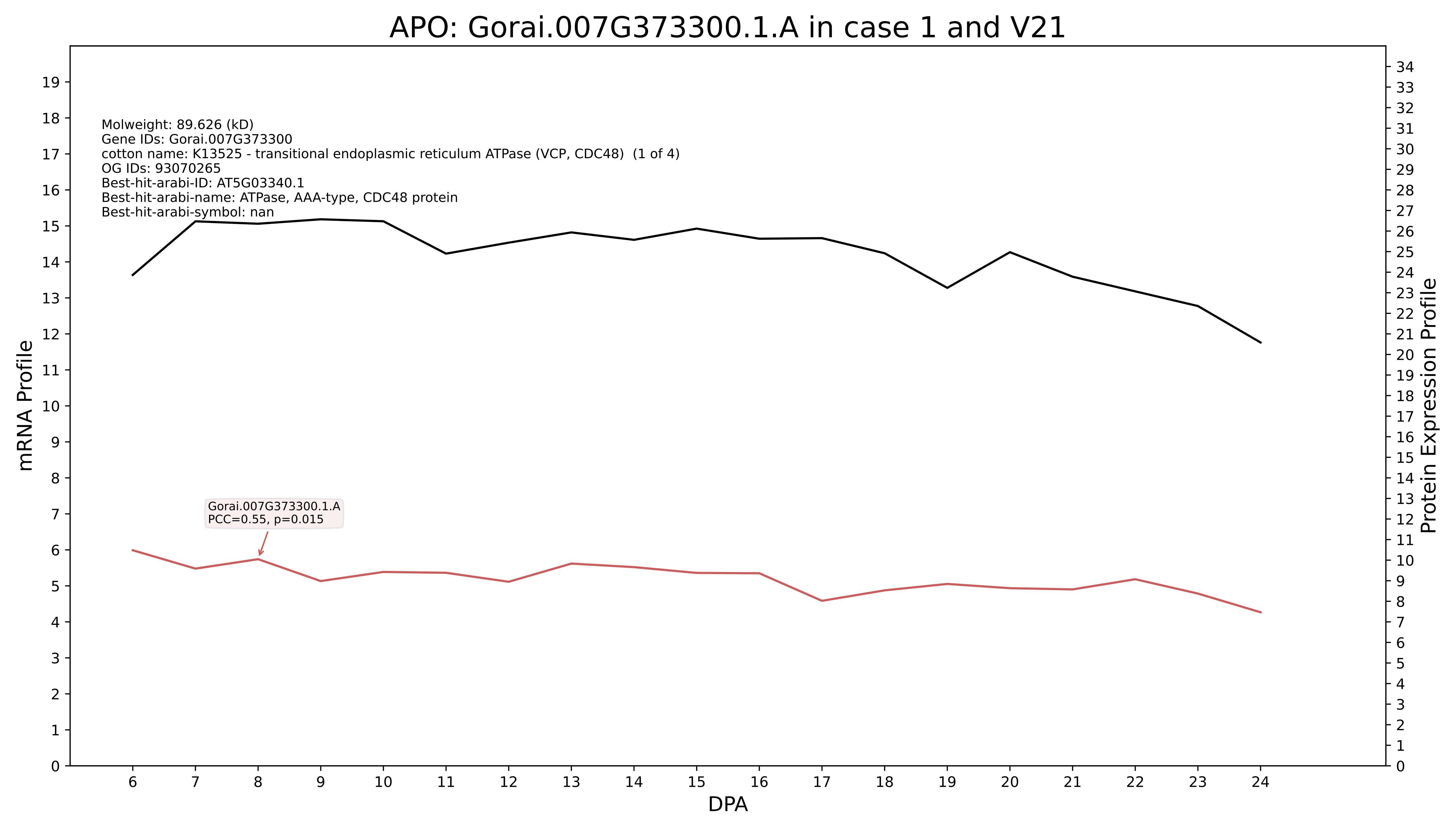

APO: Gorai.007G373300.1.A in case 1 and V21
Molweight: 89.626 (kD)
Gene IDs: Gorai.007G373300
cotton name: K13525 - transitional endoplasmic reticulum ATPase (VCP, CDC48) (1 of 4)
OG IDs: 93070265
Best-hit-arabi-ID: AT5G03340.1
Best-hit-arabi-name: ATPase, AAA-type, CDC48 protein
Best-hit-arabi-symbol: nan
Gorai.007G373300.1.A
PCC=0.55, p=0.015
mRNA Profile
Protein Expression Profile
DPA

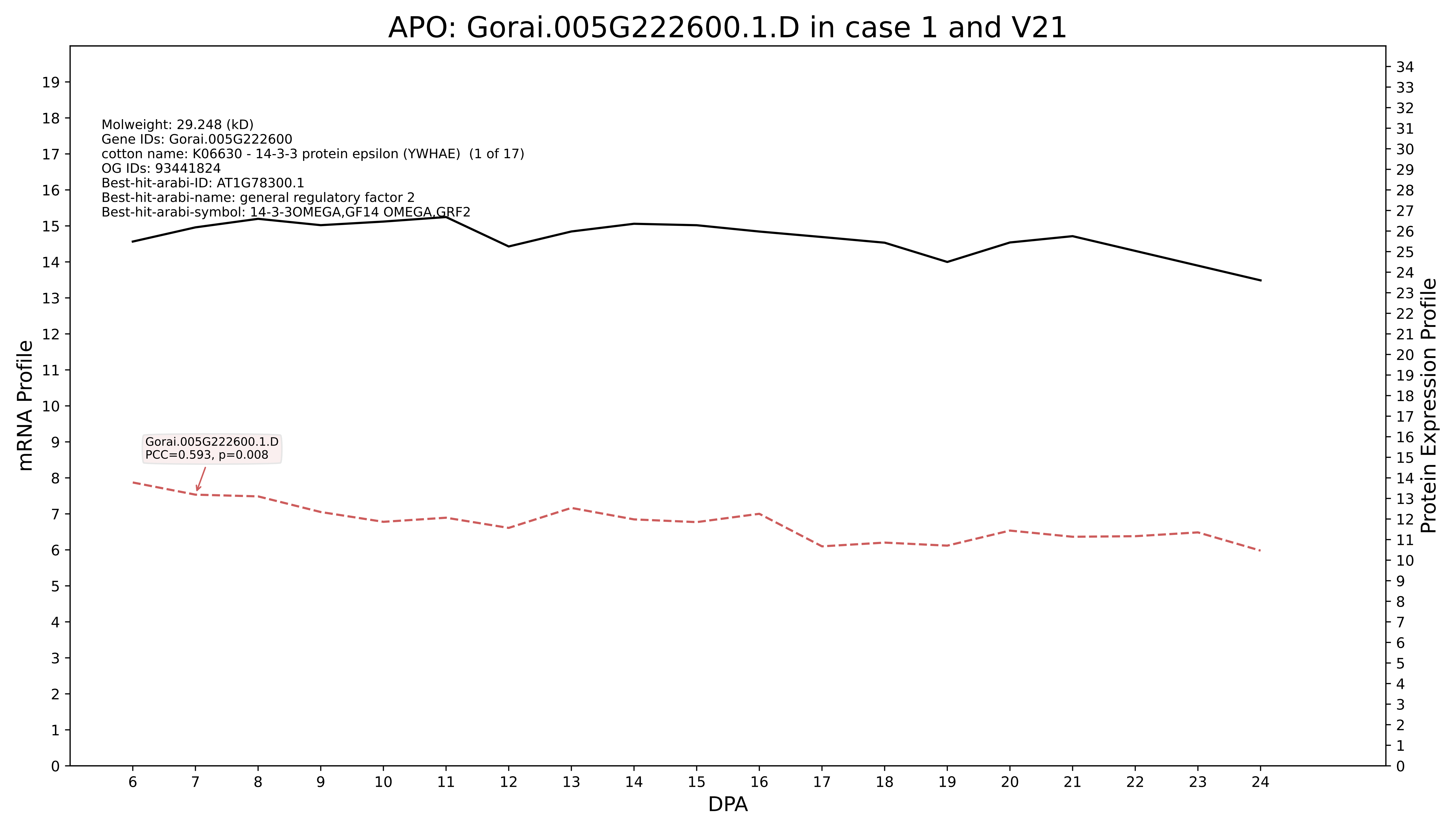

APO: Gorai.005G222600.1.D in case 1 and V21
Molweight: 29.248 (kD)
Gene IDs: Gorai.005G222600
cotton name: K06630 - 14-3-3 protein epsilon (YWHAE) (1 of 17)
OG IDs: 93441824
Best-hit-arabi-ID: AT1G78300.1
Best-hit-arabi-name: general regulatory factor 2
Best-hit-arabi-symbol: 14-3-3OMEGA,GF14 OMEGA,GRF2
Gorai.005G222600.1.D
PCC=0.593, p=0.008
mRNA Profile
Protein Expression Profile
DPA

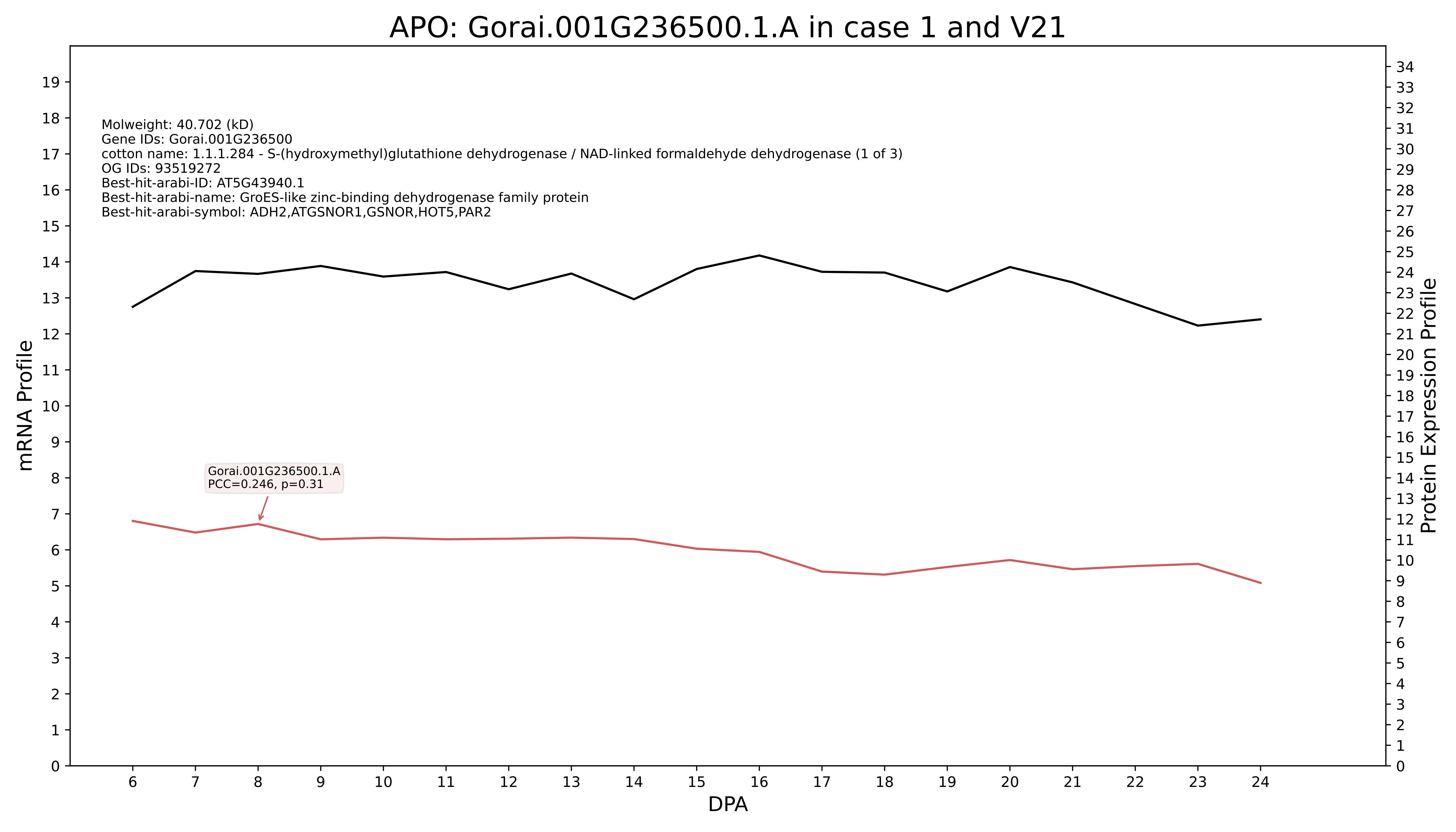
APO: Gorai.001G236500.1.A in case 1 and V21
Molweight: 40.702 (kD)
Gene IDs: Gorai.001G236500
cotton name: 1.1.1.284 - S-(hydroxymethyl)glutathione dehydrogenase / NAD-linked formaldehyde dehydrogenase (1 of 3)
OG IDs: 93519272
Best-hit-arabi-ID: AT5G43940.1
Best-hit-arabi-name: GroES-like zinc-binding dehydrogenase family protein
Best-hit-arabi-symbol: ADH2,ATGSNOR1,GSNOR,HOT5,PAR2
Gorai.001G236500.1.A
PCC=0.246, p=0.31
mRNA Profile
Protein Expression Profile
DPA

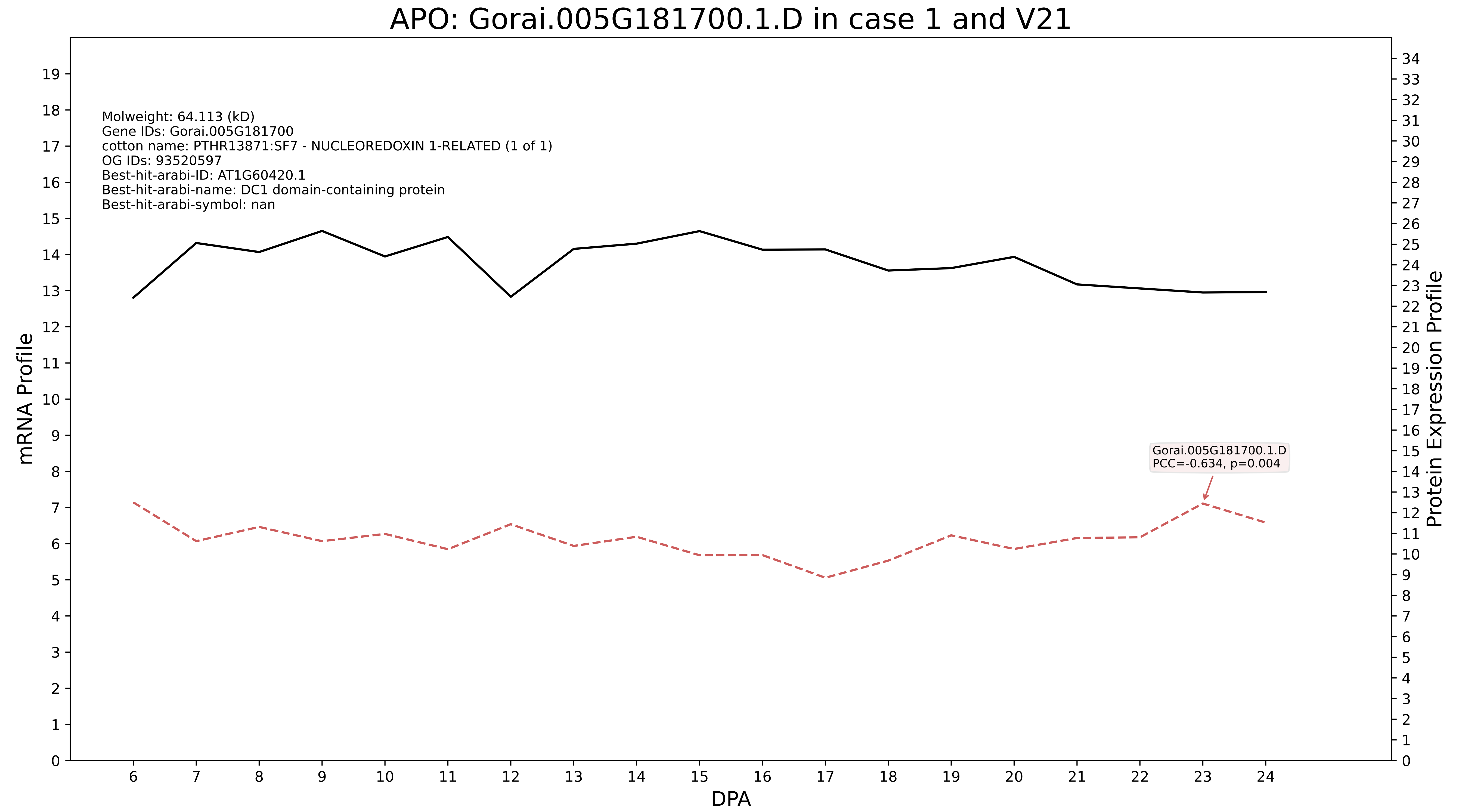

APO: Gorai.005G181700.1.D in case 1 and V21
Molweight: 64.113 (kD)
Gene IDs: Gorai.005G181700
cotton name: PTHR13871:SF7 - NUCLEOREDOXIN 1-RELATED (1 of 1)
OG IDs: 93520597
Best-hit-arabi-ID: AT1G60420.1
Best-hit-arabi-name: DC1 domain-containing protein
Best-hit-arabi-symbol: nan
Gorai.005G181700.1.D
PCC=-0.634, p=0.004
mRNA Profile
Protein Expression Profile
DPA
0 1 2 3 4 5 6 7 8 9 10 11 12 13 14 15 16 17 18 19
0 1 2 3 4 5 6 7 8 9 10 11 12 13 14 15 16 17 18 19 20 21 22 23 24 25 26 27 28 29 30 31 32 33 34
6 7 8 9 10 11 12 13 14 15 16 17 18 19 20 21 22 23 24

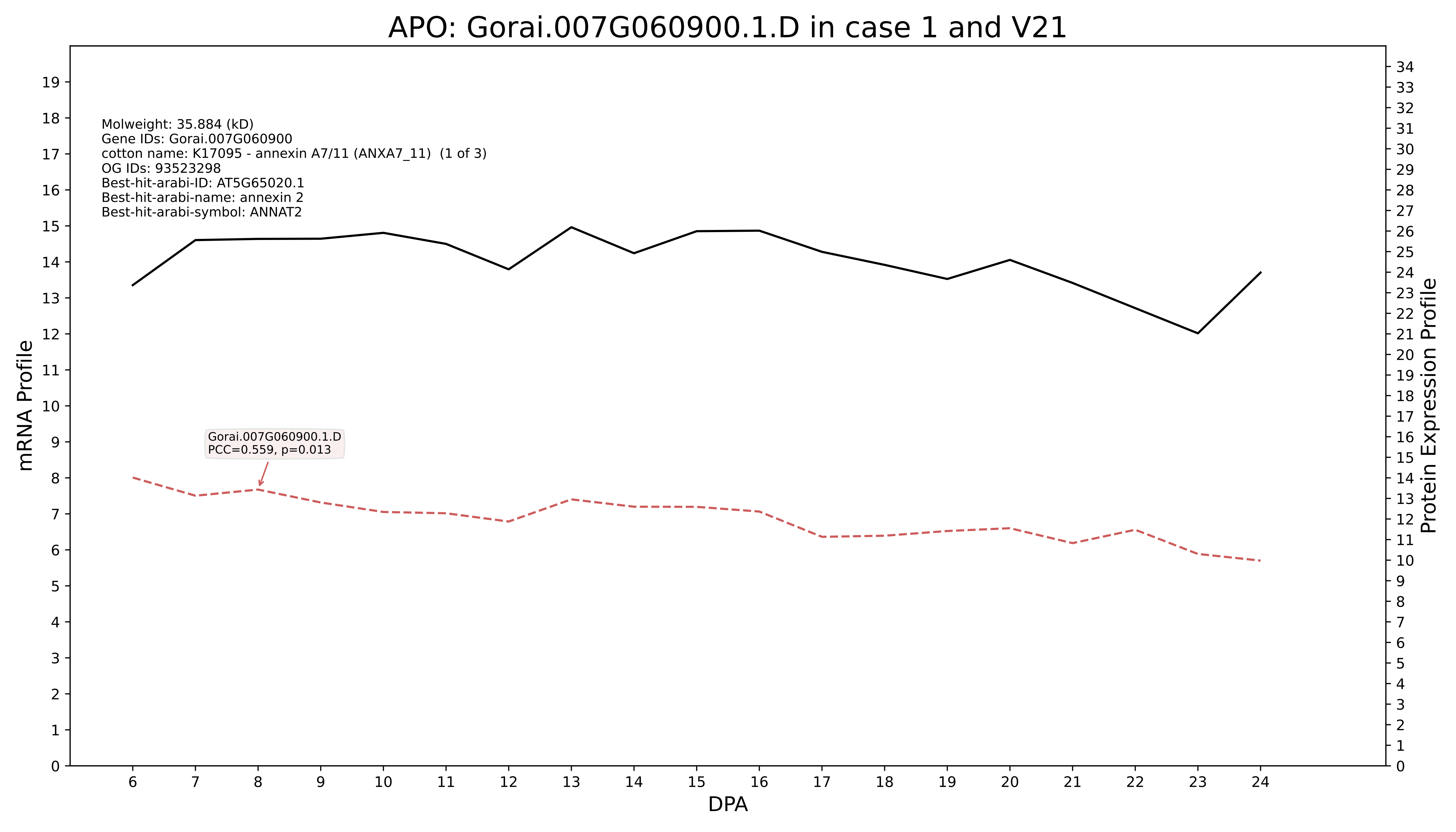
APO: Gorai.007G060900.1.D in case 1 and V21
Molweight: 35.884 (kD)
Gene IDs: Gorai.007G060900
cotton name: K17095 - annexin A7/11 (ANXA7_11) (1 of 3)
OG IDs: 93523298
Best-hit-arabi-ID: AT5G65020.1
Best-hit-arabi-name: annexin 2
Best-hit-arabi-symbol: ANNAT2
Gorai.007G060900.1.D
PCC=0.559, p=0.013
mRNA Profile
Protein Expression Profile
DPA

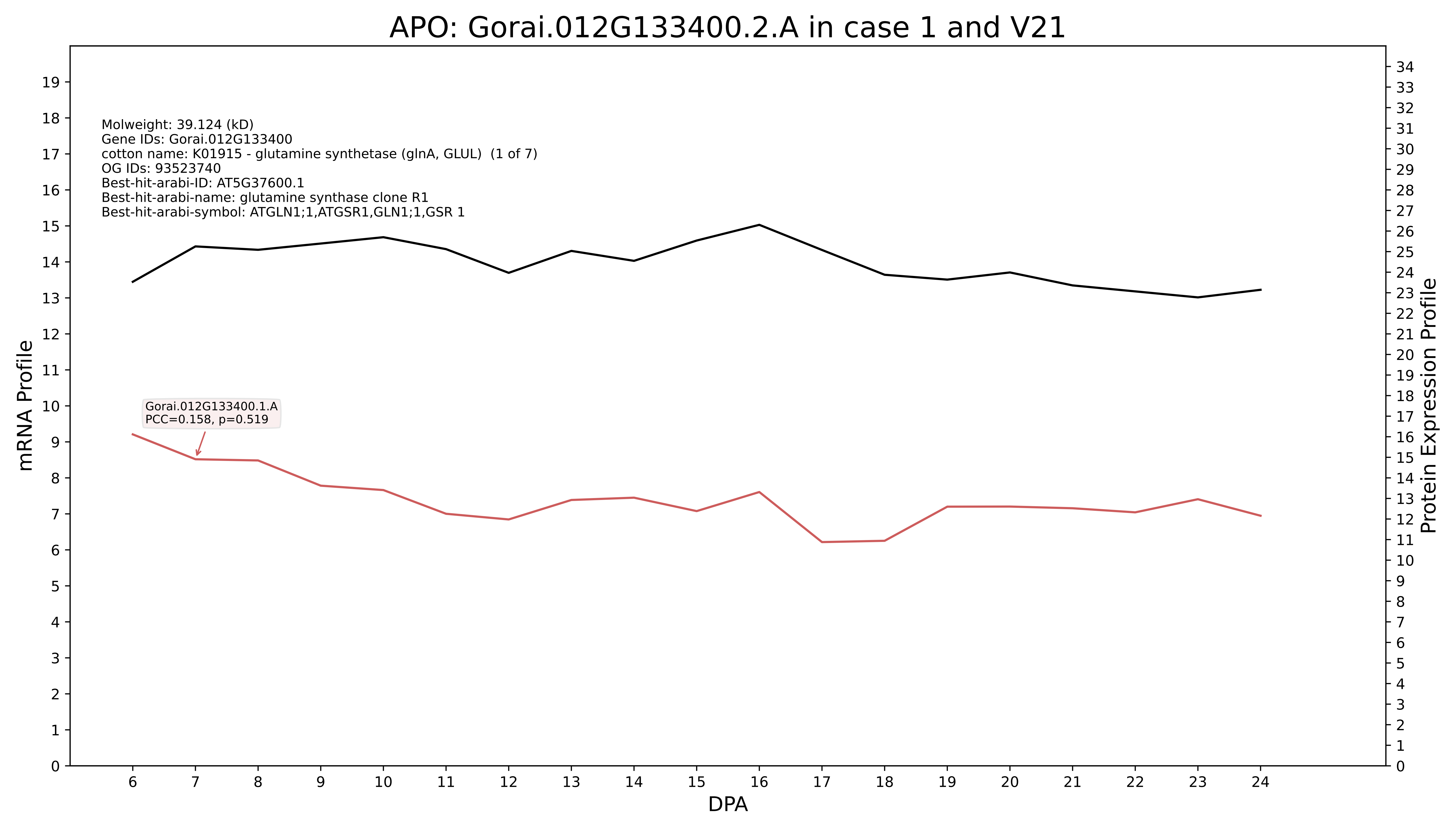
APO: Gorai.012G133400.2.A in case 1 and V21
Molweight: 39.124 (kD)
Gene IDs: Gorai.012G133400
cotton name: K01915 - glutamine synthetase (glnA, GLUL) (1 of 7)
OG IDs: 93523740
Best-hit-arabi-ID: AT5G37600.1
Best-hit-arabi-name: glutamine synthase clone R1
Best-hit-arabi-symbol: ATGLN1;1,ATGSR1,GLN1;1,GSR 1
Gorai.012G133400.1.A
PCC=0.158, p=0.519
mRNA Profile
Protein Expression Profile
DPA

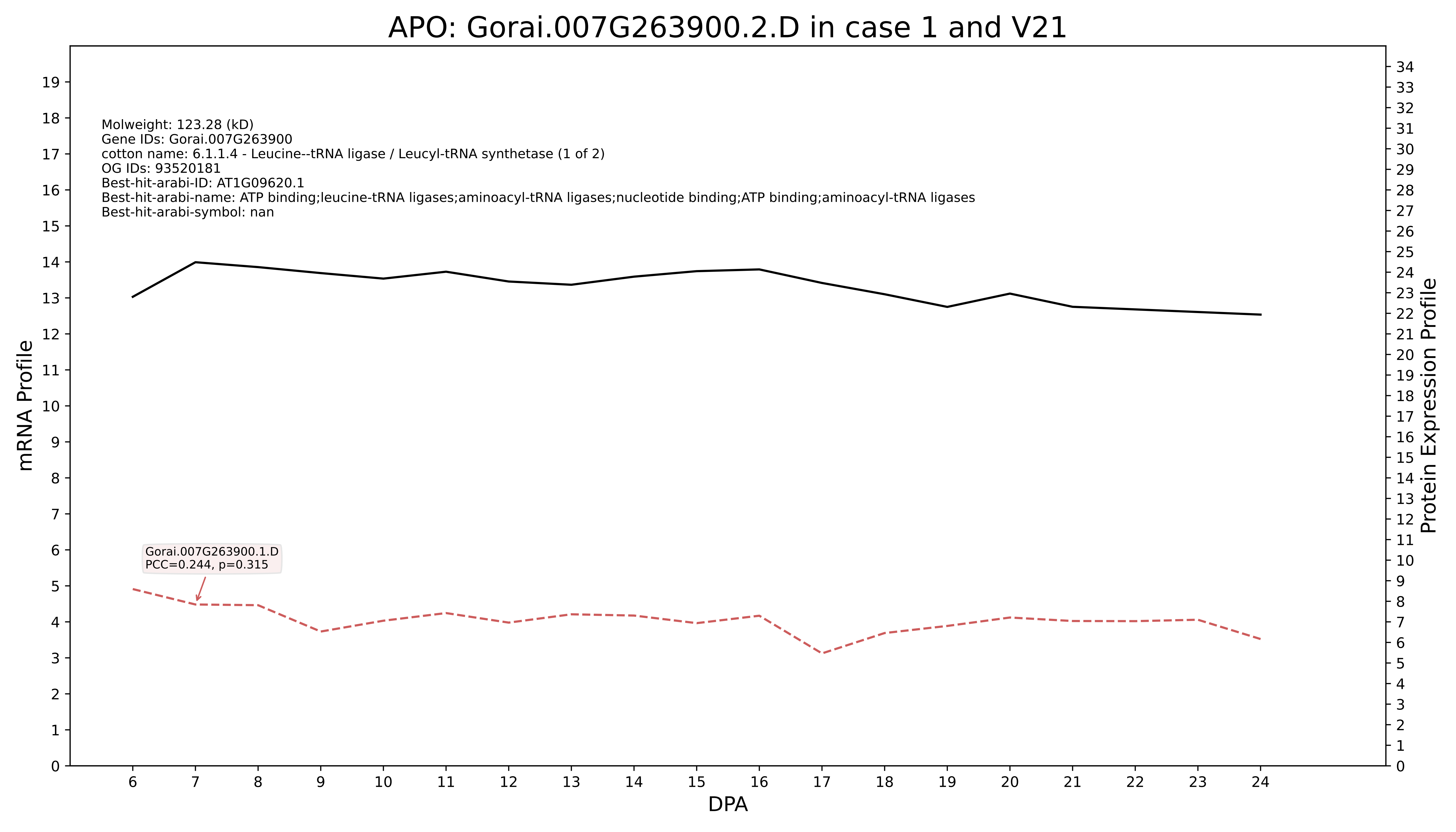
APO: Gorai.007G263900.2.D in case 1 and V21
Molweight: 123.28 (kD)
Gene IDs: Gorai.007G263900
cotton name: 6.1.1.4 - Leucine--tRNA ligase / Leucyl-tRNA synthetase (1 of 2)
OG IDs: 93520181
Best-hit-arabi-ID: AT1G09620.1
Best-hit-arabi-name: ATP binding;leucine-tRNA ligases;aminoacyl-tRNA ligases;nucleotide binding;ATP binding;aminoacyl-tRNA ligases
Best-hit-arabi-symbol: nan
Gorai.007G263900.1.D
PCC=0.244, p=0.315
mRNA Profile
Protein Expression Profile
DPA

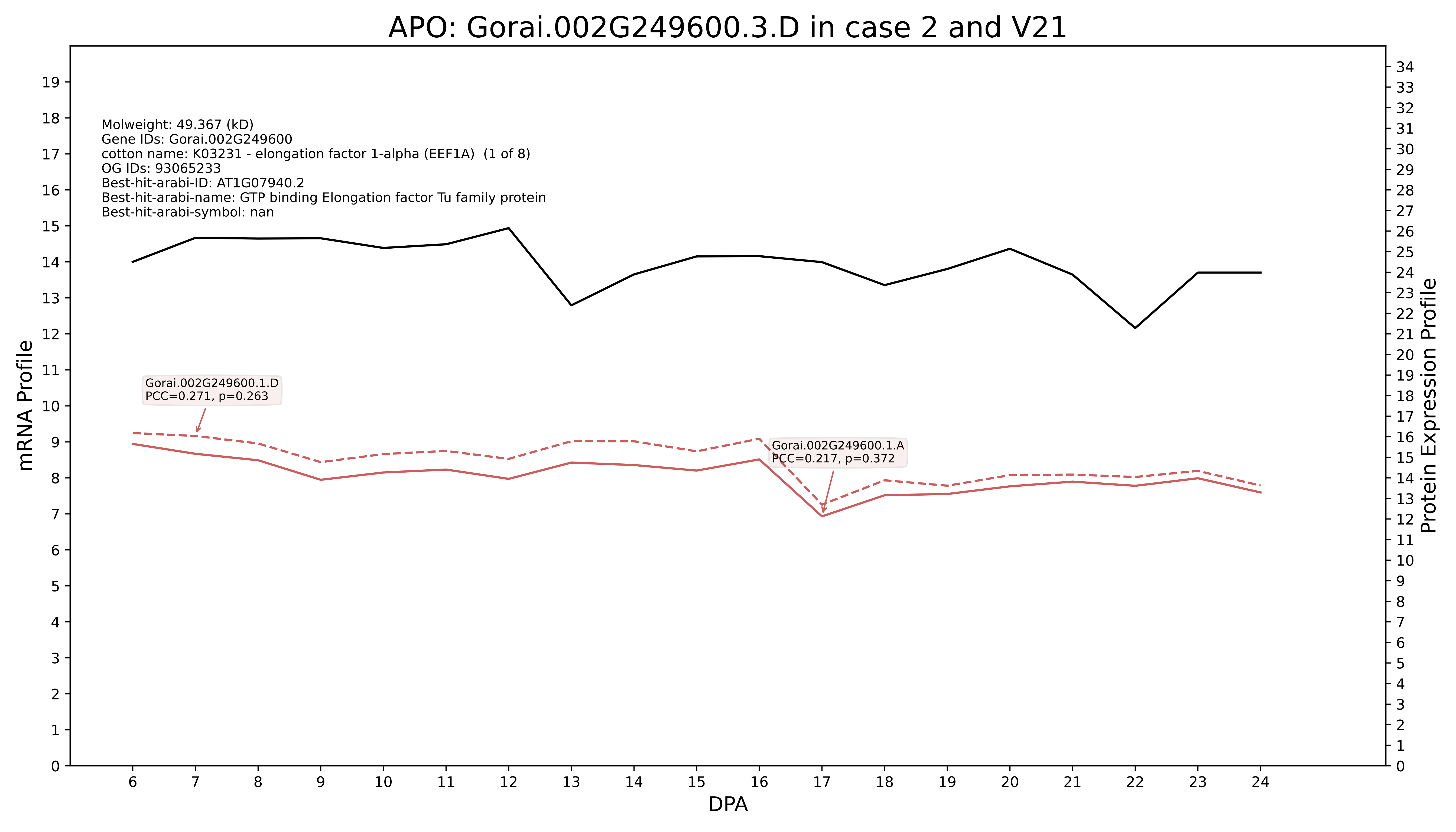

APO: Gorai.002G249600.3.D in case 2 and V21
Molweight: 49.367 (kD)
Gene IDs: Gorai.002G249600
cotton name: K03231 - elongation factor 1-alpha (EEF1A) (1 of 8)
OG IDs: 93065233
Best-hit-arabi-ID: AT1G07940.2
Best-hit-arabi-name: GTP binding Elongation factor Tu family protein
Best-hit-arabi-symbol: nan
Gorai.002G249600.1.D
PCC=0.271, p=0.263
Gorai.002G249600.1.A
PCC=0.217, p=0.372
mRNA Profile
Protein Expression Profile
DPA

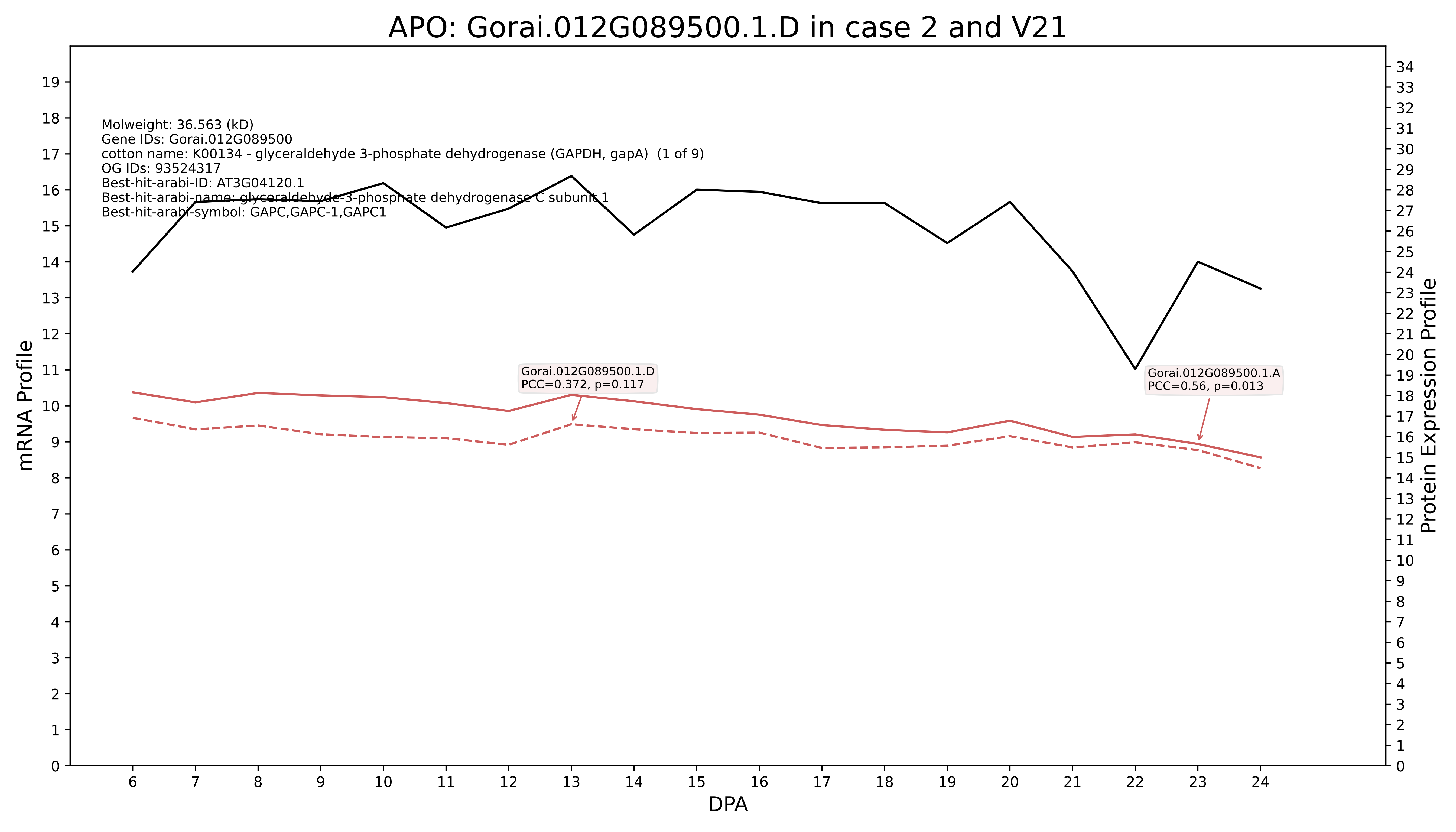

APO: Gorai.012G089500.1.D in case 2 and V21
Molweight: 36.563 (kD)
Gene IDs: Gorai.012G089500
cotton name: K00134 - glyceraldehyde 3-phosphate dehydrogenase (GAPDH, gapA) (1 of 9)
OG IDs: 93524317
Best-hit-arabi-ID: AT3G04120.1
Best-hit-arabi-name: glyceraldehyde-3-phosphate dehydrogenase C subunit 1
Best-hit-arabi-symbol: GAPC,GAPC-1,GAPC1
Gorai.012G089500.1.D
PCC=0.372, p=0.117
Gorai.012G089500.1.A
PCC=0.56, p=0.013
mRNA Profile
Protein Expression Profile
DPA

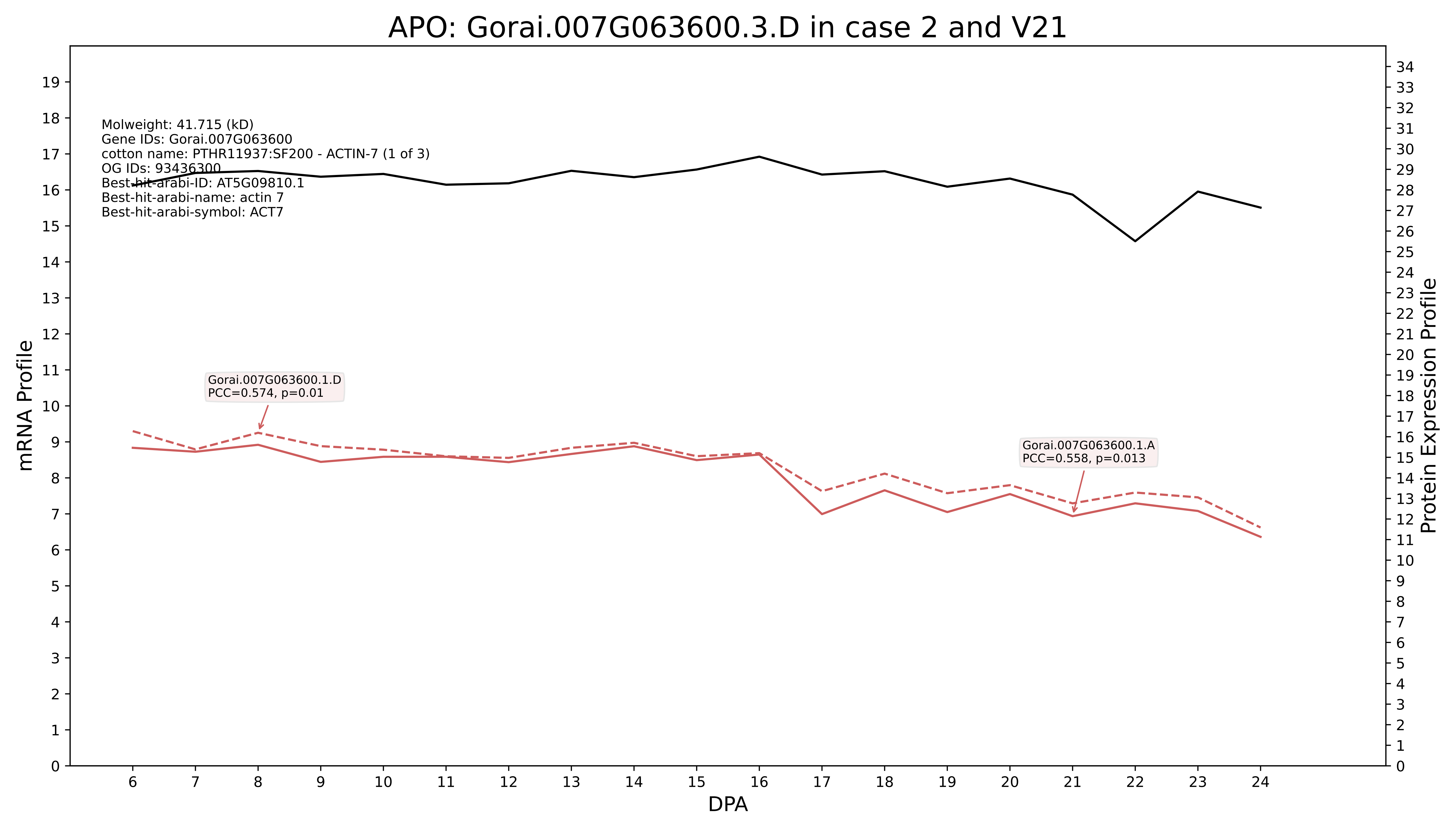

APO: Gorai.007G063600.3.D in case 2 and V21
Molweight: 41.715 (kD)
Gene IDs: Gorai.007G063600
cotton name: PTHR11937:SF200 - ACTIN-7 (1 of 3)
OG IDs: 93436300
Best-hit-arabi-ID: AT5G09810.1
Best-hit-arabi-name: actin 7
Best-hit-arabi-symbol: ACT7
Gorai.007G063600.1.D
PCC=0.574, p=0.01
Gorai.007G063600.1.A
PCC=0.558, p=0.013
mRNA Profile
Protein Expression Profile
DPA

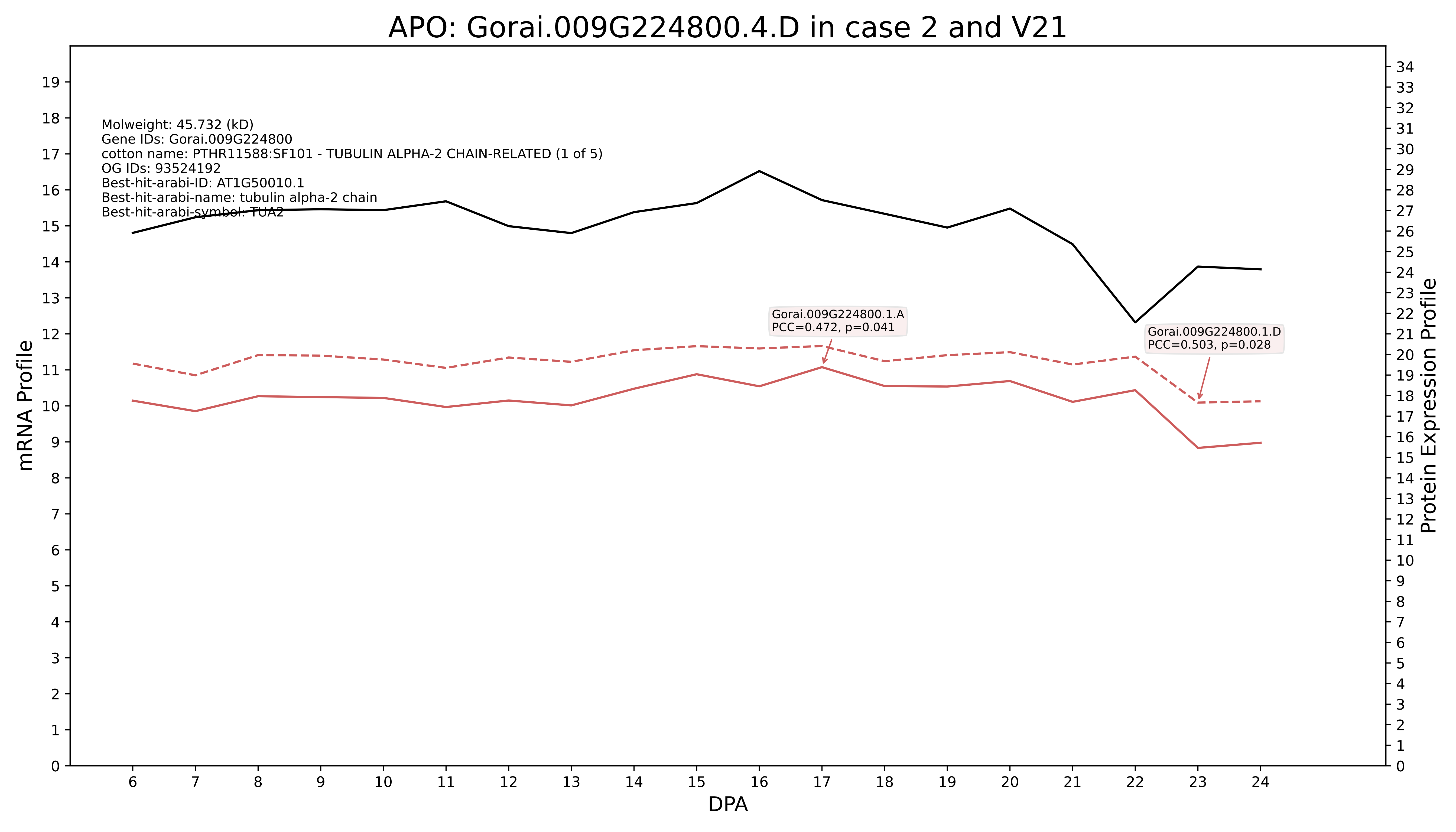

APO: Gorai.009G224800.4.D in case 2 and V21
Molweight: 45.732 (kD)
Gene IDs: Gorai.009G224800
cotton name: PTHR11588:SF101 - TUBULIN ALPHA-2 CHAIN-RELATED (1 of 5)
OG IDs: 93524192
Best-hit-arabi-ID: AT1G50010.1
Best-hit-arabi-name: tubulin alpha-2 chain
Best-hit-arabi-symbol: TUA2
Gorai.009G224800.1.A
PCC=0.472, p=0.041
Gorai.009G224800.1.D
PCC=0.503, p=0.028
mRNA Profile
Protein Expression Profile
DPA

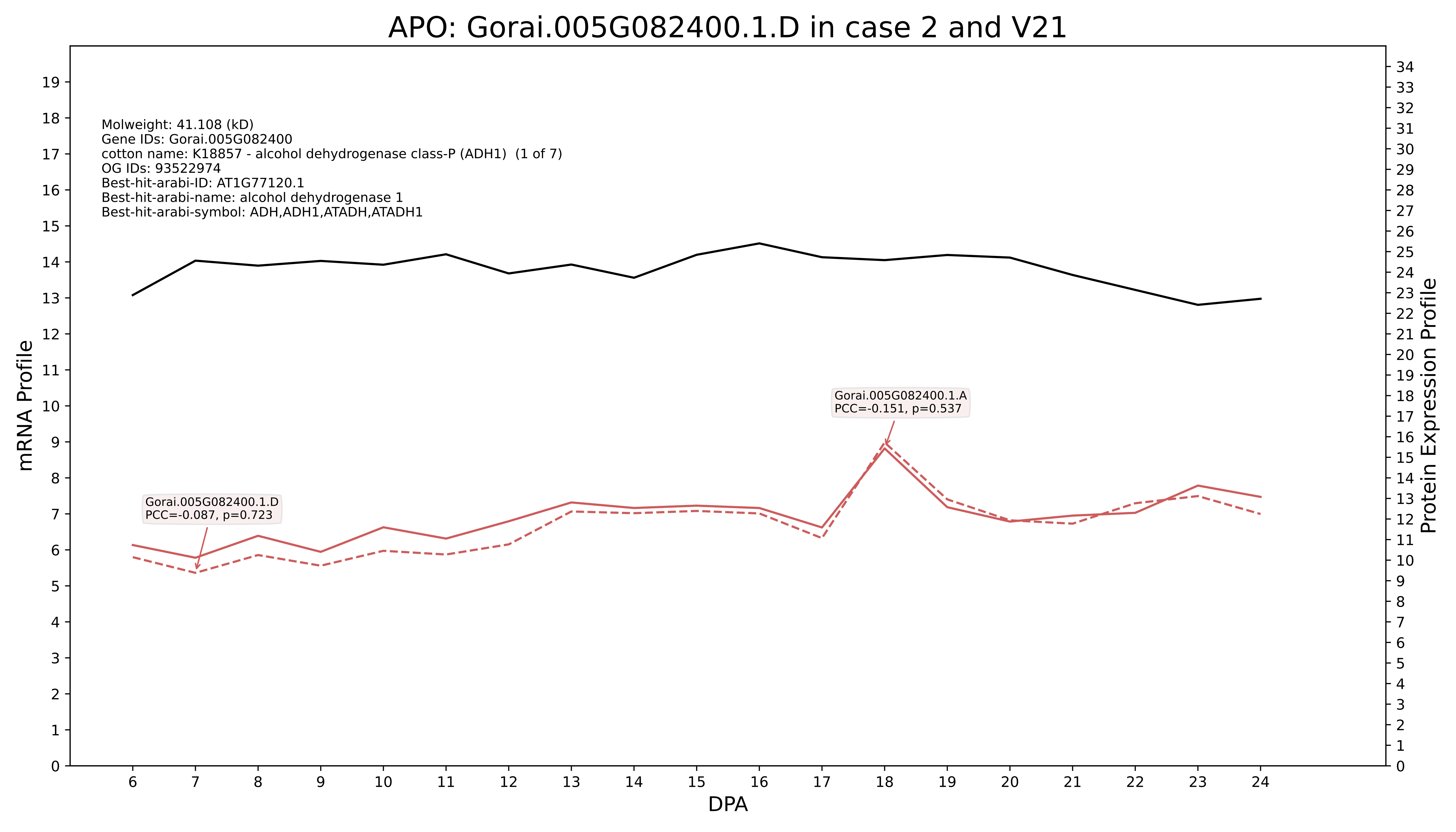
APO: Gorai.005G082400.1.D in case 2 and V21
Molweight: 41.108 (kD)
Gene IDs: Gorai.005G082400
cotton name: K18857 - alcohol dehydrogenase class-P (ADH1) (1 of 7)
OG IDs: 93522974
Best-hit-arabi-ID: AT1G77120.1
Best-hit-arabi-name: alcohol dehydrogenase 1
Best-hit-arabi-symbol: ADH,ADH1,ATADH,ATADH1
Gorai.005G082400.1.A
PCC=-0.151, p=0.537
Gorai.005G082400.1.D
PCC=-0.087, p=0.723
mRNA Profile
Protein Expression Profile
DPA

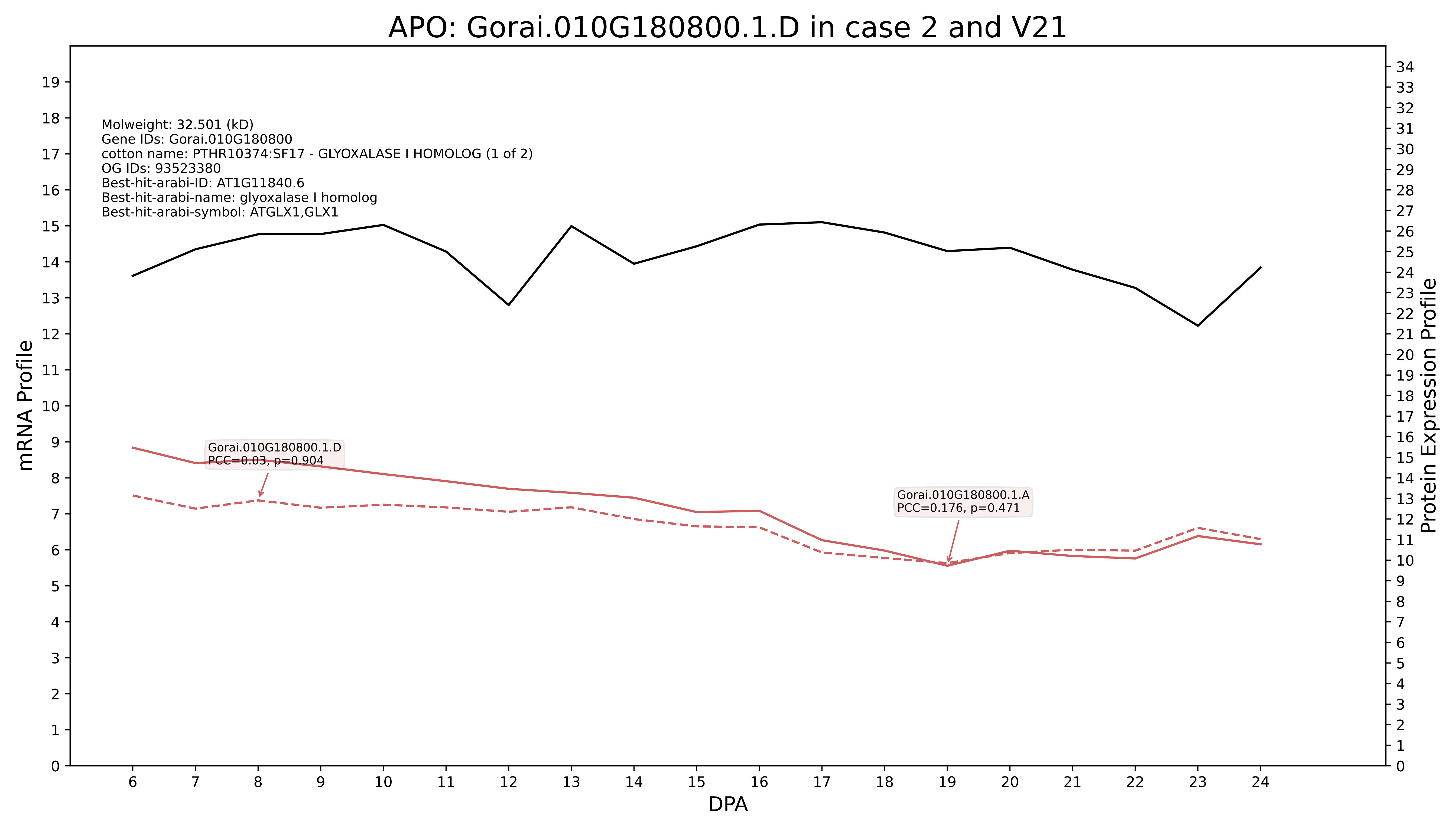
APO: Gorai.010G180800.1.D in case 2 and V21
Molweight: 32.501 (kD)
Gene IDs: Gorai.010G180800
cotton name: PTHR10374:SF17 - GLYOXALASE I HOMOLOG (1 of 2)
OG IDs: 93523380
Best-hit-arabi-ID: AT1G11840.6
Best-hit-arabi-name: glyoxalase I homolog
Best-hit-arabi-symbol: ATGLX1,GLX1
Gorai.010G180800.1.D
PCC=0.03, p=0.904
Gorai.010G180800.1.A
PCC=0.176, p=0.471
mRNA Profile
Protein Expression Profile
DPA

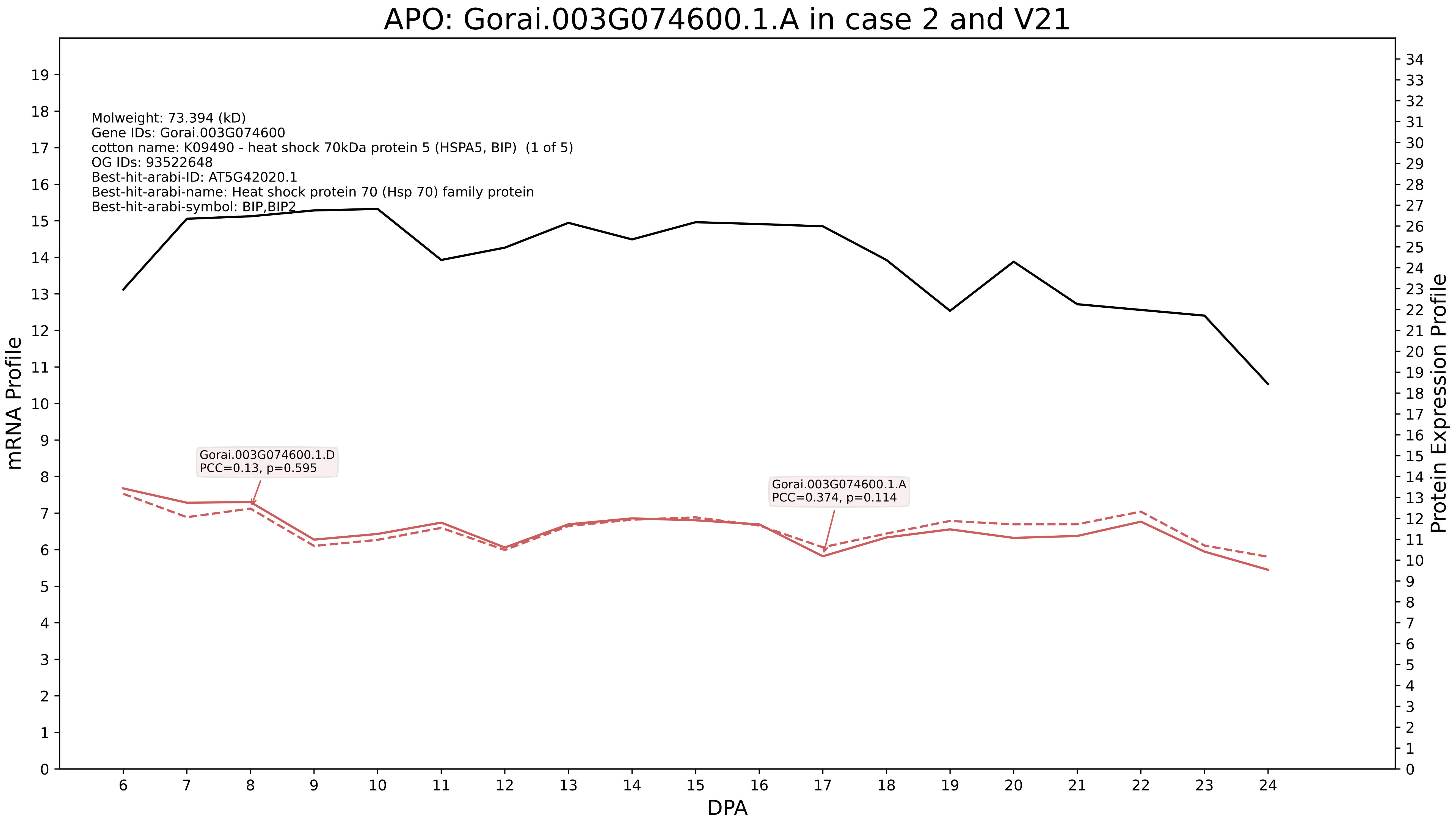
APO: Gorai.003G074600.1.A in case 2 and V21
Molweight: 73.394 (kD)
Gene IDs: Gorai.003G074600
cotton name: K09490 - heat shock 70kDa protein 5 (HSPA5, BIP) (1 of 5)
OG IDs: 93522648
Best-hit-arabi-ID: AT5G42020.1
Best-hit-arabi-name: Heat shock protein 70 (Hsp 70) family protein
Best-hit-arabi-symbol: BIP,BIP2
Gorai.003G074600.1.D
PCC=0.13, p=0.595
Gorai.003G074600.1.A
PCC=0.374, p=0.114
mRNA Profile
Protein Expression Profile
DPA

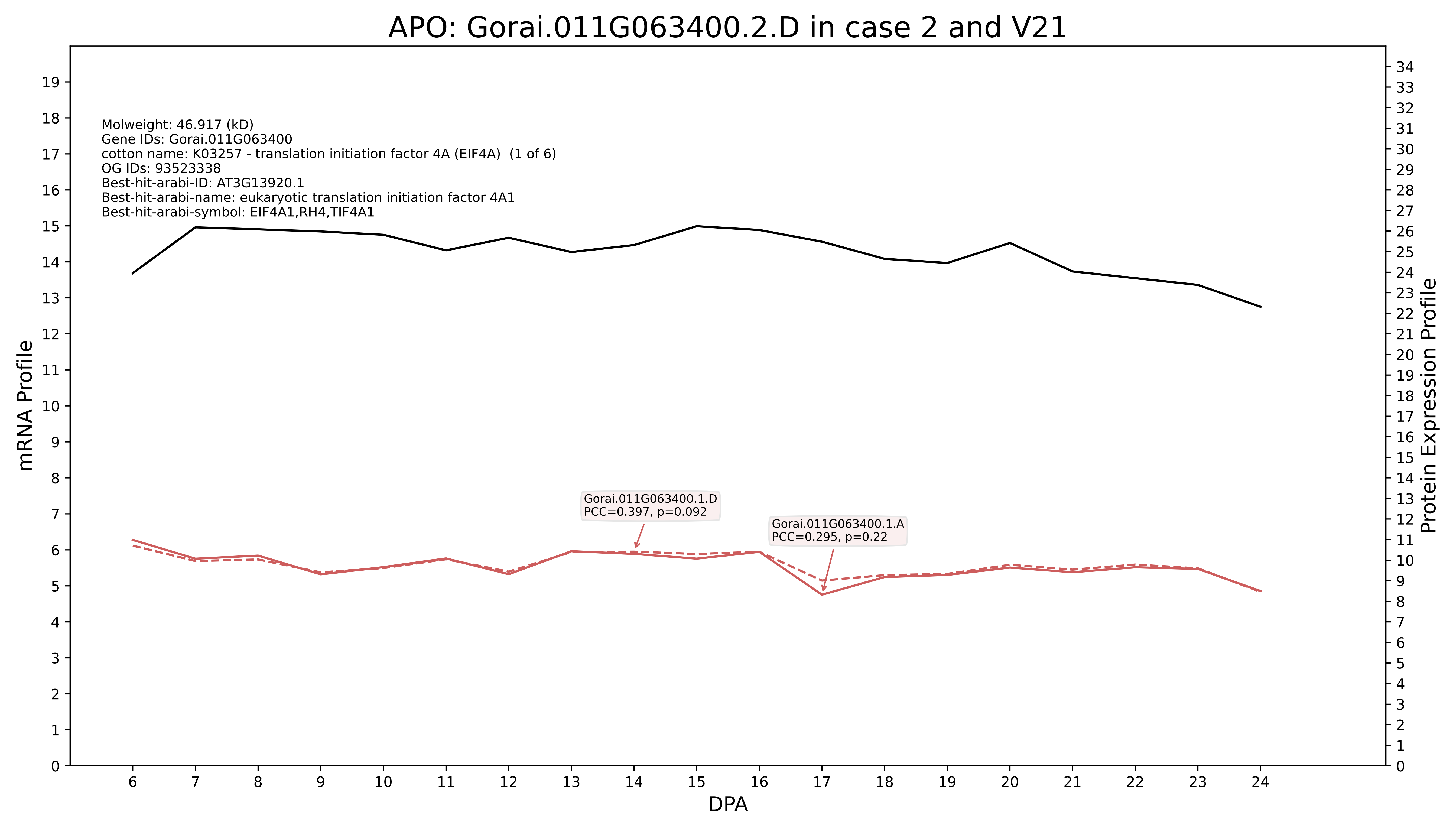

APO: Gorai.011G063400.2.D in case 2 and V21
Molweight: 46.917 (kD)
Gene IDs: Gorai.011G063400
cotton name: K03257 - translation initiation factor 4A (EIF4A) (1 of 6)
OG IDs: 93523338
Best-hit-arabi-ID: AT3G13920.1
Best-hit-arabi-name: eukaryotic translation initiation factor 4A1
Best-hit-arabi-symbol: EIF4A1,RH4,TIF4A1
Gorai.011G063400.1.D
PCC=0.397, p=0.092
Gorai.011G063400.1.A
PCC=0.295, p=0.22
mRNA Profile
Protein Expression Profile
DPA

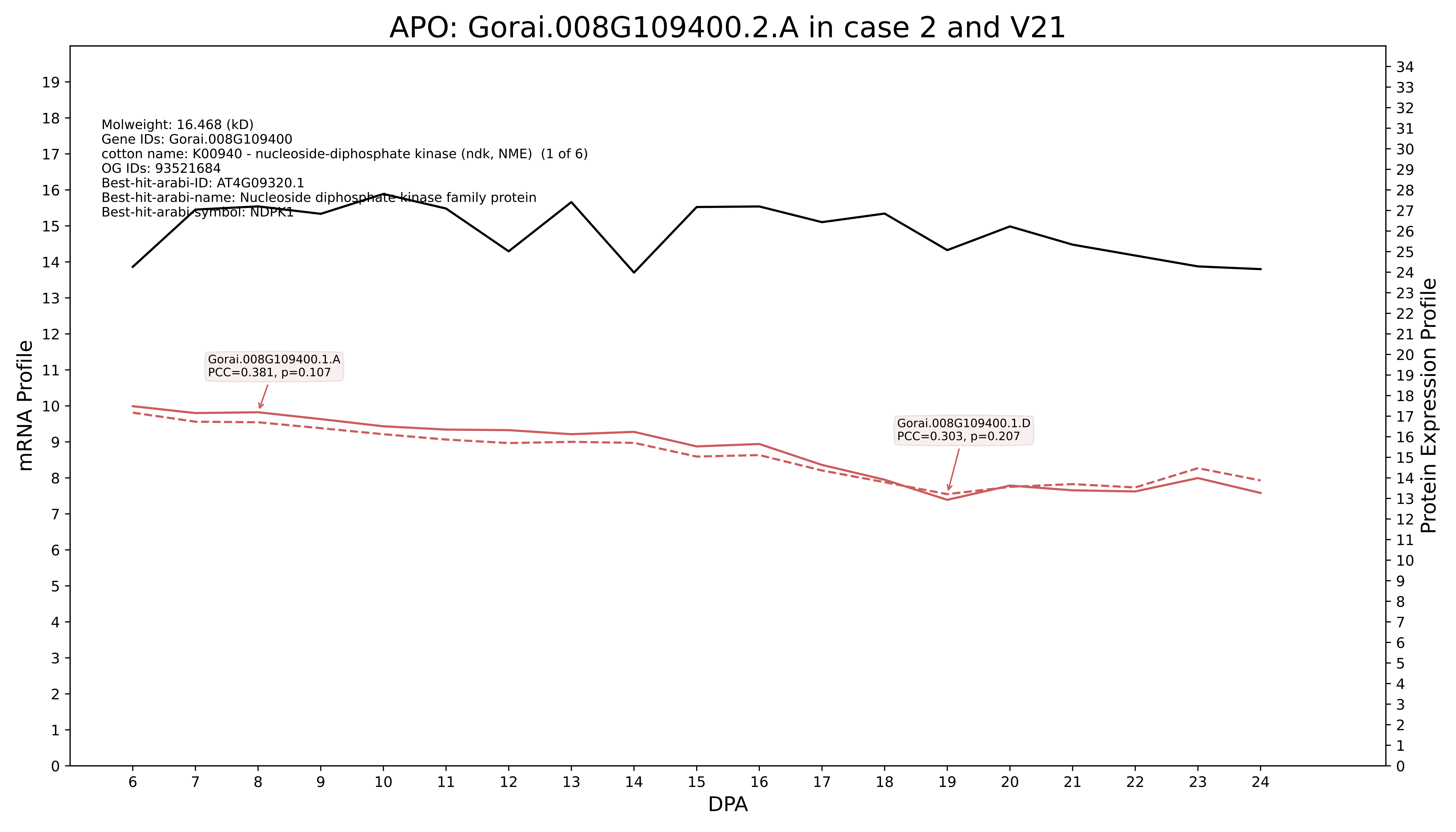
APO: Gorai.008G109400.2.A in case 2 and V21
Molweight: 16.468 (kD)
Gene IDs: Gorai.008G109400
cotton name: K00940 - nucleoside-diphosphate kinase (ndk, NME) (1 of 6)
OG IDs: 93521684
Best-hit-arabi-ID: AT4G09320.1
Best-hit-arabi-name: Nucleoside diphosphate kinase family protein
Best-hit-arabi-symbol: NDPK1
Gorai.008G109400.1.A
PCC=0.381, p=0.107
Gorai.008G109400.1.D
PCC=0.303, p=0.207
mRNA Profile
Protein Expression Profile
DPA

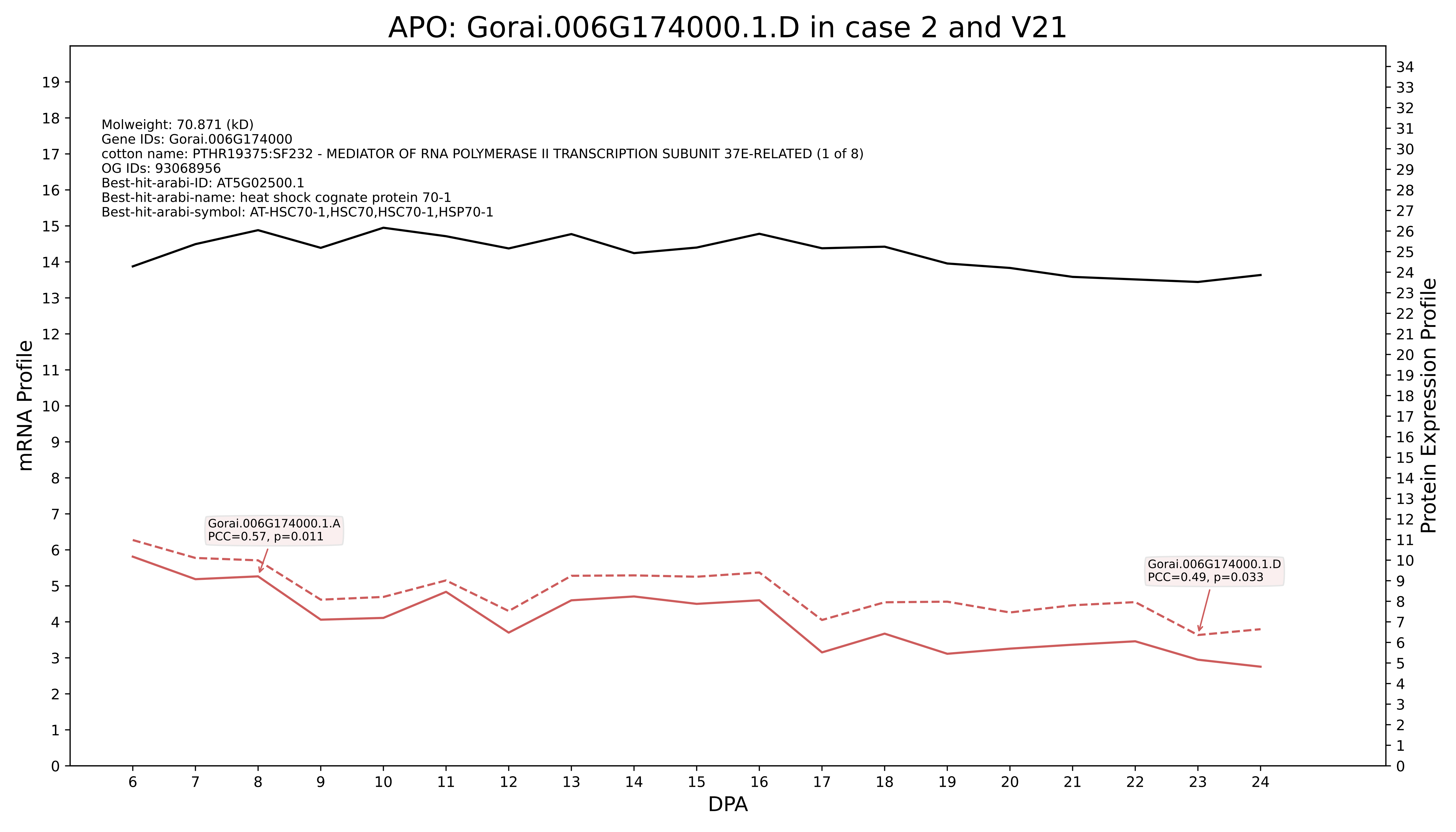
APO: Gorai.006G174000.1.D in case 2 and V21
Molweight: 70.871 (kD)
Gene IDs: Gorai.006G174000
cotton name: PTHR19375:SF232 - MEDIATOR OF RNA POLYMERASE II TRANSCRIPTION SUBUNIT 37E-RELATED (1 of 8)
OG IDs: 93068956
Best-hit-arabi-ID: AT5G02500.1
Best-hit-arabi-name: heat shock cognate protein 70-1
Best-hit-arabi-symbol: AT-HSC70-1,HSC70,HSC70-1,HSP70-1
Gorai.006G174000.1.A
PCC=0.57, p=0.011
Gorai.006G174000.1.D
PCC=0.49, p=0.033
mRNA Profile
Protein Expression Profile
DPA

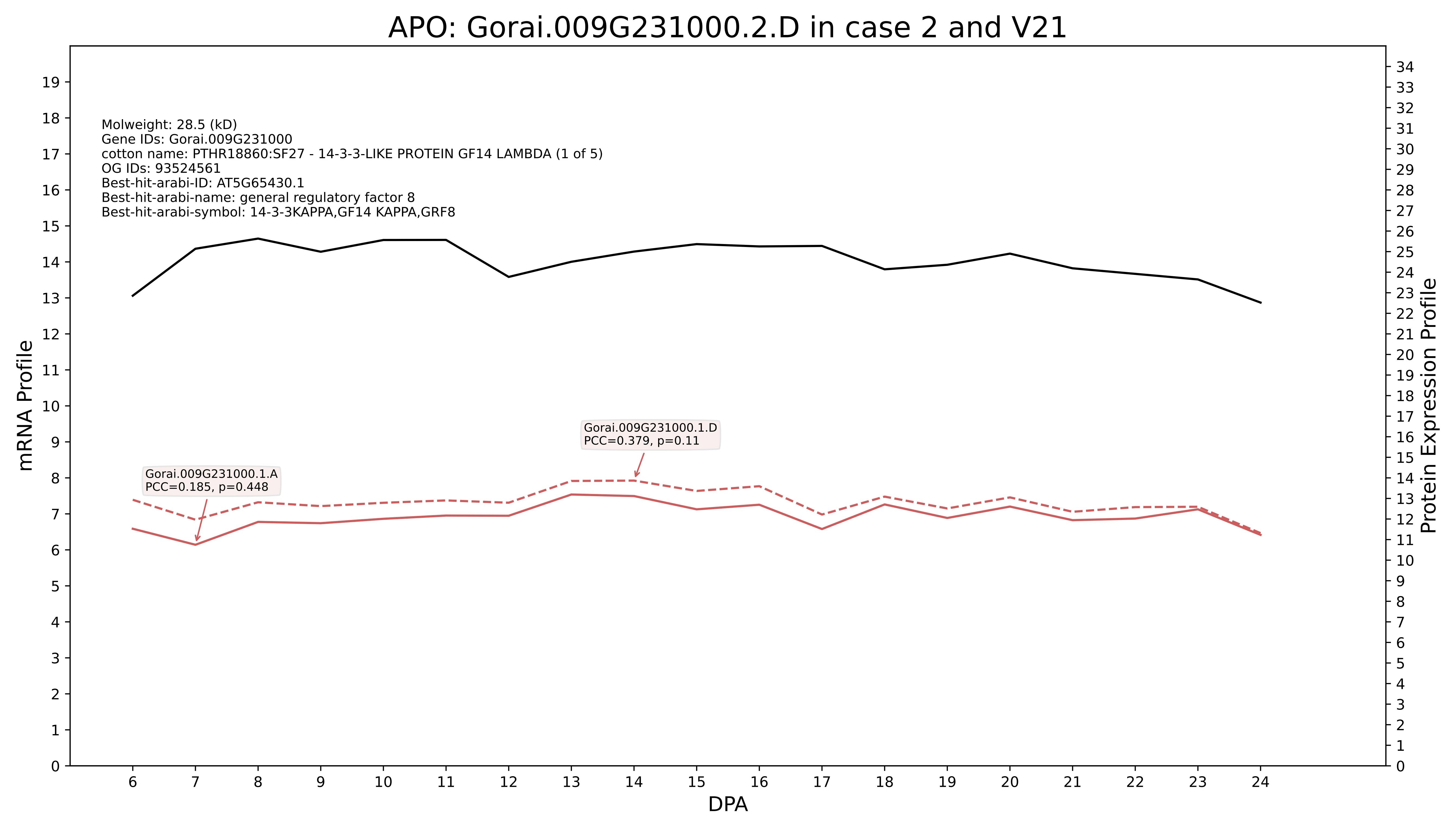
APO: Gorai.009G231000.2.D in case 2 and V21
Molweight: 28.5 (kD)
Gene IDs: Gorai.009G231000
cotton name: PTHR18860:SF27 - 14-3-3-LIKE PROTEIN GF14 LAMBDA (1 of 5)
OG IDs: 93524561
Best-hit-arabi-ID: AT5G65430.1
Best-hit-arabi-name: general regulatory factor 8
Best-hit-arabi-symbol: 14-3-3KAPPA,GF14 KAPPA,GRF8
Gorai.009G231000.1.D
PCC=0.379, p=0.11
Gorai.009G231000.1.A
PCC=0.185, p=0.448
mRNA Profile
Protein Expression Profile
DPA

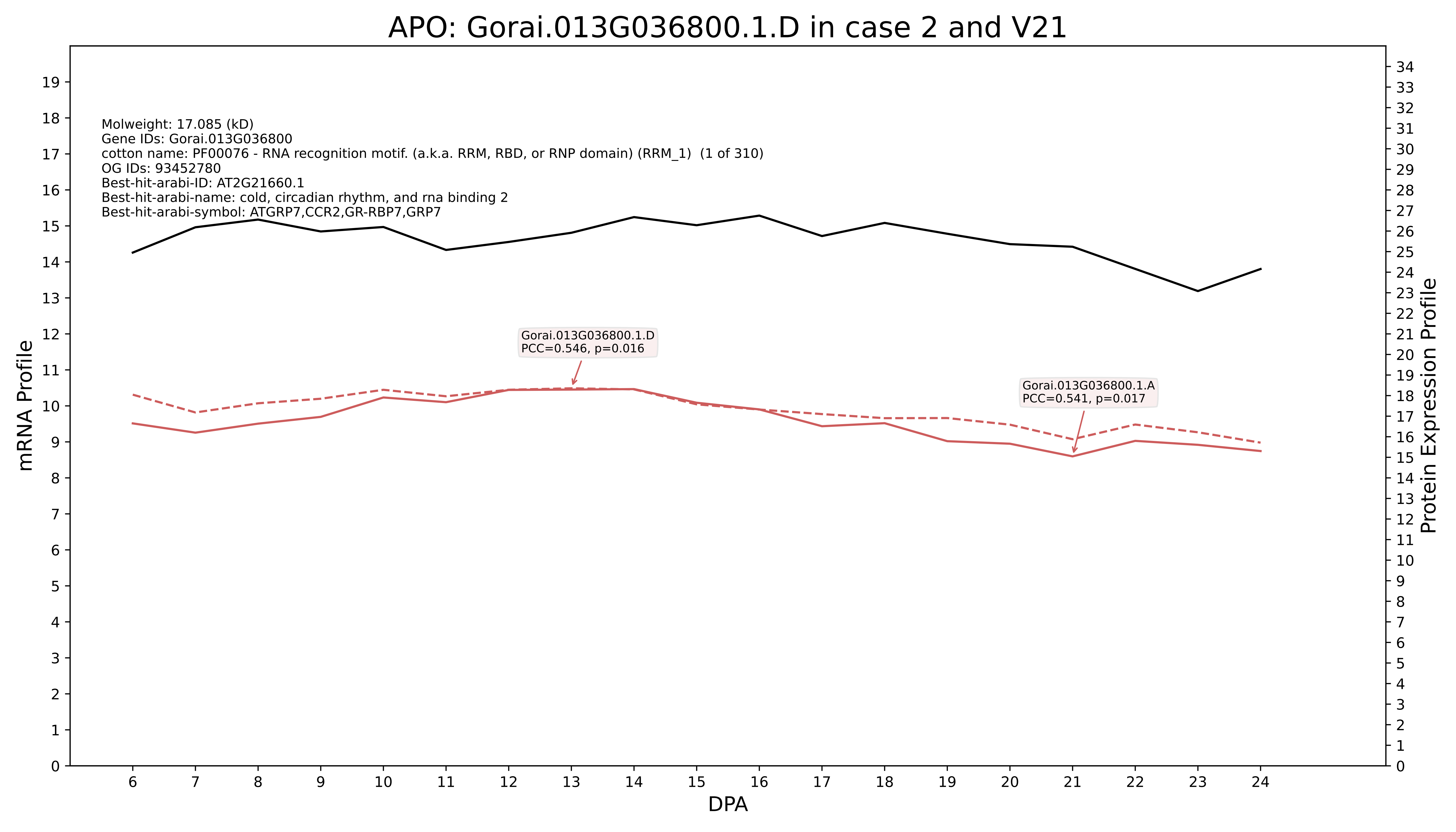
APO: Gorai.013G036800.1.D in case 2 and V21
Molweight: 17.085 (kD)
Gene IDs: Gorai.013G036800
cotton name: PF00076 - RNA recognition motif. (a.k.a. RRM, RBD, or RNP domain) (RRM_1) (1 of 310)
OG IDs: 93452780
Best-hit-arabi-ID: AT2G21660.1
Best-hit-arabi-name: cold, circadian rhythm, and rna binding 2
Best-hit-arabi-symbol: ATGRP7,CCR2,GR-RBP7,GRP7
Gorai.013G036800.1.D
PCC=0.546, p=0.016
Gorai.013G036800.1.A
PCC=0.541, p=0.017
mRNA Profile
Protein Expression Profile
DPA

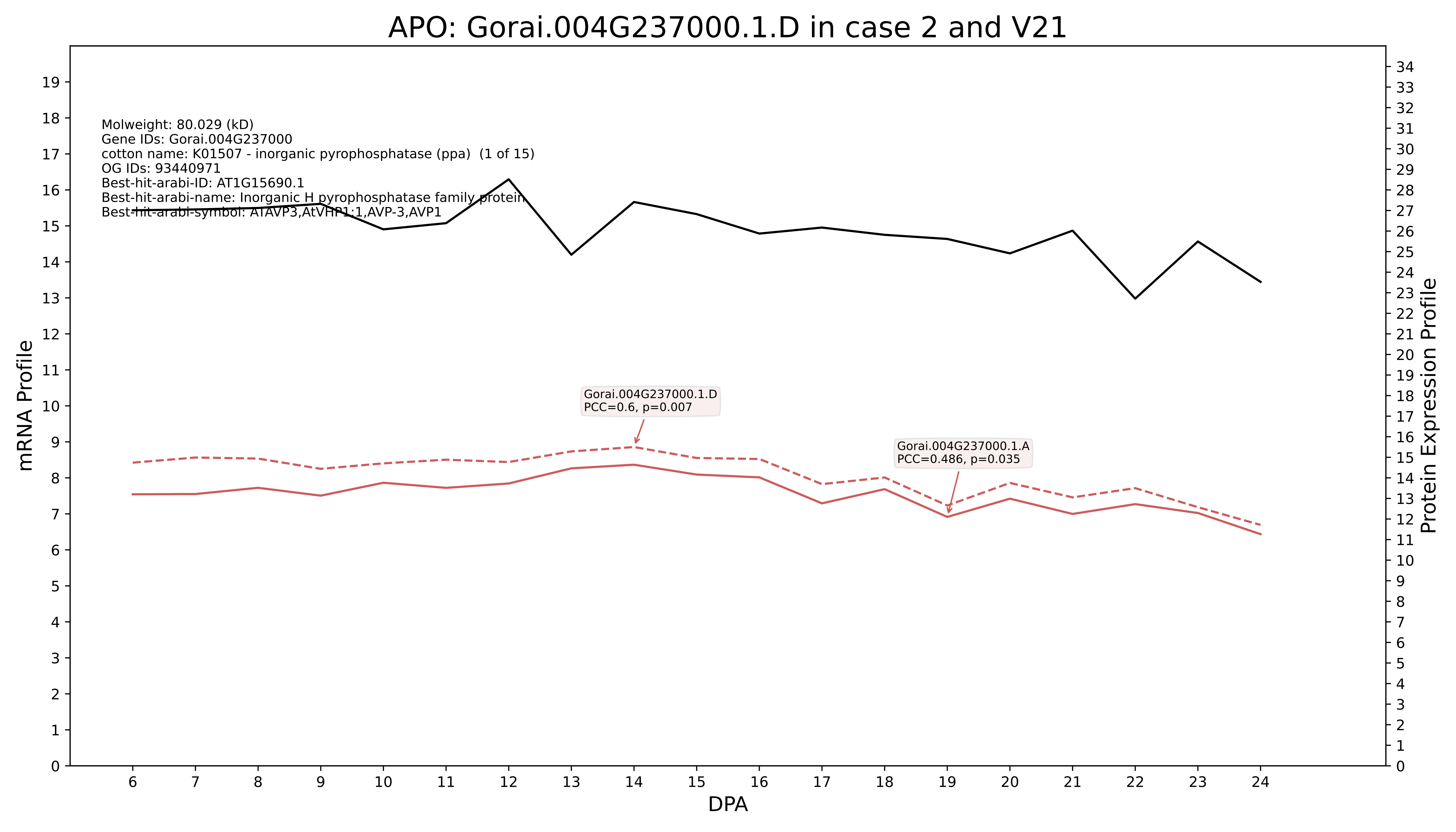
APO: Gorai.004G237000.1.D in case 2 and V21
Molweight: 80.029 (kD)
Gene IDs: Gorai.004G237000
cotton name: K01507 - inorganic pyrophosphatase (ppa) (1 of 15)
OG IDs: 93440971
Best-hit-arabi-ID: AT1G15690.1
Best-hit-arabi-name: Inorganic H pyrophosphatase family protein
Best-hit-arabi-symbol: ATAVP3,AtVHP1;1,AVP-3,AVP1
Gorai.004G237000.1.D
PCC=0.6, p=0.007
Gorai.004G237000.1.A
PCC=0.486, p=0.035
mRNA Profile
Protein Expression Profile
DPA

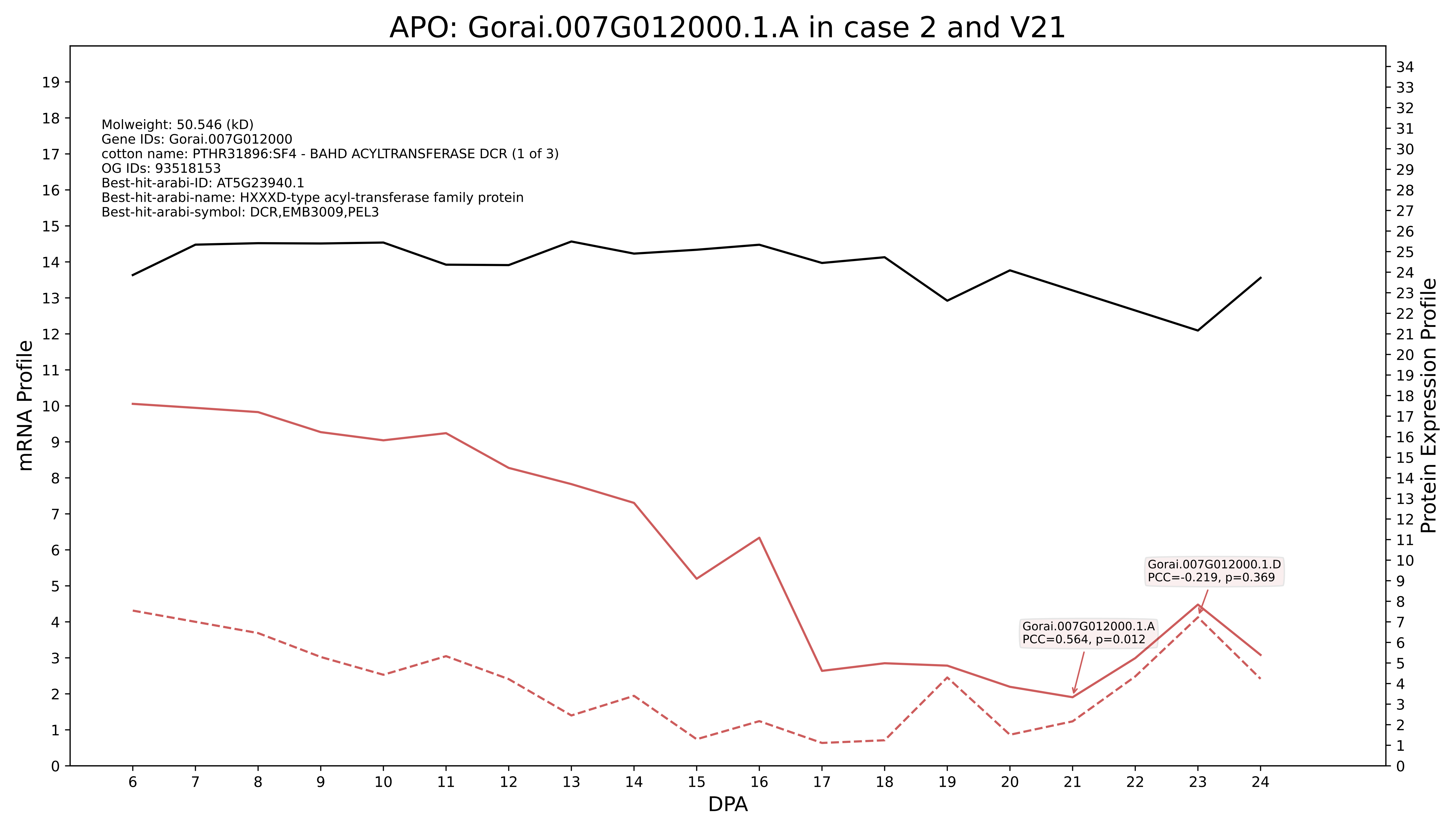

APO: Gorai.007G012000.1.A in case 2 and V21
Molweight: 50.546 (kD)
Gene IDs: Gorai.007G012000
cotton name: PTHR31896:SF4 - BAHD ACYLTRANSFERASE DCR (1 of 3)
OG IDs: 93518153
Best-hit-arabi-ID: AT5G23940.1
Best-hit-arabi-name: HXXXD-type acyl-transferase family protein
Best-hit-arabi-symbol: DCR,EMB3009,PEL3
Gorai.007G012000.1.D
PCC=-0.219, p=0.369
Gorai.007G012000.1.A
PCC=0.564, p=0.012
mRNA Profile
Protein Expression Profile
DPA

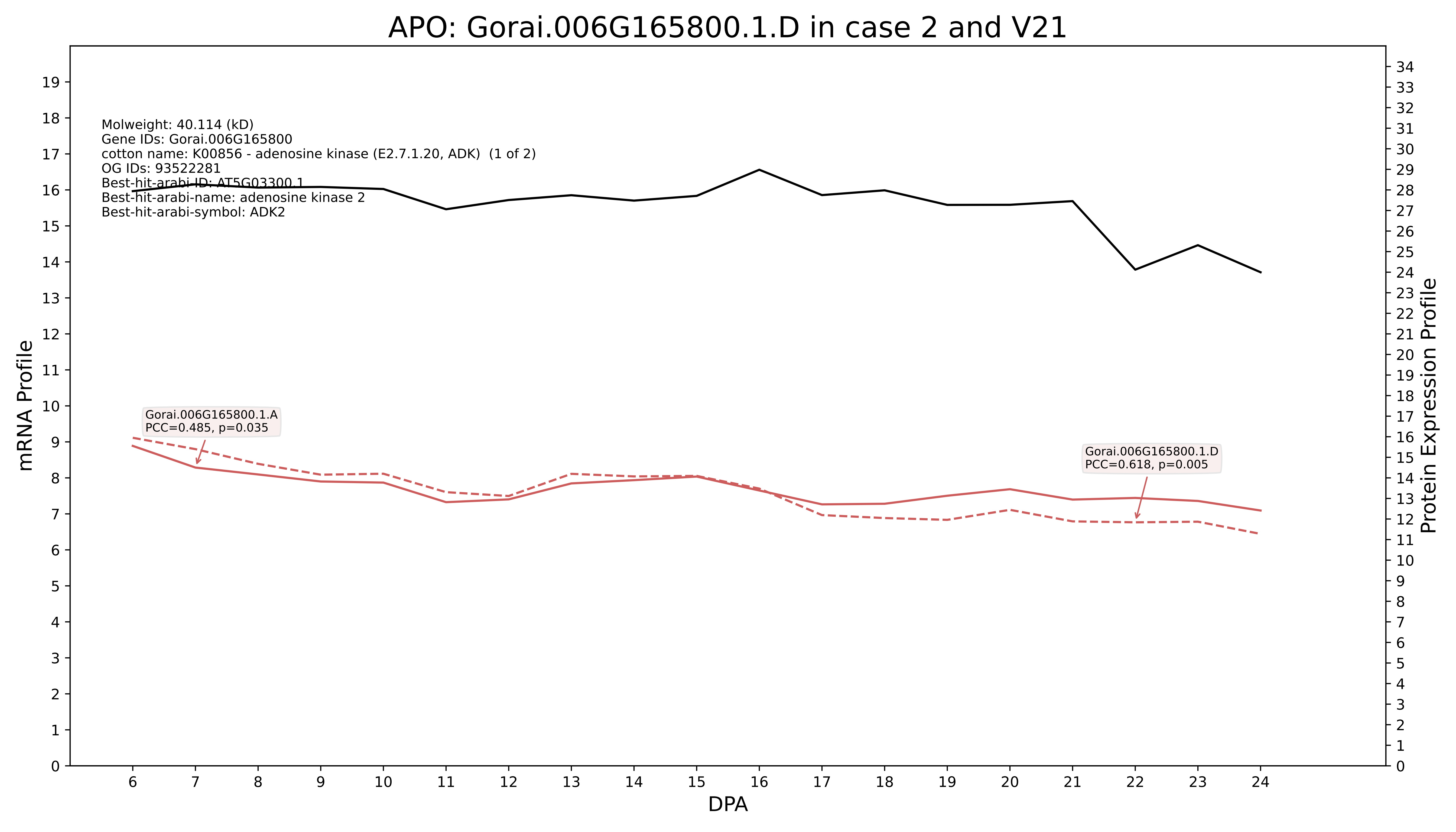
APO: Gorai.006G165800.1.D in case 2 and V21
Molweight: 40.114 (kD)
Gene IDs: Gorai.006G165800
cotton name: K00856 - adenosine kinase (E2.7.1.20, ADK) (1 of 2)
OG IDs: 93522281
Best-hit-arabi-ID: AT5G03300.1
Best-hit-arabi-name: adenosine kinase 2
Best-hit-arabi-symbol: ADK2
Gorai.006G165800.1.A
PCC=0.485, p=0.035
Gorai.006G165800.1.D
PCC=0.618, p=0.005
mRNA Profile
Protein Expression Profile
DPA

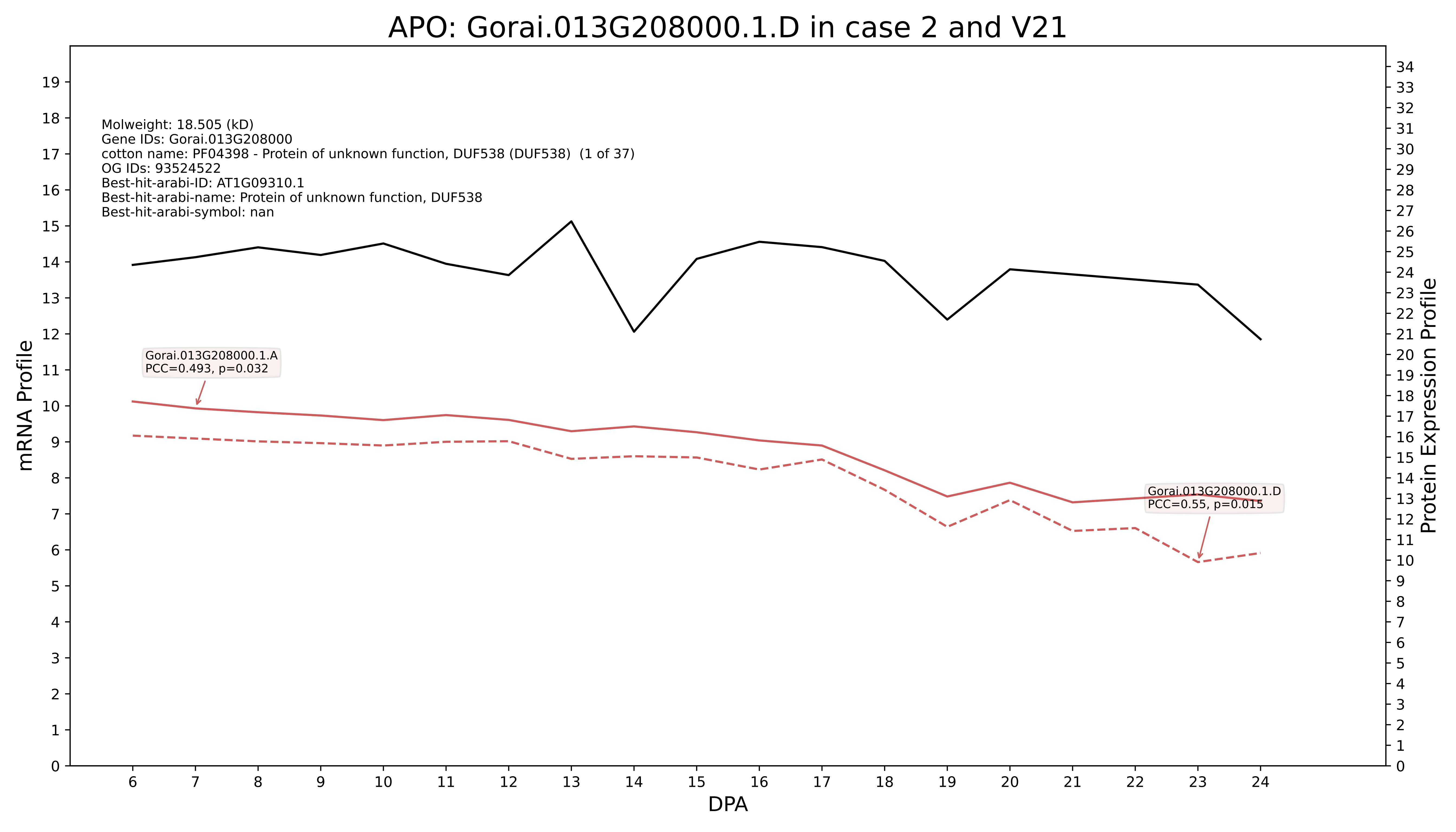

APO: Gorai.013G208000.1.D in case 2 and V21
Molweight: 18.505 (kD)
Gene IDs: Gorai.013G208000
cotton name: PF04398 - Protein of unknown function, DUF538 (DUF538) (1 of 37)
OG IDs: 93524522
Best-hit-arabi-ID: AT1G09310.1
Best-hit-arabi-name: Protein of unknown function, DUF538
Best-hit-arabi-symbol: nan
Gorai.013G208000.1.A
PCC=0.493, p=0.032
Gorai.013G208000.1.D
PCC=0.55, p=0.015
mRNA Profile
Protein Expression Profile
DPA

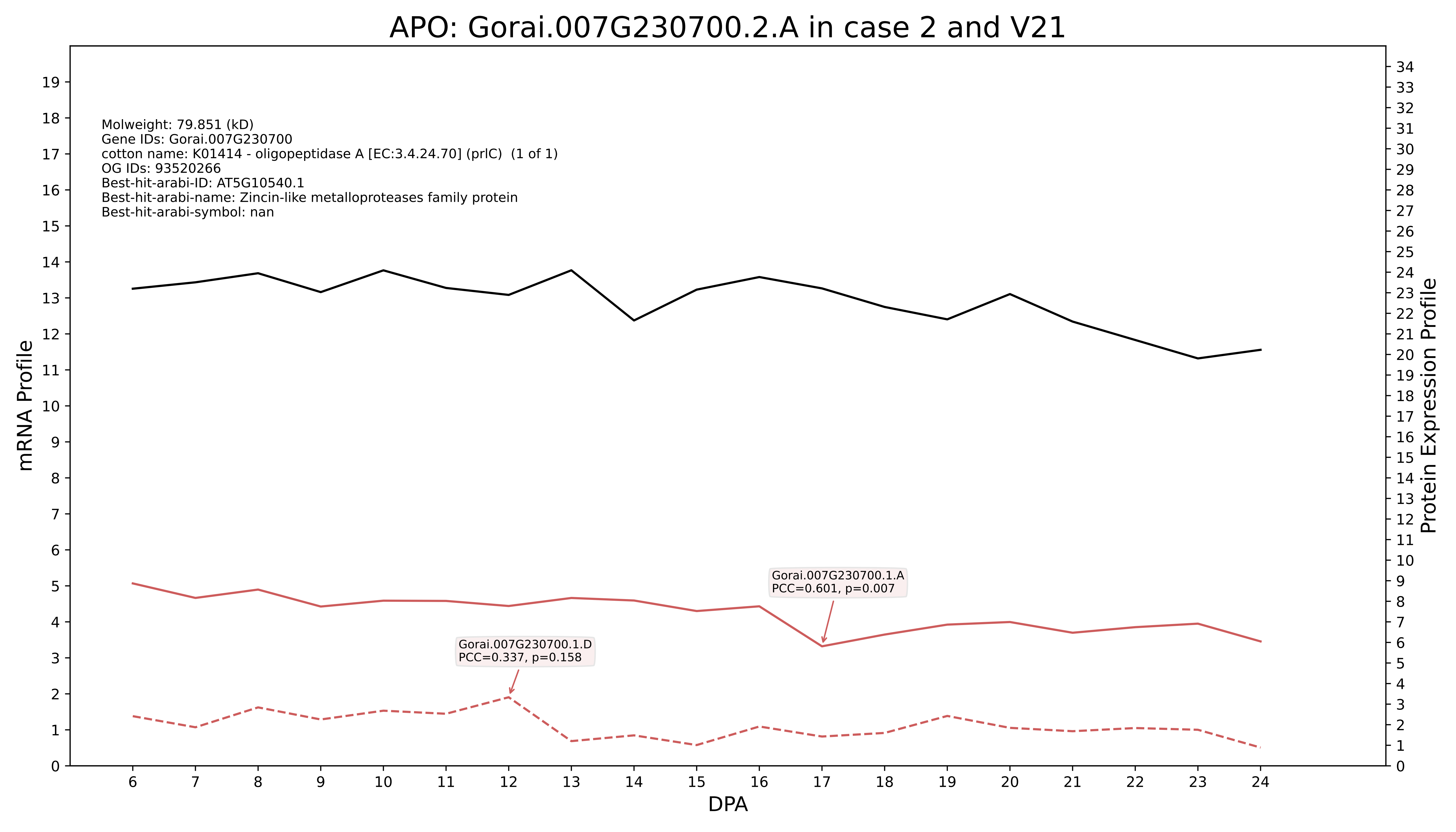
APO: Gorai.007G230700.2.A in case 2 and V21
Molweight: 79.851 (kD)
Gene IDs: Gorai.007G230700
cotton name: K01414 - oligopeptidase A [EC:3.4.24.70] (prlC) (1 of 1)
OG IDs: 93520266
Best-hit-arabi-ID: AT5G10540.1
Best-hit-arabi-name: Zincin-like metalloproteases family protein
Best-hit-arabi-symbol: nan
Gorai.007G230700.1.A
PCC=0.601, p=0.007
Gorai.007G230700.1.D
PCC=0.337, p=0.158
mRNA Profile
Protein Expression Profile
DPA

### APO: Gorai.006G223900.1.D in case 2 and V21

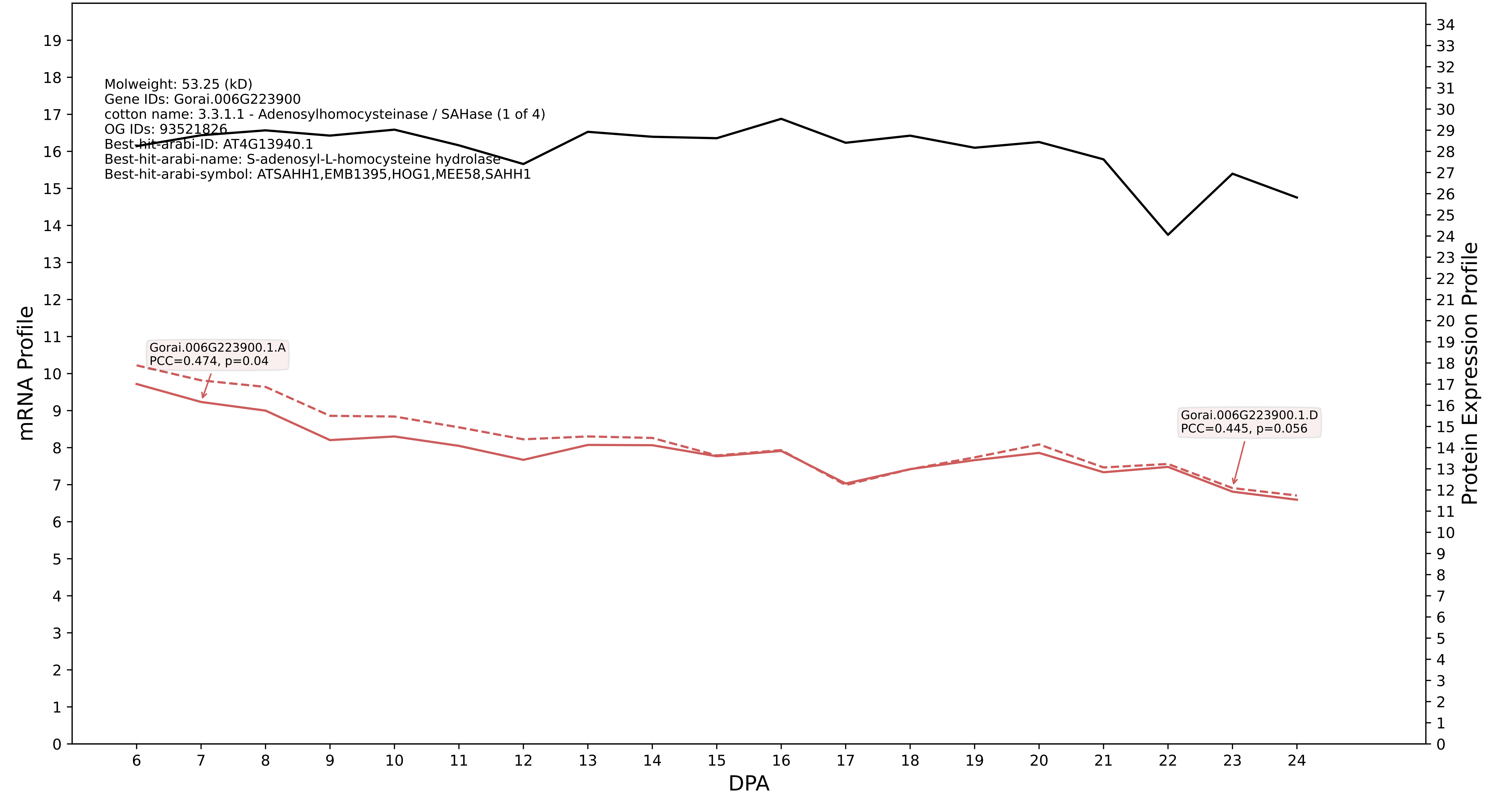

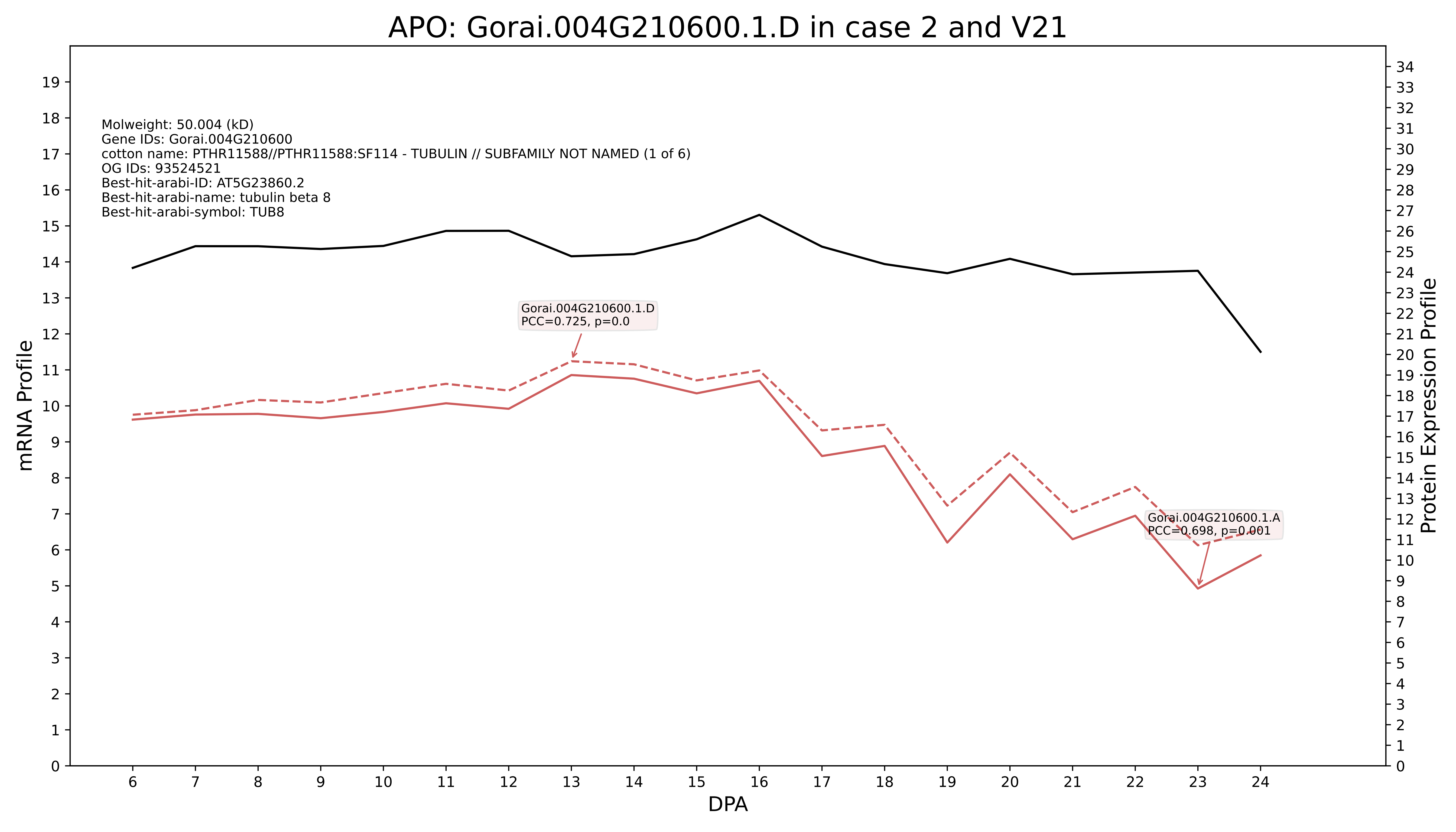
APO: Gorai.004G210600.1.D in case 2 and V21
Molweight: 50.004 (kD)
Gene IDs: Gorai.004G210600
cotton name: PTHR11588//PTHR11588:SF114 - TUBULIN // SUBFAMILY NOT NAMED (1 of 6)
OG IDs: 93524521
Best-hit-arabi-ID: AT5G23860.2
Best-hit-arabi-name: tubulin beta 8
Best-hit-arabi-symbol: TUB8
Gorai.004G210600.1.D
PCC=0.725, p=0.0
Gorai.004G210600.1.A
PCC=0.698, p=0.001
mRNA Profile
Protein Expression Profile
DPA

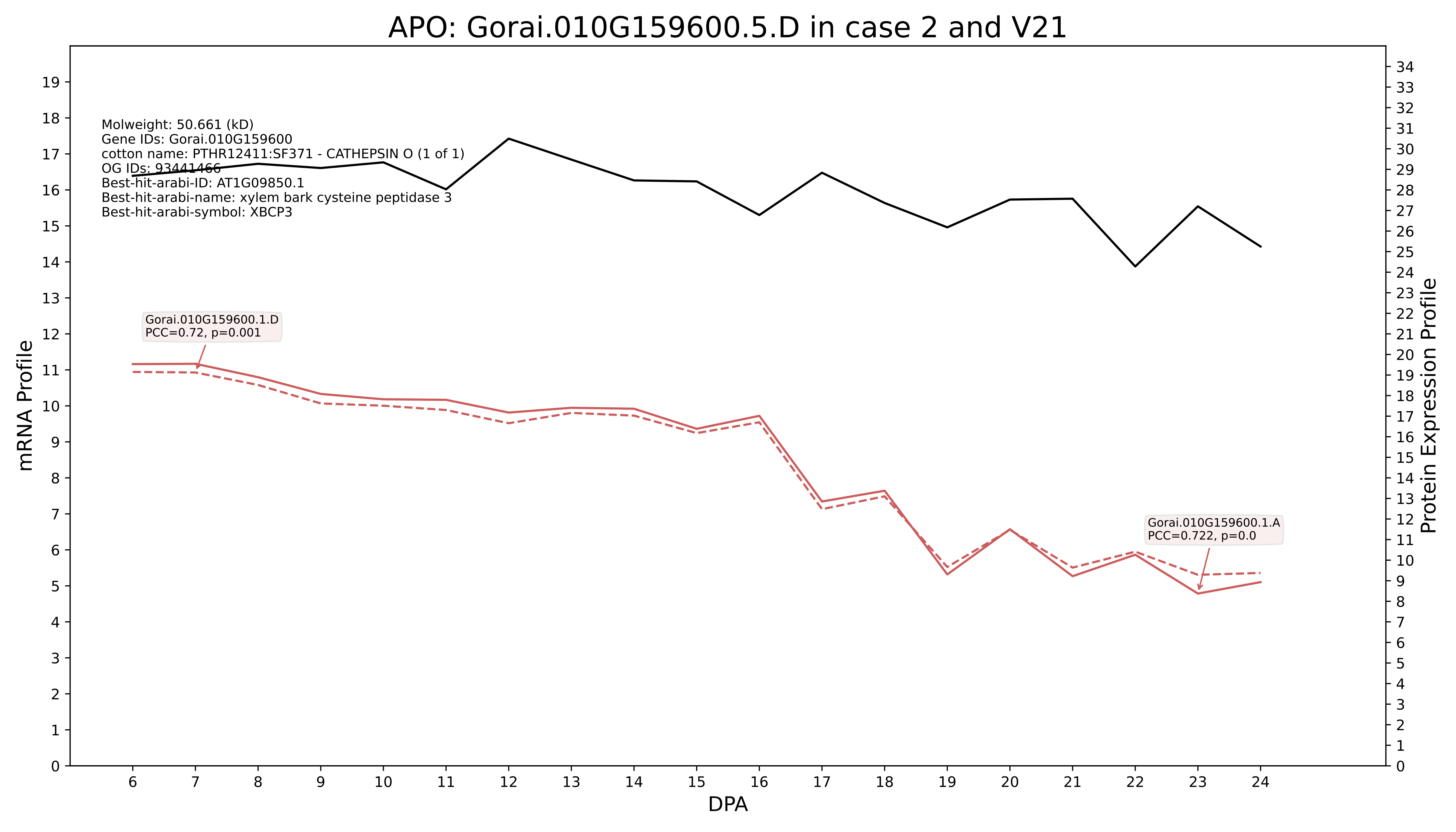

APO: Gorai.010G159600.5.D in case 2 and V21
Molweight: 50.661 (kD)
Gene IDs: Gorai.010G159600
cotton name: PTHR12411:SF371 - CATHEPSIN O (1 of 1)
OG IDs: 93441466
Best-hit-arabi-ID: AT1G09850.1
Best-hit-arabi-name: xylem bark cysteine peptidase 3
Best-hit-arabi-symbol: XBCP3
Gorai.010G159600.1.D
PCC=0.72, p=0.001
Gorai.010G159600.1.A
PCC=0.722, p=0.0
mRNA Profile
Protein Expression Profile
DPA

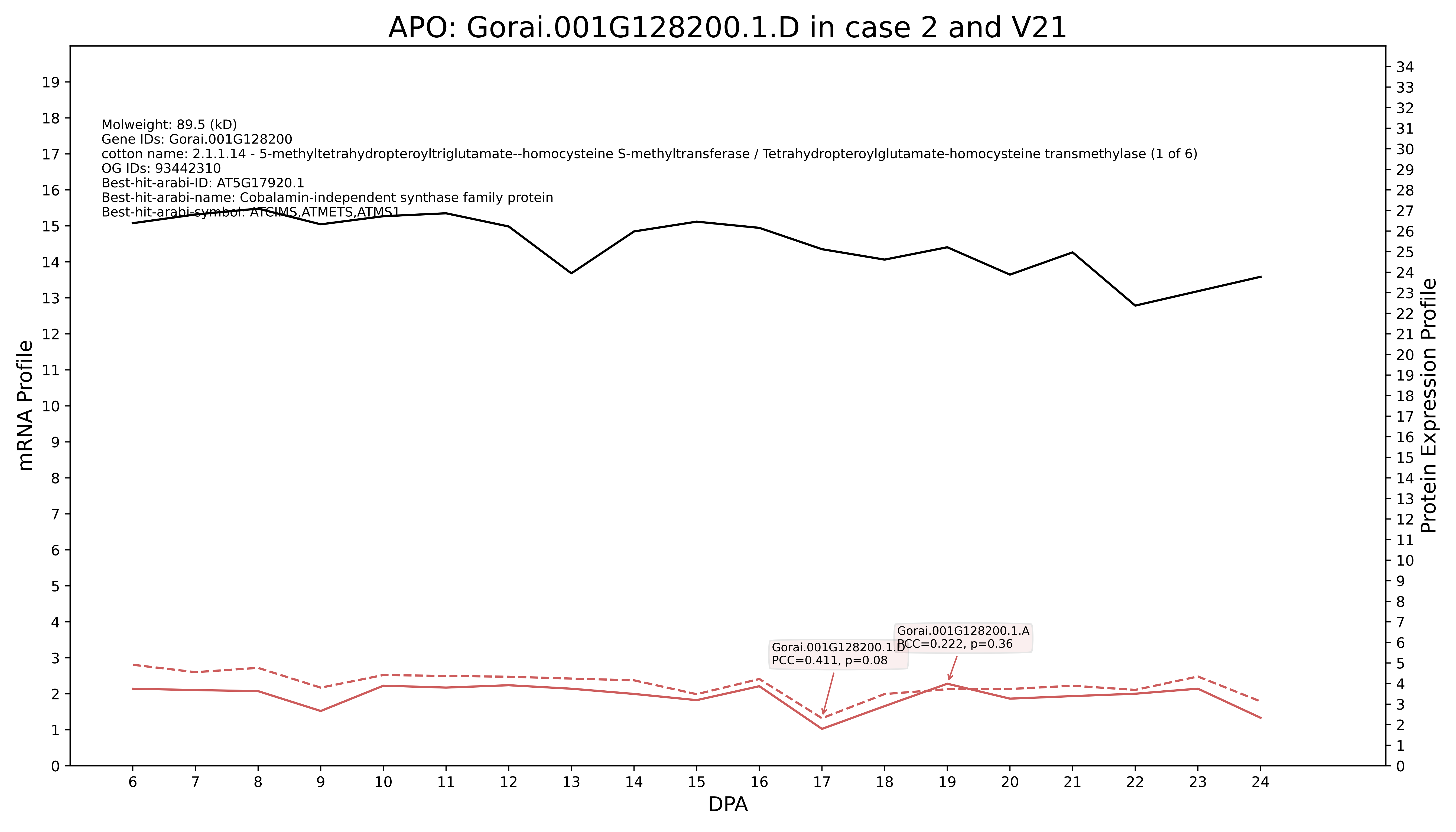
APO: Gorai.001G128200.1.D in case 2 and V21
Molweight: 89.5 (kD)
Gene IDs: Gorai.001G128200
cotton name: 2.1.1.14 - 5-methyltetrahydropteroyltriglutamate--homocysteine S-methyltransferase / Tetrahydropteroylglutamate-homocysteine transmethylase (1 of 6)
OG IDs: 93442310
Best-hit-arabi-ID: AT5G17920.1
Best-hit-arabi-name: Cobalamin-independent synthase family protein
Best-hit-arabi-symbol: ATCIMS,ATMETS,ATMS1
Gorai.001G128200.1.D
PCC=0.411, p=0.08
Gorai.001G128200.1.A
PCC=0.222, p=0.36
mRNA Profile
Protein Expression Profile
DPA

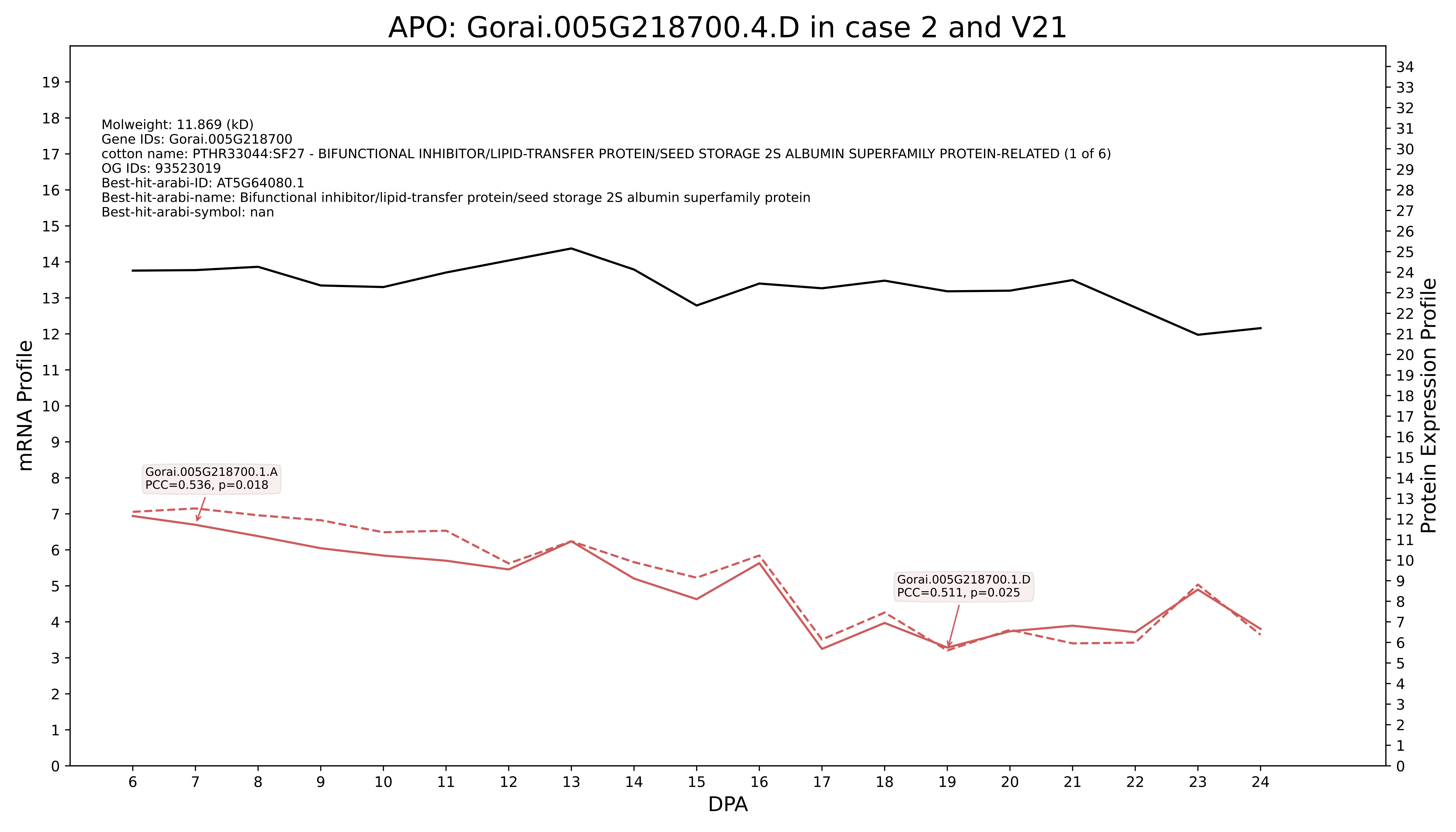

APO: Gorai.005G218700.4.D in case 2 and V21
Molweight: 11.869 (kD)
Gene IDs: Gorai.005G218700
cotton name: PTHR33044:SF27 - BIFUNCTIONAL INHIBITOR/LIPID-TRANSFER PROTEIN/SEED STORAGE 2S ALBUMIN SUPERFAMILY PROTEIN-RELATED (1 of 6)
OG IDs: 93523019
Best-hit-arabi-ID: AT5G64080.1
Best-hit-arabi-name: Bifunctional inhibitor/lipid-transfer protein/seed storage 2S albumin superfamily protein
Best-hit-arabi-symbol: nan
Gorai.005G218700.1.A
PCC=0.536, p=0.018
Gorai.005G218700.1.D
PCC=0.511, p=0.025
mRNA Profile
Protein Expression Profile
DPA

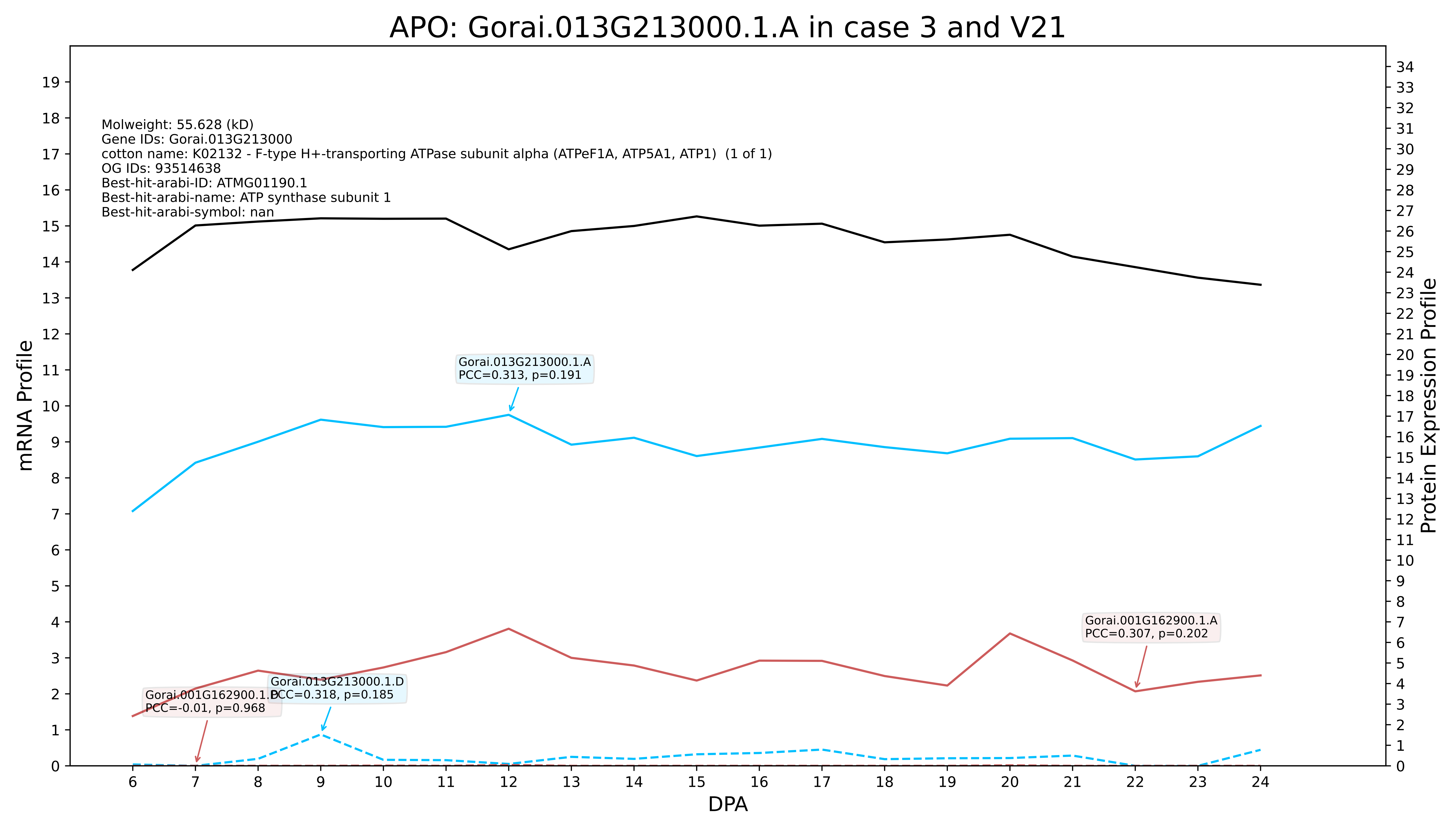

APO: Gorai.013G213000.1.A in case 3 and V21
Molweight: 55.628 (kD)
Gene IDs: Gorai.013G213000
cotton name: K02132 - F-type H+-transporting ATPase subunit alpha (ATPeF1A, ATP5A1, ATP1) (1 of 1)
OG IDs: 93514638
Best-hit-arabi-ID: ATMG01190.1
Best-hit-arabi-name: ATP synthase subunit 1
Best-hit-arabi-symbol: nan
Gorai.013G213000.1.A
PCC=0.313, p=0.191
Gorai.001G162900.1.A
PCC=0.307, p=0.202
Gorai.013G213000.1.D
PCC=0.318, p=0.185
Gorai.001G162900.1.D
PCC=-0.01, p=0.968
mRNA Profile
Protein Expression Profile
DPA

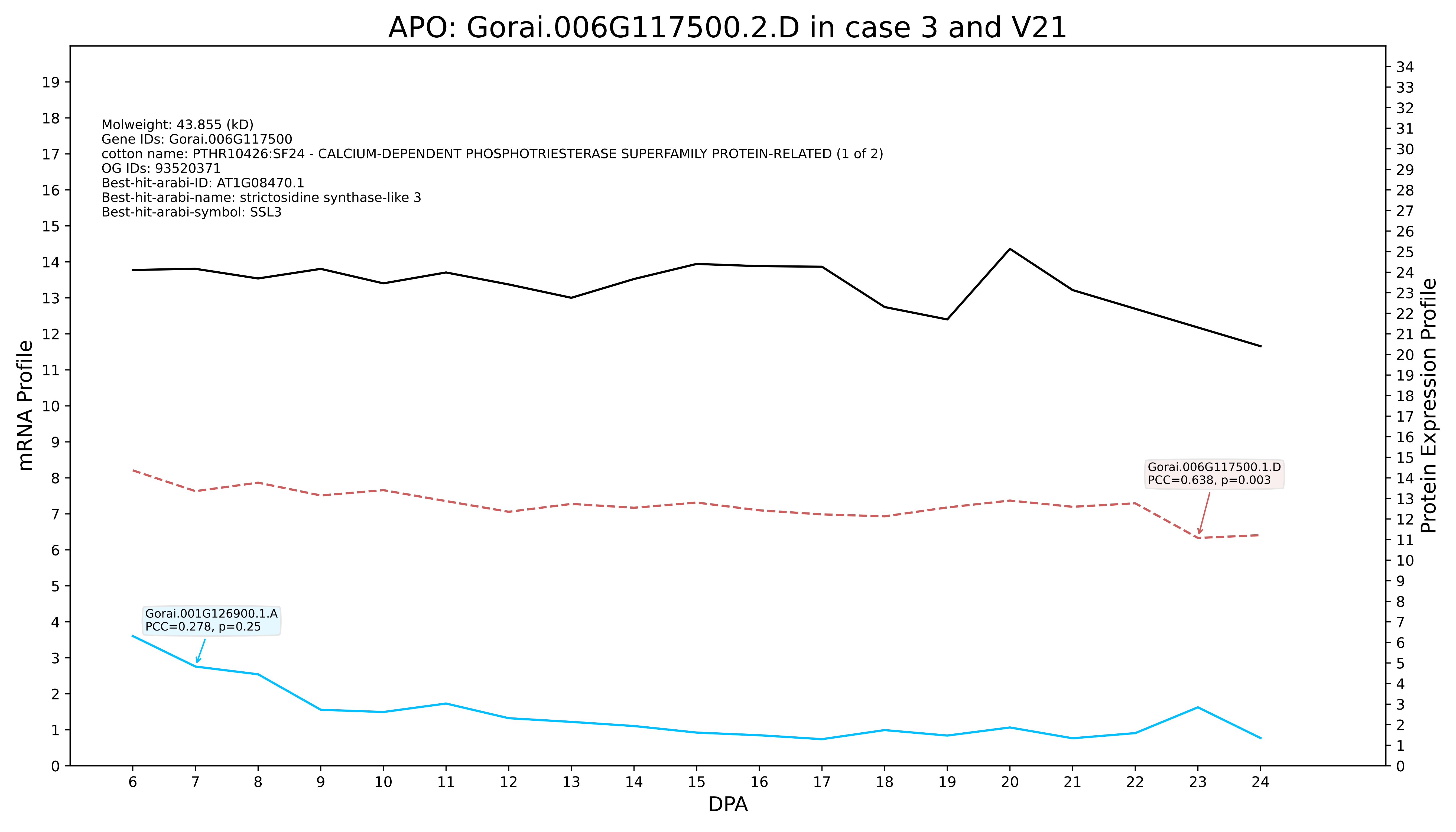

APO: Gorai.006G117500.2.D in case 3 and V21
Molweight: 43.855 (kD)
Gene IDs: Gorai.006G117500
cotton name: PTHR10426:SF24 - CALCIUM-DEPENDENT PHOSPHOTRIESTERASE SUPERFAMILY PROTEIN-RELATED (1 of 2)
OG IDs: 93520371
Best-hit-arabi-ID: AT1G08470.1
Best-hit-arabi-name: strictosidine synthase-like 3
Best-hit-arabi-symbol: SSL3
Gorai.006G117500.1.D
PCC=0.638, p=0.003
Gorai.001G126900.1.A
PCC=0.278, p=0.25
mRNA Profile
Protein Expression Profile
DPA

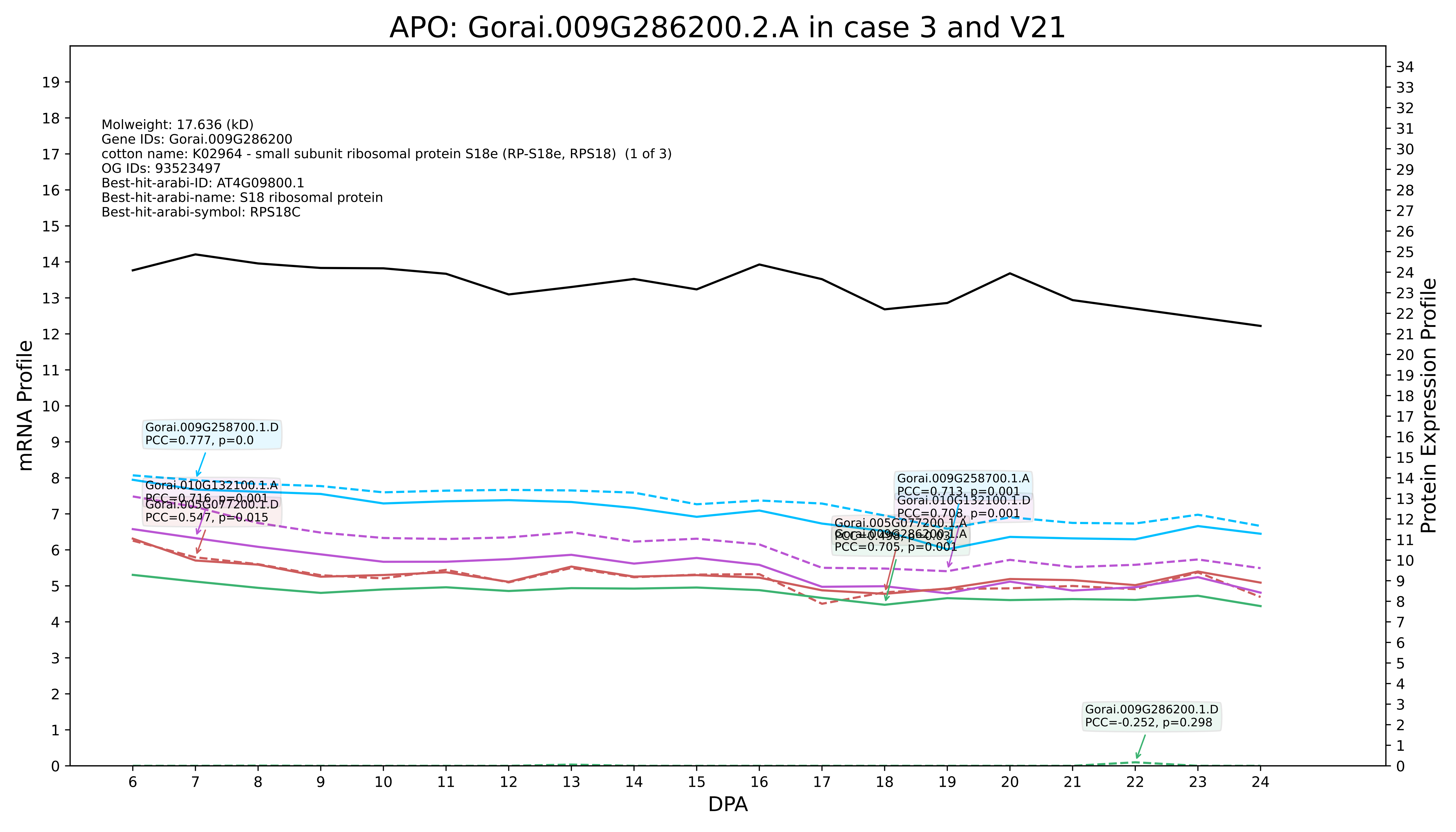

APO: Gorai.009G286200.2.A in case 3 and V21
Molweight: 17.636 (kD)
Gene IDs: Gorai.009G286200
cotton name: K02964 - small subunit ribosomal protein S18e (RP-S18e, RPS18) (1 of 3)
OG IDs: 93523497
Best-hit-arabi-ID: AT4G09800.1
Best-hit-arabi-name: S18 ribosomal protein
Best-hit-arabi-symbol: RPS18C
Gorai.009G258700.1.D
PCC=0.777, p=0.0
Gorai.010G132100.1.A
PCC=0.716, p=0.001
Gorai.005G077200.1.D
PCC=0.547, p=0.015
Gorai.009G258700.1.A
PCC=0.713, p=0.001
Gorai.010G132100.1.D
PCC=0.708, p=0.001
Gorai.005G077200.1.A
Gorai.009G286200.1.A
PCC=0.705, p=0.001
Gorai.009G286200.1.D
PCC=-0.252, p=0.298
mRNA Profile
Protein Expression Profile
DPA

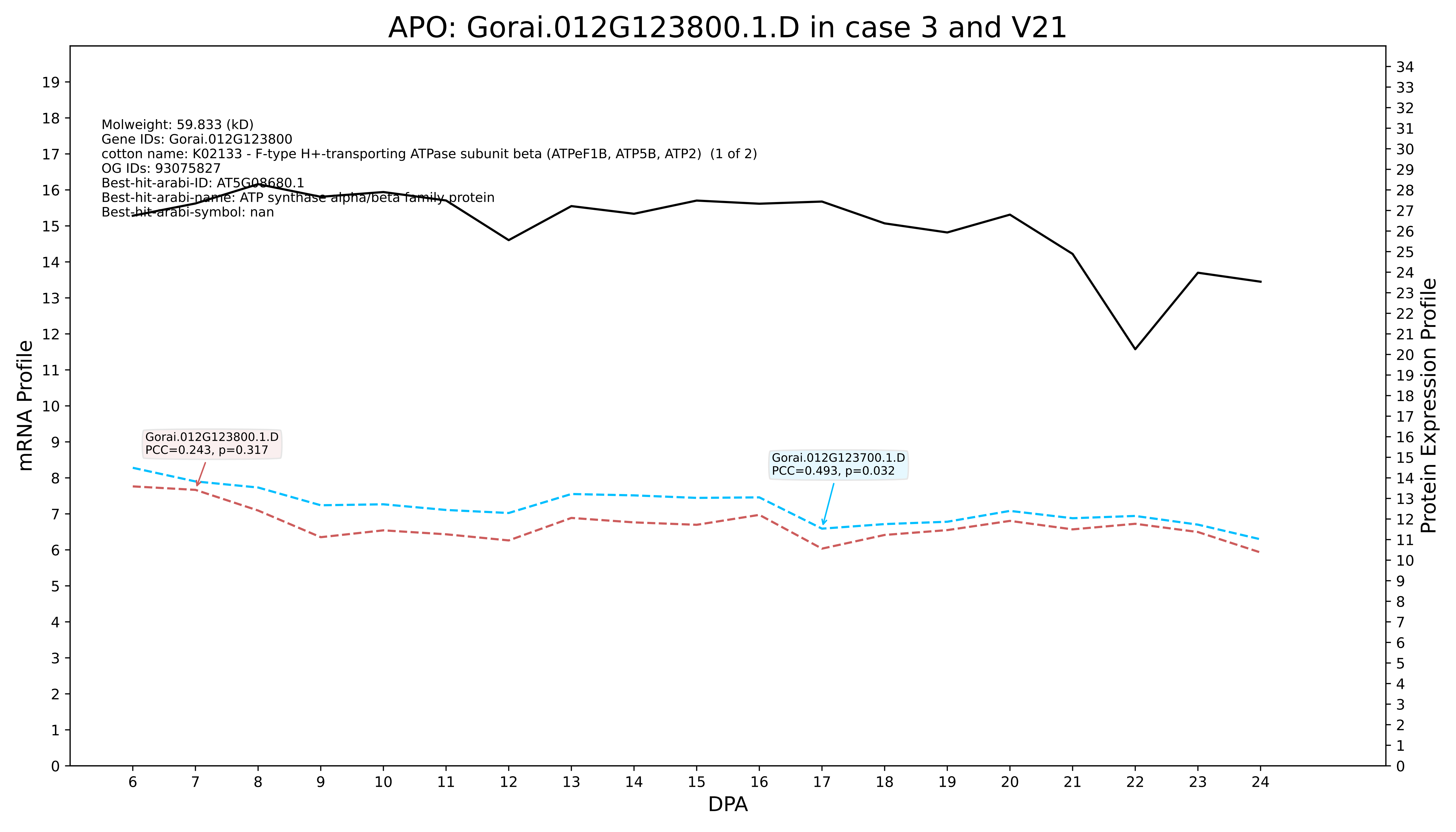
APO: Gorai.012G123800.1.D in case 3 and V21
Molweight: 59.833 (kD)
Gene IDs: Gorai.012G123800
cotton name: K02133 - F-type H+-transporting ATPase subunit beta (ATPeF1B, ATP5B, ATP2) (1 of 2)
OG IDs: 93075827
Best-hit-arabi-ID: AT5G08680.1
Best-hit-arabi-name: ATP synthase alpha/beta family protein
Best-hit-arabi-symbol: nan
Gorai.012G123800.1.D
PCC=0.243, p=0.317
Gorai.012G123700.1.D
PCC=0.493, p=0.032
mRNA Profile
Protein Expression Profile
DPA

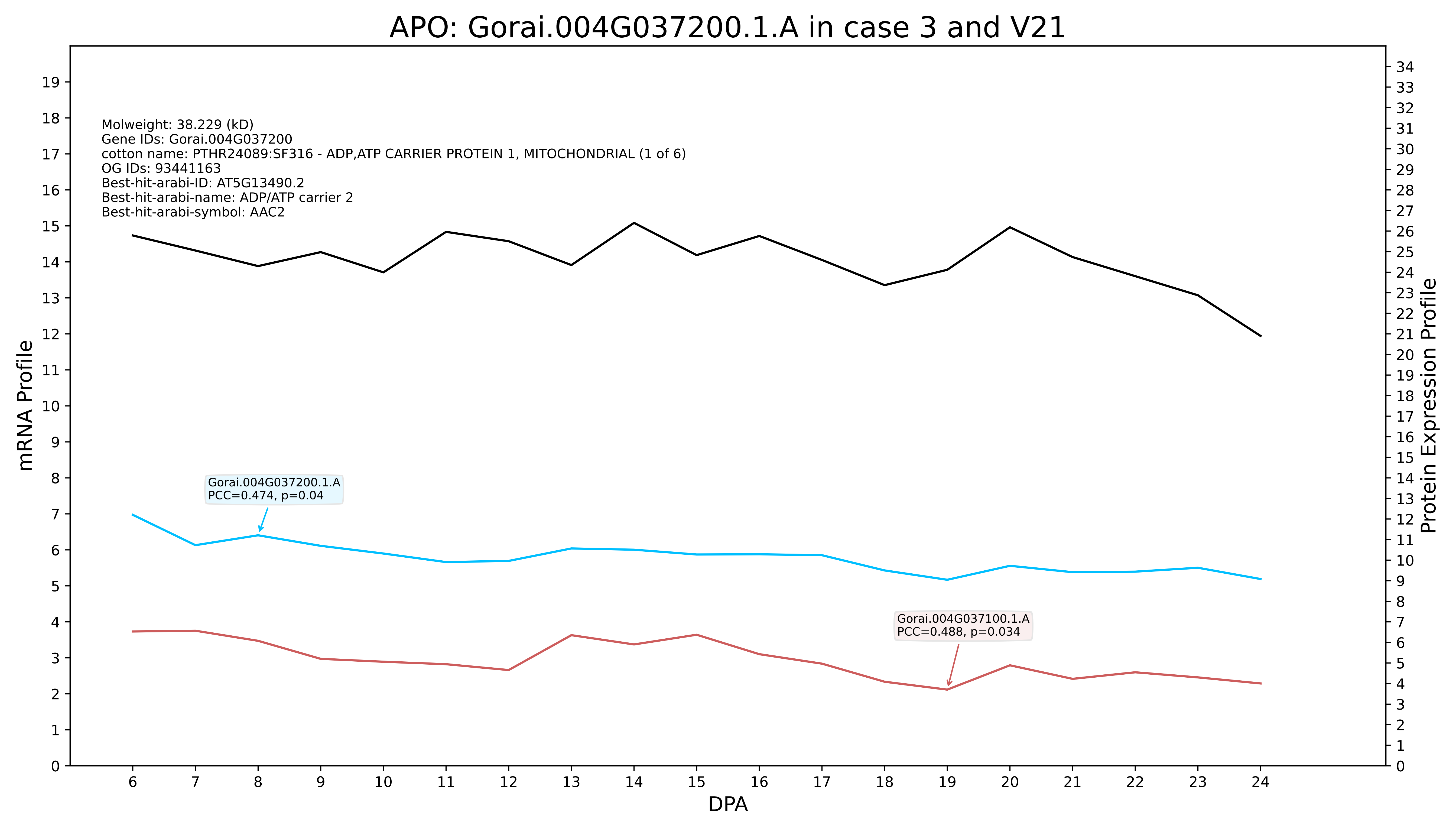
APO: Gorai.004G037200.1.A in case 3 and V21
Molweight: 38.229 (kD)
Gene IDs: Gorai.004G037200
cotton name: PTHR24089:SF316 - ADP,ATP CARRIER PROTEIN 1, MITOCHONDRIAL (1 of 6)
OG IDs: 93441163
Best-hit-arabi-ID: AT5G13490.2
Best-hit-arabi-name: ADP/ATP carrier 2
Best-hit-arabi-symbol: AAC2
Gorai.004G037200.1.A
PCC=0.474, p=0.04
Gorai.004G037100.1.A
PCC=0.488, p=0.034
mRNA Profile
Protein Expression Profile
DPA

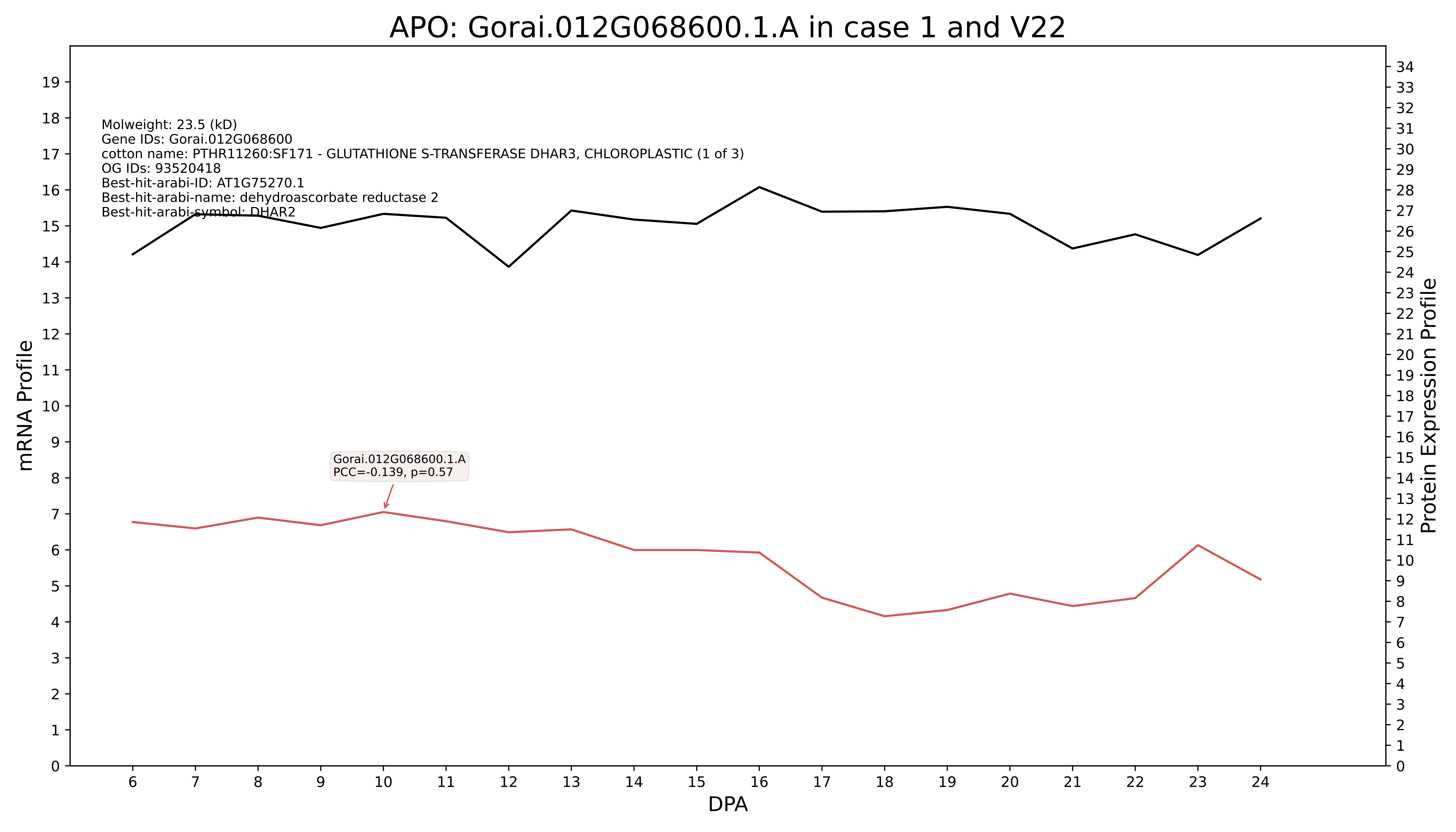

APO: Gorai.012G068600.1.A in case 1 and V22
Molweight: 23.5 (kD)
Gene IDs: Gorai.012G068600
cotton name: PTHR11260:SF171 - GLUTATHIONE S-TRANSFERASE DHAR3, CHLOROPLASTIC (1 of 3)
OG IDs: 93520418
Best-hit-arabi-ID: AT1G75270.1
Best-hit-arabi-name: dehydroascorbate reductase 2
Best-hit-arabi-symbol: DHAR2
Gorai.012G068600.1.A
PCC=-0.139, p=0.57
mRNA Profile
Protein Expression Profile
DPA

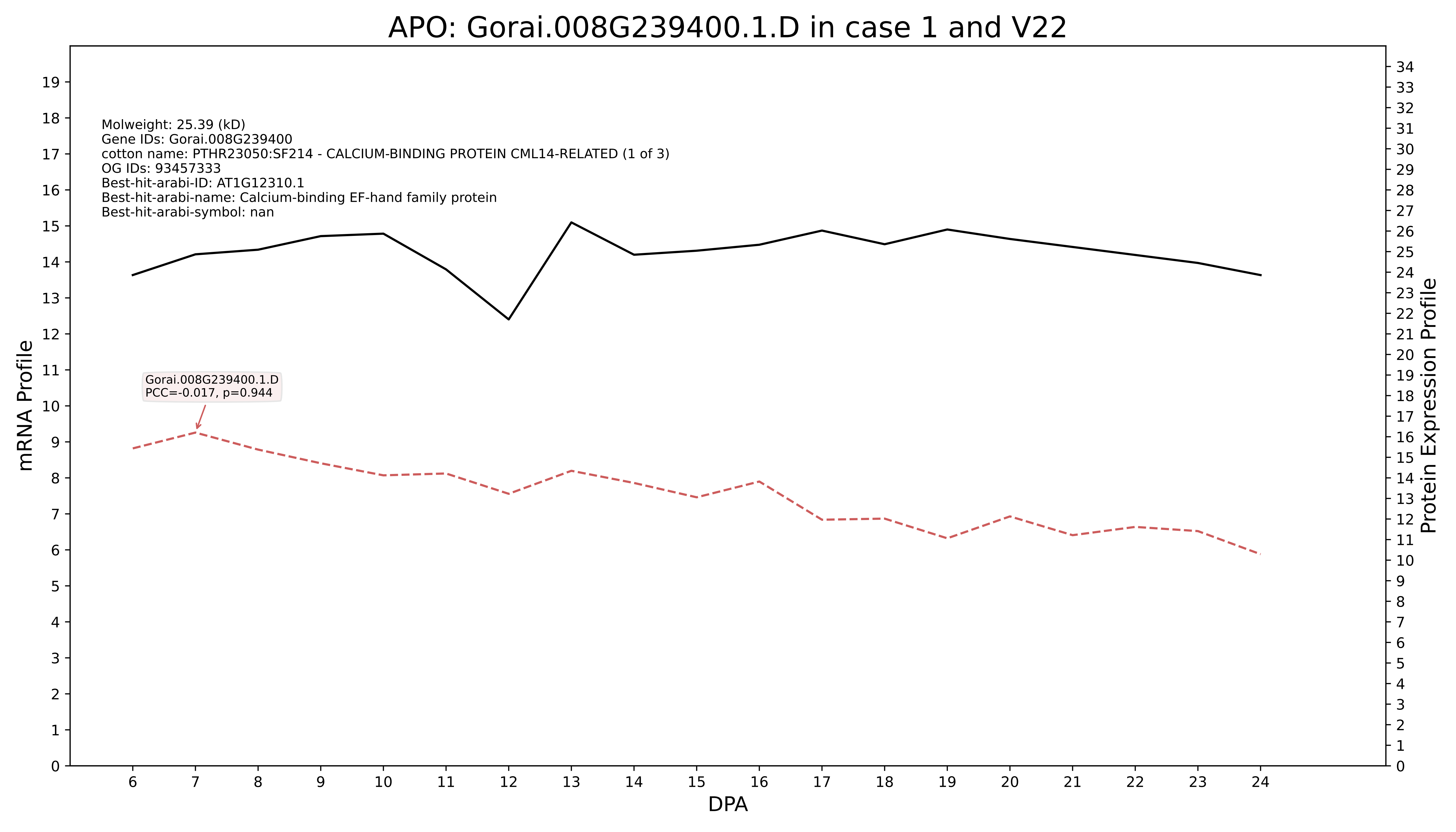

APO: Gorai.008G239400.1.D in case 1 and V22
Molweight: 25.39 (kD)
Gene IDs: Gorai.008G239400
cotton name: PTHR23050:SF214 - CALCIUM-BINDING PROTEIN CML14-RELATED (1 of 3)
OG IDs: 93457333
Best-hit-arabi-ID: AT1G12310.1
Best-hit-arabi-name: Calcium-binding EF-hand family protein
Best-hit-arabi-symbol: nan
Gorai.008G239400.1.D
PCC=-0.017, p=0.944
mRNA Profile
Protein Expression Profile
DPA

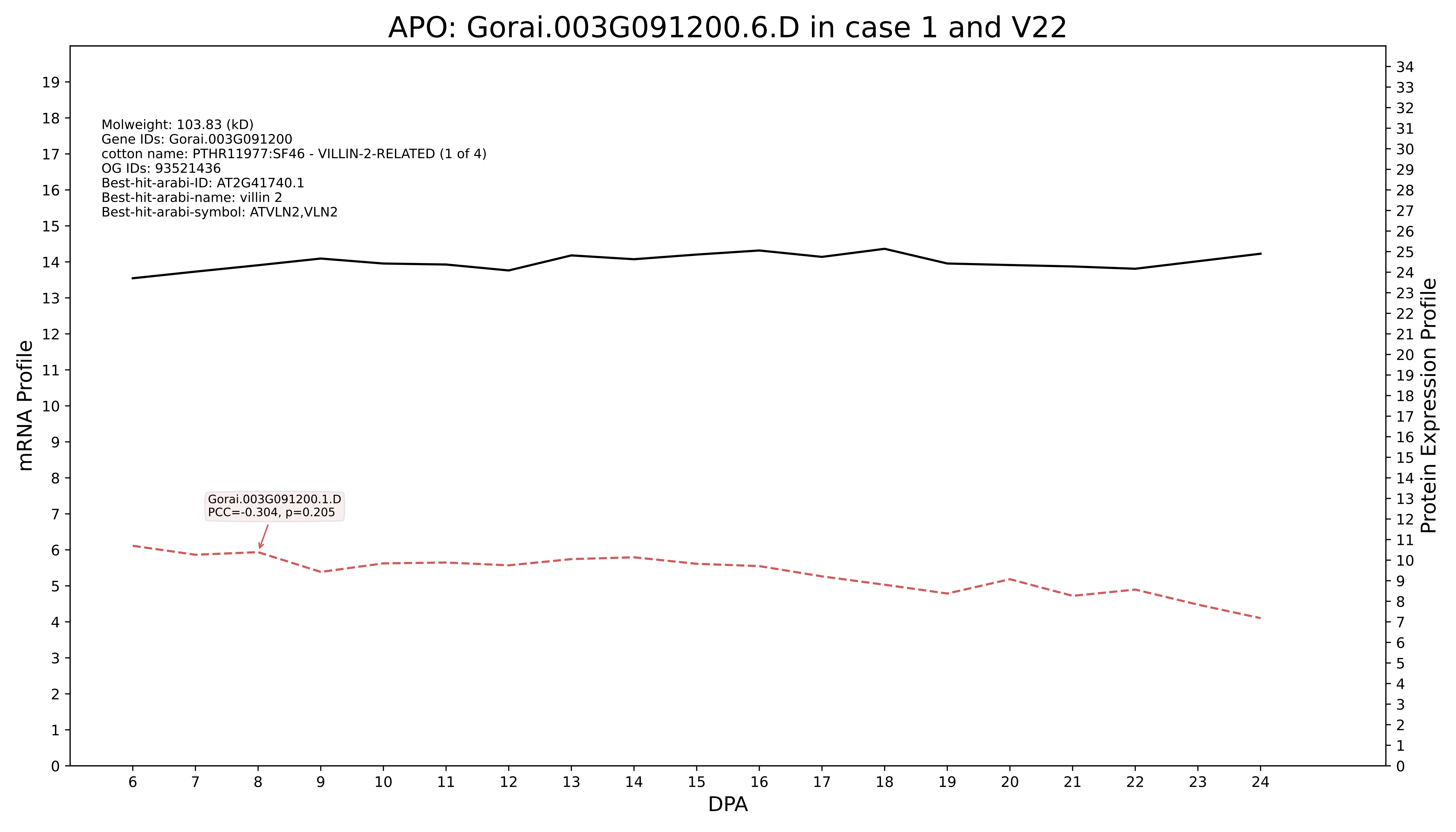

APO: Gorai.003G091200.6.D in case 1 and V22
Molweight: 103.83 (kD)
Gene IDs: Gorai.003G091200
cotton name: PTHR11977:SF46 - VILLIN-2-RELATED (1 of 4)
OG IDs: 93521436
Best-hit-arabi-ID: AT2G41740.1
Best-hit-arabi-name: villin 2
Best-hit-arabi-symbol: ATVLN2,VLN2
Gorai.003G091200.1.D
PCC=-0.304, p=0.205
mRNA Profile
Protein Expression Profile
DPA

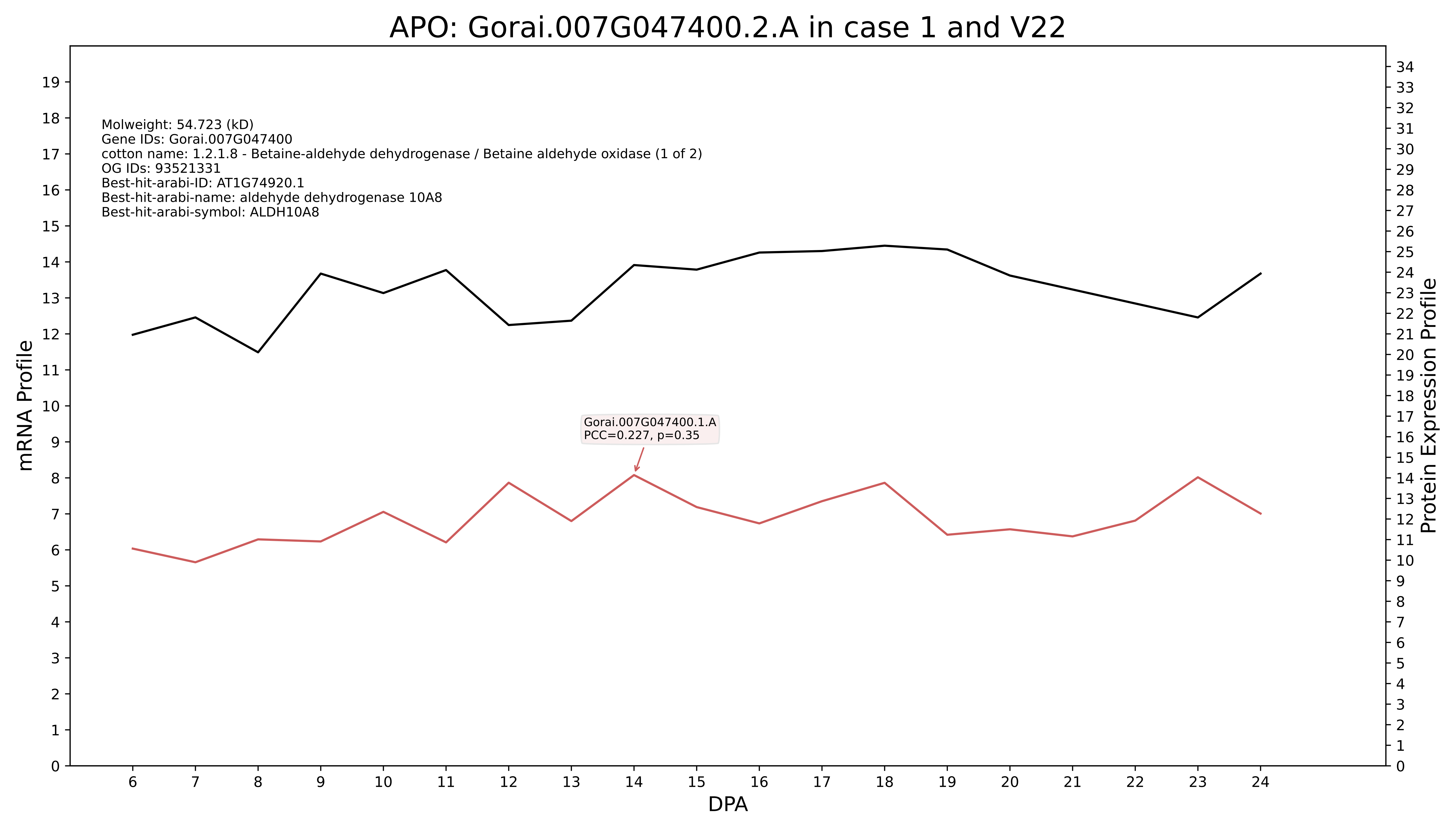

APO: Gorai.007G047400.2.A in case 1 and V22
Molweight: 54.723 (kD)
Gene IDs: Gorai.007G047400
cotton name: 1.2.1.8 - Betaine-aldehyde dehydrogenase / Betaine aldehyde oxidase (1 of 2)
OG IDs: 93521331
Best-hit-arabi-ID: AT1G74920.1
Best-hit-arabi-name: aldehyde dehydrogenase 10A8
Best-hit-arabi-symbol: ALDH10A8
Gorai.007G047400.1.A
PCC=0.227, p=0.35
mRNA Profile
Protein Expression Profile
DPA

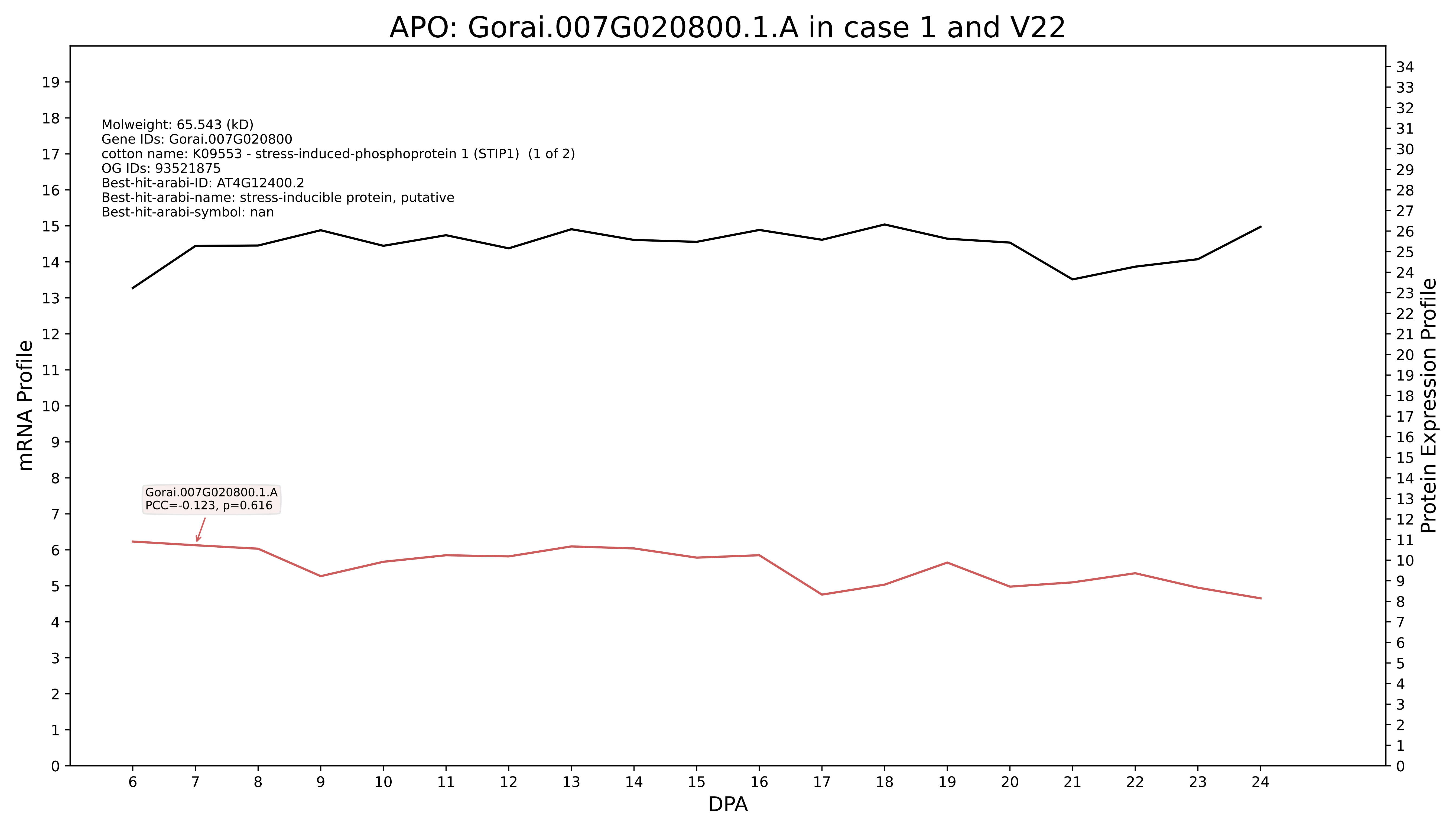

APO: Gorai.007G020800.1.A in case 1 and V22
Molweight: 65.543 (kD)
Gene IDs: Gorai.007G020800
cotton name: K09553 - stress-induced-phosphoprotein 1 (STIP1) (1 of 2)
OG IDs: 93521875
Best-hit-arabi-ID: AT4G12400.2
Best-hit-arabi-name: stress-inducible protein, putative
Best-hit-arabi-symbol: nan
Gorai.007G020800.1.A
PCC=-0.123, p=0.616
mRNA Profile
Protein Expression Profile
DPA

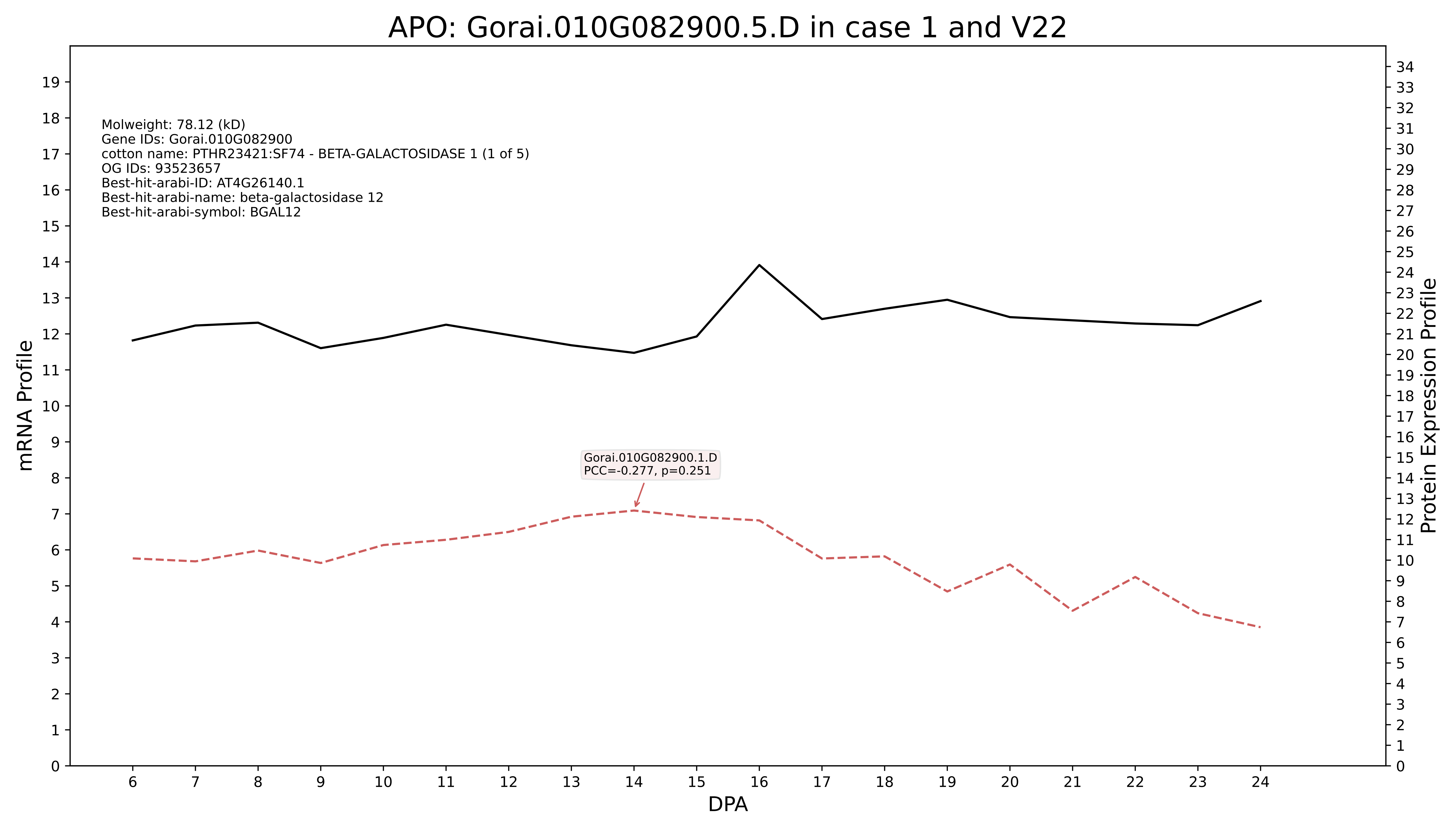

APO: Gorai.010G082900.5.D in case 1 and V22
Molweight: 78.12 (kD)
Gene IDs: Gorai.010G082900
cotton name: PTHR23421:SF74 - BETA-GALACTOSIDASE 1 (1 of 5)
OG IDs: 93523657
Best-hit-arabi-ID: AT4G26140.1
Best-hit-arabi-name: beta-galactosidase 12
Best-hit-arabi-symbol: BGAL12
Gorai.010G082900.1.D
PCC=-0.277, p=0.251
mRNA Profile
Protein Expression Profile
DPA

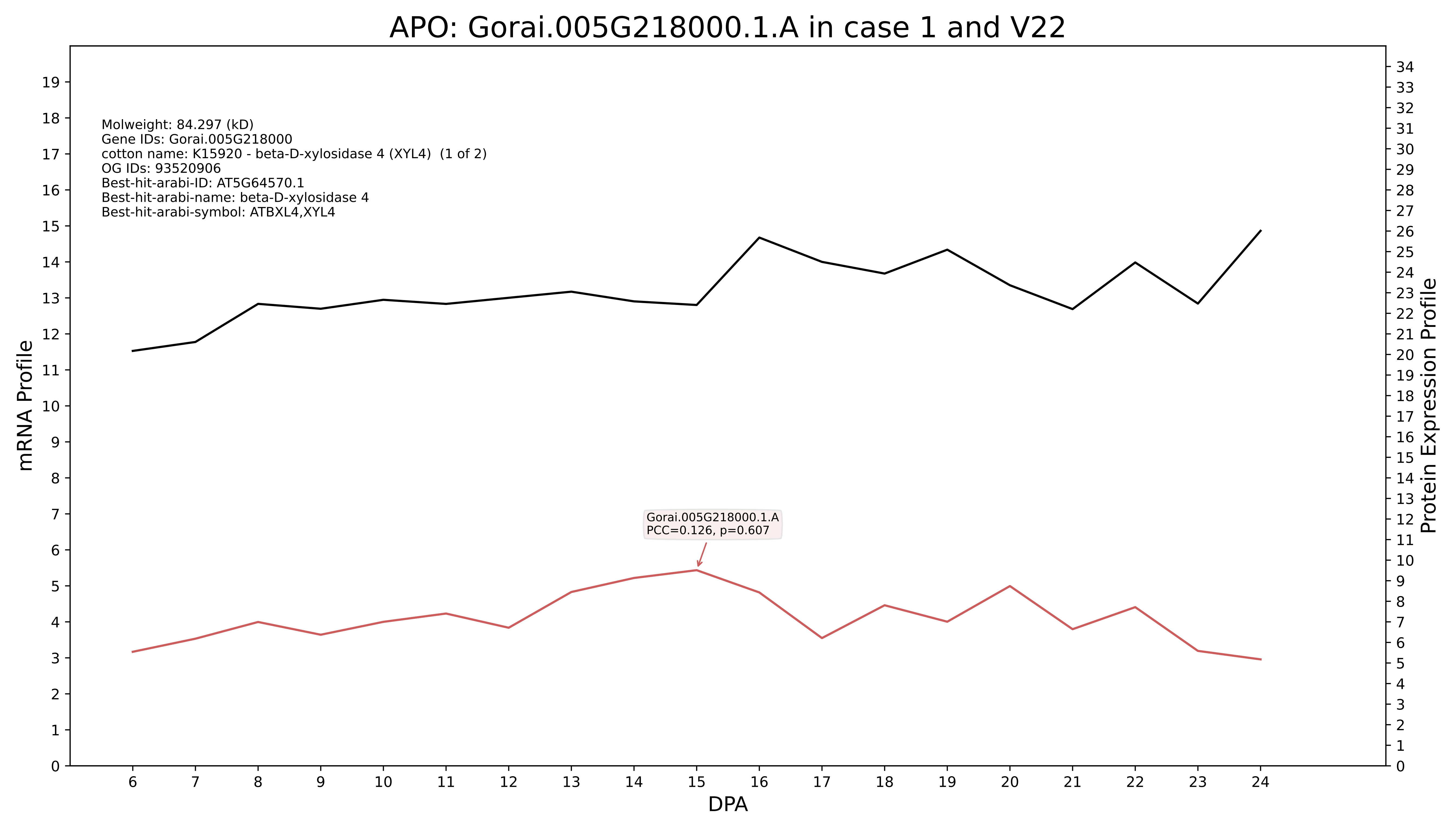
APO: Gorai.005G218000.1.A in case 1 and V22
Molweight: 84.297 (kD)
Gene IDs: Gorai.005G218000
cotton name: K15920 - beta-D-xylosidase 4 (XYL4) (1 of 2)
OG IDs: 93520906
Best-hit-arabi-ID: AT5G64570.1
Best-hit-arabi-name: beta-D-xylosidase 4
Best-hit-arabi-symbol: ATBXL4,XYL4
Gorai.005G218000.1.A
PCC=0.126, p=0.607
mRNA Profile
Protein Expression Profile
DPA

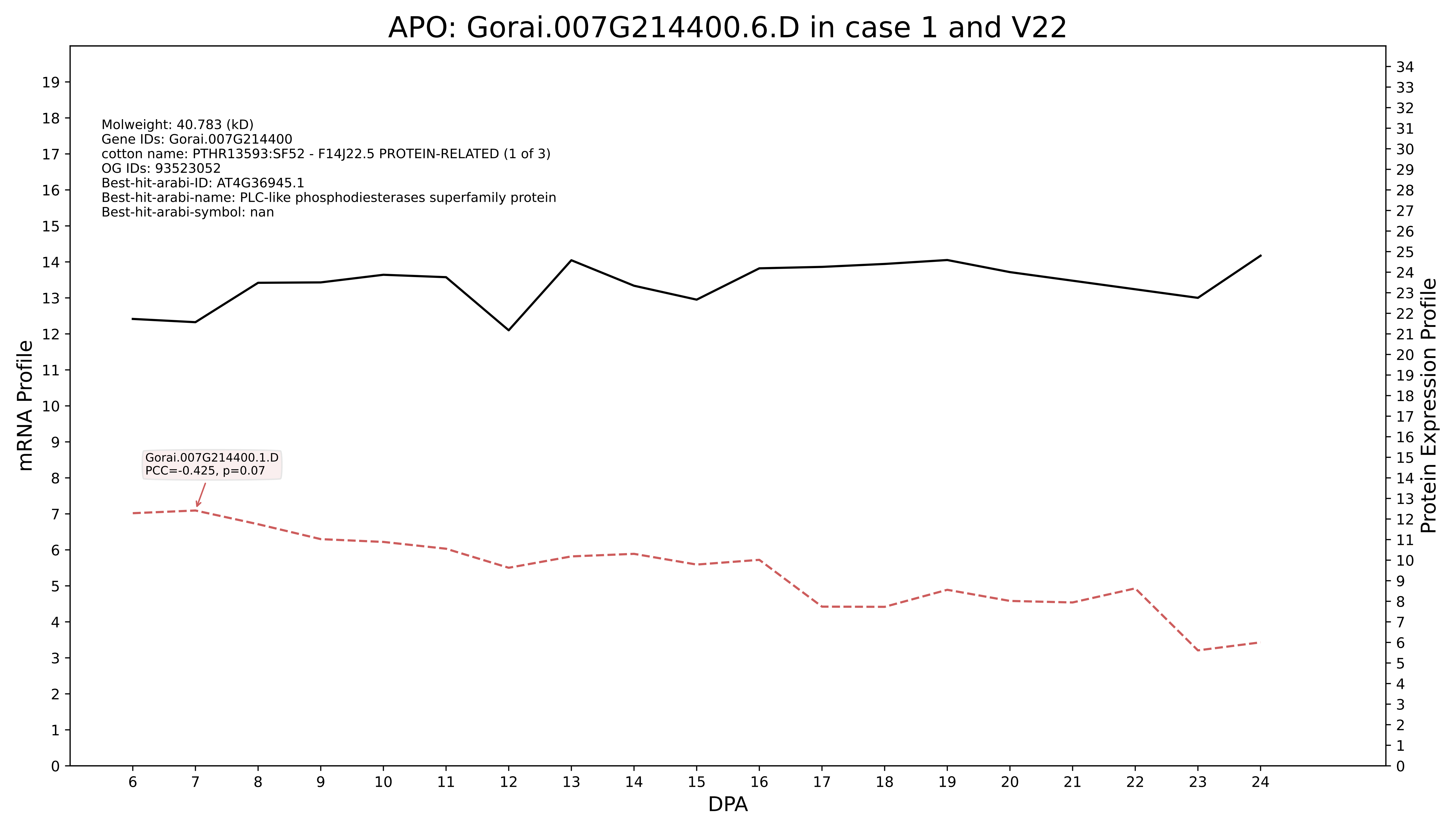
APO: Gorai.007G214400.6.D in case 1 and V22
Molweight: 40.783 (kD)
Gene IDs: Gorai.007G214400
cotton name: PTHR13593:SF52 - F14J22.5 PROTEIN-RELATED (1 of 3)
OG IDs: 93523052
Best-hit-arabi-ID: AT4G36945.1
Best-hit-arabi-name: PLC-like phosphodiesterases superfamily protein
Best-hit-arabi-symbol: nan
Gorai.007G214400.1.D
PCC=-0.425, p=0.07
mRNA Profile
Protein Expression Profile
DPA

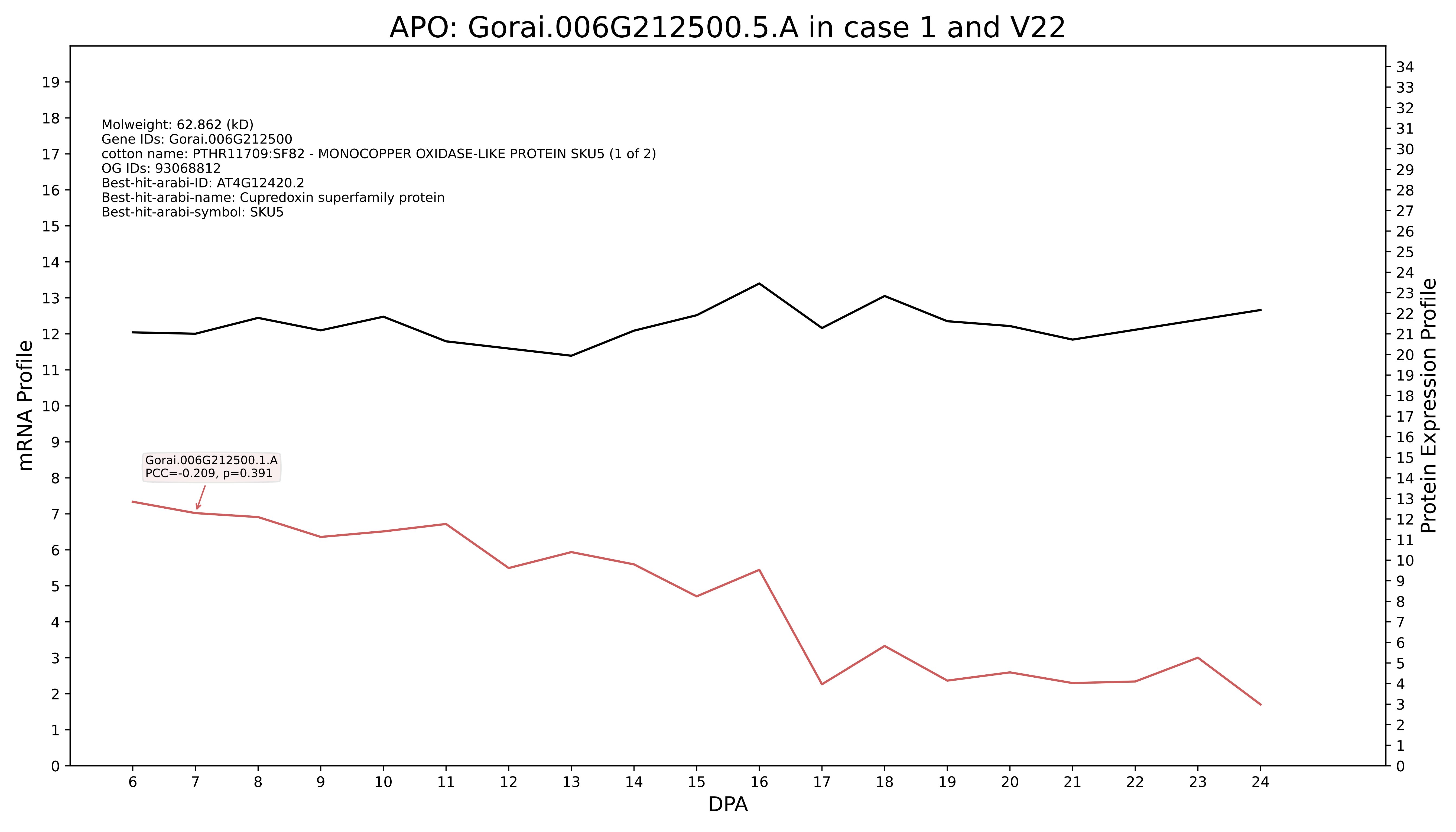
APO: Gorai.006G212500.5.A in case 1 and V22
Molweight: 62.862 (kD)
Gene IDs: Gorai.006G212500
cotton name: PTHR11709:SF82 - MONOCOPPER OXIDASE-LIKE PROTEIN SKU5 (1 of 2)
OG IDs: 93068812
Best-hit-arabi-ID: AT4G12420.2
Best-hit-arabi-name: Cupredoxin superfamily protein
Best-hit-arabi-symbol: SKU5
Gorai.006G212500.1.A
PCC=-0.209, p=0.391
mRNA Profile
Protein Expression Profile
DPA

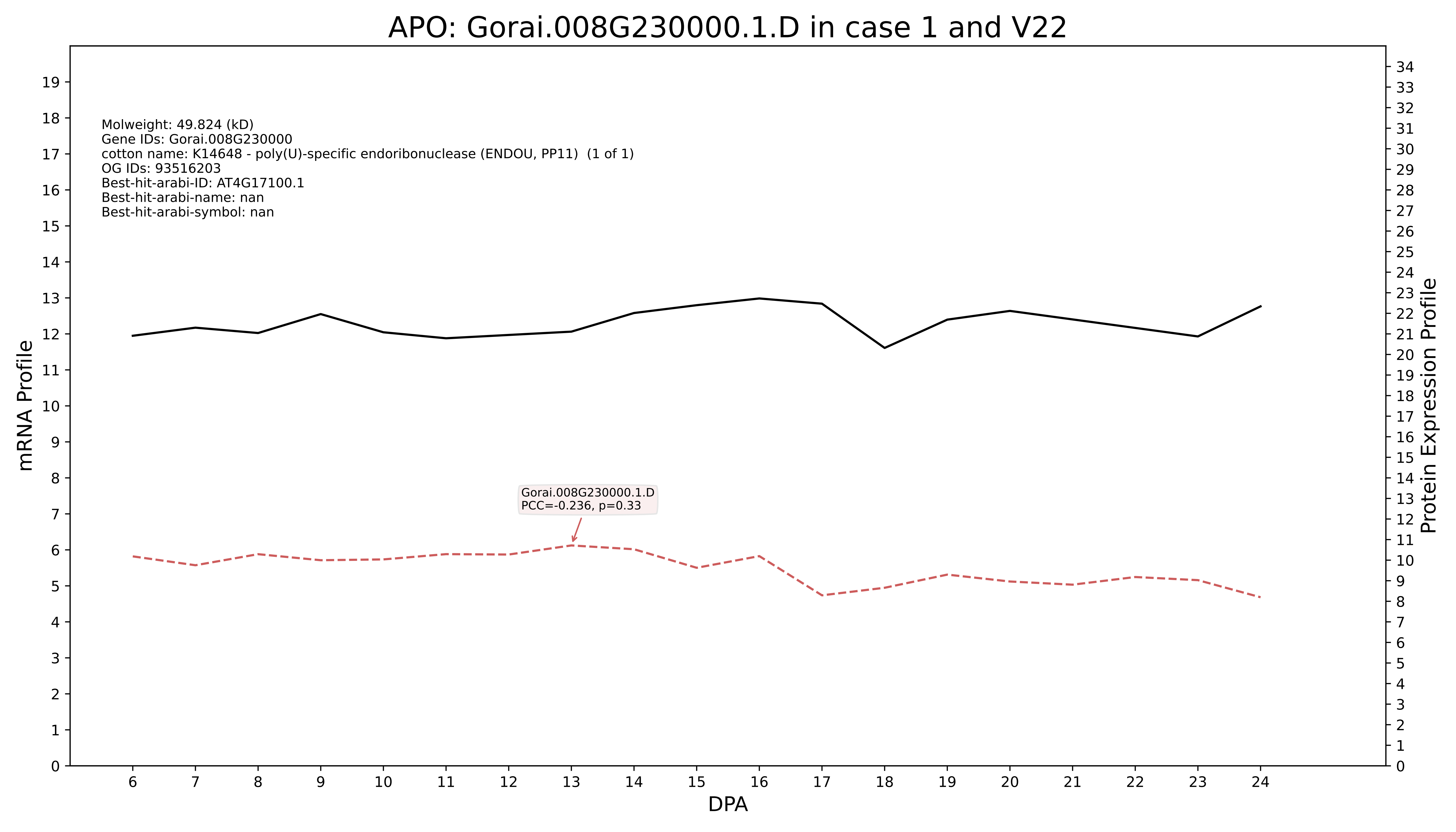

APO: Gorai.008G230000.1.D in case 1 and V22
Molweight: 49.824 (kD)
Gene IDs: Gorai.008G230000
cotton name: K14648 - poly(U)-specific endoribonuclease (ENDOU, PP11) (1 of 1)
OG IDs: 93516203
Best-hit-arabi-ID: AT4G17100.1
Best-hit-arabi-name: nan
Best-hit-arabi-symbol: nan
Gorai.008G230000.1.D
PCC=-0.236, p=0.33
mRNA Profile
Protein Expression Profile
DPA

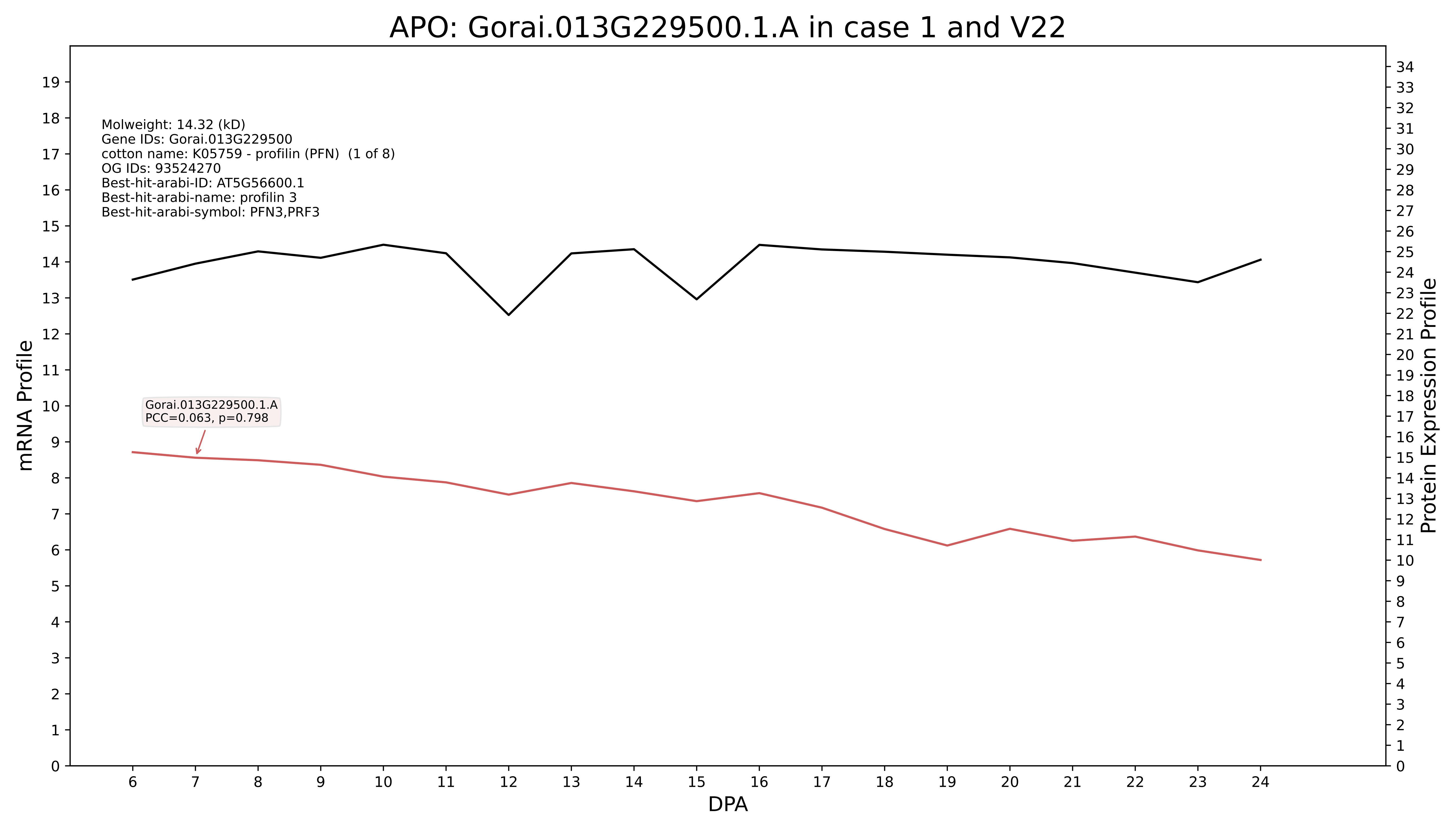

APO: Gorai.013G229500.1.A in case 1 and V22
Molweight: 14.32 (kD)
Gene IDs: Gorai.013G229500
cotton name: K05759 - profilin (PFN) (1 of 8)
OG IDs: 93524270
Best-hit-arabi-ID: AT5G56600.1
Best-hit-arabi-name: profilin 3
Best-hit-arabi-symbol: PFN3,PRF3
Gorai.013G229500.1.A
PCC=0.063, p=0.798
mRNA Profile
Protein Expression Profile
DPA

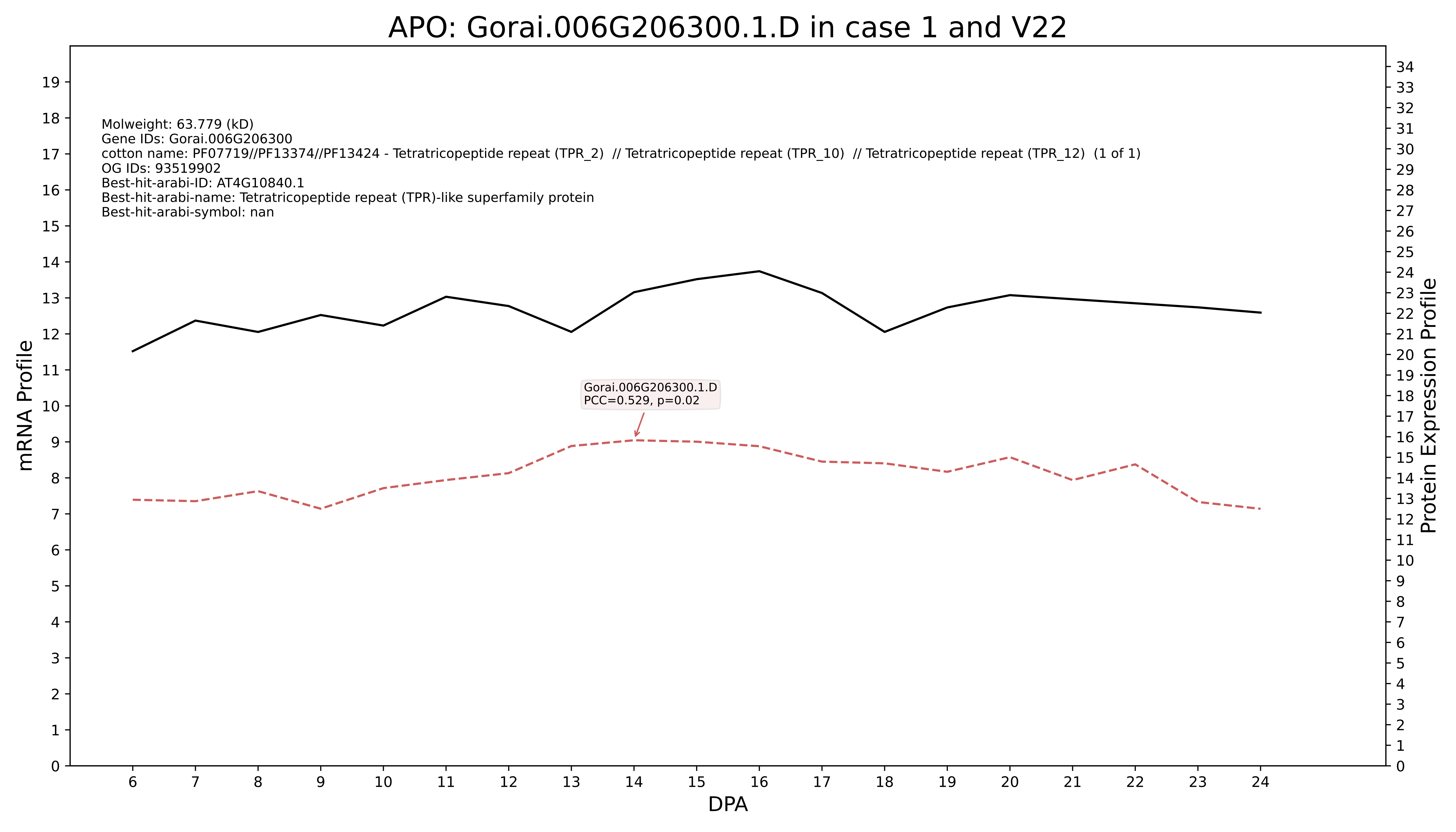

APO: Gorai.006G206300.1.D in case 1 and V22
Molweight: 63.779 (kD)
Gene IDs: Gorai.006G206300
cotton name: PF07719//PF13374//PF13424 - Tetratricopeptide repeat (TPR_2) // Tetratricopeptide repeat (TPR_10) // Tetratricopeptide repeat (TPR_12) (1 of 1)
OG IDs: 93519902
Best-hit-arabi-ID: AT4G10840.1
Best-hit-arabi-name: Tetratricopeptide repeat (TPR)-like superfamily protein
Best-hit-arabi-symbol: nan
Gorai.006G206300.1.D
PCC=0.529, p=0.02
mRNA Profile
Protein Expression Profile
DPA

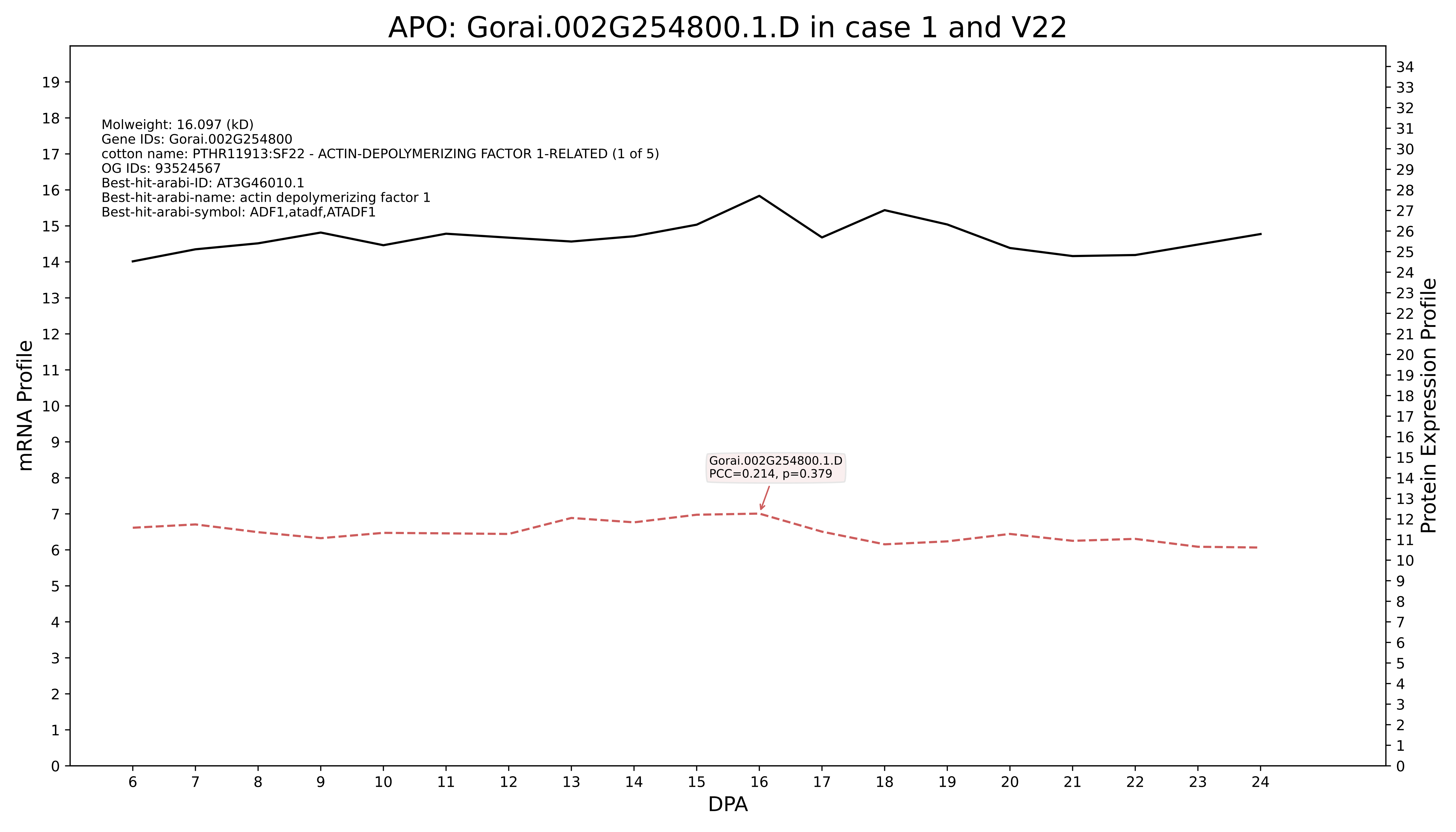
APO: Gorai.002G254800.1.D in case 1 and V22
Molweight: 16.097 (kD)
Gene IDs: Gorai.002G254800
cotton name: PTHR11913:SF22 - ACTIN-DEPOLYMERIZING FACTOR 1-RELATED (1 of 5)
OG IDs: 93524567
Best-hit-arabi-ID: AT3G46010.1
Best-hit-arabi-name: actin depolymerizing factor 1
Best-hit-arabi-symbol: ADF1,atadf,ATADF1
Gorai.002G254800.1.D
PCC=0.214, p=0.379
mRNA Profile
Protein Expression Profile
DPA
0 1 2 3 4 5 6 7 8 9 10 11 12 13 14 15 16 17 18 19
0 1 2 3 4 5 6 7 8 9 10 11 12 13 14 15 16 17 18 19 20 21 22 23 24 25 26 27 28 29 30 31 32 33 34
6 7 8 9 10 11 12 13 14 15 16 17 18 19 20 21 22 23 24

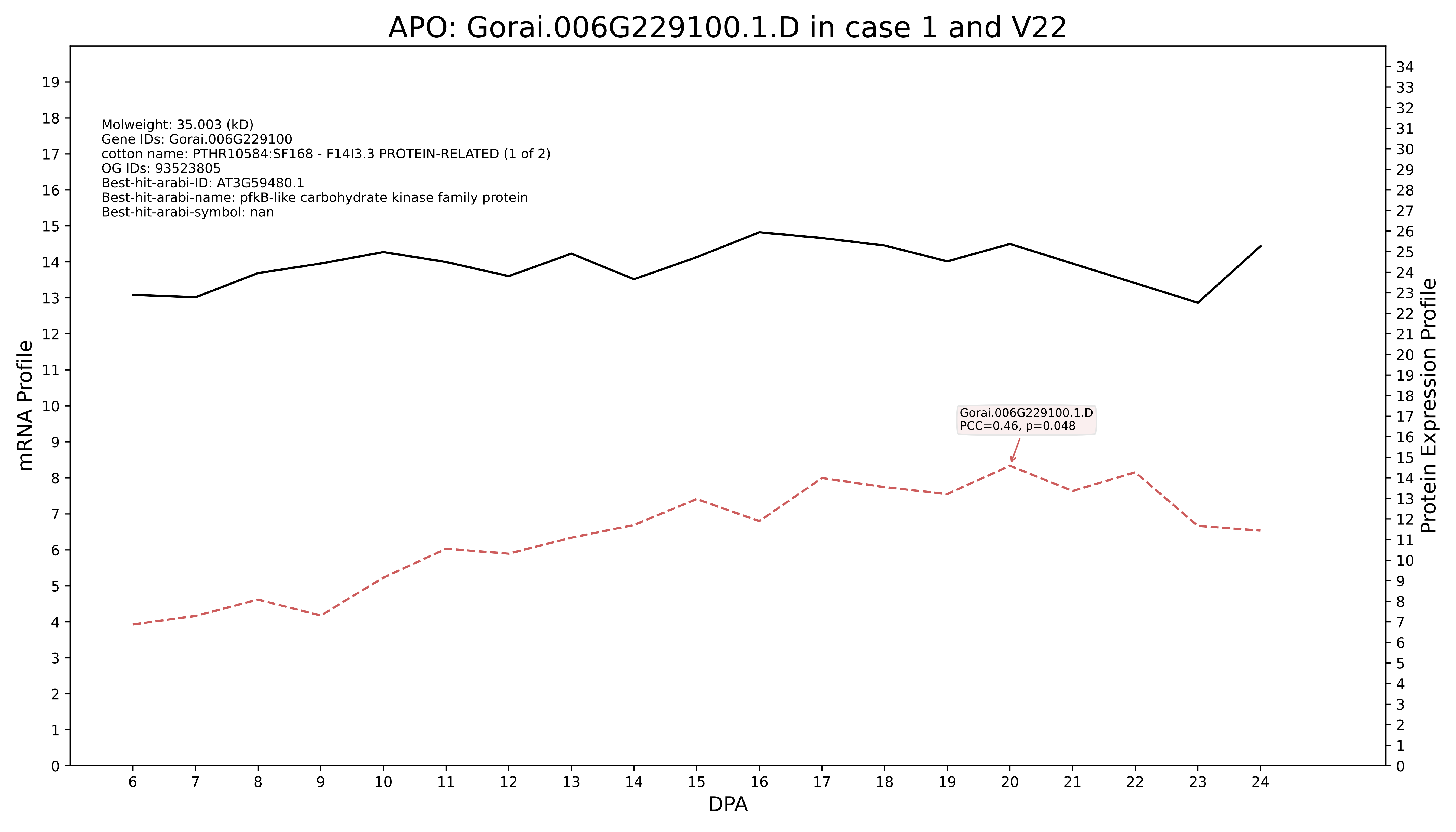
APO: Gorai.006G229100.1.D in case 1 and V22
Molweight: 35.003 (kD)
Gene IDs: Gorai.006G229100
cotton name: PTHR10584:SF168 - F14I3.3 PROTEIN-RELATED (1 of 2)
OG IDs: 93523805
Best-hit-arabi-ID: AT3G59480.1
Best-hit-arabi-name: pfkB-like carbohydrate kinase family protein
Best-hit-arabi-symbol: nan
Gorai.006G229100.1.D
PCC=0.46, p=0.048
mRNA Profile
Protein Expression Profile
DPA

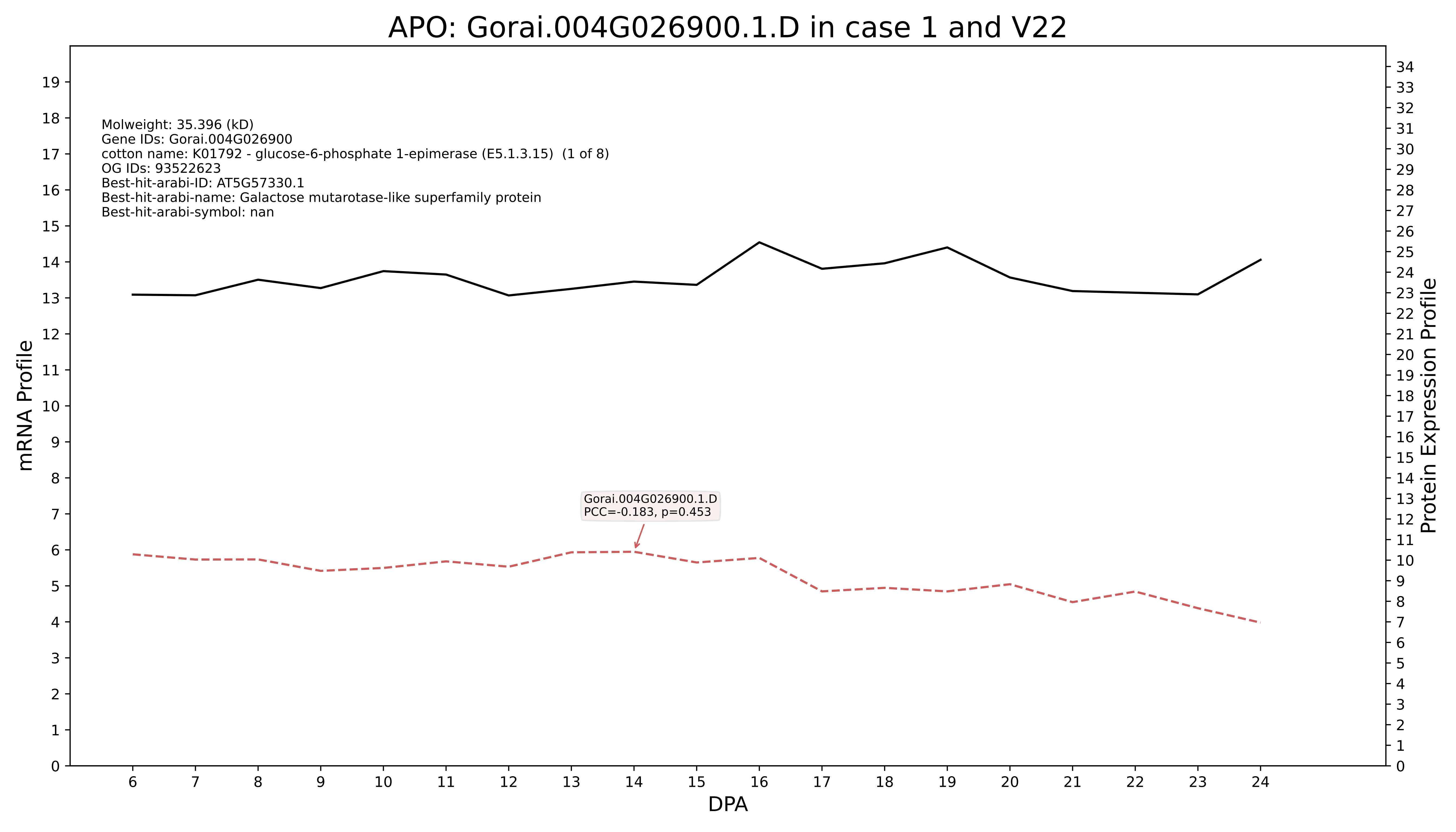
APO: Gorai.004G026900.1.D in case 1 and V22
Molweight: 35.396 (kD)
Gene IDs: Gorai.004G026900
cotton name: K01792 - glucose-6-phosphate 1-epimerase (E5.1.3.15) (1 of 8)
OG IDs: 93522623
Best-hit-arabi-ID: AT5G57330.1
Best-hit-arabi-name: Galactose mutarotase-like superfamily protein
Best-hit-arabi-symbol: nan
Gorai.004G026900.1.D
PCC=-0.183, p=0.453
mRNA Profile
Protein Expression Profile
DPA

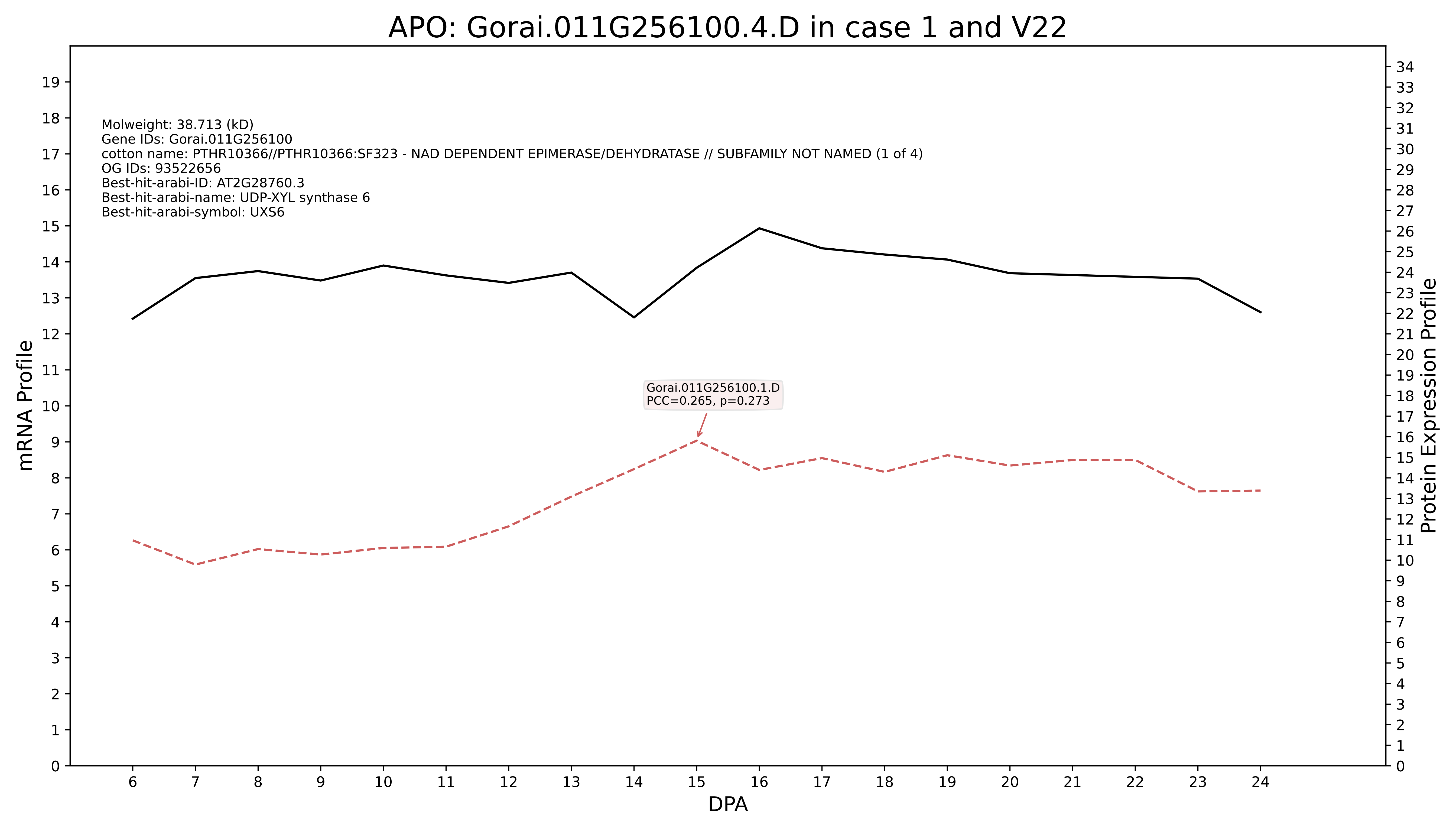
APO: Gorai.011G256100.4.D in case 1 and V22
Molweight: 38.713 (kD)
Gene IDs: Gorai.011G256100
cotton name: PTHR10366//PTHR10366:SF323 - NAD DEPENDENT EPIMERASE/DEHYDRATASE // SUBFAMILY NOT NAMED (1 of 4)
OG IDs: 93522656
Best-hit-arabi-ID: AT2G28760.3
Best-hit-arabi-name: UDP-XYL synthase 6
Best-hit-arabi-symbol: UXS6
Gorai.011G256100.1.D
PCC=0.265, p=0.273
mRNA Profile
Protein Expression Profile
DPA

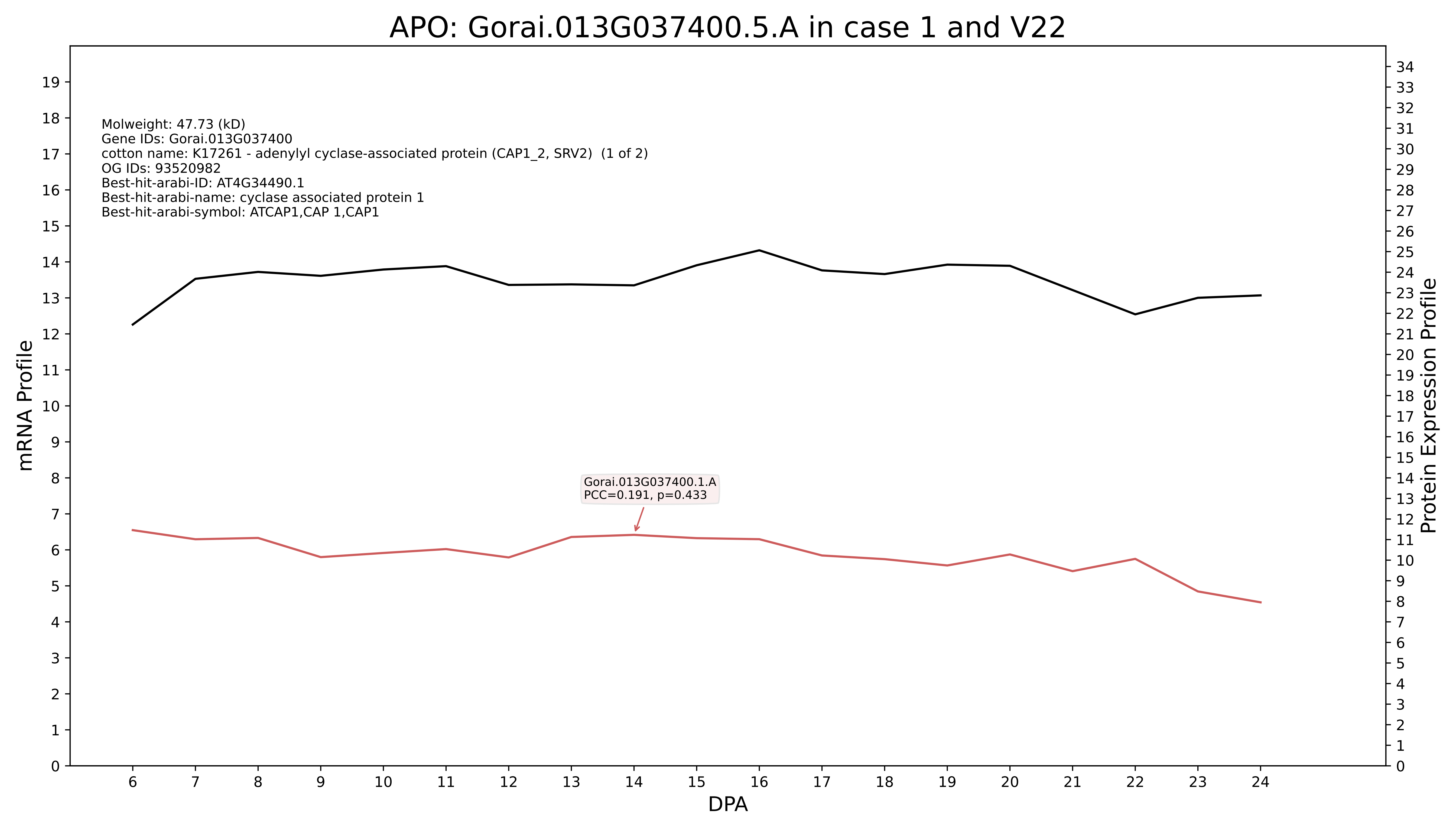

APO: Gorai.013G037400.5.A in case 1 and V22
Molweight: 47.73 (kD)
Gene IDs: Gorai.013G037400
cotton name: K17261 - adenylyl cyclase-associated protein (CAP1_2, SRV2) (1 of 2)
OG IDs: 93520982
Best-hit-arabi-ID: AT4G34490.1
Best-hit-arabi-name: cyclase associated protein 1
Best-hit-arabi-symbol: ATCAP1,CAP 1,CAP1
Gorai.013G037400.1.A
PCC=0.191, p=0.433
mRNA Profile
Protein Expression Profile
DPA

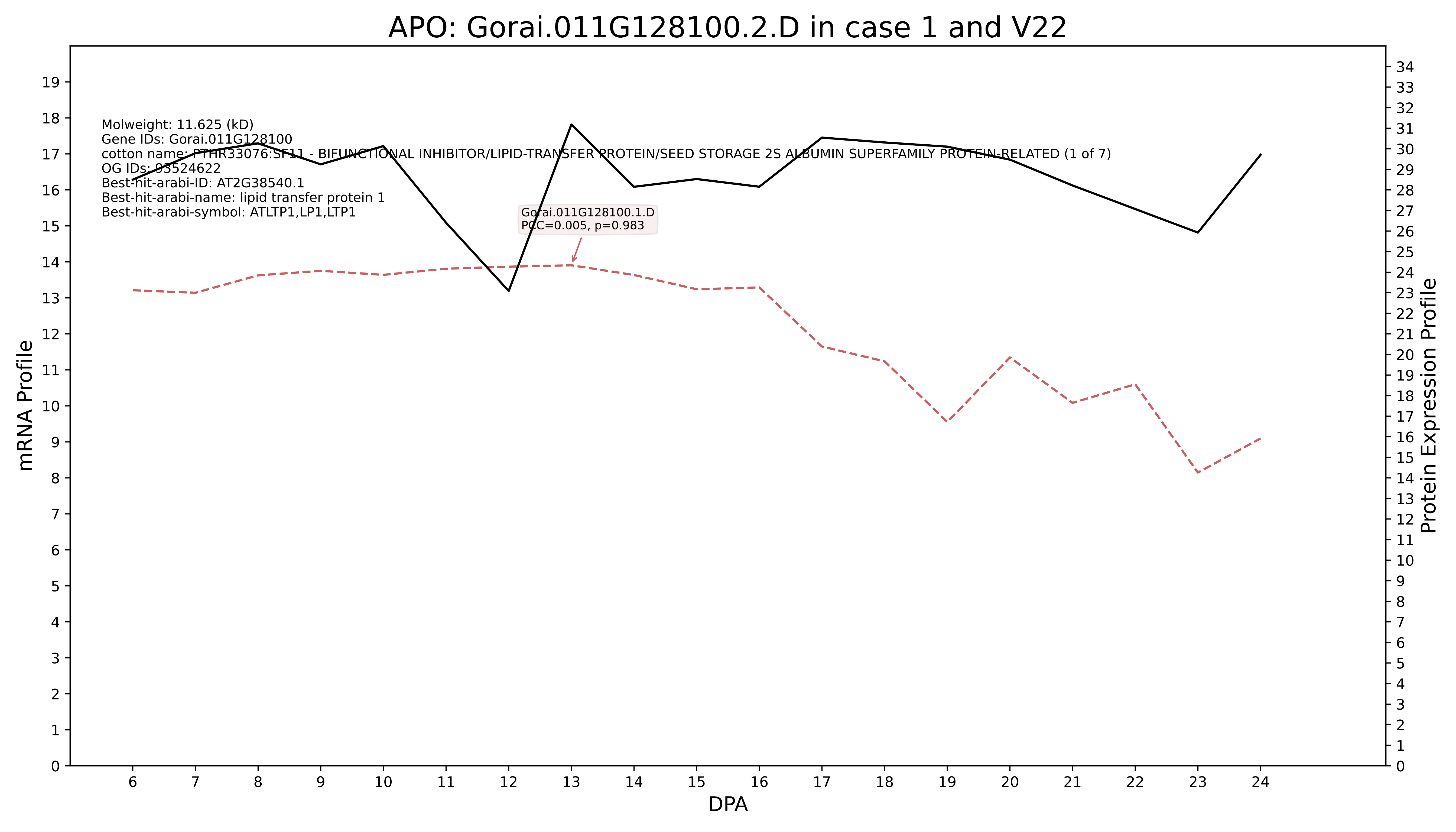
APO: Gorai.011G128100.2.D in case 1 and V22
Molweight: 11.625 (kD)
Gene IDs: Gorai.011G128100
cotton name: PTHR33076:SF11 - BIFUNCTIONAL INHIBITOR/LIPID-TRANSFER PROTEIN/SEED STORAGE 2S ALBUMIN SUPERFAMILY PROTEIN-RELATED (1 of 7)
OG IDs: 93524622
Best-hit-arabi-ID: AT2G38540.1
Best-hit-arabi-name: lipid transfer protein 1
Best-hit-arabi-symbol: ATLTP1,LP1,LTP1
Gorai.011G128100.1.D
PCC=0.005, p=0.983
mRNA Profile
Protein Expression Profile
DPA

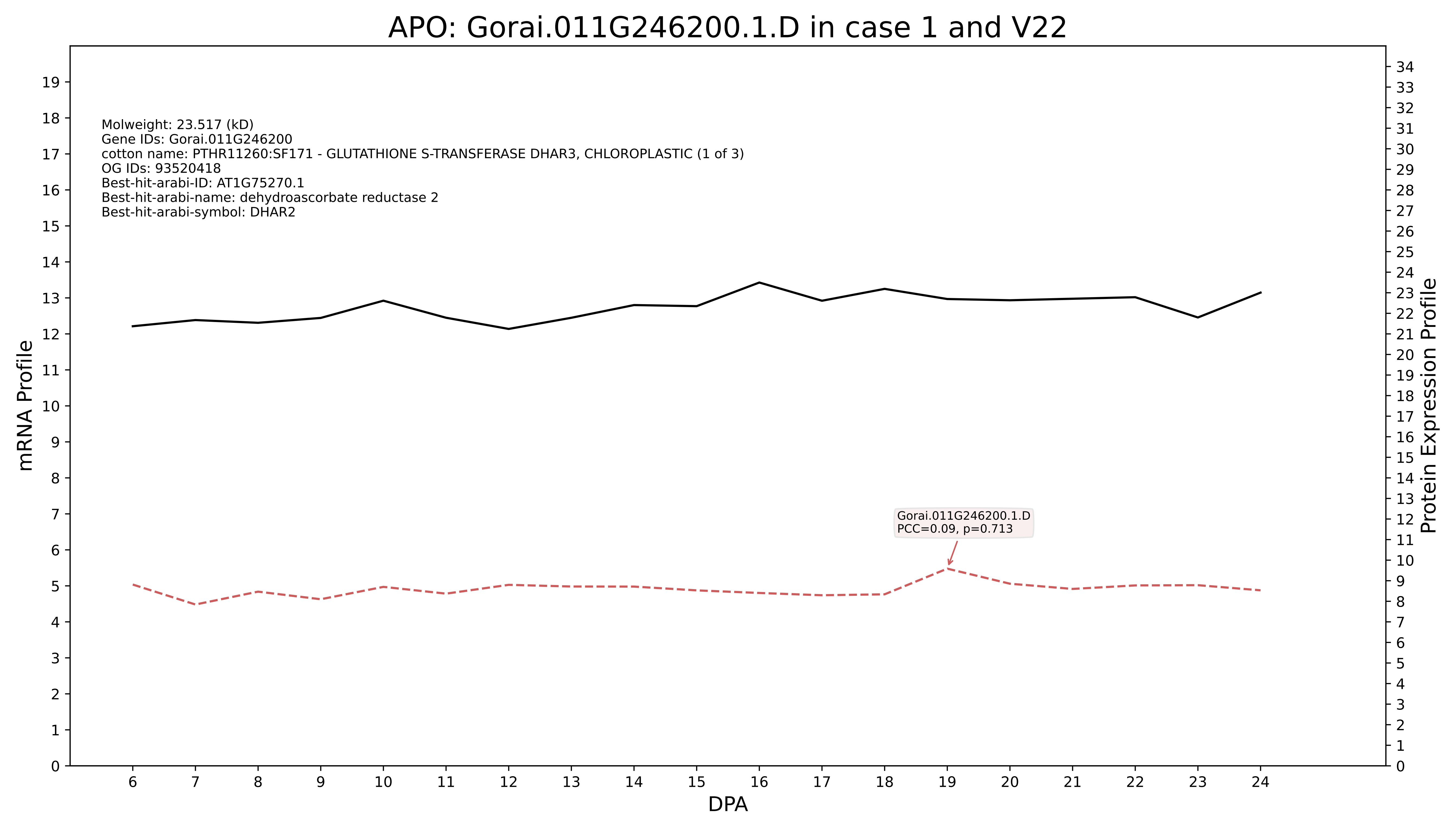

APO: Gorai.011G246200.1.D in case 1 and V22
Molweight: 23.517 (kD)
Gene IDs: Gorai.011G246200
cotton name: PTHR11260:SF171 - GLUTATHIONE S-TRANSFERASE DHAR3, CHLOROPLASTIC (1 of 3)
OG IDs: 93520418
Best-hit-arabi-ID: AT1G75270.1
Best-hit-arabi-name: dehydroascorbate reductase 2
Best-hit-arabi-symbol: DHAR2
Gorai.011G246200.1.D
PCC=0.09, p=0.713
mRNA Profile
Protein Expression Profile
DPA

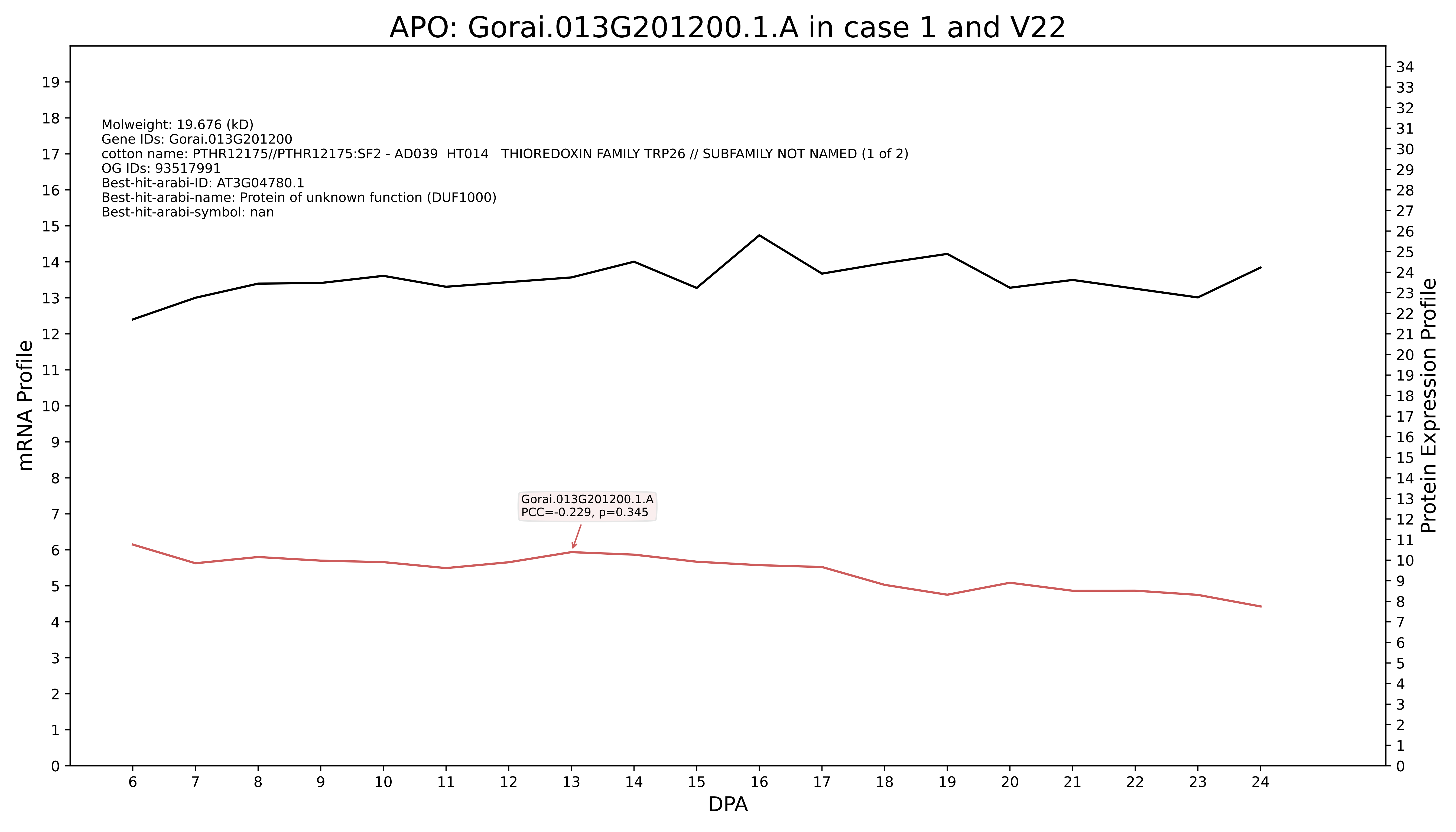

APO: Gorai.013G201200.1.A in case 1 and V22
Molweight: 19.676 (kD)
Gene IDs: Gorai.013G201200
cotton name: PTHR12175//PTHR12175:SF2 - AD039 HT014 THIOREDOXIN FAMILY TRP26 // SUBFAMILY NOT NAMED (1 of 2)
OG IDs: 93517991
Best-hit-arabi-ID: AT3G04780.1
Best-hit-arabi-name: Protein of unknown function (DUF1000)
Best-hit-arabi-symbol: nan
Gorai.013G201200.1.A
PCC=-0.229, p=0.345
mRNA Profile
Protein Expression Profile
DPA

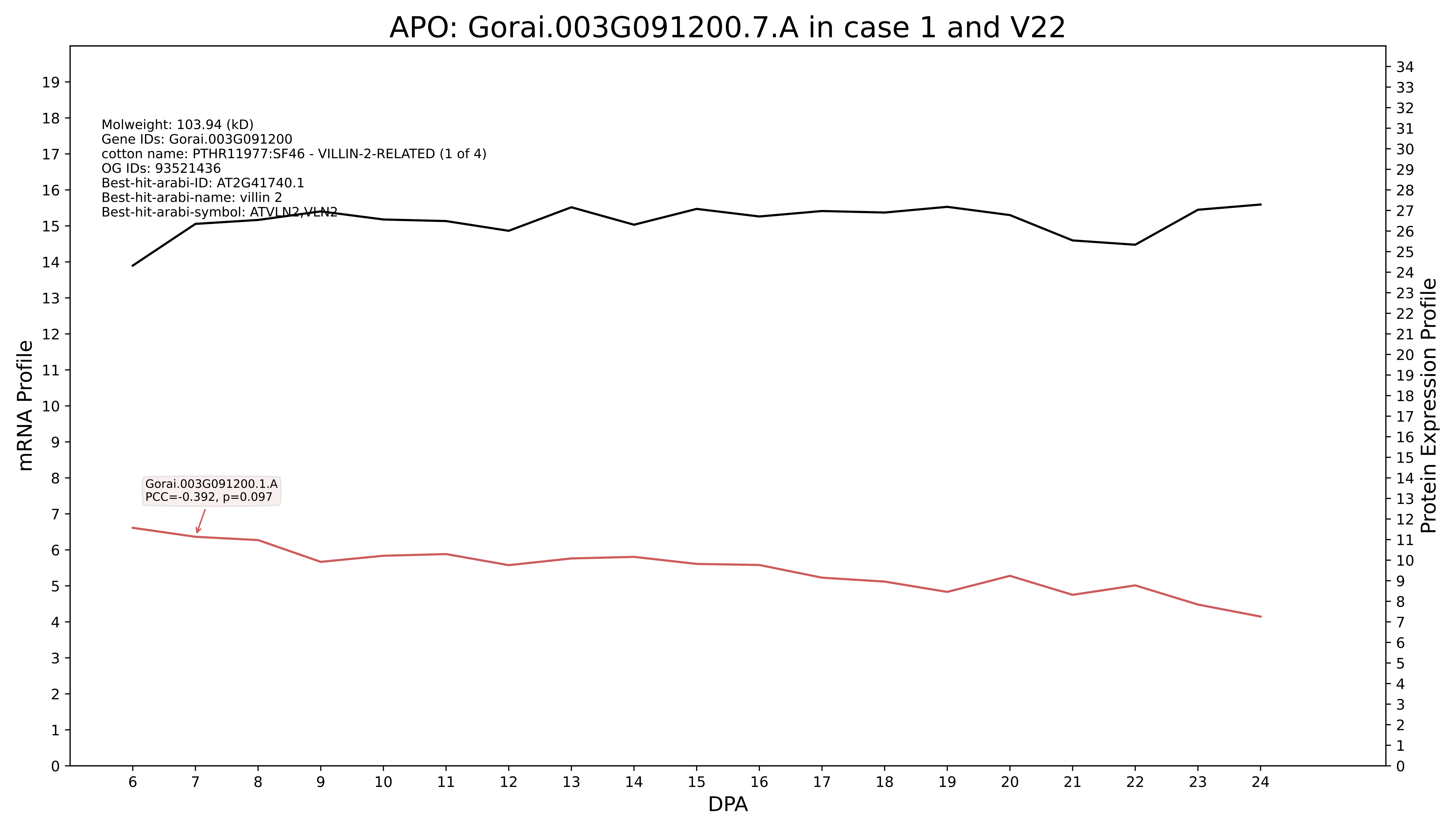
APO: Gorai.003G091200.7.A in case 1 and V22
Molweight: 103.94 (kD)
Gene IDs: Gorai.003G091200
cotton name: PTHR11977:SF46 - VILLIN-2-RELATED (1 of 4)
OG IDs: 93521436
Best-hit-arabi-ID: AT2G41740.1
Best-hit-arabi-name: villin 2
Best-hit-arabi-symbol: ATVLN2,VLN2
Gorai.003G091200.1.A
PCC=-0.392, p=0.097
mRNA Profile
Protein Expression Profile
DPA

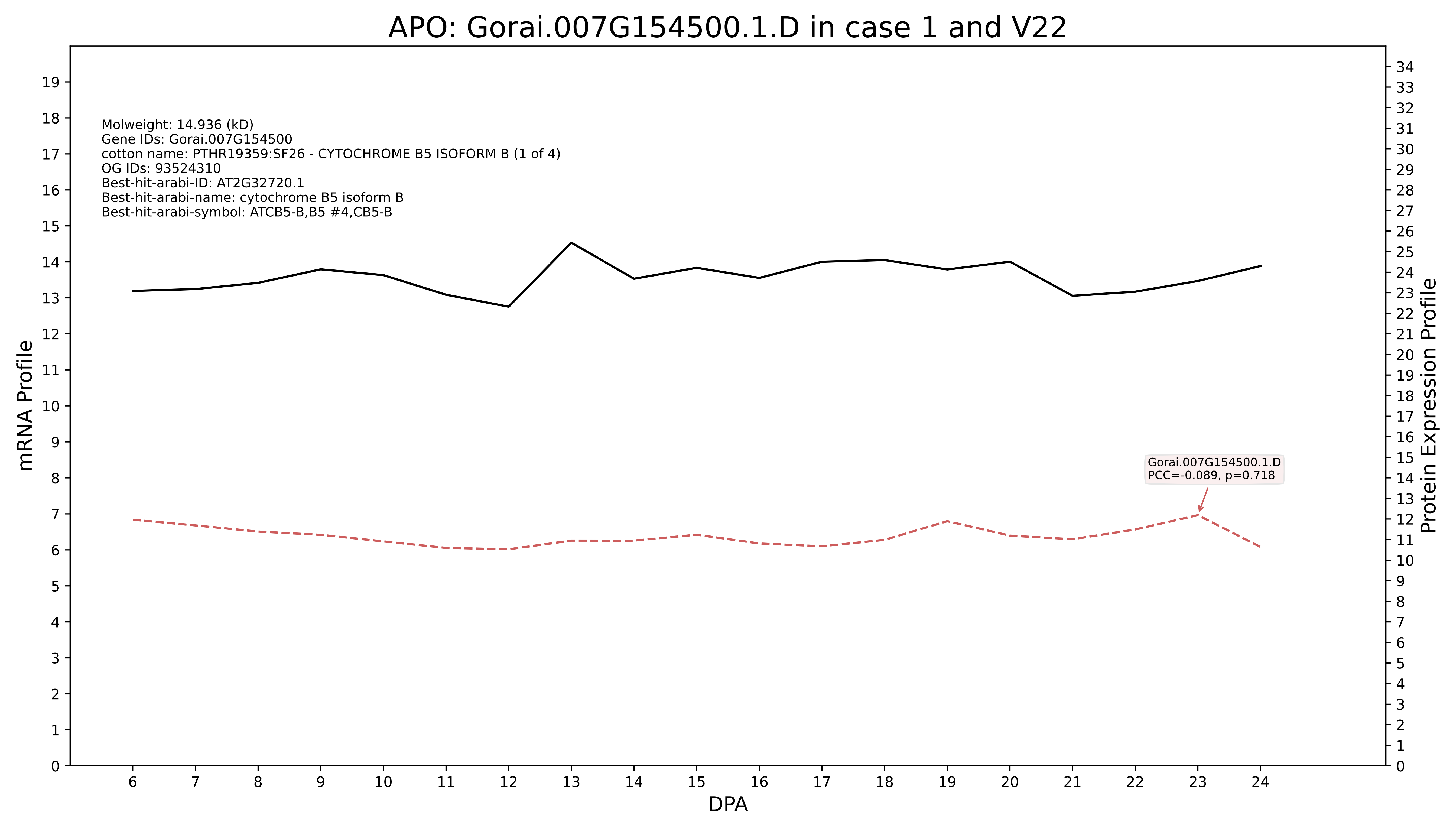
APO: Gorai.007G154500.1.D in case 1 and V22
Molweight: 14.936 (kD)
Gene IDs: Gorai.007G154500
cotton name: PTHR19359:SF26 - CYTOCHROME B5 ISOFORM B (1 of 4)
OG IDs: 93524310
Best-hit-arabi-ID: AT2G32720.1
Best-hit-arabi-name: cytochrome B5 isoform B
Best-hit-arabi-symbol: ATCB5-B,B5 #4,CB5-B
Gorai.007G154500.1.D
PCC=-0.089, p=0.718
mRNA Profile
Protein Expression Profile
DPA

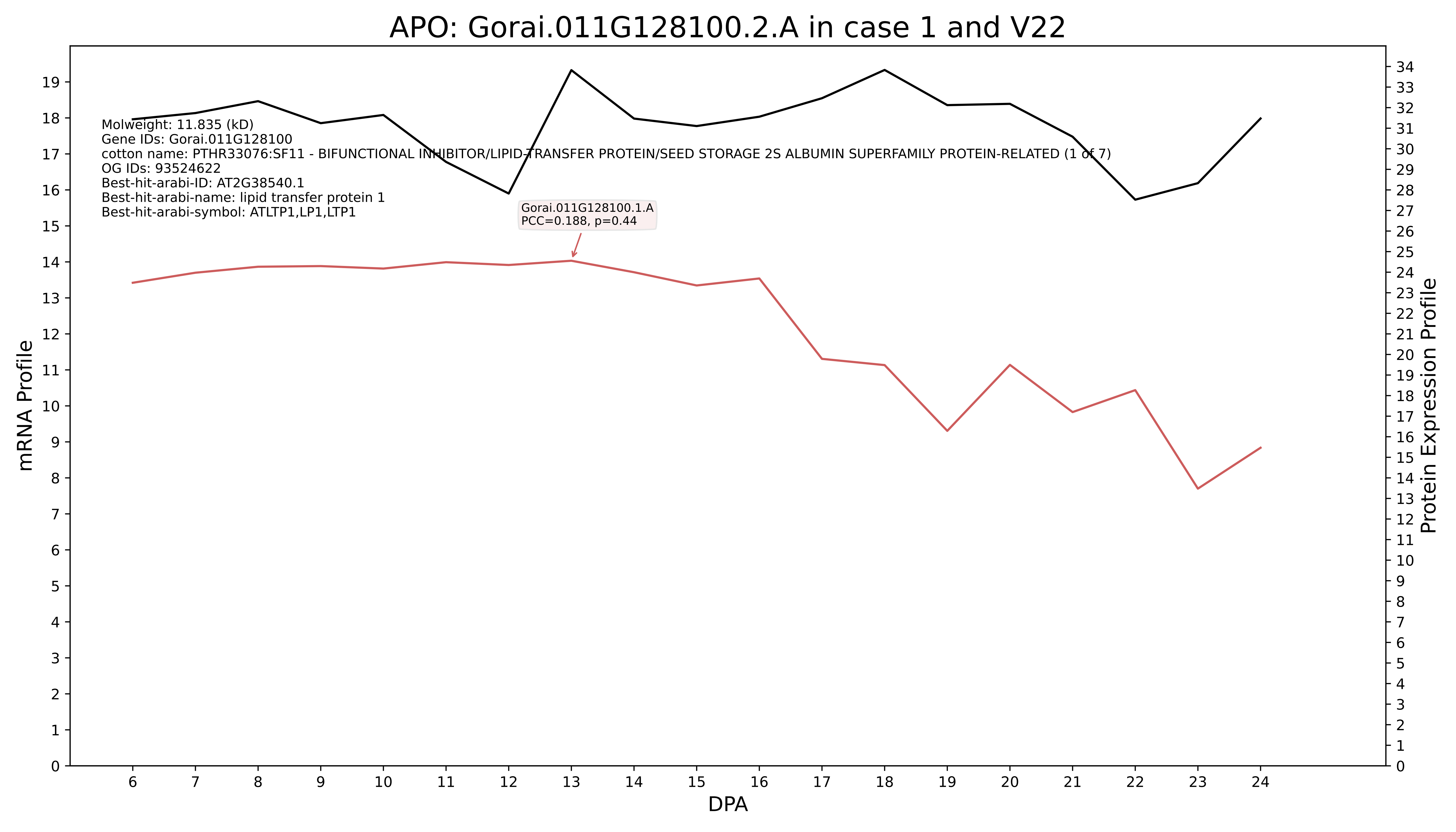

APO: Gorai.011G128100.2.A in case 1 and V22
Molweight: 11.835 (kD)
Gene IDs: Gorai.011G128100
cotton name: PTHR33076:SF11 - BIFUNCTIONAL INHIBITOR/LIPID-TRANSFER PROTEIN/SEED STORAGE 2S ALBUMIN SUPERFAMILY PROTEIN-RELATED (1 of 7)
OG IDs: 93524622
Best-hit-arabi-ID: AT2G38540.1
Best-hit-arabi-name: lipid transfer protein 1
Best-hit-arabi-symbol: ATLTP1,LP1,LTP1
Gorai.011G128100.1.A
PCC=0.188, p=0.44
mRNA Profile
Protein Expression Profile
DPA

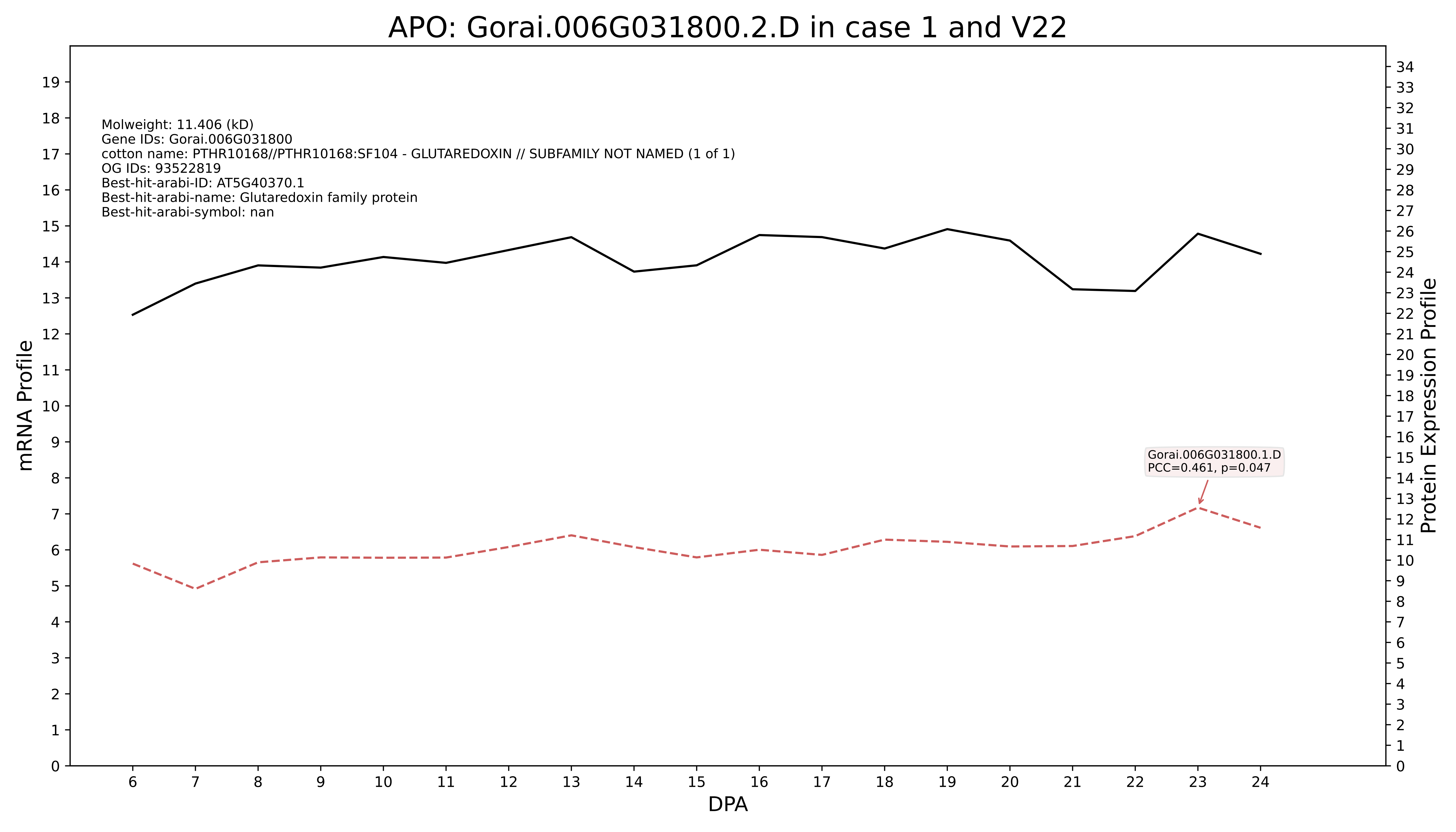
APO: Gorai.006G031800.2.D in case 1 and V22
Molweight: 11.406 (kD)
Gene IDs: Gorai.006G031800
cotton name: PTHR10168//PTHR10168:SF104 - GLUTAREDOXIN // SUBFAMILY NOT NAMED (1 of 1)
OG IDs: 93522819
Best-hit-arabi-ID: AT5G40370.1
Best-hit-arabi-name: Glutaredoxin family protein
Best-hit-arabi-symbol: nan
Gorai.006G031800.1.D
PCC=0.461, p=0.047
mRNA Profile
Protein Expression Profile
DPA

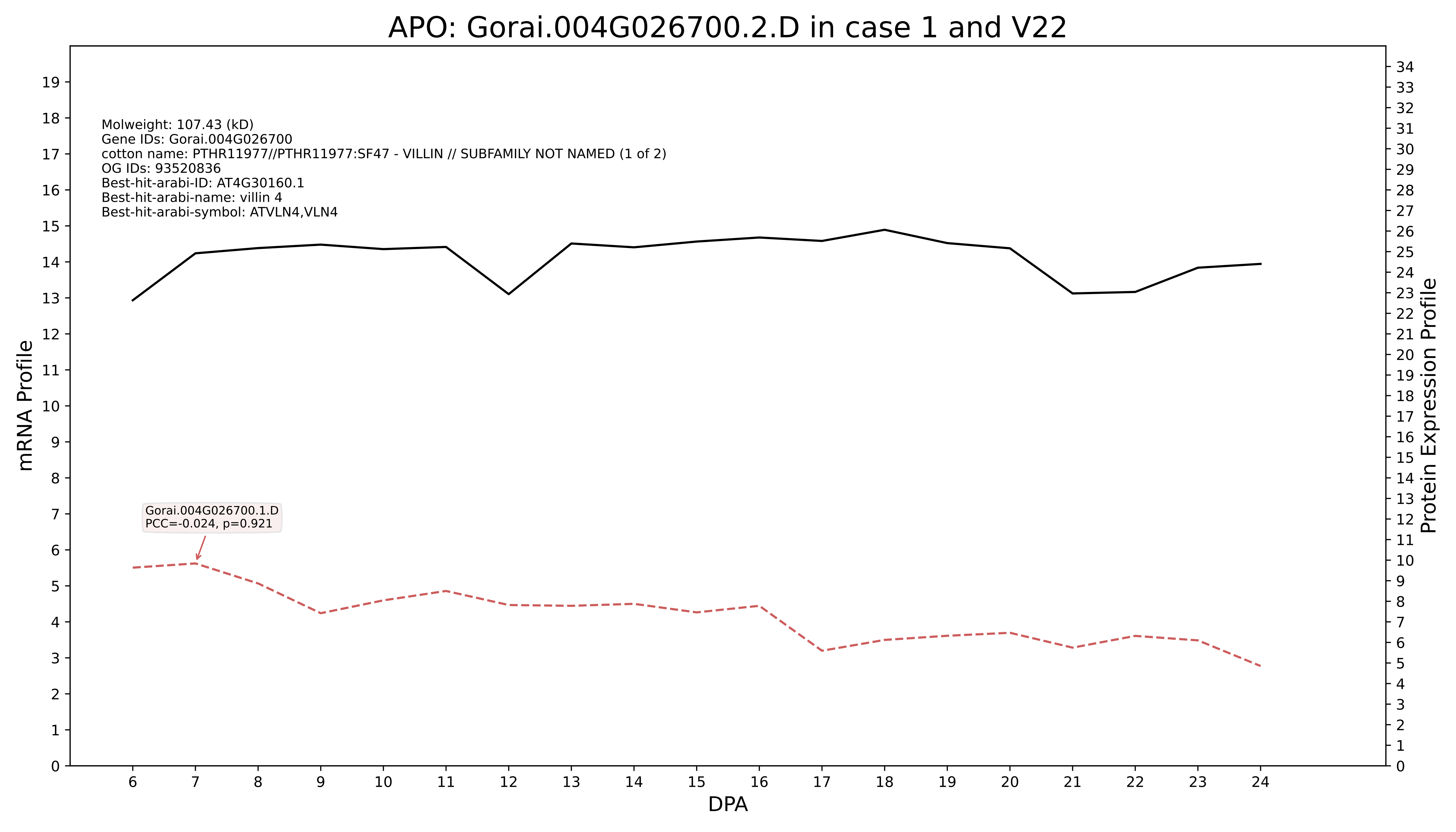

APO: Gorai.004G026700.2.D in case 1 and V22
Molweight: 107.43 (kD)
Gene IDs: Gorai.004G026700
cotton name: PTHR11977//PTHR11977:SF47 - VILLIN // SUBFAMILY NOT NAMED (1 of 2)
OG IDs: 93520836
Best-hit-arabi-ID: AT4G30160.1
Best-hit-arabi-name: villin 4
Best-hit-arabi-symbol: ATVLN4,VLN4
Gorai.004G026700.1.D
PCC=-0.024, p=0.921
mRNA Profile
Protein Expression Profile
DPA

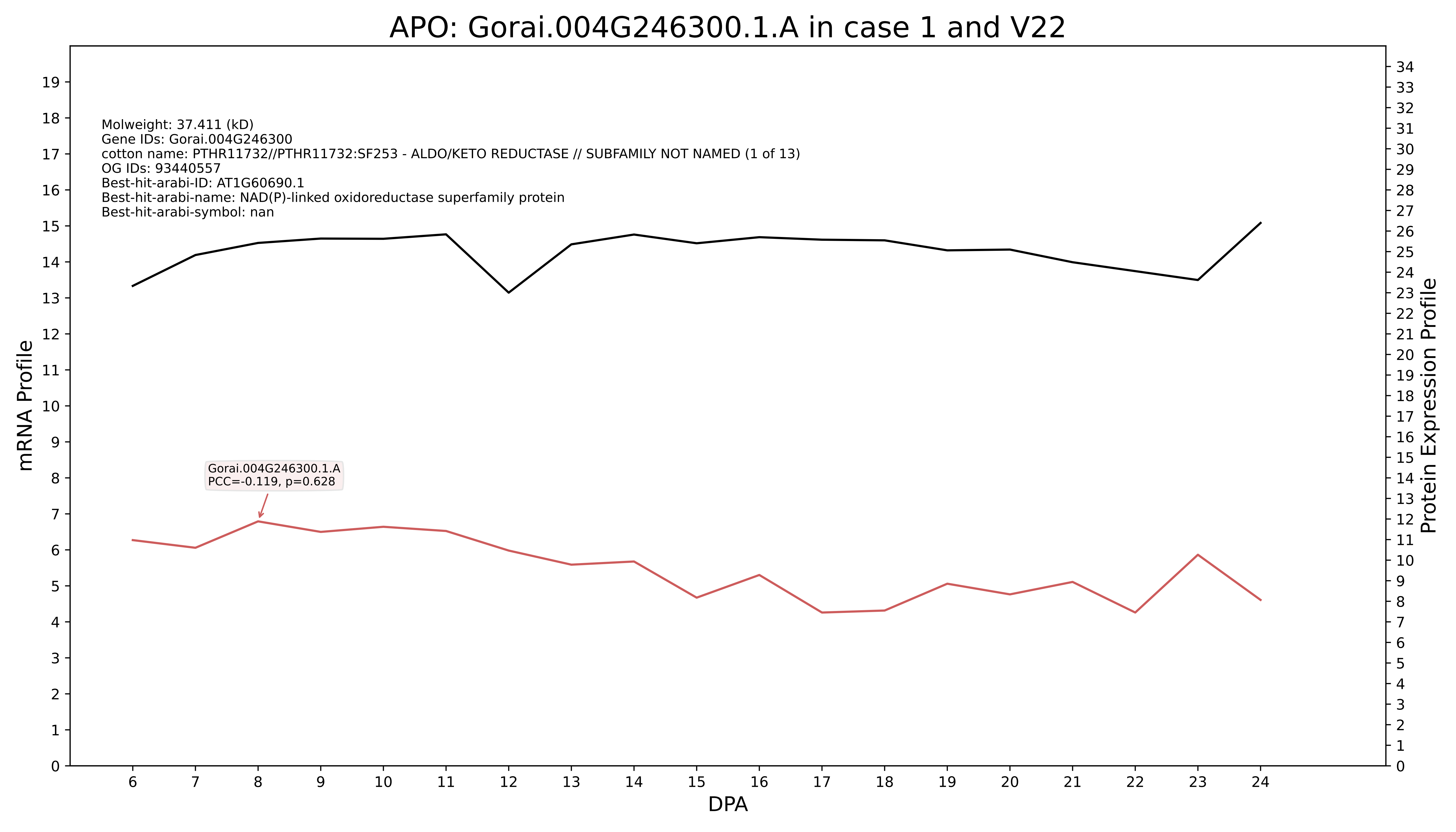

APO: Gorai.004G246300.1.A in case 1 and V22
Molweight: 37.411 (kD)
Gene IDs: Gorai.004G246300
cotton name: PTHR11732//PTHR11732:SF253 - ALDO/KETO REDUCTASE // SUBFAMILY NOT NAMED (1 of 13)
OG IDs: 93440557
Best-hit-arabi-ID: AT1G60690.1
Best-hit-arabi-name: NAD(P)-linked oxidoreductase superfamily protein
Best-hit-arabi-symbol: nan
Gorai.004G246300.1.A
PCC=-0.119, p=0.628
mRNA Profile
Protein Expression Profile
DPA

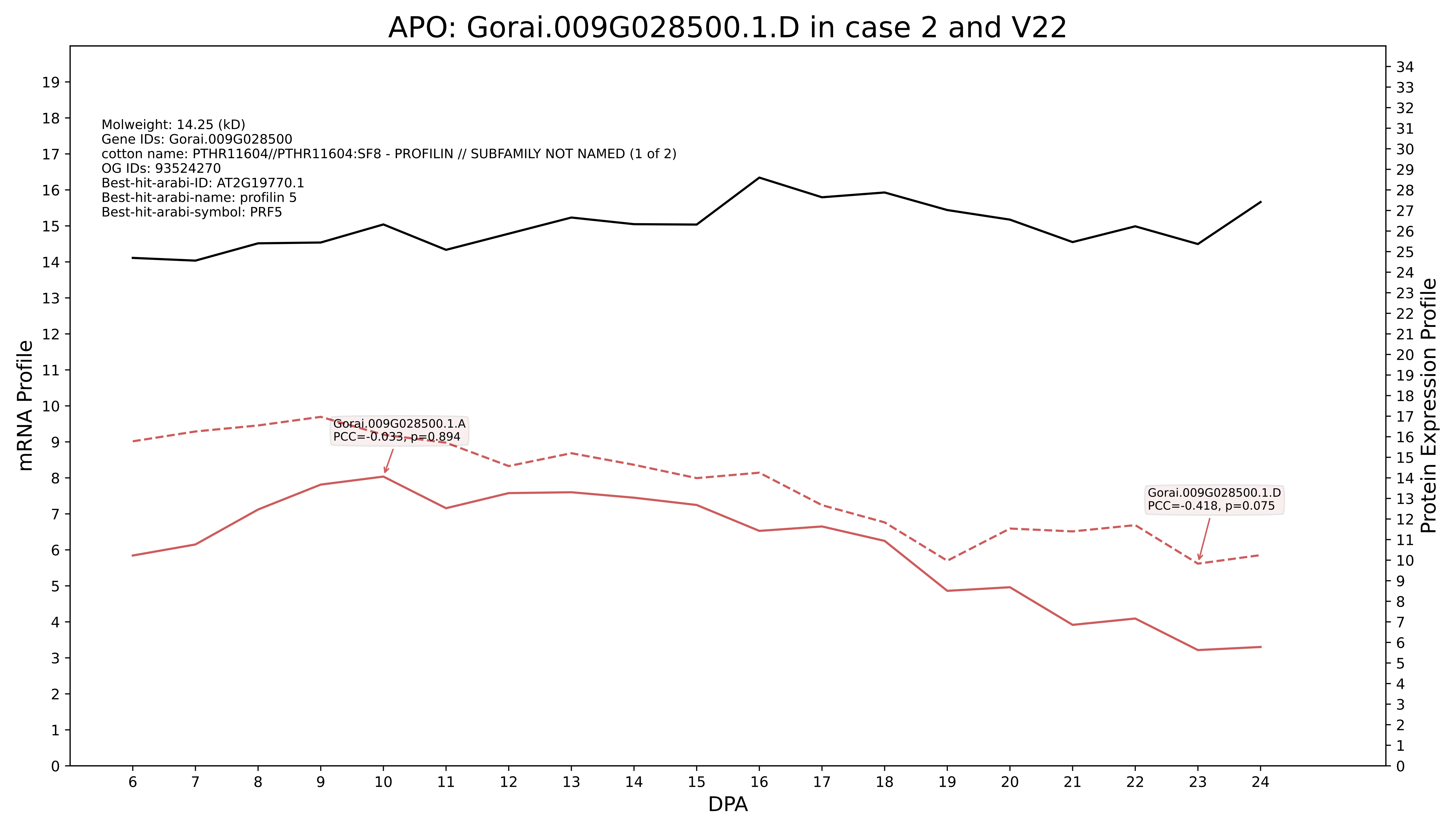

APO: Gorai.009G028500.1.D in case 2 and V22
Molweight: 14.25 (kD)
Gene IDs: Gorai.009G028500
cotton name: PTHR11604//PTHR11604:SF8 - PROFILIN // SUBFAMILY NOT NAMED (1 of 2)
OG IDs: 93524270
Best-hit-arabi-ID: AT2G19770.1
Best-hit-arabi-name: profilin 5
Best-hit-arabi-symbol: PRF5
Gorai.009G028500.1.A
PCC=-0.033, p=0.894
Gorai.009G028500.1.D
PCC=-0.418, p=0.075
mRNA Profile
Protein Expression Profile
DPA

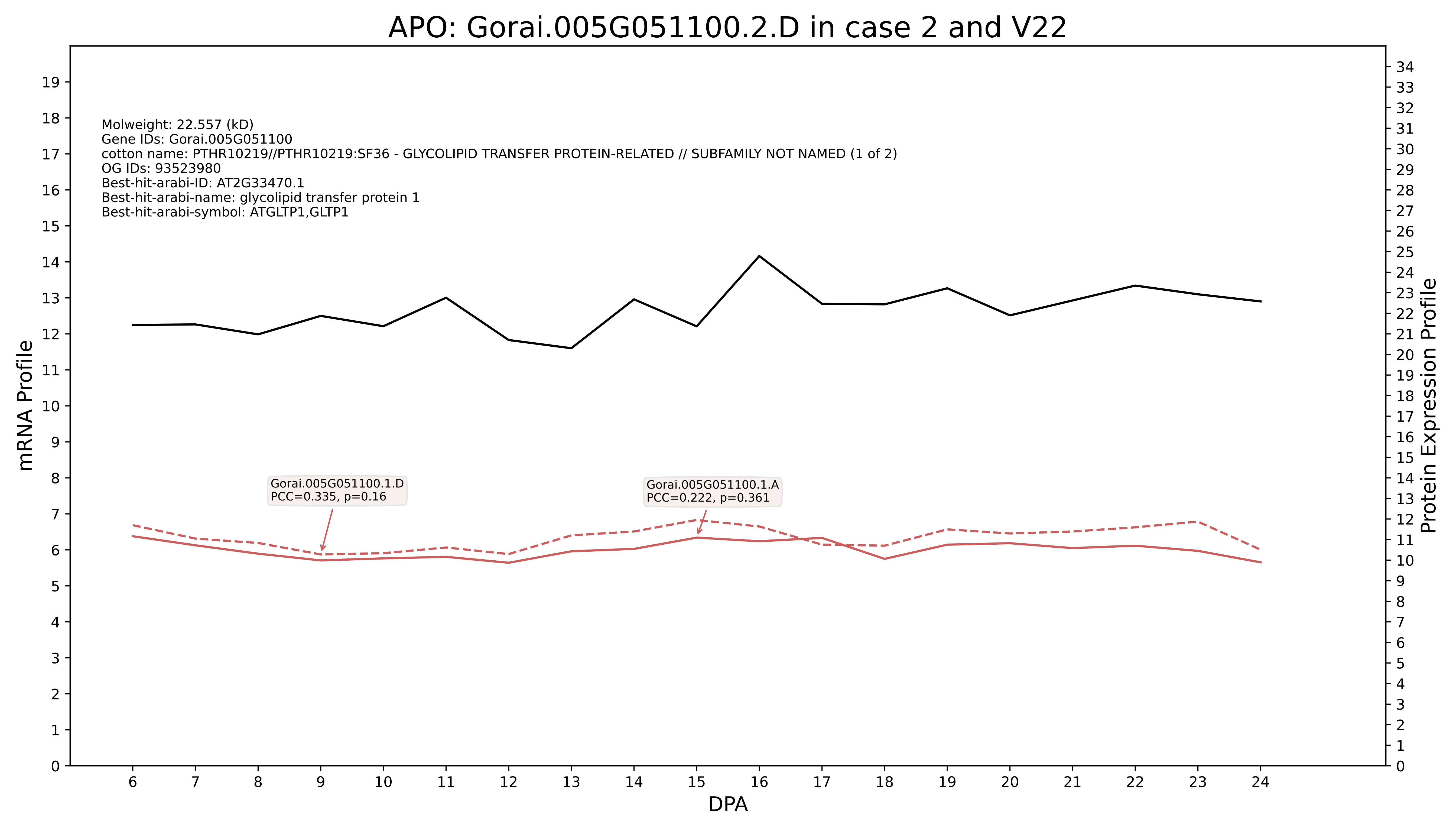
APO: Gorai.005G051100.2.D in case 2 and V22
Molweight: 22.557 (kD)
Gene IDs: Gorai.005G051100
cotton name: PTHR10219//PTHR10219:SF36 - GLYCOLIPID TRANSFER PROTEIN-RELATED // SUBFAMILY NOT NAMED (1 of 2)
OG IDs: 93523980
Best-hit-arabi-ID: AT2G33470.1
Best-hit-arabi-name: glycolipid transfer protein 1
Best-hit-arabi-symbol: ATGLTP1,GLTP1
Gorai.005G051100.1.D
PCC=0.335, p=0.16
Gorai.005G051100.1.A
PCC=0.222, p=0.361
mRNA Profile
Protein Expression Profile
DPA

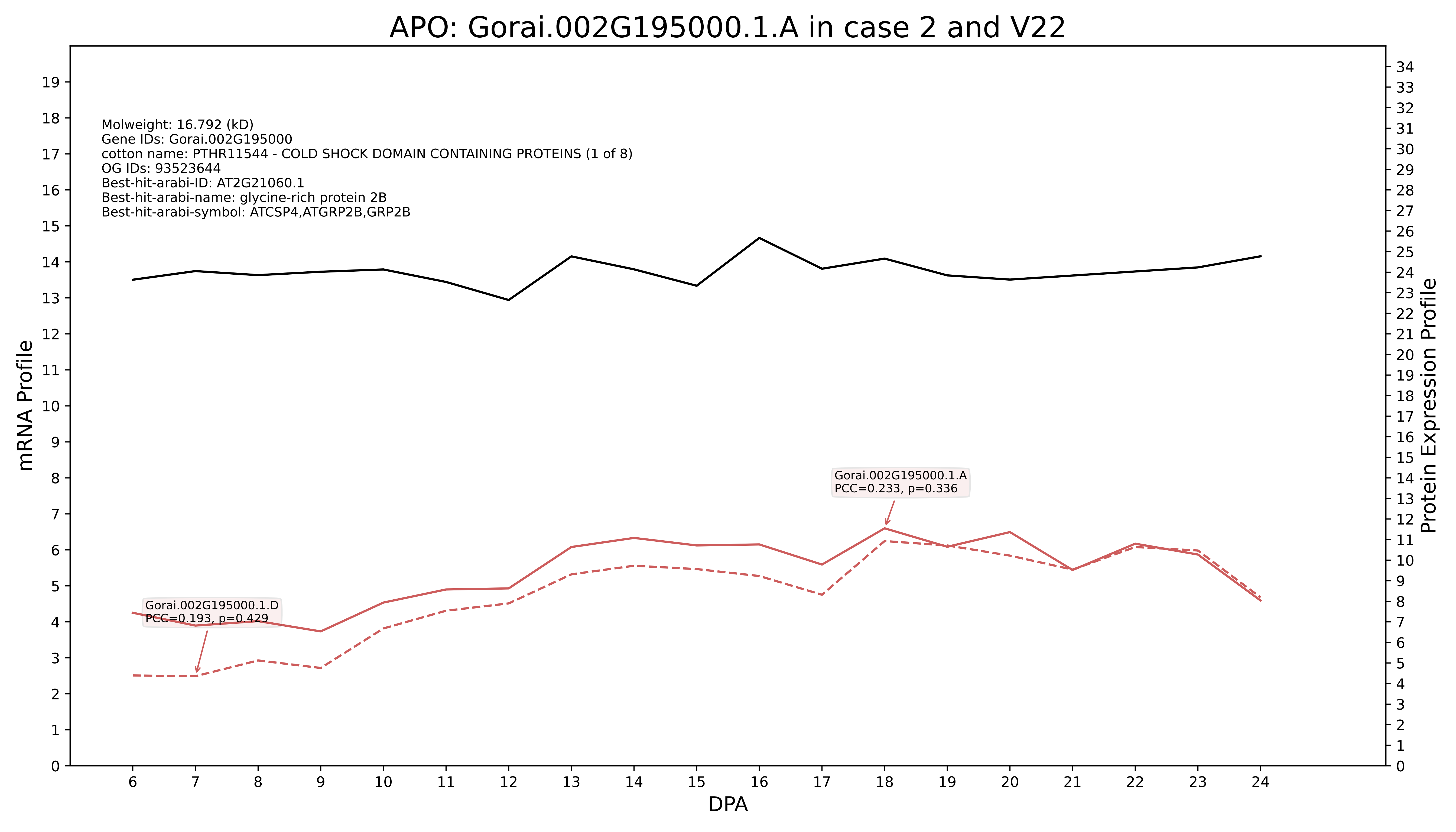

APO: Gorai.002G195000.1.A in case 2 and V22
Molweight: 16.792 (kD)
Gene IDs: Gorai.002G195000
cotton name: PTHR11544 - COLD SHOCK DOMAIN CONTAINING PROTEINS (1 of 8)
OG IDs: 93523644
Best-hit-arabi-ID: AT2G21060.1
Best-hit-arabi-name: glycine-rich protein 2B
Best-hit-arabi-symbol: ATCSP4,ATGRP2B,GRP2B
Gorai.002G195000.1.A
PCC=0.233, p=0.336
Gorai.002G195000.1.D
PCC=0.193, p=0.429
mRNA Profile
Protein Expression Profile
DPA

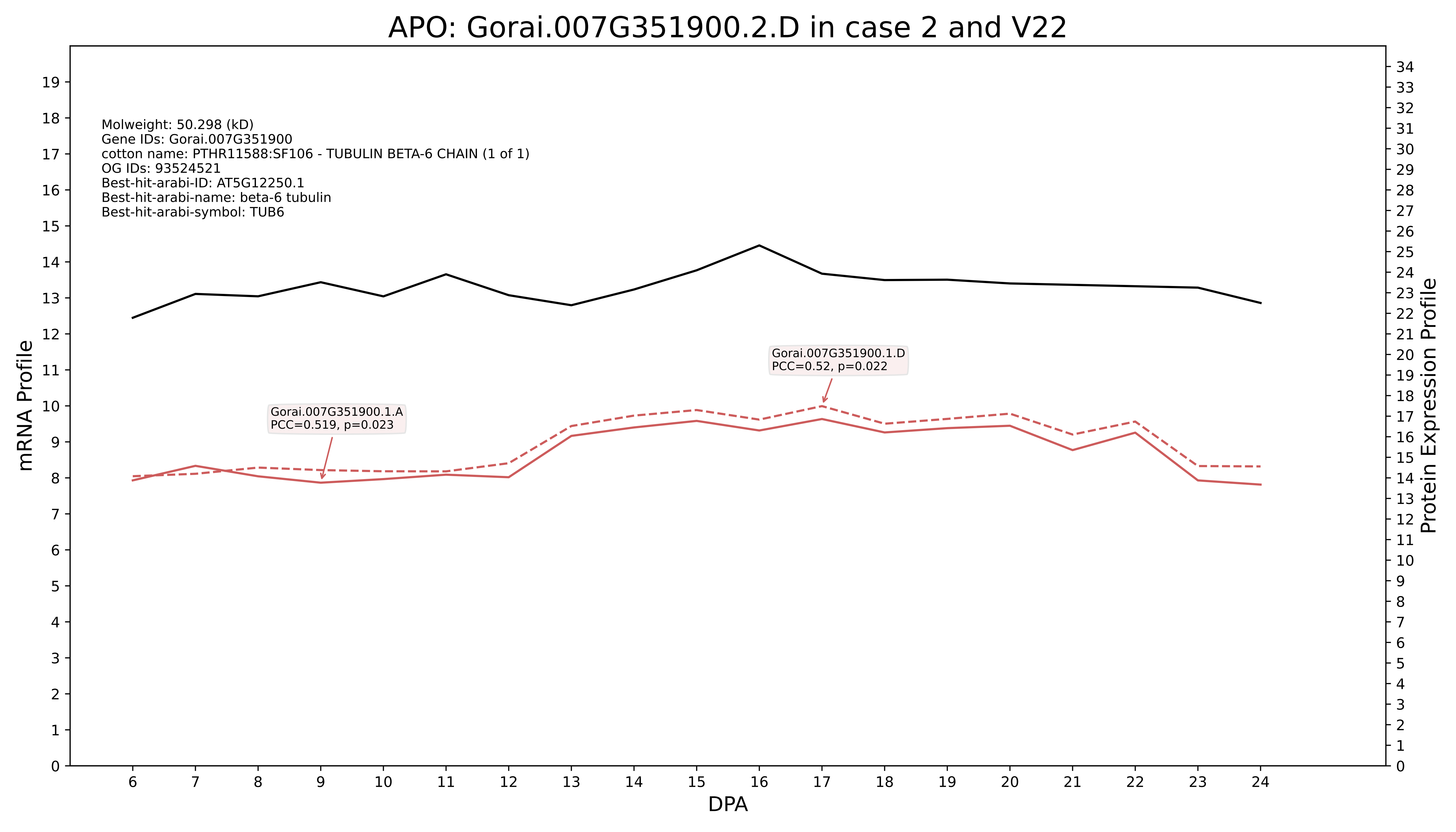
APO: Gorai.007G351900.2.D in case 2 and V22
Molweight: 50.298 (kD)
Gene IDs: Gorai.007G351900
cotton name: PTHR11588:SF106 - TUBULIN BETA-6 CHAIN (1 of 1)
OG IDs: 93524521
Best-hit-arabi-ID: AT5G12250.1
Best-hit-arabi-name: beta-6 tubulin
Best-hit-arabi-symbol: TUB6
Gorai.007G351900.1.D
PCC=0.52, p=0.022
Gorai.007G351900.1.A
PCC=0.519, p=0.023
mRNA Profile
Protein Expression Profile
DPA

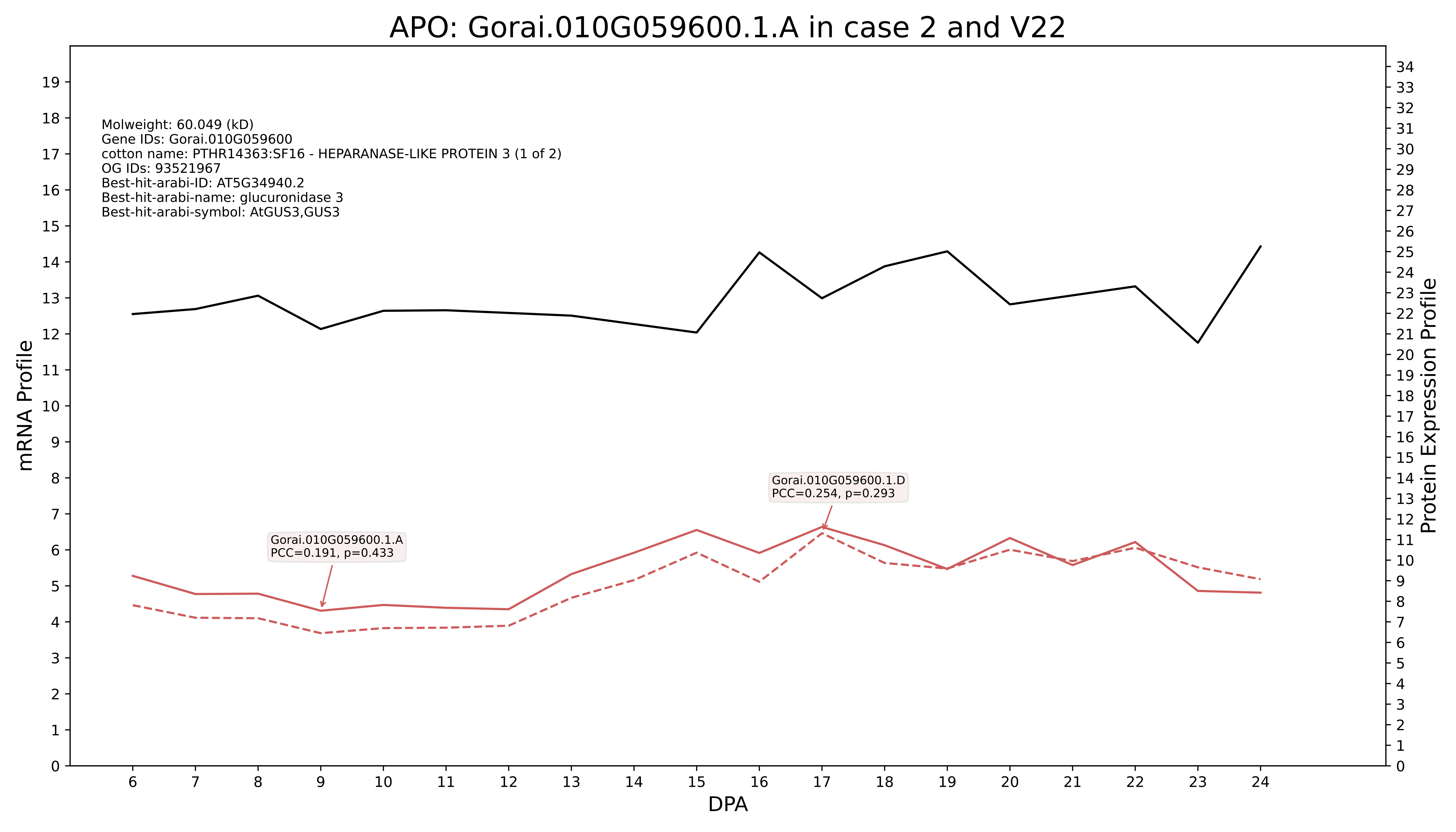

APO: Gorai.010G059600.1.A in case 2 and V22
Molweight: 60.049 (kD)
Gene IDs: Gorai.010G059600
cotton name: PTHR14363:SF16 - HEPARANASE-LIKE PROTEIN 3 (1 of 2)
OG IDs: 93521967
Best-hit-arabi-ID: AT5G34940.2
Best-hit-arabi-name: glucuronidase 3
Best-hit-arabi-symbol: AtGUS3,GUS3
Gorai.010G059600.1.D
PCC=0.254, p=0.293
Gorai.010G059600.1.A
PCC=0.191, p=0.433
mRNA Profile
Protein Expression Profile
DPA

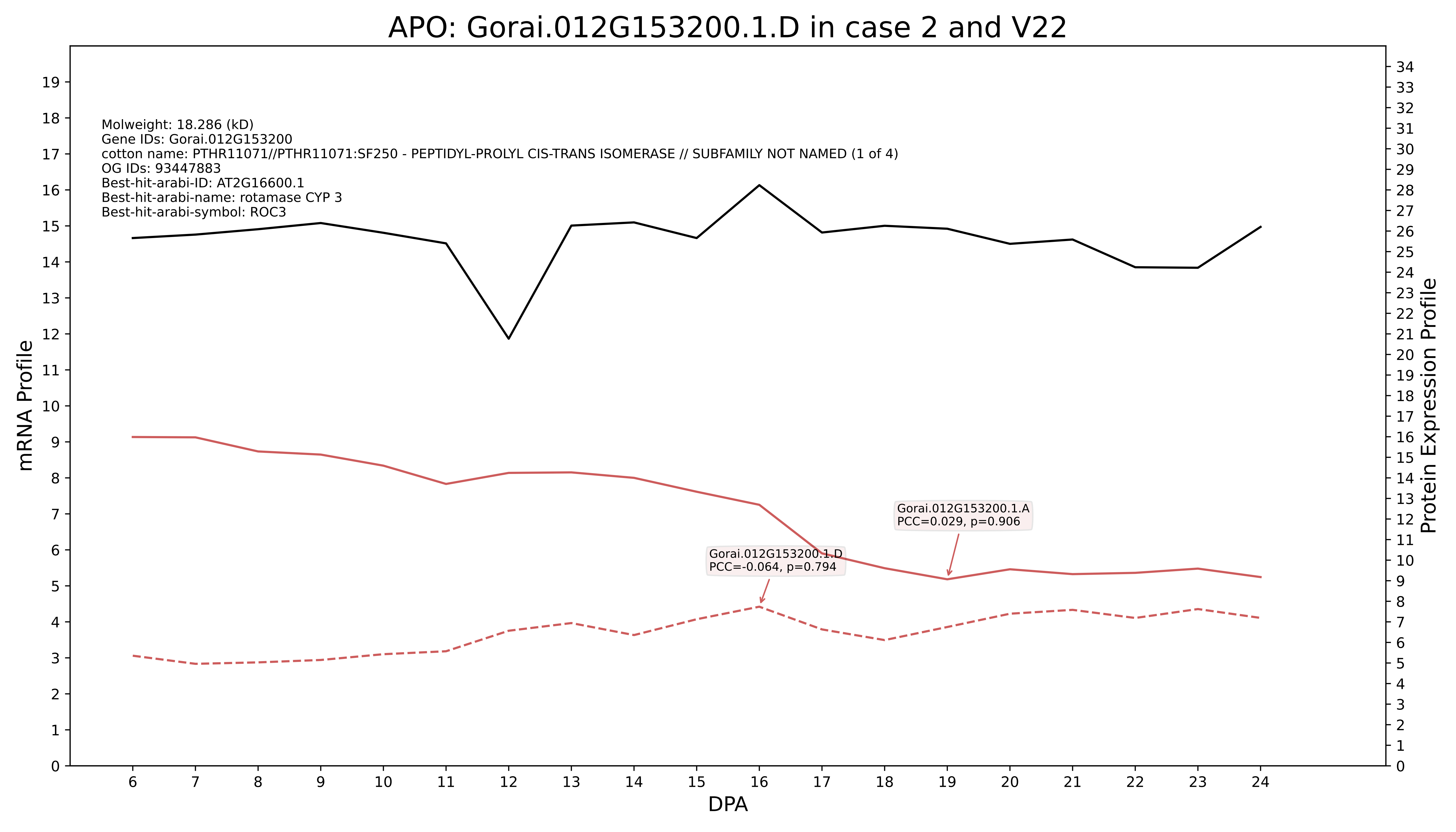
APO: Gorai.012G153200.1.D in case 2 and V22
Molweight: 18.286 (kD)
Gene IDs: Gorai.012G153200
cotton name: PTHR11071//PTHR11071:SF250 - PEPTIDYL-PROLYL CIS-TRANS ISOMERASE // SUBFAMILY NOT NAMED (1 of 4)
OG IDs: 93447883
Best-hit-arabi-ID: AT2G16600.1
Best-hit-arabi-name: rotamase CYP 3
Best-hit-arabi-symbol: ROC3
Gorai.012G153200.1.A
PCC=0.029, p=0.906
Gorai.012G153200.1.D
PCC=-0.064, p=0.794
mRNA Profile
Protein Expression Profile
DPA

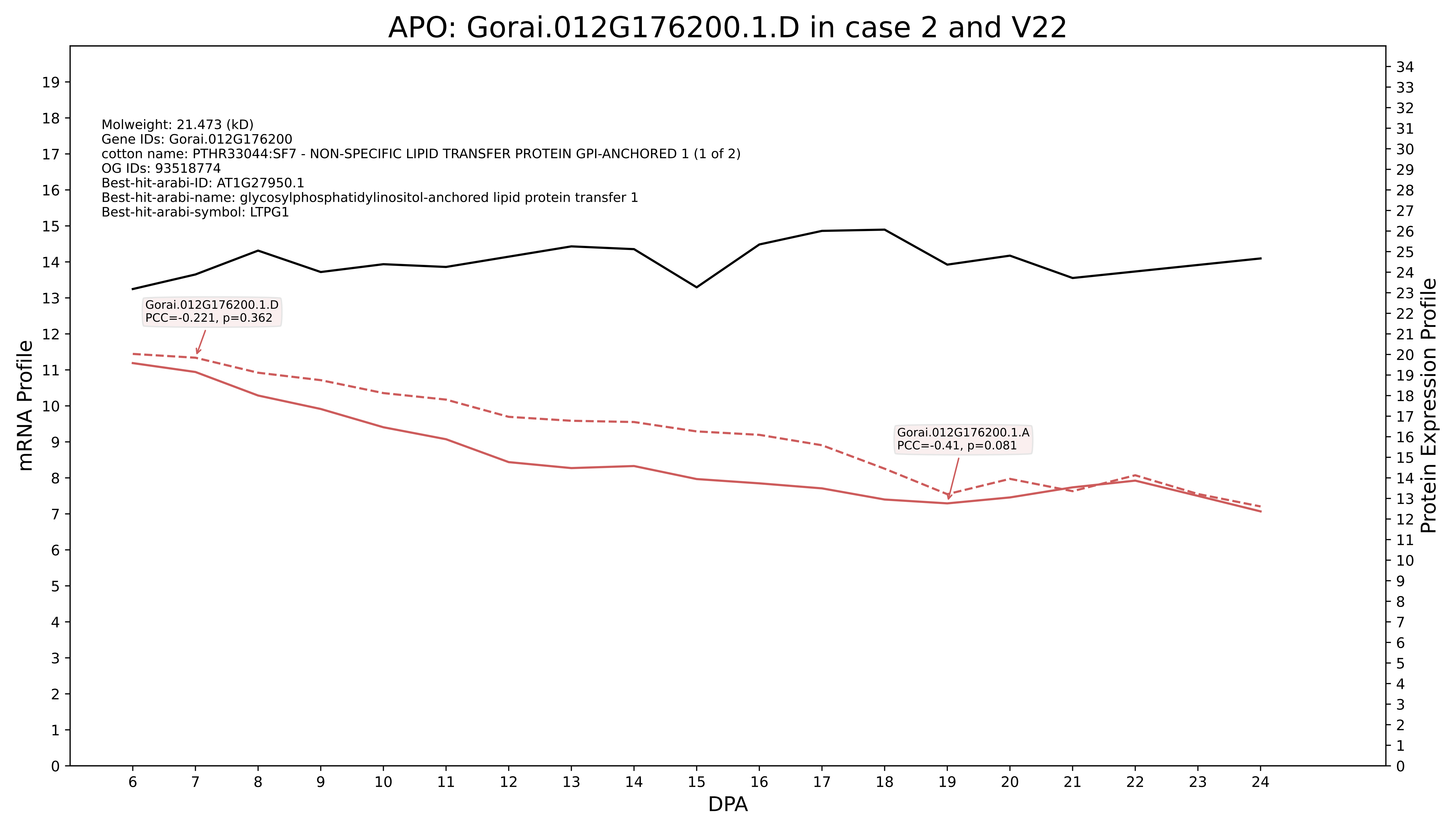
APO: Gorai.012G176200.1.D in case 2 and V22
Molweight: 21.473 (kD)
Gene IDs: Gorai.012G176200
cotton name: PTHR33044:SF7 - NON-SPECIFIC LIPID TRANSFER PROTEIN GPI-ANCHORED 1 (1 of 2)
OG IDs: 93518774
Best-hit-arabi-ID: AT1G27950.1
Best-hit-arabi-name: glycosylphosphatidylinositol-anchored lipid protein transfer 1
Best-hit-arabi-symbol: LTPG1
Gorai.012G176200.1.D
PCC=-0.221, p=0.362
Gorai.012G176200.1.A
PCC=-0.41, p=0.081
mRNA Profile
Protein Expression Profile
DPA

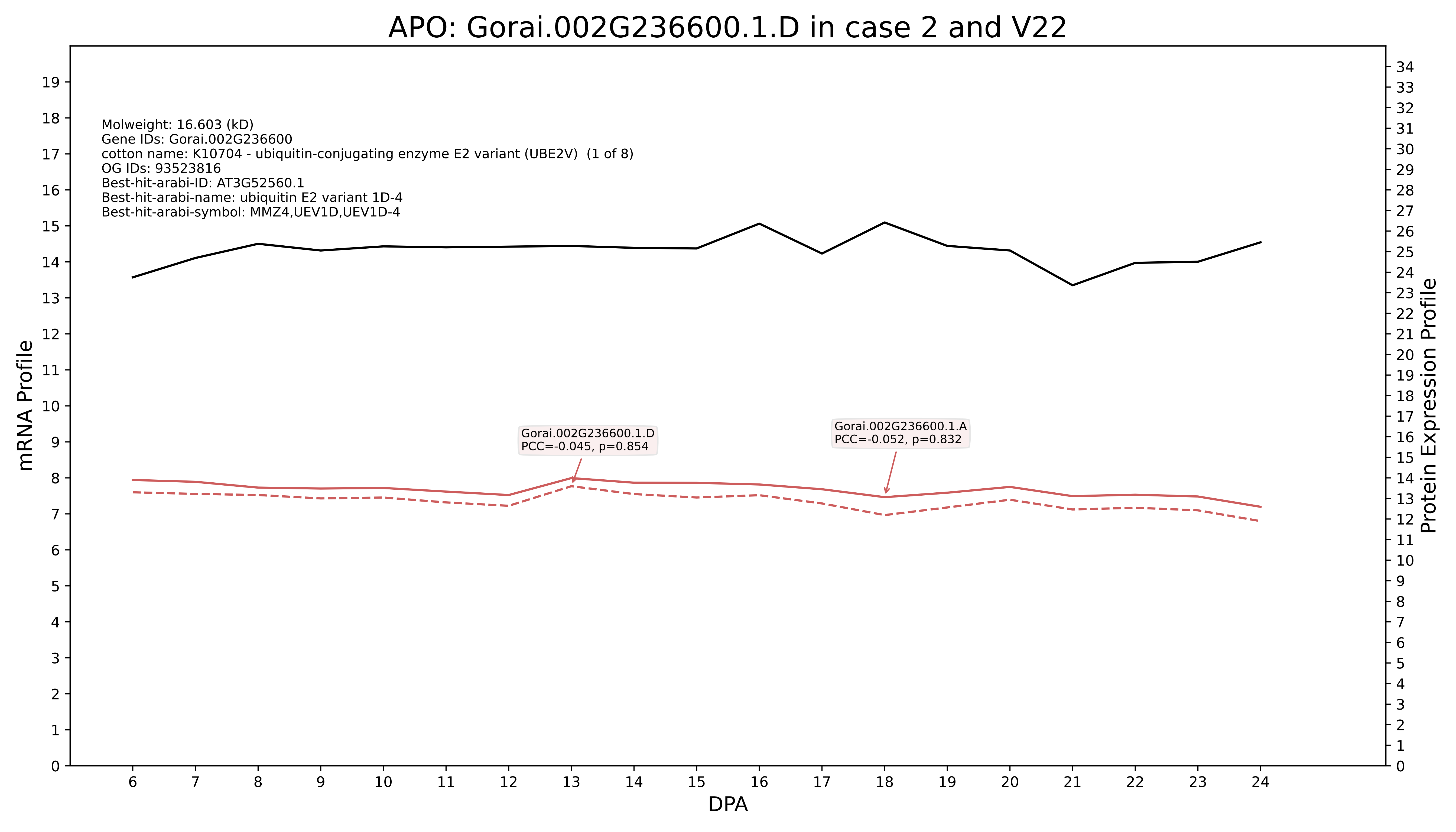
APO: Gorai.002G236600.1.D in case 2 and V22
Molweight: 16.603 (kD)
Gene IDs: Gorai.002G236600
cotton name: K10704 - ubiquitin-conjugating enzyme E2 variant (UBE2V) (1 of 8)
OG IDs: 93523816
Best-hit-arabi-ID: AT3G52560.1
Best-hit-arabi-name: ubiquitin E2 variant 1D-4
Best-hit-arabi-symbol: MMZ4,UEV1D,UEV1D-4
Gorai.002G236600.1.D
PCC=-0.045, p=0.854
Gorai.002G236600.1.A
PCC=-0.052, p=0.832
mRNA Profile
Protein Expression Profile
DPA

### APO: Gorai.002G177600.1.D in case 2 and V22

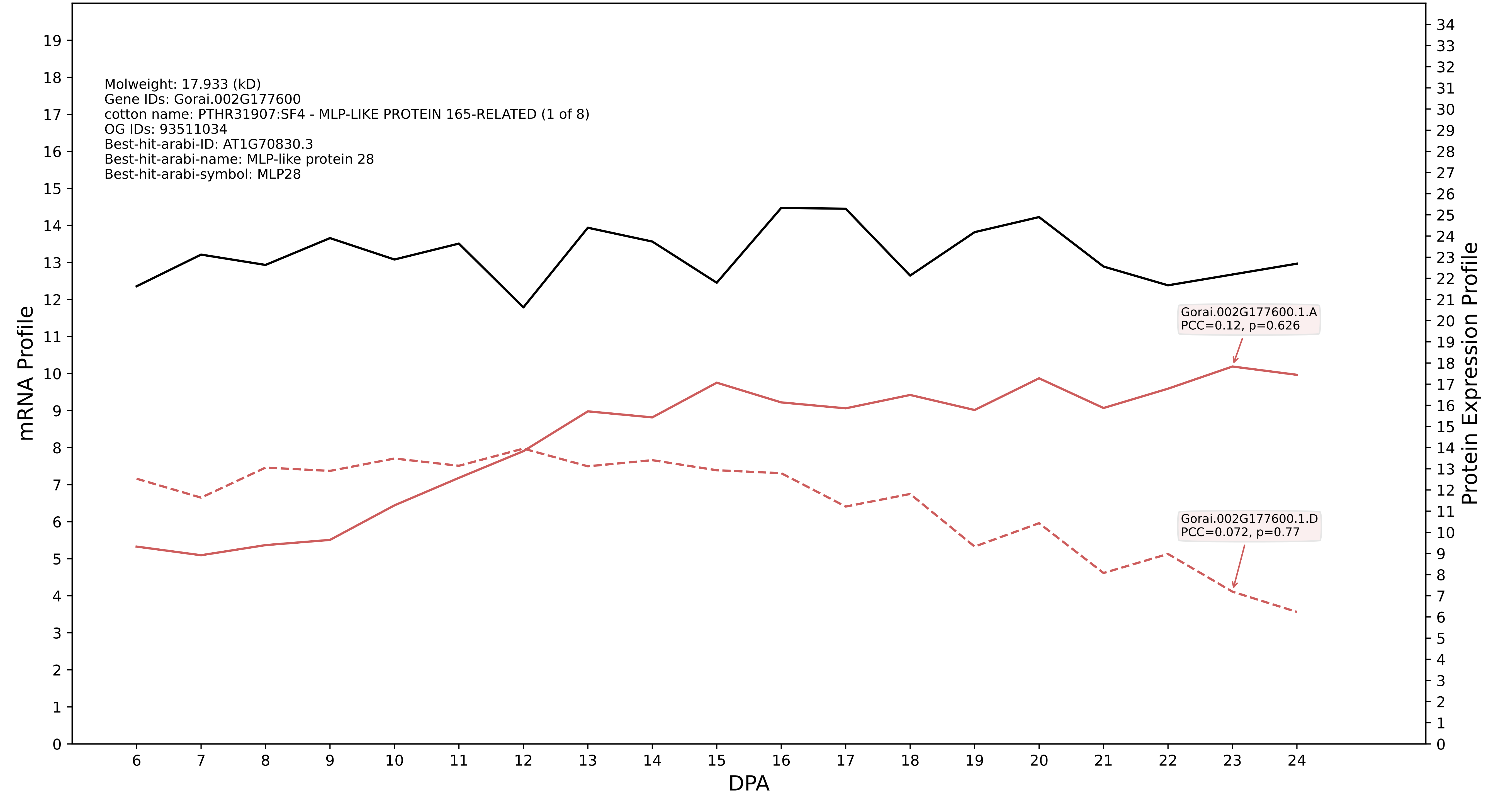

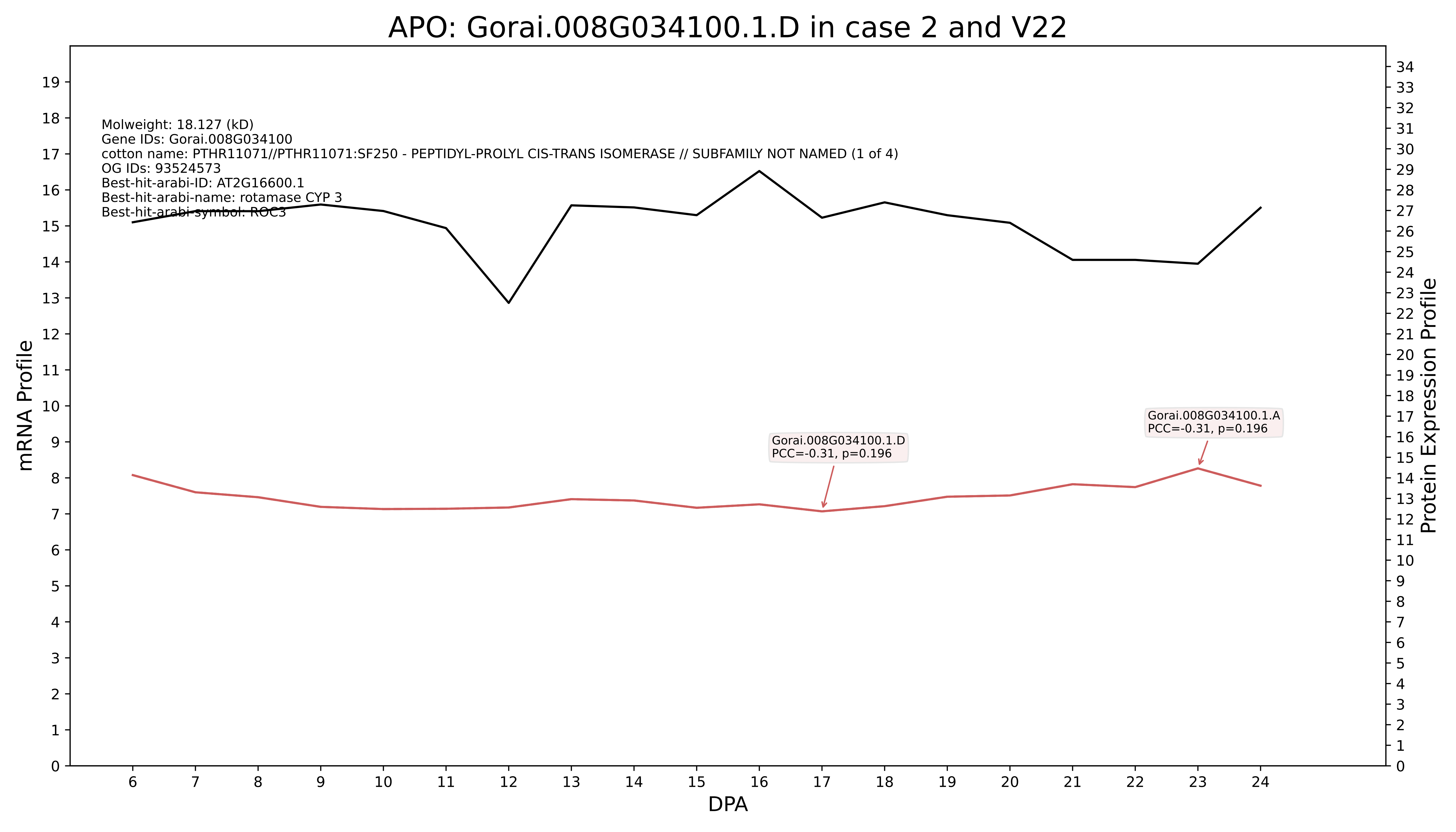
APO: Gorai.008G034100.1.D in case 2 and V22
Molweight: 18.127 (kD)
Gene IDs: Gorai.008G034100
cotton name: PTHR11071//PTHR11071:SF250 - PEPTIDYL-PROLYL CIS-TRANS ISOMERASE // SUBFAMILY NOT NAMED (1 of 4)
OG IDs: 93524573
Best-hit-arabi-ID: AT2G16600.1
Best-hit-arabi-name: rotamase CYP 3
Best-hit-arabi-symbol: ROC3
Gorai.008G034100.1.D
PCC=-0.31, p=0.196
Gorai.008G034100.1.A
PCC=-0.31, p=0.196
mRNA Profile
Protein Expression Profile
DPA

### APO: Gorai.009G080400.1.D in case 3 and V22

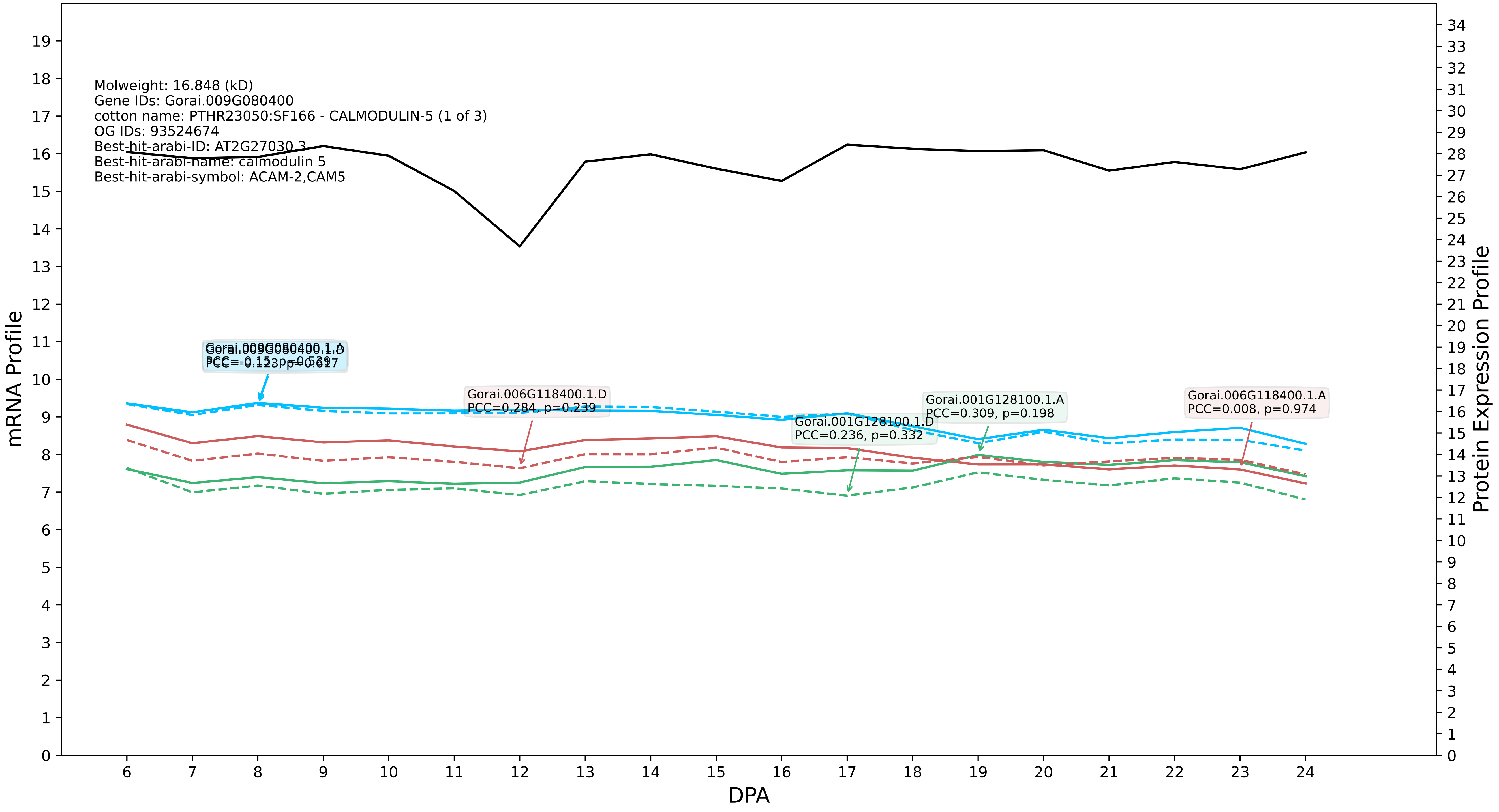

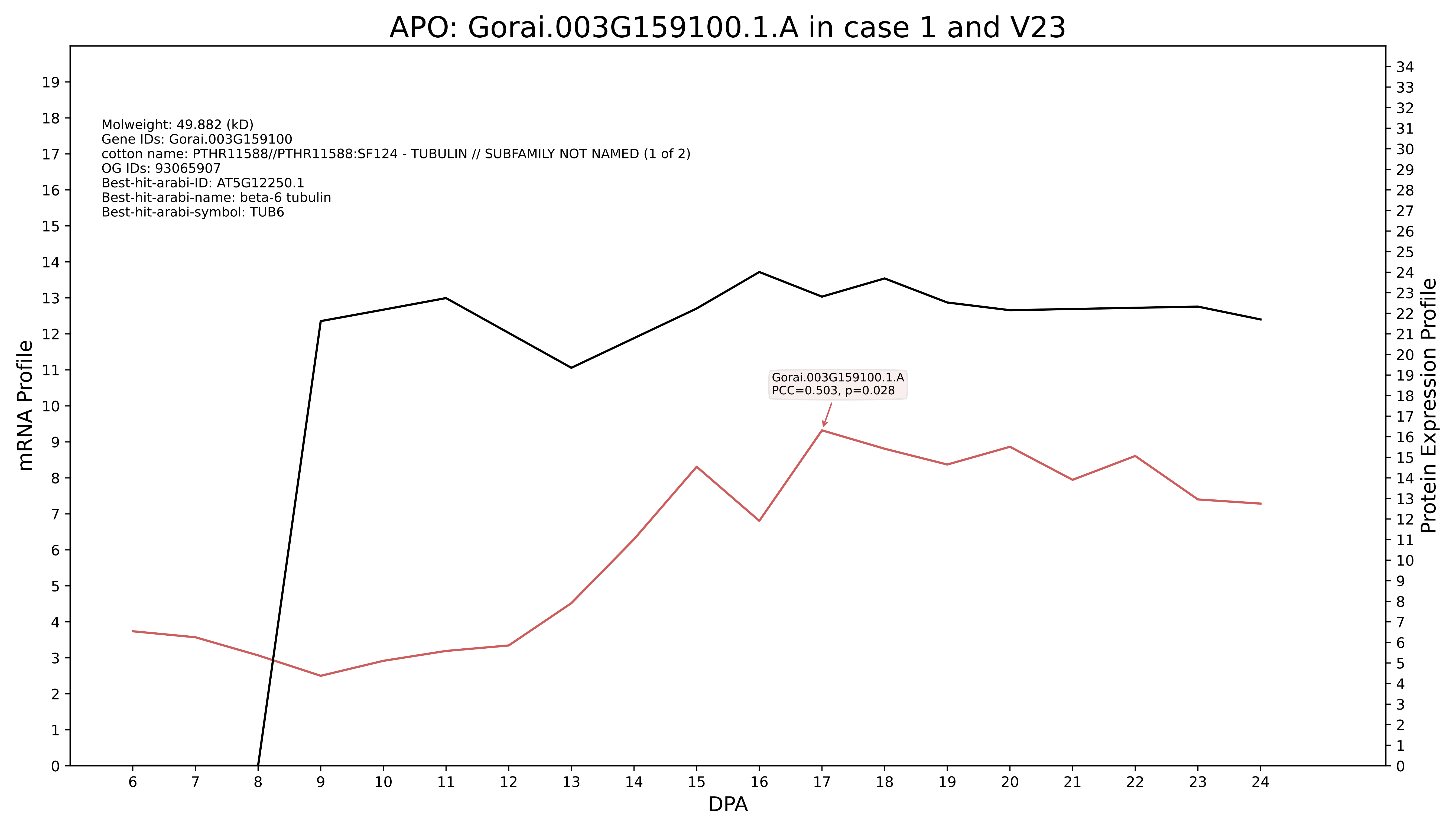
APO: Gorai.003G159100.1.A in case 1 and V23
Molweight: 49.882 (kD)
Gene IDs: Gorai.003G159100
cotton name: PTHR11588//PTHR11588:SF124 - TUBULIN // SUBFAMILY NOT NAMED (1 of 2)
OG IDs: 93065907
Best-hit-arabi-ID: AT5G12250.1
Best-hit-arabi-name: beta-6 tubulin
Best-hit-arabi-symbol: TUB6
Gorai.003G159100.1.A
PCC=0.503, p=0.028
mRNA Profile
Protein Expression Profile
DPA

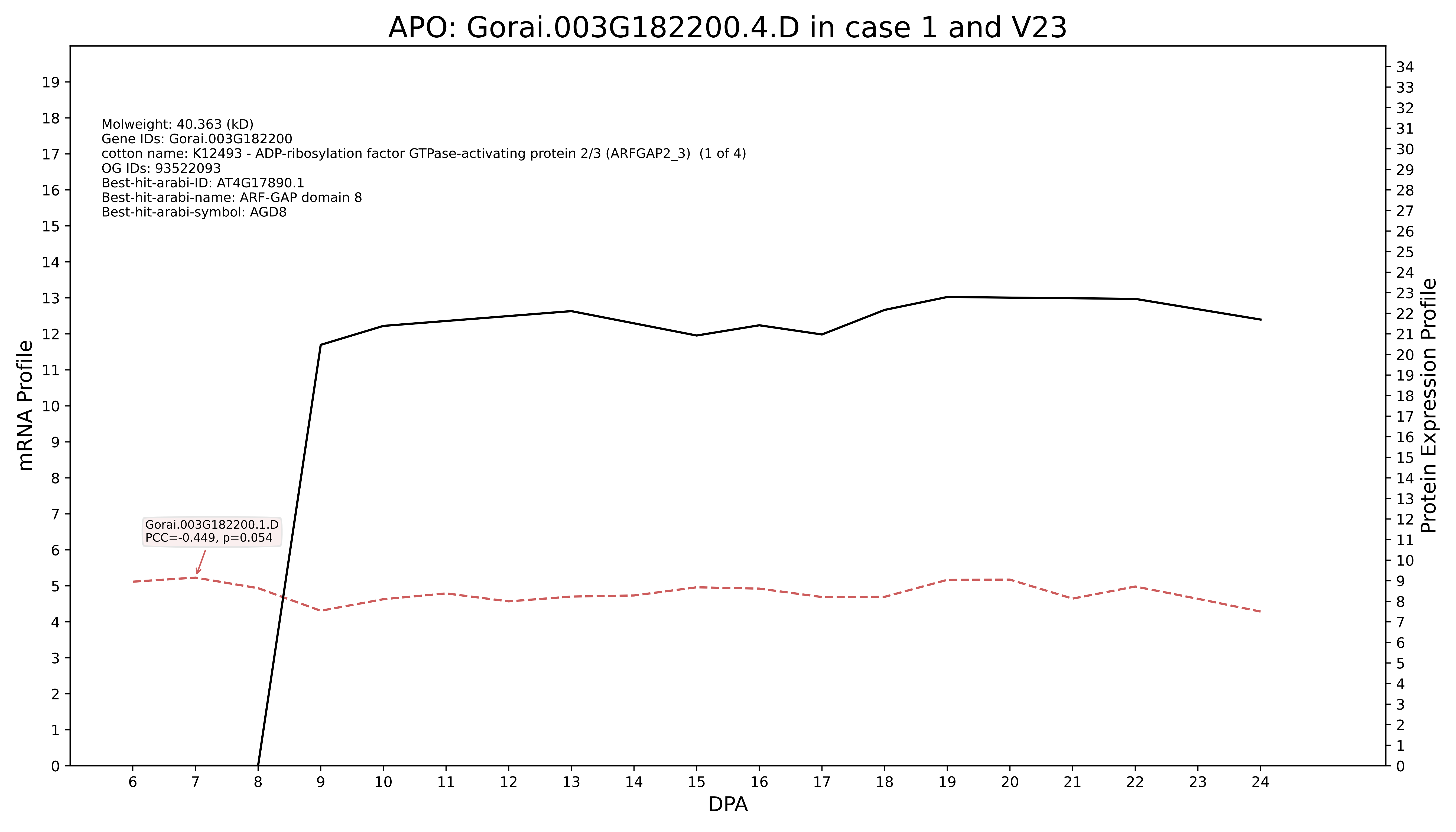
APO: Gorai.003G182200.4.D in case 1 and V23
Molweight: 40.363 (kD)
Gene IDs: Gorai.003G182200
cotton name: K12493 - ADP-ribosylation factor GTPase-activating protein 2/3 (ARFGAP2_3) (1 of 4)
OG IDs: 93522093
Best-hit-arabi-ID: AT4G17890.1
Best-hit-arabi-name: ARF-GAP domain 8
Best-hit-arabi-symbol: AGD8
Gorai.003G182200.1.D
PCC=-0.449, p=0.054
mRNA Profile
Protein Expression Profile
DPA

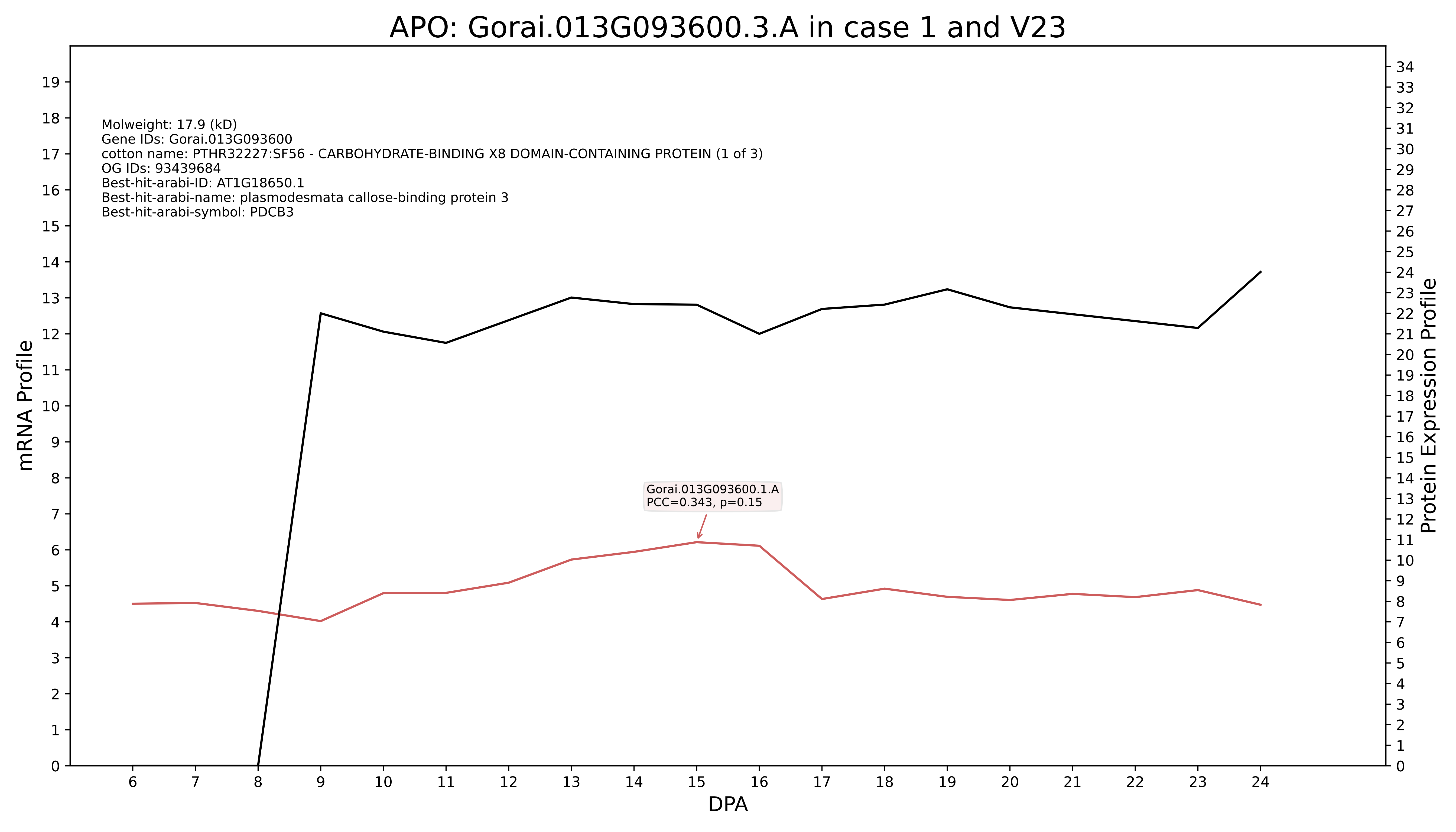
APO: Gorai.013G093600.3.A in case 1 and V23
Molweight: 17.9 (kD)
Gene IDs: Gorai.013G093600
cotton name: PTHR32227:SF56 - CARBOHYDRATE-BINDING X8 DOMAIN-CONTAINING PROTEIN (1 of 3)
OG IDs: 93439684
Best-hit-arabi-ID: AT1G18650.1
Best-hit-arabi-name: plasmodesmata callose-binding protein 3
Best-hit-arabi-symbol: PDCB3
Gorai.013G093600.1.A
PCC=0.343, p=0.15
mRNA Profile
Protein Expression Profile
DPA

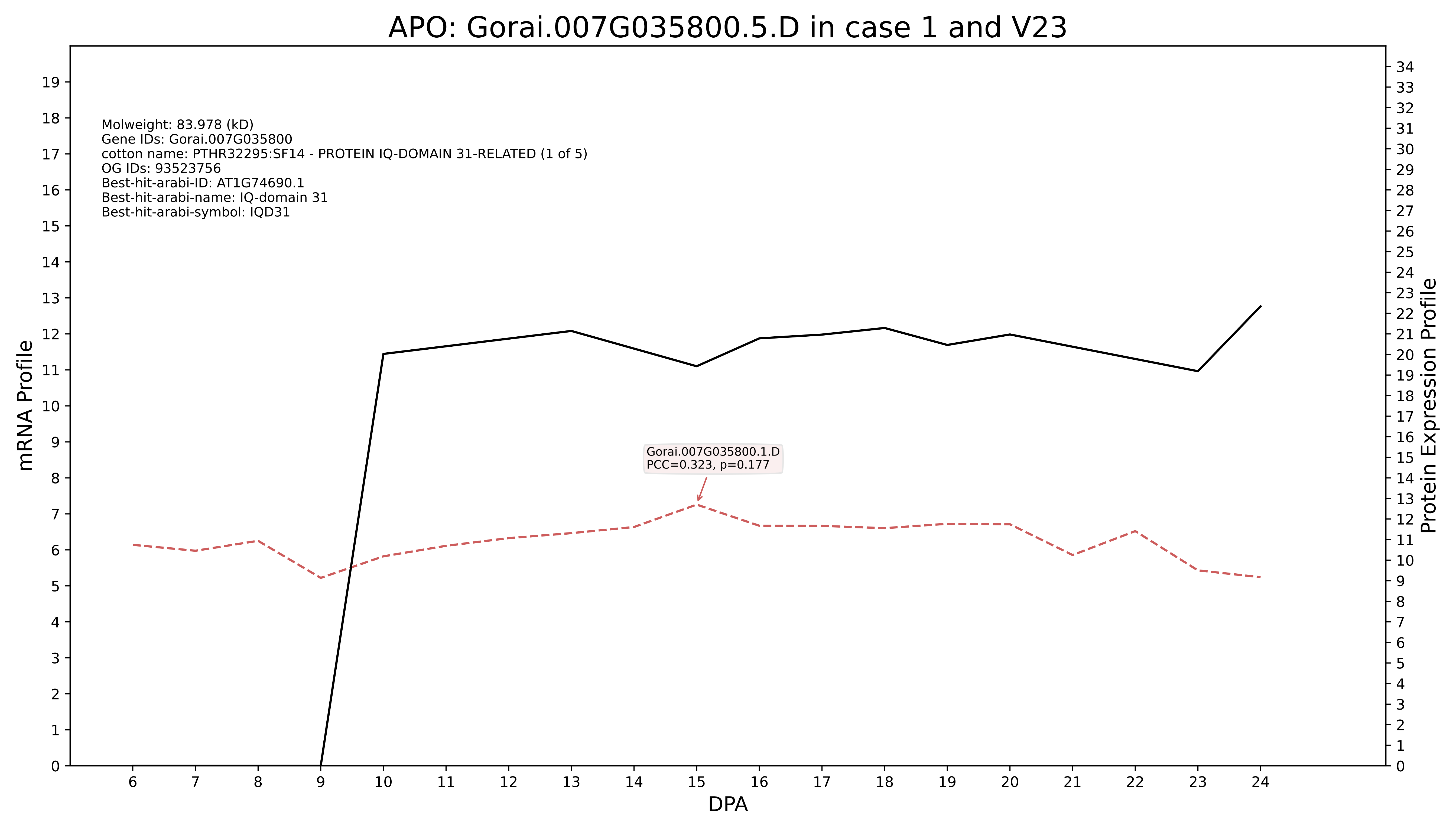

APO: Gorai.007G035800.5.D in case 1 and V23
Molweight: 83.978 (kD)
Gene IDs: Gorai.007G035800
cotton name: PTHR32295:SF14 - PROTEIN IQ-DOMAIN 31-RELATED (1 of 5)
OG IDs: 93523756
Best-hit-arabi-ID: AT1G74690.1
Best-hit-arabi-name: IQ-domain 31
Best-hit-arabi-symbol: IQD31
Gorai.007G035800.1.D
PCC=0.323, p=0.177
mRNA Profile
Protein Expression Profile
DPA

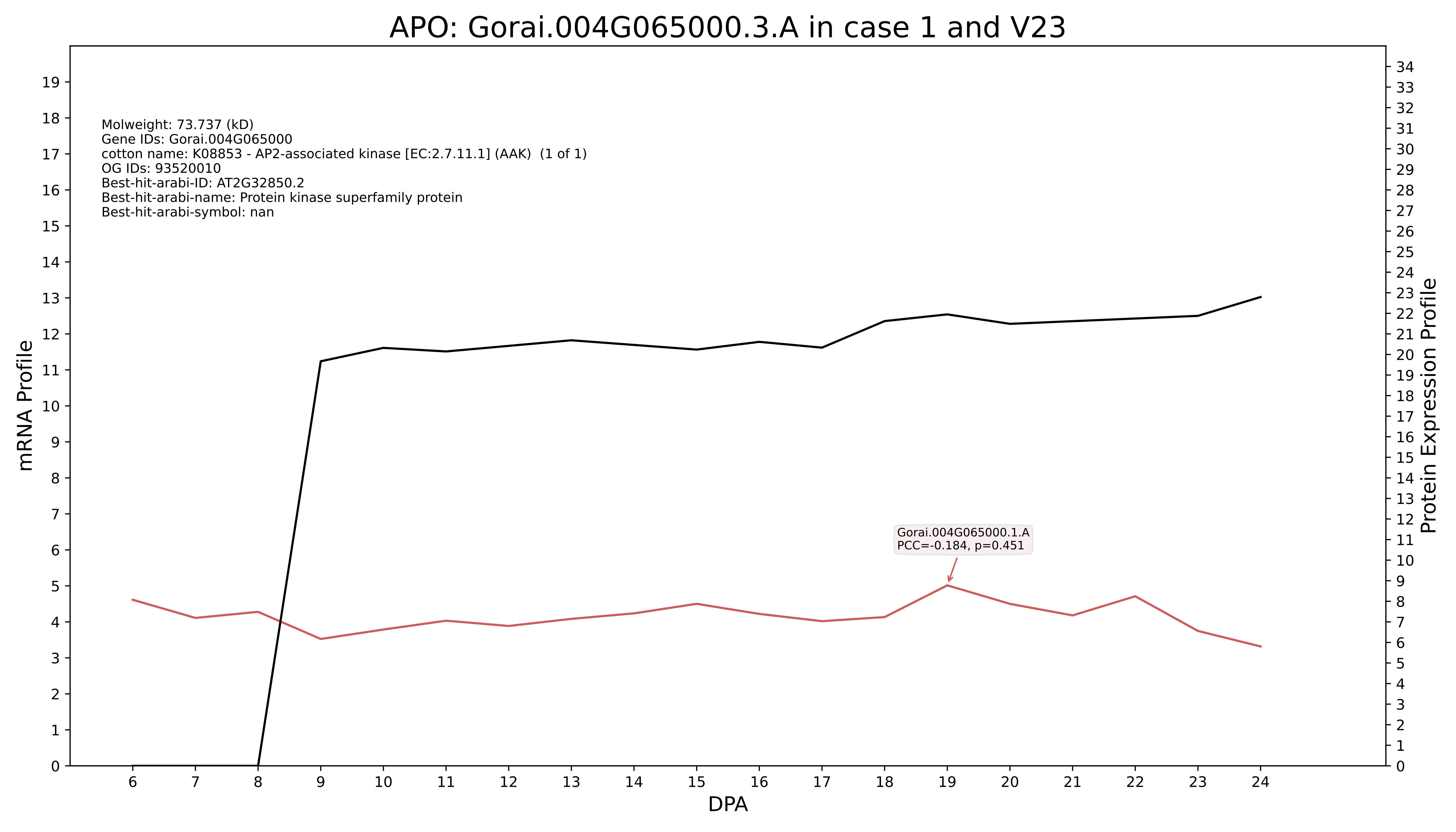
APO: Gorai.004G065000.3.A in case 1 and V23
Molweight: 73.737 (kD)
Gene IDs: Gorai.004G065000
cotton name: K08853 - AP2-associated kinase [EC:2.7.11.1] (AAK) (1 of 1)
OG IDs: 93520010
Best-hit-arabi-ID: AT2G32850.2
Best-hit-arabi-name: Protein kinase superfamily protein
Best-hit-arabi-symbol: nan
Gorai.004G065000.1.A
PCC=-0.184, p=0.451
mRNA Profile
Protein Expression Profile
DPA

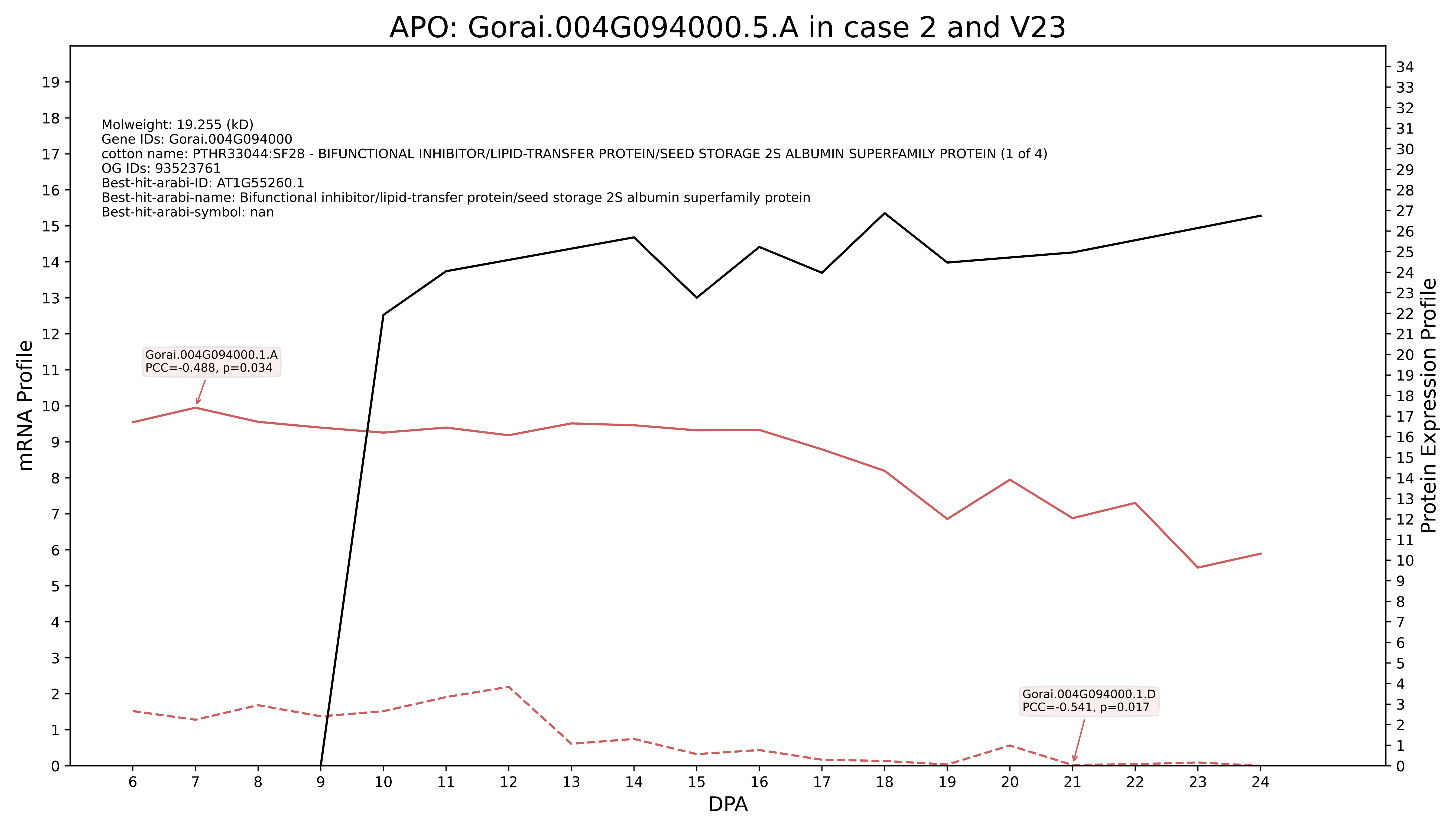
APO: Gorai.004G094000.5.A in case 2 and V23
Molweight: 19.255 (kD)
Gene IDs: Gorai.004G094000
cotton name: PTHR33044:SF28 - BIFUNCTIONAL INHIBITOR/LIPID-TRANSFER PROTEIN/SEED STORAGE 2S ALBUMIN SUPERFAMILY PROTEIN (1 of 4)
OG IDs: 93523761
Best-hit-arabi-ID: AT1G55260.1
Best-hit-arabi-name: Bifunctional inhibitor/lipid-transfer protein/seed storage 2S albumin superfamily protein
Best-hit-arabi-symbol: nan
Gorai.004G094000.1.A
PCC=-0.488, p=0.034
Gorai.004G094000.1.D
PCC=-0.541, p=0.017
mRNA Profile
Protein Expression Profile
DPA

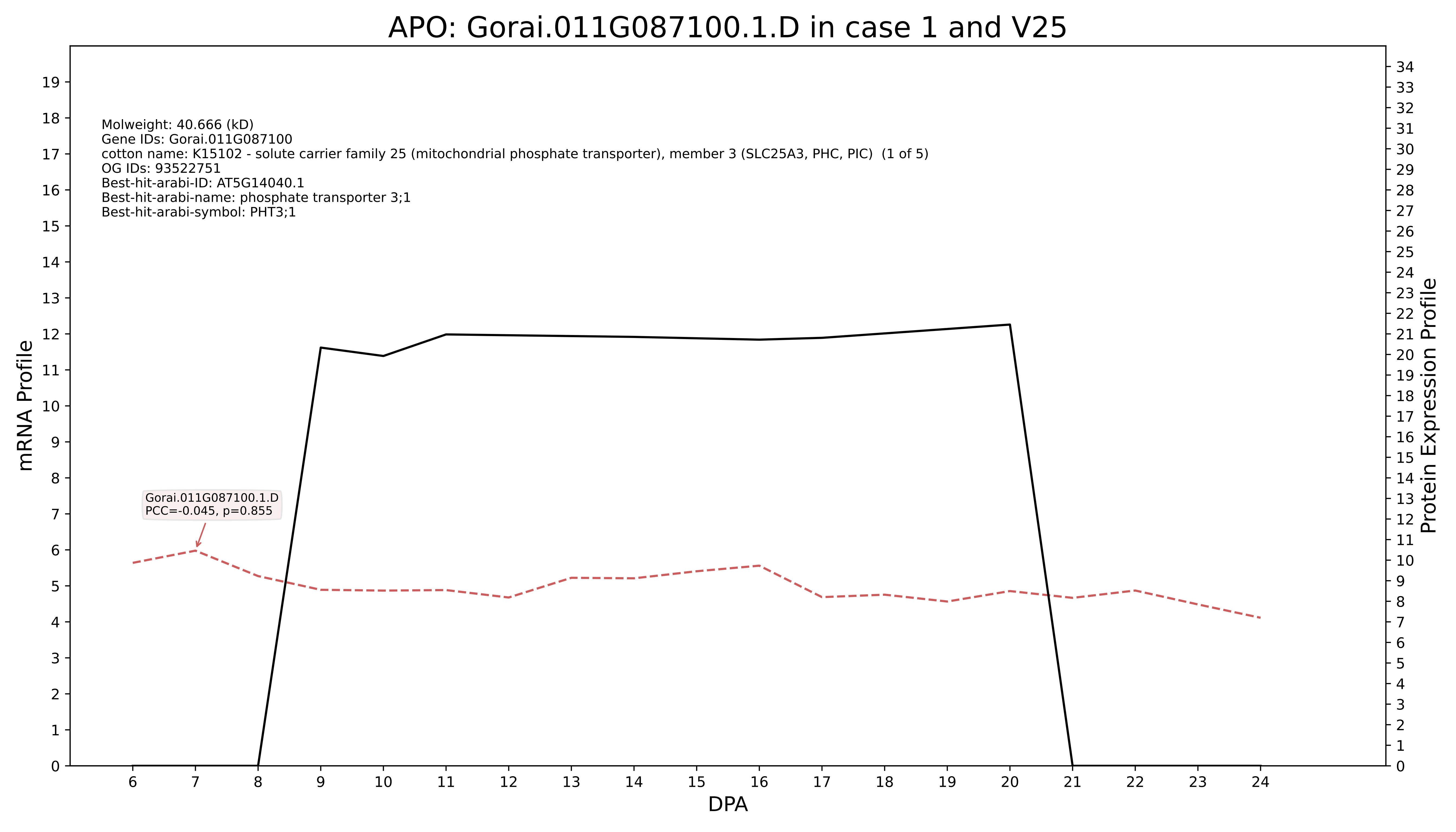

APO: Gorai.011G087100.1.D in case 1 and V25
Molweight: 40.666 (kD)
Gene IDs: Gorai.011G087100
cotton name: K15102 - solute carrier family 25 (mitochondrial phosphate transporter), member 3 (SLC25A3, PHC, PIC) (1 of 5)
OG IDs: 93522751
Best-hit-arabi-ID: AT5G14040.1
Best-hit-arabi-name: phosphate transporter 3;1
Best-hit-arabi-symbol: PHT3;1
Gorai.011G087100.1.D
PCC=-0.045, p=0.855
mRNA Profile
Protein Expression Profile
DPA

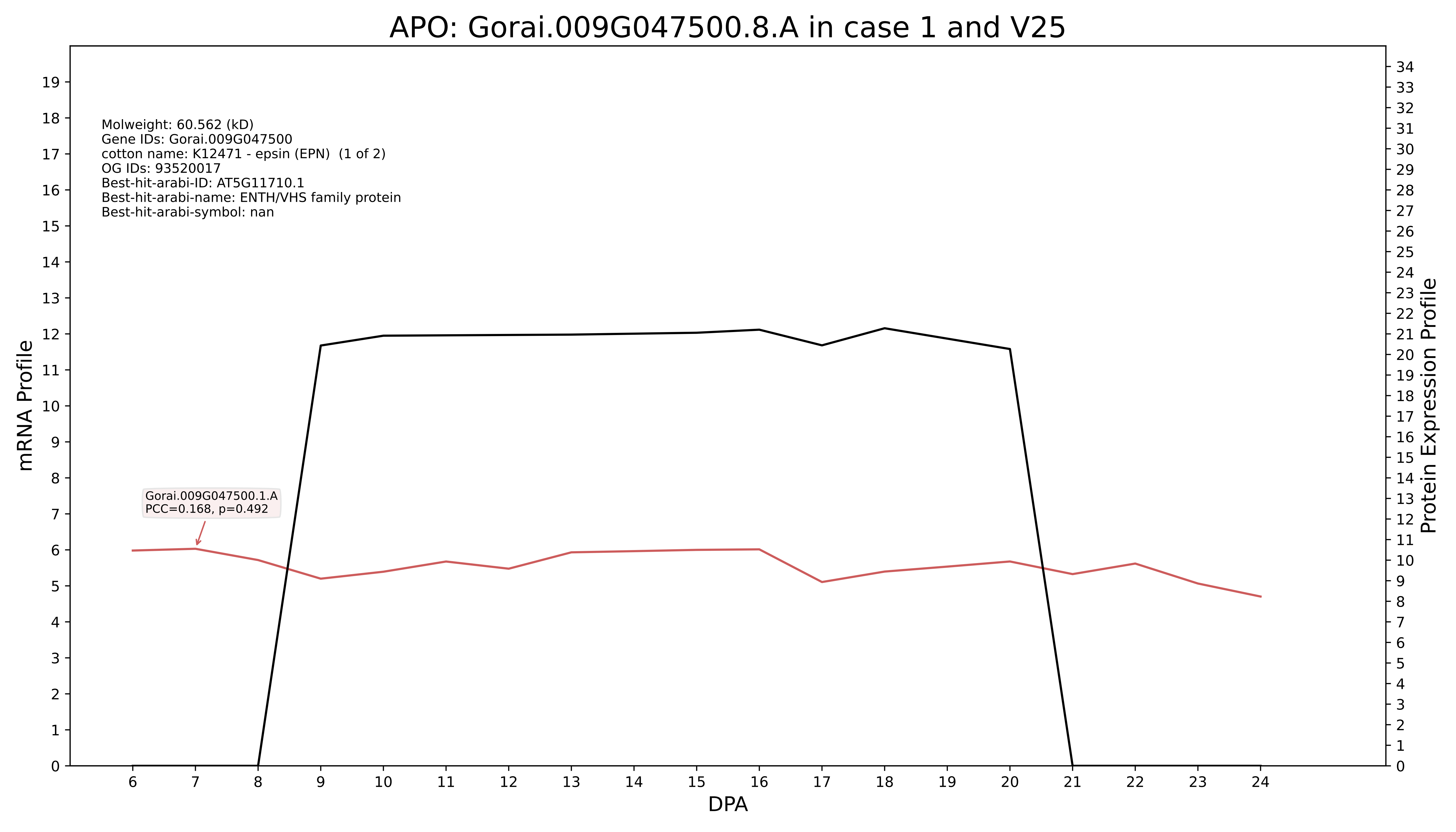
APO: Gorai.009G047500.8.A in case 1 and V25
Molweight: 60.562 (kD)
Gene IDs: Gorai.009G047500
cotton name: K12471 - epsin (EPN) (1 of 2)
OG IDs: 93520017
Best-hit-arabi-ID: AT5G11710.1
Best-hit-arabi-name: ENTH/VHS family protein
Best-hit-arabi-symbol: nan
Gorai.009G047500.1.A
PCC=0.168, p=0.492
mRNA Profile
Protein Expression Profile
DPA

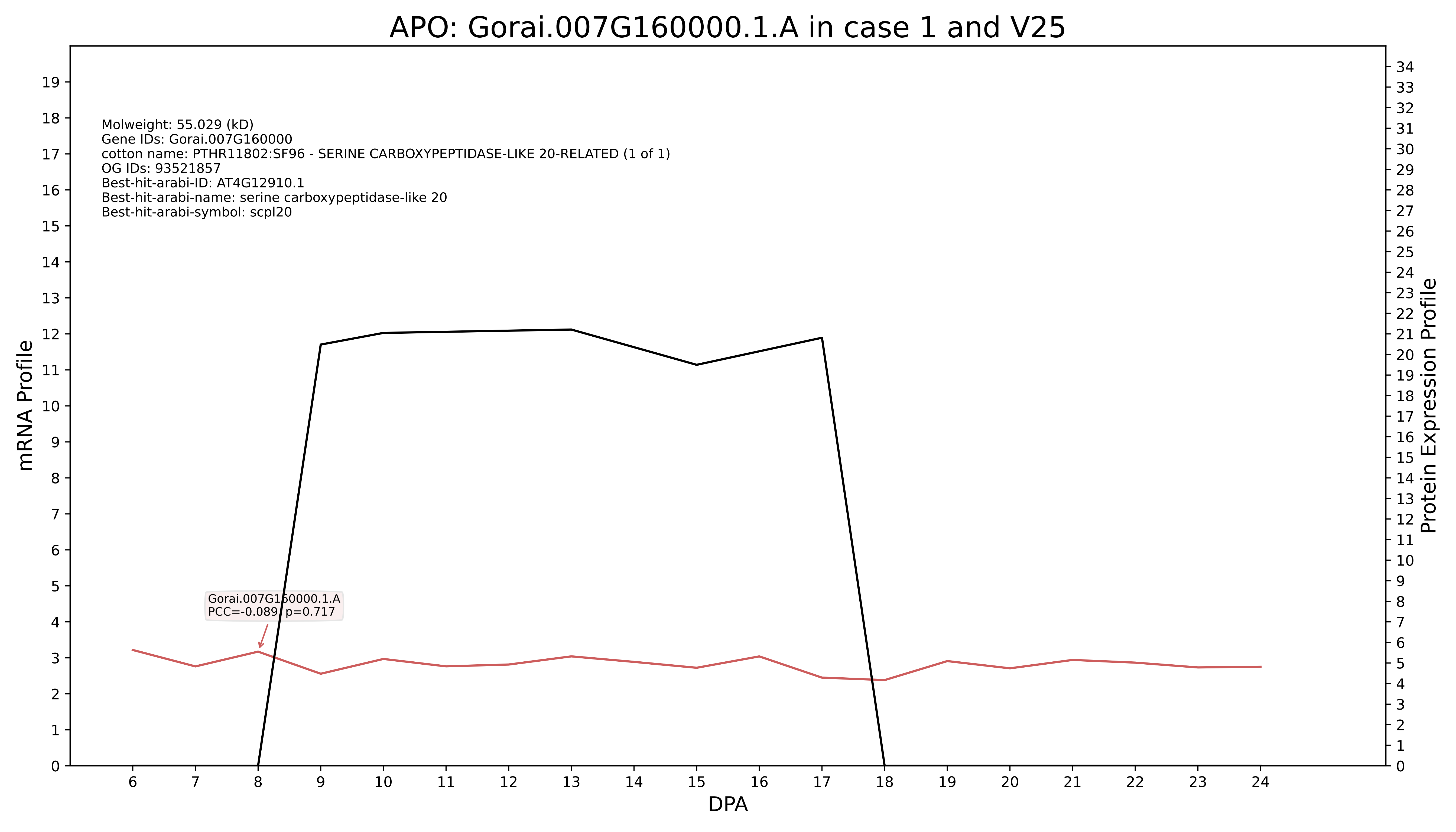

APO: Gorai.007G160000.1.A in case 1 and V25
Molweight: 55.029 (kD)
Gene IDs: Gorai.007G160000
cotton name: PTHR11802:SF96 - SERINE CARBOXYPEPTIDASE-LIKE 20-RELATED (1 of 1)
OG IDs: 93521857
Best-hit-arabi-ID: AT4G12910.1
Best-hit-arabi-name: serine carboxypeptidase-like 20
Best-hit-arabi-symbol: scpl20
Gorai.007G160000.1.A
PCC=-0.089, p=0.717
mRNA Profile
Protein Expression Profile
DPA

### APO: Gorai.008G175300.4.A in case 1 and V25

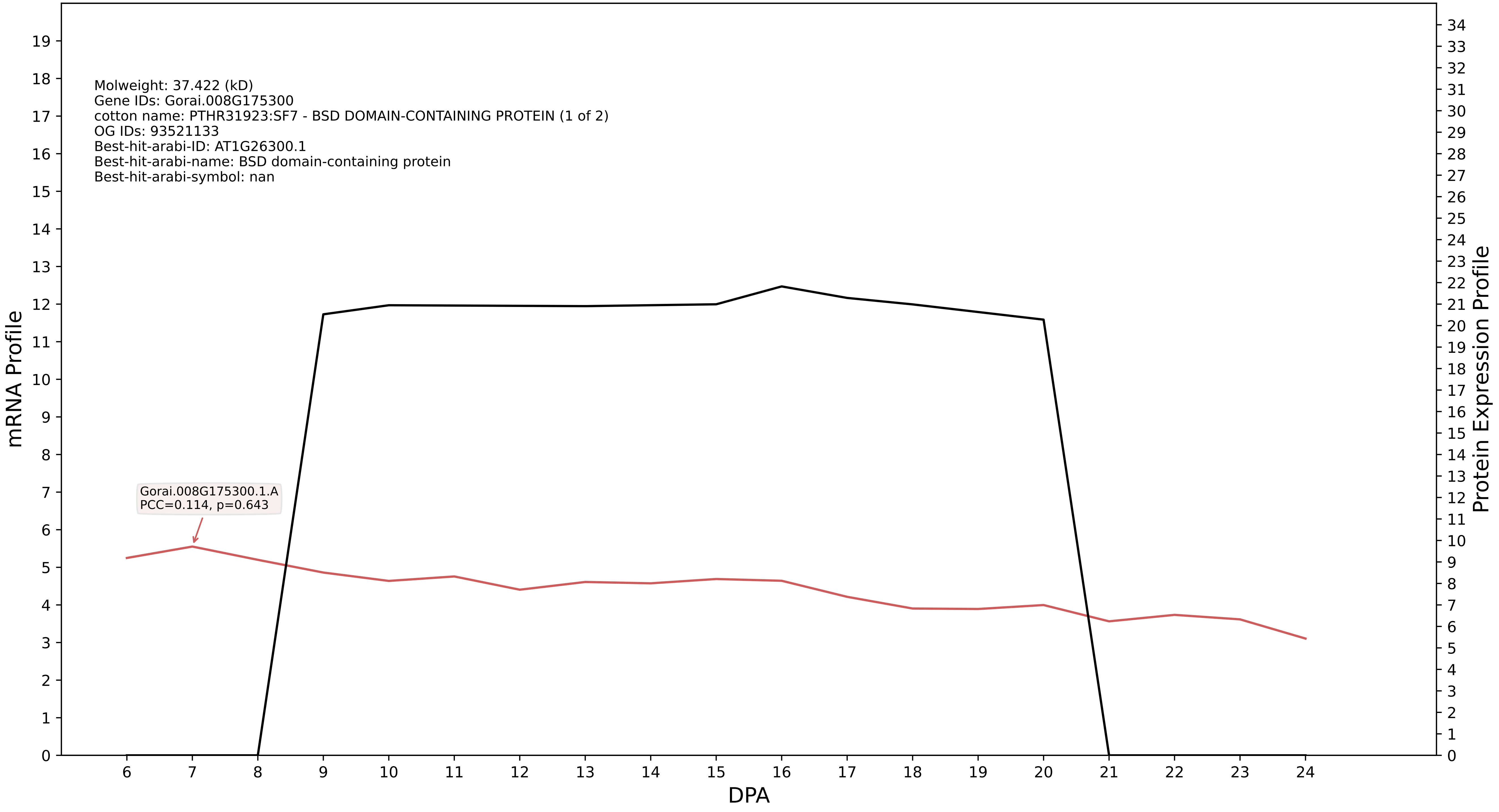

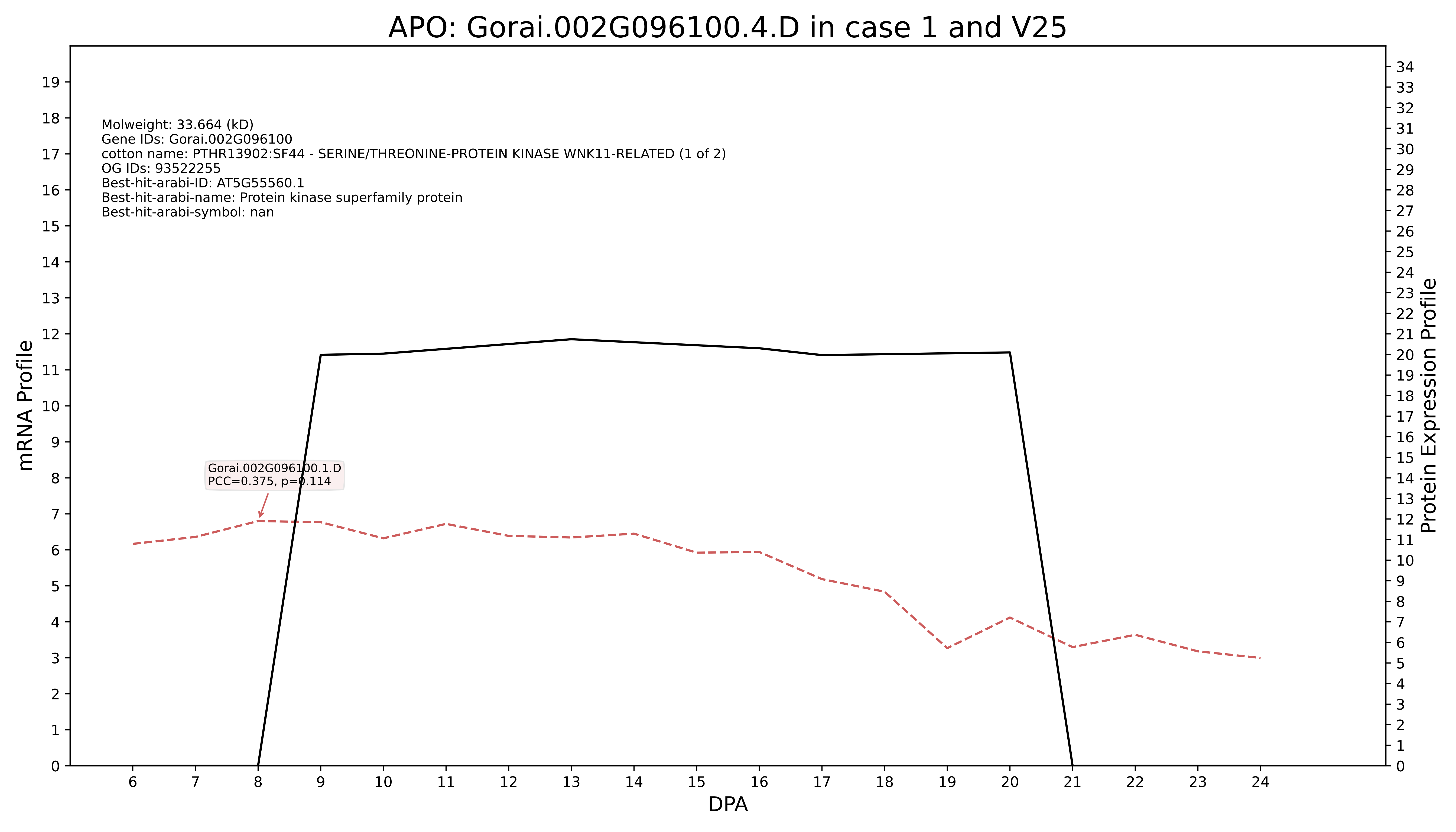

APO: Gorai.002G096100.4.D in case 1 and V25
Molweight: 33.664 (kD)
Gene IDs: Gorai.002G096100
cotton name: PTHR13902:SF44 - SERINE/THREONINE-PROTEIN KINASE WNK11-RELATED (1 of 2)
OG IDs: 93522255
Best-hit-arabi-ID: AT5G55560.1
Best-hit-arabi-name: Protein kinase superfamily protein
Best-hit-arabi-symbol: nan
Gorai.002G096100.1.D
PCC=0.375, p=0.114
mRNA Profile
Protein Expression Profile
DPA

### APO: Gorai.010G186700.4.D in case 1 and V25

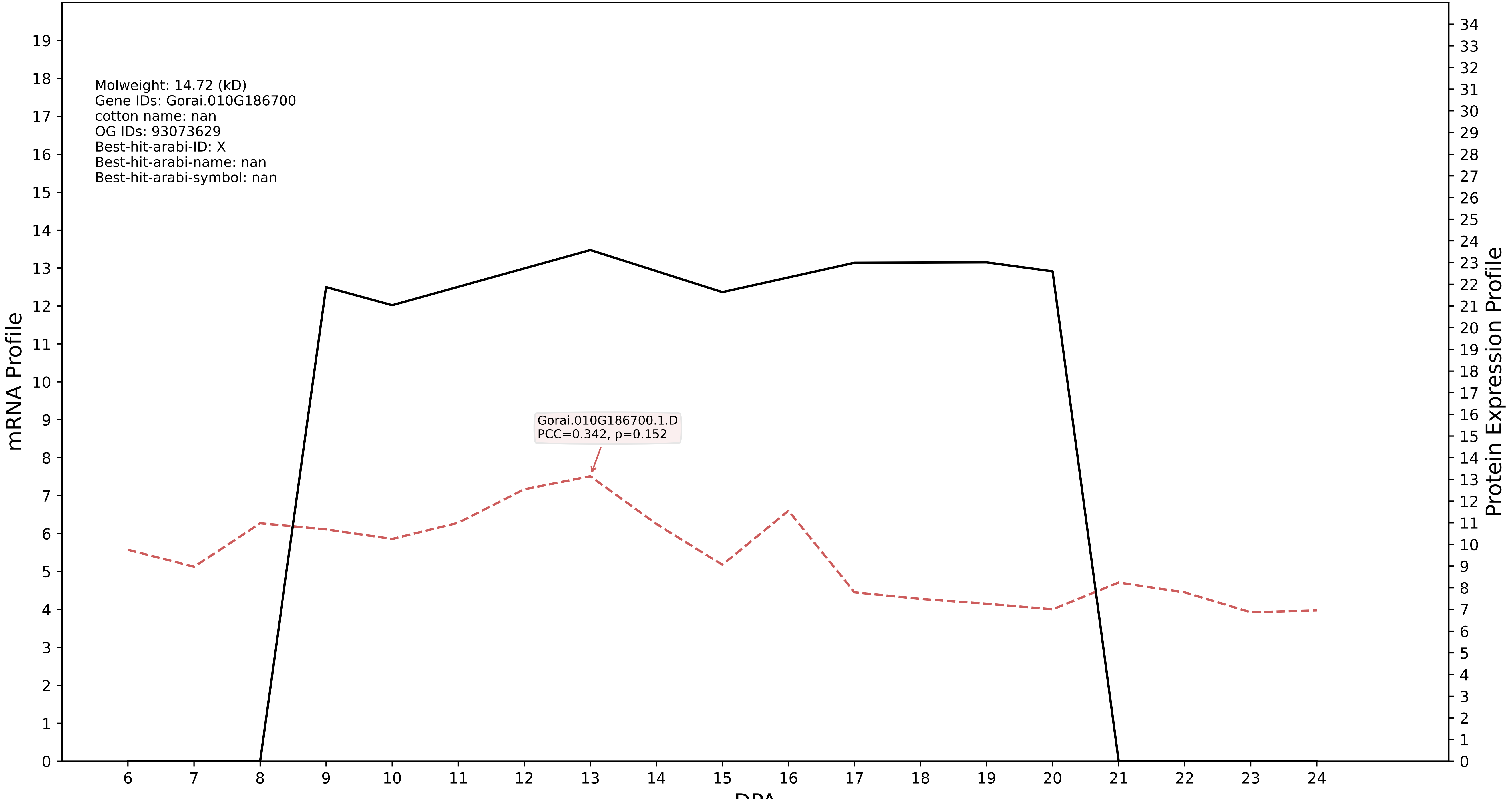

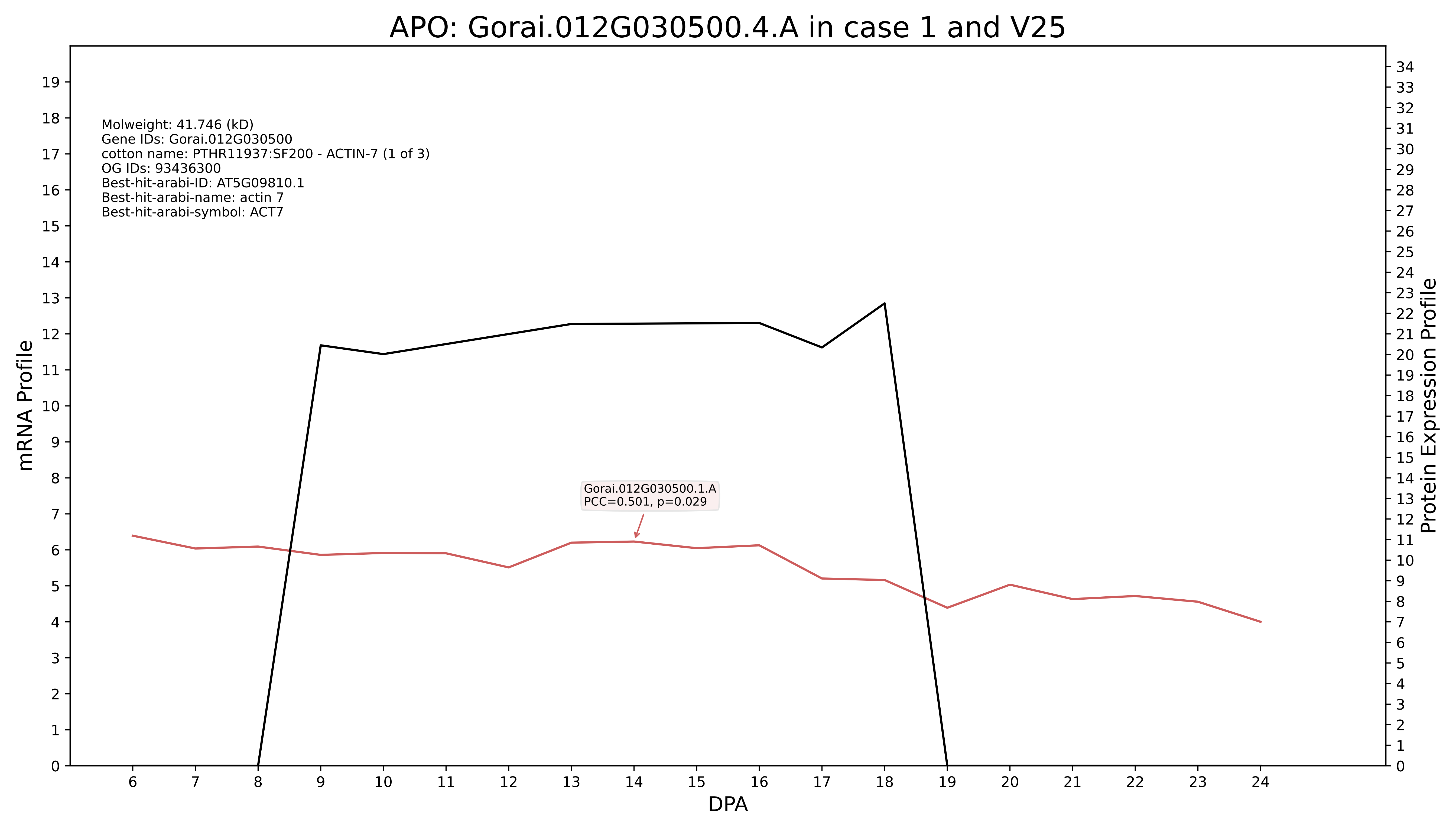
APO: Gorai.012G030500.4.A in case 1 and V25
Molweight: 41.746 (kD)
Gene IDs: Gorai.012G030500
cotton name: PTHR11937:SF200 - ACTIN-7 (1 of 3)
OG IDs: 93436300
Best-hit-arabi-ID: AT5G09810.1
Best-hit-arabi-name: actin 7
Best-hit-arabi-symbol: ACT7
Gorai.012G030500.1.A
PCC=0.501, p=0.029
mRNA Profile
Protein Expression Profile
DPA

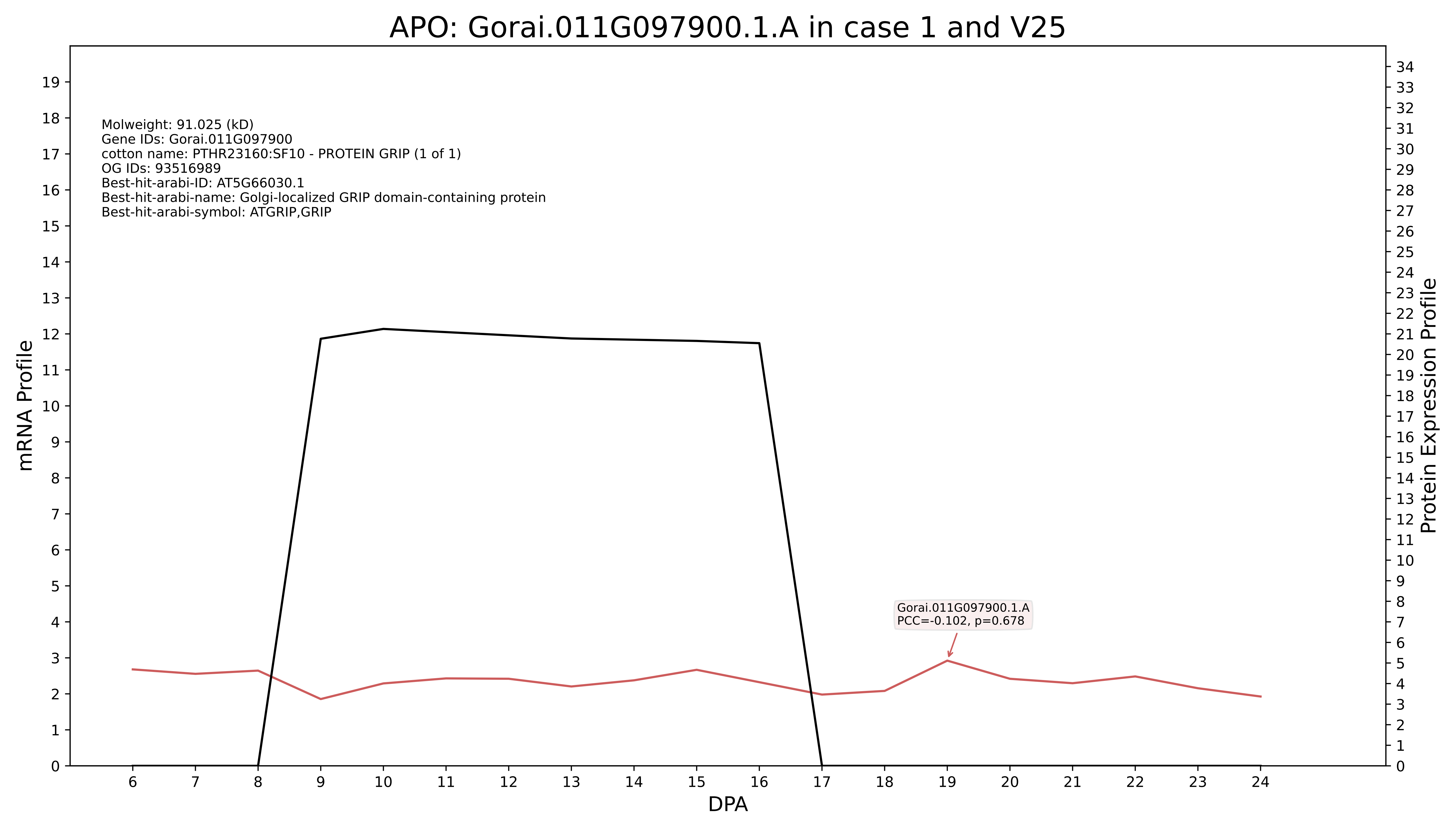

APO: Gorai.011G097900.1.A in case 1 and V25
Molweight: 91.025 (kD)
Gene IDs: Gorai.011G097900
cotton name: PTHR23160:SF10 - PROTEIN GRIP (1 of 1)
OG IDs: 93516989
Best-hit-arabi-ID: AT5G66030.1
Best-hit-arabi-name: Golgi-localized GRIP domain-containing protein
Best-hit-arabi-symbol: ATGRIP,GRIP
Gorai.011G097900.1.A
PCC=-0.102, p=0.678
mRNA Profile
Protein Expression Profile
DPA

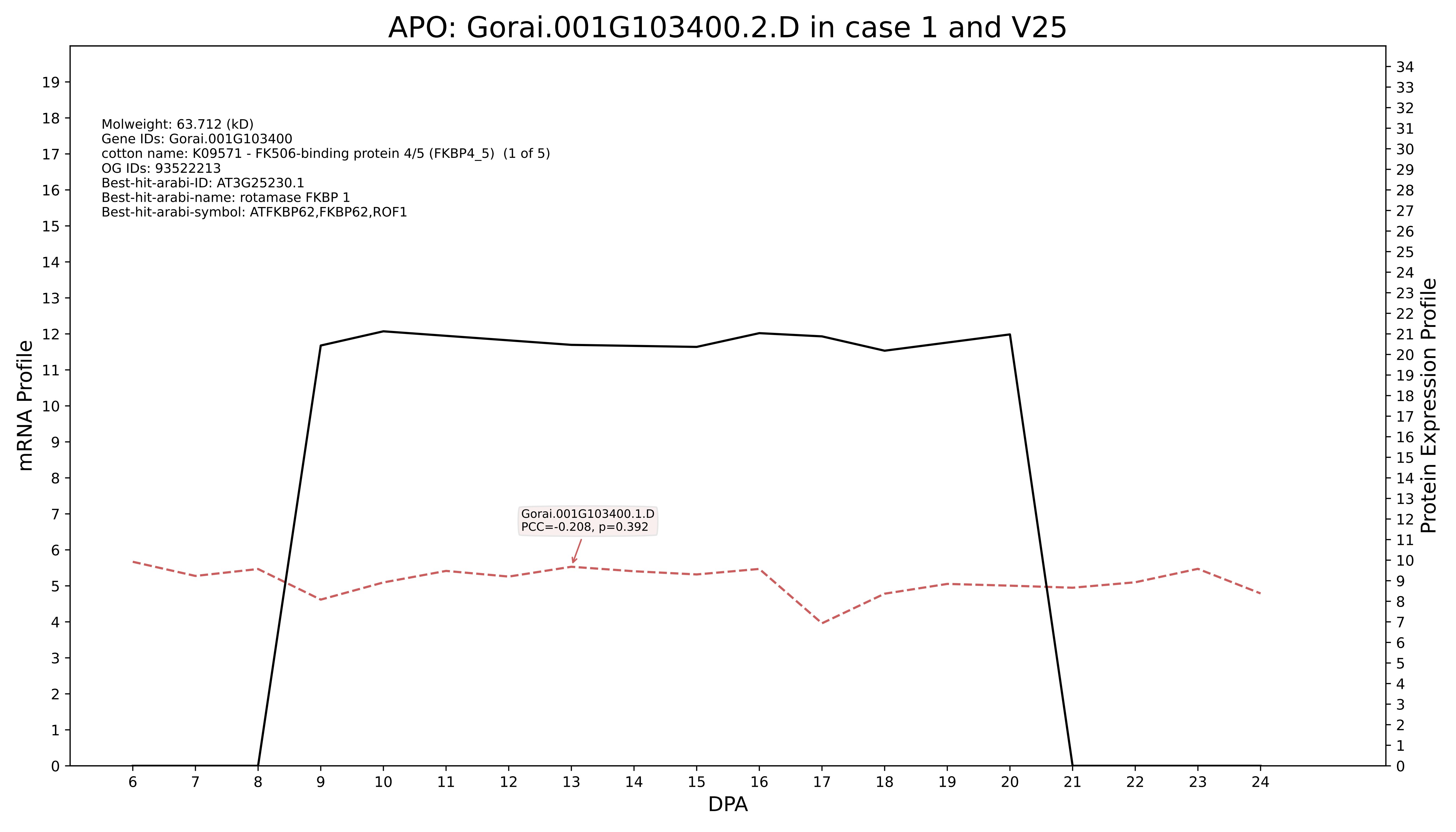
APO: Gorai.001G103400.2.D in case 1 and V25
Molweight: 63.712 (kD)
Gene IDs: Gorai.001G103400
cotton name: K09571 - FK506-binding protein 4/5 (FKBP4_5) (1 of 5)
OG IDs: 93522213
Best-hit-arabi-ID: AT3G25230.1
Best-hit-arabi-name: rotamase FKBP 1
Best-hit-arabi-symbol: ATFKBP62,FKBP62,ROF1
Gorai.001G103400.1.D
PCC=-0.208, p=0.392
mRNA Profile
Protein Expression Profile
DPA

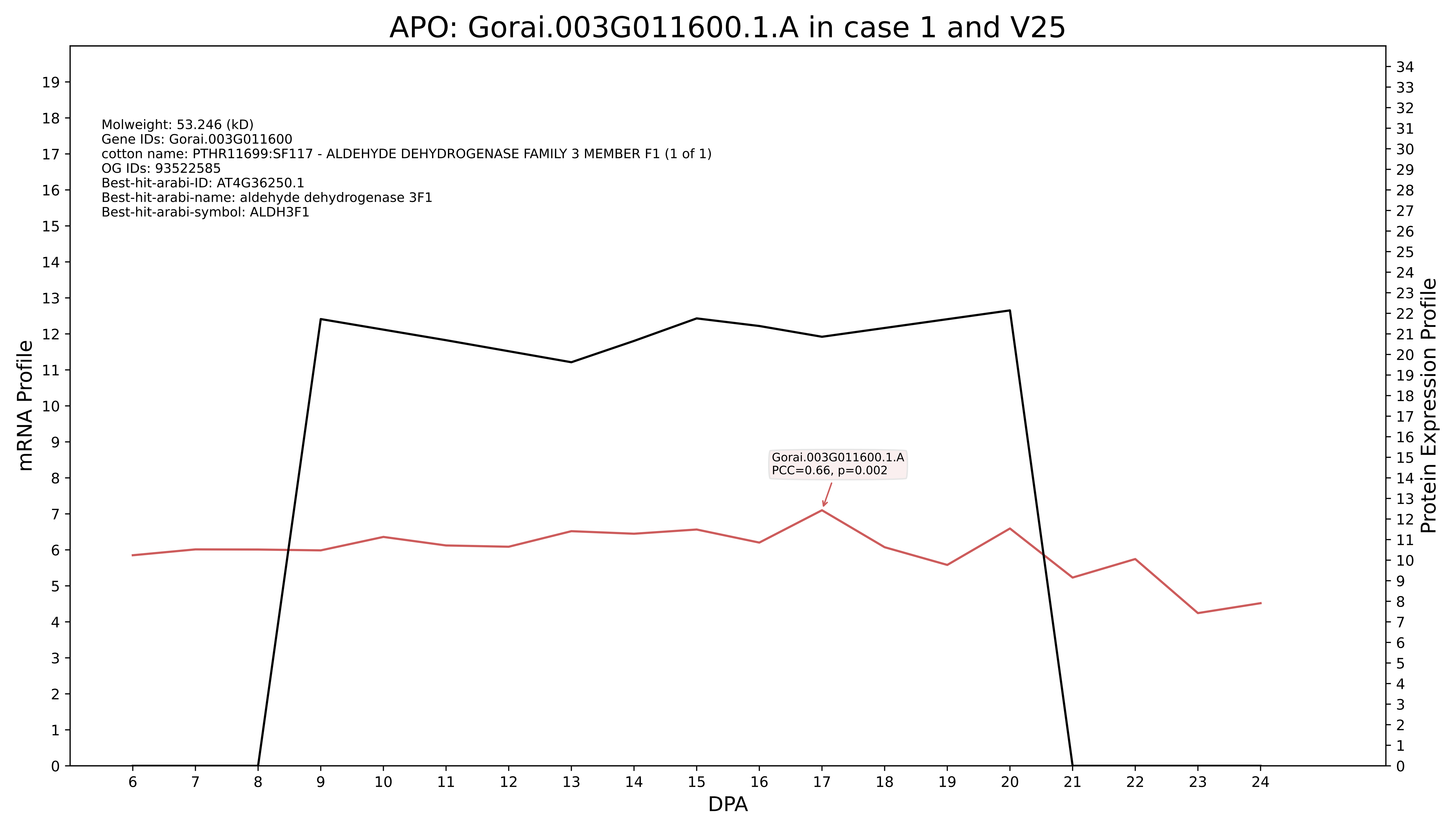

APO: Gorai.003G011600.1.A in case 1 and V25
Molweight: 53.246 (kD)
Gene IDs: Gorai.003G011600
cotton name: PTHR11699:SF117 - ALDEHYDE DEHYDROGENASE FAMILY 3 MEMBER F1 (1 of 1)
OG IDs: 93522585
Best-hit-arabi-ID: AT4G36250.1
Best-hit-arabi-name: aldehyde dehydrogenase 3F1
Best-hit-arabi-symbol: ALDH3F1
Gorai.003G011600.1.A
PCC=0.66, p=0.002
mRNA Profile
Protein Expression Profile
DPA

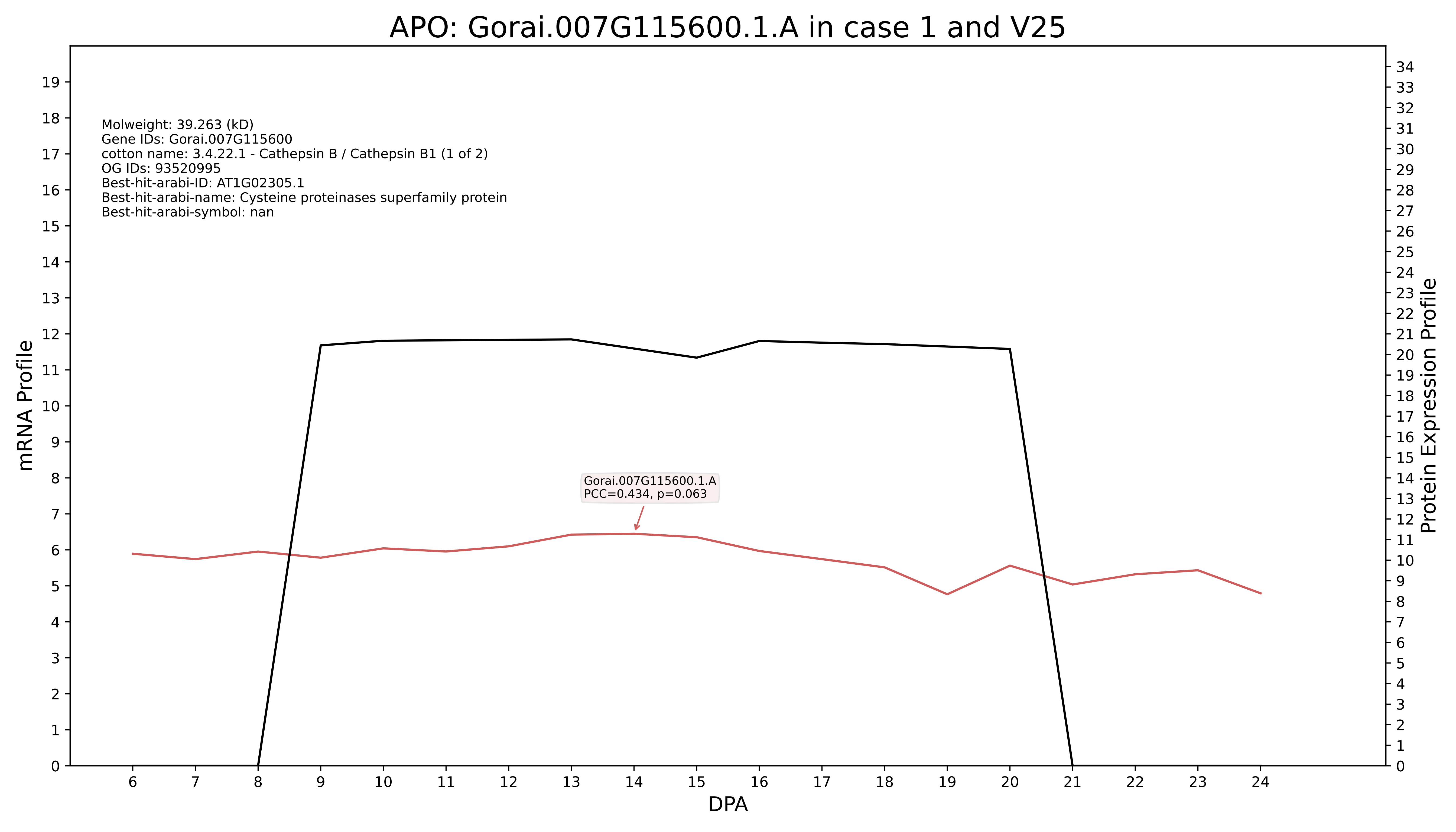
APO: Gorai.007G115600.1.A in case 1 and V25
Molweight: 39.263 (kD)
Gene IDs: Gorai.007G115600
cotton name: 3.4.22.1 - Cathepsin B / Cathepsin B1 (1 of 2)
OG IDs: 93520995
Best-hit-arabi-ID: AT1G02305.1
Best-hit-arabi-name: Cysteine proteinases superfamily protein
Best-hit-arabi-symbol: nan
Gorai.007G115600.1.A
PCC=0.434, p=0.063
mRNA Profile
Protein Expression Profile
DPA

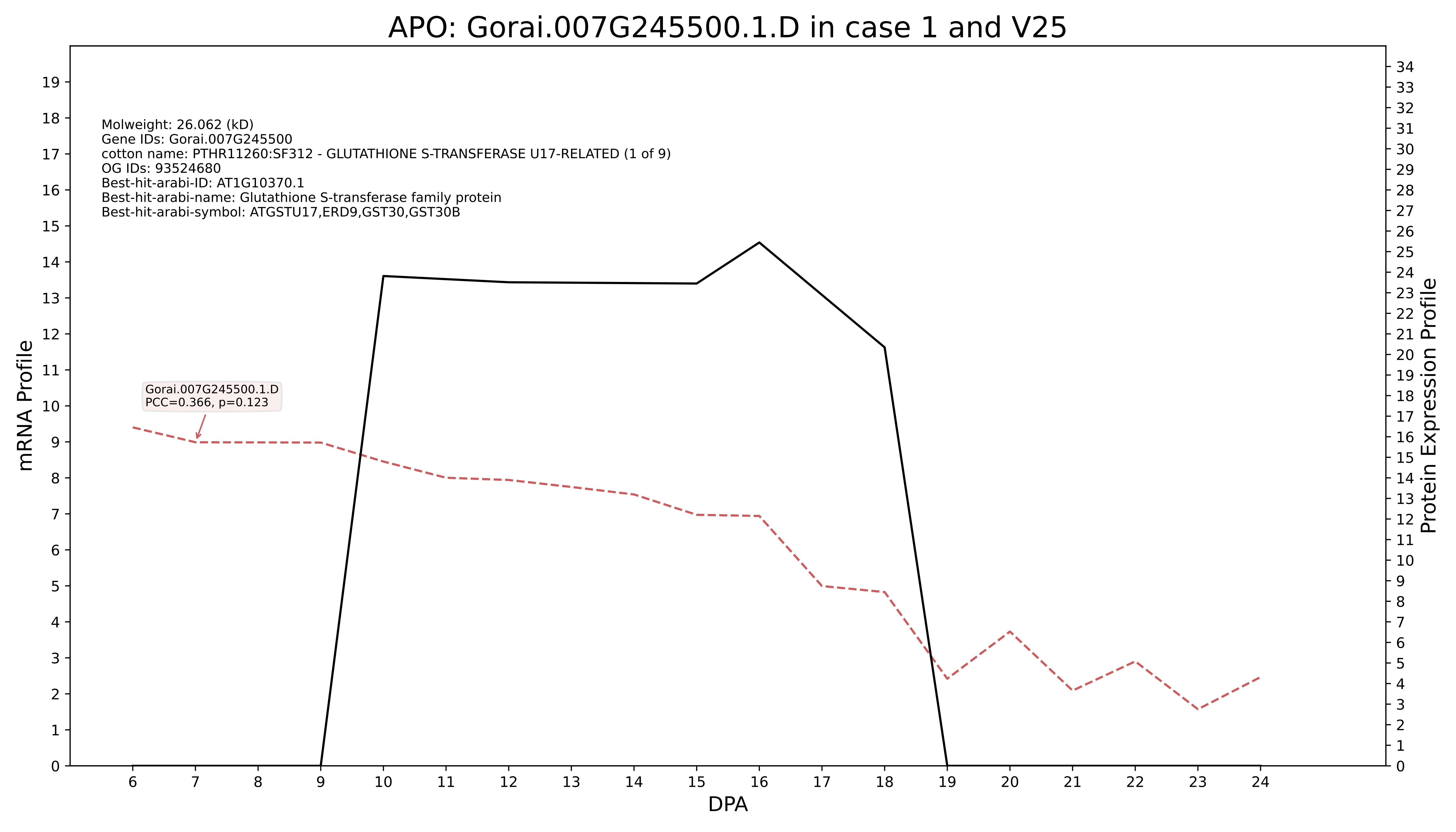
APO: Gorai.007G245500.1.D in case 1 and V25
Molweight: 26.062 (kD)
Gene IDs: Gorai.007G245500
cotton name: PTHR11260:SF312 - GLUTATHIONE S-TRANSFERASE U17-RELATED (1 of 9)
OG IDs: 93524680
Best-hit-arabi-ID: AT1G10370.1
Best-hit-arabi-name: Glutathione S-transferase family protein
Best-hit-arabi-symbol: ATGSTU17,ERD9,GST30,GST30B
Gorai.007G245500.1.D
PCC=0.366, p=0.123
mRNA Profile
Protein Expression Profile
DPA

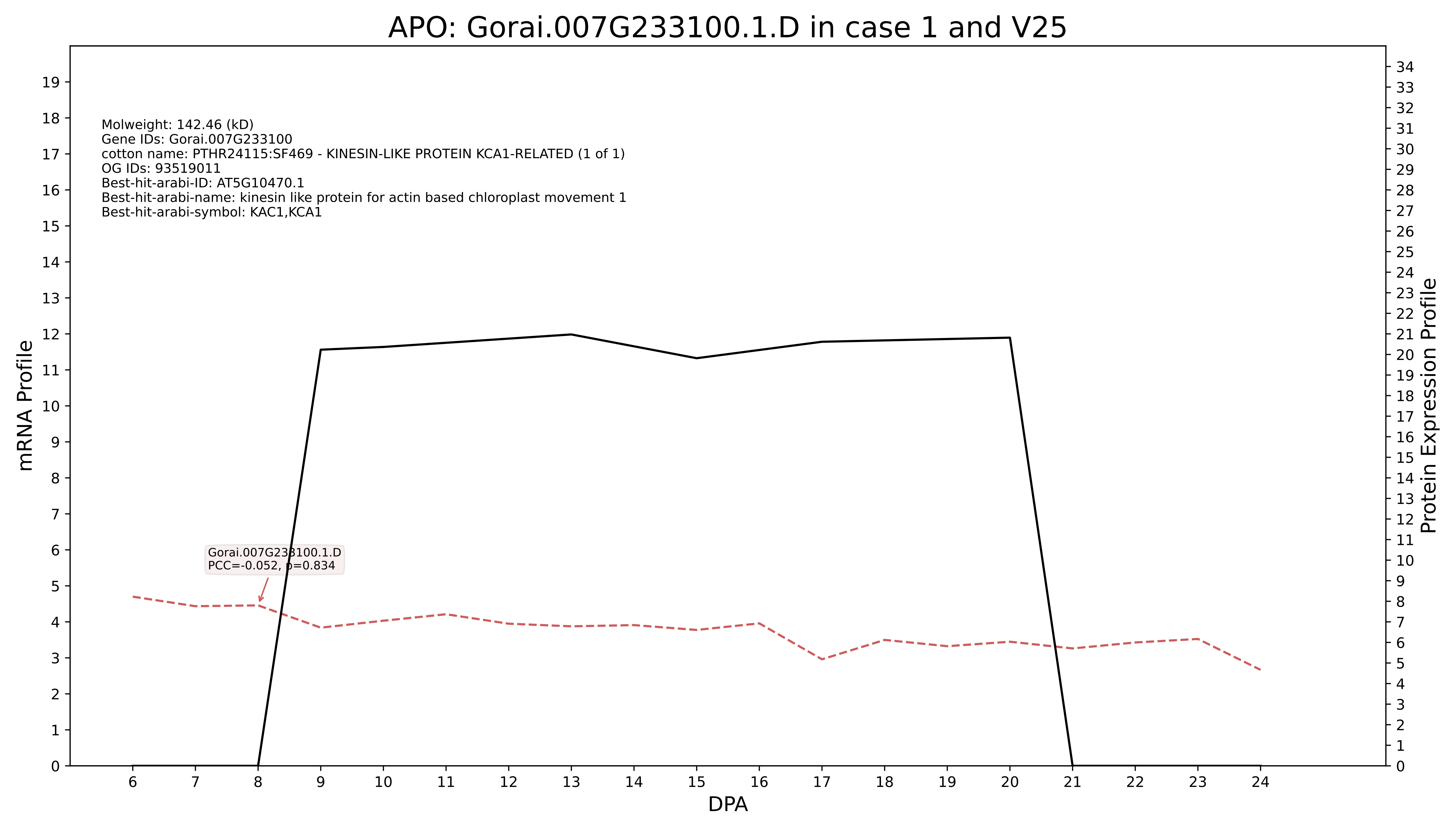
APO: Gorai.007G233100.1.D in case 1 and V25
Molweight: 142.46 (kD)
Gene IDs: Gorai.007G233100
cotton name: PTHR24115:SF469 - KINESIN-LIKE PROTEIN KCA1-RELATED (1 of 1)
OG IDs: 93519011
Best-hit-arabi-ID: AT5G10470.1
Best-hit-arabi-name: kinesin like protein for actin based chloroplast movement 1
Best-hit-arabi-symbol: KAC1,KCA1
Gorai.007G233100.1.D
PCC=-0.052, p=0.834
mRNA Profile
Protein Expression Profile
DPA

### APO: Gorai.001G230800.1.D in case 1 and V25

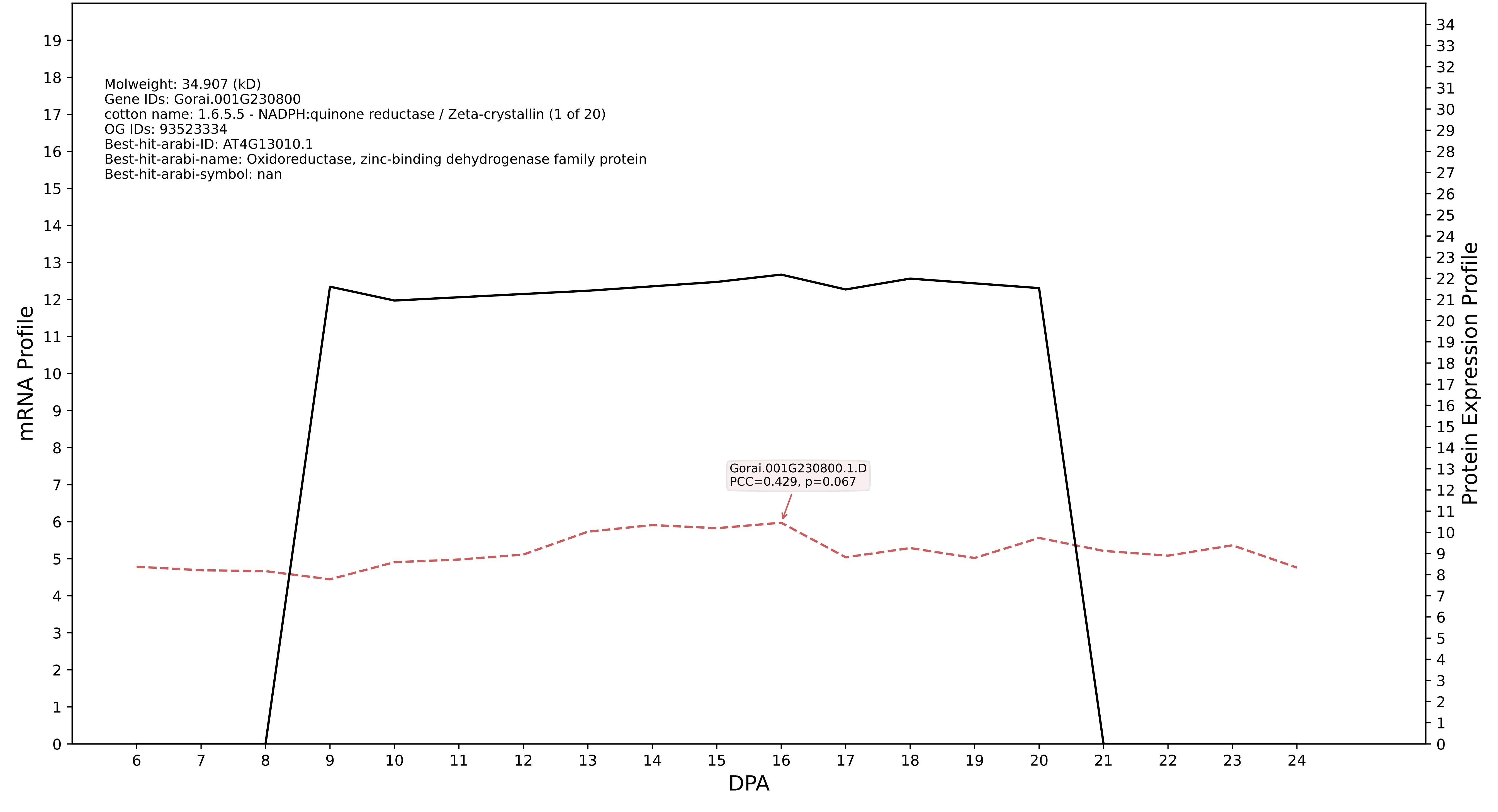

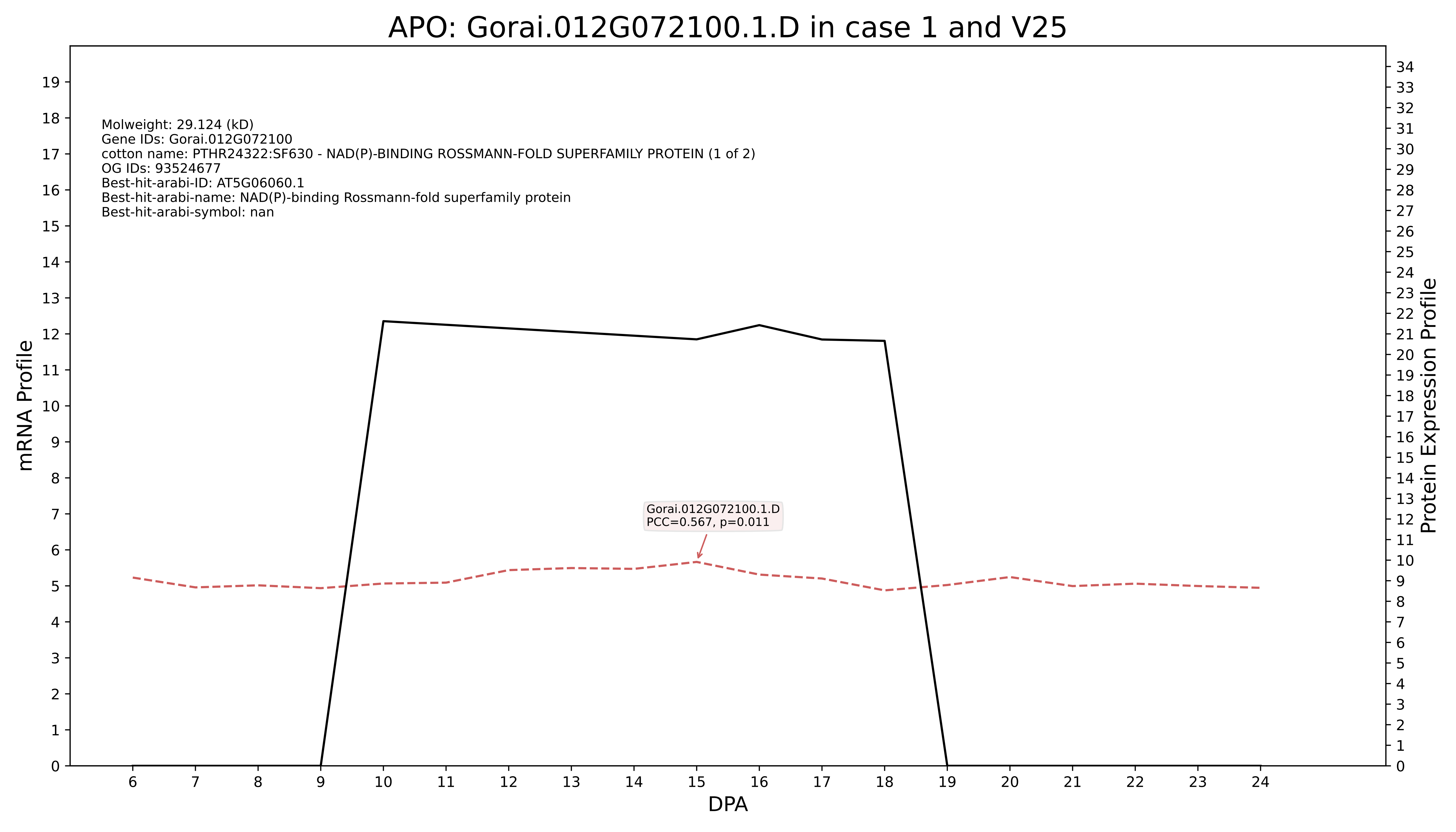
APO: Gorai.012G072100.1.D in case 1 and V25
Molweight: 29.124 (kD)
Gene IDs: Gorai.012G072100
cotton name: PTHR24322:SF630 - NAD(P)-BINDING ROSSMANN-FOLD SUPERFAMILY PROTEIN (1 of 2)
OG IDs: 93524677
Best-hit-arabi-ID: AT5G06060.1
Best-hit-arabi-name: NAD(P)-binding Rossmann-fold superfamily protein
Best-hit-arabi-symbol: nan
Gorai.012G072100.1.D
PCC=0.567, p=0.011
mRNA Profile
Protein Expression Profile
DPA

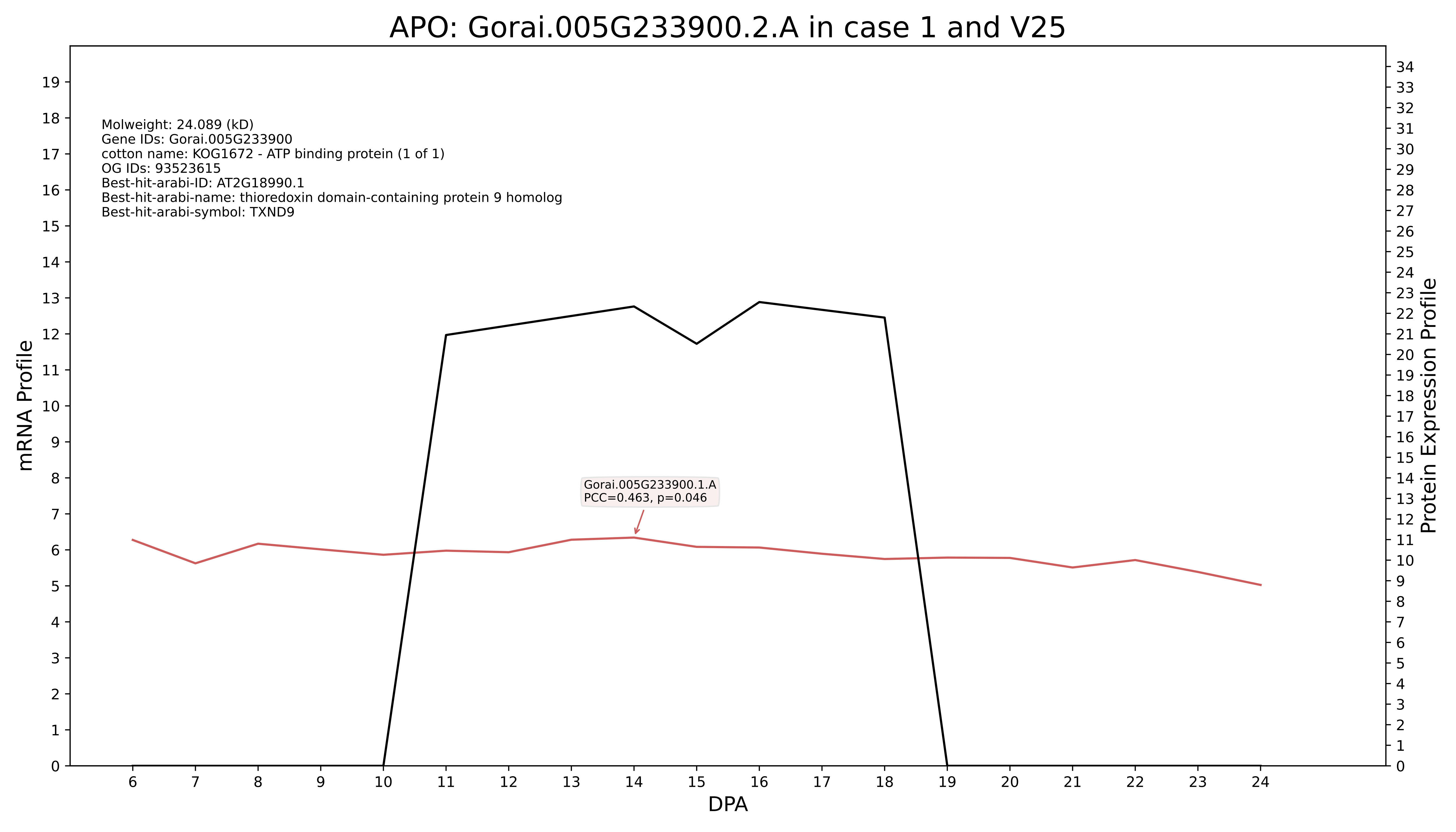
APO: Gorai.005G233900.2.A in case 1 and V25
Molweight: 24.089 (kD)
Gene IDs: Gorai.005G233900
cotton name: KOG1672 - ATP binding protein (1 of 1)
OG IDs: 93523615
Best-hit-arabi-ID: AT2G18990.1
Best-hit-arabi-name: thioredoxin domain-containing protein 9 homolog
Best-hit-arabi-symbol: TXND9
Gorai.005G233900.1.A
PCC=0.463, p=0.046
mRNA Profile
Protein Expression Profile
DPA

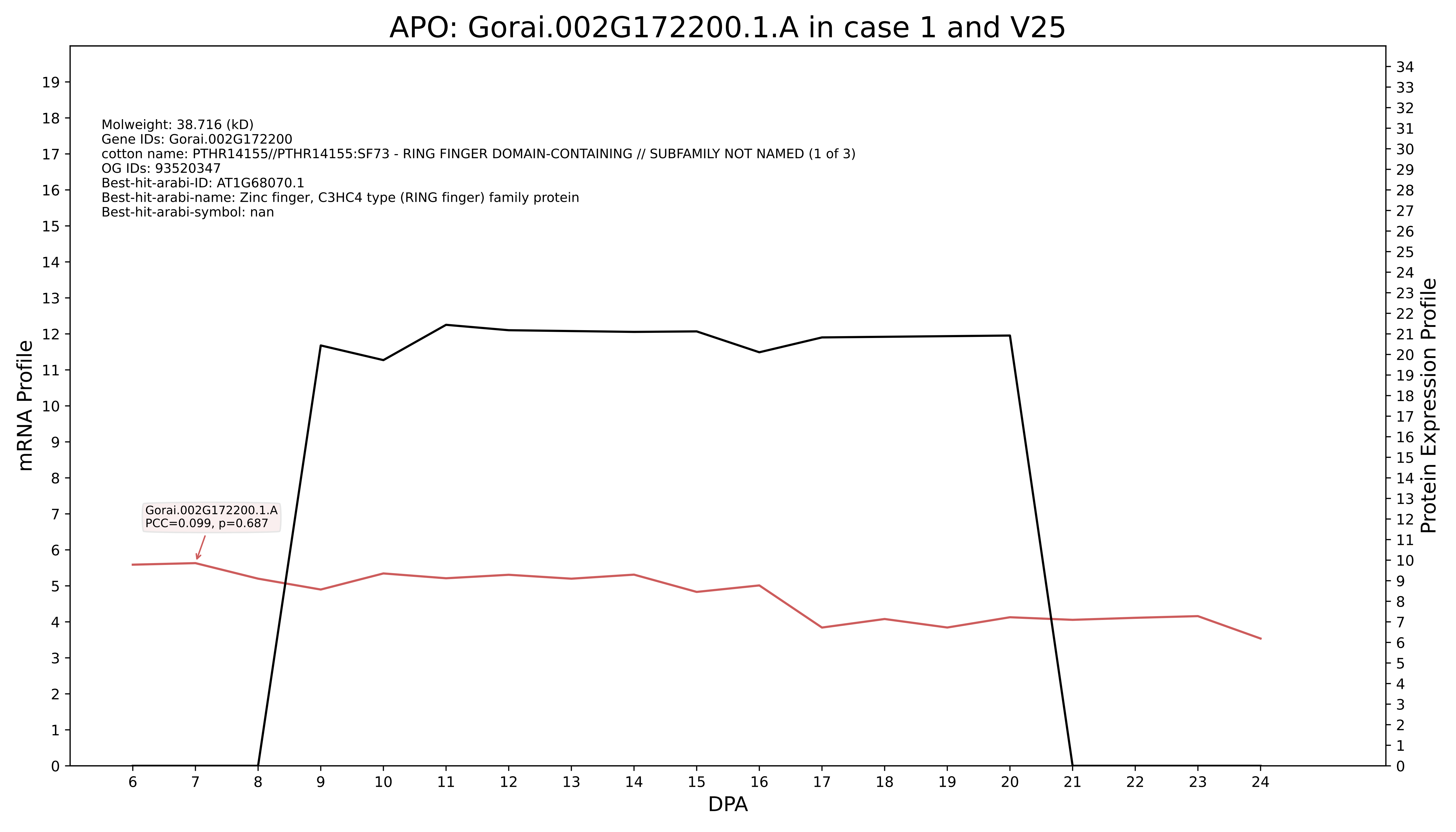
APO: Gorai.002G172200.1.A in case 1 and V25
Molweight: 38.716 (kD)
Gene IDs: Gorai.002G172200
cotton name: PTHR14155//PTHR14155:SF73 - RING FINGER DOMAIN-CONTAINING // SUBFAMILY NOT NAMED (1 of 3)
OG IDs: 93520347
Best-hit-arabi-ID: AT1G68070.1
Best-hit-arabi-name: Zinc finger, C3HC4 type (RING finger) family protein
Best-hit-arabi-symbol: nan
Gorai.002G172200.1.A
PCC=0.099, p=0.687
mRNA Profile
Protein Expression Profile
DPA

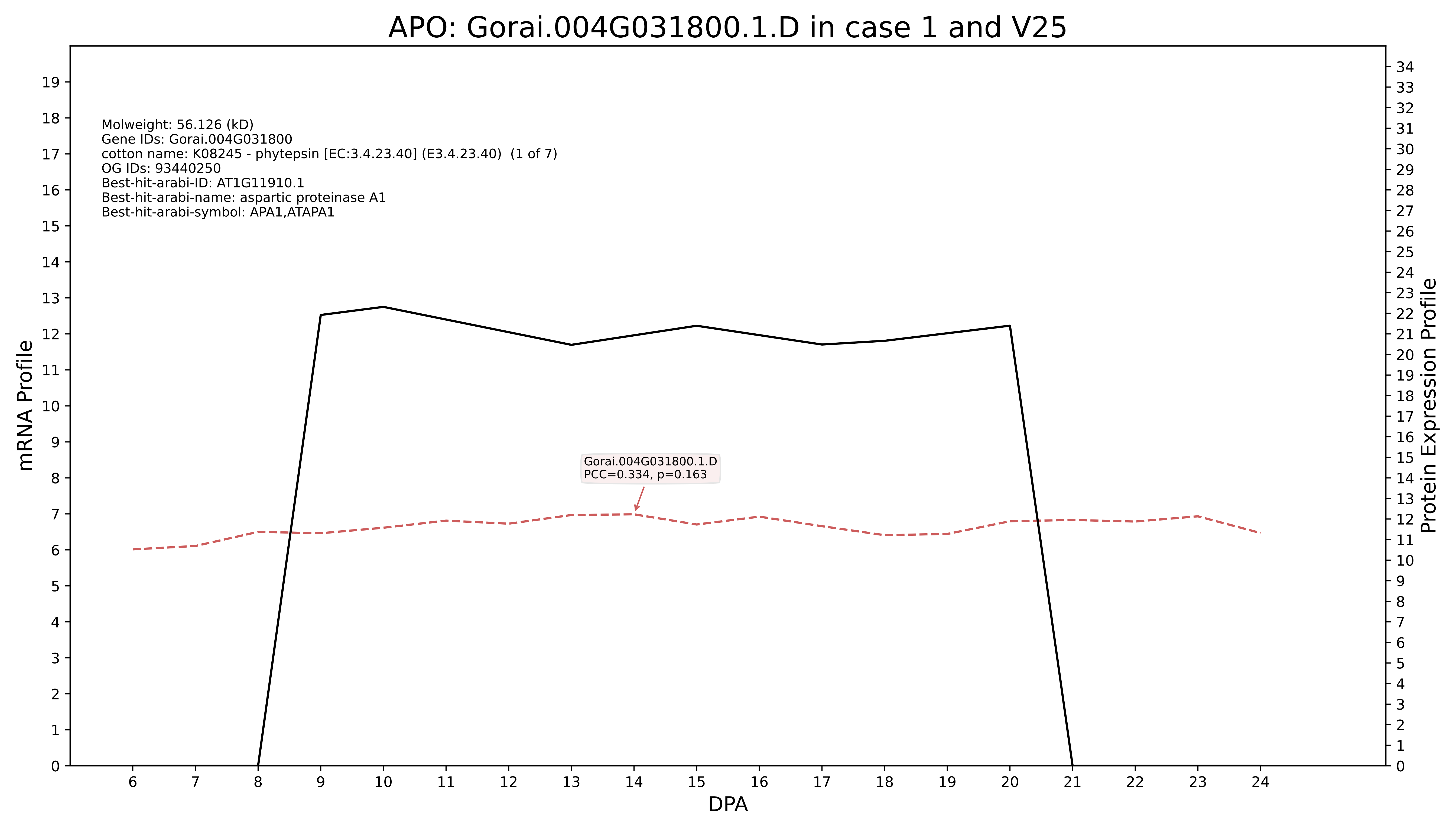

APO: Gorai.004G031800.1.D in case 1 and V25
Molweight: 56.126 (kD)
Gene IDs: Gorai.004G031800
cotton name: K08245 - phytepsin [EC:3.4.23.40] (E3.4.23.40) (1 of 7)
OG IDs: 93440250
Best-hit-arabi-ID: AT1G11910.1
Best-hit-arabi-name: aspartic proteinase A1
Best-hit-arabi-symbol: APA1,ATAPA1
Gorai.004G031800.1.D
PCC=0.334, p=0.163
mRNA Profile
Protein Expression Profile
DPA

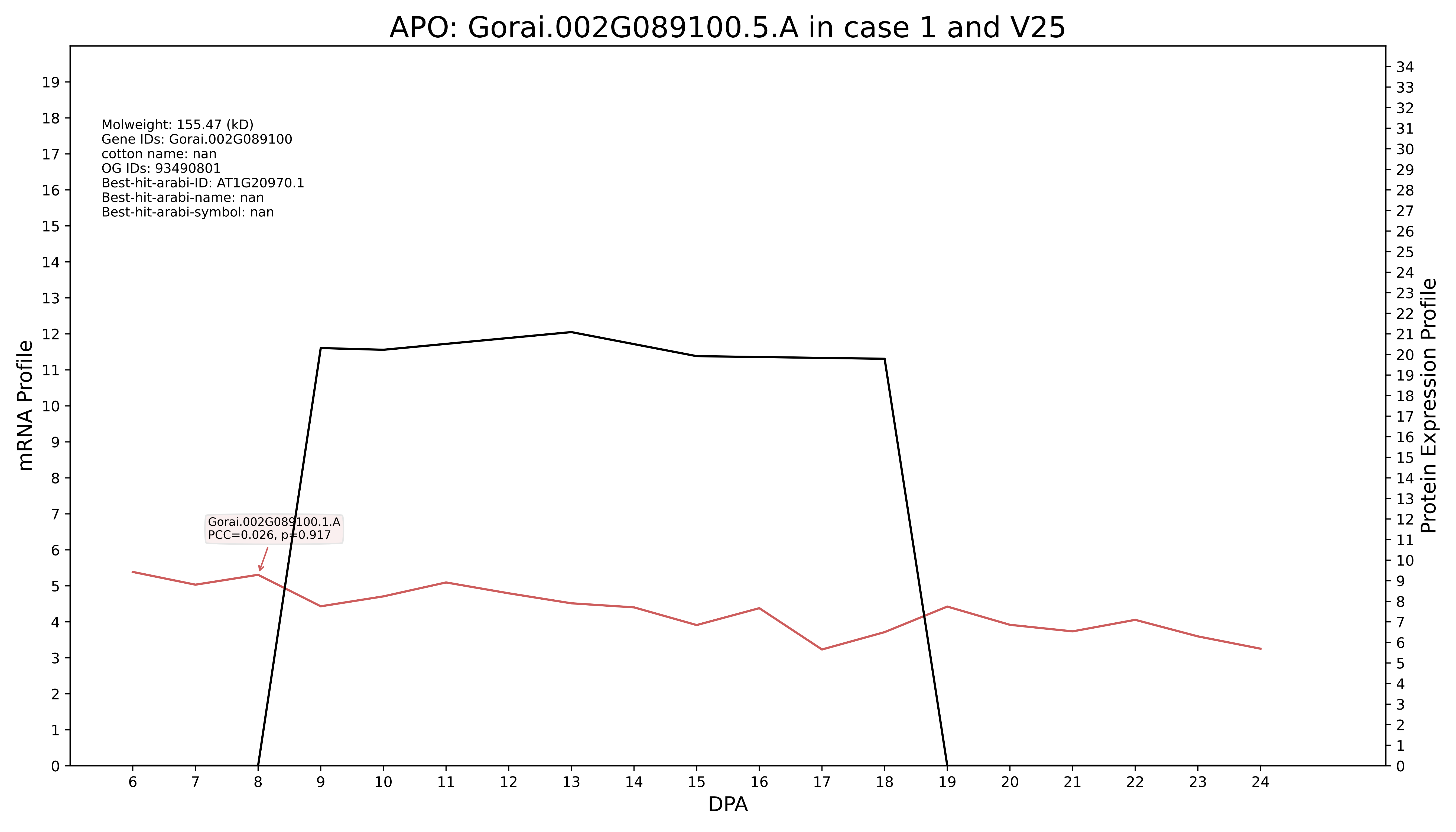
APO: Gorai.002G089100.5.A in case 1 and V25
Molweight: 155.47 (kD)
Gene IDs: Gorai.002G089100
cotton name: nan
OG IDs: 93490801
Best-hit-arabi-ID: AT1G20970.1
Best-hit-arabi-name: nan
Best-hit-arabi-symbol: nan
Gorai.002G089100.1.A
PCC=0.026, p=0.917
mRNA Profile
Protein Expression Profile
DPA

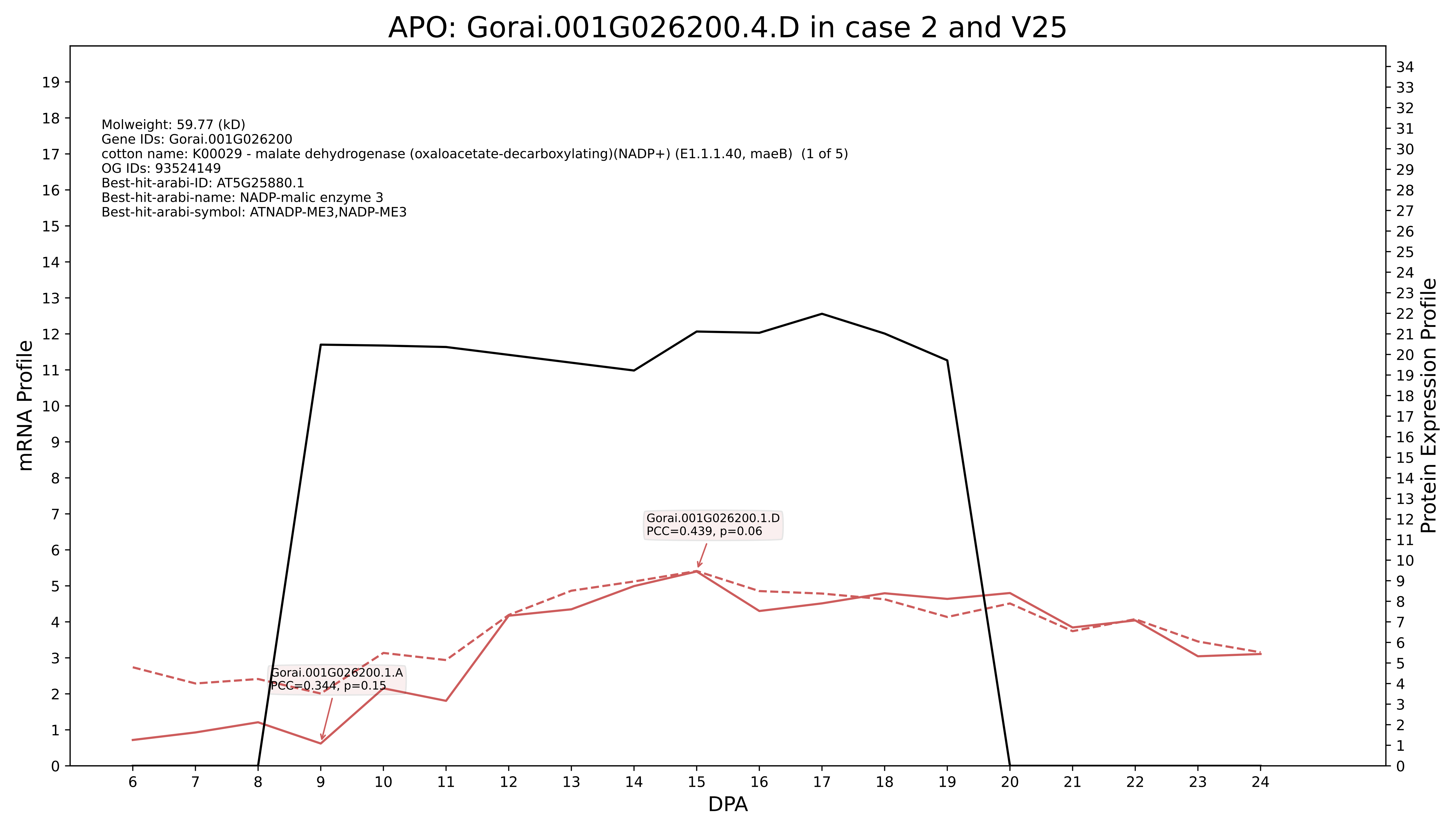
APO: Gorai.001G026200.4.D in case 2 and V25
Molweight: 59.77 (kD)
Gene IDs: Gorai.001G026200
cotton name: K00029 - malate dehydrogenase (oxaloacetate-decarboxylating)(NADP+) (E1.1.1.40, maeB) (1 of 5)
OG IDs: 93524149
Best-hit-arabi-ID: AT5G25880.1
Best-hit-arabi-name: NADP-malic enzyme 3
Best-hit-arabi-symbol: ATNADP-ME3,NADP-ME3
Gorai.001G026200.1.D
PCC=0.439, p=0.06
Gorai.001G026200.1.A
PCC=0.344, p=0.15
mRNA Profile
Protein Expression Profile
DPA

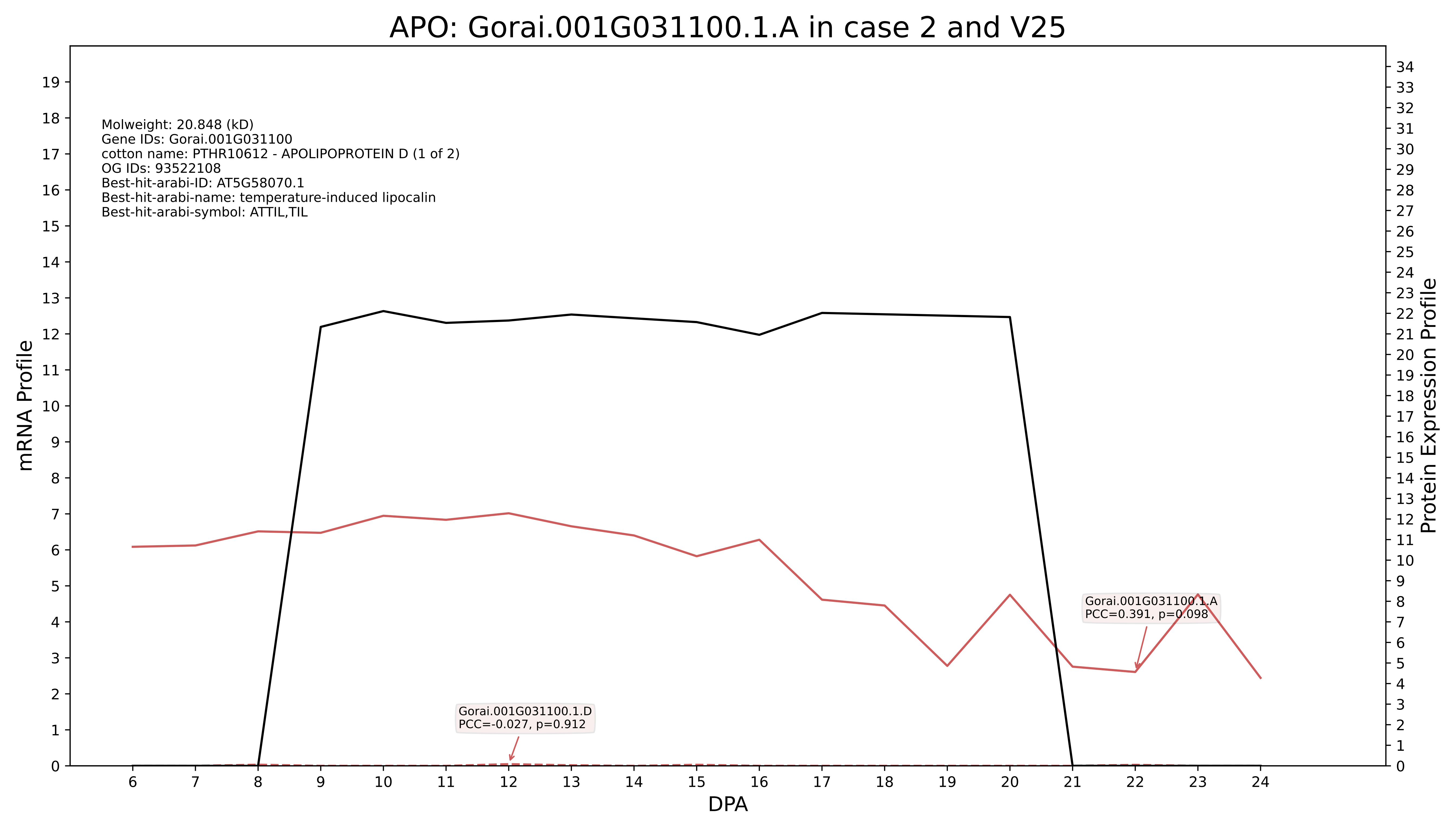

APO: Gorai.001G031100.1.A in case 2 and V25
Molweight: 20.848 (kD)
Gene IDs: Gorai.001G031100
cotton name: PTHR10612 - APOLIPOPROTEIN D (1 of 2)
OG IDs: 93522108
Best-hit-arabi-ID: AT5G58070.1
Best-hit-arabi-name: temperature-induced lipocalin
Best-hit-arabi-symbol: ATTIL,TIL
Gorai.001G031100.1.A
PCC=0.391, p=0.098
Gorai.001G031100.1.D
PCC=-0.027, p=0.912
mRNA Profile
Protein Expression Profile
DPA

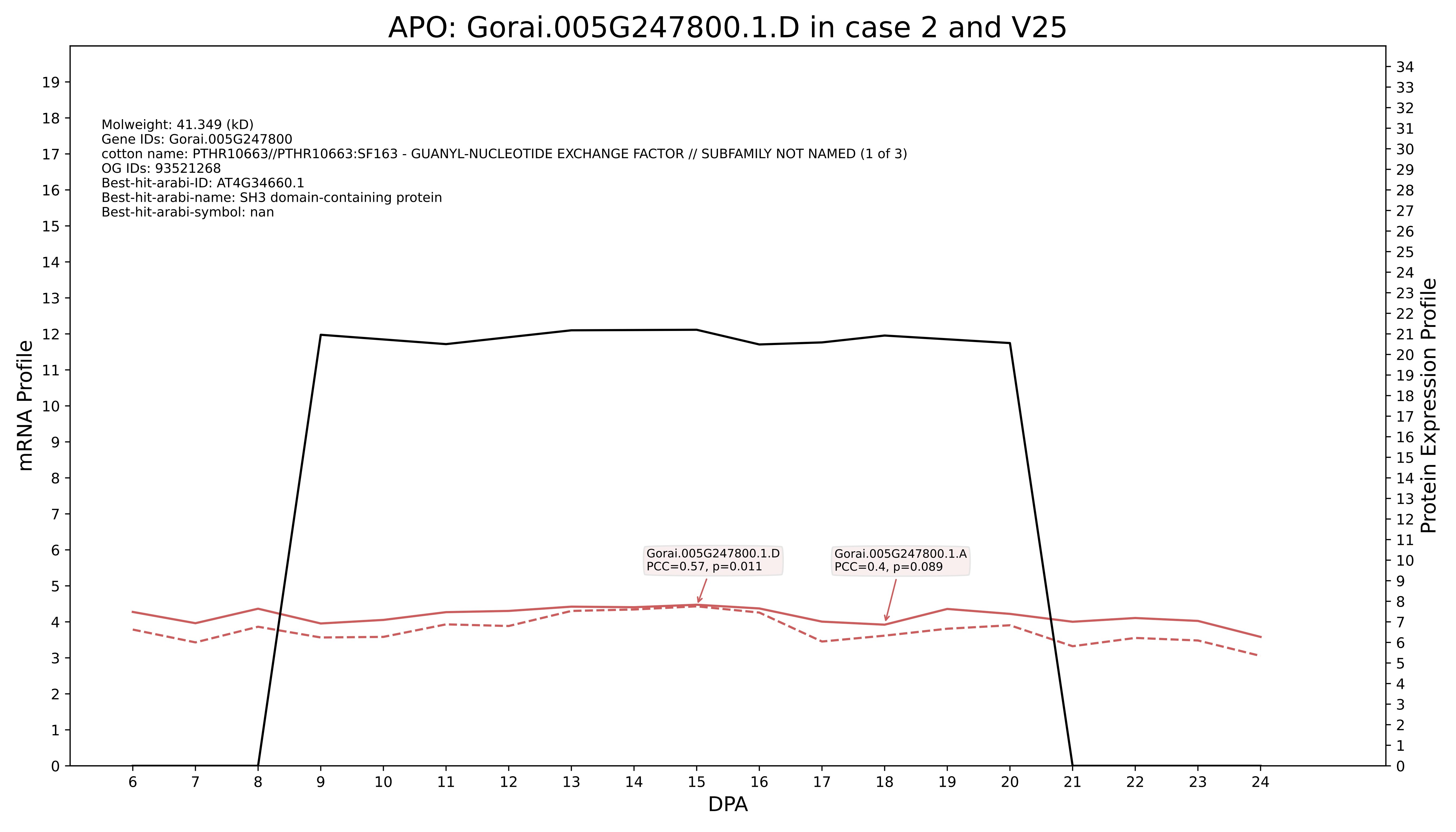

APO: Gorai.005G247800.1.D in case 2 and V25
Molweight: 41.349 (kD)
Gene IDs: Gorai.005G247800
cotton name: PTHR10663//PTHR10663:SF163 - GUANYL-NUCLEOTIDE EXCHANGE FACTOR // SUBFAMILY NOT NAMED (1 of 3)
OG IDs: 93521268
Best-hit-arabi-ID: AT4G34660.1
Best-hit-arabi-name: SH3 domain-containing protein
Best-hit-arabi-symbol: nan
Gorai.005G247800.1.D
PCC=0.57, p=0.011
Gorai.005G247800.1.A
PCC=0.4, p=0.089
mRNA Profile
Protein Expression Profile
DPA

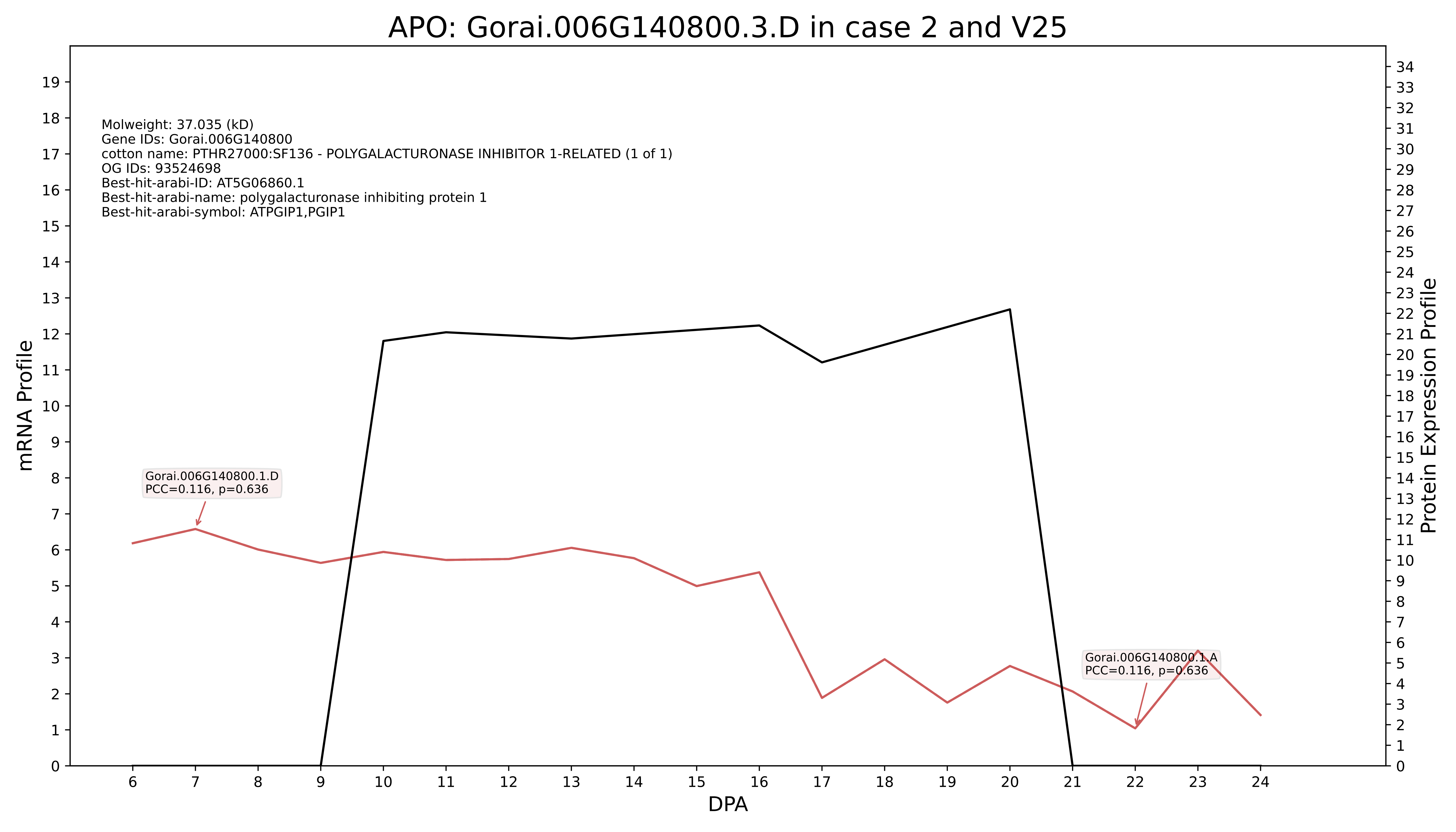

APO: Gorai.006G140800.3.D in case 2 and V25
Molweight: 37.035 (kD)
Gene IDs: Gorai.006G140800
cotton name: PTHR27000:SF136 - POLYGALACTURONASE INHIBITOR 1-RELATED (1 of 1)
OG IDs: 93524698
Best-hit-arabi-ID: AT5G06860.1
Best-hit-arabi-name: polygalacturonase inhibiting protein 1
Best-hit-arabi-symbol: ATPGIP1,PGIP1
Gorai.006G140800.1.D
PCC=0.116, p=0.636
Gorai.006G140800.1.A
PCC=0.116, p=0.636
mRNA Profile
Protein Expression Profile
DPA

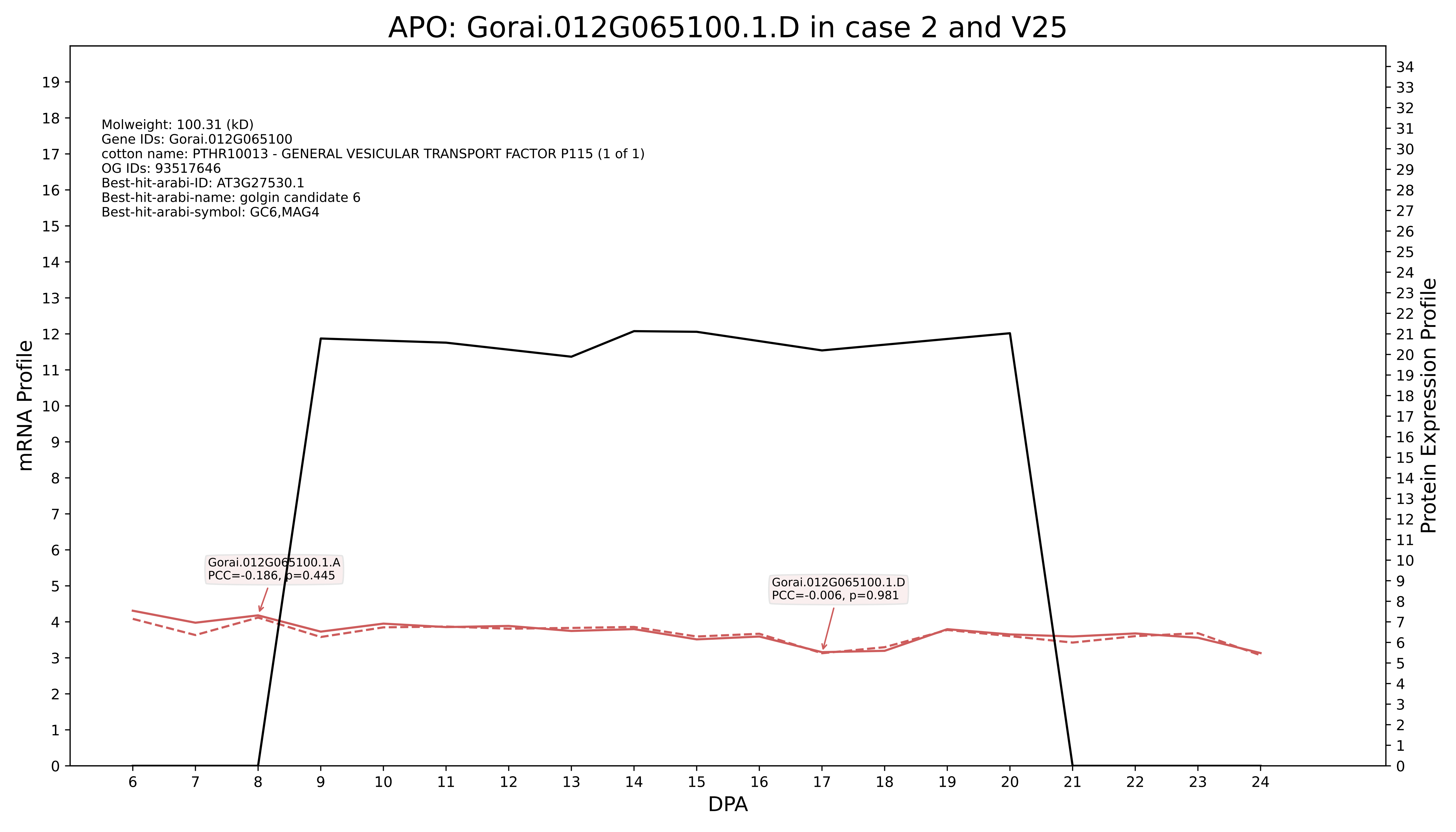

APO: Gorai.012G065100.1.D in case 2 and V25
Molweight: 100.31 (kD)
Gene IDs: Gorai.012G065100
cotton name: PTHR10013 - GENERAL VESICULAR TRANSPORT FACTOR P115 (1 of 1)
OG IDs: 93517646
Best-hit-arabi-ID: AT3G27530.1
Best-hit-arabi-name: golgin candidate 6
Best-hit-arabi-symbol: GC6,MAG4
Gorai.012G065100.1.A
PCC=-0.186, p=0.445
Gorai.012G065100.1.D
PCC=-0.006, p=0.981
mRNA Profile
Protein Expression Profile
DPA

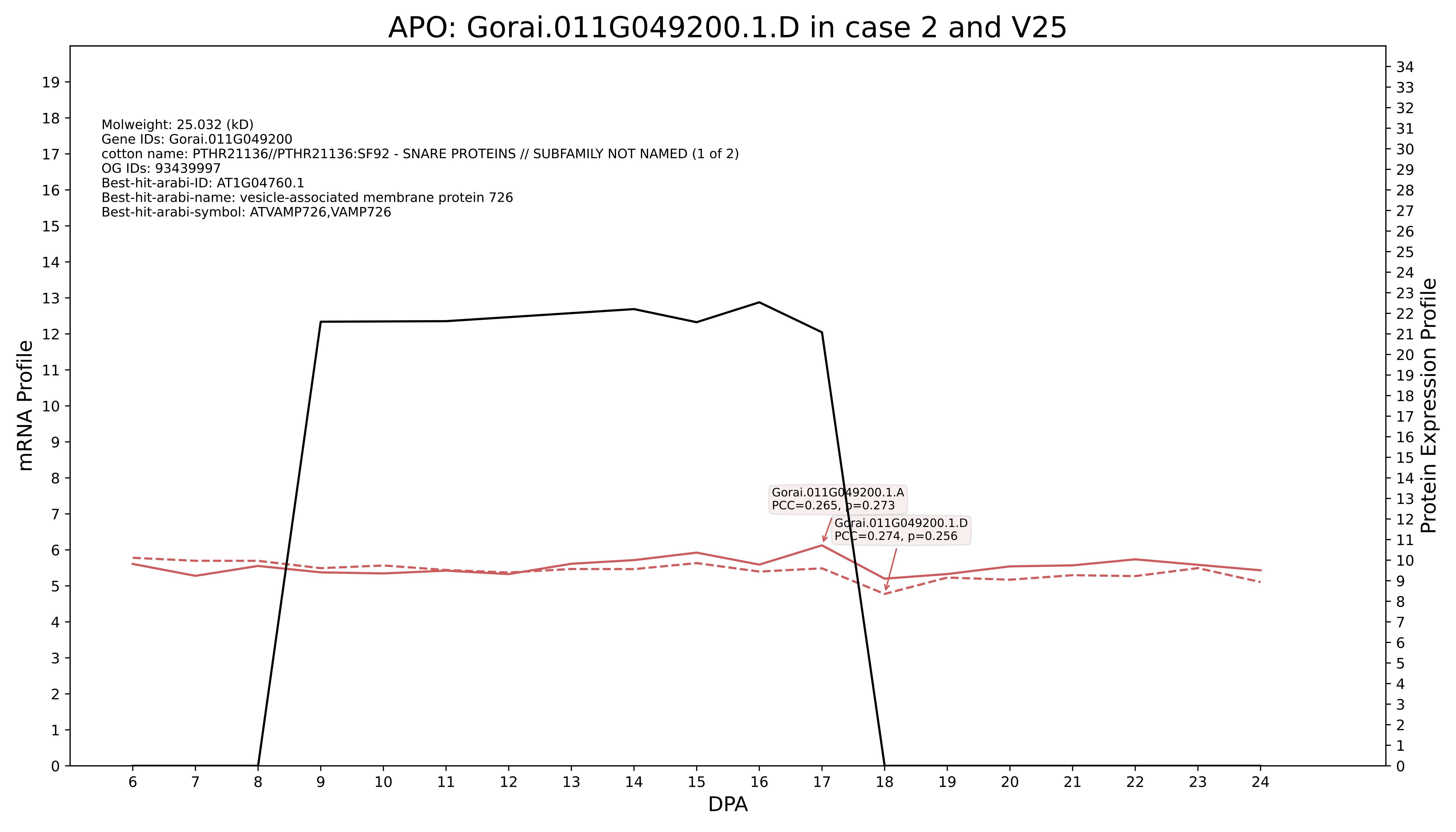

APO: Gorai.011G049200.1.D in case 2 and V25
Molweight: 25.032 (kD)
Gene IDs: Gorai.011G049200
cotton name: PTHR21136//PTHR21136:SF92 - SNARE PROTEINS // SUBFAMILY NOT NAMED (1 of 2)
OG IDs: 93439997
Best-hit-arabi-ID: AT1G04760.1
Best-hit-arabi-name: vesicle-associated membrane protein 726
Best-hit-arabi-symbol: ATVAMP726,VAMP726
Gorai.011G049200.1.A
PCC=0.265, p=0.273
Gorai.011G049200.1.D
PCC=0.274, p=0.256
mRNA Profile
Protein Expression Profile
DPA

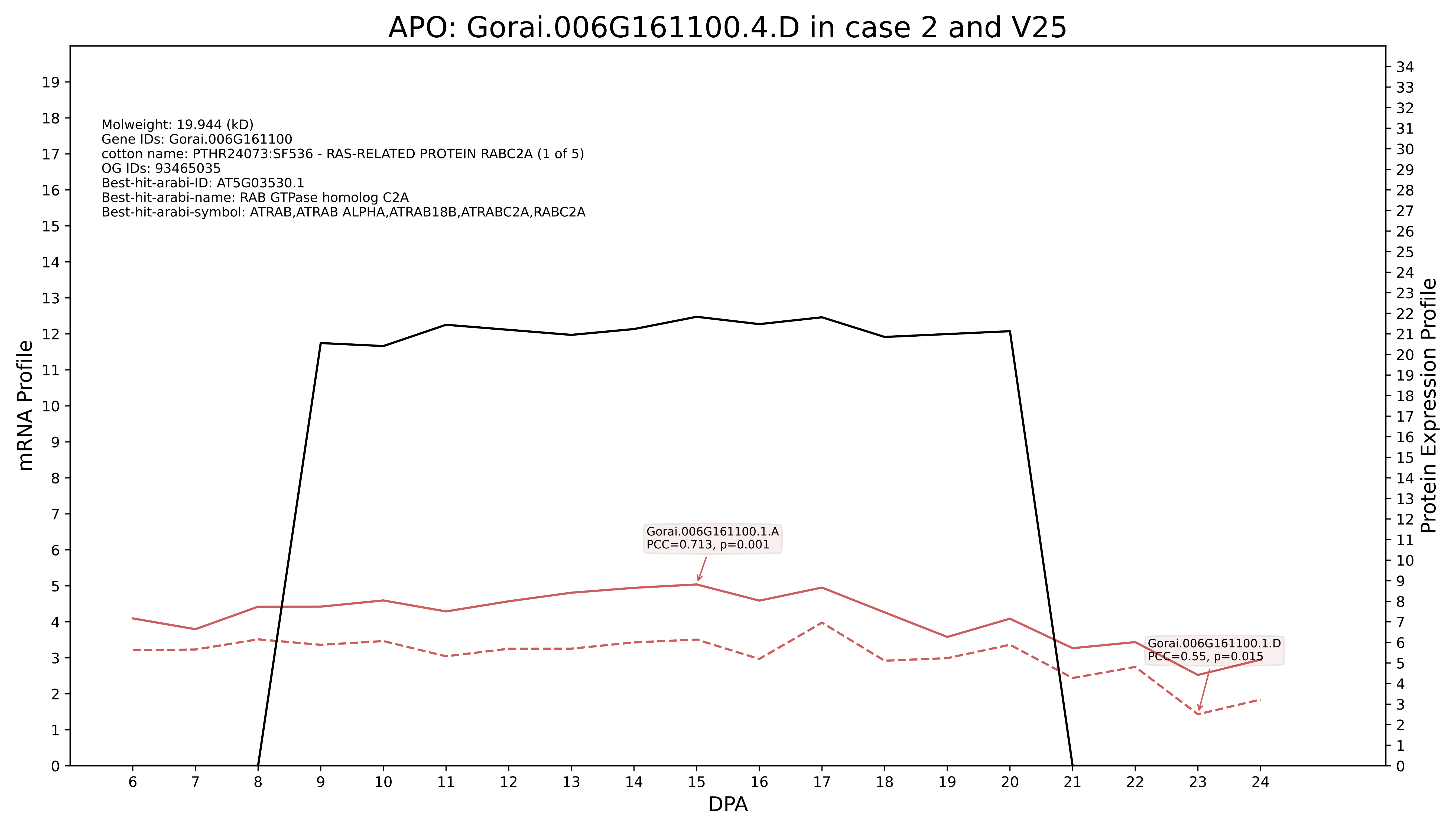

APO: Gorai.006G161100.4.D in case 2 and V25
Molweight: 19.944 (kD)
Gene IDs: Gorai.006G161100
cotton name: PTHR24073:SF536 - RAS-RELATED PROTEIN RABC2A (1 of 5)
OG IDs: 93465035
Best-hit-arabi-ID: AT5G03530.1
Best-hit-arabi-name: RAB GTPase homolog C2A
Best-hit-arabi-symbol: ATRAB,ATRAB ALPHA,ATRAB18B,ATRABC2A,RABC2A
Gorai.006G161100.1.A
PCC=0.713, p=0.001
Gorai.006G161100.1.D
PCC=0.55, p=0.015
mRNA Profile
Protein Expression Profile
DPA

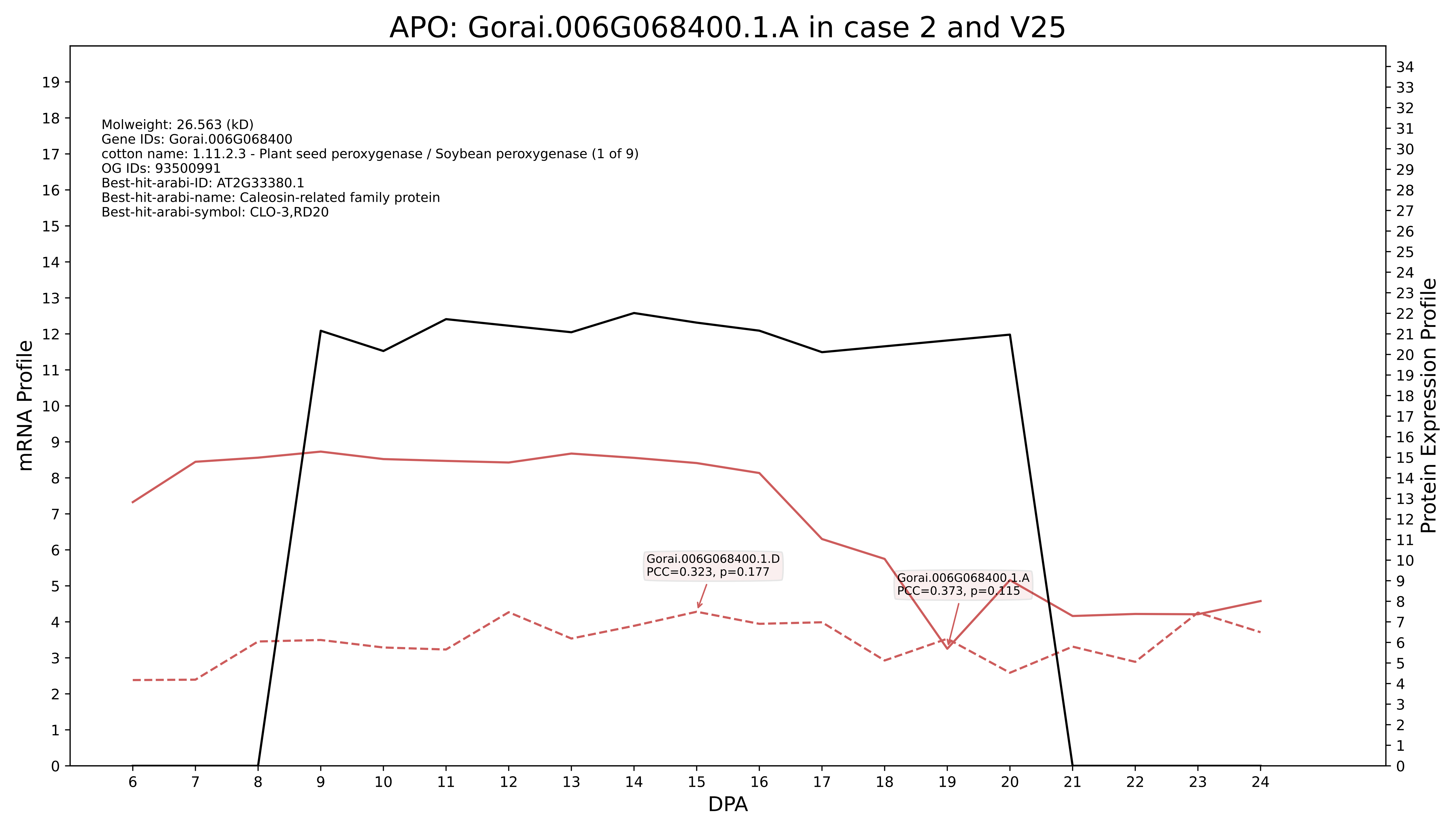

APO: Gorai.006G068400.1.A in case 2 and V25
Molweight: 26.563 (kD)
Gene IDs: Gorai.006G068400
cotton name: 1.11.2.3 - Plant seed peroxygenase / Soybean peroxygenase (1 of 9)
OG IDs: 93500991
Best-hit-arabi-ID: AT2G33380.1
Best-hit-arabi-name: Caleosin-related family protein
Best-hit-arabi-symbol: CLO-3,RD20
Gorai.006G068400.1.D
PCC=0.323, p=0.177
Gorai.006G068400.1.A
PCC=0.373, p=0.115
mRNA Profile
Protein Expression Profile
DPA

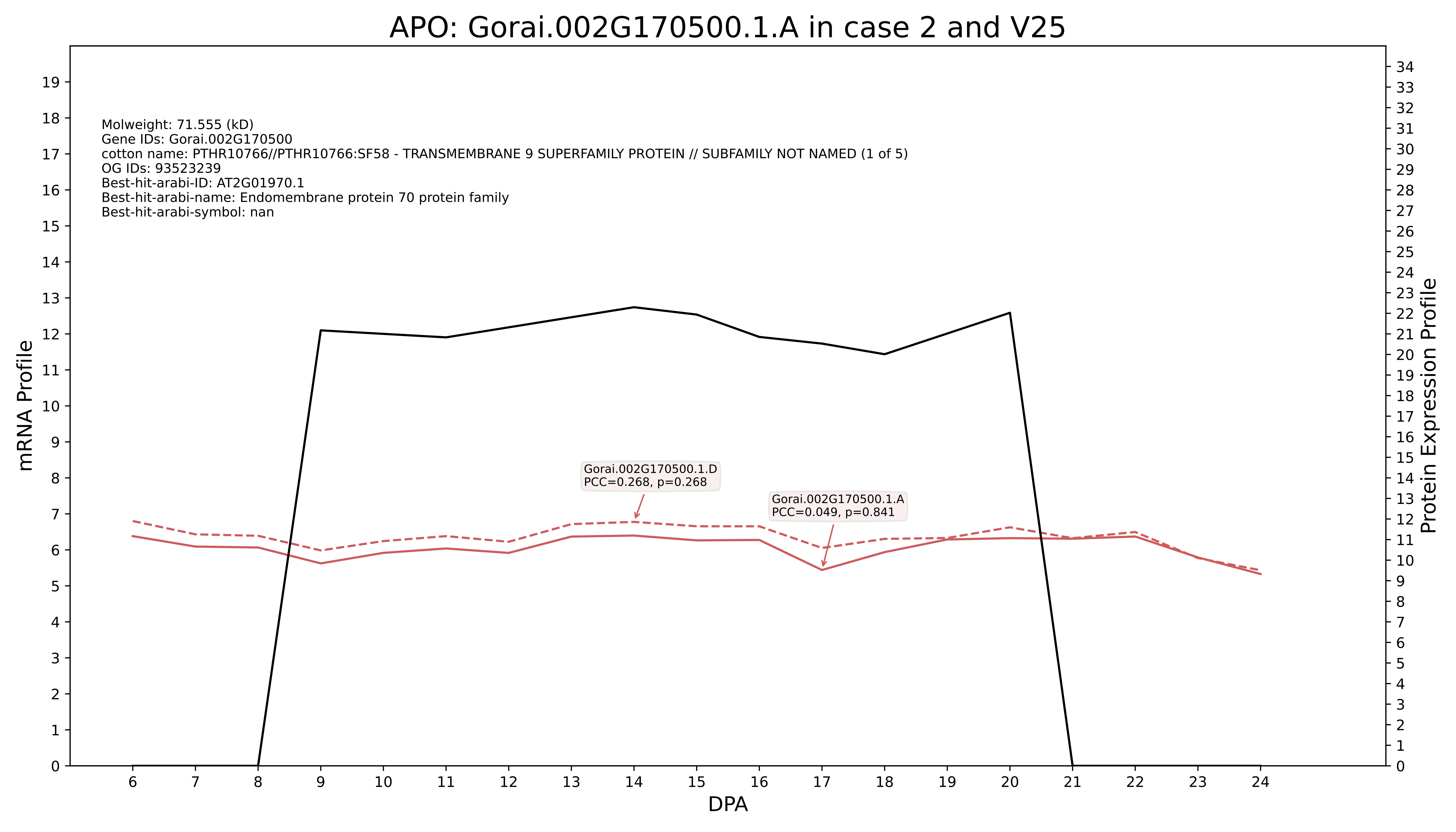
APO: Gorai.002G170500.1.A in case 2 and V25
Molweight: 71.555 (kD)
Gene IDs: Gorai.002G170500
cotton name: PTHR10766//PTHR10766:SF58 - TRANSMEMBRANE 9 SUPERFAMILY PROTEIN // SUBFAMILY NOT NAMED (1 of 5)
OG IDs: 93523239
Best-hit-arabi-ID: AT2G01970.1
Best-hit-arabi-name: Endomembrane protein 70 protein family
Best-hit-arabi-symbol: nan
Gorai.002G170500.1.D
PCC=0.268, p=0.268
Gorai.002G170500.1.A
PCC=0.049, p=0.841
mRNA Profile
Protein Expression Profile
DPA

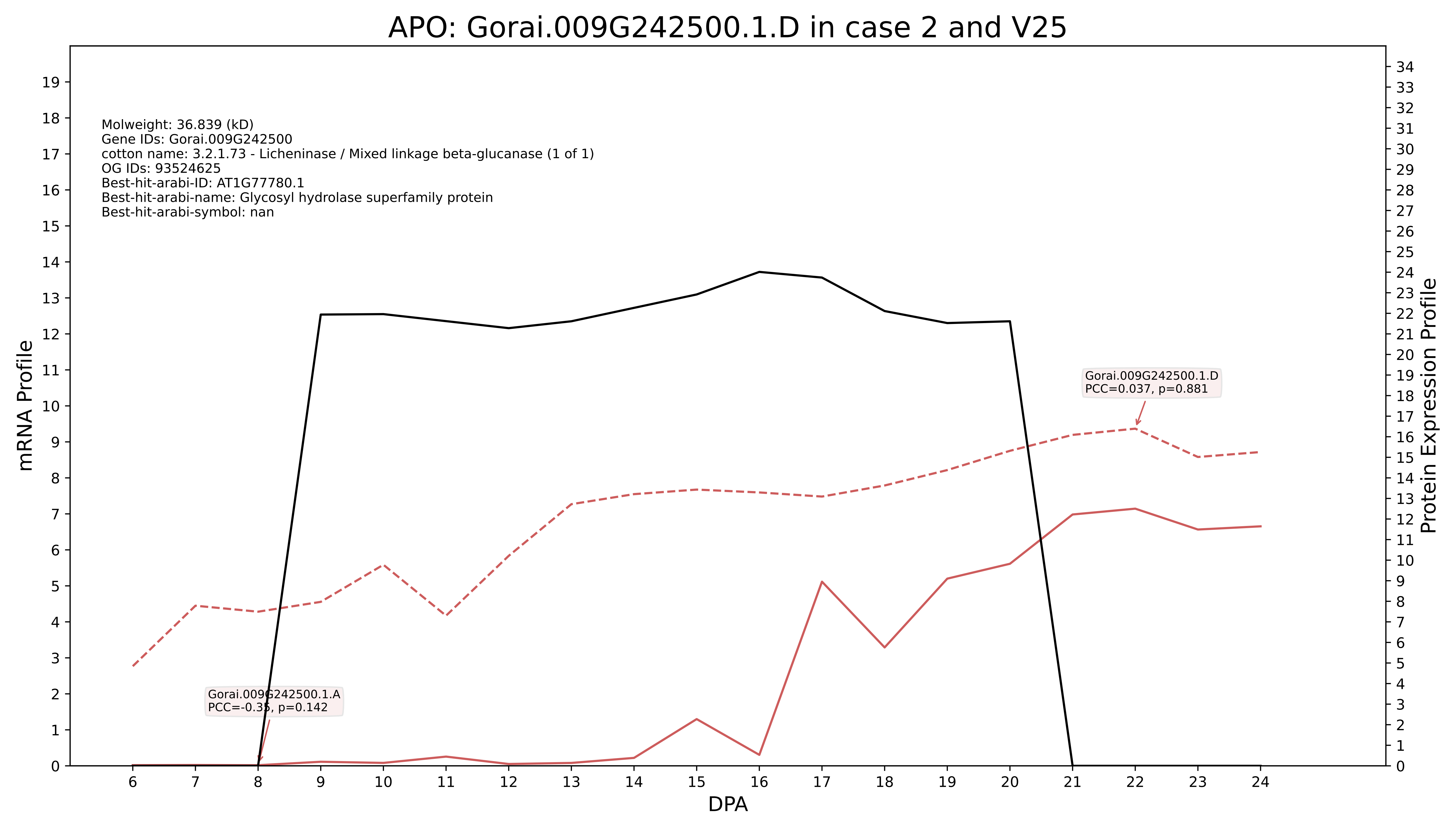

APO: Gorai.009G242500.1.D in case 2 and V25
Molweight: 36.839 (kD)
Gene IDs: Gorai.009G242500
cotton name: 3.2.1.73 - Licheninase / Mixed linkage beta-glucanase (1 of 1)
OG IDs: 93524625
Best-hit-arabi-ID: AT1G77780.1
Best-hit-arabi-name: Glycosyl hydrolase superfamily protein
Best-hit-arabi-symbol: nan
Gorai.009G242500.1.D
PCC=0.037, p=0.881
Gorai.009G242500.1.A
PCC=-0.35, p=0.142
mRNA Profile
Protein Expression Profile
DPA

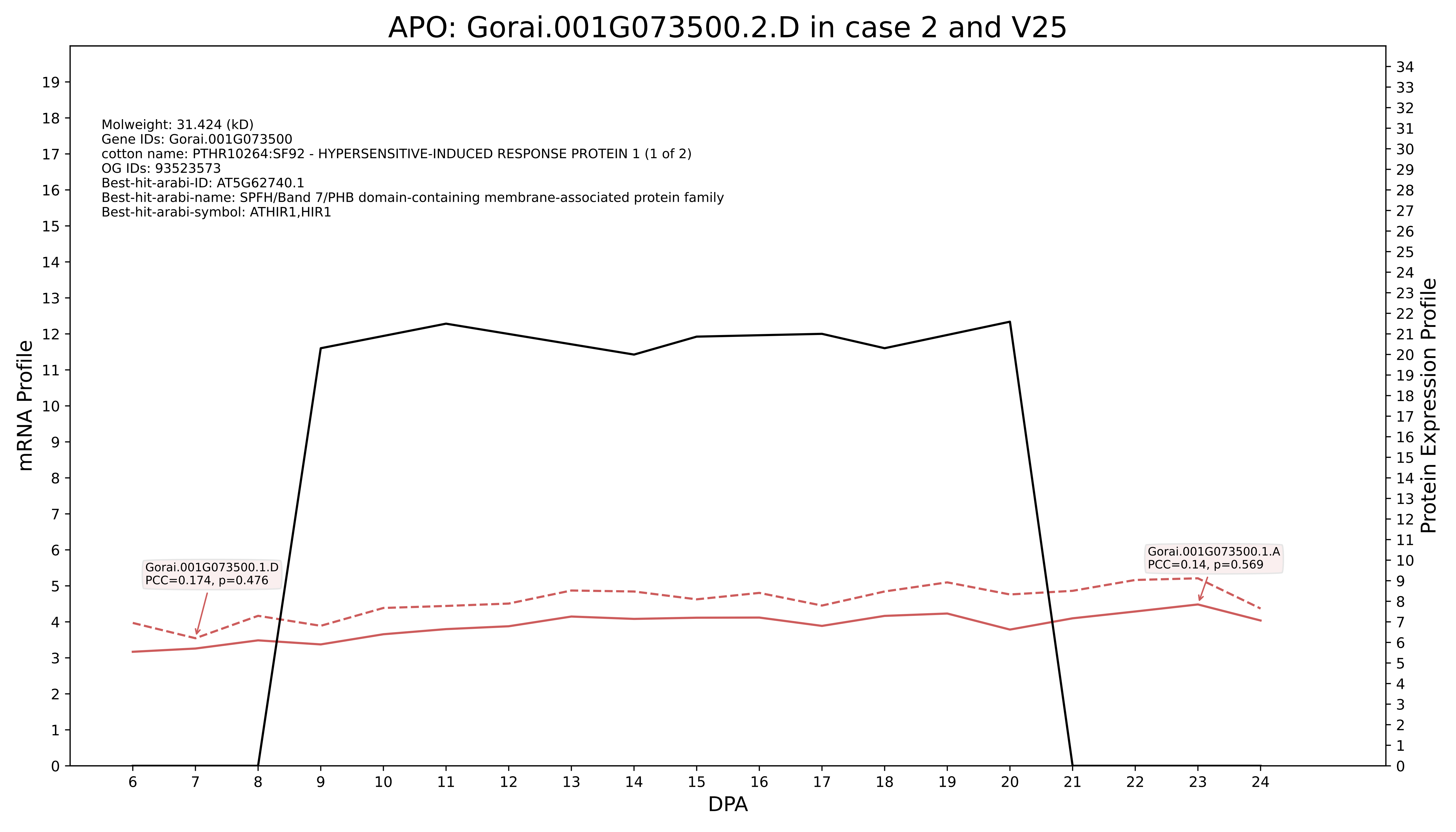

APO: Gorai.001G073500.2.D in case 2 and V25
Molweight: 31.424 (kD)
Gene IDs: Gorai.001G073500
cotton name: PTHR10264:SF92 - HYPERSENSITIVE-INDUCED RESPONSE PROTEIN 1 (1 of 2)
OG IDs: 93523573
Best-hit-arabi-ID: AT5G62740.1
Best-hit-arabi-name: SPFH/Band 7/PHB domain-containing membrane-associated protein family
Best-hit-arabi-symbol: ATHIR1,HIR1
Gorai.001G073500.1.D
PCC=0.174, p=0.476
Gorai.001G073500.1.A
PCC=0.14, p=0.569
mRNA Profile
Protein Expression Profile
DPA

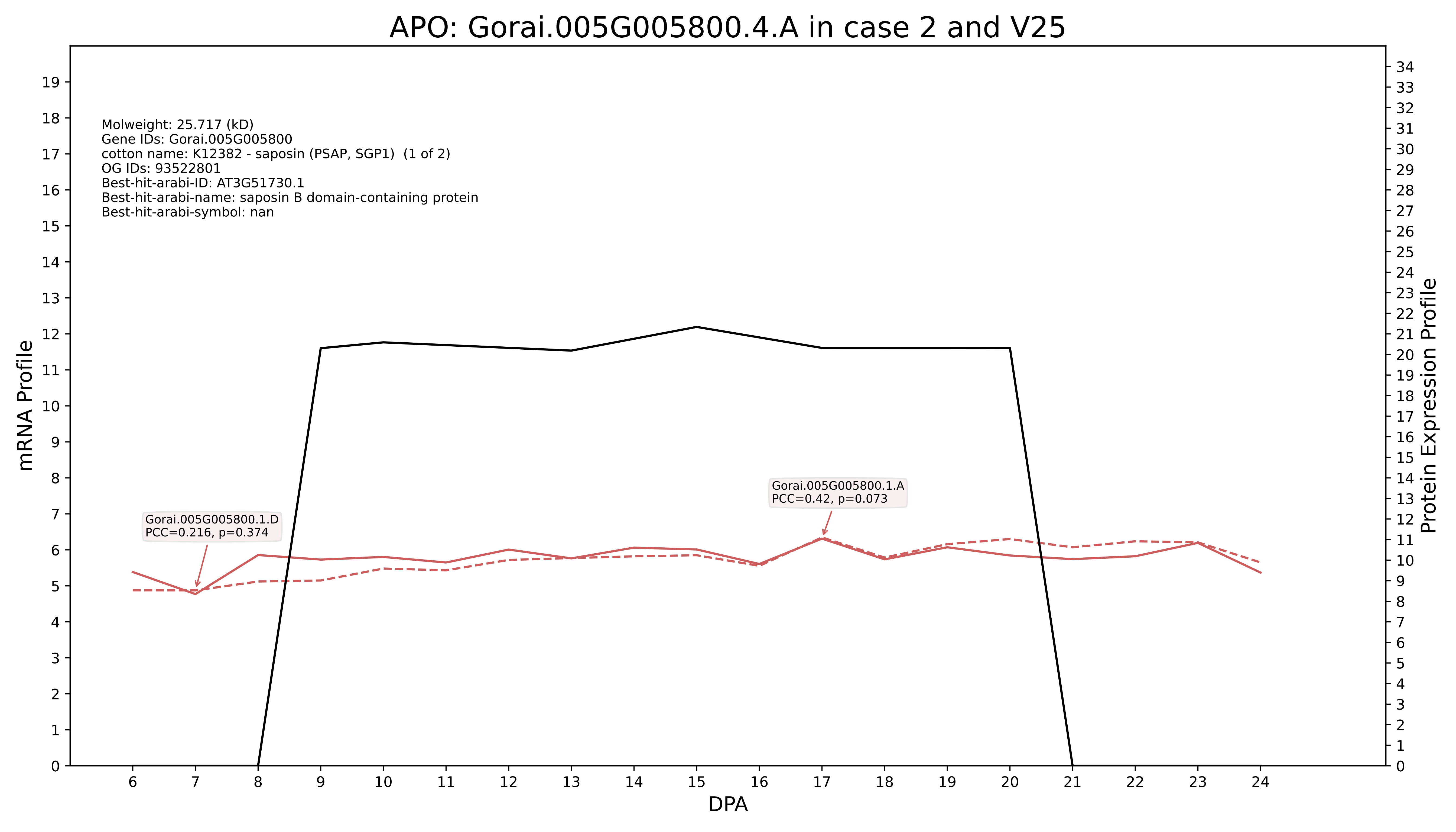
APO: Gorai.005G005800.4.A in case 2 and V25
Molweight: 25.717 (kD)
Gene IDs: Gorai.005G005800
cotton name: K12382 - saposin (PSAP, SGP1) (1 of 2)
OG IDs: 93522801
Best-hit-arabi-ID: AT3G51730.1
Best-hit-arabi-name: saposin B domain-containing protein
Best-hit-arabi-symbol: nan
Gorai.005G005800.1.D
PCC=0.216, p=0.374
Gorai.005G005800.1.A
PCC=0.42, p=0.073
mRNA Profile
Protein Expression Profile
DPA

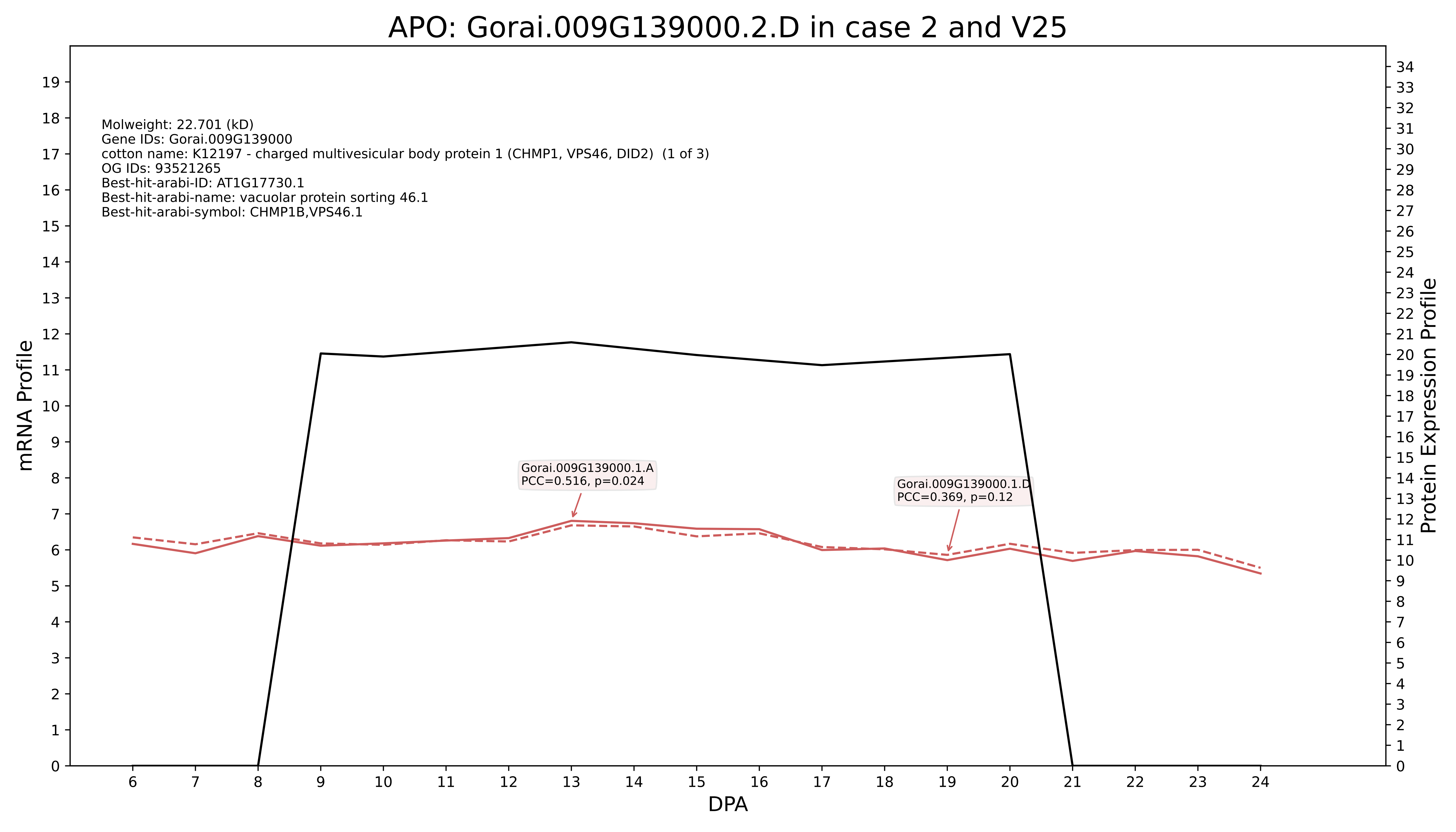
APO: Gorai.009G139000.2.D in case 2 and V25
Molweight: 22.701 (kD)
Gene IDs: Gorai.009G139000
cotton name: K12197 - charged multivesicular body protein 1 (CHMP1, VPS46, DID2) (1 of 3)
OG IDs: 93521265
Best-hit-arabi-ID: AT1G17730.1
Best-hit-arabi-name: vacuolar protein sorting 46.1
Best-hit-arabi-symbol: CHMP1B,VPS46.1
Gorai.009G139000.1.A
PCC=0.516, p=0.024
Gorai.009G139000.1.D
PCC=0.369, p=0.12
mRNA Profile
Protein Expression Profile
DPA

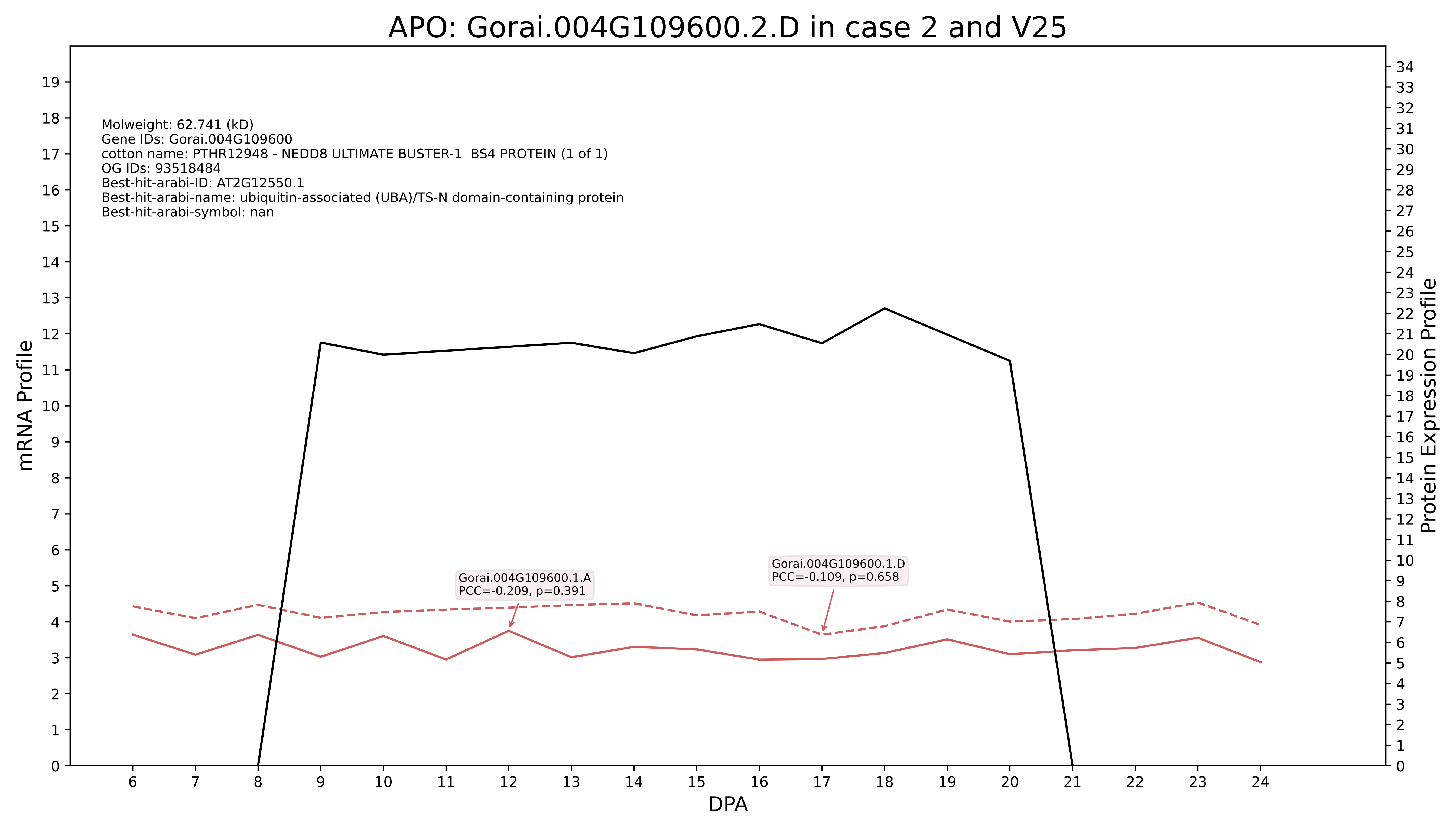
APO: Gorai.004G109600.2.D in case 2 and V25
Molweight: 62.741 (kD)
Gene IDs: Gorai.004G109600
cotton name: PTHR12948 - NEDD8 ULTIMATE BUSTER-1 BS4 PROTEIN (1 of 1)
OG IDs: 93518484
Best-hit-arabi-ID: AT2G12550.1
Best-hit-arabi-name: ubiquitin-associated (UBA)/TS-N domain-containing protein
Best-hit-arabi-symbol: nan
Gorai.004G109600.1.A
PCC=-0.209, p=0.391
Gorai.004G109600.1.D
PCC=-0.109, p=0.658
mRNA Profile
Protein Expression Profile
DPA

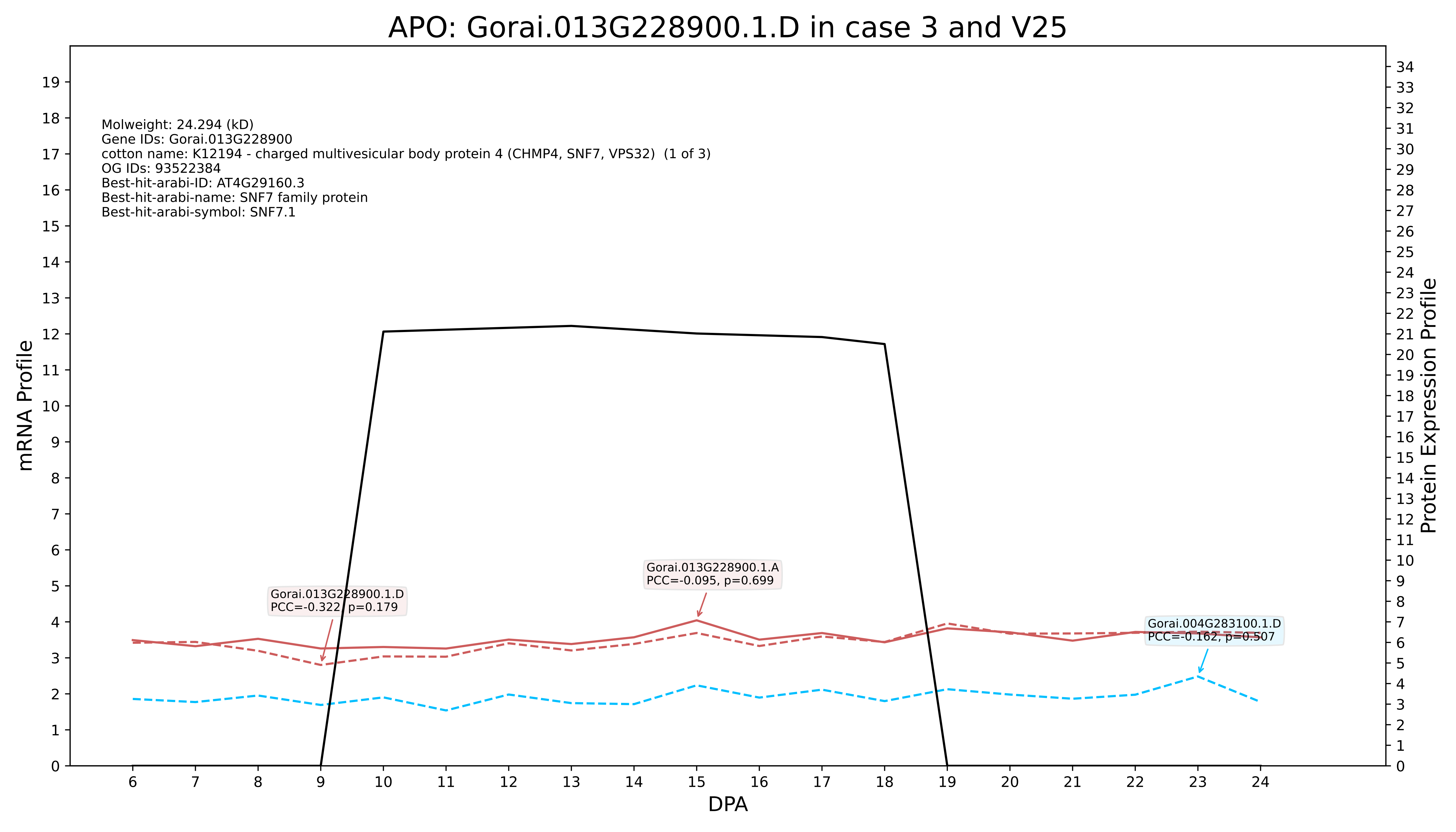

APO: Gorai.013G228900.1.D in case 3 and V25
Molweight: 24.294 (kD)
Gene IDs: Gorai.013G228900
cotton name: K12194 - charged multivesicular body protein 4 (CHMP4, SNF7, VPS32) (1 of 3)
OG IDs: 93522384
Best-hit-arabi-ID: AT4G29160.3
Best-hit-arabi-name: SNF7 family protein
Best-hit-arabi-symbol: SNF7.1
Gorai.013G228900.1.D
PCC=-0.322, p=0.179
Gorai.013G228900.1.A
PCC=-0.095, p=0.699
Gorai.004G283100.1.D
PCC=-0.162, p=0.507
mRNA Profile
Protein Expression Profile
DPA

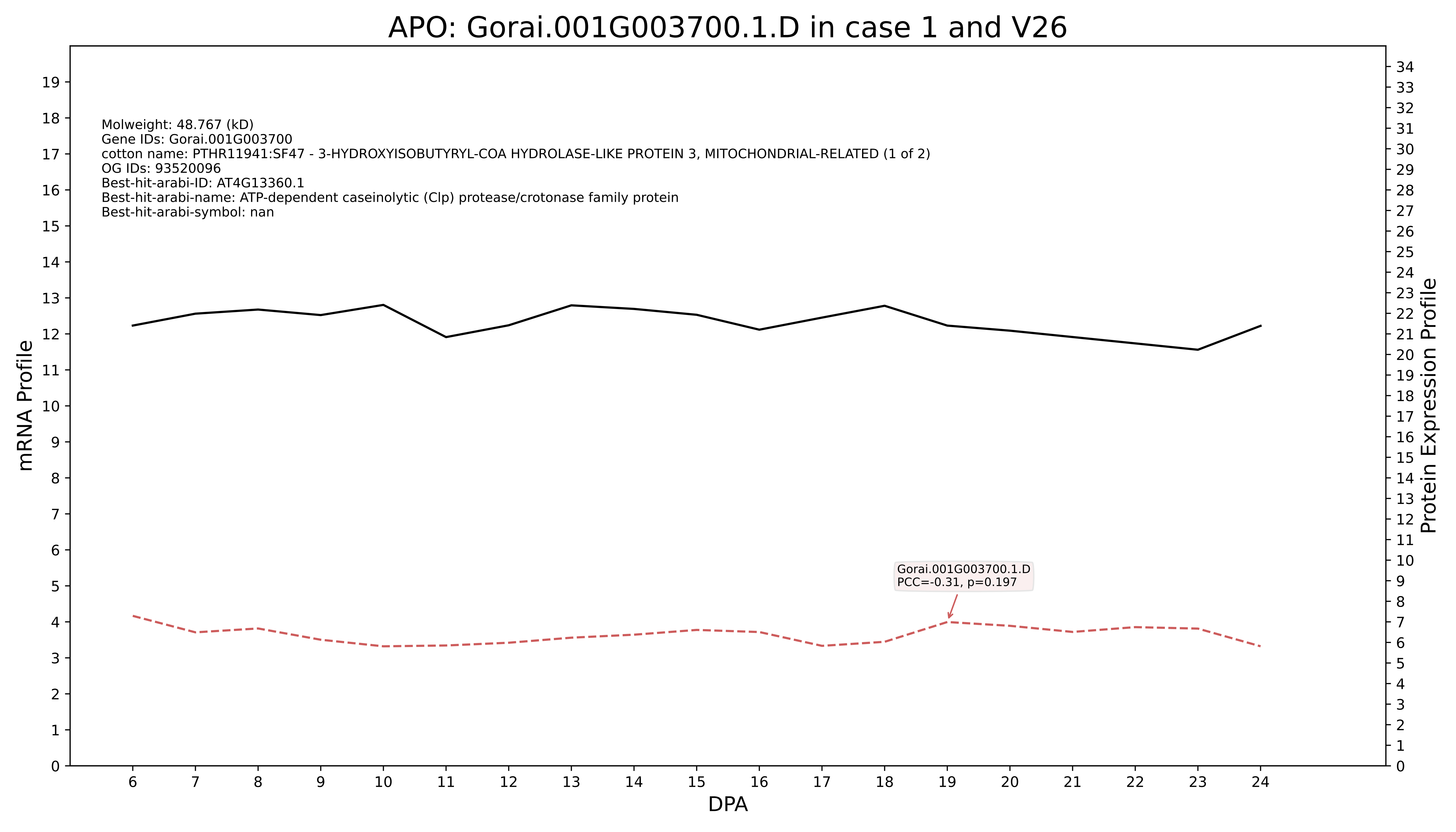

APO: Gorai.001G003700.1.D in case 1 and V26
Molweight: 48.767 (kD)
Gene IDs: Gorai.001G003700
cotton name: PTHR11941:SF47 - 3-HYDROXYISOBUTYRYL-COA HYDROLASE-LIKE PROTEIN 3, MITOCHONDRIAL-RELATED (1 of 2)
OG IDs: 93520096
Best-hit-arabi-ID: AT4G13360.1
Best-hit-arabi-name: ATP-dependent caseinolytic (Clp) protease/crotonase family protein
Best-hit-arabi-symbol: nan
Gorai.001G003700.1.D
PCC=-0.31, p=0.197
mRNA Profile
Protein Expression Profile
DPA

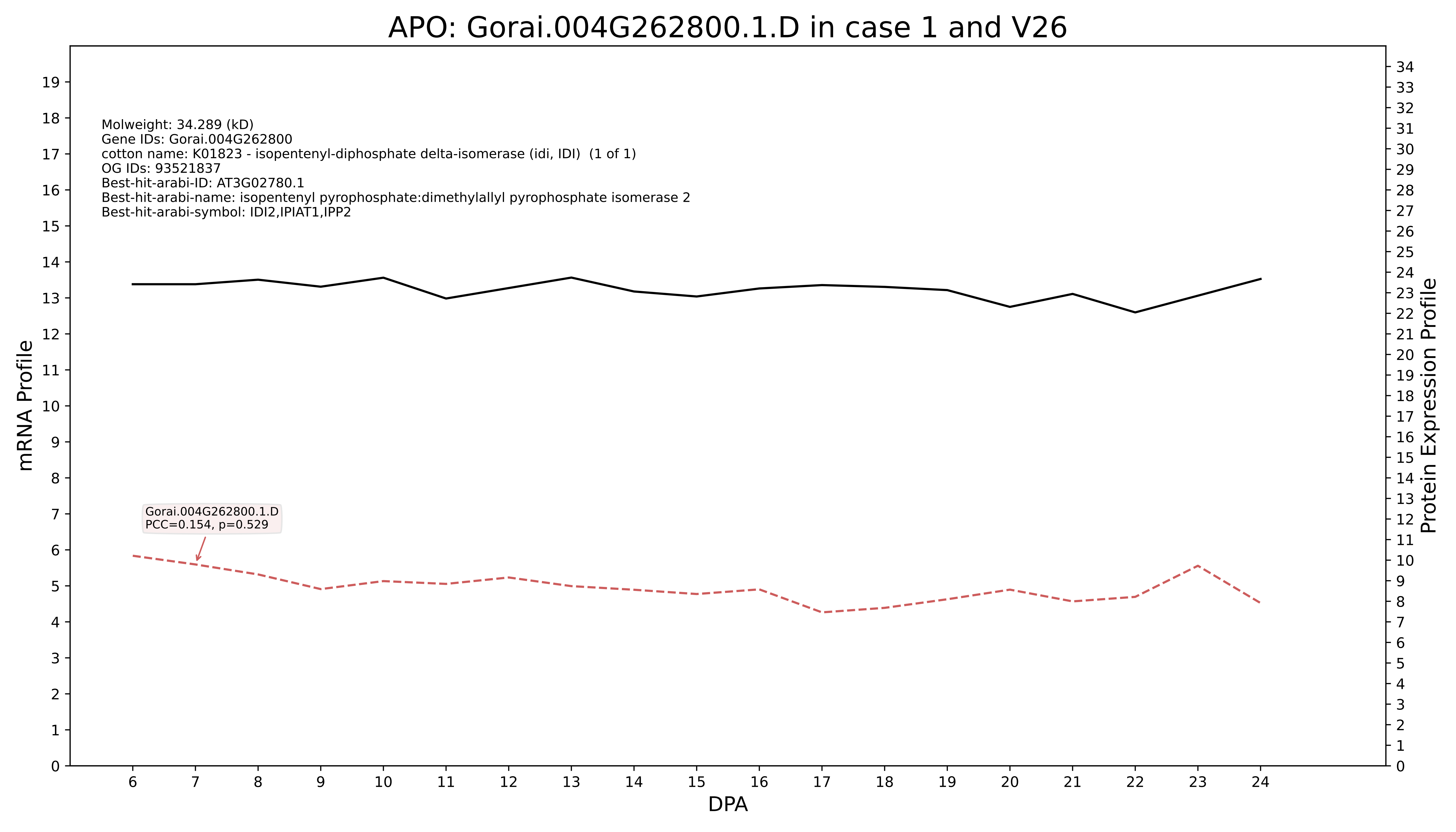
APO: Gorai.004G262800.1.D in case 1 and V26
Molweight: 34.289 (kD)
Gene IDs: Gorai.004G262800
cotton name: K01823 - isopentenyl-diphosphate delta-isomerase (idi, IDI) (1 of 1)
OG IDs: 93521837
Best-hit-arabi-ID: AT3G02780.1
Best-hit-arabi-name: isopentenyl pyrophosphate:dimethylallyl pyrophosphate isomerase 2
Best-hit-arabi-symbol: IDI2,IPIAT1,IPP2
Gorai.004G262800.1.D
PCC=0.154, p=0.529
mRNA Profile
Protein Expression Profile
DPA

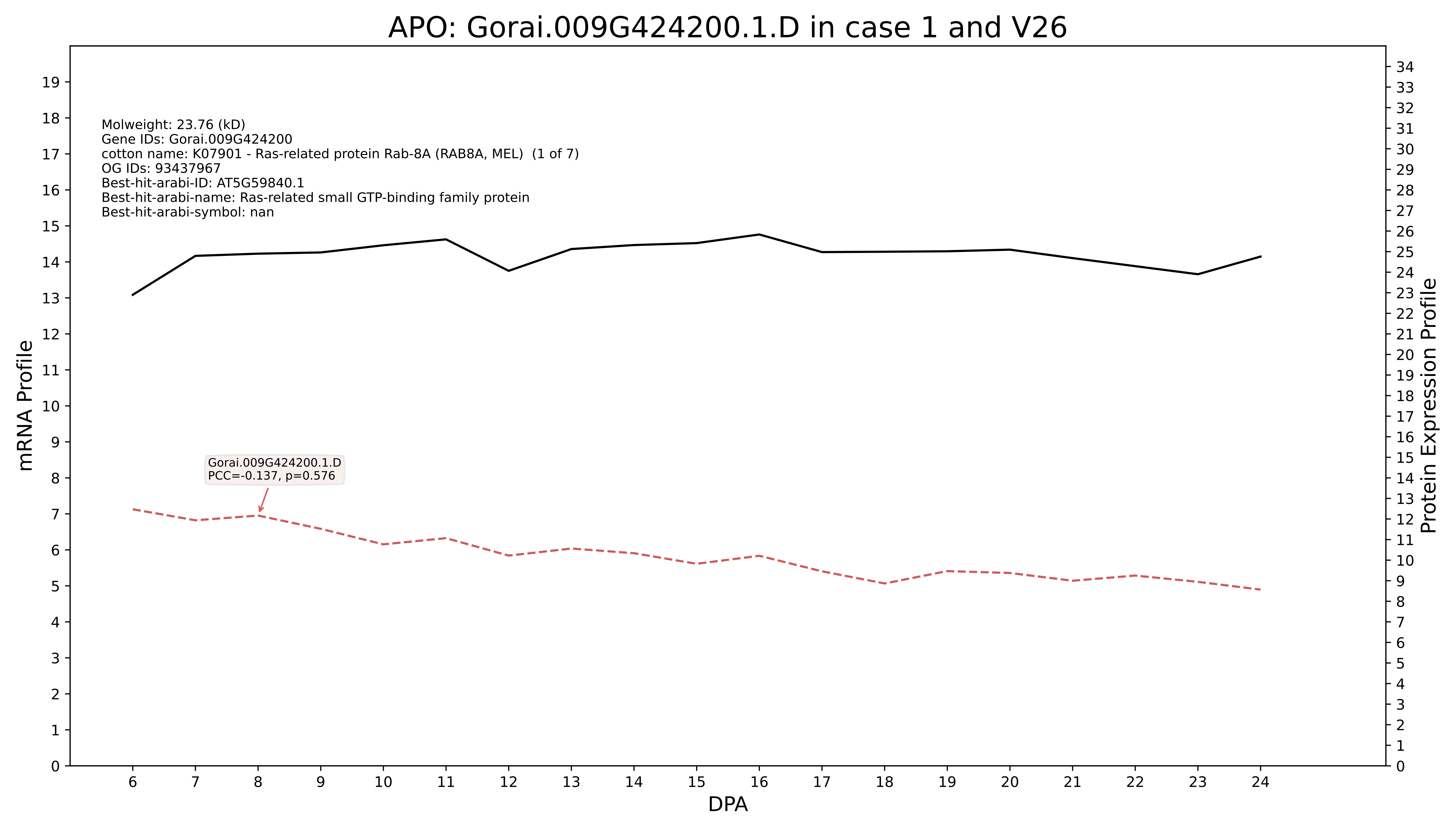

APO: Gorai.009G424200.1.D in case 1 and V26
Molweight: 23.76 (kD)
Gene IDs: Gorai.009G424200
cotton name: K07901 - Ras-related protein Rab-8A (RAB8A, MEL) (1 of 7)
OG IDs: 93437967
Best-hit-arabi-ID: AT5G59840.1
Best-hit-arabi-name: Ras-related small GTP-binding family protein
Best-hit-arabi-symbol: nan
Gorai.009G424200.1.D
PCC=-0.137, p=0.576
mRNA Profile
Protein Expression Profile
DPA

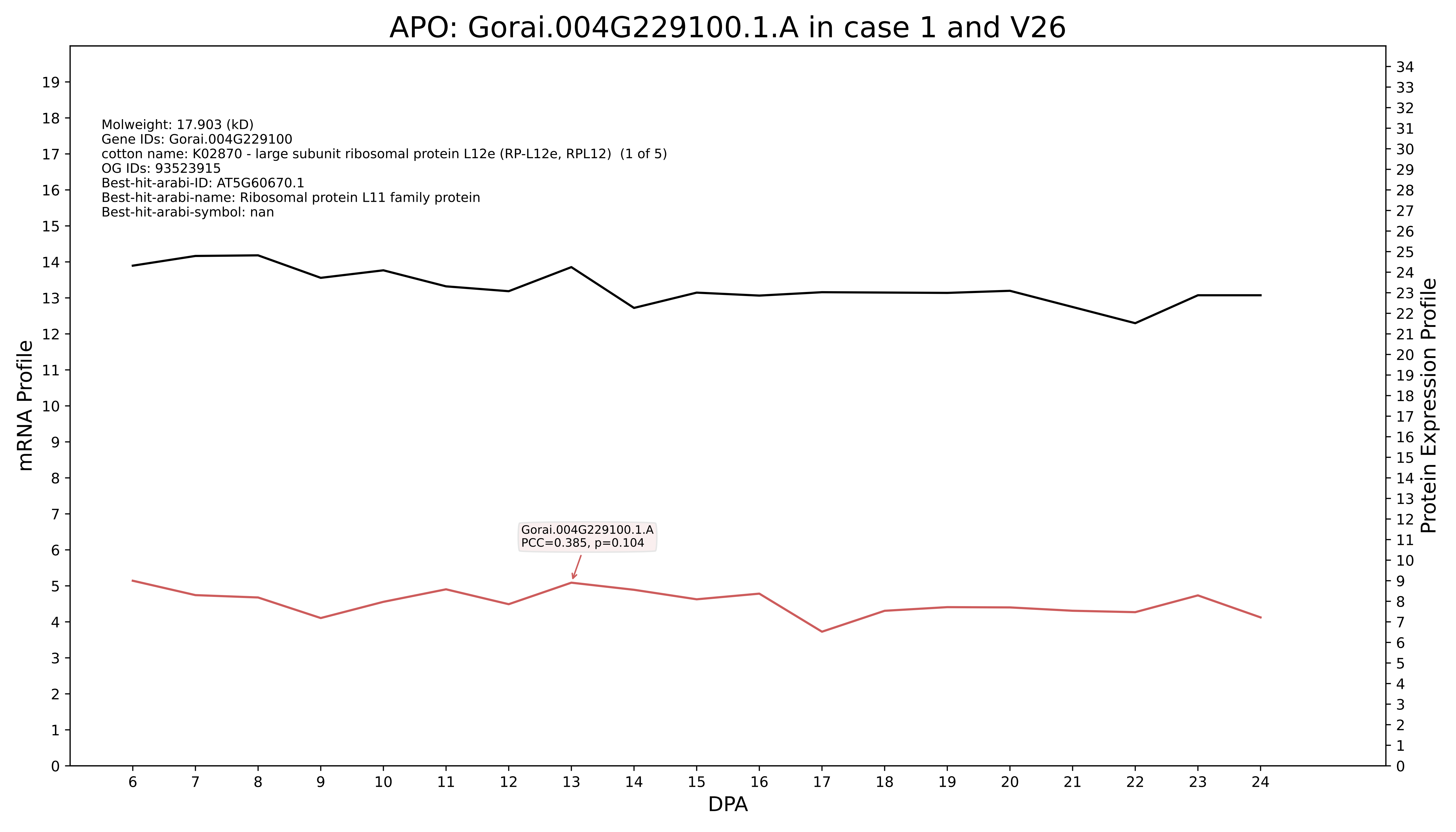
APO: Gorai.004G229100.1.A in case 1 and V26
Molweight: 17.903 (kD)
Gene IDs: Gorai.004G229100
cotton name: K02870 - large subunit ribosomal protein L12e (RP-L12e, RPL12) (1 of 5)
OG IDs: 93523915
Best-hit-arabi-ID: AT5G60670.1
Best-hit-arabi-name: Ribosomal protein L11 family protein
Best-hit-arabi-symbol: nan
Gorai.004G229100.1.A
PCC=0.385, p=0.104
mRNA Profile
Protein Expression Profile
DPA

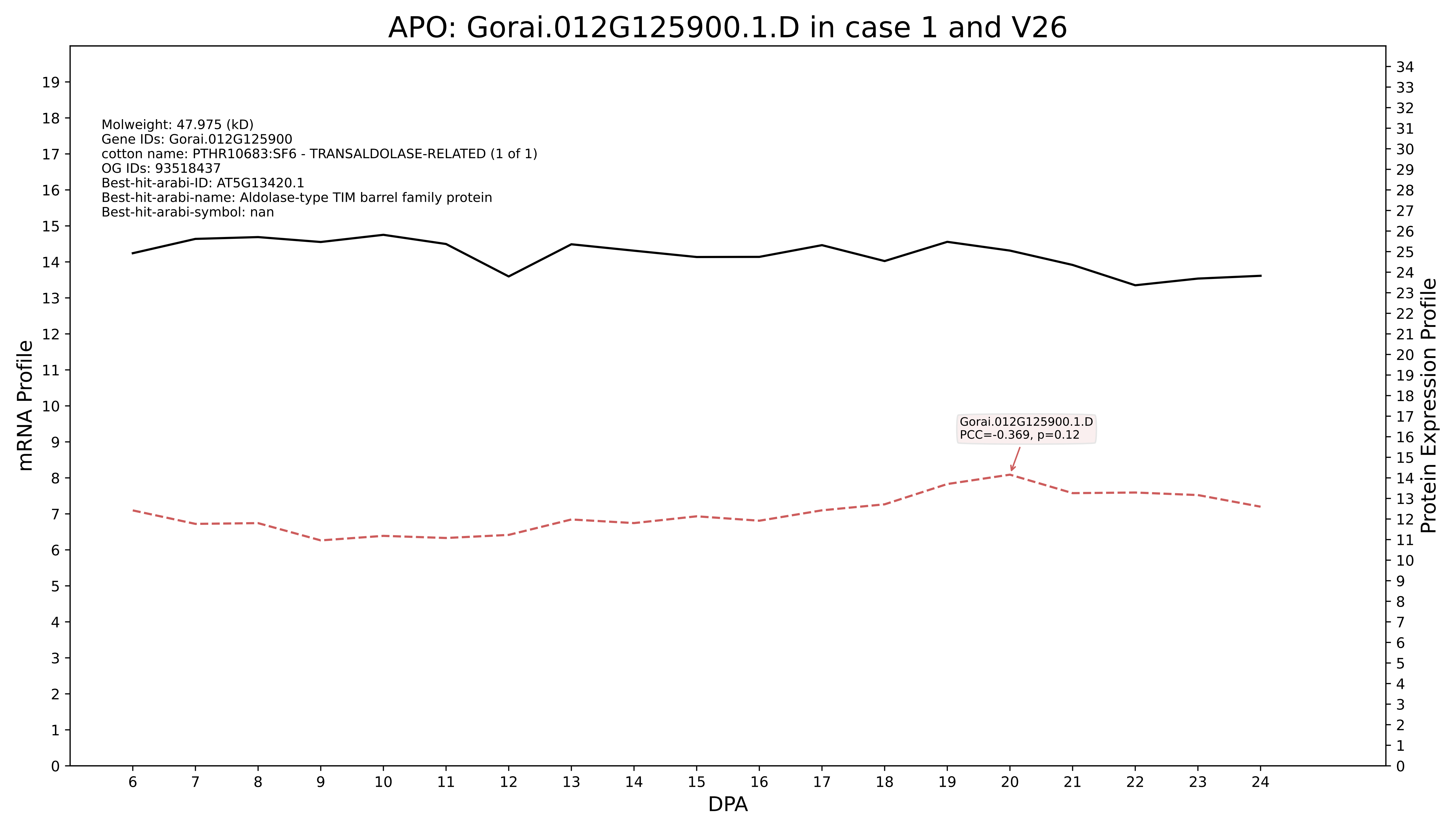

APO: Gorai.012G125900.1.D in case 1 and V26
Molweight: 47.975 (kD)
Gene IDs: Gorai.012G125900
cotton name: PTHR10683:SF6 - TRANSALDOLASE-RELATED (1 of 1)
OG IDs: 93518437
Best-hit-arabi-ID: AT5G13420.1
Best-hit-arabi-name: Aldolase-type TIM barrel family protein
Best-hit-arabi-symbol: nan
Gorai.012G125900.1.D
PCC=-0.369, p=0.12
mRNA Profile
Protein Expression Profile
DPA

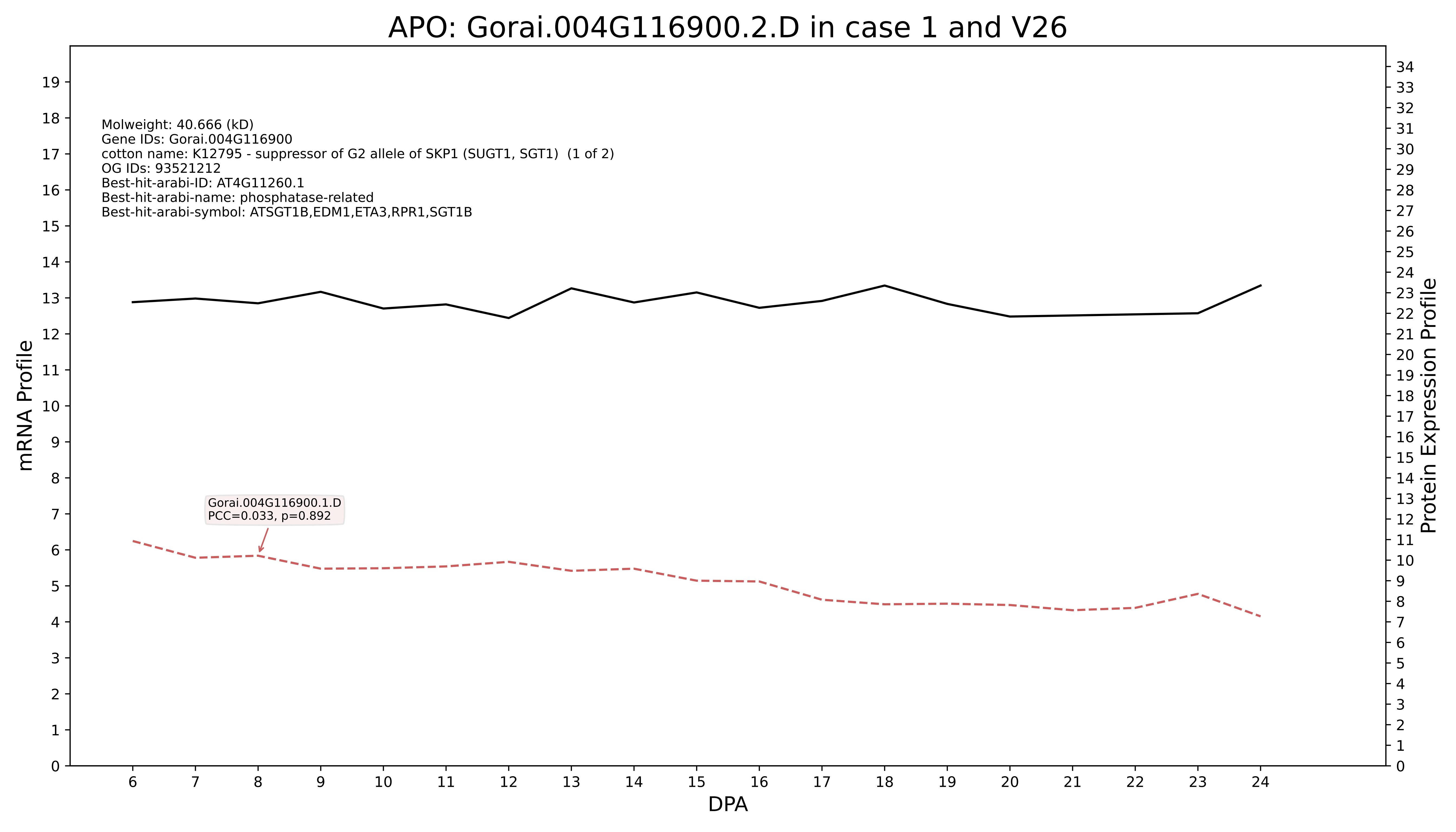
APO: Gorai.004G116900.2.D in case 1 and V26
Molweight: 40.666 (kD)
Gene IDs: Gorai.004G116900
cotton name: K12795 - suppressor of G2 allele of SKP1 (SUGT1, SGT1) (1 of 2)
OG IDs: 93521212
Best-hit-arabi-ID: AT4G11260.1
Best-hit-arabi-name: phosphatase-related
Best-hit-arabi-symbol: ATSGT1B,EDM1,ETA3,RPR1,SGT1B
Gorai.004G116900.1.D
PCC=0.033, p=0.892
mRNA Profile
Protein Expression Profile
DPA

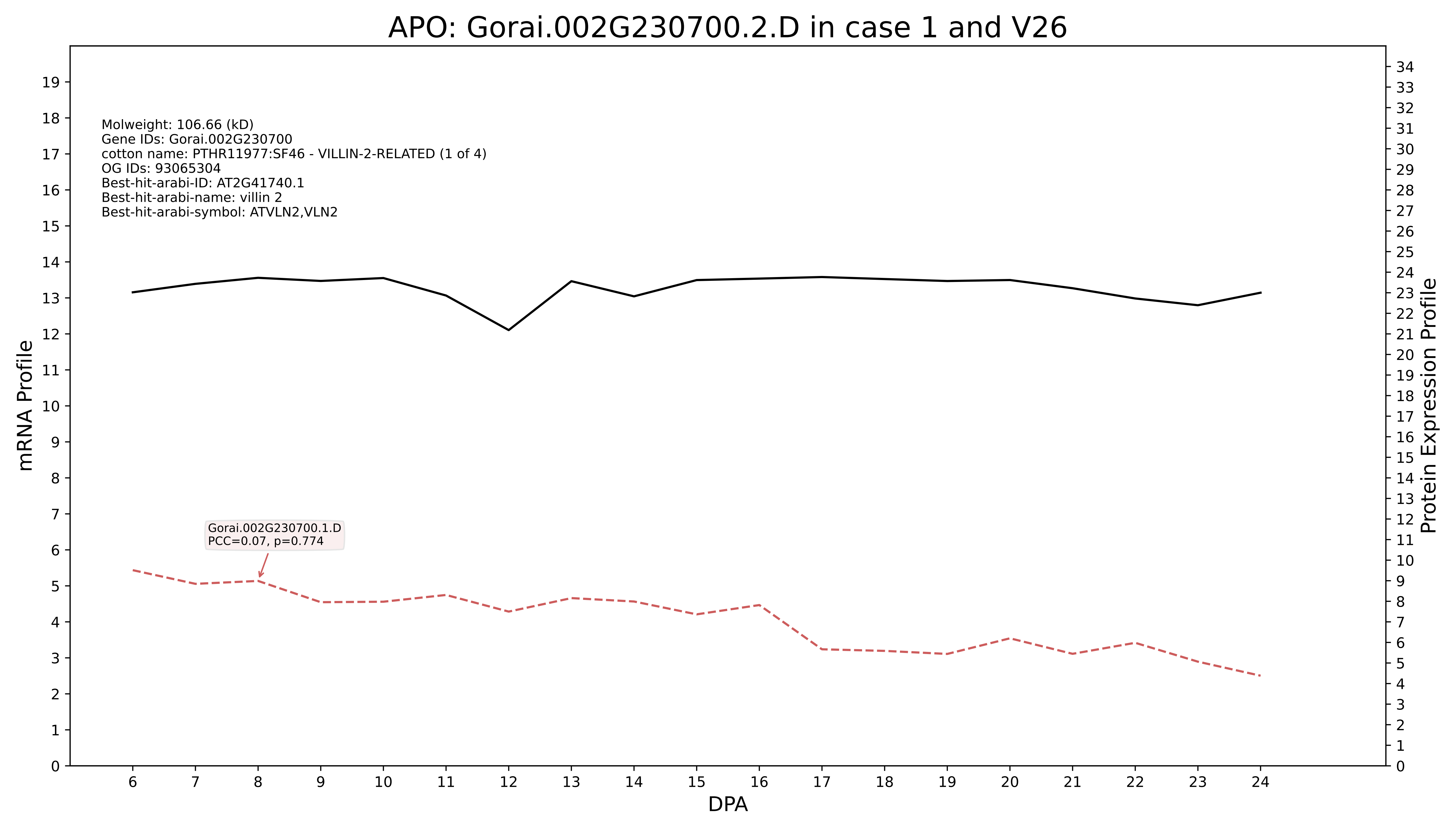

APO: Gorai.002G230700.2.D in case 1 and V26
Molweight: 106.66 (kD)
Gene IDs: Gorai.002G230700
cotton name: PTHR11977:SF46 - VILLIN-2-RELATED (1 of 4)
OG IDs: 93065304
Best-hit-arabi-ID: AT2G41740.1
Best-hit-arabi-name: villin 2
Best-hit-arabi-symbol: ATVLN2,VLN2
Gorai.002G230700.1.D
PCC=0.07, p=0.774
mRNA Profile
Protein Expression Profile
DPA

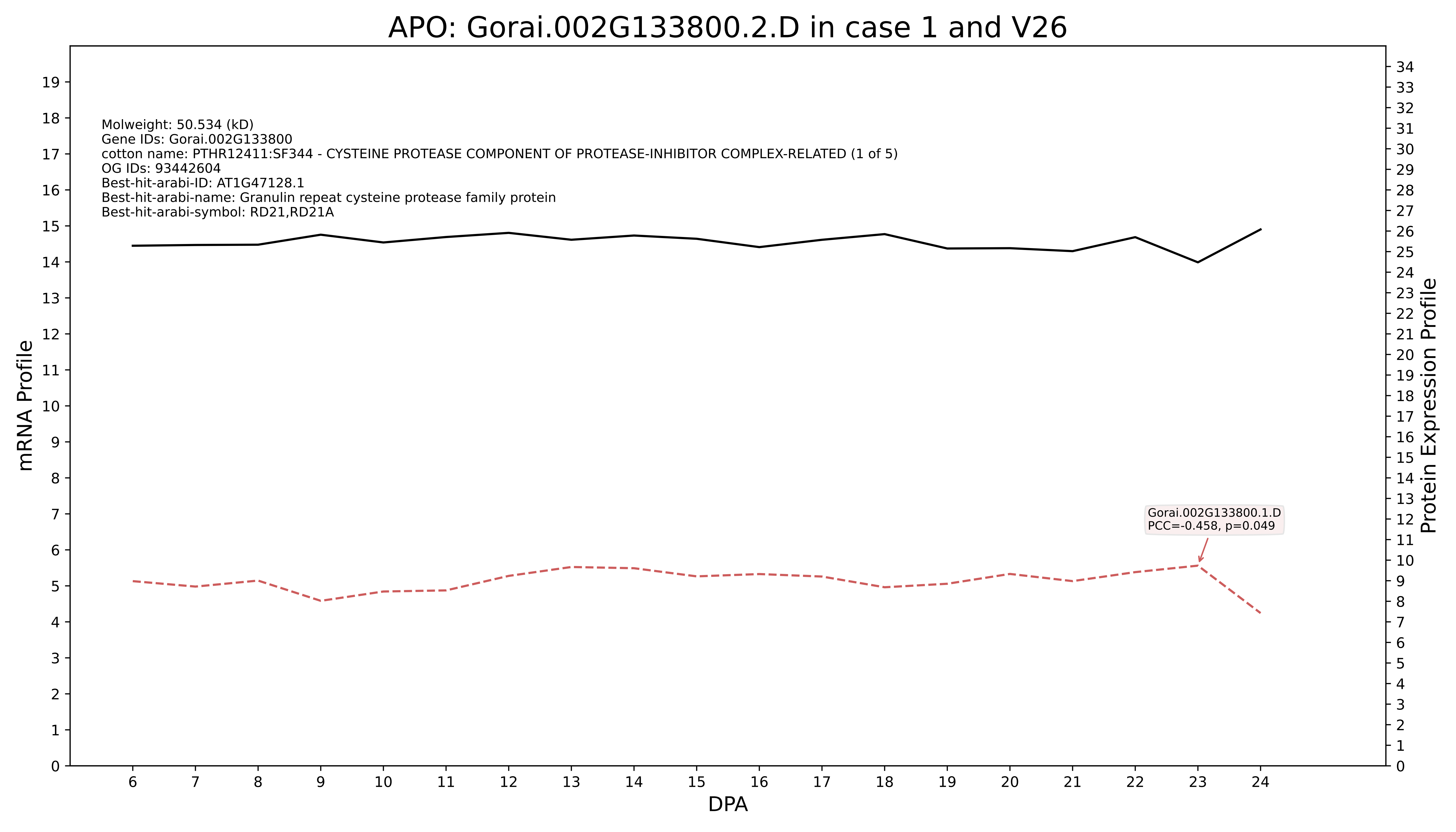
APO: Gorai.002G133800.2.D in case 1 and V26
Molweight: 50.534 (kD)
Gene IDs: Gorai.002G133800
cotton name: PTHR12411:SF344 - CYSTEINE PROTEASE COMPONENT OF PROTEASE-INHIBITOR COMPLEX-RELATED (1 of 5)
OG IDs: 93442604
Best-hit-arabi-ID: AT1G47128.1
Best-hit-arabi-name: Granulin repeat cysteine protease family protein
Best-hit-arabi-symbol: RD21,RD21A
Gorai.002G133800.1.D
PCC=-0.458, p=0.049
mRNA Profile
Protein Expression Profile
DPA

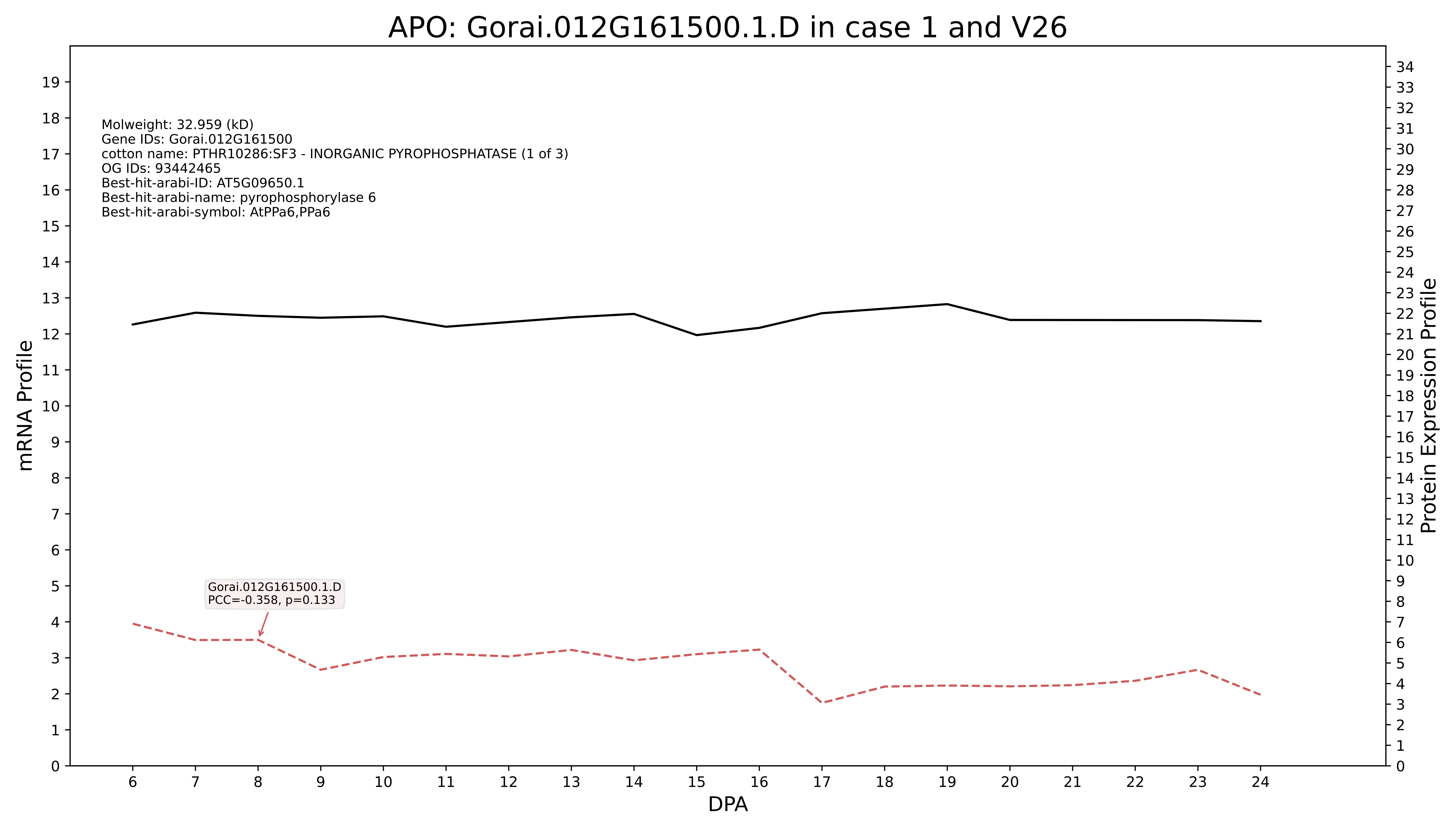
APO: Gorai.012G161500.1.D in case 1 and V26
Molweight: 32.959 (kD)
Gene IDs: Gorai.012G161500
cotton name: PTHR10286:SF3 - INORGANIC PYROPHOSPHATASE (1 of 3)
OG IDs: 93442465
Best-hit-arabi-ID: AT5G09650.1
Best-hit-arabi-name: pyrophosphorylase 6
Best-hit-arabi-symbol: AtPPa6,PPa6
Gorai.012G161500.1.D
PCC=-0.358, p=0.133
mRNA Profile
Protein Expression Profile
DPA

### APO: Gorai.005G060700.1.D in case 1 and V26

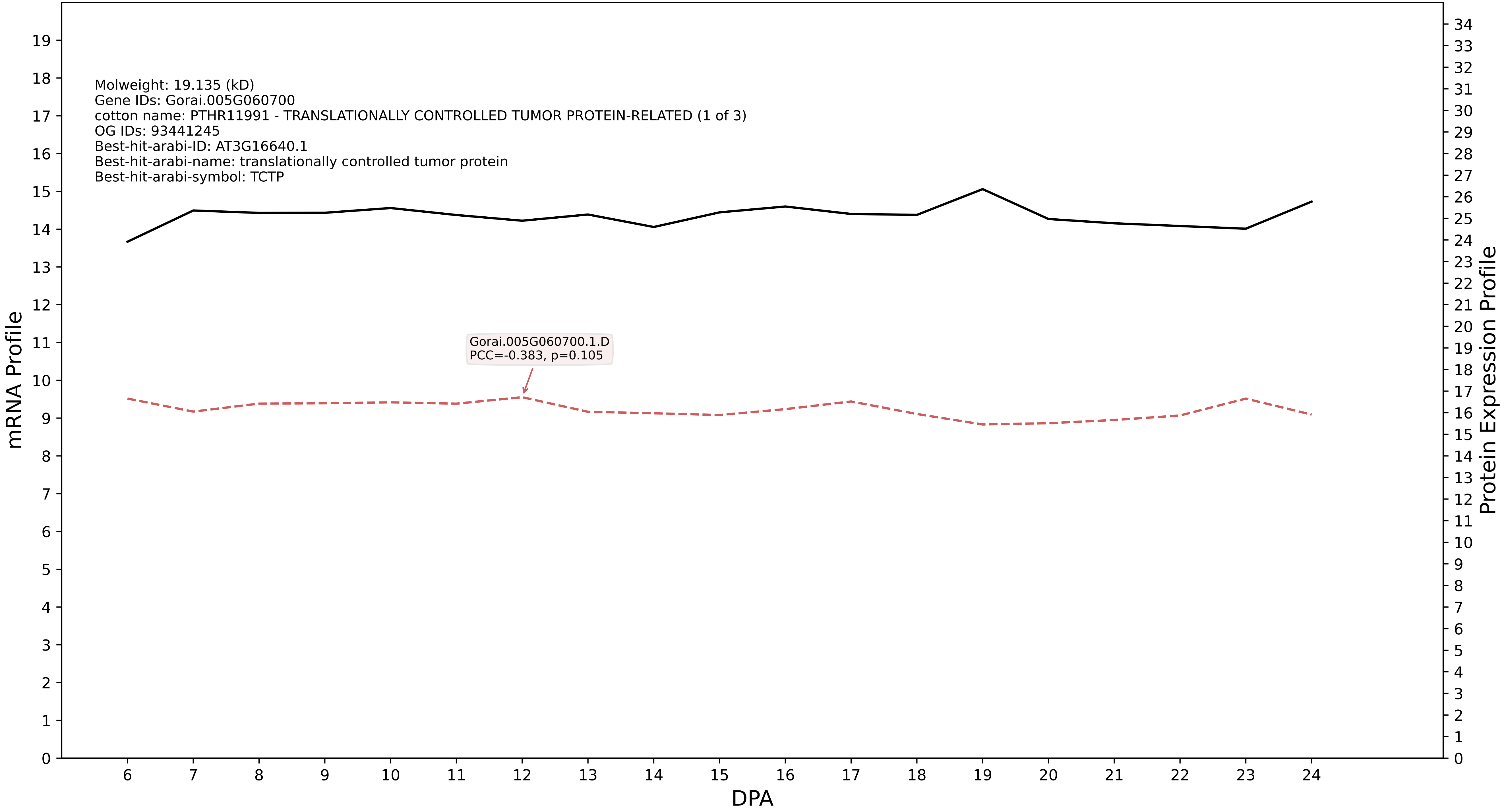

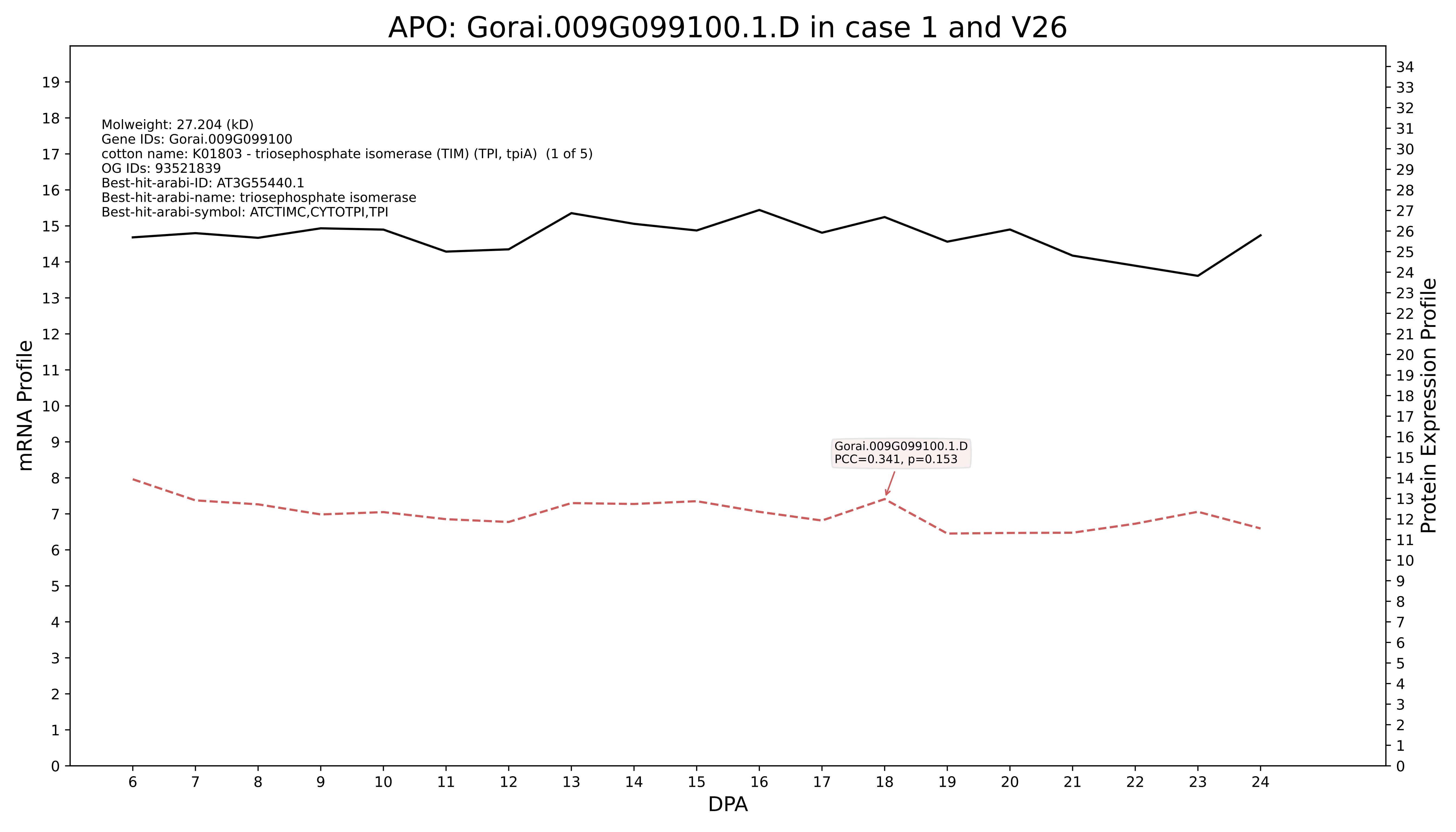
APO: Gorai.009G099100.1.D in case 1 and V26
Molweight: 27.204 (kD)
Gene IDs: Gorai.009G099100
cotton name: K01803 - triosephosphate isomerase (TIM) (TPI, tpiA) (1 of 5)
OG IDs: 93521839
Best-hit-arabi-ID: AT3G55440.1
Best-hit-arabi-name: triosephosphate isomerase
Best-hit-arabi-symbol: ATCTIMC,CYTOTPI,TPI
Gorai.009G099100.1.D
PCC=0.341, p=0.153
mRNA Profile
Protein Expression Profile
DPA

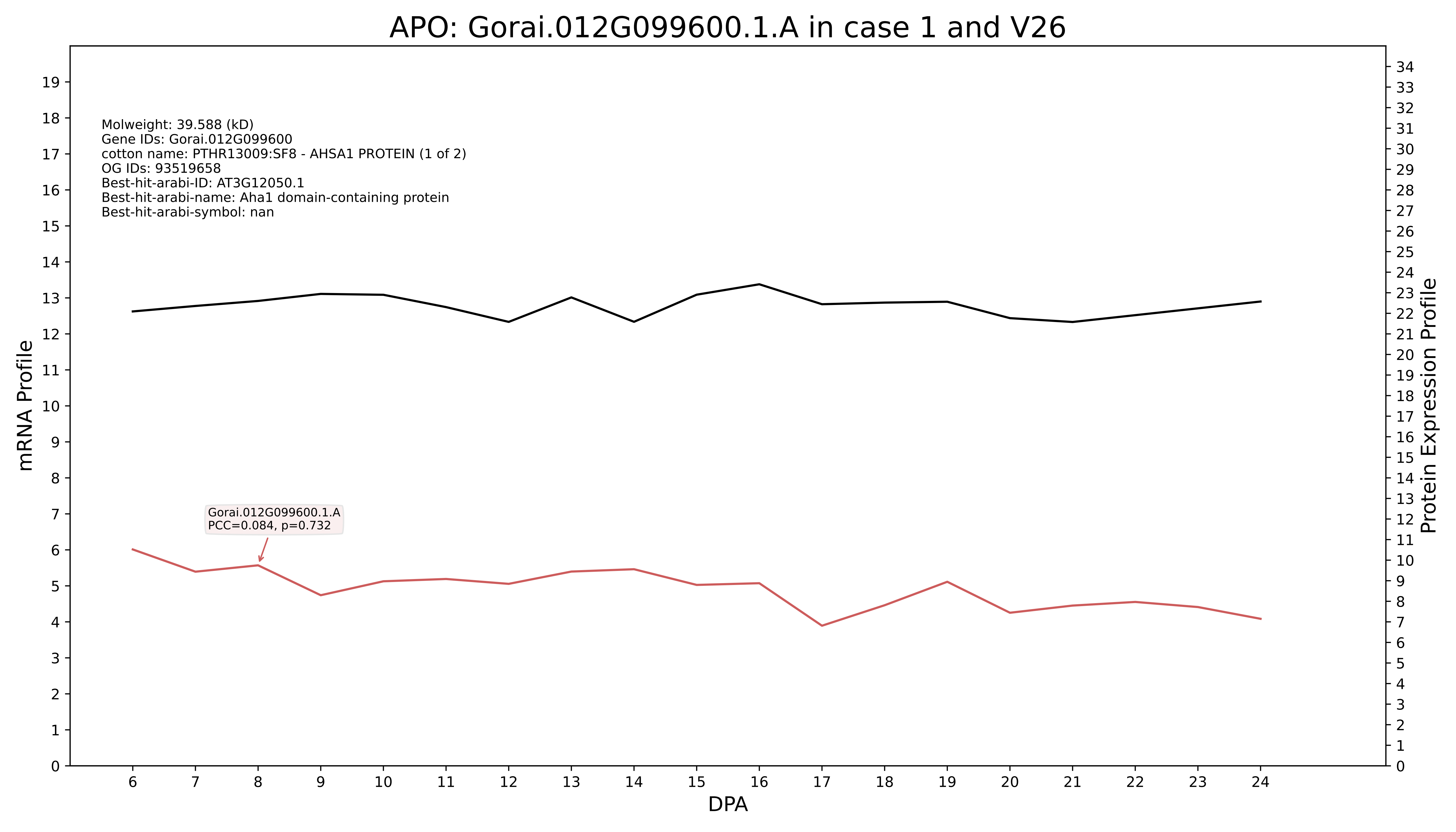
APO: Gorai.012G099600.1.A in case 1 and V26
Molweight: 39.588 (kD)
Gene IDs: Gorai.012G099600
cotton name: PTHR13009:SF8 - AHSA1 PROTEIN (1 of 2)
OG IDs: 93519658
Best-hit-arabi-ID: AT3G12050.1
Best-hit-arabi-name: Aha1 domain-containing protein
Best-hit-arabi-symbol: nan
Gorai.012G099600.1.A
PCC=0.084, p=0.732
mRNA Profile
Protein Expression Profile
DPA

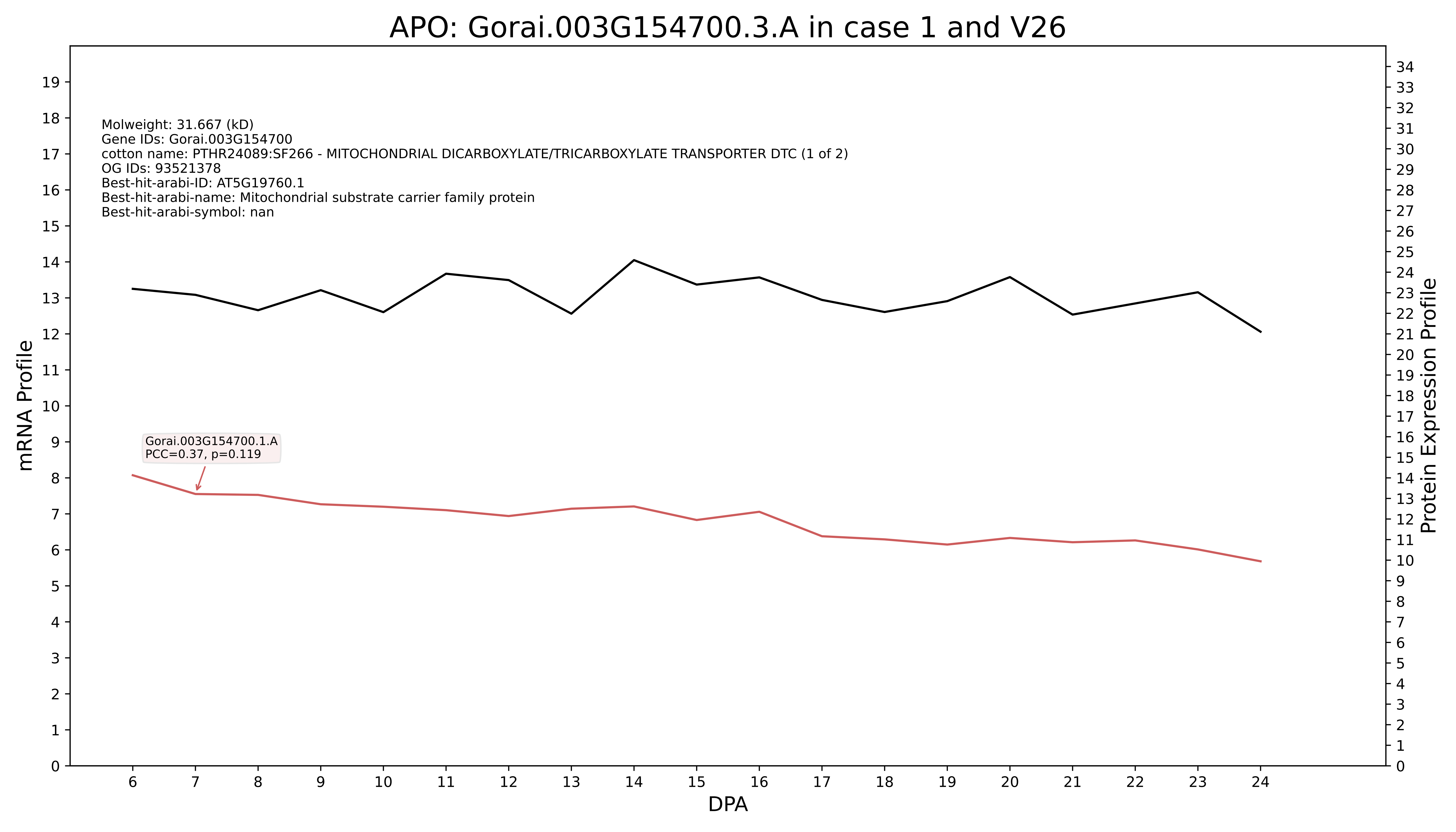
APO: Gorai.003G154700.3.A in case 1 and V26
Molweight: 31.667 (kD)
Gene IDs: Gorai.003G154700
cotton name: PTHR24089:SF266 - MITOCHONDRIAL DICARBOXYLATE/TRICARBOXYLATE TRANSPORTER DTC (1 of 2)
OG IDs: 93521378
Best-hit-arabi-ID: AT5G19760.1
Best-hit-arabi-name: Mitochondrial substrate carrier family protein
Best-hit-arabi-symbol: nan
Gorai.003G154700.1.A
PCC=0.37, p=0.119
mRNA Profile
Protein Expression Profile
DPA

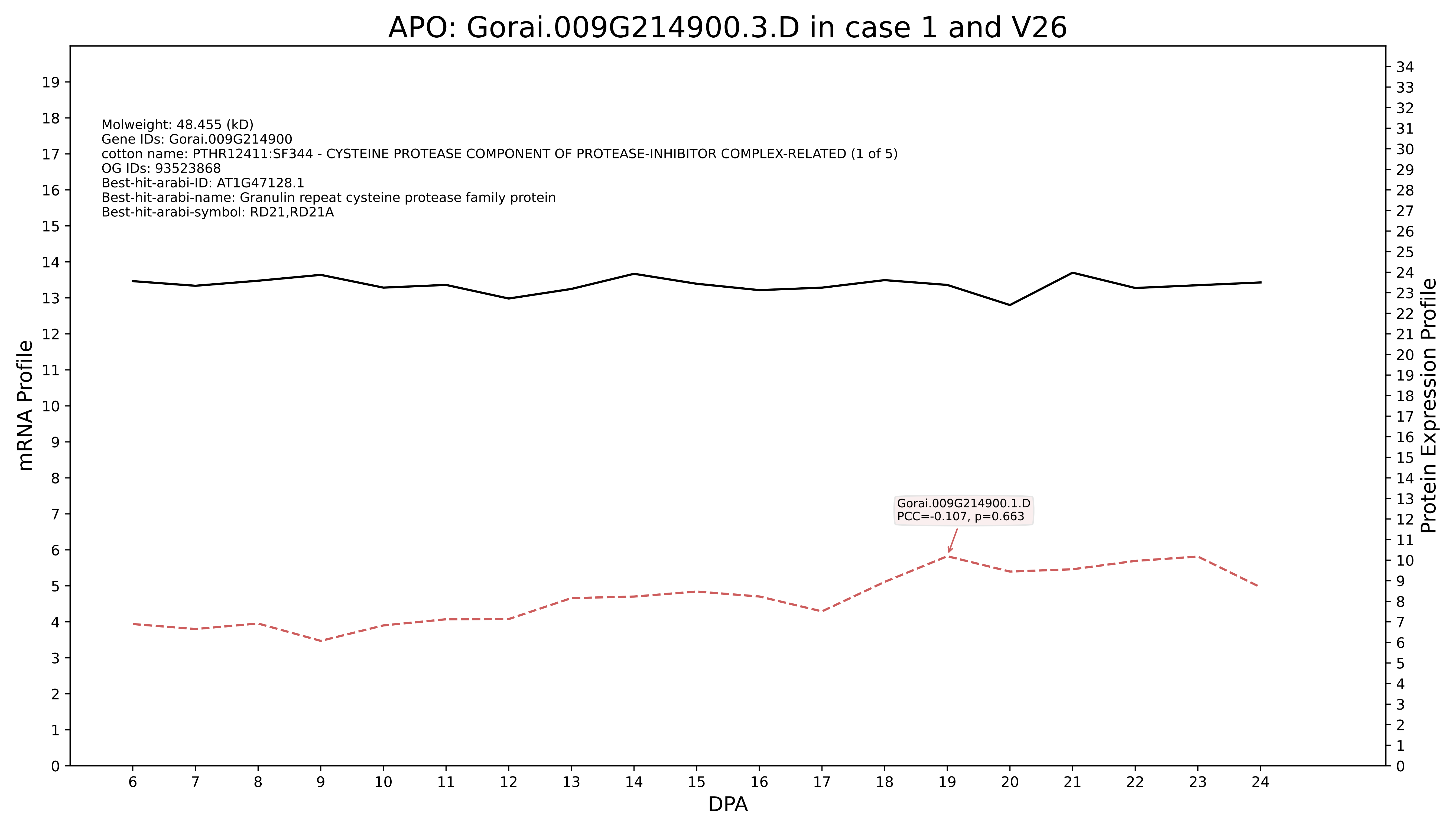
APO: Gorai.009G214900.3.D in case 1 and V26
Molweight: 48.455 (kD)
Gene IDs: Gorai.009G214900
cotton name: PTHR12411:SF344 - CYSTEINE PROTEASE COMPONENT OF PROTEASE-INHIBITOR COMPLEX-RELATED (1 of 5)
OG IDs: 93523868
Best-hit-arabi-ID: AT1G47128.1
Best-hit-arabi-name: Granulin repeat cysteine protease family protein
Best-hit-arabi-symbol: RD21,RD21A
Gorai.009G214900.1.D
PCC=-0.107, p=0.663
mRNA Profile
Protein Expression Profile
DPA

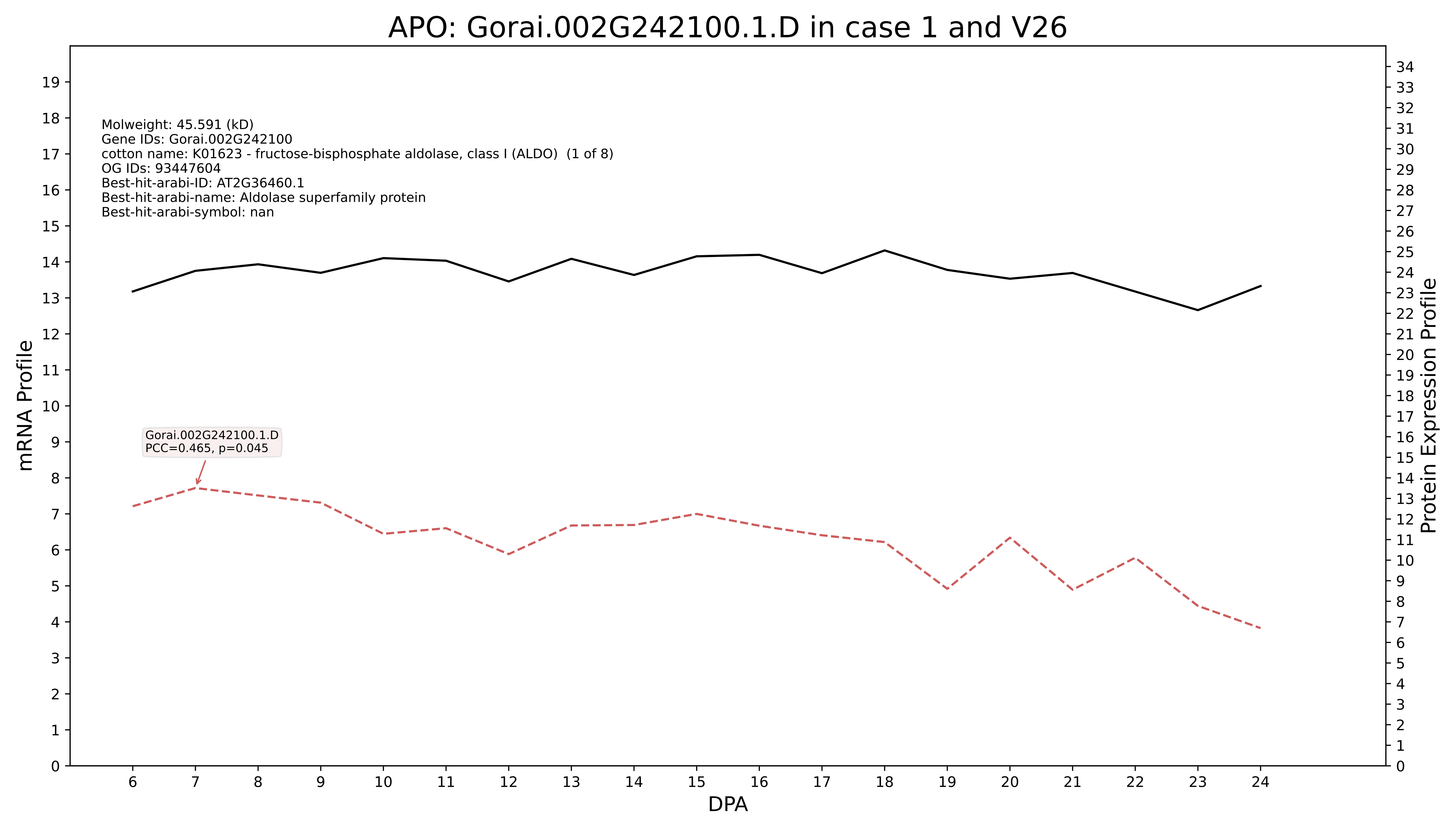

APO: Gorai.002G242100.1.D in case 1 and V26
Molweight: 45.591 (kD)
Gene IDs: Gorai.002G242100
cotton name: K01623 - fructose-bisphosphate aldolase, class I (ALDO) (1 of 8)
OG IDs: 93447604
Best-hit-arabi-ID: AT2G36460.1
Best-hit-arabi-name: Aldolase superfamily protein
Best-hit-arabi-symbol: nan
Gorai.002G242100.1.D
PCC=0.465, p=0.045
mRNA Profile
Protein Expression Profile
DPA

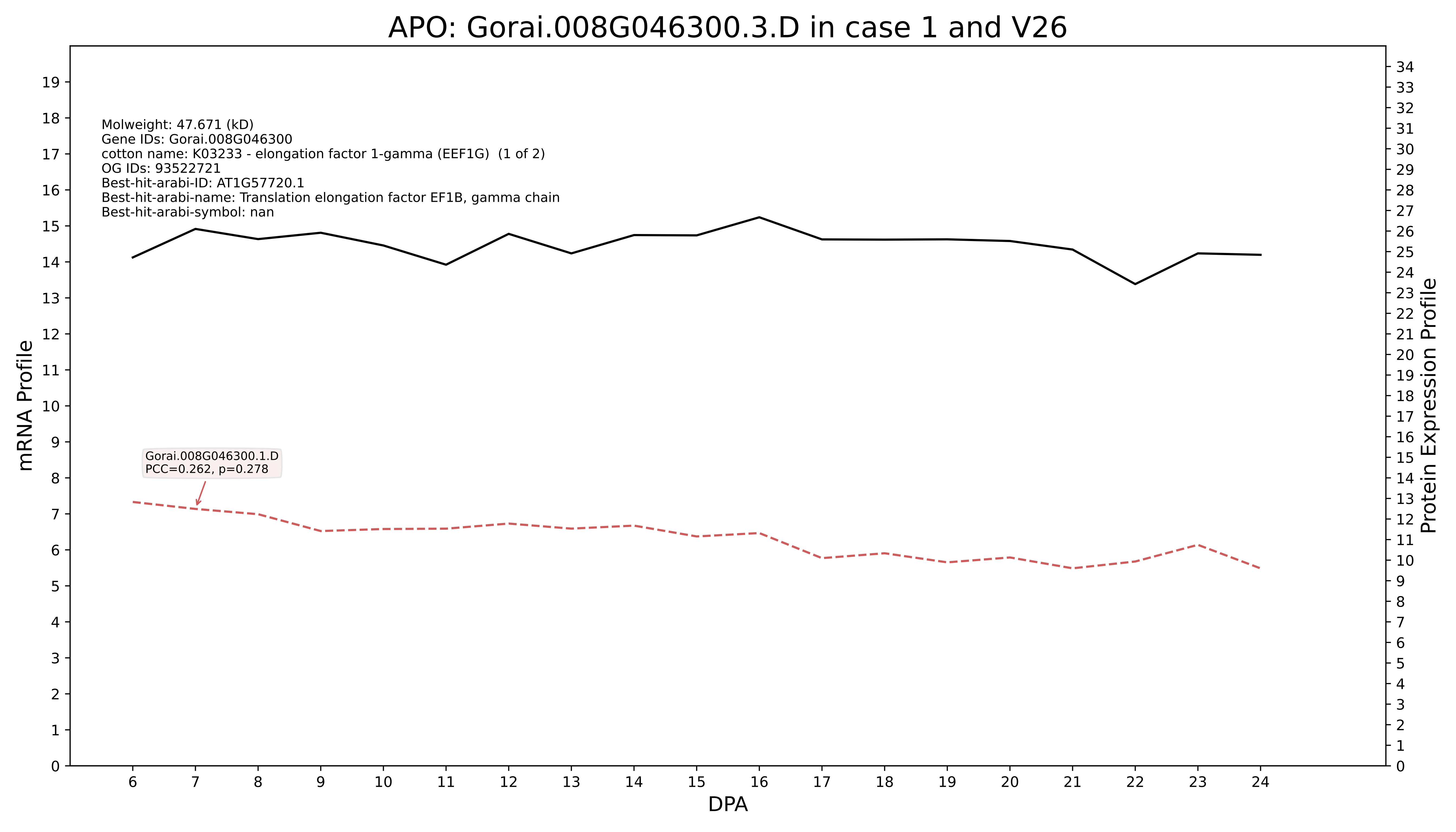

APO: Gorai.008G046300.3.D in case 1 and V26
Molweight: 47.671 (kD)
Gene IDs: Gorai.008G046300
cotton name: K03233 - elongation factor 1-gamma (EEF1G) (1 of 2)
OG IDs: 93522721
Best-hit-arabi-ID: AT1G57720.1
Best-hit-arabi-name: Translation elongation factor EF1B, gamma chain
Best-hit-arabi-symbol: nan
Gorai.008G046300.1.D
PCC=0.262, p=0.278
mRNA Profile
Protein Expression Profile
DPA

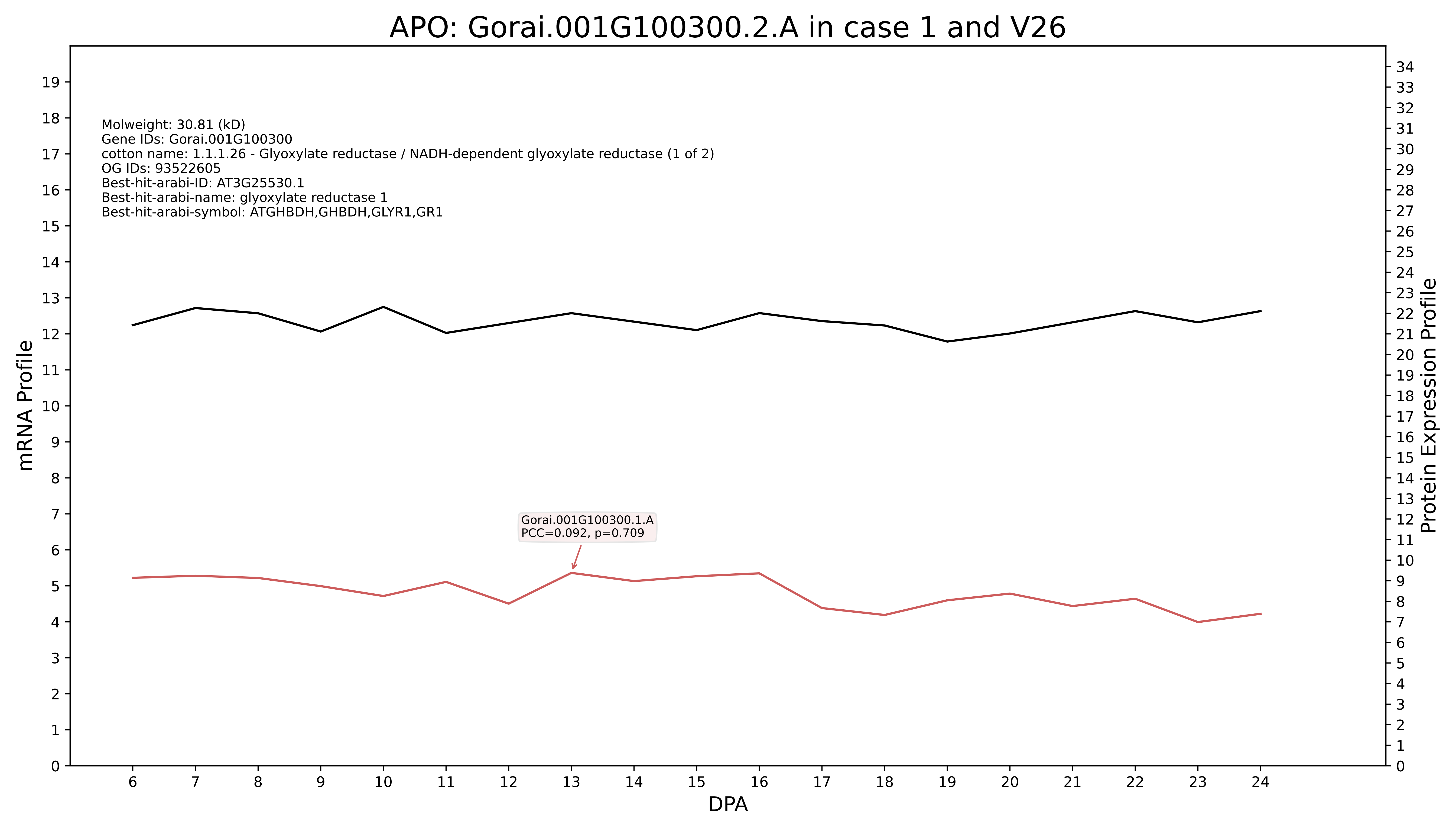
APO: Gorai.001G100300.2.A in case 1 and V26
Molweight: 30.81 (kD)
Gene IDs: Gorai.001G100300
cotton name: 1.1.1.26 - Glyoxylate reductase / NADH-dependent glyoxylate reductase (1 of 2)
OG IDs: 93522605
Best-hit-arabi-ID: AT3G25530.1
Best-hit-arabi-name: glyoxylate reductase 1
Best-hit-arabi-symbol: ATGHBDH,GHBDH,GLYR1,GR1
Gorai.001G100300.1.A
PCC=0.092, p=0.709
mRNA Profile
Protein Expression Profile
DPA

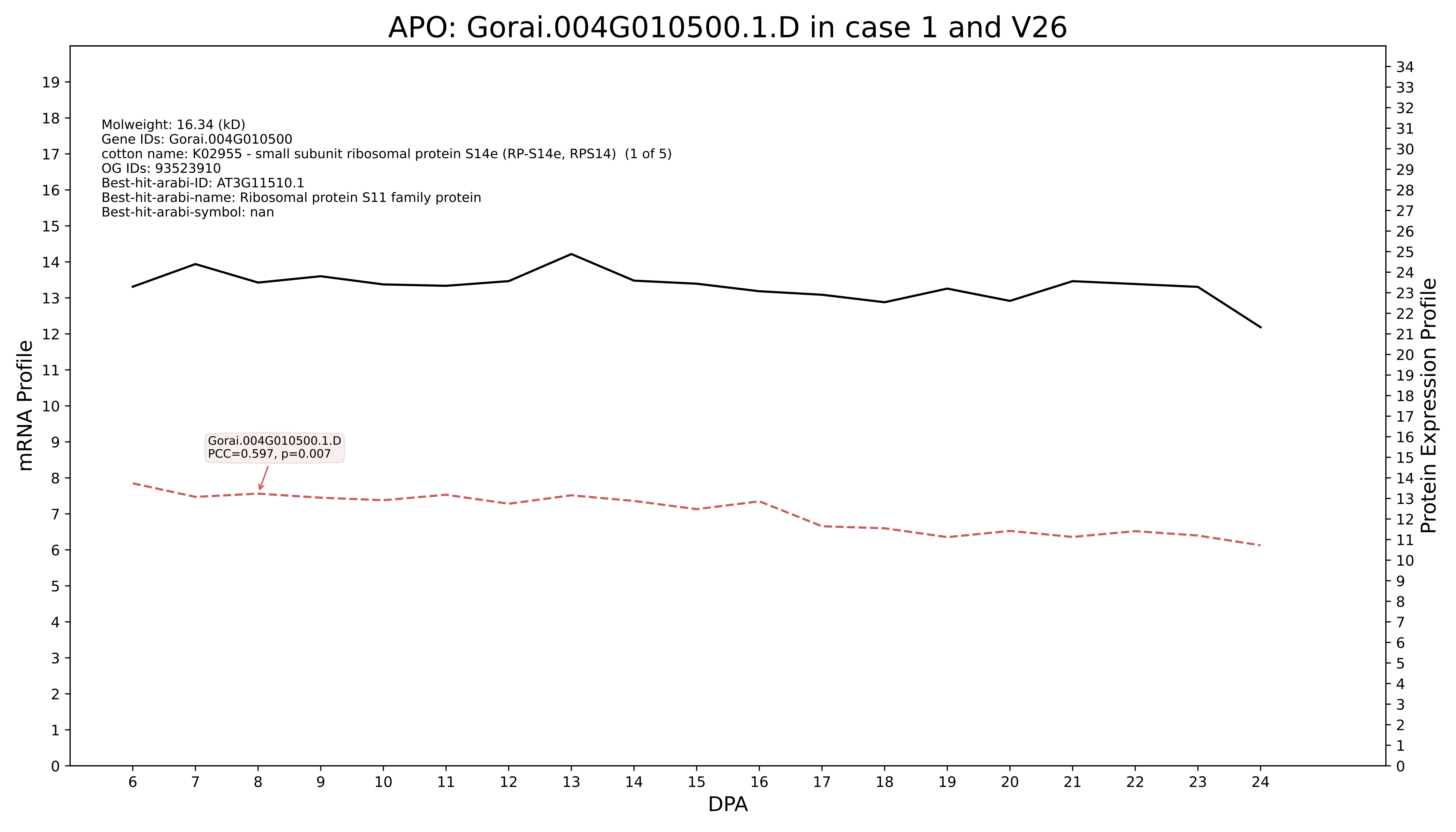

APO: Gorai.004G010500.1.D in case 1 and V26
Molweight: 16.34 (kD)
Gene IDs: Gorai.004G010500
cotton name: K02955 - small subunit ribosomal protein S14e (RP-S14e, RPS14) (1 of 5)
OG IDs: 93523910
Best-hit-arabi-ID: AT3G11510.1
Best-hit-arabi-name: Ribosomal protein S11 family protein
Best-hit-arabi-symbol: nan
Gorai.004G010500.1.D
PCC=0.597, p=0.007
mRNA Profile
Protein Expression Profile
DPA

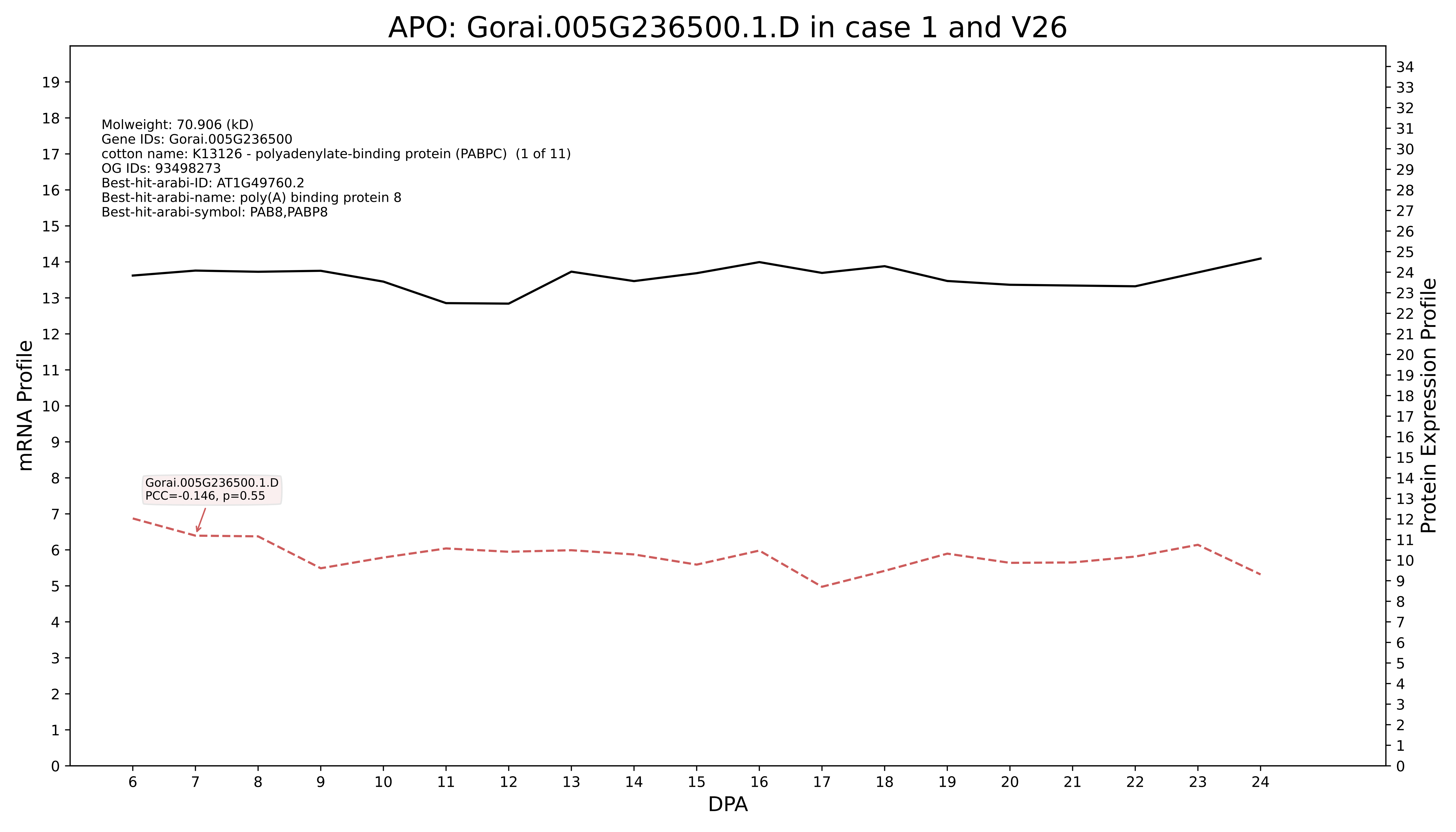

APO: Gorai.005G236500.1.D in case 1 and V26
Molweight: 70.906 (kD)
Gene IDs: Gorai.005G236500
cotton name: K13126 - polyadenylate-binding protein (PABPC) (1 of 11)
OG IDs: 93498273
Best-hit-arabi-ID: AT1G49760.2
Best-hit-arabi-name: poly(A) binding protein 8
Best-hit-arabi-symbol: PAB8,PABP8
Gorai.005G236500.1.D
PCC=-0.146, p=0.55
mRNA Profile
Protein Expression Profile
DPA

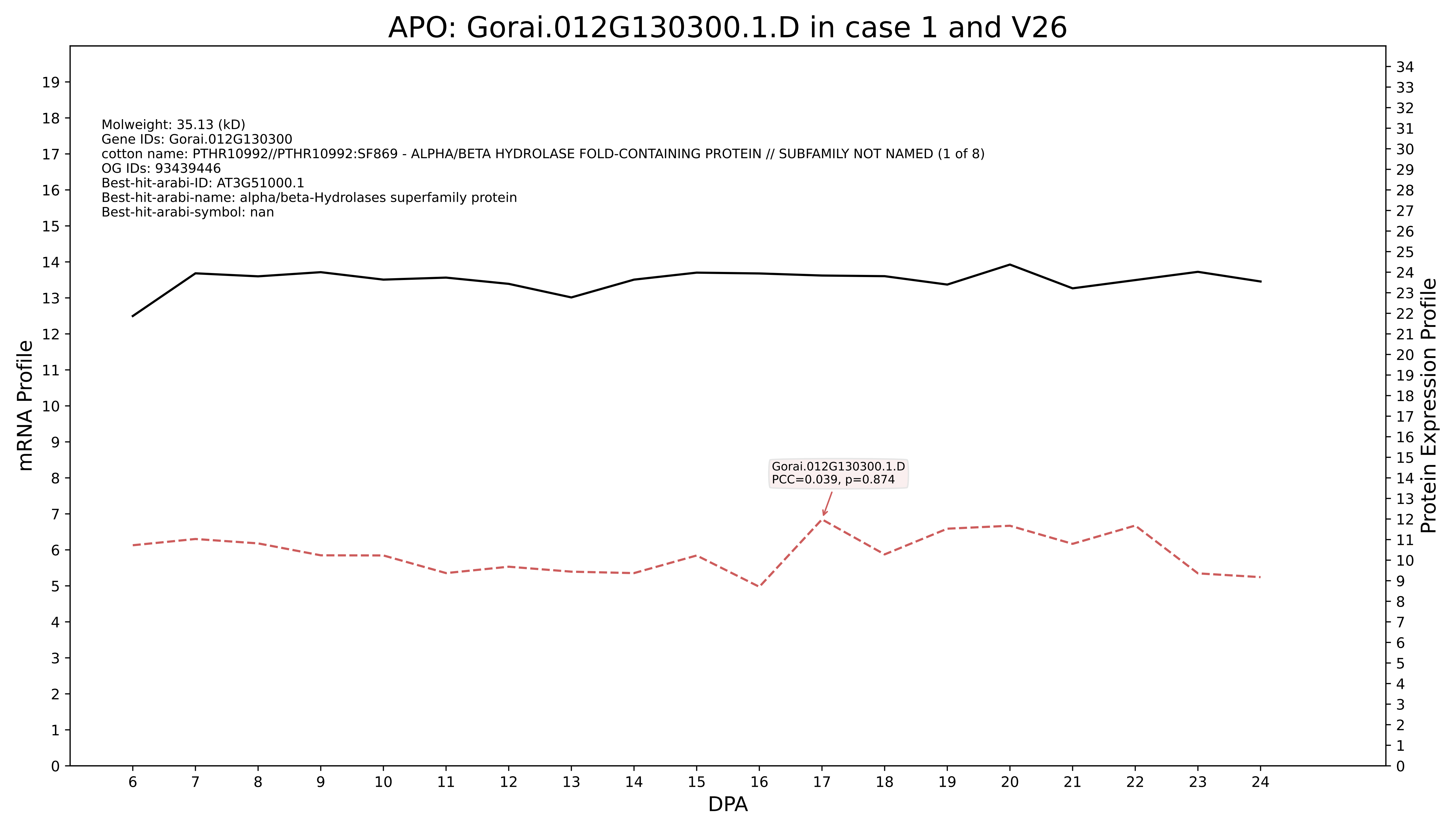
APO: Gorai.012G130300.1.D in case 1 and V26
Molweight: 35.13 (kD)
Gene IDs: Gorai.012G130300
cotton name: PTHR10992//PTHR10992:SF869 - ALPHA/BETA HYDROLASE FOLD-CONTAINING PROTEIN // SUBFAMILY NOT NAMED (1 of 8)
OG IDs: 93439446
Best-hit-arabi-ID: AT3G51000.1
Best-hit-arabi-name: alpha/beta-Hydrolases superfamily protein
Best-hit-arabi-symbol: nan
Gorai.012G130300.1.D
PCC=0.039, p=0.874
mRNA Profile
Protein Expression Profile
DPA

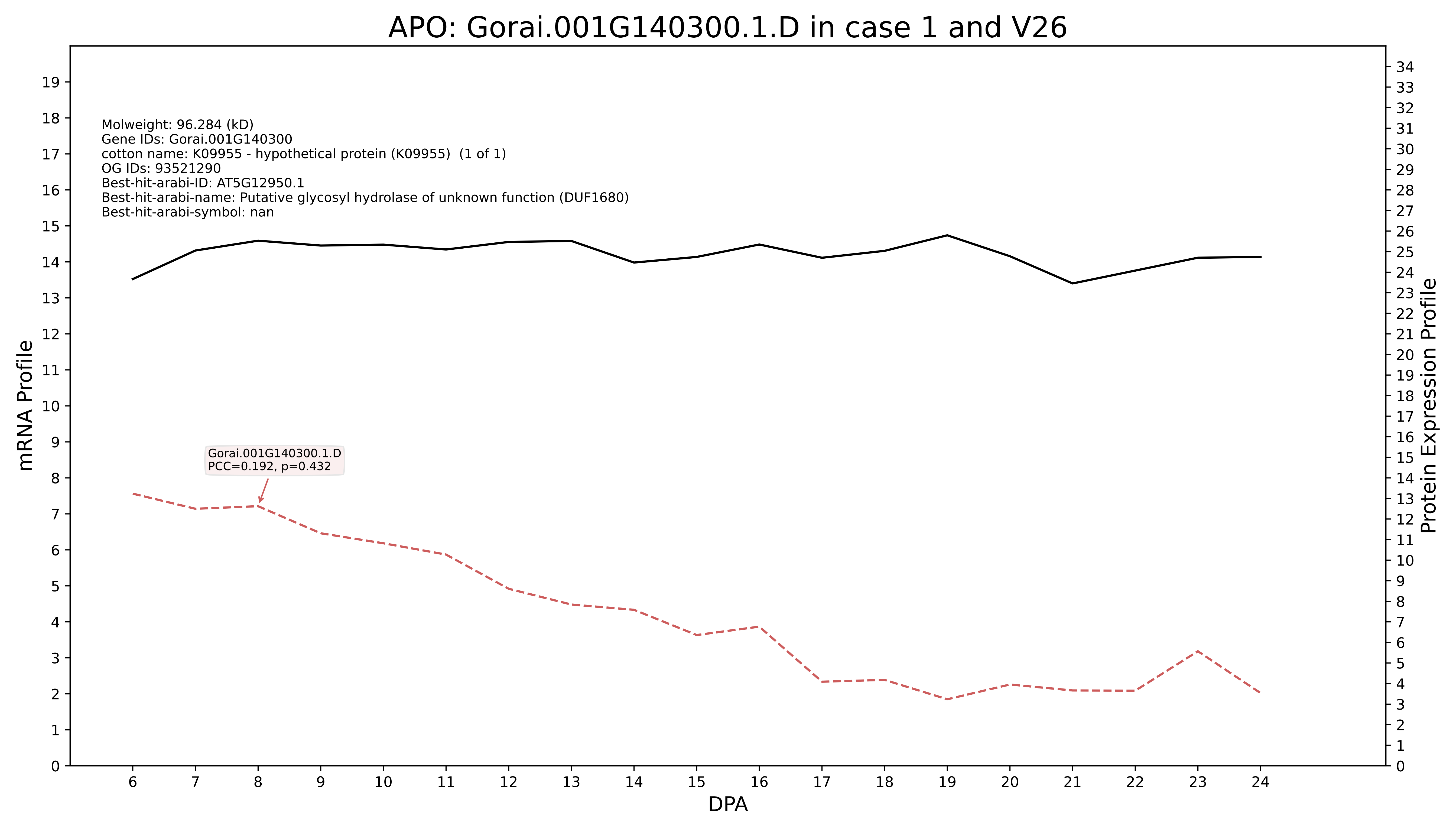
APO: Gorai.001G140300.1.D in case 1 and V26
Molweight: 96.284 (kD)
Gene IDs: Gorai.001G140300
cotton name: K09955 - hypothetical protein (K09955) (1 of 1)
OG IDs: 93521290
Best-hit-arabi-ID: AT5G12950.1
Best-hit-arabi-name: Putative glycosyl hydrolase of unknown function (DUF1680)
Best-hit-arabi-symbol: nan
Gorai.001G140300.1.D
PCC=0.192, p=0.432
mRNA Profile
Protein Expression Profile
DPA

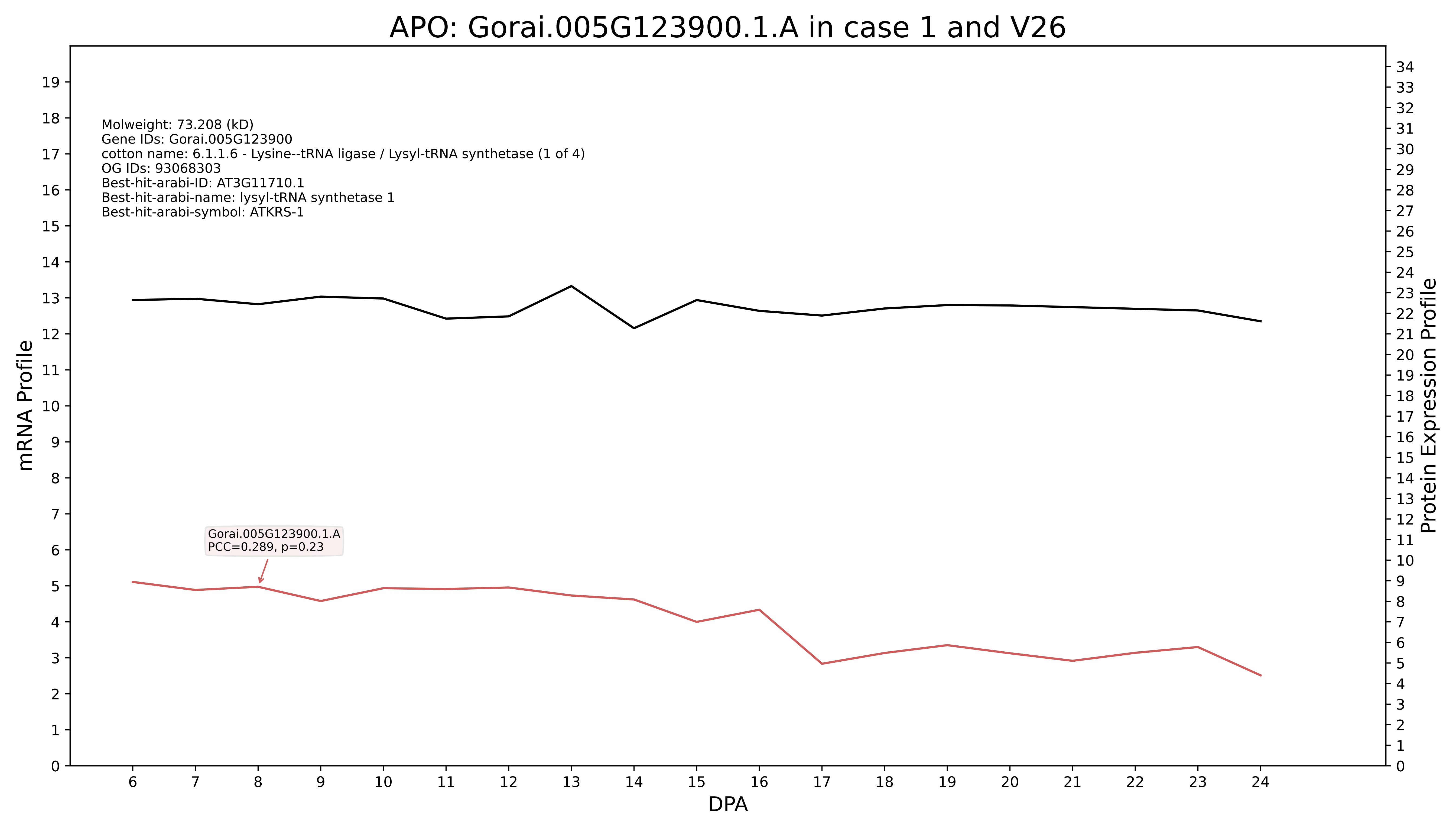
APO: Gorai.005G123900.1.A in case 1 and V26
Molweight: 73.208 (kD)
Gene IDs: Gorai.005G123900
cotton name: 6.1.1.6 - Lysine--tRNA ligase / Lysyl-tRNA synthetase (1 of 4)
OG IDs: 93068303
Best-hit-arabi-ID: AT3G11710.1
Best-hit-arabi-name: lysyl-tRNA synthetase 1
Best-hit-arabi-symbol: ATKRS-1
Gorai.005G123900.1.A
PCC=0.289, p=0.23
mRNA Profile
Protein Expression Profile
DPA

#### APO: Gorai.010G161100.1.D in case 1 and V26

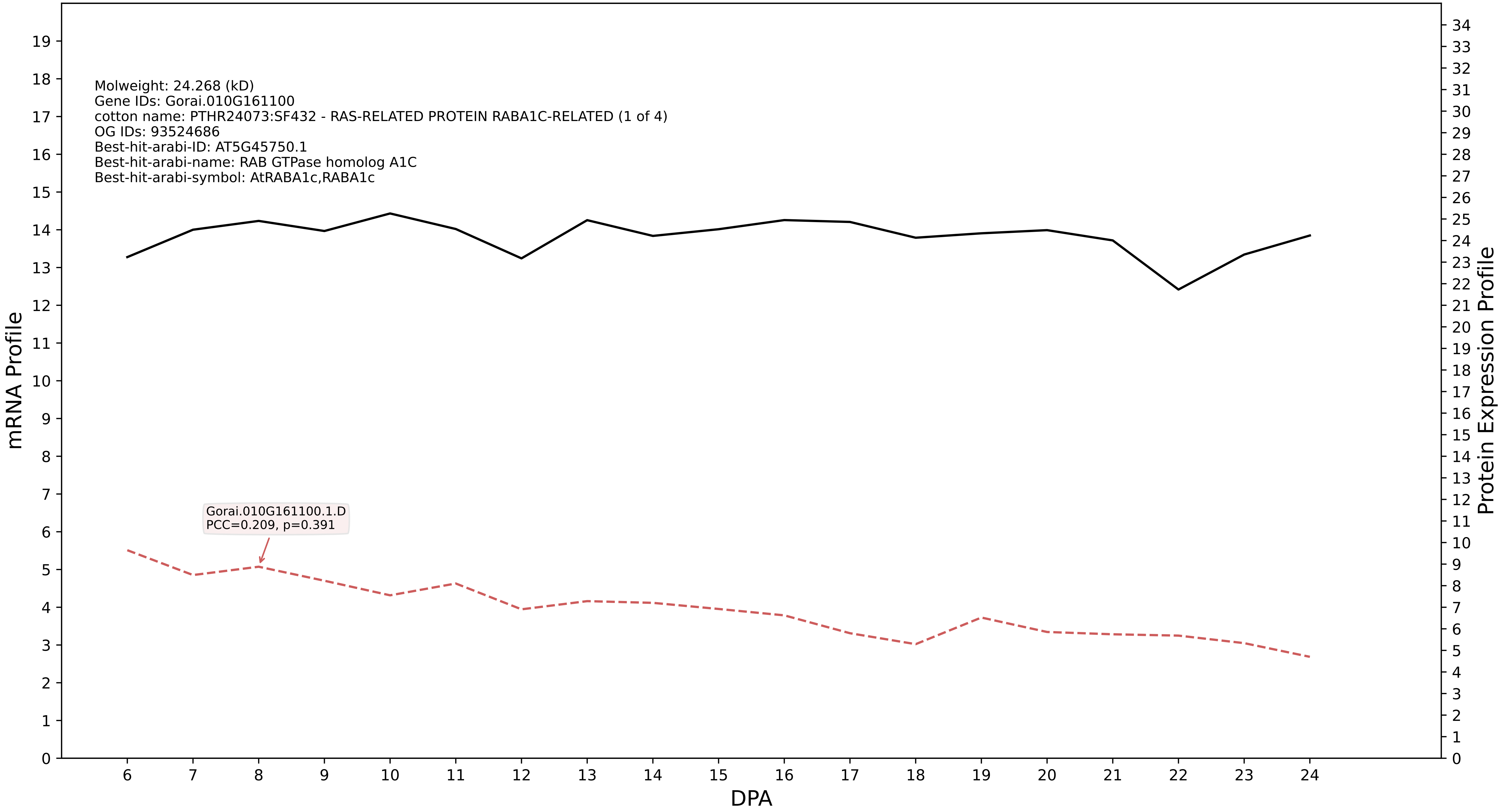

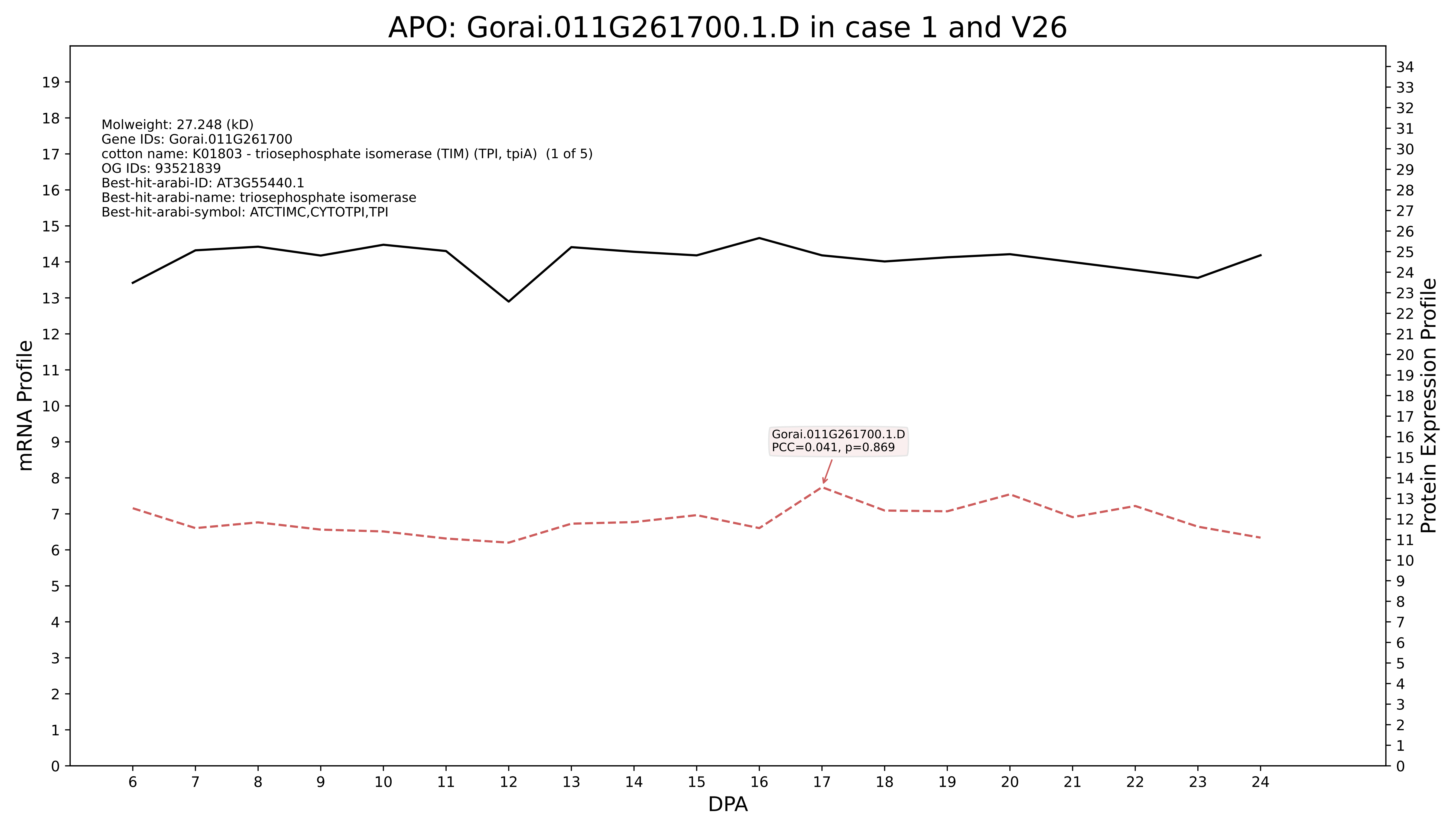
APO: Gorai.011G261700.1.D in case 1 and V26
Molweight: 27.248 (kD)
Gene IDs: Gorai.011G261700
cotton name: K01803 - triosephosphate isomerase (TIM) (TPI, tpiA) (1 of 5)
OG IDs: 93521839
Best-hit-arabi-ID: AT3G55440.1
Best-hit-arabi-name: triosephosphate isomerase
Best-hit-arabi-symbol: ATCTIMC,CYTOTPI,TPI
Gorai.011G261700.1.D
PCC=0.041, p=0.869
mRNA Profile
Protein Expression Profile
DPA

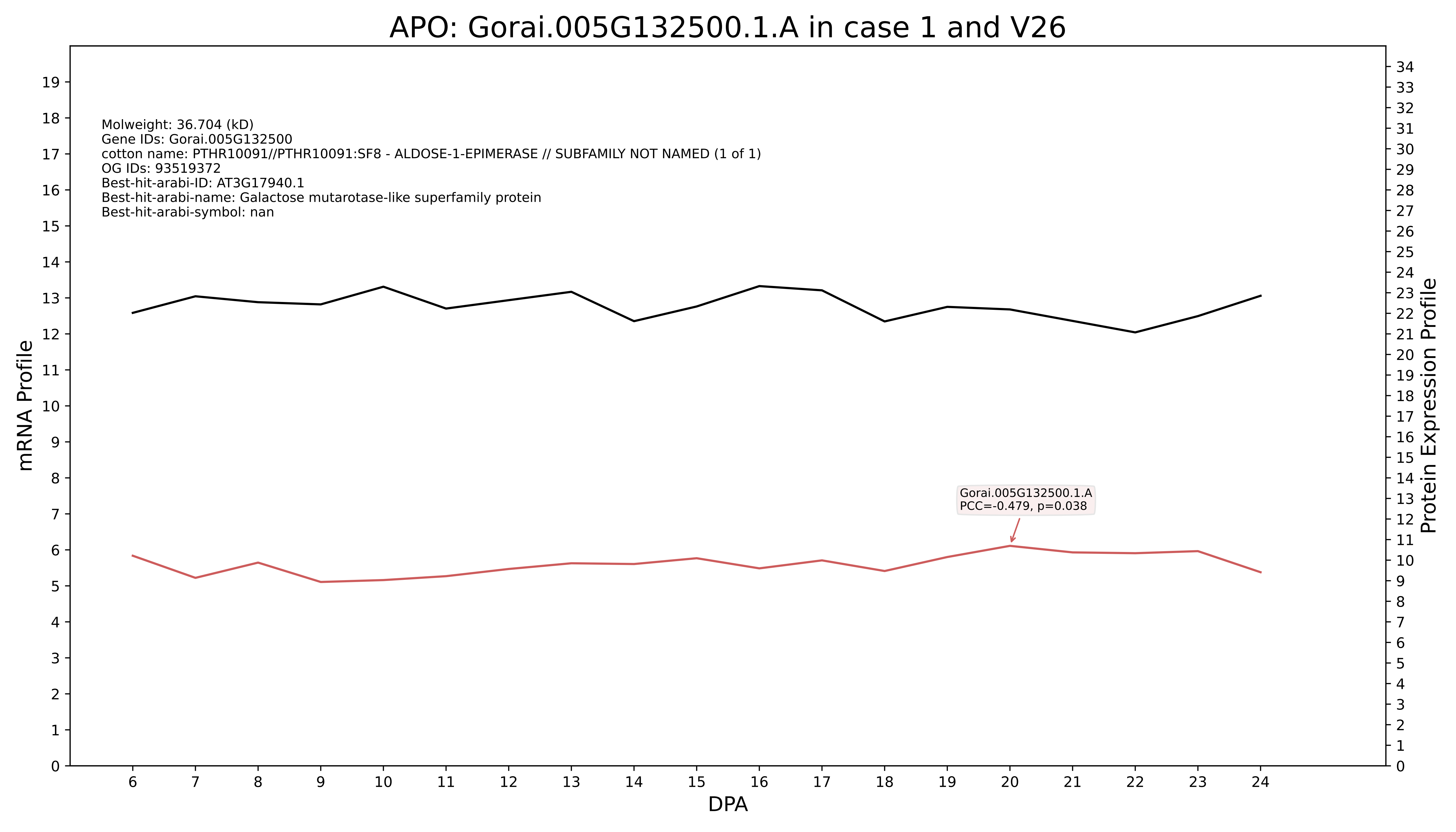

APO: Gorai.005G132500.1.A in case 1 and V26
Molweight: 36.704 (kD)
Gene IDs: Gorai.005G132500
cotton name: PTHR10091//PTHR10091:SF8 - ALDOSE-1-EPIMERASE // SUBFAMILY NOT NAMED (1 of 1)
OG IDs: 93519372
Best-hit-arabi-ID: AT3G17940.1
Best-hit-arabi-name: Galactose mutarotase-like superfamily protein
Best-hit-arabi-symbol: nan
Gorai.005G132500.1.A
PCC=-0.479, p=0.038
mRNA Profile
Protein Expression Profile
DPA

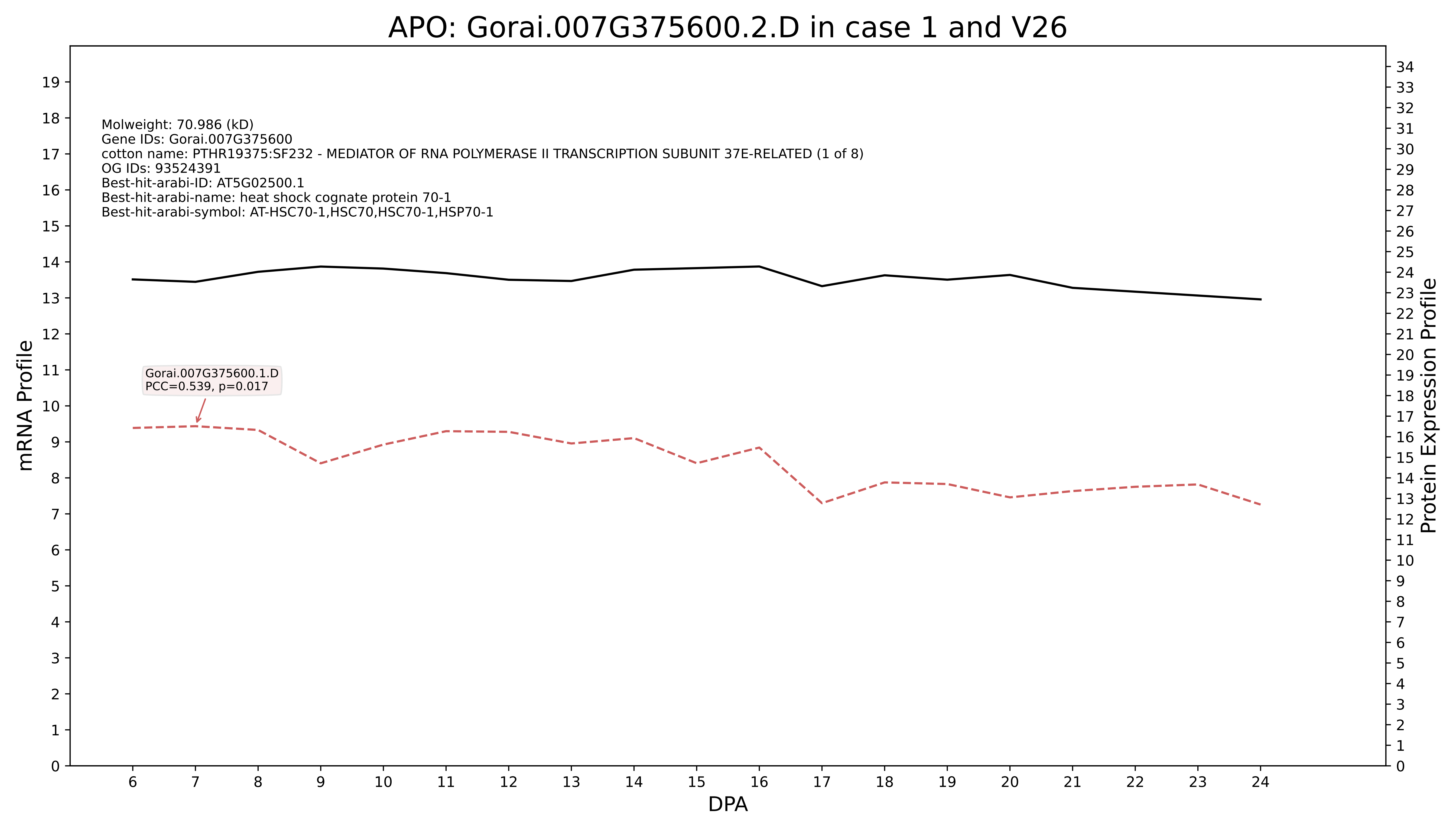
APO: Gorai.007G375600.2.D in case 1 and V26
Molweight: 70.986 (kD)
Gene IDs: Gorai.007G375600
cotton name: PTHR19375:SF232 - MEDIATOR OF RNA POLYMERASE II TRANSCRIPTION SUBUNIT 37E-RELATED (1 of 8)
OG IDs: 93524391
Best-hit-arabi-ID: AT5G02500.1
Best-hit-arabi-name: heat shock cognate protein 70-1
Best-hit-arabi-symbol: AT-HSC70-1,HSC70,HSC70-1,HSP70-1
Gorai.007G375600.1.D
PCC=0.539, p=0.017
mRNA Profile
Protein Expression Profile
DPA

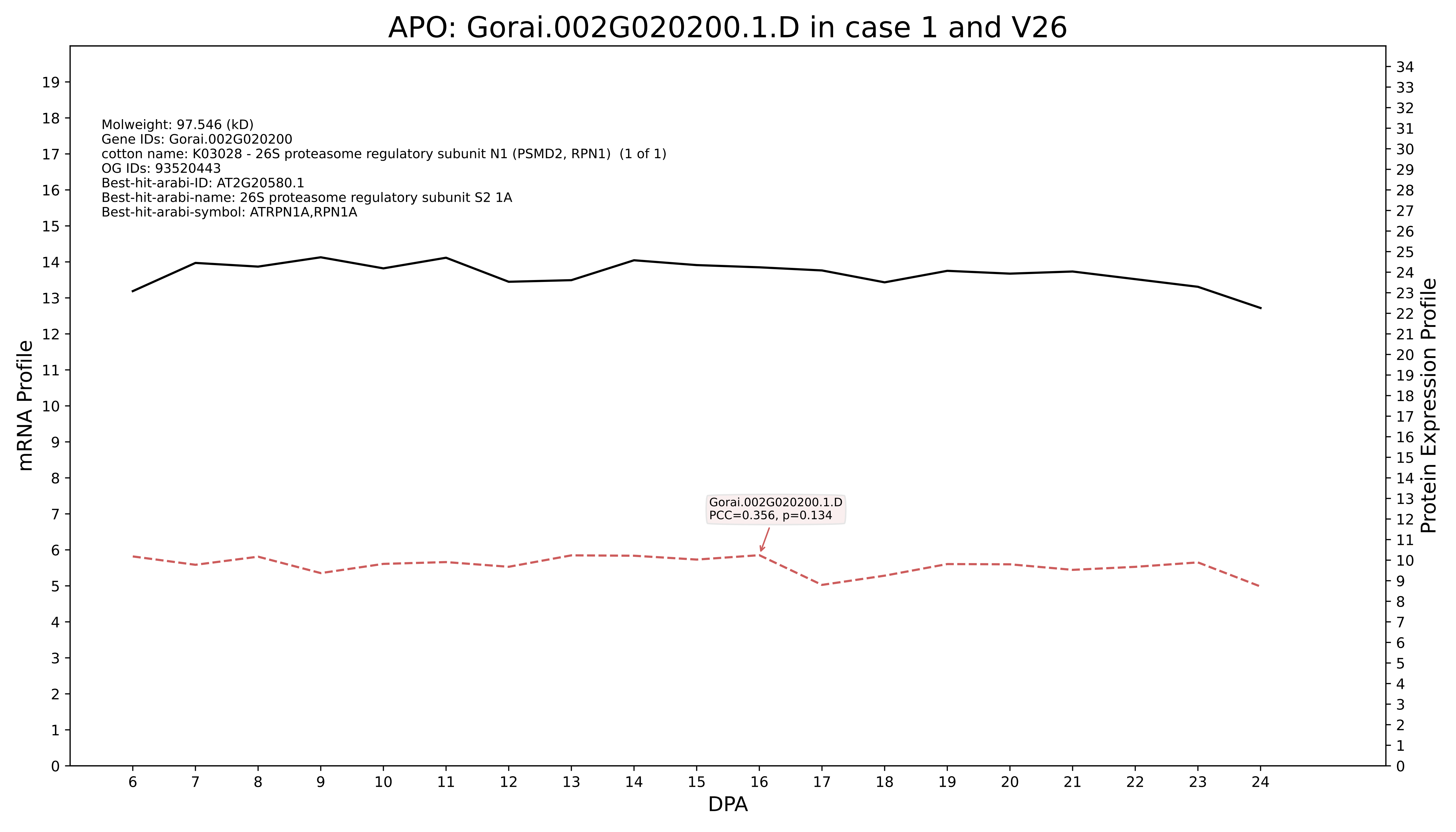
APO: Gorai.002G020200.1.D in case 1 and V26
Molweight: 97.546 (kD)
Gene IDs: Gorai.002G020200
cotton name: K03028 - 26S proteasome regulatory subunit N1 (PSMD2, RPN1) (1 of 1)
OG IDs: 93520443
Best-hit-arabi-ID: AT2G20580.1
Best-hit-arabi-name: 26S proteasome regulatory subunit S2 1A
Best-hit-arabi-symbol: ATRPN1A,RPN1A
Gorai.002G020200.1.D
PCC=0.356, p=0.134
mRNA Profile
Protein Expression Profile
DPA

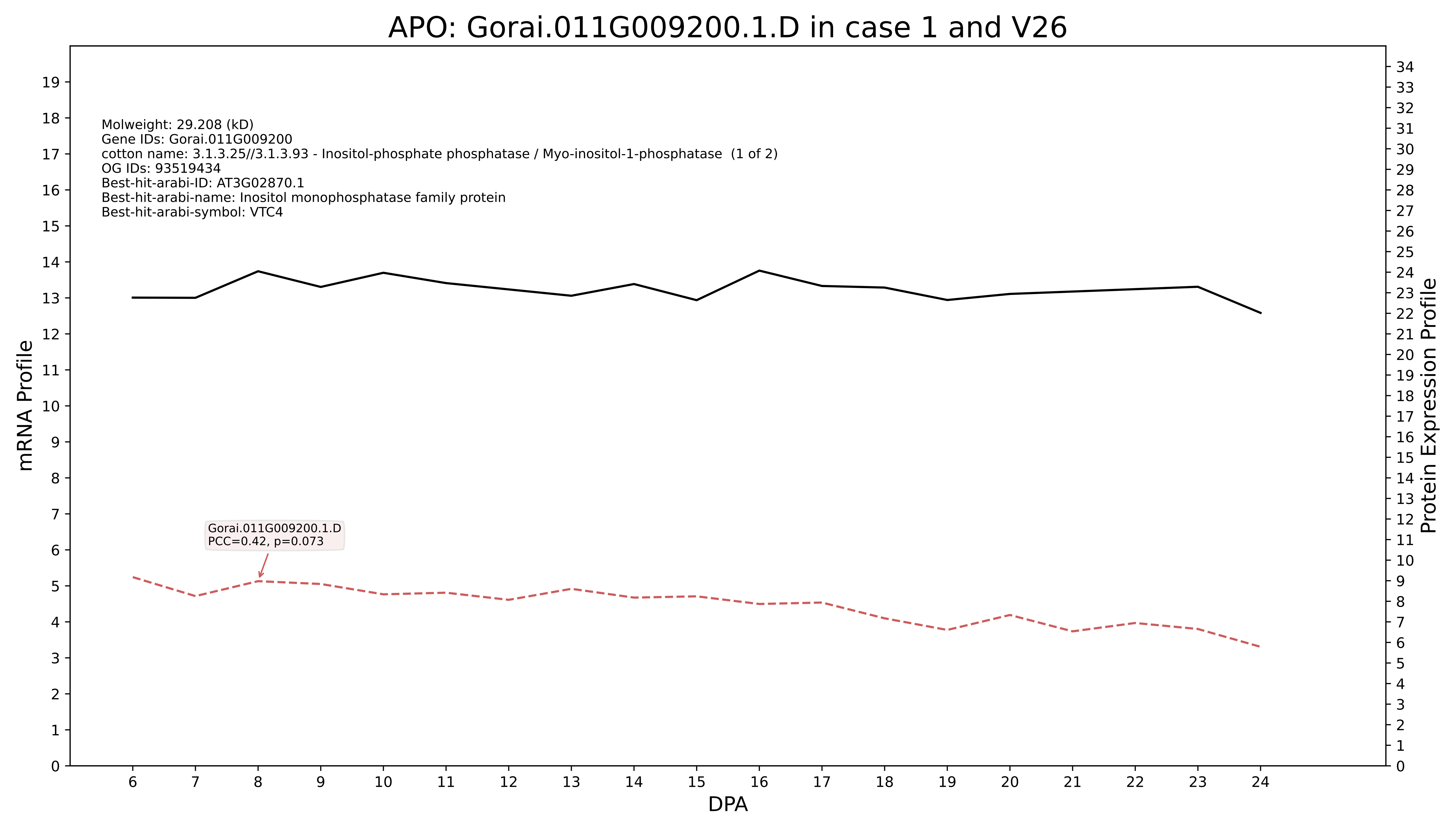
APO: Gorai.011G009200.1.D in case 1 and V26
Molweight: 29.208 (kD)
Gene IDs: Gorai.011G009200
cotton name: 3.1.3.25//3.1.3.93 - Inositol-phosphate phosphatase / Myo-inositol-1-phosphatase (1 of 2)
OG IDs: 93519434
Best-hit-arabi-ID: AT3G02870.1
Best-hit-arabi-name: Inositol monophosphatase family protein
Best-hit-arabi-symbol: VTC4
Gorai.011G009200.1.D
PCC=0.42, p=0.073
mRNA Profile
Protein Expression Profile
DPA

### APO: Gorai.006G214200.2.D in case 1 and V26

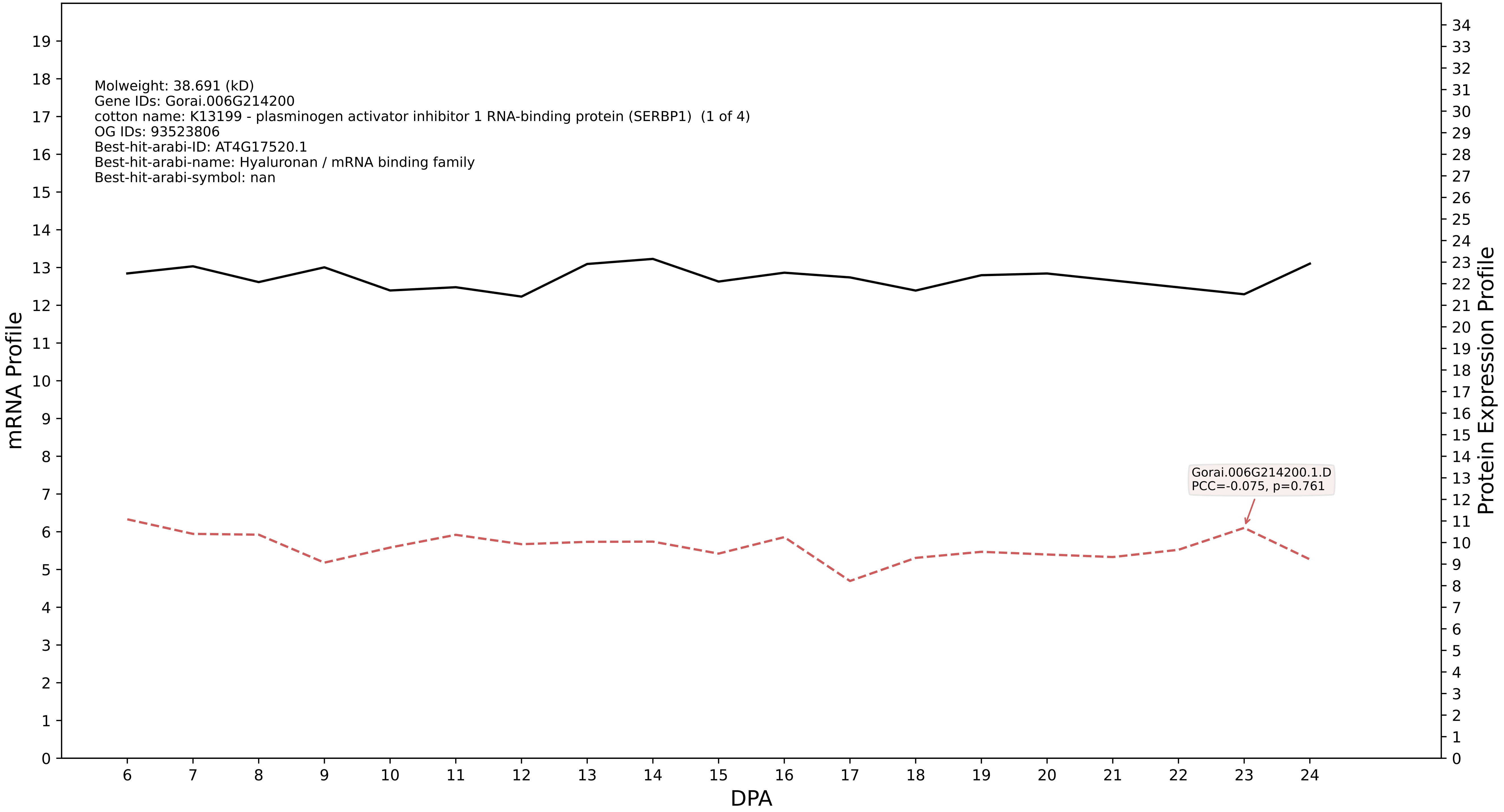

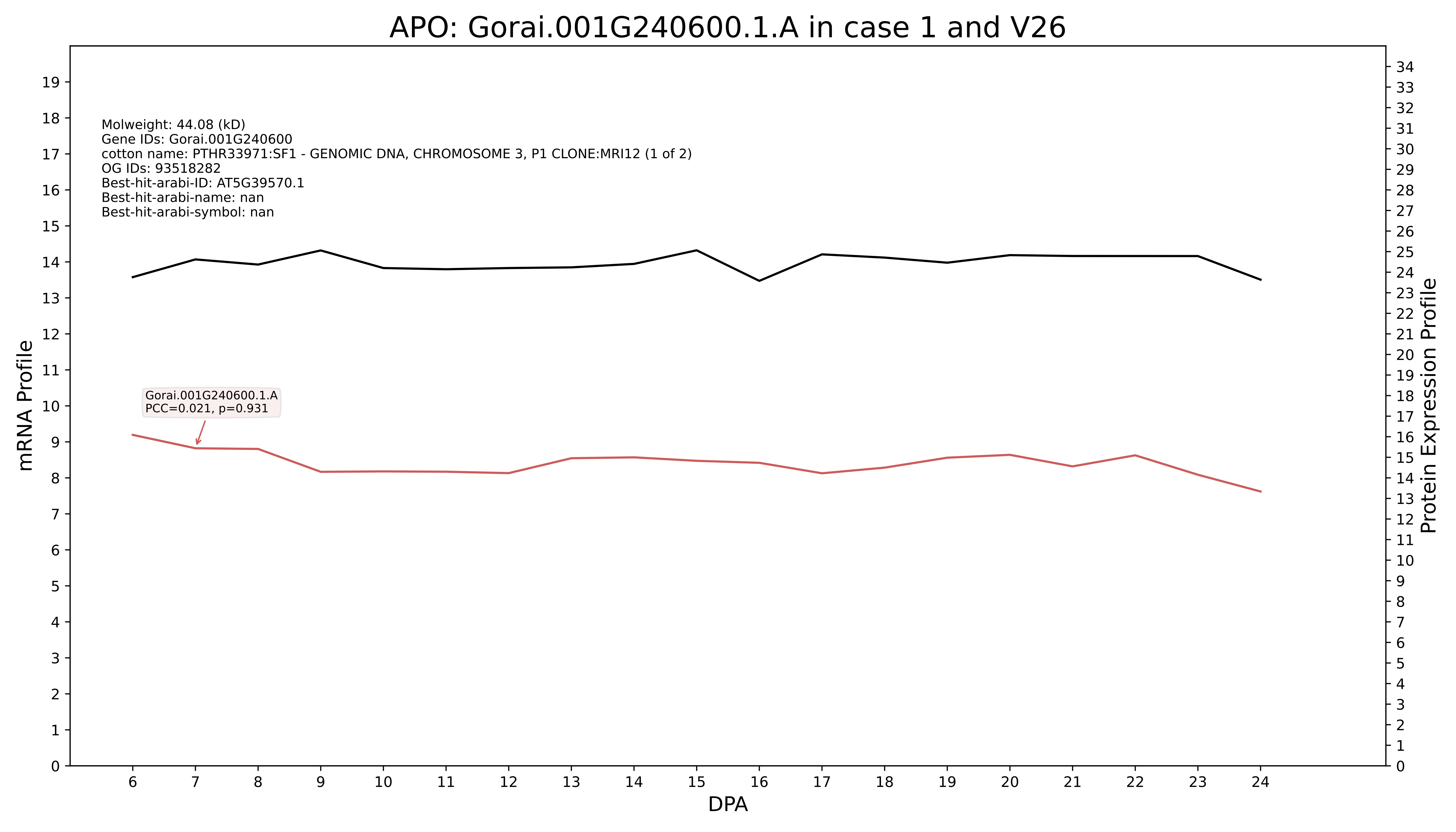
APO: Gorai.001G240600.1.A in case 1 and V26
Molweight: 44.08 (kD)
Gene IDs: Gorai.001G240600
cotton name: PTHR33971:SF1 - GENOMIC DNA, CHROMOSOME 3, P1 CLONE:MRI12 (1 of 2)
OG IDs: 93518282
Best-hit-arabi-ID: AT5G39570.1
Best-hit-arabi-name: nan
Best-hit-arabi-symbol: nan
Gorai.001G240600.1.A
PCC=0.021, p=0.931
mRNA Profile
Protein Expression Profile
DPA

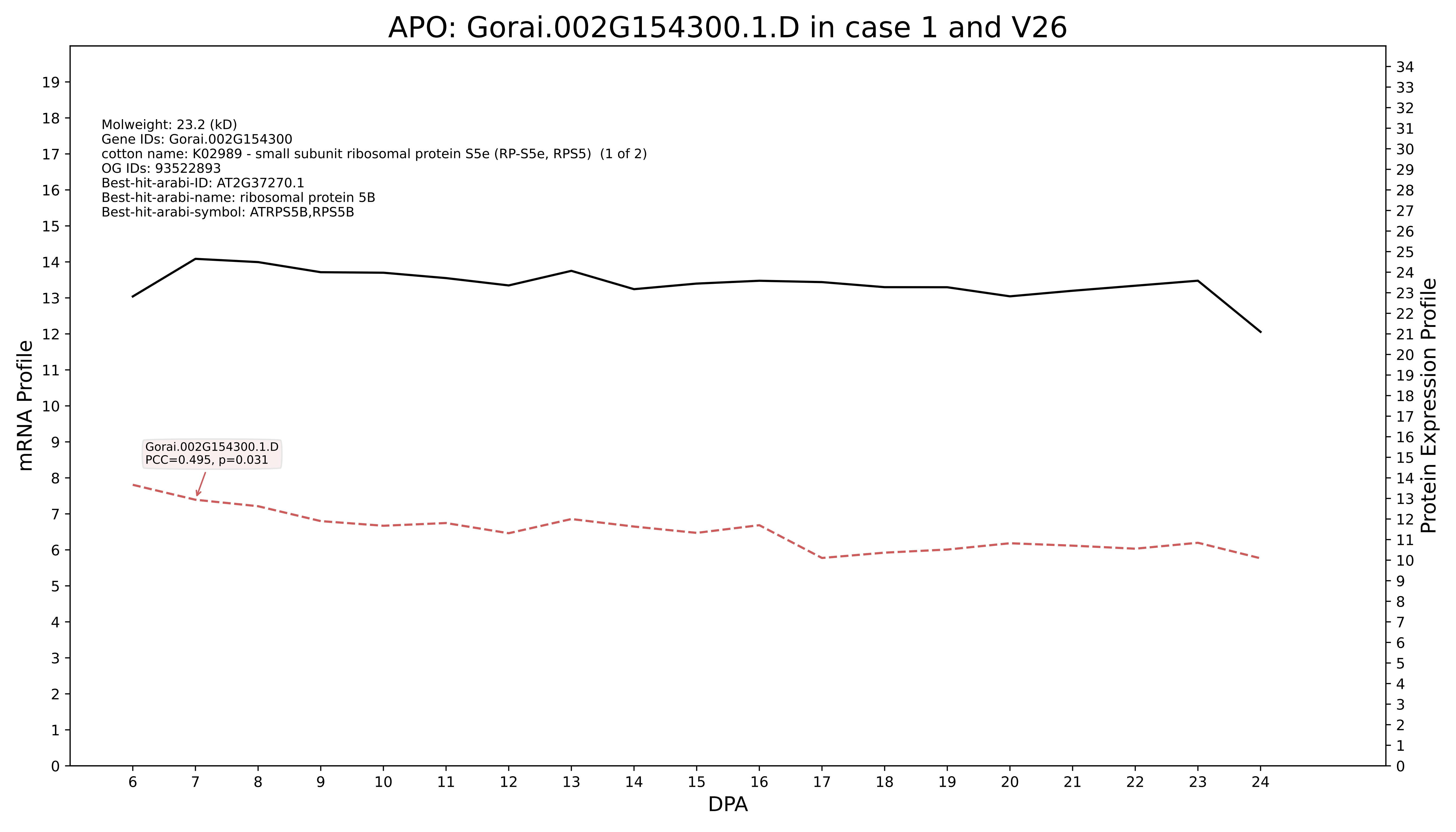
APO: Gorai.002G154300.1.D in case 1 and V26
Molweight: 23.2 (kD)
Gene IDs: Gorai.002G154300
cotton name: K02989 - small subunit ribosomal protein S5e (RP-S5e, RPS5) (1 of 2)
OG IDs: 93522893
Best-hit-arabi-ID: AT2G37270.1
Best-hit-arabi-name: ribosomal protein 5B
Best-hit-arabi-symbol: ATRPS5B,RPS5B
Gorai.002G154300.1.D
PCC=0.495, p=0.031
mRNA Profile
Protein Expression Profile
DPA

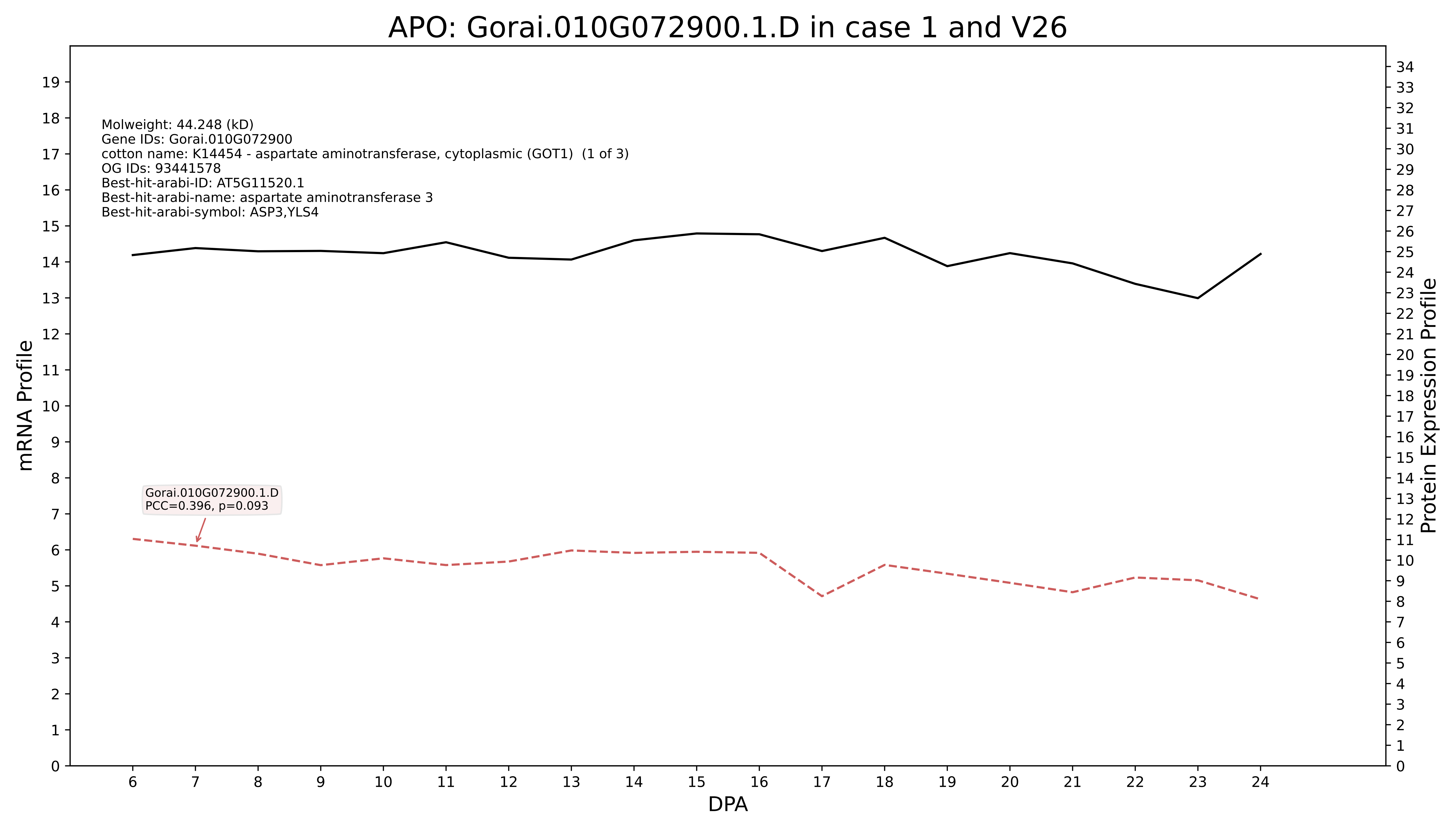
APO: Gorai.010G072900.1.D in case 1 and V26
Molweight: 44.248 (kD)
Gene IDs: Gorai.010G072900
cotton name: K14454 - aspartate aminotransferase, cytoplasmic (GOT1) (1 of 3)
OG IDs: 93441578
Best-hit-arabi-ID: AT5G11520.1
Best-hit-arabi-name: aspartate aminotransferase 3
Best-hit-arabi-symbol: ASP3,YLS4
Gorai.010G072900.1.D
PCC=0.396, p=0.093
mRNA Profile
Protein Expression Profile
DPA

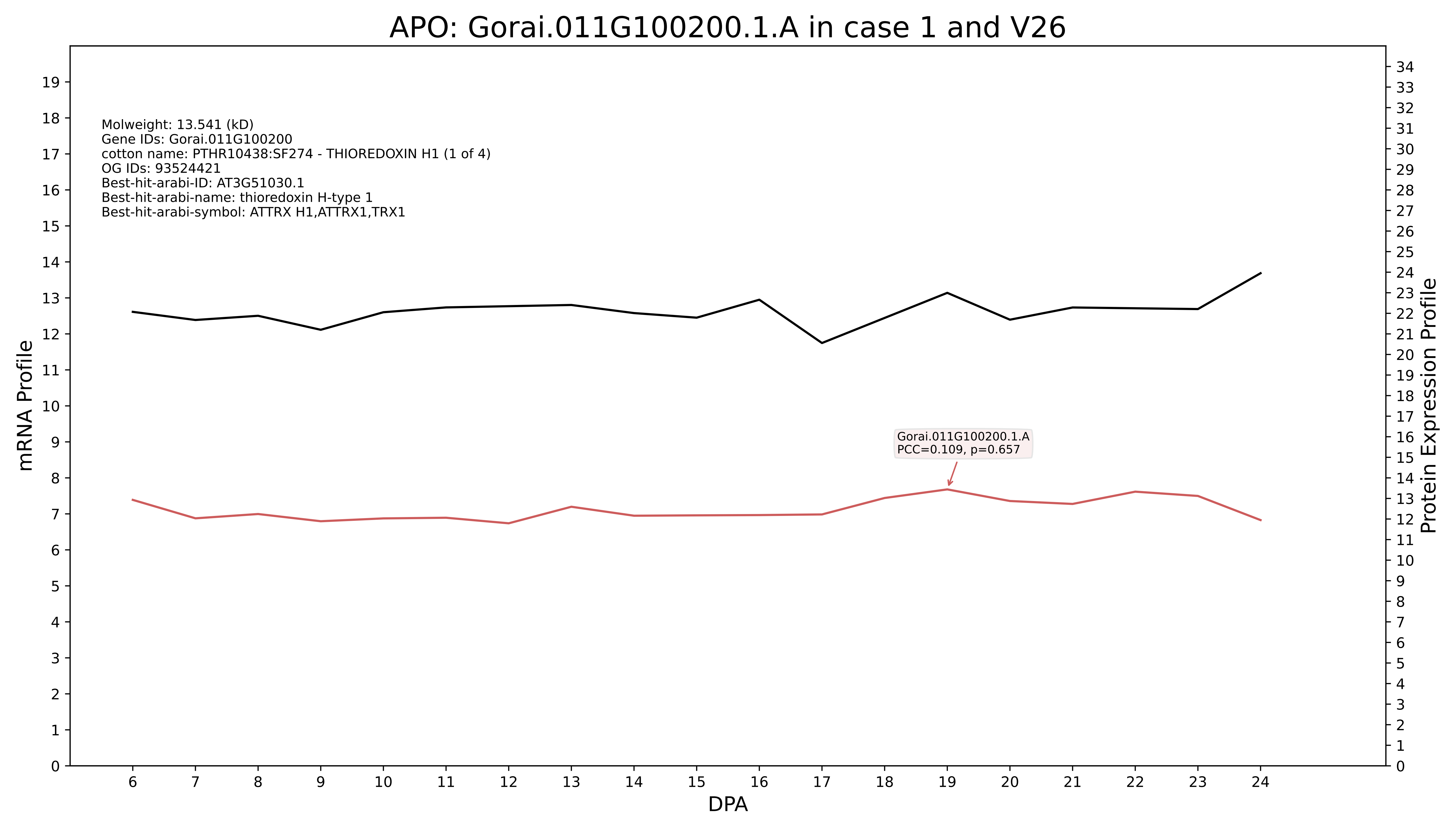
APO: Gorai.011G100200.1.A in case 1 and V26
Molweight: 13.541 (kD)
Gene IDs: Gorai.011G100200
cotton name: PTHR10438:SF274 - THIOREDOXIN H1 (1 of 4)
OG IDs: 93524421
Best-hit-arabi-ID: AT3G51030.1
Best-hit-arabi-name: thioredoxin H-type 1
Best-hit-arabi-symbol: ATTRX H1,ATTRX1,TRX1
Gorai.011G100200.1.A
PCC=0.109, p=0.657
mRNA Profile
Protein Expression Profile
DPA

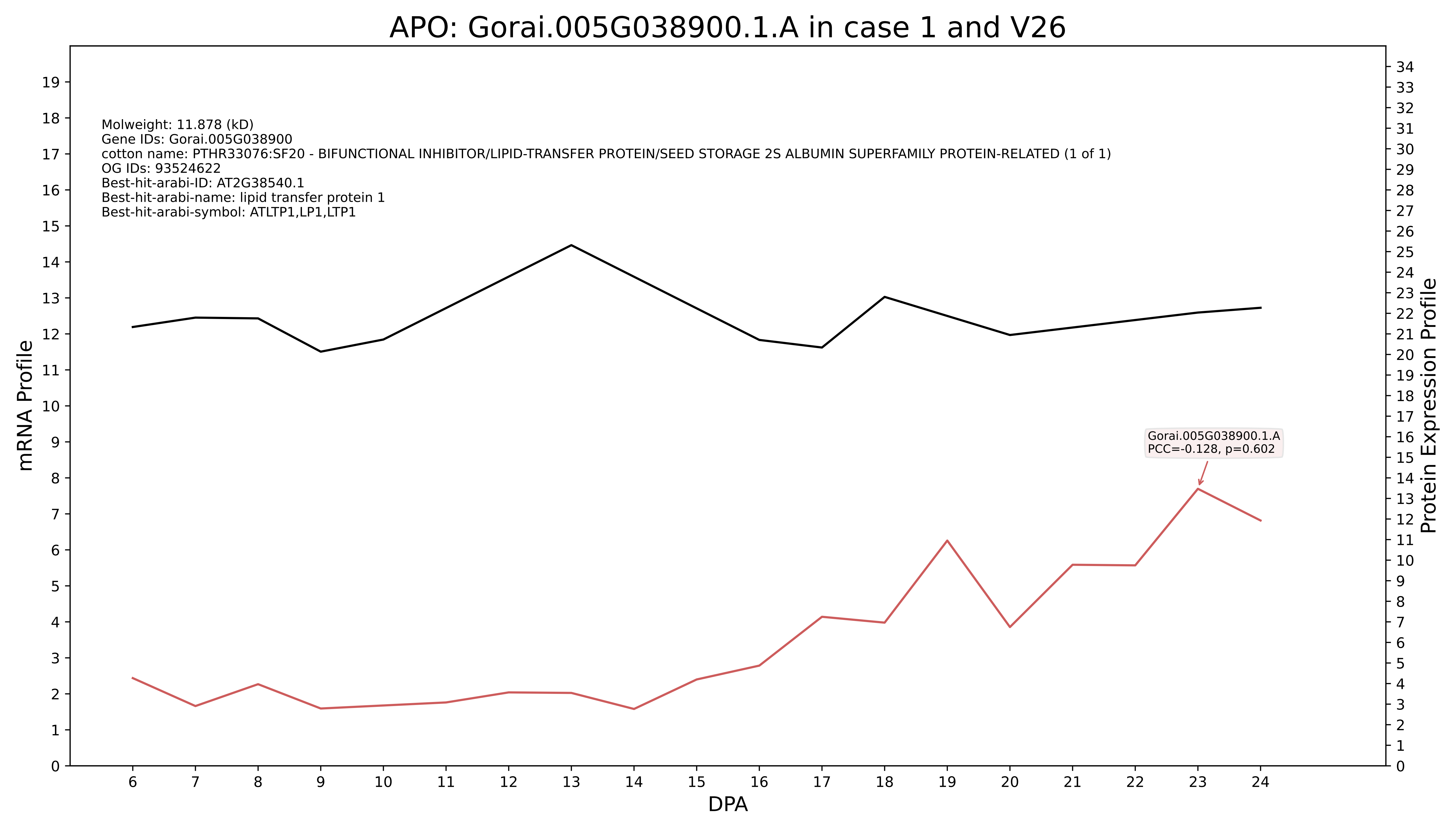
APO: Gorai.005G038900.1.A in case 1 and V26
Molweight: 11.878 (kD)
Gene IDs: Gorai.005G038900
cotton name: PTHR33076:SF20 - BIFUNCTIONAL INHIBITOR/LIPID-TRANSFER PROTEIN/SEED STORAGE 2S ALBUMIN SUPERFAMILY PROTEIN-RELATED (1 of 1)
OG IDs: 93524622
Best-hit-arabi-ID: AT2G38540.1
Best-hit-arabi-name: lipid transfer protein 1
Best-hit-arabi-symbol: ATLTP1,LP1,LTP1
Gorai.005G038900.1.A
PCC=-0.128, p=0.602
mRNA Profile
Protein Expression Profile
DPA

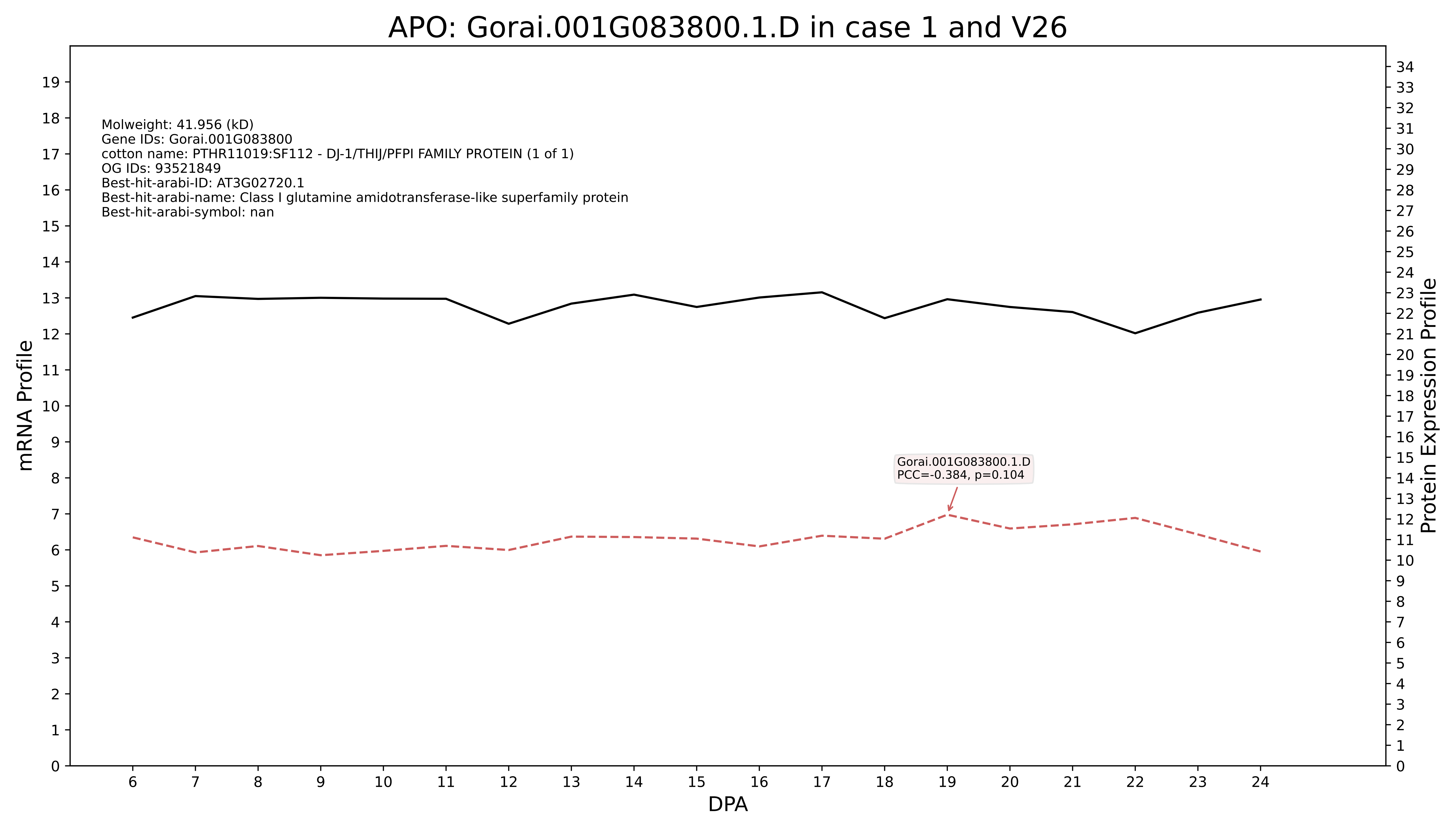
APO: Gorai.001G083800.1.D in case 1 and V26
Molweight: 41.956 (kD)
Gene IDs: Gorai.001G083800
cotton name: PTHR11019:SF112 - DJ-1/THIJ/PFPI FAMILY PROTEIN (1 of 1)
OG IDs: 93521849
Best-hit-arabi-ID: AT3G02720.1
Best-hit-arabi-name: Class I glutamine amidotransferase-like superfamily protein
Best-hit-arabi-symbol: nan
Gorai.001G083800.1.D
PCC=-0.384, p=0.104
mRNA Profile
Protein Expression Profile
DPA

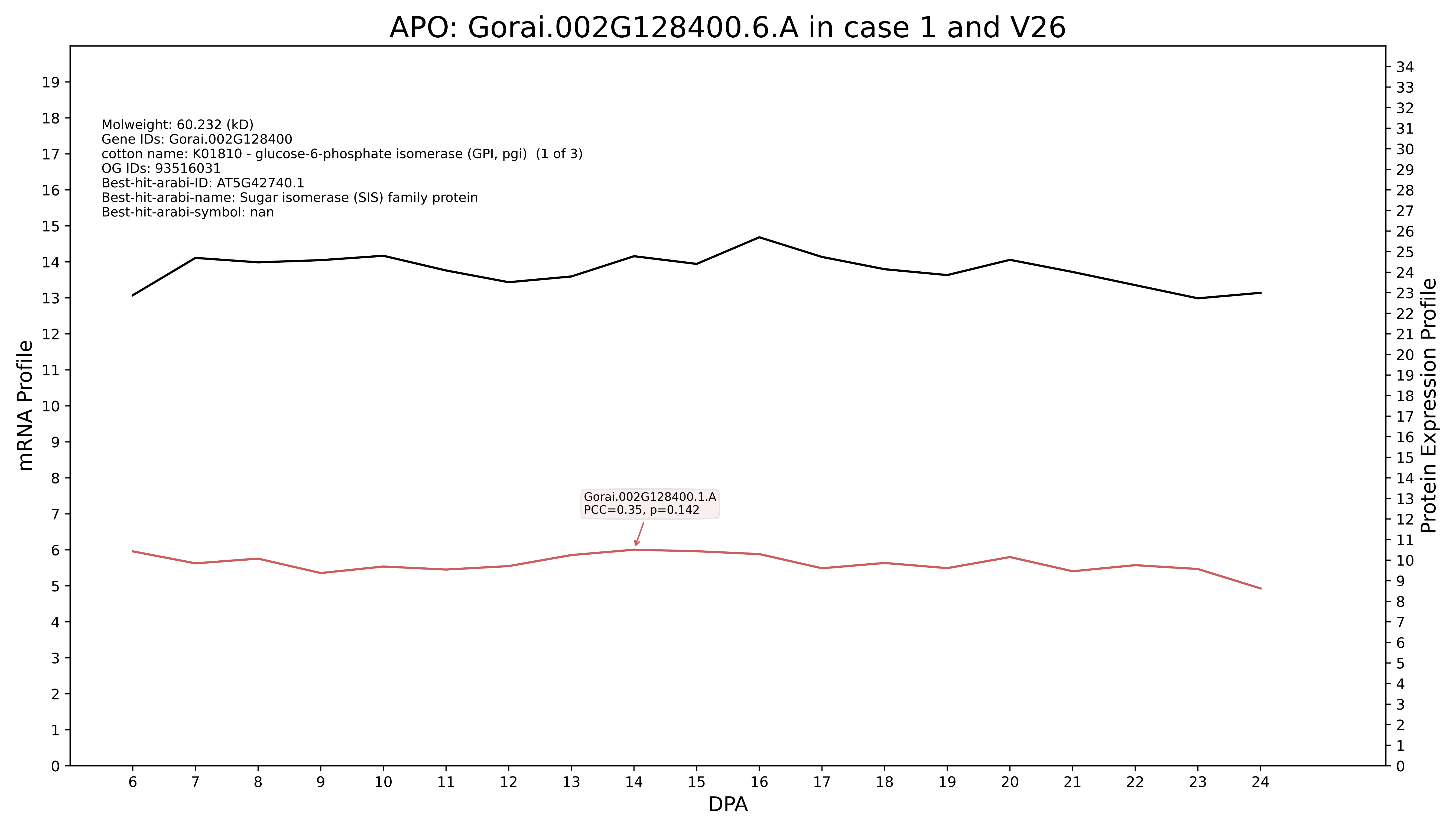
APO: Gorai.002G128400.6.A in case 1 and V26
Molweight: 60.232 (kD)
Gene IDs: Gorai.002G128400
cotton name: K01810 - glucose-6-phosphate isomerase (GPI, pgi) (1 of 3)
OG IDs: 93516031
Best-hit-arabi-ID: AT5G42740.1
Best-hit-arabi-name: Sugar isomerase (SIS) family protein
Best-hit-arabi-symbol: nan
Gorai.002G128400.1.A
PCC=0.35, p=0.142
mRNA Profile
Protein Expression Profile
DPA

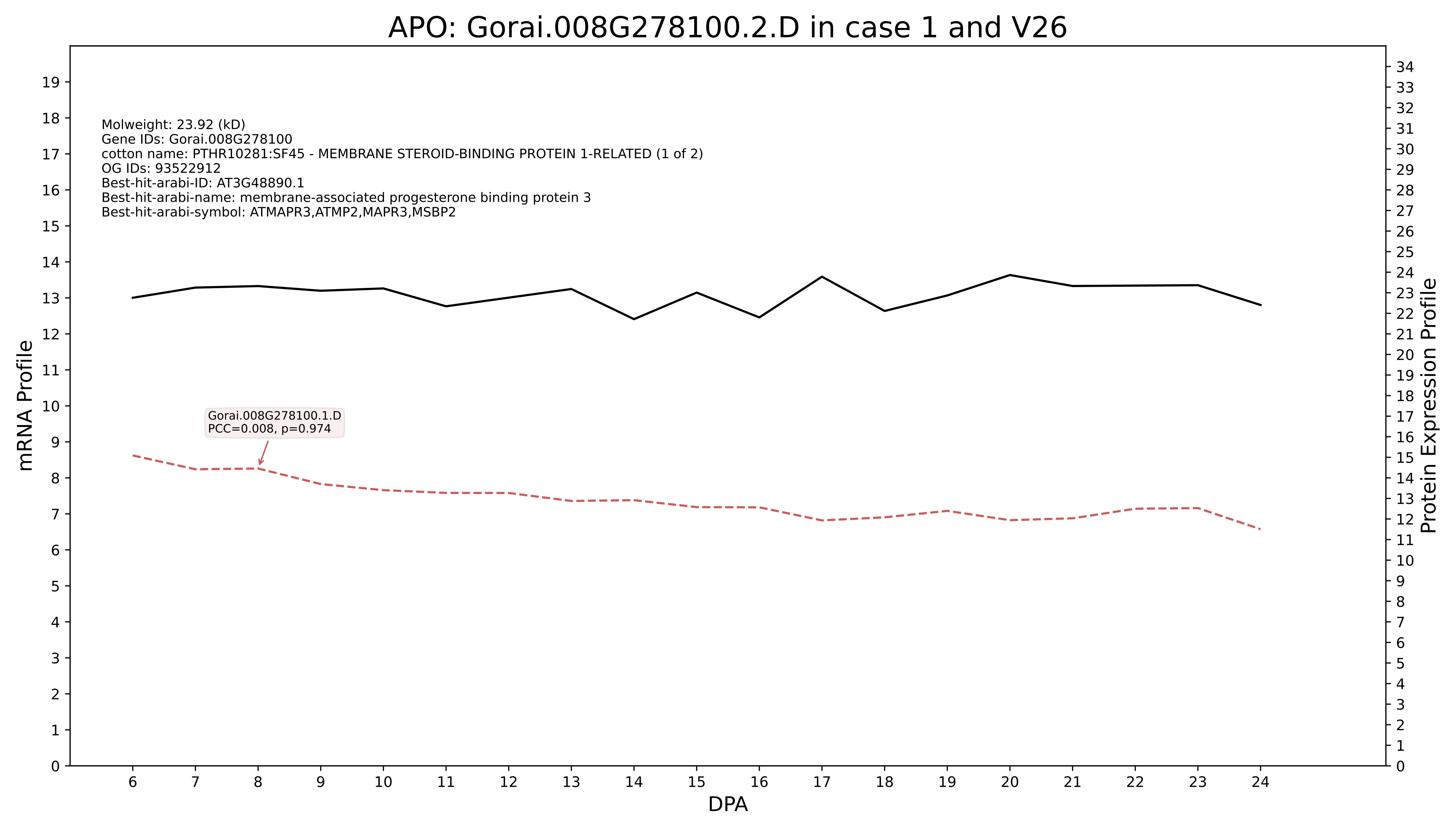
APO: Gorai.008G278100.2.D in case 1 and V26
Molweight: 23.92 (kD)
Gene IDs: Gorai.008G278100
cotton name: PTHR10281:SF45 - MEMBRANE STEROID-BINDING PROTEIN 1-RELATED (1 of 2)
OG IDs: 93522912
Best-hit-arabi-ID: AT3G48890.1
Best-hit-arabi-name: membrane-associated progesterone binding protein 3
Best-hit-arabi-symbol: ATMAPR3,ATMP2,MAPR3,MSBP2
Gorai.008G278100.1.D
PCC=0.008, p=0.974
mRNA Profile
Protein Expression Profile
DPA

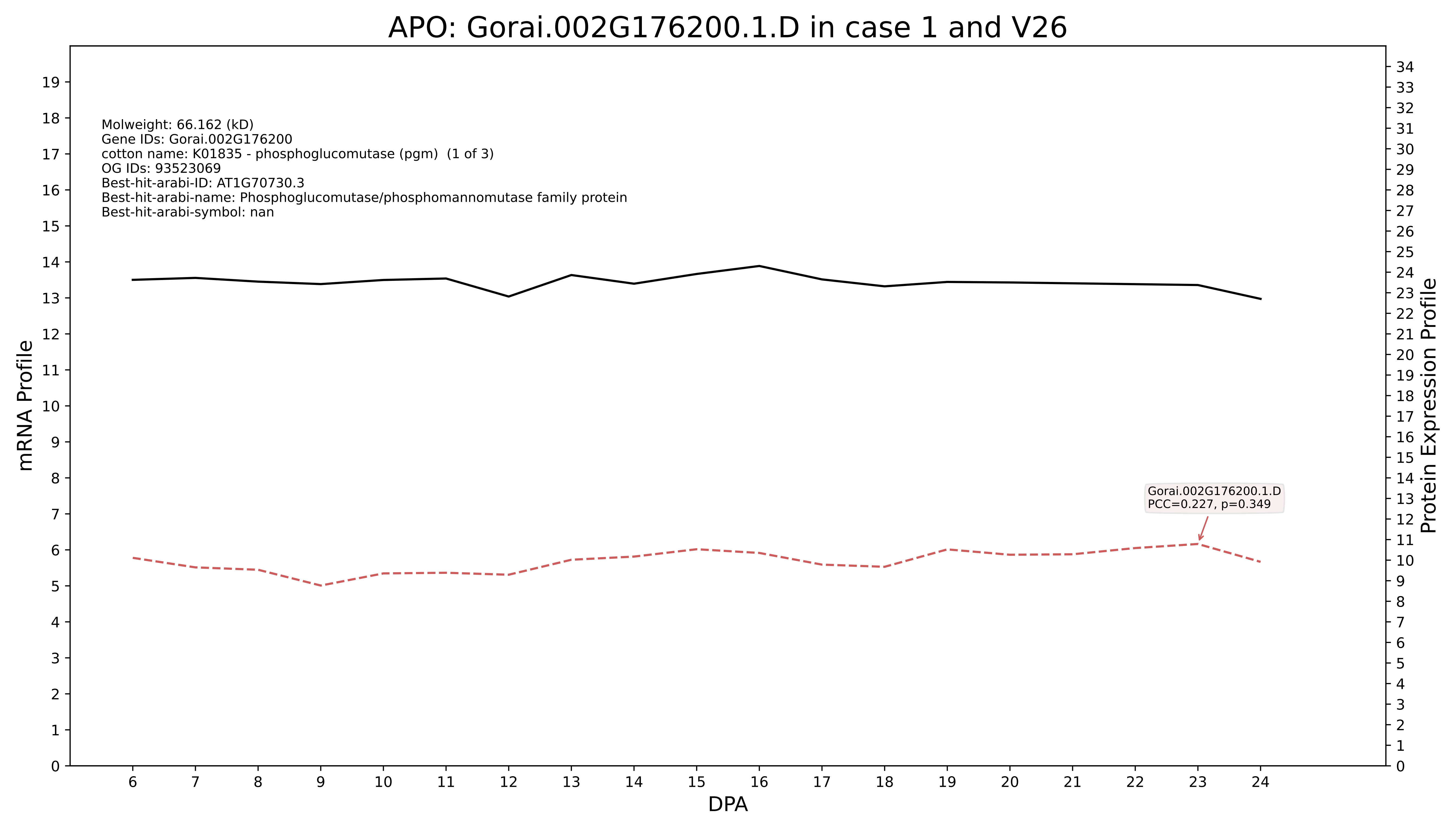

APO: Gorai.002G176200.1.D in case 1 and V26
Molweight: 66.162 (kD)
Gene IDs: Gorai.002G176200
cotton name: K01835 - phosphoglucomutase (pgm) (1 of 3)
OG IDs: 93523069
Best-hit-arabi-ID: AT1G70730.3
Best-hit-arabi-name: Phosphoglucomutase/phosphomannomutase family protein
Best-hit-arabi-symbol: nan
Gorai.002G176200.1.D
PCC=0.227, p=0.349
mRNA Profile
Protein Expression Profile
DPA

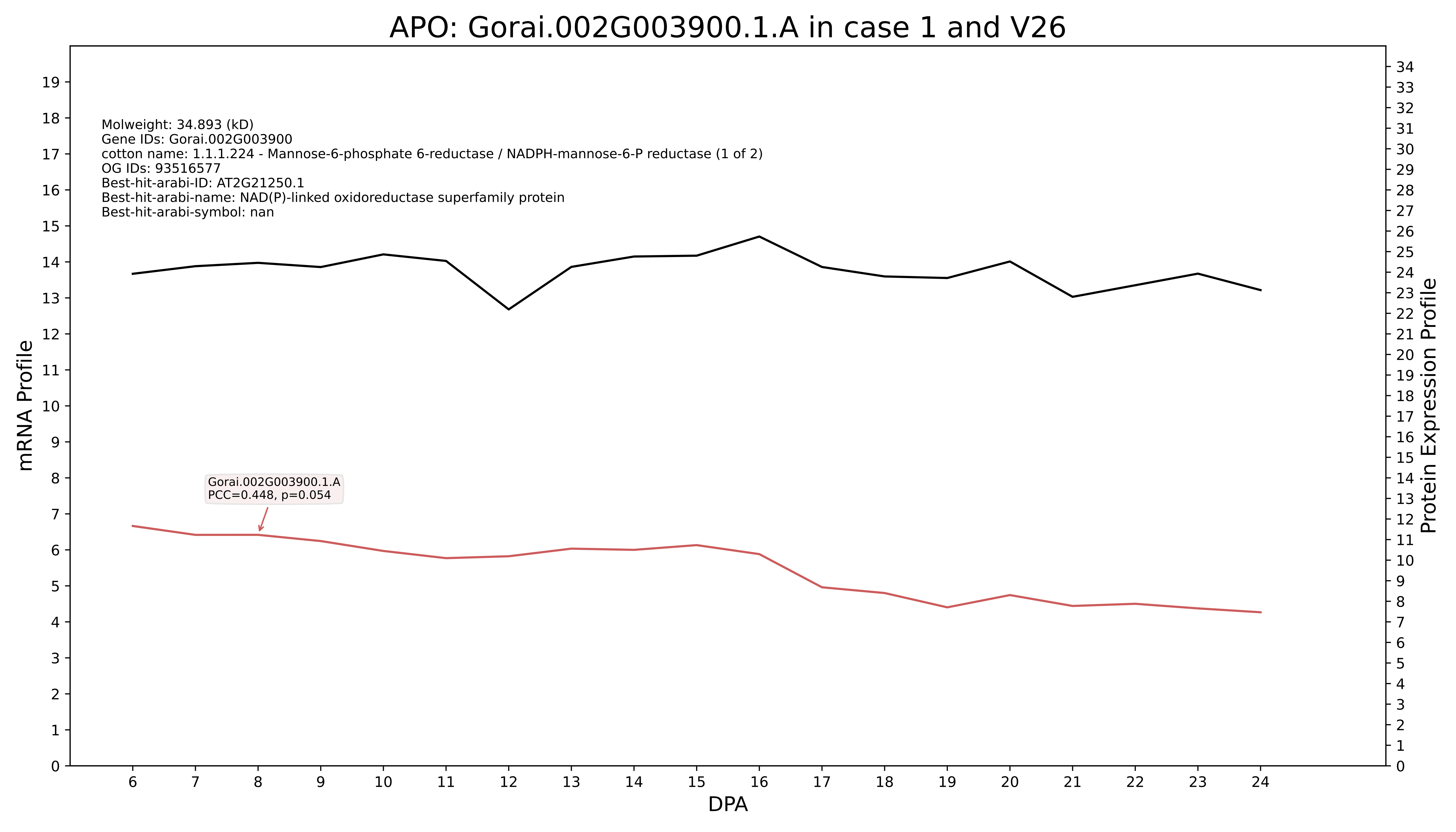
APO: Gorai.002G003900.1.A in case 1 and V26
Molweight: 34.893 (kD)
Gene IDs: Gorai.002G003900
cotton name: 1.1.1.224 - Mannose-6-phosphate 6-reductase / NADPH-mannose-6-P reductase (1 of 2)
OG IDs: 93516577
Best-hit-arabi-ID: AT2G21250.1
Best-hit-arabi-name: NAD(P)-linked oxidoreductase superfamily protein
Best-hit-arabi-symbol: nan
Gorai.002G003900.1.A
PCC=0.448, p=0.054
mRNA Profile
Protein Expression Profile
DPA

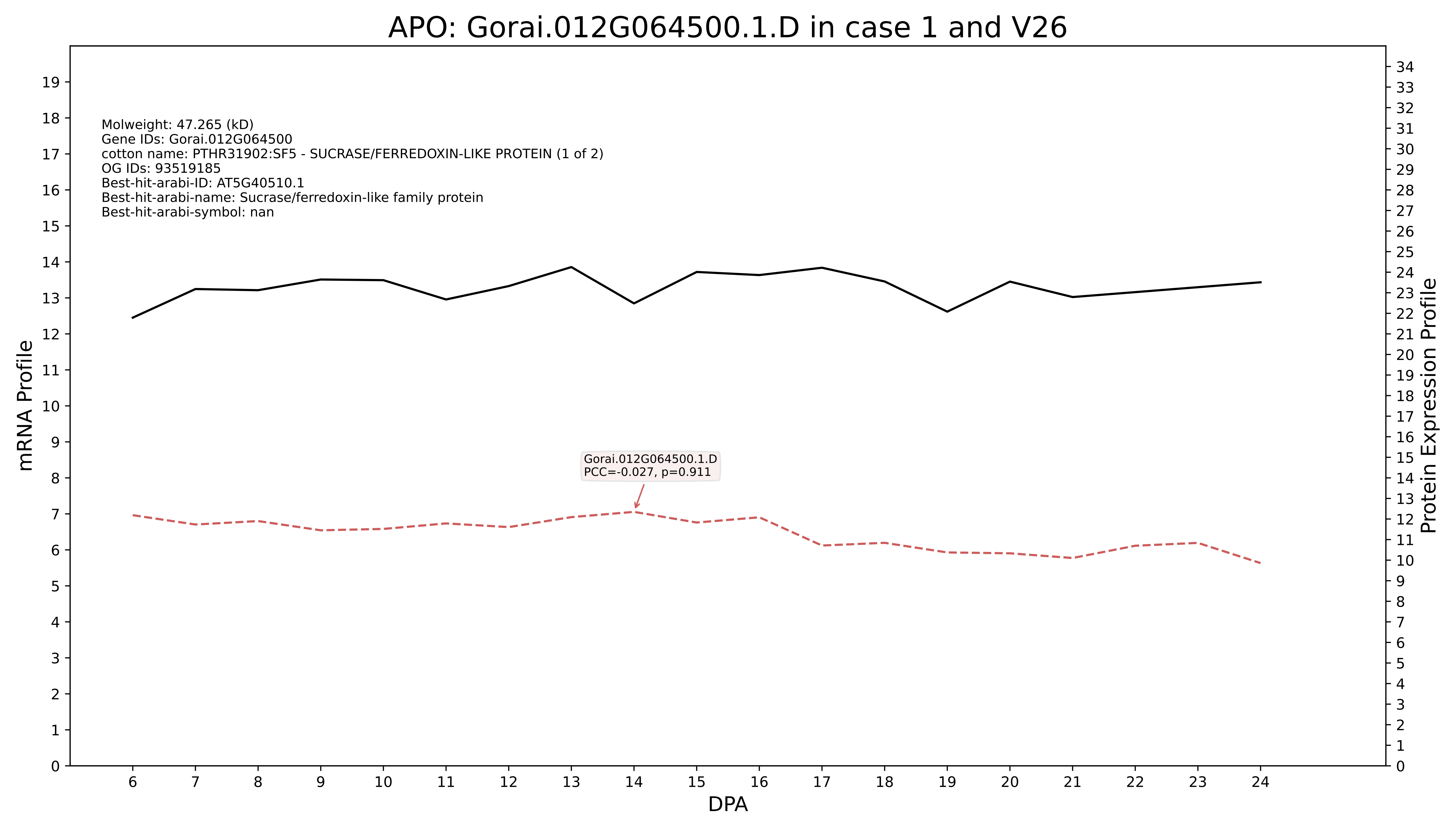
APO: Gorai.012G064500.1.D in case 1 and V26
Molweight: 47.265 (kD)
Gene IDs: Gorai.012G064500
cotton name: PTHR31902:SF5 - SUCRASE/FERREDOXIN-LIKE PROTEIN (1 of 2)
OG IDs: 93519185
Best-hit-arabi-ID: AT5G40510.1
Best-hit-arabi-name: Sucrase/ferredoxin-like family protein
Best-hit-arabi-symbol: nan
Gorai.012G064500.1.D
PCC=-0.027, p=0.911
mRNA Profile
Protein Expression Profile
DPA

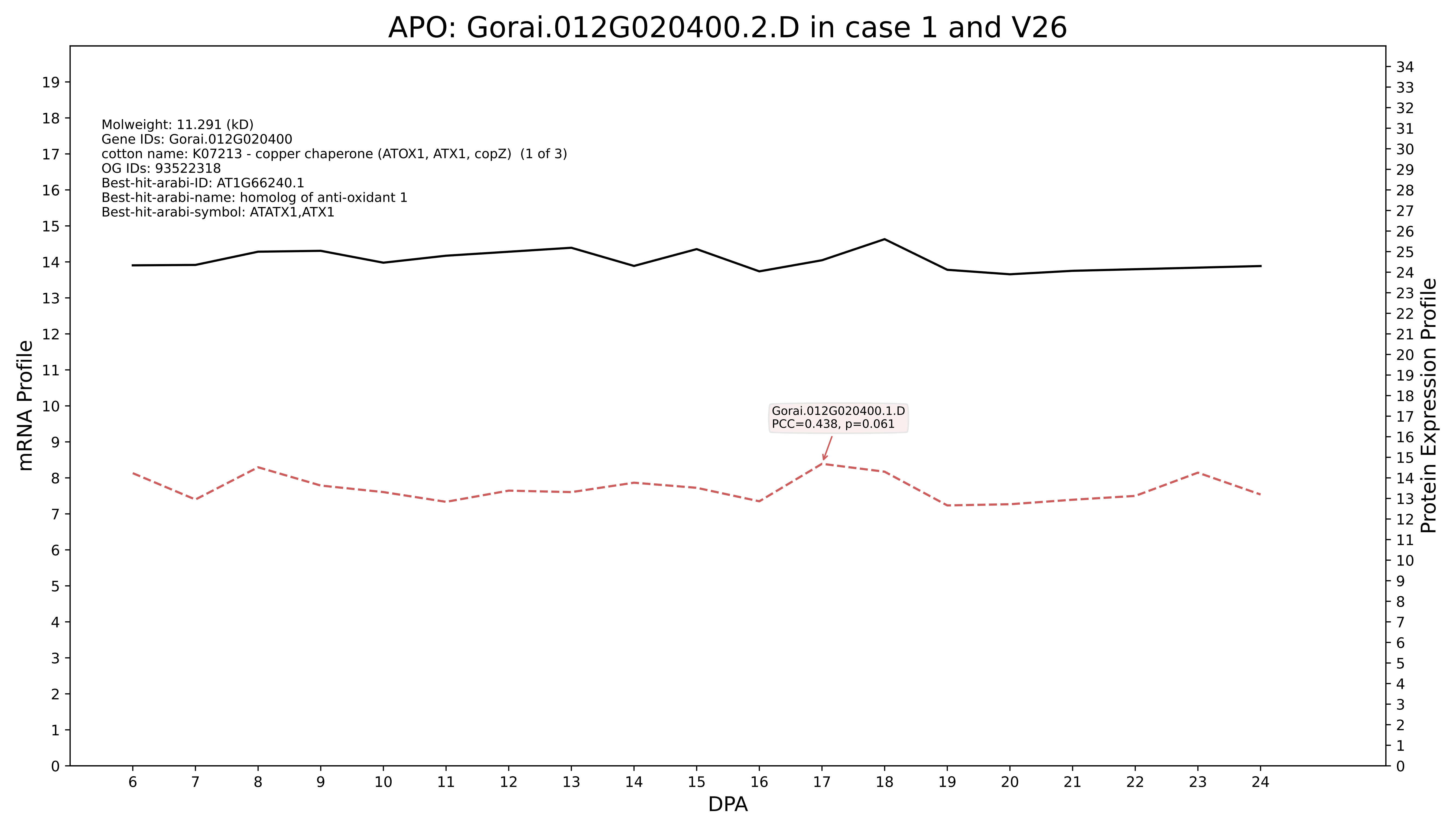

APO: Gorai.012G020400.2.D in case 1 and V26
Molweight: 11.291 (kD)
Gene IDs: Gorai.012G020400
cotton name: K07213 - copper chaperone (ATOX1, ATX1, copZ) (1 of 3)
OG IDs: 93522318
Best-hit-arabi-ID: AT1G66240.1
Best-hit-arabi-name: homolog of anti-oxidant 1
Best-hit-arabi-symbol: ATATX1,ATX1
Gorai.012G020400.1.D
PCC=0.438, p=0.061
mRNA Profile
Protein Expression Profile
DPA

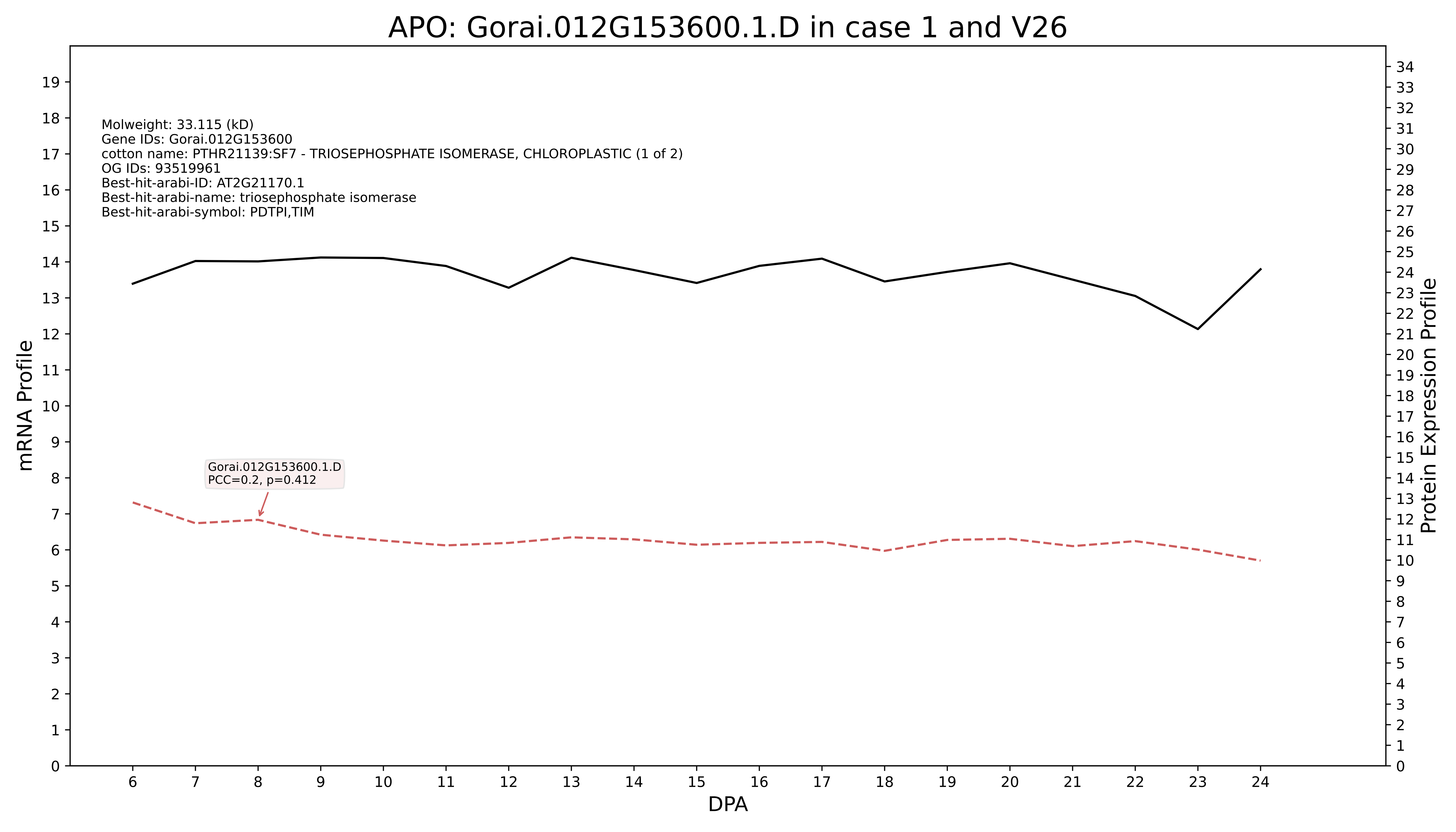

APO: Gorai.012G153600.1.D in case 1 and V26
Molweight: 33.115 (kD)
Gene IDs: Gorai.012G153600
cotton name: PTHR21139:SF7 - TRIOSEPHOSPHATE ISOMERASE, CHLOROPLASTIC (1 of 2)
OG IDs: 93519961
Best-hit-arabi-ID: AT2G21170.1
Best-hit-arabi-name: triosephosphate isomerase
Best-hit-arabi-symbol: PDTPI,TIM
Gorai.012G153600.1.D
PCC=0.2, p=0.412
mRNA Profile
Protein Expression Profile
DPA

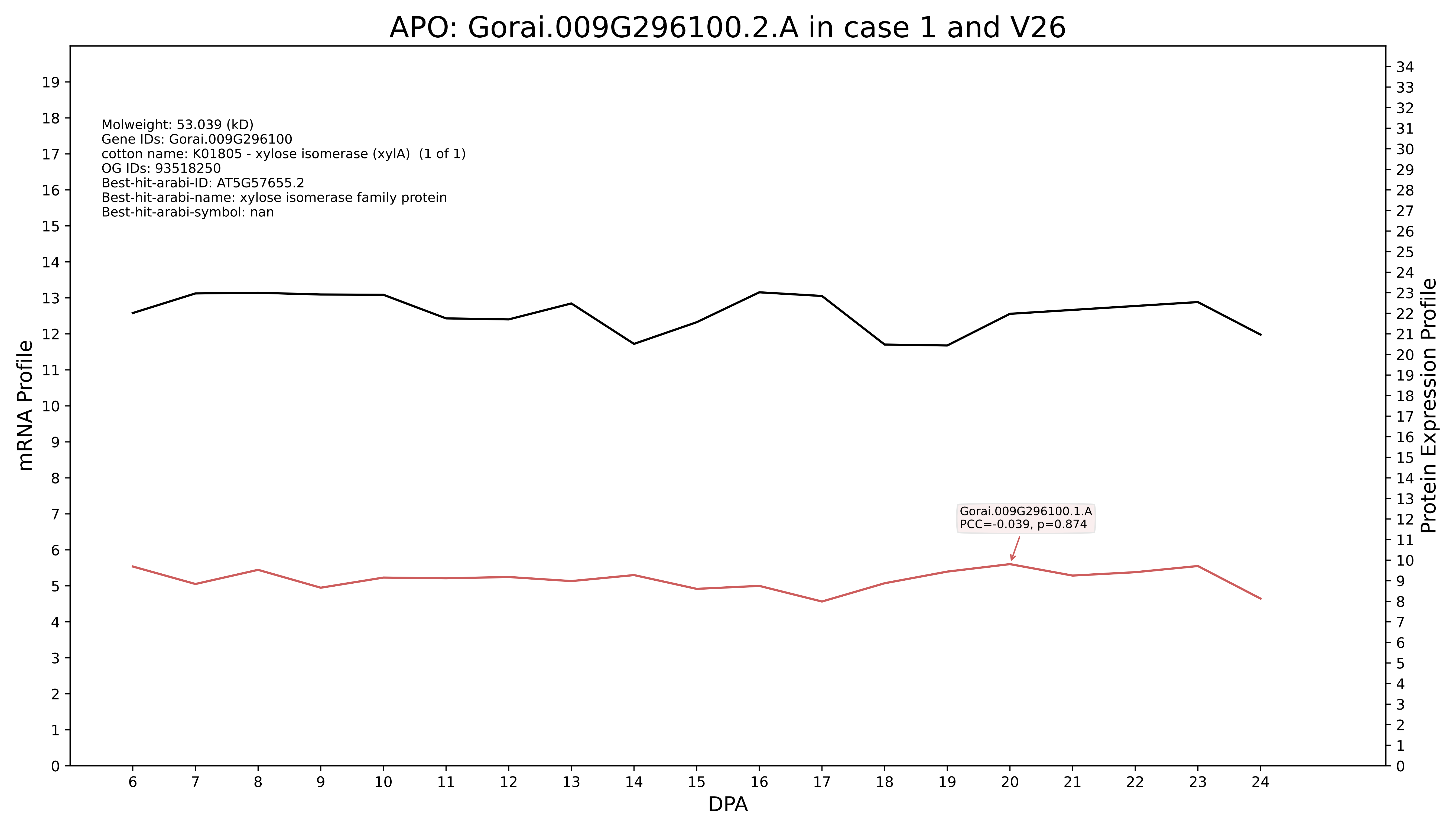
APO: Gorai.009G296100.2.A in case 1 and V26
Molweight: 53.039 (kD)
Gene IDs: Gorai.009G296100
cotton name: K01805 - xylose isomerase (xylA) (1 of 1)
OG IDs: 93518250
Best-hit-arabi-ID: AT5G57655.2
Best-hit-arabi-name: xylose isomerase family protein
Best-hit-arabi-symbol: nan
Gorai.009G296100.1.A
PCC=-0.039, p=0.874
mRNA Profile
Protein Expression Profile
DPA

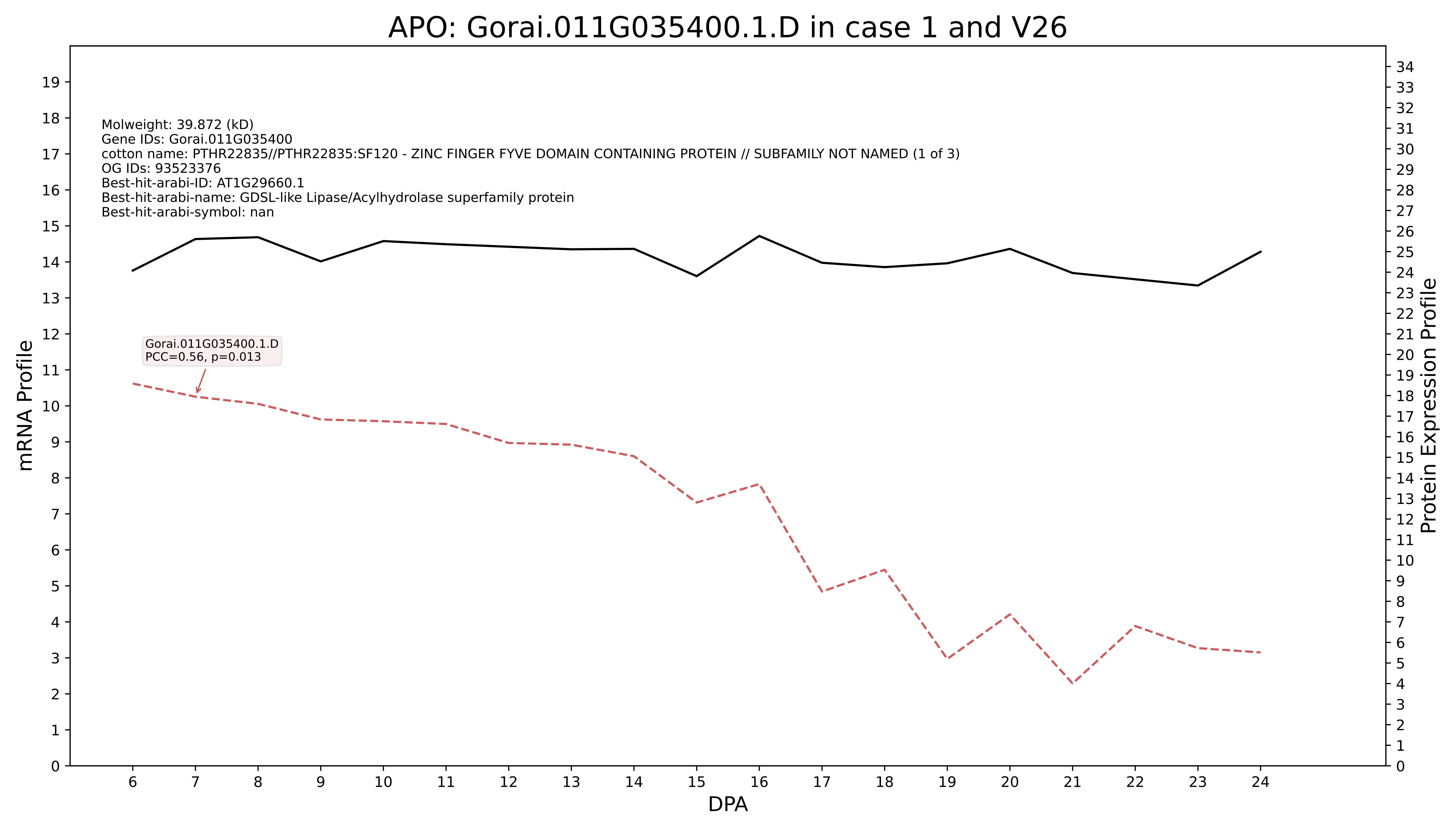
APO: Gorai.011G035400.1.D in case 1 and V26
Molweight: 39.872 (kD)
Gene IDs: Gorai.011G035400
cotton name: PTHR22835//PTHR22835:SF120 - ZINC FINGER FYVE DOMAIN CONTAINING PROTEIN // SUBFAMILY NOT NAMED (1 of 3)
OG IDs: 93523376
Best-hit-arabi-ID: AT1G29660.1
Best-hit-arabi-name: GDSL-like Lipase/Acylhydrolase superfamily protein
Best-hit-arabi-symbol: nan
Gorai.011G035400.1.D
PCC=0.56, p=0.013
mRNA Profile
Protein Expression Profile
DPA

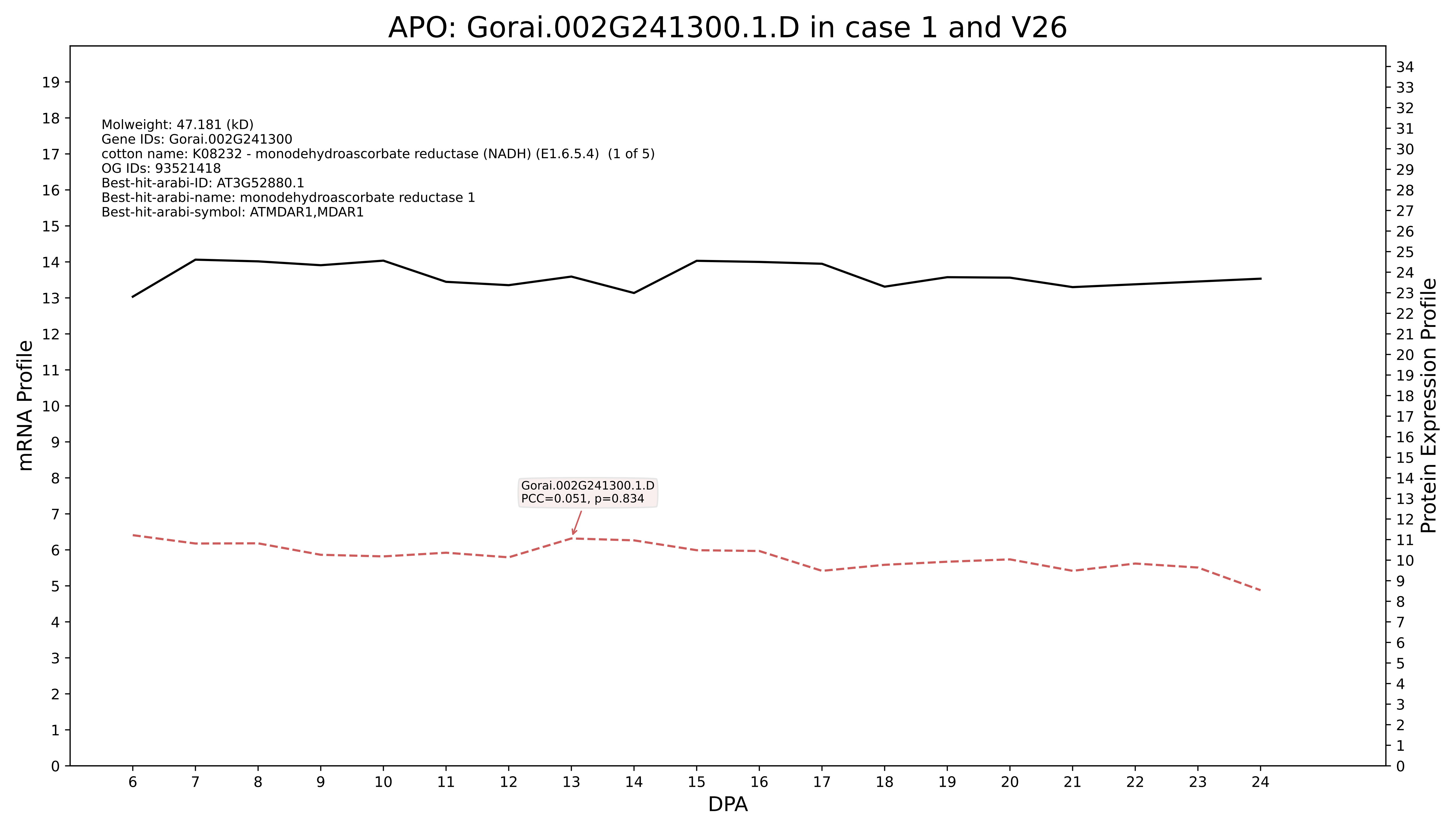
APO: Gorai.002G241300.1.D in case 1 and V26
Molweight: 47.181 (kD)
Gene IDs: Gorai.002G241300
cotton name: K08232 - monodehydroascorbate reductase (NADH) (E1.6.5.4) (1 of 5)
OG IDs: 93521418
Best-hit-arabi-ID: AT3G52880.1
Best-hit-arabi-name: monodehydroascorbate reductase 1
Best-hit-arabi-symbol: ATMDAR1,MDAR1
Gorai.002G241300.1.D
PCC=0.051, p=0.834
mRNA Profile
Protein Expression Profile
DPA

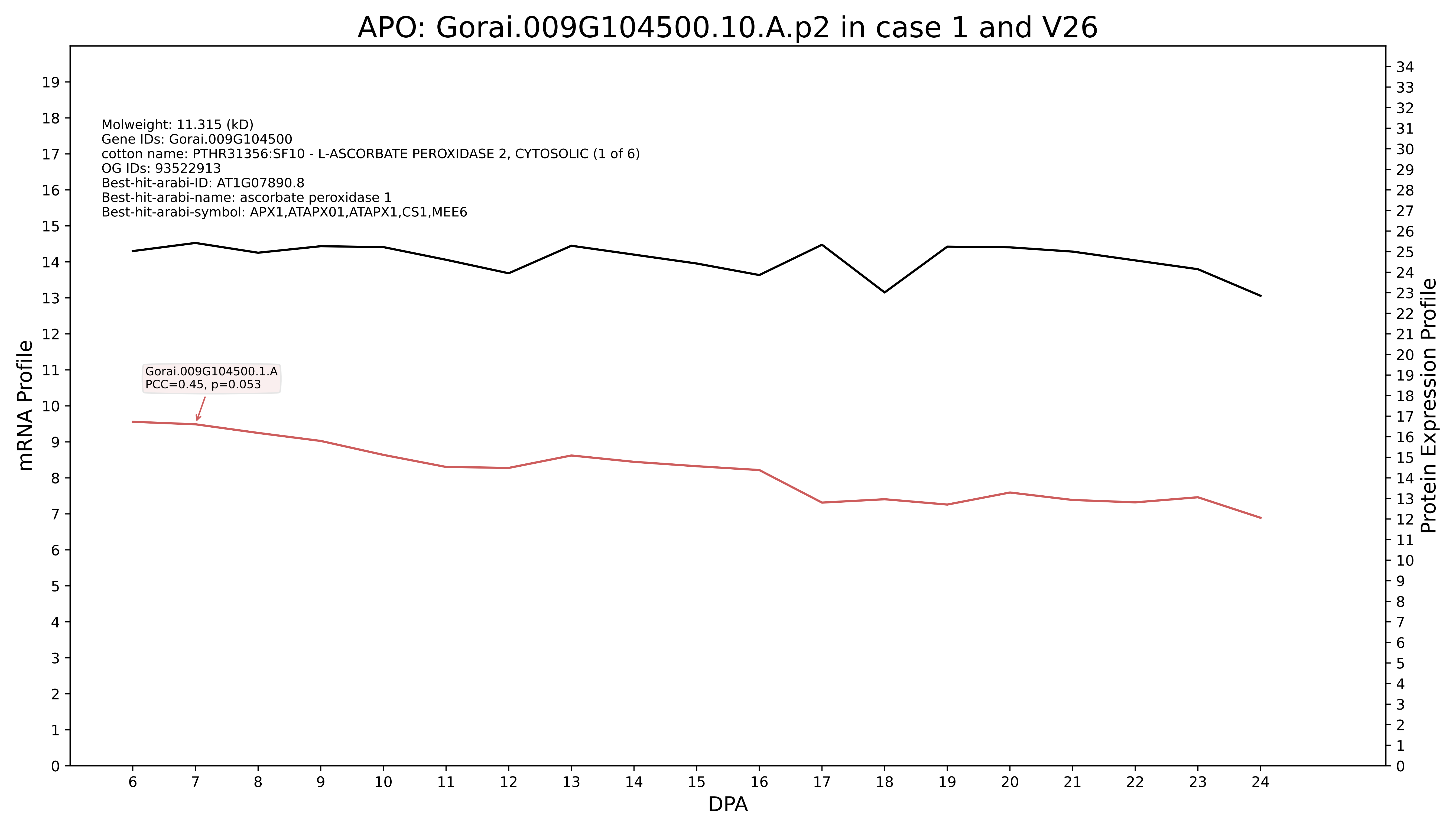
APO: Gorai.009G104500.10.A.p2 in case 1 and V26
Molweight: 11.315 (kD)
Gene IDs: Gorai.009G104500
cotton name: PTHR31356:SF10 - L-ASCORBATE PEROXIDASE 2, CYTOSOLIC (1 of 6)
OG IDs: 93522913
Best-hit-arabi-ID: AT1G07890.8
Best-hit-arabi-name: ascorbate peroxidase 1
Best-hit-arabi-symbol: APX1,ATAPX01,ATAPX1,CS1,MEE6
Gorai.009G104500.1.A
PCC=0.45, p=0.053
mRNA Profile
Protein Expression Profile
DPA

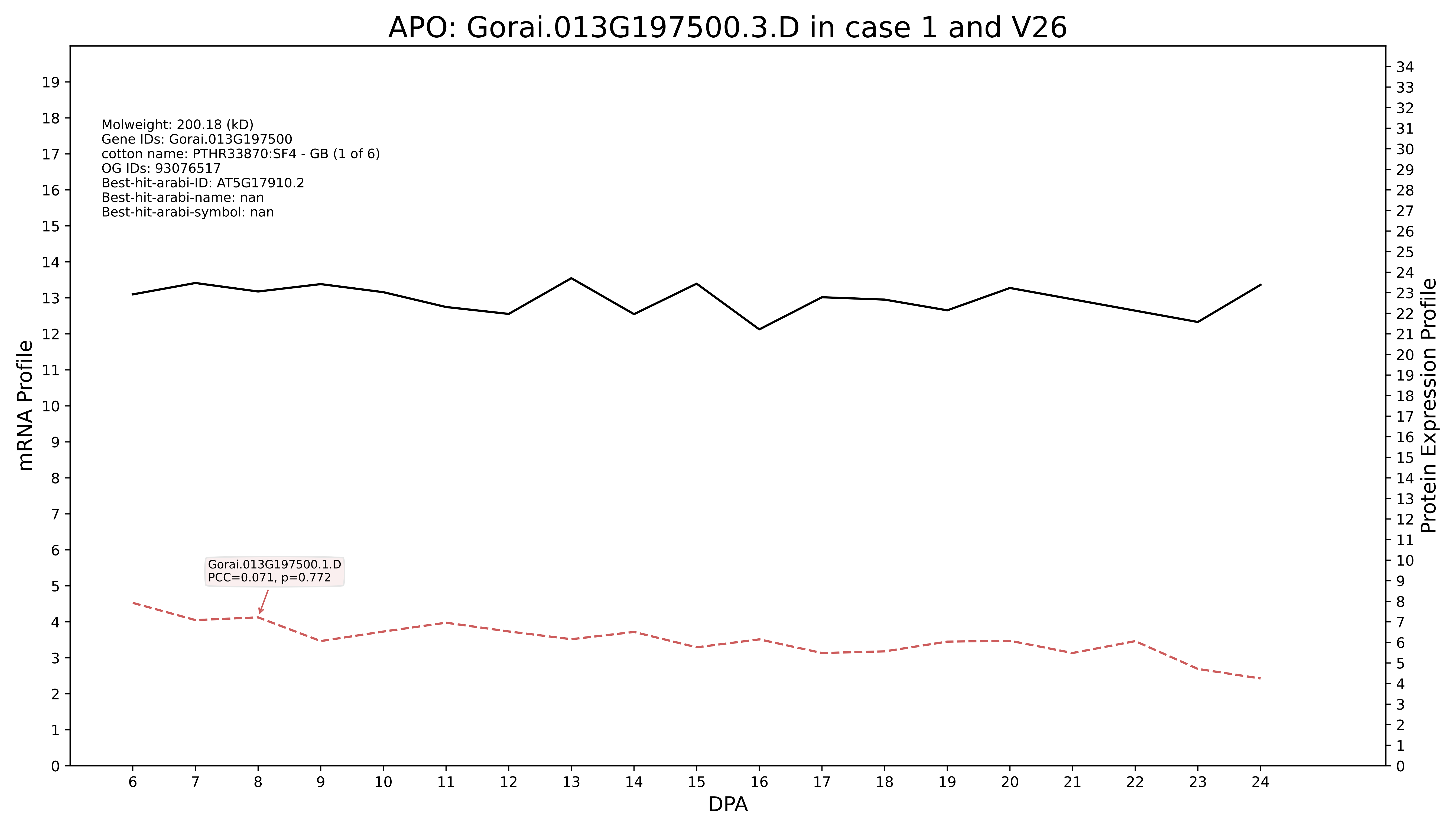

APO: Gorai.013G197500.3.D in case 1 and V26
Molweight: 200.18 (kD)
Gene IDs: Gorai.013G197500
cotton name: PTHR33870:SF4 - GB (1 of 6)
OG IDs: 93076517
Best-hit-arabi-ID: AT5G17910.2
Best-hit-arabi-name: nan
Best-hit-arabi-symbol: nan
Gorai.013G197500.1.D
PCC=0.071, p=0.772
mRNA Profile
Protein Expression Profile
DPA

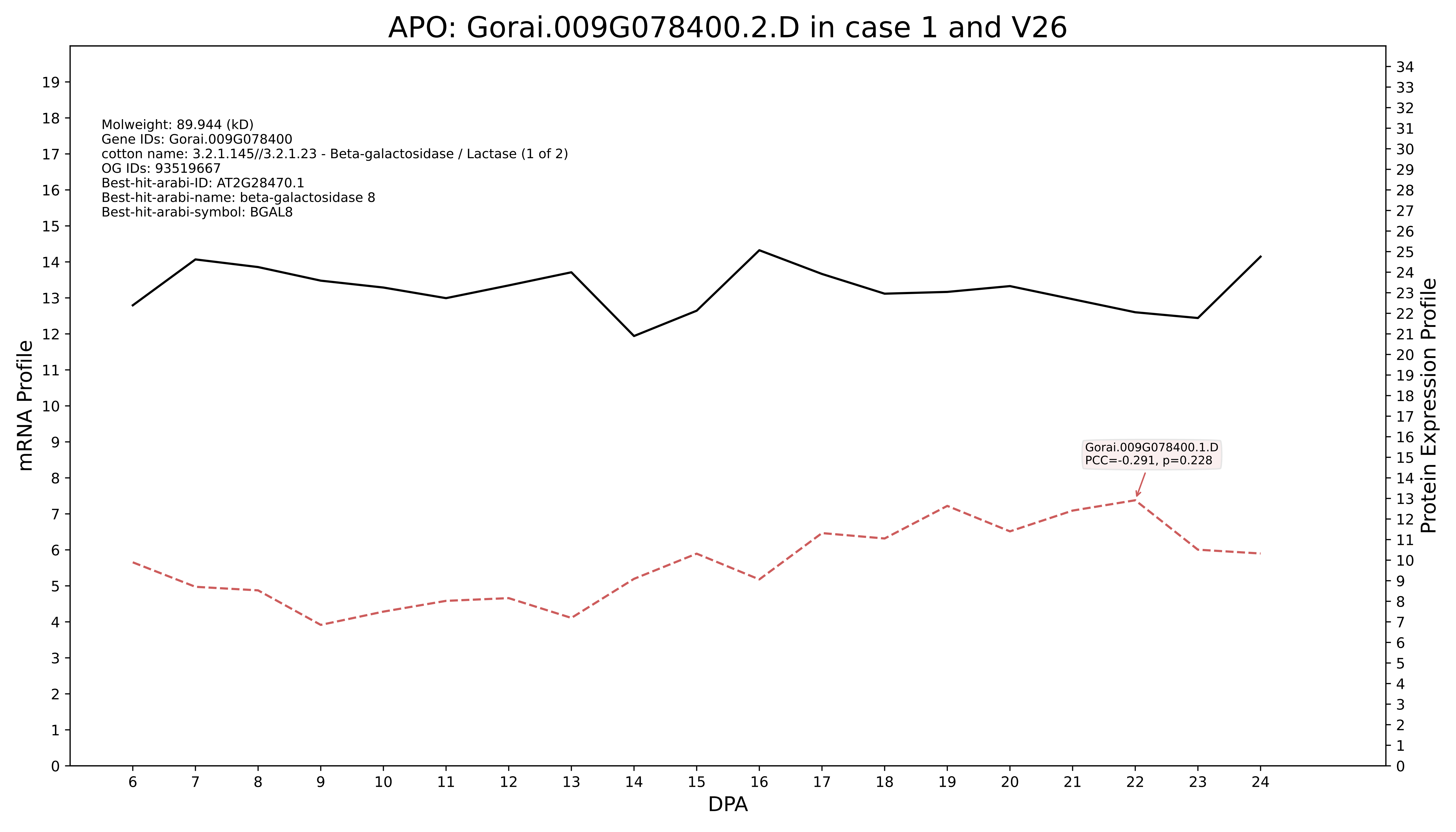
APO: Gorai.009G078400.2.D in case 1 and V26
Molweight: 89.944 (kD)
Gene IDs: Gorai.009G078400
cotton name: 3.2.1.145//3.2.1.23 - Beta-galactosidase / Lactase (1 of 2)
OG IDs: 93519667
Best-hit-arabi-ID: AT2G28470.1
Best-hit-arabi-name: beta-galactosidase 8
Best-hit-arabi-symbol: BGAL8
Gorai.009G078400.1.D
PCC=-0.291, p=0.228
mRNA Profile
Protein Expression Profile
DPA

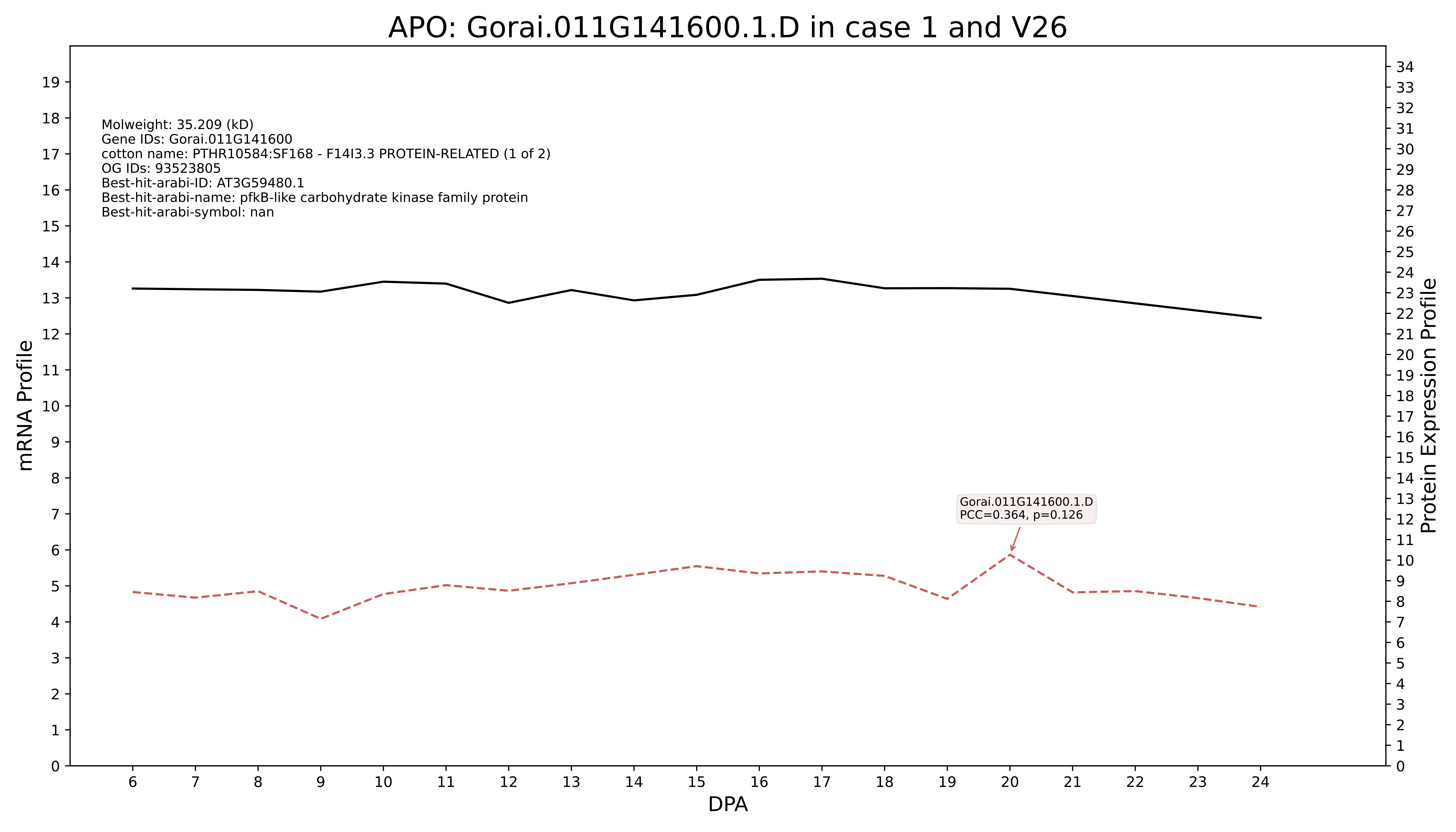

APO: Gorai.011G141600.1.D in case 1 and V26
Molweight: 35.209 (kD)
Gene IDs: Gorai.011G141600
cotton name: PTHR10584:SF168 - F14I3.3 PROTEIN-RELATED (1 of 2)
OG IDs: 93523805
Best-hit-arabi-ID: AT3G59480.1
Best-hit-arabi-name: pfkB-like carbohydrate kinase family protein
Best-hit-arabi-symbol: nan
Gorai.011G141600.1.D
PCC=0.364, p=0.126
mRNA Profile
Protein Expression Profile
DPA

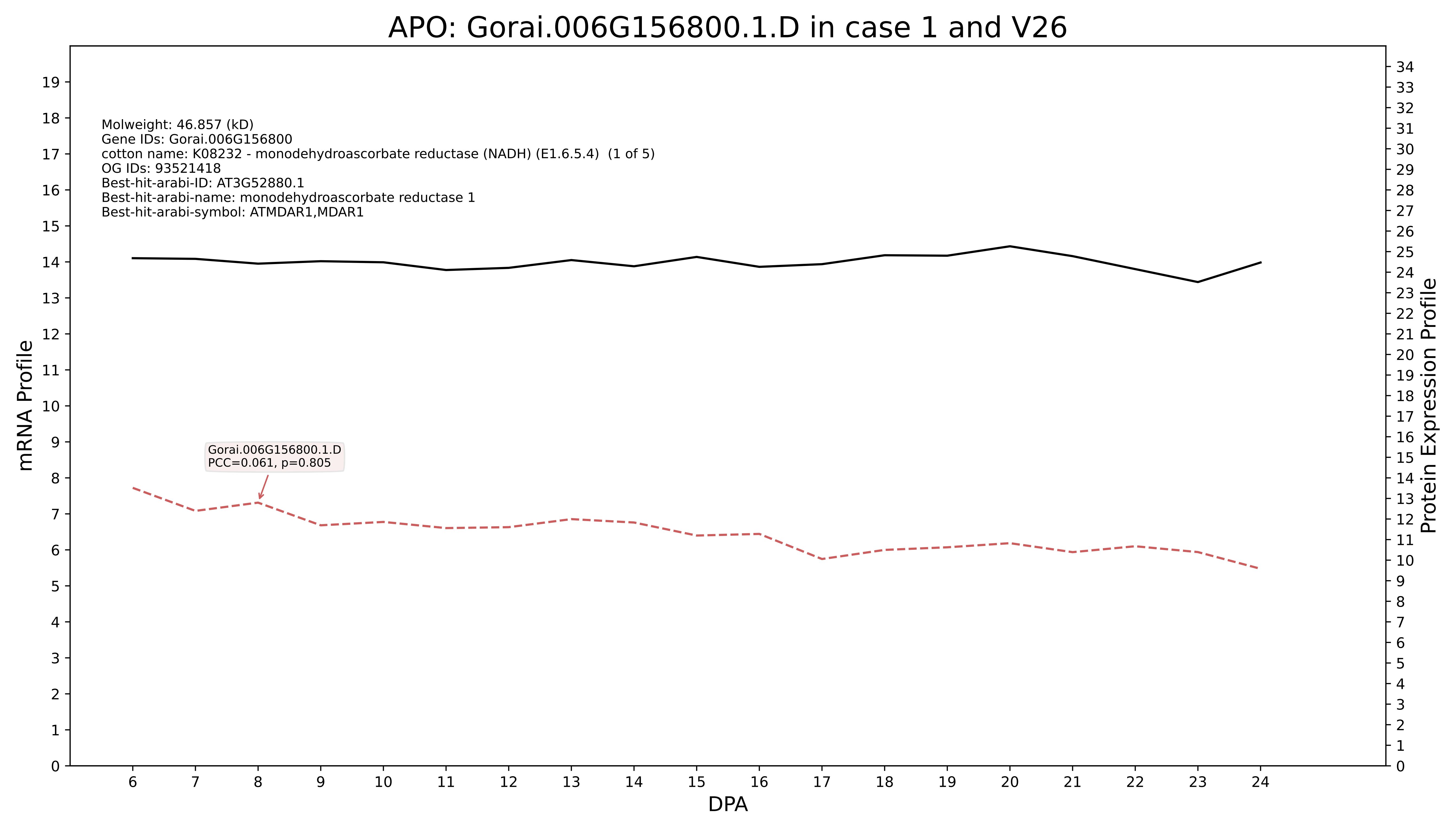
APO: Gorai.006G156800.1.D in case 1 and V26
Molweight: 46.857 (kD)
Gene IDs: Gorai.006G156800
cotton name: K08232 - monodehydroascorbate reductase (NADH) (E1.6.5.4) (1 of 5)
OG IDs: 93521418
Best-hit-arabi-ID: AT3G52880.1
Best-hit-arabi-name: monodehydroascorbate reductase 1
Best-hit-arabi-symbol: ATMDAR1,MDAR1
Gorai.006G156800.1.D
PCC=0.061, p=0.805
mRNA Profile
Protein Expression Profile
DPA

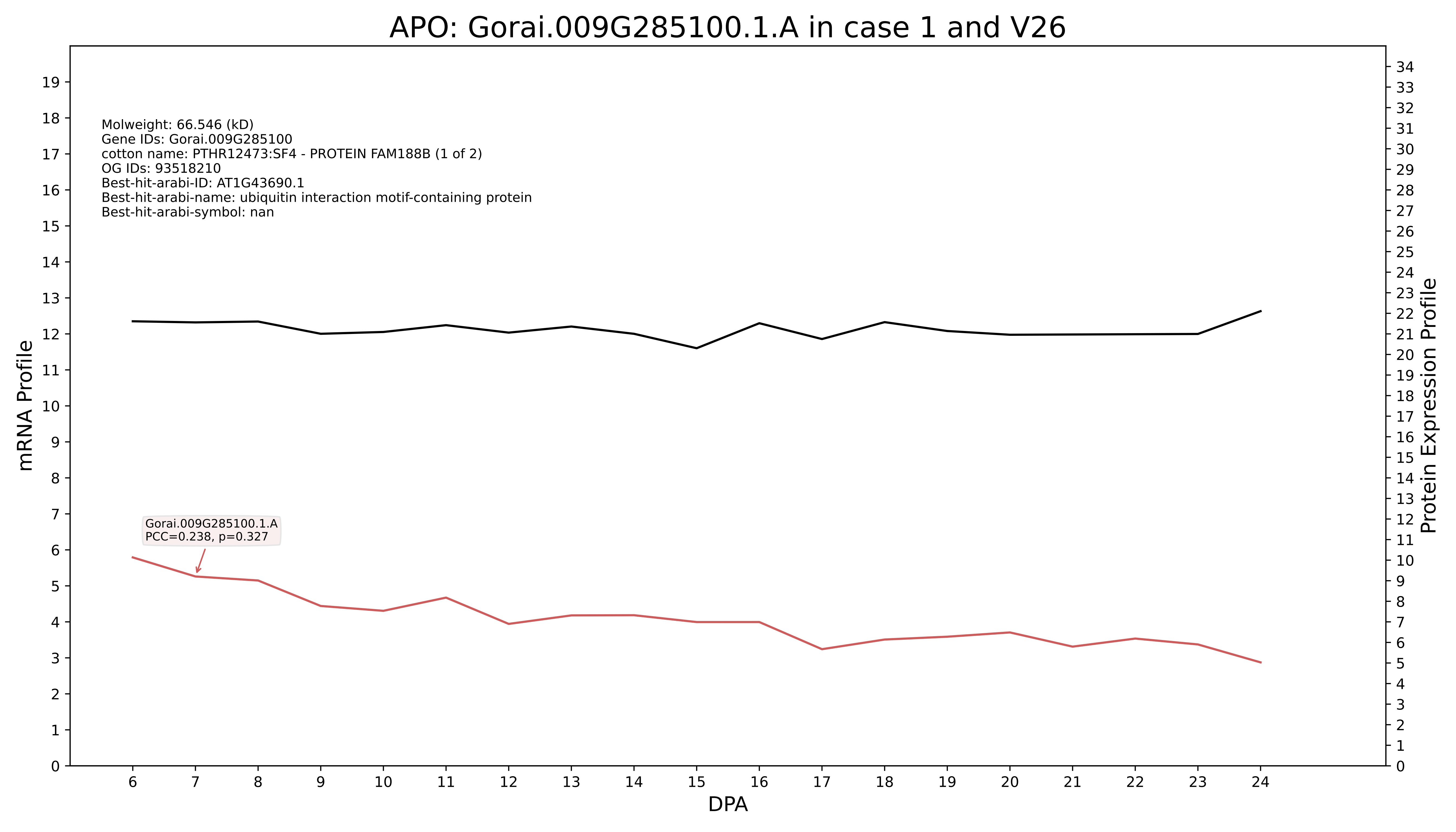
APO: Gorai.009G285100.1.A in case 1 and V26
Molweight: 66.546 (kD)
Gene IDs: Gorai.009G285100
cotton name: PTHR12473:SF4 - PROTEIN FAM188B (1 of 2)
OG IDs: 93518210
Best-hit-arabi-ID: AT1G43690.1
Best-hit-arabi-name: ubiquitin interaction motif-containing protein
Best-hit-arabi-symbol: nan
Gorai.009G285100.1.A
PCC=0.238, p=0.327
mRNA Profile
Protein Expression Profile
DPA

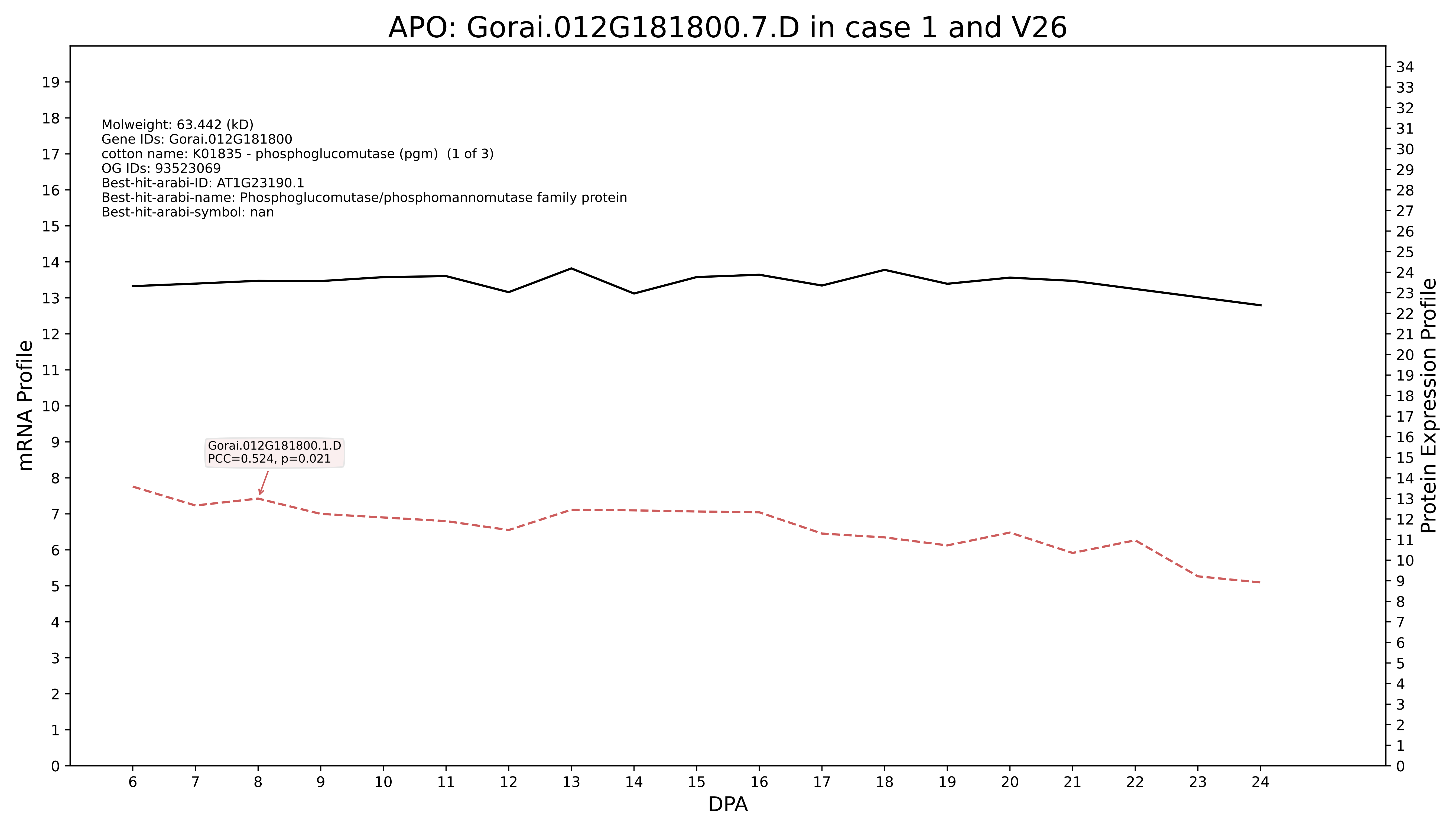
APO: Gorai.012G181800.7.D in case 1 and V26
Molweight: 63.442 (kD)
Gene IDs: Gorai.012G181800
cotton name: K01835 - phosphoglucomutase (pgm) (1 of 3)
OG IDs: 93523069
Best-hit-arabi-ID: AT1G23190.1
Best-hit-arabi-name: Phosphoglucomutase/phosphomannomutase family protein
Best-hit-arabi-symbol: nan
Gorai.012G181800.1.D
PCC=0.524, p=0.021
mRNA Profile
Protein Expression Profile
DPA

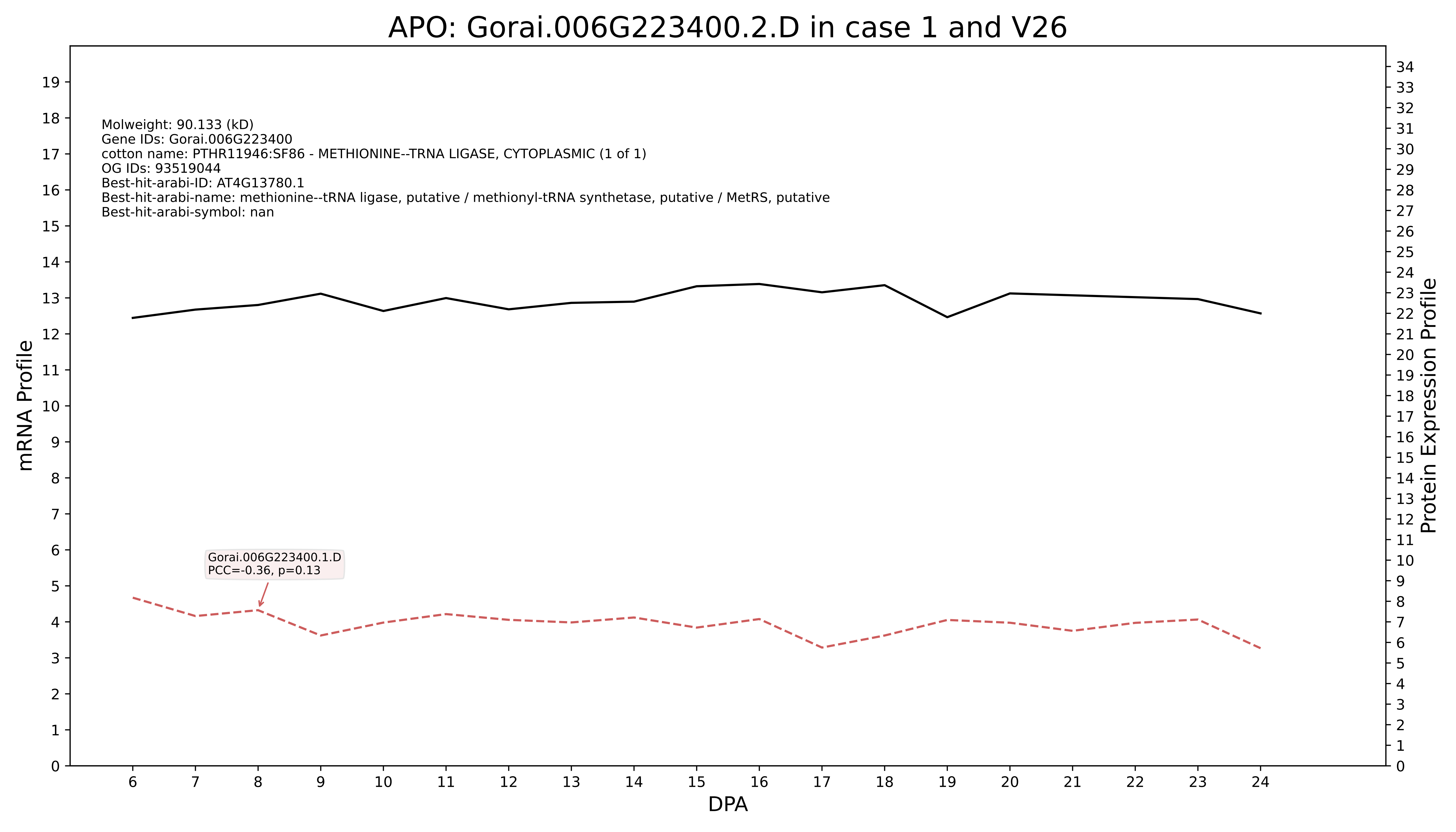
APO: Gorai.006G223400.2.D in case 1 and V26
Molweight: 90.133 (kD)
Gene IDs: Gorai.006G223400
cotton name: PTHR11946:SF86 - METHIONINE--TRNA LIGASE, CYTOPLASMIC (1 of 1)
OG IDs: 93519044
Best-hit-arabi-ID: AT4G13780.1
Best-hit-arabi-name: methionine--tRNA ligase, putative / methionyl-tRNA synthetase, putative / MetRS, putative
Best-hit-arabi-symbol: nan
Gorai.006G223400.1.D
PCC=-0.36, p=0.13
mRNA Profile
Protein Expression Profile
DPA

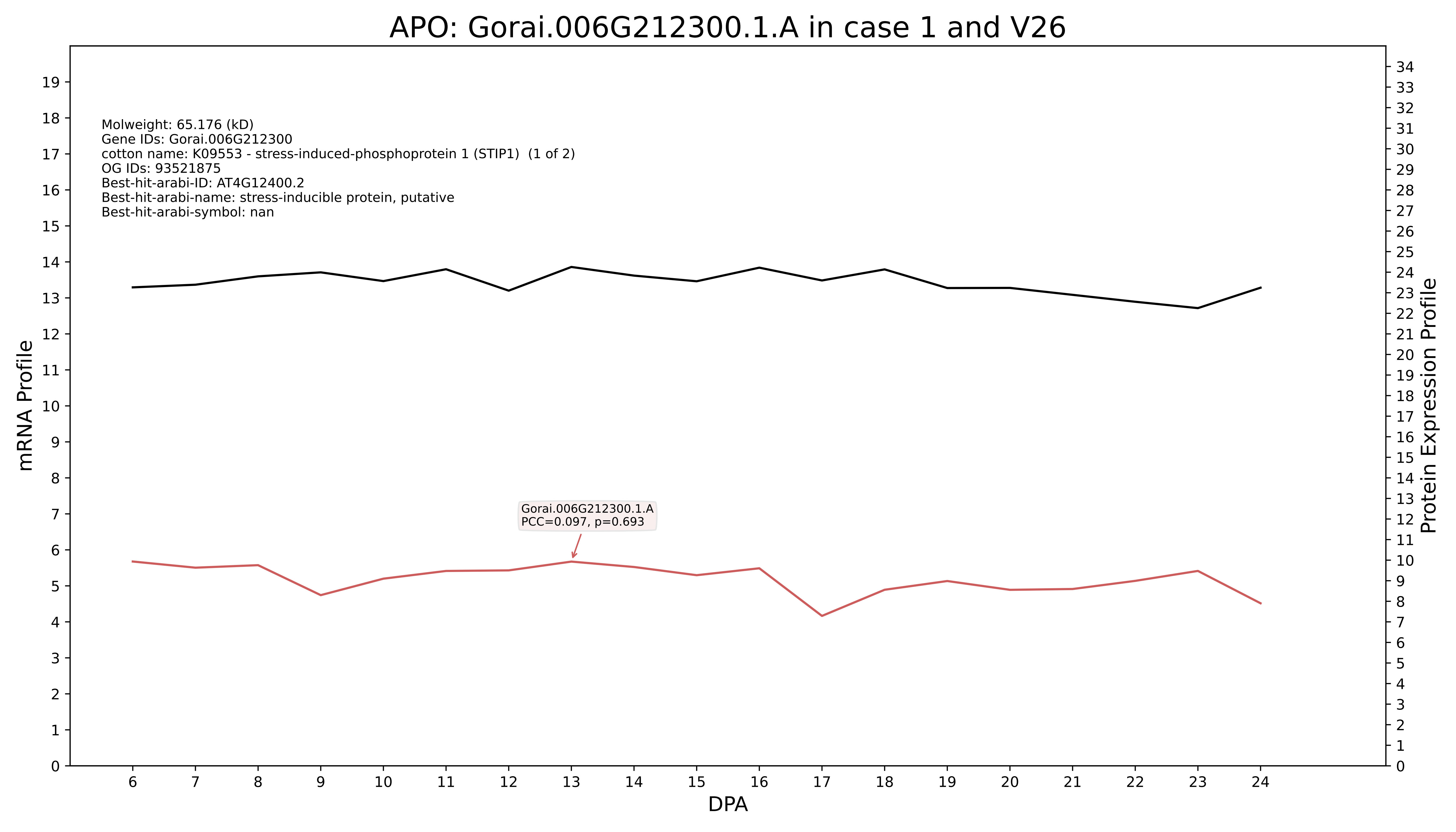

APO: Gorai.006G212300.1.A in case 1 and V26
Molweight: 65.176 (kD)
Gene IDs: Gorai.006G212300
cotton name: K09553 - stress-induced-phosphoprotein 1 (STIP1) (1 of 2)
OG IDs: 93521875
Best-hit-arabi-ID: AT4G12400.2
Best-hit-arabi-name: stress-inducible protein, putative
Best-hit-arabi-symbol: nan
Gorai.006G212300.1.A
PCC=0.097, p=0.693
mRNA Profile
Protein Expression Profile
DPA

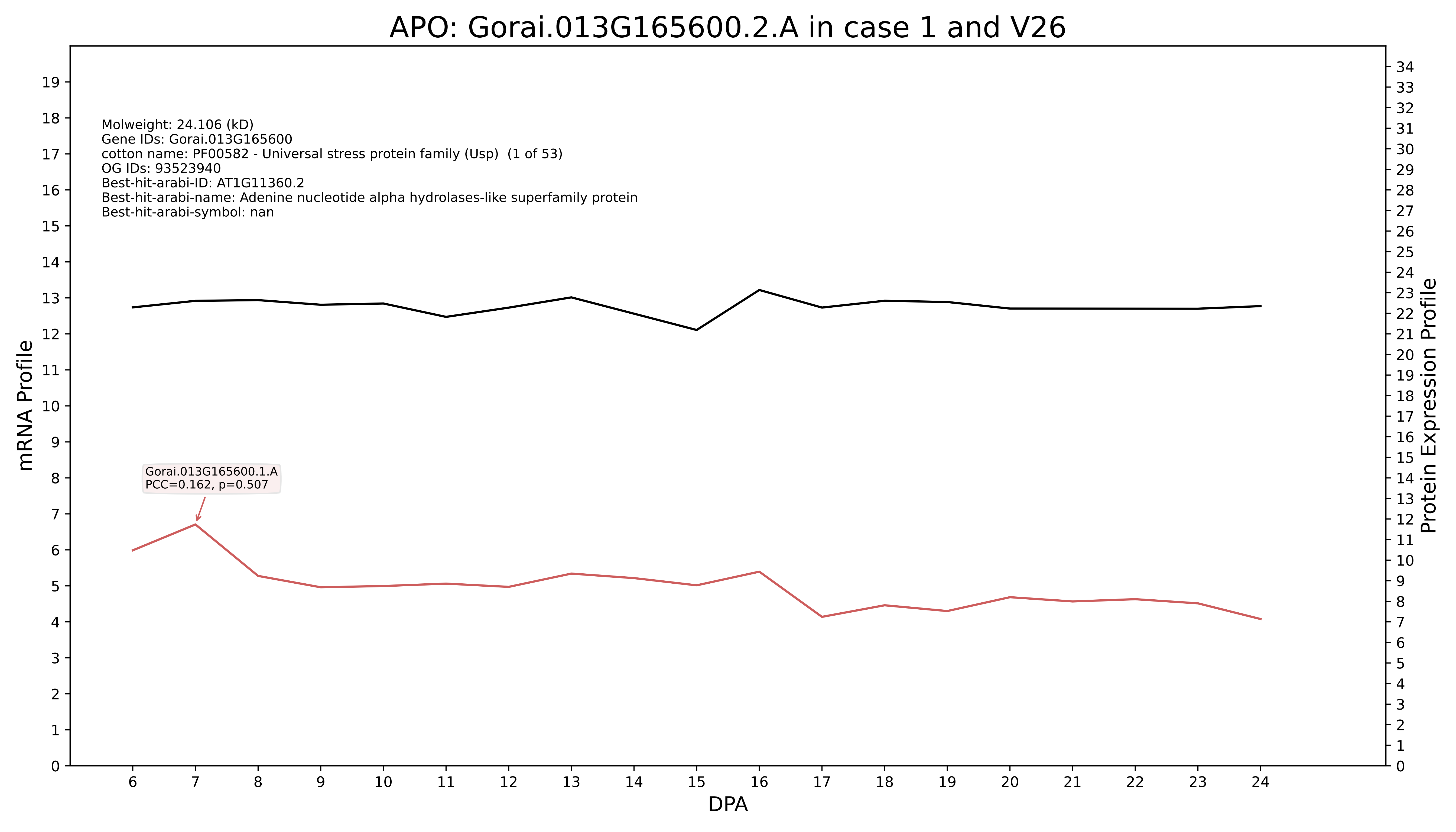
APO: Gorai.013G165600.2.A in case 1 and V26
Molweight: 24.106 (kD)
Gene IDs: Gorai.013G165600
cotton name: PF00582 - Universal stress protein family (Usp) (1 of 53)
OG IDs: 93523940
Best-hit-arabi-ID: AT1G11360.2
Best-hit-arabi-name: Adenine nucleotide alpha hydrolases-like superfamily protein
Best-hit-arabi-symbol: nan
Gorai.013G165600.1.A
PCC=0.162, p=0.507
mRNA Profile
Protein Expression Profile
DPA

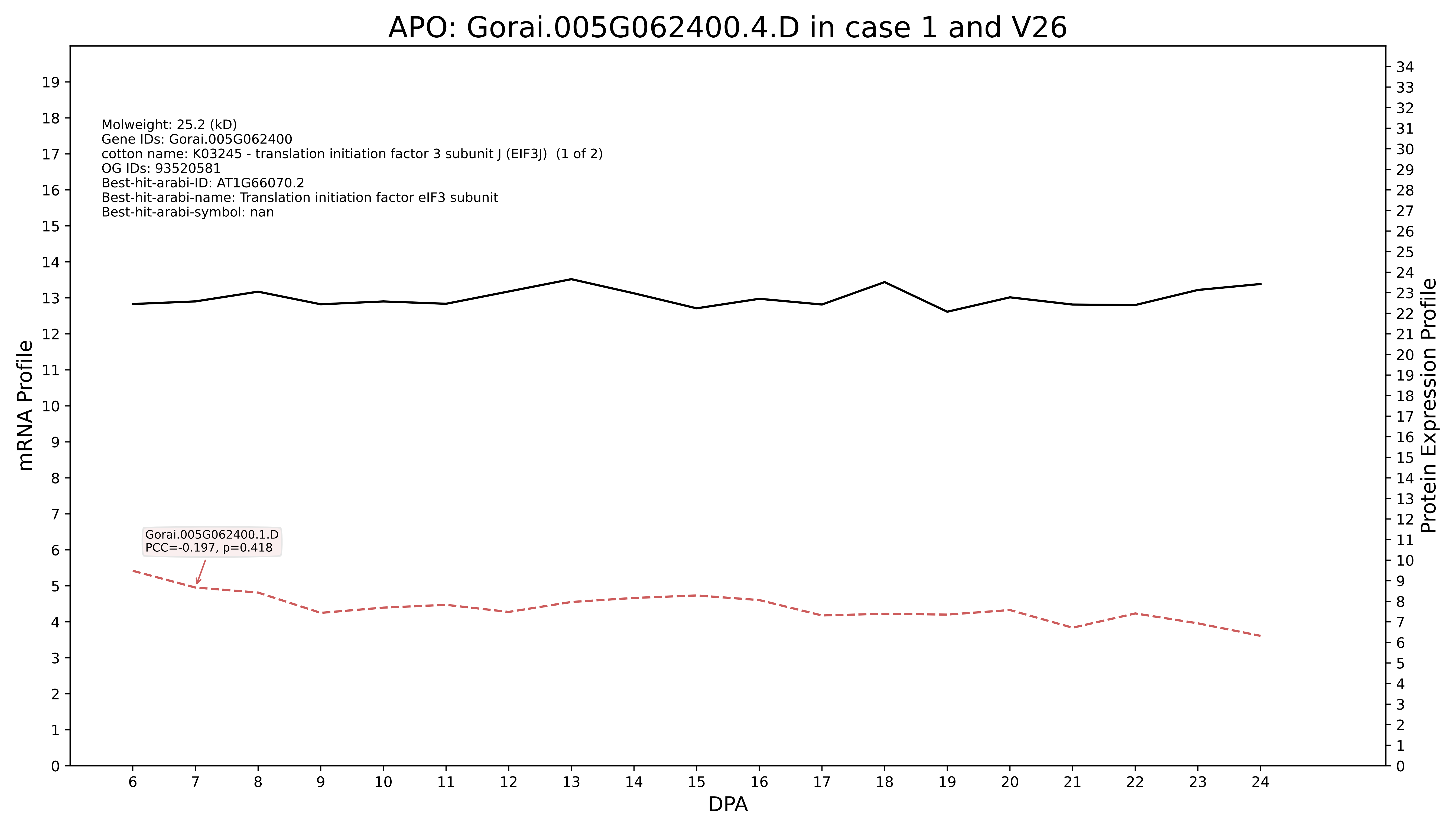
APO: Gorai.005G062400.4.D in case 1 and V26
Molweight: 25.2 (kD)
Gene IDs: Gorai.005G062400
cotton name: K03245 - translation initiation factor 3 subunit J (EIF3J) (1 of 2)
OG IDs: 93520581
Best-hit-arabi-ID: AT1G66070.2
Best-hit-arabi-name: Translation initiation factor eIF3 subunit
Best-hit-arabi-symbol: nan
Gorai.005G062400.1.D
PCC=-0.197, p=0.418
mRNA Profile
Protein Expression Profile
DPA

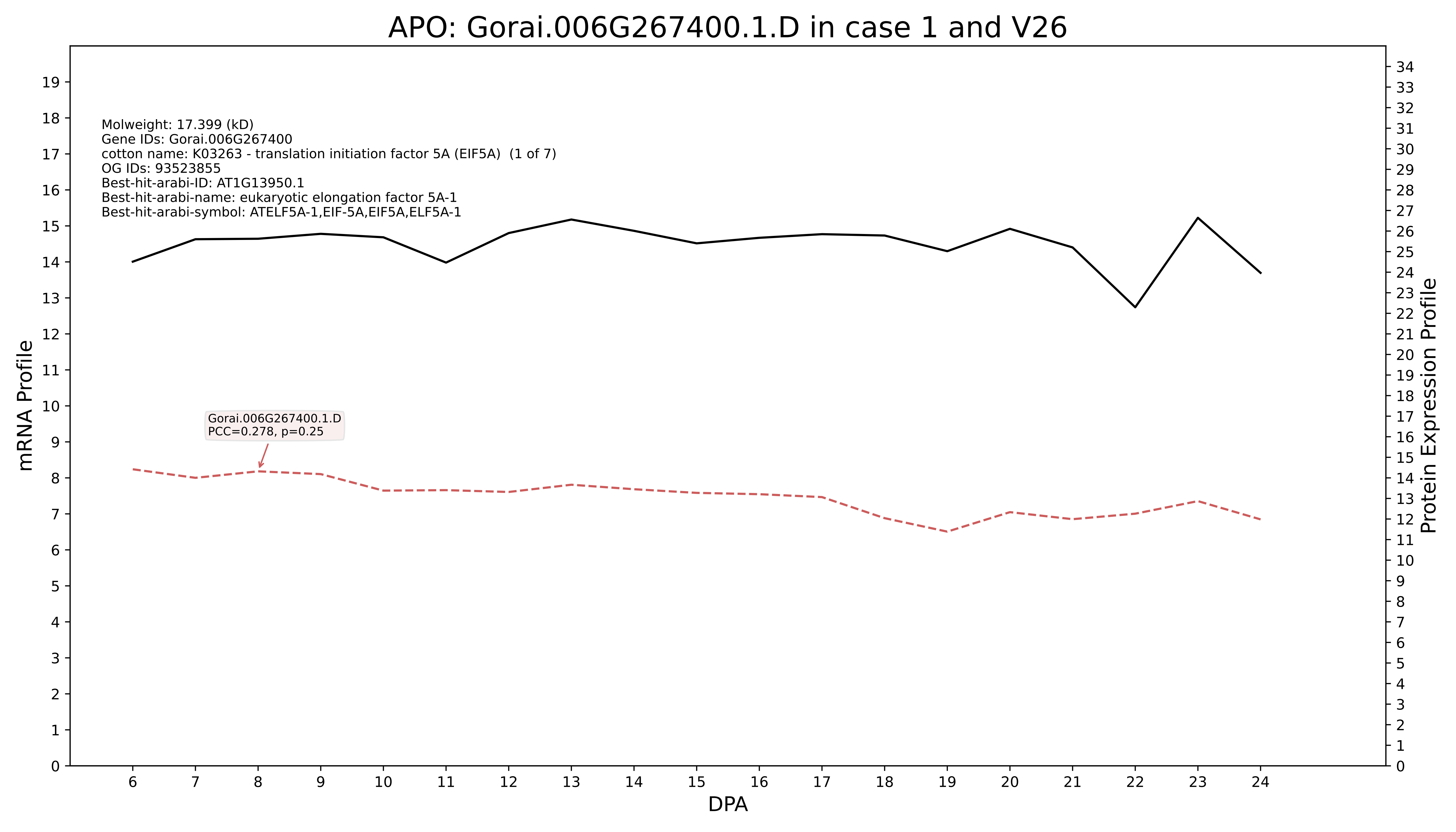
APO: Gorai.006G267400.1.D in case 1 and V26
Molweight: 17.399 (kD)
Gene IDs: Gorai.006G267400
cotton name: K03263 - translation initiation factor 5A (EIF5A) (1 of 7)
OG IDs: 93523855
Best-hit-arabi-ID: AT1G13950.1
Best-hit-arabi-name: eukaryotic elongation factor 5A-1
Best-hit-arabi-symbol: ATELF5A-1,EIF-5A,EIF5A,ELF5A-1
Gorai.006G267400.1.D
PCC=0.278, p=0.25
mRNA Profile
Protein Expression Profile
DPA

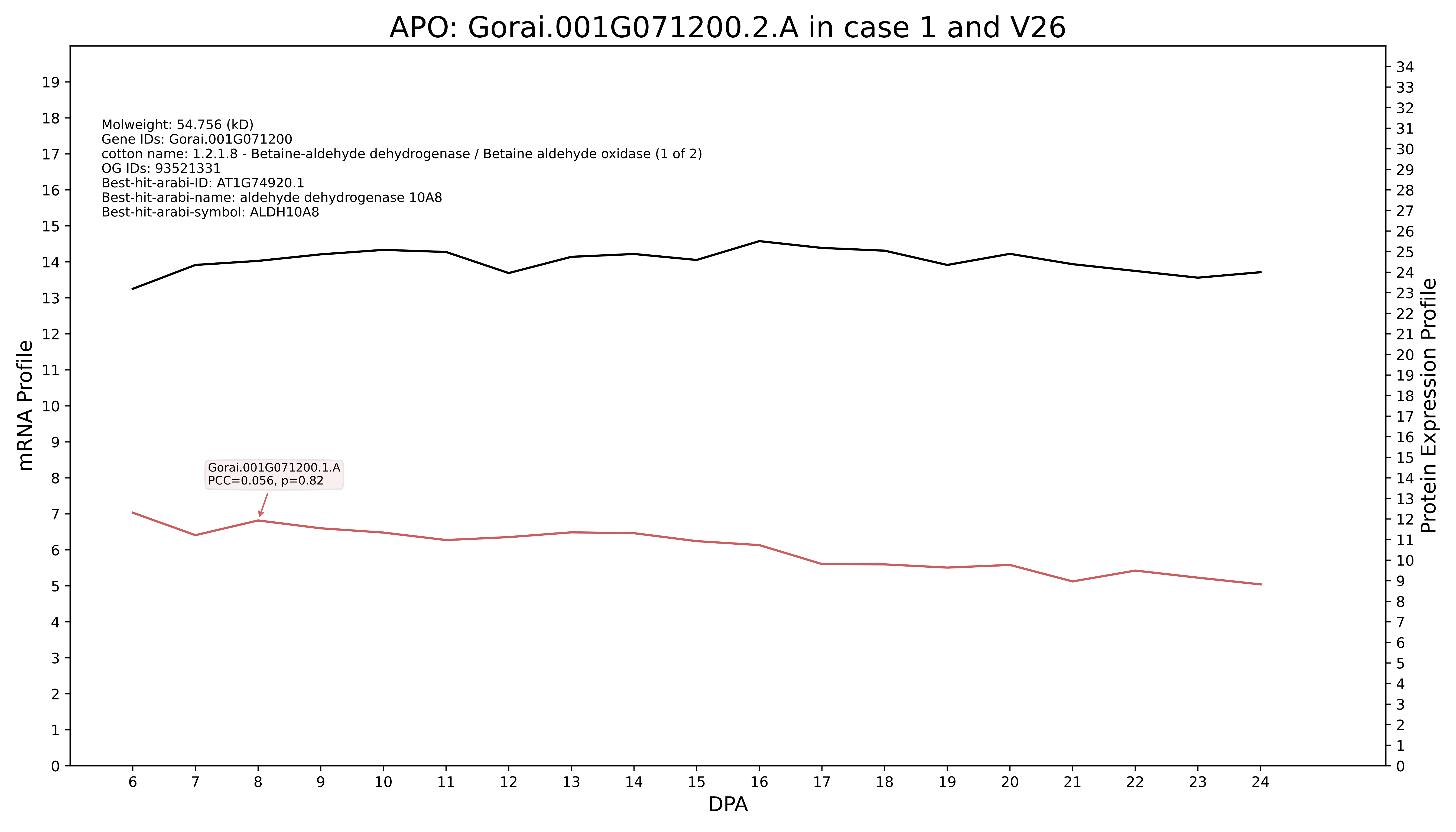
APO: Gorai.001G071200.2.A in case 1 and V26
Molweight: 54.756 (kD)
Gene IDs: Gorai.001G071200
cotton name: 1.2.1.8 - Betaine-aldehyde dehydrogenase / Betaine aldehyde oxidase (1 of 2)
OG IDs: 93521331
Best-hit-arabi-ID: AT1G74920.1
Best-hit-arabi-name: aldehyde dehydrogenase 10A8
Best-hit-arabi-symbol: ALDH10A8
Gorai.001G071200.1.A
PCC=0.056, p=0.82
mRNA Profile
Protein Expression Profile
DPA

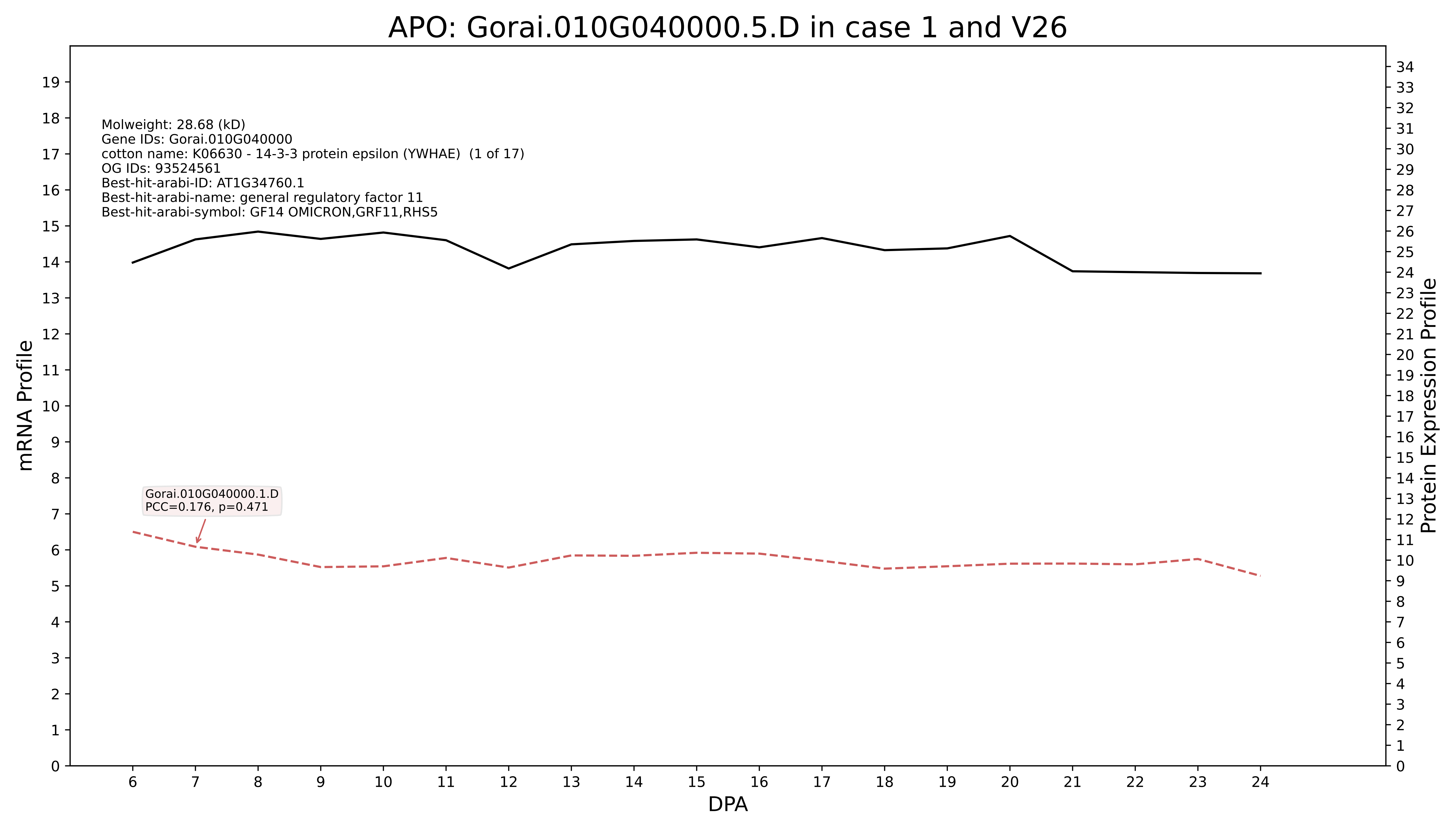
APO: Gorai.010G040000.5.D in case 1 and V26
Molweight: 28.68 (kD)
Gene IDs: Gorai.010G040000
cotton name: K06630 - 14-3-3 protein epsilon (YWHAE) (1 of 17)
OG IDs: 93524561
Best-hit-arabi-ID: AT1G34760.1
Best-hit-arabi-name: general regulatory factor 11
Best-hit-arabi-symbol: GF14 OMICRON,GRF11,RHS5
Gorai.010G040000.1.D
PCC=0.176, p=0.471
mRNA Profile
Protein Expression Profile
DPA

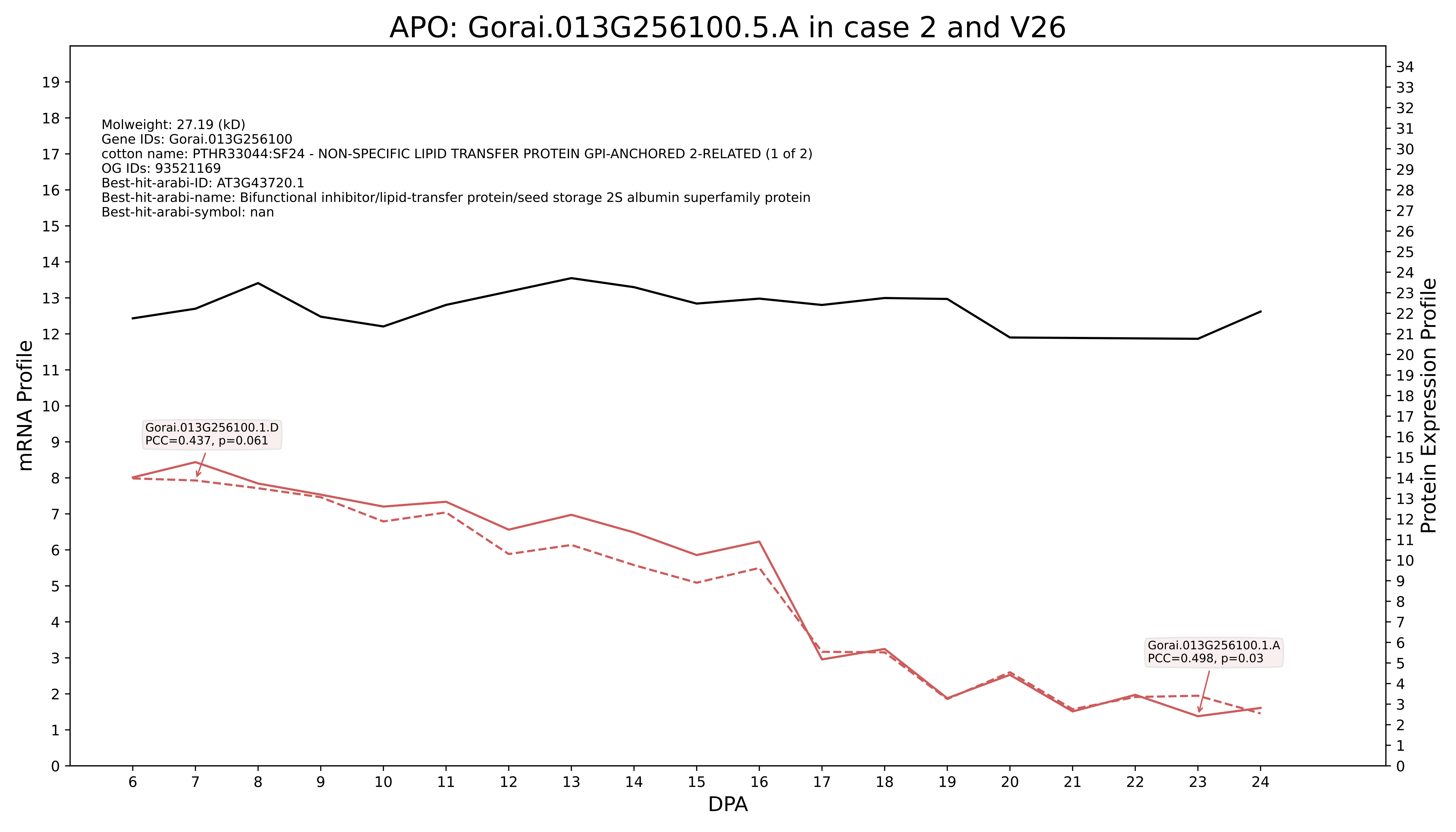
APO: Gorai.013G256100.5.A in case 2 and V26
Molweight: 27.19 (kD)
Gene IDs: Gorai.013G256100
cotton name: PTHR33044:SF24 - NON-SPECIFIC LIPID TRANSFER PROTEIN GPI-ANCHORED 2-RELATED (1 of 2)
OG IDs: 93521169
Best-hit-arabi-ID: AT3G43720.1
Best-hit-arabi-name: Bifunctional inhibitor/lipid-transfer protein/seed storage 2S albumin superfamily protein
Best-hit-arabi-symbol: nan
Gorai.013G256100.1.D
PCC=0.437, p=0.061
Gorai.013G256100.1.A
PCC=0.498, p=0.03
mRNA Profile
Protein Expression Profile
DPA

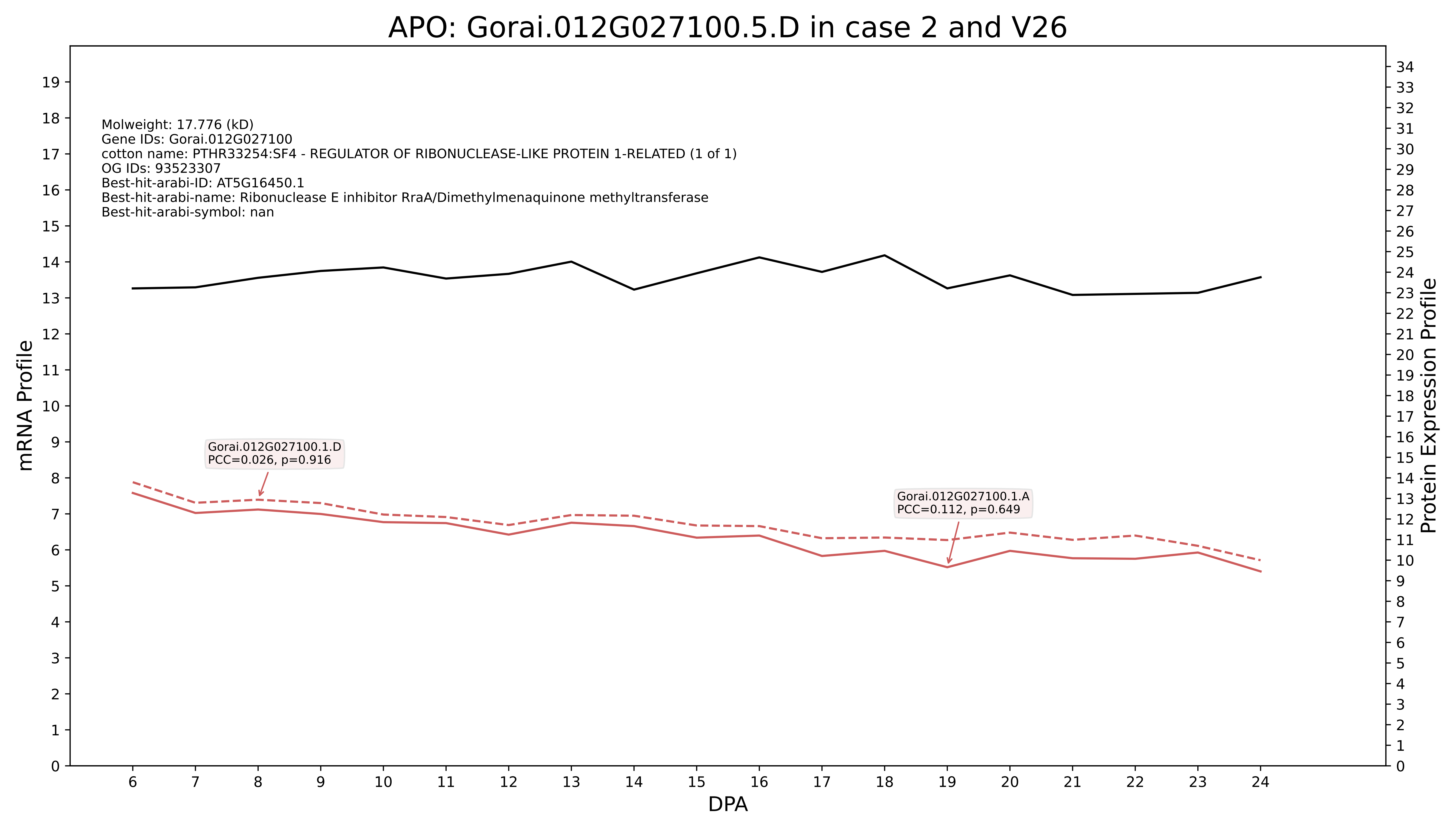

APO: Gorai.012G027100.5.D in case 2 and V26
Molweight: 17.776 (kD)
Gene IDs: Gorai.012G027100
cotton name: PTHR33254:SF4 - REGULATOR OF RIBONUCLEASE-LIKE PROTEIN 1-RELATED (1 of 1)
OG IDs: 93523307
Best-hit-arabi-ID: AT5G16450.1
Best-hit-arabi-name: Ribonuclease E inhibitor RraA/Dimethylmenaquinone methyltransferase
Best-hit-arabi-symbol: nan
Gorai.012G027100.1.D
PCC=0.026, p=0.916
Gorai.012G027100.1.A
PCC=0.112, p=0.649
mRNA Profile
Protein Expression Profile
DPA

### APO: Gorai.004G212800.3.D in case 2 and V26

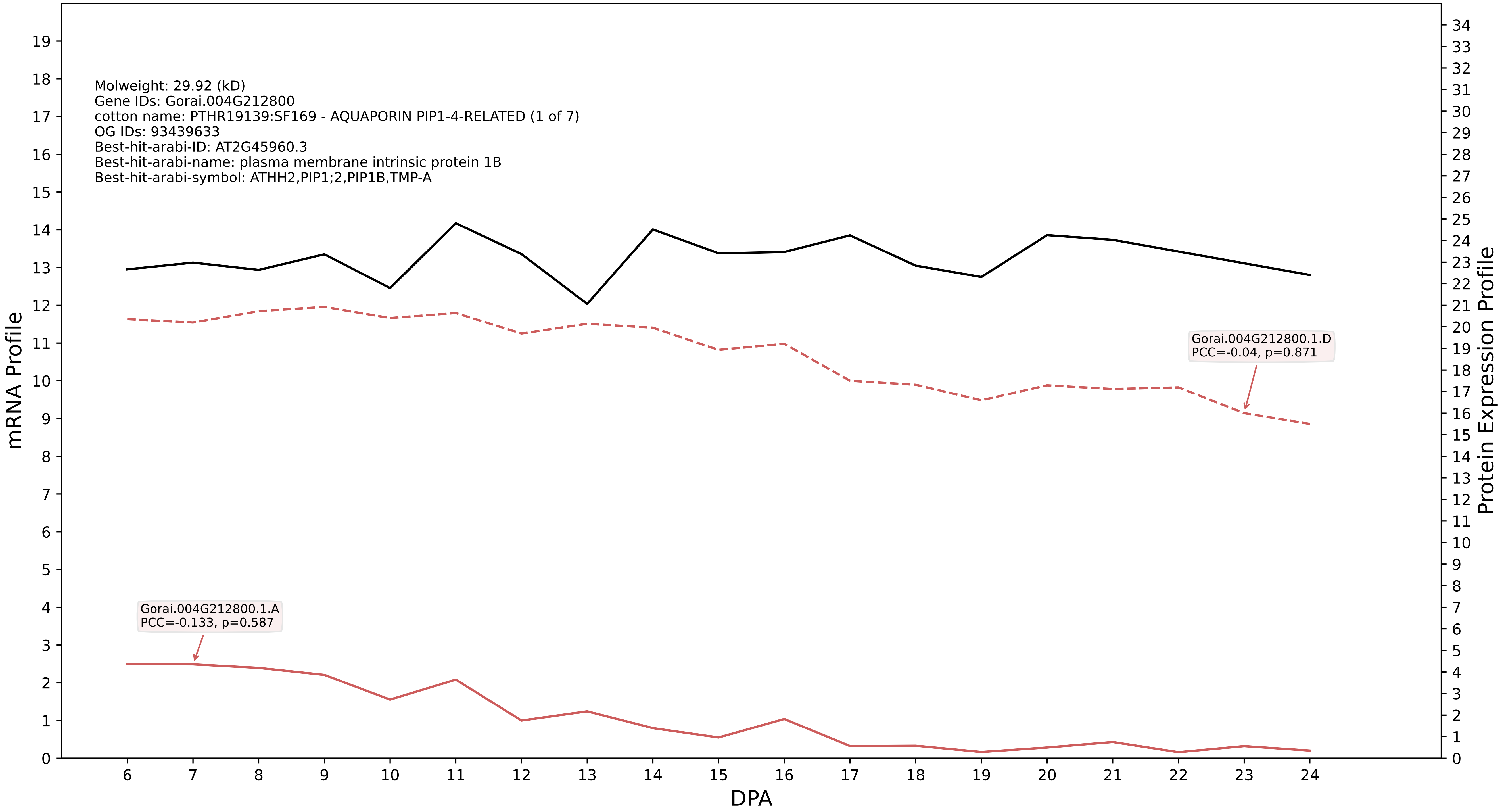

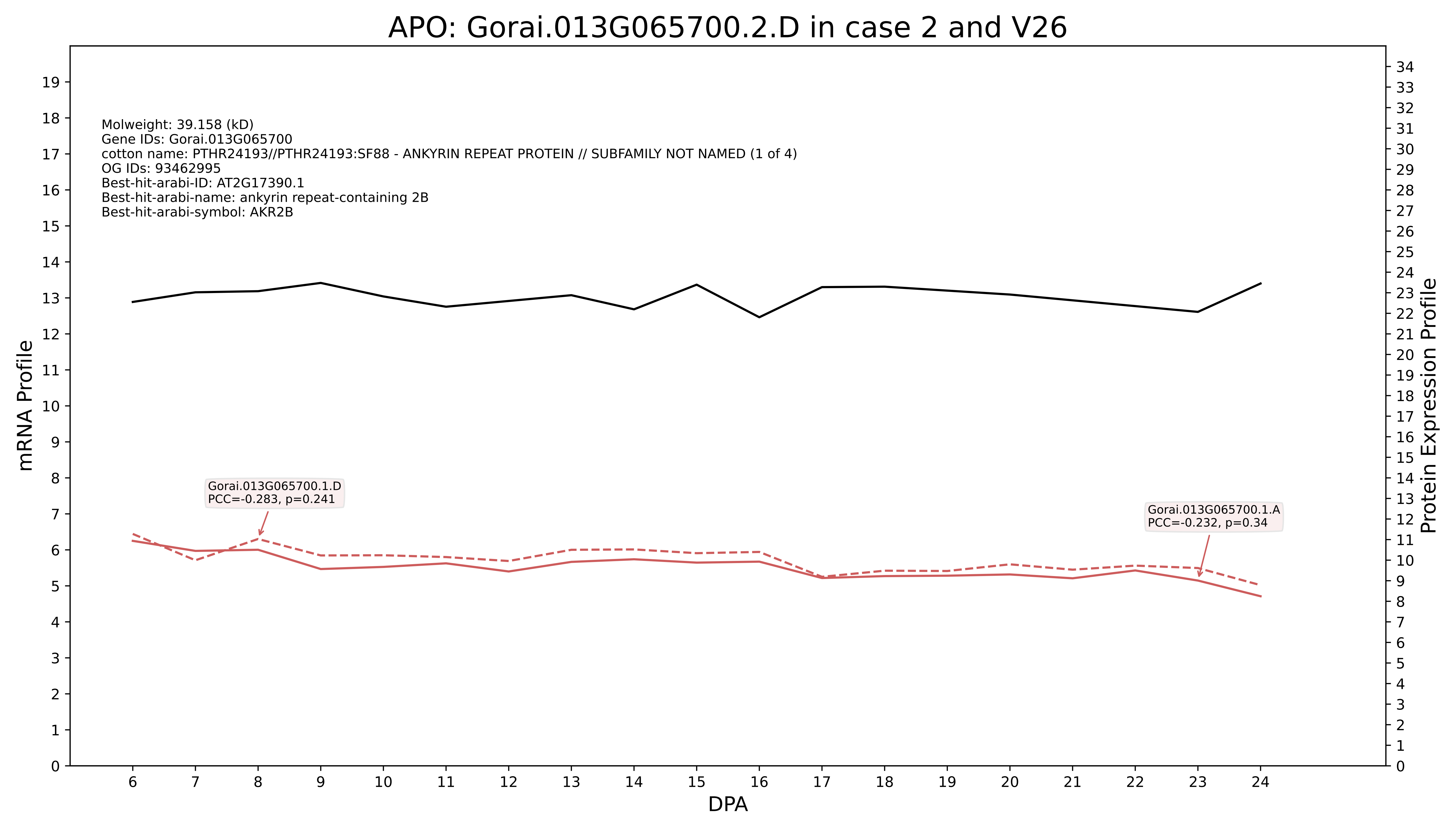
APO: Gorai.013G065700.2.D in case 2 and V26
Molweight: 39.158 (kD)
Gene IDs: Gorai.013G065700
cotton name: PTHR24193//PTHR24193:SF88 - ANKYRIN REPEAT PROTEIN // SUBFAMILY NOT NAMED (1 of 4)
OG IDs: 93462995
Best-hit-arabi-ID: AT2G17390.1
Best-hit-arabi-name: ankyrin repeat-containing 2B
Best-hit-arabi-symbol: AKR2B
Gorai.013G065700.1.D
PCC=-0.283, p=0.241
Gorai.013G065700.1.A
PCC=-0.232, p=0.34
mRNA Profile
Protein Expression Profile
DPA

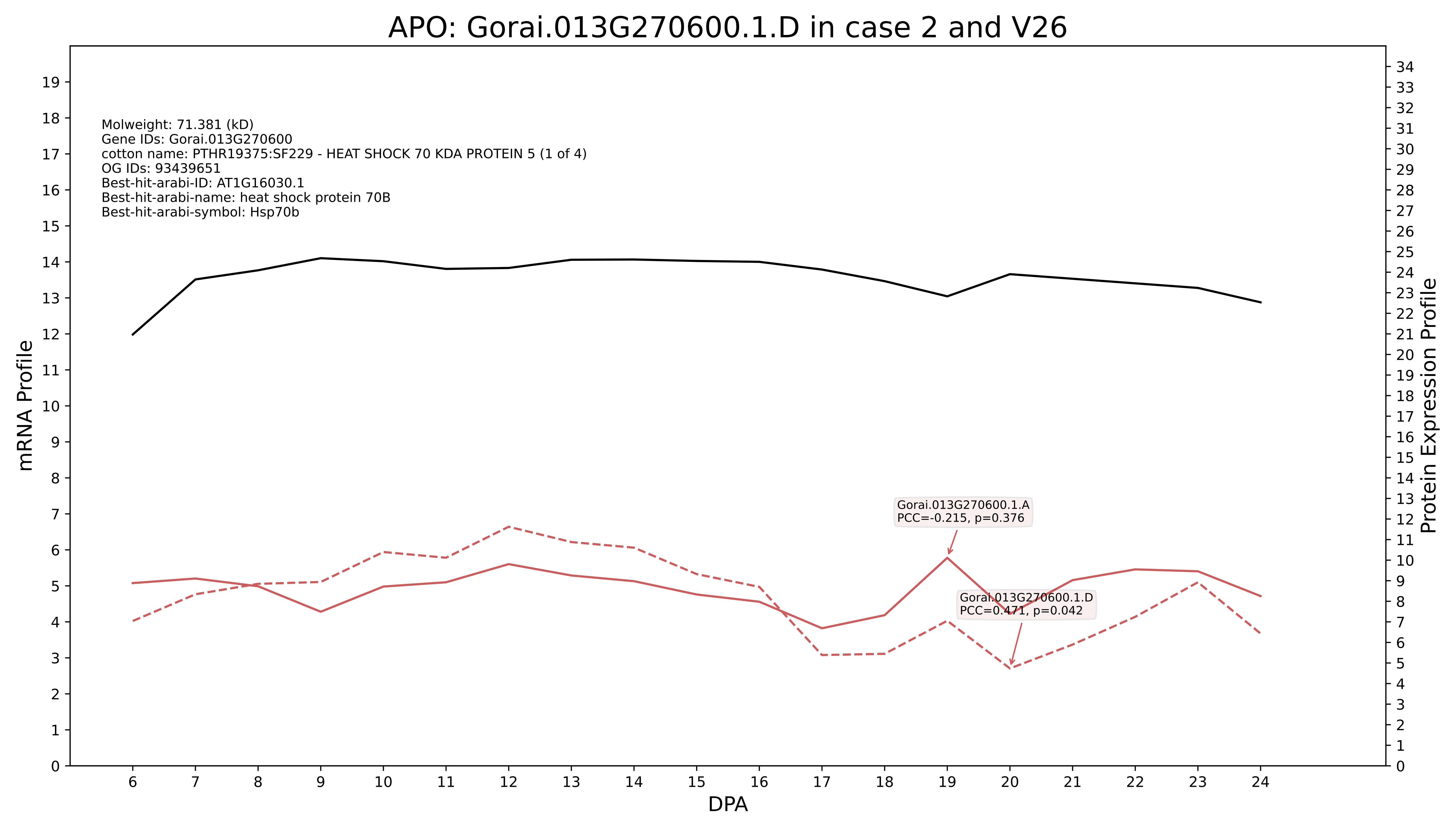

APO: Gorai.013G270600.1.D in case 2 and V26
Molweight: 71.381 (kD)
Gene IDs: Gorai.013G270600
cotton name: PTHR19375:SF229 - HEAT SHOCK 70 KDA PROTEIN 5 (1 of 4)
OG IDs: 93439651
Best-hit-arabi-ID: AT1G16030.1
Best-hit-arabi-name: heat shock protein 70B
Best-hit-arabi-symbol: Hsp70b
Gorai.013G270600.1.A
PCC=-0.215, p=0.376
Gorai.013G270600.1.D
PCC=0.471, p=0.042
mRNA Profile
Protein Expression Profile
DPA
0 1 2 3 4 5 6 7 8 9 10 11 12 13 14 15 16 17 18 19
0 1 2 3 4 5 6 7 8 9 10 11 12 13 14 15 16 17 18 19 20 21 22 23 24 25 26 27 28 29 30 31 32 33 34
6 7 8 9 10 11 12 13 14 15 16 17 18 19 20 21 22 23 24

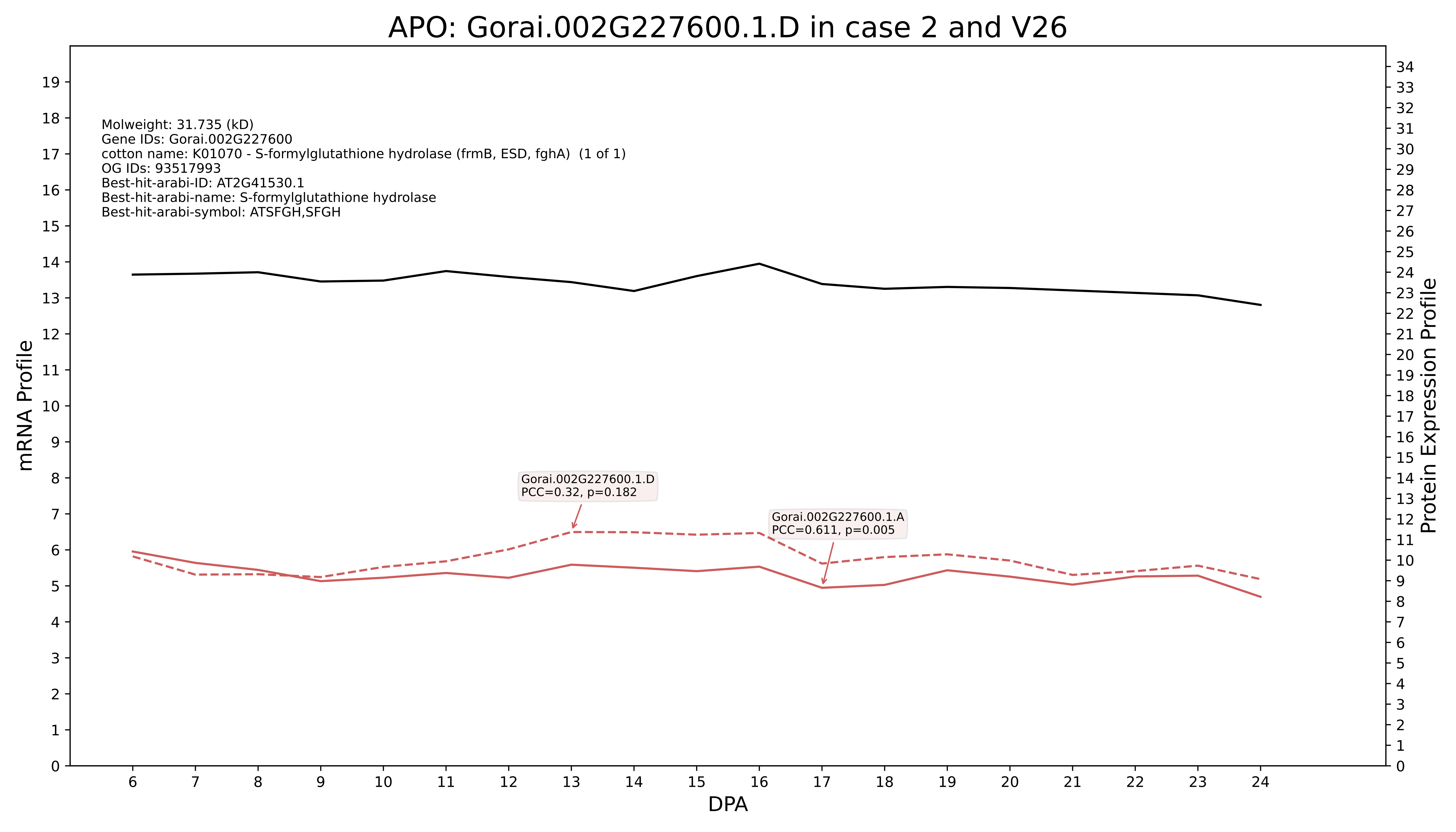
APO: Gorai.002G227600.1.D in case 2 and V26
Molweight: 31.735 (kD)
Gene IDs: Gorai.002G227600
cotton name: K01070 - S-formylglutathione hydrolase (frmB, ESD, fghA) (1 of 1)
OG IDs: 93517993
Best-hit-arabi-ID: AT2G41530.1
Best-hit-arabi-name: S-formylglutathione hydrolase
Best-hit-arabi-symbol: ATSFGH,SFGH
Gorai.002G227600.1.D
PCC=0.32, p=0.182
Gorai.002G227600.1.A
PCC=0.611, p=0.005
mRNA Profile
Protein Expression Profile
DPA

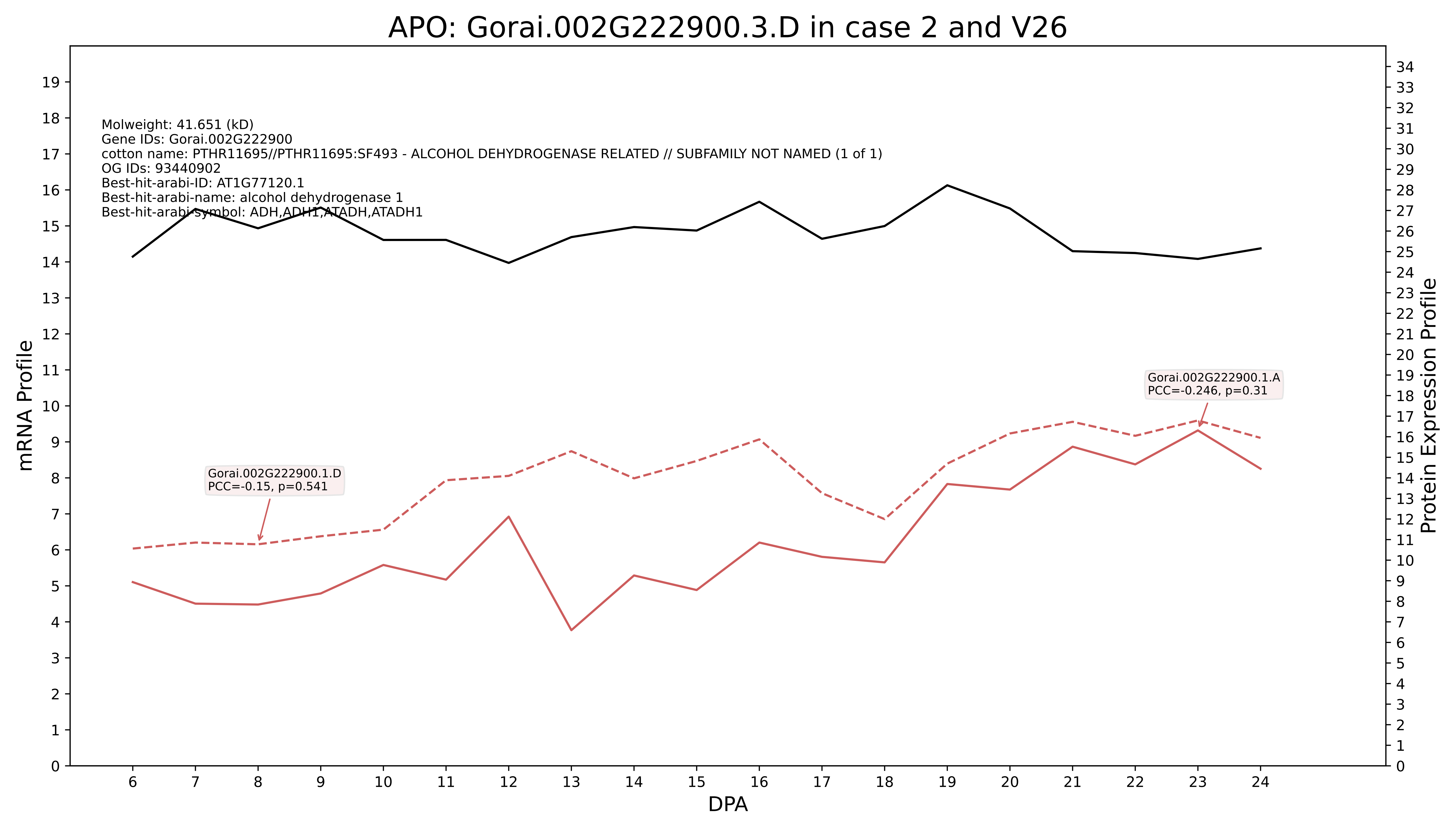
APO: Gorai.002G222900.3.D in case 2 and V26
Molweight: 41.651 (kD)
Gene IDs: Gorai.002G222900
cotton name: PTHR11695//PTHR11695:SF493 - ALCOHOL DEHYDROGENASE RELATED // SUBFAMILY NOT NAMED (1 of 1)
OG IDs: 93440902
Best-hit-arabi-ID: AT1G77120.1
Best-hit-arabi-name: alcohol dehydrogenase 1
Best-hit-arabi-symbol: ADH,ADH1,ATADH,ATADH1
Gorai.002G222900.1.A
PCC=-0.246, p=0.31
Gorai.002G222900.1.D
PCC=-0.15, p=0.541
mRNA Profile
Protein Expression Profile
DPA

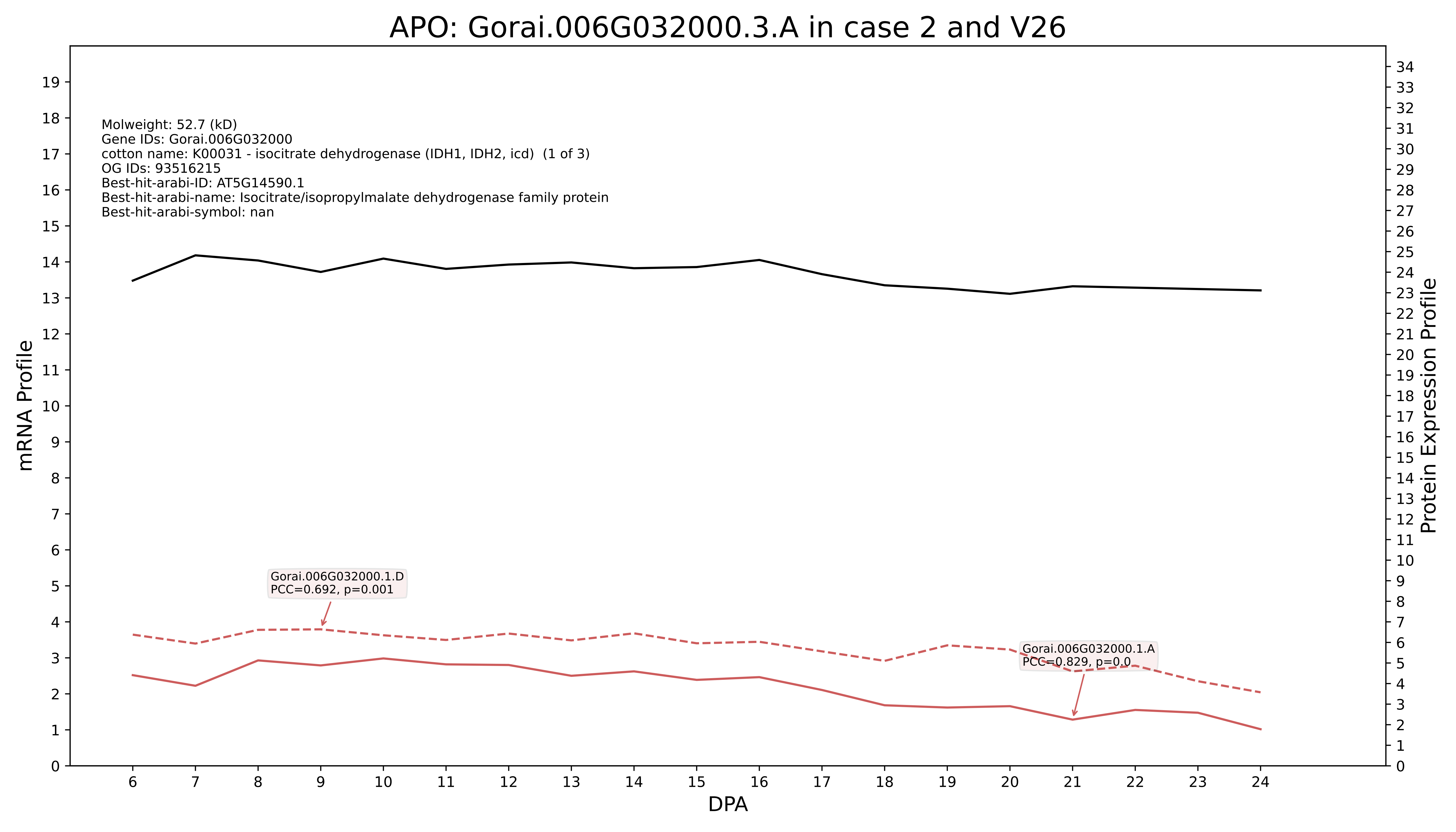
APO: Gorai.006G032000.3.A in case 2 and V26
Molweight: 52.7 (kD)
Gene IDs: Gorai.006G032000
cotton name: K00031 - isocitrate dehydrogenase (IDH1, IDH2, icd) (1 of 3)
OG IDs: 93516215
Best-hit-arabi-ID: AT5G14590.1
Best-hit-arabi-name: Isocitrate/isopropylmalate dehydrogenase family protein
Best-hit-arabi-symbol: nan
Gorai.006G032000.1.D
PCC=0.692, p=0.001
Gorai.006G032000.1.A
PCC=0.829, p=0.0
mRNA Profile
Protein Expression Profile
DPA

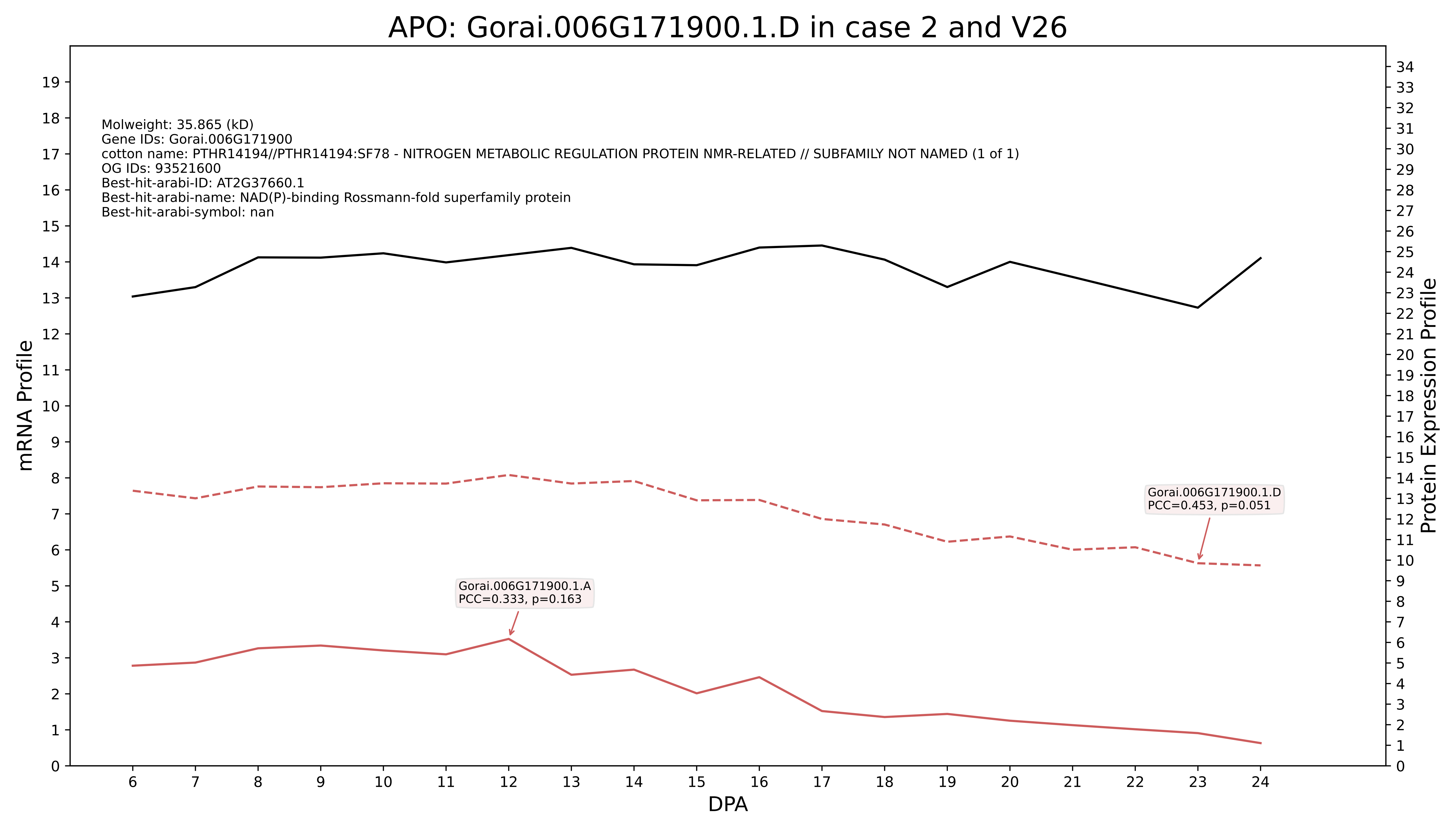
APO: Gorai.006G171900.1.D in case 2 and V26
Molweight: 35.865 (kD)
Gene IDs: Gorai.006G171900
cotton name: PTHR14194//PTHR14194:SF78 - NITROGEN METABOLIC REGULATION PROTEIN NMR-RELATED // SUBFAMILY NOT NAMED (1 of 1)
OG IDs: 93521600
Best-hit-arabi-ID: AT2G37660.1
Best-hit-arabi-name: NAD(P)-binding Rossmann-fold superfamily protein
Best-hit-arabi-symbol: nan
Gorai.006G171900.1.D
PCC=0.453, p=0.051
Gorai.006G171900.1.A
PCC=0.333, p=0.163
mRNA Profile
Protein Expression Profile
DPA

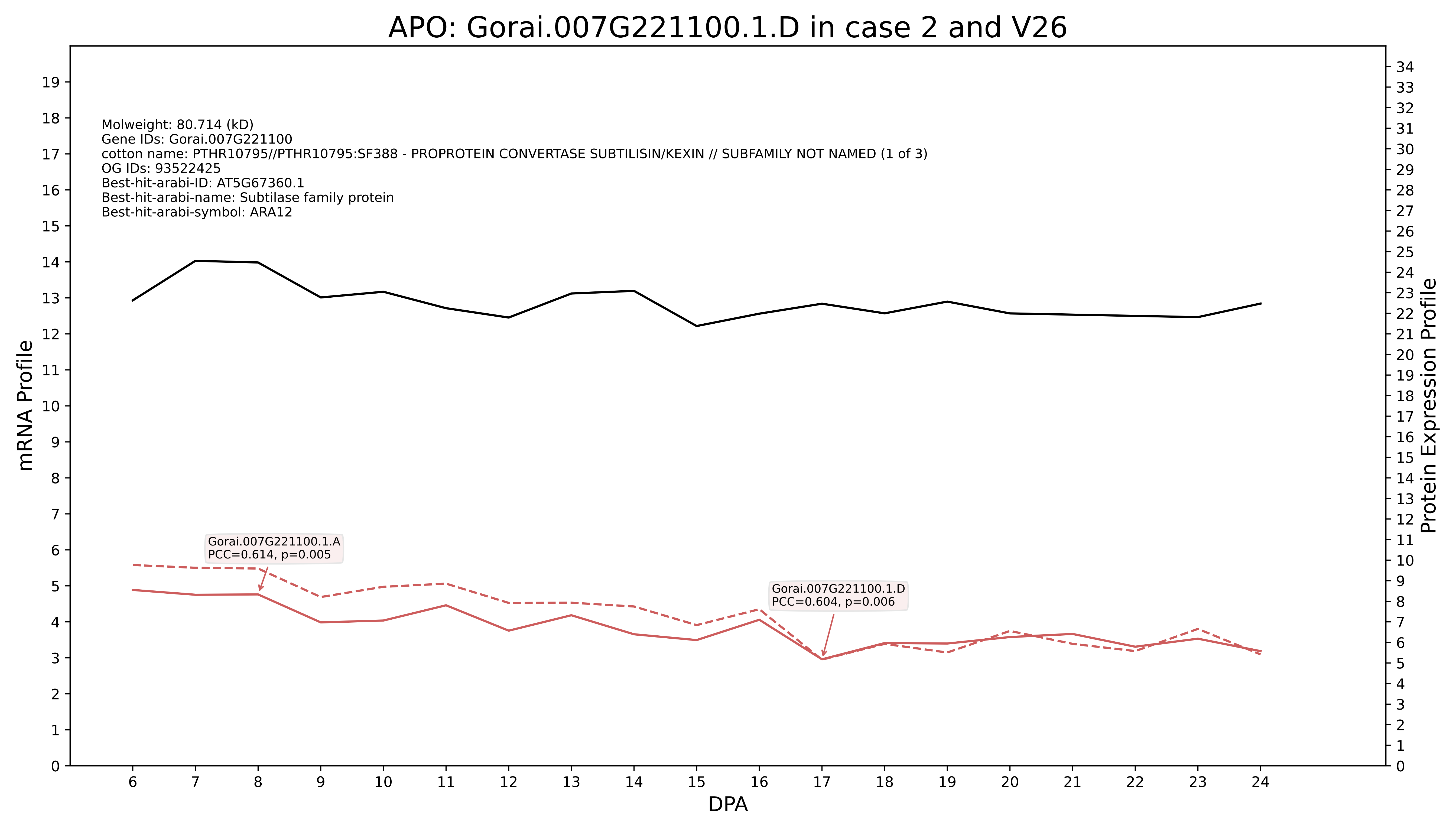

APO: Gorai.007G221100.1.D in case 2 and V26
Molweight: 80.714 (kD)
Gene IDs: Gorai.007G221100
cotton name: PTHR10795//PTHR10795:SF388 - PROPROTEIN CONVERTASE SUBTILISIN/KEXIN // SUBFAMILY NOT NAMED (1 of 3)
OG IDs: 93522425
Best-hit-arabi-ID: AT5G67360.1
Best-hit-arabi-name: Subtilase family protein
Best-hit-arabi-symbol: ARA12
Gorai.007G221100.1.A
PCC=0.614, p=0.005
Gorai.007G221100.1.D
PCC=0.604, p=0.006
mRNA Profile
Protein Expression Profile
DPA

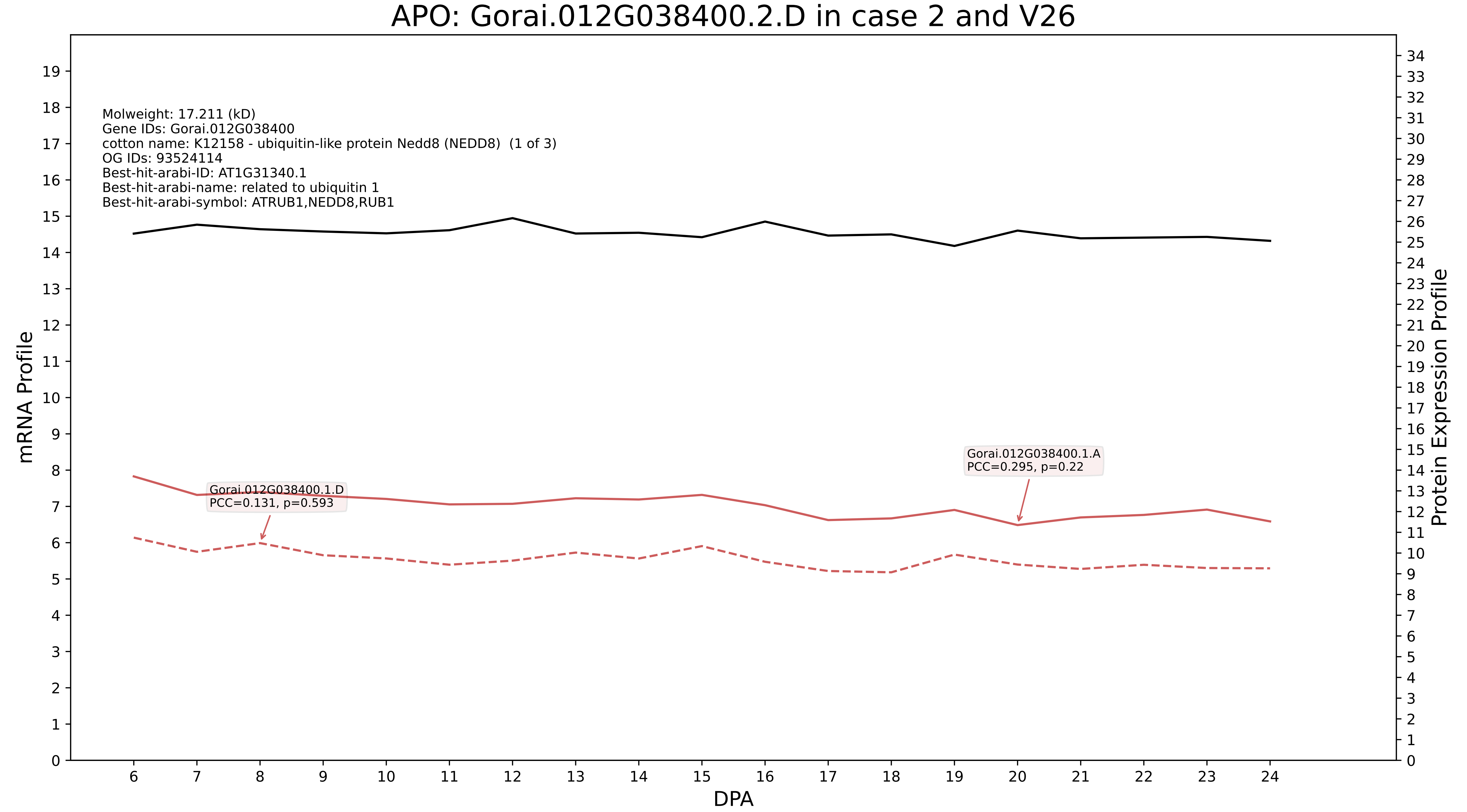

APO: Gorai.012G038400.2.D in case 2 and V26
Molweight: 17.211 (kD)
Gene IDs: Gorai.012G038400
cotton name: K12158 - ubiquitin-like protein Nedd8 (NEDD8) (1 of 3)
OG IDs: 93524114
Best-hit-arabi-ID: AT1G31340.1
Best-hit-arabi-name: related to ubiquitin 1
Best-hit-arabi-symbol: ATRUB1,NEDD8,RUB1
Gorai.012G038400.1.D
PCC=0.131, p=0.593
Gorai.012G038400.1.A
PCC=0.295, p=0.22
mRNA Profile
Protein Expression Profile
DPA

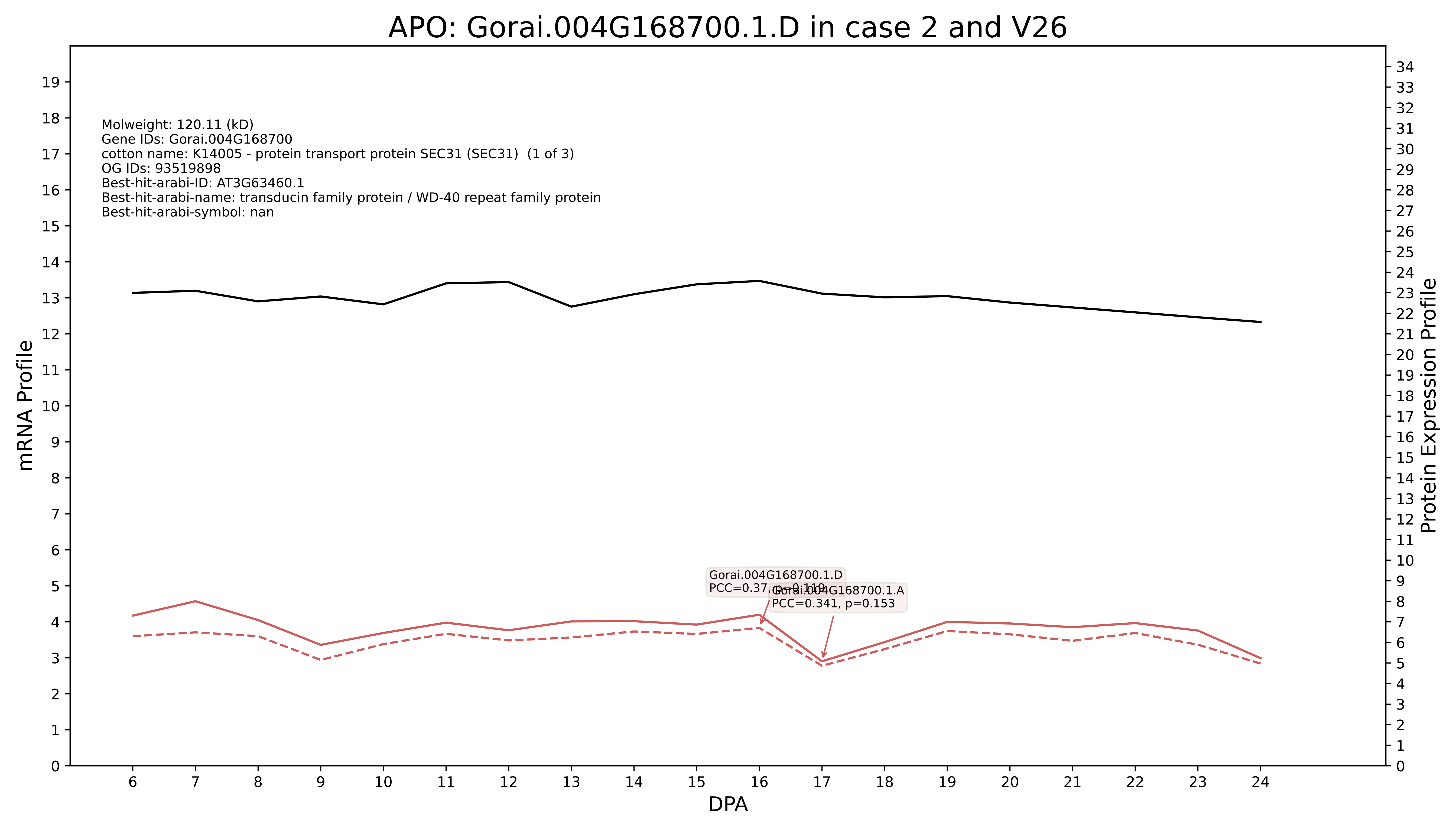
APO: Gorai.004G168700.1.D in case 2 and V26
Molweight: 120.11 (kD)
Gene IDs: Gorai.004G168700
cotton name: K14005 - protein transport protein SEC31 (SEC31) (1 of 3)
OG IDs: 93519898
Best-hit-arabi-ID: AT3G63460.1
Best-hit-arabi-name: transducin family protein / WD-40 repeat family protein
Best-hit-arabi-symbol: nan
Gorai.004G168700.1.D
PCC=0.37, p=0.119
Gorai.004G168700.1.A
PCC=0.341, p=0.153
mRNA Profile
Protein Expression Profile
DPA

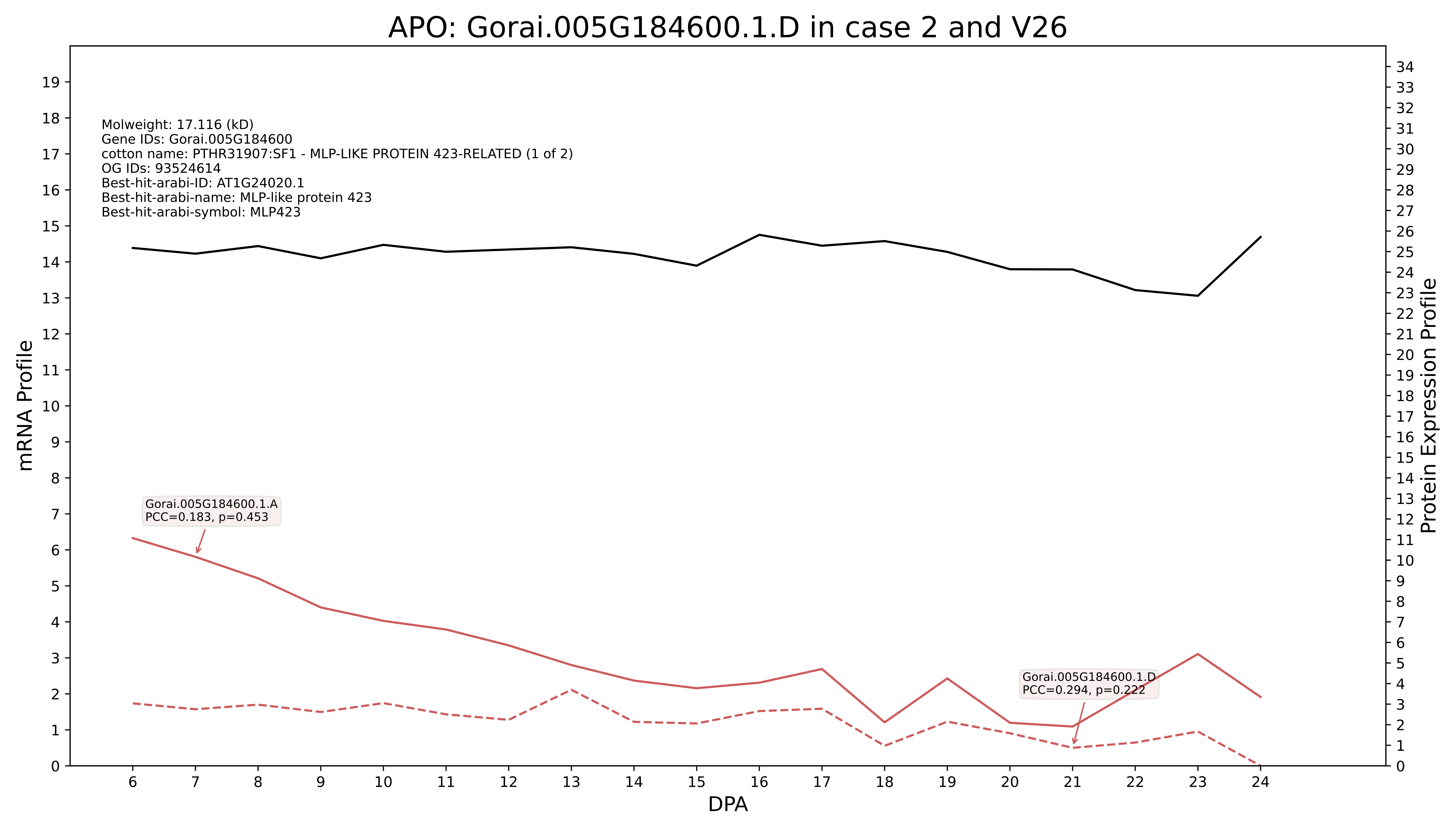
APO: Gorai.005G184600.1.D in case 2 and V26
Molweight: 17.116 (kD)
Gene IDs: Gorai.005G184600
cotton name: PTHR31907:SF1 - MLP-LIKE PROTEIN 423-RELATED (1 of 2)
OG IDs: 93524614
Best-hit-arabi-ID: AT1G24020.1
Best-hit-arabi-name: MLP-like protein 423
Best-hit-arabi-symbol: MLP423
Gorai.005G184600.1.A
PCC=0.183, p=0.453
Gorai.005G184600.1.D
PCC=0.294, p=0.222
mRNA Profile
Protein Expression Profile
DPA

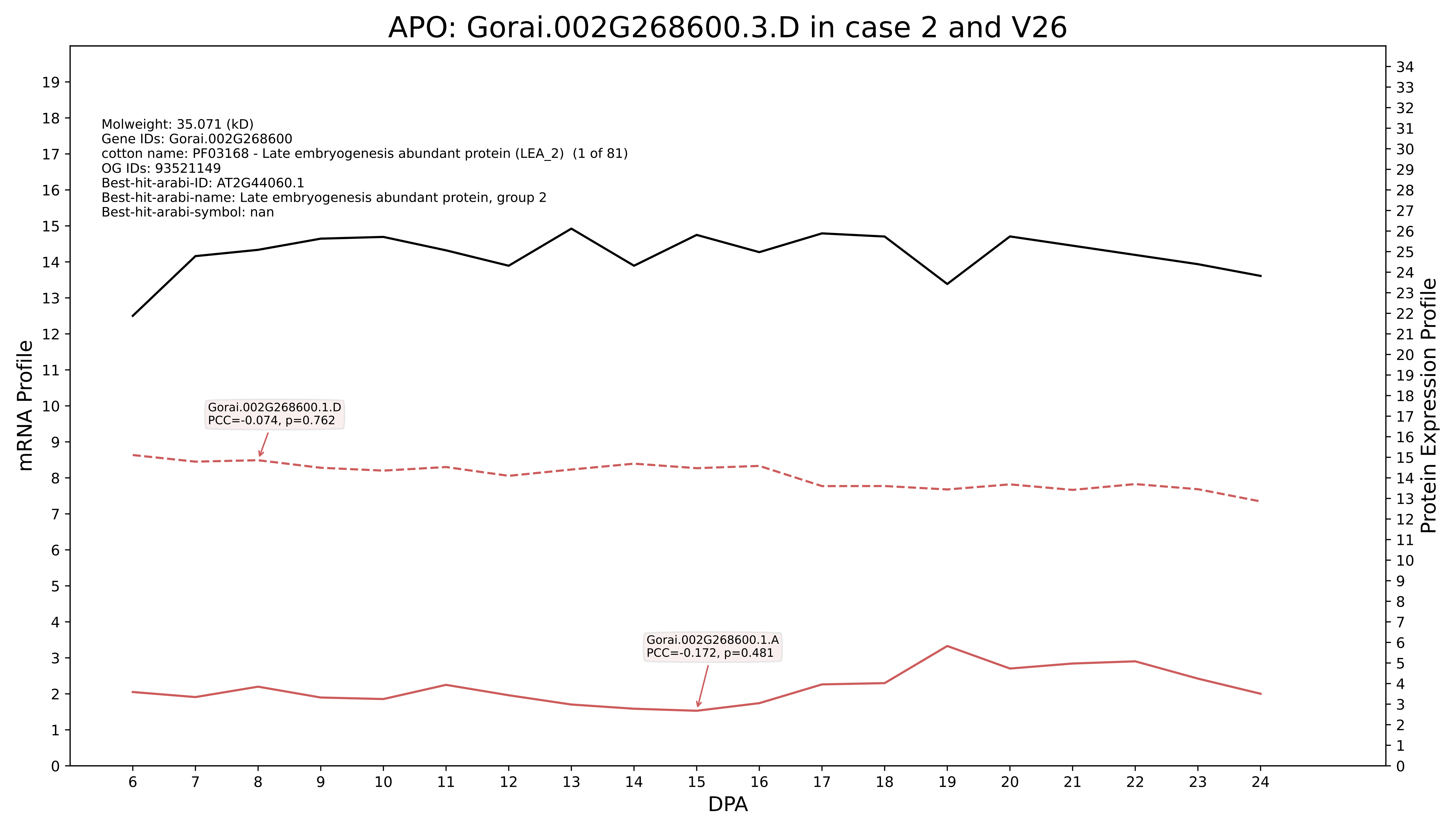
APO: Gorai.002G268600.3.D in case 2 and V26
Molweight: 35.071 (kD)
Gene IDs: Gorai.002G268600
cotton name: PF03168 - Late embryogenesis abundant protein (LEA_2) (1 of 81)
OG IDs: 93521149
Best-hit-arabi-ID: AT2G44060.1
Best-hit-arabi-name: Late embryogenesis abundant protein, group 2
Best-hit-arabi-symbol: nan
Gorai.002G268600.1.D
PCC=-0.074, p=0.762
Gorai.002G268600.1.A
PCC=-0.172, p=0.481
mRNA Profile
Protein Expression Profile
DPA

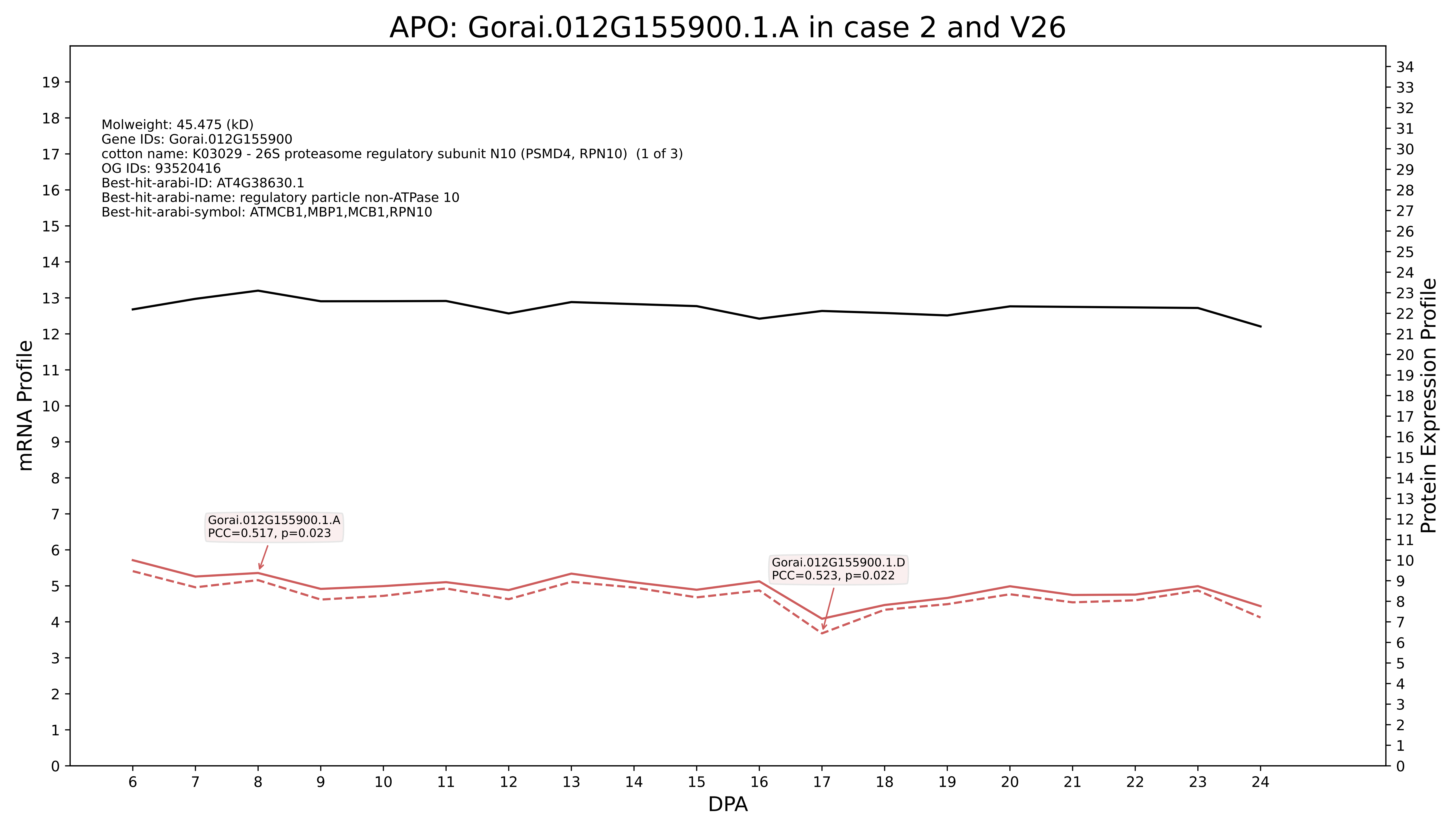
APO: Gorai.012G155900.1.A in case 2 and V26
Molweight: 45.475 (kD)
Gene IDs: Gorai.012G155900
cotton name: K03029 - 26S proteasome regulatory subunit N10 (PSMD4, RPN10) (1 of 3)
OG IDs: 93520416
Best-hit-arabi-ID: AT4G38630.1
Best-hit-arabi-name: regulatory particle non-ATPase 10
Best-hit-arabi-symbol: ATMCB1,MBP1,MCB1,RPN10
Gorai.012G155900.1.A
PCC=0.517, p=0.023
Gorai.012G155900.1.D
PCC=0.523, p=0.022
mRNA Profile
Protein Expression Profile
DPA

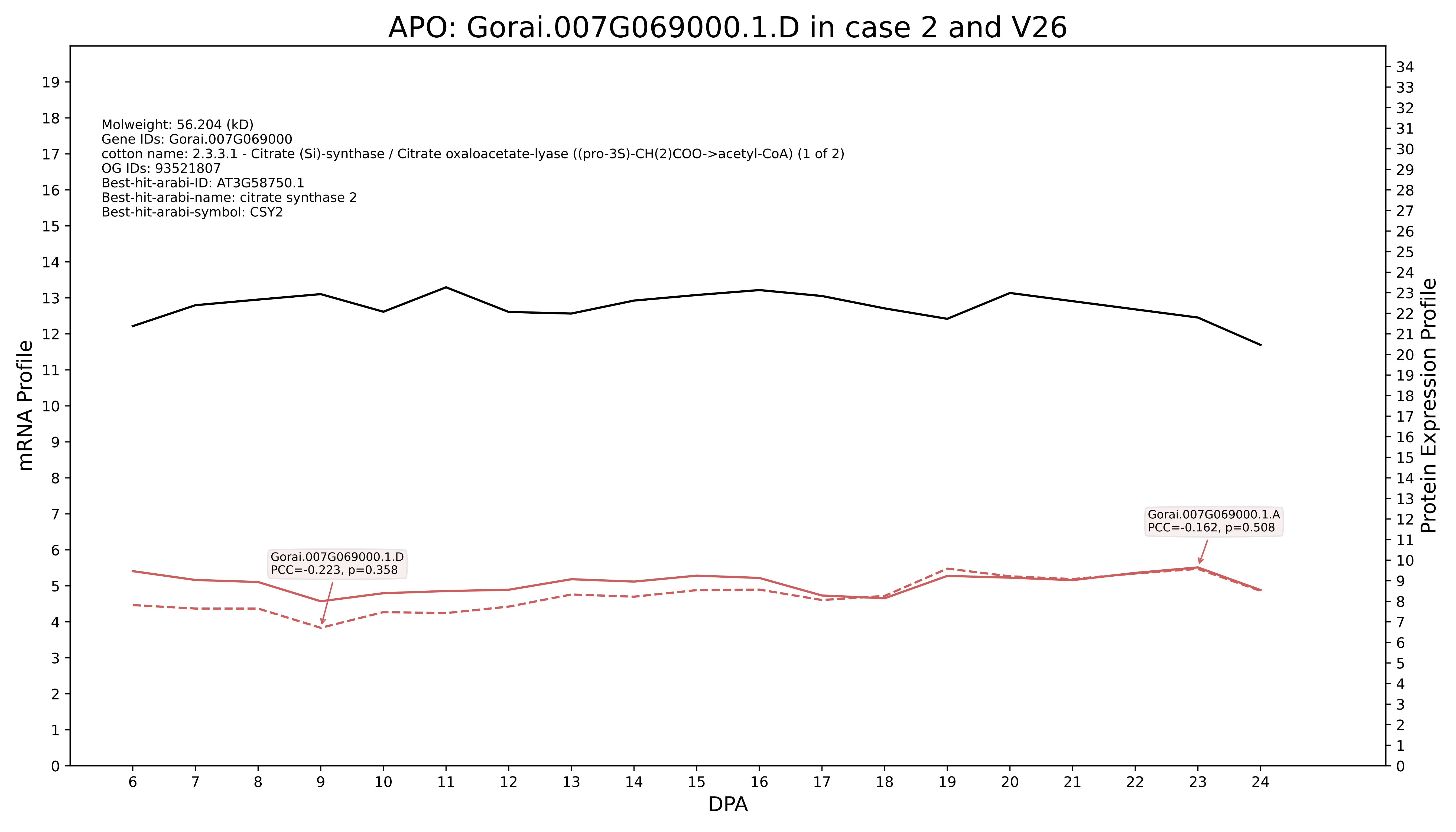

APO: Gorai.007G069000.1.D in case 2 and V26
Molweight: 56.204 (kD)
Gene IDs: Gorai.007G069000
cotton name: 2.3.3.1 - Citrate (Si)-synthase / Citrate oxaloacetate-lyase ((pro-3S)-CH(2)COO->acetyl-CoA) (1 of 2)
OG IDs: 93521807
Best-hit-arabi-ID: AT3G58750.1
Best-hit-arabi-name: citrate synthase 2
Best-hit-arabi-symbol: CSY2
Gorai.007G069000.1.D
PCC=-0.223, p=0.358
Gorai.007G069000.1.A
PCC=-0.162, p=0.508
mRNA Profile
Protein Expression Profile
DPA

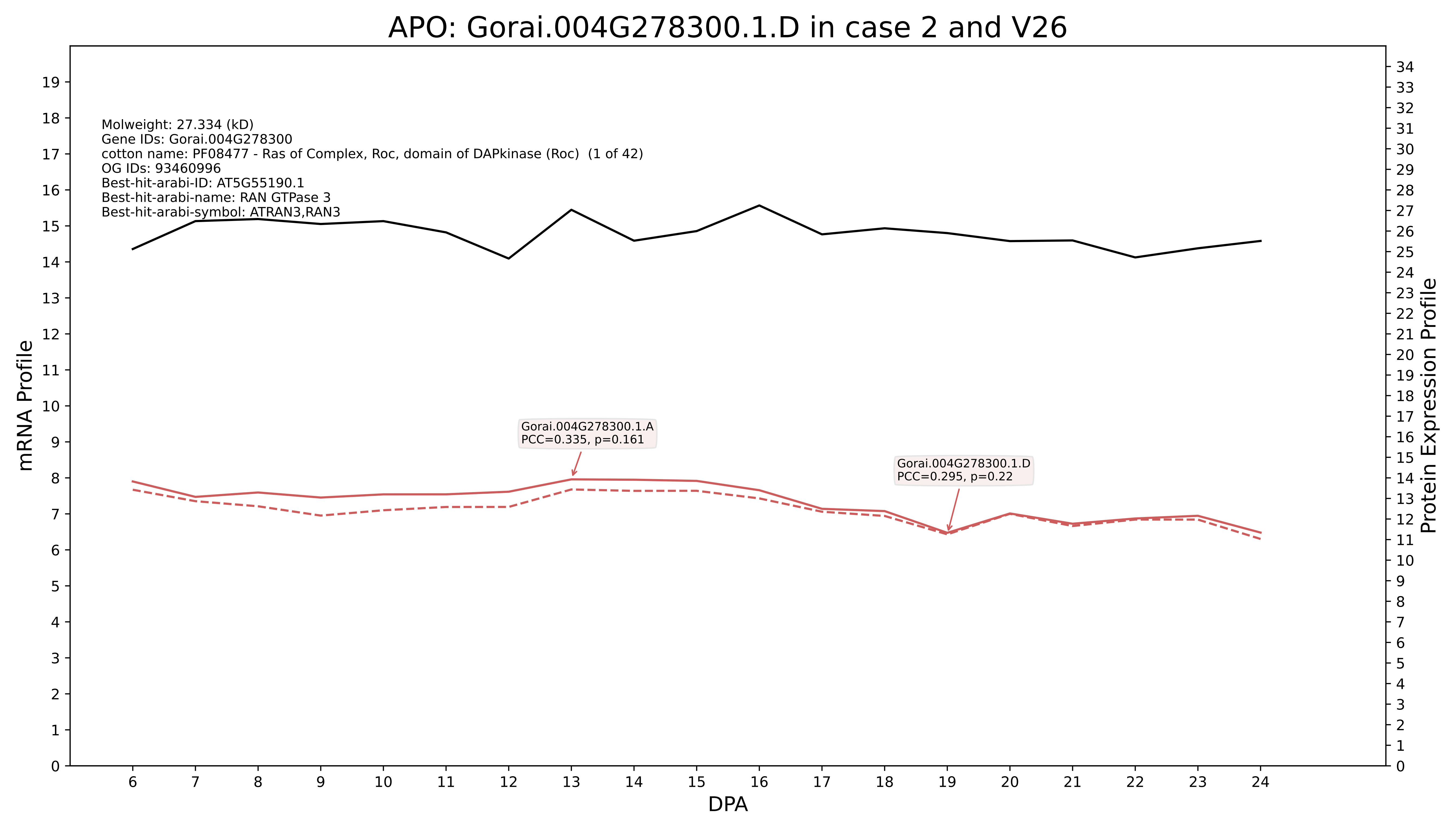
APO: Gorai.004G278300.1.D in case 2 and V26
Molweight: 27.334 (kD)
Gene IDs: Gorai.004G278300
cotton name: PF08477 - Ras of Complex, Roc, domain of DAPkinase (Roc) (1 of 42)
OG IDs: 93460996
Best-hit-arabi-ID: AT5G55190.1
Best-hit-arabi-name: RAN GTPase 3
Best-hit-arabi-symbol: ATRAN3,RAN3
Gorai.004G278300.1.A
PCC=0.335, p=0.161
Gorai.004G278300.1.D
PCC=0.295, p=0.22
mRNA Profile
Protein Expression Profile
DPA

### APO: Gorai.013G264400.1.D in case 3 and V26

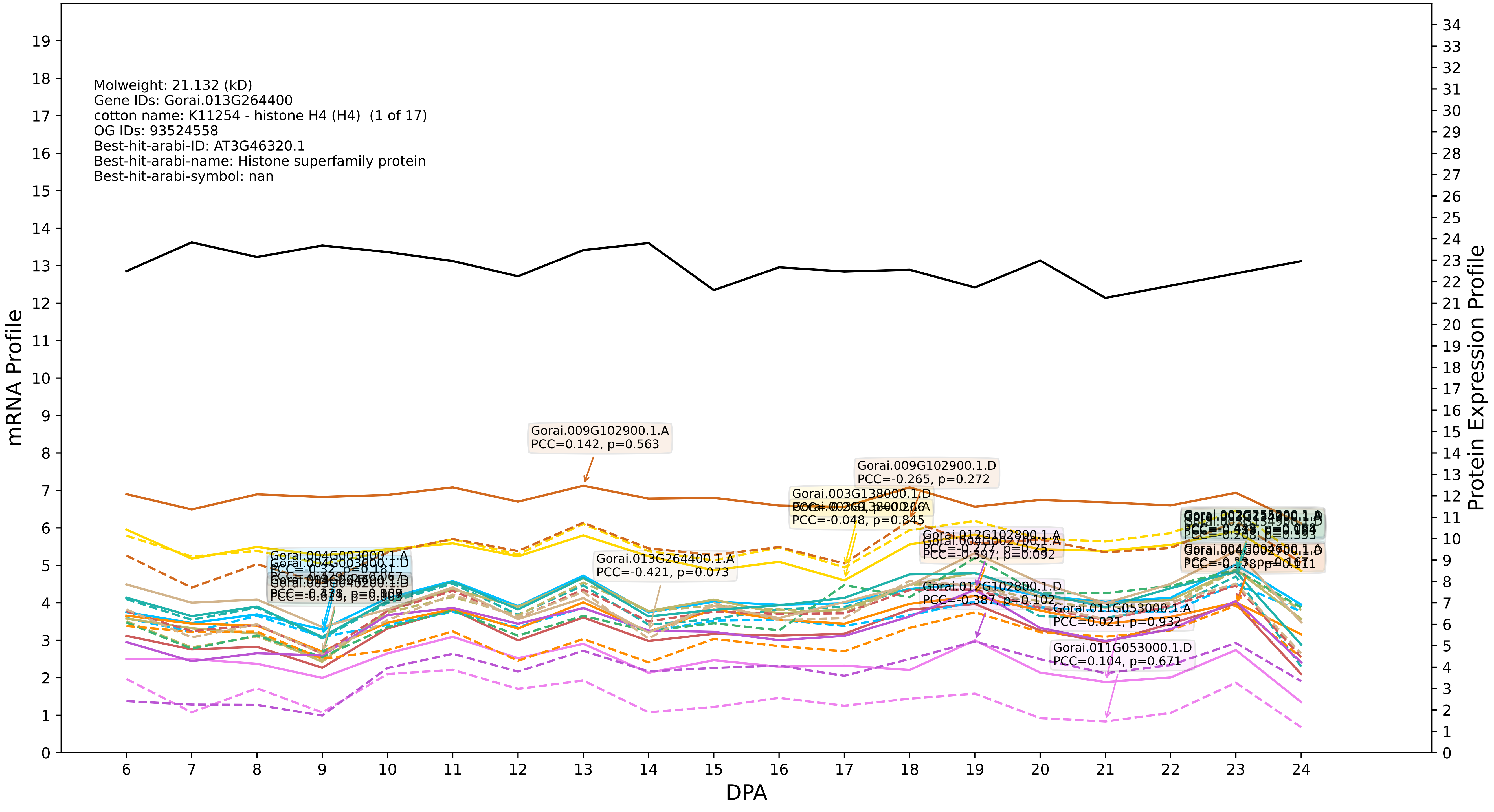

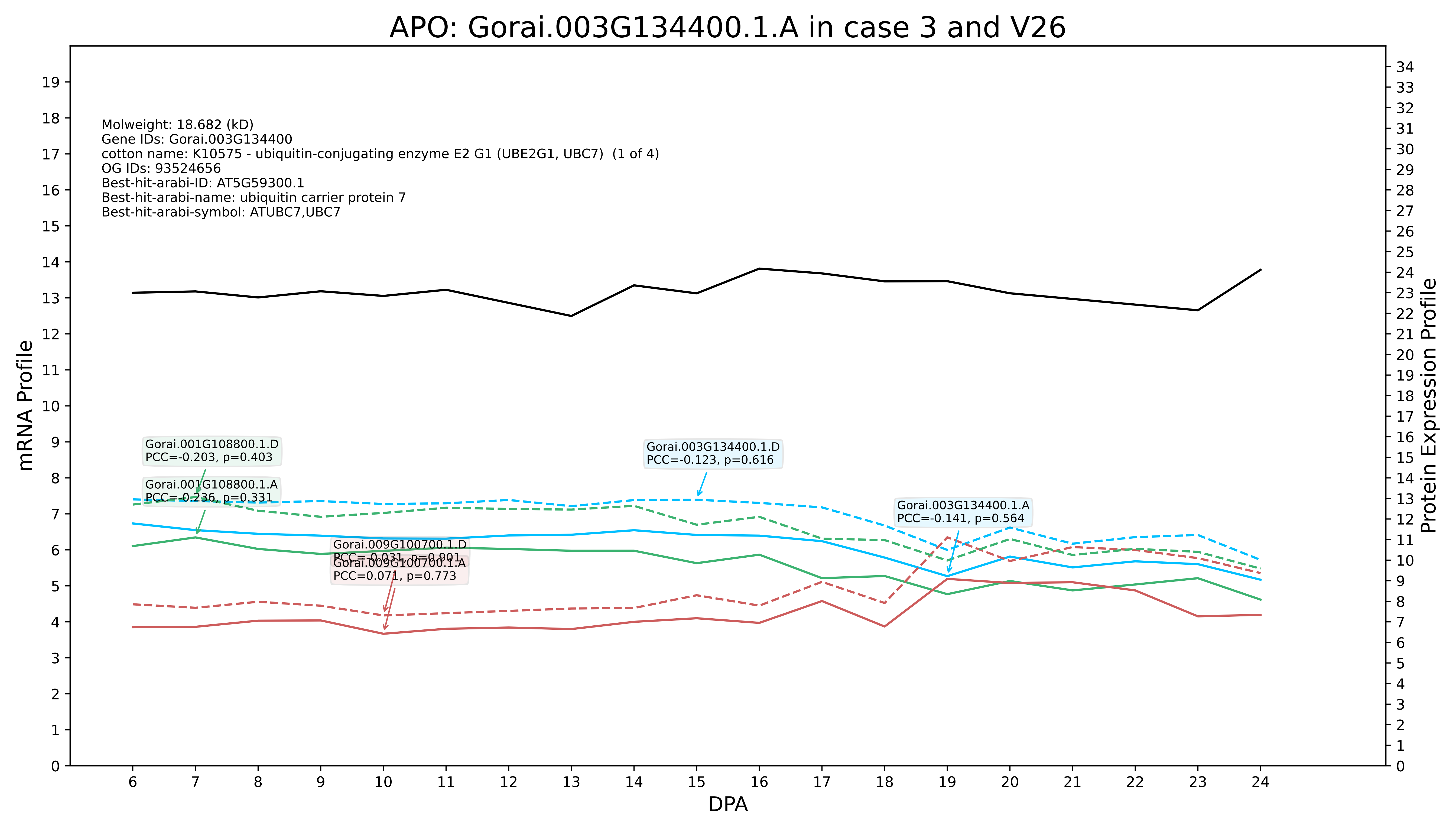

APO: Gorai.003G134400.1.A in case 3 and V26
Molweight: 18.682 (kD)
Gene IDs: Gorai.003G134400
cotton name: K10575 - ubiquitin-conjugating enzyme E2 G1 (UBE2G1, UBC7) (1 of 4)
OG IDs: 93524656
Best-hit-arabi-ID: AT5G59300.1
Best-hit-arabi-name: ubiquitin carrier protein 7
Best-hit-arabi-symbol: ATUBC7,UBC7
Gorai.001G108800.1.D
PCC=-0.203, p=0.403
Gorai.001G108800.1.A
PCC=-0.236, p=0.331
Gorai.003G134400.1.D
PCC=-0.123, p=0.616
Gorai.003G134400.1.A
PCC=-0.141, p=0.564
Gorai.009G100700.1.D
PCC=-0.031, p=0.901
Gorai.009G100700.1.A
PCC=0.071, p=0.773
mRNA Profile
Protein Expression Profile
DPA

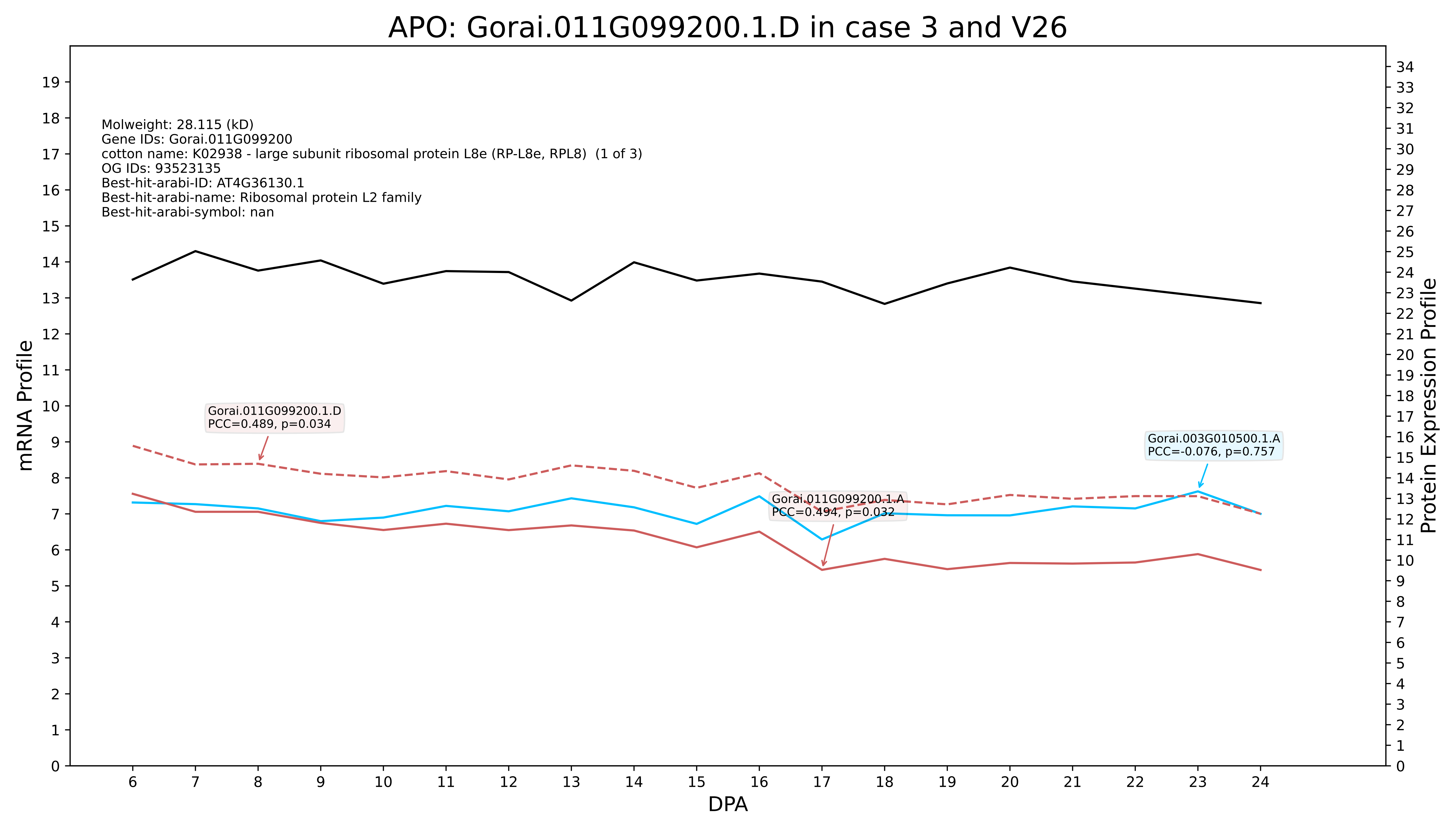

APO: Gorai.011G099200.1.D in case 3 and V26
Molweight: 28.115 (kD)
Gene IDs: Gorai.011G099200
cotton name: K02938 - large subunit ribosomal protein L8e (RP-L8e, RPL8) (1 of 3)
OG IDs: 93523135
Best-hit-arabi-ID: AT4G36130.1
Best-hit-arabi-name: Ribosomal protein L2 family
Best-hit-arabi-symbol: nan
Gorai.011G099200.1.D
PCC=0.489, p=0.034
Gorai.003G010500.1.A
PCC=-0.076, p=0.757
Gorai.011G099200.1.A
PCC=0.494, p=0.032
mRNA Profile
Protein Expression Profile
DPA

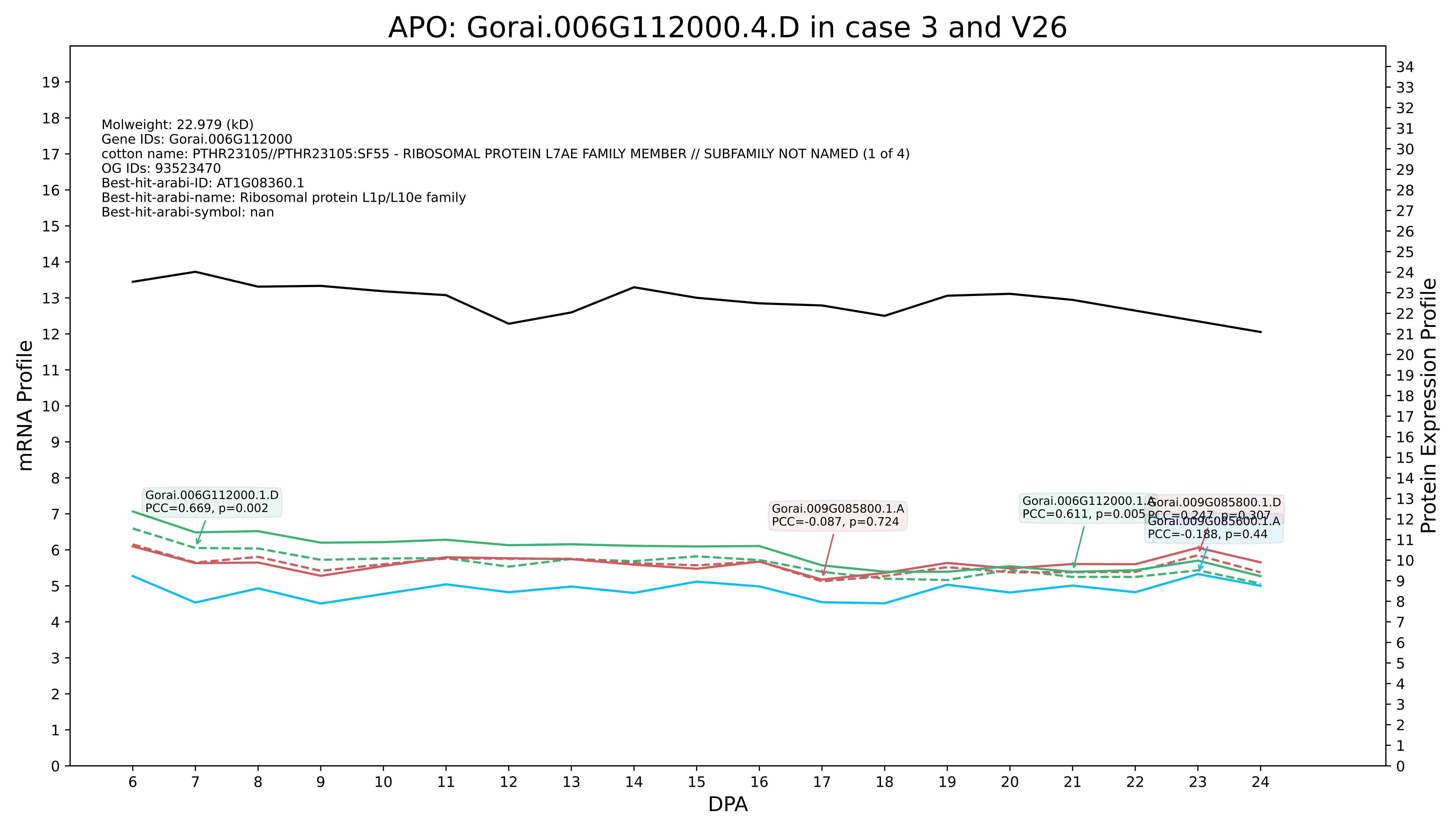

APO: Gorai.006G112000.4.D in case 3 and V26
Molweight: 22.979 (kD)
Gene IDs: Gorai.006G112000
cotton name: PTHR23105//PTHR23105:SF55 - RIBOSOMAL PROTEIN L7AE FAMILY MEMBER // SUBFAMILY NOT NAMED (1 of 4)
OG IDs: 93523470
Best-hit-arabi-ID: AT1G08360.1
Best-hit-arabi-name: Ribosomal protein L1p/L10e family
Best-hit-arabi-symbol: nan
Gorai.006G112000.1.D
PCC=0.669, p=0.002
Gorai.009G085800.1.A
PCC=-0.087, p=0.724
Gorai.006G112000.1.A
PCC=0.611, p=0.005
Gorai.009G085800.1.D
PCC=0.247, p=0.307
Gorai.009G085600.1.A
PCC=-0.188, p=0.44
mRNA Profile
Protein Expression Profile
DPA

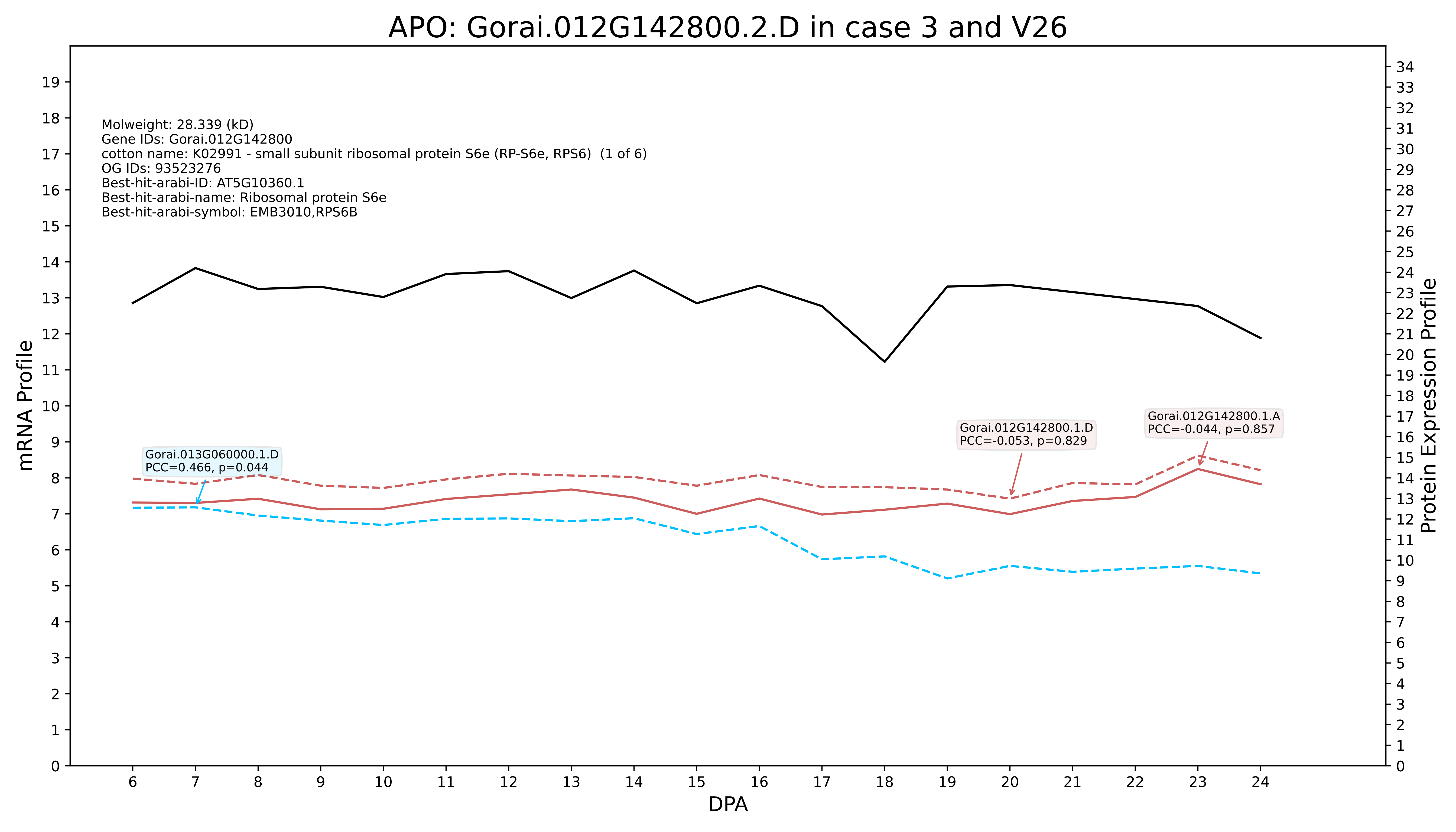
APO: Gorai.012G142800.2.D in case 3 and V26
Molweight: 28.339 (kD)
Gene IDs: Gorai.012G142800
cotton name: K02991 - small subunit ribosomal protein S6e (RP-S6e, RPS6) (1 of 6)
OG IDs: 93523276
Best-hit-arabi-ID: AT5G10360.1
Best-hit-arabi-name: Ribosomal protein S6e
Best-hit-arabi-symbol: EMB3010,RPS6B
Gorai.013G060000.1.D
PCC=0.466, p=0.044
Gorai.012G142800.1.D
PCC=-0.053, p=0.829
Gorai.012G142800.1.A
PCC=-0.044, p=0.857
mRNA Profile
Protein Expression Profile
DPA

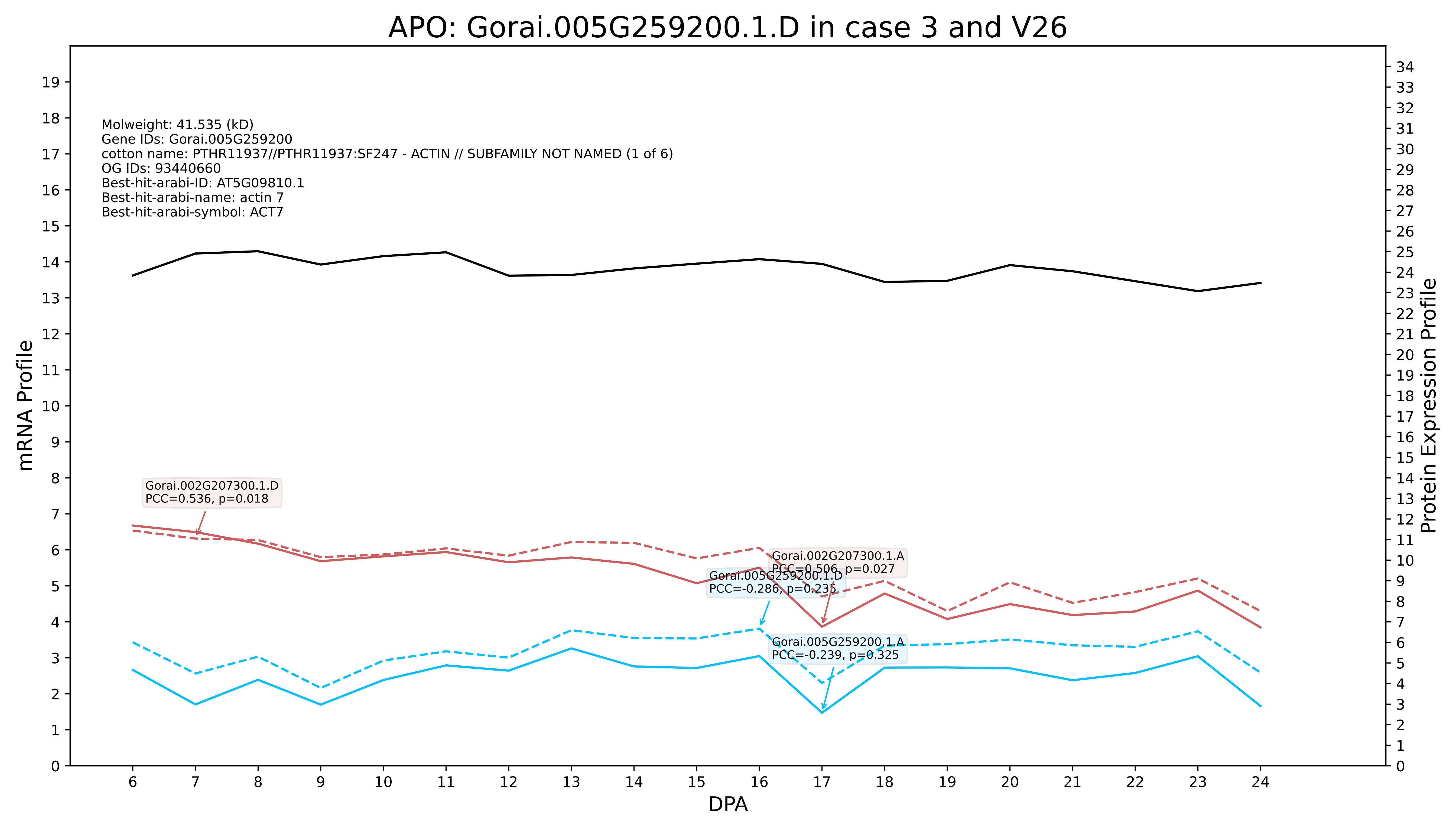

APO: Gorai.005G259200.1.D in case 3 and V26
Molweight: 41.535 (kD)
Gene IDs: Gorai.005G259200
cotton name: PTHR11937//PTHR11937:SF247 - ACTIN // SUBFAMILY NOT NAMED (1 of 6)
OG IDs: 93440660
Best-hit-arabi-ID: AT5G09810.1
Best-hit-arabi-name: actin 7
Best-hit-arabi-symbol: ACT7
Gorai.002G207300.1.D
PCC=0.536, p=0.018
Gorai.002G207300.1.A
PCC=0.506, p=0.027
Gorai.005G259200.1.D
PCC=-0.286, p=0.235
Gorai.005G259200.1.A
PCC=-0.239, p=0.325
mRNA Profile
Protein Expression Profile
DPA

### APO: Gorai.013G269700.3.A in case 1 and V27

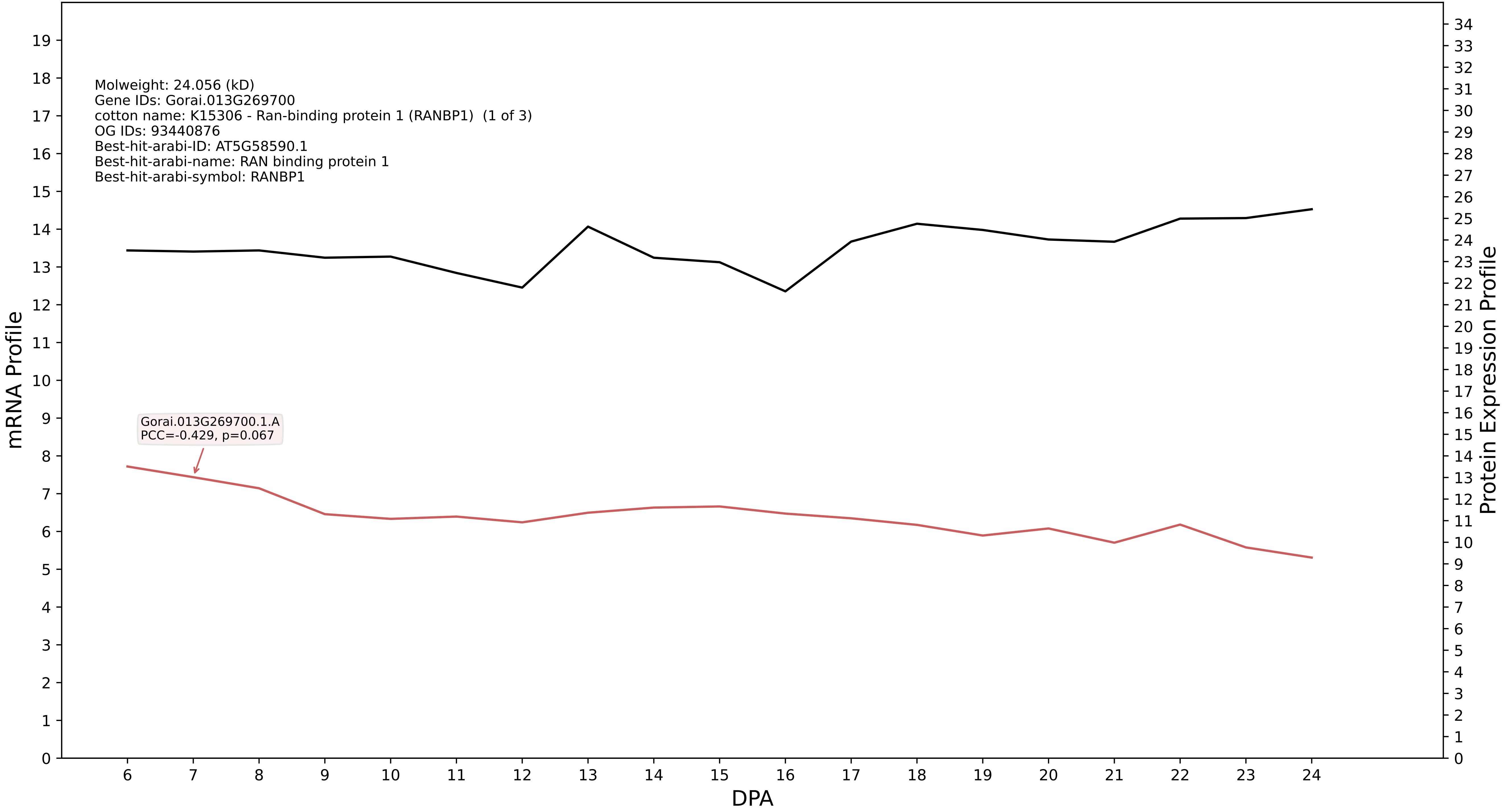

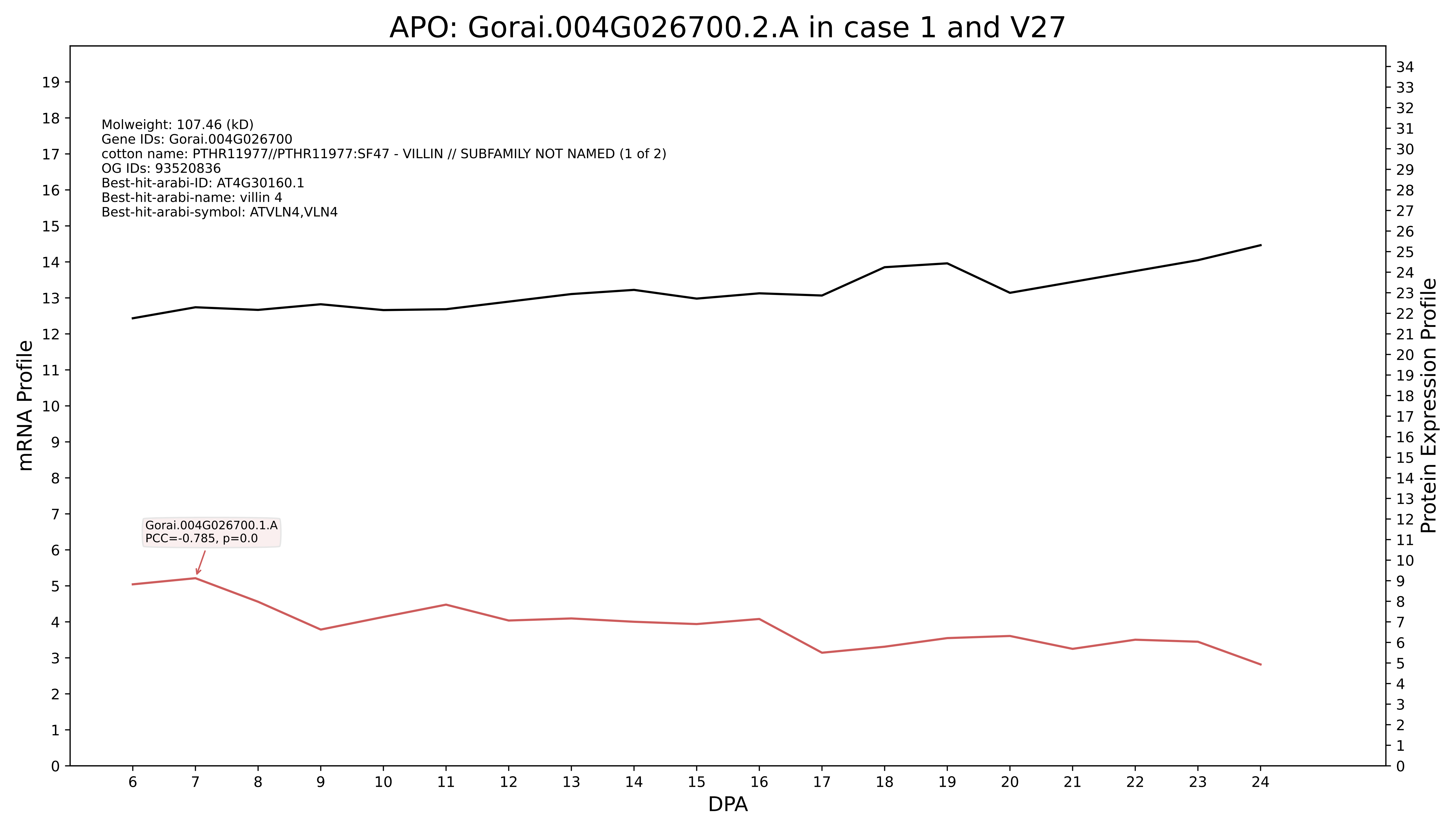
APO: Gorai.004G026700.2.A in case 1 and V27
Molweight: 107.46 (kD)
Gene IDs: Gorai.004G026700
cotton name: PTHR11977//PTHR11977:SF47 - VILLIN // SUBFAMILY NOT NAMED (1 of 2)
OG IDs: 93520836
Best-hit-arabi-ID: AT4G30160.1
Best-hit-arabi-name: villin 4
Best-hit-arabi-symbol: ATVLN4,VLN4
Gorai.004G026700.1.A
PCC=-0.785, p=0.0
mRNA Profile
Protein Expression Profile
DPA

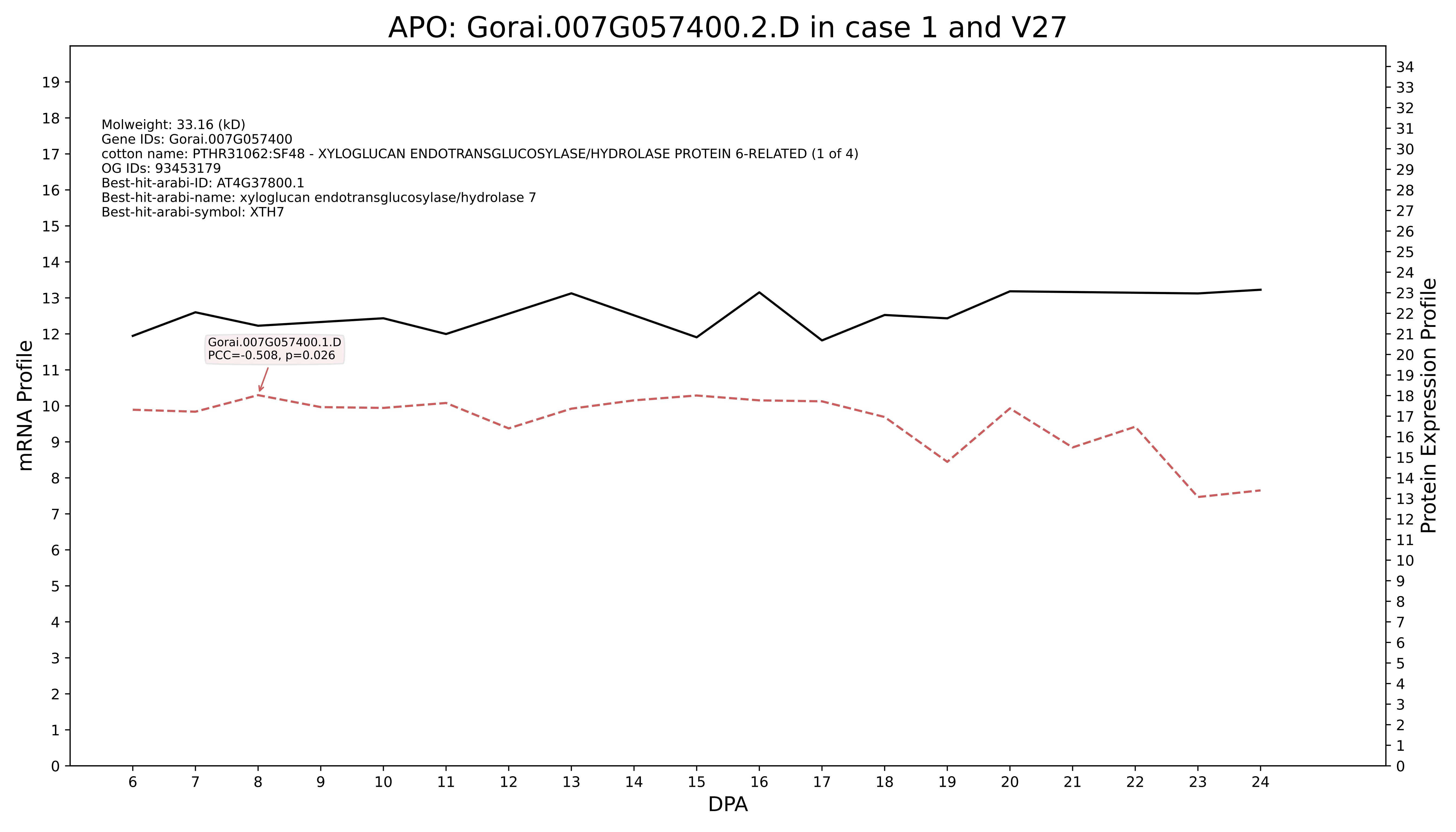

APO: Gorai.007G057400.2.D in case 1 and V27
Molweight: 33.16 (kD)
Gene IDs: Gorai.007G057400
cotton name: PTHR31062:SF48 - XYLOGLUCAN ENDOTRANSGLUCOSYLASE/HYDROLASE PROTEIN 6-RELATED (1 of 4)
OG IDs: 93453179
Best-hit-arabi-ID: AT4G37800.1
Best-hit-arabi-name: xyloglucan endotransglucosylase/hydrolase 7
Best-hit-arabi-symbol: XTH7
Gorai.007G057400.1.D
PCC=-0.508, p=0.026
mRNA Profile
Protein Expression Profile
DPA

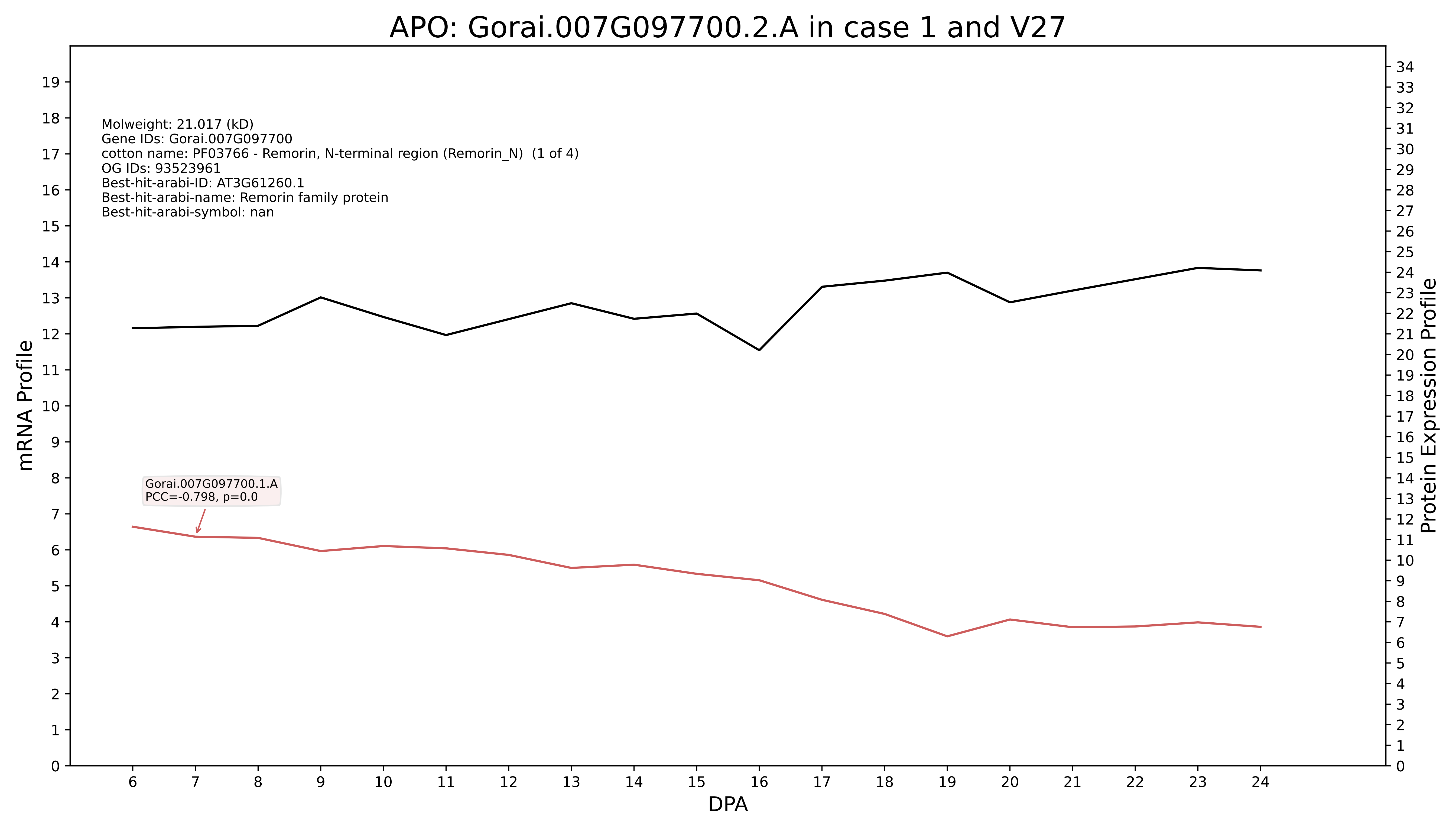

APO: Gorai.007G097700.2.A in case 1 and V27
Molweight: 21.017 (kD)
Gene IDs: Gorai.007G097700
cotton name: PF03766 - Remorin, N-terminal region (Remorin_N) (1 of 4)
OG IDs: 93523961
Best-hit-arabi-ID: AT3G61260.1
Best-hit-arabi-name: Remorin family protein
Best-hit-arabi-symbol: nan
Gorai.007G097700.1.A
PCC=-0.798, p=0.0
mRNA Profile
Protein Expression Profile
DPA

### APO: Gorai.009G189500.3.D in case 1 and V27

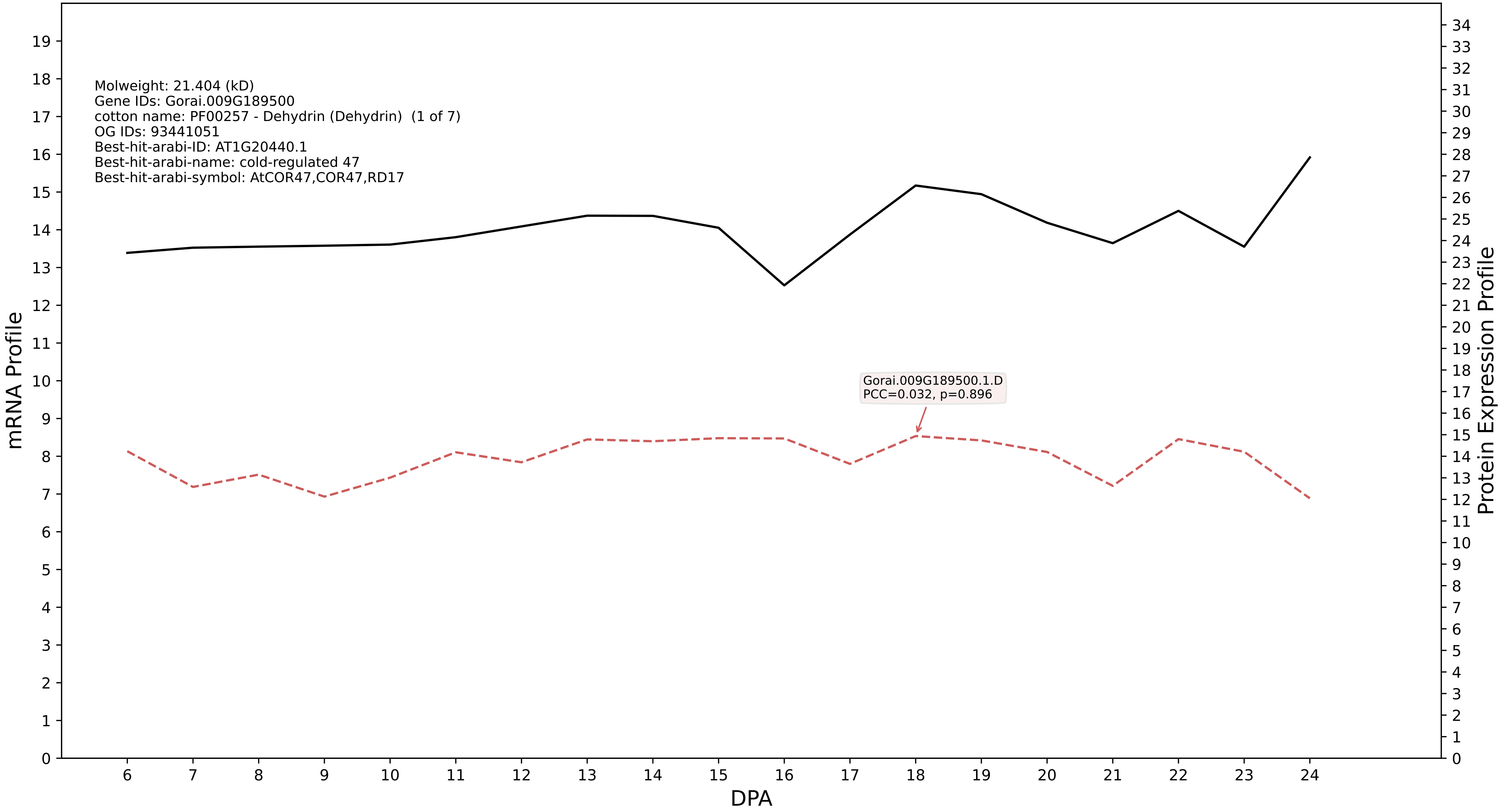

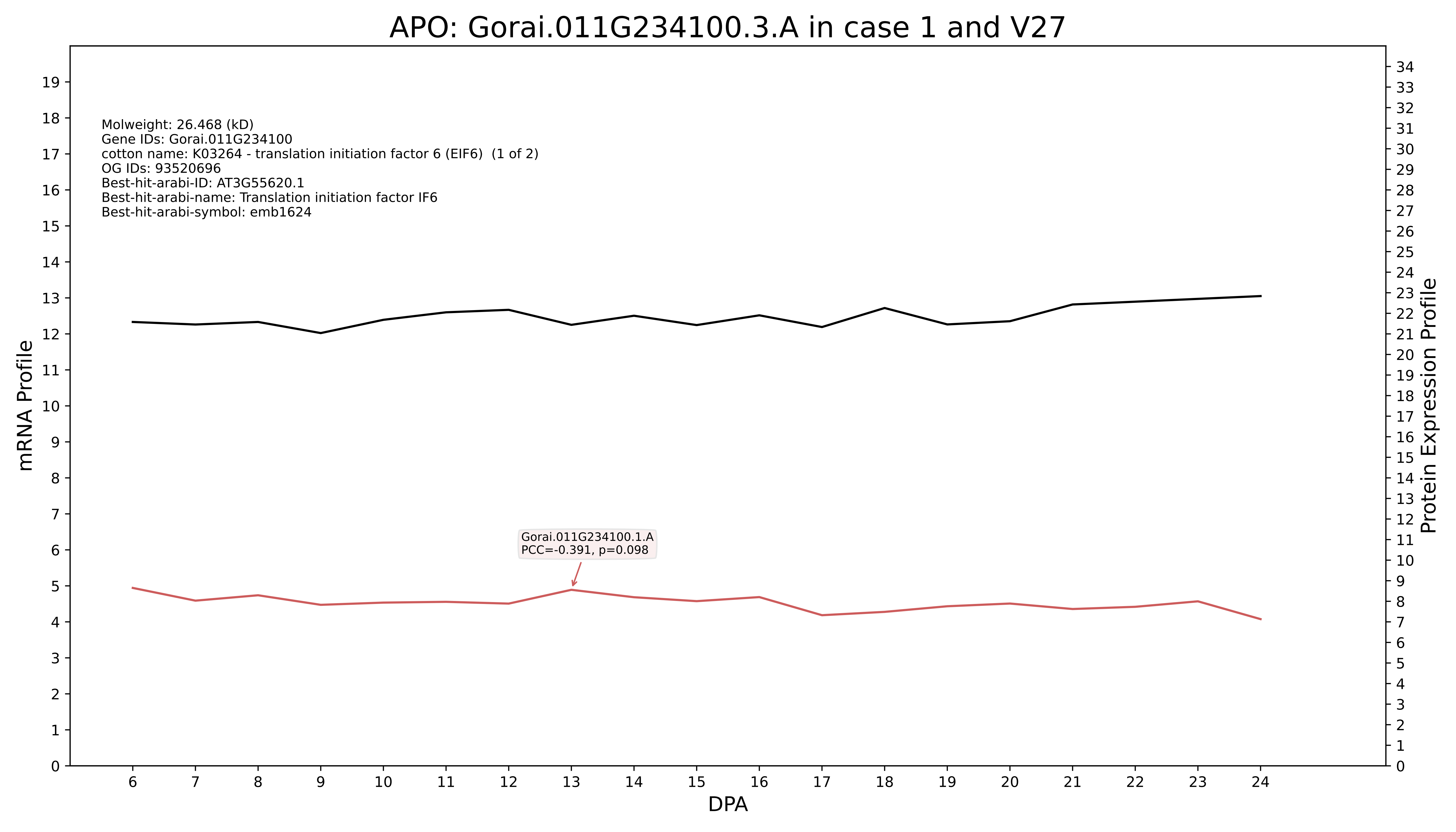

APO: Gorai.011G234100.3.A in case 1 and V27
Molweight: 26.468 (kD)
Gene IDs: Gorai.011G234100
cotton name: K03264 - translation initiation factor 6 (EIF6) (1 of 2)
OG IDs: 93520696
Best-hit-arabi-ID: AT3G55620.1
Best-hit-arabi-name: Translation initiation factor IF6
Best-hit-arabi-symbol: emb1624
Gorai.011G234100.1.A
PCC=-0.391, p=0.098
mRNA Profile
Protein Expression Profile
DPA

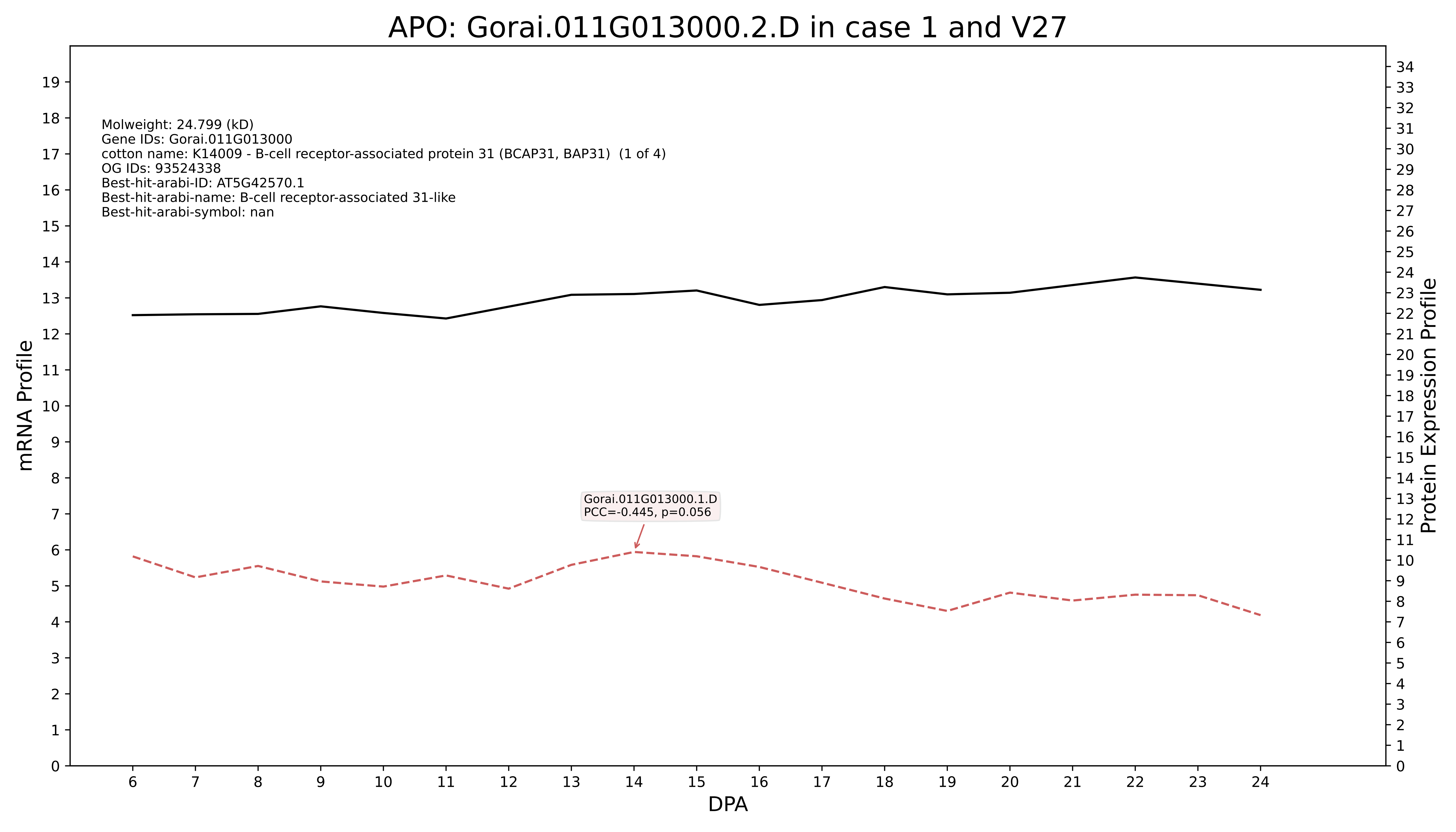

APO: Gorai.011G013000.2.D in case 1 and V27
Molweight: 24.799 (kD)
Gene IDs: Gorai.011G013000
cotton name: K14009 - B-cell receptor-associated protein 31 (BCAP31, BAP31) (1 of 4)
OG IDs: 93524338
Best-hit-arabi-ID: AT5G42570.1
Best-hit-arabi-name: B-cell receptor-associated 31-like
Best-hit-arabi-symbol: nan
Gorai.011G013000.1.D
PCC=-0.445, p=0.056
mRNA Profile
Protein Expression Profile
DPA

### APO: Gorai.010G186700.4.A in case 1 and V27

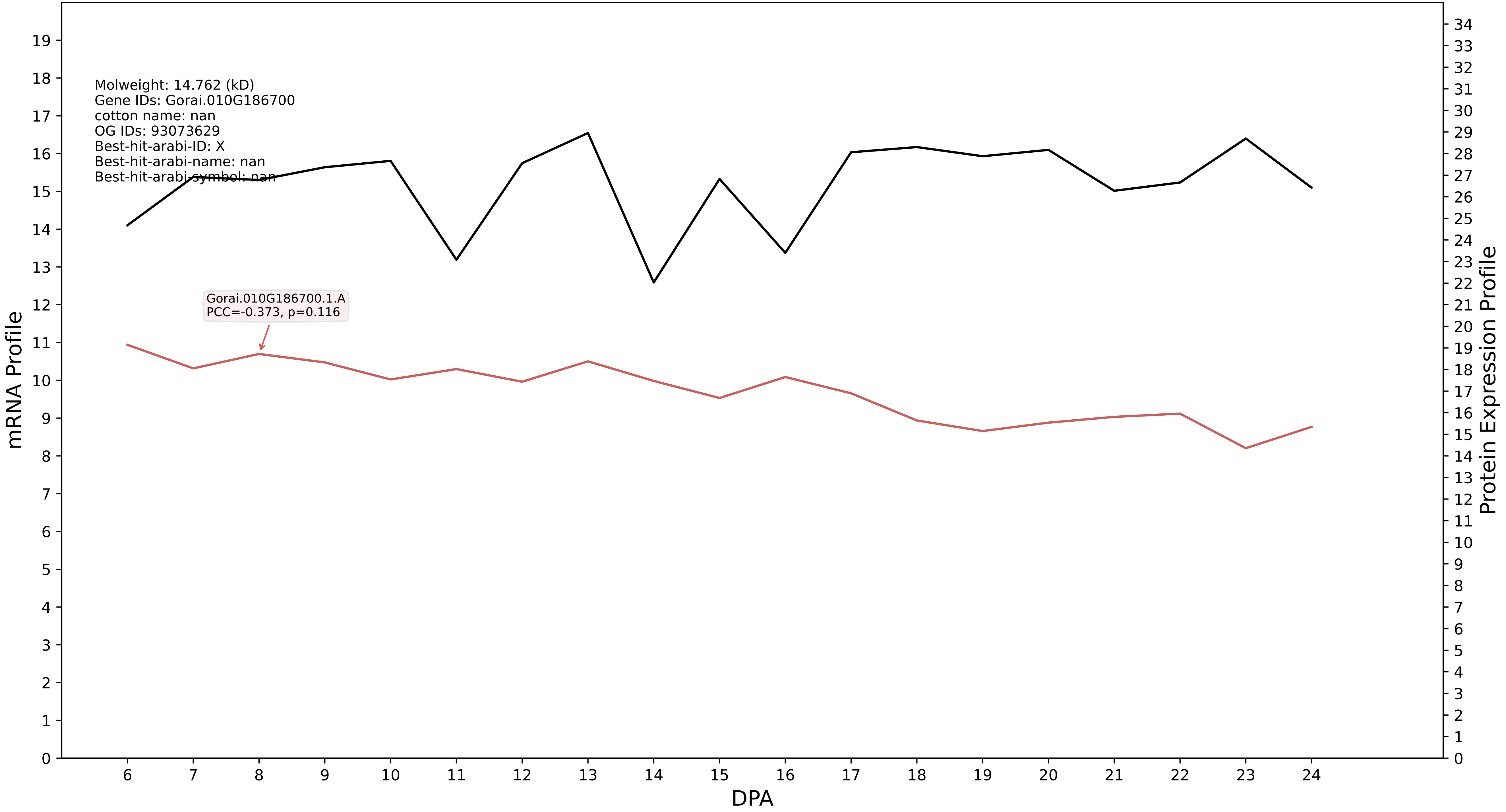

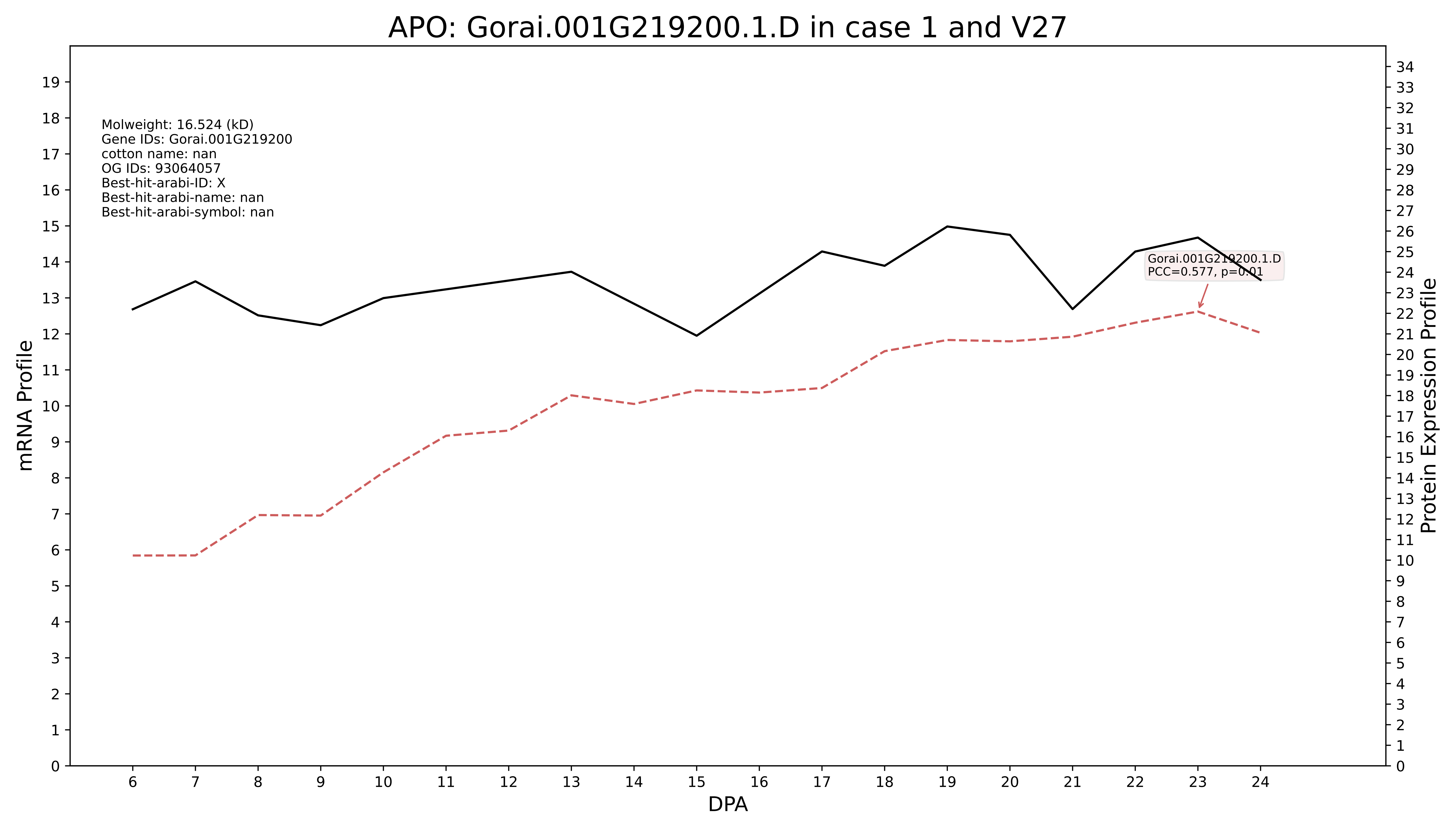
APO: Gorai.001G219200.1.D in case 1 and V27
Molweight: 16.524 (kD)
Gene IDs: Gorai.001G219200
cotton name: nan
OG IDs: 93064057
Best-hit-arabi-ID: X
Best-hit-arabi-name: nan
Best-hit-arabi-symbol: nan
Gorai.001G219200.1.D
PCC=0.577, p=0.01
mRNA Profile
Protein Expression Profile
DPA

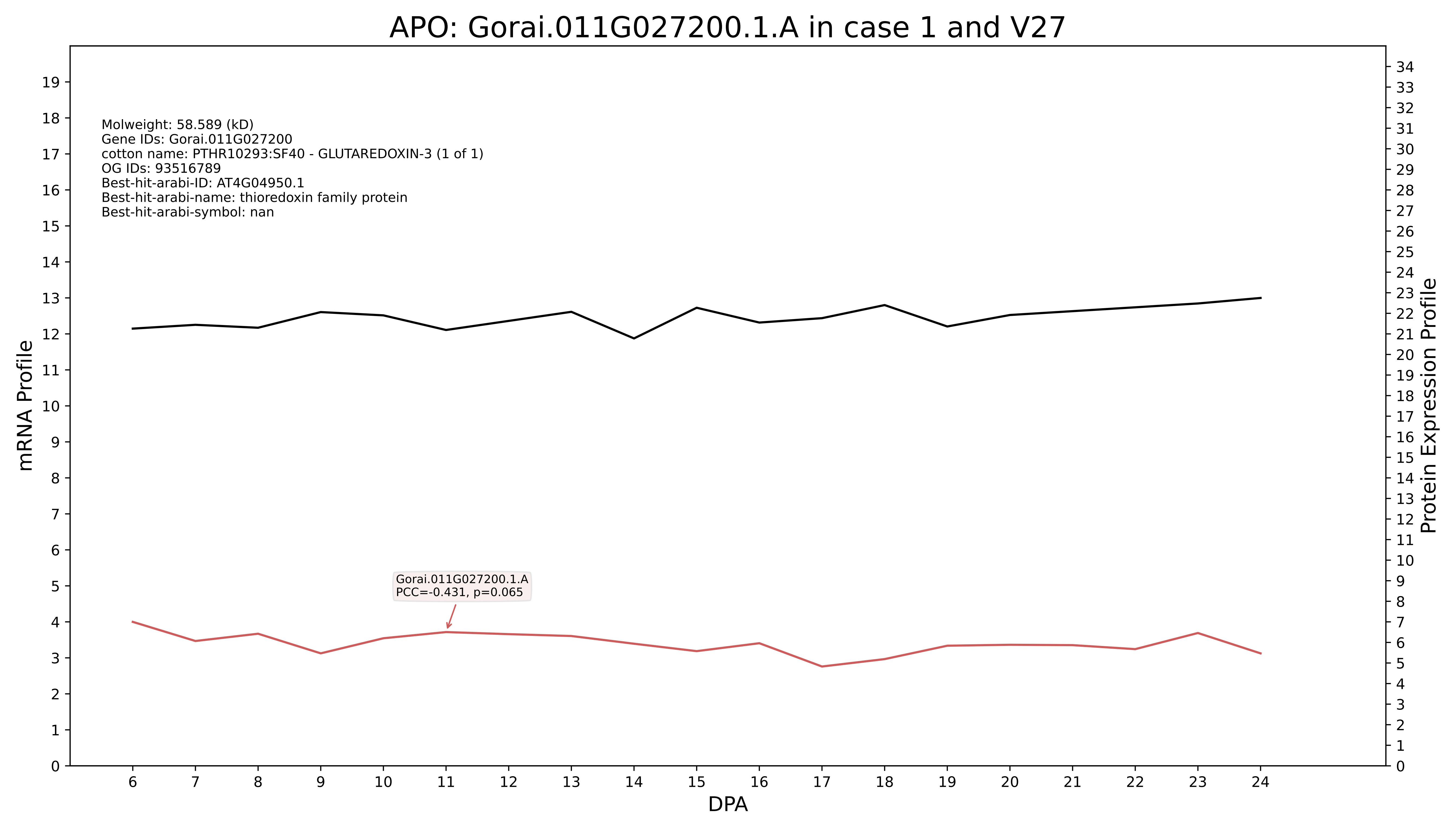

APO: Gorai.011G027200.1.A in case 1 and V27
Molweight: 58.589 (kD)
Gene IDs: Gorai.011G027200
cotton name: PTHR10293:SF40 - GLUTAREDOXIN-3 (1 of 1)
OG IDs: 93516789
Best-hit-arabi-ID: AT4G04950.1
Best-hit-arabi-name: thioredoxin family protein
Best-hit-arabi-symbol: nan
Gorai.011G027200.1.A
PCC=-0.431, p=0.065
mRNA Profile
Protein Expression Profile
DPA

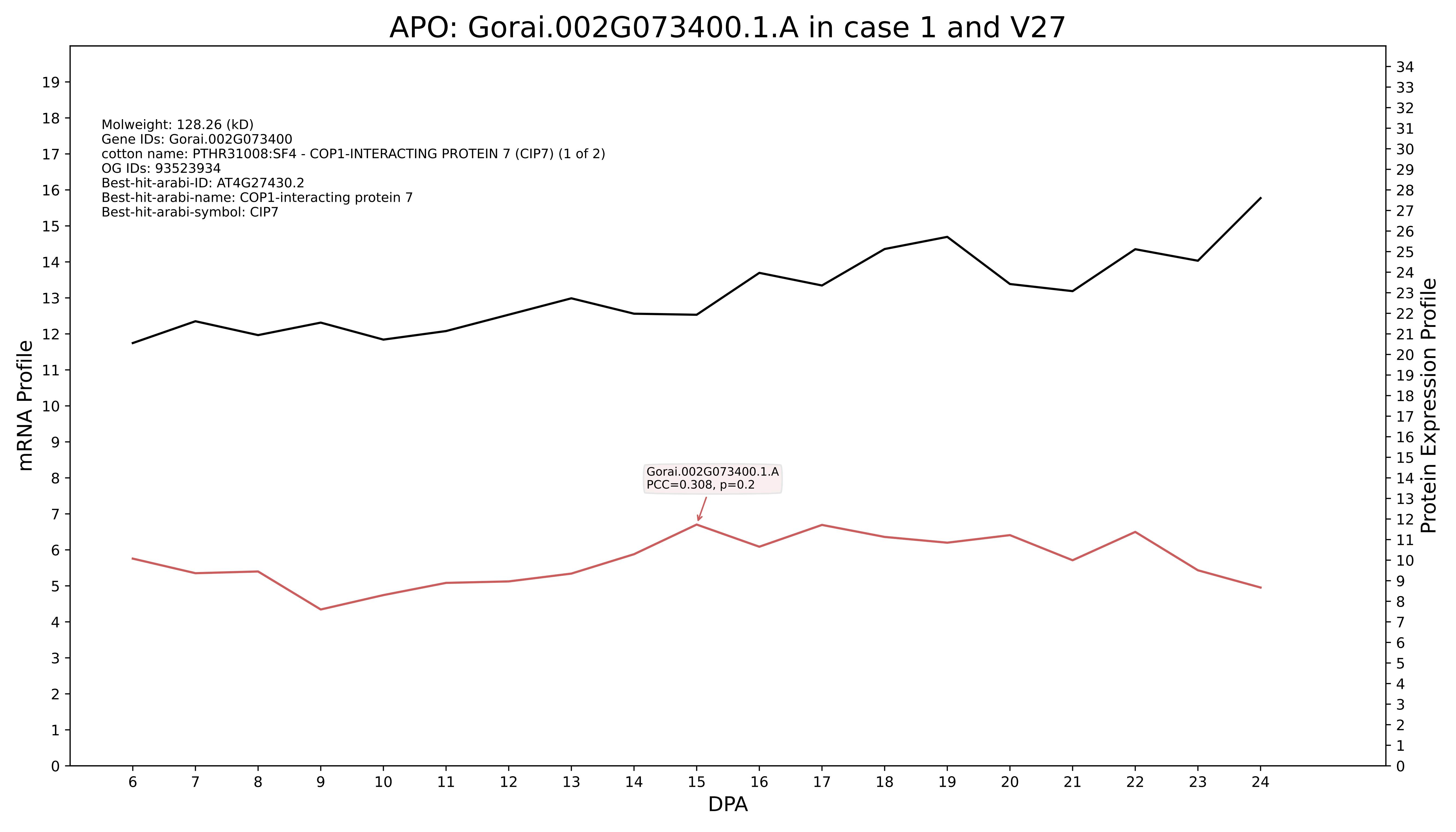

APO: Gorai.002G073400.1.A in case 1 and V27
Molweight: 128.26 (kD)
Gene IDs: Gorai.002G073400
cotton name: PTHR31008:SF4 - COP1-INTERACTING PROTEIN 7 (CIP7) (1 of 2)
OG IDs: 93523934
Best-hit-arabi-ID: AT4G27430.2
Best-hit-arabi-name: COP1-interacting protein 7
Best-hit-arabi-symbol: CIP7
Gorai.002G073400.1.A
PCC=0.308, p=0.2
mRNA Profile
Protein Expression Profile
DPA

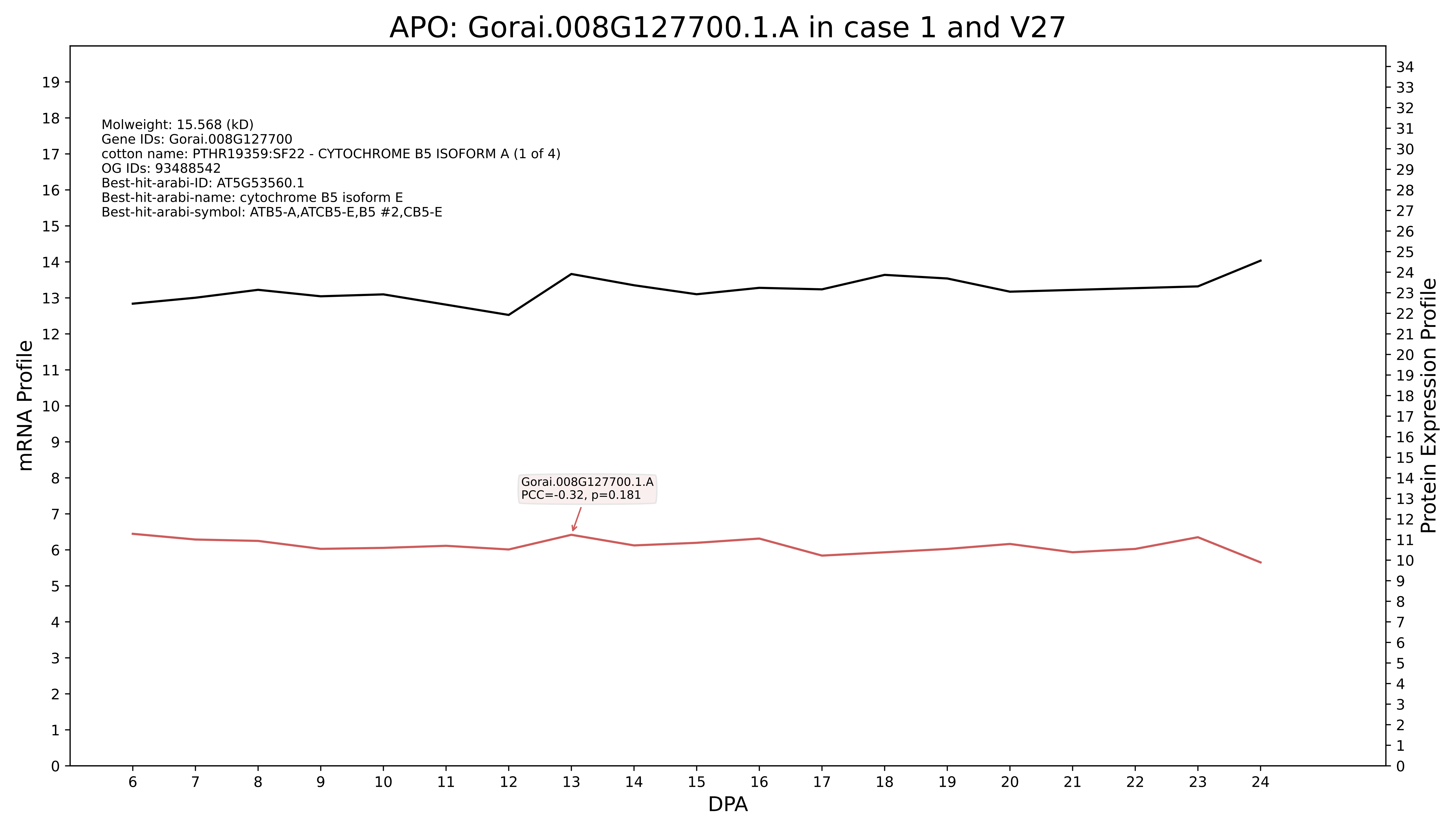

APO: Gorai.008G127700.1.A in case 1 and V27
Molweight: 15.568 (kD)
Gene IDs: Gorai.008G127700
cotton name: PTHR19359:SF22 - CYTOCHROME B5 ISOFORM A (1 of 4)
OG IDs: 93488542
Best-hit-arabi-ID: AT5G53560.1
Best-hit-arabi-name: cytochrome B5 isoform E
Best-hit-arabi-symbol: ATB5-A,ATCB5-E,B5 #2,CB5-E
Gorai.008G127700.1.A
PCC=-0.32, p=0.181
mRNA Profile
Protein Expression Profile
DPA

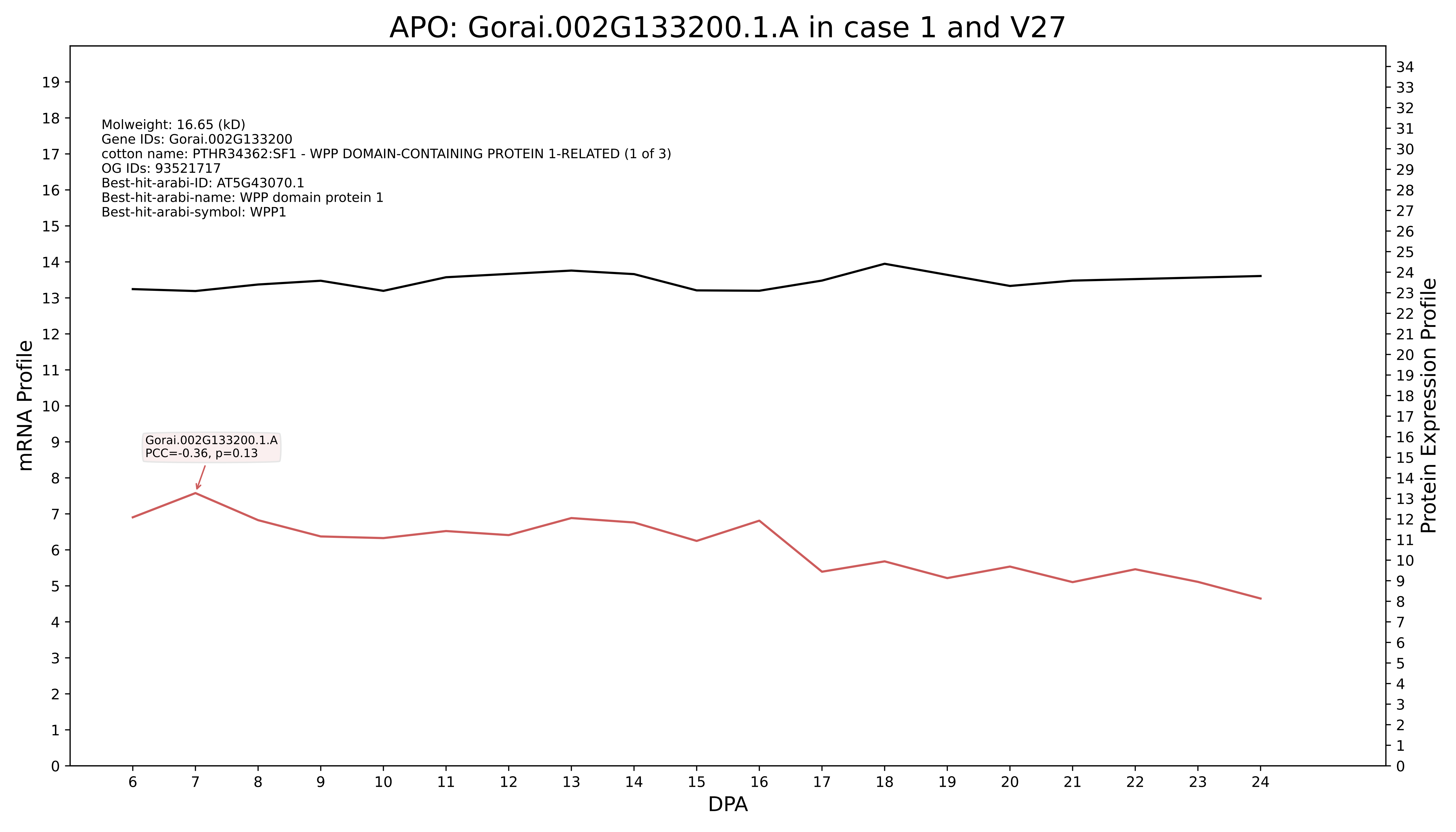

APO: Gorai.002G133200.1.A in case 1 and V27
Molweight: 16.65 (kD)
Gene IDs: Gorai.002G133200
cotton name: PTHR34362:SF1 - WPP DOMAIN-CONTAINING PROTEIN 1-RELATED (1 of 3)
OG IDs: 93521717
Best-hit-arabi-ID: AT5G43070.1
Best-hit-arabi-name: WPP domain protein 1
Best-hit-arabi-symbol: WPP1
Gorai.002G133200.1.A
PCC=-0.36, p=0.13
mRNA Profile
Protein Expression Profile
DPA

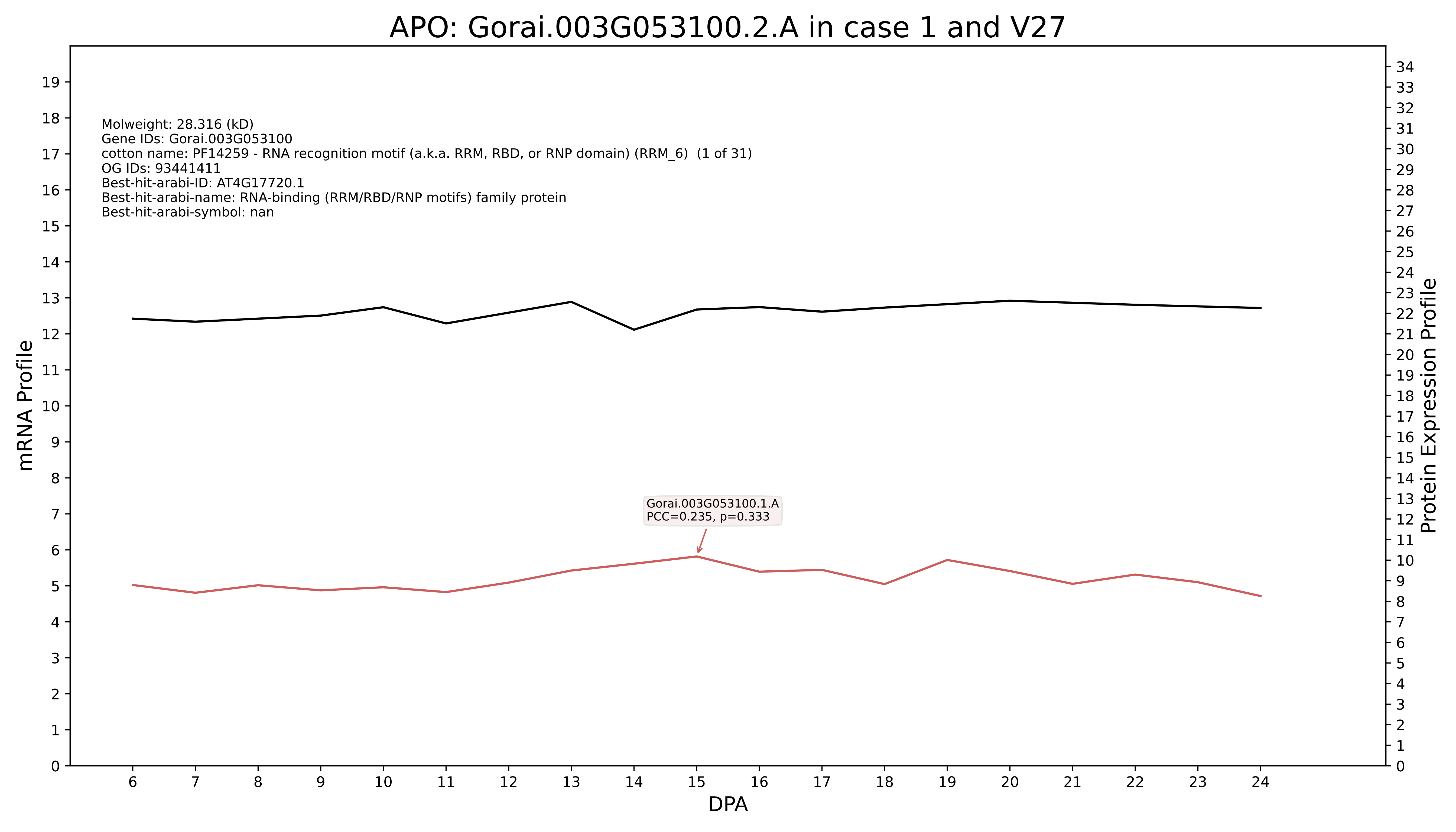
APO: Gorai.003G053100.2.A in case 1 and V27
Molweight: 28.316 (kD)
Gene IDs: Gorai.003G053100
cotton name: PF14259 - RNA recognition motif (a.k.a. RRM, RBD, or RNP domain) (RRM_6) (1 of 31)
OG IDs: 93441411
Best-hit-arabi-ID: AT4G17720.1
Best-hit-arabi-name: RNA-binding (RRM/RBD/RNP motifs) family protein
Best-hit-arabi-symbol: nan
Gorai.003G053100.1.A
PCC=0.235, p=0.333
mRNA Profile
Protein Expression Profile
DPA

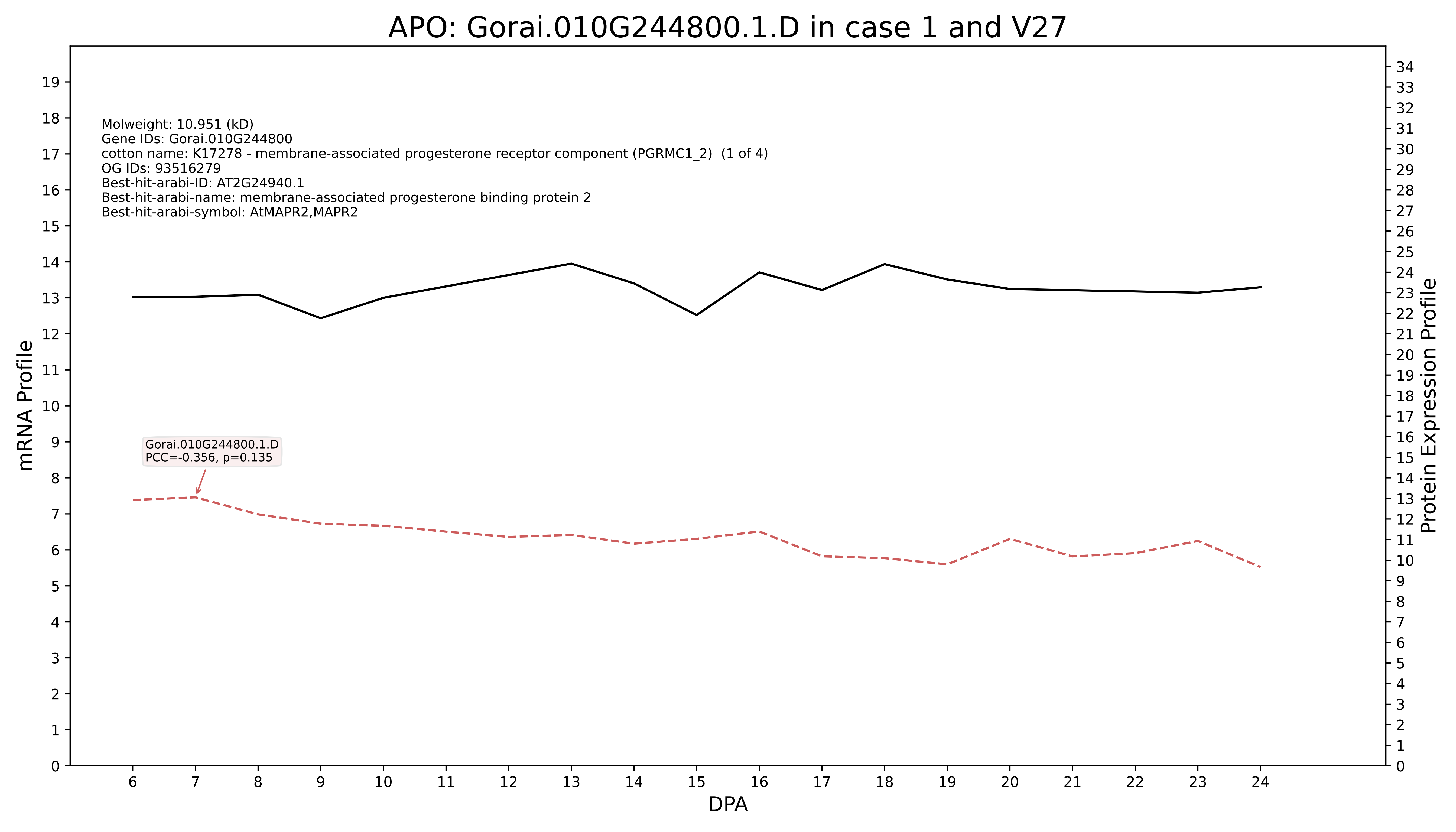

APO: Gorai.010G244800.1.D in case 1 and V27
Molweight: 10.951 (kD)
Gene IDs: Gorai.010G244800
cotton name: K17278 - membrane-associated progesterone receptor component (PGRMC1_2) (1 of 4)
OG IDs: 93516279
Best-hit-arabi-ID: AT2G24940.1
Best-hit-arabi-name: membrane-associated progesterone binding protein 2
Best-hit-arabi-symbol: AtMAPR2,MAPR2
Gorai.010G244800.1.D
PCC=-0.356, p=0.135
mRNA Profile
Protein Expression Profile
DPA

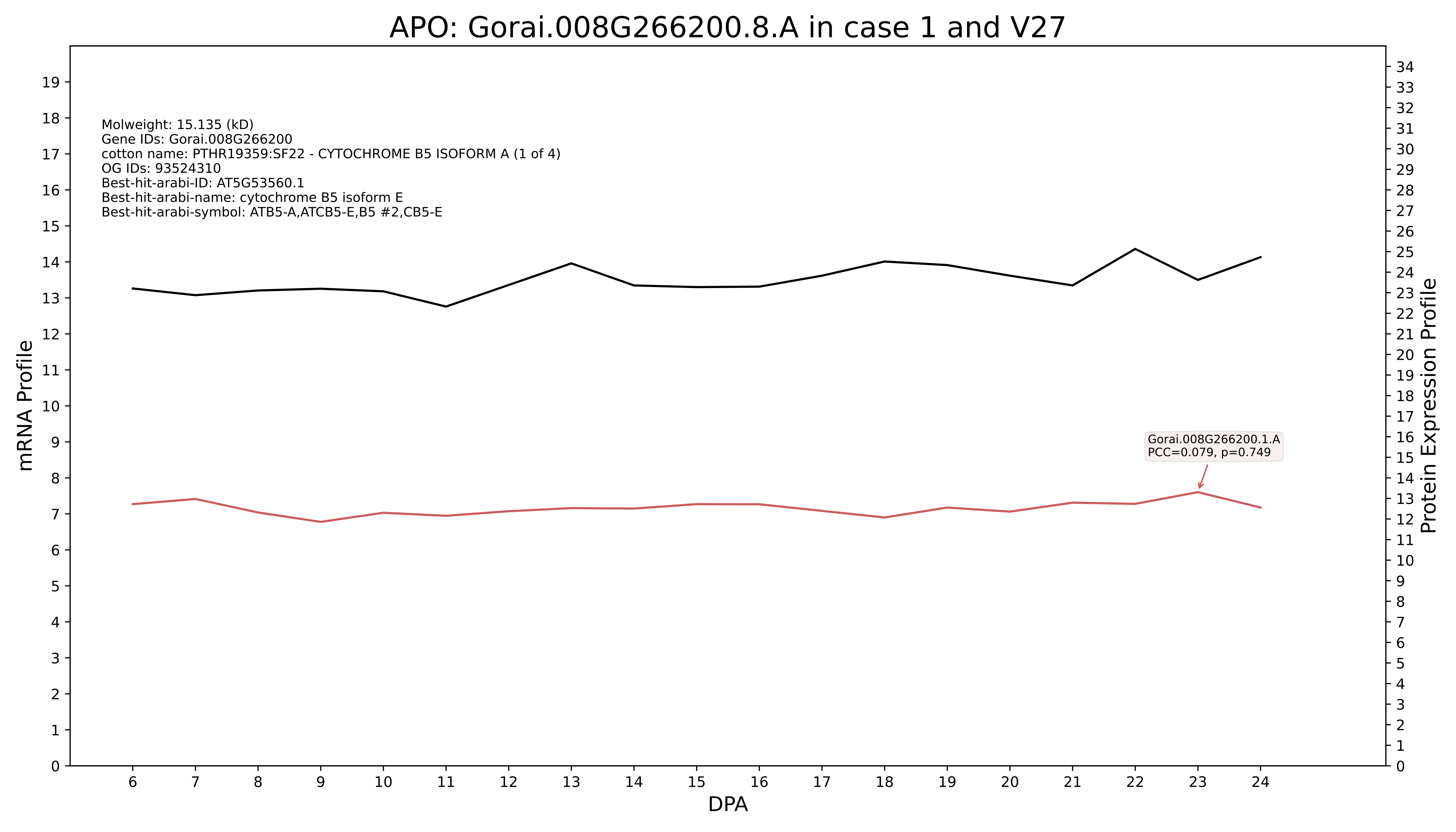
APO: Gorai.008G266200.8.A in case 1 and V27
Molweight: 15.135 (kD)
Gene IDs: Gorai.008G266200
cotton name: PTHR19359:SF22 - CYTOCHROME B5 ISOFORM A (1 of 4)
OG IDs: 93524310
Best-hit-arabi-ID: AT5G53560.1
Best-hit-arabi-name: cytochrome B5 isoform E
Best-hit-arabi-symbol: ATB5-A,ATCB5-E,B5 #2,CB5-E
Gorai.008G266200.1.A
PCC=0.079, p=0.749
mRNA Profile
Protein Expression Profile
DPA

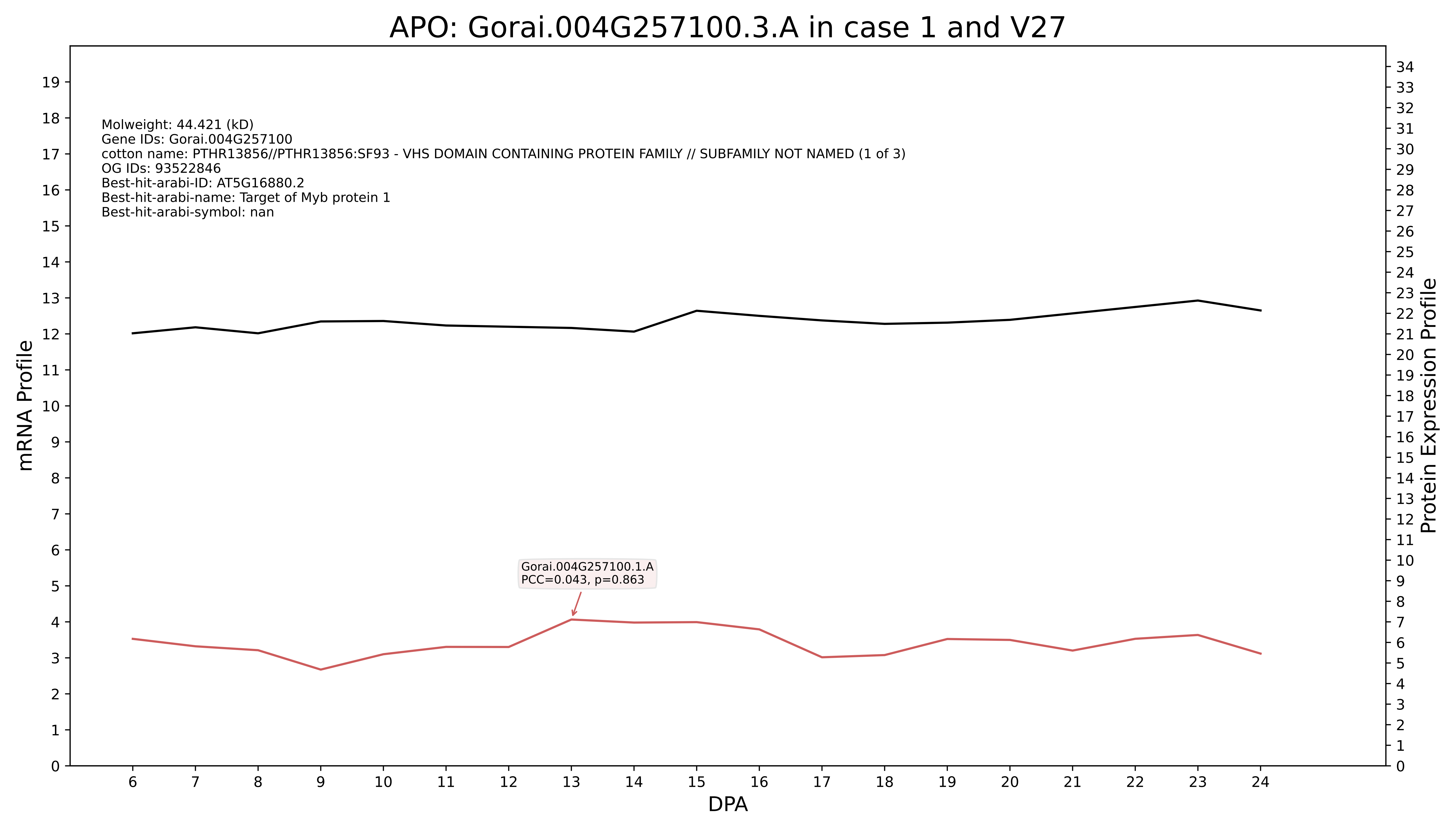

APO: Gorai.004G257100.3.A in case 1 and V27
Molweight: 44.421 (kD)
Gene IDs: Gorai.004G257100
cotton name: PTHR13856//PTHR13856:SF93 - VHS DOMAIN CONTAINING PROTEIN FAMILY // SUBFAMILY NOT NAMED (1 of 3)
OG IDs: 93522846
Best-hit-arabi-ID: AT5G16880.2
Best-hit-arabi-name: Target of Myb protein 1
Best-hit-arabi-symbol: nan
Gorai.004G257100.1.A
PCC=0.043, p=0.863
mRNA Profile
Protein Expression Profile
DPA

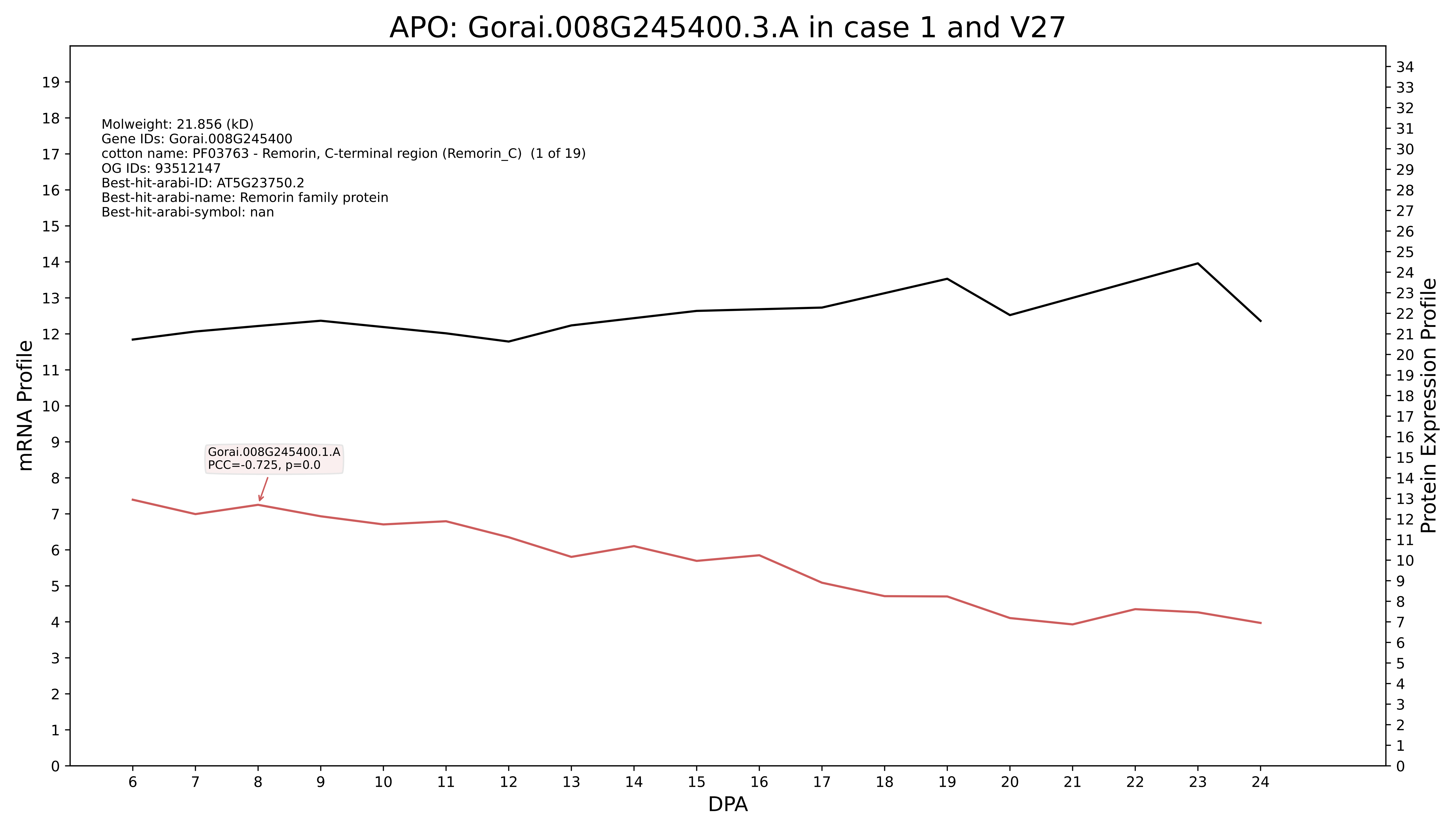
APO: Gorai.008G245400.3.A in case 1 and V27
Molweight: 21.856 (kD)
Gene IDs: Gorai.008G245400
cotton name: PF03763 - Remorin, C-terminal region (Remorin_C) (1 of 19)
OG IDs: 93512147
Best-hit-arabi-ID: AT5G23750.2
Best-hit-arabi-name: Remorin family protein
Best-hit-arabi-symbol: nan
Gorai.008G245400.1.A
PCC=-0.725, p=0.0
mRNA Profile
Protein Expression Profile
DPA

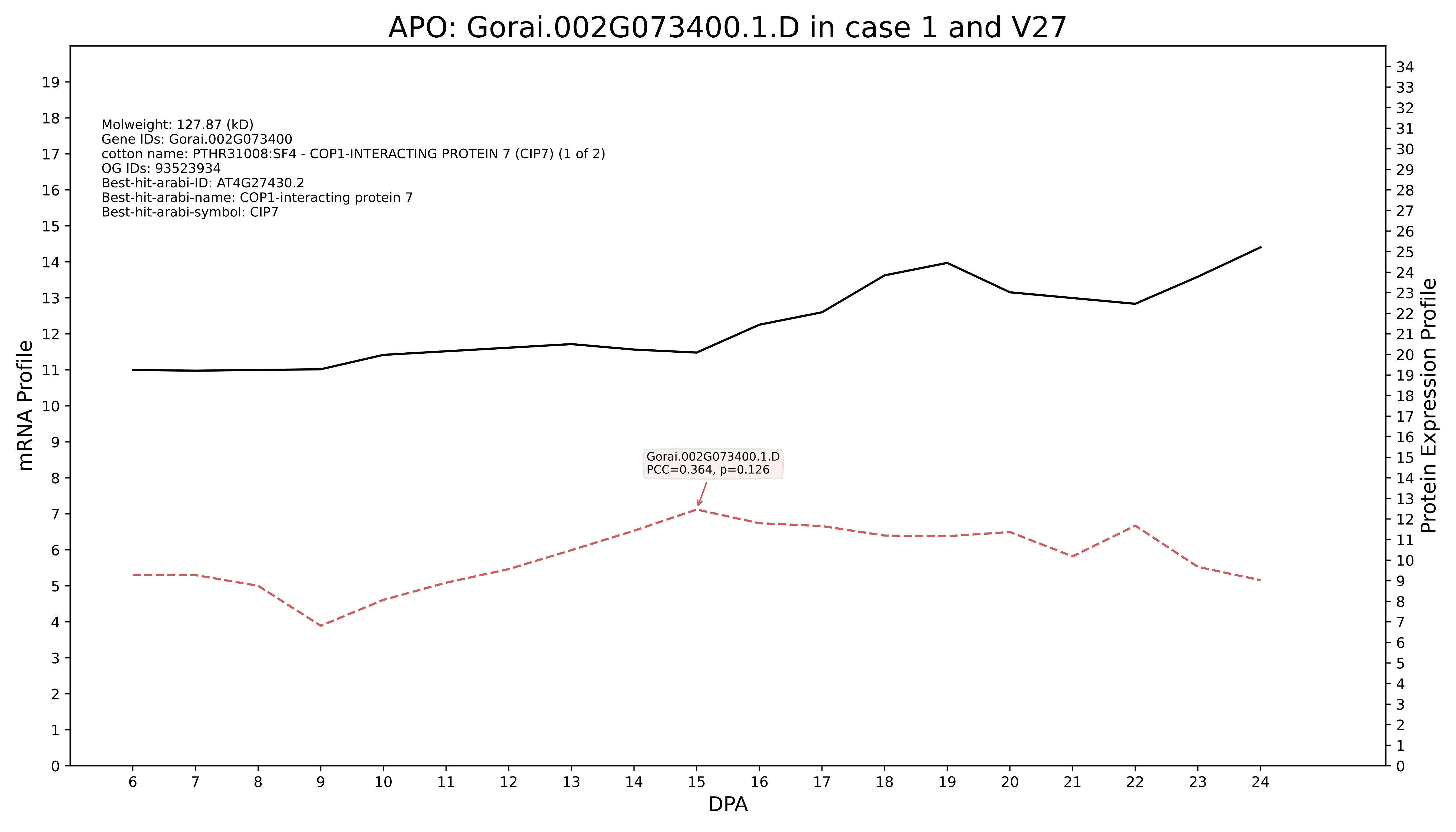

APO: Gorai.002G073400.1.D in case 1 and V27
Molweight: 127.87 (kD)
Gene IDs: Gorai.002G073400
cotton name: PTHR31008:SF4 - COP1-INTERACTING PROTEIN 7 (CIP7) (1 of 2)
OG IDs: 93523934
Best-hit-arabi-ID: AT4G27430.2
Best-hit-arabi-name: COP1-interacting protein 7
Best-hit-arabi-symbol: CIP7
Gorai.002G073400.1.D
PCC=0.364, p=0.126
mRNA Profile
Protein Expression Profile
DPA

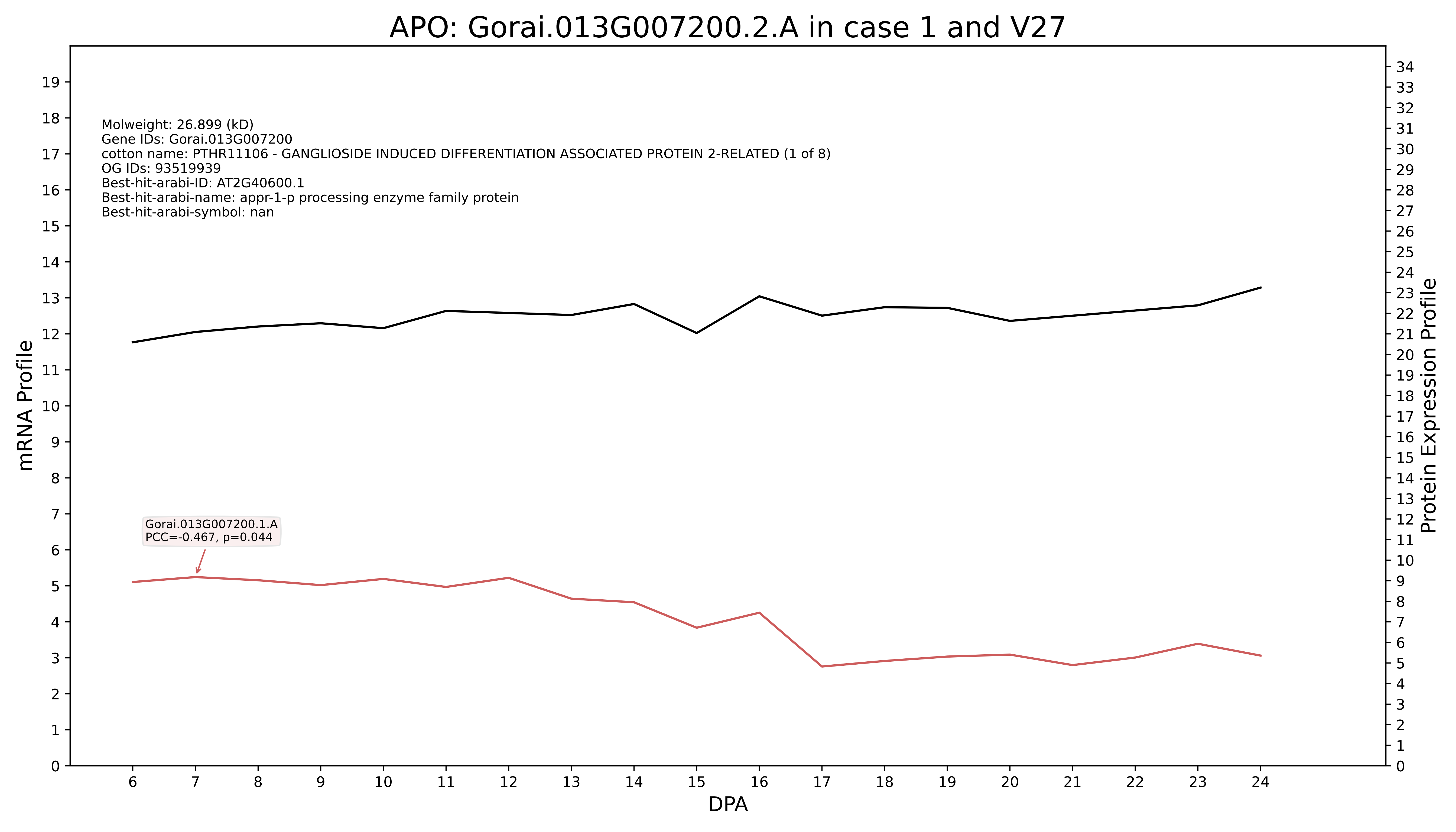
APO: Gorai.013G007200.2.A in case 1 and V27
Molweight: 26.899 (kD)
Gene IDs: Gorai.013G007200
cotton name: PTHR11106 - GANGLIOSIDE INDUCED DIFFERENTIATION ASSOCIATED PROTEIN 2-RELATED (1 of 8)
OG IDs: 93519939
Best-hit-arabi-ID: AT2G40600.1
Best-hit-arabi-name: appr-1-p processing enzyme family protein
Best-hit-arabi-symbol: nan
Gorai.013G007200.1.A
PCC=-0.467, p=0.044
mRNA Profile
Protein Expression Profile
DPA

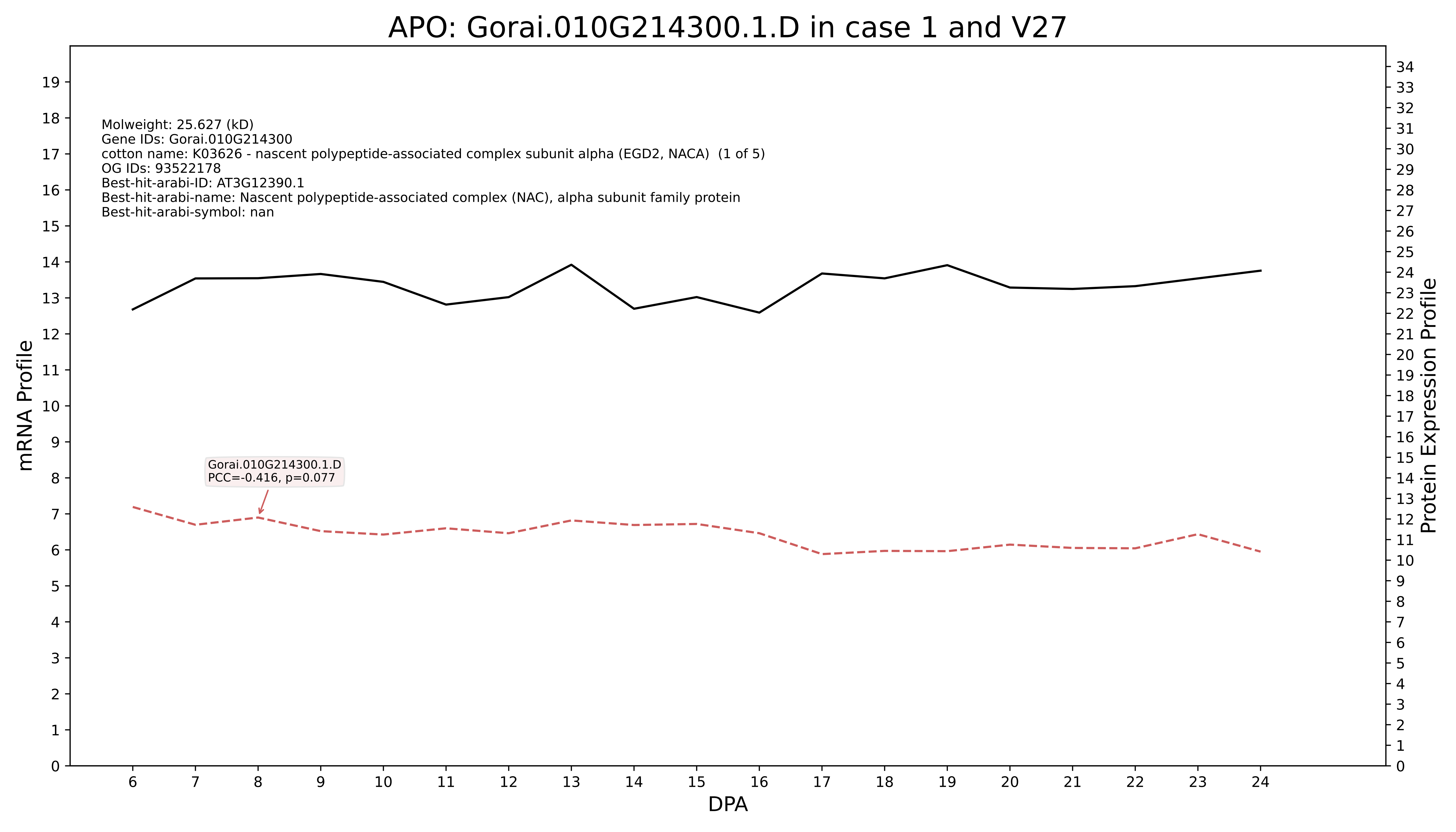

APO: Gorai.010G214300.1.D in case 1 and V27
Molweight: 25.627 (kD)
Gene IDs: Gorai.010G214300
cotton name: K03626 - nascent polypeptide-associated complex subunit alpha (EGD2, NACA) (1 of 5)
OG IDs: 93522178
Best-hit-arabi-ID: AT3G12390.1
Best-hit-arabi-name: Nascent polypeptide-associated complex (NAC), alpha subunit family protein
Best-hit-arabi-symbol: nan
Gorai.010G214300.1.D
PCC=-0.416, p=0.077
mRNA Profile
Protein Expression Profile
DPA

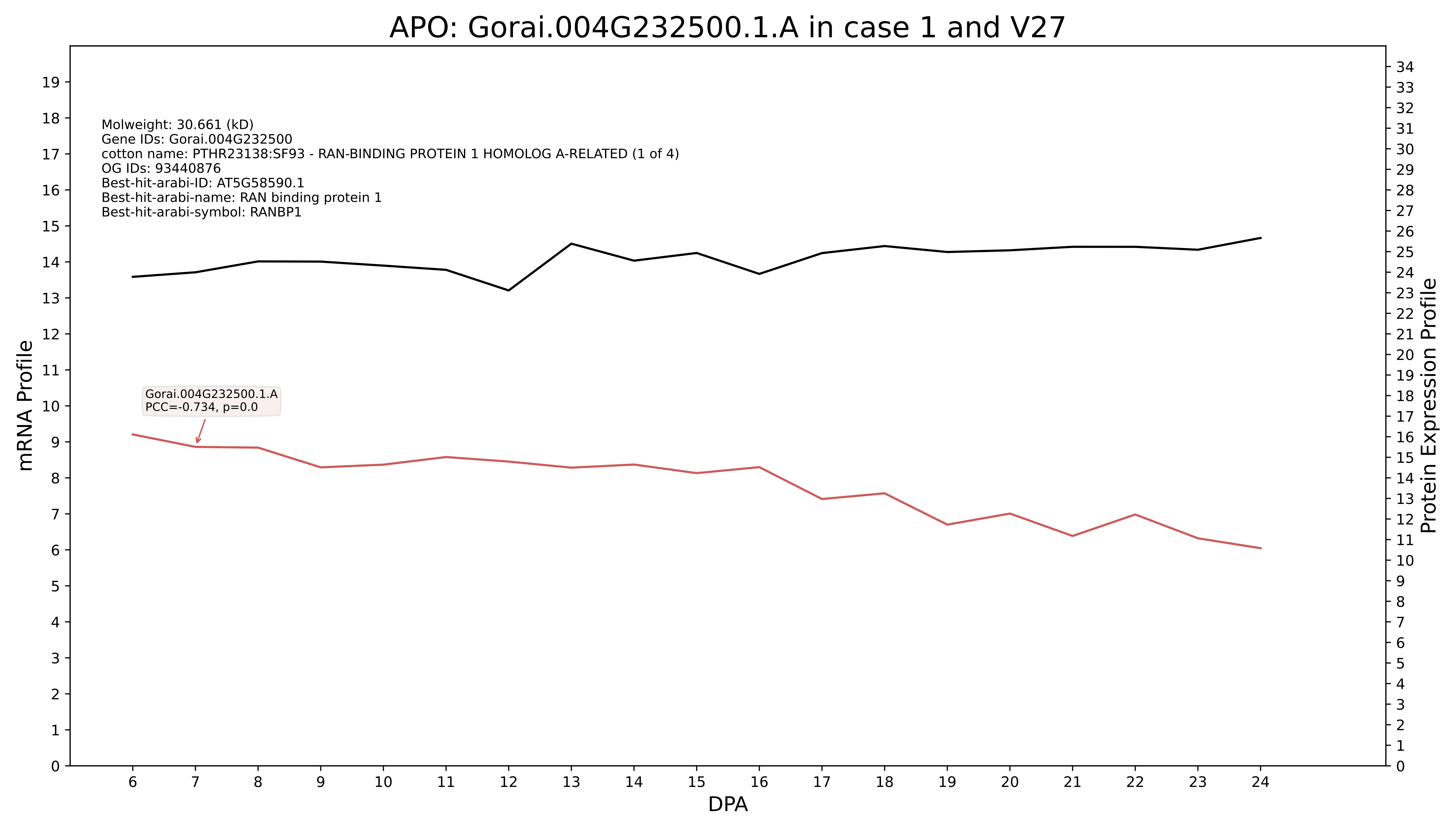
APO: Gorai.004G232500.1.A in case 1 and V27
Molweight: 30.661 (kD)
Gene IDs: Gorai.004G232500
cotton name: PTHR23138:SF93 - RAN-BINDING PROTEIN 1 HOMOLOG A-RELATED (1 of 4)
OG IDs: 93440876
Best-hit-arabi-ID: AT5G58590.1
Best-hit-arabi-name: RAN binding protein 1
Best-hit-arabi-symbol: RANBP1
Gorai.004G232500.1.A
PCC=-0.734, p=0.0
mRNA Profile
Protein Expression Profile
DPA

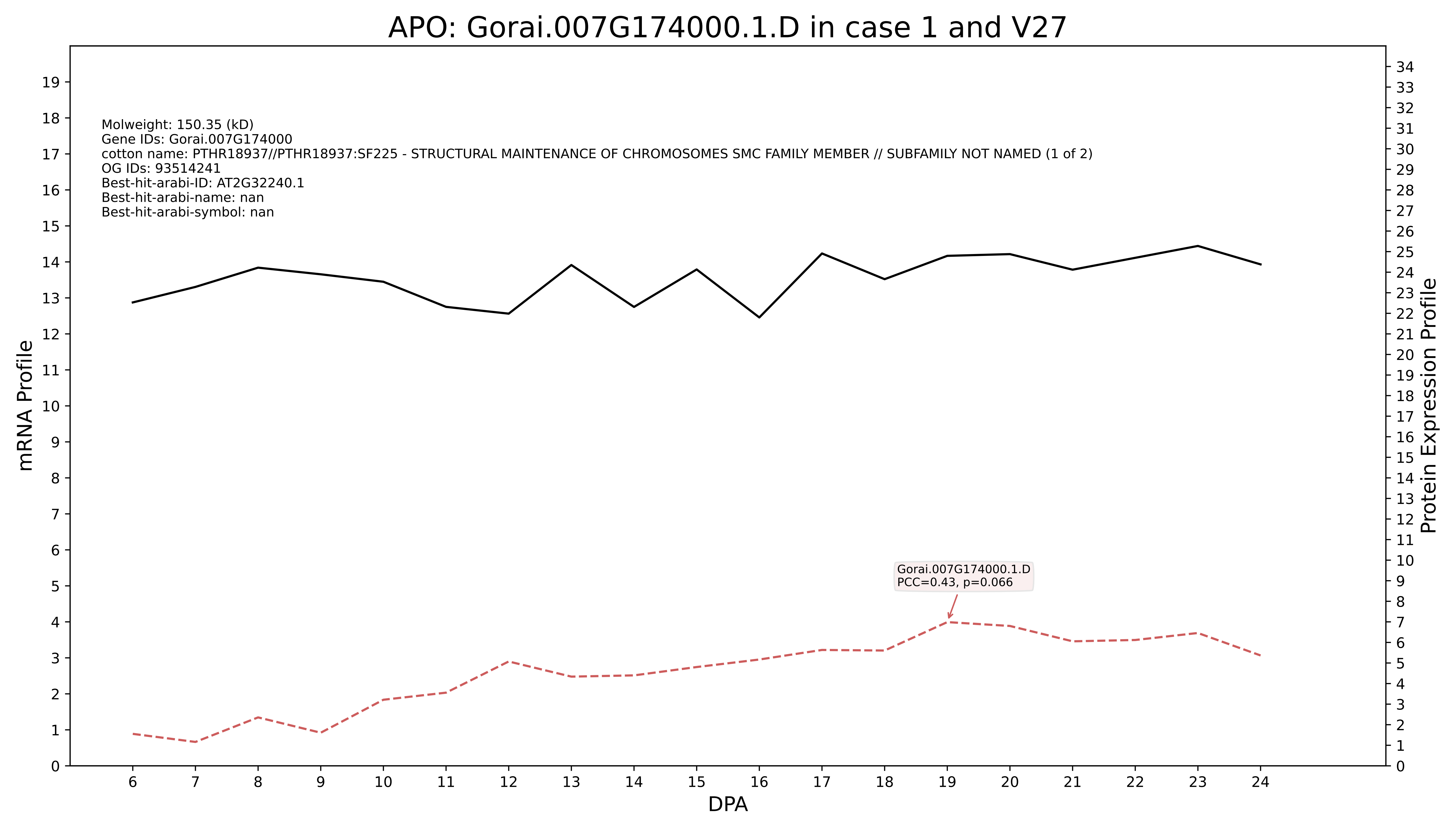
APO: Gorai.007G174000.1.D in case 1 and V27
Molweight: 150.35 (kD)
Gene IDs: Gorai.007G174000
cotton name: PTHR18937//PTHR18937:SF225 - STRUCTURAL MAINTENANCE OF CHROMOSOMES SMC FAMILY MEMBER // SUBFAMILY NOT NAMED (1 of 2)
OG IDs: 93514241
Best-hit-arabi-ID: AT2G32240.1
Best-hit-arabi-name: nan
Best-hit-arabi-symbol: nan
Gorai.007G174000.1.D
PCC=0.43, p=0.066
mRNA Profile
Protein Expression Profile
DPA

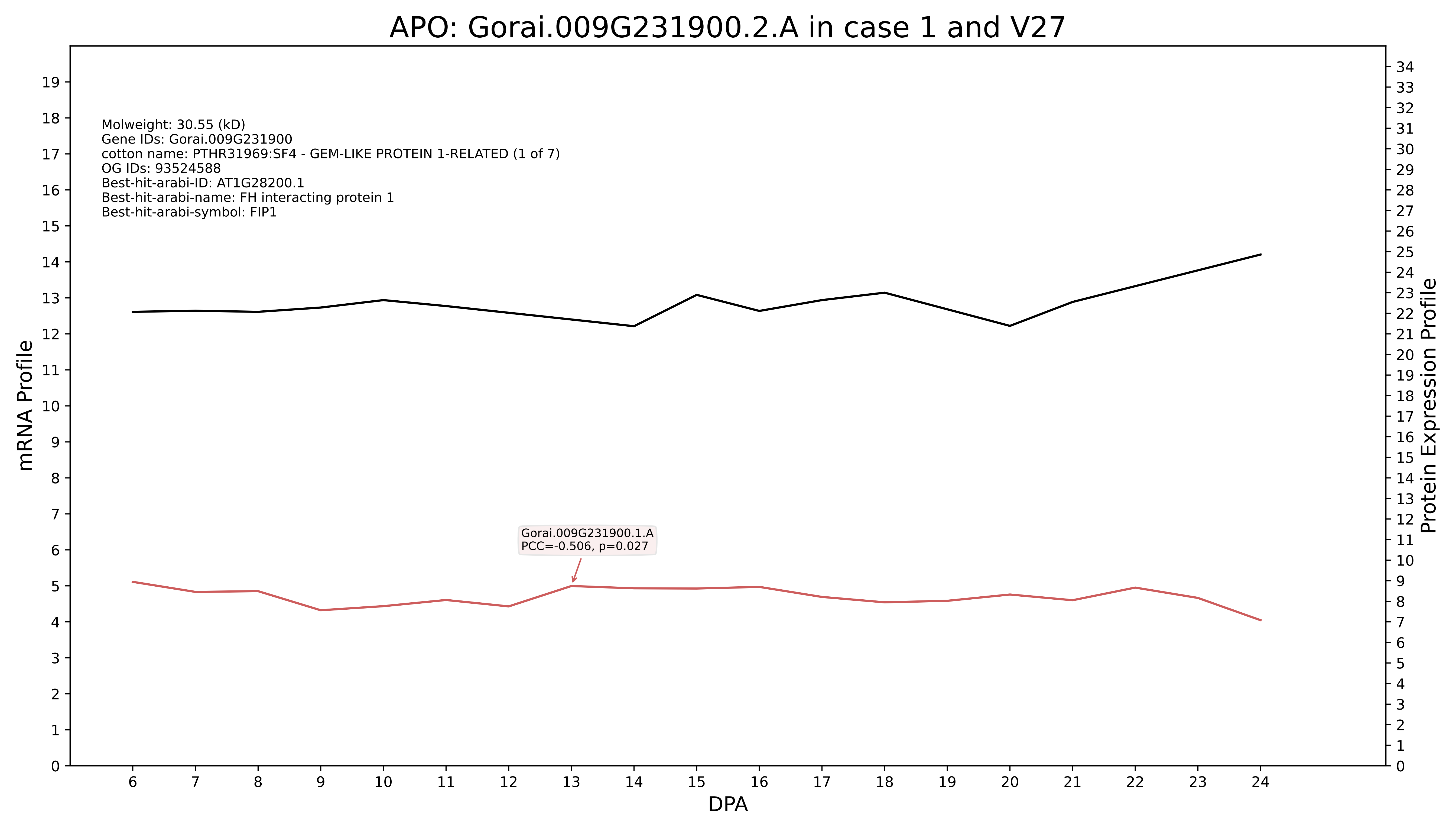
APO: Gorai.009G231900.2.A in case 1 and V27
Molweight: 30.55 (kD)
Gene IDs: Gorai.009G231900
cotton name: PTHR31969:SF4 - GEM-LIKE PROTEIN 1-RELATED (1 of 7)
OG IDs: 93524588
Best-hit-arabi-ID: AT1G28200.1
Best-hit-arabi-name: FH interacting protein 1
Best-hit-arabi-symbol: FIP1
Gorai.009G231900.1.A
PCC=-0.506, p=0.027
mRNA Profile
Protein Expression Profile
DPA

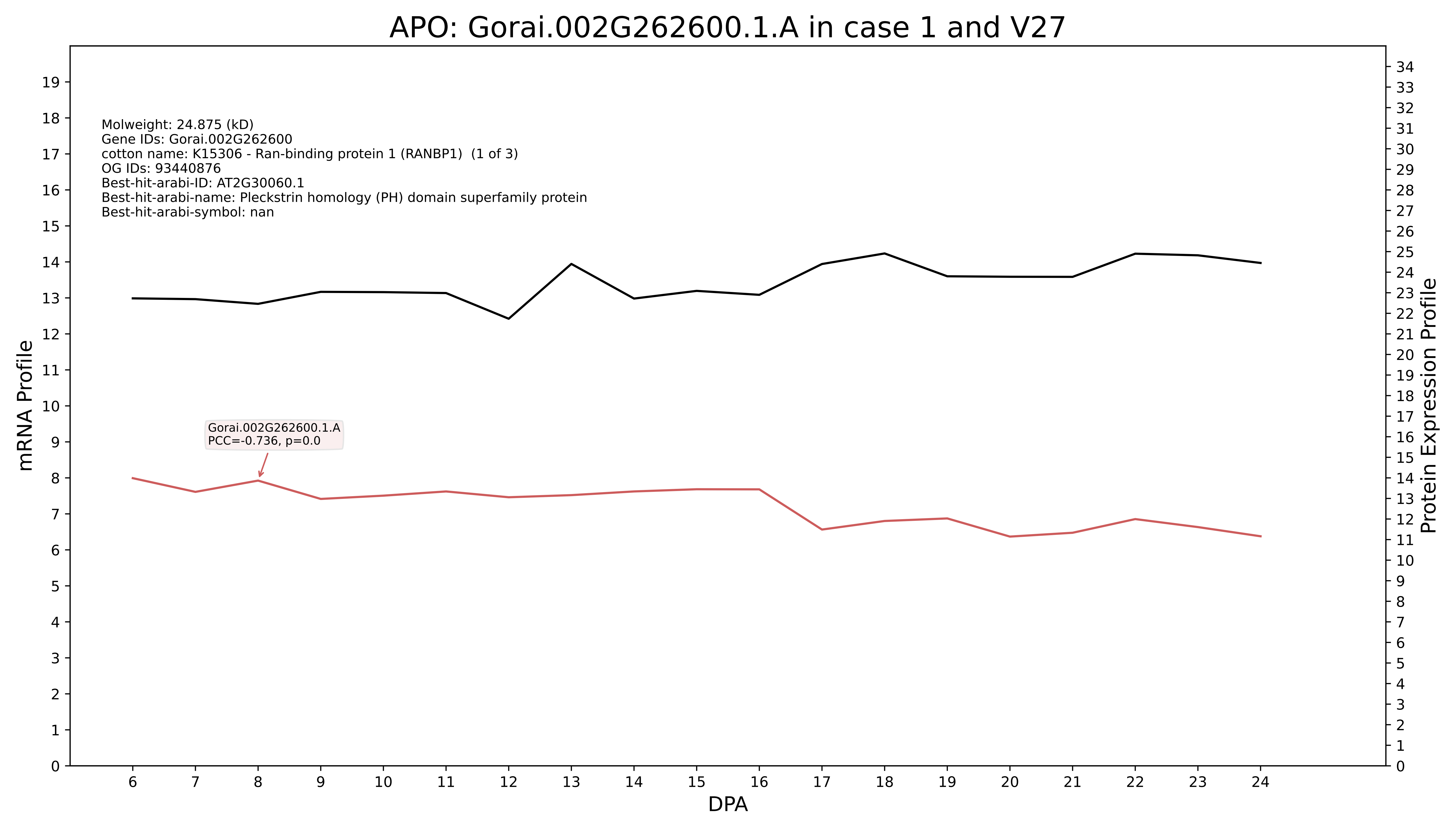
APO: Gorai.002G262600.1.A in case 1 and V27
Molweight: 24.875 (kD)
Gene IDs: Gorai.002G262600
cotton name: K15306 - Ran-binding protein 1 (RANBP1) (1 of 3)
OG IDs: 93440876
Best-hit-arabi-ID: AT2G30060.1
Best-hit-arabi-name: Pleckstrin homology (PH) domain superfamily protein
Best-hit-arabi-symbol: nan
Gorai.002G262600.1.A
PCC=-0.736, p=0.0
mRNA Profile
Protein Expression Profile
DPA

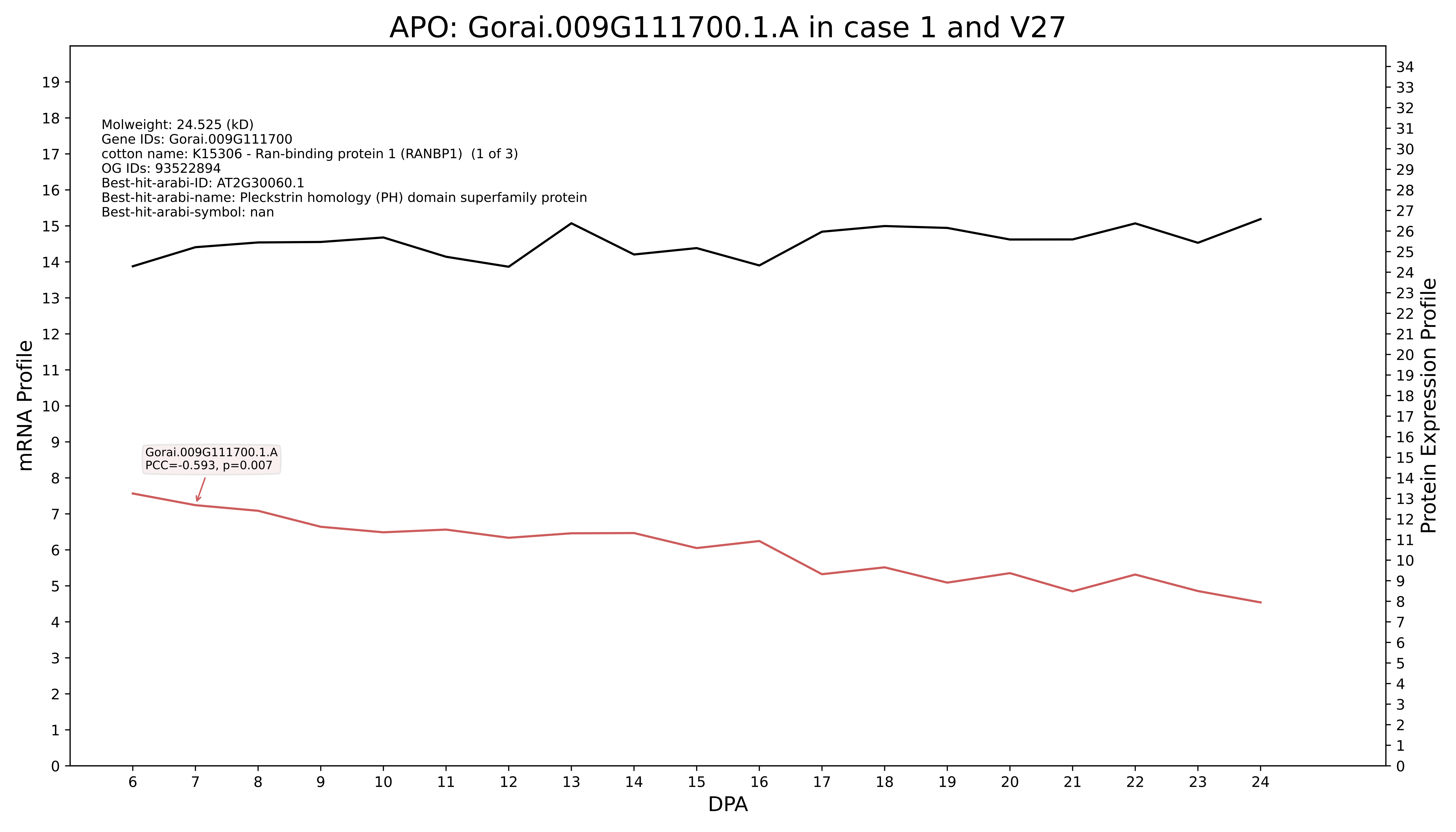
APO: Gorai.009G111700.1.A in case 1 and V27
Molweight: 24.525 (kD)
Gene IDs: Gorai.009G111700
cotton name: K15306 - Ran-binding protein 1 (RANBP1) (1 of 3)
OG IDs: 93522894
Best-hit-arabi-ID: AT2G30060.1
Best-hit-arabi-name: Pleckstrin homology (PH) domain superfamily protein
Best-hit-arabi-symbol: nan
Gorai.009G111700.1.A
PCC=-0.593, p=0.007
mRNA Profile
Protein Expression Profile
DPA

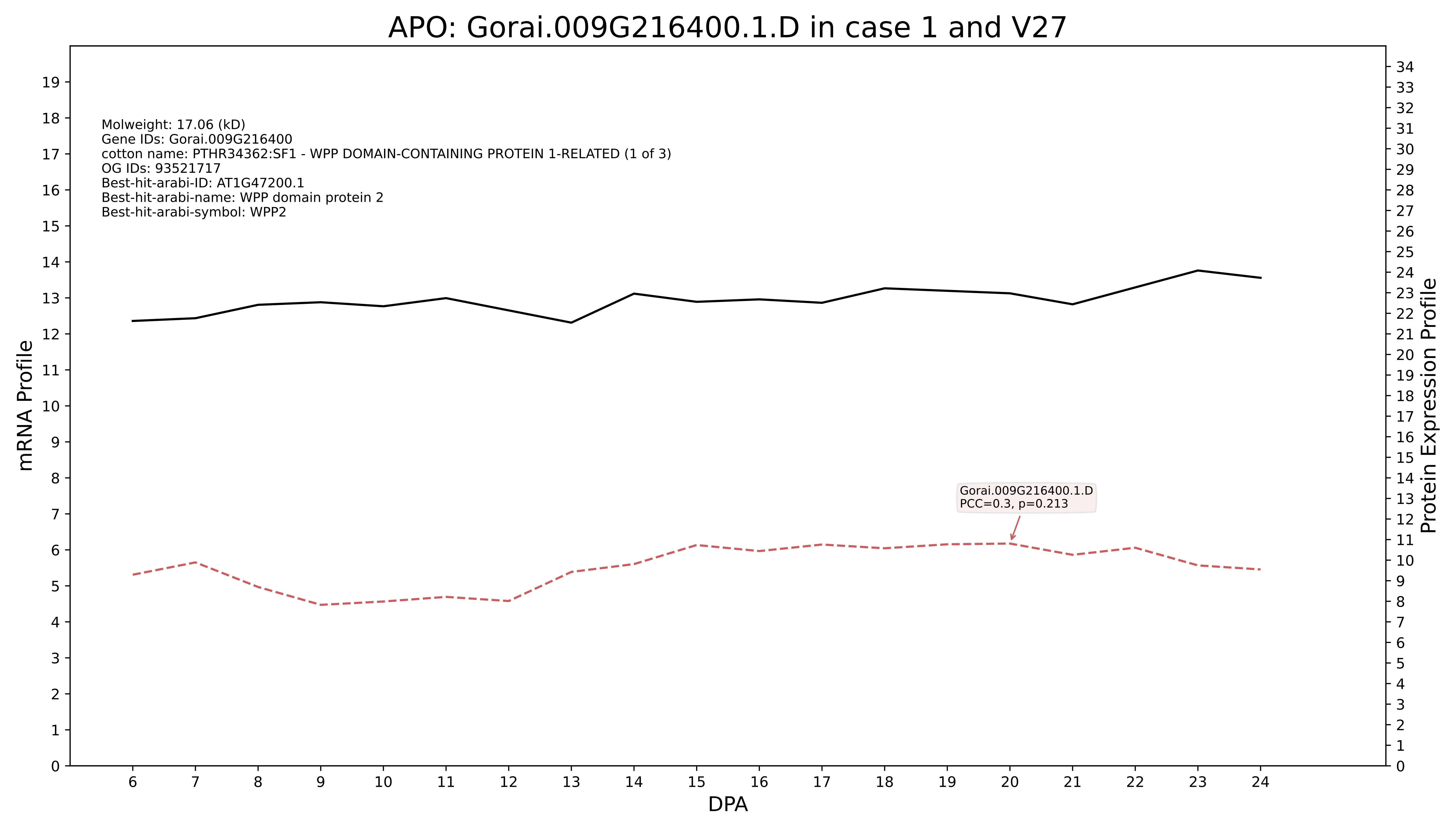

APO: Gorai.009G216400.1.D in case 1 and V27
Molweight: 17.06 (kD)
Gene IDs: Gorai.009G216400
cotton name: PTHR34362:SF1 - WPP DOMAIN-CONTAINING PROTEIN 1-RELATED (1 of 3)
OG IDs: 93521717
Best-hit-arabi-ID: AT1G47200.1
Best-hit-arabi-name: WPP domain protein 2
Best-hit-arabi-symbol: WPP2
Gorai.009G216400.1.D
PCC=0.3, p=0.213
mRNA Profile
Protein Expression Profile
DPA

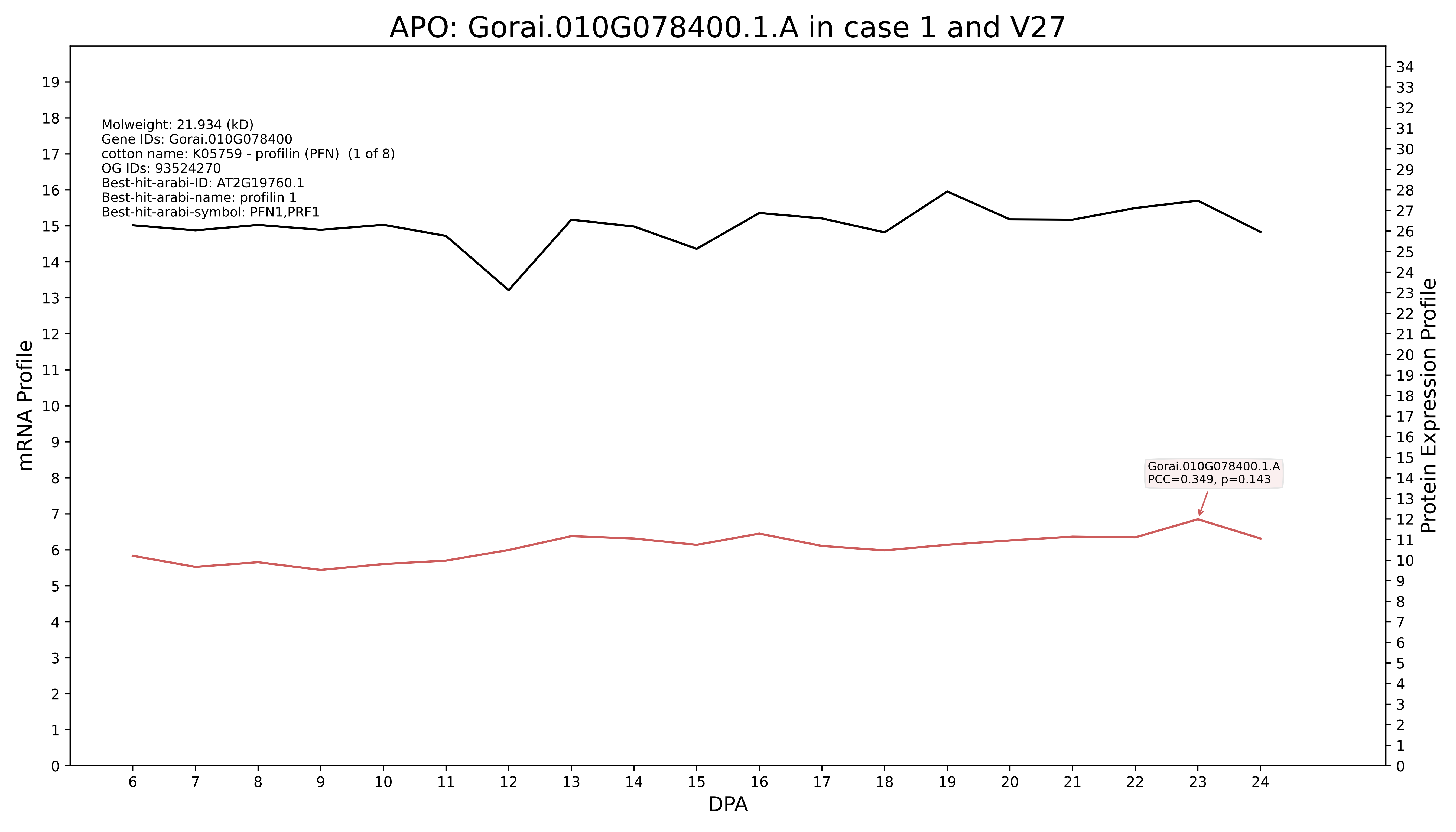

APO: Gorai.010G078400.1.A in case 1 and V27
Molweight: 21.934 (kD)
Gene IDs: Gorai.010G078400
cotton name: K05759 - profilin (PFN) (1 of 8)
OG IDs: 93524270
Best-hit-arabi-ID: AT2G19760.1
Best-hit-arabi-name: profilin 1
Best-hit-arabi-symbol: PFN1,PRF1
Gorai.010G078400.1.A
PCC=0.349, p=0.143
mRNA Profile
Protein Expression Profile
DPA

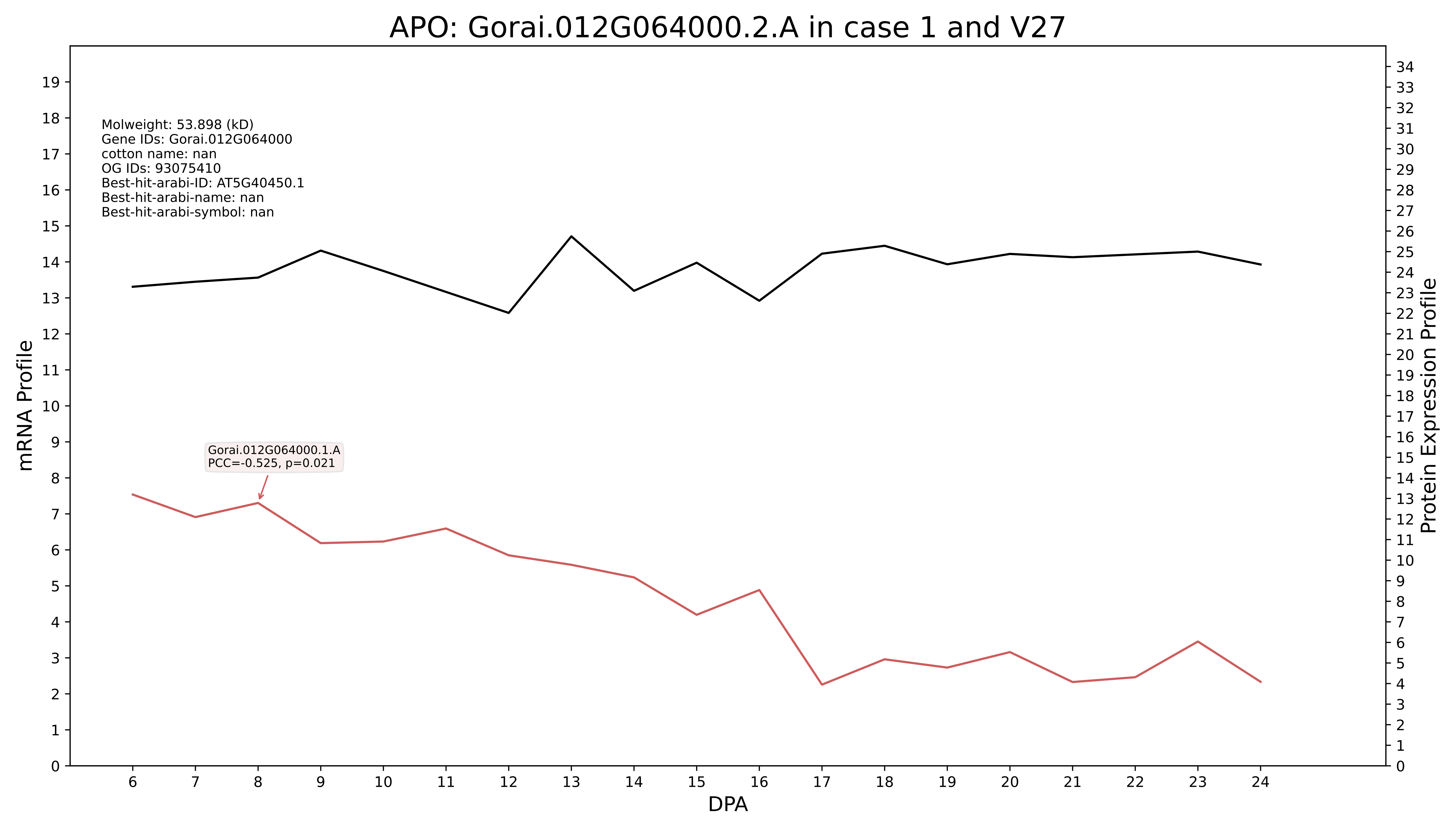

APO: Gorai.012G064000.2.A in case 1 and V27
Molweight: 53.898 (kD)
Gene IDs: Gorai.012G064000
cotton name: nan
OG IDs: 93075410
Best-hit-arabi-ID: AT5G40450.1
Best-hit-arabi-name: nan
Best-hit-arabi-symbol: nan
Gorai.012G064000.1.A
PCC=-0.525, p=0.021
mRNA Profile
Protein Expression Profile
DPA

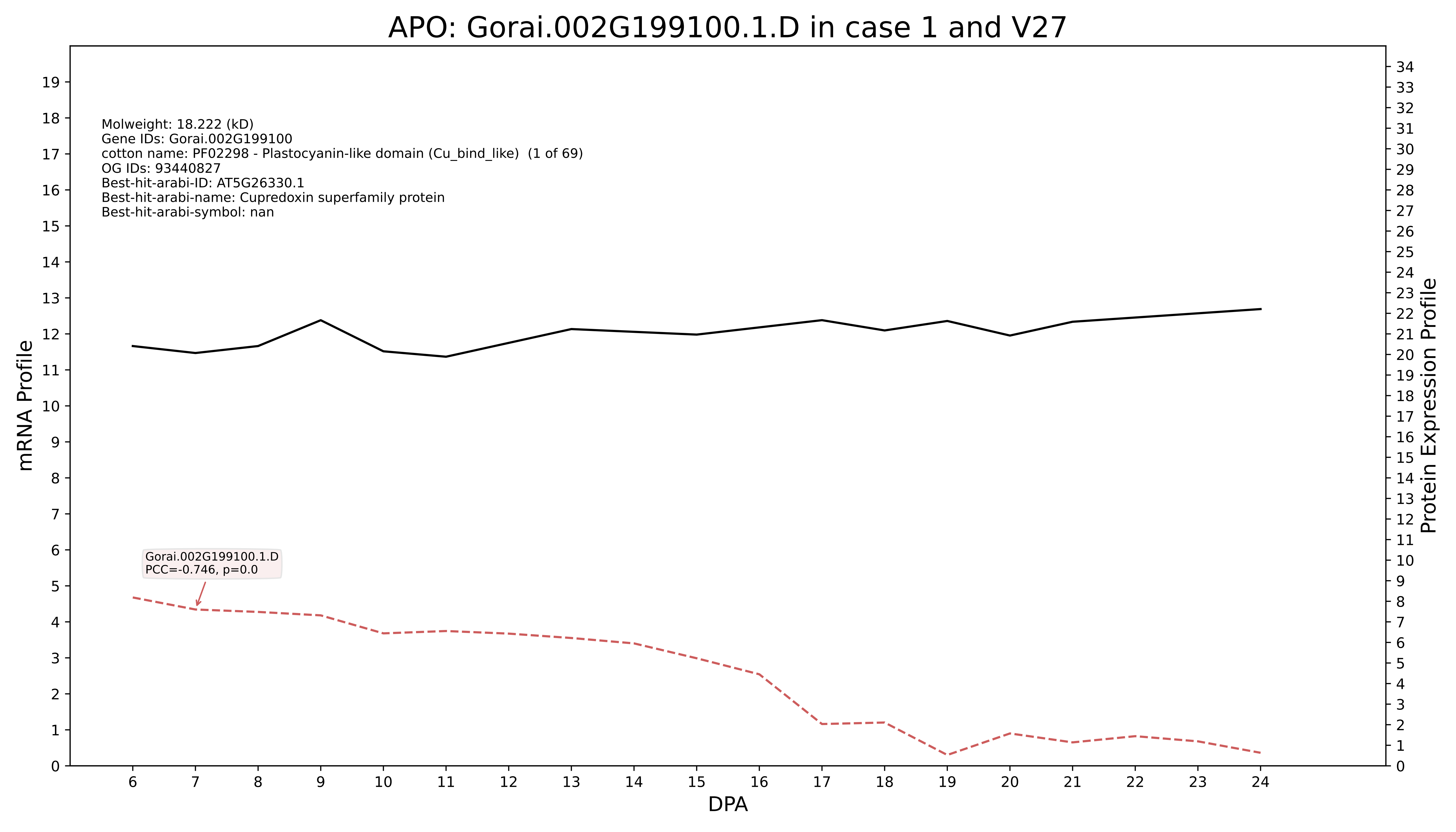
APO: Gorai.002G199100.1.D in case 1 and V27
Molweight: 18.222 (kD)
Gene IDs: Gorai.002G199100
cotton name: PF02298 - Plastocyanin-like domain (Cu_bind_like) (1 of 69)
OG IDs: 93440827
Best-hit-arabi-ID: AT5G26330.1
Best-hit-arabi-name: Cupredoxin superfamily protein
Best-hit-arabi-symbol: nan
Gorai.002G199100.1.D
PCC=-0.746, p=0.0
mRNA Profile
Protein Expression Profile
DPA

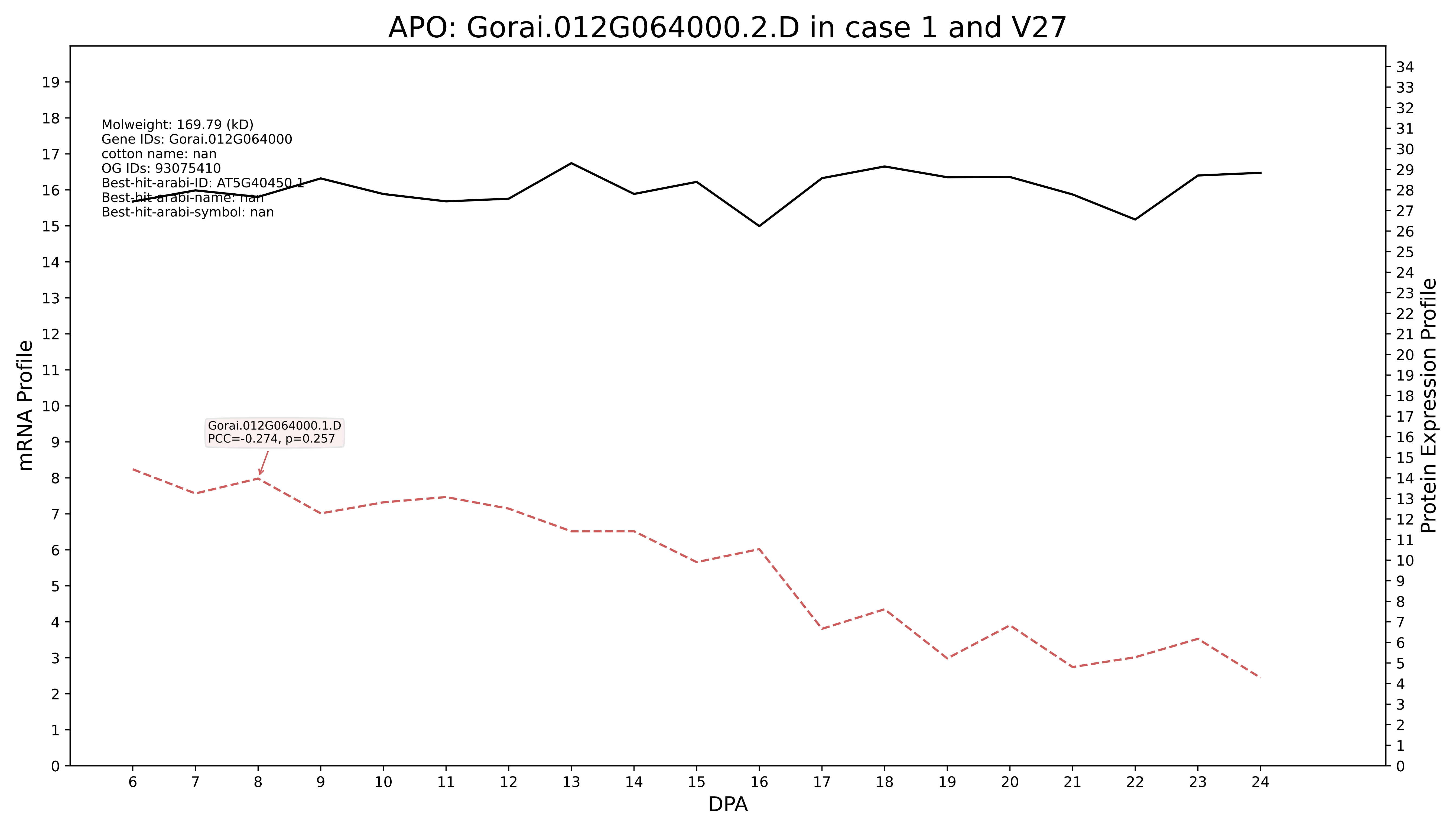

APO: Gorai.012G064000.2.D in case 1 and V27
Molweight: 169.79 (kD)
Gene IDs: Gorai.012G064000
cotton name: nan
OG IDs: 93075410
Best-hit-arabi-ID: AT5G40450.1
Best-hit-arabi-name: nan
Best-hit-arabi-symbol: nan
Gorai.012G064000.1.D
PCC=-0.274, p=0.257
mRNA Profile
Protein Expression Profile
DPA

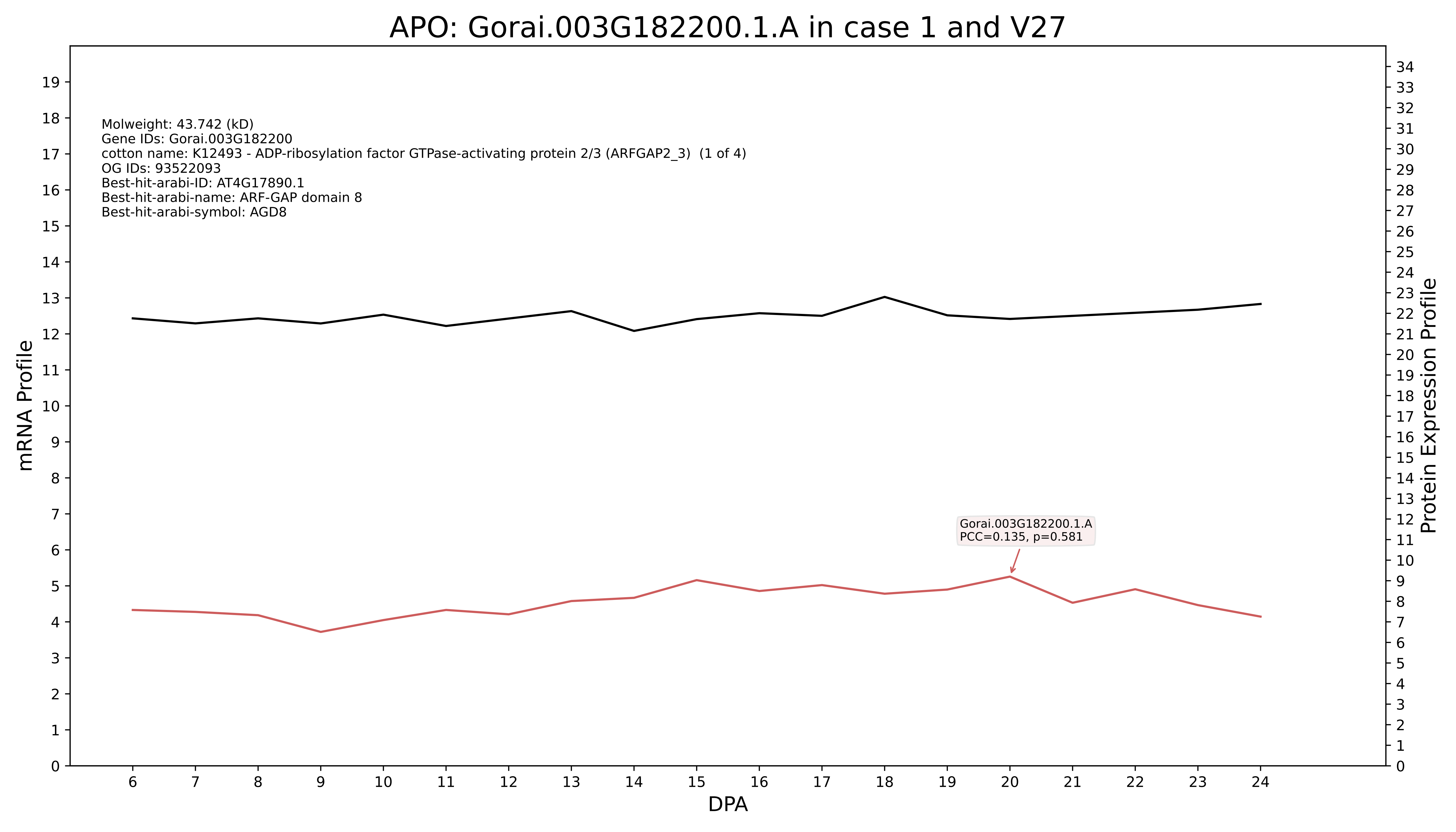
APO: Gorai.003G182200.1.A in case 1 and V27
Molweight: 43.742 (kD)
Gene IDs: Gorai.003G182200
cotton name: K12493 - ADP-ribosylation factor GTPase-activating protein 2/3 (ARFGAP2_3) (1 of 4)
OG IDs: 93522093
Best-hit-arabi-ID: AT4G17890.1
Best-hit-arabi-name: ARF-GAP domain 8
Best-hit-arabi-symbol: AGD8
Gorai.003G182200.1.A
PCC=0.135, p=0.581
mRNA Profile
Protein Expression Profile
DPA

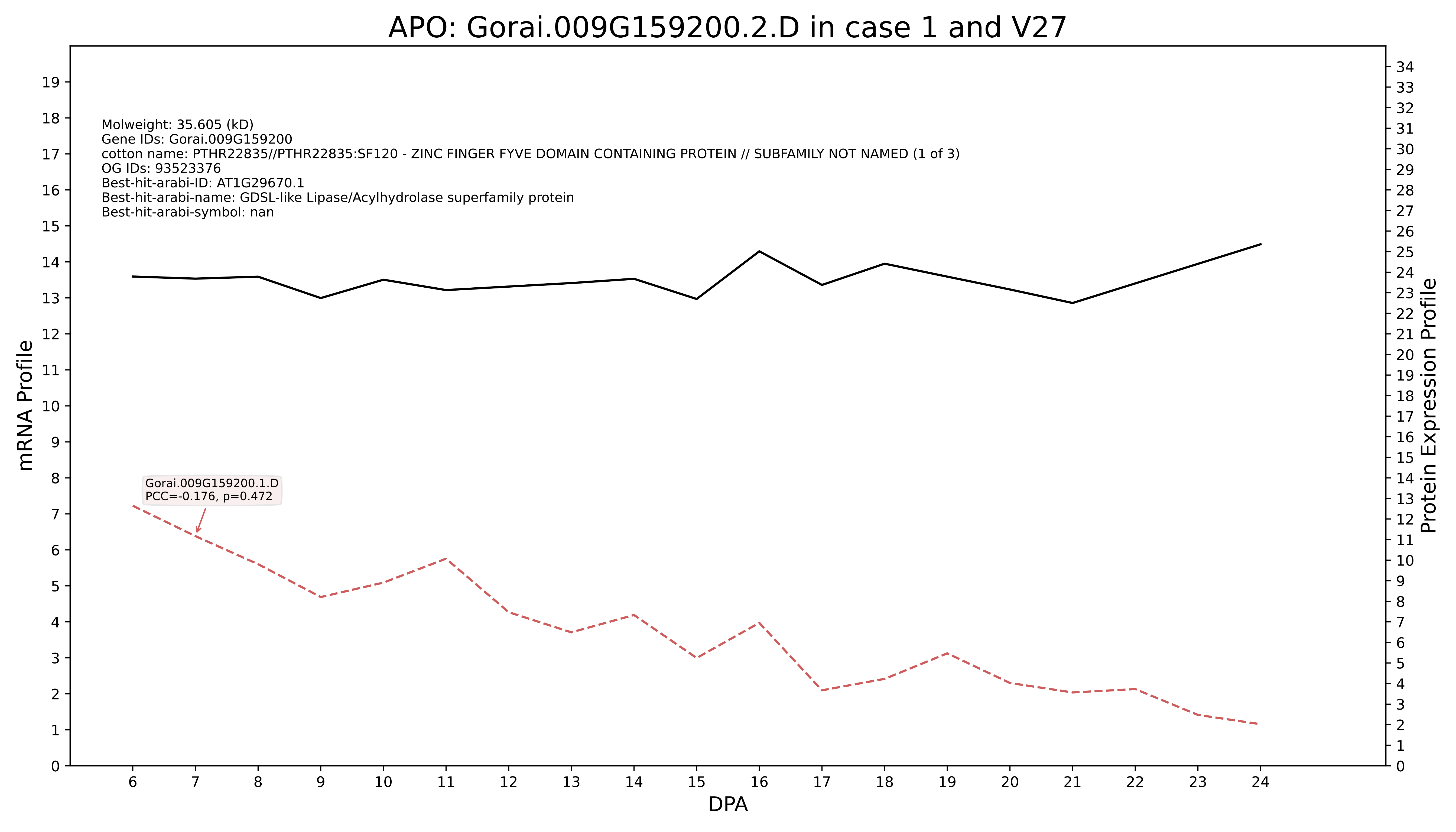
APO: Gorai.009G159200.2.D in case 1 and V27
Molweight: 35.605 (kD)
Gene IDs: Gorai.009G159200
cotton name: PTHR22835//PTHR22835:SF120 - ZINC FINGER FYVE DOMAIN CONTAINING PROTEIN // SUBFAMILY NOT NAMED (1 of 3)
OG IDs: 93523376
Best-hit-arabi-ID: AT1G29670.1
Best-hit-arabi-name: GDSL-like Lipase/Acylhydrolase superfamily protein
Best-hit-arabi-symbol: nan
Gorai.009G159200.1.D
PCC=-0.176, p=0.472
mRNA Profile
Protein Expression Profile
DPA

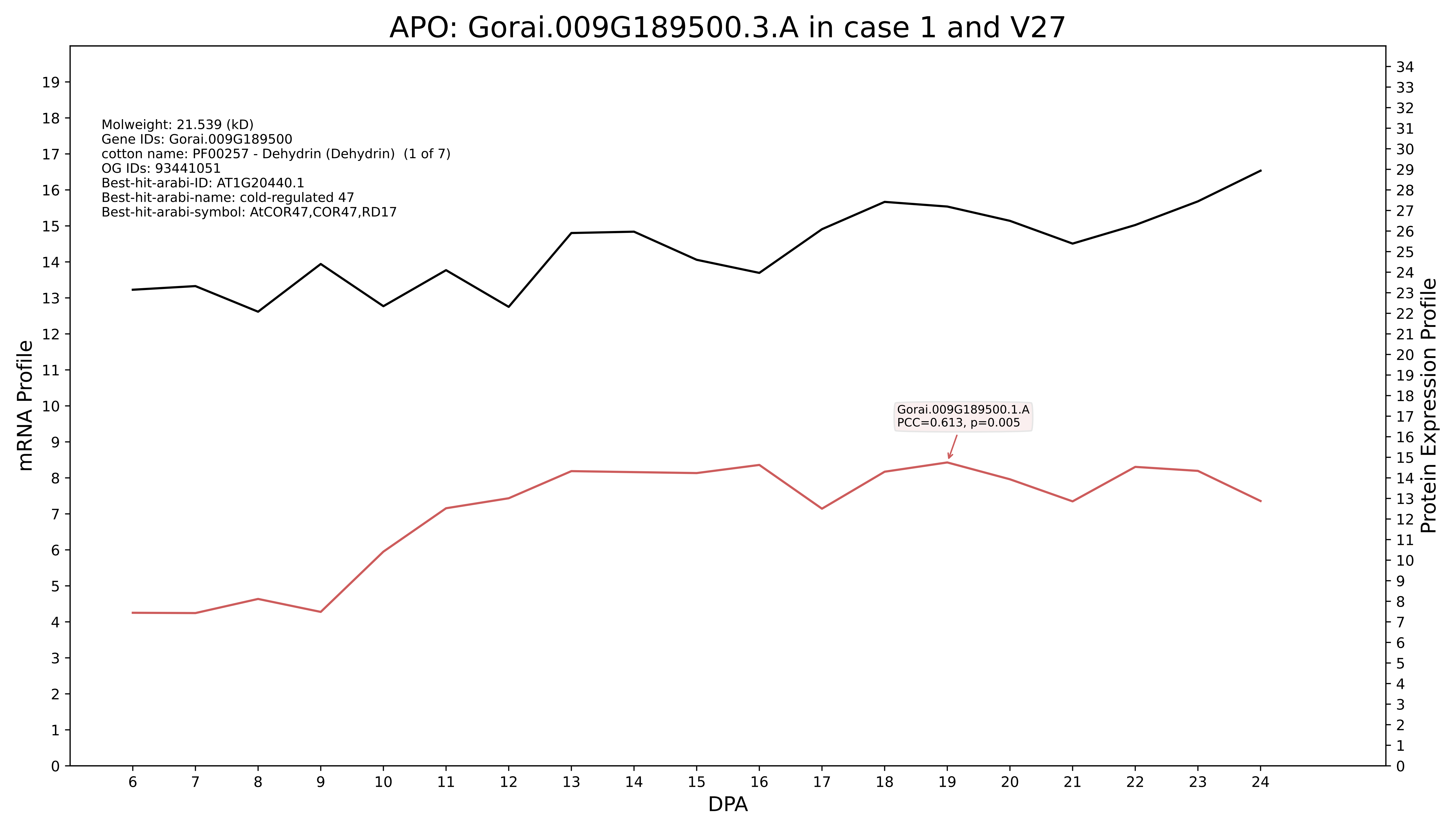
APO: Gorai.009G189500.3.A in case 1 and V27
Molweight: 21.539 (kD)
Gene IDs: Gorai.009G189500
cotton name: PF00257 - Dehydrin (Dehydrin) (1 of 7)
OG IDs: 93441051
Best-hit-arabi-ID: AT1G20440.1
Best-hit-arabi-name: cold-regulated 47
Best-hit-arabi-symbol: AtCOR47,COR47,RD17
Gorai.009G189500.1.A
PCC=0.613, p=0.005
mRNA Profile
Protein Expression Profile
DPA

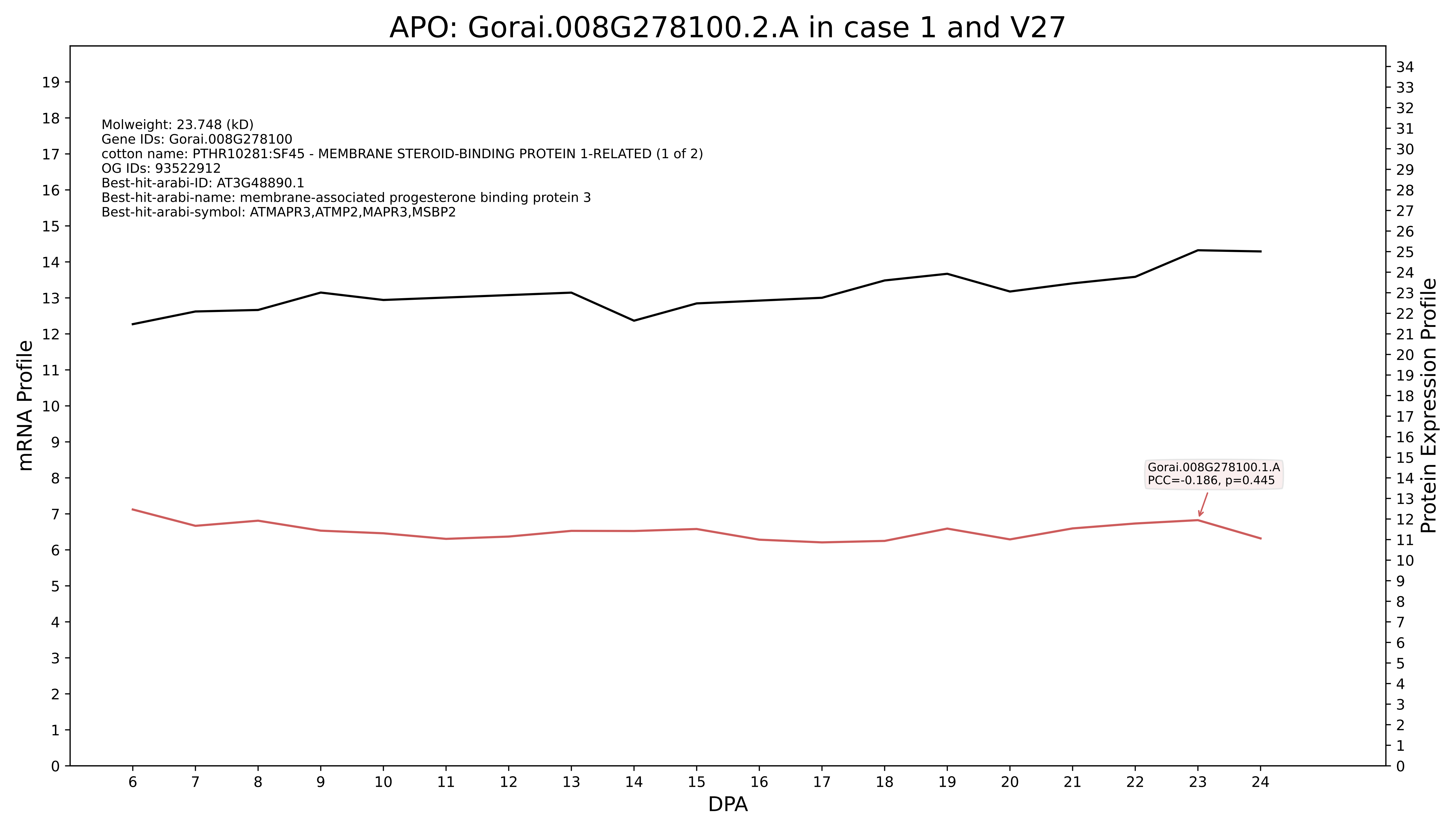

APO: Gorai.008G278100.2.A in case 1 and V27
Molweight: 23.748 (kD)
Gene IDs: Gorai.008G278100
cotton name: PTHR10281:SF45 - MEMBRANE STEROID-BINDING PROTEIN 1-RELATED (1 of 2)
OG IDs: 93522912
Best-hit-arabi-ID: AT3G48890.1
Best-hit-arabi-name: membrane-associated progesterone binding protein 3
Best-hit-arabi-symbol: ATMAPR3,ATMP2,MAPR3,MSBP2
Gorai.008G278100.1.A
PCC=-0.186, p=0.445
mRNA Profile
Protein Expression Profile
DPA

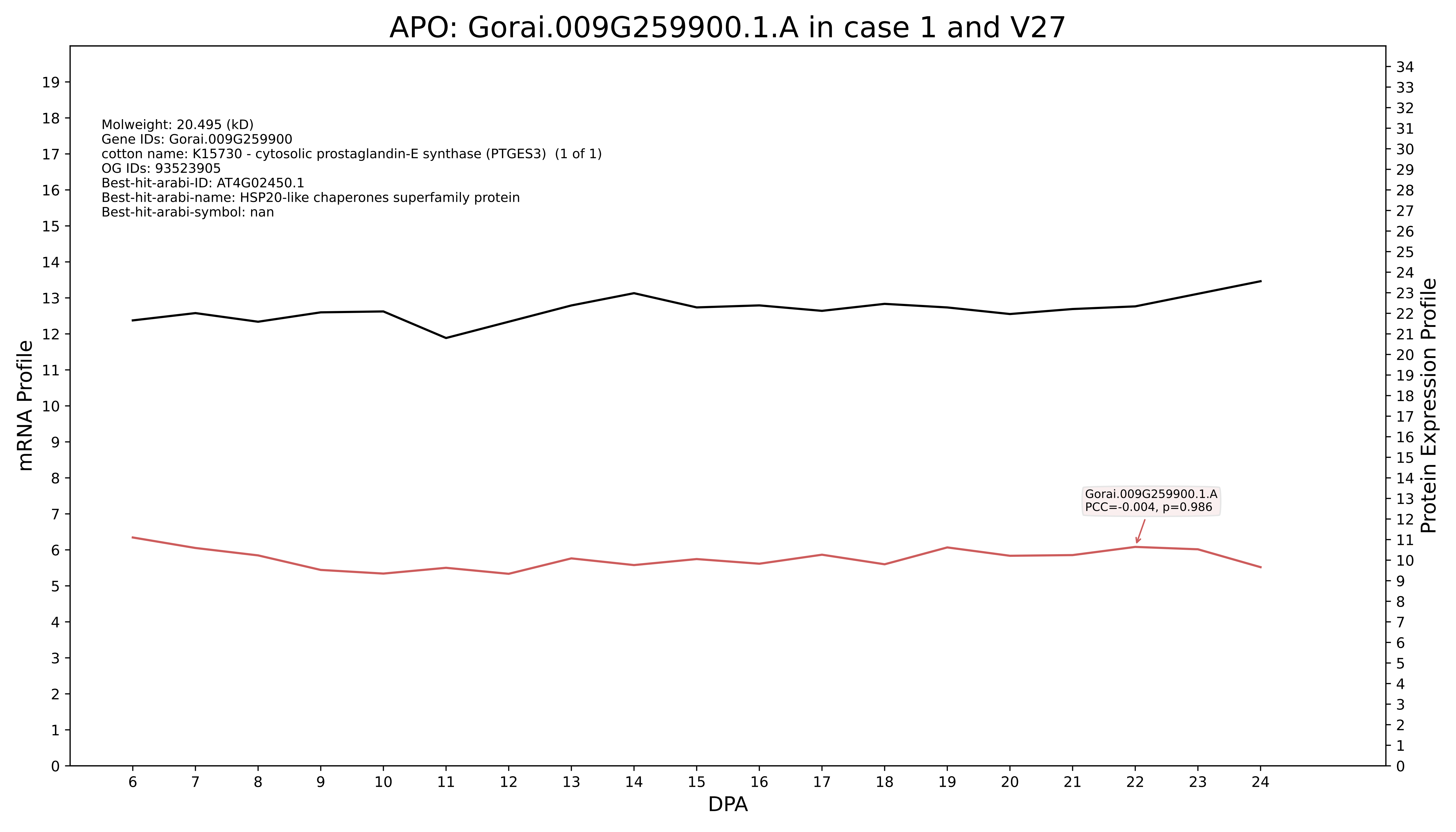
APO: Gorai.009G259900.1.A in case 1 and V27
Molweight: 20.495 (kD)
Gene IDs: Gorai.009G259900
cotton name: K15730 - cytosolic prostaglandin-E synthase (PTGES3) (1 of 1)
OG IDs: 93523905
Best-hit-arabi-ID: AT4G02450.1
Best-hit-arabi-name: HSP20-like chaperones superfamily protein
Best-hit-arabi-symbol: nan
Gorai.009G259900.1.A
PCC=-0.004, p=0.986
mRNA Profile
Protein Expression Profile
DPA

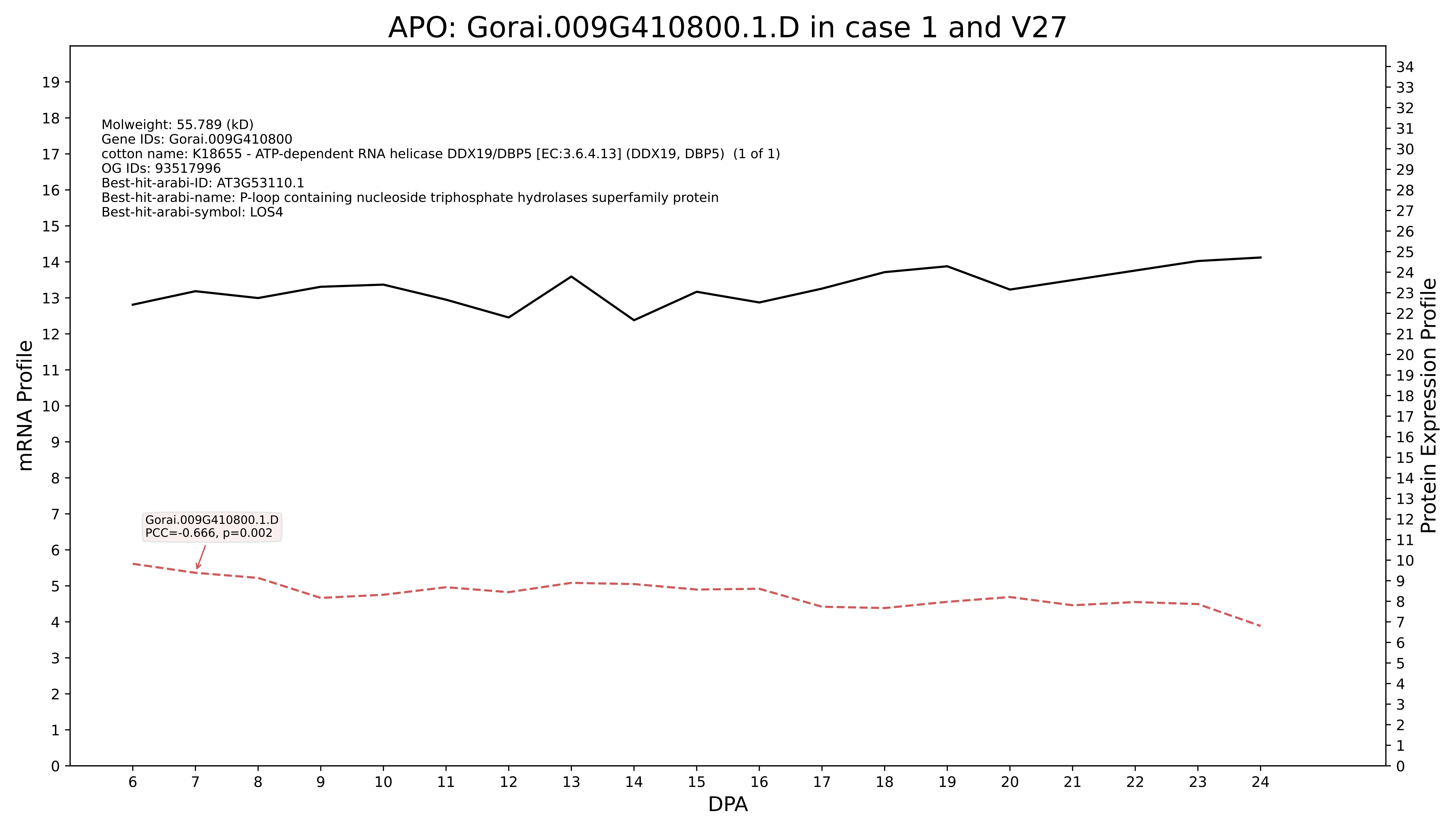

APO: Gorai.009G410800.1.D in case 1 and V27
Molweight: 55.789 (kD)
Gene IDs: Gorai.009G410800
cotton name: K18655 - ATP-dependent RNA helicase DDX19/DBP5 [EC:3.6.4.13] (DDX19, DBP5) (1 of 1)
OG IDs: 93517996
Best-hit-arabi-ID: AT3G53110.1
Best-hit-arabi-name: P-loop containing nucleoside triphosphate hydrolases superfamily protein
Best-hit-arabi-symbol: LOS4
Gorai.009G410800.1.D
PCC=-0.666, p=0.002
mRNA Profile
Protein Expression Profile
DPA

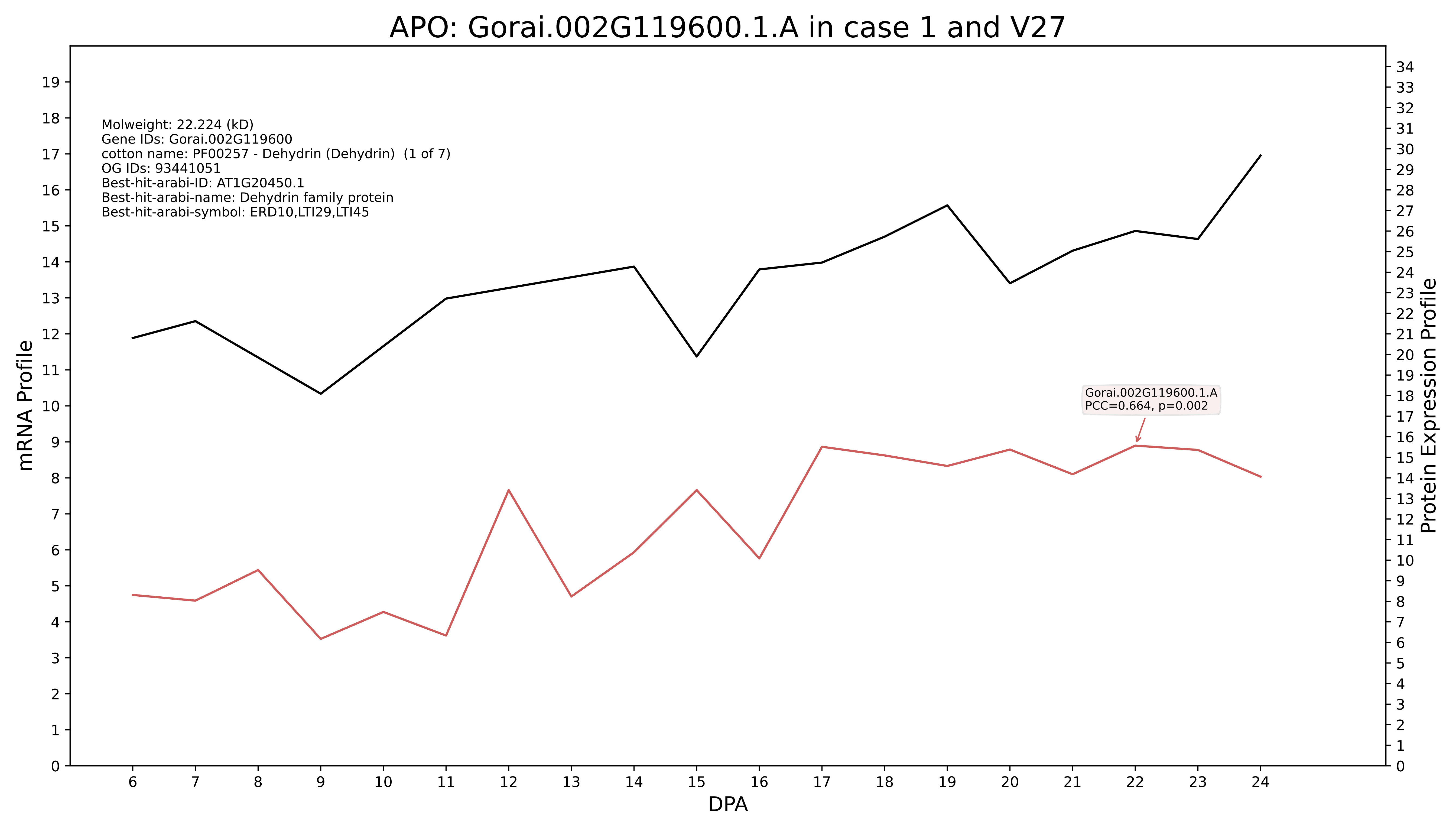
APO: Gorai.002G119600.1.A in case 1 and V27
Molweight: 22.224 (kD)
Gene IDs: Gorai.002G119600
cotton name: PF00257 - Dehydrin (Dehydrin) (1 of 7)
OG IDs: 93441051
Best-hit-arabi-ID: AT1G20450.1
Best-hit-arabi-name: Dehydrin family protein
Best-hit-arabi-symbol: ERD10,LTI29,LTI45
Gorai.002G119600.1.A
PCC=0.664, p=0.002
mRNA Profile
Protein Expression Profile
DPA

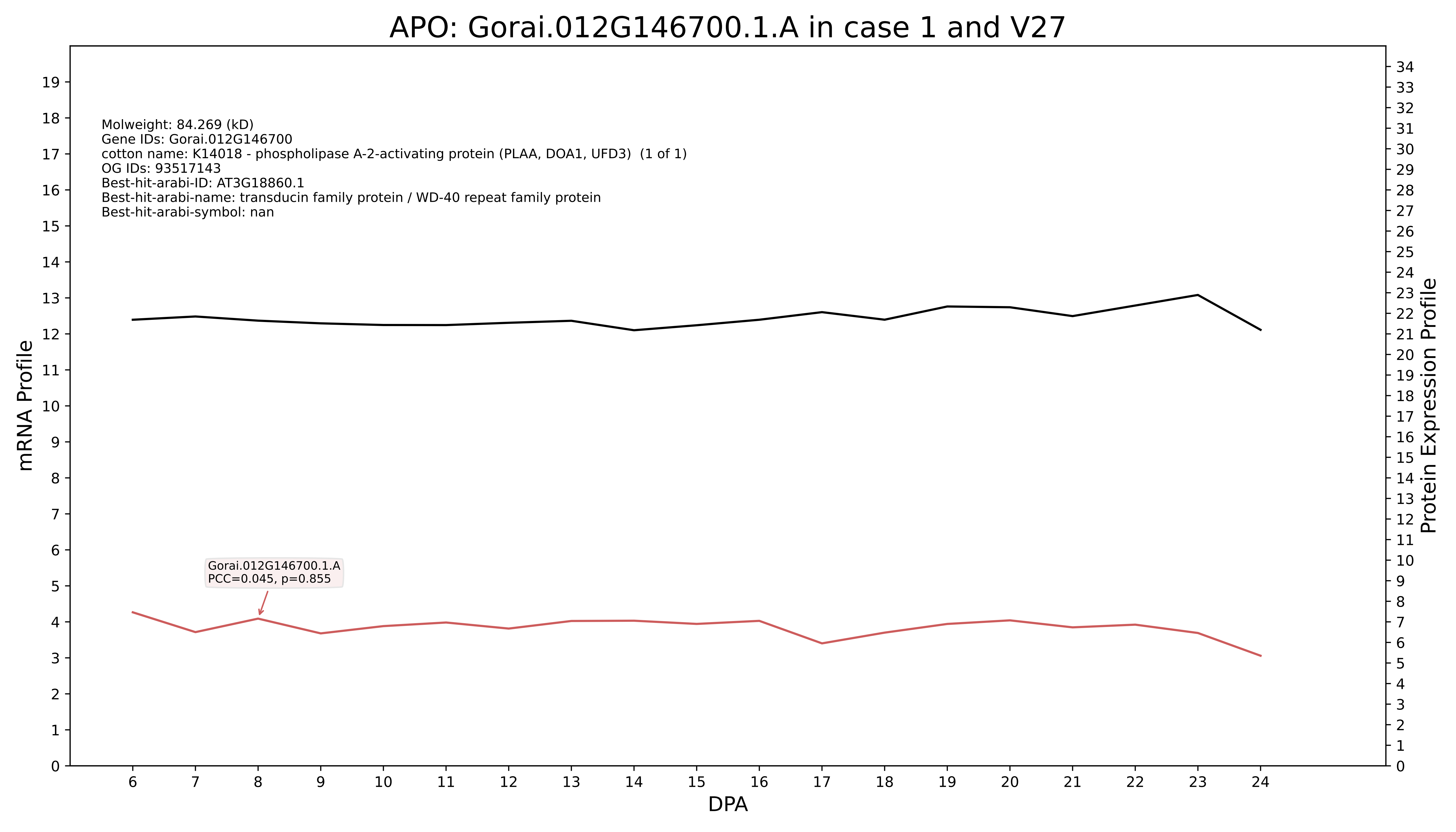

APO: Gorai.012G146700.1.A in case 1 and V27
Molweight: 84.269 (kD)
Gene IDs: Gorai.012G146700
cotton name: K14018 - phospholipase A-2-activating protein (PLAA, DOA1, UFD3) (1 of 1)
OG IDs: 93517143
Best-hit-arabi-ID: AT3G18860.1
Best-hit-arabi-name: transducin family protein / WD-40 repeat family protein
Best-hit-arabi-symbol: nan
Gorai.012G146700.1.A
PCC=0.045, p=0.855
mRNA Profile
Protein Expression Profile
DPA

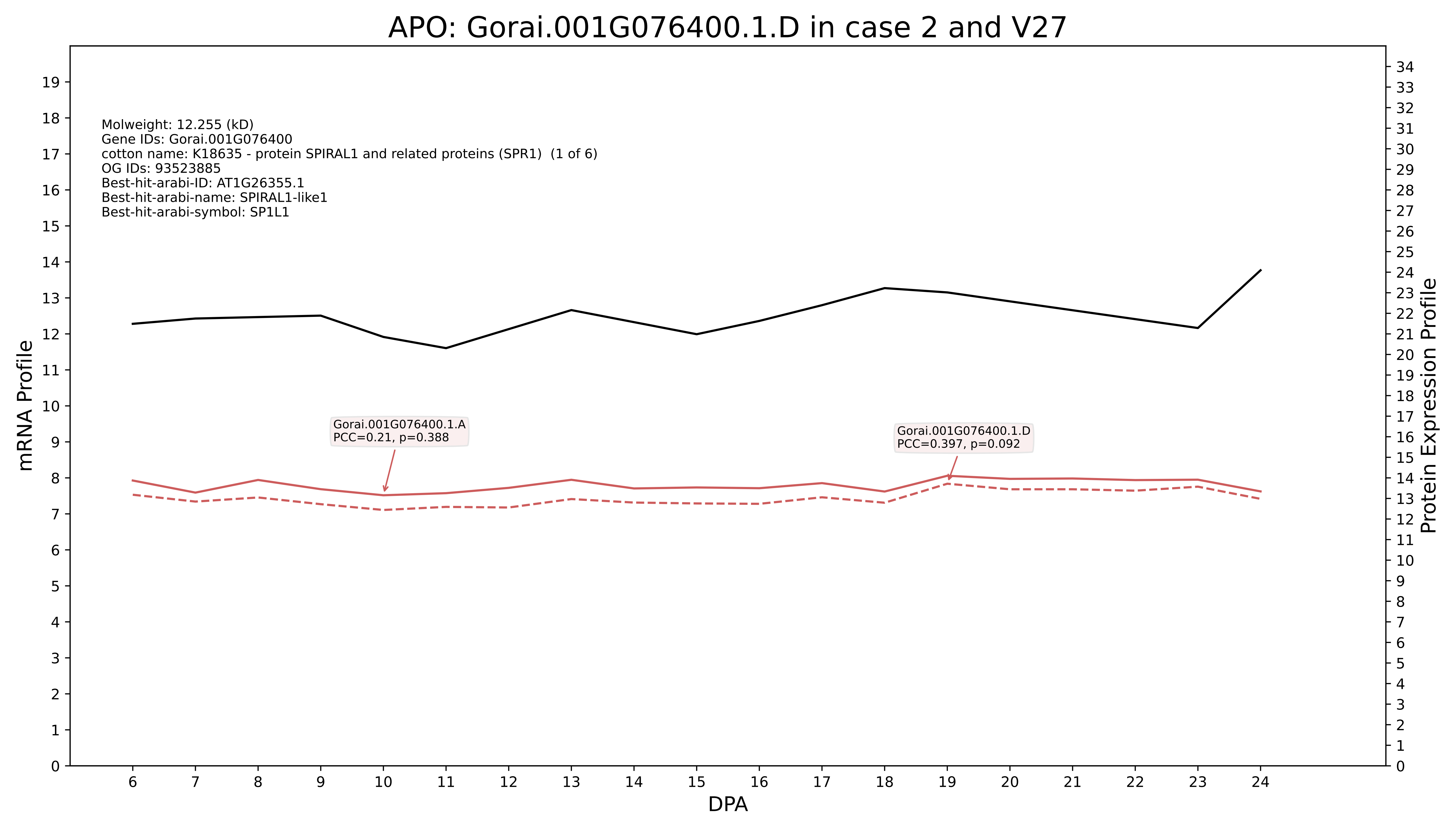
APO: Gorai.001G076400.1.D in case 2 and V27
Molweight: 12.255 (kD)
Gene IDs: Gorai.001G076400
cotton name: K18635 - protein SPIRAL1 and related proteins (SPR1) (1 of 6)
OG IDs: 93523885
Best-hit-arabi-ID: AT1G26355.1
Best-hit-arabi-name: SPIRAL1-like1
Best-hit-arabi-symbol: SP1L1
Gorai.001G076400.1.A
PCC=0.21, p=0.388
Gorai.001G076400.1.D
PCC=0.397, p=0.092
mRNA Profile
Protein Expression Profile
DPA

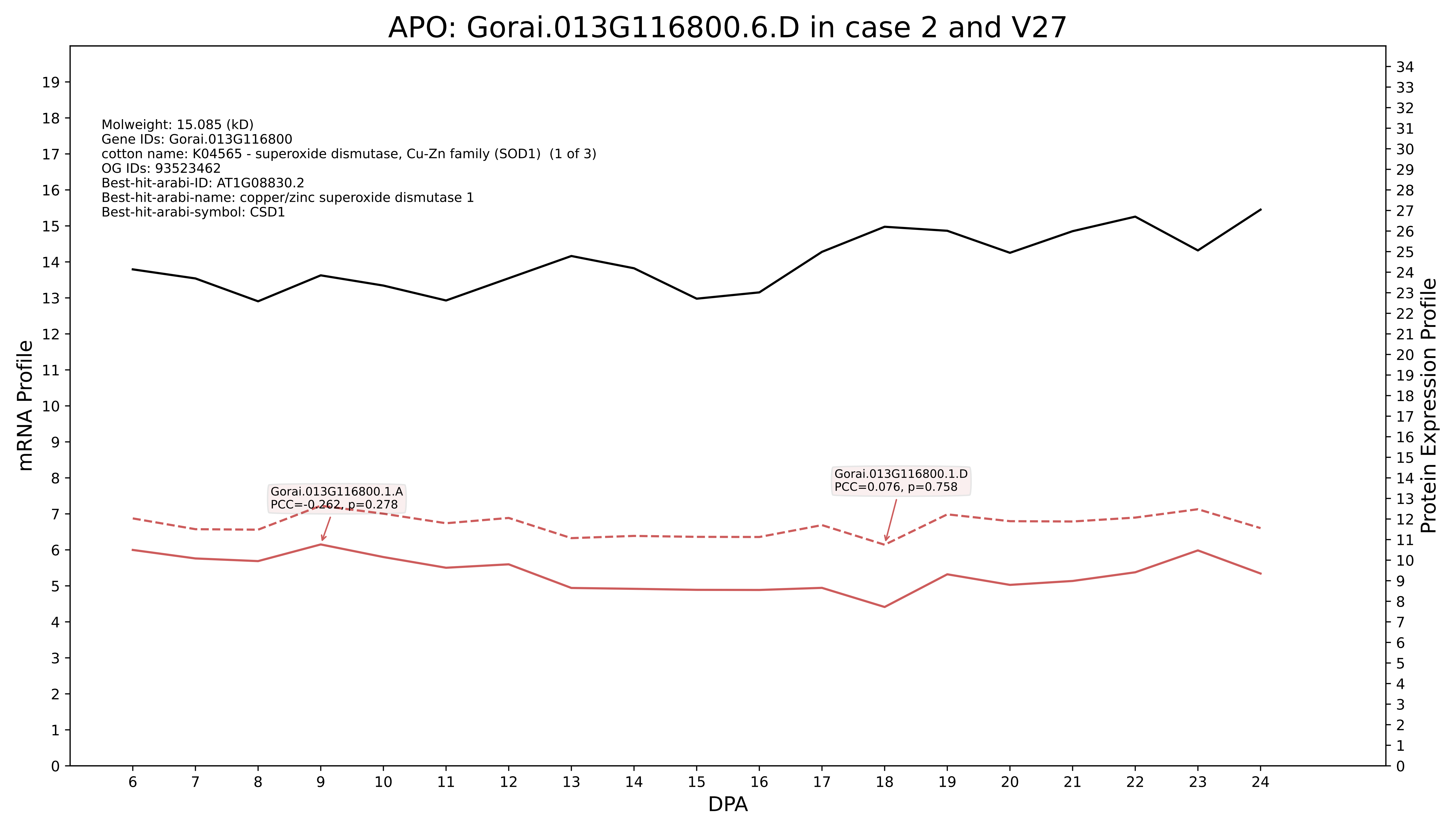
APO: Gorai.013G116800.6.D in case 2 and V27
Molweight: 15.085 (kD)
Gene IDs: Gorai.013G116800
cotton name: K04565 - superoxide dismutase, Cu-Zn family (SOD1) (1 of 3)
OG IDs: 93523462
Best-hit-arabi-ID: AT1G08830.2
Best-hit-arabi-name: copper/zinc superoxide dismutase 1
Best-hit-arabi-symbol: CSD1
Gorai.013G116800.1.A
PCC=-0.262, p=0.278
Gorai.013G116800.1.D
PCC=0.076, p=0.758
mRNA Profile
Protein Expression Profile
DPA

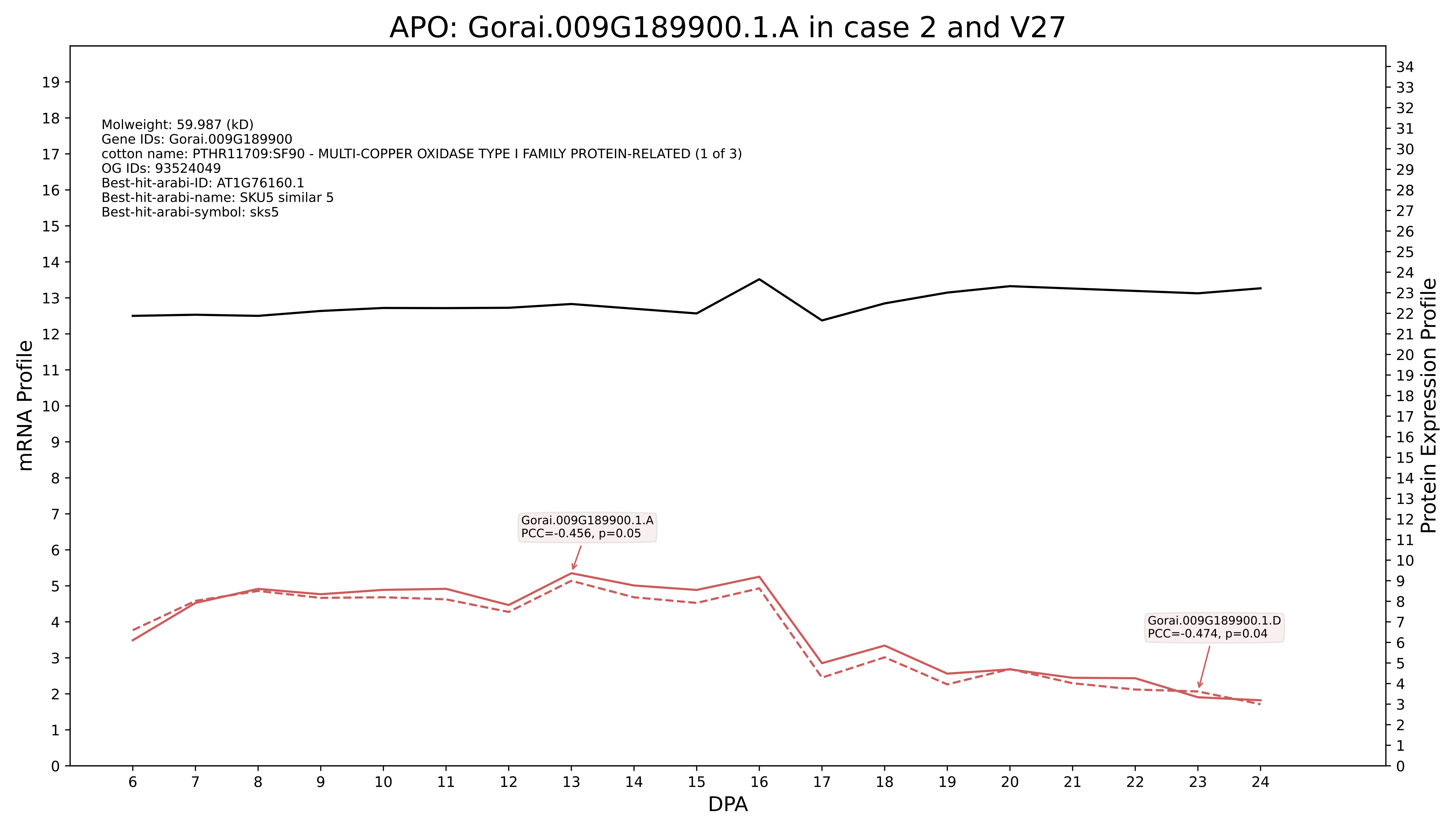
APO: Gorai.009G189900.1.A in case 2 and V27
Molweight: 59.987 (kD)
Gene IDs: Gorai.009G189900
cotton name: PTHR11709:SF90 - MULTI-COPPER OXIDASE TYPE I FAMILY PROTEIN-RELATED (1 of 3)
OG IDs: 93524049
Best-hit-arabi-ID: AT1G76160.1
Best-hit-arabi-name: SKU5 similar 5
Best-hit-arabi-symbol: sks5
Gorai.009G189900.1.A
PCC=-0.456, p=0.05
Gorai.009G189900.1.D
PCC=-0.474, p=0.04
mRNA Profile
Protein Expression Profile
DPA

### APO: Gorai.001G233900.2.D in case 2 and V27

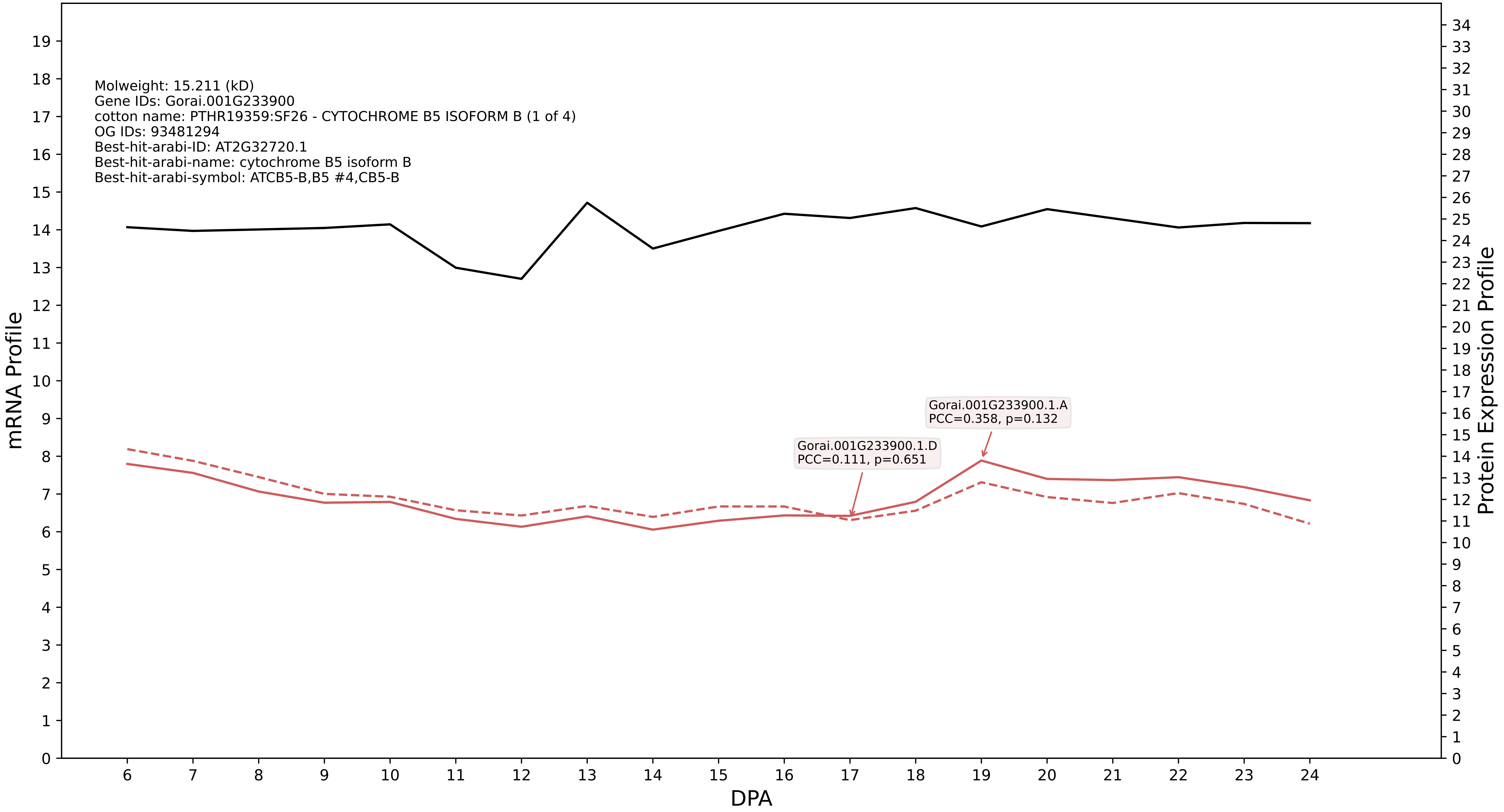

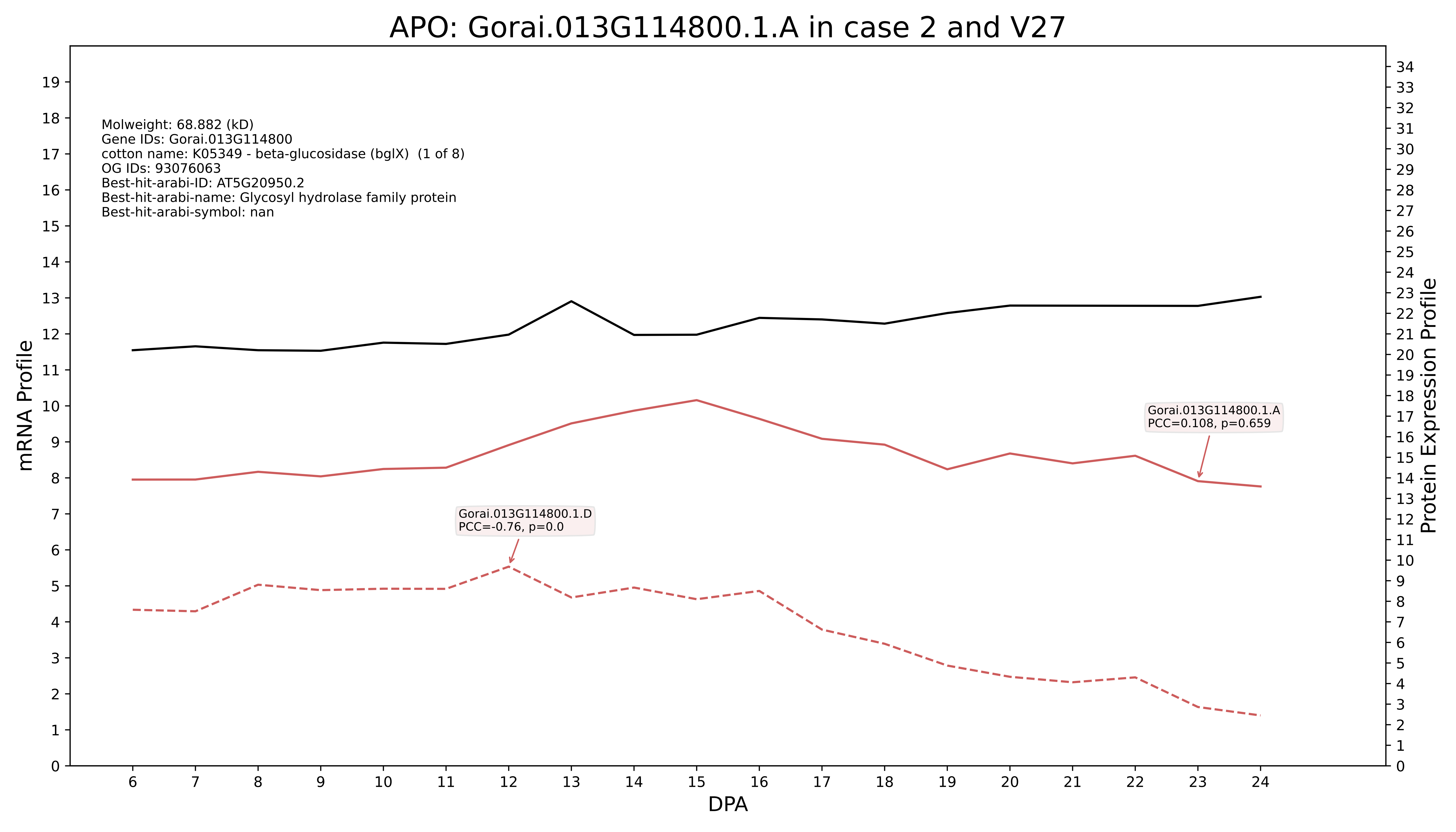

APO: Gorai.013G114800.1.A in case 2 and V27
Molweight: 68.882 (kD)
Gene IDs: Gorai.013G114800
cotton name: K05349 - beta-glucosidase (bglX) (1 of 8)
OG IDs: 93076063
Best-hit-arabi-ID: AT5G20950.2
Best-hit-arabi-name: Glycosyl hydrolase family protein
Best-hit-arabi-symbol: nan
Gorai.013G114800.1.A
PCC=0.108, p=0.659
Gorai.013G114800.1.D
PCC=-0.76, p=0.0
mRNA Profile
Protein Expression Profile
DPA

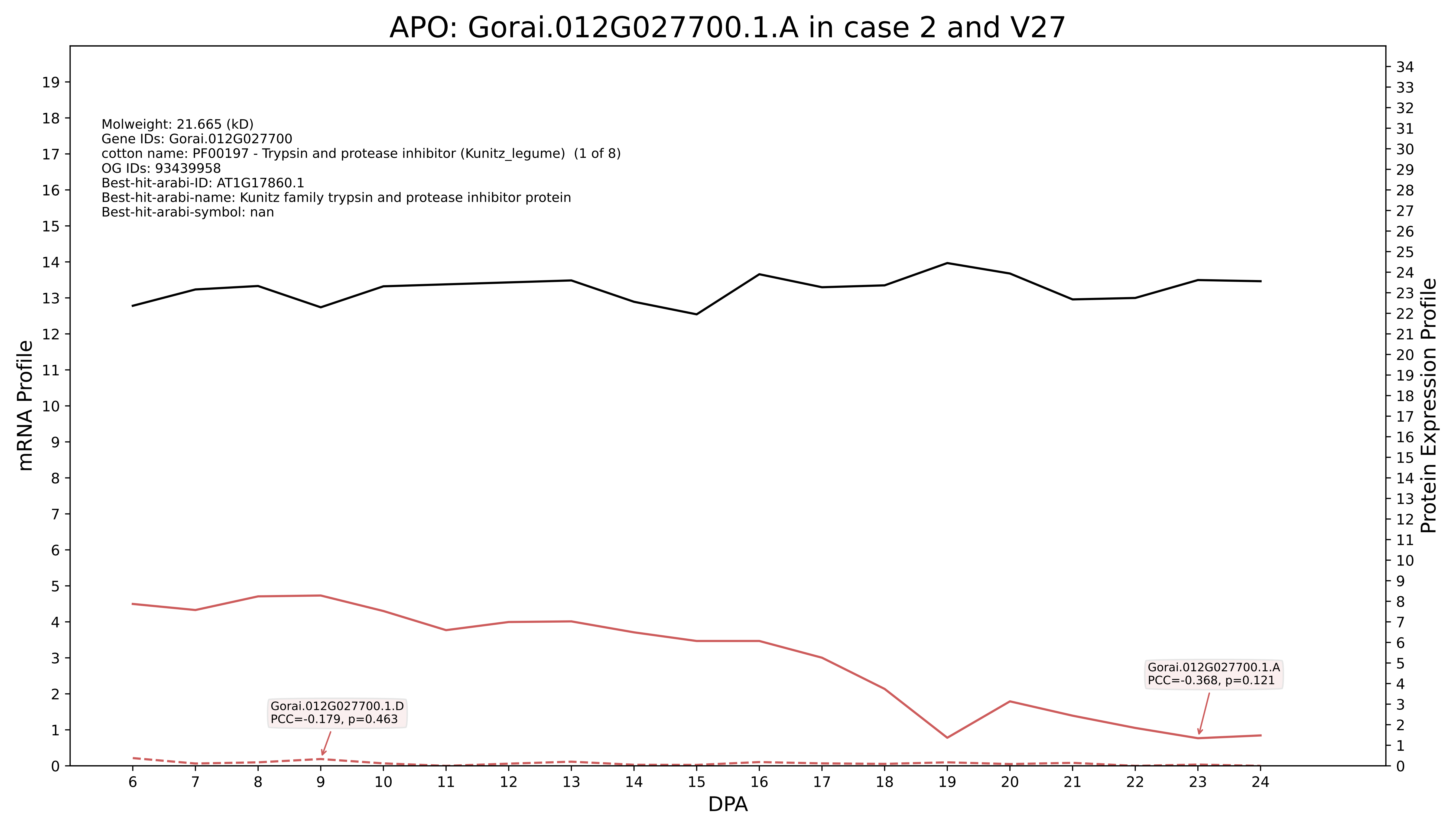
APO: Gorai.012G027700.1.A in case 2 and V27
Molweight: 21.665 (kD)
Gene IDs: Gorai.012G027700
cotton name: PF00197 - Trypsin and protease inhibitor (Kunitz_legume) (1 of 8)
OG IDs: 93439958
Best-hit-arabi-ID: AT1G17860.1
Best-hit-arabi-name: Kunitz family trypsin and protease inhibitor protein
Best-hit-arabi-symbol: nan
Gorai.012G027700.1.D
PCC=-0.179, p=0.463
Gorai.012G027700.1.A
PCC=-0.368, p=0.121
mRNA Profile
Protein Expression Profile
DPA

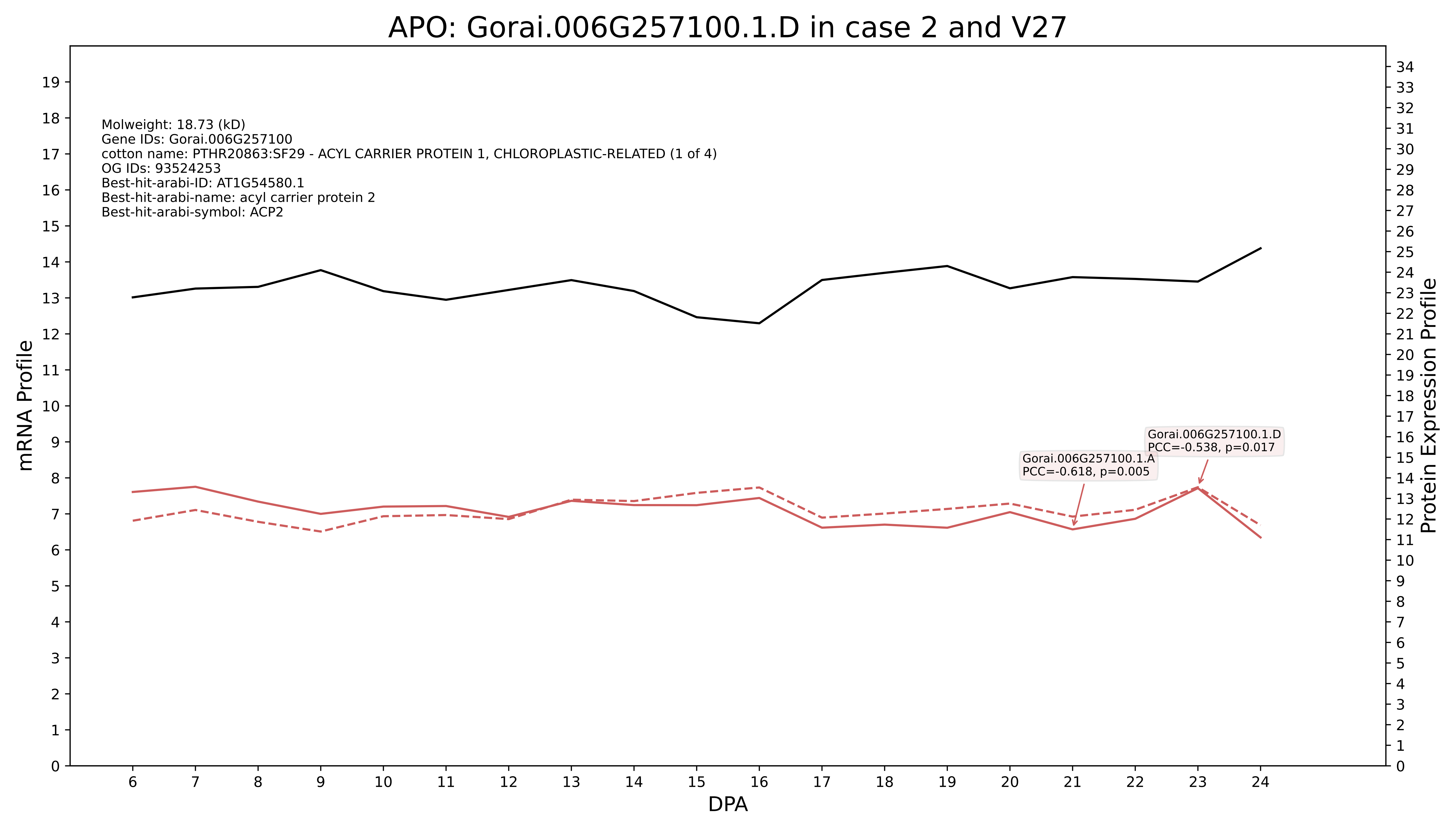
APO: Gorai.006G257100.1.D in case 2 and V27
Molweight: 18.73 (kD)
Gene IDs: Gorai.006G257100
cotton name: PTHR20863:SF29 - ACYL CARRIER PROTEIN 1, CHLOROPLASTIC-RELATED (1 of 4)
OG IDs: 93524253
Best-hit-arabi-ID: AT1G54580.1
Best-hit-arabi-name: acyl carrier protein 2
Best-hit-arabi-symbol: ACP2
Gorai.006G257100.1.A
PCC=-0.618, p=0.005
Gorai.006G257100.1.D
PCC=-0.538, p=0.017
mRNA Profile
Protein Expression Profile
DPA

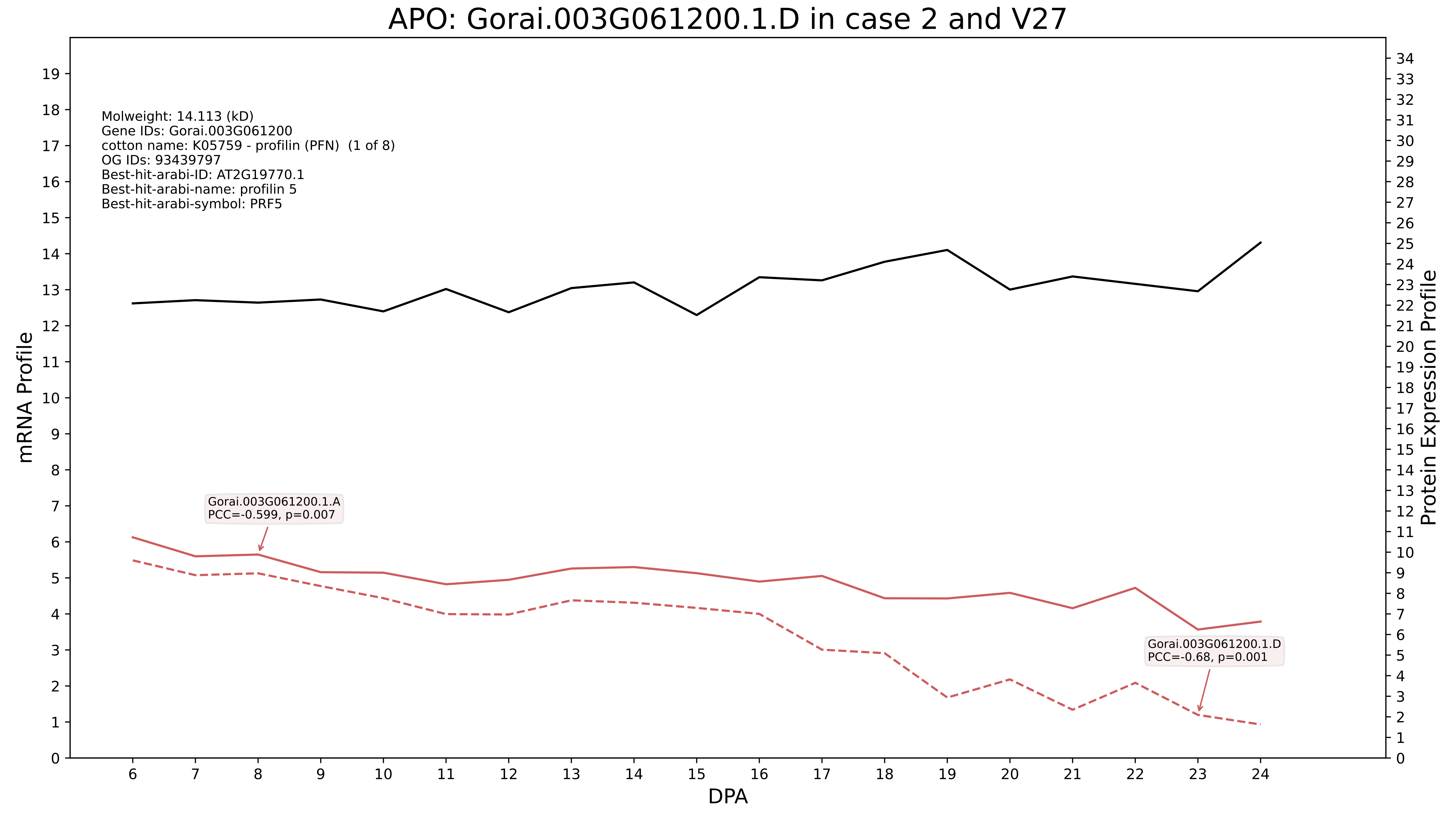

APO: Gorai.003G061200.1.D in case 2 and V27
Molweight: 14.113 (kD)
Gene IDs: Gorai.003G061200
cotton name: K05759 - profilin (PFN) (1 of 8)
OG IDs: 93439797
Best-hit-arabi-ID: AT2G19770.1
Best-hit-arabi-name: profilin 5
Best-hit-arabi-symbol: PRF5
Gorai.003G061200.1.A
PCC=-0.599, p=0.007
Gorai.003G061200.1.D
PCC=-0.68, p=0.001
mRNA Profile
Protein Expression Profile
DPA

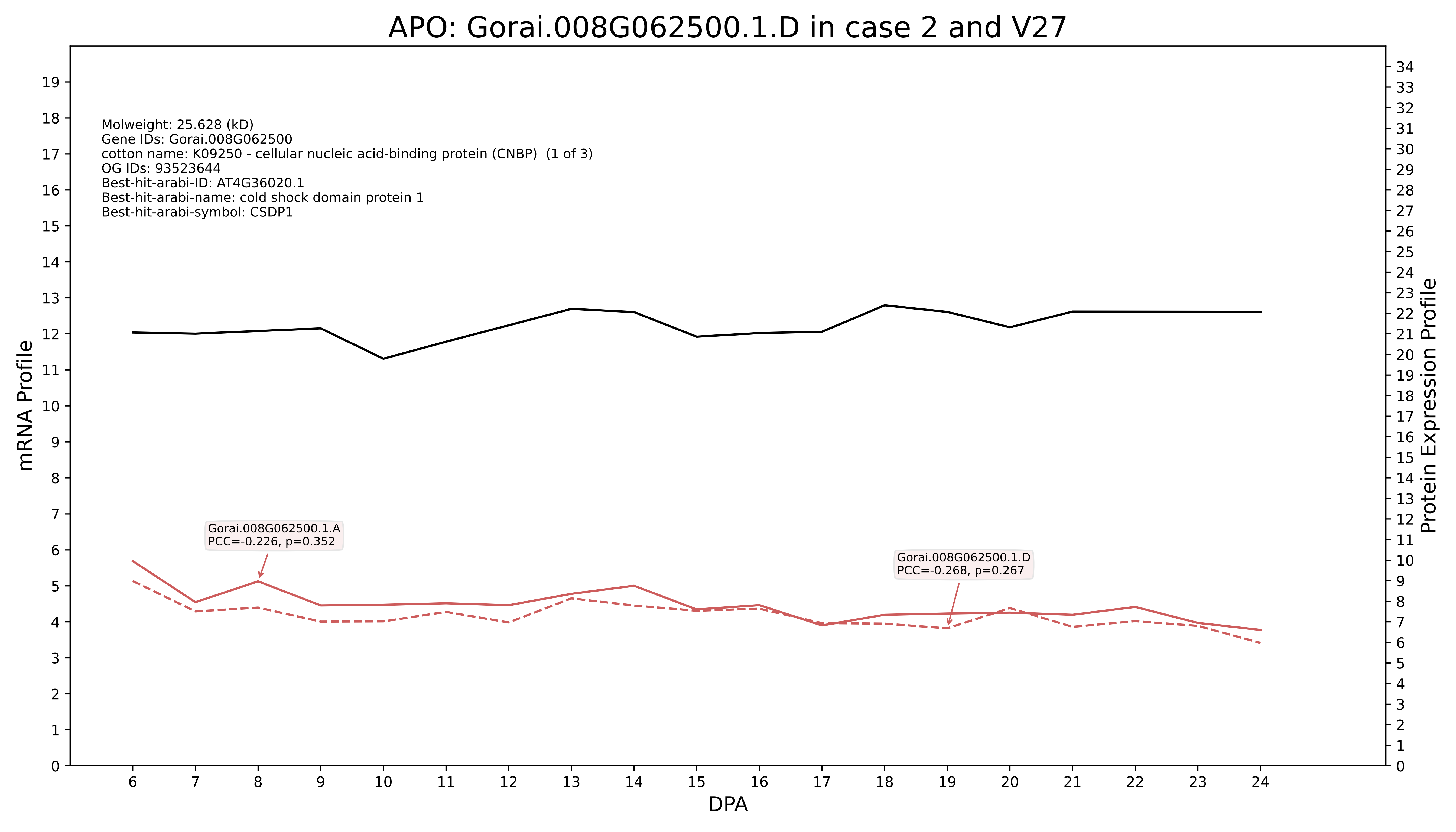
APO: Gorai.008G062500.1.D in case 2 and V27
Molweight: 25.628 (kD)
Gene IDs: Gorai.008G062500
cotton name: K09250 - cellular nucleic acid-binding protein (CNBP) (1 of 3)
OG IDs: 93523644
Best-hit-arabi-ID: AT4G36020.1
Best-hit-arabi-name: cold shock domain protein 1
Best-hit-arabi-symbol: CSDP1
Gorai.008G062500.1.A
PCC=-0.226, p=0.352
Gorai.008G062500.1.D
PCC=-0.268, p=0.267
mRNA Profile
Protein Expression Profile
DPA

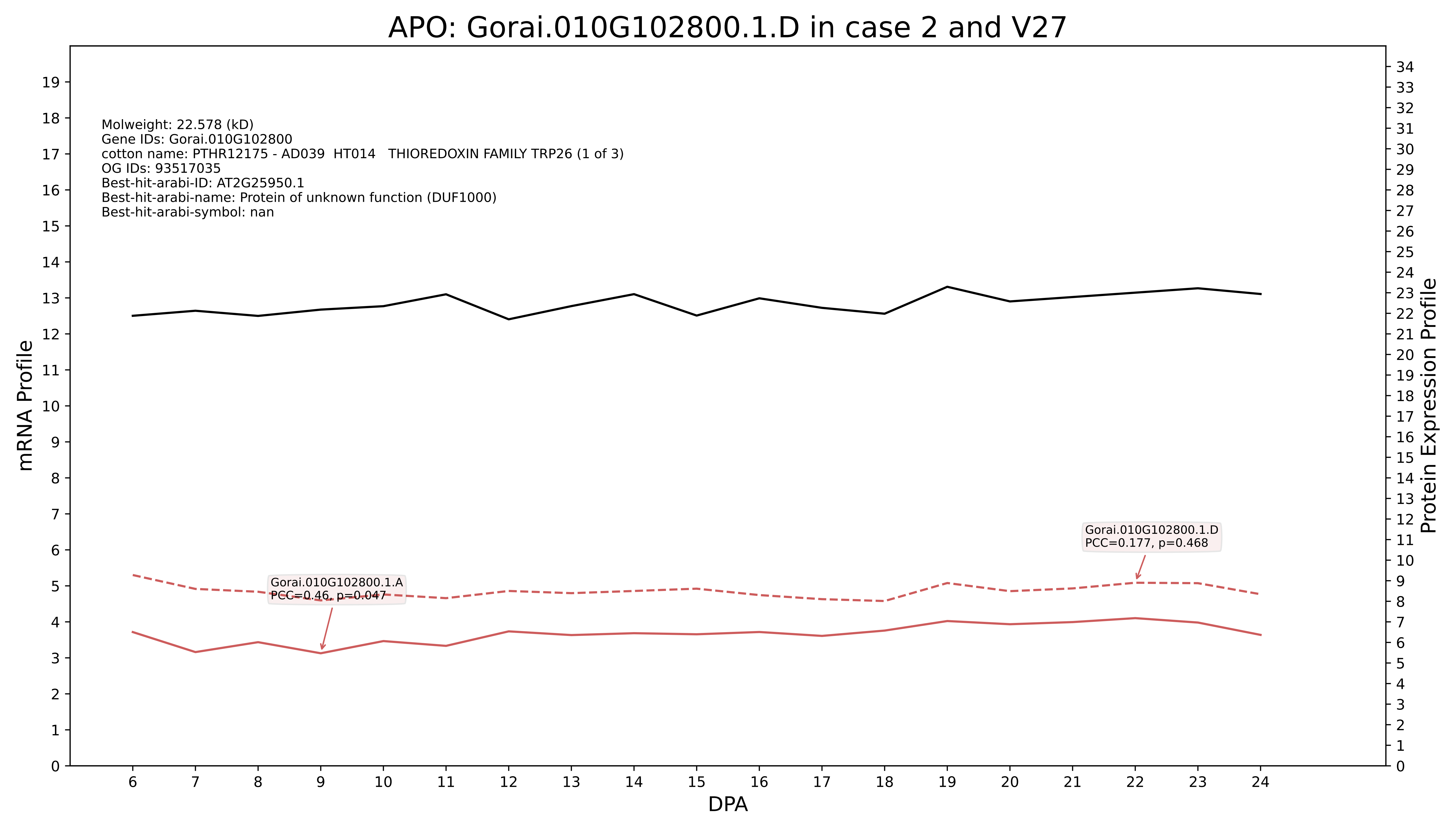

APO: Gorai.010G102800.1.D in case 2 and V27
Molweight: 22.578 (kD)
Gene IDs: Gorai.010G102800
cotton name: PTHR12175 - AD039 HT014 THIOREDOXIN FAMILY TRP26 (1 of 3)
OG IDs: 93517035
Best-hit-arabi-ID: AT2G25950.1
Best-hit-arabi-name: Protein of unknown function (DUF1000)
Best-hit-arabi-symbol: nan
Gorai.010G102800.1.A
PCC=0.46, p=0.047
Gorai.010G102800.1.D
PCC=0.177, p=0.468
mRNA Profile
Protein Expression Profile
DPA

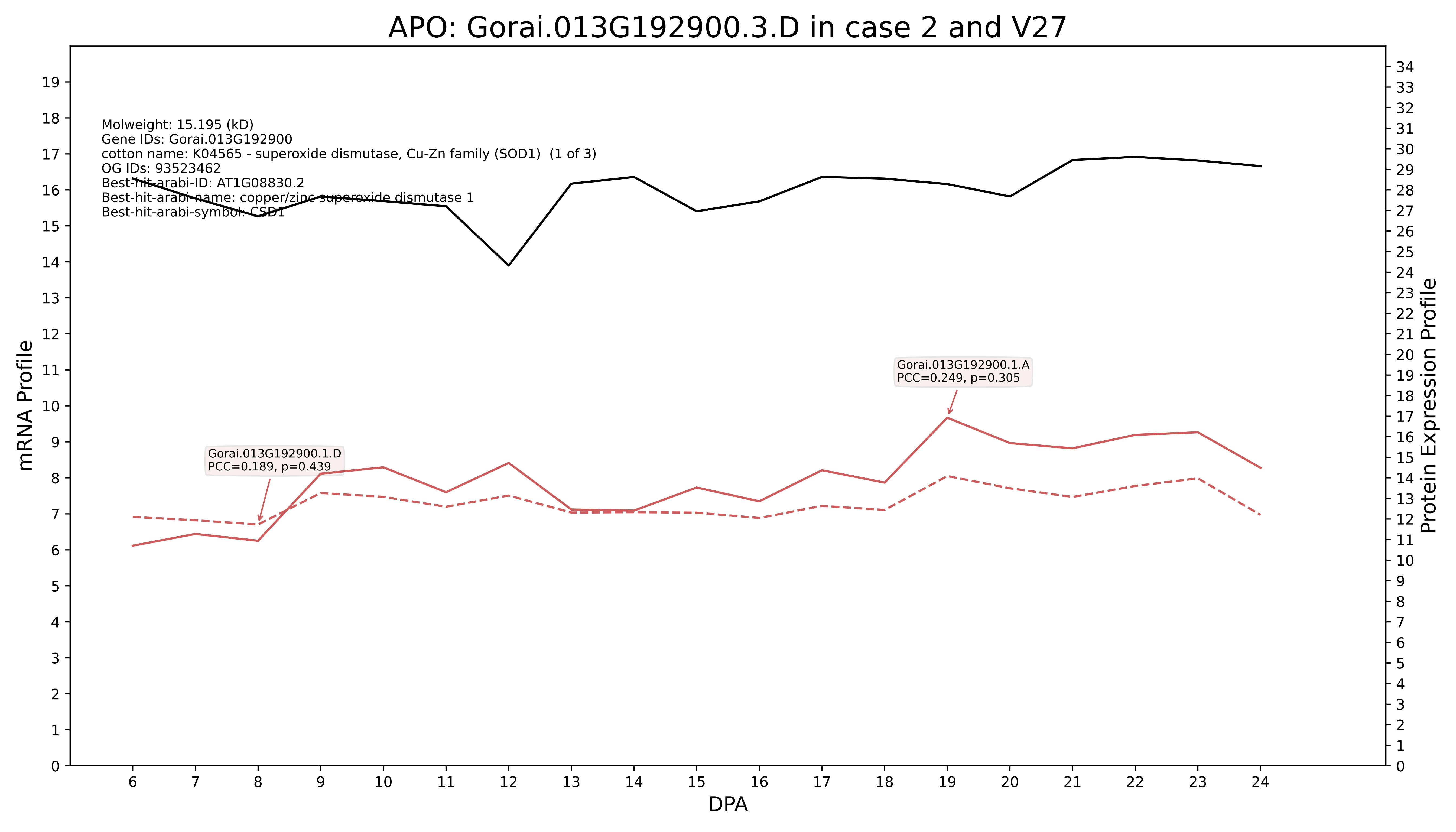
APO: Gorai.013G192900.3.D in case 2 and V27
Molweight: 15.195 (kD)
Gene IDs: Gorai.013G192900
cotton name: K04565 - superoxide dismutase, Cu-Zn family (SOD1) (1 of 3)
OG IDs: 93523462
Best-hit-arabi-ID: AT1G08830.2
Best-hit-arabi-name: copper/zinc superoxide dismutase 1
Best-hit-arabi-symbol: CSD1
Gorai.013G192900.1.A
PCC=0.249, p=0.305
Gorai.013G192900.1.D
PCC=0.189, p=0.439
mRNA Profile
Protein Expression Profile
DPA

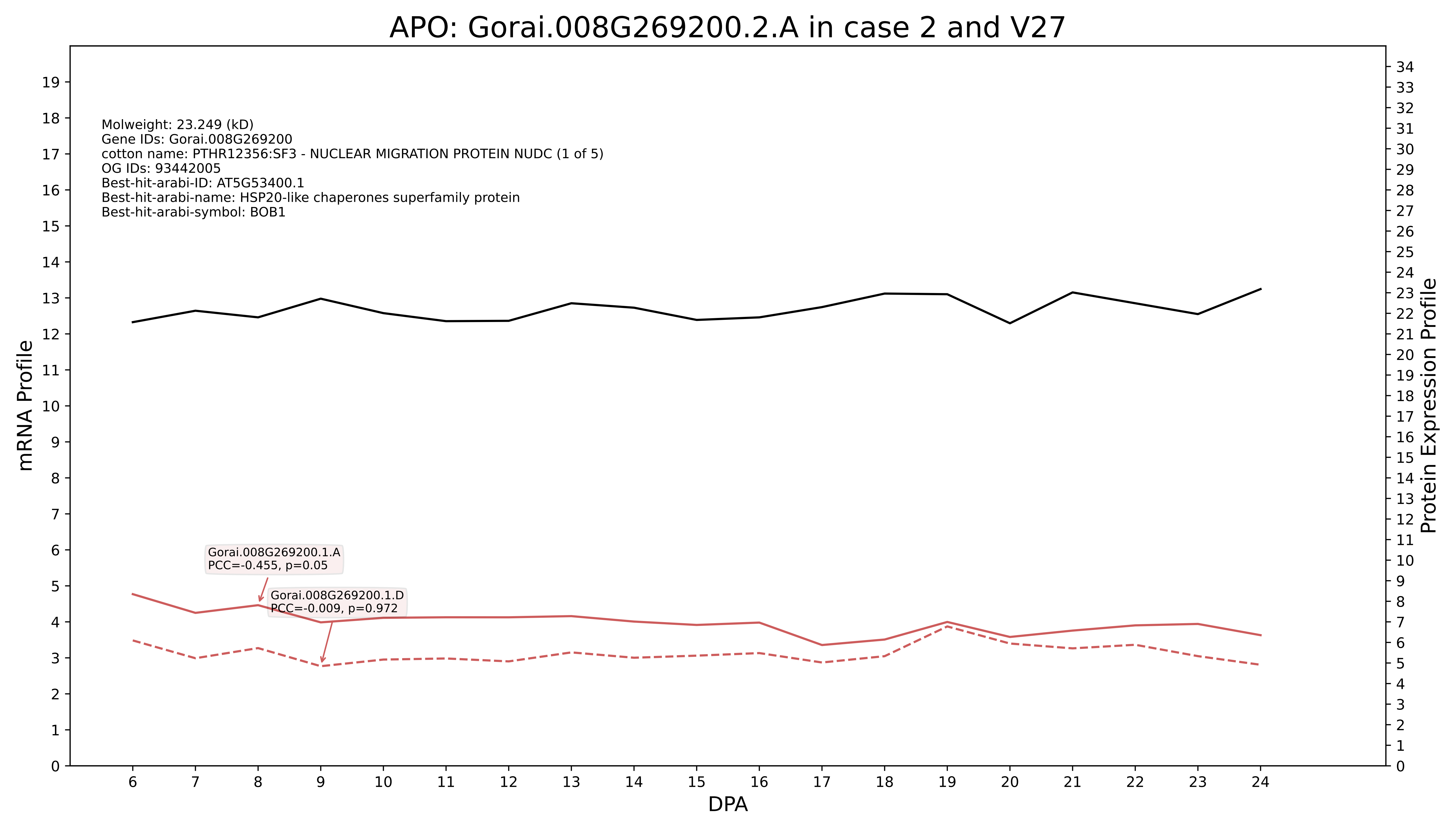

APO: Gorai.008G269200.2.A in case 2 and V27
Molweight: 23.249 (kD)
Gene IDs: Gorai.008G269200
cotton name: PTHR12356:SF3 - NUCLEAR MIGRATION PROTEIN NUDC (1 of 5)
OG IDs: 93442005
Best-hit-arabi-ID: AT5G53400.1
Best-hit-arabi-name: HSP20-like chaperones superfamily protein
Best-hit-arabi-symbol: BOB1
Gorai.008G269200.1.A
PCC=-0.455, p=0.05
Gorai.008G269200.1.D
PCC=-0.009, p=0.972
mRNA Profile
Protein Expression Profile
DPA

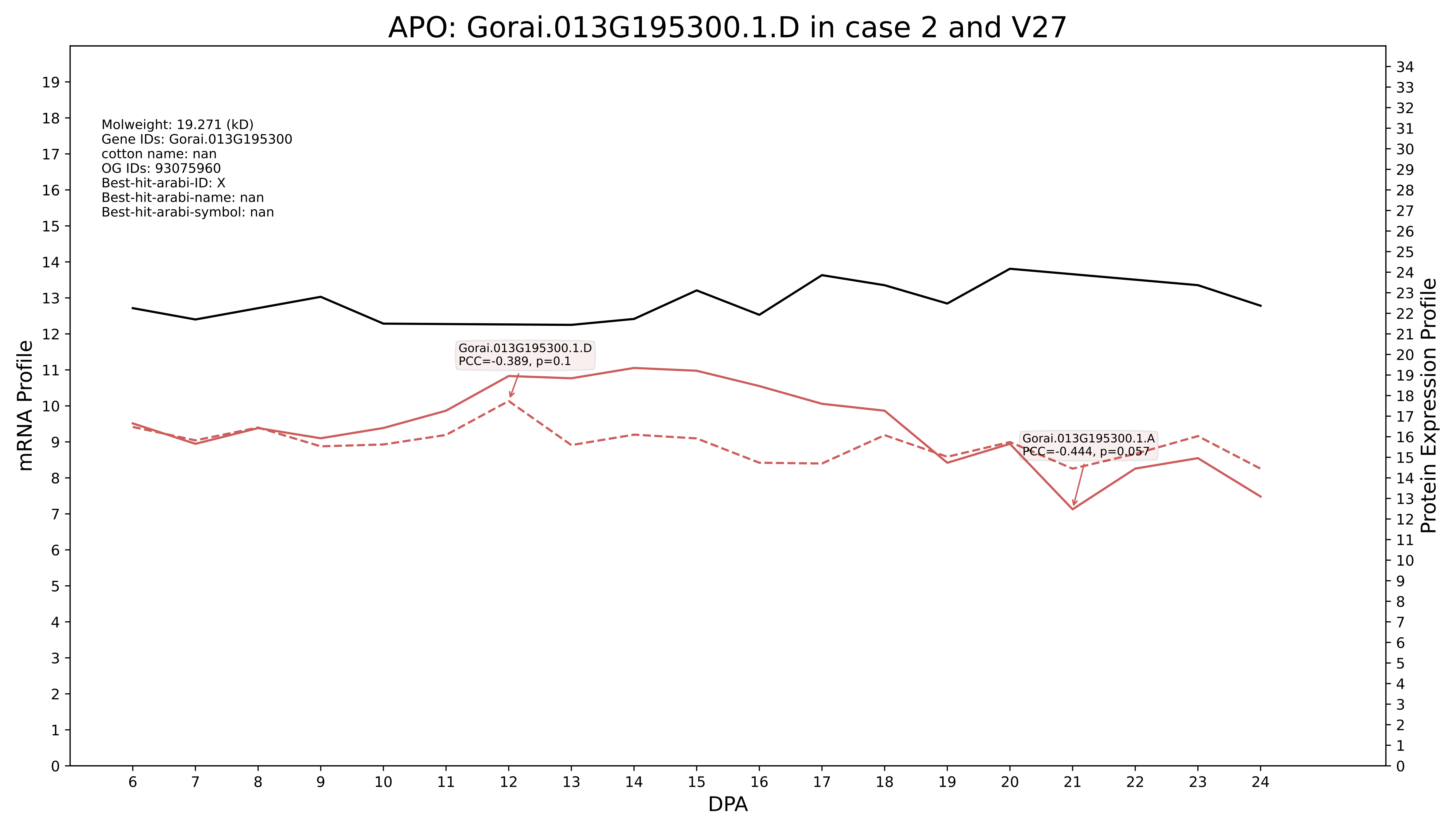

APO: Gorai.013G195300.1.D in case 2 and V27
Molweight: 19.271 (kD)
Gene IDs: Gorai.013G195300
cotton name: nan
OG IDs: 93075960
Best-hit-arabi-ID: X
Best-hit-arabi-name: nan
Best-hit-arabi-symbol: nan
Gorai.013G195300.1.D
PCC=-0.389, p=0.1
Gorai.013G195300.1.A
PCC=-0.444, p=0.057
mRNA Profile
Protein Expression Profile
DPA

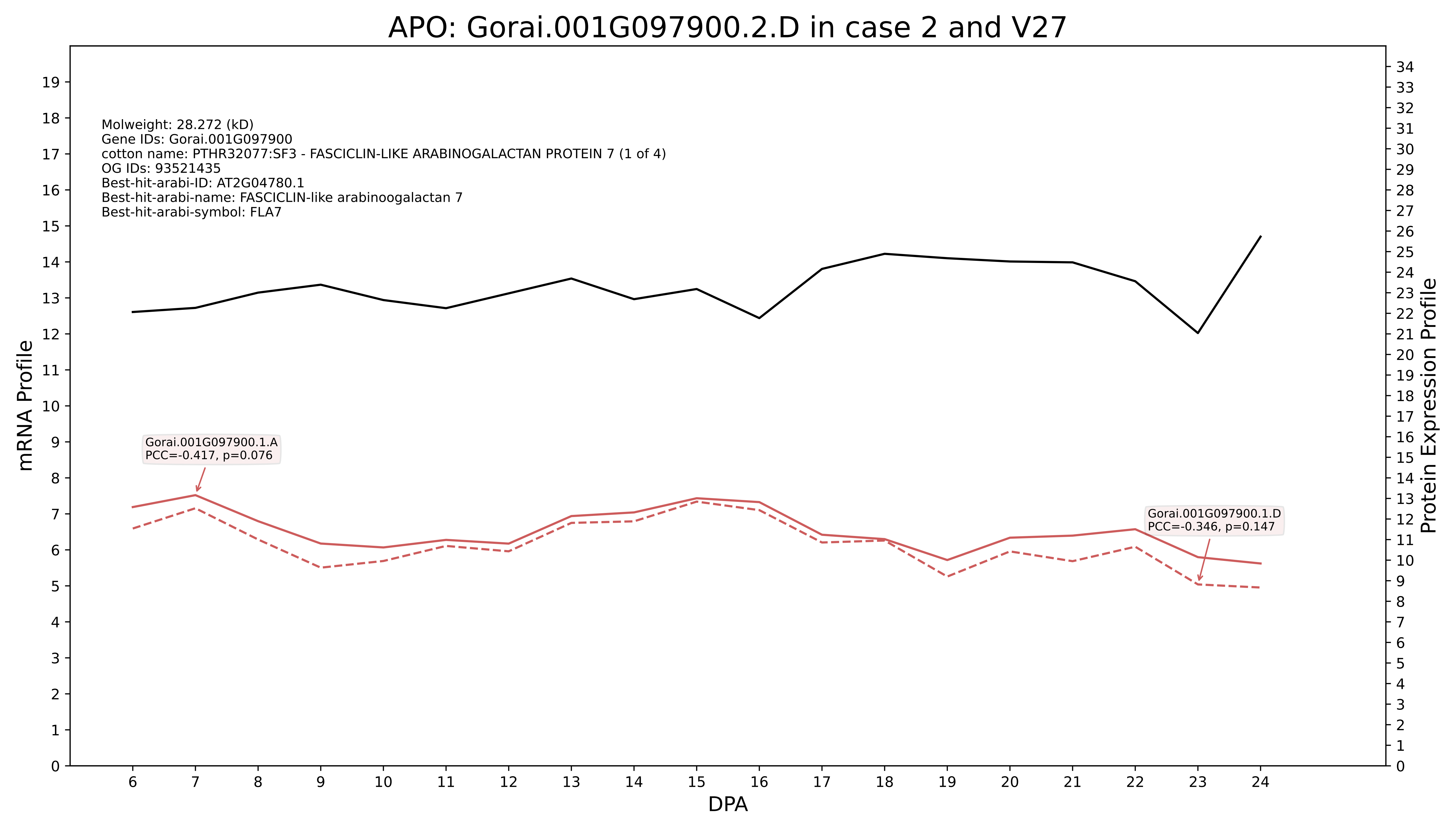

APO: Gorai.001G097900.2.D in case 2 and V27
Molweight: 28.272 (kD)
Gene IDs: Gorai.001G097900
cotton name: PTHR32077:SF3 - FASCICLIN-LIKE ARABINOGALACTAN PROTEIN 7 (1 of 4)
OG IDs: 93521435
Best-hit-arabi-ID: AT2G04780.1
Best-hit-arabi-name: FASCICLIN-like arabinoogalactan 7
Best-hit-arabi-symbol: FLA7
Gorai.001G097900.1.A
PCC=-0.417, p=0.076
Gorai.001G097900.1.D
PCC=-0.346, p=0.147
mRNA Profile
Protein Expression Profile
DPA

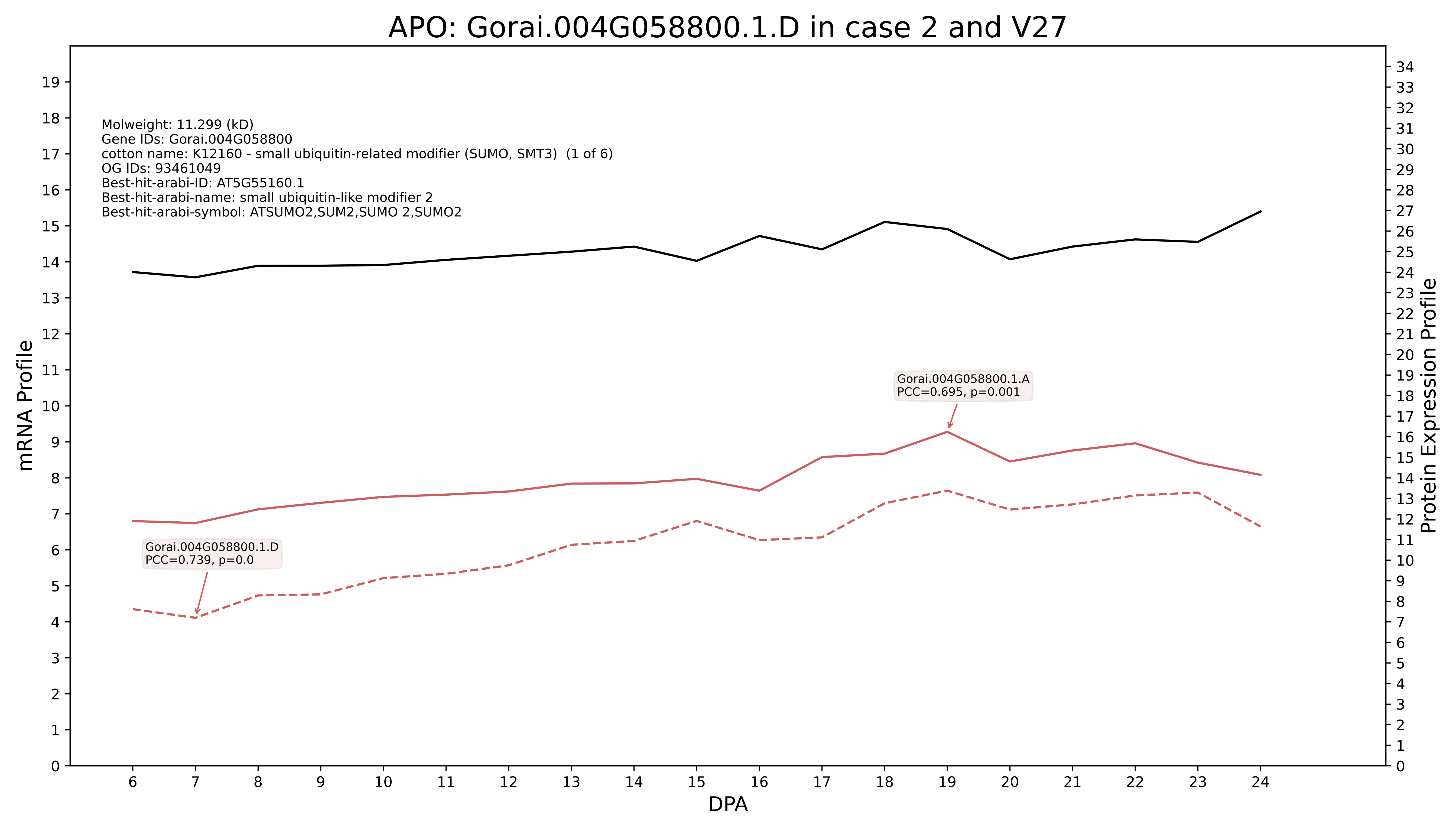
APO: Gorai.004G058800.1.D in case 2 and V27
Molweight: 11.299 (kD)
Gene IDs: Gorai.004G058800
cotton name: K12160 - small ubiquitin-related modifier (SUMO, SMT3) (1 of 6)
OG IDs: 93461049
Best-hit-arabi-ID: AT5G55160.1
Best-hit-arabi-name: small ubiquitin-like modifier 2
Best-hit-arabi-symbol: ATSUMO2,SUM2,SUMO 2,SUMO2
Gorai.004G058800.1.A
PCC=0.695, p=0.001
Gorai.004G058800.1.D
PCC=0.739, p=0.0
mRNA Profile
Protein Expression Profile
DPA

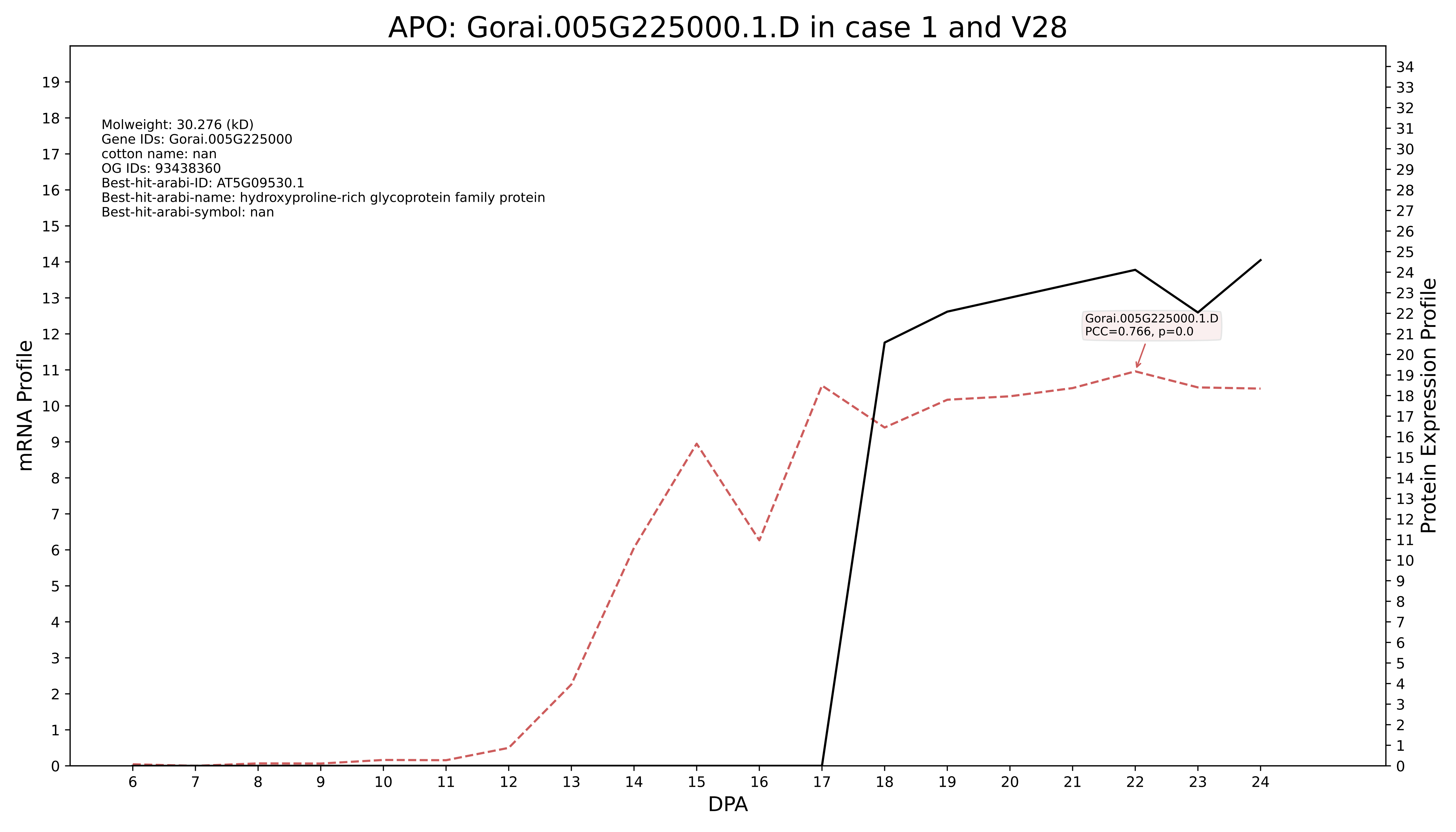
APO: Gorai.005G225000.1.D in case 1 and V28
Molweight: 30.276 (kD)
Gene IDs: Gorai.005G225000
cotton name: nan
OG IDs: 93438360
Best-hit-arabi-ID: AT5G09530.1
Best-hit-arabi-name: hydroxyproline-rich glycoprotein family protein
Best-hit-arabi-symbol: nan
Gorai.005G225000.1.D
PCC=0.766, p=0.0
mRNA Profile
Protein Expression Profile
DPA

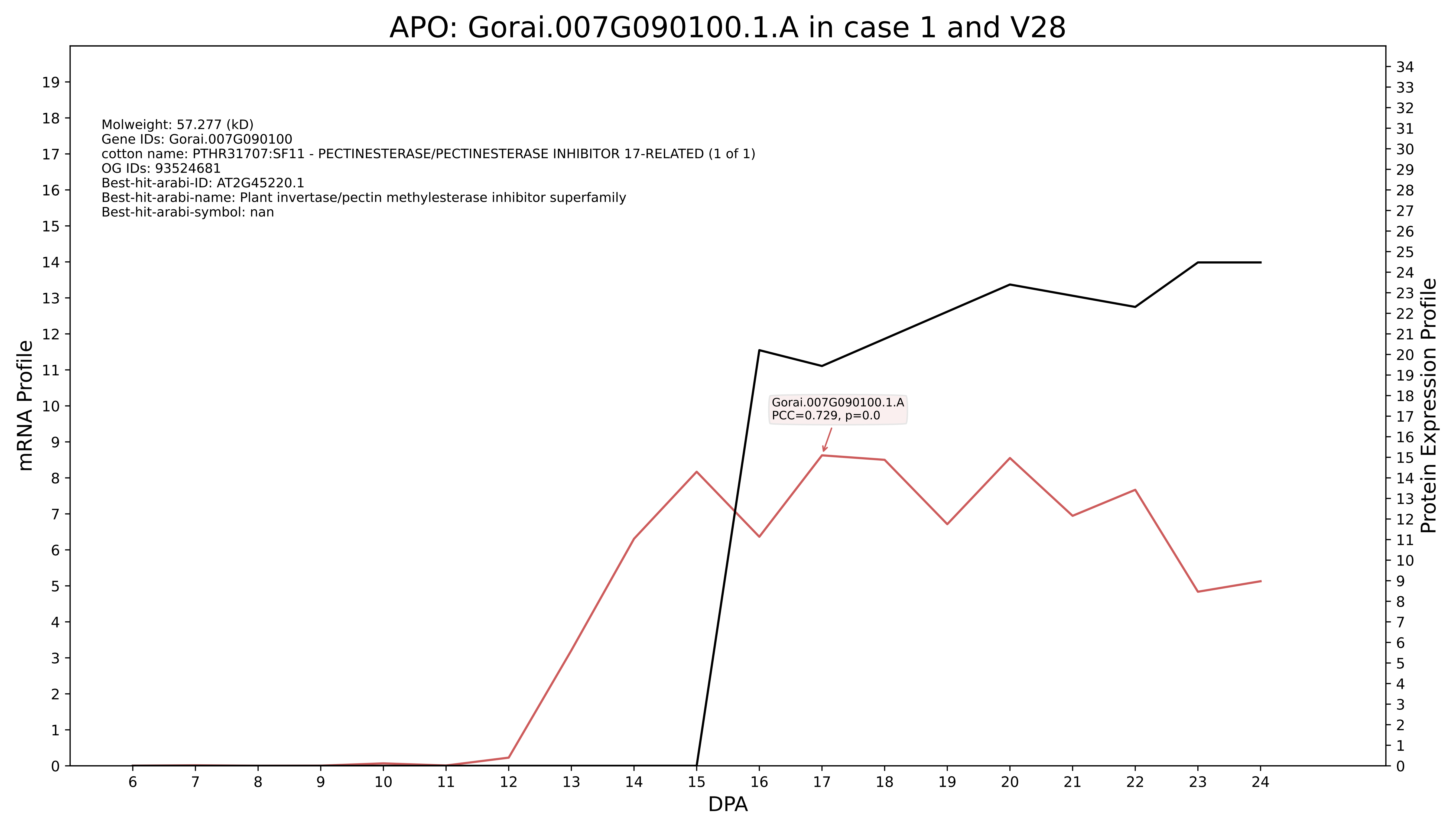

APO: Gorai.007G090100.1.A in case 1 and V28
Molweight: 57.277 (kD)
Gene IDs: Gorai.007G090100
cotton name: PTHR31707:SF11 - PECTINESTERASE/PECTINESTERASE INHIBITOR 17-RELATED (1 of 1)
OG IDs: 93524681
Best-hit-arabi-ID: AT2G45220.1
Best-hit-arabi-name: Plant invertase/pectin methylesterase inhibitor superfamily
Best-hit-arabi-symbol: nan
Gorai.007G090100.1.A
PCC=0.729, p=0.0
mRNA Profile
Protein Expression Profile
DPA

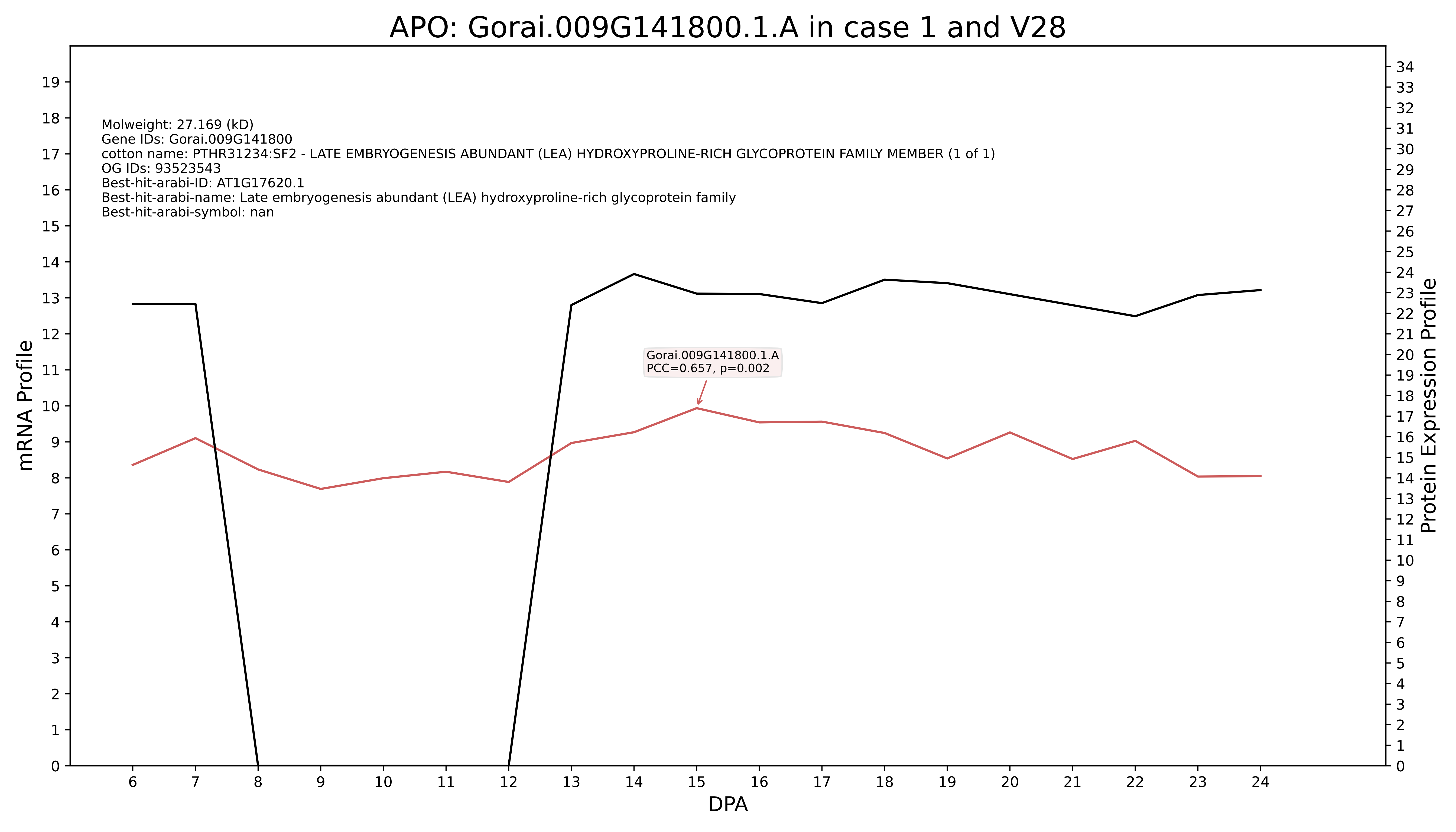

APO: Gorai.009G141800.1.A in case 1 and V28
Molweight: 27.169 (kD)
Gene IDs: Gorai.009G141800
cotton name: PTHR31234:SF2 - LATE EMBRYOGENESIS ABUNDANT (LEA) HYDROXYPROLINE-RICH GLYCOPROTEIN FAMILY MEMBER (1 of 1)
OG IDs: 93523543
Best-hit-arabi-ID: AT1G17620.1
Best-hit-arabi-name: Late embryogenesis abundant (LEA) hydroxyproline-rich glycoprotein family
Best-hit-arabi-symbol: nan
Gorai.009G141800.1.A
PCC=0.657, p=0.002
mRNA Profile
Protein Expression Profile
DPA

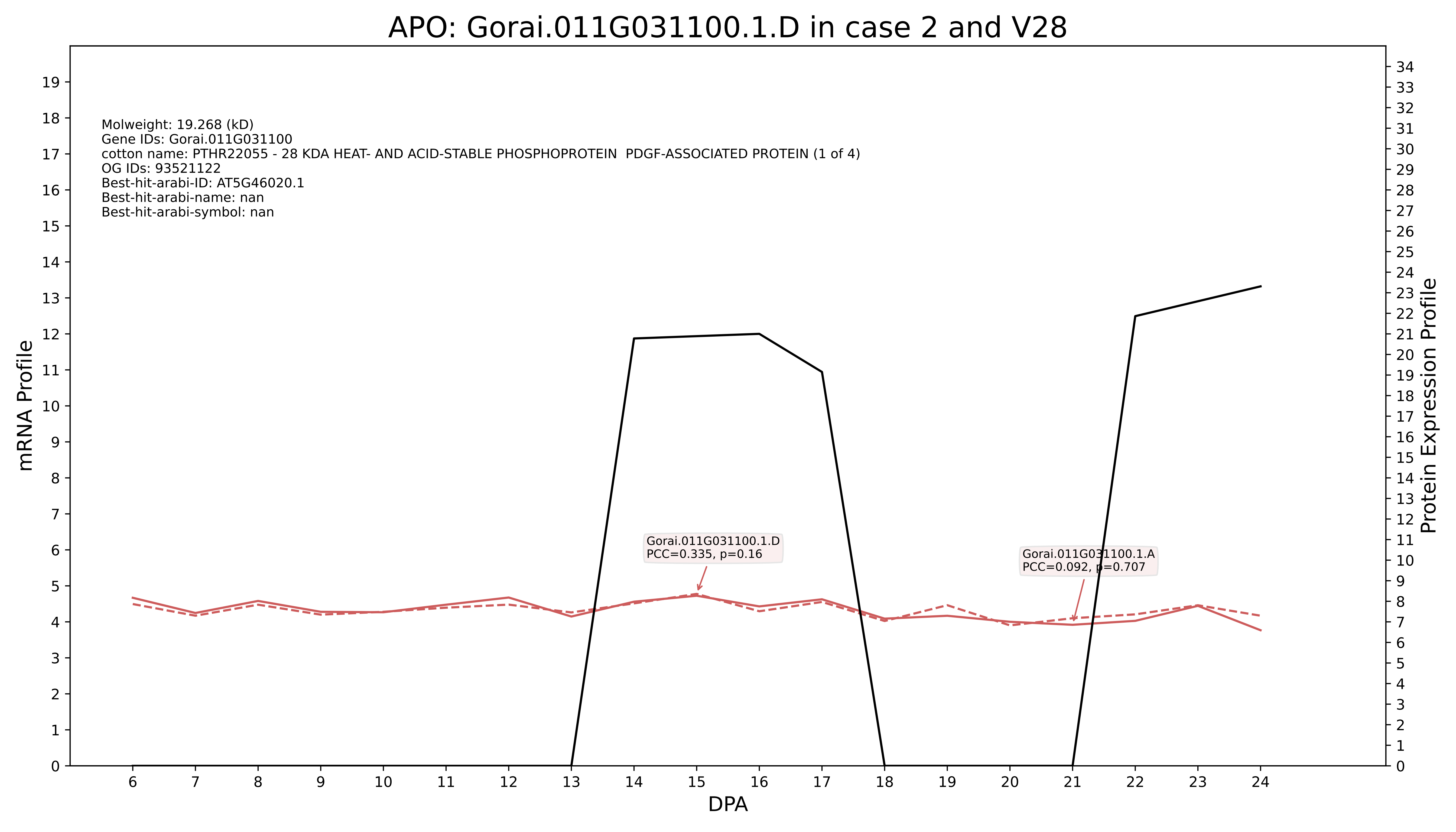
APO: Gorai.011G031100.1.D in case 2 and V28
Molweight: 19.268 (kD)
Gene IDs: Gorai.011G031100
cotton name: PTHR22055 - 28 KDA HEAT- AND ACID-STABLE PHOSPHOPROTEIN PDGF-ASSOCIATED PROTEIN (1 of 4)
OG IDs: 93521122
Best-hit-arabi-ID: AT5G46020.1
Best-hit-arabi-name: nan
Best-hit-arabi-symbol: nan
Gorai.011G031100.1.D
PCC=0.335, p=0.16
Gorai.011G031100.1.A
PCC=0.092, p=0.707
mRNA Profile
Protein Expression Profile
DPA
