## Supplementary Figure S11 for "Systems-level proteomic models of cotton fiber development: A high-resolution resource to analyze cell dynamics and trait engineering"

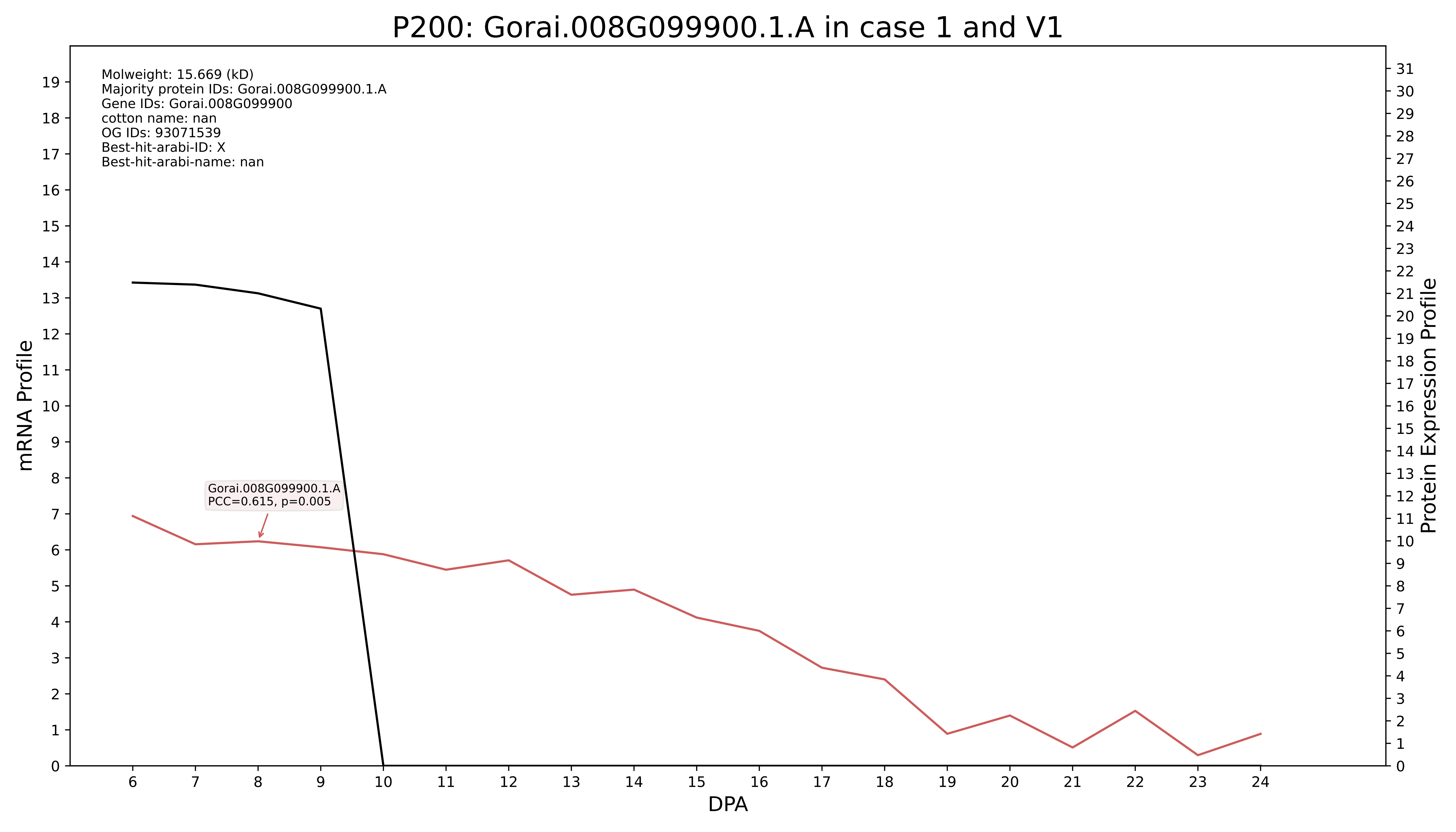
P200: Gorai.008G099900.1.A in case 1 and V1
Molweight: 15.669 (kD)
Majority protein IDs: Gorai.008G099900.1.A
Gene IDs: Gorai.008G099900
cotton name: nan
OG IDs: 93071539
Best-hit-arabi-ID: X
Best-hit-arabi-name: nan
Gorai.008G099900.1.A
PCC=0.615, p=0.005
mRNA Profile
Protein Expression Profile
DPA

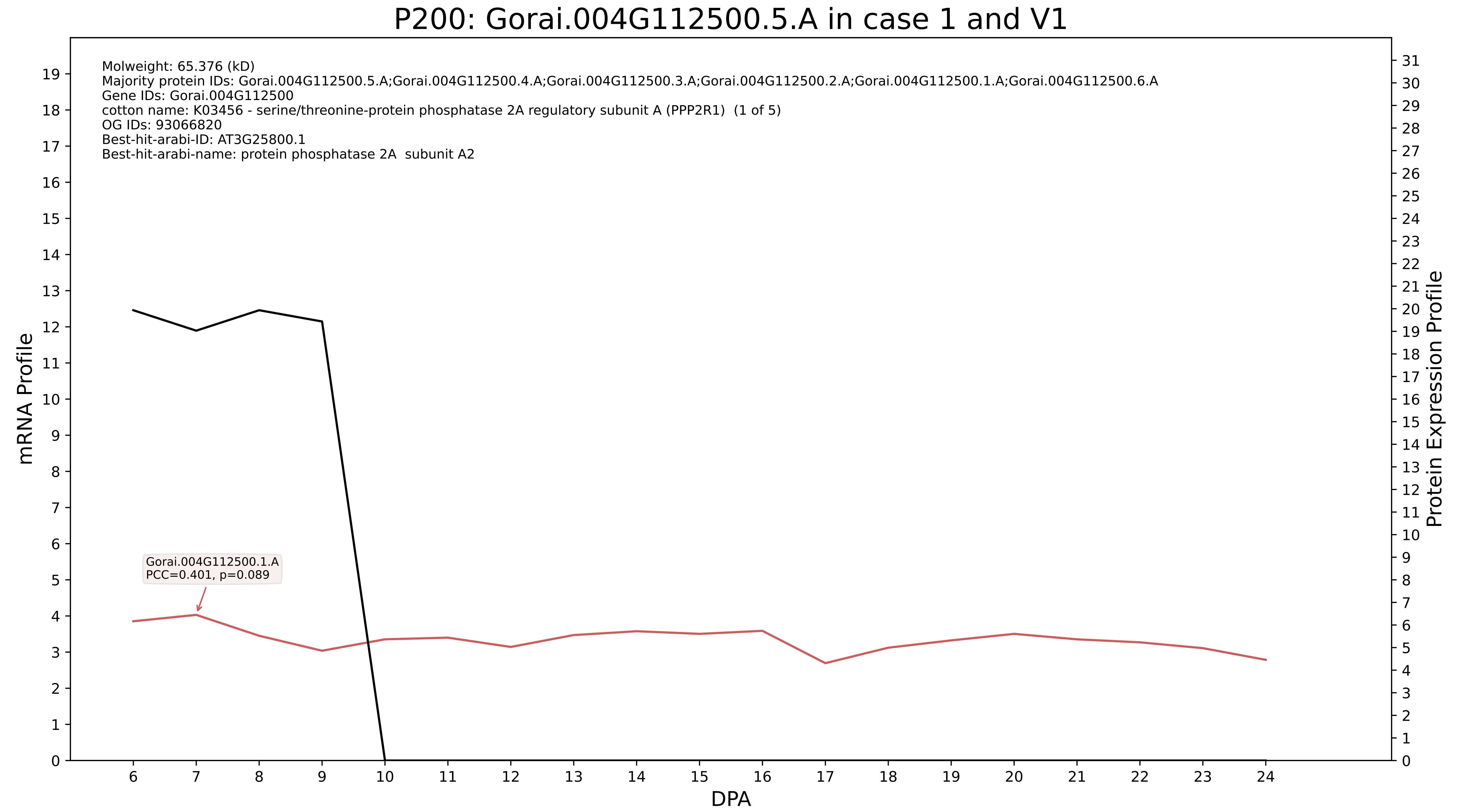
P200: Gorai.004G112500.5.A in case 1 and V1
Molweight: 65.376 (kD)
Majority protein IDs: Gorai.004G112500.5.A;Gorai.004G112500.4.A;Gorai.004G112500.3.A;Gorai.004G112500.2.A;Gorai.004G112500.1.A;Gorai.004G112500.6.A
Gene IDs: Gorai.004G112500
cotton name: K03456 - serine/threonine-protein phosphatase 2A regulatory subunit A (PPP2R1) (1 of 5)
OG IDs: 93066820
Best-hit-arabi-ID: AT3G25800.1
Best-hit-arabi-name: protein phosphatase 2A subunit A2
Gorai.004G112500.1.A
PCC=0.401, p=0.089
mRNA Profile
Protein Expression Profile
DPA

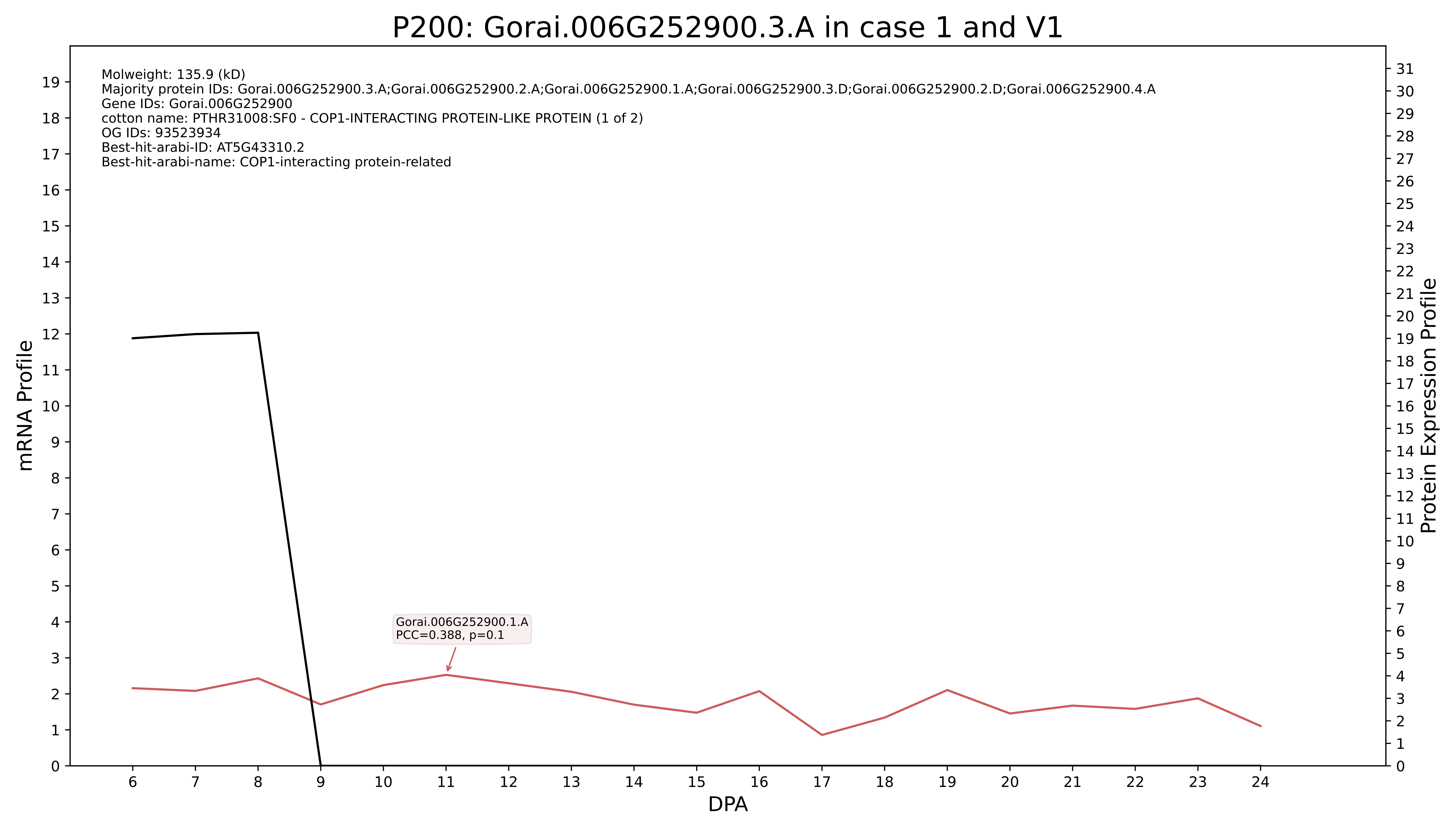
P200: Gorai.006G252900.3.A in case 1 and V1
Molweight: 135.9 (kD)
Majority protein IDs: Gorai.006G252900.3.A;Gorai.006G252900.2.A;Gorai.006G252900.1.A;Gorai.006G252900.3.D;Gorai.006G252900.2.D;Gorai.006G252900.4.A
Gene IDs: Gorai.006G252900
cotton name: PTHR31008:SF0 - COP1-INTERACTING PROTEIN-LIKE PROTEIN (1 of 2)
OG IDs: 93523934
Best-hit-arabi-ID: AT5G43310.2
Best-hit-arabi-name: COP1-interacting protein-related
Gorai.006G252900.1.A
PCC=0.388, p=0.1
mRNA Profile
Protein Expression Profile
DPA

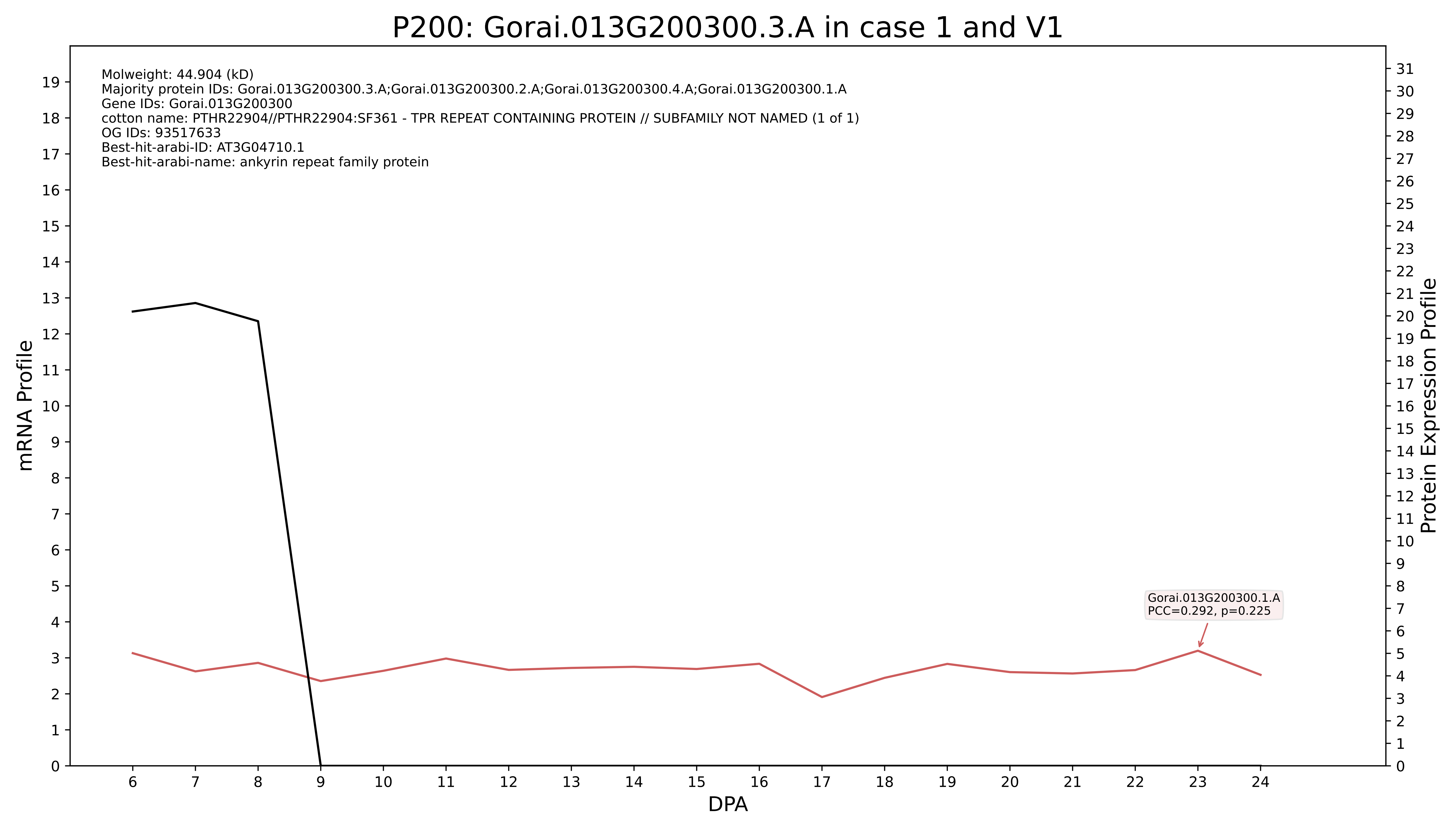

P200: Gorai.013G200300.3.A in case 1 and V1
Molweight: 44.904 (kD)
Majority protein IDs: Gorai.013G200300.3.A;Gorai.013G200300.2.A;Gorai.013G200300.4.A;Gorai.013G200300.1.A
Gene IDs: Gorai.013G200300
cotton name: PTHR22904//PTHR22904:SF361 - TPR REPEAT CONTAINING PROTEIN // SUBFAMILY NOT NAMED (1 of 1)
OG IDs: 93517633
Best-hit-arabi-ID: AT3G04710.1
Best-hit-arabi-name: ankyrin repeat family protein
Gorai.013G200300.1.A
PCC=0.292, p=0.225
mRNA Profile
Protein Expression Profile
DPA

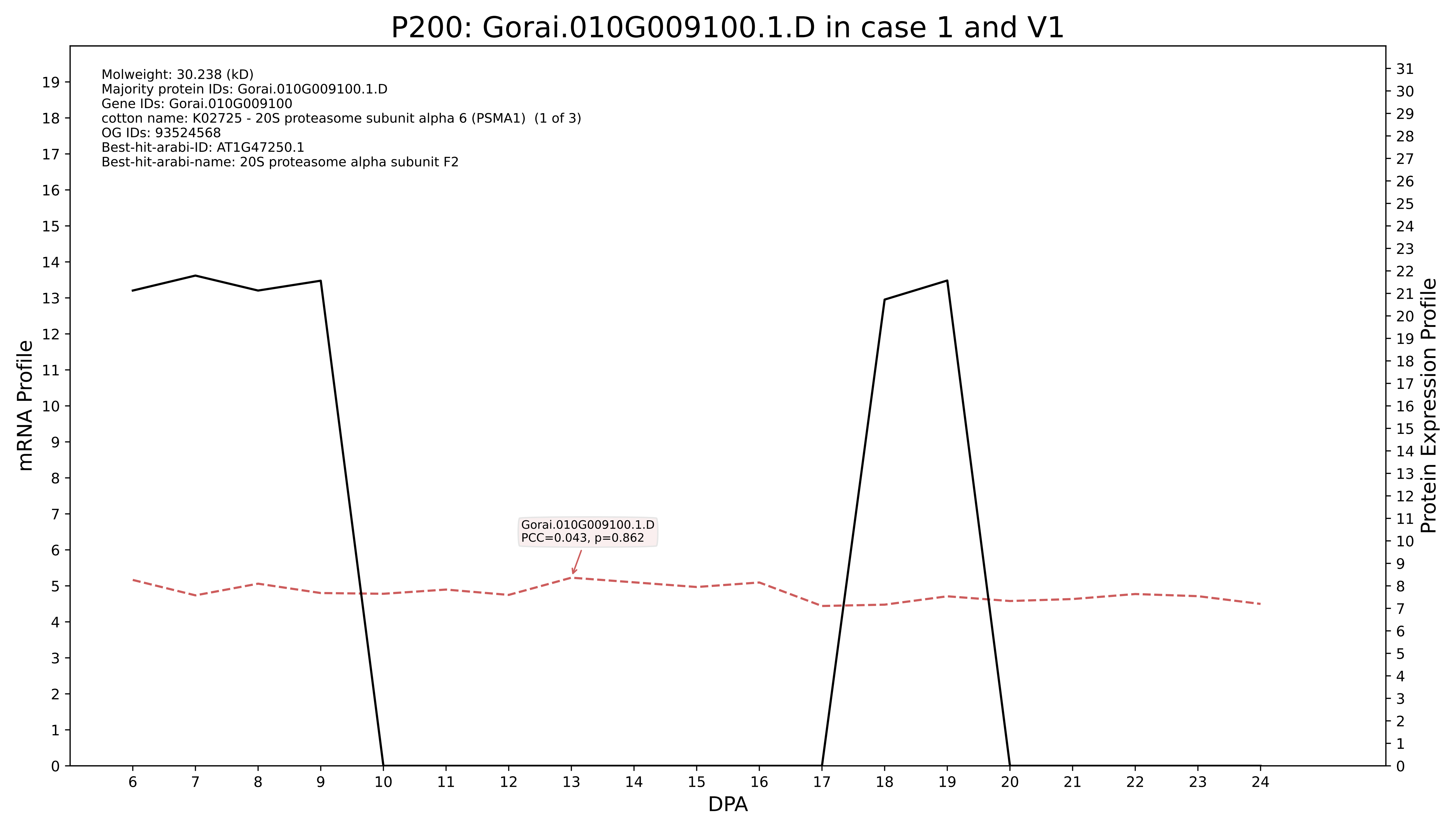
P200: Gorai.010G009100.1.D in case 1 and V1
Molweight: 30.238 (kD)
Majority protein IDs: Gorai.010G009100.1.D
Gene IDs: Gorai.010G009100
cotton name: K02725 - 20S proteasome subunit alpha 6 (PSMA1) (1 of 3)
OG IDs: 93524568
Best-hit-arabi-ID: AT1G47250.1
Best-hit-arabi-name: 20S proteasome alpha subunit F2
Gorai.010G009100.1.D
PCC=0.043, p=0.862
mRNA Profile
Protein Expression Profile
DPA

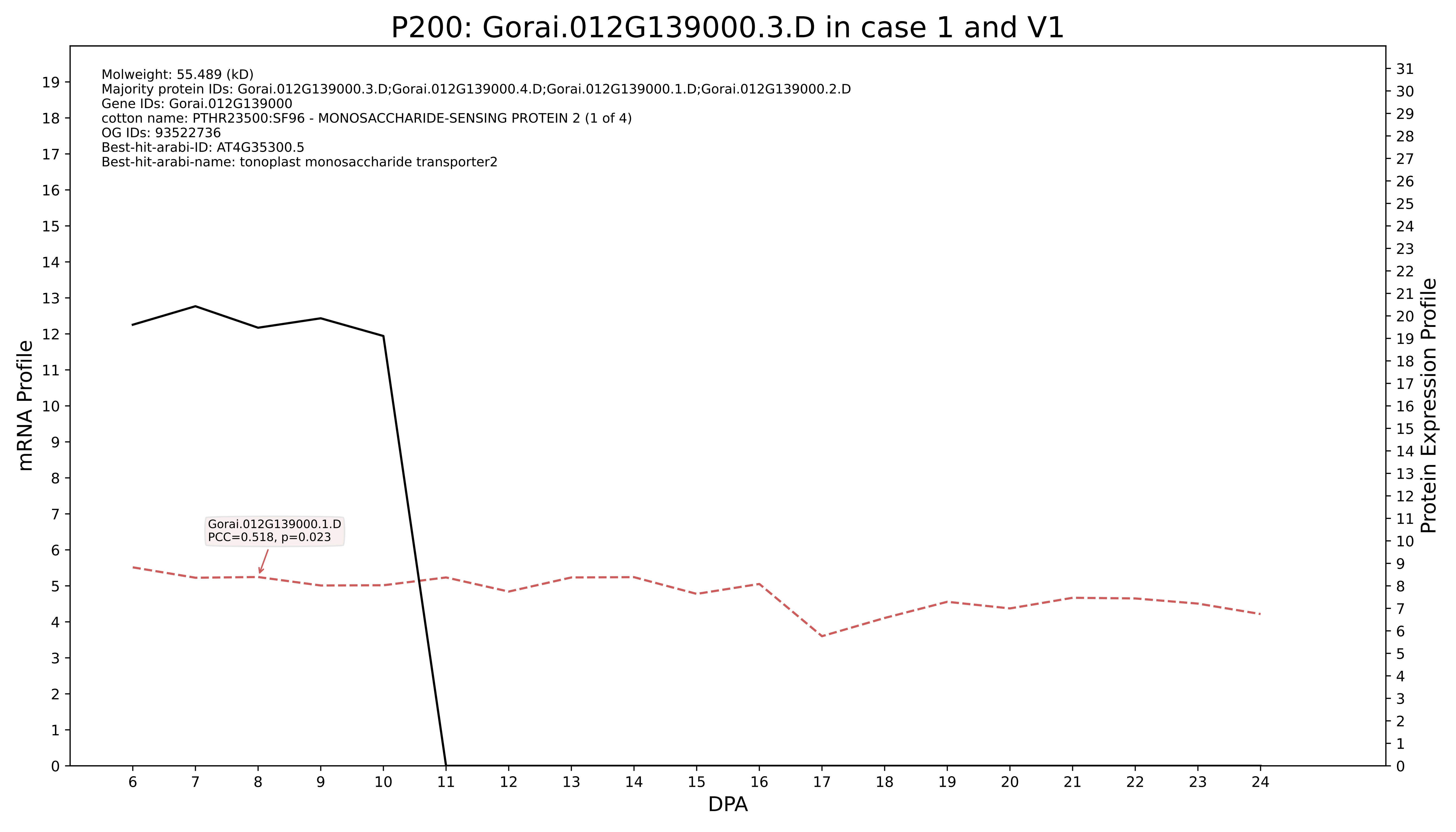
P200: Gorai.012G139000.3.D in case 1 and V1
Molweight: 55.489 (kD)
Majority protein IDs: Gorai.012G139000.3.D;Gorai.012G139000.4.D;Gorai.012G139000.1.D;Gorai.012G139000.2.D
Gene IDs: Gorai.012G139000
cotton name: PTHR23500:SF96 - MONOSACCHARIDE-SENSING PROTEIN 2 (1 of 4)
OG IDs: 93522736
Best-hit-arabi-ID: AT4G35300.5
Best-hit-arabi-name: tonoplast monosaccharide transporter2
Gorai.012G139000.1.D
PCC=0.518, p=0.023
mRNA Profile
Protein Expression Profile
DPA

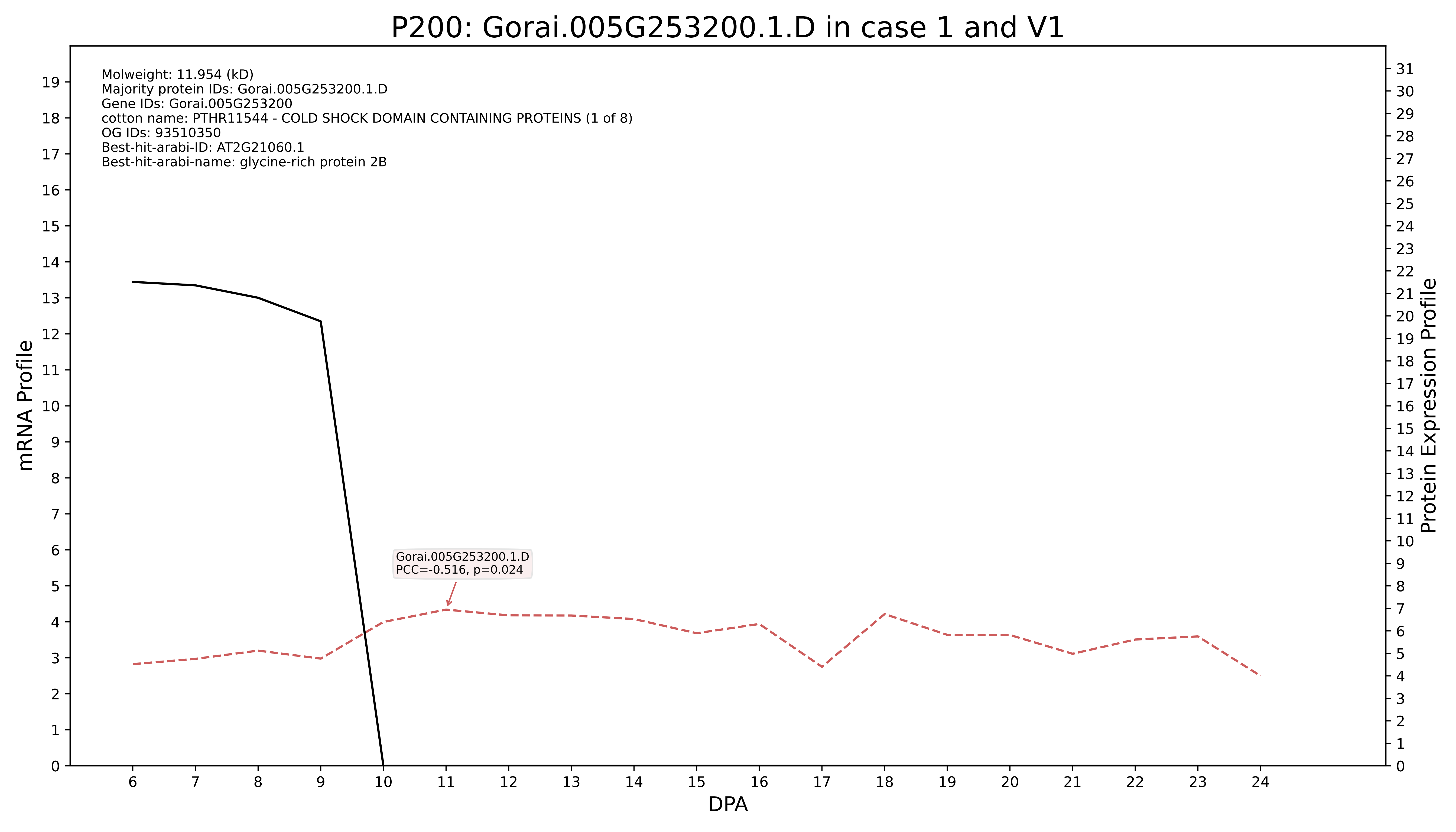
P200: Gorai.005G253200.1.D in case 1 and V1
Molweight: 11.954 (kD)
Majority protein IDs: Gorai.005G253200.1.D
Gene IDs: Gorai.005G253200
cotton name: PTHR11544 - COLD SHOCK DOMAIN CONTAINING PROTEINS (1 of 8)
OG IDs: 93510350
Best-hit-arabi-ID: AT2G21060.1
Best-hit-arabi-name: glycine-rich protein 2B
Gorai.005G253200.1.D
PCC=-0.516, p=0.024
mRNA Profile
Protein Expression Profile
DPA

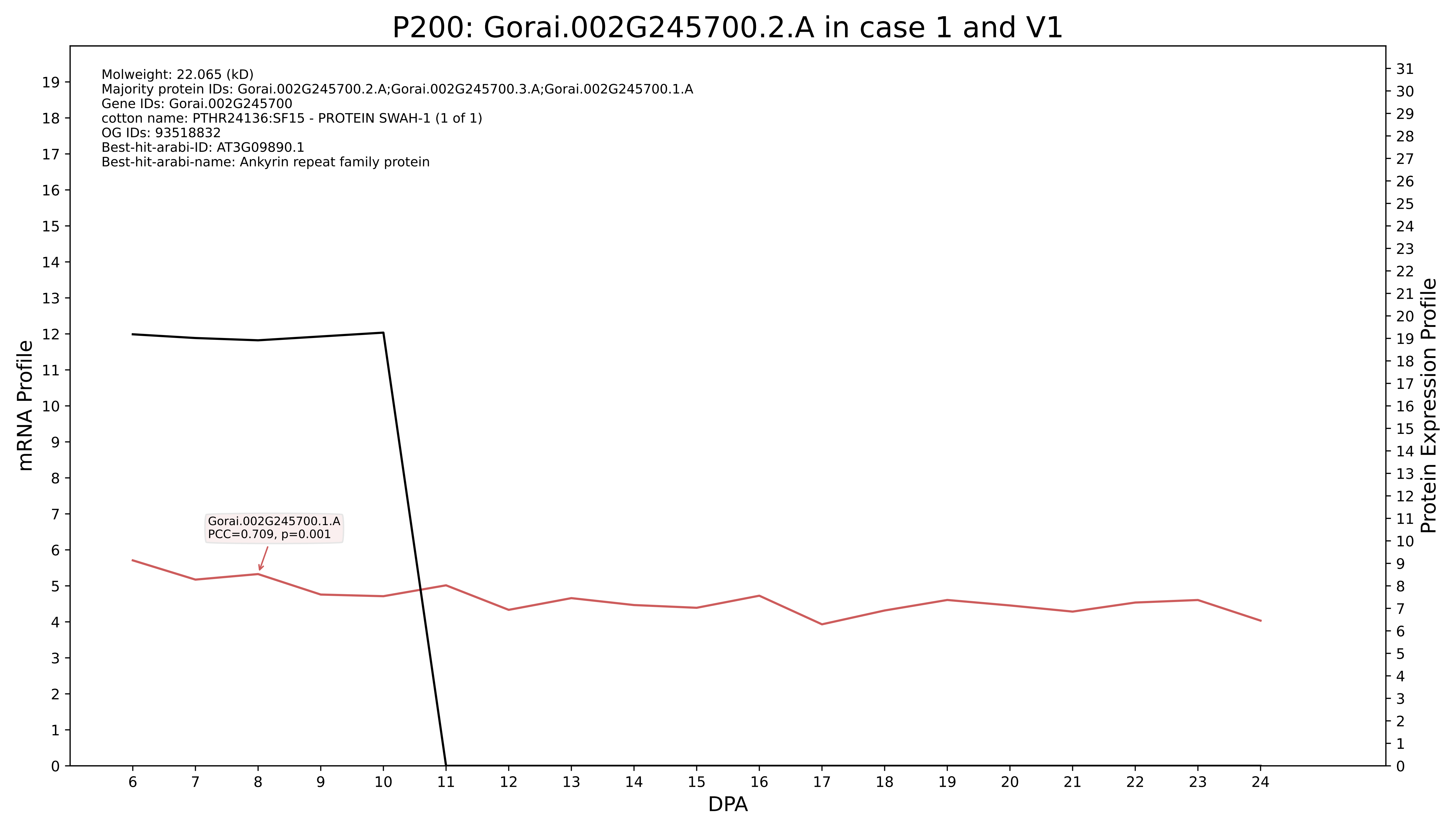

P200: Gorai.002G245700.2.A in case 1 and V1
Molweight: 22.065 (kD)
Majority protein IDs: Gorai.002G245700.2.A;Gorai.002G245700.3.A;Gorai.002G245700.1.A
Gene IDs: Gorai.002G245700
cotton name: PTHR24136:SF15 - PROTEIN SWAH-1 (1 of 1)
OG IDs: 93518832
Best-hit-arabi-ID: AT3G09890.1
Best-hit-arabi-name: Ankyrin repeat family protein
Gorai.002G245700.1.A
PCC=0.709, p=0.001
mRNA Profile
Protein Expression Profile
DPA
0 1 2 3 4 5 6 7 8 9 10 11 12 13 14 15 16 17 18 19
0 1 2 3 4 5 6 7 8 9 10 11 12 13 14 15 16 17 18 19 20 21 22 23 24 25 26 27 28 29 30 31
6 7 8 9 10 11 12 13 14 15 16 17 18 19 20 21 22 23 24

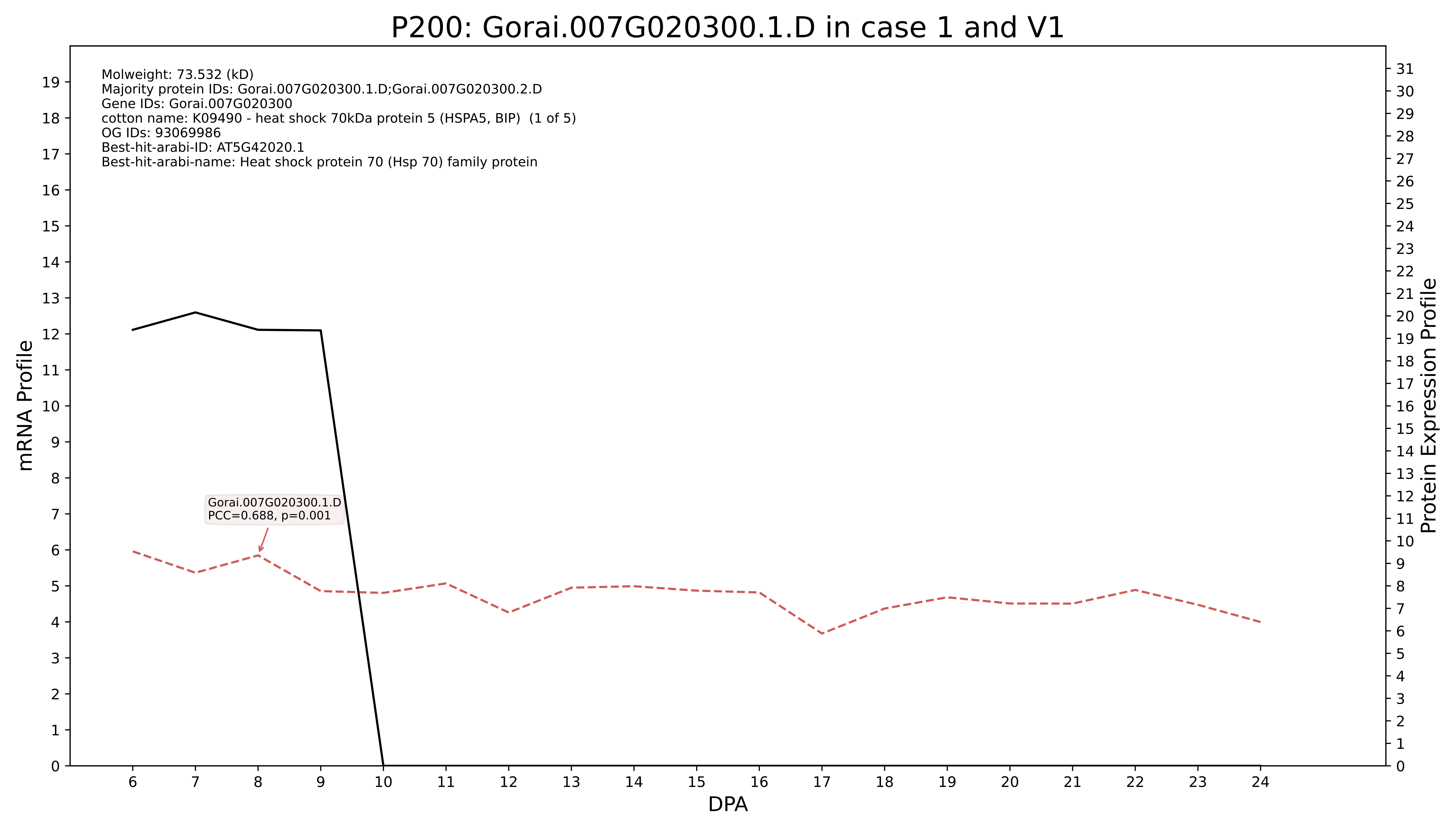
P200: Gorai.007G020300.1.D in case 1 and V1
Molweight: 73.532 (kD)
Majority protein IDs: Gorai.007G020300.1.D;Gorai.007G020300.2.D
Gene IDs: Gorai.007G020300
cotton name: K09490 - heat shock 70kDa protein 5 (HSPA5, BIP) (1 of 5)
OG IDs: 93069986
Best-hit-arabi-ID: AT5G42020.1
Best-hit-arabi-name: Heat shock protein 70 (Hsp 70) family protein
Gorai.007G020300.1.D
PCC=0.688, p=0.001
mRNA Profile
Protein Expression Profile
DPA

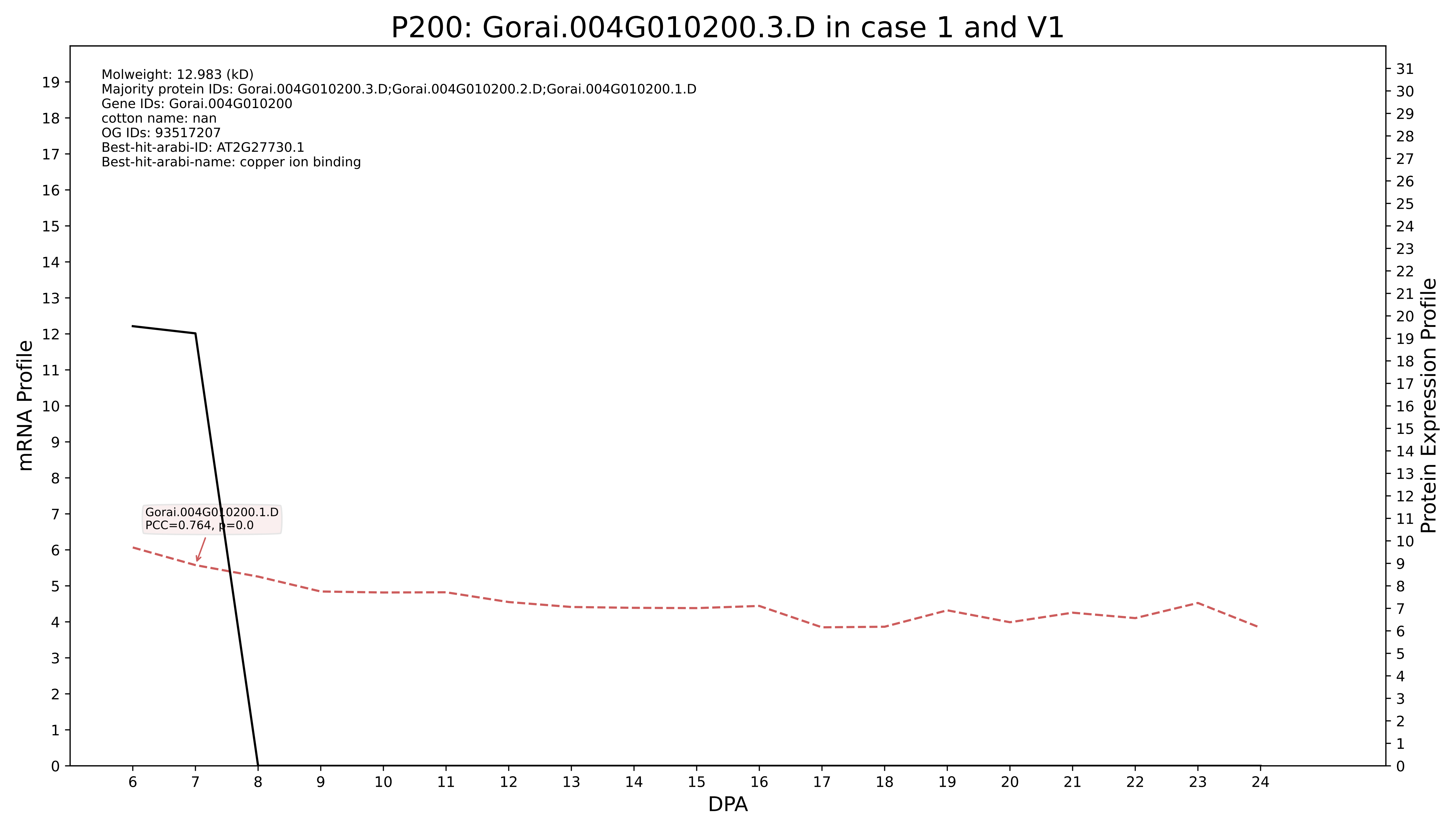
P200: Gorai.004G010200.3.D in case 1 and V1
Molweight: 12.983 (kD)
Majority protein IDs: Gorai.004G010200.3.D;Gorai.004G010200.2.D;Gorai.004G010200.1.D
Gene IDs: Gorai.004G010200
cotton name: nan
OG IDs: 93517207
Best-hit-arabi-ID: AT2G27730.1
Best-hit-arabi-name: copper ion binding
Gorai.004G010200.1.D
PCC=0.764, p=0.0
mRNA Profile
Protein Expression Profile
DPA

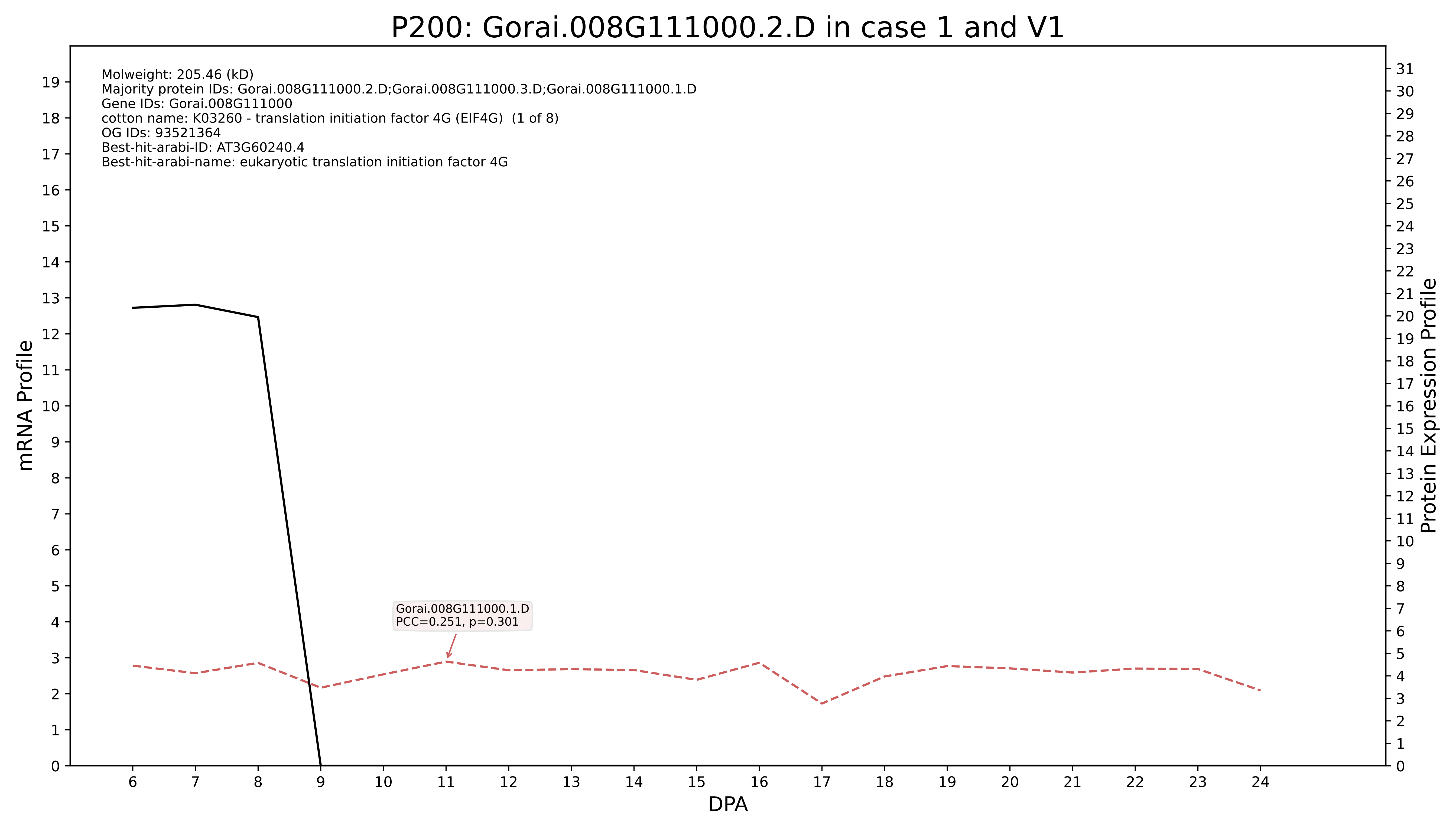
P200: Gorai.008G111000.2.D in case 1 and V1
Molweight: 205.46 (kD)
Majority protein IDs: Gorai.008G111000.2.D;Gorai.008G111000.3.D;Gorai.008G111000.1.D
Gene IDs: Gorai.008G111000
cotton name: K03260 - translation initiation factor 4G (EIF4G) (1 of 8)
OG IDs: 93521364
Best-hit-arabi-ID: AT3G60240.4
Best-hit-arabi-name: eukaryotic translation initiation factor 4G
Gorai.008G111000.1.D
PCC=0.251, p=0.301
mRNA Profile
Protein Expression Profile
DPA

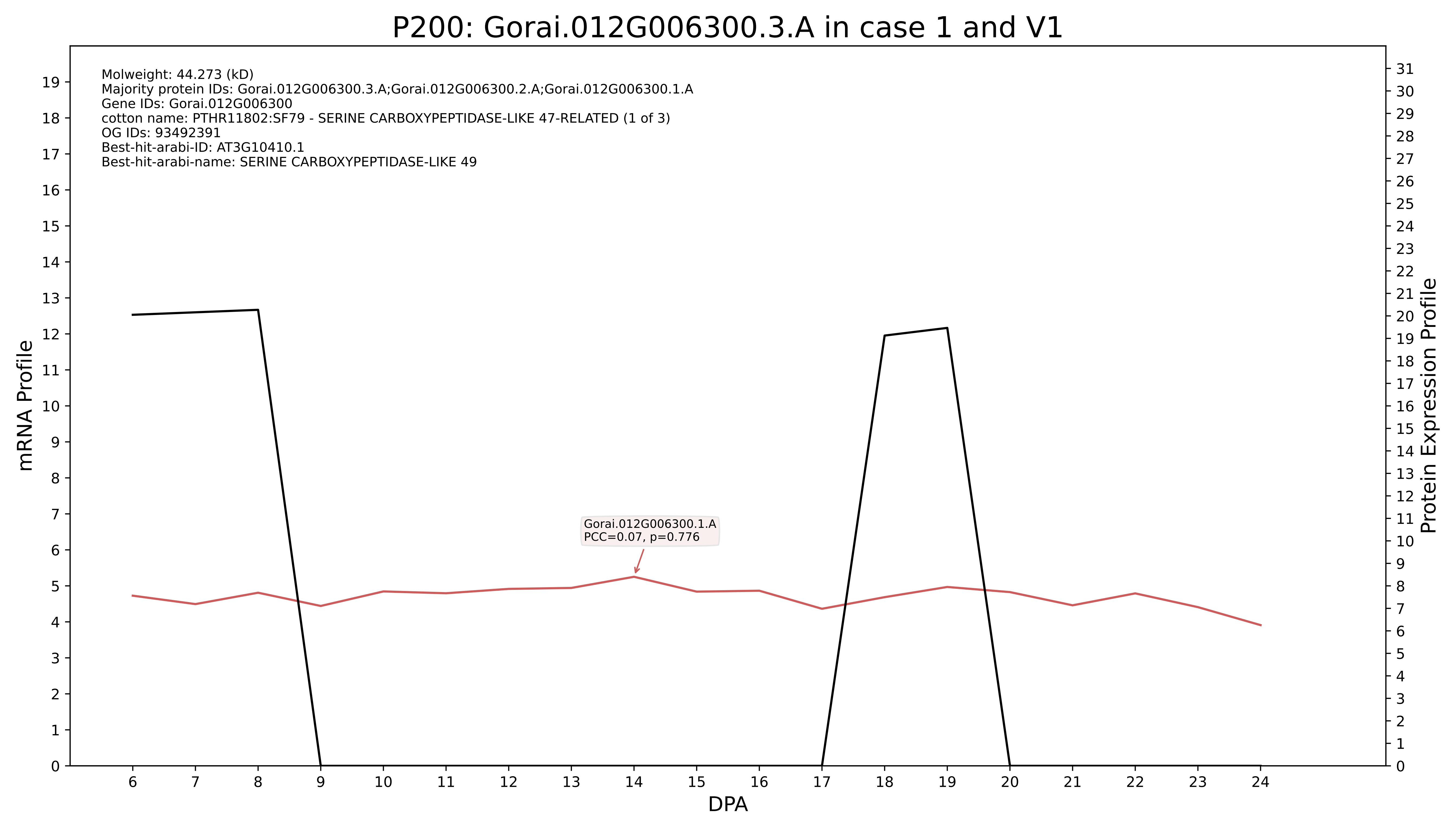
P200: Gorai.012G006300.3.A in case 1 and V1
Molweight: 44.273 (kD)
Majority protein IDs: Gorai.012G006300.3.A;Gorai.012G006300.2.A;Gorai.012G006300.1.A
Gene IDs: Gorai.012G006300
cotton name: PTHR11802:SF79 - SERINE CARBOXYPEPTIDASE-LIKE 47-RELATED (1 of 3)
OG IDs: 93492391
Best-hit-arabi-ID: AT3G10410.1
Best-hit-arabi-name: SERINE CARBOXYPEPTIDASE-LIKE 49
Gorai.012G006300.1.A
PCC=0.07, p=0.776
mRNA Profile
Protein Expression Profile
DPA

### P200: Gorai.007G049000.2.D in case 1 and V1

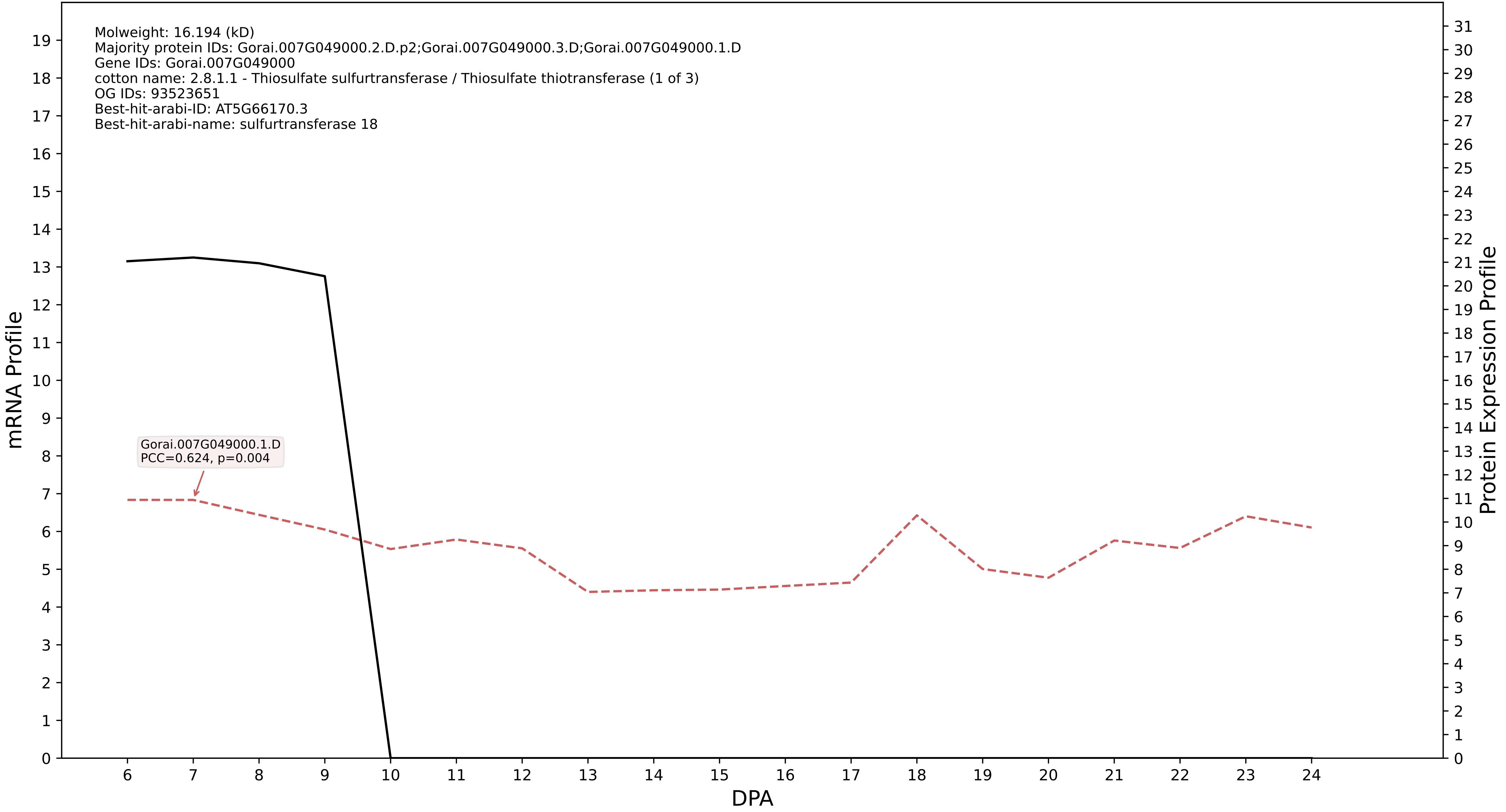

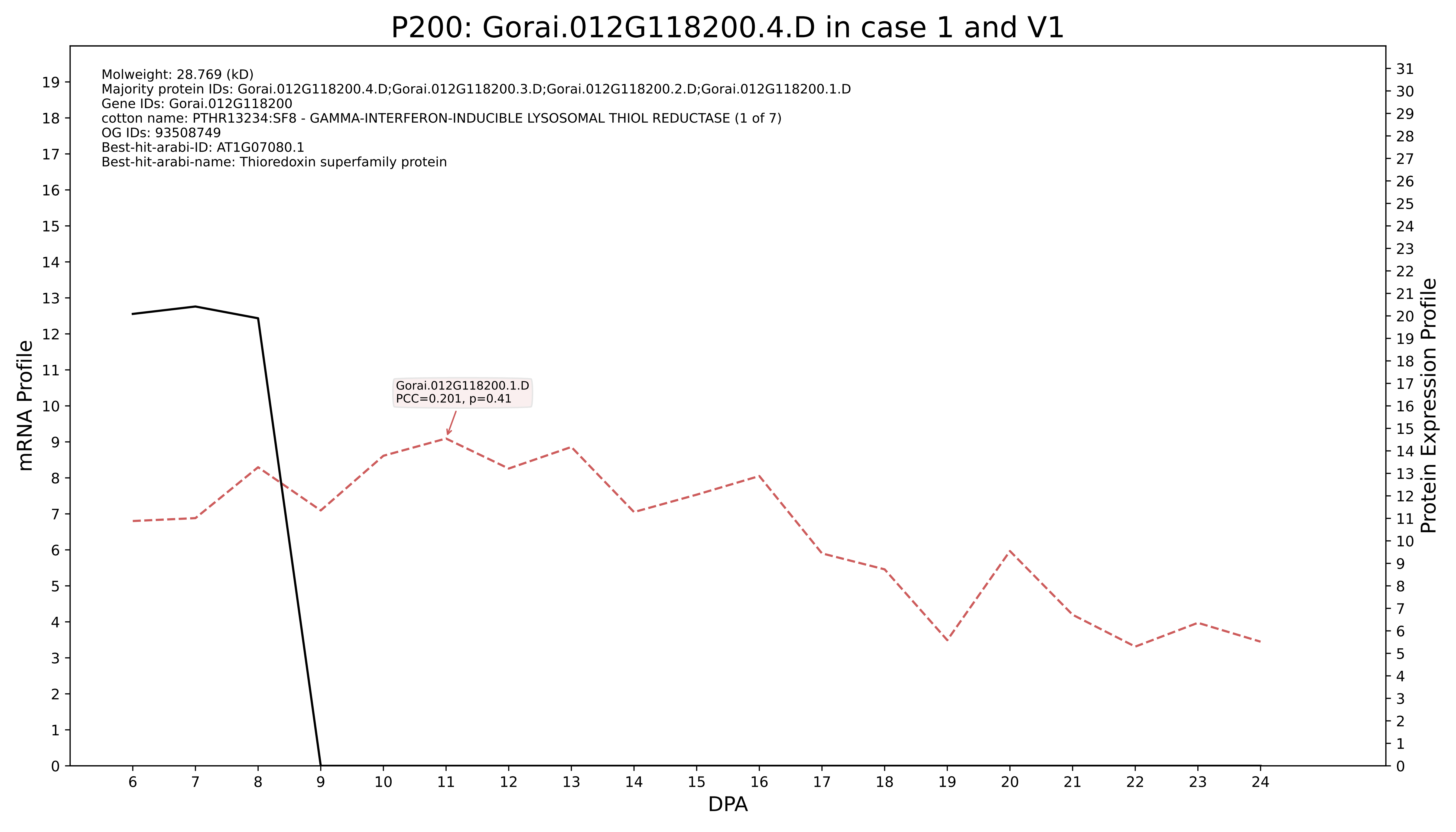
P200: Gorai.012G118200.4.D in case 1 and V1
Molweight: 28.769 (kD)
Majority protein IDs: Gorai.012G118200.4.D;Gorai.012G118200.3.D;Gorai.012G118200.2.D;Gorai.012G118200.1.D
Gene IDs: Gorai.012G118200
cotton name: PTHR13234:SF8 - GAMMA-INTERFERON-INDUCIBLE LYSOSOMAL THIOL REDUCTASE (1 of 7)
OG IDs: 93508749
Best-hit-arabi-ID: AT1G07080.1
Best-hit-arabi-name: Thioredoxin superfamily protein
Gorai.012G118200.1.D
PCC=0.201, p=0.41
mRNA Profile
Protein Expression Profile
DPA

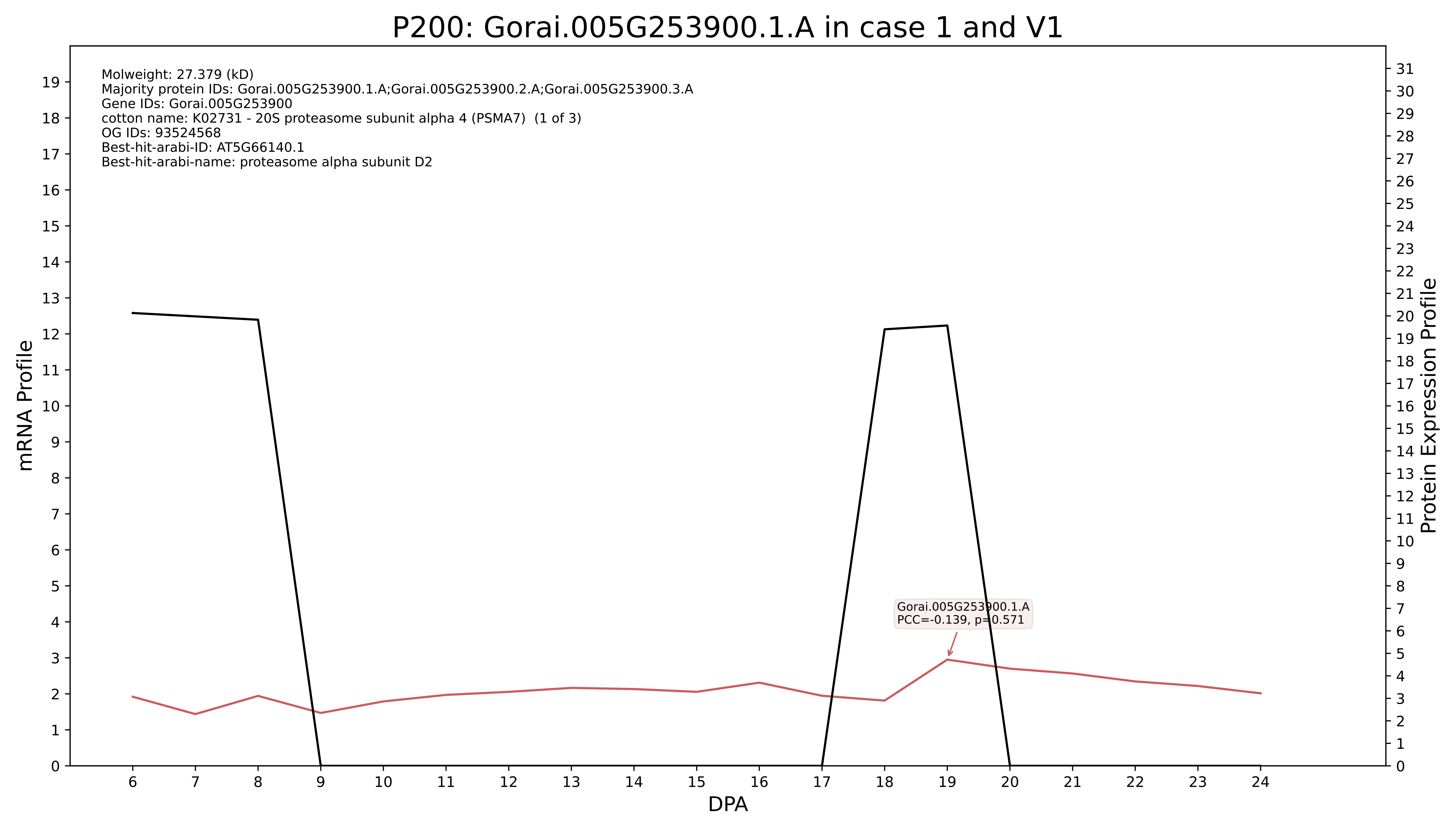
P200: Gorai.005G253900.1.A in case 1 and V1
Molweight: 27.379 (kD)
Majority protein IDs: Gorai.005G253900.1.A;Gorai.005G253900.2.A;Gorai.005G253900.3.A
Gene IDs: Gorai.005G253900
cotton name: K02731 - 20S proteasome subunit alpha 4 (PSMA7) (1 of 3)
OG IDs: 93524568
Best-hit-arabi-ID: AT5G66140.1
Best-hit-arabi-name: proteasome alpha subunit D2
Gorai.005G253900.1.A
PCC=-0.139, p=0.571
mRNA Profile
Protein Expression Profile
DPA

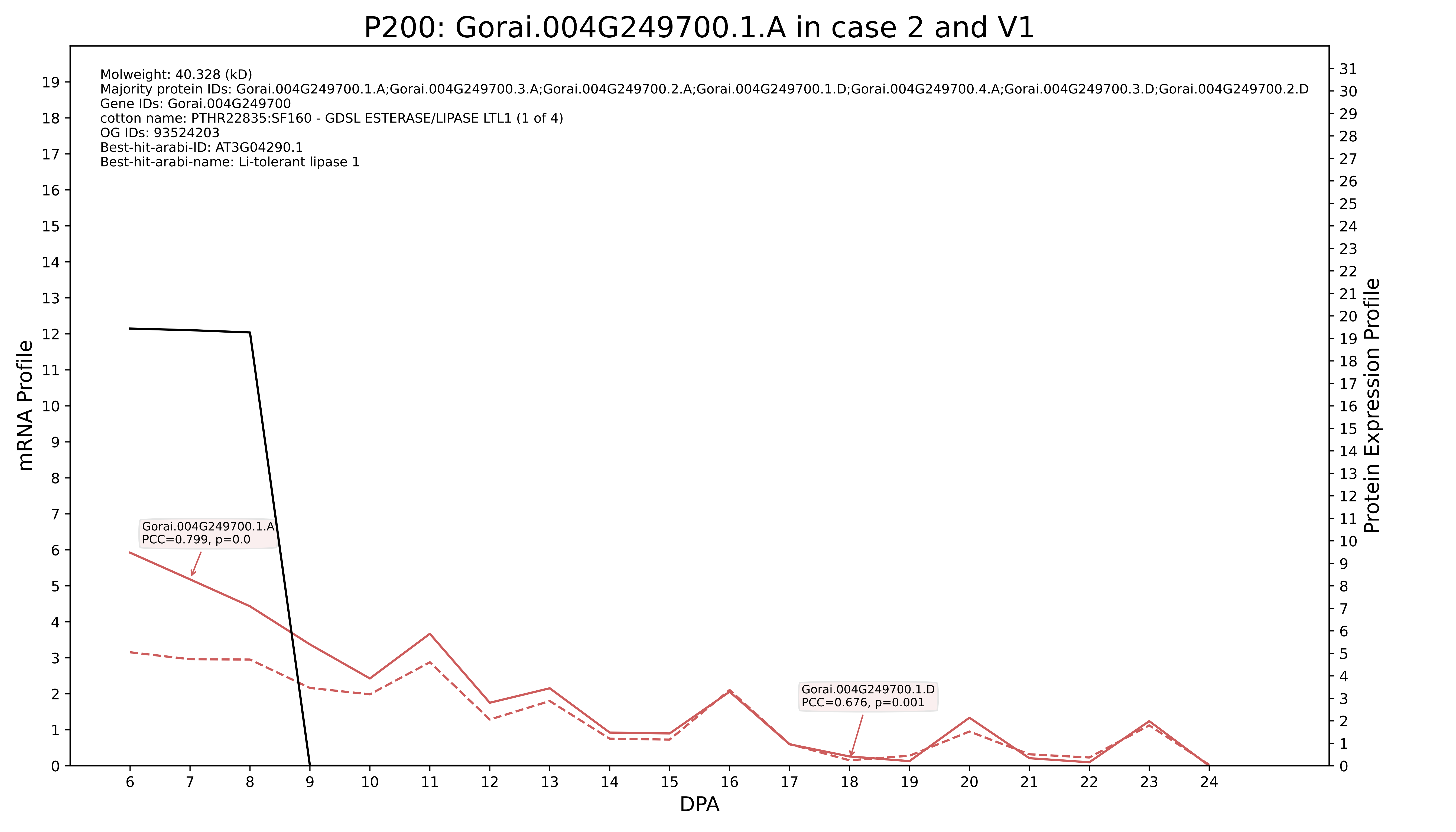

P200: Gorai.004G249700.1.A in case 2 and V1
Molweight: 40.328 (kD)
Majority protein IDs: Gorai.004G249700.1.A;Gorai.004G249700.3.A;Gorai.004G249700.2.A;Gorai.004G249700.1.D;Gorai.004G249700.4.A;Gorai.004G249700.3.D;Gorai.004G249700.2.D
Gene IDs: Gorai.004G249700
cotton name: PTHR22835:SF160 - GDSL ESTERASE/LIPASE LTL1 (1 of 4)
OG IDs: 93524203
Best-hit-arabi-ID: AT3G04290.1
Best-hit-arabi-name: Li-tolerant lipase 1
Gorai.004G249700.1.A
PCC=0.799, p=0.0
Gorai.004G249700.1.D
PCC=0.676, p=0.001
mRNA Profile
Protein Expression Profile
DPA

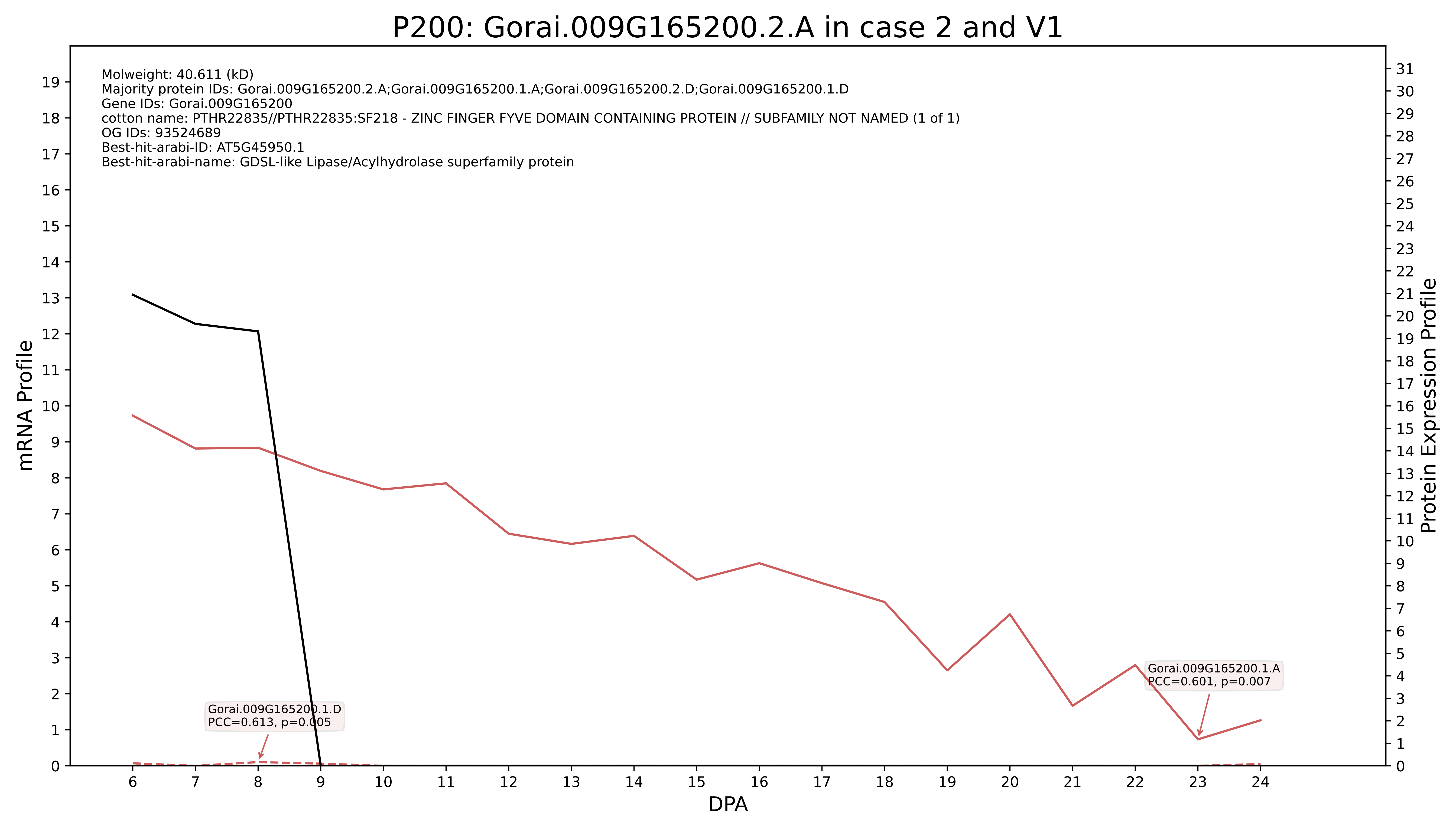
P200: Gorai.009G165200.2.A in case 2 and V1
Molweight: 40.611 (kD)
Majority protein IDs: Gorai.009G165200.2.A;Gorai.009G165200.1.A;Gorai.009G165200.2.D;Gorai.009G165200.1.D
Gene IDs: Gorai.009G165200
cotton name: PTHR22835//PTHR22835:SF218 - ZINC FINGER FYVE DOMAIN CONTAINING PROTEIN // SUBFAMILY NOT NAMED (1 of 1)
OG IDs: 93524689
Best-hit-arabi-ID: AT5G45950.1
Best-hit-arabi-name: GDSL-like Lipase/Acylhydrolase superfamily protein
Gorai.009G165200.1.D
PCC=0.613, p=0.005
Gorai.009G165200.1.A
PCC=0.601, p=0.007
mRNA Profile
Protein Expression Profile
DPA

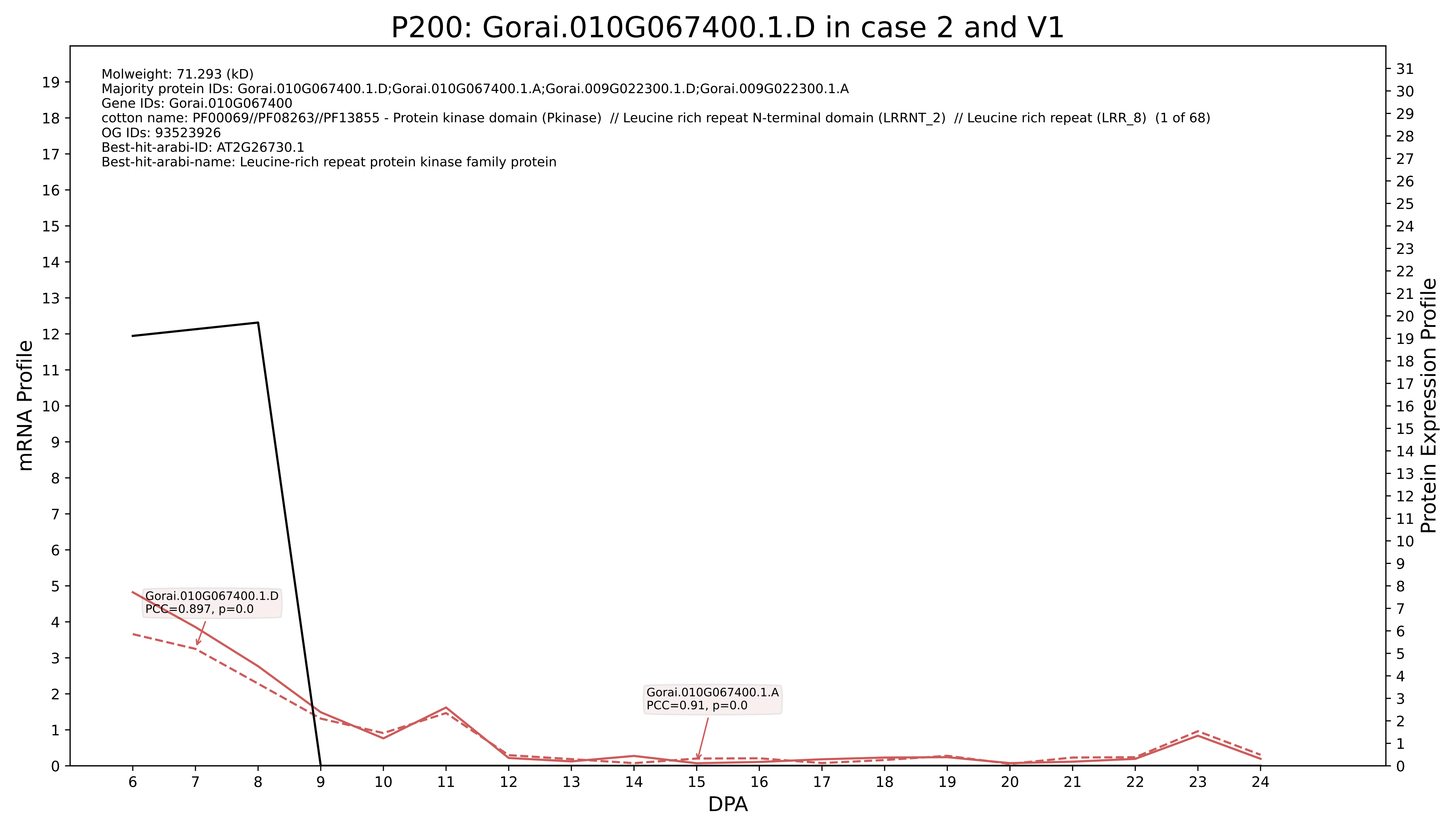
P200: Gorai.010G067400.1.D in case 2 and V1
Molweight: 71.293 (kD)
Majority protein IDs: Gorai.010G067400.1.D;Gorai.010G067400.1.A;Gorai.009G022300.1.D;Gorai.009G022300.1.A
Gene IDs: Gorai.010G067400
cotton name: PF00069//PF08263//PF13855 - Protein kinase domain (Pkinase) // Leucine rich repeat N-terminal domain (LRRNT_2) // Leucine rich repeat (LRR_8) (1 of 68)
OG IDs: 93523926
Best-hit-arabi-ID: AT2G26730.1
Best-hit-arabi-name: Leucine-rich repeat protein kinase family protein
Gorai.010G067400.1.D
PCC=0.897, p=0.0
Gorai.010G067400.1.A
PCC=0.91, p=0.0
mRNA Profile
Protein Expression Profile
DPA

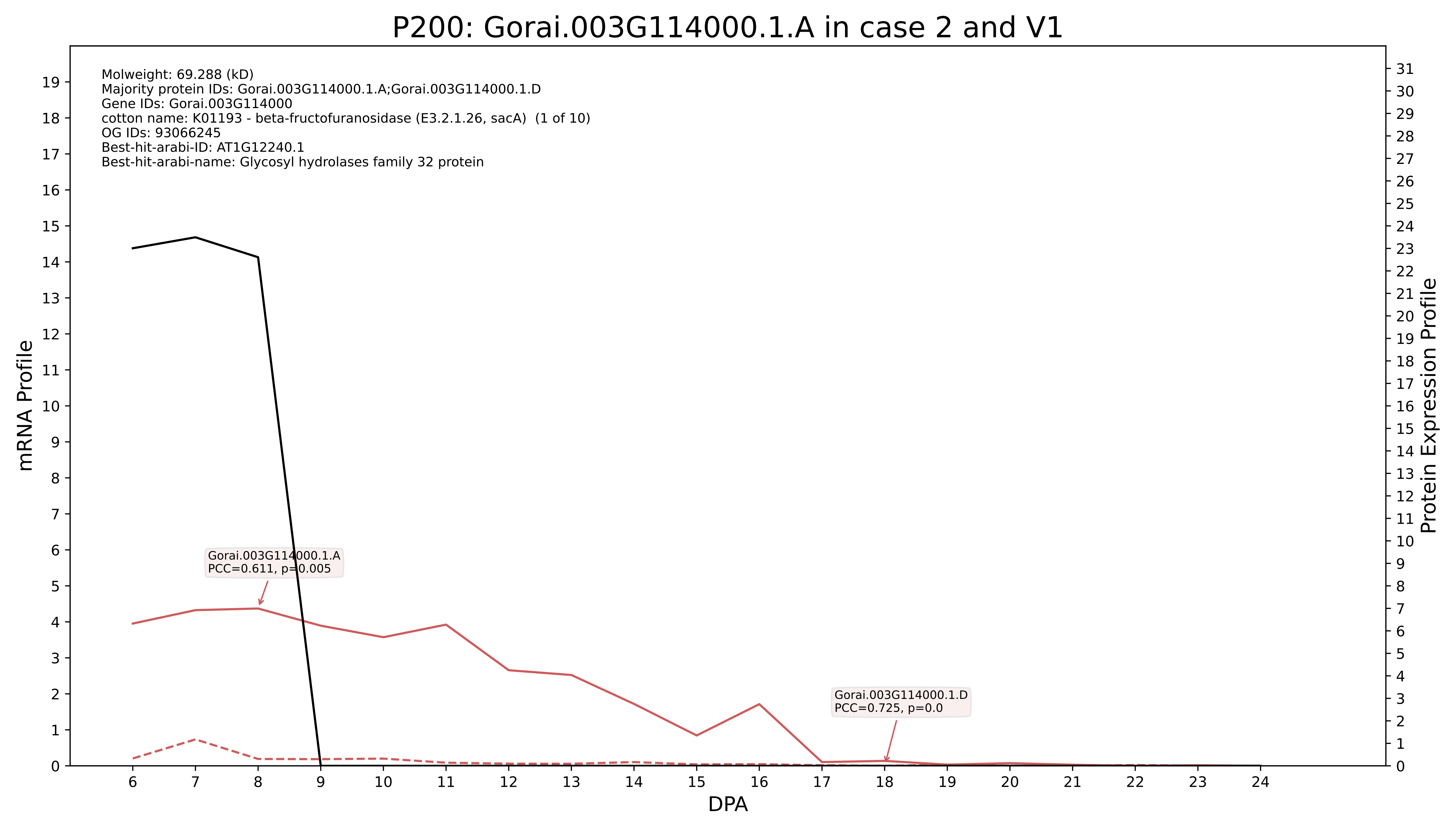

P200: Gorai.003G114000.1.A in case 2 and V1
Molweight: 69.288 (kD)
Majority protein IDs: Gorai.003G114000.1.A;Gorai.003G114000.1.D
Gene IDs: Gorai.003G114000
cotton name: K01193 - beta-fructofuranosidase (E3.2.1.26, sacA) (1 of 10)
OG IDs: 93066245
Best-hit-arabi-ID: AT1G12240.1
Best-hit-arabi-name: Glycosyl hydrolases family 32 protein
Gorai.003G114000.1.A
PCC=0.611, p=0.005
Gorai.003G114000.1.D
PCC=0.725, p=0.0
mRNA Profile
Protein Expression Profile
DPA

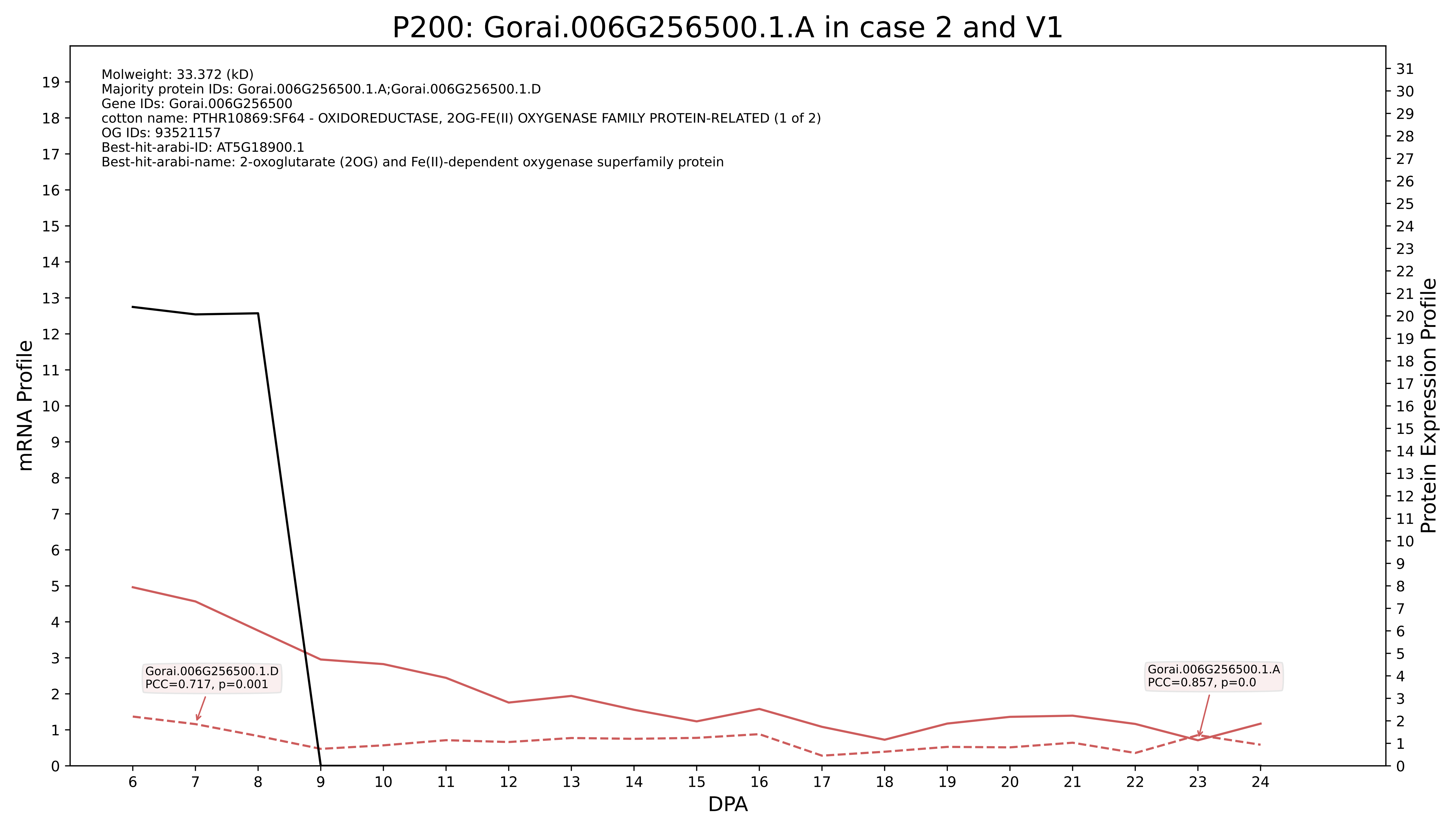

P200: Gorai.006G256500.1.A in case 2 and V1
Molweight: 33.372 (kD)
Majority protein IDs: Gorai.006G256500.1.A;Gorai.006G256500.1.D
Gene IDs: Gorai.006G256500
cotton name: PTHR10869:SF64 - OXIDOREDUCTASE, 2OG-FE(II) OXYGENASE FAMILY PROTEIN-RELATED (1 of 2)
OG IDs: 93521157
Best-hit-arabi-ID: AT5G18900.1
Best-hit-arabi-name: 2-oxoglutarate (2OG) and Fe(II)-dependent oxygenase superfamily protein
mRNA Profile
Protein Expression Profile
DPA
Gorai.006G256500.1.D
PCC=0.717, p=0.001
Gorai.006G256500.1.A
PCC=0.857, p=0.0

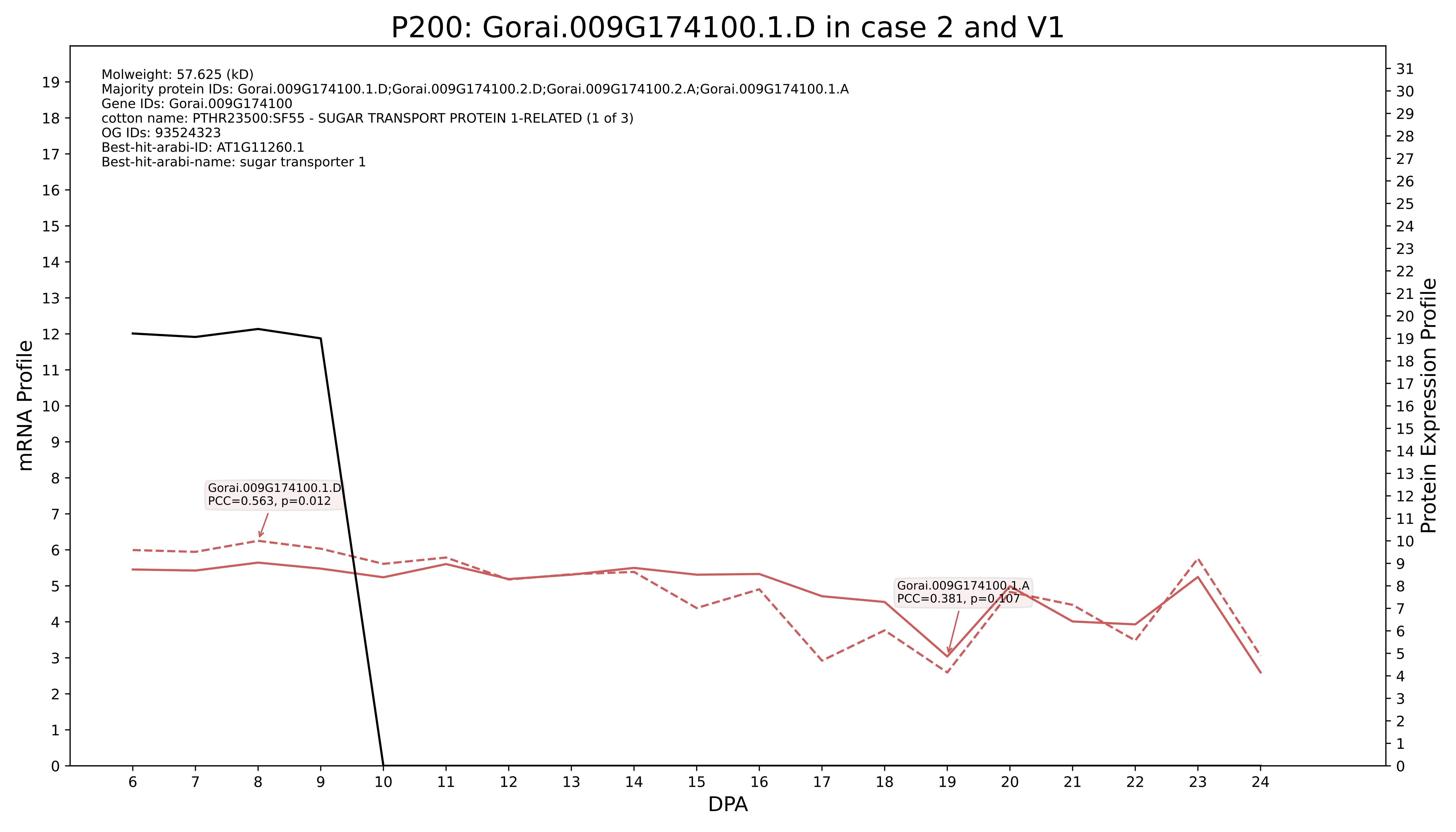
P200: Gorai.009G174100.1.D in case 2 and V1
Molweight: 57.625 (kD)
Majority protein IDs: Gorai.009G174100.1.D;Gorai.009G174100.2.D;Gorai.009G174100.2.A;Gorai.009G174100.1.A
Gene IDs: Gorai.009G174100
cotton name: PTHR23500:SF55 - SUGAR TRANSPORT PROTEIN 1-RELATED (1 of 3)
OG IDs: 93524323
Best-hit-arabi-ID: AT1G11260.1
Best-hit-arabi-name: sugar transporter 1
Gorai.009G174100.1.D
PCC=0.563, p=0.012
Gorai.009G174100.1.A
PCC=0.381, p=0.107
mRNA Profile
Protein Expression Profile
DPA

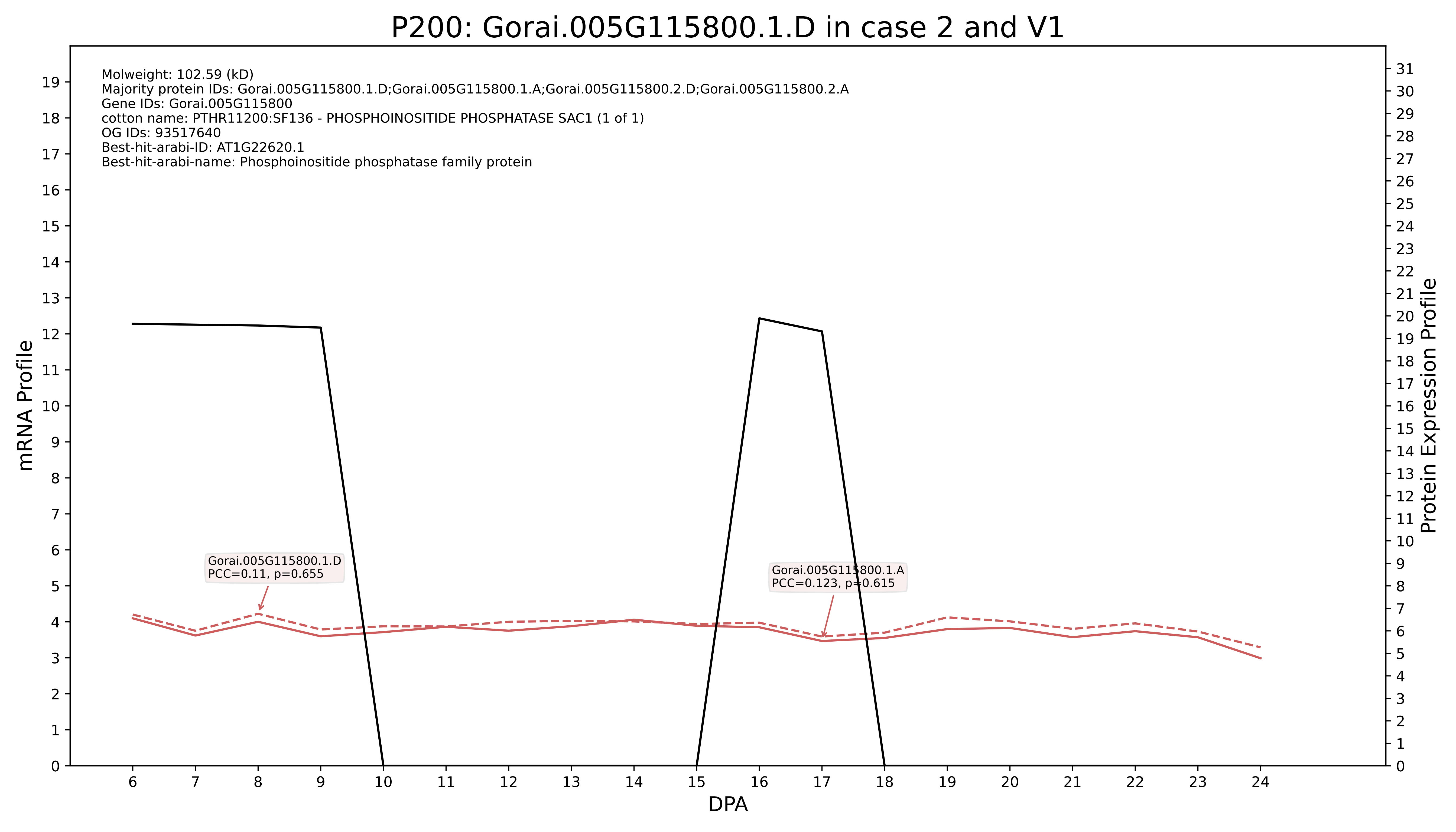
P200: Gorai.005G115800.1.D in case 2 and V1
Molweight: 102.59 (kD)
Majority protein IDs: Gorai.005G115800.1.D;Gorai.005G115800.1.A;Gorai.005G115800.2.D;Gorai.005G115800.2.A
Gene IDs: Gorai.005G115800
cotton name: PTHR11200:SF136 - PHOSPHOINOSITIDE PHOSPHATASE SAC1 (1 of 1)
OG IDs: 93517640
Best-hit-arabi-ID: AT1G22620.1
Best-hit-arabi-name: Phosphoinositide phosphatase family protein
Gorai.005G115800.1.D
PCC=0.11, p=0.655
Gorai.005G115800.1.A
PCC=0.123, p=0.615
mRNA Profile
Protein Expression Profile
DPA

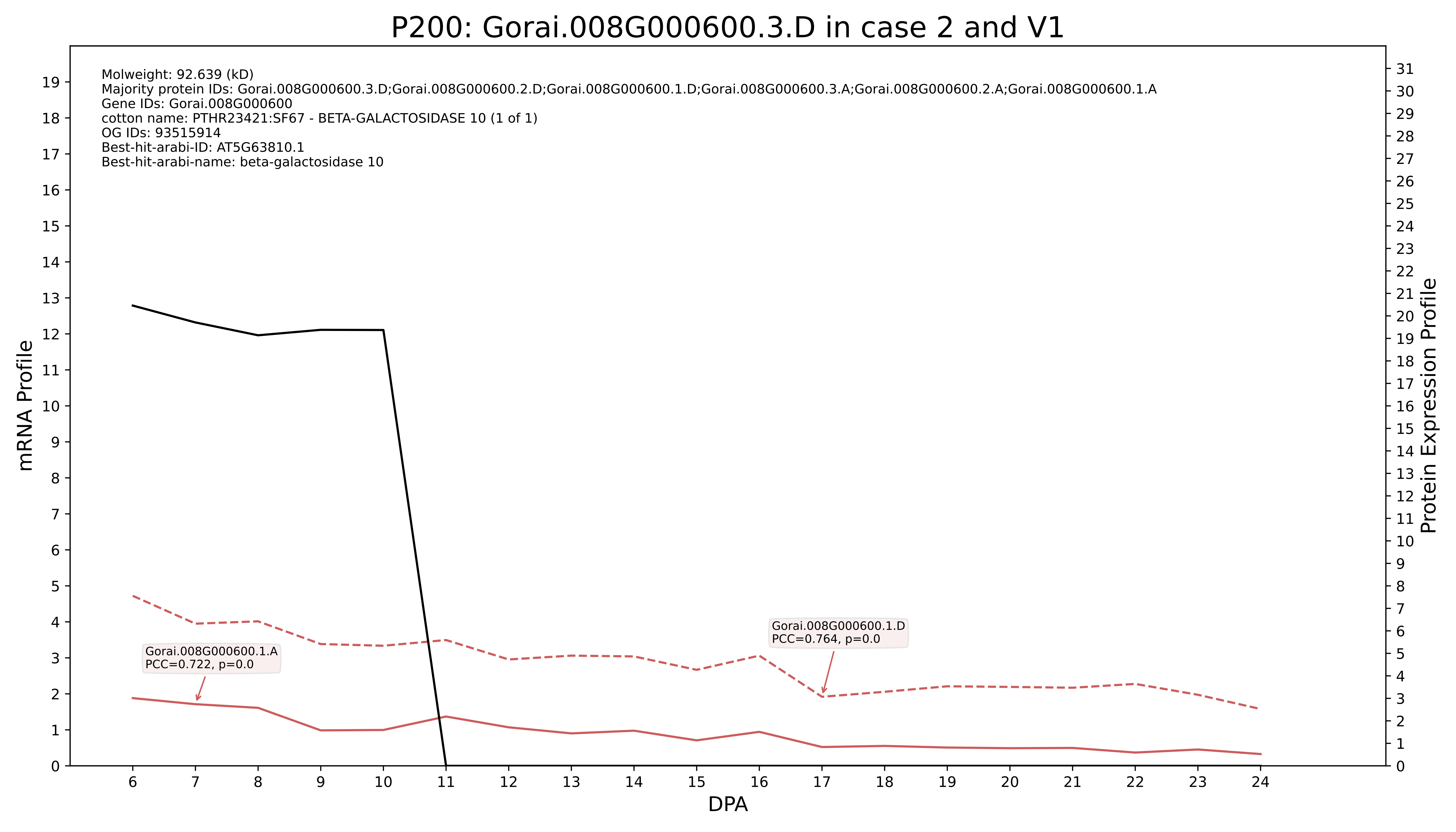
P200: Gorai.008G000600.3.D in case 2 and V1
Molweight: 92.639 (kD)
Majority protein IDs: Gorai.008G000600.3.D;Gorai.008G000600.2.D;Gorai.008G000600.1.D;Gorai.008G000600.3.A;Gorai.008G000600.2.A;Gorai.008G000600.1.A
Gene IDs: Gorai.008G000600
cotton name: PTHR23421:SF67 - BETA-GALACTOSIDASE 10 (1 of 1)
OG IDs: 93515914
Best-hit-arabi-ID: AT5G63810.1
Best-hit-arabi-name: beta-galactosidase 10
Gorai.008G000600.1.A
PCC=0.722, p=0.0
Gorai.008G000600.1.D
PCC=0.764, p=0.0
mRNA Profile
Protein Expression Profile
DPA

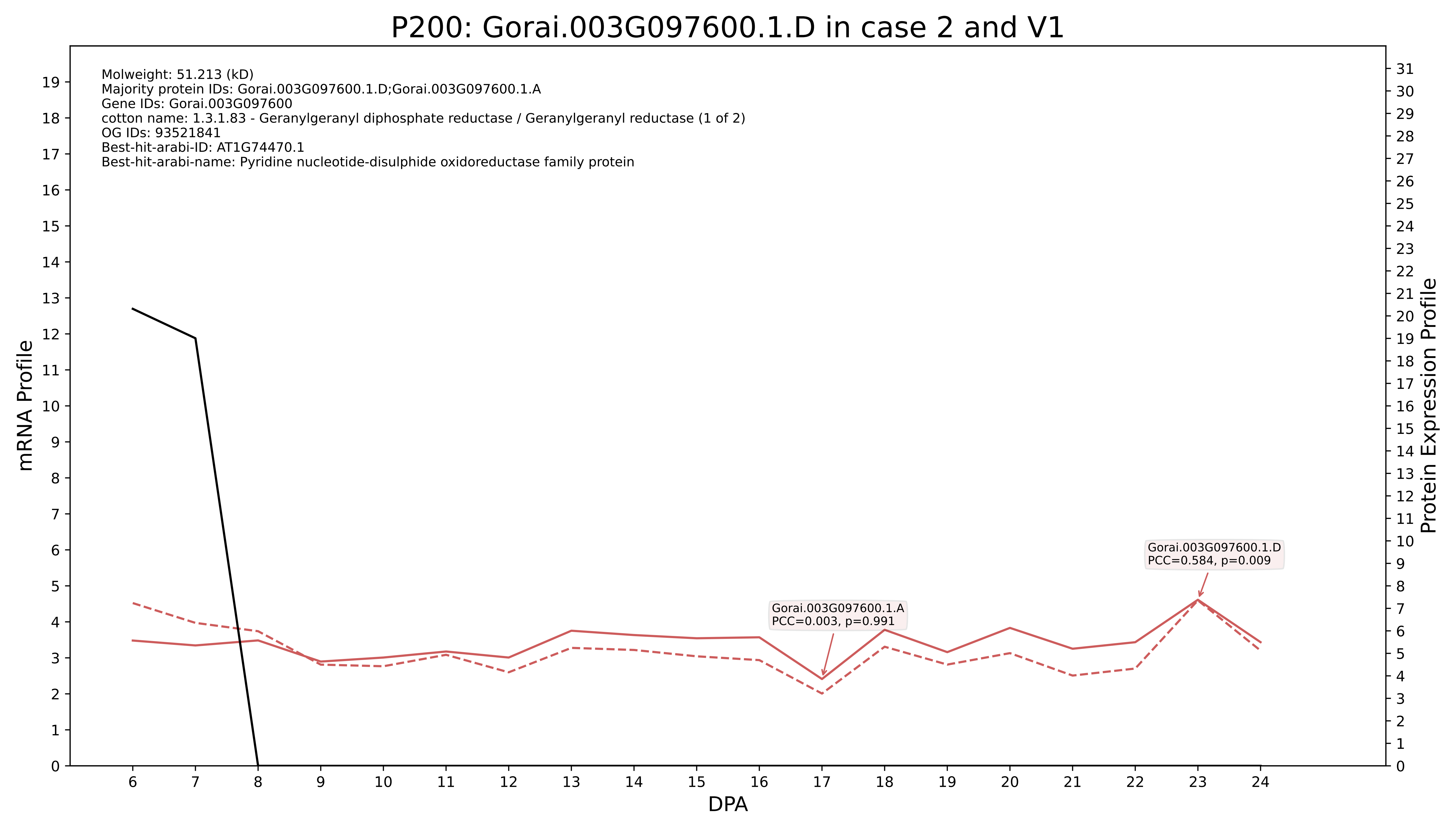
P200: Gorai.003G097600.1.D in case 2 and V1
Molweight: 51.213 (kD)
Majority protein IDs: Gorai.003G097600.1.D;Gorai.003G097600.1.A
Gene IDs: Gorai.003G097600
cotton name: 1.3.1.83 - Geranylgeranyl diphosphate reductase / Geranylgeranyl reductase (1 of 2)
OG IDs: 93521841
Best-hit-arabi-ID: AT1G74470.1
Best-hit-arabi-name: Pyridine nucleotide-disulphide oxidoreductase family protein
Gorai.003G097600.1.A
PCC=0.003, p=0.991
Gorai.003G097600.1.D
PCC=0.584, p=0.009
mRNA Profile
Protein Expression Profile
DPA

### P200: Gorai.003G125400.2.D in case 2 and V1

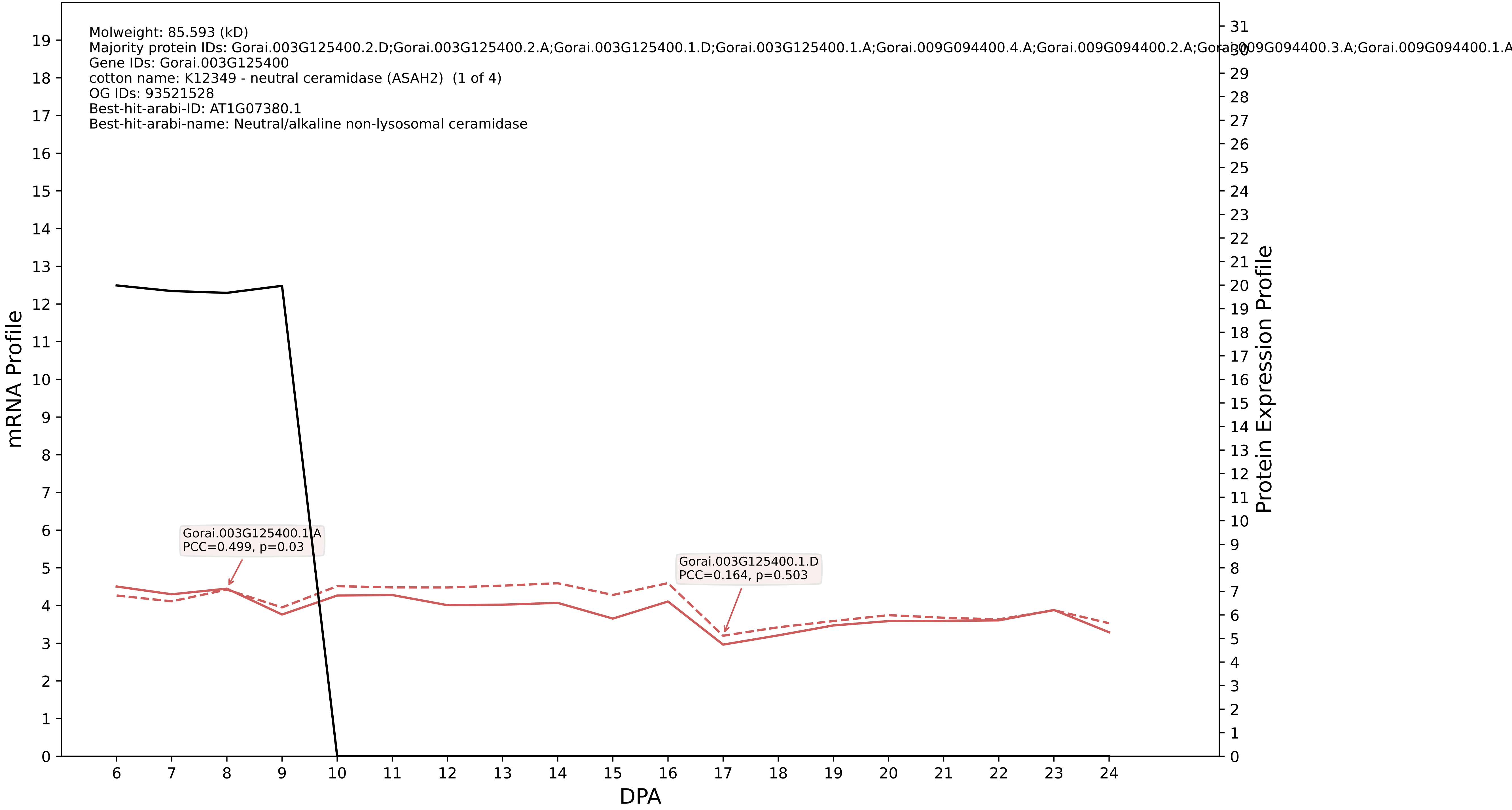

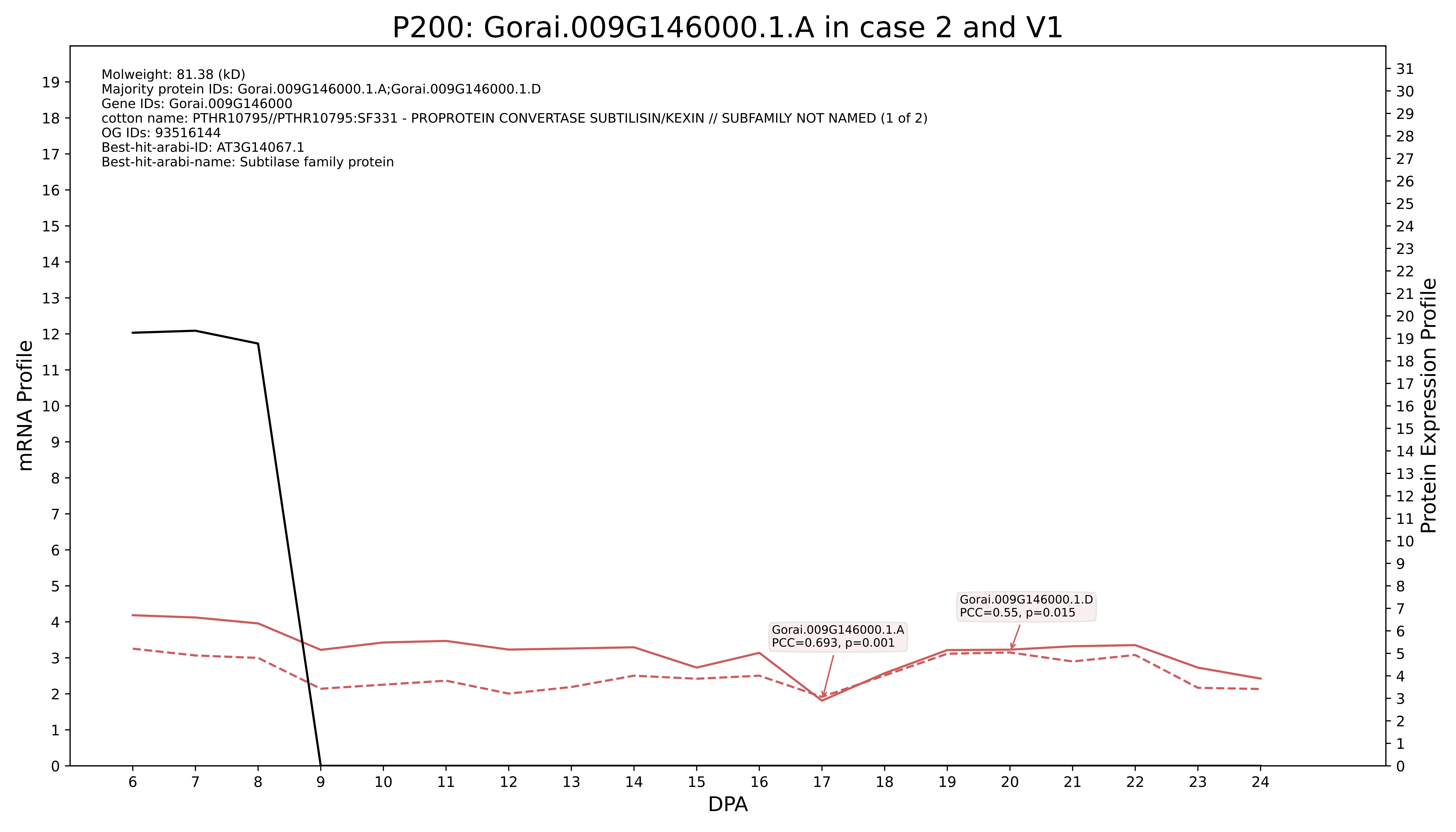
P200: Gorai.009G146000.1.A in case 2 and V1
Molweight: 81.38 (kD)
Majority protein IDs: Gorai.009G146000.1.A;Gorai.009G146000.1.D
Gene IDs: Gorai.009G146000
cotton name: PTHR10795//PTHR10795:SF331 - PROPROTEIN CONVERTASE SUBTILISIN/KEXIN // SUBFAMILY NOT NAMED (1 of 2)
OG IDs: 93516144
Best-hit-arabi-ID: AT3G14067.1
Best-hit-arabi-name: Subtilase family protein
Gorai.009G146000.1.A
PCC=0.693, p=0.001
Gorai.009G146000.1.D
PCC=0.55, p=0.015
mRNA Profile
Protein Expression Profile
DPA

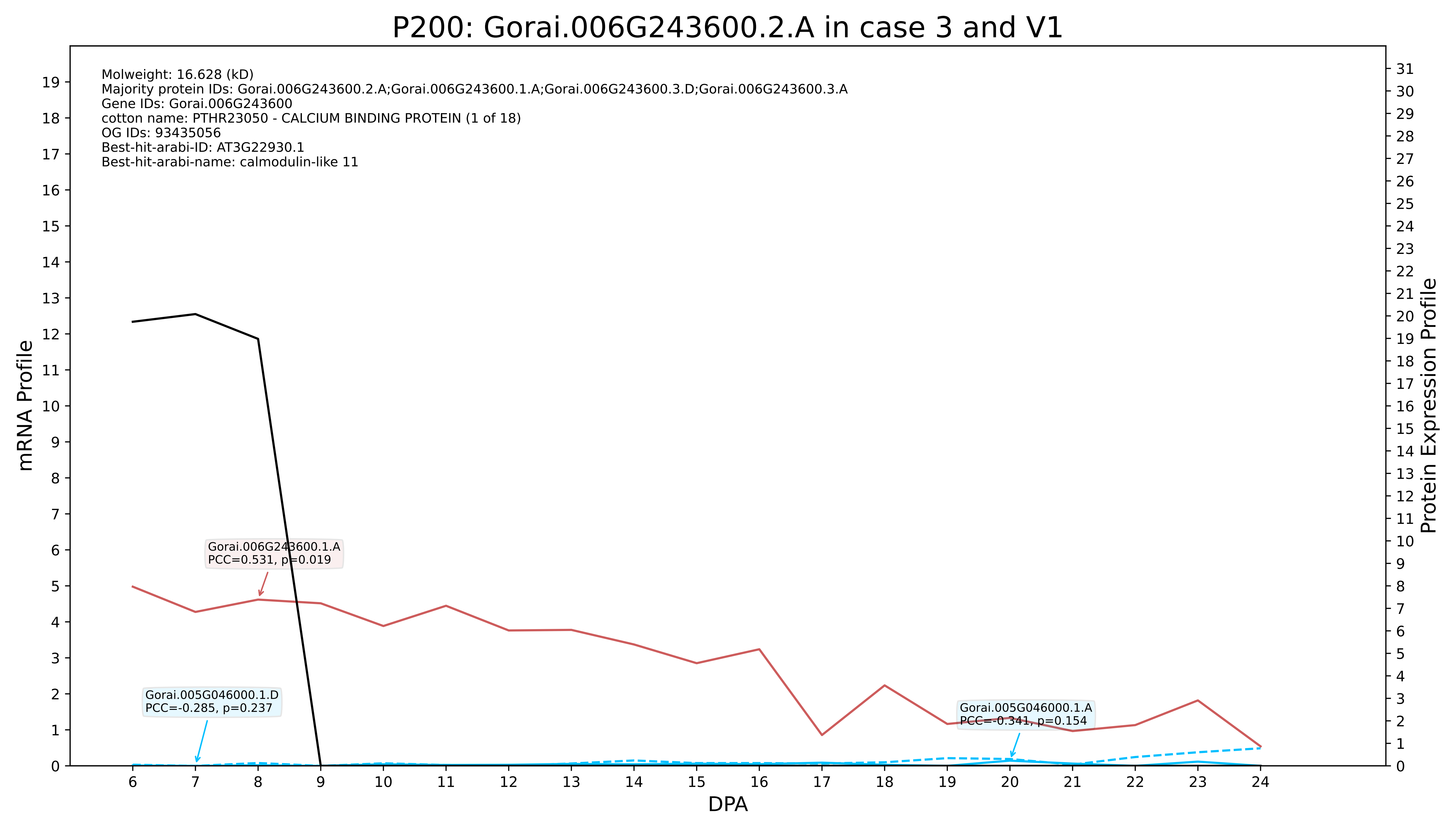
P200: Gorai.006G243600.2.A in case 3 and V1
Molweight: 16.628 (kD)
Majority protein IDs: Gorai.006G243600.2.A;Gorai.006G243600.1.A;Gorai.006G243600.3.D;Gorai.006G243600.3.A
Gene IDs: Gorai.006G243600
cotton name: PTHR23050 - CALCIUM BINDING PROTEIN (1 of 18)
OG IDs: 93435056
Best-hit-arabi-ID: AT3G22930.1
Best-hit-arabi-name: calmodulin-like 11
Gorai.006G243600.1.A
PCC=0.531, p=0.019
Gorai.005G046000.1.D
PCC=-0.285, p=0.237
Gorai.005G046000.1.A
PCC=-0.341, p=0.154
mRNA Profile
Protein Expression Profile
DPA

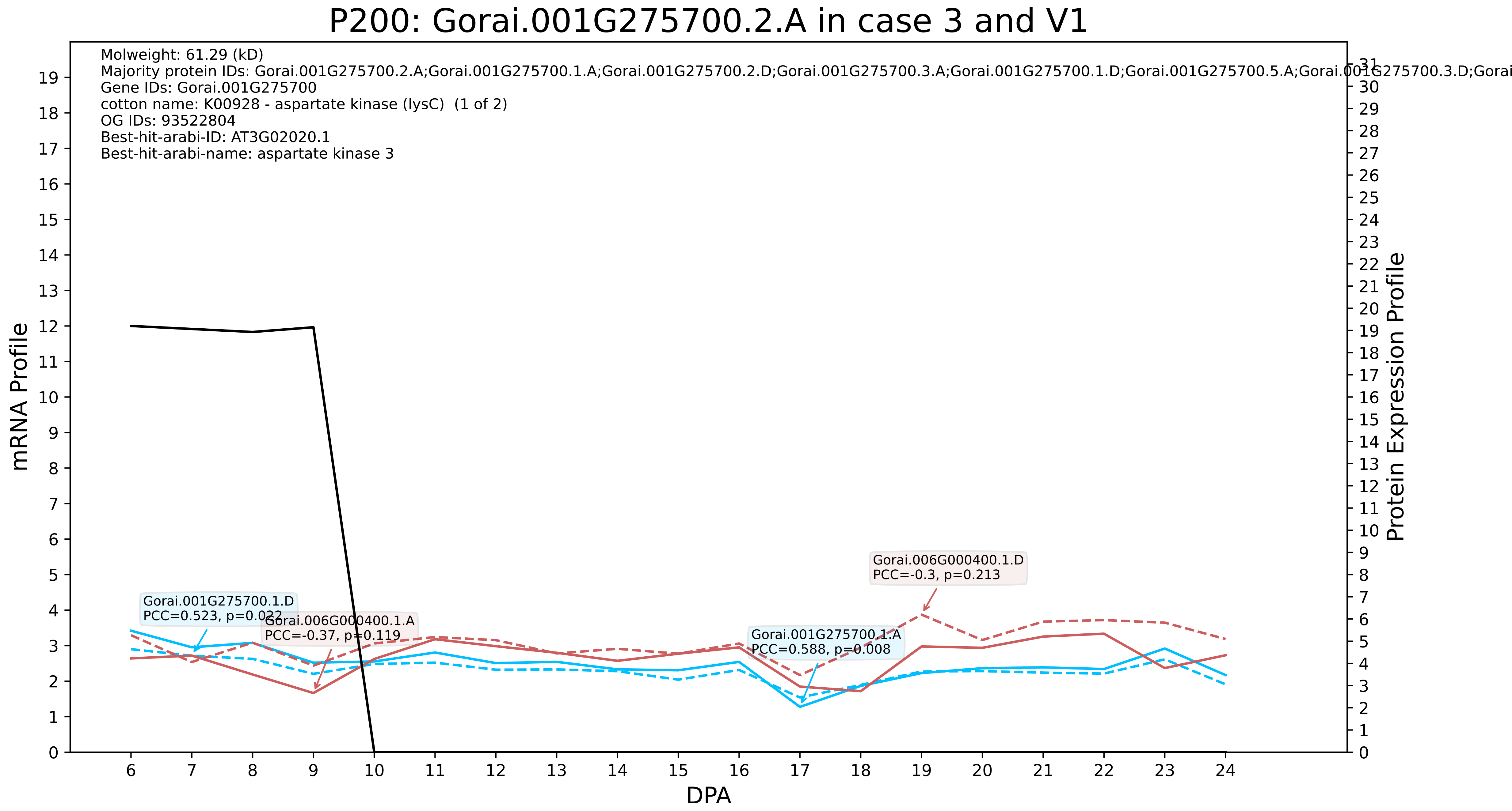
P200: Gorai.001G275700.2.A in case 3 and V1
Molweight: 61.29 (kD)
Majority protein IDs: Gorai.001G275700.2.A;Gorai.001G275700.1.A;Gorai.001G275700.2.D;Gorai.001G275700.3.A;Gorai.001G275700.1.D;Gorai.001G275700.5.A;Gorai.001G275700.3.D;Gorai
Gene IDs: Gorai.001G275700
cotton name: K00928 - aspartate kinase (lysC) (1 of 2)
OG IDs: 93522804
Best-hit-arabi-ID: AT3G02020.1
Best-hit-arabi-name: aspartate kinase 3
Gorai.001G275700.1.D
PCC=0.523, p=0.022
Gorai.006G000400.1.A
PCC=-0.37, p=0.119
Gorai.001G275700.1.A
PCC=0.588, p=0.008
Gorai.006G000400.1.D
PCC=-0.3, p=0.213
mRNA Profile
Protein Expression Profile
DPA

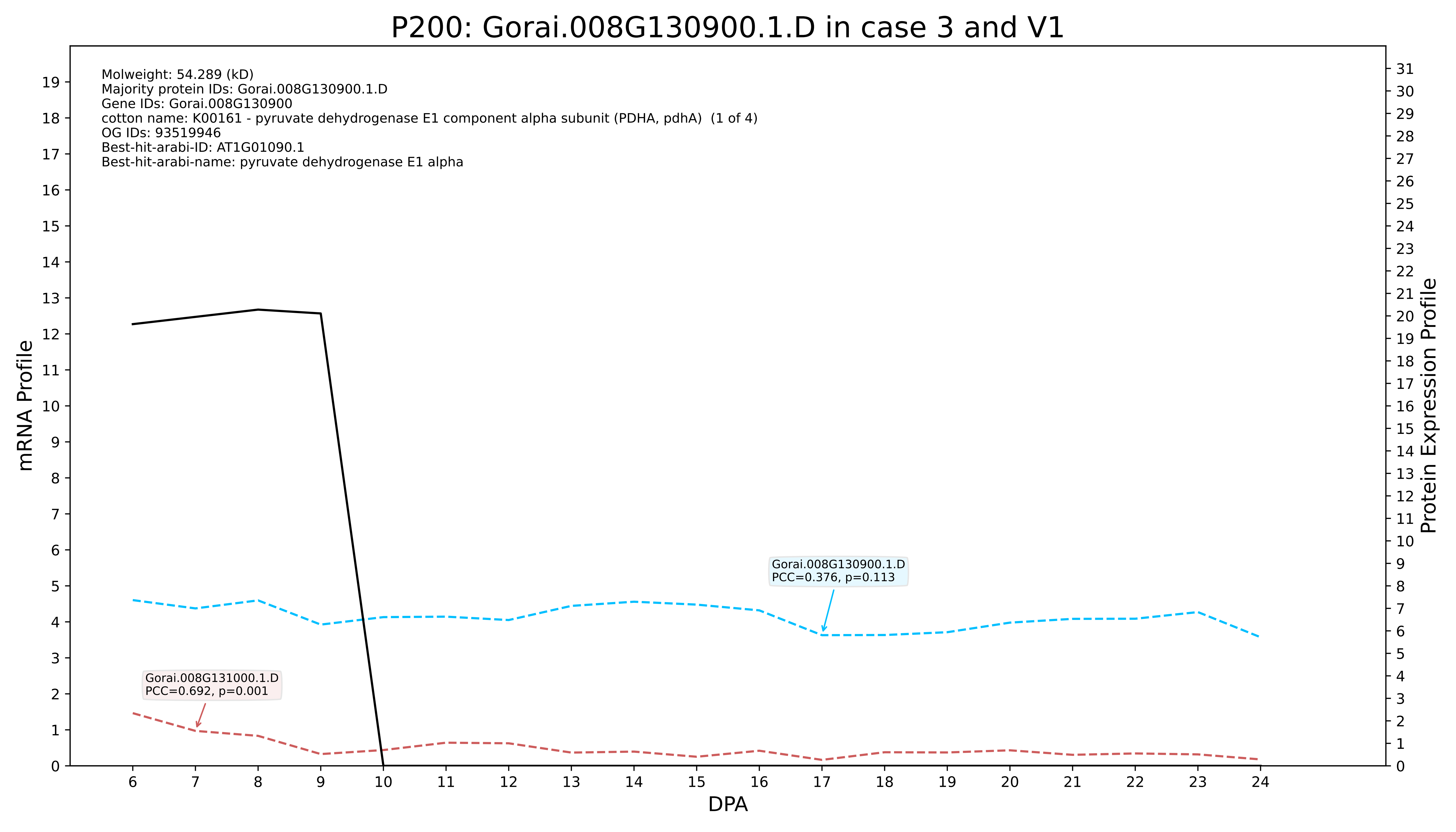

P200: Gorai.008G130900.1.D in case 3 and V1
Molweight: 54.289 (kD)
Majority protein IDs: Gorai.008G130900.1.D
Gene IDs: Gorai.008G130900
cotton name: K00161 - pyruvate dehydrogenase E1 component alpha subunit (PDHA, pdhA) (1 of 4)
OG IDs: 93519946
Best-hit-arabi-ID: AT1G01090.1
Best-hit-arabi-name: pyruvate dehydrogenase E1 alpha
Gorai.008G130900.1.D
PCC=0.376, p=0.113
Gorai.008G131000.1.D
PCC=0.692, p=0.001
mRNA Profile
Protein Expression Profile
DPA

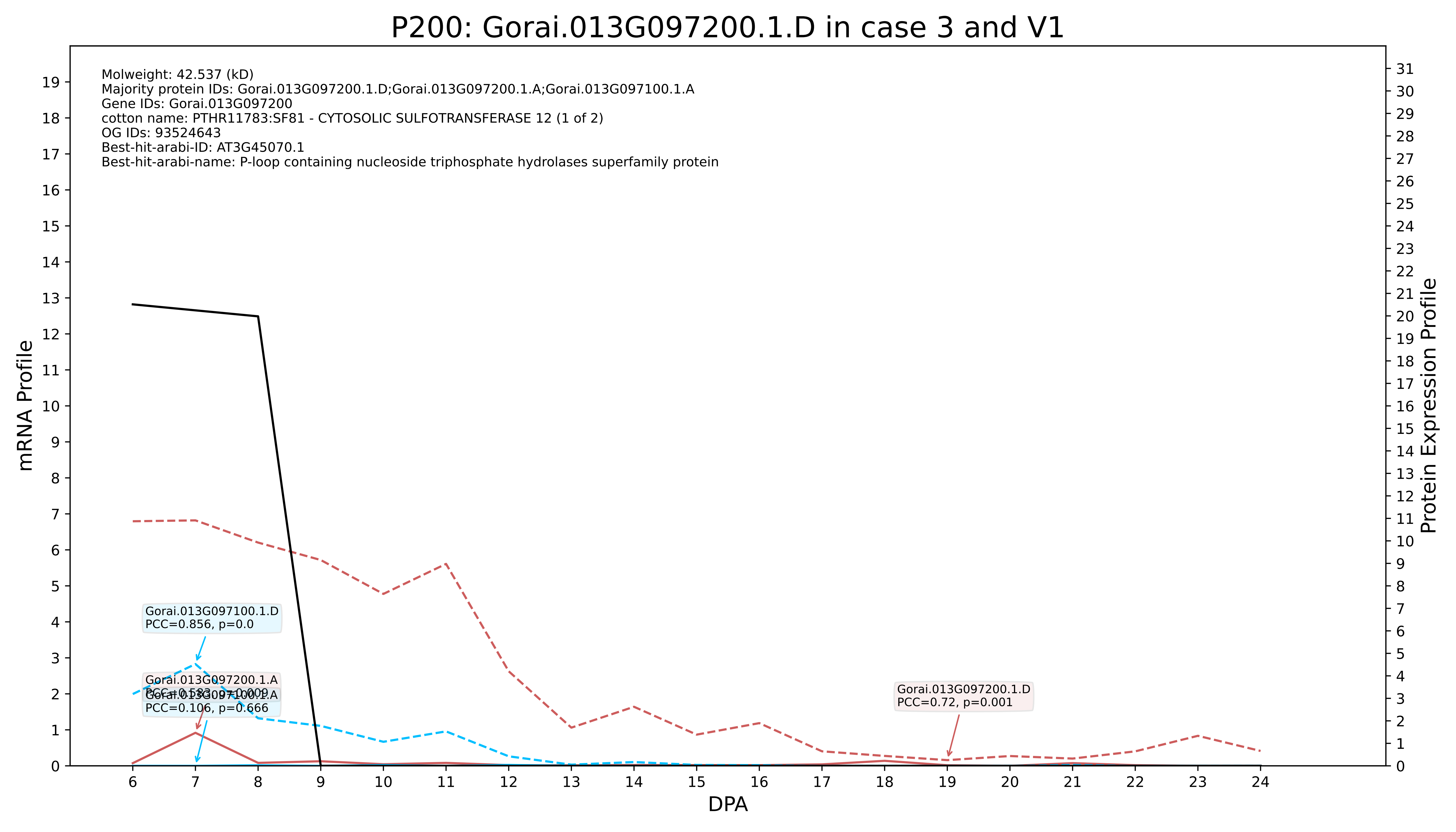
P200: Gorai.013G097200.1.D in case 3 and V1
Molweight: 42.537 (kD)
Majority protein IDs: Gorai.013G097200.1.D;Gorai.013G097200.1.A;Gorai.013G097100.1.A
Gene IDs: Gorai.013G097200
cotton name: PTHR11783:SF81 - CYTOSOLIC SULFOTRANSFERASE 12 (1 of 2)
OG IDs: 93524643
Best-hit-arabi-ID: AT3G45070.1
Best-hit-arabi-name: P-loop containing nucleoside triphosphate hydrolases superfamily protein
Gorai.013G097100.1.D
PCC=0.856, p=0.0
Gorai.013G097200.1.A
Gorai.013G097100.1.A
PCC=0.106, p=0.666
Gorai.013G097200.1.D
PCC=0.72, p=0.001
mRNA Profile
Protein Expression Profile
DPA

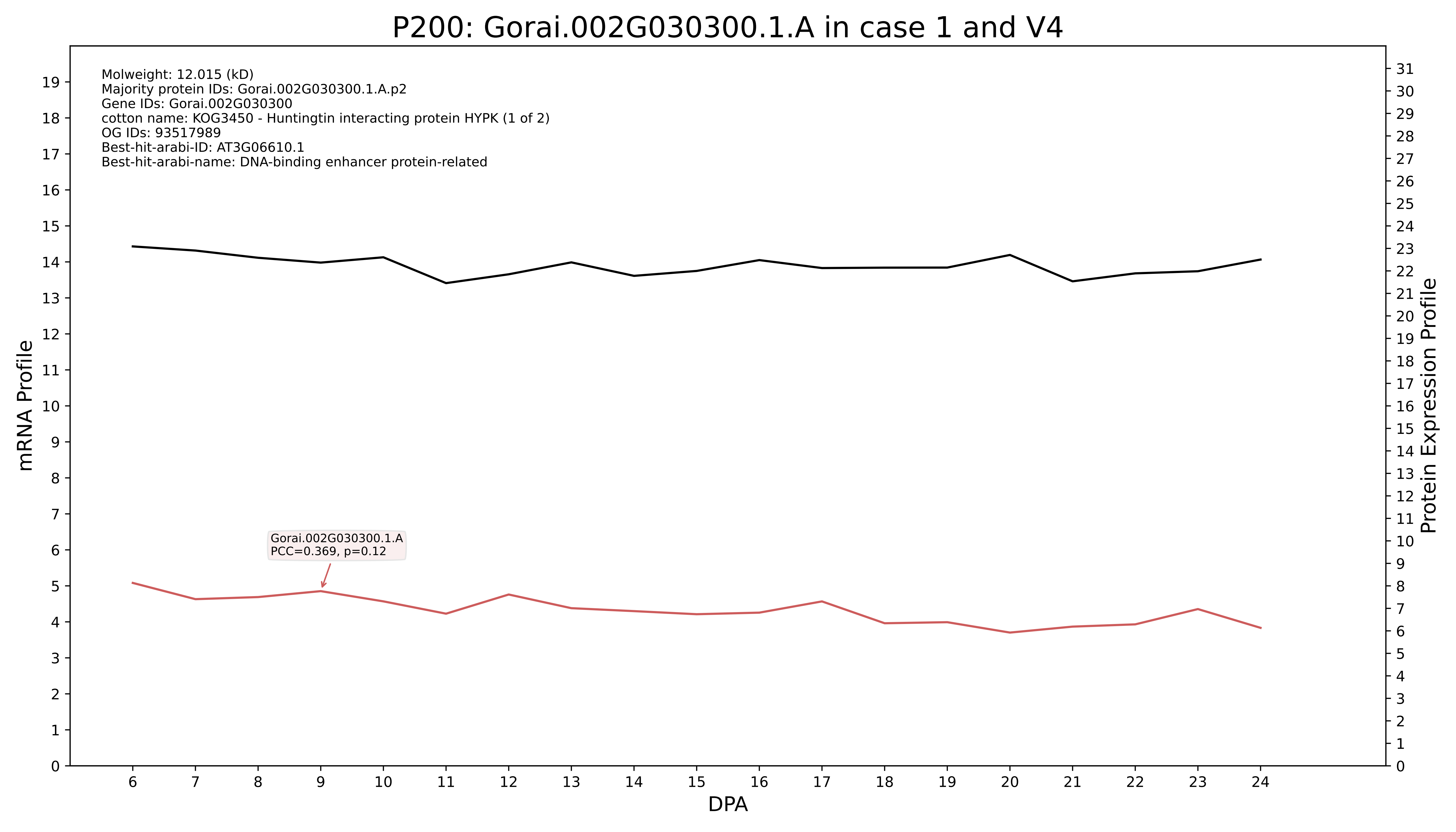

P200: Gorai.002G030300.1.A in case 1 and V4
Molweight: 12.015 (kD)
Majority protein IDs: Gorai.002G030300.1.A.p2
Gene IDs: Gorai.002G030300
cotton name: KOG3450 - Huntingtin interacting protein HYPK (1 of 2)
OG IDs: 93517989
Best-hit-arabi-ID: AT3G06610.1
Best-hit-arabi-name: DNA-binding enhancer protein-related
Gorai.002G030300.1.A
PCC=0.369, p=0.12
mRNA Profile
Protein Expression Profile
DPA

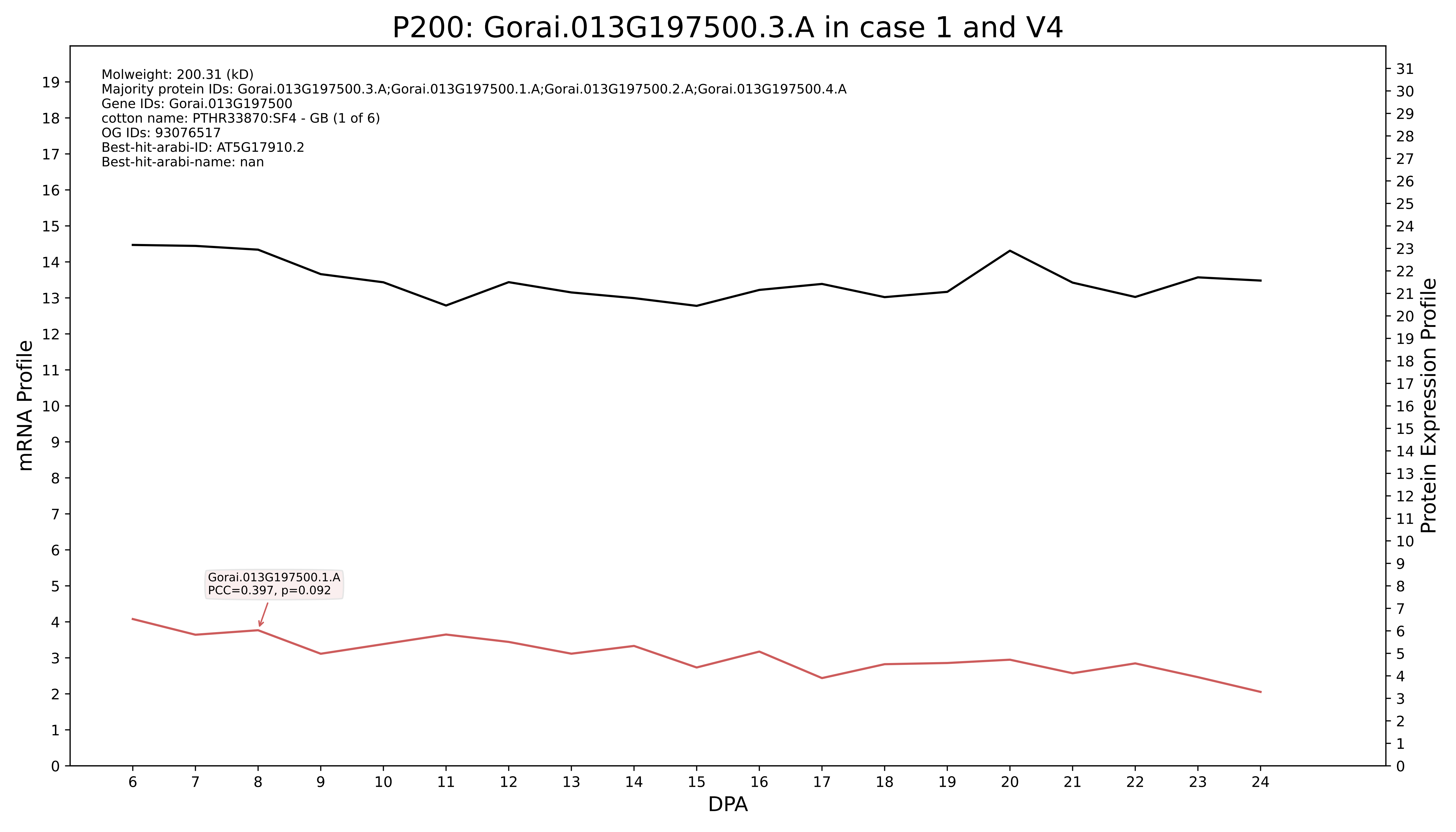
P200: Gorai.013G197500.3.A in case 1 and V4
Molweight: 200.31 (kD)
Majority protein IDs: Gorai.013G197500.3.A;Gorai.013G197500.1.A;Gorai.013G197500.2.A;Gorai.013G197500.4.A
Gene IDs: Gorai.013G197500
cotton name: PTHR33870:SF4 - GB (1 of 6)
OG IDs: 93076517
Best-hit-arabi-ID: AT5G17910.2
Best-hit-arabi-name: nan
Gorai.013G197500.1.A
PCC=0.397, p=0.092
mRNA Profile
Protein Expression Profile
DPA

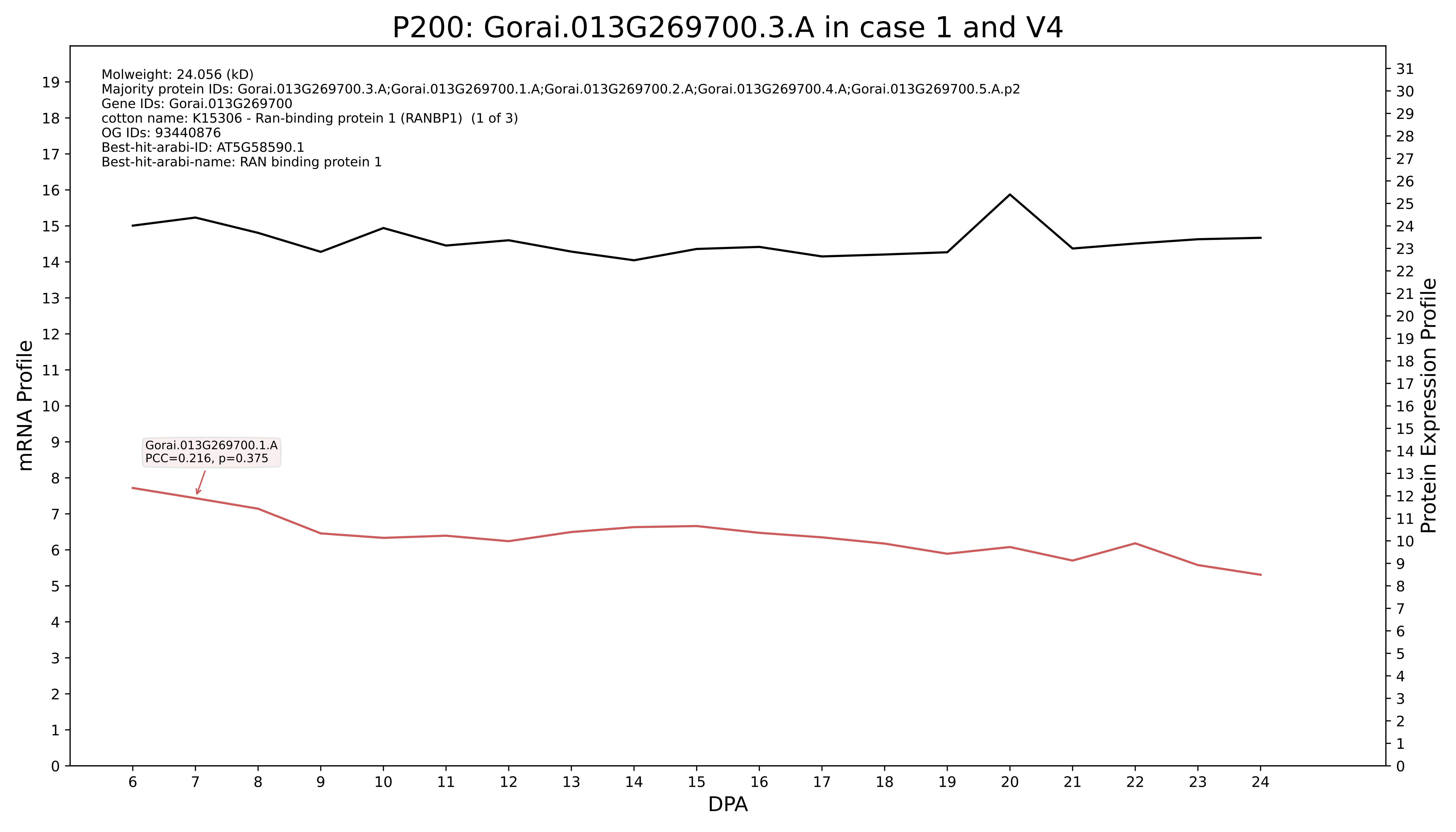
P200: Gorai.013G269700.3.A in case 1 and V4
Molweight: 24.056 (kD)
Majority protein IDs: Gorai.013G269700.3.A;Gorai.013G269700.1.A;Gorai.013G269700.2.A;Gorai.013G269700.4.A;Gorai.013G269700.5.A.p2
Gene IDs: Gorai.013G269700
cotton name: K15306 - Ran-binding protein 1 (RANBP1) (1 of 3)
OG IDs: 93440876
Best-hit-arabi-ID: AT5G58590.1
Best-hit-arabi-name: RAN binding protein 1
Gorai.013G269700.1.A
PCC=0.216, p=0.375
mRNA Profile
Protein Expression Profile
DPA

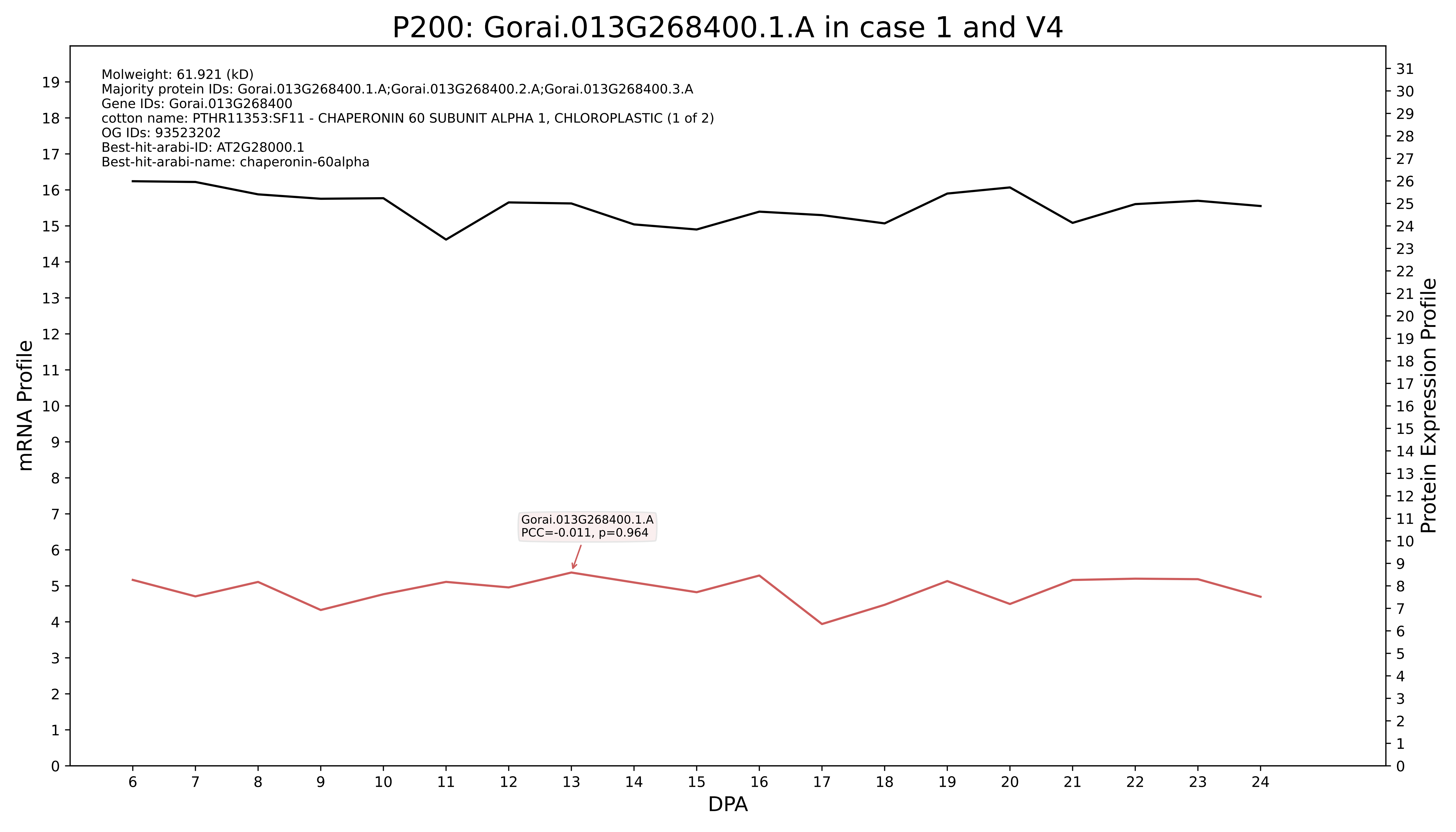
P200: Gorai.013G268400.1.A in case 1 and V4
Molweight: 61.921 (kD)
Majority protein IDs: Gorai.013G268400.1.A;Gorai.013G268400.2.A;Gorai.013G268400.3.A
Gene IDs: Gorai.013G268400
cotton name: PTHR11353:SF11 - CHAPERONIN 60 SUBUNIT ALPHA 1, CHLOROPLASTIC (1 of 2)
OG IDs: 93523202
Best-hit-arabi-ID: AT2G28000.1
Best-hit-arabi-name: chaperonin-60alpha
Gorai.013G268400.1.A
PCC=-0.011, p=0.964
mRNA Profile
Protein Expression Profile
DPA

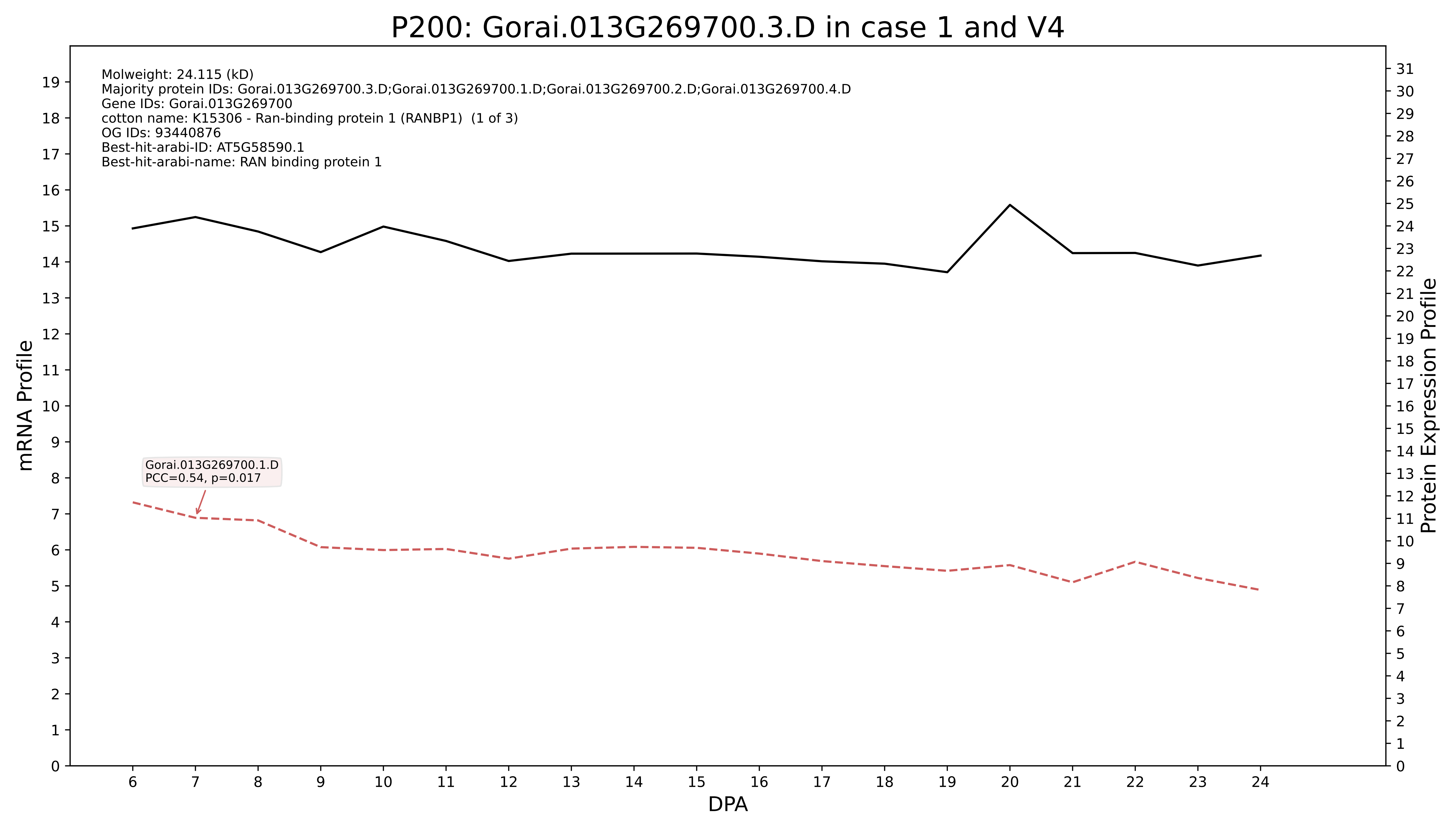
P200: Gorai.013G269700.3.D in case 1 and V4
Molweight: 24.115 (kD)
Majority protein IDs: Gorai.013G269700.3.D;Gorai.013G269700.1.D;Gorai.013G269700.2.D;Gorai.013G269700.4.D
Gene IDs: Gorai.013G269700
cotton name: K15306 - Ran-binding protein 1 (RANBP1) (1 of 3)
OG IDs: 93440876
Best-hit-arabi-ID: AT5G58590.1
Best-hit-arabi-name: RAN binding protein 1
Gorai.013G269700.1.D
PCC=0.54, p=0.017
mRNA Profile
Protein Expression Profile
DPA

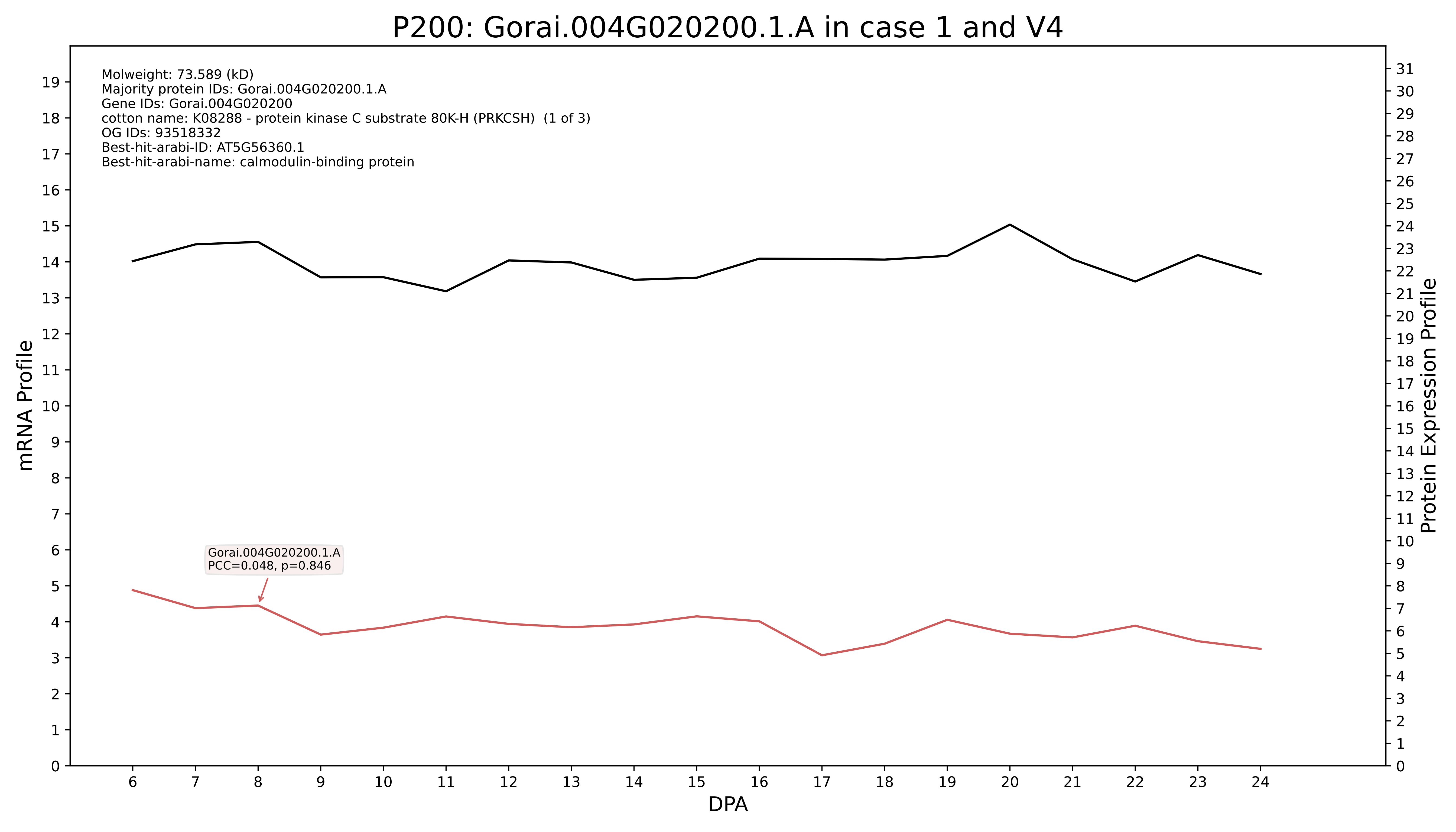

P200: Gorai.004G020200.1.A in case 1 and V4
Molweight: 73.589 (kD)
Majority protein IDs: Gorai.004G020200.1.A
Gene IDs: Gorai.004G020200
cotton name: K08288 - protein kinase C substrate 80K-H (PRKCSH) (1 of 3)
OG IDs: 93518332
Best-hit-arabi-ID: AT5G56360.1
Best-hit-arabi-name: calmodulin-binding protein
Gorai.004G020200.1.A
PCC=0.048, p=0.846
mRNA Profile
Protein Expression Profile
DPA

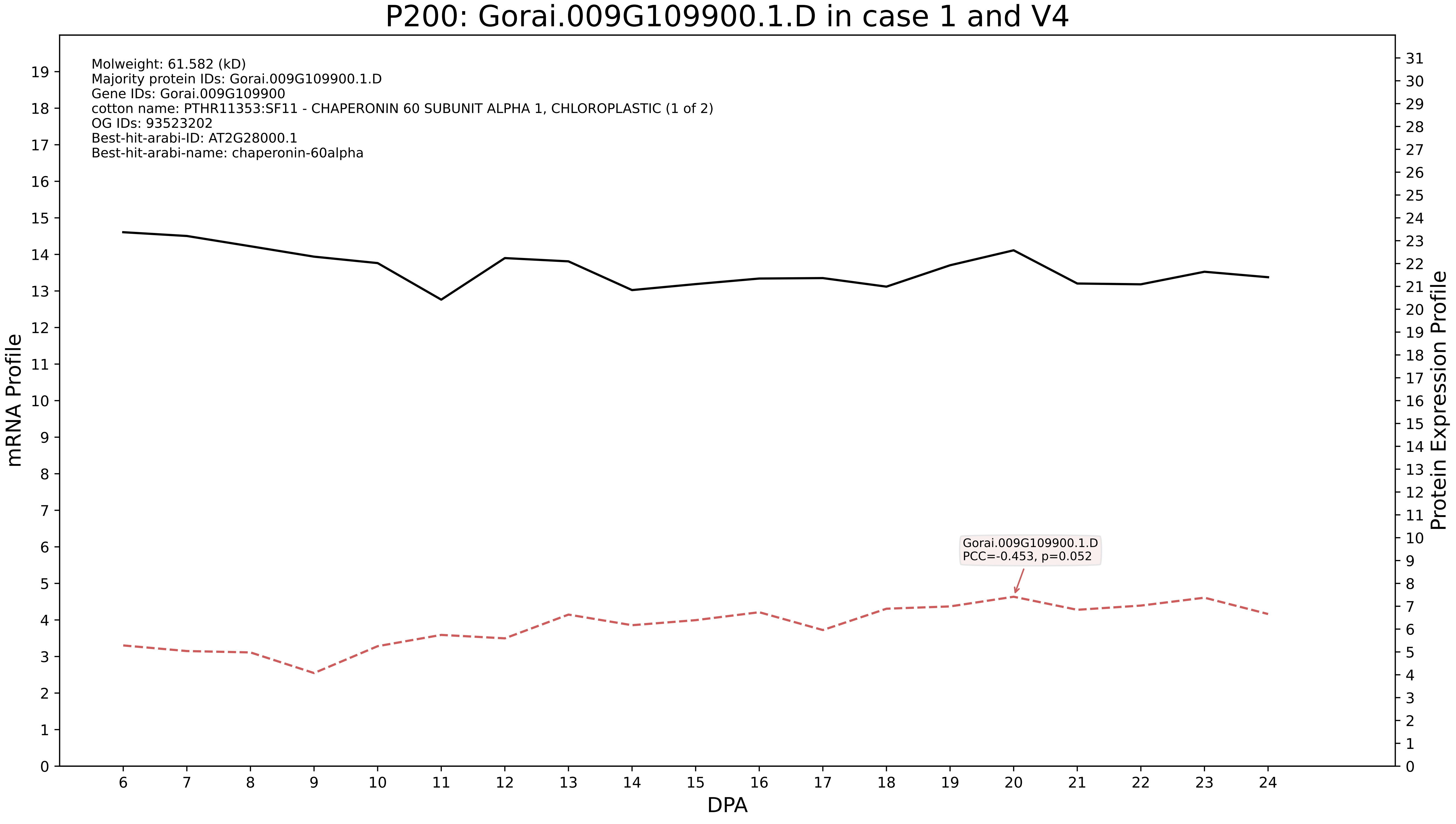
P200: Gorai.009G109900.1.D in case 1 and V4
Molweight: 61.582 (kD)
Majority protein IDs: Gorai.009G109900.1.D
Gene IDs: Gorai.009G109900
cotton name: PTHR11353:SF11 - CHAPERONIN 60 SUBUNIT ALPHA 1, CHLOROPLASTIC (1 of 2)
OG IDs: 93523202
Best-hit-arabi-ID: AT2G28000.1
Best-hit-arabi-name: chaperonin-60alpha
Gorai.009G109900.1.D
PCC=-0.453, p=0.052
mRNA Profile
Protein Expression Profile
DPA

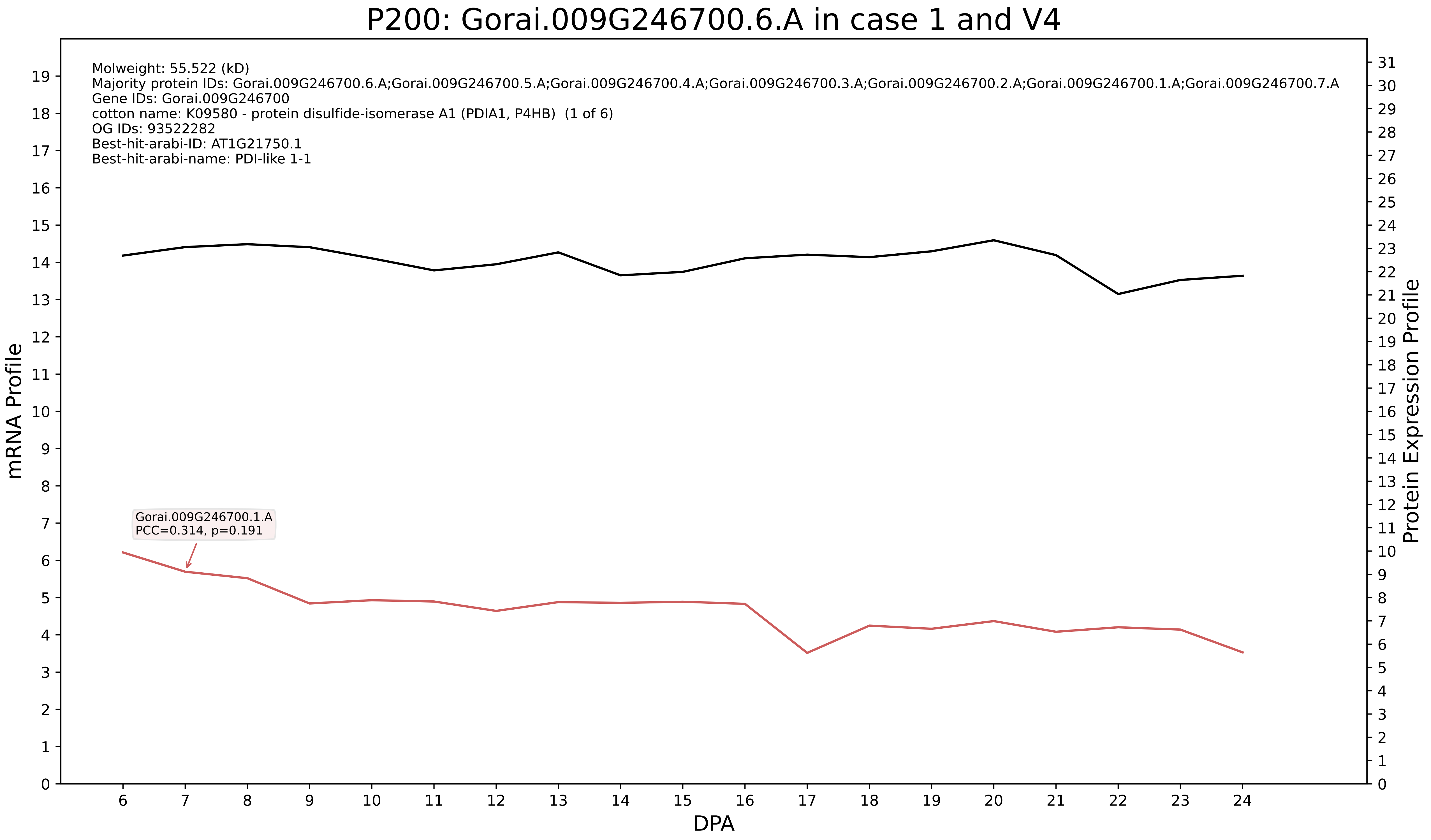

P200: Gorai.009G246700.6.A in case 1 and V4
Molweight: 55.522 (kD)
Majority protein IDs: Gorai.009G246700.6.A;Gorai.009G246700.5.A;Gorai.009G246700.4.A;Gorai.009G246700.3.A;Gorai.009G246700.2.A;Gorai.009G246700.1.A;Gorai.009G246700.7.A
Gene IDs: Gorai.009G246700
cotton name: K09580 - protein disulfide-isomerase A1 (PDIA1, P4HB) (1 of 6)
OG IDs: 93522282
Best-hit-arabi-ID: AT1G21750.1
Best-hit-arabi-name: PDI-like 1-1
Gorai.009G246700.1.A
PCC=0.314, p=0.191
mRNA Profile
Protein Expression Profile
DPA

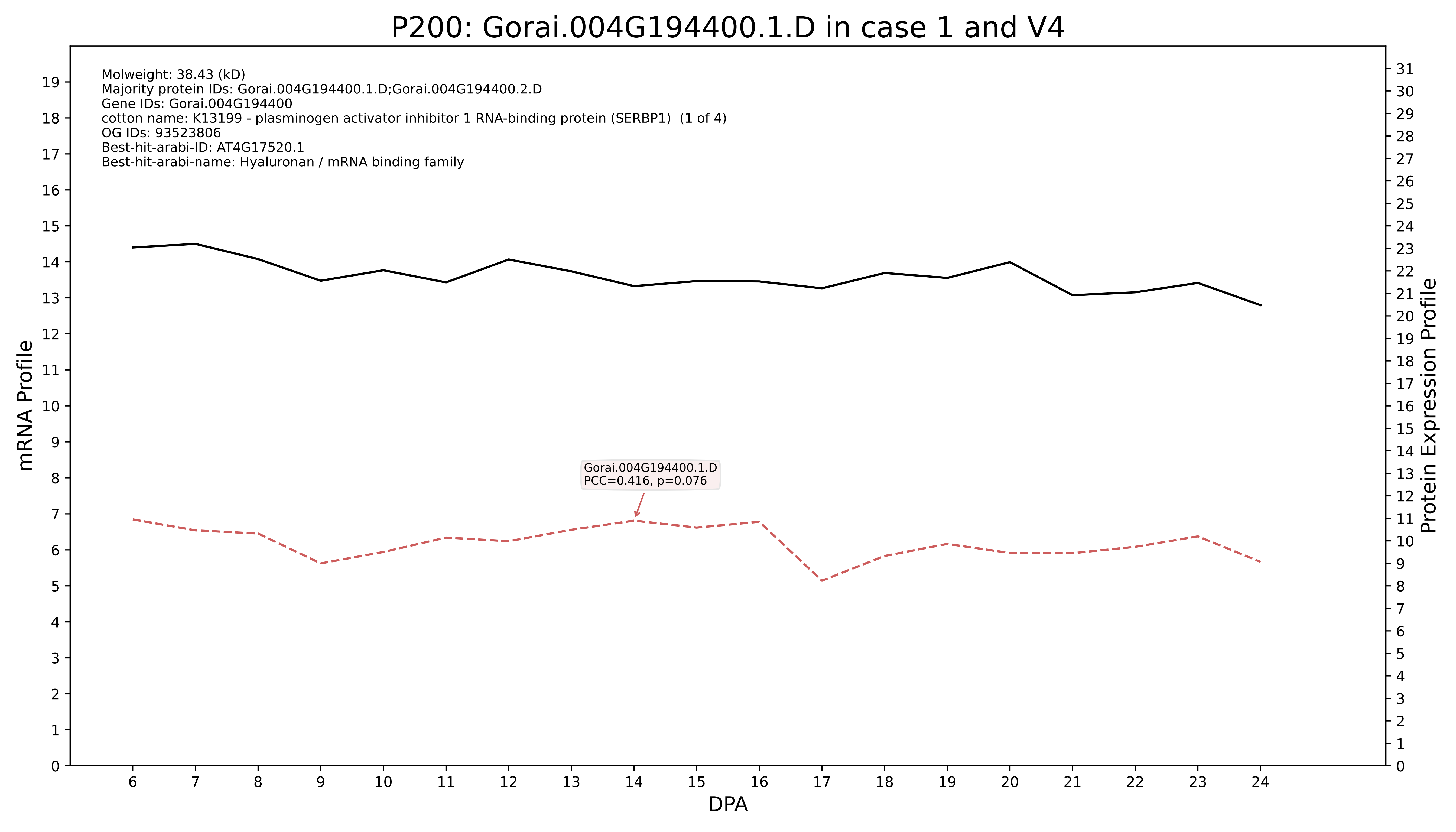
P200: Gorai.004G194400.1.D in case 1 and V4
Molweight: 38.43 (kD)
Majority protein IDs: Gorai.004G194400.1.D;Gorai.004G194400.2.D
Gene IDs: Gorai.004G194400
cotton name: K13199 - plasminogen activator inhibitor 1 RNA-binding protein (SERBP1) (1 of 4)
OG IDs: 93523806
Best-hit-arabi-ID: AT4G17520.1
Best-hit-arabi-name: Hyaluronan / mRNA binding family
Gorai.004G194400.1.D
PCC=0.416, p=0.076
mRNA Profile
Protein Expression Profile
DPA

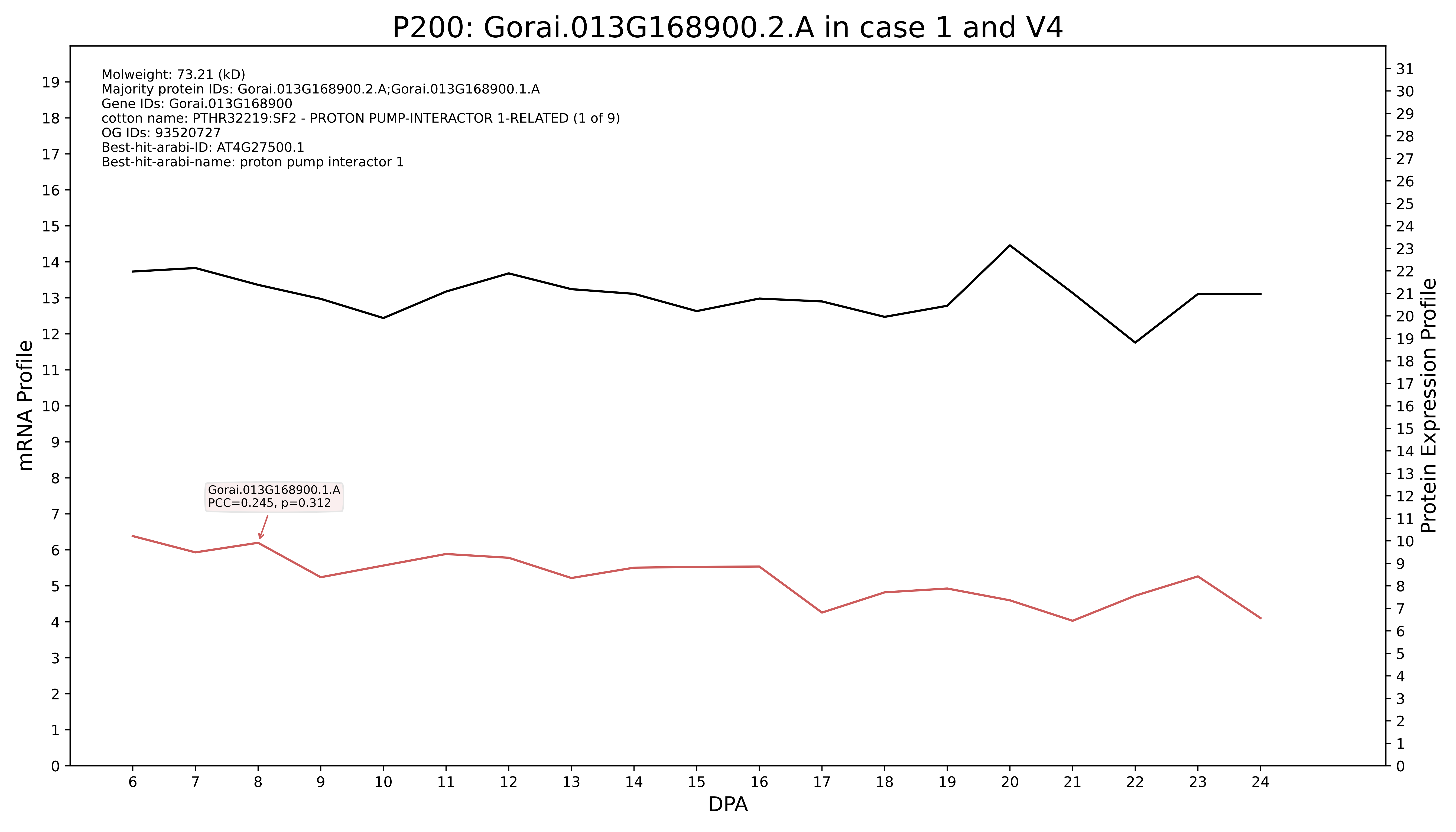
P200: Gorai.013G168900.2.A in case 1 and V4
Molweight: 73.21 (kD)
Majority protein IDs: Gorai.013G168900.2.A;Gorai.013G168900.1.A
Gene IDs: Gorai.013G168900
cotton name: PTHR32219:SF2 - PROTON PUMP-INTERACTOR 1-RELATED (1 of 9)
OG IDs: 93520727
Best-hit-arabi-ID: AT4G27500.1
Best-hit-arabi-name: proton pump interactor 1
Gorai.013G168900.1.A
PCC=0.245, p=0.312
mRNA Profile
Protein Expression Profile
DPA

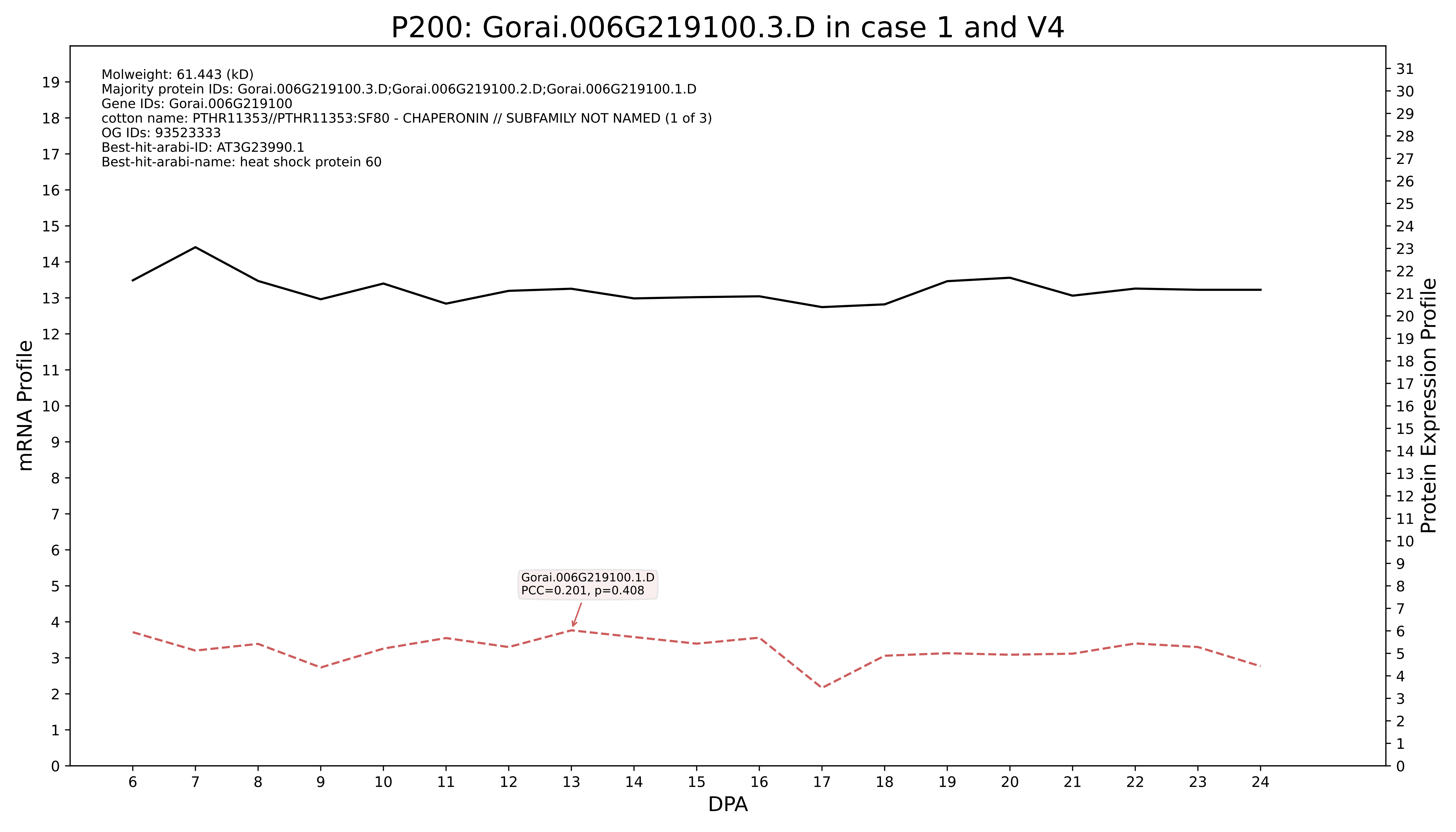
P200: Gorai.006G219100.3.D in case 1 and V4
Molweight: 61.443 (kD)
Majority protein IDs: Gorai.006G219100.3.D;Gorai.006G219100.2.D;Gorai.006G219100.1.D
Gene IDs: Gorai.006G219100
cotton name: PTHR11353//PTHR11353:SF80 - CHAPERONIN // SUBFAMILY NOT NAMED (1 of 3)
OG IDs: 93523333
Best-hit-arabi-ID: AT3G23990.1
Best-hit-arabi-name: heat shock protein 60
Gorai.006G219100.1.D
PCC=0.201, p=0.408
mRNA Profile
Protein Expression Profile
DPA

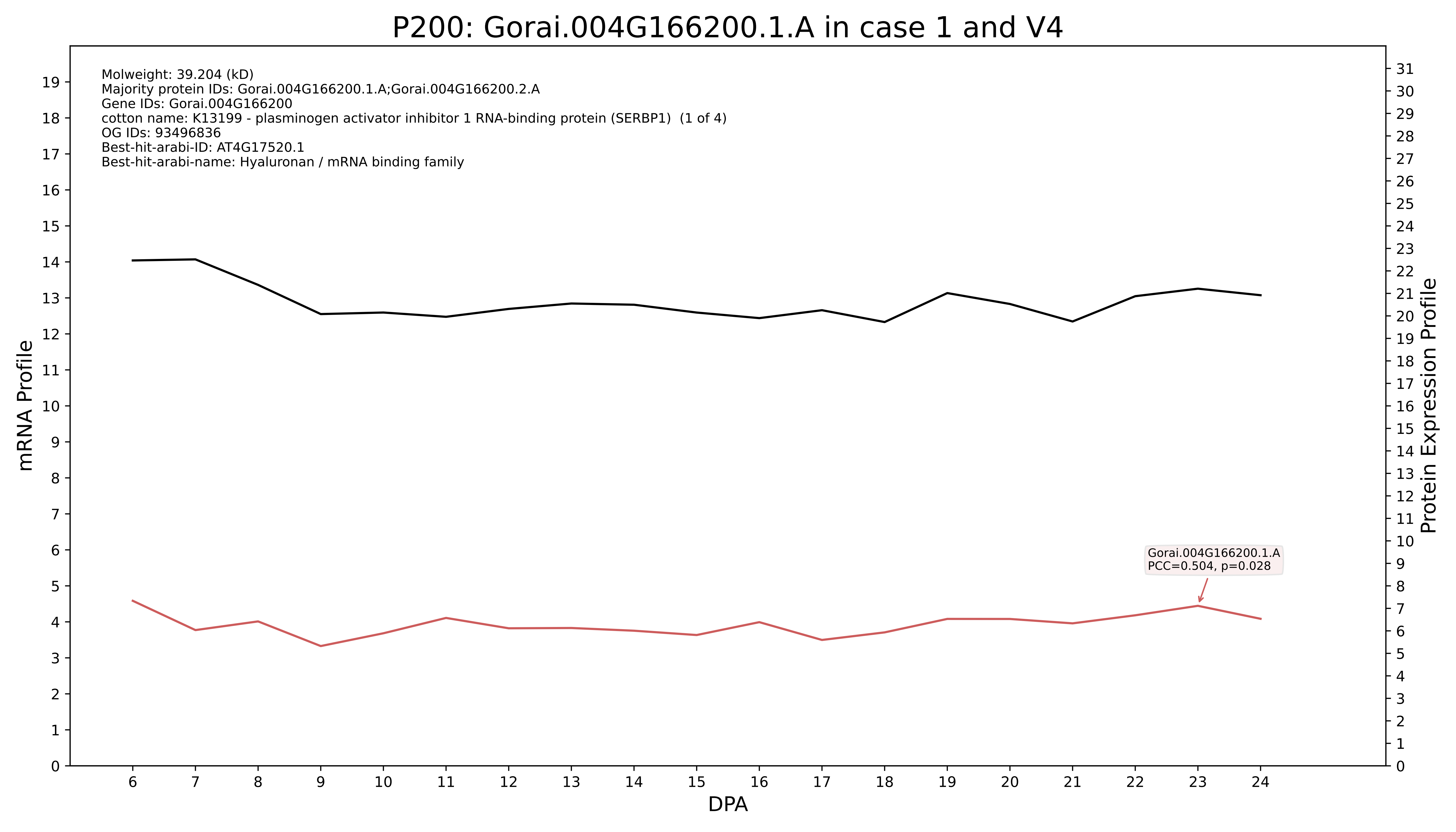
P200: Gorai.004G166200.1.A in case 1 and V4
Molweight: 39.204 (kD)
Majority protein IDs: Gorai.004G166200.1.A;Gorai.004G166200.2.A
Gene IDs: Gorai.004G166200
cotton name: K13199 - plasminogen activator inhibitor 1 RNA-binding protein (SERBP1) (1 of 4)
OG IDs: 93496836
Best-hit-arabi-ID: AT4G17520.1
Best-hit-arabi-name: Hyaluronan / mRNA binding family
Gorai.004G166200.1.A
PCC=0.504, p=0.028
mRNA Profile
Protein Expression Profile
DPA

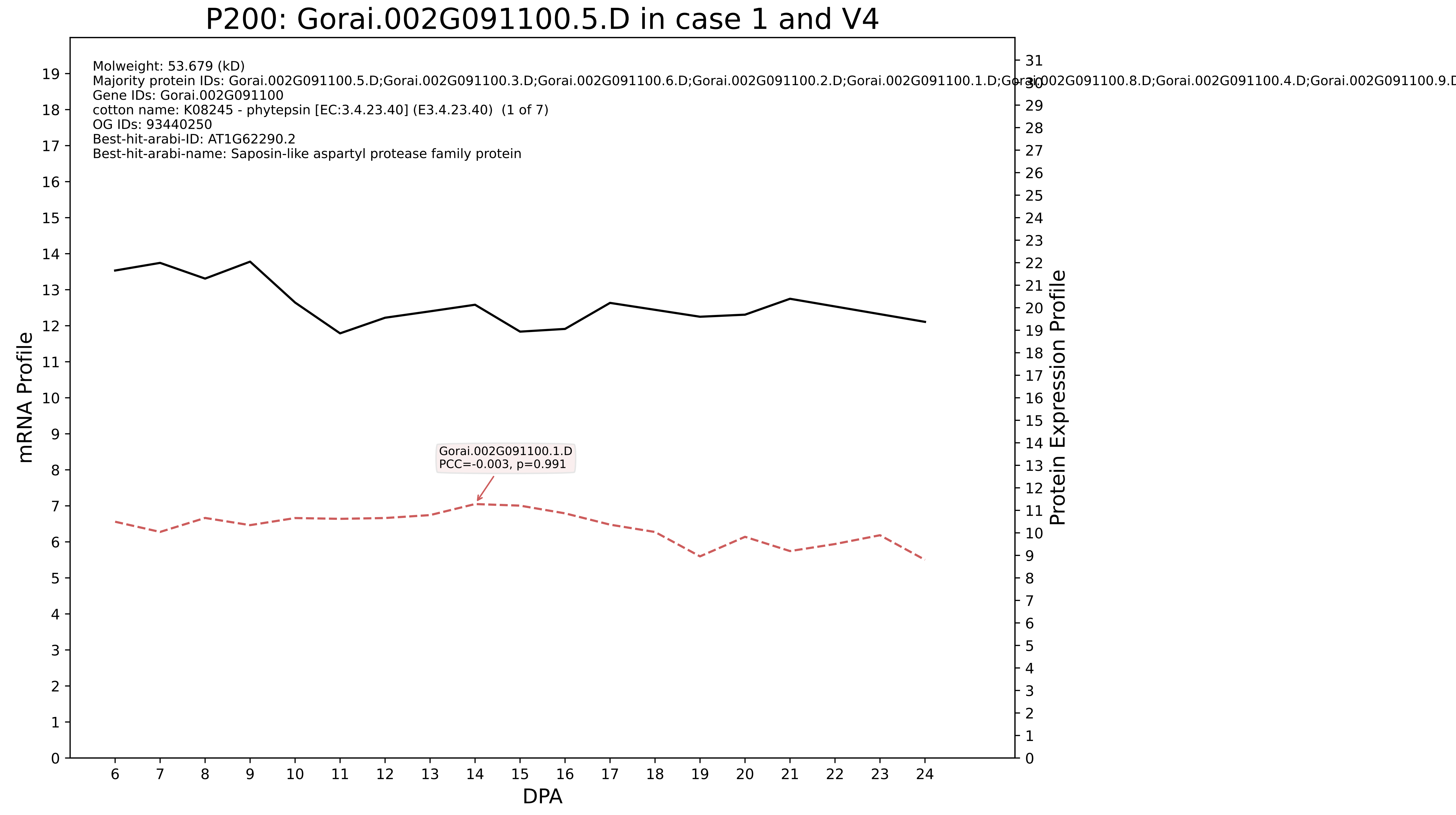

P200: Gorai.002G091100.5.D in case 1 and V4
Molweight: 53.679 (kD)
Majority protein IDs: Gorai.002G091100.5.D;Gorai.002G091100.3.D;Gorai.002G091100.6.D;Gorai.002G091100.2.D;Gorai.002G091100.1.D;Gorai.002G091100.8.D;Gorai.002G091100.4.D;Gorai.002G091100.9.D
Gene IDs: Gorai.002G091100
cotton name: K08245 - phytepsin [EC:3.4.23.40] (E3.4.23.40) (1 of 7)
OG IDs: 93440250
Best-hit-arabi-ID: AT1G62290.2
Best-hit-arabi-name: Saposin-like aspartyl protease family protein
Gorai.002G091100.1.D
PCC=-0.003, p=0.991
mRNA Profile
Protein Expression Profile
DPA

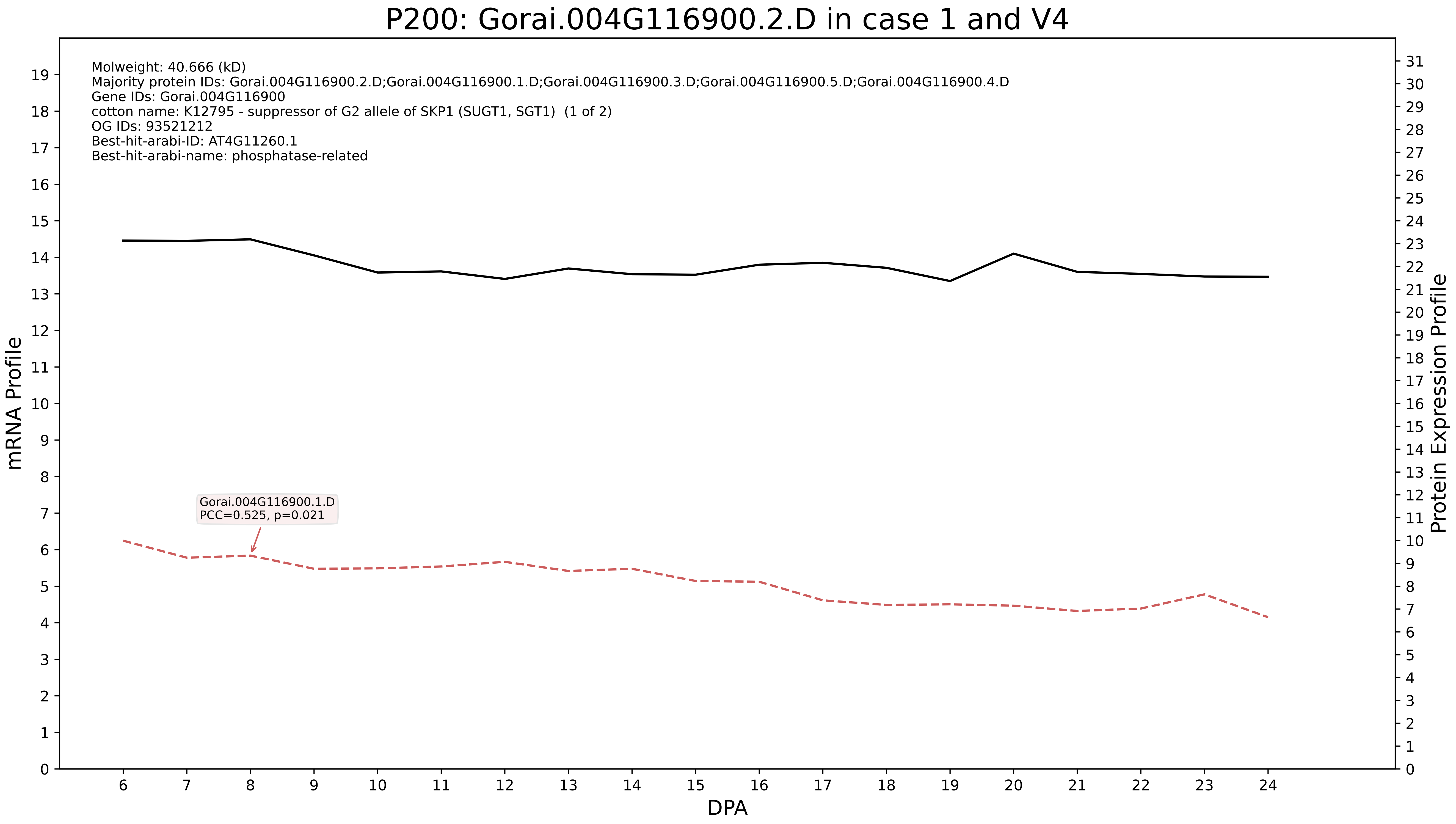
P200: Gorai.004G116900.2.D in case 1 and V4
Molweight: 40.666 (kD)
Majority protein IDs: Gorai.004G116900.2.D;Gorai.004G116900.1.D;Gorai.004G116900.3.D;Gorai.004G116900.5.D;Gorai.004G116900.4.D
Gene IDs: Gorai.004G116900
cotton name: K12795 - suppressor of G2 allele of SKP1 (SUGT1, SGT1) (1 of 2)
OG IDs: 93521212
Best-hit-arabi-ID: AT4G11260.1
Best-hit-arabi-name: phosphatase-related
Gorai.004G116900.1.D
PCC=0.525, p=0.021
mRNA Profile
Protein Expression Profile
DPA

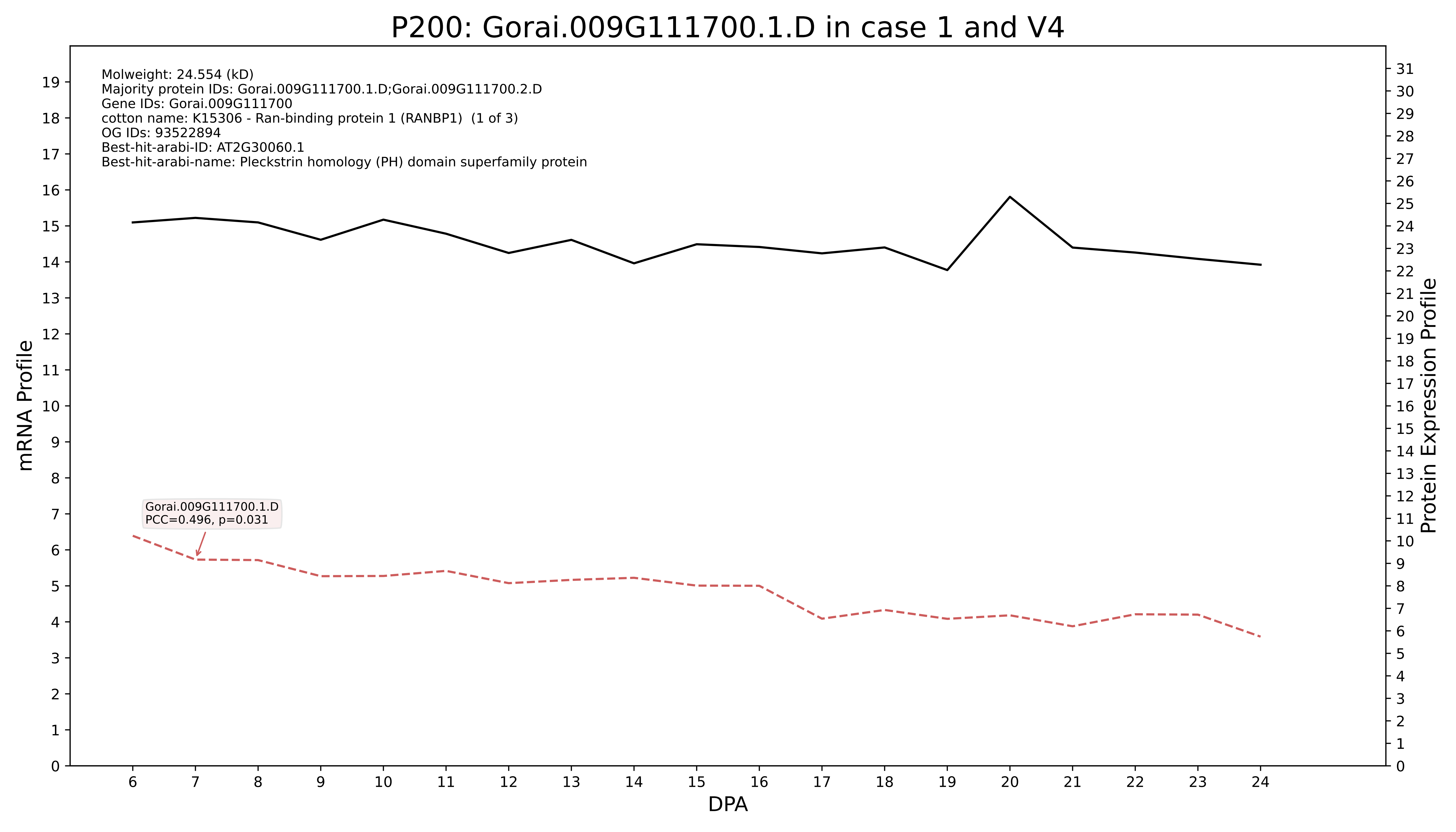
P200: Gorai.009G111700.1.D in case 1 and V4
Molweight: 24.554 (kD)
Majority protein IDs: Gorai.009G111700.1.D;Gorai.009G111700.2.D
Gene IDs: Gorai.009G111700
cotton name: K15306 - Ran-binding protein 1 (RANBP1) (1 of 3)
OG IDs: 93522894
Best-hit-arabi-ID: AT2G30060.1
Best-hit-arabi-name: Pleckstrin homology (PH) domain superfamily protein
Gorai.009G111700.1.D
PCC=0.496, p=0.031
mRNA Profile
Protein Expression Profile
DPA

### P200: Gorai.012G082700.1.D in case 1 and V4

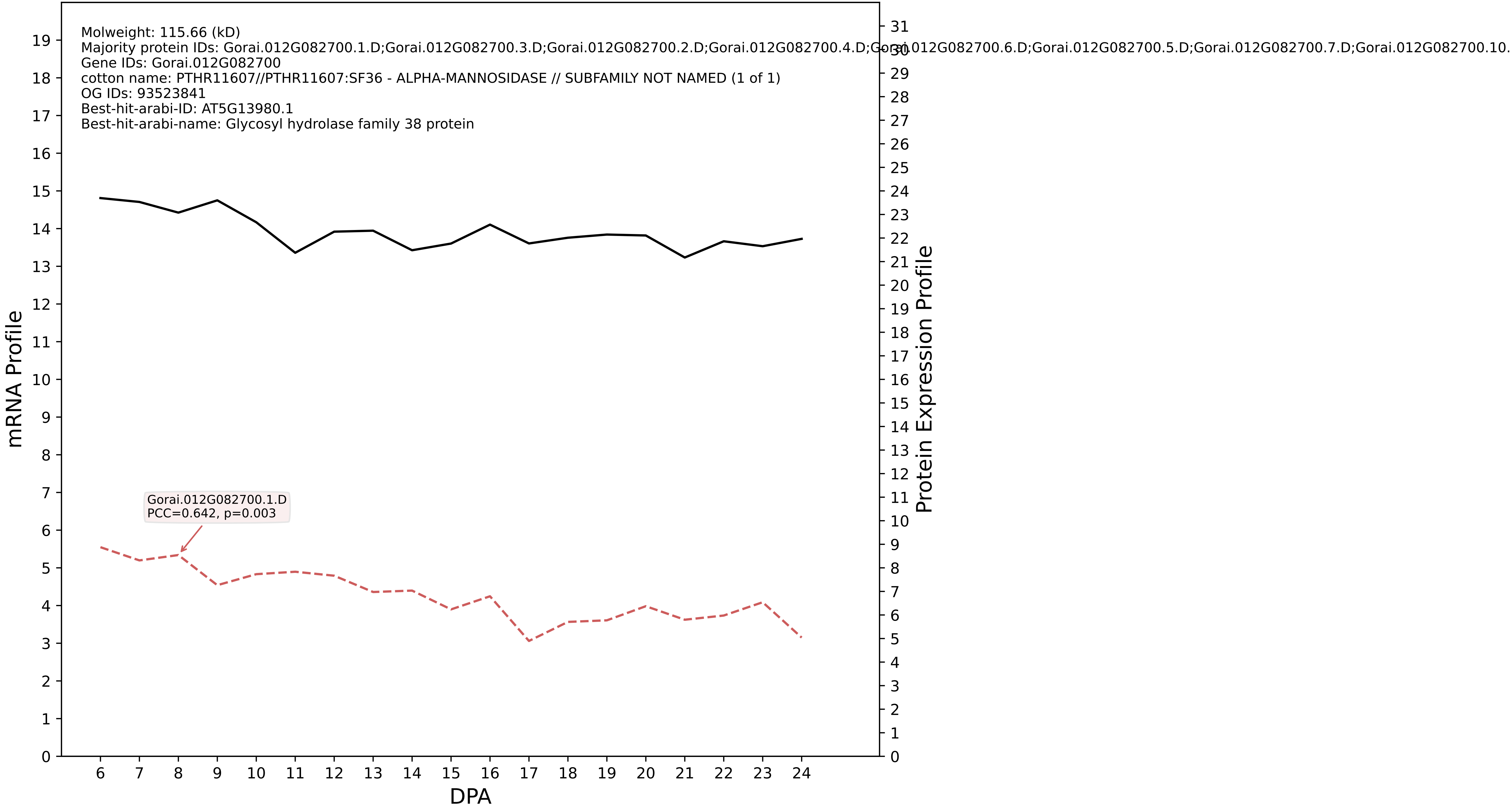

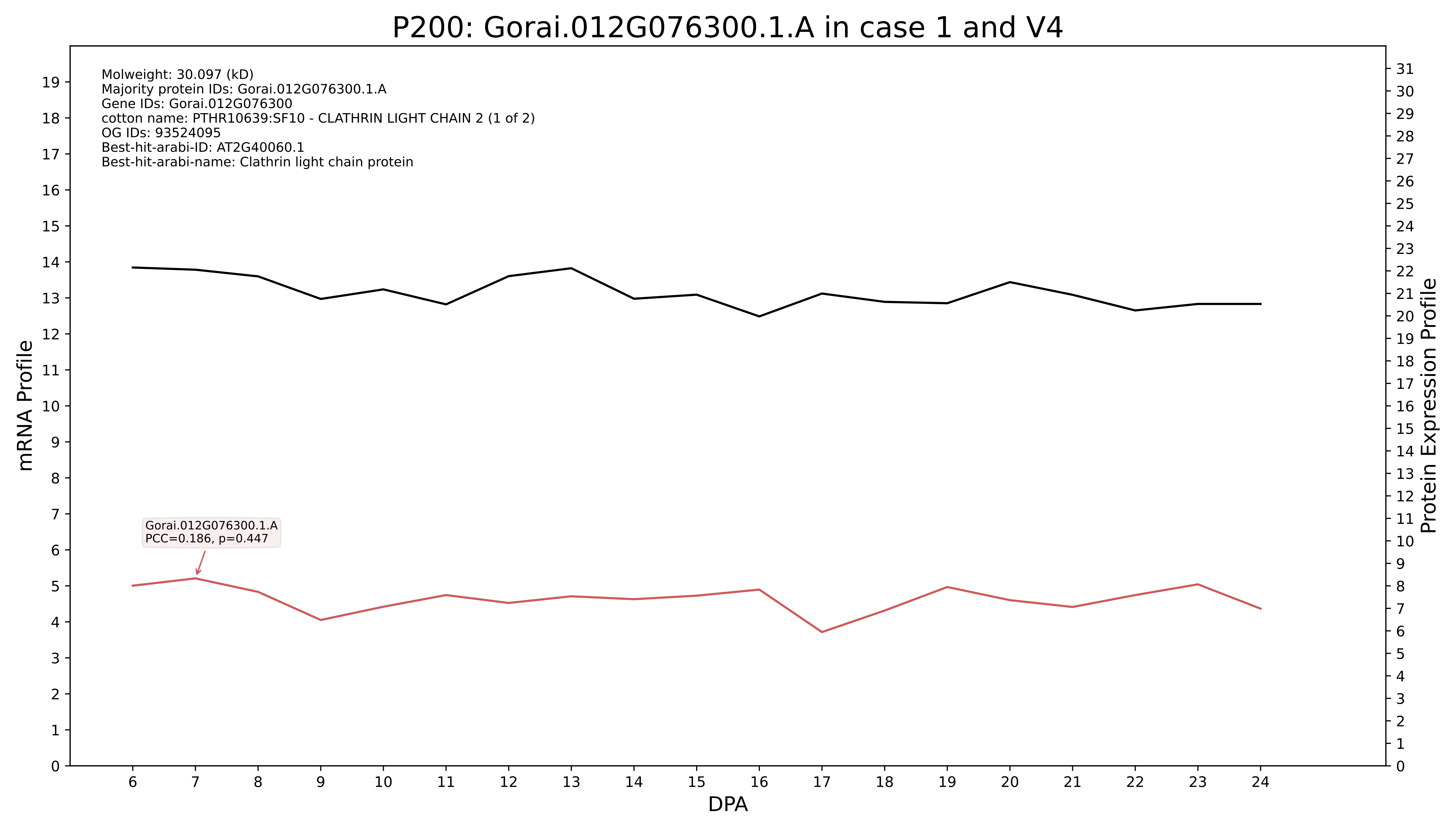
P200: Gorai.012G076300.1.A in case 1 and V4
Molweight: 30.097 (kD)
Majority protein IDs: Gorai.012G076300.1.A
Gene IDs: Gorai.012G076300
cotton name: PTHR10639:SF10 - CLATHRIN LIGHT CHAIN 2 (1 of 2)
OG IDs: 93524095
Best-hit-arabi-ID: AT2G40060.1
Best-hit-arabi-name: Clathrin light chain protein
Gorai.012G076300.1.A
PCC=0.186, p=0.447
mRNA Profile
Protein Expression Profile
DPA

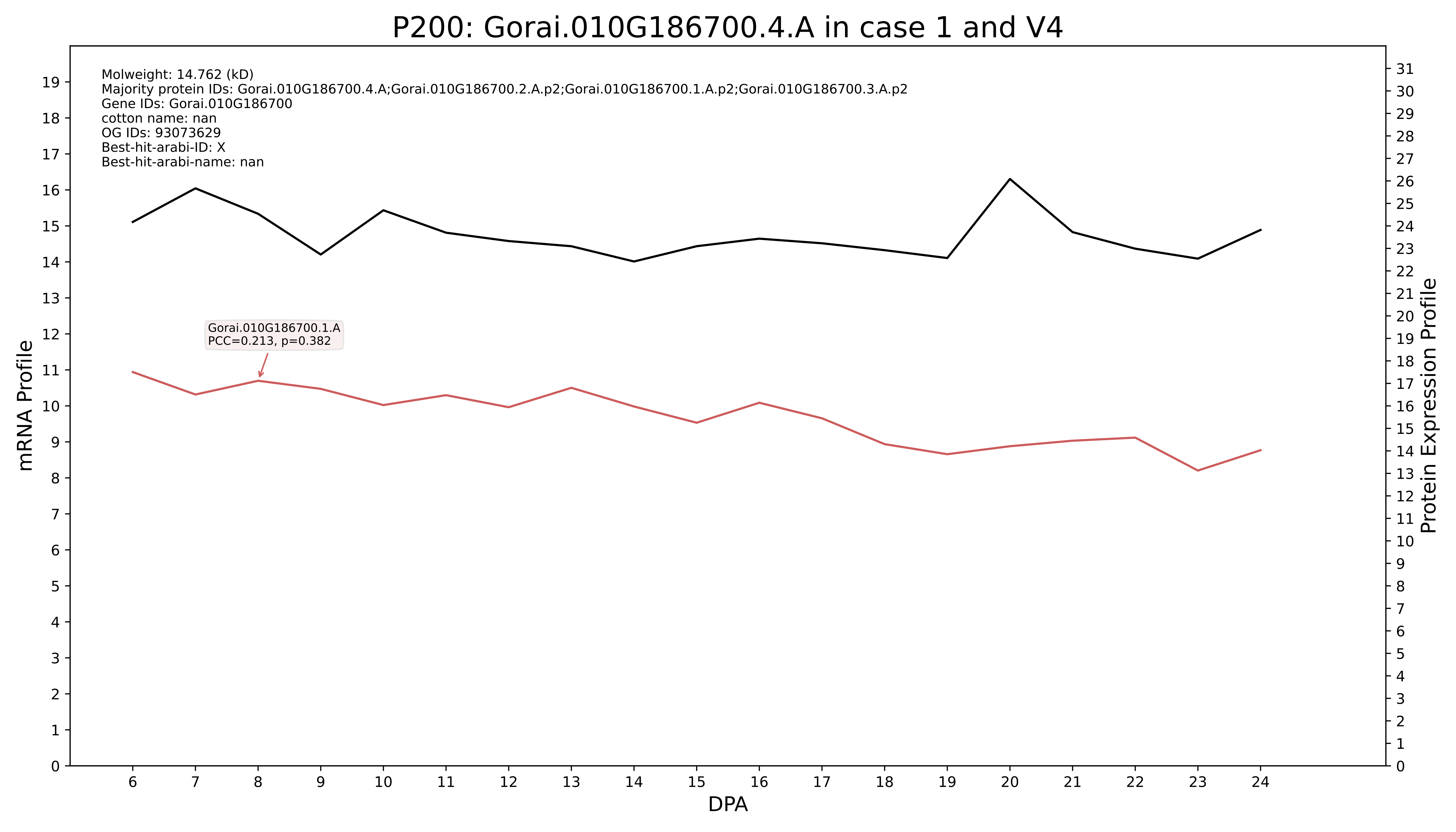
P200: Gorai.010G186700.4.A in case 1 and V4
Molweight: 14.762 (kD)
Majority protein IDs: Gorai.010G186700.4.A;Gorai.010G186700.2.A.p2;Gorai.010G186700.1.A.p2;Gorai.010G186700.3.A.p2
Gene IDs: Gorai.010G186700
cotton name: nan
OG IDs: 93073629
Best-hit-arabi-ID: X
Best-hit-arabi-name: nan
Gorai.010G186700.1.A
PCC=0.213, p=0.382
mRNA Profile
Protein Expression Profile
DPA

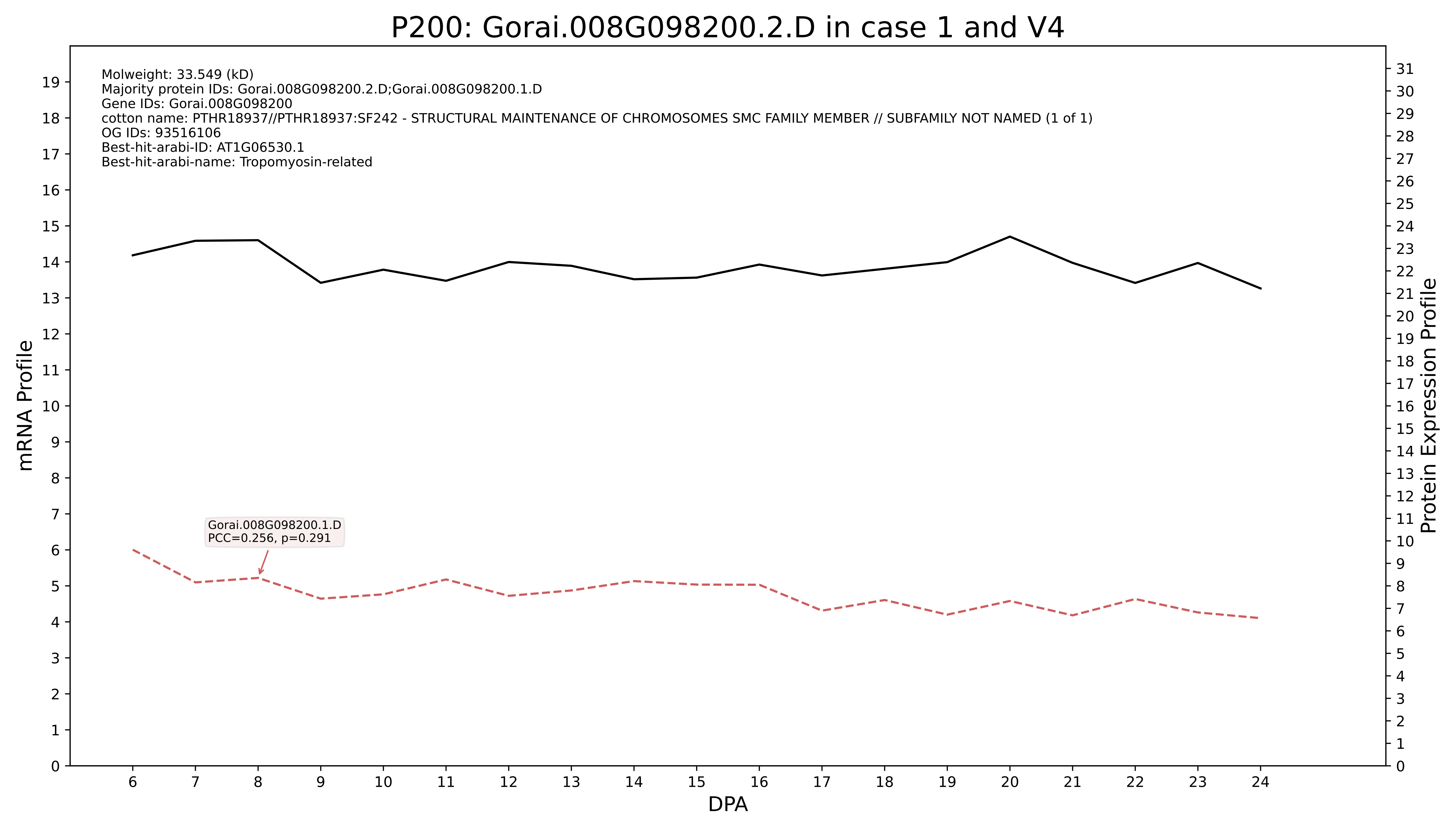

P200: Gorai.008G098200.2.D in case 1 and V4
Molweight: 33.549 (kD)
Majority protein IDs: Gorai.008G098200.2.D;Gorai.008G098200.1.D
Gene IDs: Gorai.008G098200
cotton name: PTHR18937//PTHR18937:SF242 - STRUCTURAL MAINTENANCE OF CHROMOSOMES SMC FAMILY MEMBER // SUBFAMILY NOT NAMED (1 of 1)
OG IDs: 93516106
Best-hit-arabi-ID: AT1G06530.1
Best-hit-arabi-name: Tropomyosin-related
Gorai.008G098200.1.D
PCC=0.256, p=0.291
mRNA Profile
Protein Expression Profile
DPA

### P200: Gorai.013G077100.1.D in case 1 and V4

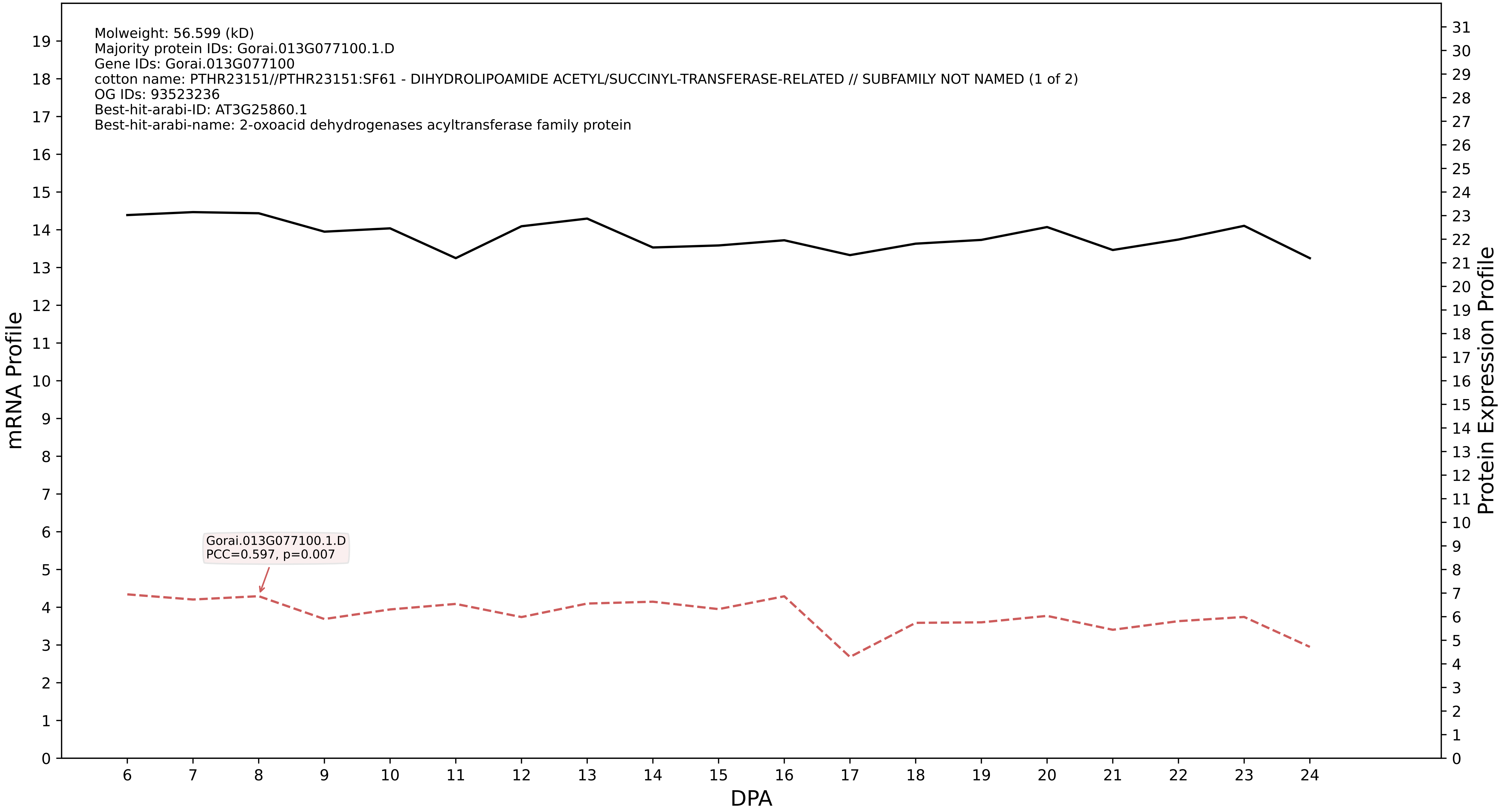

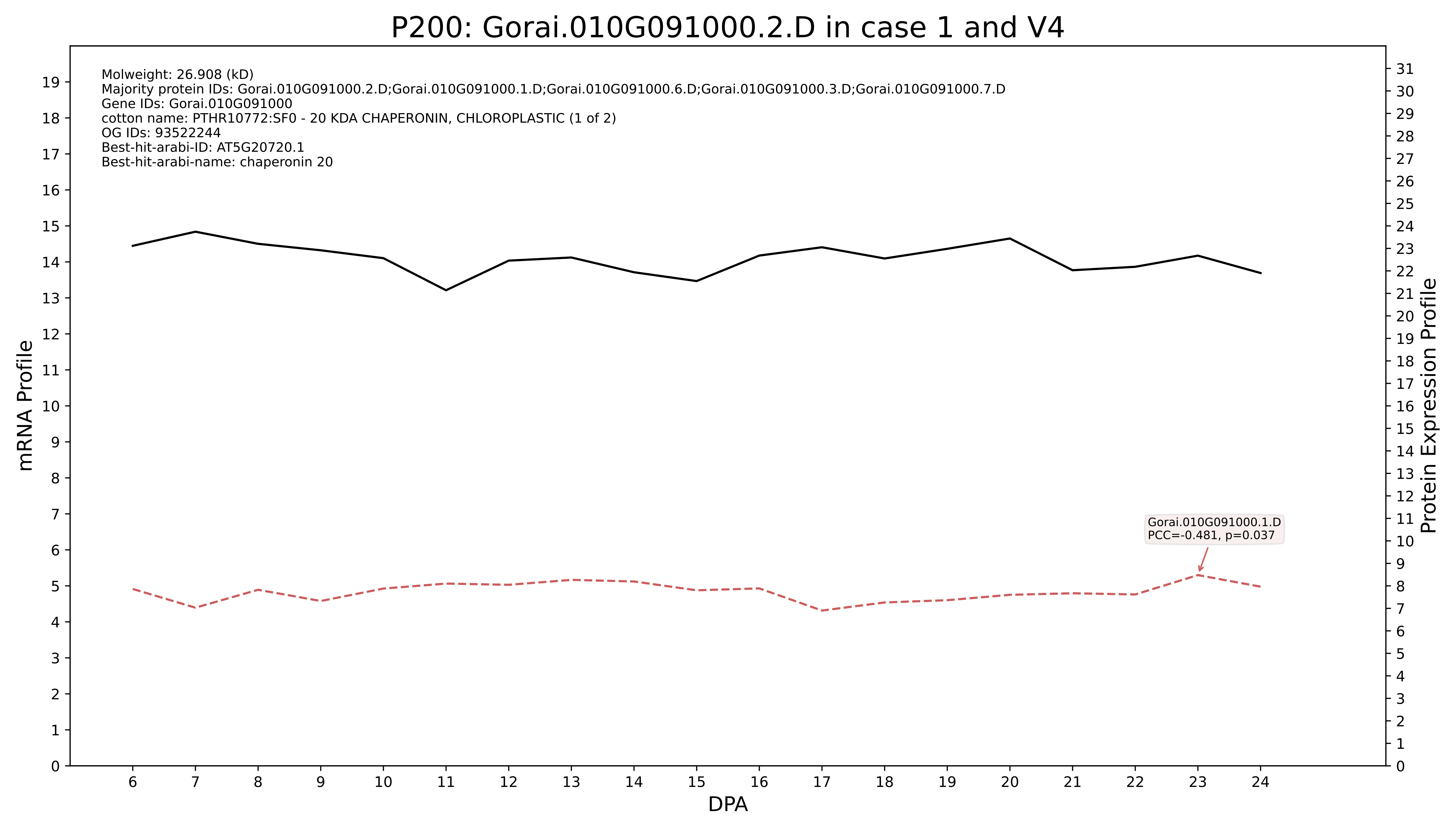

P200: Gorai.010G091000.2.D in case 1 and V4
Molweight: 26.908 (kD)
Majority protein IDs: Gorai.010G091000.2.D;Gorai.010G091000.1.D;Gorai.010G091000.6.D;Gorai.010G091000.3.D;Gorai.010G091000.7.D
Gene IDs: Gorai.010G091000
cotton name: PTHR10772:SF0 - 20 KDA CHAPERONIN, CHLOROPLASTIC (1 of 2)
OG IDs: 93522244
Best-hit-arabi-ID: AT5G20720.1
Best-hit-arabi-name: chaperonin 20
Gorai.010G091000.1.D
PCC=-0.481, p=0.037
mRNA Profile
Protein Expression Profile
DPA

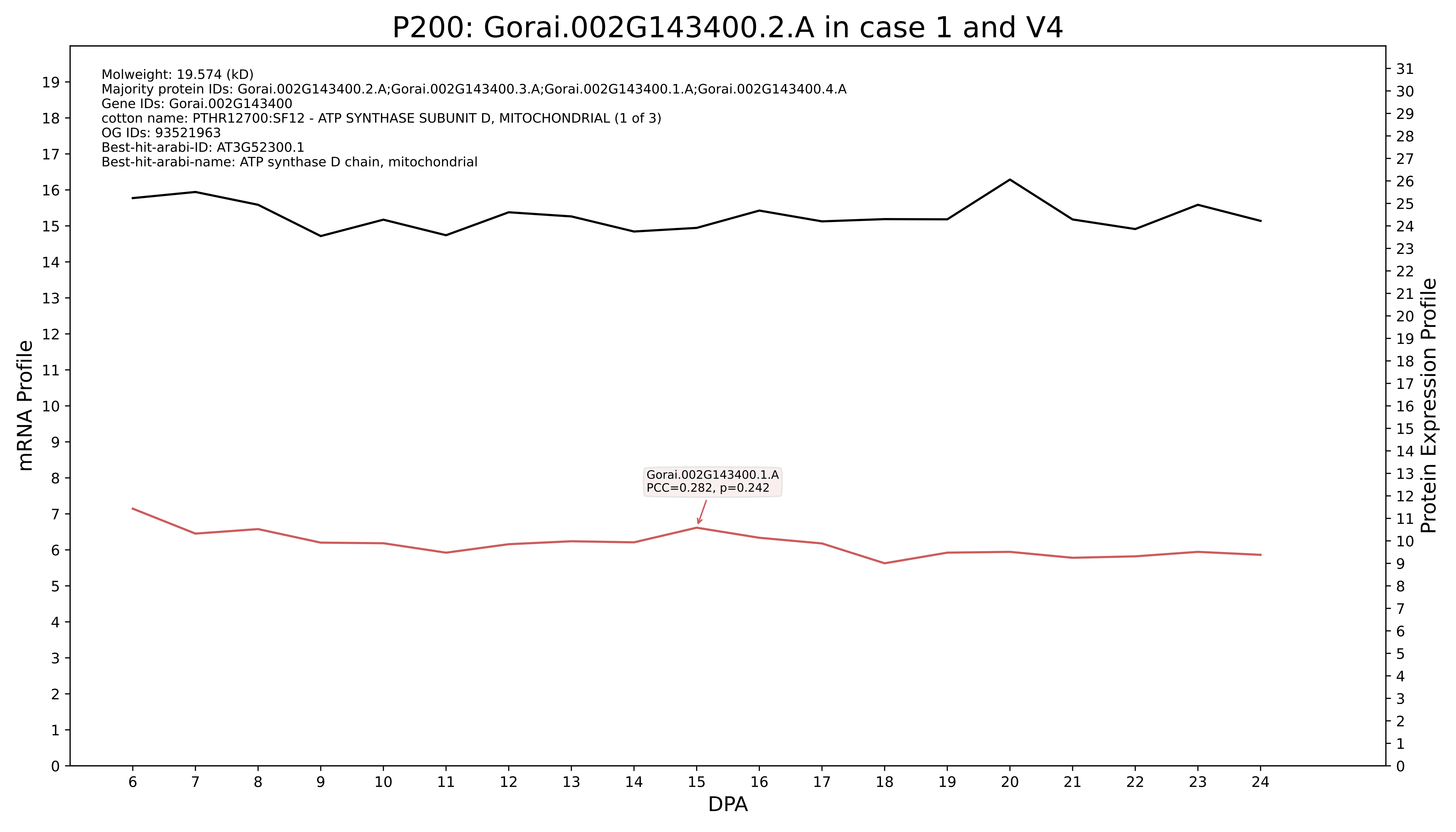
P200: Gorai.002G143400.2.A in case 1 and V4
Molweight: 19.574 (kD)
Majority protein IDs: Gorai.002G143400.2.A;Gorai.002G143400.3.A;Gorai.002G143400.1.A;Gorai.002G143400.4.A
Gene IDs: Gorai.002G143400
cotton name: PTHR12700:SF12 - ATP SYNTHASE SUBUNIT D, MITOCHONDRIAL (1 of 3)
OG IDs: 93521963
Best-hit-arabi-ID: AT3G52300.1
Best-hit-arabi-name: ATP synthase D chain, mitochondrial
Gorai.002G143400.1.A
PCC=0.282, p=0.242
mRNA Profile
Protein Expression Profile
DPA

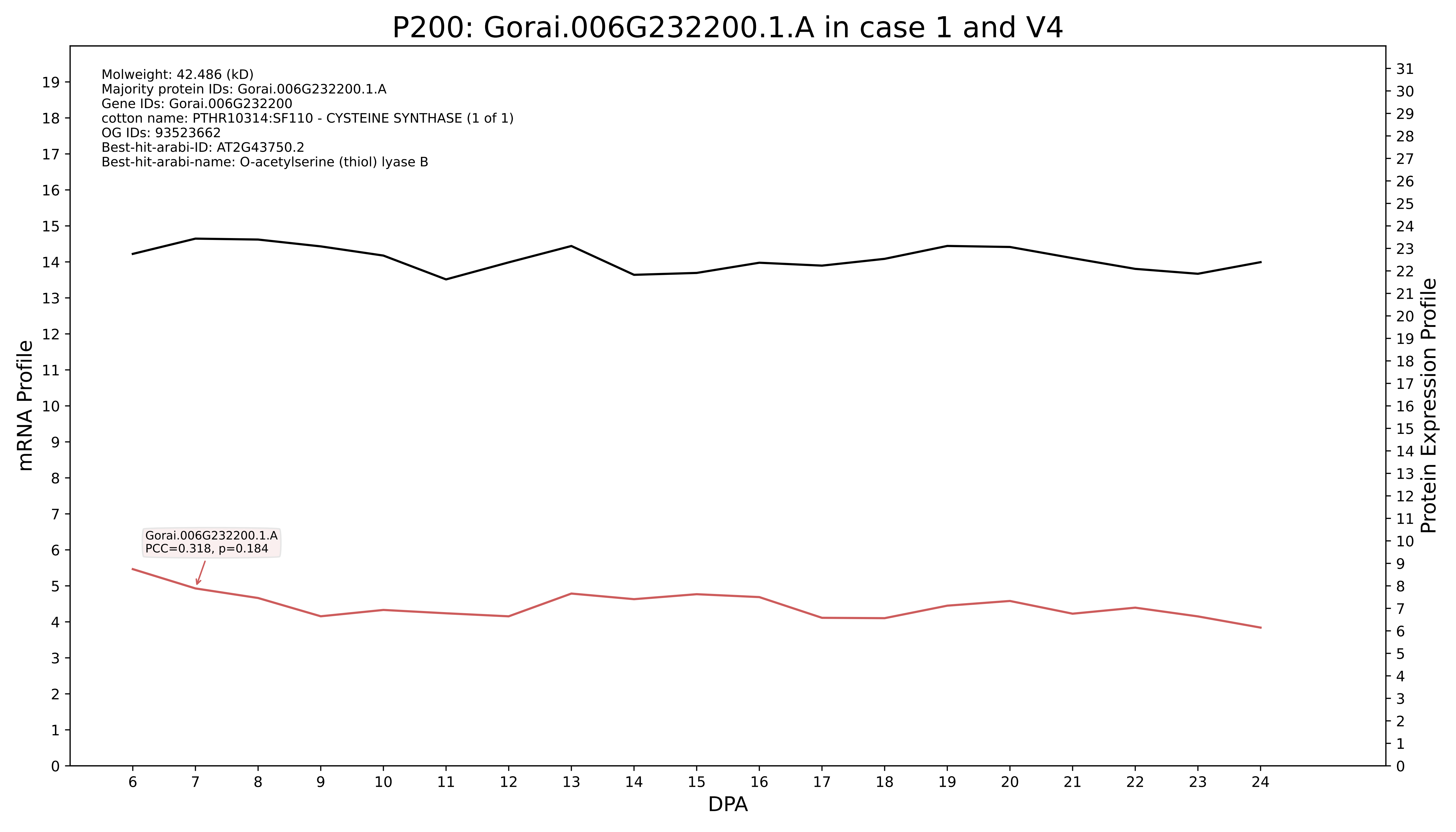

P200: Gorai.006G232200.1.A in case 1 and V4
Molweight: 42.486 (kD)
Majority protein IDs: Gorai.006G232200.1.A
Gene IDs: Gorai.006G232200
cotton name: PTHR10314:SF110 - CYSTEINE SYNTHASE (1 of 1)
OG IDs: 93523662
Best-hit-arabi-ID: AT2G43750.2
Best-hit-arabi-name: O-acetylserine (thiol) lyase B
Gorai.006G232200.1.A
PCC=0.318, p=0.184
mRNA Profile
Protein Expression Profile
DPA

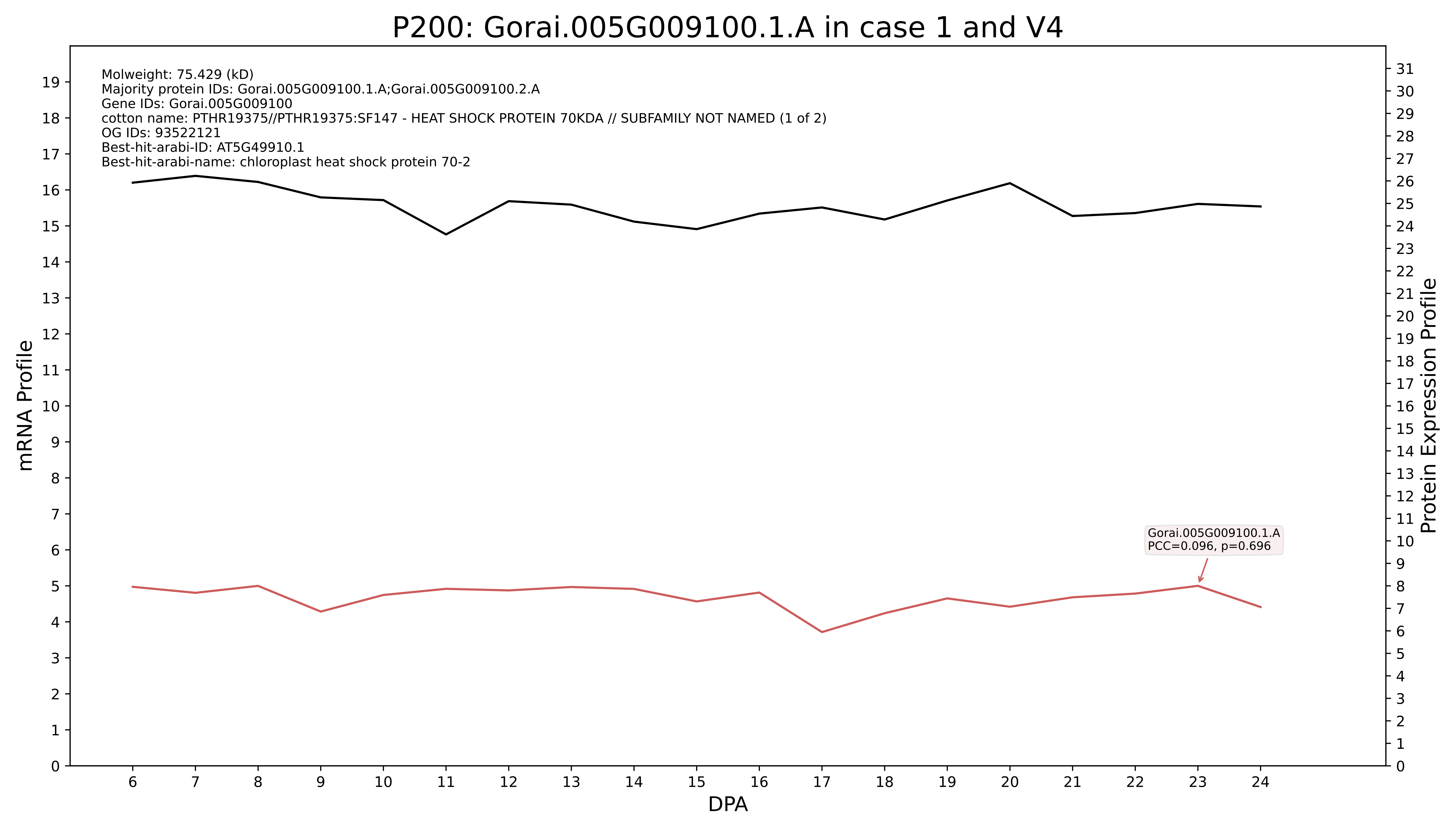

P200: Gorai.005G009100.1.A in case 1 and V4
Molweight: 75.429 (kD)
Majority protein IDs: Gorai.005G009100.1.A;Gorai.005G009100.2.A
Gene IDs: Gorai.005G009100
cotton name: PTHR19375//PTHR19375:SF147 - HEAT SHOCK PROTEIN 70KDA // SUBFAMILY NOT NAMED (1 of 2)
OG IDs: 93522121
Best-hit-arabi-ID: AT5G49910.1
Best-hit-arabi-name: chloroplast heat shock protein 70-2
Gorai.005G009100.1.A
PCC=0.096, p=0.696
mRNA Profile
Protein Expression Profile
DPA

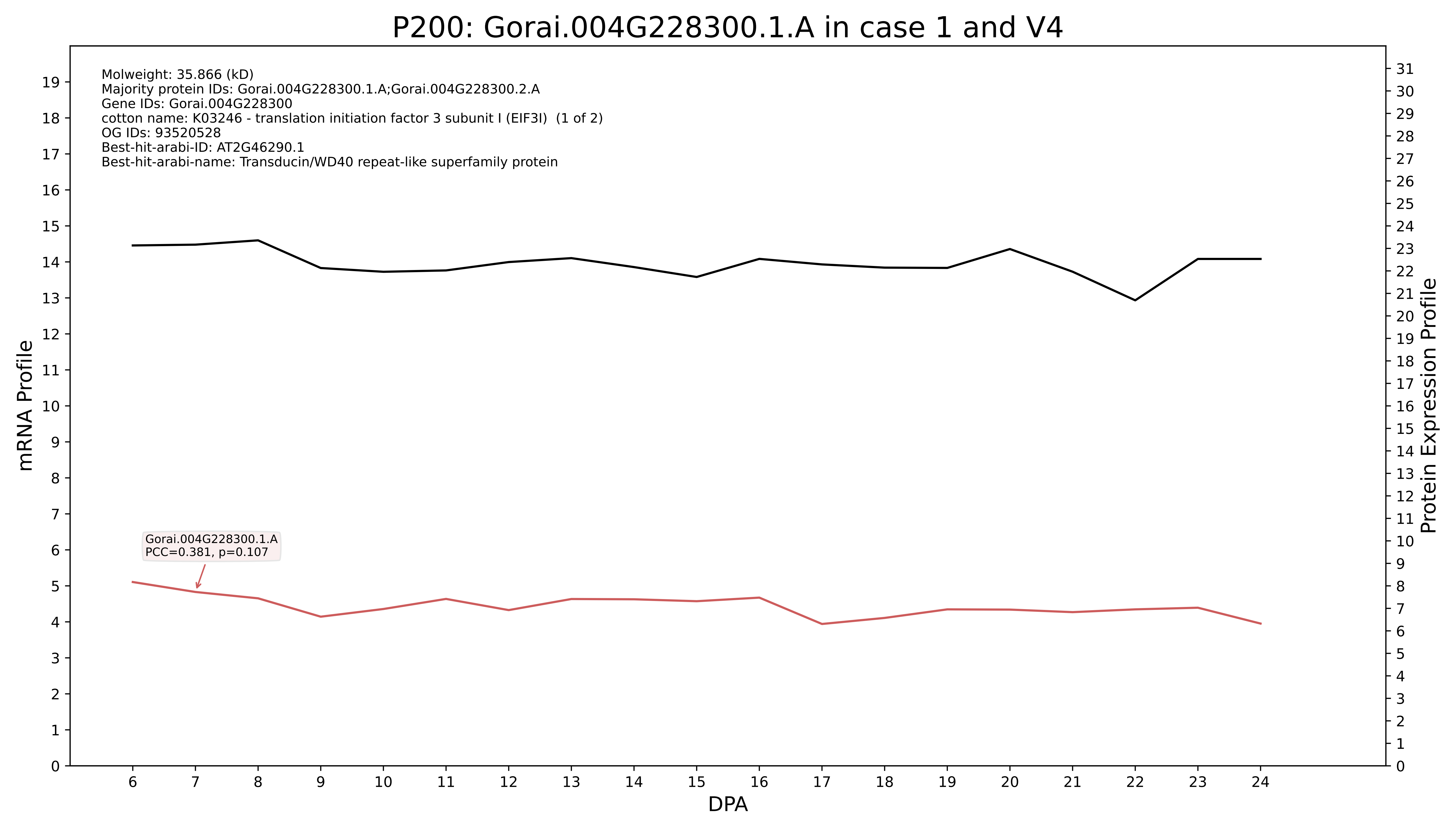
P200: Gorai.004G228300.1.A in case 1 and V4
Molweight: 35.866 (kD)
Majority protein IDs: Gorai.004G228300.1.A;Gorai.004G228300.2.A
Gene IDs: Gorai.004G228300
cotton name: K03246 - translation initiation factor 3 subunit I (EIF3I) (1 of 2)
OG IDs: 93520528
Best-hit-arabi-ID: AT2G46290.1
Best-hit-arabi-name: Transducin/WD40 repeat-like superfamily protein
Gorai.004G228300.1.A
PCC=0.381, p=0.107
mRNA Profile
Protein Expression Profile
DPA

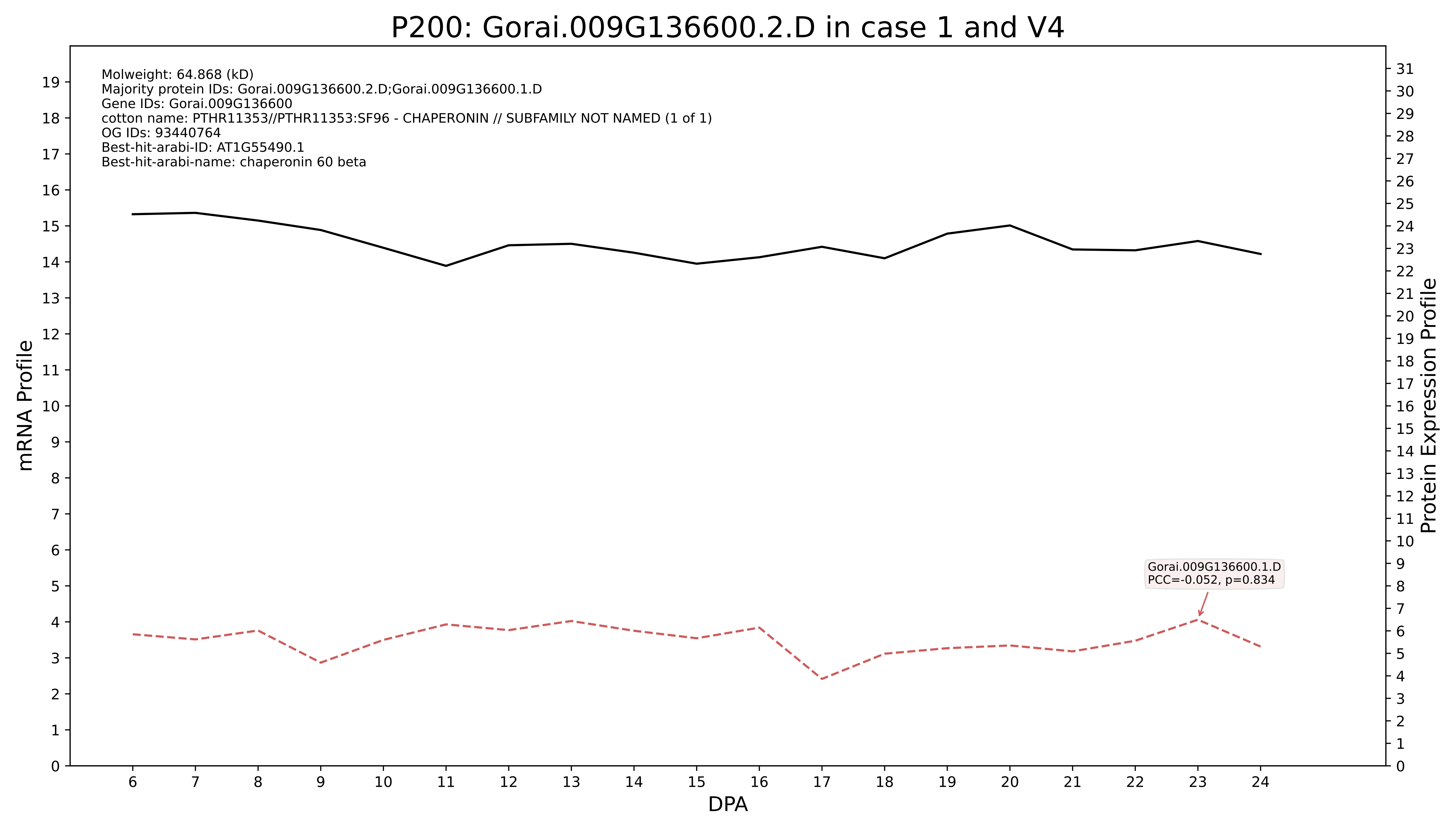
P200: Gorai.009G136600.2.D in case 1 and V4
Molweight: 64.868 (kD)
Majority protein IDs: Gorai.009G136600.2.D;Gorai.009G136600.1.D
Gene IDs: Gorai.009G136600
cotton name: PTHR11353//PTHR11353:SF96 - CHAPERONIN // SUBFAMILY NOT NAMED (1 of 1)
OG IDs: 93440764
Best-hit-arabi-ID: AT1G55490.1
Best-hit-arabi-name: chaperonin 60 beta
Gorai.009G136600.1.D
PCC=-0.052, p=0.834
mRNA Profile
Protein Expression Profile
DPA

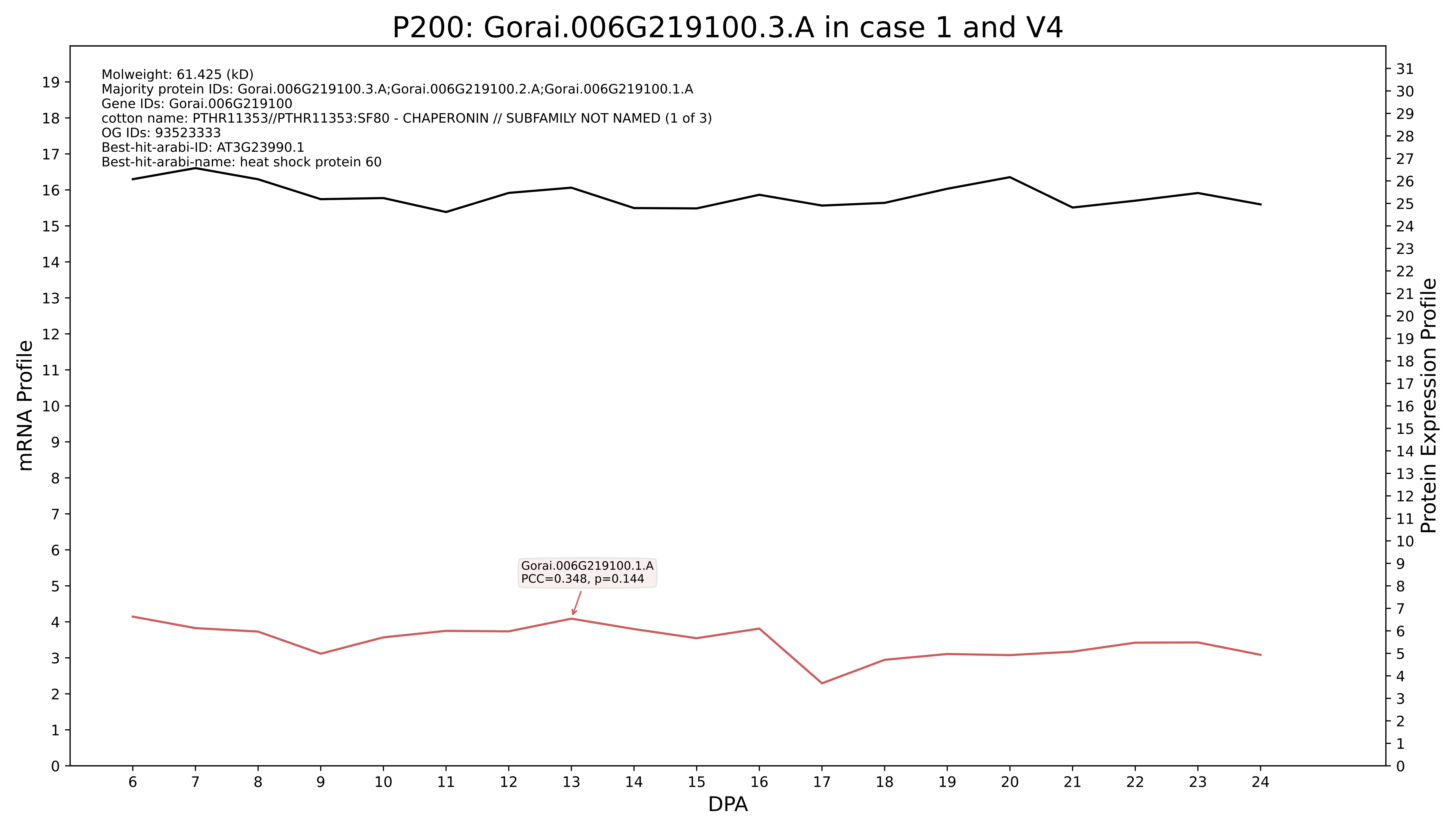

P200: Gorai.006G219100.3.A in case 1 and V4
Molweight: 61.425 (kD)
Majority protein IDs: Gorai.006G219100.3.A;Gorai.006G219100.2.A;Gorai.006G219100.1.A
Gene IDs: Gorai.006G219100
cotton name: PTHR11353//PTHR11353:SF80 - CHAPERONIN // SUBFAMILY NOT NAMED (1 of 3)
OG IDs: 93523333
Best-hit-arabi-ID: AT3G23990.1
Best-hit-arabi-name: heat shock protein 60
Gorai.006G219100.1.A
PCC=0.348, p=0.144
mRNA Profile
Protein Expression Profile
DPA

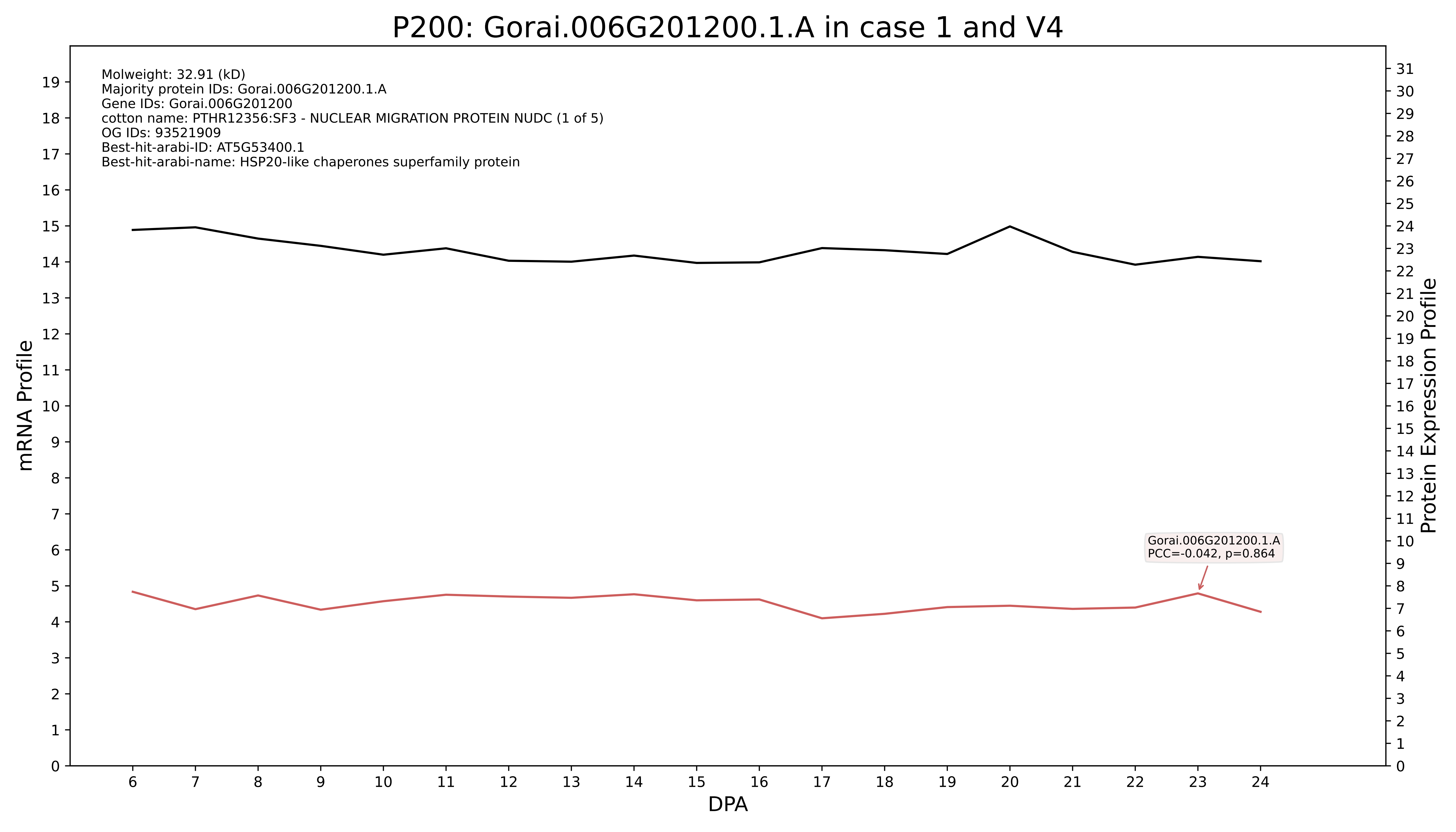
P200: Gorai.006G201200.1.A in case 1 and V4
Molweight: 32.91 (kD)
Majority protein IDs: Gorai.006G201200.1.A
Gene IDs: Gorai.006G201200
cotton name: PTHR12356:SF3 - NUCLEAR MIGRATION PROTEIN NUDC (1 of 5)
OG IDs: 93521909
Best-hit-arabi-ID: AT5G53400.1
Best-hit-arabi-name: HSP20-like chaperones superfamily protein
Gorai.006G201200.1.A
PCC=-0.042, p=0.864
mRNA Profile
Protein Expression Profile
DPA

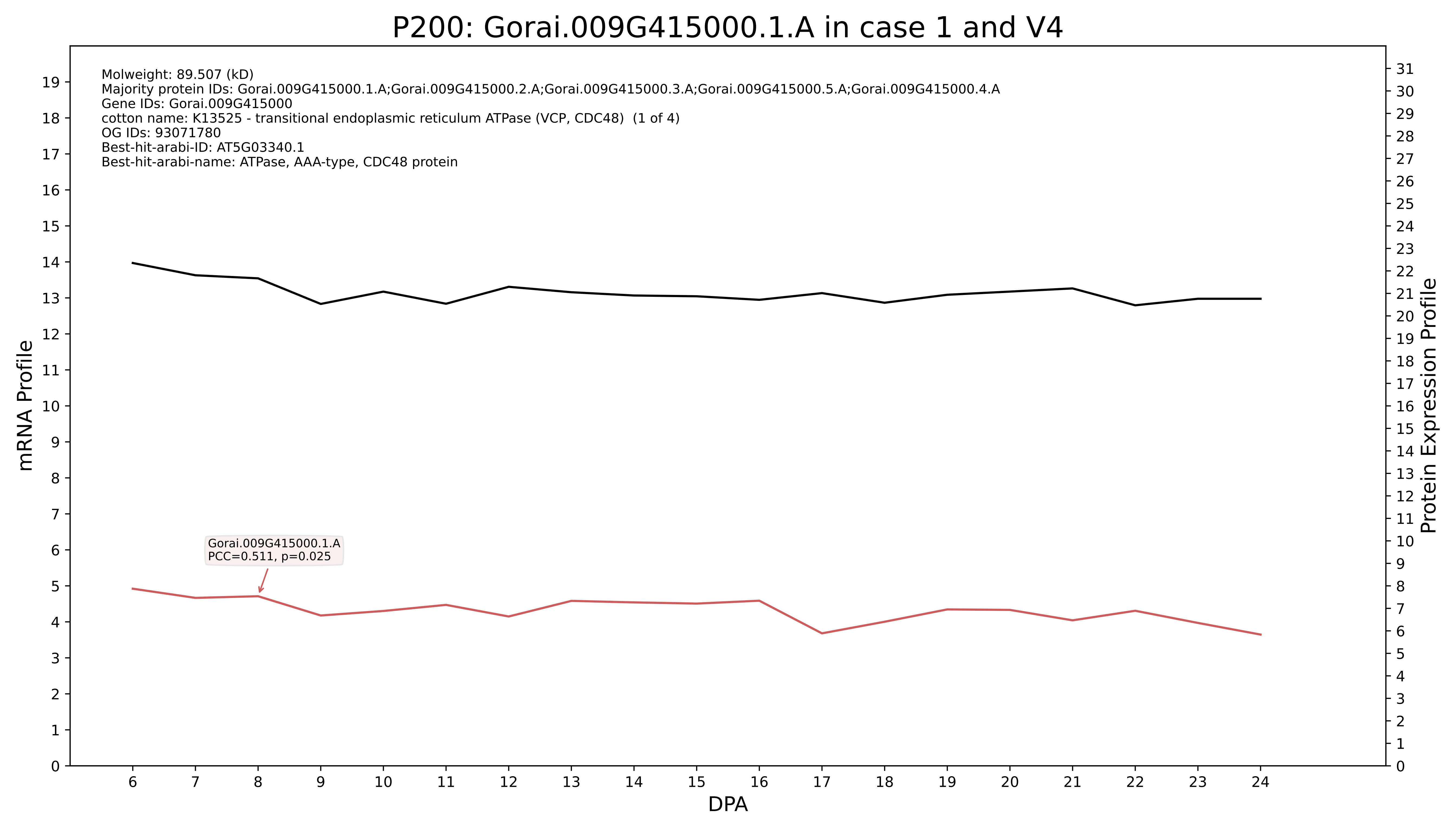

P200: Gorai.009G415000.1.A in case 1 and V4
Molweight: 89.507 (kD)
Majority protein IDs: Gorai.009G415000.1.A;Gorai.009G415000.2.A;Gorai.009G415000.3.A;Gorai.009G415000.5.A;Gorai.009G415000.4.A
Gene IDs: Gorai.009G415000
cotton name: K13525 - transitional endoplasmic reticulum ATPase (VCP, CDC48) (1 of 4)
OG IDs: 93071780
Best-hit-arabi-ID: AT5G03340.1
Best-hit-arabi-name: ATPase, AAA-type, CDC48 protein
Gorai.009G415000.1.A
PCC=0.511, p=0.025
mRNA Profile
Protein Expression Profile
DPA

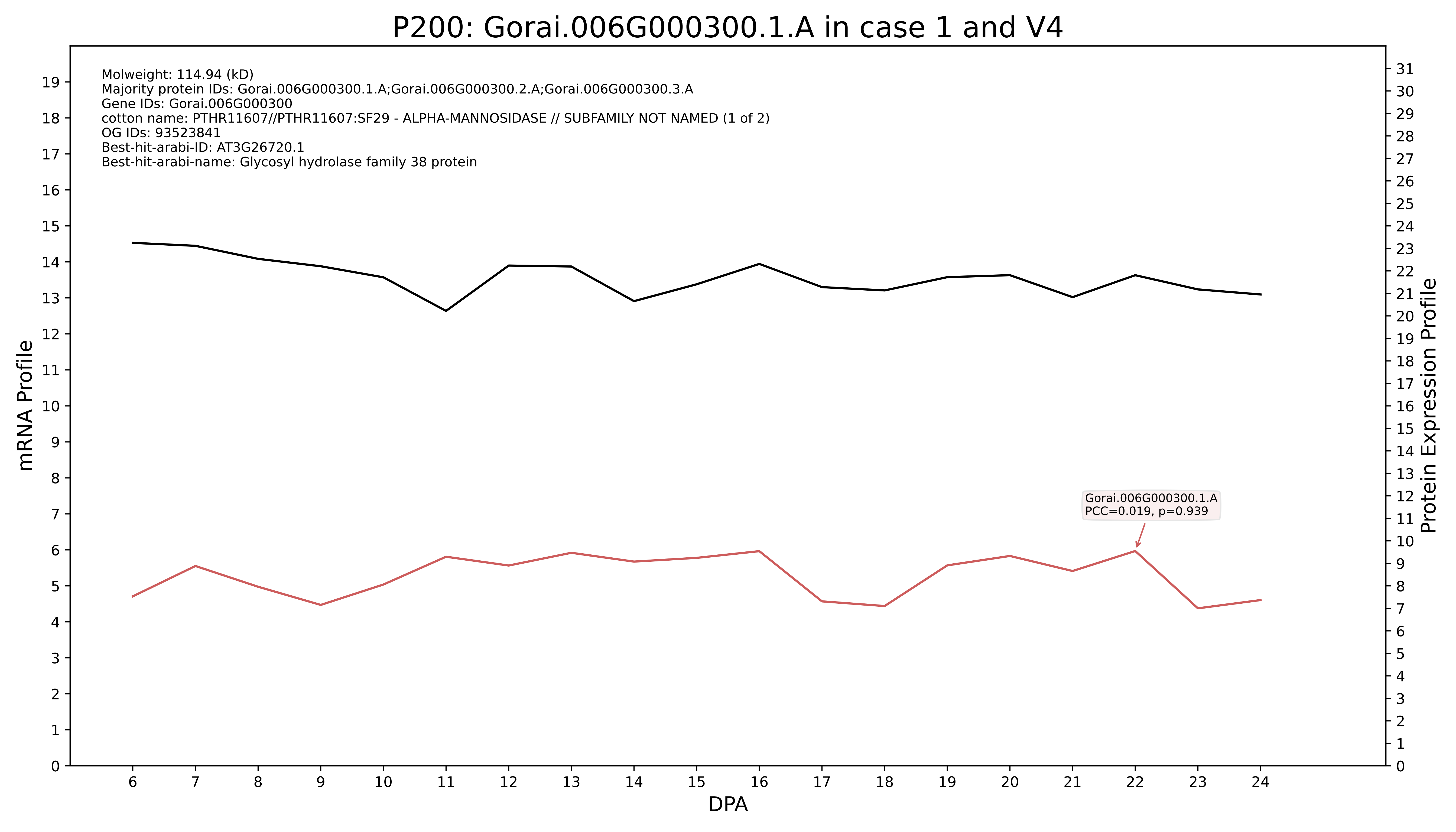
P200: Gorai.006G000300.1.A in case 1 and V4
Molweight: 114.94 (kD)
Majority protein IDs: Gorai.006G000300.1.A;Gorai.006G000300.2.A;Gorai.006G000300.3.A
Gene IDs: Gorai.006G000300
cotton name: PTHR11607//PTHR11607:SF29 - ALPHA-MANNOSIDASE // SUBFAMILY NOT NAMED (1 of 2)
OG IDs: 93523841
Best-hit-arabi-ID: AT3G26720.1
Best-hit-arabi-name: Glycosyl hydrolase family 38 protein
Gorai.006G000300.1.A
PCC=0.019, p=0.939
mRNA Profile
Protein Expression Profile
DPA

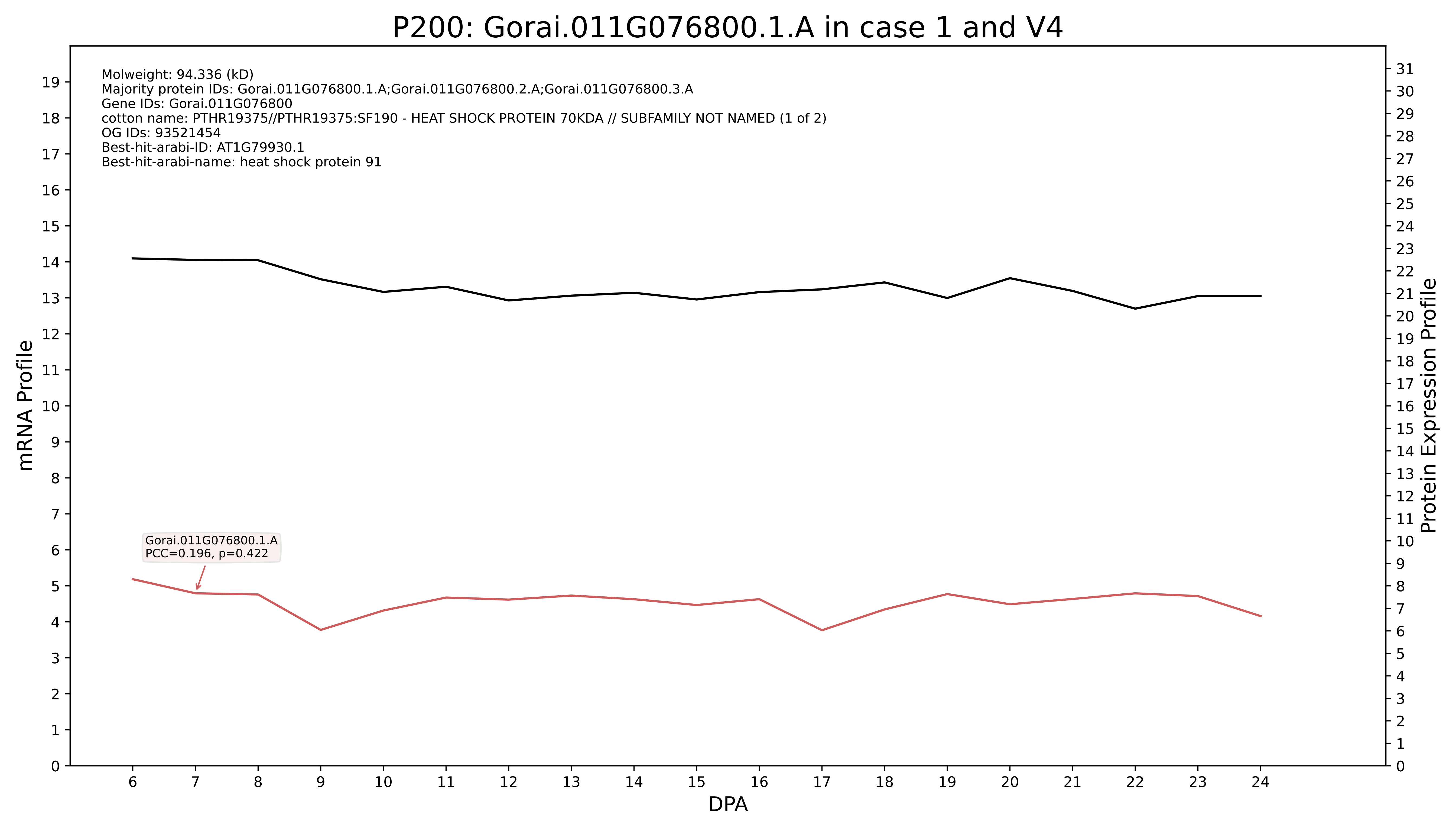
P200: Gorai.011G076800.1.A in case 1 and V4
Molweight: 94.336 (kD)
Majority protein IDs: Gorai.011G076800.1.A;Gorai.011G076800.2.A;Gorai.011G076800.3.A
Gene IDs: Gorai.011G076800
cotton name: PTHR19375//PTHR19375:SF190 - HEAT SHOCK PROTEIN 70KDA // SUBFAMILY NOT NAMED (1 of 2)
OG IDs: 93521454
Best-hit-arabi-ID: AT1G79930.1
Best-hit-arabi-name: heat shock protein 91
Gorai.011G076800.1.A
PCC=0.196, p=0.422
mRNA Profile
Protein Expression Profile
DPA

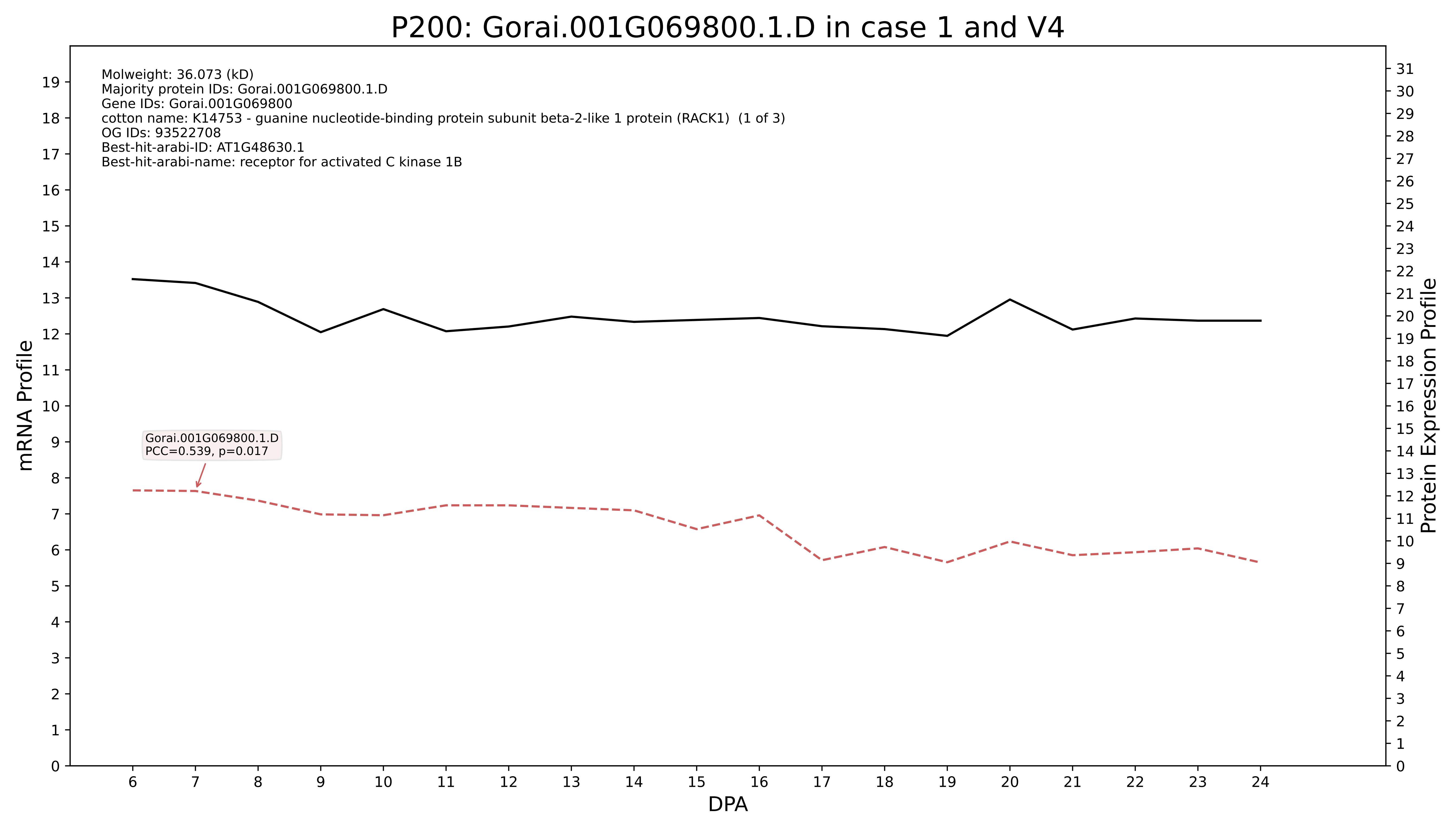
P200: Gorai.001G069800.1.D in case 1 and V4
Molweight: 36.073 (kD)
Majority protein IDs: Gorai.001G069800.1.D
Gene IDs: Gorai.001G069800
cotton name: K14753 - guanine nucleotide-binding protein subunit beta-2-like 1 protein (RACK1) (1 of 3)
OG IDs: 93522708
Best-hit-arabi-ID: AT1G48630.1
Best-hit-arabi-name: receptor for activated C kinase 1B
Gorai.001G069800.1.D
PCC=0.539, p=0.017
mRNA Profile
Protein Expression Profile
DPA

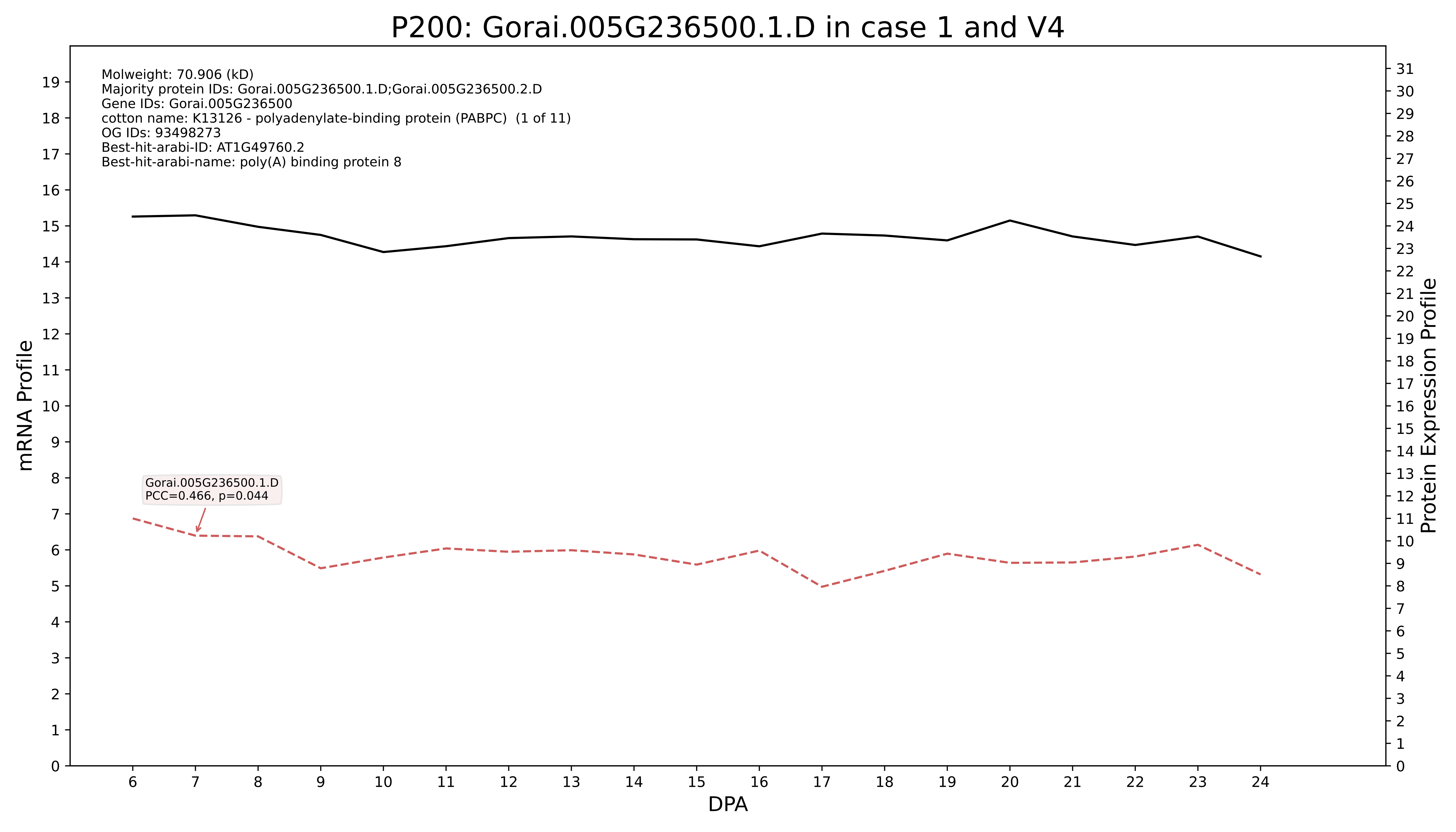

P200: Gorai.005G236500.1.D in case 1 and V4
Molweight: 70.906 (kD)
Majority protein IDs: Gorai.005G236500.1.D;Gorai.005G236500.2.D
Gene IDs: Gorai.005G236500
cotton name: K13126 - polyadenylate-binding protein (PABPC) (1 of 11)
OG IDs: 93498273
Best-hit-arabi-ID: AT1G49760.2
Best-hit-arabi-name: poly(A) binding protein 8
Gorai.005G236500.1.D
PCC=0.466, p=0.044
mRNA Profile
Protein Expression Profile
DPA

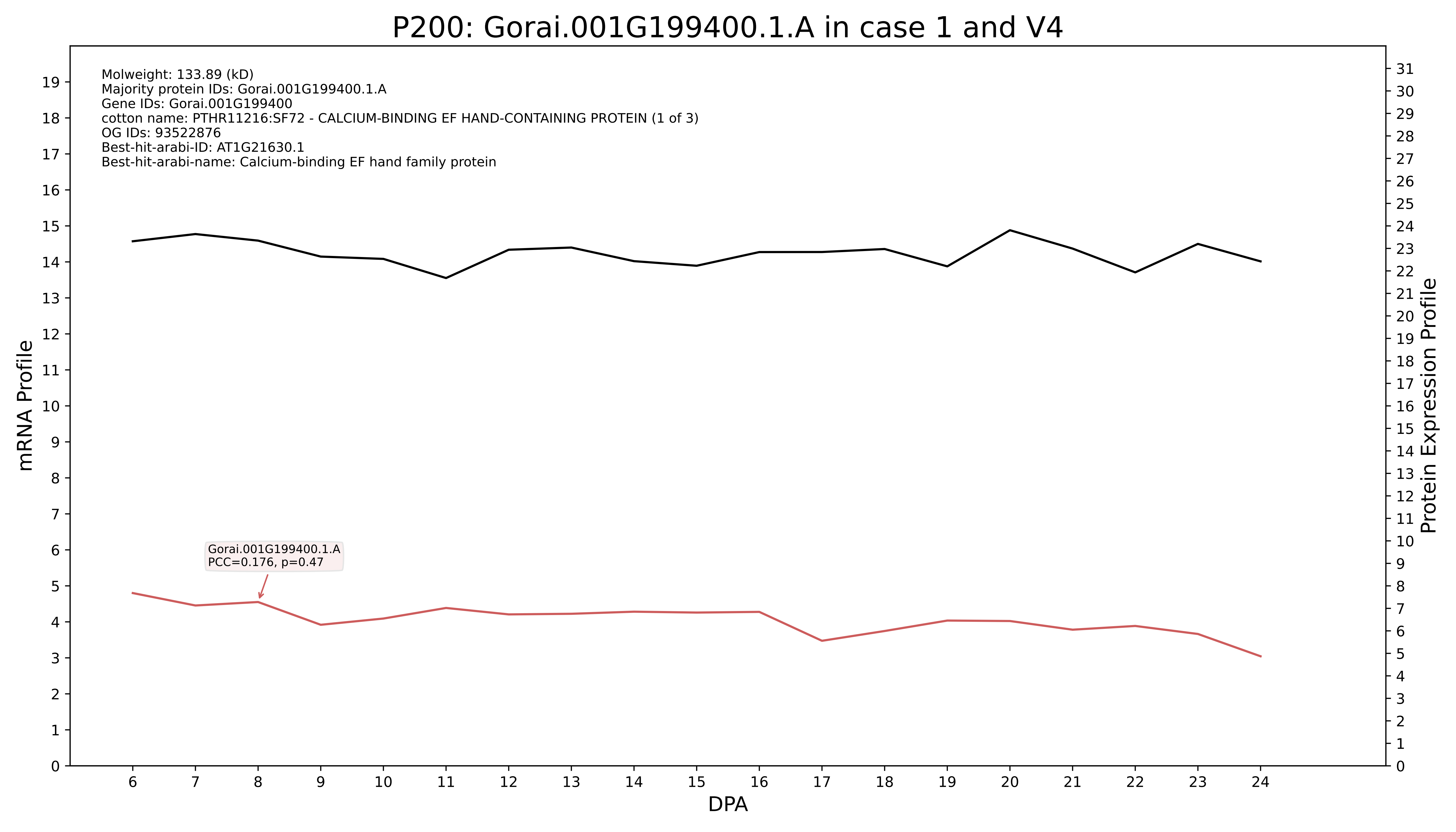
P200: Gorai.001G199400.1.A in case 1 and V4
Molweight: 133.89 (kD)
Majority protein IDs: Gorai.001G199400.1.A
Gene IDs: Gorai.001G199400
cotton name: PTHR11216:SF72 - CALCIUM-BINDING EF HAND-CONTAINING PROTEIN (1 of 3)
OG IDs: 93522876
Best-hit-arabi-ID: AT1G21630.1
Best-hit-arabi-name: Calcium-binding EF hand family protein
Gorai.001G199400.1.A
PCC=0.176, p=0.47
mRNA Profile
Protein Expression Profile
DPA

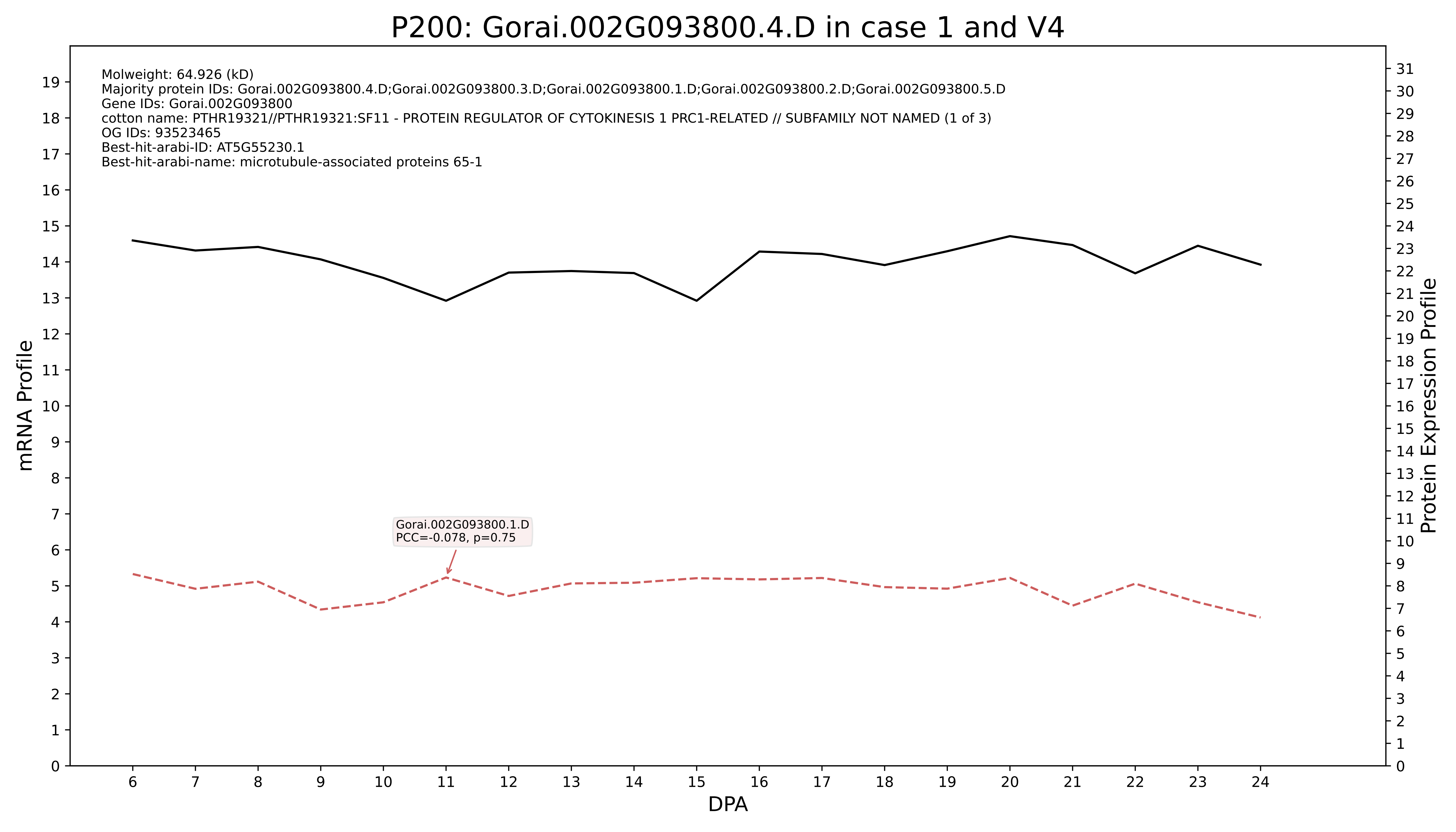

P200: Gorai.002G093800.4.D in case 1 and V4
Molweight: 64.926 (kD)
Majority protein IDs: Gorai.002G093800.4.D;Gorai.002G093800.3.D;Gorai.002G093800.1.D;Gorai.002G093800.2.D;Gorai.002G093800.5.D
Gene IDs: Gorai.002G093800
cotton name: PTHR19321//PTHR19321:SF11 - PROTEIN REGULATOR OF CYTOKINESIS 1 PRC1-RELATED // SUBFAMILY NOT NAMED (1 of 3)
OG IDs: 93523465
Best-hit-arabi-ID: AT5G55230.1
Best-hit-arabi-name: microtubule-associated proteins 65-1
Gorai.002G093800.1.D
PCC=-0.078, p=0.75
mRNA Profile
Protein Expression Profile
DPA

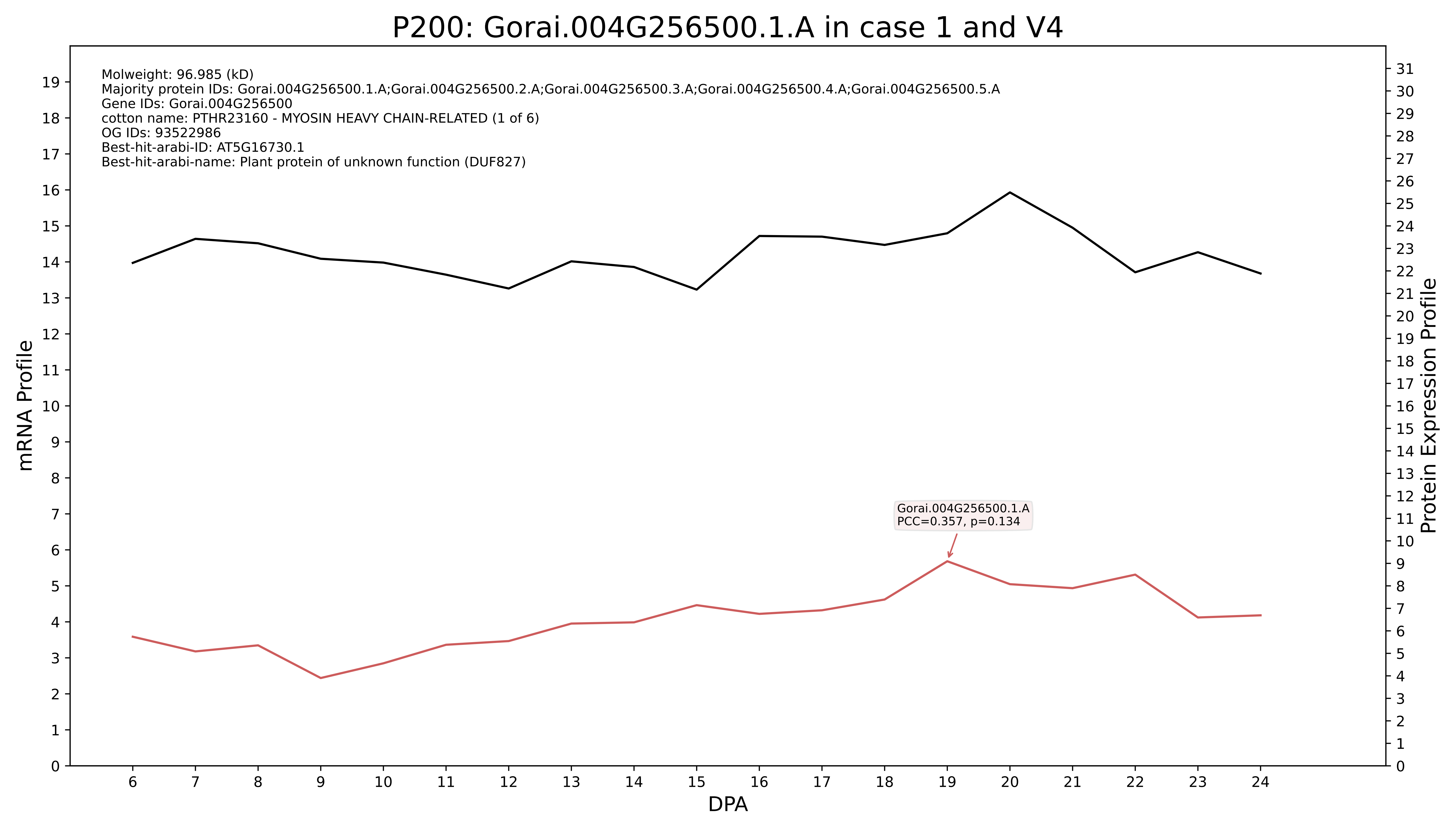
P200: Gorai.004G256500.1.A in case 1 and V4
Molweight: 96.985 (kD)
Majority protein IDs: Gorai.004G256500.1.A;Gorai.004G256500.2.A;Gorai.004G256500.3.A;Gorai.004G256500.4.A;Gorai.004G256500.5.A
Gene IDs: Gorai.004G256500
cotton name: PTHR23160 - MYOSIN HEAVY CHAIN-RELATED (1 of 6)
OG IDs: 93522986
Best-hit-arabi-ID: AT5G16730.1
Best-hit-arabi-name: Plant protein of unknown function (DUF827)
Gorai.004G256500.1.A
PCC=0.357, p=0.134
mRNA Profile
Protein Expression Profile
DPA

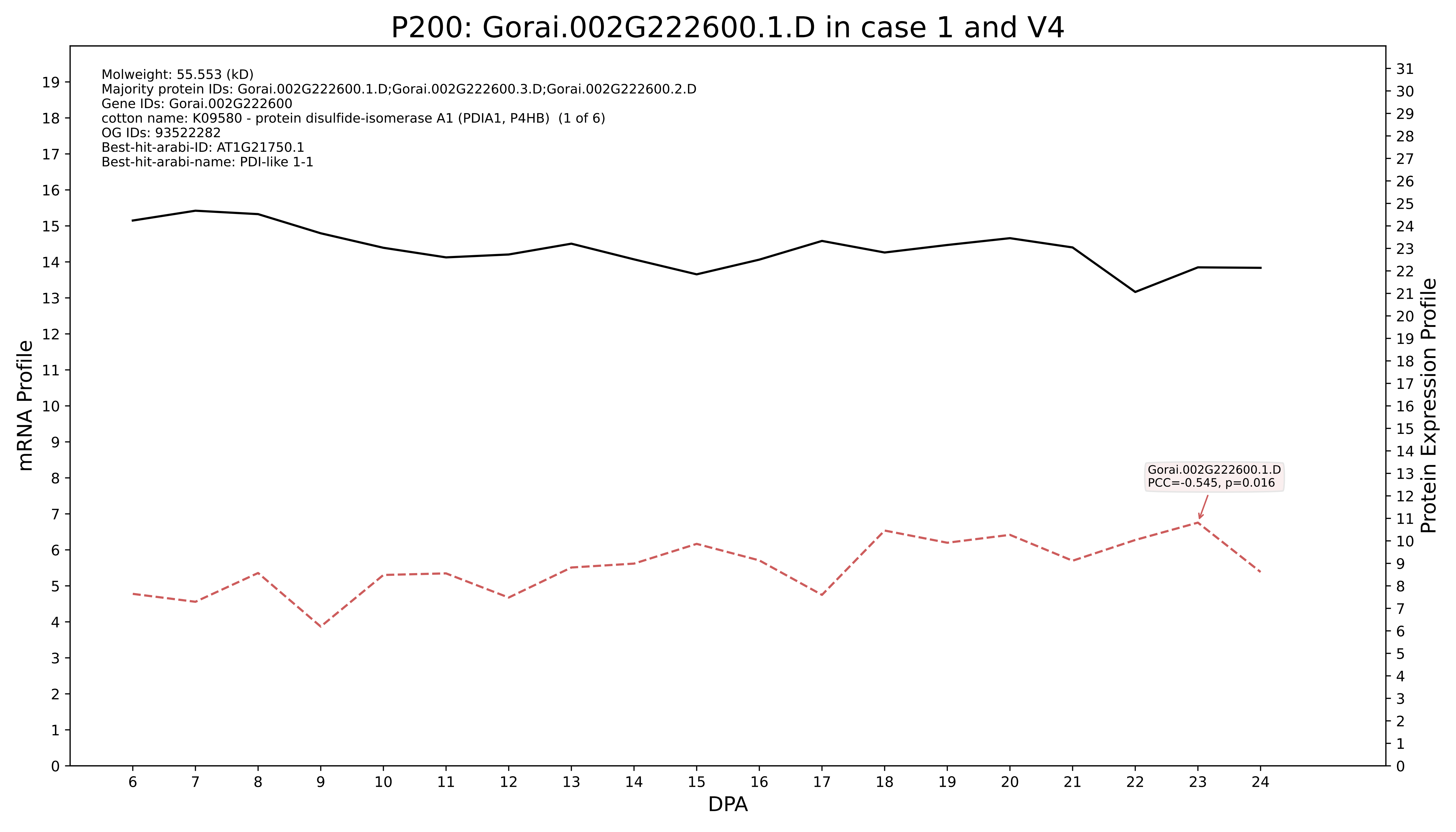
P200: Gorai.002G222600.1.D in case 1 and V4
Molweight: 55.553 (kD)
Majority protein IDs: Gorai.002G222600.1.D;Gorai.002G222600.3.D;Gorai.002G222600.2.D
Gene IDs: Gorai.002G222600
cotton name: K09580 - protein disulfide-isomerase A1 (PDIA1, P4HB) (1 of 6)
OG IDs: 93522282
Best-hit-arabi-ID: AT1G21750.1
Best-hit-arabi-name: PDI-like 1-1
Gorai.002G222600.1.D
PCC=-0.545, p=0.016
mRNA Profile
Protein Expression Profile
DPA

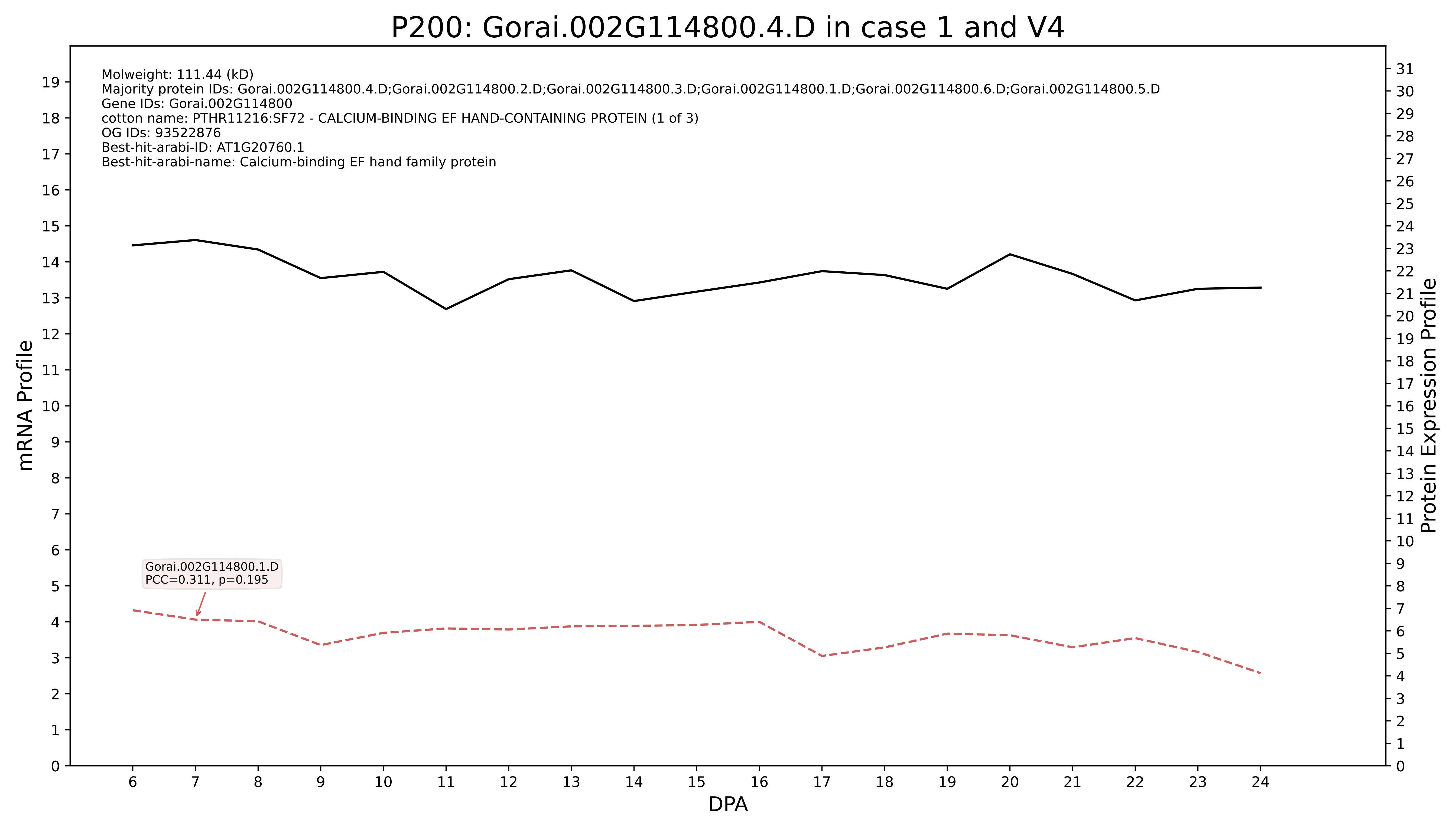
P200: Gorai.002G114800.4.D in case 1 and V4
Molweight: 111.44 (kD)
Majority protein IDs: Gorai.002G114800.4.D;Gorai.002G114800.2.D;Gorai.002G114800.3.D;Gorai.002G114800.1.D;Gorai.002G114800.6.D;Gorai.002G114800.5.D
Gene IDs: Gorai.002G114800
cotton name: PTHR11216:SF72 - CALCIUM-BINDING EF HAND-CONTAINING PROTEIN (1 of 3)
OG IDs: 93522876
Best-hit-arabi-ID: AT1G20760.1
Best-hit-arabi-name: Calcium-binding EF hand family protein
Gorai.002G114800.1.D
PCC=0.311, p=0.195
mRNA Profile
Protein Expression Profile
DPA

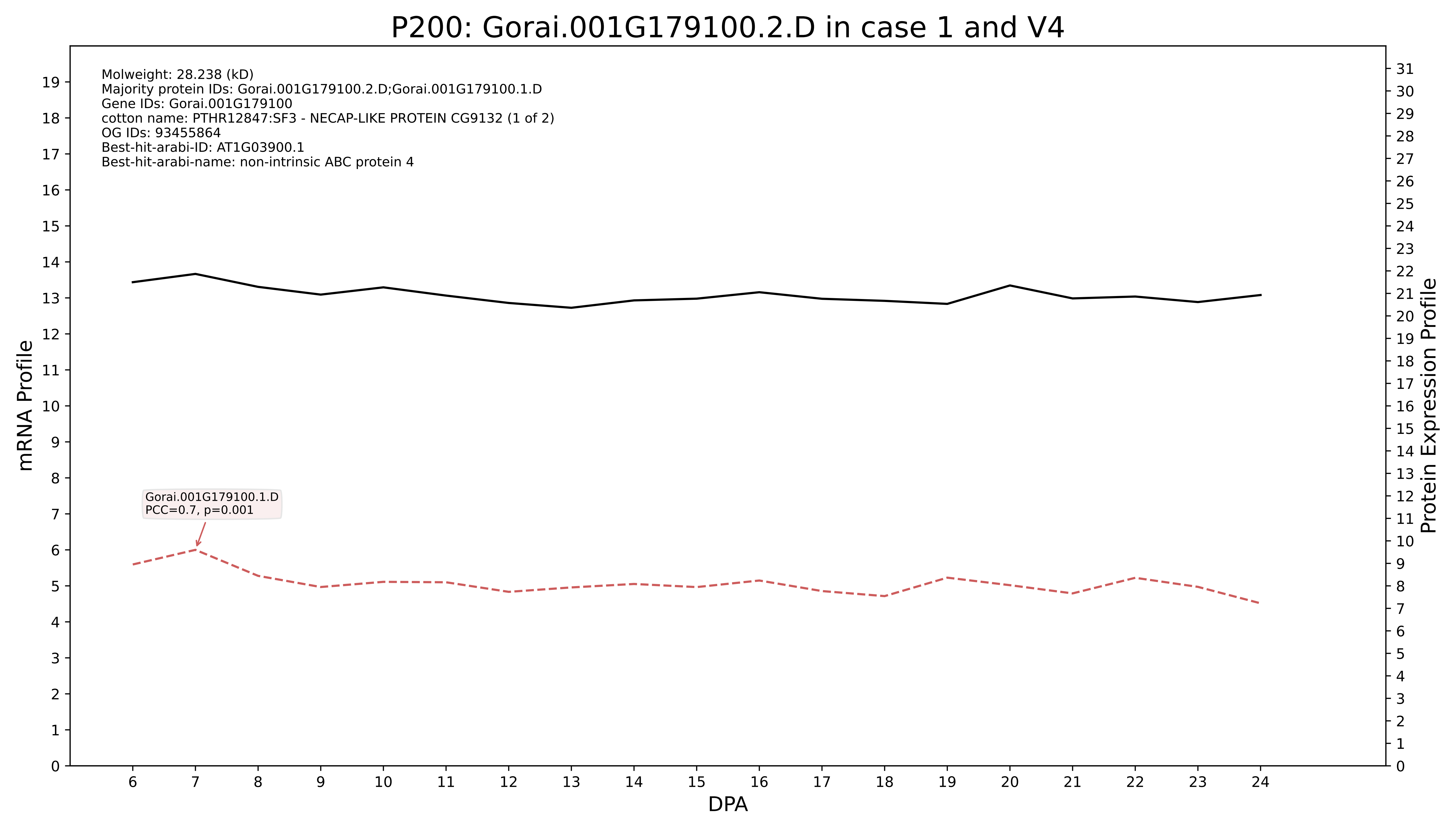

P200: Gorai.001G179100.2.D in case 1 and V4
Molweight: 28.238 (kD)
Majority protein IDs: Gorai.001G179100.2.D;Gorai.001G179100.1.D
Gene IDs: Gorai.001G179100
cotton name: PTHR12847:SF3 - NECAP-LIKE PROTEIN CG9132 (1 of 2)
OG IDs: 93455864
Best-hit-arabi-ID: AT1G03900.1
Best-hit-arabi-name: non-intrinsic ABC protein 4
Gorai.001G179100.1.D
PCC=0.7, p=0.001
mRNA Profile
Protein Expression Profile
DPA

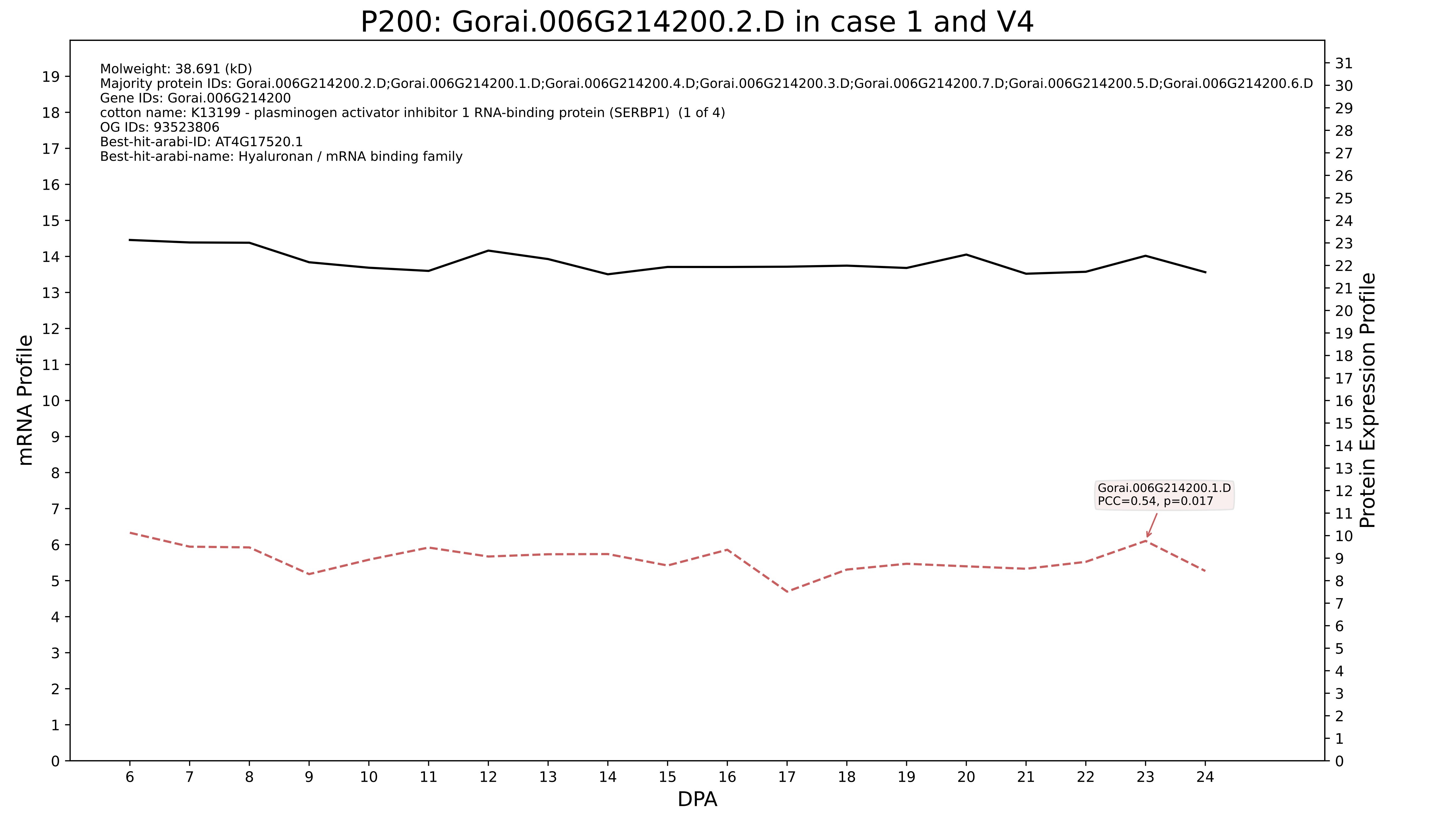

P200: Gorai.006G214200.2.D in case 1 and V4
Molweight: 38.691 (kD)
Majority protein IDs: Gorai.006G214200.2.D;Gorai.006G214200.1.D;Gorai.006G214200.4.D;Gorai.006G214200.3.D;Gorai.006G214200.7.D;Gorai.006G214200.5.D;Gorai.006G214200.6.D
Gene IDs: Gorai.006G214200
cotton name: K13199 - plasminogen activator inhibitor 1 RNA-binding protein (SERBP1) (1 of 4)
OG IDs: 93523806
Best-hit-arabi-ID: AT4G17520.1
Best-hit-arabi-name: Hyaluronan / mRNA binding family
Gorai.006G214200.1.D
PCC=0.54, p=0.017
mRNA Profile
Protein Expression Profile
DPA

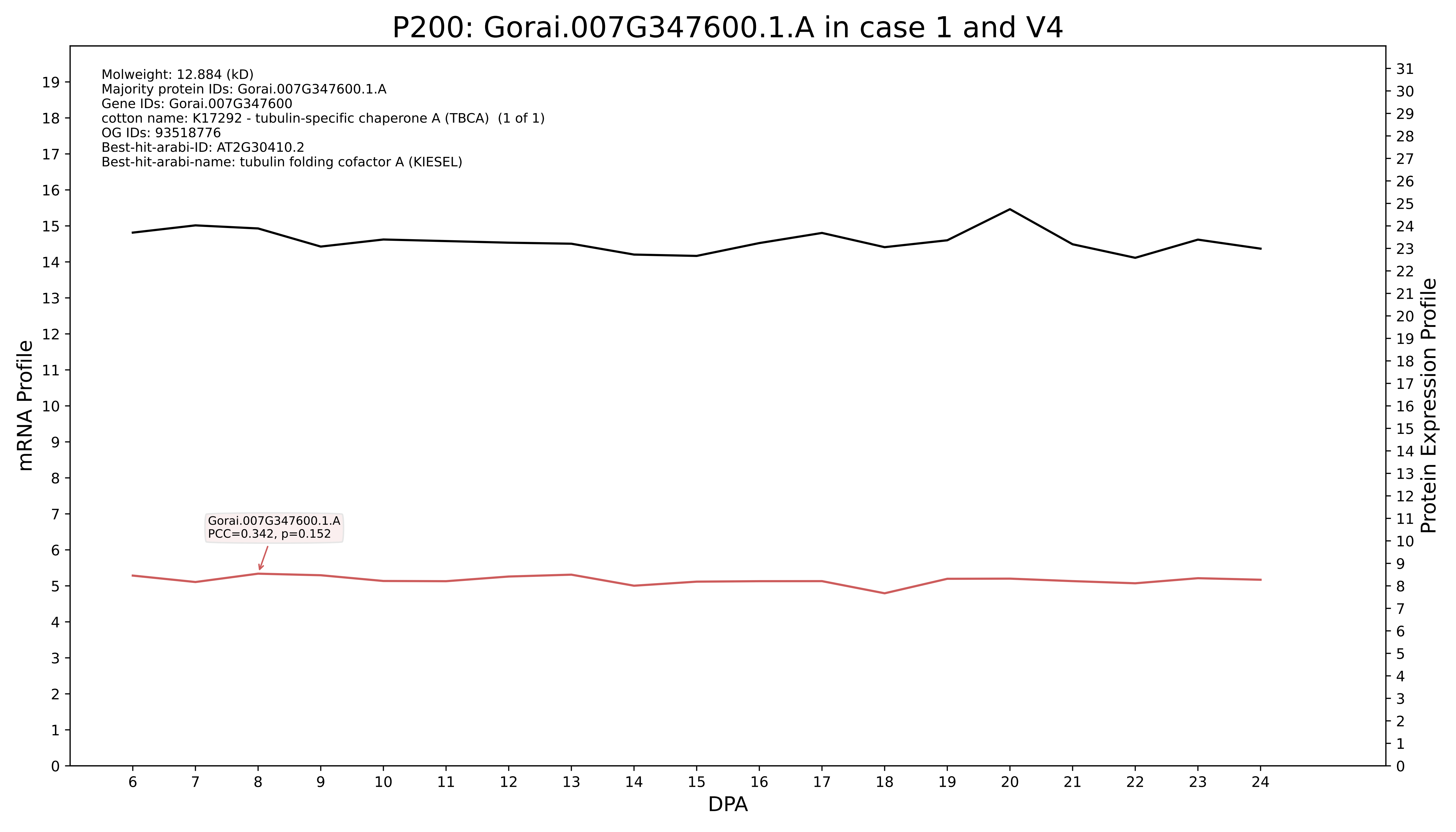
P200: Gorai.007G347600.1.A in case 1 and V4
Molweight: 12.884 (kD)
Majority protein IDs: Gorai.007G347600.1.A
Gene IDs: Gorai.007G347600
cotton name: K17292 - tubulin-specific chaperone A (TBCA) (1 of 1)
OG IDs: 93518776
Best-hit-arabi-ID: AT2G30410.2
Best-hit-arabi-name: tubulin folding cofactor A (KIESEL)
Gorai.007G347600.1.A
PCC=0.342, p=0.152
mRNA Profile
Protein Expression Profile
DPA

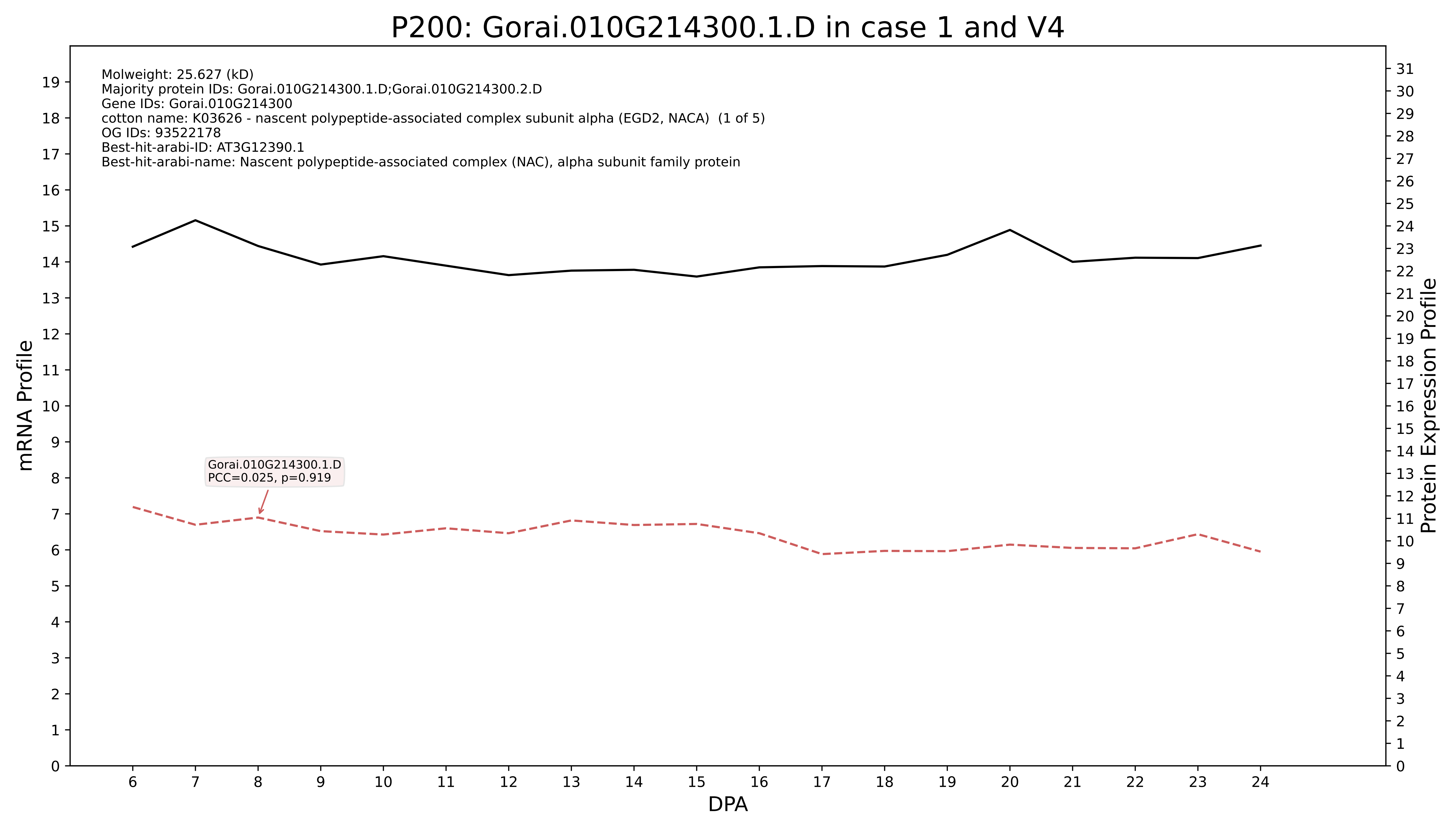

P200: Gorai.010G214300.1.D in case 1 and V4
Molweight: 25.627 (kD)
Majority protein IDs: Gorai.010G214300.1.D;Gorai.010G214300.2.D
Gene IDs: Gorai.010G214300
cotton name: K03626 - nascent polypeptide-associated complex subunit alpha (EGD2, NACA) (1 of 5)
OG IDs: 93522178
Best-hit-arabi-ID: AT3G12390.1
Best-hit-arabi-name: Nascent polypeptide-associated complex (NAC), alpha subunit family protein
Gorai.010G214300.1.D
PCC=0.025, p=0.919
mRNA Profile
Protein Expression Profile
DPA

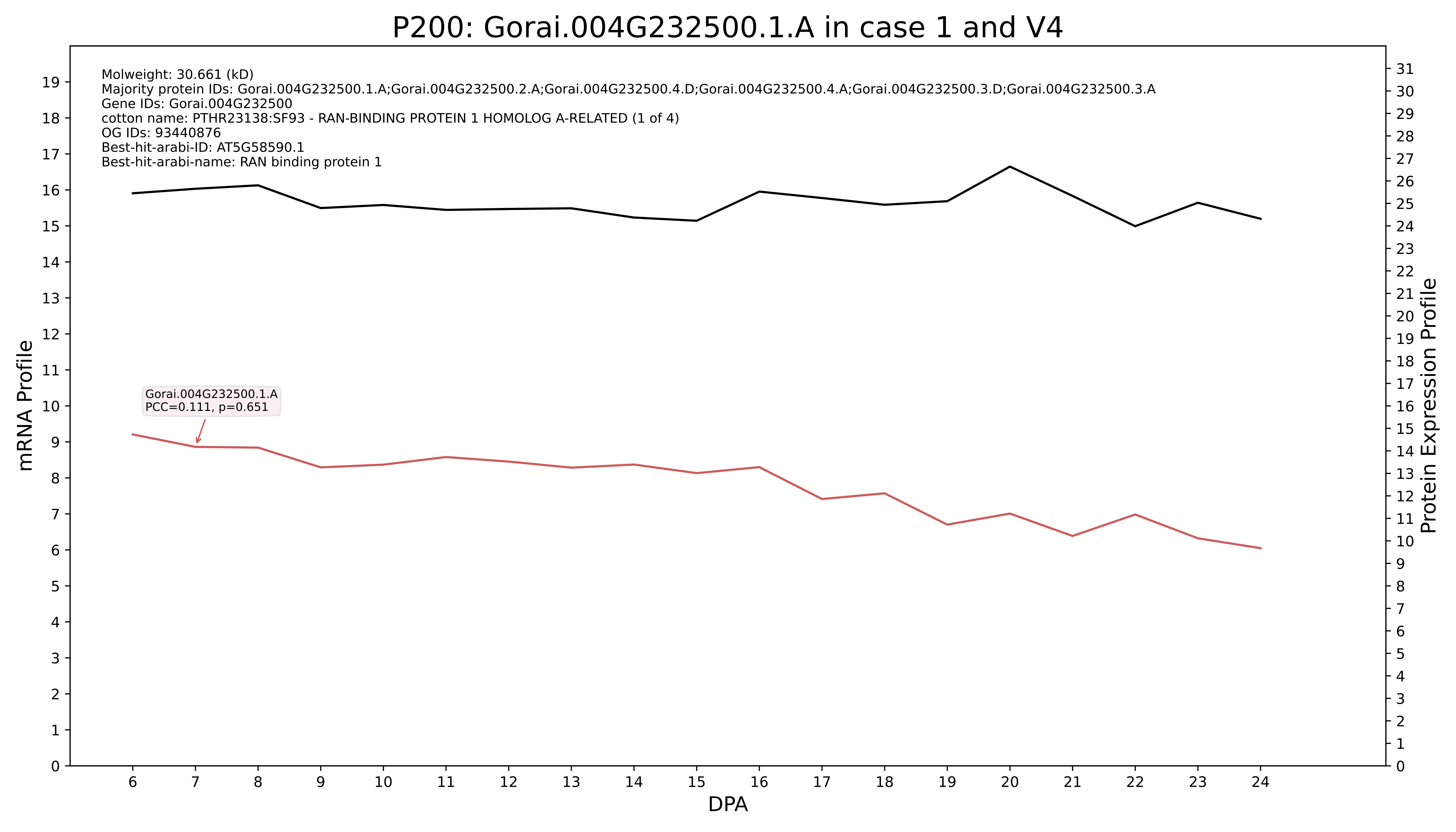
P200: Gorai.004G232500.1.A in case 1 and V4
Molweight: 30.661 (kD)
Majority protein IDs: Gorai.004G232500.1.A;Gorai.004G232500.2.A;Gorai.004G232500.4.D;Gorai.004G232500.4.A;Gorai.004G232500.3.D;Gorai.004G232500.3.A
Gene IDs: Gorai.004G232500
cotton name: PTHR23138:SF93 - RAN-BINDING PROTEIN 1 HOMOLOG A-RELATED (1 of 4)
OG IDs: 93440876
Best-hit-arabi-ID: AT5G58590.1
Best-hit-arabi-name: RAN binding protein 1
Gorai.004G232500.1.A
PCC=0.111, p=0.651
mRNA Profile
Protein Expression Profile
DPA

### P200: Gorai.009G418800.2.A in case 1 and V4

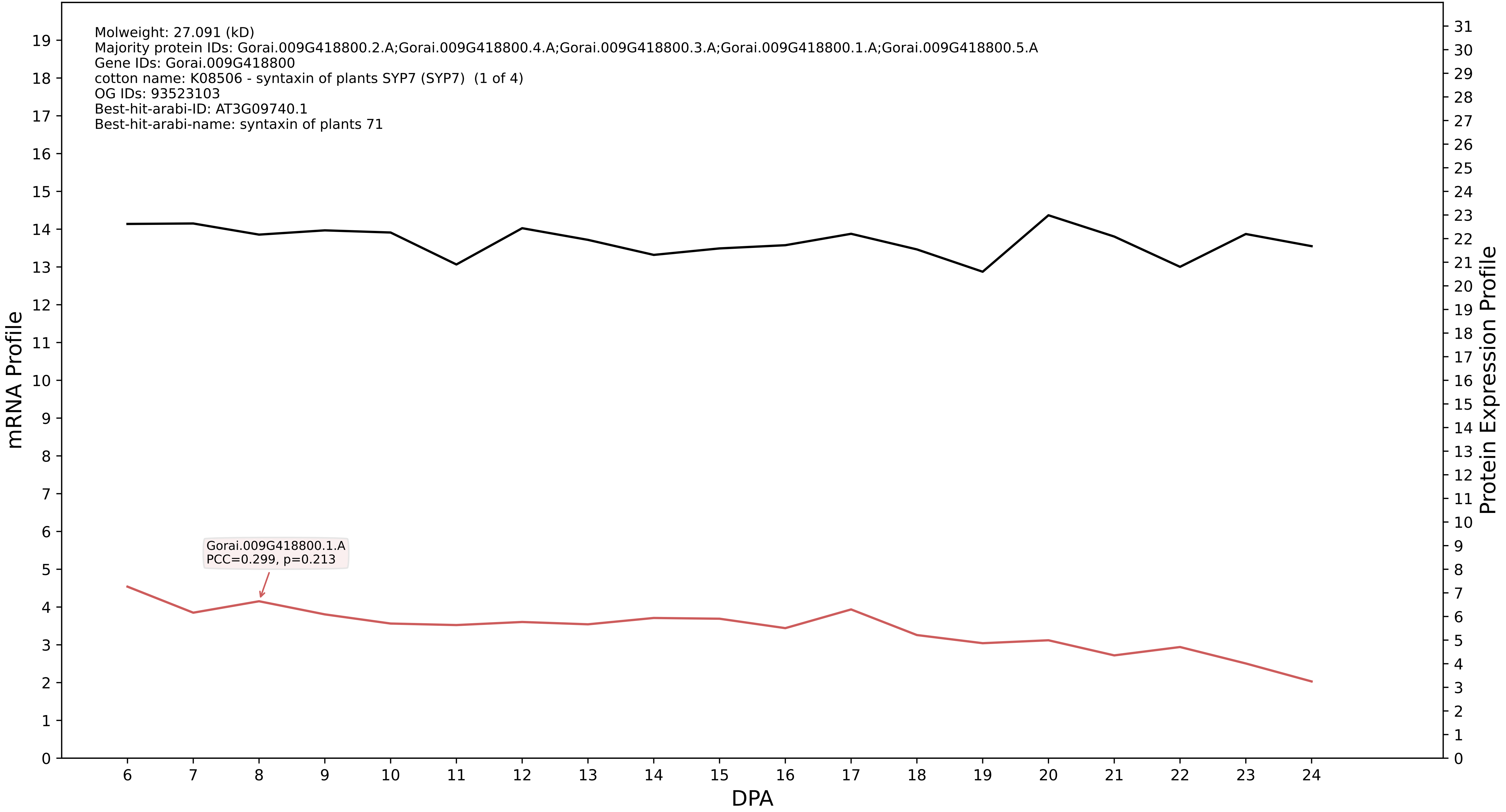

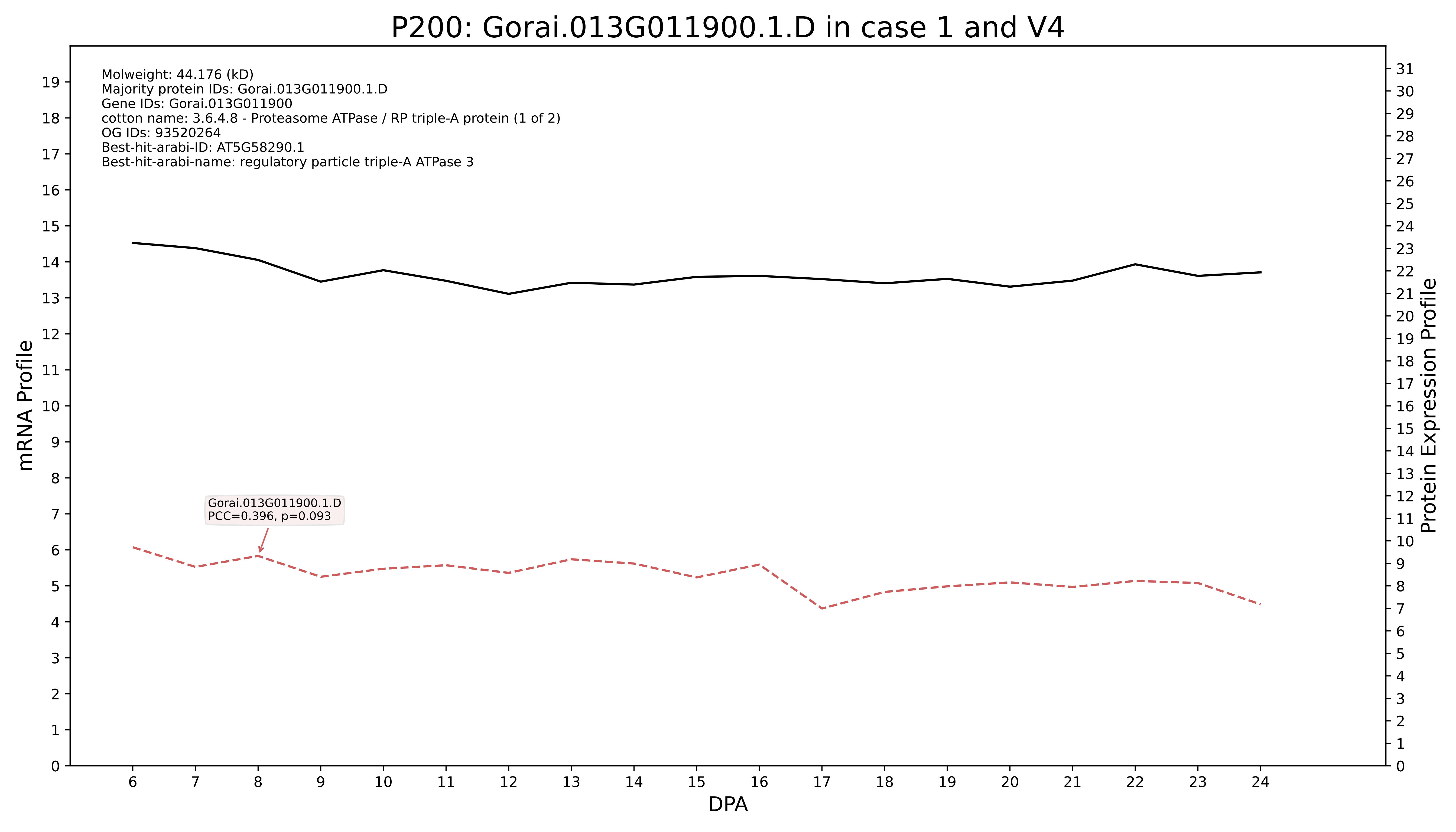
P200: Gorai.013G011900.1.D in case 1 and V4
Molweight: 44.176 (kD)
Majority protein IDs: Gorai.013G011900.1.D
Gene IDs: Gorai.013G011900
cotton name: 3.6.4.8 - Proteasome ATPase / RP triple-A protein (1 of 2)
OG IDs: 93520264
Best-hit-arabi-ID: AT5G58290.1
Best-hit-arabi-name: regulatory particle triple-A ATPase 3
Gorai.013G011900.1.D
PCC=0.396, p=0.093
mRNA Profile
Protein Expression Profile
DPA

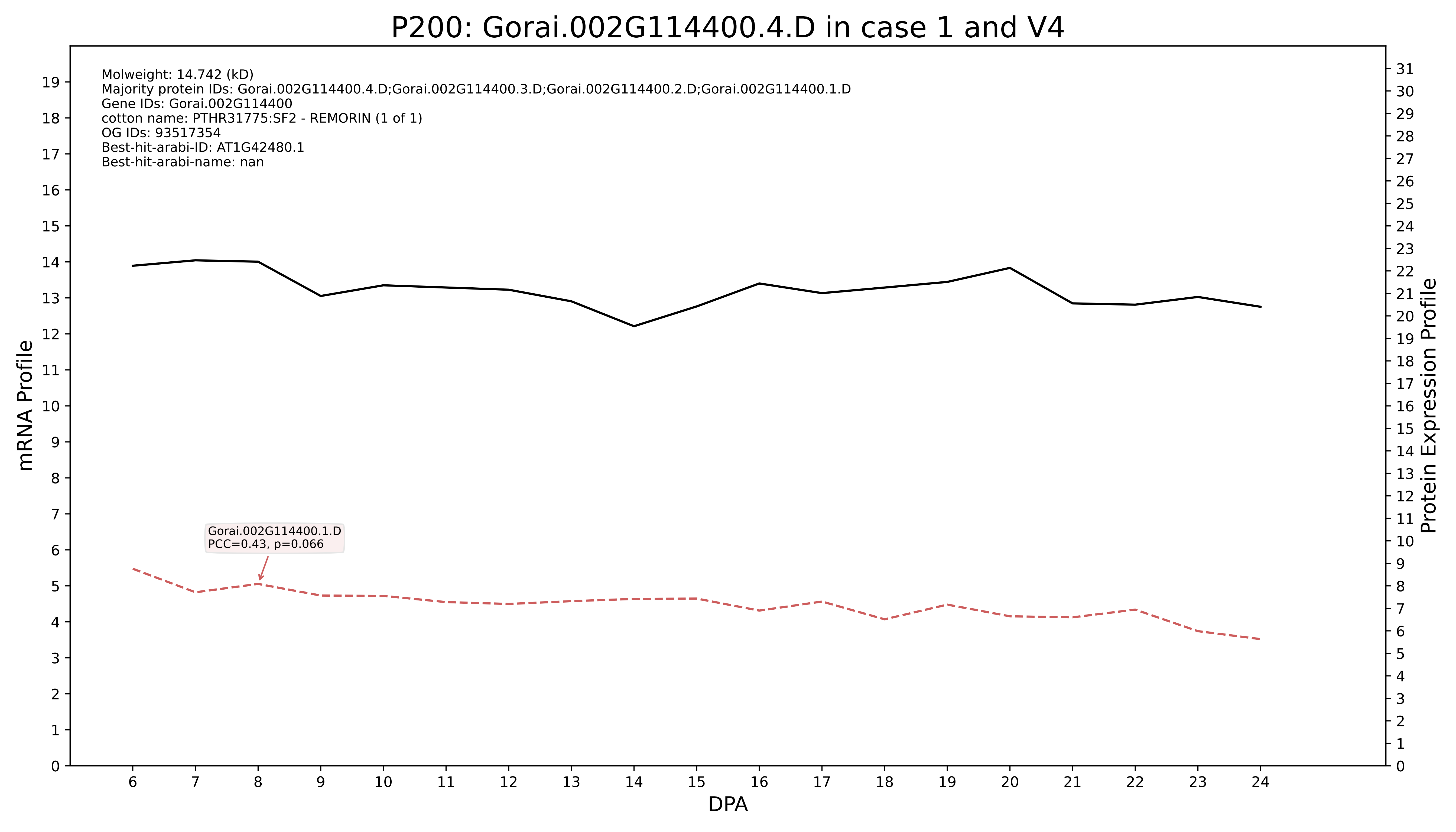

P200: Gorai.002G114400.4.D in case 1 and V4
Molweight: 14.742 (kD)
Majority protein IDs: Gorai.002G114400.4.D;Gorai.002G114400.3.D;Gorai.002G114400.2.D;Gorai.002G114400.1.D
Gene IDs: Gorai.002G114400
cotton name: PTHR31775:SF2 - REMORIN (1 of 1)
OG IDs: 93517354
Best-hit-arabi-ID: AT1G42480.1
Best-hit-arabi-name: nan
Gorai.002G114400.1.D
PCC=0.43, p=0.066
mRNA Profile
Protein Expression Profile
DPA

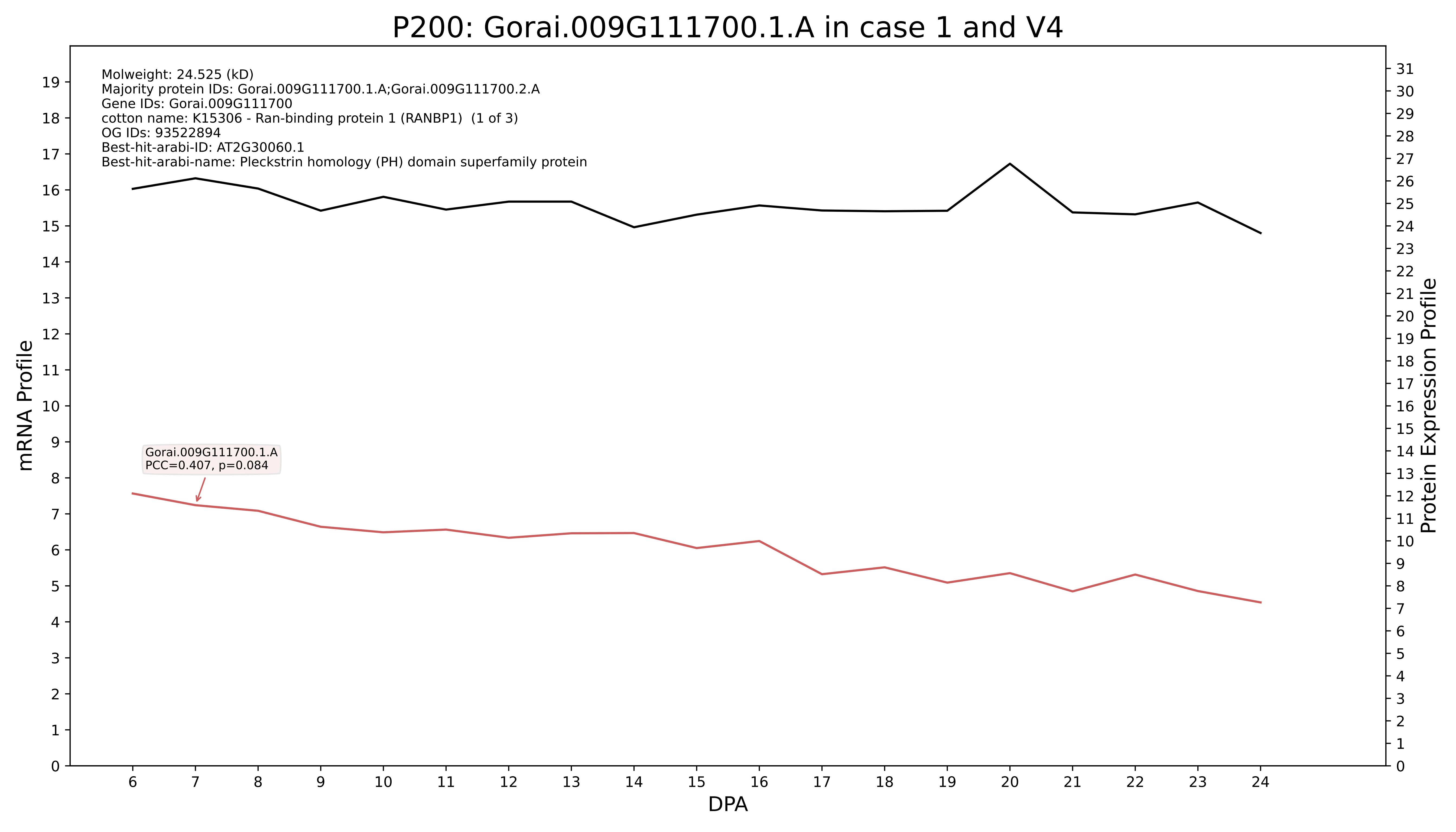
P200: Gorai.009G111700.1.A in case 1 and V4
Molweight: 24.525 (kD)
Majority protein IDs: Gorai.009G111700.1.A;Gorai.009G111700.2.A
Gene IDs: Gorai.009G111700
cotton name: K15306 - Ran-binding protein 1 (RANBP1) (1 of 3)
OG IDs: 93522894
Best-hit-arabi-ID: AT2G30060.1
Best-hit-arabi-name: Pleckstrin homology (PH) domain superfamily protein
Gorai.009G111700.1.A
PCC=0.407, p=0.084
mRNA Profile
Protein Expression Profile
DPA

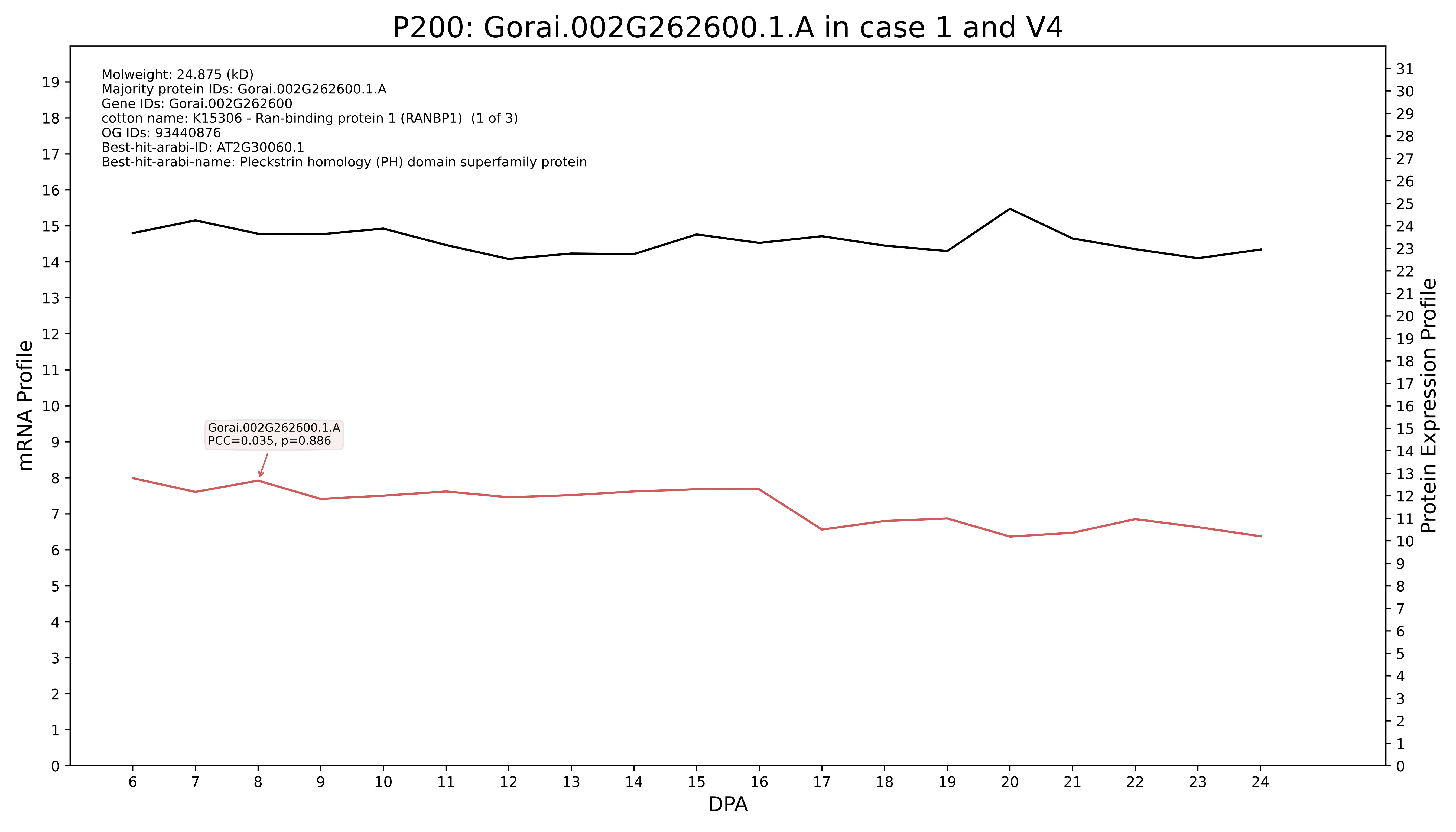
P200: Gorai.002G262600.1.A in case 1 and V4
Molweight: 24.875 (kD)
Majority protein IDs: Gorai.002G262600.1.A
Gene IDs: Gorai.002G262600
cotton name: K15306 - Ran-binding protein 1 (RANBP1) (1 of 3)
OG IDs: 93440876
Best-hit-arabi-ID: AT2G30060.1
Best-hit-arabi-name: Pleckstrin homology (PH) domain superfamily protein
Gorai.002G262600.1.A
PCC=0.035, p=0.886
mRNA Profile
Protein Expression Profile
DPA

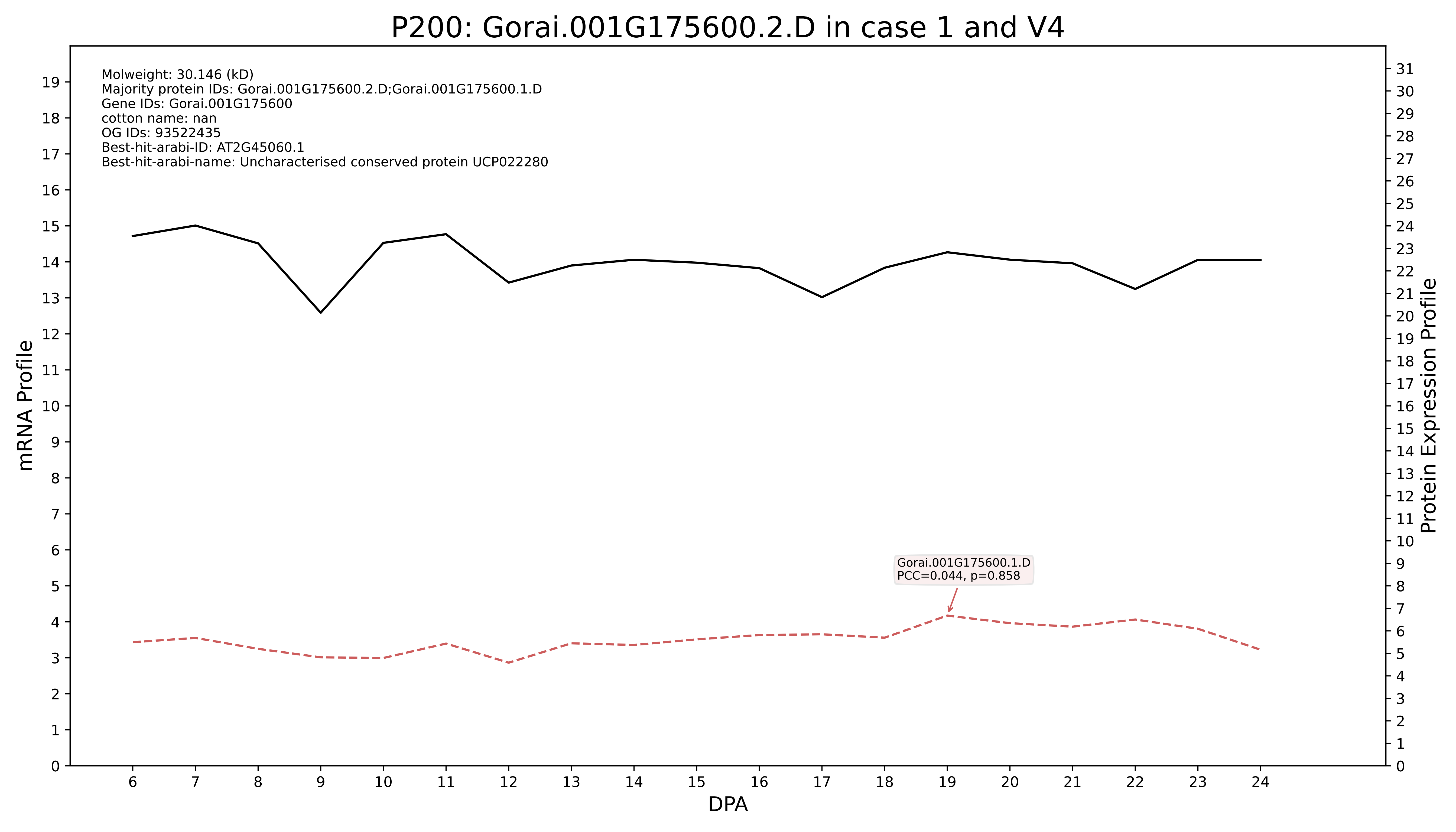

P200: Gorai.001G175600.2.D in case 1 and V4
Molweight: 30.146 (kD)
Majority protein IDs: Gorai.001G175600.2.D;Gorai.001G175600.1.D
Gene IDs: Gorai.001G175600
cotton name: nan
OG IDs: 93522435
Best-hit-arabi-ID: AT2G45060.1
Best-hit-arabi-name: Uncharacterised conserved protein UCP022280
Gorai.001G175600.1.D
PCC=0.044, p=0.858
mRNA Profile
Protein Expression Profile
DPA

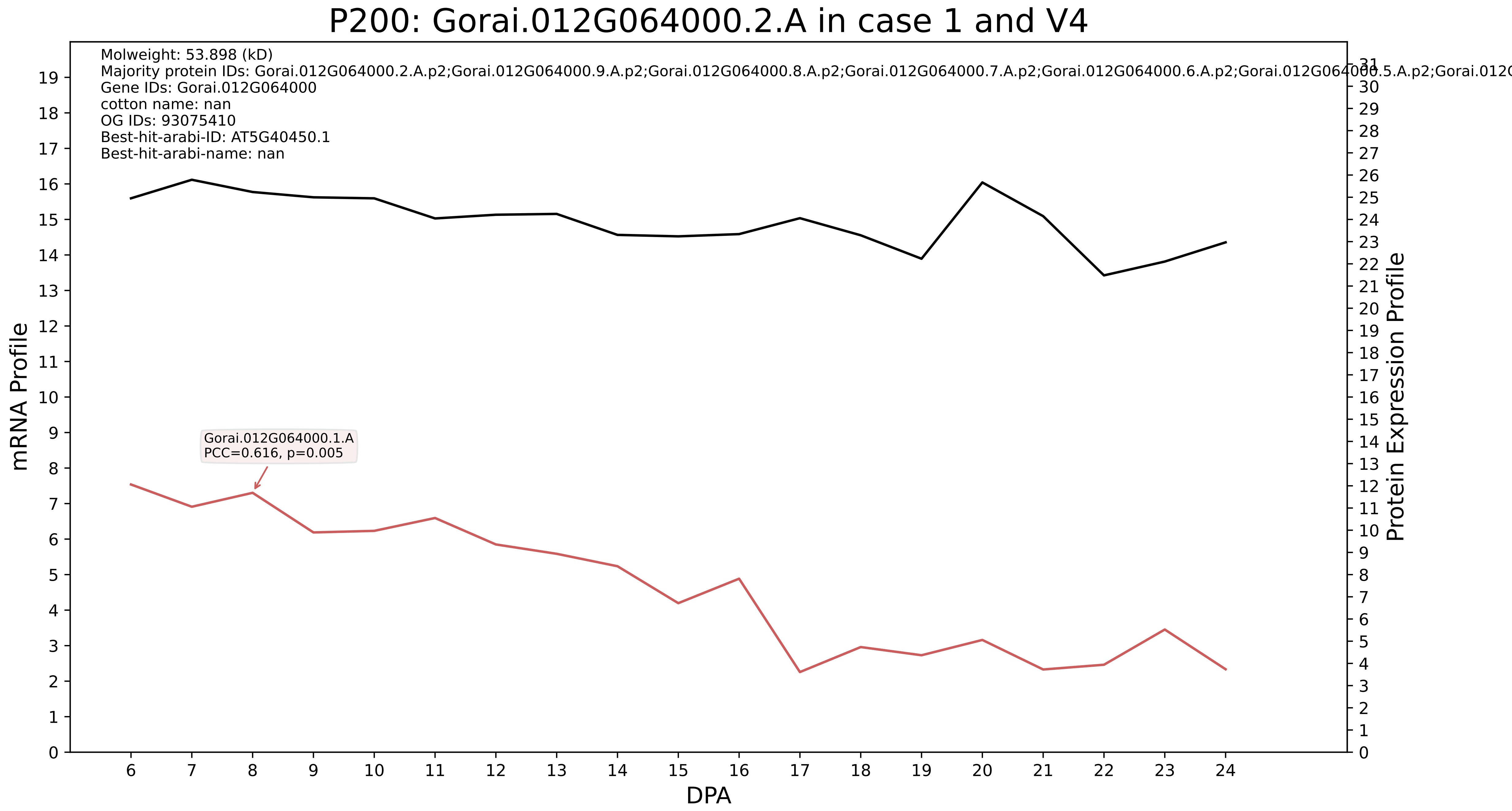

P200: Gorai.012G064000.2.A in case 1 and V4
Molweight: 53.898 (kD)
Majority protein IDs: Gorai.012G064000.2.A.p2;Gorai.012G064000.9.A.p2;Gorai.012G064000.8.A.p2;Gorai.012G064000.7.A.p2;Gorai.012G064000.6.A.p2;Gorai.012G064000.5.A.p2;Gorai.012G
Gene IDs: Gorai.012G064000
cotton name: nan
OG IDs: 93075410
Best-hit-arabi-ID: AT5G40450.1
Best-hit-arabi-name: nan
Gorai.012G064000.1.A
PCC=0.616, p=0.005
mRNA Profile
Protein Expression Profile
DPA

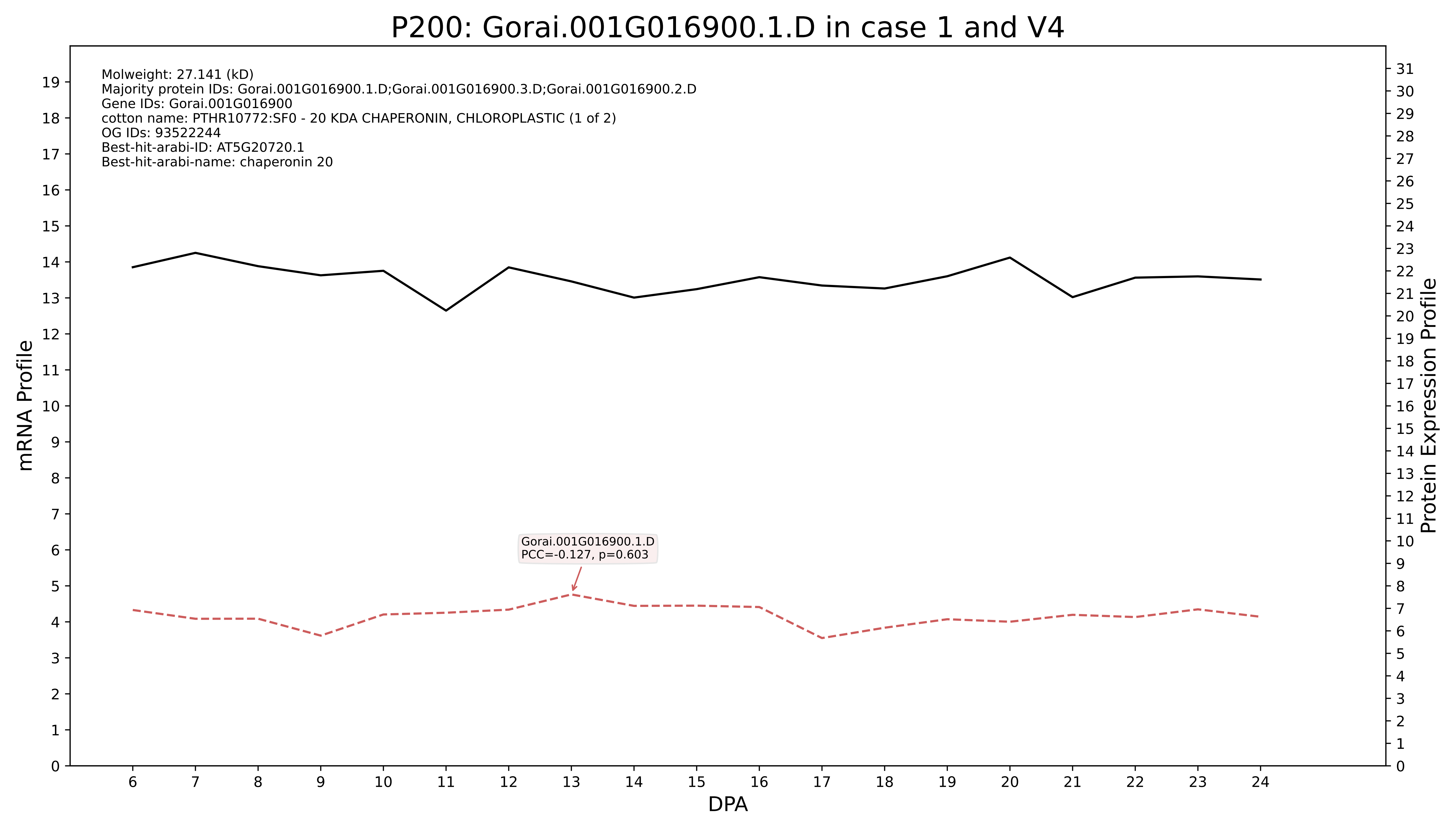

P200: Gorai.001G016900.1.D in case 1 and V4
Molweight: 27.141 (kD)
Majority protein IDs: Gorai.001G016900.1.D;Gorai.001G016900.3.D;Gorai.001G016900.2.D
Gene IDs: Gorai.001G016900
cotton name: PTHR10772:SF0 - 20 KDA CHAPERONIN, CHLOROPLASTIC (1 of 2)
OG IDs: 93522244
Best-hit-arabi-ID: AT5G20720.1
Best-hit-arabi-name: chaperonin 20
Gorai.001G016900.1.D
PCC=-0.127, p=0.603
mRNA Profile
Protein Expression Profile
DPA

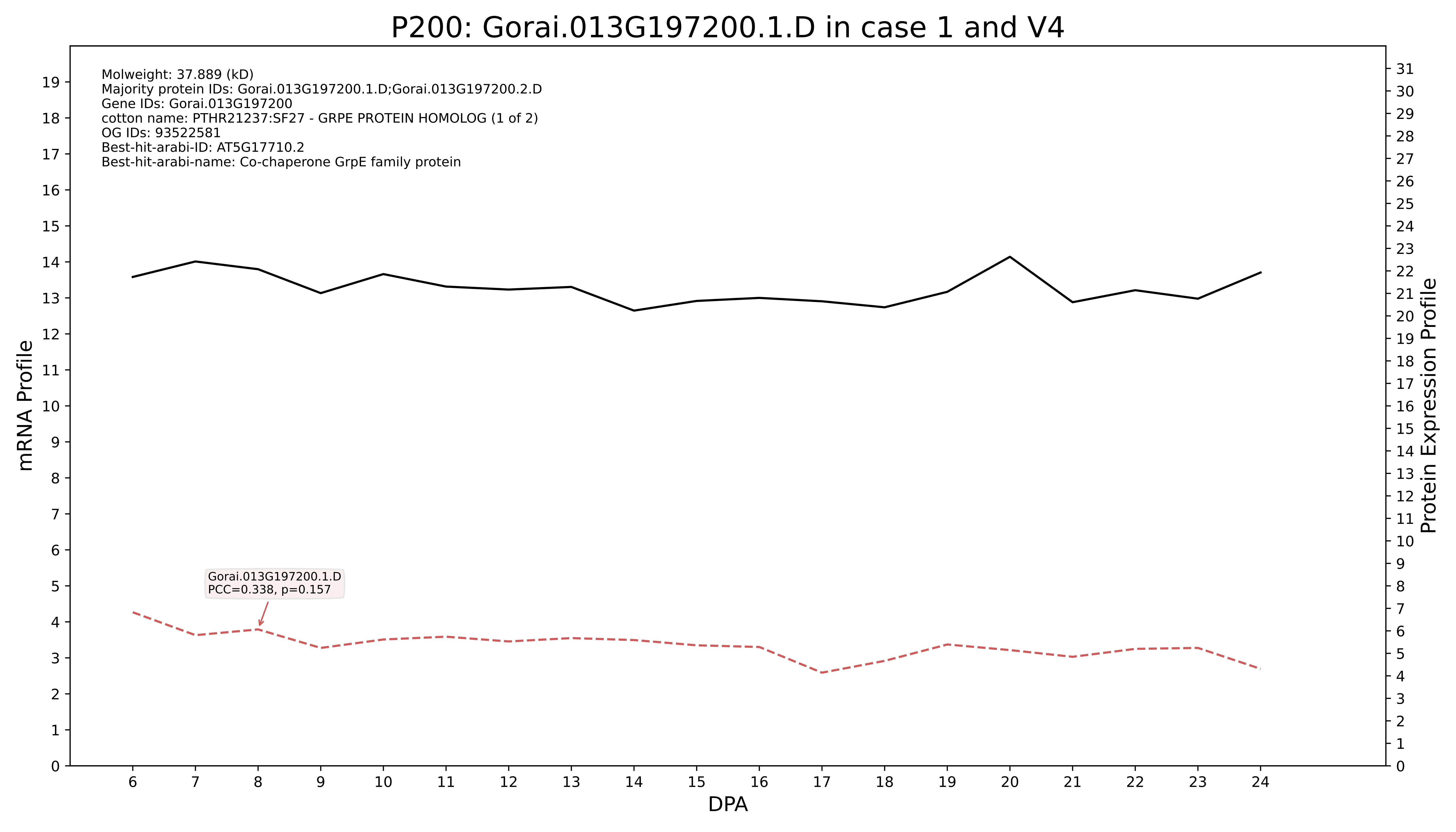
P200: Gorai.013G197200.1.D in case 1 and V4
Molweight: 37.889 (kD)
Majority protein IDs: Gorai.013G197200.1.D;Gorai.013G197200.2.D
Gene IDs: Gorai.013G197200
cotton name: PTHR21237:SF27 - GRPE PROTEIN HOMOLOG (1 of 2)
OG IDs: 93522581
Best-hit-arabi-ID: AT5G17710.2
Best-hit-arabi-name: Co-chaperone GrpE family protein
Gorai.013G197200.1.D
PCC=0.338, p=0.157
mRNA Profile
Protein Expression Profile
DPA

### P200: Gorai.006G156700.1.A in case 1 and V4

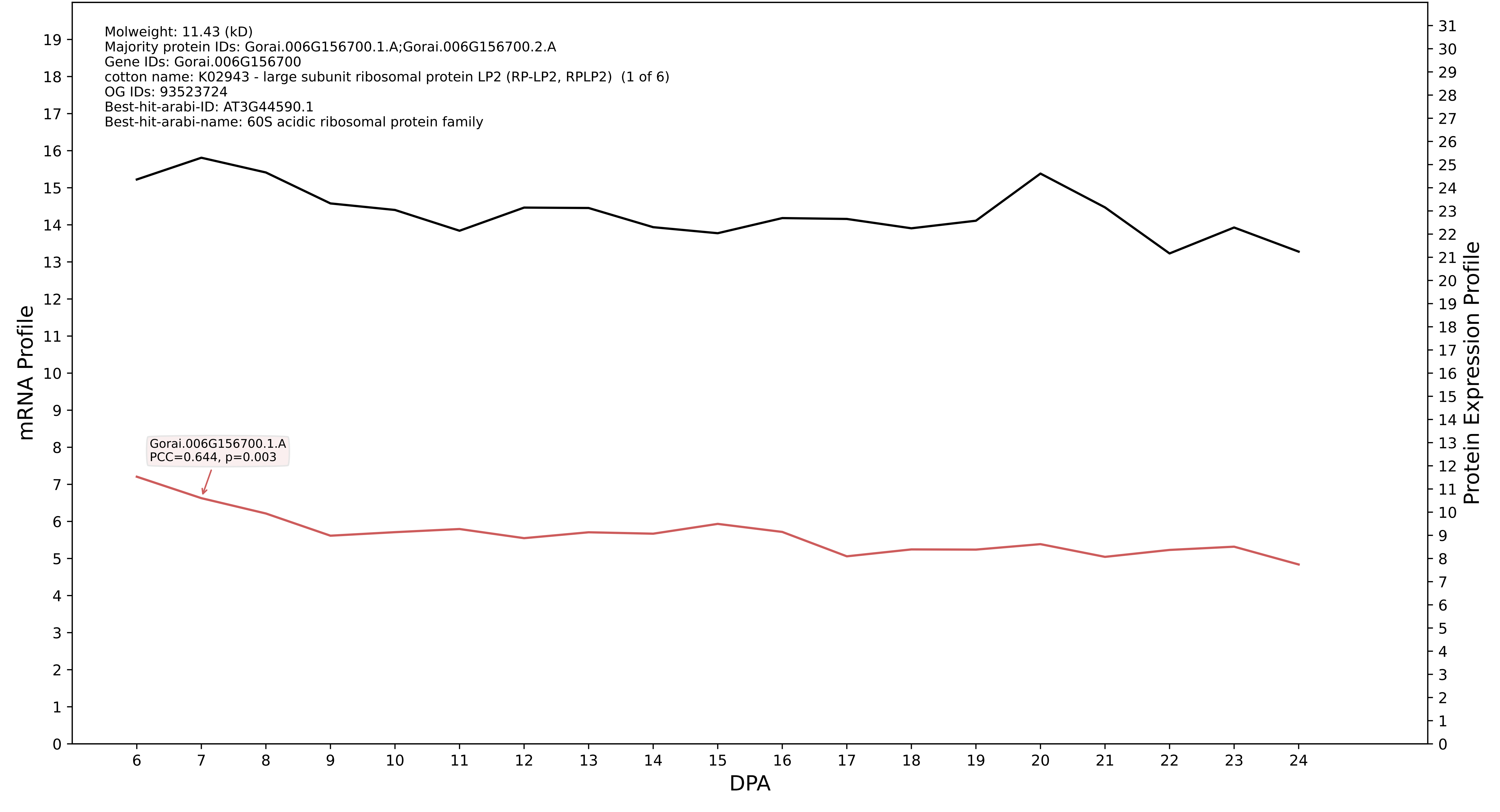

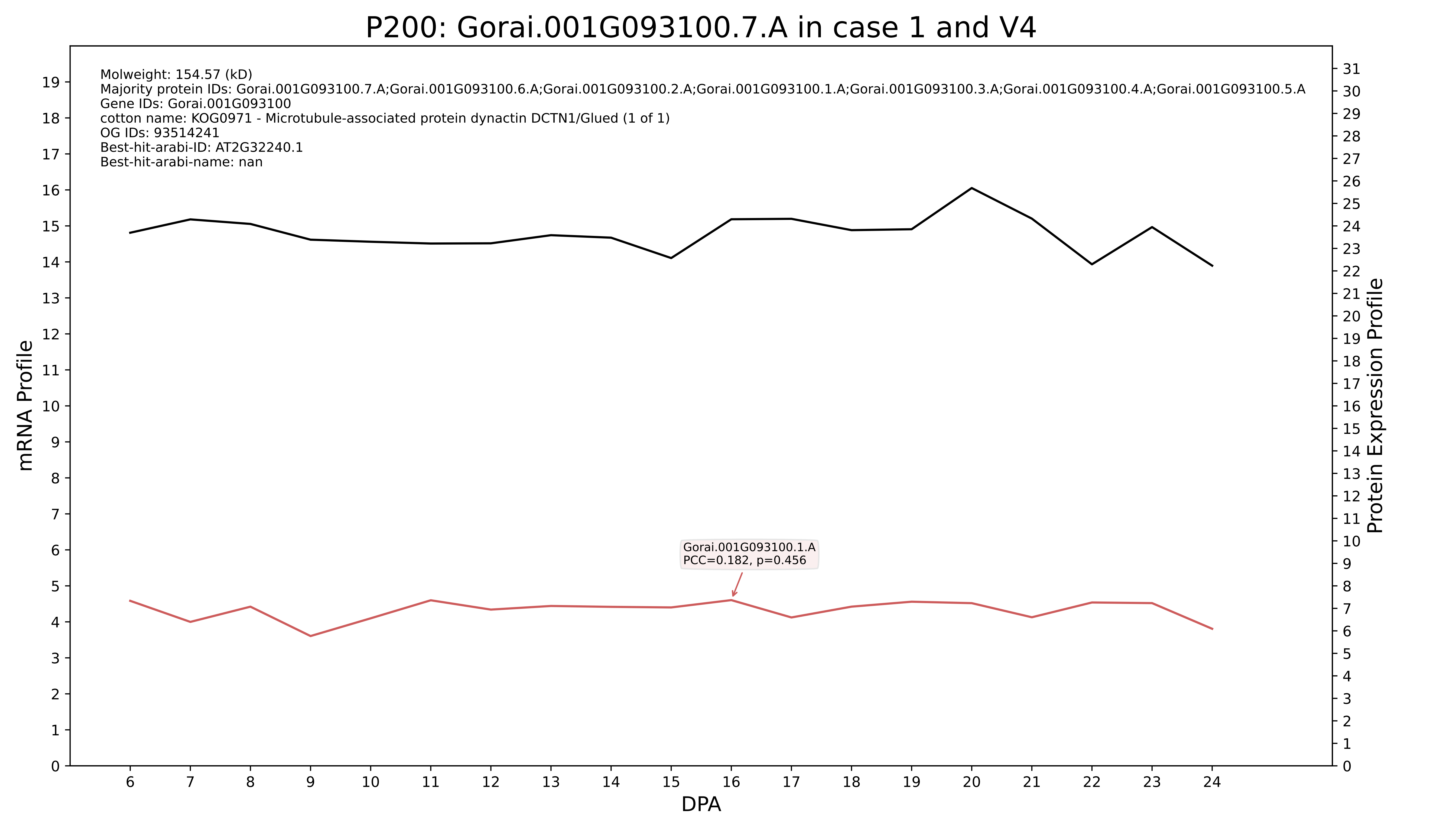
P200: Gorai.001G093100.7.A in case 1 and V4
Molweight: 154.57 (kD)
Majority protein IDs: Gorai.001G093100.7.A;Gorai.001G093100.6.A;Gorai.001G093100.2.A;Gorai.001G093100.1.A;Gorai.001G093100.3.A;Gorai.001G093100.4.A;Gorai.001G093100.5.A
Gene IDs: Gorai.001G093100
cotton name: KOG0971 - Microtubule-associated protein dynactin DCTN1/Glued (1 of 1)
OG IDs: 93514241
Best-hit-arabi-ID: AT2G32240.1
Best-hit-arabi-name: nan
Gorai.001G093100.1.A
PCC=0.182, p=0.456
mRNA Profile
Protein Expression Profile
DPA
0 1 2 3 4 5 6 7 8 9 10 11 12 13 14 15 16 17 18 19
0 1 2 3 4 5 6 7 8 9 10 11 12 13 14 15 16 17 18 19 20 21 22 23 24 25 26 27 28 29 30 31
6 7 8 9 10 11 12 13 14 15 16 17 18 19 20 21 22 23 24

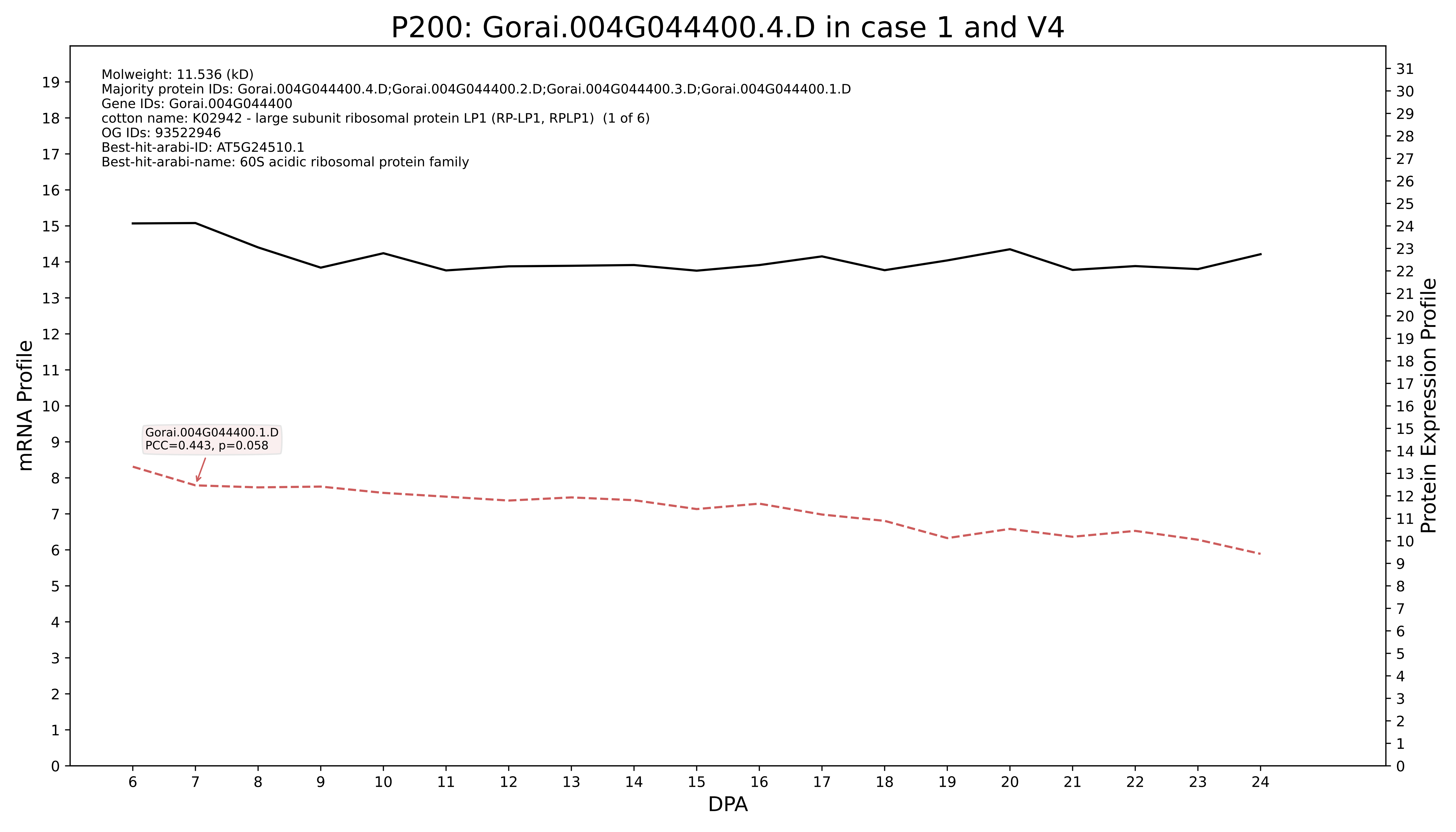
P200: Gorai.004G044400.4.D in case 1 and V4
Molweight: 11.536 (kD)
Majority protein IDs: Gorai.004G044400.4.D;Gorai.004G044400.2.D;Gorai.004G044400.3.D;Gorai.004G044400.1.D
Gene IDs: Gorai.004G044400
cotton name: K02942 - large subunit ribosomal protein LP1 (RP-LP1, RPLP1) (1 of 6)
OG IDs: 93522946
Best-hit-arabi-ID: AT5G24510.1
Best-hit-arabi-name: 60S acidic ribosomal protein family
Gorai.004G044400.1.D
PCC=0.443, p=0.058
mRNA Profile
Protein Expression Profile
DPA

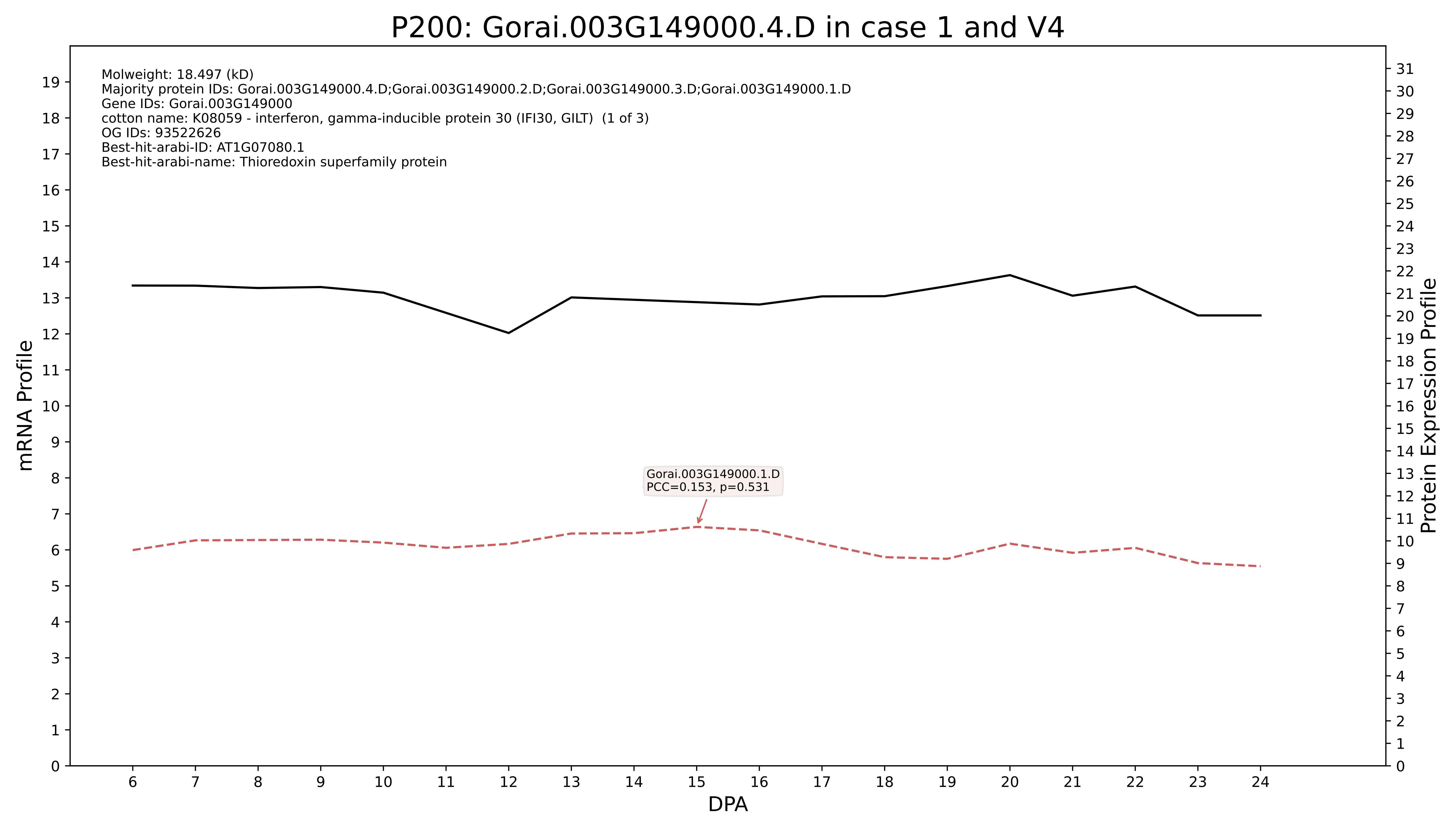

P200: Gorai.003G149000.4.D in case 1 and V4
Molweight: 18.497 (kD)
Majority protein IDs: Gorai.003G149000.4.D;Gorai.003G149000.2.D;Gorai.003G149000.3.D;Gorai.003G149000.1.D
Gene IDs: Gorai.003G149000
cotton name: K08059 - interferon, gamma-inducible protein 30 (IFI30, GILT) (1 of 3)
OG IDs: 93522626
Best-hit-arabi-ID: AT1G07080.1
Best-hit-arabi-name: Thioredoxin superfamily protein
Gorai.003G149000.1.D
PCC=0.153, p=0.531
mRNA Profile
Protein Expression Profile
DPA

00: Gorai.012G064000.2.D in case 1 and V4

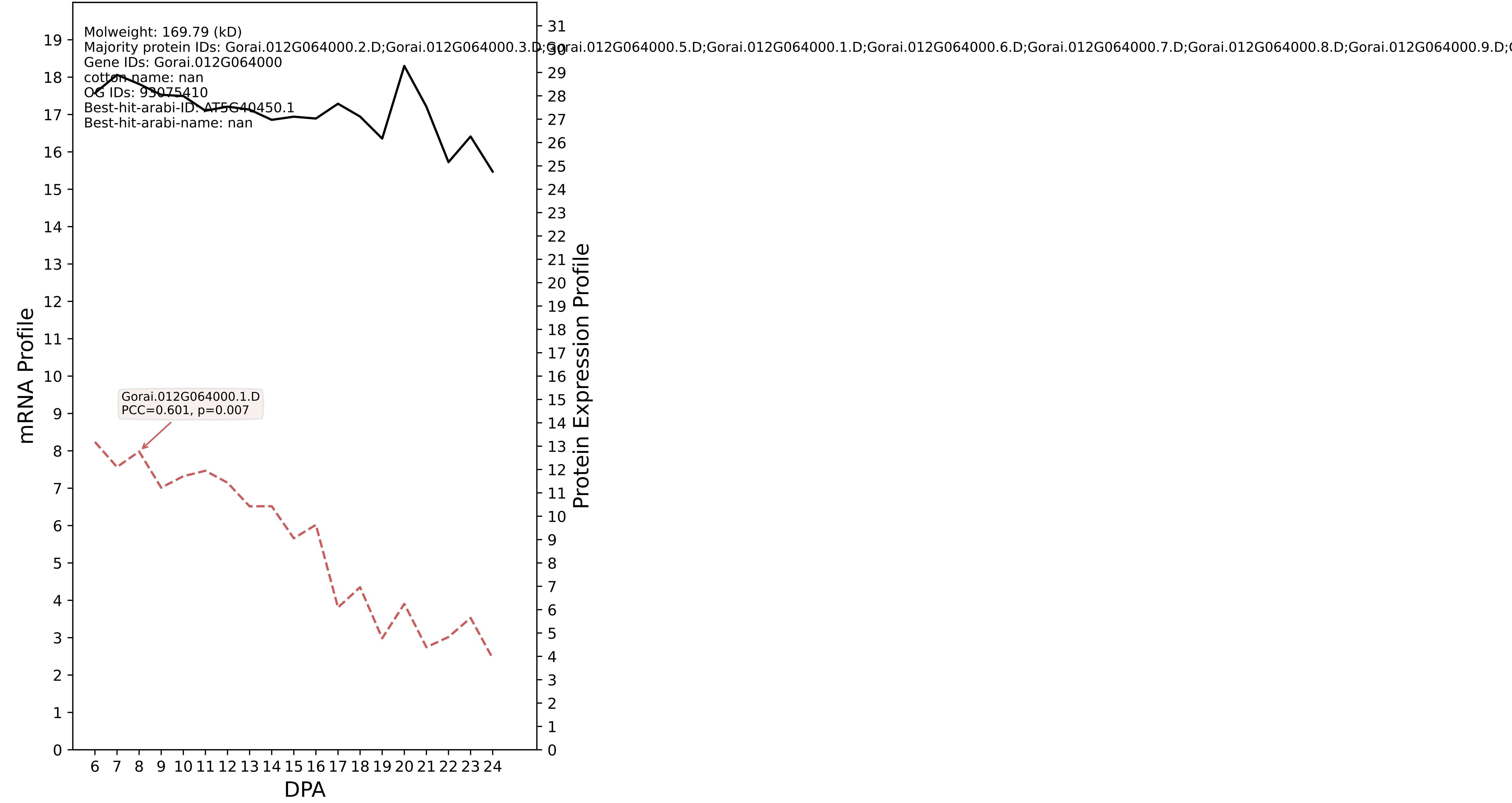

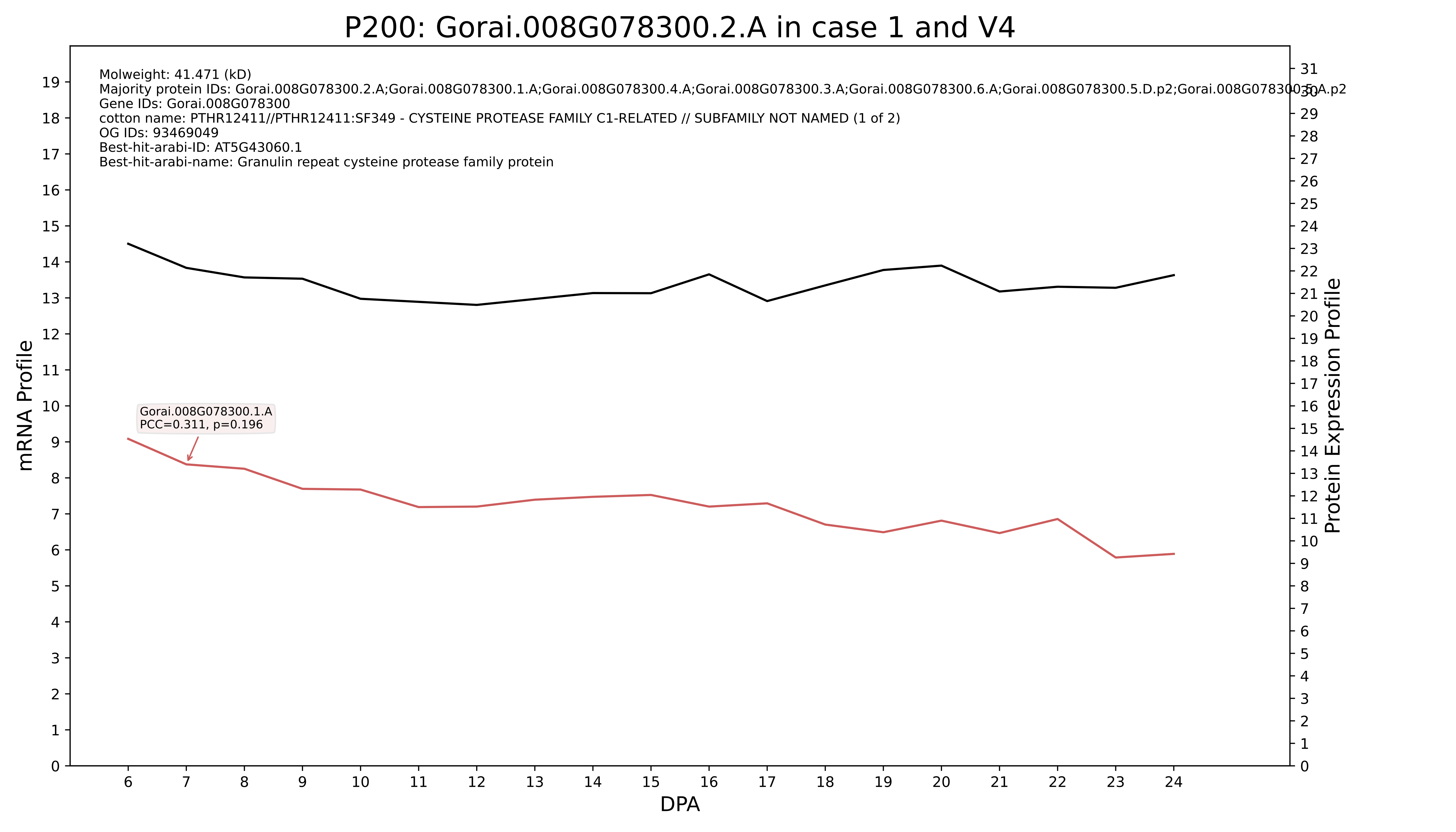
P200: Gorai.008G078300.2.A in case 1 and V4
Molweight: 41.471 (kD)
Majority protein IDs: Gorai.008G078300.2.A;Gorai.008G078300.1.A;Gorai.008G078300.4.A;Gorai.008G078300.3.A;Gorai.008G078300.6.A;Gorai.008G078300.5.D.p2;Gorai.008G078300.5.A.p2
Gene IDs: Gorai.008G078300
cotton name: PTHR12411//PTHR12411:SF349 - CYSTEINE PROTEASE FAMILY C1-RELATED // SUBFAMILY NOT NAMED (1 of 2)
OG IDs: 93469049
Best-hit-arabi-ID: AT5G43060.1
Best-hit-arabi-name: Granulin repeat cysteine protease family protein
Gorai.008G078300.1.A
PCC=0.311, p=0.196
mRNA Profile
Protein Expression Profile
DPA

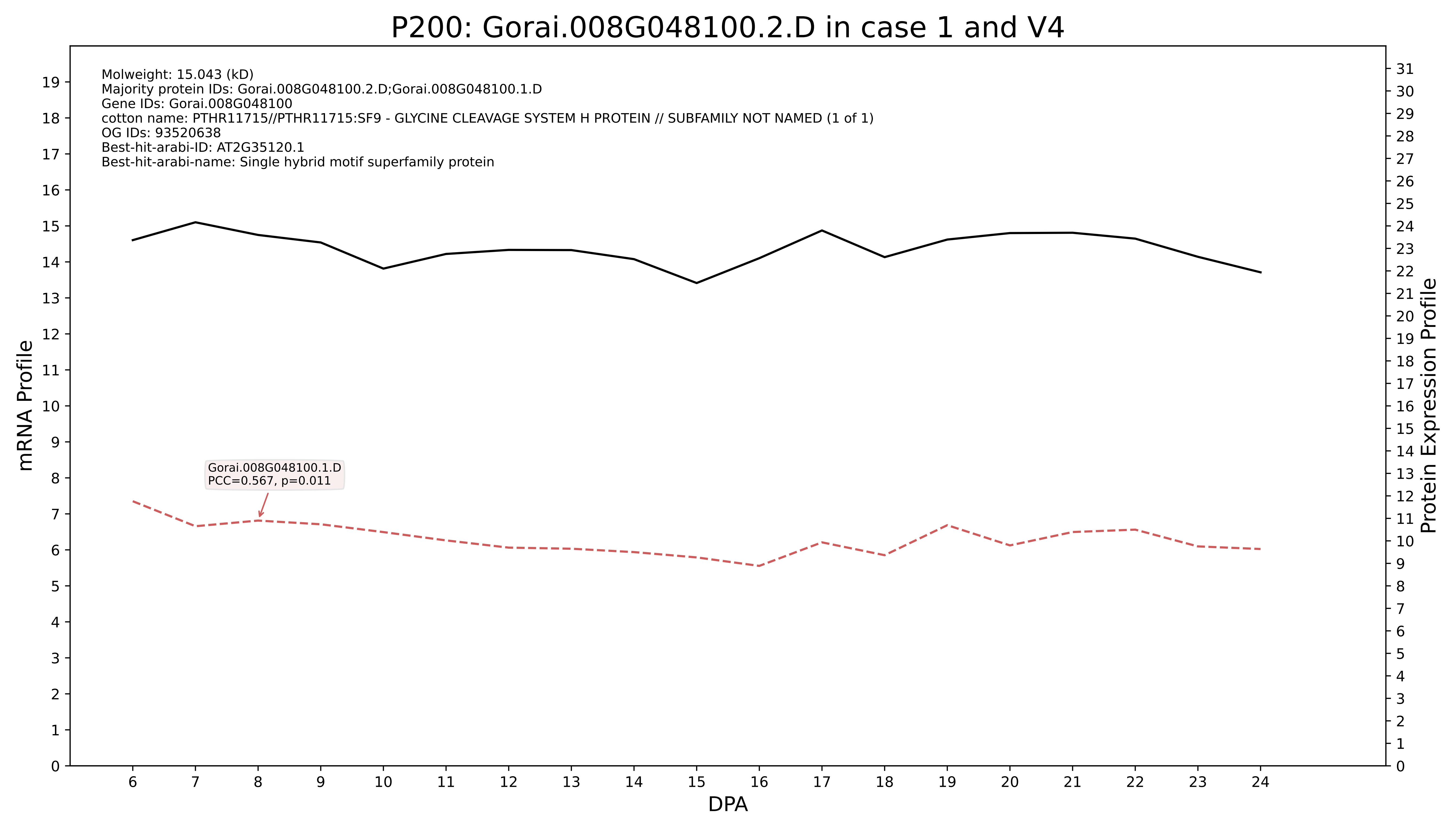

P200: Gorai.008G048100.2.D in case 1 and V4
Molweight: 15.043 (kD)
Majority protein IDs: Gorai.008G048100.2.D;Gorai.008G048100.1.D
Gene IDs: Gorai.008G048100
cotton name: PTHR11715//PTHR11715:SF9 - GLYCINE CLEAVAGE SYSTEM H PROTEIN // SUBFAMILY NOT NAMED (1 of 1)
OG IDs: 93520638
Best-hit-arabi-ID: AT2G35120.1
Best-hit-arabi-name: Single hybrid motif superfamily protein
Gorai.008G048100.1.D
PCC=0.567, p=0.011
mRNA Profile
Protein Expression Profile
DPA

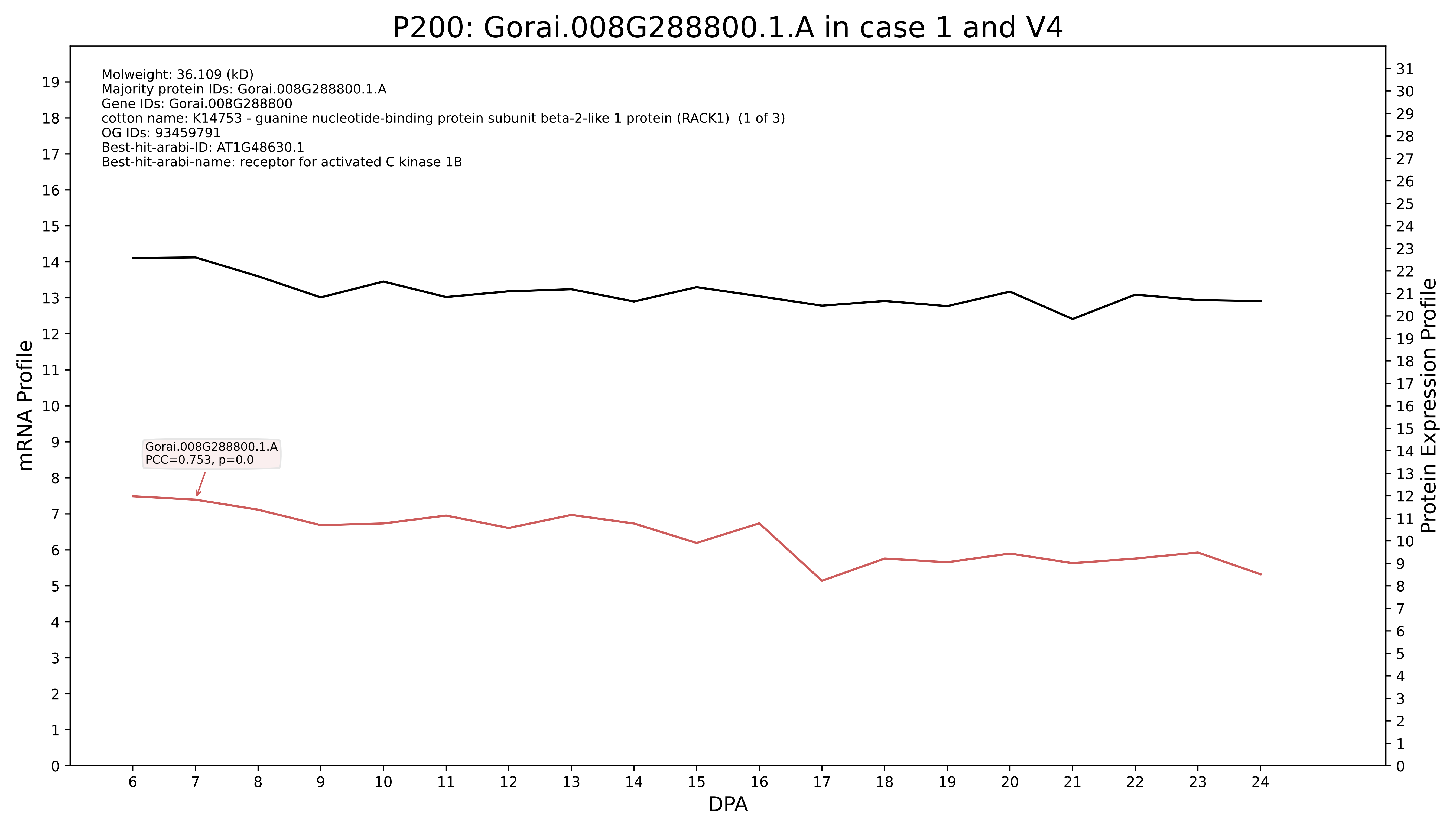
P200: Gorai.008G288800.1.A in case 1 and V4
Molweight: 36.109 (kD)
Majority protein IDs: Gorai.008G288800.1.A
Gene IDs: Gorai.008G288800
cotton name: K14753 - guanine nucleotide-binding protein subunit beta-2-like 1 protein (RACK1) (1 of 3)
OG IDs: 93459791
Best-hit-arabi-ID: AT1G48630.1
Best-hit-arabi-name: receptor for activated C kinase 1B
Gorai.008G288800.1.A
PCC=0.753, p=0.0
mRNA Profile
Protein Expression Profile
DPA

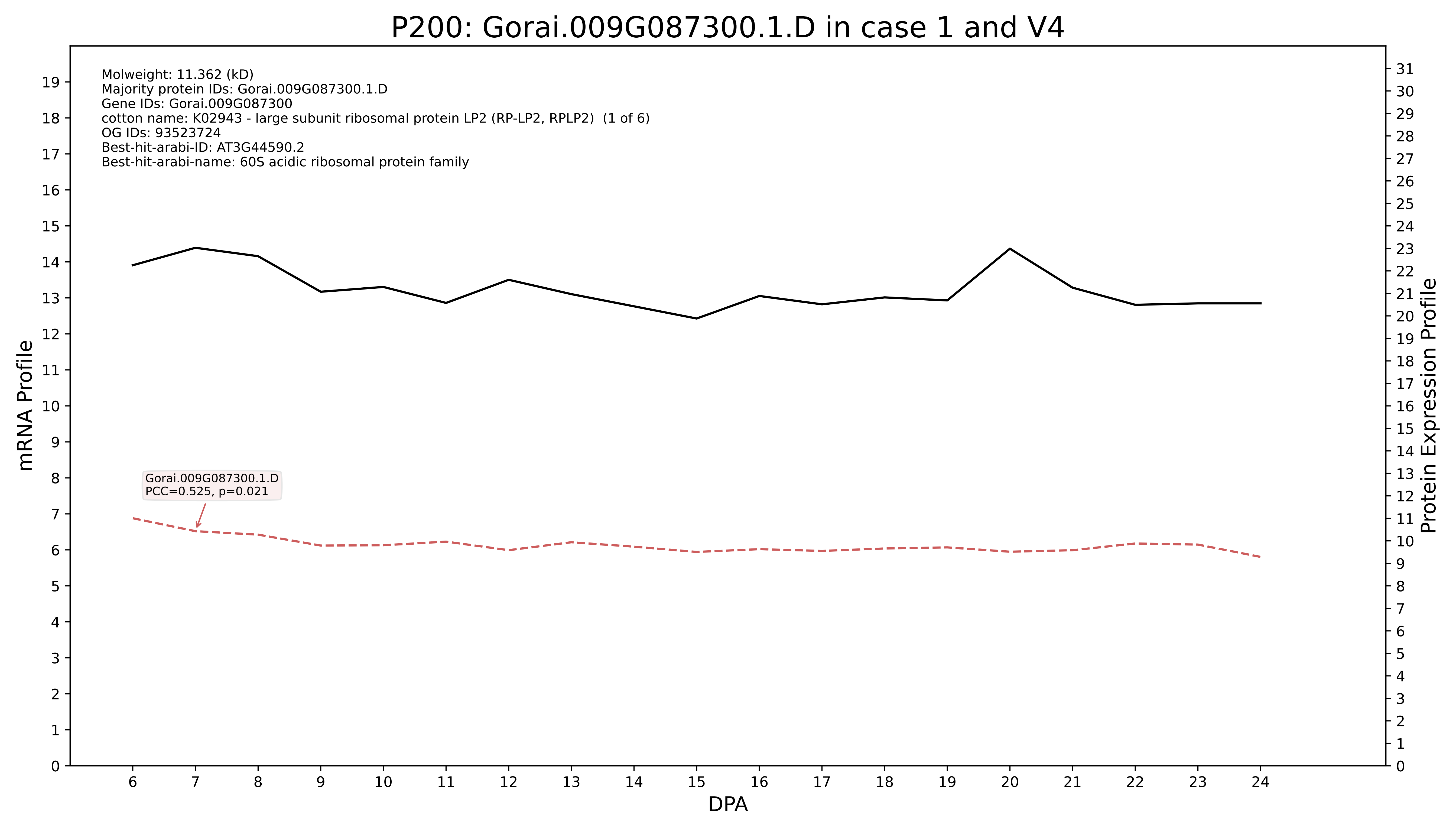
P200: Gorai.009G087300.1.D in case 1 and V4
Molweight: 11.362 (kD)
Majority protein IDs: Gorai.009G087300.1.D
Gene IDs: Gorai.009G087300
cotton name: K02943 - large subunit ribosomal protein LP2 (RP-LP2, RPLP2) (1 of 6)
OG IDs: 93523724
Best-hit-arabi-ID: AT3G44590.2
Best-hit-arabi-name: 60S acidic ribosomal protein family
Gorai.009G087300.1.D
PCC=0.525, p=0.021
mRNA Profile
Protein Expression Profile
DPA

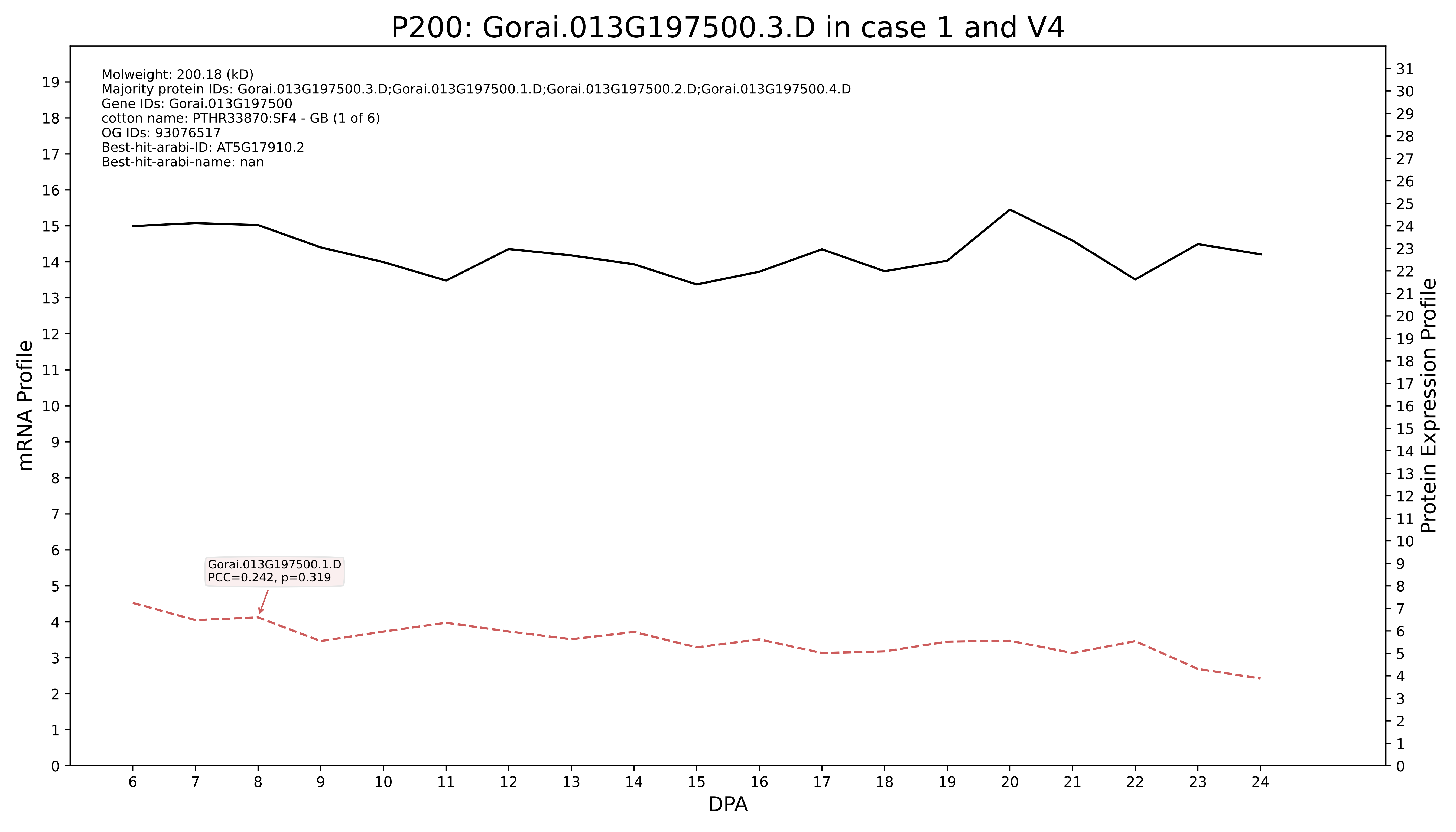
P200: Gorai.013G197500.3.D in case 1 and V4
Molweight: 200.18 (kD)
Majority protein IDs: Gorai.013G197500.3.D;Gorai.013G197500.1.D;Gorai.013G197500.2.D;Gorai.013G197500.4.D
Gene IDs: Gorai.013G197500
cotton name: PTHR33870:SF4 - GB (1 of 6)
OG IDs: 93076517
Best-hit-arabi-ID: AT5G17910.2
Best-hit-arabi-name: nan
Gorai.013G197500.1.D
PCC=0.242, p=0.319
mRNA Profile
Protein Expression Profile
DPA

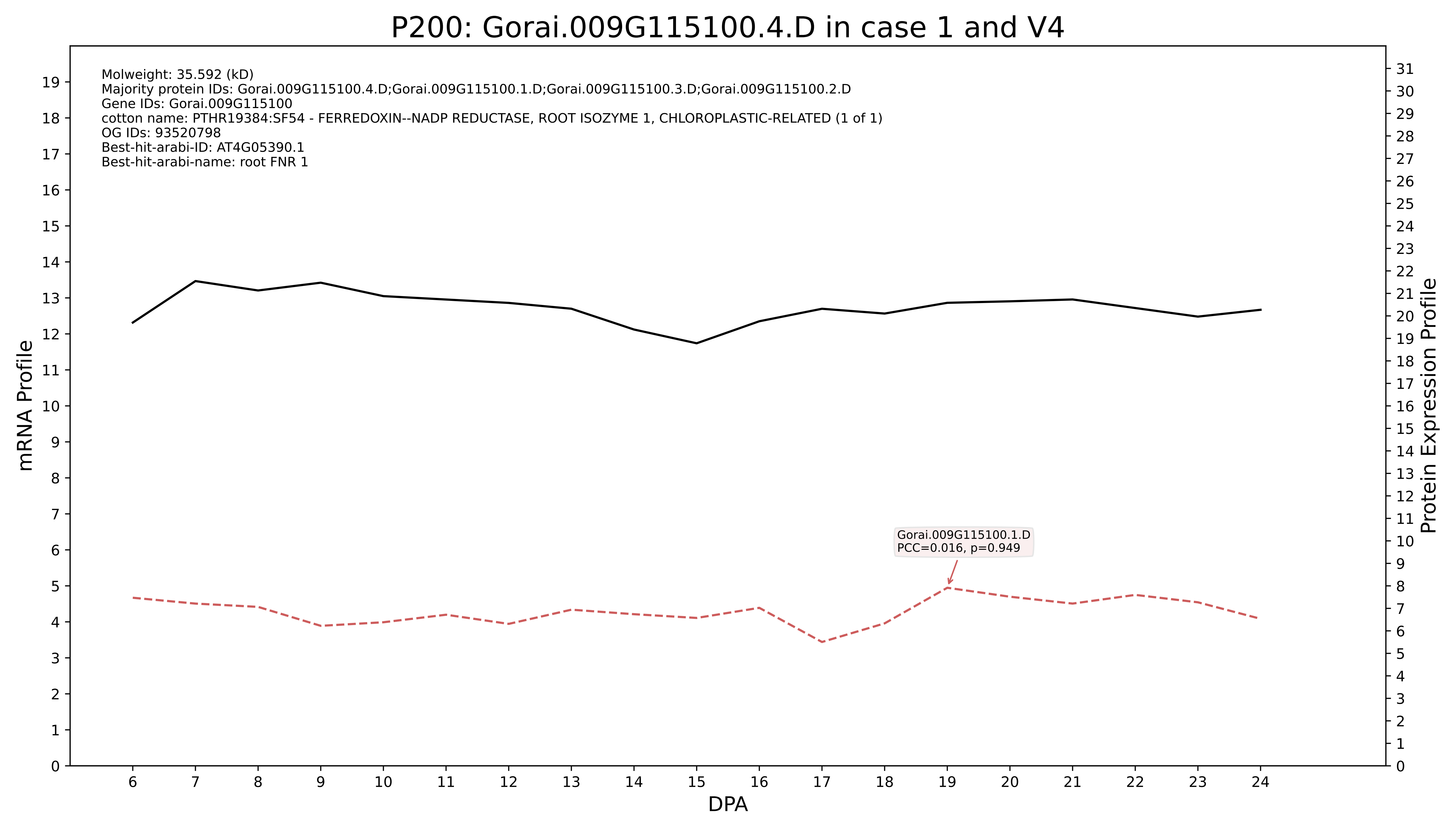
P200: Gorai.009G115100.4.D in case 1 and V4
Molweight: 35.592 (kD)
Majority protein IDs: Gorai.009G115100.4.D;Gorai.009G115100.1.D;Gorai.009G115100.3.D;Gorai.009G115100.2.D
Gene IDs: Gorai.009G115100
cotton name: PTHR19384:SF54 - FERREDOXIN--NADP REDUCTASE, ROOT ISOZYME 1, CHLOROPLASTIC-RELATED (1 of 1)
OG IDs: 93520798
Best-hit-arabi-ID: AT4G05390.1
Best-hit-arabi-name: root FNR 1
Gorai.009G115100.1.D
PCC=0.016, p=0.949
mRNA Profile
Protein Expression Profile
DPA

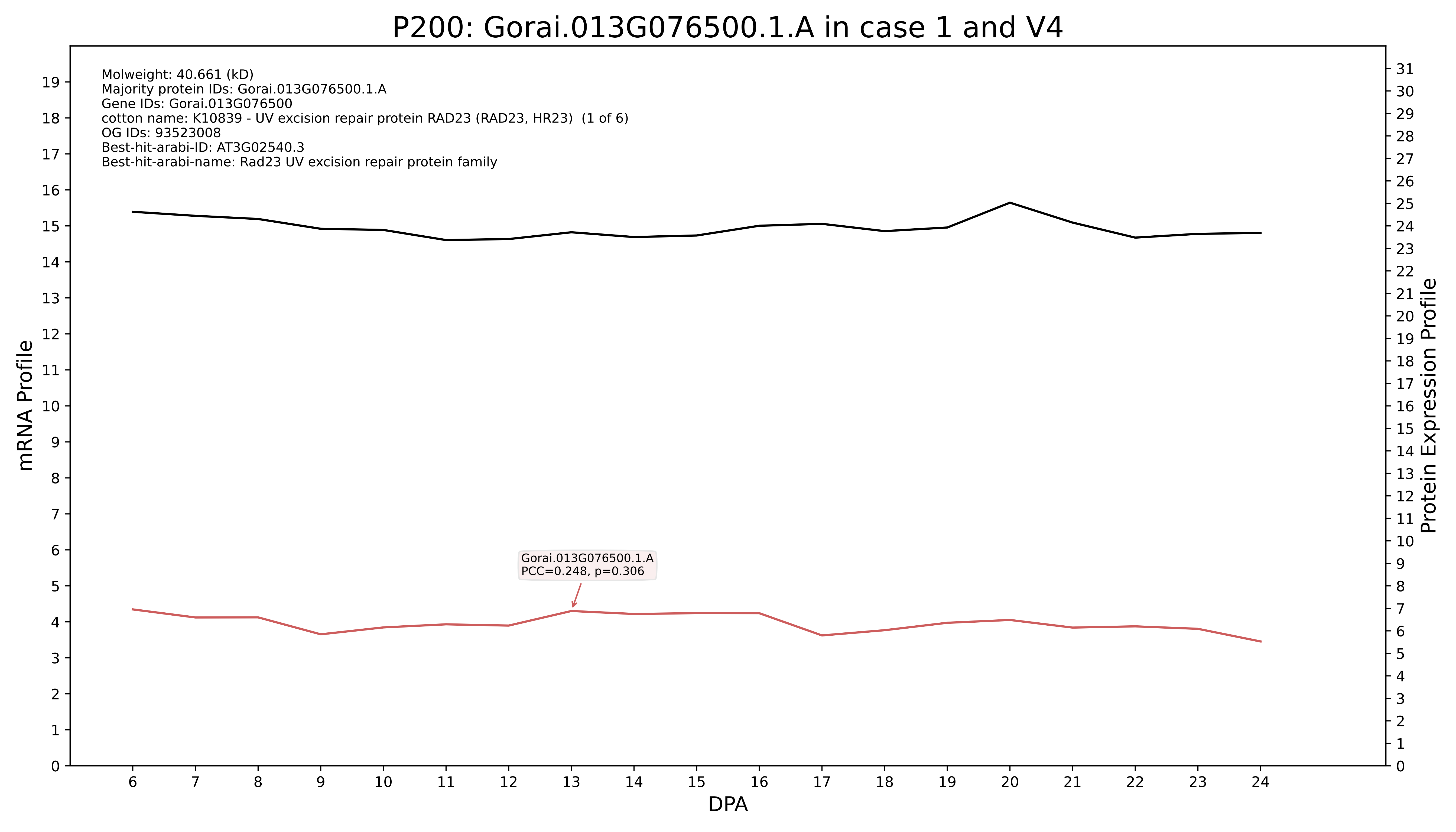

P200: Gorai.013G076500.1.A in case 1 and V4
Molweight: 40.661 (kD)
Majority protein IDs: Gorai.013G076500.1.A
Gene IDs: Gorai.013G076500
cotton name: K10839 - UV excision repair protein RAD23 (RAD23, HR23) (1 of 6)
OG IDs: 93523008
Best-hit-arabi-ID: AT3G02540.3
Best-hit-arabi-name: Rad23 UV excision repair protein family
Gorai.013G076500.1.A
PCC=0.248, p=0.306
mRNA Profile
Protein Expression Profile
DPA

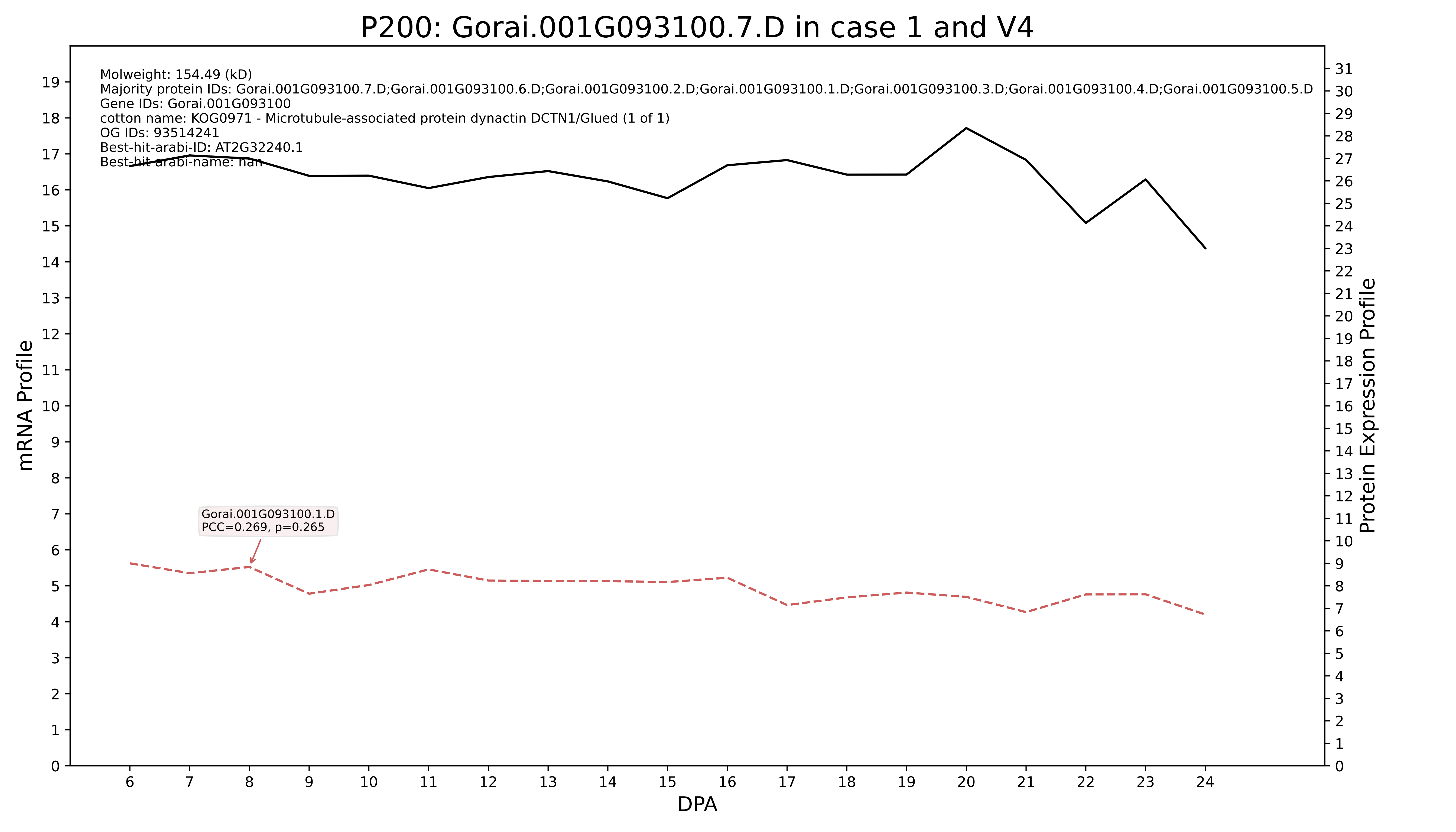
P200: Gorai.001G093100.7.D in case 1 and V4
Molweight: 154.49 (kD)
Majority protein IDs: Gorai.001G093100.7.D;Gorai.001G093100.6.D;Gorai.001G093100.2.D;Gorai.001G093100.1.D;Gorai.001G093100.3.D;Gorai.001G093100.4.D;Gorai.001G093100.5.D
Gene IDs: Gorai.001G093100
cotton name: KOG0971 - Microtubule-associated protein dynactin DCTN1/Glued (1 of 1)
OG IDs: 93514241
Best-hit-arabi-ID: AT2G32240.1
Best-hit-arabi-name: nan
Gorai.001G093100.1.D
PCC=0.269, p=0.265
mRNA Profile
Protein Expression Profile
DPA

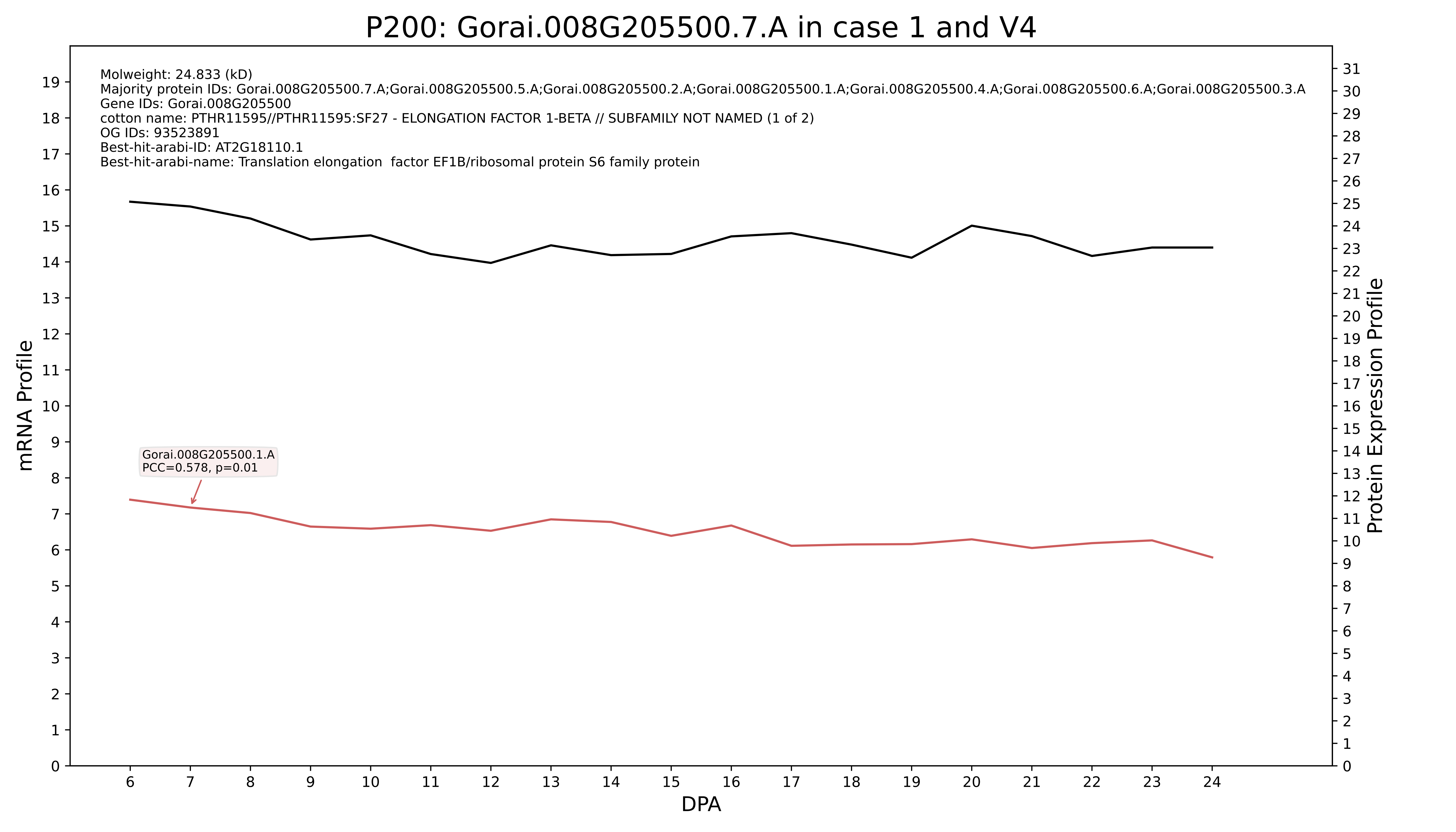

P200: Gorai.008G205500.7.A in case 1 and V4
Molweight: 24.833 (kD)
Majority protein IDs: Gorai.008G205500.7.A;Gorai.008G205500.5.A;Gorai.008G205500.2.A;Gorai.008G205500.1.A;Gorai.008G205500.4.A;Gorai.008G205500.6.A;Gorai.008G205500.3.A
Gene IDs: Gorai.008G205500
cotton name: PTHR11595//PTHR11595:SF27 - ELONGATION FACTOR 1-BETA // SUBFAMILY NOT NAMED (1 of 2)
OG IDs: 93523891
Best-hit-arabi-ID: AT2G18110.1
Best-hit-arabi-name: Translation elongation factor EF1B/ribosomal protein S6 family protein
Gorai.008G205500.1.A
PCC=0.578, p=0.01
mRNA Profile
Protein Expression Profile
DPA

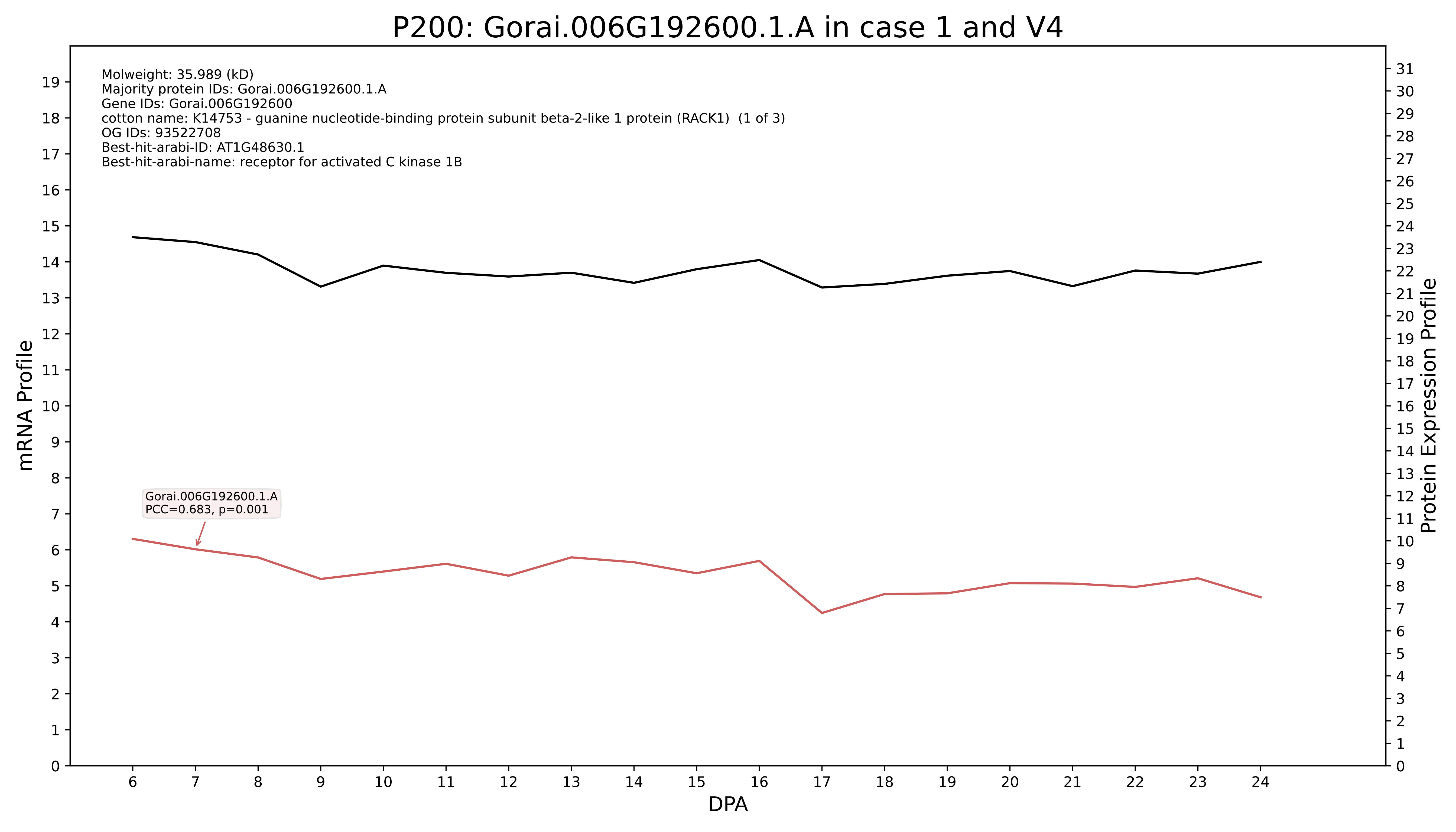
P200: Gorai.006G192600.1.A in case 1 and V4
Molweight: 35.989 (kD)
Majority protein IDs: Gorai.006G192600.1.A
Gene IDs: Gorai.006G192600
cotton name: K14753 - guanine nucleotide-binding protein subunit beta-2-like 1 protein (RACK1) (1 of 3)
OG IDs: 93522708
Best-hit-arabi-ID: AT1G48630.1
Best-hit-arabi-name: receptor for activated C kinase 1B
Gorai.006G192600.1.A
PCC=0.683, p=0.001
mRNA Profile
Protein Expression Profile
DPA

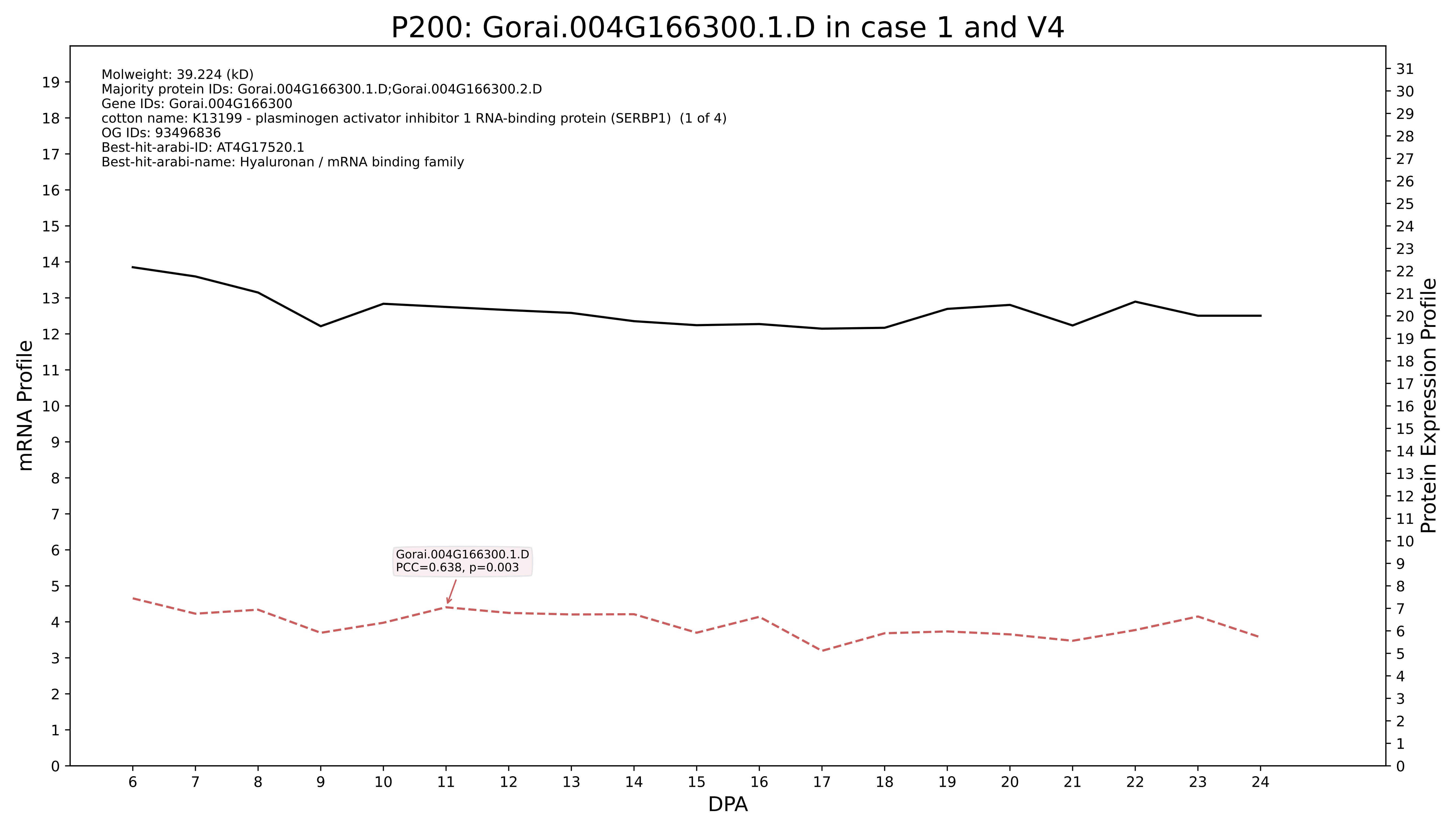

P200: Gorai.004G166300.1.D in case 1 and V4
Molweight: 39.224 (kD)
Majority protein IDs: Gorai.004G166300.1.D;Gorai.004G166300.2.D
Gene IDs: Gorai.004G166300
cotton name: K13199 - plasminogen activator inhibitor 1 RNA-binding protein (SERBP1) (1 of 4)
OG IDs: 93496836
Best-hit-arabi-ID: AT4G17520.1
Best-hit-arabi-name: Hyaluronan / mRNA binding family
Gorai.004G166300.1.D
PCC=0.638, p=0.003
mRNA Profile
Protein Expression Profile
DPA

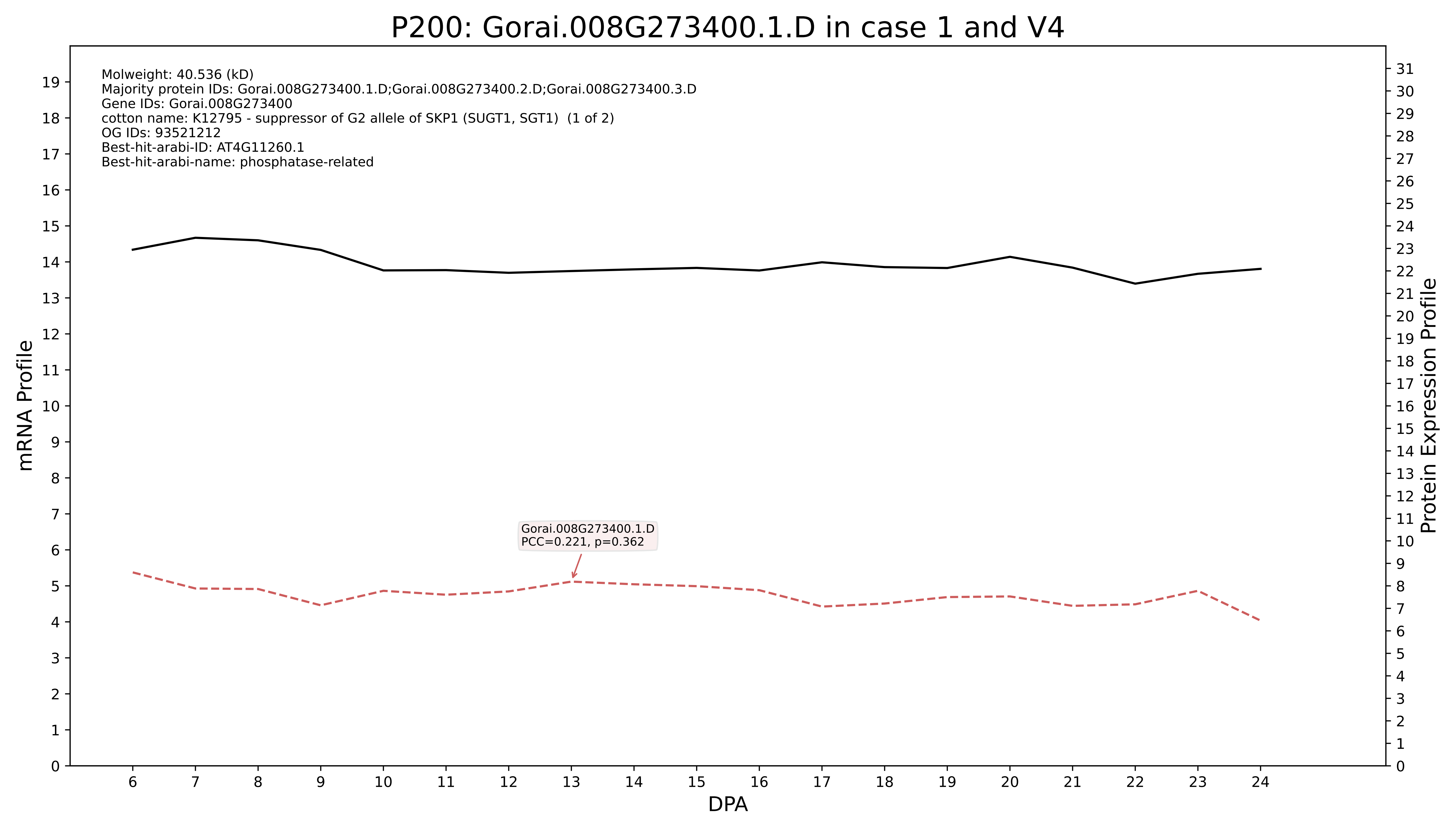
P200: Gorai.008G273400.1.D in case 1 and V4
Molweight: 40.536 (kD)
Majority protein IDs: Gorai.008G273400.1.D;Gorai.008G273400.2.D;Gorai.008G273400.3.D
Gene IDs: Gorai.008G273400
cotton name: K12795 - suppressor of G2 allele of SKP1 (SUGT1, SGT1) (1 of 2)
OG IDs: 93521212
Best-hit-arabi-ID: AT4G11260.1
Best-hit-arabi-name: phosphatase-related
Gorai.008G273400.1.D
PCC=0.221, p=0.362
mRNA Profile
Protein Expression Profile
DPA

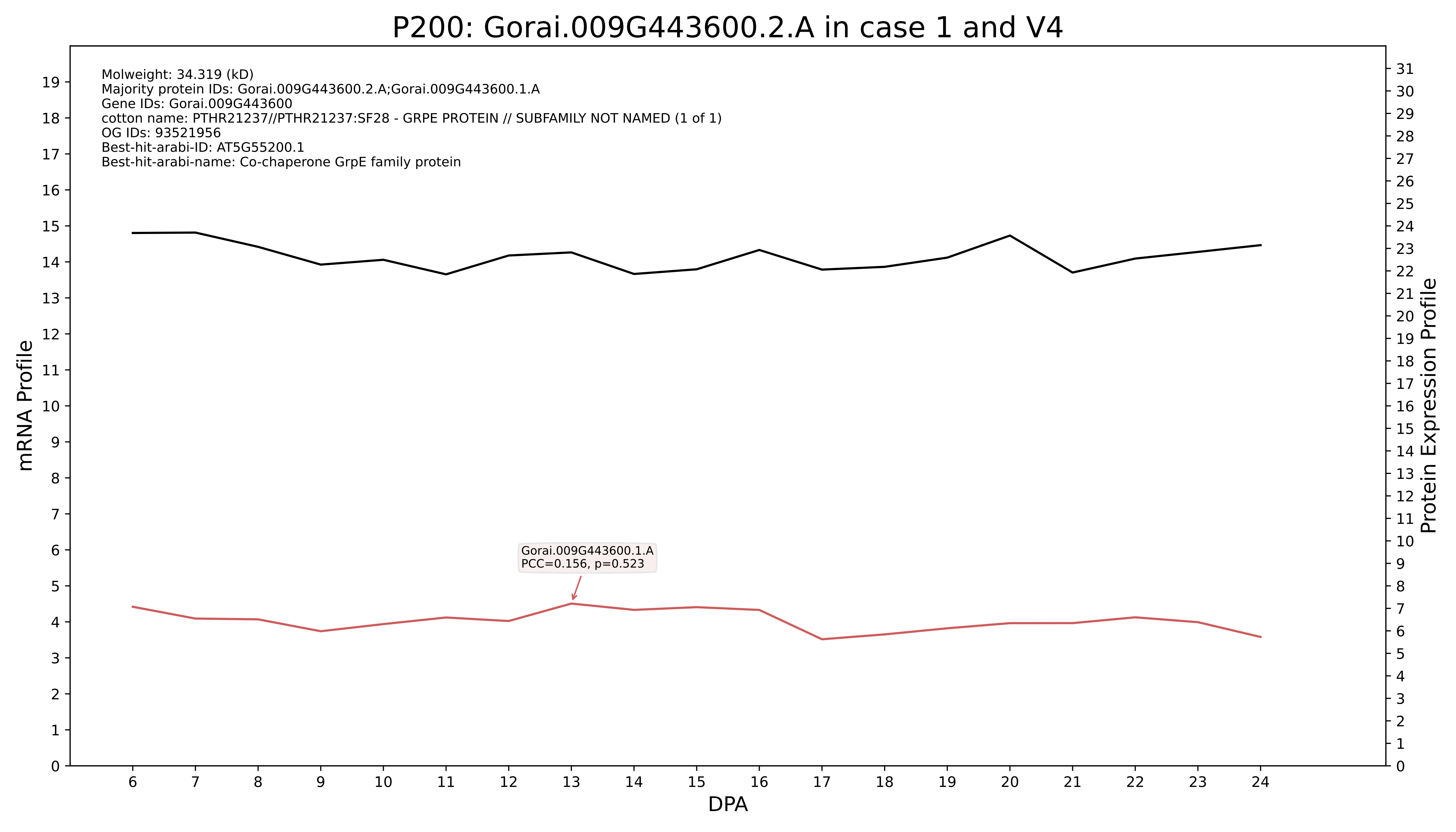

P200: Gorai.009G443600.2.A in case 1 and V4
Molweight: 34.319 (kD)
Majority protein IDs: Gorai.009G443600.2.A;Gorai.009G443600.1.A
Gene IDs: Gorai.009G443600
cotton name: PTHR21237//PTHR21237:SF28 - GRPE PROTEIN // SUBFAMILY NOT NAMED (1 of 1)
OG IDs: 93521956
Best-hit-arabi-ID: AT5G55200.1
Best-hit-arabi-name: Co-chaperone GrpE family protein
Gorai.009G443600.1.A
PCC=0.156, p=0.523
mRNA Profile
Protein Expression Profile
DPA

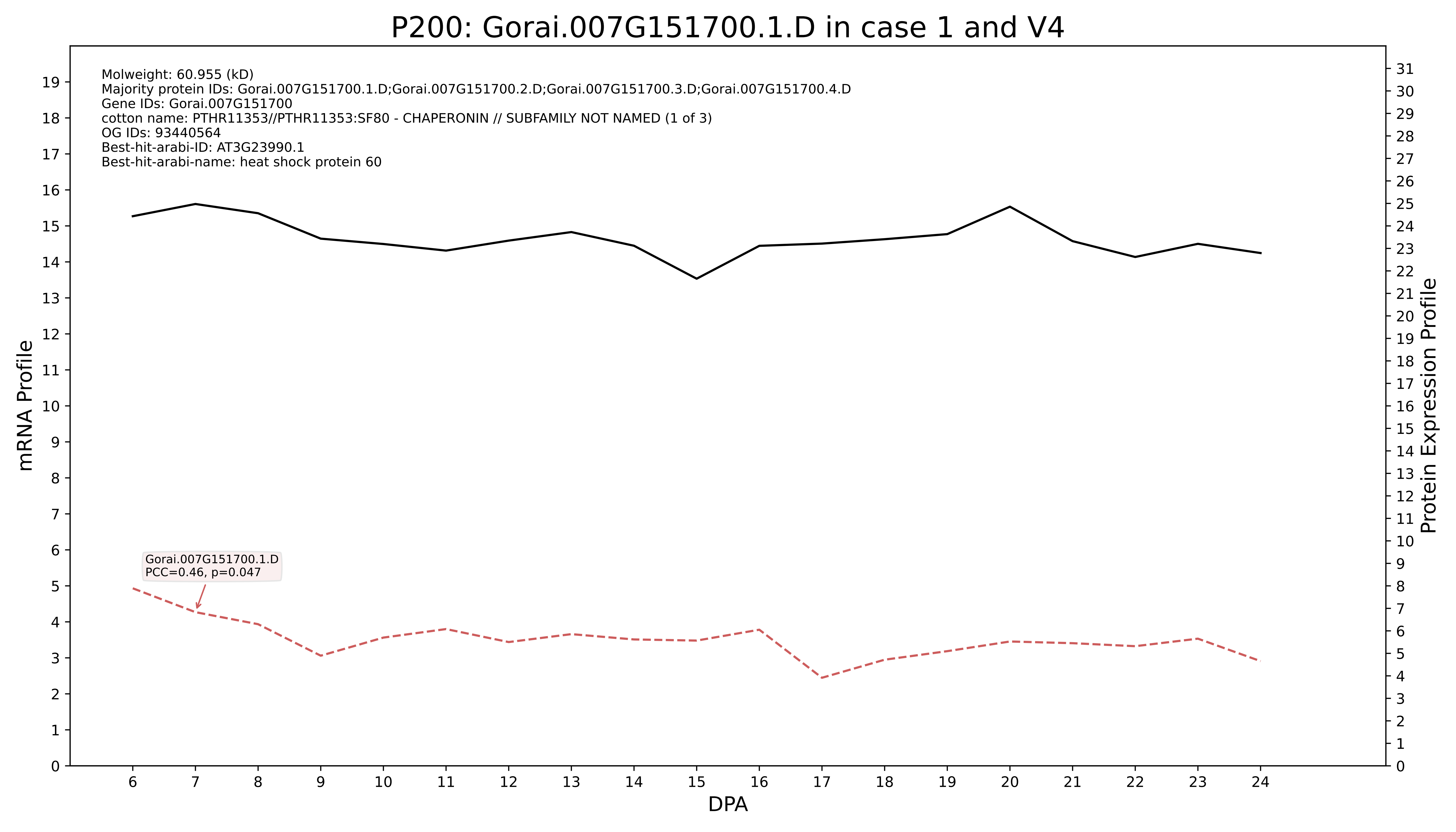
P200: Gorai.007G151700.1.D in case 1 and V4
Molweight: 60.955 (kD)
Majority protein IDs: Gorai.007G151700.1.D;Gorai.007G151700.2.D;Gorai.007G151700.3.D;Gorai.007G151700.4.D
Gene IDs: Gorai.007G151700
cotton name: PTHR11353//PTHR11353:SF80 - CHAPERONIN // SUBFAMILY NOT NAMED (1 of 3)
OG IDs: 93440564
Best-hit-arabi-ID: AT3G23990.1
Best-hit-arabi-name: heat shock protein 60
Gorai.007G151700.1.D
PCC=0.46, p=0.047
mRNA Profile
Protein Expression Profile
DPA

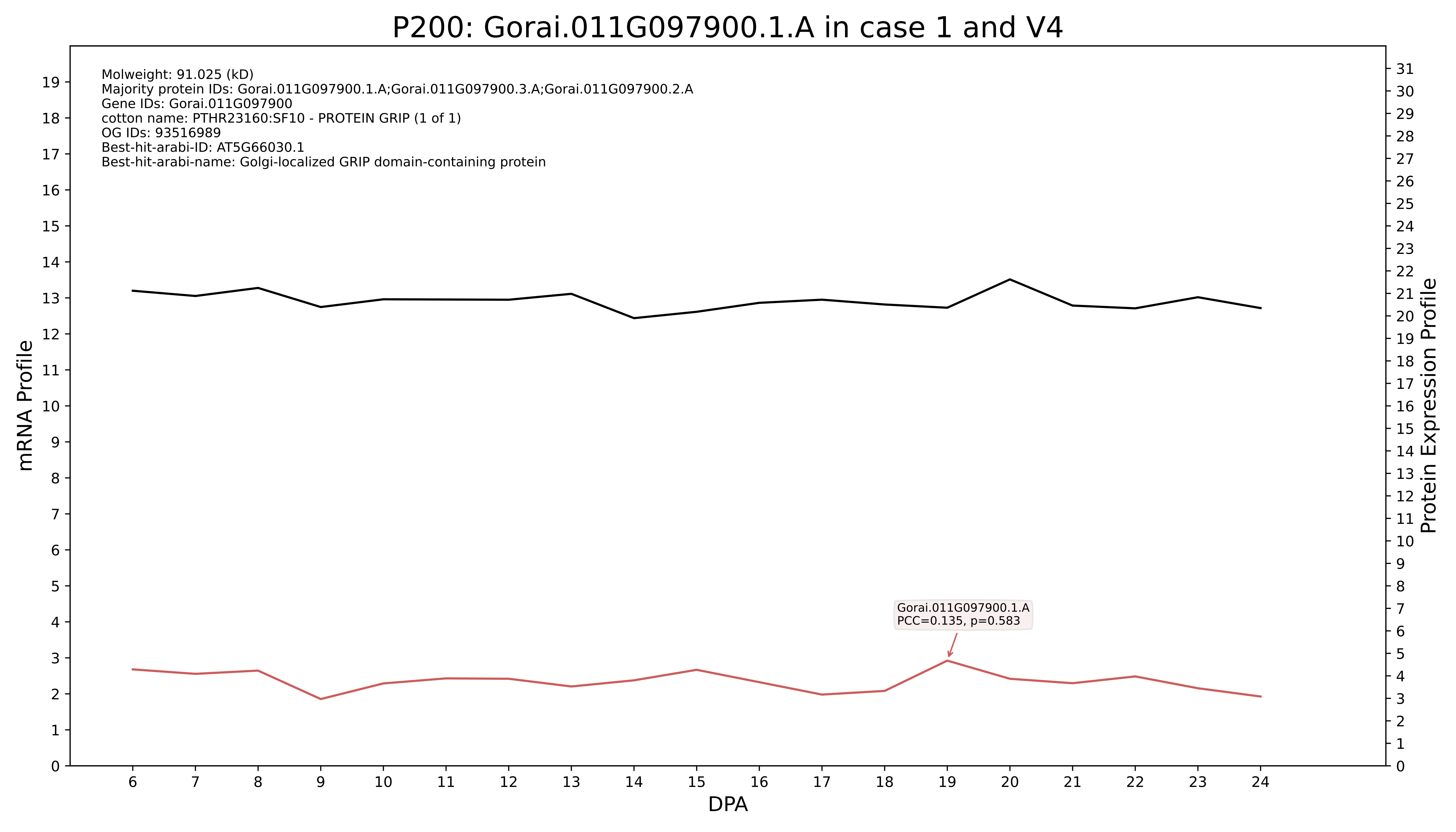

P200: Gorai.011G097900.1.A in case 1 and V4
Molweight: 91.025 (kD)
Majority protein IDs: Gorai.011G097900.1.A;Gorai.011G097900.3.A;Gorai.011G097900.2.A
Gene IDs: Gorai.011G097900
cotton name: PTHR23160:SF10 - PROTEIN GRIP (1 of 1)
OG IDs: 93516989
Best-hit-arabi-ID: AT5G66030.1
Best-hit-arabi-name: Golgi-localized GRIP domain-containing protein
Gorai.011G097900.1.A
PCC=0.135, p=0.583
mRNA Profile
Protein Expression Profile
DPA

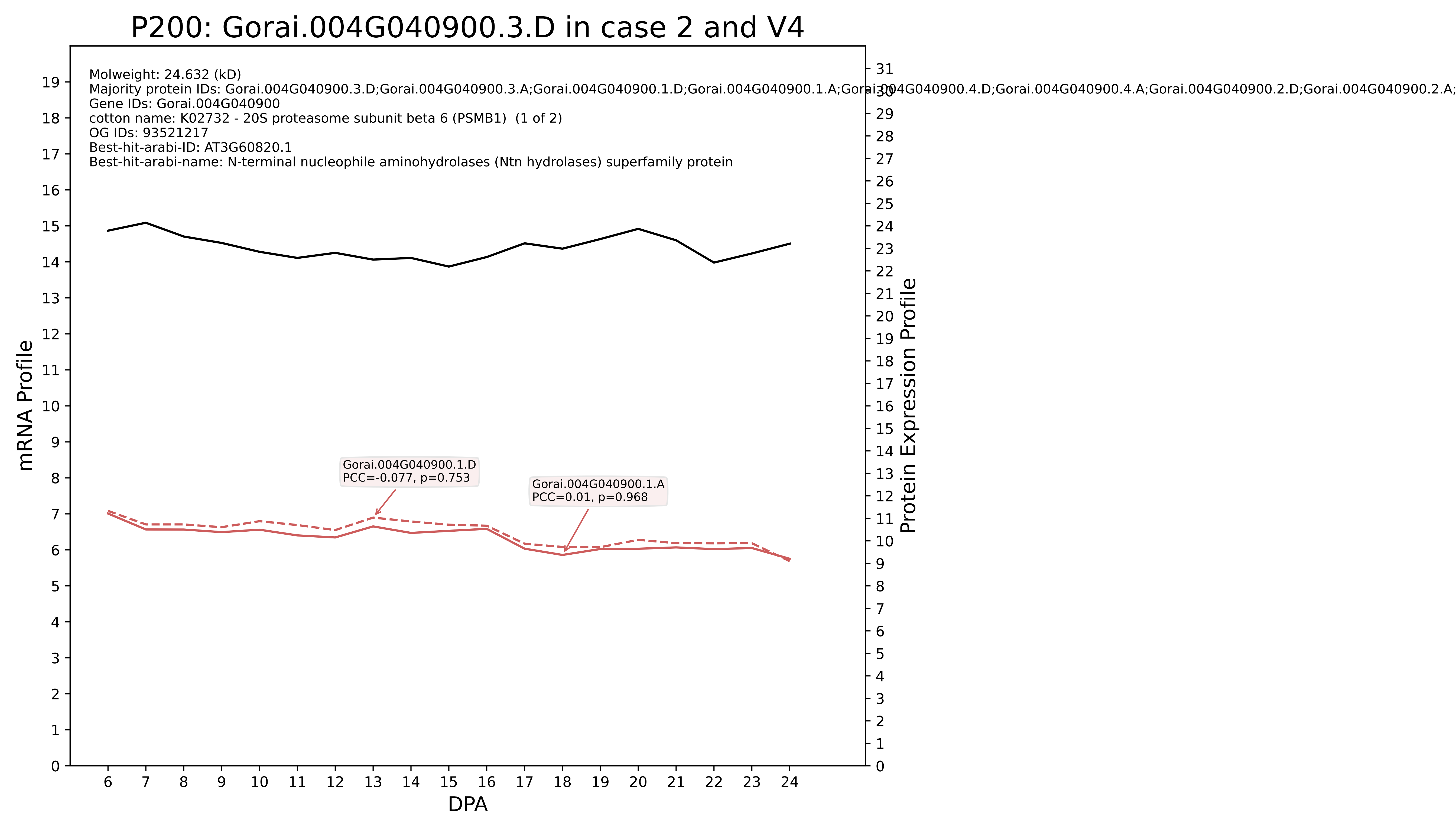
P200: Gorai.004G040900.3.D in case 2 and V4
Molweight: 24.632 (kD)
Majority protein IDs: Gorai.004G040900.3.D;Gorai.004G040900.3.A;Gorai.004G040900.1.D;Gorai.004G040900.1.A;Gorai.004G040900.4.D;Gorai.004G040900.4.A;Gorai.004G040900.2.D;Gorai.004G040900.2.A;
Gene IDs: Gorai.004G040900
cotton name: K02732 - 20S proteasome subunit beta 6 (PSMB1) (1 of 2)
OG IDs: 93521217
Best-hit-arabi-ID: AT3G60820.1
Best-hit-arabi-name: N-terminal nucleophile aminohydrolases (Ntn hydrolases) superfamily protein
Gorai.004G040900.1.D
PCC=-0.077, p=0.753
Gorai.004G040900.1.A
PCC=0.01, p=0.968
mRNA Profile
Protein Expression Profile
DPA

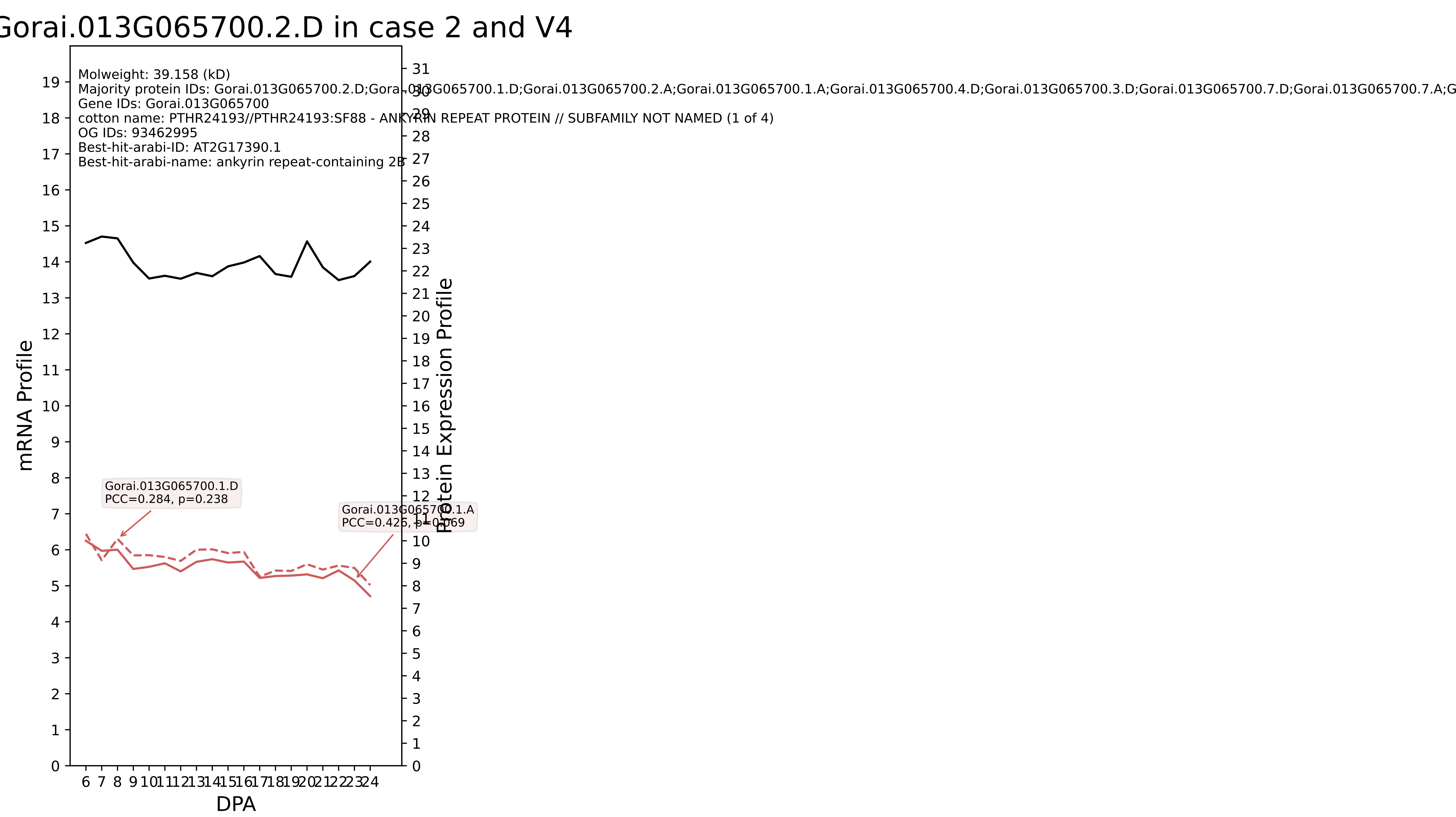
Gorai.013G065700.2.D in case 2 and V4
Molweight: 39.158 (kD)
Majority protein IDs: Gorai.013G065700.2.D;Gorai.013G065700.1.D;Gorai.013G065700.2.A;Gorai.013G065700.1.A;Gorai.013G065700.4.D;Gorai.013G065700.3.D;Gorai.013G065700.7.D;Gorai.013G065700.7.A;G
Gene IDs: Gorai.013G065700
cotton name: PTHR24193//PTHR24193:SF88 - ANKYRIN REPEAT PROTEIN // SUBFAMILY NOT NAMED (1 of 4)
OG IDs: 93462995
Best-hit-arabi-ID: AT2G17390.1
Best-hit-arabi-name: ankyrin repeat-containing 2B
Gorai.013G065700.1.D
PCC=0.284, p=0.238
Gorai.013G065700.1.A
PCC=0.426, p=0.069
mRNA Profile
Protein Expression Profile
DPA
6 7 8 9 10 11 12 13 14 15 16 17 18 19 20 21 22 23 24
0 1 2 3 4 5 6 7 8 9 10 11 12 13 14 15 16 17 18 19
0 1 2 3 4 5 6 7 8 9 10 11 12 13 14 15 16 17 18 19 20 21 22 23 24 25 26 27 28 29 30 31

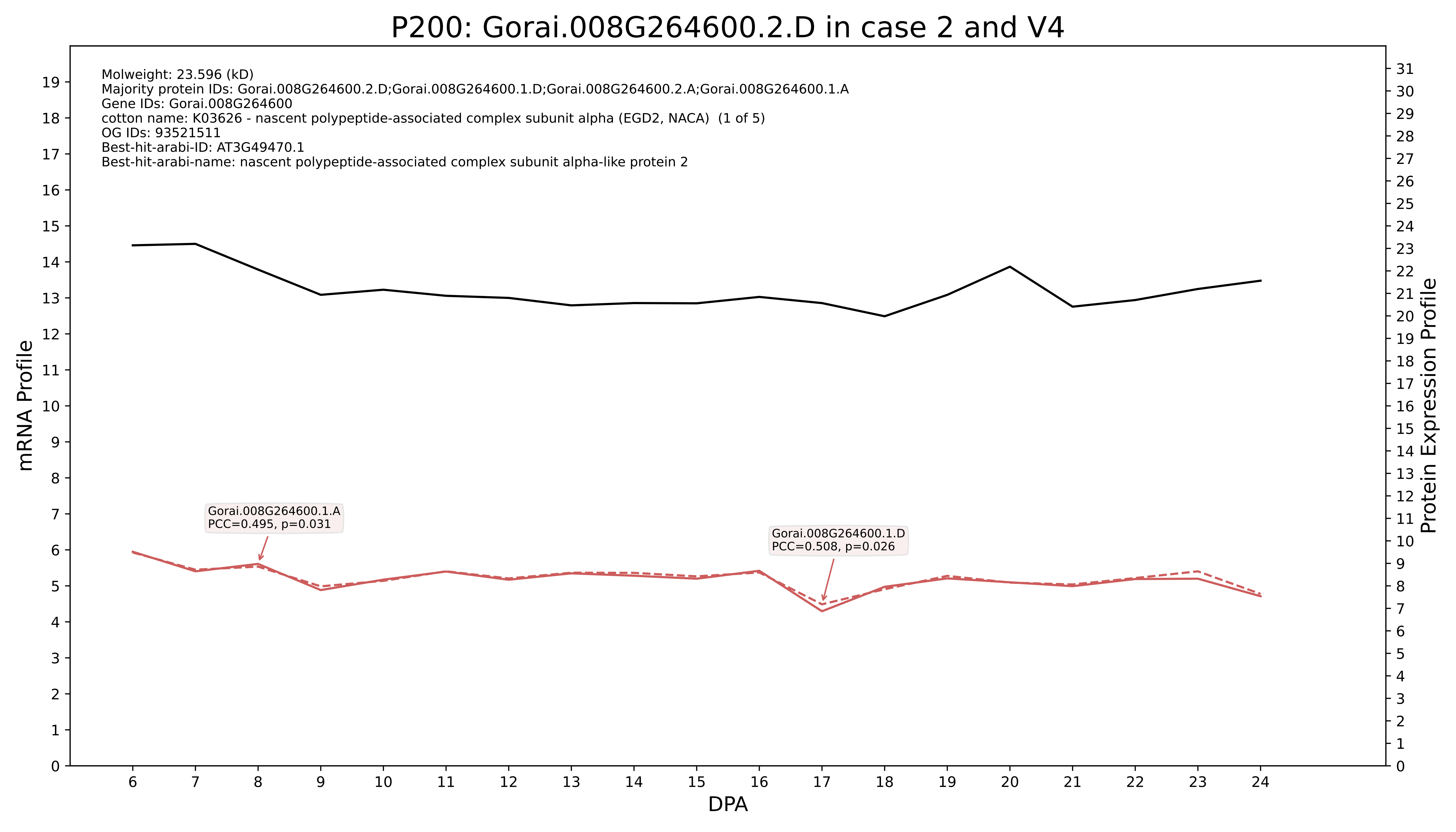
P200: Gorai.008G264600.2.D in case 2 and V4
Molweight: 23.596 (kD)
Majority protein IDs: Gorai.008G264600.2.D;Gorai.008G264600.1.D;Gorai.008G264600.2.A;Gorai.008G264600.1.A
Gene IDs: Gorai.008G264600
cotton name: K03626 - nascent polypeptide-associated complex subunit alpha (EGD2, NACA) (1 of 5)
OG IDs: 93521511
Best-hit-arabi-ID: AT3G49470.1
Best-hit-arabi-name: nascent polypeptide-associated complex subunit alpha-like protein 2
Gorai.008G264600.1.A
PCC=0.495, p=0.031
Gorai.008G264600.1.D
PCC=0.508, p=0.026
mRNA Profile
Protein Expression Profile
DPA

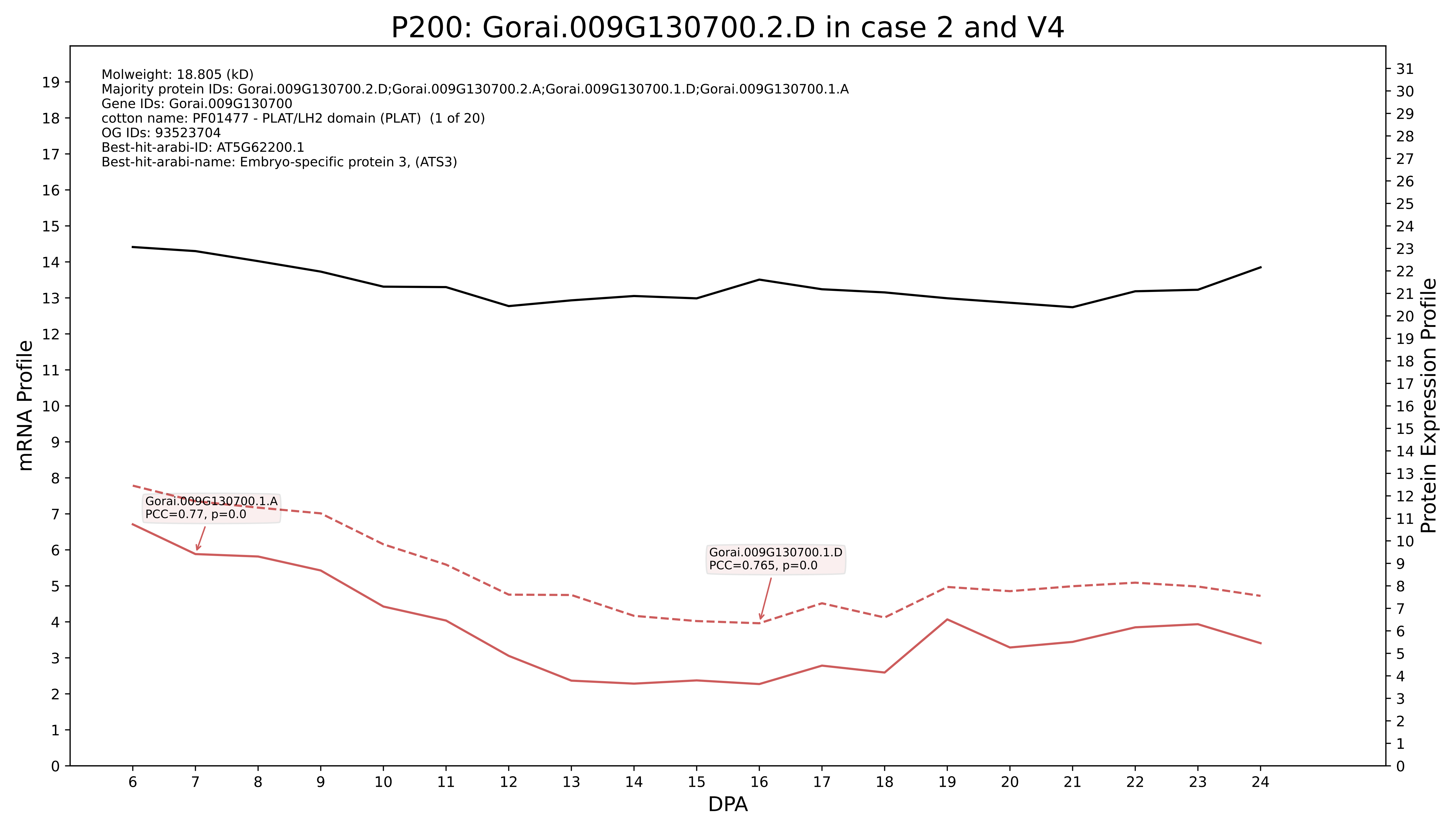

P200: Gorai.009G130700.2.D in case 2 and V4
Molweight: 18.805 (kD)
Majority protein IDs: Gorai.009G130700.2.D;Gorai.009G130700.2.A;Gorai.009G130700.1.D;Gorai.009G130700.1.A
Gene IDs: Gorai.009G130700
cotton name: PF01477 - PLAT/LH2 domain (PLAT) (1 of 20)
OG IDs: 93523704
Best-hit-arabi-ID: AT5G62200.1
Best-hit-arabi-name: Embryo-specific protein 3, (ATS3)
Gorai.009G130700.1.A
PCC=0.77, p=0.0
Gorai.009G130700.1.D
PCC=0.765, p=0.0
mRNA Profile
Protein Expression Profile
DPA

### P200: Gorai.009G182500.1.D in case 2 and V4

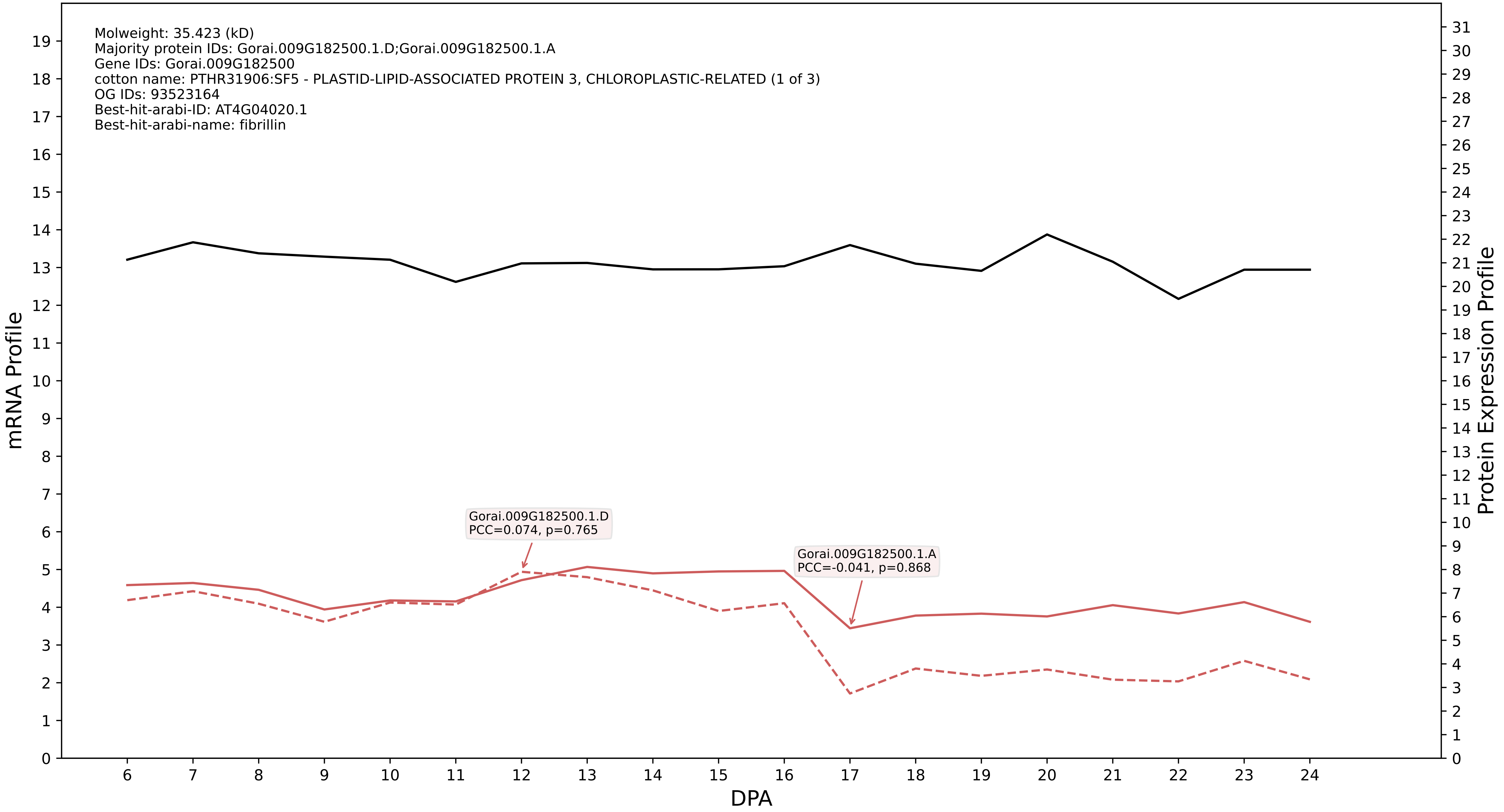

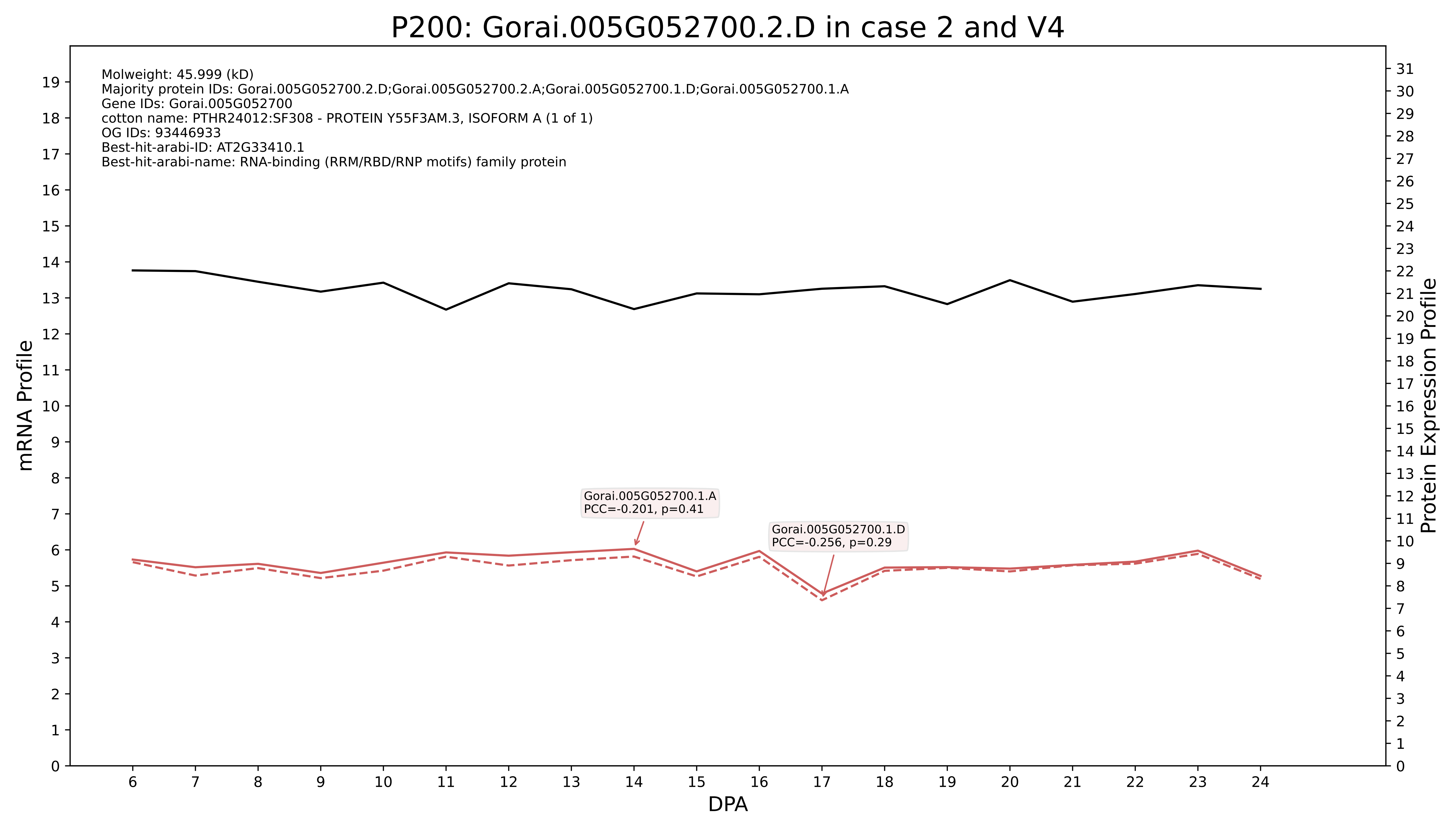
P200: Gorai.005G052700.2.D in case 2 and V4
Molweight: 45.999 (kD)
Majority protein IDs: Gorai.005G052700.2.D;Gorai.005G052700.2.A;Gorai.005G052700.1.D;Gorai.005G052700.1.A
Gene IDs: Gorai.005G052700
cotton name: PTHR24012:SF308 - PROTEIN Y55F3AM.3, ISOFORM A (1 of 1)
OG IDs: 93446933
Best-hit-arabi-ID: AT2G33410.1
Best-hit-arabi-name: RNA-binding (RRM/RBD/RNP motifs) family protein
Gorai.005G052700.1.A
PCC=-0.201, p=0.41
Gorai.005G052700.1.D
PCC=-0.256, p=0.29
mRNA Profile
Protein Expression Profile
DPA

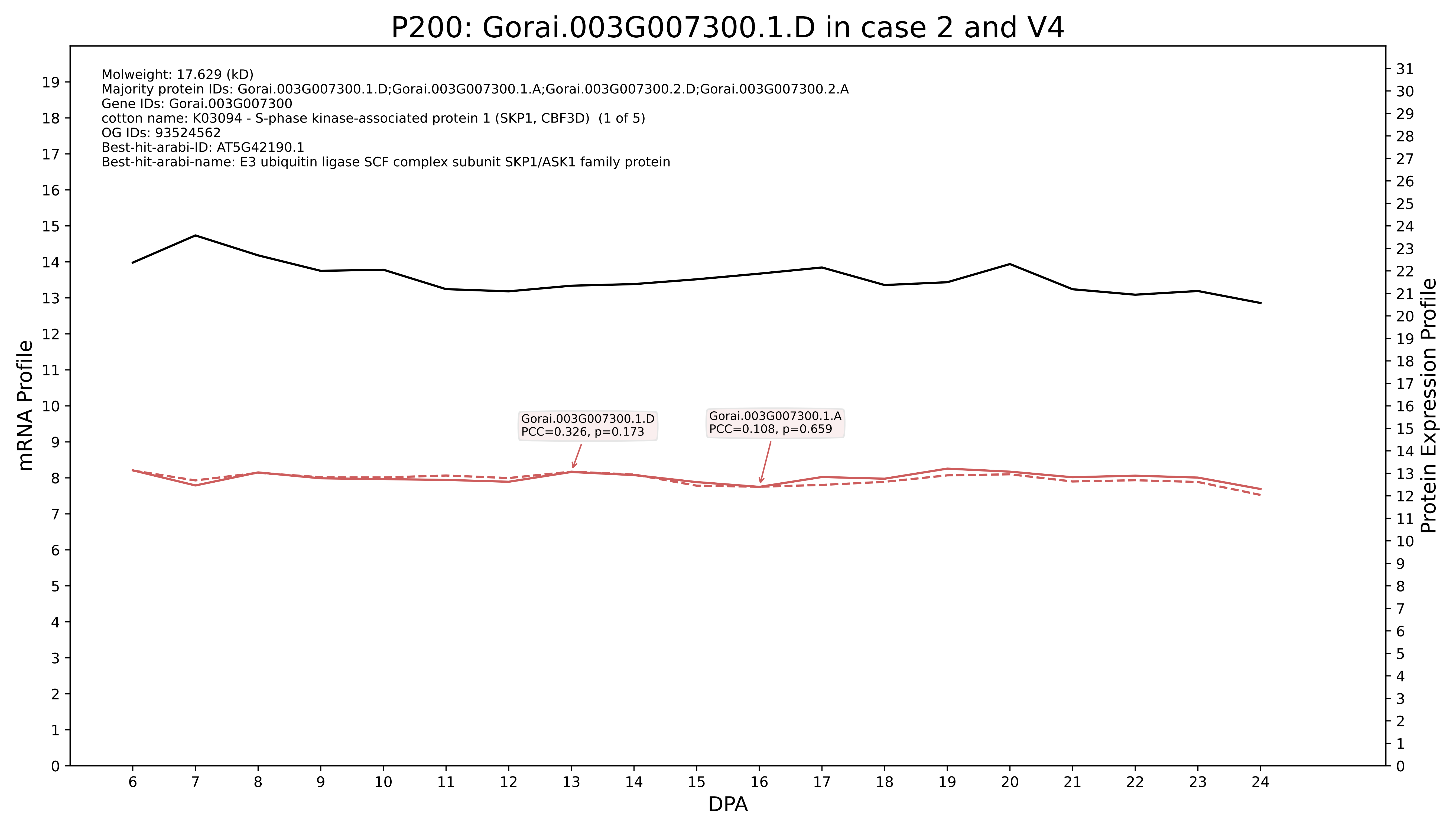
P200: Gorai.003G007300.1.D in case 2 and V4
Molweight: 17.629 (kD)
Majority protein IDs: Gorai.003G007300.1.D;Gorai.003G007300.1.A;Gorai.003G007300.2.D;Gorai.003G007300.2.A
Gene IDs: Gorai.003G007300
cotton name: K03094 - S-phase kinase-associated protein 1 (SKP1, CBF3D) (1 of 5)
OG IDs: 93524562
Best-hit-arabi-ID: AT5G42190.1
Best-hit-arabi-name: E3 ubiquitin ligase SCF complex subunit SKP1/ASK1 family protein
Gorai.003G007300.1.D
PCC=0.326, p=0.173
Gorai.003G007300.1.A
PCC=0.108, p=0.659
mRNA Profile
Protein Expression Profile
DPA

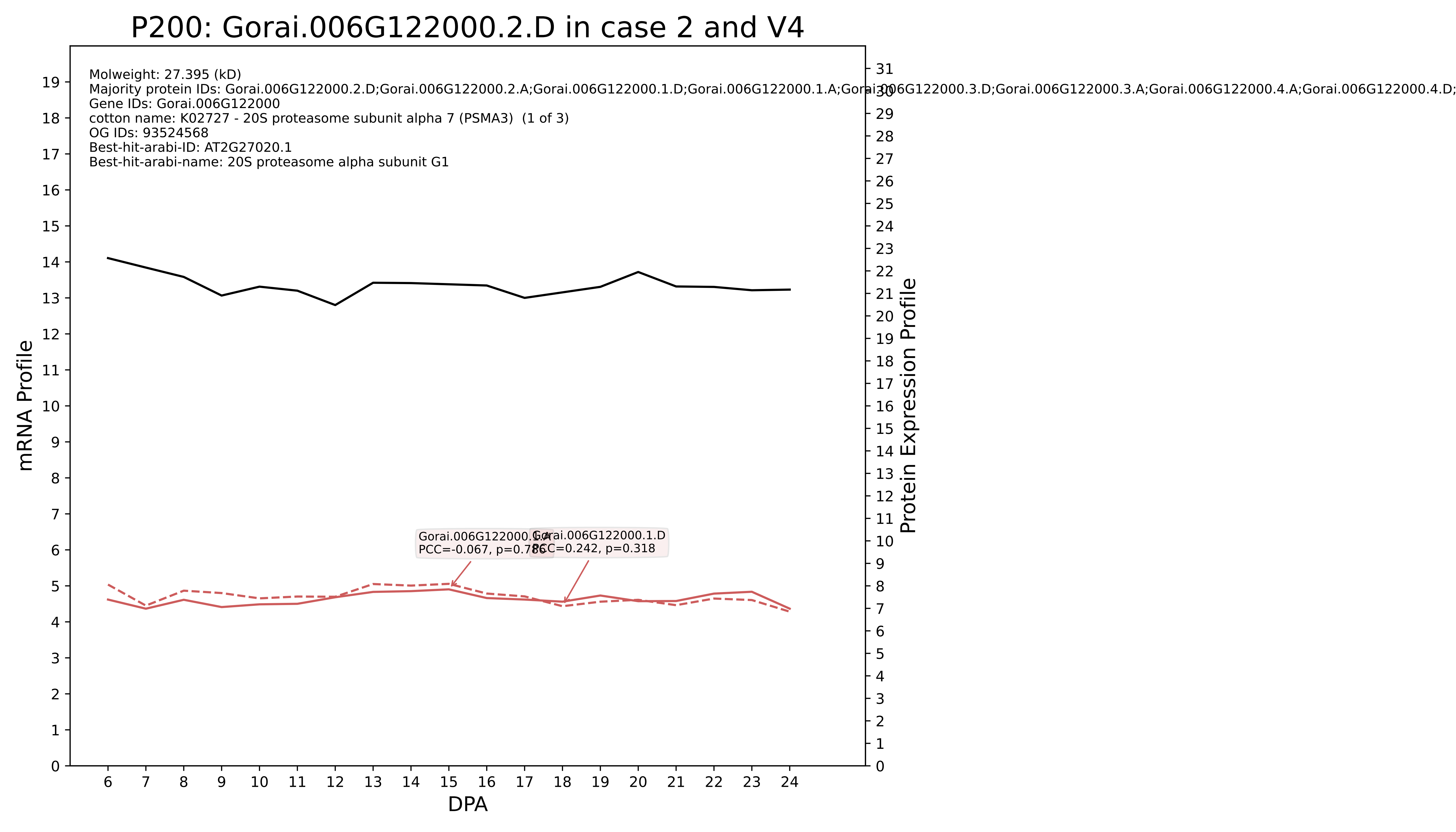
P200: Gorai.006G122000.2.D in case 2 and V4
Molweight: 27.395 (kD)
Majority protein IDs: Gorai.006G122000.2.D;Gorai.006G122000.2.A;Gorai.006G122000.1.D;Gorai.006G122000.1.A;Gorai.006G122000.3.D;Gorai.006G122000.3.A;Gorai.006G122000.4.A;Gorai.006G122000.4.D;
Gene IDs: Gorai.006G122000
cotton name: K02727 - 20S proteasome subunit alpha 7 (PSMA3) (1 of 3)
OG IDs: 93524568
Best-hit-arabi-ID: AT2G27020.1
Best-hit-arabi-name: 20S proteasome alpha subunit G1
Gorai.006G122000.1.A
PCC=-0.067, p=0.786
Gorai.006G122000.1.D
PCC=0.242, p=0.318
mRNA Profile
Protein Expression Profile
DPA

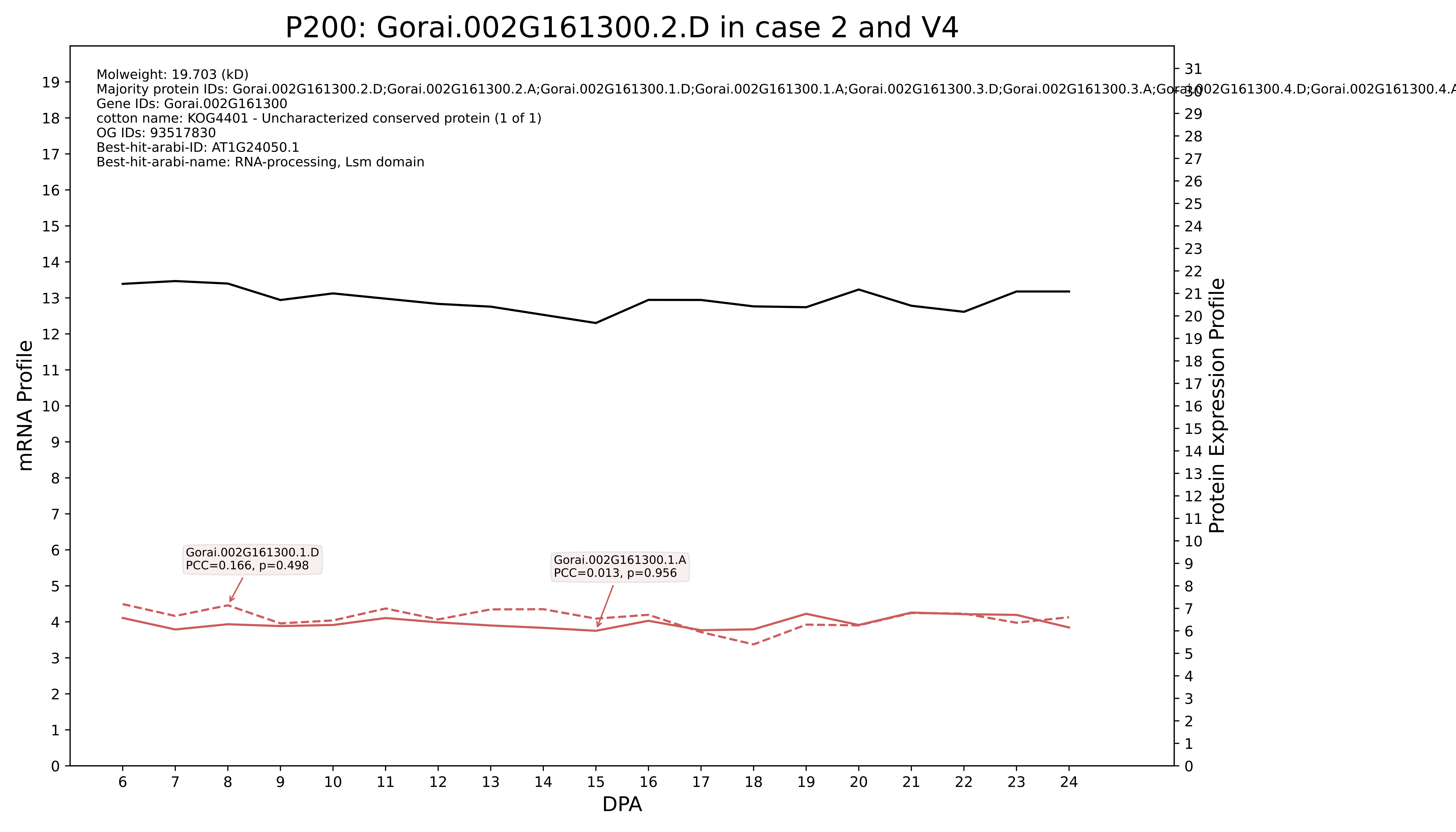

P200: Gorai.002G161300.2.D in case 2 and V4
Molweight: 19.703 (kD)
Majority protein IDs: Gorai.002G161300.2.D;Gorai.002G161300.2.A;Gorai.002G161300.1.D;Gorai.002G161300.1.A;Gorai.002G161300.3.D;Gorai.002G161300.3.A;Gorai.002G161300.4.D;Gorai.002G161300.4.A
Gene IDs: Gorai.002G161300
cotton name: KOG4401 - Uncharacterized conserved protein (1 of 1)
OG IDs: 93517830
Best-hit-arabi-ID: AT1G24050.1
Best-hit-arabi-name: RNA-processing, Lsm domain
Gorai.002G161300.1.D
PCC=0.166, p=0.498
Gorai.002G161300.1.A
PCC=0.013, p=0.956
mRNA Profile
Protein Expression Profile
DPA

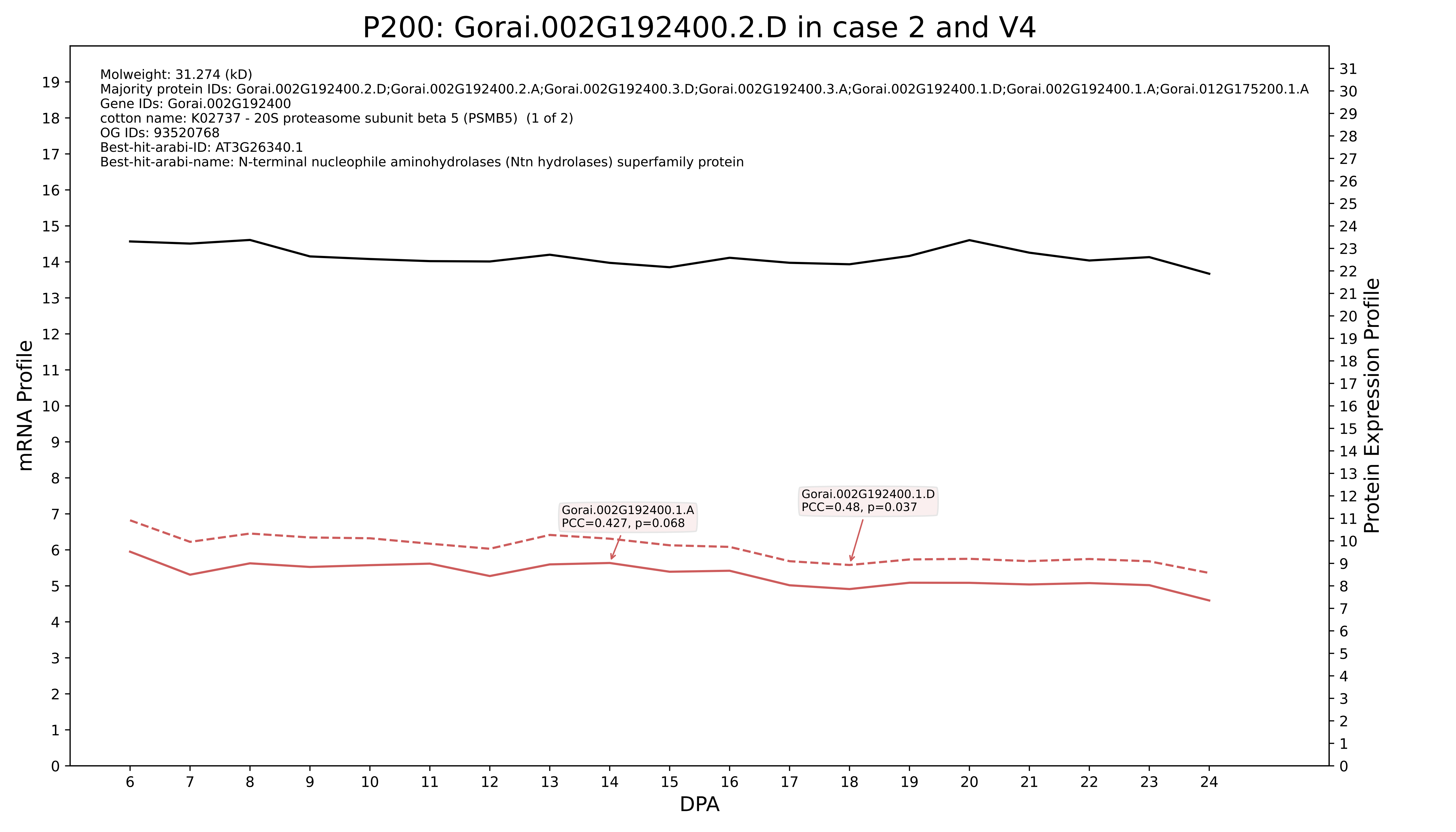
P200: Gorai.002G192400.2.D in case 2 and V4
Molweight: 31.274 (kD)
Majority protein IDs: Gorai.002G192400.2.D;Gorai.002G192400.2.A;Gorai.002G192400.3.D;Gorai.002G192400.3.A;Gorai.002G192400.1.D;Gorai.002G192400.1.A;Gorai.012G175200.1.A
Gene IDs: Gorai.002G192400
cotton name: K02737 - 20S proteasome subunit beta 5 (PSMB5) (1 of 2)
OG IDs: 93520768
Best-hit-arabi-ID: AT3G26340.1
Best-hit-arabi-name: N-terminal nucleophile aminohydrolases (Ntn hydrolases) superfamily protein
Gorai.002G192400.1.A
PCC=0.427, p=0.068
Gorai.002G192400.1.D
PCC=0.48, p=0.037
mRNA Profile
Protein Expression Profile
DPA

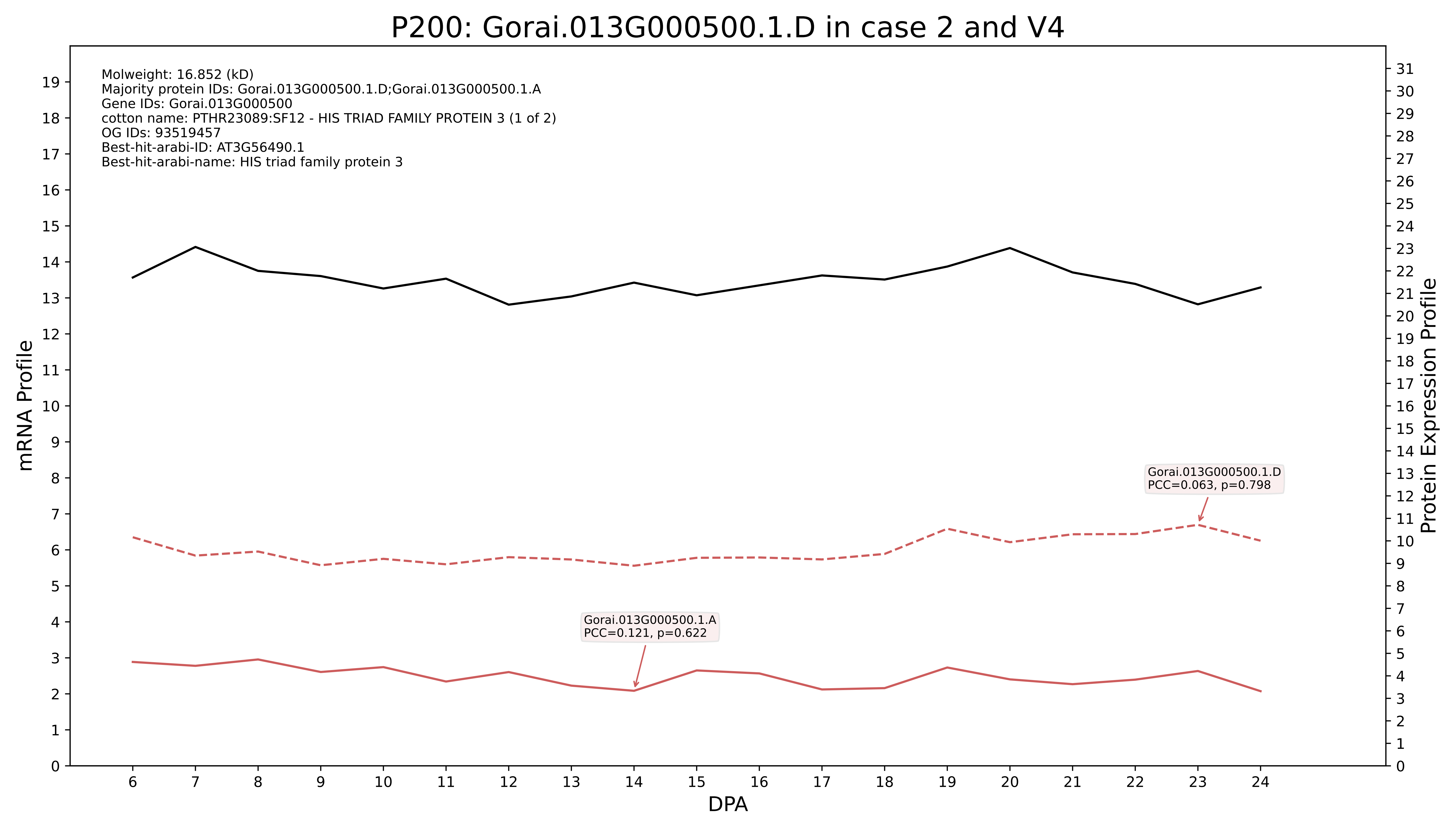
P200: Gorai.013G000500.1.D in case 2 and V4
Molweight: 16.852 (kD)
Majority protein IDs: Gorai.013G000500.1.D;Gorai.013G000500.1.A
Gene IDs: Gorai.013G000500
cotton name: PTHR23089:SF12 - HIS TRIAD FAMILY PROTEIN 3 (1 of 2)
OG IDs: 93519457
Best-hit-arabi-ID: AT3G56490.1
Best-hit-arabi-name: HIS triad family protein 3
Gorai.013G000500.1.D
PCC=0.063, p=0.798
Gorai.013G000500.1.A
PCC=0.121, p=0.622
mRNA Profile
Protein Expression Profile
DPA

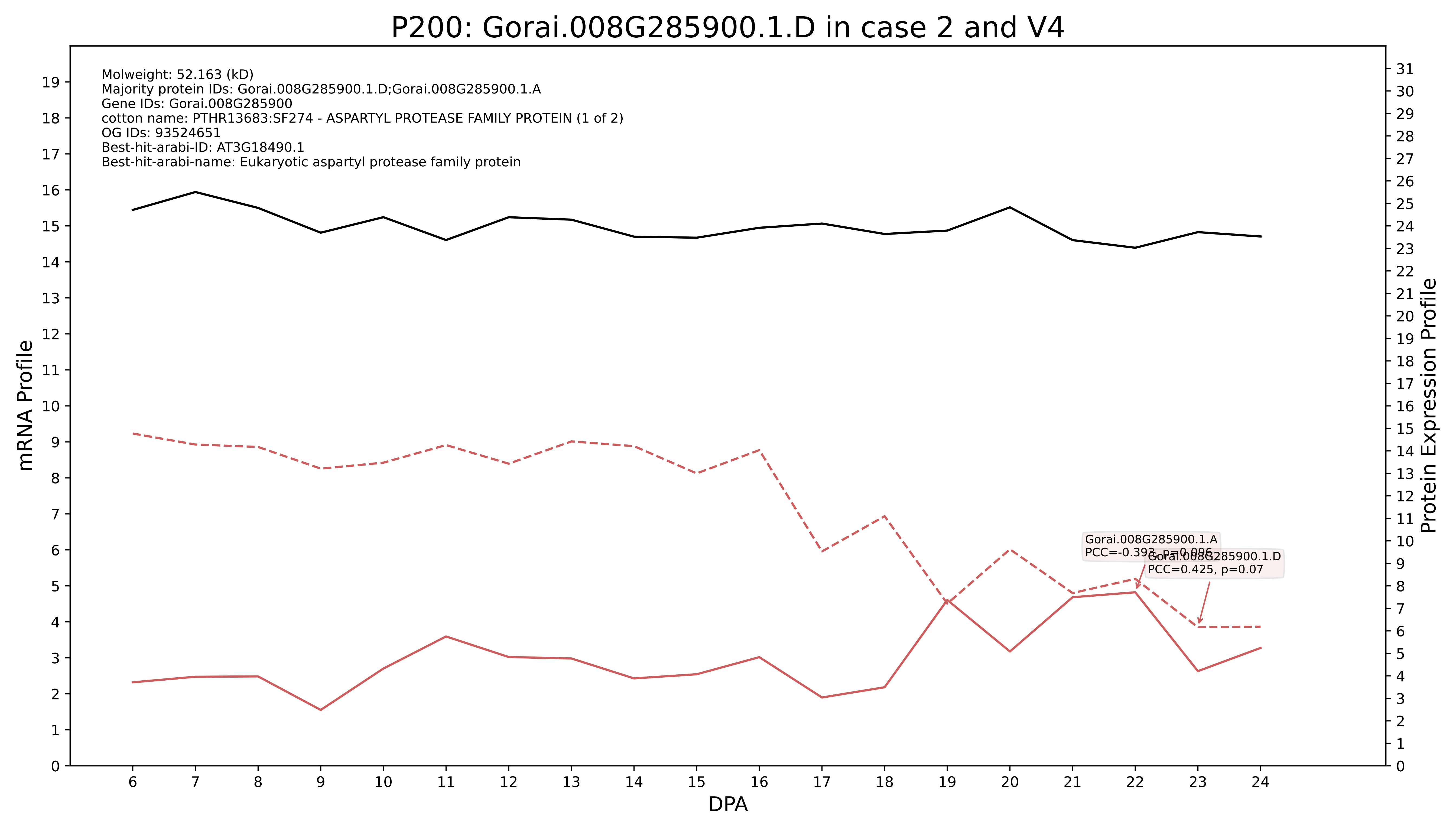
P200: Gorai.008G285900.1.D in case 2 and V4
Molweight: 52.163 (kD)
Majority protein IDs: Gorai.008G285900.1.D;Gorai.008G285900.1.A
Gene IDs: Gorai.008G285900
cotton name: PTHR13683:SF274 - ASPARTYL PROTEASE FAMILY PROTEIN (1 of 2)
OG IDs: 93524651
Best-hit-arabi-ID: AT3G18490.1
Best-hit-arabi-name: Eukaryotic aspartyl protease family protein
Gorai.008G285900.1.A
PCC=-0.393, p=0.096
Gorai.008G285900.1.D
PCC=0.425, p=0.07
mRNA Profile
Protein Expression Profile
DPA

200: Gorai.009G329800.1.A in case 2 and V4

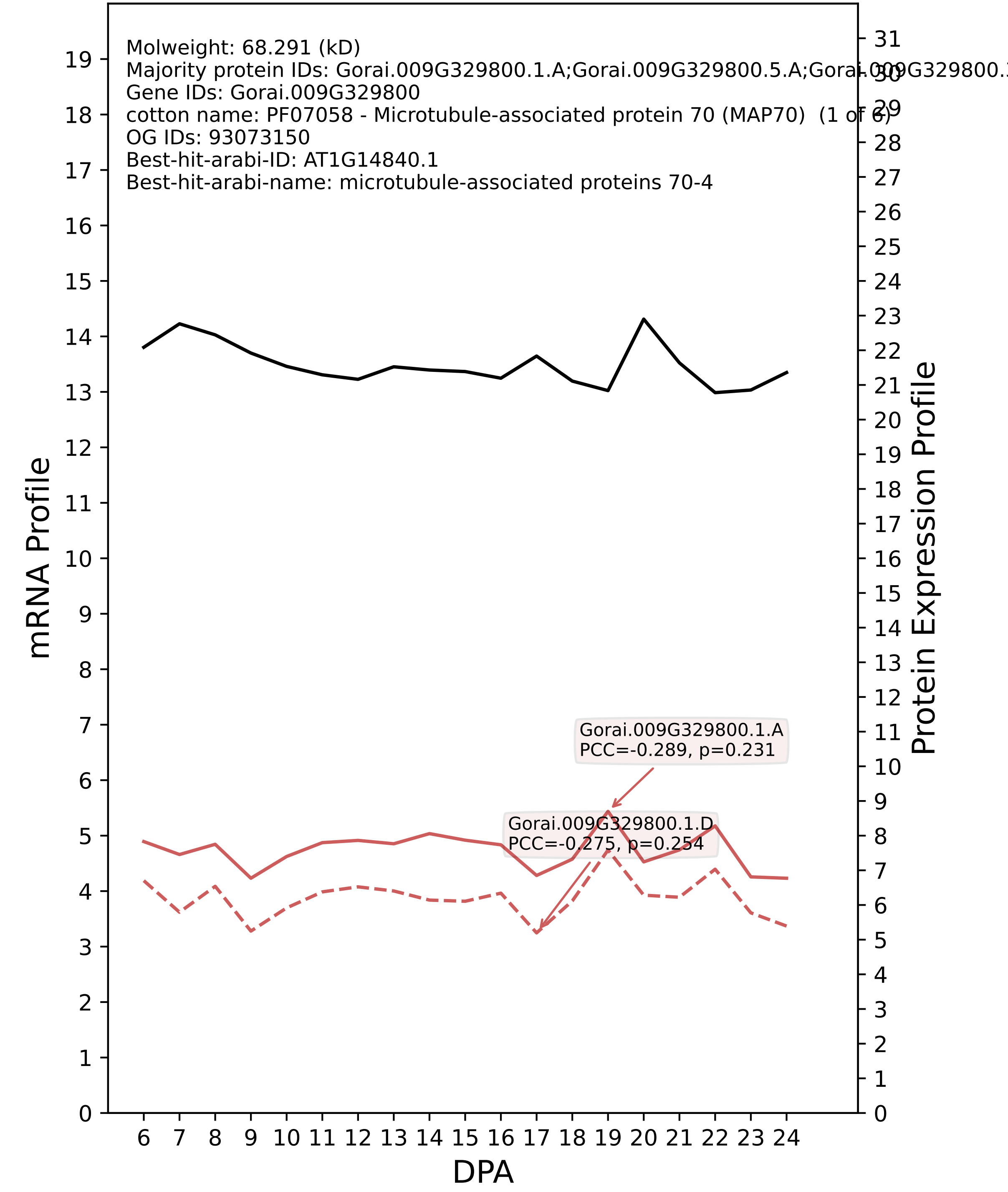

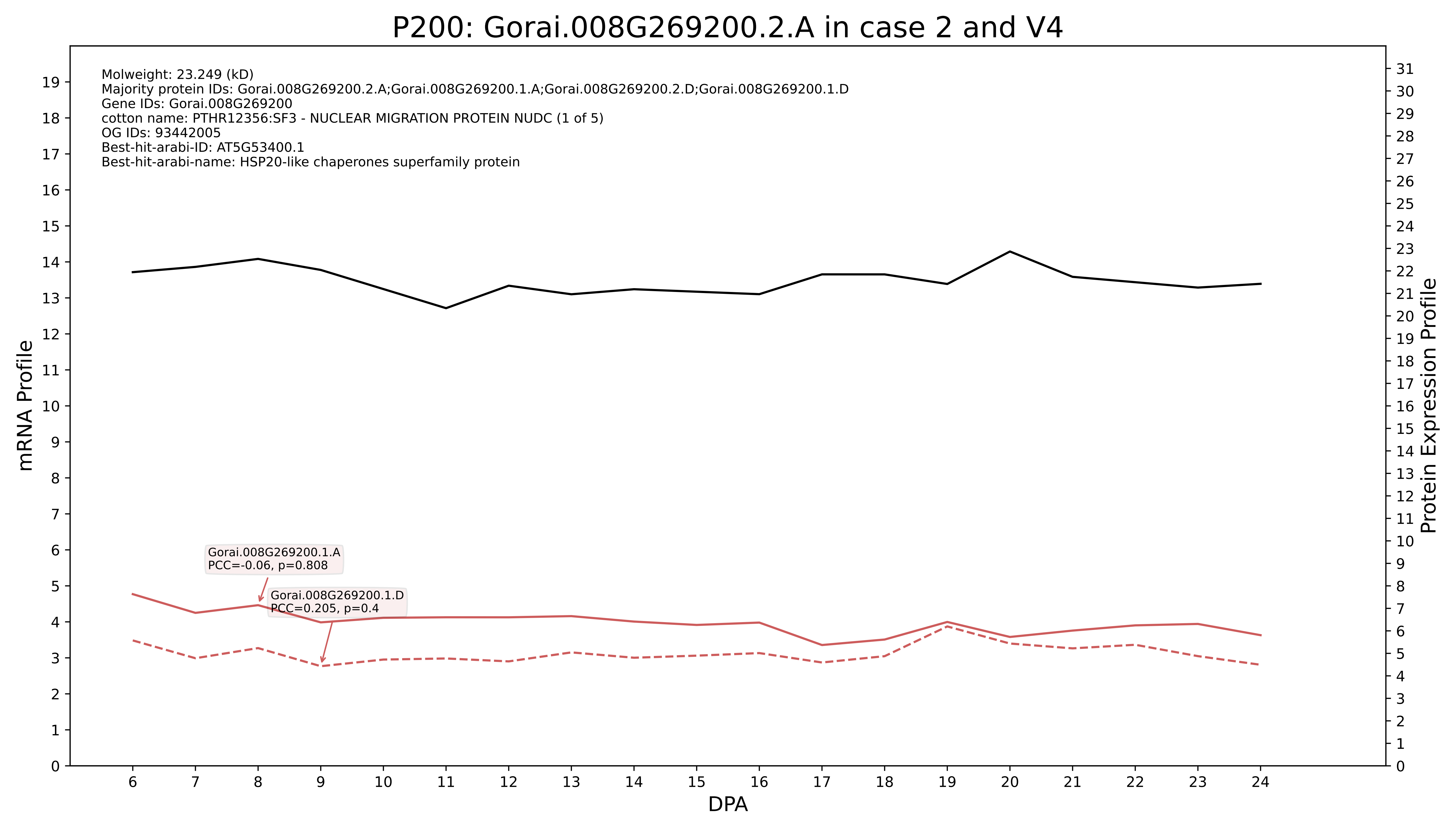

P200: Gorai.008G269200.2.A in case 2 and V4
Molweight: 23.249 (kD)
Majority protein IDs: Gorai.008G269200.2.A;Gorai.008G269200.1.A;Gorai.008G269200.2.D;Gorai.008G269200.1.D
Gene IDs: Gorai.008G269200
cotton name: PTHR12356:SF3 - NUCLEAR MIGRATION PROTEIN NUDC (1 of 5)
OG IDs: 93442005
Best-hit-arabi-ID: AT5G53400.1
Best-hit-arabi-name: HSP20-like chaperones superfamily protein
Gorai.008G269200.1.A
PCC=-0.06, p=0.808
Gorai.008G269200.1.D
PCC=0.205, p=0.4
mRNA Profile
Protein Expression Profile
DPA

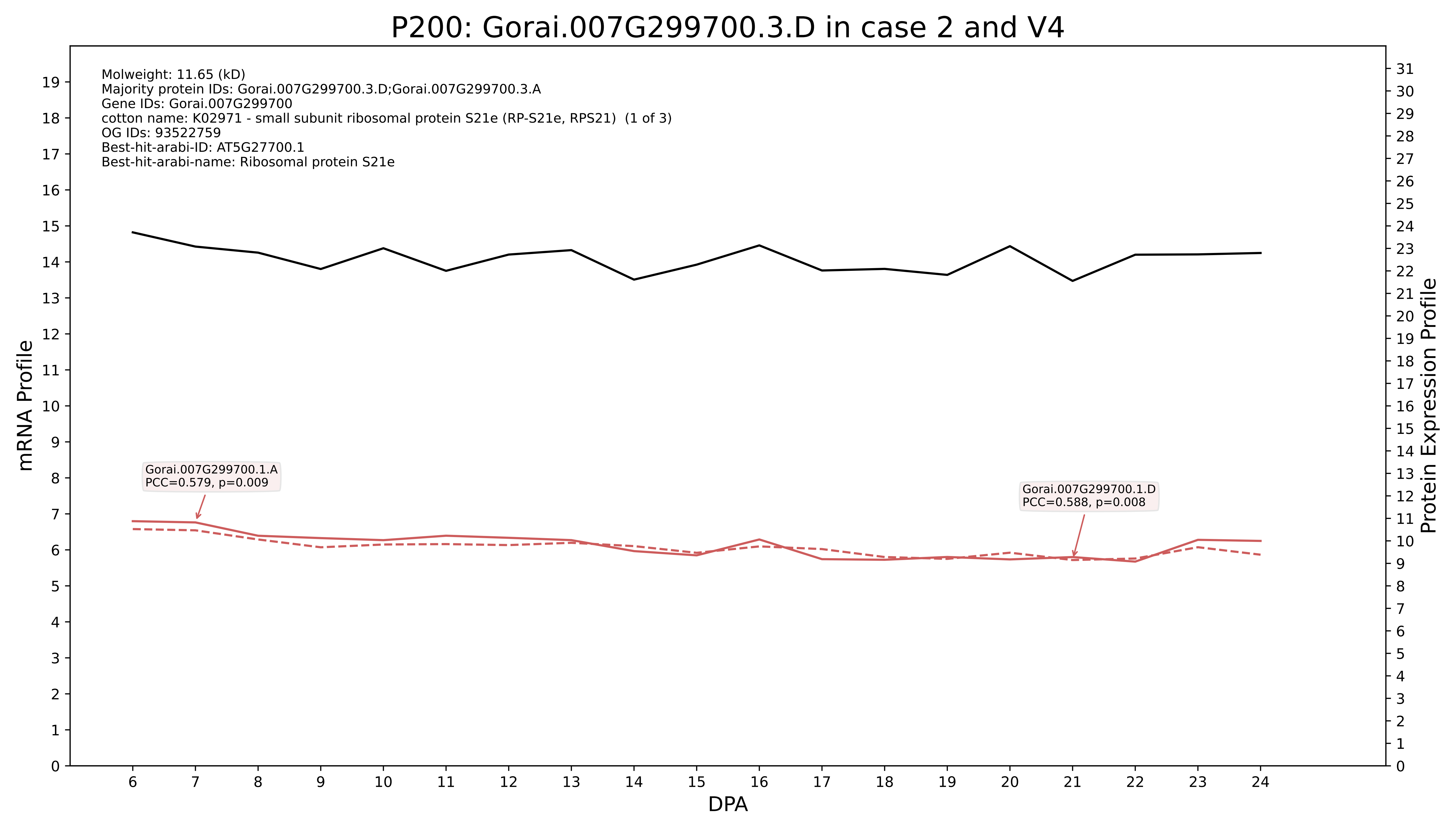
P200: Gorai.007G299700.3.D in case 2 and V4
Molweight: 11.65 (kD)
Majority protein IDs: Gorai.007G299700.3.D;Gorai.007G299700.3.A
Gene IDs: Gorai.007G299700
cotton name: K02971 - small subunit ribosomal protein S21e (RP-S21e, RPS21) (1 of 3)
OG IDs: 93522759
Best-hit-arabi-ID: AT5G27700.1
Best-hit-arabi-name: Ribosomal protein S21e
Gorai.007G299700.1.A
PCC=0.579, p=0.009
Gorai.007G299700.1.D
PCC=0.588, p=0.008
mRNA Profile
Protein Expression Profile
DPA

### P200: Gorai.010G159600.5.D in case 2 and V4

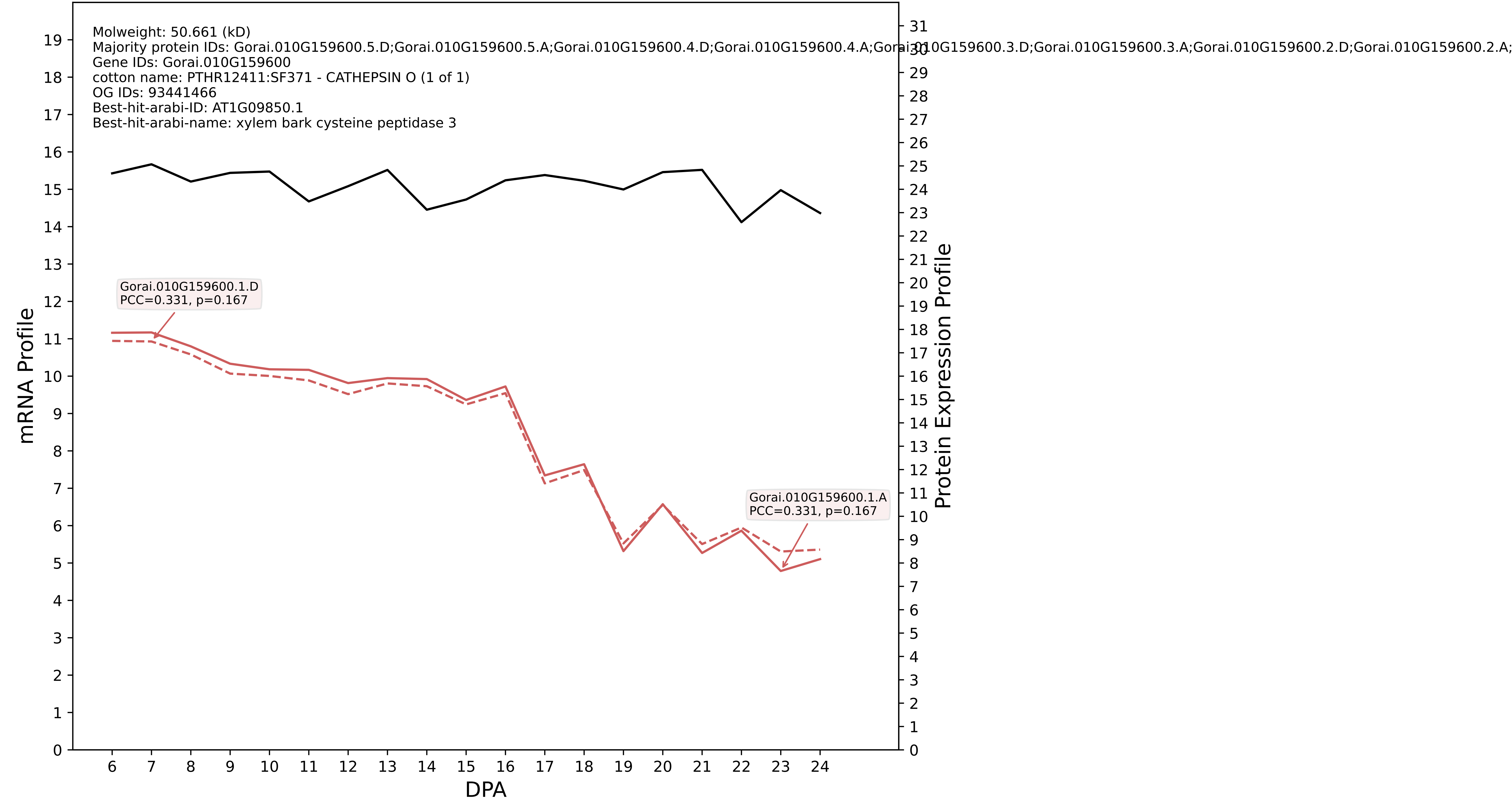

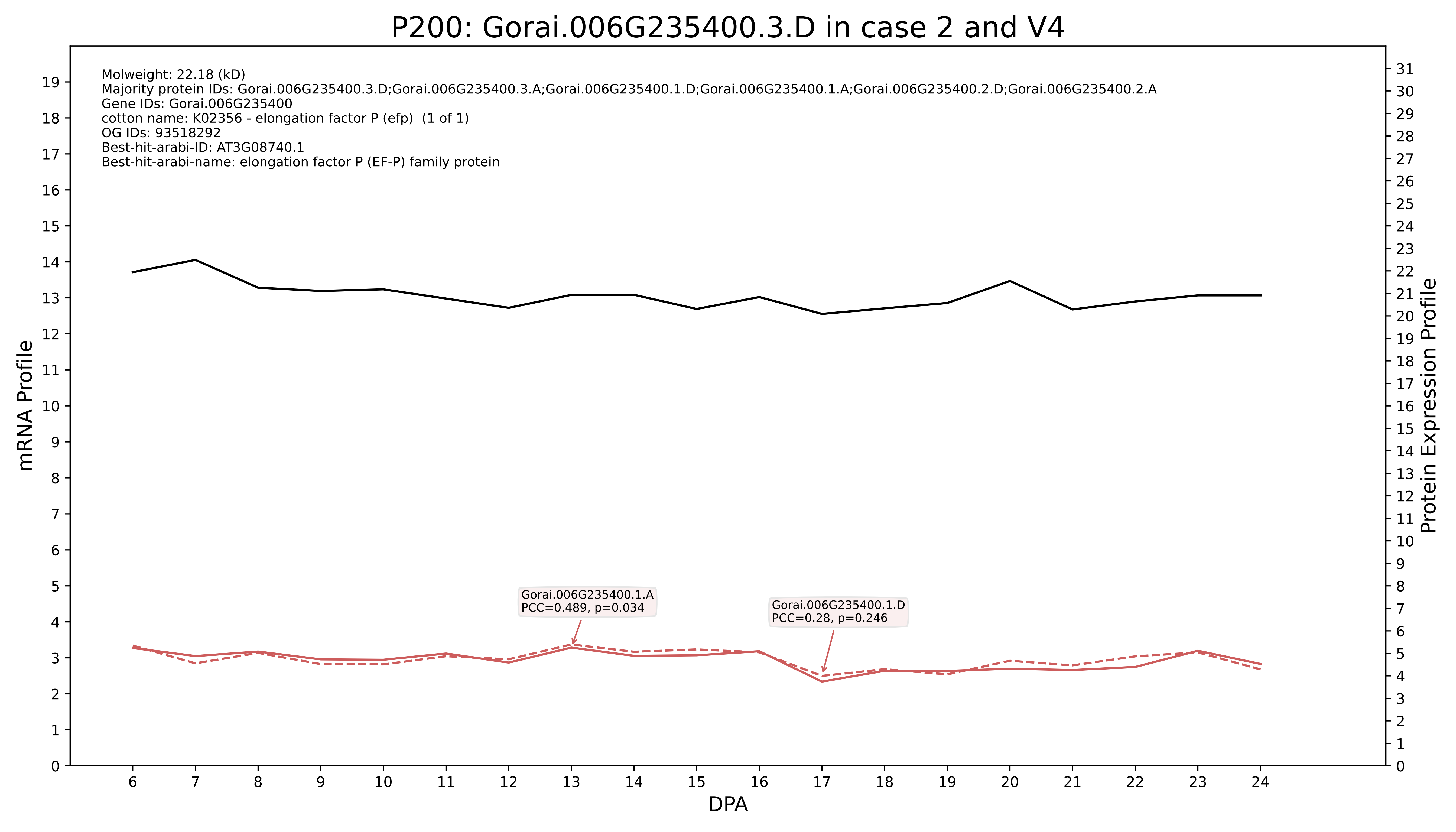
P200: Gorai.006G235400.3.D in case 2 and V4
Molweight: 22.18 (kD)
Majority protein IDs: Gorai.006G235400.3.D;Gorai.006G235400.3.A;Gorai.006G235400.1.D;Gorai.006G235400.1.A;Gorai.006G235400.2.D;Gorai.006G235400.2.A
Gene IDs: Gorai.006G235400
cotton name: K02356 - elongation factor P (efp) (1 of 1)
OG IDs: 93518292
Best-hit-arabi-ID: AT3G08740.1
Best-hit-arabi-name: elongation factor P (EF-P) family protein
Gorai.006G235400.1.A
PCC=0.489, p=0.034
Gorai.006G235400.1.D
PCC=0.28, p=0.246
mRNA Profile
Protein Expression Profile
DPA

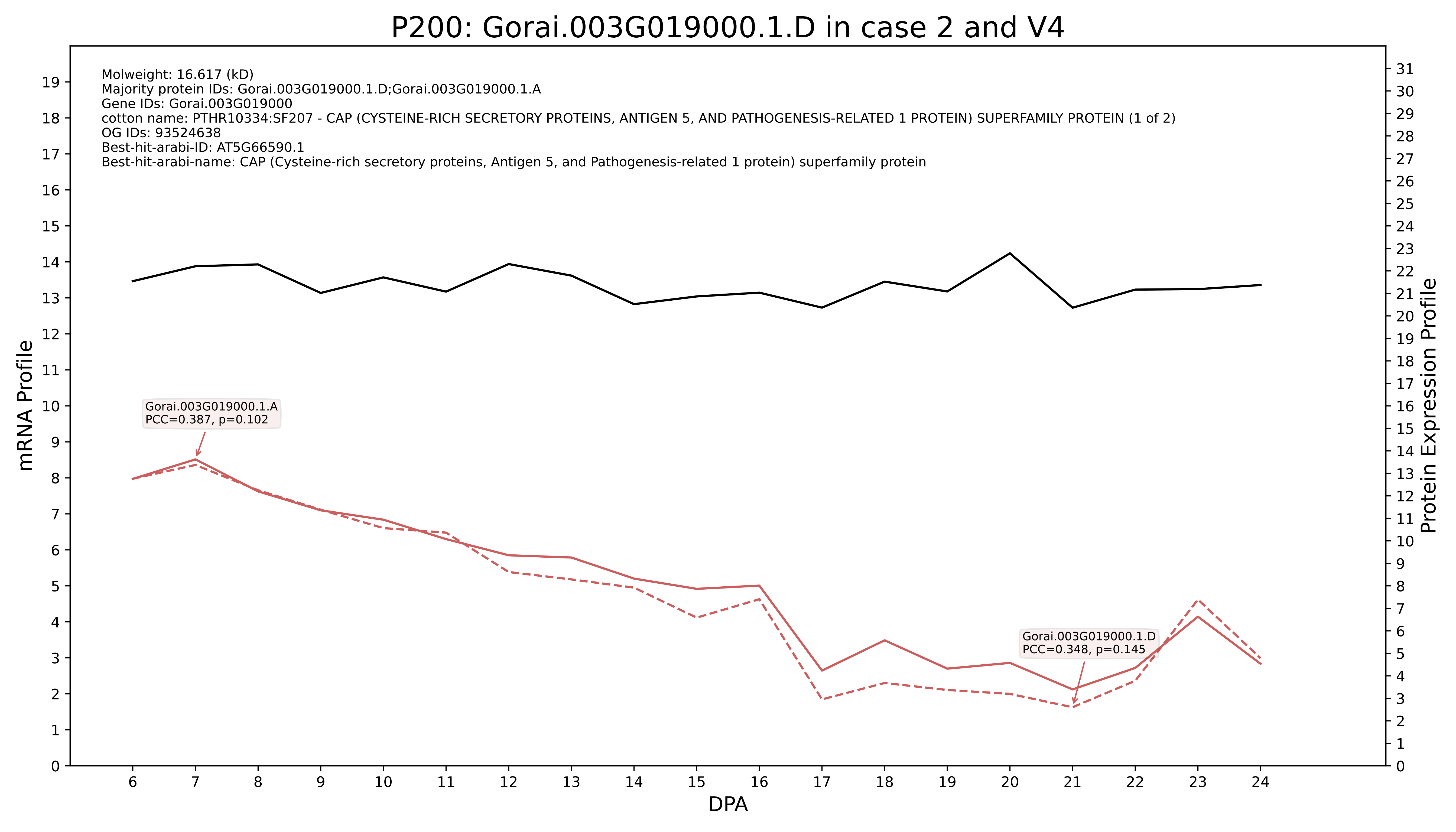
P200: Gorai.003G019000.1.D in case 2 and V4
Molweight: 16.617 (kD)
Majority protein IDs: Gorai.003G019000.1.D;Gorai.003G019000.1.A
Gene IDs: Gorai.003G019000
cotton name: PTHR10334:SF207 - CAP (CYSTEINE-RICH SECRETORY PROTEINS, ANTIGEN 5, AND PATHOGENESIS-RELATED 1 PROTEIN) SUPERFAMILY PROTEIN (1 of 2)
OG IDs: 93524638
Best-hit-arabi-ID: AT5G66590.1
Best-hit-arabi-name: CAP (Cysteine-rich secretory proteins, Antigen 5, and Pathogenesis-related 1 protein) superfamily protein
Gorai.003G019000.1.A
PCC=0.387, p=0.102
Gorai.003G019000.1.D
PCC=0.348, p=0.145
mRNA Profile
Protein Expression Profile
DPA

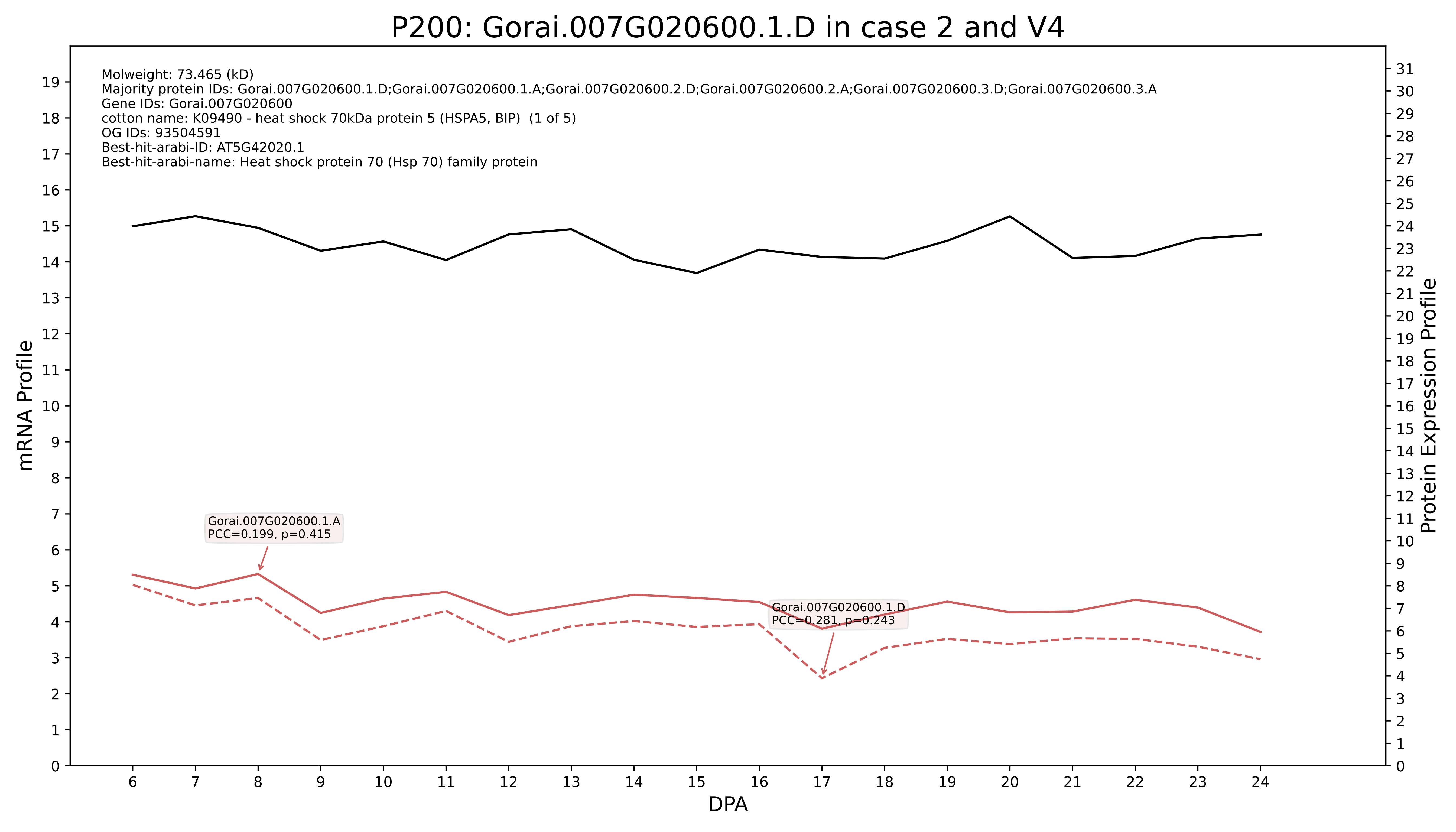

P200: Gorai.007G020600.1.D in case 2 and V4
Molweight: 73.465 (kD)
Majority protein IDs: Gorai.007G020600.1.D;Gorai.007G020600.1.A;Gorai.007G020600.2.D;Gorai.007G020600.2.A;Gorai.007G020600.3.D;Gorai.007G020600.3.A
Gene IDs: Gorai.007G020600
cotton name: K09490 - heat shock 70kDa protein 5 (HSPA5, BIP) (1 of 5)
OG IDs: 93504591
Best-hit-arabi-ID: AT5G42020.1
Best-hit-arabi-name: Heat shock protein 70 (Hsp 70) family protein
Gorai.007G020600.1.A
PCC=0.199, p=0.415
Gorai.007G020600.1.D
PCC=0.281, p=0.243
mRNA Profile
Protein Expression Profile
DPA

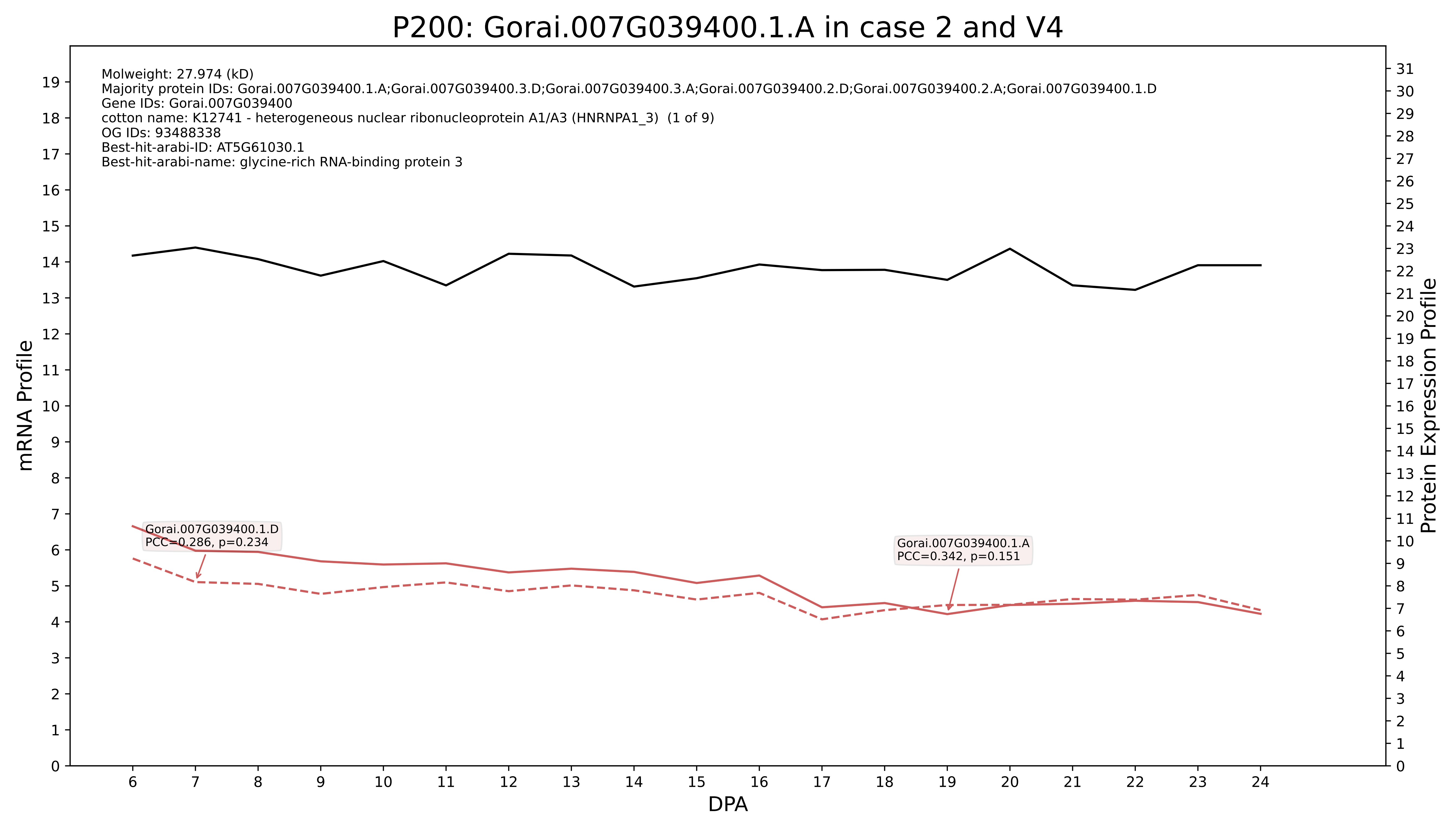
P200: Gorai.007G039400.1.A in case 2 and V4
Molweight: 27.974 (kD)
Majority protein IDs: Gorai.007G039400.1.A;Gorai.007G039400.3.D;Gorai.007G039400.3.A;Gorai.007G039400.2.D;Gorai.007G039400.2.A;Gorai.007G039400.1.D
Gene IDs: Gorai.007G039400
cotton name: K12741 - heterogeneous nuclear ribonucleoprotein A1/A3 (HNRNPA1_3) (1 of 9)
OG IDs: 93488338
Best-hit-arabi-ID: AT5G61030.1
Best-hit-arabi-name: glycine-rich RNA-binding protein 3
Gorai.007G039400.1.D
PCC=0.286, p=0.234
Gorai.007G039400.1.A
PCC=0.342, p=0.151
mRNA Profile
Protein Expression Profile
DPA

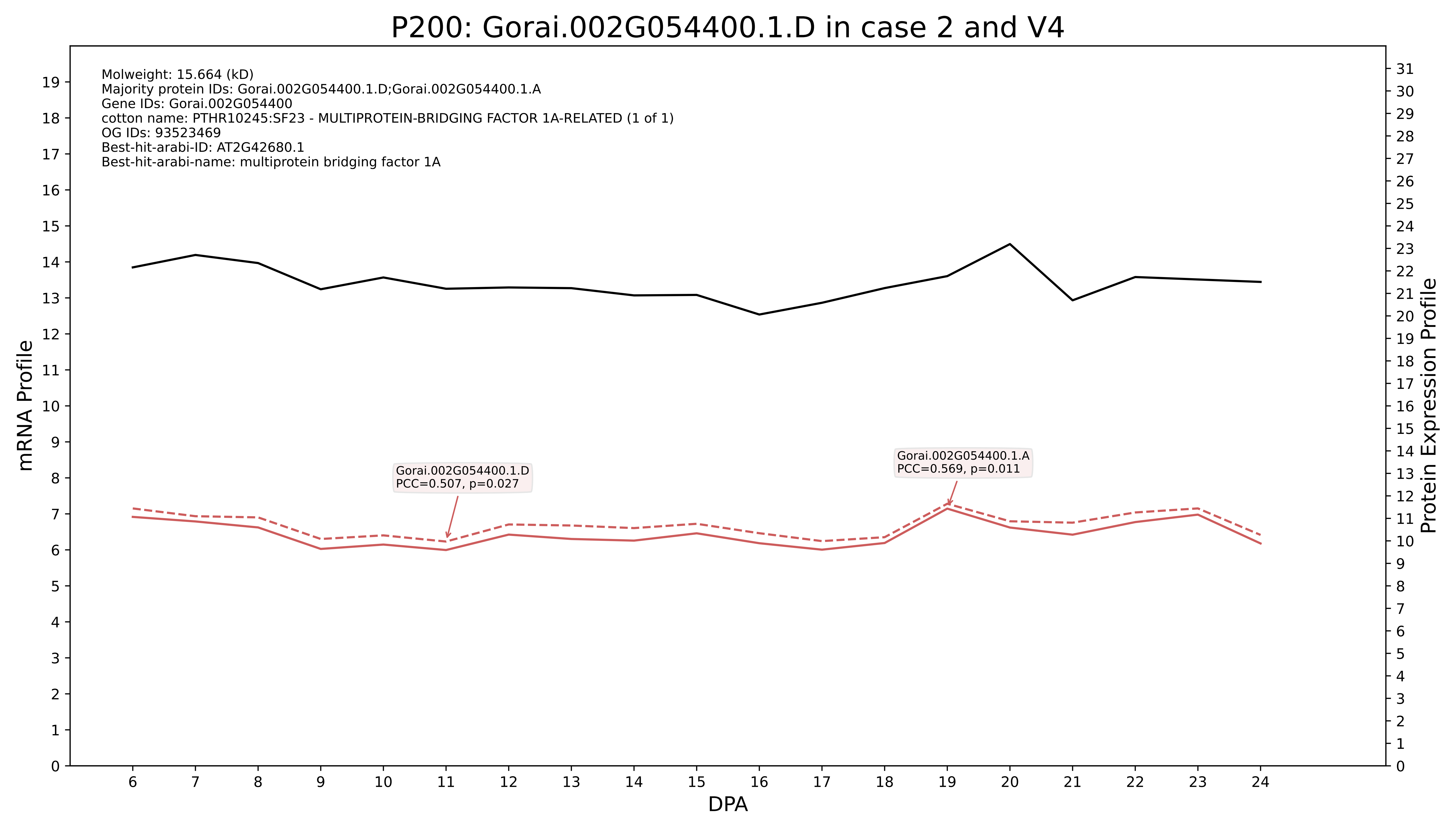

P200: Gorai.002G054400.1.D in case 2 and V4
Molweight: 15.664 (kD)
Majority protein IDs: Gorai.002G054400.1.D;Gorai.002G054400.1.A
Gene IDs: Gorai.002G054400
cotton name: PTHR10245:SF23 - MULTIPROTEIN-BRIDGING FACTOR 1A-RELATED (1 of 1)
OG IDs: 93523469
Best-hit-arabi-ID: AT2G42680.1
Best-hit-arabi-name: multiprotein bridging factor 1A
Gorai.002G054400.1.D
PCC=0.507, p=0.027
Gorai.002G054400.1.A
PCC=0.569, p=0.011
mRNA Profile
Protein Expression Profile
DPA

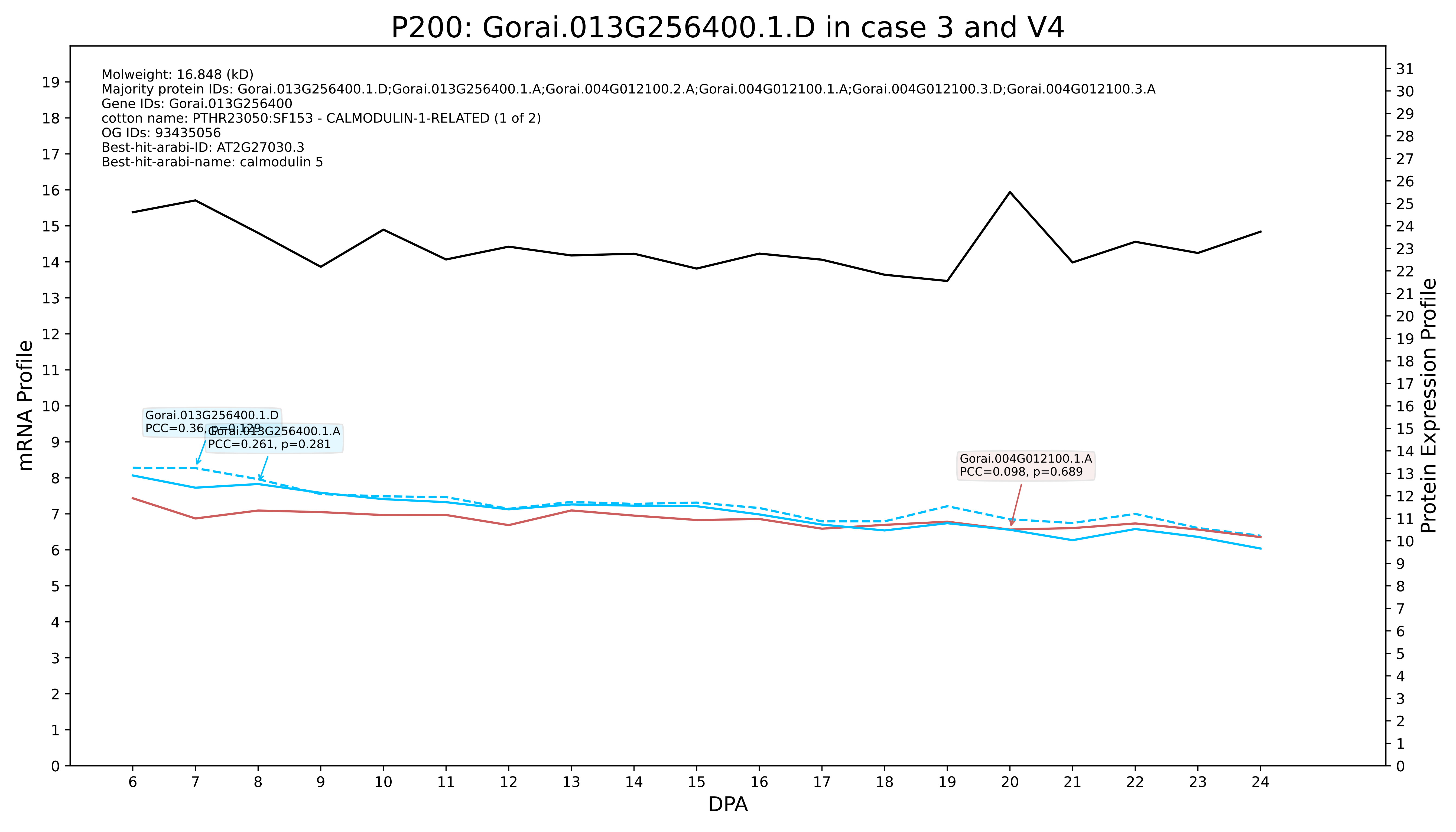
P200: Gorai.013G256400.1.D in case 3 and V4
Molweight: 16.848 (kD)
Majority protein IDs: Gorai.013G256400.1.D;Gorai.013G256400.1.A;Gorai.004G012100.2.A;Gorai.004G012100.1.A;Gorai.004G012100.3.D;Gorai.004G012100.3.A
Gene IDs: Gorai.013G256400
cotton name: PTHR23050:SF153 - CALMODULIN-1-RELATED (1 of 2)
OG IDs: 93435056
Best-hit-arabi-ID: AT2G27030.3
Best-hit-arabi-name: calmodulin 5
Gorai.013G256400.1.D
PCC=0.36, p=0.129
Gorai.013G256400.1.A
PCC=0.261, p=0.281
Gorai.004G012100.1.A
PCC=0.098, p=0.689
mRNA Profile
Protein Expression Profile
DPA

### P200: Gorai.009G080400.1.D in case 3 and V4

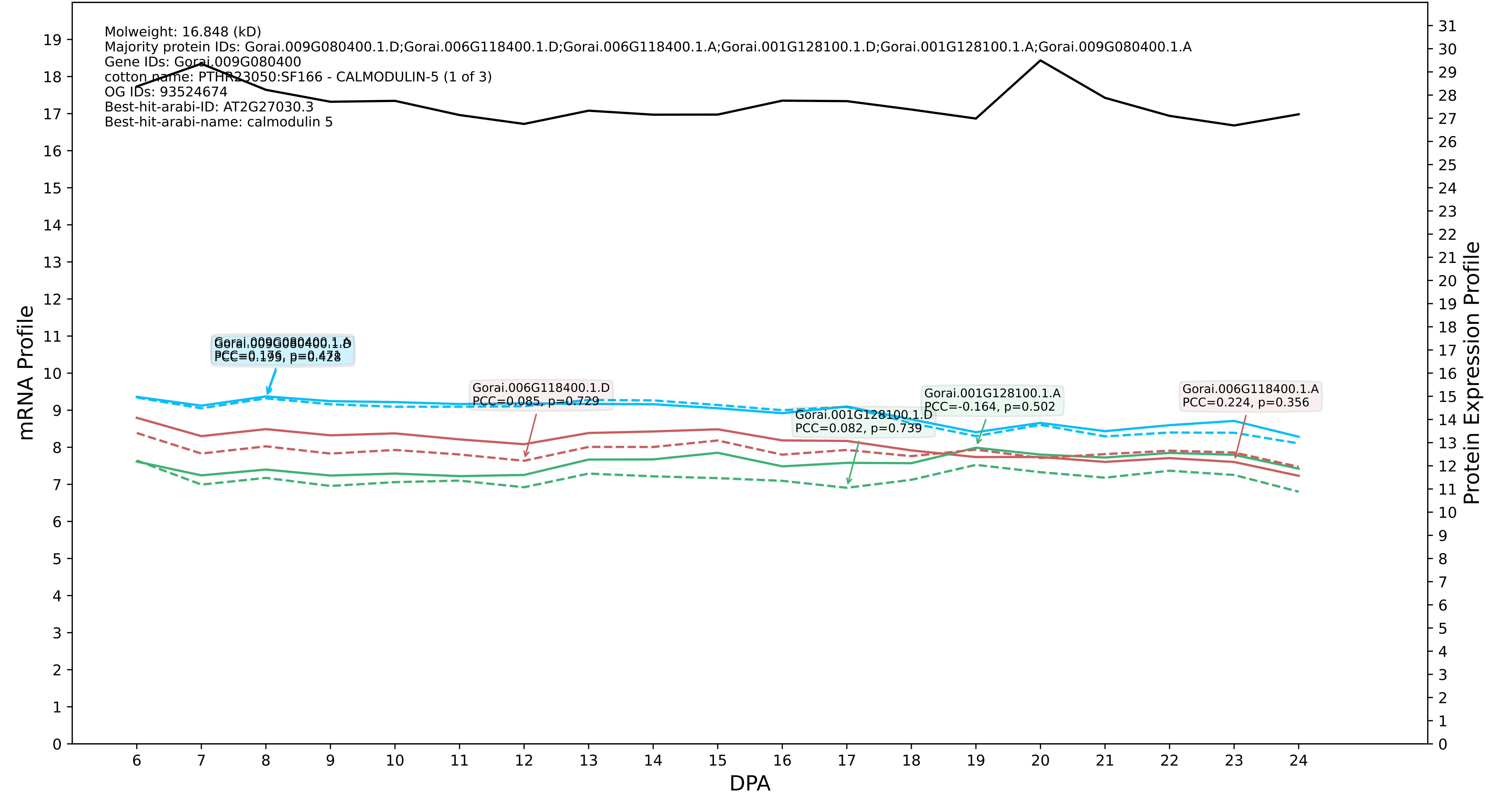

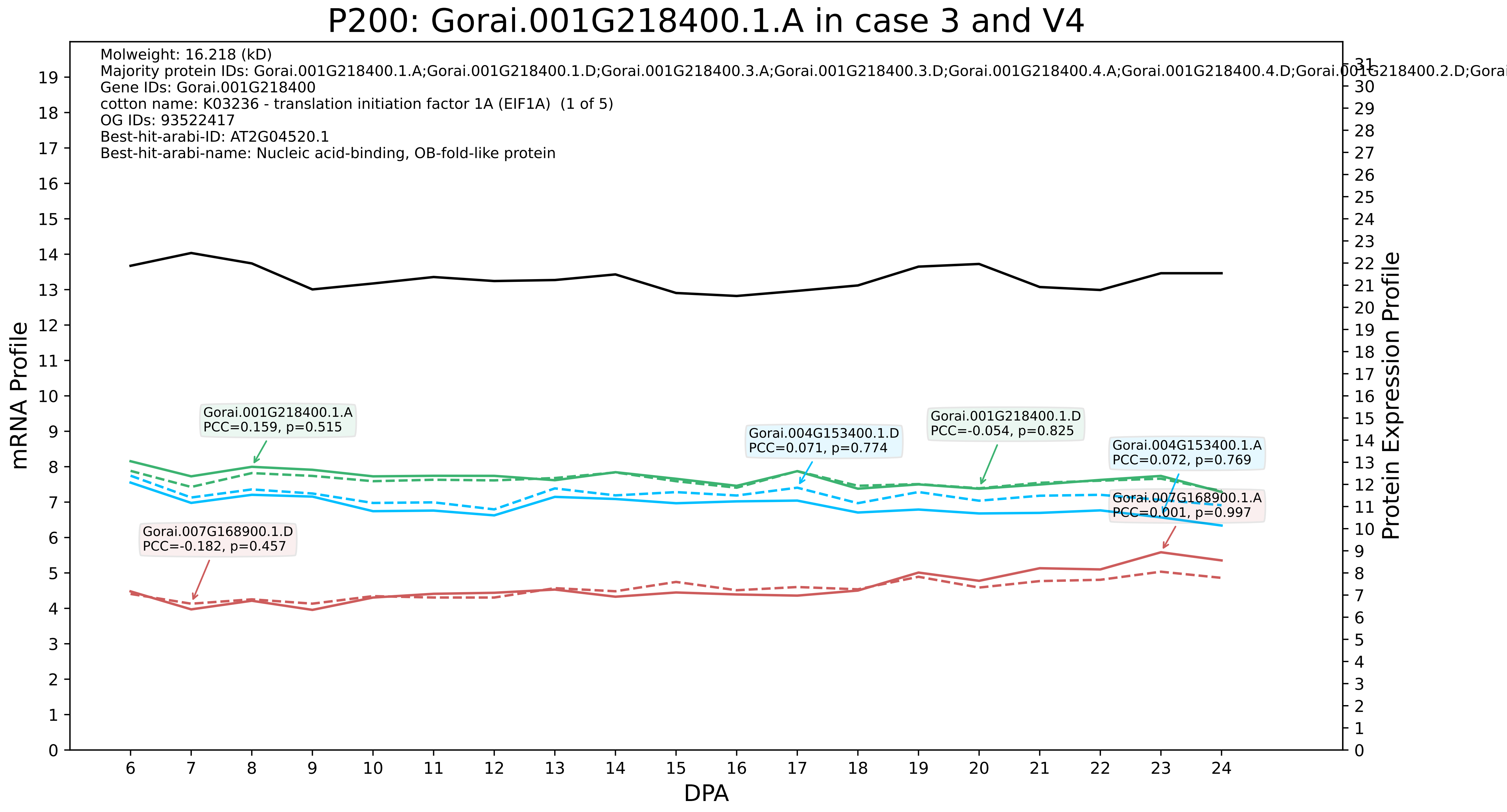

P200: Gorai.001G218400.1.A in case 3 and V4
Molweight: 16.218 (kD)
Majority protein IDs: Gorai.001G218400.1.A;Gorai.001G218400.1.D;Gorai.001G218400.3.A;Gorai.001G218400.3.D;Gorai.001G218400.4.A;Gorai.001G218400.4.D;Gorai.001G218400.2.D;Gorai
Gene IDs: Gorai.001G218400
cotton name: K03236 - translation initiation factor 1A (EIF1A) (1 of 5)
OG IDs: 93522417
Best-hit-arabi-ID: AT2G04520.1
Best-hit-arabi-name: Nucleic acid-binding, OB-fold-like protein
Gorai.001G218400.1.A
PCC=0.159, p=0.515
Gorai.004G153400.1.D
PCC=0.071, p=0.774
Gorai.001G218400.1.D
PCC=-0.054, p=0.825
Gorai.004G153400.1.A
PCC=0.072, p=0.769
Gorai.007G168900.1.A
PCC=0.001, p=0.997
Gorai.007G168900.1.D
PCC=-0.182, p=0.457
mRNA Profile
Protein Expression Profile
DPA

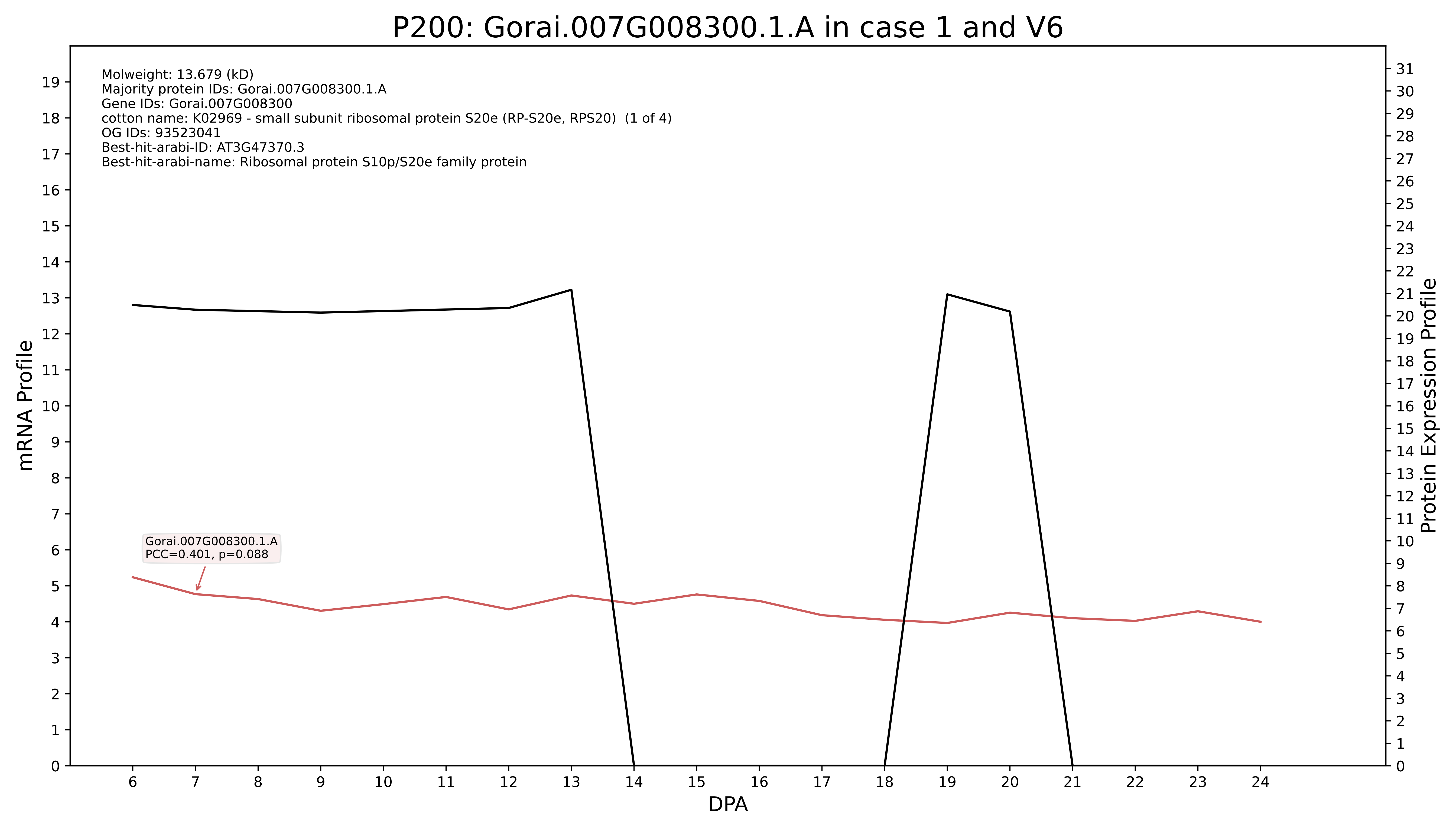
P200: Gorai.007G008300.1.A in case 1 and V6
Molweight: 13.679 (kD)
Majority protein IDs: Gorai.007G008300.1.A
Gene IDs: Gorai.007G008300
cotton name: K02969 - small subunit ribosomal protein S20e (RP-S20e, RPS20) (1 of 4)
OG IDs: 93523041
Best-hit-arabi-ID: AT3G47370.3
Best-hit-arabi-name: Ribosomal protein S10p/S20e family protein
Gorai.007G008300.1.A
PCC=0.401, p=0.088
mRNA Profile
Protein Expression Profile
DPA

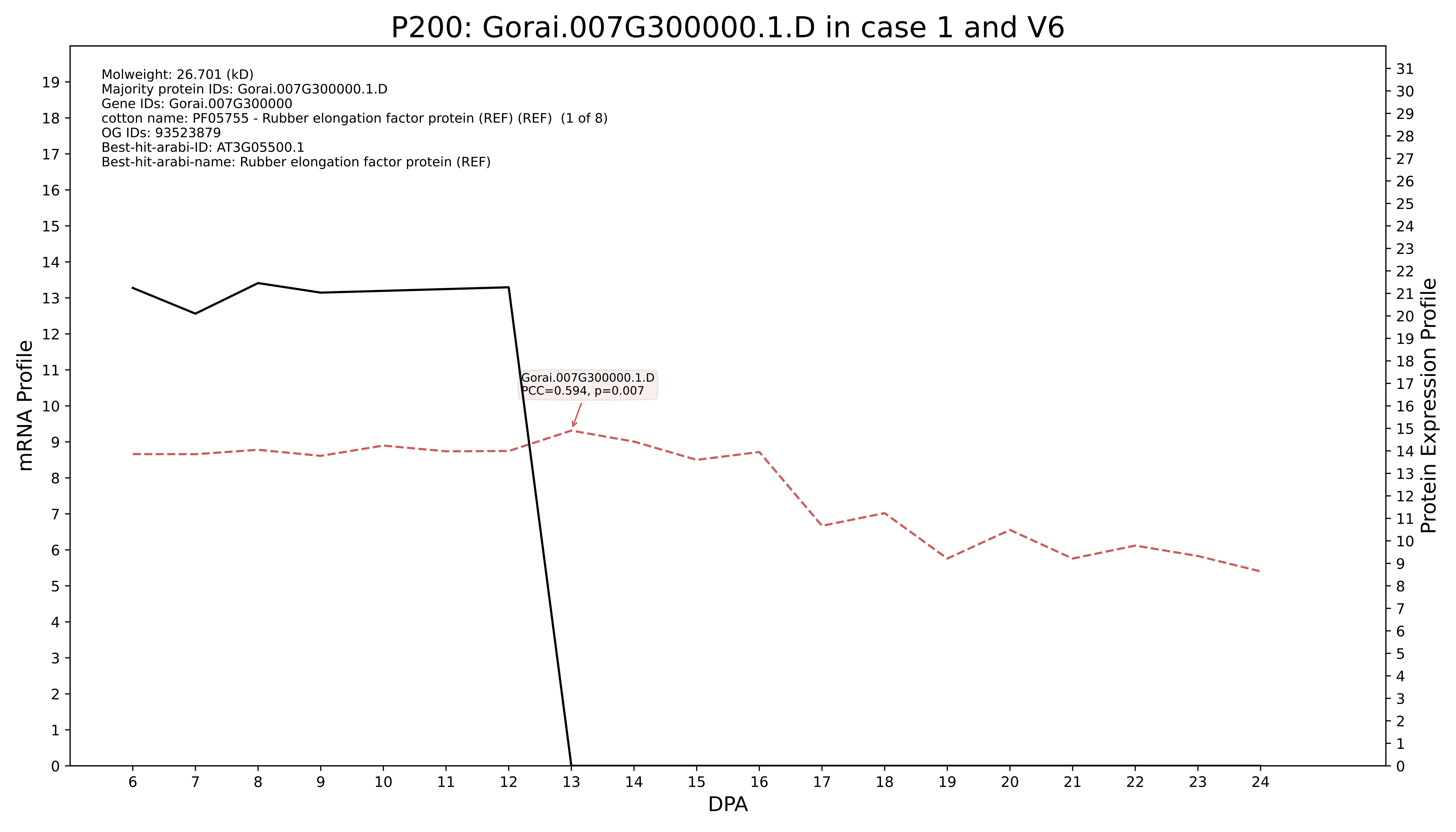

P200: Gorai.007G300000.1.D in case 1 and V6
Molweight: 26.701 (kD)
Majority protein IDs: Gorai.007G300000.1.D
Gene IDs: Gorai.007G300000
cotton name: PF05755 - Rubber elongation factor protein (REF) (REF) (1 of 8)
OG IDs: 93523879
Best-hit-arabi-ID: AT3G05500.1
Best-hit-arabi-name: Rubber elongation factor protein (REF)
Gorai.007G300000.1.D
PCC=0.594, p=0.007
mRNA Profile
Protein Expression Profile
DPA

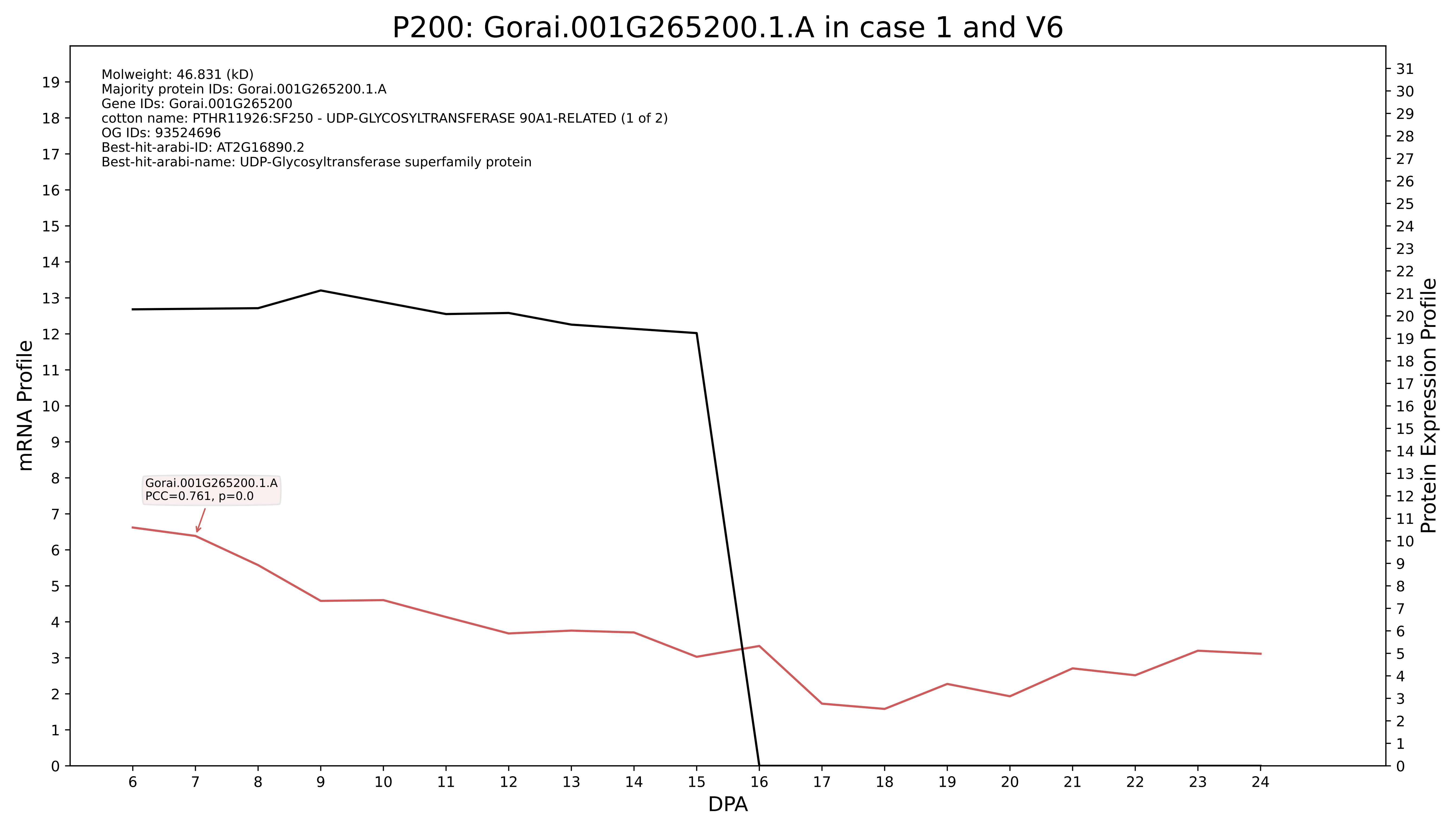
P200: Gorai.001G265200.1.A in case 1 and V6
Molweight: 46.831 (kD)
Majority protein IDs: Gorai.001G265200.1.A
Gene IDs: Gorai.001G265200
cotton name: PTHR11926:SF250 - UDP-GLYCOSYLTRANSFERASE 90A1-RELATED (1 of 2)
OG IDs: 93524696
Best-hit-arabi-ID: AT2G16890.2
Best-hit-arabi-name: UDP-Glycosyltransferase superfamily protein
Gorai.001G265200.1.A
PCC=0.761, p=0.0
mRNA Profile
Protein Expression Profile
DPA

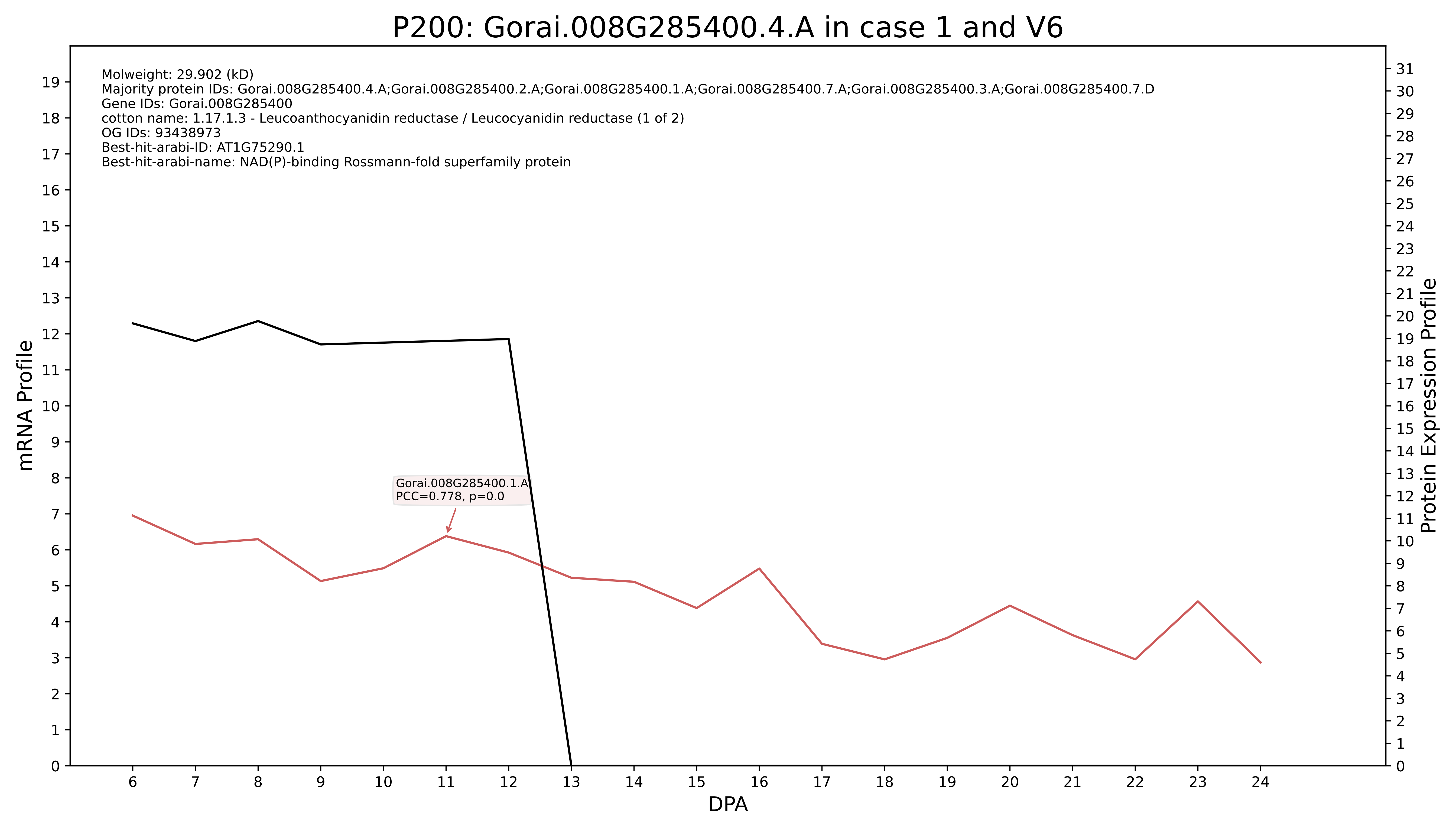

P200: Gorai.008G285400.4.A in case 1 and V6
Molweight: 29.902 (kD)
Majority protein IDs: Gorai.008G285400.4.A;Gorai.008G285400.2.A;Gorai.008G285400.1.A;Gorai.008G285400.7.A;Gorai.008G285400.3.A;Gorai.008G285400.7.D
Gene IDs: Gorai.008G285400
cotton name: 1.17.1.3 - Leucoanthocyanidin reductase / Leucocyanidin reductase (1 of 2)
OG IDs: 93438973
Best-hit-arabi-ID: AT1G75290.1
Best-hit-arabi-name: NAD(P)-binding Rossmann-fold superfamily protein
Gorai.008G285400.1.A
PCC=0.778, p=0.0
mRNA Profile
Protein Expression Profile
DPA

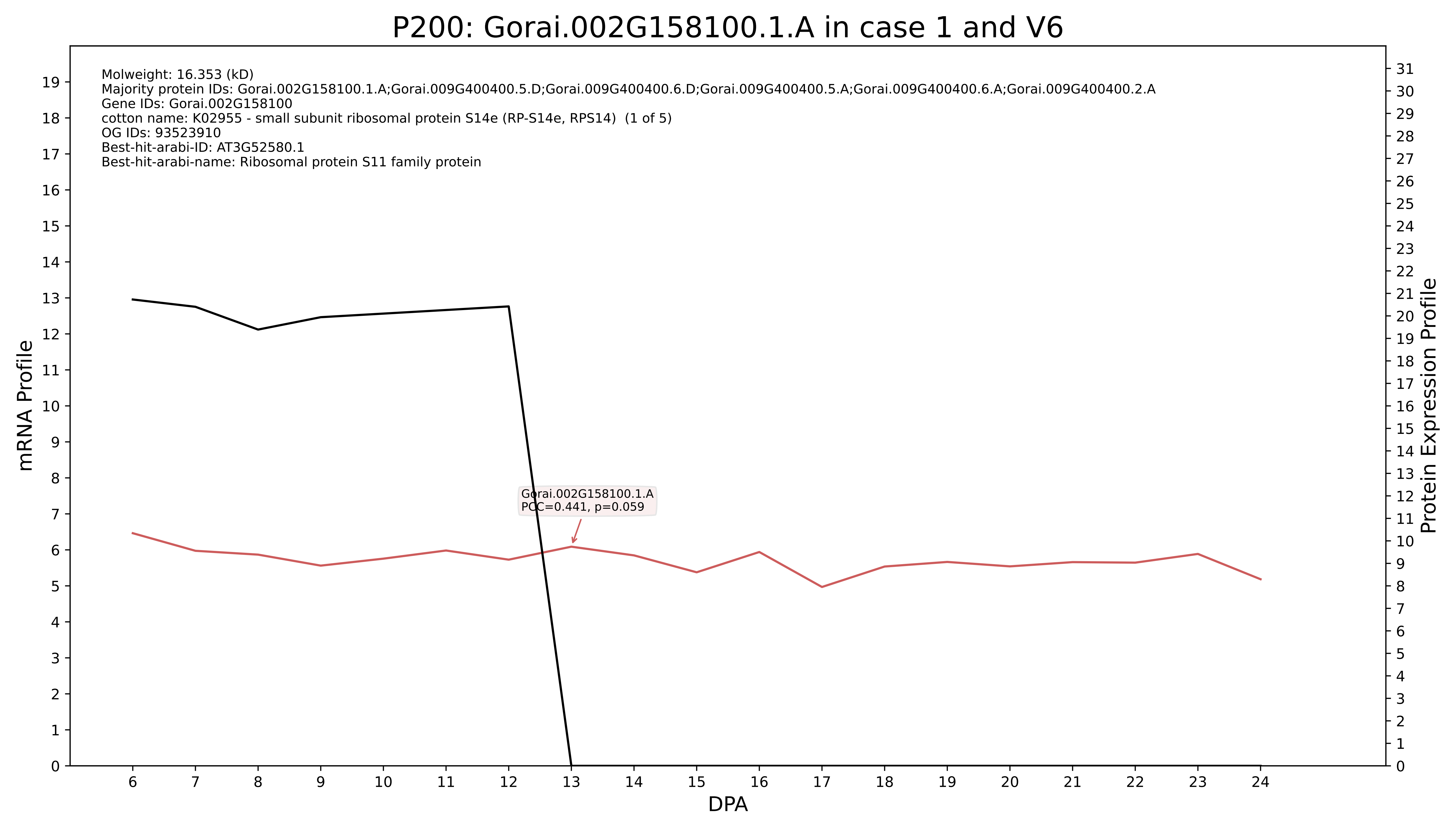

P200: Gorai.002G158100.1.A in case 1 and V6
Molweight: 16.353 (kD)
Majority protein IDs: Gorai.002G158100.1.A;Gorai.009G400400.5.D;Gorai.009G400400.6.D;Gorai.009G400400.5.A;Gorai.009G400400.6.A;Gorai.009G400400.2.A
Gene IDs: Gorai.002G158100
cotton name: K02955 - small subunit ribosomal protein S14e (RP-S14e, RPS14) (1 of 5)
OG IDs: 93523910
Best-hit-arabi-ID: AT3G52580.1
Best-hit-arabi-name: Ribosomal protein S11 family protein
Gorai.002G158100.1.A
PCC=0.441, p=0.059
mRNA Profile
Protein Expression Profile
DPA

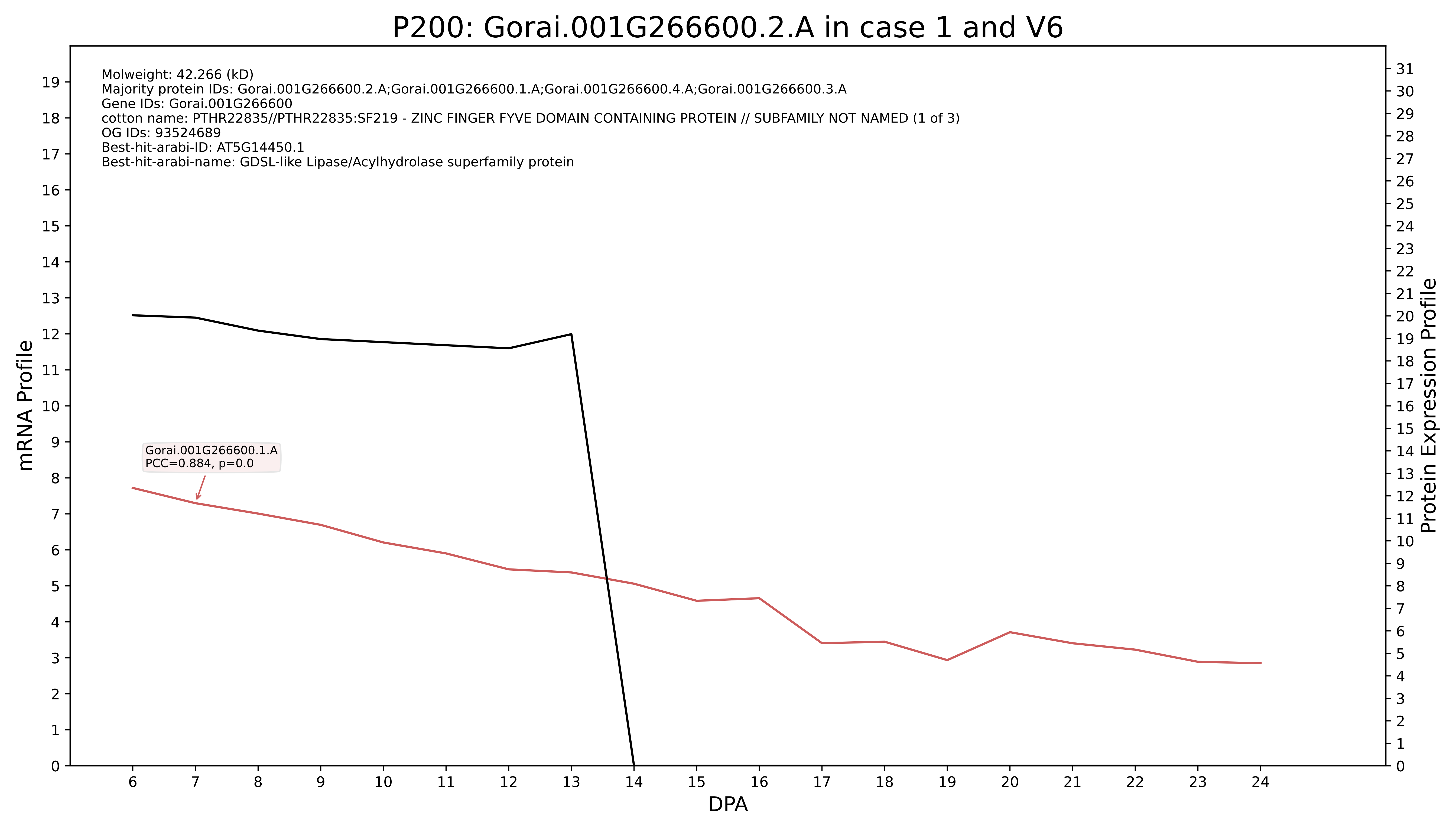
P200: Gorai.001G266600.2.A in case 1 and V6
Molweight: 42.266 (kD)
Majority protein IDs: Gorai.001G266600.2.A;Gorai.001G266600.1.A;Gorai.001G266600.4.A;Gorai.001G266600.3.A
Gene IDs: Gorai.001G266600
cotton name: PTHR22835//PTHR22835:SF219 - ZINC FINGER FYVE DOMAIN CONTAINING PROTEIN // SUBFAMILY NOT NAMED (1 of 3)
OG IDs: 93524689
Best-hit-arabi-ID: AT5G14450.1
Best-hit-arabi-name: GDSL-like Lipase/Acylhydrolase superfamily protein
Gorai.001G266600.1.A
PCC=0.884, p=0.0
mRNA Profile
Protein Expression Profile
DPA

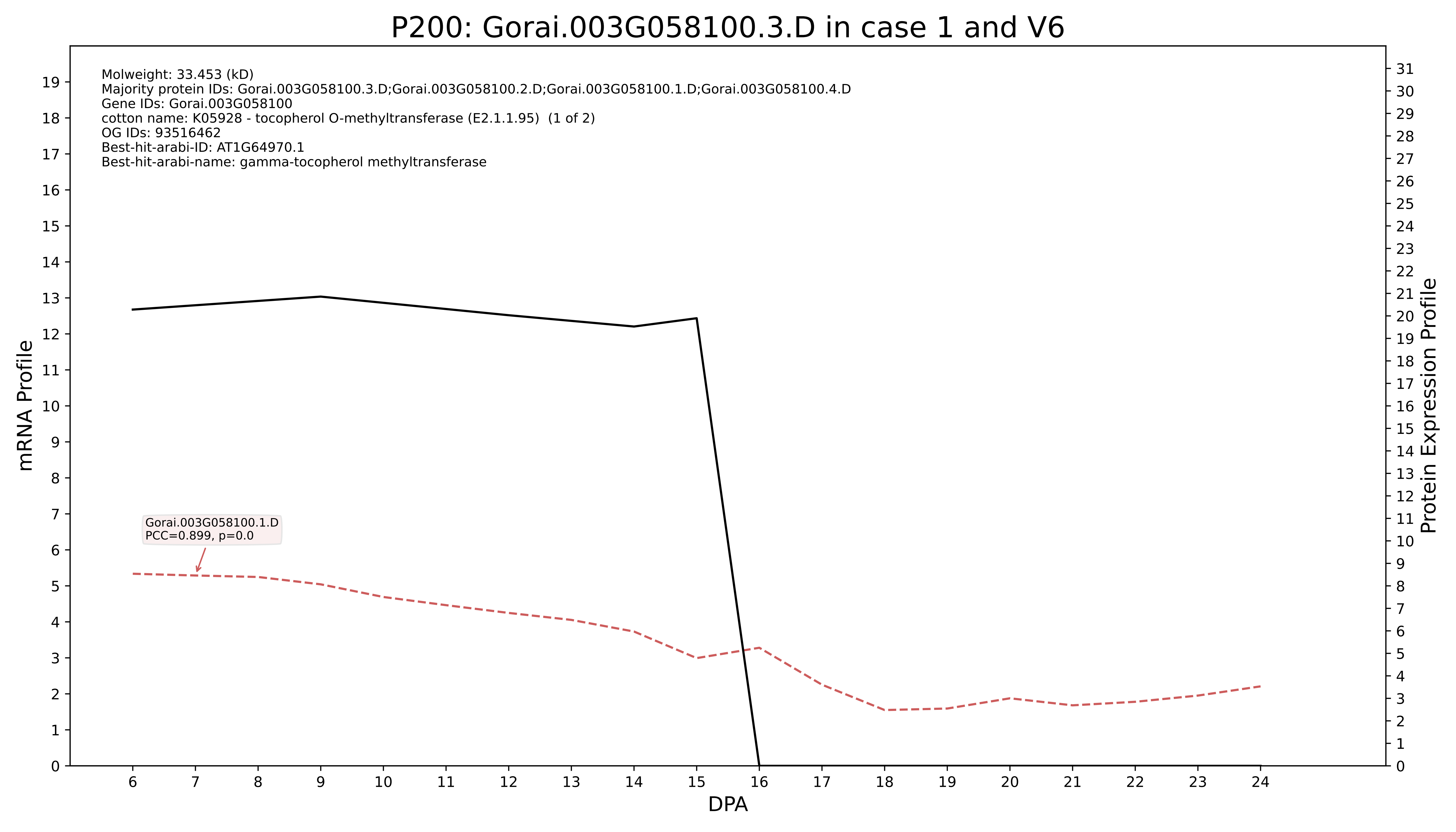
P200: Gorai.003G058100.3.D in case 1 and V6
Molweight: 33.453 (kD)
Majority protein IDs: Gorai.003G058100.3.D;Gorai.003G058100.2.D;Gorai.003G058100.1.D;Gorai.003G058100.4.D
Gene IDs: Gorai.003G058100
cotton name: K05928 - tocopherol O-methyltransferase (E2.1.1.95) (1 of 2)
OG IDs: 93516462
Best-hit-arabi-ID: AT1G64970.1
Best-hit-arabi-name: gamma-tocopherol methyltransferase
Gorai.003G058100.1.D
PCC=0.899, p=0.0
mRNA Profile
Protein Expression Profile
DPA

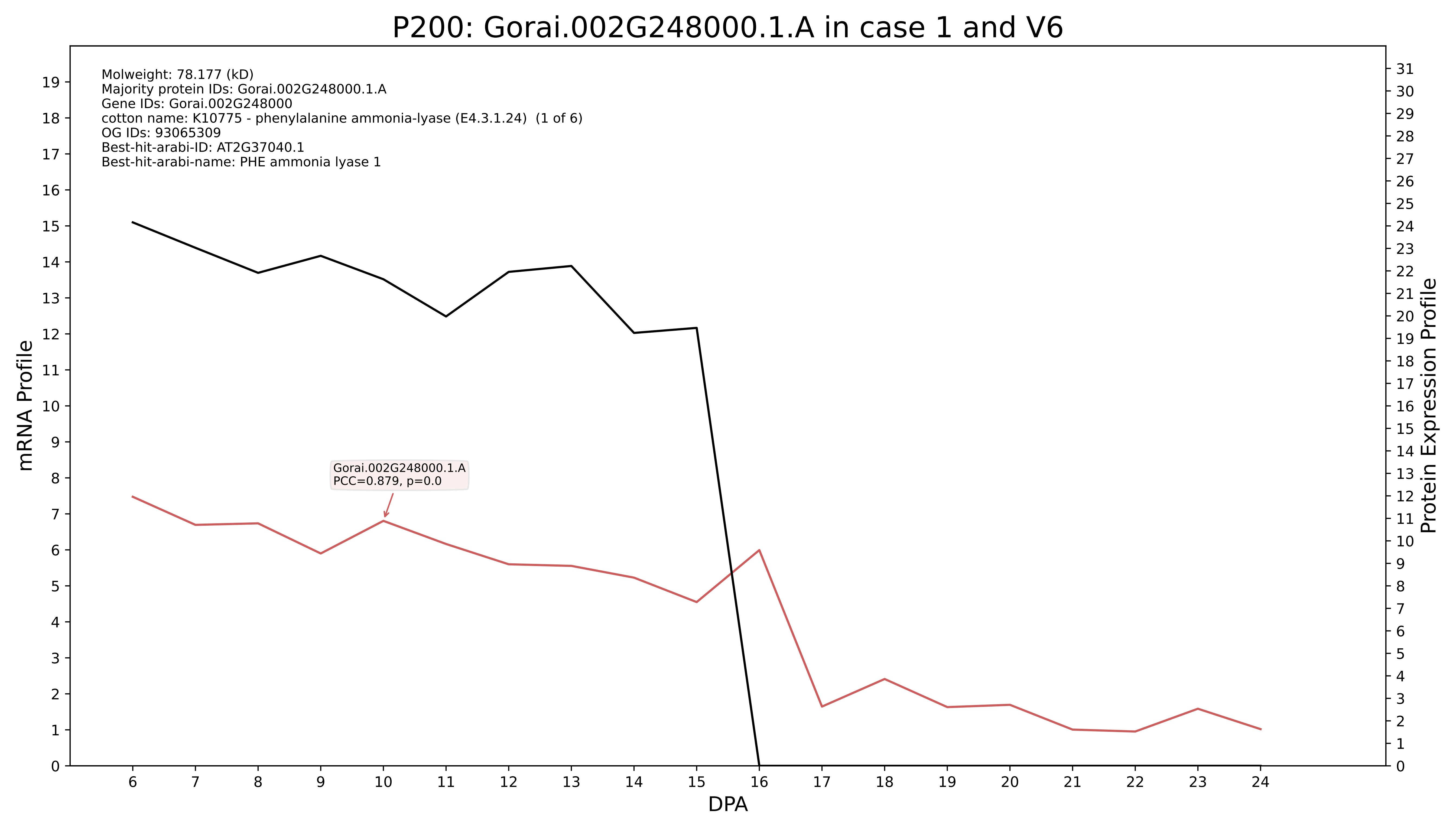
P200: Gorai.002G248000.1.A in case 1 and V6
Molweight: 78.177 (kD)
Majority protein IDs: Gorai.002G248000.1.A
Gene IDs: Gorai.002G248000
cotton name: K10775 - phenylalanine ammonia-lyase (E4.3.1.24) (1 of 6)
OG IDs: 93065309
Best-hit-arabi-ID: AT2G37040.1
Best-hit-arabi-name: PHE ammonia lyase 1
Gorai.002G248000.1.A
PCC=0.879, p=0.0
mRNA Profile
Protein Expression Profile
DPA

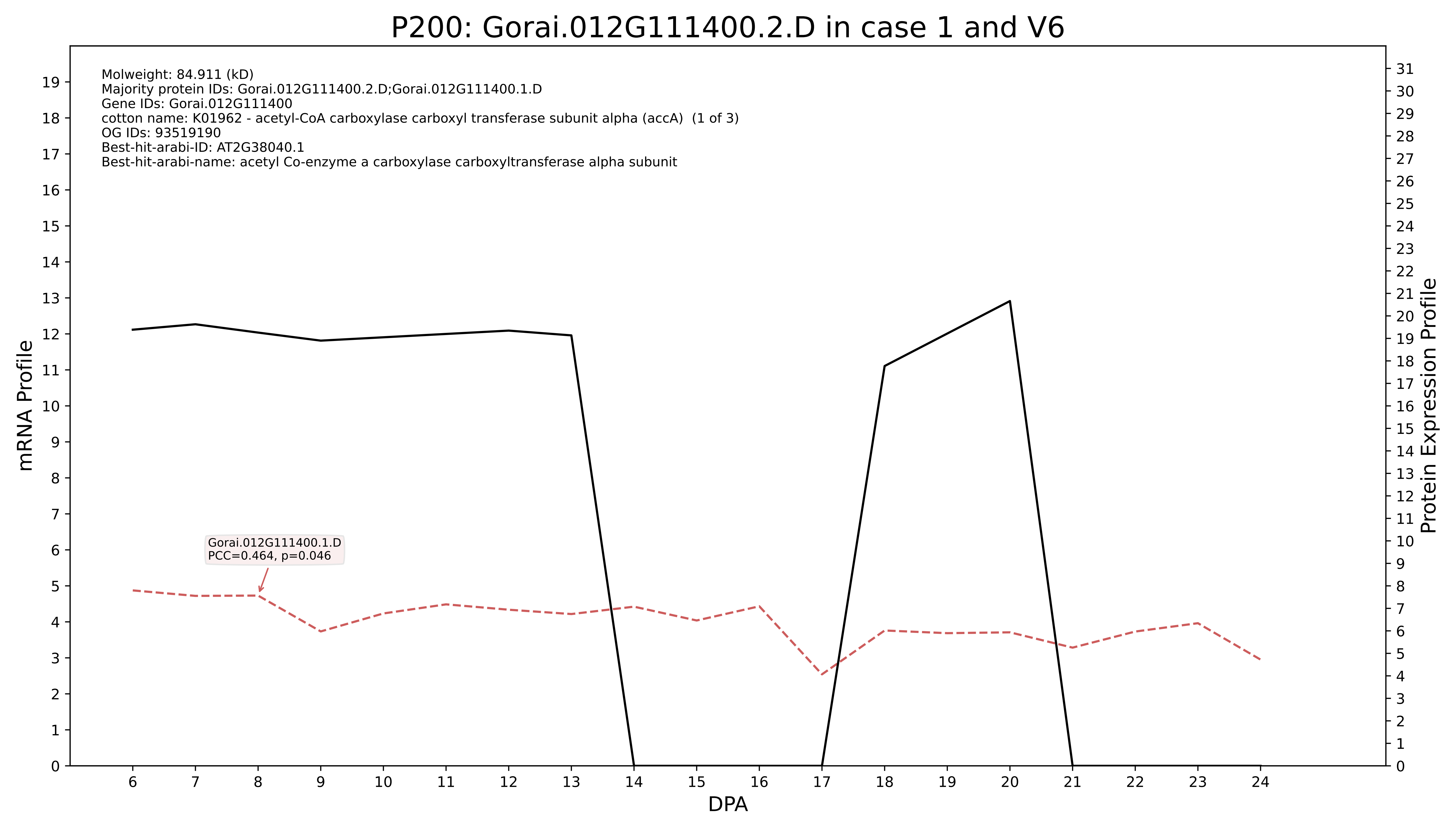

P200: Gorai.012G111400.2.D in case 1 and V6
Molweight: 84.911 (kD)
Majority protein IDs: Gorai.012G111400.2.D;Gorai.012G111400.1.D
Gene IDs: Gorai.012G111400
cotton name: K01962 - acetyl-CoA carboxylase carboxyl transferase subunit alpha (accA) (1 of 3)
OG IDs: 93519190
Best-hit-arabi-ID: AT2G38040.1
Best-hit-arabi-name: acetyl Co-enzyme a carboxylase carboxyltransferase alpha subunit
Gorai.012G111400.1.D
PCC=0.464, p=0.046
mRNA Profile
Protein Expression Profile
DPA

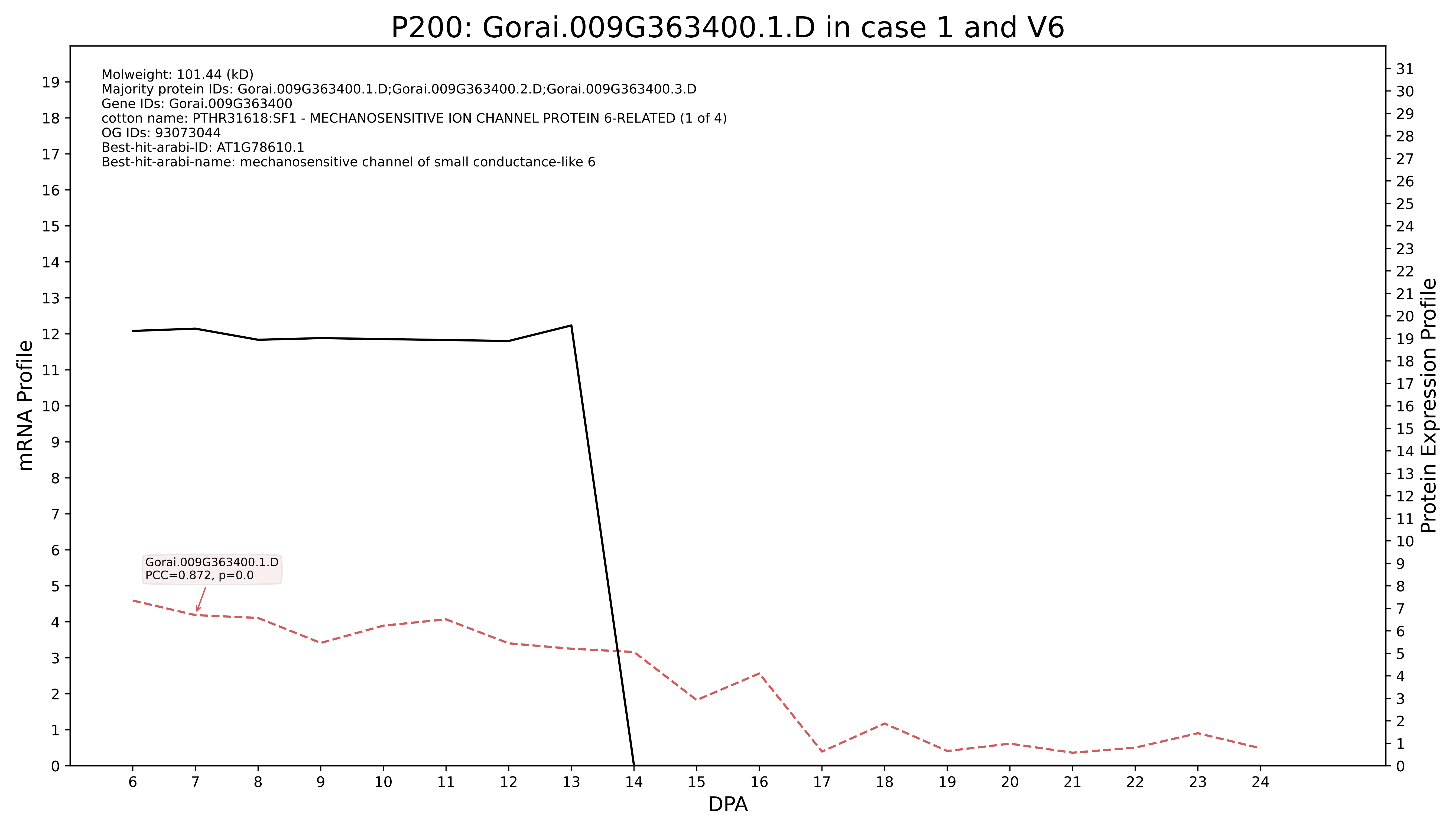
P200: Gorai.009G363400.1.D in case 1 and V6
Molweight: 101.44 (kD)
Majority protein IDs: Gorai.009G363400.1.D;Gorai.009G363400.2.D;Gorai.009G363400.3.D
Gene IDs: Gorai.009G363400
cotton name: PTHR31618:SF1 - MECHANOSENSITIVE ION CHANNEL PROTEIN 6-RELATED (1 of 4)
OG IDs: 93073044
Best-hit-arabi-ID: AT1G78610.1
Best-hit-arabi-name: mechanosensitive channel of small conductance-like 6
Gorai.009G363400.1.D
PCC=0.872, p=0.0
mRNA Profile
Protein Expression Profile
DPA

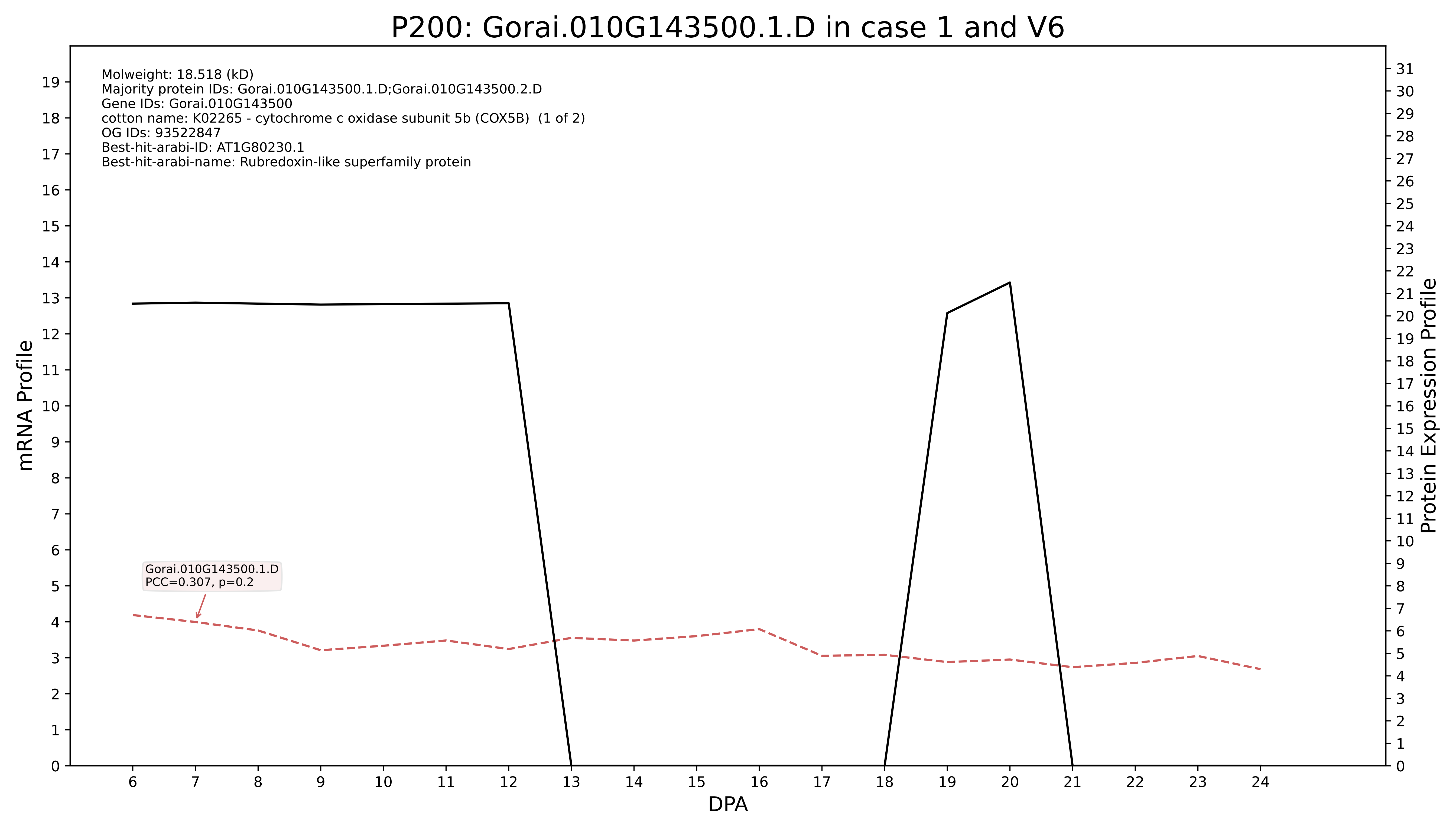
P200: Gorai.010G143500.1.D in case 1 and V6
Molweight: 18.518 (kD)
Majority protein IDs: Gorai.010G143500.1.D;Gorai.010G143500.2.D
Gene IDs: Gorai.010G143500
cotton name: K02265 - cytochrome c oxidase subunit 5b (COX5B) (1 of 2)
OG IDs: 93522847
Best-hit-arabi-ID: AT1G80230.1
Best-hit-arabi-name: Rubredoxin-like superfamily protein
Gorai.010G143500.1.D
PCC=0.307, p=0.2
mRNA Profile
Protein Expression Profile
DPA

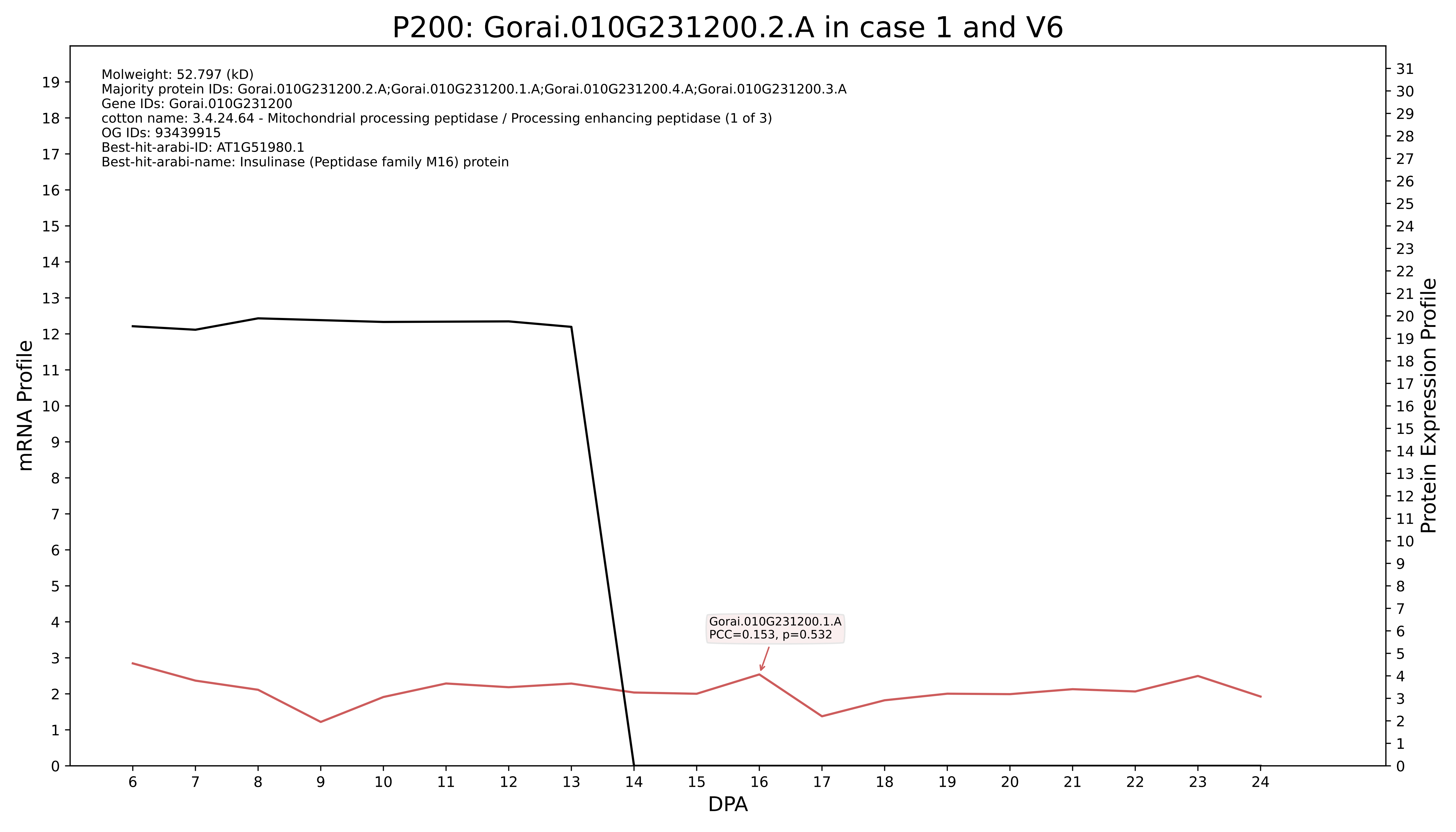

P200: Gorai.010G231200.2.A in case 1 and V6
Molweight: 52.797 (kD)
Majority protein IDs: Gorai.010G231200.2.A;Gorai.010G231200.1.A;Gorai.010G231200.4.A;Gorai.010G231200.3.A
Gene IDs: Gorai.010G231200
cotton name: 3.4.24.64 - Mitochondrial processing peptidase / Processing enhancing peptidase (1 of 3)
OG IDs: 93439915
Best-hit-arabi-ID: AT1G51980.1
Best-hit-arabi-name: Insulinase (Peptidase family M16) protein
Gorai.010G231200.1.A
PCC=0.153, p=0.532
mRNA Profile
Protein Expression Profile
DPA

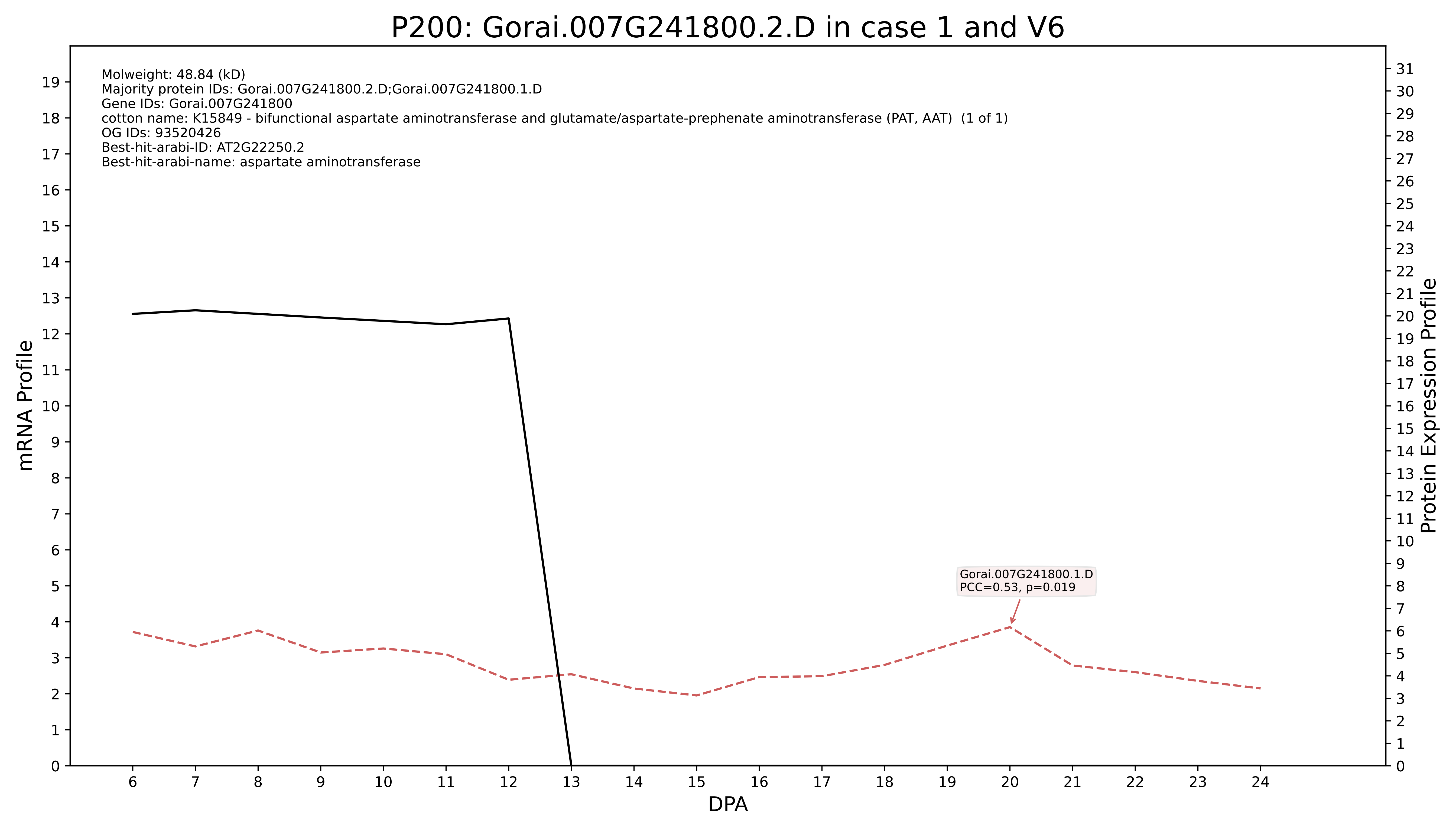
P200: Gorai.007G241800.2.D in case 1 and V6
Molweight: 48.84 (kD)
Majority protein IDs: Gorai.007G241800.2.D;Gorai.007G241800.1.D
Gene IDs: Gorai.007G241800
cotton name: K15849 - bifunctional aspartate aminotransferase and glutamate/aspartate-prephenate aminotransferase (PAT, AAT) (1 of 1)
OG IDs: 93520426
Best-hit-arabi-ID: AT2G22250.2
Best-hit-arabi-name: aspartate aminotransferase
Gorai.007G241800.1.D
PCC=0.53, p=0.019
mRNA Profile
Protein Expression Profile
DPA

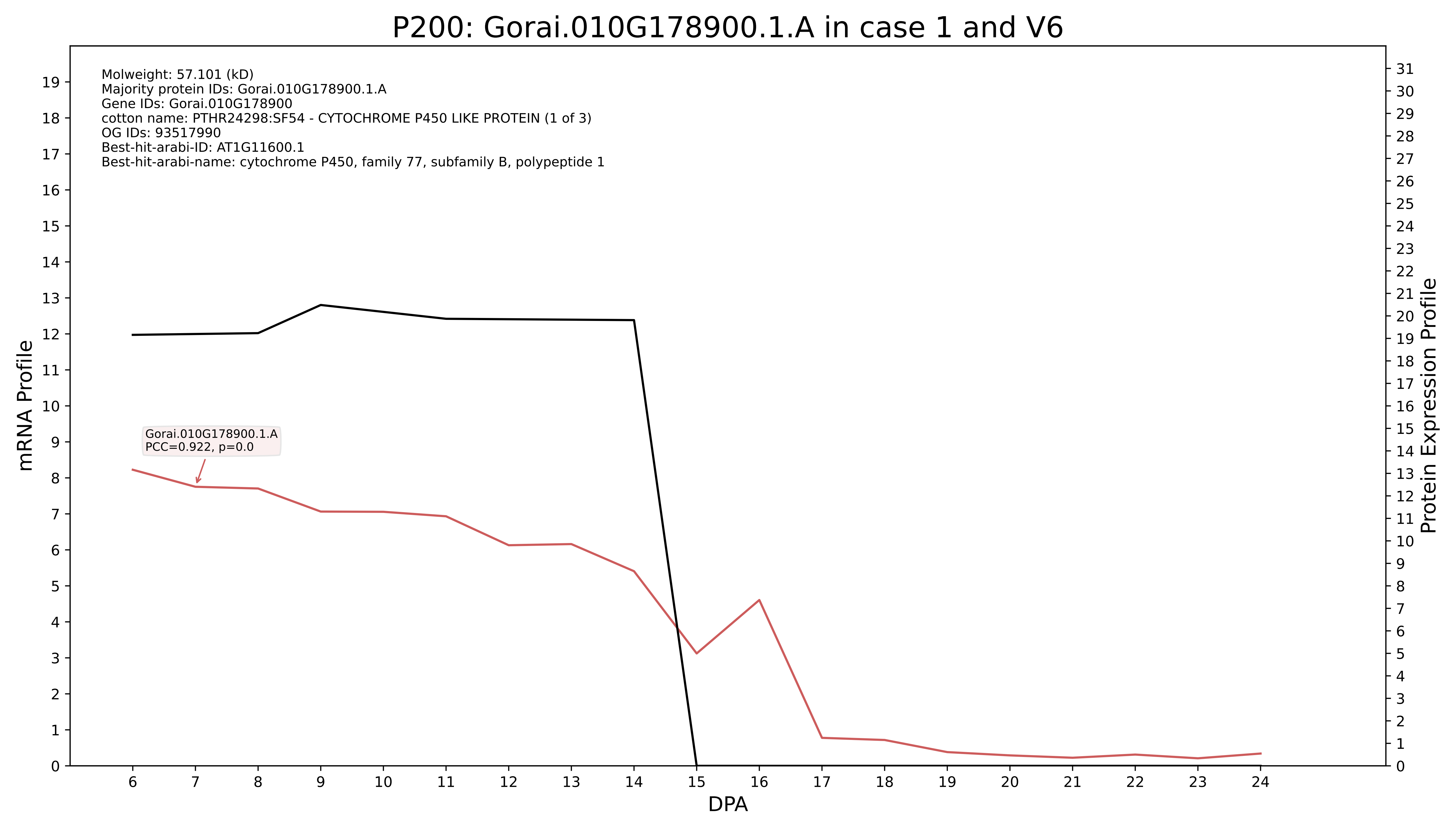
P200: Gorai.010G178900.1.A in case 1 and V6
Molweight: 57.101 (kD)
Majority protein IDs: Gorai.010G178900.1.A
Gene IDs: Gorai.010G178900
cotton name: PTHR24298:SF54 - CYTOCHROME P450 LIKE PROTEIN (1 of 3)
OG IDs: 93517990
Best-hit-arabi-ID: AT1G11600.1
Best-hit-arabi-name: cytochrome P450, family 77, subfamily B, polypeptide 1
Gorai.010G178900.1.A
PCC=0.922, p=0.0
mRNA Profile
Protein Expression Profile
DPA

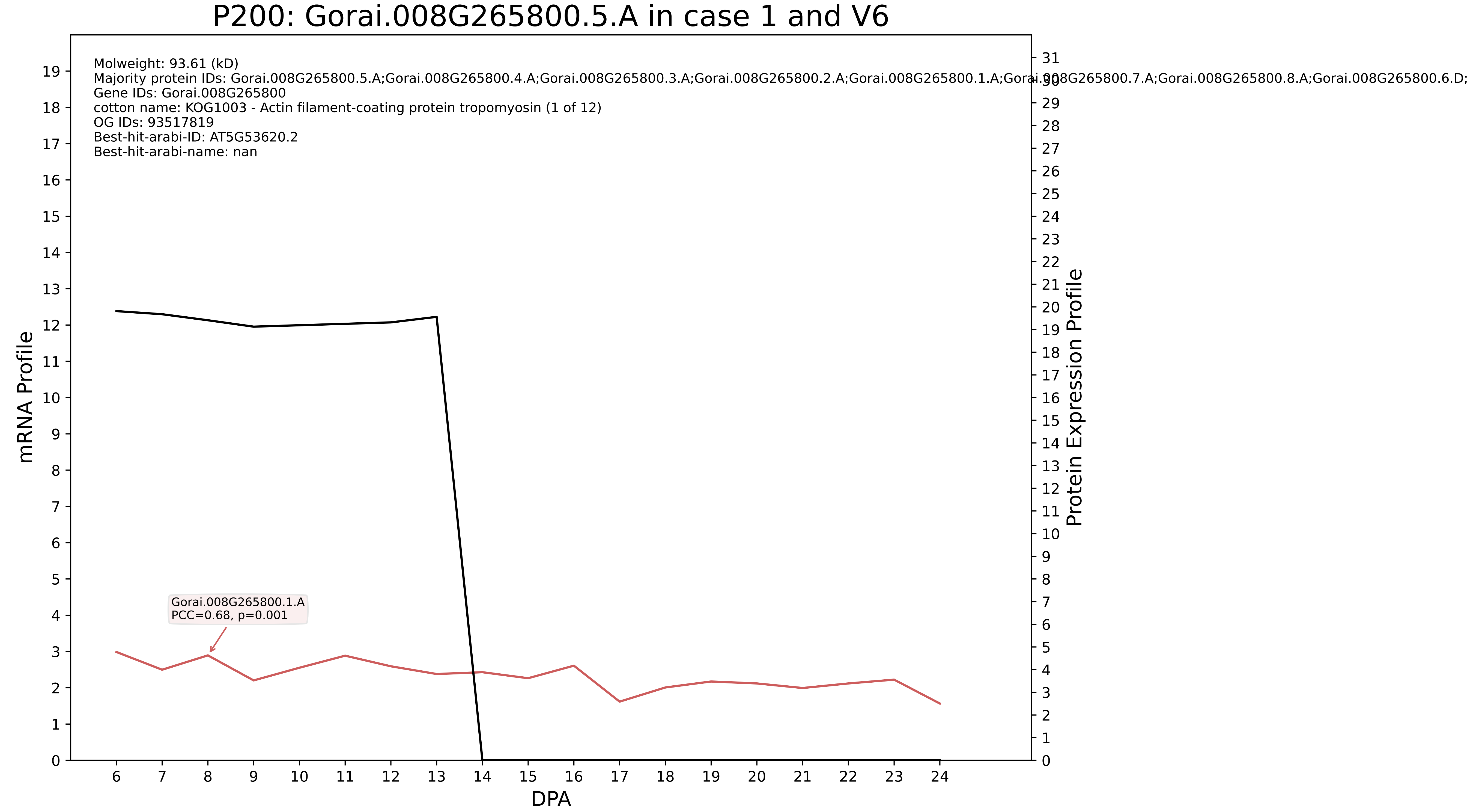
P200: Gorai.008G265800.5.A in case 1 and V6
Molweight: 93.61 (kD)
Majority protein IDs: Gorai.008G265800.5.A;Gorai.008G265800.4.A;Gorai.008G265800.3.A;Gorai.008G265800.2.A;Gorai.008G265800.1.A;Gorai.008G265800.7.A;Gorai.008G265800.8.A;Gorai.008G265800.6.D;
Gene IDs: Gorai.008G265800
cotton name: KOG1003 - Actin filament-coating protein tropomyosin (1 of 12)
OG IDs: 93517819
Best-hit-arabi-ID: AT5G53620.2
Best-hit-arabi-name: nan
Gorai.008G265800.1.A
PCC=0.68, p=0.001
mRNA Profile
Protein Expression Profile
DPA

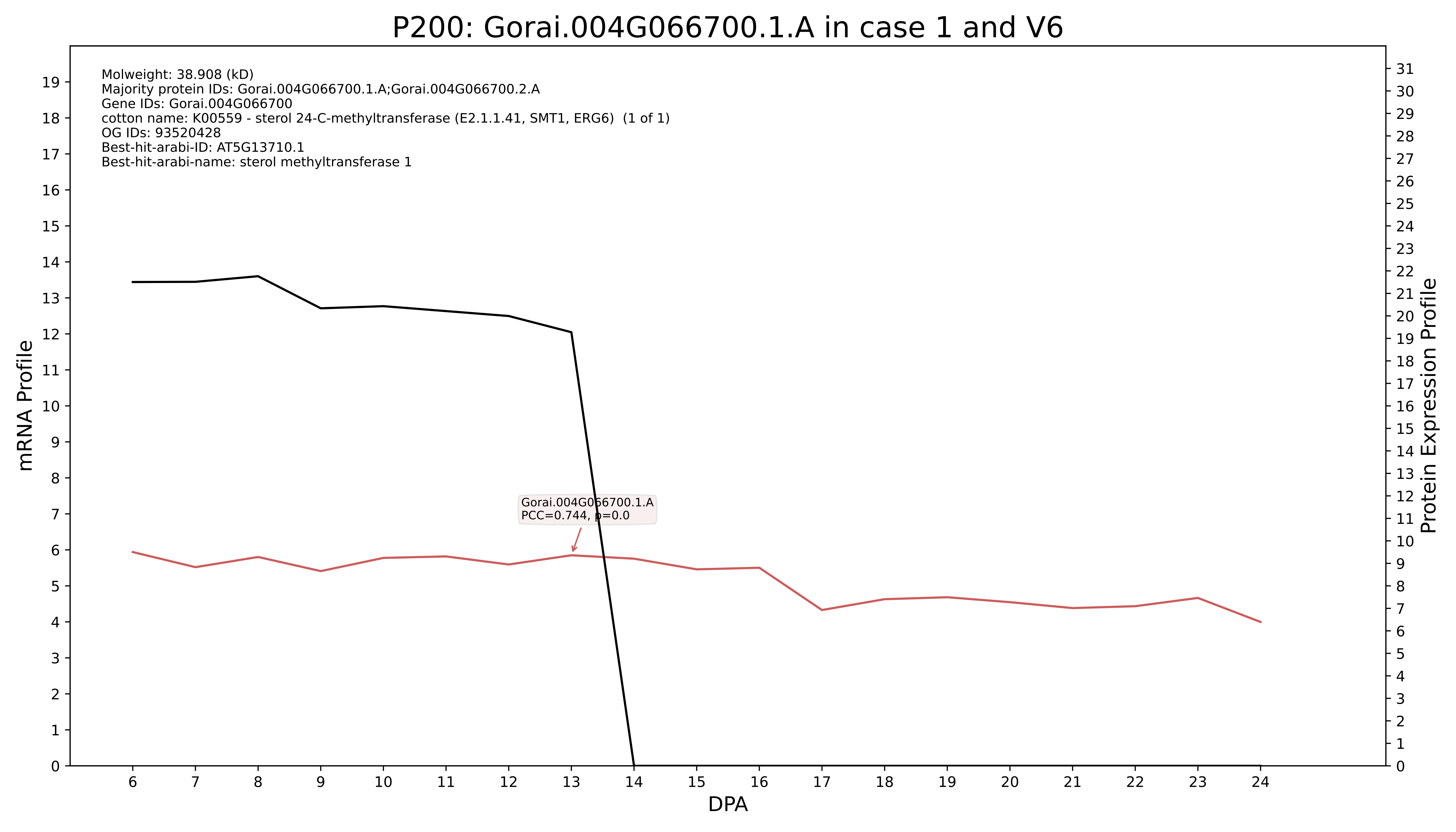
P200: Gorai.004G066700.1.A in case 1 and V6
Molweight: 38.908 (kD)
Majority protein IDs: Gorai.004G066700.1.A;Gorai.004G066700.2.A
Gene IDs: Gorai.004G066700
cotton name: K00559 - sterol 24-C-methyltransferase (E2.1.1.41, SMT1, ERG6) (1 of 1)
OG IDs: 93520428
Best-hit-arabi-ID: AT5G13710.1
Best-hit-arabi-name: sterol methyltransferase 1
Gorai.004G066700.1.A
PCC=0.744, p=0.0
mRNA Profile
Protein Expression Profile
DPA

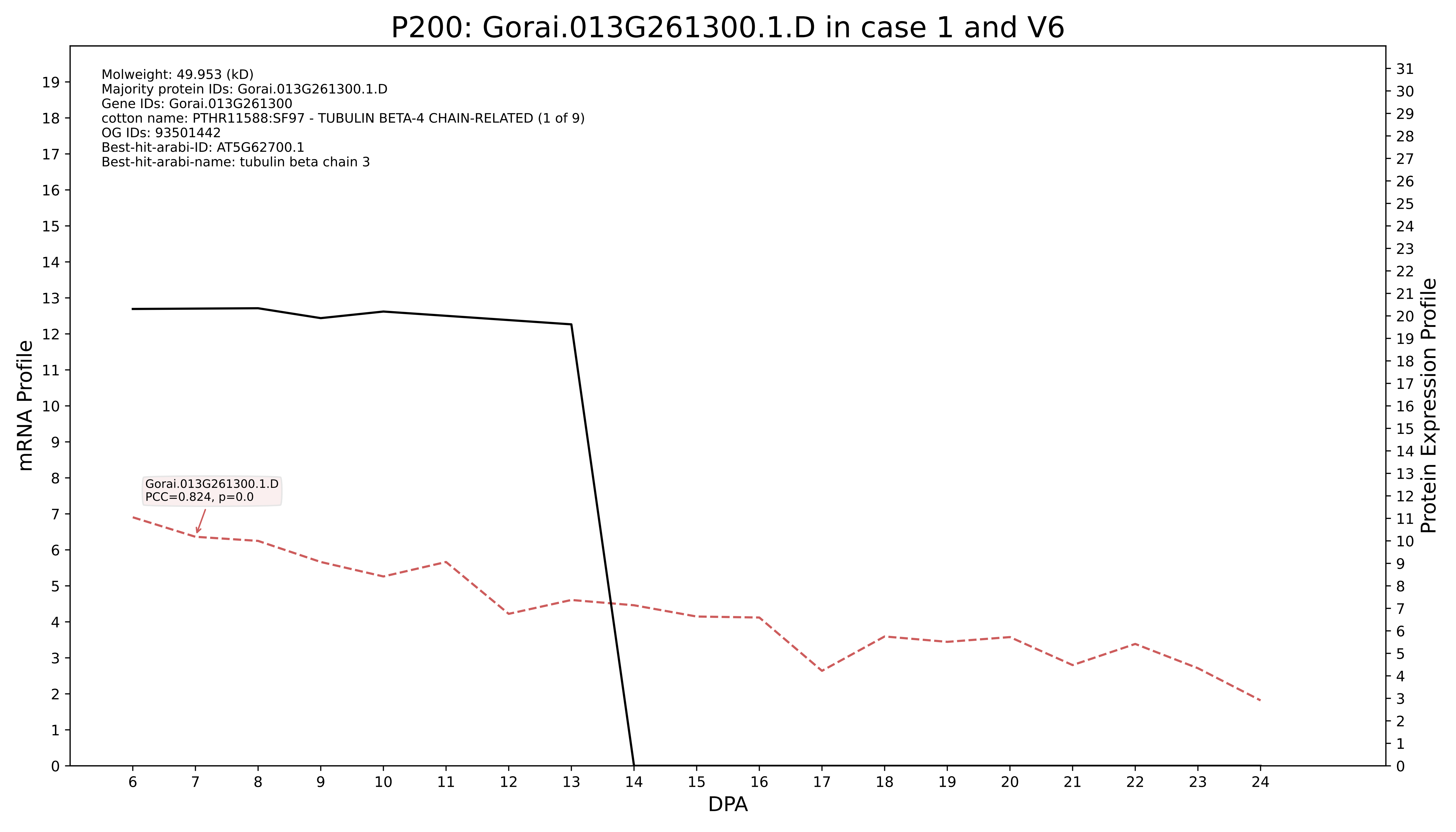
P200: Gorai.013G261300.1.D in case 1 and V6
Molweight: 49.953 (kD)
Majority protein IDs: Gorai.013G261300.1.D
Gene IDs: Gorai.013G261300
cotton name: PTHR11588:SF97 - TUBULIN BETA-4 CHAIN-RELATED (1 of 9)
OG IDs: 93501442
Best-hit-arabi-ID: AT5G62700.1
Best-hit-arabi-name: tubulin beta chain 3
Gorai.013G261300.1.D
PCC=0.824, p=0.0
mRNA Profile
Protein Expression Profile
DPA

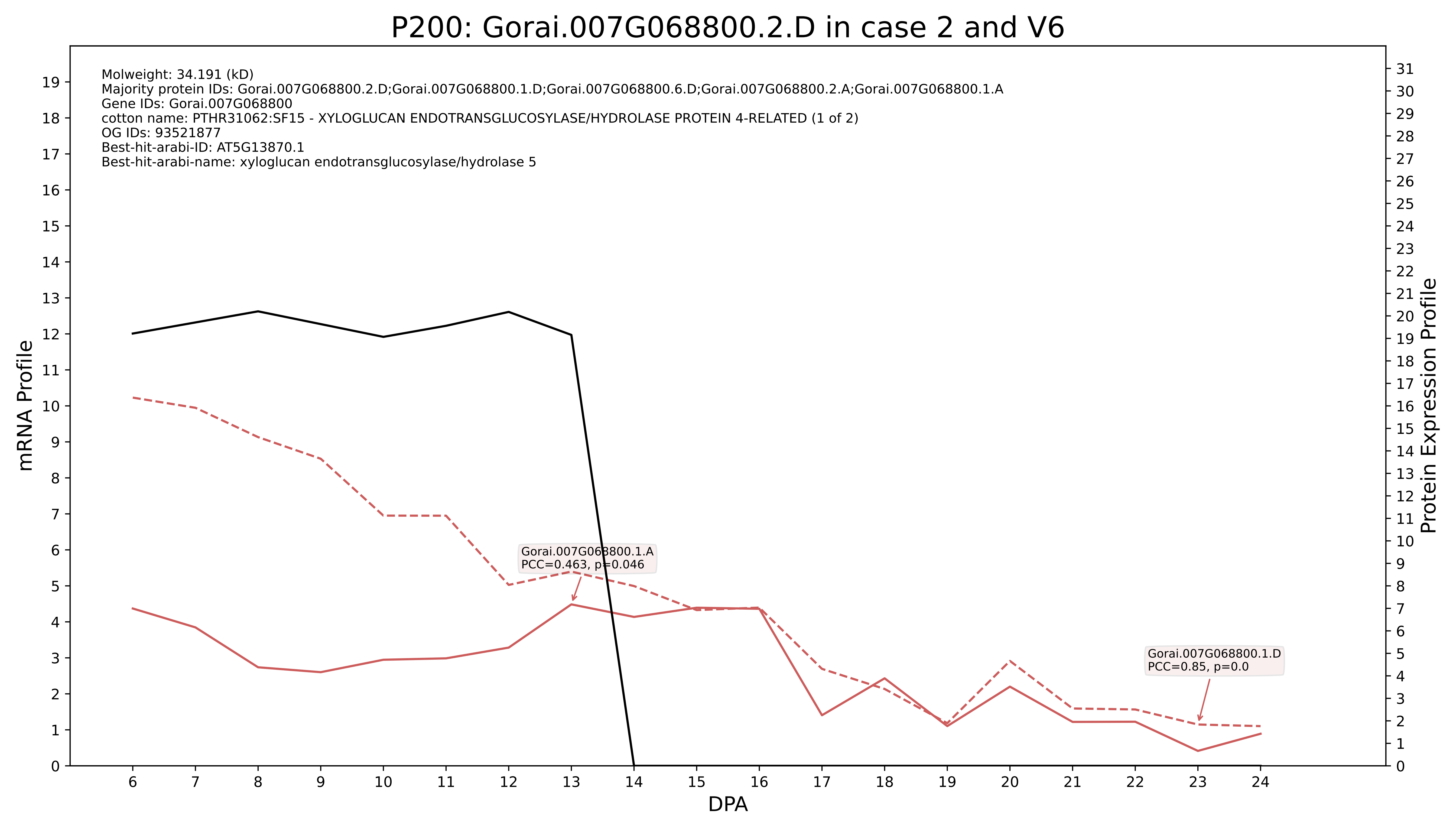
P200: Gorai.007G068800.2.D in case 2 and V6
Molweight: 34.191 (kD)
Majority protein IDs: Gorai.007G068800.2.D;Gorai.007G068800.1.D;Gorai.007G068800.6.D;Gorai.007G068800.2.A;Gorai.007G068800.1.A
Gene IDs: Gorai.007G068800
cotton name: PTHR31062:SF15 - XYLOGLUCAN ENDOTRANSGLUCOSYLASE/HYDROLASE PROTEIN 4-RELATED (1 of 2)
OG IDs: 93521877
Best-hit-arabi-ID: AT5G13870.1
Best-hit-arabi-name: xyloglucan endotransglucosylase/hydrolase 5
Gorai.007G068800.1.A
PCC=0.463, p=0.046
Gorai.007G068800.1.D
PCC=0.85, p=0.0
mRNA Profile
Protein Expression Profile
DPA

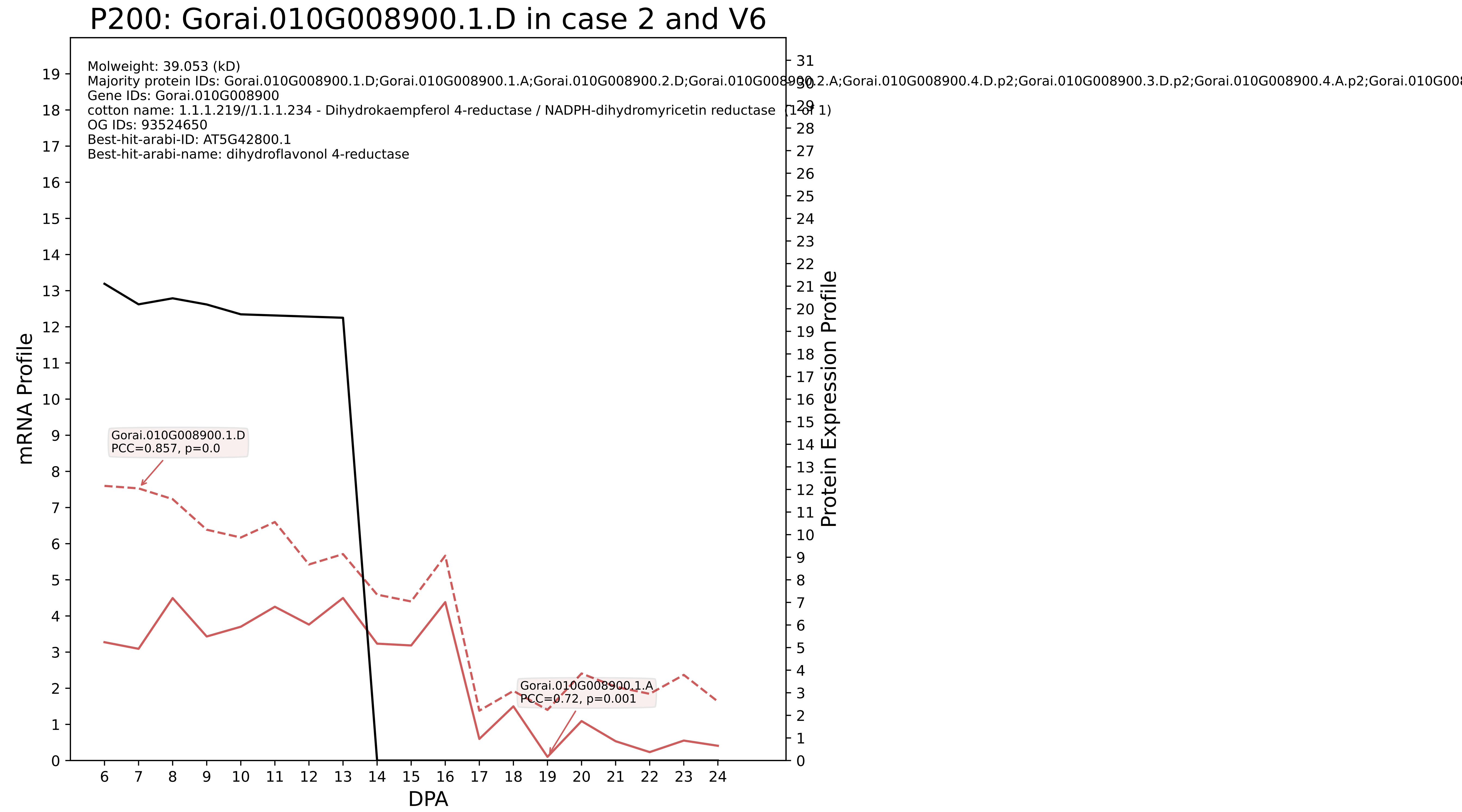

P200: Gorai.010G008900.1.D in case 2 and V6
Molweight: 39.053 (kD)
Majority protein IDs: Gorai.010G008900.1.D;Gorai.010G008900.1.A;Gorai.010G008900.2.D;Gorai.010G008900.2.A;Gorai.010G008900.4.D.p2;Gorai.010G008900.3.D.p2;Gorai.010G008900.4.A.p2;Gorai.010G008
Gene IDs: Gorai.010G008900
cotton name: 1.1.1.219//1.1.1.234 - Dihydrokaempferol 4-reductase / NADPH-dihydromyricetin reductase (1 of 1)
OG IDs: 93524650
Best-hit-arabi-ID: AT5G42800.1
Best-hit-arabi-name: dihydroflavonol 4-reductase
Gorai.010G008900.1.D
PCC=0.857, p=0.0
Gorai.010G008900.1.A
PCC=0.72, p=0.001
mRNA Profile
Protein Expression Profile
DPA

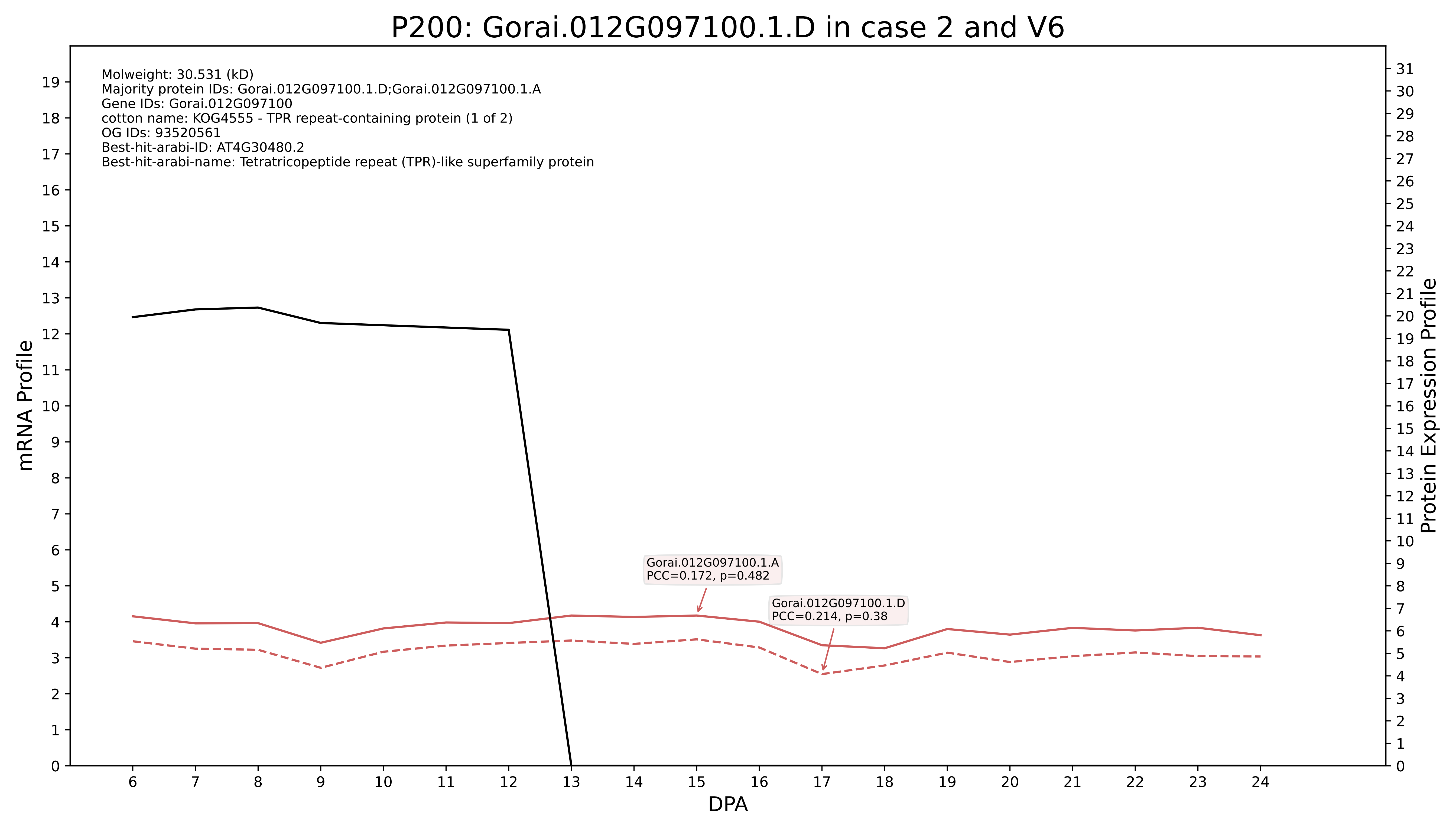

P200: Gorai.012G097100.1.D in case 2 and V6
Molweight: 30.531 (kD)
Majority protein IDs: Gorai.012G097100.1.D;Gorai.012G097100.1.A
Gene IDs: Gorai.012G097100
cotton name: KOG4555 - TPR repeat-containing protein (1 of 2)
OG IDs: 93520561
Best-hit-arabi-ID: AT4G30480.2
Best-hit-arabi-name: Tetratricopeptide repeat (TPR)-like superfamily protein
Gorai.012G097100.1.A
PCC=0.172, p=0.482
Gorai.012G097100.1.D
PCC=0.214, p=0.38
mRNA Profile
Protein Expression Profile
DPA

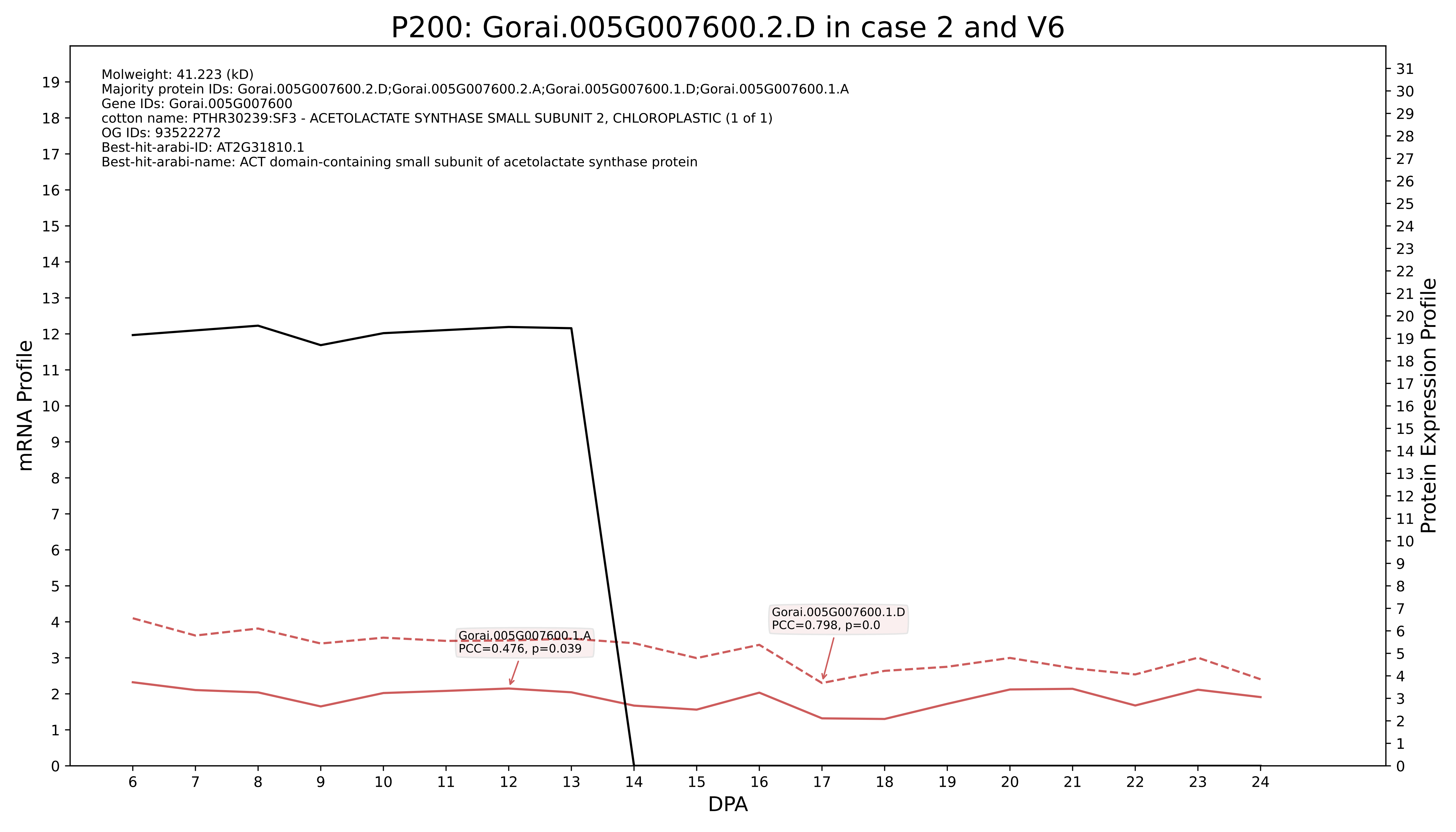
P200: Gorai.005G007600.2.D in case 2 and V6
Molweight: 41.223 (kD)
Majority protein IDs: Gorai.005G007600.2.D;Gorai.005G007600.2.A;Gorai.005G007600.1.D;Gorai.005G007600.1.A
Gene IDs: Gorai.005G007600
cotton name: PTHR30239:SF3 - ACETOLACTATE SYNTHASE SMALL SUBUNIT 2, CHLOROPLASTIC (1 of 1)
OG IDs: 93522272
Best-hit-arabi-ID: AT2G31810.1
Best-hit-arabi-name: ACT domain-containing small subunit of acetolactate synthase protein
Gorai.005G007600.1.A
PCC=0.476, p=0.039
Gorai.005G007600.1.D
PCC=0.798, p=0.0
mRNA Profile
Protein Expression Profile
DPA

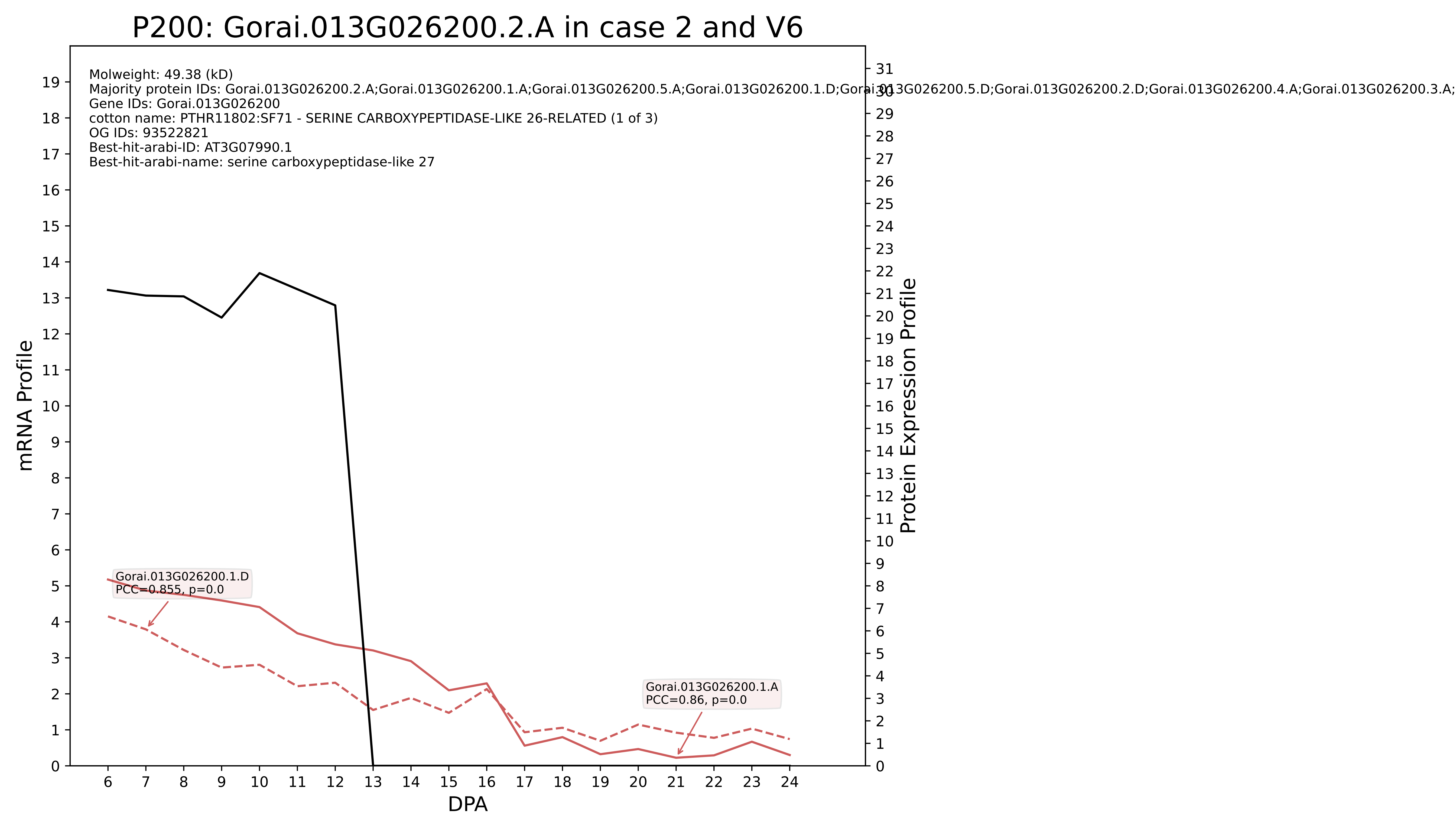
P200: Gorai.013G026200.2.A in case 2 and V6
Molweight: 49.38 (kD)
Majority protein IDs: Gorai.013G026200.2.A;Gorai.013G026200.1.A;Gorai.013G026200.5.A;Gorai.013G026200.1.D;Gorai.013G026200.5.D;Gorai.013G026200.2.D;Gorai.013G026200.4.A;Gorai.013G026200.3.A;
Gene IDs: Gorai.013G026200
cotton name: PTHR11802:SF71 - SERINE CARBOXYPEPTIDASE-LIKE 26-RELATED (1 of 3)
OG IDs: 93522821
Best-hit-arabi-ID: AT3G07990.1
Best-hit-arabi-name: serine carboxypeptidase-like 27
Gorai.013G026200.1.D
PCC=0.855, p=0.0
Gorai.013G026200.1.A
PCC=0.86, p=0.0
mRNA Profile
Protein Expression Profile
DPA

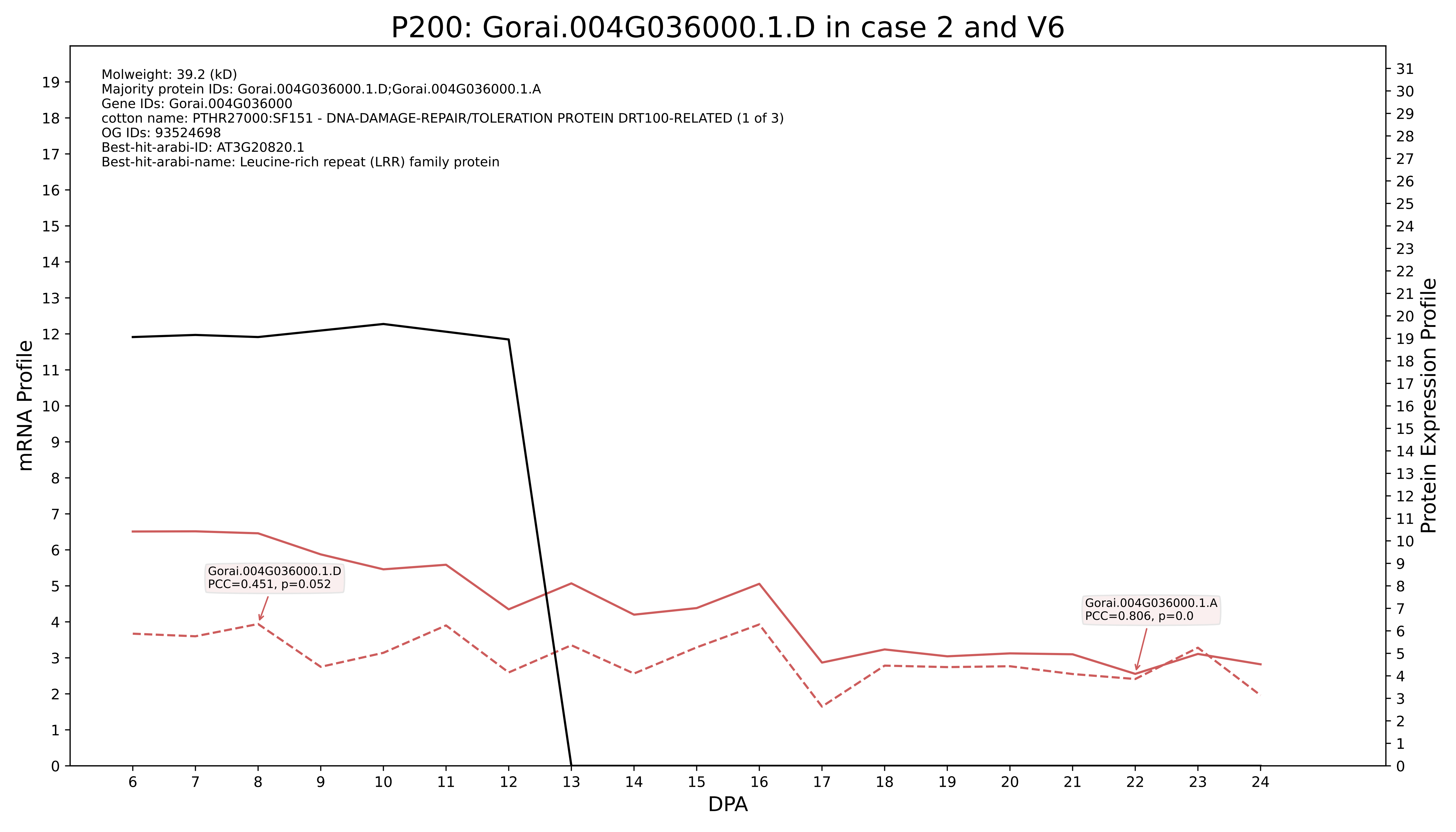

P200: Gorai.004G036000.1.D in case 2 and V6
Molweight: 39.2 (kD)
Majority protein IDs: Gorai.004G036000.1.D;Gorai.004G036000.1.A
Gene IDs: Gorai.004G036000
cotton name: PTHR27000:SF151 - DNA-DAMAGE-REPAIR/TOLERATION PROTEIN DRT100-RELATED (1 of 3)
OG IDs: 93524698
Best-hit-arabi-ID: AT3G20820.1
Best-hit-arabi-name: Leucine-rich repeat (LRR) family protein
Gorai.004G036000.1.D
PCC=0.451, p=0.052
Gorai.004G036000.1.A
PCC=0.806, p=0.0
mRNA Profile
Protein Expression Profile
DPA

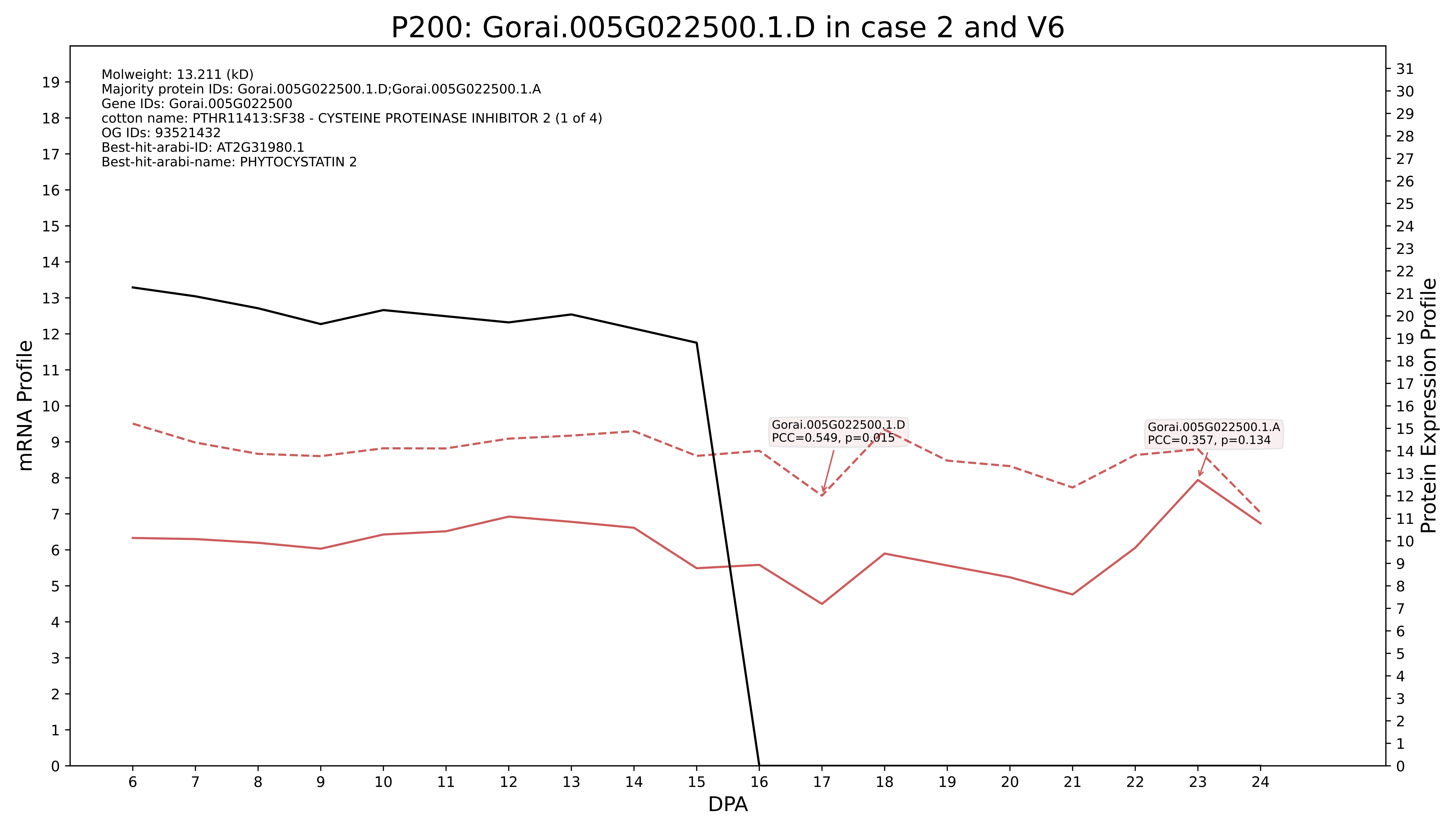
P200: Gorai.005G022500.1.D in case 2 and V6
Molweight: 13.211 (kD)
Majority protein IDs: Gorai.005G022500.1.D;Gorai.005G022500.1.A
Gene IDs: Gorai.005G022500
cotton name: PTHR11413:SF38 - CYSTEINE PROTEINASE INHIBITOR 2 (1 of 4)
OG IDs: 93521432
Best-hit-arabi-ID: AT2G31980.1
Best-hit-arabi-name: PHYTOCYSTATIN 2
Gorai.005G022500.1.D
PCC=0.549, p=0.015
Gorai.005G022500.1.A
PCC=0.357, p=0.134
mRNA Profile
Protein Expression Profile
DPA

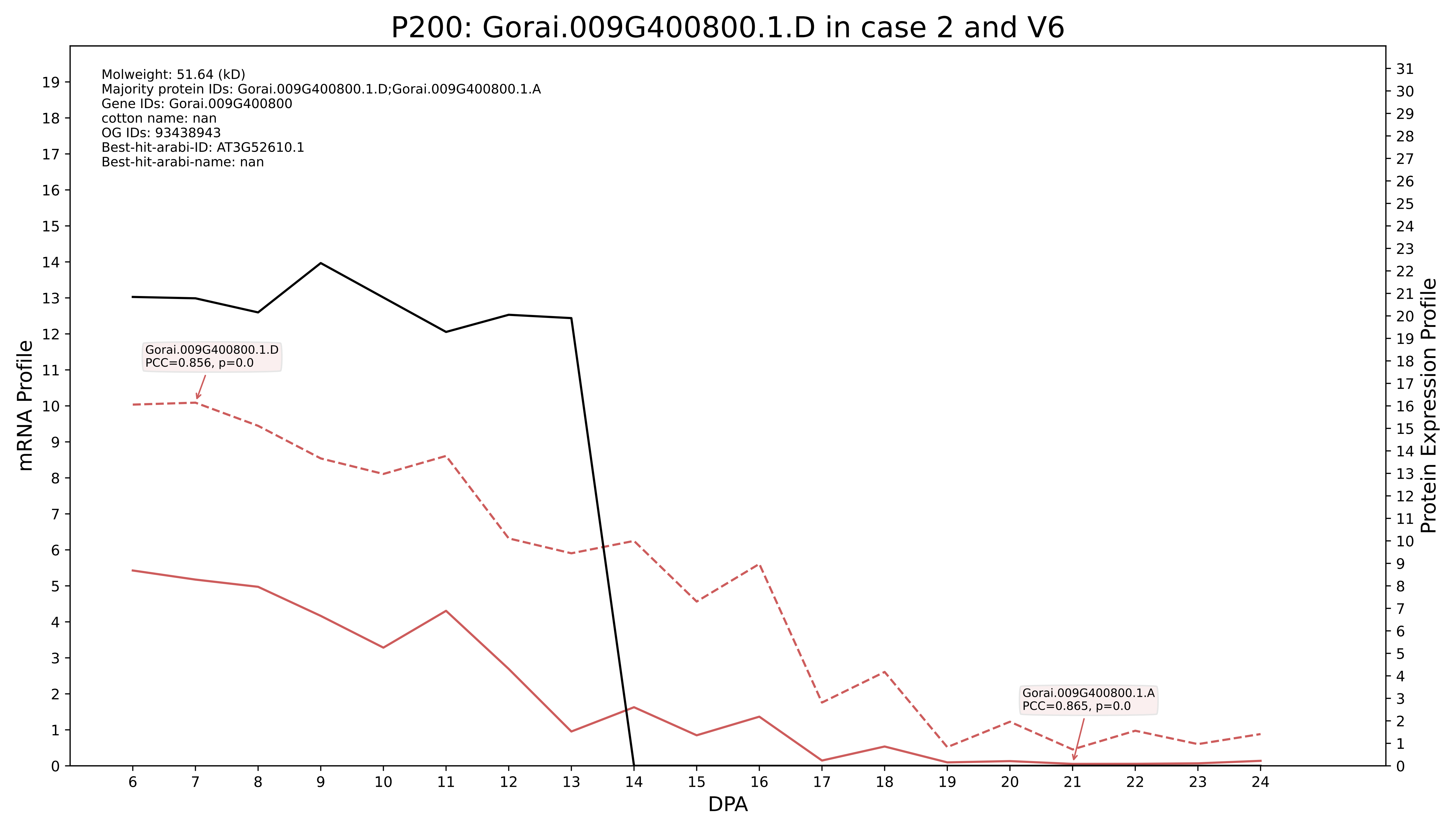
P200: Gorai.009G400800.1.D in case 2 and V6
Molweight: 51.64 (kD)
Majority protein IDs: Gorai.009G400800.1.D;Gorai.009G400800.1.A
Gene IDs: Gorai.009G400800
cotton name: nan
OG IDs: 93438943
Best-hit-arabi-ID: AT3G52610.1
Best-hit-arabi-name: nan
Gorai.009G400800.1.D
PCC=0.856, p=0.0
Gorai.009G400800.1.A
PCC=0.865, p=0.0
mRNA Profile
Protein Expression Profile
DPA

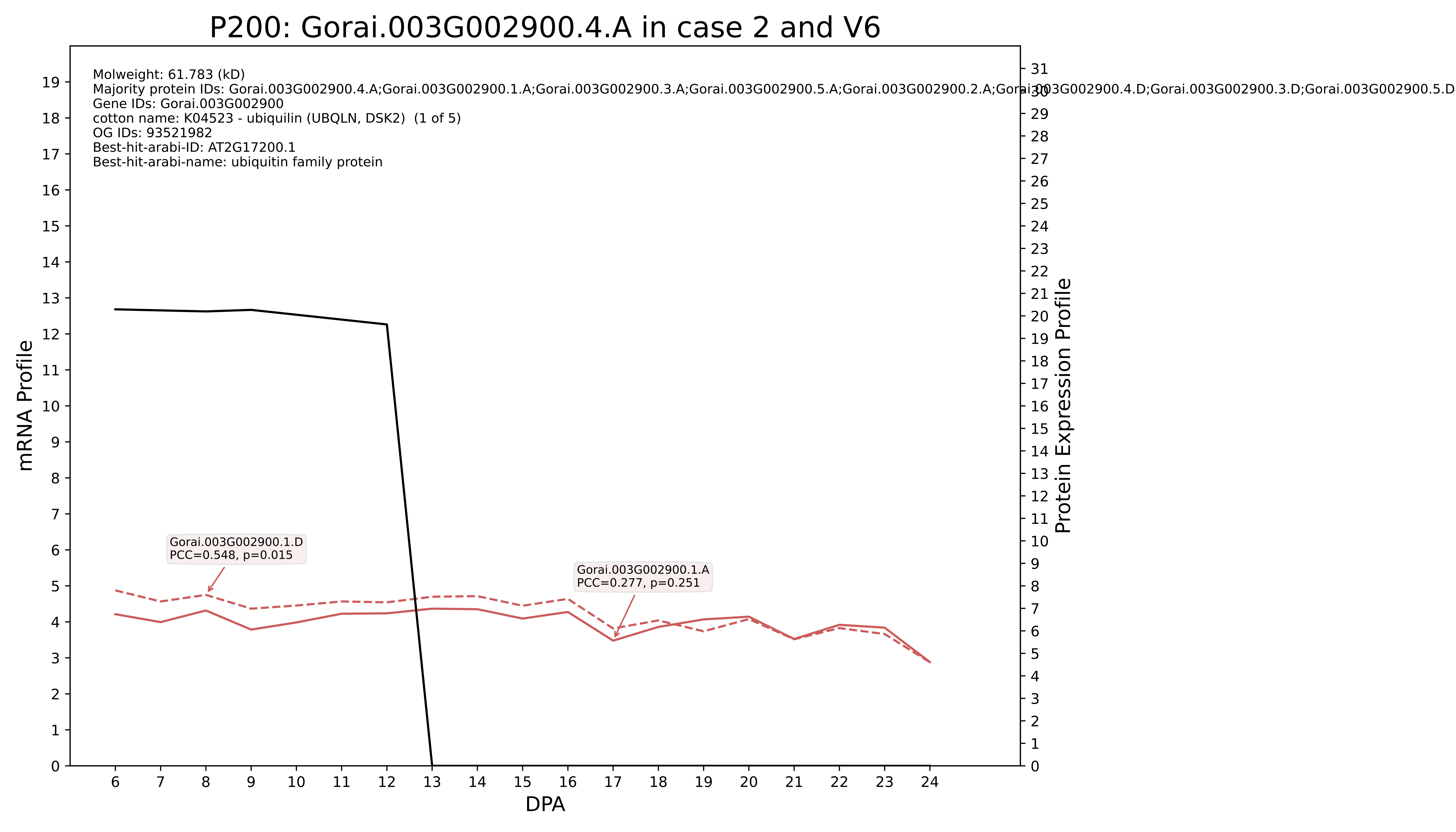
P200: Gorai.003G002900.4.A in case 2 and V6
Molweight: 61.783 (kD)
Majority protein IDs: Gorai.003G002900.4.A;Gorai.003G002900.1.A;Gorai.003G002900.3.A;Gorai.003G002900.5.A;Gorai.003G002900.2.A;Gorai.003G002900.4.D;Gorai.003G002900.3.D;Gorai.003G002900.5.D
Gene IDs: Gorai.003G002900
cotton name: K04523 - ubiquilin (UBQLN, DSK2) (1 of 5)
OG IDs: 93521982
Best-hit-arabi-ID: AT2G17200.1
Best-hit-arabi-name: ubiquitin family protein
Gorai.003G002900.1.D
PCC=0.548, p=0.015
Gorai.003G002900.1.A
PCC=0.277, p=0.251
mRNA Profile
Protein Expression Profile
DPA

### P200: Gorai.004G013500.1.D in case 2 and V6

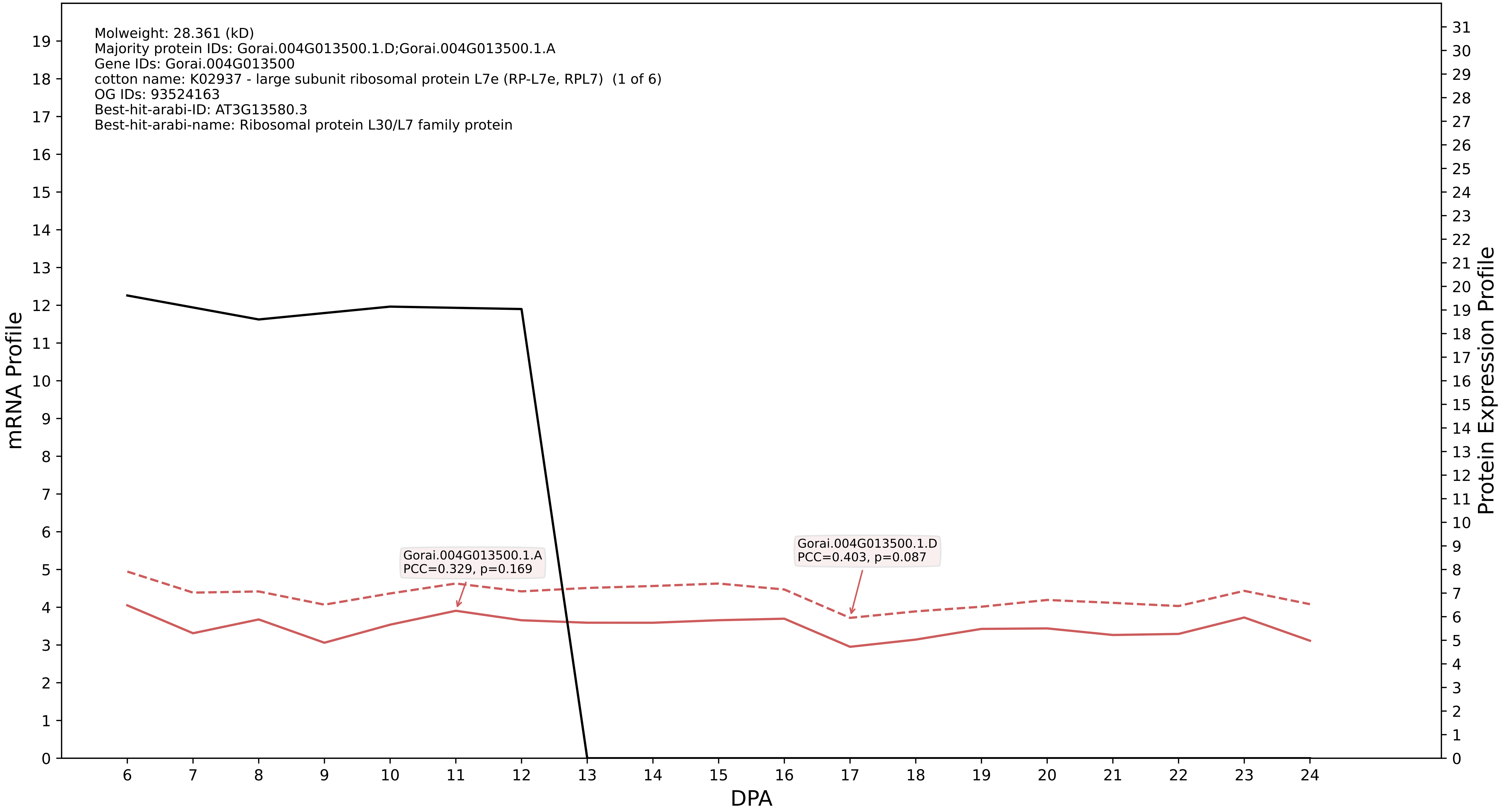

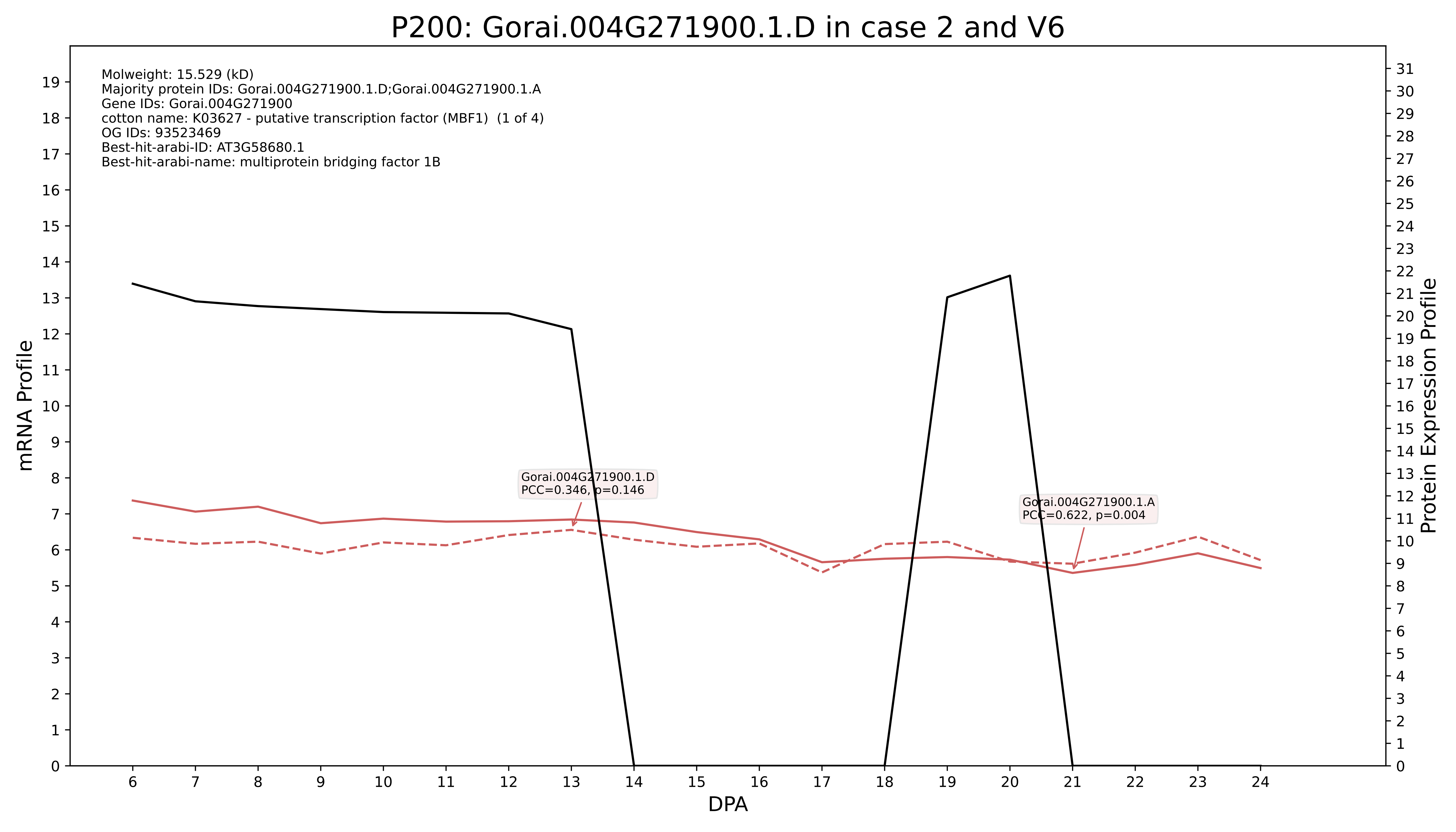
P200: Gorai.004G271900.1.D in case 2 and V6
Molweight: 15.529 (kD)
Majority protein IDs: Gorai.004G271900.1.D;Gorai.004G271900.1.A
Gene IDs: Gorai.004G271900
cotton name: K03627 - putative transcription factor (MBF1) (1 of 4)
OG IDs: 93523469
Best-hit-arabi-ID: AT3G58680.1
Best-hit-arabi-name: multiprotein bridging factor 1B
Gorai.004G271900.1.D
PCC=0.346, p=0.146
Gorai.004G271900.1.A
PCC=0.622, p=0.004
mRNA Profile
Protein Expression Profile
DPA

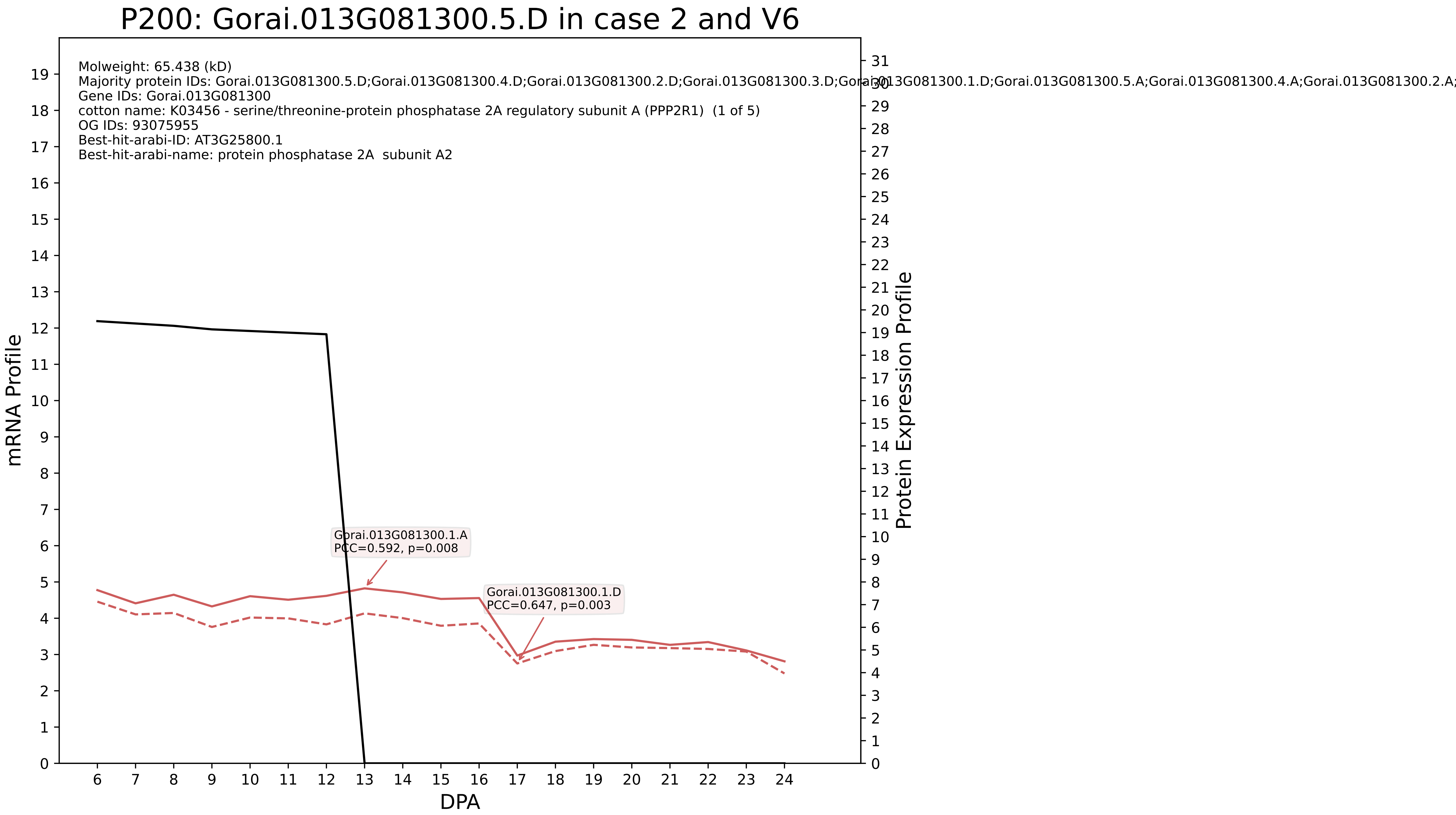
P200: Gorai.013G081300.5.D in case 2 and V6
Molweight: 65.438 (kD)
Majority protein IDs: Gorai.013G081300.5.D;Gorai.013G081300.4.D;Gorai.013G081300.2.D;Gorai.013G081300.3.D;Gorai.013G081300.1.D;Gorai.013G081300.5.A;Gorai.013G081300.4.A;Gorai.013G081300.2.A;
Gene IDs: Gorai.013G081300
cotton name: K03456 - serine/threonine-protein phosphatase 2A regulatory subunit A (PPP2R1) (1 of 5)
OG IDs: 93075955
Best-hit-arabi-ID: AT3G25800.1
Best-hit-arabi-name: protein phosphatase 2A subunit A2
Gorai.013G081300.1.A
PCC=0.592, p=0.008
Gorai.013G081300.1.D
PCC=0.647, p=0.003
mRNA Profile
Protein Expression Profile
DPA

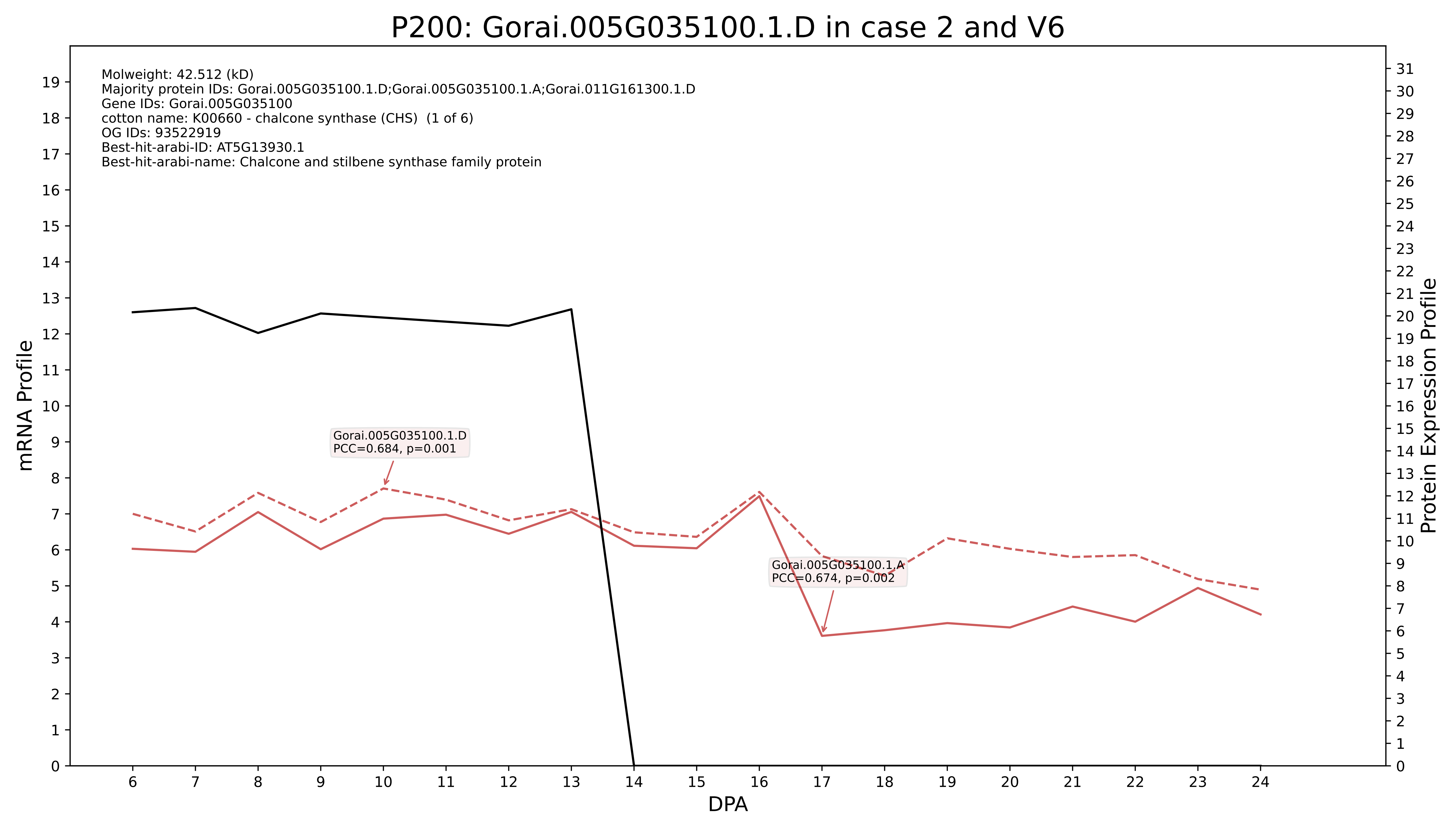
P200: Gorai.005G035100.1.D in case 2 and V6
Molweight: 42.512 (kD)
Majority protein IDs: Gorai.005G035100.1.D;Gorai.005G035100.1.A;Gorai.011G161300.1.D
Gene IDs: Gorai.005G035100
cotton name: K00660 - chalcone synthase (CHS) (1 of 6)
OG IDs: 93522919
Best-hit-arabi-ID: AT5G13930.1
Best-hit-arabi-name: Chalcone and stilbene synthase family protein
Gorai.005G035100.1.D
PCC=0.684, p=0.001
Gorai.005G035100.1.A
PCC=0.674, p=0.002
mRNA Profile
Protein Expression Profile
DPA

200: Gorai.002G103600.2.D in case 2 and V6

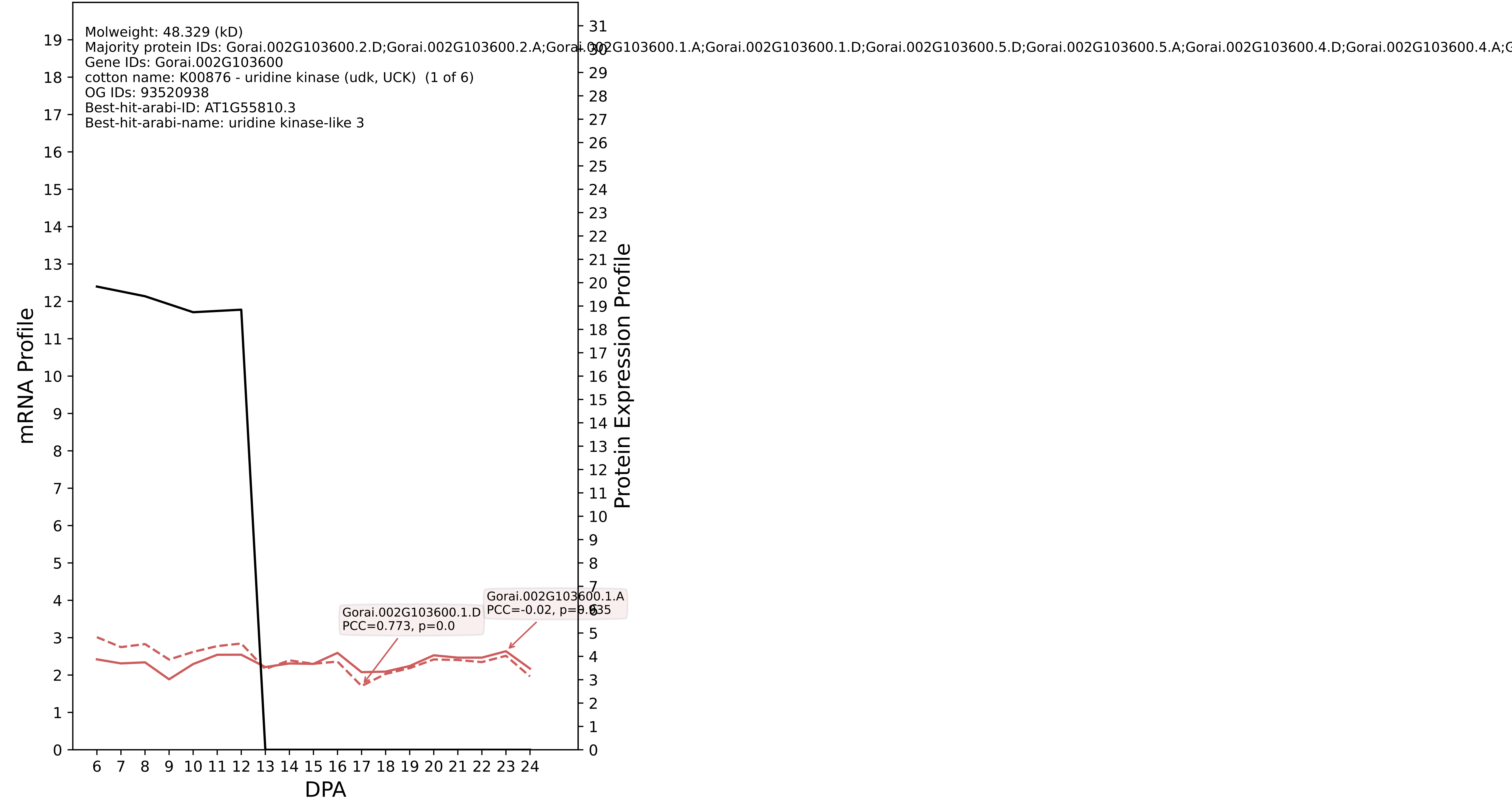

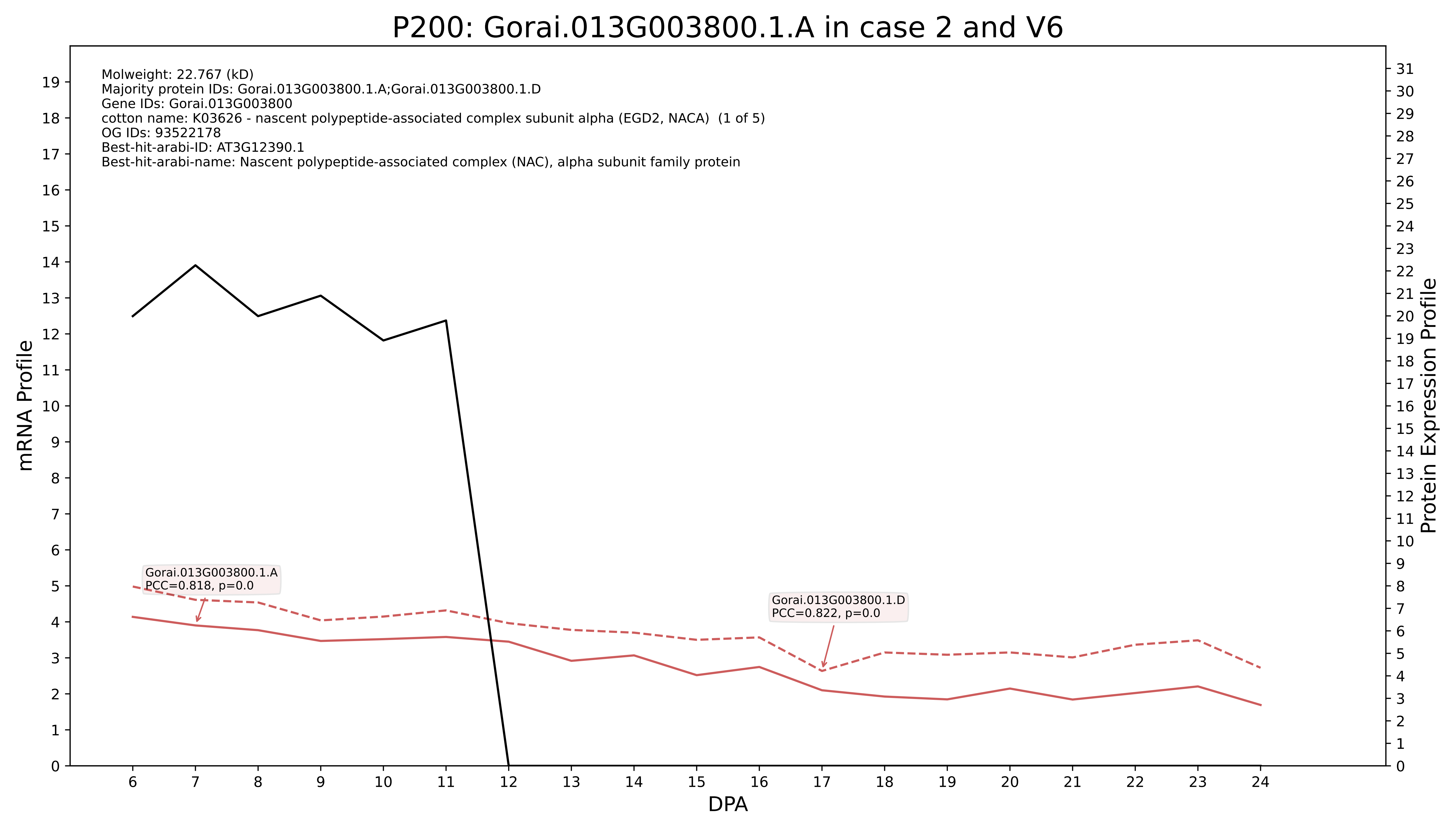
P200: Gorai.013G003800.1.A in case 2 and V6
Molweight: 22.767 (kD)
Majority protein IDs: Gorai.013G003800.1.A;Gorai.013G003800.1.D
Gene IDs: Gorai.013G003800
cotton name: K03626 - nascent polypeptide-associated complex subunit alpha (EGD2, NACA) (1 of 5)
OG IDs: 93522178
Best-hit-arabi-ID: AT3G12390.1
Best-hit-arabi-name: Nascent polypeptide-associated complex (NAC), alpha subunit family protein
Gorai.013G003800.1.A
PCC=0.818, p=0.0
Gorai.013G003800.1.D
PCC=0.822, p=0.0
mRNA Profile
Protein Expression Profile
DPA

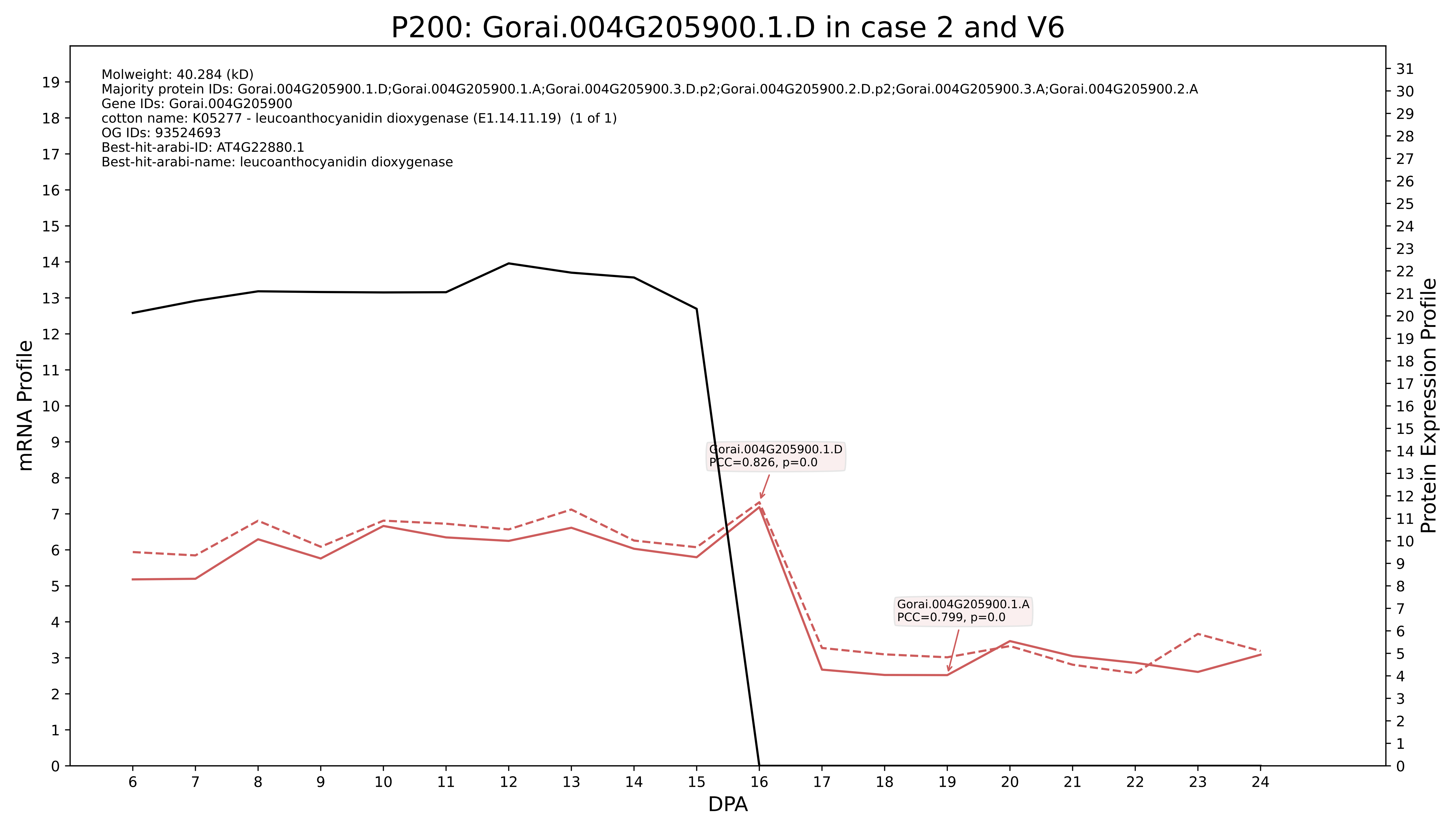

P200: Gorai.004G205900.1.D in case 2 and V6
Molweight: 40.284 (kD)
Majority protein IDs: Gorai.004G205900.1.D;Gorai.004G205900.1.A;Gorai.004G205900.3.D.p2;Gorai.004G205900.2.D.p2;Gorai.004G205900.3.A;Gorai.004G205900.2.A
Gene IDs: Gorai.004G205900
cotton name: K05277 - leucoanthocyanidin dioxygenase (E1.14.11.19) (1 of 1)
OG IDs: 93524693
Best-hit-arabi-ID: AT4G22880.1
Best-hit-arabi-name: leucoanthocyanidin dioxygenase
Gorai.004G205900.1.D
PCC=0.826, p=0.0
Gorai.004G205900.1.A
PCC=0.799, p=0.0
mRNA Profile
Protein Expression Profile
DPA

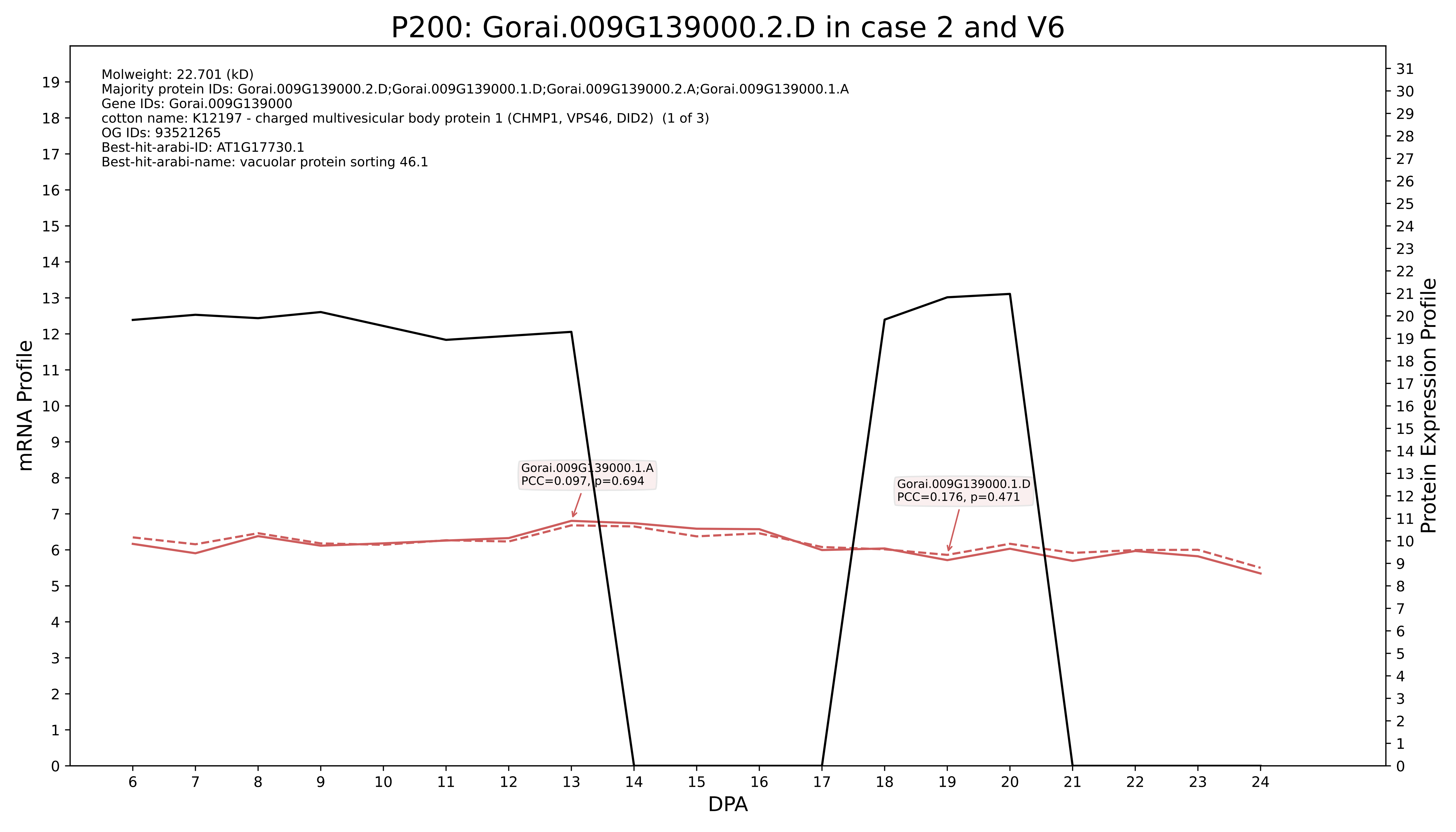
P200: Gorai.009G139000.2.D in case 2 and V6
Molweight: 22.701 (kD)
Majority protein IDs: Gorai.009G139000.2.D;Gorai.009G139000.1.D;Gorai.009G139000.2.A;Gorai.009G139000.1.A
Gene IDs: Gorai.009G139000
cotton name: K12197 - charged multivesicular body protein 1 (CHMP1, VPS46, DID2) (1 of 3)
OG IDs: 93521265
Best-hit-arabi-ID: AT1G17730.1
Best-hit-arabi-name: vacuolar protein sorting 46.1
Gorai.009G139000.1.A
PCC=0.097, p=0.694
Gorai.009G139000.1.D
PCC=0.176, p=0.471
mRNA Profile
Protein Expression Profile
DPA

### P200: Gorai.007G367100.1.D in case 3 and V6

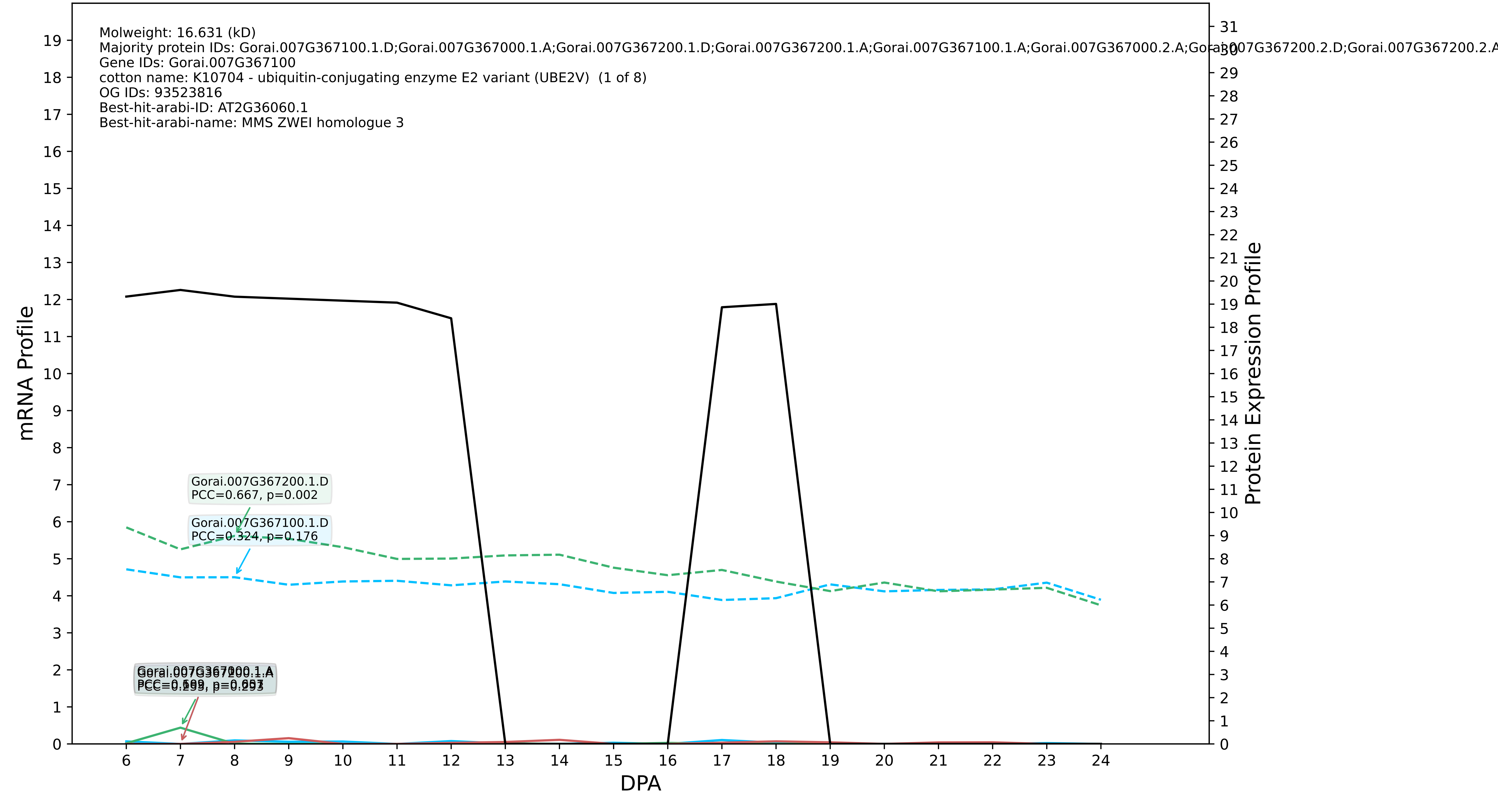

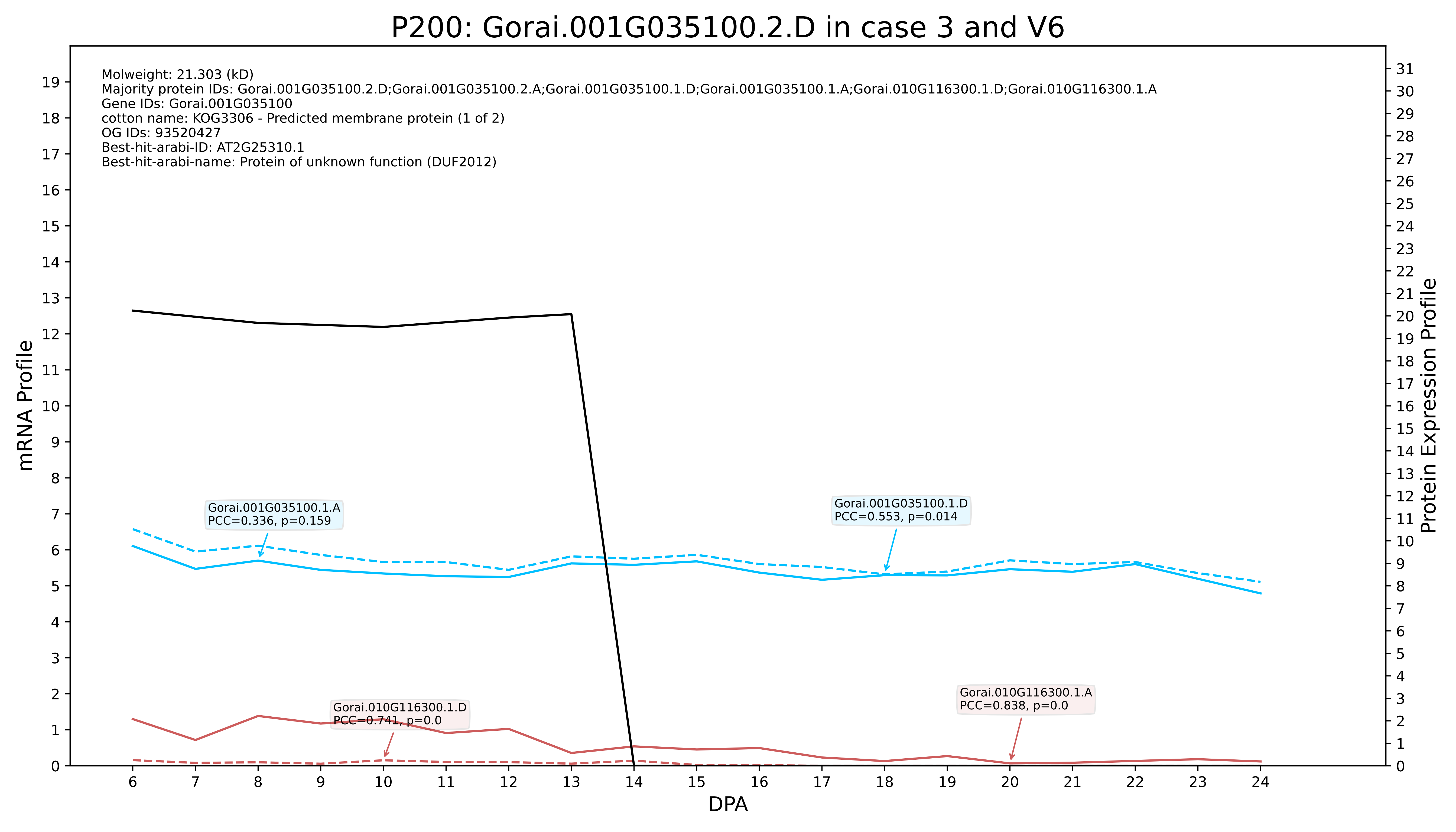
P200: Gorai.001G035100.2.D in case 3 and V6
Molweight: 21.303 (kD)
Majority protein IDs: Gorai.001G035100.2.D;Gorai.001G035100.2.A;Gorai.001G035100.1.D;Gorai.001G035100.1.A;Gorai.010G116300.1.D;Gorai.010G116300.1.A
Gene IDs: Gorai.001G035100
cotton name: KOG3306 - Predicted membrane protein (1 of 2)
OG IDs: 93520427
Best-hit-arabi-ID: AT2G25310.1
Best-hit-arabi-name: Protein of unknown function (DUF2012)
Gorai.001G035100.1.A
PCC=0.336, p=0.159
Gorai.001G035100.1.D
PCC=0.553, p=0.014
Gorai.010G116300.1.D
PCC=0.741, p=0.0
Gorai.010G116300.1.A
PCC=0.838, p=0.0
mRNA Profile
Protein Expression Profile
DPA

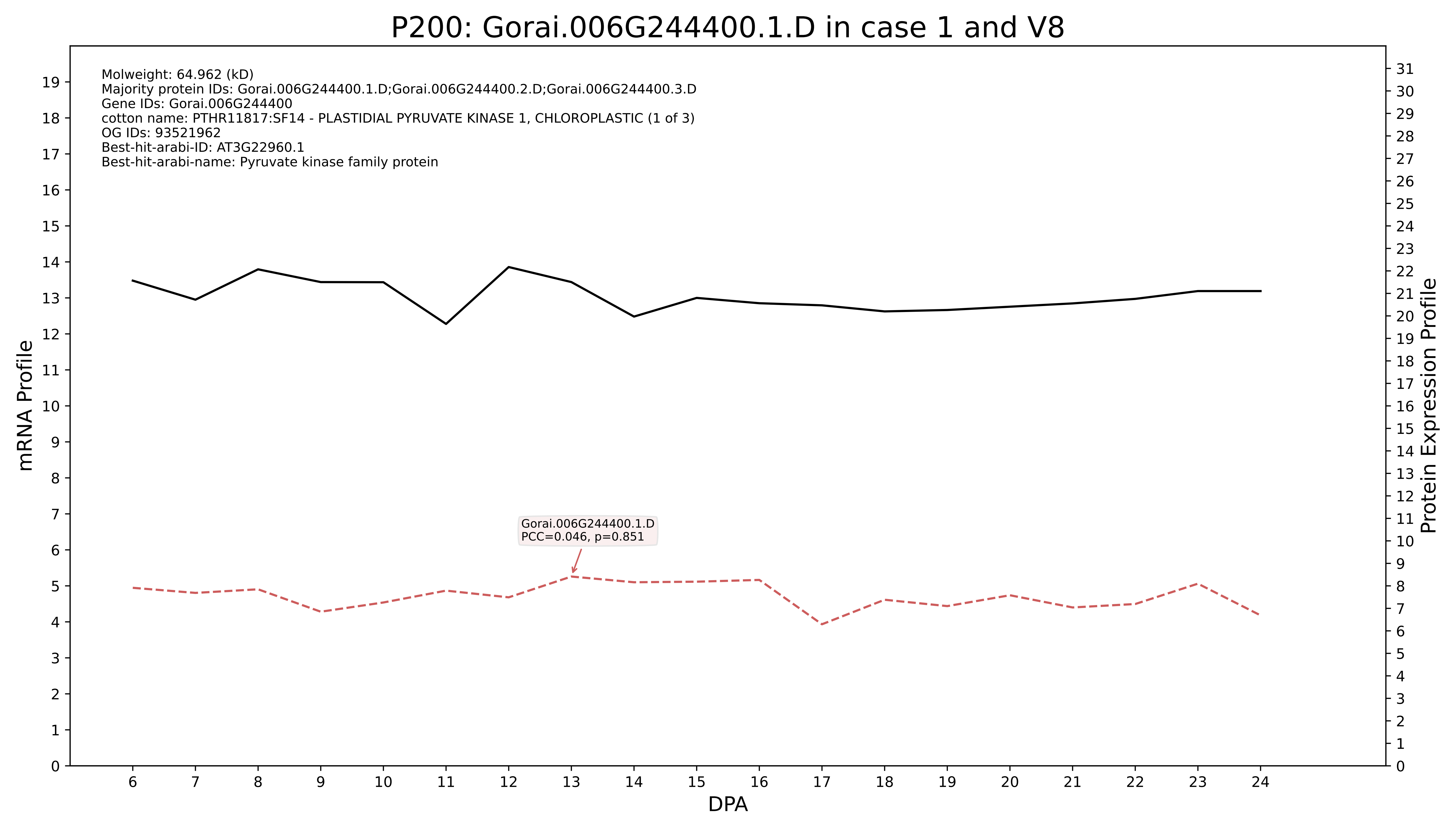
P200: Gorai.006G244400.1.D in case 1 and V8
Molweight: 64.962 (kD)
Majority protein IDs: Gorai.006G244400.1.D;Gorai.006G244400.2.D;Gorai.006G244400.3.D
Gene IDs: Gorai.006G244400
cotton name: PTHR11817:SF14 - PLASTIDIAL PYRUVATE KINASE 1, CHLOROPLASTIC (1 of 3)
OG IDs: 93521962
Best-hit-arabi-ID: AT3G22960.1
Best-hit-arabi-name: Pyruvate kinase family protein
Gorai.006G244400.1.D
PCC=0.046, p=0.851
mRNA Profile
Protein Expression Profile
DPA

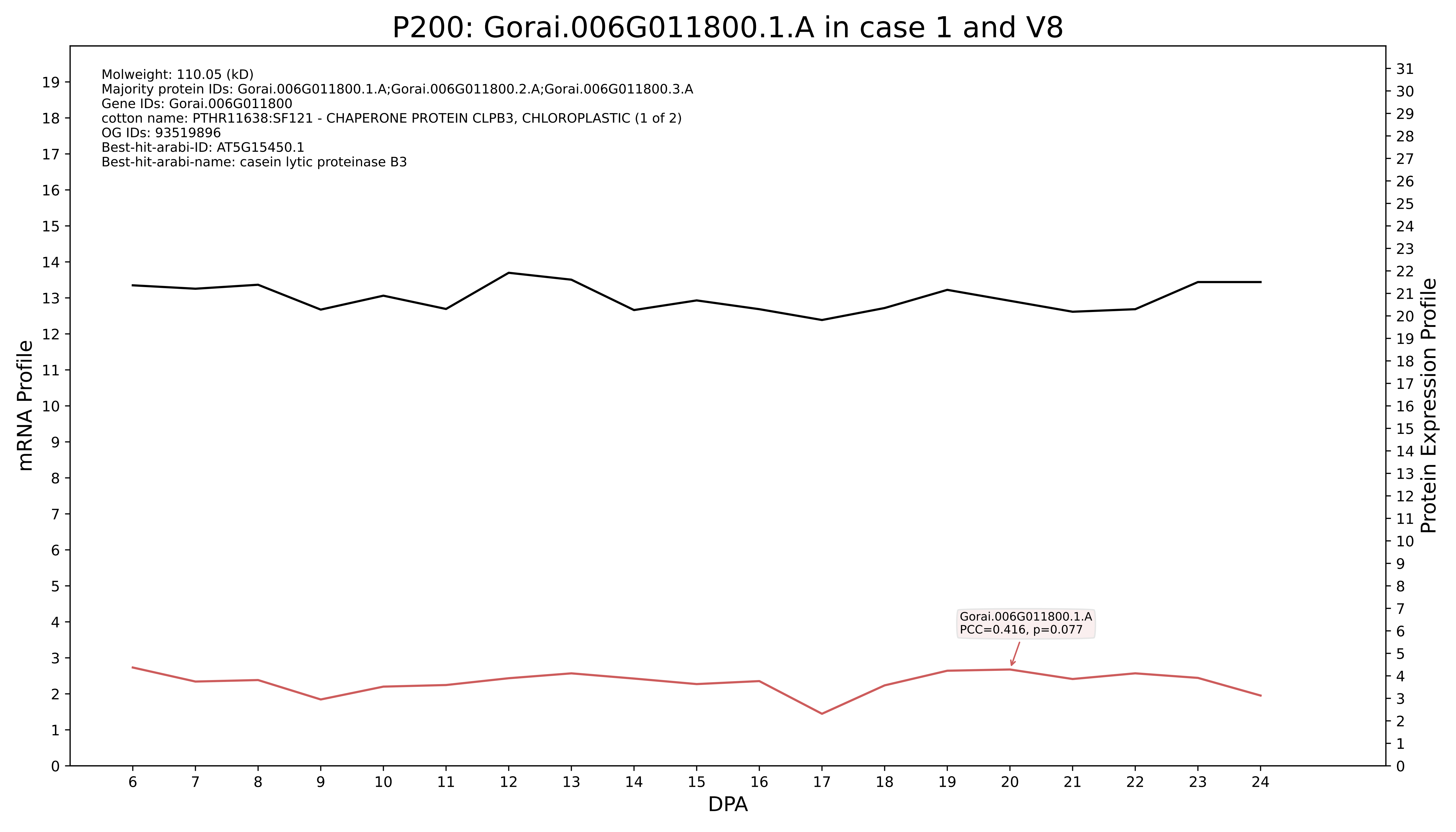

P200: Gorai.006G011800.1.A in case 1 and V8
Molweight: 110.05 (kD)
Majority protein IDs: Gorai.006G011800.1.A;Gorai.006G011800.2.A;Gorai.006G011800.3.A
Gene IDs: Gorai.006G011800
cotton name: PTHR11638:SF121 - CHAPERONE PROTEIN CLPB3, CHLOROPLASTIC (1 of 2)
OG IDs: 93519896
Best-hit-arabi-ID: AT5G15450.1
Best-hit-arabi-name: casein lytic proteinase B3
Gorai.006G011800.1.A
PCC=0.416, p=0.077
mRNA Profile
Protein Expression Profile
DPA

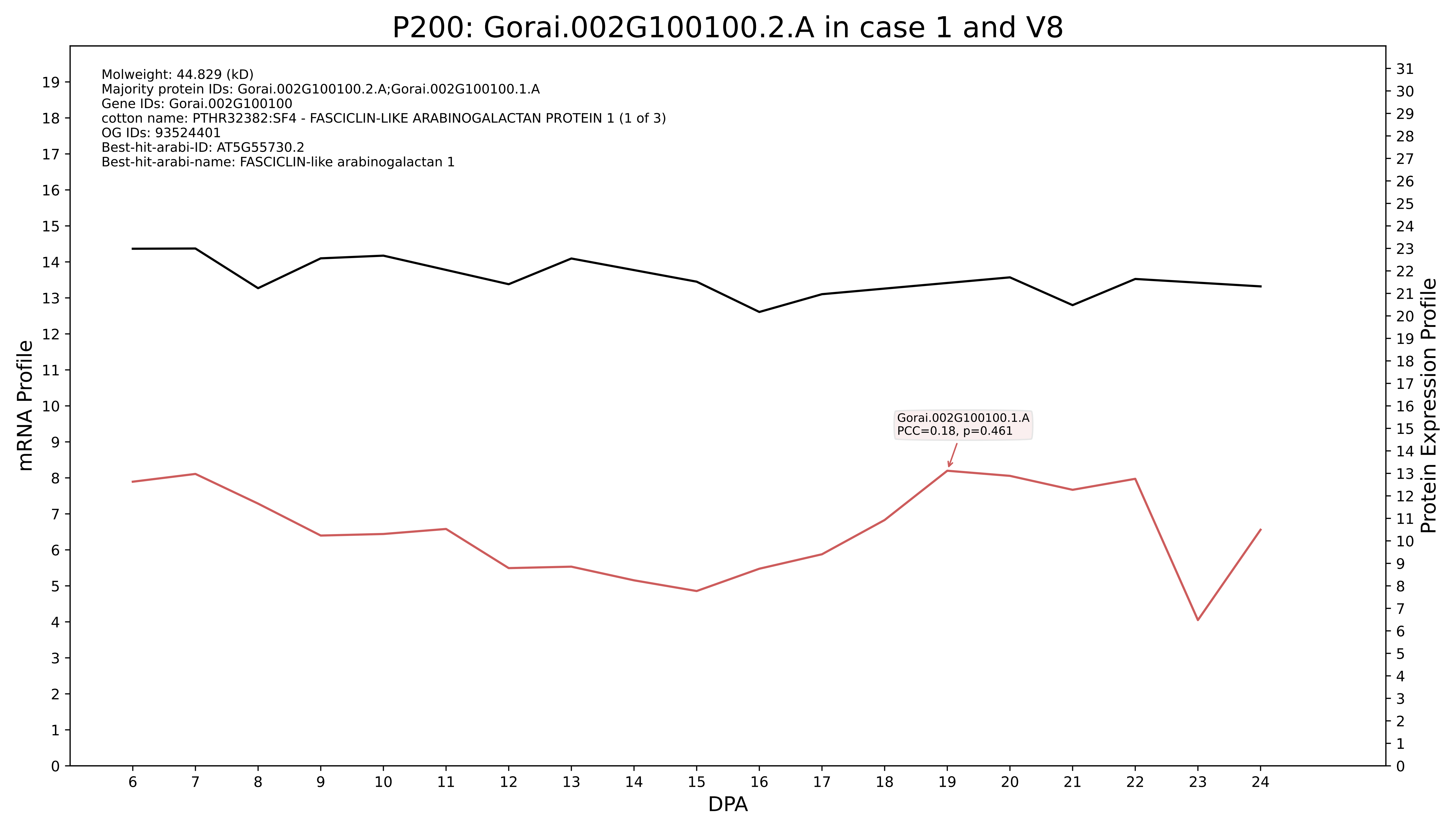

P200: Gorai.002G100100.2.A in case 1 and V8
Molweight: 44.829 (kD)
Majority protein IDs: Gorai.002G100100.2.A;Gorai.002G100100.1.A
Gene IDs: Gorai.002G100100
cotton name: PTHR32382:SF4 - FASCICLIN-LIKE ARABINOGALACTAN PROTEIN 1 (1 of 3)
OG IDs: 93524401
Best-hit-arabi-ID: AT5G55730.2
Best-hit-arabi-name: FASCICLIN-like arabinogalactan 1
Gorai.002G100100.1.A
PCC=0.18, p=0.461
mRNA Profile
Protein Expression Profile
DPA

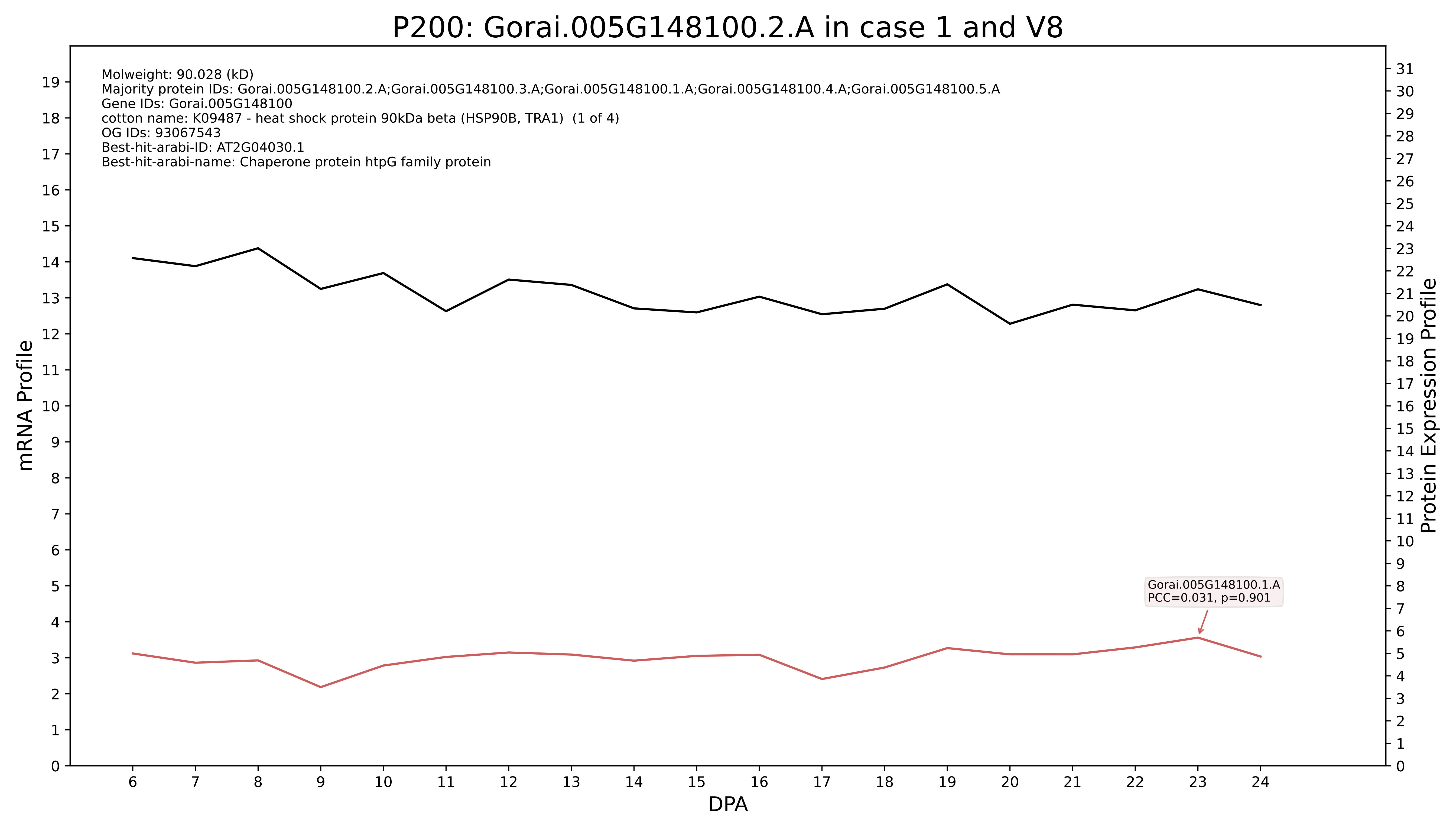
P200: Gorai.005G148100.2.A in case 1 and V8
Molweight: 90.028 (kD)
Majority protein IDs: Gorai.005G148100.2.A;Gorai.005G148100.3.A;Gorai.005G148100.1.A;Gorai.005G148100.4.A;Gorai.005G148100.5.A
Gene IDs: Gorai.005G148100
cotton name: K09487 - heat shock protein 90kDa beta (HSP90B, TRA1) (1 of 4)
OG IDs: 93067543
Best-hit-arabi-ID: AT2G04030.1
Best-hit-arabi-name: Chaperone protein htpG family protein
Gorai.005G148100.1.A
PCC=0.031, p=0.901
mRNA Profile
Protein Expression Profile
DPA

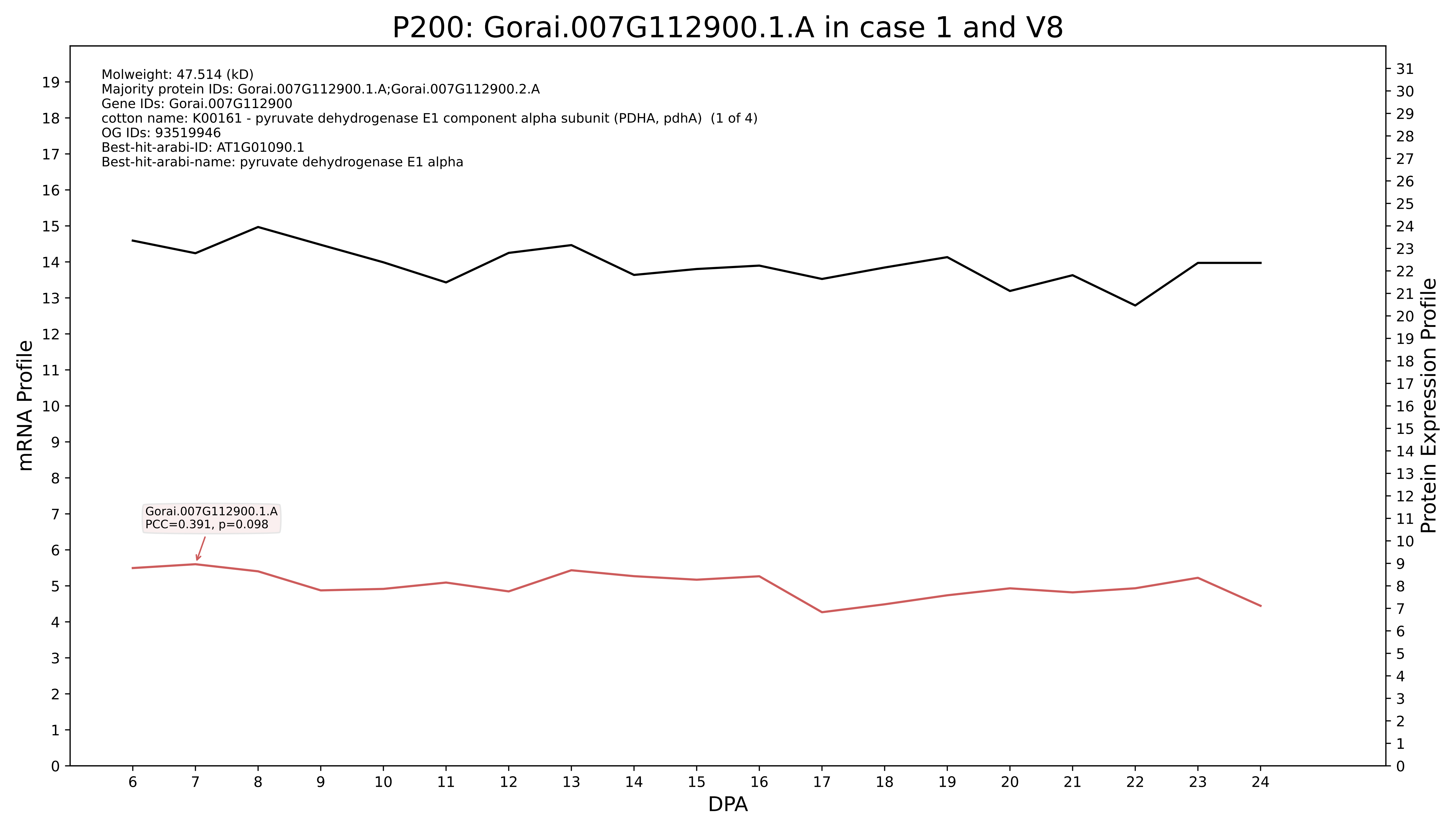
P200: Gorai.007G112900.1.A in case 1 and V8
Molweight: 47.514 (kD)
Majority protein IDs: Gorai.007G112900.1.A;Gorai.007G112900.2.A
Gene IDs: Gorai.007G112900
cotton name: K00161 - pyruvate dehydrogenase E1 component alpha subunit (PDHA, pdhA) (1 of 4)
OG IDs: 93519946
Best-hit-arabi-ID: AT1G01090.1
Best-hit-arabi-name: pyruvate dehydrogenase E1 alpha
Gorai.007G112900.1.A
PCC=0.391, p=0.098
mRNA Profile
Protein Expression Profile
DPA

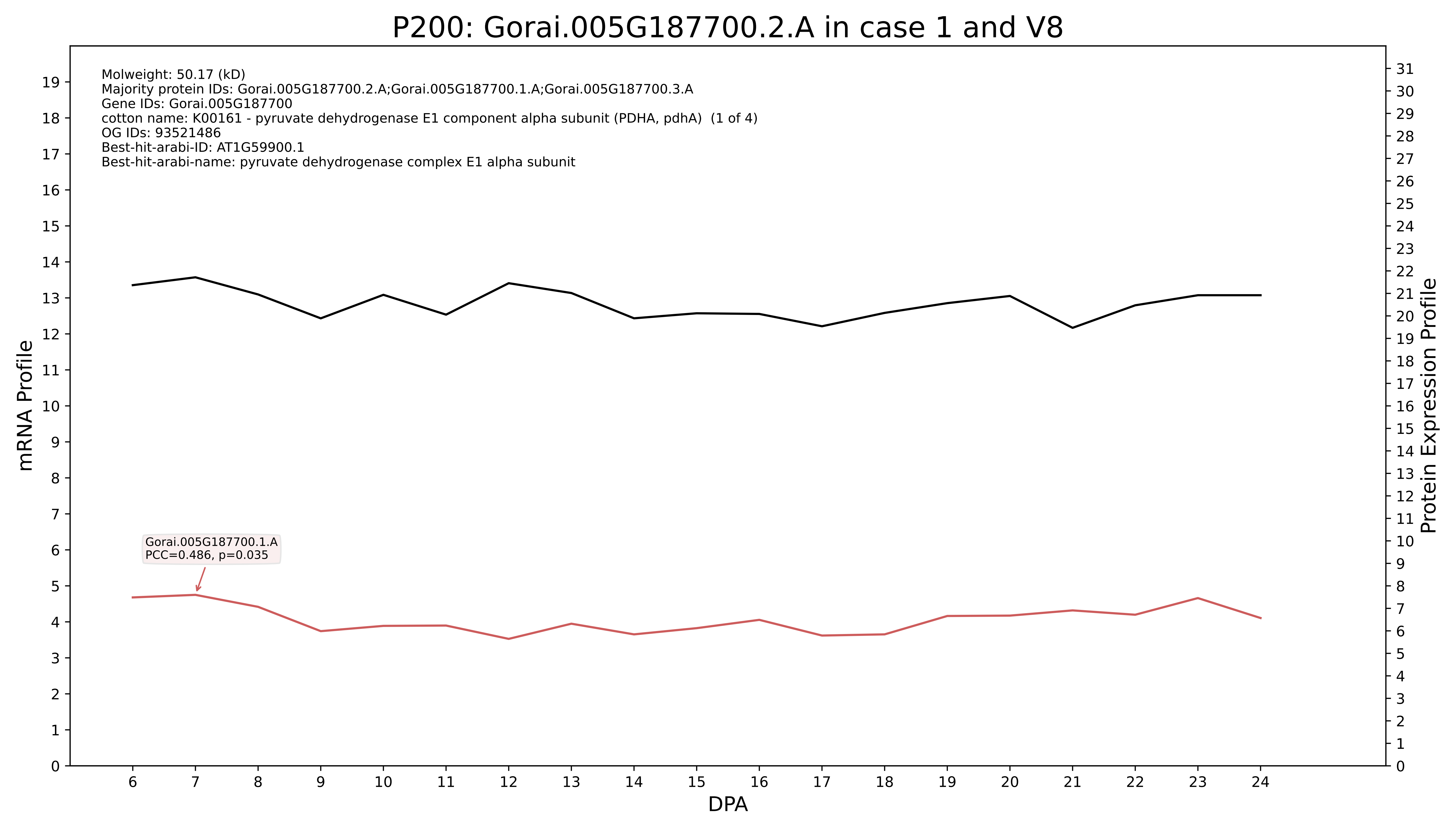
P200: Gorai.005G187700.2.A in case 1 and V8
Molweight: 50.17 (kD)
Majority protein IDs: Gorai.005G187700.2.A;Gorai.005G187700.1.A;Gorai.005G187700.3.A
Gene IDs: Gorai.005G187700
cotton name: K00161 - pyruvate dehydrogenase E1 component alpha subunit (PDHA, pdhA) (1 of 4)
OG IDs: 93521486
Best-hit-arabi-ID: AT1G59900.1
Best-hit-arabi-name: pyruvate dehydrogenase complex E1 alpha subunit
Gorai.005G187700.1.A
PCC=0.486, p=0.035
mRNA Profile
Protein Expression Profile
DPA

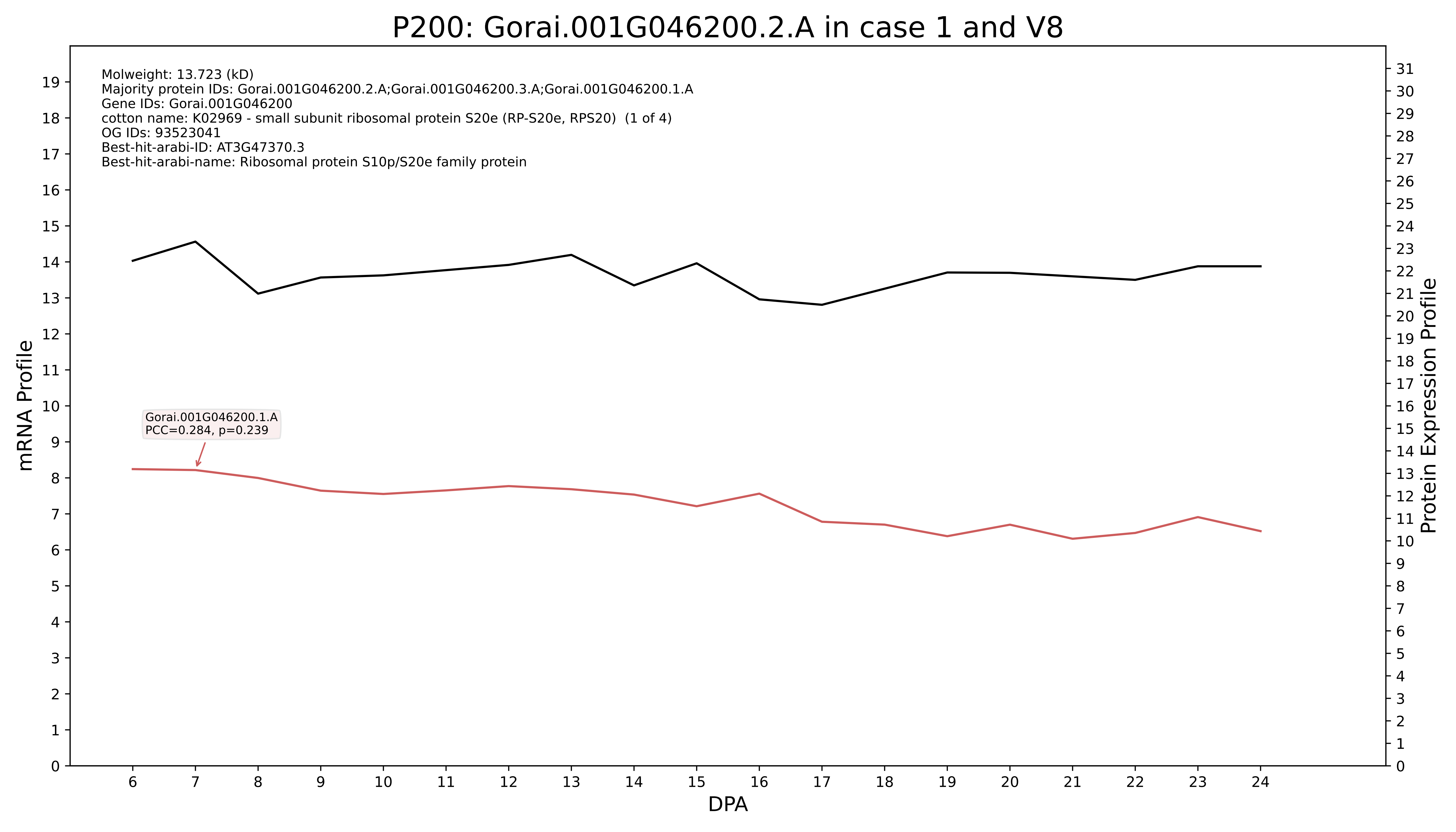
P200: Gorai.001G046200.2.A in case 1 and V8
Molweight: 13.723 (kD)
Majority protein IDs: Gorai.001G046200.2.A;Gorai.001G046200.3.A;Gorai.001G046200.1.A
Gene IDs: Gorai.001G046200
cotton name: K02969 - small subunit ribosomal protein S20e (RP-S20e, RPS20) (1 of 4)
OG IDs: 93523041
Best-hit-arabi-ID: AT3G47370.3
Best-hit-arabi-name: Ribosomal protein S10p/S20e family protein
Gorai.001G046200.1.A
PCC=0.284, p=0.239
mRNA Profile
Protein Expression Profile
DPA

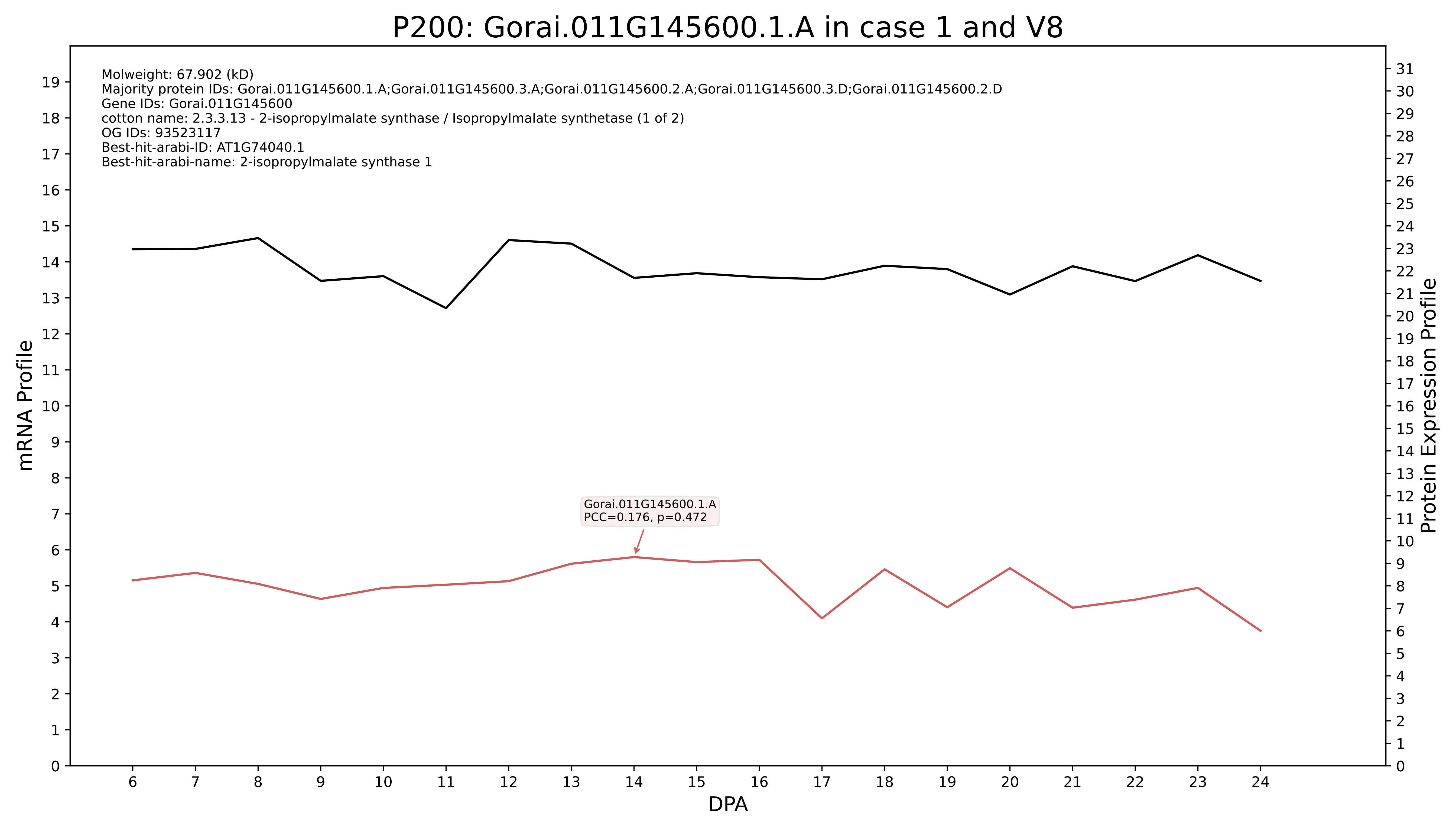

P200: Gorai.011G145600.1.A in case 1 and V8
Molweight: 67.902 (kD)
Majority protein IDs: Gorai.011G145600.1.A;Gorai.011G145600.3.A;Gorai.011G145600.2.A;Gorai.011G145600.3.D;Gorai.011G145600.2.D
Gene IDs: Gorai.011G145600
cotton name: 2.3.3.13 - 2-isopropylmalate synthase / Isopropylmalate synthetase (1 of 2)
OG IDs: 93523117
Best-hit-arabi-ID: AT1G74040.1
Best-hit-arabi-name: 2-isopropylmalate synthase 1
Gorai.011G145600.1.A
PCC=0.176, p=0.472
mRNA Profile
Protein Expression Profile
DPA

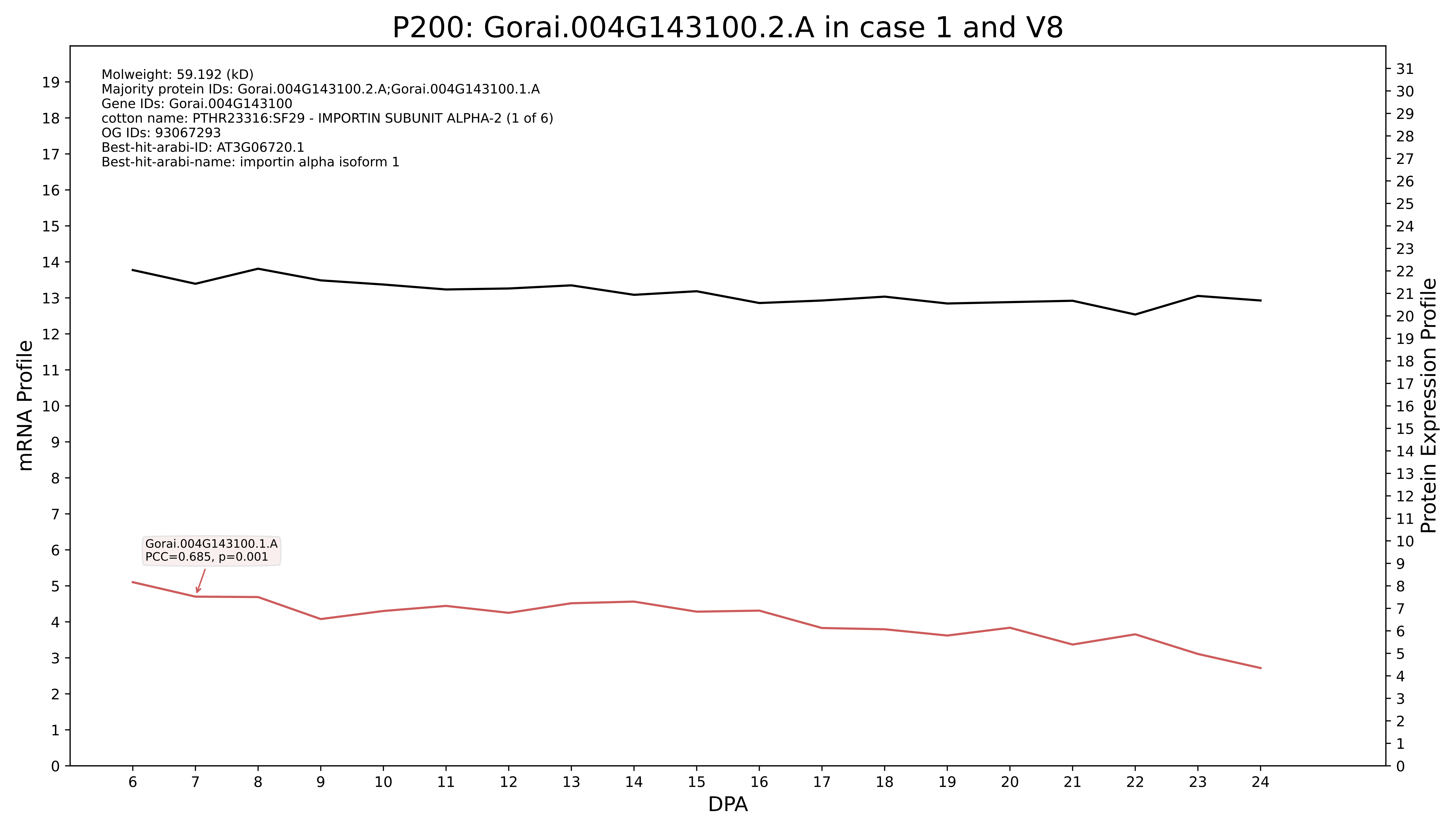
P200: Gorai.004G143100.2.A in case 1 and V8
Molweight: 59.192 (kD)
Majority protein IDs: Gorai.004G143100.2.A;Gorai.004G143100.1.A
Gene IDs: Gorai.004G143100
cotton name: PTHR23316:SF29 - IMPORTIN SUBUNIT ALPHA-2 (1 of 6)
OG IDs: 93067293
Best-hit-arabi-ID: AT3G06720.1
Best-hit-arabi-name: importin alpha isoform 1
Gorai.004G143100.1.A
PCC=0.685, p=0.001
mRNA Profile
Protein Expression Profile
DPA

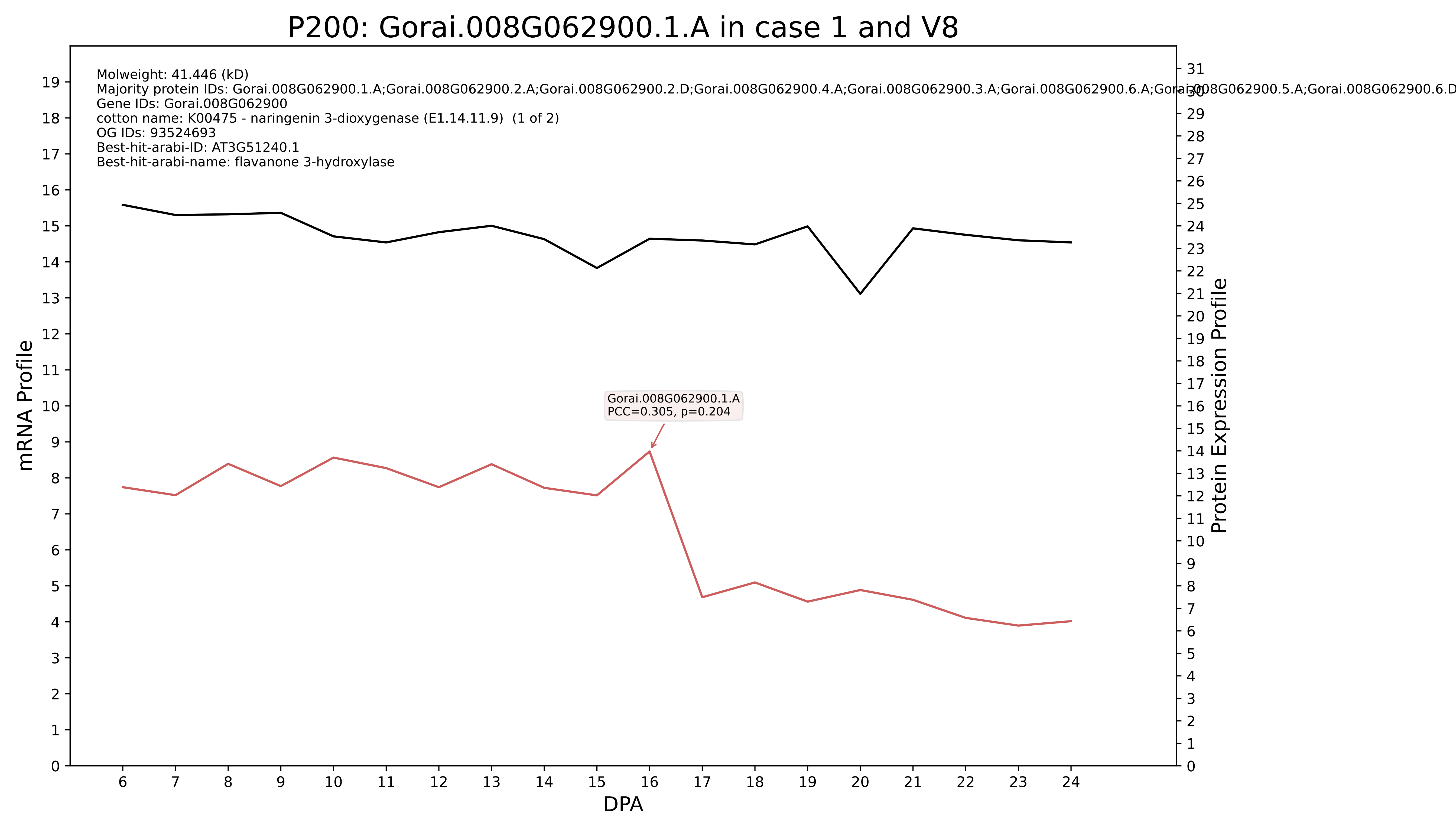

P200: Gorai.008G062900.1.A in case 1 and V8
Molweight: 41.446 (kD)
Majority protein IDs: Gorai.008G062900.1.A;Gorai.008G062900.2.A;Gorai.008G062900.2.D;Gorai.008G062900.4.A;Gorai.008G062900.3.A;Gorai.008G062900.6.A;Gorai.008G062900.5.A;Gorai.008G062900.6.D
Gene IDs: Gorai.008G062900
cotton name: K00475 - naringenin 3-dioxygenase (E1.14.11.9) (1 of 2)
OG IDs: 93524693
Best-hit-arabi-ID: AT3G51240.1
Best-hit-arabi-name: flavanone 3-hydroxylase
Gorai.008G062900.1.A
PCC=0.305, p=0.204
mRNA Profile
Protein Expression Profile
DPA

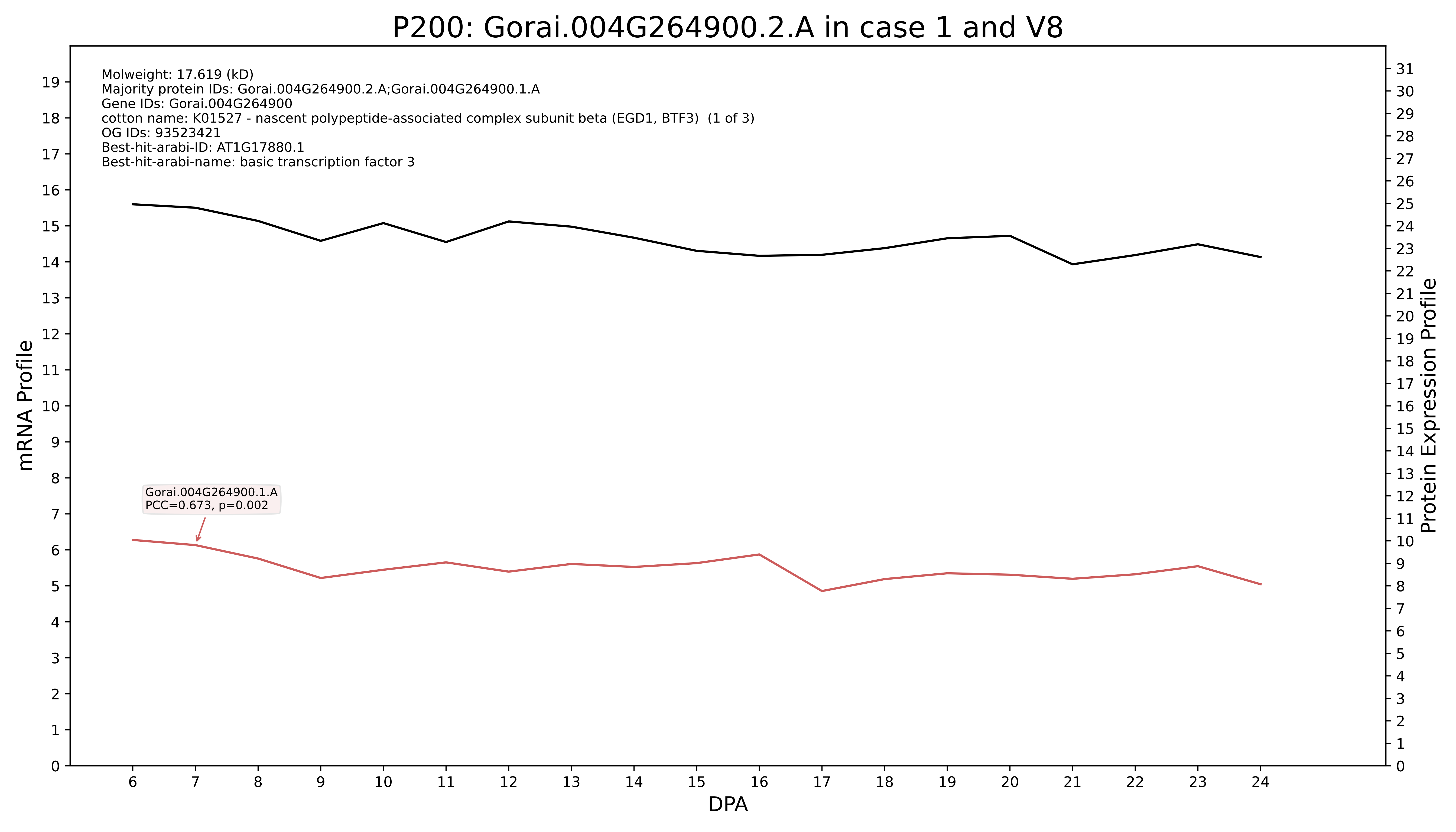
P200: Gorai.004G264900.2.A in case 1 and V8
Molweight: 17.619 (kD)
Majority protein IDs: Gorai.004G264900.2.A;Gorai.004G264900.1.A
Gene IDs: Gorai.004G264900
cotton name: K01527 - nascent polypeptide-associated complex subunit beta (EGD1, BTF3) (1 of 3)
OG IDs: 93523421
Best-hit-arabi-ID: AT1G17880.1
Best-hit-arabi-name: basic transcription factor 3
Gorai.004G264900.1.A
PCC=0.673, p=0.002
mRNA Profile
Protein Expression Profile
DPA

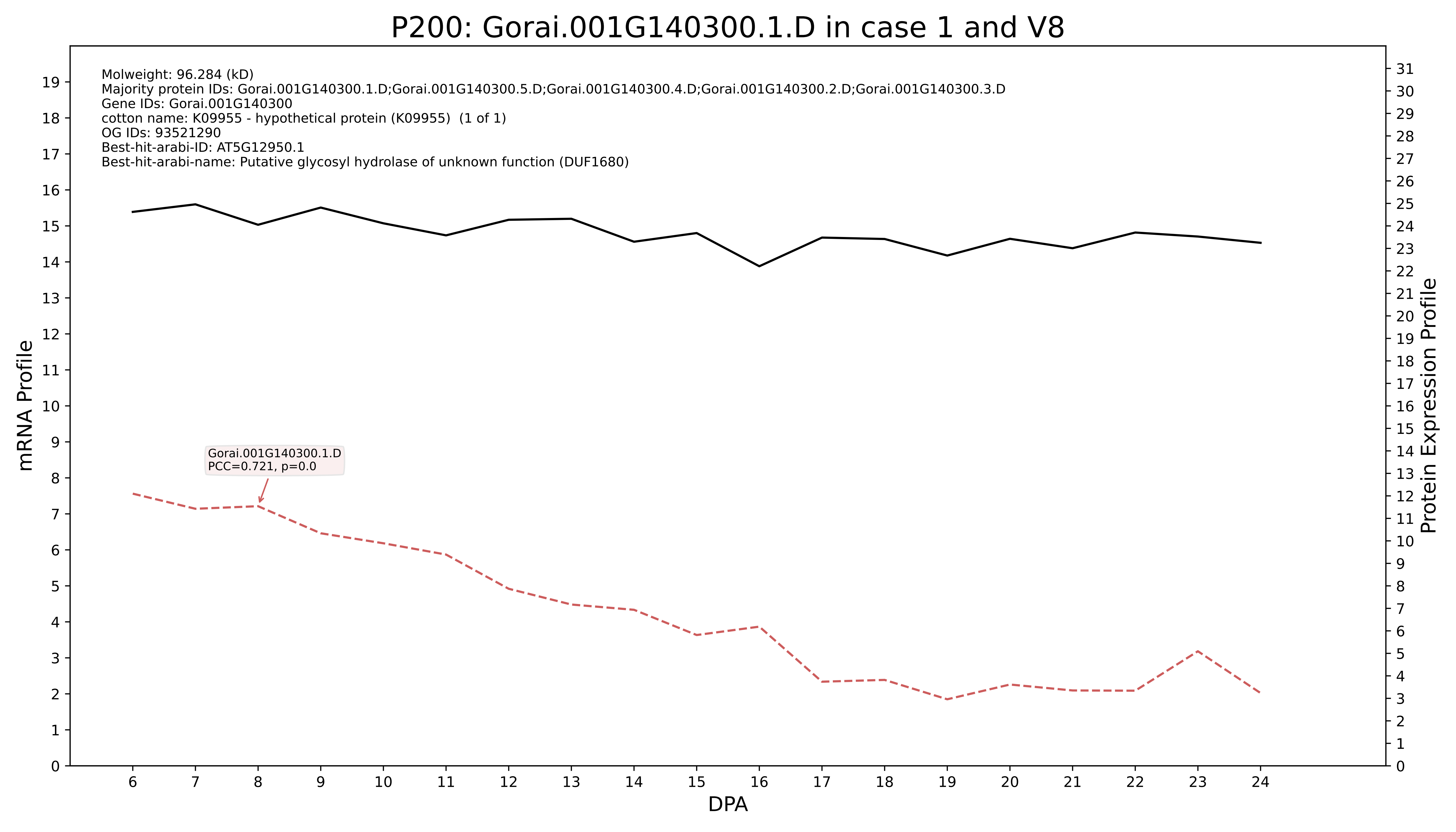
P200: Gorai.001G140300.1.D in case 1 and V8
Molweight: 96.284 (kD)
Majority protein IDs: Gorai.001G140300.1.D;Gorai.001G140300.5.D;Gorai.001G140300.4.D;Gorai.001G140300.2.D;Gorai.001G140300.3.D
Gene IDs: Gorai.001G140300
cotton name: K09955 - hypothetical protein (K09955) (1 of 1)
OG IDs: 93521290
Best-hit-arabi-ID: AT5G12950.1
Best-hit-arabi-name: Putative glycosyl hydrolase of unknown function (DUF1680)
Gorai.001G140300.1.D
PCC=0.721, p=0.0
mRNA Profile
Protein Expression Profile
DPA

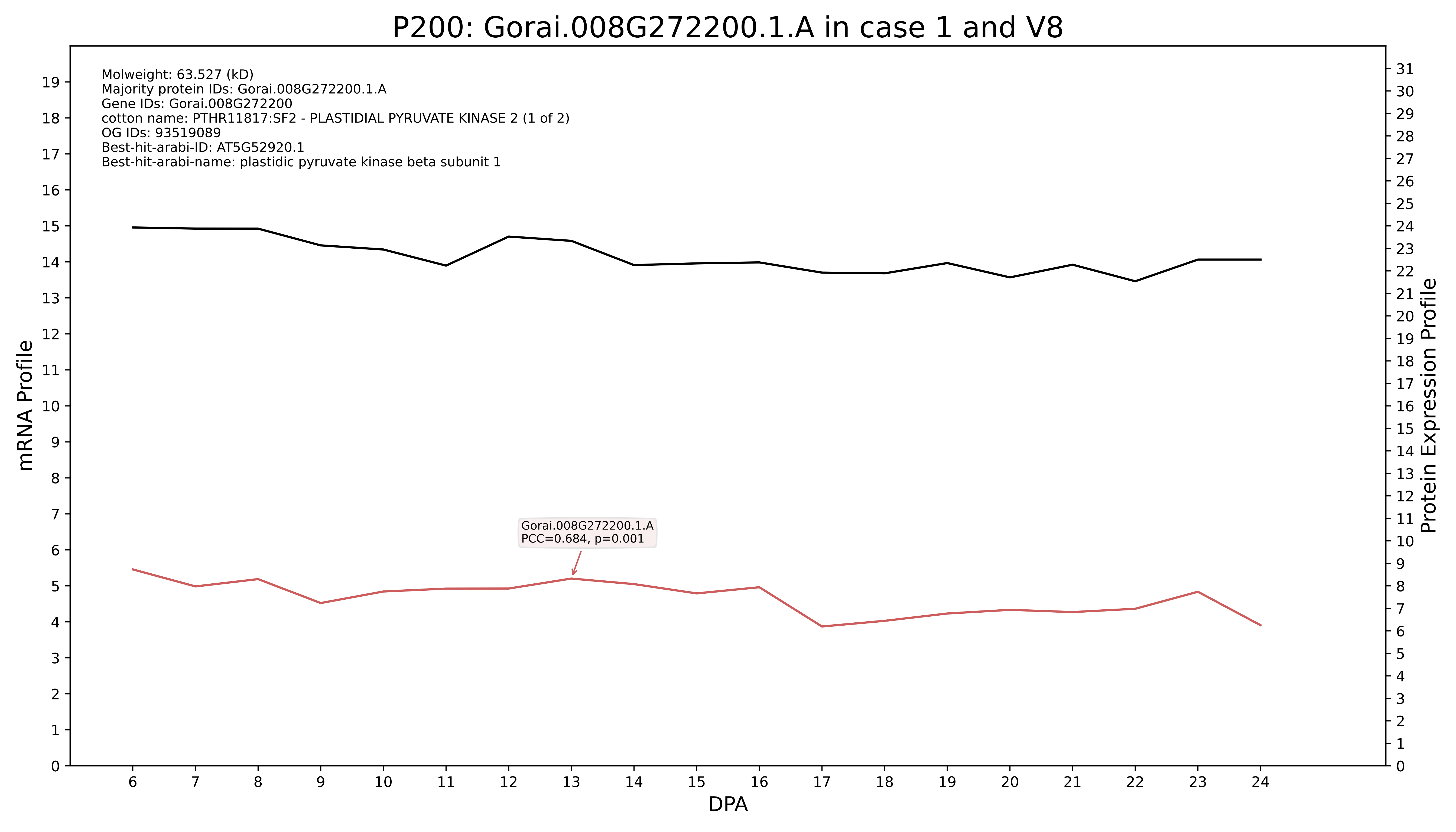
P200: Gorai.008G272200.1.A in case 1 and V8
Molweight: 63.527 (kD)
Majority protein IDs: Gorai.008G272200.1.A
Gene IDs: Gorai.008G272200
cotton name: PTHR11817:SF2 - PLASTIDIAL PYRUVATE KINASE 2 (1 of 2)
OG IDs: 93519089
Best-hit-arabi-ID: AT5G52920.1
Best-hit-arabi-name: plastidic pyruvate kinase beta subunit 1
Gorai.008G272200.1.A
PCC=0.684, p=0.001
mRNA Profile
Protein Expression Profile
DPA
0 1 2 3 4 5 6 7 8 9 10 11 12 13 14 15 16 17 18 19
0 1 2 3 4 5 6 7 8 9 10 11 12 13 14 15 16 17 18 19 20 21 22 23 24 25 26 27 28 29 30 31
6 7 8 9 10 11 12 13 14 15 16 17 18 19 20 21 22 23 24

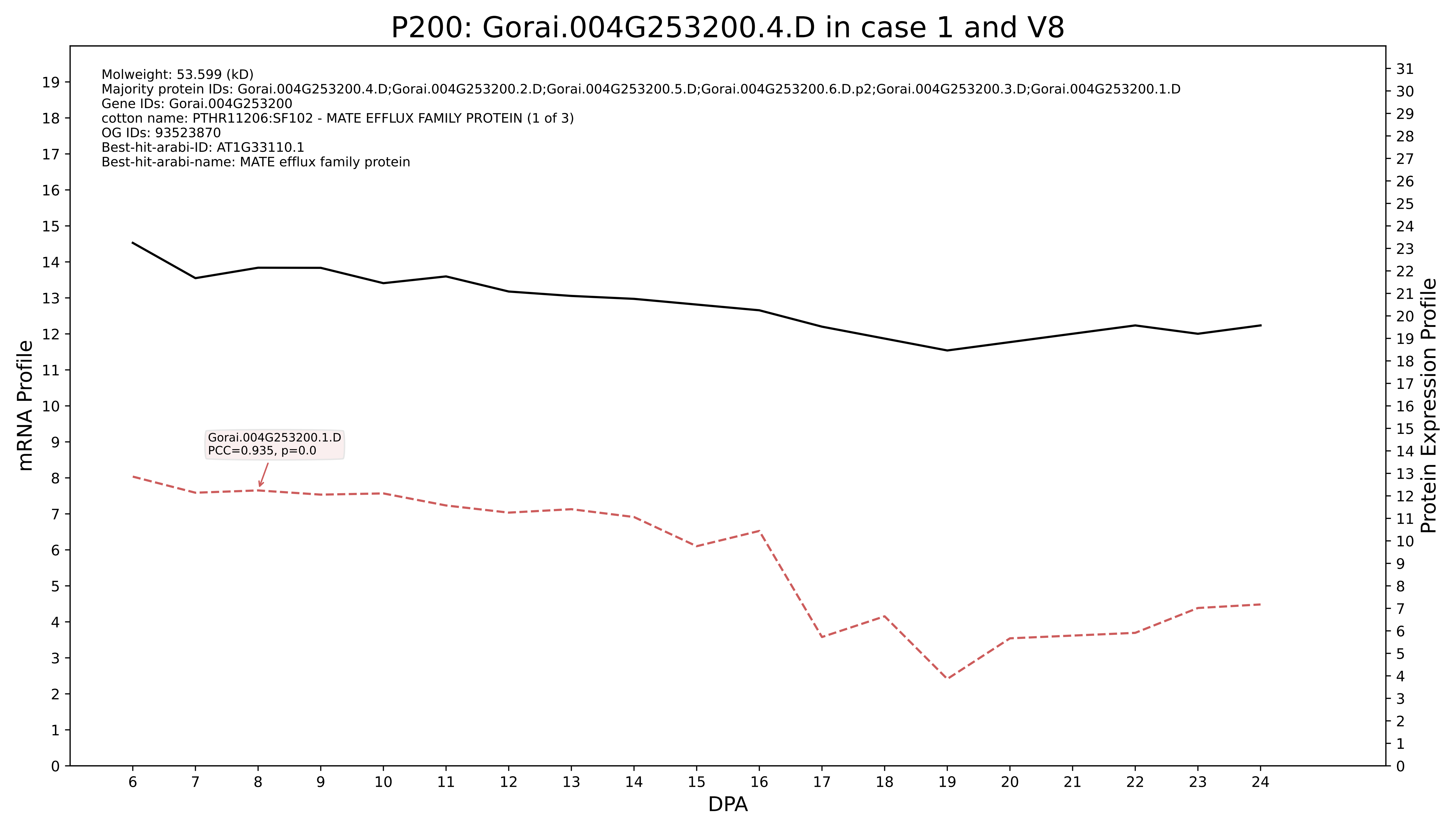
P200: Gorai.004G253200.4.D in case 1 and V8
Molweight: 53.599 (kD)
Majority protein IDs: Gorai.004G253200.4.D;Gorai.004G253200.2.D;Gorai.004G253200.5.D;Gorai.004G253200.6.D.p2;Gorai.004G253200.3.D;Gorai.004G253200.1.D
Gene IDs: Gorai.004G253200
cotton name: PTHR11206:SF102 - MATE EFFLUX FAMILY PROTEIN (1 of 3)
OG IDs: 93523870
Best-hit-arabi-ID: AT1G33110.1
Best-hit-arabi-name: MATE efflux family protein
Gorai.004G253200.1.D
PCC=0.935, p=0.0
mRNA Profile
Protein Expression Profile
DPA

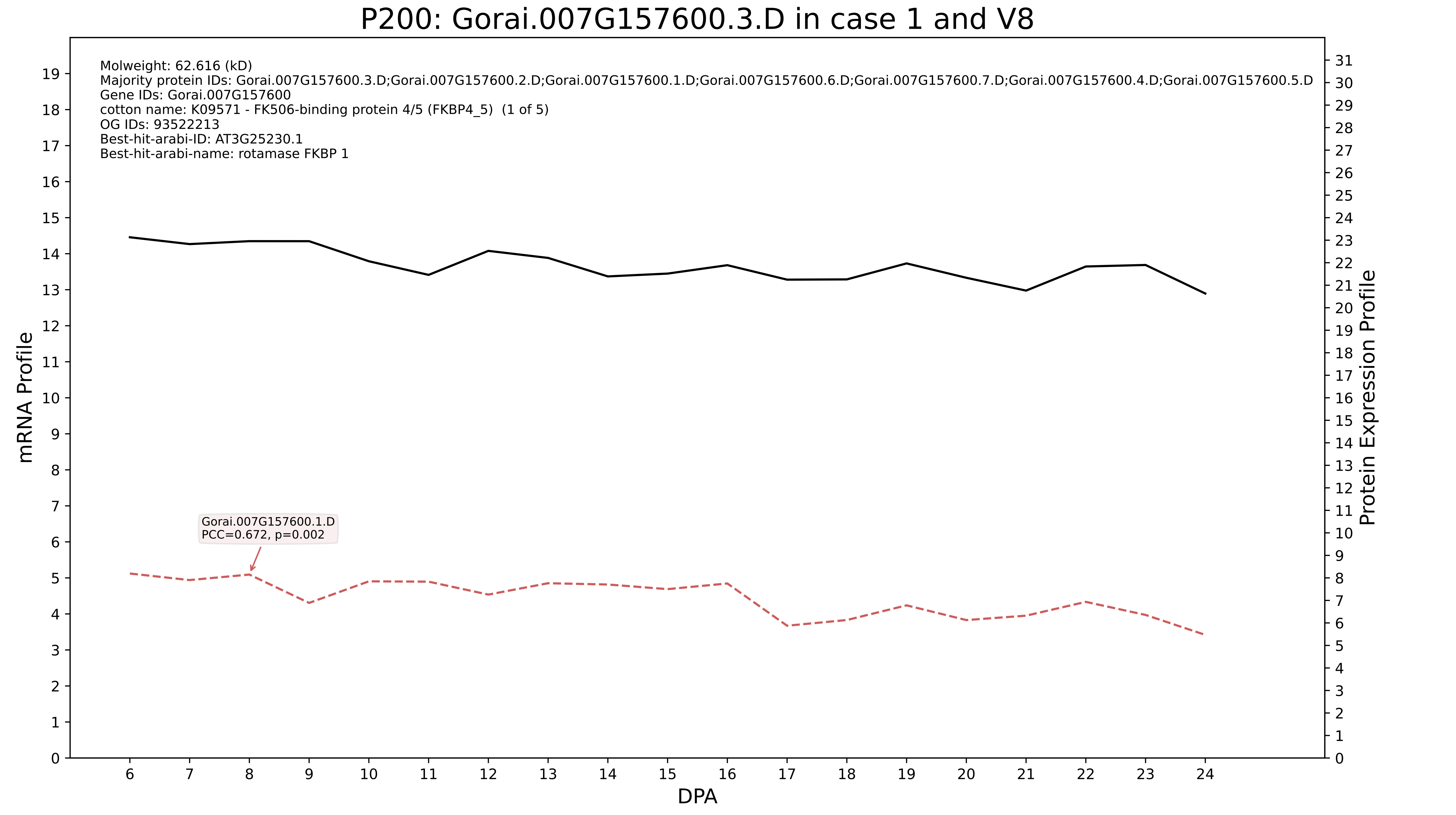
P200: Gorai.007G157600.3.D in case 1 and V8
Molweight: 62.616 (kD)
Majority protein IDs: Gorai.007G157600.3.D;Gorai.007G157600.2.D;Gorai.007G157600.1.D;Gorai.007G157600.6.D;Gorai.007G157600.7.D;Gorai.007G157600.4.D;Gorai.007G157600.5.D
Gene IDs: Gorai.007G157600
cotton name: K09571 - FK506-binding protein 4/5 (FKBP4_5) (1 of 5)
OG IDs: 93522213
Best-hit-arabi-ID: AT3G25230.1
Best-hit-arabi-name: rotamase FKBP 1
Gorai.007G157600.1.D
PCC=0.672, p=0.002
mRNA Profile
Protein Expression Profile
DPA

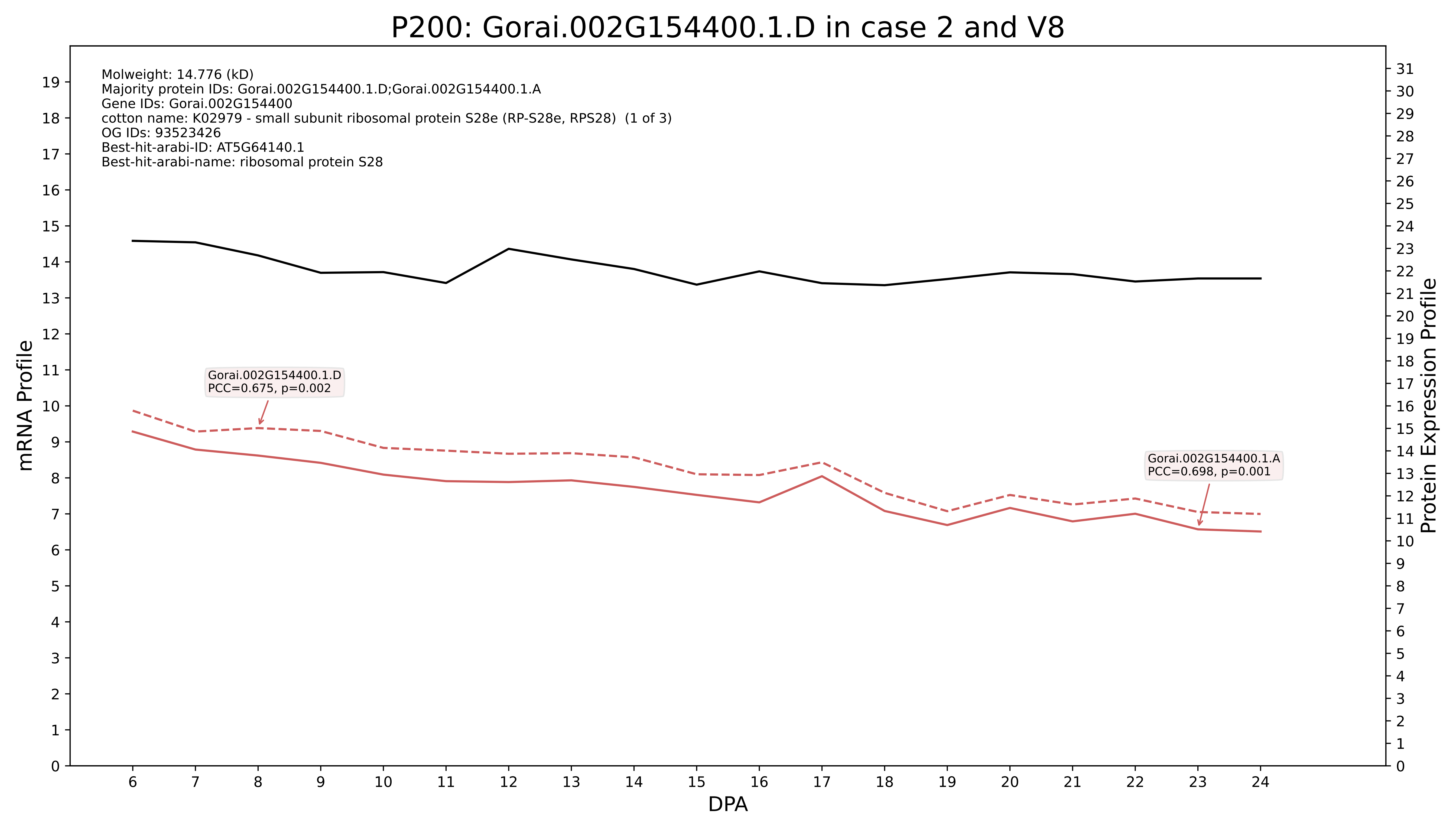
P200: Gorai.002G154400.1.D in case 2 and V8
Molweight: 14.776 (kD)
Majority protein IDs: Gorai.002G154400.1.D;Gorai.002G154400.1.A
Gene IDs: Gorai.002G154400
cotton name: K02979 - small subunit ribosomal protein S28e (RP-S28e, RPS28) (1 of 3)
OG IDs: 93523426
Best-hit-arabi-ID: AT5G64140.1
Best-hit-arabi-name: ribosomal protein S28
Gorai.002G154400.1.D
PCC=0.675, p=0.002
Gorai.002G154400.1.A
PCC=0.698, p=0.001
mRNA Profile
Protein Expression Profile
DPA

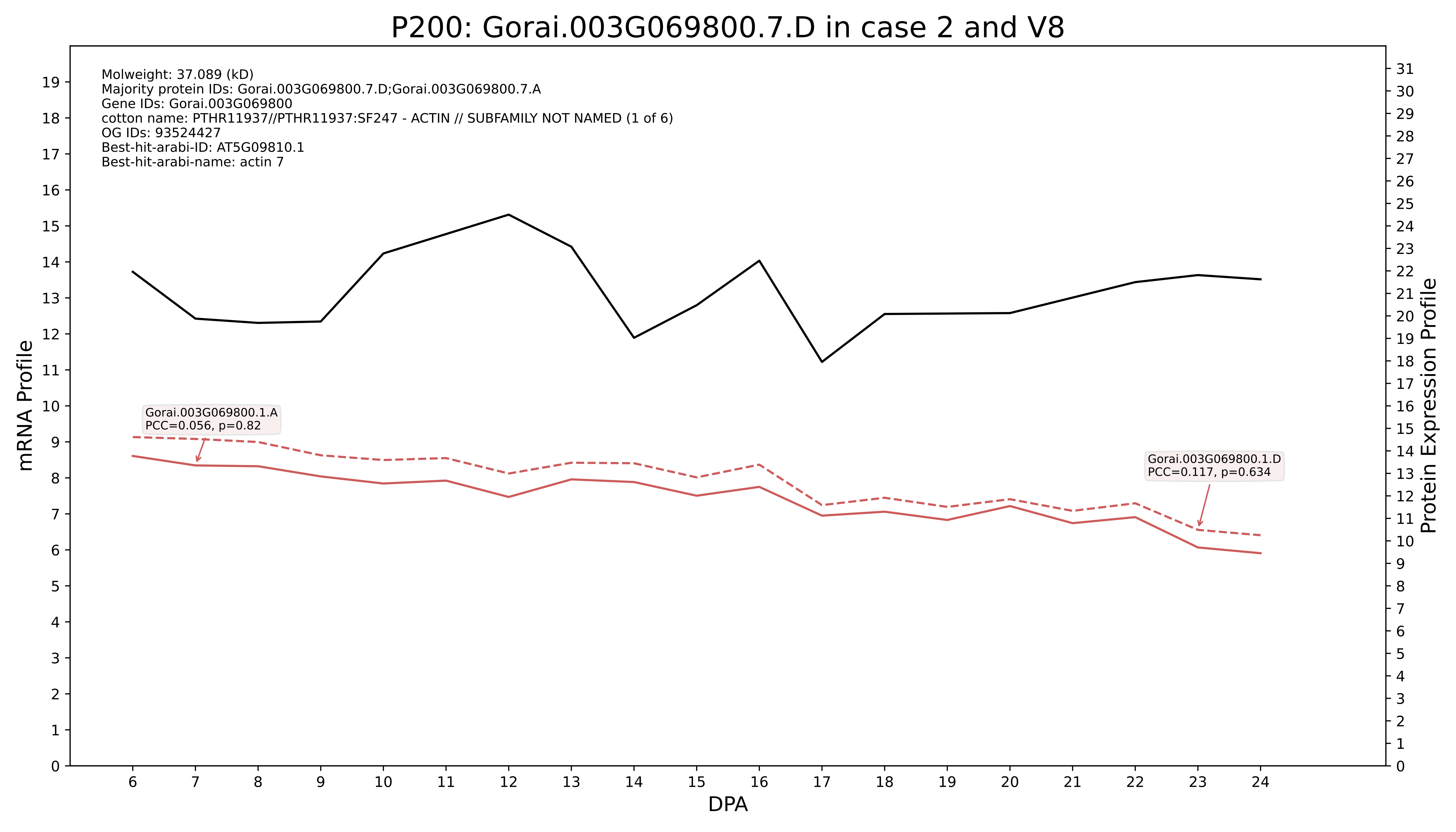

P200: Gorai.003G069800.7.D in case 2 and V8
Molweight: 37.089 (kD)
Majority protein IDs: Gorai.003G069800.7.D;Gorai.003G069800.7.A
Gene IDs: Gorai.003G069800
cotton name: PTHR11937//PTHR11937:SF247 - ACTIN // SUBFAMILY NOT NAMED (1 of 6)
OG IDs: 93524427
Best-hit-arabi-ID: AT5G09810.1
Best-hit-arabi-name: actin 7
Gorai.003G069800.1.A
PCC=0.056, p=0.82
Gorai.003G069800.1.D
PCC=0.117, p=0.634
mRNA Profile
Protein Expression Profile
DPA

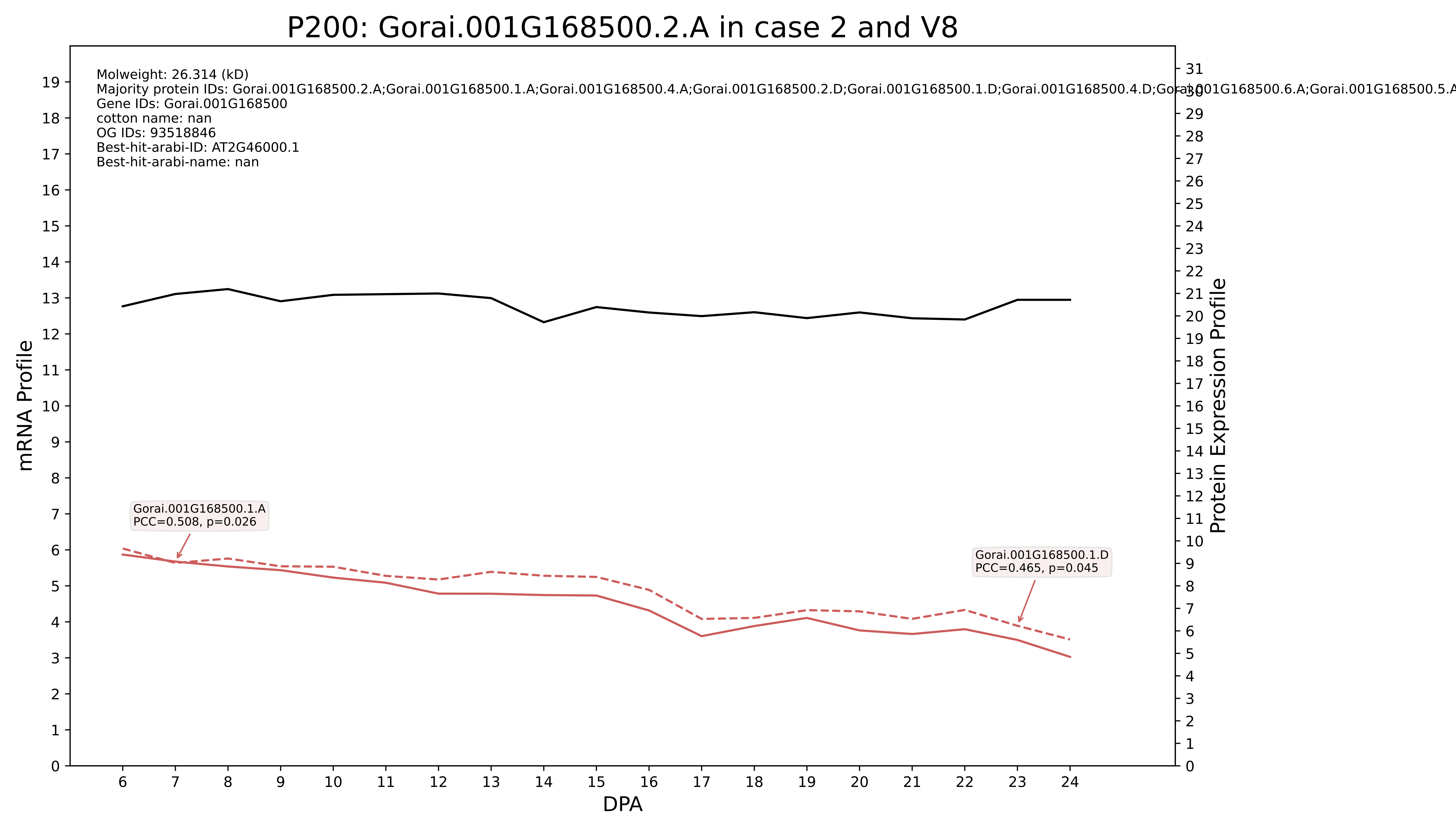

P200: Gorai.001G168500.2.A in case 2 and V8
Molweight: 26.314 (kD)
Majority protein IDs: Gorai.001G168500.2.A;Gorai.001G168500.1.A;Gorai.001G168500.4.A;Gorai.001G168500.2.D;Gorai.001G168500.1.D;Gorai.001G168500.4.D;Gorai.001G168500.6.A;Gorai.001G168500.5.A
Gene IDs: Gorai.001G168500
cotton name: nan
OG IDs: 93518846
Best-hit-arabi-ID: AT2G46000.1
Best-hit-arabi-name: nan
Gorai.001G168500.1.A
PCC=0.508, p=0.026
Gorai.001G168500.1.D
PCC=0.465, p=0.045
mRNA Profile
Protein Expression Profile
DPA

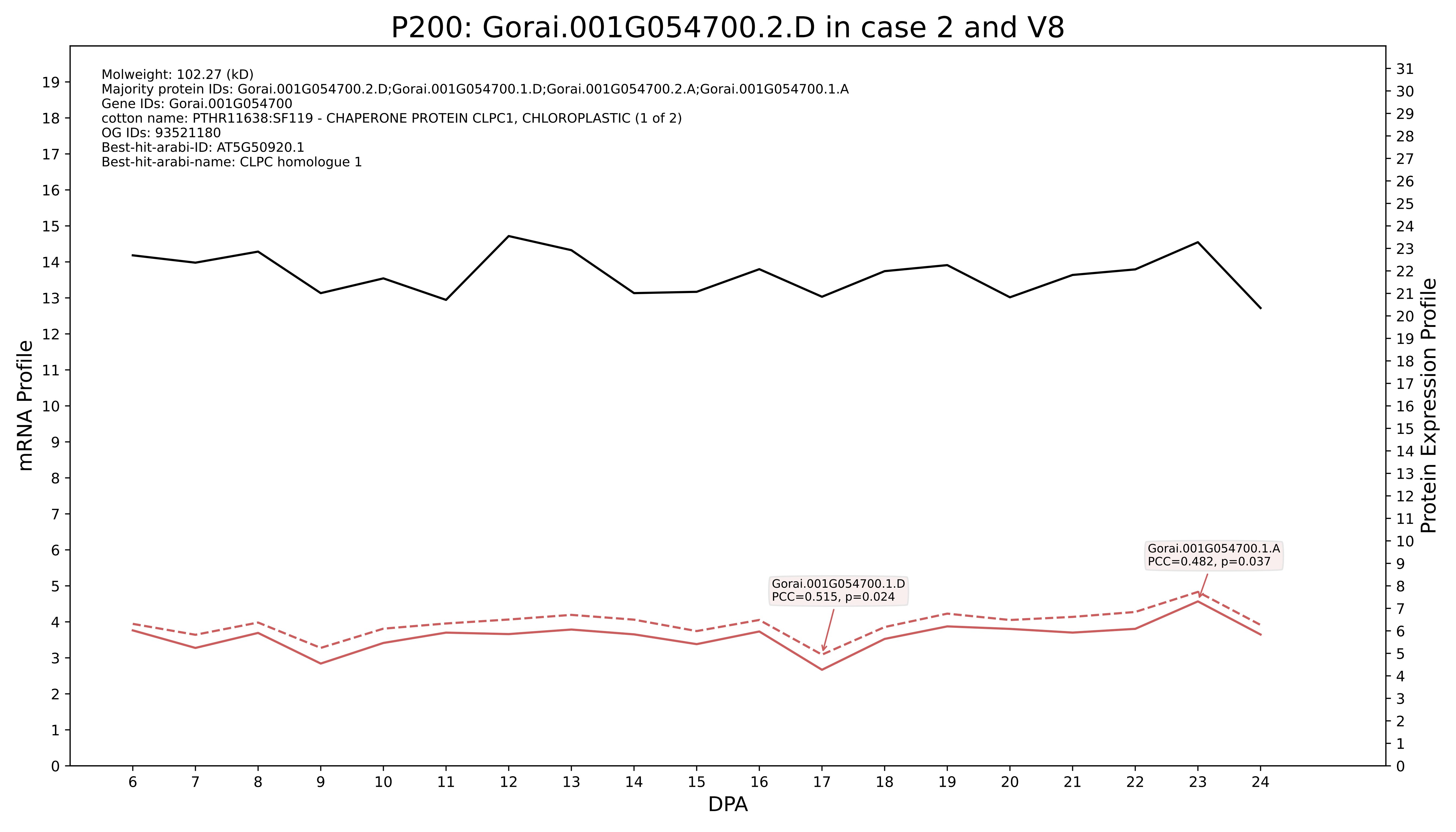
P200: Gorai.001G054700.2.D in case 2 and V8
Molweight: 102.27 (kD)
Majority protein IDs: Gorai.001G054700.2.D;Gorai.001G054700.1.D;Gorai.001G054700.2.A;Gorai.001G054700.1.A
Gene IDs: Gorai.001G054700
cotton name: PTHR11638:SF119 - CHAPERONE PROTEIN CLPC1, CHLOROPLASTIC (1 of 2)
OG IDs: 93521180
Best-hit-arabi-ID: AT5G50920.1
Best-hit-arabi-name: CLPC homologue 1
Gorai.001G054700.1.D
PCC=0.515, p=0.024
Gorai.001G054700.1.A
PCC=0.482, p=0.037
mRNA Profile
Protein Expression Profile
DPA

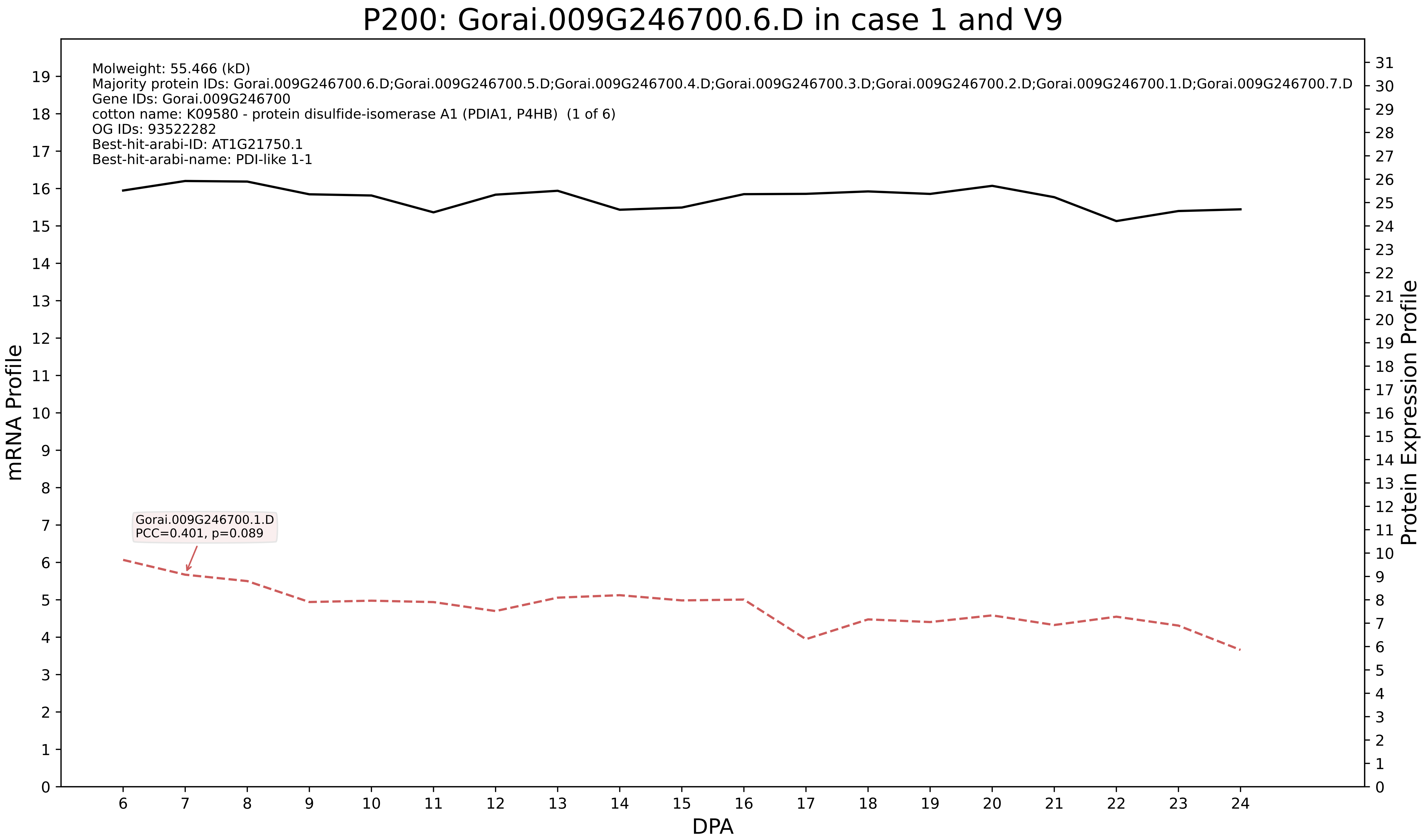
P200: Gorai.009G246700.6.D in case 1 and V9
Molweight: 55.466 (kD)
Majority protein IDs: Gorai.009G246700.6.D;Gorai.009G246700.5.D;Gorai.009G246700.4.D;Gorai.009G246700.3.D;Gorai.009G246700.2.D;Gorai.009G246700.1.D;Gorai.009G246700.7.D
Gene IDs: Gorai.009G246700
cotton name: K09580 - protein disulfide-isomerase A1 (PDIA1, P4HB) (1 of 6)
OG IDs: 93522282
Best-hit-arabi-ID: AT1G21750.1
Best-hit-arabi-name: PDI-like 1-1
Gorai.009G246700.1.D
PCC=0.401, p=0.089
mRNA Profile
Protein Expression Profile
DPA

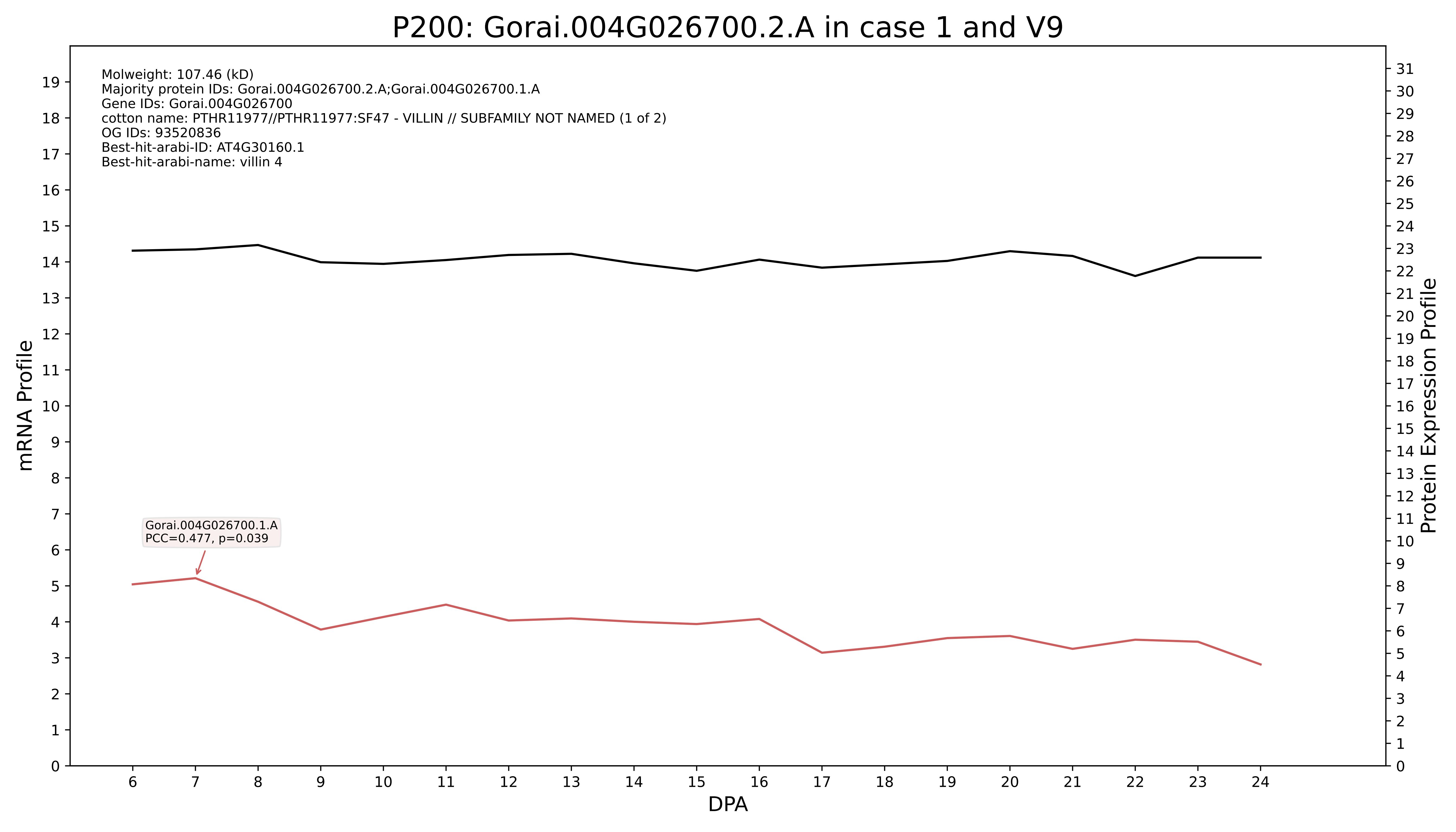

P200: Gorai.004G026700.2.A in case 1 and V9
Molweight: 107.46 (kD)
Majority protein IDs: Gorai.004G026700.2.A;Gorai.004G026700.1.A
Gene IDs: Gorai.004G026700
cotton name: PTHR11977//PTHR11977:SF47 - VILLIN // SUBFAMILY NOT NAMED (1 of 2)
OG IDs: 93520836
Best-hit-arabi-ID: AT4G30160.1
Best-hit-arabi-name: villin 4
Gorai.004G026700.1.A
PCC=0.477, p=0.039
mRNA Profile
Protein Expression Profile
DPA

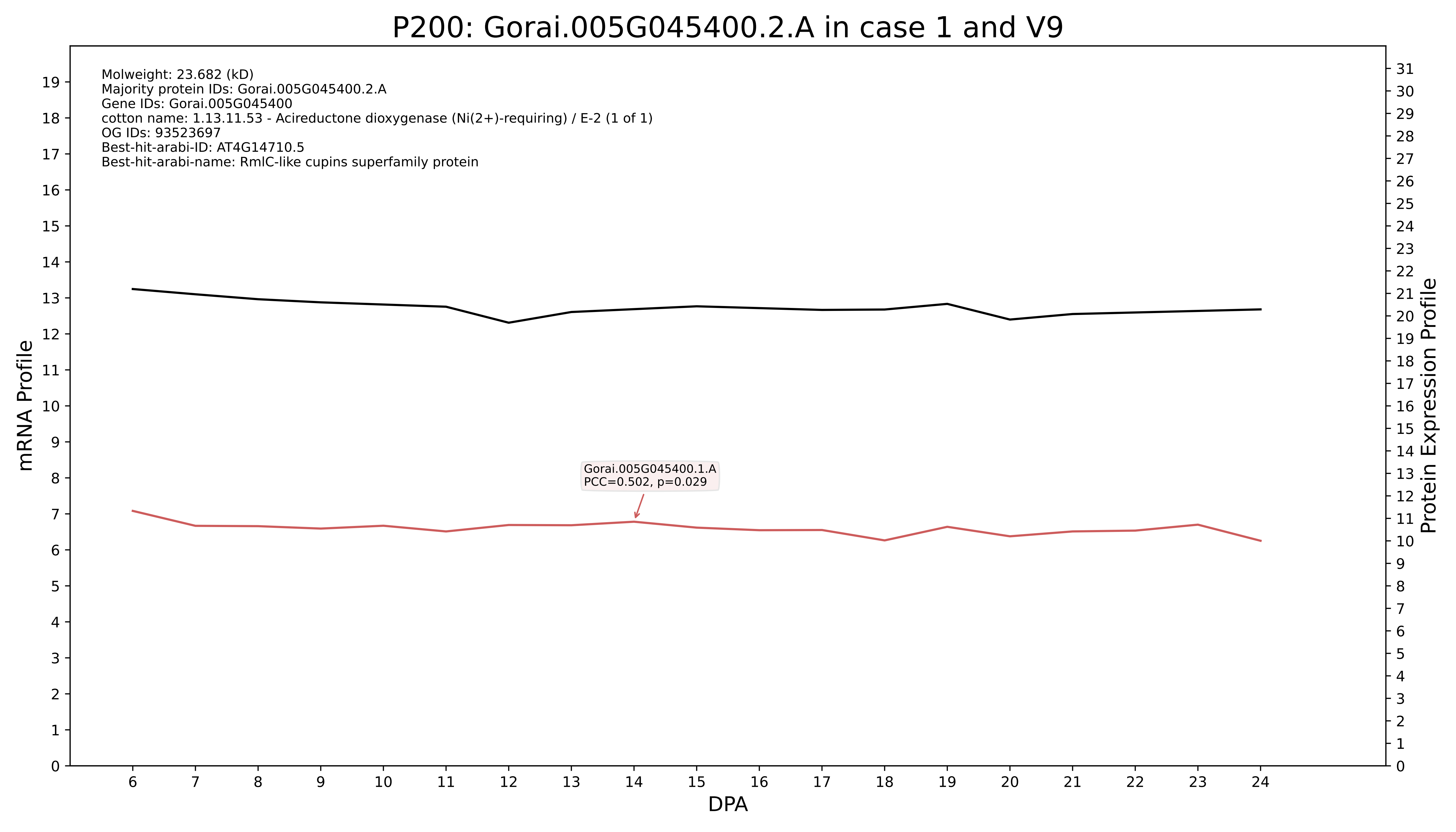
P200: Gorai.005G045400.2.A in case 1 and V9
Molweight: 23.682 (kD)
Majority protein IDs: Gorai.005G045400.2.A
Gene IDs: Gorai.005G045400
cotton name: 1.13.11.53 - Acireductone dioxygenase (Ni(2+)-requiring) / E-2 (1 of 1)
OG IDs: 93523697
Best-hit-arabi-ID: AT4G14710.5
Best-hit-arabi-name: RmlC-like cupins superfamily protein
Gorai.005G045400.1.A
PCC=0.502, p=0.029
mRNA Profile
Protein Expression Profile
DPA

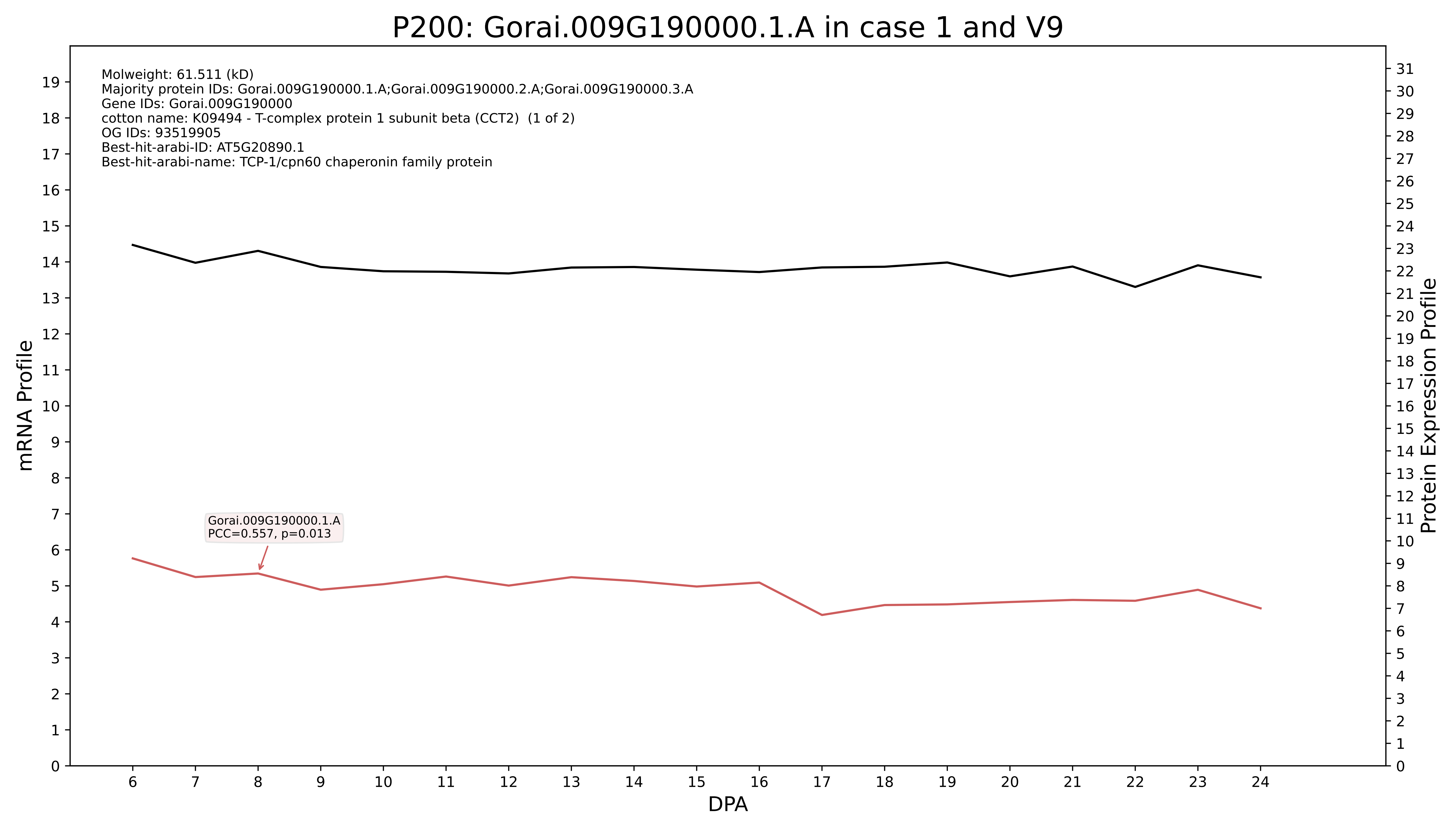
P200: Gorai.009G190000.1.A in case 1 and V9
Molweight: 61.511 (kD)
Majority protein IDs: Gorai.009G190000.1.A;Gorai.009G190000.2.A;Gorai.009G190000.3.A
Gene IDs: Gorai.009G190000
cotton name: K09494 - T-complex protein 1 subunit beta (CCT2) (1 of 2)
OG IDs: 93519905
Best-hit-arabi-ID: AT5G20890.1
Best-hit-arabi-name: TCP-1/cpn60 chaperonin family protein
Gorai.009G190000.1.A
PCC=0.557, p=0.013
mRNA Profile
Protein Expression Profile
DPA

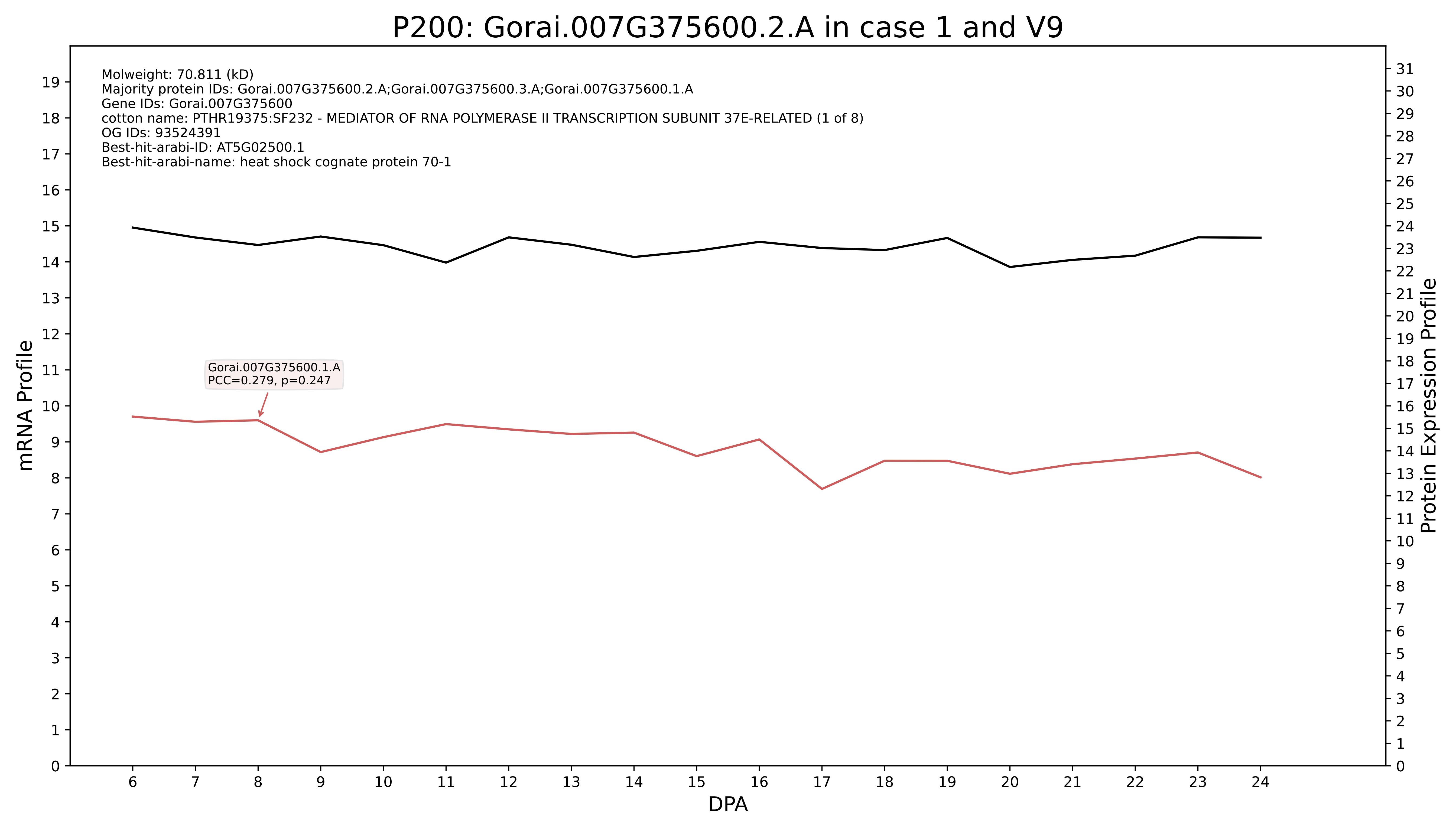
P200: Gorai.007G375600.2.A in case 1 and V9
Molweight: 70.811 (kD)
Majority protein IDs: Gorai.007G375600.2.A;Gorai.007G375600.3.A;Gorai.007G375600.1.A
Gene IDs: Gorai.007G375600
cotton name: PTHR19375:SF232 - MEDIATOR OF RNA POLYMERASE II TRANSCRIPTION SUBUNIT 37E-RELATED (1 of 8)
OG IDs: 93524391
Best-hit-arabi-ID: AT5G02500.1
Best-hit-arabi-name: heat shock cognate protein 70-1
Gorai.007G375600.1.A
PCC=0.279, p=0.247
mRNA Profile
Protein Expression Profile
DPA

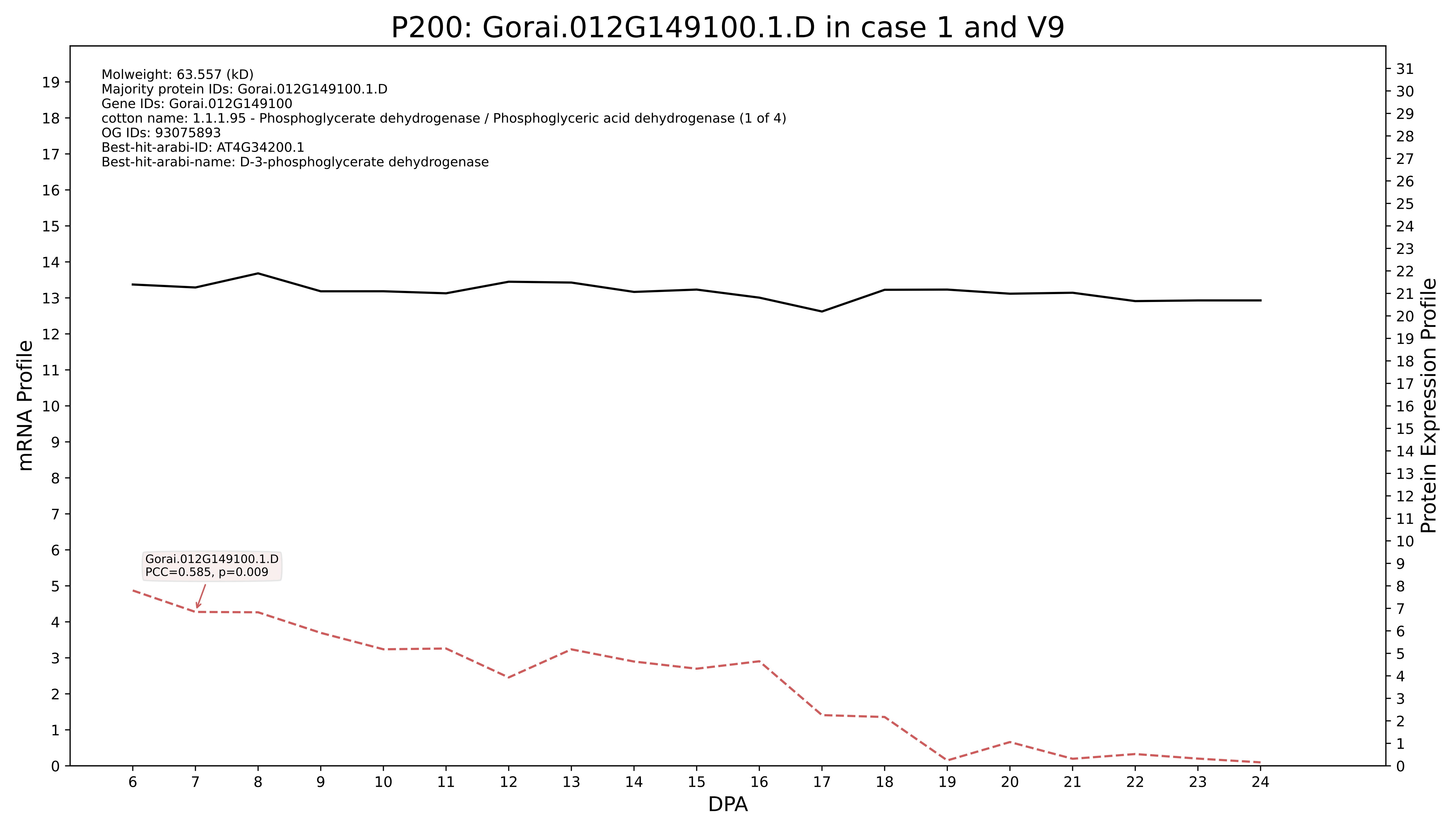
P200: Gorai.012G149100.1.D in case 1 and V9
Molweight: 63.557 (kD)
Majority protein IDs: Gorai.012G149100.1.D
Gene IDs: Gorai.012G149100
cotton name: 1.1.1.95 - Phosphoglycerate dehydrogenase / Phosphoglyceric acid dehydrogenase (1 of 4)
OG IDs: 93075893
Best-hit-arabi-ID: AT4G34200.1
Best-hit-arabi-name: D-3-phosphoglycerate dehydrogenase
Gorai.012G149100.1.D
PCC=0.585, p=0.009
mRNA Profile
Protein Expression Profile
DPA

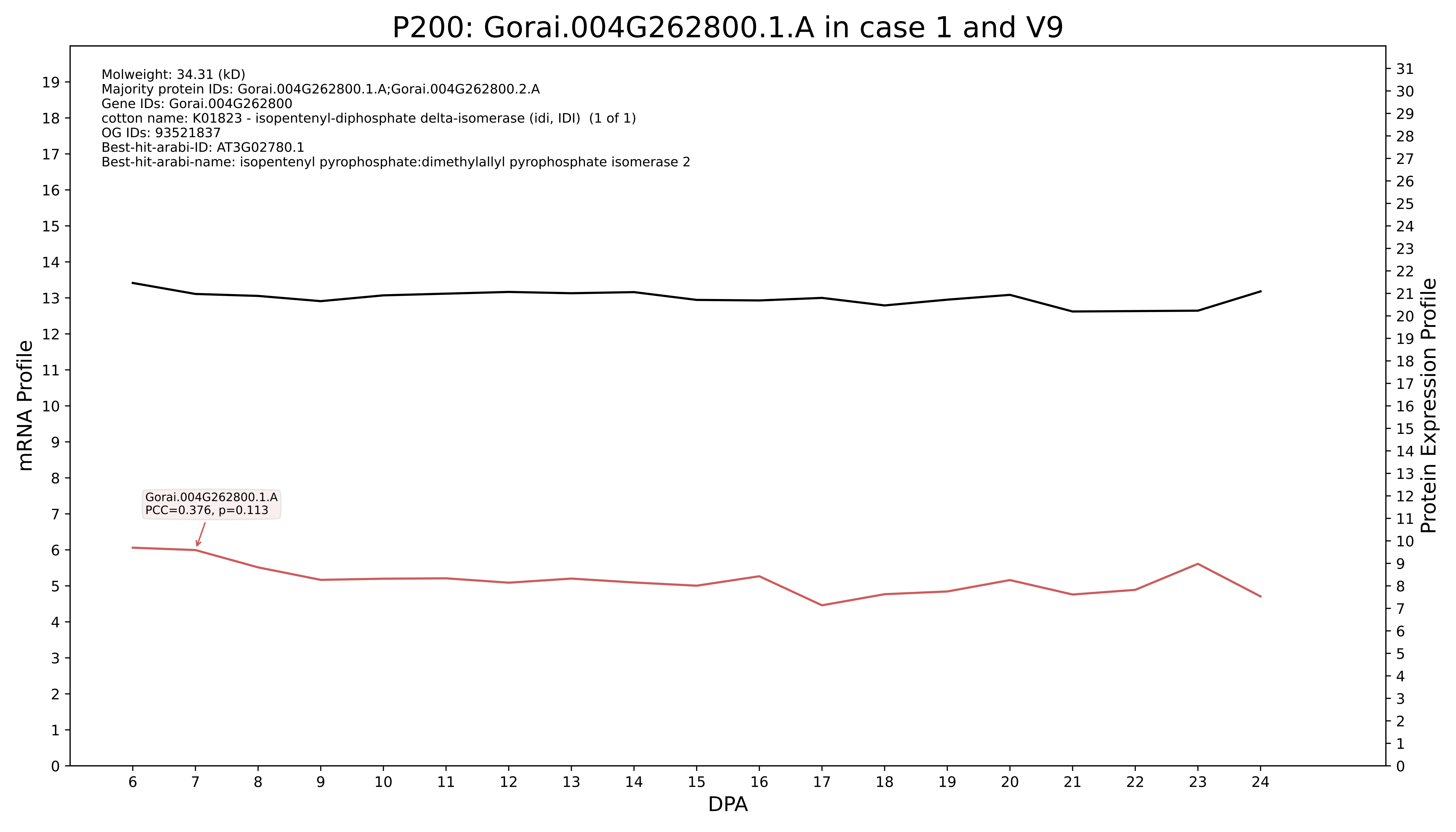
P200: Gorai.004G262800.1.A in case 1 and V9
Molweight: 34.31 (kD)
Majority protein IDs: Gorai.004G262800.1.A;Gorai.004G262800.2.A
Gene IDs: Gorai.004G262800
cotton name: K01823 - isopentenyl-diphosphate delta-isomerase (idi, IDI) (1 of 1)
OG IDs: 93521837
Best-hit-arabi-ID: AT3G02780.1
Best-hit-arabi-name: isopentenyl pyrophosphate:dimethylallyl pyrophosphate isomerase 2
Gorai.004G262800.1.A
PCC=0.376, p=0.113
mRNA Profile
Protein Expression Profile
DPA

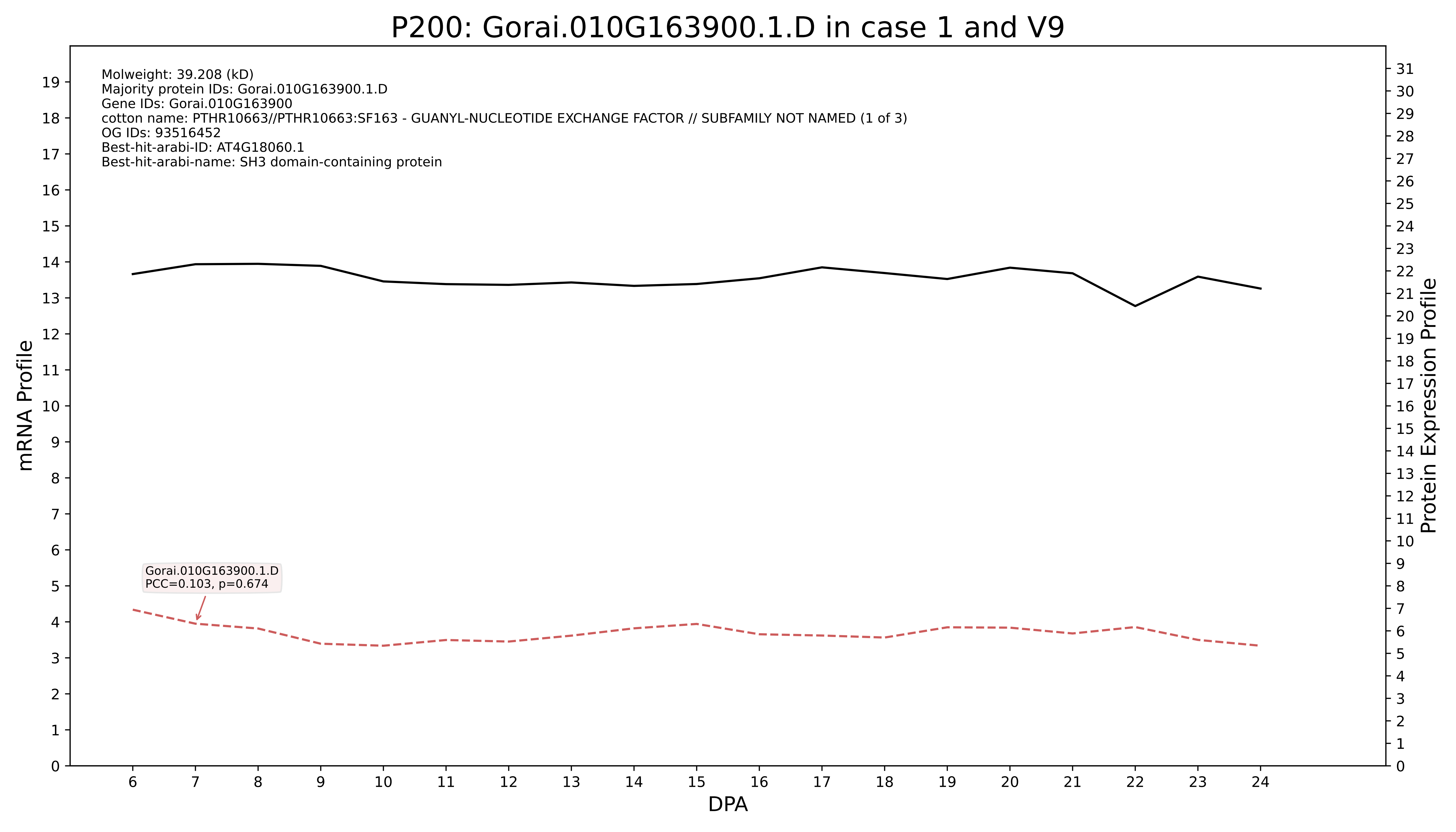
P200: Gorai.010G163900.1.D in case 1 and V9
Molweight: 39.208 (kD)
Majority protein IDs: Gorai.010G163900.1.D
Gene IDs: Gorai.010G163900
cotton name: PTHR10663//PTHR10663:SF163 - GUANYL-NUCLEOTIDE EXCHANGE FACTOR // SUBFAMILY NOT NAMED (1 of 3)
OG IDs: 93516452
Best-hit-arabi-ID: AT4G18060.1
Best-hit-arabi-name: SH3 domain-containing protein
Gorai.010G163900.1.D
PCC=0.103, p=0.674
mRNA Profile
Protein Expression Profile
DPA

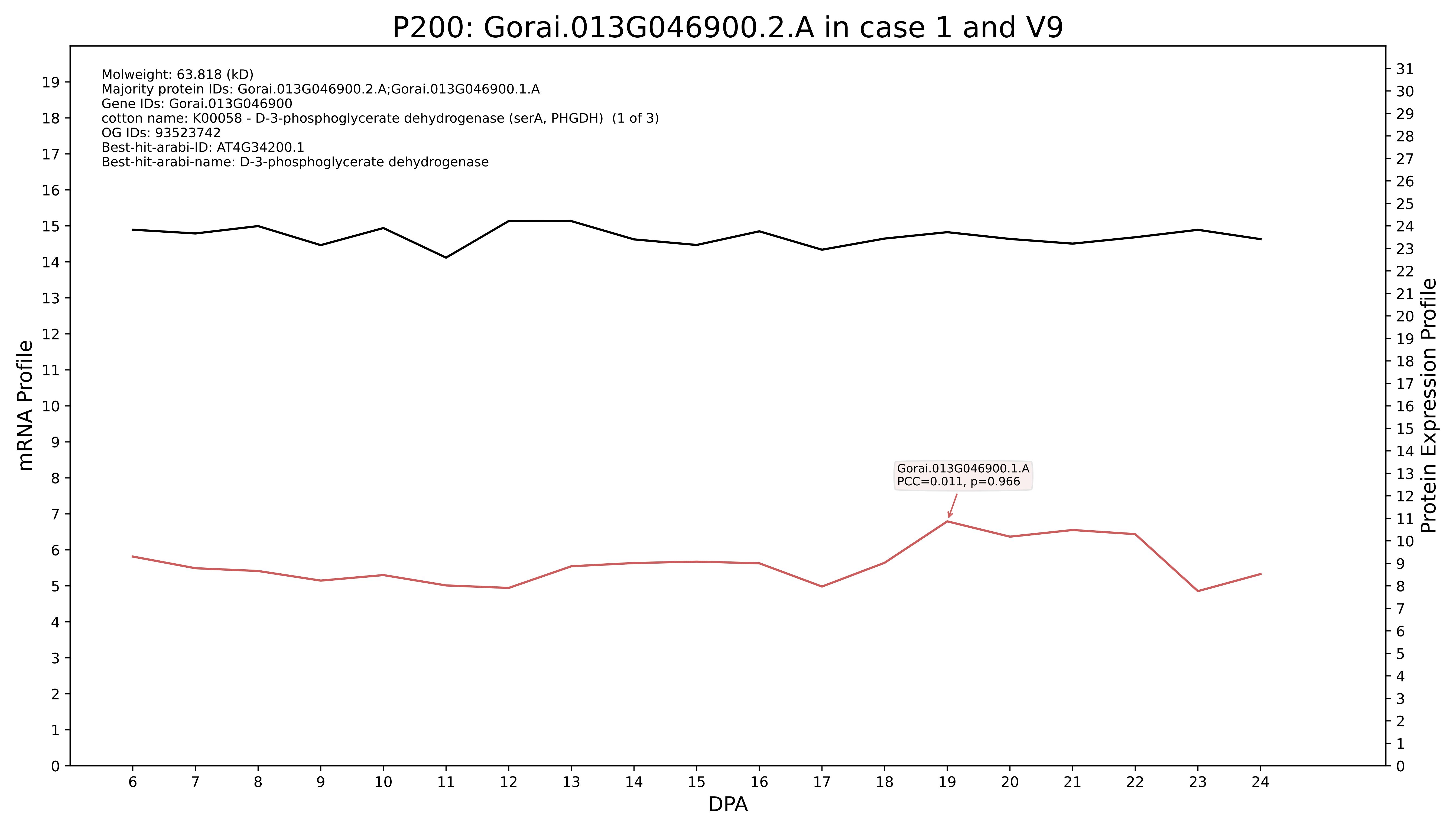
P200: Gorai.013G046900.2.A in case 1 and V9
Molweight: 63.818 (kD)
Majority protein IDs: Gorai.013G046900.2.A;Gorai.013G046900.1.A
Gene IDs: Gorai.013G046900
cotton name: K00058 - D-3-phosphoglycerate dehydrogenase (serA, PHGDH) (1 of 3)
OG IDs: 93523742
Best-hit-arabi-ID: AT4G34200.1
Best-hit-arabi-name: D-3-phosphoglycerate dehydrogenase
Gorai.013G046900.1.A
PCC=0.011, p=0.966
mRNA Profile
Protein Expression Profile
DPA

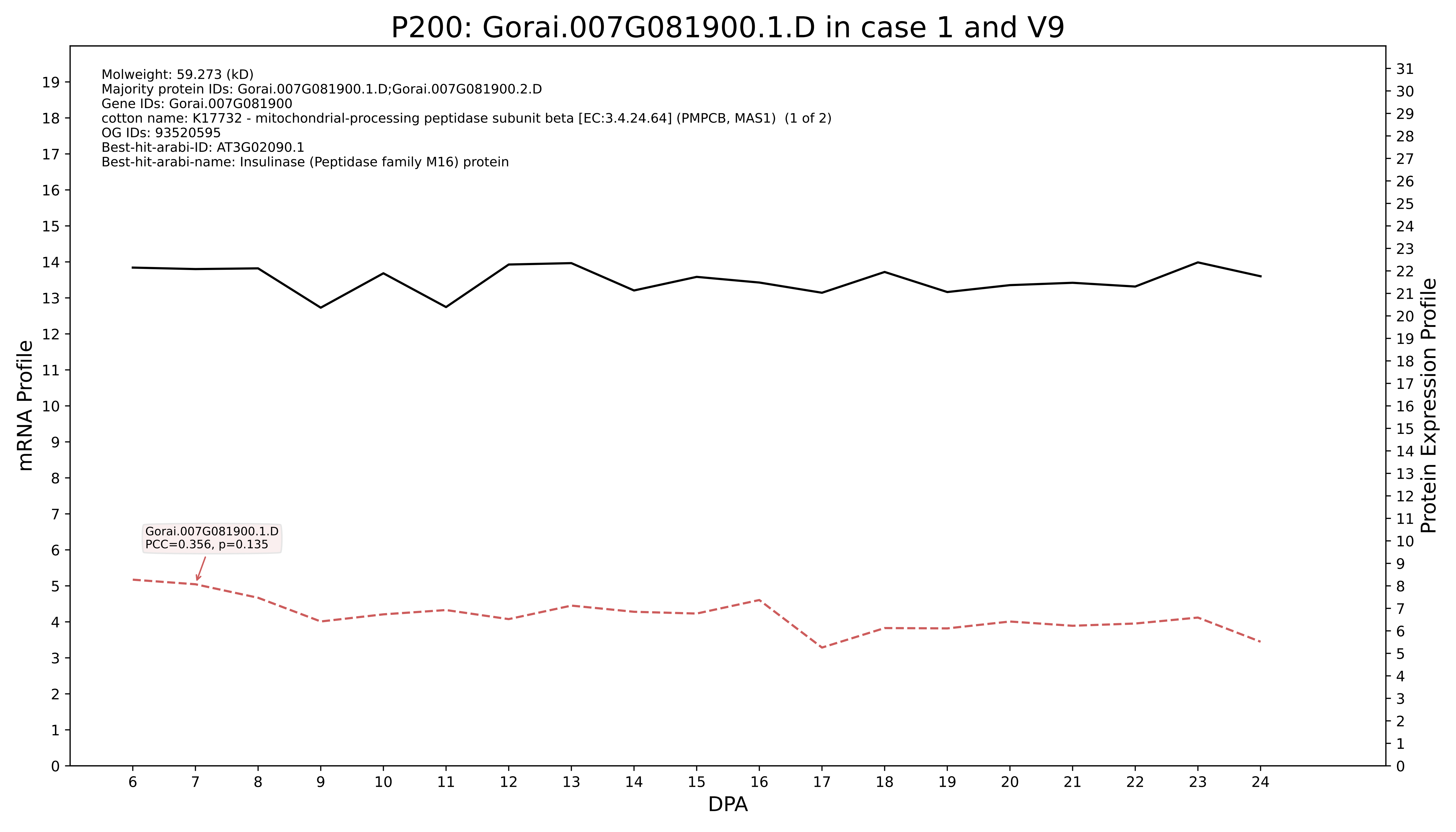
P200: Gorai.007G081900.1.D in case 1 and V9
Molweight: 59.273 (kD)
Majority protein IDs: Gorai.007G081900.1.D;Gorai.007G081900.2.D
Gene IDs: Gorai.007G081900
cotton name: K17732 - mitochondrial-processing peptidase subunit beta [EC:3.4.24.64] (PMPCB, MAS1) (1 of 2)
OG IDs: 93520595
Best-hit-arabi-ID: AT3G02090.1
Best-hit-arabi-name: Insulinase (Peptidase family M16) protein
Gorai.007G081900.1.D
PCC=0.356, p=0.135
mRNA Profile
Protein Expression Profile
DPA

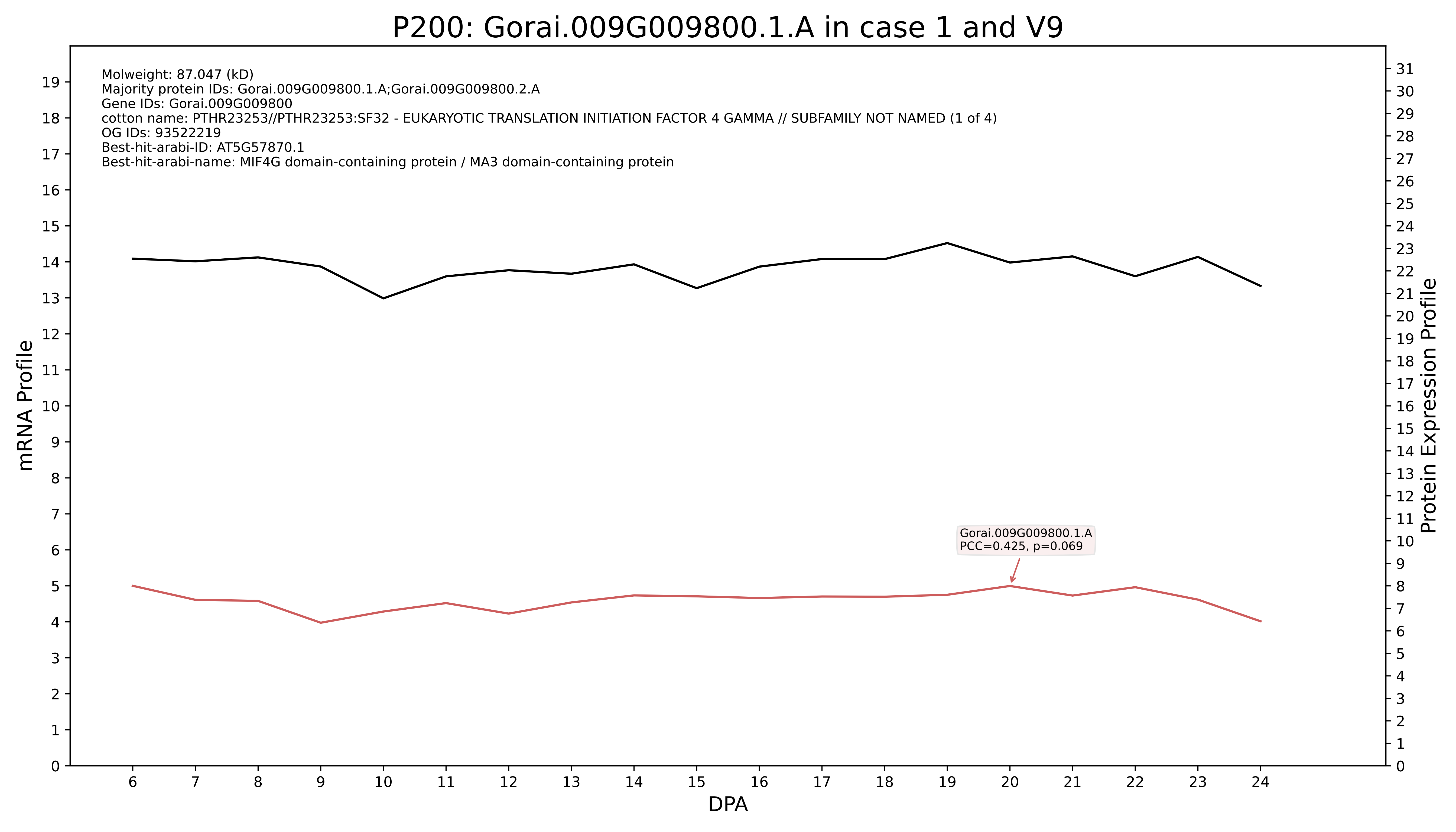
P200: Gorai.009G009800.1.A in case 1 and V9
Molweight: 87.047 (kD)
Majority protein IDs: Gorai.009G009800.1.A;Gorai.009G009800.2.A
Gene IDs: Gorai.009G009800
cotton name: PTHR23253//PTHR23253:SF32 - EUKARYOTIC TRANSLATION INITIATION FACTOR 4 GAMMA // SUBFAMILY NOT NAMED (1 of 4)
OG IDs: 93522219
Best-hit-arabi-ID: AT5G57870.1
Best-hit-arabi-name: MIF4G domain-containing protein / MA3 domain-containing protein
Gorai.009G009800.1.A
PCC=0.425, p=0.069
mRNA Profile
Protein Expression Profile
DPA

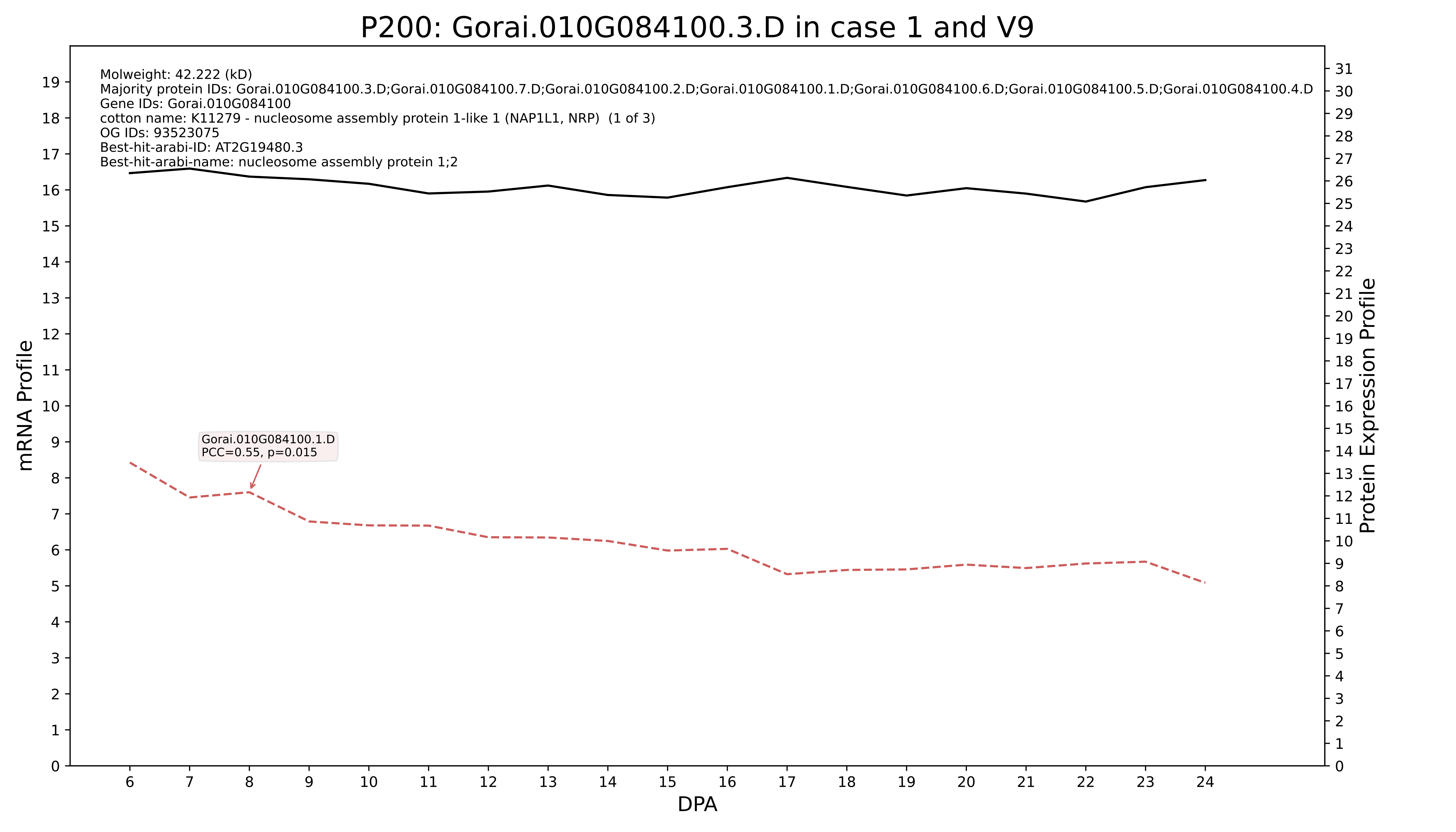
P200: Gorai.010G084100.3.D in case 1 and V9
Molweight: 42.222 (kD)
Majority protein IDs: Gorai.010G084100.3.D;Gorai.010G084100.7.D;Gorai.010G084100.2.D;Gorai.010G084100.1.D;Gorai.010G084100.6.D;Gorai.010G084100.5.D;Gorai.010G084100.4.D
Gene IDs: Gorai.010G084100
cotton name: K11279 - nucleosome assembly protein 1-like 1 (NAP1L1, NRP) (1 of 3)
OG IDs: 93523075
Best-hit-arabi-ID: AT2G19480.3
Best-hit-arabi-name: nucleosome assembly protein 1;2
Gorai.010G084100.1.D
PCC=0.55, p=0.015
mRNA Profile
Protein Expression Profile
DPA

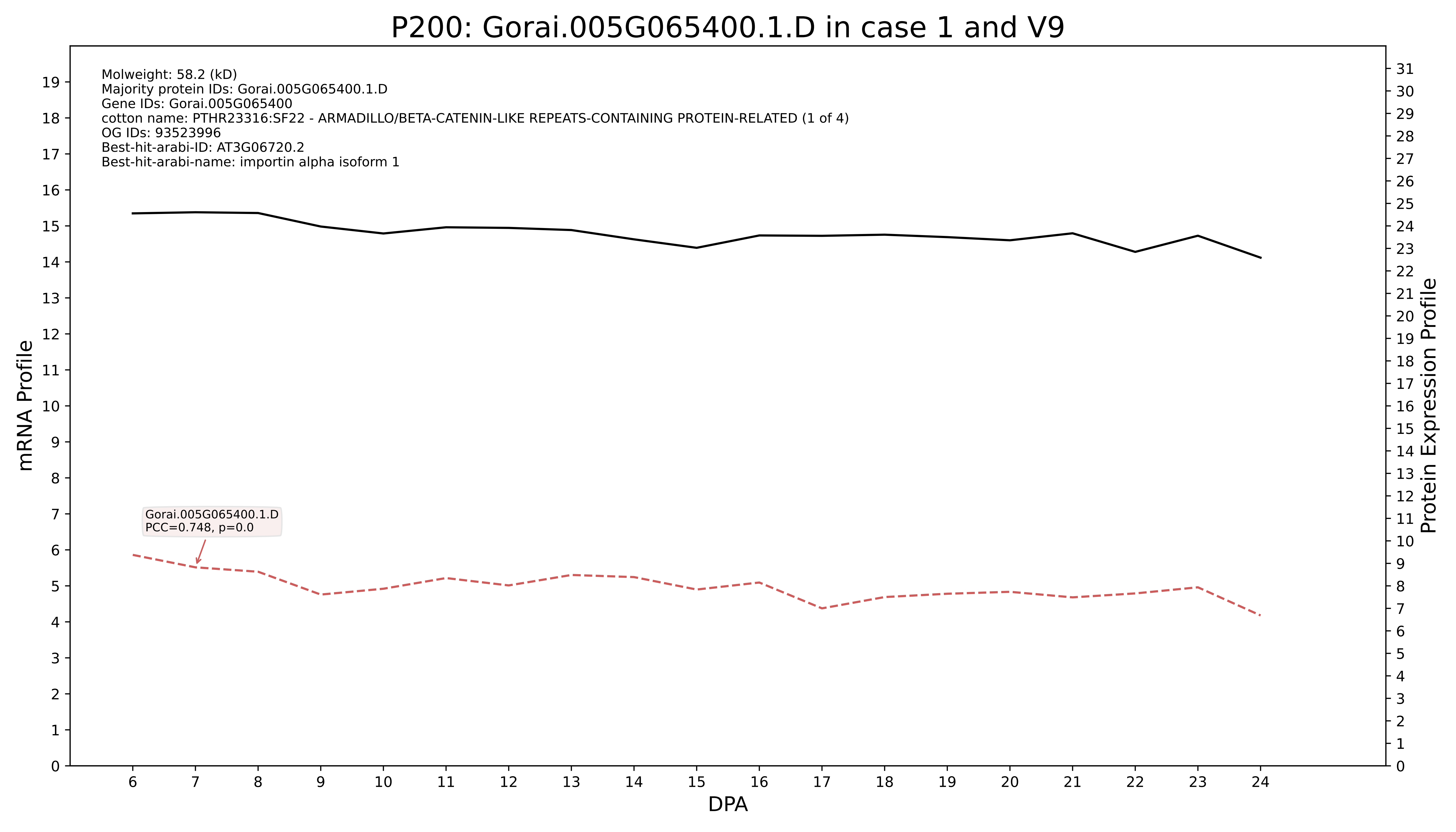
P200: Gorai.005G065400.1.D in case 1 and V9
Molweight: 58.2 (kD)
Majority protein IDs: Gorai.005G065400.1.D
Gene IDs: Gorai.005G065400
cotton name: PTHR23316:SF22 - ARMADILLO/BETA-CATENIN-LIKE REPEATS-CONTAINING PROTEIN-RELATED (1 of 4)
OG IDs: 93523996
Best-hit-arabi-ID: AT3G06720.2
Best-hit-arabi-name: importin alpha isoform 1
Gorai.005G065400.1.D
PCC=0.748, p=0.0
mRNA Profile
Protein Expression Profile
DPA

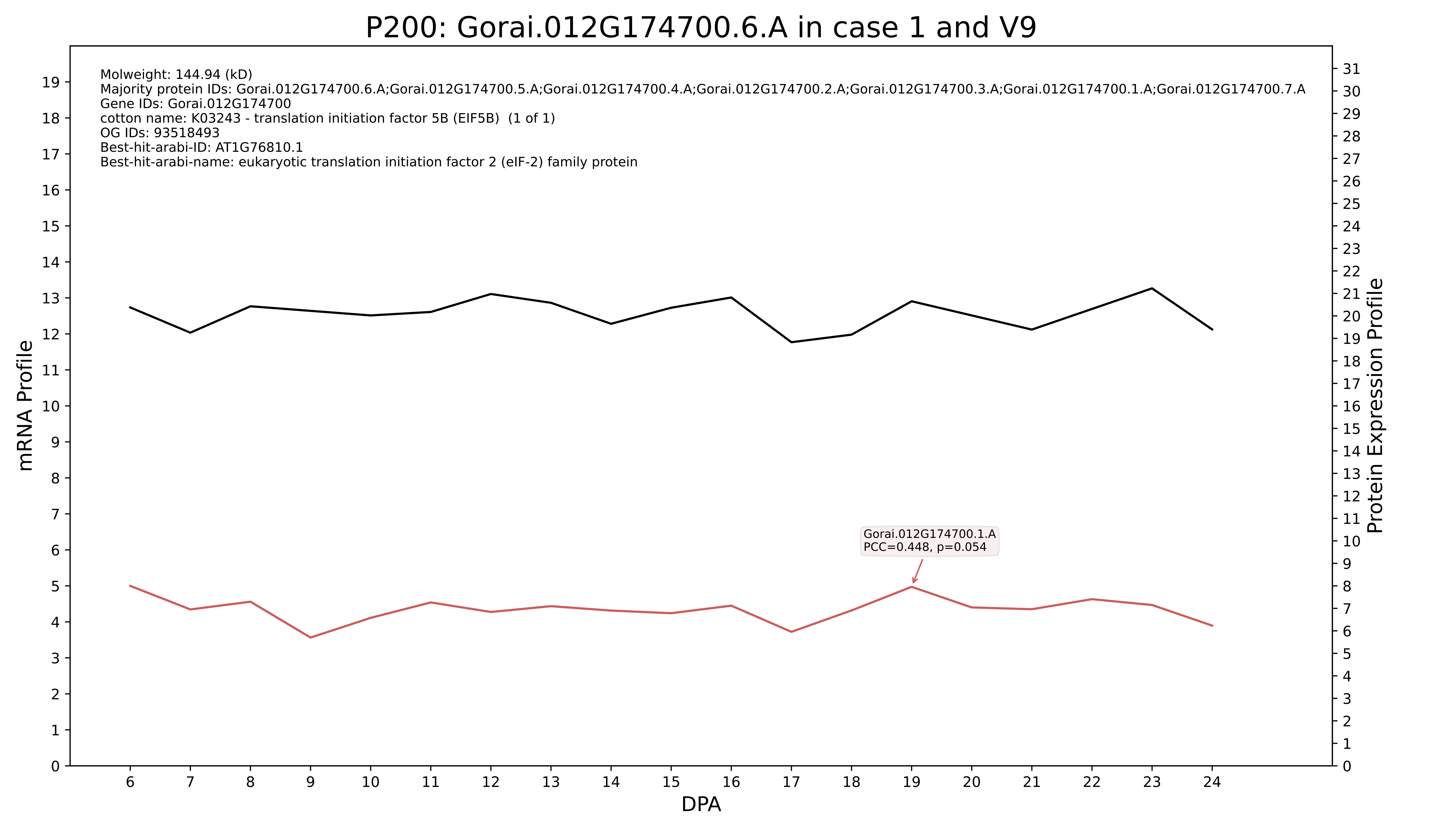
P200: Gorai.012G174700.6.A in case 1 and V9
Molweight: 144.94 (kD)
Majority protein IDs: Gorai.012G174700.6.A;Gorai.012G174700.5.A;Gorai.012G174700.4.A;Gorai.012G174700.2.A;Gorai.012G174700.3.A;Gorai.012G174700.1.A;Gorai.012G174700.7.A
Gene IDs: Gorai.012G174700
cotton name: K03243 - translation initiation factor 5B (EIF5B) (1 of 1)
OG IDs: 93518493
Best-hit-arabi-ID: AT1G76810.1
Best-hit-arabi-name: eukaryotic translation initiation factor 2 (eIF-2) family protein
Gorai.012G174700.1.A
PCC=0.448, p=0.054
mRNA Profile
Protein Expression Profile
DPA

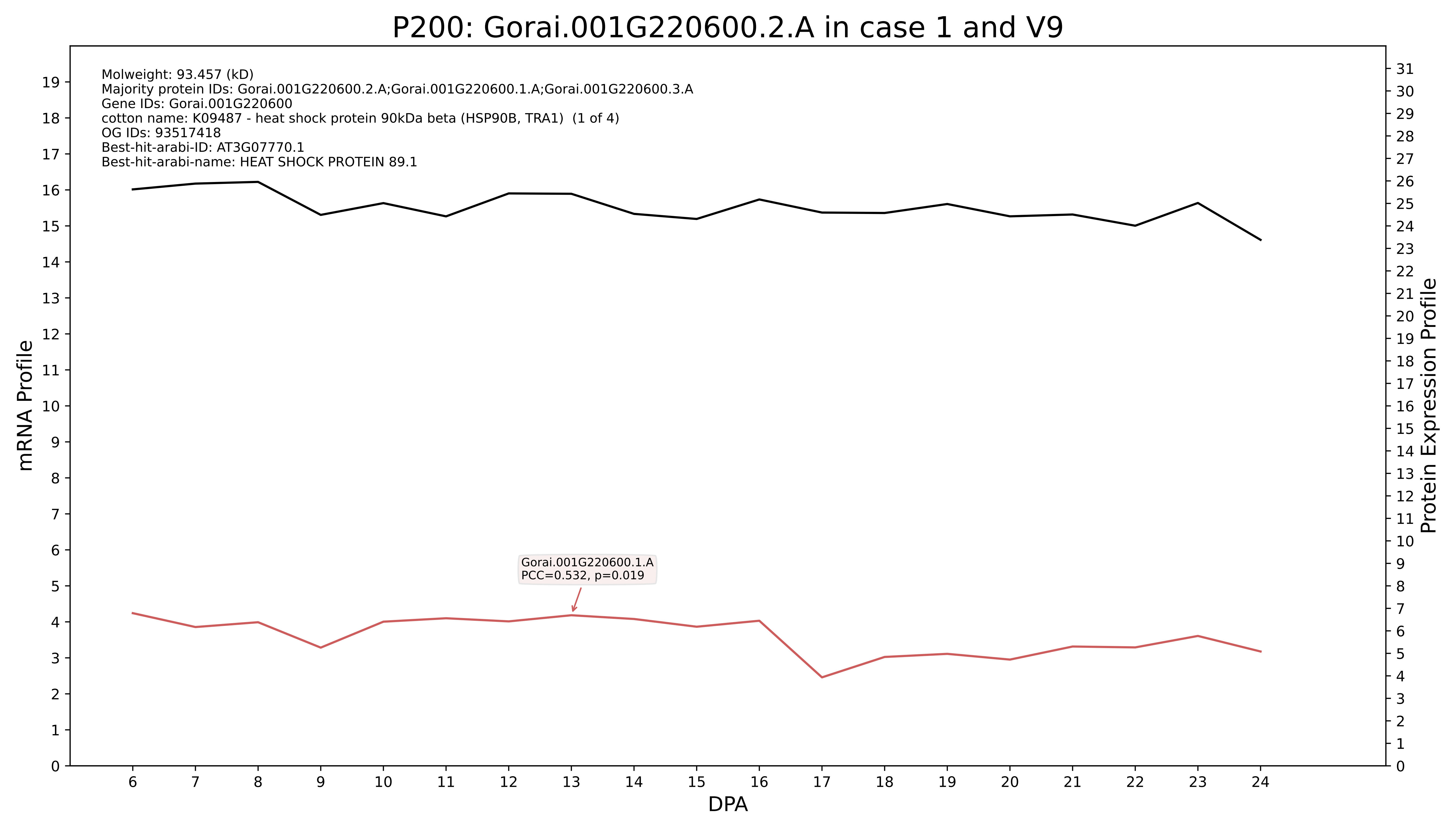
P200: Gorai.001G220600.2.A in case 1 and V9
Molweight: 93.457 (kD)
Majority protein IDs: Gorai.001G220600.2.A;Gorai.001G220600.1.A;Gorai.001G220600.3.A
Gene IDs: Gorai.001G220600
cotton name: K09487 - heat shock protein 90kDa beta (HSP90B, TRA1) (1 of 4)
OG IDs: 93517418
Best-hit-arabi-ID: AT3G07770.1
Best-hit-arabi-name: HEAT SHOCK PROTEIN 89.1
Gorai.001G220600.1.A
PCC=0.532, p=0.019
mRNA Profile
Protein Expression Profile
DPA

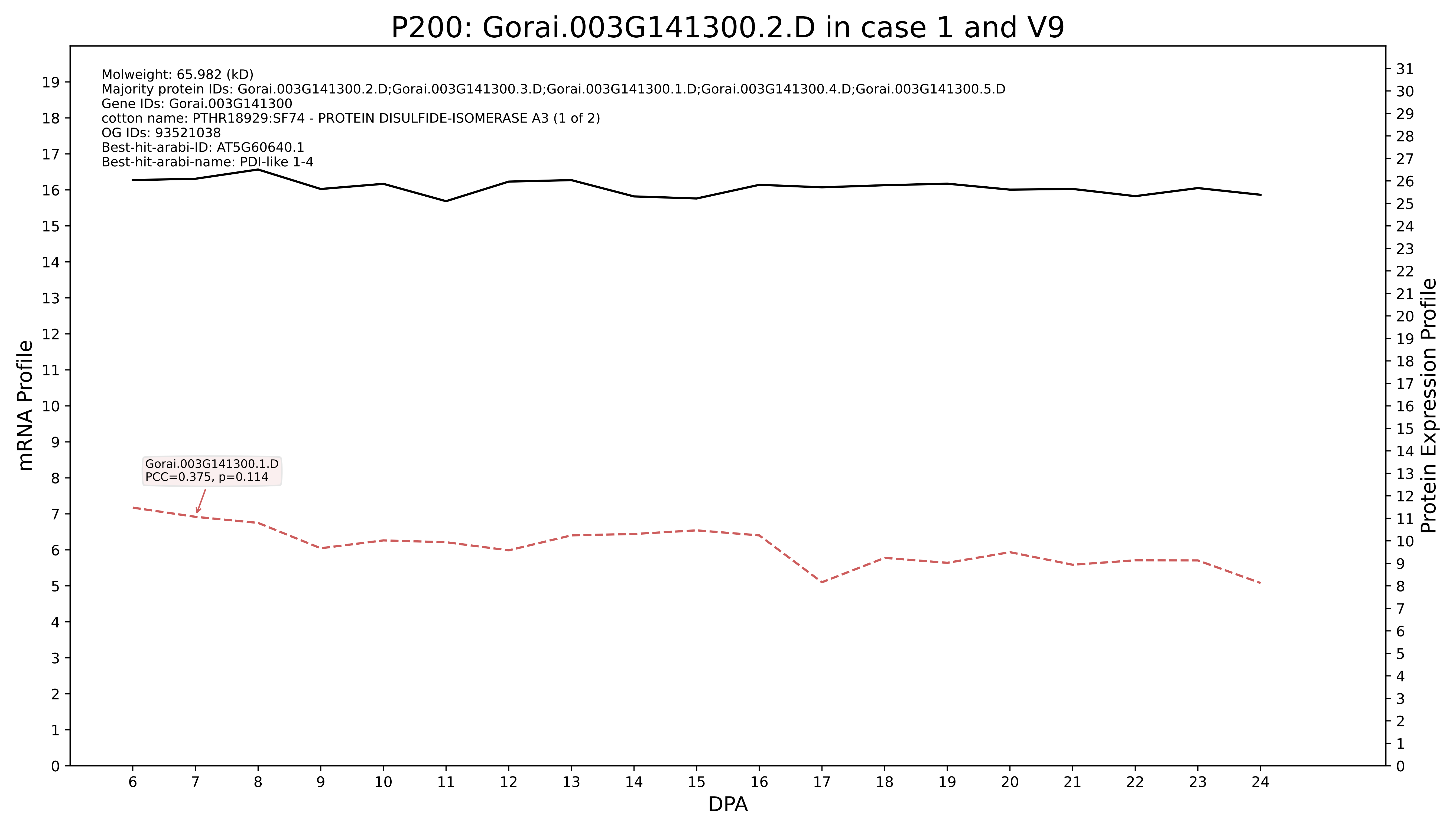

P200: Gorai.003G141300.2.D in case 1 and V9
Molweight: 65.982 (kD)
Majority protein IDs: Gorai.003G141300.2.D;Gorai.003G141300.3.D;Gorai.003G141300.1.D;Gorai.003G141300.4.D;Gorai.003G141300.5.D
Gene IDs: Gorai.003G141300
cotton name: PTHR18929:SF74 - PROTEIN DISULFIDE-ISOMERASE A3 (1 of 2)
OG IDs: 93521038
Best-hit-arabi-ID: AT5G60640.1
Best-hit-arabi-name: PDI-like 1-4
Gorai.003G141300.1.D
PCC=0.375, p=0.114
mRNA Profile
Protein Expression Profile
DPA

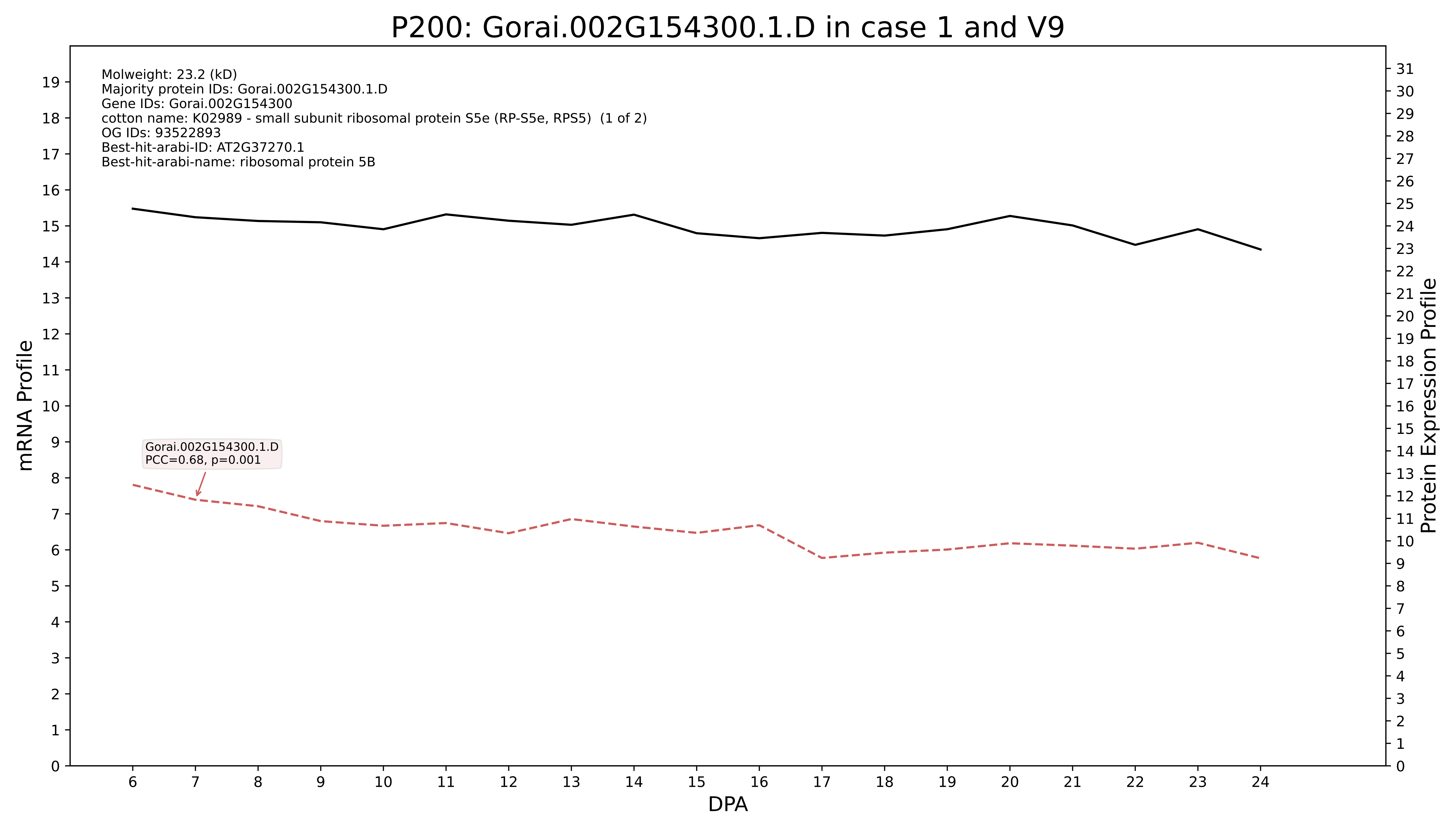
P200: Gorai.002G154300.1.D in case 1 and V9
Molweight: 23.2 (kD)
Majority protein IDs: Gorai.002G154300.1.D
Gene IDs: Gorai.002G154300
cotton name: K02989 - small subunit ribosomal protein S5e (RP-S5e, RPS5) (1 of 2)
OG IDs: 93522893
Best-hit-arabi-ID: AT2G37270.1
Best-hit-arabi-name: ribosomal protein 5B
Gorai.002G154300.1.D
PCC=0.68, p=0.001
mRNA Profile
Protein Expression Profile
DPA

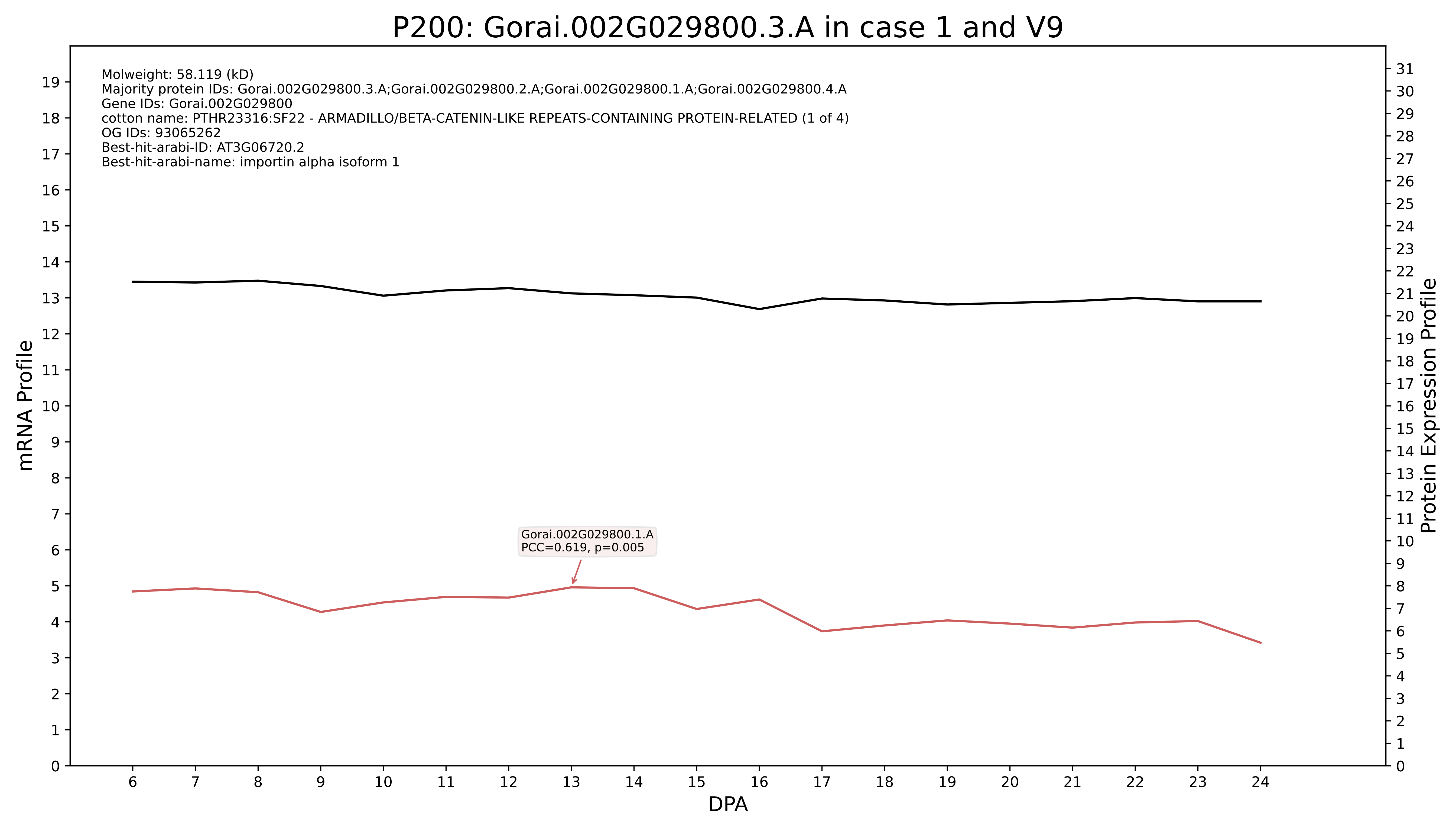
P200: Gorai.002G029800.3.A in case 1 and V9
Molweight: 58.119 (kD)
Majority protein IDs: Gorai.002G029800.3.A;Gorai.002G029800.2.A;Gorai.002G029800.1.A;Gorai.002G029800.4.A
Gene IDs: Gorai.002G029800
cotton name: PTHR23316:SF22 - ARMADILLO/BETA-CATENIN-LIKE REPEATS-CONTAINING PROTEIN-RELATED (1 of 4)
OG IDs: 93065262
Best-hit-arabi-ID: AT3G06720.2
Best-hit-arabi-name: importin alpha isoform 1
Gorai.002G029800.1.A
PCC=0.619, p=0.005
mRNA Profile
Protein Expression Profile
DPA

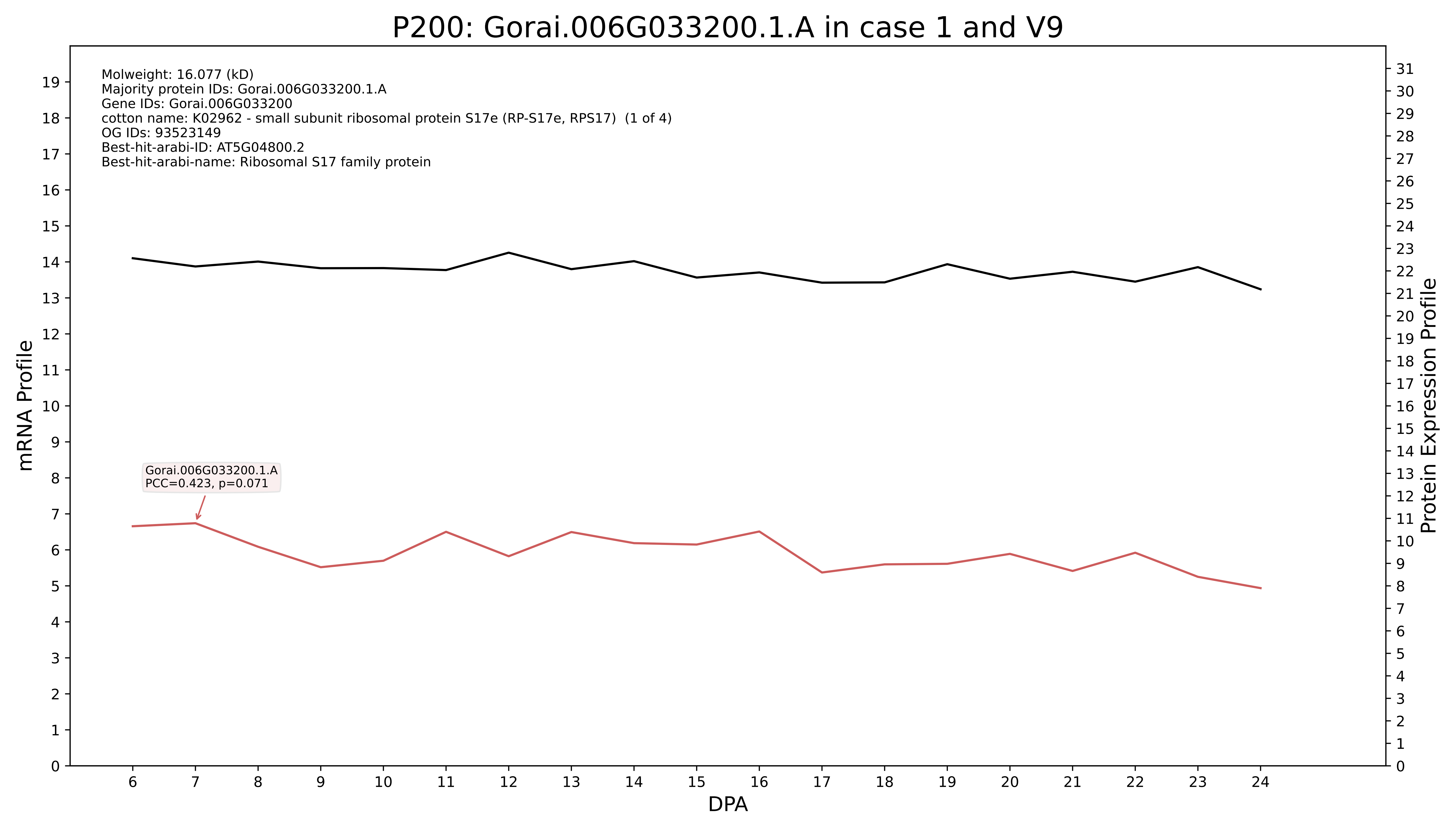
P200: Gorai.006G033200.1.A in case 1 and V9
Molweight: 16.077 (kD)
Majority protein IDs: Gorai.006G033200.1.A
Gene IDs: Gorai.006G033200
cotton name: K02962 - small subunit ribosomal protein S17e (RP-S17e, RPS17) (1 of 4)
OG IDs: 93523149
Best-hit-arabi-ID: AT5G04800.2
Best-hit-arabi-name: Ribosomal S17 family protein
Gorai.006G033200.1.A
PCC=0.423, p=0.071
mRNA Profile
Protein Expression Profile
DPA

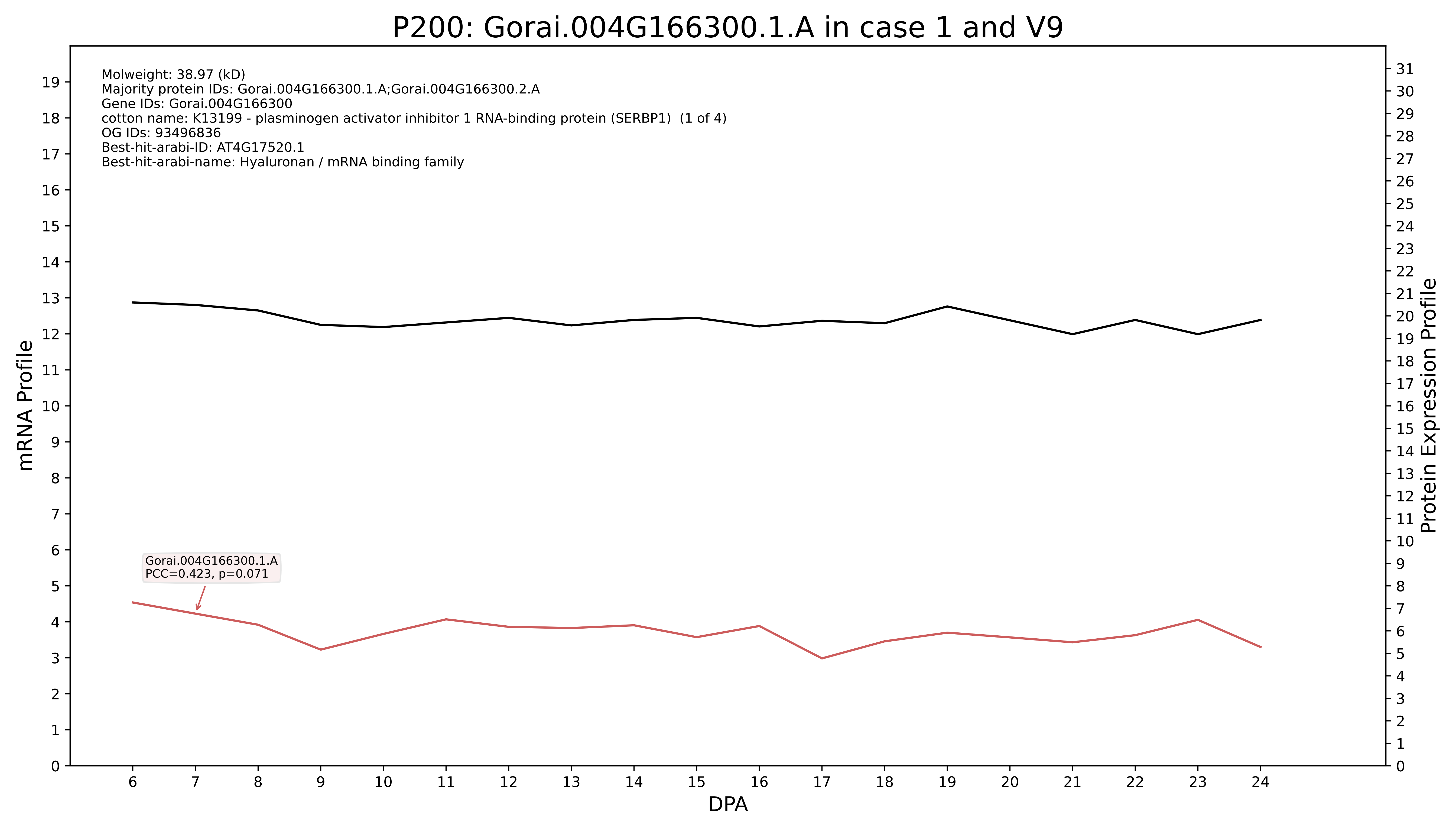

P200: Gorai.004G166300.1.A in case 1 and V9
Molweight: 38.97 (kD)
Majority protein IDs: Gorai.004G166300.1.A;Gorai.004G166300.2.A
Gene IDs: Gorai.004G166300
cotton name: K13199 - plasminogen activator inhibitor 1 RNA-binding protein (SERBP1) (1 of 4)
OG IDs: 93496836
Best-hit-arabi-ID: AT4G17520.1
Best-hit-arabi-name: Hyaluronan / mRNA binding family
Gorai.004G166300.1.A
PCC=0.423, p=0.071
mRNA Profile
Protein Expression Profile
DPA

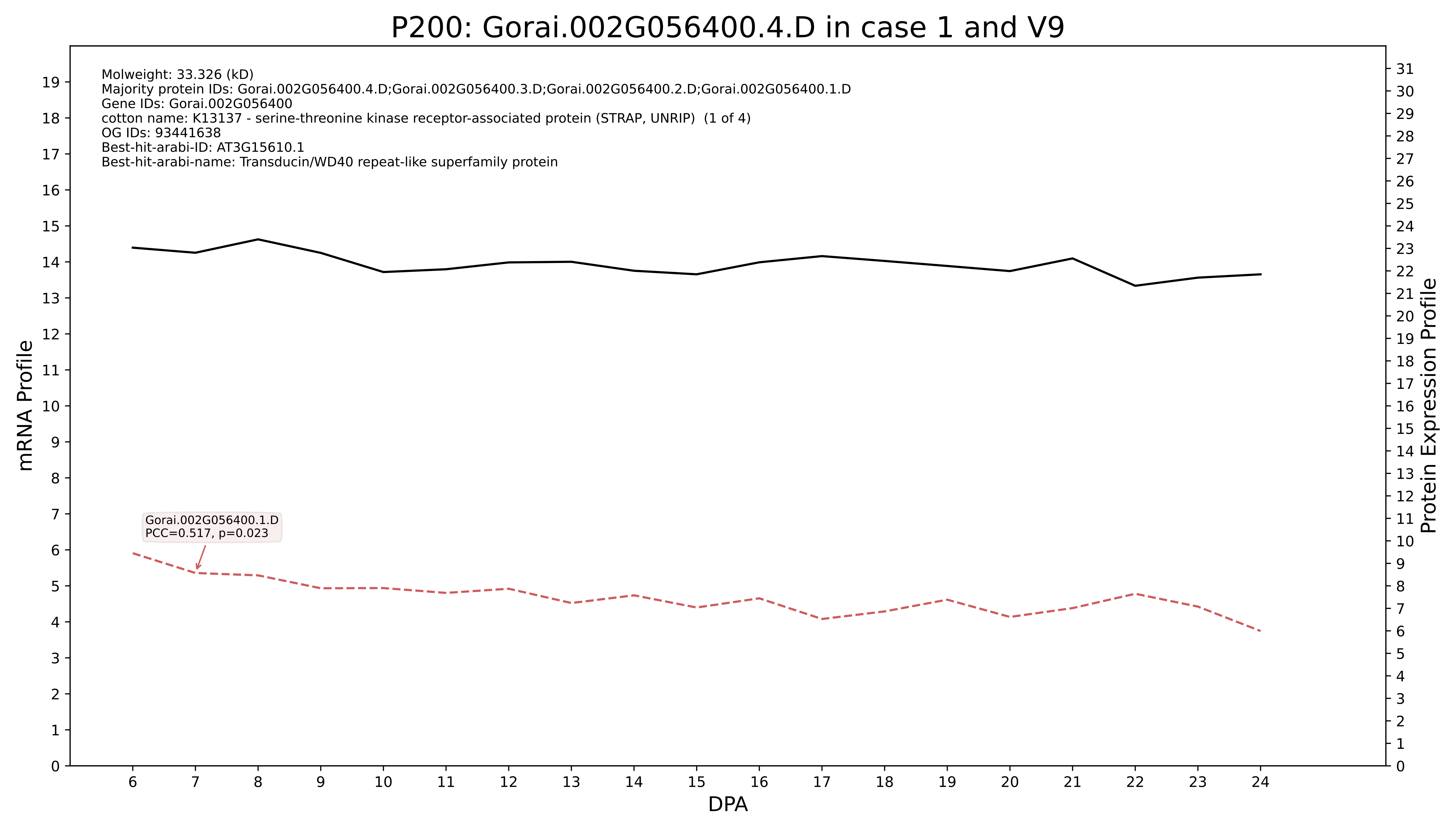

P200: Gorai.002G056400.4.D in case 1 and V9
Molweight: 33.326 (kD)
Majority protein IDs: Gorai.002G056400.4.D;Gorai.002G056400.3.D;Gorai.002G056400.2.D;Gorai.002G056400.1.D
Gene IDs: Gorai.002G056400
cotton name: K13137 - serine-threonine kinase receptor-associated protein (STRAP, UNRIP) (1 of 4)
OG IDs: 93441638
Best-hit-arabi-ID: AT3G15610.1
Best-hit-arabi-name: Transducin/WD40 repeat-like superfamily protein
Gorai.002G056400.1.D
PCC=0.517, p=0.023
mRNA Profile
Protein Expression Profile
DPA

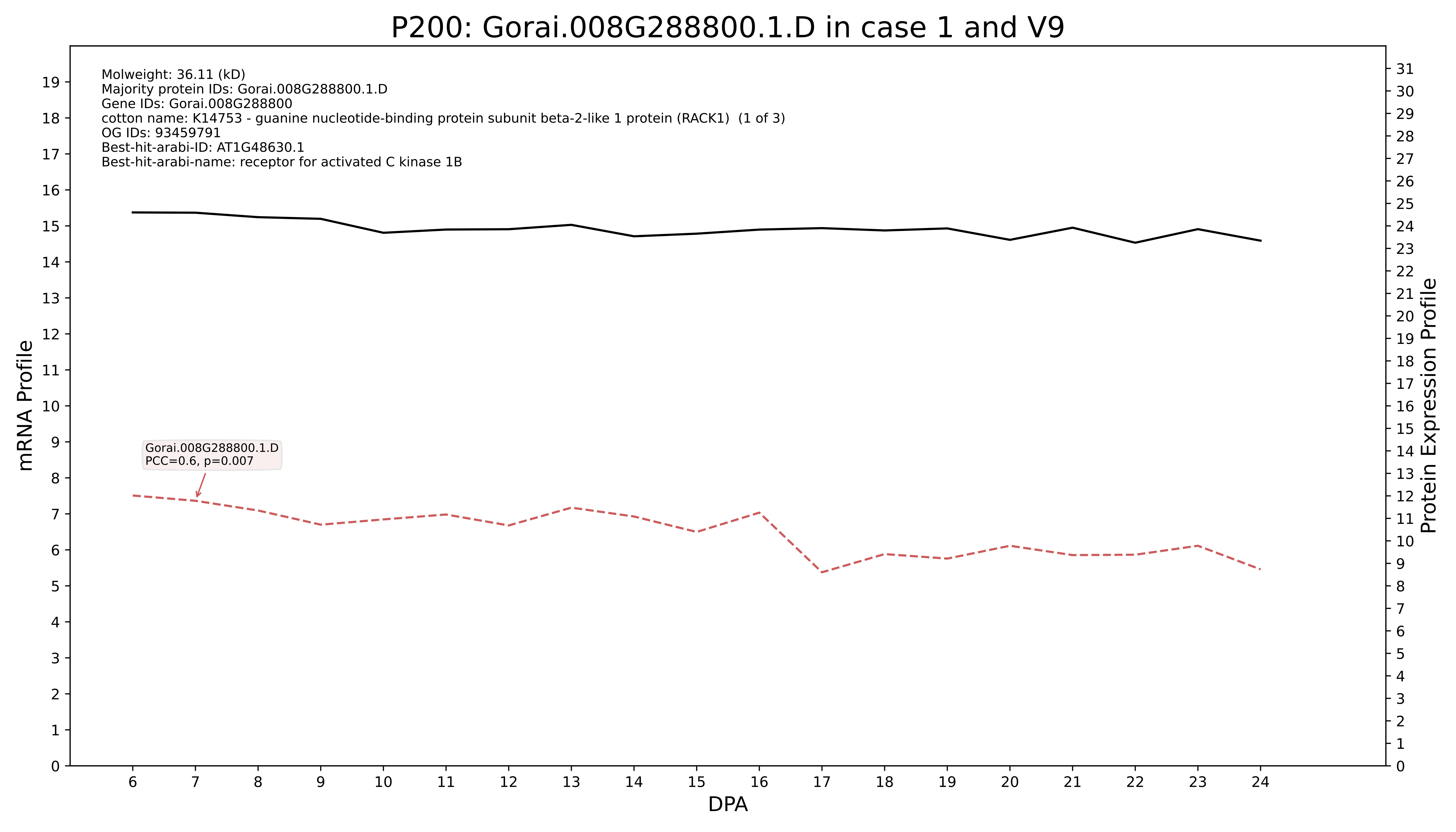

P200: Gorai.008G288800.1.D in case 1 and V9
Molweight: 36.11 (kD)
Majority protein IDs: Gorai.008G288800.1.D
Gene IDs: Gorai.008G288800
cotton name: K14753 - guanine nucleotide-binding protein subunit beta-2-like 1 protein (RACK1) (1 of 3)
OG IDs: 93459791
Best-hit-arabi-ID: AT1G48630.1
Best-hit-arabi-name: receptor for activated C kinase 1B
Gorai.008G288800.1.D
PCC=0.6, p=0.007
mRNA Profile
Protein Expression Profile
DPA

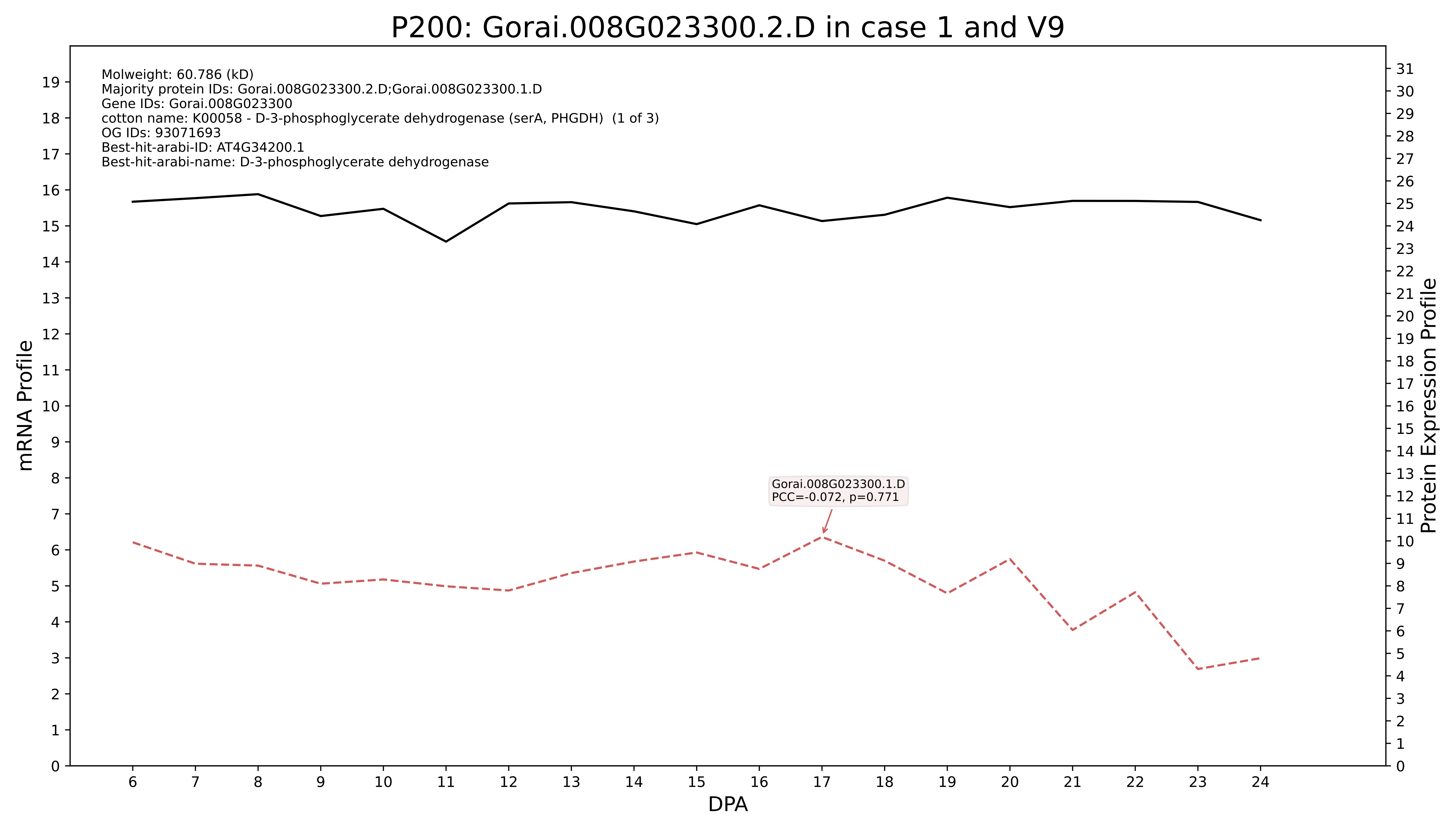

P200: Gorai.008G023300.2.D in case 1 and V9
Molweight: 60.786 (kD)
Majority protein IDs: Gorai.008G023300.2.D;Gorai.008G023300.1.D
Gene IDs: Gorai.008G023300
cotton name: K00058 - D-3-phosphoglycerate dehydrogenase (serA, PHGDH) (1 of 3)
OG IDs: 93071693
Best-hit-arabi-ID: AT4G34200.1
Best-hit-arabi-name: D-3-phosphoglycerate dehydrogenase
Gorai.008G023300.1.D
PCC=-0.072, p=0.771
mRNA Profile
Protein Expression Profile
DPA

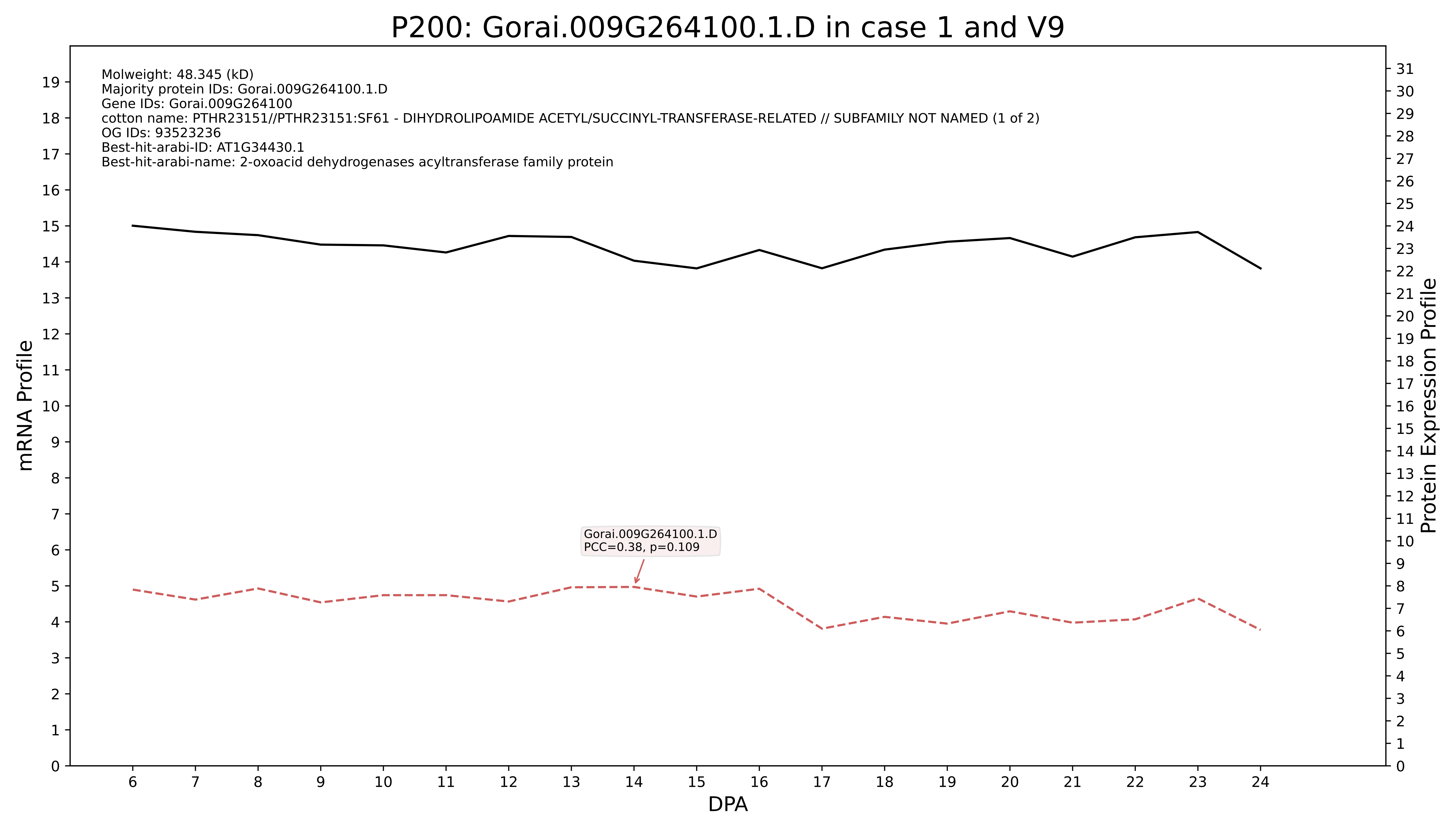
P200: Gorai.009G264100.1.D in case 1 and V9
Molweight: 48.345 (kD)
Majority protein IDs: Gorai.009G264100.1.D
Gene IDs: Gorai.009G264100
cotton name: PTHR23151//PTHR23151:SF61 - DIHYDROLIPOAMIDE ACETYL/SUCCINYL-TRANSFERASE-RELATED // SUBFAMILY NOT NAMED (1 of 2)
OG IDs: 93523236
Best-hit-arabi-ID: AT1G34430.1
Best-hit-arabi-name: 2-oxoacid dehydrogenases acyltransferase family protein
Gorai.009G264100.1.D
PCC=0.38, p=0.109
mRNA Profile
Protein Expression Profile
DPA

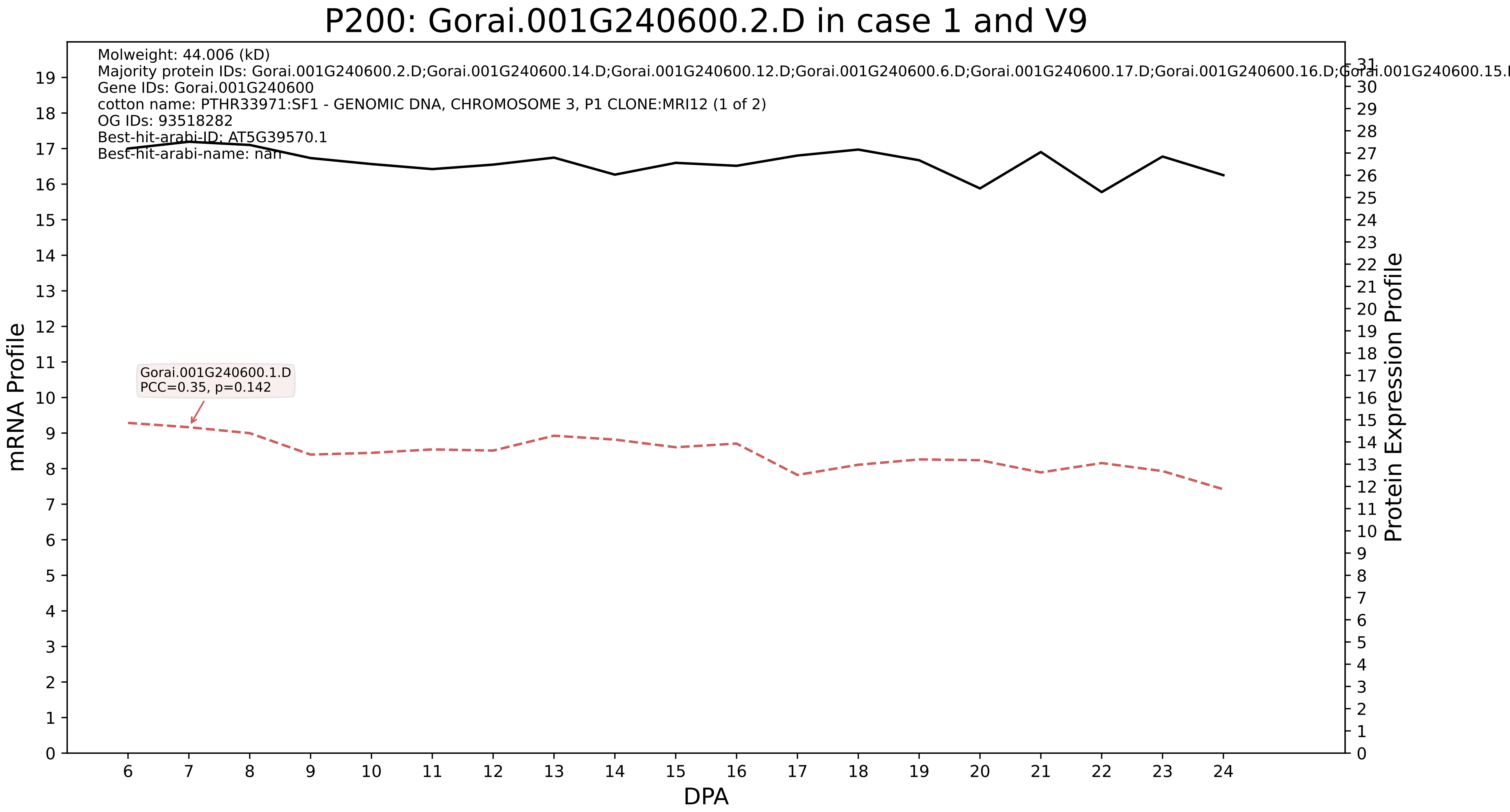

P200: Gorai.001G240600.2.D in case 1 and V9
Molweight: 44.006 (kD)
Majority protein IDs: Gorai.001G240600.2.D;Gorai.001G240600.14.D;Gorai.001G240600.12.D;Gorai.001G240600.6.D;Gorai.001G240600.17.D;Gorai.001G240600.16.D;Gorai.001G240600.15.
Gene IDs: Gorai.001G240600
cotton name: PTHR33971:SF1 - GENOMIC DNA, CHROMOSOME 3, P1 CLONE:MRI12 (1 of 2)
OG IDs: 93518282
Best-hit-arabi-ID: AT5G39570.1
Best-hit-arabi-name: nan
Gorai.001G240600.1.D
PCC=0.35, p=0.142
mRNA Profile
Protein Expression Profile
DPA

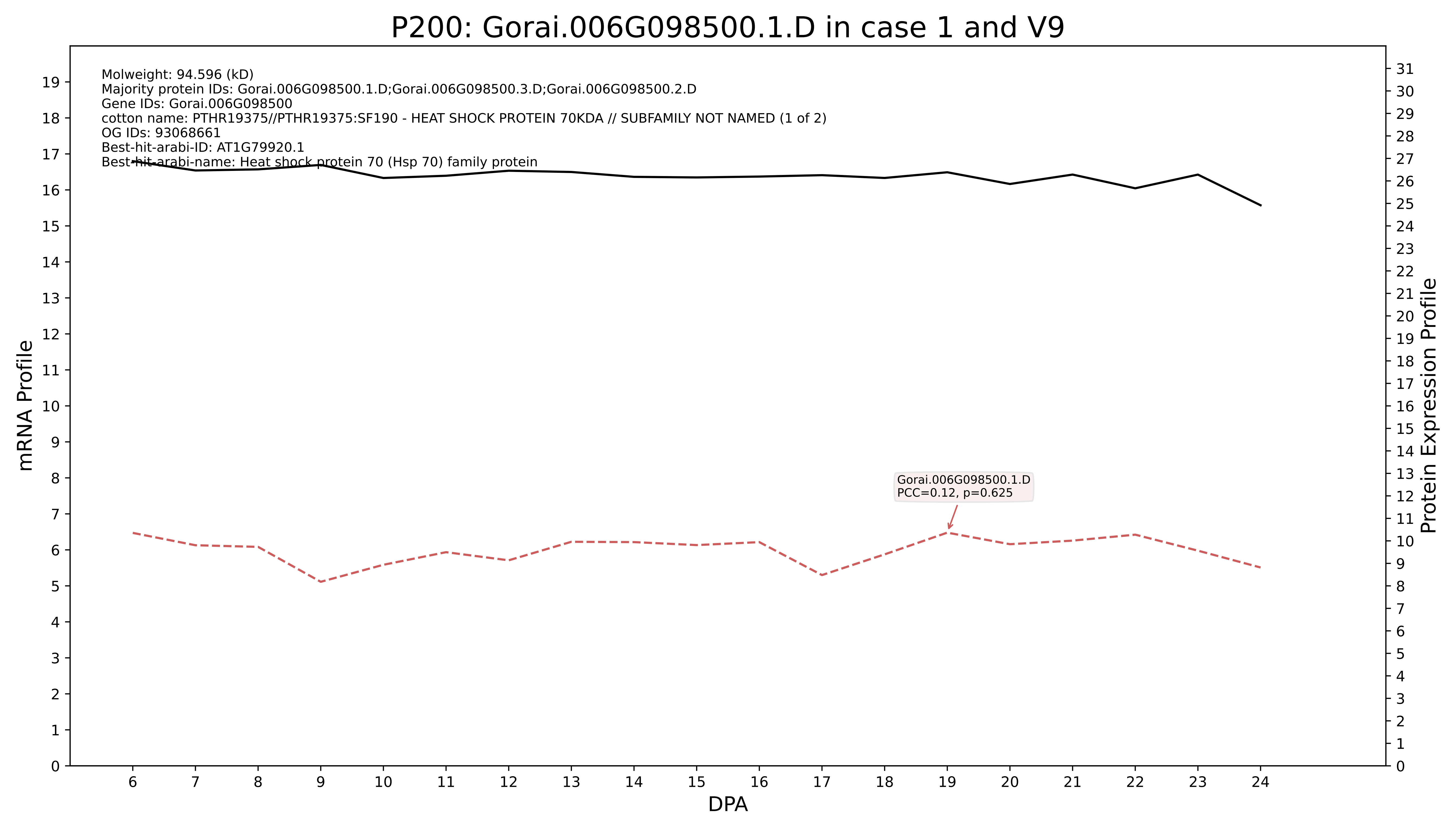
P200: Gorai.006G098500.1.D in case 1 and V9
Molweight: 94.596 (kD)
Majority protein IDs: Gorai.006G098500.1.D;Gorai.006G098500.3.D;Gorai.006G098500.2.D
Gene IDs: Gorai.006G098500
cotton name: PTHR19375//PTHR19375:SF190 - HEAT SHOCK PROTEIN 70KDA // SUBFAMILY NOT NAMED (1 of 2)
OG IDs: 93068661
Best-hit-arabi-ID: AT1G79920.1
Best-hit-arabi-name: Heat shock protein 70 (Hsp 70) family protein
Gorai.006G098500.1.D
PCC=0.12, p=0.625
mRNA Profile
Protein Expression Profile
DPA

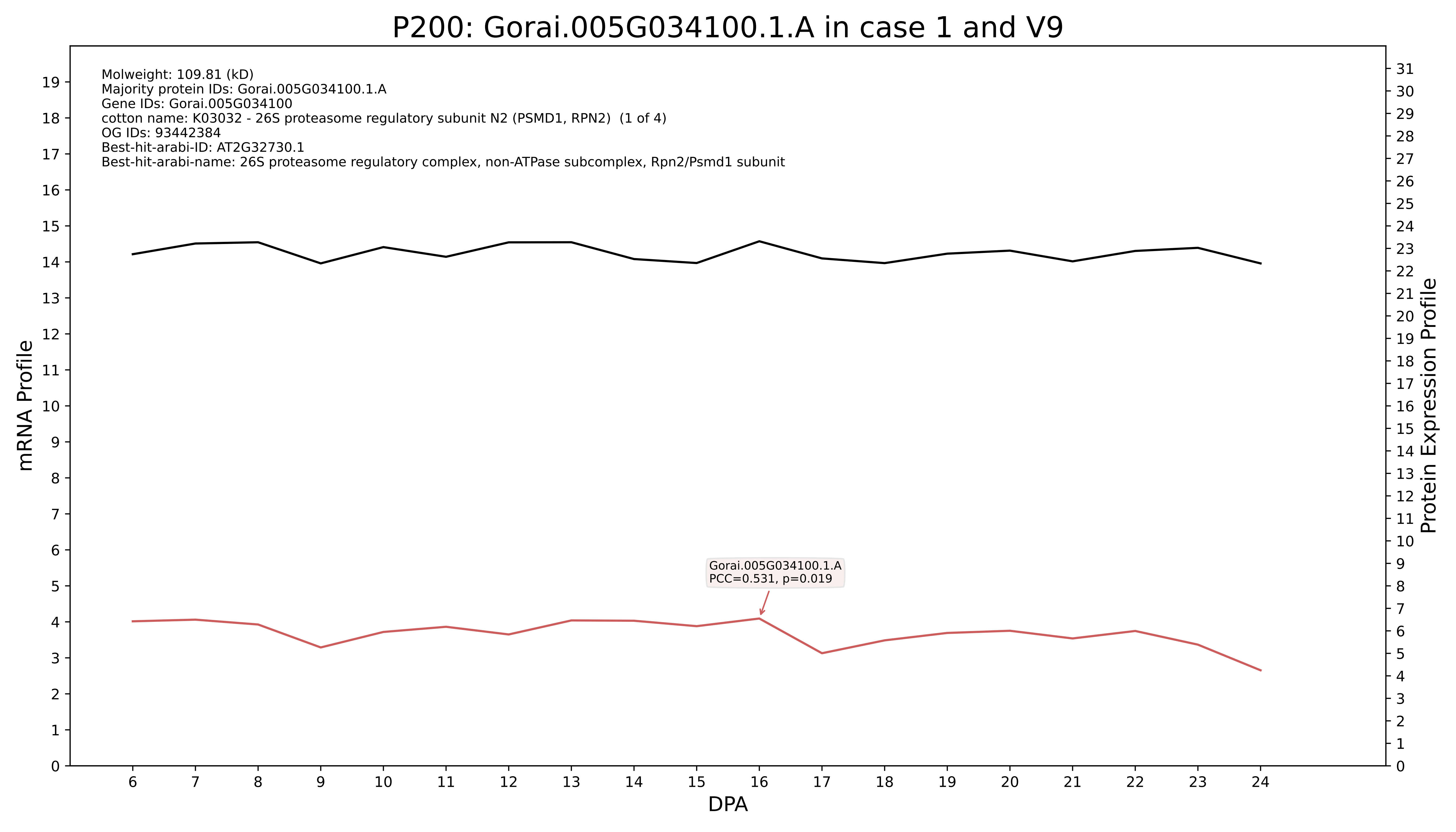

P200: Gorai.005G034100.1.A in case 1 and V9
Molweight: 109.81 (kD)
Majority protein IDs: Gorai.005G034100.1.A
Gene IDs: Gorai.005G034100
cotton name: K03032 - 26S proteasome regulatory subunit N2 (PSMD1, RPN2) (1 of 4)
OG IDs: 93442384
Best-hit-arabi-ID: AT2G32730.1
Best-hit-arabi-name: 26S proteasome regulatory complex, non-ATPase subcomplex, Rpn2/Psmd1 subunit
Gorai.005G034100.1.A
PCC=0.531, p=0.019
mRNA Profile
Protein Expression Profile
DPA
6 7 8 9 10 11 12 13 14 15 16 17 18 19 20 21 22 23 24
0 1 2 3 4 5 6 7 8 9 10 11 12 13 14 15 16 17 18 19
0 1 2 3 4 5 6 7 8 9 10 11 12 13 14 15 16 17 18 19 20 21 22 23 24 25 26 27 28 29 30 31

### P200: Gorai.007G336900.1.D in case 1 and V9

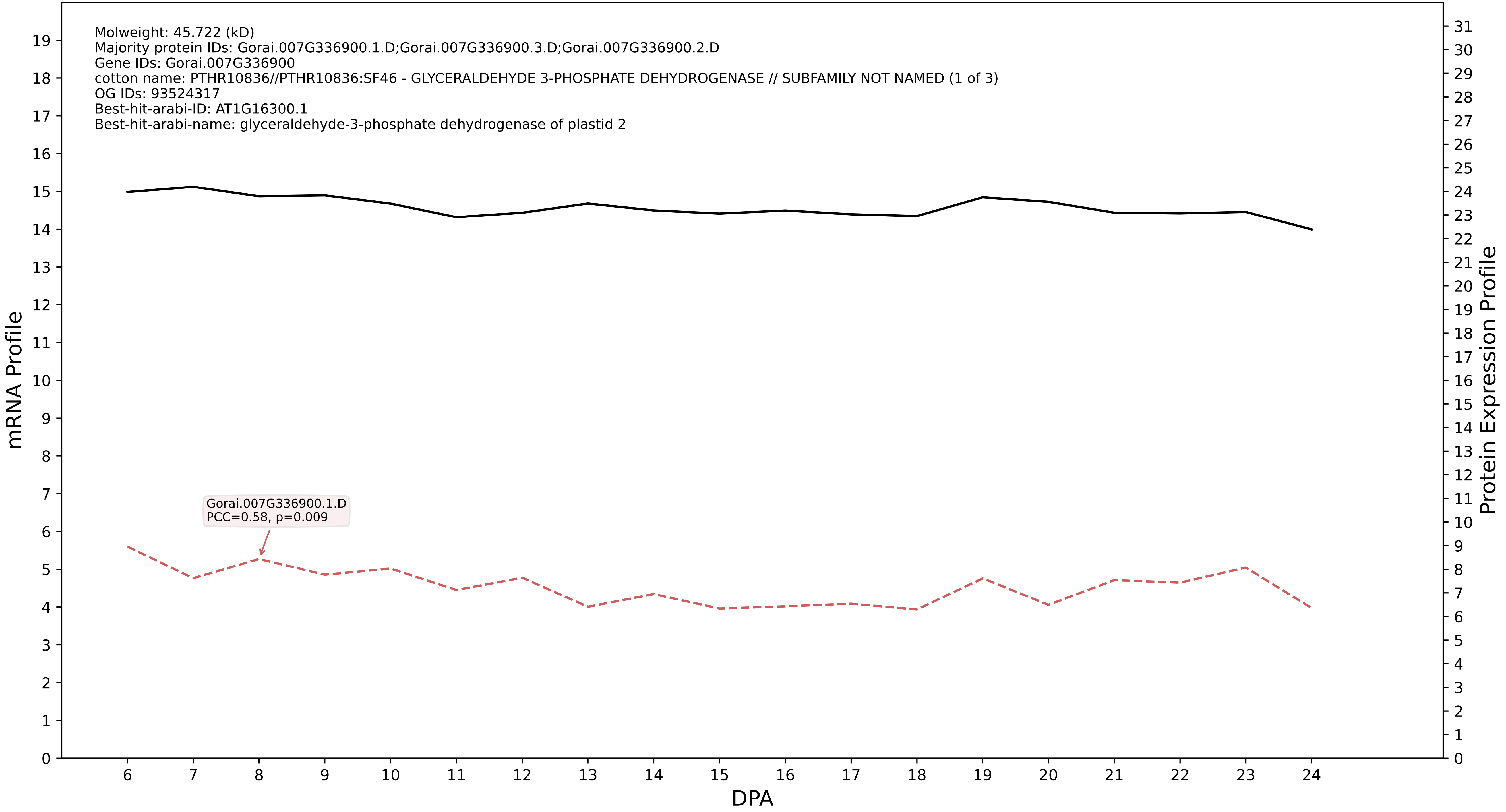

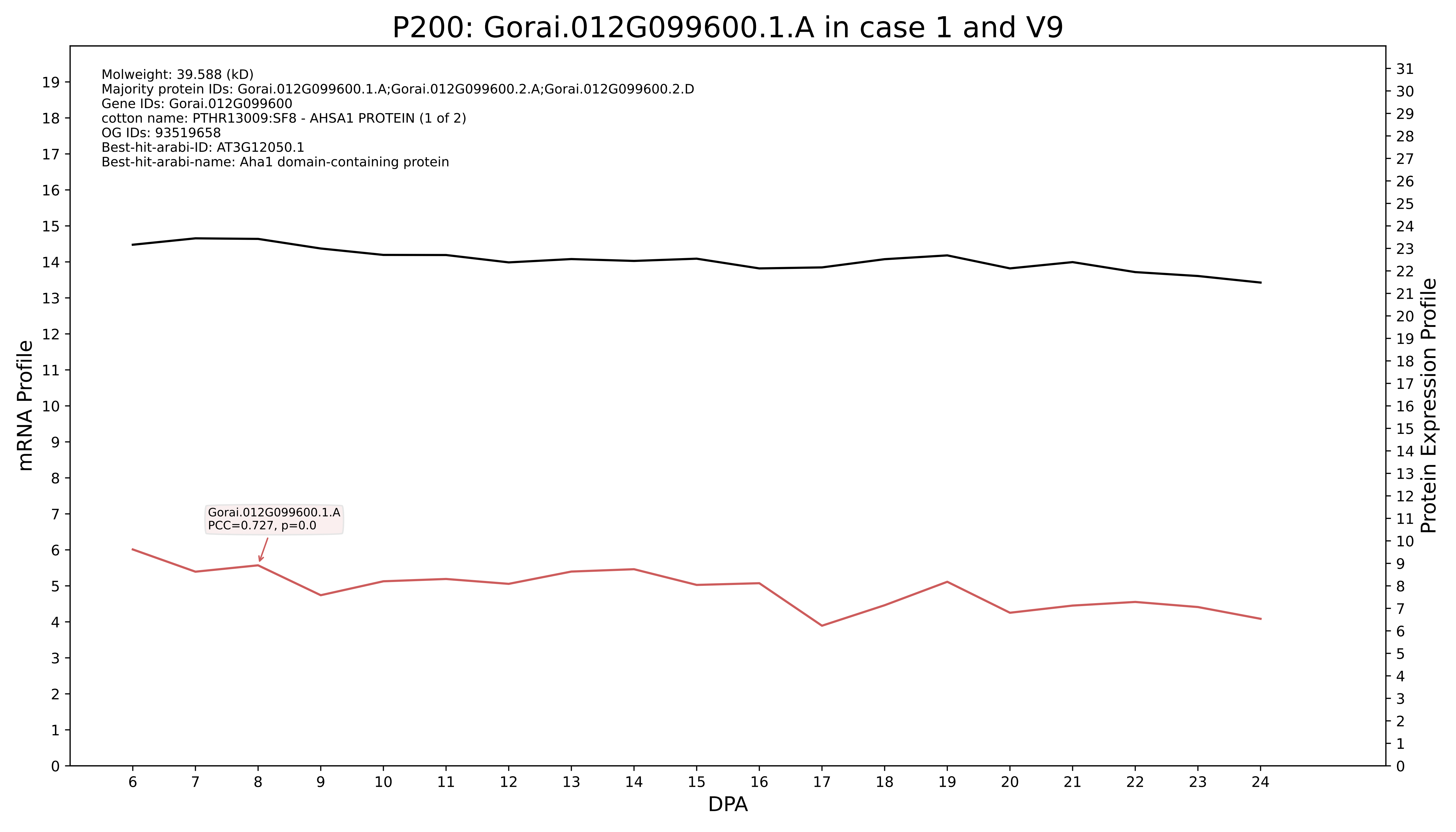
P200: Gorai.012G099600.1.A in case 1 and V9
Molweight: 39.588 (kD)
Majority protein IDs: Gorai.012G099600.1.A;Gorai.012G099600.2.A;Gorai.012G099600.2.D
Gene IDs: Gorai.012G099600
cotton name: PTHR13009:SF8 - AHSA1 PROTEIN (1 of 2)
OG IDs: 93519658
Best-hit-arabi-ID: AT3G12050.1
Best-hit-arabi-name: Aha1 domain-containing protein
Gorai.012G099600.1.A
PCC=0.727, p=0.0
mRNA Profile
Protein Expression Profile
DPA

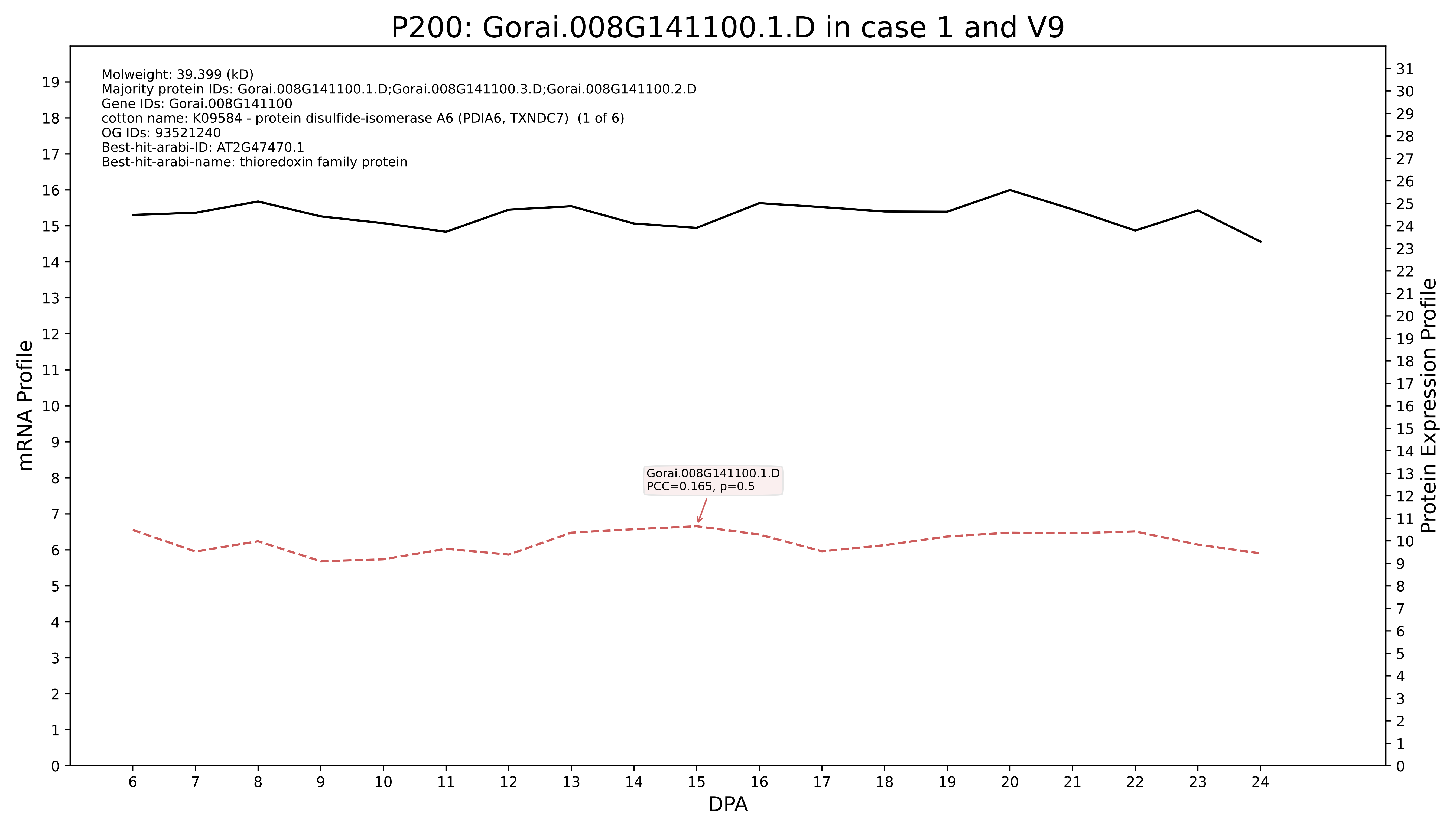
P200: Gorai.008G141100.1.D in case 1 and V9
Molweight: 39.399 (kD)
Majority protein IDs: Gorai.008G141100.1.D;Gorai.008G141100.3.D;Gorai.008G141100.2.D
Gene IDs: Gorai.008G141100
cotton name: K09584 - protein disulfide-isomerase A6 (PDIA6, TXNDC7) (1 of 6)
OG IDs: 93521240
Best-hit-arabi-ID: AT2G47470.1
Best-hit-arabi-name: thioredoxin family protein
Gorai.008G141100.1.D
PCC=0.165, p=0.5
mRNA Profile
Protein Expression Profile
DPA

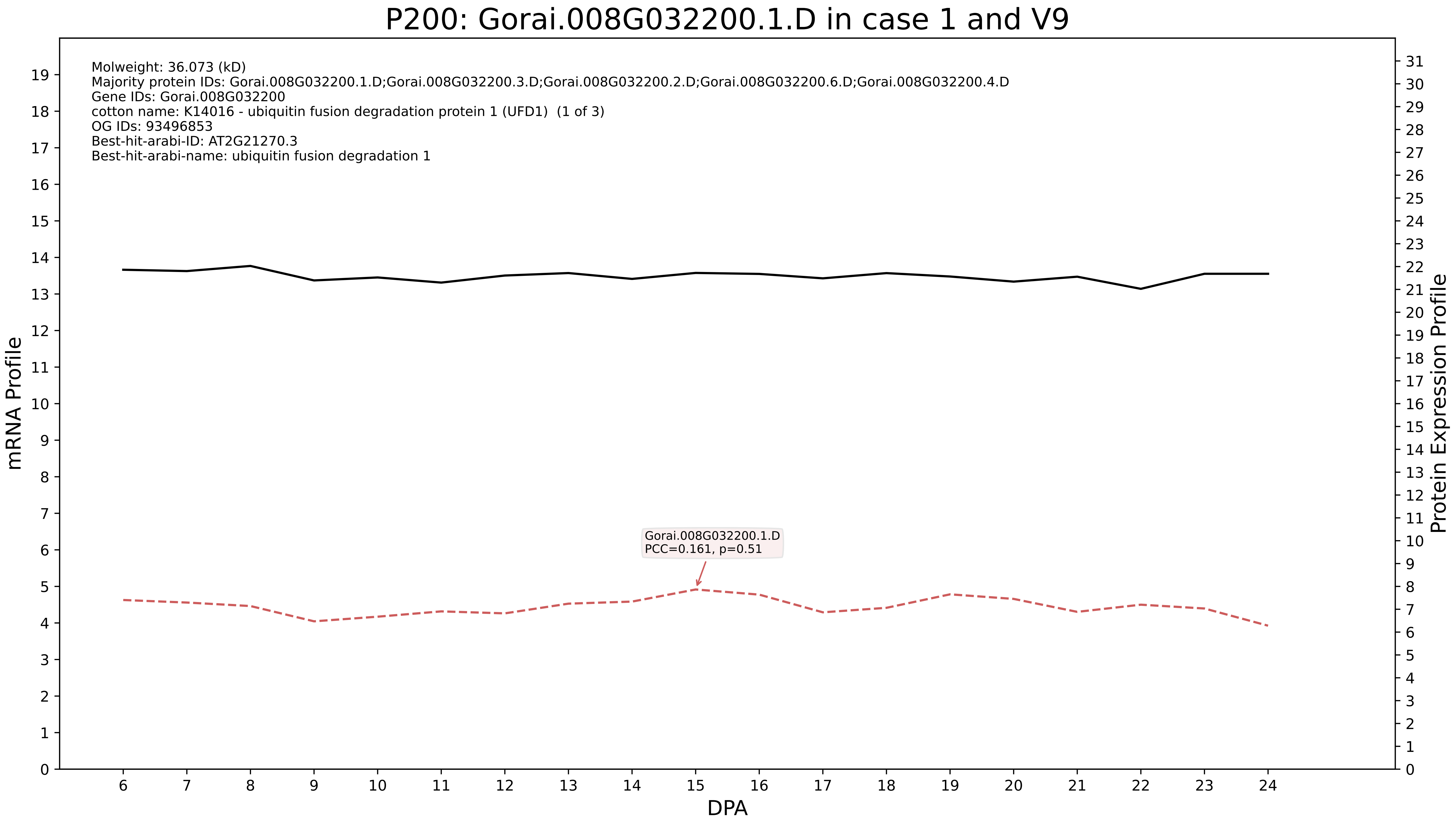
P200: Gorai.008G032200.1.D in case 1 and V9
Molweight: 36.073 (kD)
Majority protein IDs: Gorai.008G032200.1.D;Gorai.008G032200.3.D;Gorai.008G032200.2.D;Gorai.008G032200.6.D;Gorai.008G032200.4.D
Gene IDs: Gorai.008G032200
cotton name: K14016 - ubiquitin fusion degradation protein 1 (UFD1) (1 of 3)
OG IDs: 93496853
Best-hit-arabi-ID: AT2G21270.3
Best-hit-arabi-name: ubiquitin fusion degradation 1
Gorai.008G032200.1.D
PCC=0.161, p=0.51
mRNA Profile
Protein Expression Profile
DPA

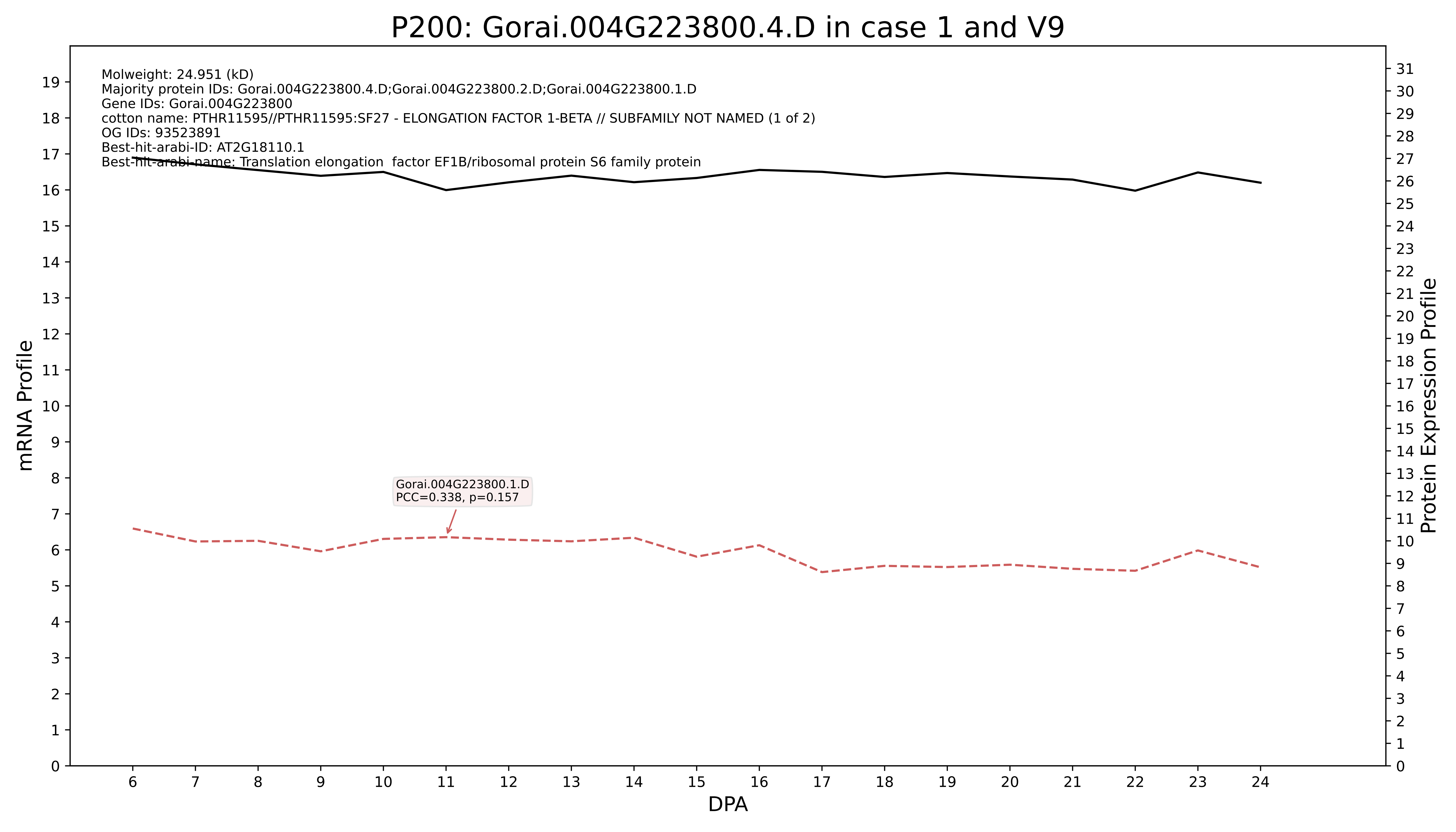
P200: Gorai.004G223800.4.D in case 1 and V9
Molweight: 24.951 (kD)
Majority protein IDs: Gorai.004G223800.4.D;Gorai.004G223800.2.D;Gorai.004G223800.1.D
Gene IDs: Gorai.004G223800
cotton name: PTHR11595//PTHR11595:SF27 - ELONGATION FACTOR 1-BETA // SUBFAMILY NOT NAMED (1 of 2)
OG IDs: 93523891
Best-hit-arabi-ID: AT2G18110.1
Best-hit-arabi-name: Translation elongation factor EF1B/ribosomal protein S6 family protein
Gorai.004G223800.1.D
PCC=0.338, p=0.157
mRNA Profile
Protein Expression Profile
DPA

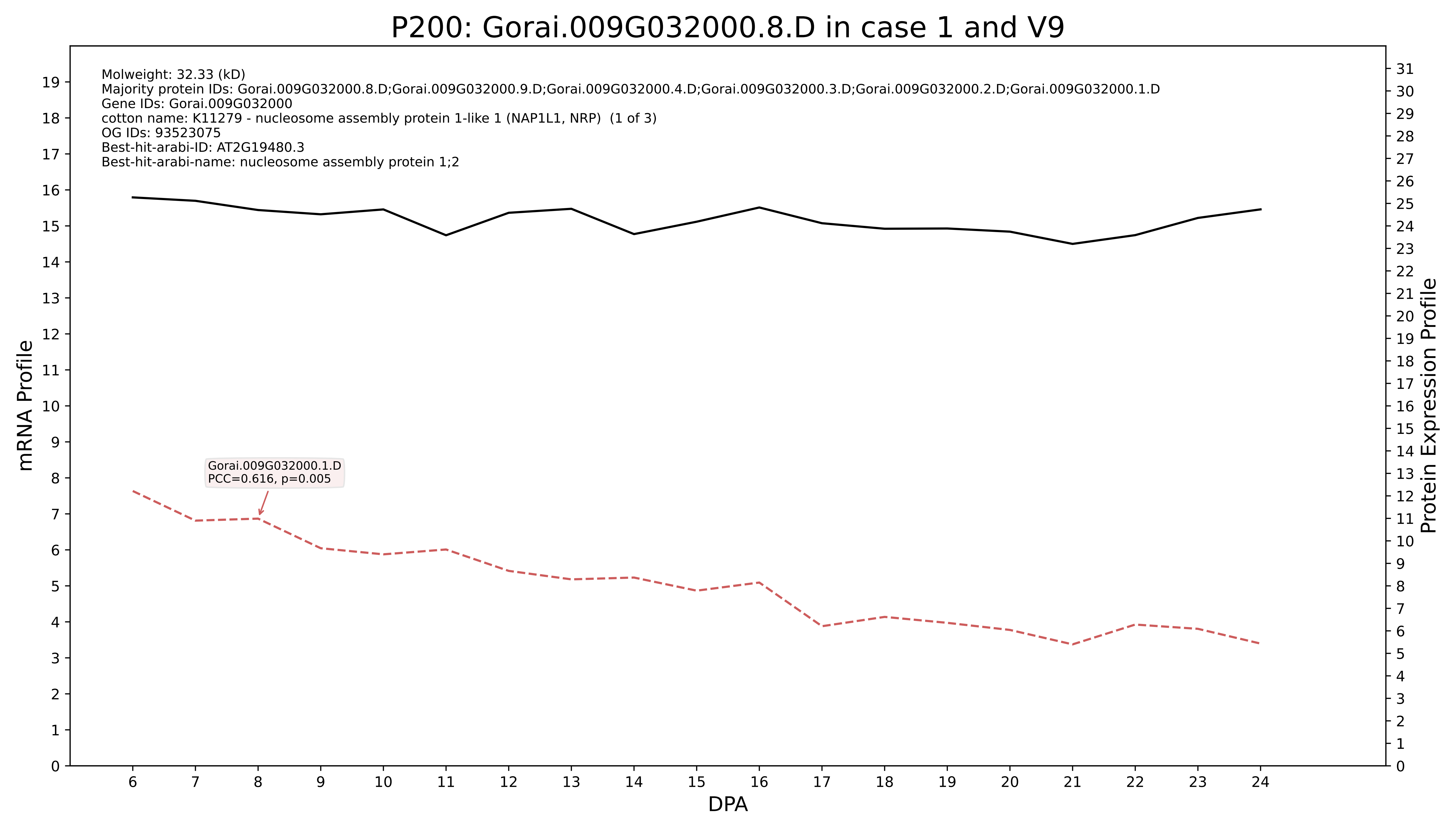
P200: Gorai.009G032000.8.D in case 1 and V9
Molweight: 32.33 (kD)
Majority protein IDs: Gorai.009G032000.8.D;Gorai.009G032000.9.D;Gorai.009G032000.4.D;Gorai.009G032000.3.D;Gorai.009G032000.2.D;Gorai.009G032000.1.D
Gene IDs: Gorai.009G032000
cotton name: K11279 - nucleosome assembly protein 1-like 1 (NAP1L1, NRP) (1 of 3)
OG IDs: 93523075
Best-hit-arabi-ID: AT2G19480.3
Best-hit-arabi-name: nucleosome assembly protein 1;2
Gorai.009G032000.1.D
PCC=0.616, p=0.005
mRNA Profile
Protein Expression Profile
DPA

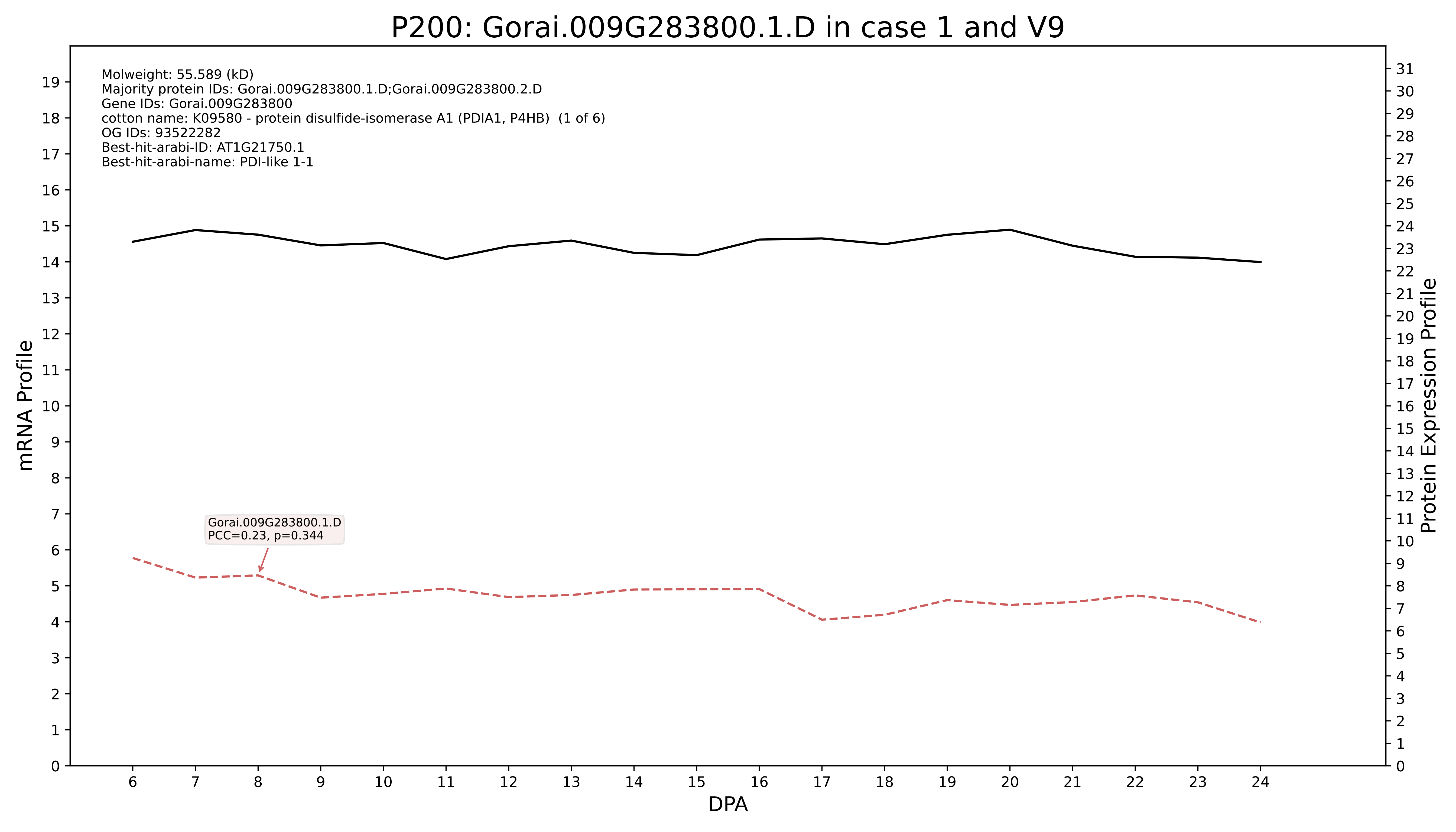
P200: Gorai.009G283800.1.D in case 1 and V9
Molweight: 55.589 (kD)
Majority protein IDs: Gorai.009G283800.1.D;Gorai.009G283800.2.D
Gene IDs: Gorai.009G283800
cotton name: K09580 - protein disulfide-isomerase A1 (PDIA1, P4HB) (1 of 6)
OG IDs: 93522282
Best-hit-arabi-ID: AT1G21750.1
Best-hit-arabi-name: PDI-like 1-1
Gorai.009G283800.1.D
PCC=0.23, p=0.344
mRNA Profile
Protein Expression Profile
DPA

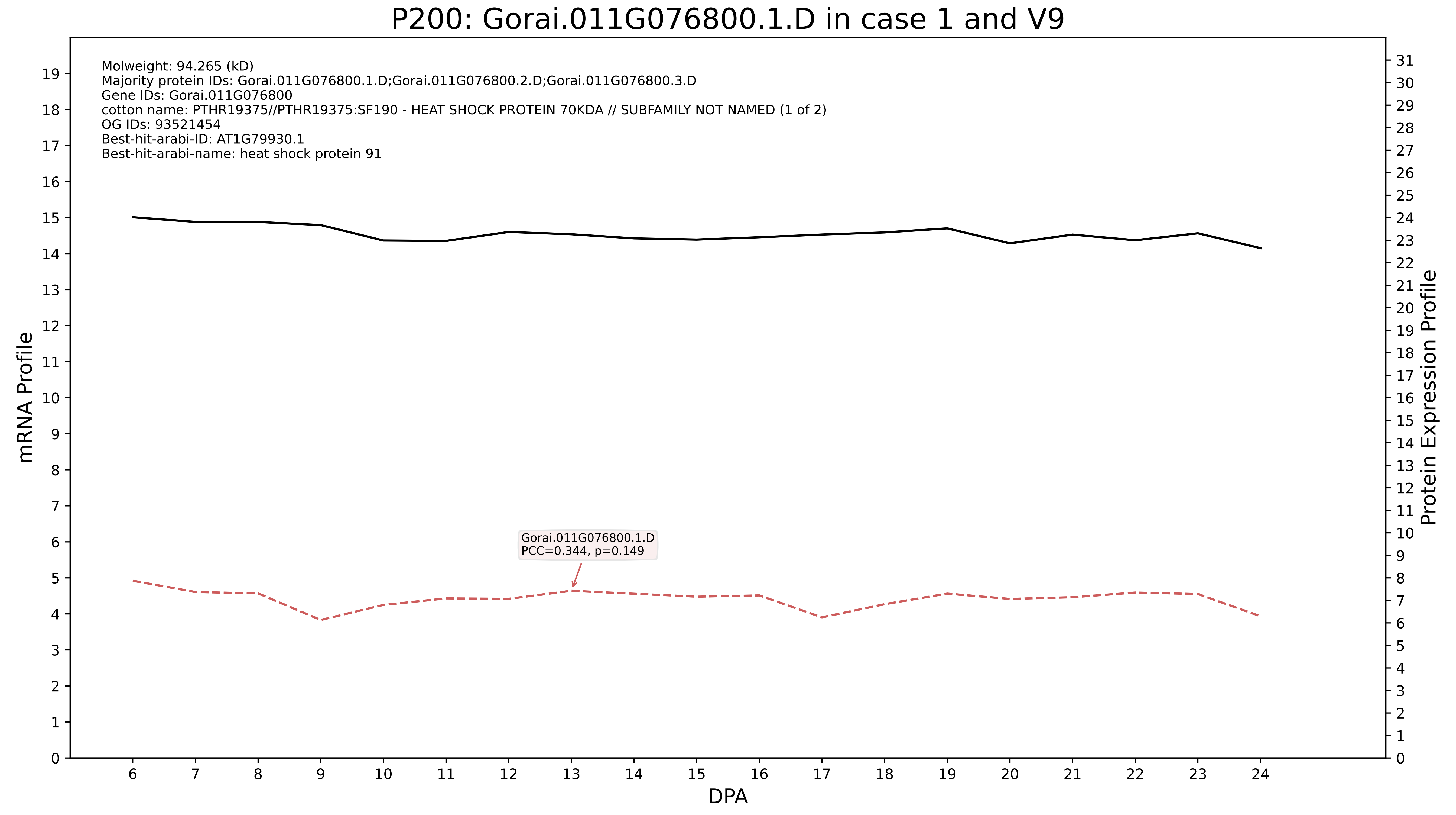
P200: Gorai.011G076800.1.D in case 1 and V9
Molweight: 94.265 (kD)
Majority protein IDs: Gorai.011G076800.1.D;Gorai.011G076800.2.D;Gorai.011G076800.3.D
Gene IDs: Gorai.011G076800
cotton name: PTHR19375//PTHR19375:SF190 - HEAT SHOCK PROTEIN 70KDA // SUBFAMILY NOT NAMED (1 of 2)
OG IDs: 93521454
Best-hit-arabi-ID: AT1G79930.1
Best-hit-arabi-name: heat shock protein 91
Gorai.011G076800.1.D
PCC=0.344, p=0.149
mRNA Profile
Protein Expression Profile
DPA

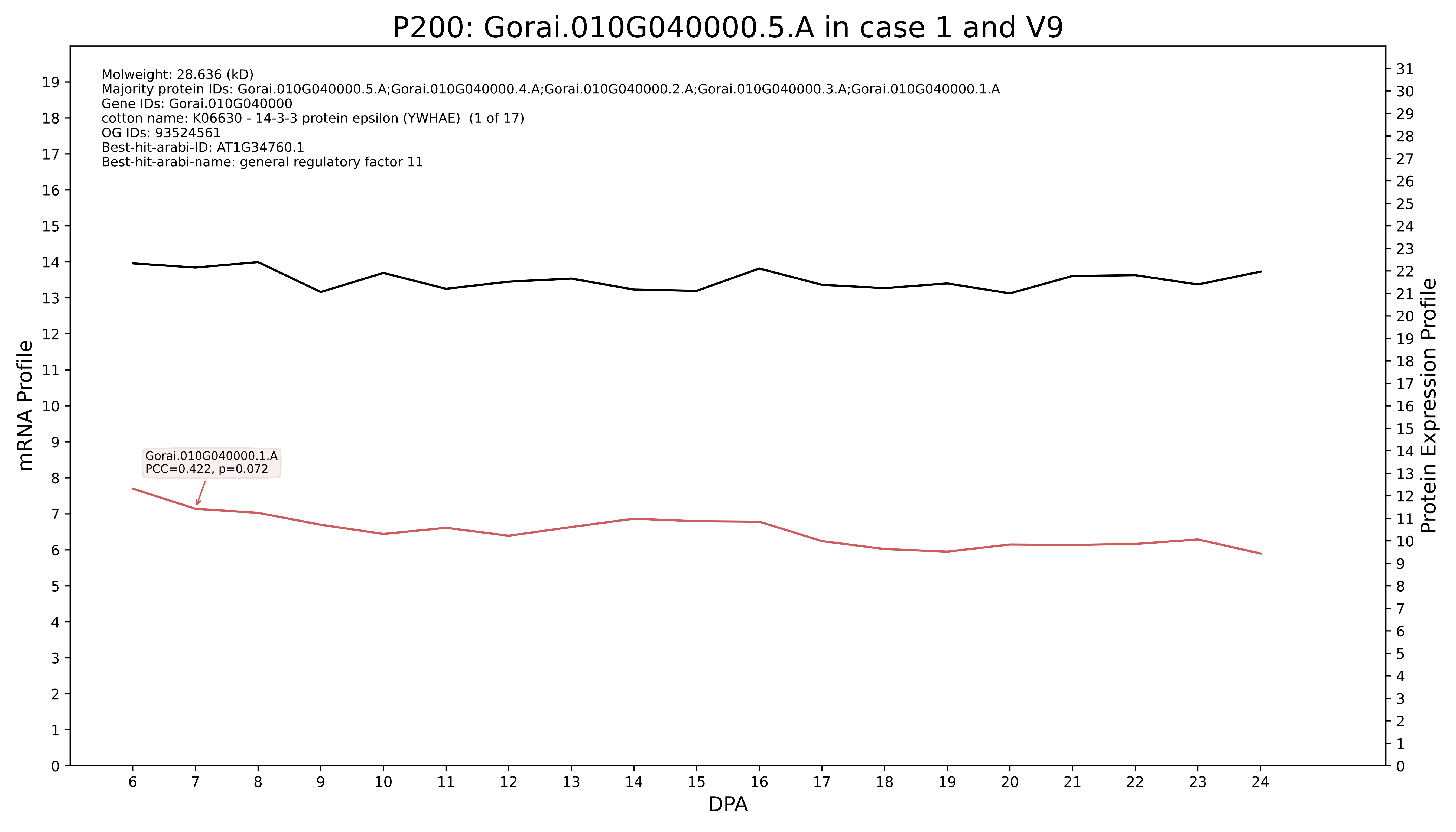
P200: Gorai.010G040000.5.A in case 1 and V9
Molweight: 28.636 (kD)
Majority protein IDs: Gorai.010G040000.5.A;Gorai.010G040000.4.A;Gorai.010G040000.2.A;Gorai.010G040000.3.A;Gorai.010G040000.1.A
Gene IDs: Gorai.010G040000
cotton name: K06630 - 14-3-3 protein epsilon (YWHAE) (1 of 17)
OG IDs: 93524561
Best-hit-arabi-ID: AT1G34760.1
Best-hit-arabi-name: general regulatory factor 11
Gorai.010G040000.1.A
PCC=0.422, p=0.072
mRNA Profile
Protein Expression Profile
DPA

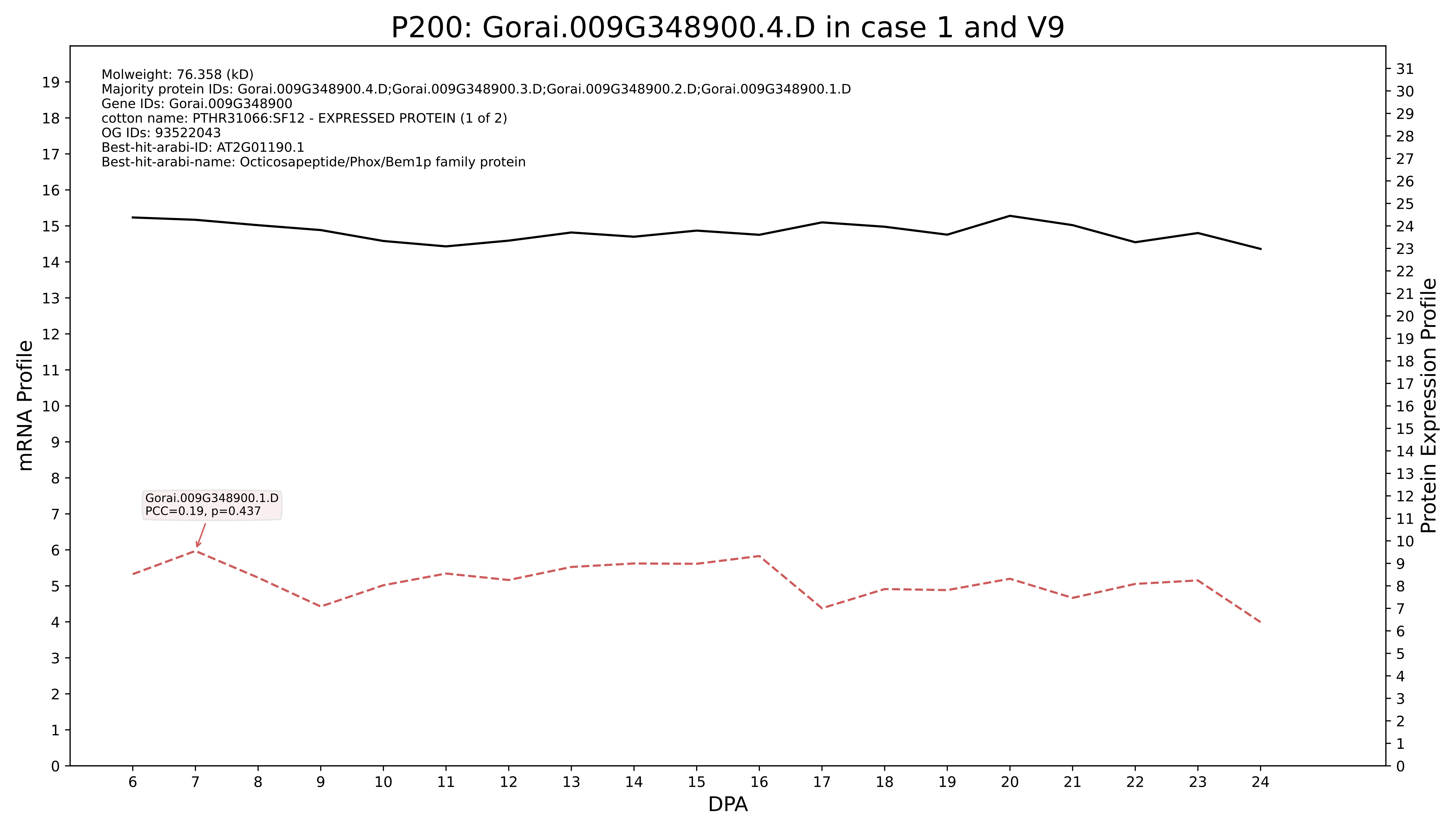
P200: Gorai.009G348900.4.D in case 1 and V9
Molweight: 76.358 (kD)
Majority protein IDs: Gorai.009G348900.4.D;Gorai.009G348900.3.D;Gorai.009G348900.2.D;Gorai.009G348900.1.D
Gene IDs: Gorai.009G348900
cotton name: PTHR31066:SF12 - EXPRESSED PROTEIN (1 of 2)
OG IDs: 93522043
Best-hit-arabi-ID: AT2G01190.1
Best-hit-arabi-name: Octicosapeptide/Phox/Bem1p family protein
Gorai.009G348900.1.D
PCC=0.19, p=0.437
mRNA Profile
Protein Expression Profile
DPA

### P200: Gorai.005G183500.2.D in case 1 and V9

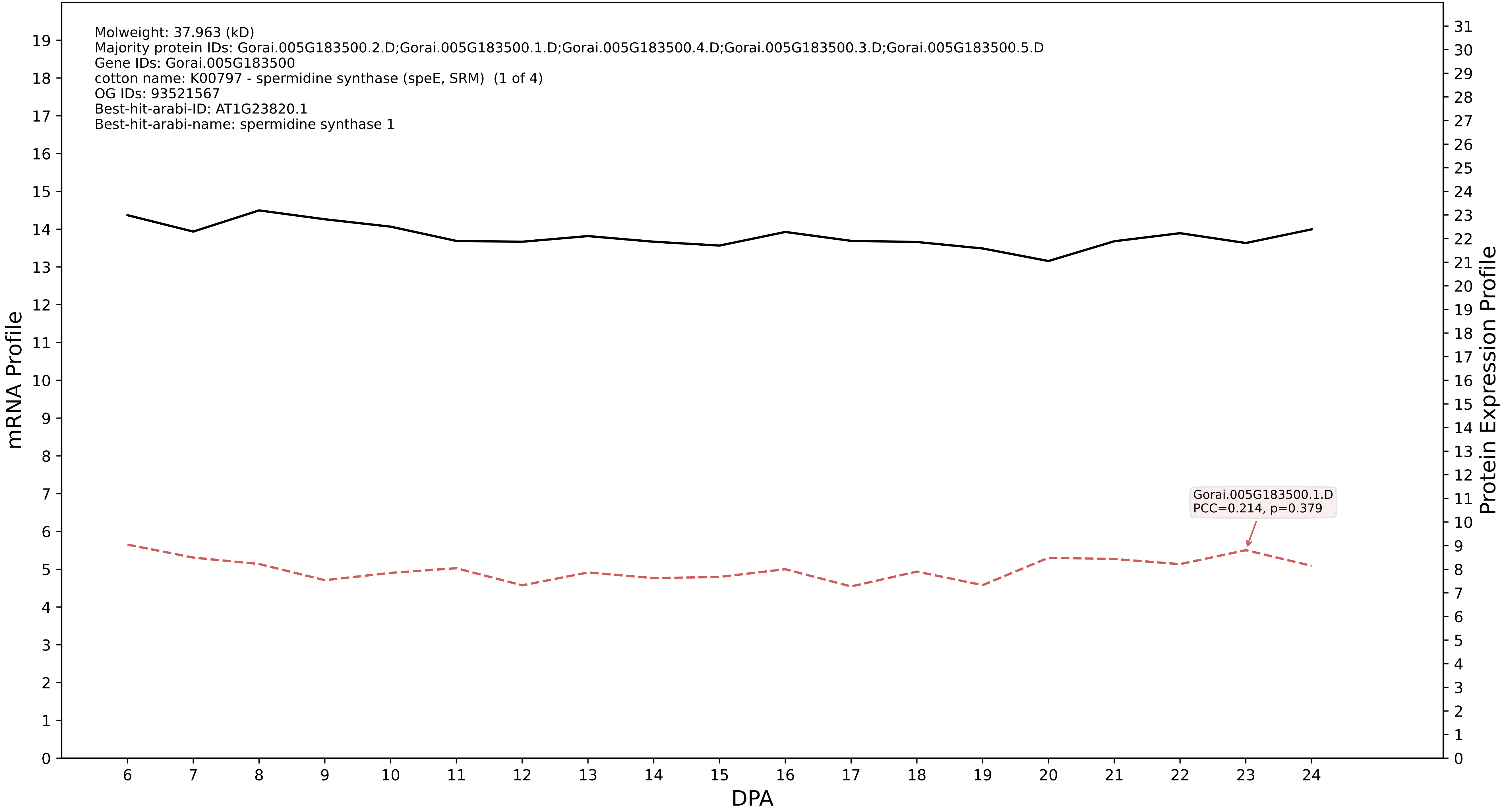

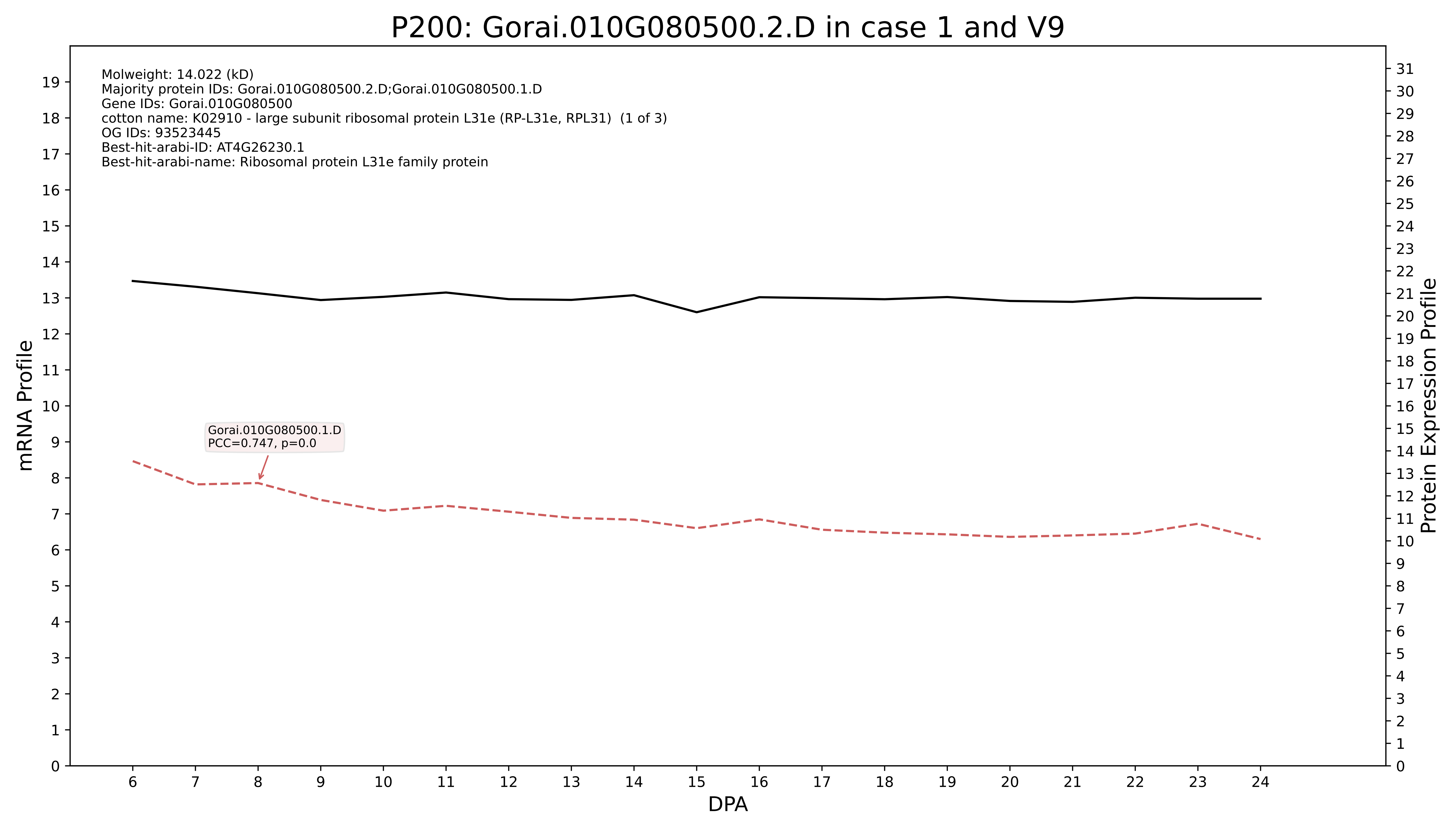
P200: Gorai.010G080500.2.D in case 1 and V9
Molweight: 14.022 (kD)
Majority protein IDs: Gorai.010G080500.2.D;Gorai.010G080500.1.D
Gene IDs: Gorai.010G080500
cotton name: K02910 - large subunit ribosomal protein L31e (RP-L31e, RPL31) (1 of 3)
OG IDs: 93523445
Best-hit-arabi-ID: AT4G26230.1
Best-hit-arabi-name: Ribosomal protein L31e family protein
Gorai.010G080500.1.D
PCC=0.747, p=0.0
mRNA Profile
Protein Expression Profile
DPA

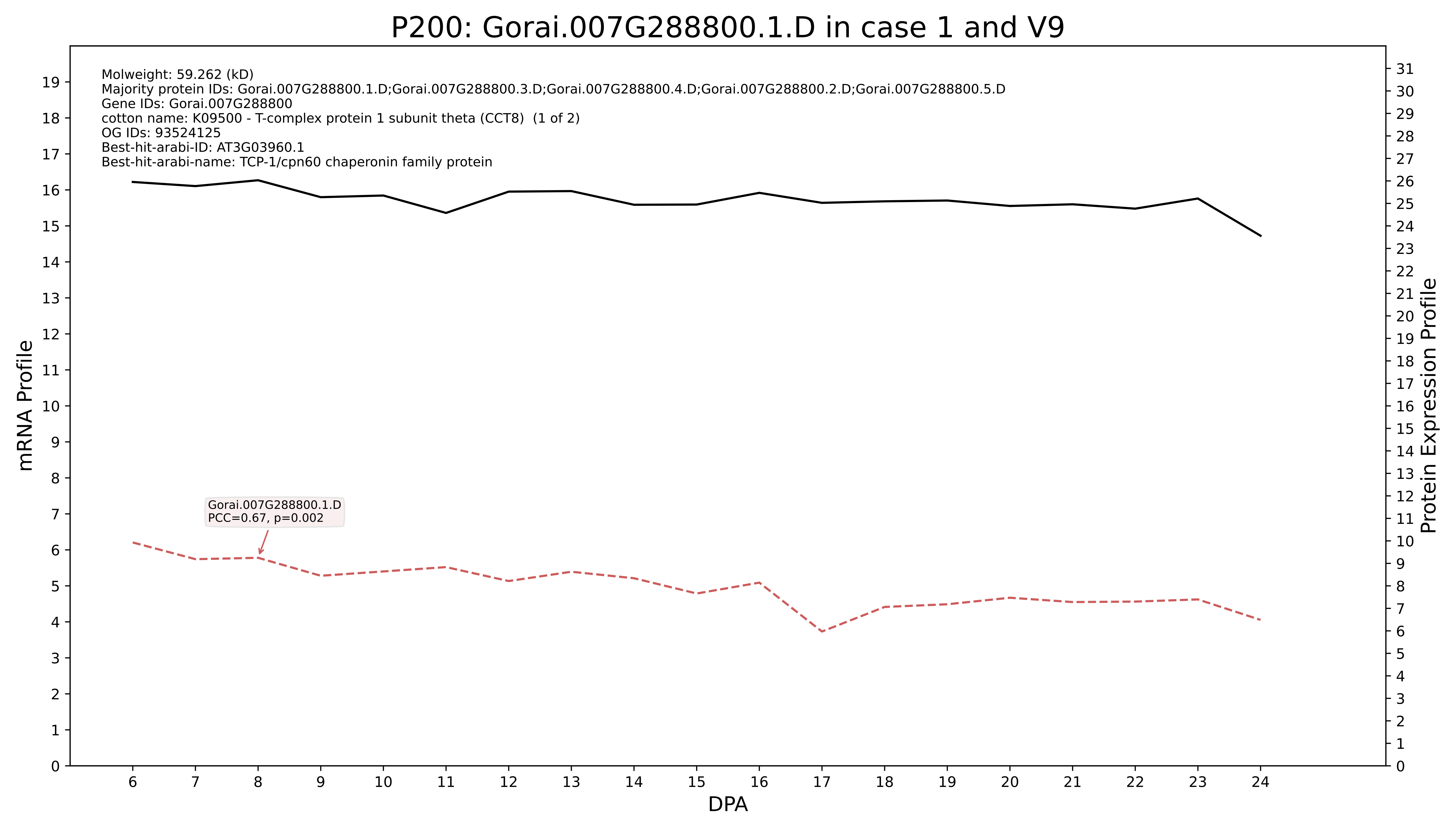
P200: Gorai.007G288800.1.D in case 1 and V9
Molweight: 59.262 (kD)
Majority protein IDs: Gorai.007G288800.1.D;Gorai.007G288800.3.D;Gorai.007G288800.4.D;Gorai.007G288800.2.D;Gorai.007G288800.5.D
Gene IDs: Gorai.007G288800
cotton name: K09500 - T-complex protein 1 subunit theta (CCT8) (1 of 2)
OG IDs: 93524125
Best-hit-arabi-ID: AT3G03960.1
Best-hit-arabi-name: TCP-1/cpn60 chaperonin family protein
Gorai.007G288800.1.D
PCC=0.67, p=0.002
mRNA Profile
Protein Expression Profile
DPA

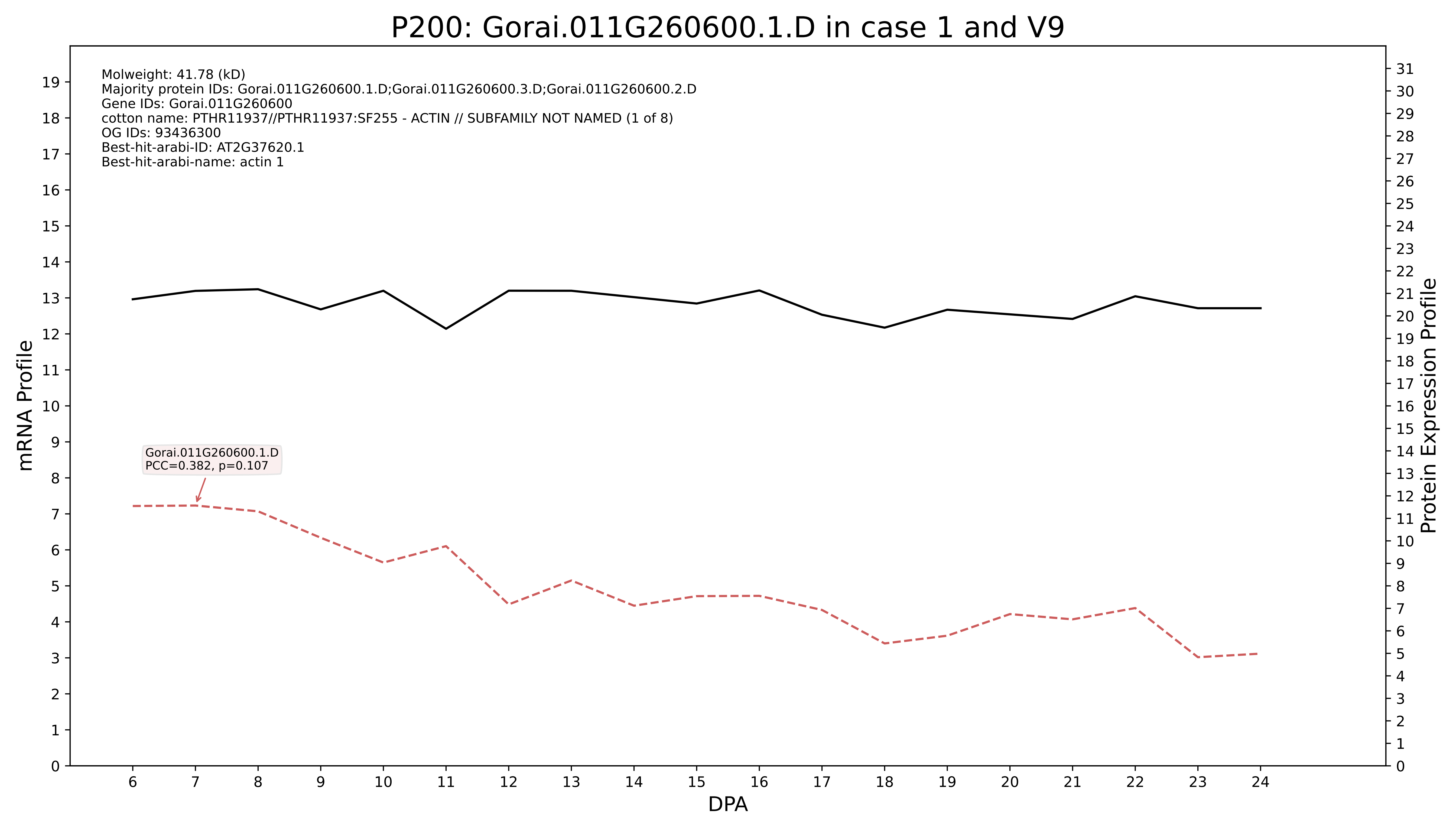
P200: Gorai.011G260600.1.D in case 1 and V9
Molweight: 41.78 (kD)
Majority protein IDs: Gorai.011G260600.1.D;Gorai.011G260600.3.D;Gorai.011G260600.2.D
Gene IDs: Gorai.011G260600
cotton name: PTHR11937//PTHR11937:SF255 - ACTIN // SUBFAMILY NOT NAMED (1 of 8)
OG IDs: 93436300
Best-hit-arabi-ID: AT2G37620.1
Best-hit-arabi-name: actin 1
Gorai.011G260600.1.D
PCC=0.382, p=0.107
mRNA Profile
Protein Expression Profile
DPA

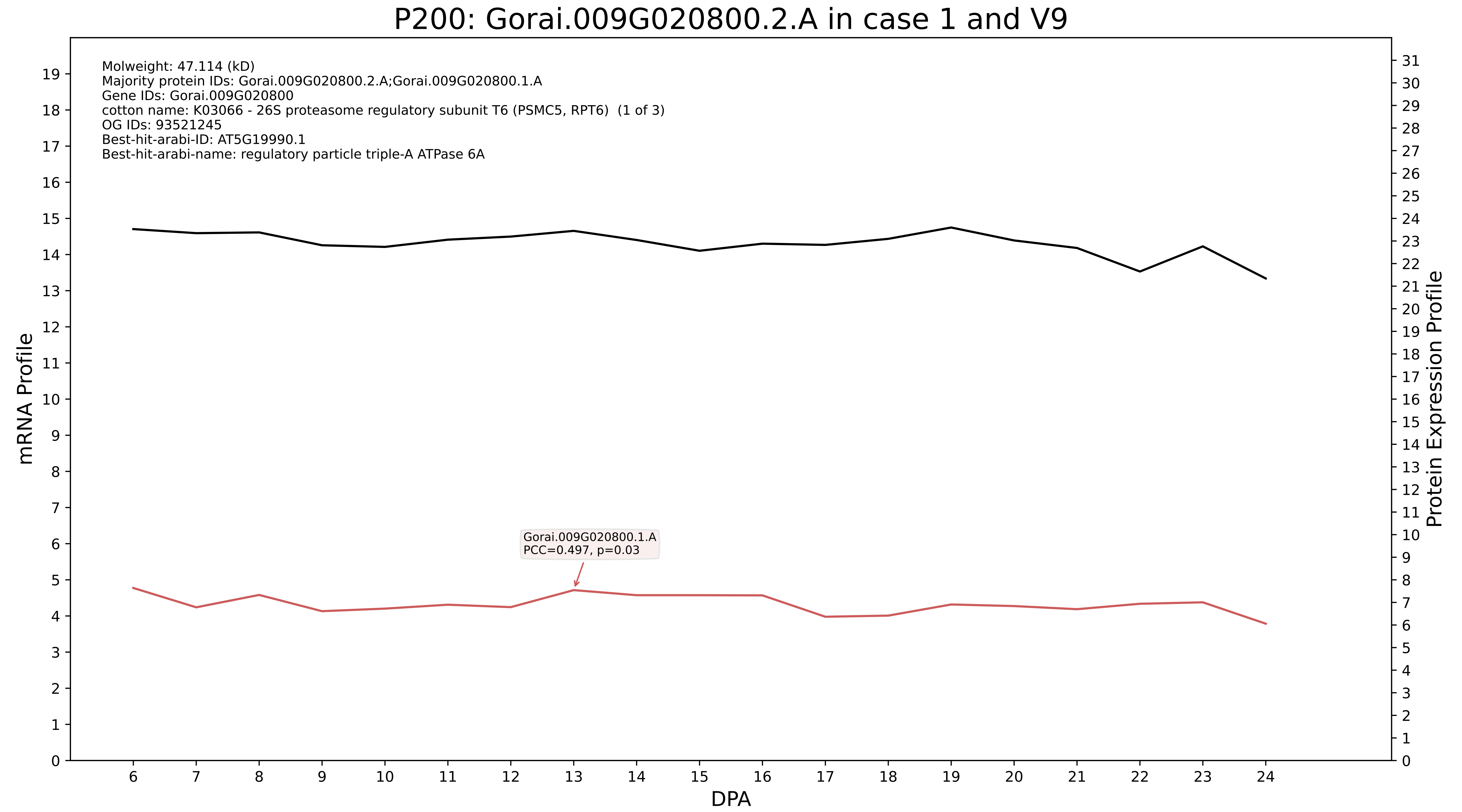
P200: Gorai.009G020800.2.A in case 1 and V9
Molweight: 47.114 (kD)
Majority protein IDs: Gorai.009G020800.2.A;Gorai.009G020800.1.A
Gene IDs: Gorai.009G020800
cotton name: K03066 - 26S proteasome regulatory subunit T6 (PSMC5, RPT6) (1 of 3)
OG IDs: 93521245
Best-hit-arabi-ID: AT5G19990.1
Best-hit-arabi-name: regulatory particle triple-A ATPase 6A
Gorai.009G020800.1.A
PCC=0.497, p=0.03
mRNA Profile
Protein Expression Profile
DPA
0 1 2 3 4 5 6 7 8 9 10 11 12 13 14 15 16 17 18 19
0 1 2 3 4 5 6 7 8 9 10 11 12 13 14 15 16 17 18 19 20 21 22 23 24 25 26 27 28 29 30 31
6 7 8 9 10 11 12 13 14 15 16 17 18 19 20 21 22 23 24

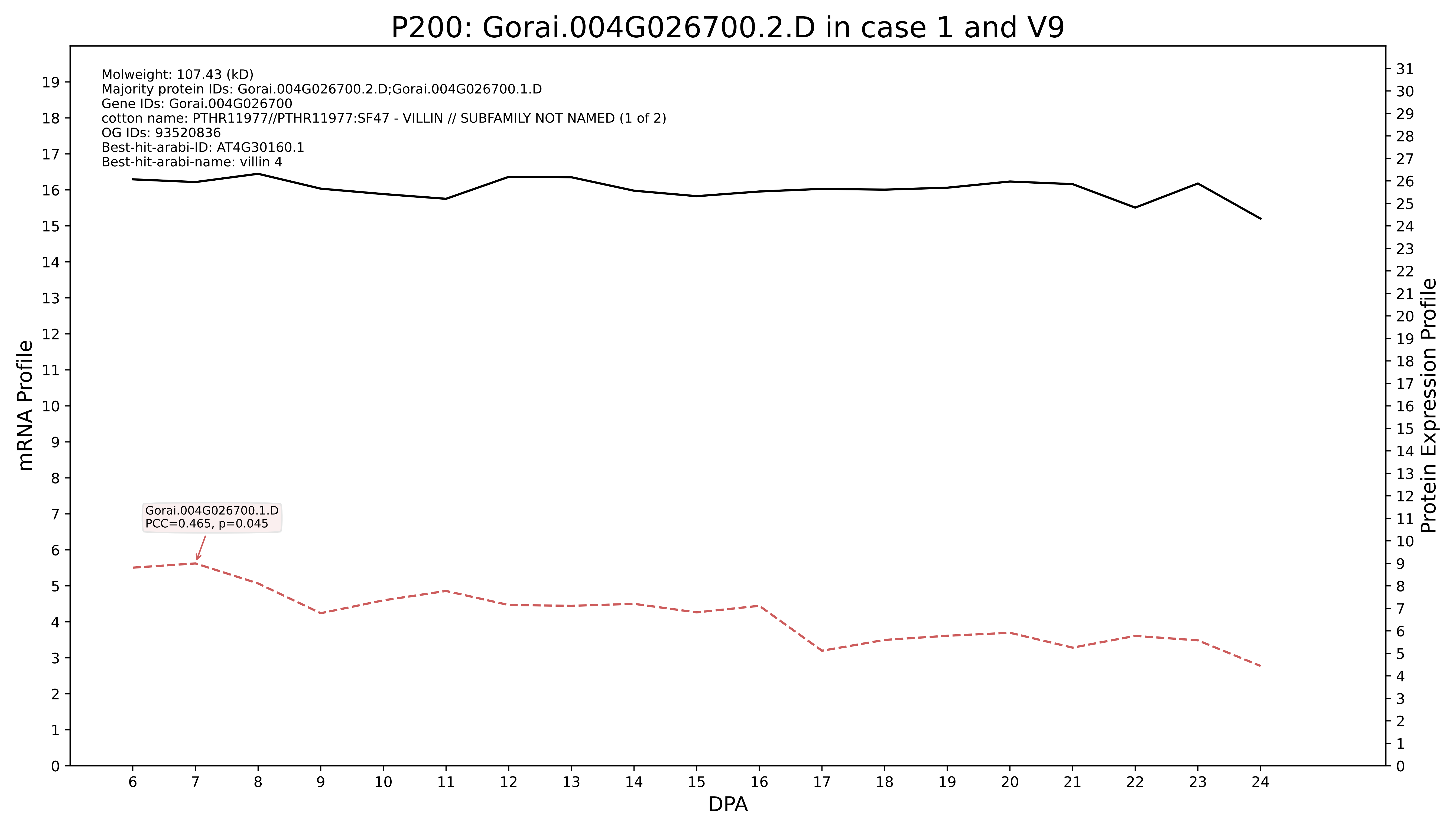

P200: Gorai.004G026700.2.D in case 1 and V9
Molweight: 107.43 (kD)
Majority protein IDs: Gorai.004G026700.2.D;Gorai.004G026700.1.D
Gene IDs: Gorai.004G026700
cotton name: PTHR11977//PTHR11977:SF47 - VILLIN // SUBFAMILY NOT NAMED (1 of 2)
OG IDs: 93520836
Best-hit-arabi-ID: AT4G30160.1
Best-hit-arabi-name: villin 4
Gorai.004G026700.1.D
PCC=0.465, p=0.045
mRNA Profile
Protein Expression Profile
DPA

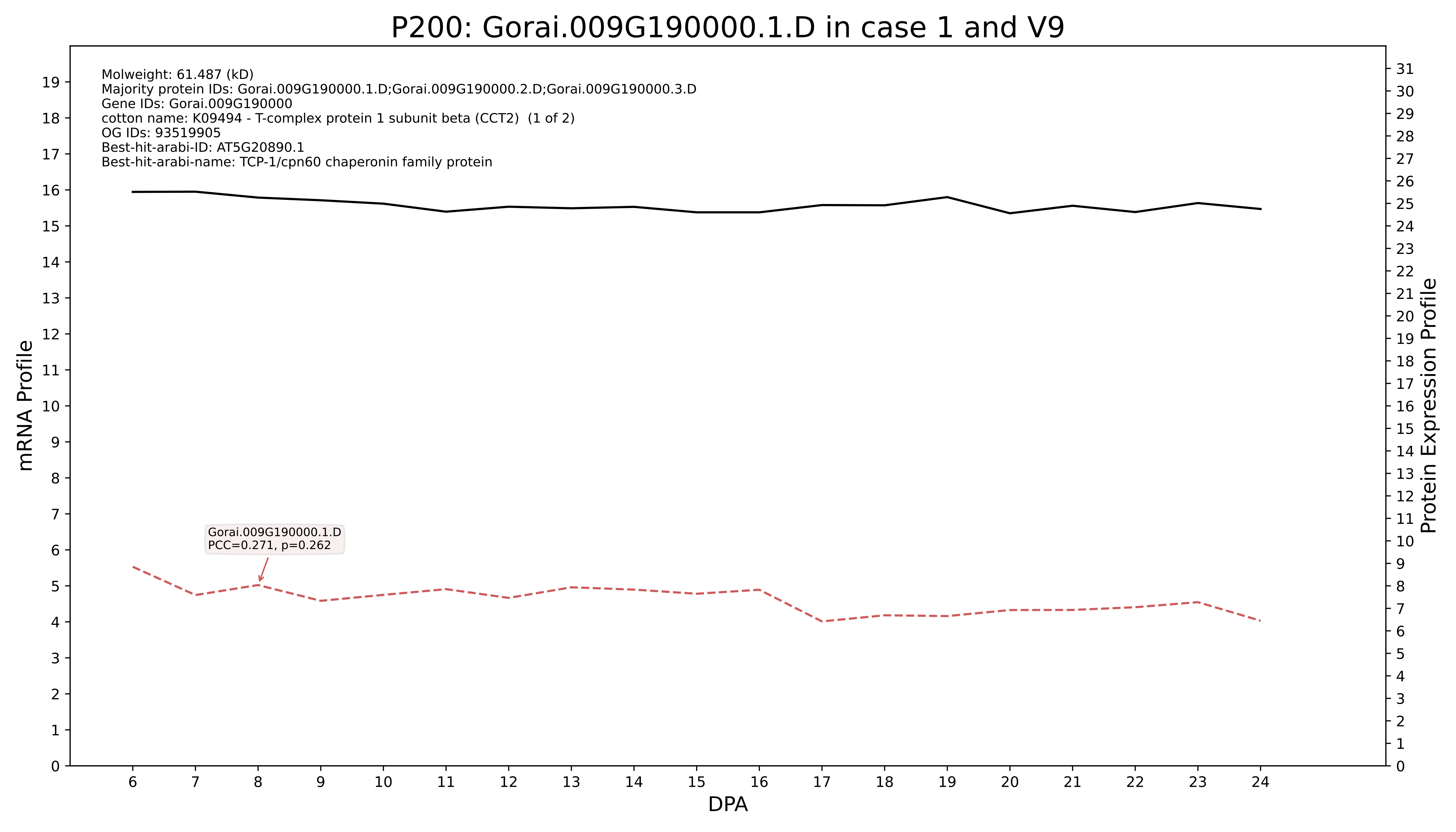
P200: Gorai.009G190000.1.D in case 1 and V9
Molweight: 61.487 (kD)
Majority protein IDs: Gorai.009G190000.1.D;Gorai.009G190000.2.D;Gorai.009G190000.3.D
Gene IDs: Gorai.009G190000
cotton name: K09494 - T-complex protein 1 subunit beta (CCT2) (1 of 2)
OG IDs: 93519905
Best-hit-arabi-ID: AT5G20890.1
Best-hit-arabi-name: TCP-1/cpn60 chaperonin family protein
Gorai.009G190000.1.D
PCC=0.271, p=0.262
mRNA Profile
Protein Expression Profile
DPA

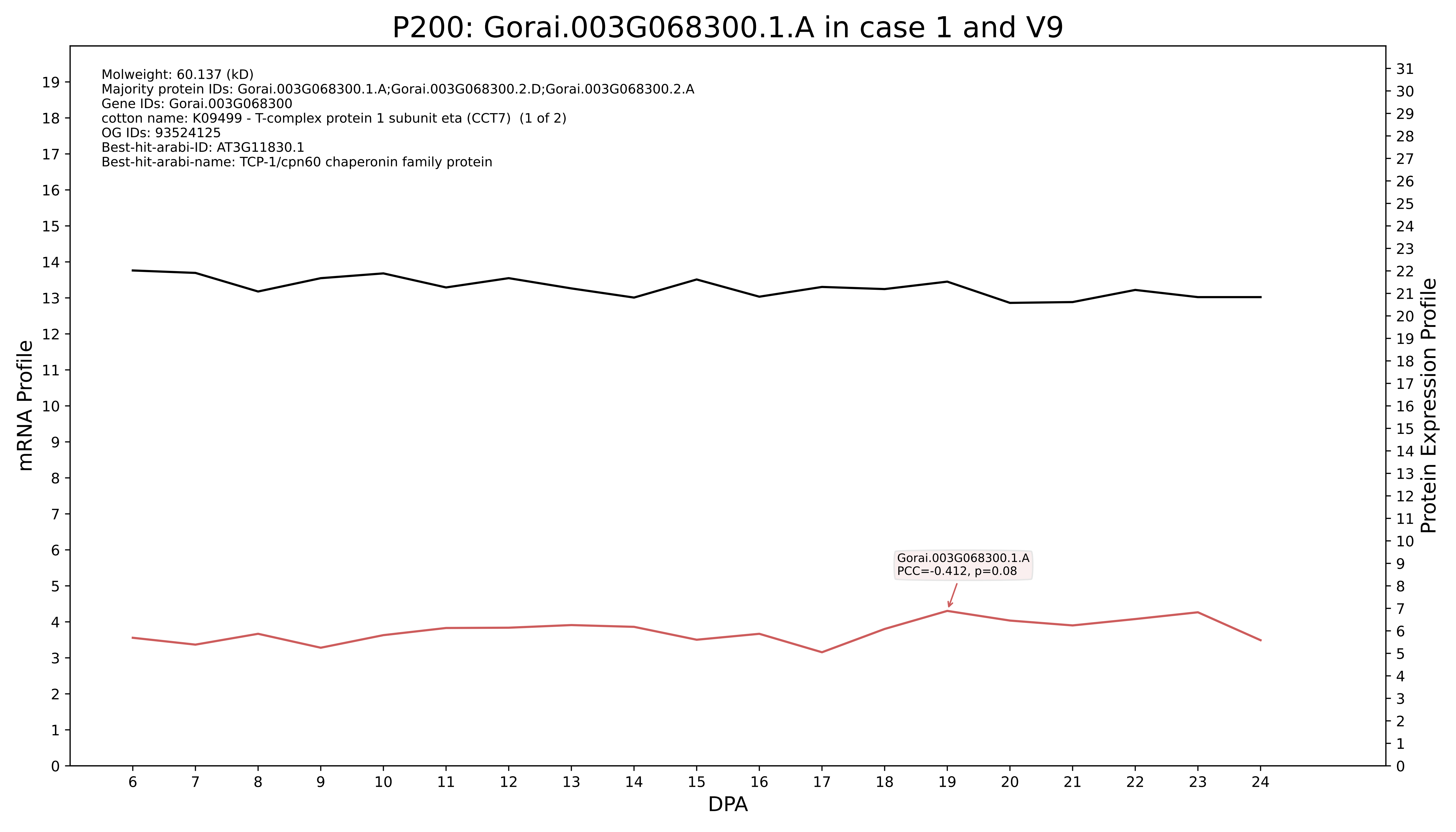

P200: Gorai.003G068300.1.A in case 1 and V9
Molweight: 60.137 (kD)
Majority protein IDs: Gorai.003G068300.1.A;Gorai.003G068300.2.D;Gorai.003G068300.2.A
Gene IDs: Gorai.003G068300
cotton name: K09499 - T-complex protein 1 subunit eta (CCT7) (1 of 2)
OG IDs: 93524125
Best-hit-arabi-ID: AT3G11830.1
Best-hit-arabi-name: TCP-1/cpn60 chaperonin family protein
Gorai.003G068300.1.A
PCC=-0.412, p=0.08
mRNA Profile
Protein Expression Profile
DPA

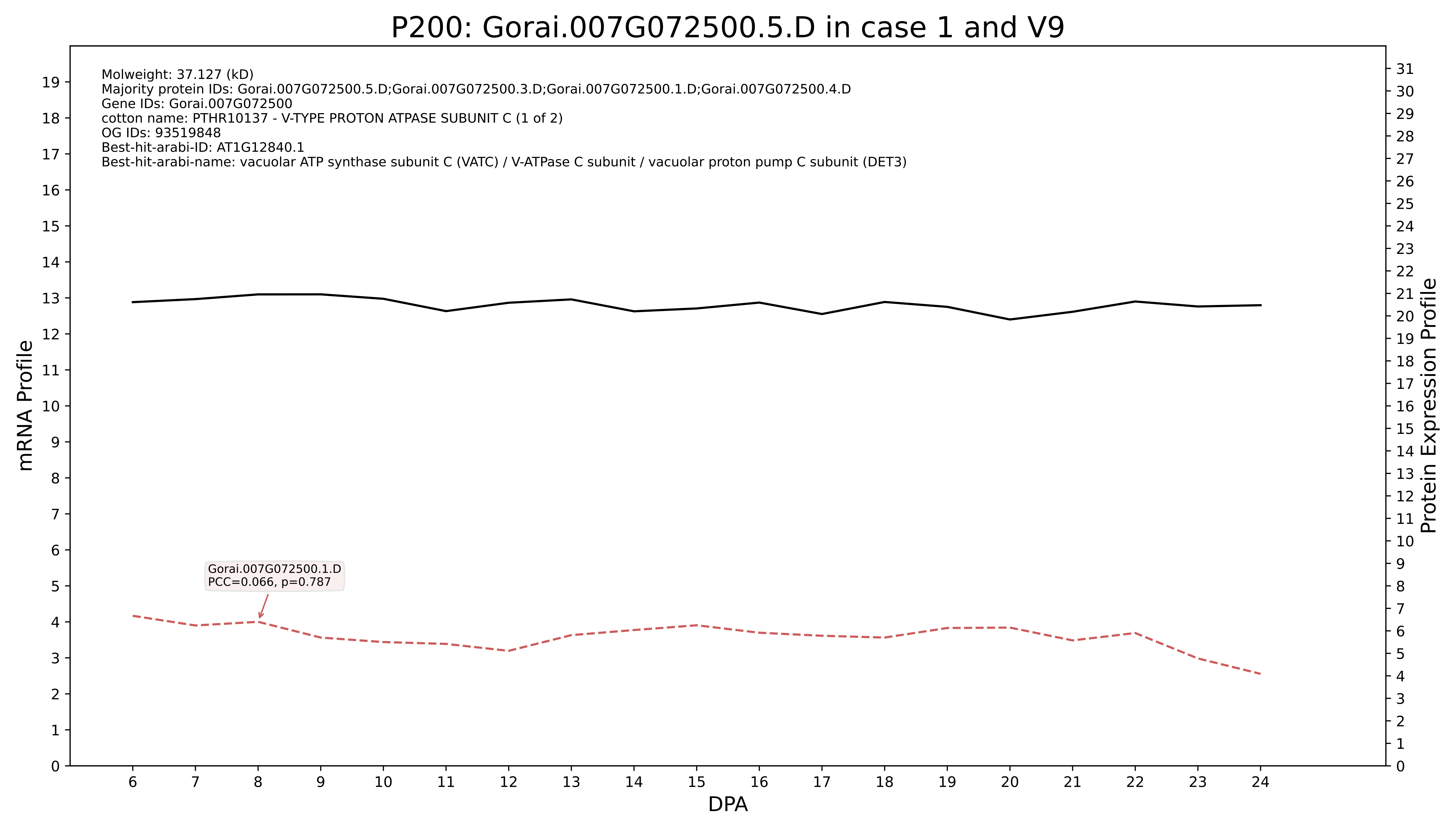

P200: Gorai.007G072500.5.D in case 1 and V9
Molweight: 37.127 (kD)
Majority protein IDs: Gorai.007G072500.5.D;Gorai.007G072500.3.D;Gorai.007G072500.1.D;Gorai.007G072500.4.D
Gene IDs: Gorai.007G072500
cotton name: PTHR10137 - V-TYPE PROTON ATPASE SUBUNIT C (1 of 2)
OG IDs: 93519848
Best-hit-arabi-ID: AT1G12840.1
Best-hit-arabi-name: vacuolar ATP synthase subunit C (VATC) / V-ATPase C subunit / vacuolar proton pump C subunit (DET3)
Gorai.007G072500.1.D
PCC=0.066, p=0.787
mRNA Profile
Protein Expression Profile
DPA

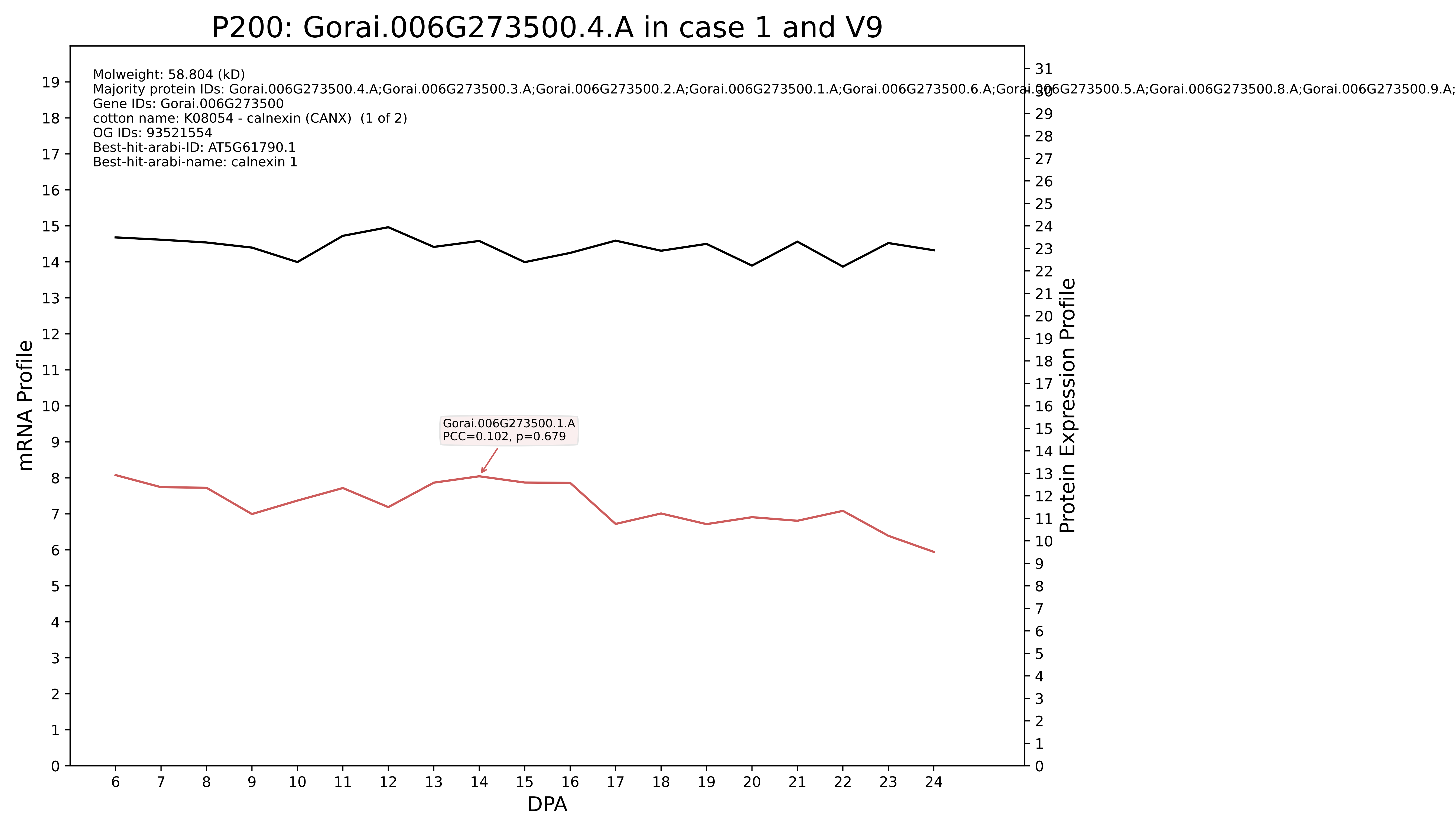

P200: Gorai.006G273500.4.A in case 1 and V9
Molweight: 58.804 (kD)
Majority protein IDs: Gorai.006G273500.4.A;Gorai.006G273500.3.A;Gorai.006G273500.2.A;Gorai.006G273500.1.A;Gorai.006G273500.6.A;Gorai.006G273500.5.A;Gorai.006G273500.8.A;Gorai.006G273500.9.A;
Gene IDs: Gorai.006G273500
cotton name: K08054 - calnexin (CANX) (1 of 2)
OG IDs: 93521554
Best-hit-arabi-ID: AT5G61790.1
Best-hit-arabi-name: calnexin 1
Gorai.006G273500.1.A
PCC=0.102, p=0.679
mRNA Profile
Protein Expression Profile
DPA

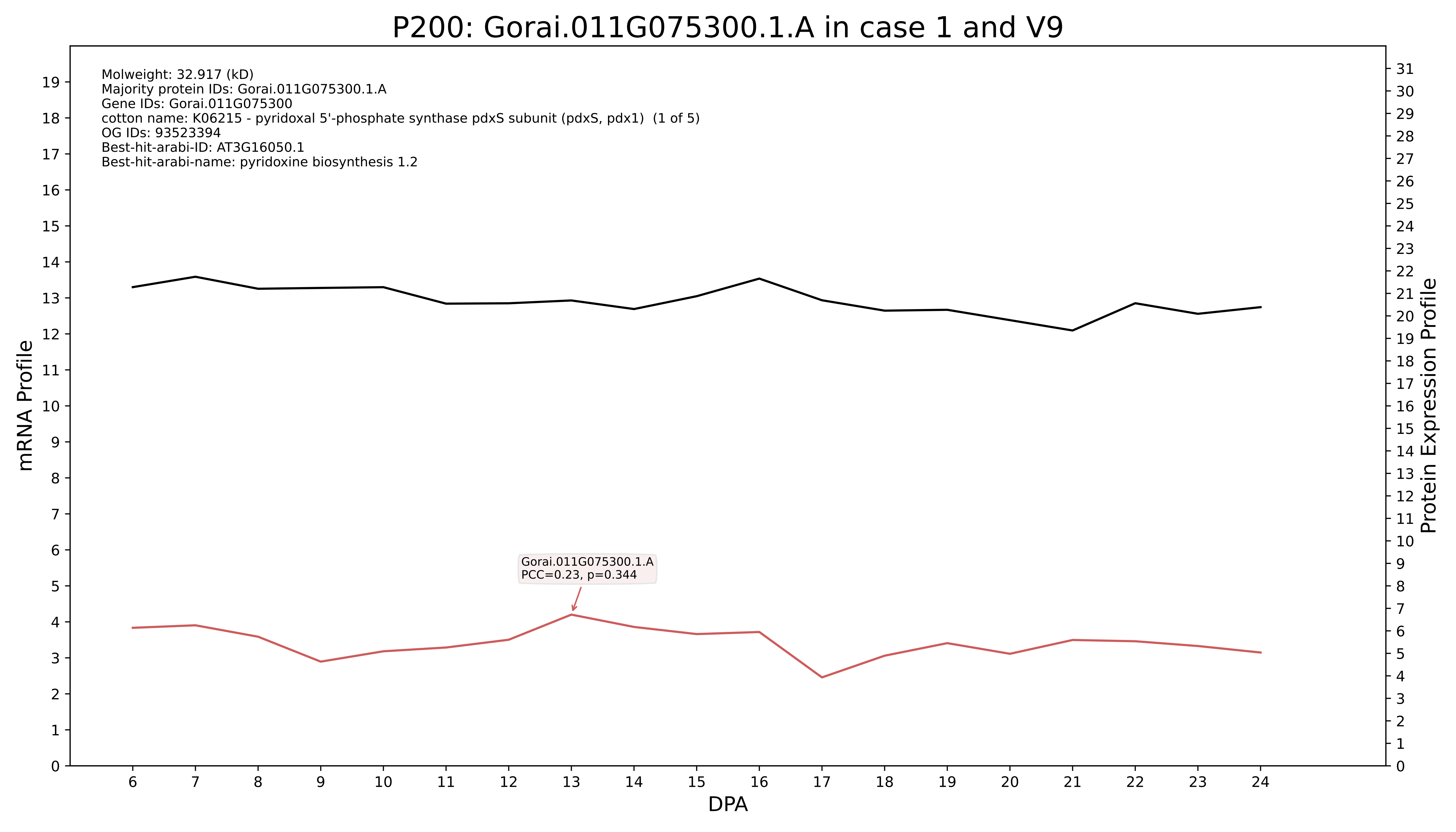

P200: Gorai.011G075300.1.A in case 1 and V9
Molweight: 32.917 (kD)
Majority protein IDs: Gorai.011G075300.1.A
Gene IDs: Gorai.011G075300
cotton name: K06215 - pyridoxal 5'-phosphate synthase pdxS subunit (pdxS, pdx1) (1 of 5)
OG IDs: 93523394
Best-hit-arabi-ID: AT3G16050.1
Best-hit-arabi-name: pyridoxine biosynthesis 1.2
Gorai.011G075300.1.A
PCC=0.23, p=0.344
mRNA Profile
Protein Expression Profile
DPA

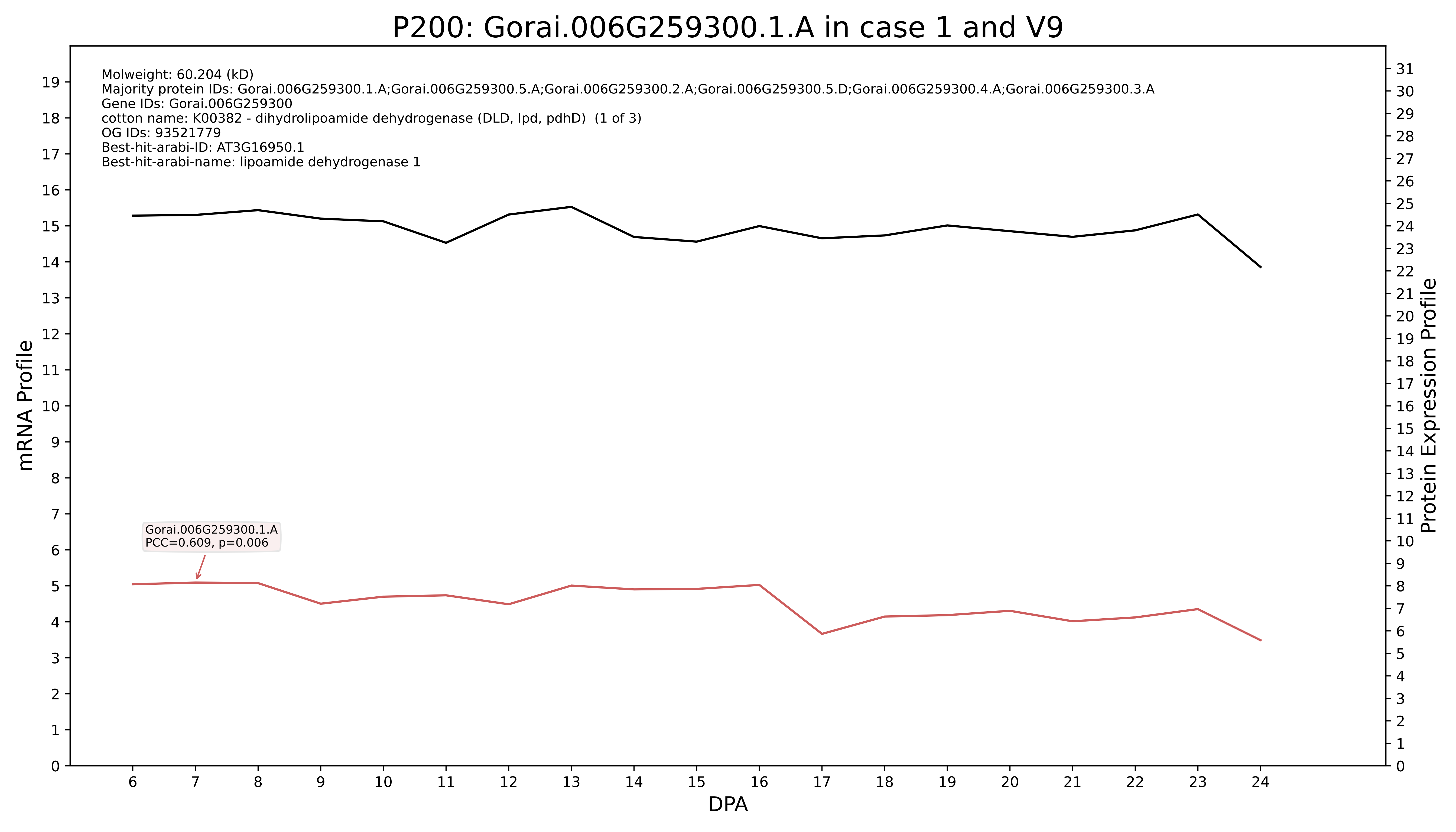

P200: Gorai.006G259300.1.A in case 1 and V9
Molweight: 60.204 (kD)
Majority protein IDs: Gorai.006G259300.1.A;Gorai.006G259300.5.A;Gorai.006G259300.2.A;Gorai.006G259300.5.D;Gorai.006G259300.4.A;Gorai.006G259300.3.A
Gene IDs: Gorai.006G259300
cotton name: K00382 - dihydrolipoamide dehydrogenase (DLD, lpd, pdhD) (1 of 3)
OG IDs: 93521779
Best-hit-arabi-ID: AT3G16950.1
Best-hit-arabi-name: lipoamide dehydrogenase 1
Gorai.006G259300.1.A
PCC=0.609, p=0.006
mRNA Profile
Protein Expression Profile
DPA

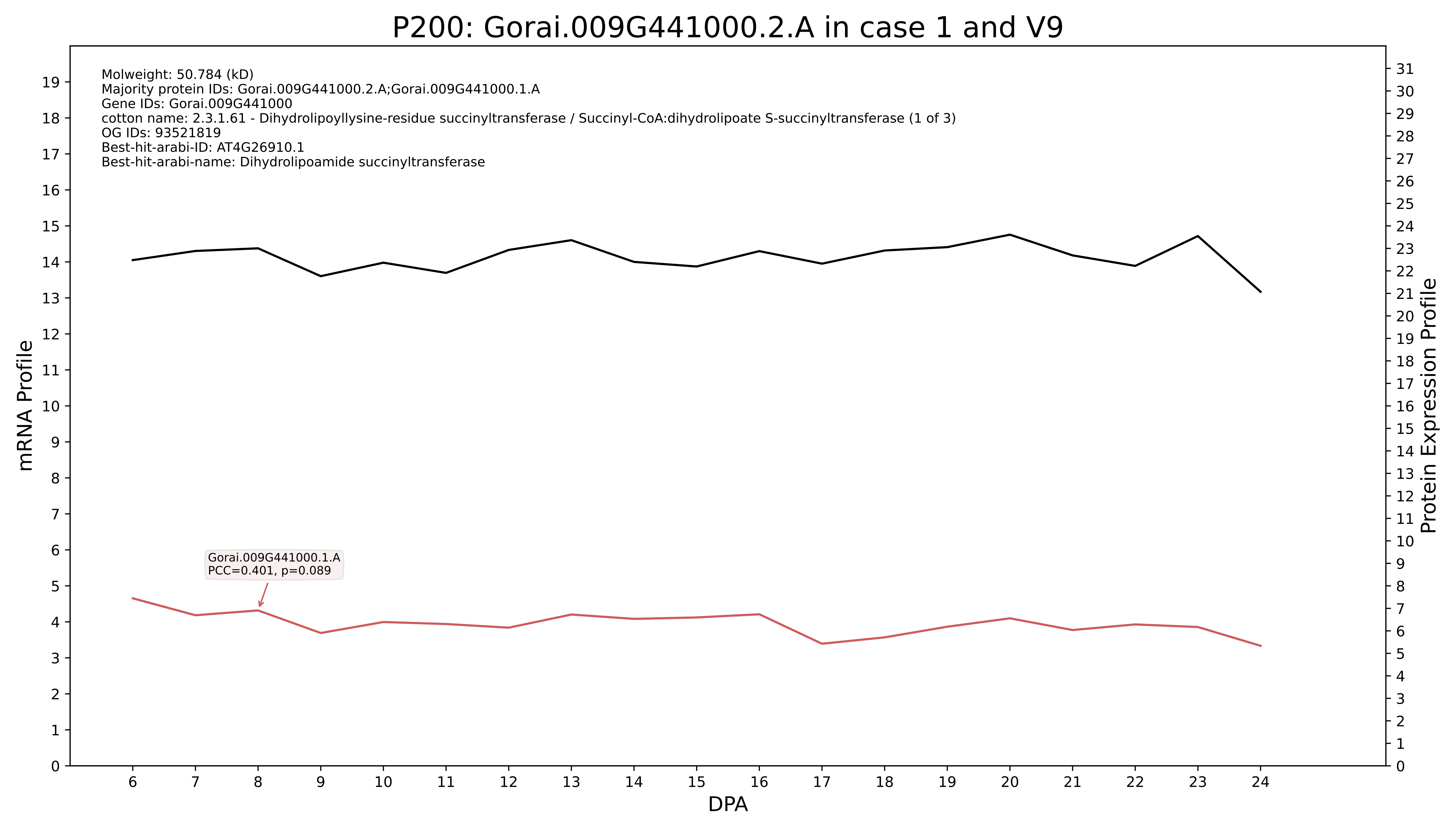
P200: Gorai.009G441000.2.A in case 1 and V9
Molweight: 50.784 (kD)
Majority protein IDs: Gorai.009G441000.2.A;Gorai.009G441000.1.A
Gene IDs: Gorai.009G441000
cotton name: 2.3.1.61 - Dihydrolipoyllysine-residue succinyltransferase / Succinyl-CoA:dihydrolipoate S-succinyltransferase (1 of 3)
OG IDs: 93521819
Best-hit-arabi-ID: AT4G26910.1
Best-hit-arabi-name: Dihydrolipoamide succinyltransferase
Gorai.009G441000.1.A
PCC=0.401, p=0.089
mRNA Profile
Protein Expression Profile
DPA

### P200: Gorai.002G126800.1.A in case 1 and V9

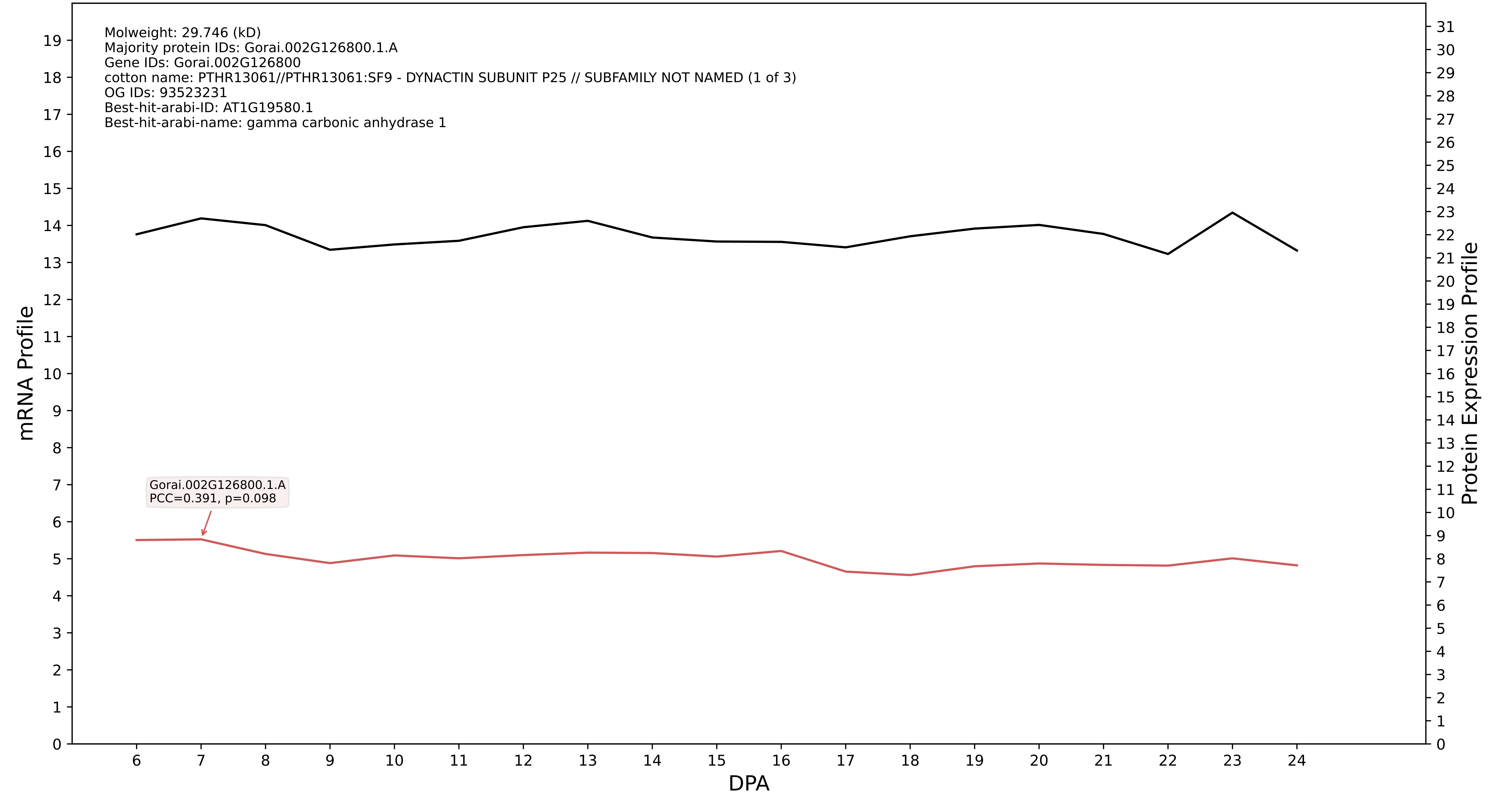

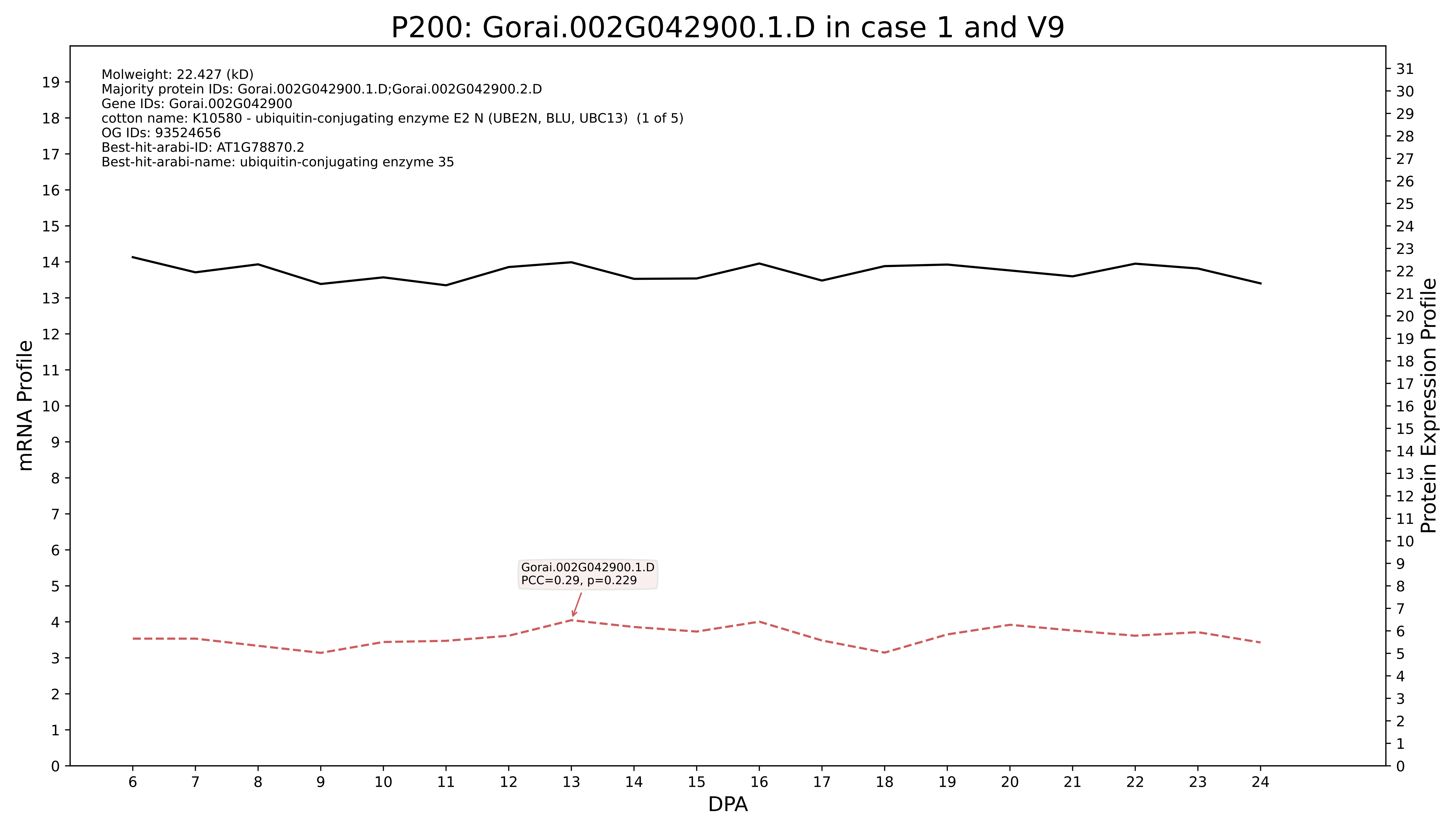
P200: Gorai.002G042900.1.D in case 1 and V9
Molweight: 22.427 (kD)
Majority protein IDs: Gorai.002G042900.1.D;Gorai.002G042900.2.D
Gene IDs: Gorai.002G042900
cotton name: K10580 - ubiquitin-conjugating enzyme E2 N (UBE2N, BLU, UBC13) (1 of 5)
OG IDs: 93524656
Best-hit-arabi-ID: AT1G78870.2
Best-hit-arabi-name: ubiquitin-conjugating enzyme 35
Gorai.002G042900.1.D
PCC=0.29, p=0.229
mRNA Profile
Protein Expression Profile
DPA

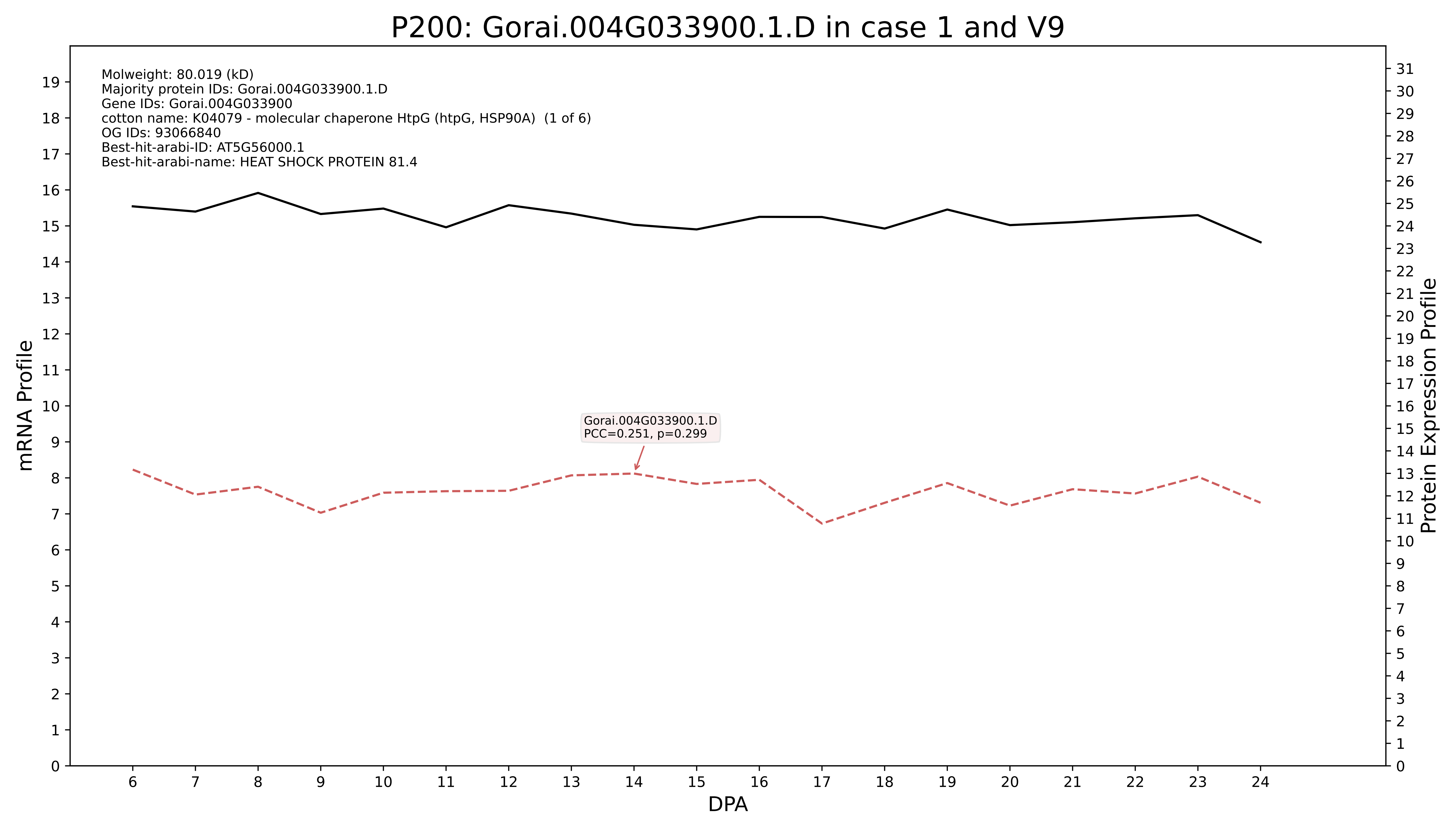
P200: Gorai.004G033900.1.D in case 1 and V9
Molweight: 80.019 (kD)
Majority protein IDs: Gorai.004G033900.1.D
Gene IDs: Gorai.004G033900
cotton name: K04079 - molecular chaperone HtpG (htpG, HSP90A) (1 of 6)
OG IDs: 93066840
Best-hit-arabi-ID: AT5G56000.1
Best-hit-arabi-name: HEAT SHOCK PROTEIN 81.4
Gorai.004G033900.1.D
PCC=0.251, p=0.299
mRNA Profile
Protein Expression Profile
DPA

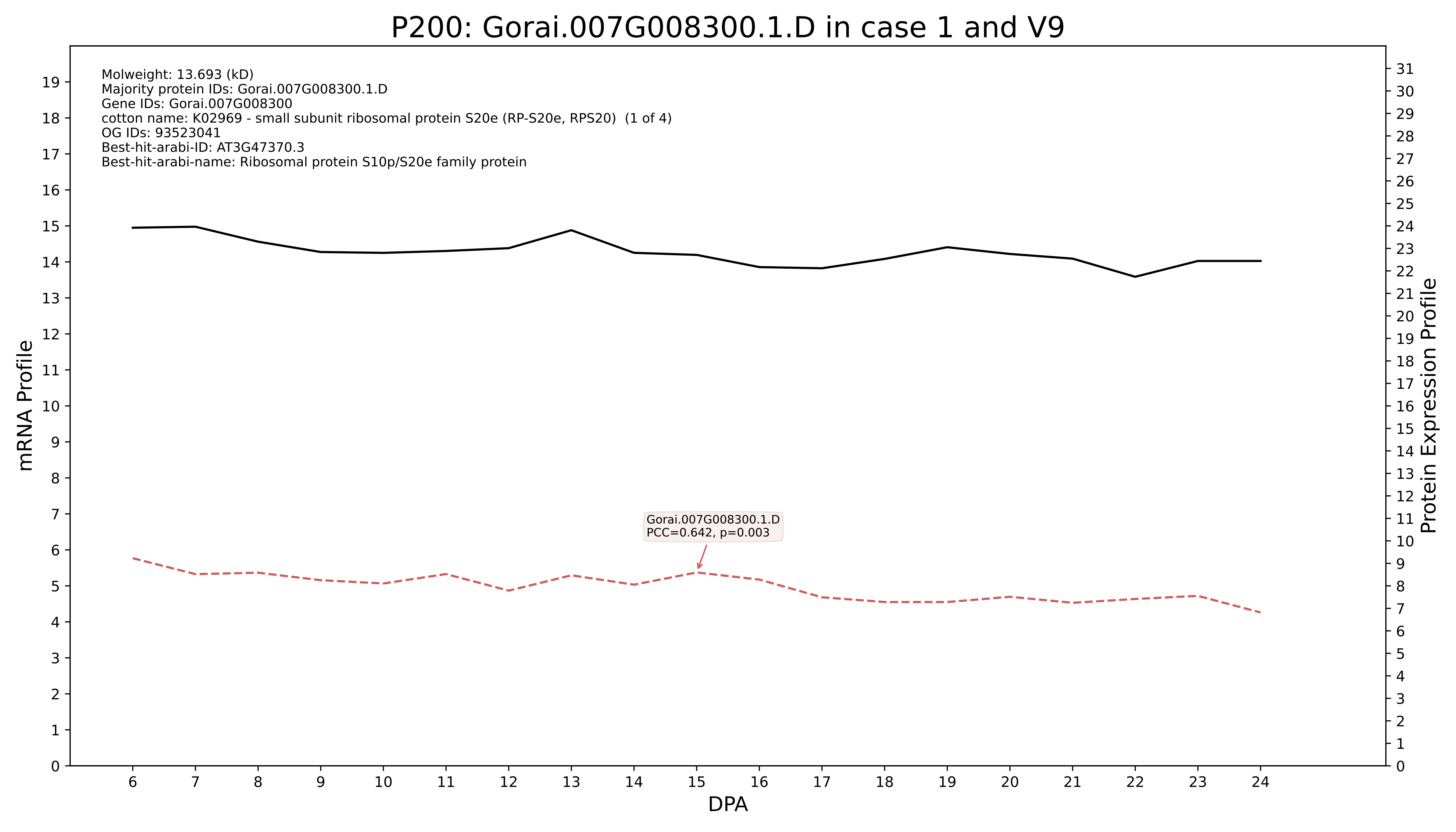
P200: Gorai.007G008300.1.D in case 1 and V9
Molweight: 13.693 (kD)
Majority protein IDs: Gorai.007G008300.1.D
Gene IDs: Gorai.007G008300
cotton name: K02969 - small subunit ribosomal protein S20e (RP-S20e, RPS20) (1 of 4)
OG IDs: 93523041
Best-hit-arabi-ID: AT3G47370.3
Best-hit-arabi-name: Ribosomal protein S10p/S20e family protein
Gorai.007G008300.1.D
PCC=0.642, p=0.003
mRNA Profile
Protein Expression Profile
DPA

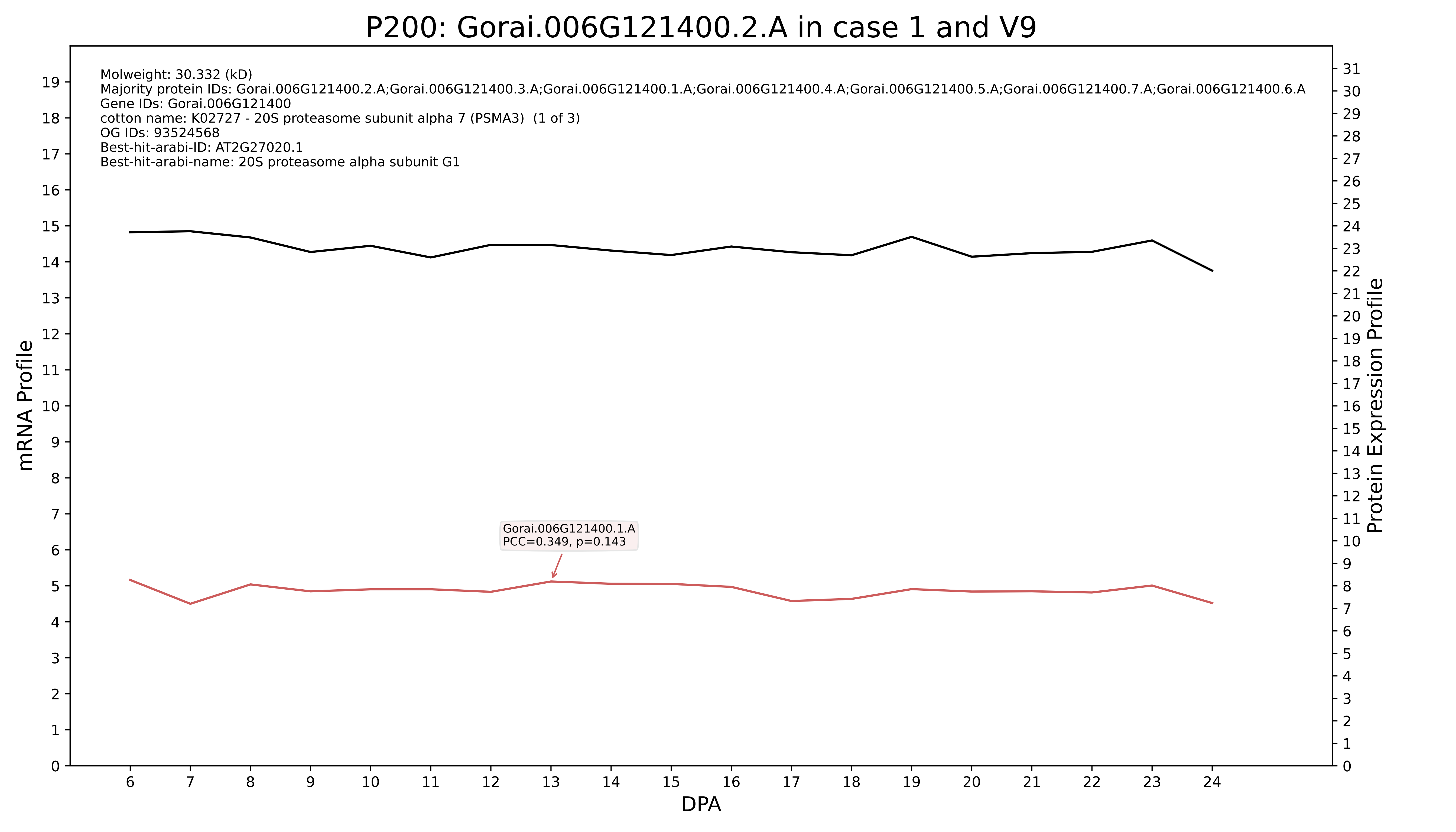

P200: Gorai.006G121400.2.A in case 1 and V9
Molweight: 30.332 (kD)
Majority protein IDs: Gorai.006G121400.2.A;Gorai.006G121400.3.A;Gorai.006G121400.1.A;Gorai.006G121400.4.A;Gorai.006G121400.5.A;Gorai.006G121400.7.A;Gorai.006G121400.6.A
Gene IDs: Gorai.006G121400
cotton name: K02727 - 20S proteasome subunit alpha 7 (PSMA3) (1 of 3)
OG IDs: 93524568
Best-hit-arabi-ID: AT2G27020.1
Best-hit-arabi-name: 20S proteasome alpha subunit G1
Gorai.006G121400.1.A
PCC=0.349, p=0.143
mRNA Profile
Protein Expression Profile
DPA

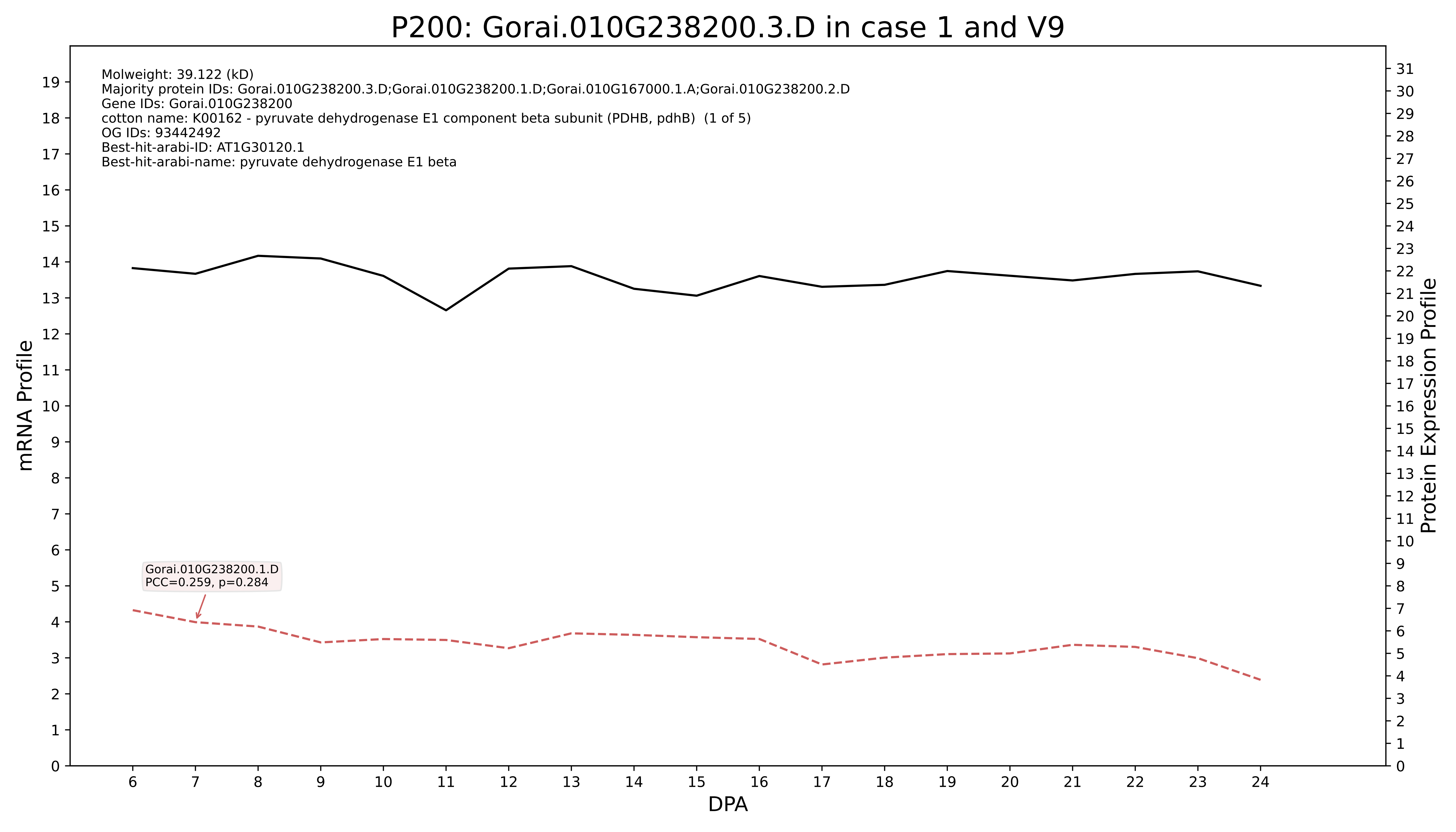
P200: Gorai.010G238200.3.D in case 1 and V9
Molweight: 39.122 (kD)
Majority protein IDs: Gorai.010G238200.3.D;Gorai.010G238200.1.D;Gorai.010G167000.1.A;Gorai.010G238200.2.D
Gene IDs: Gorai.010G238200
cotton name: K00162 - pyruvate dehydrogenase E1 component beta subunit (PDHB, pdhB) (1 of 5)
OG IDs: 93442492
Best-hit-arabi-ID: AT1G30120.1
Best-hit-arabi-name: pyruvate dehydrogenase E1 beta
Gorai.010G238200.1.D
PCC=0.259, p=0.284
mRNA Profile
Protein Expression Profile
DPA

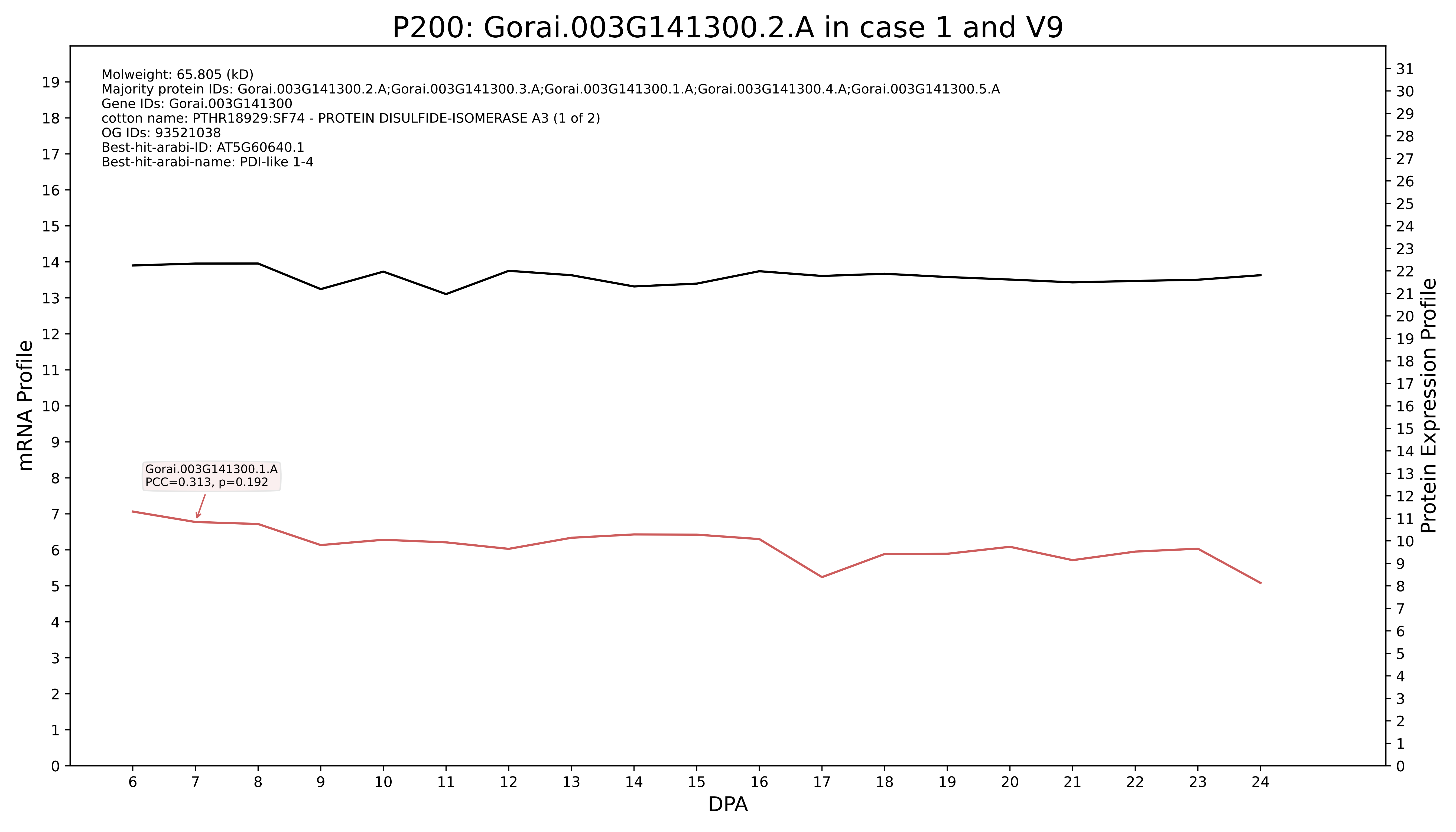

P200: Gorai.003G141300.2.A in case 1 and V9
Molweight: 65.805 (kD)
Majority protein IDs: Gorai.003G141300.2.A;Gorai.003G141300.3.A;Gorai.003G141300.1.A;Gorai.003G141300.4.A;Gorai.003G141300.5.A
Gene IDs: Gorai.003G141300
cotton name: PTHR18929:SF74 - PROTEIN DISULFIDE-ISOMERASE A3 (1 of 2)
OG IDs: 93521038
Best-hit-arabi-ID: AT5G60640.1
Best-hit-arabi-name: PDI-like 1-4
Gorai.003G141300.1.A
PCC=0.313, p=0.192
mRNA Profile
Protein Expression Profile
DPA

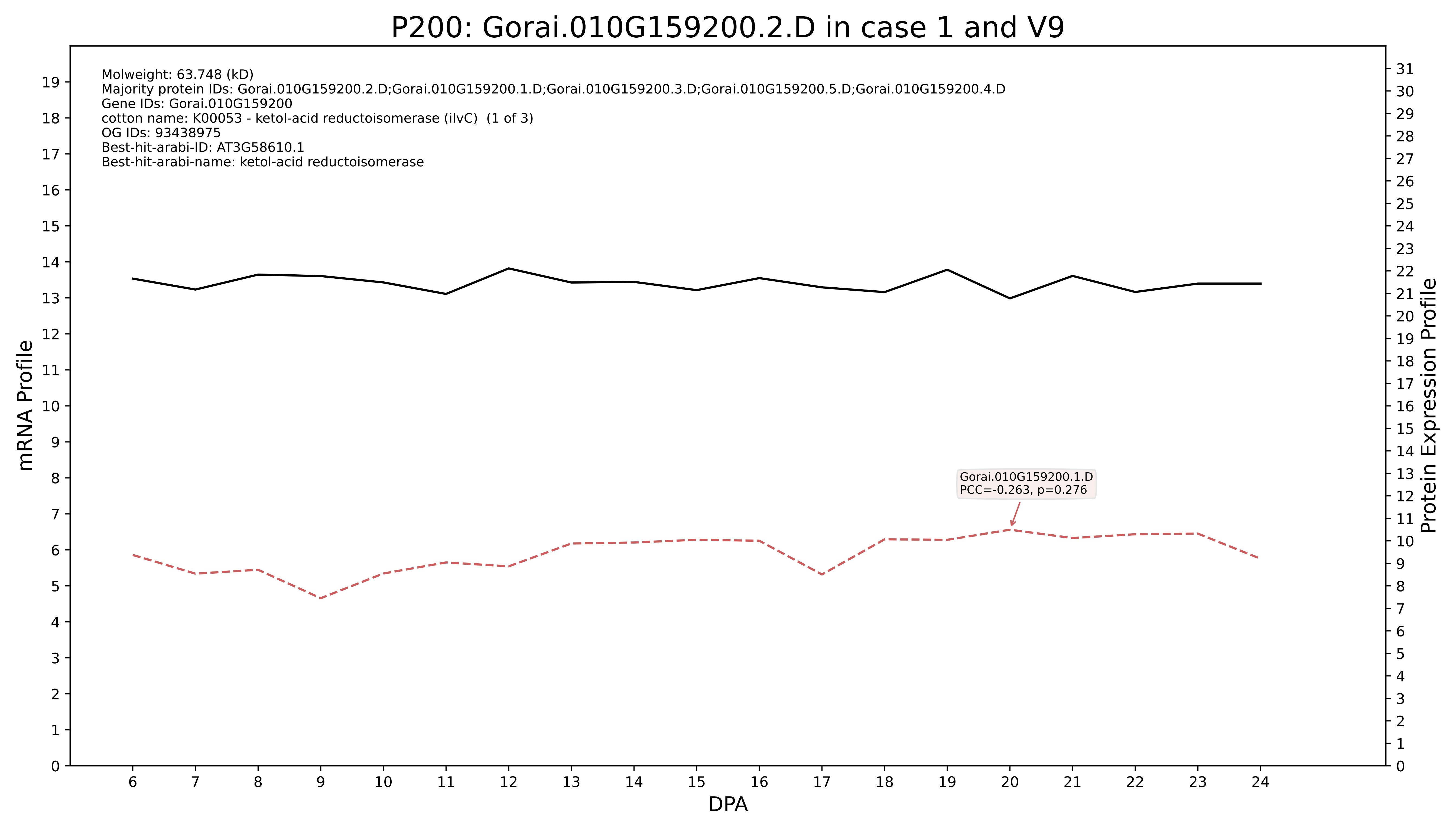

P200: Gorai.010G159200.2.D in case 1 and V9
Molweight: 63.748 (kD)
Majority protein IDs: Gorai.010G159200.2.D;Gorai.010G159200.1.D;Gorai.010G159200.3.D;Gorai.010G159200.5.D;Gorai.010G159200.4.D
Gene IDs: Gorai.010G159200
cotton name: K00053 - ketol-acid reductoisomerase (ilvC) (1 of 3)
OG IDs: 93438975
Best-hit-arabi-ID: AT3G58610.1
Best-hit-arabi-name: ketol-acid reductoisomerase
Gorai.010G159200.1.D
PCC=-0.263, p=0.276
mRNA Profile
Protein Expression Profile
DPA

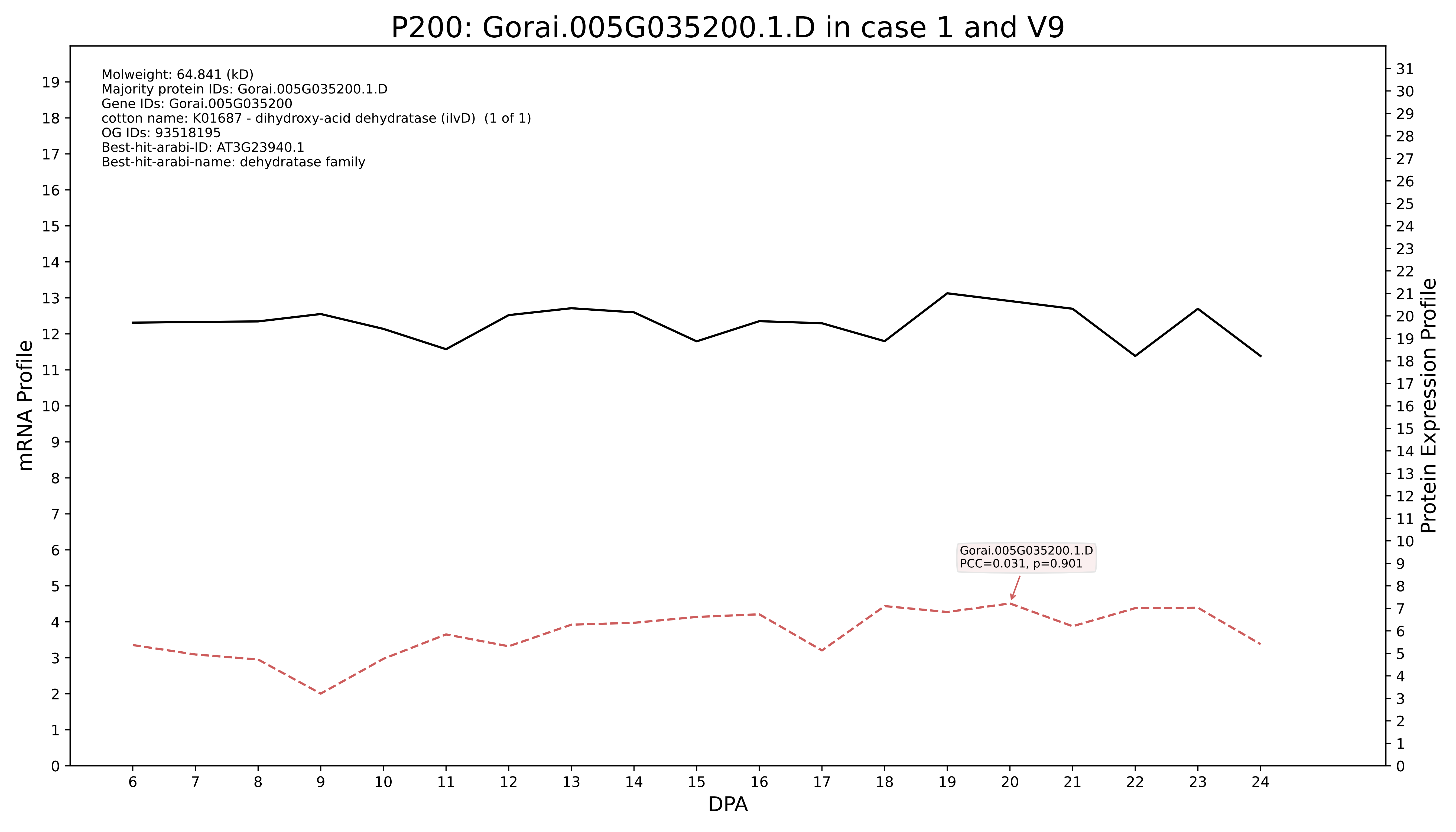
P200: Gorai.005G035200.1.D in case 1 and V9
Molweight: 64.841 (kD)
Majority protein IDs: Gorai.005G035200.1.D
Gene IDs: Gorai.005G035200
cotton name: K01687 - dihydroxy-acid dehydratase (ilvD) (1 of 1)
OG IDs: 93518195
Best-hit-arabi-ID: AT3G23940.1
Best-hit-arabi-name: dehydratase family
Gorai.005G035200.1.D
PCC=0.031, p=0.901
mRNA Profile
Protein Expression Profile
DPA

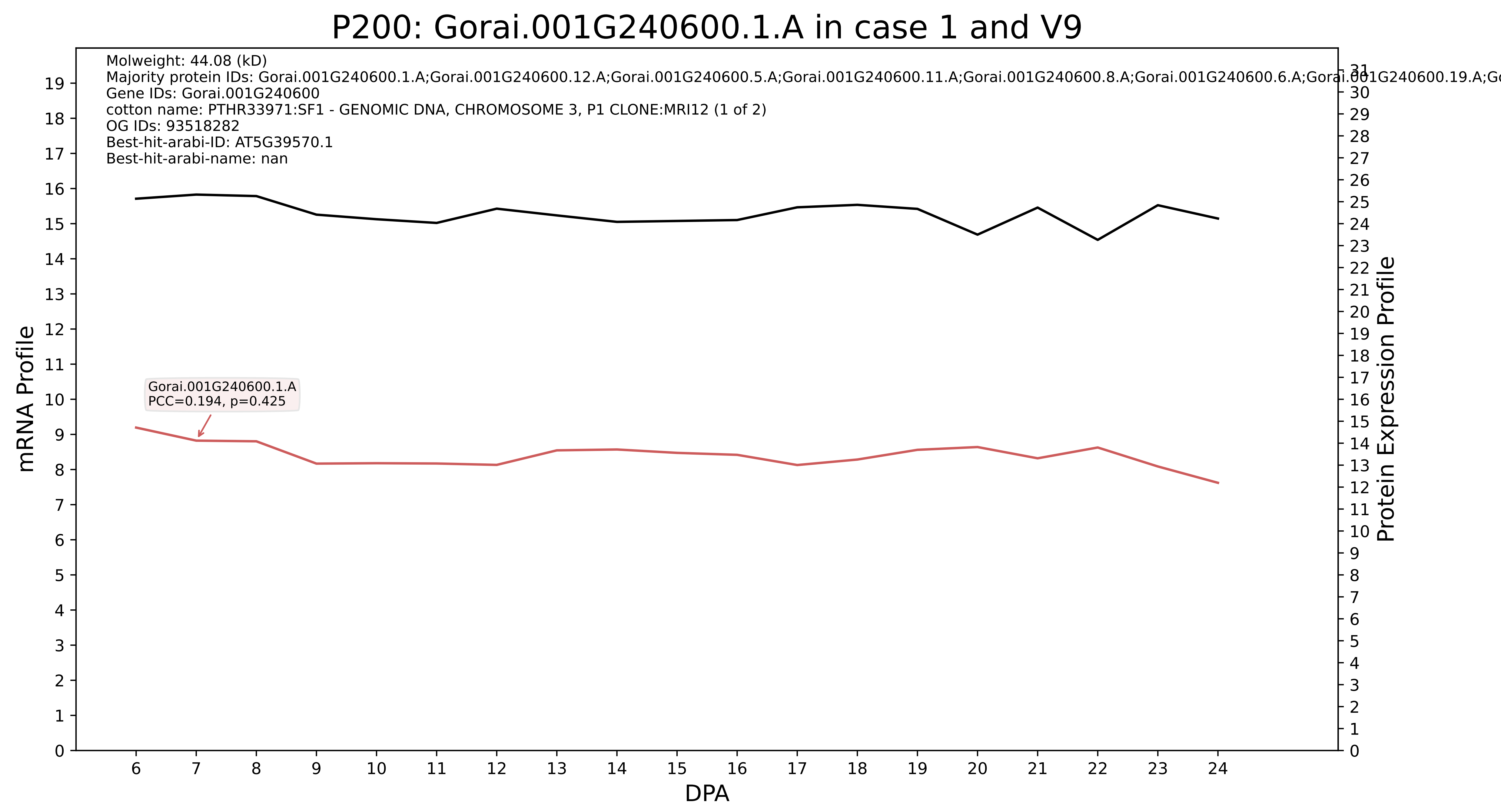
P200: Gorai.001G240600.1.A in case 1 and V9
Molweight: 44.08 (kD)
Majority protein IDs: Gorai.001G240600.1.A;Gorai.001G240600.12.A;Gorai.001G240600.5.A;Gorai.001G240600.11.A;Gorai.001G240600.8.A;Gorai.001G240600.6.A;Gorai.001G240600.19.A;Go
Gene IDs: Gorai.001G240600
cotton name: PTHR33971:SF1 - GENOMIC DNA, CHROMOSOME 3, P1 CLONE:MRI12 (1 of 2)
OG IDs: 93518282
Best-hit-arabi-ID: AT5G39570.1
Best-hit-arabi-name: nan
Gorai.001G240600.1.A
PCC=0.194, p=0.425
mRNA Profile
Protein Expression Profile
DPA

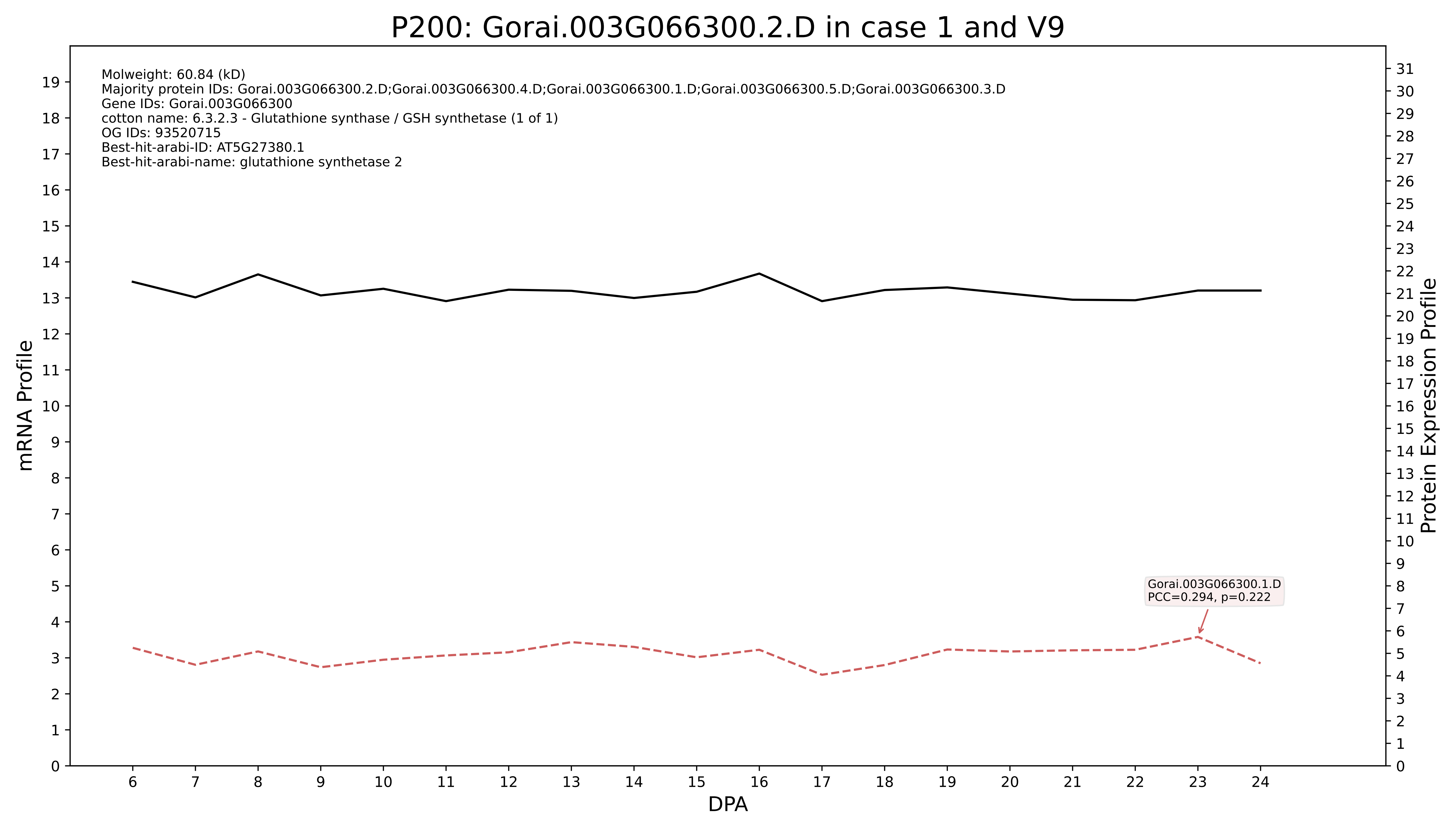
P200: Gorai.003G066300.2.D in case 1 and V9
Molweight: 60.84 (kD)
Majority protein IDs: Gorai.003G066300.2.D;Gorai.003G066300.4.D;Gorai.003G066300.1.D;Gorai.003G066300.5.D;Gorai.003G066300.3.D
Gene IDs: Gorai.003G066300
cotton name: 6.3.2.3 - Glutathione synthase / GSH synthetase (1 of 1)
OG IDs: 93520715
Best-hit-arabi-ID: AT5G27380.1
Best-hit-arabi-name: glutathione synthetase 2
Gorai.003G066300.1.D
PCC=0.294, p=0.222
mRNA Profile
Protein Expression Profile
DPA

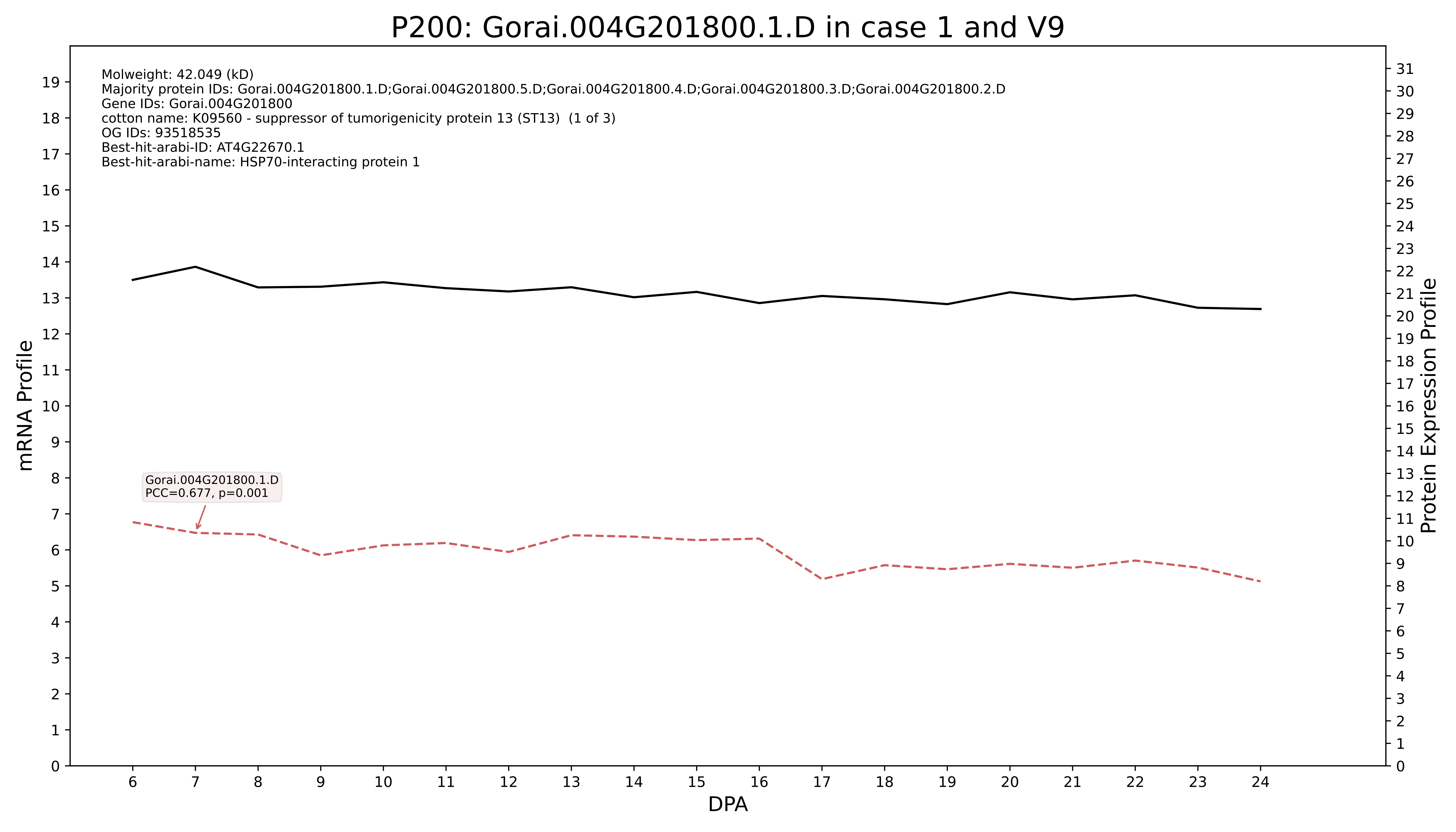
P200: Gorai.004G201800.1.D in case 1 and V9
Molweight: 42.049 (kD)
Majority protein IDs: Gorai.004G201800.1.D;Gorai.004G201800.5.D;Gorai.004G201800.4.D;Gorai.004G201800.3.D;Gorai.004G201800.2.D
Gene IDs: Gorai.004G201800
cotton name: K09560 - suppressor of tumorigenicity protein 13 (ST13) (1 of 3)
OG IDs: 93518535
Best-hit-arabi-ID: AT4G22670.1
Best-hit-arabi-name: HSP70-interacting protein 1
Gorai.004G201800.1.D
PCC=0.677, p=0.001
mRNA Profile
Protein Expression Profile
DPA

### P200: Gorai.011G044400.1.D in case 1 and V9

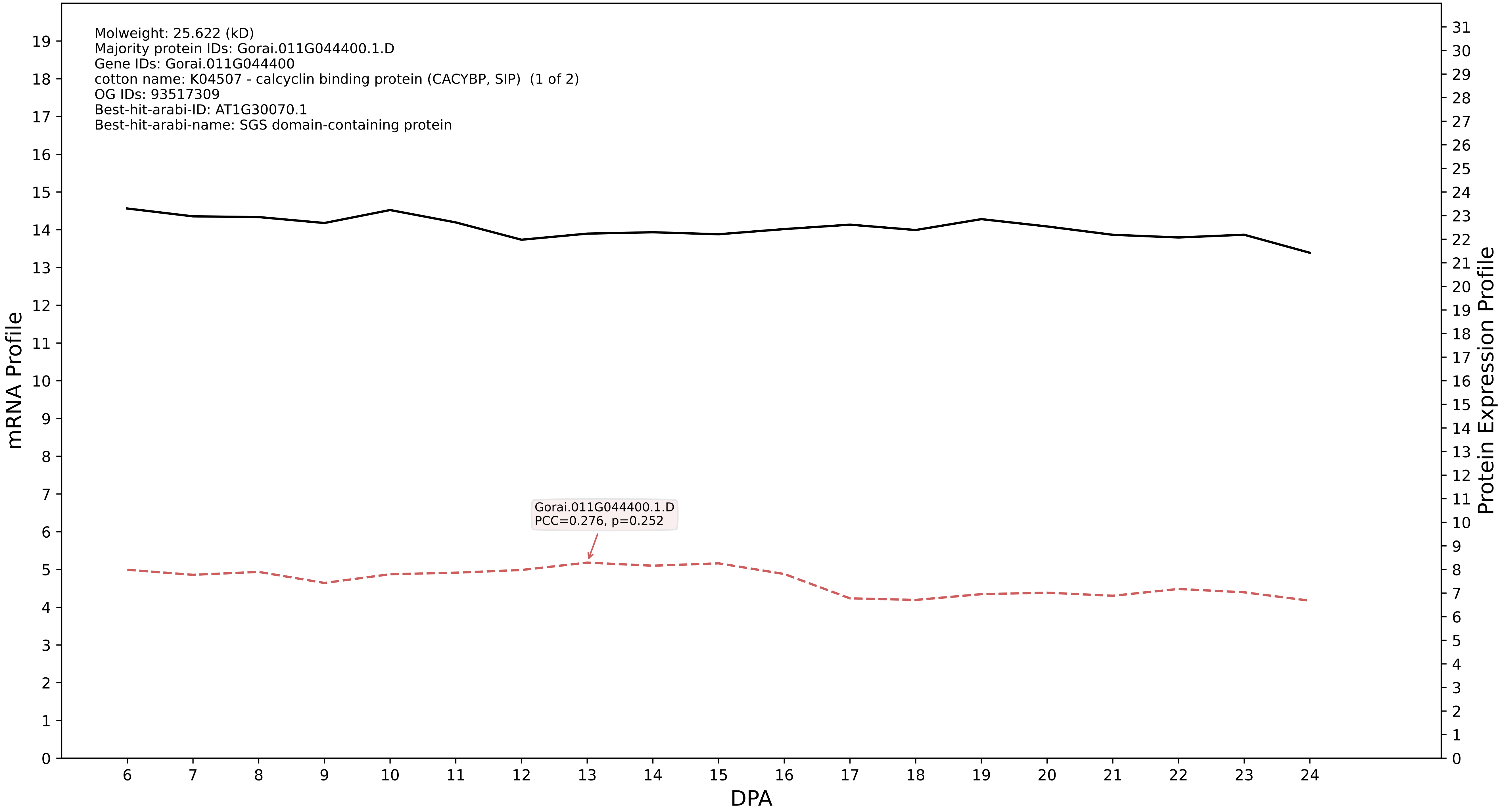

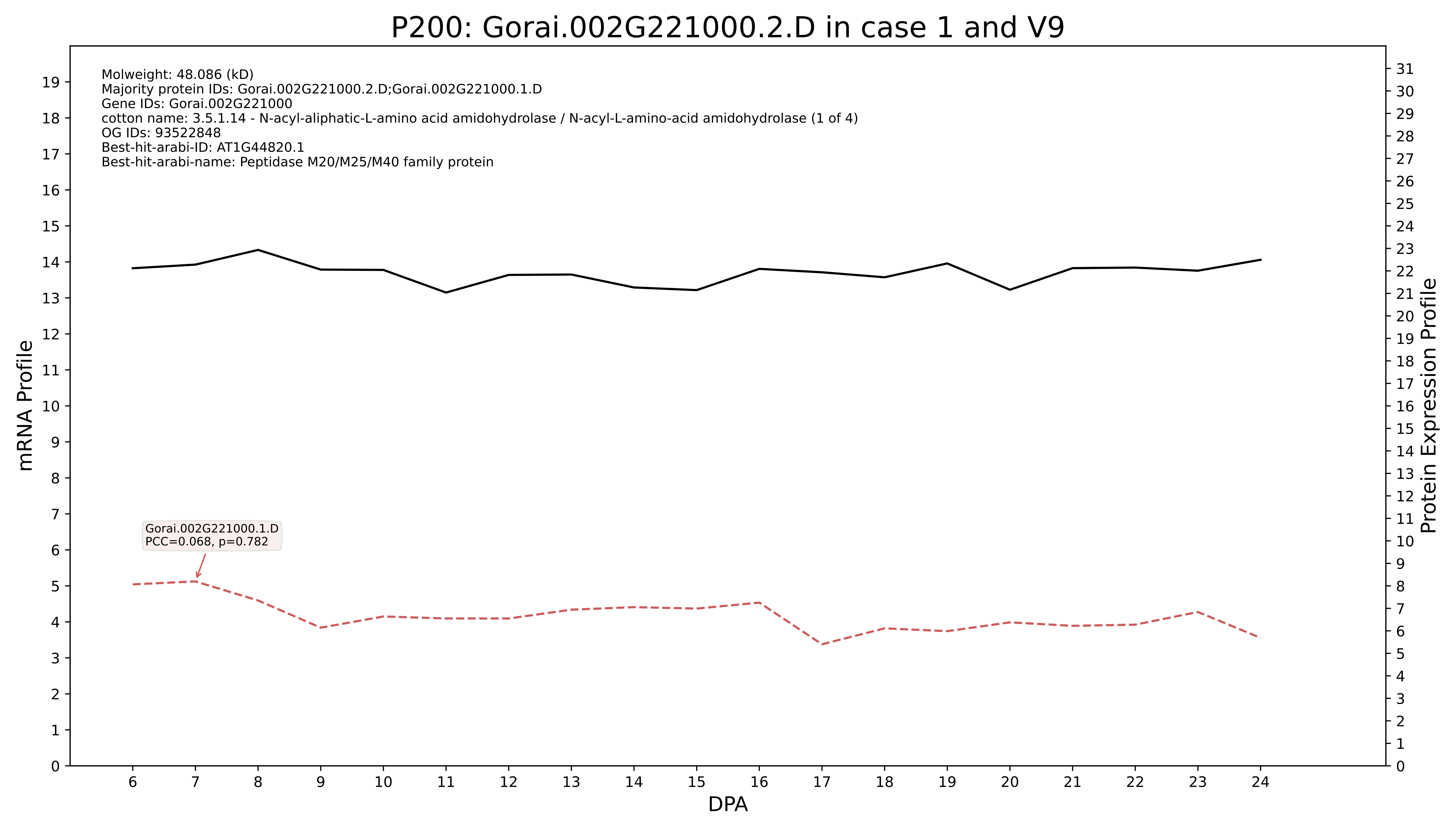
P200: Gorai.002G221000.2.D in case 1 and V9
Molweight: 48.086 (kD)
Majority protein IDs: Gorai.002G221000.2.D;Gorai.002G221000.1.D
Gene IDs: Gorai.002G221000
cotton name: 3.5.1.14 - N-acyl-aliphatic-L-amino acid amidohydrolase / N-acyl-L-amino-acid amidohydrolase (1 of 4)
OG IDs: 93522848
Best-hit-arabi-ID: AT1G44820.1
Best-hit-arabi-name: Peptidase M20/M25/M40 family protein
Gorai.002G221000.1.D
PCC=0.068, p=0.782
mRNA Profile
Protein Expression Profile
DPA

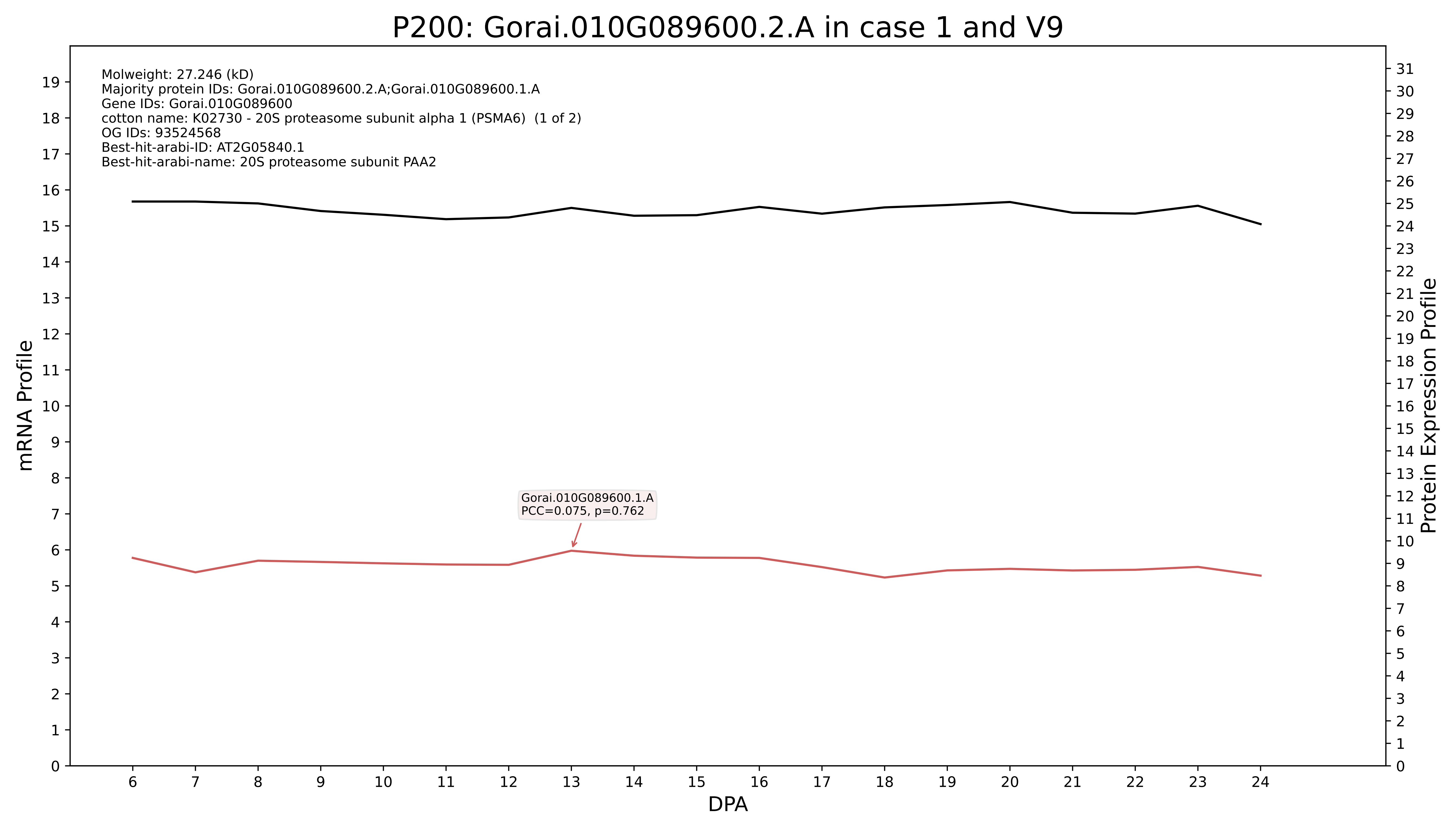
P200: Gorai.010G089600.2.A in case 1 and V9
Molweight: 27.246 (kD)
Majority protein IDs: Gorai.010G089600.2.A;Gorai.010G089600.1.A
Gene IDs: Gorai.010G089600
cotton name: K02730 - 20S proteasome subunit alpha 1 (PSMA6) (1 of 2)
OG IDs: 93524568
Best-hit-arabi-ID: AT2G05840.1
Best-hit-arabi-name: 20S proteasome subunit PAA2
Gorai.010G089600.1.A
PCC=0.075, p=0.762
mRNA Profile
Protein Expression Profile
DPA

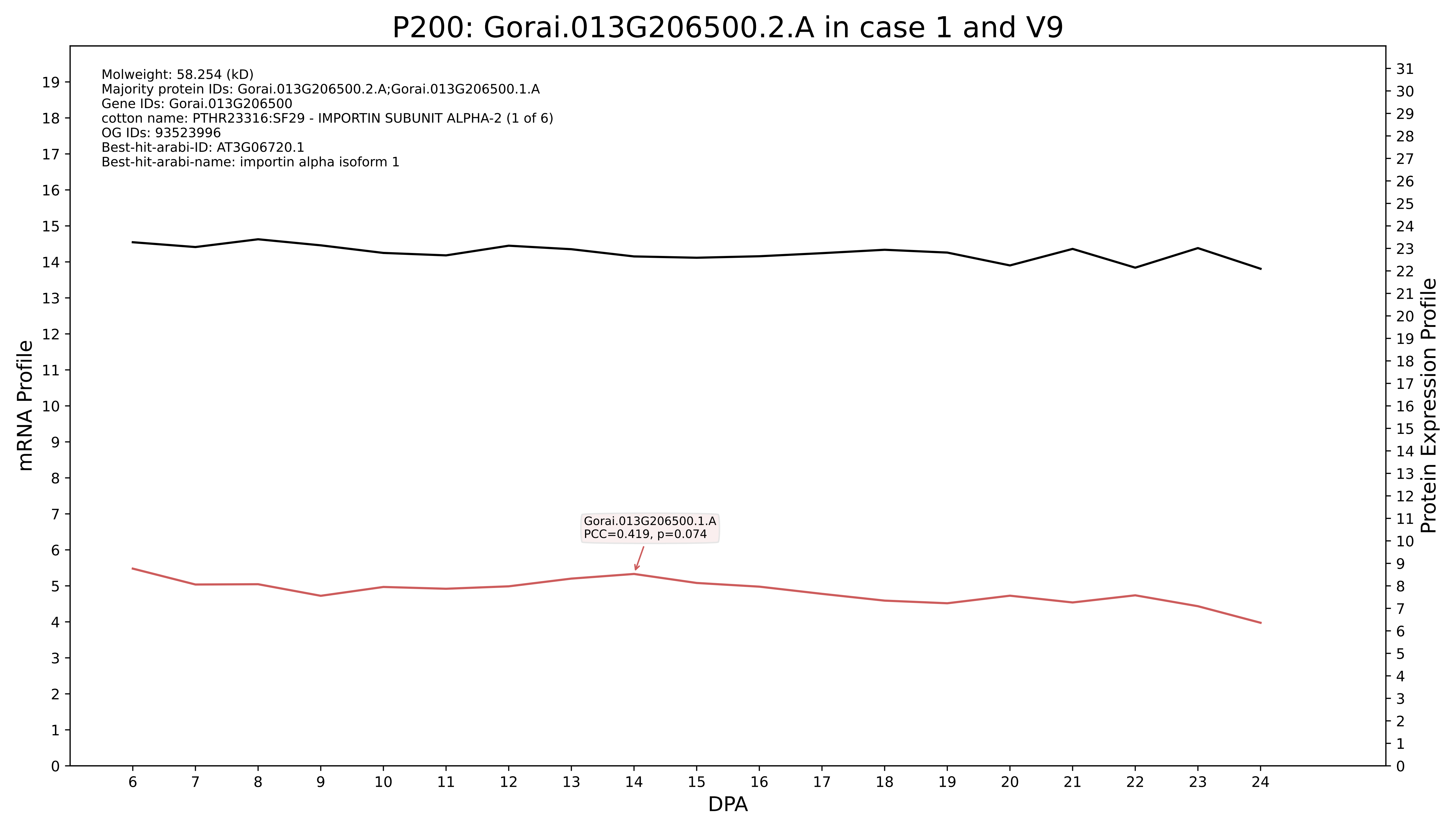
P200: Gorai.013G206500.2.A in case 1 and V9
Molweight: 58.254 (kD)
Majority protein IDs: Gorai.013G206500.2.A;Gorai.013G206500.1.A
Gene IDs: Gorai.013G206500
cotton name: PTHR23316:SF29 - IMPORTIN SUBUNIT ALPHA-2 (1 of 6)
OG IDs: 93523996
Best-hit-arabi-ID: AT3G06720.1
Best-hit-arabi-name: importin alpha isoform 1
Gorai.013G206500.1.A
PCC=0.419, p=0.074
mRNA Profile
Protein Expression Profile
DPA

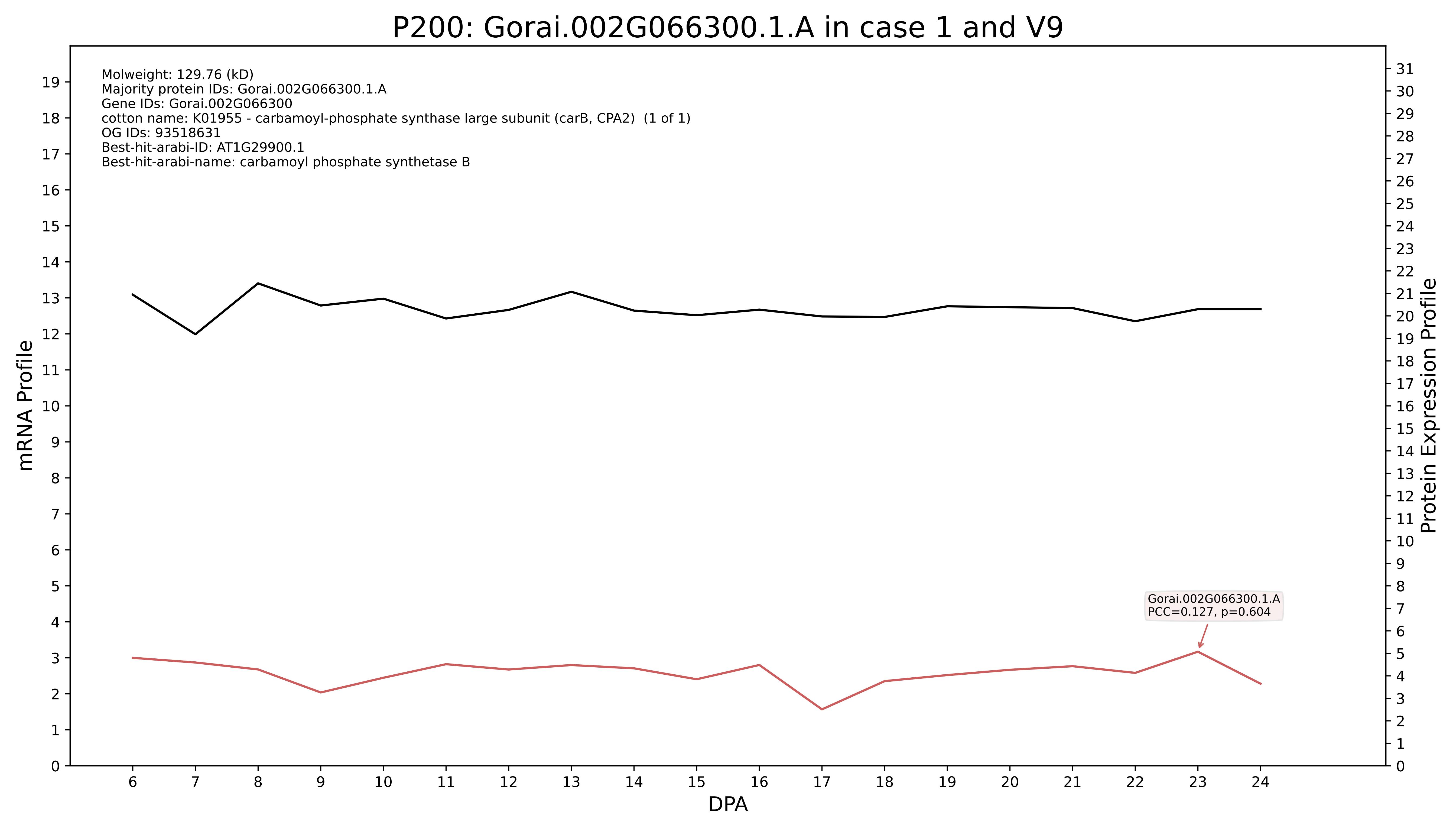

P200: Gorai.002G066300.1.A in case 1 and V9
Molweight: 129.76 (kD)
Majority protein IDs: Gorai.002G066300.1.A
Gene IDs: Gorai.002G066300
cotton name: K01955 - carbamoyl-phosphate synthase large subunit (carB, CPA2) (1 of 1)
OG IDs: 93518631
Best-hit-arabi-ID: AT1G29900.1
Best-hit-arabi-name: carbamoyl phosphate synthetase B
Gorai.002G066300.1.A
PCC=0.127, p=0.604
mRNA Profile
Protein Expression Profile
DPA

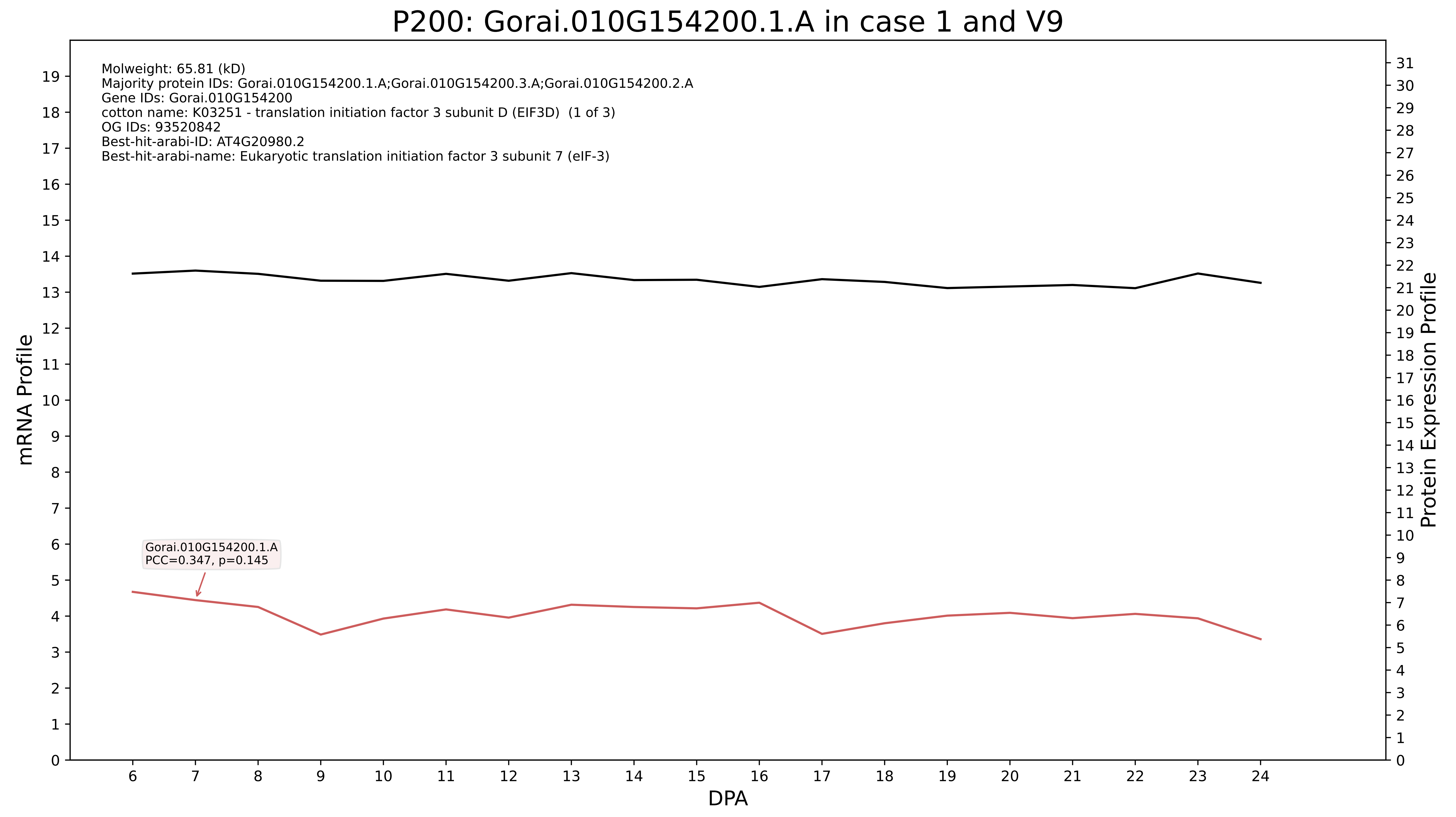
P200: Gorai.010G154200.1.A in case 1 and V9
Molweight: 65.81 (kD)
Majority protein IDs: Gorai.010G154200.1.A;Gorai.010G154200.3.A;Gorai.010G154200.2.A
Gene IDs: Gorai.010G154200
cotton name: K03251 - translation initiation factor 3 subunit D (EIF3D) (1 of 3)
OG IDs: 93520842
Best-hit-arabi-ID: AT4G20980.2
Best-hit-arabi-name: Eukaryotic translation initiation factor 3 subunit 7 (eIF-3)
Gorai.010G154200.1.A
PCC=0.347, p=0.145
mRNA Profile
Protein Expression Profile
DPA

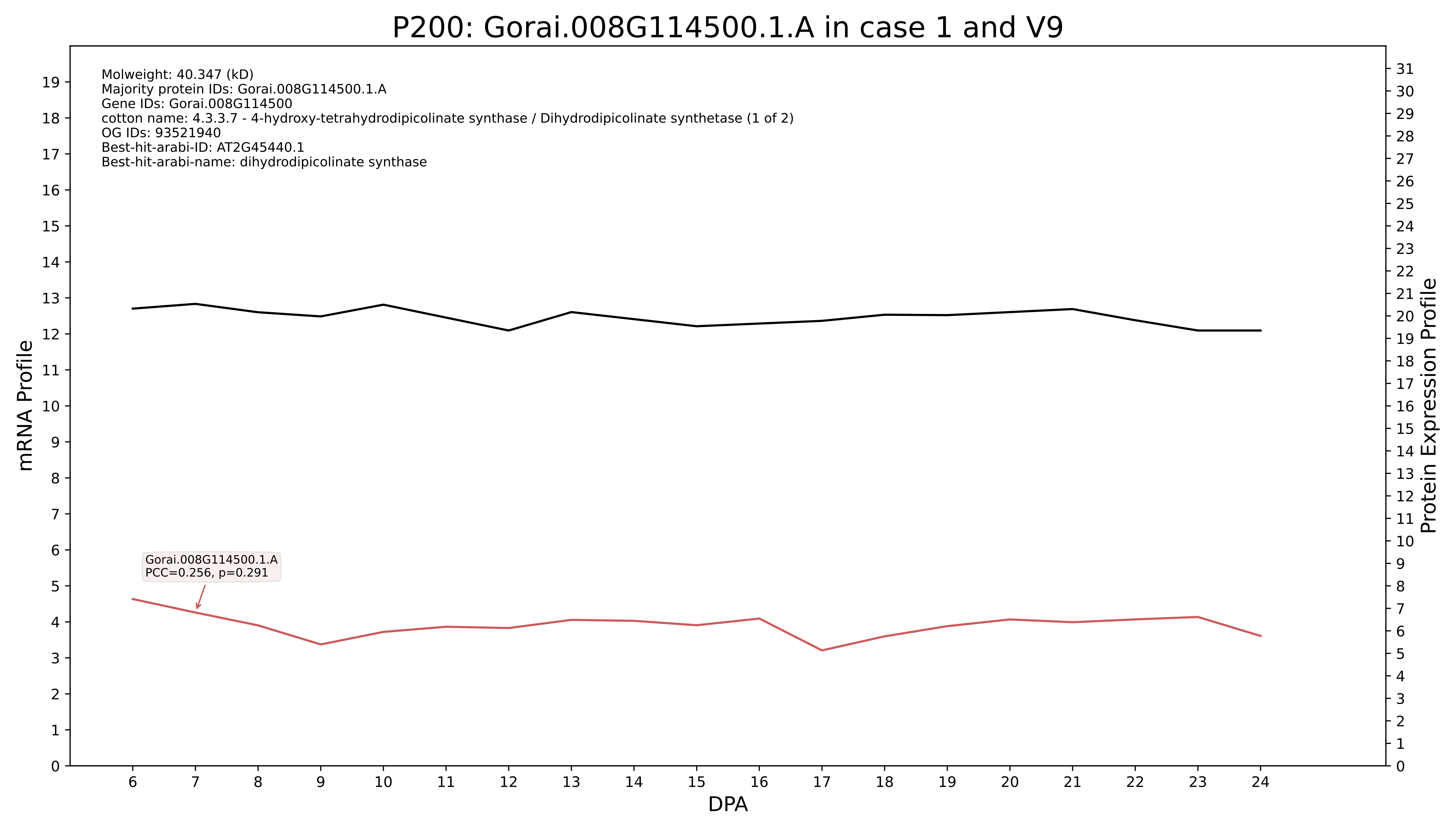
P200: Gorai.008G114500.1.A in case 1 and V9
Molweight: 40.347 (kD)
Majority protein IDs: Gorai.008G114500.1.A
Gene IDs: Gorai.008G114500
cotton name: 4.3.3.7 - 4-hydroxy-tetrahydrodipicolinate synthase / Dihydrodipicolinate synthetase (1 of 2)
OG IDs: 93521940
Best-hit-arabi-ID: AT2G45440.1
Best-hit-arabi-name: dihydrodipicolinate synthase
Gorai.008G114500.1.A
PCC=0.256, p=0.291
mRNA Profile
Protein Expression Profile
DPA

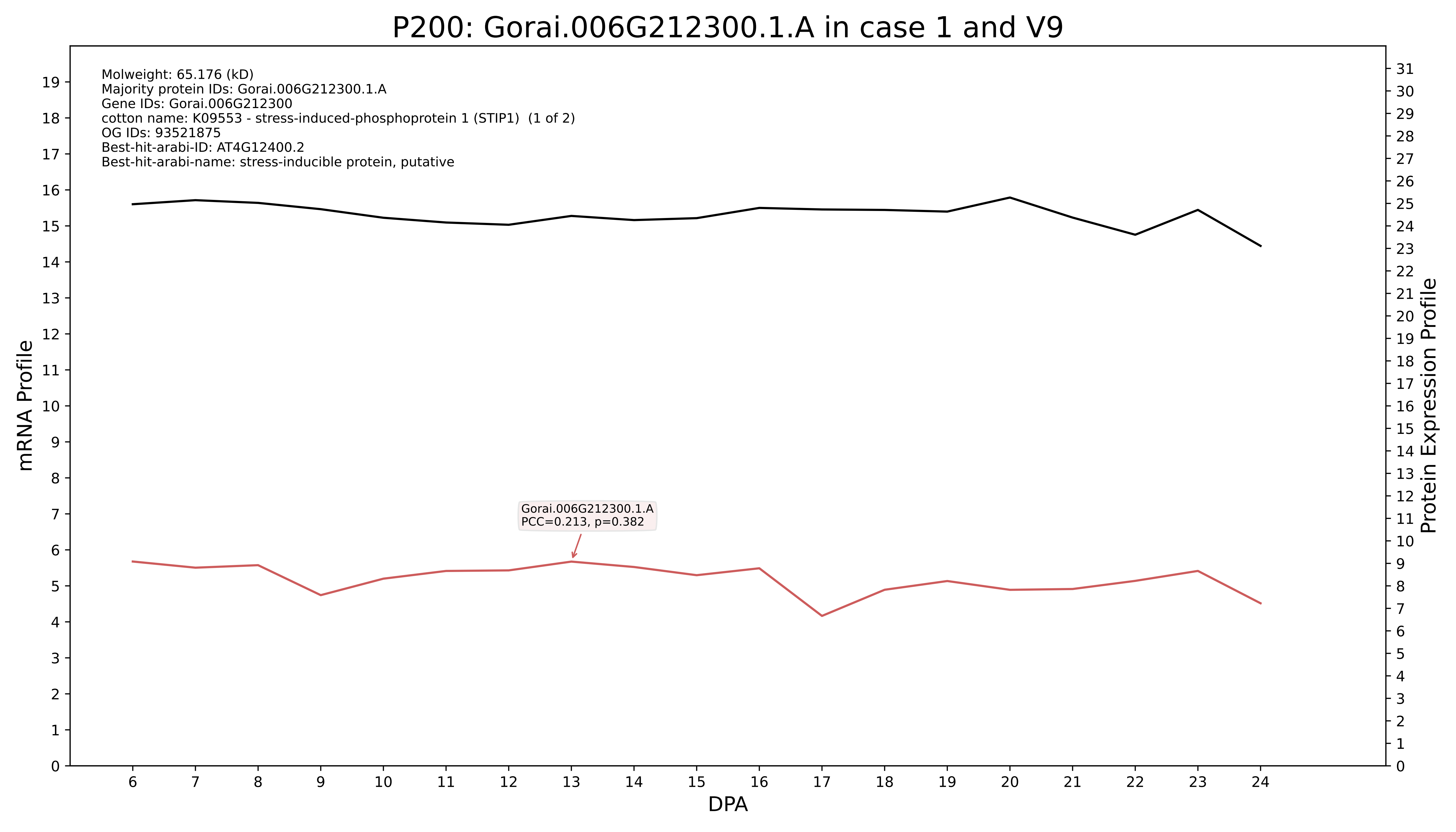
P200: Gorai.006G212300.1.A in case 1 and V9
Molweight: 65.176 (kD)
Majority protein IDs: Gorai.006G212300.1.A
Gene IDs: Gorai.006G212300
cotton name: K09553 - stress-induced-phosphoprotein 1 (STIP1) (1 of 2)
OG IDs: 93521875
Best-hit-arabi-ID: AT4G12400.2
Best-hit-arabi-name: stress-inducible protein, putative
Gorai.006G212300.1.A
PCC=0.213, p=0.382
mRNA Profile
Protein Expression Profile
DPA

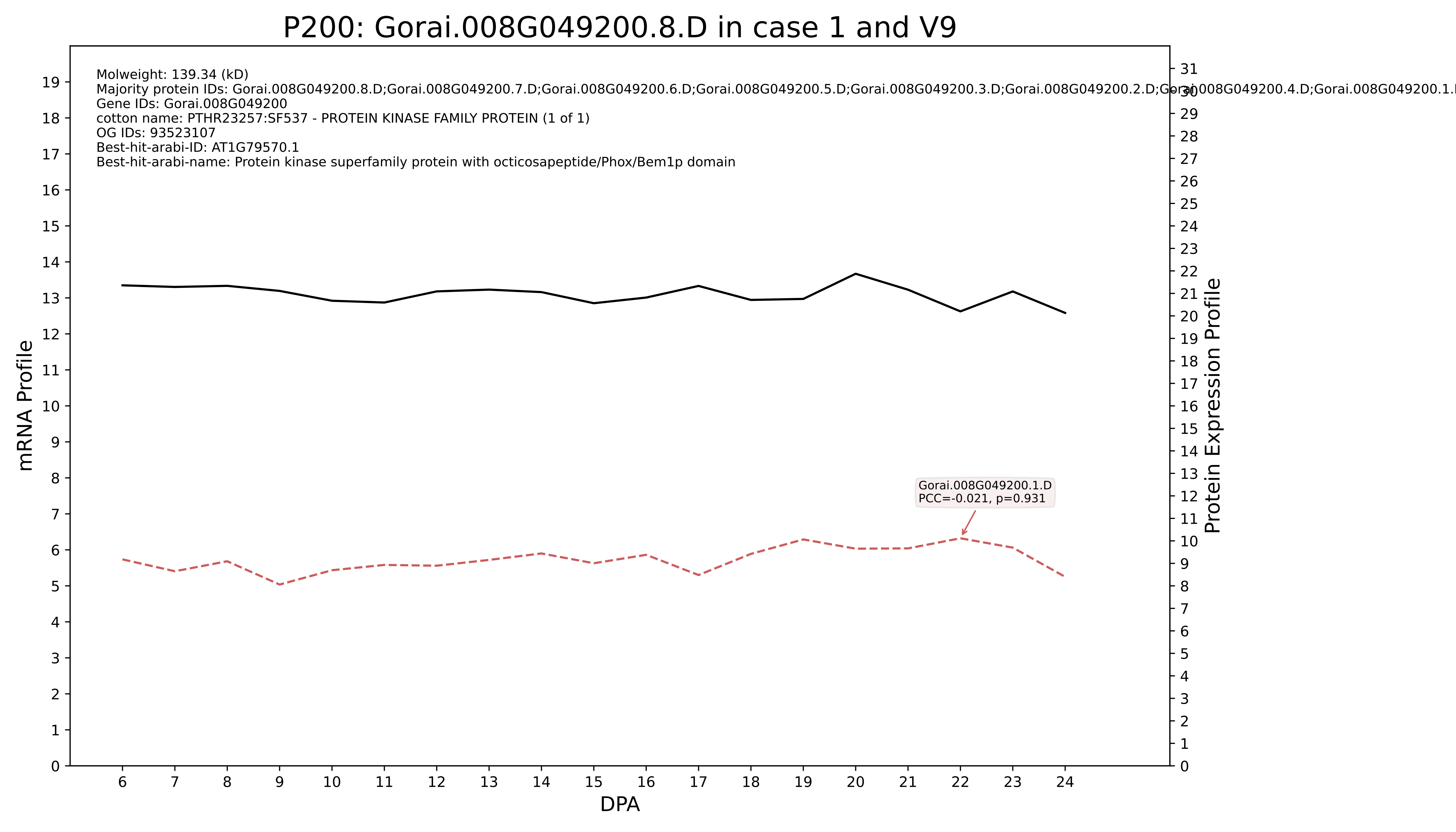
P200: Gorai.008G049200.8.D in case 1 and V9
Molweight: 139.34 (kD)
Majority protein IDs: Gorai.008G049200.8.D;Gorai.008G049200.7.D;Gorai.008G049200.6.D;Gorai.008G049200.5.D;Gorai.008G049200.3.D;Gorai.008G049200.2.D;Gorai.008G049200.4.D;Gorai.008G049200.1.D
Gene IDs: Gorai.008G049200
cotton name: PTHR23257:SF537 - PROTEIN KINASE FAMILY PROTEIN (1 of 1)
OG IDs: 93523107
Best-hit-arabi-ID: AT1G79570.1
Best-hit-arabi-name: Protein kinase superfamily protein with octicosapeptide/Phox/Bem1p domain
Gorai.008G049200.1.D
PCC=-0.021, p=0.931
mRNA Profile
Protein Expression Profile
DPA

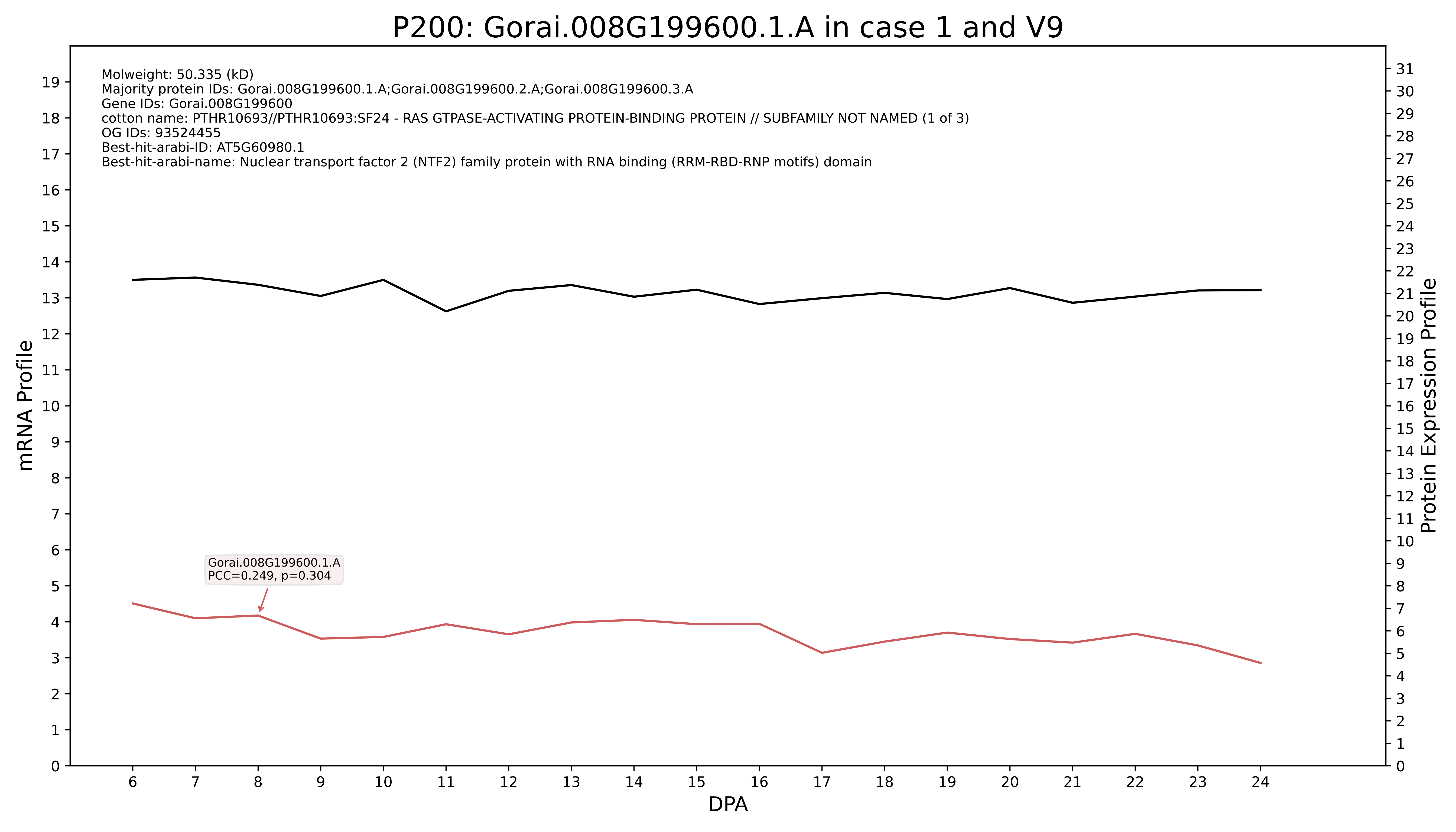
P200: Gorai.008G199600.1.A in case 1 and V9
Molweight: 50.335 (kD)
Majority protein IDs: Gorai.008G199600.1.A;Gorai.008G199600.2.A;Gorai.008G199600.3.A
Gene IDs: Gorai.008G199600
cotton name: PTHR10693//PTHR10693:SF24 - RAS GTPASE-ACTIVATING PROTEIN-BINDING PROTEIN // SUBFAMILY NOT NAMED (1 of 3)
OG IDs: 93524455
Best-hit-arabi-ID: AT5G60980.1
Best-hit-arabi-name: Nuclear transport factor 2 (NTF2) family protein with RNA binding (RRM-RBD-RNP motifs) domain
Gorai.008G199600.1.A
PCC=0.249, p=0.304
mRNA Profile
Protein Expression Profile
DPA

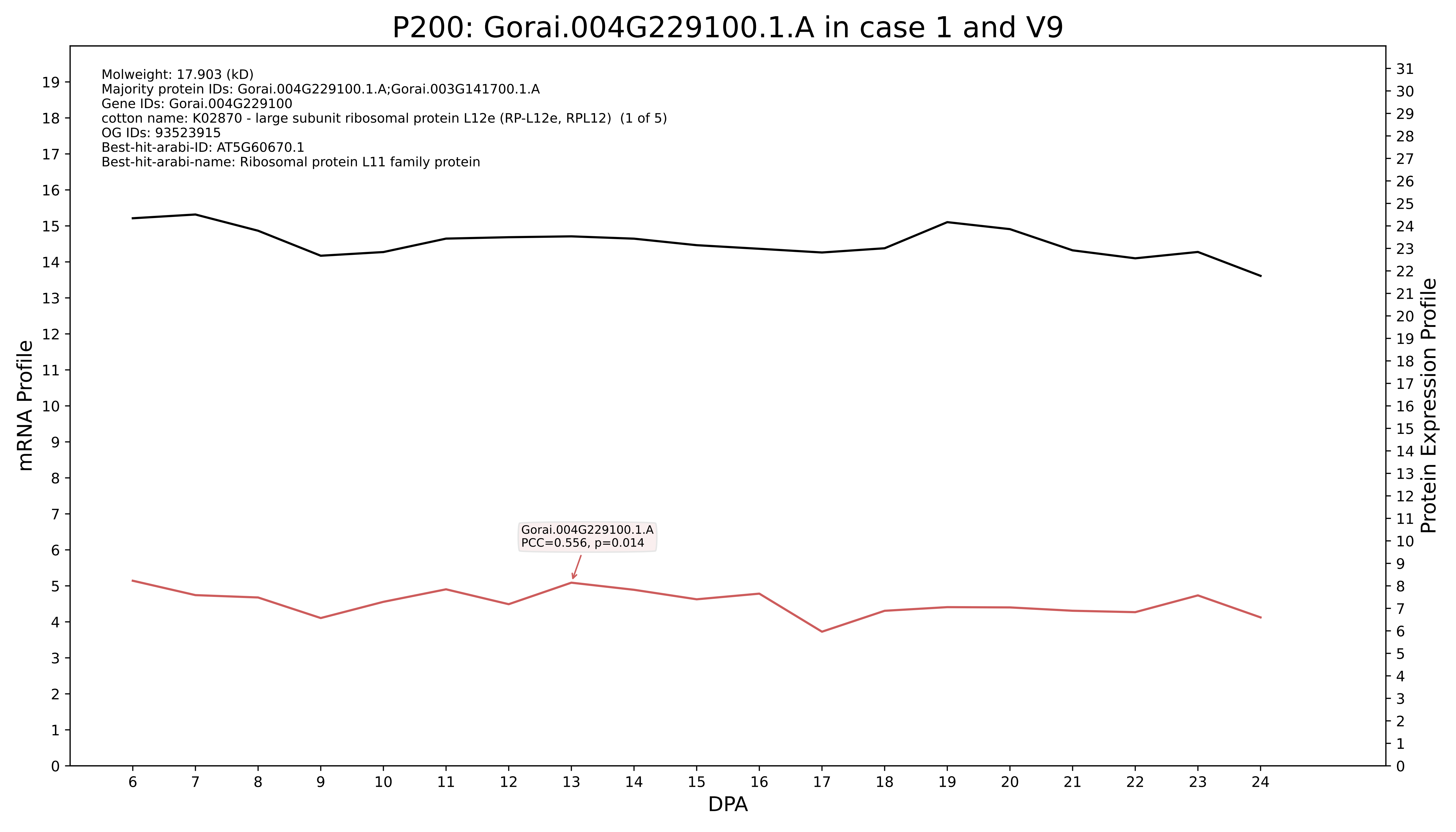
P200: Gorai.004G229100.1.A in case 1 and V9
Molweight: 17.903 (kD)
Majority protein IDs: Gorai.004G229100.1.A;Gorai.003G141700.1.A
Gene IDs: Gorai.004G229100
cotton name: K02870 - large subunit ribosomal protein L12e (RP-L12e, RPL12) (1 of 5)
OG IDs: 93523915
Best-hit-arabi-ID: AT5G60670.1
Best-hit-arabi-name: Ribosomal protein L11 family protein
Gorai.004G229100.1.A
PCC=0.556, p=0.014
mRNA Profile
Protein Expression Profile
DPA

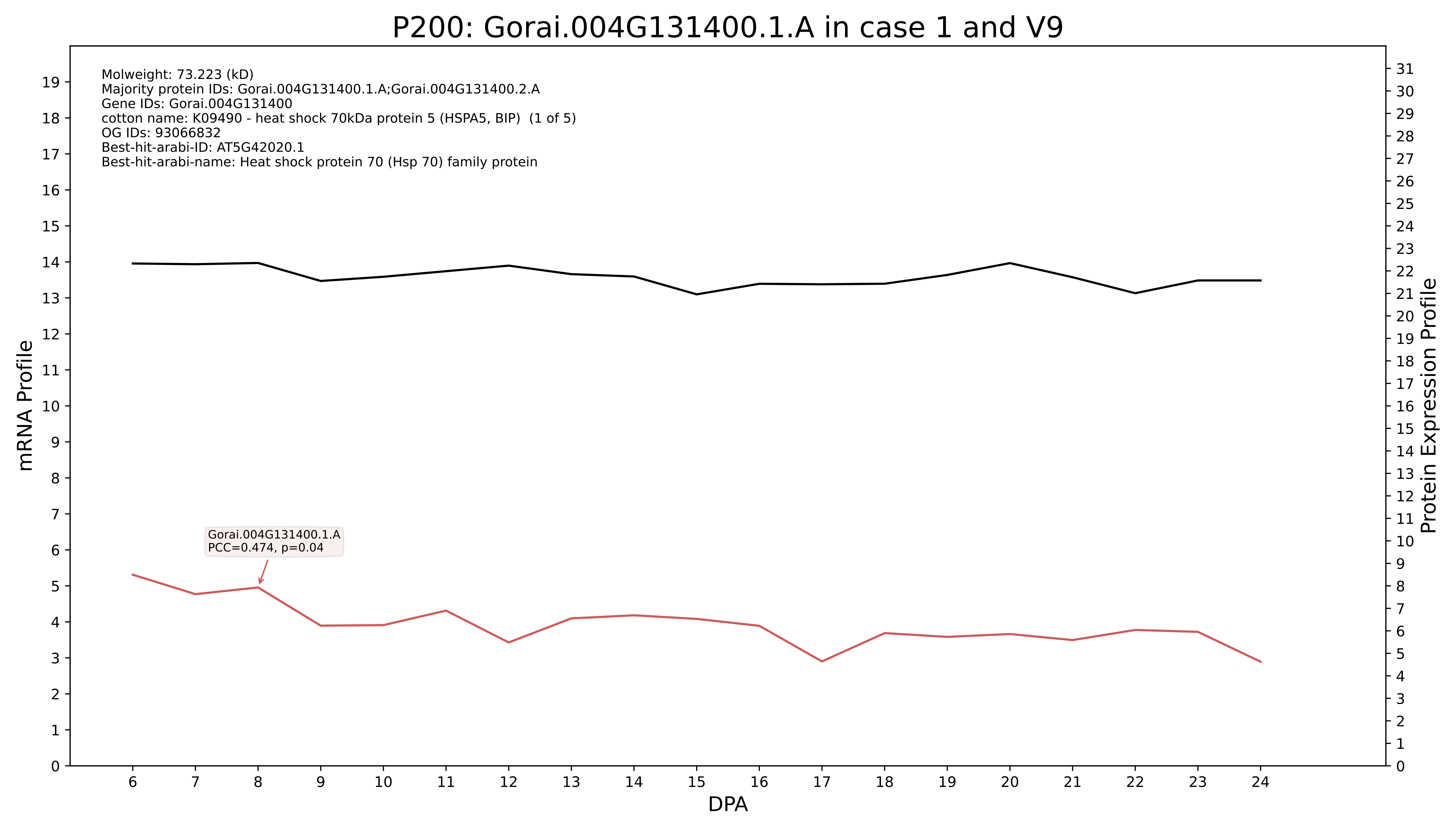
P200: Gorai.004G131400.1.A in case 1 and V9
Molweight: 73.223 (kD)
Majority protein IDs: Gorai.004G131400.1.A;Gorai.004G131400.2.A
Gene IDs: Gorai.004G131400
cotton name: K09490 - heat shock 70kDa protein 5 (HSPA5, BIP) (1 of 5)
OG IDs: 93066832
Best-hit-arabi-ID: AT5G42020.1
Best-hit-arabi-name: Heat shock protein 70 (Hsp 70) family protein
Gorai.004G131400.1.A
PCC=0.474, p=0.04
mRNA Profile
Protein Expression Profile
DPA

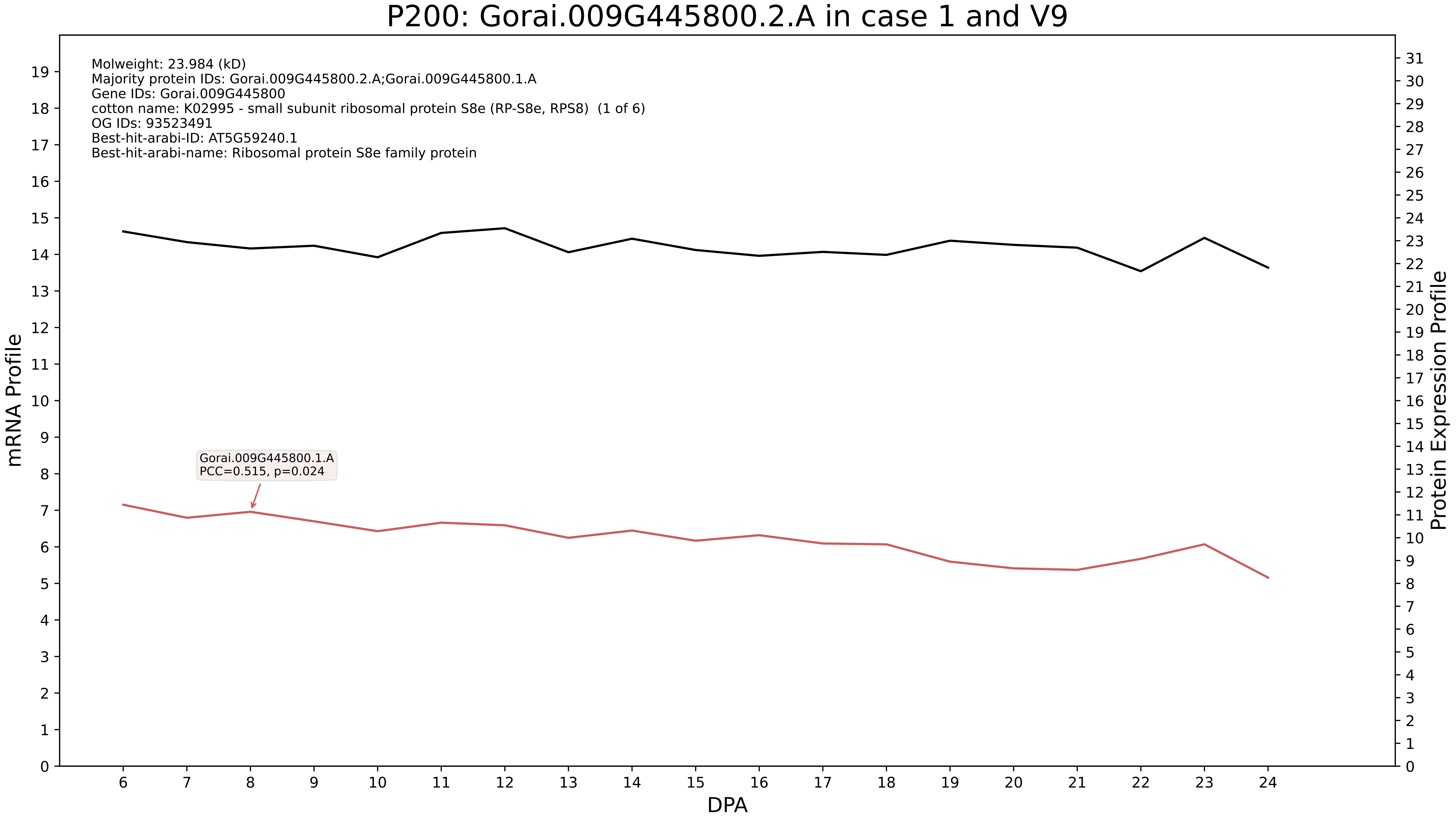

P200: Gorai.009G445800.2.A in case 1 and V9
Molweight: 23.984 (kD)
Majority protein IDs: Gorai.009G445800.2.A;Gorai.009G445800.1.A
Gene IDs: Gorai.009G445800
cotton name: K02995 - small subunit ribosomal protein S8e (RP-S8e, RPS8) (1 of 6)
OG IDs: 93523491
Best-hit-arabi-ID: AT5G59240.1
Best-hit-arabi-name: Ribosomal protein S8e family protein
Gorai.009G445800.1.A
PCC=0.515, p=0.024
mRNA Profile
Protein Expression Profile
DPA

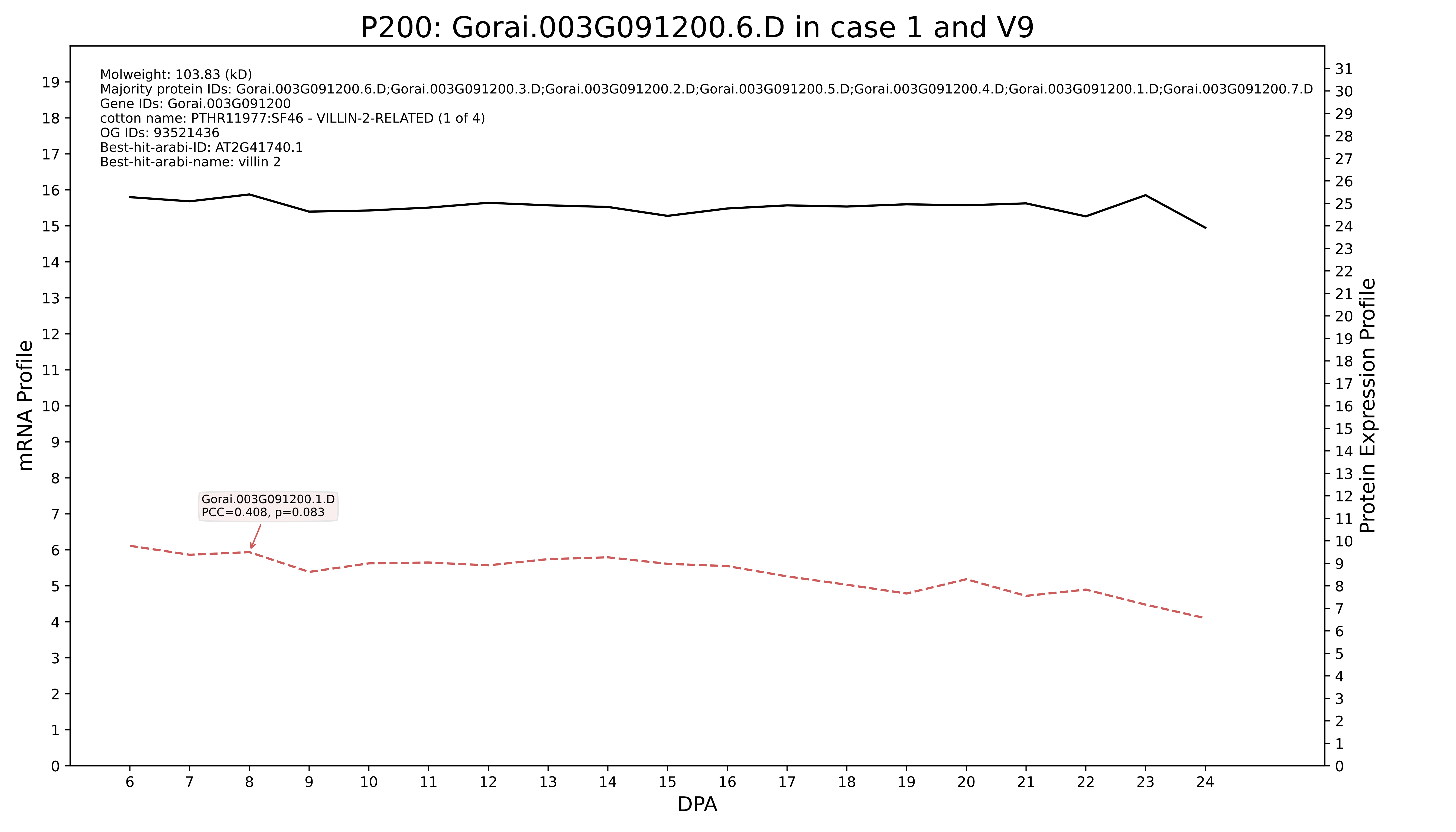
P200: Gorai.003G091200.6.D in case 1 and V9
Molweight: 103.83 (kD)
Majority protein IDs: Gorai.003G091200.6.D;Gorai.003G091200.3.D;Gorai.003G091200.2.D;Gorai.003G091200.5.D;Gorai.003G091200.4.D;Gorai.003G091200.1.D;Gorai.003G091200.7.D
Gene IDs: Gorai.003G091200
cotton name: PTHR11977:SF46 - VILLIN-2-RELATED (1 of 4)
OG IDs: 93521436
Best-hit-arabi-ID: AT2G41740.1
Best-hit-arabi-name: villin 2
Gorai.003G091200.1.D
PCC=0.408, p=0.083
mRNA Profile
Protein Expression Profile
DPA

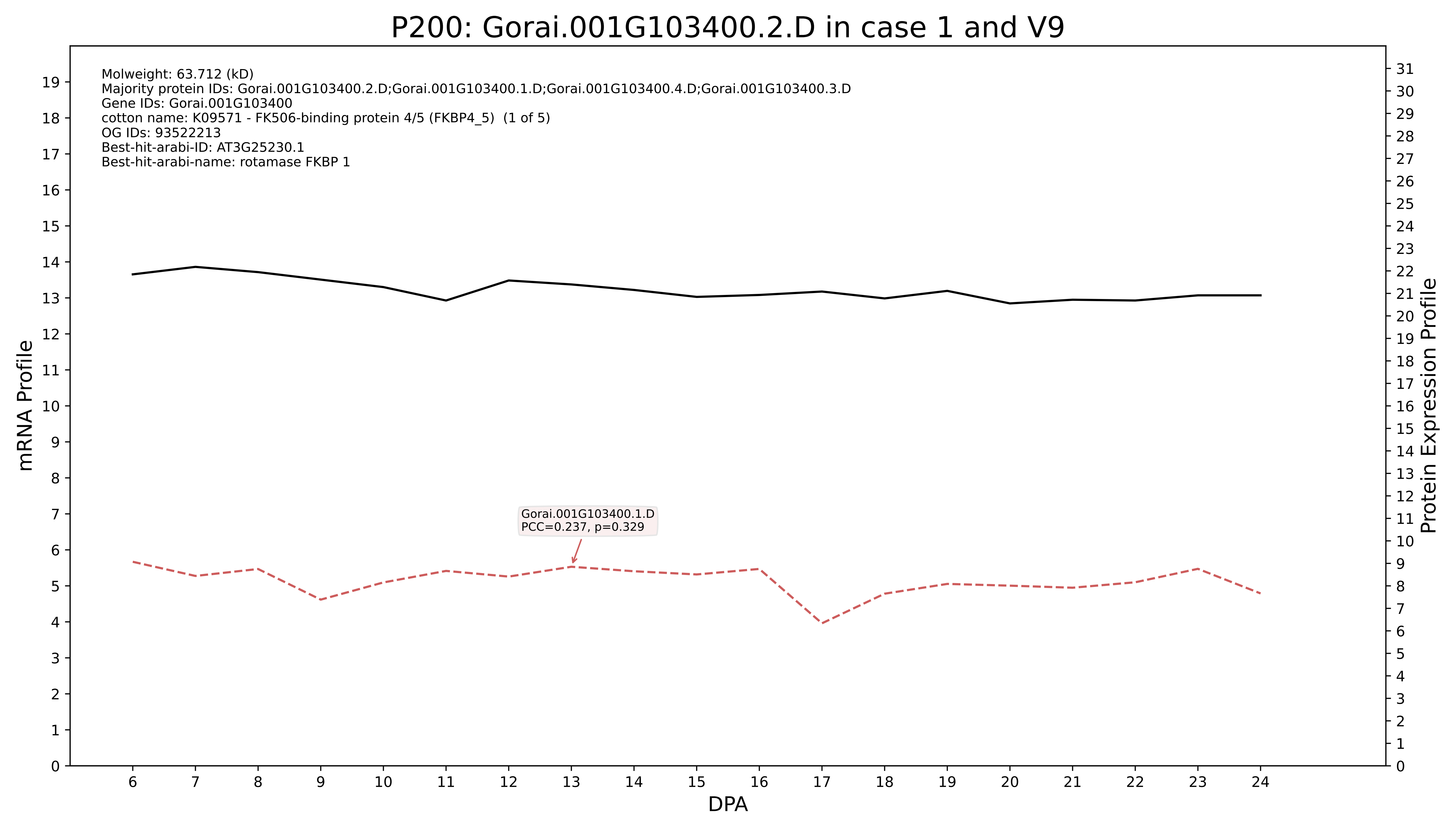
P200: Gorai.001G103400.2.D in case 1 and V9
Molweight: 63.712 (kD)
Majority protein IDs: Gorai.001G103400.2.D;Gorai.001G103400.1.D;Gorai.001G103400.4.D;Gorai.001G103400.3.D
Gene IDs: Gorai.001G103400
cotton name: K09571 - FK506-binding protein 4/5 (FKBP4_5) (1 of 5)
OG IDs: 93522213
Best-hit-arabi-ID: AT3G25230.1
Best-hit-arabi-name: rotamase FKBP 1
Gorai.001G103400.1.D
PCC=0.237, p=0.329
mRNA Profile
Protein Expression Profile
DPA

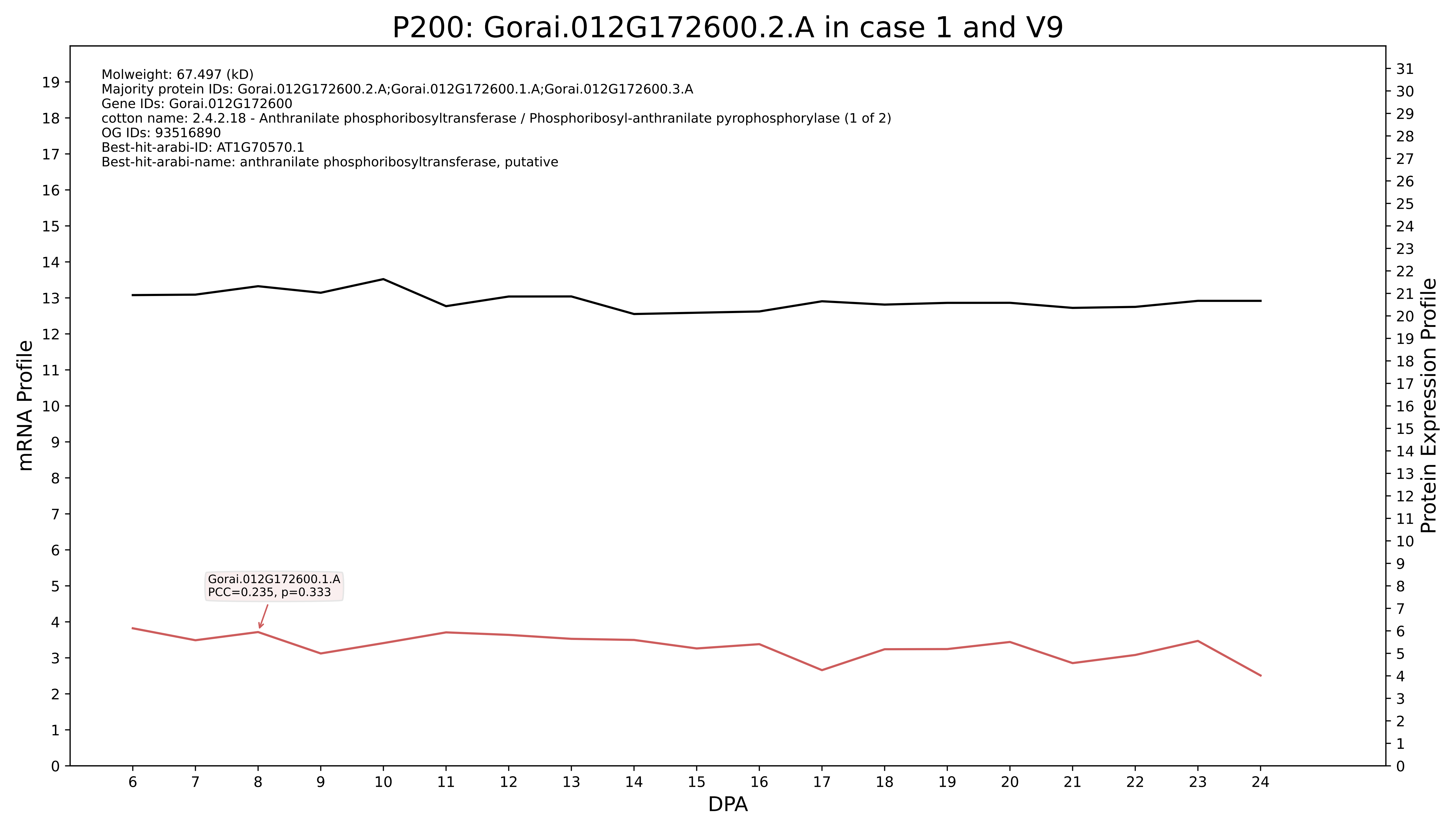
P200: Gorai.012G172600.2.A in case 1 and V9
Molweight: 67.497 (kD)
Majority protein IDs: Gorai.012G172600.2.A;Gorai.012G172600.1.A;Gorai.012G172600.3.A
Gene IDs: Gorai.012G172600
cotton name: 2.4.2.18 - Anthranilate phosphoribosyltransferase / Phosphoribosyl-anthranilate pyrophosphorylase (1 of 2)
OG IDs: 93516890
Best-hit-arabi-ID: AT1G70570.1
Best-hit-arabi-name: anthranilate phosphoribosyltransferase, putative
Gorai.012G172600.1.A
PCC=0.235, p=0.333
mRNA Profile
Protein Expression Profile
DPA

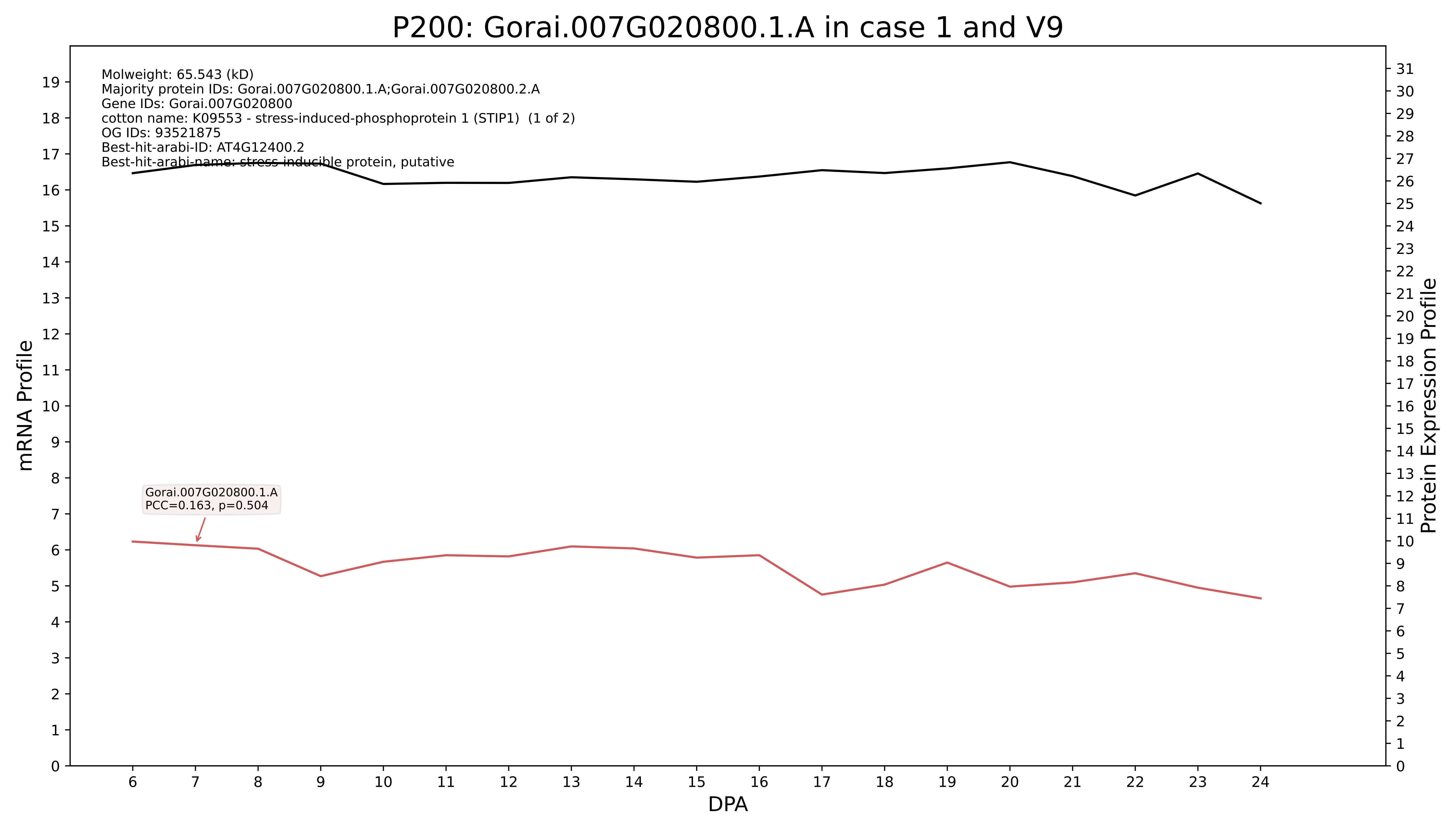
P200: Gorai.007G020800.1.A in case 1 and V9
Molweight: 65.543 (kD)
Majority protein IDs: Gorai.007G020800.1.A;Gorai.007G020800.2.A
Gene IDs: Gorai.007G020800
cotton name: K09553 - stress-induced-phosphoprotein 1 (STIP1) (1 of 2)
OG IDs: 93521875
Best-hit-arabi-ID: AT4G12400.2
Best-hit-arabi-name: stress-inducible protein, putative
Gorai.007G020800.1.A
PCC=0.163, p=0.504
mRNA Profile
Protein Expression Profile
DPA

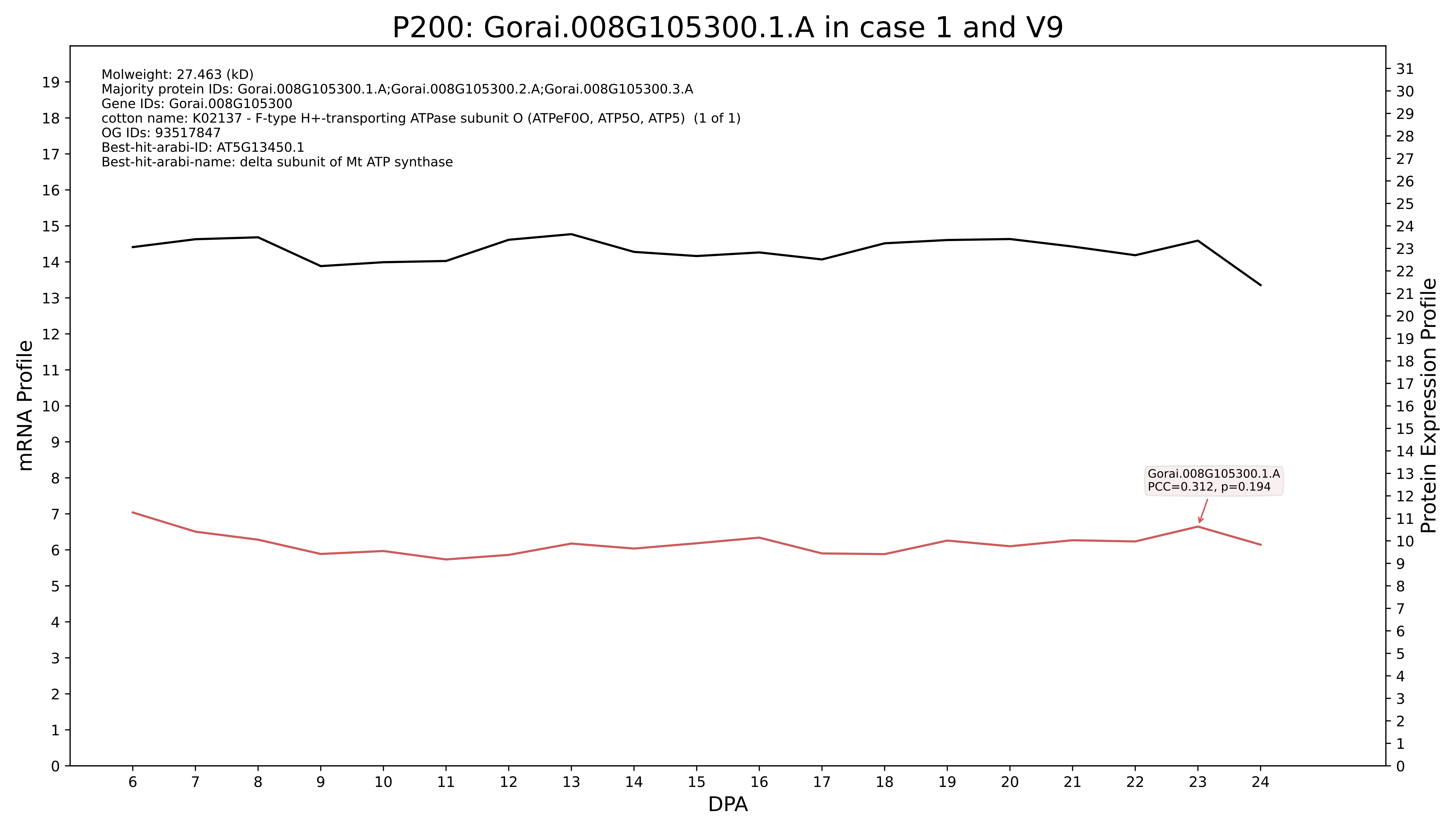
P200: Gorai.008G105300.1.A in case 1 and V9
Molweight: 27.463 (kD)
Majority protein IDs: Gorai.008G105300.1.A;Gorai.008G105300.2.A;Gorai.008G105300.3.A
Gene IDs: Gorai.008G105300
cotton name: K02137 - F-type H+-transporting ATPase subunit O (ATPeF0O, ATP5O, ATP5) (1 of 1)
OG IDs: 93517847
Best-hit-arabi-ID: AT5G13450.1
Best-hit-arabi-name: delta subunit of Mt ATP synthase
Gorai.008G105300.1.A
PCC=0.312, p=0.194
mRNA Profile
Protein Expression Profile
DPA

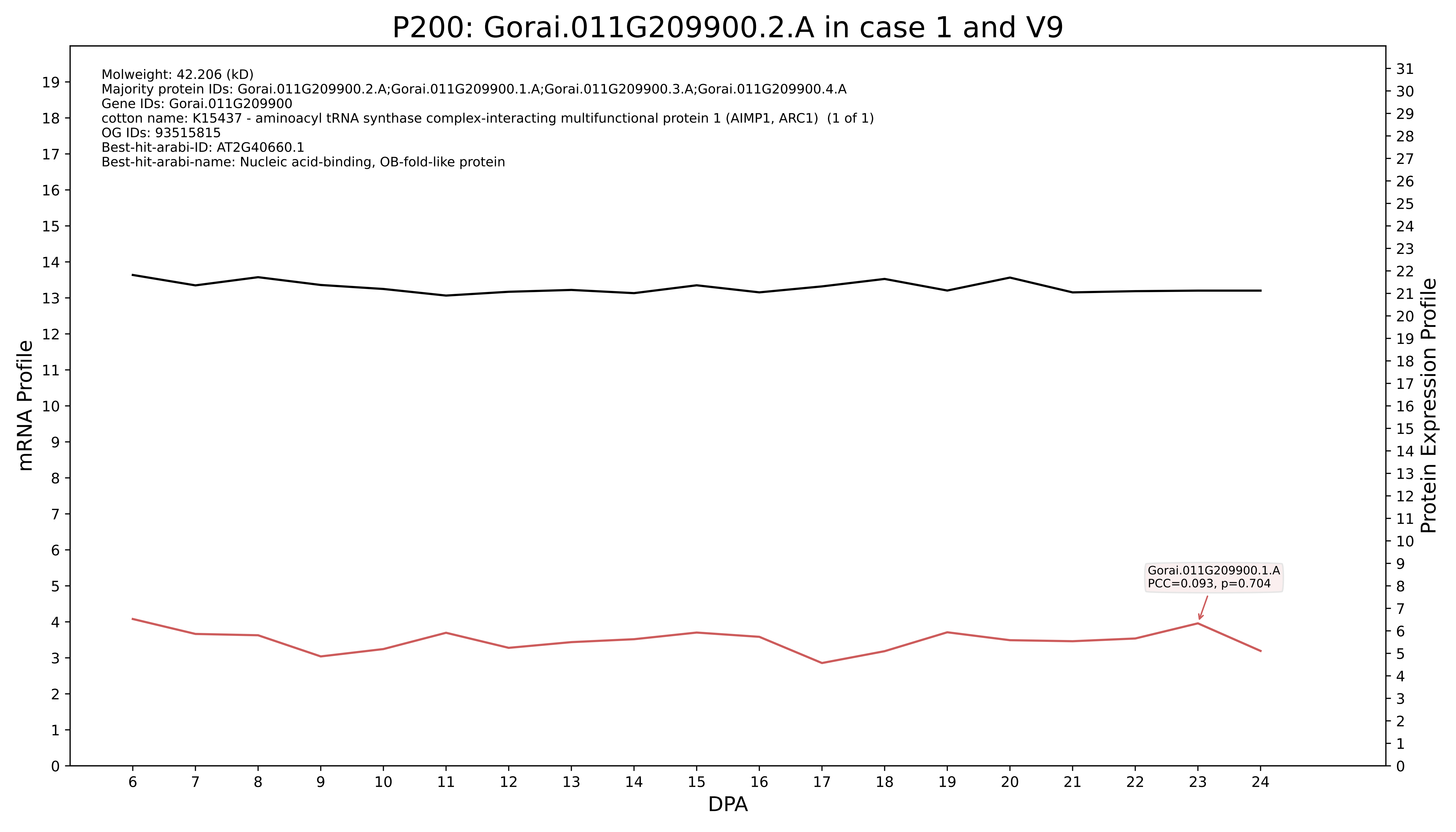

P200: Gorai.011G209900.2.A in case 1 and V9
Molweight: 42.206 (kD)
Majority protein IDs: Gorai.011G209900.2.A;Gorai.011G209900.1.A;Gorai.011G209900.3.A;Gorai.011G209900.4.A
Gene IDs: Gorai.011G209900
cotton name: K15437 - aminoacyl tRNA synthase complex-interacting multifunctional protein 1 (AIMP1, ARC1) (1 of 1)
OG IDs: 93515815
Best-hit-arabi-ID: AT2G40660.1
Best-hit-arabi-name: Nucleic acid-binding, OB-fold-like protein
Gorai.011G209900.1.A
PCC=0.093, p=0.704
mRNA Profile
Protein Expression Profile
DPA

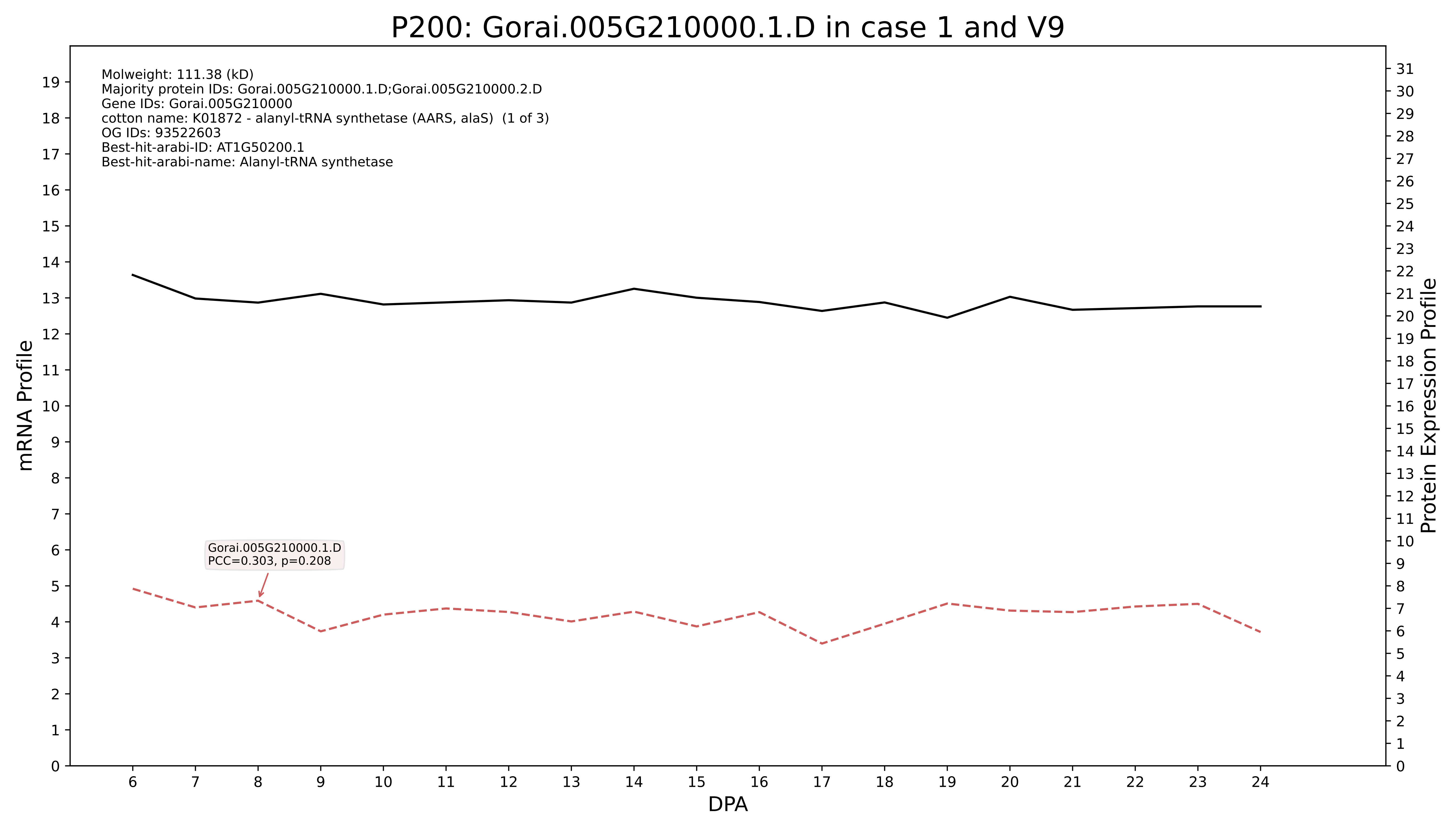
P200: Gorai.005G210000.1.D in case 1 and V9
Molweight: 111.38 (kD)
Majority protein IDs: Gorai.005G210000.1.D;Gorai.005G210000.2.D
Gene IDs: Gorai.005G210000
cotton name: K01872 - alanyl-tRNA synthetase (AARS, alaS) (1 of 3)
OG IDs: 93522603
Best-hit-arabi-ID: AT1G50200.1
Best-hit-arabi-name: Alanyl-tRNA synthetase
Gorai.005G210000.1.D
PCC=0.303, p=0.208
mRNA Profile
Protein Expression Profile
DPA

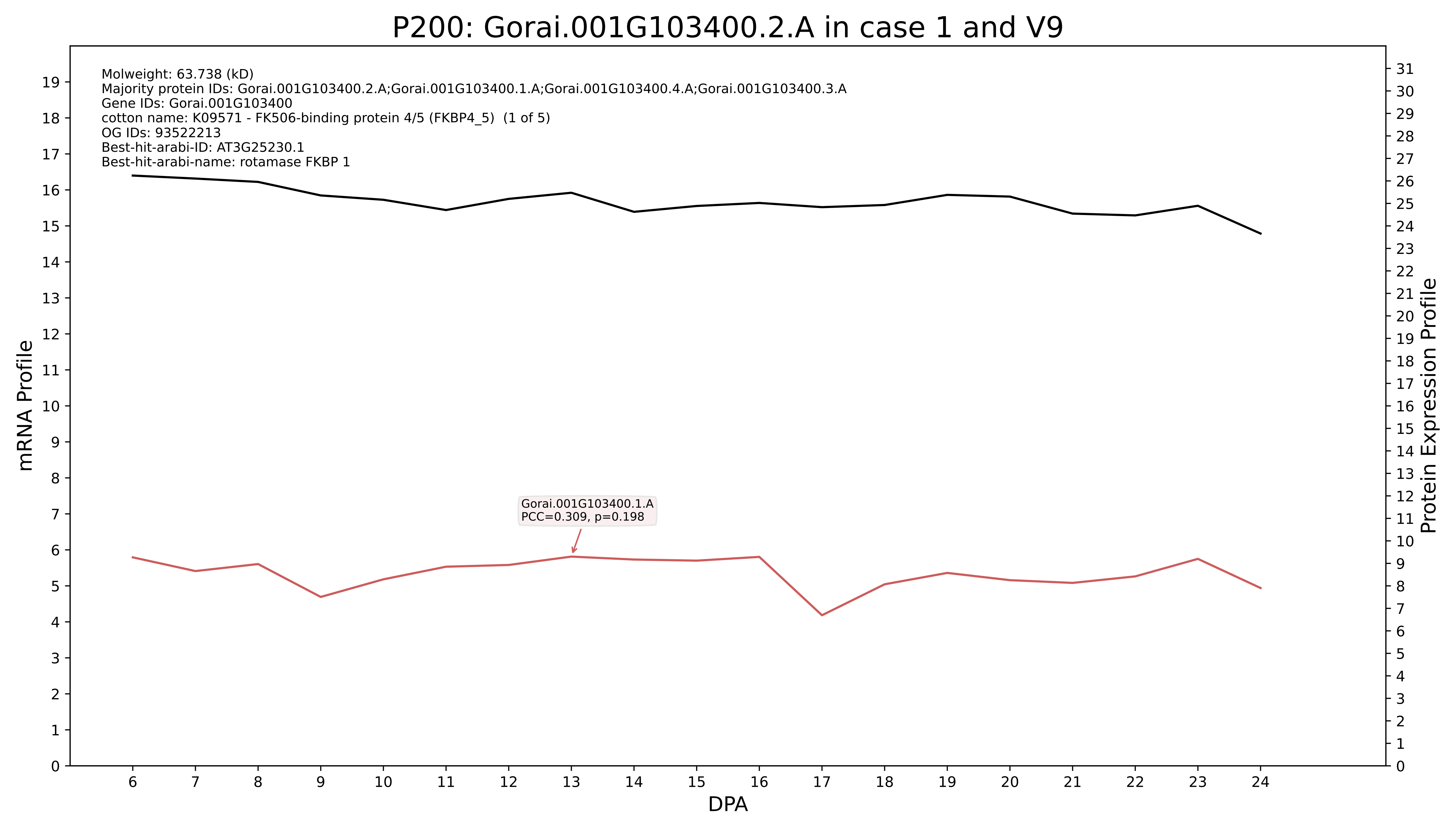
P200: Gorai.001G103400.2.A in case 1 and V9
Molweight: 63.738 (kD)
Majority protein IDs: Gorai.001G103400.2.A;Gorai.001G103400.1.A;Gorai.001G103400.4.A;Gorai.001G103400.3.A
Gene IDs: Gorai.001G103400
cotton name: K09571 - FK506-binding protein 4/5 (FKBP4_5) (1 of 5)
OG IDs: 93522213
Best-hit-arabi-ID: AT3G25230.1
Best-hit-arabi-name: rotamase FKBP 1
Gorai.001G103400.1.A
PCC=0.309, p=0.198
mRNA Profile
Protein Expression Profile
DPA

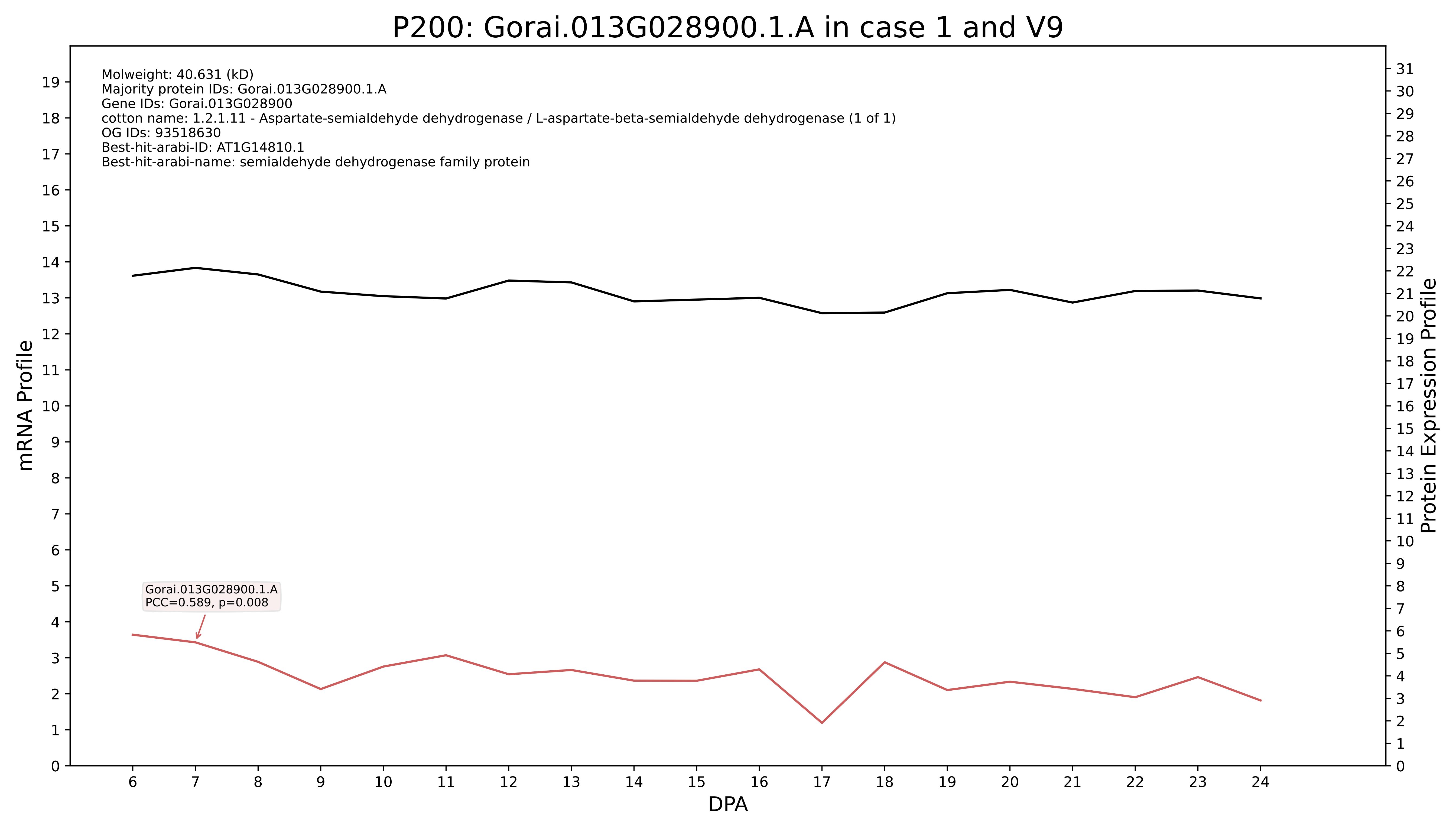
P200: Gorai.013G028900.1.A in case 1 and V9
Molweight: 40.631 (kD)
Majority protein IDs: Gorai.013G028900.1.A
Gene IDs: Gorai.013G028900
cotton name: 1.2.1.11 - Aspartate-semialdehyde dehydrogenase / L-aspartate-beta-semialdehyde dehydrogenase (1 of 1)
OG IDs: 93518630
Best-hit-arabi-ID: AT1G14810.1
Best-hit-arabi-name: semialdehyde dehydrogenase family protein
Gorai.013G028900.1.A
PCC=0.589, p=0.008
mRNA Profile
Protein Expression Profile
DPA

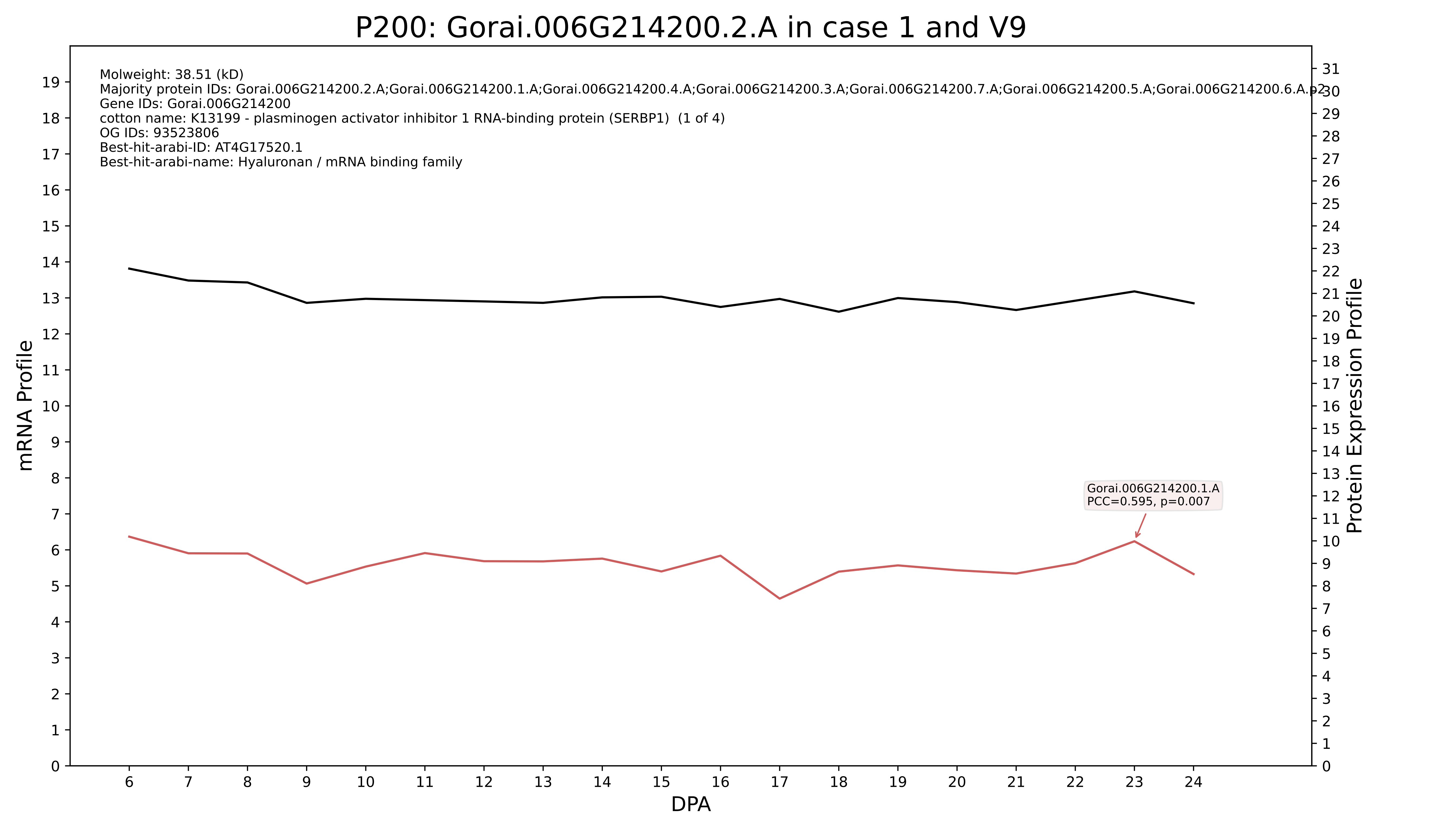
P200: Gorai.006G214200.2.A in case 1 and V9
Molweight: 38.51 (kD)
Majority protein IDs: Gorai.006G214200.2.A;Gorai.006G214200.1.A;Gorai.006G214200.4.A;Gorai.006G214200.3.A;Gorai.006G214200.7.A;Gorai.006G214200.5.A;Gorai.006G214200.6.A;p2
Gene IDs: Gorai.006G214200
cotton name: K13199 - plasminogen activator inhibitor 1 RNA-binding protein (SERBP1) (1 of 4)
OG IDs: 93523806
Best-hit-arabi-ID: AT4G17520.1
Best-hit-arabi-name: Hyaluronan / mRNA binding family
Gorai.006G214200.1.A
PCC=0.595, p=0.007
mRNA Profile
Protein Expression Profile
DPA

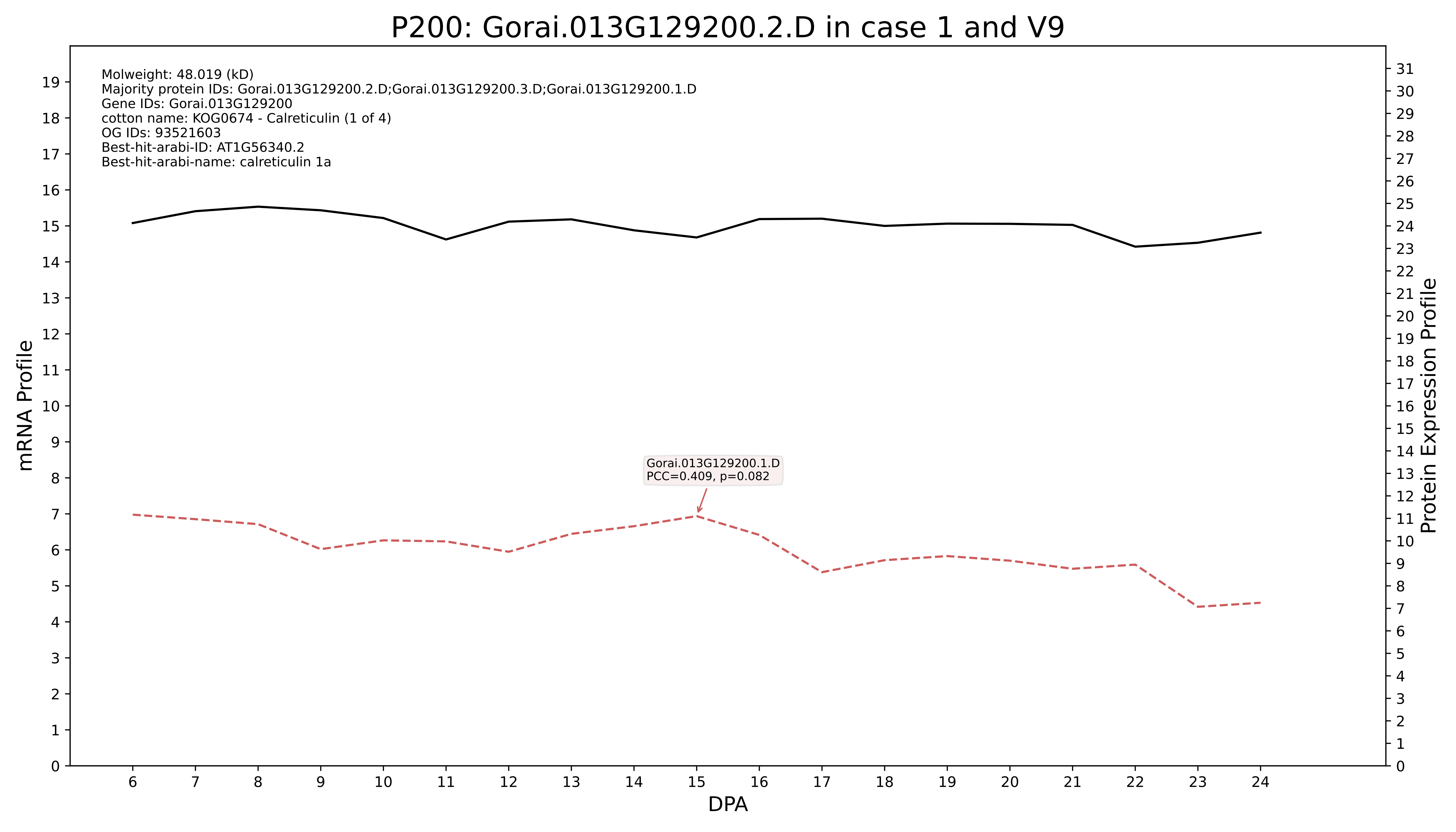
P200: Gorai.013G129200.2.D in case 1 and V9
Molweight: 48.019 (kD)
Majority protein IDs: Gorai.013G129200.2.D;Gorai.013G129200.3.D;Gorai.013G129200.1.D
Gene IDs: Gorai.013G129200
cotton name: KOG0674 - Calreticulin (1 of 4)
OG IDs: 93521603
Best-hit-arabi-ID: AT1G56340.2
Best-hit-arabi-name: calreticulin 1a
Gorai.013G129200.1.D
PCC=0.409, p=0.082
mRNA Profile
Protein Expression Profile
DPA

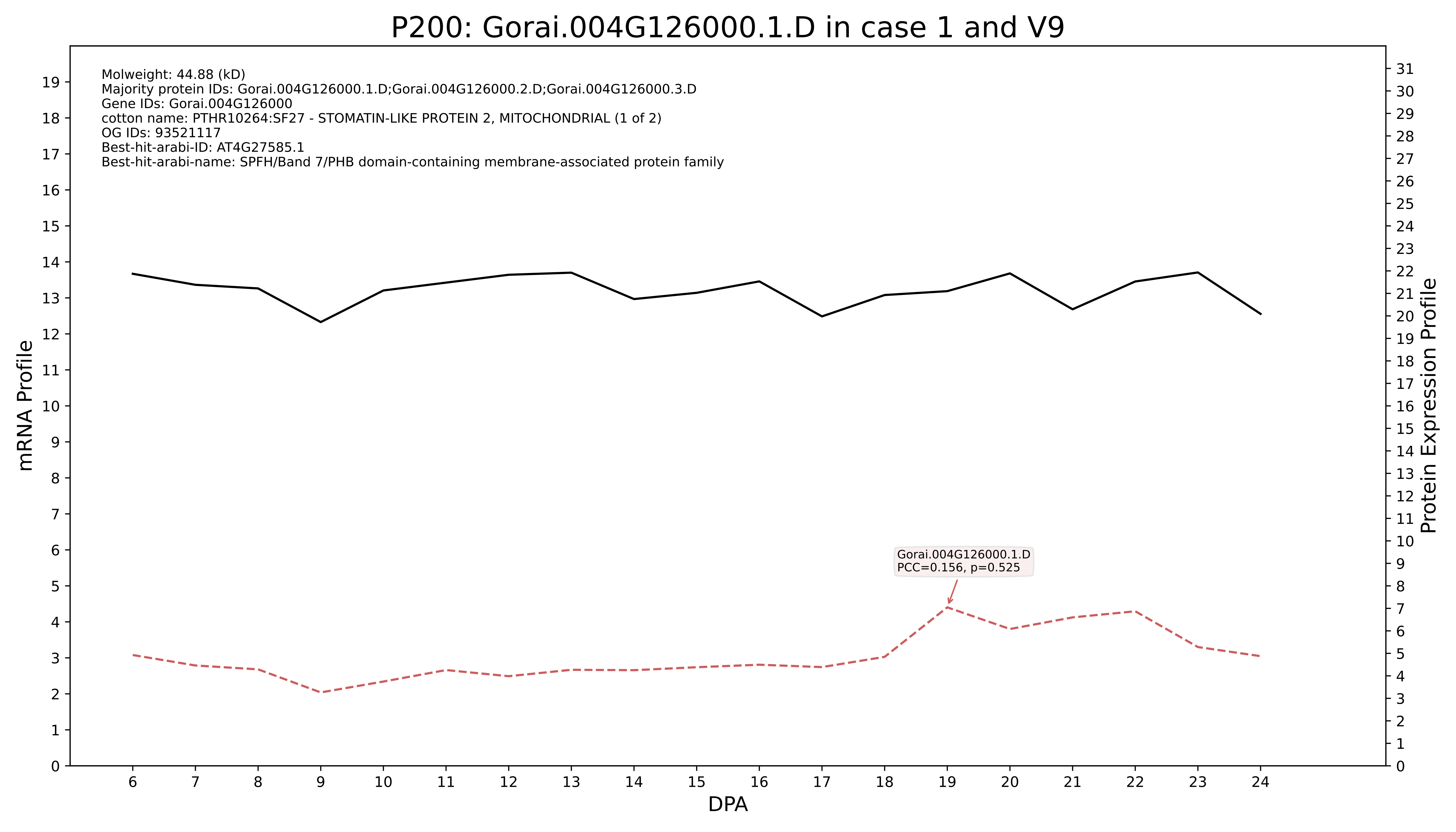
P200: Gorai.004G126000.1.D in case 1 and V9
Molweight: 44.88 (kD)
Majority protein IDs: Gorai.004G126000.1.D;Gorai.004G126000.2.D;Gorai.004G126000.3.D
Gene IDs: Gorai.004G126000
cotton name: PTHR10264:SF27 - STOMATIN-LIKE PROTEIN 2, MITOCHONDRIAL (1 of 2)
OG IDs: 93521117
Best-hit-arabi-ID: AT4G27585.1
Best-hit-arabi-name: SPFH/Band 7/PHB domain-containing membrane-associated protein family
Gorai.004G126000.1.D
PCC=0.156, p=0.525
mRNA Profile
Protein Expression Profile
DPA

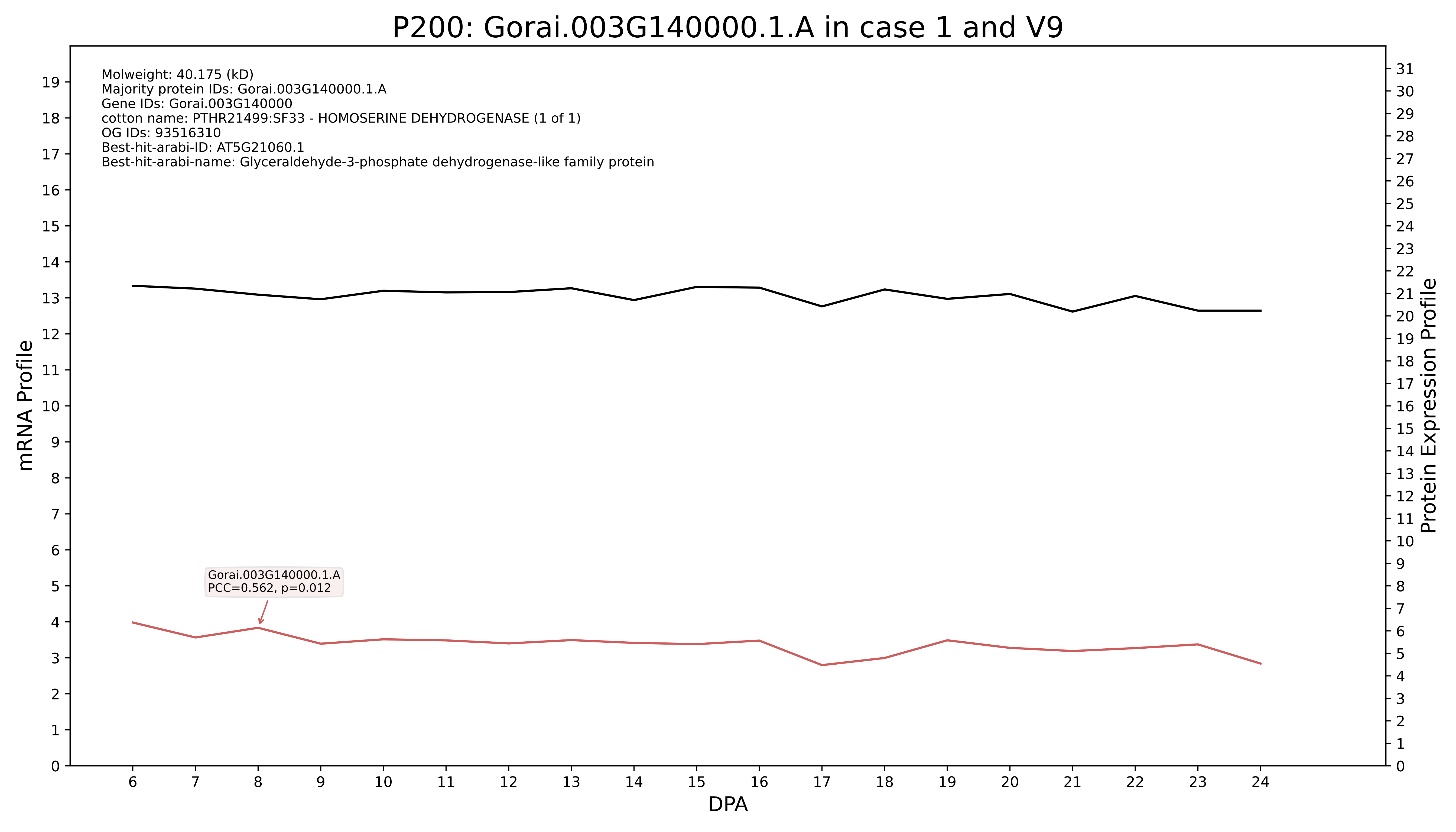
P200: Gorai.003G140000.1.A in case 1 and V9
Molweight: 40.175 (kD)
Majority protein IDs: Gorai.003G140000.1.A
Gene IDs: Gorai.003G140000
cotton name: PTHR21499:SF33 - HOMOSERINE DEHYDROGENASE (1 of 1)
OG IDs: 93516310
Best-hit-arabi-ID: AT5G21060.1
Best-hit-arabi-name: Glyceraldehyde-3-phosphate dehydrogenase-like family protein
Gorai.003G140000.1.A
PCC=0.562, p=0.012
mRNA Profile
Protein Expression Profile
DPA

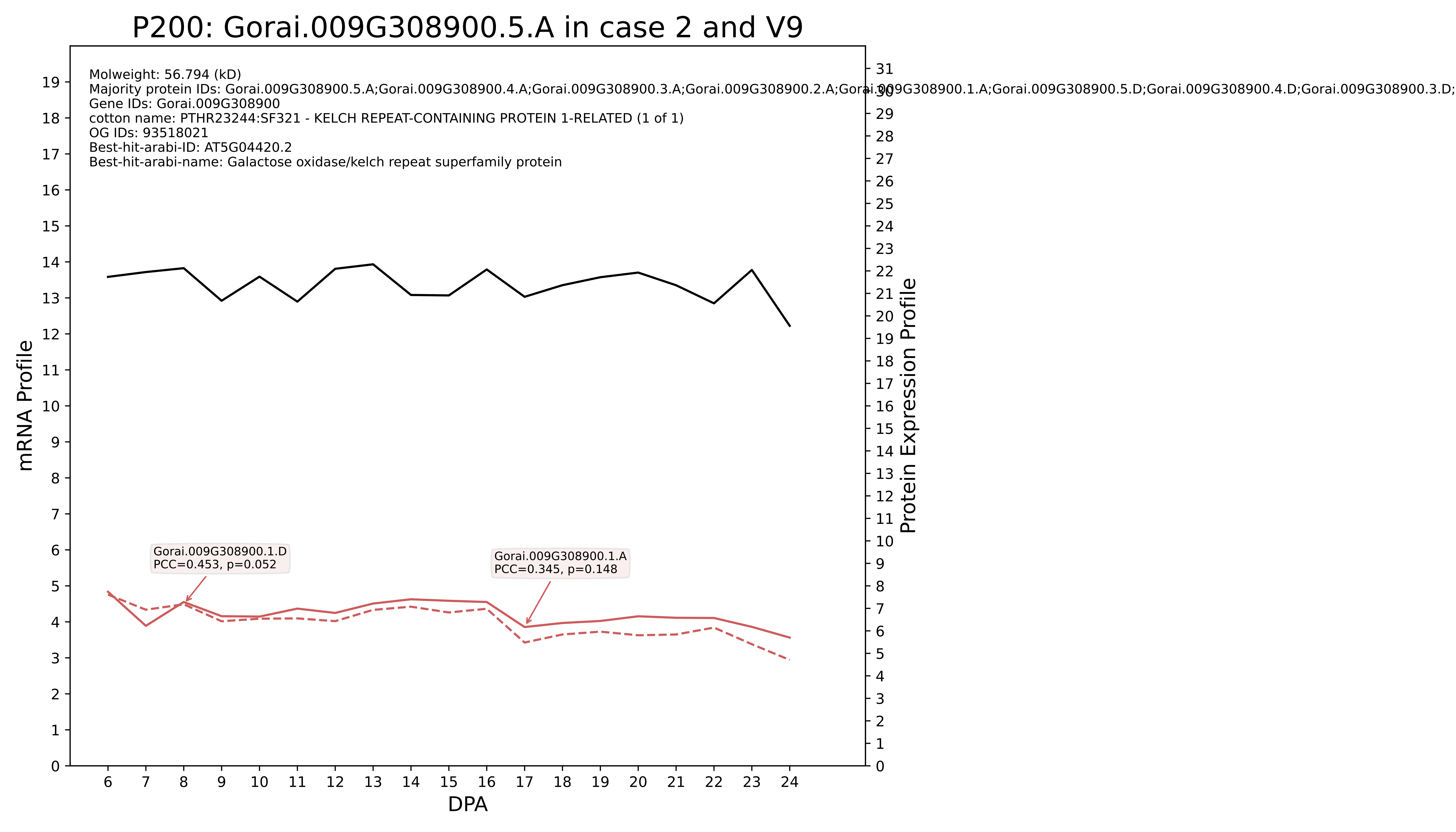
P200: Gorai.009G308900.5.A in case 2 and V9
Molweight: 56.794 (kD)
Majority protein IDs: Gorai.009G308900.5.A;Gorai.009G308900.4.A;Gorai.009G308900.3.A;Gorai.009G308900.2.A;Gorai.009G308900.1.A;Gorai.009G308900.5.D;Gorai.009G308900.4.D;Gorai.009G308900.3.D;
Gene IDs: Gorai.009G308900
cotton name: PTHR23244:SF321 - KELCH REPEAT-CONTAINING PROTEIN 1-RELATED (1 of 1)
OG IDs: 93518021
Best-hit-arabi-ID: AT5G04420.2
Best-hit-arabi-name: Galactose oxidase/kelch repeat superfamily protein
Gorai.009G308900.1.D
PCC=0.453, p=0.052
Gorai.009G308900.1.A
PCC=0.345, p=0.148
mRNA Profile
Protein Expression Profile
DPA

### P200: Gorai.009G044500.1.D in case 2 and V9

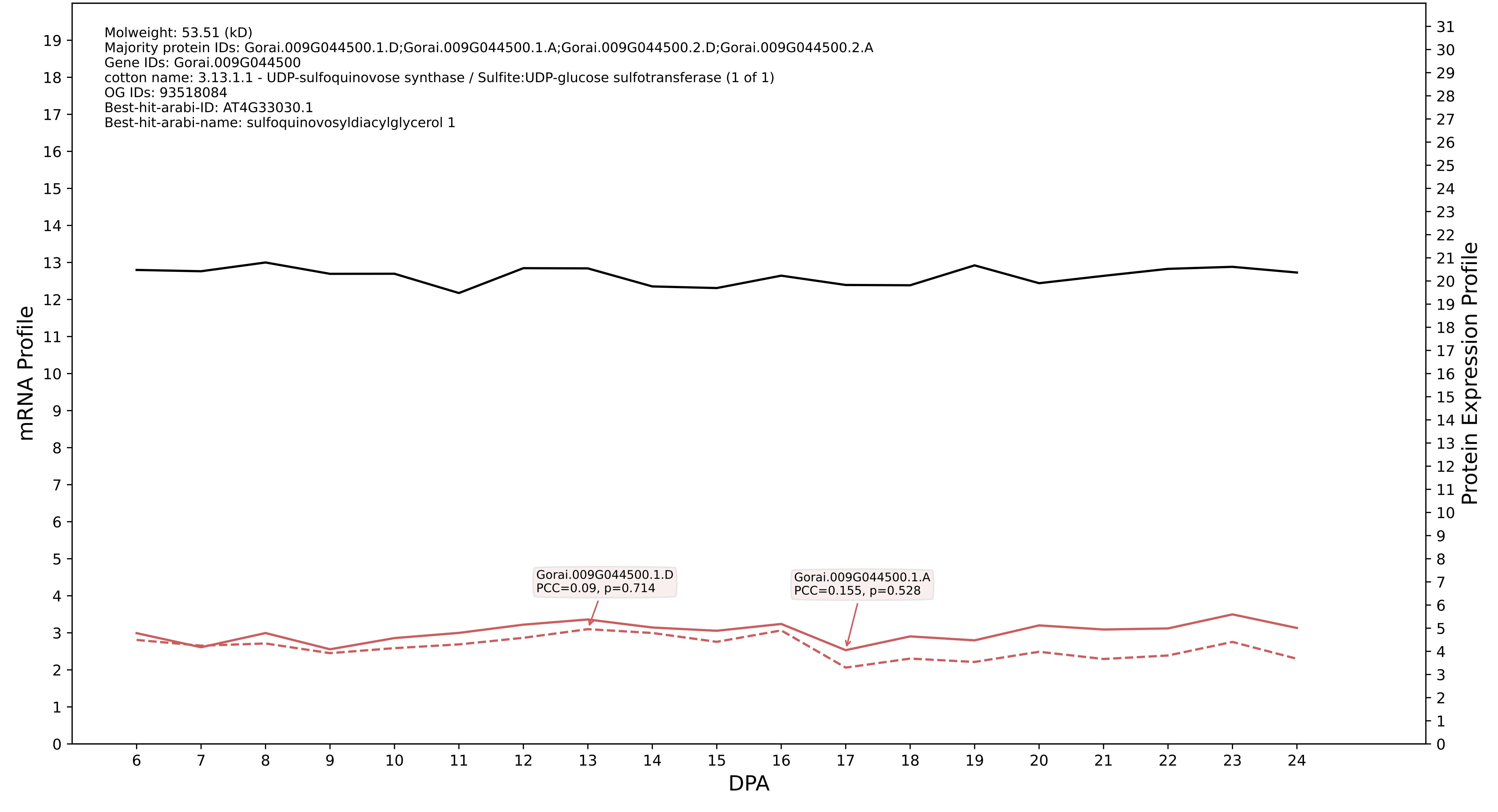

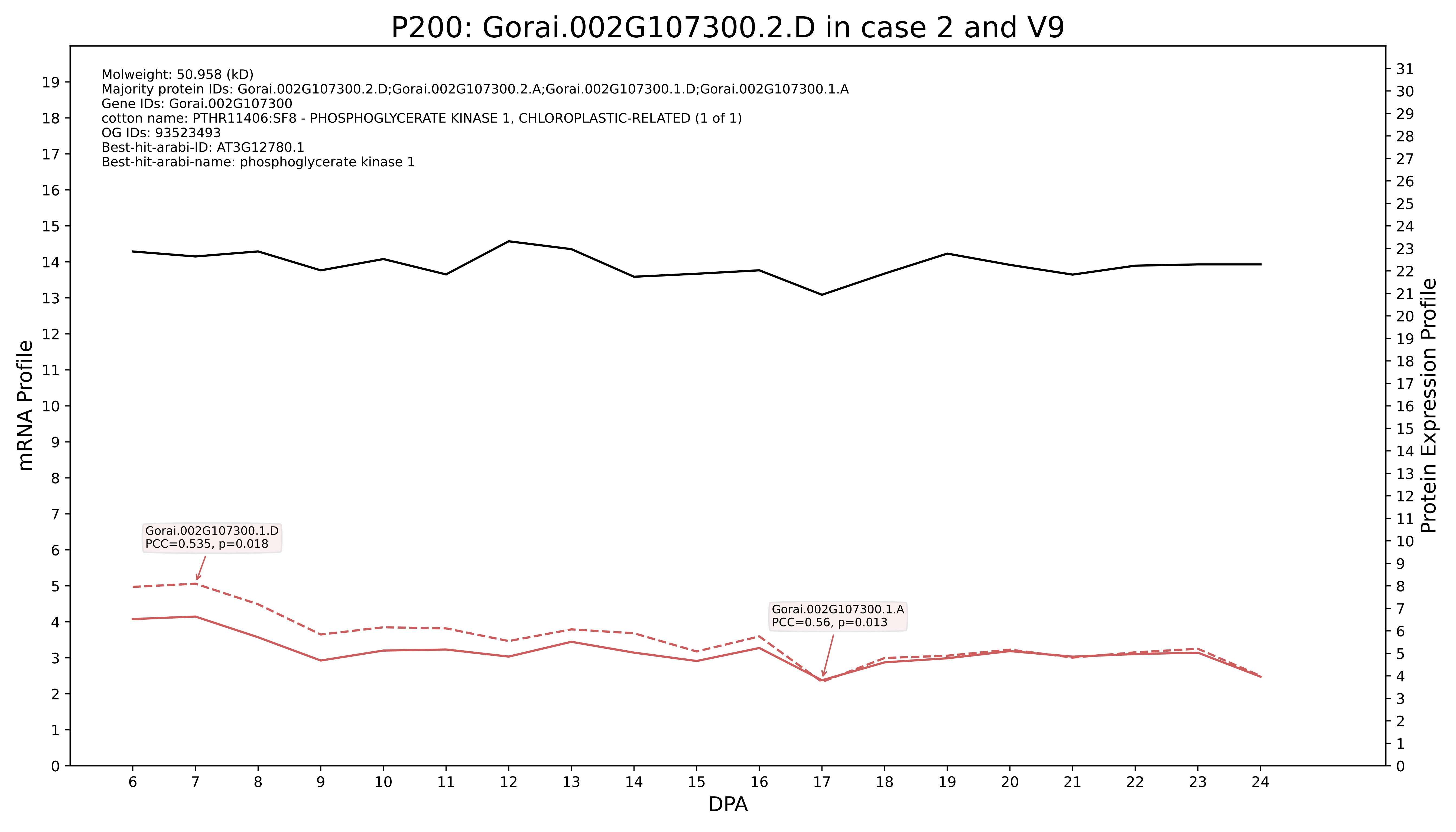
P200: Gorai.002G107300.2.D in case 2 and V9
Molweight: 50.958 (kD)
Majority protein IDs: Gorai.002G107300.2.D;Gorai.002G107300.2.A;Gorai.002G107300.1.D;Gorai.002G107300.1.A
Gene IDs: Gorai.002G107300
cotton name: PTHR11406:SF8 - PHOSPHOGLYCERATE KINASE 1, CHLOROPLASTIC-RELATED (1 of 1)
OG IDs: 93523493
Best-hit-arabi-ID: AT3G12780.1
Best-hit-arabi-name: phosphoglycerate kinase 1
Gorai.002G107300.1.D
PCC=0.535, p=0.018
Gorai.002G107300.1.A
PCC=0.56, p=0.013
mRNA Profile
Protein Expression Profile
DPA

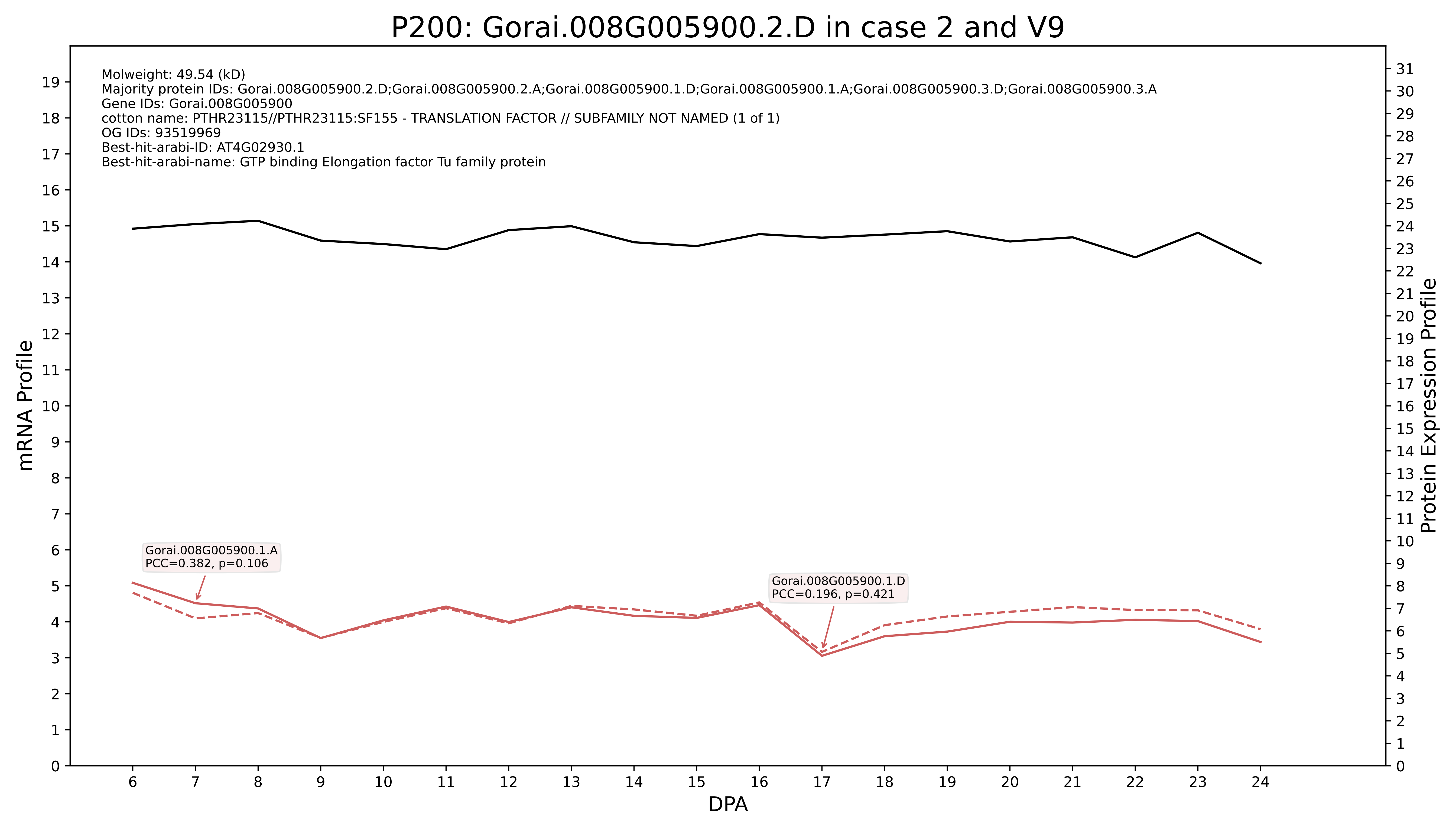

P200: Gorai.008G005900.2.D in case 2 and V9
Molweight: 49.54 (kD)
Majority protein IDs: Gorai.008G005900.2.D;Gorai.008G005900.2.A;Gorai.008G005900.1.D;Gorai.008G005900.1.A;Gorai.008G005900.3.D;Gorai.008G005900.3.A
Gene IDs: Gorai.008G005900
cotton name: PTHR23115//PTHR23115:SF155 - TRANSLATION FACTOR // SUBFAMILY NOT NAMED (1 of 1)
OG IDs: 93519969
Best-hit-arabi-ID: AT4G02930.1
Best-hit-arabi-name: GTP binding Elongation factor Tu family protein
Gorai.008G005900.1.A
PCC=0.382, p=0.106
Gorai.008G005900.1.D
PCC=0.196, p=0.421
mRNA Profile
Protein Expression Profile
DPA

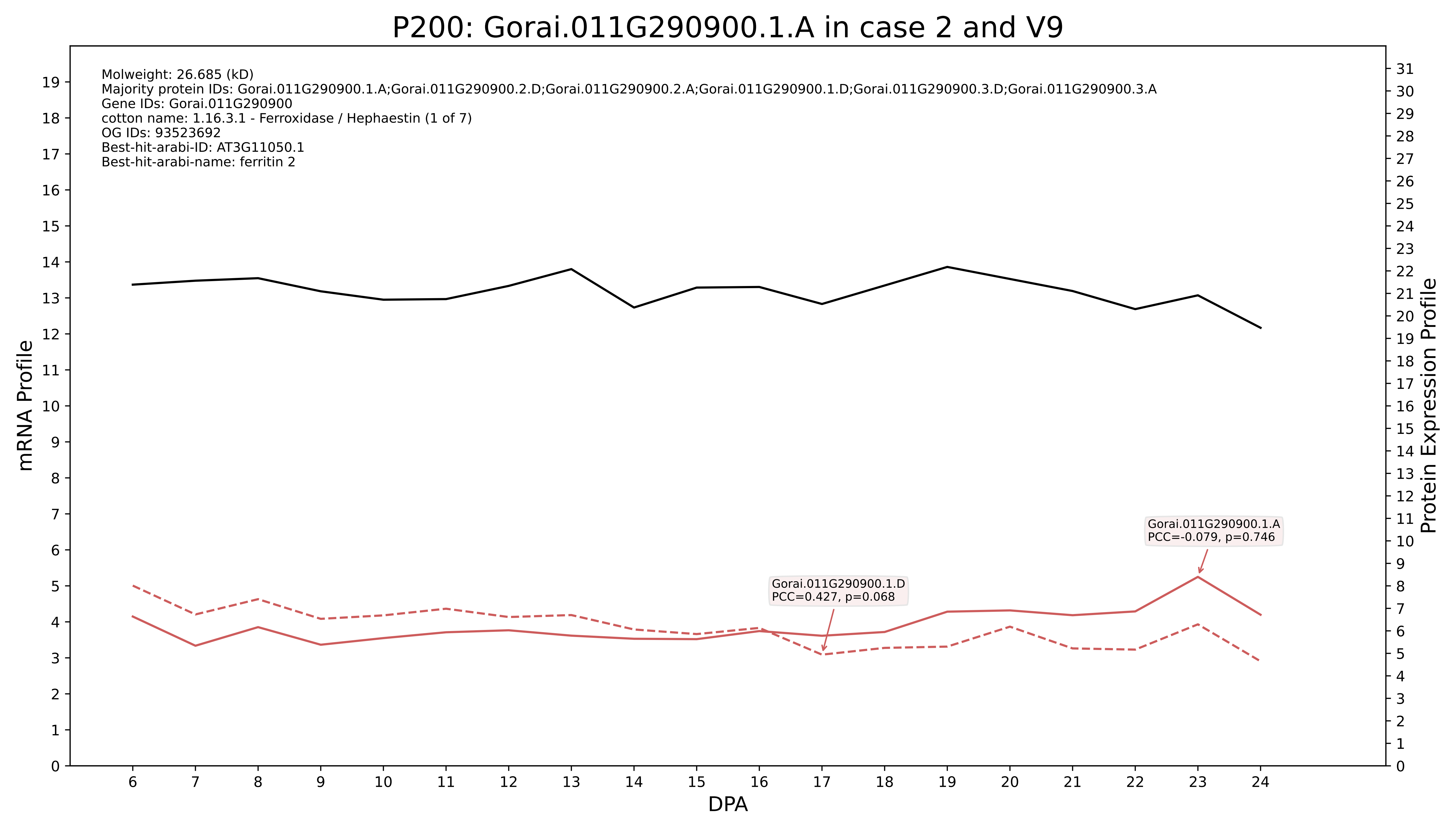
P200: Gorai.011G290900.1.A in case 2 and V9
Molweight: 26.685 (kD)
Majority protein IDs: Gorai.011G290900.1.A;Gorai.011G290900.2.D;Gorai.011G290900.2.A;Gorai.011G290900.1.D;Gorai.011G290900.3.D;Gorai.011G290900.3.A
Gene IDs: Gorai.011G290900
cotton name: 1.16.3.1 - Ferroxidase / Hephaestin (1 of 7)
OG IDs: 93523692
Best-hit-arabi-ID: AT3G11050.1
Best-hit-arabi-name: ferritin 2
Gorai.011G290900.1.A
PCC=-0.079, p=0.746
Gorai.011G290900.1.D
PCC=0.427, p=0.068
mRNA Profile
Protein Expression Profile
DPA

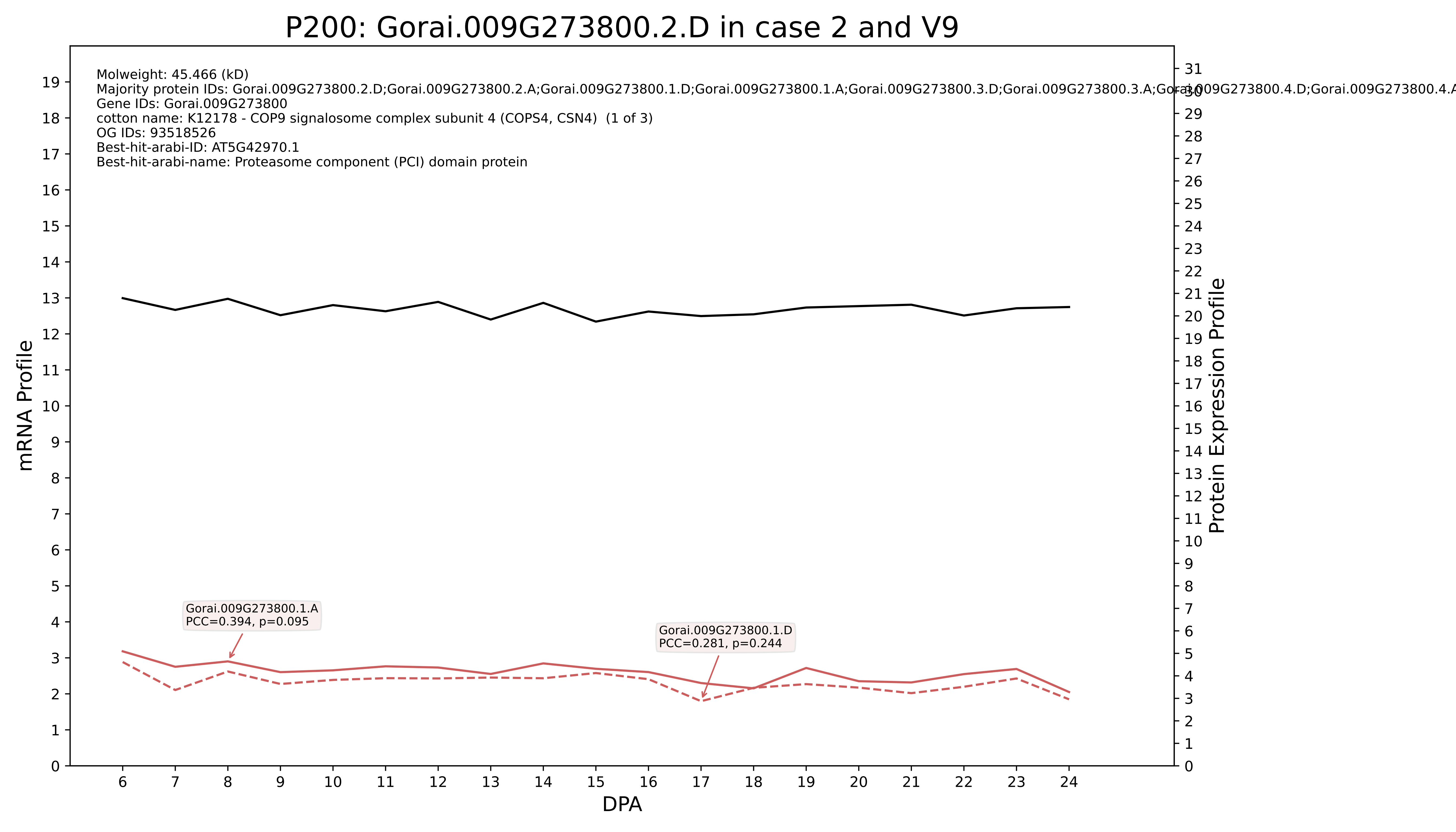
P200: Gorai.009G273800.2.D in case 2 and V9
Molweight: 45.466 (kD)
Majority protein IDs: Gorai.009G273800.2.D;Gorai.009G273800.2.A;Gorai.009G273800.1.D;Gorai.009G273800.1.A;Gorai.009G273800.3.D;Gorai.009G273800.3.A;Gorai.009G273800.4.D;Gorai.009G273800.4.A
Gene IDs: Gorai.009G273800
cotton name: K12178 - COP9 signalosome complex subunit 4 (COPS4, CSN4) (1 of 3)
OG IDs: 93518526
Best-hit-arabi-ID: AT5G42970.1
Best-hit-arabi-name: Proteasome component (PCI) domain protein
Gorai.009G273800.1.A
PCC=0.394, p=0.095
Gorai.009G273800.1.D
PCC=0.281, p=0.244
mRNA Profile
Protein Expression Profile
DPA

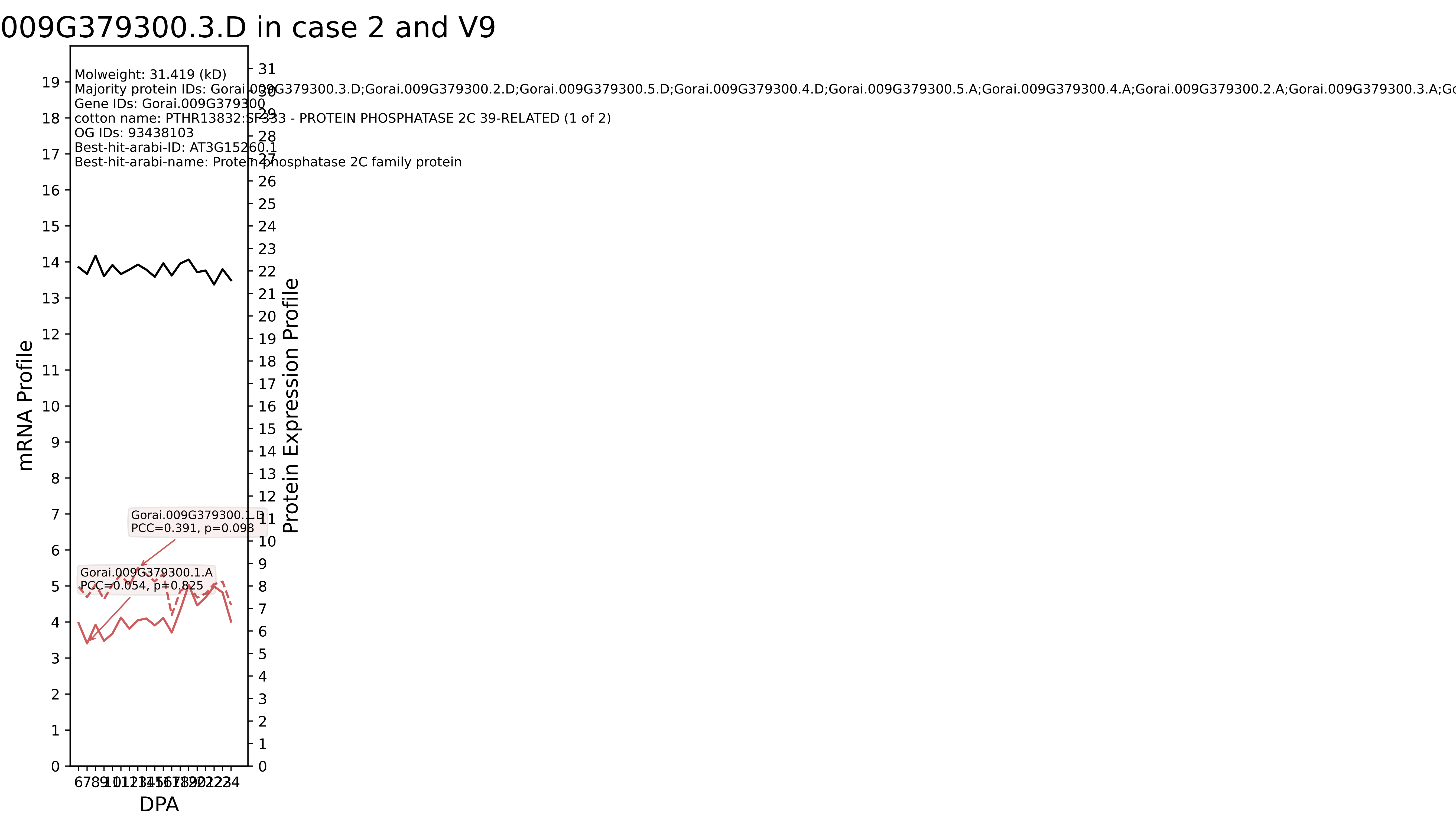
009G379300.3.D in case 2 and V9
Molweight: 31.419 (kD)
Majority protein IDs: Gorai.009G379300.3.D;Gorai.009G379300.2.D;Gorai.009G379300.5.D;Gorai.009G379300.4.D;Gorai.009G379300.5.A;Gorai.009G379300.4.A;Gorai.009G379300.2.A;Gorai.009G379300.3.A;Go
Gene IDs: Gorai.009G379300
cotton name: PTHR13832:SF333 - PROTEIN PHOSPHATASE 2C 39-RELATED (1 of 2)
OG IDs: 93438103
Best-hit-arabi-ID: AT3G15260.1
Best-hit-arabi-name: Protein phosphatase 2C family protein
Gorai.009G379300.1.D
PCC=0.391, p=0.098
Gorai.009G379300.1.A
PCC=0.054, p=0.825
mRNA Profile
Protein Expression Profile
DPA

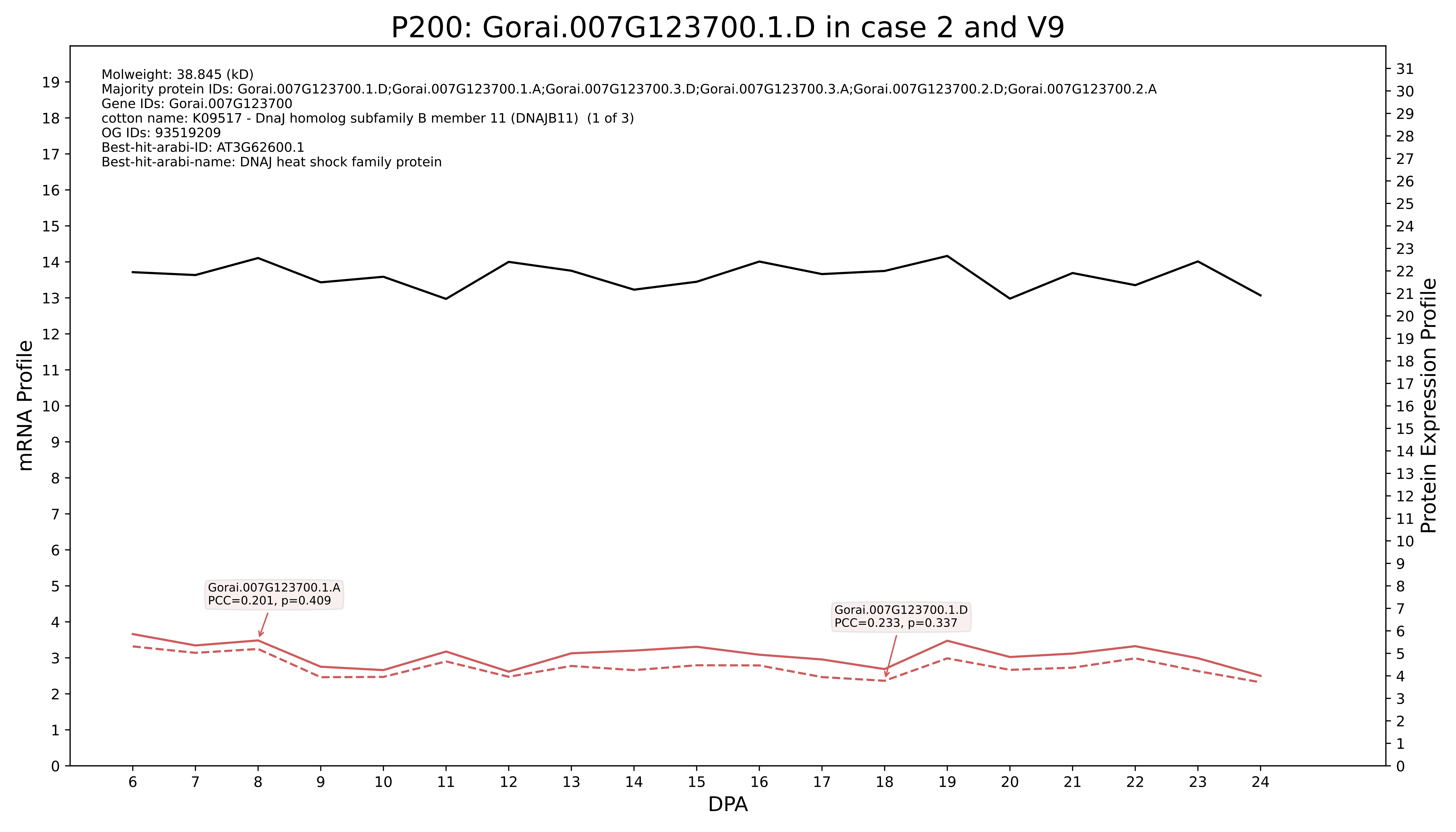
P200: Gorai.007G123700.1.D in case 2 and V9
Molweight: 38.845 (kD)
Majority protein IDs: Gorai.007G123700.1.D;Gorai.007G123700.1.A;Gorai.007G123700.3.D;Gorai.007G123700.3.A;Gorai.007G123700.2.D;Gorai.007G123700.2.A
Gene IDs: Gorai.007G123700
cotton name: K09517 - DnaJ homolog subfamily B member 11 (DNAJB11) (1 of 3)
OG IDs: 93519209
Best-hit-arabi-ID: AT3G62600.1
Best-hit-arabi-name: DNAJ heat shock family protein
Gorai.007G123700.1.A
PCC=0.201, p=0.409
Gorai.007G123700.1.D
PCC=0.233, p=0.337
mRNA Profile
Protein Expression Profile
DPA

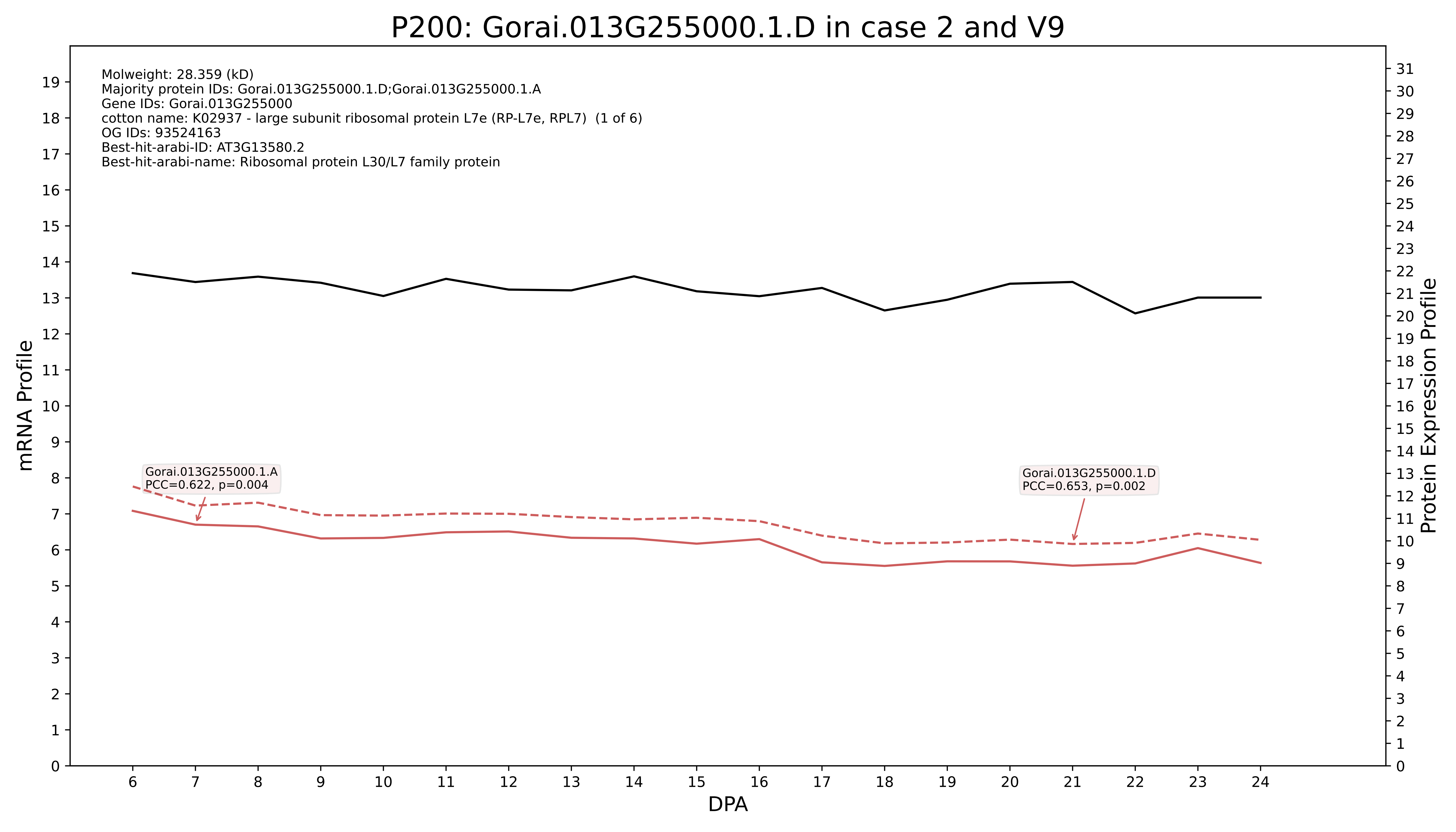
P200: Gorai.013G255000.1.D in case 2 and V9
Molweight: 28.359 (kD)
Majority protein IDs: Gorai.013G255000.1.D;Gorai.013G255000.1.A
Gene IDs: Gorai.013G255000
cotton name: K02937 - large subunit ribosomal protein L7e (RP-L7e, RPL7) (1 of 6)
OG IDs: 93524163
Best-hit-arabi-ID: AT3G13580.2
Best-hit-arabi-name: Ribosomal protein L30/L7 family protein
Gorai.013G255000.1.A
PCC=0.622, p=0.004
Gorai.013G255000.1.D
PCC=0.653, p=0.002
mRNA Profile
Protein Expression Profile
DPA

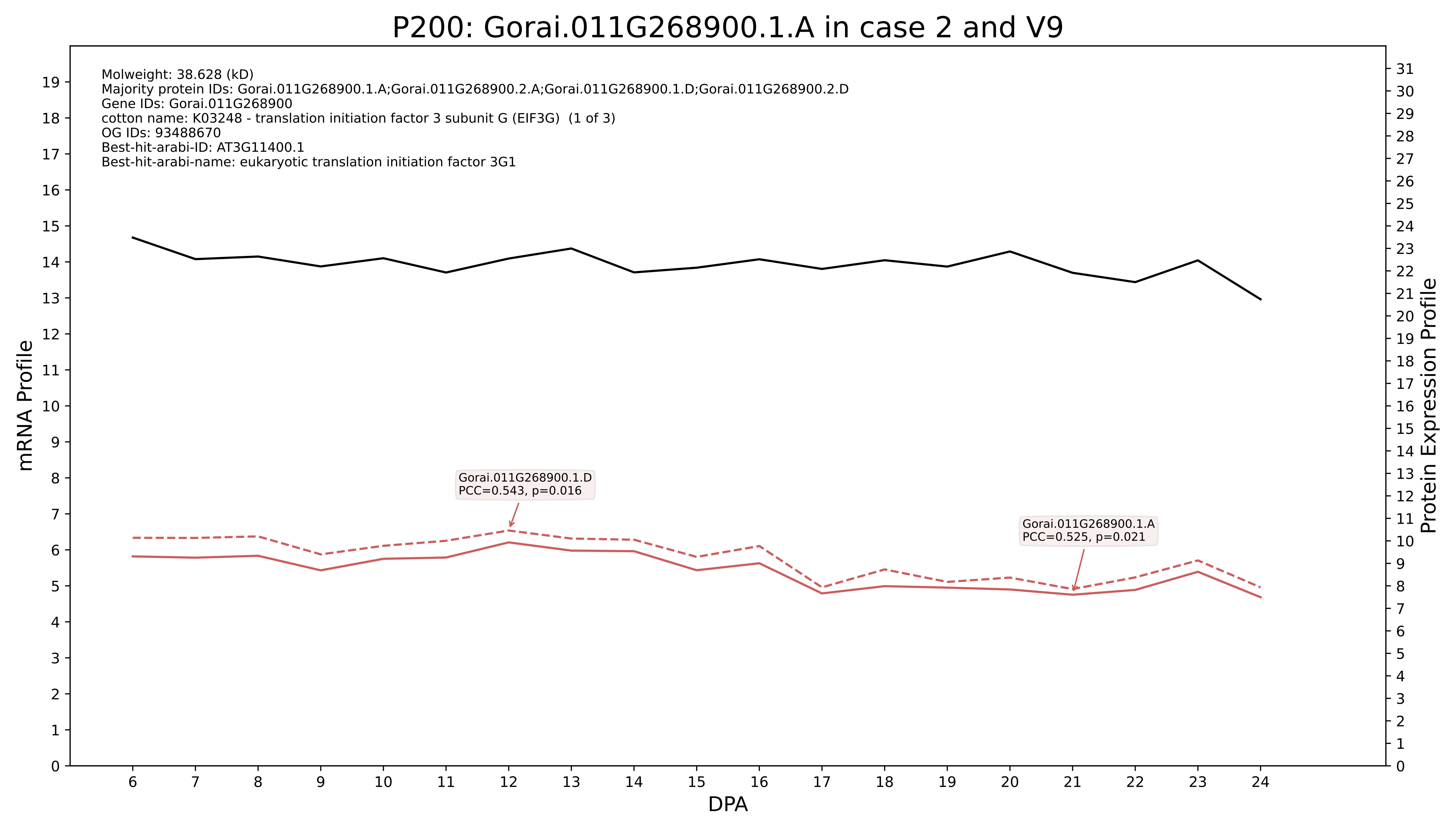
P200: Gorai.011G268900.1.A in case 2 and V9
Molweight: 38.628 (kD)
Majority protein IDs: Gorai.011G268900.1.A;Gorai.011G268900.2.A;Gorai.011G268900.1.D;Gorai.011G268900.2.D
Gene IDs: Gorai.011G268900
cotton name: K03248 - translation initiation factor 3 subunit G (EIF3G) (1 of 3)
OG IDs: 93488670
Best-hit-arabi-ID: AT3G11400.1
Best-hit-arabi-name: eukaryotic translation initiation factor 3G1
Gorai.011G268900.1.D
PCC=0.543, p=0.016
Gorai.011G268900.1.A
PCC=0.525, p=0.021
mRNA Profile
Protein Expression Profile
DPA

200: Gorai.002G115100.6.D in case 2 and V9

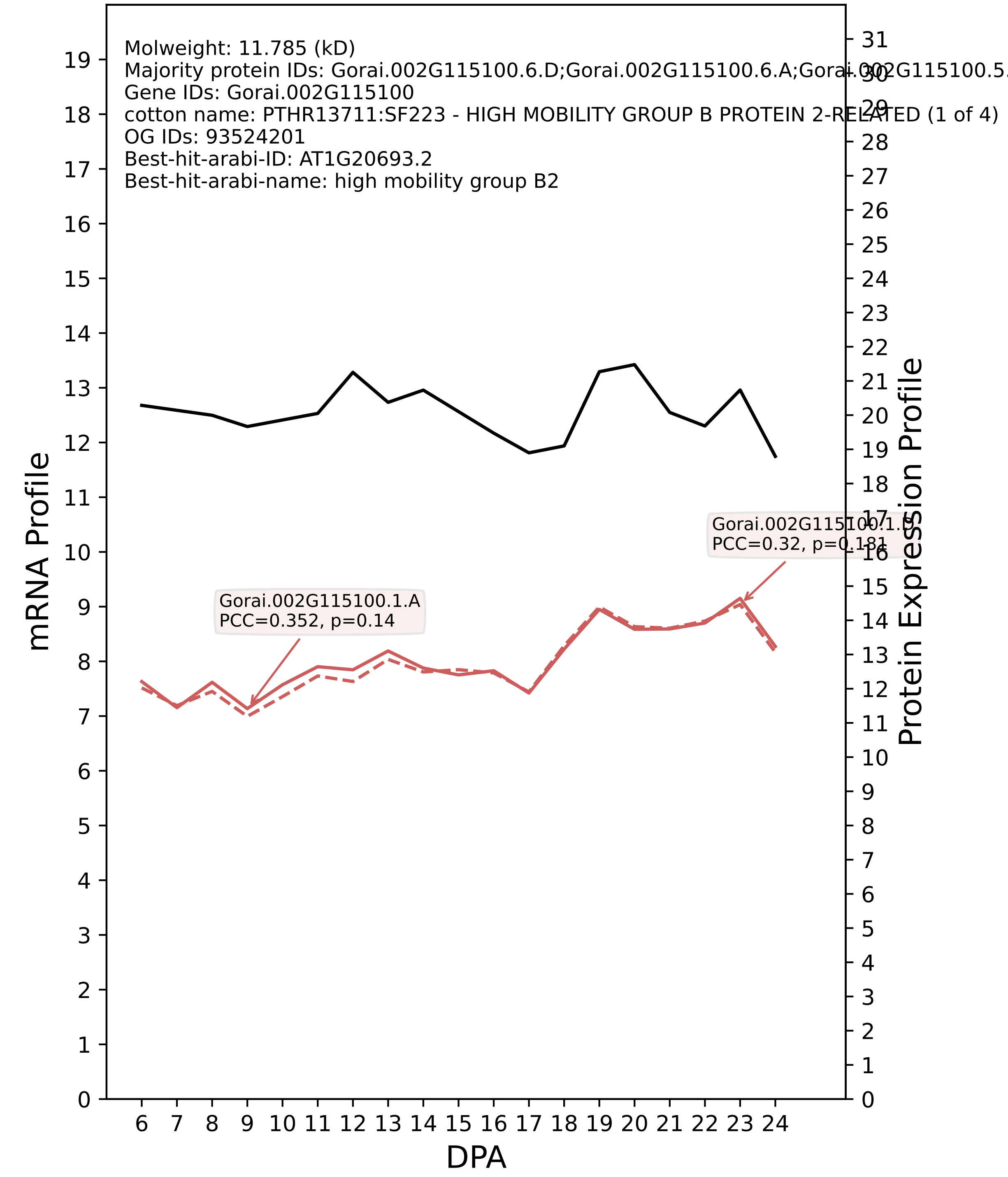

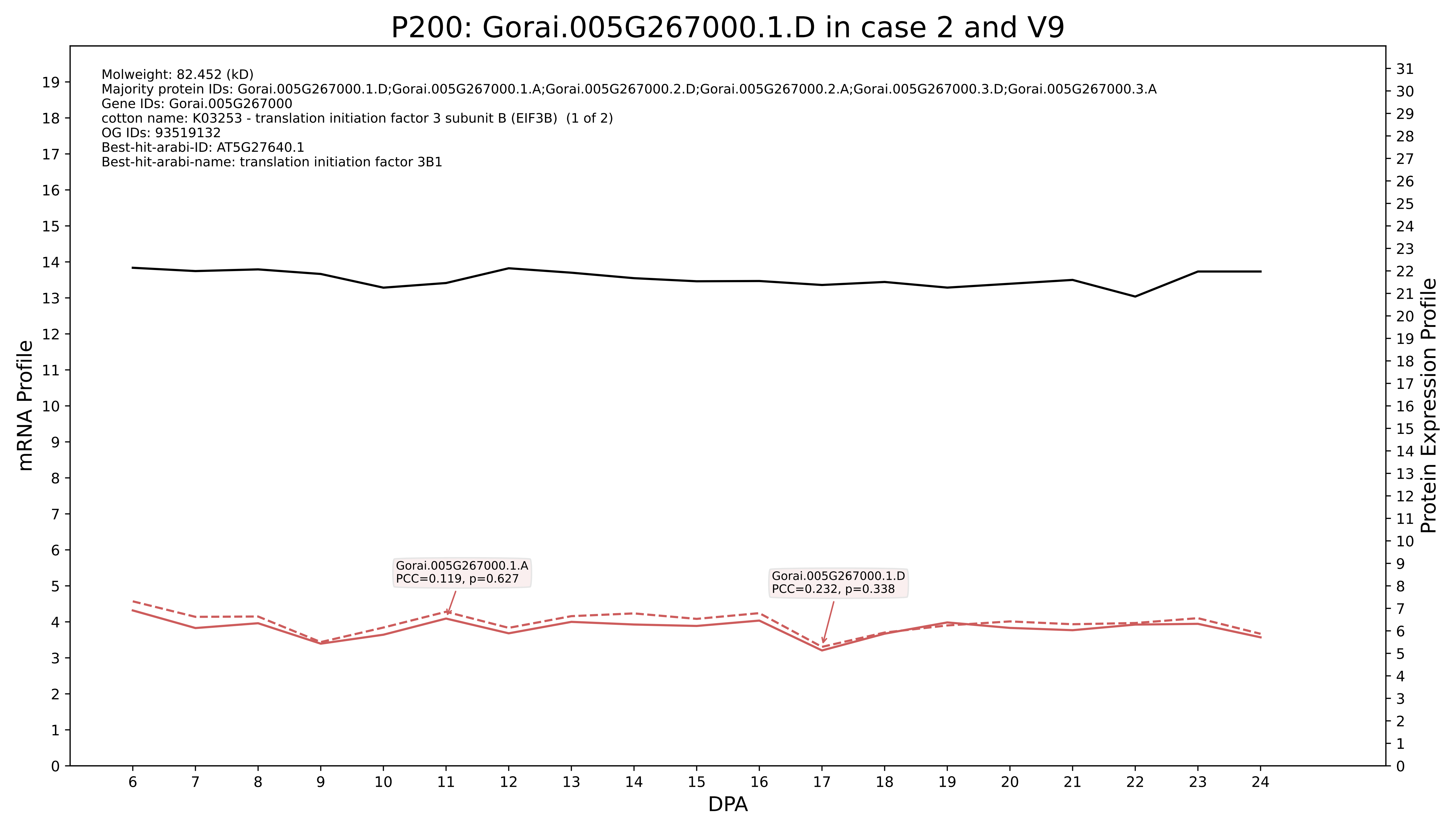
P200: Gorai.005G267000.1.D in case 2 and V9
Molweight: 82.452 (kD)
Majority protein IDs: Gorai.005G267000.1.D;Gorai.005G267000.1.A;Gorai.005G267000.2.D;Gorai.005G267000.2.A;Gorai.005G267000.3.D;Gorai.005G267000.3.A
Gene IDs: Gorai.005G267000
cotton name: K03253 - translation initiation factor 3 subunit B (EIF3B) (1 of 2)
OG IDs: 93519132
Best-hit-arabi-ID: AT5G27640.1
Best-hit-arabi-name: translation initiation factor 3B1
Gorai.005G267000.1.A
PCC=0.119, p=0.627
Gorai.005G267000.1.D
PCC=0.232, p=0.338
mRNA Profile
Protein Expression Profile
DPA

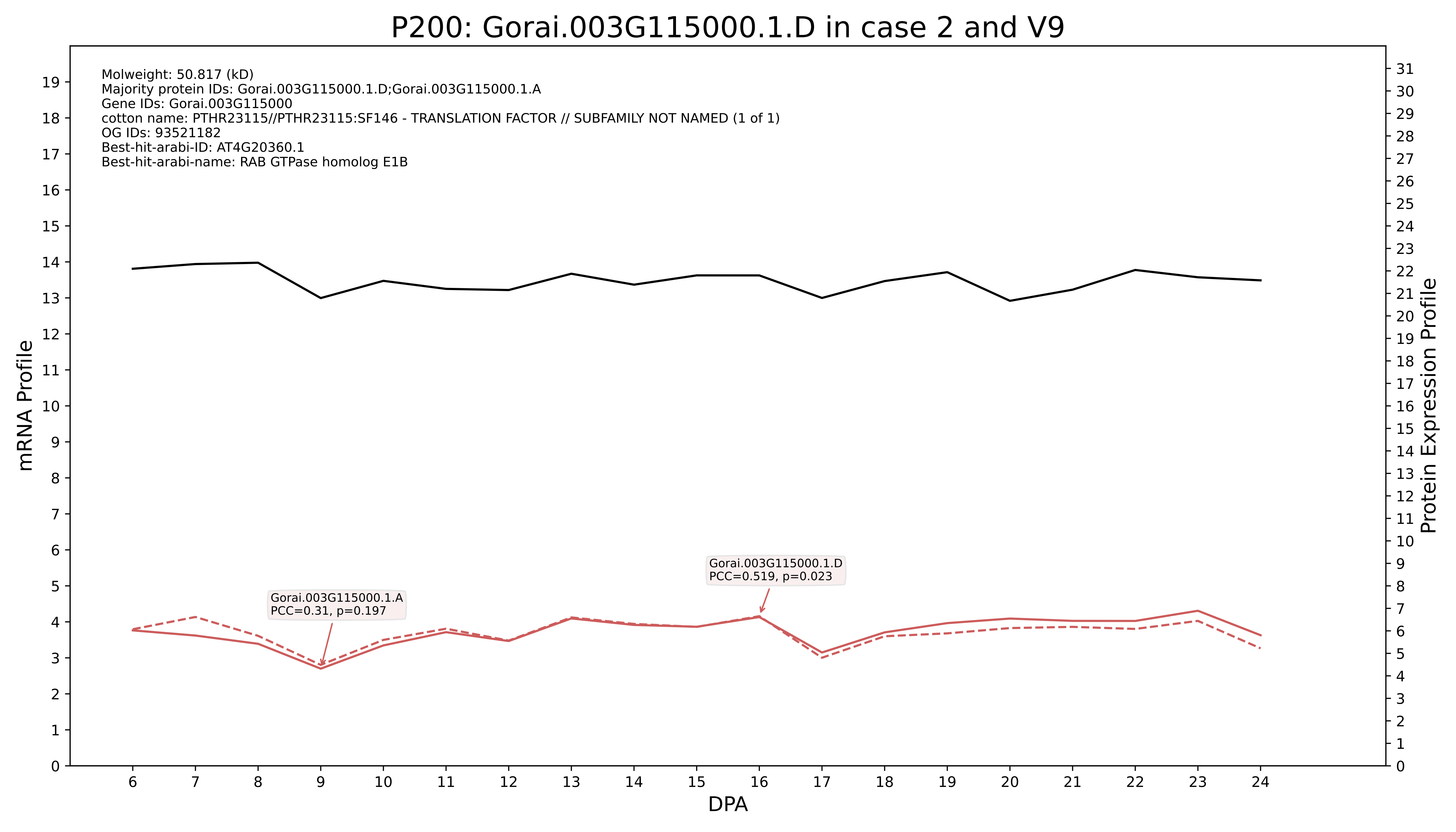

P200: Gorai.003G115000.1.D in case 2 and V9
Molweight: 50.817 (kD)
Majority protein IDs: Gorai.003G115000.1.D;Gorai.003G115000.1.A
Gene IDs: Gorai.003G115000
cotton name: PTHR23115//PTHR23115:SF146 - TRANSLATION FACTOR // SUBFAMILY NOT NAMED (1 of 1)
OG IDs: 93521182
Best-hit-arabi-ID: AT4G20360.1
Best-hit-arabi-name: RAB GTPase homolog E1B
Gorai.003G115000.1.A
PCC=0.31, p=0.197
Gorai.003G115000.1.D
PCC=0.519, p=0.023
mRNA Profile
Protein Expression Profile
DPA

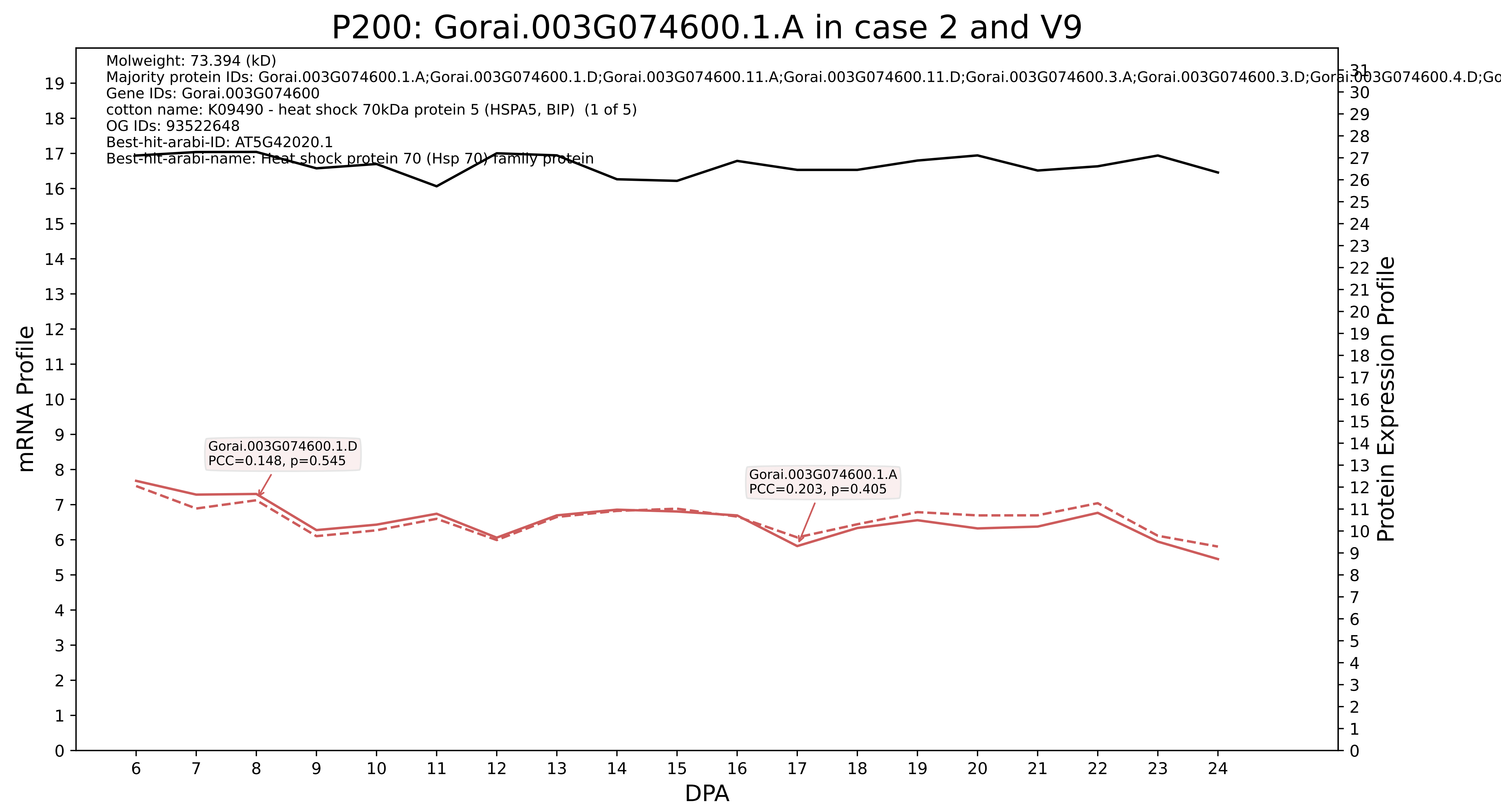
P200: Gorai.003G074600.1.A in case 2 and V9
Molweight: 73.394 (kD)
Majority protein IDs: Gorai.003G074600.1.A;Gorai.003G074600.1.D;Gorai.003G074600.11.A;Gorai.003G074600.11.D;Gorai.003G074600.3.A;Gorai.003G074600.3.D;Gorai.003G074600.4.D;Go
Gene IDs: Gorai.003G074600
cotton name: K09490 - heat shock 70kDa protein 5 (HSPA5, BIP) (1 of 5)
OG IDs: 93522648
Best-hit-arabi-ID: AT5G42020.1
Best-hit-arabi-name: Heat shock protein 70 (Hsp 70) family protein
Gorai.003G074600.1.D
PCC=0.148, p=0.545
Gorai.003G074600.1.A
PCC=0.203, p=0.405
mRNA Profile
Protein Expression Profile
DPA

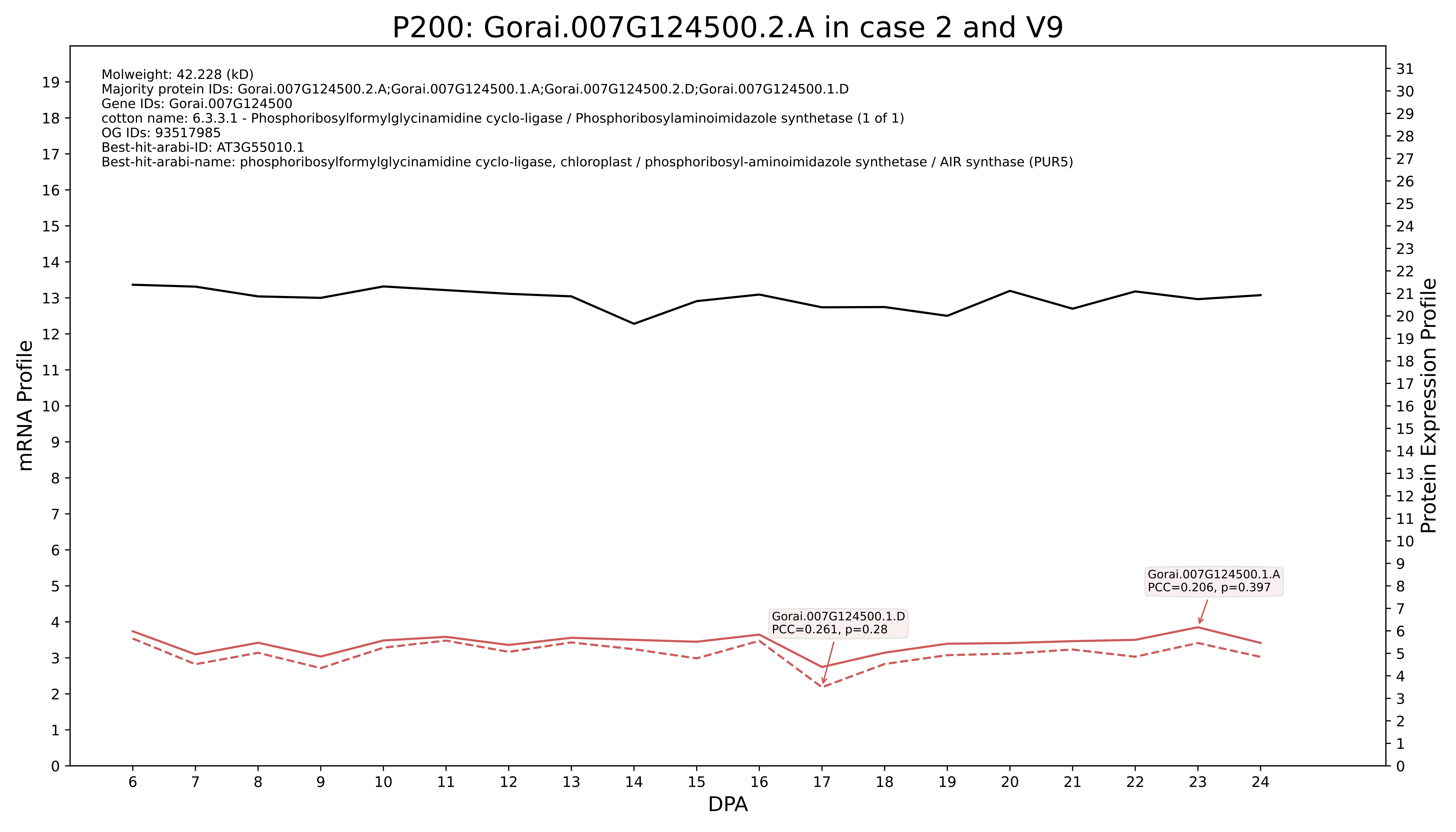

P200: Gorai.007G124500.2.A in case 2 and V9
Molweight: 42.228 (kD)
Majority protein IDs: Gorai.007G124500.2.A;Gorai.007G124500.1.A;Gorai.007G124500.2.D;Gorai.007G124500.1.D
Gene IDs: Gorai.007G124500
cotton name: 6.3.3.1 - Phosphoribosylformylglycinamidine cyclo-ligase / Phosphoribosylaminoimidazole synthetase (1 of 1)
OG IDs: 93517985
Best-hit-arabi-ID: AT3G55010.1
Best-hit-arabi-name: phosphoribosylformylglycinamidine cyclo-ligase, chloroplast / phosphoribosyl-aminoimidazole synthetase / AIR synthase (PUR5)
Gorai.007G124500.1.A
PCC=0.206, p=0.397
Gorai.007G124500.1.D
PCC=0.261, p=0.28
mRNA Profile
Protein Expression Profile
DPA

200: Gorai.008G030600.3.D in case 2 and V9

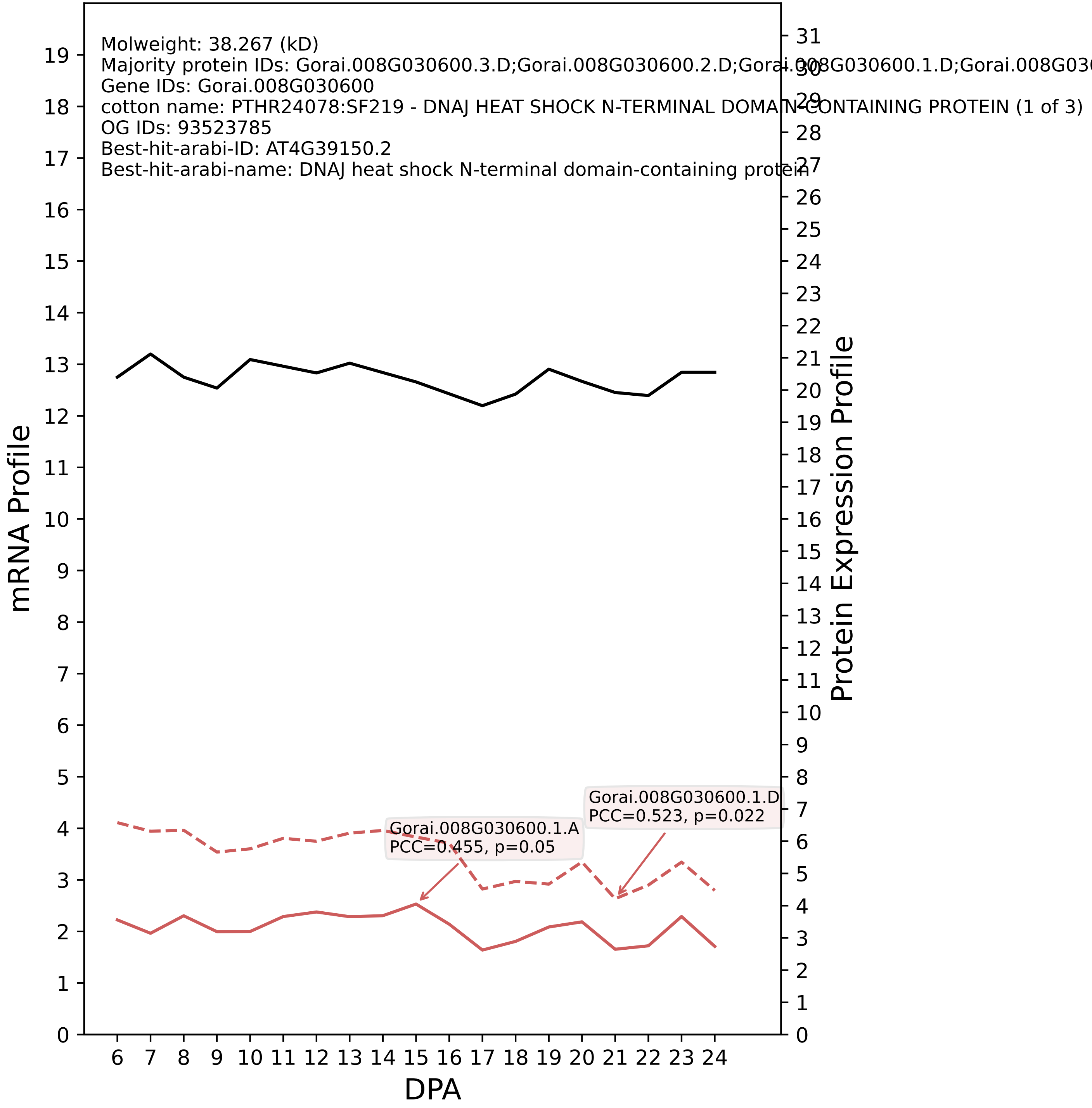

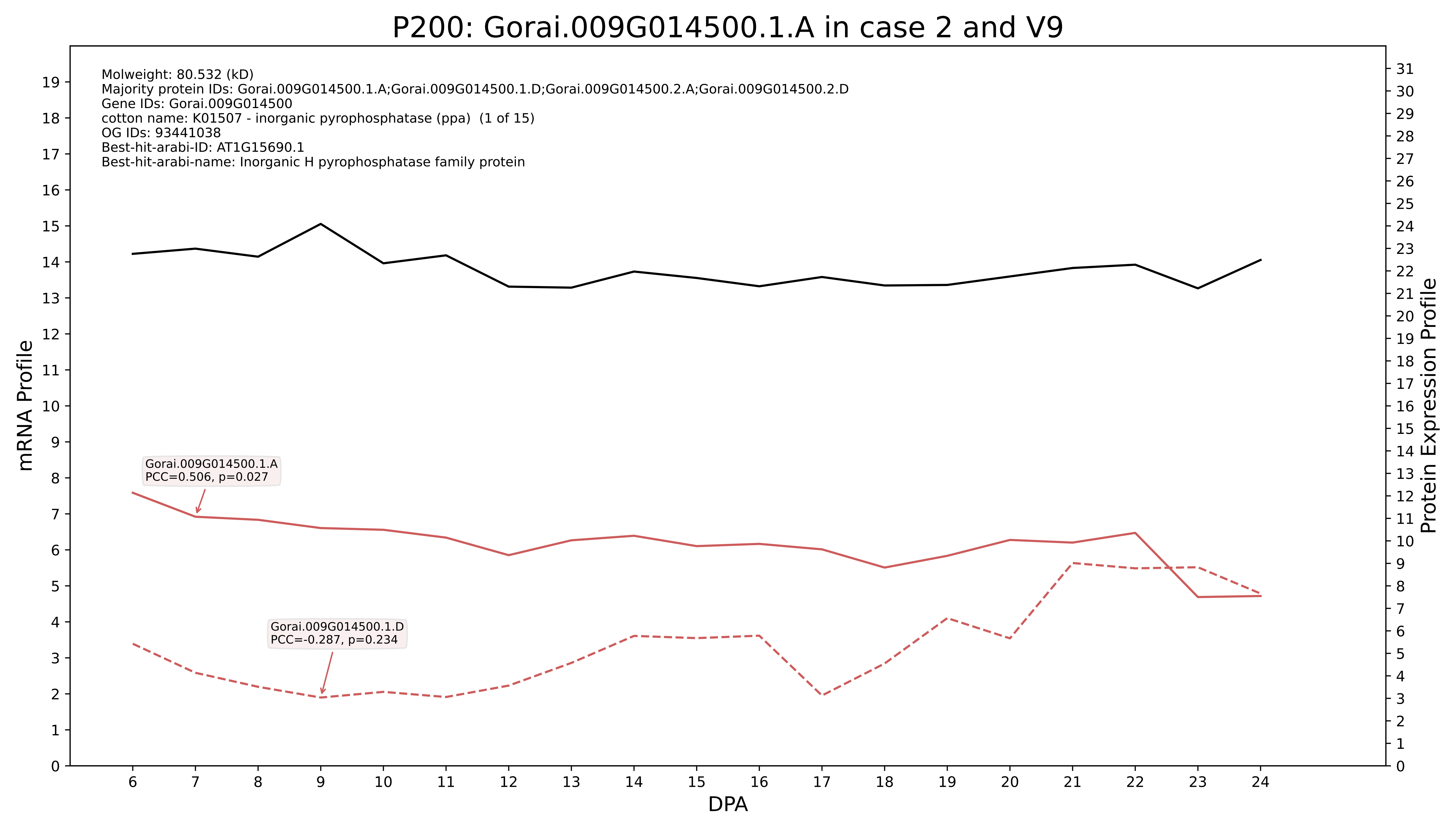
P200: Gorai.009G014500.1.A in case 2 and V9
Molweight: 80.532 (kD)
Majority protein IDs: Gorai.009G014500.1.A;Gorai.009G014500.1.D;Gorai.009G014500.2.A;Gorai.009G014500.2.D
Gene IDs: Gorai.009G014500
cotton name: K01507 - inorganic pyrophosphatase (ppa) (1 of 15)
OG IDs: 93441038
Best-hit-arabi-ID: AT1G15690.1
Best-hit-arabi-name: Inorganic H pyrophosphatase family protein
Gorai.009G014500.1.A
PCC=0.506, p=0.027
Gorai.009G014500.1.D
PCC=-0.287, p=0.234
mRNA Profile
Protein Expression Profile
DPA

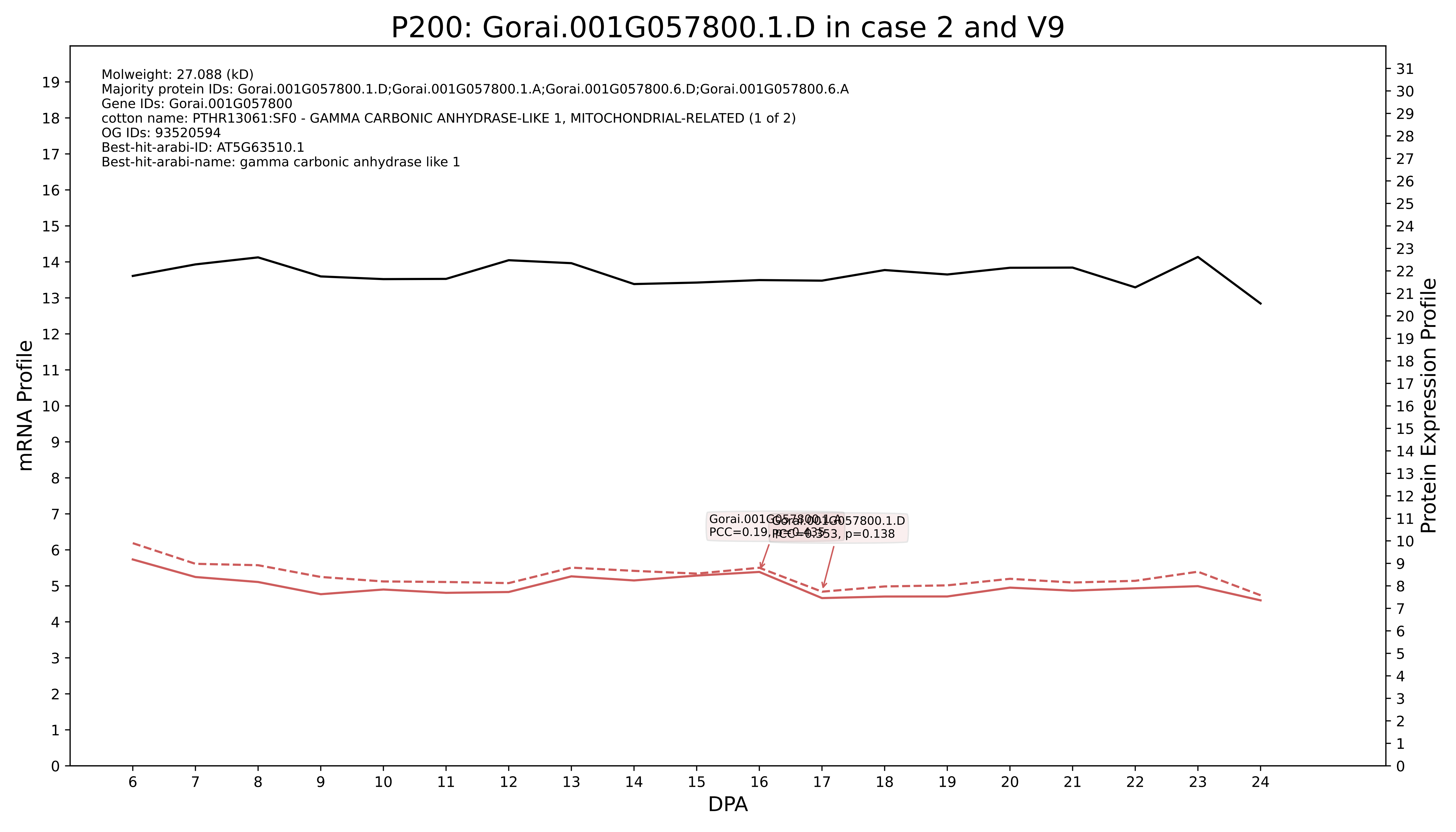

P200: Gorai.001G057800.1.D in case 2 and V9
Molweight: 27.088 (kD)
Majority protein IDs: Gorai.001G057800.1.D;Gorai.001G057800.1.A;Gorai.001G057800.6.D;Gorai.001G057800.6.A
Gene IDs: Gorai.001G057800
cotton name: PTHR13061:SF0 - GAMMA CARBONIC ANHYDRASE-LIKE 1, MITOCHONDRIAL-RELATED (1 of 2)
OG IDs: 93520594
Best-hit-arabi-ID: AT5G63510.1
Best-hit-arabi-name: gamma carbonic anhydrase like 1
Gorai.001G057800.1.A
Gorai.001G057800.1.D
PCC=0.353, p=0.138
mRNA Profile
Protein Expression Profile
DPA

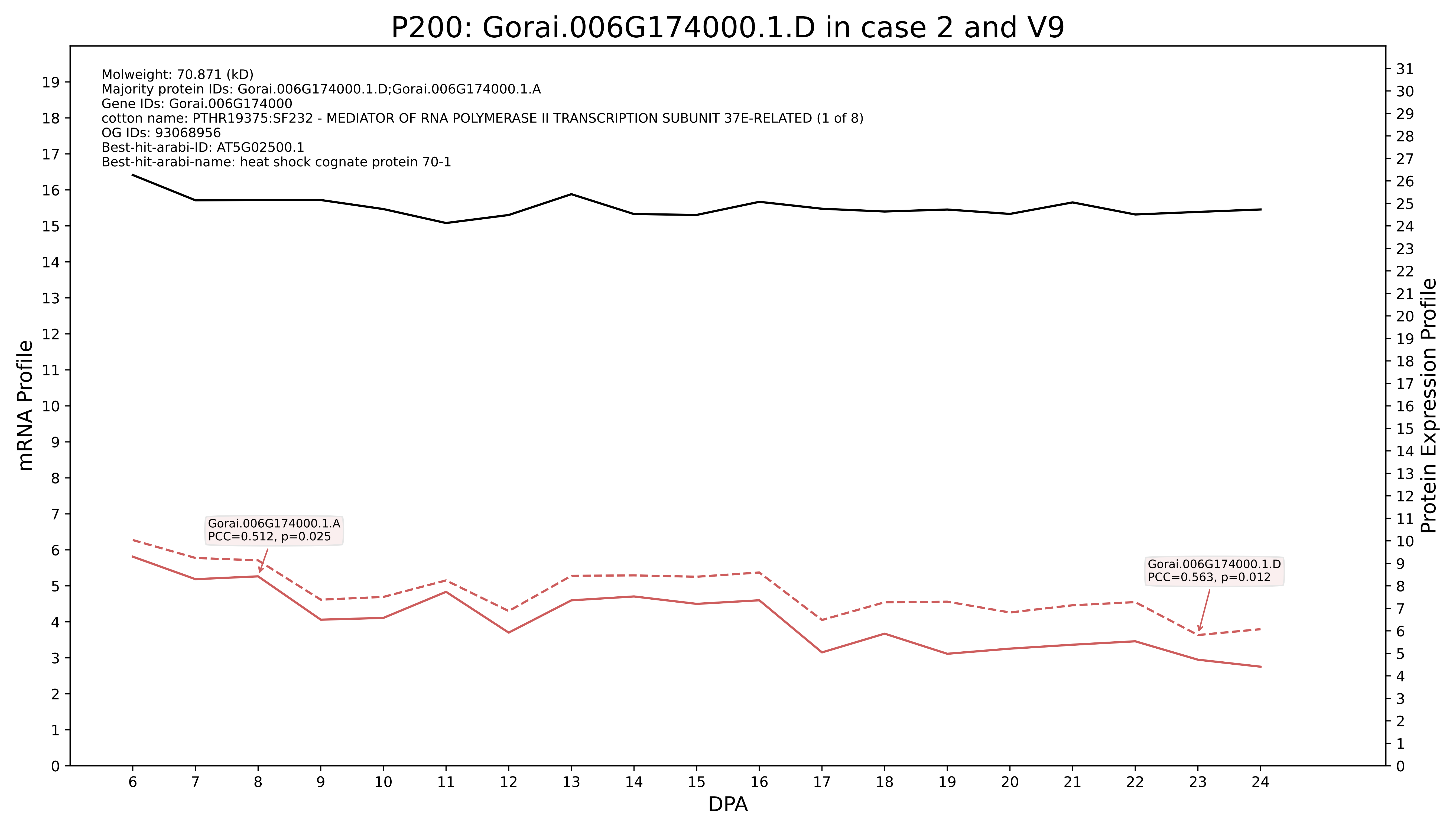

P200: Gorai.006G174000.1.D in case 2 and V9
Molweight: 70.871 (kD)
Majority protein IDs: Gorai.006G174000.1.D;Gorai.006G174000.1.A
Gene IDs: Gorai.006G174000
cotton name: PTHR19375:SF232 - MEDIATOR OF RNA POLYMERASE II TRANSCRIPTION SUBUNIT 37E-RELATED (1 of 8)
OG IDs: 93068956
Best-hit-arabi-ID: AT5G02500.1
Best-hit-arabi-name: heat shock cognate protein 70-1
Gorai.006G174000.1.A
PCC=0.512, p=0.025
Gorai.006G174000.1.D
PCC=0.563, p=0.012
mRNA Profile
Protein Expression Profile
DPA

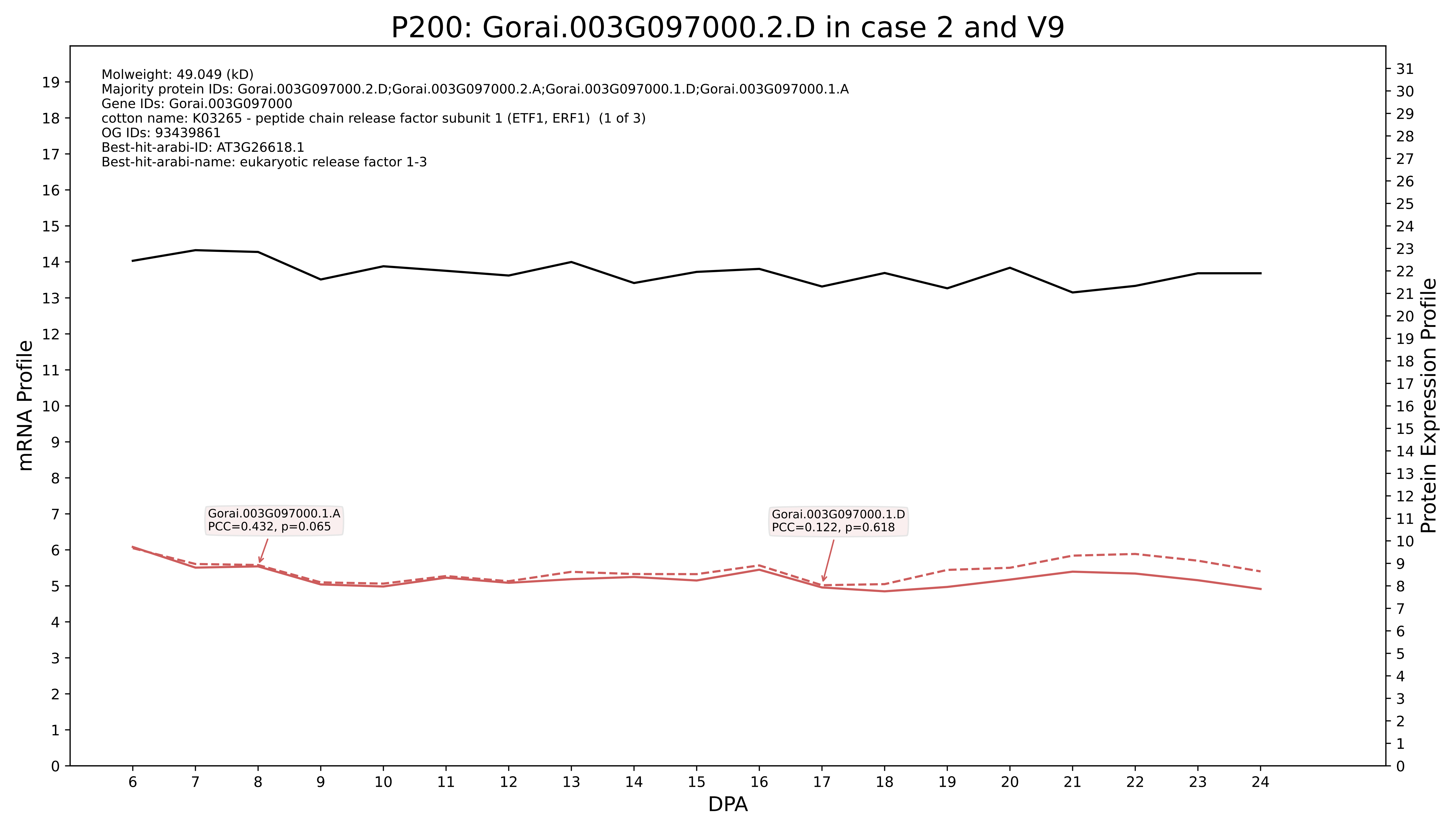
P200: Gorai.003G097000.2.D in case 2 and V9
Molweight: 49.049 (kD)
Majority protein IDs: Gorai.003G097000.2.D;Gorai.003G097000.2.A;Gorai.003G097000.1.D;Gorai.003G097000.1.A
Gene IDs: Gorai.003G097000
cotton name: K03265 - peptide chain release factor subunit 1 (ETF1, ERF1) (1 of 3)
OG IDs: 93439861
Best-hit-arabi-ID: AT3G26618.1
Best-hit-arabi-name: eukaryotic release factor 1-3
Gorai.003G097000.1.A
PCC=0.432, p=0.065
Gorai.003G097000.1.D
PCC=0.122, p=0.618
mRNA Profile
Protein Expression Profile
DPA

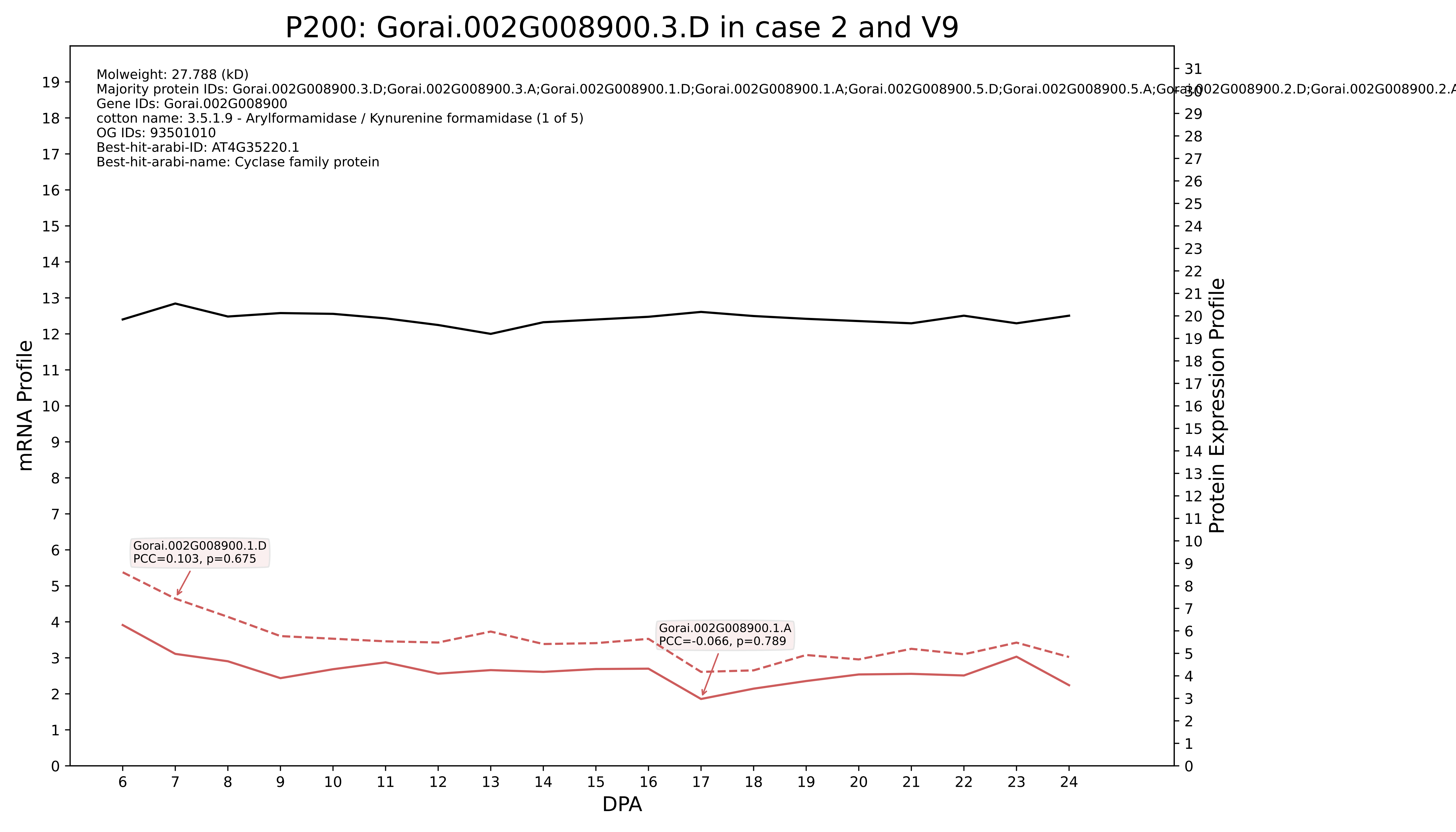
P200: Gorai.002G008900.3.D in case 2 and V9
Molweight: 27.788 (kD)
Majority protein IDs: Gorai.002G008900.3.D;Gorai.002G008900.3.A;Gorai.002G008900.1.D;Gorai.002G008900.1.A;Gorai.002G008900.5.D;Gorai.002G008900.5.A;Gorai.002G008900.2.D;Gorai.002G008900.2.A
Gene IDs: Gorai.002G008900
cotton name: 3.5.1.9 - Arylformamidase / Kynurenine formamidase (1 of 5)
OG IDs: 93501010
Best-hit-arabi-ID: AT4G35220.1
Best-hit-arabi-name: Cyclase family protein
Gorai.002G008900.1.D
PCC=0.103, p=0.675
Gorai.002G008900.1.A
PCC=-0.066, p=0.789
mRNA Profile
Protein Expression Profile
DPA

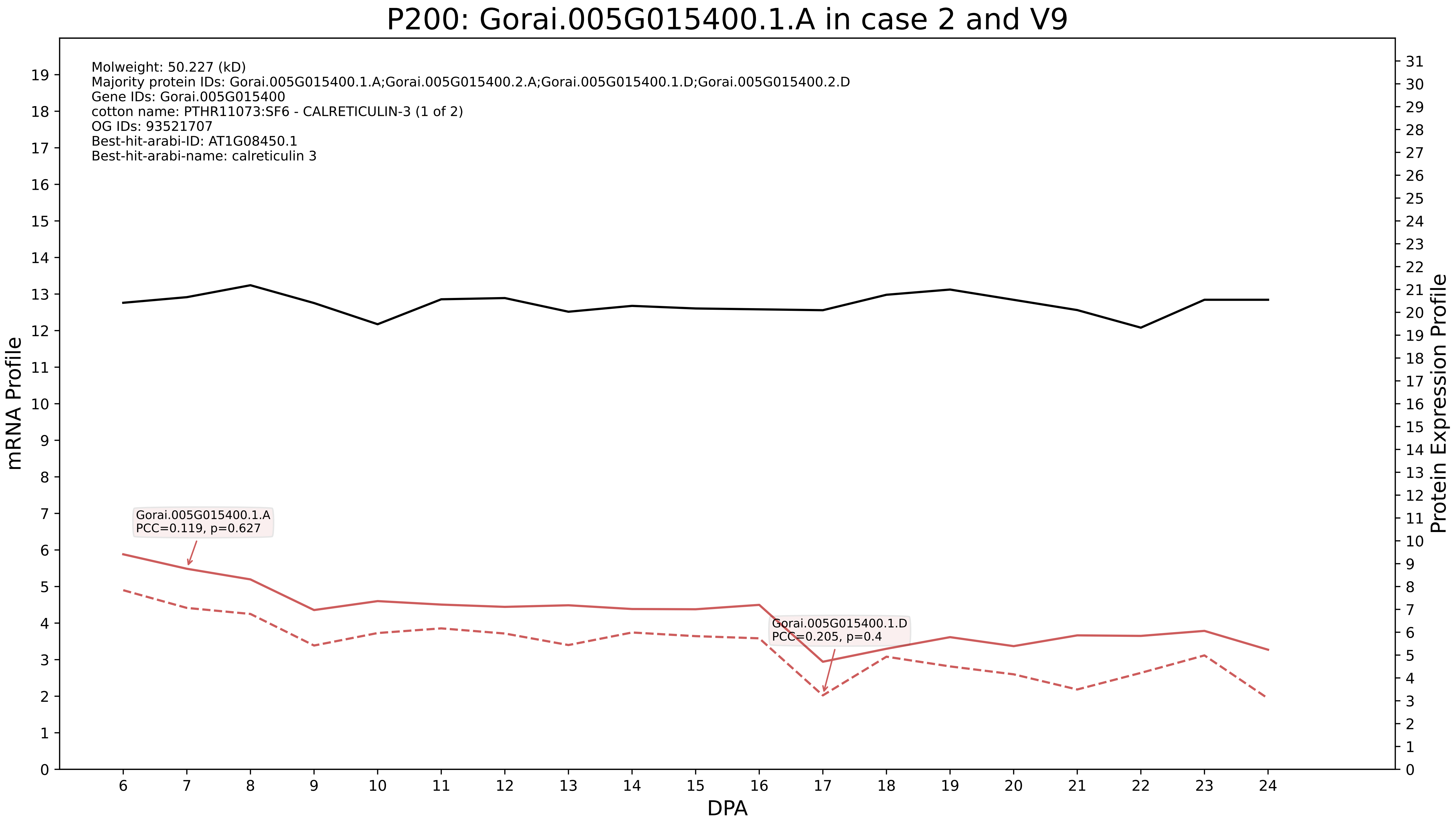
P200: Gorai.005G015400.1.A in case 2 and V9
Molweight: 50.227 (kD)
Majority protein IDs: Gorai.005G015400.1.A;Gorai.005G015400.2.A;Gorai.005G015400.1.D;Gorai.005G015400.2.D
Gene IDs: Gorai.005G015400
cotton name: PTHR11073:SF6 - CALRETICULIN-3 (1 of 2)
OG IDs: 93521707
Best-hit-arabi-ID: AT1G08450.1
Best-hit-arabi-name: calreticulin 3
Gorai.005G015400.1.A
PCC=0.119, p=0.627
Gorai.005G015400.1.D
PCC=0.205, p=0.4
mRNA Profile
Protein Expression Profile
DPA

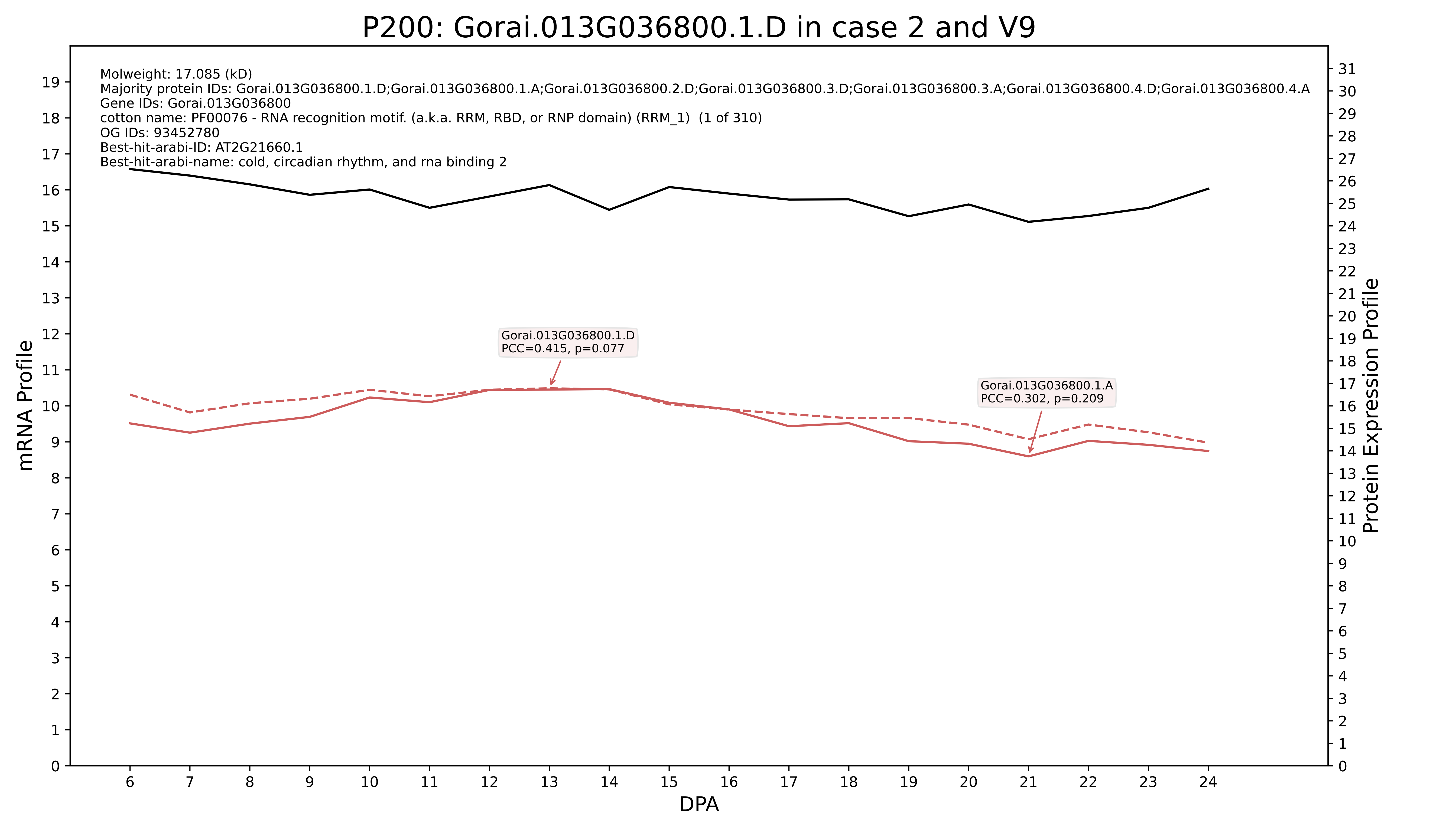
P200: Gorai.013G036800.1.D in case 2 and V9
Molweight: 17.085 (kD)
Majority protein IDs: Gorai.013G036800.1.D;Gorai.013G036800.1.A;Gorai.013G036800.2.D;Gorai.013G036800.3.D;Gorai.013G036800.3.A;Gorai.013G036800.4.D;Gorai.013G036800.4.A
Gene IDs: Gorai.013G036800
cotton name: PF00076 - RNA recognition motif. (a.k.a. RRM, RBD, or RNP domain) (RRM_1) (1 of 310)
OG IDs: 93452780
Best-hit-arabi-ID: AT2G21660.1
Best-hit-arabi-name: cold, circadian rhythm, and rna binding 2
Gorai.013G036800.1.D
PCC=0.415, p=0.077
Gorai.013G036800.1.A
PCC=0.302, p=0.209
mRNA Profile
Protein Expression Profile
DPA

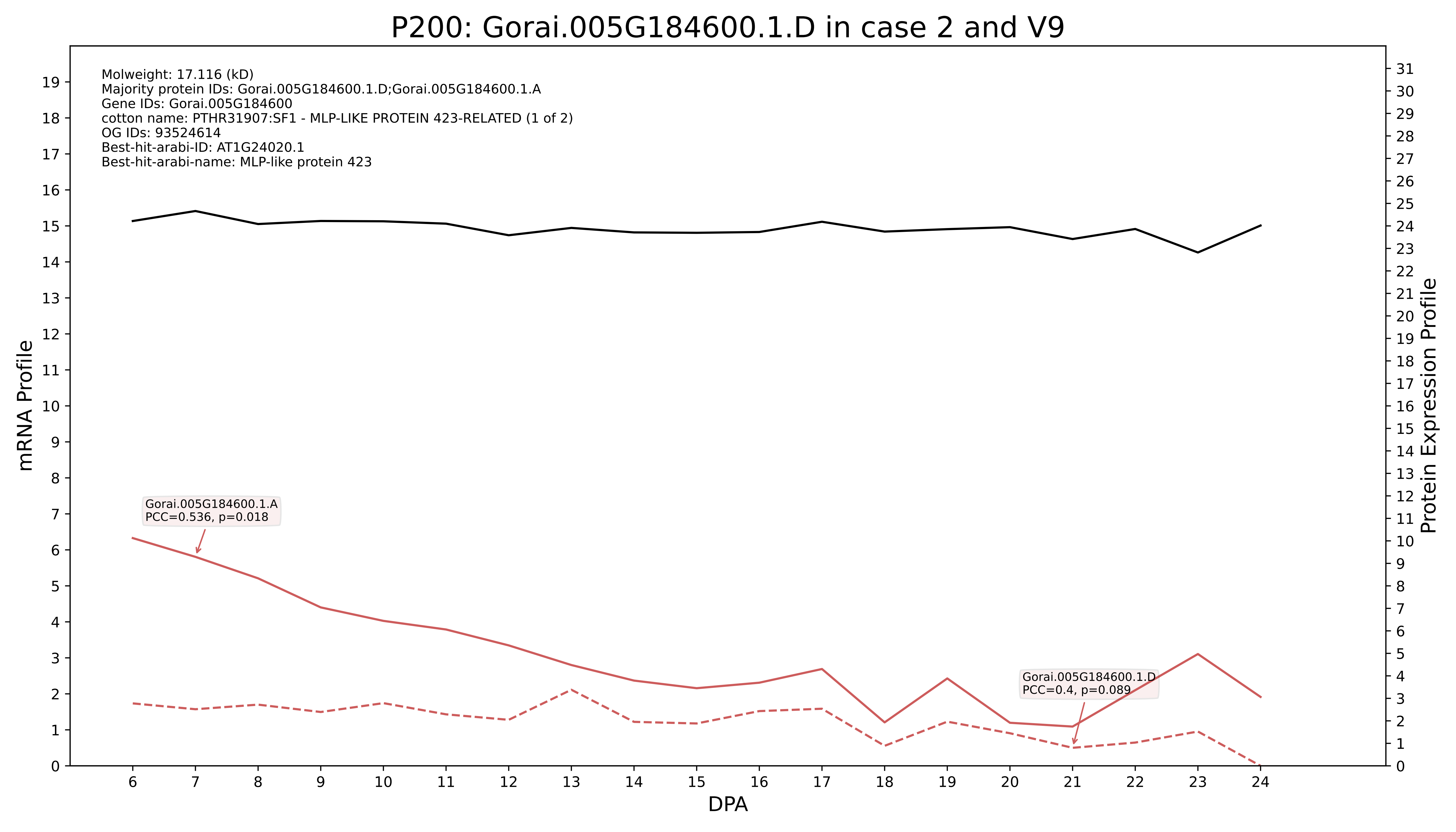

P200: Gorai.005G184600.1.D in case 2 and V9
Molweight: 17.116 (kD)
Majority protein IDs: Gorai.005G184600.1.D;Gorai.005G184600.1.A
Gene IDs: Gorai.005G184600
cotton name: PTHR31907:SF1 - MLP-LIKE PROTEIN 423-RELATED (1 of 2)
OG IDs: 93524614
Best-hit-arabi-ID: AT1G24020.1
Best-hit-arabi-name: MLP-like protein 423
Gorai.005G184600.1.A
PCC=0.536, p=0.018
Gorai.005G184600.1.D
PCC=0.4, p=0.089
mRNA Profile
Protein Expression Profile
DPA

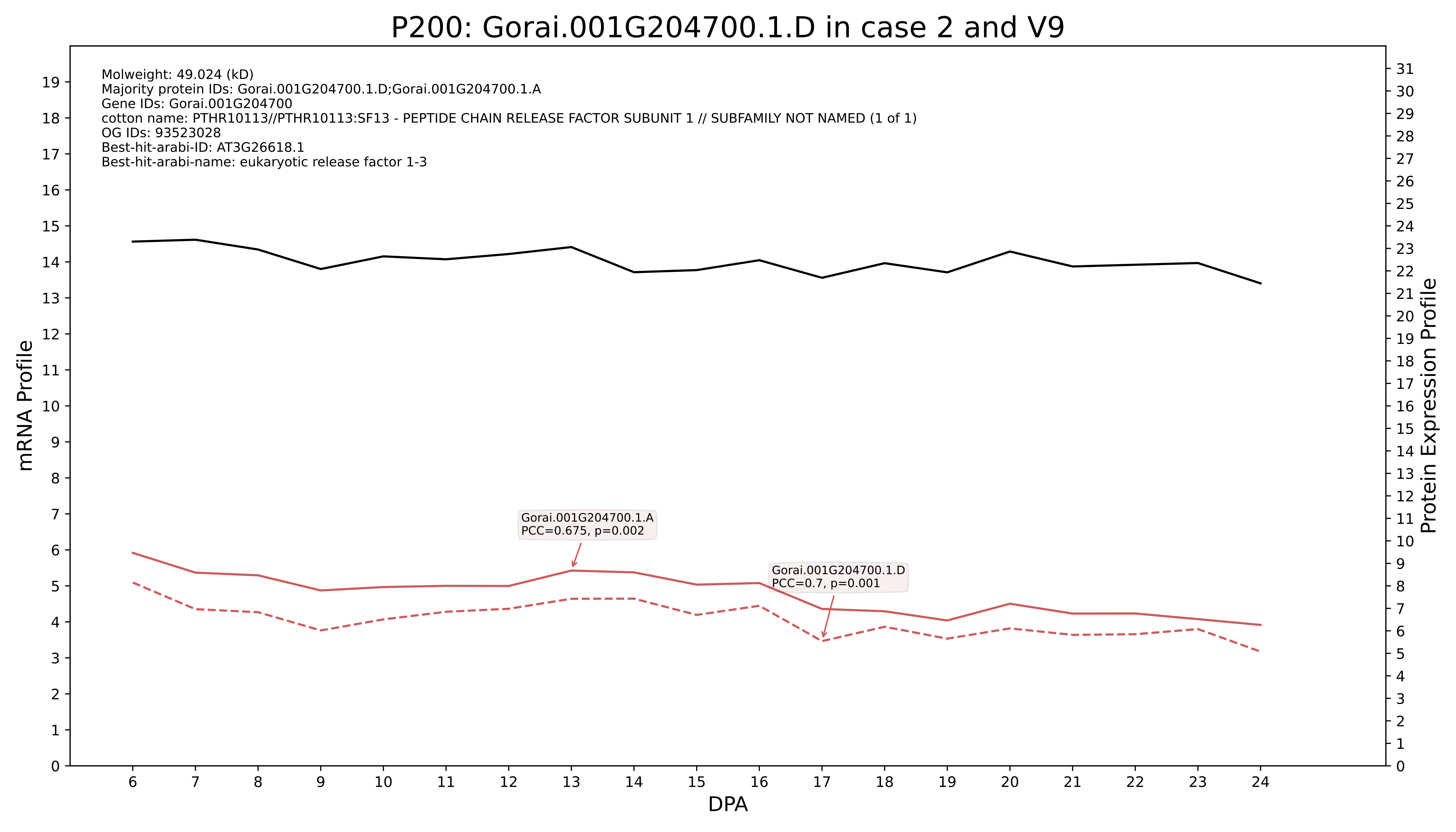
P200: Gorai.001G204700.1.D in case 2 and V9
Molweight: 49.024 (kD)
Majority protein IDs: Gorai.001G204700.1.D;Gorai.001G204700.1.A
Gene IDs: Gorai.001G204700
cotton name: PTHR10113//PTHR10113:SF13 - PEPTIDE CHAIN RELEASE FACTOR SUBUNIT 1 // SUBFAMILY NOT NAMED (1 of 1)
OG IDs: 93523028
Best-hit-arabi-ID: AT3G26618.1
Best-hit-arabi-name: eukaryotic release factor 1-3
Gorai.001G204700.1.A
PCC=0.675, p=0.002
Gorai.001G204700.1.D
PCC=0.7, p=0.001
mRNA Profile
Protein Expression Profile
DPA

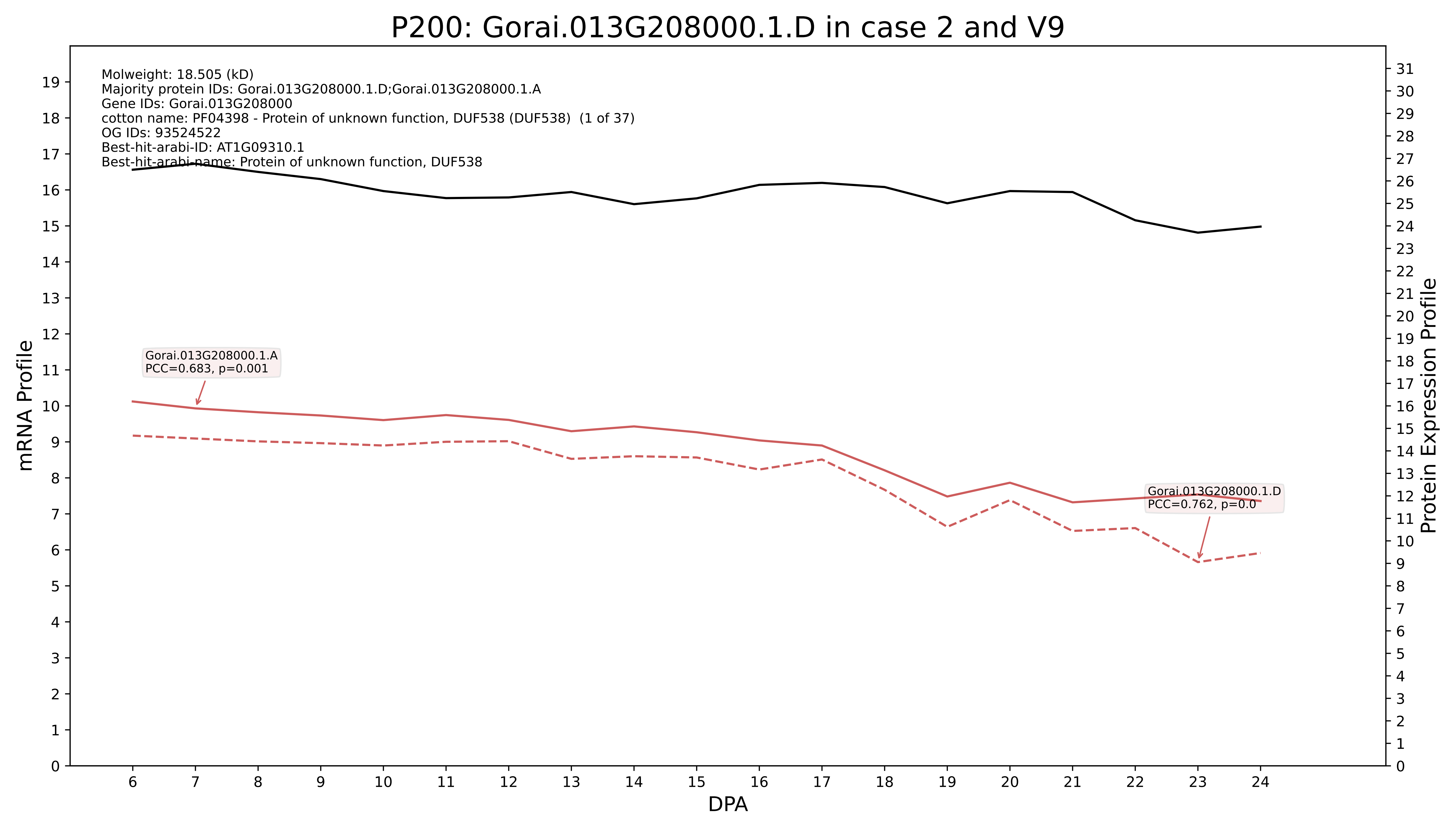
P200: Gorai.013G208000.1.D in case 2 and V9
Molweight: 18.505 (kD)
Majority protein IDs: Gorai.013G208000.1.D;Gorai.013G208000.1.A
Gene IDs: Gorai.013G208000
cotton name: PF04398 - Protein of unknown function, DUF538 (DUF538) (1 of 37)
OG IDs: 93524522
Best-hit-arabi-ID: AT1G09310.1
Best-hit-arabi-name: Protein of unknown function, DUF538
Gorai.013G208000.1.A
PCC=0.683, p=0.001
Gorai.013G208000.1.D
PCC=0.762, p=0.0
mRNA Profile
Protein Expression Profile
DPA

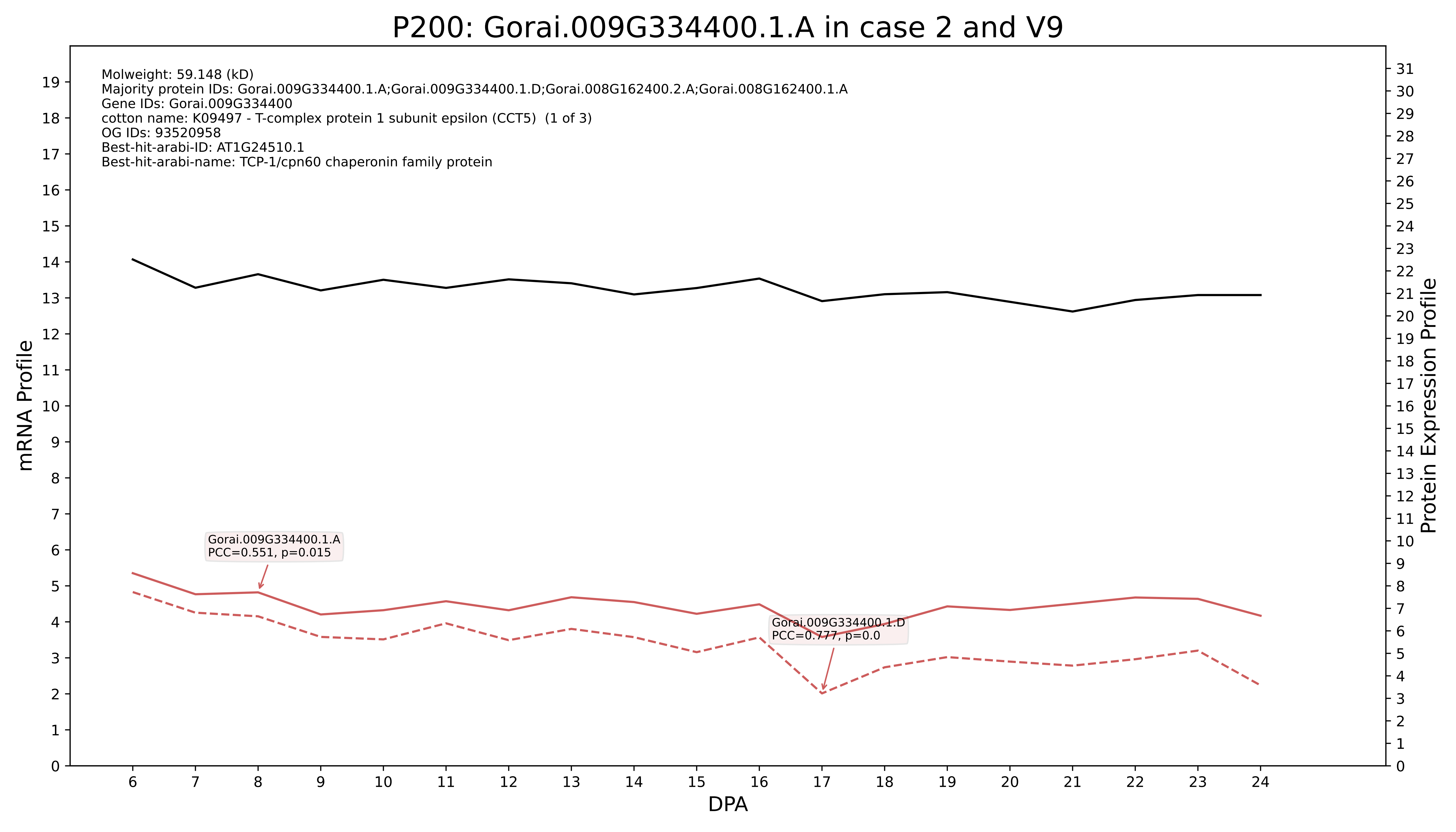
P200: Gorai.009G334400.1.A in case 2 and V9
Molweight: 59.148 (kD)
Majority protein IDs: Gorai.009G334400.1.A;Gorai.009G334400.1.D;Gorai.008G162400.2.A;Gorai.008G162400.1.A
Gene IDs: Gorai.009G334400
cotton name: K09497 - T-complex protein 1 subunit epsilon (CCT5) (1 of 3)
OG IDs: 93520958
Best-hit-arabi-ID: AT1G24510.1
Best-hit-arabi-name: TCP-1/cpn60 chaperonin family protein
Gorai.009G334400.1.A
PCC=0.551, p=0.015
Gorai.009G334400.1.D
PCC=0.777, p=0.0
mRNA Profile
Protein Expression Profile
DPA

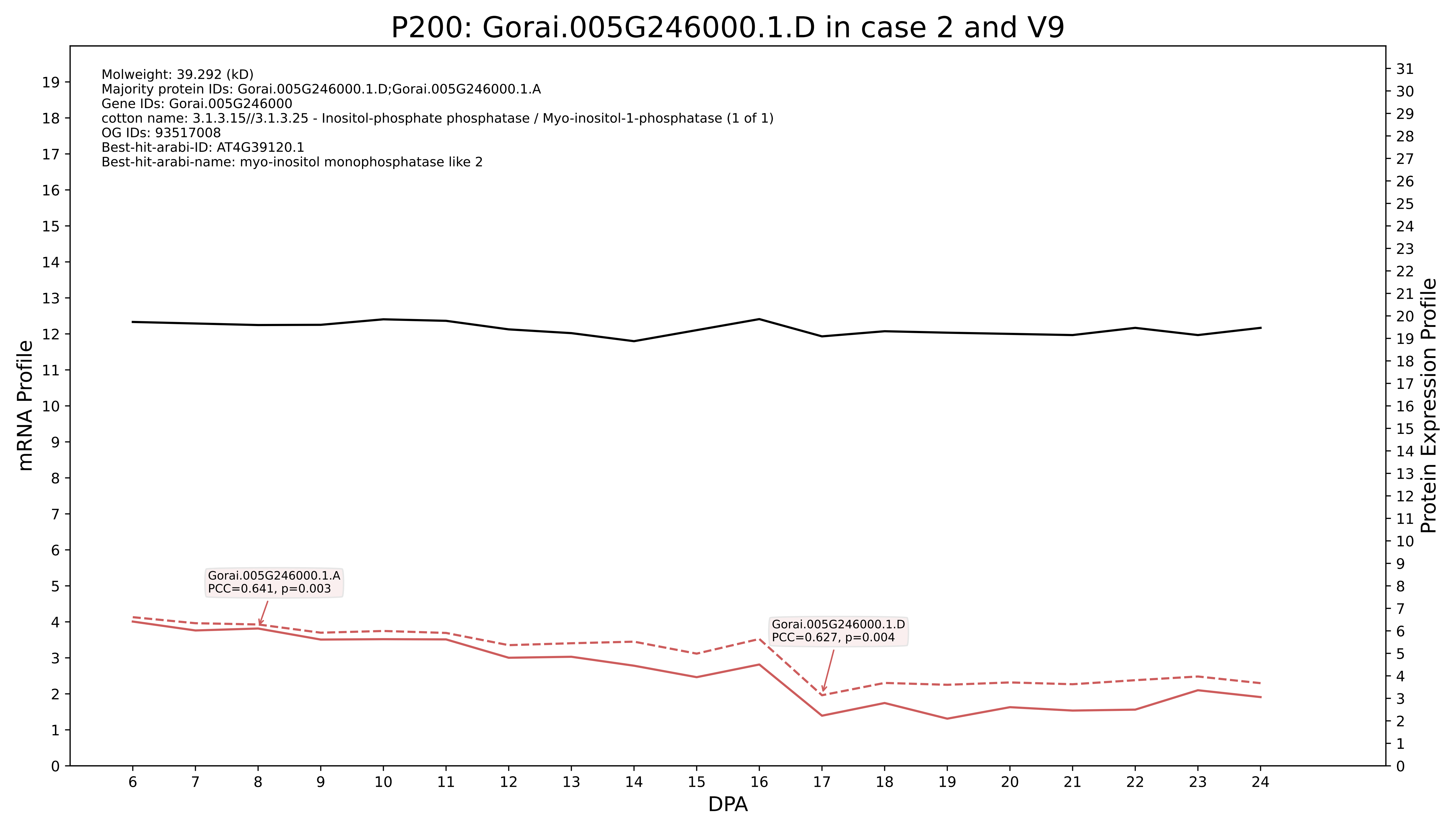
P200: Gorai.005G246000.1.D in case 2 and V9
Molweight: 39.292 (kD)
Majority protein IDs: Gorai.005G246000.1.D;Gorai.005G246000.1.A
Gene IDs: Gorai.005G246000
cotton name: 3.1.3.15//3.1.3.25 - Inositol-phosphate phosphatase / Myo-inositol-1-phosphatase (1 of 1)
OG IDs: 93517008
Best-hit-arabi-ID: AT4G39120.1
Best-hit-arabi-name: myo-inositol monophosphatase like 2
Gorai.005G246000.1.A
PCC=0.641, p=0.003
Gorai.005G246000.1.D
PCC=0.627, p=0.004
mRNA Profile
Protein Expression Profile
DPA

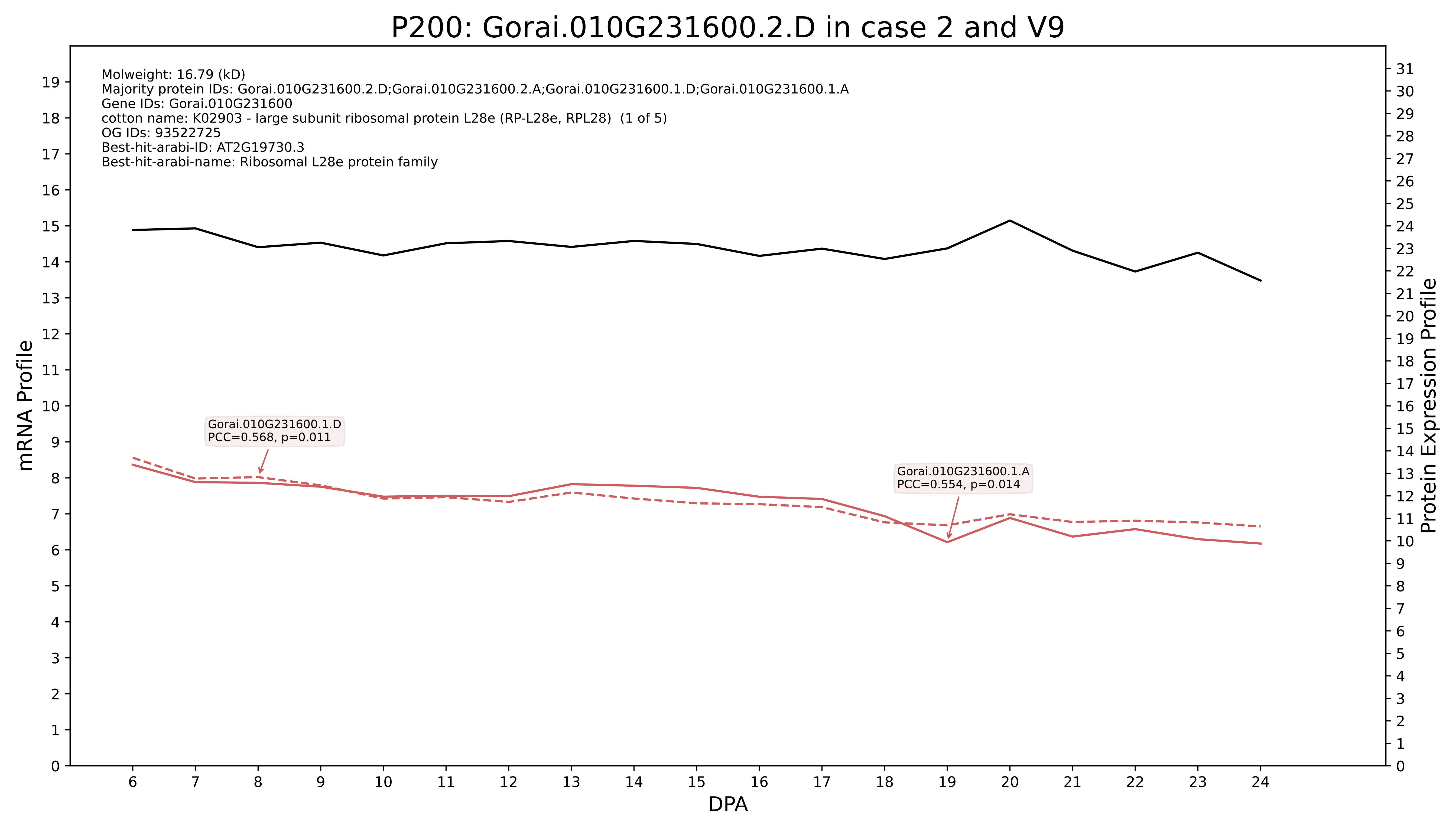
P200: Gorai.010G231600.2.D in case 2 and V9
Molweight: 16.79 (kD)
Majority protein IDs: Gorai.010G231600.2.D;Gorai.010G231600.2.A;Gorai.010G231600.1.D;Gorai.010G231600.1.A
Gene IDs: Gorai.010G231600
cotton name: K02903 - large subunit ribosomal protein L28e (RP-L28e, RPL28) (1 of 5)
OG IDs: 93522725
Best-hit-arabi-ID: AT2G19730.3
Best-hit-arabi-name: Ribosomal L28e protein family
Gorai.010G231600.1.D
PCC=0.568, p=0.011
Gorai.010G231600.1.A
PCC=0.554, p=0.014
mRNA Profile
Protein Expression Profile
DPA

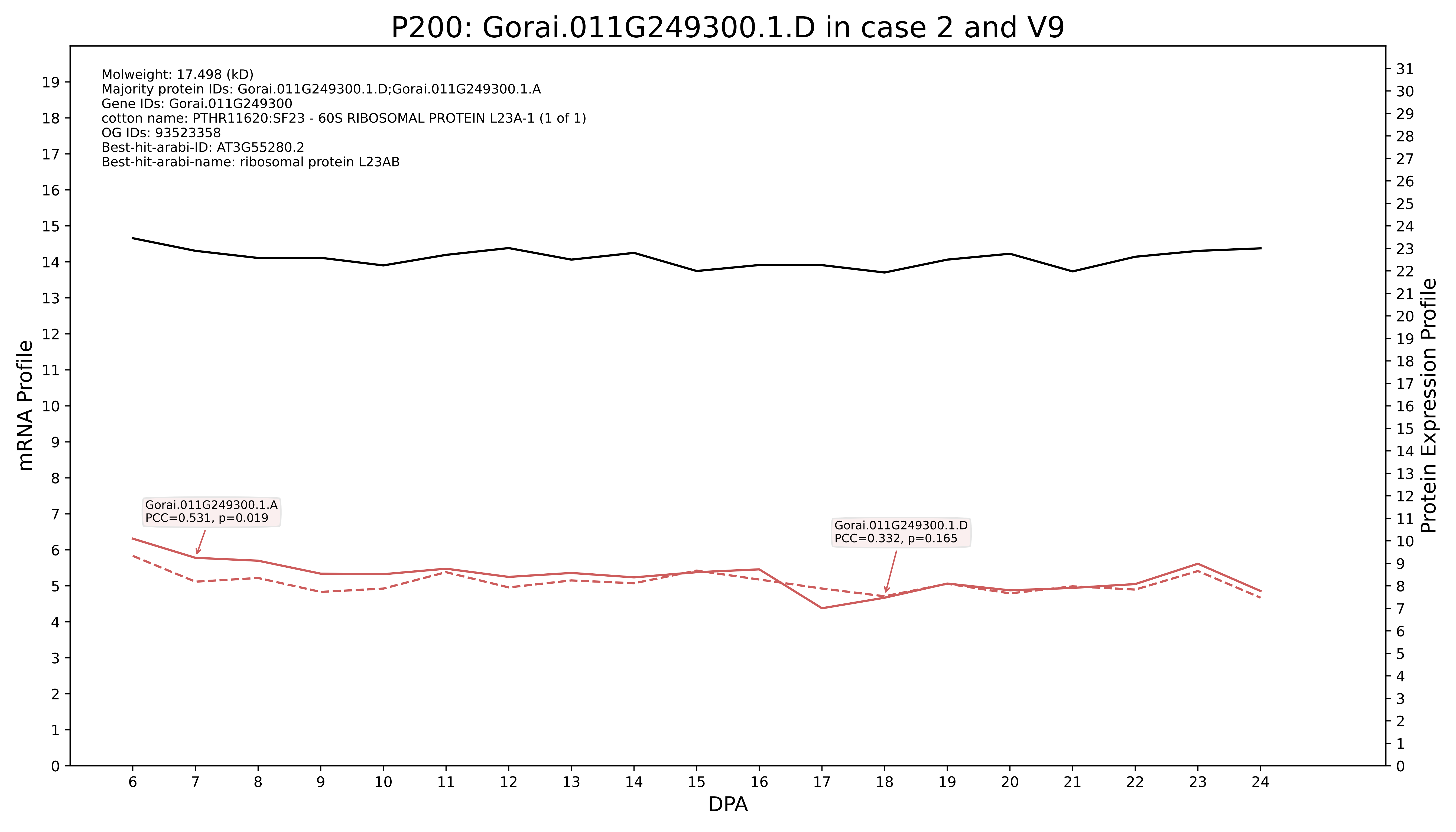
P200: Gorai.011G249300.1.D in case 2 and V9
Molweight: 17.498 (kD)
Majority protein IDs: Gorai.011G249300.1.D;Gorai.011G249300.1.A
Gene IDs: Gorai.011G249300
cotton name: PTHR11620:SF23 - 60S RIBOSOMAL PROTEIN L23A-1 (1 of 1)
OG IDs: 93523358
Best-hit-arabi-ID: AT3G55280.2
Best-hit-arabi-name: ribosomal protein L23AB
Gorai.011G249300.1.A
PCC=0.531, p=0.019
Gorai.011G249300.1.D
PCC=0.332, p=0.165
mRNA Profile
Protein Expression Profile
DPA

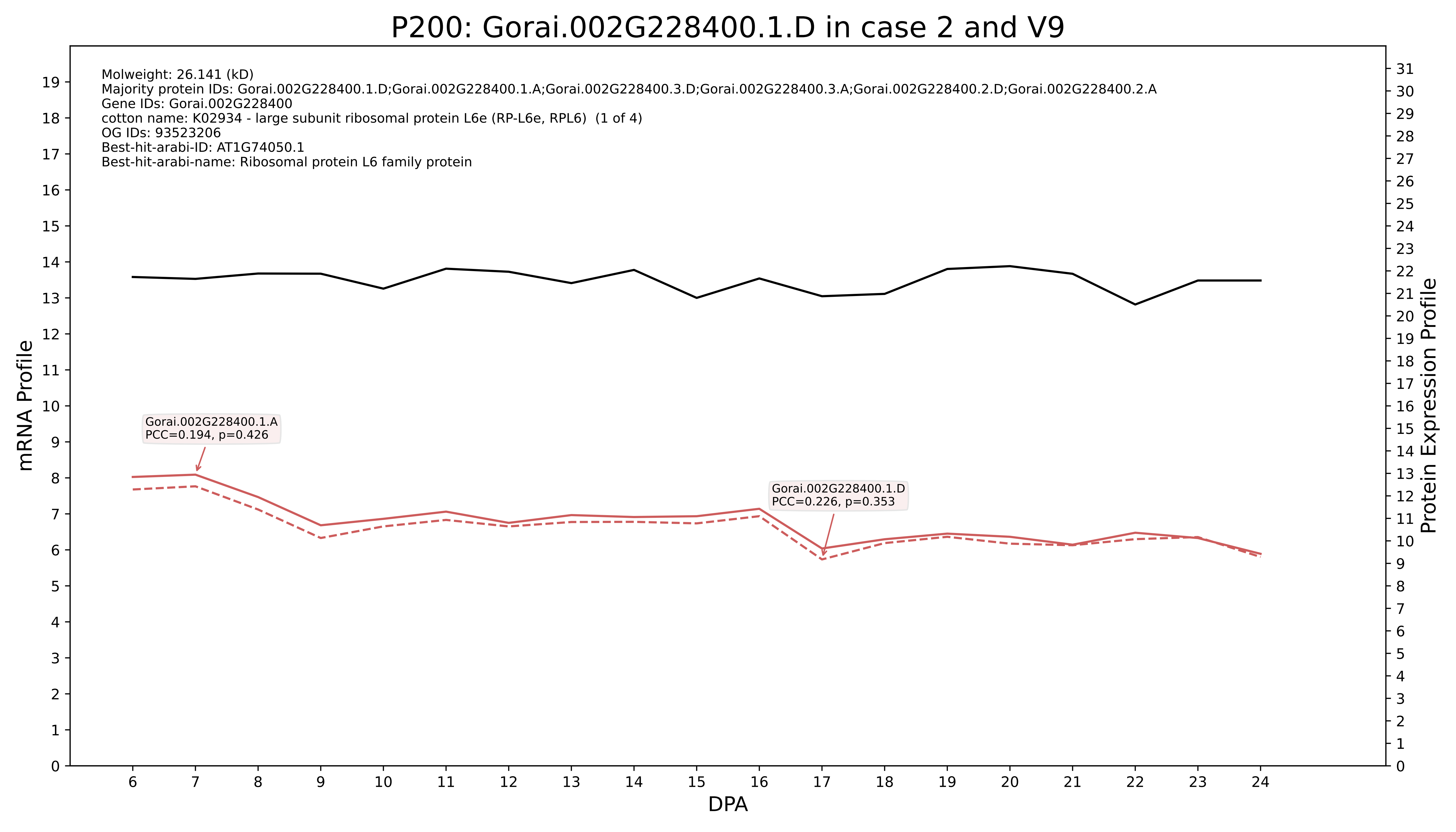
P200: Gorai.002G228400.1.D in case 2 and V9
Molweight: 26.141 (kD)
Majority protein IDs: Gorai.002G228400.1.D;Gorai.002G228400.1.A;Gorai.002G228400.3.D;Gorai.002G228400.3.A;Gorai.002G228400.2.D;Gorai.002G228400.2.A
Gene IDs: Gorai.002G228400
cotton name: K02934 - large subunit ribosomal protein L6e (RP-L6e, RPL6) (1 of 4)
OG IDs: 93523206
Best-hit-arabi-ID: AT1G74050.1
Best-hit-arabi-name: Ribosomal protein L6 family protein
Gorai.002G228400.1.A
PCC=0.194, p=0.426
Gorai.002G228400.1.D
PCC=0.226, p=0.353
mRNA Profile
Protein Expression Profile
DPA

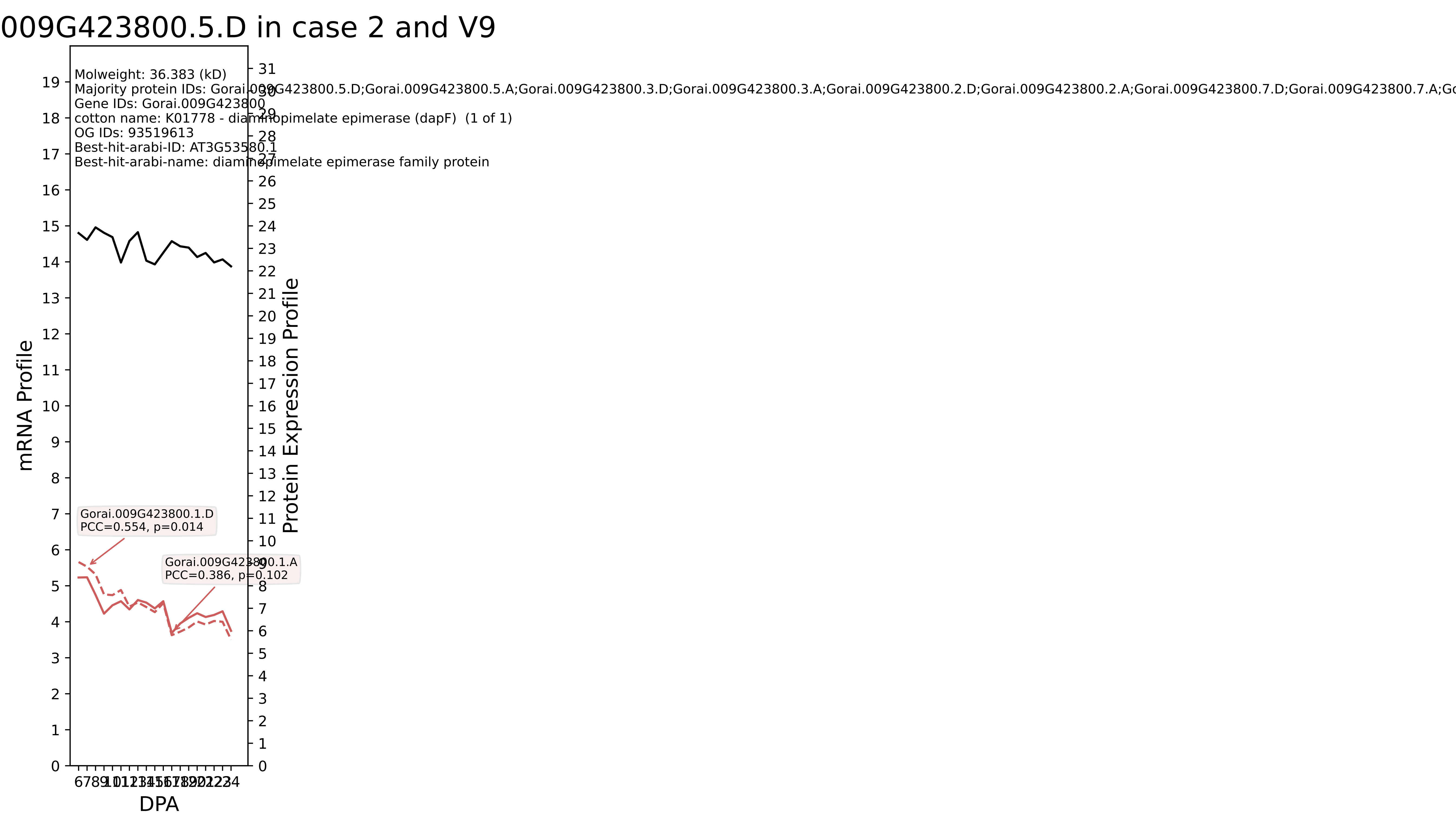
009G423800.5.D in case 2 and V9
Molweight: 36.383 (kD)
Majority protein IDs: Gorai.009G423800.5.D;Gorai.009G423800.5.A;Gorai.009G423800.3.D;Gorai.009G423800.3.A;Gorai.009G423800.2.D;Gorai.009G423800.2.A;Gorai.009G423800.7.D;Gorai.009G423800.7.A;Go
Gene IDs: Gorai.009G423800
cotton name: K01778 - diaminopimelate epimerase (dapF) (1 of 1)
OG IDs: 93519613
Best-hit-arabi-ID: AT3G53580.1
Best-hit-arabi-name: diaminopimelate epimerase family protein
Gorai.009G423800.1.D
PCC=0.554, p=0.014
Gorai.009G423800.1.A
PCC=0.386, p=0.102
mRNA Profile
Protein Expression Profile
DPA

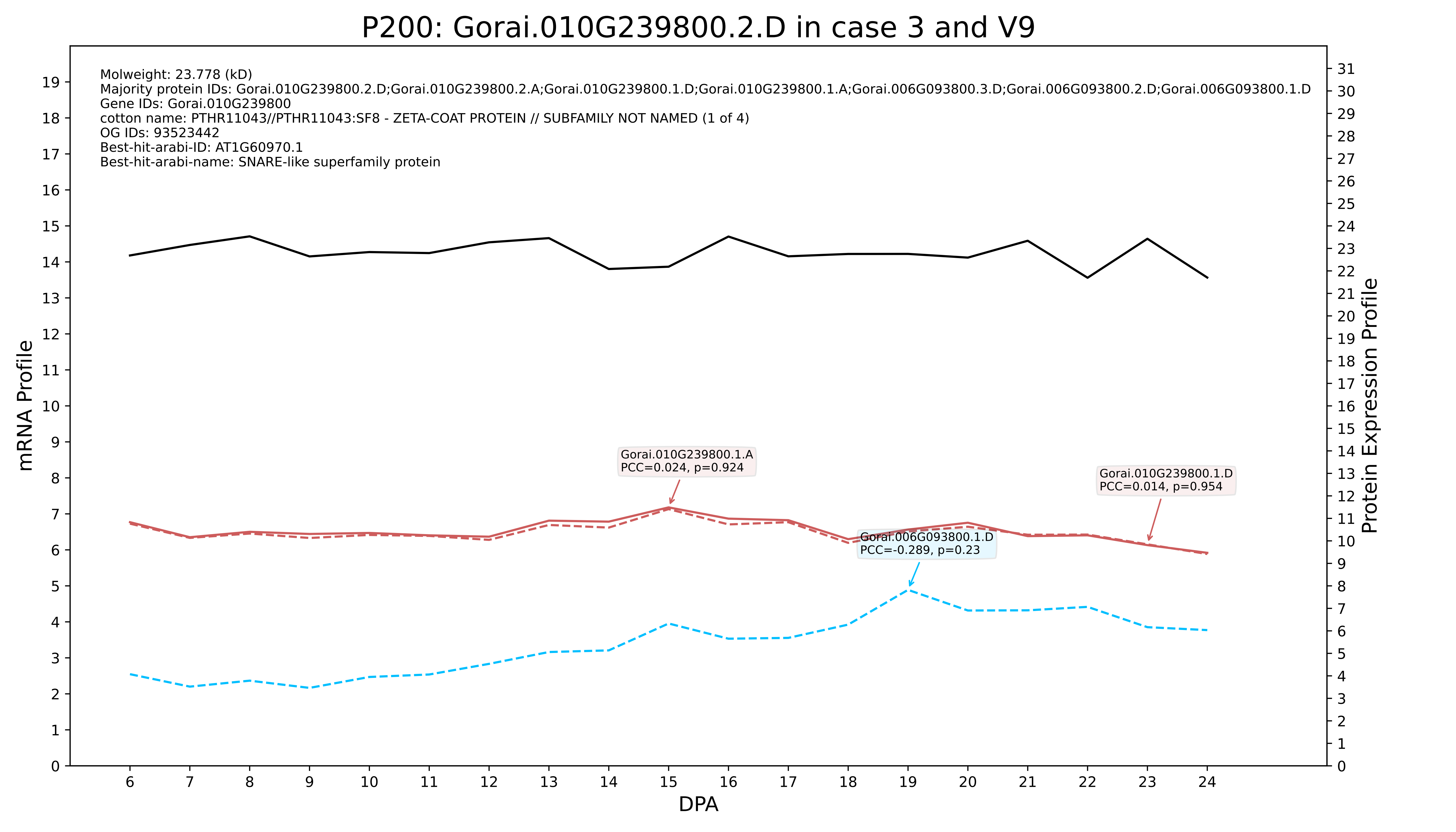

P200: Gorai.010G239800.2.D in case 3 and V9
Molweight: 23.778 (kD)
Majority protein IDs: Gorai.010G239800.2.D;Gorai.010G239800.2.A;Gorai.010G239800.1.D;Gorai.010G239800.1.A;Gorai.006G093800.3.D;Gorai.006G093800.2.D;Gorai.006G093800.1.D
Gene IDs: Gorai.010G239800
cotton name: PTHR11043//PTHR11043:SF8 - ZETA-COAT PROTEIN // SUBFAMILY NOT NAMED (1 of 4)
OG IDs: 93523442
Best-hit-arabi-ID: AT1G60970.1
Best-hit-arabi-name: SNARE-like superfamily protein
Gorai.010G239800.1.A
PCC=0.024, p=0.924
Gorai.010G239800.1.D
PCC=0.014, p=0.954
Gorai.006G093800.1.D
PCC=-0.289, p=0.23
mRNA Profile
Protein Expression Profile
DPA

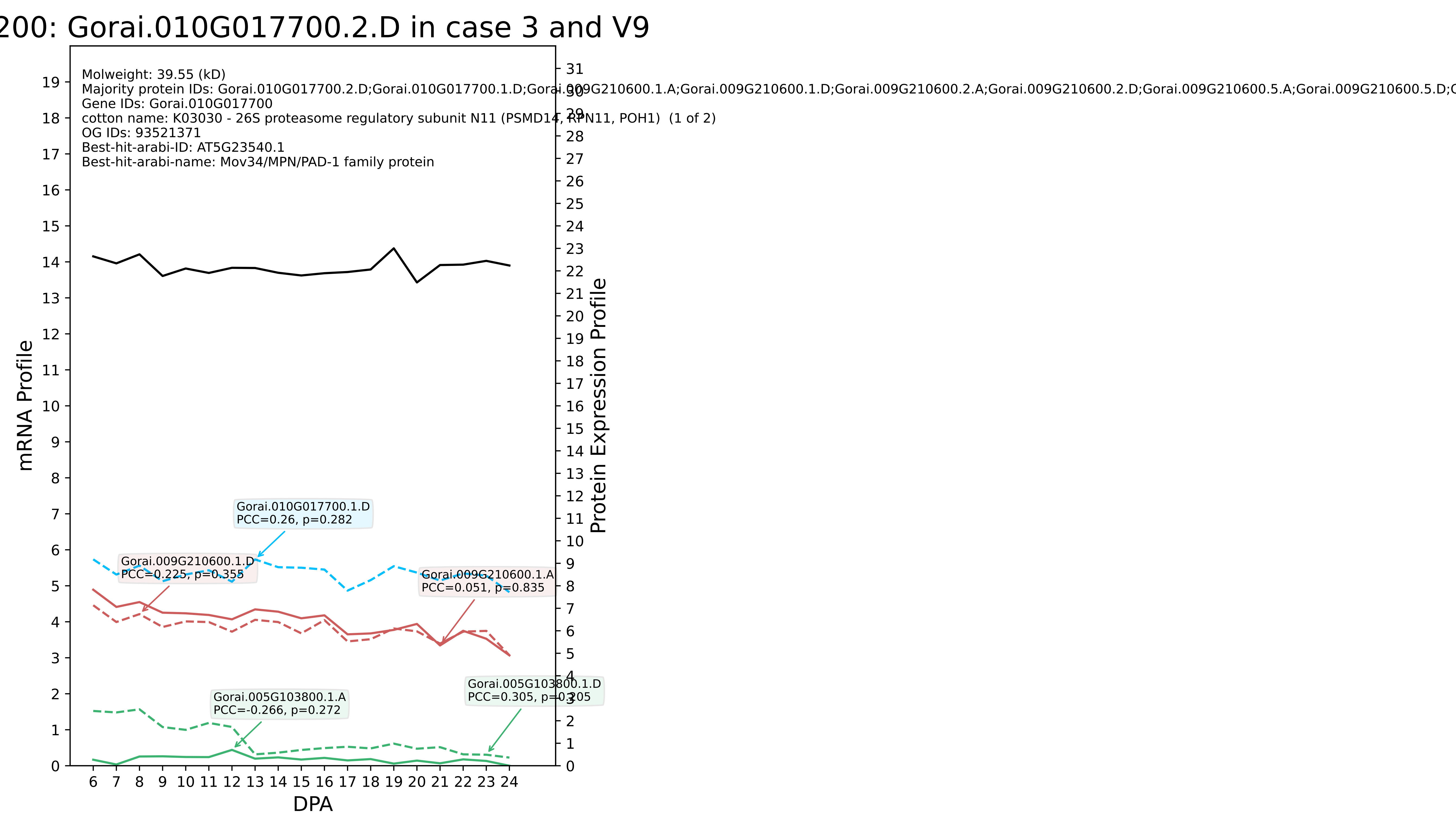
200: Gorai.010G017700.2.D in case 3 and V9
Molweight: 39.55 (kD)
Majority protein IDs: Gorai.010G017700.2.D;Gorai.010G017700.1.D;Gorai.009G210600.1.A;Gorai.009G210600.1.D;Gorai.009G210600.2.A;Gorai.009G210600.2.D;Gorai.009G210600.5.A;Gorai.009G210600.5.D;G
Gene IDs: Gorai.010G017700
cotton name: K03030 - 26S proteasome regulatory subunit N11 (PSMD14, RPN11, POH1) (1 of 2)
OG IDs: 93521371
Best-hit-arabi-ID: AT5G23540.1
Best-hit-arabi-name: Mov34/MPN/PAD-1 family protein
Gorai.010G017700.1.D
PCC=0.26, p=0.282
Gorai.009G210600.1.D
PCC=0.225, p=0.355
Gorai.009G210600.1.A
PCC=0.051, p=0.835
Gorai.005G103800.1.A
PCC=-0.266, p=0.272
Gorai.005G103800.1.D
PCC=0.305, p=0.205
mRNA Profile
Protein Expression Profile
DPA

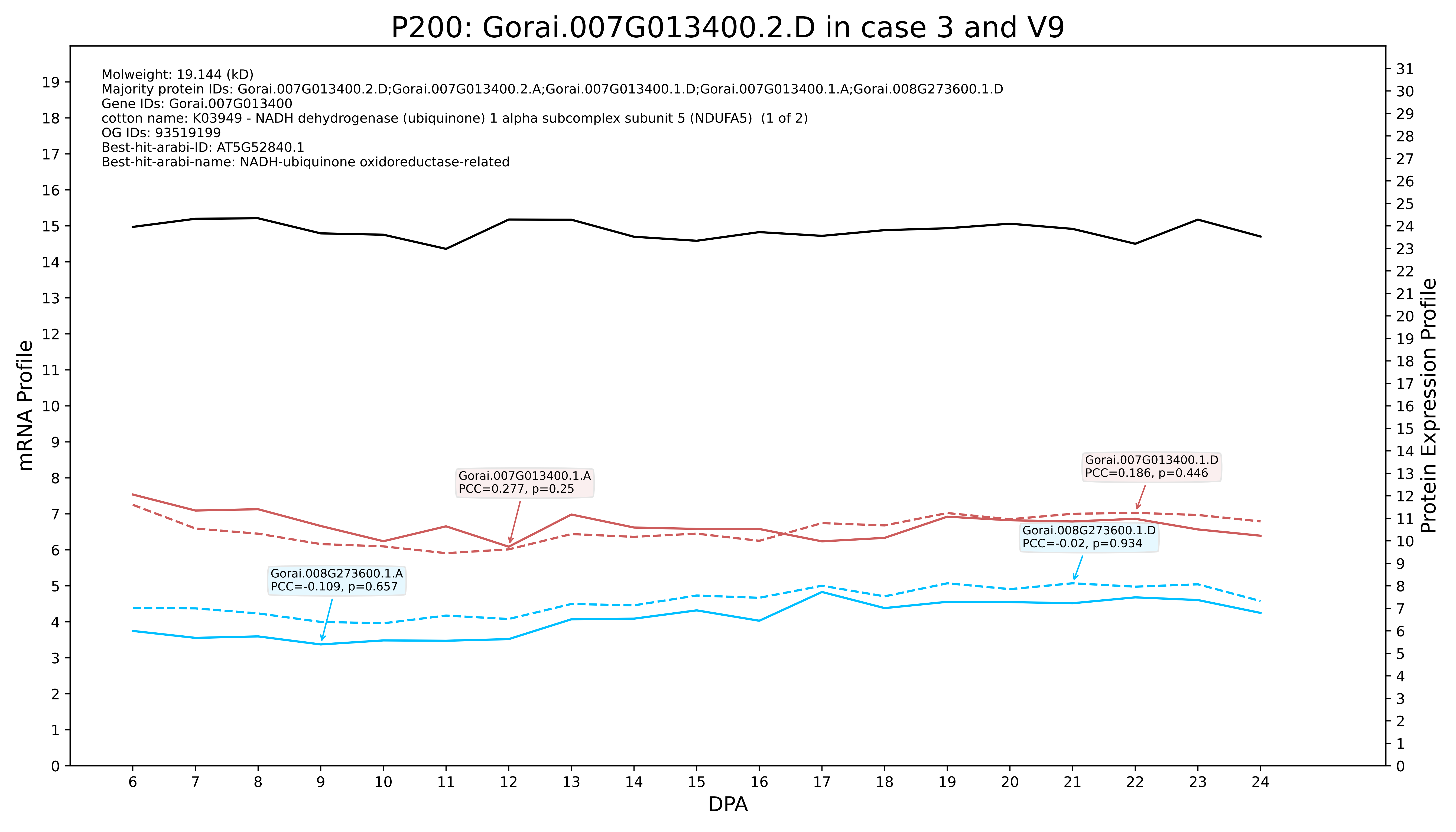

P200: Gorai.007G013400.2.D in case 3 and V9
Molweight: 19.144 (kD)
Majority protein IDs: Gorai.007G013400.2.D;Gorai.007G013400.2.A;Gorai.007G013400.1.D;Gorai.007G013400.1.A;Gorai.008G273600.1.D
Gene IDs: Gorai.007G013400
cotton name: K03949 - NADH dehydrogenase (ubiquinone) 1 alpha subcomplex subunit 5 (NDUFA5) (1 of 2)
OG IDs: 93519199
Best-hit-arabi-ID: AT5G52840.1
Best-hit-arabi-name: NADH-ubiquinone oxidoreductase-related
Gorai.007G013400.1.A
PCC=0.277, p=0.25
Gorai.007G013400.1.D
PCC=0.186, p=0.446
Gorai.008G273600.1.D
PCC=-0.02, p=0.934
Gorai.008G273600.1.A
PCC=-0.109, p=0.657
mRNA Profile
Protein Expression Profile
DPA

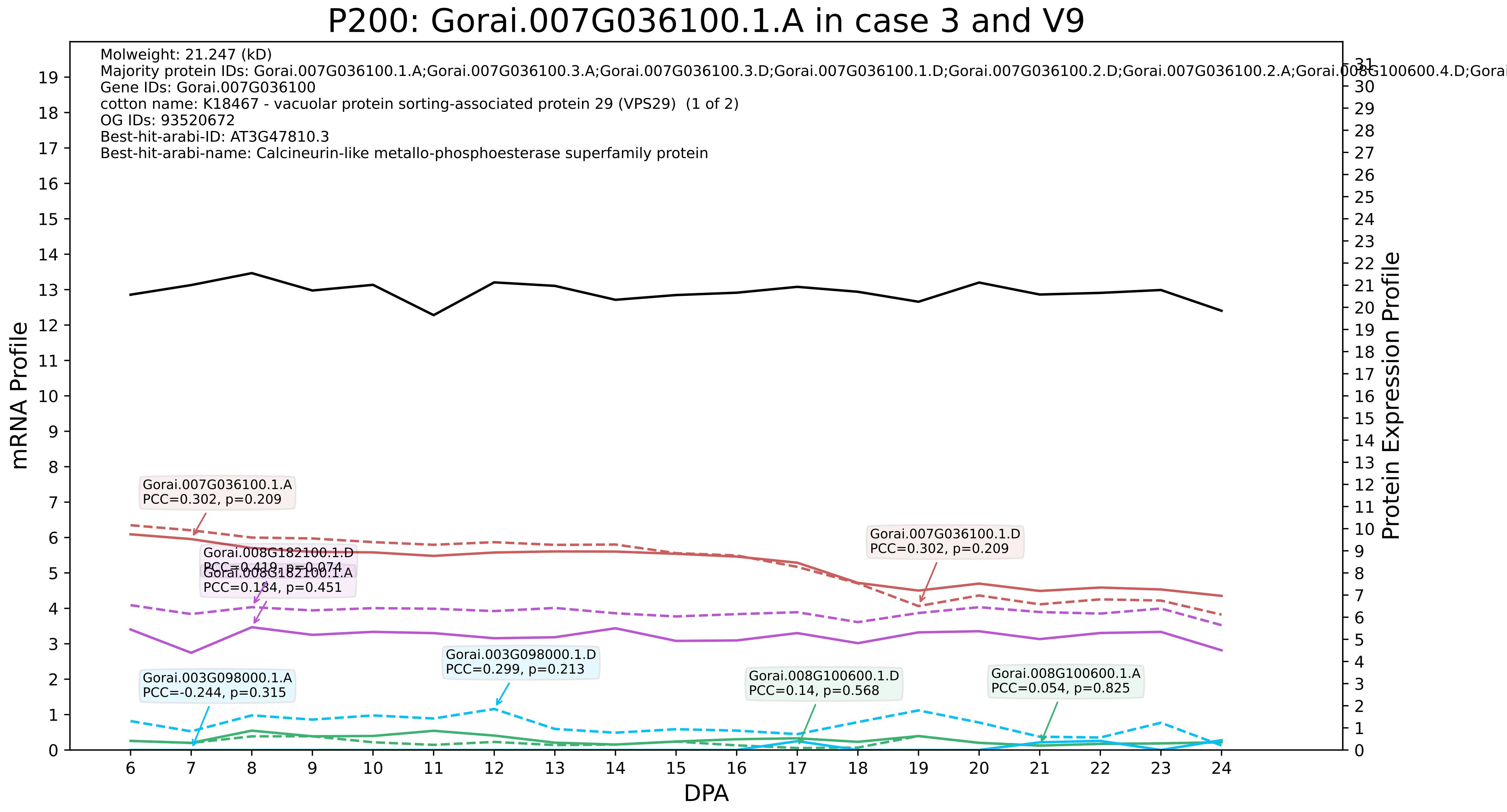
P200: Gorai.007G036100.1.A in case 3 and V9
Molweight: 21.247 (kD)
Majority protein IDs: Gorai.007G036100.1.A;Gorai.007G036100.3.A;Gorai.007G036100.3.D;Gorai.007G036100.1.D;Gorai.007G036100.2.D;Gorai.007G036100.2.A;Gorai.008G100600.4.D;Gorai
Gene IDs: Gorai.007G036100
cotton name: K18467 - vacuolar protein sorting-associated protein 29 (VPS29) (1 of 2)
OG IDs: 93520672
Best-hit-arabi-ID: AT3G47810.3
Best-hit-arabi-name: Calcineurin-like metallo-phosphoesterase superfamily protein
mRNA Profile
Protein Expression Profile
DPA
Gorai.007G036100.1.A
PCC=0.302, p=0.209
Gorai.007G036100.1.D
PCC=0.302, p=0.209
Gorai.008G182100.1.D
PCC=0.419, p=0.074
Gorai.008G182100.1.A
PCC=0.184, p=0.451
Gorai.003G098000.1.D
PCC=0.299, p=0.213
Gorai.003G098000.1.A
PCC=-0.244, p=0.315
Gorai.008G100600.1.D
PCC=0.14, p=0.568
Gorai.008G100600.1.A
PCC=0.054, p=0.825

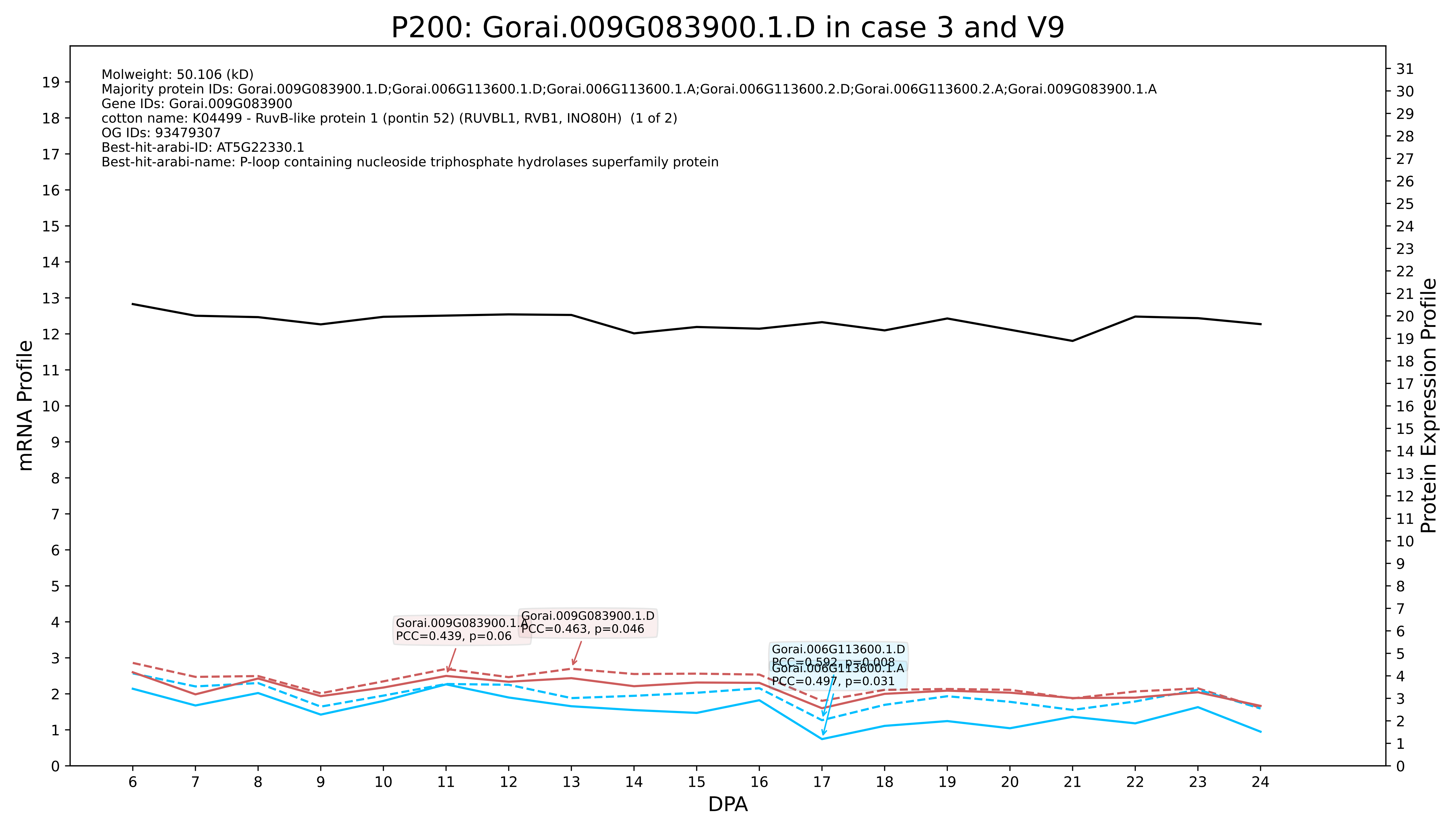

P200: Gorai.009G083900.1.D in case 3 and V9
Molweight: 50.106 (kD)
Majority protein IDs: Gorai.009G083900.1.D;Gorai.006G113600.1.D;Gorai.006G113600.1.A;Gorai.006G113600.2.D;Gorai.006G113600.2.A;Gorai.009G083900.1.A
Gene IDs: Gorai.009G083900
cotton name: K04499 - RuvB-like protein 1 (pontin 52) (RUVBL1, RVB1, INO80H) (1 of 2)
OG IDs: 93479307
Best-hit-arabi-ID: AT5G22330.1
Best-hit-arabi-name: P-loop containing nucleoside triphosphate hydrolases superfamily protein
Gorai.009G083900.1.A
PCC=0.439, p=0.06
Gorai.009G083900.1.D
PCC=0.463, p=0.046
Gorai.006G113600.1.D
PCC=0.592, p=0.008
Gorai.006G113600.1.A
PCC=0.497, p=0.031
mRNA Profile
Protein Expression Profile
DPA

### P200: Gorai.006G246500.1.A in case 3 and V9

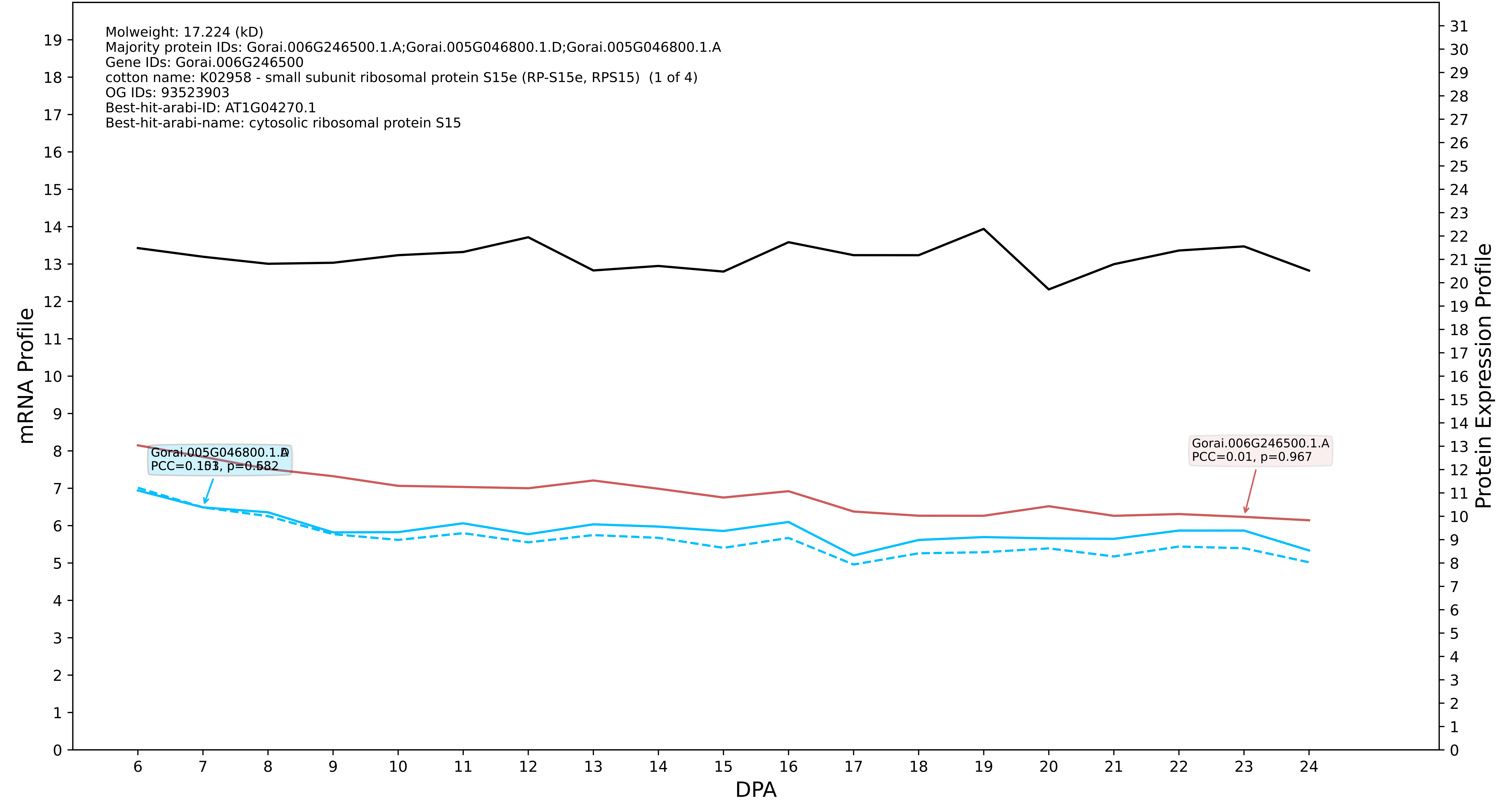

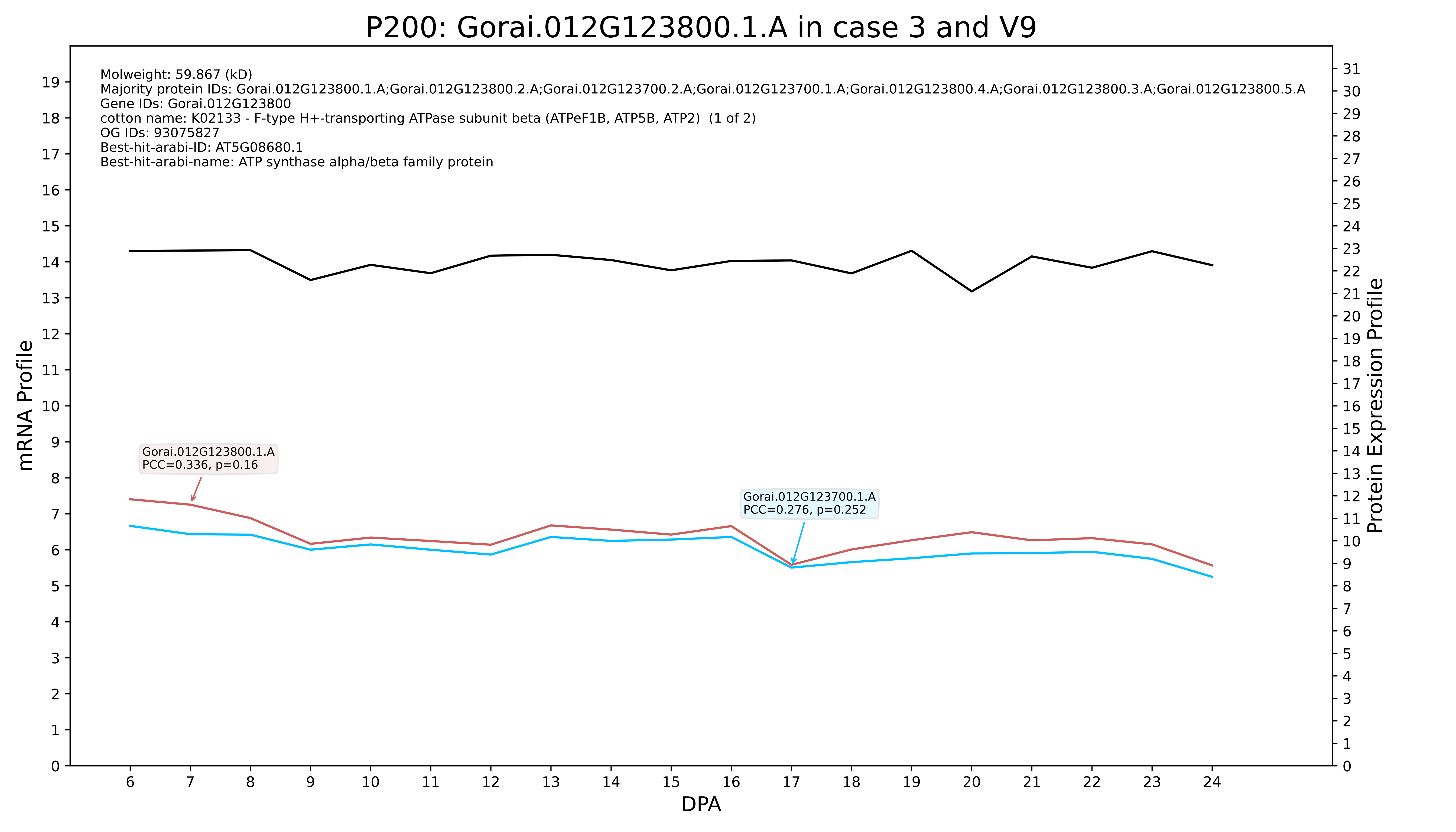
P200: Gorai.012G123800.1.A in case 3 and V9
Molweight: 59.867 (kD)
Majority protein IDs: Gorai.012G123800.1.A;Gorai.012G123800.2.A;Gorai.012G123700.2.A;Gorai.012G123700.1.A;Gorai.012G123800.4.A;Gorai.012G123800.3.A;Gorai.012G123800.5.A
Gene IDs: Gorai.012G123800
cotton name: K02133 - F-type H+-transporting ATPase subunit beta (ATPeF1B, ATP5B, ATP2) (1 of 2)
OG IDs: 93075827
Best-hit-arabi-ID: AT5G08680.1
Best-hit-arabi-name: ATP synthase alpha/beta family protein
Gorai.012G123800.1.A
PCC=0.336, p=0.16
Gorai.012G123700.1.A
PCC=0.276, p=0.252
mRNA Profile
Protein Expression Profile
DPA

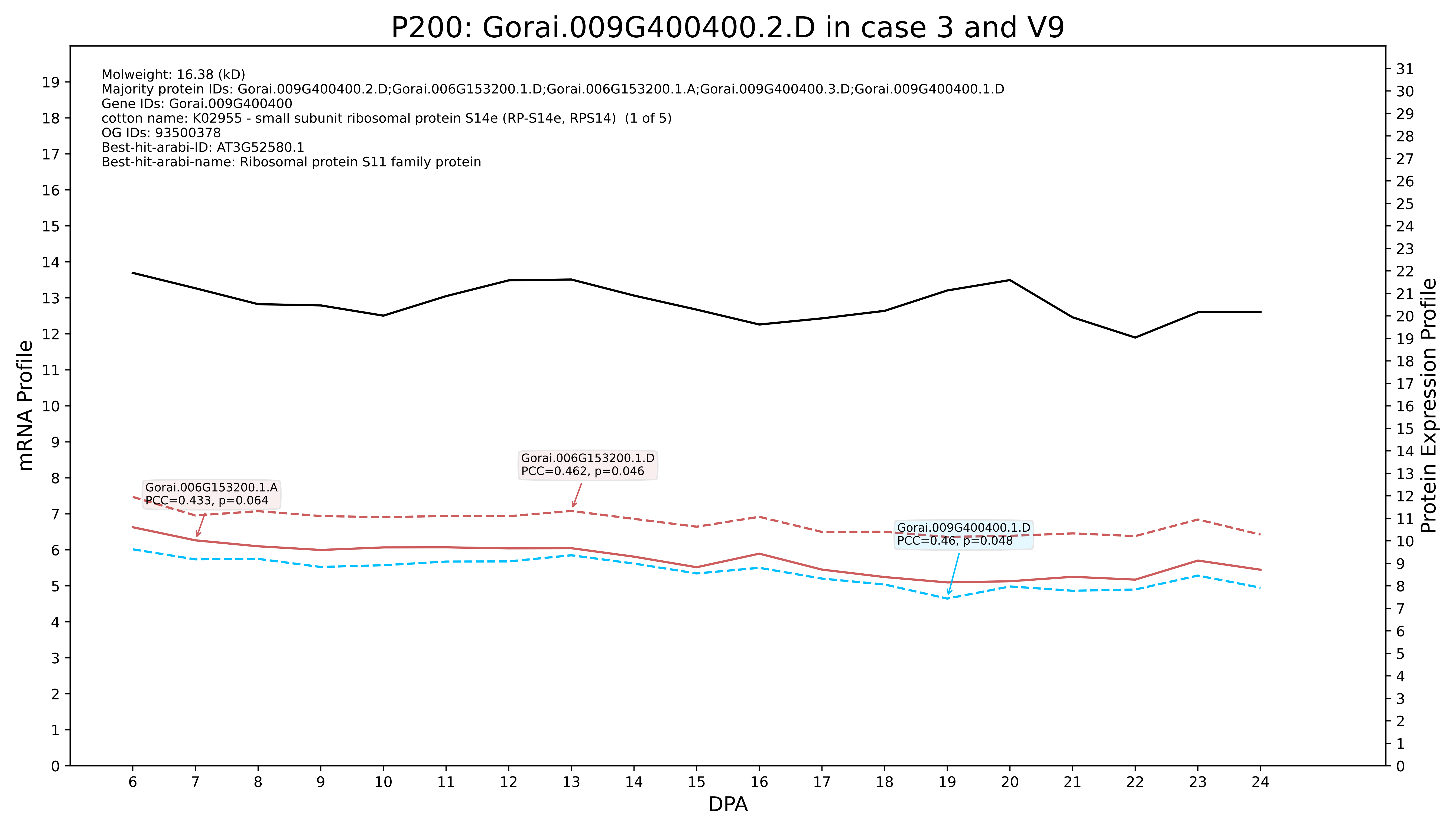
P200: Gorai.009G400400.2.D in case 3 and V9
Molweight: 16.38 (kD)
Majority protein IDs: Gorai.009G400400.2.D;Gorai.006G153200.1.D;Gorai.006G153200.1.A;Gorai.009G400400.3.D;Gorai.009G400400.1.D
Gene IDs: Gorai.009G400400
cotton name: K02955 - small subunit ribosomal protein S14e (RP-S14e, RPS14) (1 of 5)
OG IDs: 93500378
Best-hit-arabi-ID: AT3G52580.1
Best-hit-arabi-name: Ribosomal protein S11 family protein
Gorai.006G153200.1.D
PCC=0.462, p=0.046
Gorai.006G153200.1.A
PCC=0.433, p=0.064
Gorai.009G400400.1.D
PCC=0.46, p=0.048
mRNA Profile
Protein Expression Profile
DPA

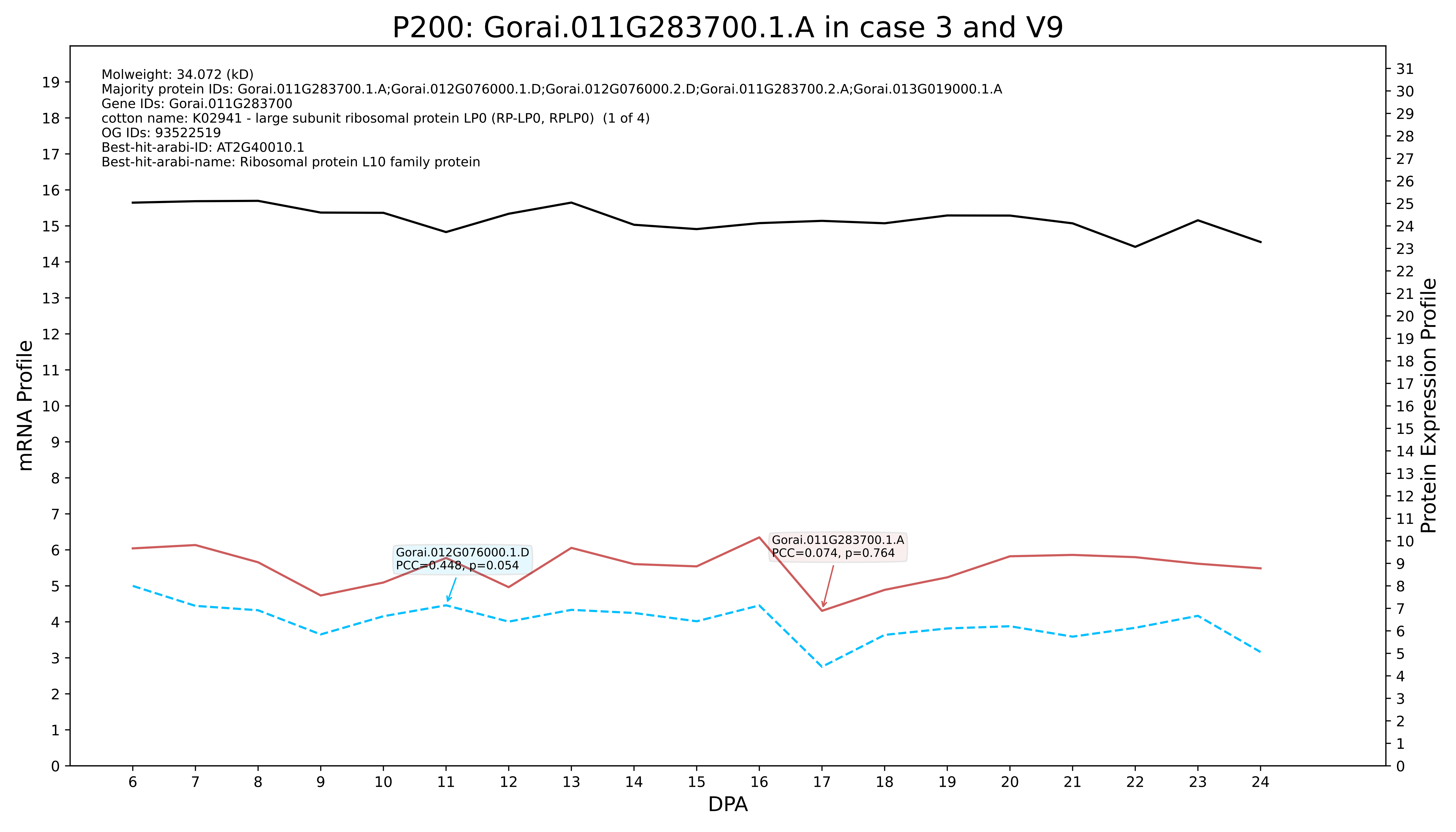
P200: Gorai.011G283700.1.A in case 3 and V9
Molweight: 34.072 (kD)
Majority protein IDs: Gorai.011G283700.1.A;Gorai.012G076000.1.D;Gorai.012G076000.2.D;Gorai.011G283700.2.A;Gorai.013G019000.1.A
Gene IDs: Gorai.011G283700
cotton name: K02941 - large subunit ribosomal protein LP0 (RP-LP0, RPLP0) (1 of 4)
OG IDs: 93522519
Best-hit-arabi-ID: AT2G40010.1
Best-hit-arabi-name: Ribosomal protein L10 family protein
Gorai.012G076000.1.D
PCC=0.448, p=0.054
Gorai.011G283700.1.A
PCC=0.074, p=0.764
mRNA Profile
Protein Expression Profile
DPA

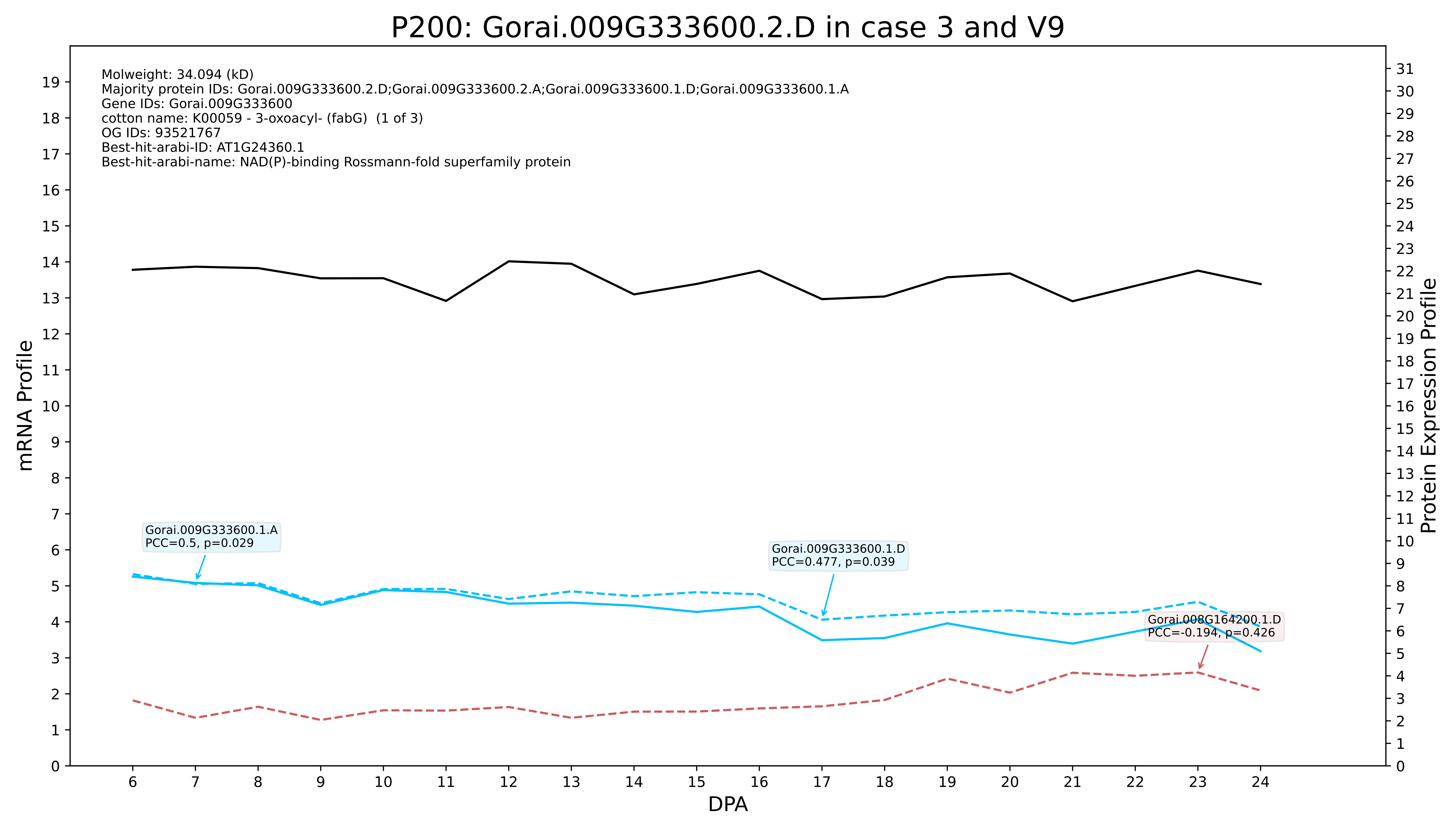

P200: Gorai.009G333600.2.D in case 3 and V9
Molweight: 34.094 (kD)
Majority protein IDs: Gorai.009G333600.2.D;Gorai.009G333600.2.A;Gorai.009G333600.1.D;Gorai.009G333600.1.A
Gene IDs: Gorai.009G333600
cotton name: K00059 - 3-oxoacyl- (fabG) (1 of 3)
OG IDs: 93521767
Best-hit-arabi-ID: AT1G24360.1
Best-hit-arabi-name: NAD(P)-binding Rossmann-fold superfamily protein
Gorai.009G333600.1.A
PCC=0.5, p=0.029
Gorai.009G333600.1.D
PCC=0.477, p=0.039
Gorai.008G164200.1.D
PCC=-0.194, p=0.426
mRNA Profile
Protein Expression Profile
DPA

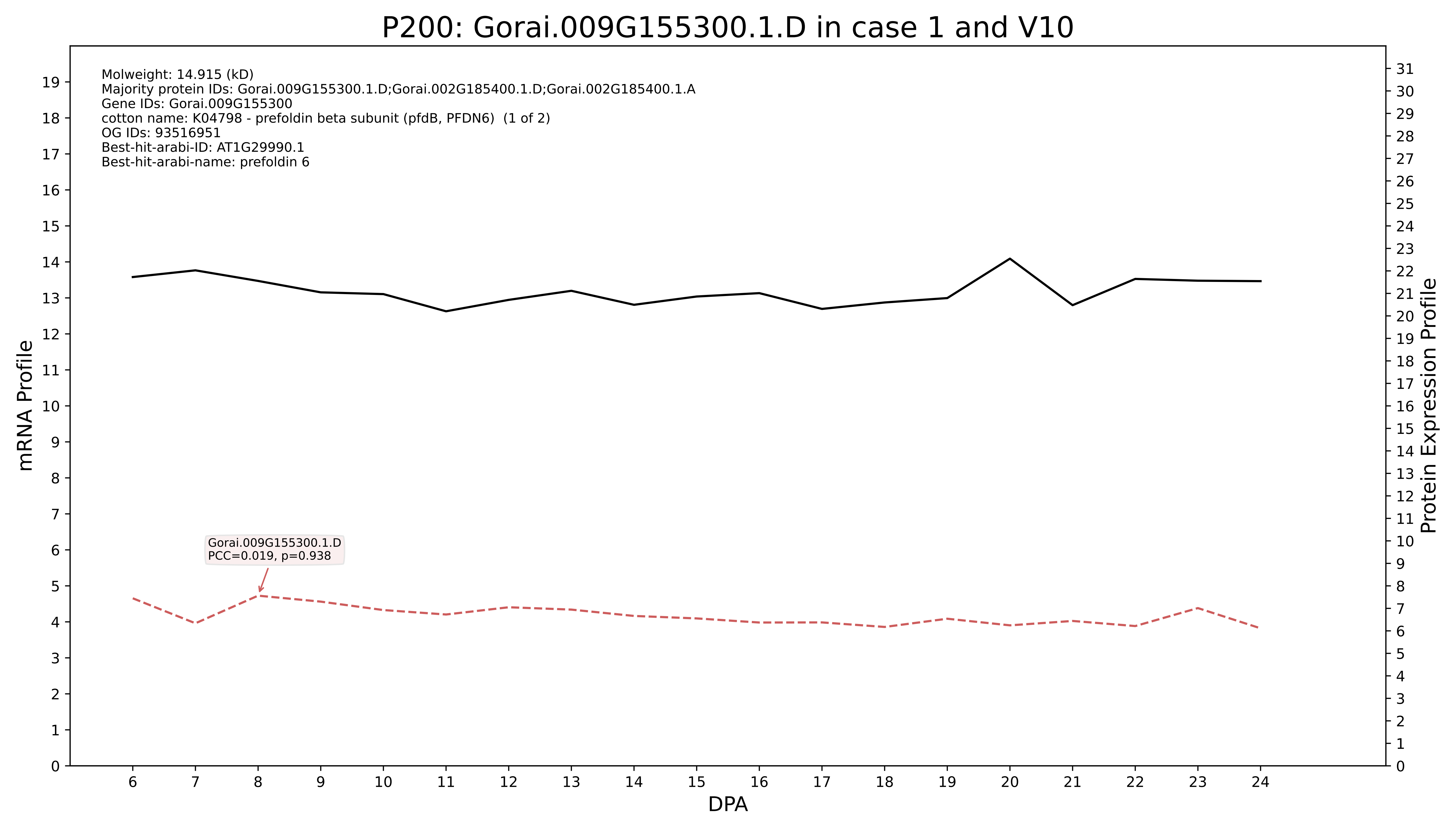
P200: Gorai.009G155300.1.D in case 1 and V10
Molweight: 14.915 (kD)
Majority protein IDs: Gorai.009G155300.1.D;Gorai.002G185400.1.D;Gorai.002G185400.1.A
Gene IDs: Gorai.009G155300
cotton name: K04798 - prefoldin beta subunit (pfdB, PFDN6) (1 of 2)
OG IDs: 93516951
Best-hit-arabi-ID: AT1G29990.1
Best-hit-arabi-name: prefoldin 6
Gorai.009G155300.1.D
PCC=0.019, p=0.938
mRNA Profile
Protein Expression Profile
DPA

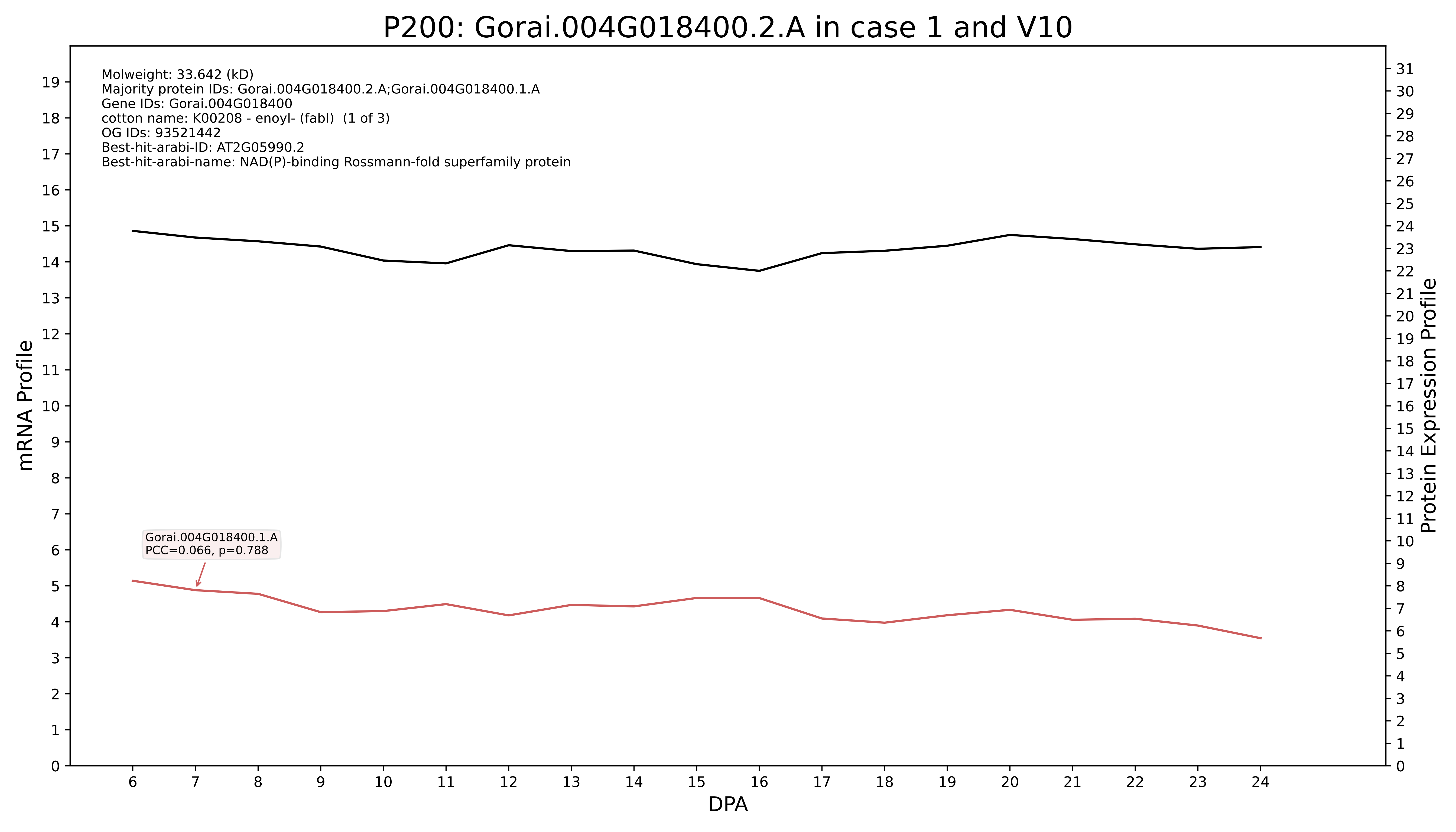
P200: Gorai.004G018400.2.A in case 1 and V10
Molweight: 33.642 (kD)
Majority protein IDs: Gorai.004G018400.2.A;Gorai.004G018400.1.A
Gene IDs: Gorai.004G018400
cotton name: K00208 - enoyl- (fabI) (1 of 3)
OG IDs: 93521442
Best-hit-arabi-ID: AT2G05990.2
Best-hit-arabi-name: NAD(P)-binding Rossmann-fold superfamily protein
Gorai.004G018400.1.A
PCC=0.066, p=0.788
mRNA Profile
Protein Expression Profile
DPA

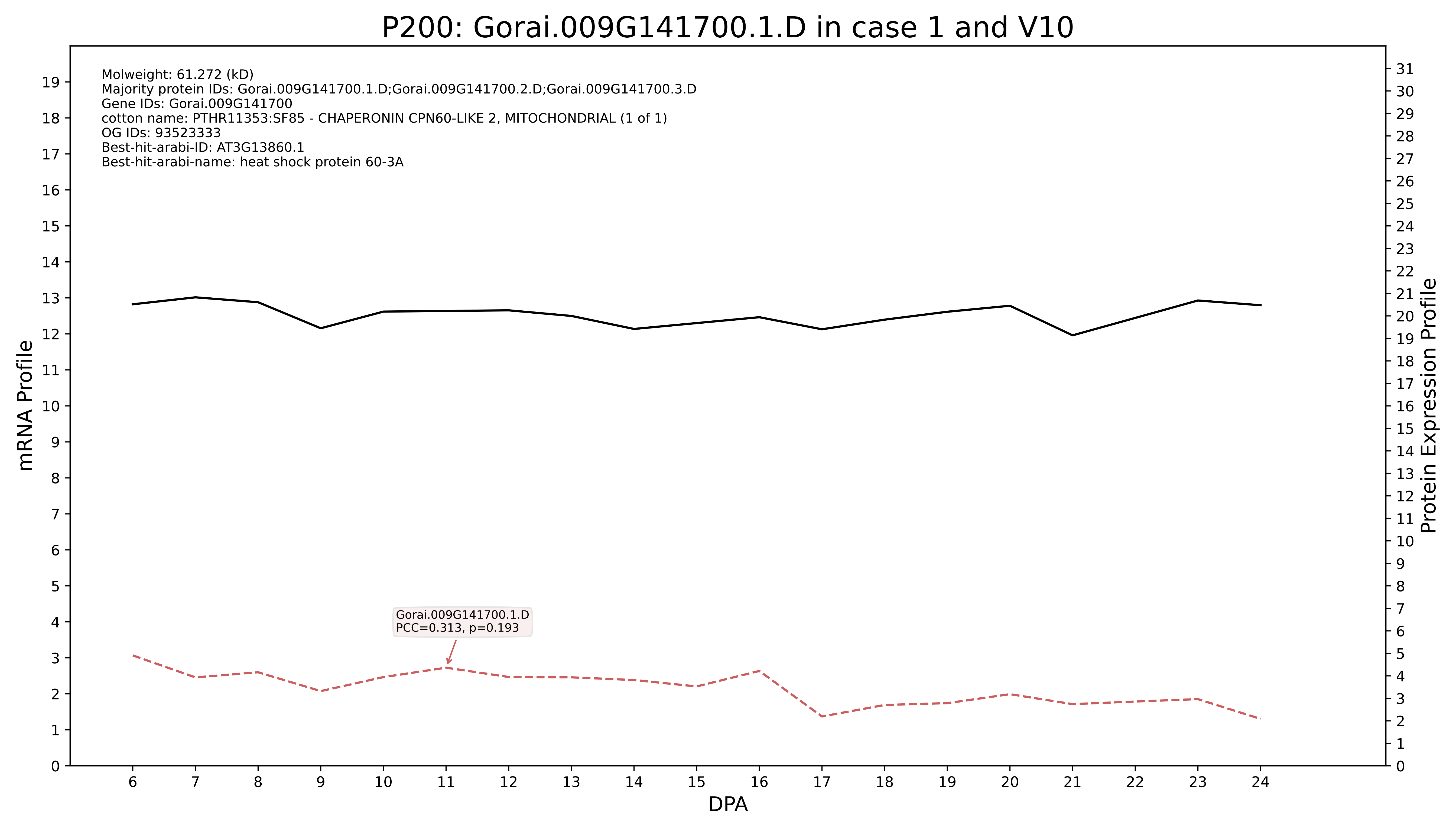
P200: Gorai.009G141700.1.D in case 1 and V10
Molweight: 61.272 (kD)
Majority protein IDs: Gorai.009G141700.1.D;Gorai.009G141700.2.D;Gorai.009G141700.3.D
Gene IDs: Gorai.009G141700
cotton name: PTHR11353:SF85 - CHAPERONIN CPN60-LIKE 2, MITOCHONDRIAL (1 of 1)
OG IDs: 93523333
Best-hit-arabi-ID: AT3G13860.1
Best-hit-arabi-name: heat shock protein 60-3A
Gorai.009G141700.1.D
PCC=0.313, p=0.193
mRNA Profile
Protein Expression Profile
DPA

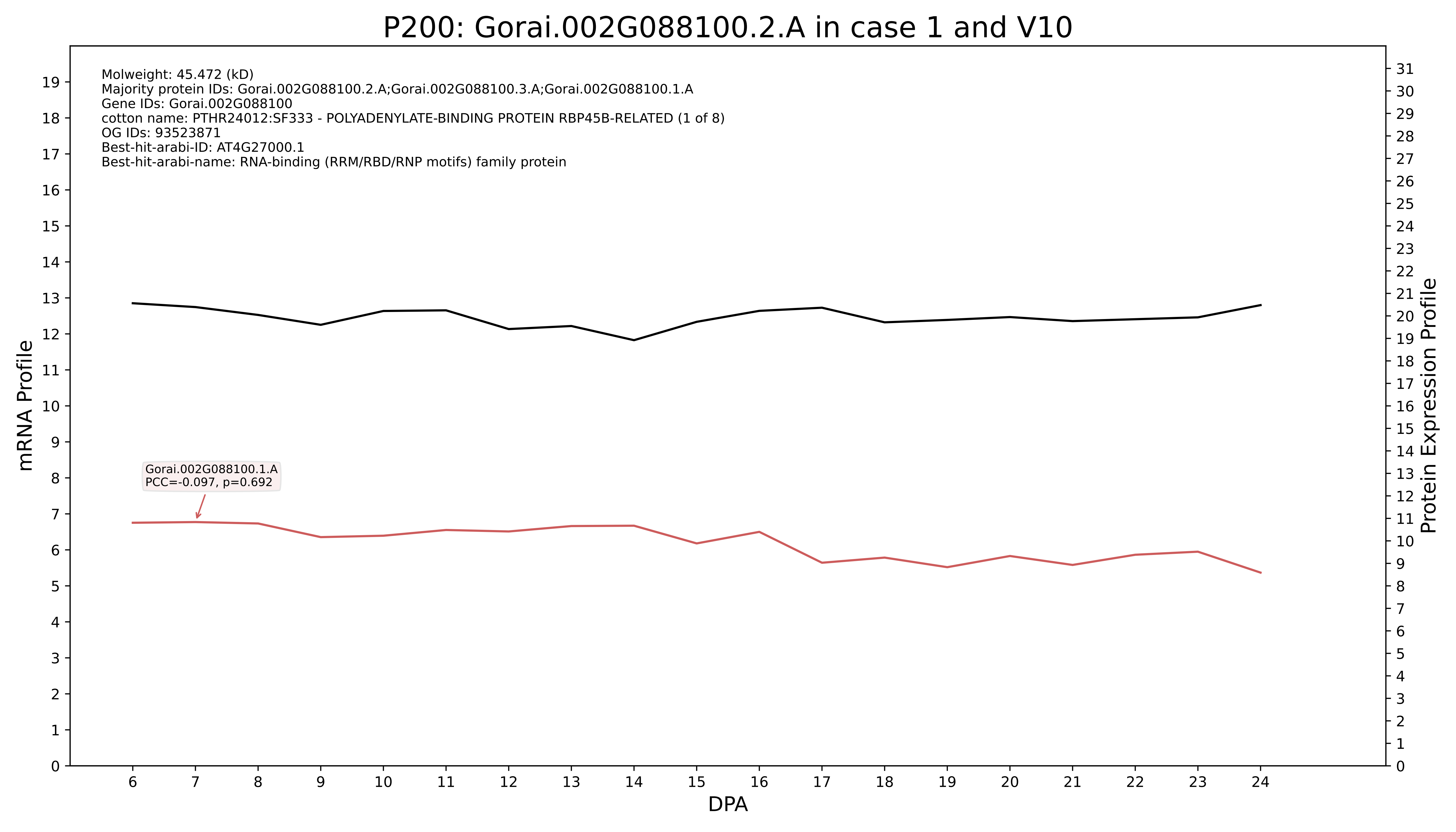
P200: Gorai.002G088100.2.A in case 1 and V10
Molweight: 45.472 (kD)
Majority protein IDs: Gorai.002G088100.2.A;Gorai.002G088100.3.A;Gorai.002G088100.1.A
Gene IDs: Gorai.002G088100
cotton name: PTHR24012:SF333 - POLYADENYLATE-BINDING PROTEIN RBP45B-RELATED (1 of 8)
OG IDs: 93523871
Best-hit-arabi-ID: AT4G27000.1
Best-hit-arabi-name: RNA-binding (RRM/RBD/RNP motifs) family protein
Gorai.002G088100.1.A
PCC=-0.097, p=0.692
mRNA Profile
Protein Expression Profile
DPA

#### P200: Gorai.004G096800.2.D in case 1 and V10

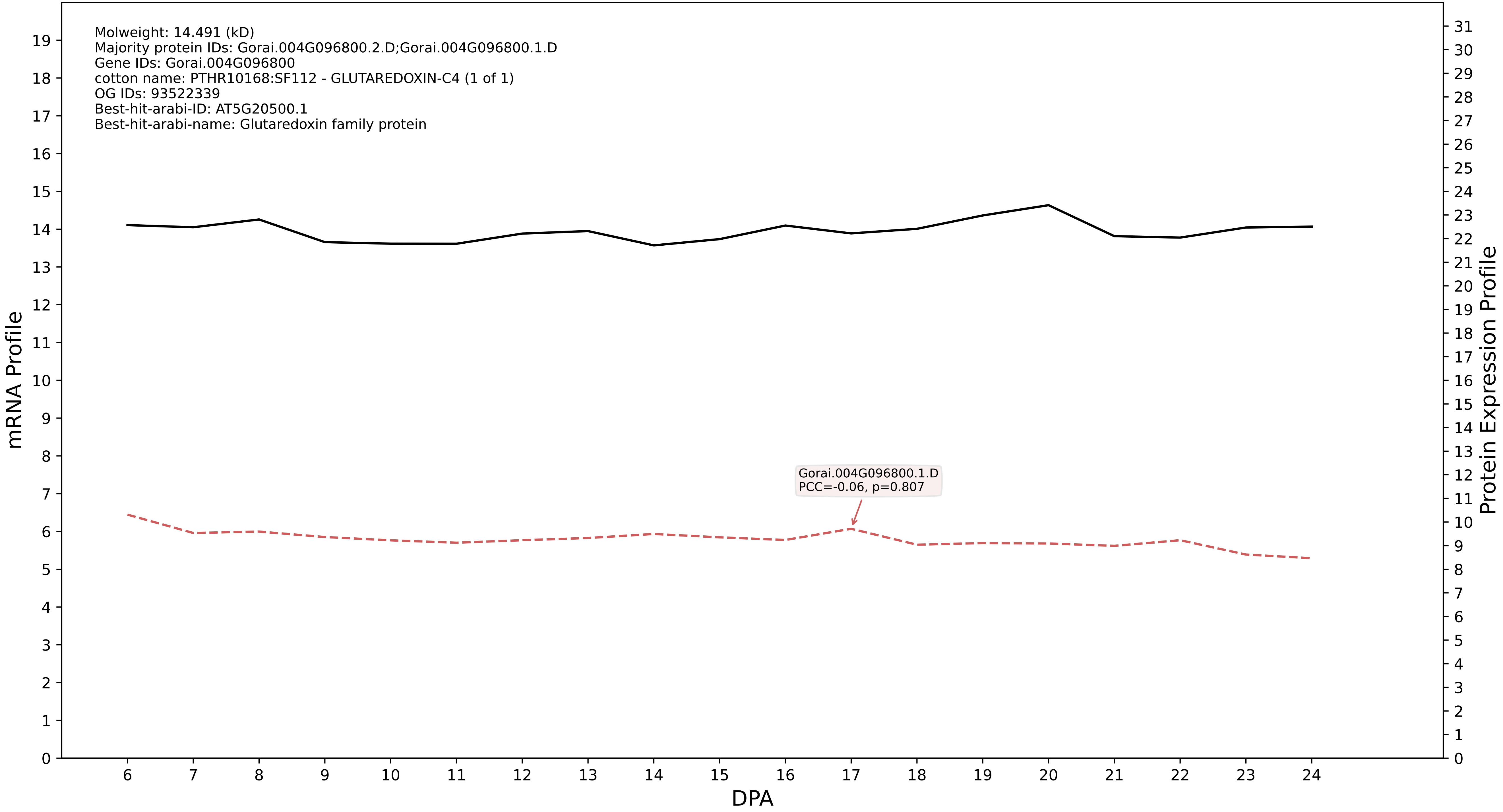

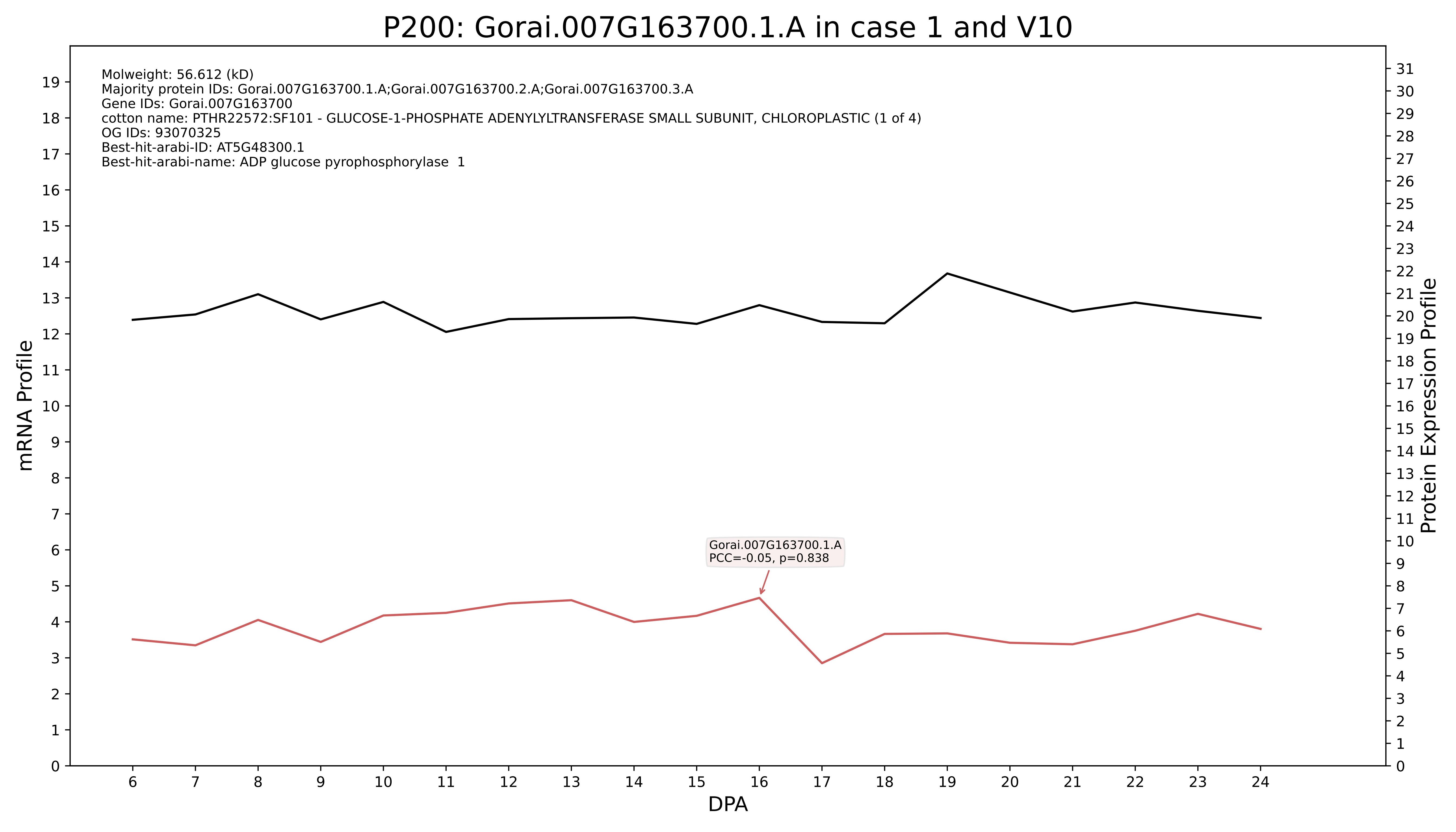
P200: Gorai.007G163700.1.A in case 1 and V10
Molweight: 56.612 (kD)
Majority protein IDs: Gorai.007G163700.1.A;Gorai.007G163700.2.A;Gorai.007G163700.3.A
Gene IDs: Gorai.007G163700
cotton name: PTHR22572:SF101 - GLUCOSE-1-PHOSPHATE ADENYLYLTRANSFERASE SMALL SUBUNIT, CHLOROPLASTIC (1 of 4)
OG IDs: 93070325
Best-hit-arabi-ID: AT5G48300.1
Best-hit-arabi-name: ADP glucose pyrophosphorylase 1
Gorai.007G163700.1.A
PCC=-0.05, p=0.838
mRNA Profile
Protein Expression Profile
DPA

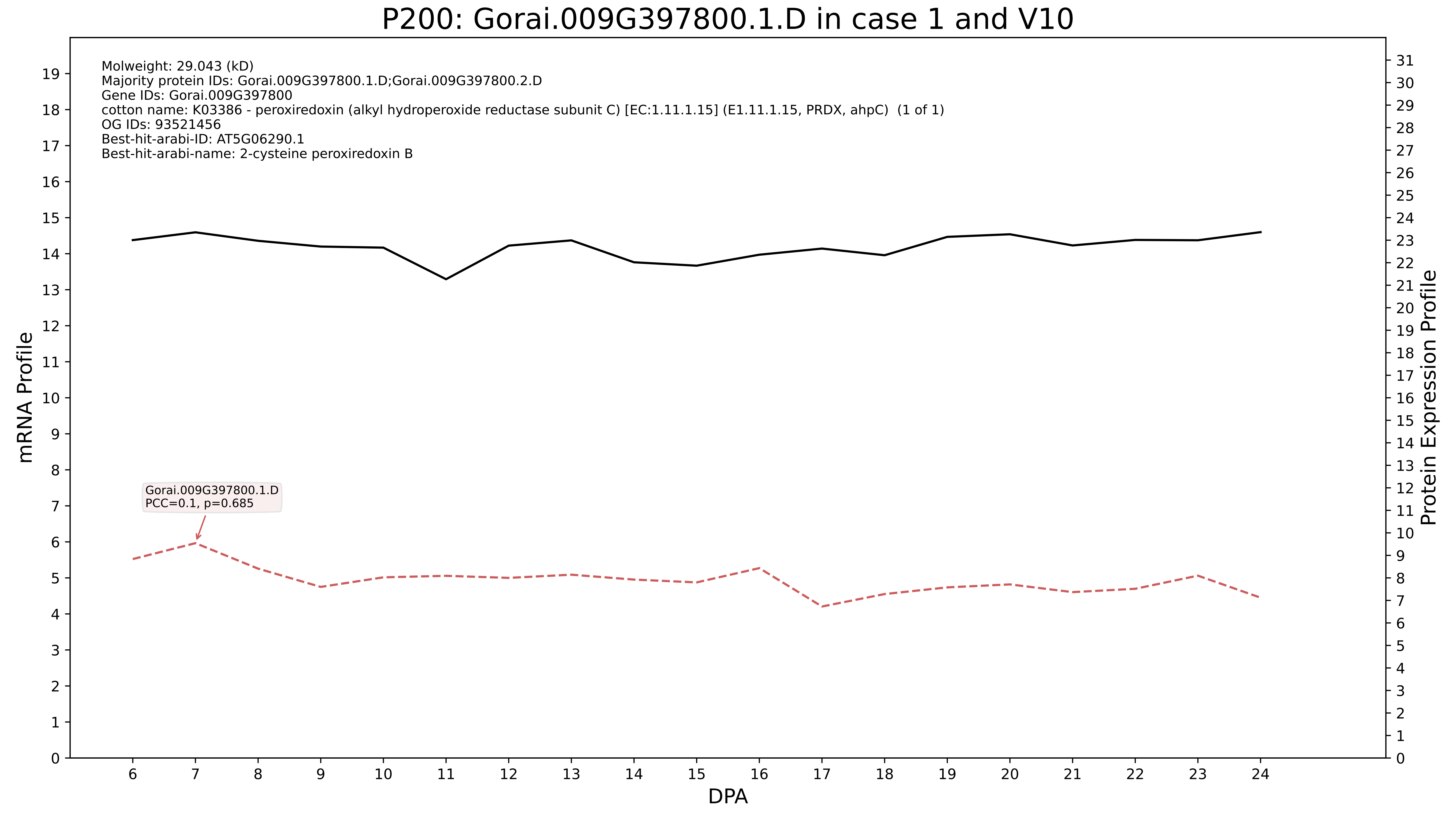
P200: Gorai.009G397800.1.D in case 1 and V10
Molweight: 29.043 (kD)
Majority protein IDs: Gorai.009G397800.1.D;Gorai.009G397800.2.D
Gene IDs: Gorai.009G397800
cotton name: K03386 - peroxiredoxin (alkyl hydroperoxide reductase subunit C) [EC:1.11.1.15] (E1.11.1.15, PRDX, ahpC) (1 of 1)
OG IDs: 93521456
Best-hit-arabi-ID: AT5G06290.1
Best-hit-arabi-name: 2-cysteine peroxiredoxin B
Gorai.009G397800.1.D
PCC=0.1, p=0.685
mRNA Profile
Protein Expression Profile
DPA

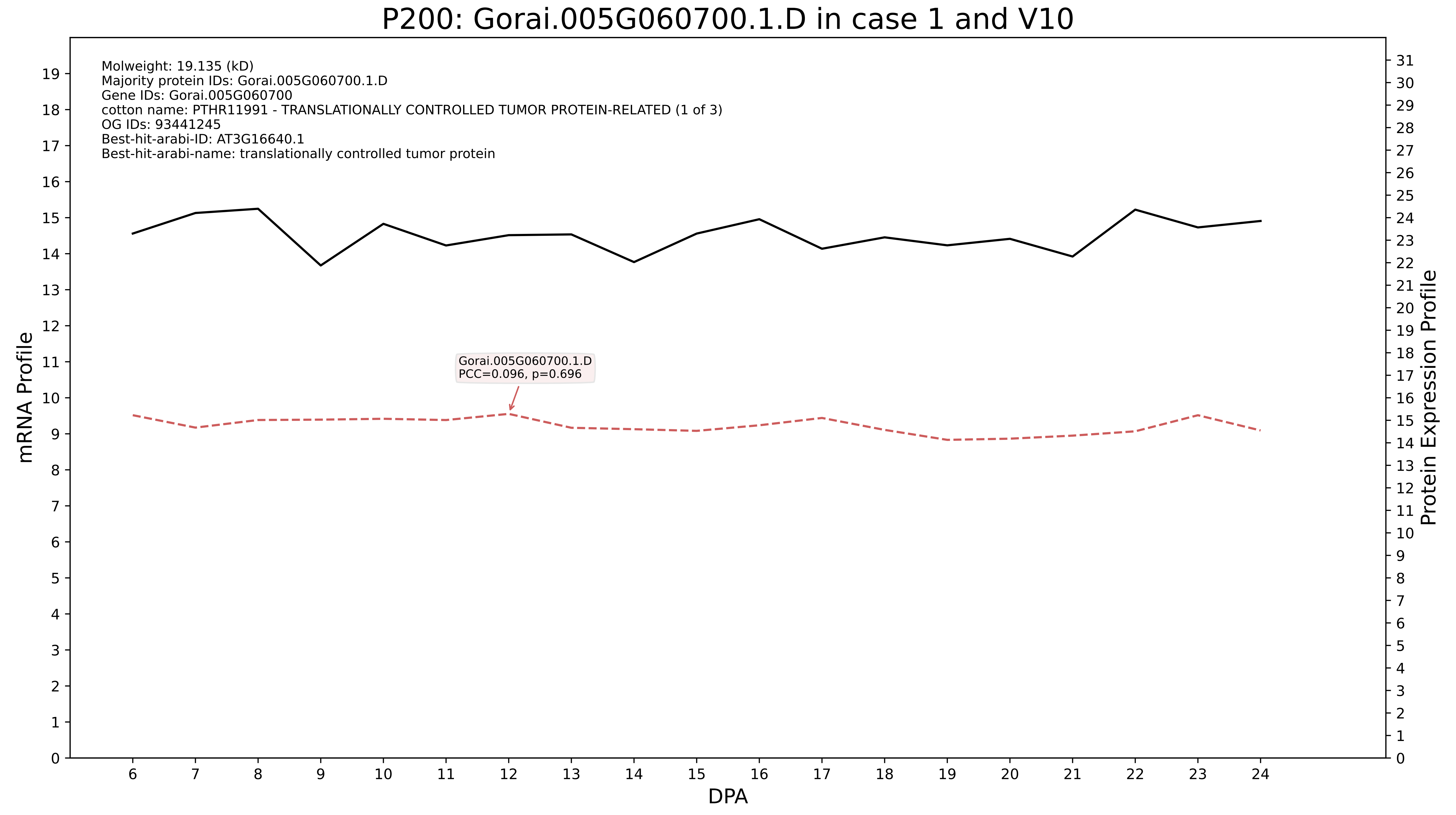
P200: Gorai.005G060700.1.D in case 1 and V10
Molweight: 19.135 (kD)
Majority protein IDs: Gorai.005G060700.1.D
Gene IDs: Gorai.005G060700
cotton name: PTHR11991 - TRANSLATIONALLY CONTROLLED TUMOR PROTEIN-RELATED (1 of 3)
OG IDs: 93441245
Best-hit-arabi-ID: AT3G16640.1
Best-hit-arabi-name: translationally controlled tumor protein
Gorai.005G060700.1.D
PCC=0.096, p=0.696
mRNA Profile
Protein Expression Profile
DPA

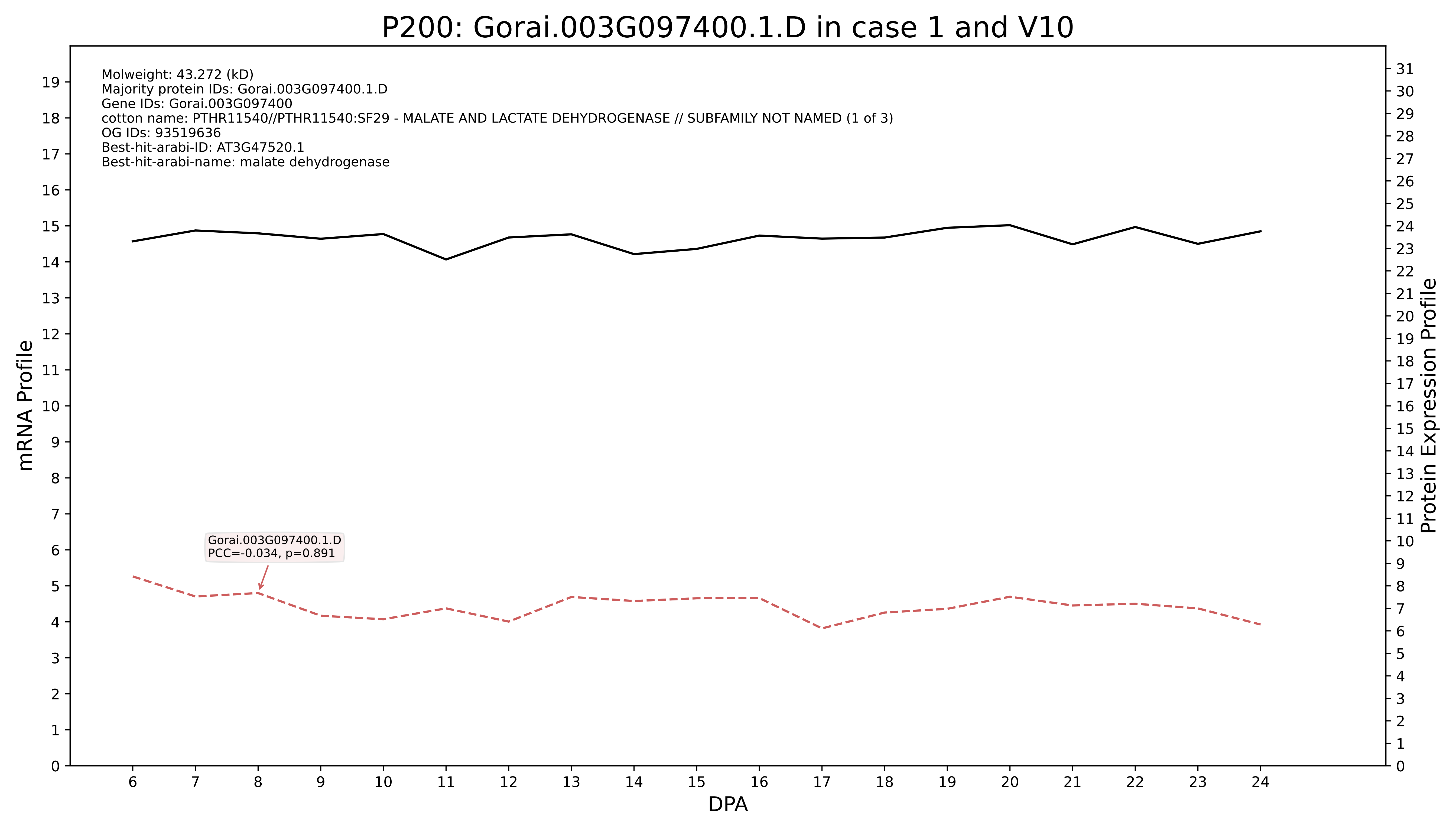
P200: Gorai.003G097400.1.D in case 1 and V10
Molweight: 43.272 (kD)
Majority protein IDs: Gorai.003G097400.1.D
Gene IDs: Gorai.003G097400
cotton name: PTHR11540//PTHR11540:SF29 - MALATE AND LACTATE DEHYDROGENASE // SUBFAMILY NOT NAMED (1 of 3)
OG IDs: 93519636
Best-hit-arabi-ID: AT3G47520.1
Best-hit-arabi-name: malate dehydrogenase
Gorai.003G097400.1.D
PCC=-0.034, p=0.891
mRNA Profile
Protein Expression Profile
DPA

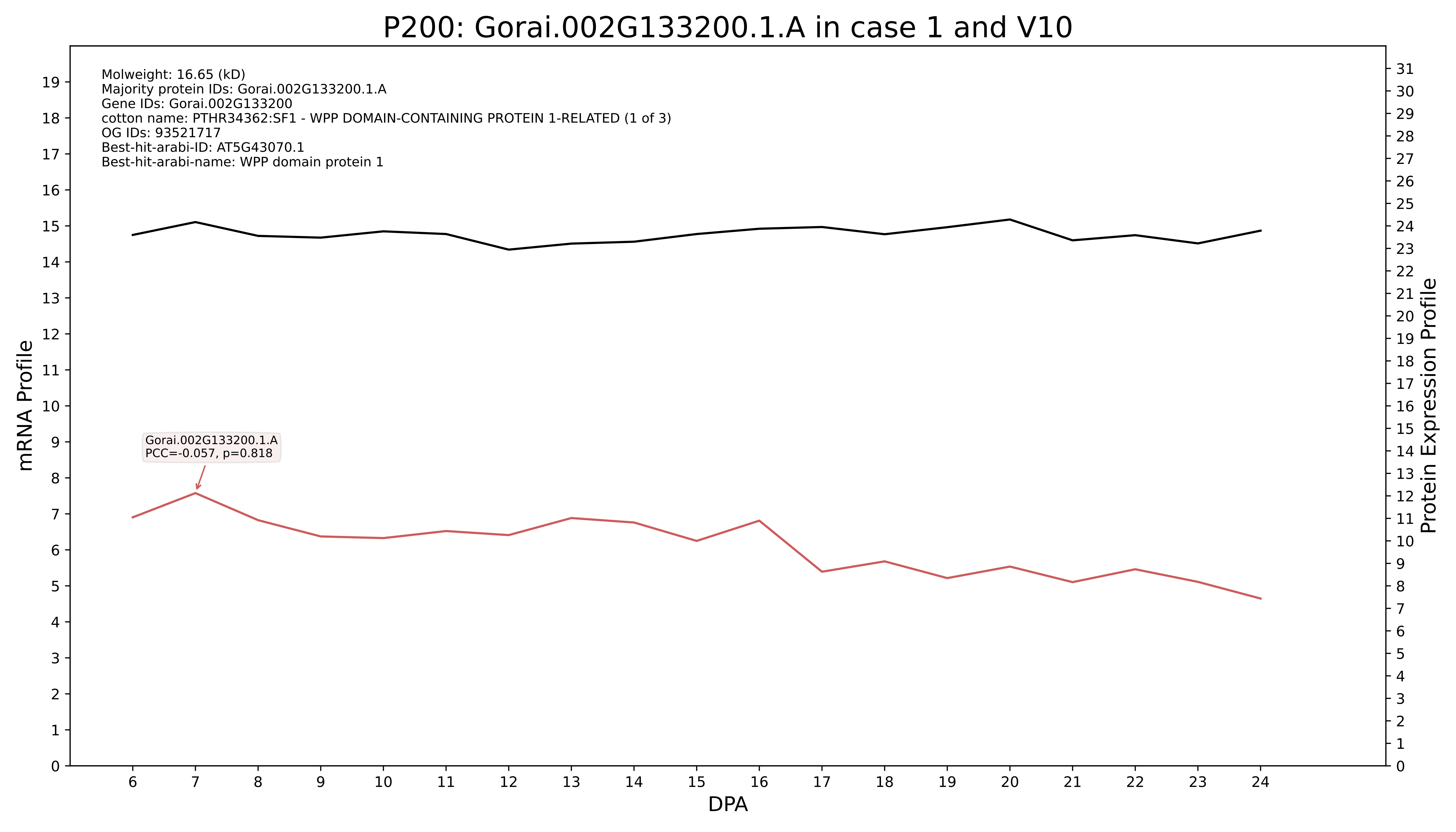

P200: Gorai.002G133200.1.A in case 1 and V10
Molweight: 16.65 (kD)
Majority protein IDs: Gorai.002G133200.1.A
Gene IDs: Gorai.002G133200
cotton name: PTHR34362:SF1 - WPP DOMAIN-CONTAINING PROTEIN 1-RELATED (1 of 3)
OG IDs: 93521717
Best-hit-arabi-ID: AT5G43070.1
Best-hit-arabi-name: WPP domain protein 1
Gorai.002G133200.1.A
PCC=-0.057, p=0.818
mRNA Profile
Protein Expression Profile
DPA

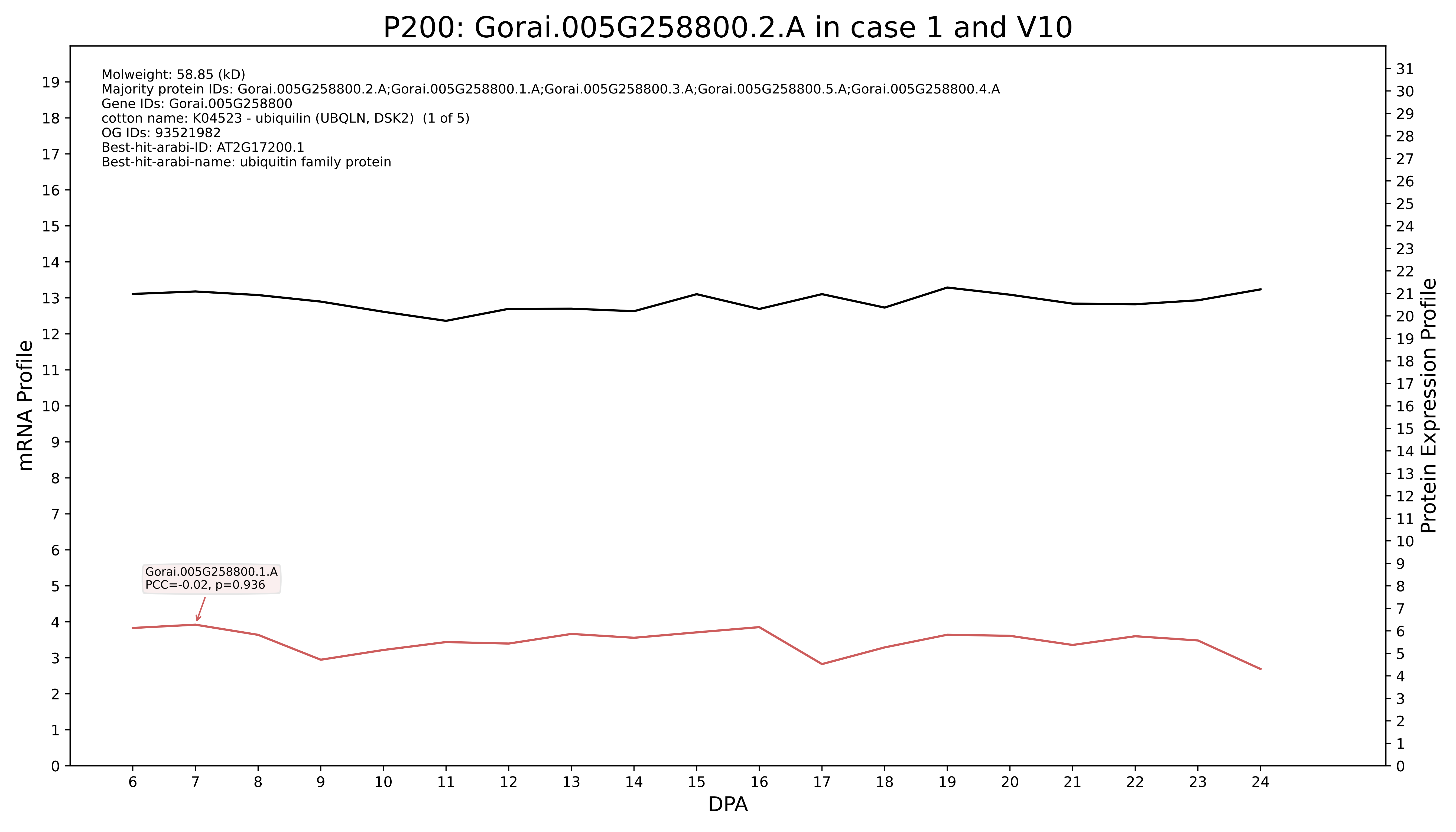
P200: Gorai.005G258800.2.A in case 1 and V10
Molweight: 58.85 (kD)
Majority protein IDs: Gorai.005G258800.2.A;Gorai.005G258800.1.A;Gorai.005G258800.3.A;Gorai.005G258800.5.A;Gorai.005G258800.4.A
Gene IDs: Gorai.005G258800
cotton name: K04523 - ubiquilin (UBQLN, DSK2) (1 of 5)
OG IDs: 93521982
Best-hit-arabi-ID: AT2G17200.1
Best-hit-arabi-name: ubiquitin family protein
Gorai.005G258800.1.A
PCC=-0.02, p=0.936
mRNA Profile
Protein Expression Profile
DPA

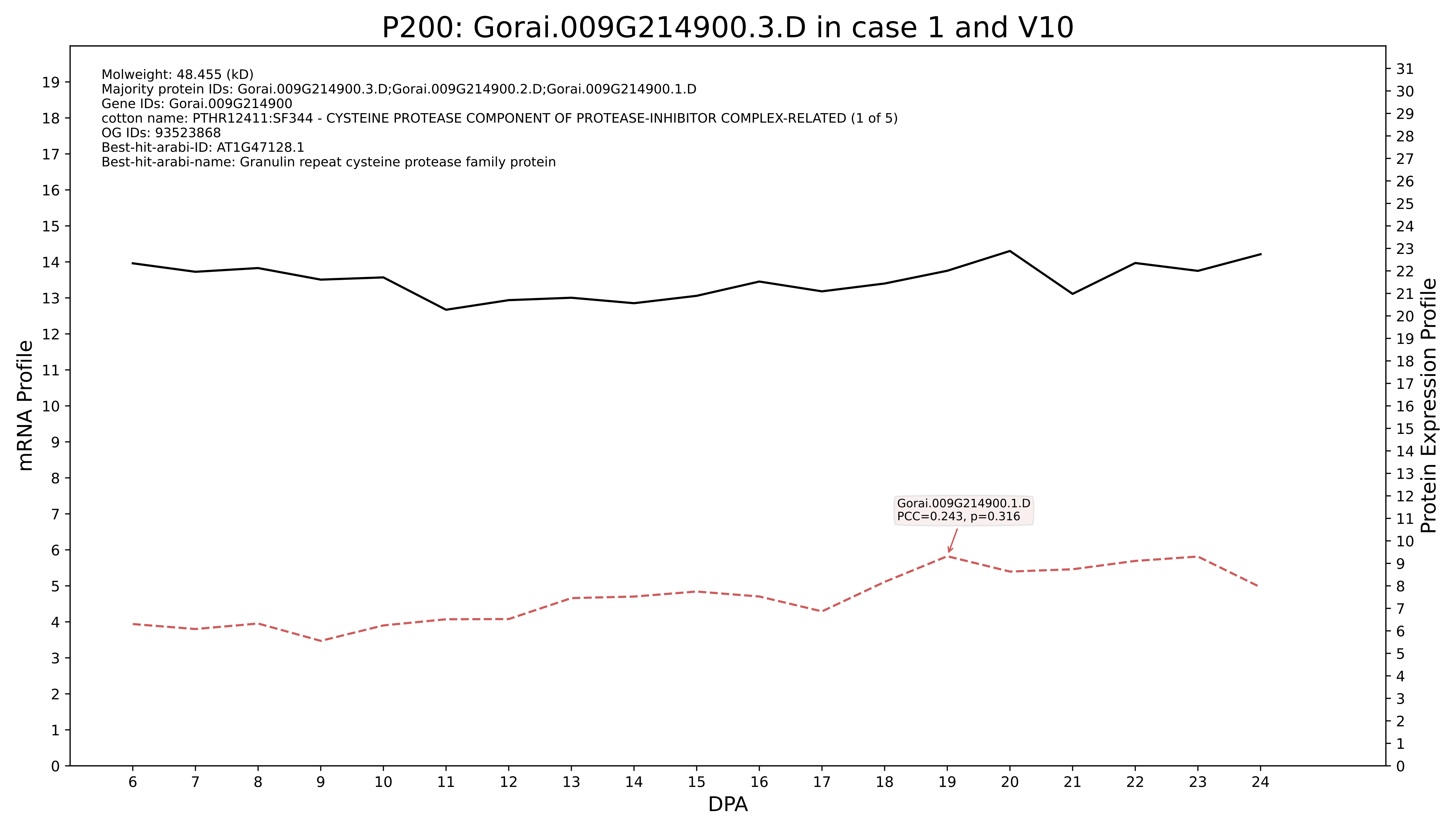
P200: Gorai.009G214900.3.D in case 1 and V10
Molweight: 48.455 (kD)
Majority protein IDs: Gorai.009G214900.3.D;Gorai.009G214900.2.D;Gorai.009G214900.1.D
Gene IDs: Gorai.009G214900
cotton name: PTHR12411:SF344 - CYSTEINE PROTEASE COMPONENT OF PROTEASE-INHIBITOR COMPLEX-RELATED (1 of 5)
OG IDs: 93523868
Best-hit-arabi-ID: AT1G47128.1
Best-hit-arabi-name: Granulin repeat cysteine protease family protein
Gorai.009G214900.1.D
PCC=0.243, p=0.316
mRNA Profile
Protein Expression Profile
DPA

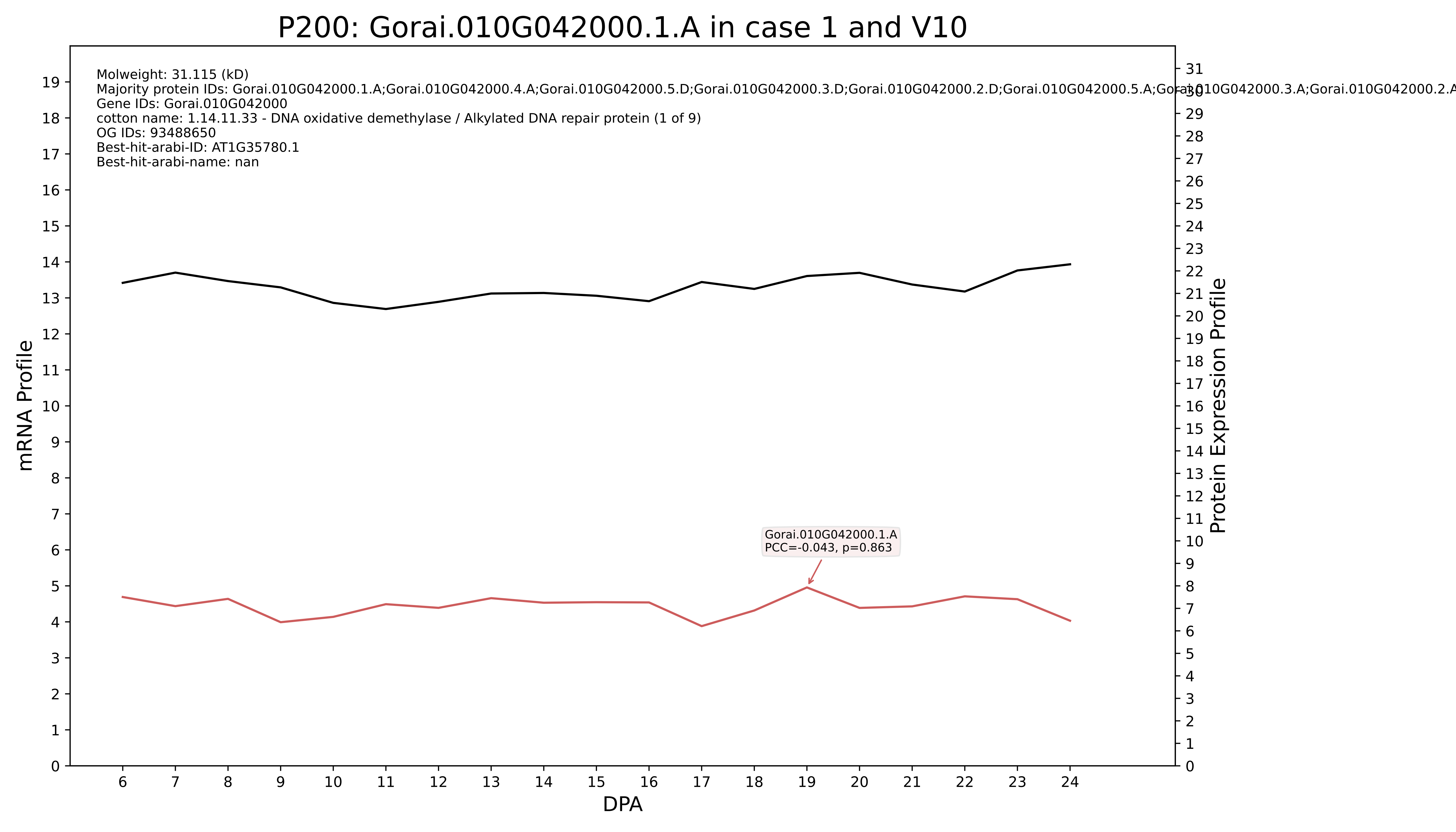

P200: Gorai.010G042000.1.A in case 1 and V10
Molweight: 31.115 (kD)
Majority protein IDs: Gorai.010G042000.1.A;Gorai.010G042000.4.A;Gorai.010G042000.5.D;Gorai.010G042000.3.D;Gorai.010G042000.2.D;Gorai.010G042000.5.A;Gorai.010G042000.3.A;Gorai.010G042000.2.A
Gene IDs: Gorai.010G042000
cotton name: 1.14.11.33 - DNA oxidative demethylase / Alkylated DNA repair protein (1 of 9)
OG IDs: 93488650
Best-hit-arabi-ID: AT1G35780.1
Best-hit-arabi-name: nan
Gorai.010G042000.1.A
PCC=-0.043, p=0.863
mRNA Profile
Protein Expression Profile
DPA

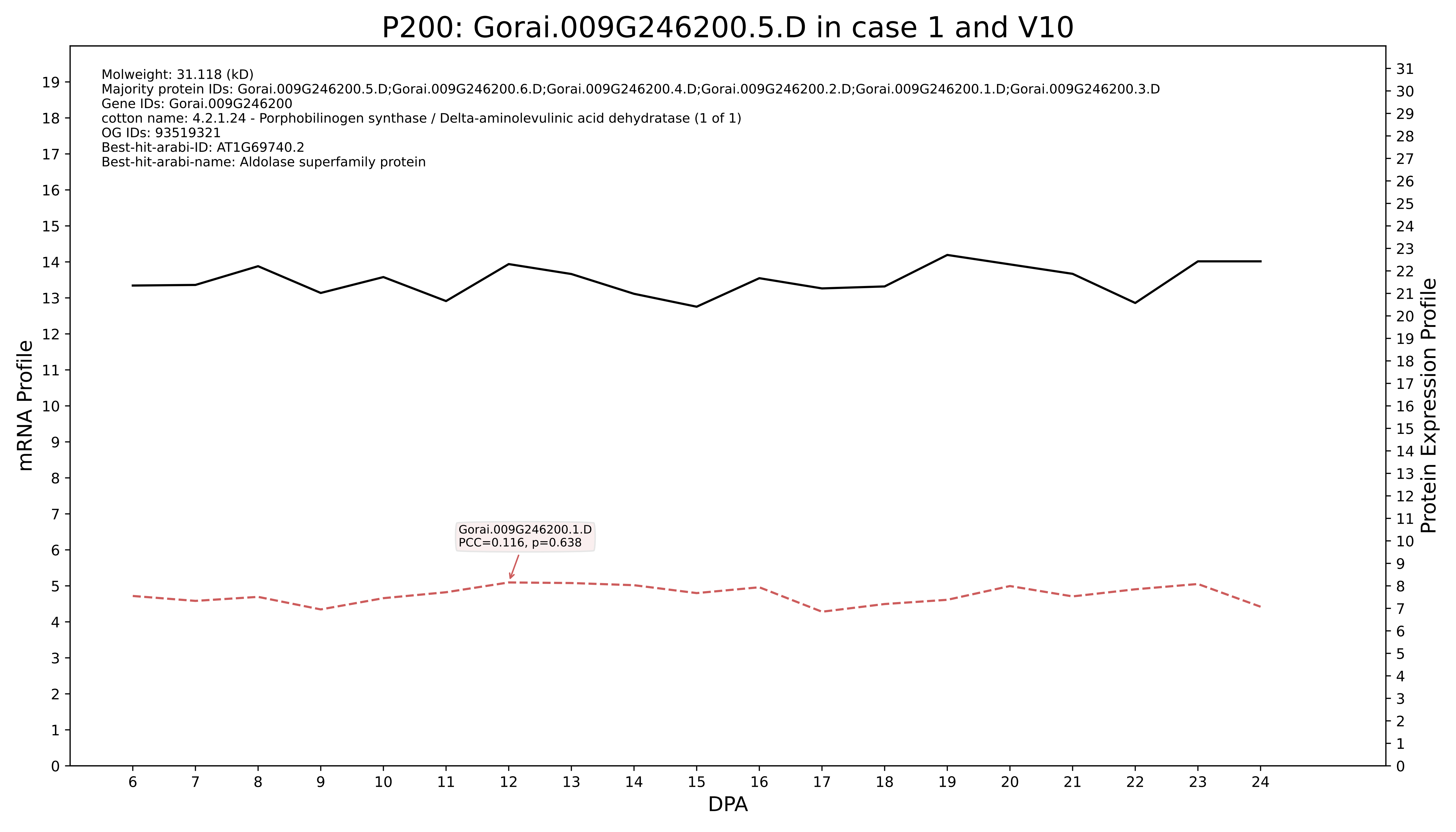
P200: Gorai.009G246200.5.D in case 1 and V10
Molweight: 31.118 (kD)
Majority protein IDs: Gorai.009G246200.5.D;Gorai.009G246200.6.D;Gorai.009G246200.4.D;Gorai.009G246200.2.D;Gorai.009G246200.1.D;Gorai.009G246200.3.D
Gene IDs: Gorai.009G246200
cotton name: 4.2.1.24 - Porphobilinogen synthase / Delta-aminolevulinic acid dehydratase (1 of 1)
OG IDs: 93519321
Best-hit-arabi-ID: AT1G69740.2
Best-hit-arabi-name: Aldolase superfamily protein
Gorai.009G246200.1.D
PCC=0.116, p=0.638
mRNA Profile
Protein Expression Profile
DPA

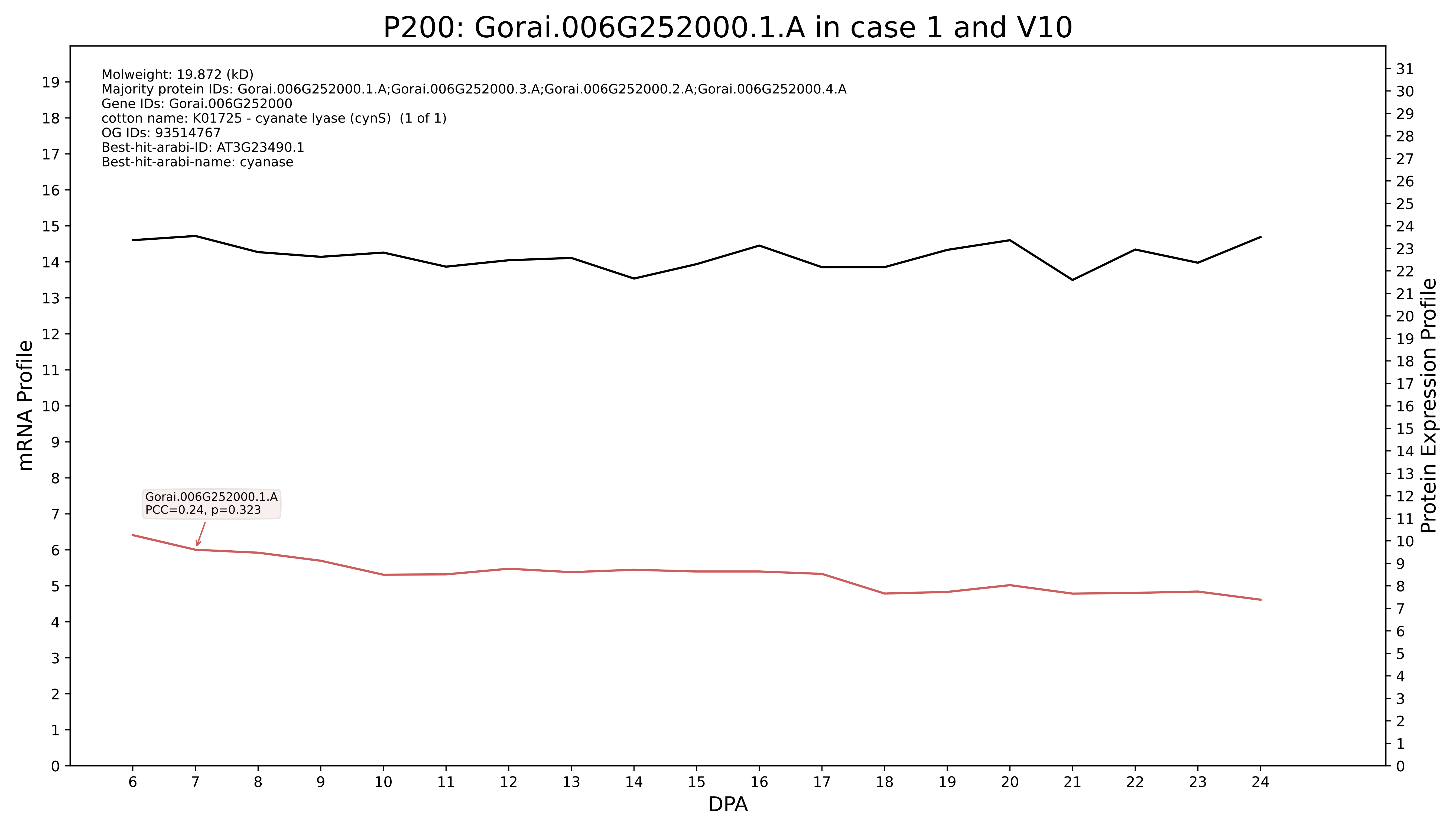
P200: Gorai.006G252000.1.A in case 1 and V10
Molweight: 19.872 (kD)
Majority protein IDs: Gorai.006G252000.1.A;Gorai.006G252000.3.A;Gorai.006G252000.2.A;Gorai.006G252000.4.A
Gene IDs: Gorai.006G252000
cotton name: K01725 - cyanate lyase (cynS) (1 of 1)
OG IDs: 93514767
Best-hit-arabi-ID: AT3G23490.1
Best-hit-arabi-name: cyanase
Gorai.006G252000.1.A
PCC=0.24, p=0.323
mRNA Profile
Protein Expression Profile
DPA

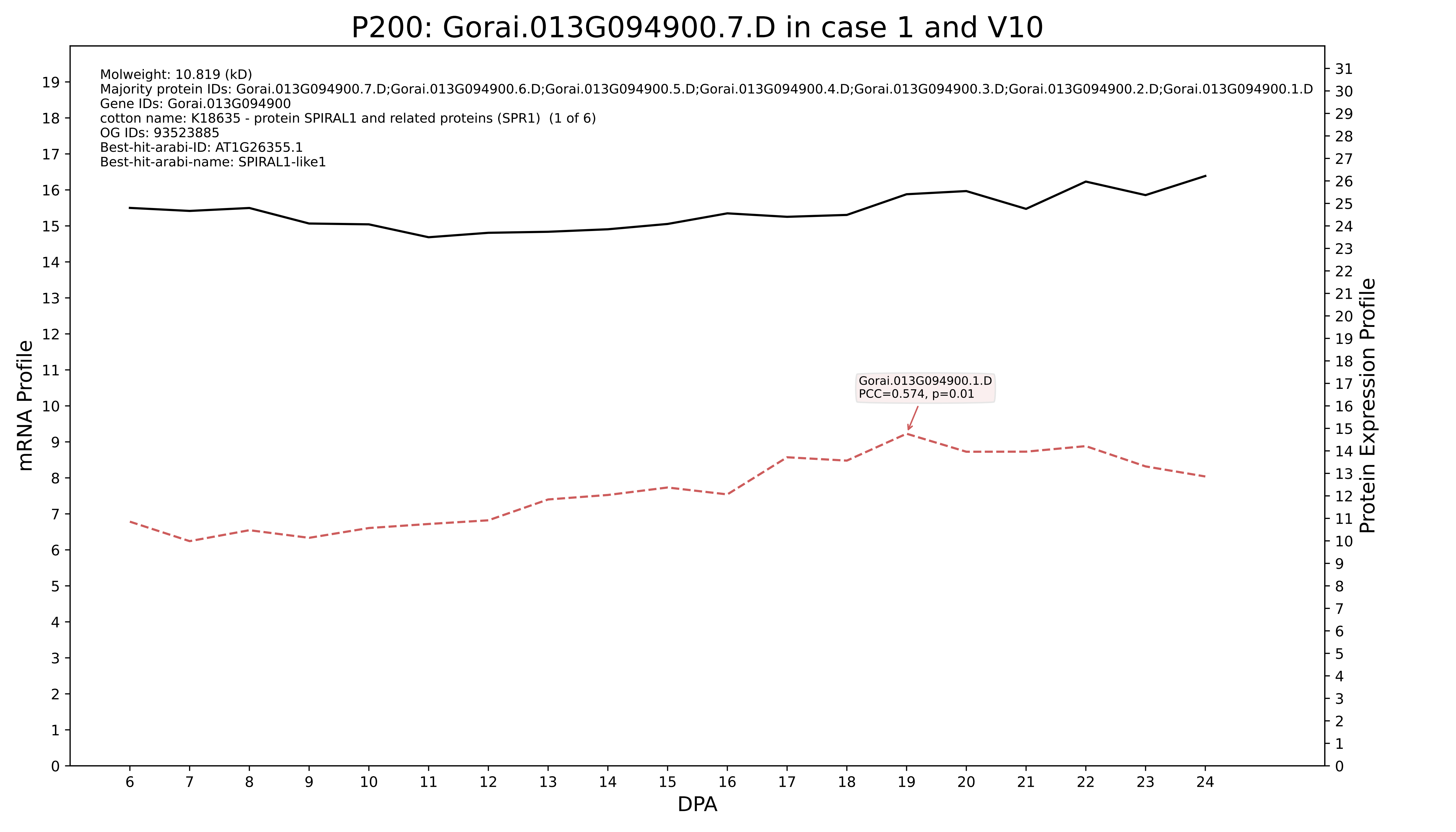
P200: Gorai.013G094900.7.D in case 1 and V10
Molweight: 10.819 (kD)
Majority protein IDs: Gorai.013G094900.7.D;Gorai.013G094900.6.D;Gorai.013G094900.5.D;Gorai.013G094900.4.D;Gorai.013G094900.3.D;Gorai.013G094900.2.D;Gorai.013G094900.1.D
Gene IDs: Gorai.013G094900
cotton name: K18635 - protein SPIRAL1 and related proteins (SPR1) (1 of 6)
OG IDs: 93523885
Best-hit-arabi-ID: AT1G26355.1
Best-hit-arabi-name: SPIRAL1-like1
Gorai.013G094900.1.D
PCC=0.574, p=0.01
mRNA Profile
Protein Expression Profile
DPA

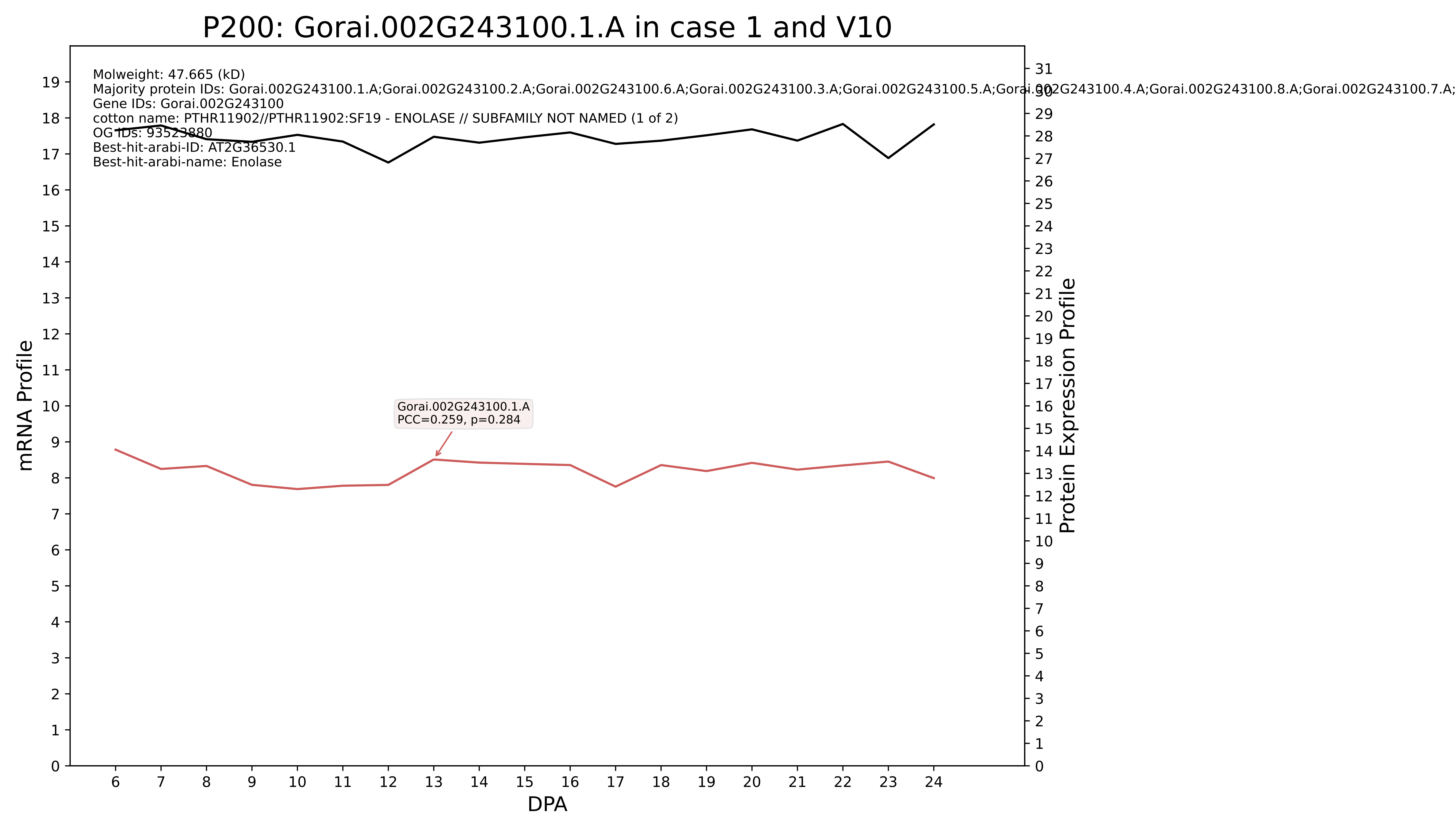
P200: Gorai.002G243100.1.A in case 1 and V10
Molweight: 47.665 (kD)
Majority protein IDs: Gorai.002G243100.1.A;Gorai.002G243100.2.A;Gorai.002G243100.6.A;Gorai.002G243100.3.A;Gorai.002G243100.5.A;Gorai.002G243100.4.A;Gorai.002G243100.8.A;Gorai.002G243100.7.A;
Gene IDs: Gorai.002G243100
cotton name: PTHR11902//PTHR11902:SF19 - ENOLASE // SUBFAMILY NOT NAMED (1 of 2)
OG IDs: 93523880
Best-hit-arabi-ID: AT2G36530.1
Best-hit-arabi-name: Enolase
Gorai.002G243100.1.A
PCC=0.259, p=0.284
mRNA Profile
Protein Expression Profile
DPA

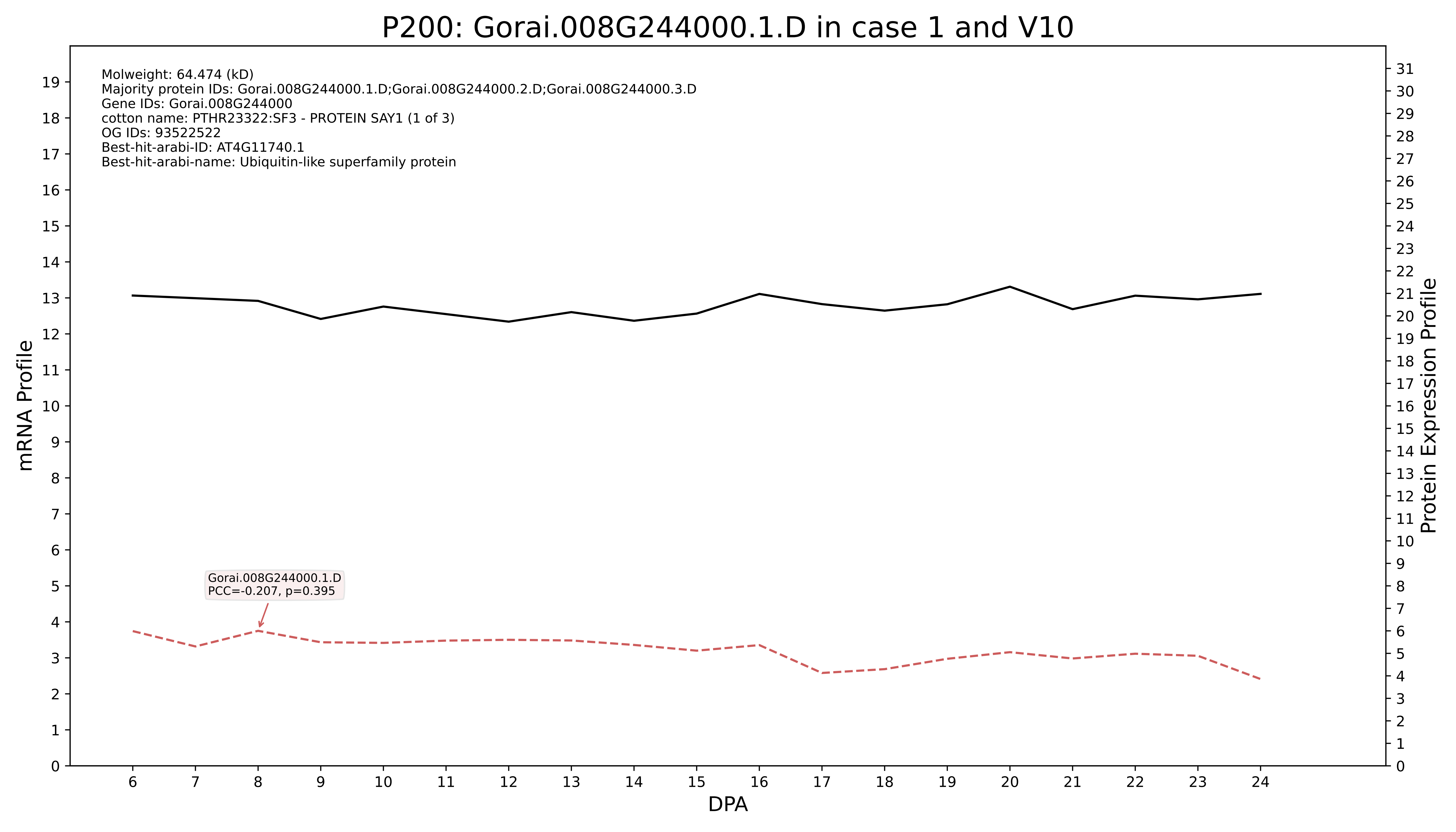
P200: Gorai.008G244000.1.D in case 1 and V10
Molweight: 64.474 (kD)
Majority protein IDs: Gorai.008G244000.1.D;Gorai.008G244000.2.D;Gorai.008G244000.3.D
Gene IDs: Gorai.008G244000
cotton name: PTHR23322:SF3 - PROTEIN SAY1 (1 of 3)
OG IDs: 93522522
Best-hit-arabi-ID: AT4G11740.1
Best-hit-arabi-name: Ubiquitin-like superfamily protein
Gorai.008G244000.1.D
PCC=-0.207, p=0.395
mRNA Profile
Protein Expression Profile
DPA

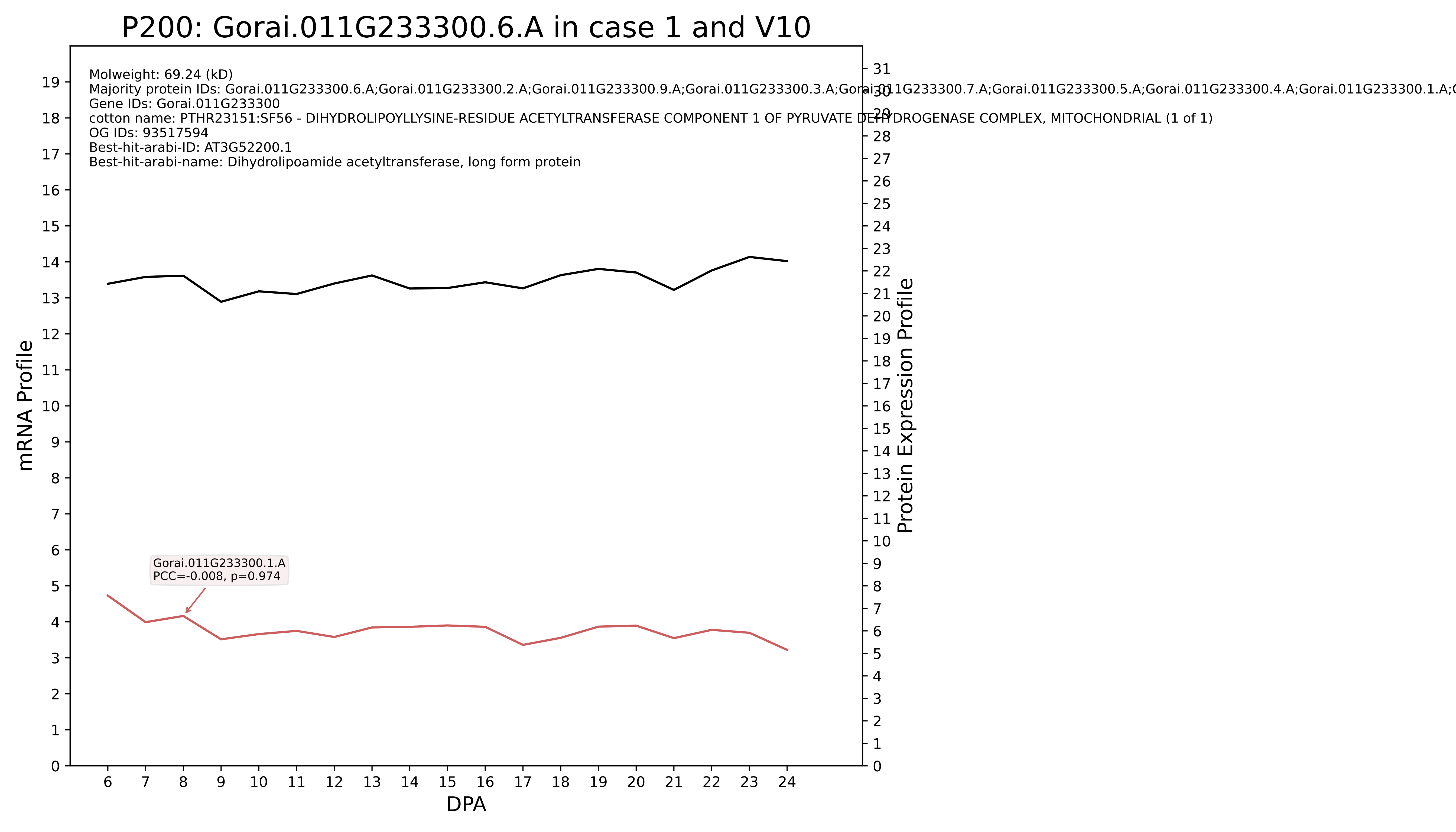
P200: Gorai.011G233300.6.A in case 1 and V10
Molweight: 69.24 (kD)
Majority protein IDs: Gorai.011G233300.6.A;Gorai.011G233300.2.A;Gorai.011G233300.9.A;Gorai.011G233300.3.A;Gorai.011G233300.7.A;Gorai.011G233300.5.A;Gorai.011G233300.4.A;Gorai.011G233300.1.A;
Gene IDs: Gorai.011G233300
cotton name: PTHR23151:SF56 - DIHYDROLIPOYLLYSINE-RESIDUE ACETYLTRANSFERASE COMPONENT 1 OF PYRUVATE DEHYDROGENASE COMPLEX, MITOCHONDRIAL (1 of 1)
OG IDs: 93517594
Best-hit-arabi-ID: AT3G52200.1
Best-hit-arabi-name: Dihydrolipoamide acetyltransferase, long form protein
Gorai.011G233300.1.A
PCC=-0.008, p=0.974
mRNA Profile
Protein Expression Profile
DPA

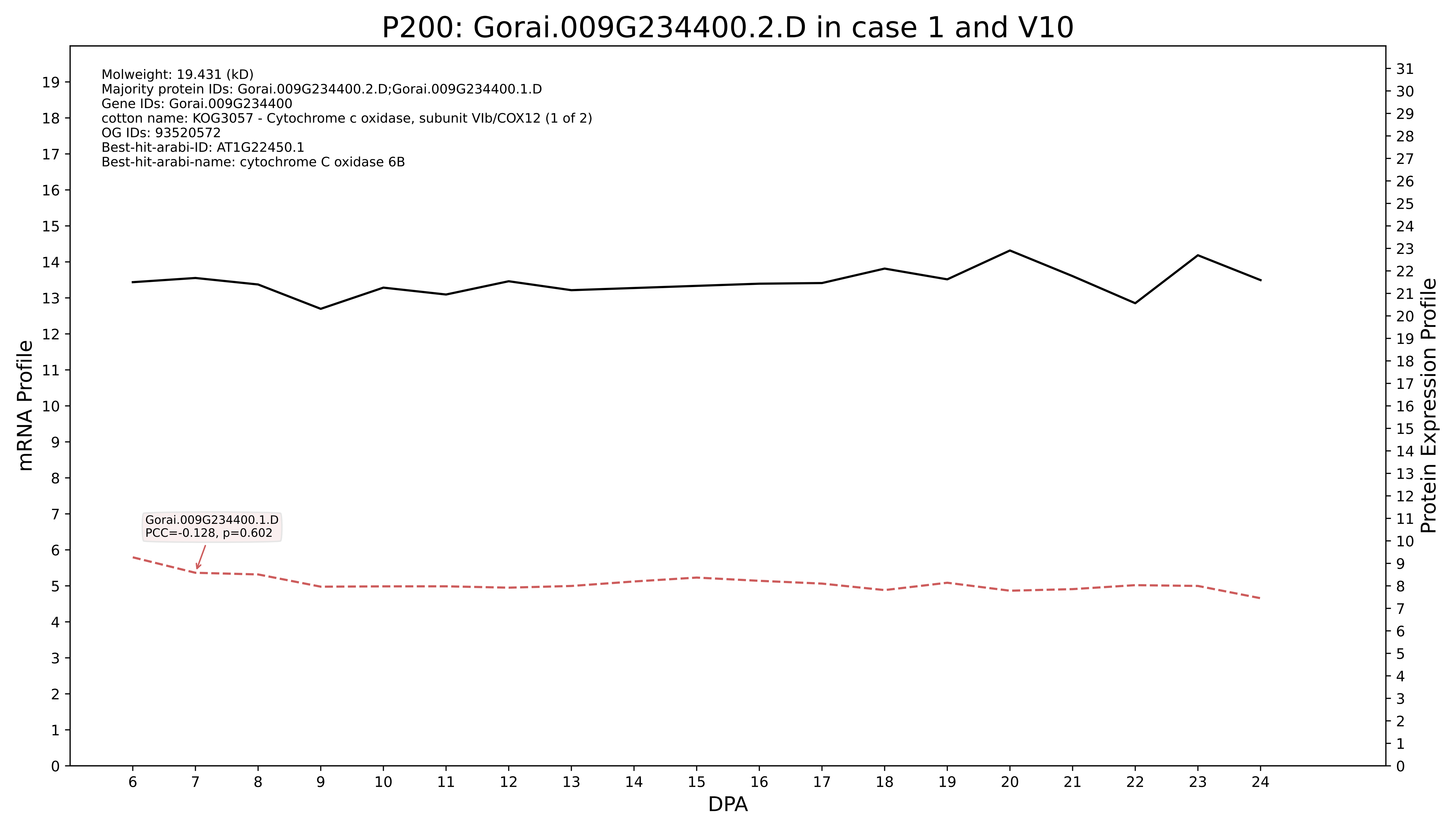
P200: Gorai.009G234400.2.D in case 1 and V10
Molweight: 19.431 (kD)
Majority protein IDs: Gorai.009G234400.2.D;Gorai.009G234400.1.D
Gene IDs: Gorai.009G234400
cotton name: KOG3057 - Cytochrome c oxidase, subunit VIb/COX12 (1 of 2)
OG IDs: 93520572
Best-hit-arabi-ID: AT1G22450.1
Best-hit-arabi-name: cytochrome C oxidase 6B
Gorai.009G234400.1.D
PCC=-0.128, p=0.602
mRNA Profile
Protein Expression Profile
DPA

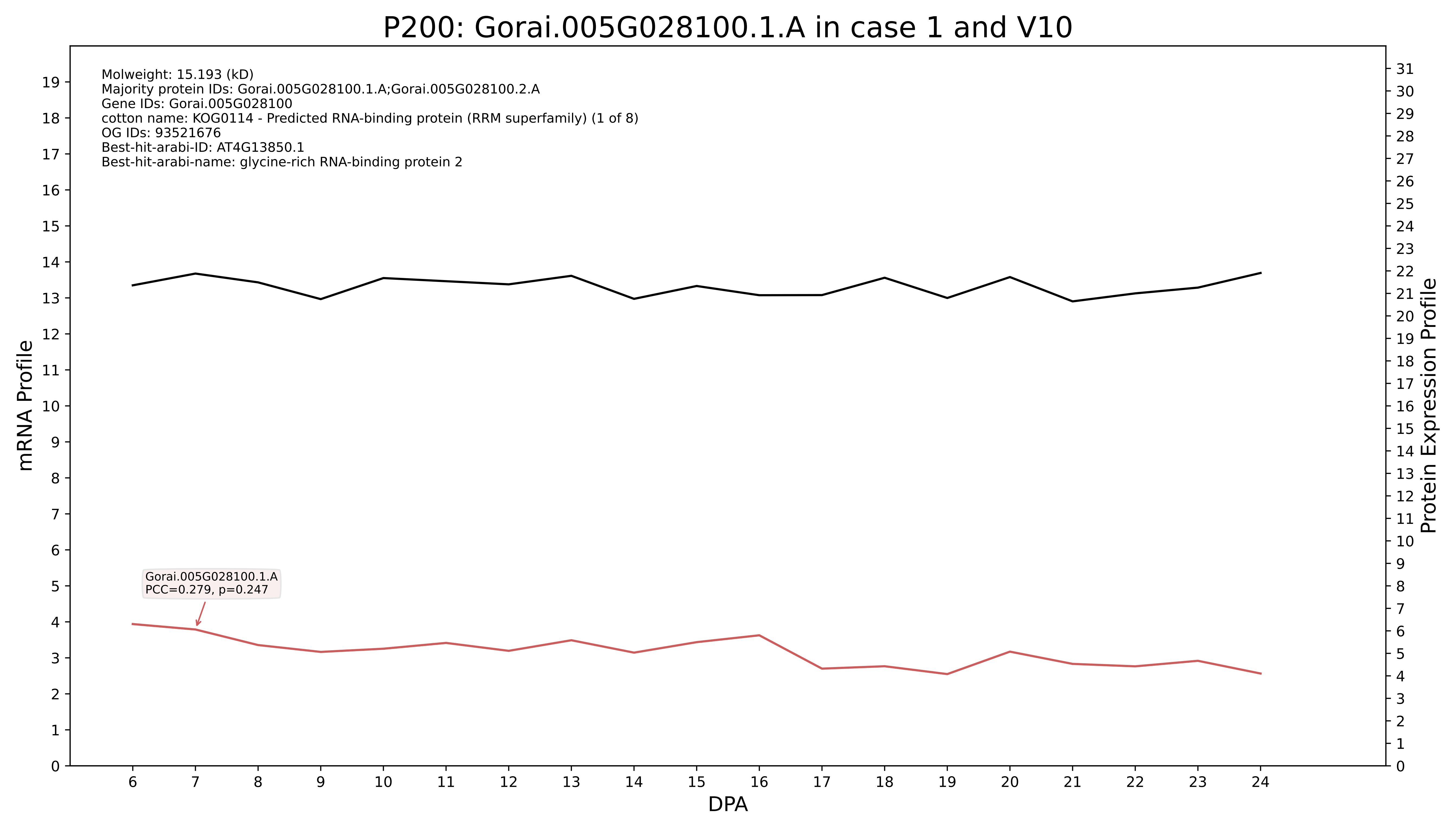
P200: Gorai.005G028100.1.A in case 1 and V10
Molweight: 15.193 (kD)
Majority protein IDs: Gorai.005G028100.1.A;Gorai.005G028100.2.A
Gene IDs: Gorai.005G028100
cotton name: KOG0114 - Predicted RNA-binding protein (RRM superfamily) (1 of 8)
OG IDs: 93521676
Best-hit-arabi-ID: AT4G13850.1
Best-hit-arabi-name: glycine-rich RNA-binding protein 2
Gorai.005G028100.1.A
PCC=0.279, p=0.247
mRNA Profile
Protein Expression Profile
DPA

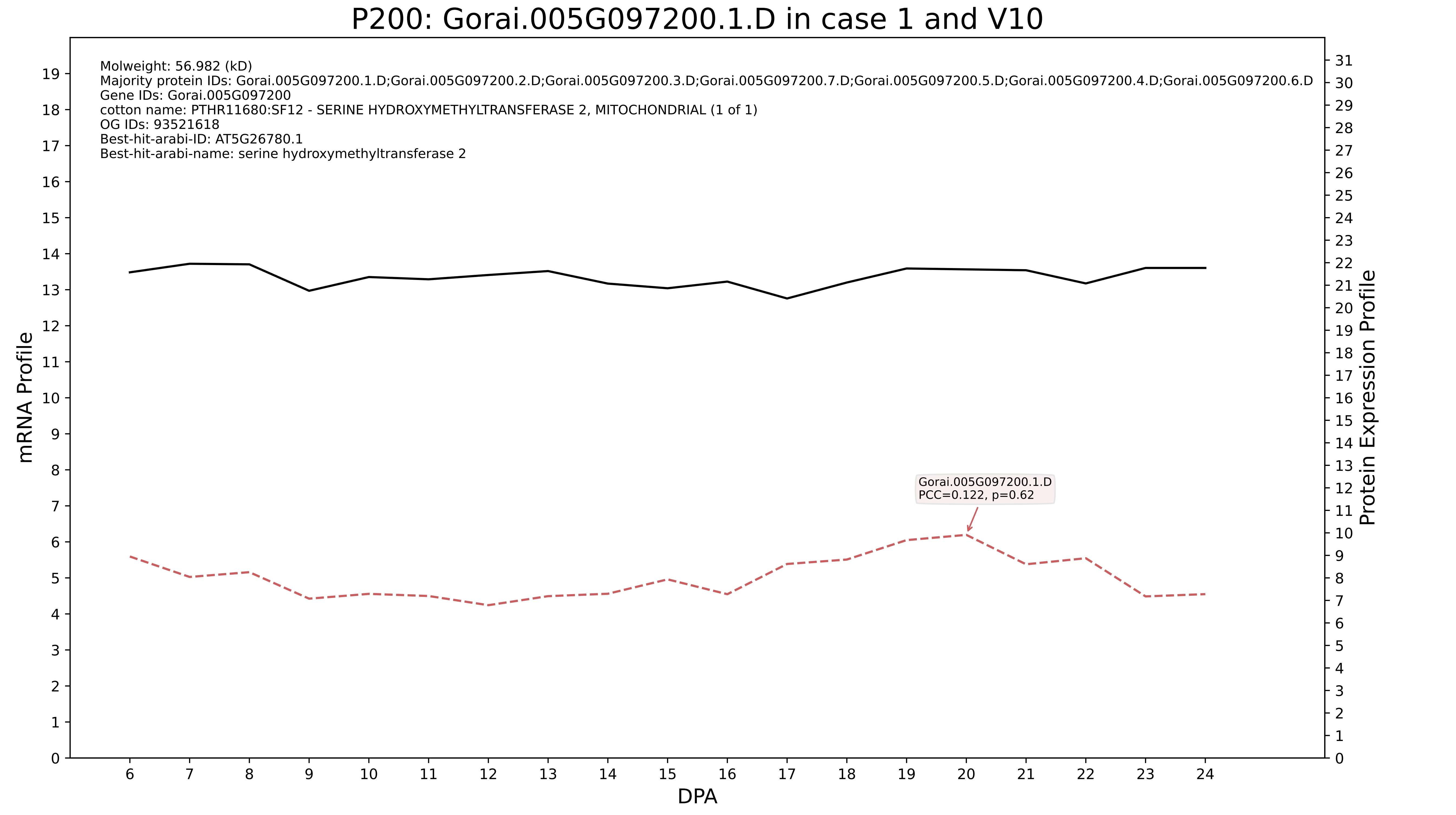

P200: Gorai.005G097200.1.D in case 1 and V10
Molweight: 56.982 (kD)
Majority protein IDs: Gorai.005G097200.1.D;Gorai.005G097200.2.D;Gorai.005G097200.3.D;Gorai.005G097200.7.D;Gorai.005G097200.5.D;Gorai.005G097200.4.D;Gorai.005G097200.6.D
Gene IDs: Gorai.005G097200
cotton name: PTHR11680:SF12 - SERINE HYDROXYMETHYLTRANSFERASE 2, MITOCHONDRIAL (1 of 1)
OG IDs: 93521618
Best-hit-arabi-ID: AT5G26780.1
Best-hit-arabi-name: serine hydroxymethyltransferase 2
Gorai.005G097200.1.D
PCC=0.122, p=0.62
mRNA Profile
Protein Expression Profile
DPA

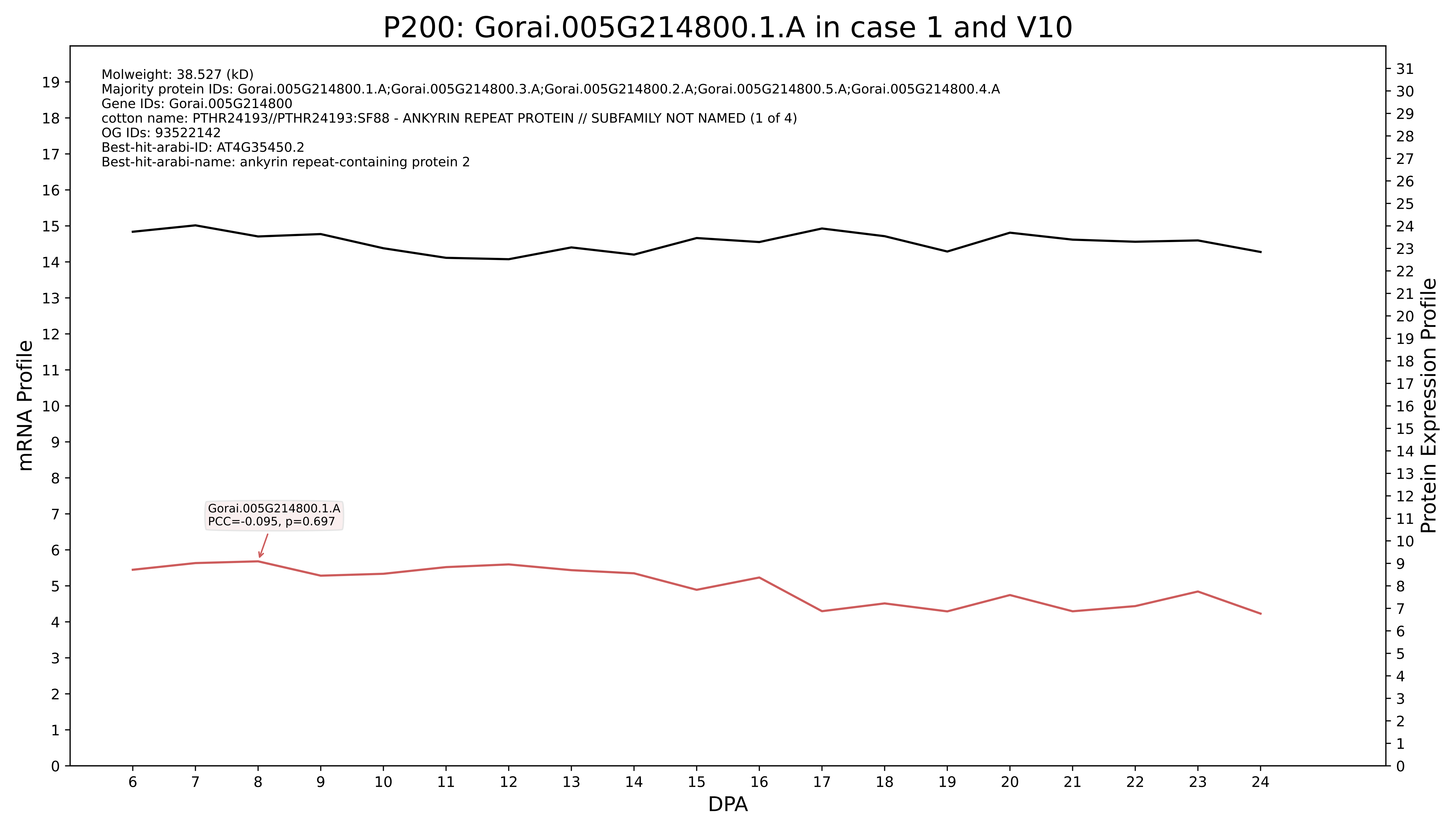
P200: Gorai.005G214800.1.A in case 1 and V10
Molweight: 38.527 (kD)
Majority protein IDs: Gorai.005G214800.1.A;Gorai.005G214800.3.A;Gorai.005G214800.2.A;Gorai.005G214800.5.A;Gorai.005G214800.4.A
Gene IDs: Gorai.005G214800
cotton name: PTHR24193//PTHR24193:SF88 - ANKYRIN REPEAT PROTEIN // SUBFAMILY NOT NAMED (1 of 4)
OG IDs: 93522142
Best-hit-arabi-ID: AT4G35450.2
Best-hit-arabi-name: ankyrin repeat-containing protein 2
Gorai.005G214800.1.A
PCC=-0.095, p=0.697
mRNA Profile
Protein Expression Profile
DPA

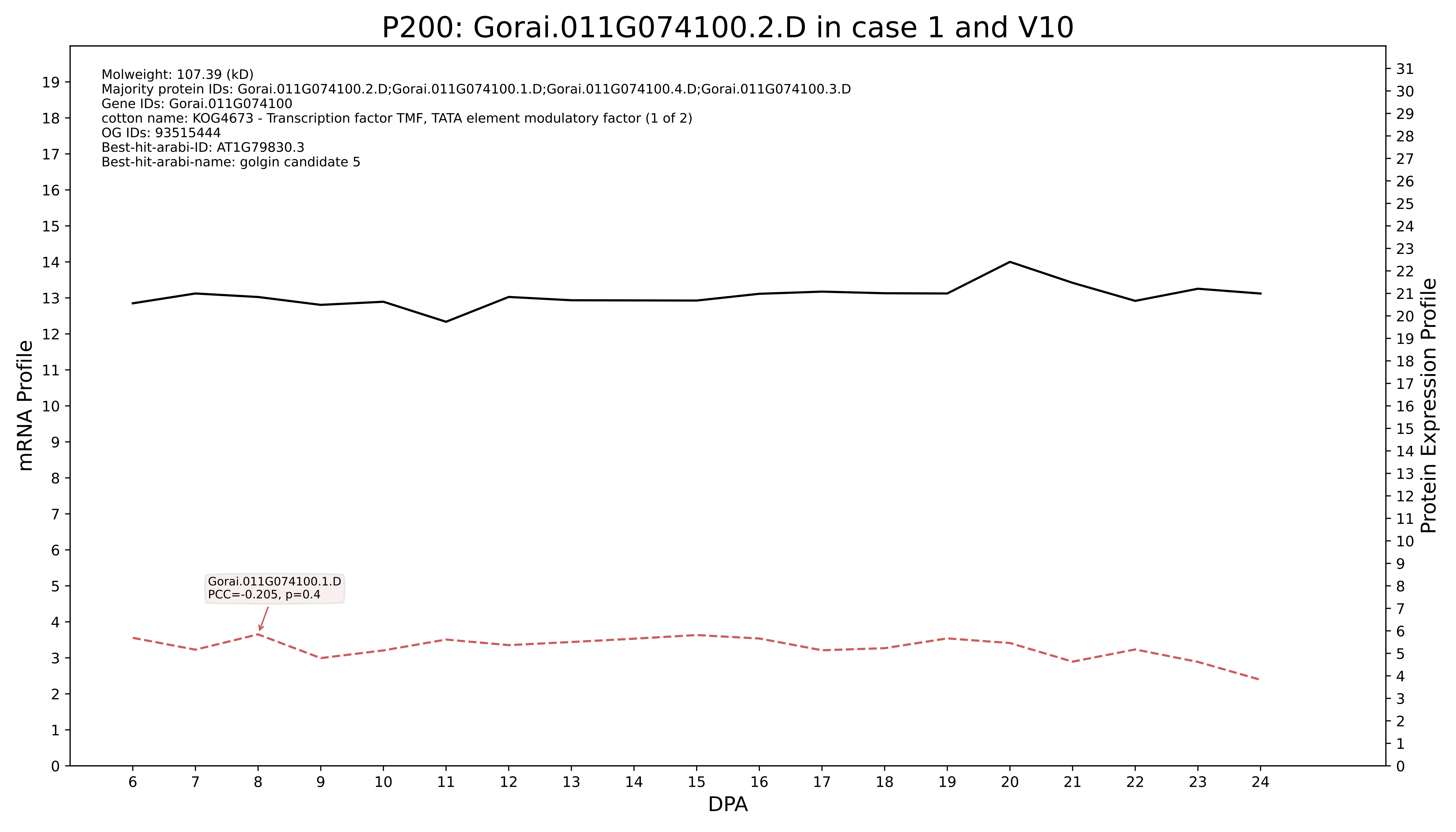
P200: Gorai.011G074100.2.D in case 1 and V10
Molweight: 107.39 (kD)
Majority protein IDs: Gorai.011G074100.2.D;Gorai.011G074100.1.D;Gorai.011G074100.4.D;Gorai.011G074100.3.D
Gene IDs: Gorai.011G074100
cotton name: KOG4673 - Transcription factor TMF, TATA element modulatory factor (1 of 2)
OG IDs: 93515444
Best-hit-arabi-ID: AT1G79830.3
Best-hit-arabi-name: golgin candidate 5
Gorai.011G074100.1.D
PCC=-0.205, p=0.4
mRNA Profile
Protein Expression Profile
DPA

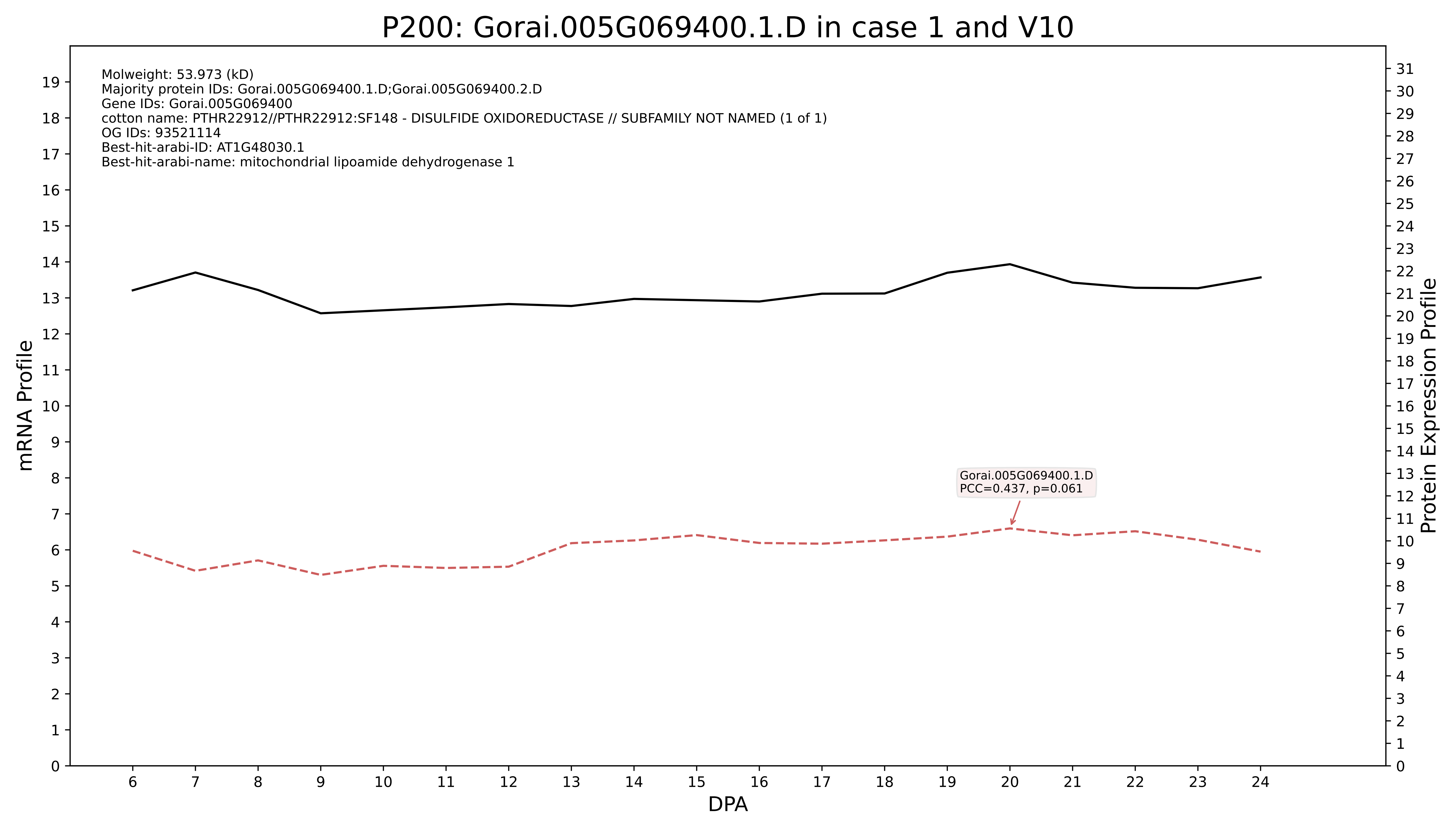
P200: Gorai.005G069400.1.D in case 1 and V10
Molweight: 53.973 (kD)
Majority protein IDs: Gorai.005G069400.1.D;Gorai.005G069400.2.D
Gene IDs: Gorai.005G069400
cotton name: PTHR22912//PTHR22912:SF148 - DISULFIDE OXIDOREDUCTASE // SUBFAMILY NOT NAMED (1 of 1)
OG IDs: 93521114
Best-hit-arabi-ID: AT1G48030.1
Best-hit-arabi-name: mitochondrial lipoamide dehydrogenase 1
Gorai.005G069400.1.D
PCC=0.437, p=0.061
mRNA Profile
Protein Expression Profile
DPA

P200: Gorai.003G182200.1.A in case 1 and V10

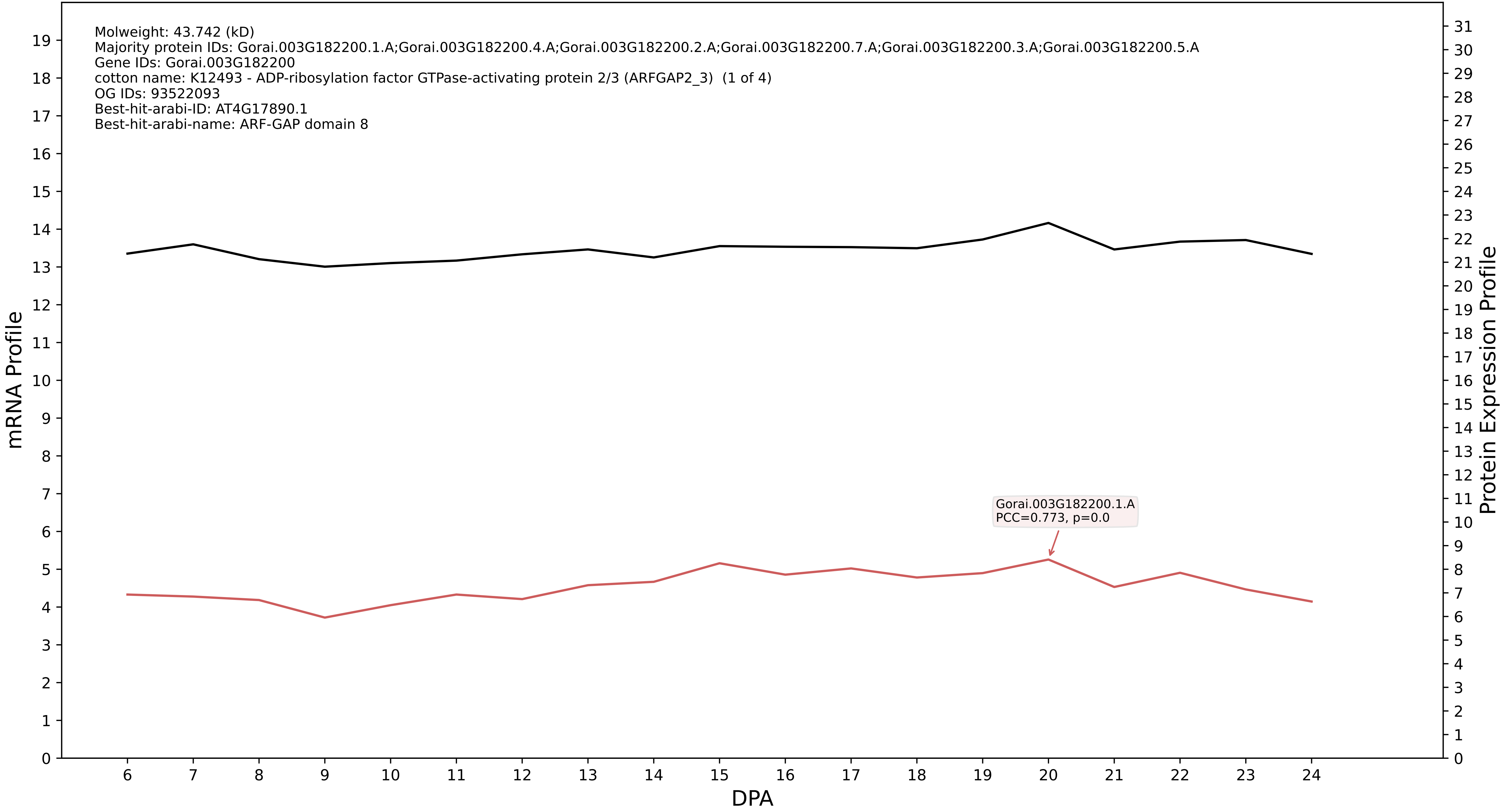

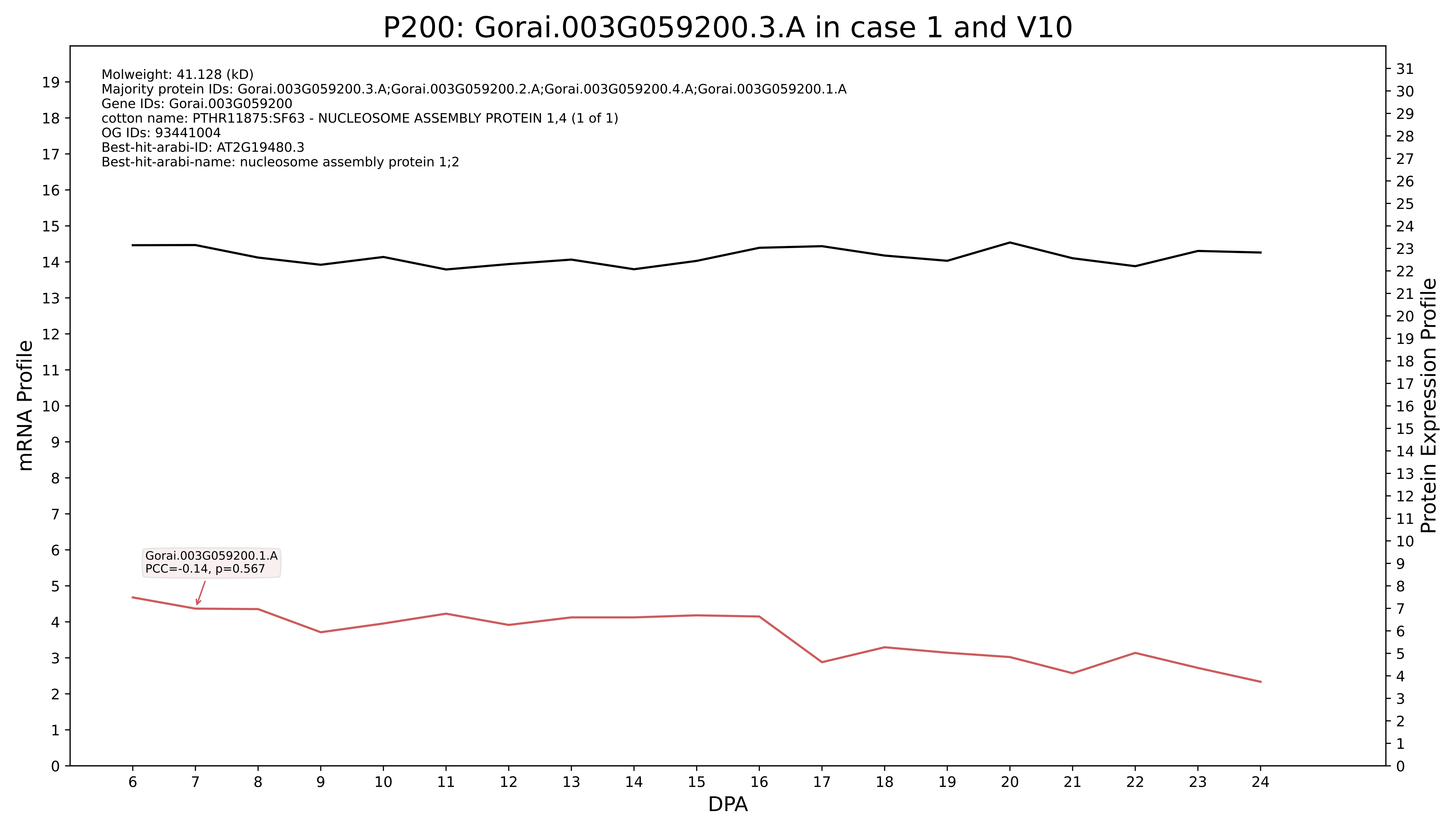

P200: Gorai.003G059200.3.A in case 1 and V10
Molweight: 41.128 (kD)
Majority protein IDs: Gorai.003G059200.3.A;Gorai.003G059200.2.A;Gorai.003G059200.4.A;Gorai.003G059200.1.A
Gene IDs: Gorai.003G059200
cotton name: PTHR11875:SF63 - NUCLEOSOME ASSEMBLY PROTEIN 1,4 (1 of 1)
OG IDs: 93441004
Best-hit-arabi-ID: AT2G19480.3
Best-hit-arabi-name: nucleosome assembly protein 1;2
Gorai.003G059200.1.A
PCC=-0.14, p=0.567
mRNA Profile
Protein Expression Profile
DPA

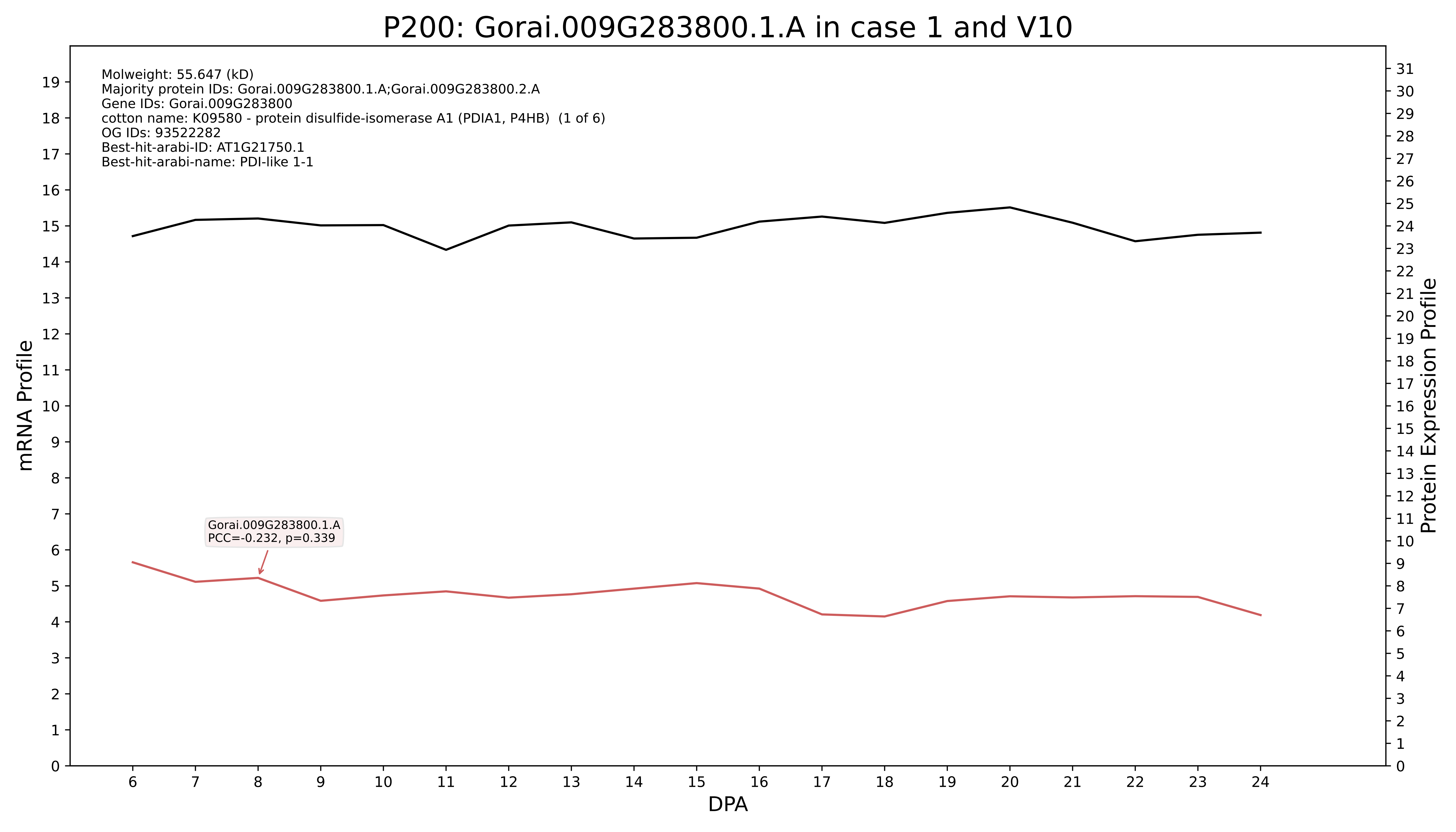

P200: Gorai.009G283800.1.A in case 1 and V10
Molweight: 55.647 (kD)
Majority protein IDs: Gorai.009G283800.1.A;Gorai.009G283800.2.A
Gene IDs: Gorai.009G283800
cotton name: K09580 - protein disulfide-isomerase A1 (PDIA1, P4HB) (1 of 6)
OG IDs: 93522282
Best-hit-arabi-ID: AT1G21750.1
Best-hit-arabi-name: PDI-like 1-1
Gorai.009G283800.1.A
PCC=-0.232, p=0.339
mRNA Profile
Protein Expression Profile
DPA

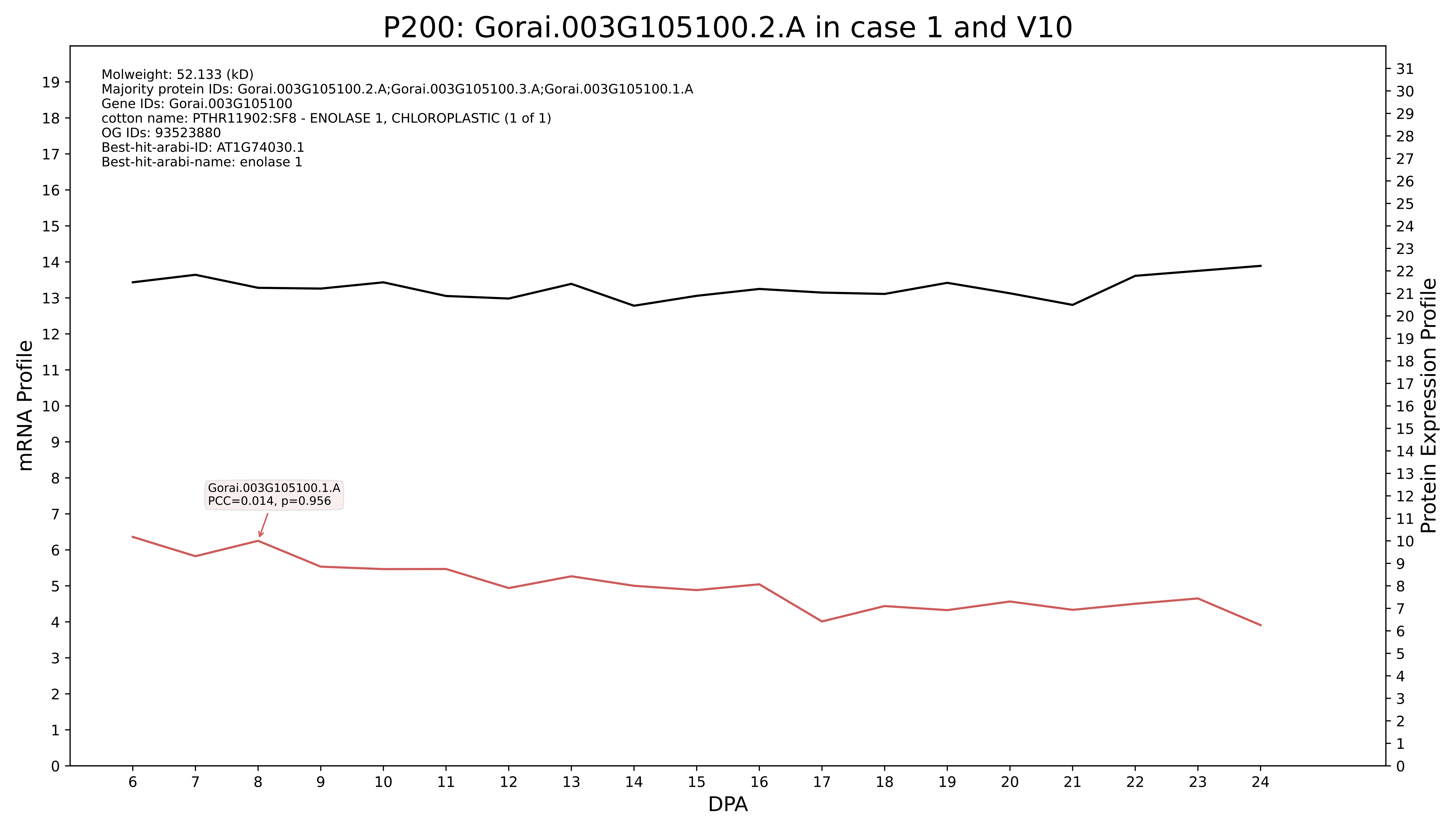
P200: Gorai.003G105100.2.A in case 1 and V10
Molweight: 52.133 (kD)
Majority protein IDs: Gorai.003G105100.2.A;Gorai.003G105100.3.A;Gorai.003G105100.1.A
Gene IDs: Gorai.003G105100
cotton name: PTHR11902:SF8 - ENOLASE 1, CHLOROPLASTIC (1 of 1)
OG IDs: 93523880
Best-hit-arabi-ID: AT1G74030.1
Best-hit-arabi-name: enolase 1
Gorai.003G105100.1.A
PCC=0.014, p=0.956
mRNA Profile
Protein Expression Profile
DPA

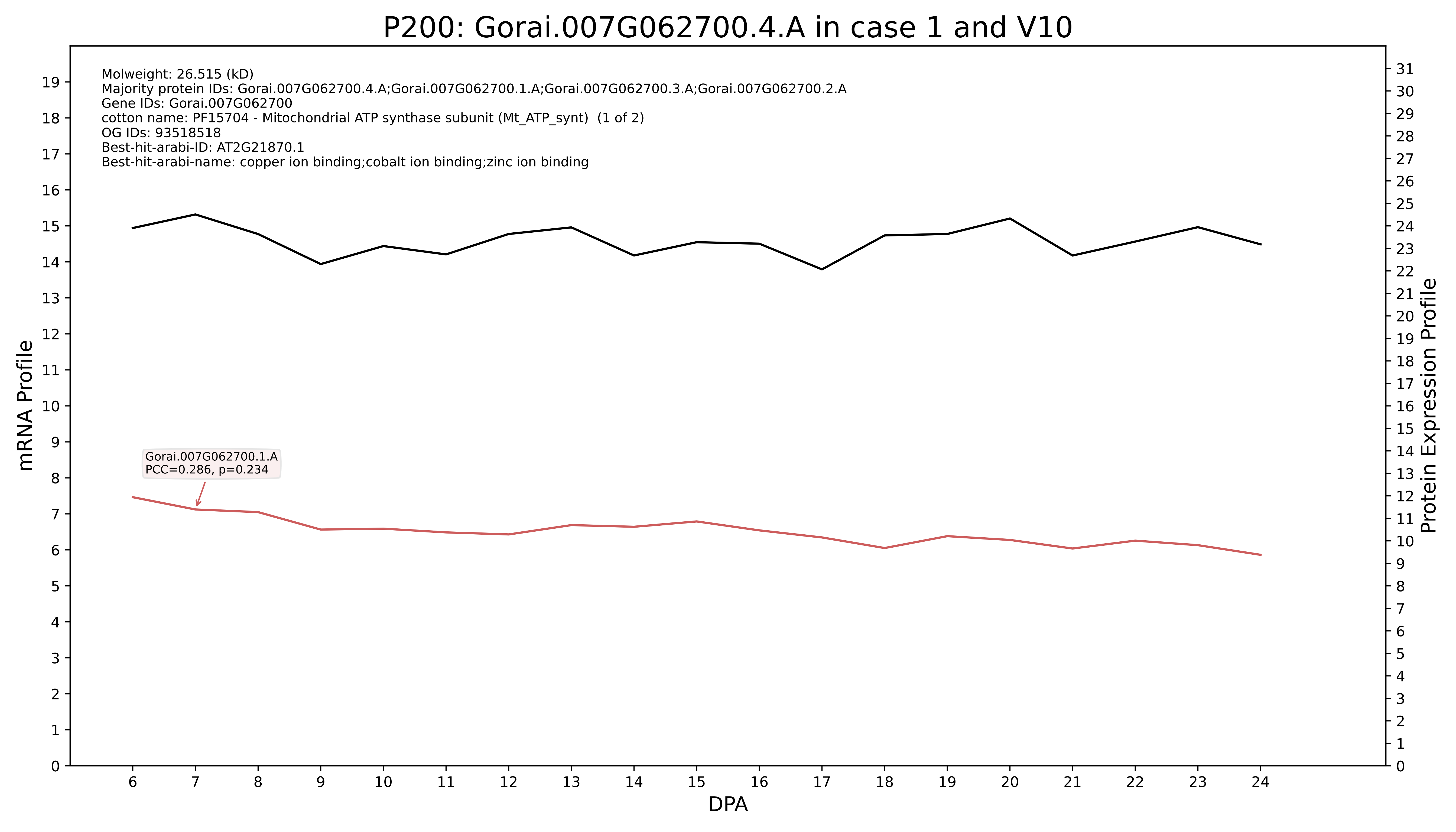
P200: Gorai.007G062700.4.A in case 1 and V10
Molweight: 26.515 (kD)
Majority protein IDs: Gorai.007G062700.4.A;Gorai.007G062700.1.A;Gorai.007G062700.3.A;Gorai.007G062700.2.A
Gene IDs: Gorai.007G062700
cotton name: PF15704 - Mitochondrial ATP synthase subunit (Mt_ATP_synt) (1 of 2)
OG IDs: 93518518
Best-hit-arabi-ID: AT2G21870.1
Best-hit-arabi-name: copper ion binding;cobalt ion binding;zinc ion binding
Gorai.007G062700.1.A
PCC=0.286, p=0.234
mRNA Profile
Protein Expression Profile
DPA

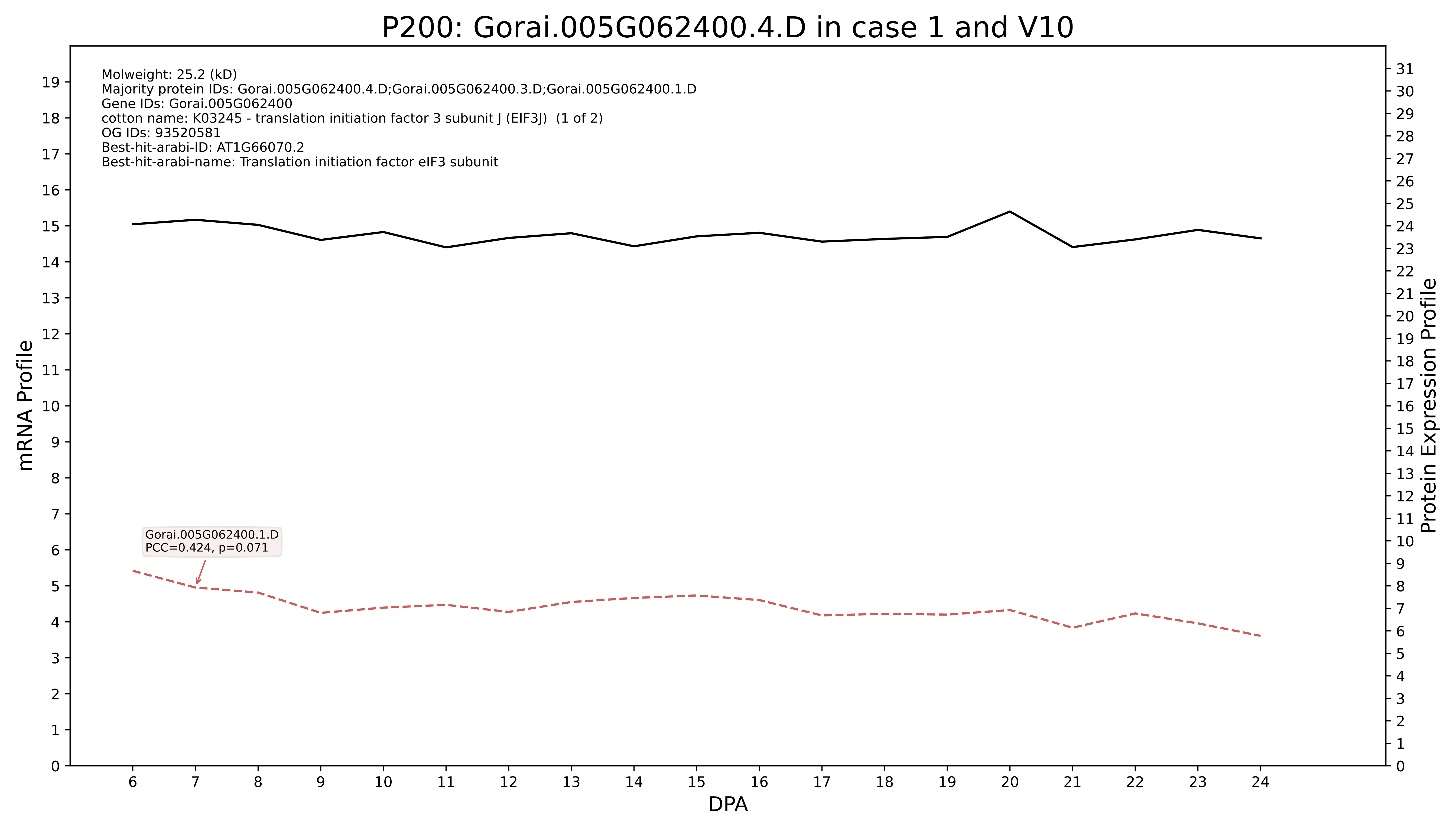
P200: Gorai.005G062400.4.D in case 1 and V10
Molweight: 25.2 (kD)
Majority protein IDs: Gorai.005G062400.4.D;Gorai.005G062400.3.D;Gorai.005G062400.1.D
Gene IDs: Gorai.005G062400
cotton name: K03245 - translation initiation factor 3 subunit J (EIF3J) (1 of 2)
OG IDs: 93520581
Best-hit-arabi-ID: AT1G66070.2
Best-hit-arabi-name: Translation initiation factor eIF3 subunit
Gorai.005G062400.1.D
PCC=0.424, p=0.071
mRNA Profile
Protein Expression Profile
DPA

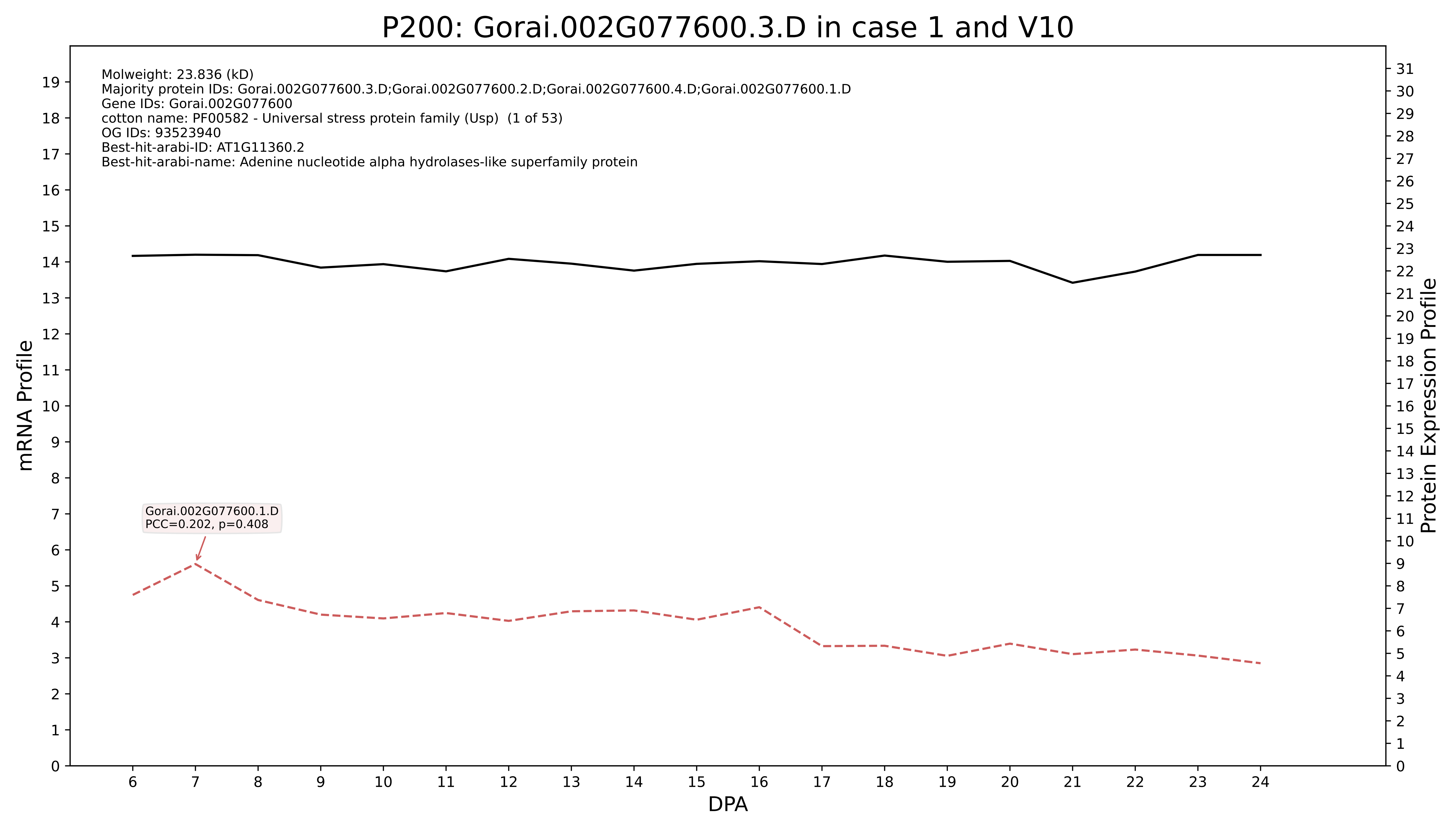
P200: Gorai.002G077600.3.D in case 1 and V10
Molweight: 23.836 (kD)
Majority protein IDs: Gorai.002G077600.3.D;Gorai.002G077600.2.D;Gorai.002G077600.4.D;Gorai.002G077600.1.D
Gene IDs: Gorai.002G077600
cotton name: PF00582 - Universal stress protein family (Usp) (1 of 53)
OG IDs: 93523940
Best-hit-arabi-ID: AT1G11360.2
Best-hit-arabi-name: Adenine nucleotide alpha hydrolases-like superfamily protein
Gorai.002G077600.1.D
PCC=0.202, p=0.408
mRNA Profile
Protein Expression Profile
DPA

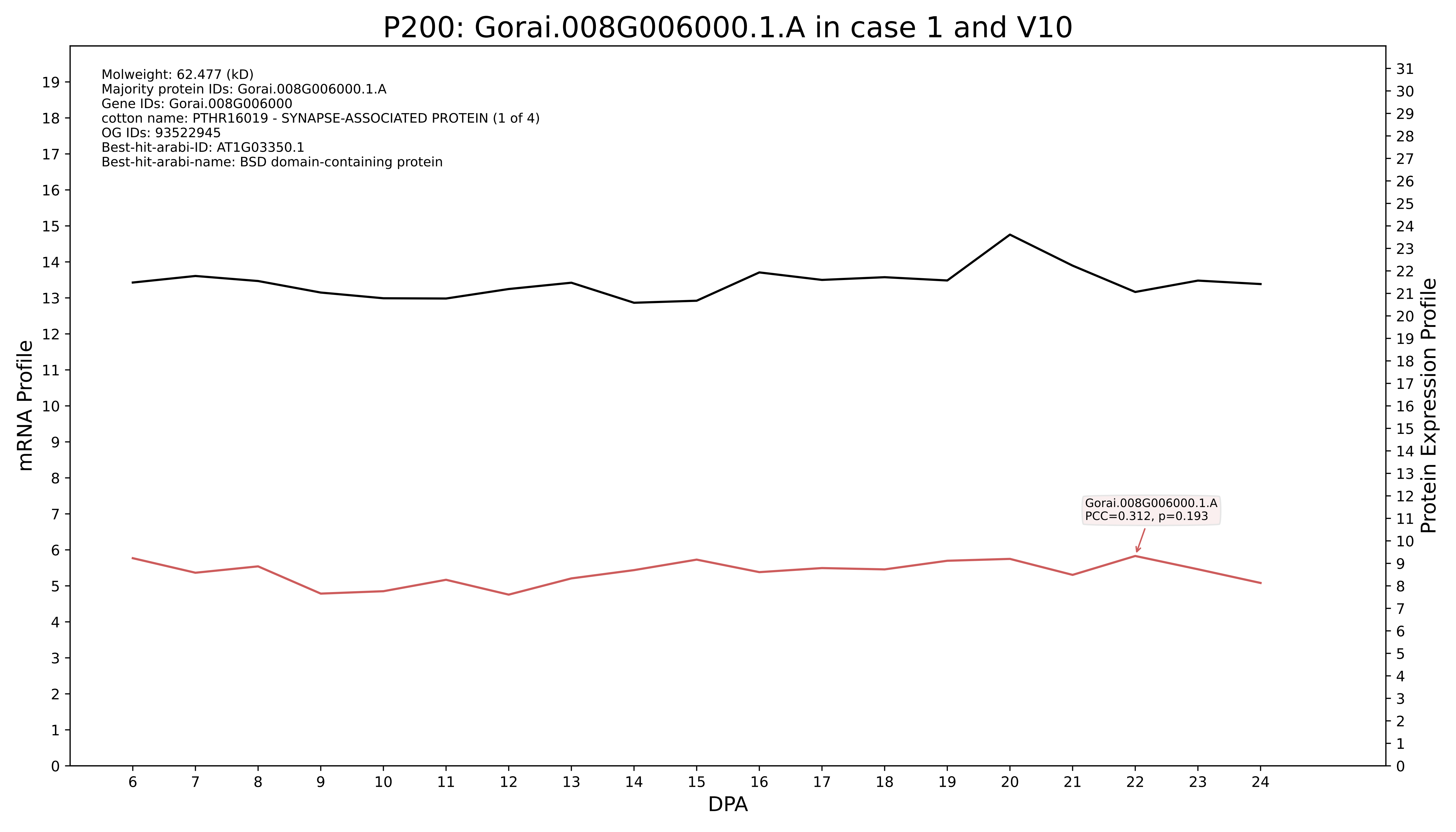
P200: Gorai.008G006000.1.A in case 1 and V10
Molweight: 62.477 (kD)
Majority protein IDs: Gorai.008G006000.1.A
Gene IDs: Gorai.008G006000
cotton name: PTHR16019 - SYNAPSE-ASSOCIATED PROTEIN (1 of 4)
OG IDs: 93522945
Best-hit-arabi-ID: AT1G03350.1
Best-hit-arabi-name: BSD domain-containing protein
Gorai.008G006000.1.A
PCC=0.312, p=0.193
mRNA Profile
Protein Expression Profile
DPA

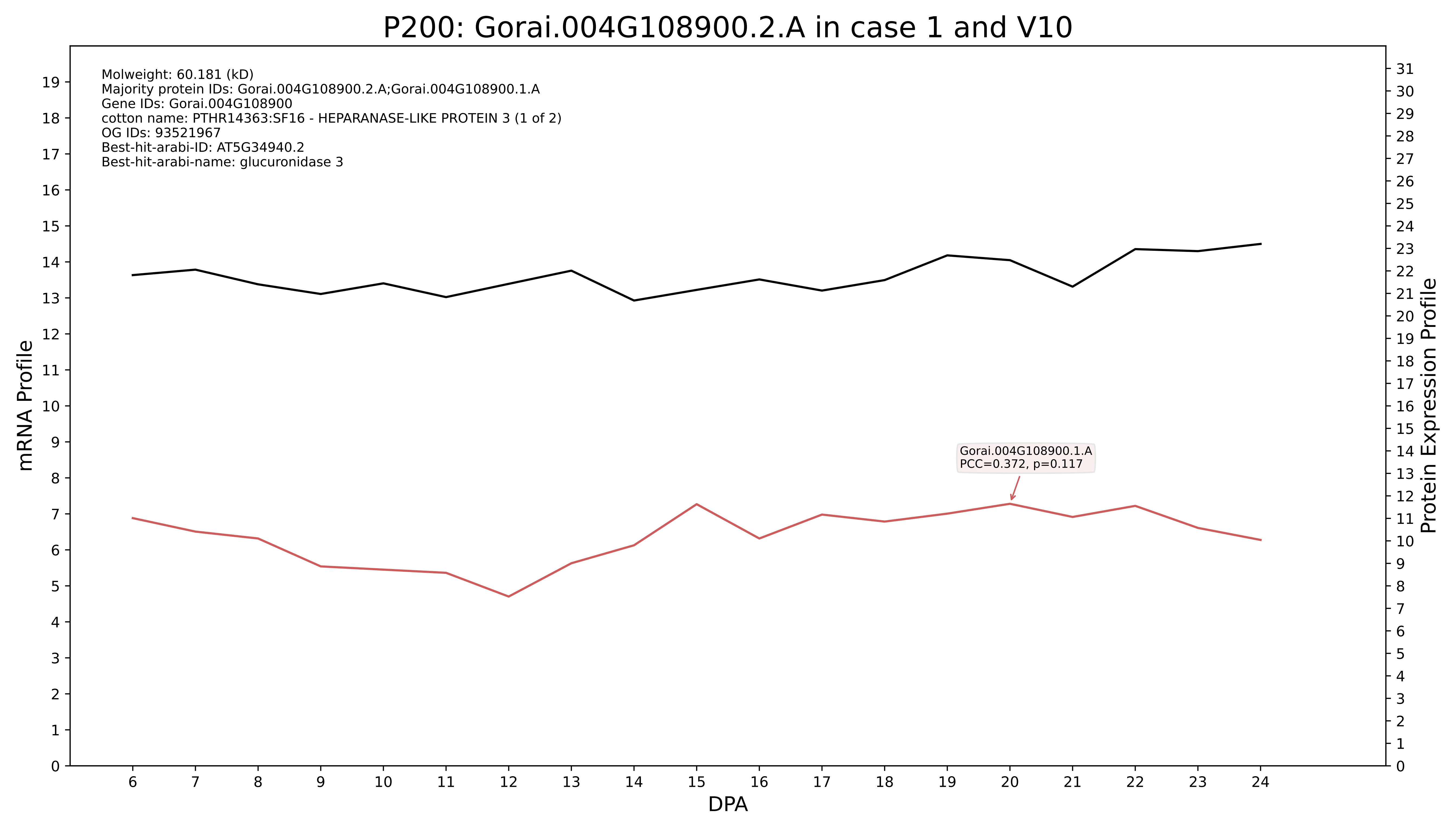

P200: Gorai.004G108900.2.A in case 1 and V10
Molweight: 60.181 (kD)
Majority protein IDs: Gorai.004G108900.2.A;Gorai.004G108900.1.A
Gene IDs: Gorai.004G108900
cotton name: PTHR14363:SF16 - HEPARANASE-LIKE PROTEIN 3 (1 of 2)
OG IDs: 93521967
Best-hit-arabi-ID: AT5G34940.2
Best-hit-arabi-name: glucuronidase 3
Gorai.004G108900.1.A
PCC=0.372, p=0.117
mRNA Profile
Protein Expression Profile
DPA

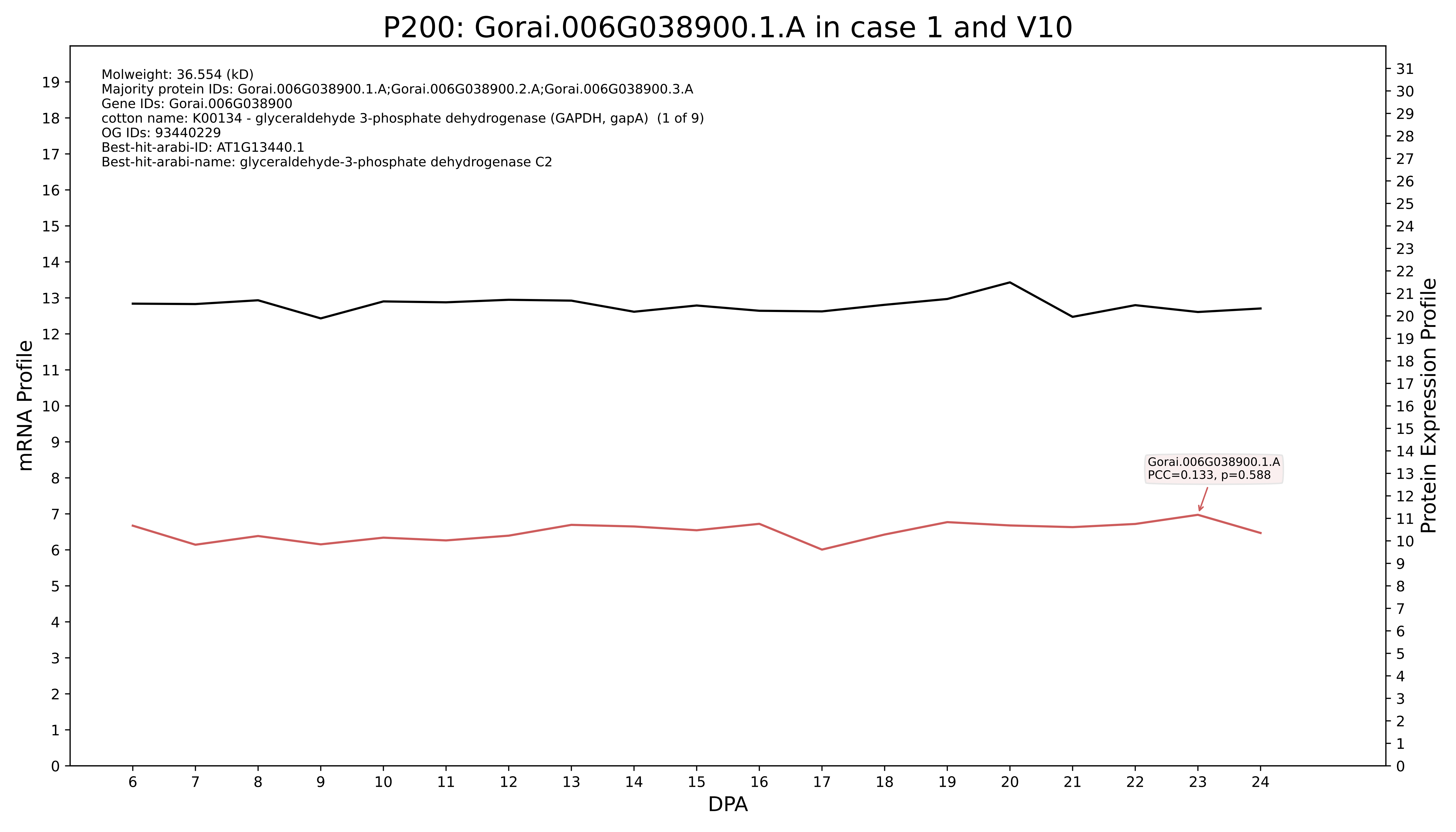
P200: Gorai.006G038900.1.A in case 1 and V10
Molweight: 36.554 (kD)
Majority protein IDs: Gorai.006G038900.1.A;Gorai.006G038900.2.A;Gorai.006G038900.3.A
Gene IDs: Gorai.006G038900
cotton name: K00134 - glyceraldehyde 3-phosphate dehydrogenase (GAPDH, gapA) (1 of 9)
OG IDs: 93440229
Best-hit-arabi-ID: AT1G13440.1
Best-hit-arabi-name: glyceraldehyde-3-phosphate dehydrogenase C2
Gorai.006G038900.1.A
PCC=0.133, p=0.588
mRNA Profile
Protein Expression Profile
DPA

### P200: Gorai.001G219200.1.D in case 1 and V10

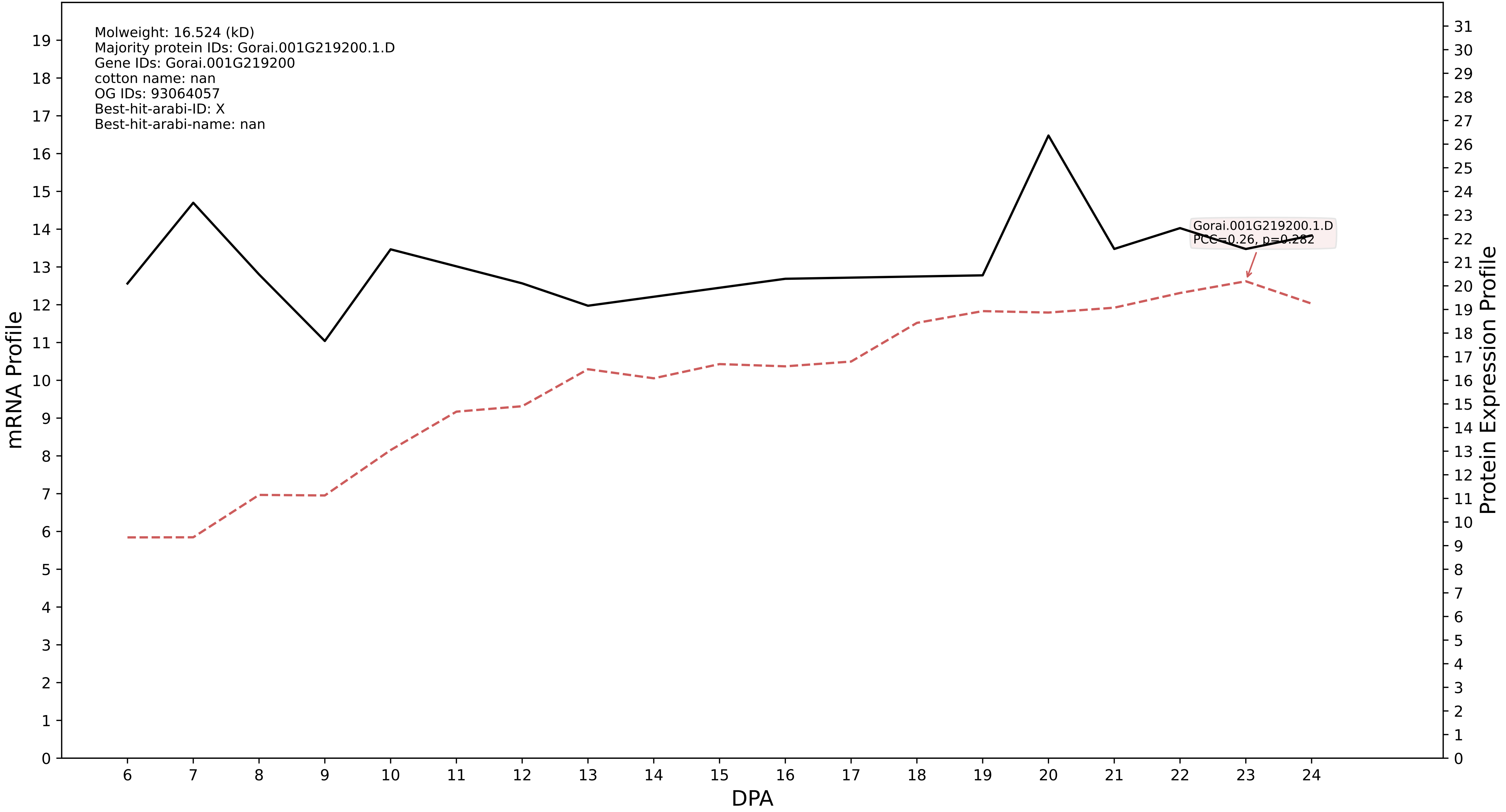

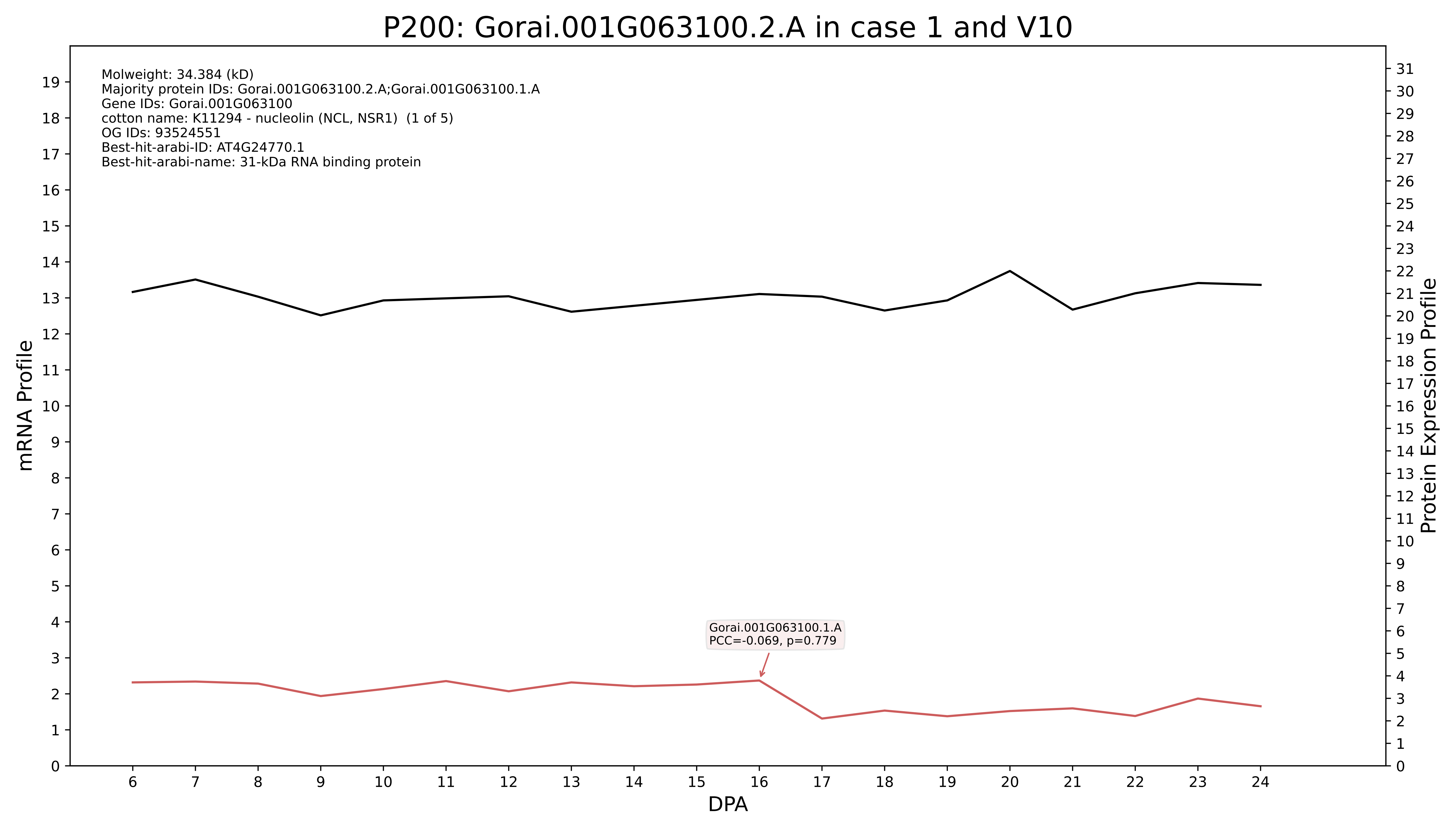
P200: Gorai.001G063100.2.A in case 1 and V10
Molweight: 34.384 (kD)
Majority protein IDs: Gorai.001G063100.2.A;Gorai.001G063100.1.A
Gene IDs: Gorai.001G063100
cotton name: K11294 - nucleolin (NCL, NSR1) (1 of 5)
OG IDs: 93524551
Best-hit-arabi-ID: AT4G24770.1
Best-hit-arabi-name: 31-kDa RNA binding protein
Gorai.001G063100.1.A
PCC=-0.069, p=0.779
mRNA Profile
Protein Expression Profile
DPA

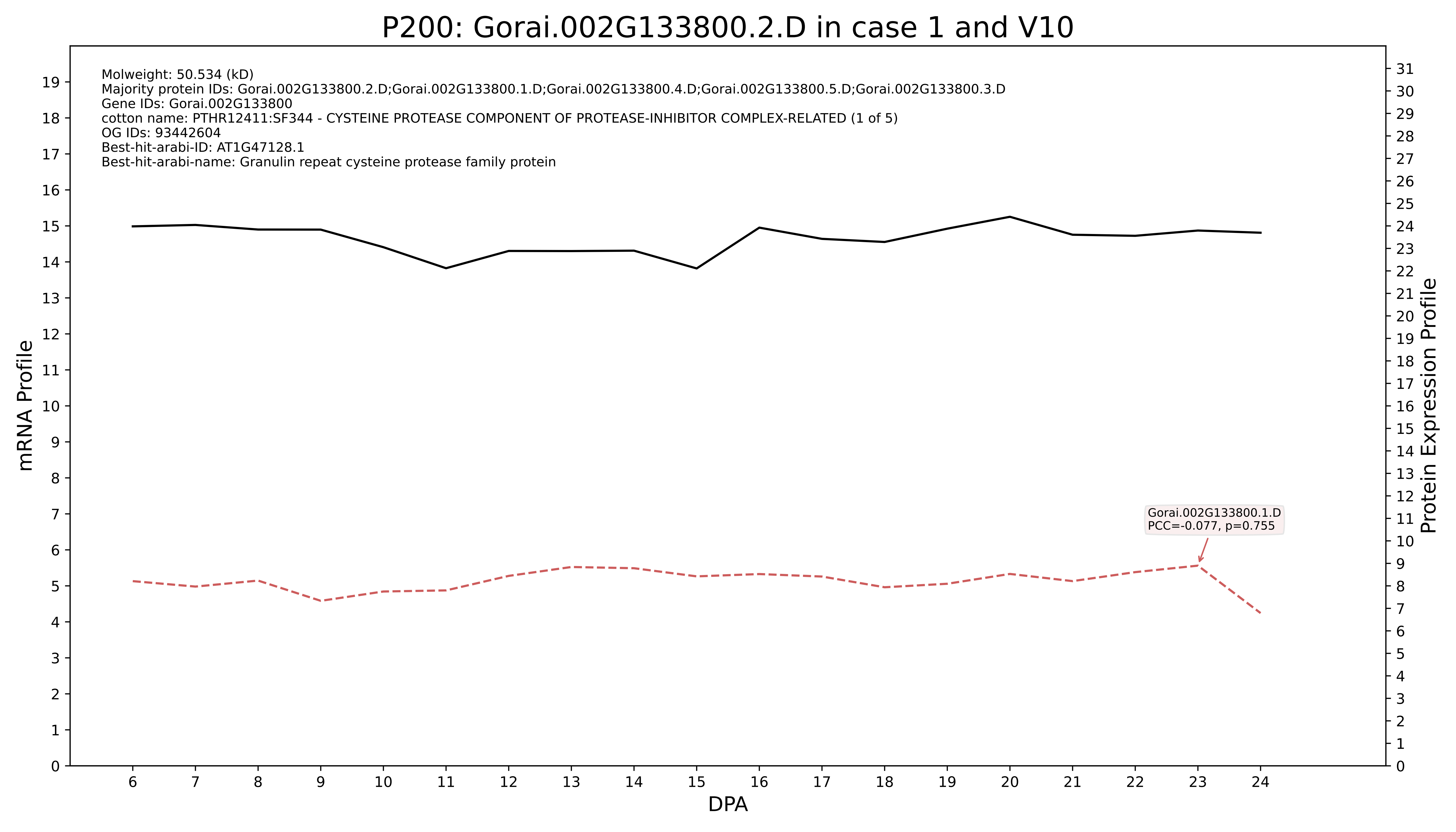

P200: Gorai.002G133800.2.D in case 1 and V10
Molweight: 50.534 (kD)
Majority protein IDs: Gorai.002G133800.2.D;Gorai.002G133800.1.D;Gorai.002G133800.4.D;Gorai.002G133800.5.D;Gorai.002G133800.3.D
Gene IDs: Gorai.002G133800
cotton name: PTHR12411:SF344 - CYSTEINE PROTEASE COMPONENT OF PROTEASE-INHIBITOR COMPLEX-RELATED (1 of 5)
OG IDs: 93442604
Best-hit-arabi-ID: AT1G47128.1
Best-hit-arabi-name: Granulin repeat cysteine protease family protein
Gorai.002G133800.1.D
PCC=-0.077, p=0.755
mRNA Profile
Protein Expression Profile
DPA

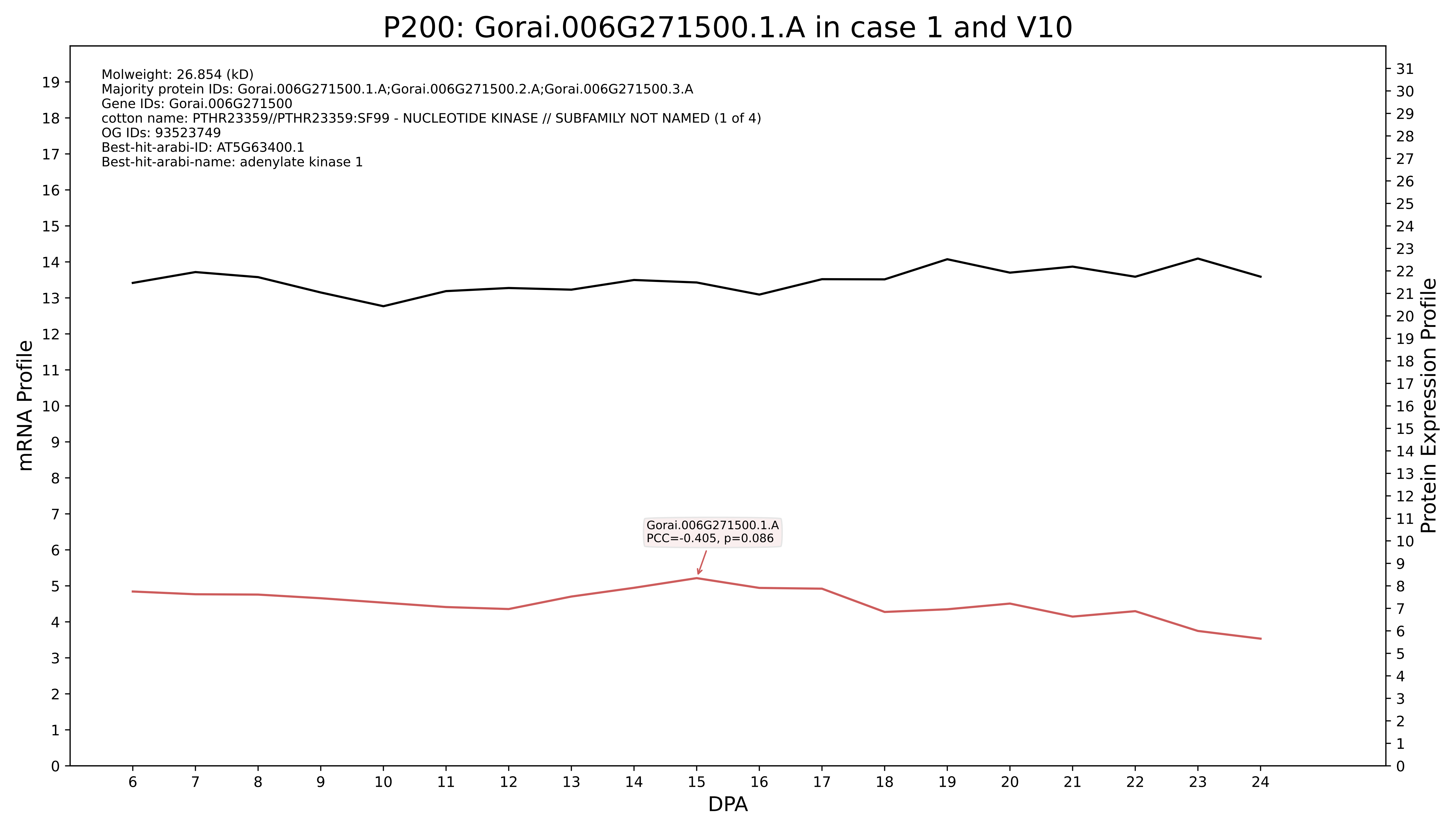
P200: Gorai.006G271500.1.A in case 1 and V10
Molweight: 26.854 (kD)
Majority protein IDs: Gorai.006G271500.1.A;Gorai.006G271500.2.A;Gorai.006G271500.3.A
Gene IDs: Gorai.006G271500
cotton name: PTHR23359//PTHR23359:SF99 - NUCLEOTIDE KINASE // SUBFAMILY NOT NAMED (1 of 4)
OG IDs: 93523749
Best-hit-arabi-ID: AT5G63400.1
Best-hit-arabi-name: adenylate kinase 1
Gorai.006G271500.1.A
PCC=-0.405, p=0.086
mRNA Profile
Protein Expression Profile
DPA

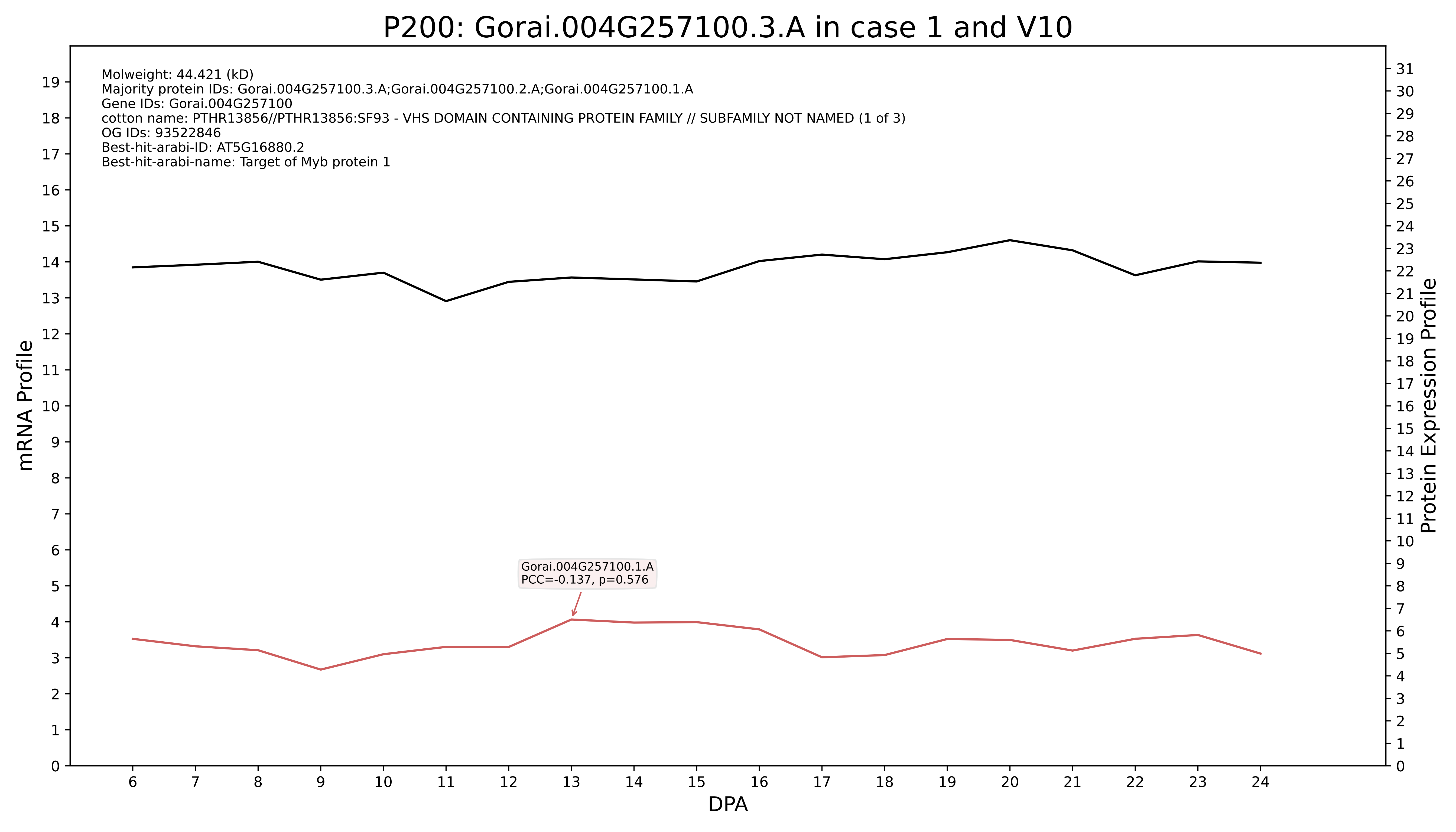
P200: Gorai.004G257100.3.A in case 1 and V10
Molweight: 44.421 (kD)
Majority protein IDs: Gorai.004G257100.3.A;Gorai.004G257100.2.A;Gorai.004G257100.1.A
Gene IDs: Gorai.004G257100
cotton name: PTHR13856//PTHR13856:SF93 - VHS DOMAIN CONTAINING PROTEIN FAMILY // SUBFAMILY NOT NAMED (1 of 3)
OG IDs: 93522846
Best-hit-arabi-ID: AT5G16880.2
Best-hit-arabi-name: Target of Myb protein 1
Gorai.004G257100.1.A
PCC=-0.137, p=0.576
mRNA Profile
Protein Expression Profile
DPA

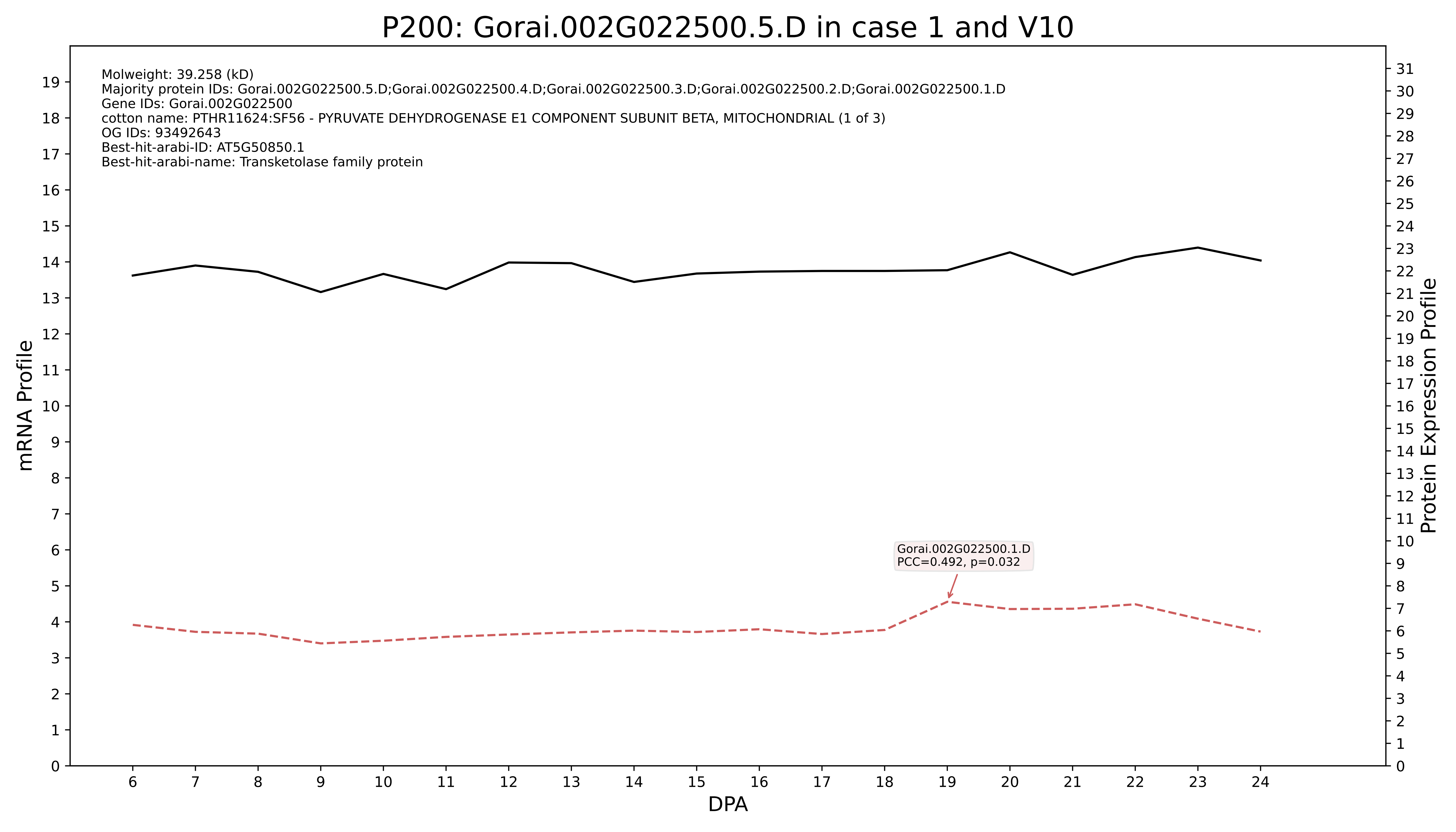

P200: Gorai.002G022500.5.D in case 1 and V10
Molweight: 39.258 (kD)
Majority protein IDs: Gorai.002G022500.5.D;Gorai.002G022500.4.D;Gorai.002G022500.3.D;Gorai.002G022500.2.D;Gorai.002G022500.1.D
Gene IDs: Gorai.002G022500
cotton name: PTHR11624:SF56 - PYRUVATE DEHYDROGENASE E1 COMPONENT SUBUNIT BETA, MITOCHONDRIAL (1 of 3)
OG IDs: 93492643
Best-hit-arabi-ID: AT5G50850.1
Best-hit-arabi-name: Transketolase family protein
Gorai.002G022500.1.D
PCC=0.492, p=0.032
mRNA Profile
Protein Expression Profile
DPA

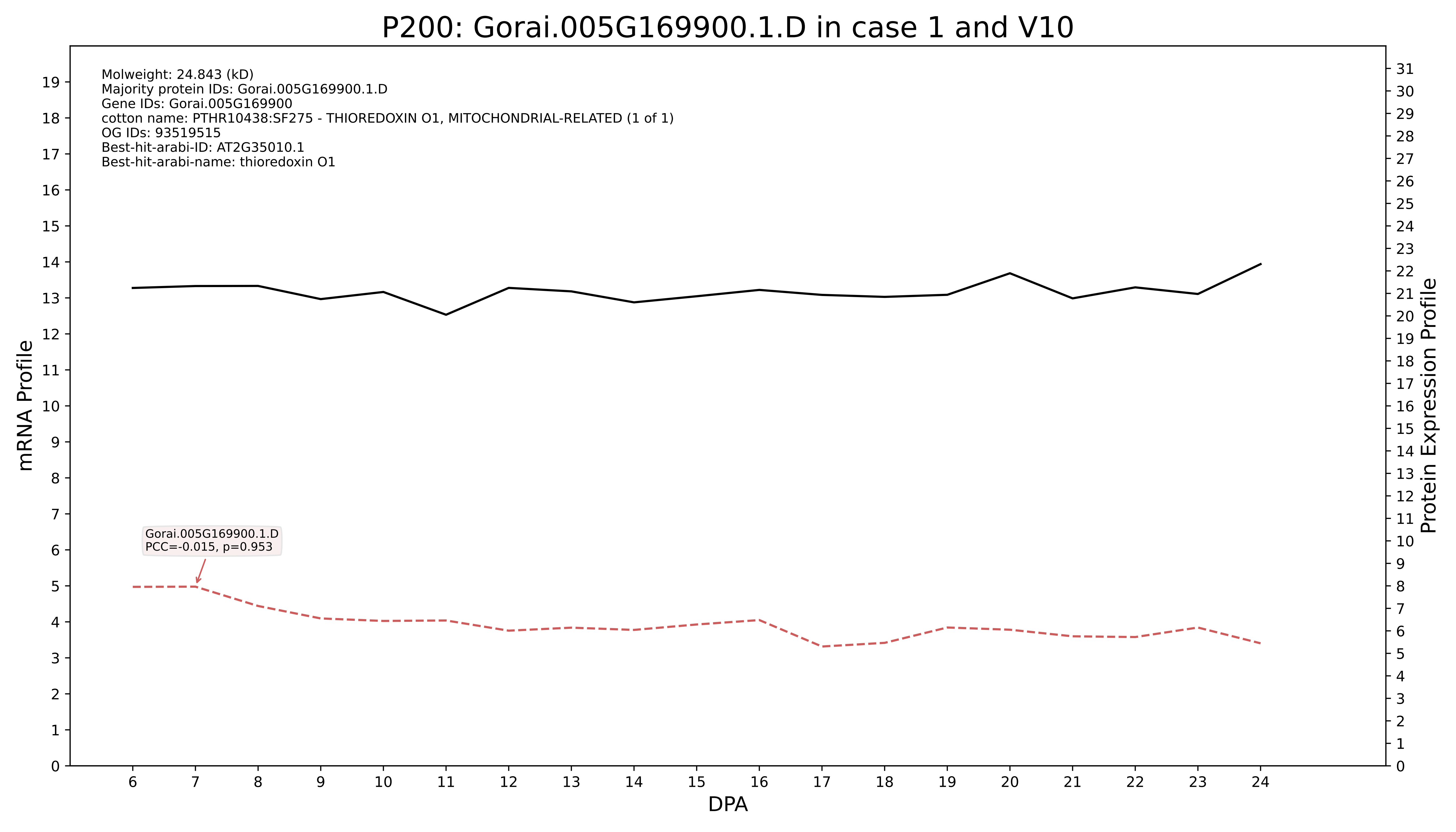
P200: Gorai.005G169900.1.D in case 1 and V10
Molweight: 24.843 (kD)
Majority protein IDs: Gorai.005G169900.1.D
Gene IDs: Gorai.005G169900
cotton name: PTHR10438:SF275 - THIOREDOXIN O1, MITOCHONDRIAL-RELATED (1 of 1)
OG IDs: 93519515
Best-hit-arabi-ID: AT2G35010.1
Best-hit-arabi-name: thioredoxin O1
Gorai.005G169900.1.D
PCC=-0.015, p=0.953
mRNA Profile
Protein Expression Profile
DPA

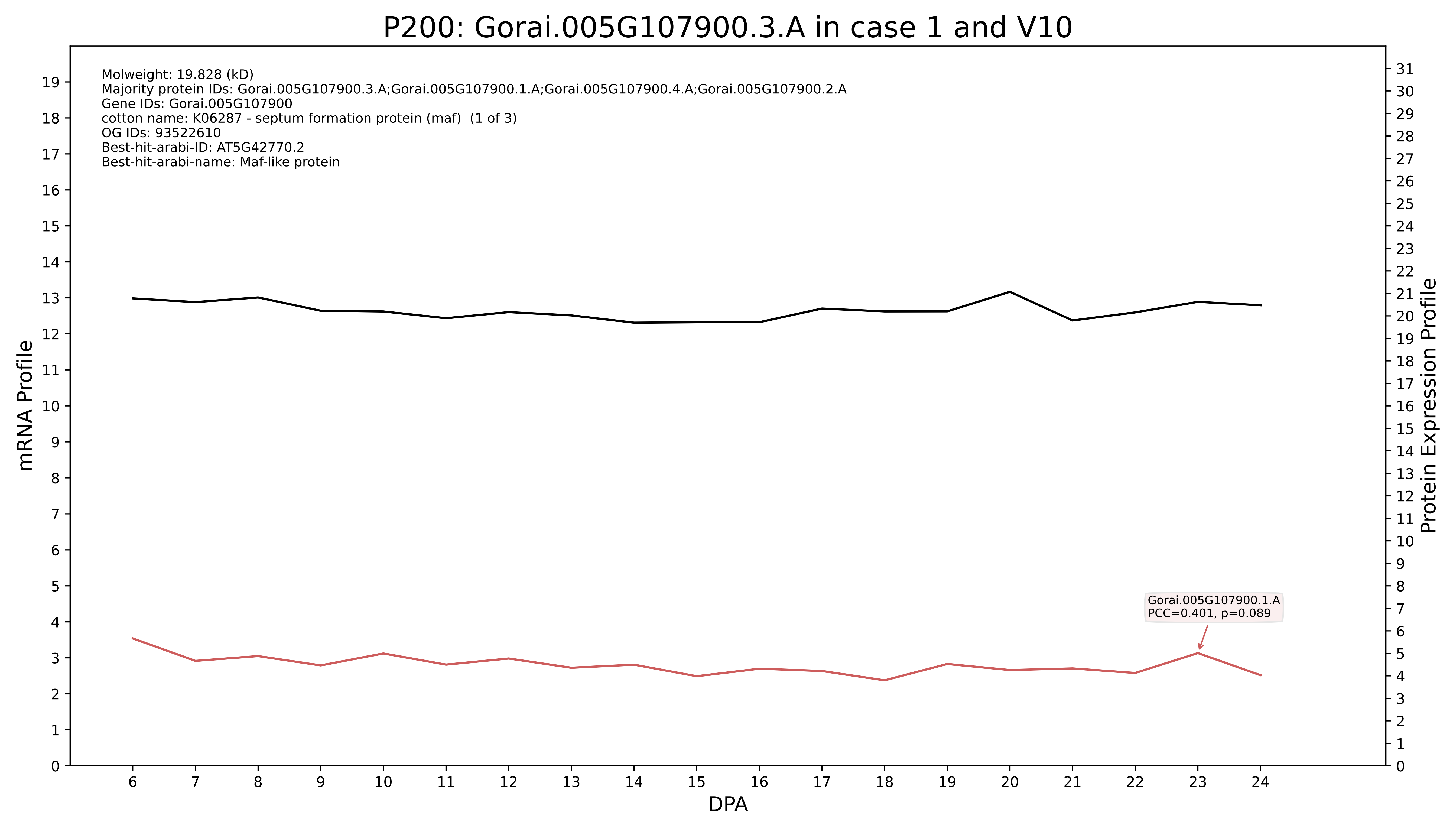

P200: Gorai.005G107900.3.A in case 1 and V10
Molweight: 19.828 (kD)
Majority protein IDs: Gorai.005G107900.3.A;Gorai.005G107900.1.A;Gorai.005G107900.4.A;Gorai.005G107900.2.A
Gene IDs: Gorai.005G107900
cotton name: K06287 - septum formation protein (maf) (1 of 3)
OG IDs: 93522610
Best-hit-arabi-ID: AT5G42770.2
Best-hit-arabi-name: Maf-like protein
Gorai.005G107900.1.A
PCC=0.401, p=0.089
mRNA Profile
Protein Expression Profile
DPA

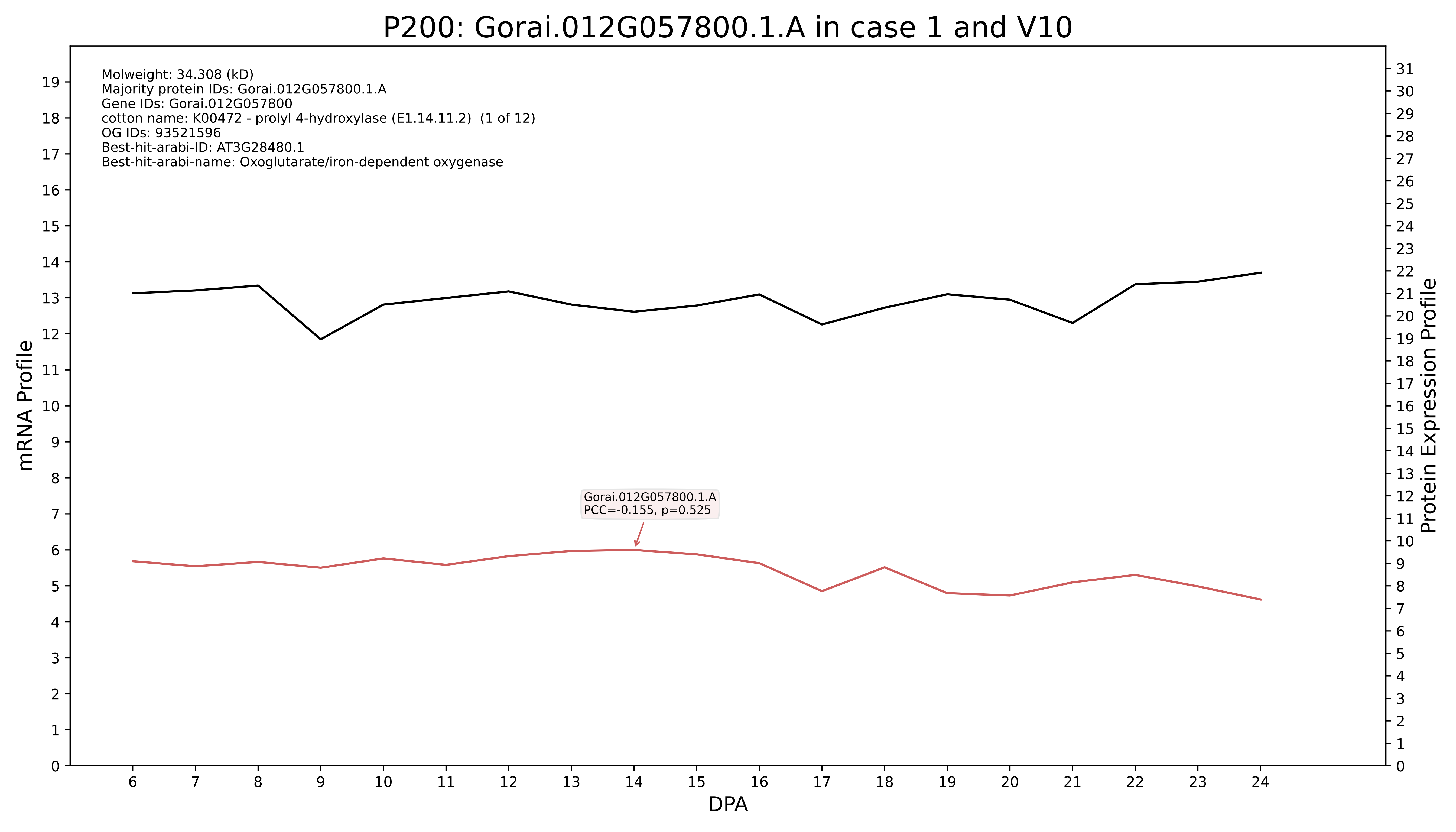
P200: Gorai.012G057800.1.A in case 1 and V10
Molweight: 34.308 (kD)
Majority protein IDs: Gorai.012G057800.1.A
Gene IDs: Gorai.012G057800
cotton name: K00472 - prolyl 4-hydroxylase (E1.14.11.2) (1 of 12)
OG IDs: 93521596
Best-hit-arabi-ID: AT3G28480.1
Best-hit-arabi-name: Oxoglutarate/iron-dependent oxygenase
Gorai.012G057800.1.A
PCC=-0.155, p=0.525
mRNA Profile
Protein Expression Profile
DPA

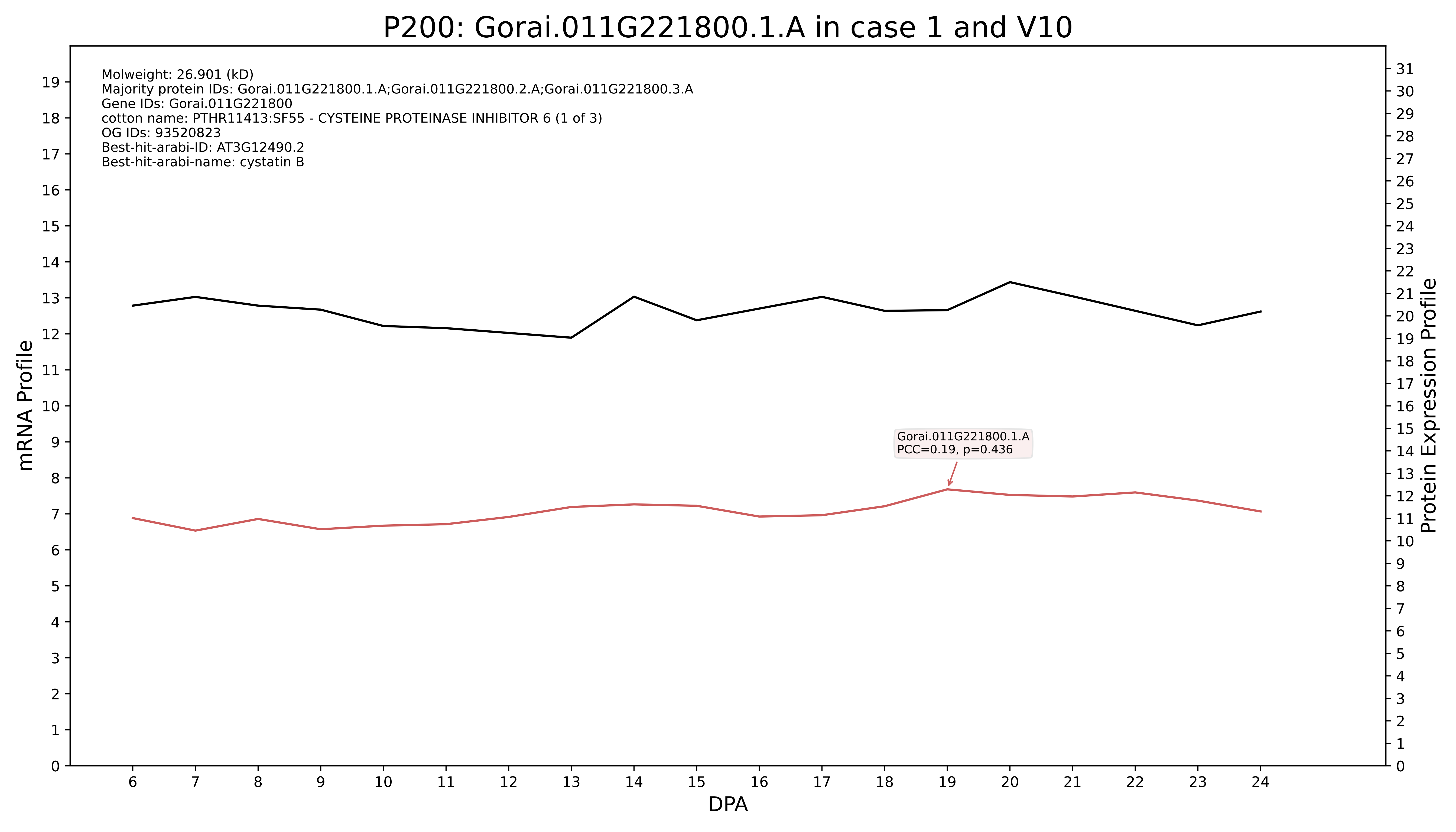
P200: Gorai.011G221800.1.A in case 1 and V10
Molweight: 26.901 (kD)
Majority protein IDs: Gorai.011G221800.1.A;Gorai.011G221800.2.A;Gorai.011G221800.3.A
Gene IDs: Gorai.011G221800
cotton name: PTHR11413:SF55 - CYSTEINE PROTEINASE INHIBITOR 6 (1 of 3)
OG IDs: 93520823
Best-hit-arabi-ID: AT3G12490.2
Best-hit-arabi-name: cystatin B
Gorai.011G221800.1.A
PCC=0.19, p=0.436
mRNA Profile
Protein Expression Profile
DPA

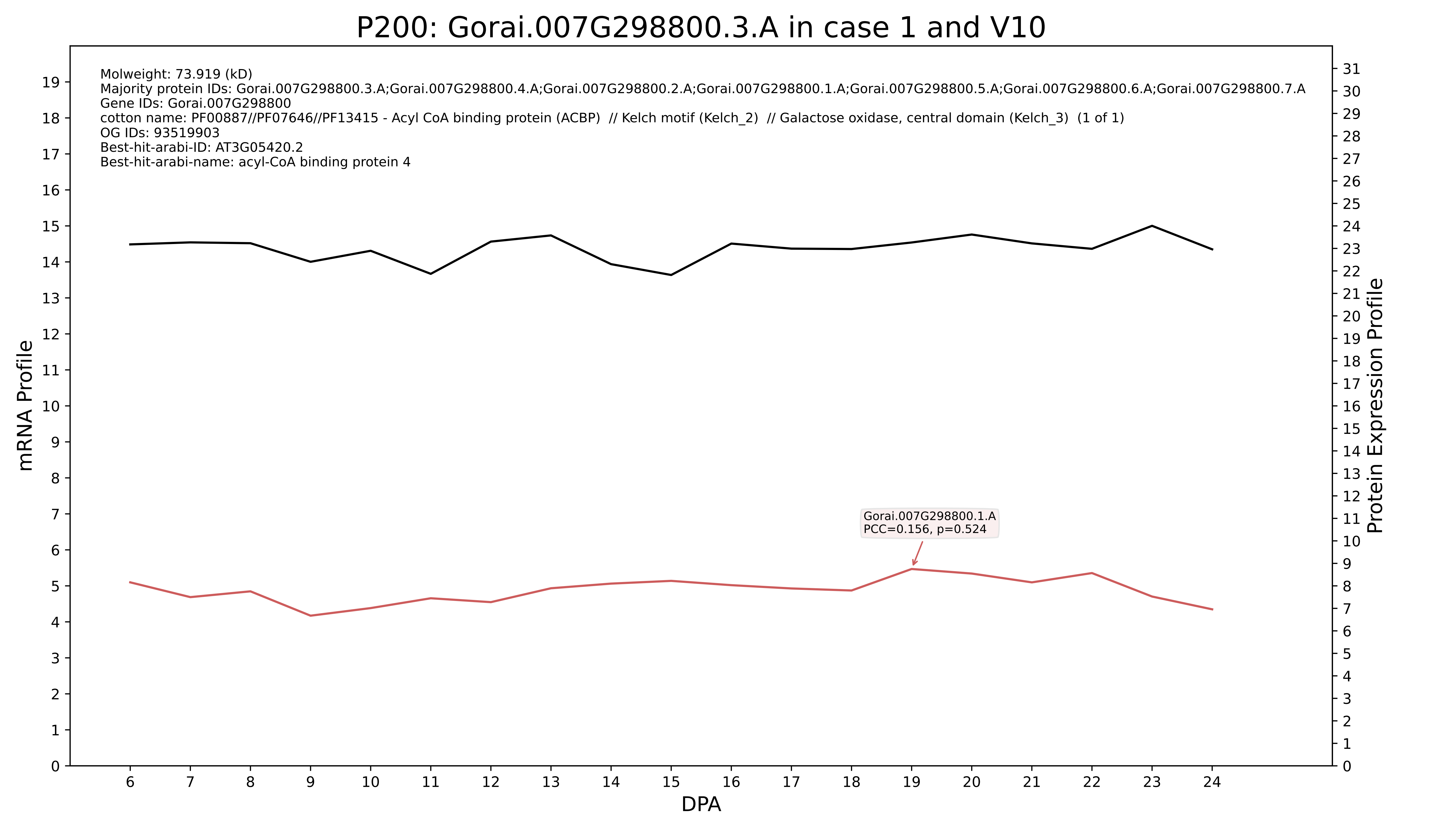
P200: Gorai.007G298800.3.A in case 1 and V10
Molweight: 73.919 (kD)
Majority protein IDs: Gorai.007G298800.3.A;Gorai.007G298800.4.A;Gorai.007G298800.2.A;Gorai.007G298800.1.A;Gorai.007G298800.5.A;Gorai.007G298800.6.A;Gorai.007G298800.7.A
Gene IDs: Gorai.007G298800
cotton name: PF00887//PF07646//PF13415 - Acyl CoA binding protein (ACBP) // Kelch motif (Kelch_2) // Galactose oxidase, central domain (Kelch_3) (1 of 1)
OG IDs: 93519903
Best-hit-arabi-ID: AT3G05420.2
Best-hit-arabi-name: acyl-CoA binding protein 4
Gorai.007G298800.1.A
PCC=0.156, p=0.524
mRNA Profile
Protein Expression Profile
DPA

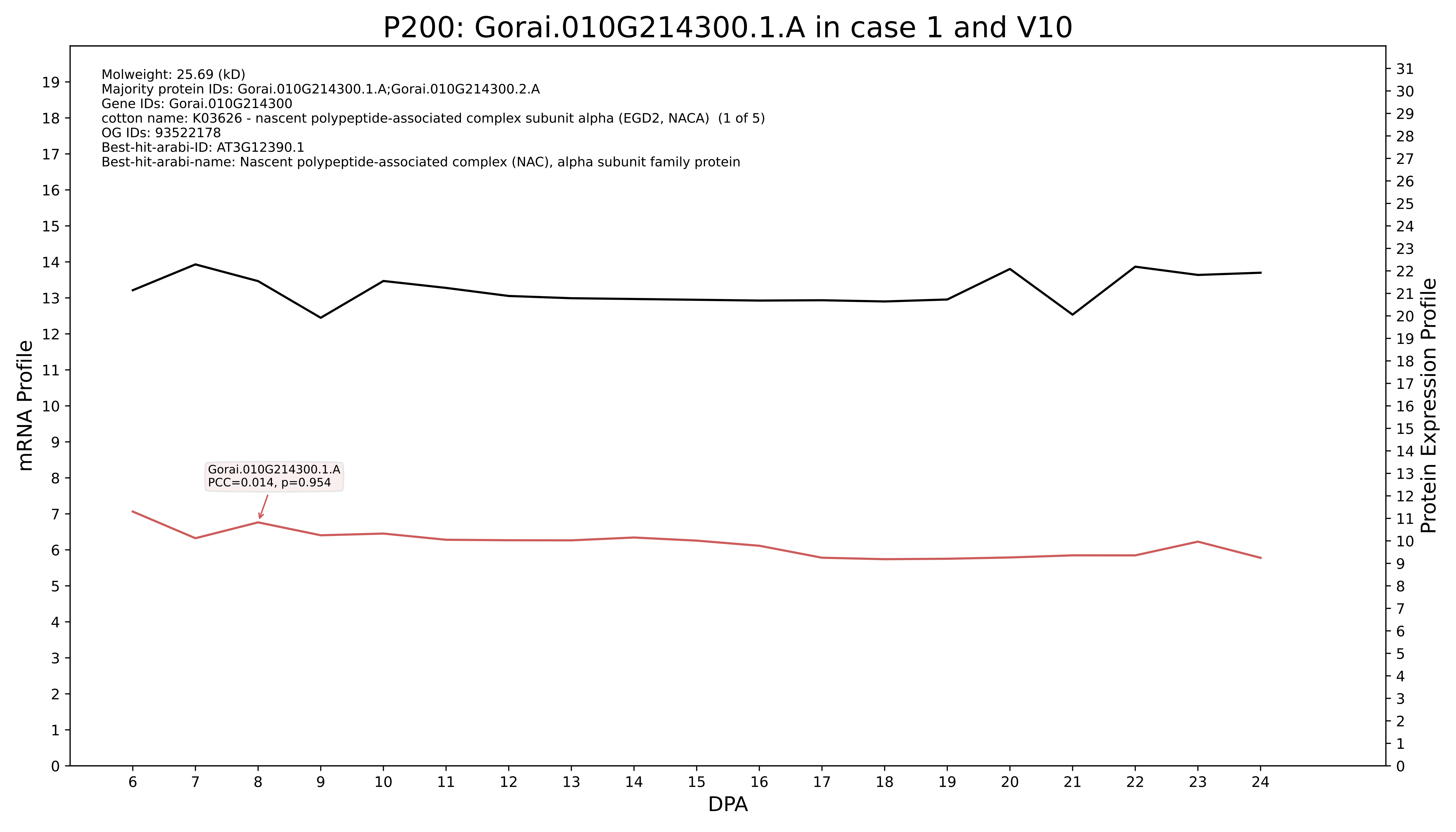
P200: Gorai.010G214300.1.A in case 1 and V10
Molweight: 25.69 (kD)
Majority protein IDs: Gorai.010G214300.1.A;Gorai.010G214300.2.A
Gene IDs: Gorai.010G214300
cotton name: K03626 - nascent polypeptide-associated complex subunit alpha (EGD2, NACA) (1 of 5)
OG IDs: 93522178
Best-hit-arabi-ID: AT3G12390.1
Best-hit-arabi-name: Nascent polypeptide-associated complex (NAC), alpha subunit family protein
Gorai.010G214300.1.A
PCC=0.014, p=0.954
mRNA Profile
Protein Expression Profile
DPA

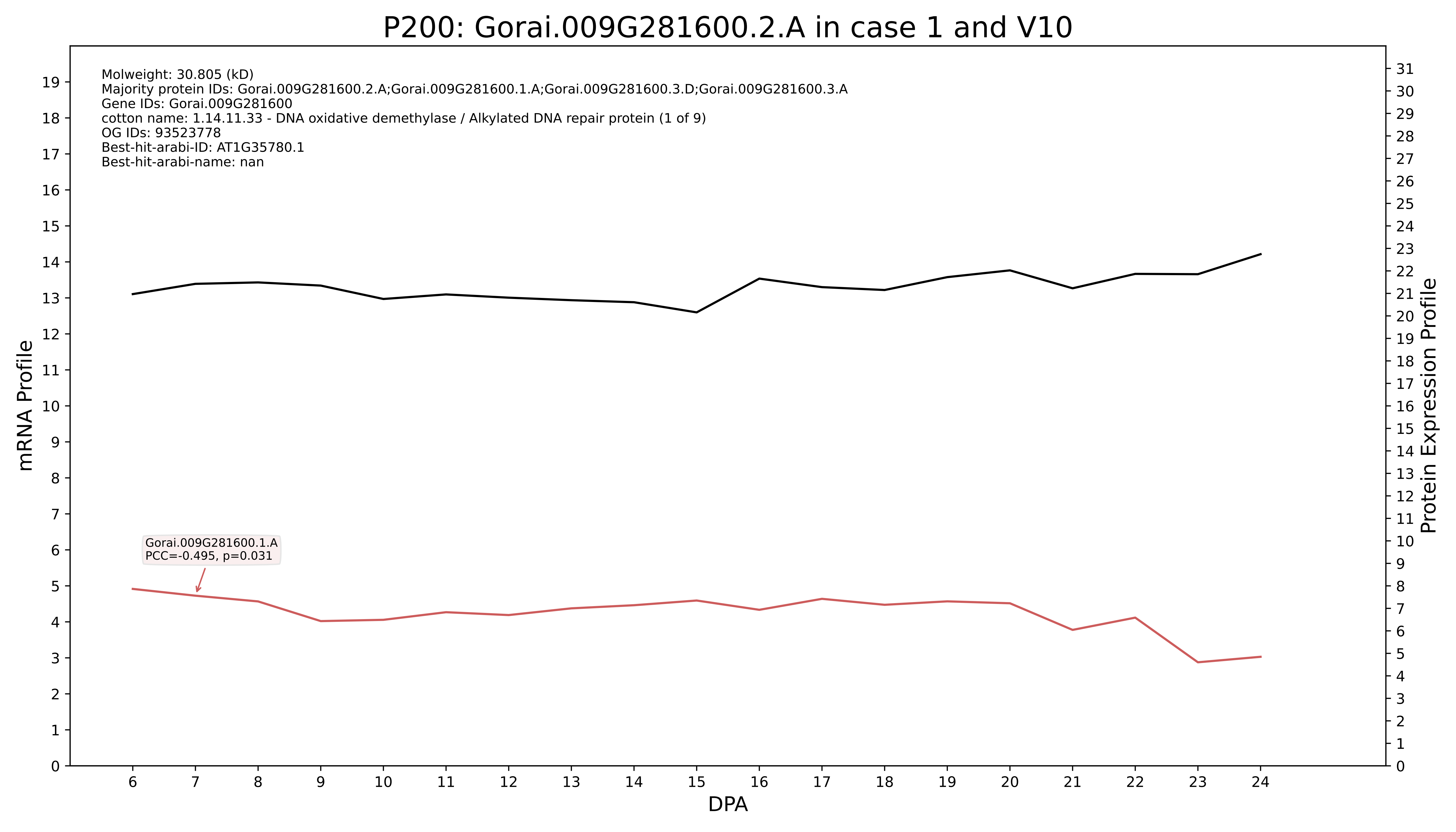
P200: Gorai.009G281600.2.A in case 1 and V10
Molweight: 30.805 (kD)
Majority protein IDs: Gorai.009G281600.2.A;Gorai.009G281600.1.A;Gorai.009G281600.3.D;Gorai.009G281600.3.A
Gene IDs: Gorai.009G281600
cotton name: 1.14.11.33 - DNA oxidative demethylase / Alkylated DNA repair protein (1 of 9)
OG IDs: 93523778
Best-hit-arabi-ID: AT1G35780.1
Best-hit-arabi-name: nan
Gorai.009G281600.1.A
PCC=-0.495, p=0.031
mRNA Profile
Protein Expression Profile
DPA

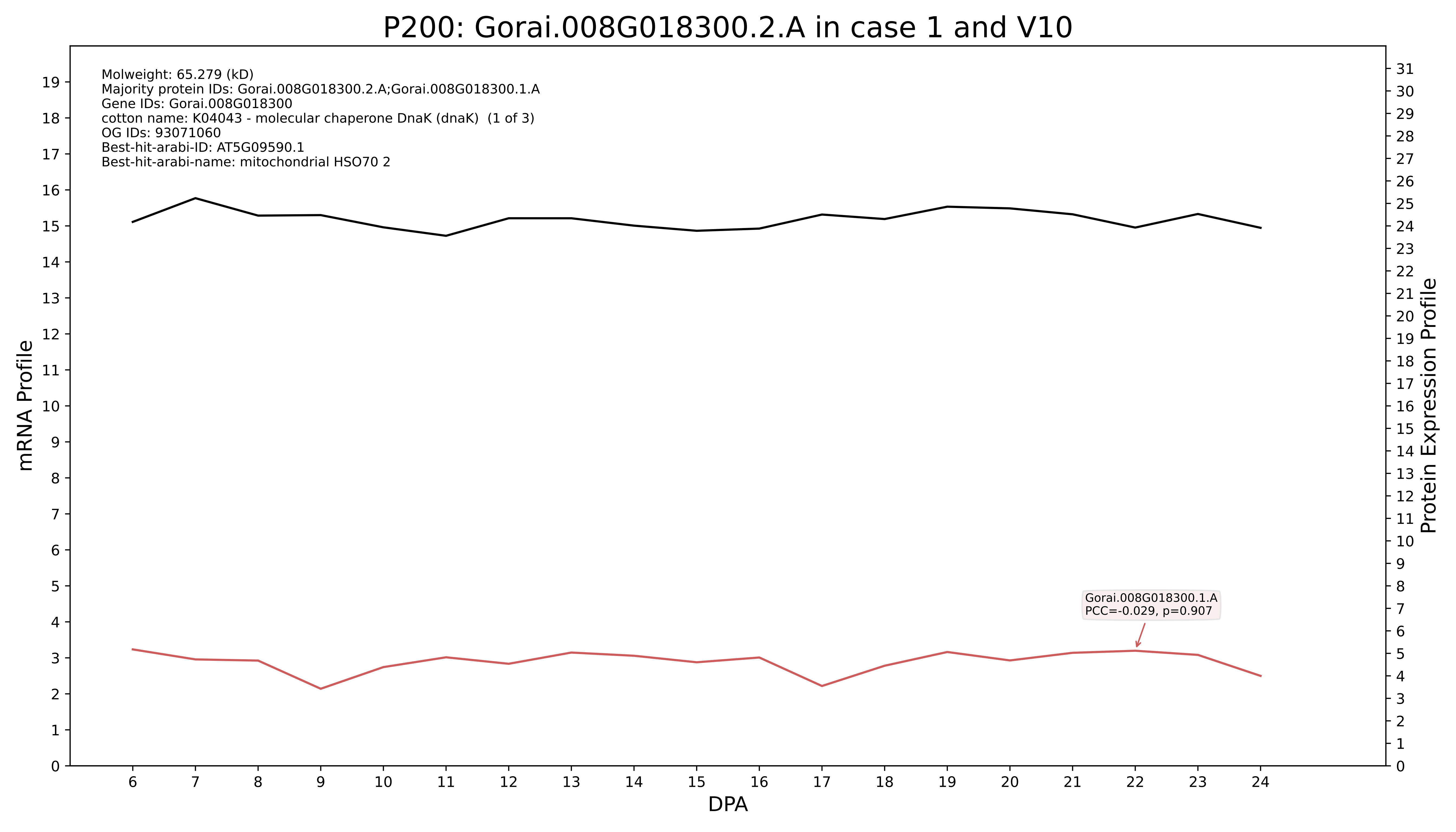
P200: Gorai.008G018300.2.A in case 1 and V10
Molweight: 65.279 (kD)
Majority protein IDs: Gorai.008G018300.2.A;Gorai.008G018300.1.A
Gene IDs: Gorai.008G018300
cotton name: K04043 - molecular chaperone DnaK (dnaK) (1 of 3)
OG IDs: 93071060
Best-hit-arabi-ID: AT5G09590.1
Best-hit-arabi-name: mitochondrial HSO70 2
Gorai.008G018300.1.A
PCC=-0.029, p=0.907
mRNA Profile
Protein Expression Profile
DPA

### P200: Gorai.013G206900.2.A in case 1 and V10

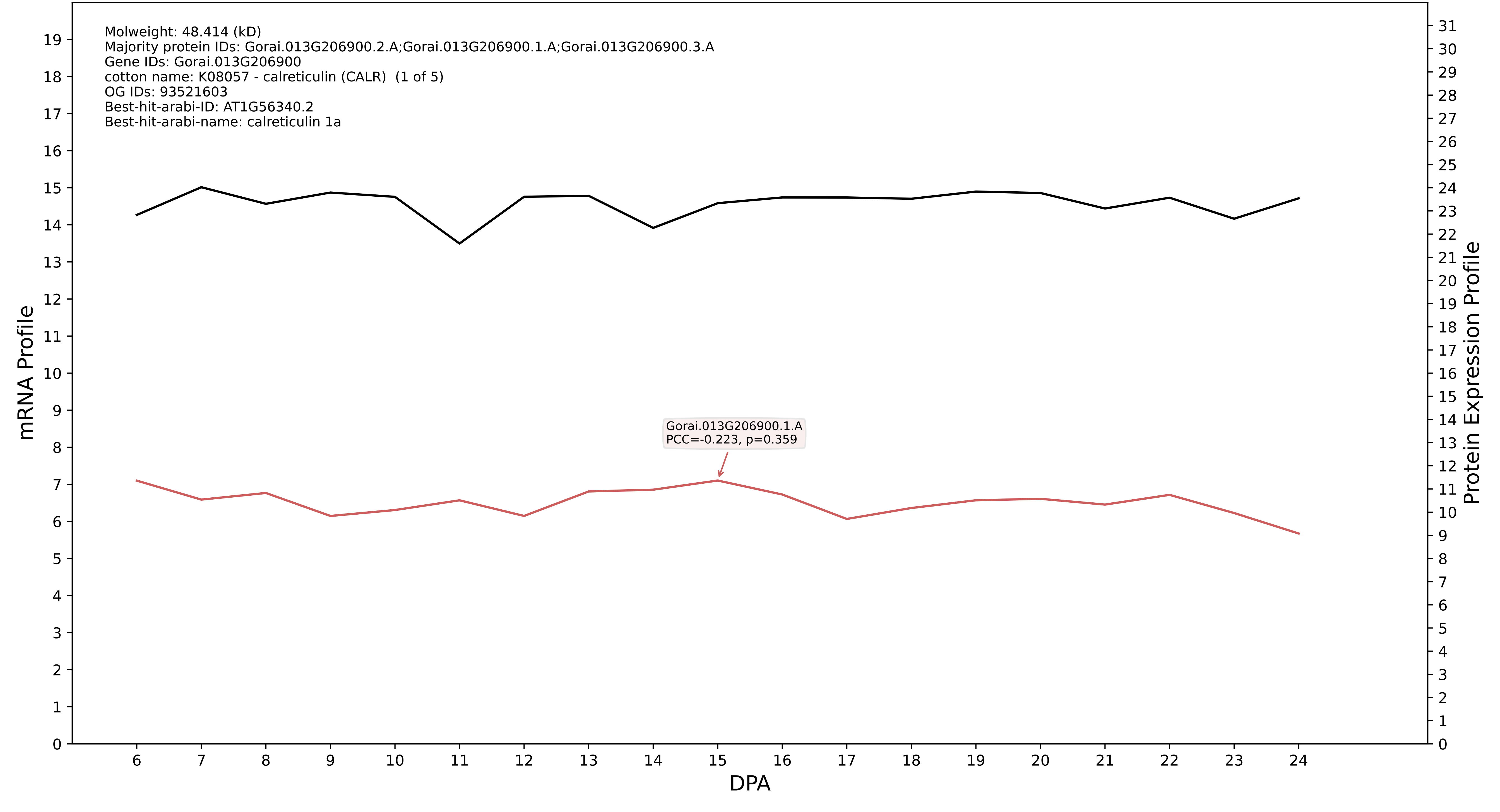

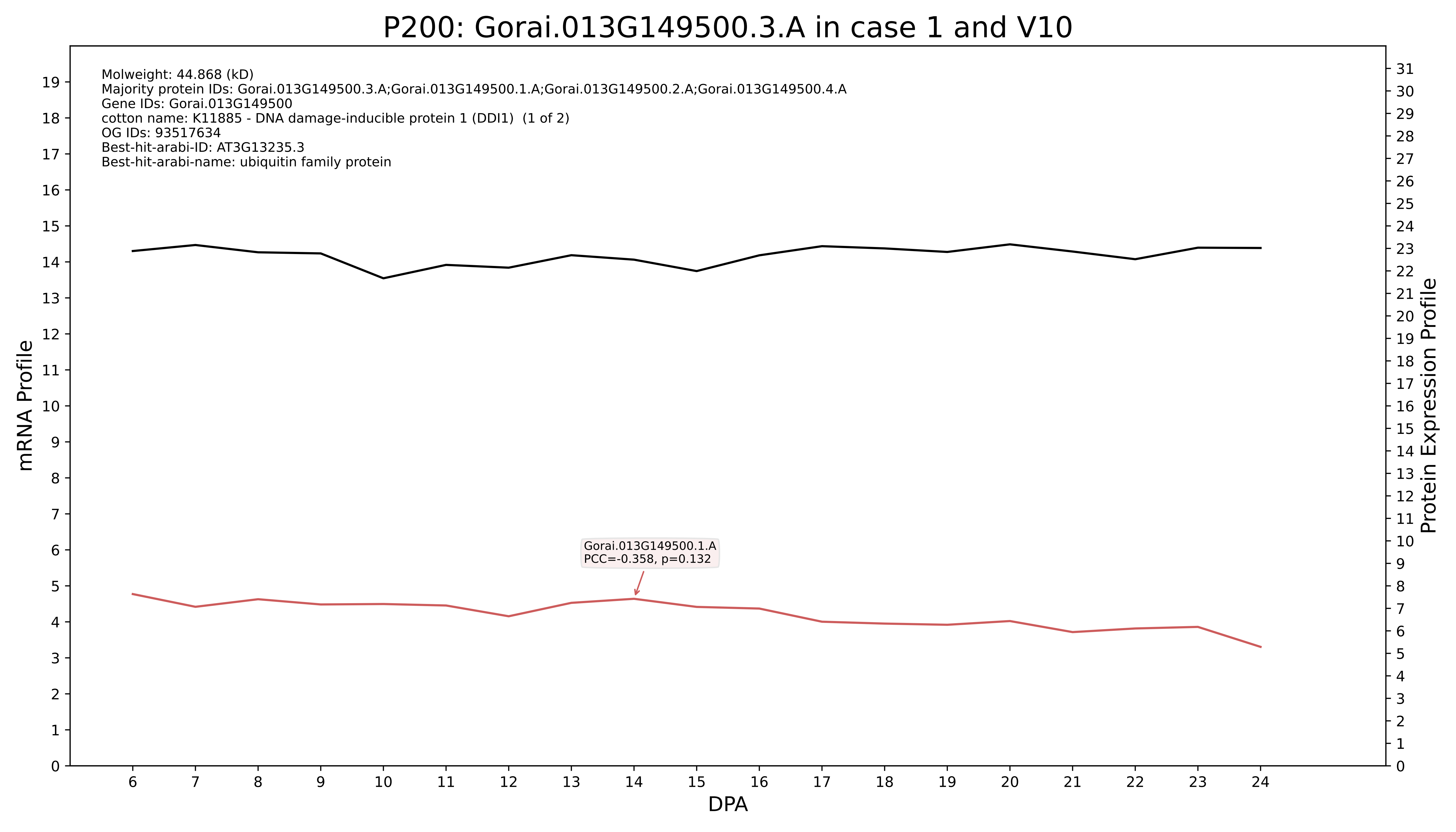

P200: Gorai.013G149500.3.A in case 1 and V10
Molweight: 44.868 (kD)
Majority protein IDs: Gorai.013G149500.3.A;Gorai.013G149500.1.A;Gorai.013G149500.2.A;Gorai.013G149500.4.A
Gene IDs: Gorai.013G149500
cotton name: K11885 - DNA damage-inducible protein 1 (DDI1) (1 of 2)
OG IDs: 93517634
Best-hit-arabi-ID: AT3G13235.3
Best-hit-arabi-name: ubiquitin family protein
Gorai.013G149500.1.A
PCC=-0.358, p=0.132
mRNA Profile
Protein Expression Profile
DPA

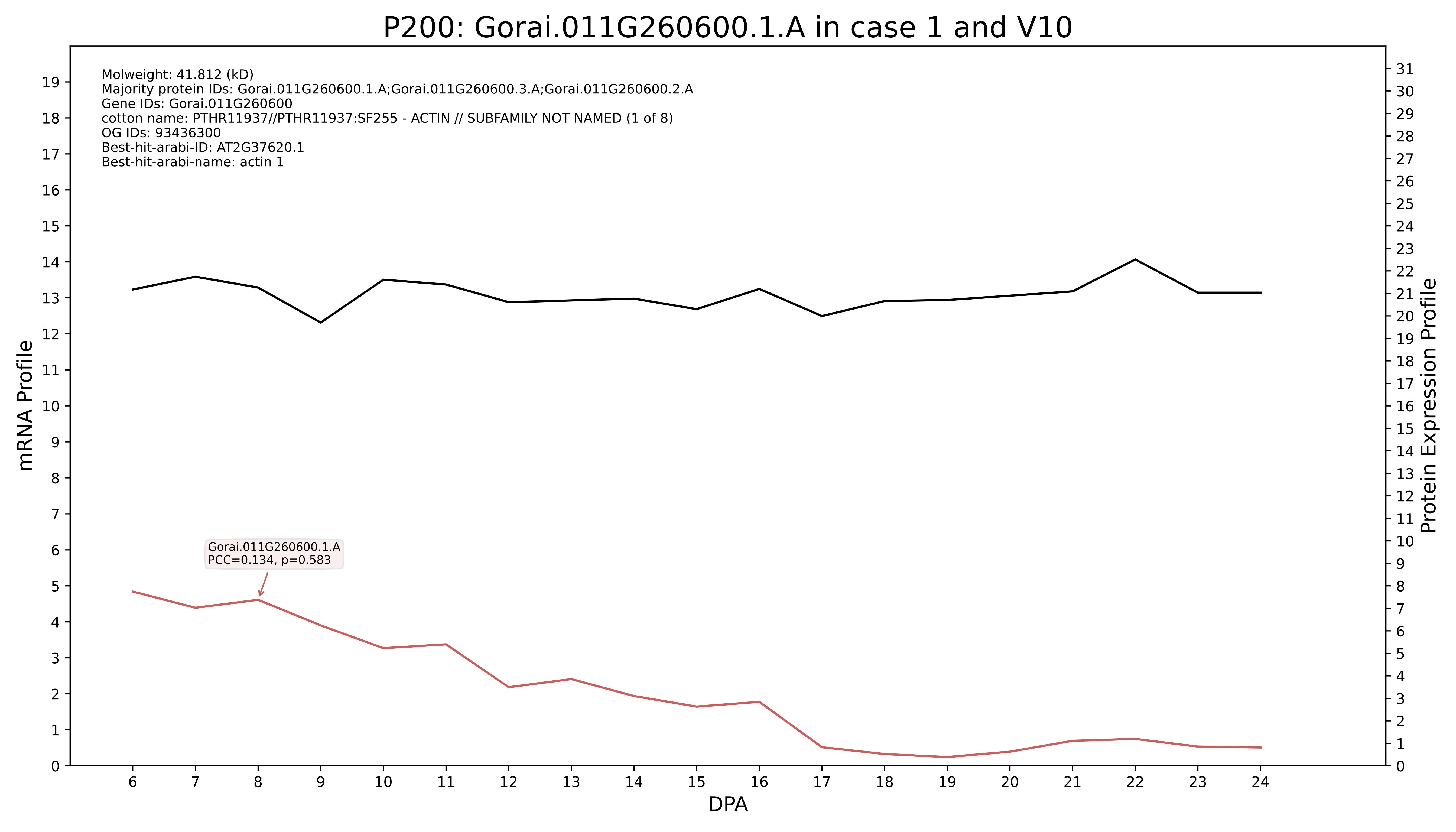
P200: Gorai.011G260600.1.A in case 1 and V10
Molweight: 41.812 (kD)
Majority protein IDs: Gorai.011G260600.1.A;Gorai.011G260600.3.A;Gorai.011G260600.2.A
Gene IDs: Gorai.011G260600
cotton name: PTHR11937//PTHR11937:SF255 - ACTIN // SUBFAMILY NOT NAMED (1 of 8)
OG IDs: 93436300
Best-hit-arabi-ID: AT2G37620.1
Best-hit-arabi-name: actin 1
Gorai.011G260600.1.A
PCC=0.134, p=0.583
mRNA Profile
Protein Expression Profile
DPA

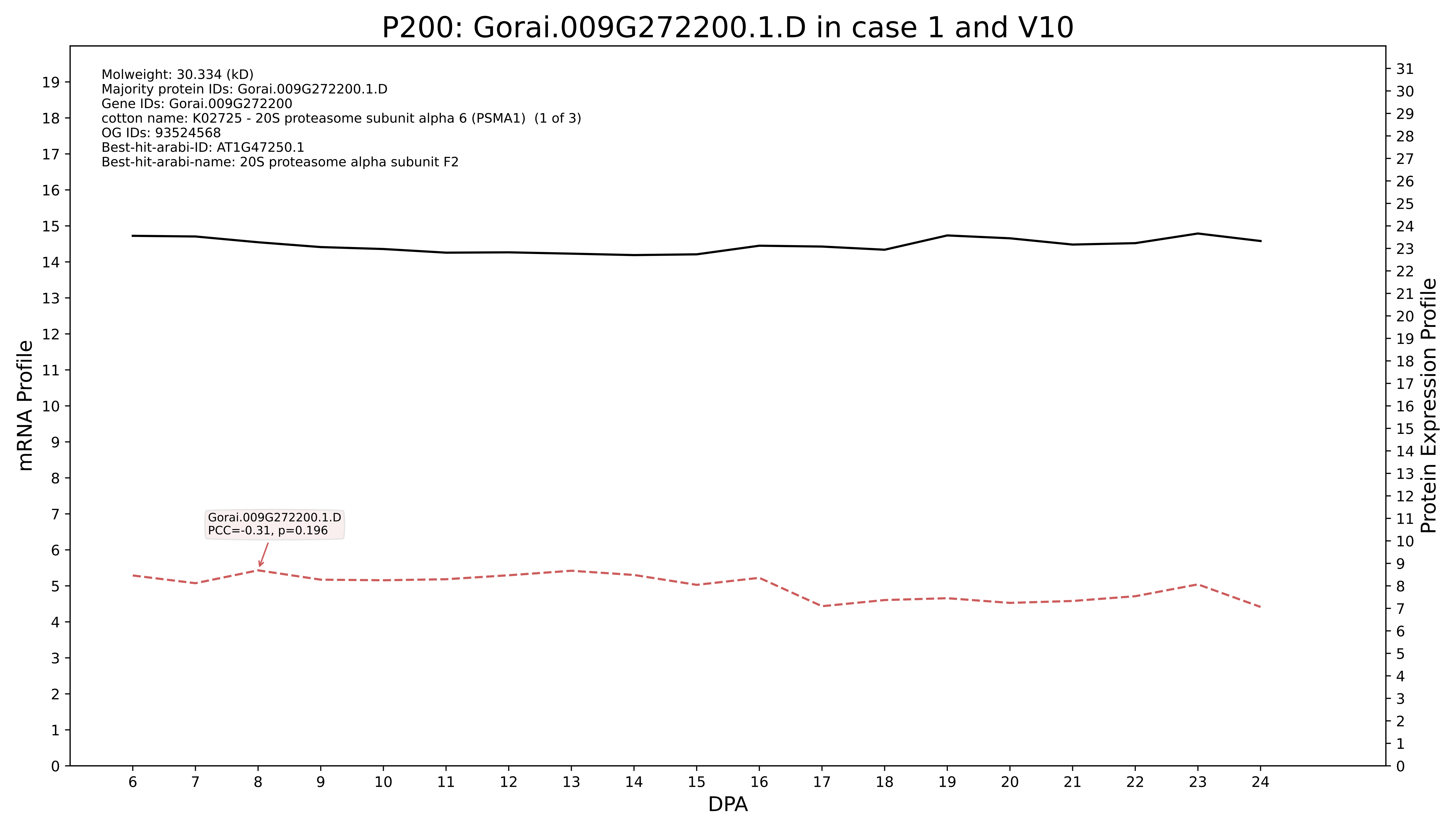
P200: Gorai.009G272200.1.D in case 1 and V10
Molweight: 30.334 (kD)
Majority protein IDs: Gorai.009G272200.1.D
Gene IDs: Gorai.009G272200
cotton name: K02725 - 20S proteasome subunit alpha 6 (PSMA1) (1 of 3)
OG IDs: 93524568
Best-hit-arabi-ID: AT1G47250.1
Best-hit-arabi-name: 20S proteasome alpha subunit F2
Gorai.009G272200.1.D
PCC=-0.31, p=0.196
mRNA Profile
Protein Expression Profile
DPA

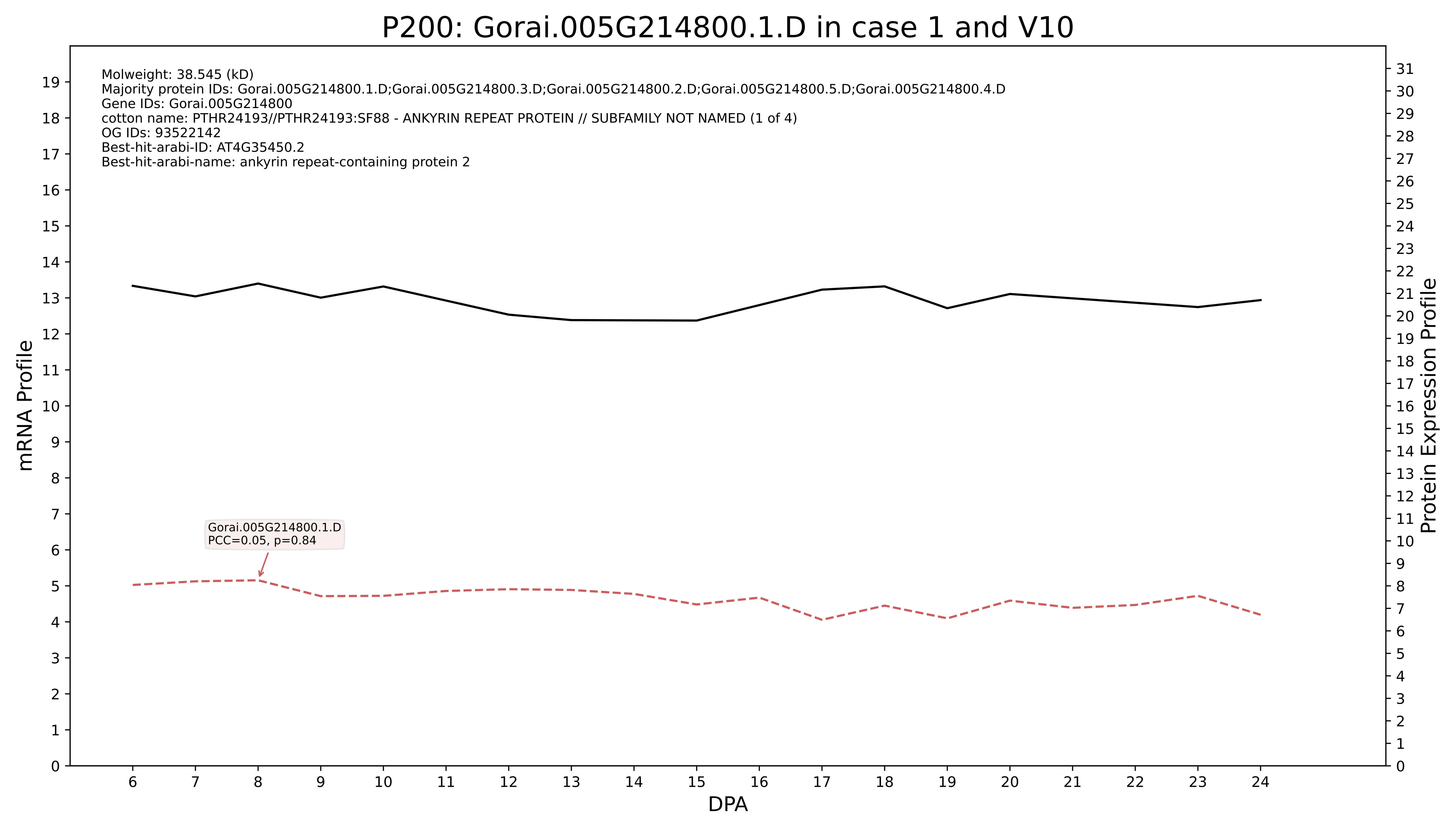
P200: Gorai.005G214800.1.D in case 1 and V10
Molweight: 38.545 (kD)
Majority protein IDs: Gorai.005G214800.1.D;Gorai.005G214800.3.D;Gorai.005G214800.2.D;Gorai.005G214800.5.D;Gorai.005G214800.4.D
Gene IDs: Gorai.005G214800
cotton name: PTHR24193//PTHR24193:SF88 - ANKYRIN REPEAT PROTEIN // SUBFAMILY NOT NAMED (1 of 4)
OG IDs: 93522142
Best-hit-arabi-ID: AT4G35450.2
Best-hit-arabi-name: ankyrin repeat-containing protein 2
Gorai.005G214800.1.D
PCC=0.05, p=0.84
mRNA Profile
Protein Expression Profile
DPA

### P200: Gorai.009G310800.1.D in case 1 and V10

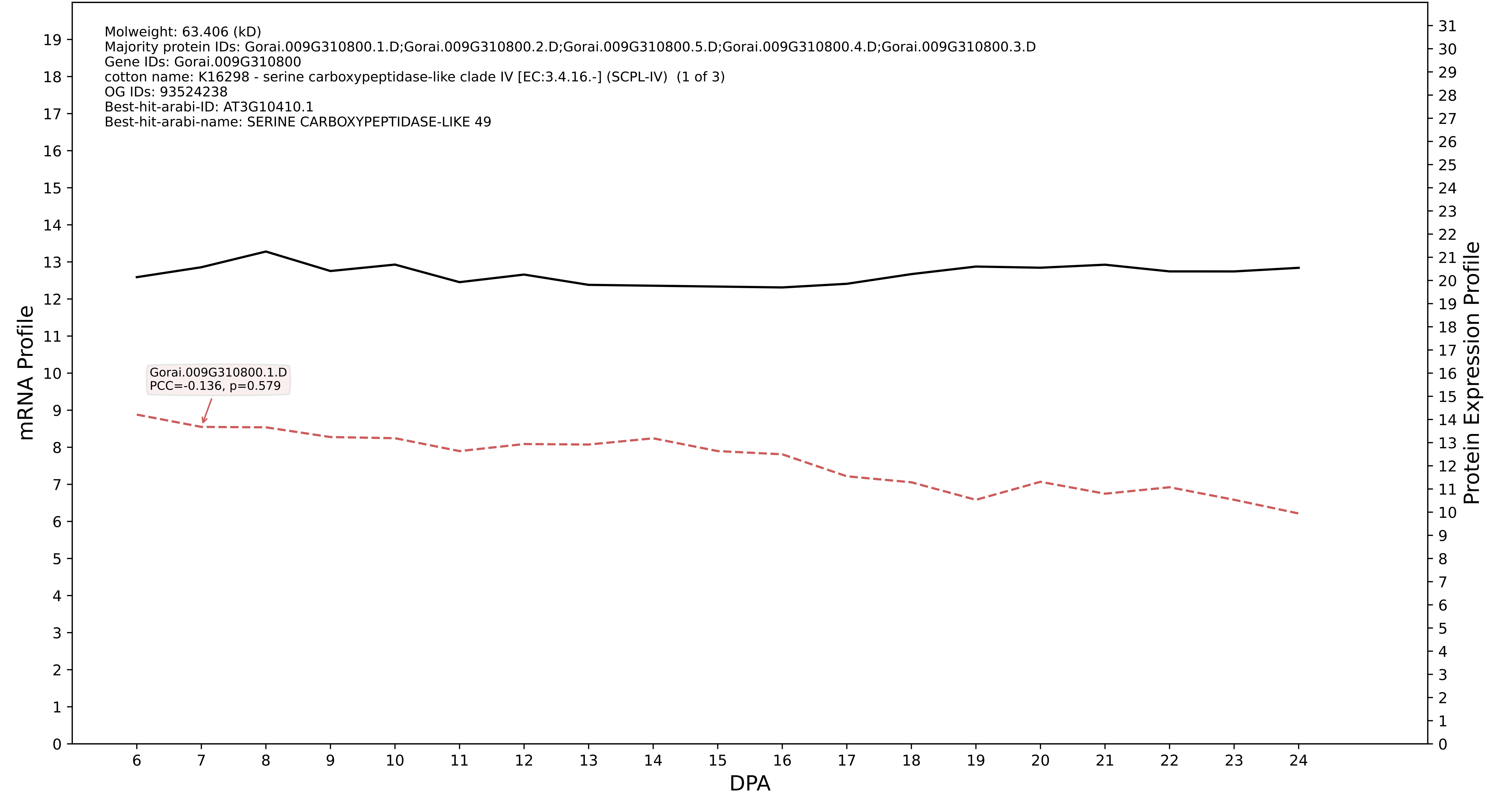

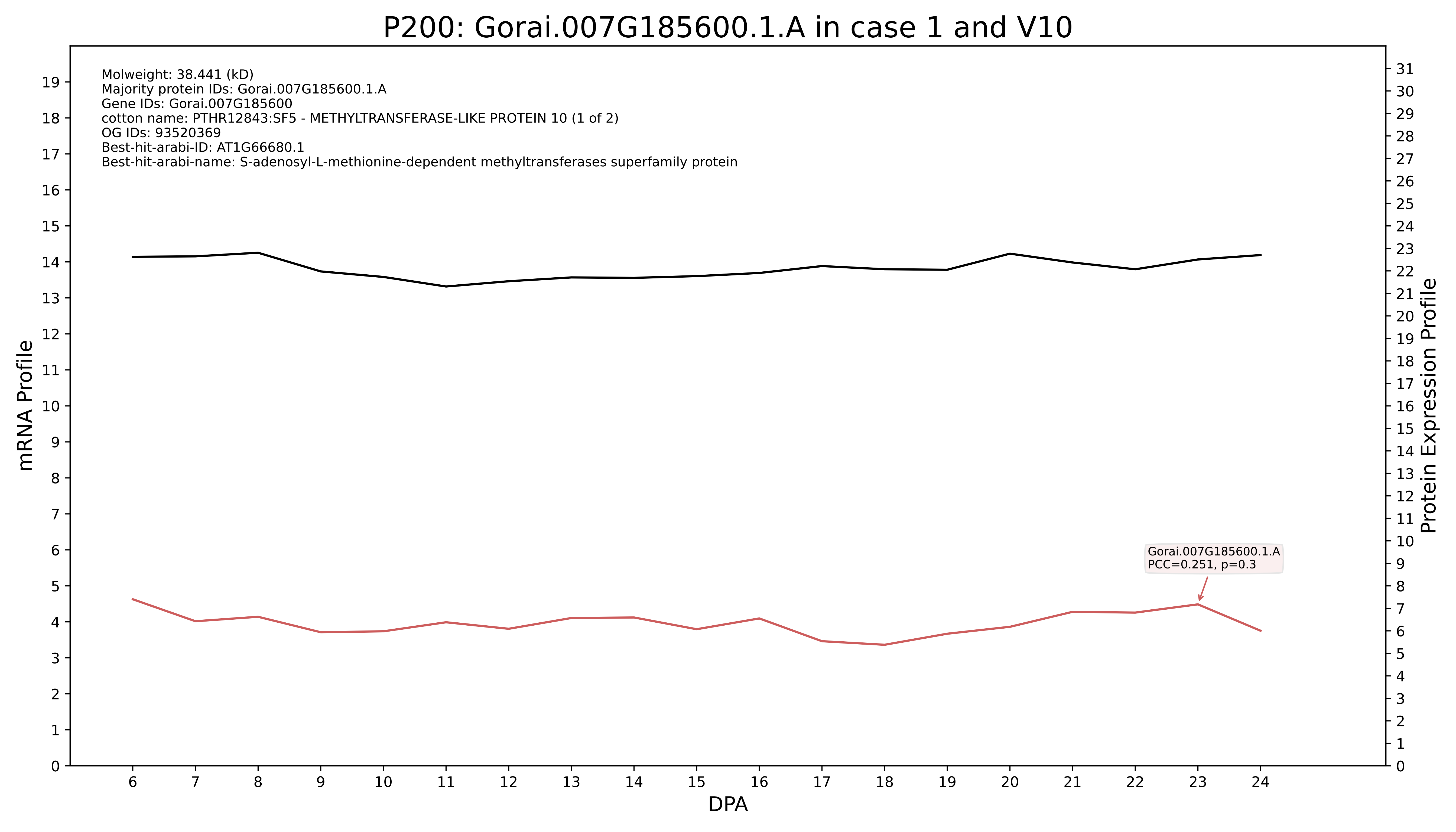
P200: Gorai.007G185600.1.A in case 1 and V10
Molweight: 38.441 (kD)
Majority protein IDs: Gorai.007G185600.1.A
Gene IDs: Gorai.007G185600
cotton name: PTHR12843:SF5 - METHYLTRANSFERASE-LIKE PROTEIN 10 (1 of 2)
OG IDs: 93520369
Best-hit-arabi-ID: AT1G66680.1
Best-hit-arabi-name: S-adenosyl-L-methionine-dependent methyltransferases superfamily protein
Gorai.007G185600.1.A
PCC=0.251, p=0.3
mRNA Profile
Protein Expression Profile
DPA

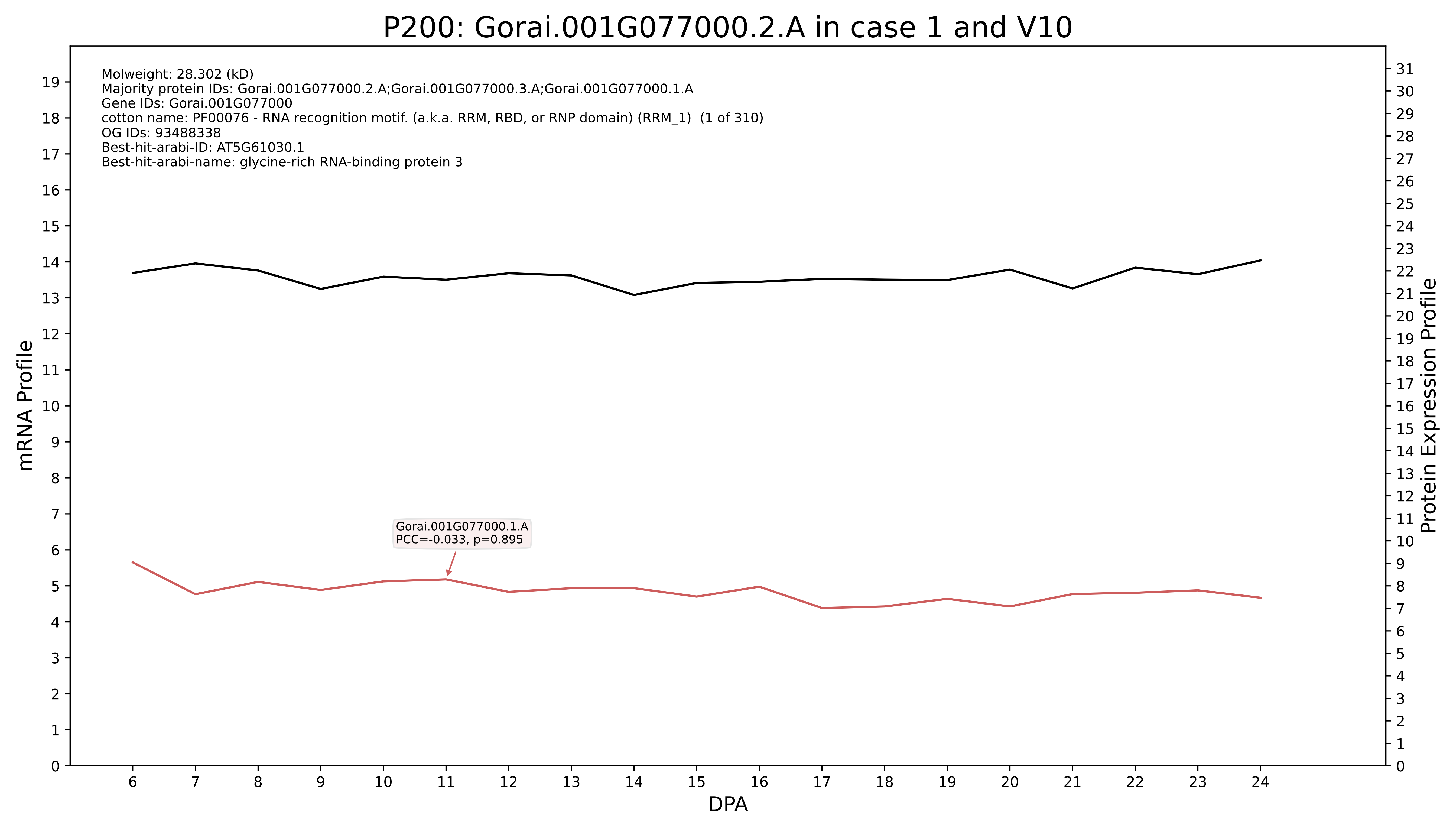
P200: Gorai.001G077000.2.A in case 1 and V10
Molweight: 28.302 (kD)
Majority protein IDs: Gorai.001G077000.2.A;Gorai.001G077000.3.A;Gorai.001G077000.1.A
Gene IDs: Gorai.001G077000
cotton name: PF00076 - RNA recognition motif. (a.k.a. RRM, RBD, or RNP domain) (RRM_1) (1 of 310)
OG IDs: 93488338
Best-hit-arabi-ID: AT5G61030.1
Best-hit-arabi-name: glycine-rich RNA-binding protein 3
Gorai.001G077000.1.A
PCC=-0.033, p=0.895
mRNA Profile
Protein Expression Profile
DPA

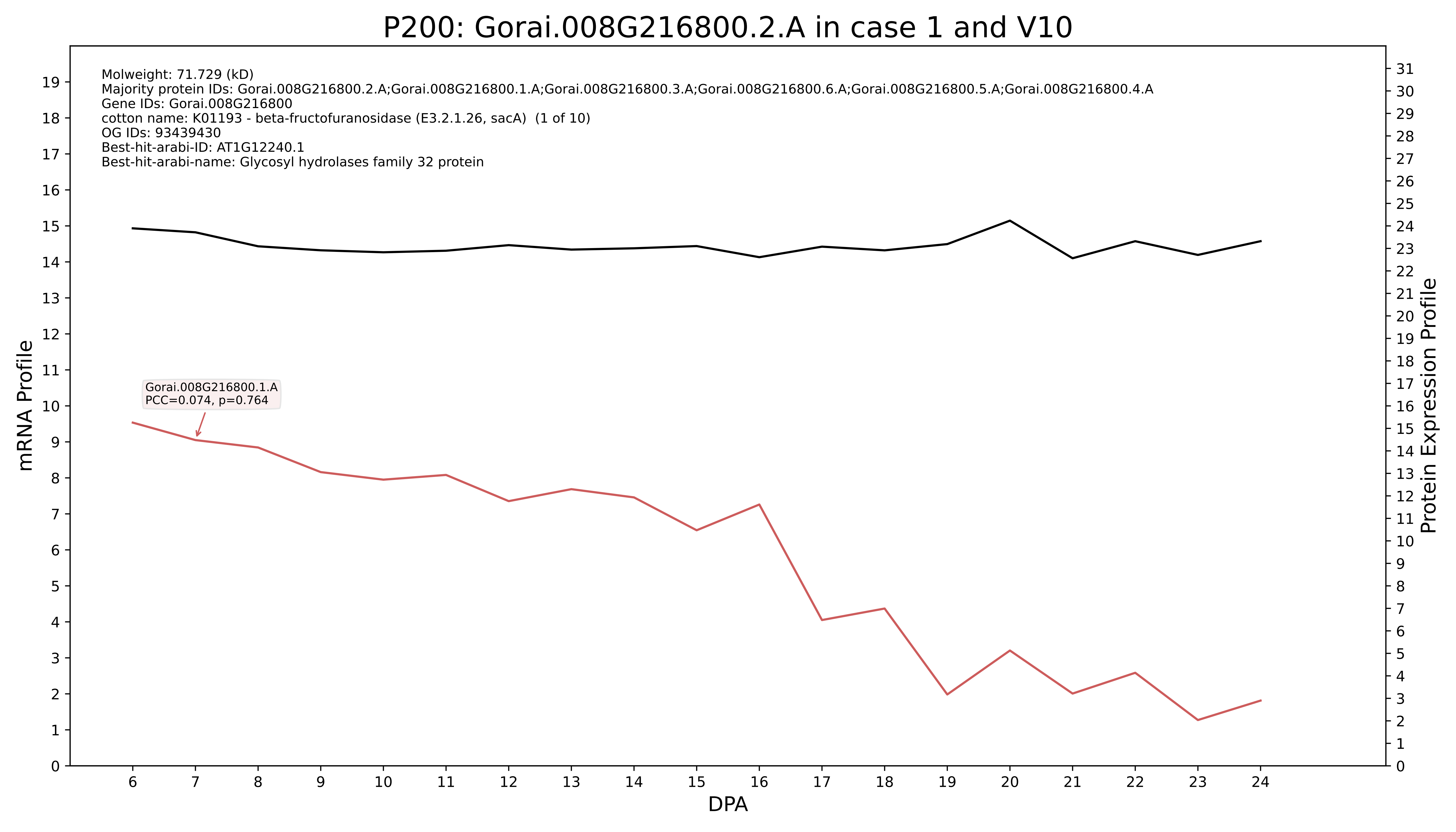
P200: Gorai.008G216800.2.A in case 1 and V10
Molweight: 71.729 (kD)
Majority protein IDs: Gorai.008G216800.2.A;Gorai.008G216800.1.A;Gorai.008G216800.3.A;Gorai.008G216800.6.A;Gorai.008G216800.5.A;Gorai.008G216800.4.A
Gene IDs: Gorai.008G216800
cotton name: K01193 - beta-fructofuranosidase (E3.2.1.26, sacA) (1 of 10)
OG IDs: 93439430
Best-hit-arabi-ID: AT1G12240.1
Best-hit-arabi-name: Glycosyl hydrolases family 32 protein
Gorai.008G216800.1.A
PCC=0.074, p=0.764
mRNA Profile
Protein Expression Profile
DPA

### P200: Gorai.002G023000.1.D in case 1 and V10

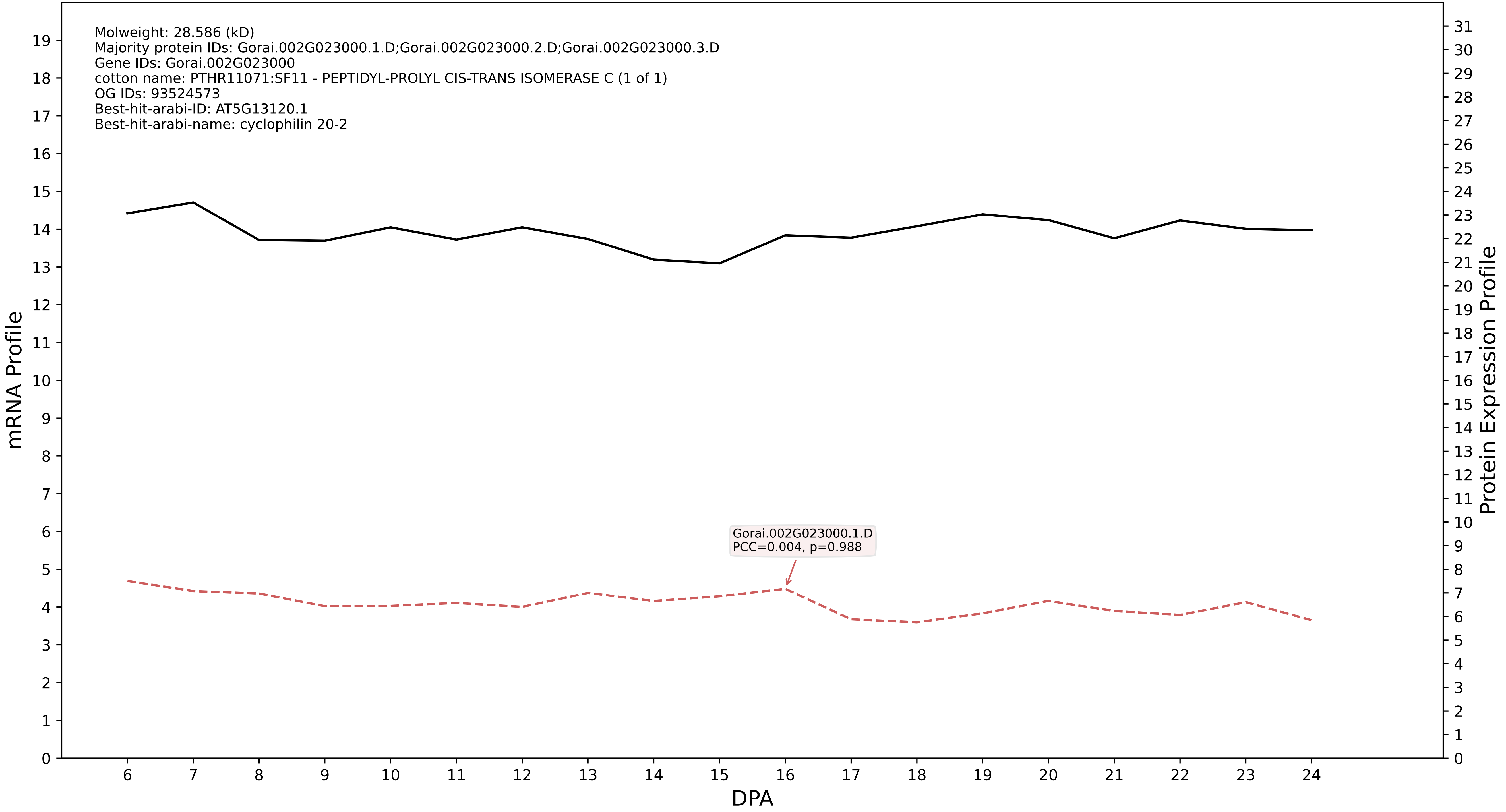

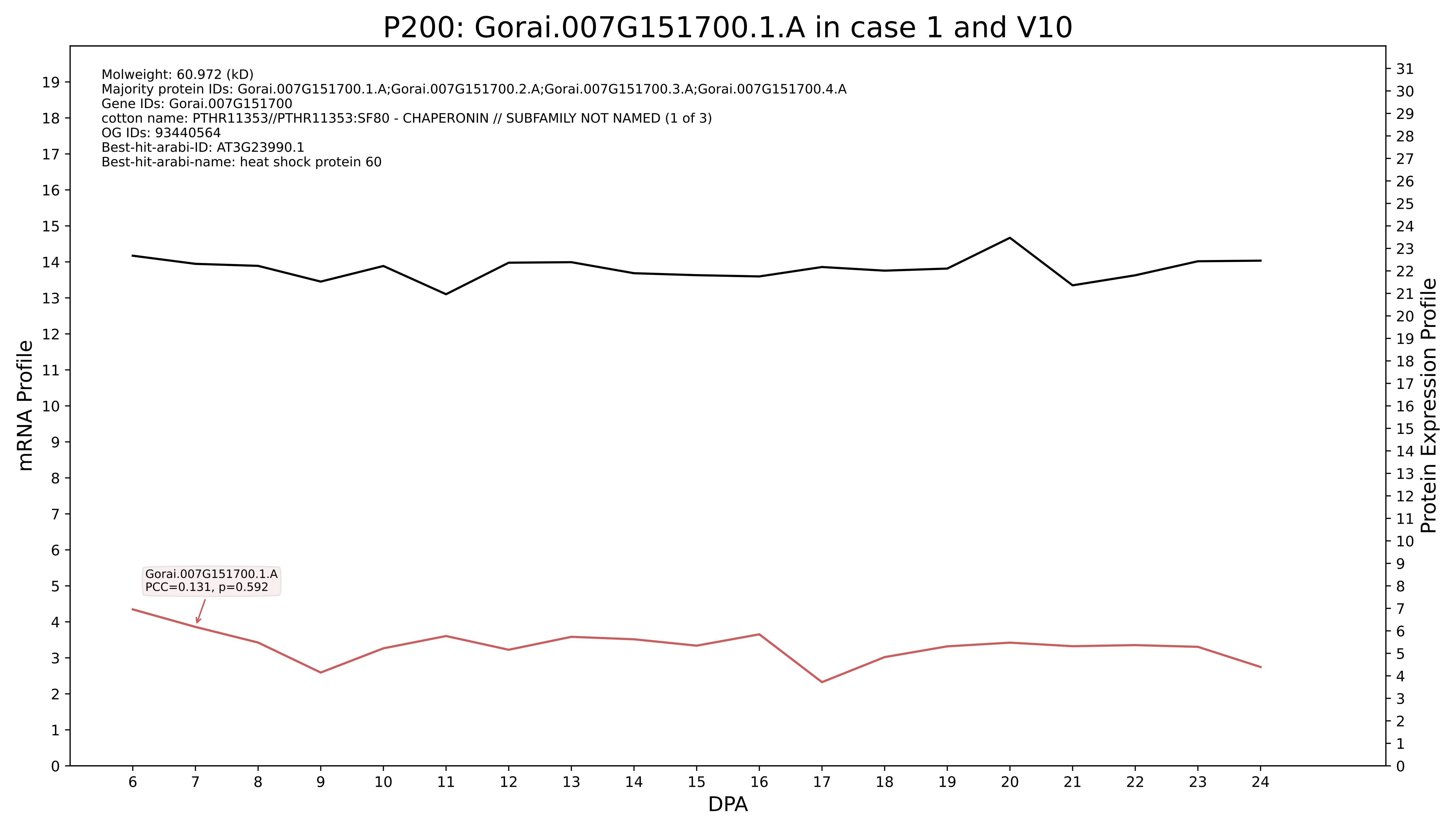
P200: Gorai.007G151700.1.A in case 1 and V10
Molweight: 60.972 (kD)
Majority protein IDs: Gorai.007G151700.1.A;Gorai.007G151700.2.A;Gorai.007G151700.3.A;Gorai.007G151700.4.A
Gene IDs: Gorai.007G151700
cotton name: PTHR11353//PTHR11353:SF80 - CHAPERONIN // SUBFAMILY NOT NAMED (1 of 3)
OG IDs: 93440564
Best-hit-arabi-ID: AT3G23990.1
Best-hit-arabi-name: heat shock protein 60
Gorai.007G151700.1.A
PCC=0.131, p=0.592
mRNA Profile
Protein Expression Profile
DPA

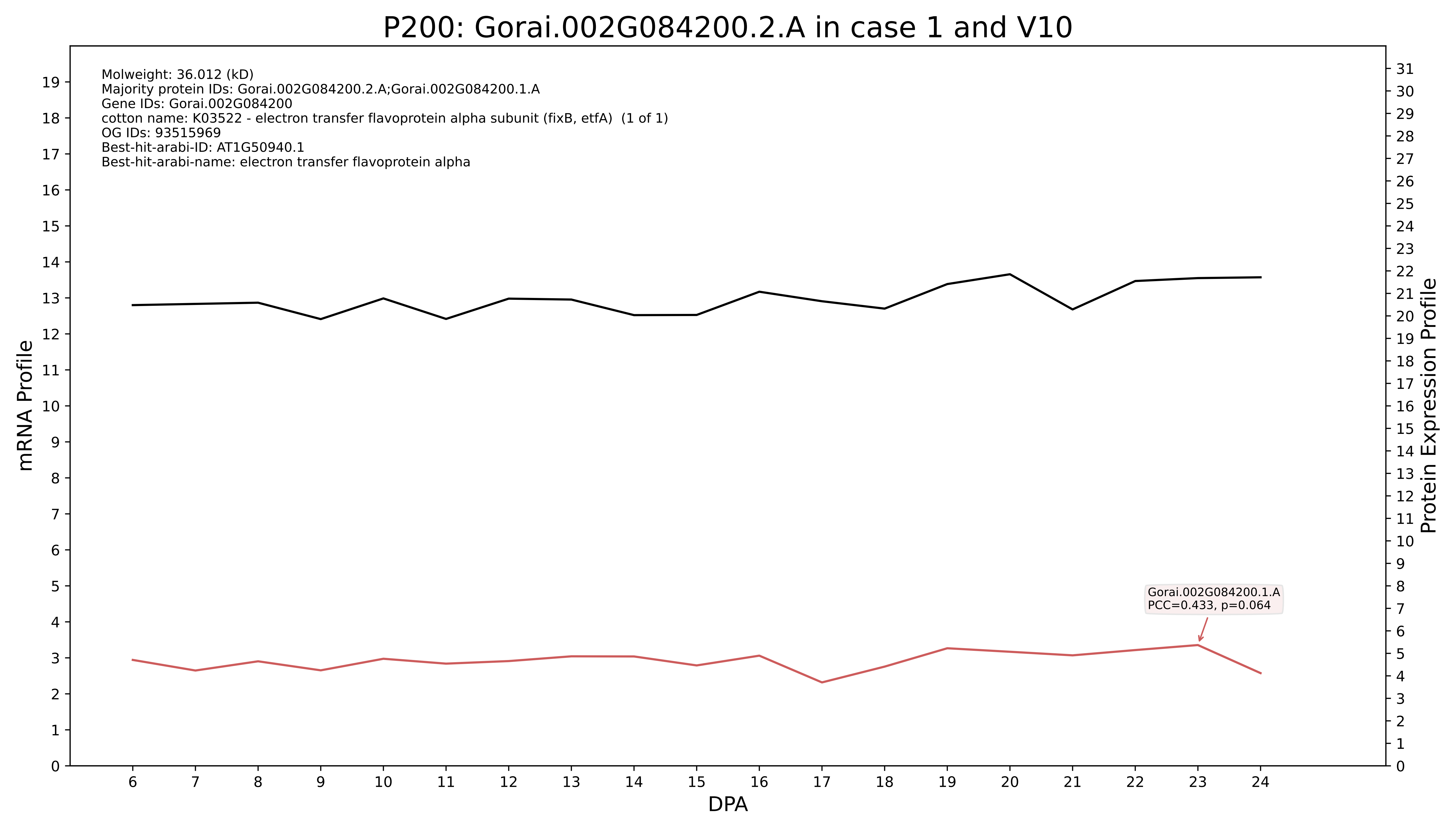
P200: Gorai.002G084200.2.A in case 1 and V10
Molweight: 36.012 (kD)
Majority protein IDs: Gorai.002G084200.2.A;Gorai.002G084200.1.A
Gene IDs: Gorai.002G084200
cotton name: K03522 - electron transfer flavoprotein alpha subunit (fixB, etfA) (1 of 1)
OG IDs: 93515969
Best-hit-arabi-ID: AT1G50940.1
Best-hit-arabi-name: electron transfer flavoprotein alpha
Gorai.002G084200.1.A
PCC=0.433, p=0.064
mRNA Profile
Protein Expression Profile
DPA

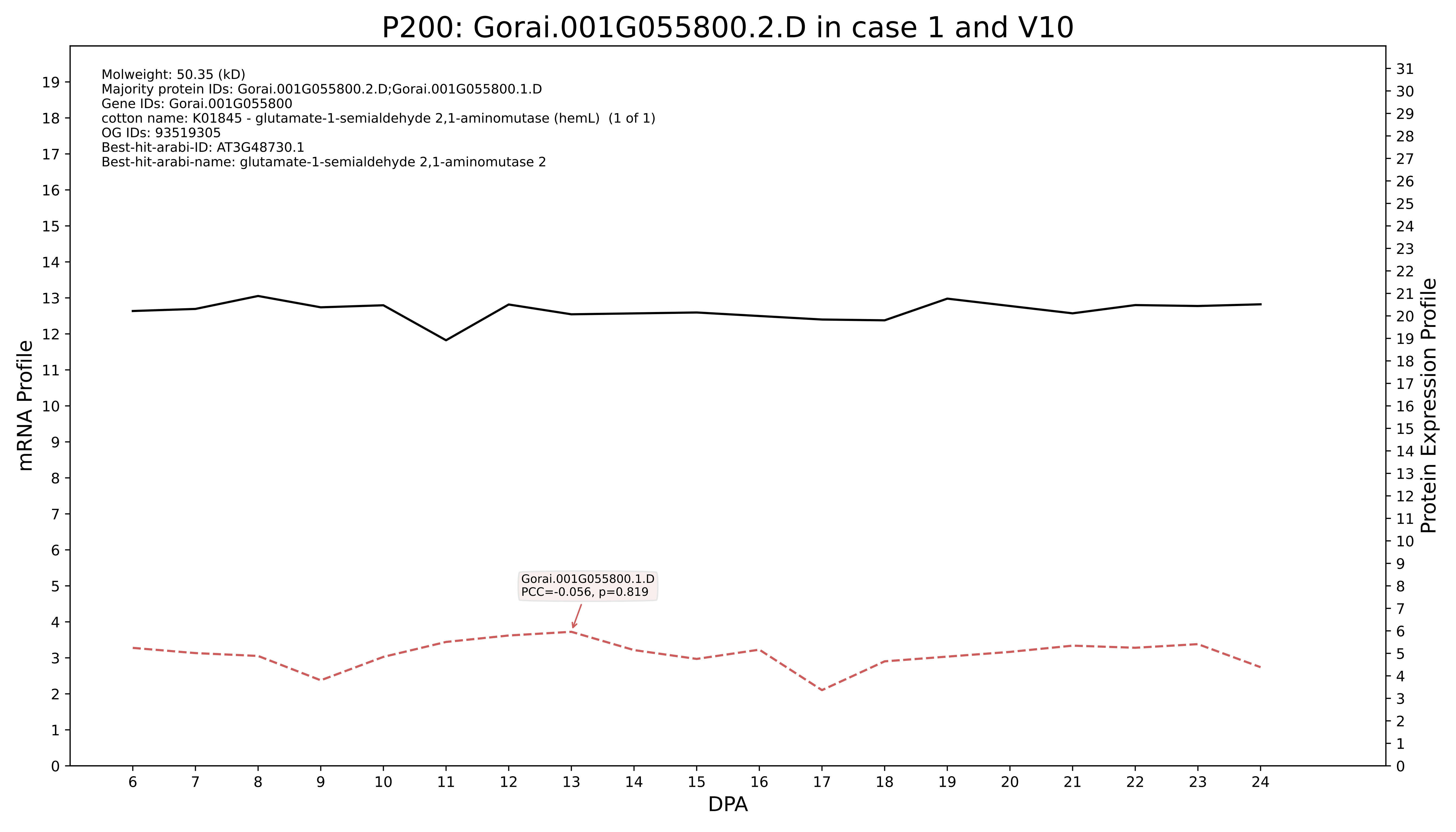

P200: Gorai.001G055800.2.D in case 1 and V10
Molweight: 50.35 (kD)
Majority protein IDs: Gorai.001G055800.2.D;Gorai.001G055800.1.D
Gene IDs: Gorai.001G055800
cotton name: K01845 - glutamate-1-semialdehyde 2,1-aminomutase (hemL) (1 of 1)
OG IDs: 93519305
Best-hit-arabi-ID: AT3G48730.1
Best-hit-arabi-name: glutamate-1-semialdehyde 2,1-aminomutase 2
Gorai.001G055800.1.D
PCC=-0.056, p=0.819
mRNA Profile
Protein Expression Profile
DPA

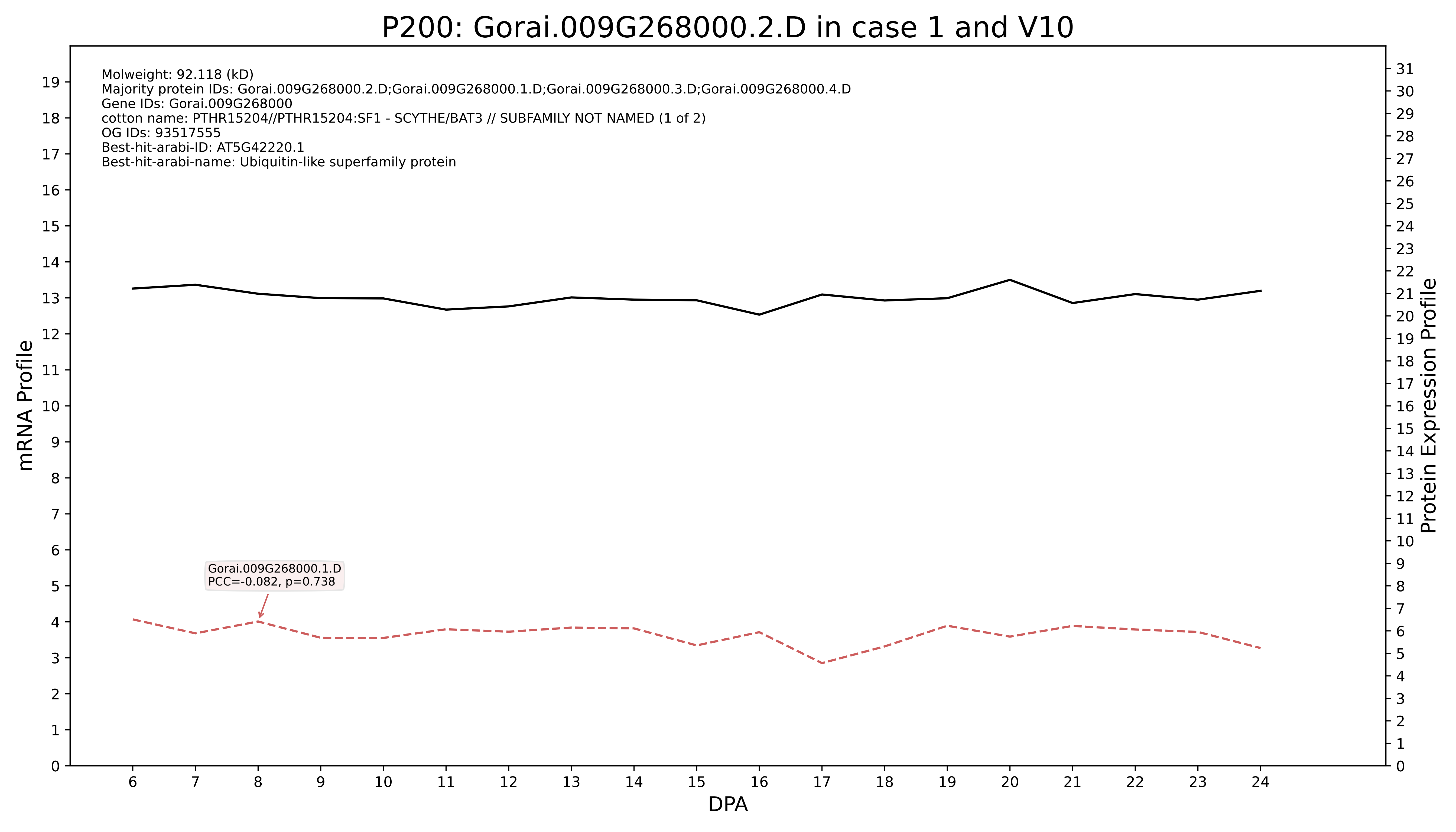

P200: Gorai.009G268000.2.D in case 1 and V10
Molweight: 92.118 (kD)
Majority protein IDs: Gorai.009G268000.2.D;Gorai.009G268000.1.D;Gorai.009G268000.3.D;Gorai.009G268000.4.D
Gene IDs: Gorai.009G268000
cotton name: PTHR15204//PTHR15204:SF1 - SCYTHE/BAT3 // SUBFAMILY NOT NAMED (1 of 2)
OG IDs: 93517555
Best-hit-arabi-ID: AT5G42220.1
Best-hit-arabi-name: Ubiquitin-like superfamily protein
Gorai.009G268000.1.D
PCC=-0.082, p=0.738
mRNA Profile
Protein Expression Profile
DPA

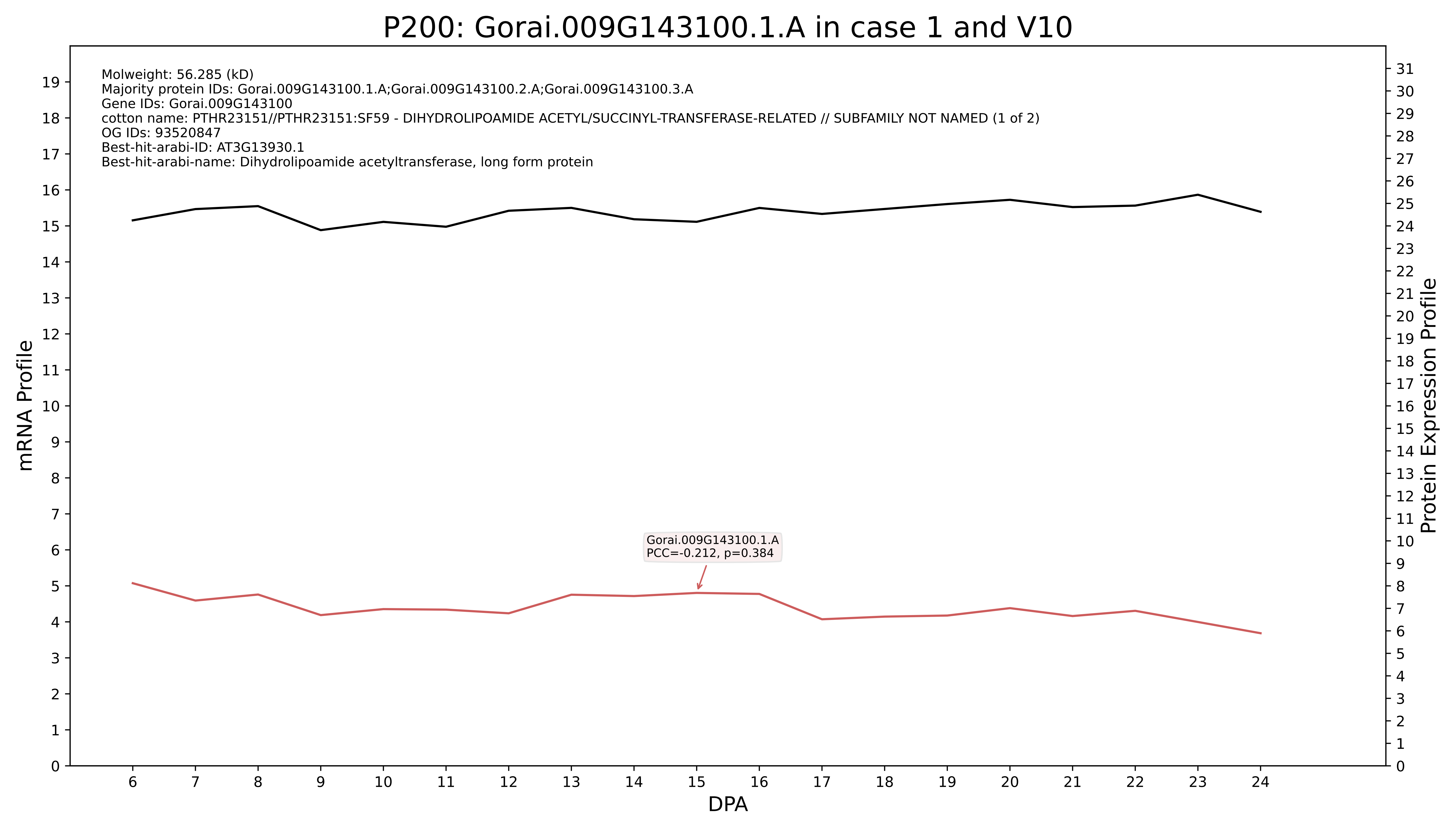
P200: Gorai.009G143100.1.A in case 1 and V10
Molweight: 56.285 (kD)
Majority protein IDs: Gorai.009G143100.1.A;Gorai.009G143100.2.A;Gorai.009G143100.3.A
Gene IDs: Gorai.009G143100
cotton name: PTHR23151//PTHR23151:SF59 - DIHYDROLIPOAMIDE ACETYL/SUCCINYL-TRANSFERASE-RELATED // SUBFAMILY NOT NAMED (1 of 2)
OG IDs: 93520847
Best-hit-arabi-ID: AT3G13930.1
Best-hit-arabi-name: Dihydrolipoamide acetyltransferase, long form protein
Gorai.009G143100.1.A
PCC=-0.212, p=0.384
mRNA Profile
Protein Expression Profile
DPA

### P200: Gorai.011G233300.6.D in case 1 and V10

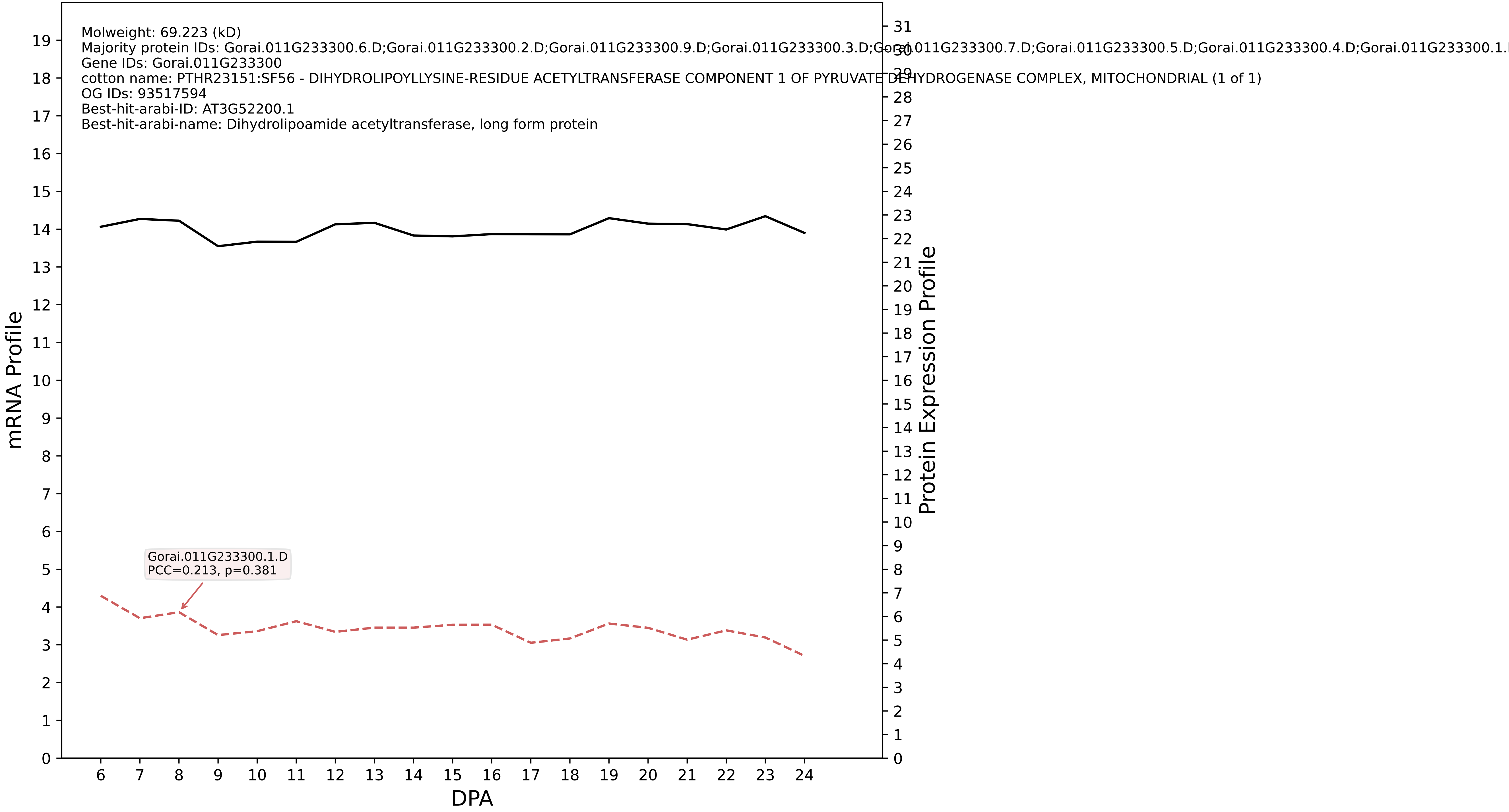

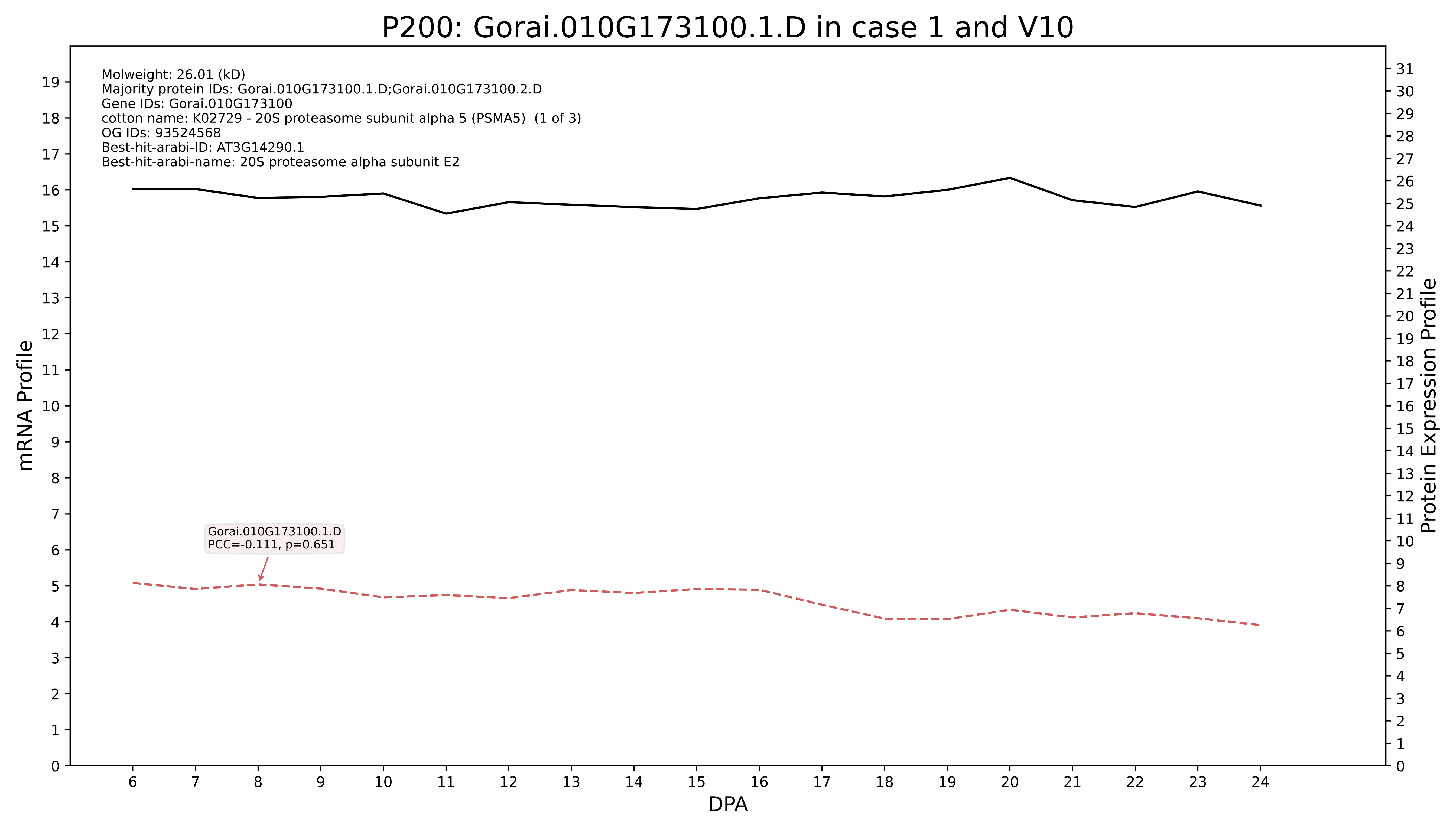
P200: Gorai.010G173100.1.D in case 1 and V10
Molweight: 26.01 (kD)
Majority protein IDs: Gorai.010G173100.1.D;Gorai.010G173100.2.D
Gene IDs: Gorai.010G173100
cotton name: K02729 - 20S proteasome subunit alpha 5 (PSMA5) (1 of 3)
OG IDs: 93524568
Best-hit-arabi-ID: AT3G14290.1
Best-hit-arabi-name: 20S proteasome alpha subunit E2
Gorai.010G173100.1.D
PCC=-0.111, p=0.651
mRNA Profile
Protein Expression Profile
DPA

### P200: Gorai.004G141000.2.D in case 1 and V10

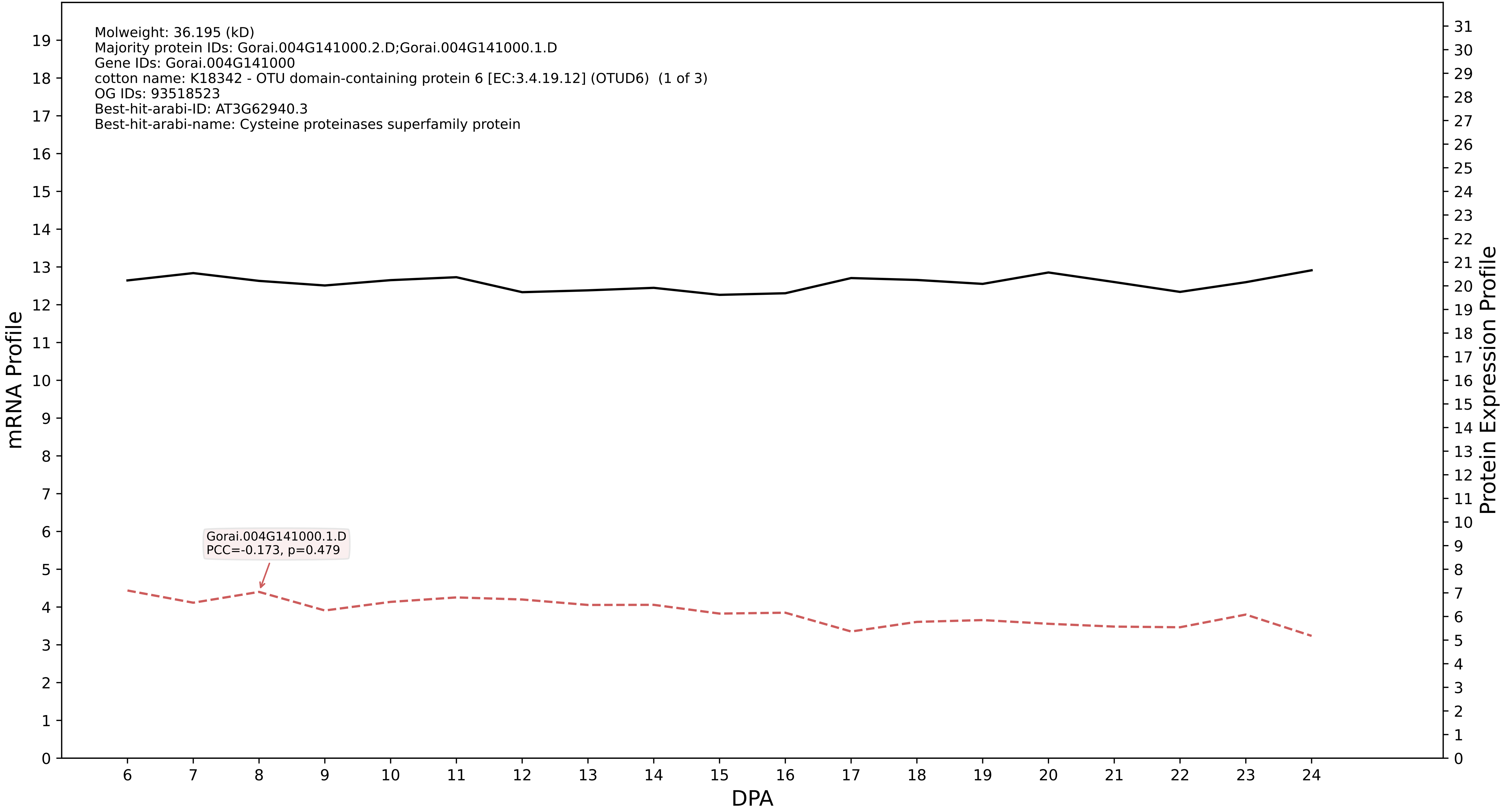

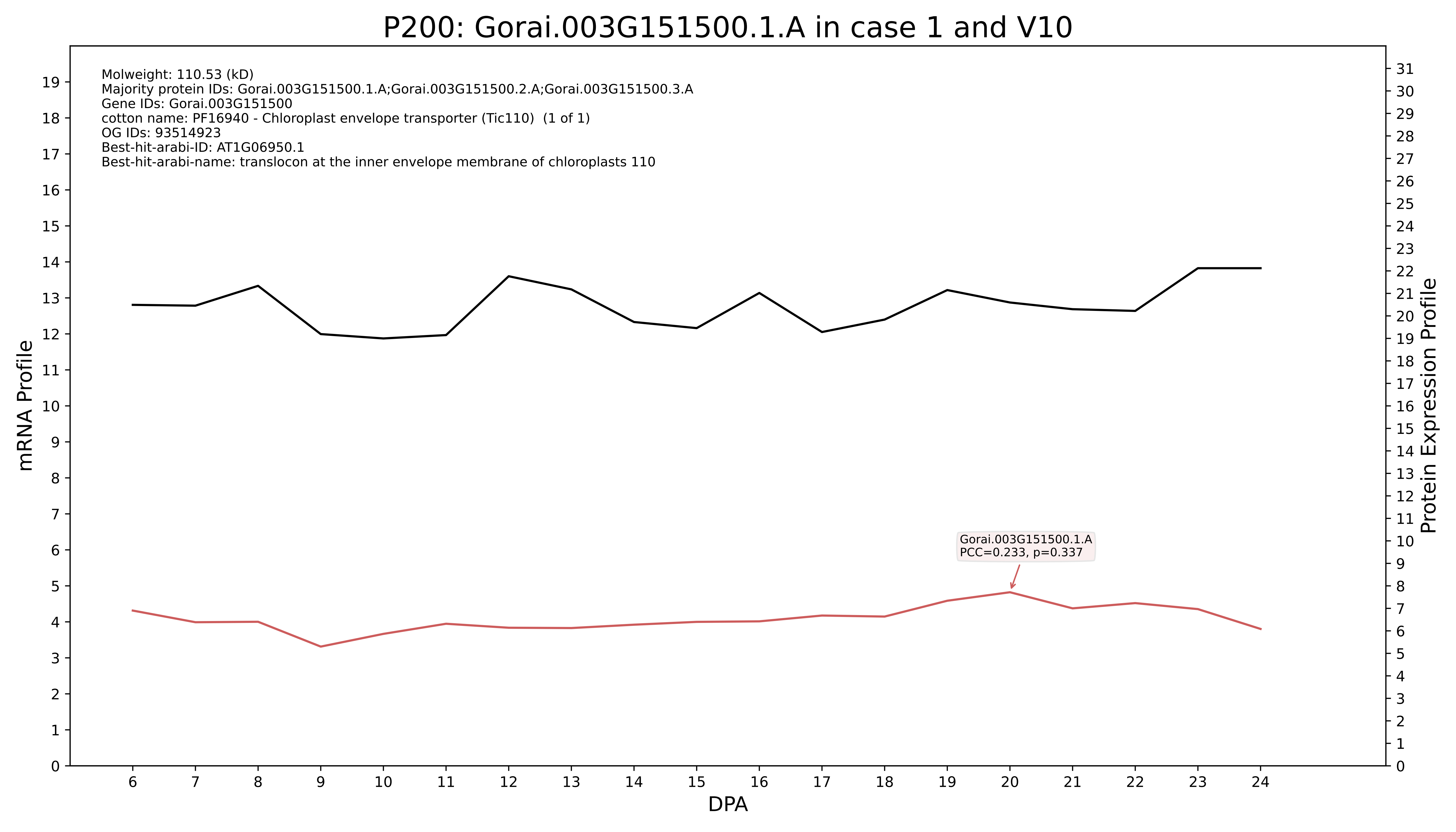
P200: Gorai.003G151500.1.A in case 1 and V10
Molweight: 110.53 (kD)
Majority protein IDs: Gorai.003G151500.1.A;Gorai.003G151500.2.A;Gorai.003G151500.3.A
Gene IDs: Gorai.003G151500
cotton name: PF16940 - Chloroplast envelope transporter (Tic110) (1 of 1)
OG IDs: 93514923
Best-hit-arabi-ID: AT1G06950.1
Best-hit-arabi-name: translocon at the inner envelope membrane of chloroplasts 110
Gorai.003G151500.1.A
PCC=0.233, p=0.337
mRNA Profile
Protein Expression Profile
DPA

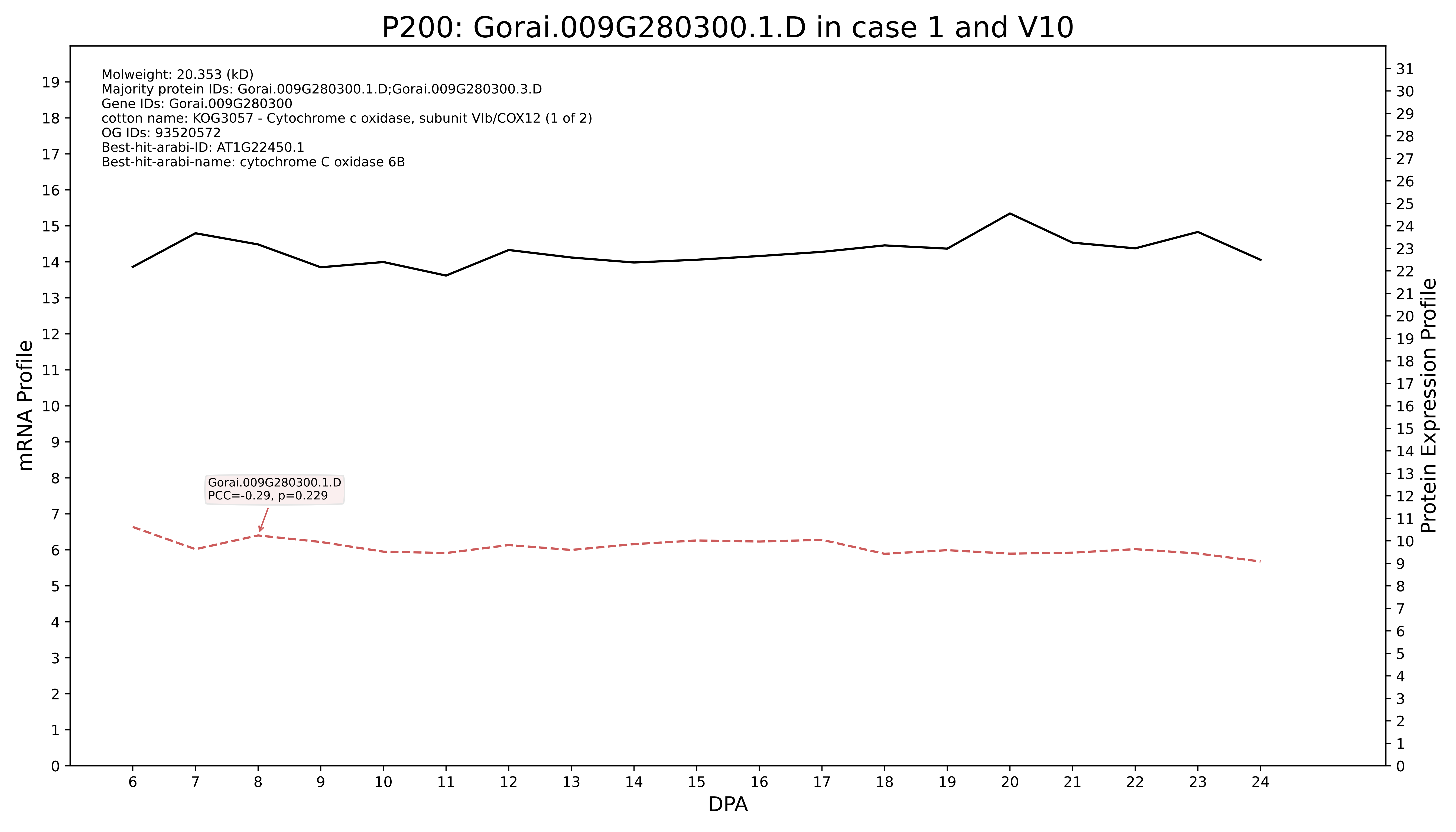
P200: Gorai.009G280300.1.D in case 1 and V10
Molweight: 20.353 (kD)
Majority protein IDs: Gorai.009G280300.1.D;Gorai.009G280300.3.D
Gene IDs: Gorai.009G280300
cotton name: KOG3057 - Cytochrome c oxidase, subunit VIb/COX12 (1 of 2)
OG IDs: 93520572
Best-hit-arabi-ID: AT1G22450.1
Best-hit-arabi-name: cytochrome C oxidase 6B
Gorai.009G280300.1.D
PCC=-0.29, p=0.229
mRNA Profile
Protein Expression Profile
DPA

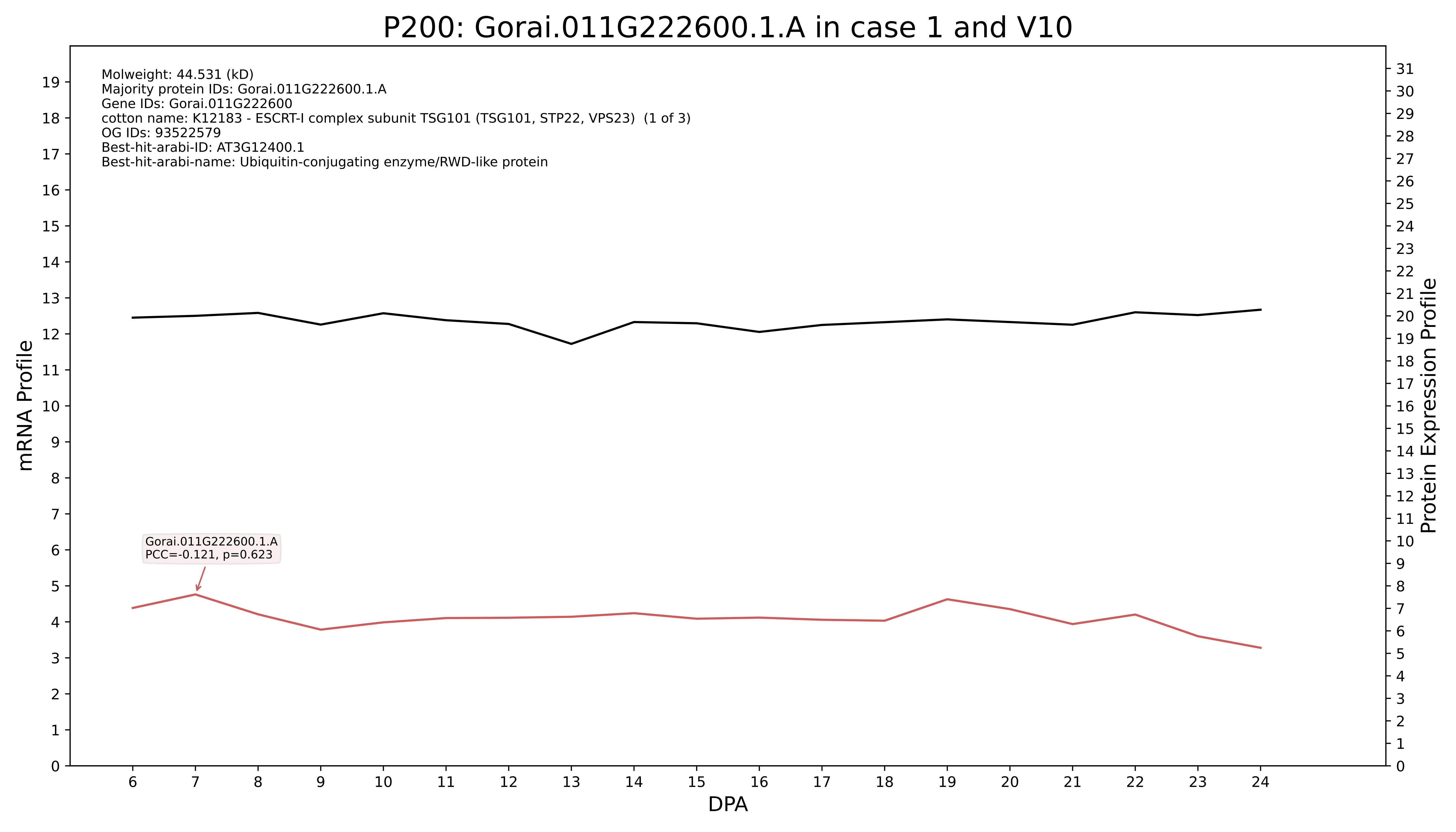
P200: Gorai.011G222600.1.A in case 1 and V10
Molweight: 44.531 (kD)
Majority protein IDs: Gorai.011G222600.1.A
Gene IDs: Gorai.011G222600
cotton name: K12183 - ESCRT-I complex subunit TSG101 (TSG101, STP22, VPS23) (1 of 3)
OG IDs: 93522579
Best-hit-arabi-ID: AT3G12400.1
Best-hit-arabi-name: Ubiquitin-conjugating enzyme/RWD-like protein
Gorai.011G222600.1.A
PCC=-0.121, p=0.623
mRNA Profile
Protein Expression Profile
DPA

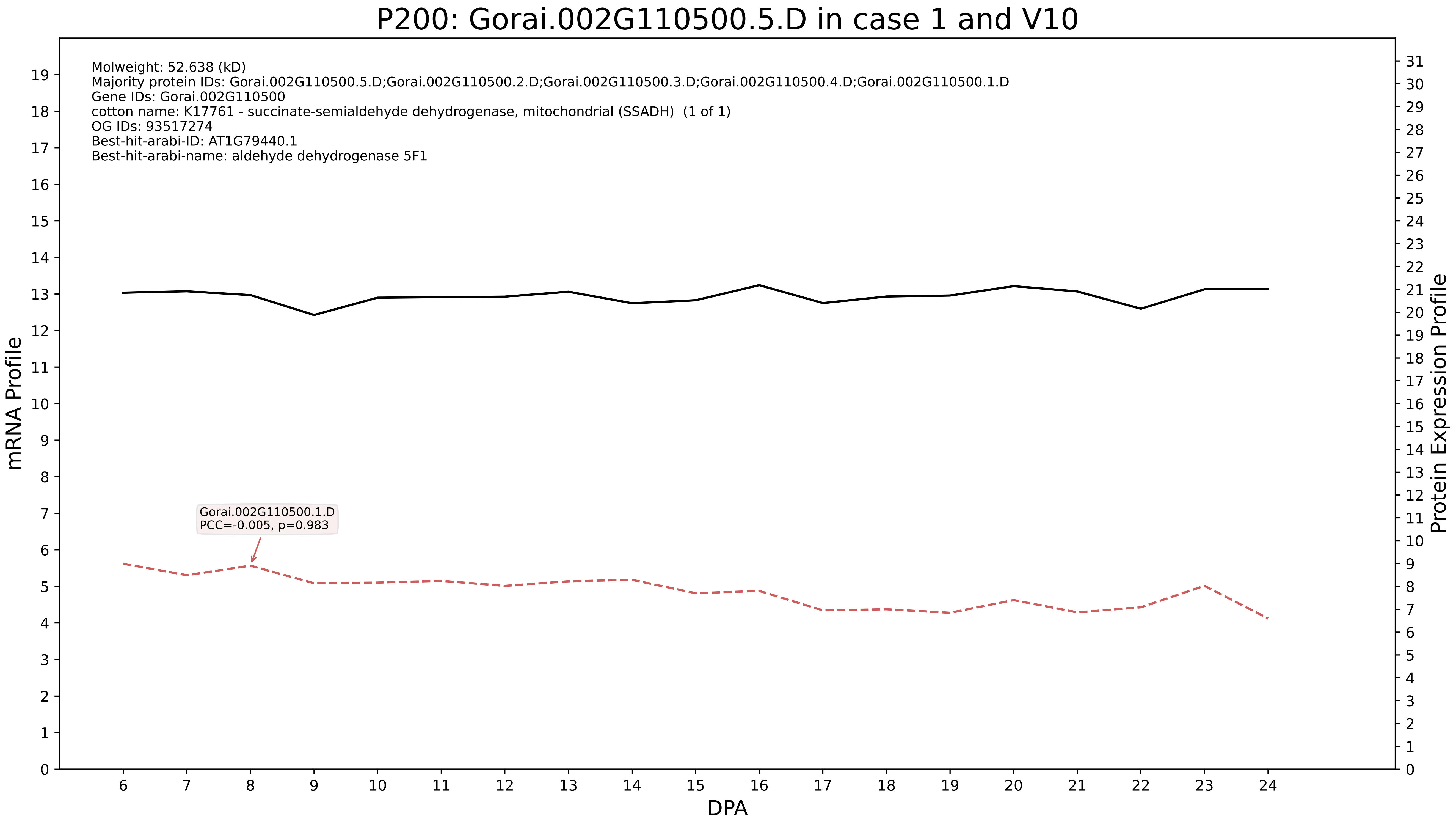
P200: Gorai.002G110500.5.D in case 1 and V10
Molweight: 52.638 (kD)
Majority protein IDs: Gorai.002G110500.5.D;Gorai.002G110500.2.D;Gorai.002G110500.3.D;Gorai.002G110500.4.D;Gorai.002G110500.1.D
Gene IDs: Gorai.002G110500
cotton name: K17761 - succinate-semialdehyde dehydrogenase, mitochondrial (SSADH) (1 of 1)
OG IDs: 93517274
Best-hit-arabi-ID: AT1G79440.1
Best-hit-arabi-name: aldehyde dehydrogenase 5F1
Gorai.002G110500.1.D
PCC=-0.005, p=0.983
mRNA Profile
Protein Expression Profile
DPA

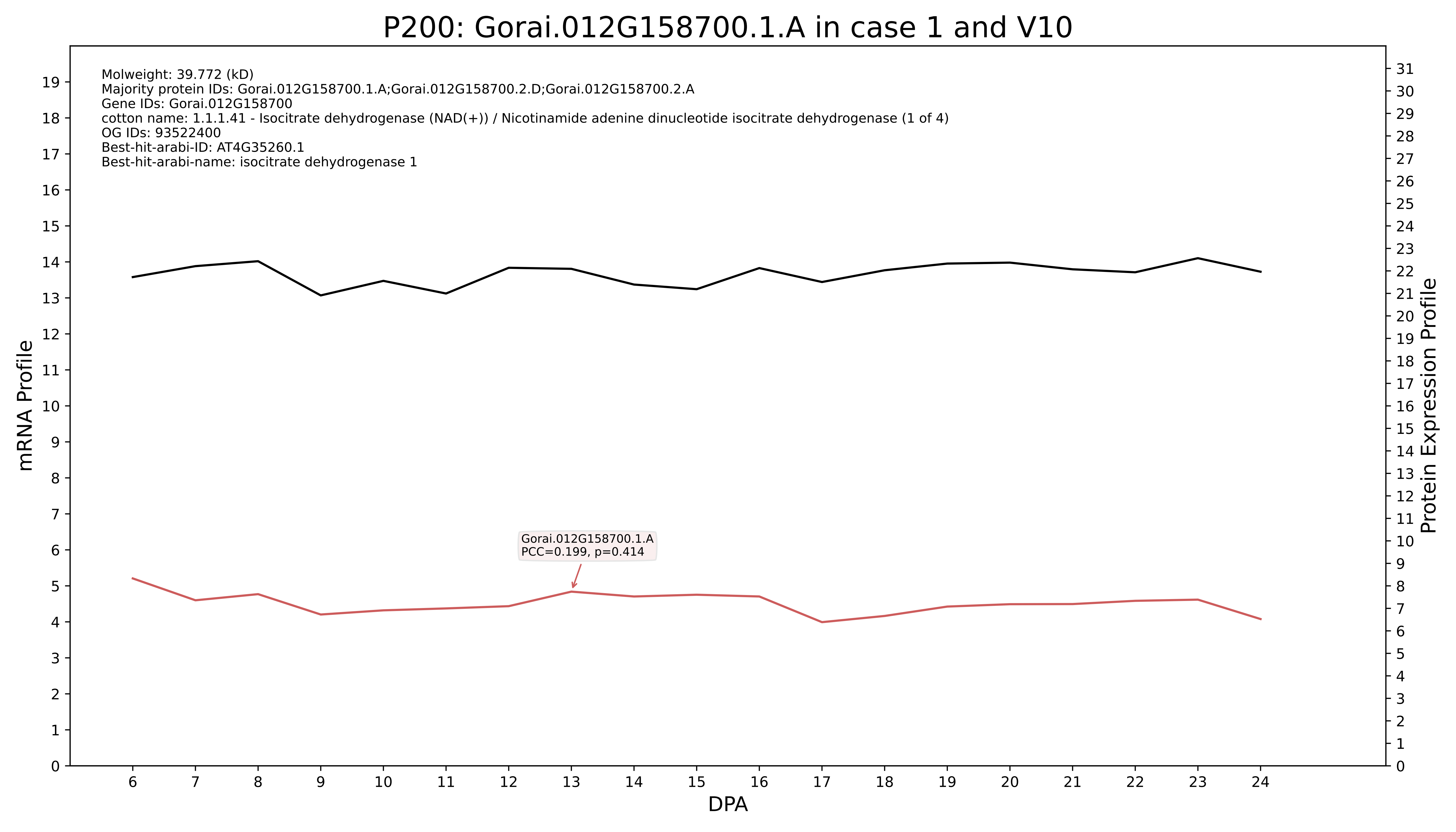
P200: Gorai.012G158700.1.A in case 1 and V10
Molweight: 39.772 (kD)
Majority protein IDs: Gorai.012G158700.1.A;Gorai.012G158700.2.D;Gorai.012G158700.2.A
Gene IDs: Gorai.012G158700
cotton name: 1.1.1.41 - Isocitrate dehydrogenase (NAD(+)) / Nicotinamide adenine dinucleotide isocitrate dehydrogenase (1 of 4)
OG IDs: 93522400
Best-hit-arabi-ID: AT4G35260.1
Best-hit-arabi-name: isocitrate dehydrogenase 1
Gorai.012G158700.1.A
PCC=0.199, p=0.414
mRNA Profile
Protein Expression Profile
DPA

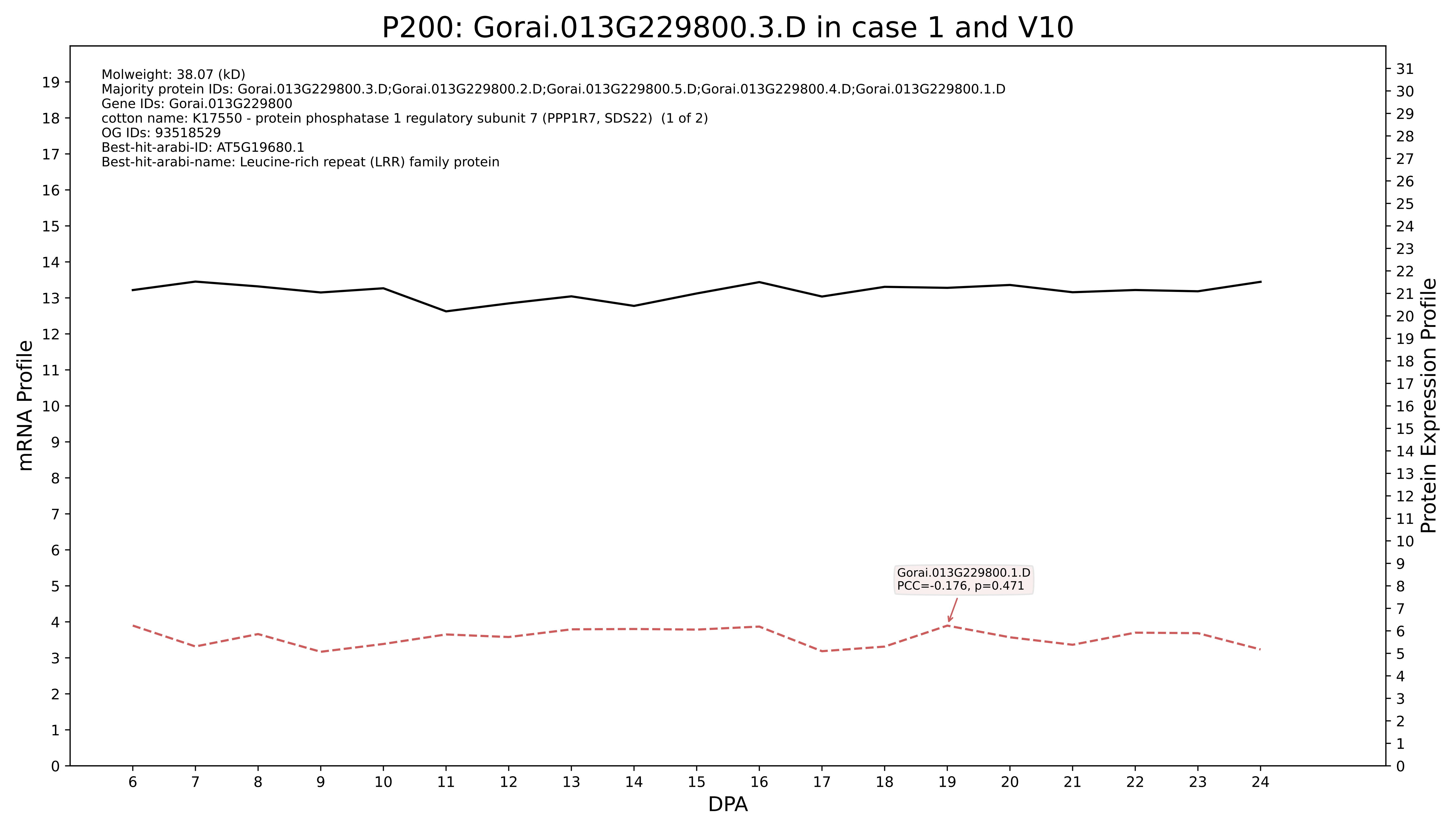
P200: Gorai.013G229800.3.D in case 1 and V10
Molweight: 38.07 (kD)
Majority protein IDs: Gorai.013G229800.3.D;Gorai.013G229800.2.D;Gorai.013G229800.5.D;Gorai.013G229800.4.D;Gorai.013G229800.1.D
Gene IDs: Gorai.013G229800
cotton name: K17550 - protein phosphatase 1 regulatory subunit 7 (PPP1R7, SDS22) (1 of 2)
OG IDs: 93518529
Best-hit-arabi-ID: AT5G19680.1
Best-hit-arabi-name: Leucine-rich repeat (LRR) family protein
Gorai.013G229800.1.D
PCC=-0.176, p=0.471
mRNA Profile
Protein Expression Profile
DPA

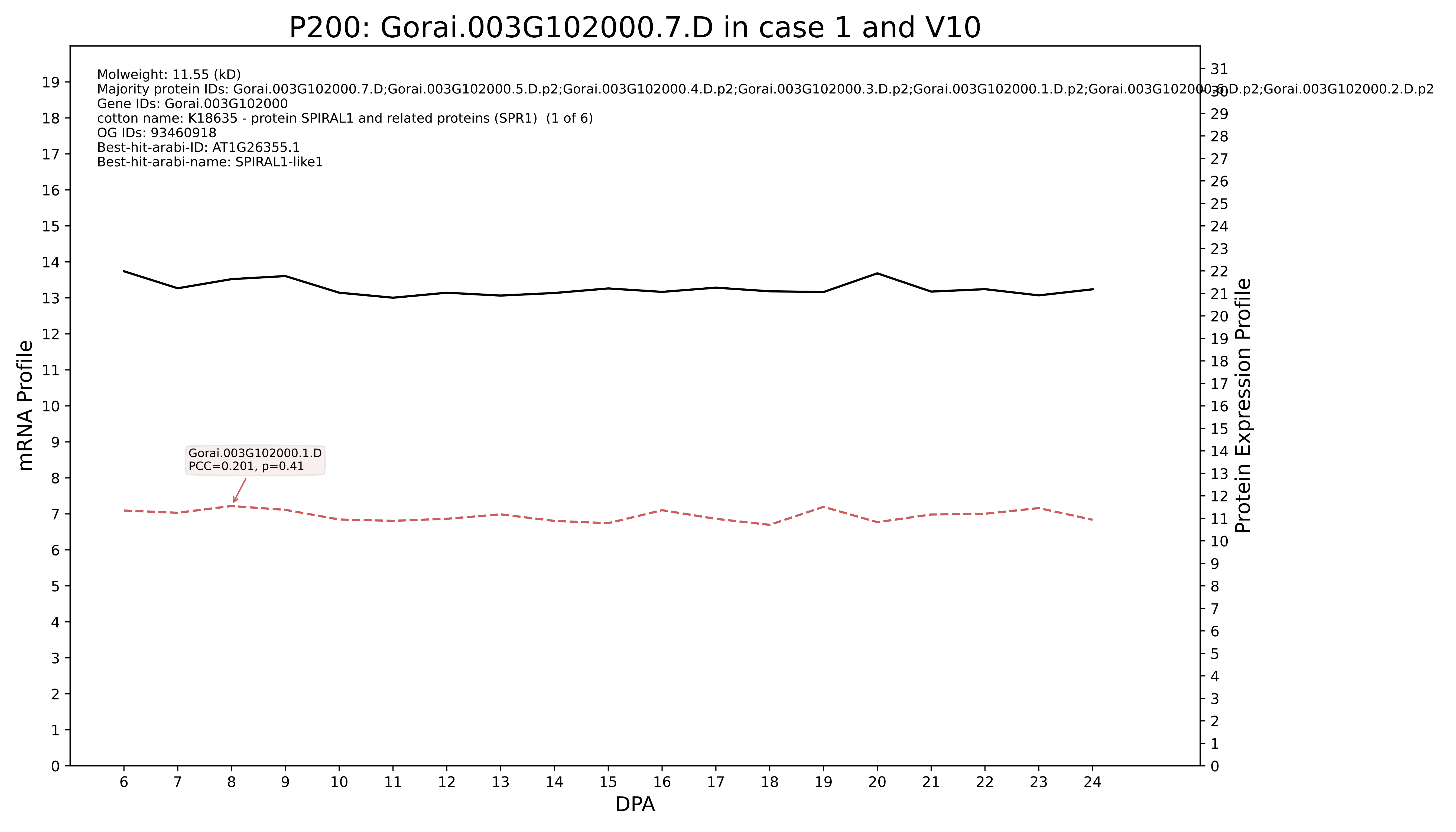
P200: Gorai.003G102000.7.D in case 1 and V10
Molweight: 11.55 (kD)
Majority protein IDs: Gorai.003G102000.7.D;Gorai.003G102000.5.D.p2;Gorai.003G102000.4.D.p2;Gorai.003G102000.3.D.p2;Gorai.003G102000.1.D.p2;Gorai.003G102000.6.D.p2;Gorai.003G102000.2.D.p2
Gene IDs: Gorai.003G102000
cotton name: K18635 - protein SPIRAL1 and related proteins (SPR1) (1 of 6)
OG IDs: 93460918
Best-hit-arabi-ID: AT1G26355.1
Best-hit-arabi-name: SPIRAL1-like1
Gorai.003G102000.1.D
PCC=0.201, p=0.41
mRNA Profile
Protein Expression Profile
DPA

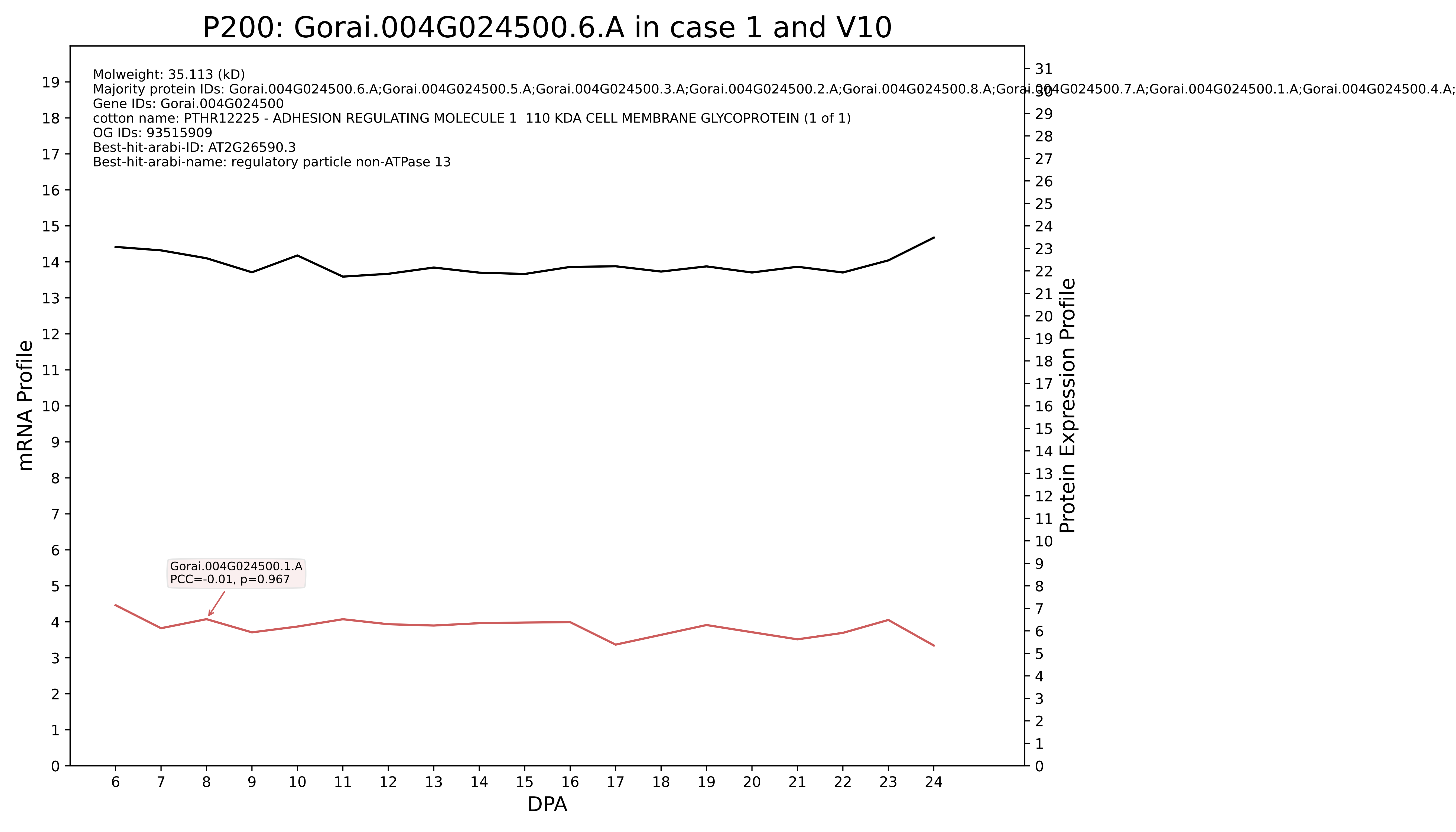
P200: Gorai.004G024500.6.A in case 1 and V10
Molweight: 35.113 (kD)
Majority protein IDs: Gorai.004G024500.6.A;Gorai.004G024500.5.A;Gorai.004G024500.3.A;Gorai.004G024500.2.A;Gorai.004G024500.8.A;Gorai.004G024500.7.A;Gorai.004G024500.1.A;Gorai.004G024500.4.A;
Gene IDs: Gorai.004G024500
cotton name: PTHR12225 - ADHESION REGULATING MOLECULE 1 110 KDA CELL MEMBRANE GLYCOPROTEIN (1 of 1)
OG IDs: 93515909
Best-hit-arabi-ID: AT2G26590.3
Best-hit-arabi-name: regulatory particle non-ATPase 13
Gorai.004G024500.1.A
PCC=-0.01, p=0.967
mRNA Profile
Protein Expression Profile
DPA

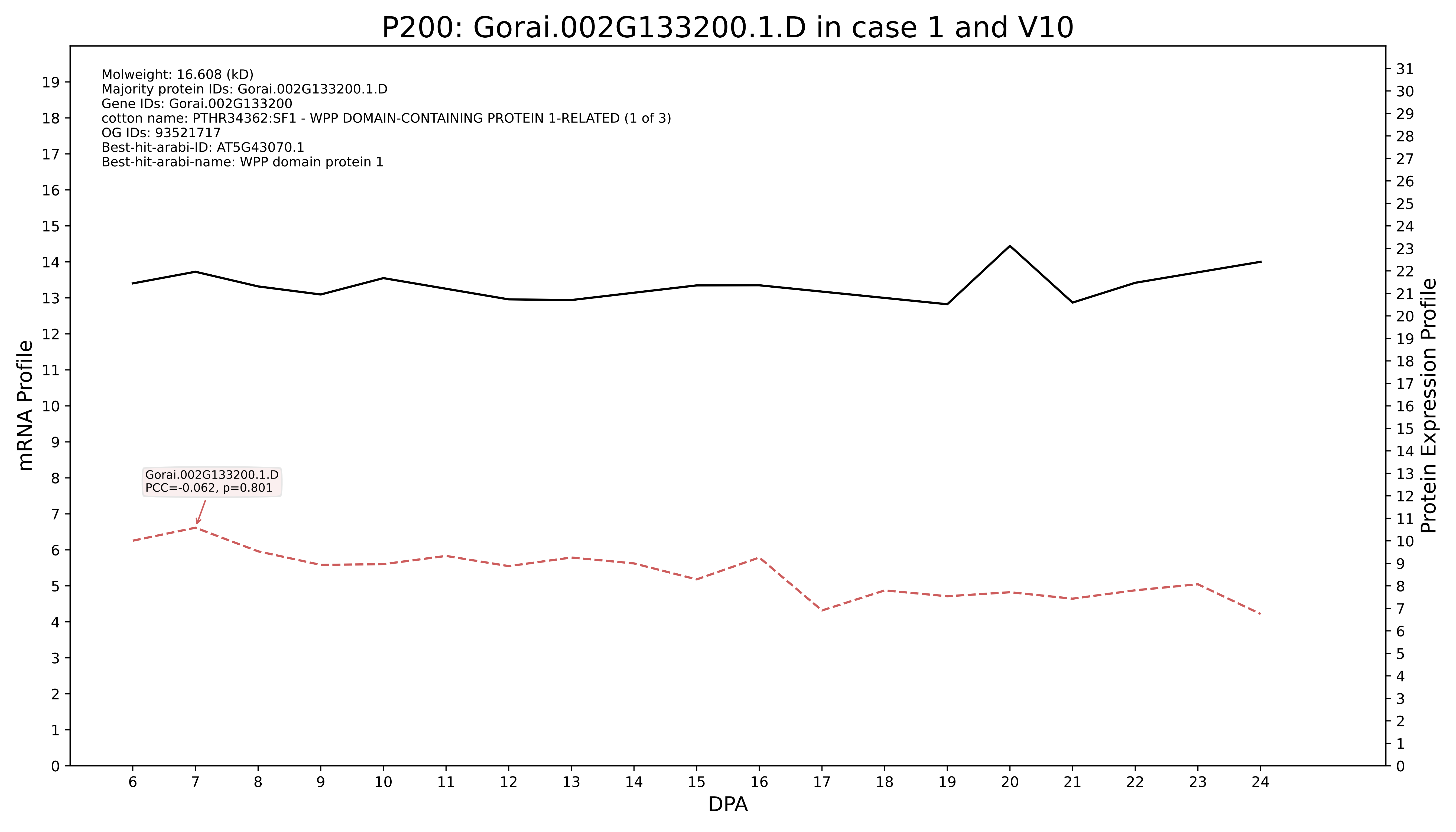

P200: Gorai.002G133200.1.D in case 1 and V10
Molweight: 16.608 (kD)
Majority protein IDs: Gorai.002G133200.1.D
Gene IDs: Gorai.002G133200
cotton name: PTHR34362:SF1 - WPP DOMAIN-CONTAINING PROTEIN 1-RELATED (1 of 3)
OG IDs: 93521717
Best-hit-arabi-ID: AT5G43070.1
Best-hit-arabi-name: WPP domain protein 1
Gorai.002G133200.1.D
PCC=-0.062, p=0.801
mRNA Profile
Protein Expression Profile
DPA

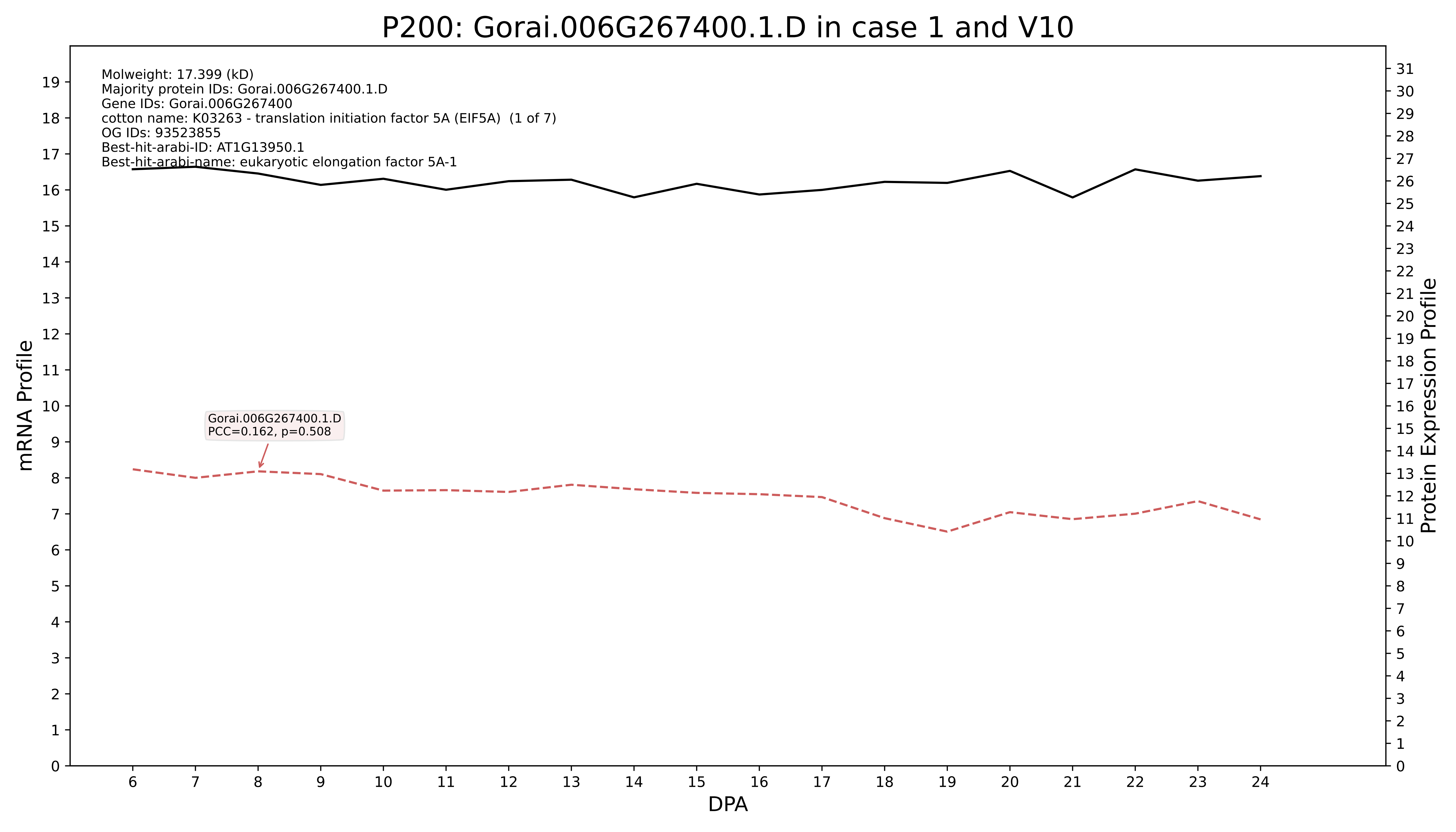

P200: Gorai.006G267400.1.D in case 1 and V10
Molweight: 17.399 (kD)
Majority protein IDs: Gorai.006G267400.1.D
Gene IDs: Gorai.006G267400
cotton name: K03263 - translation initiation factor 5A (EIF5A) (1 of 7)
OG IDs: 93523855
Best-hit-arabi-ID: AT1G13950.1
Best-hit-arabi-name: eukaryotic elongation factor 5A-1
Gorai.006G267400.1.D
PCC=0.162, p=0.508
mRNA Profile
Protein Expression Profile
DPA

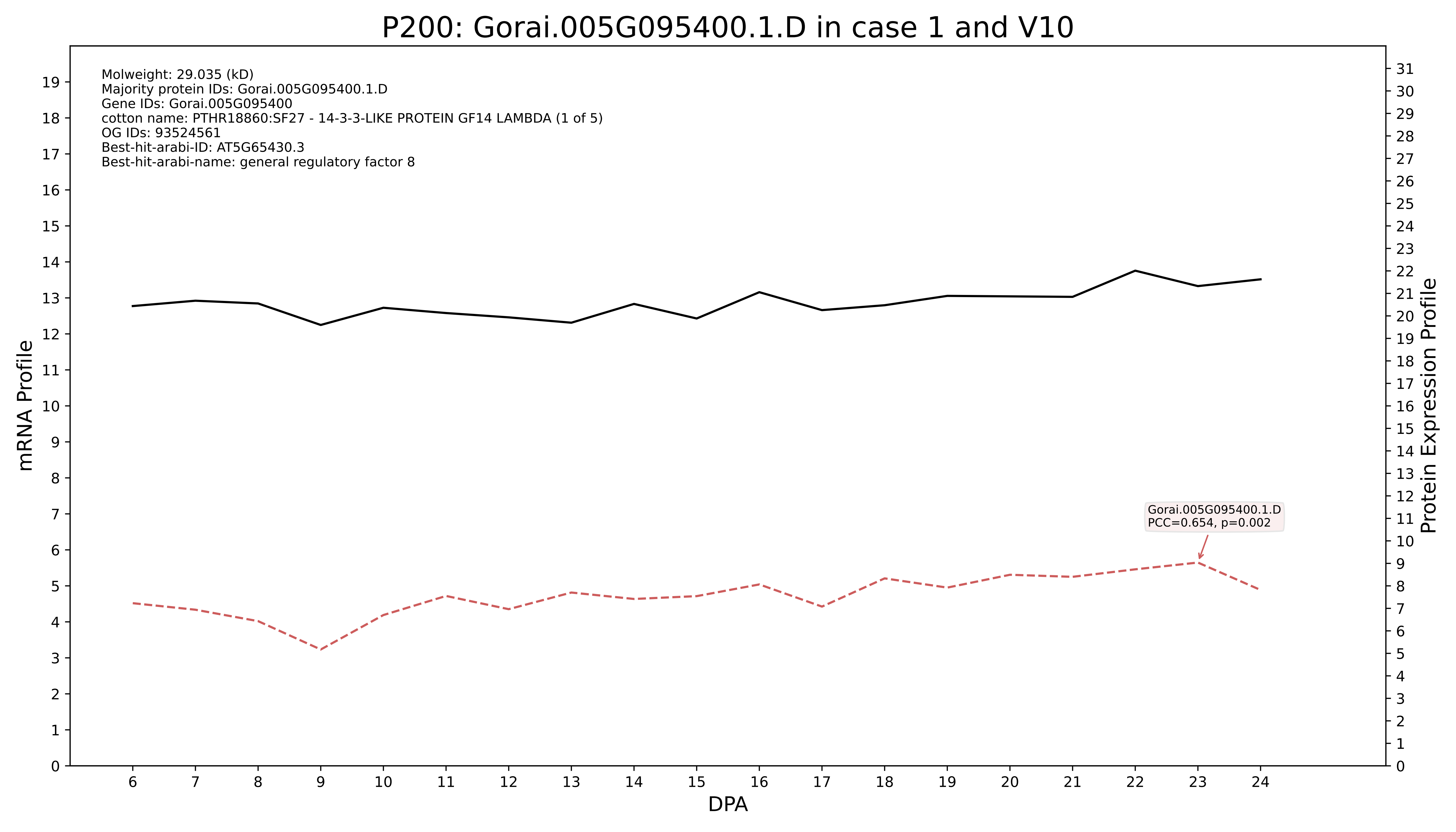
P200: Gorai.005G095400.1.D in case 1 and V10
Molweight: 29.035 (kD)
Majority protein IDs: Gorai.005G095400.1.D
Gene IDs: Gorai.005G095400
cotton name: PTHR18860:SF27 - 14-3-3-LIKE PROTEIN GF14 LAMBDA (1 of 5)
OG IDs: 93524561
Best-hit-arabi-ID: AT5G65430.3
Best-hit-arabi-name: general regulatory factor 8
Gorai.005G095400.1.D
PCC=0.654, p=0.002
mRNA Profile
Protein Expression Profile
DPA

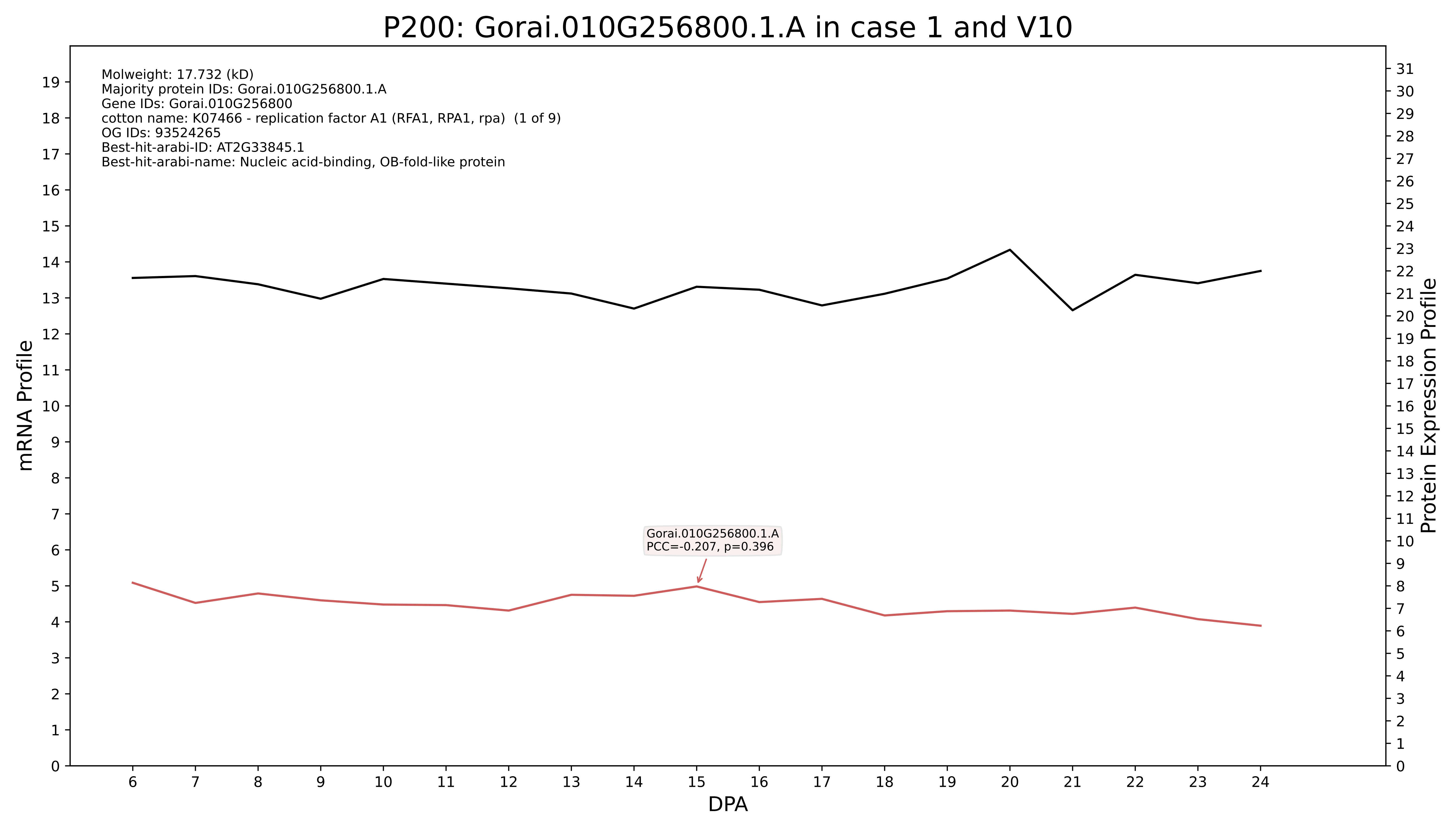
P200: Gorai.010G256800.1.A in case 1 and V10
Molweight: 17.732 (kD)
Majority protein IDs: Gorai.010G256800.1.A
Gene IDs: Gorai.010G256800
cotton name: K07466 - replication factor A1 (RFA1, RPA1, rpa) (1 of 9)
OG IDs: 93524265
Best-hit-arabi-ID: AT2G33845.1
Best-hit-arabi-name: Nucleic acid-binding, OB-fold-like protein
Gorai.010G256800.1.A
PCC=-0.207, p=0.396
mRNA Profile
Protein Expression Profile
DPA

### P200: Gorai.005G053200.2.D in case 1 and V10

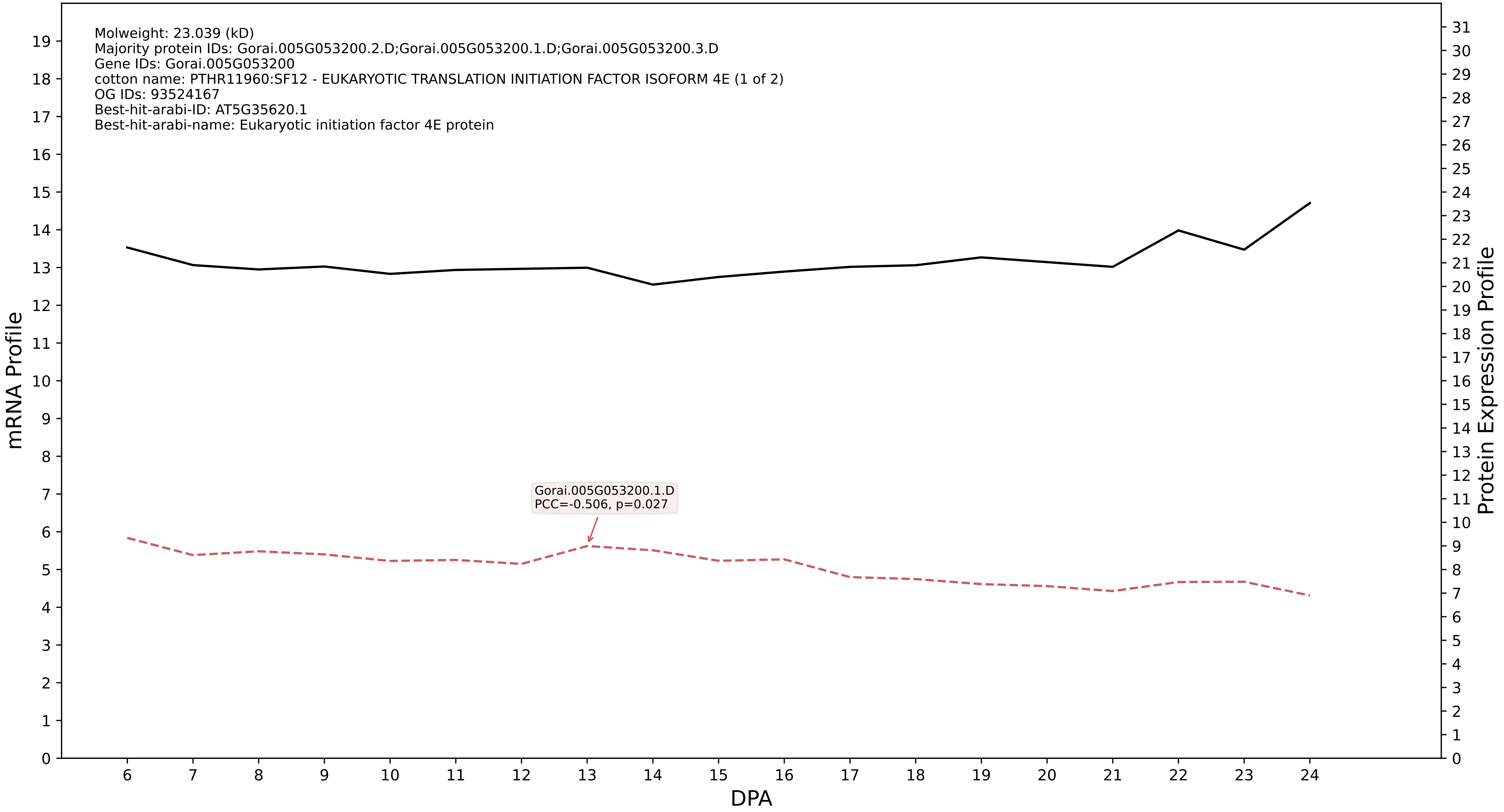

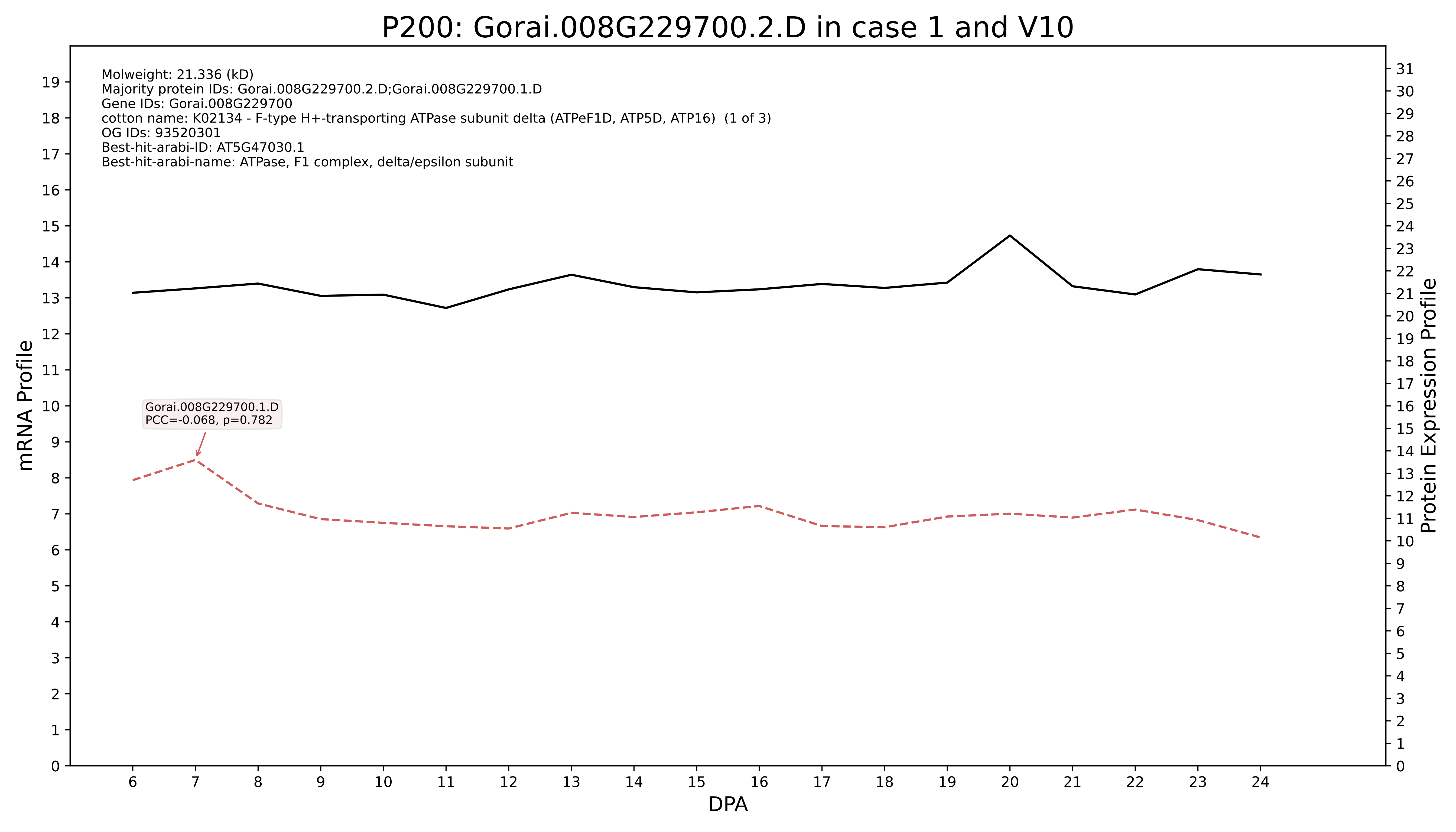
P200: Gorai.008G229700.2.D in case 1 and V10
Molweight: 21.336 (kD)
Majority protein IDs: Gorai.008G229700.2.D;Gorai.008G229700.1.D
Gene IDs: Gorai.008G229700
cotton name: K02134 - F-type H+-transporting ATPase subunit delta (ATPeF1D, ATP5D, ATP16) (1 of 3)
OG IDs: 93520301
Best-hit-arabi-ID: AT5G47030.1
Best-hit-arabi-name: ATPase, F1 complex, delta/epsilon subunit
Gorai.008G229700.1.D
PCC=-0.068, p=0.782
mRNA Profile
Protein Expression Profile
DPA

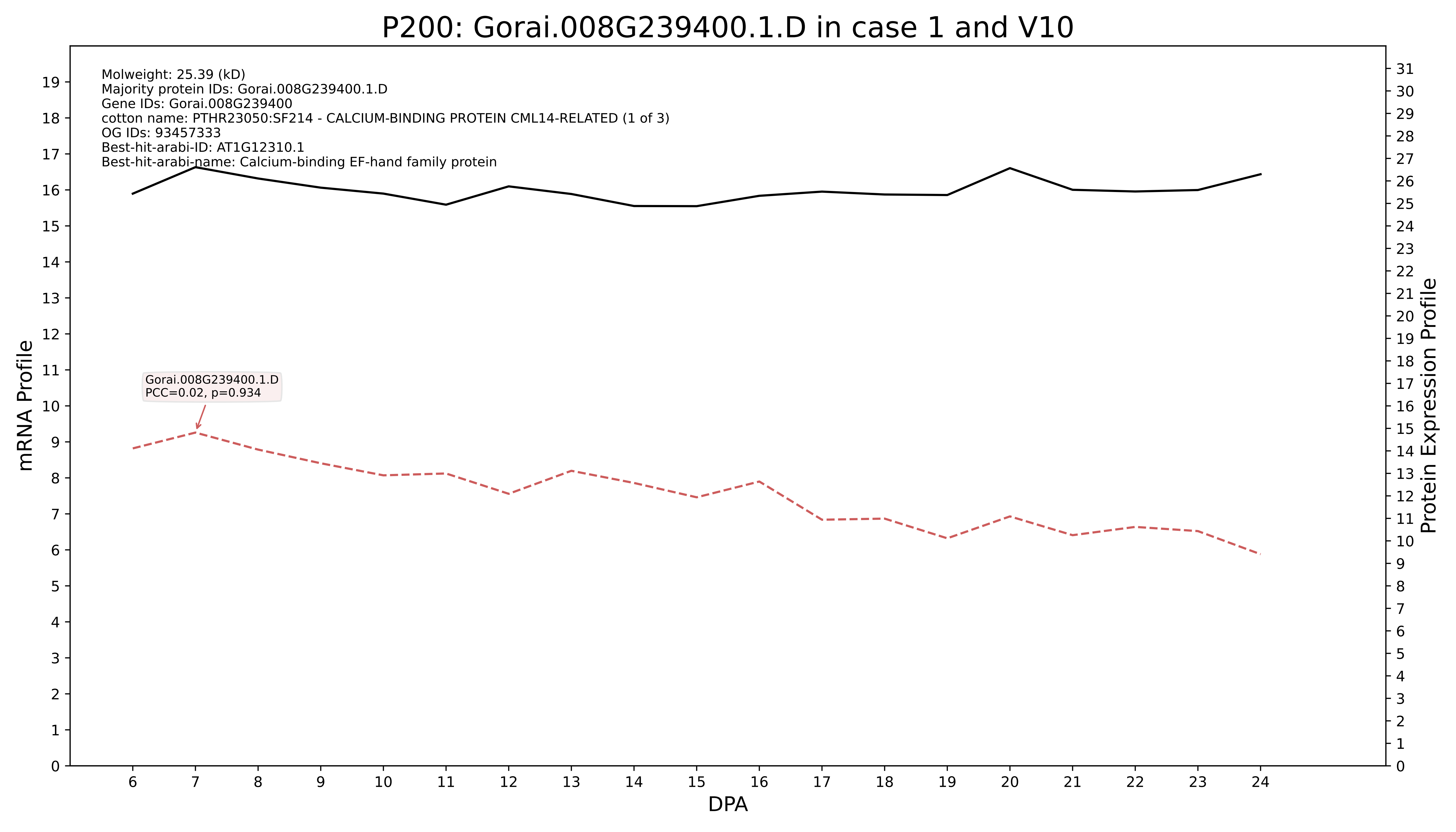
P200: Gorai.008G239400.1.D in case 1 and V10
Molweight: 25.39 (kD)
Majority protein IDs: Gorai.008G239400.1.D
Gene IDs: Gorai.008G239400
cotton name: PTHR23050:SF214 - CALCIUM-BINDING PROTEIN CML14-RELATED (1 of 3)
OG IDs: 93457333
Best-hit-arabi-ID: AT1G12310.1
Best-hit-arabi-name: Calcium-binding EF-hand family protein
Gorai.008G239400.1.D
PCC=0.02, p=0.934
mRNA Profile
Protein Expression Profile
DPA

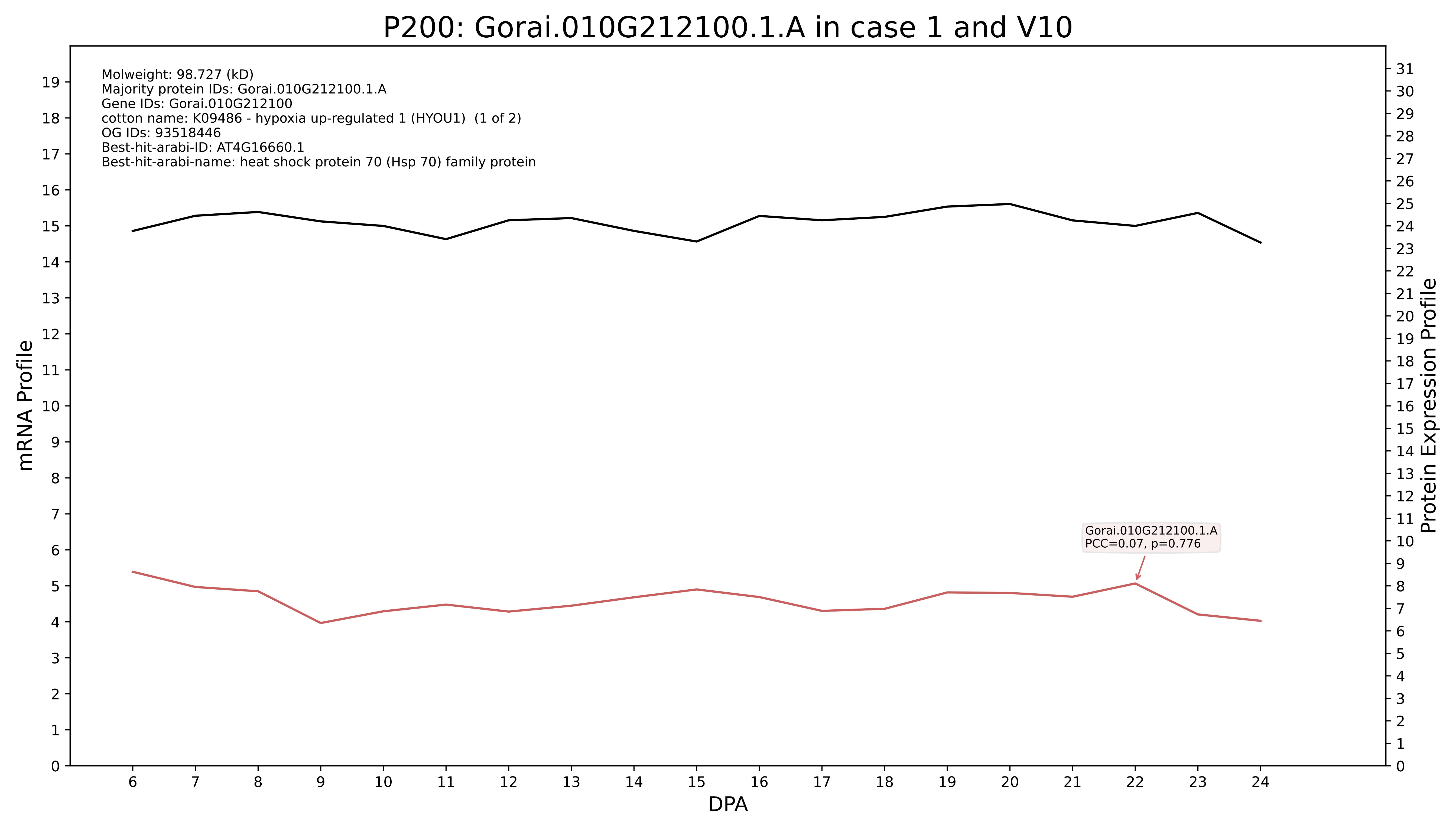

P200: Gorai.010G212100.1.A in case 1 and V10
Molweight: 98.727 (kD)
Majority protein IDs: Gorai.010G212100.1.A
Gene IDs: Gorai.010G212100
cotton name: K09486 - hypoxia up-regulated 1 (HYOU1) (1 of 2)
OG IDs: 93518446
Best-hit-arabi-ID: AT4G16660.1
Best-hit-arabi-name: heat shock protein 70 (Hsp 70) family protein
Gorai.010G212100.1.A
PCC=0.07, p=0.776
mRNA Profile
Protein Expression Profile
DPA

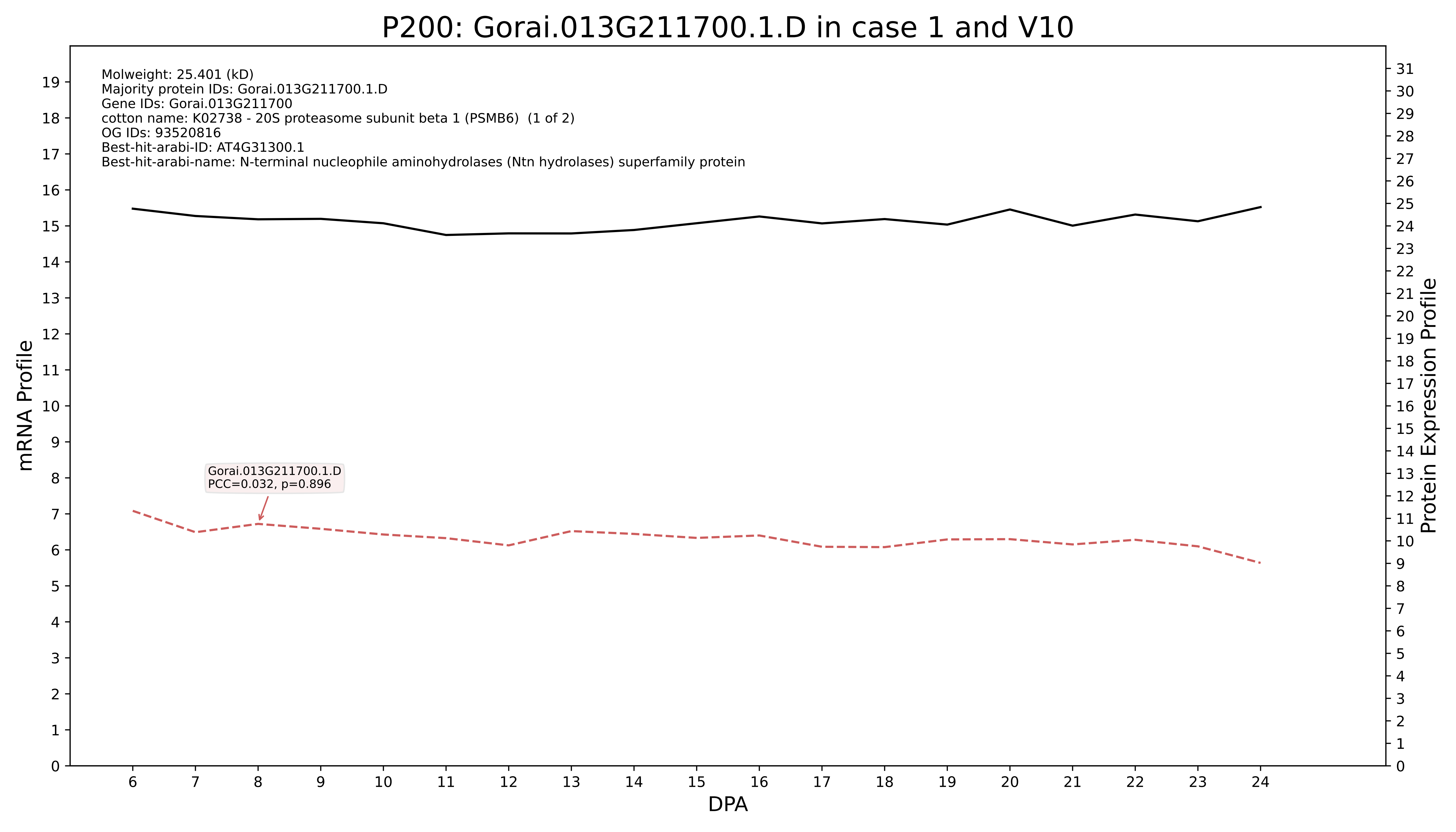

P200: Gorai.013G211700.1.D in case 1 and V10
Molweight: 25.401 (kD)
Majority protein IDs: Gorai.013G211700.1.D
Gene IDs: Gorai.013G211700
cotton name: K02738 - 20S proteasome subunit beta 1 (PSMB6) (1 of 2)
OG IDs: 93520816
Best-hit-arabi-ID: AT4G31300.1
Best-hit-arabi-name: N-terminal nucleophile aminohydrolases (Ntn hydrolases) superfamily protein
Gorai.013G211700.1.D
PCC=0.032, p=0.896
mRNA Profile
Protein Expression Profile
DPA

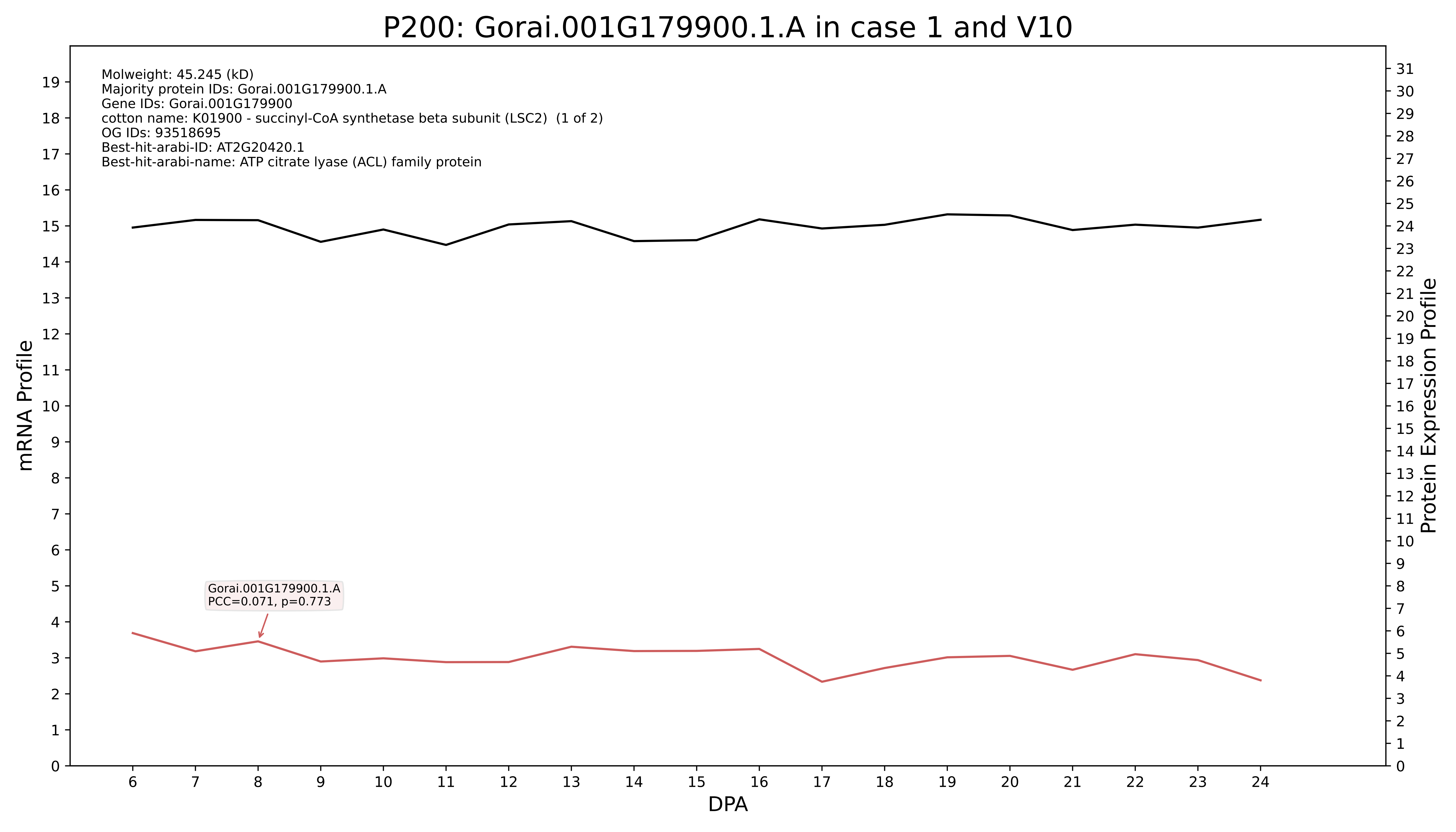

P200: Gorai.001G179900.1.A in case 1 and V10
Molweight: 45.245 (kD)
Majority protein IDs: Gorai.001G179900.1.A
Gene IDs: Gorai.001G179900
cotton name: K01900 - succinyl-CoA synthetase beta subunit (LSC2) (1 of 2)
OG IDs: 93518695
Best-hit-arabi-ID: AT2G20420.1
Best-hit-arabi-name: ATP citrate lyase (ACL) family protein
Gorai.001G179900.1.A
PCC=0.071, p=0.773
mRNA Profile
Protein Expression Profile
DPA

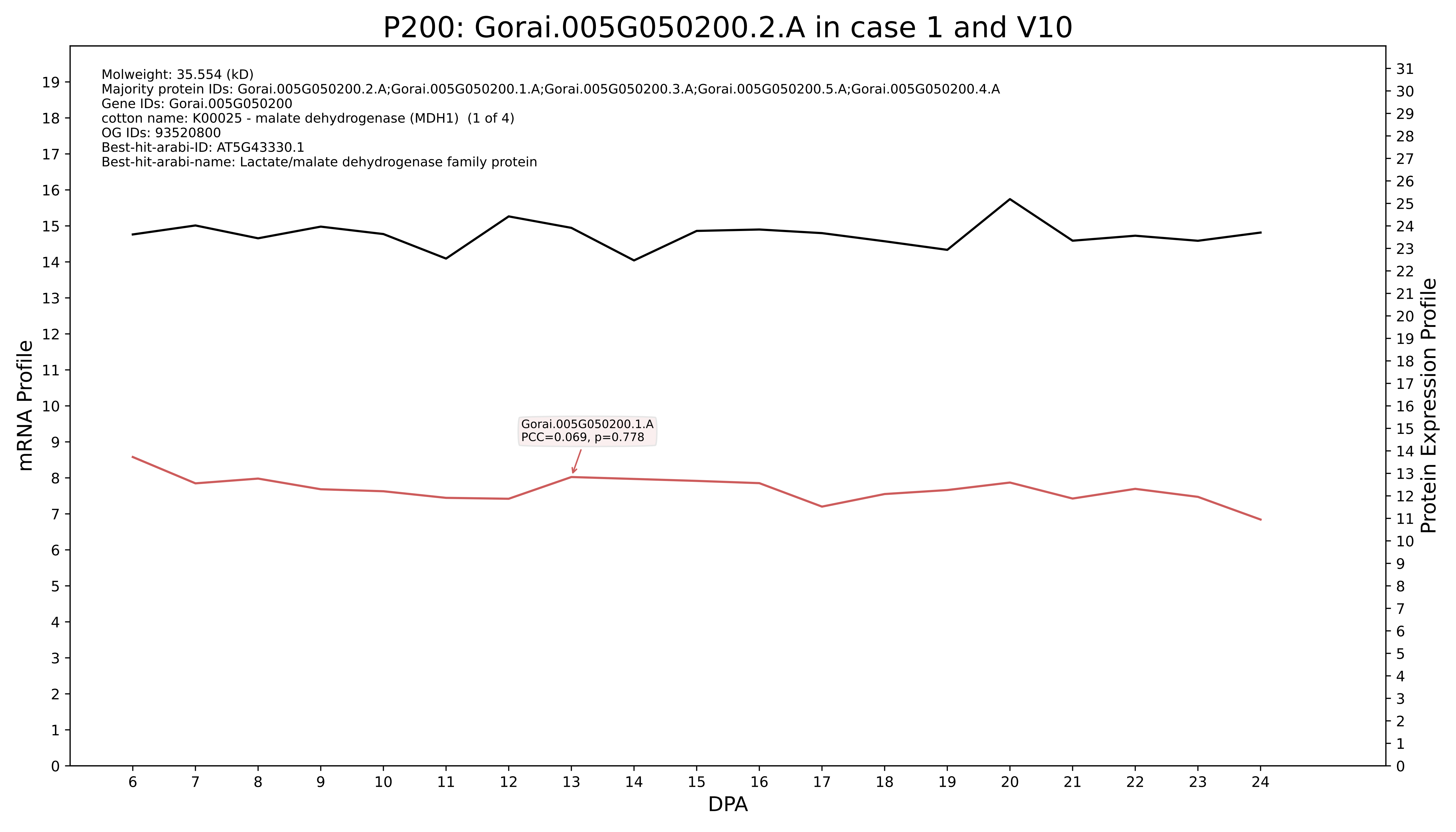
P200: Gorai.005G050200.2.A in case 1 and V10
Molweight: 35.554 (kD)
Majority protein IDs: Gorai.005G050200.2.A;Gorai.005G050200.1.A;Gorai.005G050200.3.A;Gorai.005G050200.5.A;Gorai.005G050200.4.A
Gene IDs: Gorai.005G050200
cotton name: K00025 - malate dehydrogenase (MDH1) (1 of 4)
OG IDs: 93520800
Best-hit-arabi-ID: AT5G43330.1
Best-hit-arabi-name: Lactate/malate dehydrogenase family protein
Gorai.005G050200.1.A
PCC=0.069, p=0.778
mRNA Profile
Protein Expression Profile
DPA

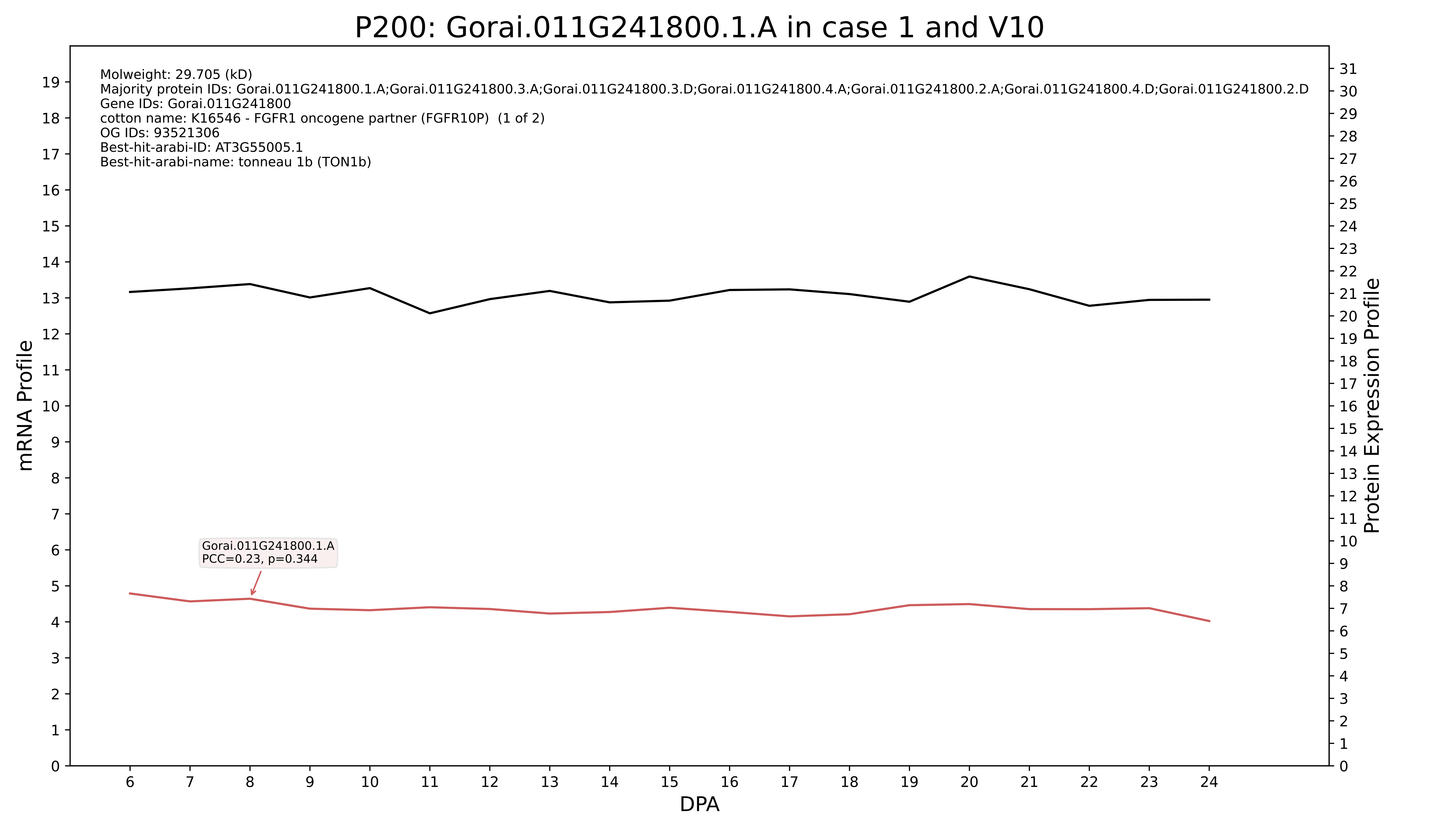
P200: Gorai.011G241800.1.A in case 1 and V10
Molweight: 29.705 (kD)
Majority protein IDs: Gorai.011G241800.1.A;Gorai.011G241800.3.A;Gorai.011G241800.3.D;Gorai.011G241800.4.A;Gorai.011G241800.2.A;Gorai.011G241800.4.D;Gorai.011G241800.2.D
Gene IDs: Gorai.011G241800
cotton name: K16546 - FGFR1 oncogene partner (FGFR10P) (1 of 2)
OG IDs: 93521306
Best-hit-arabi-ID: AT3G55005.1
Best-hit-arabi-name: tonneau 1b (TON1b)
Gorai.011G241800.1.A
PCC=0.23, p=0.344
mRNA Profile
Protein Expression Profile
DPA

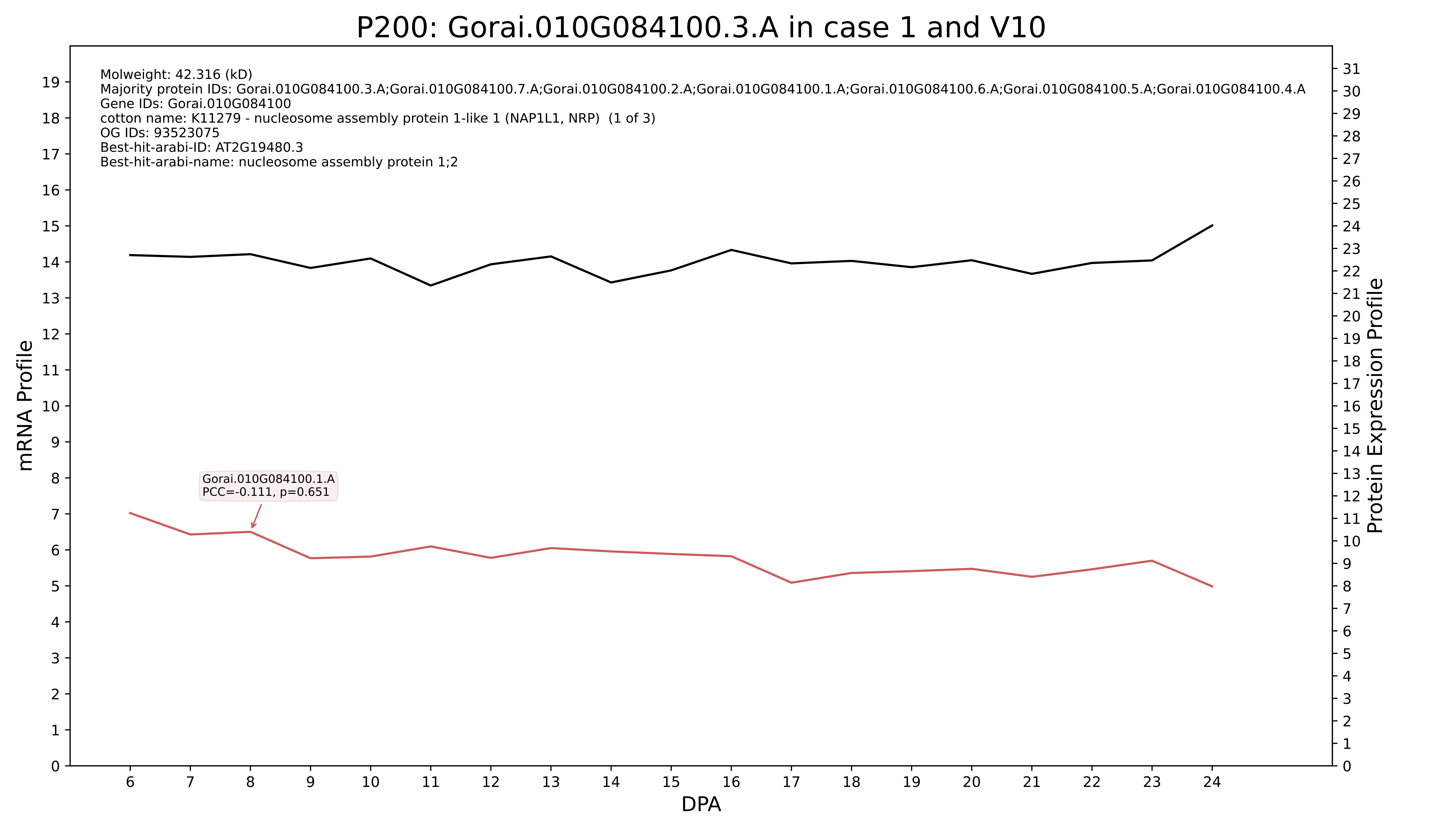

P200: Gorai.010G084100.3.A in case 1 and V10
Molweight: 42.316 (kD)
Majority protein IDs: Gorai.010G084100.3.A;Gorai.010G084100.7.A;Gorai.010G084100.2.A;Gorai.010G084100.1.A;Gorai.010G084100.6.A;Gorai.010G084100.5.A;Gorai.010G084100.4.A
Gene IDs: Gorai.010G084100
cotton name: K11279 - nucleosome assembly protein 1-like 1 (NAP1L1, NRP) (1 of 3)
OG IDs: 93523075
Best-hit-arabi-ID: AT2G19480.3
Best-hit-arabi-name: nucleosome assembly protein 1;2
Gorai.010G084100.1.A
PCC=-0.111, p=0.651
mRNA Profile
Protein Expression Profile
DPA

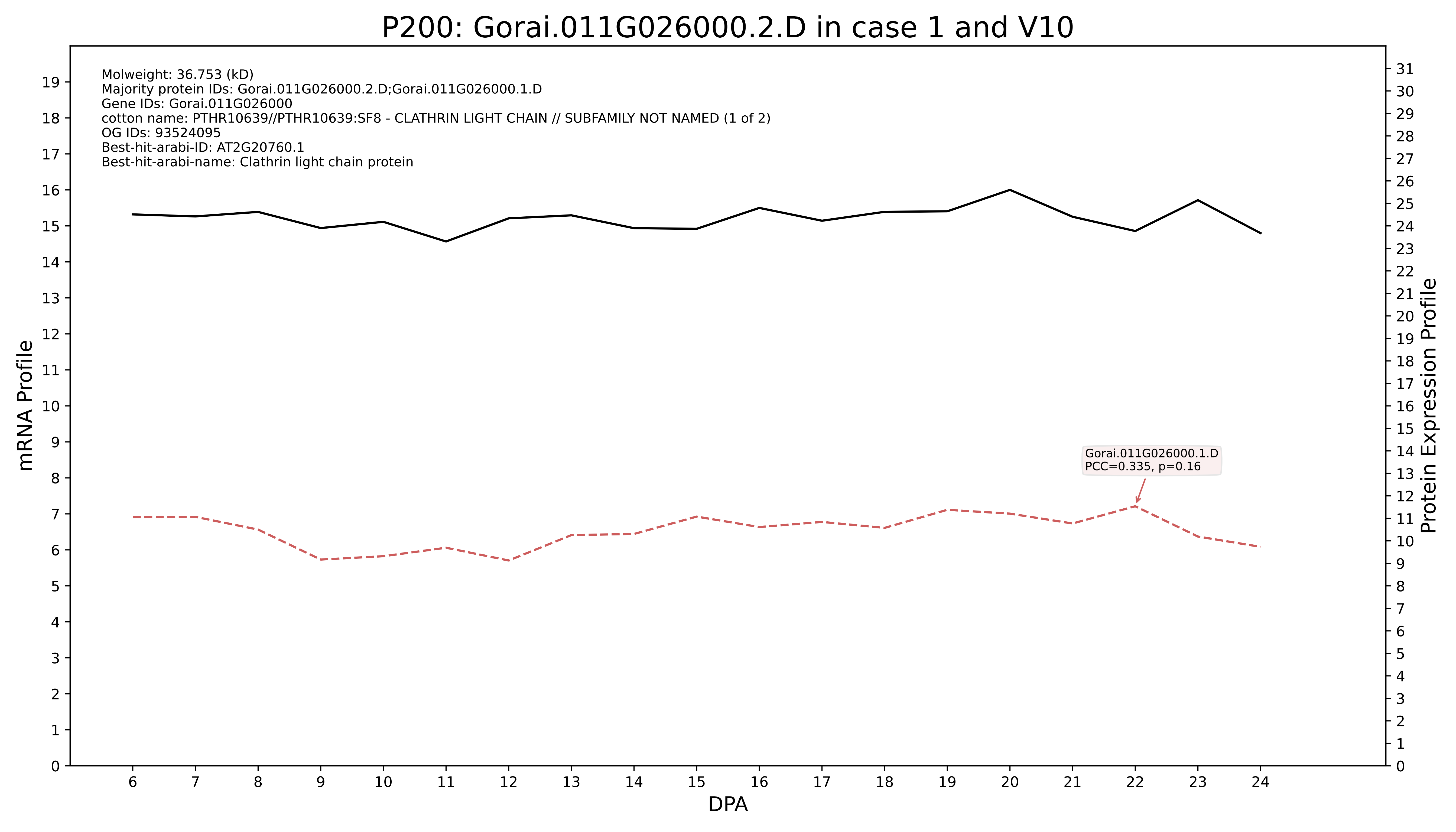
P200: Gorai.011G026000.2.D in case 1 and V10
Molweight: 36.753 (kD)
Majority protein IDs: Gorai.011G026000.2.D;Gorai.011G026000.1.D
Gene IDs: Gorai.011G026000
cotton name: PTHR10639//PTHR10639:SF8 - CLATHRIN LIGHT CHAIN // SUBFAMILY NOT NAMED (1 of 2)
OG IDs: 93524095
Best-hit-arabi-ID: AT2G20760.1
Best-hit-arabi-name: Clathrin light chain protein
Gorai.011G026000.1.D
PCC=0.335, p=0.16
mRNA Profile
Protein Expression Profile
DPA

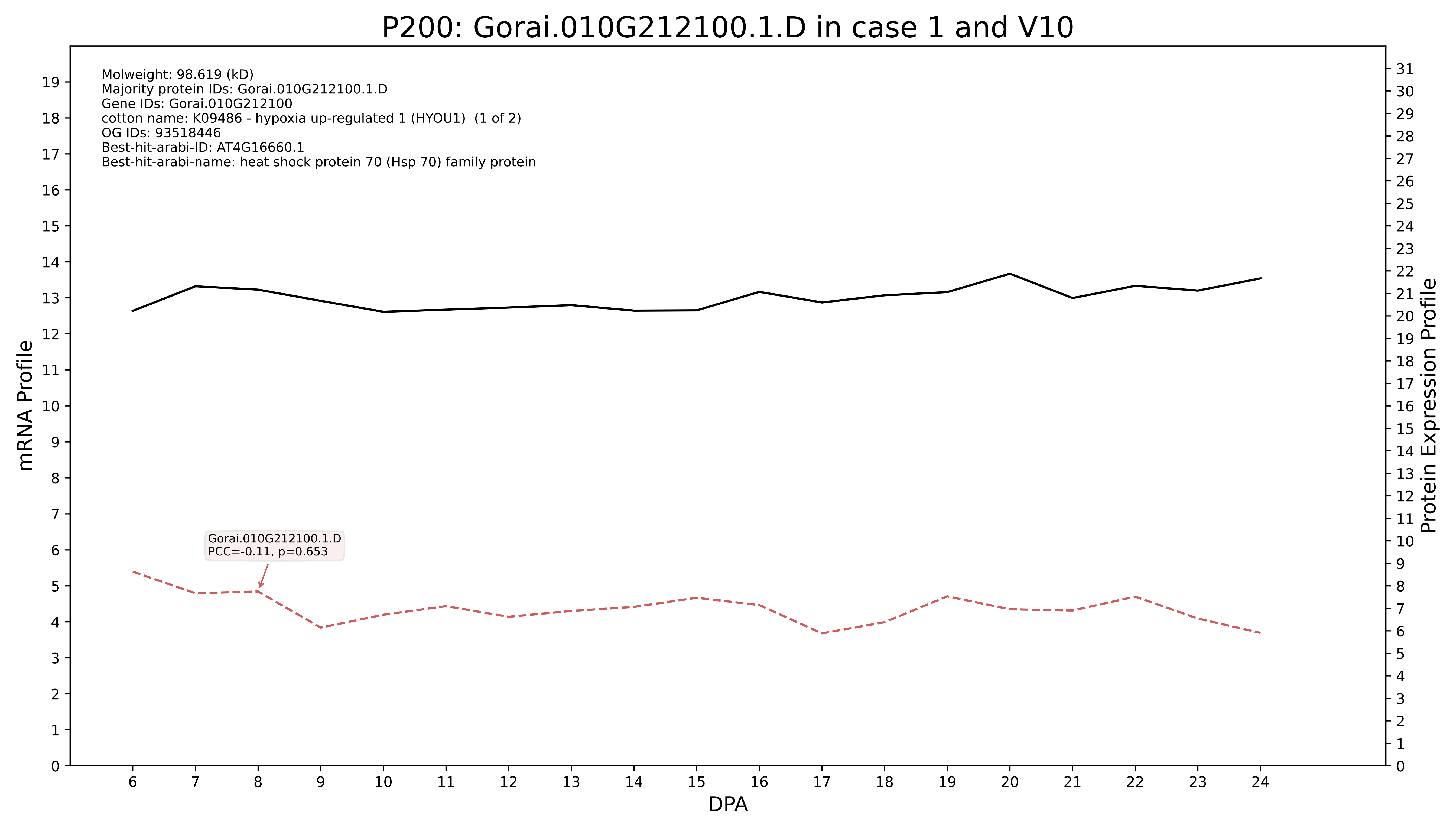

P200: Gorai.010G212100.1.D in case 1 and V10
Molweight: 98.619 (kD)
Majority protein IDs: Gorai.010G212100.1.D
Gene IDs: Gorai.010G212100
cotton name: K09486 - hypoxia up-regulated 1 (HYOU1) (1 of 2)
OG IDs: 93518446
Best-hit-arabi-ID: AT4G16660.1
Best-hit-arabi-name: heat shock protein 70 (Hsp 70) family protein
Gorai.010G212100.1.D
PCC=-0.11, p=0.653
mRNA Profile
Protein Expression Profile
DPA
0 1 2 3 4 5 6 7 8 9 10 11 12 13 14 15 16 17 18 19
0 1 2 3 4 5 6 7 8 9 10 11 12 13 14 15 16 17 18 19 20 21 22 23 24 25 26 27 28 29 30 31
6 7 8 9 10 11 12 13 14 15 16 17 18 19 20 21 22 23 24

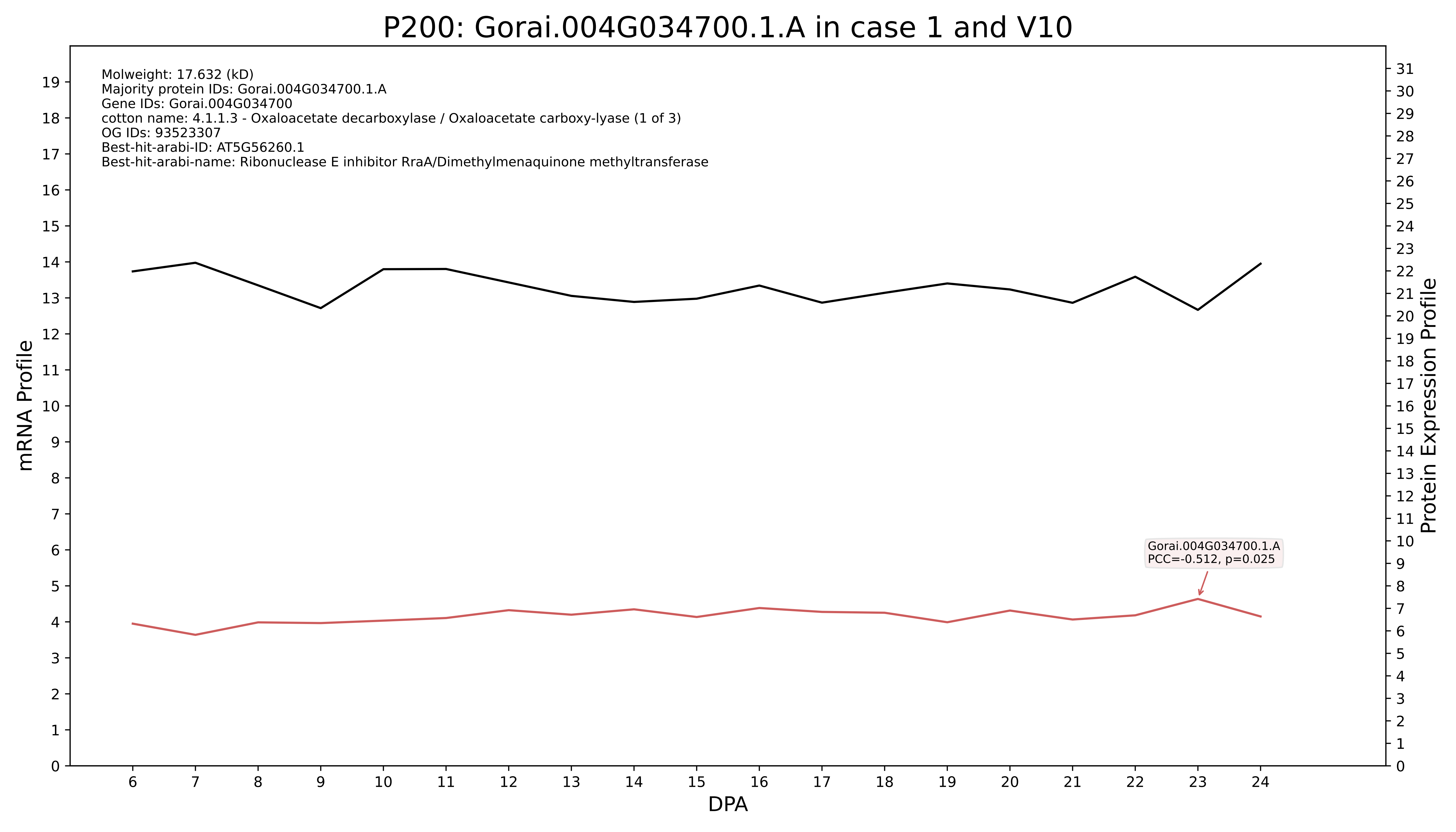

P200: Gorai.004G034700.1.A in case 1 and V10
Molweight: 17.632 (kD)
Majority protein IDs: Gorai.004G034700.1.A
Gene IDs: Gorai.004G034700
cotton name: 4.1.1.3 - Oxaloacetate decarboxylase / Oxaloacetate carboxy-lyase (1 of 3)
OG IDs: 93523307
Best-hit-arabi-ID: AT5G56260.1
Best-hit-arabi-name: Ribonuclease E inhibitor RraA/Dimethylmenaquinone methyltransferase
Gorai.004G034700.1.A
PCC=-0.512, p=0.025
mRNA Profile
Protein Expression Profile
DPA

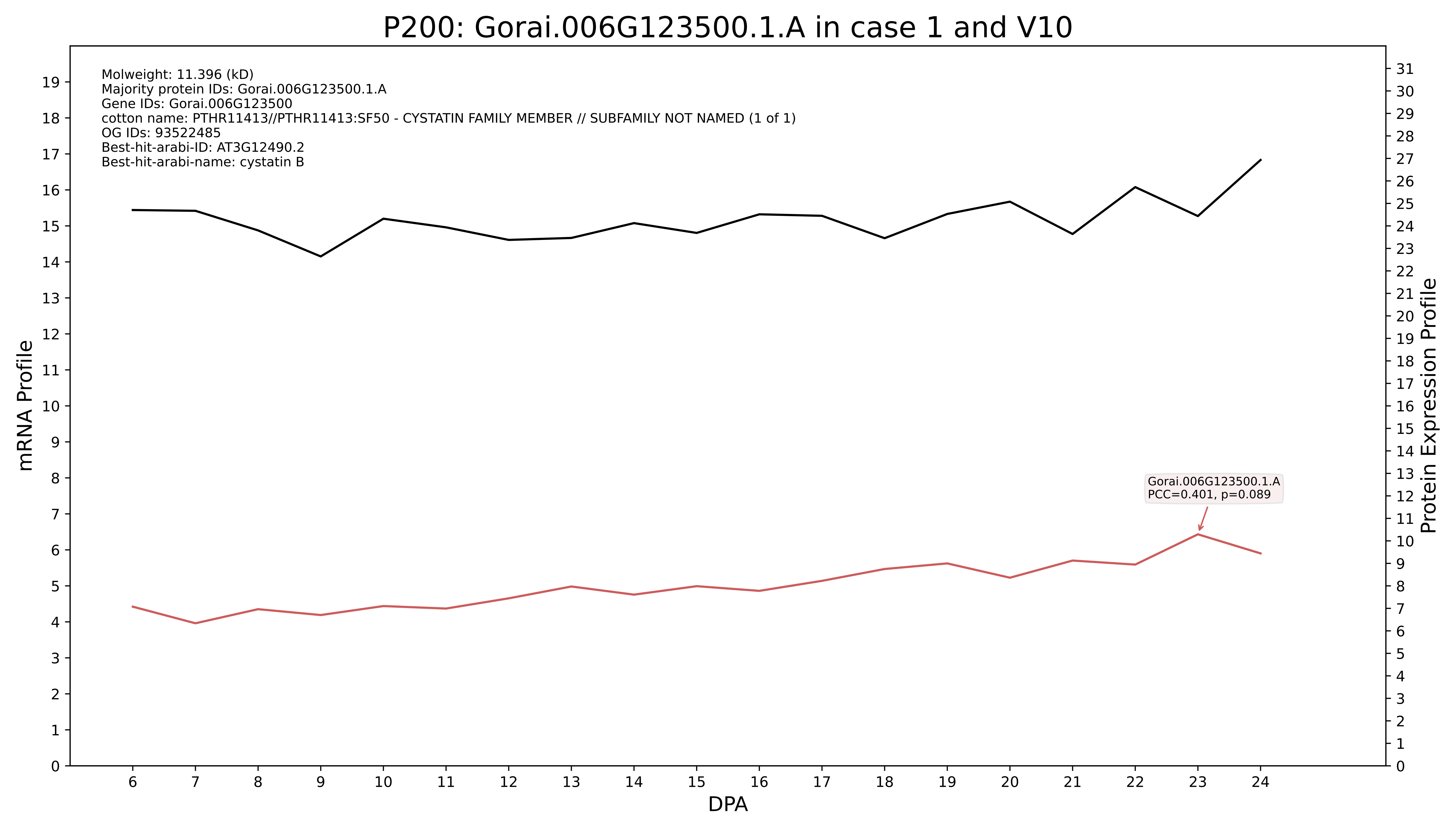
P200: Gorai.006G123500.1.A in case 1 and V10
Molweight: 11.396 (kD)
Majority protein IDs: Gorai.006G123500.1.A
Gene IDs: Gorai.006G123500
cotton name: PTHR11413//PTHR11413:SF50 - CYSTATIN FAMILY MEMBER // SUBFAMILY NOT NAMED (1 of 1)
OG IDs: 93522485
Best-hit-arabi-ID: AT3G12490.2
Best-hit-arabi-name: cystatin B
Gorai.006G123500.1.A
PCC=0.401, p=0.089
mRNA Profile
Protein Expression Profile
DPA

### P200: Gorai.009G092400.1.D in case 1 and V10

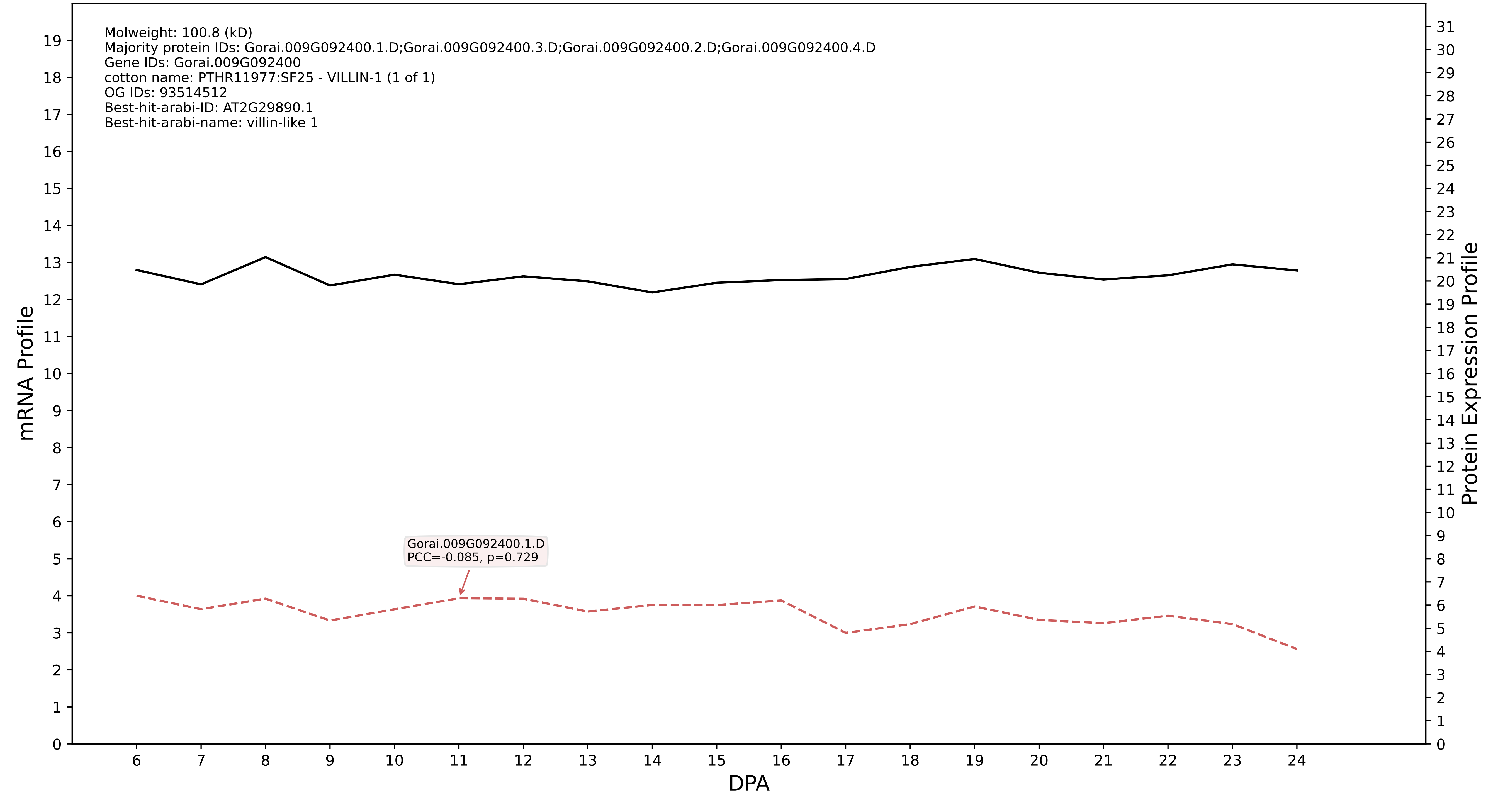

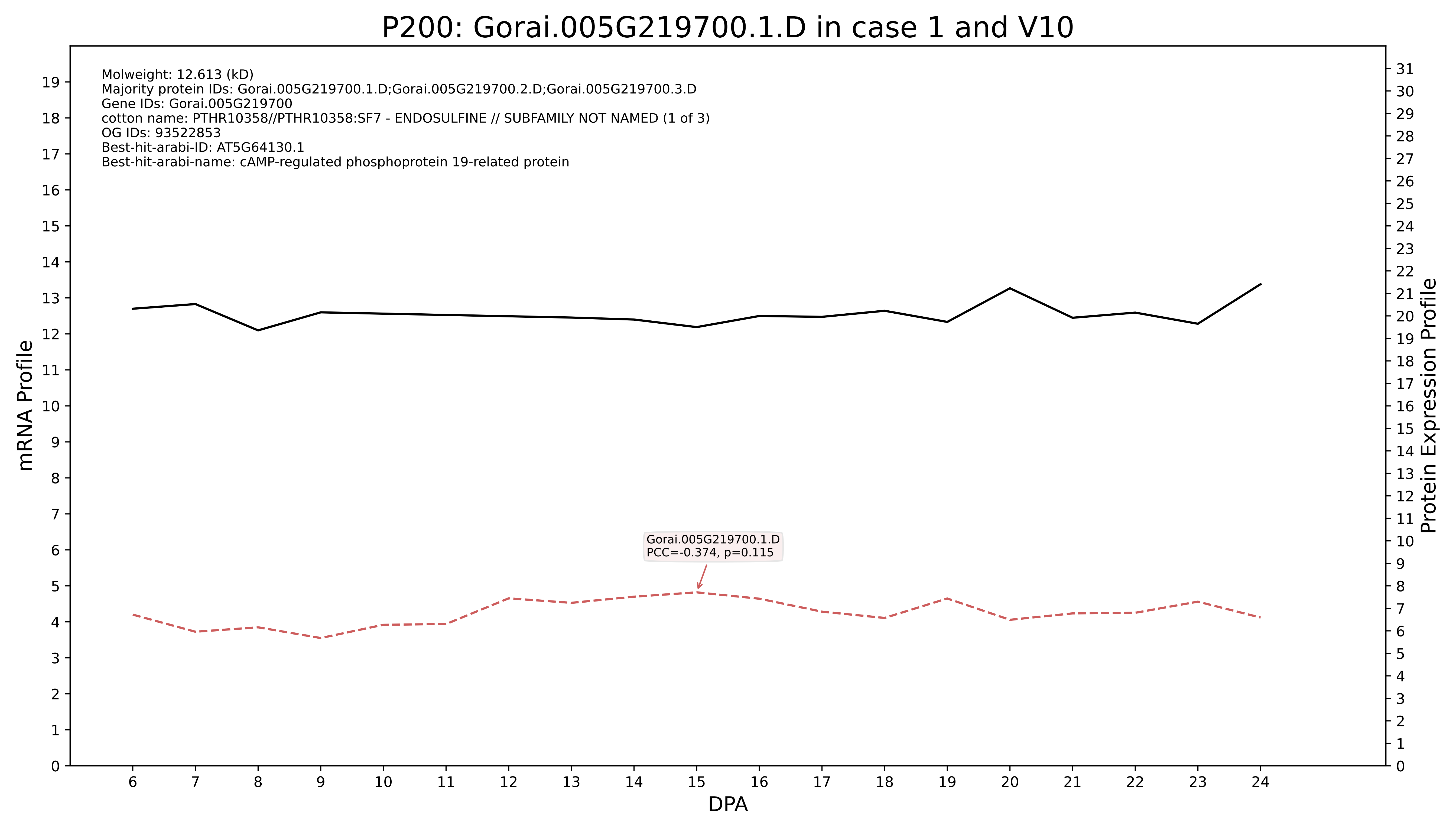

P200: Gorai.005G219700.1.D in case 1 and V10
Molweight: 12.613 (kD)
Majority protein IDs: Gorai.005G219700.1.D;Gorai.005G219700.2.D;Gorai.005G219700.3.D
Gene IDs: Gorai.005G219700
cotton name: PTHR10358//PTHR10358:SF7 - ENDOSULFINE // SUBFAMILY NOT NAMED (1 of 3)
OG IDs: 93522853
Best-hit-arabi-ID: AT5G64130.1
Best-hit-arabi-name: cAMP-regulated phosphoprotein 19-related protein
Gorai.005G219700.1.D
PCC=-0.374, p=0.115
mRNA Profile
Protein Expression Profile
DPA

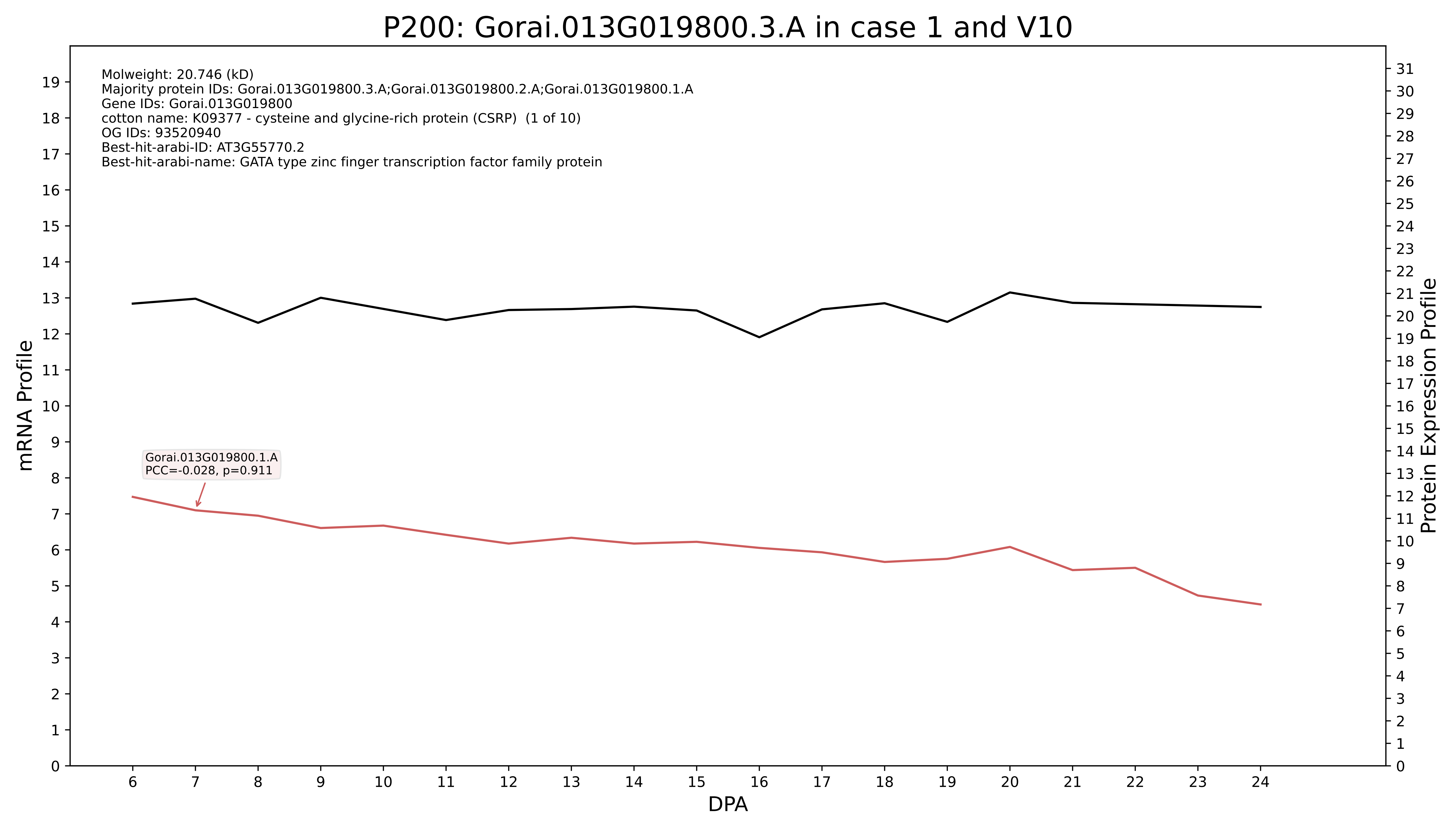
P200: Gorai.013G019800.3.A in case 1 and V10
Molweight: 20.746 (kD)
Majority protein IDs: Gorai.013G019800.3.A;Gorai.013G019800.2.A;Gorai.013G019800.1.A
Gene IDs: Gorai.013G019800
cotton name: K09377 - cysteine and glycine-rich protein (CSRP) (1 of 10)
OG IDs: 93520940
Best-hit-arabi-ID: AT3G55770.2
Best-hit-arabi-name: GATA type zinc finger transcription factor family protein
Gorai.013G019800.1.A
PCC=-0.028, p=0.911
mRNA Profile
Protein Expression Profile
DPA

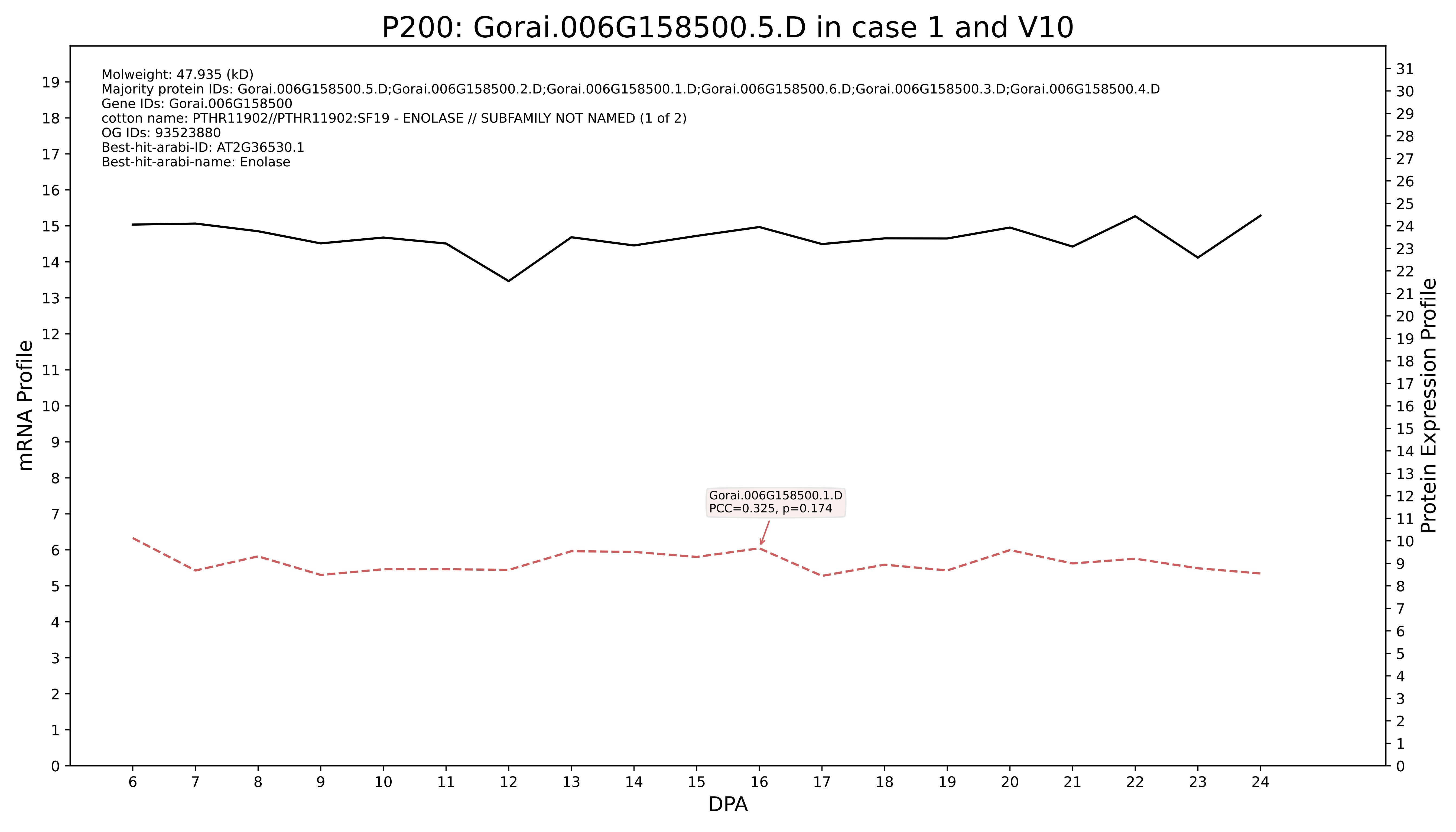
P200: Gorai.006G158500.5.D in case 1 and V10
Molweight: 47.935 (kD)
Majority protein IDs: Gorai.006G158500.5.D;Gorai.006G158500.2.D;Gorai.006G158500.1.D;Gorai.006G158500.6.D;Gorai.006G158500.3.D;Gorai.006G158500.4.D
Gene IDs: Gorai.006G158500
cotton name: PTHR11902//PTHR11902:SF19 - ENOLASE // SUBFAMILY NOT NAMED (1 of 2)
OG IDs: 93523880
Best-hit-arabi-ID: AT2G36530.1
Best-hit-arabi-name: Enolase
Gorai.006G158500.1.D
PCC=0.325, p=0.174
mRNA Profile
Protein Expression Profile
DPA

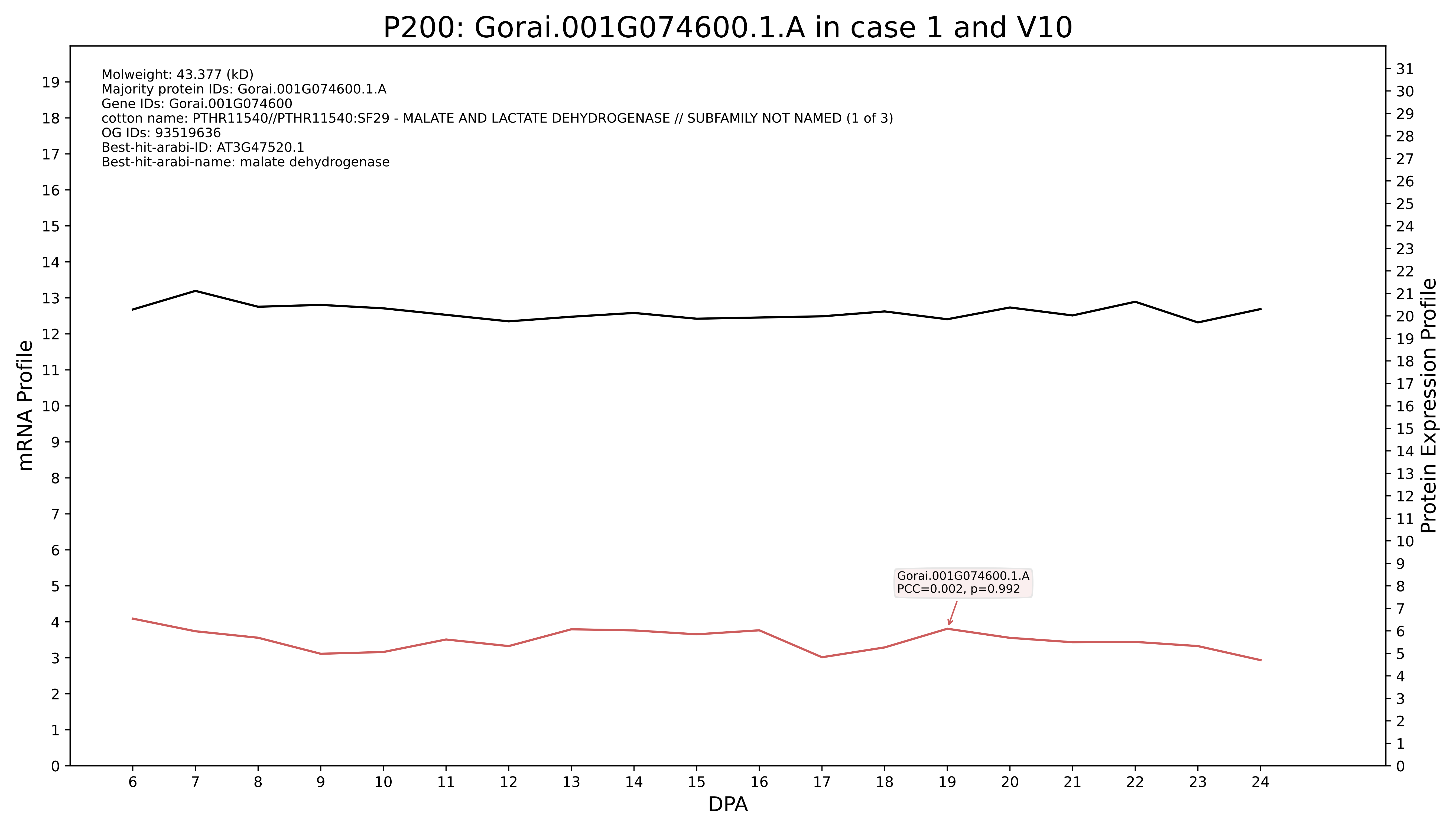
P200: Gorai.001G074600.1.A in case 1 and V10
Molweight: 43.377 (kD)
Majority protein IDs: Gorai.001G074600.1.A
Gene IDs: Gorai.001G074600
cotton name: PTHR11540//PTHR11540:SF29 - MALATE AND LACTATE DEHYDROGENASE // SUBFAMILY NOT NAMED (1 of 3)
OG IDs: 93519636
Best-hit-arabi-ID: AT3G47520.1
Best-hit-arabi-name: malate dehydrogenase
Gorai.001G074600.1.A
PCC=0.002, p=0.992
mRNA Profile
Protein Expression Profile
DPA

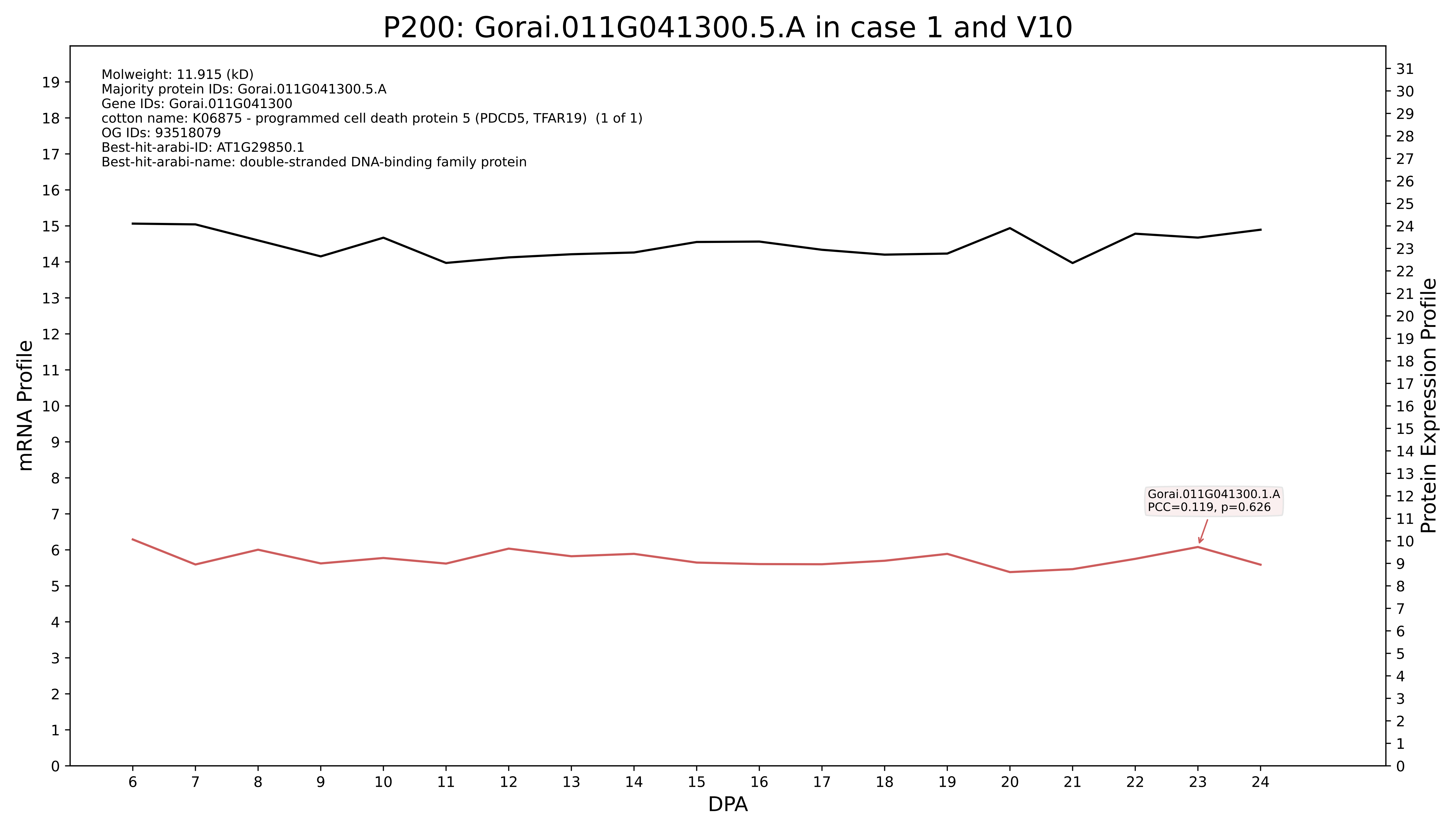
P200: Gorai.011G041300.5.A in case 1 and V10
Molweight: 11.915 (kD)
Majority protein IDs: Gorai.011G041300.5.A
Gene IDs: Gorai.011G041300
cotton name: K06875 - programmed cell death protein 5 (PDCD5, TFAR19) (1 of 1)
OG IDs: 93518079
Best-hit-arabi-ID: AT1G29850.1
Best-hit-arabi-name: double-stranded DNA-binding family protein
Gorai.011G041300.1.A
PCC=0.119, p=0.626
mRNA Profile
Protein Expression Profile
DPA

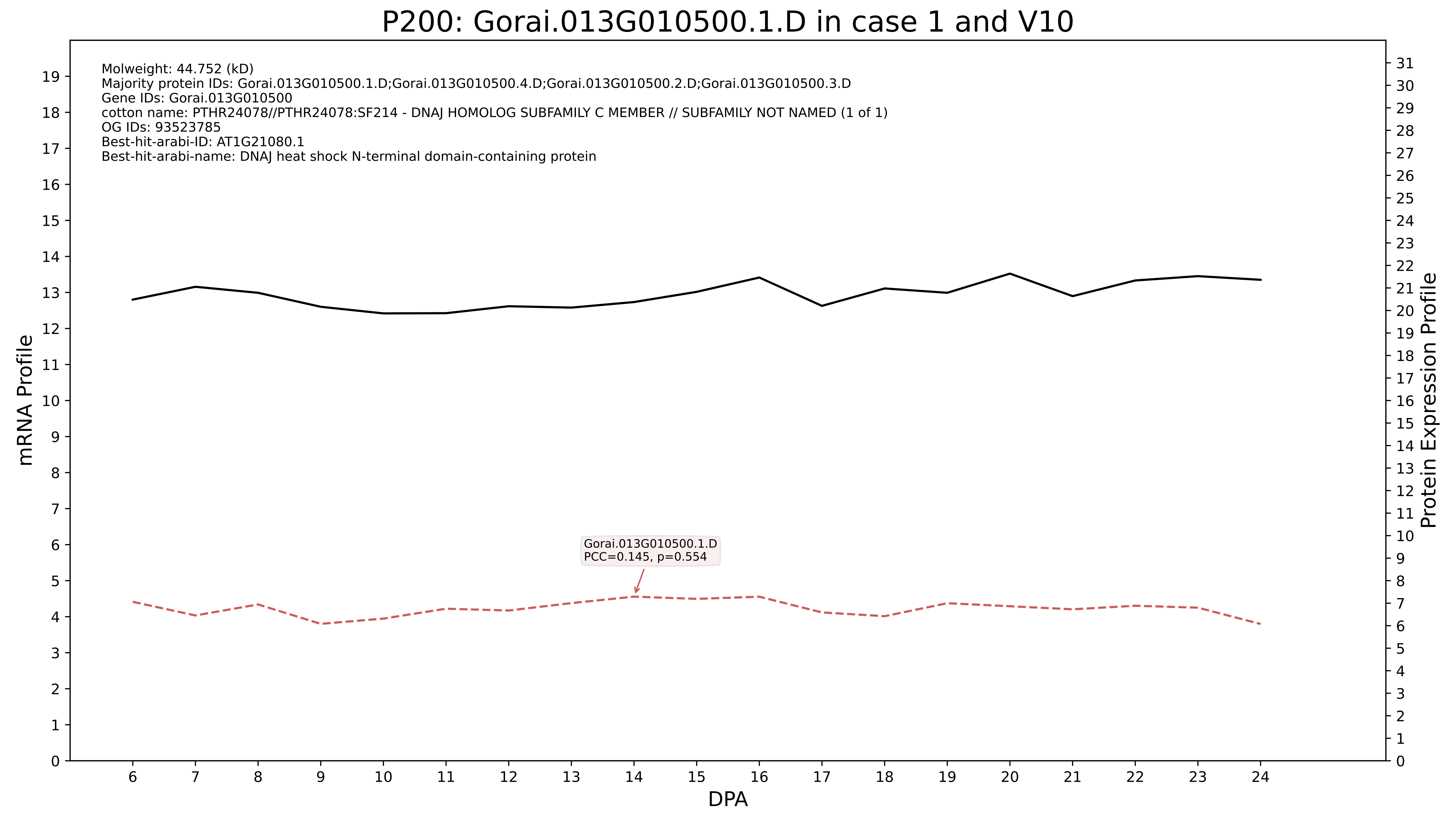

P200: Gorai.013G010500.1.D in case 1 and V10
Molweight: 44.752 (kD)
Majority protein IDs: Gorai.013G010500.1.D;Gorai.013G010500.4.D;Gorai.013G010500.2.D;Gorai.013G010500.3.D
Gene IDs: Gorai.013G010500
cotton name: PTHR24078//PTHR24078:SF214 - DNAJ HOMOLOG SUBFAMILY C MEMBER // SUBFAMILY NOT NAMED (1 of 1)
OG IDs: 93523785
Best-hit-arabi-ID: AT1G21080.1
Best-hit-arabi-name: DNAJ heat shock N-terminal domain-containing protein
Gorai.013G010500.1.D
PCC=0.145, p=0.554
mRNA Profile
Protein Expression Profile
DPA

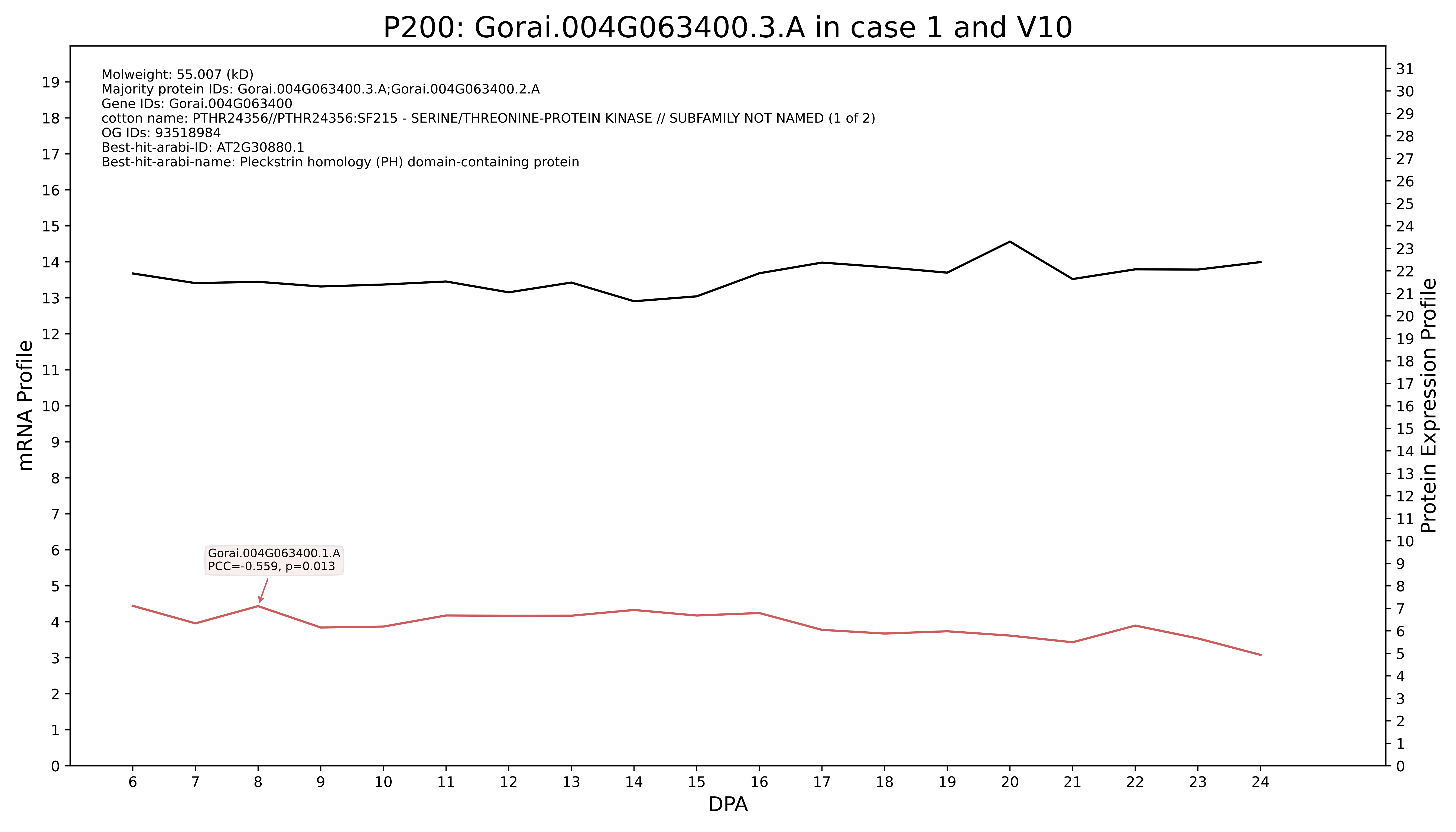
P200: Gorai.004G063400.3.A in case 1 and V10
Molweight: 55.007 (kD)
Majority protein IDs: Gorai.004G063400.3.A;Gorai.004G063400.2.A
Gene IDs: Gorai.004G063400
cotton name: PTHR24356//PTHR24356:SF215 - SERINE/THREONINE-PROTEIN KINASE // SUBFAMILY NOT NAMED (1 of 2)
OG IDs: 93518984
Best-hit-arabi-ID: AT2G30880.1
Best-hit-arabi-name: Pleckstrin homology (PH) domain-containing protein
Gorai.004G063400.1.A
PCC=-0.559, p=0.013
mRNA Profile
Protein Expression Profile
DPA

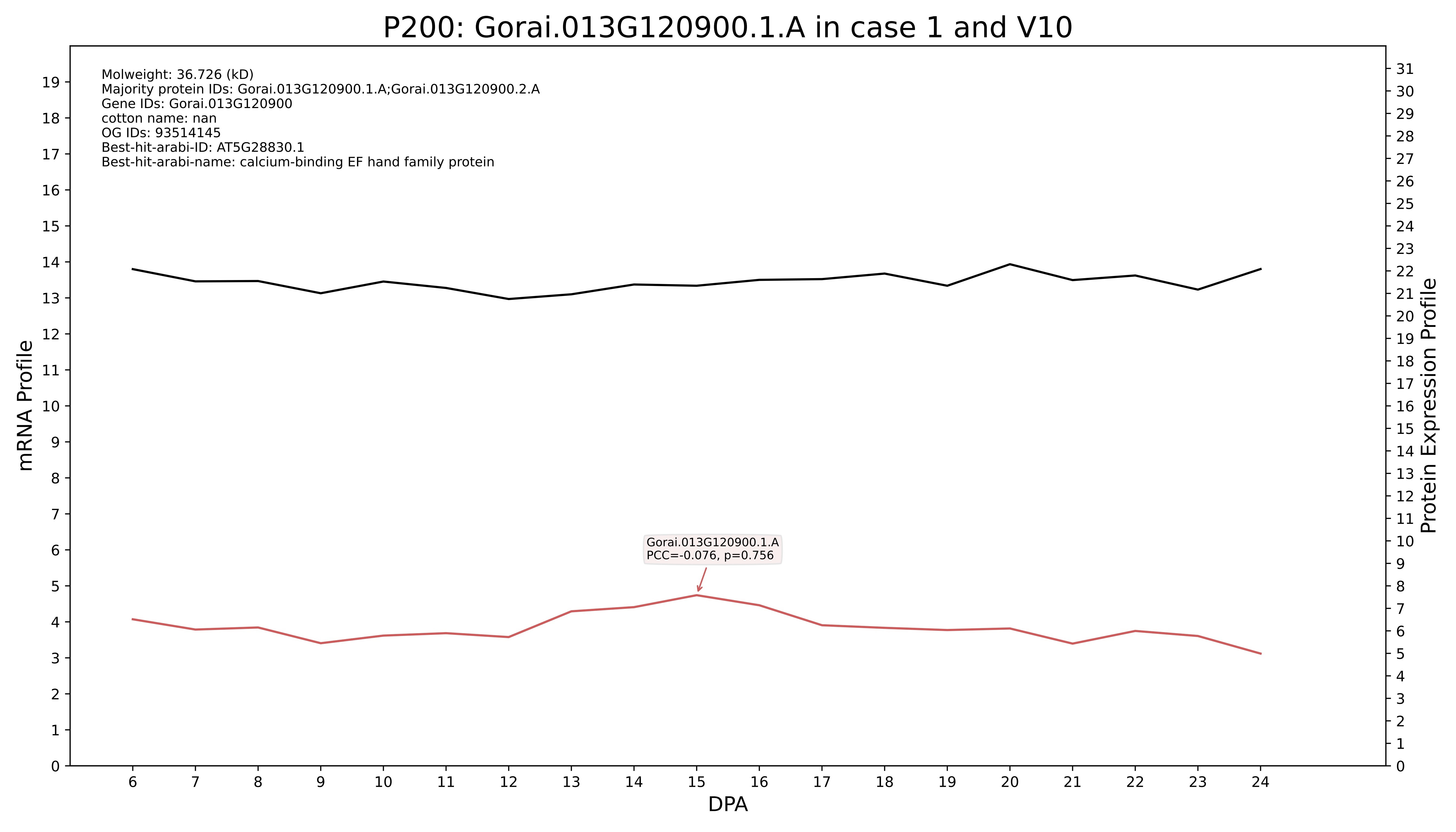
P200: Gorai.013G120900.1.A in case 1 and V10
Molweight: 36.726 (kD)
Majority protein IDs: Gorai.013G120900.1.A;Gorai.013G120900.2.A
Gene IDs: Gorai.013G120900
cotton name: nan
OG IDs: 93514145
Best-hit-arabi-ID: AT5G28830.1
Best-hit-arabi-name: calcium-binding EF hand family protein
Gorai.013G120900.1.A
PCC=-0.076, p=0.756
mRNA Profile
Protein Expression Profile
DPA

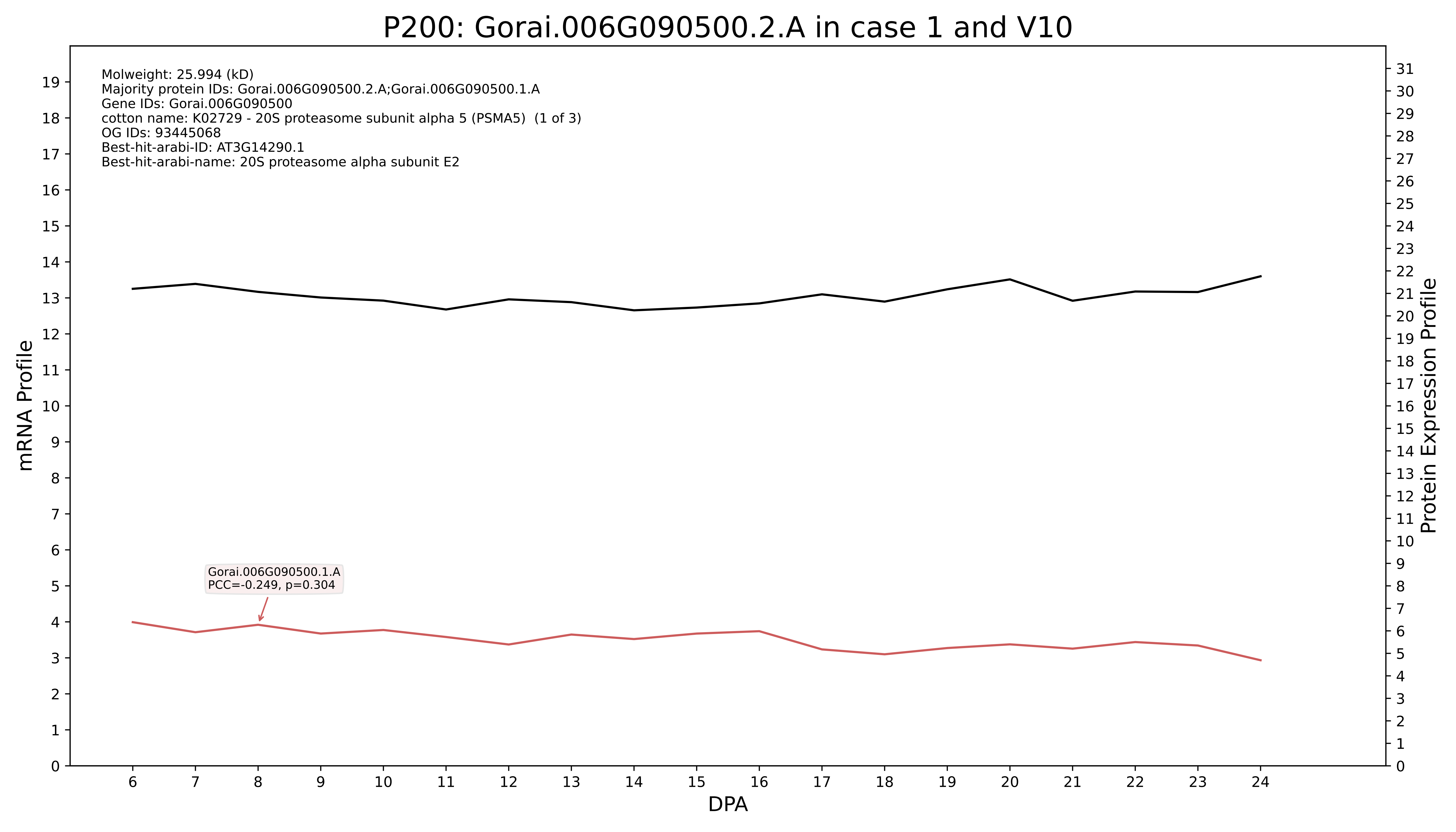

P200: Gorai.006G090500.2.A in case 1 and V10
Molweight: 25.994 (kD)
Majority protein IDs: Gorai.006G090500.2.A;Gorai.006G090500.1.A
Gene IDs: Gorai.006G090500
cotton name: K02729 - 20S proteasome subunit alpha 5 (PSMA5) (1 of 3)
OG IDs: 93445068
Best-hit-arabi-ID: AT3G14290.1
Best-hit-arabi-name: 20S proteasome alpha subunit E2
Gorai.006G090500.1.A
PCC=-0.249, p=0.304
mRNA Profile
Protein Expression Profile
DPA

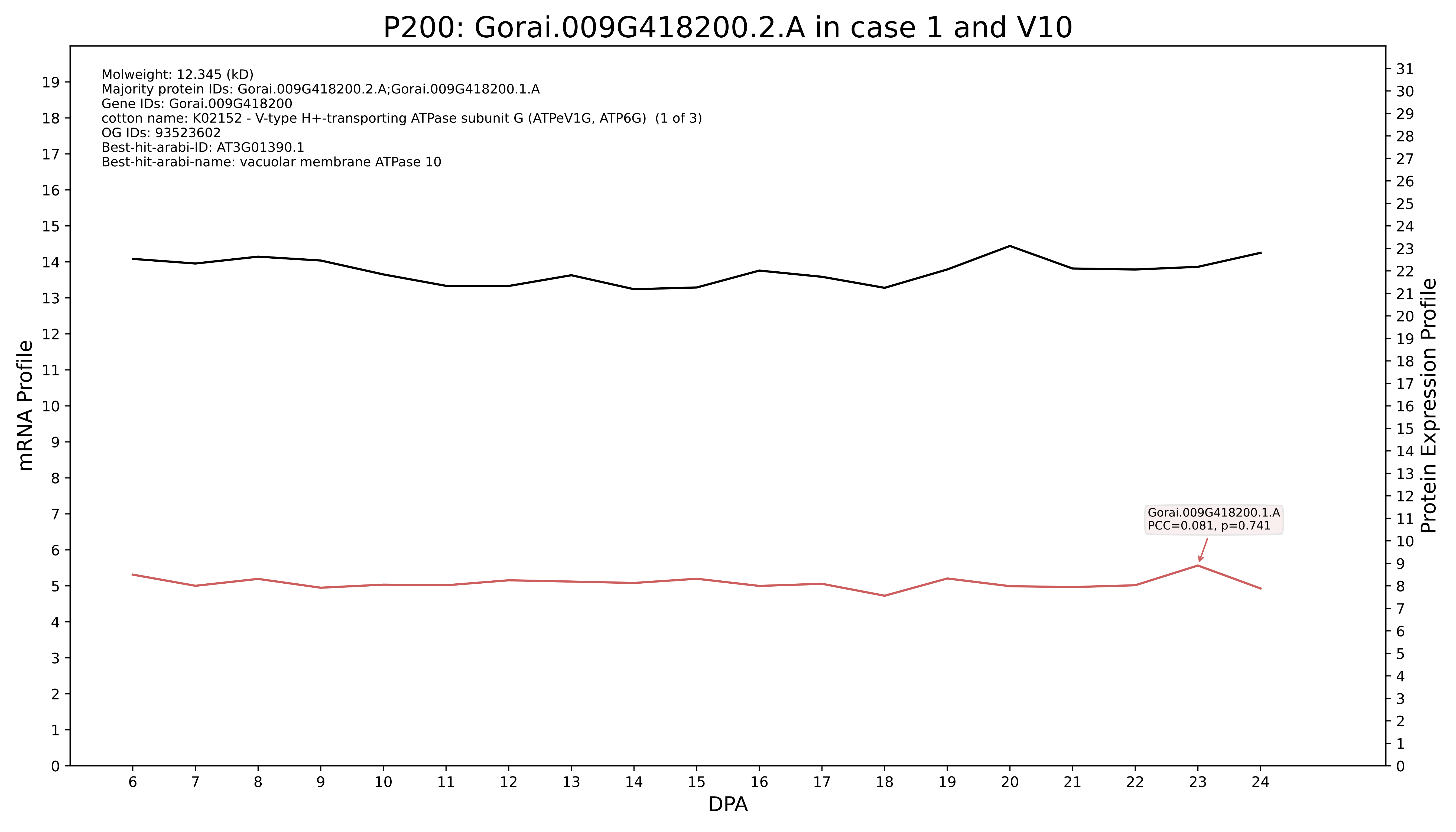
P200: Gorai.009G418200.2.A in case 1 and V10
Molweight: 12.345 (kD)
Majority protein IDs: Gorai.009G418200.2.A;Gorai.009G418200.1.A
Gene IDs: Gorai.009G418200
cotton name: K02152 - V-type H+-transporting ATPase subunit G (ATPeV1G, ATP6G) (1 of 3)
OG IDs: 93523602
Best-hit-arabi-ID: AT3G01390.1
Best-hit-arabi-name: vacuolar membrane ATPase 10
Gorai.009G418200.1.A
PCC=0.081, p=0.741
mRNA Profile
Protein Expression Profile
DPA

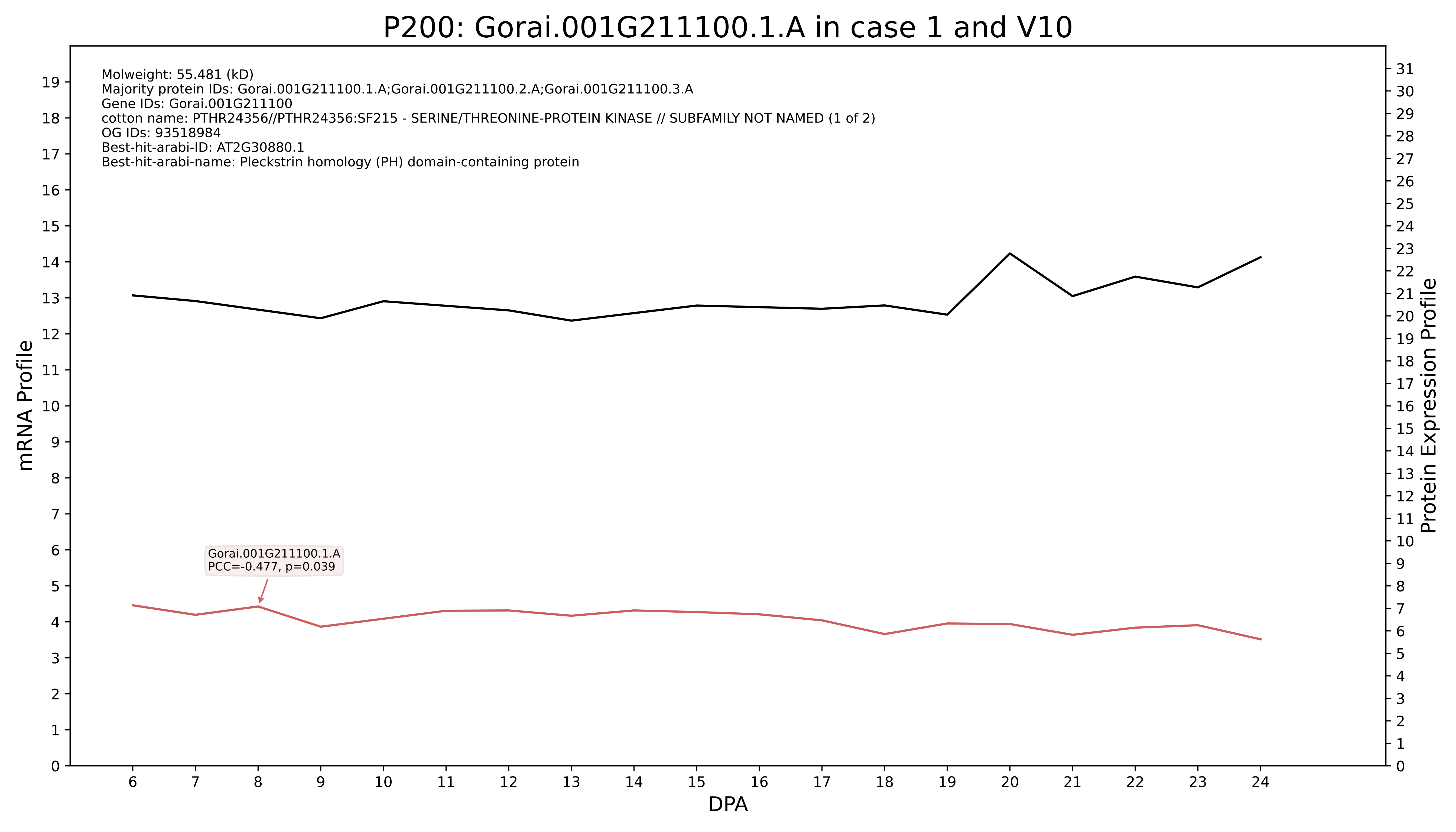
P200: Gorai.001G211100.1.A in case 1 and V10
Molweight: 55.481 (kD)
Majority protein IDs: Gorai.001G211100.1.A;Gorai.001G211100.2.A;Gorai.001G211100.3.A
Gene IDs: Gorai.001G211100
cotton name: PTHR24356//PTHR24356:SF215 - SERINE/THREONINE-PROTEIN KINASE // SUBFAMILY NOT NAMED (1 of 2)
OG IDs: 93518984
Best-hit-arabi-ID: AT2G30880.1
Best-hit-arabi-name: Pleckstrin homology (PH) domain-containing protein
Gorai.001G211100.1.A
PCC=-0.477, p=0.039
mRNA Profile
Protein Expression Profile
DPA

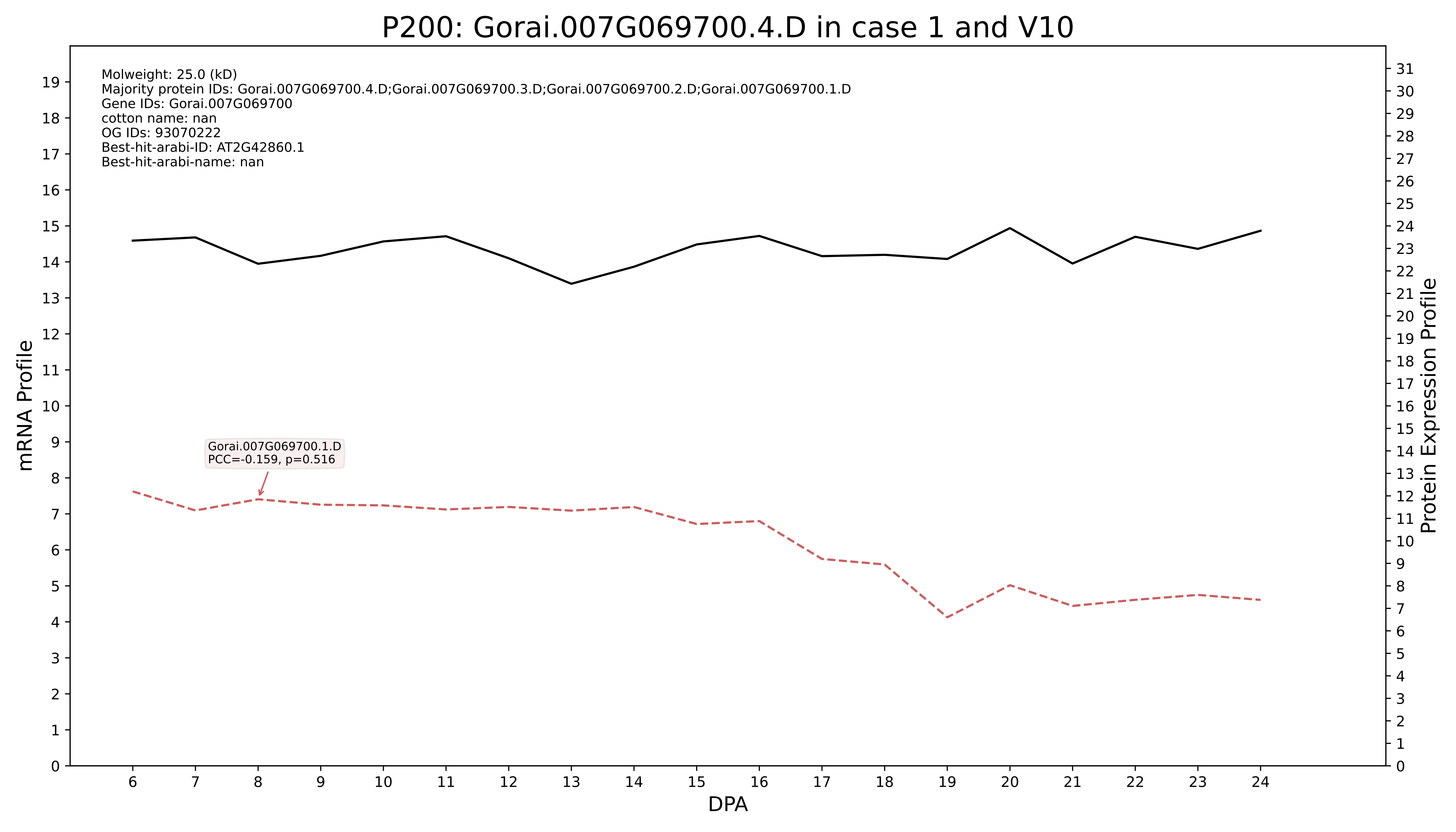

P200: Gorai.007G069700.4.D in case 1 and V10
Molweight: 25.0 (kD)
Majority protein IDs: Gorai.007G069700.4.D;Gorai.007G069700.3.D;Gorai.007G069700.2.D;Gorai.007G069700.1.D
Gene IDs: Gorai.007G069700
cotton name: nan
OG IDs: 93070222
Best-hit-arabi-ID: AT2G42860.1
Best-hit-arabi-name: nan
Gorai.007G069700.1.D
PCC=-0.159, p=0.516
mRNA Profile
Protein Expression Profile
DPA

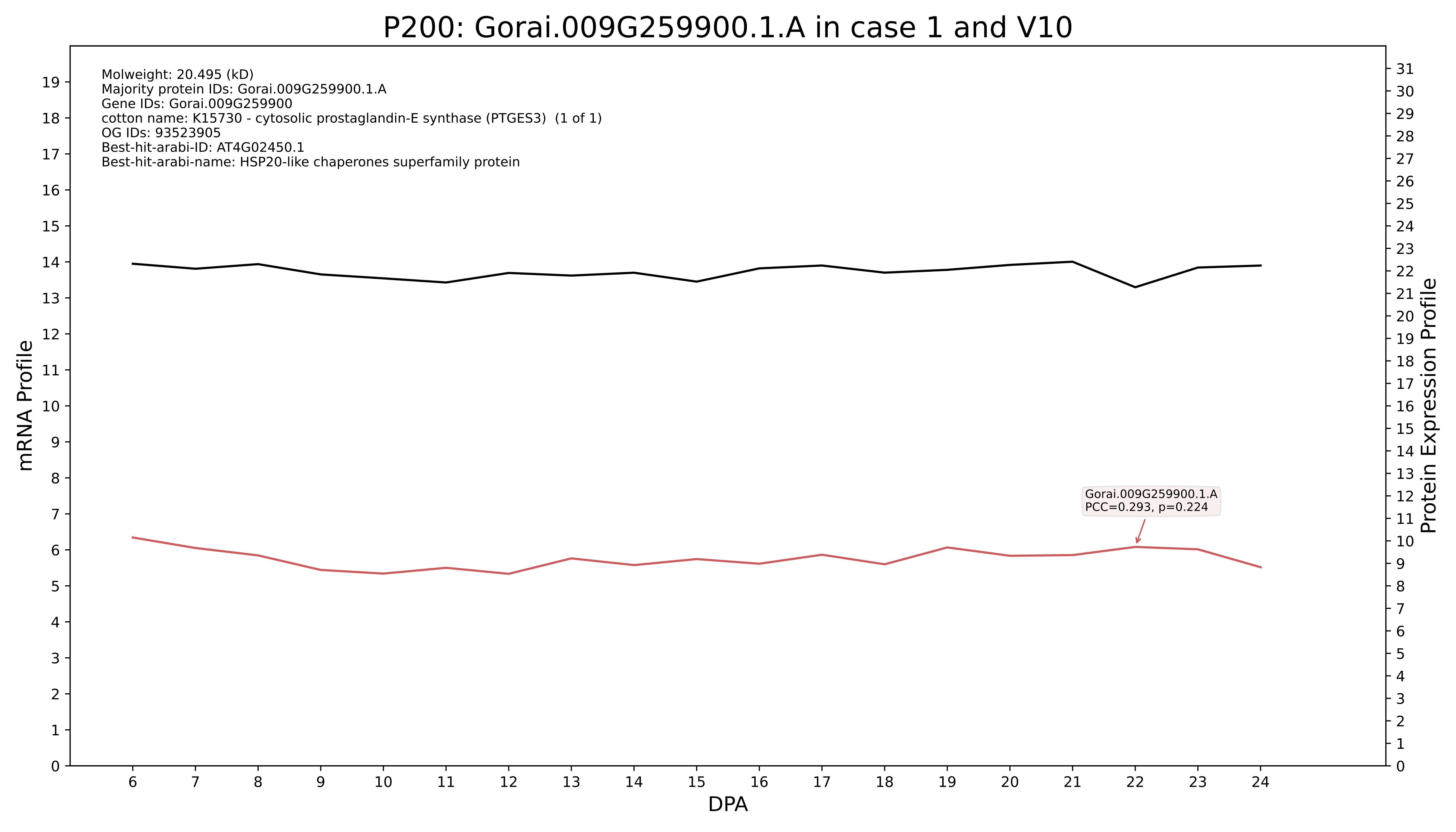
P200: Gorai.009G259900.1.A in case 1 and V10
Molweight: 20.495 (kD)
Majority protein IDs: Gorai.009G259900.1.A
Gene IDs: Gorai.009G259900
cotton name: K15730 - cytosolic prostaglandin-E synthase (PTGES3) (1 of 1)
OG IDs: 93523905
Best-hit-arabi-ID: AT4G02450.1
Best-hit-arabi-name: HSP20-like chaperones superfamily protein
Gorai.009G259900.1.A
PCC=0.293, p=0.224
mRNA Profile
Protein Expression Profile
DPA

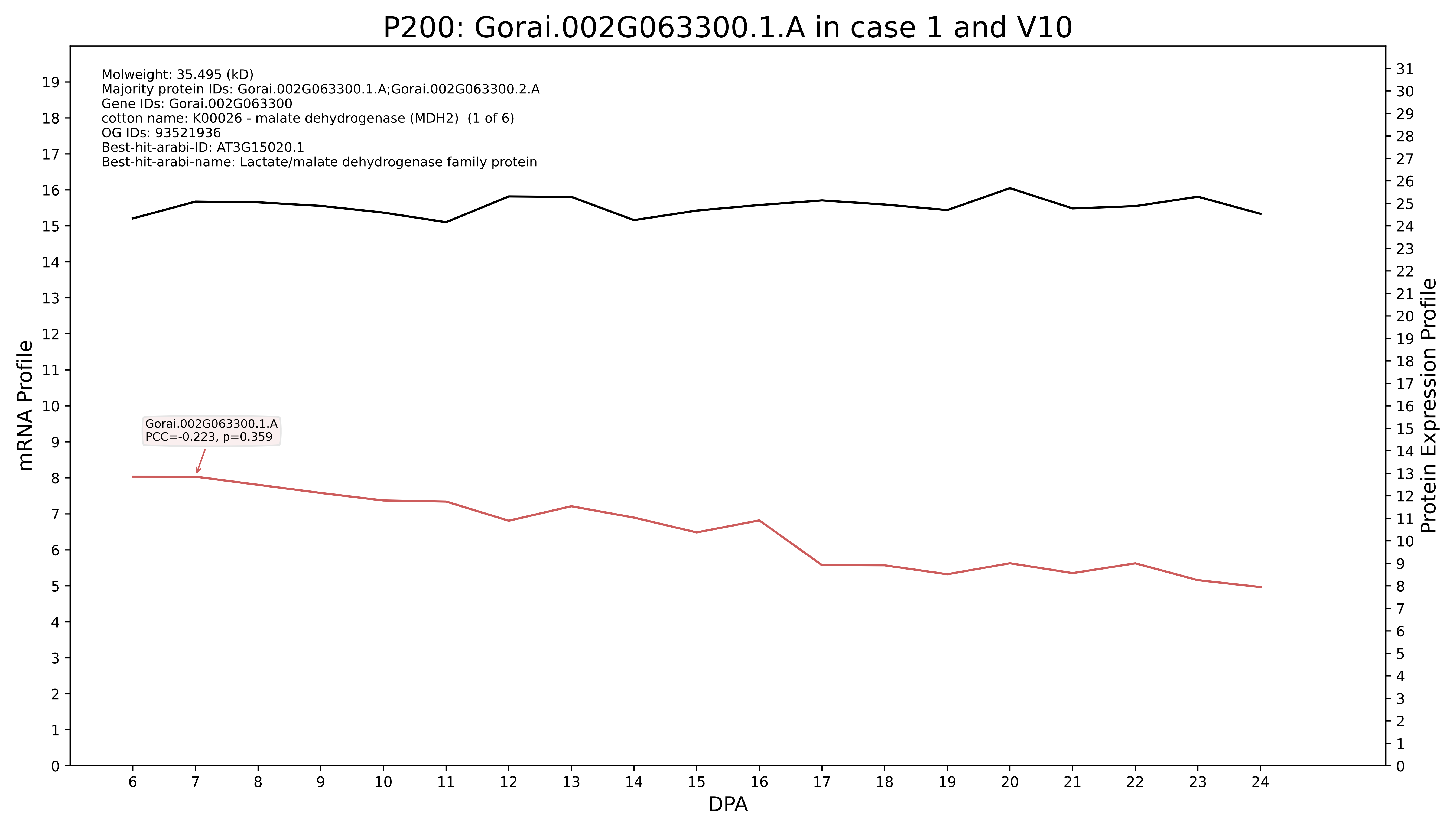
P200: Gorai.002G063300.1.A in case 1 and V10
Molweight: 35.495 (kD)
Majority protein IDs: Gorai.002G063300.1.A;Gorai.002G063300.2.A
Gene IDs: Gorai.002G063300
cotton name: K00026 - malate dehydrogenase (MDH2) (1 of 6)
OG IDs: 93521936
Best-hit-arabi-ID: AT3G15020.1
Best-hit-arabi-name: Lactate/malate dehydrogenase family protein
Gorai.002G063300.1.A
PCC=-0.223, p=0.359
mRNA Profile
Protein Expression Profile
DPA

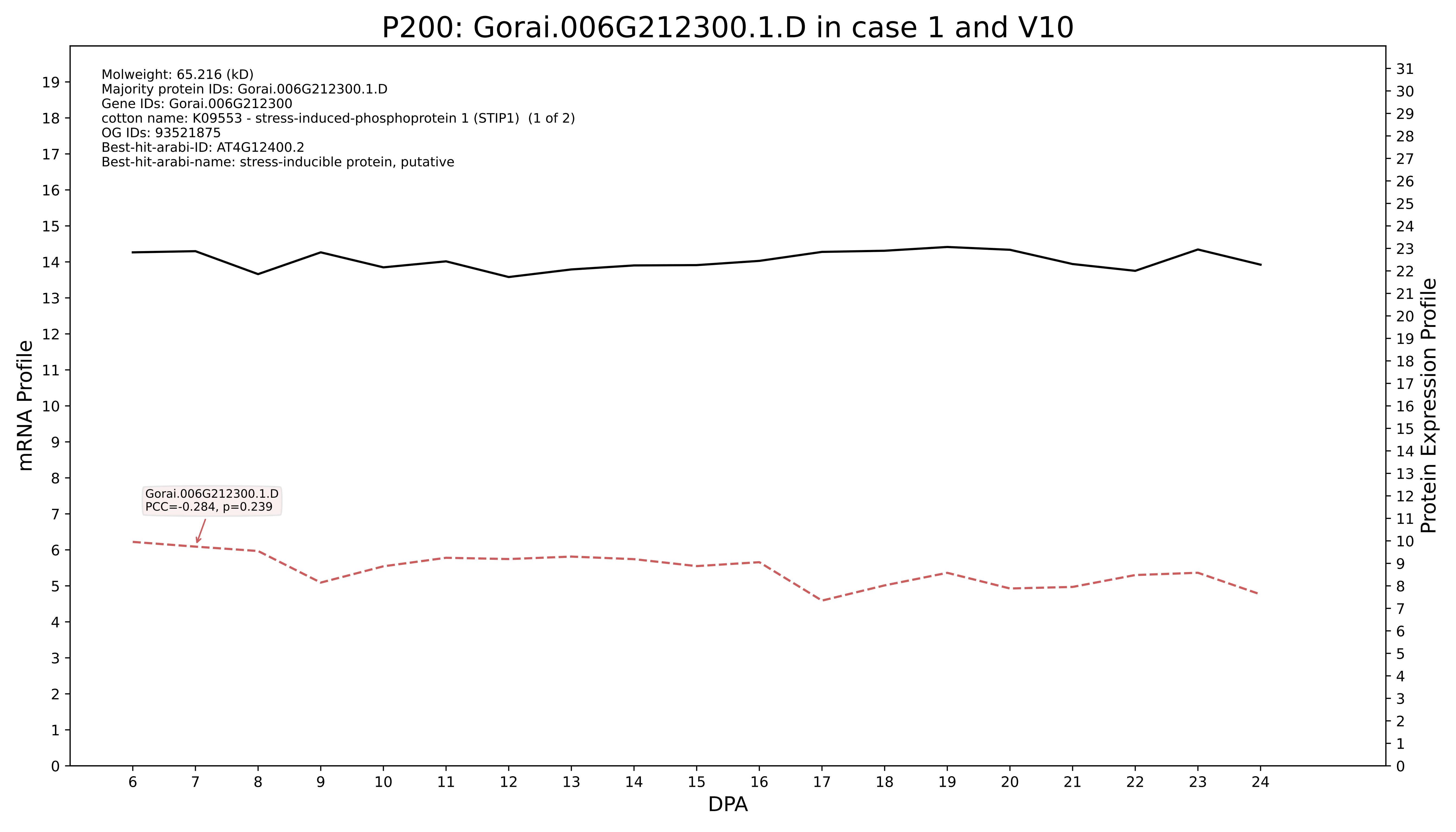
P200: Gorai.006G212300.1.D in case 1 and V10
Molweight: 65.216 (kD)
Majority protein IDs: Gorai.006G212300.1.D
Gene IDs: Gorai.006G212300
cotton name: K09553 - stress-induced-phosphoprotein 1 (STIP1) (1 of 2)
OG IDs: 93521875
Best-hit-arabi-ID: AT4G12400.2
Best-hit-arabi-name: stress-inducible protein, putative
Gorai.006G212300.1.D
PCC=-0.284, p=0.239
mRNA Profile
Protein Expression Profile
DPA

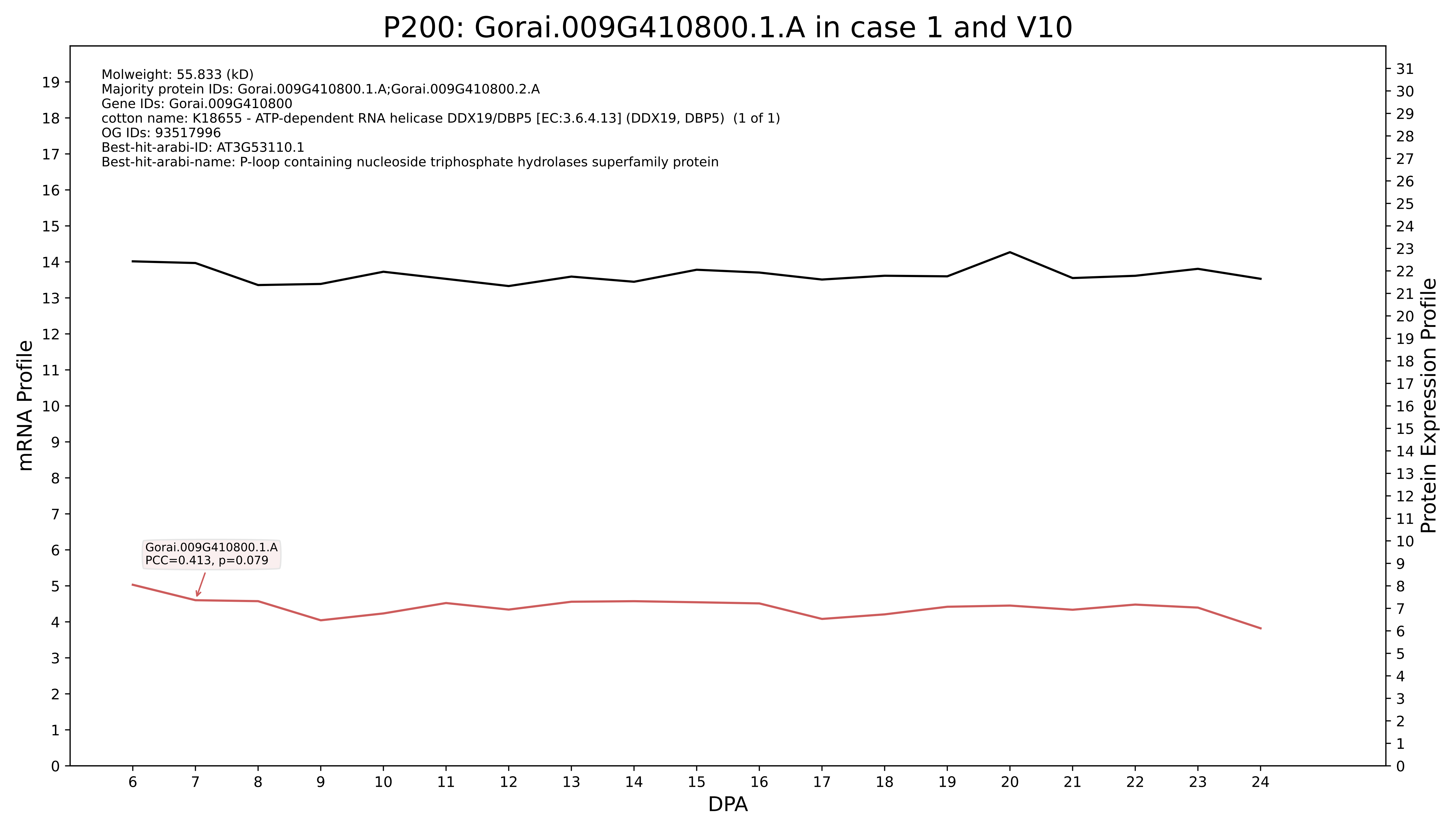

P200: Gorai.009G410800.1.A in case 1 and V10
Molweight: 55.833 (kD)
Majority protein IDs: Gorai.009G410800.1.A;Gorai.009G410800.2.A
Gene IDs: Gorai.009G410800
cotton name: K18655 - ATP-dependent RNA helicase DDX19/DBP5 [EC:3.6.4.13] (DDX19, DBP5) (1 of 1)
OG IDs: 93517996
Best-hit-arabi-ID: AT3G53110.1
Best-hit-arabi-name: P-loop containing nucleoside triphosphate hydrolases superfamily protein
Gorai.009G410800.1.A
PCC=0.413, p=0.079
mRNA Profile
Protein Expression Profile
DPA

### P200: Gorai.013G216900.2.D in case 2 and V10

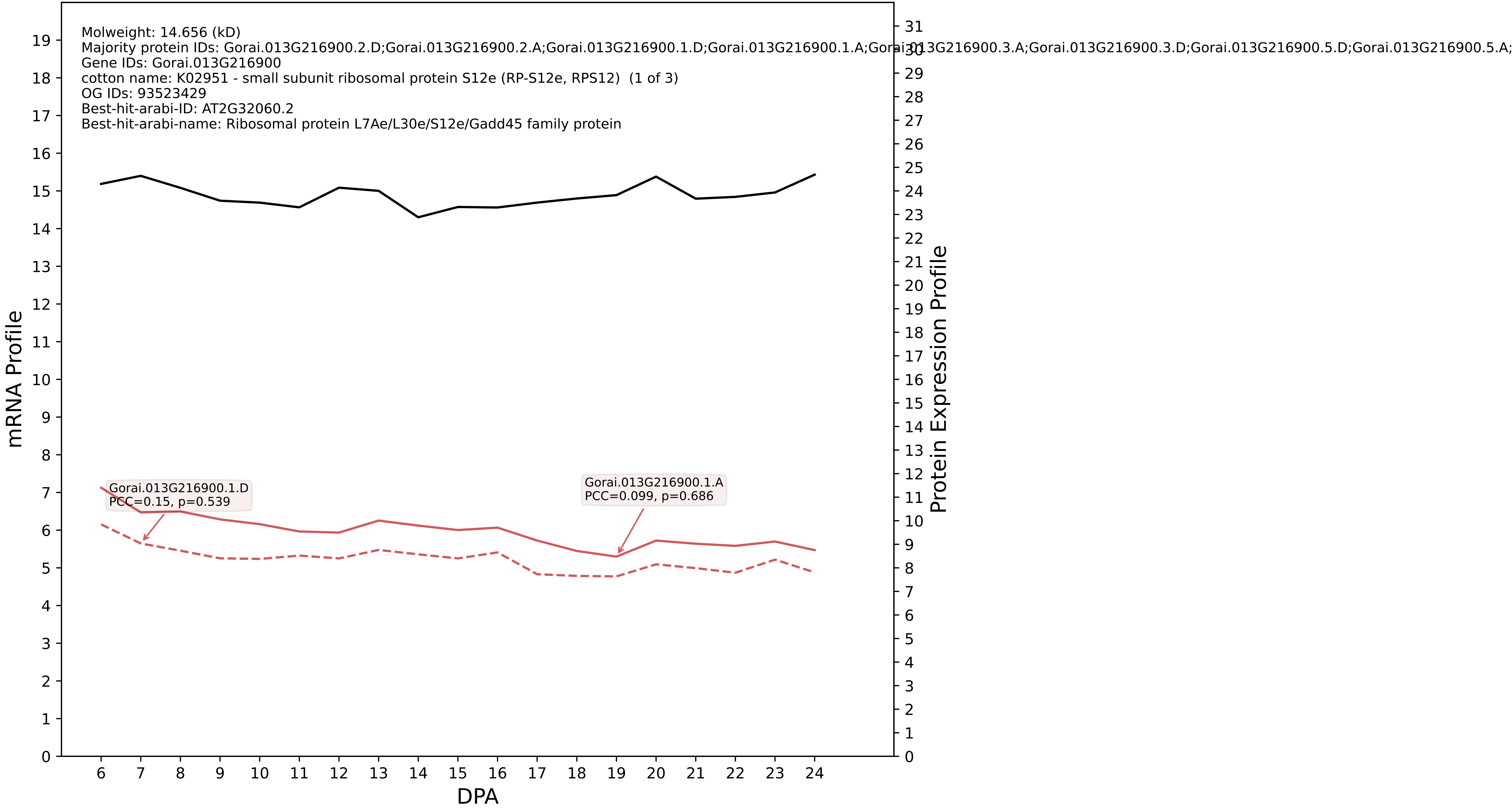

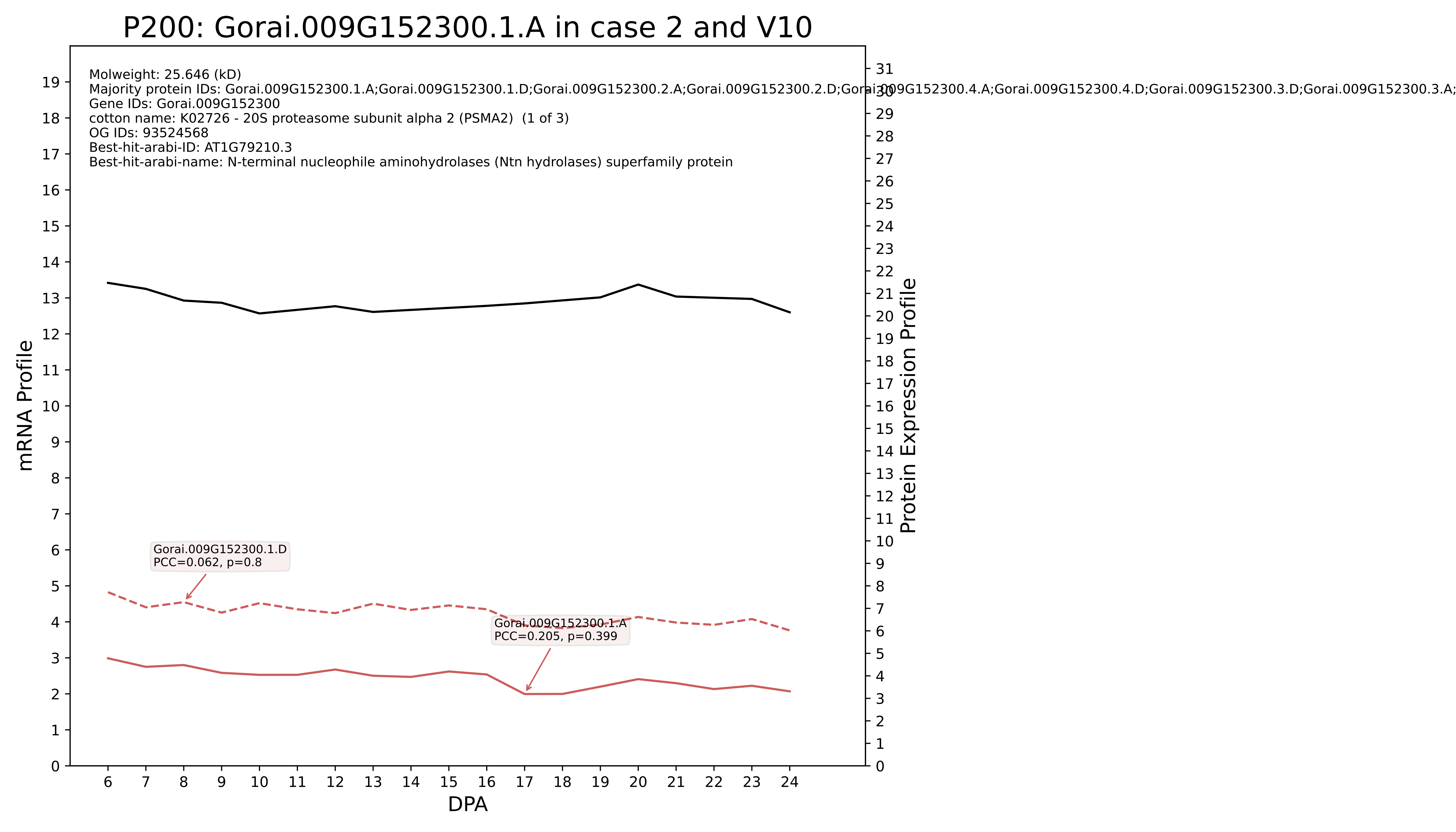

P200: Gorai.009G152300.1.A in case 2 and V10
Molweight: 25.646 (kD)
Majority protein IDs: Gorai.009G152300.1.A;Gorai.009G152300.1.D;Gorai.009G152300.2.A;Gorai.009G152300.2.D;Gorai.009G152300.4.A;Gorai.009G152300.4.D;Gorai.009G152300.3.D;Gorai.009G152300.3.A;
Gene IDs: Gorai.009G152300
cotton name: K02726 - 20S proteasome subunit alpha 2 (PSMA2) (1 of 3)
OG IDs: 93524568
Best-hit-arabi-ID: AT1G79210.3
Best-hit-arabi-name: N-terminal nucleophile aminohydrolases (Ntn hydrolases) superfamily protein
Gorai.009G152300.1.D
PCC=0.062, p=0.8
Gorai.009G152300.1.A
PCC=0.205, p=0.399
mRNA Profile
Protein Expression Profile
DPA

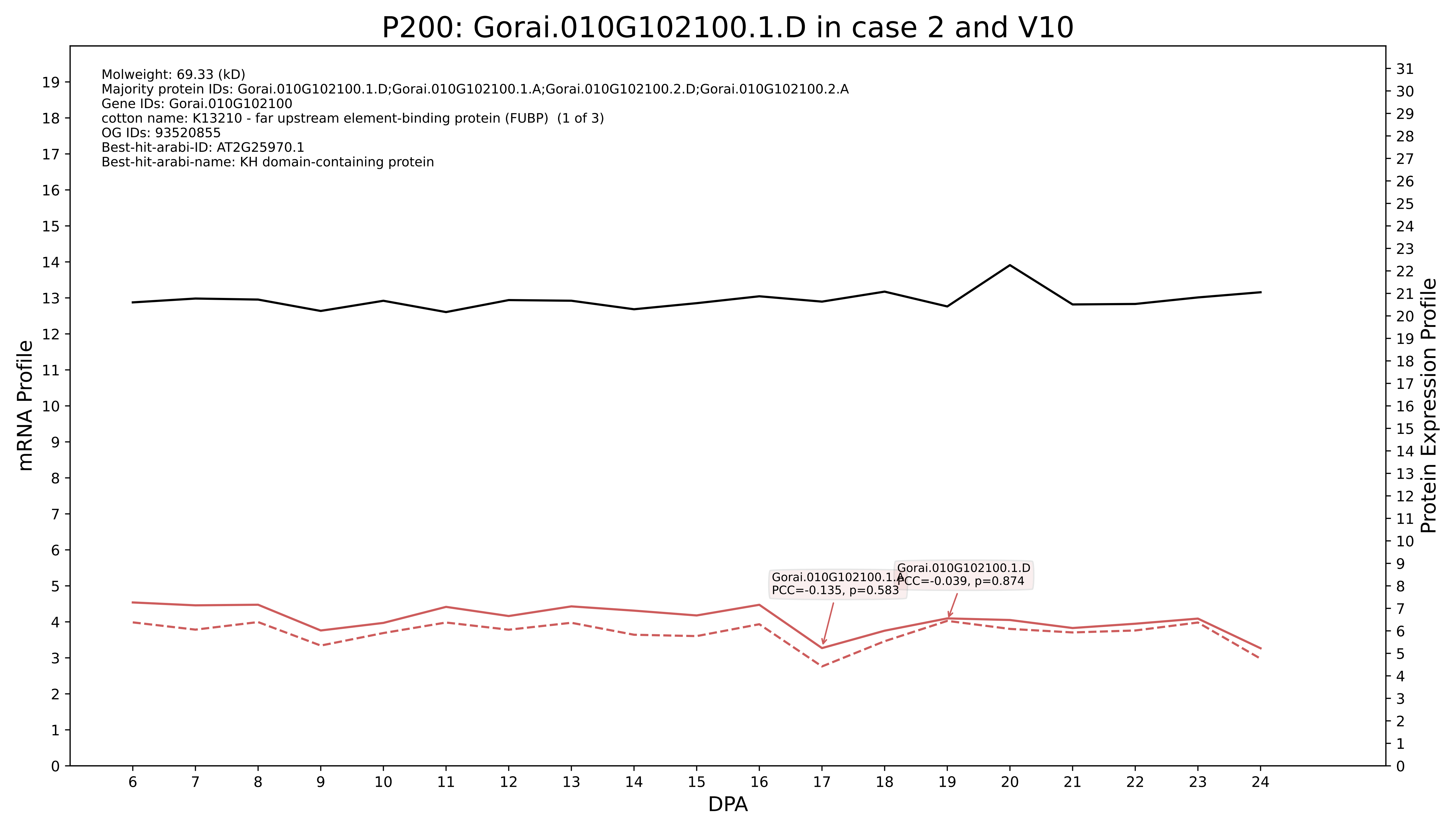
P200: Gorai.010G102100.1.D in case 2 and V10
Molweight: 69.33 (kD)
Majority protein IDs: Gorai.010G102100.1.D;Gorai.010G102100.1.A;Gorai.010G102100.2.D;Gorai.010G102100.2.A
Gene IDs: Gorai.010G102100
cotton name: K13210 - far upstream element-binding protein (FUBP) (1 of 3)
OG IDs: 93520855
Best-hit-arabi-ID: AT2G25970.1
Best-hit-arabi-name: KH domain-containing protein
Gorai.010G102100.1.A
PCC=-0.135, p=0.583
Gorai.010G102100.1.D
PCC=-0.039, p=0.874
mRNA Profile
Protein Expression Profile
DPA

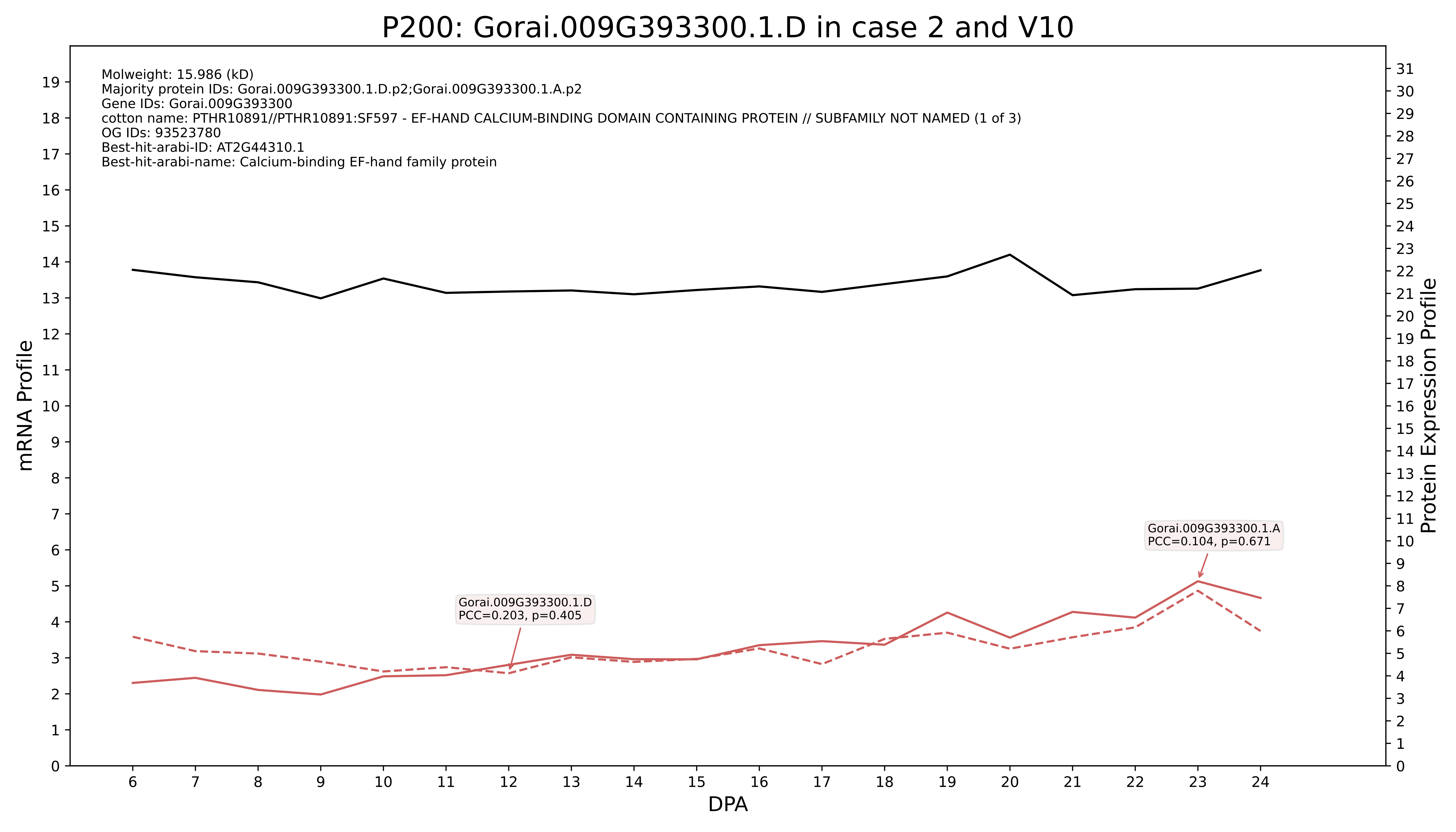

P200: Gorai.009G393300.1.D in case 2 and V10
Molweight: 15.986 (kD)
Majority protein IDs: Gorai.009G393300.1.D.p2;Gorai.009G393300.1.A.p2
Gene IDs: Gorai.009G393300
cotton name: PTHR10891//PTHR10891:SF597 - EF-HAND CALCIUM-BINDING DOMAIN CONTAINING PROTEIN // SUBFAMILY NOT NAMED (1 of 3)
OG IDs: 93523780
Best-hit-arabi-ID: AT2G44310.1
Best-hit-arabi-name: Calcium-binding EF-hand family protein
Gorai.009G393300.1.A
PCC=0.104, p=0.671
Gorai.009G393300.1.D
PCC=0.203, p=0.405
mRNA Profile
Protein Expression Profile
DPA

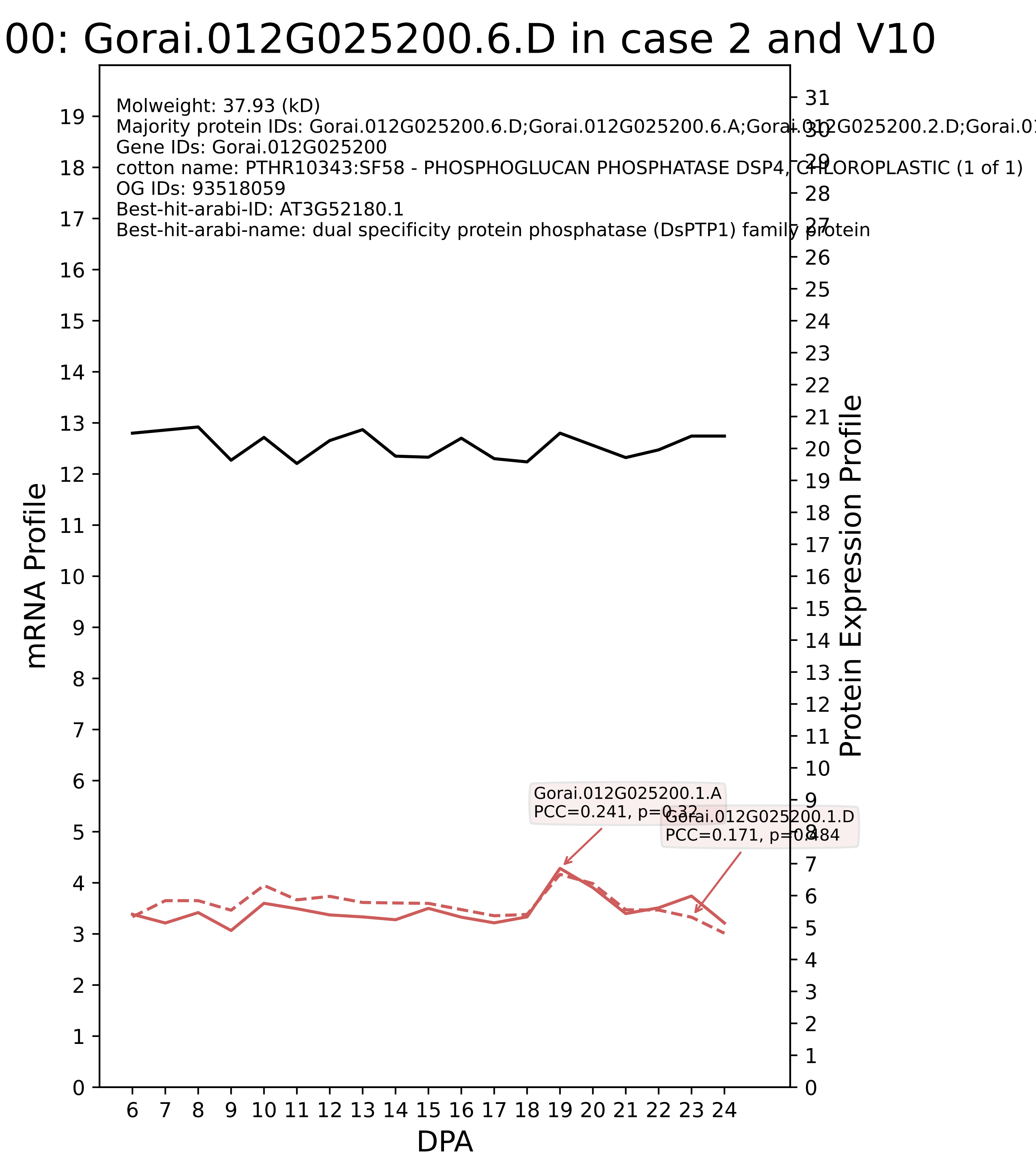

00: Gorai.012G025200.6.D in case 2 and V10
Molweight: 37.93 (kD)
Majority protein IDs: Gorai.012G025200.6.D;Gorai.012G025200.6.A;Gorai.012G025200.2.D;Gorai.012G025200.4.D;Gorai.012G025200.2.A;Gorai.012G025200.4.A;Gorai.012G025200.1.D;Gorai.012G025200.1.A;G
Gene IDs: Gorai.012G025200
cotton name: PTHR10343:SF58 - PHOSPHOGLUCAN PHOSPHATASE DSP4, CHLOROPLASTIC (1 of 1)
OG IDs: 93518059
Best-hit-arabi-ID: AT3G52180.1
Best-hit-arabi-name: dual specificity protein phosphatase (DsPTP1) family protein
Gorai.012G025200.1.A
PCC=0.241, p=0.32
Gorai.012G025200.1.D
PCC=0.171, p=0.484
mRNA Profile
Protein Expression Profile
DPA

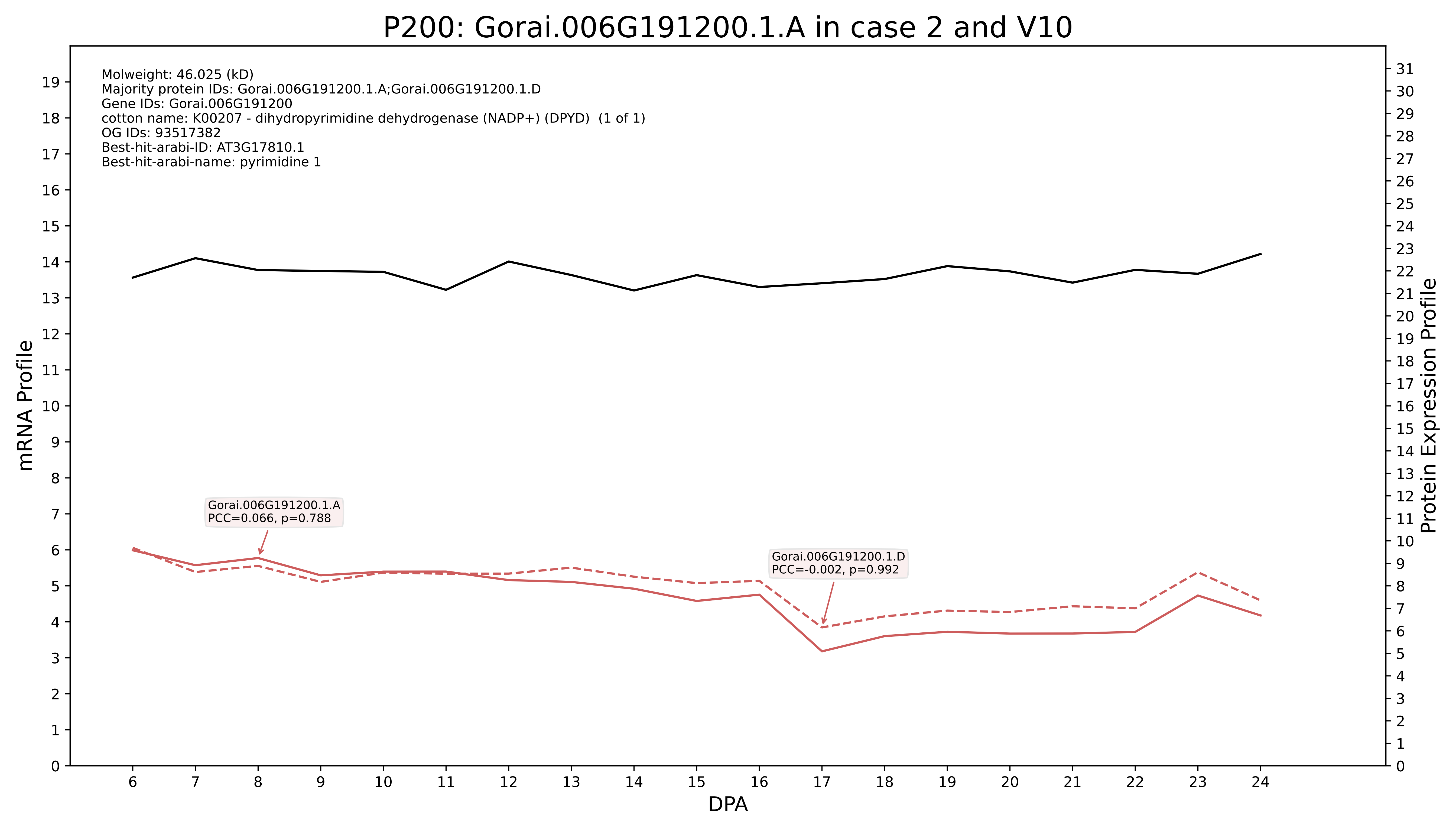
P200: Gorai.006G191200.1.A in case 2 and V10
Molweight: 46.025 (kD)
Majority protein IDs: Gorai.006G191200.1.A;Gorai.006G191200.1.D
Gene IDs: Gorai.006G191200
cotton name: K00207 - dihydropyrimidine dehydrogenase (NADP+) (DPYD) (1 of 1)
OG IDs: 93517382
Best-hit-arabi-ID: AT3G17810.1
Best-hit-arabi-name: pyrimidine 1
Gorai.006G191200.1.A
PCC=0.066, p=0.788
Gorai.006G191200.1.D
PCC=-0.002, p=0.992
mRNA Profile
Protein Expression Profile
DPA

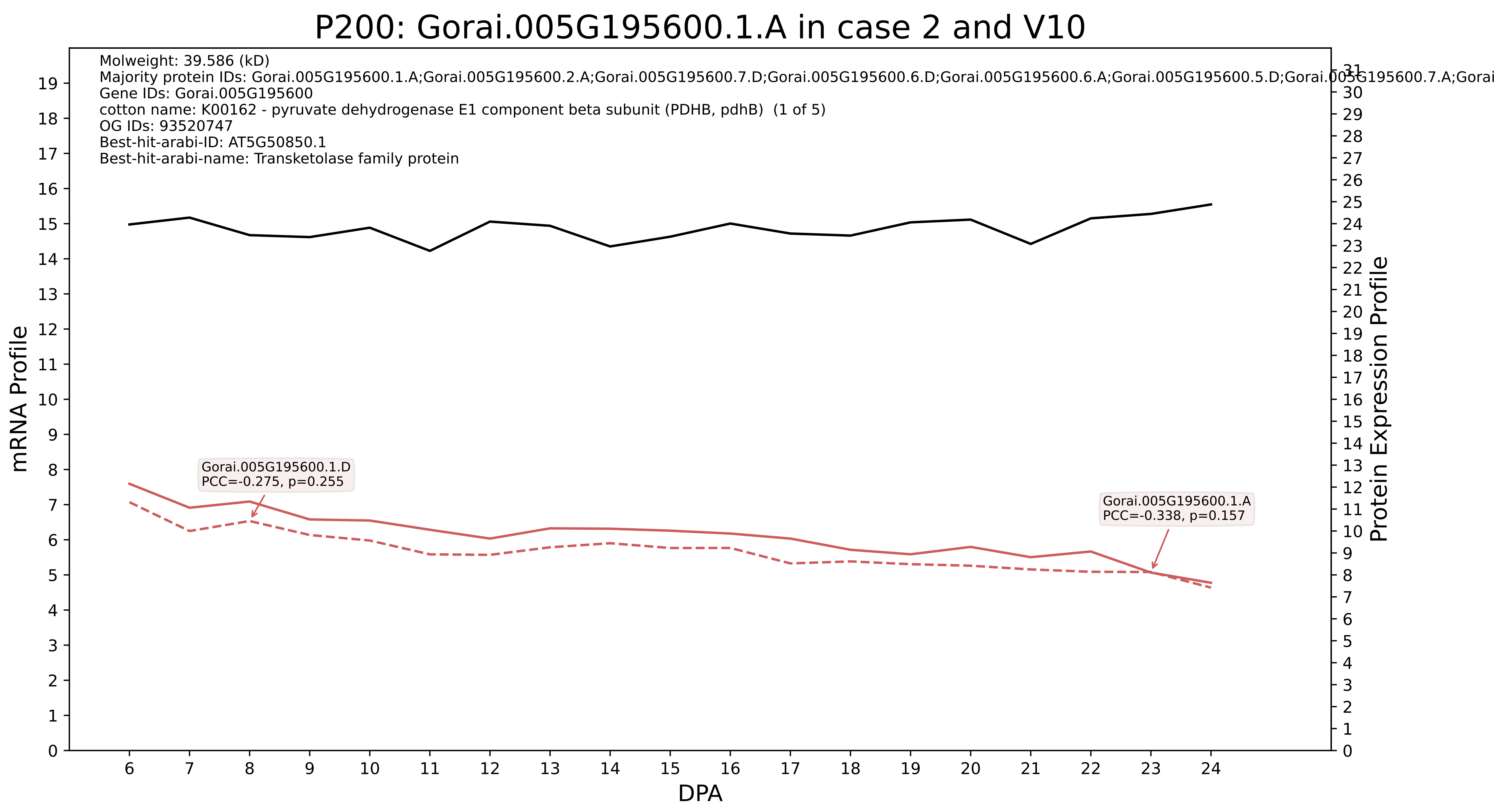

P200: Gorai.005G195600.1.A in case 2 and V10
Molweight: 39.586 (kD)
Majority protein IDs: Gorai.005G195600.1.A;Gorai.005G195600.2.A;Gorai.005G195600.7.D;Gorai.005G195600.6.D;Gorai.005G195600.6.A;Gorai.005G195600.5.D;Gorai.005G195600.7.A;Gorai
Gene IDs: Gorai.005G195600
cotton name: K00162 - pyruvate dehydrogenase E1 component beta subunit (PDHB, pdhB) (1 of 5)
OG IDs: 93520747
Best-hit-arabi-ID: AT5G50850.1
Best-hit-arabi-name: Transketolase family protein
Gorai.005G195600.1.D
PCC=-0.275, p=0.255
Gorai.005G195600.1.A
PCC=-0.338, p=0.157
mRNA Profile
Protein Expression Profile
DPA

### P200: Gorai.009G302200.2.D in case 2 and V10

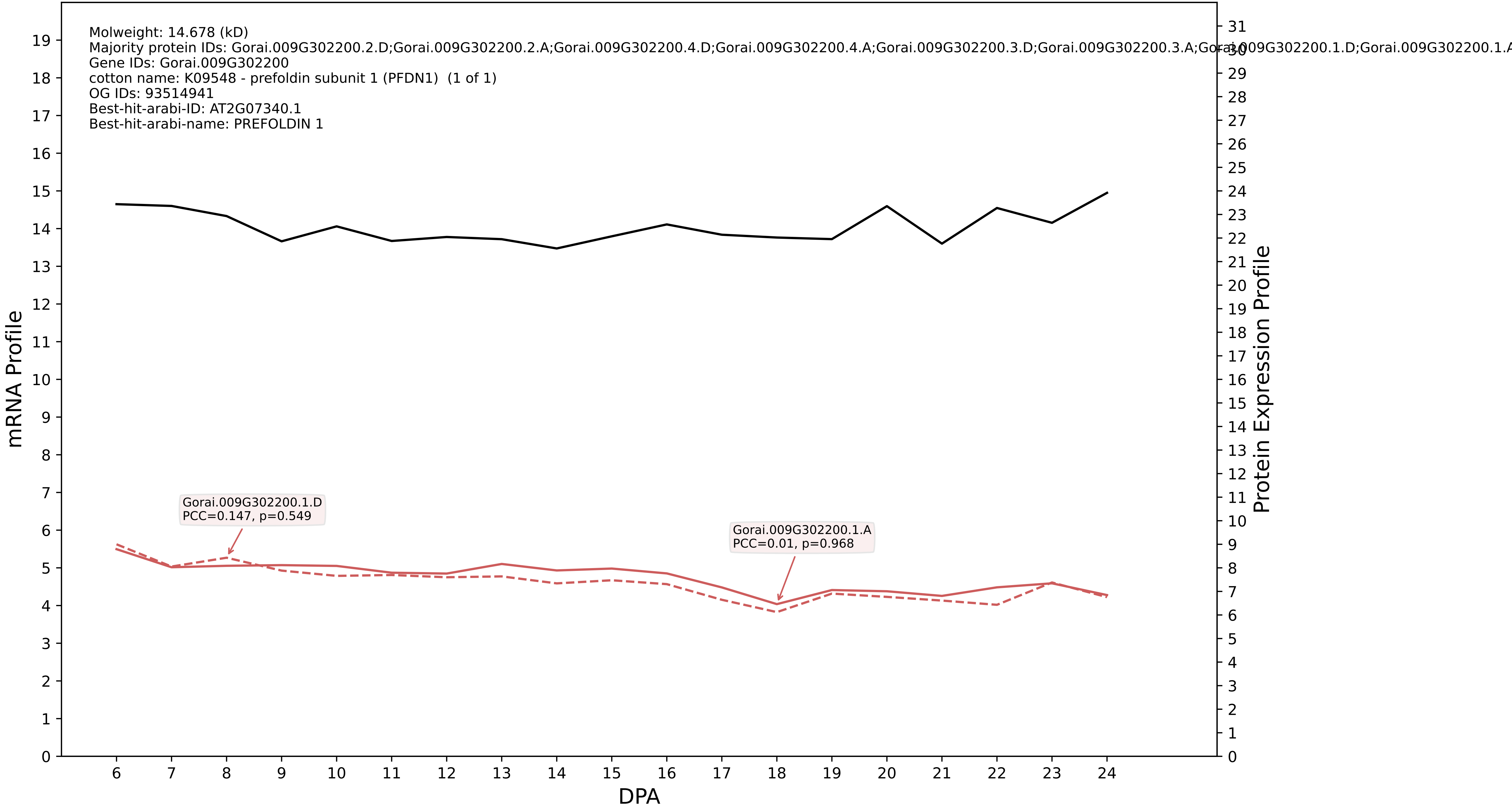

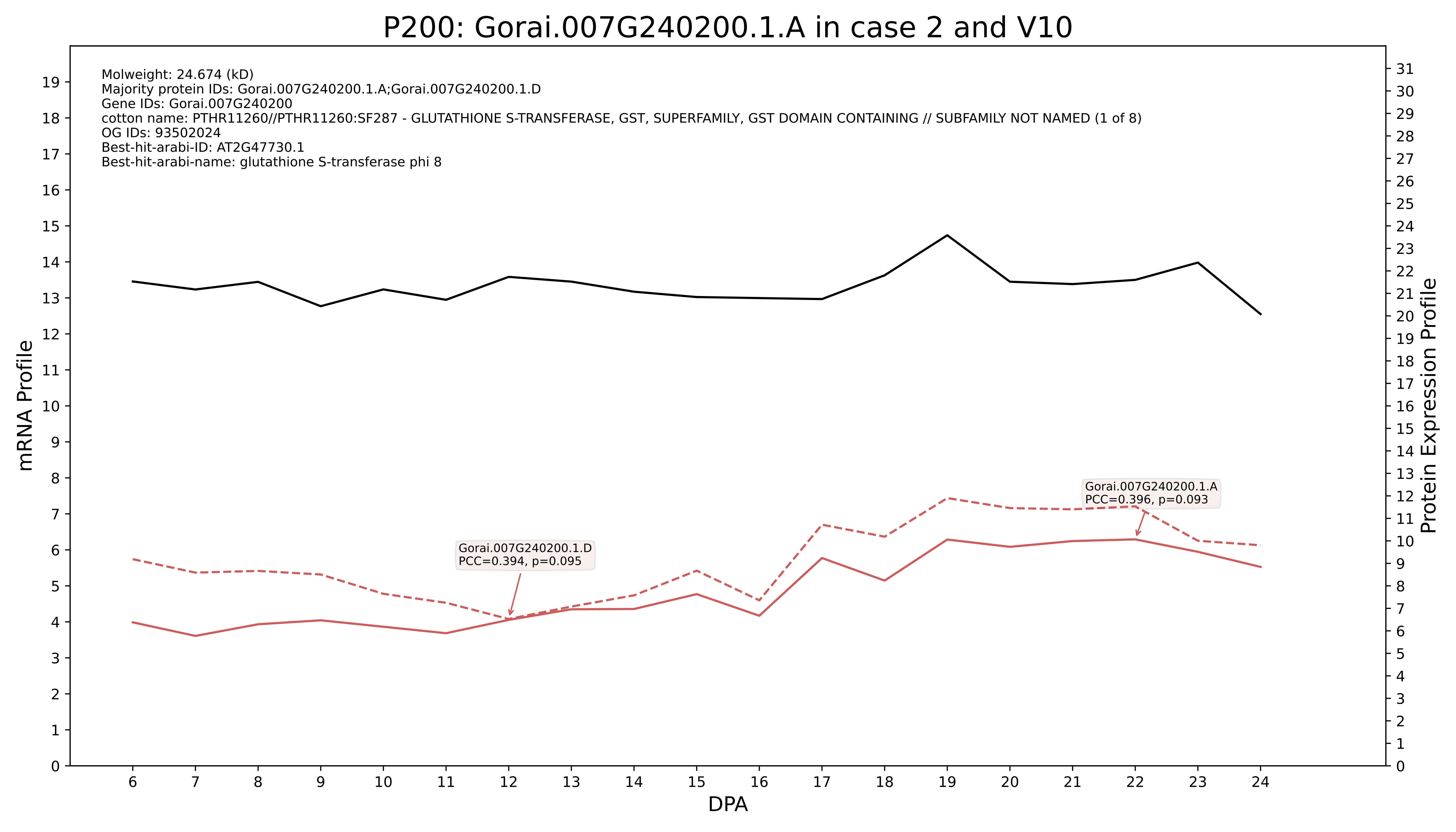
P200: Gorai.007G240200.1.A in case 2 and V10
Molweight: 24.674 (kD)
Majority protein IDs: Gorai.007G240200.1.A;Gorai.007G240200.1.D
Gene IDs: Gorai.007G240200
cotton name: PTHR11260//PTHR11260:SF287 - GLUTATHIONE S-TRANSFERASE, GST, SUPERFAMILY, GST DOMAIN CONTAINING // SUBFAMILY NOT NAMED (1 of 8)
OG IDs: 93502024
Best-hit-arabi-ID: AT2G47730.1
Best-hit-arabi-name: glutathione S-transferase phi 8
Gorai.007G240200.1.D
PCC=0.394, p=0.095
Gorai.007G240200.1.A
PCC=0.396, p=0.093
mRNA Profile
Protein Expression Profile
DPA

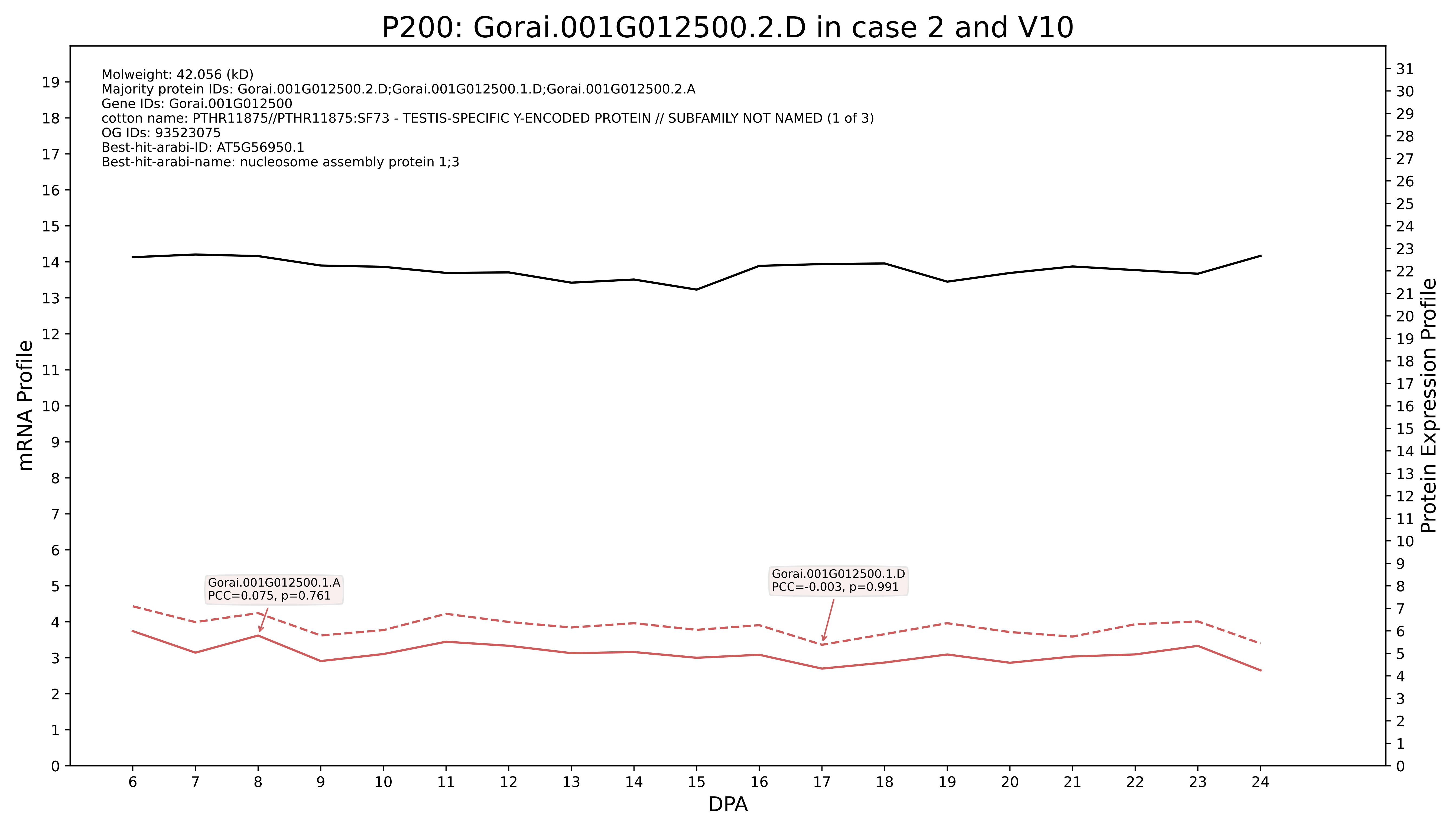

P200: Gorai.001G012500.2.D in case 2 and V10
Molweight: 42.056 (kD)
Majority protein IDs: Gorai.001G012500.2.D;Gorai.001G012500.1.D;Gorai.001G012500.2.A
Gene IDs: Gorai.001G012500
cotton name: PTHR11875//PTHR11875:SF73 - TESTIS-SPECIFIC Y-ENCODED PROTEIN // SUBFAMILY NOT NAMED (1 of 3)
OG IDs: 93523075
Best-hit-arabi-ID: AT5G56950.1
Best-hit-arabi-name: nucleosome assembly protein 1;3
Gorai.001G012500.1.A
PCC=0.075, p=0.761
Gorai.001G012500.1.D
PCC=-0.003, p=0.991
mRNA Profile
Protein Expression Profile
DPA

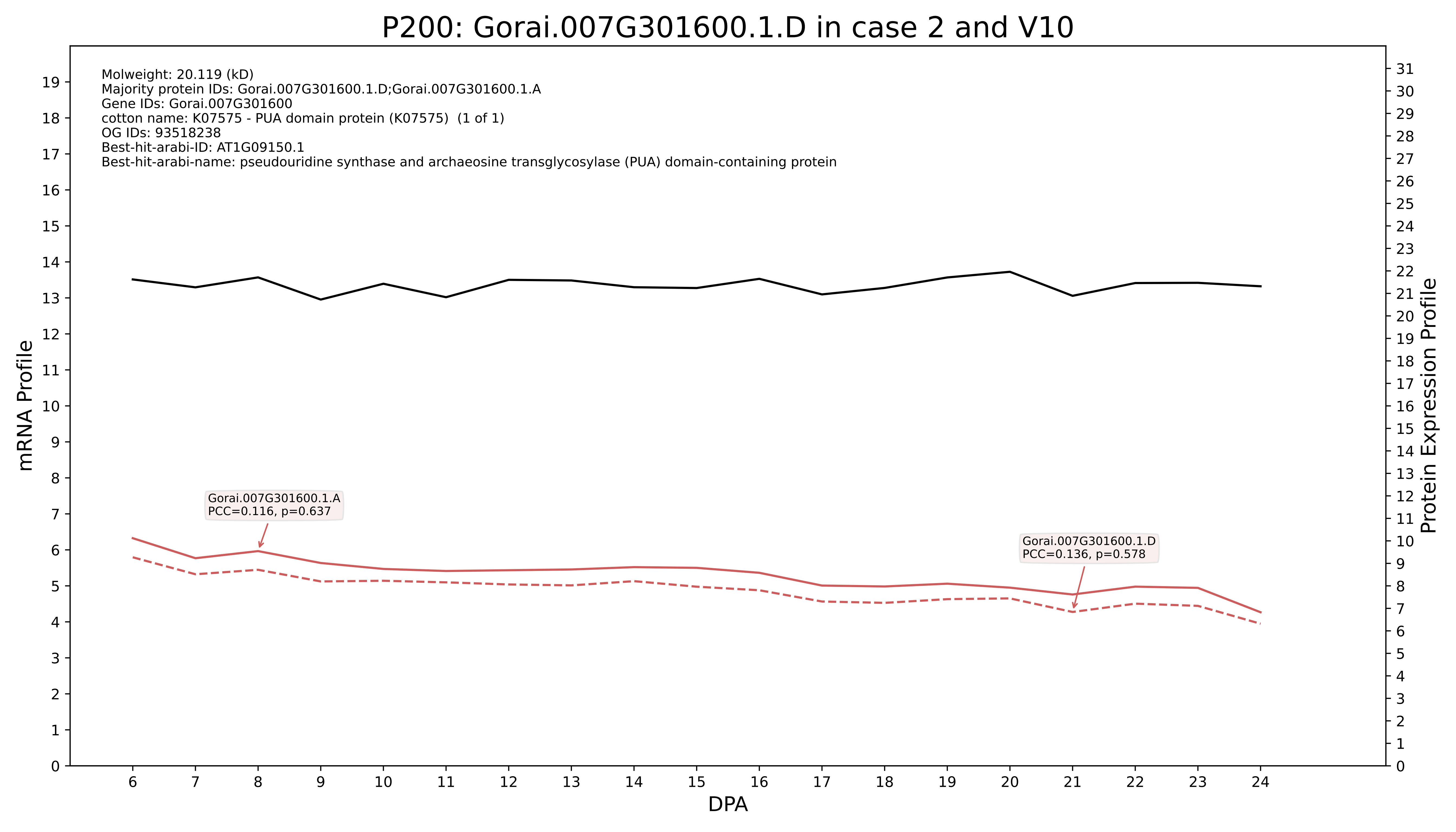
P200: Gorai.007G301600.1.D in case 2 and V10
Molweight: 20.119 (kD)
Majority protein IDs: Gorai.007G301600.1.D;Gorai.007G301600.1.A
Gene IDs: Gorai.007G301600
cotton name: K07575 - PUA domain protein (K07575) (1 of 1)
OG IDs: 93518238
Best-hit-arabi-ID: AT1G09150.1
Best-hit-arabi-name: pseudouridine synthase and archaeosine transglycosylase (PUA) domain-containing protein
Gorai.007G301600.1.A
PCC=0.116, p=0.637
Gorai.007G301600.1.D
PCC=0.136, p=0.578
mRNA Profile
Protein Expression Profile
DPA

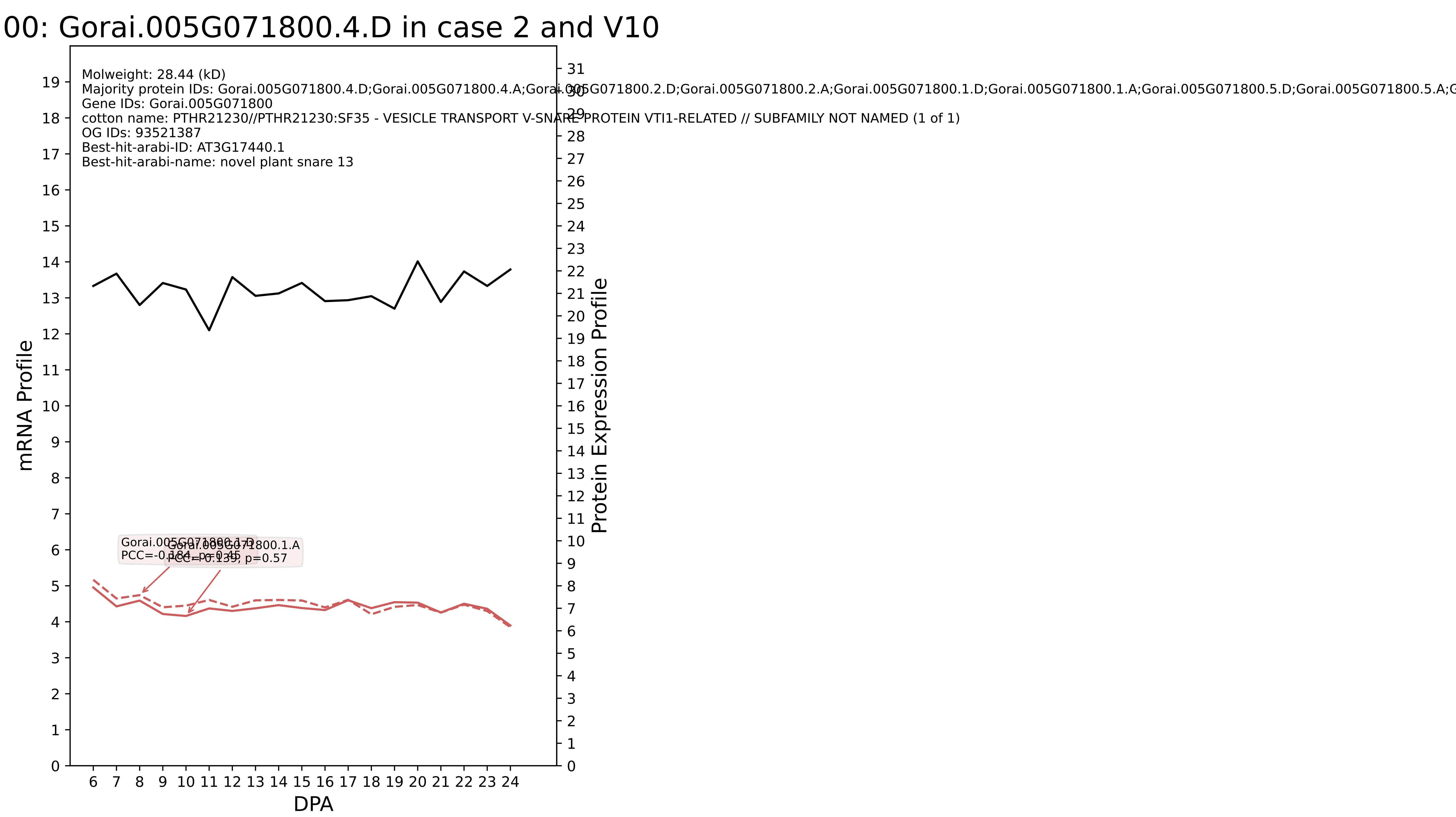
00: Gorai.005G071800.4.D in case 2 and V10
Molweight: 28.44 (kD)
Majority protein IDs: Gorai.005G071800.4.D;Gorai.005G071800.4.A;Gorai.005G071800.2.D;Gorai.005G071800.2.A;Gorai.005G071800.1.D;Gorai.005G071800.1.A;Gorai.005G071800.5.D;Gorai.005G071800.5.A;G
Gene IDs: Gorai.005G071800
cotton name: PTHR21230//PTHR21230:SF35 - VESICLE TRANSPORT V-SNARE PROTEIN VTI1-RELATED // SUBFAMILY NOT NAMED (1 of 1)
OG IDs: 93521387
Best-hit-arabi-ID: AT3G17440.1
Best-hit-arabi-name: novel plant snare 13
Gorai.005G071800.1.D
PCC=-0.184, p=0.45
Gorai.005G071800.1.A
PCC=0.139, p=0.57
mRNA Profile
Protein Expression Profile
DPA

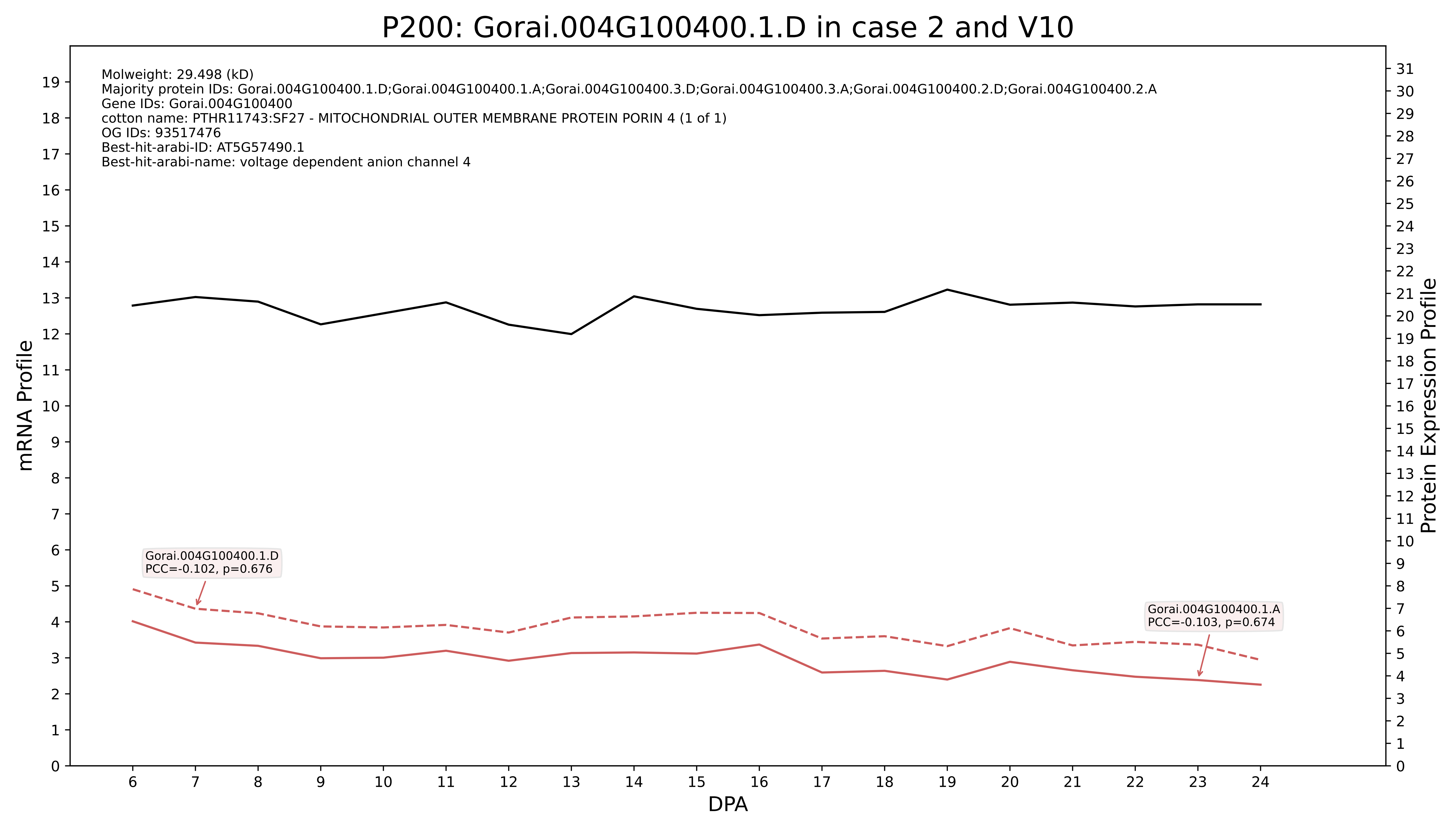
P200: Gorai.004G100400.1.D in case 2 and V10
Molweight: 29.498 (kD)
Majority protein IDs: Gorai.004G100400.1.D;Gorai.004G100400.1.A;Gorai.004G100400.3.D;Gorai.004G100400.3.A;Gorai.004G100400.2.D;Gorai.004G100400.2.A
Gene IDs: Gorai.004G100400
cotton name: PTHR11743:SF27 - MITOCHONDRIAL OUTER MEMBRANE PROTEIN PORIN 4 (1 of 1)
OG IDs: 93517476
Best-hit-arabi-ID: AT5G57490.1
Best-hit-arabi-name: voltage dependent anion channel 4
Gorai.004G100400.1.D
PCC=-0.102, p=0.676
Gorai.004G100400.1.A
PCC=-0.103, p=0.674
mRNA Profile
Protein Expression Profile
DPA

### P200: Gorai.001G095800.1.D in case 2 and V10

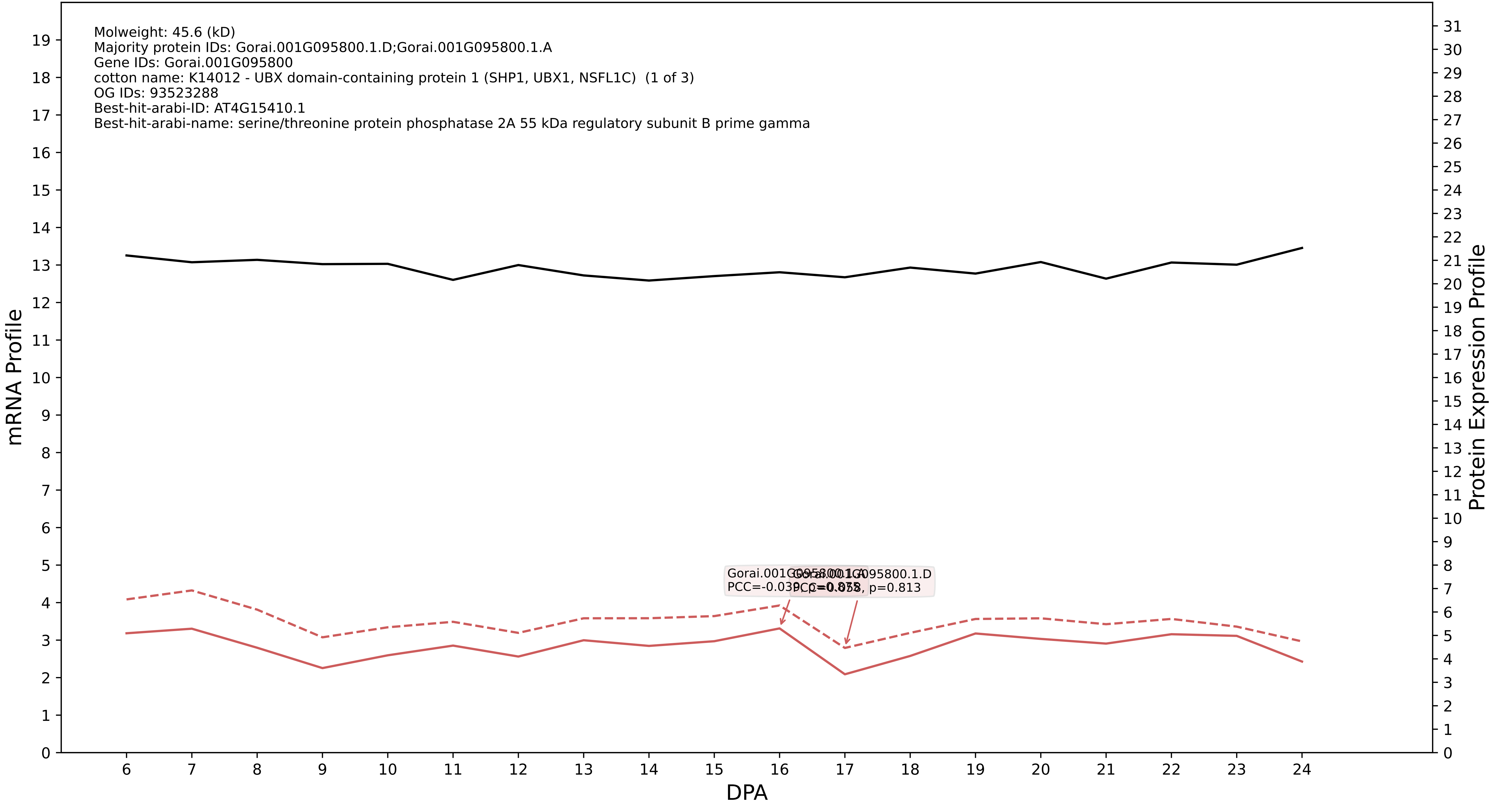

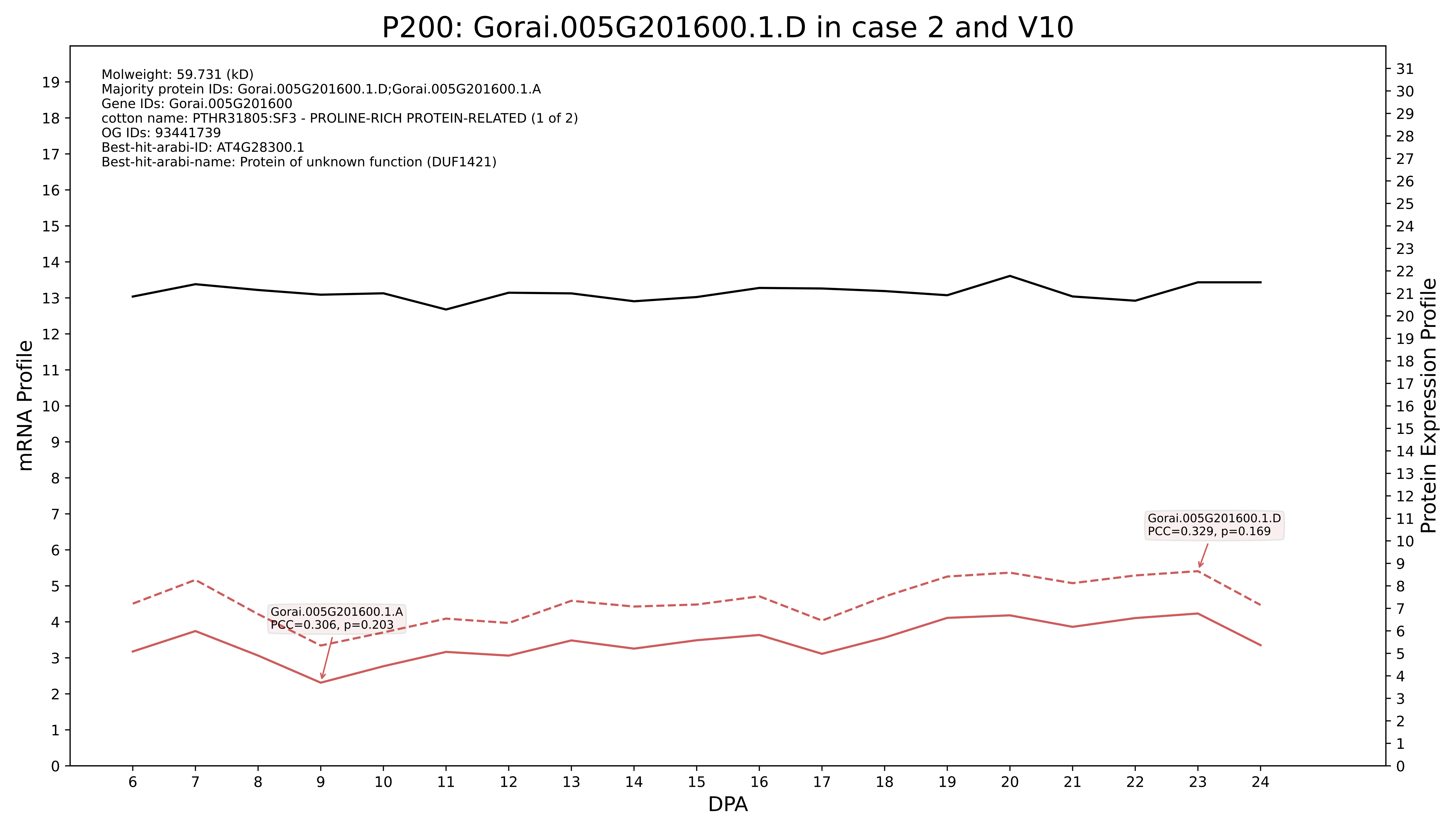
P200: Gorai.005G201600.1.D in case 2 and V10
Molweight: 59.731 (kD)
Majority protein IDs: Gorai.005G201600.1.D;Gorai.005G201600.1.A
Gene IDs: Gorai.005G201600
cotton name: PTHR31805:SF3 - PROLINE-RICH PROTEIN-RELATED (1 of 2)
OG IDs: 93441739
Best-hit-arabi-ID: AT4G28300.1
Best-hit-arabi-name: Protein of unknown function (DUF1421)
Gorai.005G201600.1.D
PCC=0.329, p=0.169
Gorai.005G201600.1.A
PCC=0.306, p=0.203
mRNA Profile
Protein Expression Profile
DPA

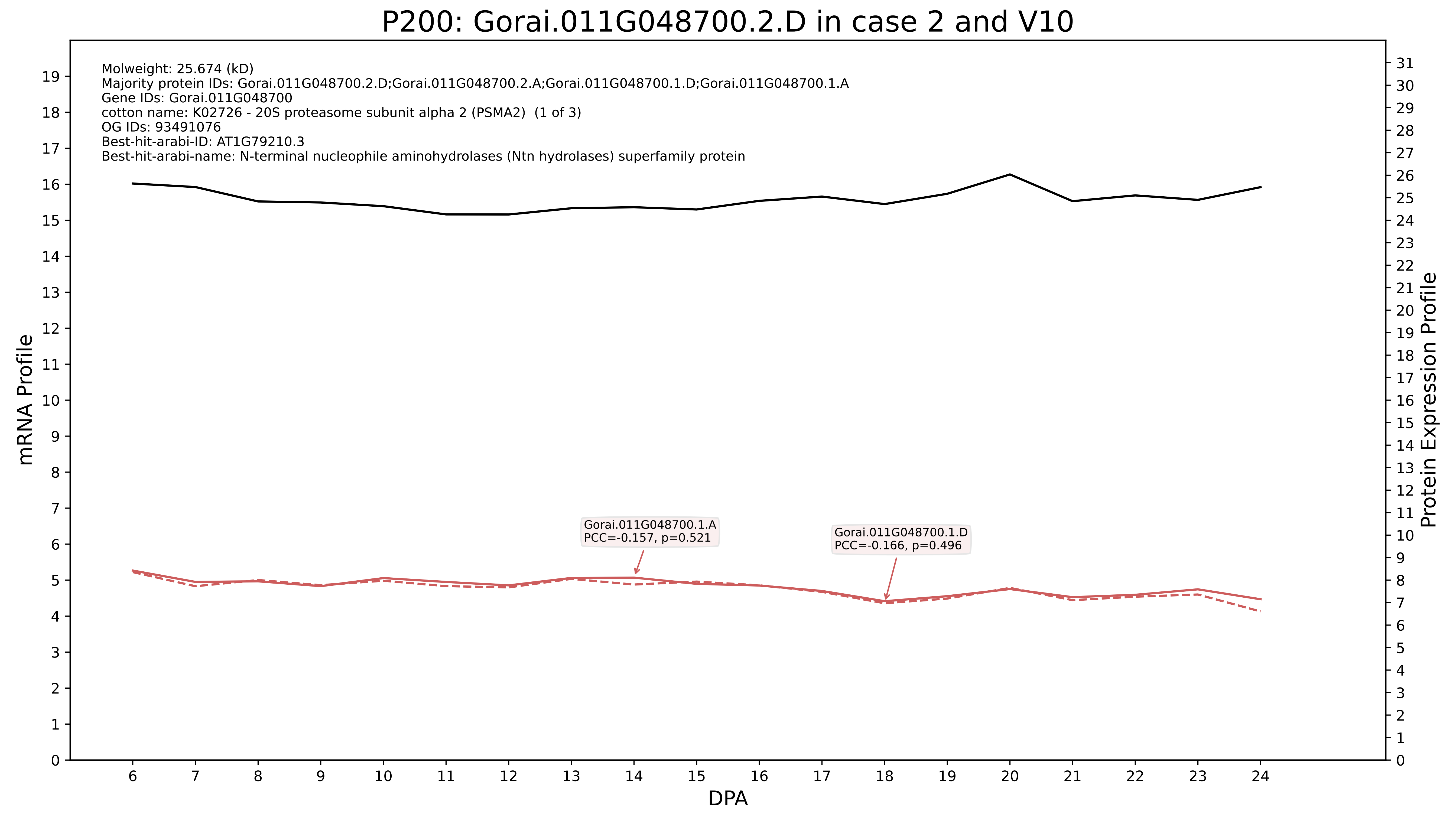

P200: Gorai.011G048700.2.D in case 2 and V10
Molweight: 25.674 (kD)
Majority protein IDs: Gorai.011G048700.2.D;Gorai.011G048700.2.A;Gorai.011G048700.1.D;Gorai.011G048700.1.A
Gene IDs: Gorai.011G048700
cotton name: K02726 - 20S proteasome subunit alpha 2 (PSMA2) (1 of 3)
OG IDs: 93491076
Best-hit-arabi-ID: AT1G79210.3
Best-hit-arabi-name: N-terminal nucleophile aminohydrolases (Ntn hydrolases) superfamily protein
Gorai.011G048700.1.A
PCC=-0.157, p=0.521
Gorai.011G048700.1.D
PCC=-0.166, p=0.496
mRNA Profile
Protein Expression Profile
DPA

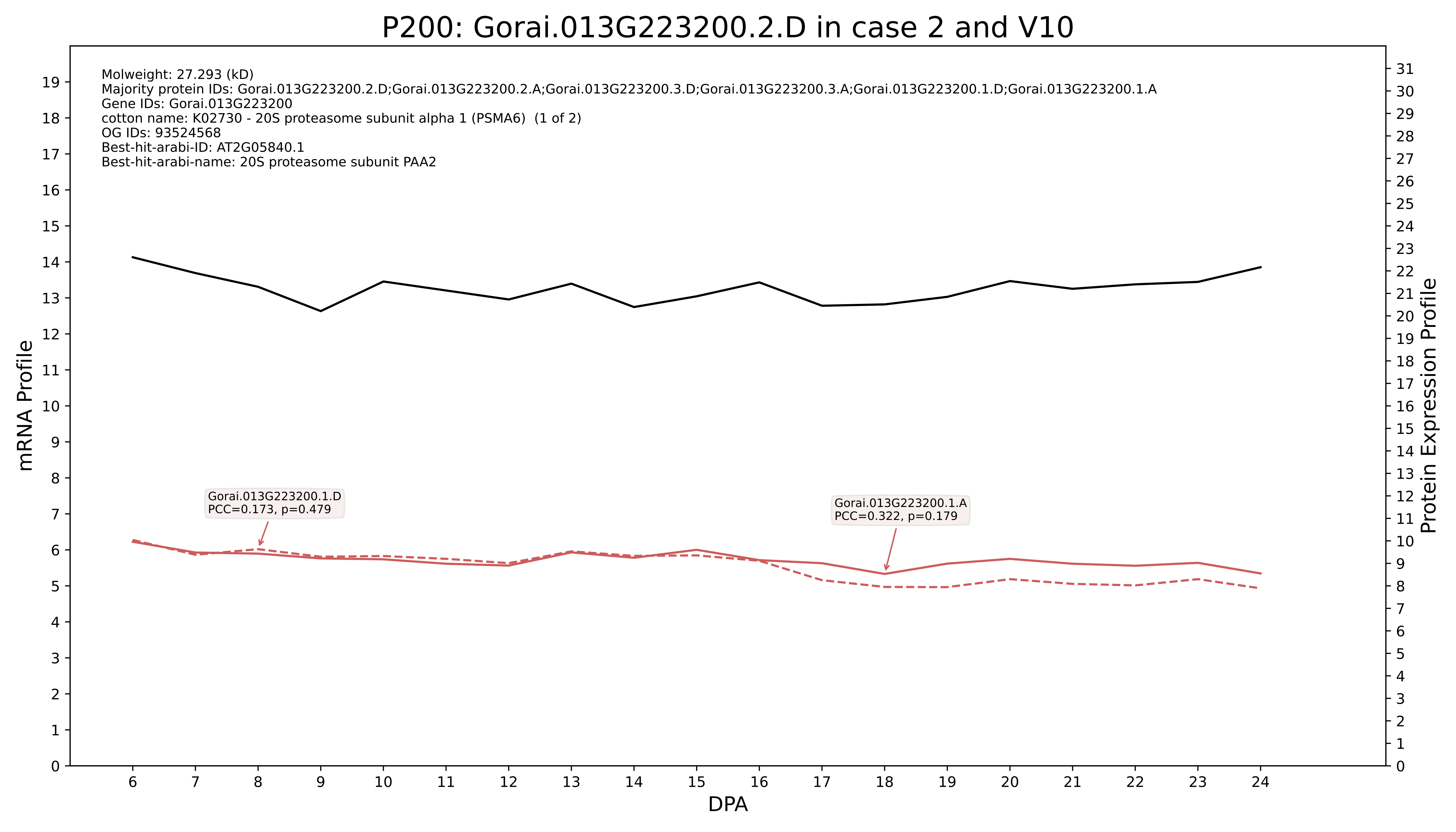

P200: Gorai.013G223200.2.D in case 2 and V10
Molweight: 27.293 (kD)
Majority protein IDs: Gorai.013G223200.2.D;Gorai.013G223200.2.A;Gorai.013G223200.3.D;Gorai.013G223200.3.A;Gorai.013G223200.1.D;Gorai.013G223200.1.A
Gene IDs: Gorai.013G223200
cotton name: K02730 - 20S proteasome subunit alpha 1 (PSMA6) (1 of 2)
OG IDs: 93524568
Best-hit-arabi-ID: AT2G05840.1
Best-hit-arabi-name: 20S proteasome subunit PAA2
Gorai.013G223200.1.D
PCC=0.173, p=0.479
Gorai.013G223200.1.A
PCC=0.322, p=0.179
mRNA Profile
Protein Expression Profile
DPA

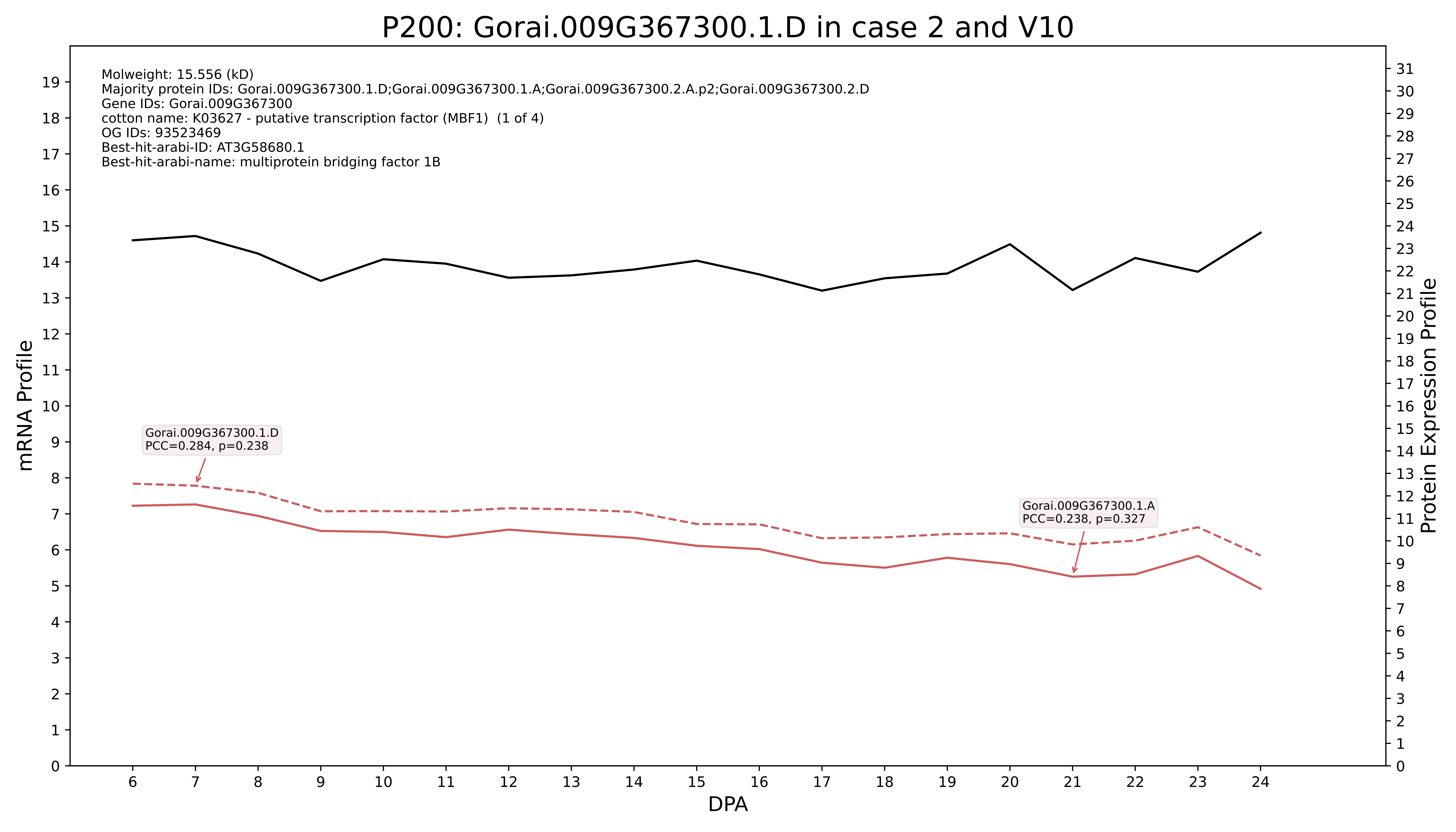
P200: Gorai.009G367300.1.D in case 2 and V10
Molweight: 15.556 (kD)
Majority protein IDs: Gorai.009G367300.1.D;Gorai.009G367300.1.A;Gorai.009G367300.2.A.p2;Gorai.009G367300.2.D
Gene IDs: Gorai.009G367300
cotton name: K03627 - putative transcription factor (MBF1) (1 of 4)
OG IDs: 93523469
Best-hit-arabi-ID: AT3G58680.1
Best-hit-arabi-name: multiprotein bridging factor 1B
Gorai.009G367300.1.D
PCC=0.284, p=0.238
Gorai.009G367300.1.A
PCC=0.238, p=0.327
mRNA Profile
Protein Expression Profile
DPA

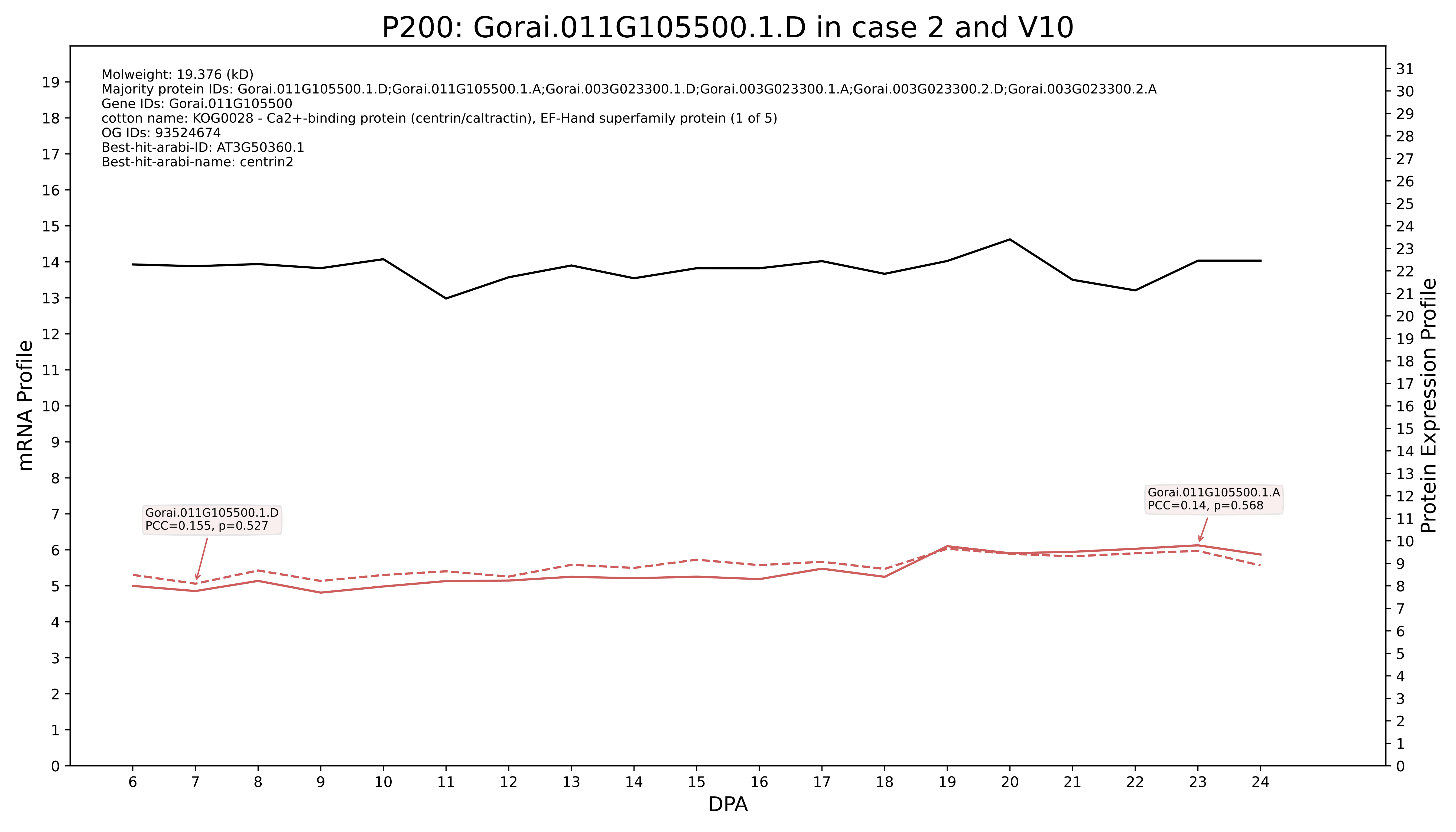
P200: Gorai.011G105500.1.D in case 2 and V10
Molweight: 19.376 (kD)
Majority protein IDs: Gorai.011G105500.1.D;Gorai.011G105500.1.A;Gorai.003G023300.1.D;Gorai.003G023300.1.A;Gorai.003G023300.2.D;Gorai.003G023300.2.A
Gene IDs: Gorai.011G105500
cotton name: KOG0028 - Ca2+-binding protein (centrin/caltractin), EF-Hand superfamily protein (1 of 5)
OG IDs: 93524674
Best-hit-arabi-ID: AT3G50360.1
Best-hit-arabi-name: centrin2
Gorai.011G105500.1.D
PCC=0.155, p=0.527
Gorai.011G105500.1.A
PCC=0.14, p=0.568
mRNA Profile
Protein Expression Profile
DPA

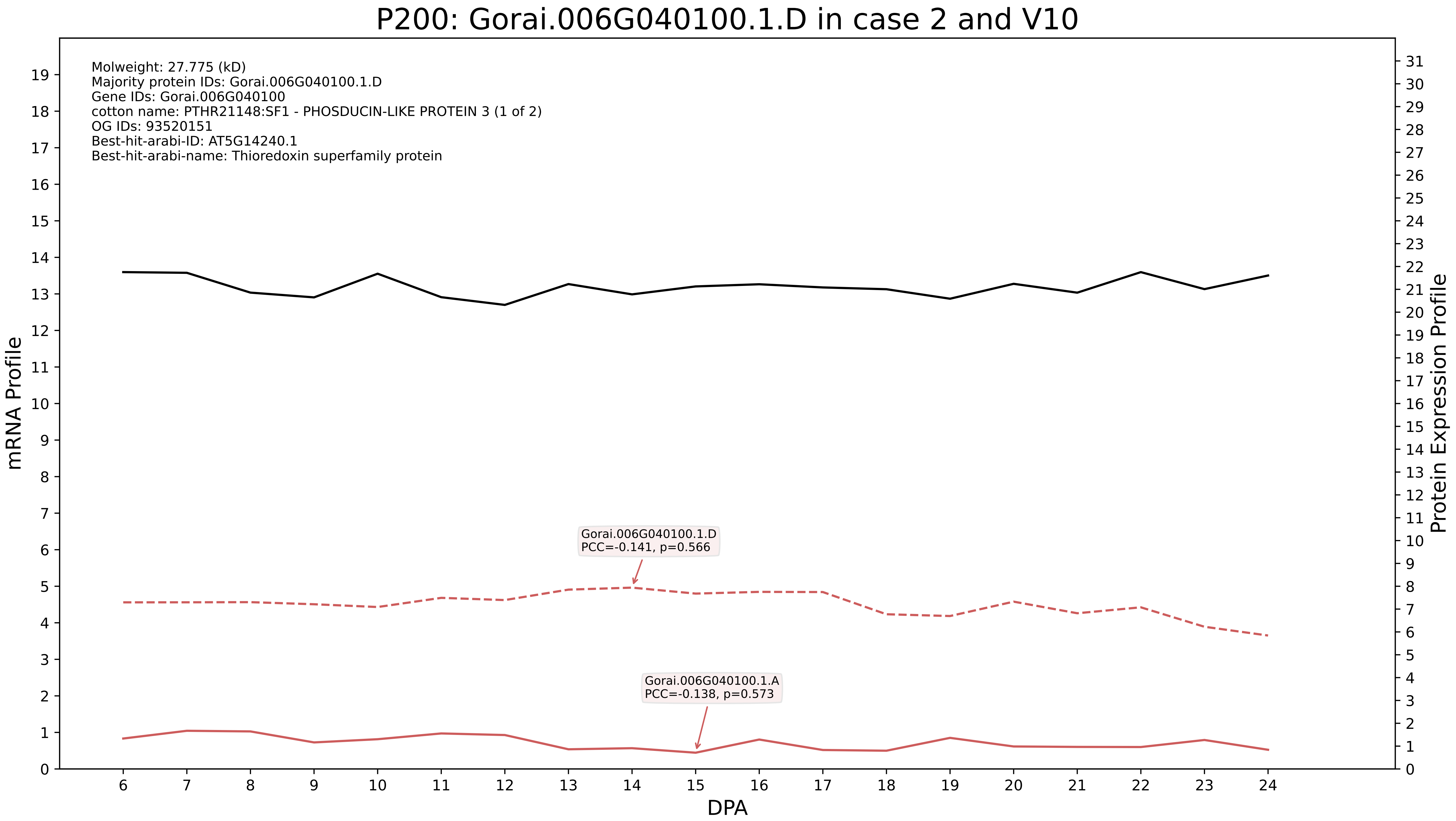
P200: Gorai.006G040100.1.D in case 2 and V10
Molweight: 27.775 (kD)
Majority protein IDs: Gorai.006G040100.1.D
Gene IDs: Gorai.006G040100
cotton name: PTHR21148:SF1 - PHOSDUCIN-LIKE PROTEIN 3 (1 of 2)
OG IDs: 93520151
Best-hit-arabi-ID: AT5G14240.1
Best-hit-arabi-name: Thioredoxin superfamily protein
Gorai.006G040100.1.D
PCC=-0.141, p=0.566
Gorai.006G040100.1.A
PCC=-0.138, p=0.573
mRNA Profile
Protein Expression Profile
DPA

### P200: Gorai.006G086200.2.D in case 2 and V10

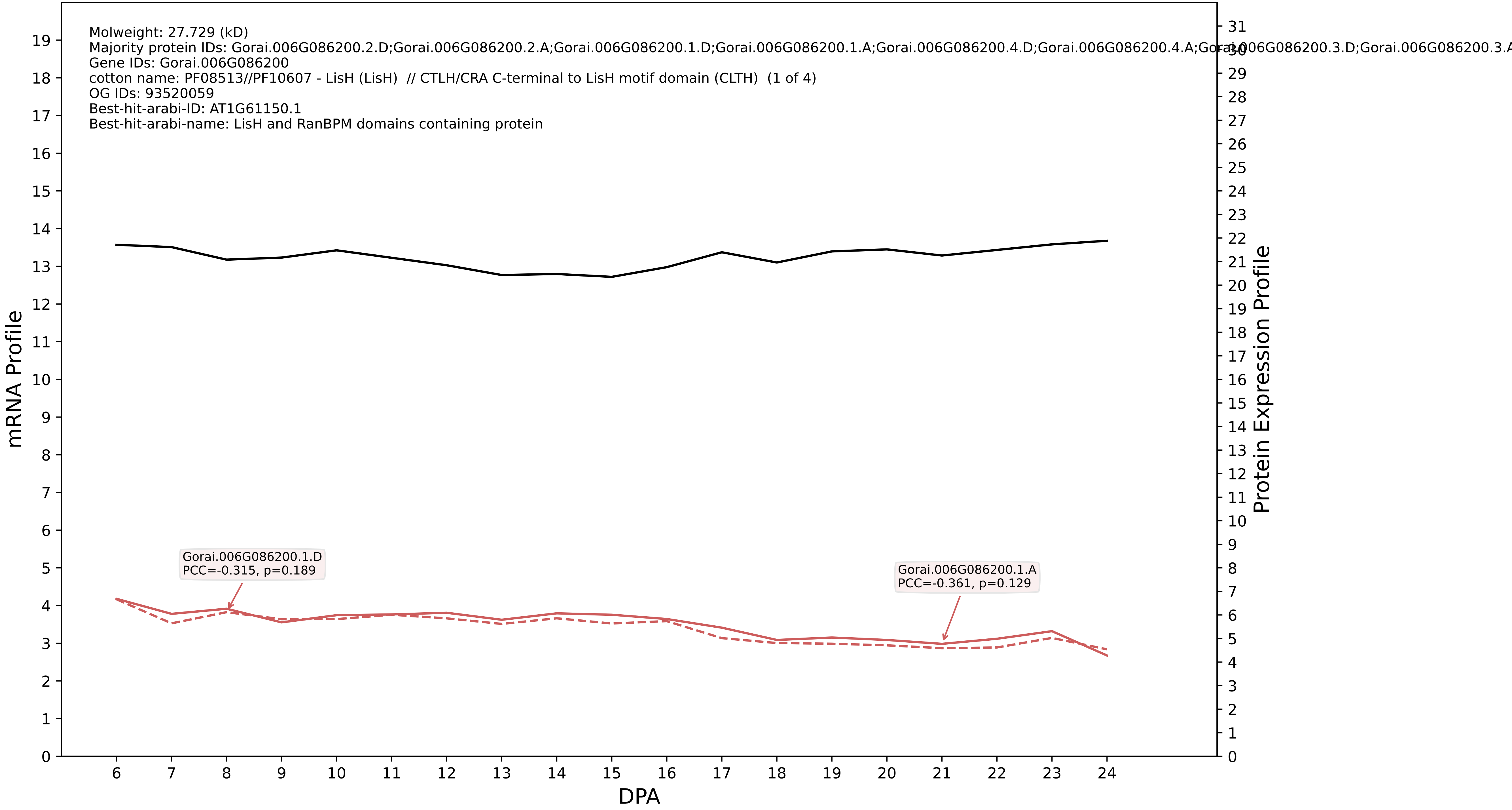

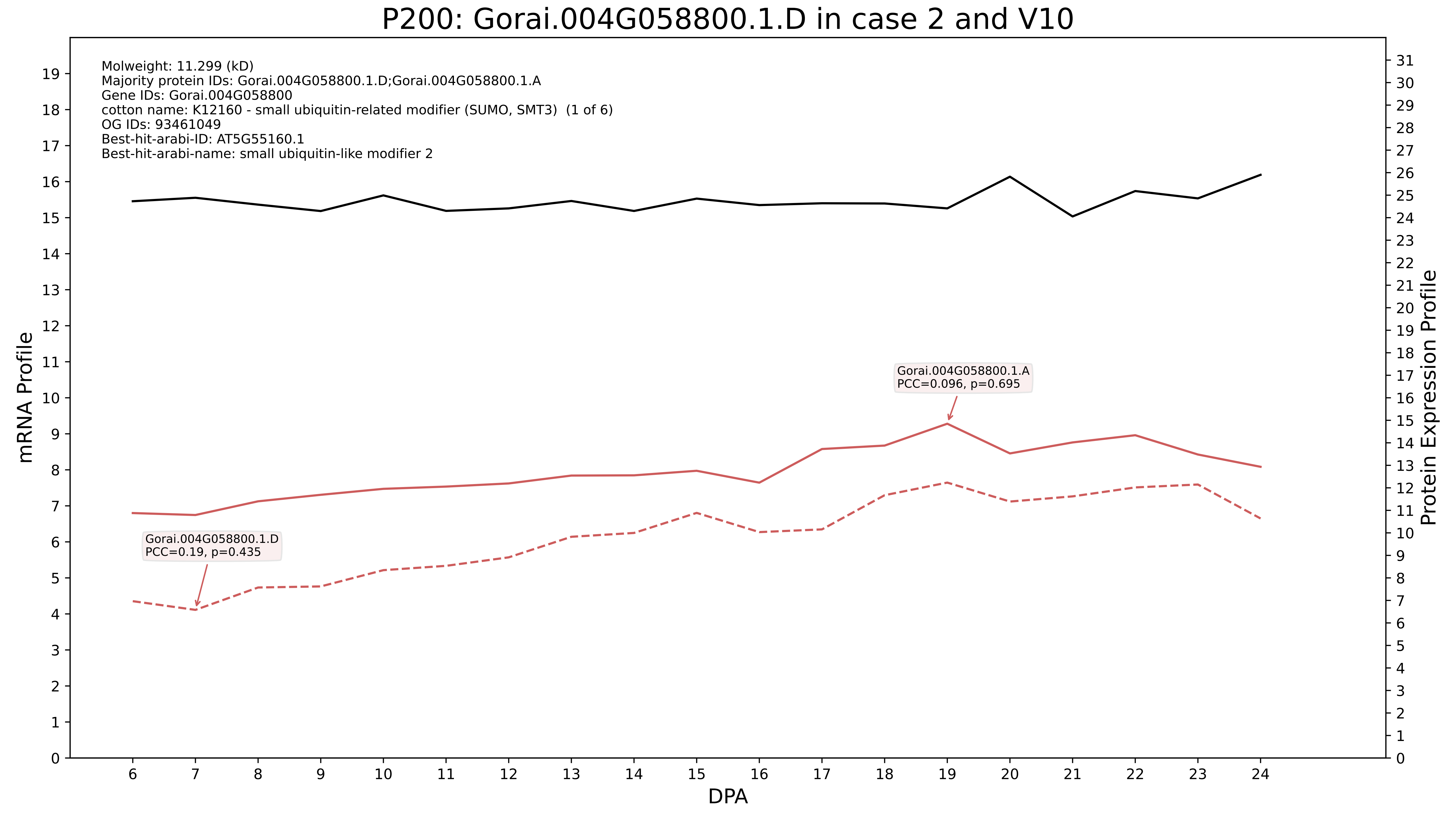
P200: Gorai.004G058800.1.D in case 2 and V10
Molweight: 11.299 (kD)
Majority protein IDs: Gorai.004G058800.1.D;Gorai.004G058800.1.A
Gene IDs: Gorai.004G058800
cotton name: K12160 - small ubiquitin-related modifier (SUMO, SMT3) (1 of 6)
OG IDs: 93461049
Best-hit-arabi-ID: AT5G55160.1
Best-hit-arabi-name: small ubiquitin-like modifier 2
Gorai.004G058800.1.A
PCC=0.096, p=0.695
Gorai.004G058800.1.D
PCC=0.19, p=0.435
mRNA Profile
Protein Expression Profile
DPA

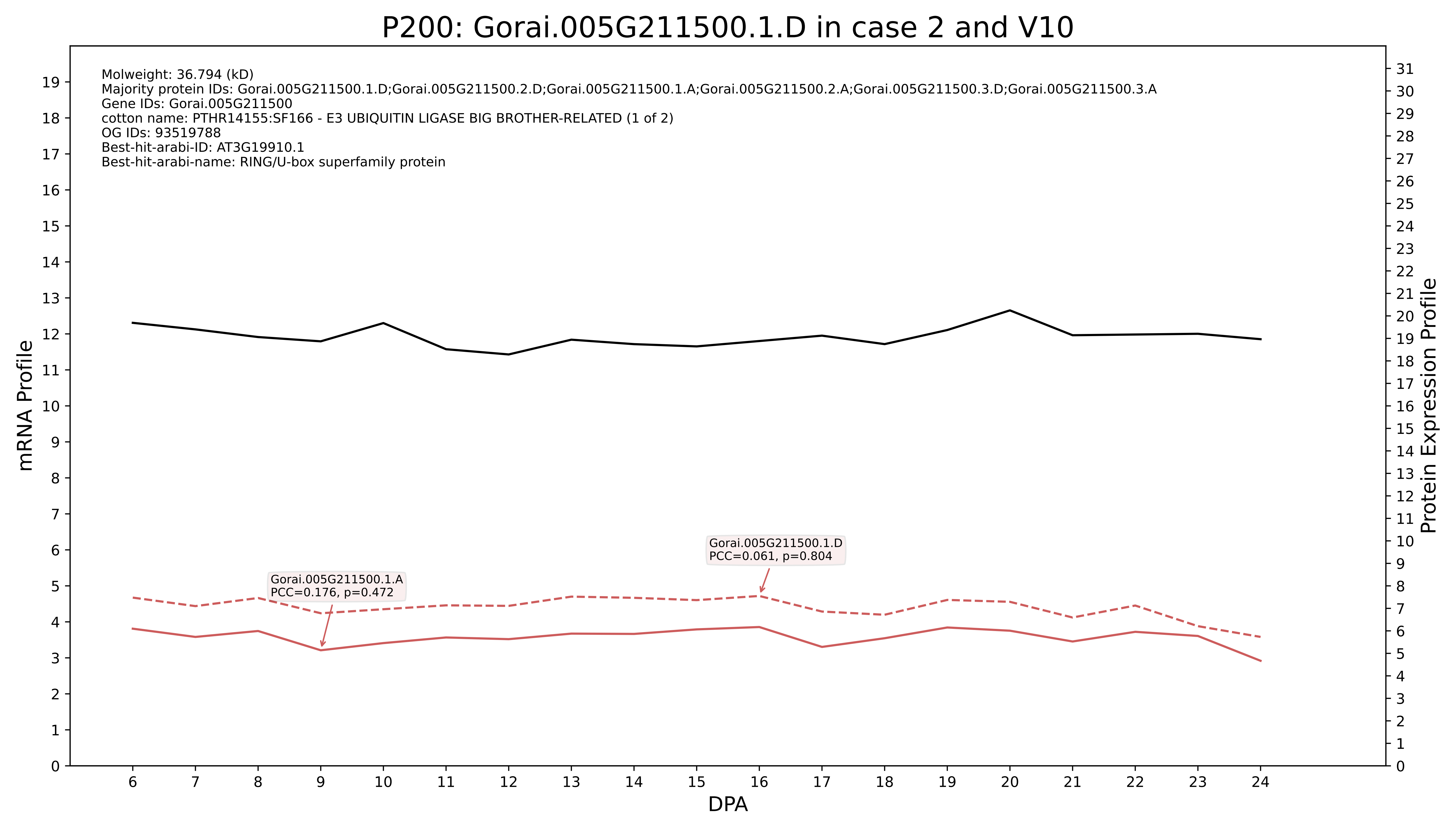
P200: Gorai.005G211500.1.D in case 2 and V10
Molweight: 36.794 (kD)
Majority protein IDs: Gorai.005G211500.1.D;Gorai.005G211500.2.D;Gorai.005G211500.1.A;Gorai.005G211500.2.A;Gorai.005G211500.3.D;Gorai.005G211500.3.A
Gene IDs: Gorai.005G211500
cotton name: PTHR14155:SF166 - E3 UBIQUITIN LIGASE BIG BROTHER-RELATED (1 of 2)
OG IDs: 93519788
Best-hit-arabi-ID: AT3G19910.1
Best-hit-arabi-name: RING/U-box superfamily protein
Gorai.005G211500.1.D
PCC=0.061, p=0.804
Gorai.005G211500.1.A
PCC=0.176, p=0.472
mRNA Profile
Protein Expression Profile
DPA

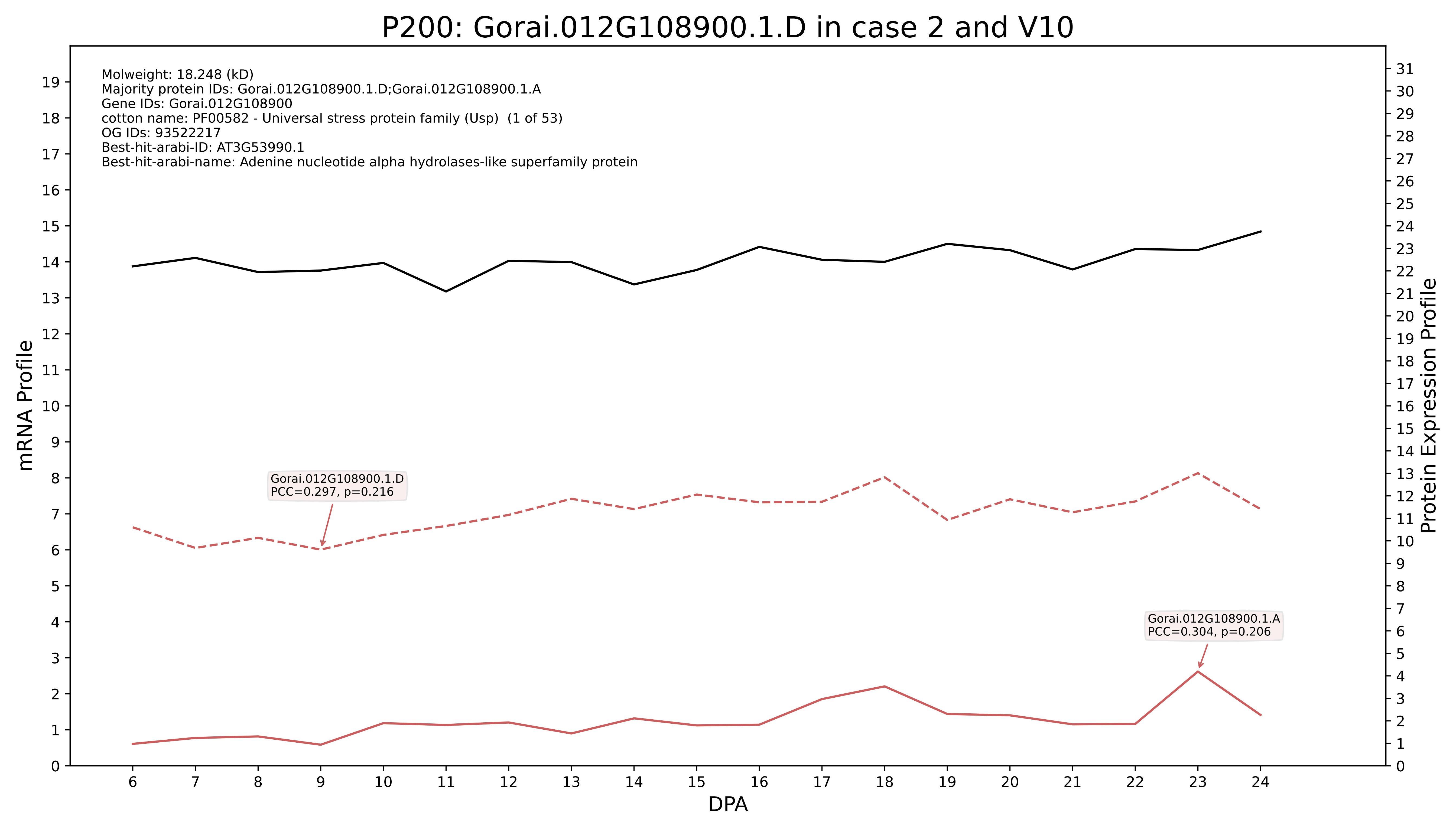
P200: Gorai.012G108900.1.D in case 2 and V10
Molweight: 18.248 (kD)
Majority protein IDs: Gorai.012G108900.1.D;Gorai.012G108900.1.A
Gene IDs: Gorai.012G108900
cotton name: PF00582 - Universal stress protein family (Usp) (1 of 53)
OG IDs: 93522217
Best-hit-arabi-ID: AT3G53990.1
Best-hit-arabi-name: Adenine nucleotide alpha hydrolases-like superfamily protein
Gorai.012G108900.1.D
PCC=0.297, p=0.216
Gorai.012G108900.1.A
PCC=0.304, p=0.206
mRNA Profile
Protein Expression Profile
DPA

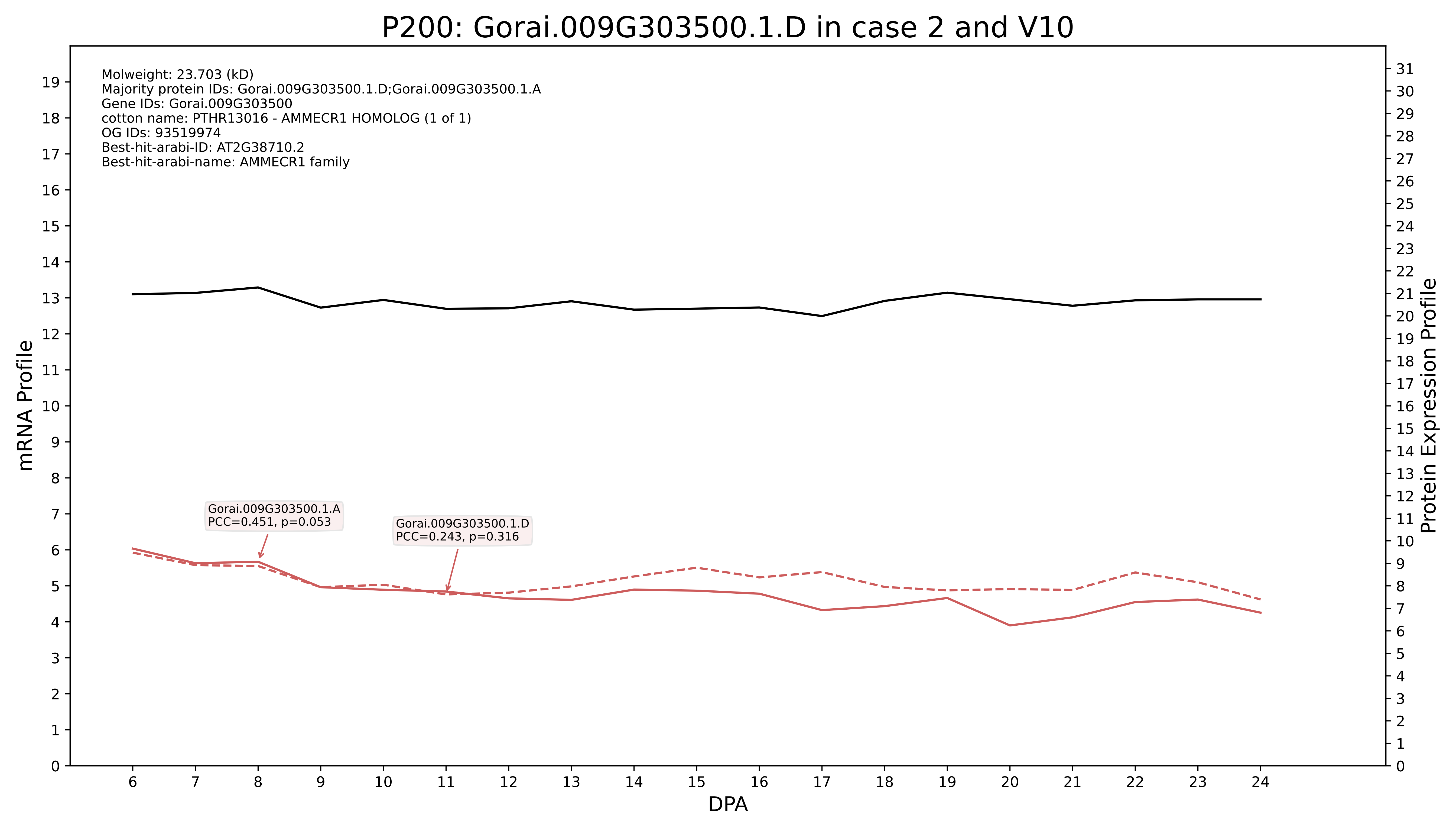

P200: Gorai.009G303500.1.D in case 2 and V10
Molweight: 23.703 (kD)
Majority protein IDs: Gorai.009G303500.1.D;Gorai.009G303500.1.A
Gene IDs: Gorai.009G303500
cotton name: PTHR13016 - AMMECR1 HOMOLOG (1 of 1)
OG IDs: 93519974
Best-hit-arabi-ID: AT2G38710.2
Best-hit-arabi-name: AMMECR1 family
Gorai.009G303500.1.A
PCC=0.451, p=0.053
Gorai.009G303500.1.D
PCC=0.243, p=0.316
mRNA Profile
Protein Expression Profile
DPA

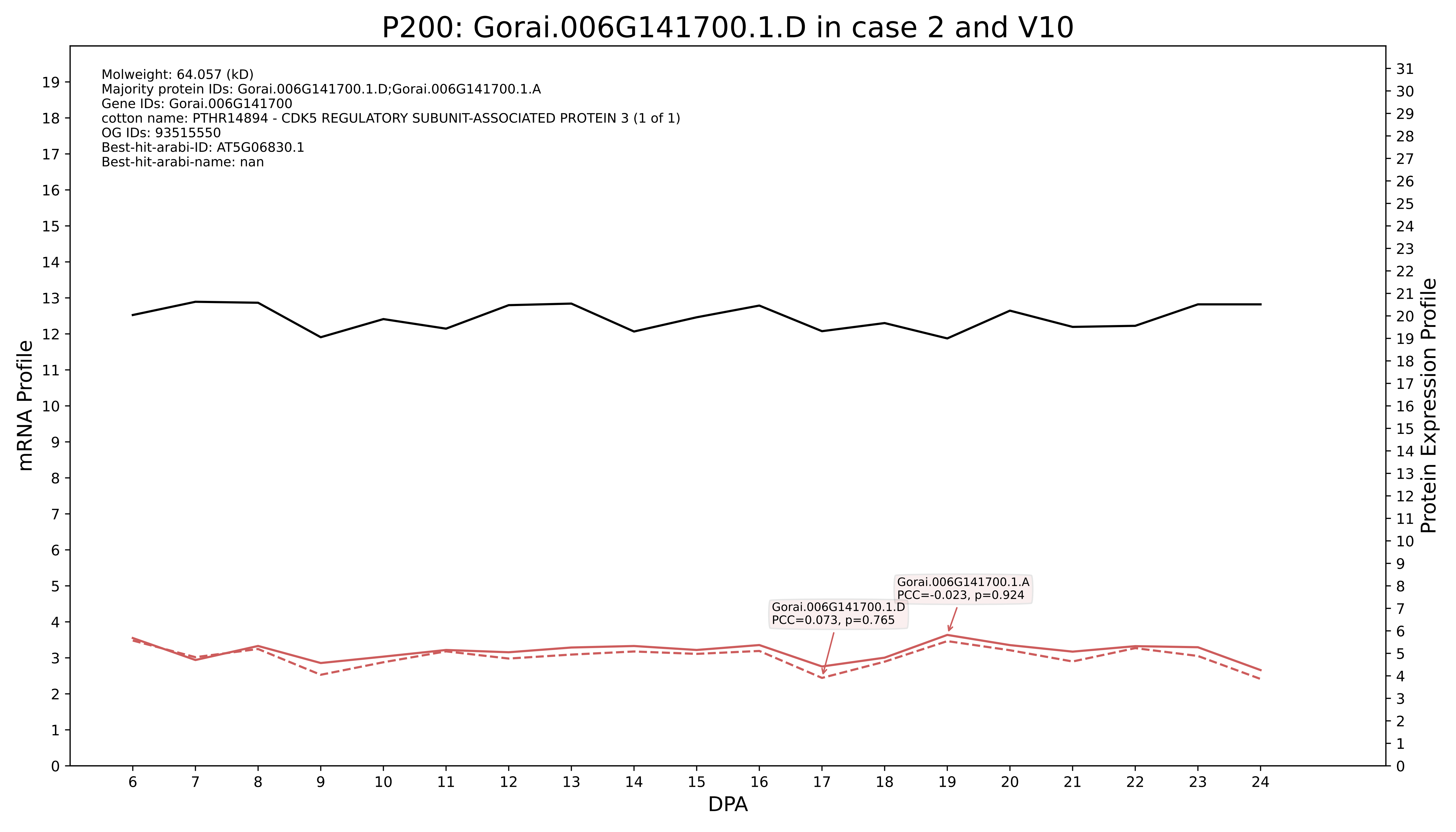
P200: Gorai.006G141700.1.D in case 2 and V10
Molweight: 64.057 (kD)
Majority protein IDs: Gorai.006G141700.1.D;Gorai.006G141700.1.A
Gene IDs: Gorai.006G141700
cotton name: PTHR14894 - CDK5 REGULATORY SUBUNIT-ASSOCIATED PROTEIN 3 (1 of 1)
OG IDs: 93515550
Best-hit-arabi-ID: AT5G06830.1
Best-hit-arabi-name: nan
Gorai.006G141700.1.A
PCC=-0.023, p=0.924
Gorai.006G141700.1.D
PCC=0.073, p=0.765
mRNA Profile
Protein Expression Profile
DPA

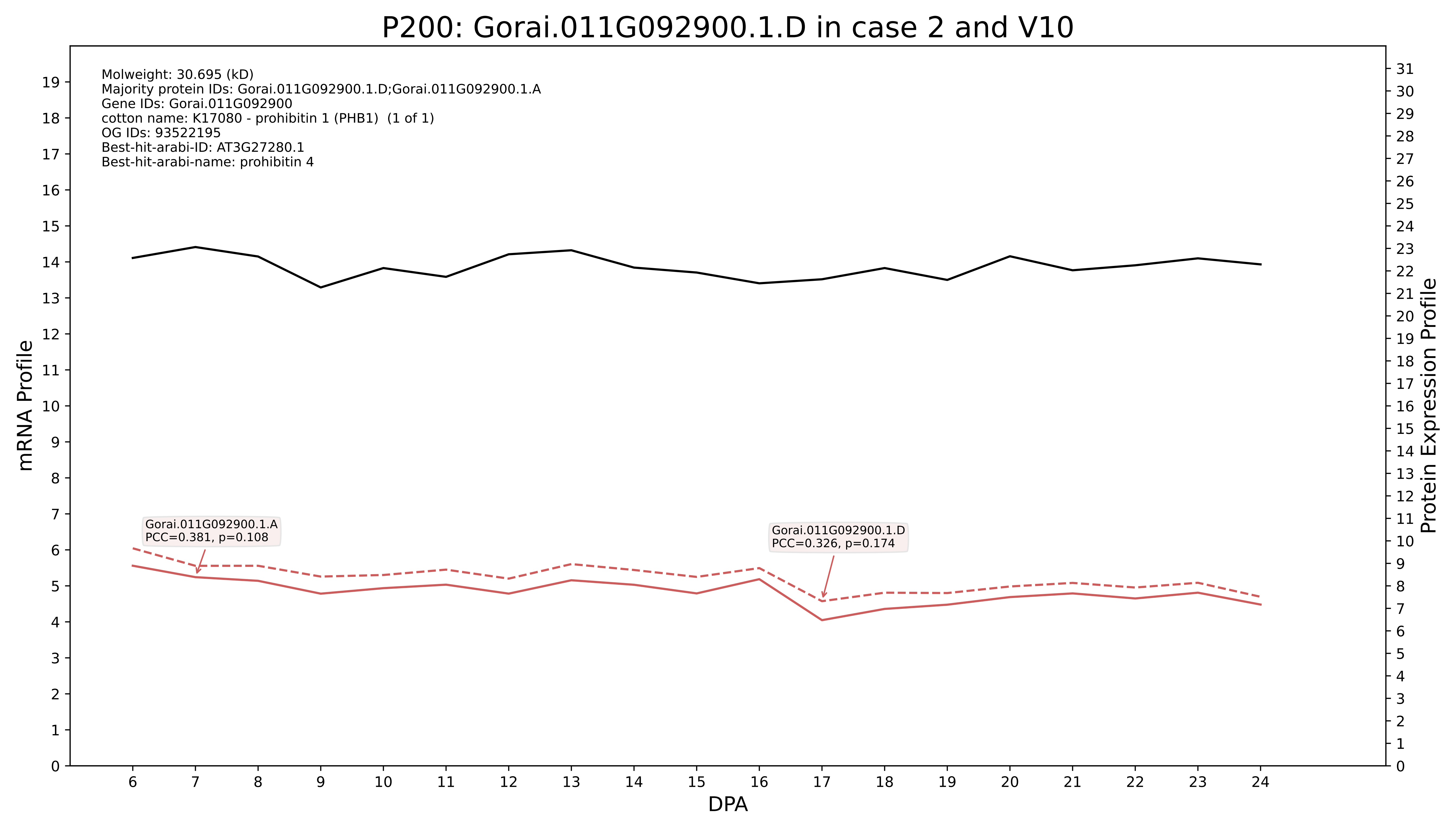

P200: Gorai.011G092900.1.D in case 2 and V10
Molweight: 30.695 (kD)
Majority protein IDs: Gorai.011G092900.1.D;Gorai.011G092900.1.A
Gene IDs: Gorai.011G092900
cotton name: K17080 - prohibitin 1 (PHB1) (1 of 1)
OG IDs: 93522195
Best-hit-arabi-ID: AT3G27280.1
Best-hit-arabi-name: prohibitin 4
Gorai.011G092900.1.A
PCC=0.381, p=0.108
Gorai.011G092900.1.D
PCC=0.326, p=0.174
mRNA Profile
Protein Expression Profile
DPA

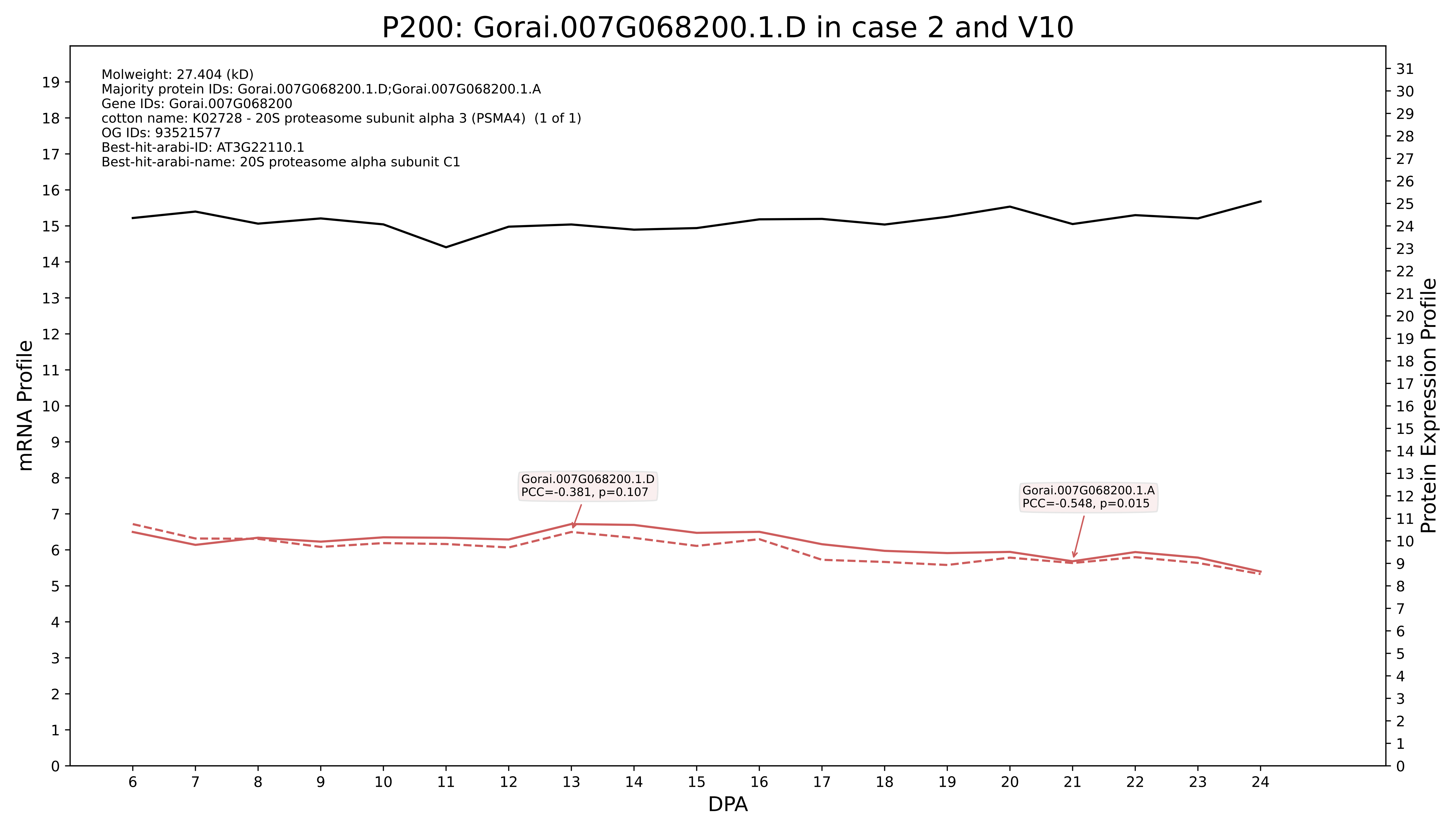
P200: Gorai.007G068200.1.D in case 2 and V10
Molweight: 27.404 (kD)
Majority protein IDs: Gorai.007G068200.1.D;Gorai.007G068200.1.A
Gene IDs: Gorai.007G068200
cotton name: K02728 - 20S proteasome subunit alpha 3 (PSMA4) (1 of 1)
OG IDs: 93521577
Best-hit-arabi-ID: AT3G22110.1
Best-hit-arabi-name: 20S proteasome alpha subunit C1
Gorai.007G068200.1.D
PCC=-0.381, p=0.107
Gorai.007G068200.1.A
PCC=-0.548, p=0.015
mRNA Profile
Protein Expression Profile
DPA

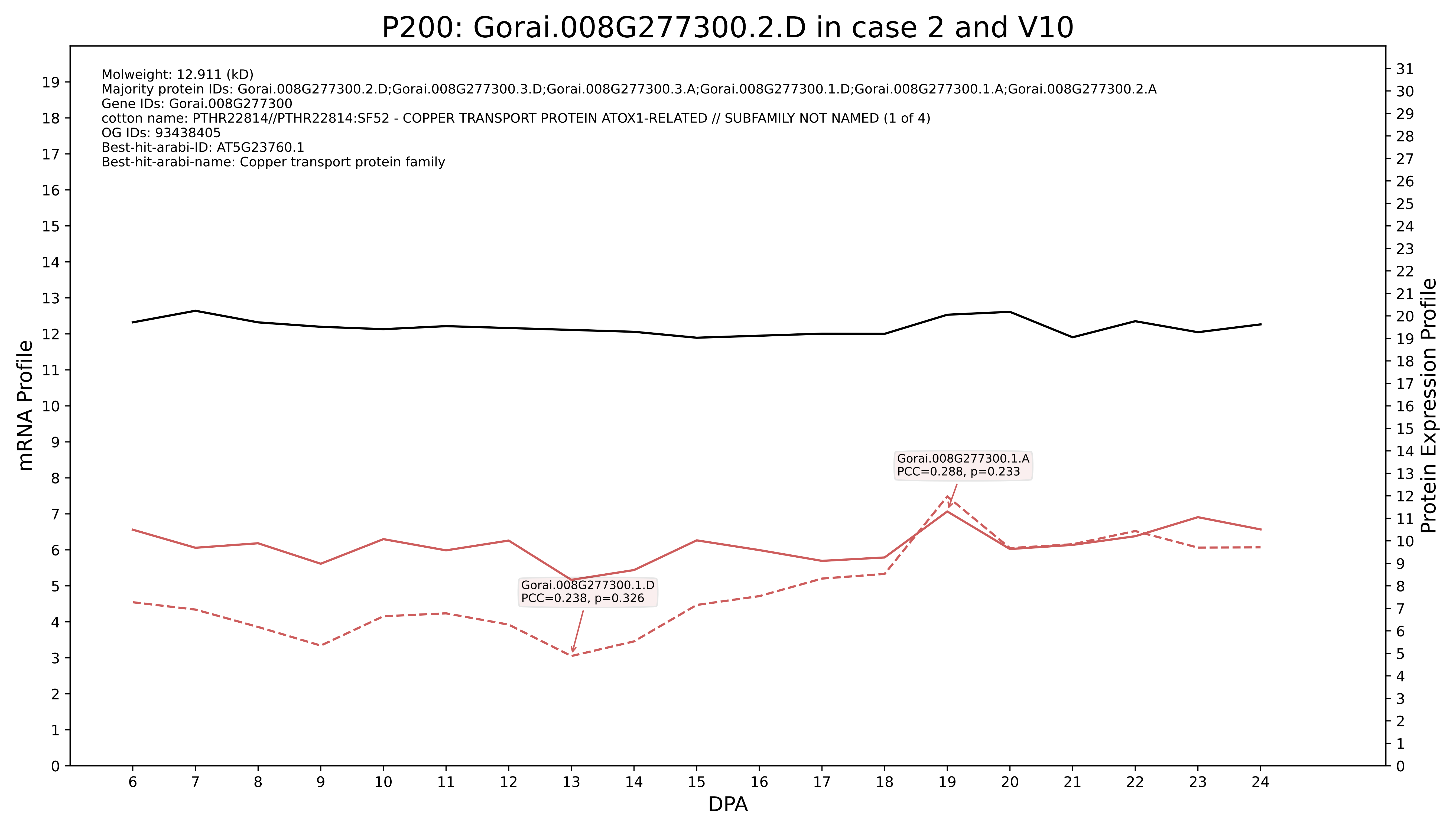
P200: Gorai.008G277300.2.D in case 2 and V10
Molweight: 12.911 (kD)
Majority protein IDs: Gorai.008G277300.2.D;Gorai.008G277300.3.D;Gorai.008G277300.3.A;Gorai.008G277300.1.D;Gorai.008G277300.1.A;Gorai.008G277300.2.A
Gene IDs: Gorai.008G277300
cotton name: PTHR22814//PTHR22814:SF52 - COPPER TRANSPORT PROTEIN ATOX1-RELATED // SUBFAMILY NOT NAMED (1 of 4)
OG IDs: 93438405
Best-hit-arabi-ID: AT5G23760.1
Best-hit-arabi-name: Copper transport protein family
Gorai.008G277300.1.A
PCC=0.288, p=0.233
Gorai.008G277300.1.D
PCC=0.238, p=0.326
mRNA Profile
Protein Expression Profile
DPA

### P200: Gorai.004G143300.1.D in case 2 and V10

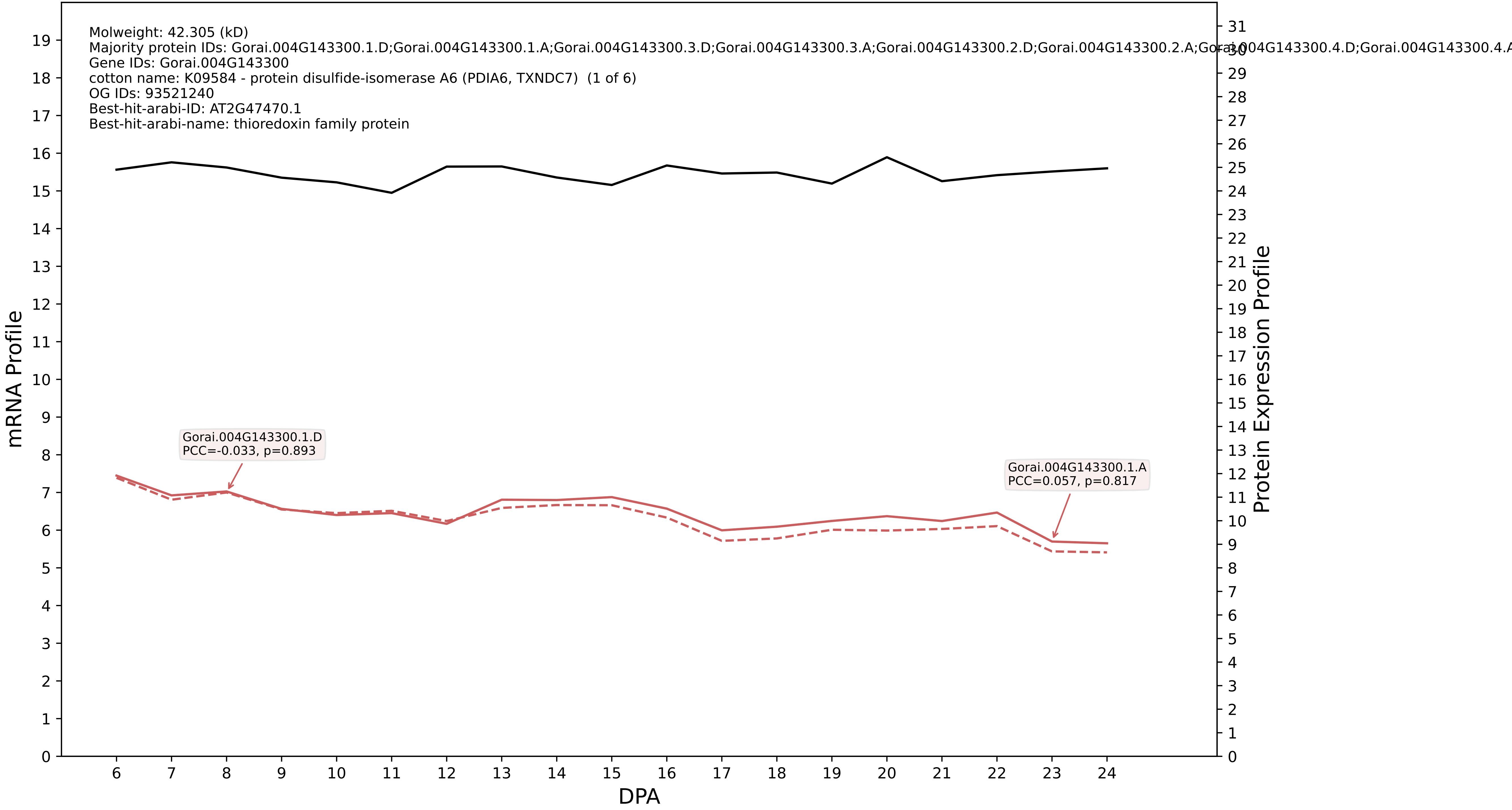

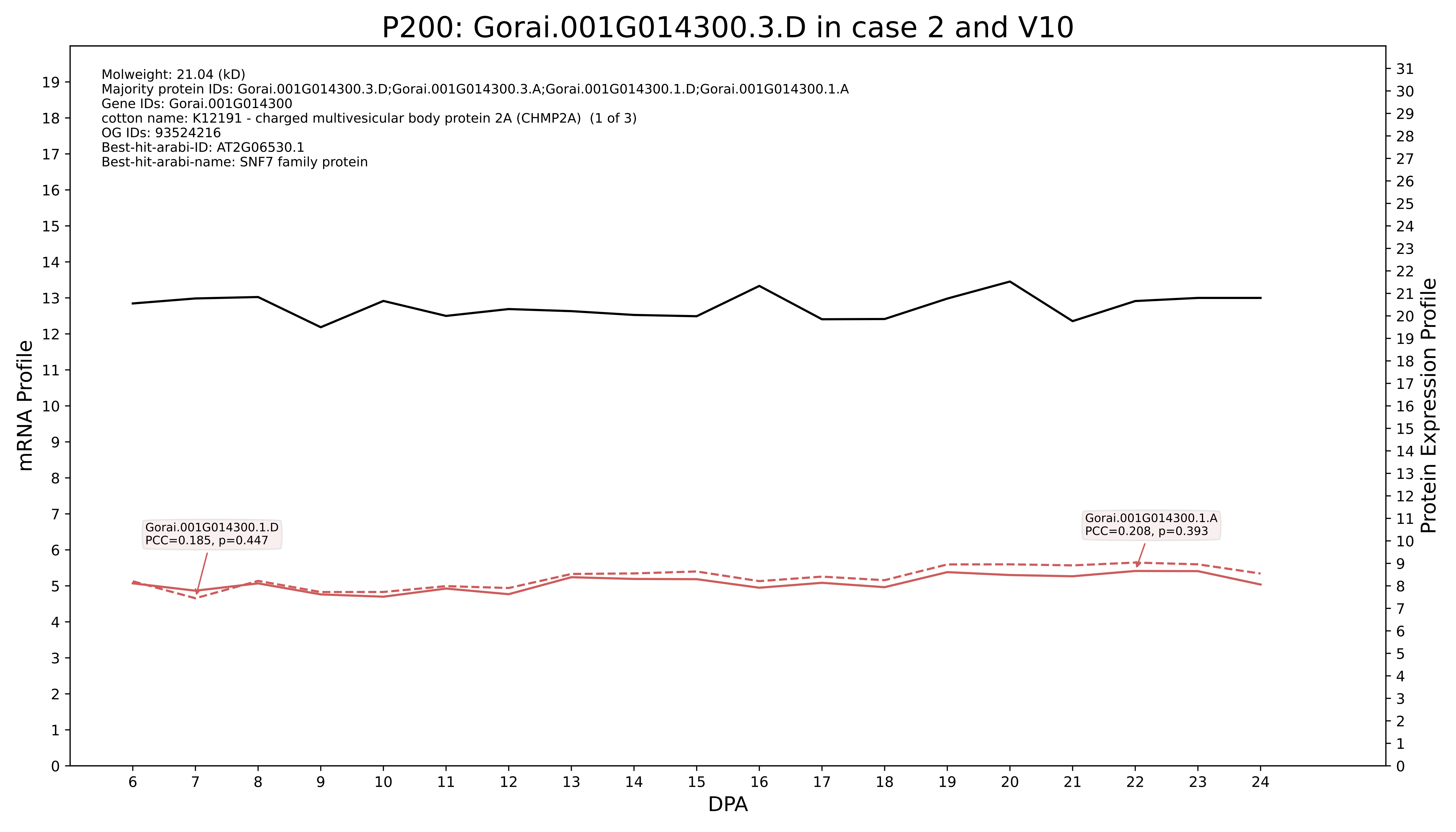

P200: Gorai.001G014300.3.D in case 2 and V10
Molweight: 21.04 (kD)
Majority protein IDs: Gorai.001G014300.3.D;Gorai.001G014300.3.A;Gorai.001G014300.1.D;Gorai.001G014300.1.A
Gene IDs: Gorai.001G014300
cotton name: K12191 - charged multivesicular body protein 2A (CHMP2A) (1 of 3)
OG IDs: 93524216
Best-hit-arabi-ID: AT2G06530.1
Best-hit-arabi-name: SNF7 family protein
Gorai.001G014300.1.D
PCC=0.185, p=0.447
Gorai.001G014300.1.A
PCC=0.208, p=0.393
mRNA Profile
Protein Expression Profile
DPA

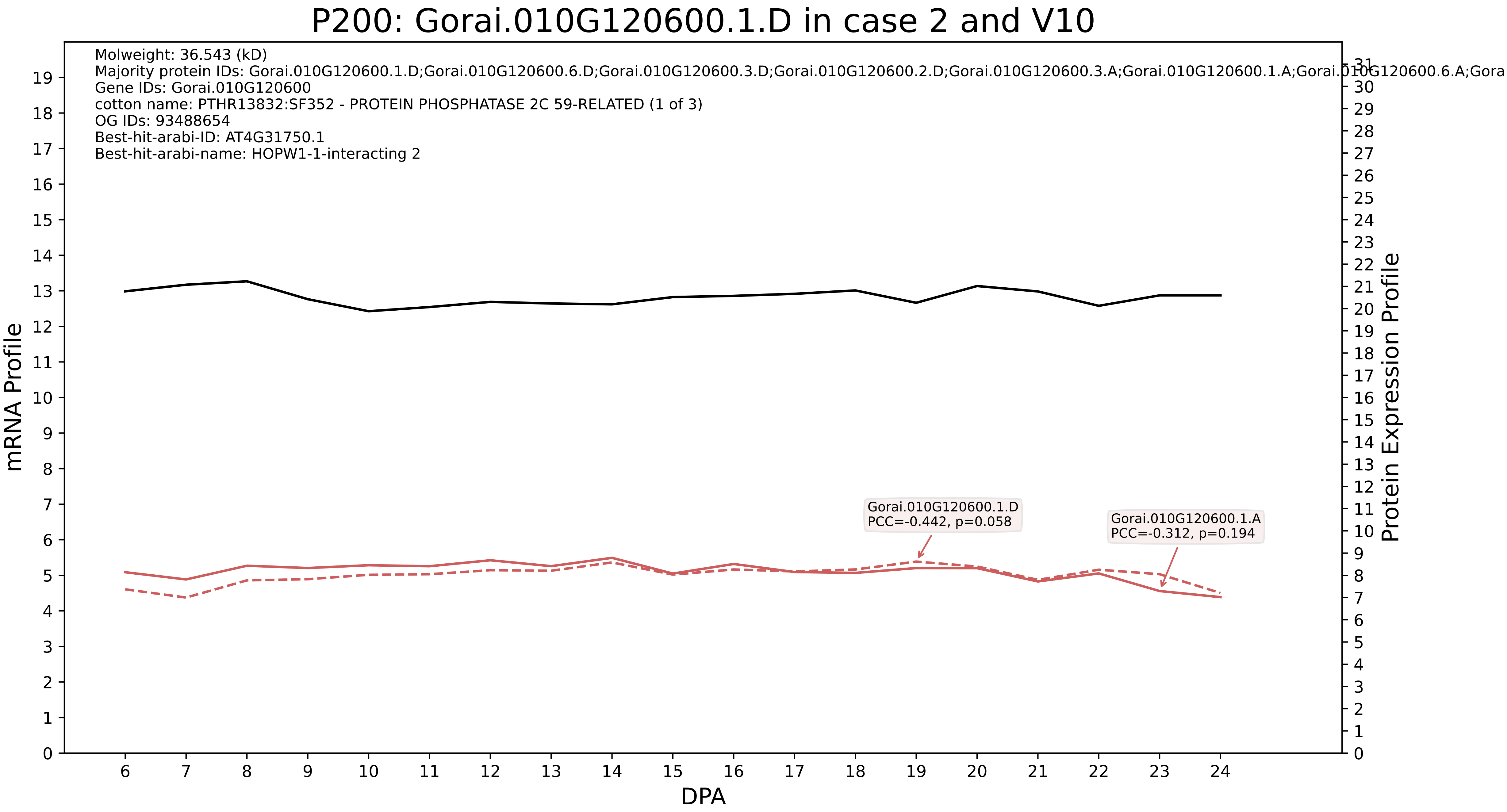
P200: Gorai.010G120600.1.D in case 2 and V10
Molweight: 36.543 (kD)
Majority protein IDs: Gorai.010G120600.1.D;Gorai.010G120600.6.D;Gorai.010G120600.3.D;Gorai.010G120600.2.D;Gorai.010G120600.3.A;Gorai.010G120600.1.A;Gorai.010G120600.6.A;Gorai
Gene IDs: Gorai.010G120600
cotton name: PTHR13832:SF352 - PROTEIN PHOSPHATASE 2C 59-RELATED (1 of 3)
OG IDs: 93488654
Best-hit-arabi-ID: AT4G31750.1
Best-hit-arabi-name: HOPW1-1-interacting 2
Gorai.010G120600.1.D
PCC=-0.442, p=0.058
Gorai.010G120600.1.A
PCC=-0.312, p=0.194
mRNA Profile
Protein Expression Profile
DPA

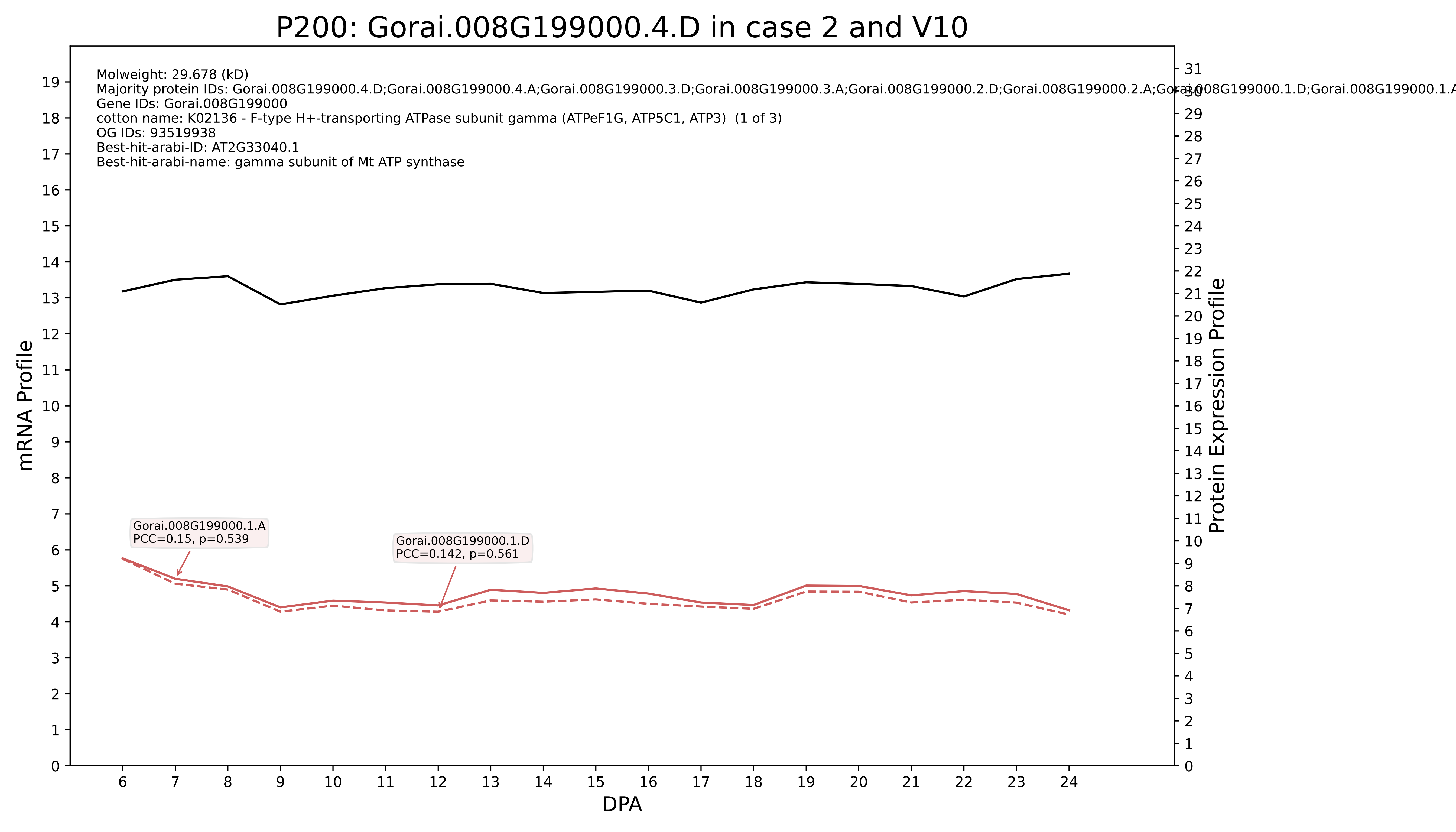
P200: Gorai.008G199000.4.D in case 2 and V10
Molweight: 29.678 (kD)
Majority protein IDs: Gorai.008G199000.4.D;Gorai.008G199000.4.A;Gorai.008G199000.3.D;Gorai.008G199000.3.A;Gorai.008G199000.2.D;Gorai.008G199000.2.A;Gorai.008G199000.1.D;Gorai.008G199000.1.A
Gene IDs: Gorai.008G199000
cotton name: K02136 - F-type H+-transporting ATPase subunit gamma (ATPeF1G, ATP5C1, ATP3) (1 of 3)
OG IDs: 93519938
Best-hit-arabi-ID: AT2G33040.1
Best-hit-arabi-name: gamma subunit of Mt ATP synthase
Gorai.008G199000.1.A
PCC=0.15, p=0.539
Gorai.008G199000.1.D
PCC=0.142, p=0.561
mRNA Profile
Protein Expression Profile
DPA

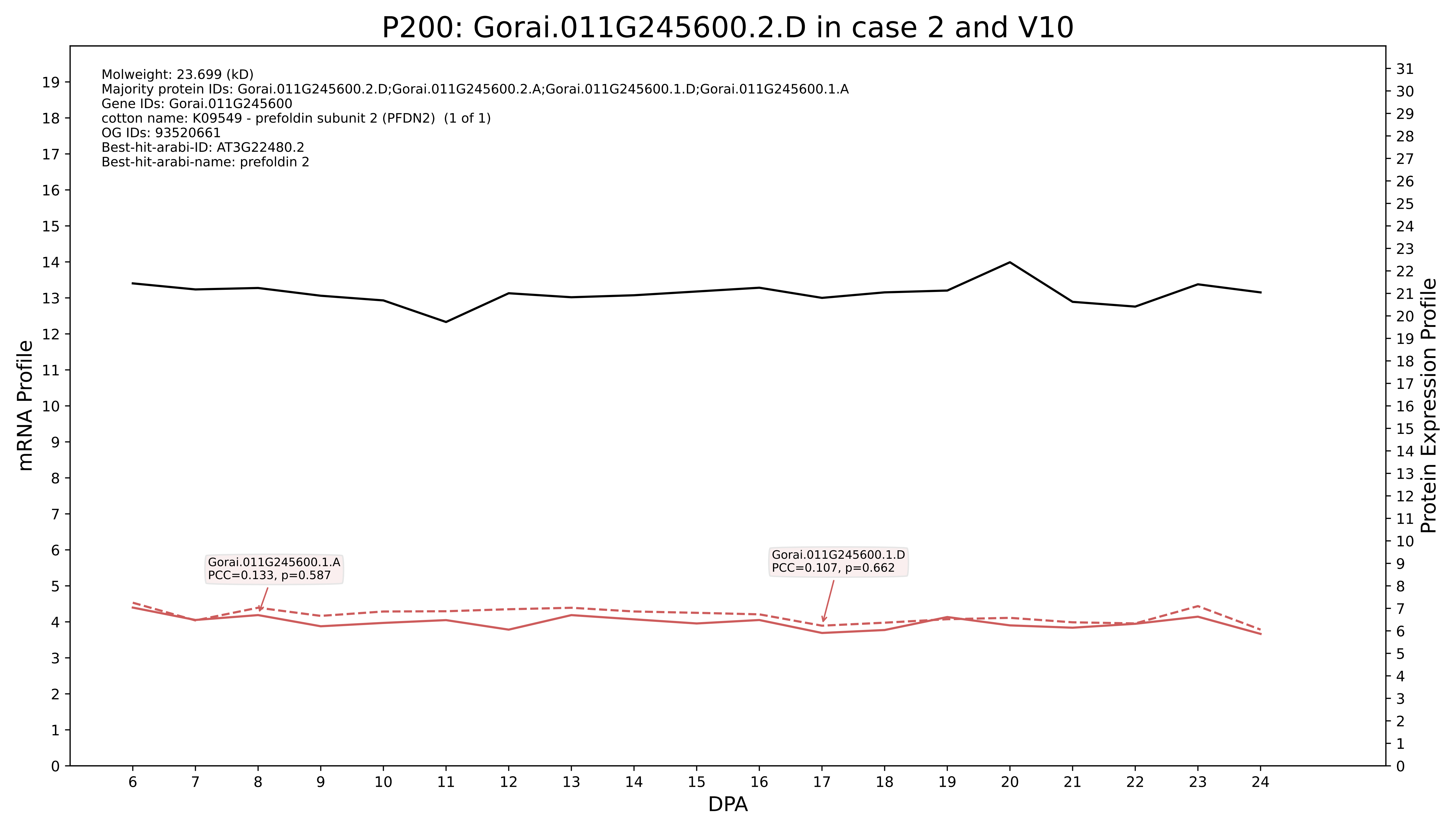
P200: Gorai.011G245600.2.D in case 2 and V10
Molweight: 23.699 (kD)
Majority protein IDs: Gorai.011G245600.2.D;Gorai.011G245600.2.A;Gorai.011G245600.1.D;Gorai.011G245600.1.A
Gene IDs: Gorai.011G245600
cotton name: K09549 - prefoldin subunit 2 (PFDN2) (1 of 1)
OG IDs: 93520661
Best-hit-arabi-ID: AT3G22480.2
Best-hit-arabi-name: prefoldin 2
Gorai.011G245600.1.A
PCC=0.133, p=0.587
Gorai.011G245600.1.D
PCC=0.107, p=0.662
mRNA Profile
Protein Expression Profile
DPA

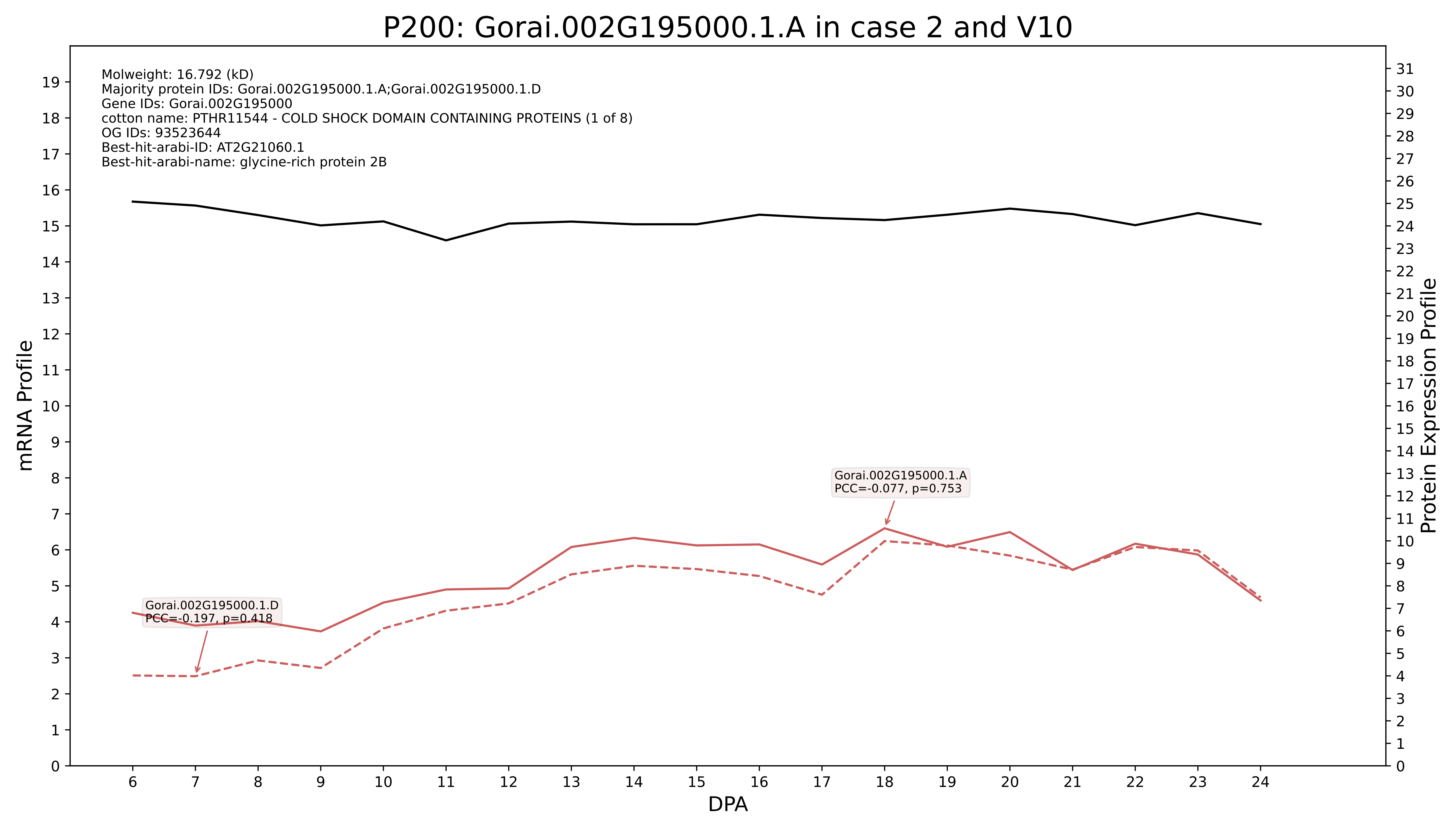

P200: Gorai.002G195000.1.A in case 2 and V10
Molweight: 16.792 (kD)
Majority protein IDs: Gorai.002G195000.1.A;Gorai.002G195000.1.D
Gene IDs: Gorai.002G195000
cotton name: PTHR11544 - COLD SHOCK DOMAIN CONTAINING PROTEINS (1 of 8)
OG IDs: 93523644
Best-hit-arabi-ID: AT2G21060.1
Best-hit-arabi-name: glycine-rich protein 2B
Gorai.002G195000.1.A
PCC=-0.077, p=0.753
Gorai.002G195000.1.D
PCC=-0.197, p=0.418
mRNA Profile
Protein Expression Profile
DPA

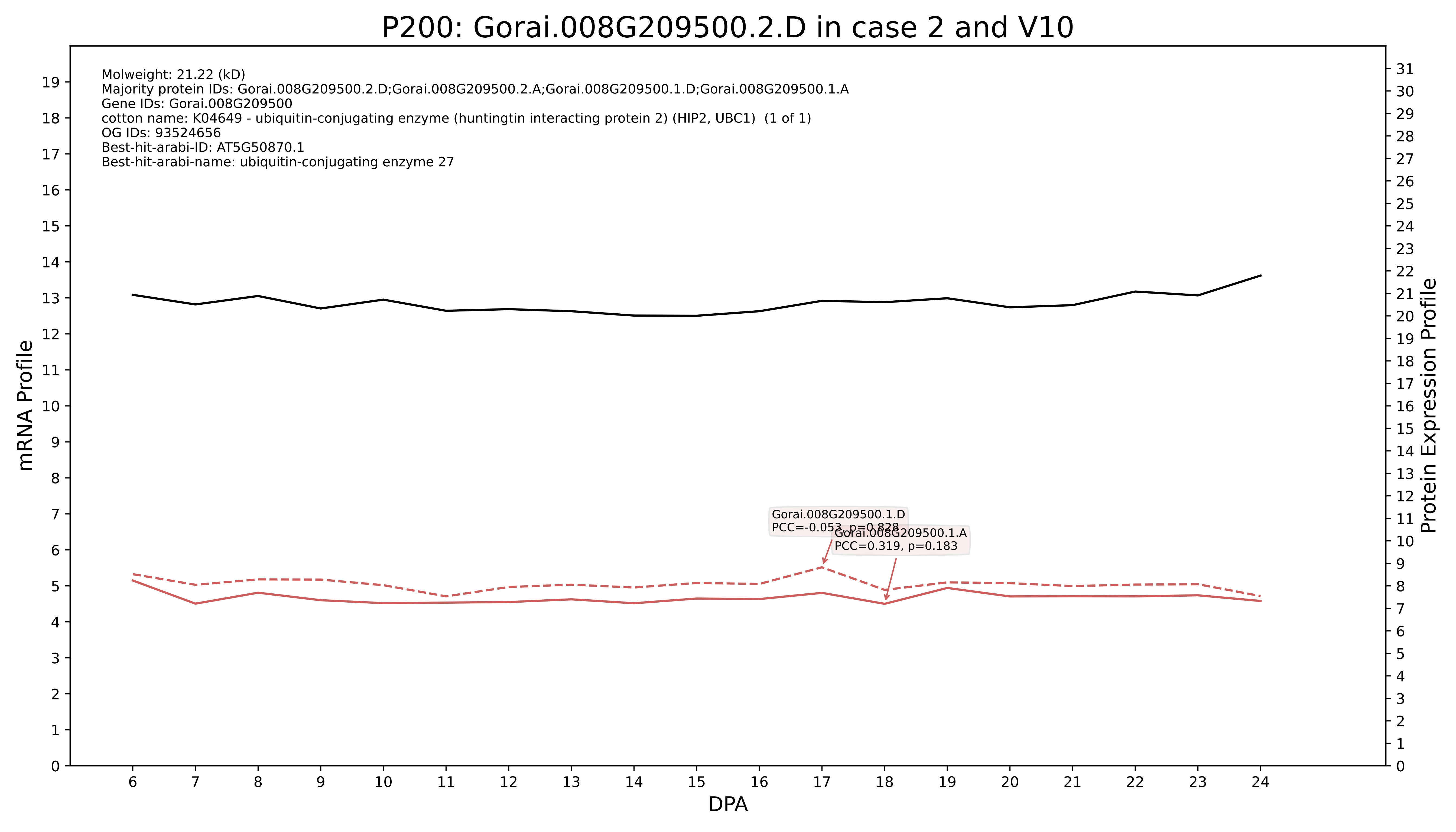

P200: Gorai.008G209500.2.D in case 2 and V10
Molweight: 21.22 (kD)
Majority protein IDs: Gorai.008G209500.2.D;Gorai.008G209500.2.A;Gorai.008G209500.1.D;Gorai.008G209500.1.A
Gene IDs: Gorai.008G209500
cotton name: K04649 - ubiquitin-conjugating enzyme (huntingtin interacting protein 2) (HIP2, UBC1) (1 of 1)
OG IDs: 93524656
Best-hit-arabi-ID: AT5G50870.1
Best-hit-arabi-name: ubiquitin-conjugating enzyme 27
Gorai.008G209500.1.D
PCC=-0.053, p=0.828
Gorai.008G209500.1.A
PCC=0.319, p=0.183
mRNA Profile
Protein Expression Profile
DPA

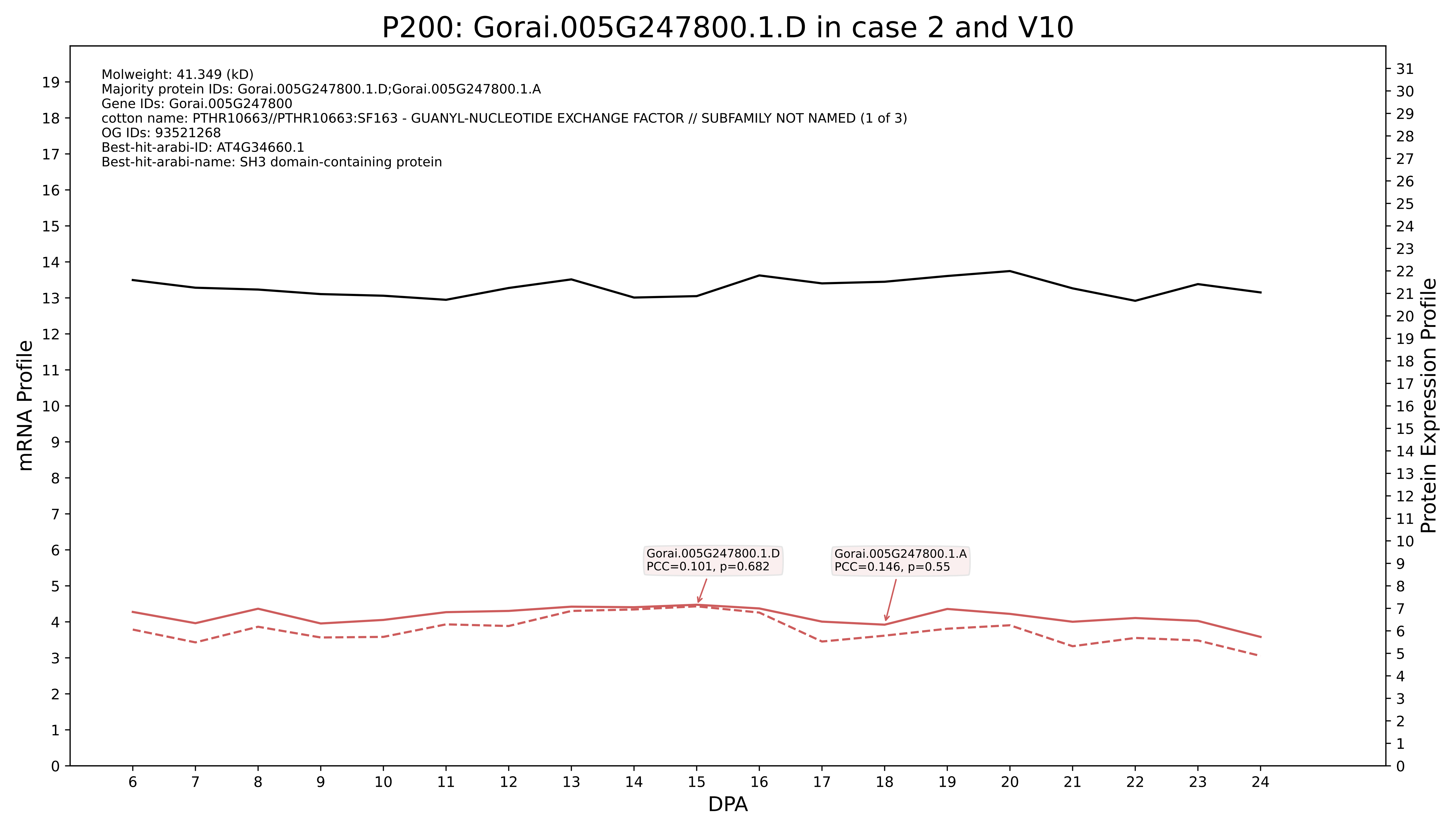
P200: Gorai.005G247800.1.D in case 2 and V10
Molweight: 41.349 (kD)
Majority protein IDs: Gorai.005G247800.1.D;Gorai.005G247800.1.A
Gene IDs: Gorai.005G247800
cotton name: PTHR10663//PTHR10663:SF163 - GUANYL-NUCLEOTIDE EXCHANGE FACTOR // SUBFAMILY NOT NAMED (1 of 3)
OG IDs: 93521268
Best-hit-arabi-ID: AT4G34660.1
Best-hit-arabi-name: SH3 domain-containing protein
Gorai.005G247800.1.D
PCC=0.101, p=0.682
Gorai.005G247800.1.A
PCC=0.146, p=0.55
mRNA Profile
Protein Expression Profile
DPA

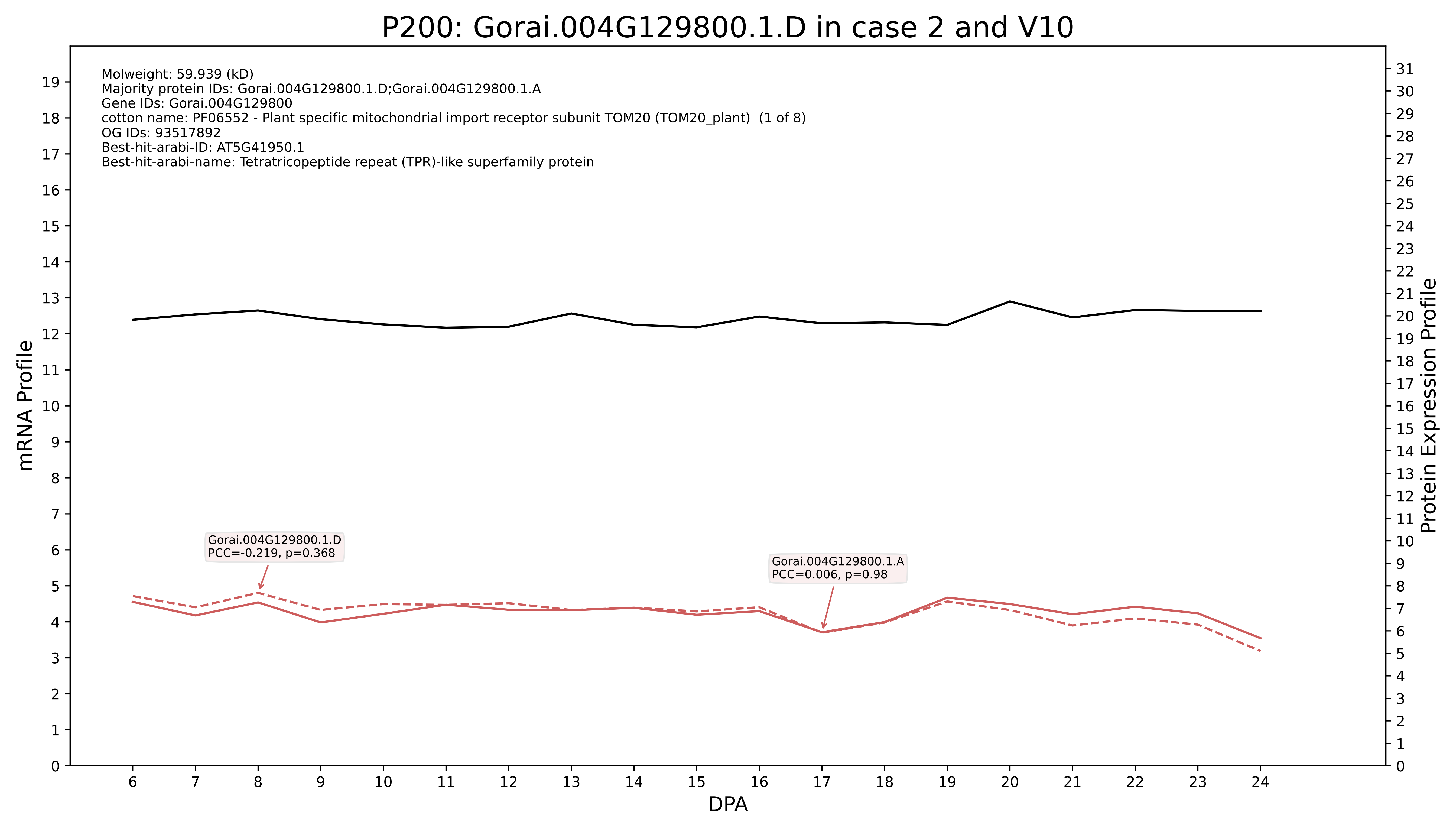

P200: Gorai.004G129800.1.D in case 2 and V10
Molweight: 59.939 (kD)
Majority protein IDs: Gorai.004G129800.1.D;Gorai.004G129800.1.A
Gene IDs: Gorai.004G129800
cotton name: PF06552 - Plant specific mitochondrial import receptor subunit TOM20 (TOM20_plant) (1 of 8)
OG IDs: 93517892
Best-hit-arabi-ID: AT5G41950.1
Best-hit-arabi-name: Tetratricopeptide repeat (TPR)-like superfamily protein
Gorai.004G129800.1.D
PCC=-0.219, p=0.368
Gorai.004G129800.1.A
PCC=0.006, p=0.98
mRNA Profile
Protein Expression Profile
DPA

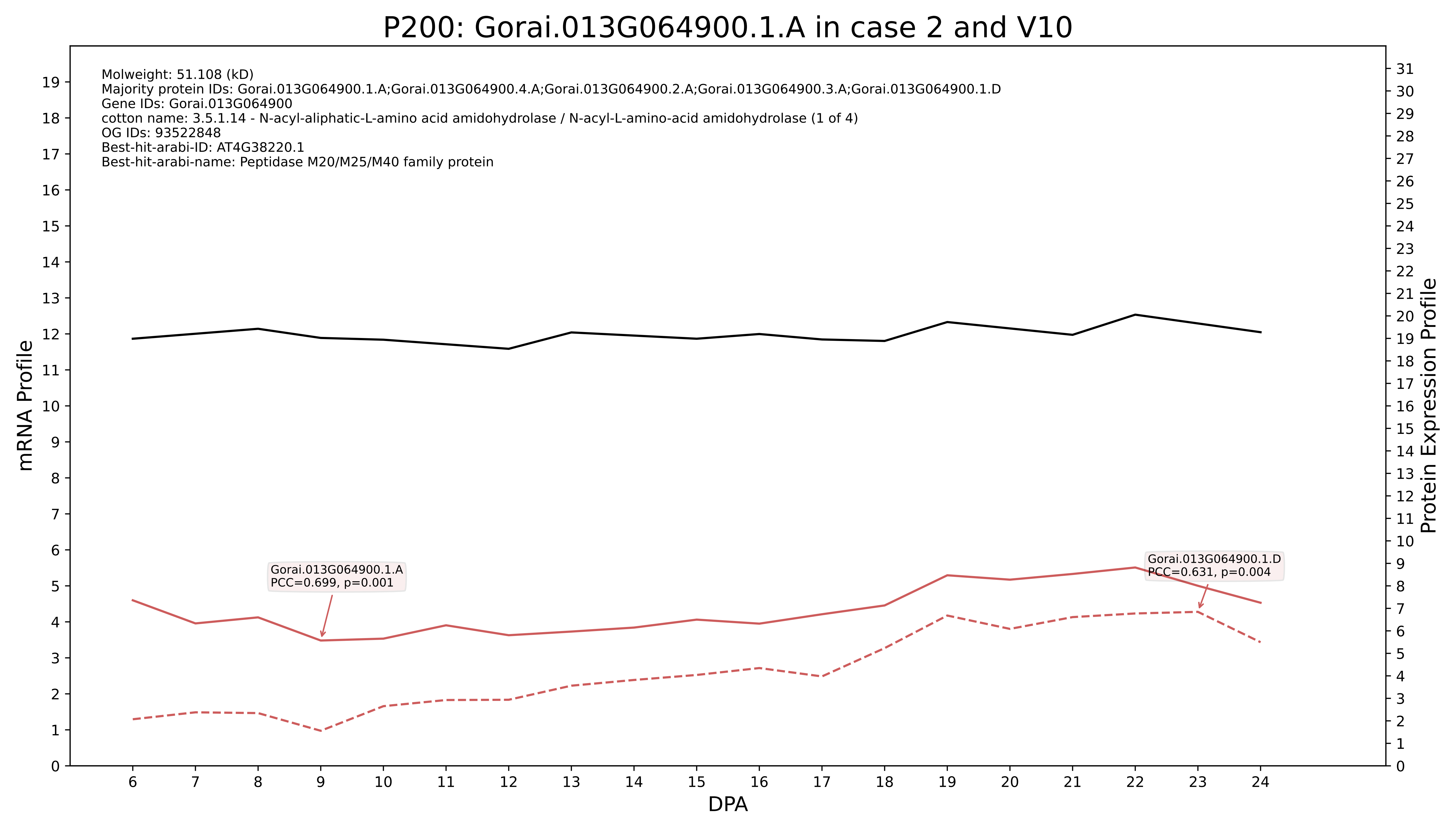
P200: Gorai.013G064900.1.A in case 2 and V10
Molweight: 51.108 (kD)
Majority protein IDs: Gorai.013G064900.1.A;Gorai.013G064900.4.A;Gorai.013G064900.2.A;Gorai.013G064900.3.A;Gorai.013G064900.1.D
Gene IDs: Gorai.013G064900
cotton name: 3.5.1.14 - N-acyl-aliphatic-L-amino acid amidohydrolase / N-acyl-L-amino-acid amidohydrolase (1 of 4)
OG IDs: 93522848
Best-hit-arabi-ID: AT4G38220.1
Best-hit-arabi-name: Peptidase M20/M25/M40 family protein
Gorai.013G064900.1.A
PCC=0.699, p=0.001
Gorai.013G064900.1.D
PCC=0.631, p=0.004
mRNA Profile
Protein Expression Profile
DPA

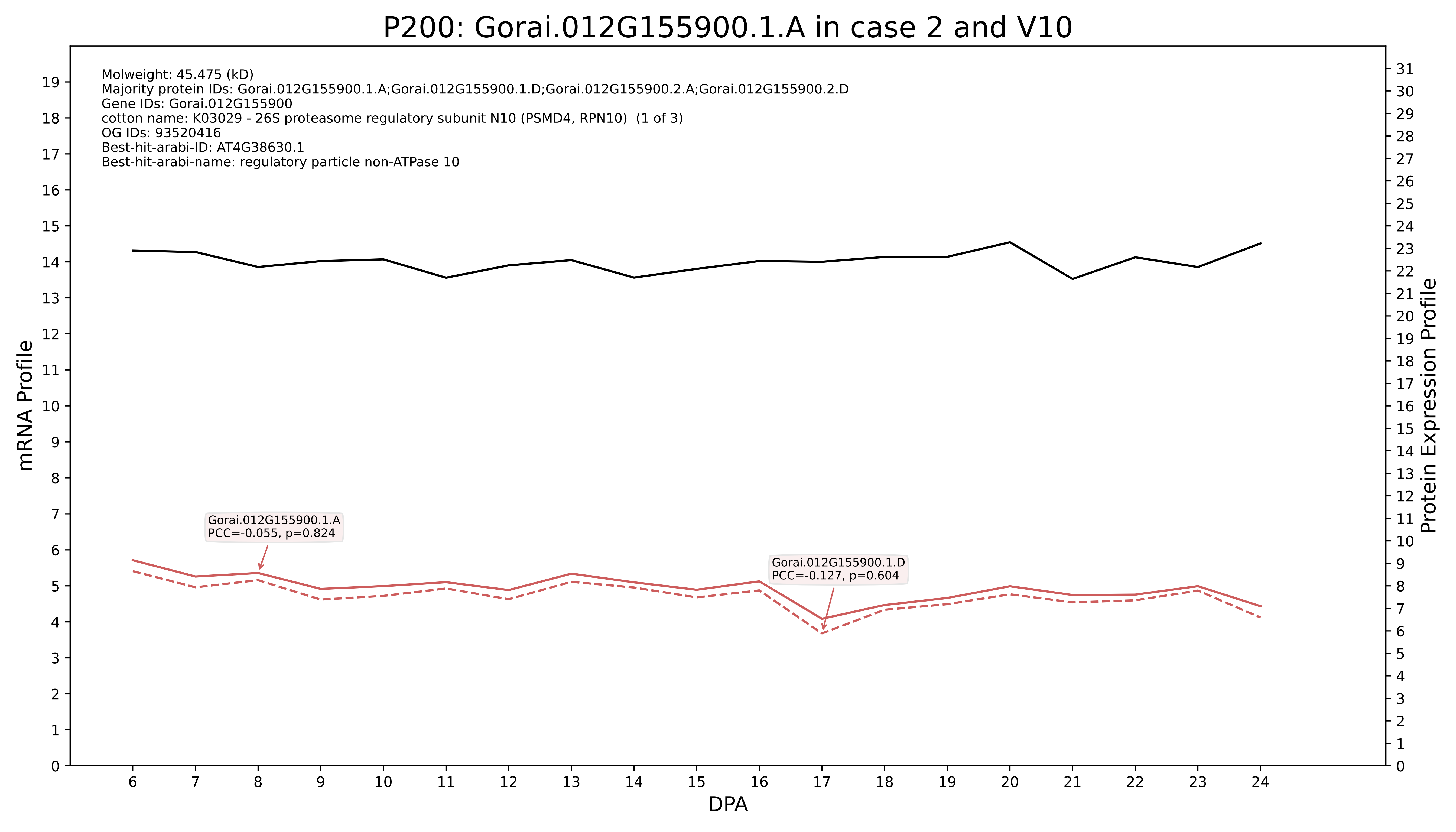
P200: Gorai.012G155900.1.A in case 2 and V10
Molweight: 45.475 (kD)
Majority protein IDs: Gorai.012G155900.1.A;Gorai.012G155900.1.D;Gorai.012G155900.2.A;Gorai.012G155900.2.D
Gene IDs: Gorai.012G155900
cotton name: K03029 - 26S proteasome regulatory subunit N10 (PSMD4, RPN10) (1 of 3)
OG IDs: 93520416
Best-hit-arabi-ID: AT4G38630.1
Best-hit-arabi-name: regulatory particle non-ATPase 10
Gorai.012G155900.1.A
PCC=-0.055, p=0.824
Gorai.012G155900.1.D
PCC=-0.127, p=0.604
mRNA Profile
Protein Expression Profile
DPA

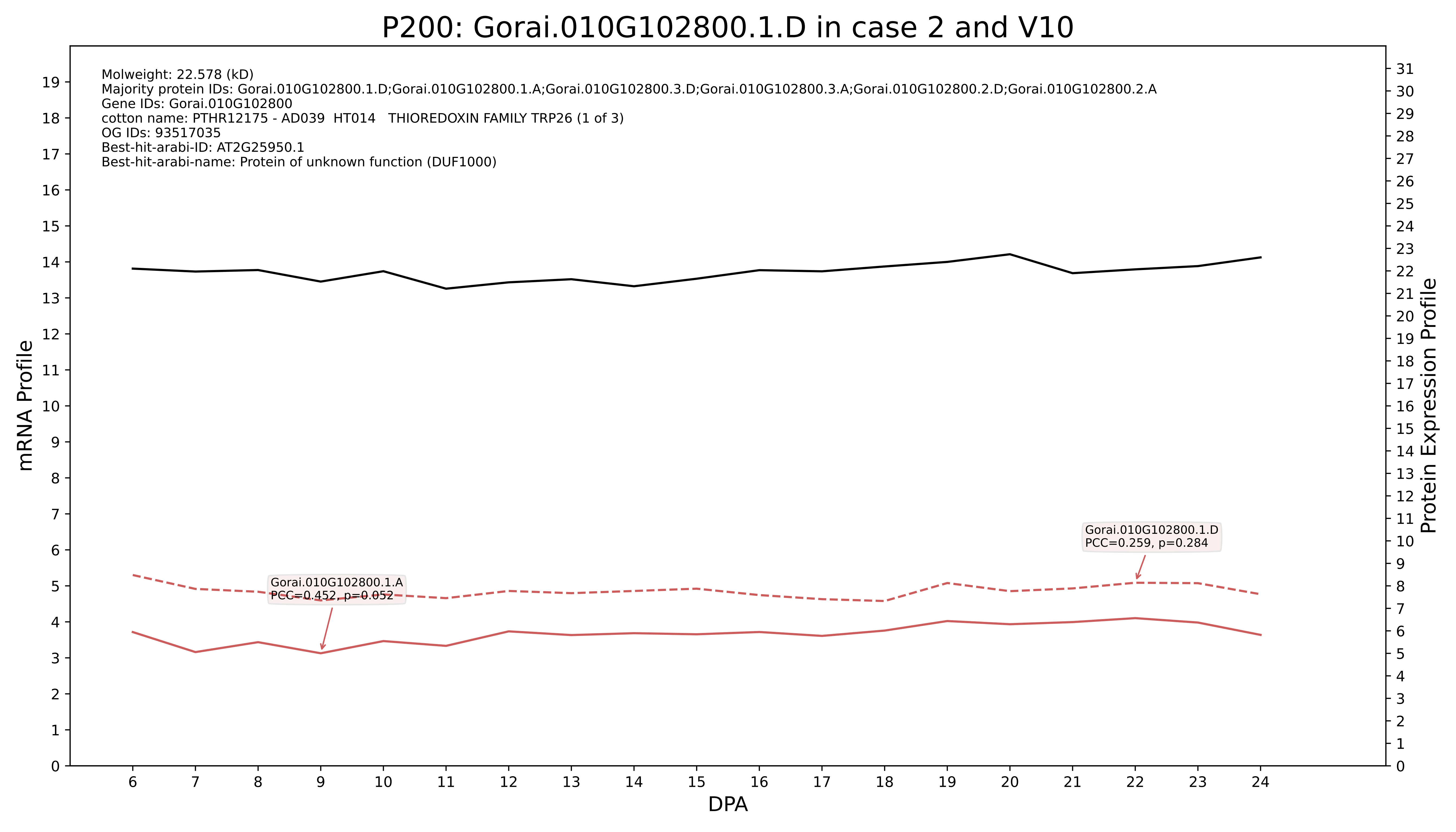
P200: Gorai.010G102800.1.D in case 2 and V10
Molweight: 22.578 (kD)
Majority protein IDs: Gorai.010G102800.1.D;Gorai.010G102800.1.A;Gorai.010G102800.3.D;Gorai.010G102800.3.A;Gorai.010G102800.2.D;Gorai.010G102800.2.A
Gene IDs: Gorai.010G102800
cotton name: PTHR12175 - AD039 HT014 THIOREDOXIN FAMILY TRP26 (1 of 3)
OG IDs: 93517035
Best-hit-arabi-ID: AT2G25950.1
Best-hit-arabi-name: Protein of unknown function (DUF1000)
Gorai.010G102800.1.A
PCC=0.452, p=0.052
Gorai.010G102800.1.D
PCC=0.259, p=0.284
mRNA Profile
Protein Expression Profile
DPA

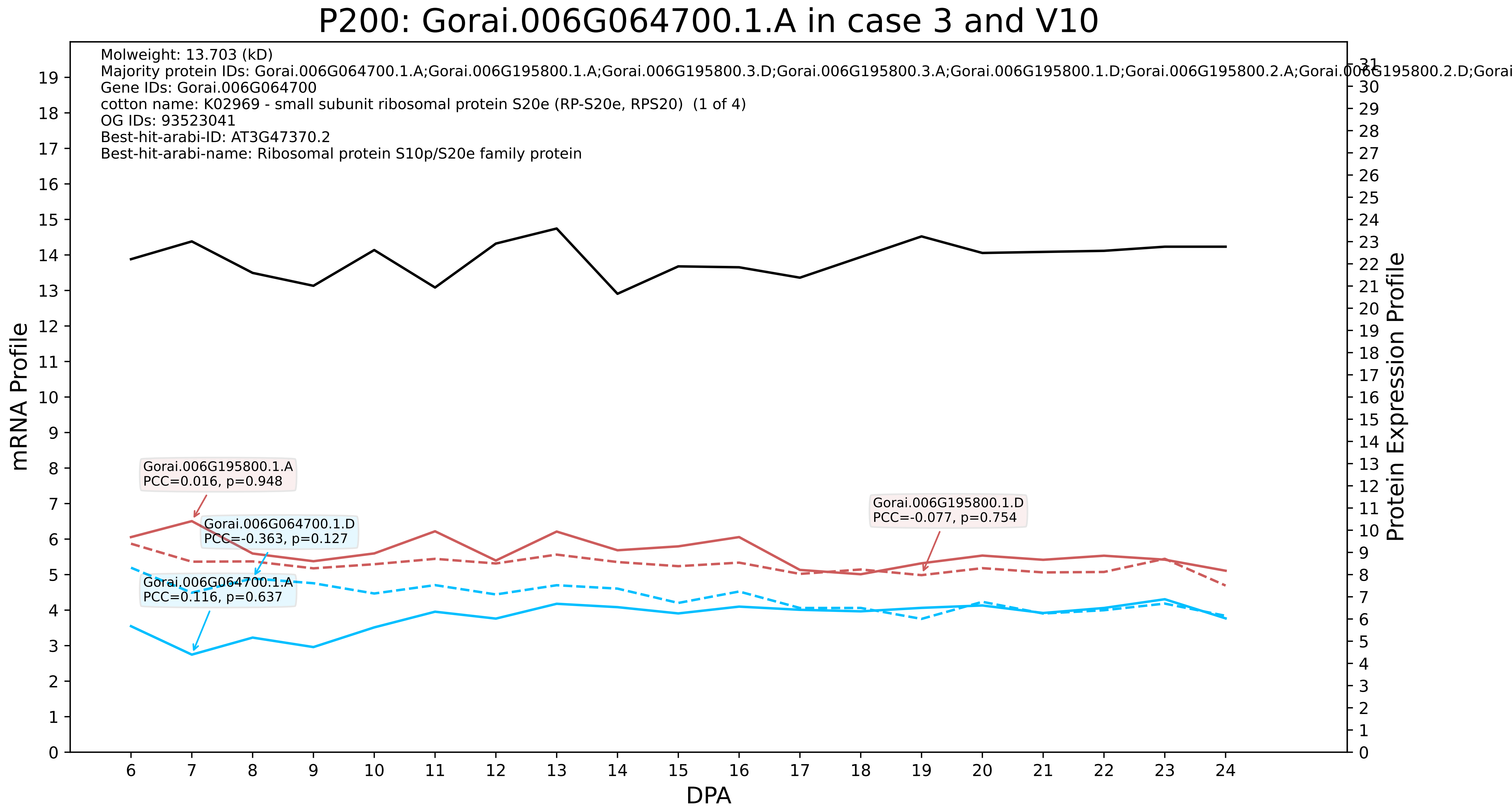

P200: Gorai.006G064700.1.A in case 3 and V10
Molweight: 13.703 (kD)
Majority protein IDs: Gorai.006G064700.1.A;Gorai.006G195800.1.A;Gorai.006G195800.3.D;Gorai.006G195800.3.A;Gorai.006G195800.1.D;Gorai.006G195800.2.A;Gorai.006G195800.2.D;Gorai
Gene IDs: Gorai.006G064700
cotton name: K02969 - small subunit ribosomal protein S20e (RP-S20e, RPS20) (1 of 4)
OG IDs: 93523041
Best-hit-arabi-ID: AT3G47370.2
Best-hit-arabi-name: Ribosomal protein S10p/S20e family protein
Gorai.006G195800.1.A
PCC=0.016, p=0.948
Gorai.006G064700.1.D
PCC=-0.363, p=0.127
Gorai.006G064700.1.A
PCC=0.116, p=0.637
Gorai.006G195800.1.D
PCC=-0.077, p=0.754
mRNA Profile
Protein Expression Profile
DPA

00: Gorai.013G163400.1.D in case 3 and V10

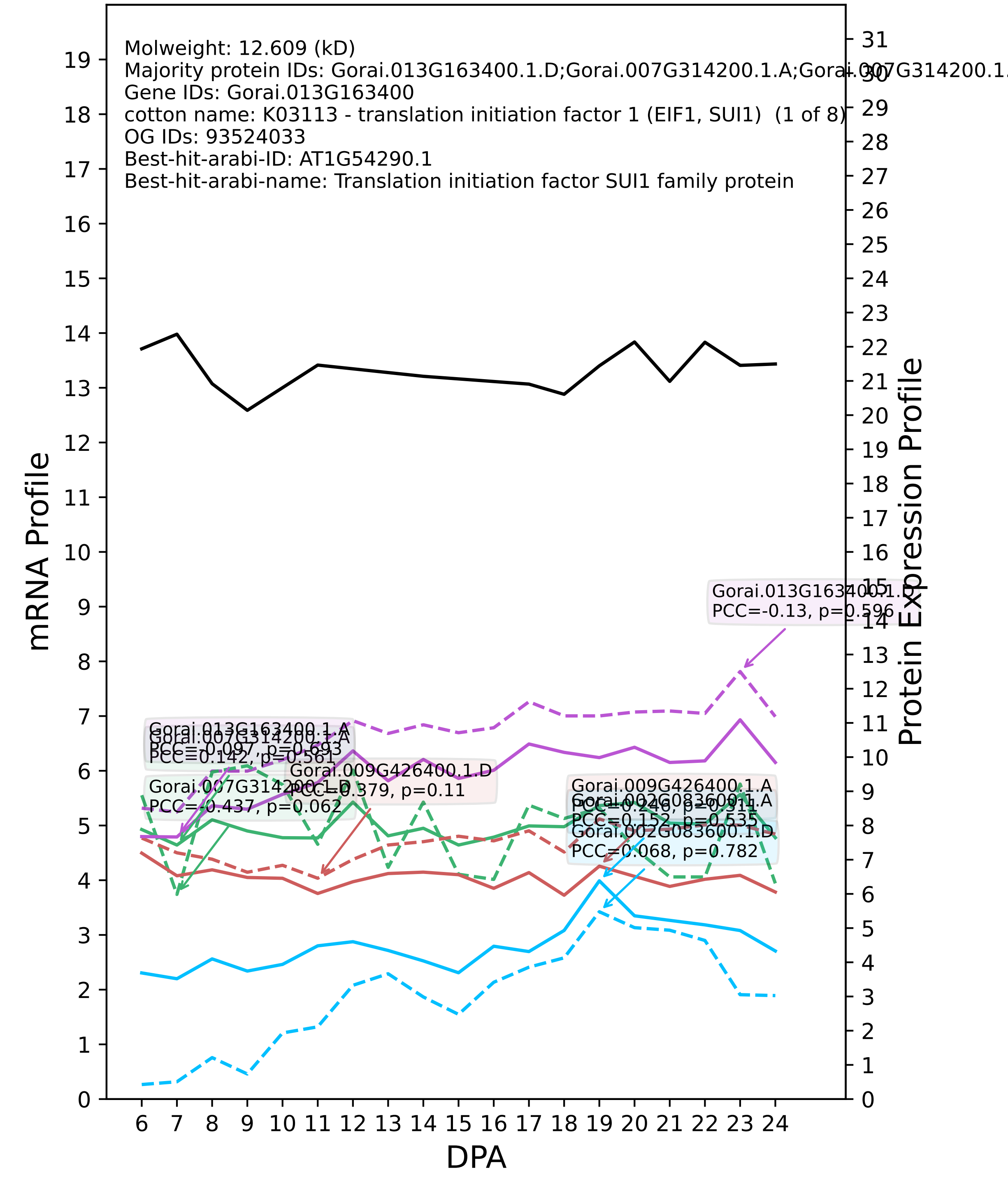

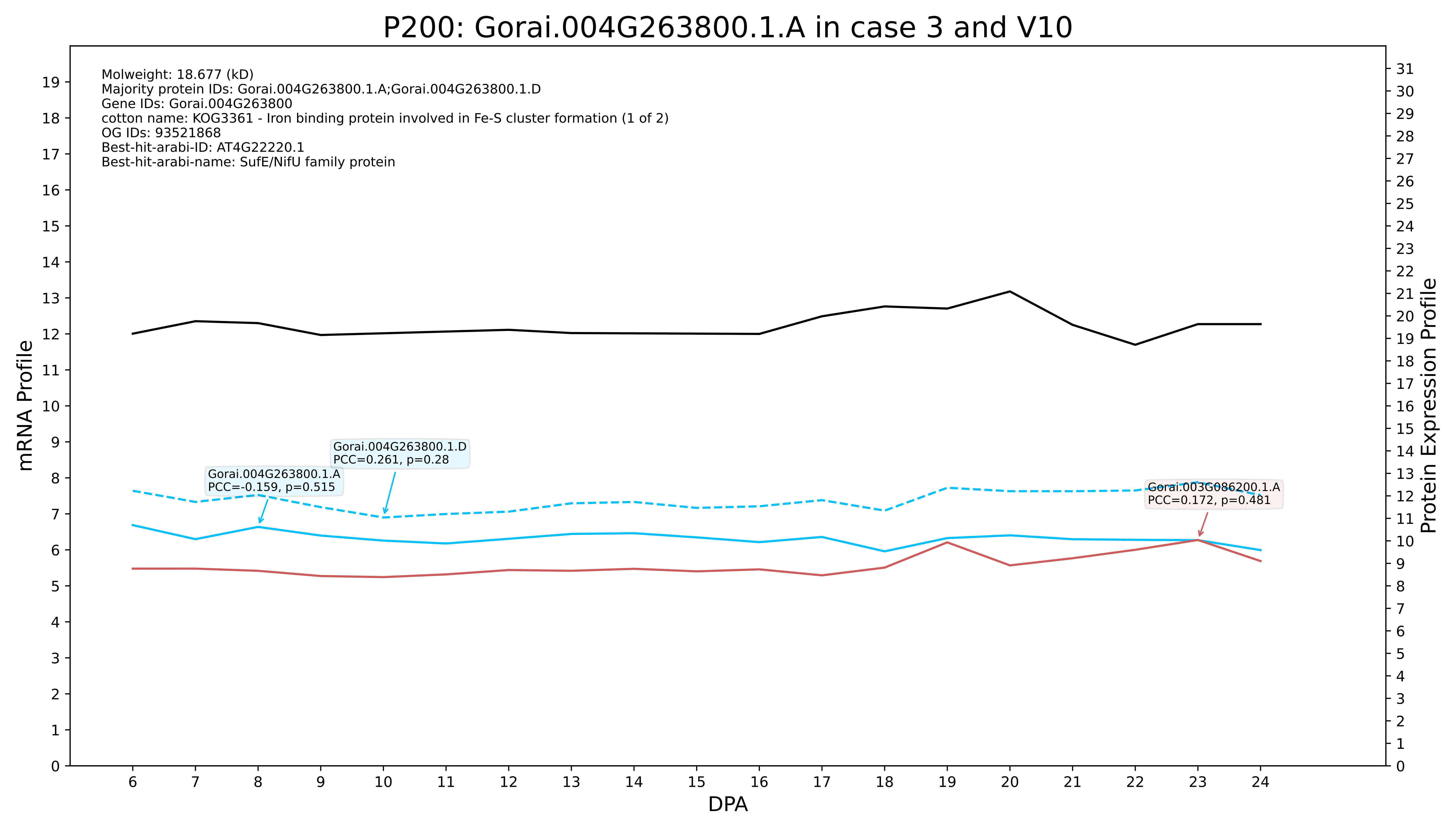

P200: Gorai.004G263800.1.A in case 3 and V10
Molweight: 18.677 (kD)
Majority protein IDs: Gorai.004G263800.1.A;Gorai.004G263800.1.D
Gene IDs: Gorai.004G263800
cotton name: KOG3361 - Iron binding protein involved in Fe-S cluster formation (1 of 2)
OG IDs: 93521868
Best-hit-arabi-ID: AT4G22220.1
Best-hit-arabi-name: SufE/NifU family protein
Gorai.004G263800.1.D
PCC=0.261, p=0.28
Gorai.004G263800.1.A
PCC=-0.159, p=0.515
Gorai.003G086200.1.A
PCC=0.172, p=0.481
mRNA Profile
Protein Expression Profile
DPA

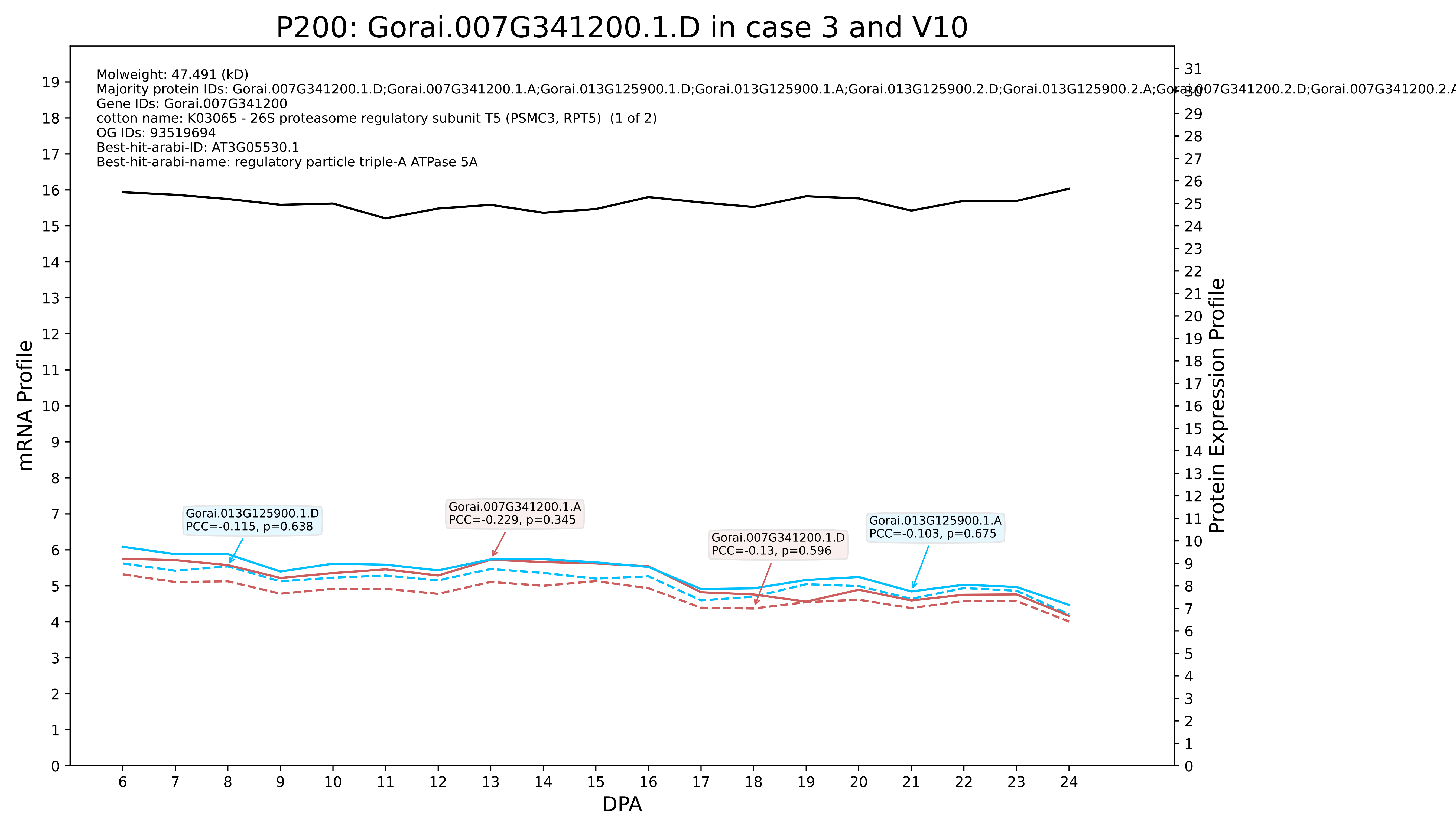

P200: Gorai.007G341200.1.D in case 3 and V10
Molweight: 47.491 (kD)
Majority protein IDs: Gorai.007G341200.1.D;Gorai.007G341200.1.A;Gorai.013G125900.1.D;Gorai.013G125900.1.A;Gorai.013G125900.2.D;Gorai.013G125900.2.A;Gorai.007G341200.2.D;Gorai.007G341200.2.A
Gene IDs: Gorai.007G341200
cotton name: K03065 - 26S proteasome regulatory subunit T5 (PSMC3, RPT5) (1 of 2)
OG IDs: 93519694
Best-hit-arabi-ID: AT3G05530.1
Best-hit-arabi-name: regulatory particle triple-A ATPase 5A
Gorai.013G125900.1.D
PCC=-0.115, p=0.638
Gorai.007G341200.1.A
PCC=-0.229, p=0.345
Gorai.007G341200.1.D
PCC=-0.13, p=0.596
Gorai.013G125900.1.A
PCC=-0.103, p=0.675
mRNA Profile
Protein Expression Profile
DPA

### P200: Gorai.008G087200.1.D in case 3 and V10

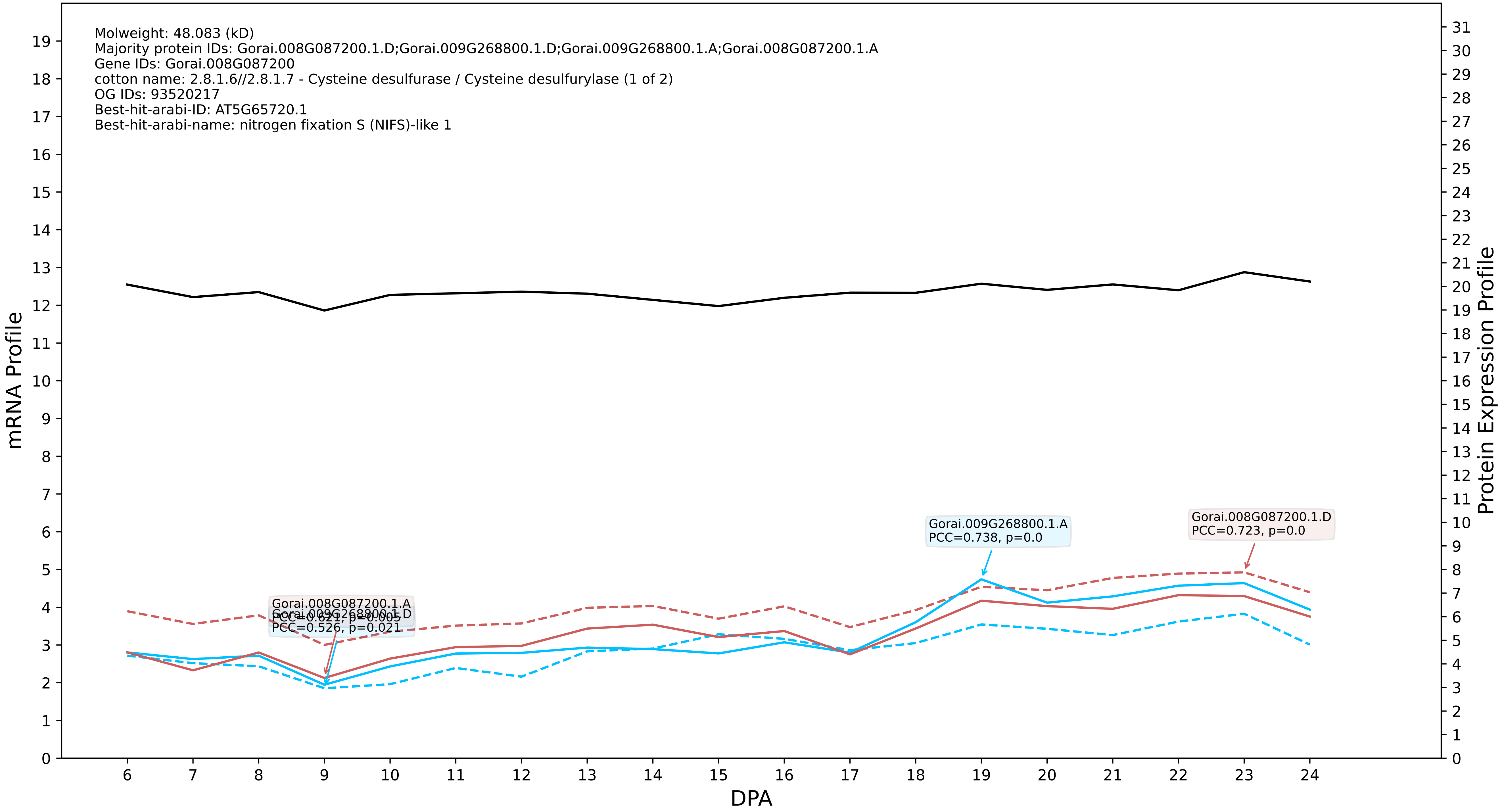

### P200: Gorai.007G256300.1.A in case 3 and V10

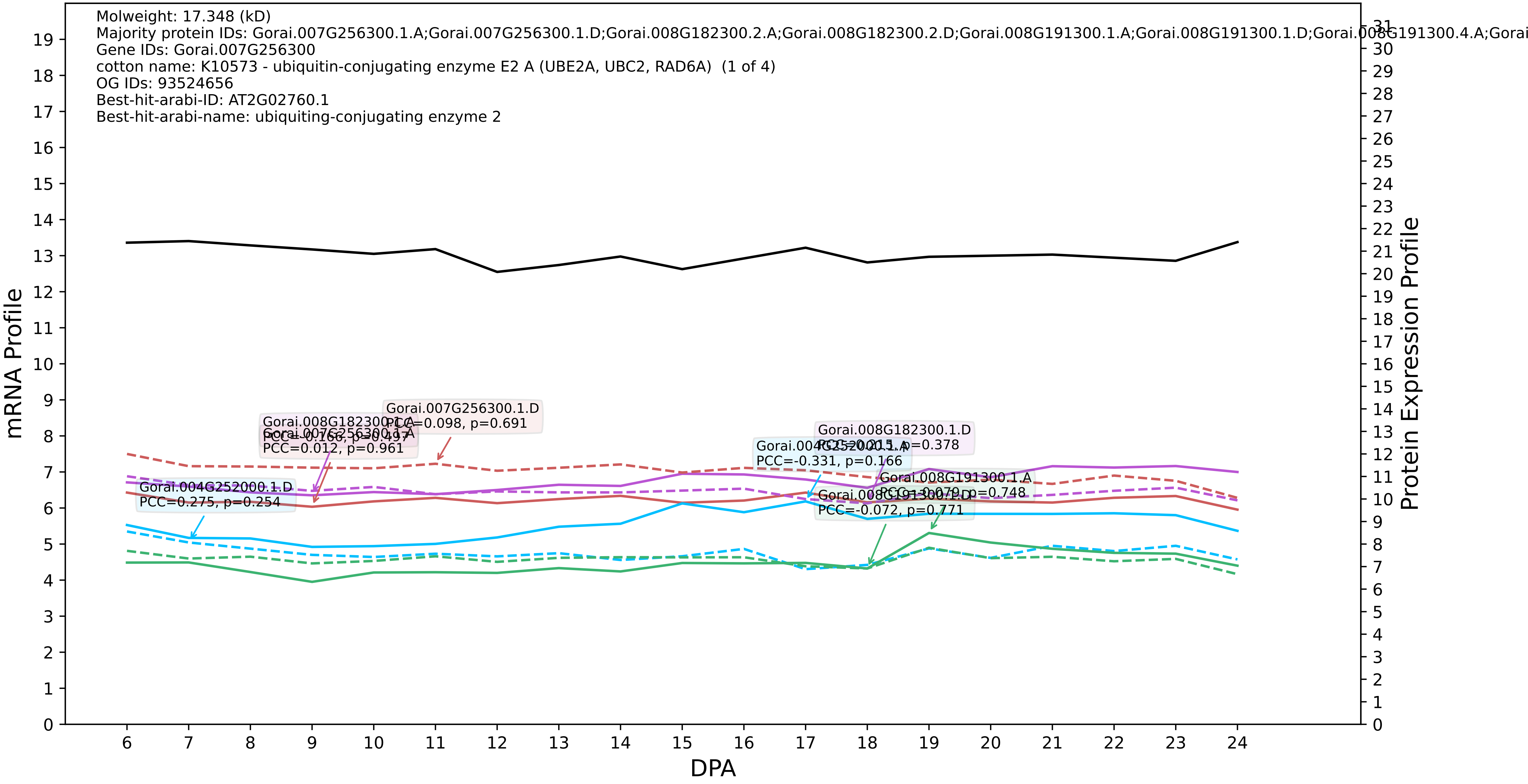

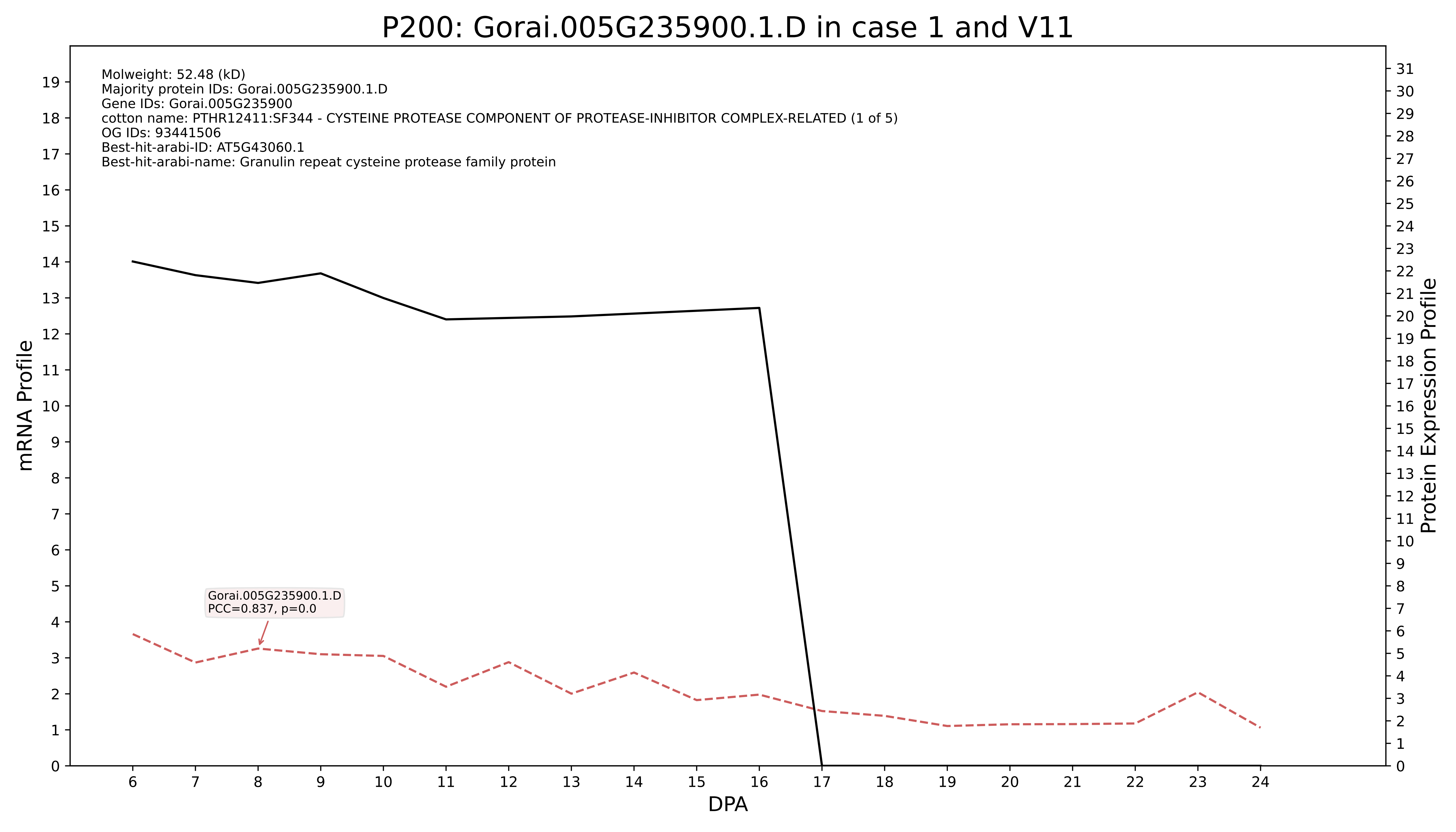

P200: Gorai.005G235900.1.D in case 1 and V11
Molweight: 52.48 (kD)
Majority protein IDs: Gorai.005G235900.1.D
Gene IDs: Gorai.005G235900
cotton name: PTHR12411:SF344 - CYSTEINE PROTEASE COMPONENT OF PROTEASE-INHIBITOR COMPLEX-RELATED (1 of 5)
OG IDs: 93441506
Best-hit-arabi-ID: AT5G43060.1
Best-hit-arabi-name: Granulin repeat cysteine protease family protein
Gorai.005G235900.1.D
PCC=0.837, p=0.0
mRNA Profile
Protein Expression Profile
DPA

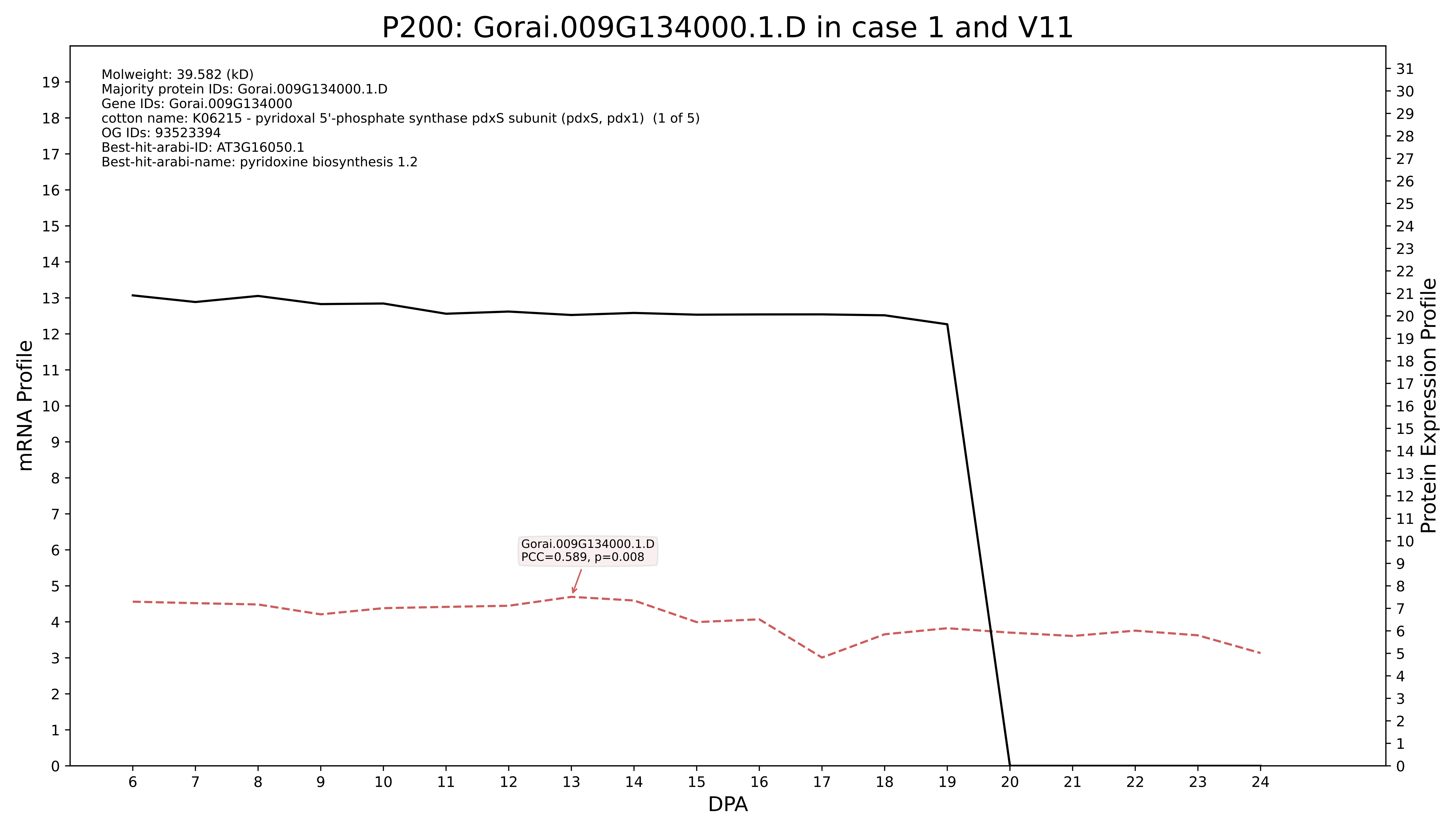
P200: Gorai.009G134000.1.D in case 1 and V11
Molweight: 39.582 (kD)
Majority protein IDs: Gorai.009G134000.1.D
Gene IDs: Gorai.009G134000
cotton name: K06215 - pyridoxal 5'-phosphate synthase pdxS subunit (pdxS, pdx1) (1 of 5)
OG IDs: 93523394
Best-hit-arabi-ID: AT3G16050.1
Best-hit-arabi-name: pyridoxine biosynthesis 1.2
Gorai.009G134000.1.D
PCC=0.589, p=0.008
mRNA Profile
Protein Expression Profile
DPA
0 1 2 3 4 5 6 7 8 9 10 11 12 13 14 15 16 17 18 19
0 1 2 3 4 5 6 7 8 9 10 11 12 13 14 15 16 17 18 19 20 21 22 23 24 25 26 27 28 29 30 31
6 7 8 9 10 11 12 13 14 15 16 17 18 19 20 21 22 23 24

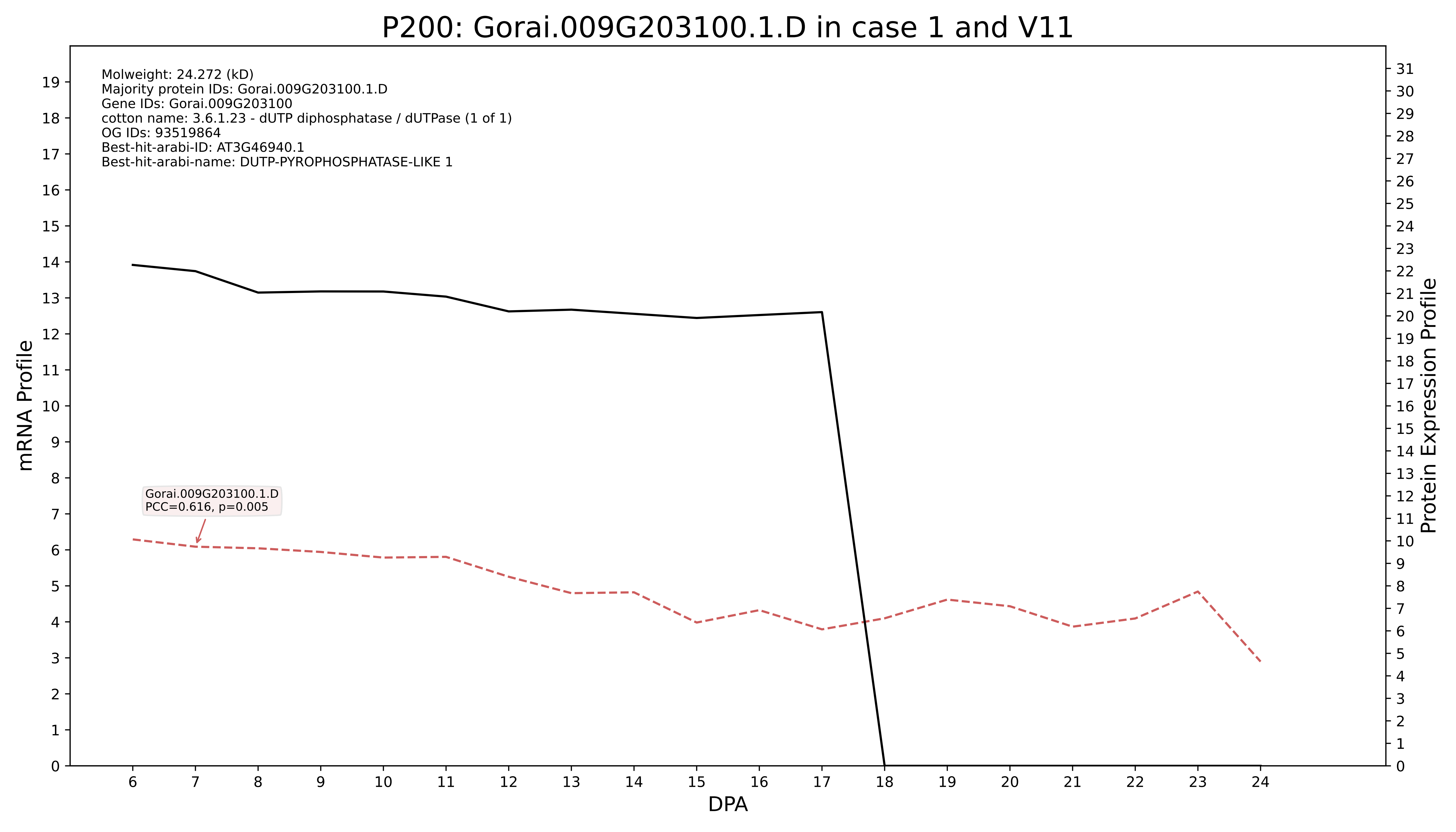
P200: Gorai.009G203100.1.D in case 1 and V11
Molweight: 24.272 (kD)
Majority protein IDs: Gorai.009G203100.1.D
Gene IDs: Gorai.009G203100
cotton name: 3.6.1.23 - dUTP diphosphatase / dUTPase (1 of 1)
OG IDs: 93519864
Best-hit-arabi-ID: AT3G46940.1
Best-hit-arabi-name: DUTP-PYROPHOSPHATASE-LIKE 1
Gorai.009G203100.1.D
PCC=0.616, p=0.005
mRNA Profile
Protein Expression Profile
DPA

P200: Gorai.001G121200.1.A in case 1 and V11

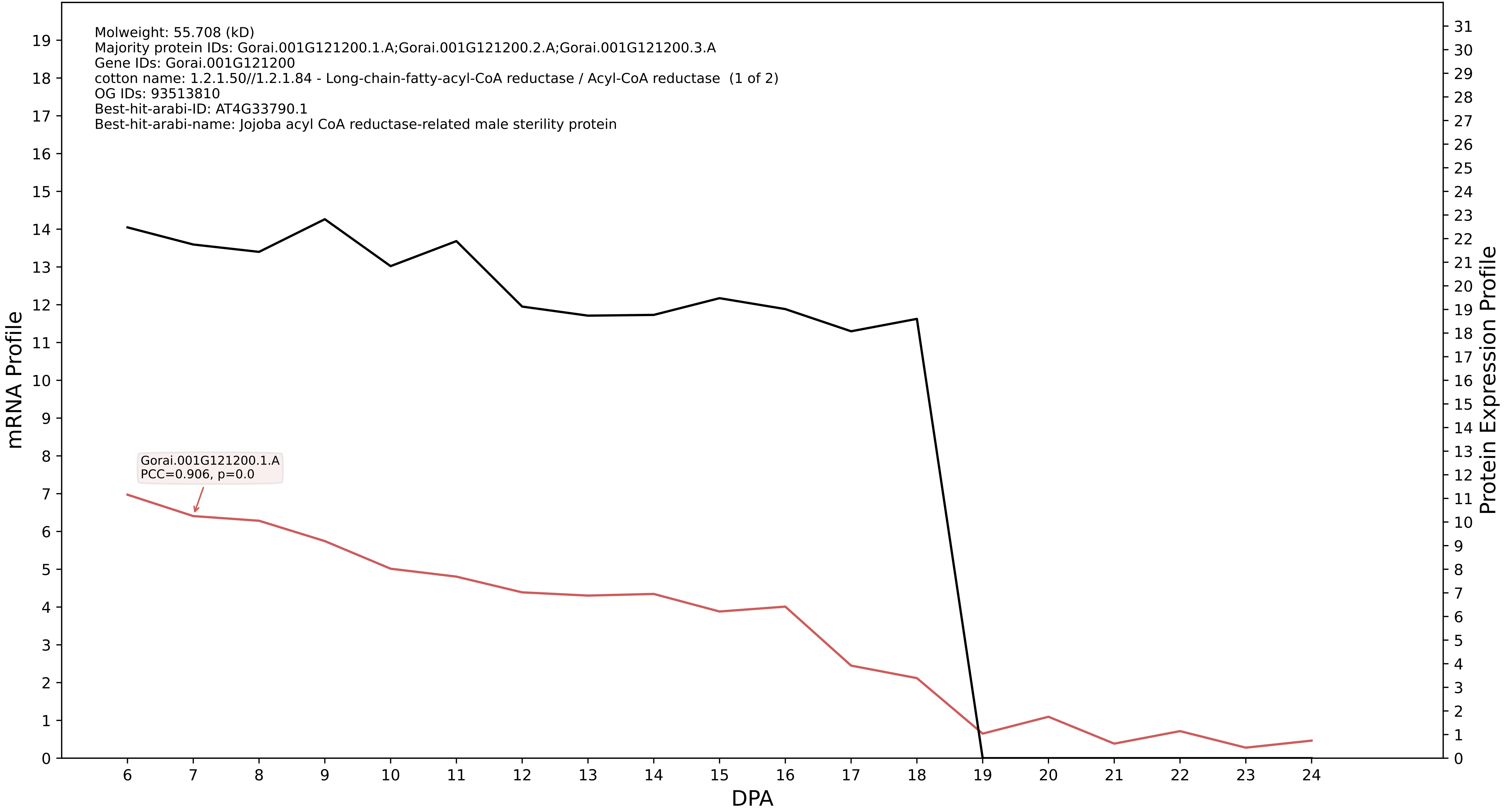

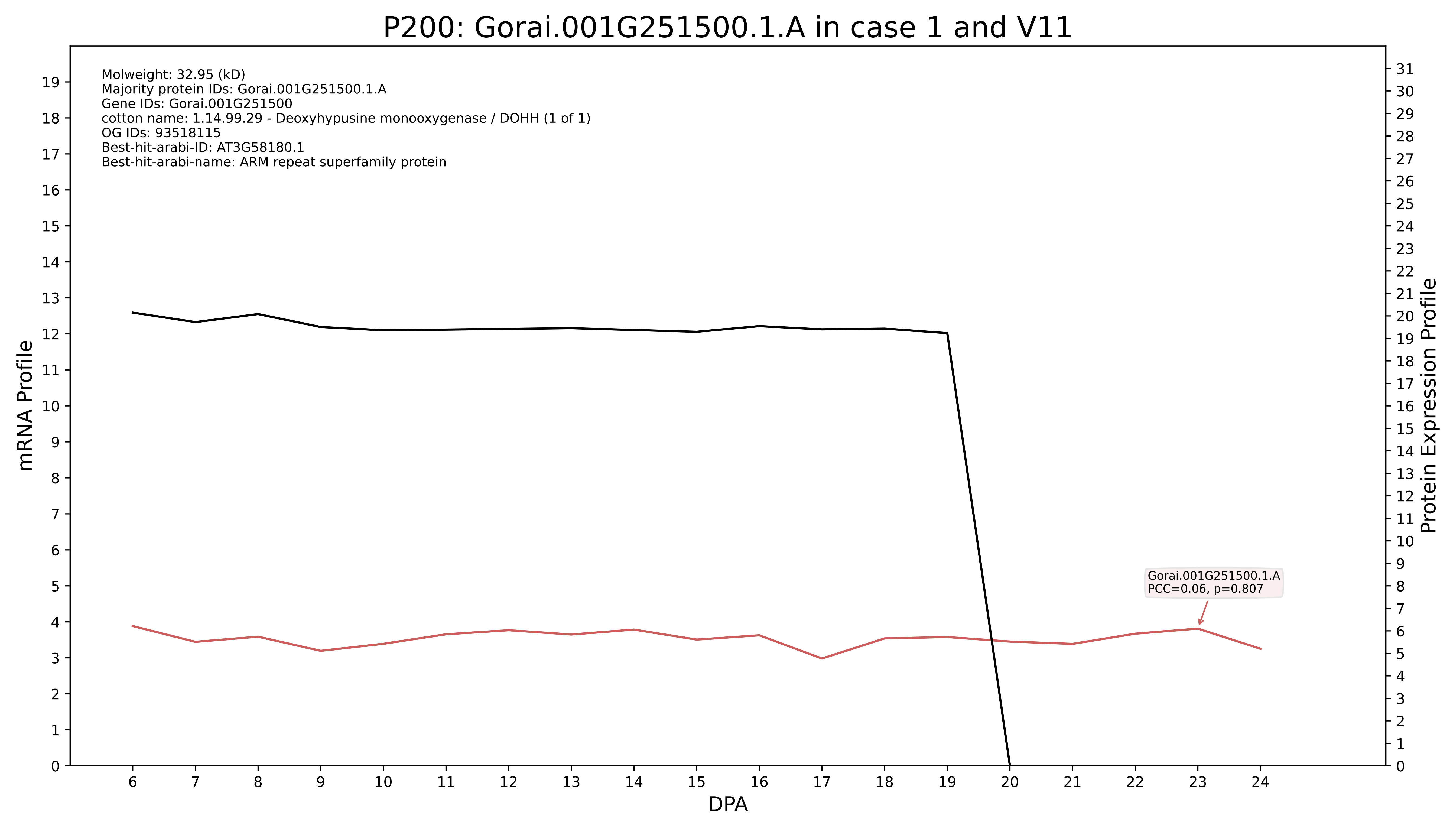
P200: Gorai.001G251500.1.A in case 1 and V11
Molweight: 32.95 (kD)
Majority protein IDs: Gorai.001G251500.1.A
Gene IDs: Gorai.001G251500
cotton name: 1.14.99.29 - Deoxyhypusine monooxygenase / DOHH (1 of 1)
OG IDs: 93518115
Best-hit-arabi-ID: AT3G58180.1
Best-hit-arabi-name: ARM repeat superfamily protein
Gorai.001G251500.1.A
PCC=0.06, p=0.807
mRNA Profile
Protein Expression Profile
DPA

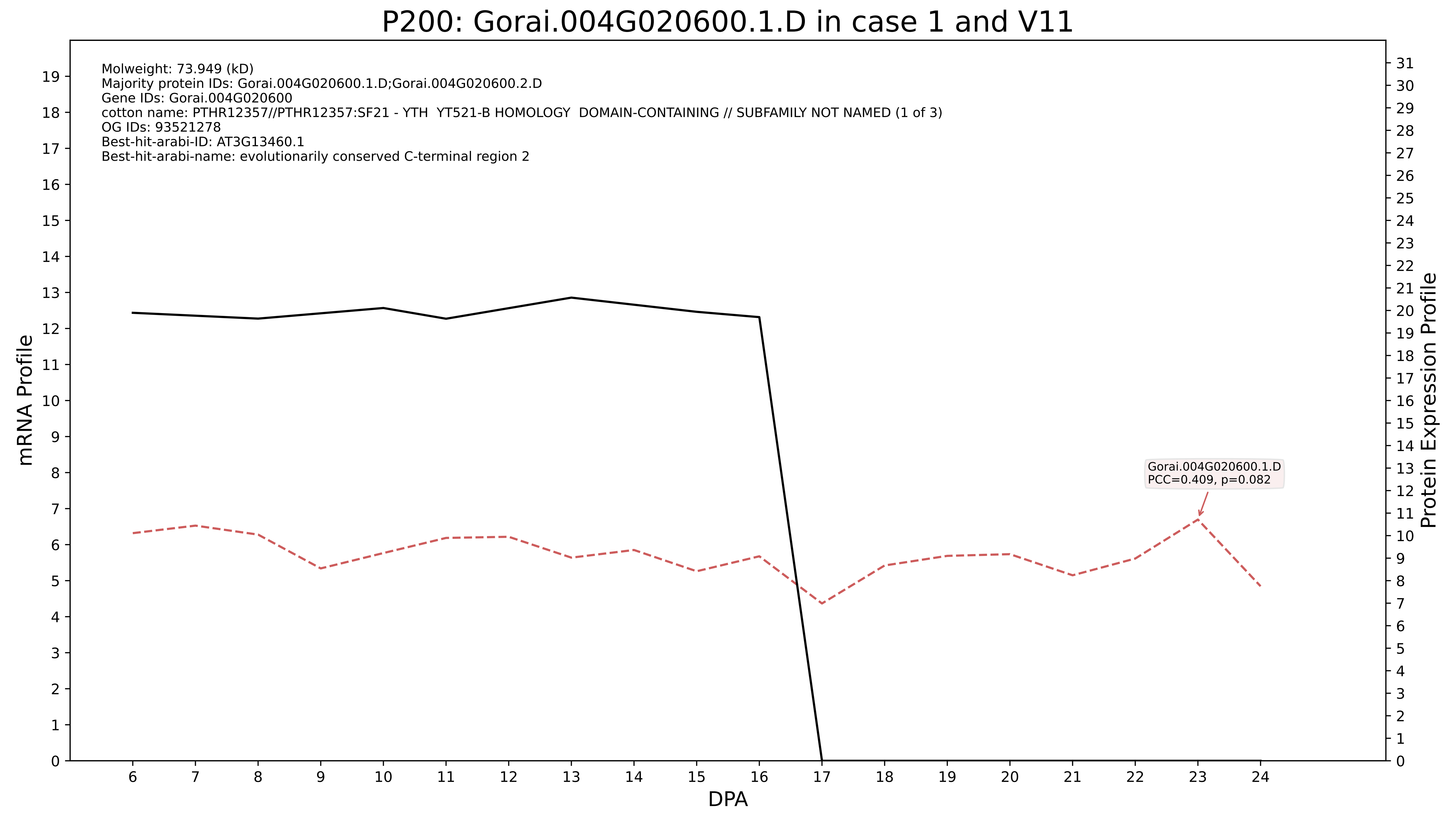
P200: Gorai.004G020600.1.D in case 1 and V11
Molweight: 73.949 (kD)
Majority protein IDs: Gorai.004G020600.1.D;Gorai.004G020600.2.D
Gene IDs: Gorai.004G020600
cotton name: PTHR12357//PTHR12357:SF21 - YTH YT521-B HOMOLOGY DOMAIN-CONTAINING // SUBFAMILY NOT NAMED (1 of 3)
OG IDs: 93521278
Best-hit-arabi-ID: AT3G13460.1
Best-hit-arabi-name: evolutionarily conserved C-terminal region 2
Gorai.004G020600.1.D
PCC=0.409, p=0.082
mRNA Profile
Protein Expression Profile
DPA

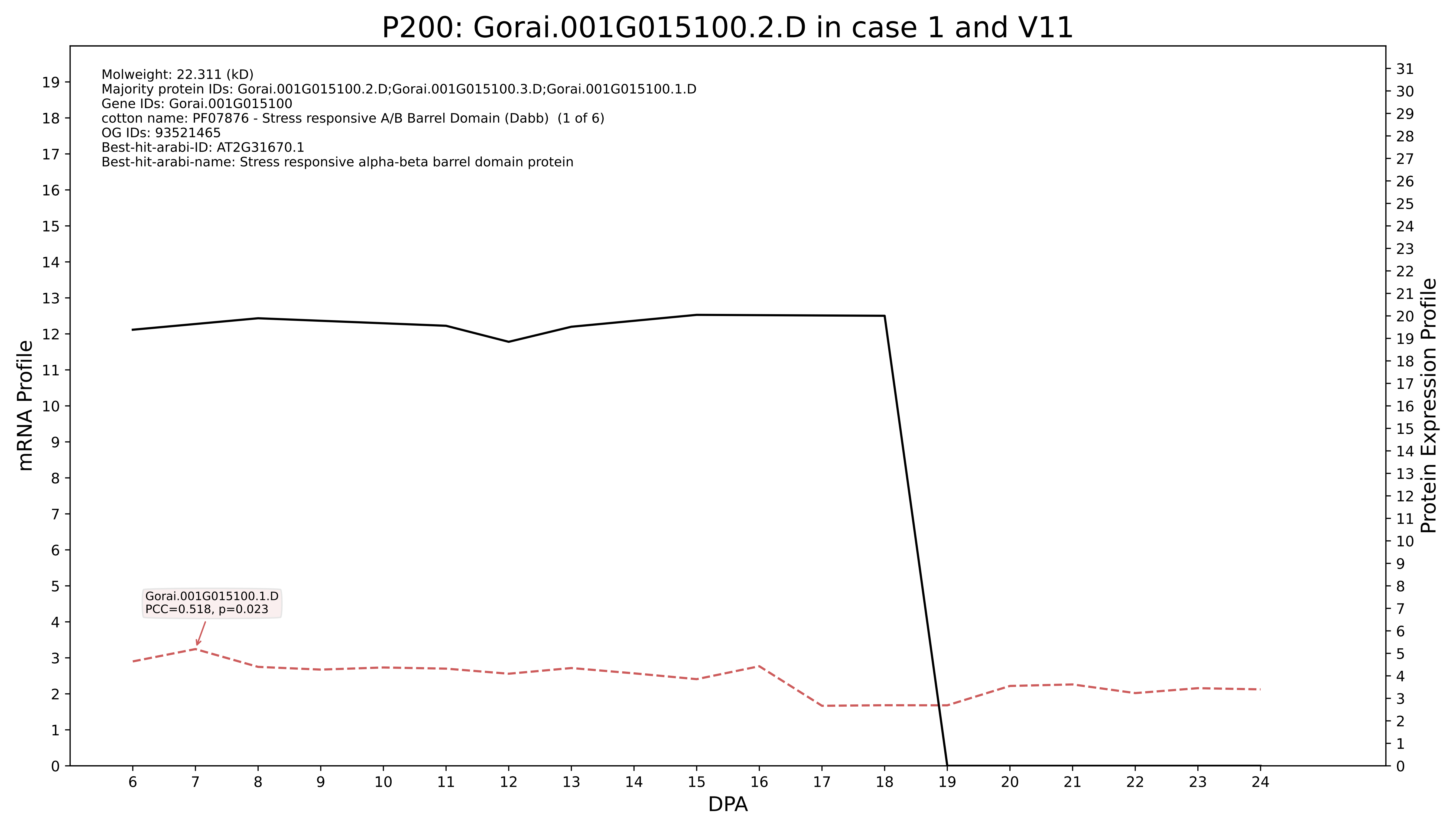
P200: Gorai.001G015100.2.D in case 1 and V11
Molweight: 22.311 (kD)
Majority protein IDs: Gorai.001G015100.2.D;Gorai.001G015100.3.D;Gorai.001G015100.1.D
Gene IDs: Gorai.001G015100
cotton name: PF07876 - Stress responsive A/B Barrel Domain (Dabb) (1 of 6)
OG IDs: 93521465
Best-hit-arabi-ID: AT2G31670.1
Best-hit-arabi-name: Stress responsive alpha-beta barrel domain protein
Gorai.001G015100.1.D
PCC=0.518, p=0.023
mRNA Profile
Protein Expression Profile
DPA

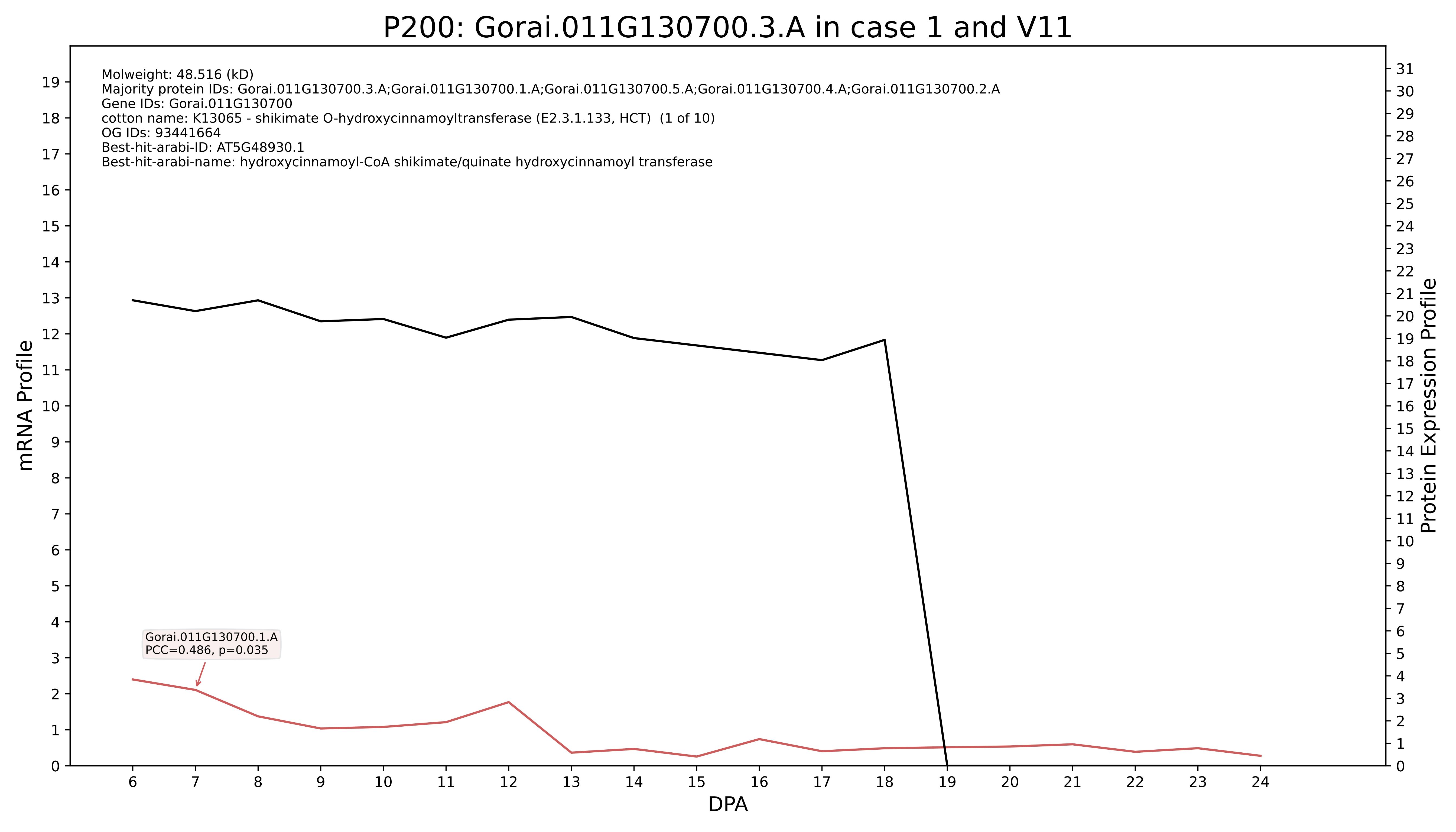

P200: Gorai.011G130700.3.A in case 1 and V11
Molweight: 48.516 (kD)
Majority protein IDs: Gorai.011G130700.3.A;Gorai.011G130700.1.A;Gorai.011G130700.5.A;Gorai.011G130700.4.A;Gorai.011G130700.2.A
Gene IDs: Gorai.011G130700
cotton name: K13065 - shikimate O-hydroxycinnamoyltransferase (E2.3.1.133, HCT) (1 of 10)
OG IDs: 93441664
Best-hit-arabi-ID: AT5G48930.1
Best-hit-arabi-name: hydroxycinnamoyl-CoA shikimate/quinate hydroxycinnamoyl transferase
Gorai.011G130700.1.A
PCC=0.486, p=0.035
mRNA Profile
Protein Expression Profile
DPA

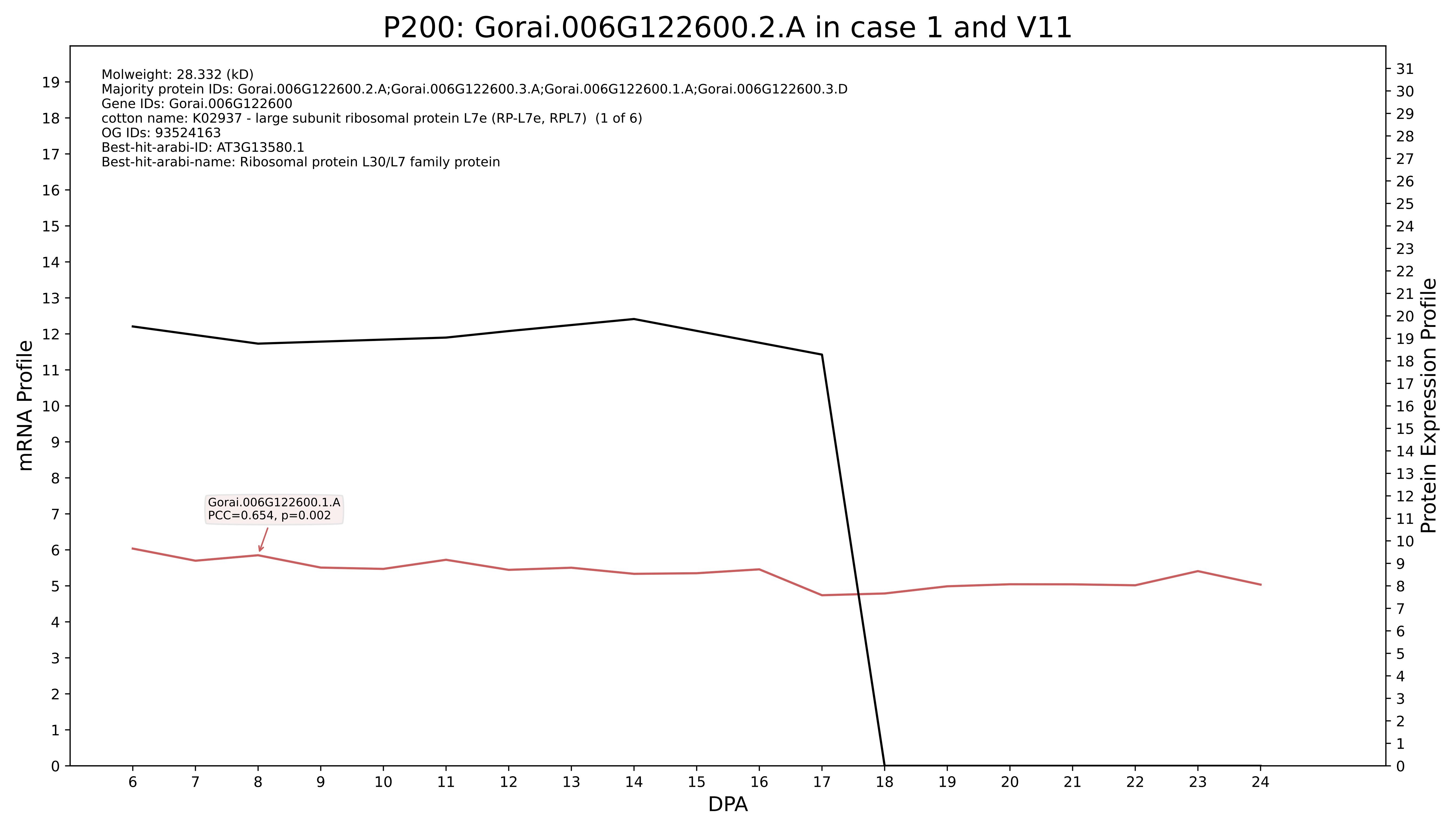
P200: Gorai.006G122600.2.A in case 1 and V11
Molweight: 28.332 (kD)
Majority protein IDs: Gorai.006G122600.2.A;Gorai.006G122600.3.A;Gorai.006G122600.1.A;Gorai.006G122600.3.D
Gene IDs: Gorai.006G122600
cotton name: K02937 - large subunit ribosomal protein L7e (RP-L7e, RPL7) (1 of 6)
OG IDs: 93524163
Best-hit-arabi-ID: AT3G13580.1
Best-hit-arabi-name: Ribosomal protein L30/L7 family protein
Gorai.006G122600.1.A
PCC=0.654, p=0.002
mRNA Profile
Protein Expression Profile
DPA

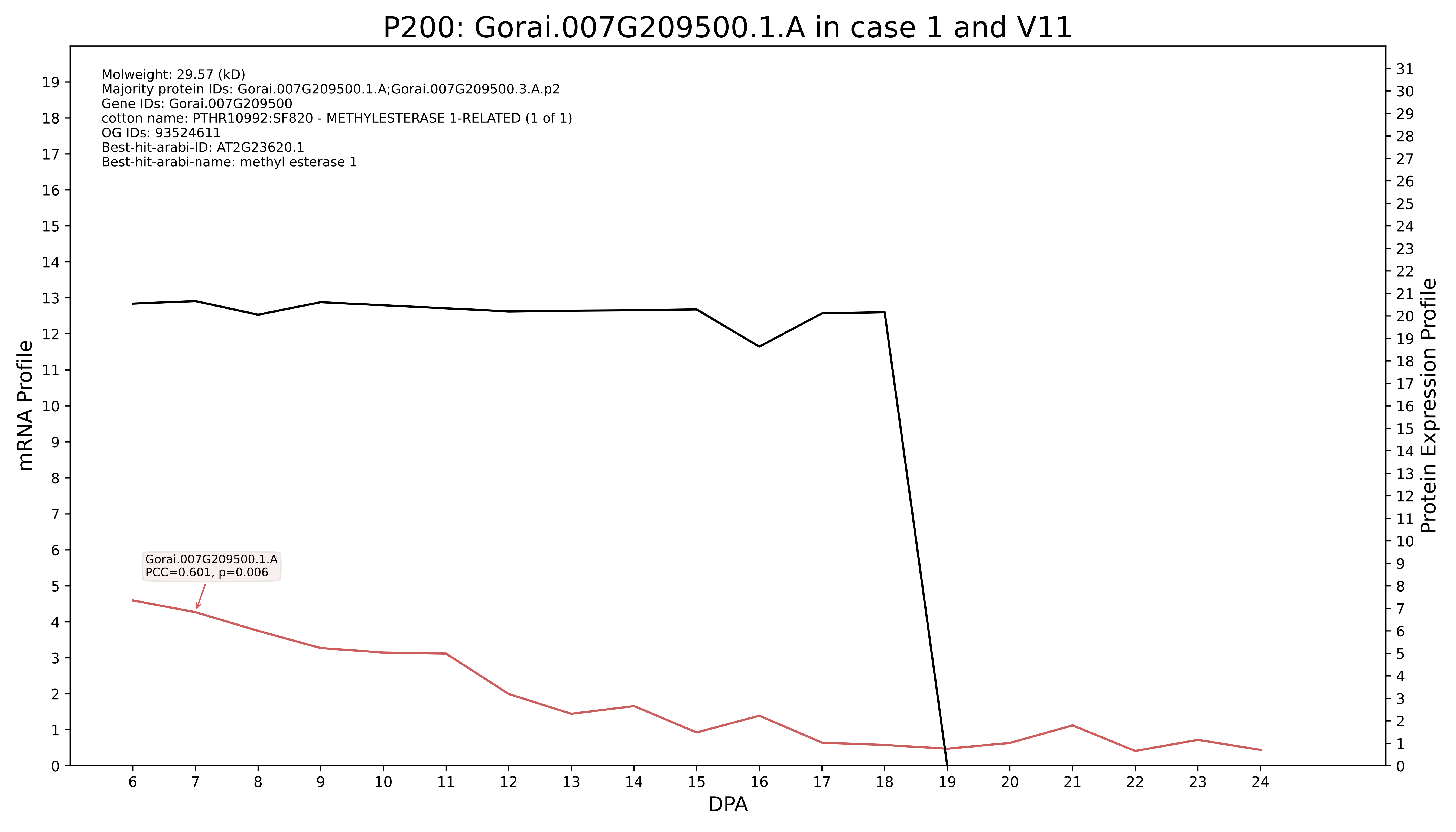
P200: Gorai.007G209500.1.A in case 1 and V11
Molweight: 29.57 (kD)
Majority protein IDs: Gorai.007G209500.1.A;Gorai.007G209500.3.A.p2
Gene IDs: Gorai.007G209500
cotton name: PTHR10992:SF820 - METHYLESTERASE 1-RELATED (1 of 1)
OG IDs: 93524611
Best-hit-arabi-ID: AT2G23620.1
Best-hit-arabi-name: methyl esterase 1
Gorai.007G209500.1.A
PCC=0.601, p=0.006
mRNA Profile
Protein Expression Profile
DPA

### P200: Gorai.008G113600.2.D in case 1 and V11

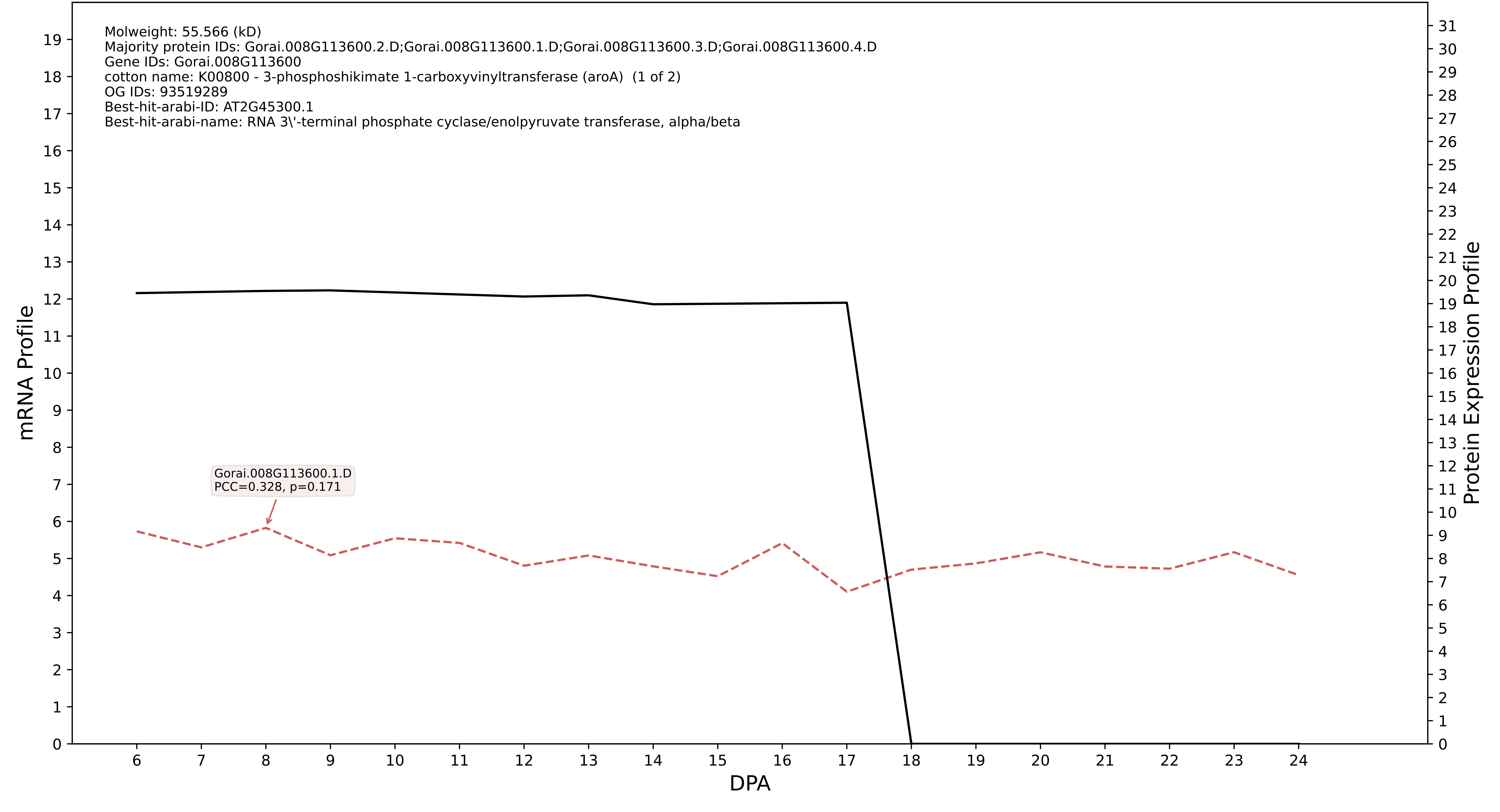

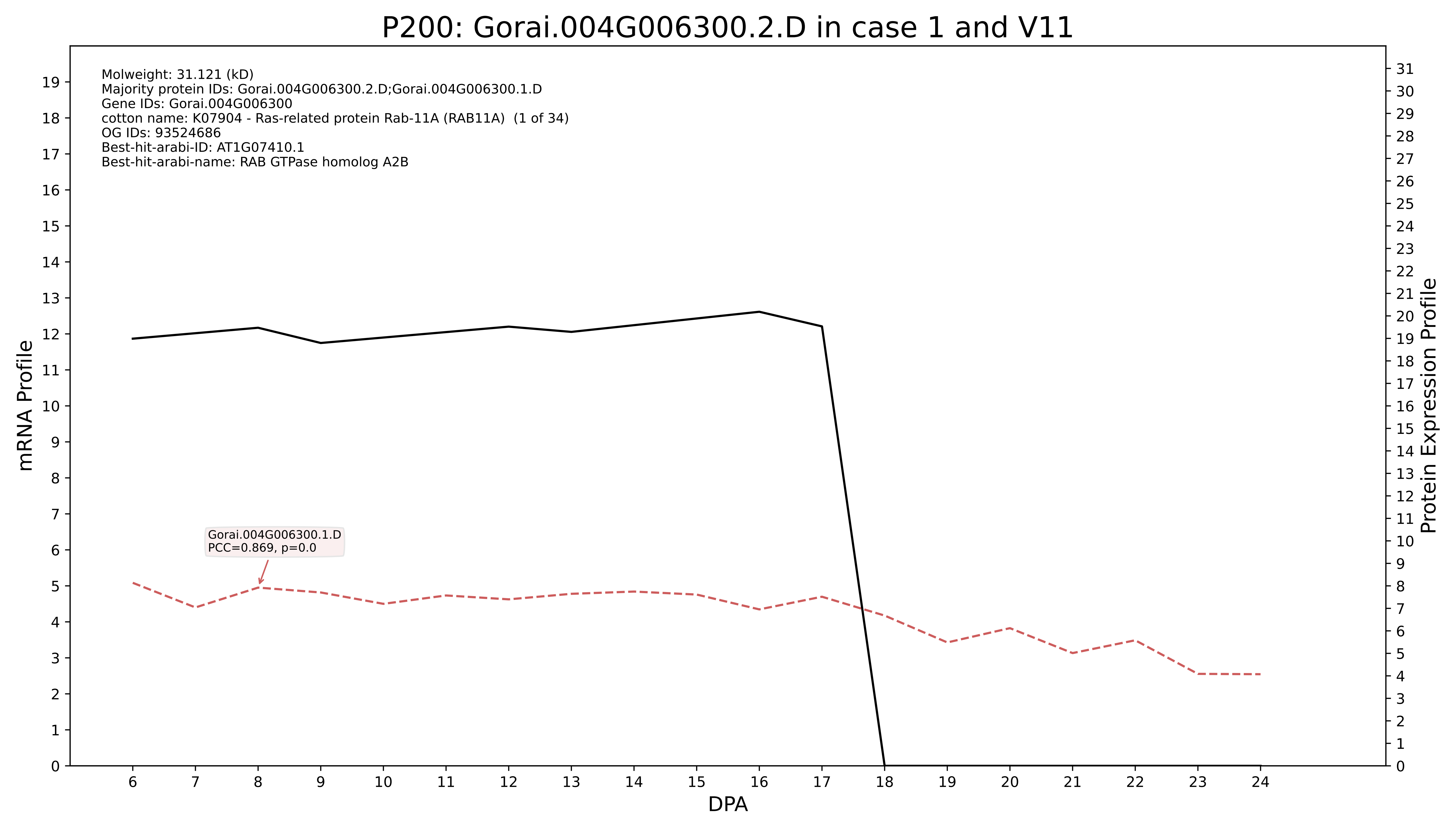
P200: Gorai.004G006300.2.D in case 1 and V11
Molweight: 31.121 (kD)
Majority protein IDs: Gorai.004G006300.2.D;Gorai.004G006300.1.D
Gene IDs: Gorai.004G006300
cotton name: K07904 - Ras-related protein Rab-11A (RAB11A) (1 of 34)
OG IDs: 93524686
Best-hit-arabi-ID: AT1G07410.1
Best-hit-arabi-name: RAB GTPase homolog A2B
Gorai.004G006300.1.D
PCC=0.869, p=0.0
mRNA Profile
Protein Expression Profile
DPA

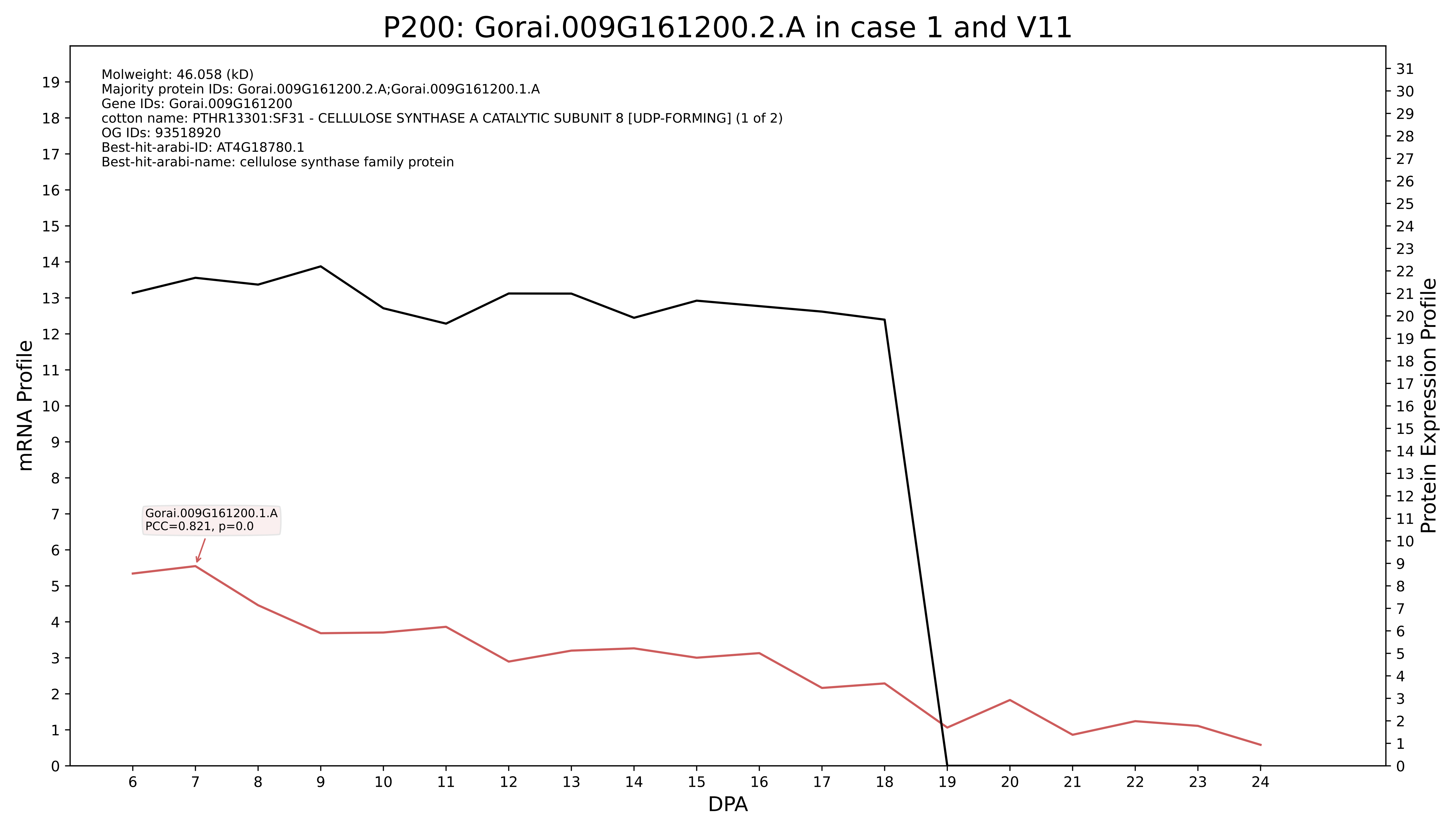

P200: Gorai.009G161200.2.A in case 1 and V11
Molweight: 46.058 (kD)
Majority protein IDs: Gorai.009G161200.2.A;Gorai.009G161200.1.A
Gene IDs: Gorai.009G161200
cotton name: PTHR13301:SF31 - CELLULOSE SYNTHASE A CATALYTIC SUBUNIT 8 [UDP-FORMING] (1 of 2)
OG IDs: 93518920
Best-hit-arabi-ID: AT4G18780.1
Best-hit-arabi-name: cellulose synthase family protein
Gorai.009G161200.1.A
PCC=0.821, p=0.0
mRNA Profile
Protein Expression Profile
DPA

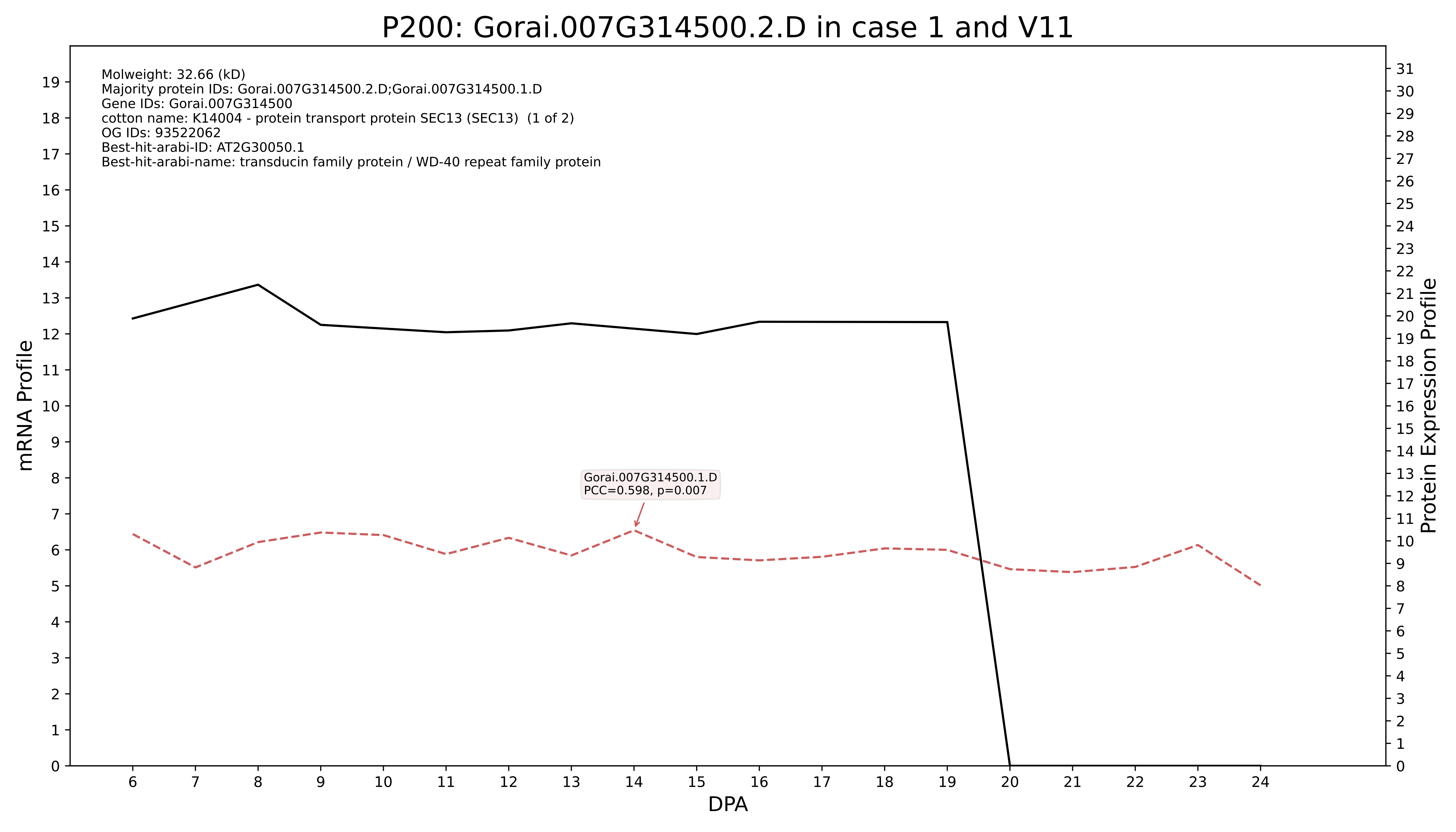
P200: Gorai.007G314500.2.D in case 1 and V11
Molweight: 32.66 (kD)
Majority protein IDs: Gorai.007G314500.2.D;Gorai.007G314500.1.D
Gene IDs: Gorai.007G314500
cotton name: K14004 - protein transport protein SEC13 (SEC13) (1 of 2)
OG IDs: 93522062
Best-hit-arabi-ID: AT2G30050.1
Best-hit-arabi-name: transducin family protein / WD-40 repeat family protein
Gorai.007G314500.1.D
PCC=0.598, p=0.007
mRNA Profile
Protein Expression Profile
DPA

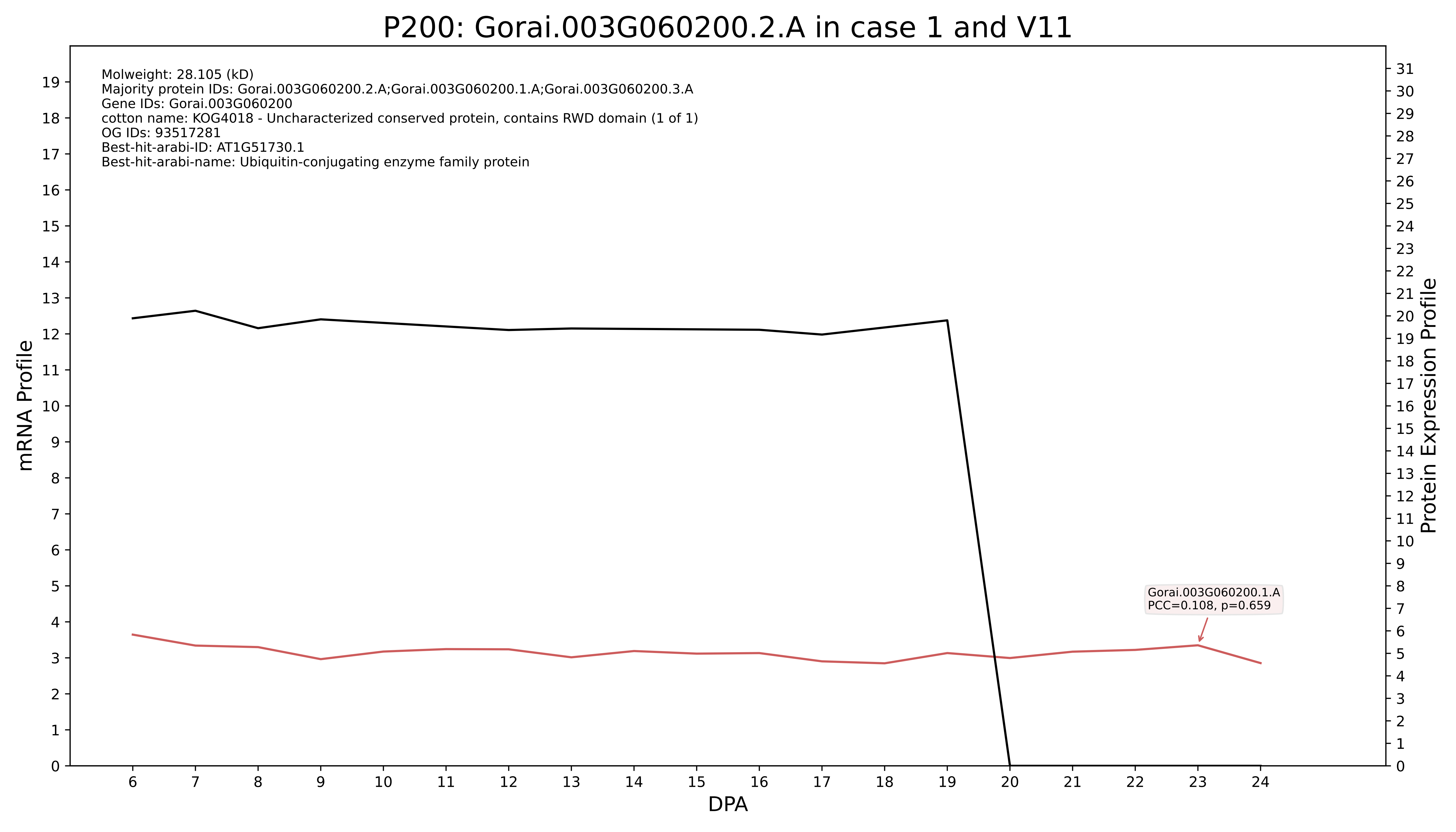
P200: Gorai.003G060200.2.A in case 1 and V11
Molweight: 28.105 (kD)
Majority protein IDs: Gorai.003G060200.2.A;Gorai.003G060200.1.A;Gorai.003G060200.3.A
Gene IDs: Gorai.003G060200
cotton name: KOG4018 - Uncharacterized conserved protein, contains RWD domain (1 of 1)
OG IDs: 93517281
Best-hit-arabi-ID: AT1G51730.1
Best-hit-arabi-name: Ubiquitin-conjugating enzyme family protein
Gorai.003G060200.1.A
PCC=0.108, p=0.659
mRNA Profile
Protein Expression Profile
DPA

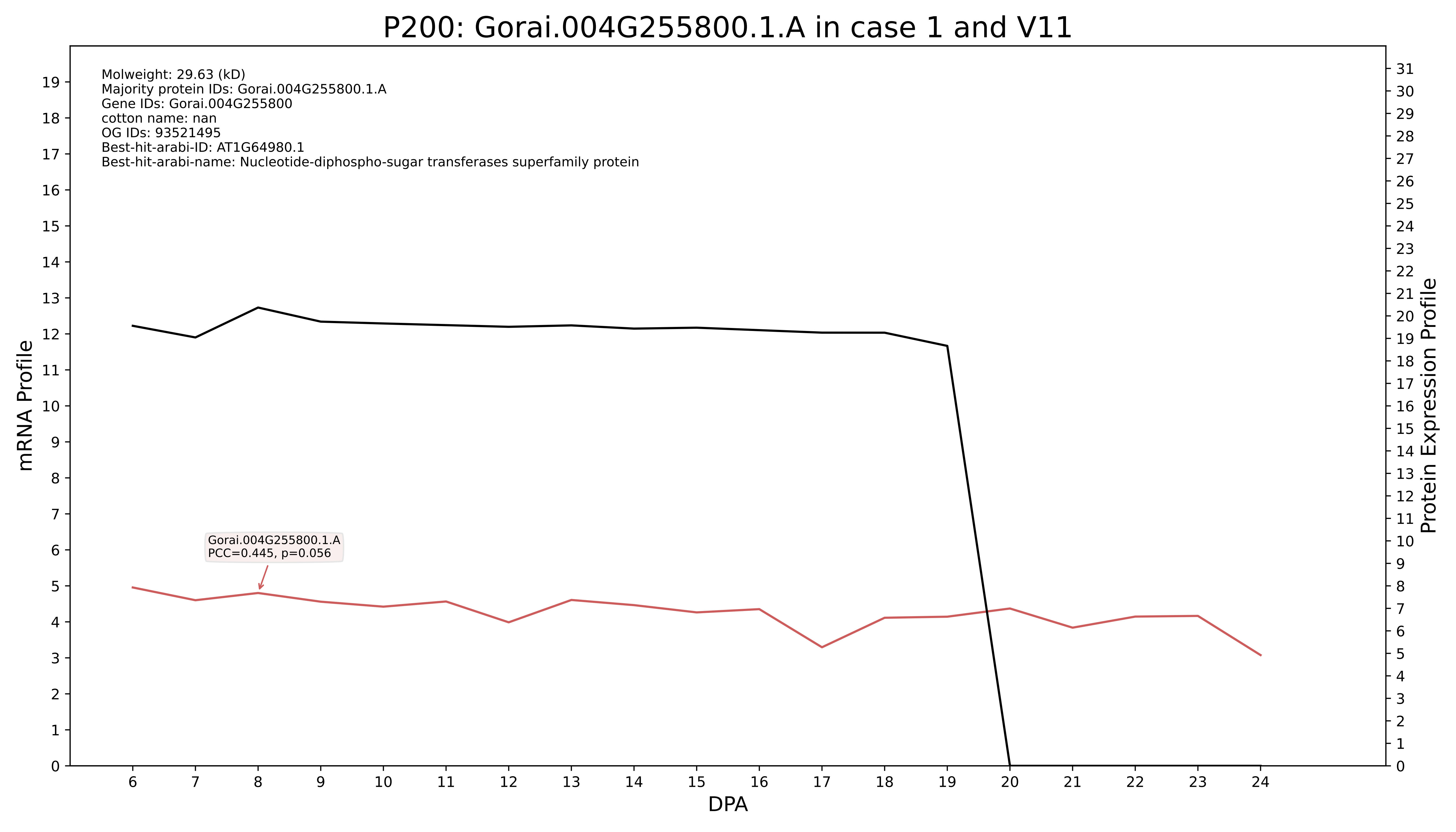

P200: Gorai.004G255800.1.A in case 1 and V11
Molweight: 29.63 (kD)
Majority protein IDs: Gorai.004G255800.1.A
Gene IDs: Gorai.004G255800
cotton name: nan
OG IDs: 93521495
Best-hit-arabi-ID: AT1G64980.1
Best-hit-arabi-name: Nucleotide-diphospho-sugar transferases superfamily protein
Gorai.004G255800.1.A
PCC=0.445, p=0.056
mRNA Profile
Protein Expression Profile
DPA

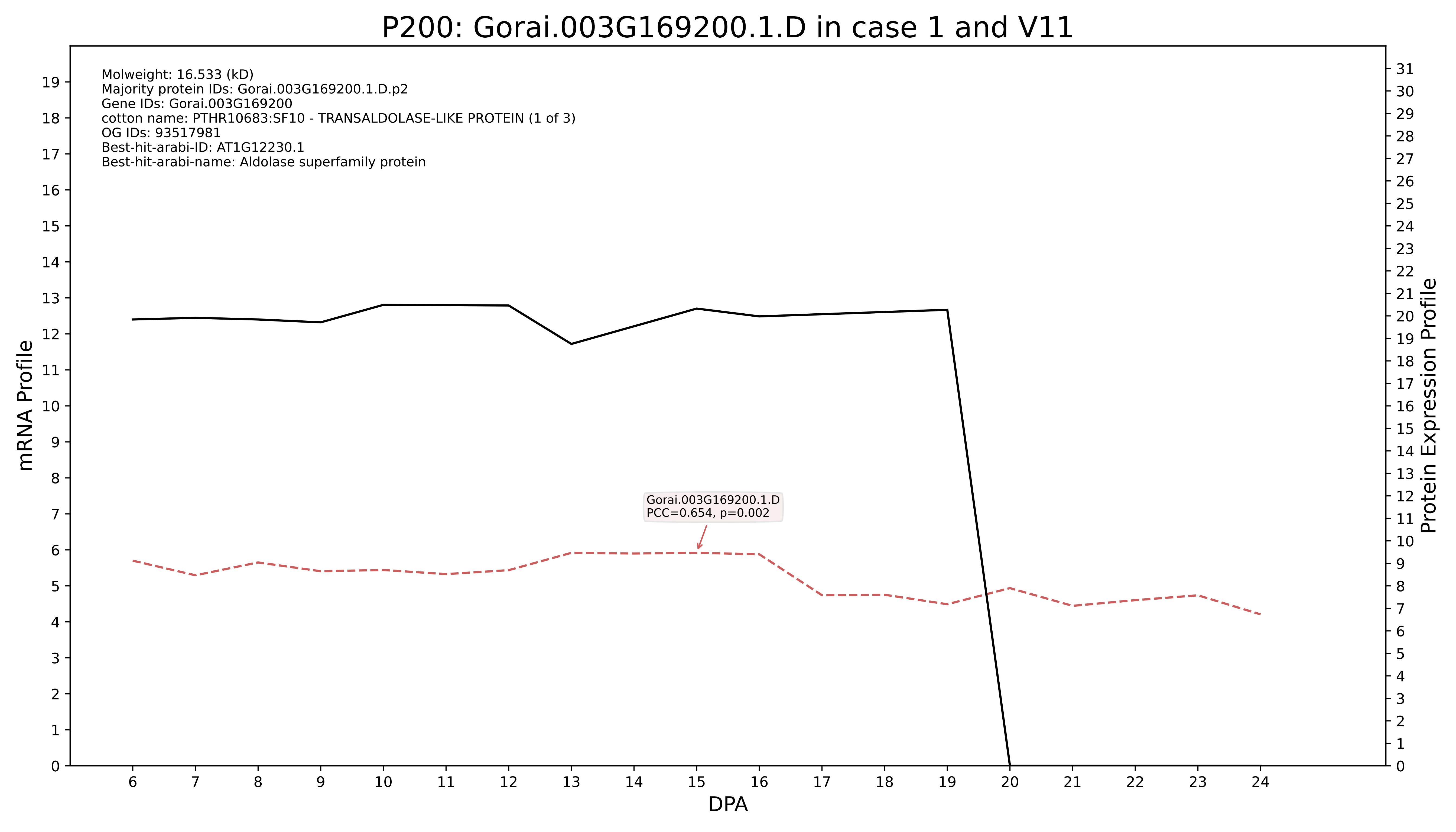

P200: Gorai.003G169200.1.D in case 1 and V11
Molweight: 16.533 (kD)
Majority protein IDs: Gorai.003G169200.1.D.p2
Gene IDs: Gorai.003G169200
cotton name: PTHR10683:SF10 - TRANSALDOLASE-LIKE PROTEIN (1 of 3)
OG IDs: 93517981
Best-hit-arabi-ID: AT1G12230.1
Best-hit-arabi-name: Aldolase superfamily protein
Gorai.003G169200.1.D
PCC=0.654, p=0.002
mRNA Profile
Protein Expression Profile
DPA

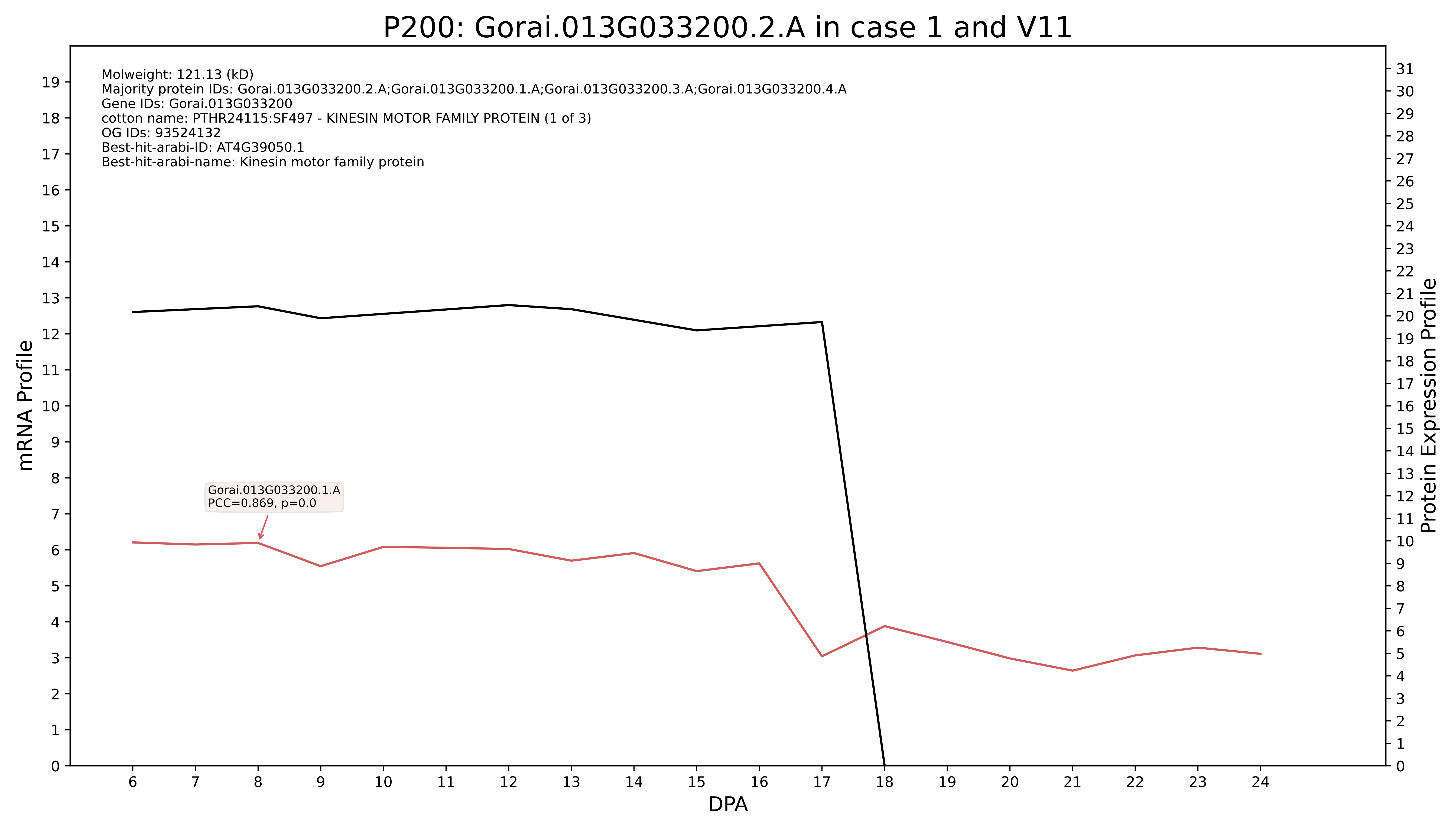

P200: Gorai.013G033200.2.A in case 1 and V11
Molweight: 121.13 (kD)
Majority protein IDs: Gorai.013G033200.2.A;Gorai.013G033200.1.A;Gorai.013G033200.3.A;Gorai.013G033200.4.A
Gene IDs: Gorai.013G033200
cotton name: PTHR24115:SF497 - KINESIN MOTOR FAMILY PROTEIN (1 of 3)
OG IDs: 93524132
Best-hit-arabi-ID: AT4G39050.1
Best-hit-arabi-name: Kinesin motor family protein
Gorai.013G033200.1.A
PCC=0.869, p=0.0
mRNA Profile
Protein Expression Profile
DPA

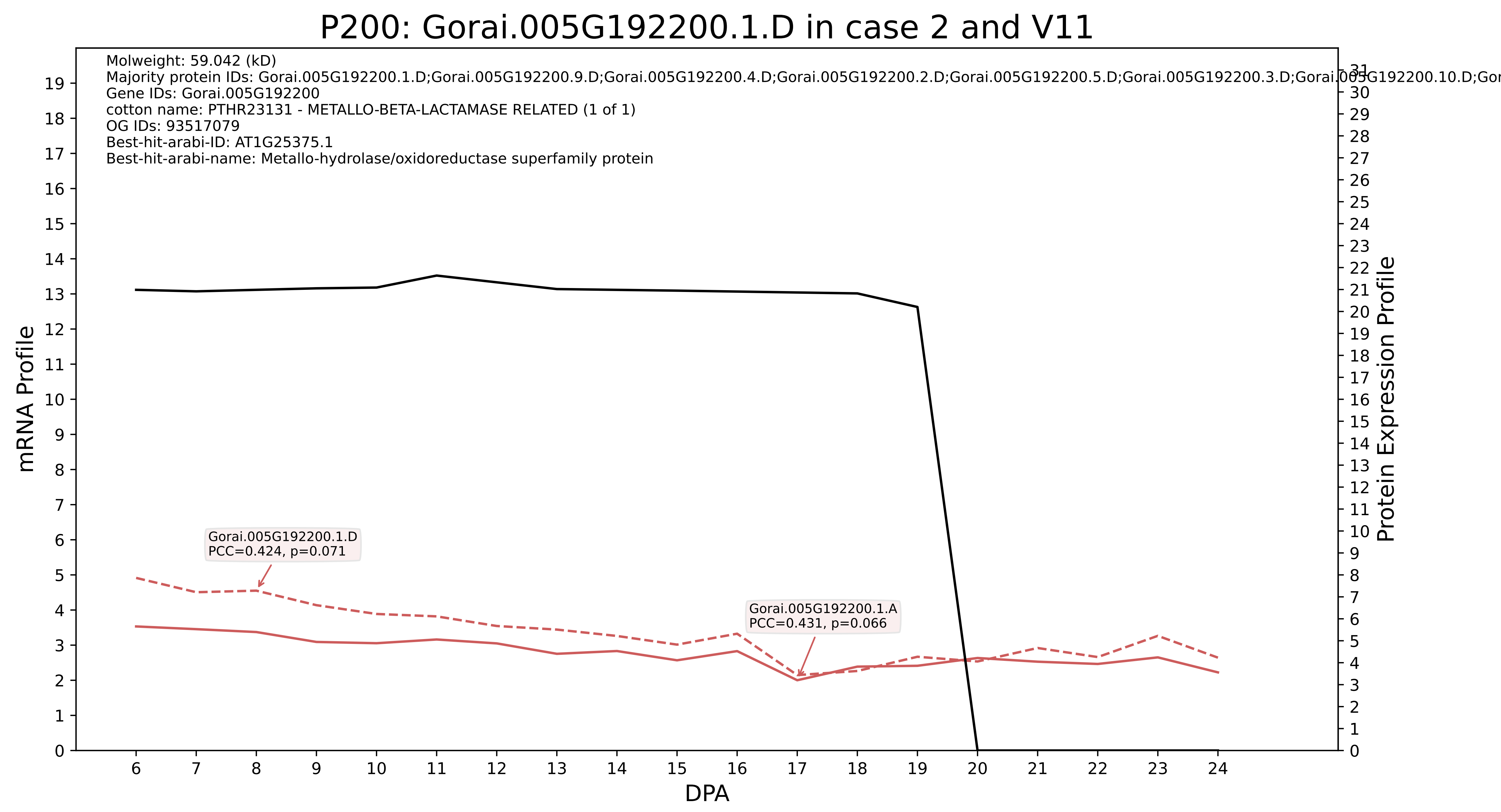
P200: Gorai.005G192200.1.D in case 2 and V11
Molweight: 59.042 (kD)
Majority protein IDs: Gorai.005G192200.1.D;Gorai.005G192200.9.D;Gorai.005G192200.4.D;Gorai.005G192200.2.D;Gorai.005G192200.5.D;Gorai.005G192200.3.D;Gorai.005G192200.10.D;Gor
Gene IDs: Gorai.005G192200
cotton name: PTHR23131 - METALLO-BETA-LACTAMASE RELATED (1 of 1)
OG IDs: 93517079
Best-hit-arabi-ID: AT1G25375.1
Best-hit-arabi-name: Metallo-hydrolase/oxidoreductase superfamily protein
Gorai.005G192200.1.D
PCC=0.424, p=0.071
Gorai.005G192200.1.A
PCC=0.431, p=0.066
mRNA Profile
Protein Expression Profile
DPA

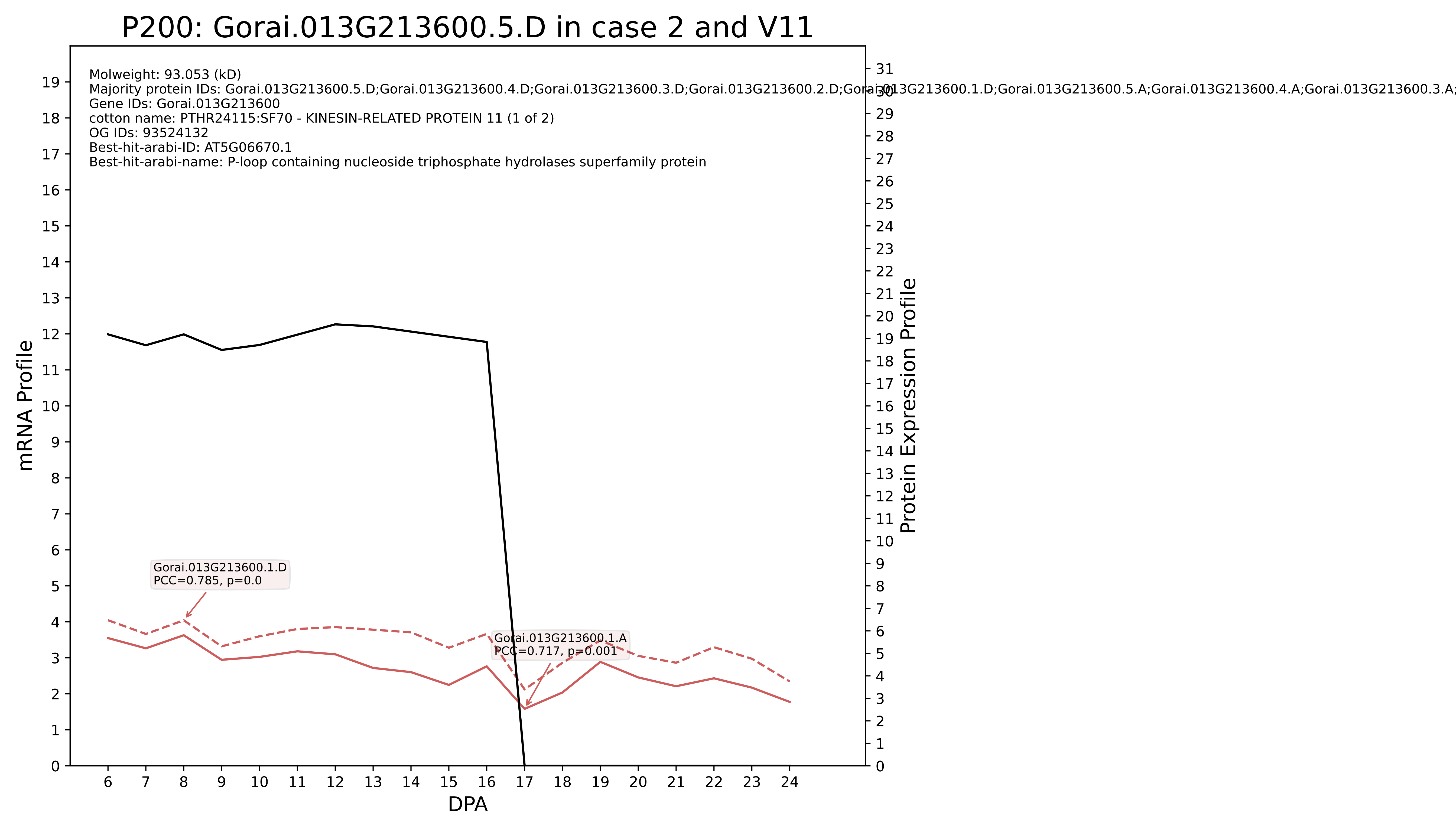

P200: Gorai.013G213600.5.D in case 2 and V11
Molweight: 93.053 (kD)
Majority protein IDs: Gorai.013G213600.5.D;Gorai.013G213600.4.D;Gorai.013G213600.3.D;Gorai.013G213600.2.D;Gorai.013G213600.1.D;Gorai.013G213600.5.A;Gorai.013G213600.4.A;Gorai.013G213600.3.A;
Gene IDs: Gorai.013G213600
cotton name: PTHR24115:SF70 - KINESIN-RELATED PROTEIN 11 (1 of 2)
OG IDs: 93524132
Best-hit-arabi-ID: AT5G06670.1
Best-hit-arabi-name: P-loop containing nucleoside triphosphate hydrolases superfamily protein
Gorai.013G213600.1.D
PCC=0.785, p=0.0
Gorai.013G213600.1.A
PCC=0.717, p=0.001
mRNA Profile
Protein Expression Profile
DPA

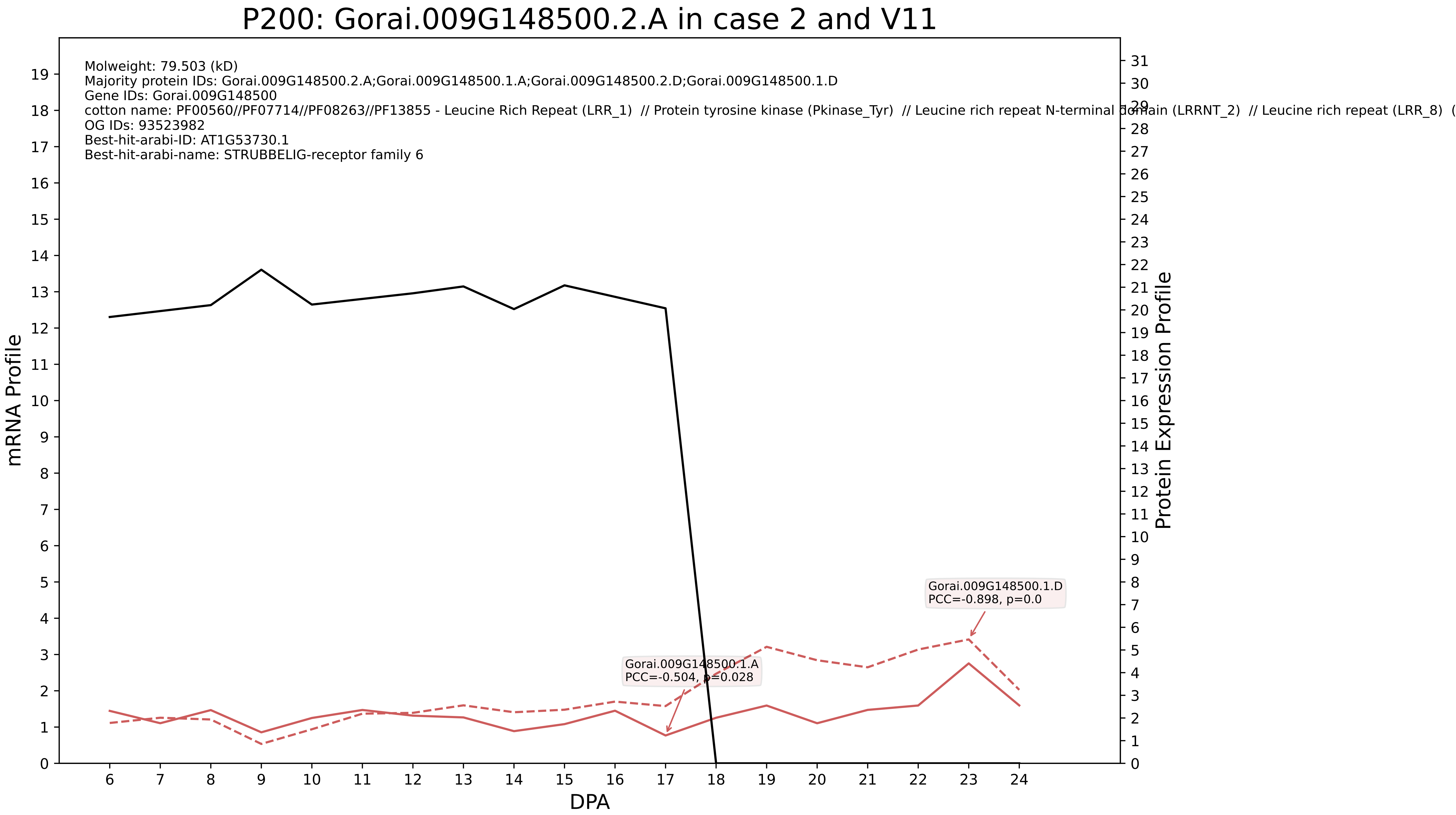

P200: Gorai.009G148500.2.A in case 2 and V11
Molweight: 79.503 (kD)
Majority protein IDs: Gorai.009G148500.2.A;Gorai.009G148500.1.A;Gorai.009G148500.2.D;Gorai.009G148500.1.D
Gene IDs: Gorai.009G148500
cotton name: PF00560//PF07714//PF08263//PF13855 - Leucine Rich Repeat (LRR_1) // Protein tyrosine kinase (Pkinase_Tyr) // Leucine rich repeat N-terminal domain (LRRNT_2) // Leucine rich repeat (LRR_8) (
OG IDs: 93523982
Best-hit-arabi-ID: AT1G53730.1
Best-hit-arabi-name: STRUBBELIG-receptor family 6
Gorai.009G148500.1.D
PCC=-0.898, p=0.0
Gorai.009G148500.1.A
PCC=-0.504, p=0.028
mRNA Profile
Protein Expression Profile
DPA

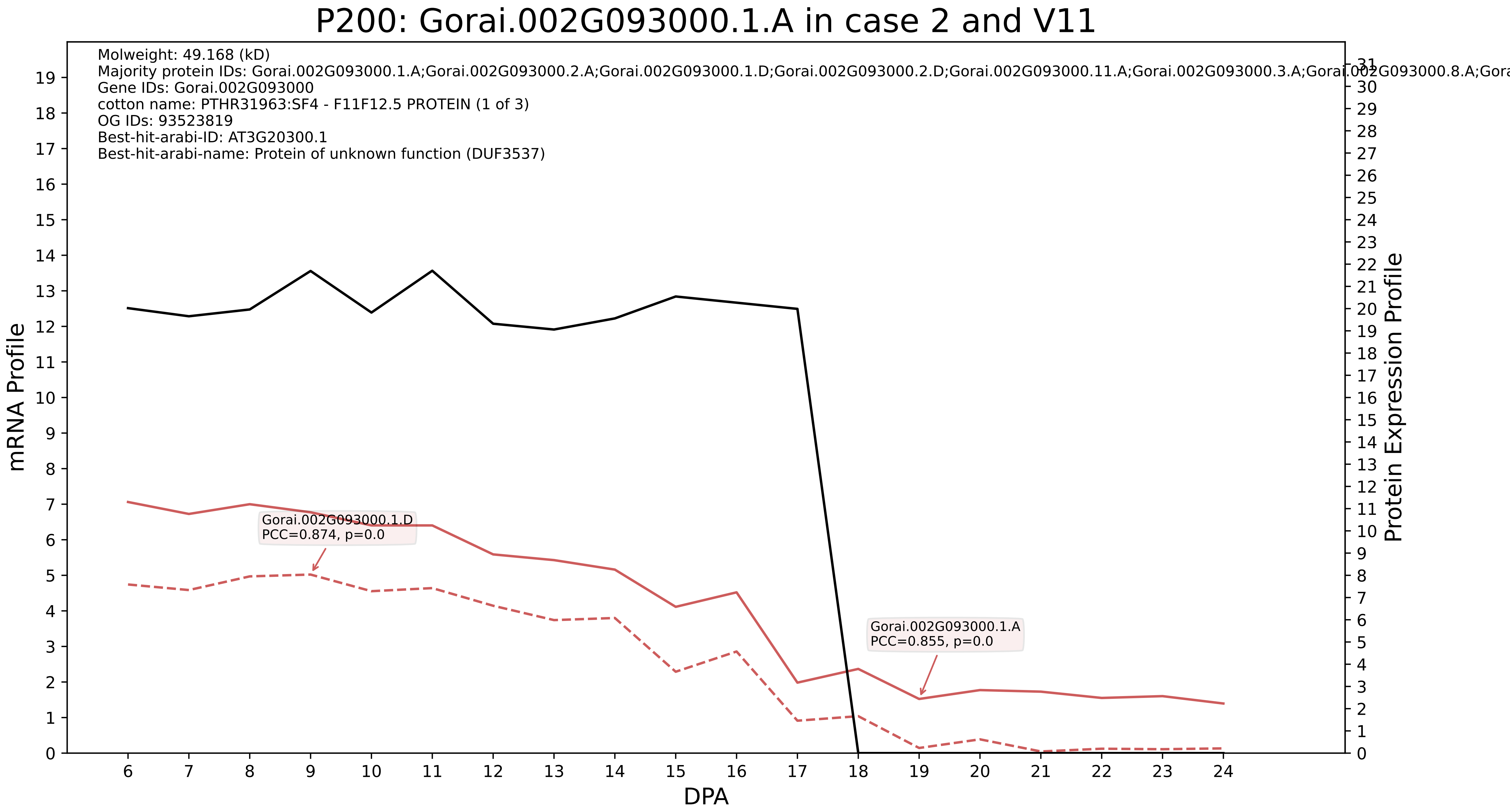

P200: Gorai.002G093000.1.A in case 2 and V11
Molweight: 49.168 (kD)
Majority protein IDs: Gorai.002G093000.1.A;Gorai.002G093000.2.A;Gorai.002G093000.1.D;Gorai.002G093000.2.D;Gorai.002G093000.11.A;Gorai.002G093000.3.A;Gorai.002G093000.8.A;Gora
Gene IDs: Gorai.002G093000
cotton name: PTHR31963:SF4 - F11F12.5 PROTEIN (1 of 3)
OG IDs: 93523819
Best-hit-arabi-ID: AT3G20300.1
Best-hit-arabi-name: Protein of unknown function (DUF3537)
Gorai.002G093000.1.D
PCC=0.874, p=0.0
Gorai.002G093000.1.A
PCC=0.855, p=0.0
mRNA Profile
Protein Expression Profile
DPA

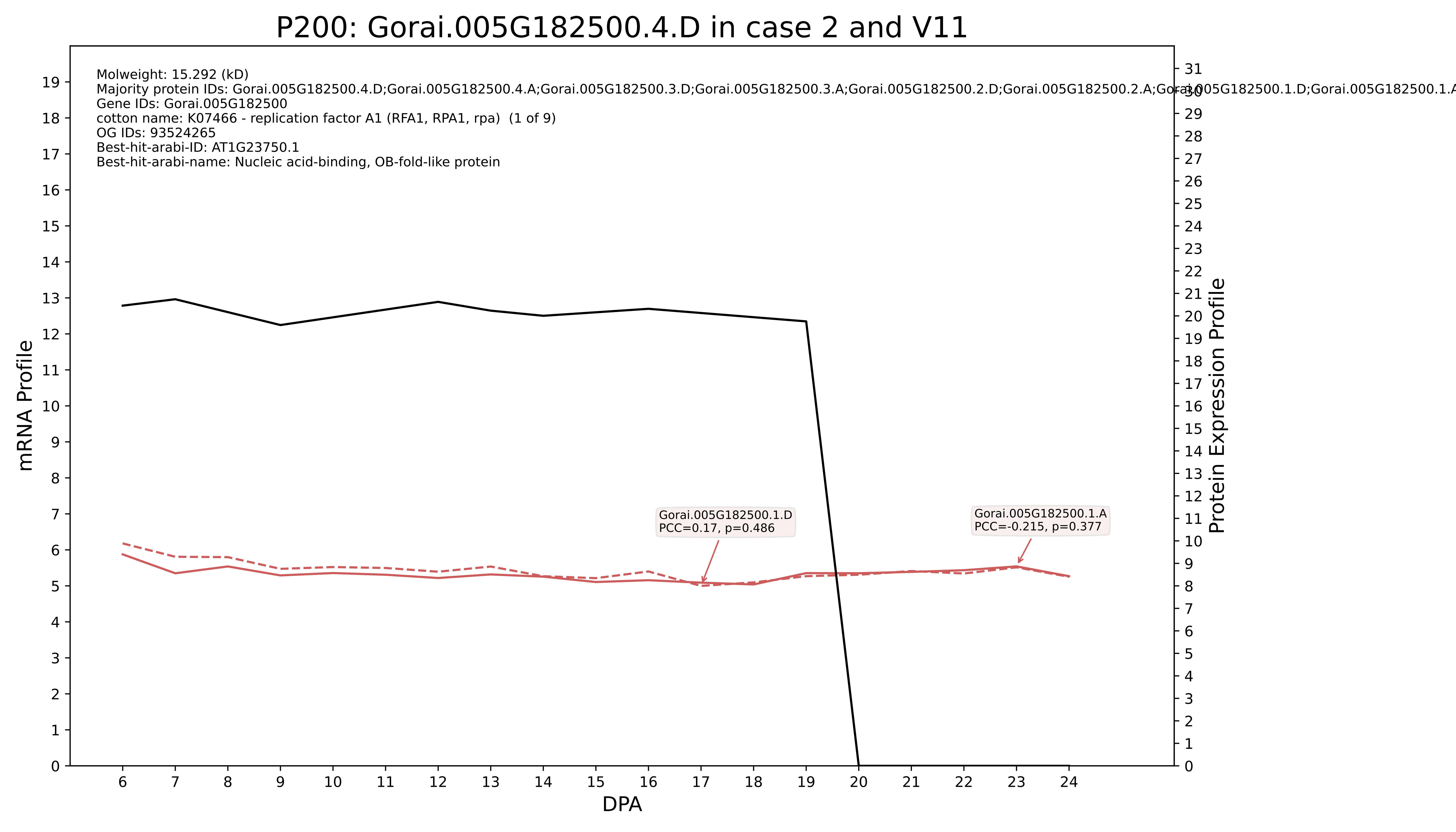

P200: Gorai.005G182500.4.D in case 2 and V11
Molweight: 15.292 (kD)
Majority protein IDs: Gorai.005G182500.4.D;Gorai.005G182500.4.A;Gorai.005G182500.3.D;Gorai.005G182500.3.A;Gorai.005G182500.2.D;Gorai.005G182500.2.A;Gorai.005G182500.1.D;Gorai.005G182500.1.A
Gene IDs: Gorai.005G182500
cotton name: K07466 - replication factor A1 (RFA1, RPA1, rpa) (1 of 9)
OG IDs: 93524265
Best-hit-arabi-ID: AT1G23750.1
Best-hit-arabi-name: Nucleic acid-binding, OB-fold-like protein
Gorai.005G182500.1.D
PCC=0.17, p=0.486
Gorai.005G182500.1.A
PCC=-0.215, p=0.377
mRNA Profile
Protein Expression Profile
DPA

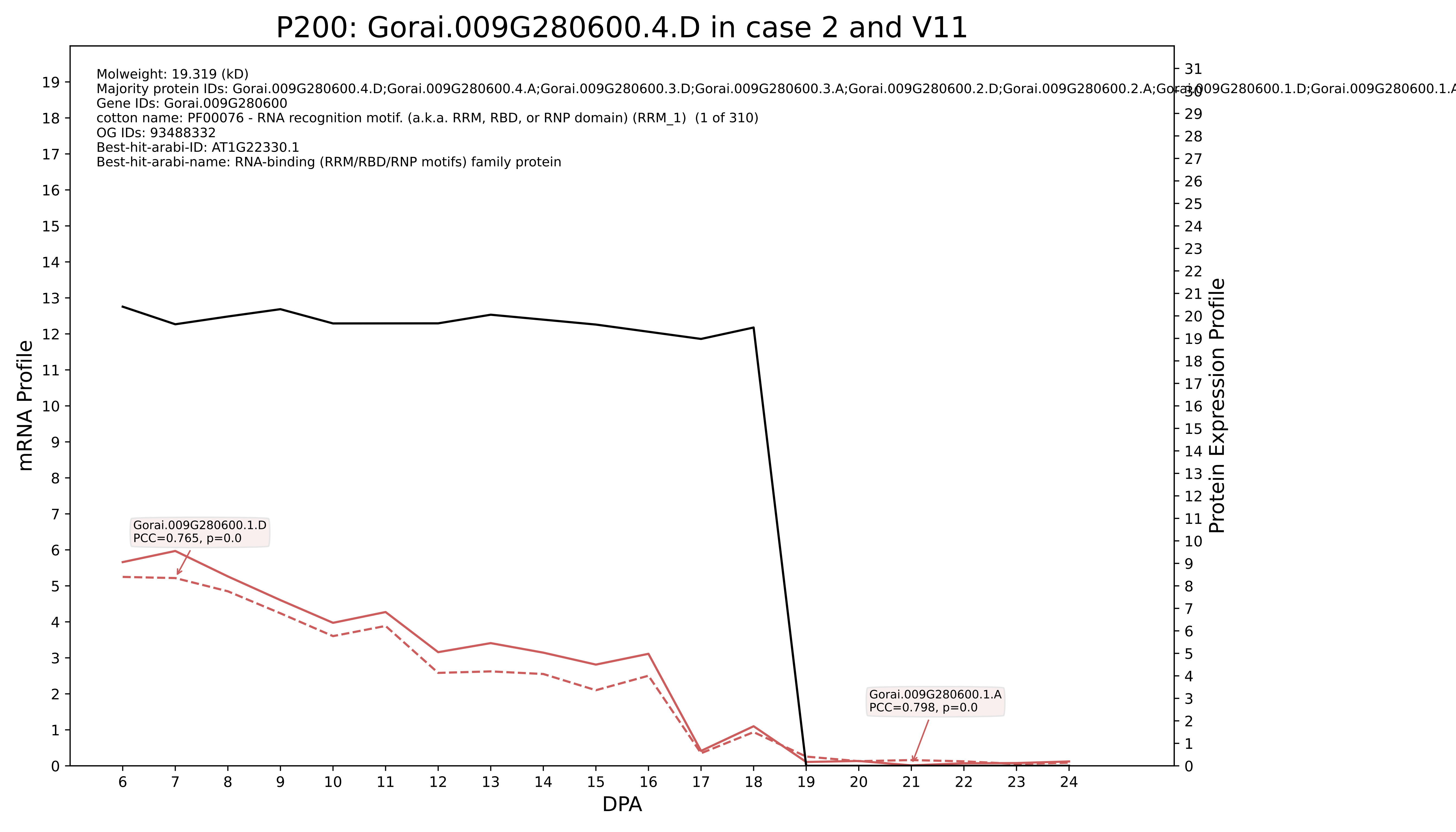

P200: Gorai.009G280600.4.D in case 2 and V11
Molweight: 19.319 (kD)
Majority protein IDs: Gorai.009G280600.4.D;Gorai.009G280600.4.A;Gorai.009G280600.3.D;Gorai.009G280600.3.A;Gorai.009G280600.2.D;Gorai.009G280600.2.A;Gorai.009G280600.1.D;Gorai.009G280600.1.A
Gene IDs: Gorai.009G280600
cotton name: PF00076 - RNA recognition motif. (a.k.a. RRM, RBD, or RNP domain) (RRM_1) (1 of 310)
OG IDs: 93488332
Best-hit-arabi-ID: AT1G22330.1
Best-hit-arabi-name: RNA-binding (RRM/RBD/RNP motifs) family protein
Gorai.009G280600.1.D
PCC=0.765, p=0.0
Gorai.009G280600.1.A
PCC=0.798, p=0.0
mRNA Profile
Protein Expression Profile
DPA

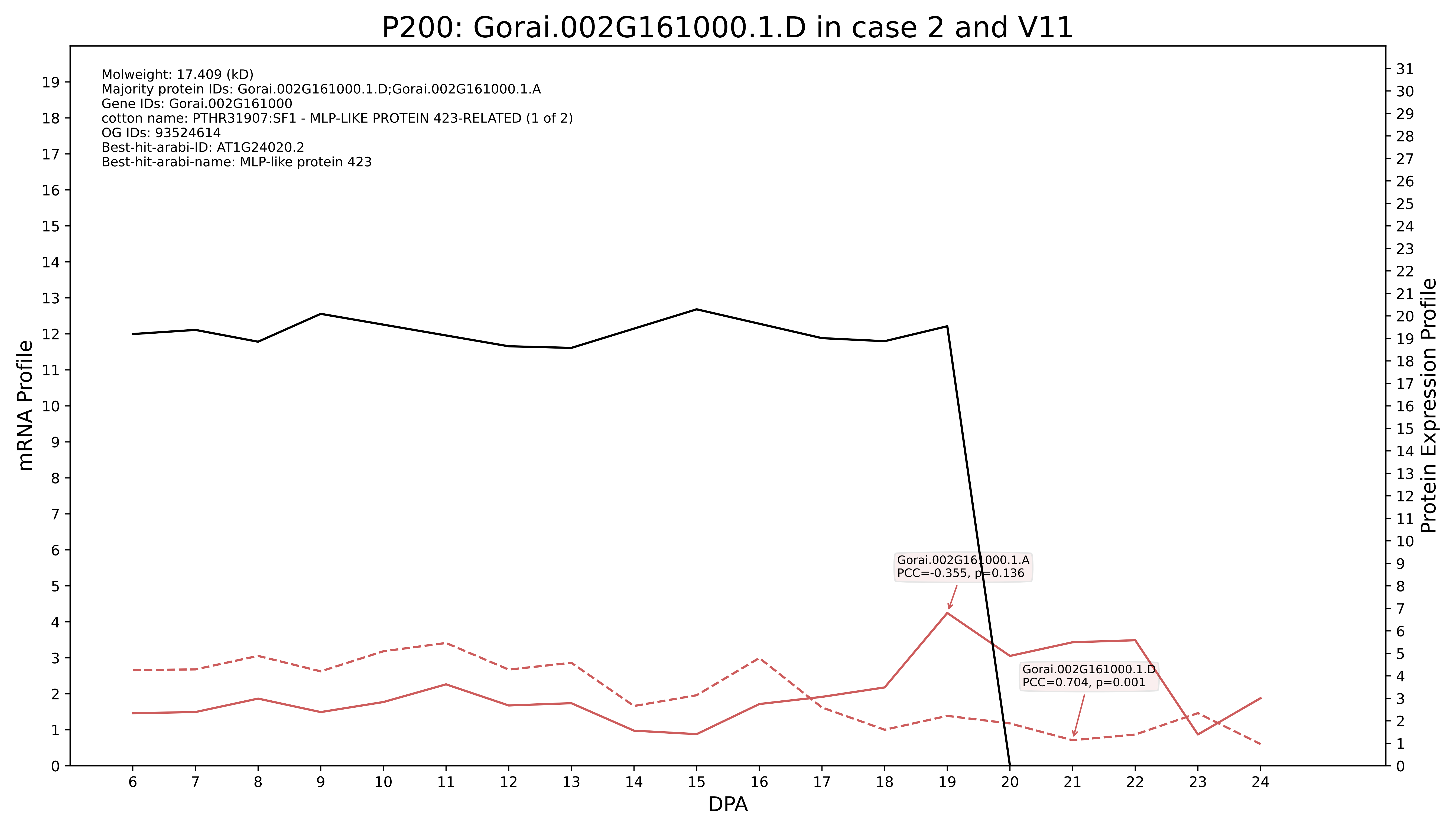

P200: Gorai.002G161000.1.D in case 2 and V11
Molweight: 17.409 (kD)
Majority protein IDs: Gorai.002G161000.1.D;Gorai.002G161000.1.A
Gene IDs: Gorai.002G161000
cotton name: PTHR31907:SF1 - MLP-LIKE PROTEIN 423-RELATED (1 of 2)
OG IDs: 93524614
Best-hit-arabi-ID: AT1G24020.2
Best-hit-arabi-name: MLP-like protein 423
Gorai.002G161000.1.A
PCC=-0.355, p=0.136
Gorai.002G161000.1.D
PCC=0.704, p=0.001
mRNA Profile
Protein Expression Profile
DPA

### P200: Gorai.001G180400.1.D in case 2 and V12

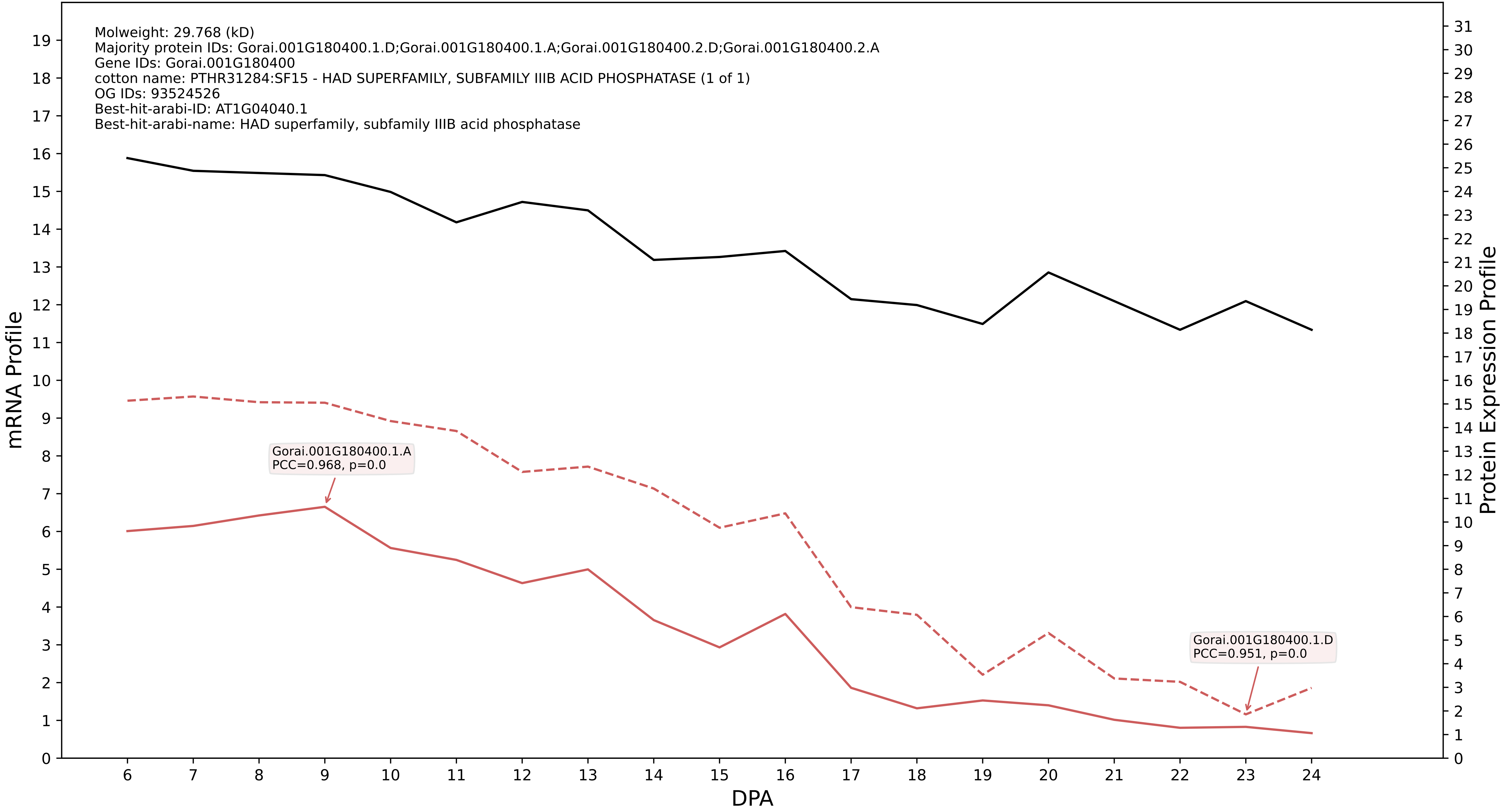

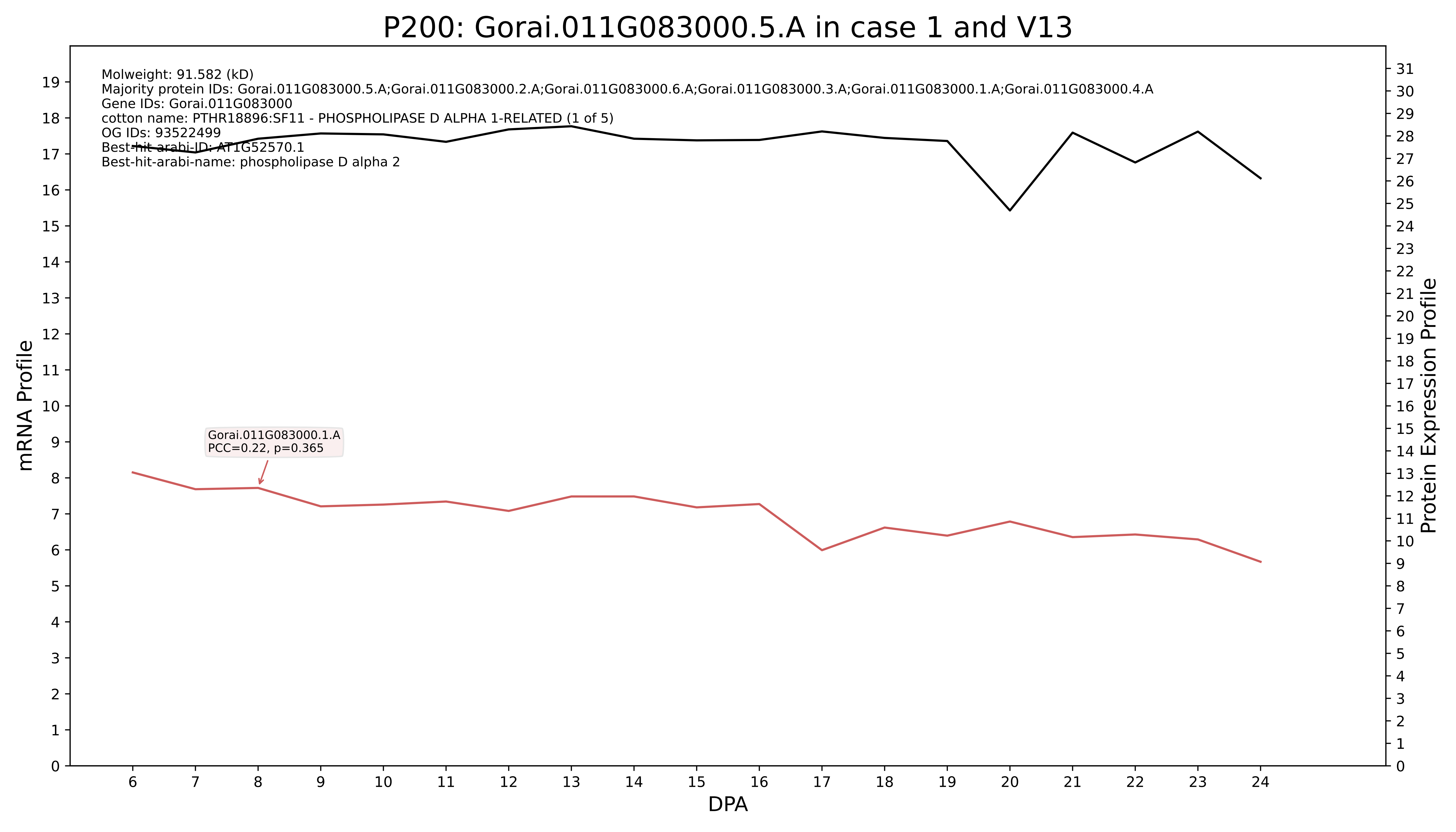
P200: Gorai.011G083000.5.A in case 1 and V13
Molweight: 91.582 (kD)
Majority protein IDs: Gorai.011G083000.5.A;Gorai.011G083000.2.A;Gorai.011G083000.6.A;Gorai.011G083000.3.A;Gorai.011G083000.1.A;Gorai.011G083000.4.A
Gene IDs: Gorai.011G083000
cotton name: PTHR18896:SF11 - PHOSPHOLIPASE D ALPHA 1-RELATED (1 of 5)
OG IDs: 93522499
Best-hit-arabi-ID: AT1G52570.1
Best-hit-arabi-name: phospholipase D alpha 2
Gorai.011G083000.1.A
PCC=0.22, p=0.365
mRNA Profile
Protein Expression Profile
DPA

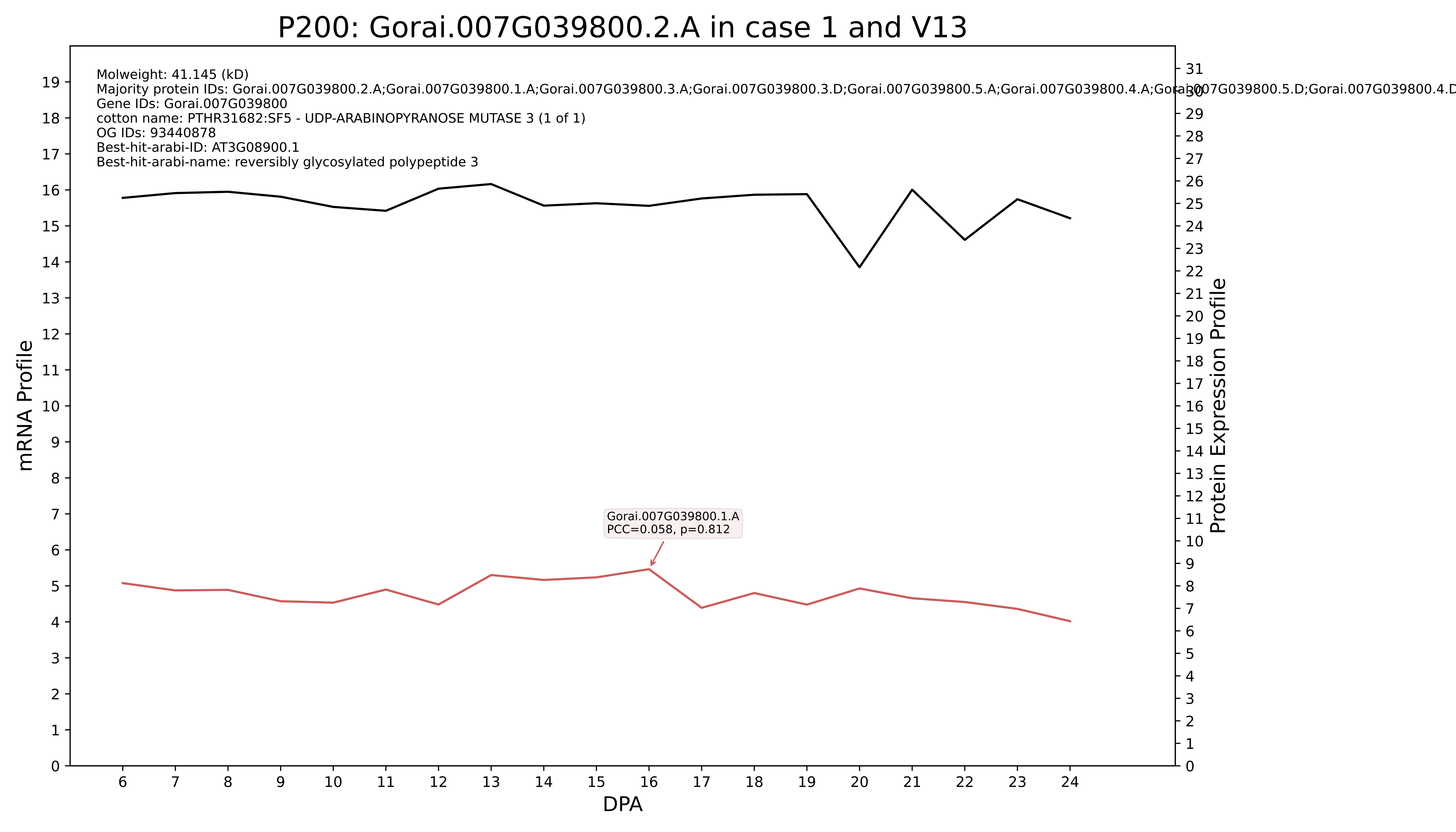

P200: Gorai.007G039800.2.A in case 1 and V13
Molweight: 41.145 (kD)
Majority protein IDs: Gorai.007G039800.2.A;Gorai.007G039800.1.A;Gorai.007G039800.3.A;Gorai.007G039800.3.D;Gorai.007G039800.5.A;Gorai.007G039800.4.A;Gorai.007G039800.5.D;Gorai.007G039800.4.D
Gene IDs: Gorai.007G039800
cotton name: PTHR31682:SF5 - UDP-ARABINOPYRANOSE MUTASE 3 (1 of 1)
OG IDs: 93440878
Best-hit-arabi-ID: AT3G08900.1
Best-hit-arabi-name: reversibly glycosylated polypeptide 3
Gorai.007G039800.1.A
PCC=0.058, p=0.812
mRNA Profile
Protein Expression Profile
DPA

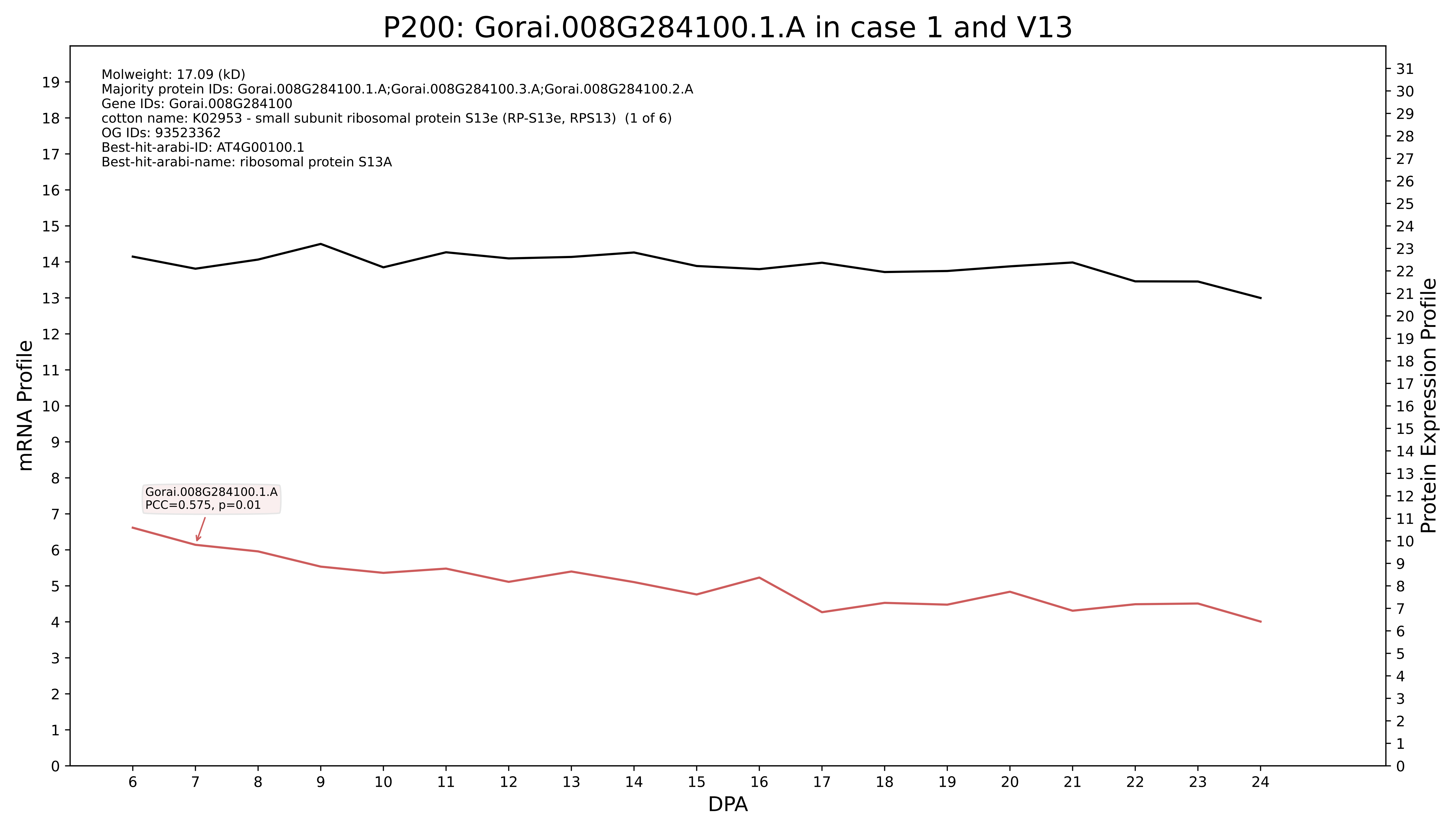

P200: Gorai.008G284100.1.A in case 1 and V13
Molweight: 17.09 (kD)
Majority protein IDs: Gorai.008G284100.1.A;Gorai.008G284100.3.A;Gorai.008G284100.2.A
Gene IDs: Gorai.008G284100
cotton name: K02953 - small subunit ribosomal protein S13e (RP-S13e, RPS13) (1 of 6)
OG IDs: 93523362
Best-hit-arabi-ID: AT4G00100.1
Best-hit-arabi-name: ribosomal protein S13A
Gorai.008G284100.1.A
PCC=0.575, p=0.01
mRNA Profile
Protein Expression Profile
DPA

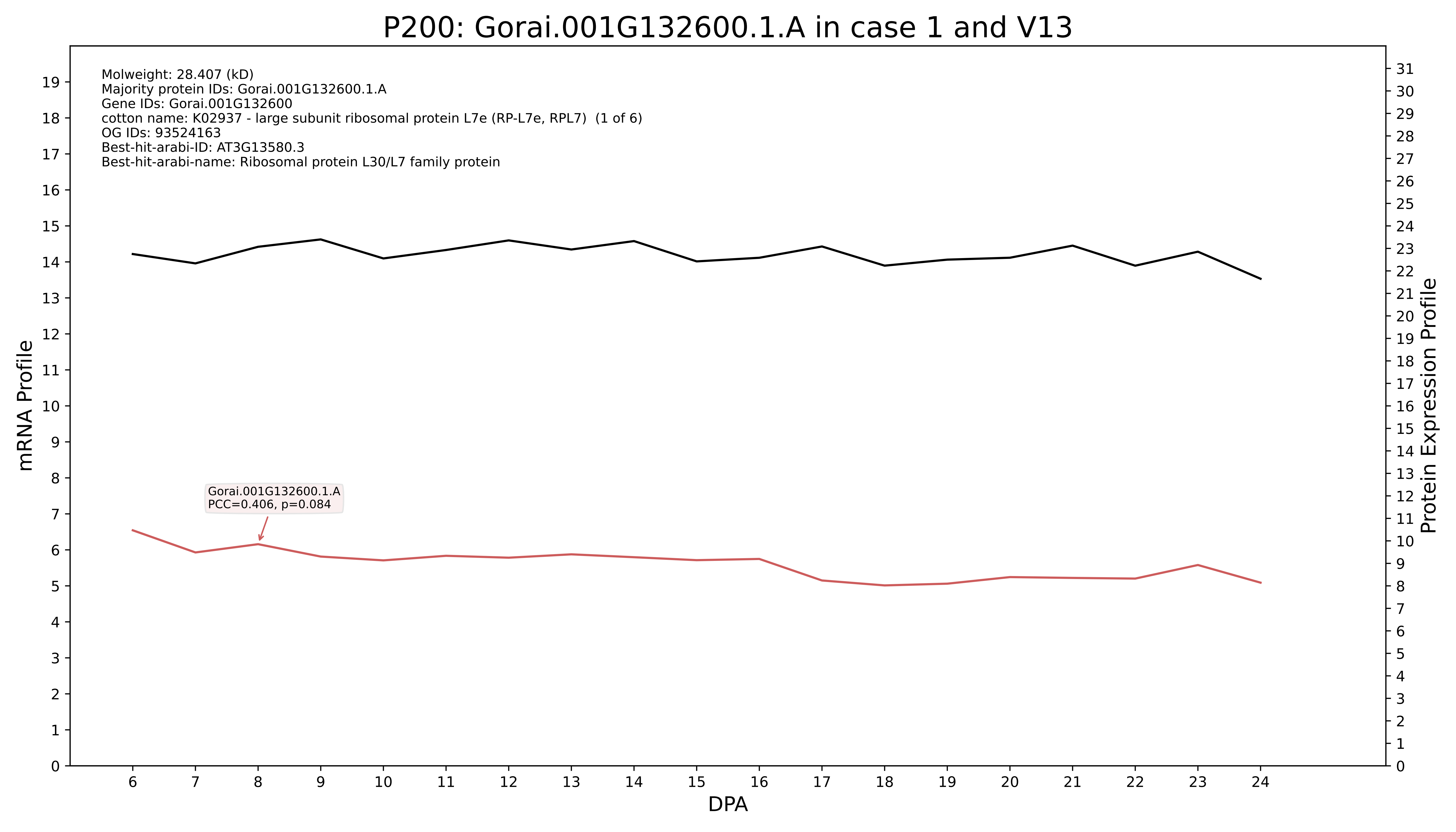
P200: Gorai.001G132600.1.A in case 1 and V13
Molweight: 28.407 (kD)
Majority protein IDs: Gorai.001G132600.1.A
Gene IDs: Gorai.001G132600
cotton name: K02937 - large subunit ribosomal protein L7e (RP-L7e, RPL7) (1 of 6)
OG IDs: 93524163
Best-hit-arabi-ID: AT3G13580.3
Best-hit-arabi-name: Ribosomal protein L30/L7 family protein
Gorai.001G132600.1.A
PCC=0.406, p=0.084
mRNA Profile
Protein Expression Profile
DPA

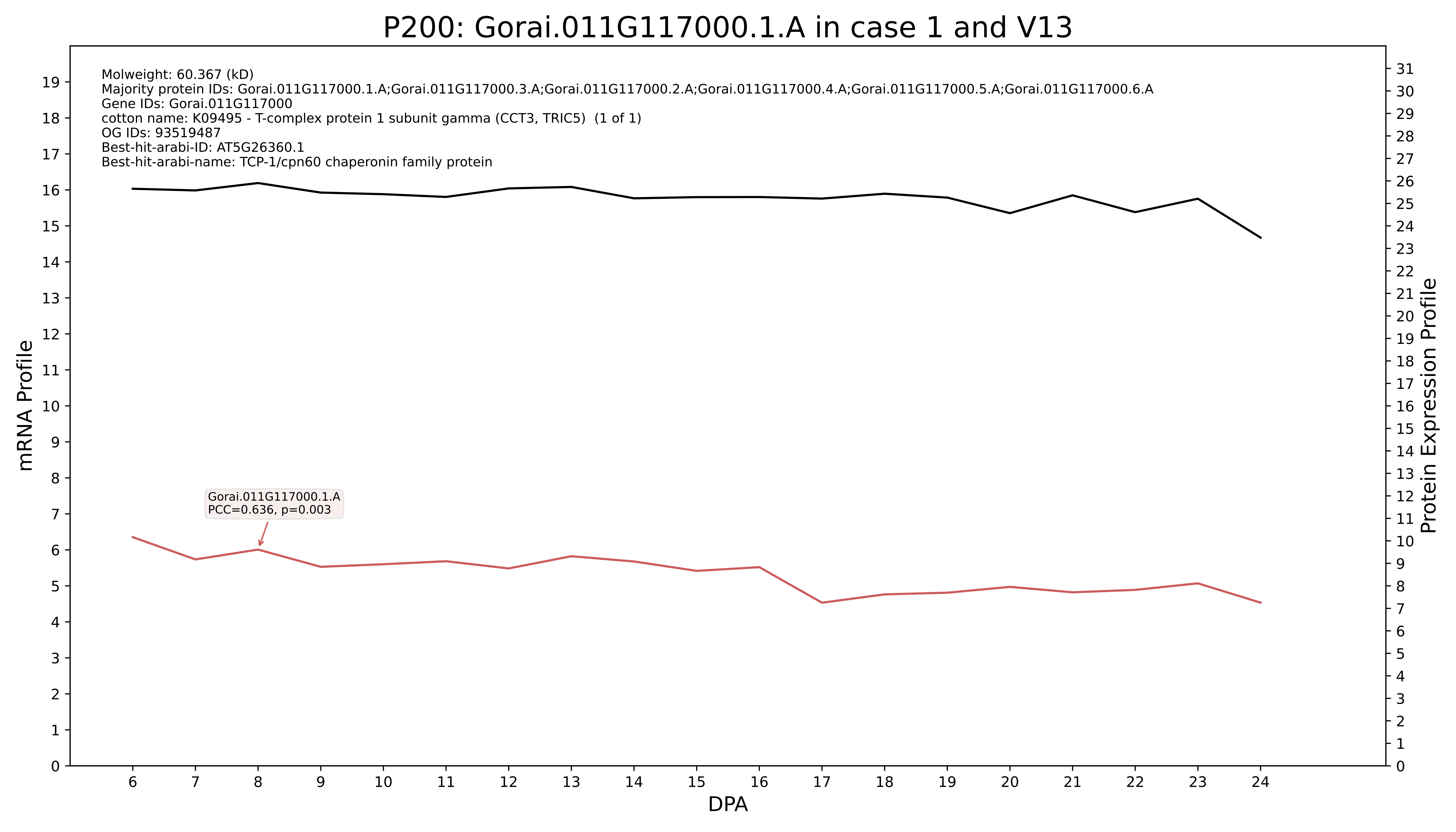
P200: Gorai.011G117000.1.A in case 1 and V13
Molweight: 60.367 (kD)
Majority protein IDs: Gorai.011G117000.1.A;Gorai.011G117000.3.A;Gorai.011G117000.2.A;Gorai.011G117000.4.A;Gorai.011G117000.5.A;Gorai.011G117000.6.A
Gene IDs: Gorai.011G117000
cotton name: K09495 - T-complex protein 1 subunit gamma (CCT3, TRIC5) (1 of 1)
OG IDs: 93519487
Best-hit-arabi-ID: AT5G26360.1
Best-hit-arabi-name: TCP-1/cpn60 chaperonin family protein
Gorai.011G117000.1.A
PCC=0.636, p=0.003
mRNA Profile
Protein Expression Profile
DPA

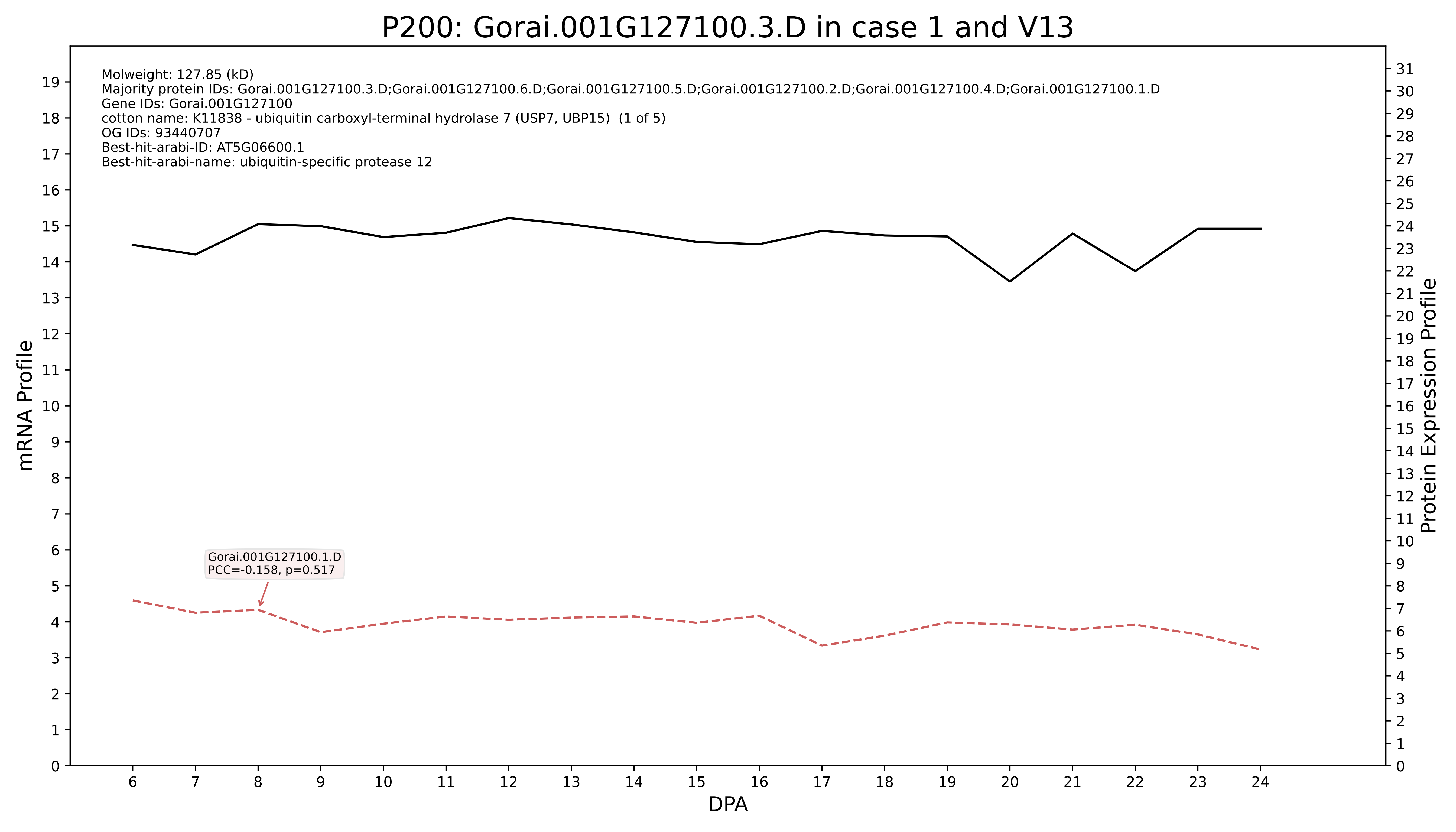
P200: Gorai.001G127100.3.D in case 1 and V13
Molweight: 127.85 (kD)
Majority protein IDs: Gorai.001G127100.3.D;Gorai.001G127100.6.D;Gorai.001G127100.5.D;Gorai.001G127100.2.D;Gorai.001G127100.4.D;Gorai.001G127100.1.D
Gene IDs: Gorai.001G127100
cotton name: K11838 - ubiquitin carboxyl-terminal hydrolase 7 (USP7, UBP15) (1 of 5)
OG IDs: 93440707
Best-hit-arabi-ID: AT5G06600.1
Best-hit-arabi-name: ubiquitin-specific protease 12
Gorai.001G127100.1.D
PCC=-0.158, p=0.517
mRNA Profile
Protein Expression Profile
DPA

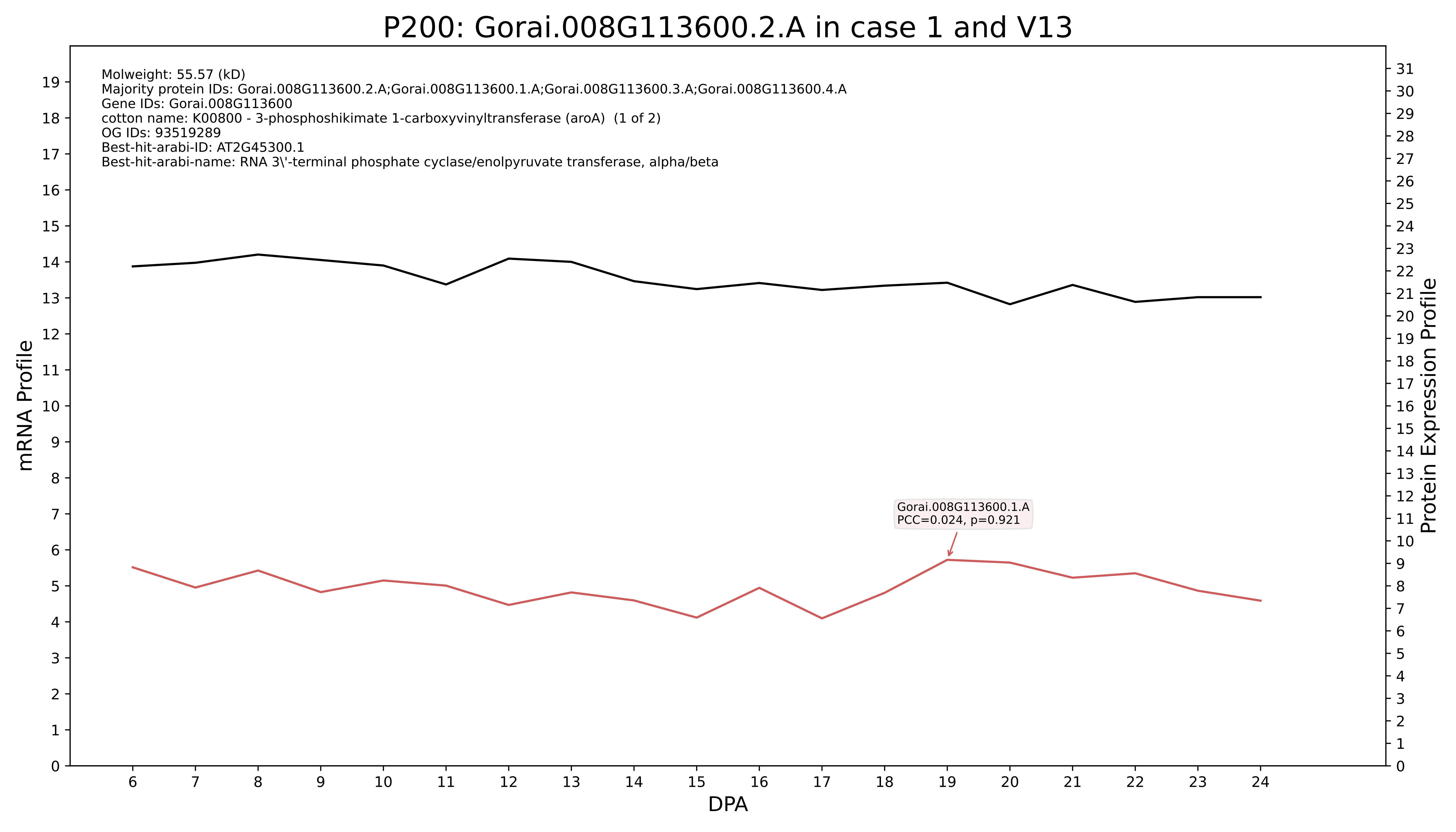
P200: Gorai.008G113600.2.A in case 1 and V13
Molweight: 55.57 (kD)
Majority protein IDs: Gorai.008G113600.2.A;Gorai.008G113600.1.A;Gorai.008G113600.3.A;Gorai.008G113600.4.A
Gene IDs: Gorai.008G113600
cotton name: K00800 - 3-phosphoshikimate 1-carboxyvinyltransferase (aroA) (1 of 2)
OG IDs: 93519289
Best-hit-arabi-ID: AT2G45300.1
Best-hit-arabi-name: RNA 3\'-terminal phosphate cyclase/enolpyruvate transferase, alpha/beta
Gorai.008G113600.1.A
PCC=0.024, p=0.921
mRNA Profile
Protein Expression Profile
DPA

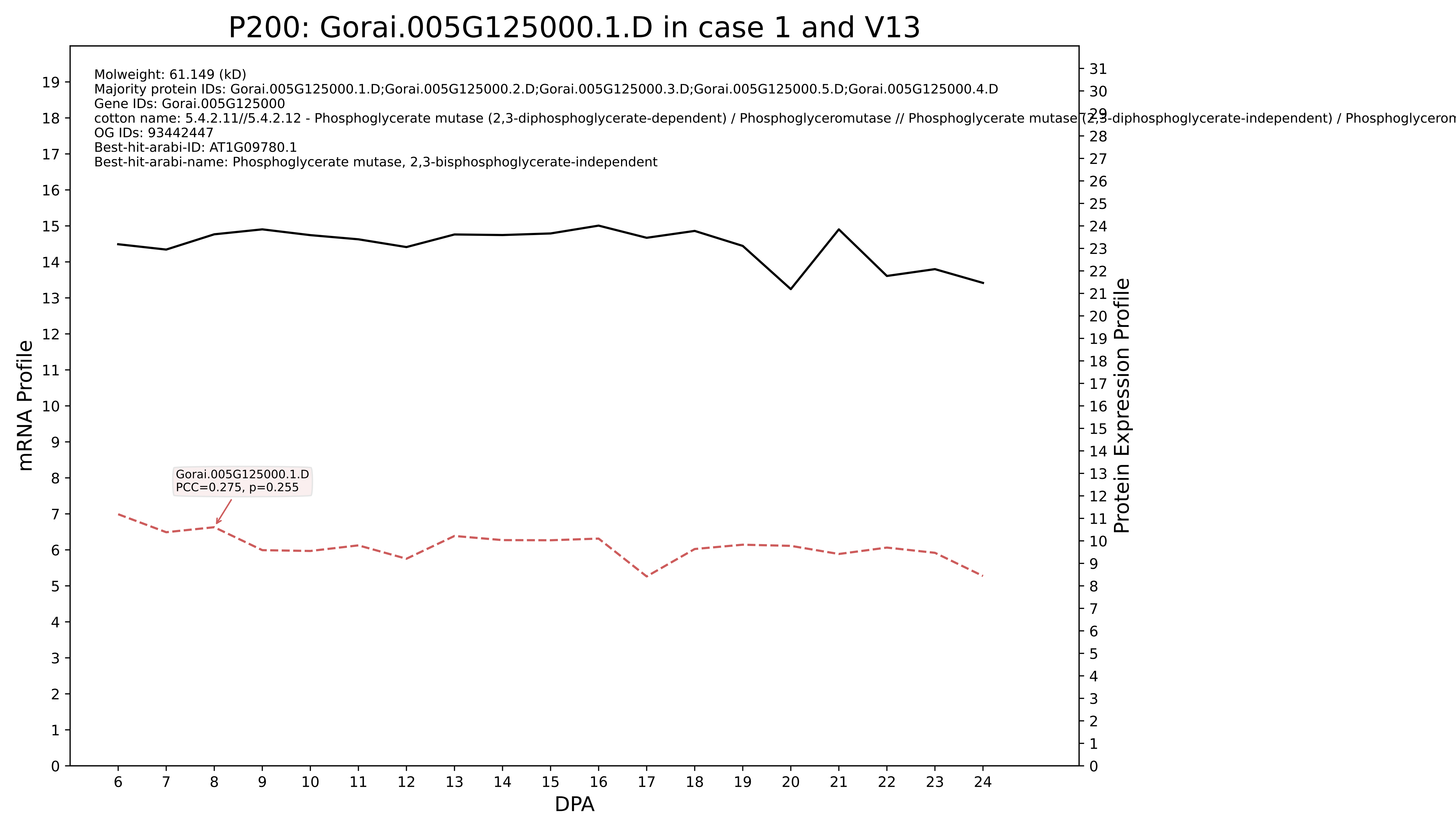
P200: Gorai.005G125000.1.D in case 1 and V13
Molweight: 61.149 (kD)
Majority protein IDs: Gorai.005G125000.1.D;Gorai.005G125000.2.D;Gorai.005G125000.3.D;Gorai.005G125000.5.D;Gorai.005G125000.4.D
Gene IDs: Gorai.005G125000
cotton name: 5.4.2.11//5.4.2.12 - Phosphoglycerate mutase (2,3-diphosphoglycerate-dependent) / Phosphoglyceromutase // Phosphoglycerate mutase (2,3-diphosphoglycerate-independent) / Phosphoglyceron
OG IDs: 93442447
Best-hit-arabi-ID: AT1G09780.1
Best-hit-arabi-name: Phosphoglycerate mutase, 2,3-bisphosphoglycerate-independent
Gorai.005G125000.1.D
PCC=0.275, p=0.255
mRNA Profile
Protein Expression Profile
DPA

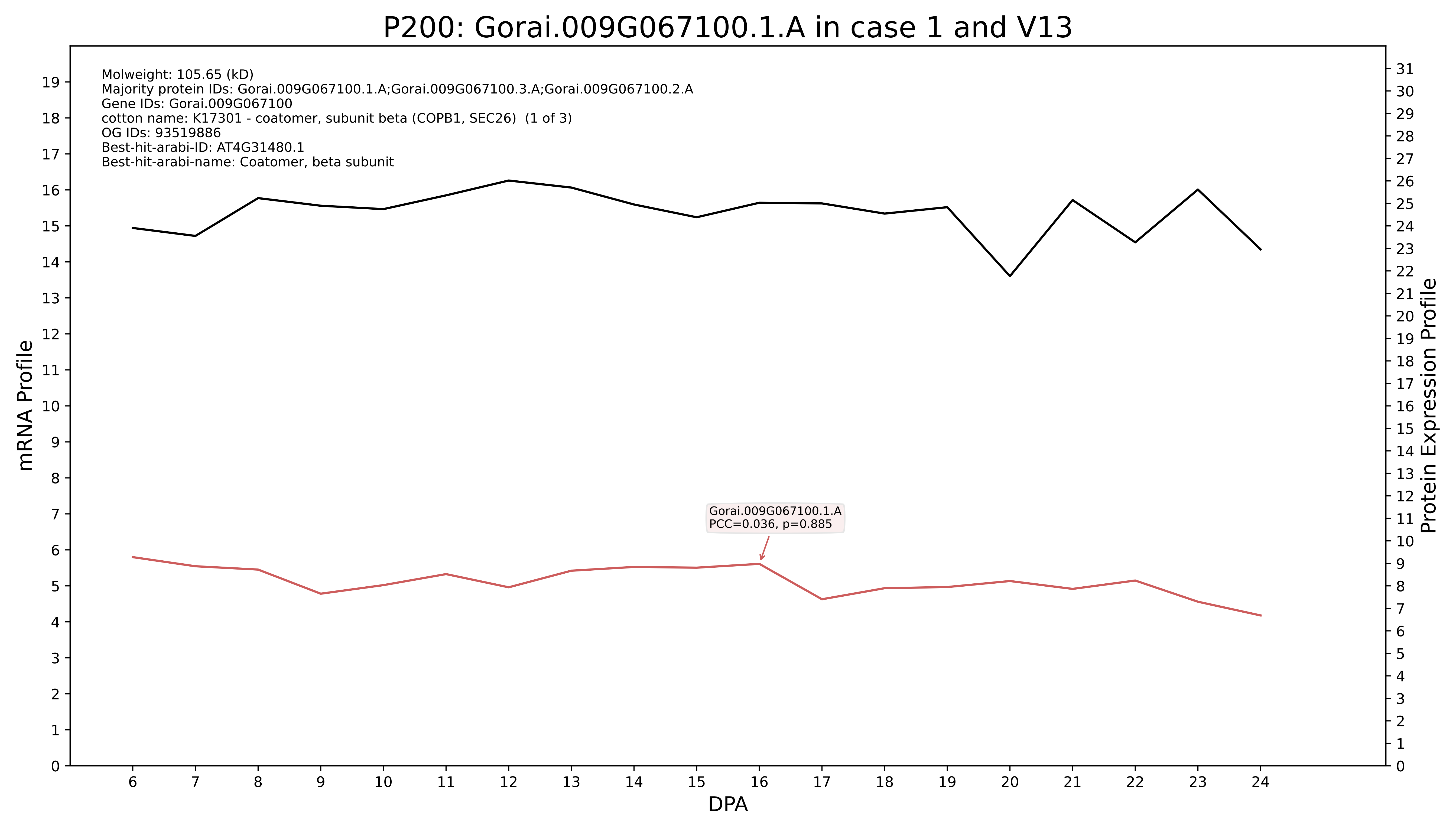

P200: Gorai.009G067100.1.A in case 1 and V13
Molweight: 105.65 (kD)
Majority protein IDs: Gorai.009G067100.1.A;Gorai.009G067100.3.A;Gorai.009G067100.2.A
Gene IDs: Gorai.009G067100
cotton name: K17301 - coatomer, subunit beta (COPB1, SEC26) (1 of 3)
OG IDs: 93519886
Best-hit-arabi-ID: AT4G31480.1
Best-hit-arabi-name: Coatomer, beta subunit
Gorai.009G067100.1.A
PCC=0.036, p=0.885
mRNA Profile
Protein Expression Profile
DPA

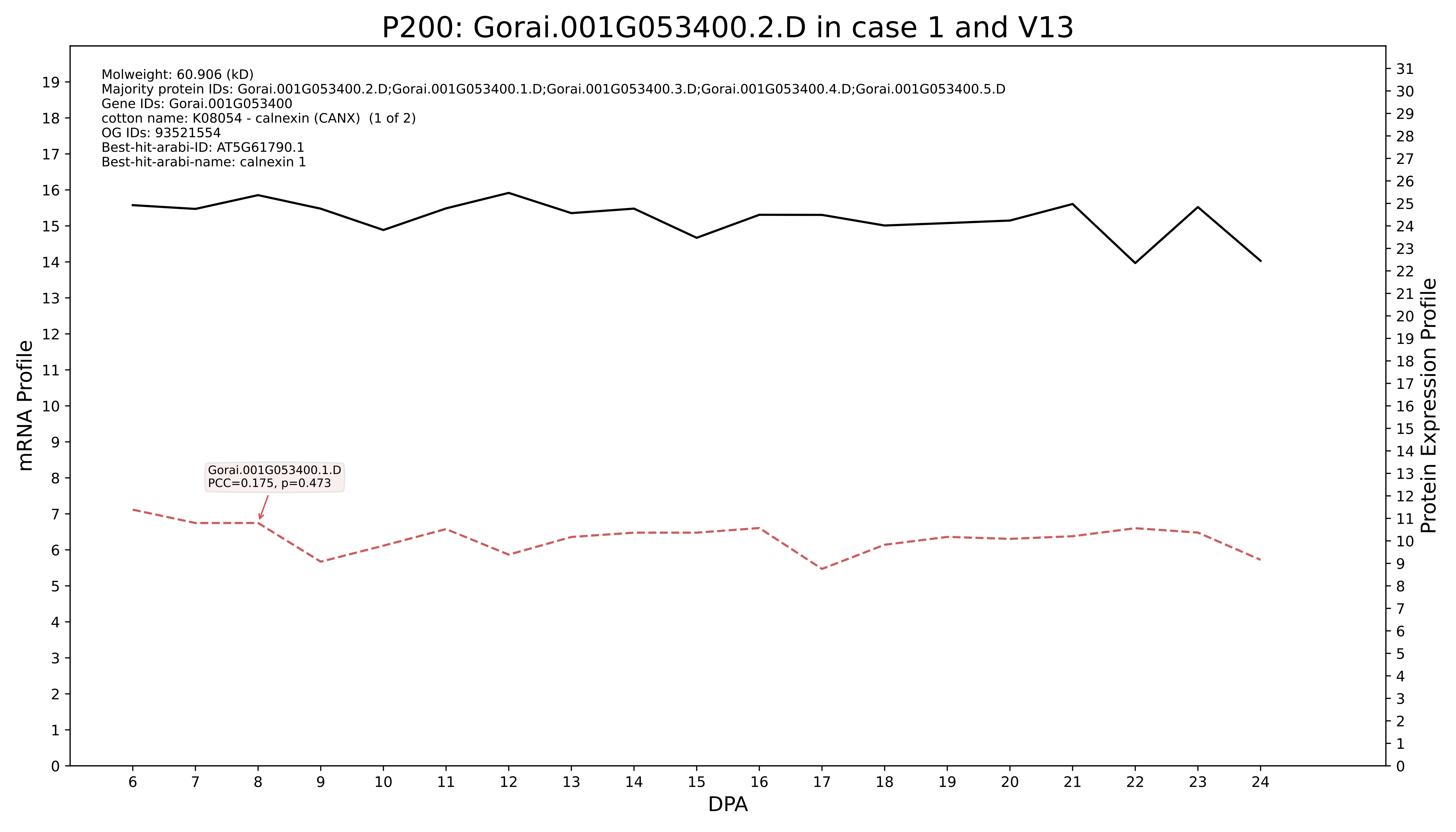
P200: Gorai.001G053400.2.D in case 1 and V13
Molweight: 60.906 (kD)
Majority protein IDs: Gorai.001G053400.2.D;Gorai.001G053400.1.D;Gorai.001G053400.3.D;Gorai.001G053400.4.D;Gorai.001G053400.5.D
Gene IDs: Gorai.001G053400
cotton name: K08054 - calnexin (CANX) (1 of 2)
OG IDs: 93521554
Best-hit-arabi-ID: AT5G61790.1
Best-hit-arabi-name: calnexin 1
Gorai.001G053400.1.D
PCC=0.175, p=0.473
mRNA Profile
Protein Expression Profile
DPA

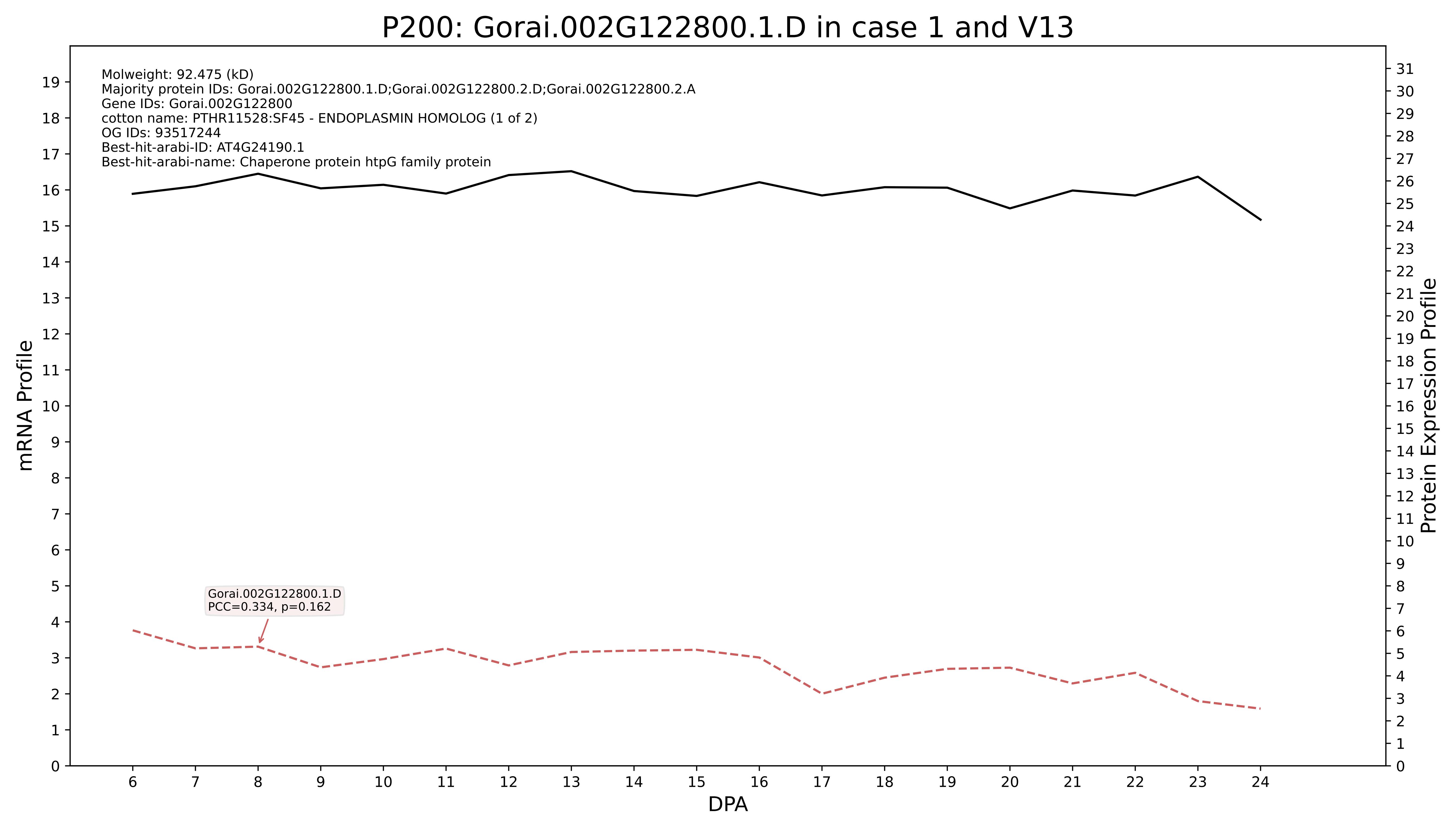

P200: Gorai.002G122800.1.D in case 1 and V13
Molweight: 92.475 (kD)
Majority protein IDs: Gorai.002G122800.1.D;Gorai.002G122800.2.D;Gorai.002G122800.2.A
Gene IDs: Gorai.002G122800
cotton name: PTHR11528:SF45 - ENDOPLASMIN HOMOLOG (1 of 2)
OG IDs: 93517244
Best-hit-arabi-ID: AT4G24190.1
Best-hit-arabi-name: Chaperone protein htpG family protein
Gorai.002G122800.1.D
PCC=0.334, p=0.162
mRNA Profile
Protein Expression Profile
DPA

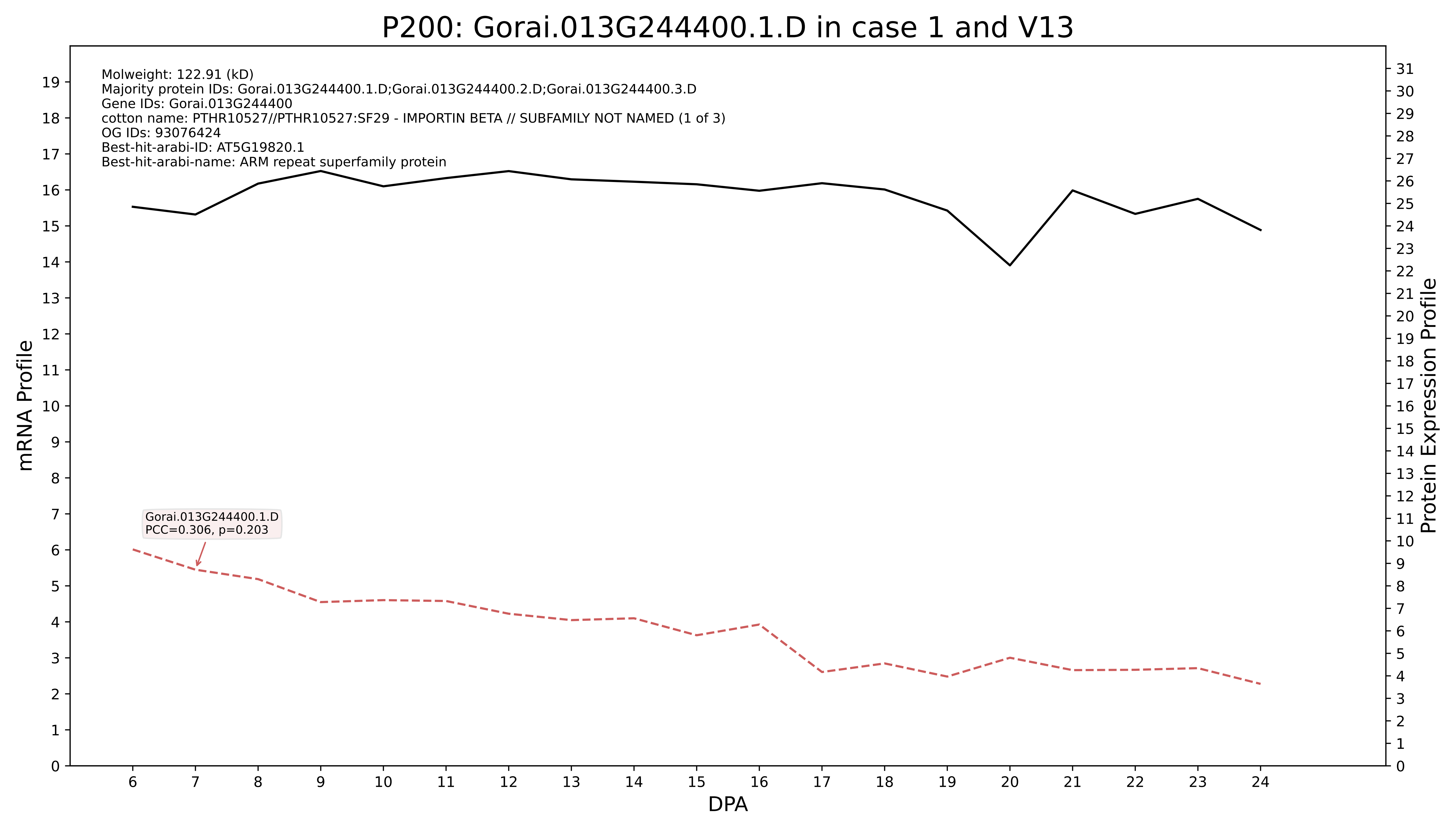
P200: Gorai.013G244400.1.D in case 1 and V13
Molweight: 122.91 (kD)
Majority protein IDs: Gorai.013G244400.1.D;Gorai.013G244400.2.D;Gorai.013G244400.3.D
Gene IDs: Gorai.013G244400
cotton name: PTHR10527//PTHR10527:SF29 - IMPORTIN BETA // SUBFAMILY NOT NAMED (1 of 3)
OG IDs: 93076424
Best-hit-arabi-ID: AT5G19820.1
Best-hit-arabi-name: ARM repeat superfamily protein
Gorai.013G244400.1.D
PCC=0.306, p=0.203
mRNA Profile
Protein Expression Profile
DPA

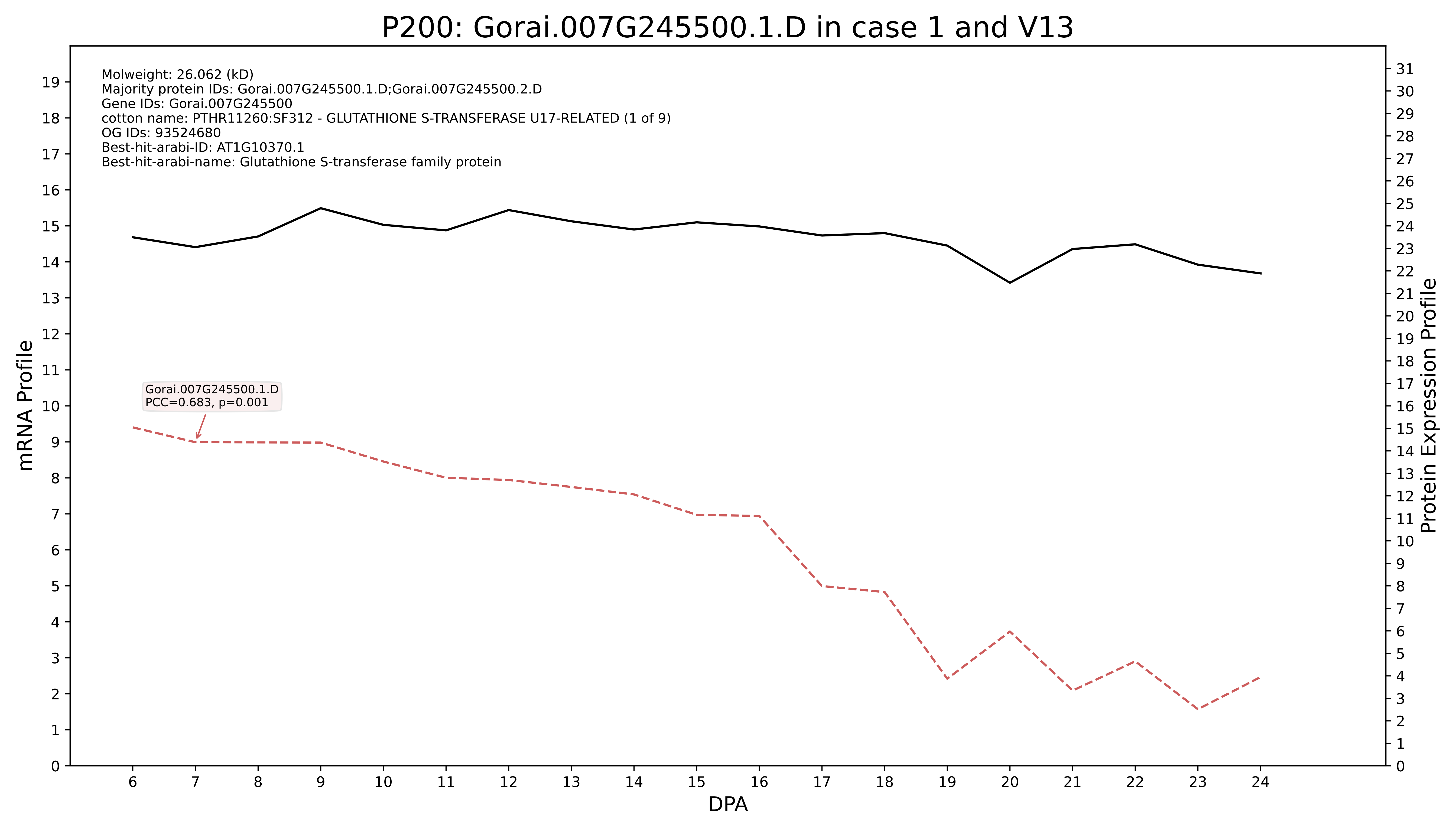

P200: Gorai.007G245500.1.D in case 1 and V13
Molweight: 26.062 (kD)
Majority protein IDs: Gorai.007G245500.1.D;Gorai.007G245500.2.D
Gene IDs: Gorai.007G245500
cotton name: PTHR11260:SF312 - GLUTATHIONE S-TRANSFERASE U17-RELATED (1 of 9)
OG IDs: 93524680
Best-hit-arabi-ID: AT1G10370.1
Best-hit-arabi-name: Glutathione S-transferase family protein
Gorai.007G245500.1.D
PCC=0.683, p=0.001
mRNA Profile
Protein Expression Profile
DPA

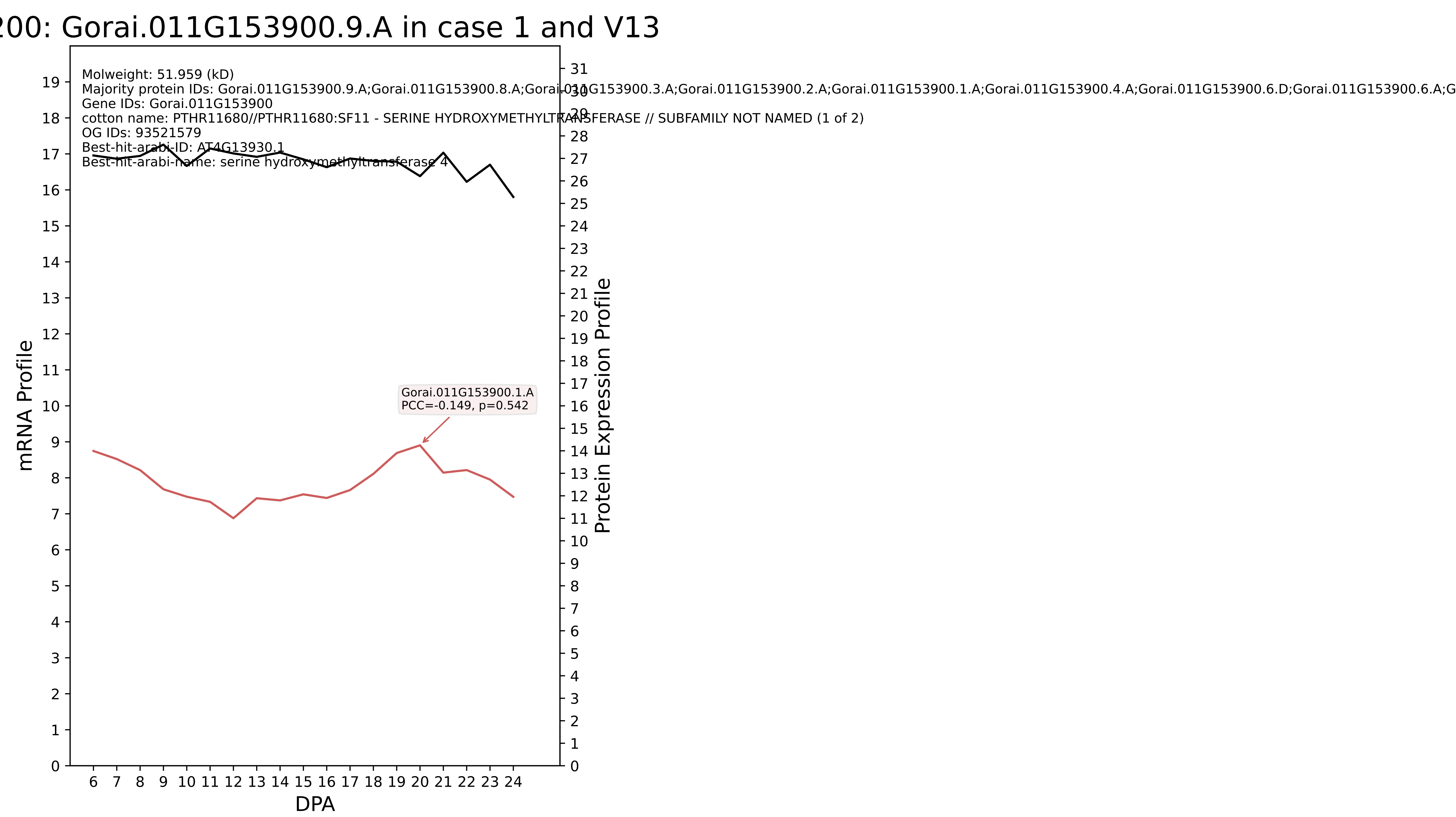
200: Gorai.011G153900.9.A in case 1 and V13
Molweight: 51.959 (kD)
Majority protein IDs: Gorai.011G153900.9.A;Gorai.011G153900.8.A;Gorai.011G153900.3.A;Gorai.011G153900.2.A;Gorai.011G153900.1.A;Gorai.011G153900.4.A;Gorai.011G153900.6.D;Gorai.011G153900.6.A;G
Gene IDs: Gorai.011G153900
cotton name: PTHR11680//PTHR11680:SF11 - SERINE HYDROXYMETHYLTRANSFERASE // SUBFAMILY NOT NAMED (1 of 2)
OG IDs: 93521579
Best-hit-arabi-ID: AT4G13930.1
Best-hit-arabi-name: serine hydroxymethyltransferase 4
Gorai.011G153900.1.A
PCC=-0.149, p=0.542
mRNA Profile
Protein Expression Profile
DPA

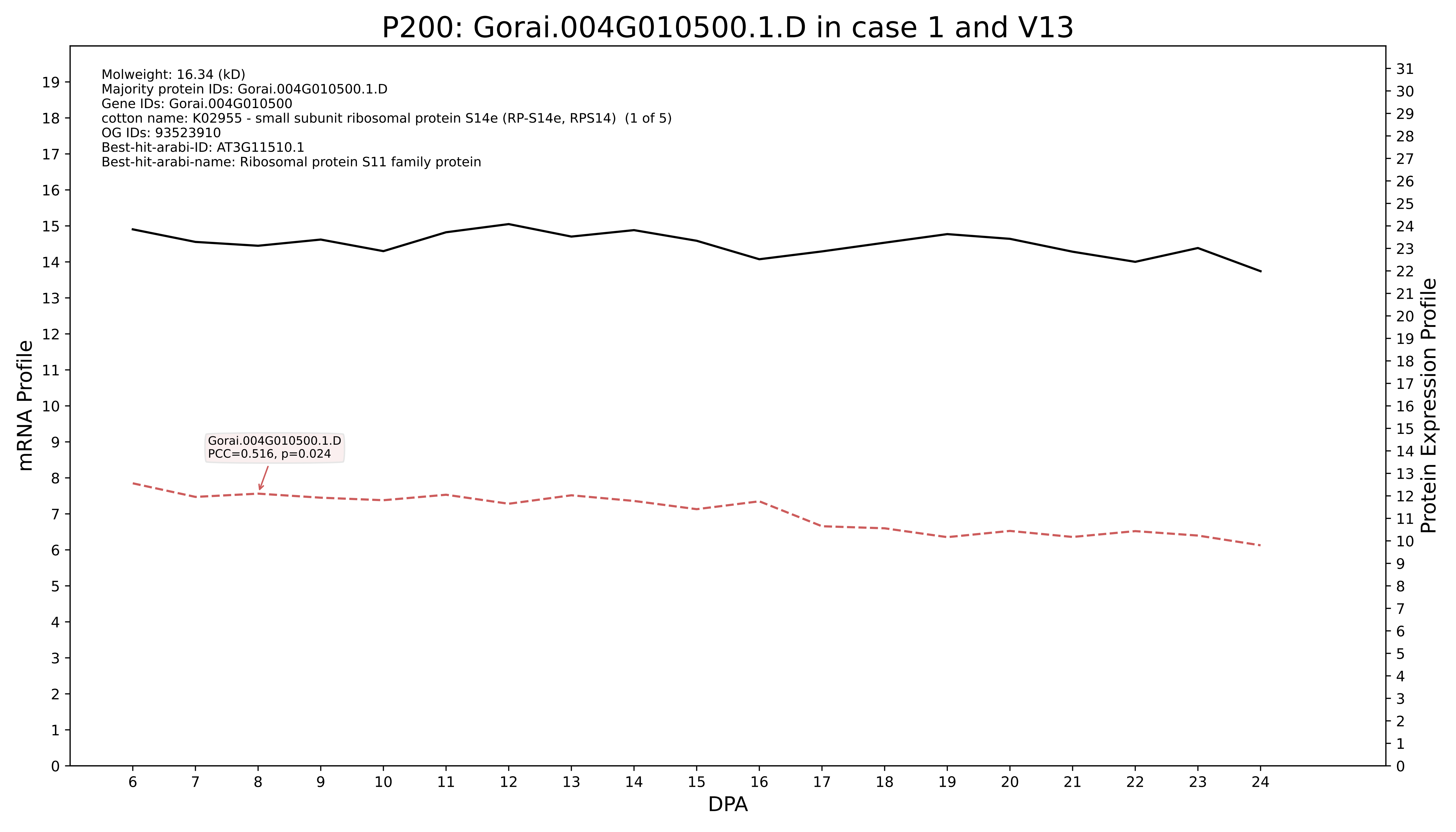
P200: Gorai.004G010500.1.D in case 1 and V13
Molweight: 16.34 (kD)
Majority protein IDs: Gorai.004G010500.1.D
Gene IDs: Gorai.004G010500
cotton name: K02955 - small subunit ribosomal protein S14e (RP-S14e, RPS14) (1 of 5)
OG IDs: 93523910
Best-hit-arabi-ID: AT3G11510.1
Best-hit-arabi-name: Ribosomal protein S11 family protein
Gorai.004G010500.1.D
PCC=0.516, p=0.024
mRNA Profile
Protein Expression Profile
DPA

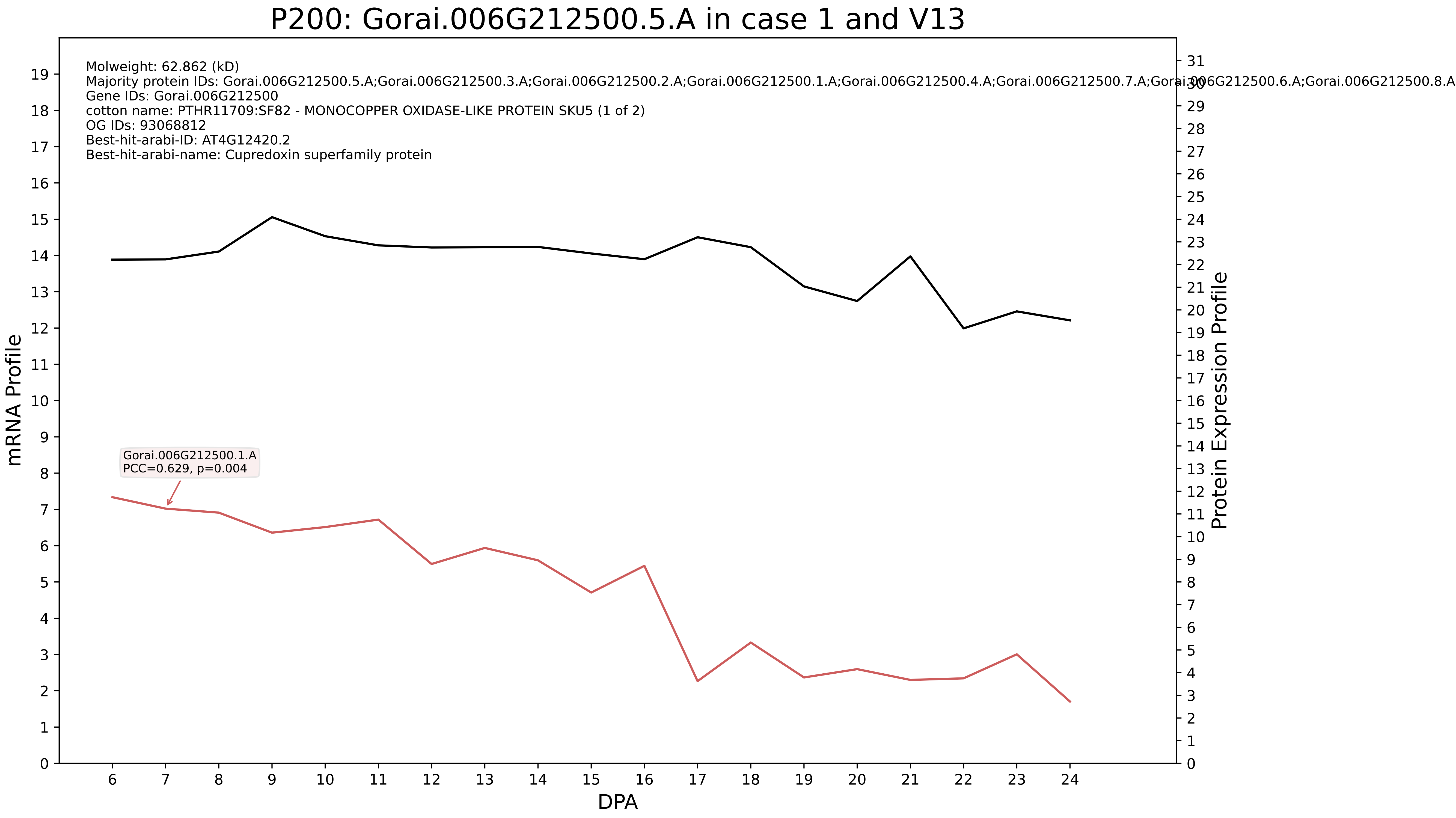

P200: Gorai.006G212500.5.A in case 1 and V13
Molweight: 62.862 (kD)
Majority protein IDs: Gorai.006G212500.5.A;Gorai.006G212500.3.A;Gorai.006G212500.2.A;Gorai.006G212500.1.A;Gorai.006G212500.4.A;Gorai.006G212500.7.A;Gorai.006G212500.6.A;Gorai.006G212500.8.A
Gene IDs: Gorai.006G212500
cotton name: PTHR11709:SF82 - MONOCOPPER OXIDASE-LIKE PROTEIN SKU5 (1 of 2)
OG IDs: 93068812
Best-hit-arabi-ID: AT4G12420.2
Best-hit-arabi-name: Cupredoxin superfamily protein
Gorai.006G212500.1.A
PCC=0.629, p=0.004
mRNA Profile
Protein Expression Profile
DPA

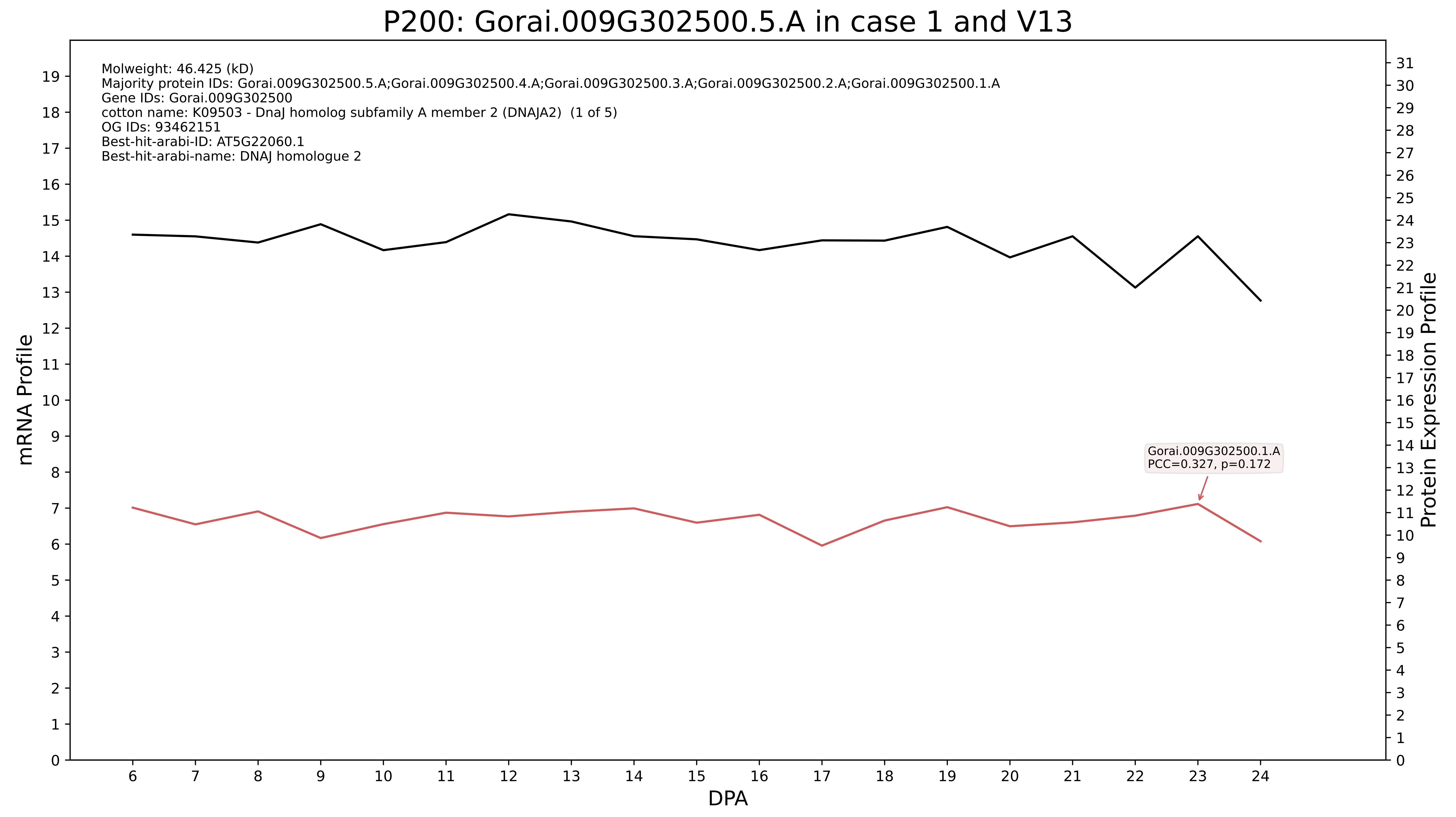
P200: Gorai.009G302500.5.A in case 1 and V13
Molweight: 46.425 (kD)
Majority protein IDs: Gorai.009G302500.5.A;Gorai.009G302500.4.A;Gorai.009G302500.3.A;Gorai.009G302500.2.A;Gorai.009G302500.1.A
Gene IDs: Gorai.009G302500
cotton name: K09503 - DnaJ homolog subfamily A member 2 (DNAJA2) (1 of 5)
OG IDs: 93462151
Best-hit-arabi-ID: AT5G22060.1
Best-hit-arabi-name: DNAJ homologue 2
Gorai.009G302500.1.A
PCC=0.327, p=0.172
mRNA Profile
Protein Expression Profile
DPA

### P200: Gorai.009G038000.6.D in case 1 and V13

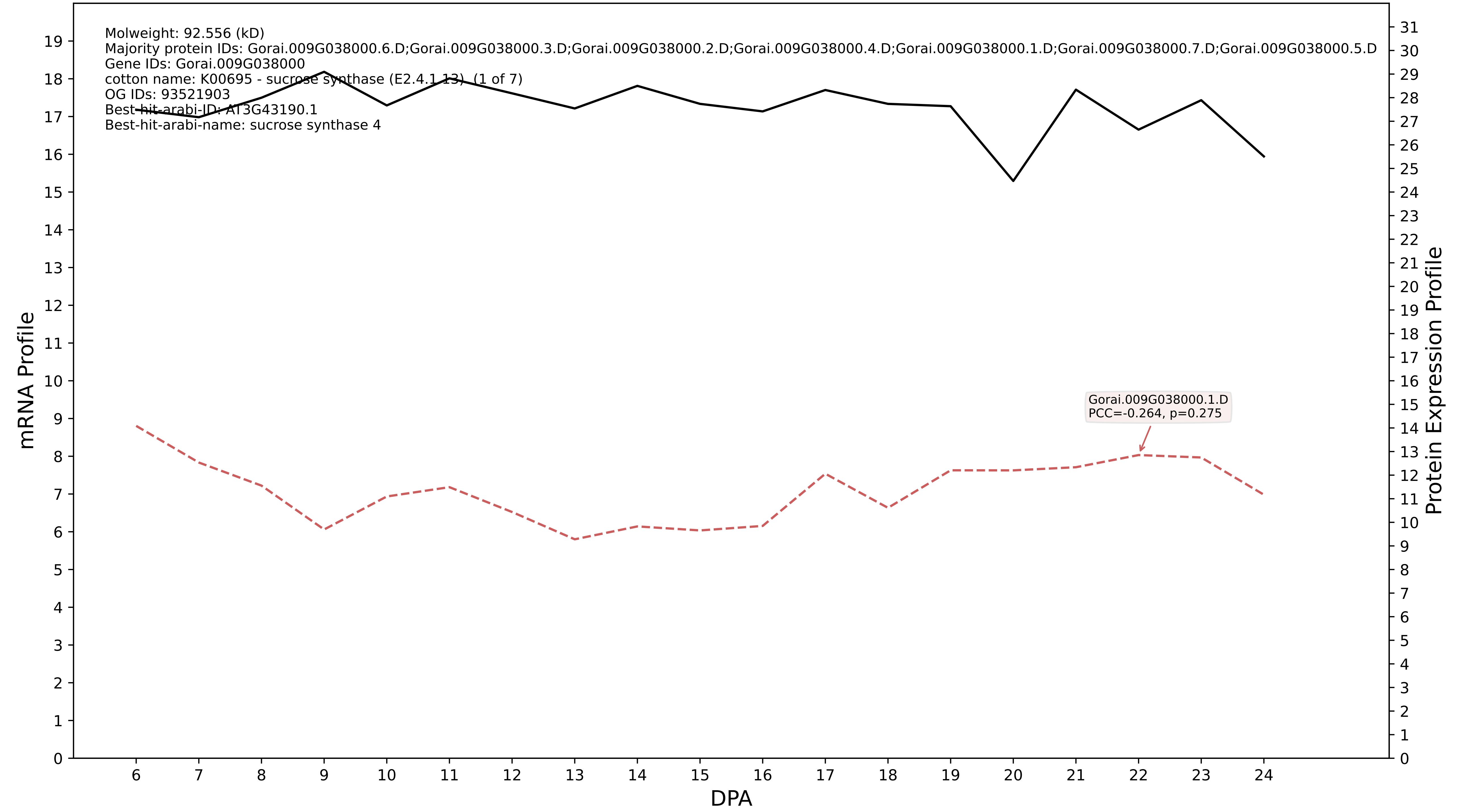

### P200: Gorai.004G095400.6.A in case 1 and V13

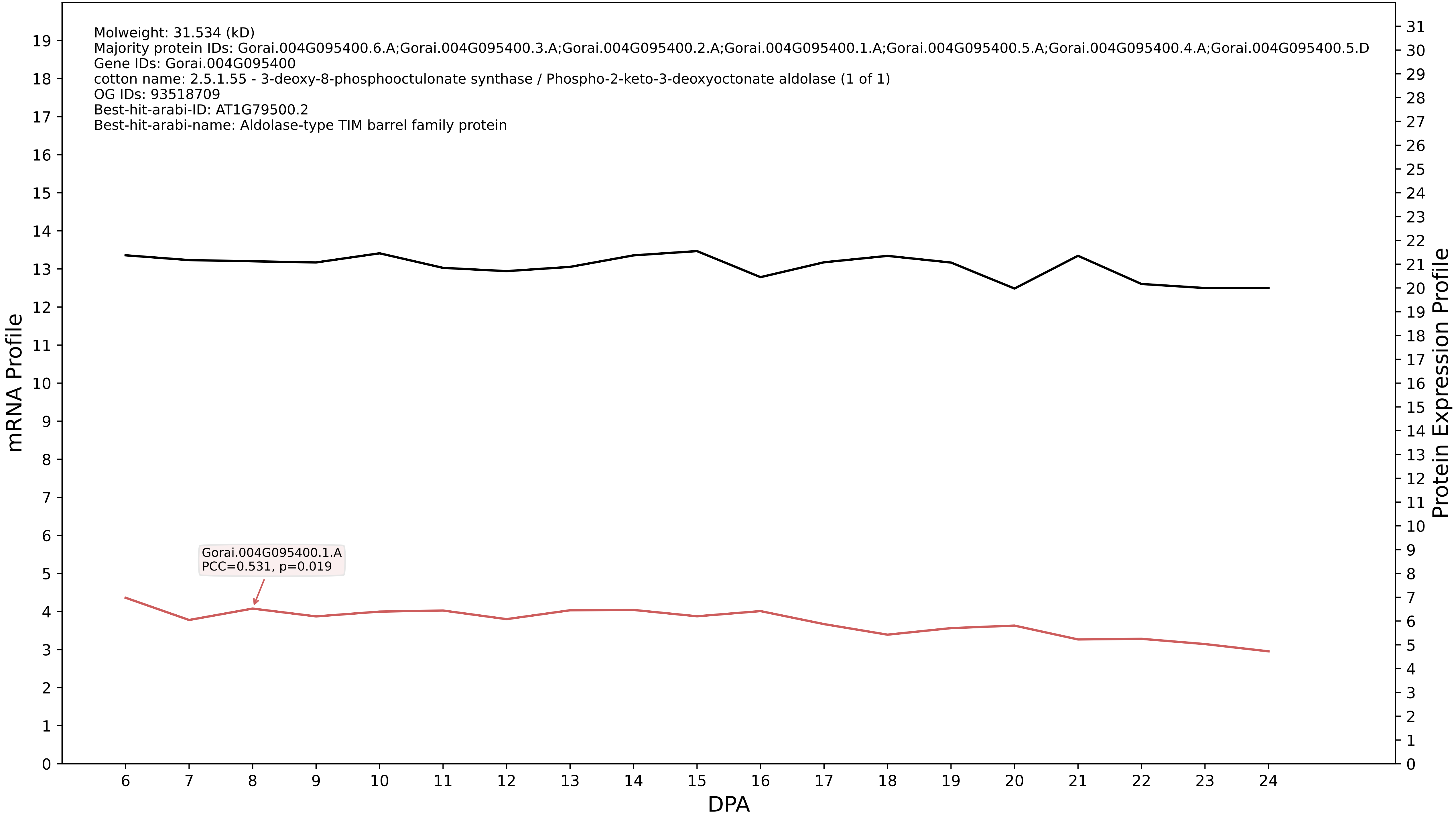

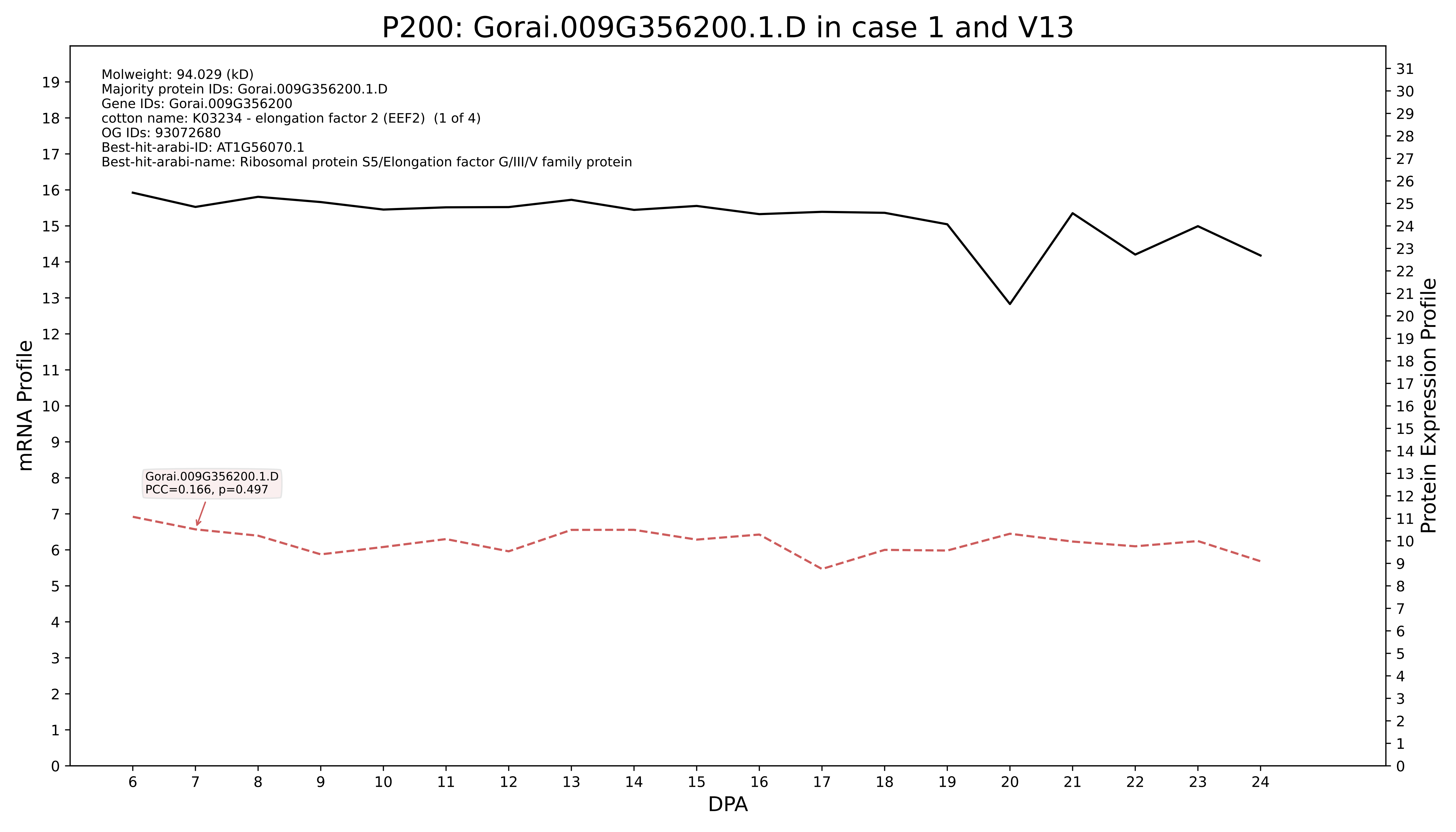
P200: Gorai.009G356200.1.D in case 1 and V13
Molweight: 94.029 (kD)
Majority protein IDs: Gorai.009G356200.1.D
Gene IDs: Gorai.009G356200
cotton name: K03234 - elongation factor 2 (EEF2) (1 of 4)
OG IDs: 93072680
Best-hit-arabi-ID: AT1G56070.1
Best-hit-arabi-name: Ribosomal protein S5/Elongation factor G/III/V family protein
Gorai.009G356200.1.D
PCC=0.166, p=0.497
mRNA Profile
Protein Expression Profile
DPA

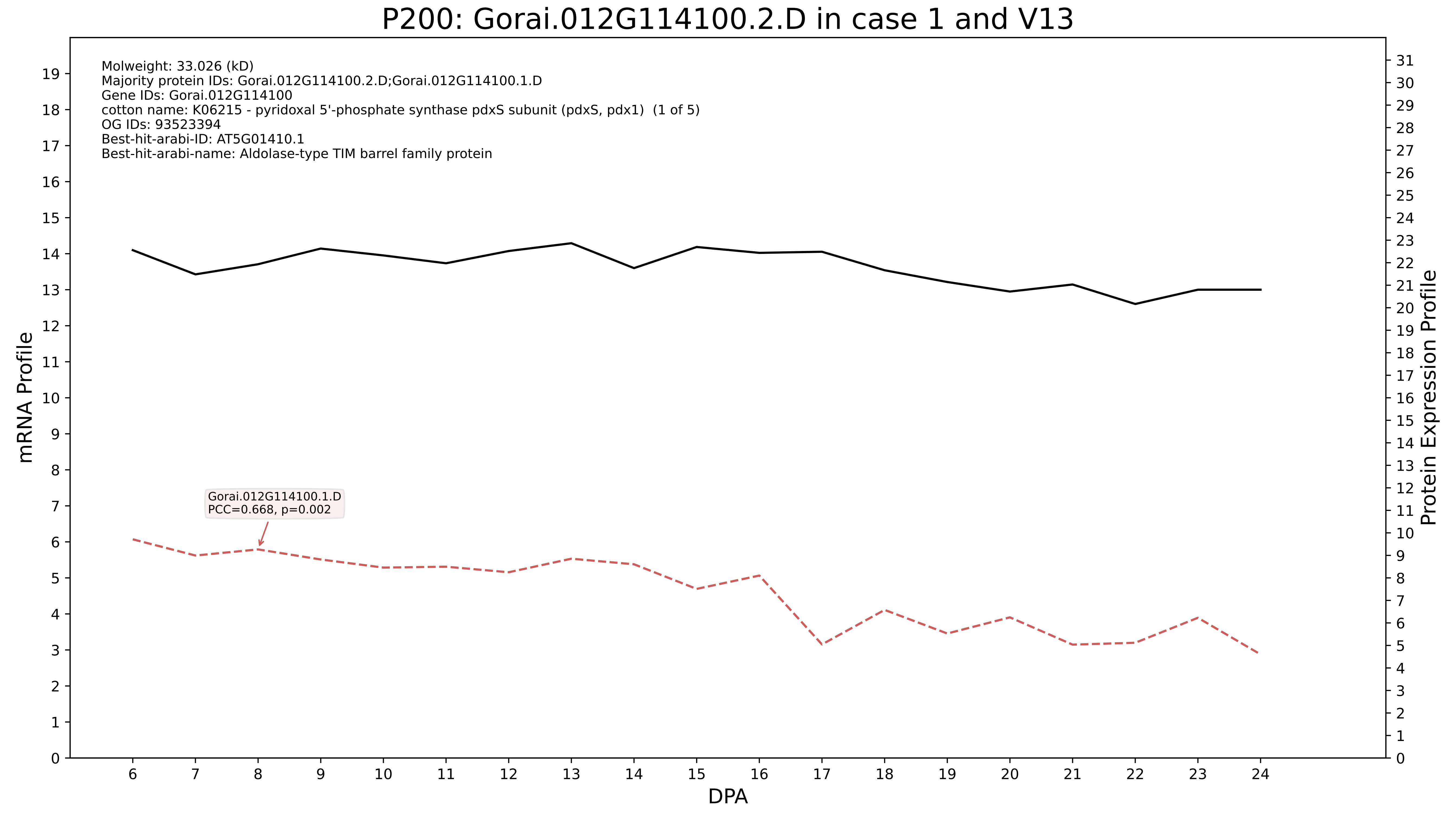

P200: Gorai.012G114100.2.D in case 1 and V13
Molweight: 33.026 (kD)
Majority protein IDs: Gorai.012G114100.2.D;Gorai.012G114100.1.D
Gene IDs: Gorai.012G114100
cotton name: K06215 - pyridoxal 5'-phosphate synthase pdxS subunit (pdxS, pdx1) (1 of 5)
OG IDs: 93523394
Best-hit-arabi-ID: AT5G01410.1
Best-hit-arabi-name: Aldolase-type TIM barrel family protein
Gorai.012G114100.1.D
PCC=0.668, p=0.002
mRNA Profile
Protein Expression Profile
DPA

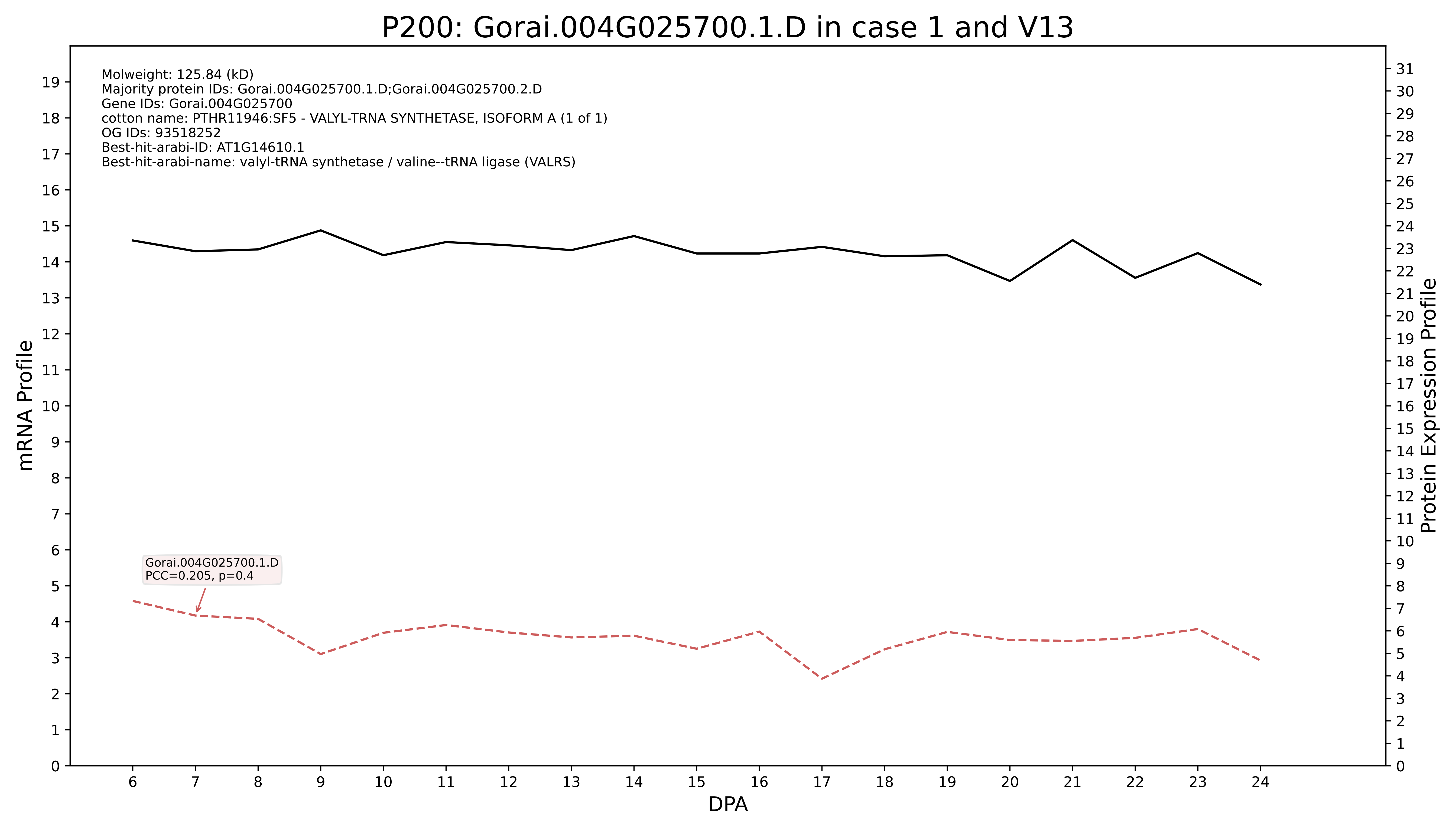

P200: Gorai.004G025700.1.D in case 1 and V13
Molweight: 125.84 (kD)
Majority protein IDs: Gorai.004G025700.1.D;Gorai.004G025700.2.D
Gene IDs: Gorai.004G025700
cotton name: PTHR11946:SF5 - VALYL-TRNA SYNTHETASE, ISOFORM A (1 of 1)
OG IDs: 93518252
Best-hit-arabi-ID: AT1G14610.1
Best-hit-arabi-name: valyl-tRNA synthetase / valine--tRNA ligase (VALRS)
Gorai.004G025700.1.D
PCC=0.205, p=0.4
mRNA Profile
Protein Expression Profile
DPA

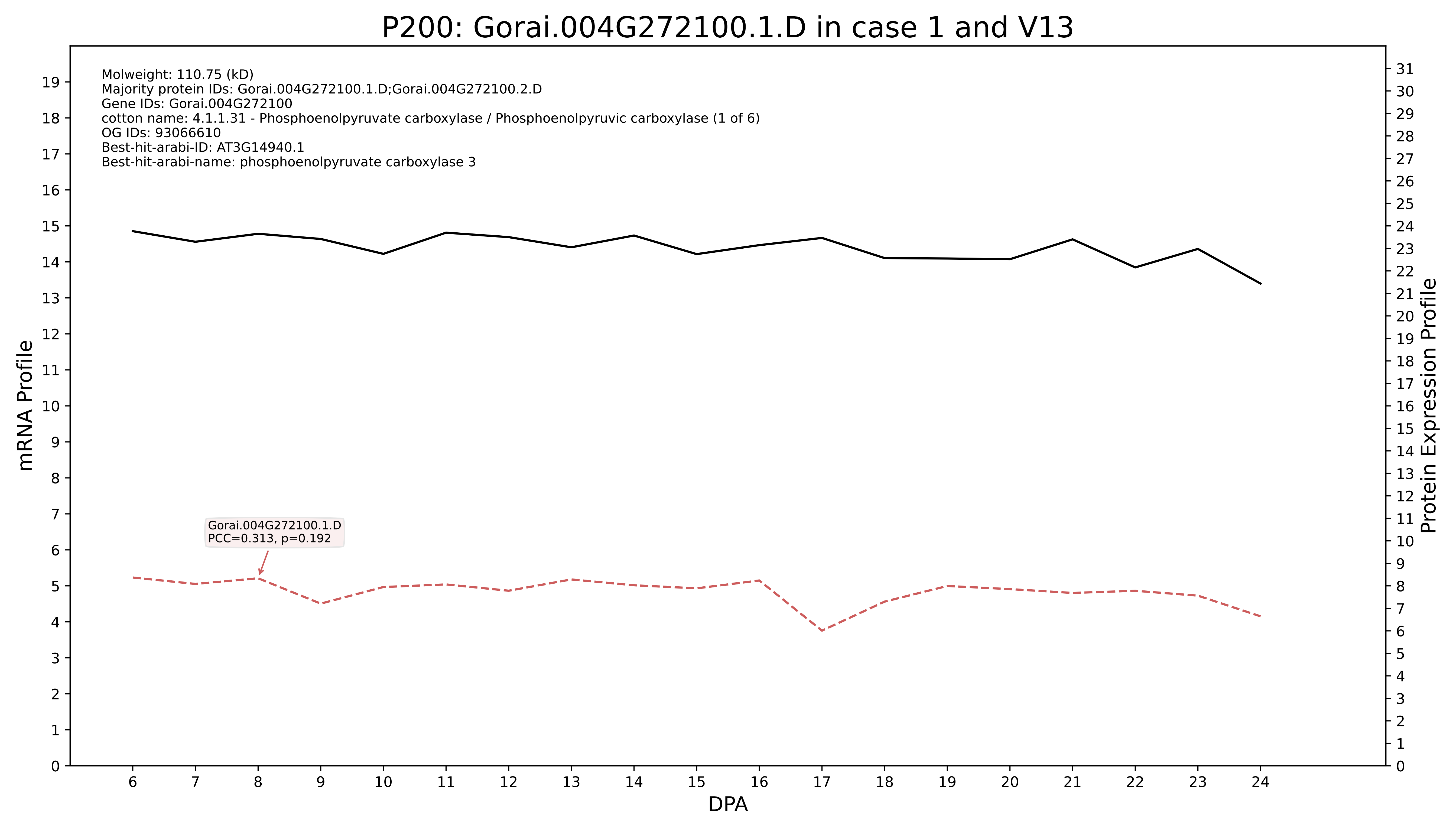

P200: Gorai.004G272100.1.D in case 1 and V13
Molweight: 110.75 (kD)
Majority protein IDs: Gorai.004G272100.1.D;Gorai.004G272100.2.D
Gene IDs: Gorai.004G272100
cotton name: 4.1.1.31 - Phosphoenolpyruvate carboxylase / Phosphoenolpyruvic carboxylase (1 of 6)
OG IDs: 93066610
Best-hit-arabi-ID: AT3G14940.1
Best-hit-arabi-name: phosphoenolpyruvate carboxylase 3
Gorai.004G272100.1.D
PCC=0.313, p=0.192
mRNA Profile
Protein Expression Profile
DPA

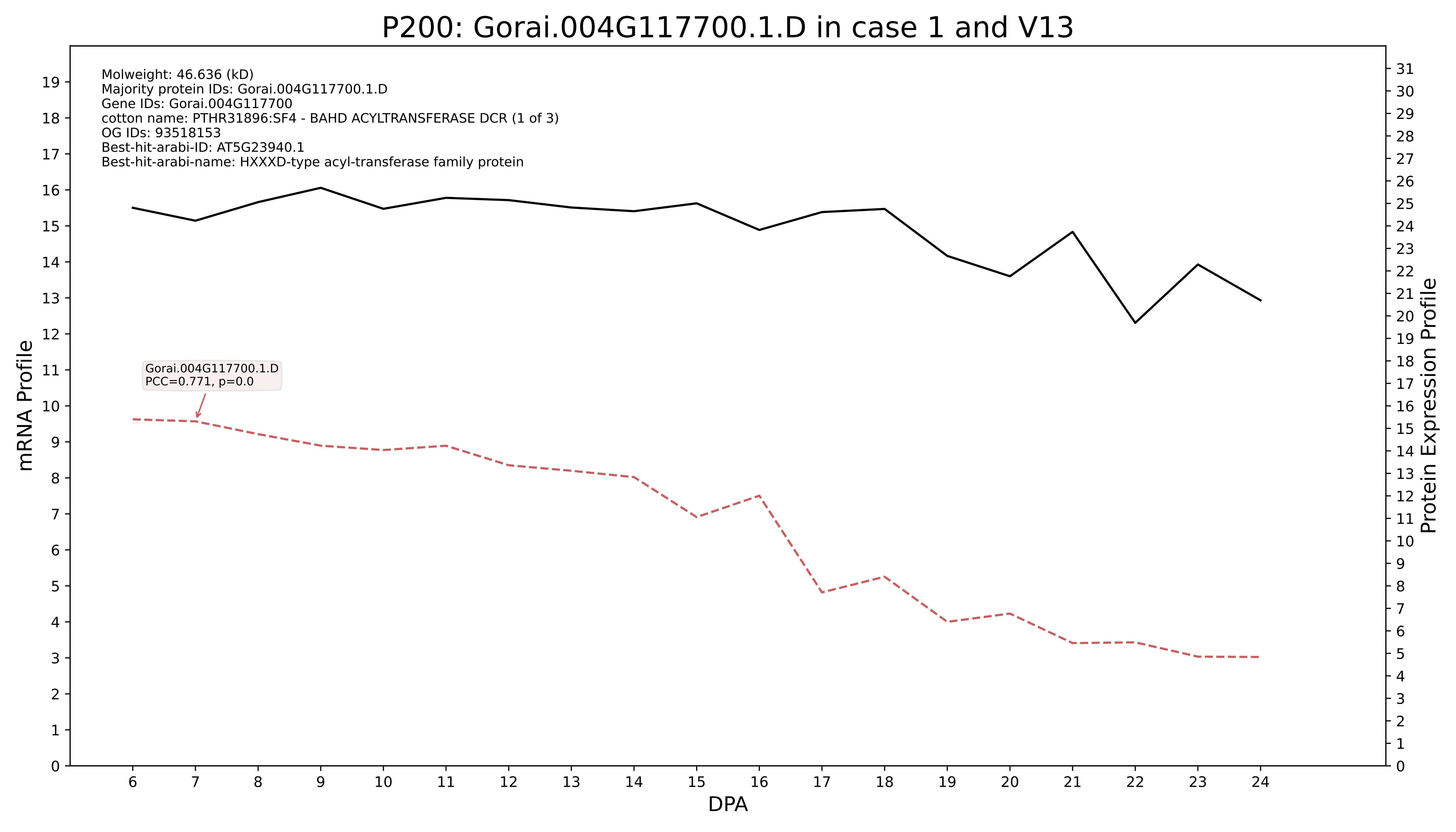

P200: Gorai.004G117700.1.D in case 1 and V13
Molweight: 46.636 (kD)
Majority protein IDs: Gorai.004G117700.1.D
Gene IDs: Gorai.004G117700
cotton name: PTHR31896:SF4 - BAHD ACYLTRANSFERASE DCR (1 of 3)
OG IDs: 93518153
Best-hit-arabi-ID: AT5G23940.1
Best-hit-arabi-name: HXXXD-type acyl-transferase family protein
Gorai.004G117700.1.D
PCC=0.771, p=0.0
mRNA Profile
Protein Expression Profile
DPA

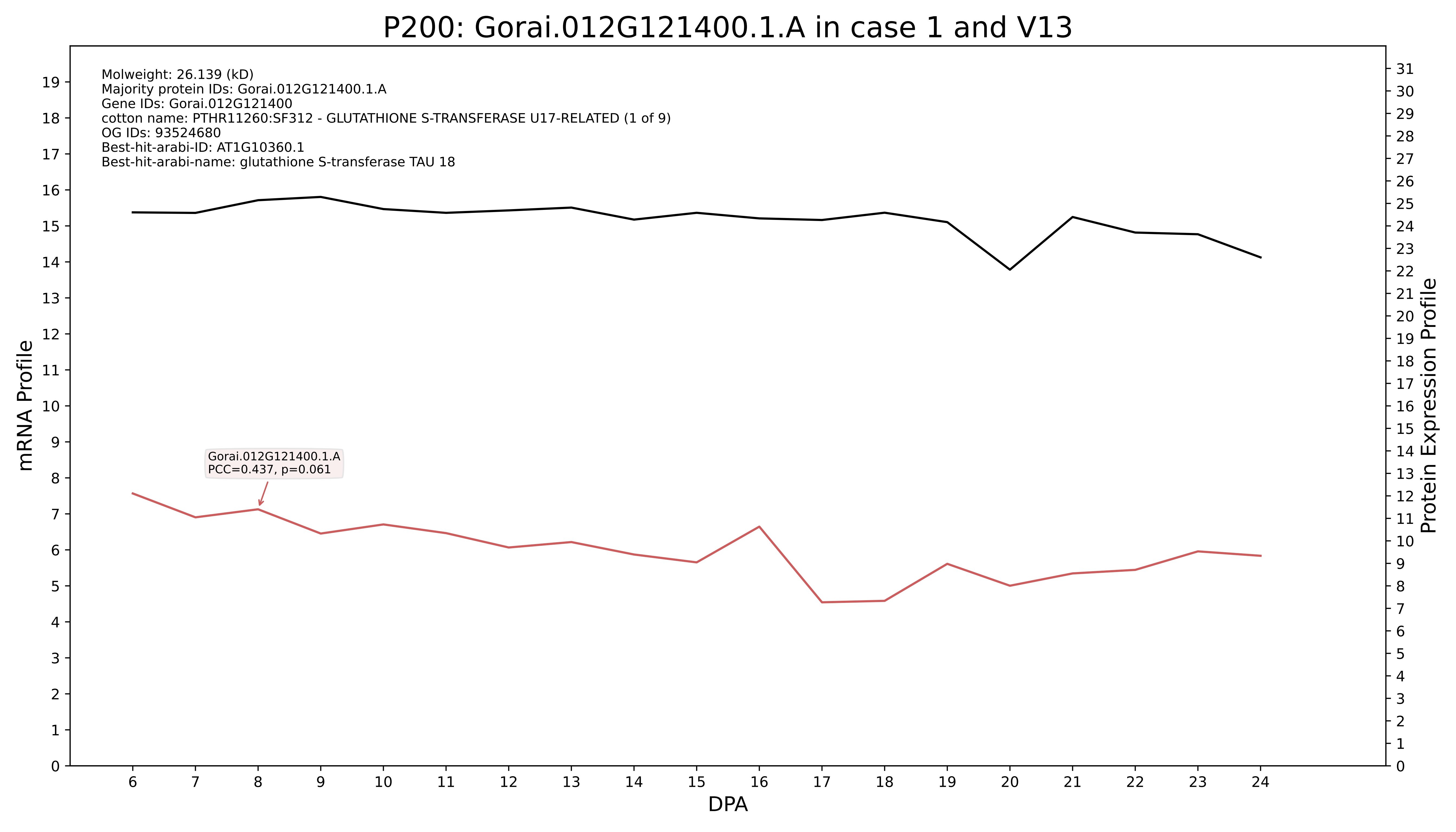
P200: Gorai.012G121400.1.A in case 1 and V13
Molweight: 26.139 (kD)
Majority protein IDs: Gorai.012G121400.1.A
Gene IDs: Gorai.012G121400
cotton name: PTHR11260:SF312 - GLUTATHIONE S-TRANSFERASE U17-RELATED (1 of 9)
OG IDs: 93524680
Best-hit-arabi-ID: AT1G10360.1
Best-hit-arabi-name: glutathione S-transferase TAU 18
Gorai.012G121400.1.A
PCC=0.437, p=0.061
mRNA Profile
Protein Expression Profile
DPA

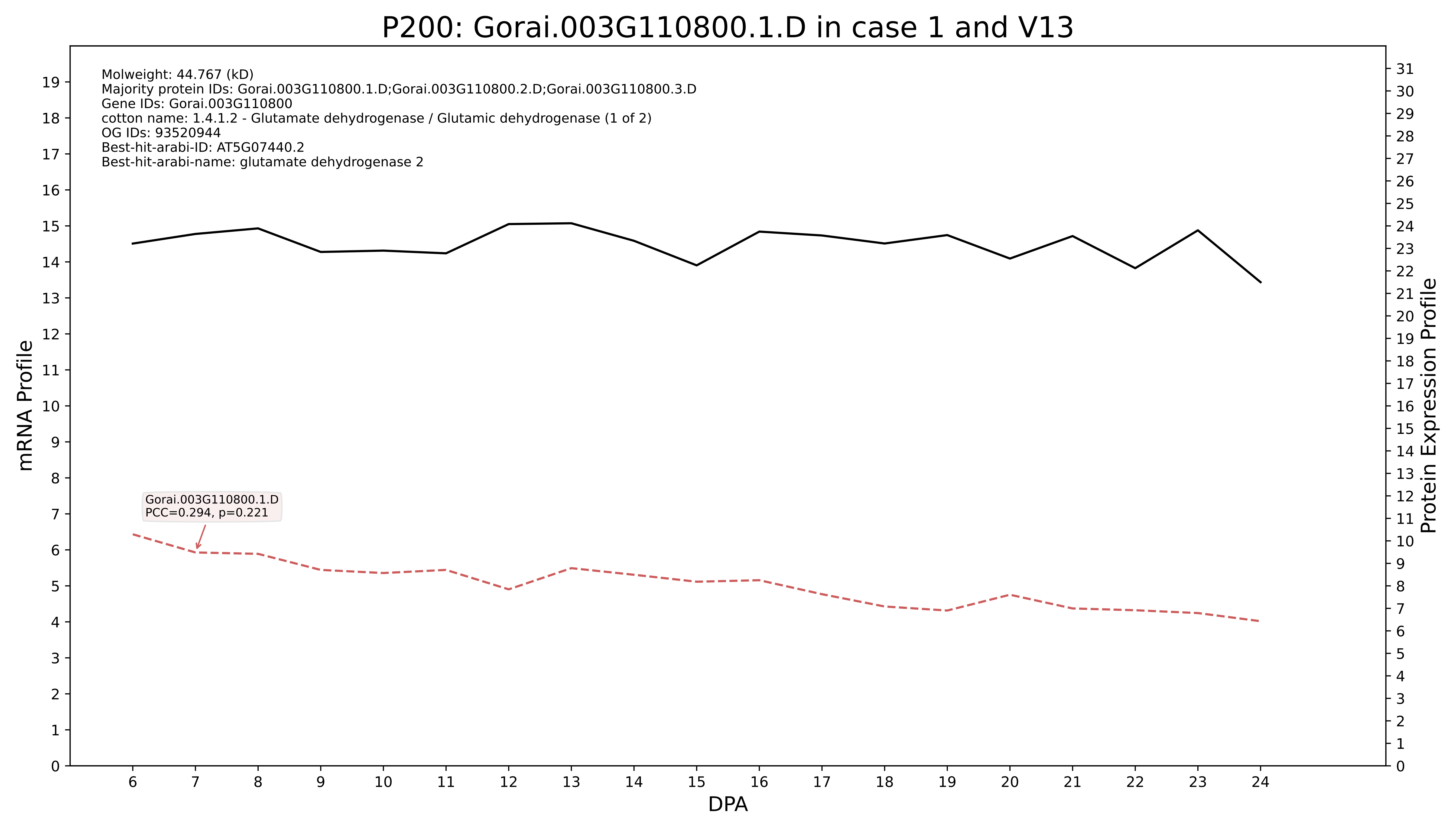
P200: Gorai.003G110800.1.D in case 1 and V13
Molweight: 44.767 (kD)
Majority protein IDs: Gorai.003G110800.1.D;Gorai.003G110800.2.D;Gorai.003G110800.3.D
Gene IDs: Gorai.003G110800
cotton name: 1.4.1.2 - Glutamate dehydrogenase / Glutamic dehydrogenase (1 of 2)
OG IDs: 93520944
Best-hit-arabi-ID: AT5G07440.2
Best-hit-arabi-name: glutamate dehydrogenase 2
Gorai.003G110800.1.D
PCC=0.294, p=0.221
mRNA Profile
Protein Expression Profile
DPA

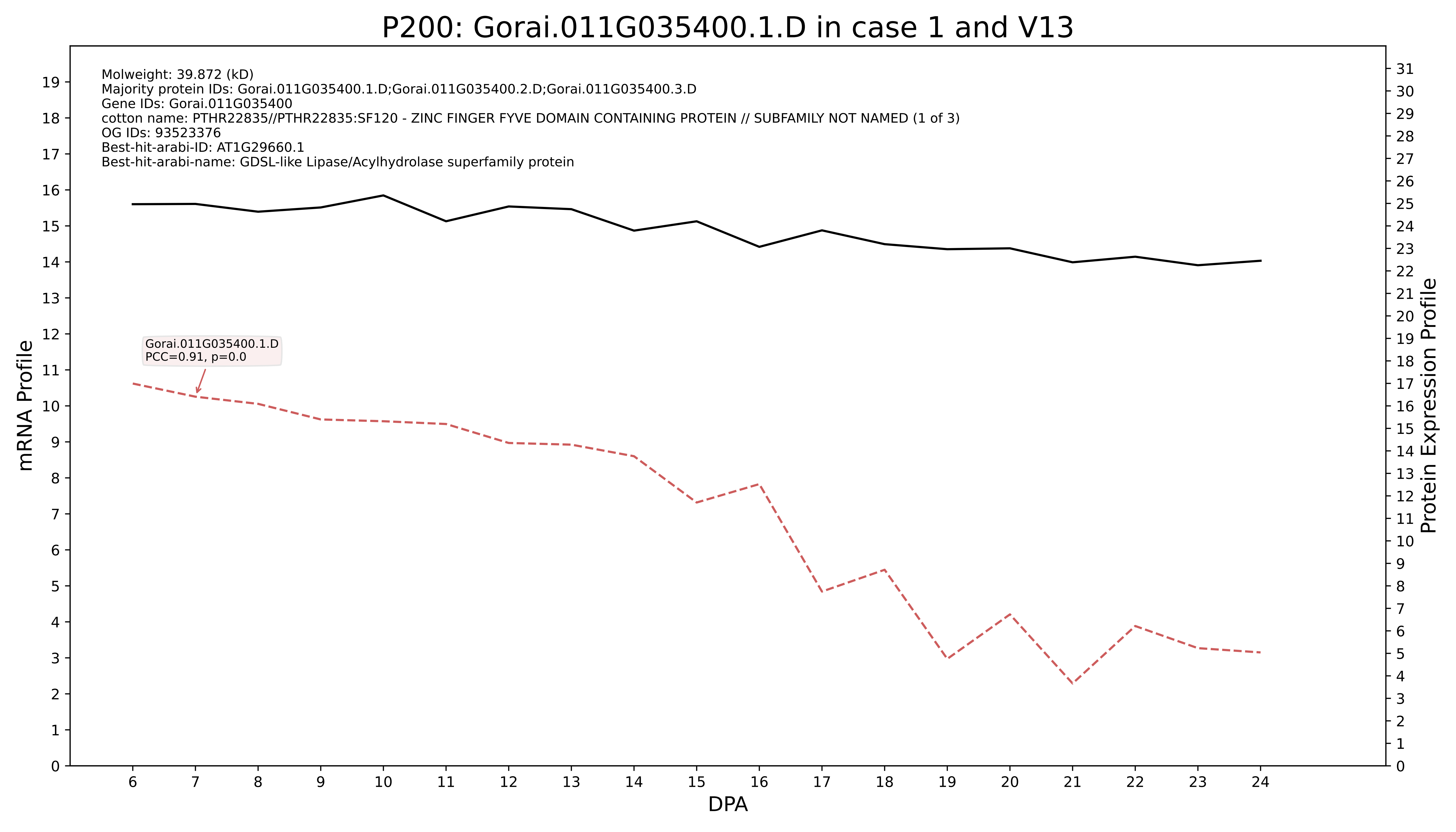
P200: Gorai.011G035400.1.D in case 1 and V13
Molweight: 39.872 (kD)
Majority protein IDs: Gorai.011G035400.1.D;Gorai.011G035400.2.D;Gorai.011G035400.3.D
Gene IDs: Gorai.011G035400
cotton name: PTHR22835//PTHR22835:SF120 - ZINC FINGER FYVE DOMAIN CONTAINING PROTEIN // SUBFAMILY NOT NAMED (1 of 3)
OG IDs: 93523376
Best-hit-arabi-ID: AT1G29660.1
Best-hit-arabi-name: GDSL-like Lipase/Acylhydrolase superfamily protein
Gorai.011G035400.1.D
PCC=0.91, p=0.0
mRNA Profile
Protein Expression Profile
DPA

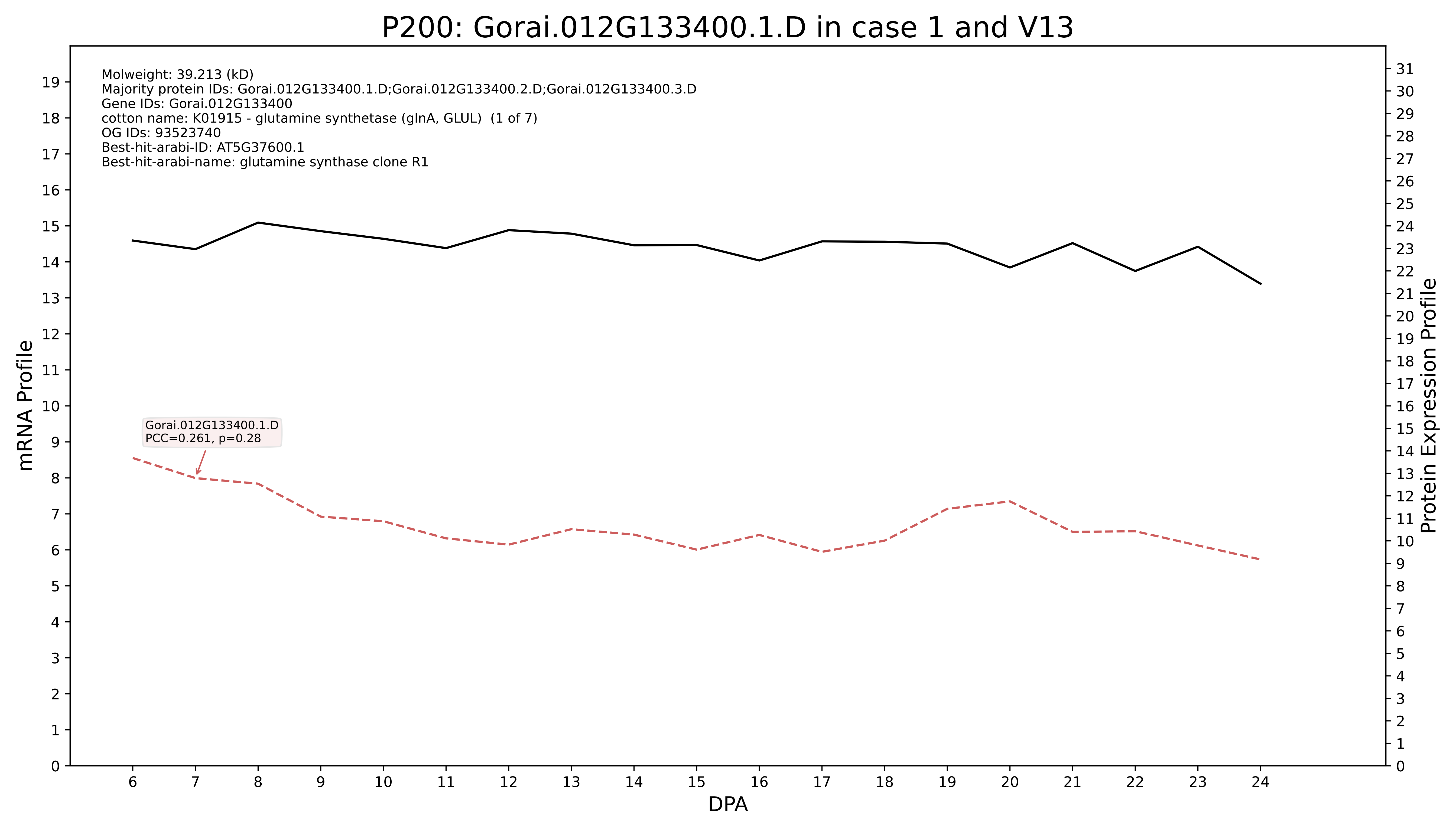

P200: Gorai.012G133400.1.D in case 1 and V13
Molweight: 39.213 (kD)
Majority protein IDs: Gorai.012G133400.1.D;Gorai.012G133400.2.D;Gorai.012G133400.3.D
Gene IDs: Gorai.012G133400
cotton name: K01915 - glutamine synthetase (glnA, GLUL) (1 of 7)
OG IDs: 93523740
Best-hit-arabi-ID: AT5G37600.1
Best-hit-arabi-name: glutamine synthase clone R1
Gorai.012G133400.1.D
PCC=0.261, p=0.28
mRNA Profile
Protein Expression Profile
DPA

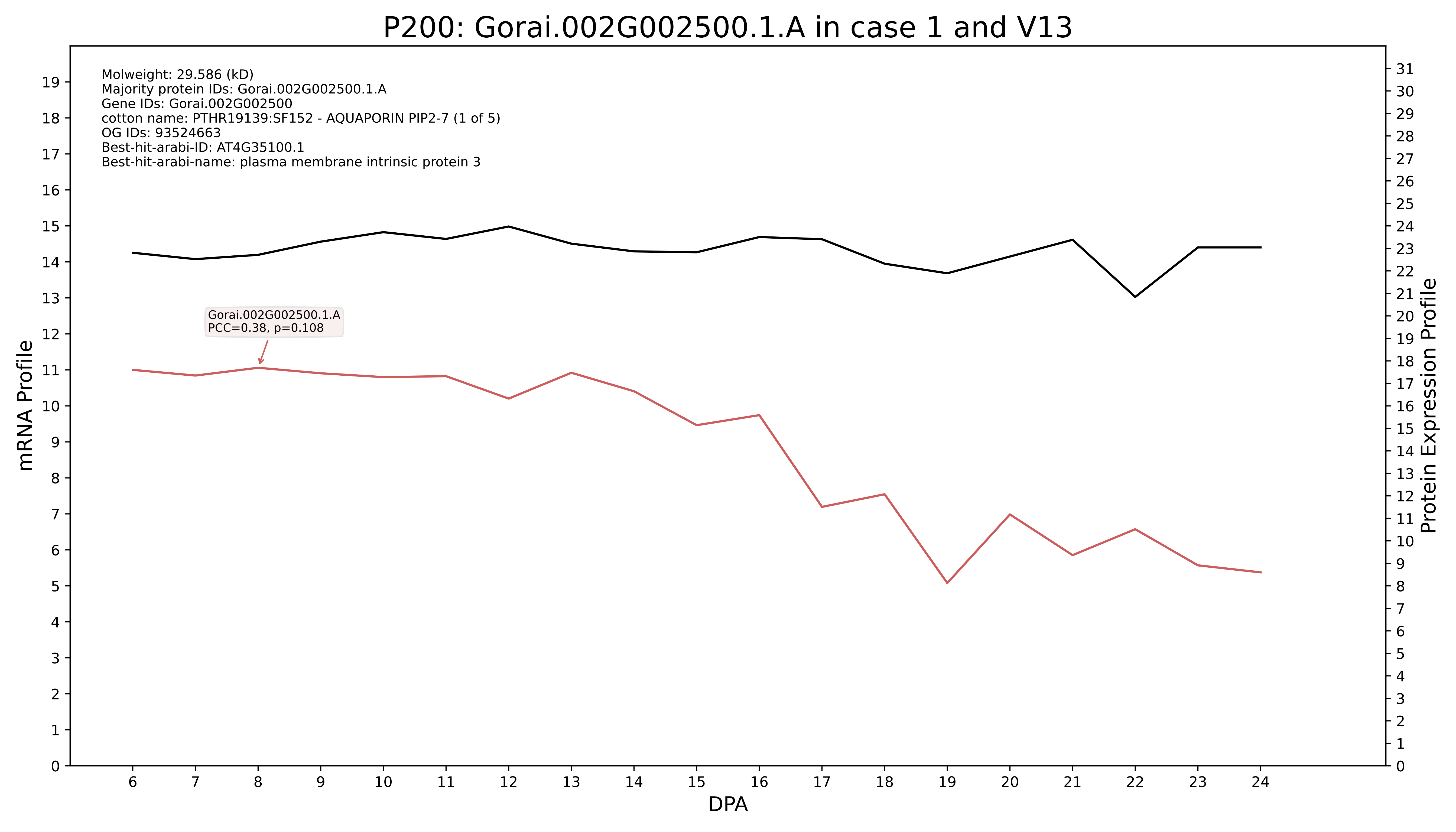
P200: Gorai.002G002500.1.A in case 1 and V13
Molweight: 29.586 (kD)
Majority protein IDs: Gorai.002G002500.1.A
Gene IDs: Gorai.002G002500
cotton name: PTHR19139:SF152 - AQUAPORIN PIP2-7 (1 of 5)
OG IDs: 93524663
Best-hit-arabi-ID: AT4G35100.1
Best-hit-arabi-name: plasma membrane intrinsic protein 3
Gorai.002G002500.1.A
PCC=0.38, p=0.108
mRNA Profile
Protein Expression Profile
DPA

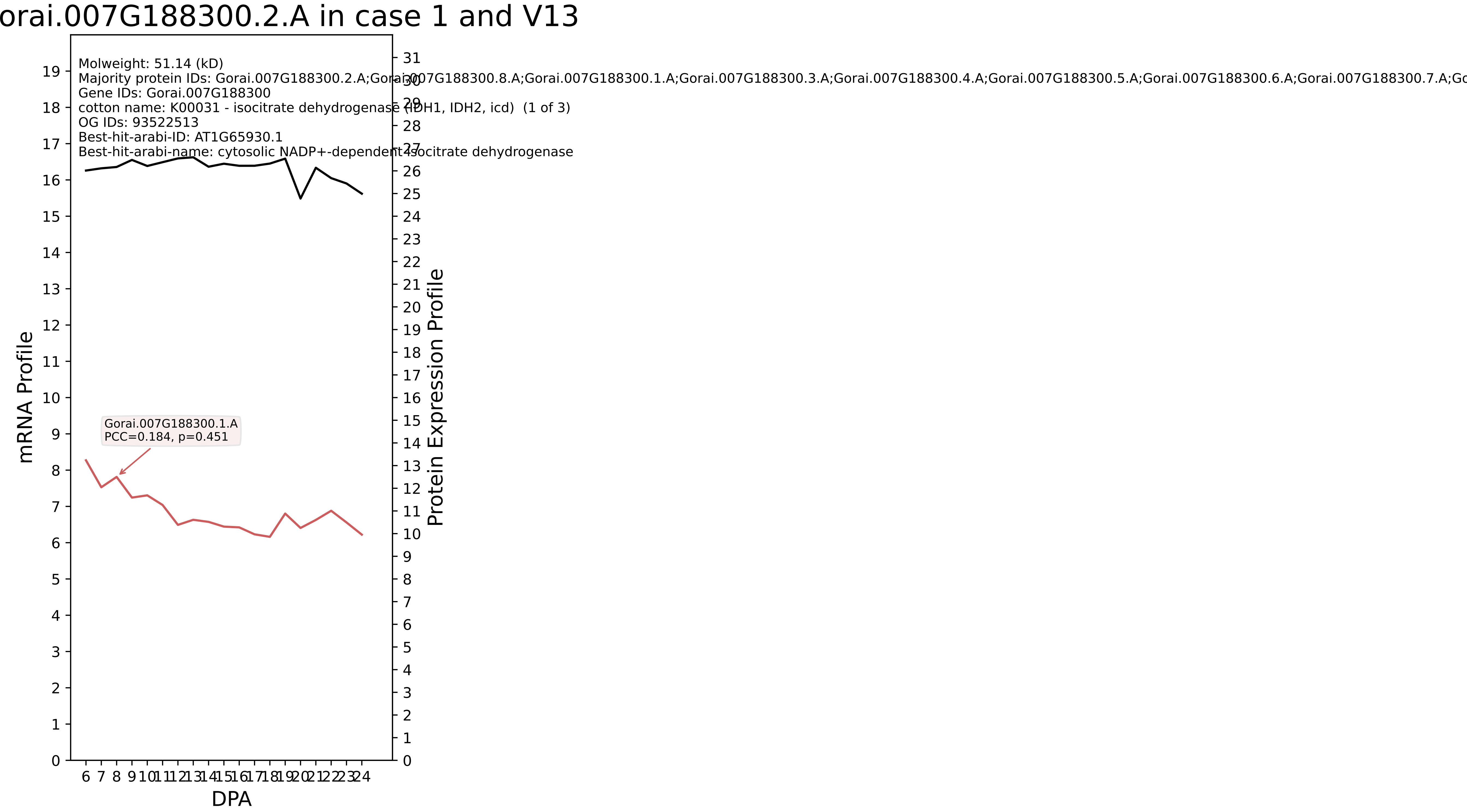
orai.007G188300.2.A in case 1 and V13
Molweight: 51.14 (kD)
Majority protein IDs: Gorai.007G188300.2.A;Gorai.007G188300.8.A;Gorai.007G188300.1.A;Gorai.007G188300.3.A;Gorai.007G188300.4.A;Gorai.007G188300.5.A;Gorai.007G188300.6.A;Gorai.007G188300.7.A;Go
Gene IDs: Gorai.007G188300
cotton name: K00031 - isocitrate dehydrogenase (IDH1, IDH2, icd) (1 of 3)
OG IDs: 93522513
Best-hit-arabi-ID: AT1G65930.1
Best-hit-arabi-name: cytosolic NADP+-dependent isocitrate dehydrogenase
Gorai.007G188300.1.A
PCC=0.184, p=0.451
mRNA Profile
Protein Expression Profile
DPA

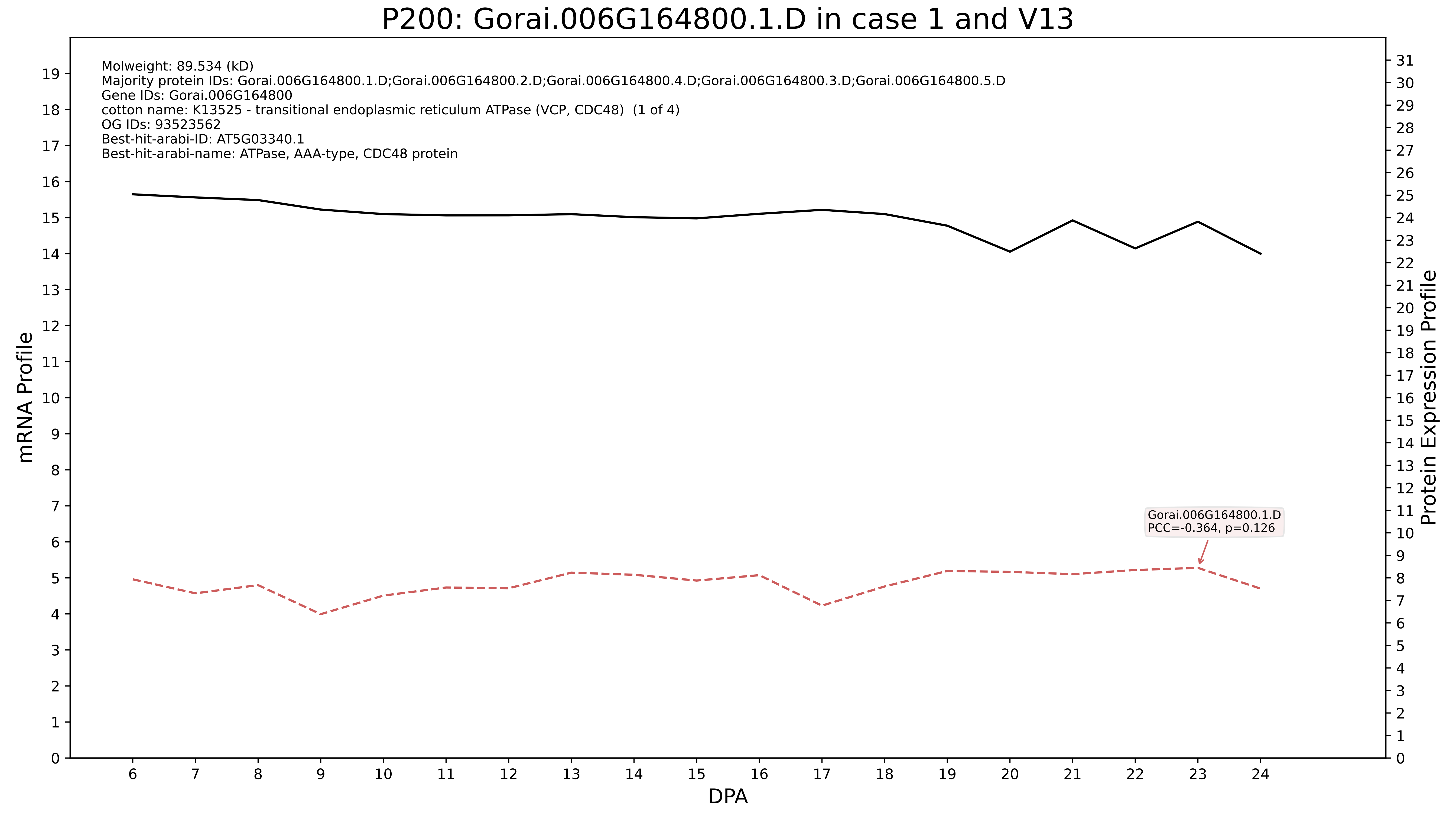
P200: Gorai.006G164800.1.D in case 1 and V13
Molweight: 89.534 (kD)
Majority protein IDs: Gorai.006G164800.1.D;Gorai.006G164800.2.D;Gorai.006G164800.4.D;Gorai.006G164800.3.D;Gorai.006G164800.5.D
Gene IDs: Gorai.006G164800
cotton name: K13525 - transitional endoplasmic reticulum ATPase (VCP, CDC48) (1 of 4)
OG IDs: 93523562
Best-hit-arabi-ID: AT5G03340.1
Best-hit-arabi-name: ATPase, AAA-type, CDC48 protein
Gorai.006G164800.1.D
PCC=-0.364, p=0.126
mRNA Profile
Protein Expression Profile
DPA

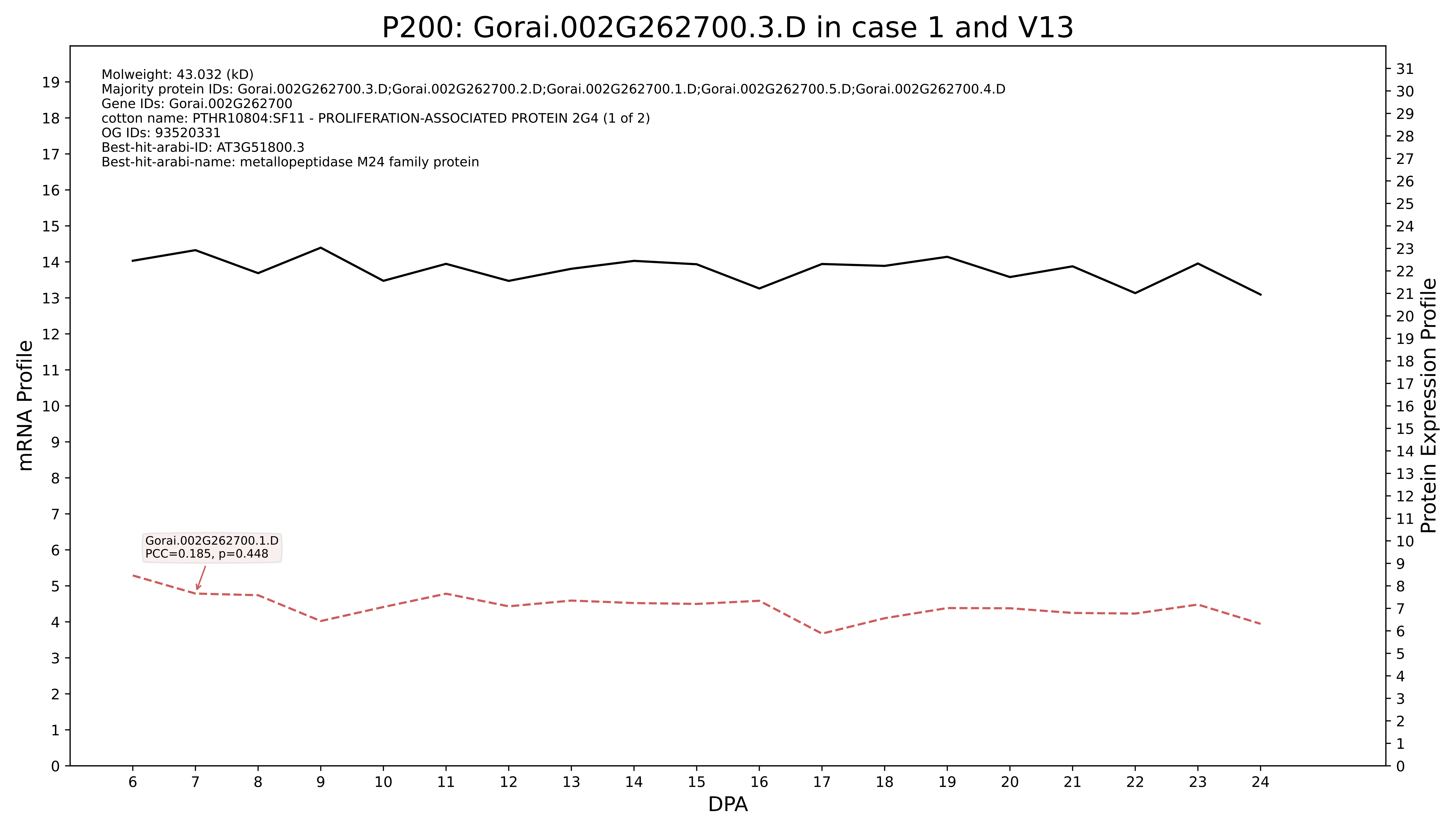
P200: Gorai.002G262700.3.D in case 1 and V13
Molweight: 43.032 (kD)
Majority protein IDs: Gorai.002G262700.3.D;Gorai.002G262700.2.D;Gorai.002G262700.1.D;Gorai.002G262700.5.D;Gorai.002G262700.4.D
Gene IDs: Gorai.002G262700
cotton name: PTHR10804:SF11 - PROLIFERATION-ASSOCIATED PROTEIN 2G4 (1 of 2)
OG IDs: 93520331
Best-hit-arabi-ID: AT3G51800.3
Best-hit-arabi-name: metallopeptidase M24 family protein
Gorai.002G262700.1.D
PCC=0.185, p=0.448
mRNA Profile
Protein Expression Profile
DPA

### P200: Gorai.005G210000.1.A in case 1 and V13

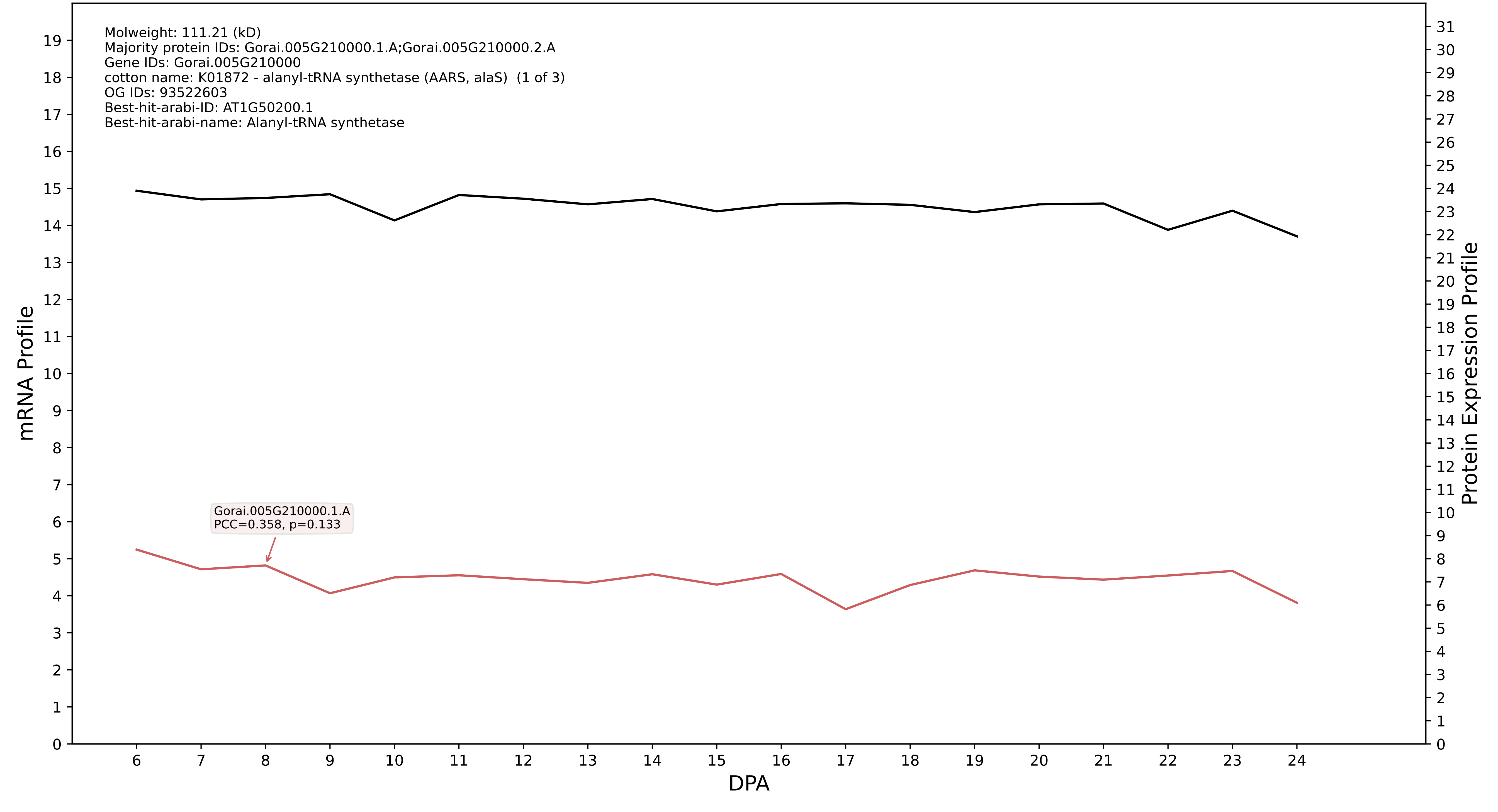

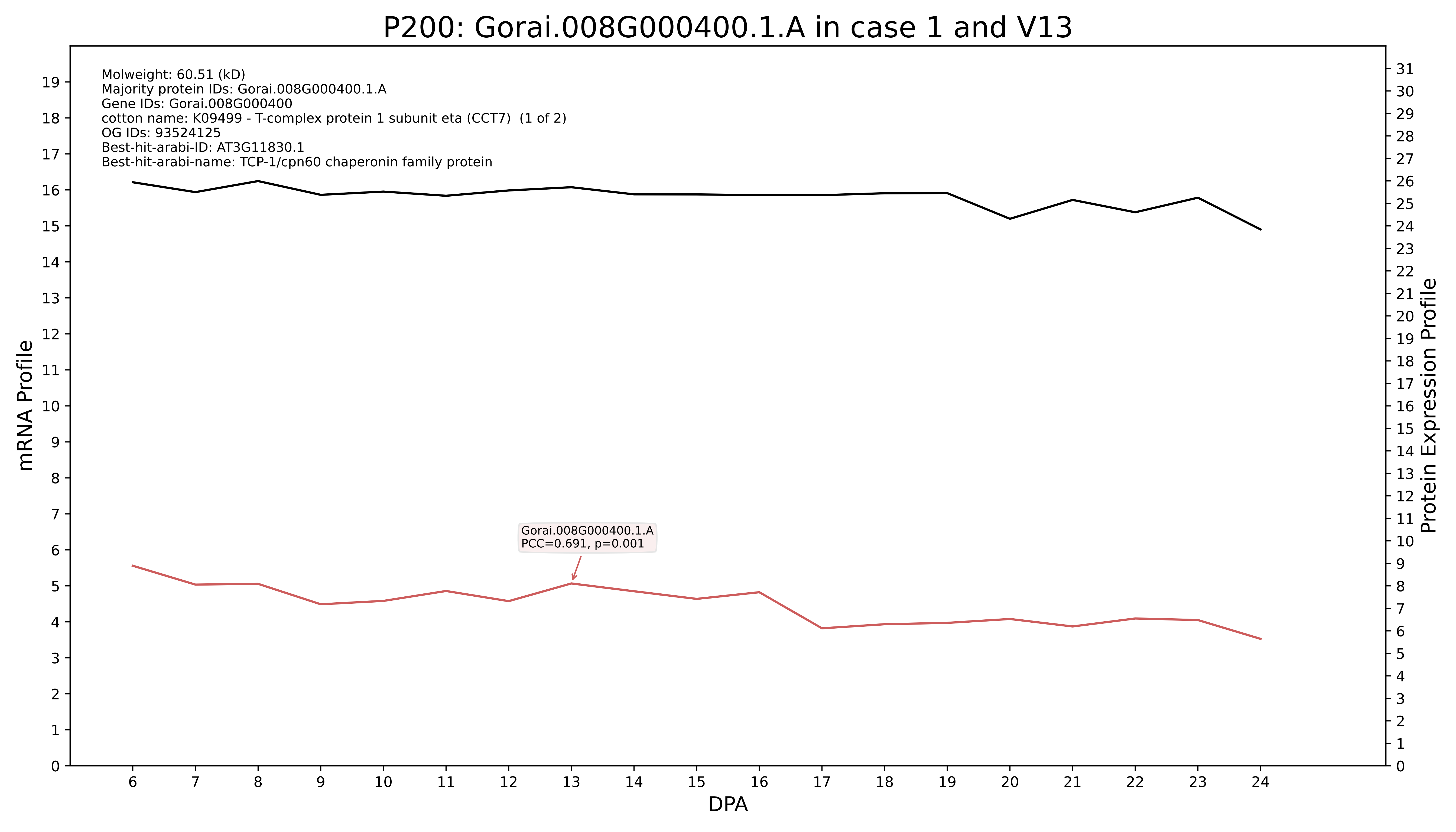

P200: Gorai.008G000400.1.A in case 1 and V13
Molweight: 60.51 (kD)
Majority protein IDs: Gorai.008G000400.1.A
Gene IDs: Gorai.008G000400
cotton name: K09499 - T-complex protein 1 subunit eta (CCT7) (1 of 2)
OG IDs: 93524125
Best-hit-arabi-ID: AT3G11830.1
Best-hit-arabi-name: TCP-1/cpn60 chaperonin family protein
Gorai.008G000400.1.A
PCC=0.691, p=0.001
mRNA Profile
Protein Expression Profile
DPA

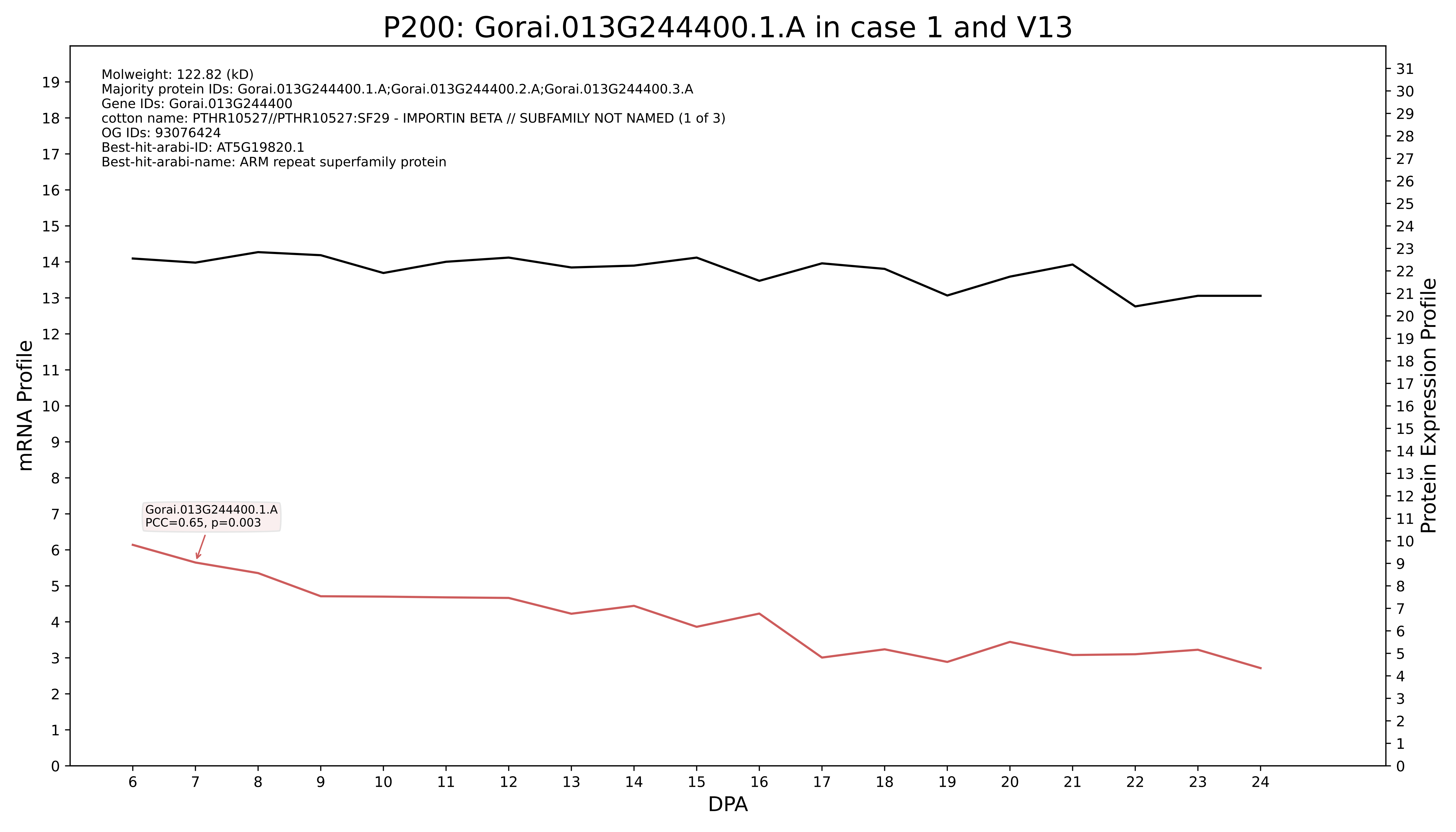

P200: Gorai.013G244400.1.A in case 1 and V13
Molweight: 122.82 (kD)
Majority protein IDs: Gorai.013G244400.1.A;Gorai.013G244400.2.A;Gorai.013G244400.3.A
Gene IDs: Gorai.013G244400
cotton name: PTHR10527//PTHR10527:SF29 - IMPORTIN BETA // SUBFAMILY NOT NAMED (1 of 3)
OG IDs: 93076424
Best-hit-arabi-ID: AT5G19820.1
Best-hit-arabi-name: ARM repeat superfamily protein
Gorai.013G244400.1.A
PCC=0.65, p=0.003
mRNA Profile
Protein Expression Profile
DPA

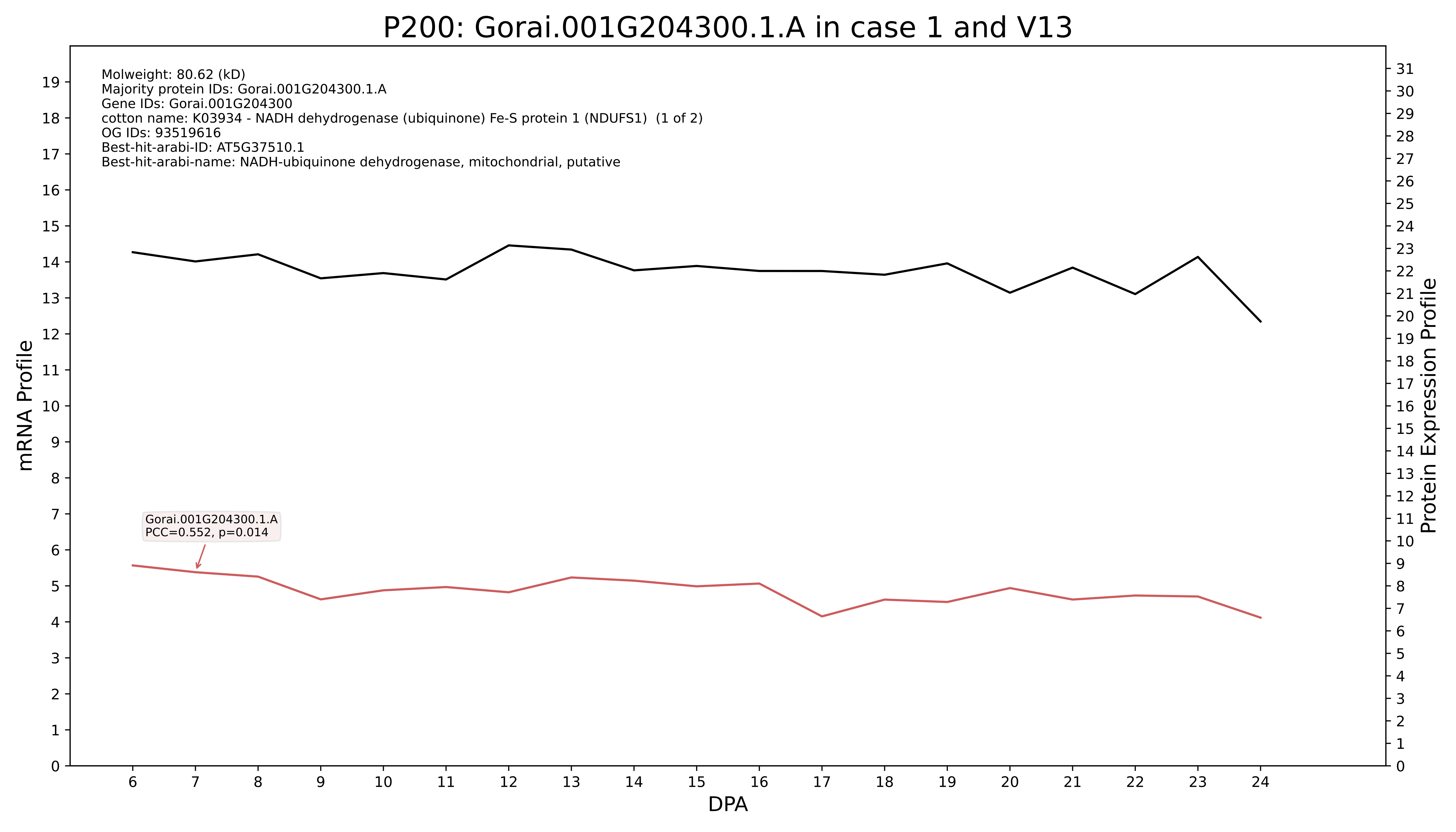

P200: Gorai.001G204300.1.A in case 1 and V13
Molweight: 80.62 (kD)
Majority protein IDs: Gorai.001G204300.1.A
Gene IDs: Gorai.001G204300
cotton name: K03934 - NADH dehydrogenase (ubiquinone) Fe-S protein 1 (NDUFS1) (1 of 2)
OG IDs: 93519616
Best-hit-arabi-ID: AT5G37510.1
Best-hit-arabi-name: NADH-ubiquinone dehydrogenase, mitochondrial, putative
Gorai.001G204300.1.A
PCC=0.552, p=0.014
mRNA Profile
Protein Expression Profile
DPA

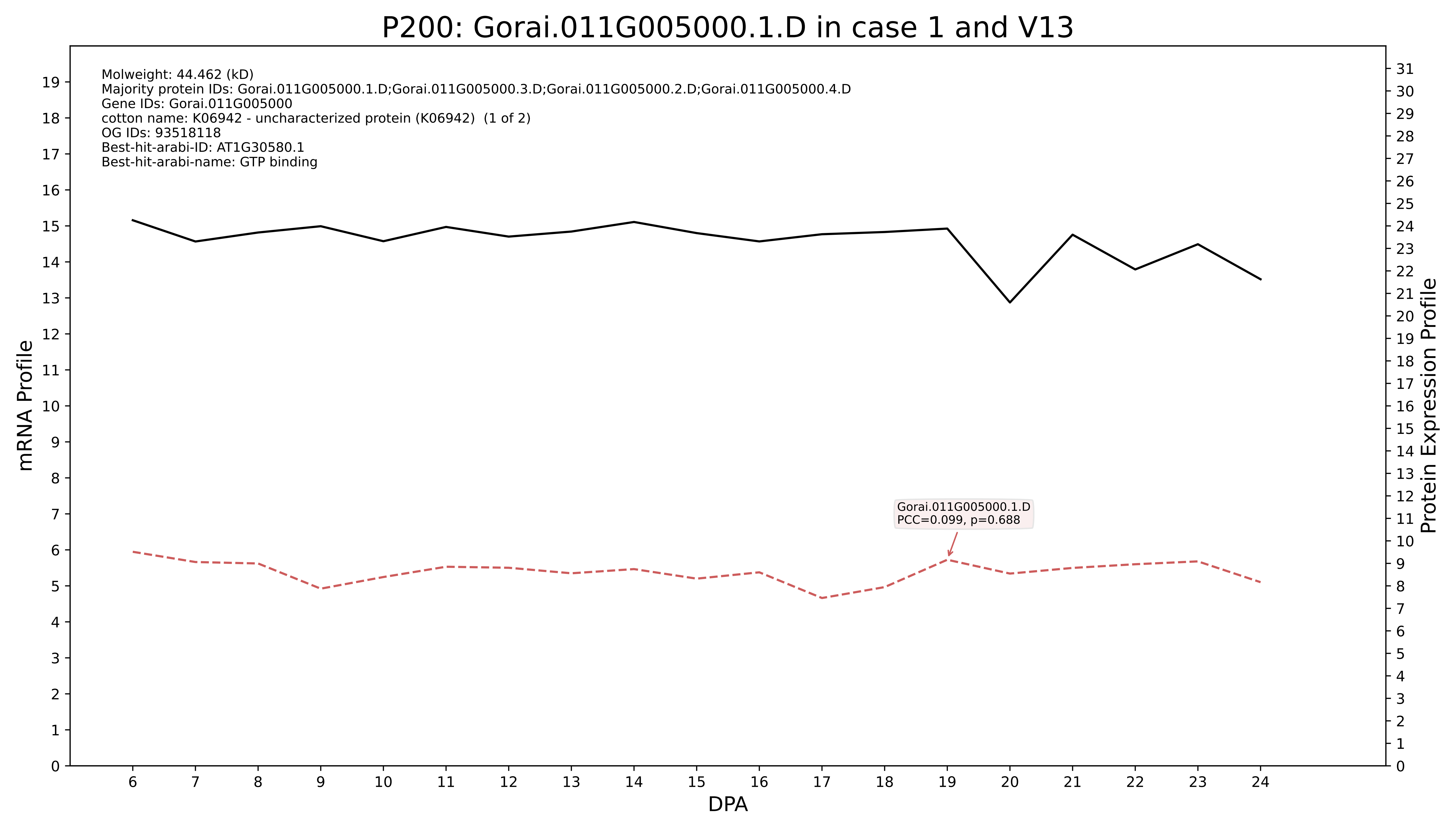

P200: Gorai.011G005000.1.D in case 1 and V13
Molweight: 44.462 (kD)
Majority protein IDs: Gorai.011G005000.1.D;Gorai.011G005000.3.D;Gorai.011G005000.2.D;Gorai.011G005000.4.D
Gene IDs: Gorai.011G005000
cotton name: K06942 - uncharacterized protein (K06942) (1 of 2)
OG IDs: 93518118
Best-hit-arabi-ID: AT1G30580.1
Best-hit-arabi-name: GTP binding
Gorai.011G005000.1.D
PCC=0.099, p=0.688
mRNA Profile
Protein Expression Profile
DPA

### P200: Gorai.013G270300.1.D in case 1 and V13

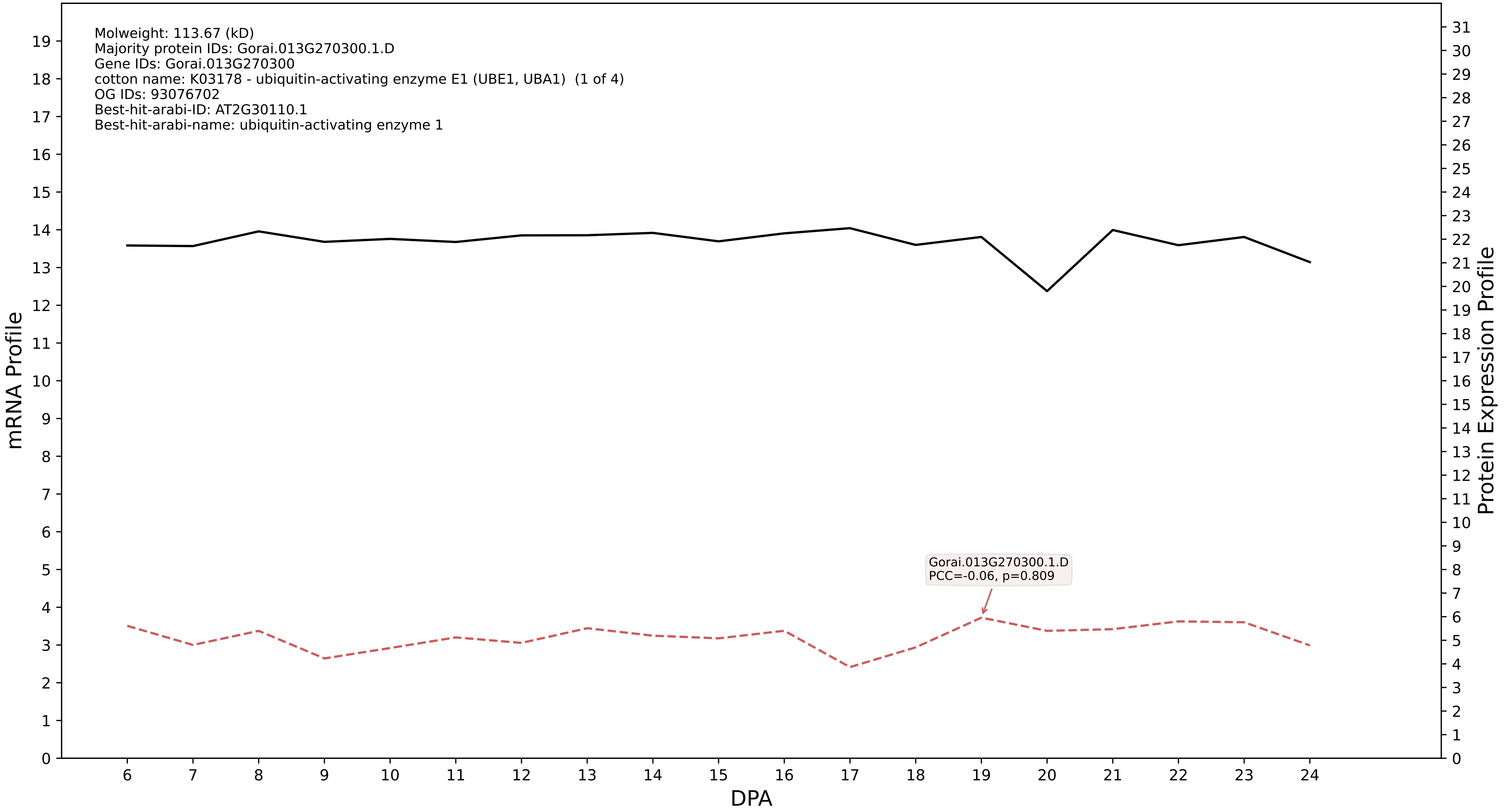

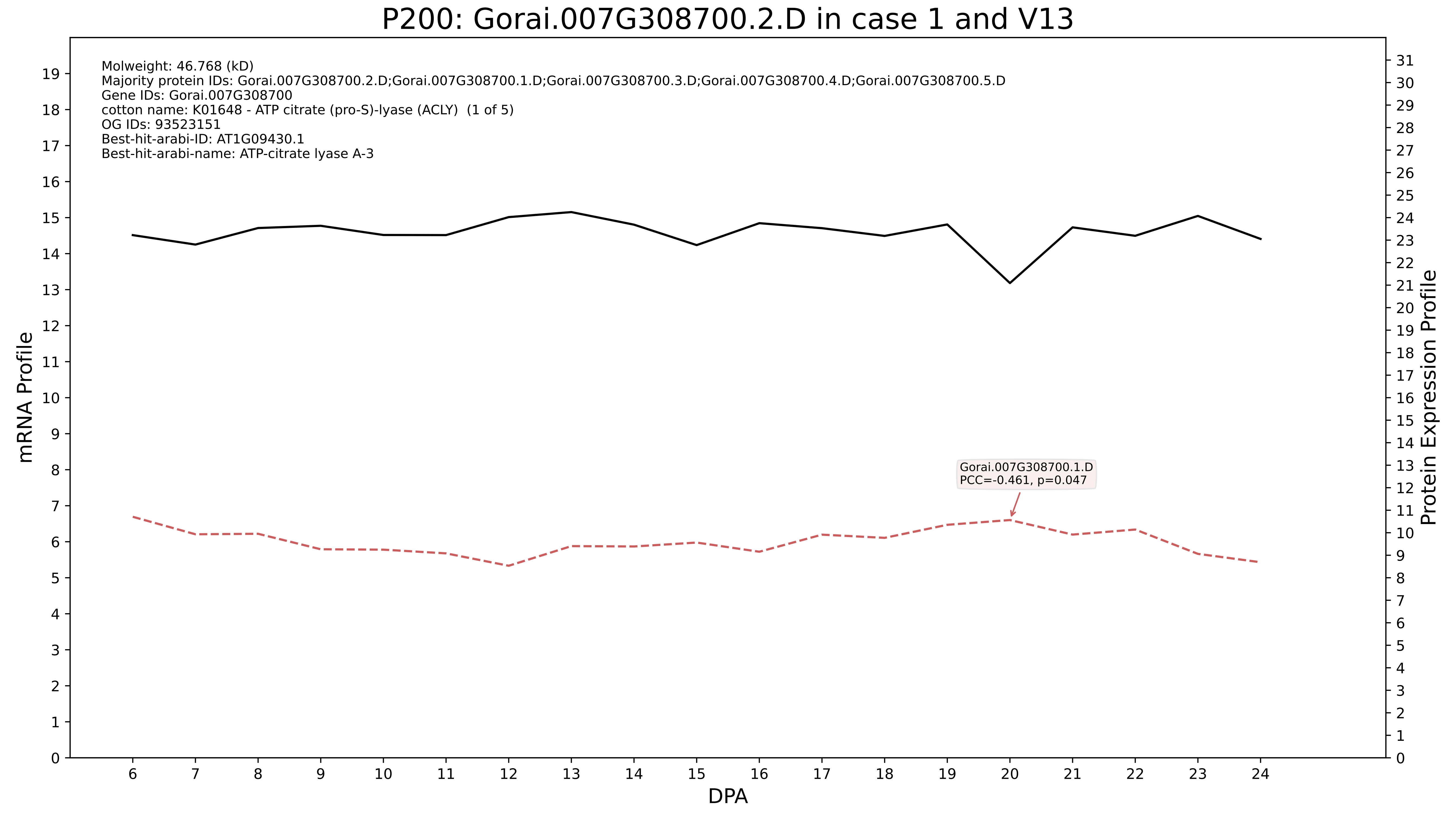

P200: Gorai.007G308700.2.D in case 1 and V13
Molweight: 46.768 (kD)
Majority protein IDs: Gorai.007G308700.2.D;Gorai.007G308700.1.D;Gorai.007G308700.3.D;Gorai.007G308700.4.D;Gorai.007G308700.5.D
Gene IDs: Gorai.007G308700
cotton name: K01648 - ATP citrate (pro-S)-lyase (ACLY) (1 of 5)
OG IDs: 93523151
Best-hit-arabi-ID: AT1G09430.1
Best-hit-arabi-name: ATP-citrate lyase A-3
Gorai.007G308700.1.D
PCC=-0.461, p=0.047
mRNA Profile
Protein Expression Profile
DPA

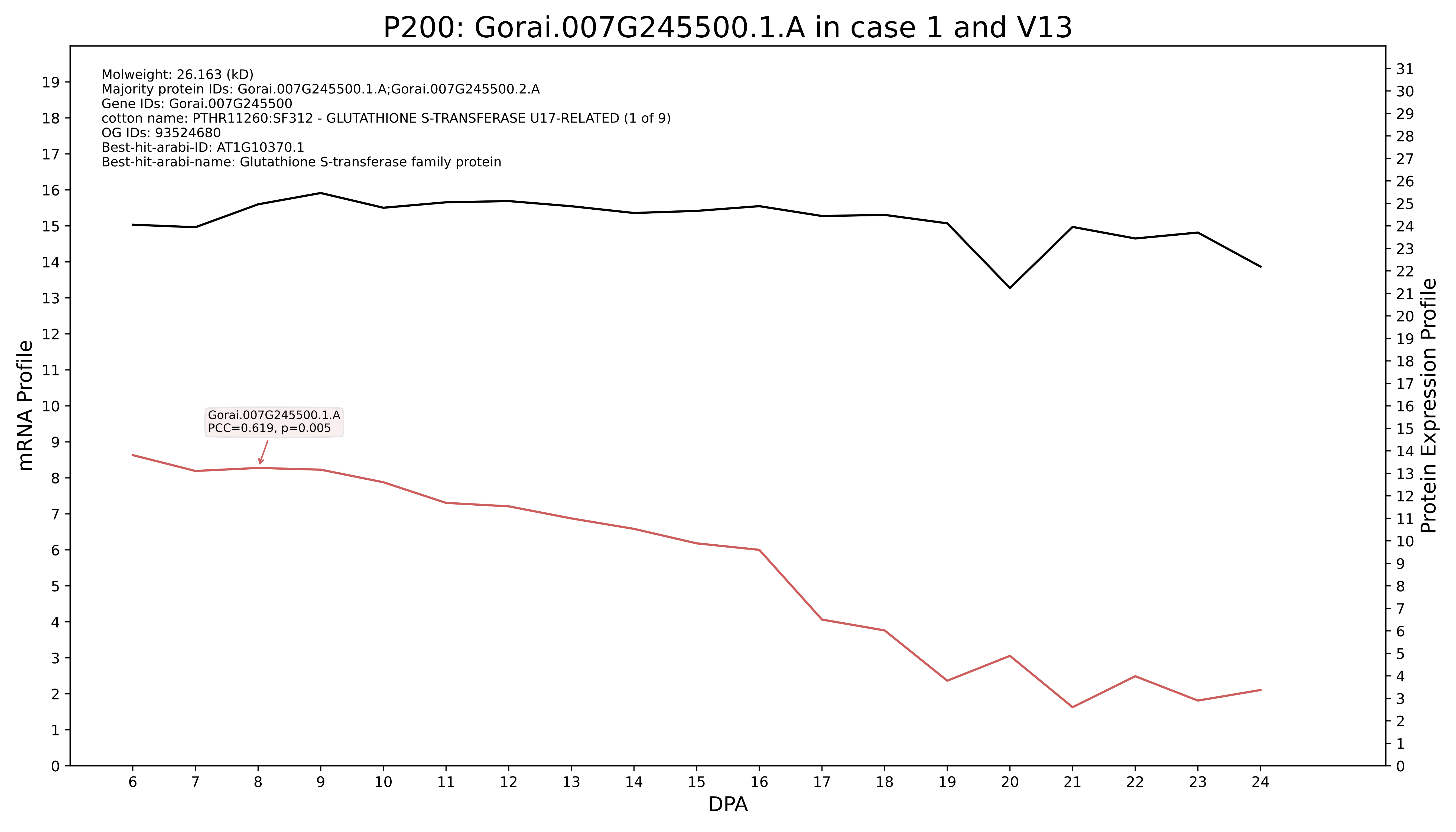
P200: Gorai.007G245500.1.A in case 1 and V13
Molweight: 26.163 (kD)
Majority protein IDs: Gorai.007G245500.1.A;Gorai.007G245500.2.A
Gene IDs: Gorai.007G245500
cotton name: PTHR11260:SF312 - GLUTATHIONE S-TRANSFERASE U17-RELATED (1 of 9)
OG IDs: 93524680
Best-hit-arabi-ID: AT1G10370.1
Best-hit-arabi-name: Glutathione S-transferase family protein
Gorai.007G245500.1.A
PCC=0.619, p=0.005
mRNA Profile
Protein Expression Profile
DPA

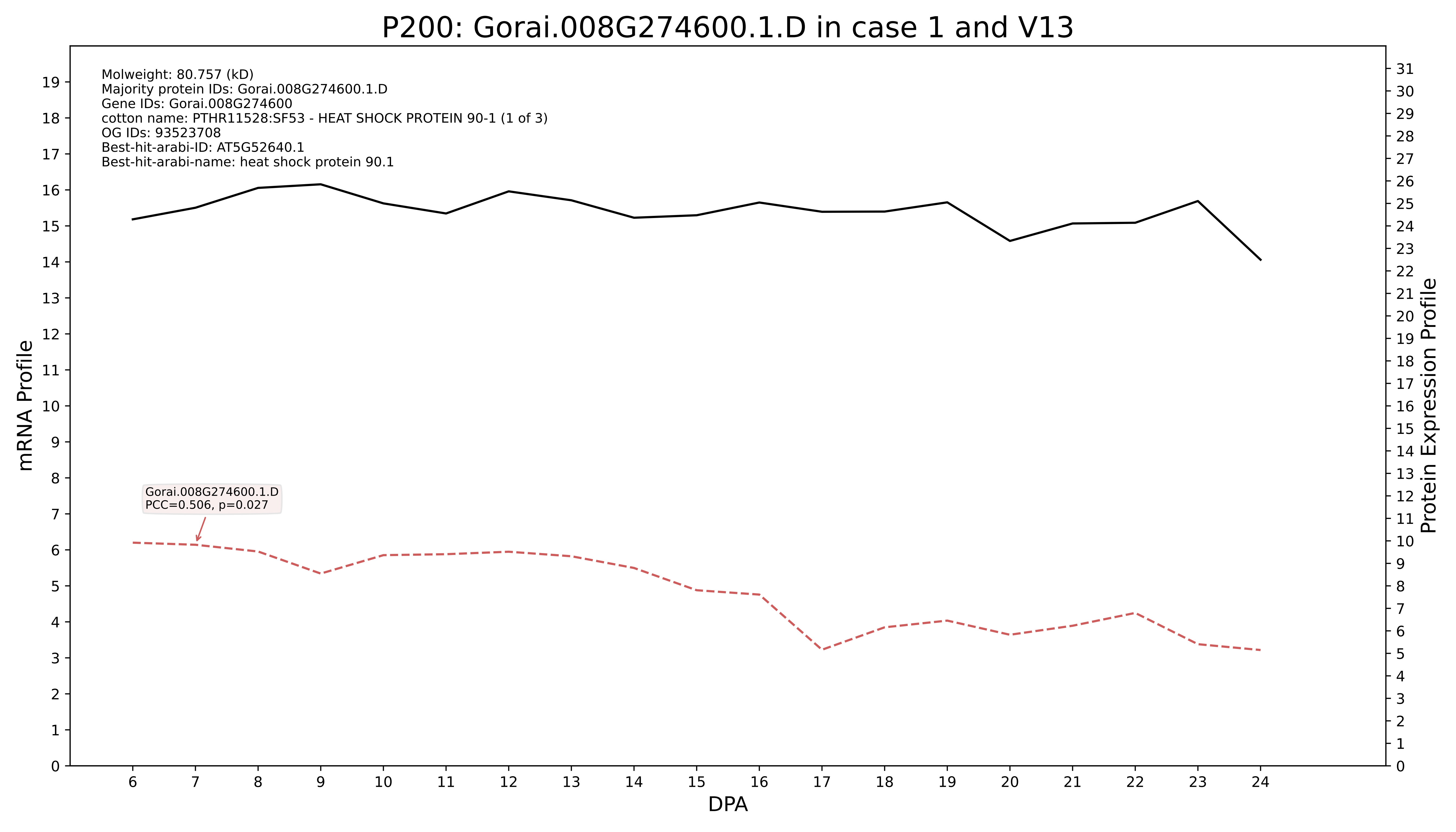

P200: Gorai.008G274600.1.D in case 1 and V13
Molweight: 80.757 (kD)
Majority protein IDs: Gorai.008G274600.1.D
Gene IDs: Gorai.008G274600
cotton name: PTHR11528:SF53 - HEAT SHOCK PROTEIN 90-1 (1 of 3)
OG IDs: 93523708
Best-hit-arabi-ID: AT5G52640.1
Best-hit-arabi-name: heat shock protein 90.1
Gorai.008G274600.1.D
PCC=0.506, p=0.027
mRNA Profile
Protein Expression Profile
DPA

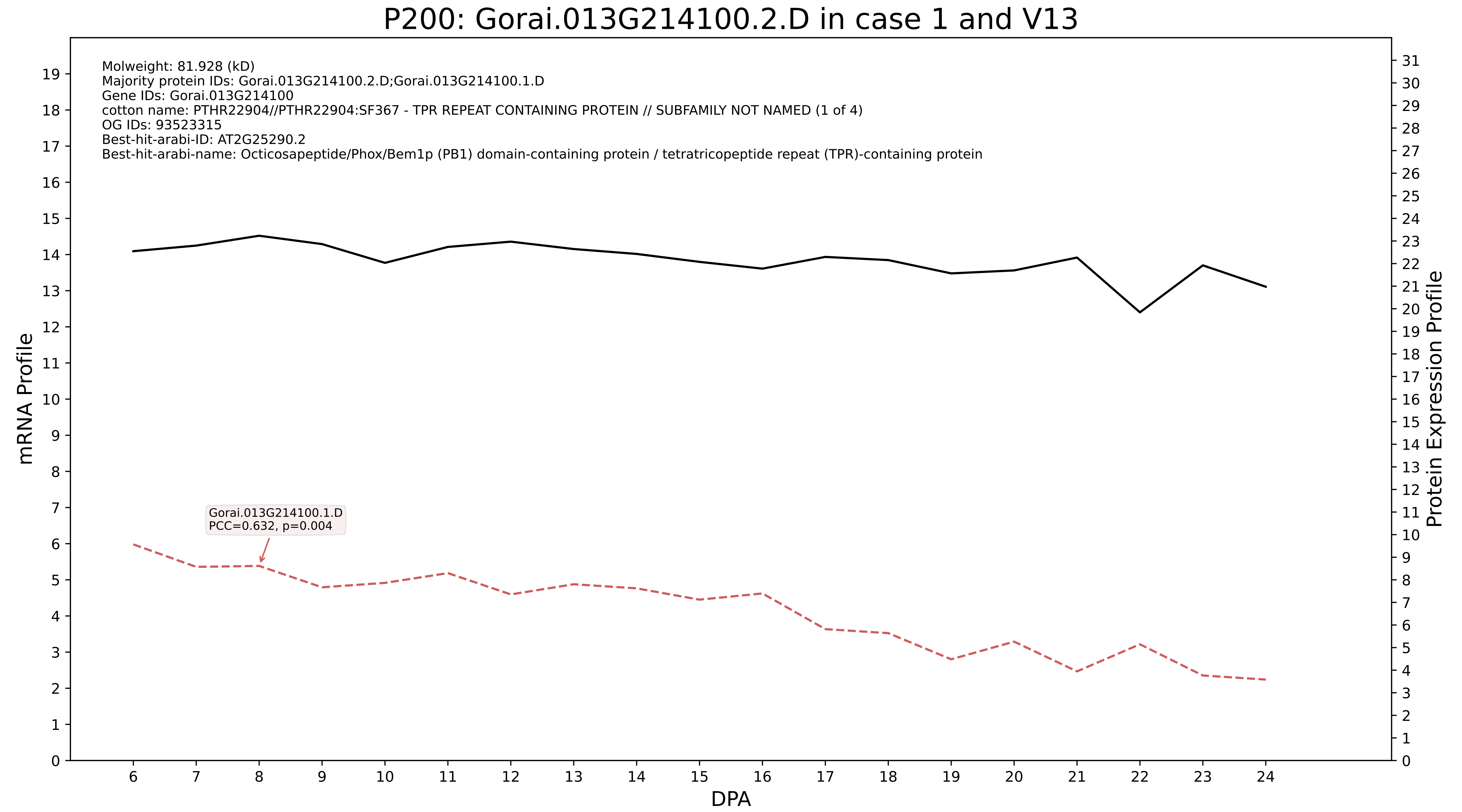

P200: Gorai.013G214100.2.D in case 1 and V13
Molweight: 81.928 (kD)
Majority protein IDs: Gorai.013G214100.2.D;Gorai.013G214100.1.D
Gene IDs: Gorai.013G214100
cotton name: PTHR22904//PTHR22904:SF367 - TPR REPEAT CONTAINING PROTEIN // SUBFAMILY NOT NAMED (1 of 4)
OG IDs: 93523315
Best-hit-arabi-ID: AT2G25290.2
Best-hit-arabi-name: Octicosapeptide/Phox/Bem1p (PB1) domain-containing protein / tetratricopeptide repeat (TPR)-containing protein
Gorai.013G214100.1.D
PCC=0.632, p=0.004
mRNA Profile
Protein Expression Profile
DPA

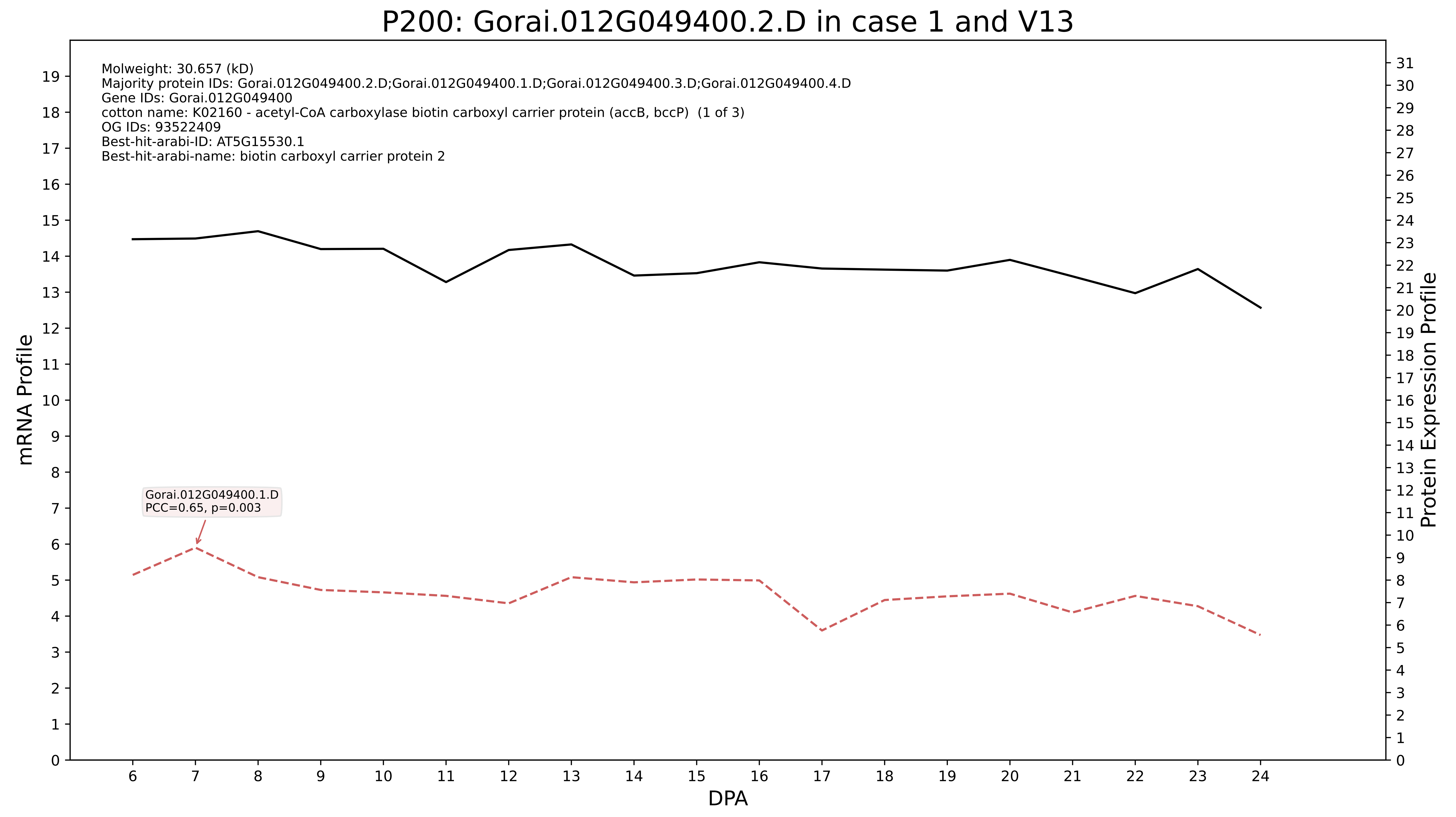

P200: Gorai.012G049400.2.D in case 1 and V13
Molweight: 30.657 (kD)
Majority protein IDs: Gorai.012G049400.2.D;Gorai.012G049400.1.D;Gorai.012G049400.3.D;Gorai.012G049400.4.D
Gene IDs: Gorai.012G049400
cotton name: K02160 - acetyl-CoA carboxylase biotin carboxyl carrier protein (accB, bccP) (1 of 3)
OG IDs: 93522409
Best-hit-arabi-ID: AT5G15530.1
Best-hit-arabi-name: biotin carboxyl carrier protein 2
Gorai.012G049400.1.D
PCC=0.65, p=0.003
mRNA Profile
Protein Expression Profile
DPA

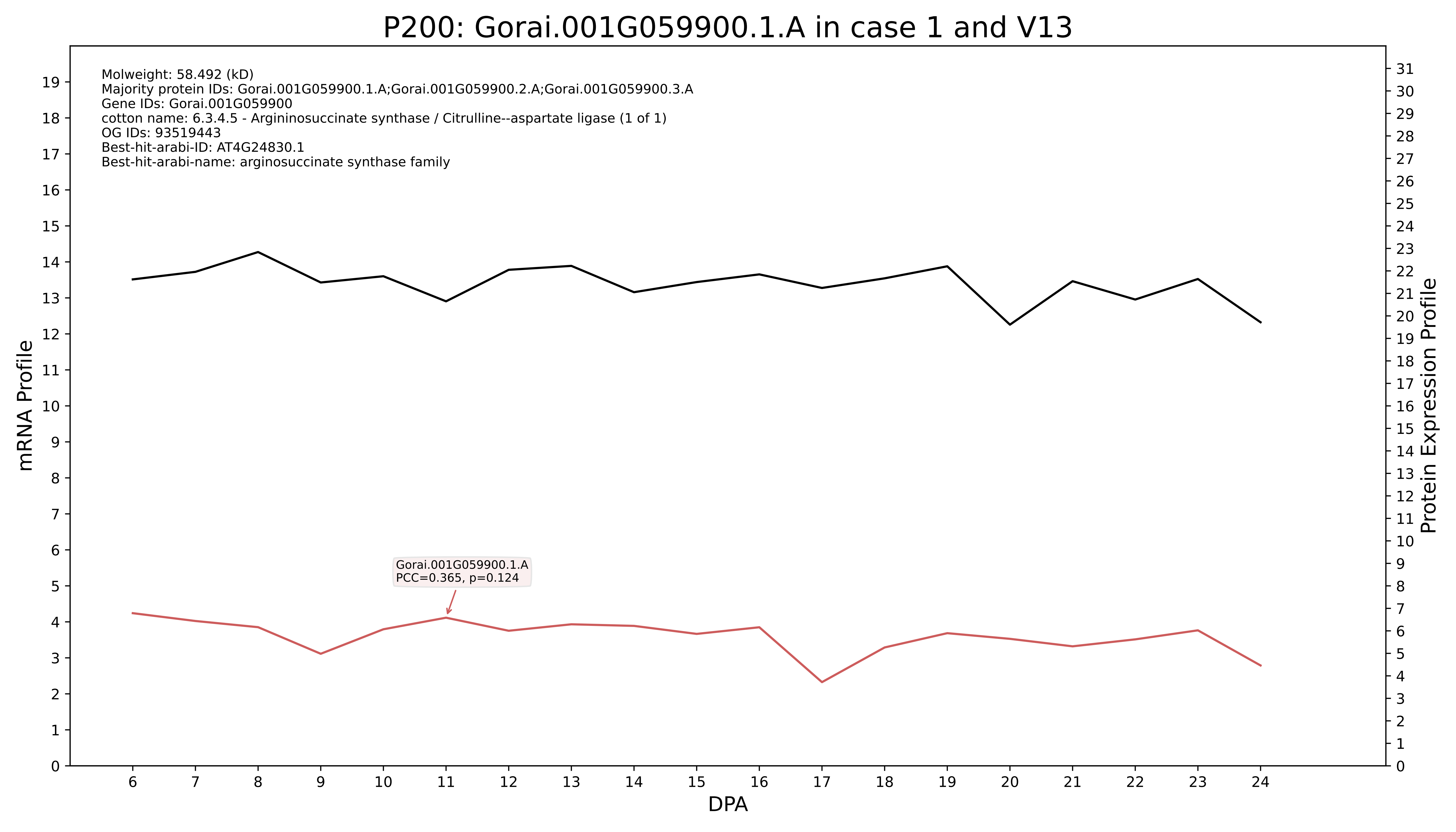

P200: Gorai.001G059900.1.A in case 1 and V13
Molweight: 58.492 (kD)
Majority protein IDs: Gorai.001G059900.1.A;Gorai.001G059900.2.A;Gorai.001G059900.3.A
Gene IDs: Gorai.001G059900
cotton name: 6.3.4.5 - Argininosuccinate synthase / Citrulline--aspartate ligase (1 of 1)
OG IDs: 93519443
Best-hit-arabi-ID: AT4G24830.1
Best-hit-arabi-name: arginosuccinate synthase family
Gorai.001G059900.1.A
PCC=0.365, p=0.124
mRNA Profile
Protein Expression Profile
DPA

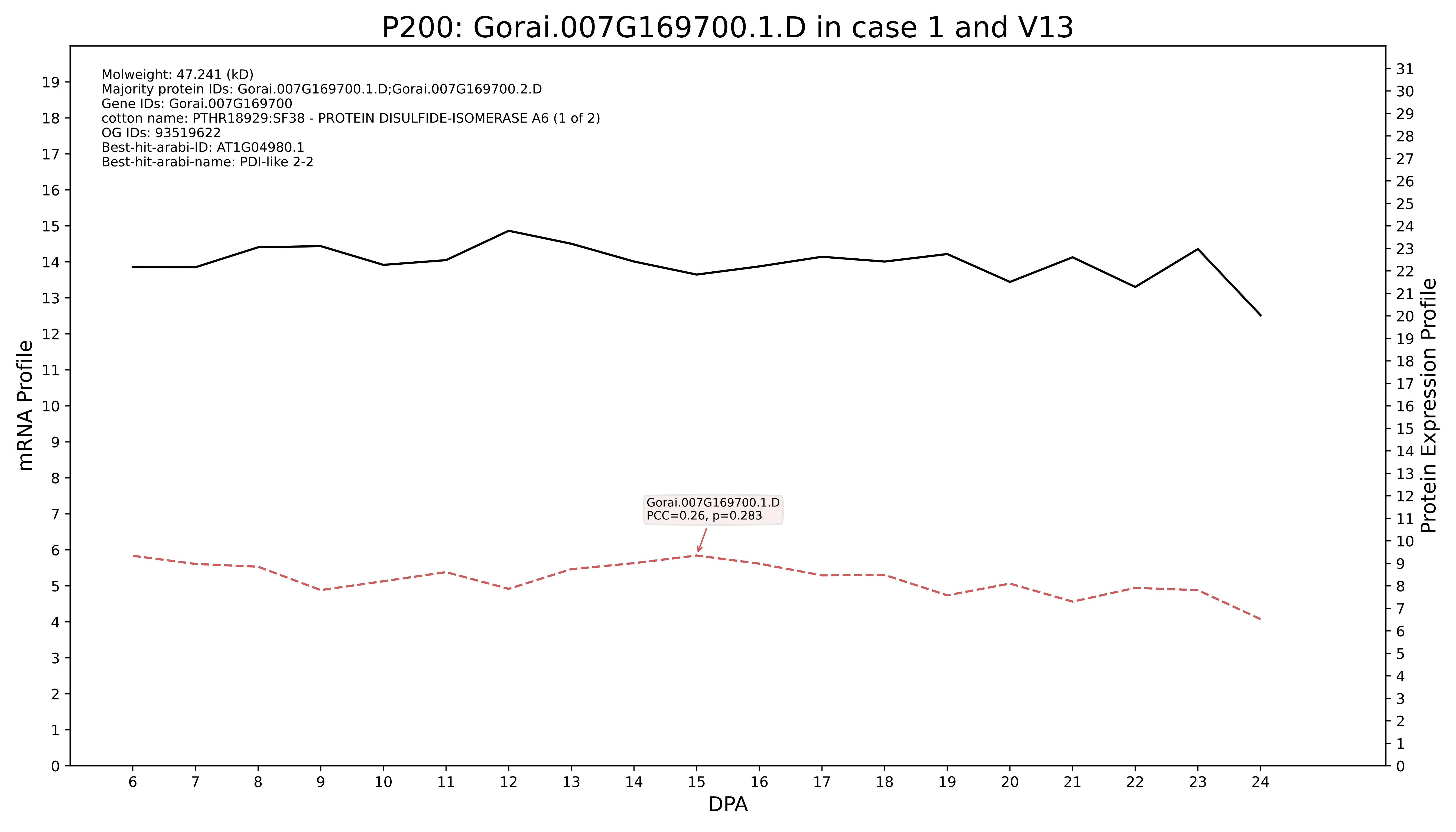
P200: Gorai.007G169700.1.D in case 1 and V13
Molweight: 47.241 (kD)
Majority protein IDs: Gorai.007G169700.1.D;Gorai.007G169700.2.D
Gene IDs: Gorai.007G169700
cotton name: PTHR18929:SF38 - PROTEIN DISULFIDE-ISOMERASE A6 (1 of 2)
OG IDs: 93519622
Best-hit-arabi-ID: AT1G04980.1
Best-hit-arabi-name: PDI-like 2-2
Gorai.007G169700.1.D
PCC=0.26, p=0.283
mRNA Profile
Protein Expression Profile
DPA

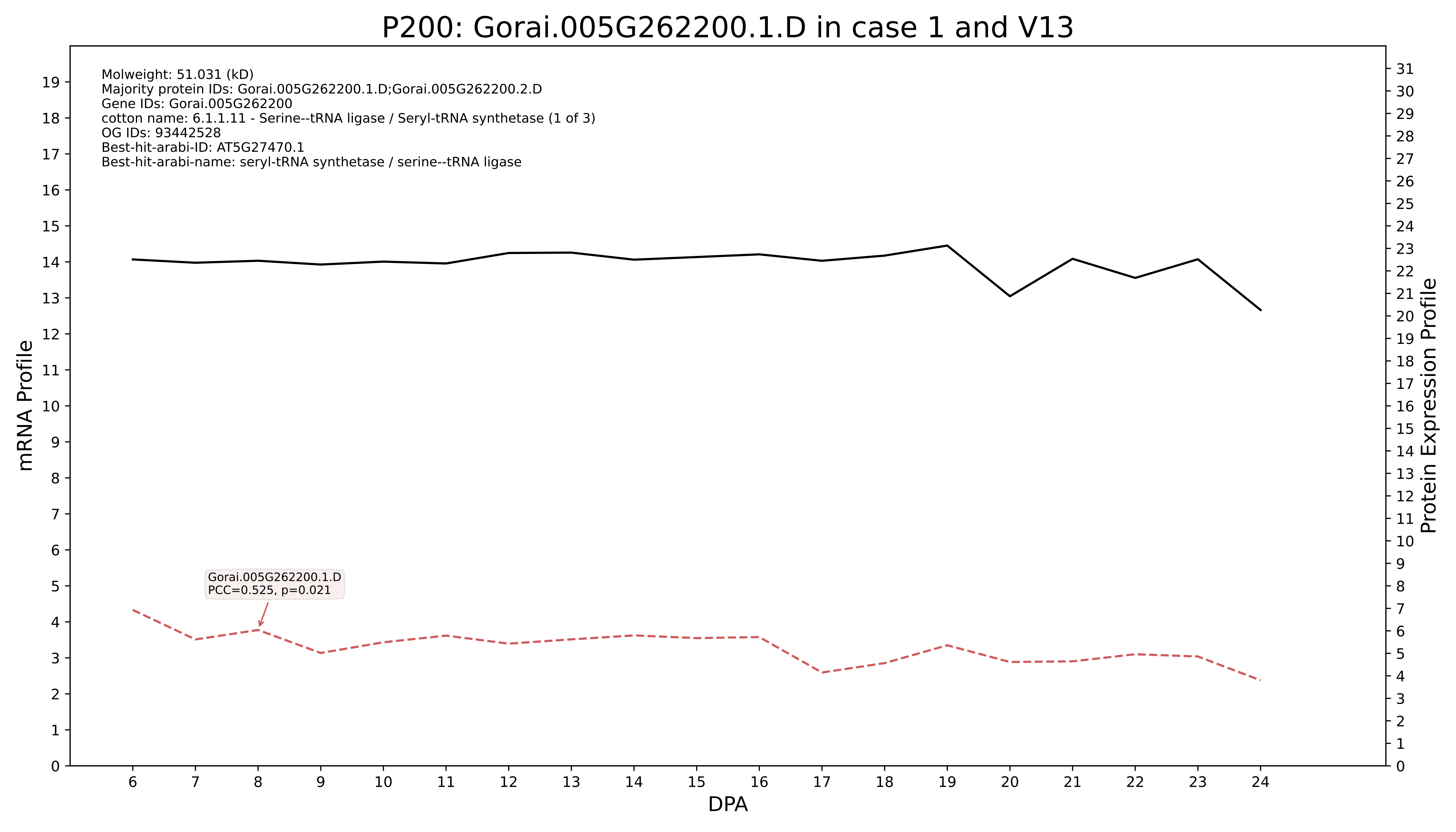

P200: Gorai.005G262200.1.D in case 1 and V13
Molweight: 51.031 (kD)
Majority protein IDs: Gorai.005G262200.1.D;Gorai.005G262200.2.D
Gene IDs: Gorai.005G262200
cotton name: 6.1.1.11 - Serine--tRNA ligase / Seryl-tRNA synthetase (1 of 3)
OG IDs: 93442528
Best-hit-arabi-ID: AT5G27470.1
Best-hit-arabi-name: seryl-tRNA synthetase / serine--tRNA ligase
Gorai.005G262200.1.D
PCC=0.525, p=0.021
mRNA Profile
Protein Expression Profile
DPA

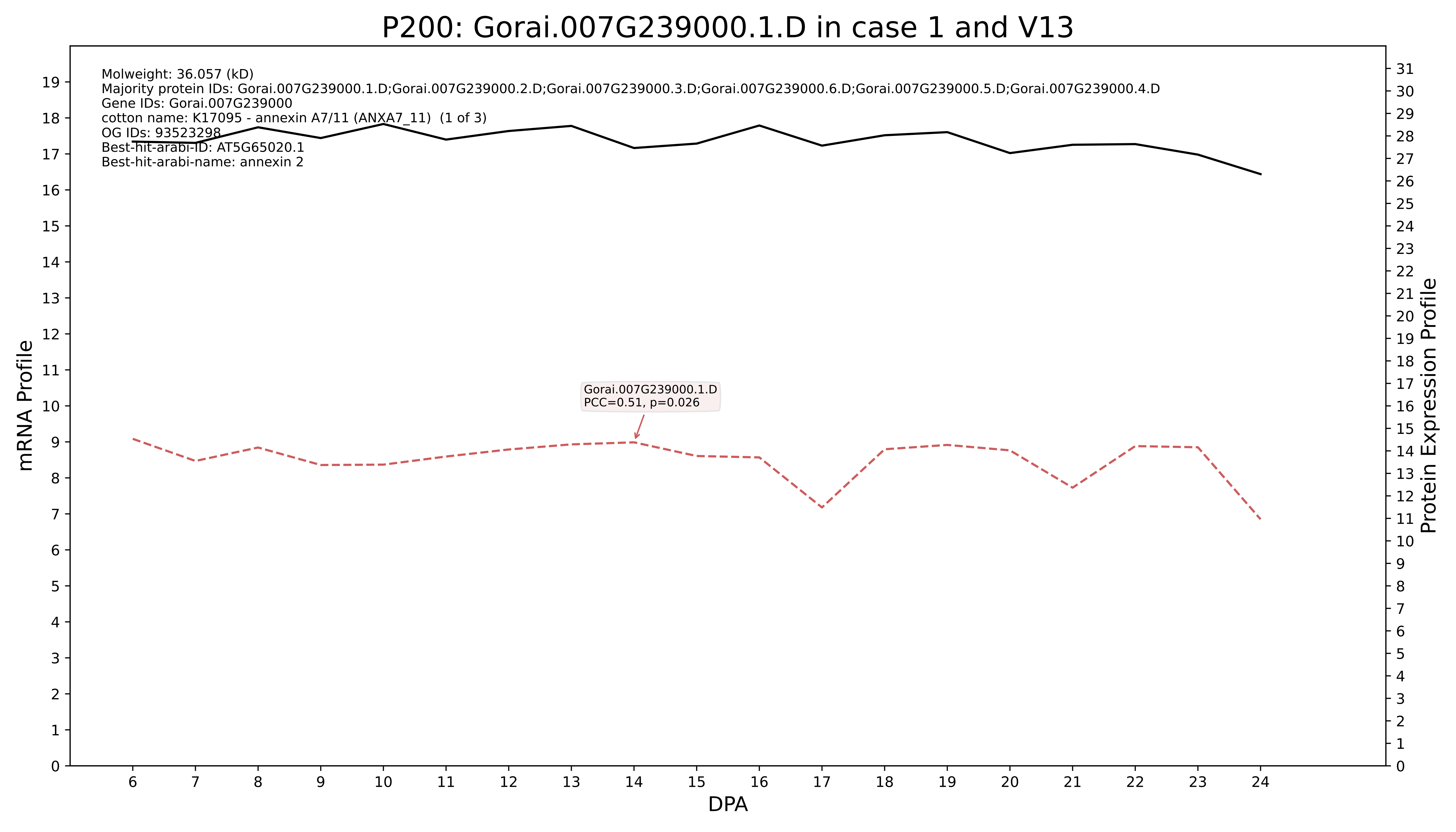

P200: Gorai.007G239000.1.D in case 1 and V13
Molweight: 36.057 (kD)
Majority protein IDs: Gorai.007G239000.1.D;Gorai.007G239000.2.D;Gorai.007G239000.3.D;Gorai.007G239000.6.D;Gorai.007G239000.5.D;Gorai.007G239000.4.D
Gene IDs: Gorai.007G239000
cotton name: K17095 - annexin A7/11 (ANXA7_11) (1 of 3)
OG IDs: 93523298
Best-hit-arabi-ID: AT5G65020.1
Best-hit-arabi-name: annexin 2
Gorai.007G239000.1.D
PCC=0.51, p=0.026
mRNA Profile
Protein Expression Profile
DPA

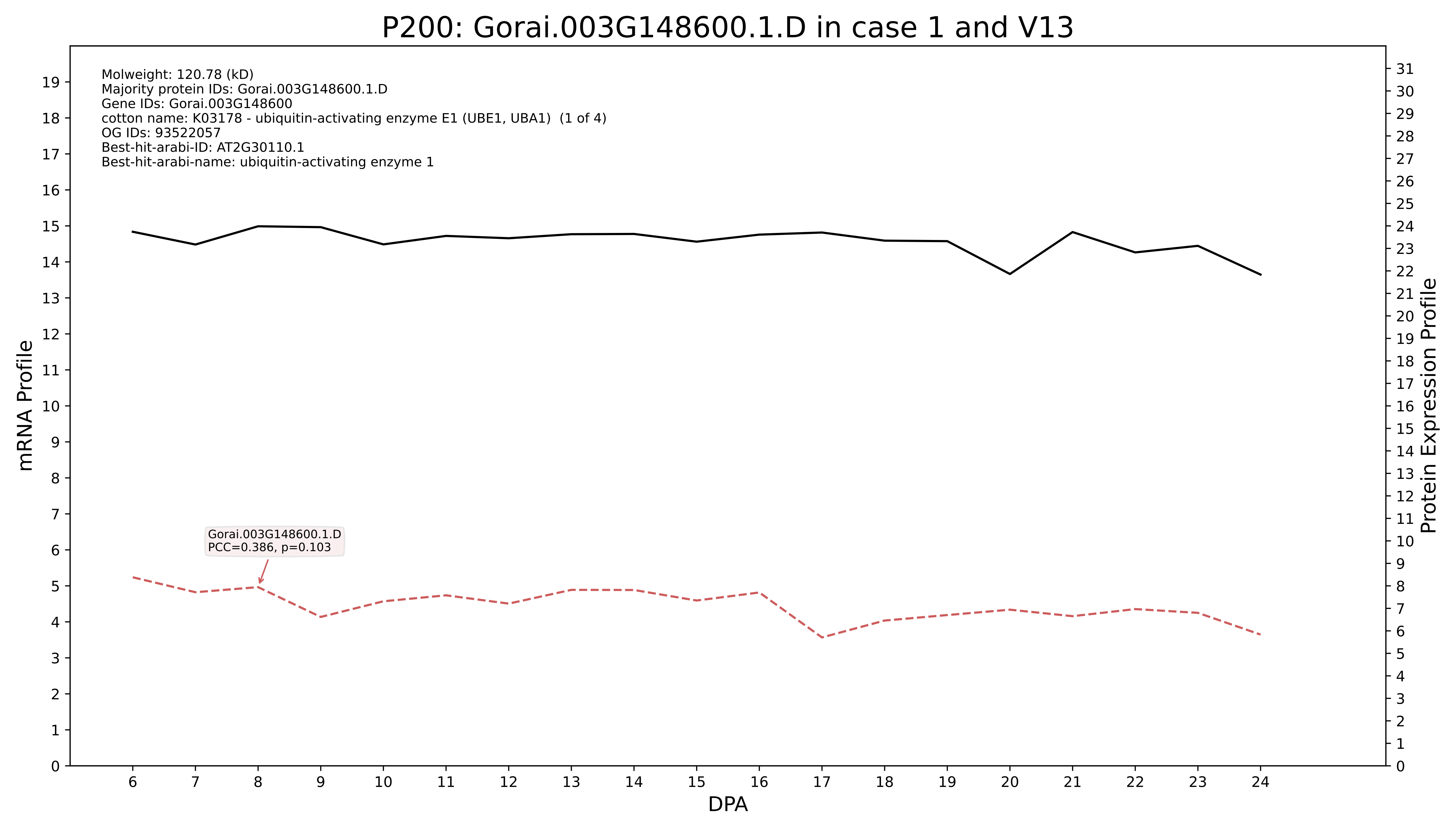
P200: Gorai.003G148600.1.D in case 1 and V13
Molweight: 120.78 (kD)
Majority protein IDs: Gorai.003G148600.1.D
Gene IDs: Gorai.003G148600
cotton name: K03178 - ubiquitin-activating enzyme E1 (UBE1, UBA1) (1 of 4)
OG IDs: 93522057
Best-hit-arabi-ID: AT2G30110.1
Best-hit-arabi-name: ubiquitin-activating enzyme 1
Gorai.003G148600.1.D
PCC=0.386, p=0.103
mRNA Profile
Protein Expression Profile
DPA

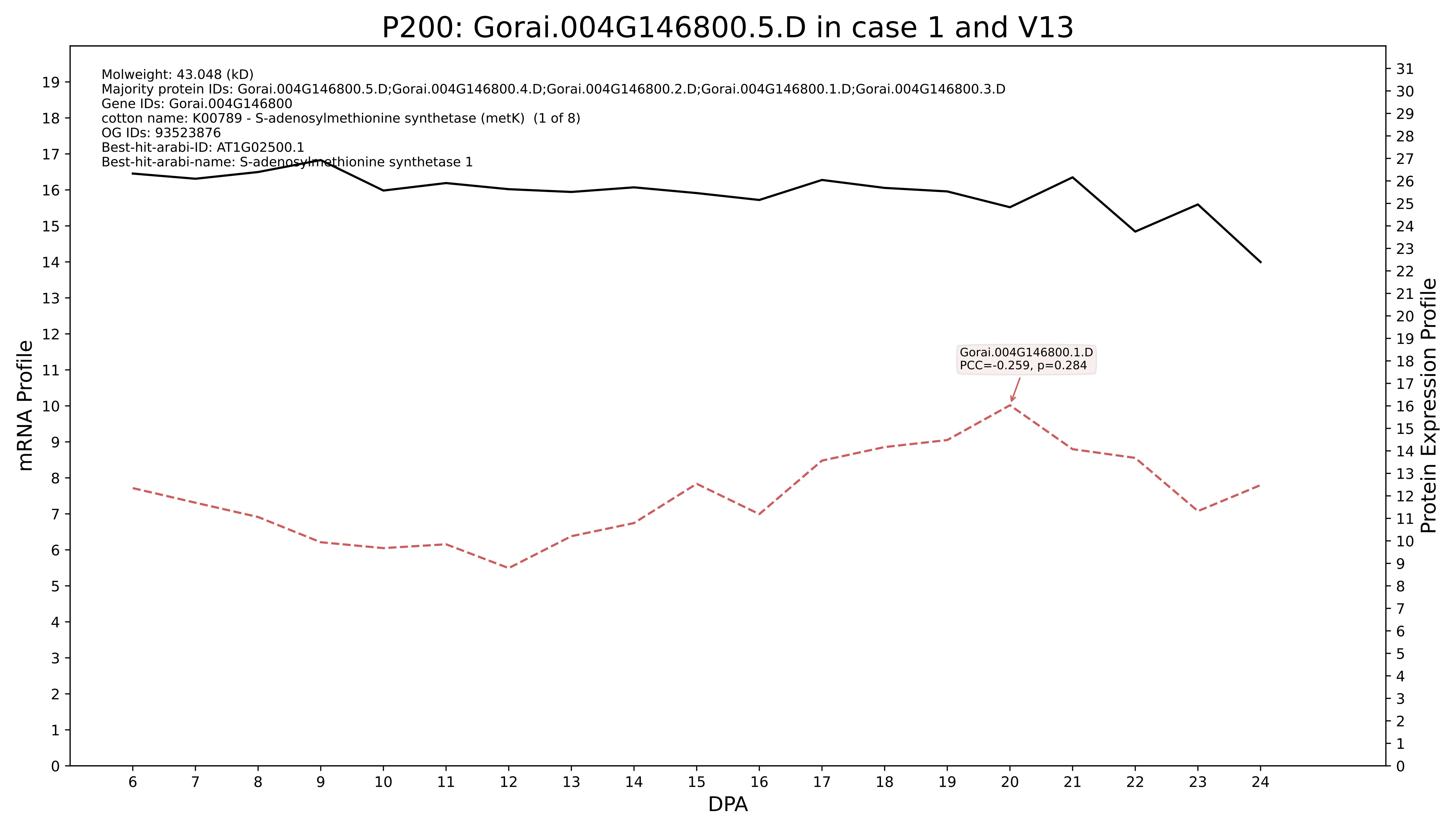
P200: Gorai.004G146800.5.D in case 1 and V13
Molweight: 43.048 (kD)
Majority protein IDs: Gorai.004G146800.5.D;Gorai.004G146800.4.D;Gorai.004G146800.2.D;Gorai.004G146800.1.D;Gorai.004G146800.3.D
Gene IDs: Gorai.004G146800
cotton name: K00789 - S-adenosylmethionine synthetase (metK) (1 of 8)
OG IDs: 93523876
Best-hit-arabi-ID: AT1G02500.1
Best-hit-arabi-name: S-adenosylmethionine synthetase 1
Gorai.004G146800.1.D
PCC=-0.259, p=0.284
mRNA Profile
Protein Expression Profile
DPA

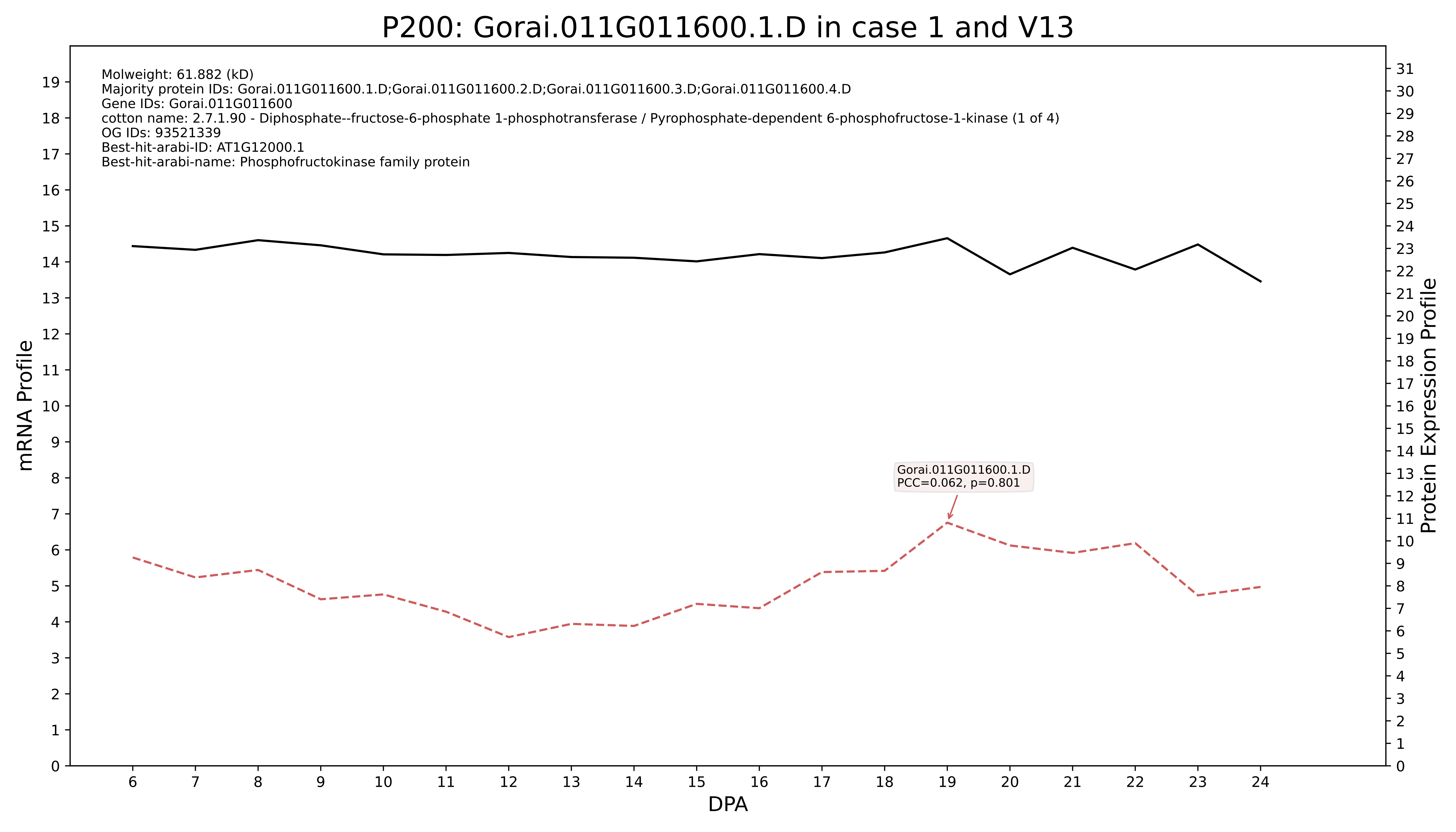
P200: Gorai.011G011600.1.D in case 1 and V13
Molweight: 61.882 (kD)
Majority protein IDs: Gorai.011G011600.1.D;Gorai.011G011600.2.D;Gorai.011G011600.3.D;Gorai.011G011600.4.D
Gene IDs: Gorai.011G011600
cotton name: 2.7.1.90 - Diphosphate--fructose-6-phosphate 1-phosphotransferase / Pyrophosphate-dependent 6-phosphofructose-1-kinase (1 of 4)
OG IDs: 93521339
Best-hit-arabi-ID: AT1G12000.1
Best-hit-arabi-name: Phosphofructokinase family protein
Gorai.011G011600.1.D
PCC=0.062, p=0.801
mRNA Profile
Protein Expression Profile
DPA

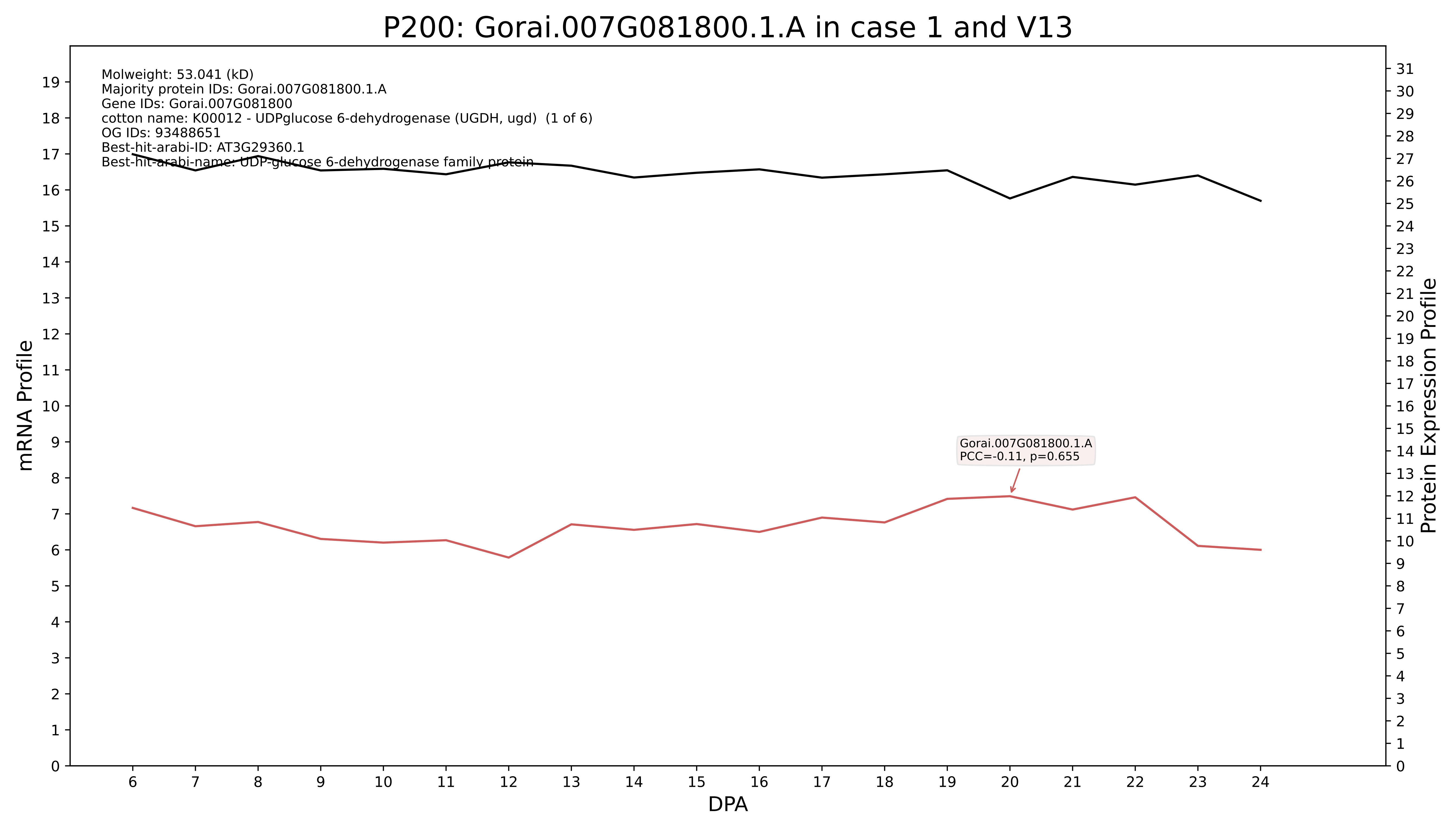
P200: Gorai.007G081800.1.A in case 1 and V13
Molweight: 53.041 (kD)
Majority protein IDs: Gorai.007G081800.1.A
Gene IDs: Gorai.007G081800
cotton name: K00012 - UDPglucose 6-dehydrogenase (UGDH, ugd) (1 of 6)
OG IDs: 93488651
Best-hit-arabi-ID: AT3G29360.1
Best-hit-arabi-name: UDP-glucose 6-dehydrogenase family protein
Gorai.007G081800.1.A
PCC=-0.11, p=0.655
mRNA Profile
Protein Expression Profile
DPA

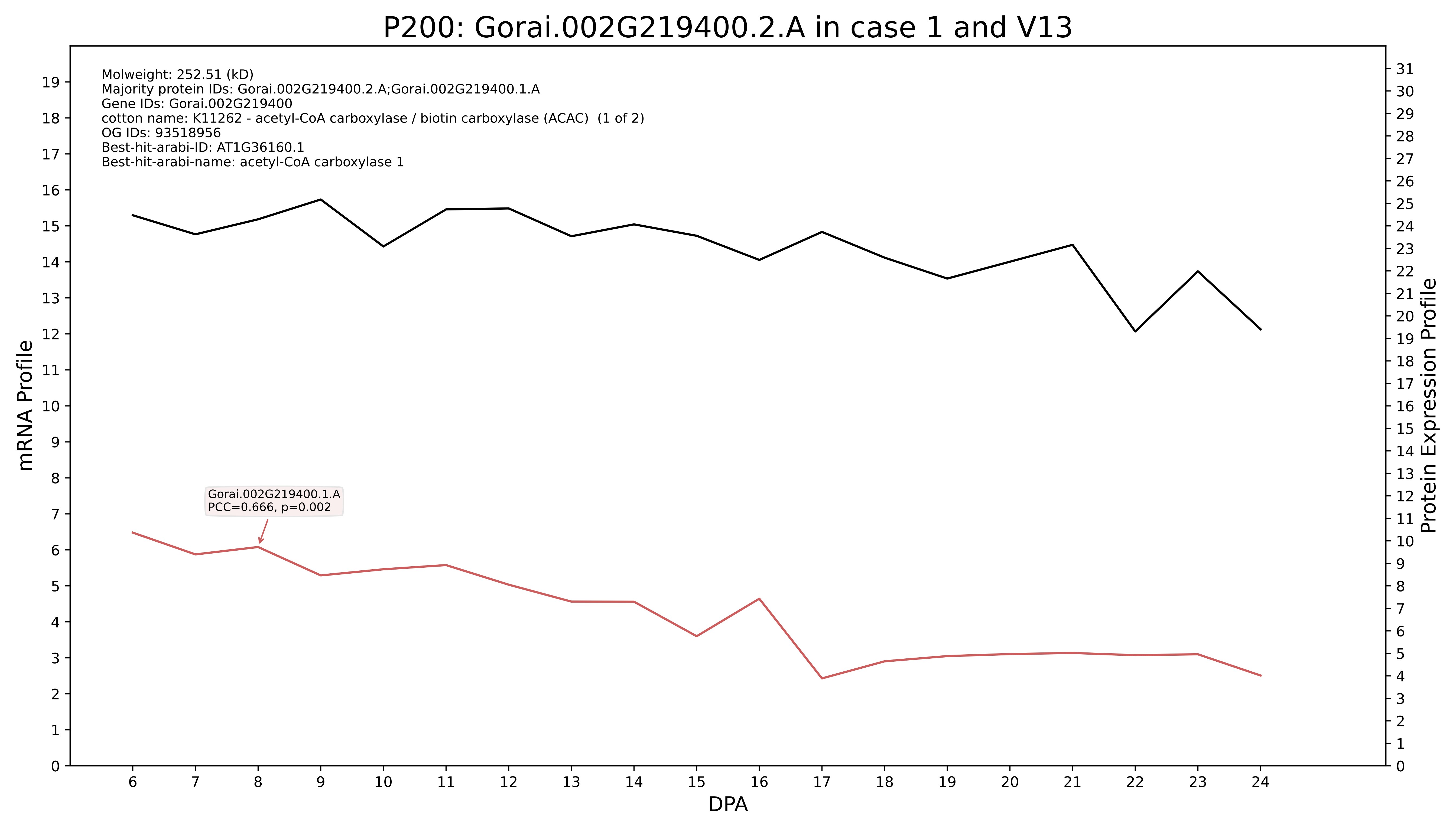
P200: Gorai.002G219400.2.A in case 1 and V13
Molweight: 252.51 (kD)
Majority protein IDs: Gorai.002G219400.2.A;Gorai.002G219400.1.A
Gene IDs: Gorai.002G219400
cotton name: K11262 - acetyl-CoA carboxylase / biotin carboxylase (ACAC) (1 of 2)
OG IDs: 93518956
Best-hit-arabi-ID: AT1G36160.1
Best-hit-arabi-name: acetyl-CoA carboxylase 1
Gorai.002G219400.1.A
PCC=0.666, p=0.002
mRNA Profile
Protein Expression Profile
DPA

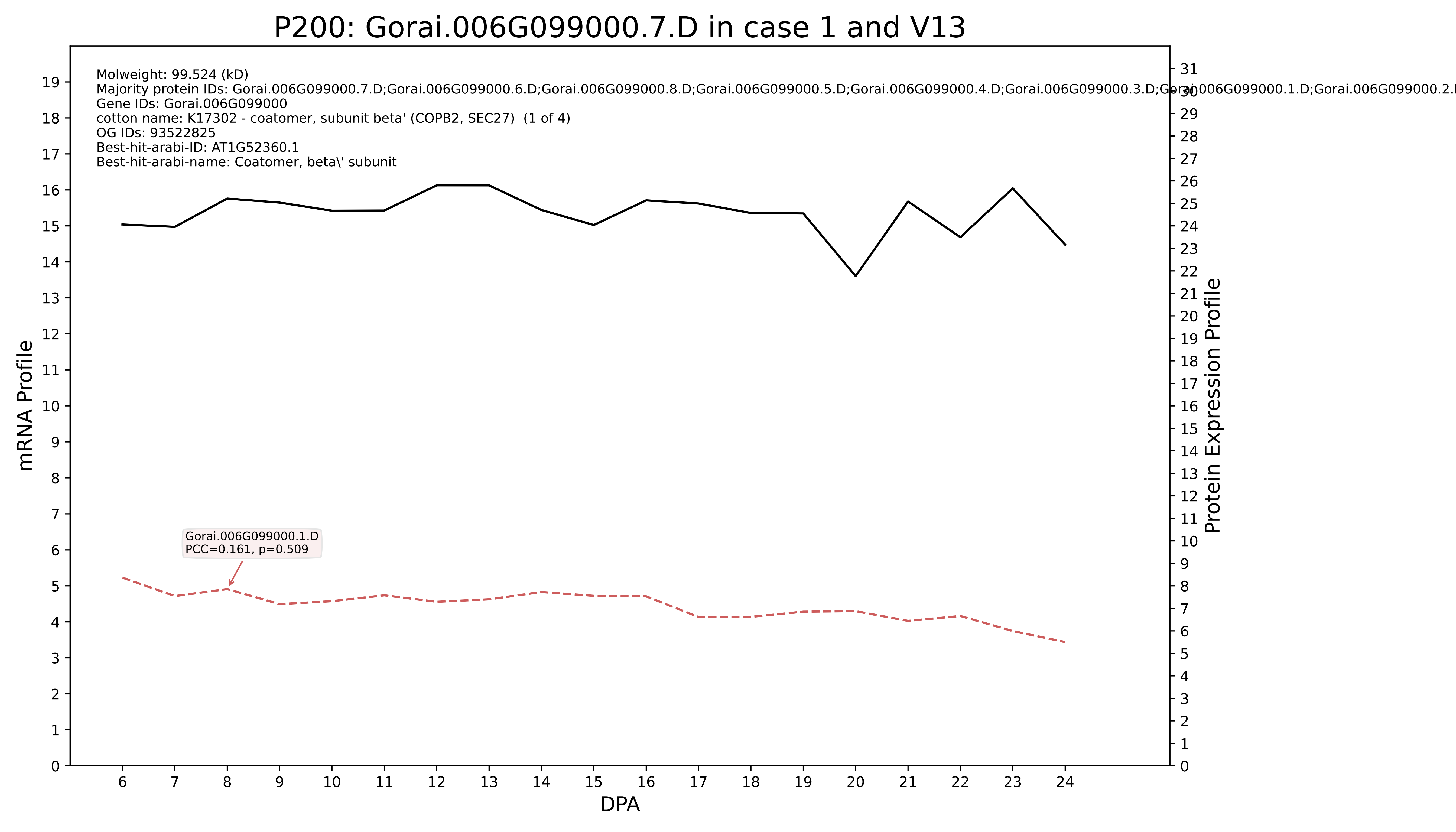
P200: Gorai.006G099000.7.D in case 1 and V13
Molweight: 99.524 (kD)
Majority protein IDs: Gorai.006G099000.7.D;Gorai.006G099000.6.D;Gorai.006G099000.8.D;Gorai.006G099000.5.D;Gorai.006G099000.4.D;Gorai.006G099000.3.D;Gorai.006G099000.1.D;Gorai.006G099000.2.D
Gene IDs: Gorai.006G099000
cotton name: K17302 - coatomer, subunit beta' (COPB2, SEC27) (1 of 4)
OG IDs: 93522825
Best-hit-arabi-ID: AT1G52360.1
Best-hit-arabi-name: Coatomer, beta\' subunit
Gorai.006G099000.1.D
PCC=0.161, p=0.509
mRNA Profile
Protein Expression Profile
DPA

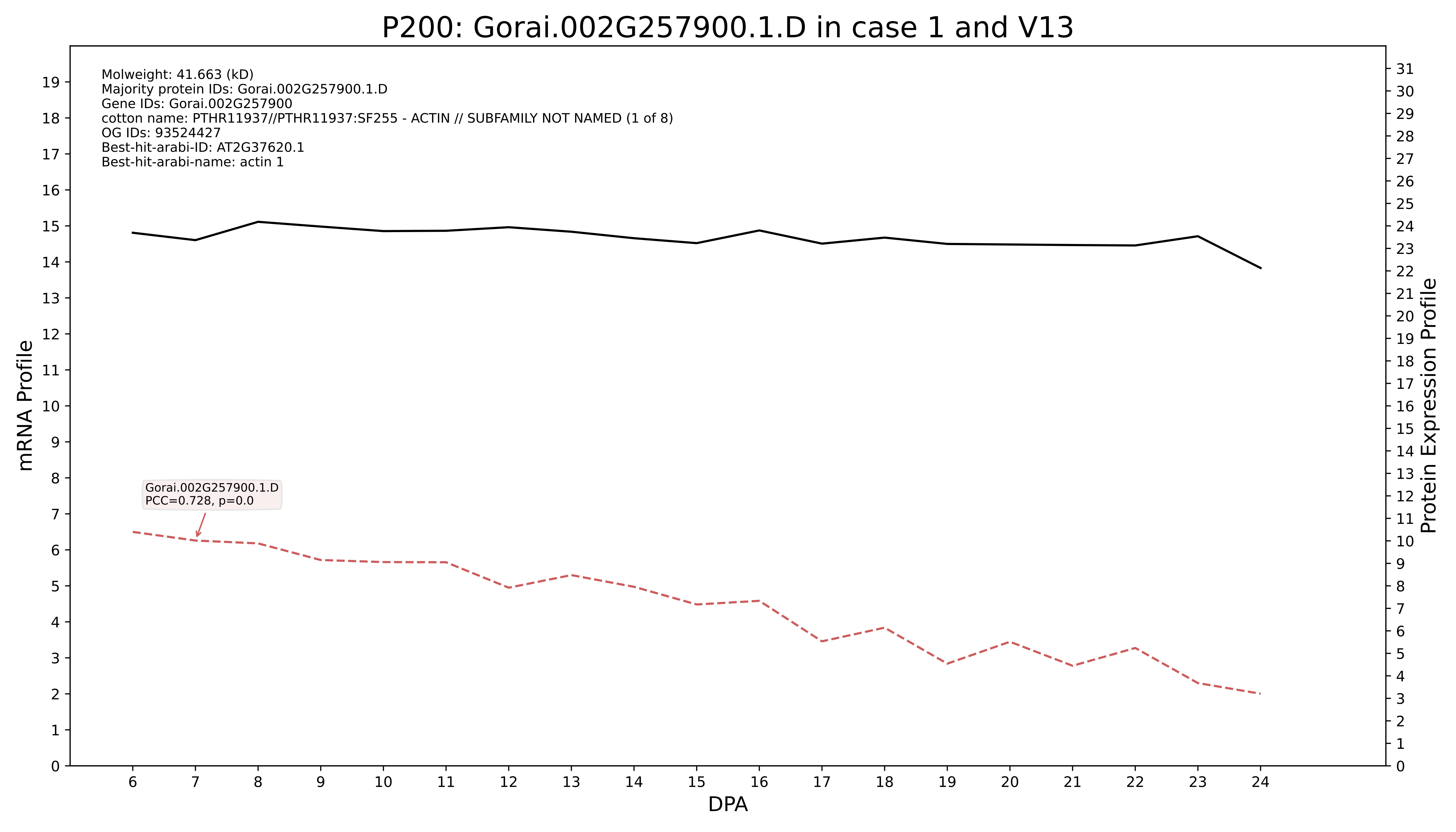

P200: Gorai.002G257900.1.D in case 1 and V13
Molweight: 41.663 (kD)
Majority protein IDs: Gorai.002G257900.1.D
Gene IDs: Gorai.002G257900
cotton name: PTHR11937//PTHR11937:SF255 - ACTIN // SUBFAMILY NOT NAMED (1 of 8)
OG IDs: 93524427
Best-hit-arabi-ID: AT2G37620.1
Best-hit-arabi-name: actin 1
Gorai.002G257900.1.D
PCC=0.728, p=0.0
mRNA Profile
Protein Expression Profile
DPA

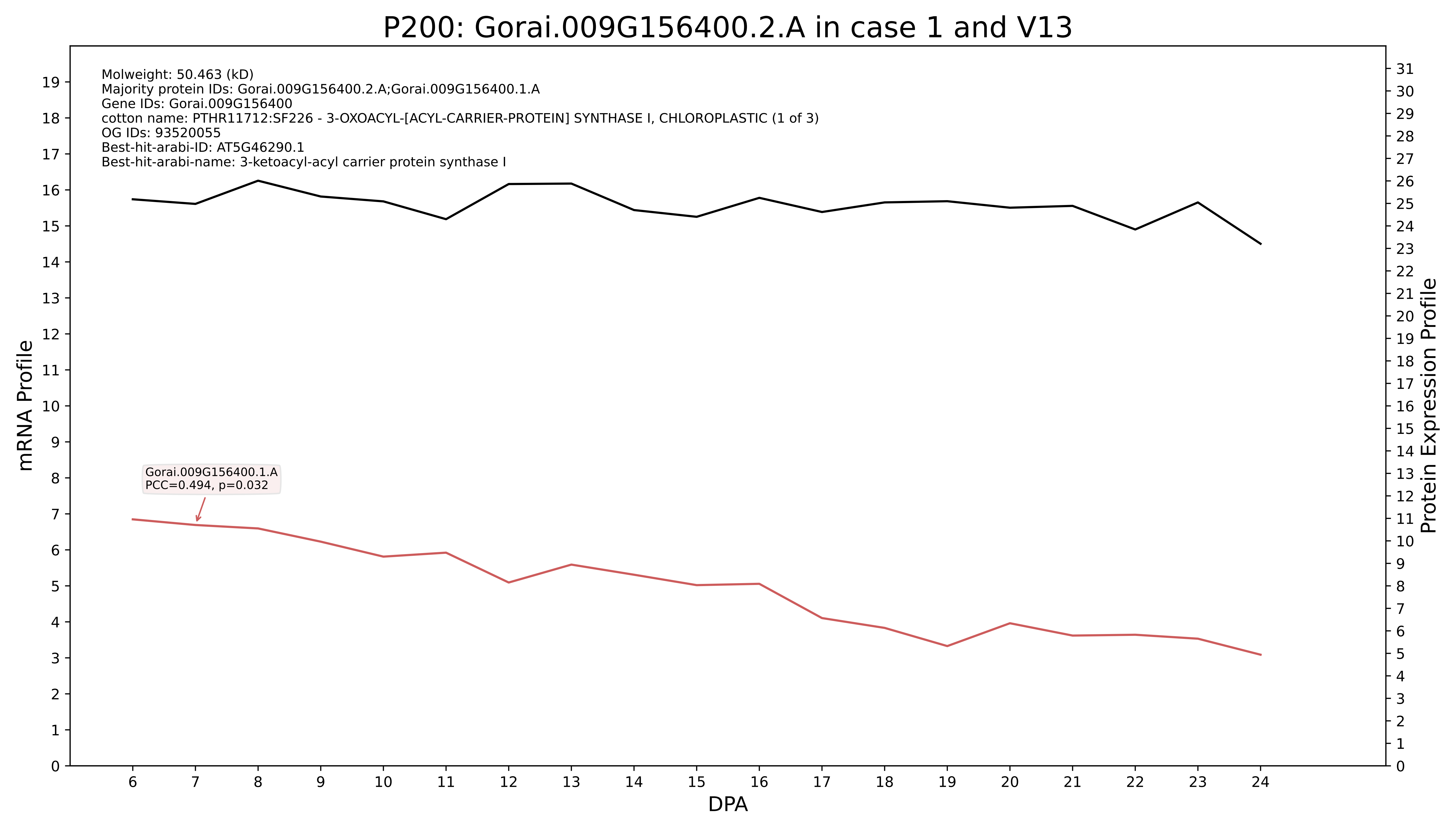
P200: Gorai.009G156400.2.A in case 1 and V13
Molweight: 50.463 (kD)
Majority protein IDs: Gorai.009G156400.2.A;Gorai.009G156400.1.A
Gene IDs: Gorai.009G156400
cotton name: PTHR11712:SF226 - 3-OXOACYL-[ACYL-CARRIER-PROTEIN] SYNTHASE I, CHLOROPLASTIC (1 of 3)
OG IDs: 93520055
Best-hit-arabi-ID: AT5G46290.1
Best-hit-arabi-name: 3-ketoacyl-acyl carrier protein synthase I
Gorai.009G156400.1.A
PCC=0.494, p=0.032
mRNA Profile
Protein Expression Profile
DPA

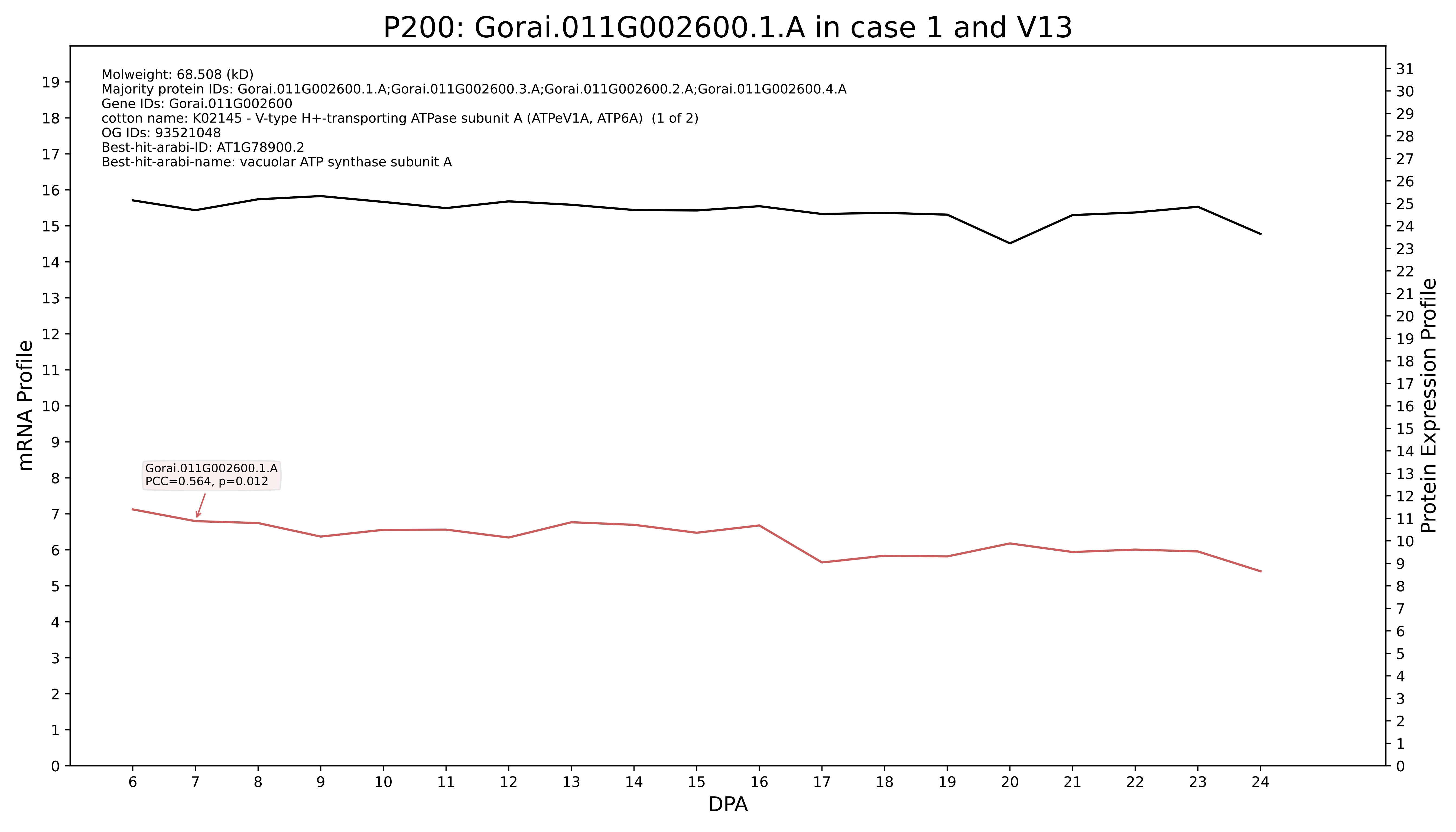
P200: Gorai.011G002600.1.A in case 1 and V13
Molweight: 68.508 (kD)
Majority protein IDs: Gorai.011G002600.1.A;Gorai.011G002600.3.A;Gorai.011G002600.2.A;Gorai.011G002600.4.A
Gene IDs: Gorai.011G002600
cotton name: K02145 - V-type H+-transporting ATPase subunit A (ATPeV1A, ATP6A) (1 of 2)
OG IDs: 93521048
Best-hit-arabi-ID: AT1G78900.2
Best-hit-arabi-name: vacuolar ATP synthase subunit A
Gorai.011G002600.1.A
PCC=0.564, p=0.012
mRNA Profile
Protein Expression Profile
DPA

### P200: Gorai.006G223600.4.D in case 1 and V13

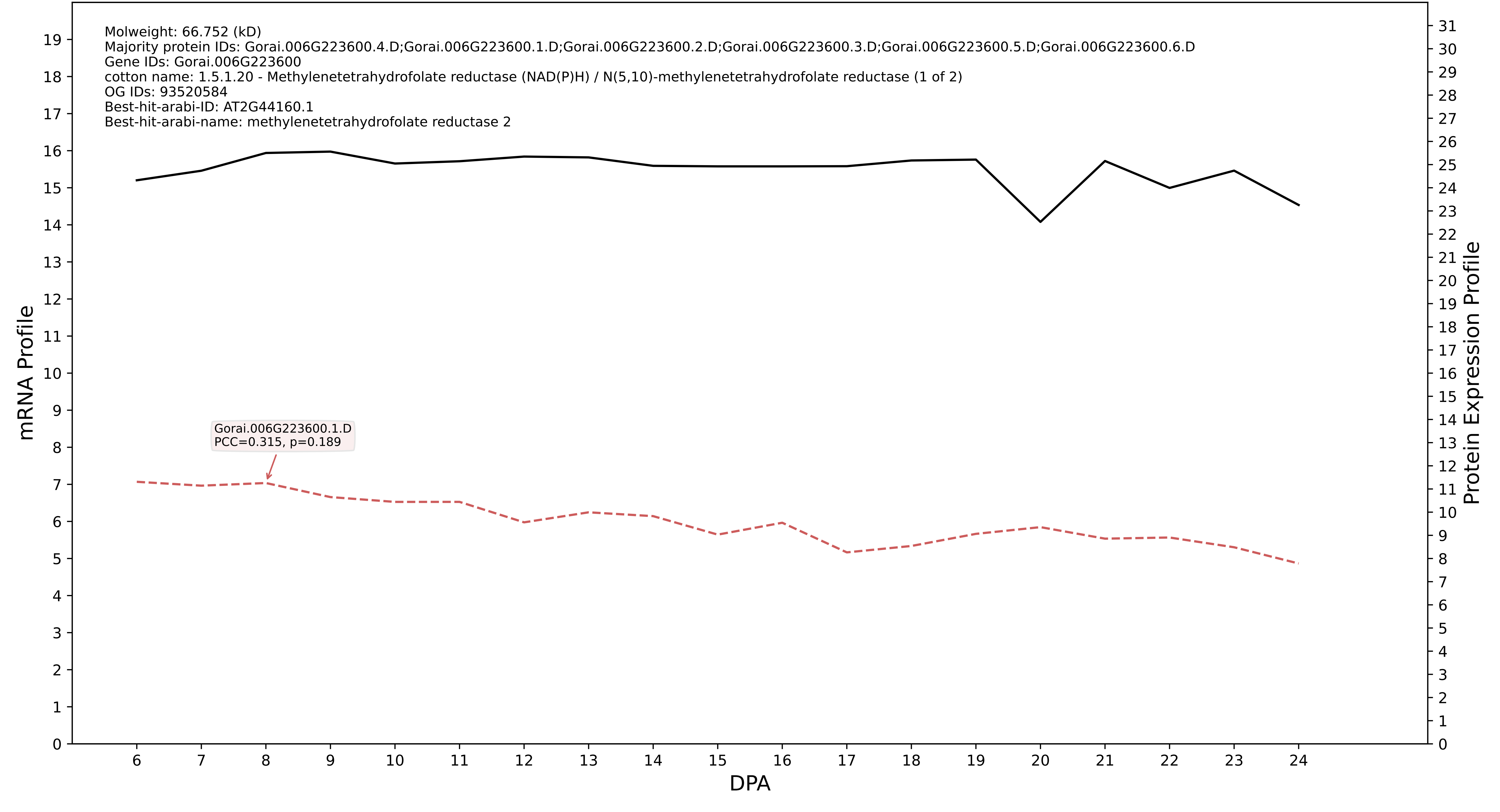

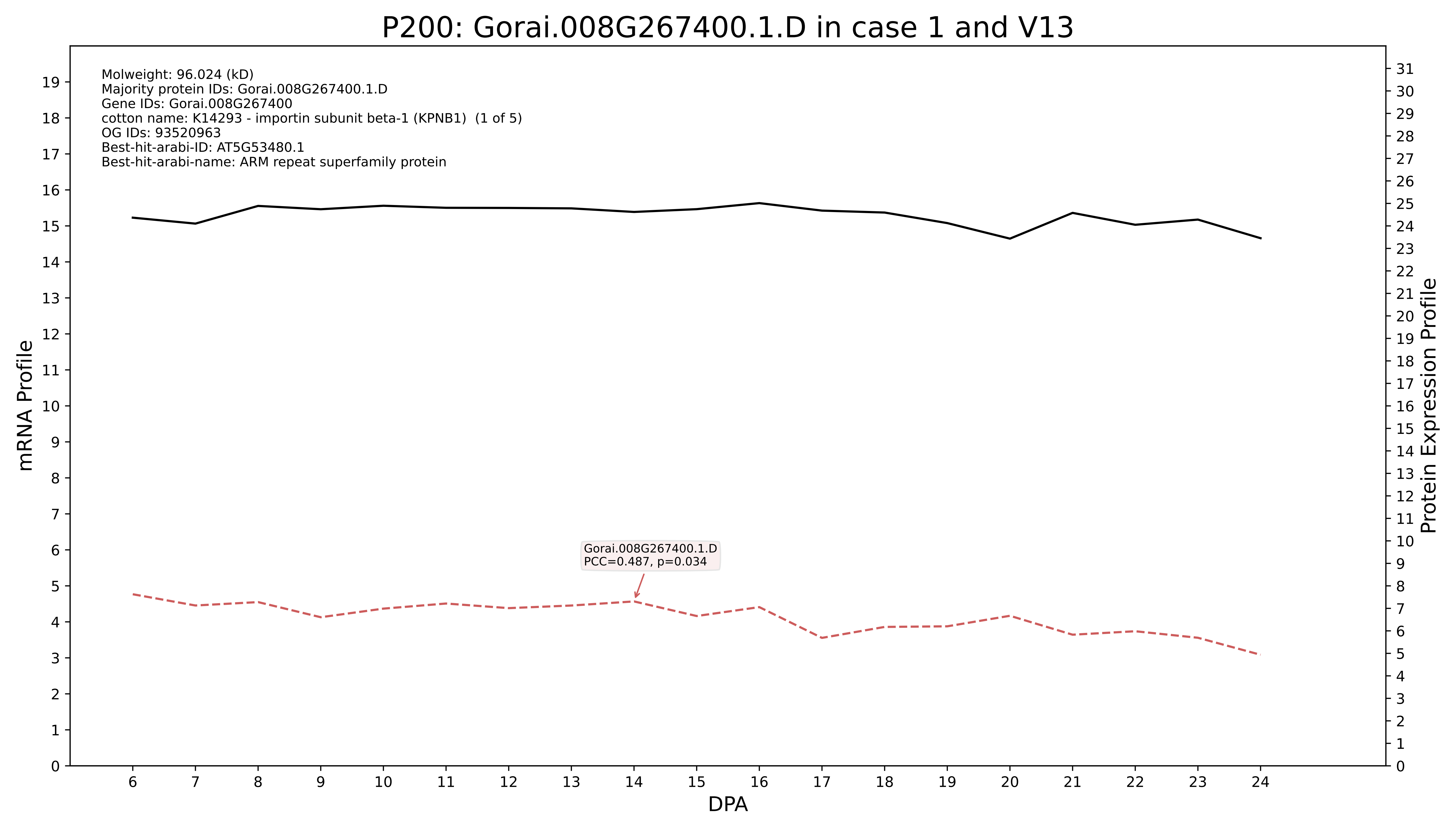
P200: Gorai.008G267400.1.D in case 1 and V13
Molweight: 96.024 (kD)
Majority protein IDs: Gorai.008G267400.1.D
Gene IDs: Gorai.008G267400
cotton name: K14293 - importin subunit beta-1 (KPNB1) (1 of 5)
OG IDs: 93520963
Best-hit-arabi-ID: AT5G53480.1
Best-hit-arabi-name: ARM repeat superfamily protein
Gorai.008G267400.1.D
PCC=0.487, p=0.034
mRNA Profile
Protein Expression Profile
DPA

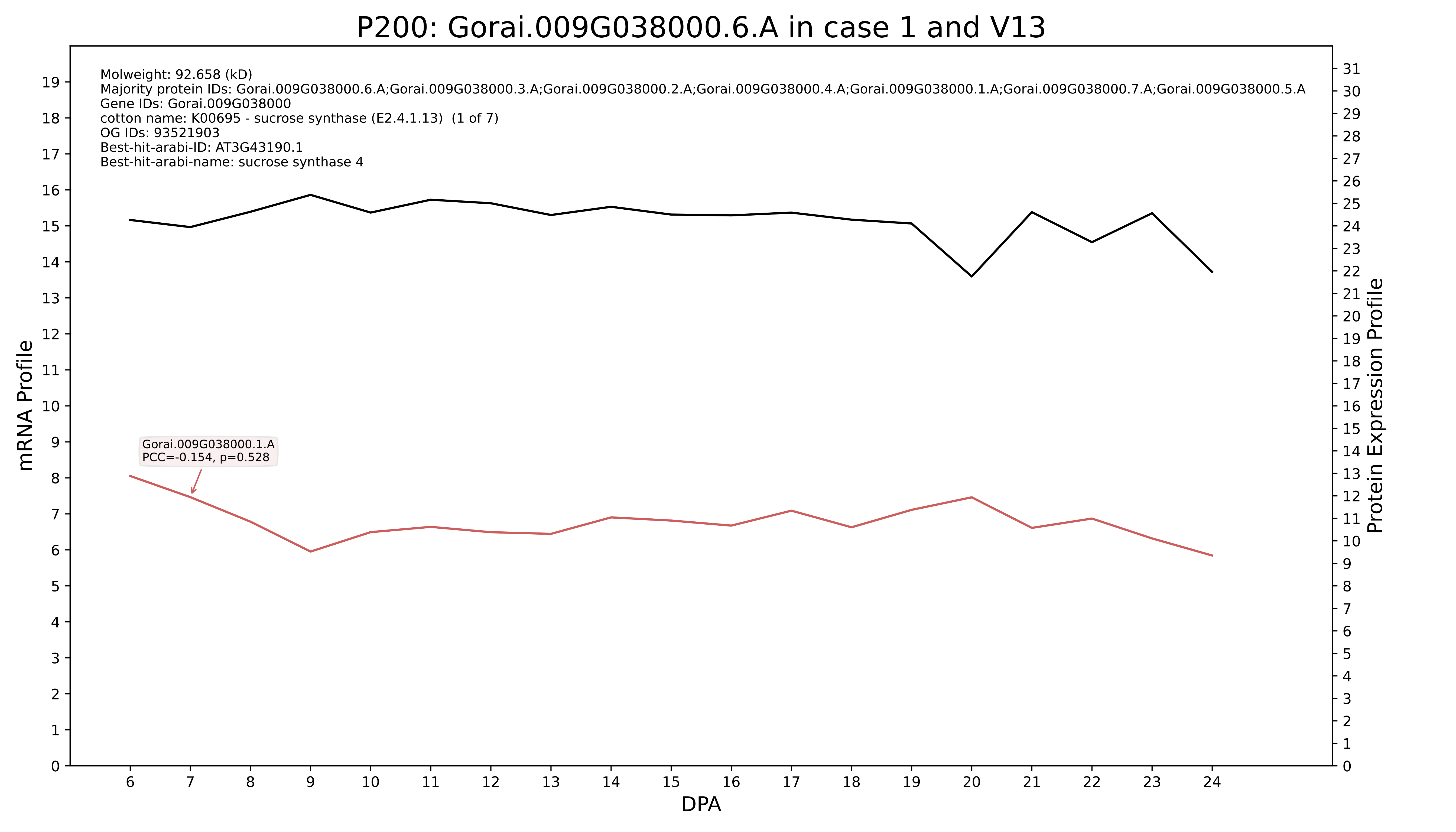
P200: Gorai.009G038000.6.A in case 1 and V13
Molweight: 92.658 (kD)
Majority protein IDs: Gorai.009G038000.6.A;Gorai.009G038000.3.A;Gorai.009G038000.2.A;Gorai.009G038000.4.A;Gorai.009G038000.1.A;Gorai.009G038000.7.A;Gorai.009G038000.5.A
Gene IDs: Gorai.009G038000
cotton name: K00695 - sucrose synthase (E2.4.1.13) (1 of 7)
OG IDs: 93521903
Best-hit-arabi-ID: AT3G43190.1
Best-hit-arabi-name: sucrose synthase 4
Gorai.009G038000.1.A
PCC=-0.154, p=0.528
mRNA Profile
Protein Expression Profile
DPA

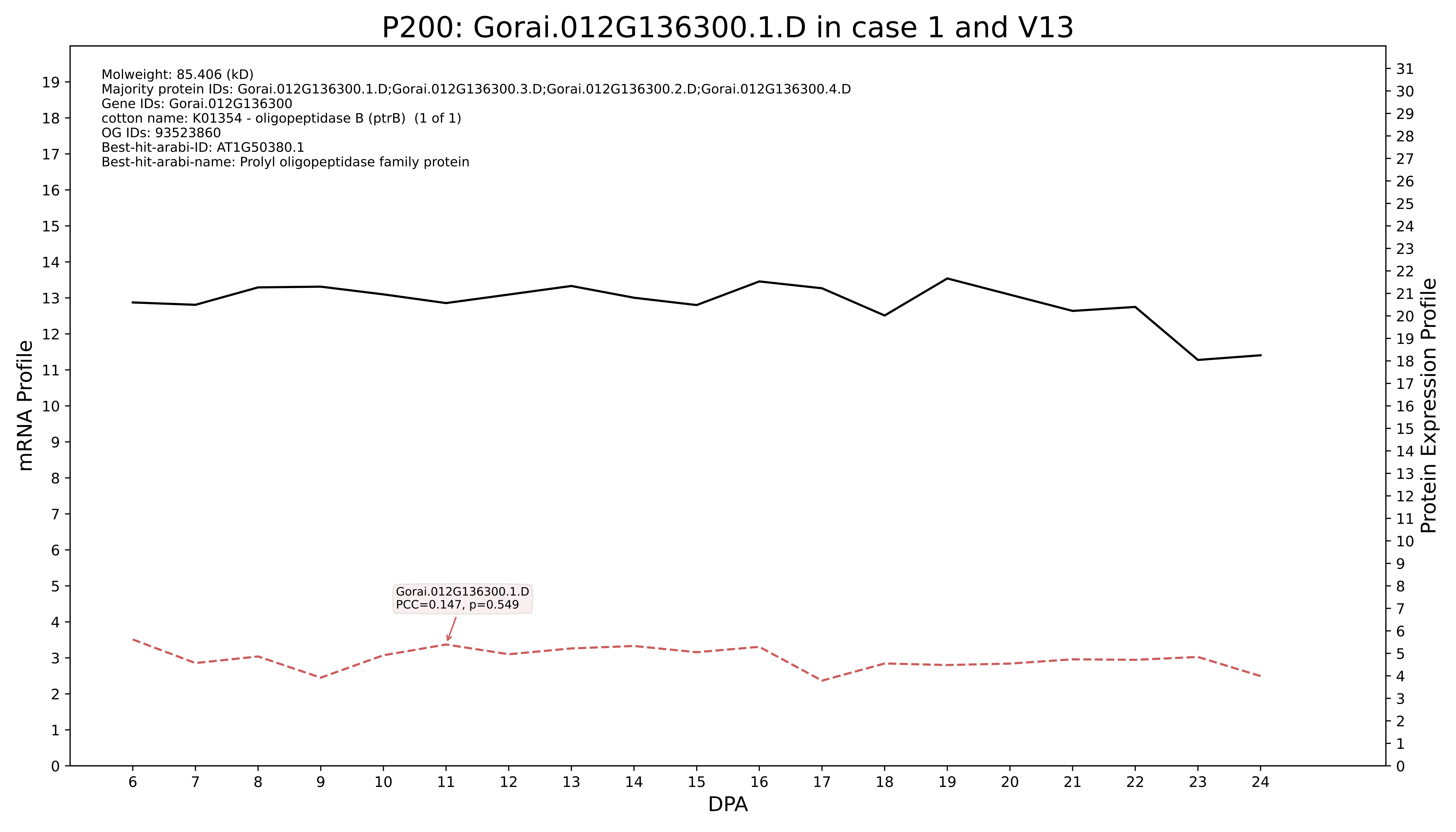
P200: Gorai.012G136300.1.D in case 1 and V13
Molweight: 85.406 (kD)
Majority protein IDs: Gorai.012G136300.1.D;Gorai.012G136300.3.D;Gorai.012G136300.2.D;Gorai.012G136300.4.D
Gene IDs: Gorai.012G136300
cotton name: K01354 - oligopeptidase B (ptrB) (1 of 1)
OG IDs: 93523860
Best-hit-arabi-ID: AT1G50380.1
Best-hit-arabi-name: Prolyl oligopeptidase family protein
Gorai.012G136300.1.D
PCC=0.147, p=0.549
mRNA Profile
Protein Expression Profile
DPA

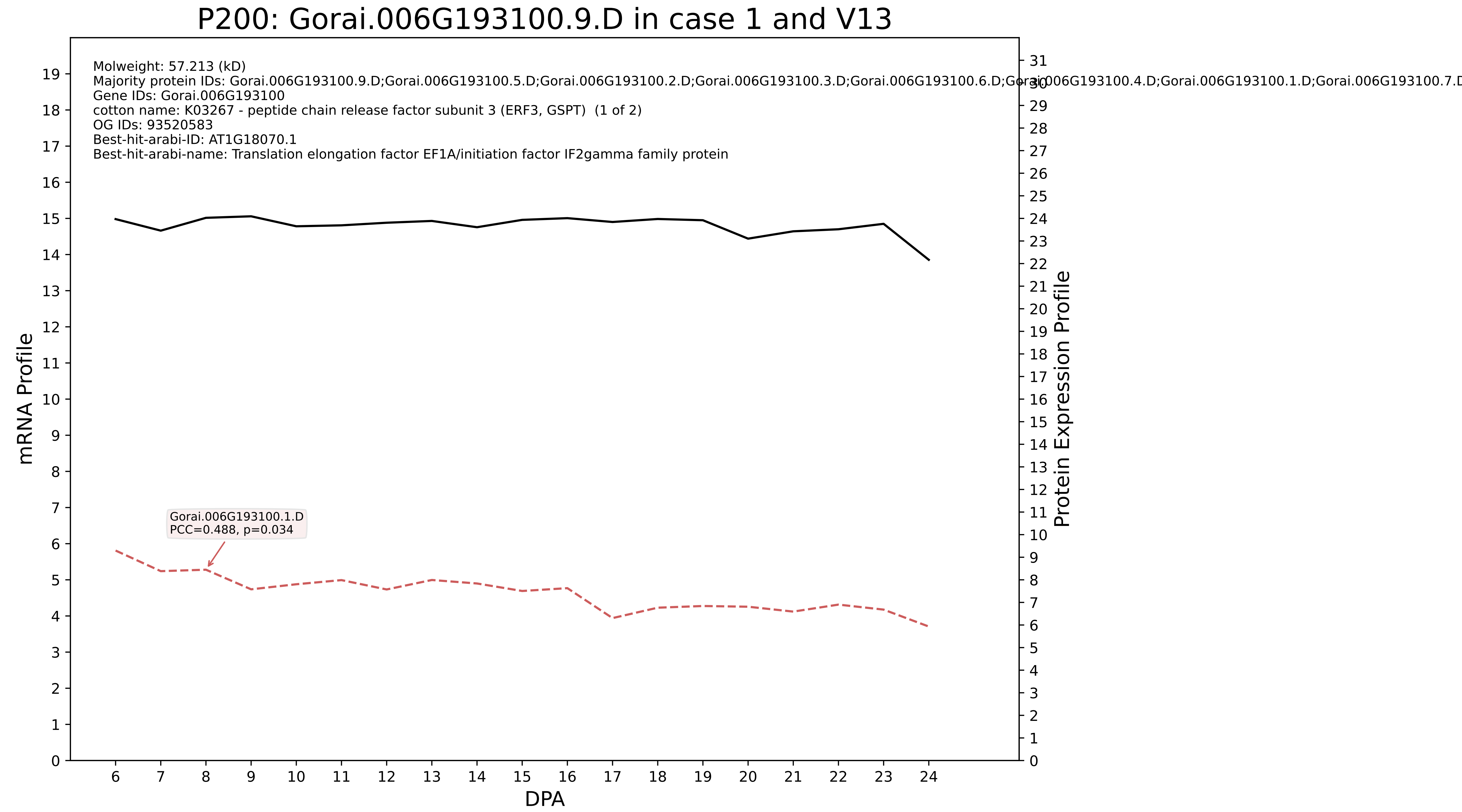
P200: Gorai.006G193100.9.D in case 1 and V13
Molweight: 57.213 (kD)
Majority protein IDs: Gorai.006G193100.9.D;Gorai.006G193100.5.D;Gorai.006G193100.2.D;Gorai.006G193100.3.D;Gorai.006G193100.6.D;Gorai.006G193100.4.D;Gorai.006G193100.1.D;Gorai.006G193100.7.D
Gene IDs: Gorai.006G193100
cotton name: K03267 - peptide chain release factor subunit 3 (ERF3, GSPT) (1 of 2)
OG IDs: 93520583
Best-hit-arabi-ID: AT1G18070.1
Best-hit-arabi-name: Translation elongation factor EF1A/initiation factor IF2gamma family protein
Gorai.006G193100.1.D
PCC=0.488, p=0.034
mRNA Profile
Protein Expression Profile
DPA

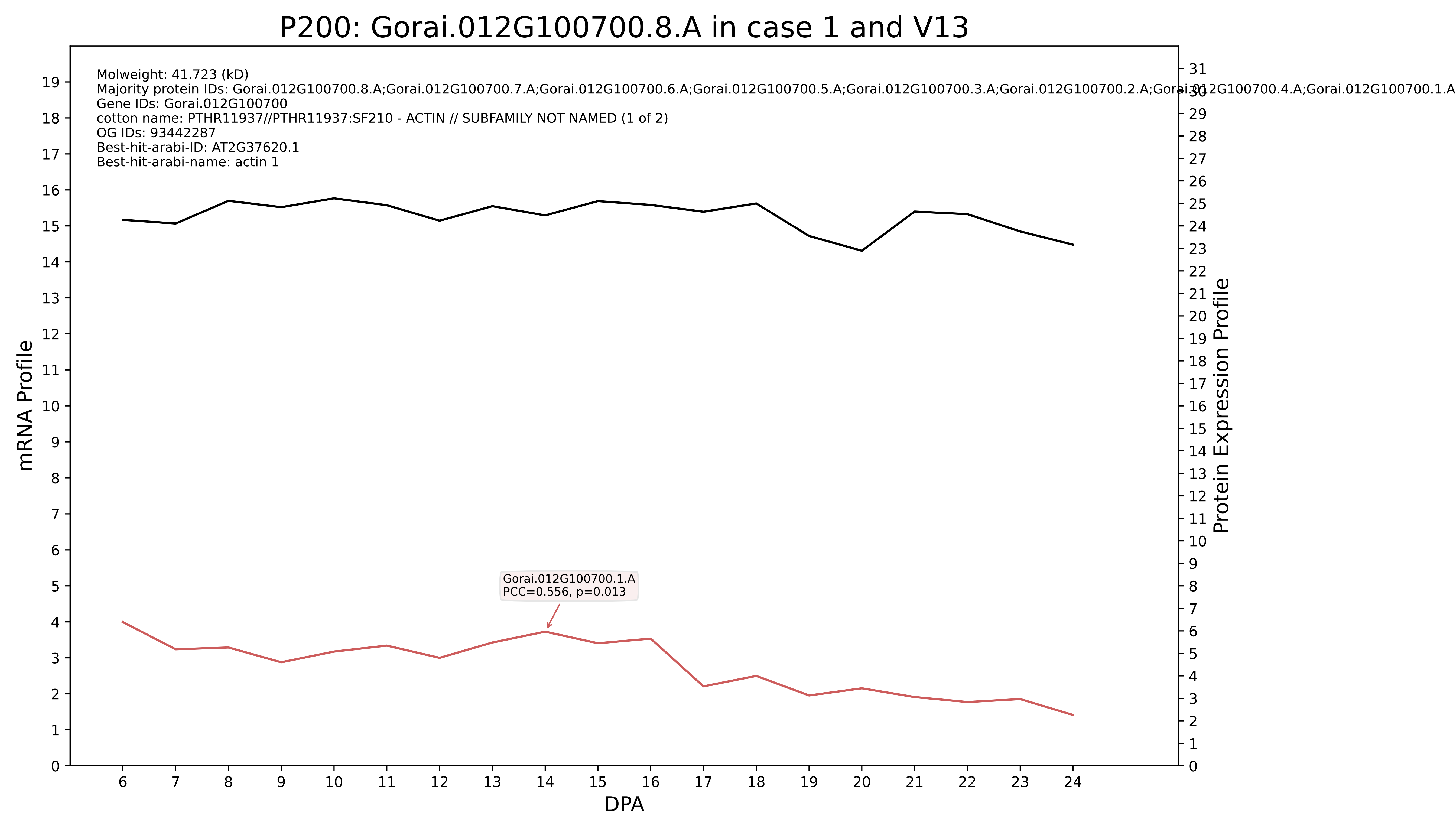
P200: Gorai.012G100700.8.A in case 1 and V13
Molweight: 41.723 (kD)
Majority protein IDs: Gorai.012G100700.8.A;Gorai.012G100700.7.A;Gorai.012G100700.6.A;Gorai.012G100700.5.A;Gorai.012G100700.3.A;Gorai.012G100700.2.A;Gorai.012G100700.4.A;Gorai.012G100700.1.A
Gene IDs: Gorai.012G100700
cotton name: PTHR11937//PTHR11937:SF210 - ACTIN // SUBFAMILY NOT NAMED (1 of 2)
OG IDs: 93442287
Best-hit-arabi-ID: AT2G37620.1
Best-hit-arabi-name: actin 1
Gorai.012G100700.1.A
PCC=0.556, p=0.013
mRNA Profile
Protein Expression Profile
DPA

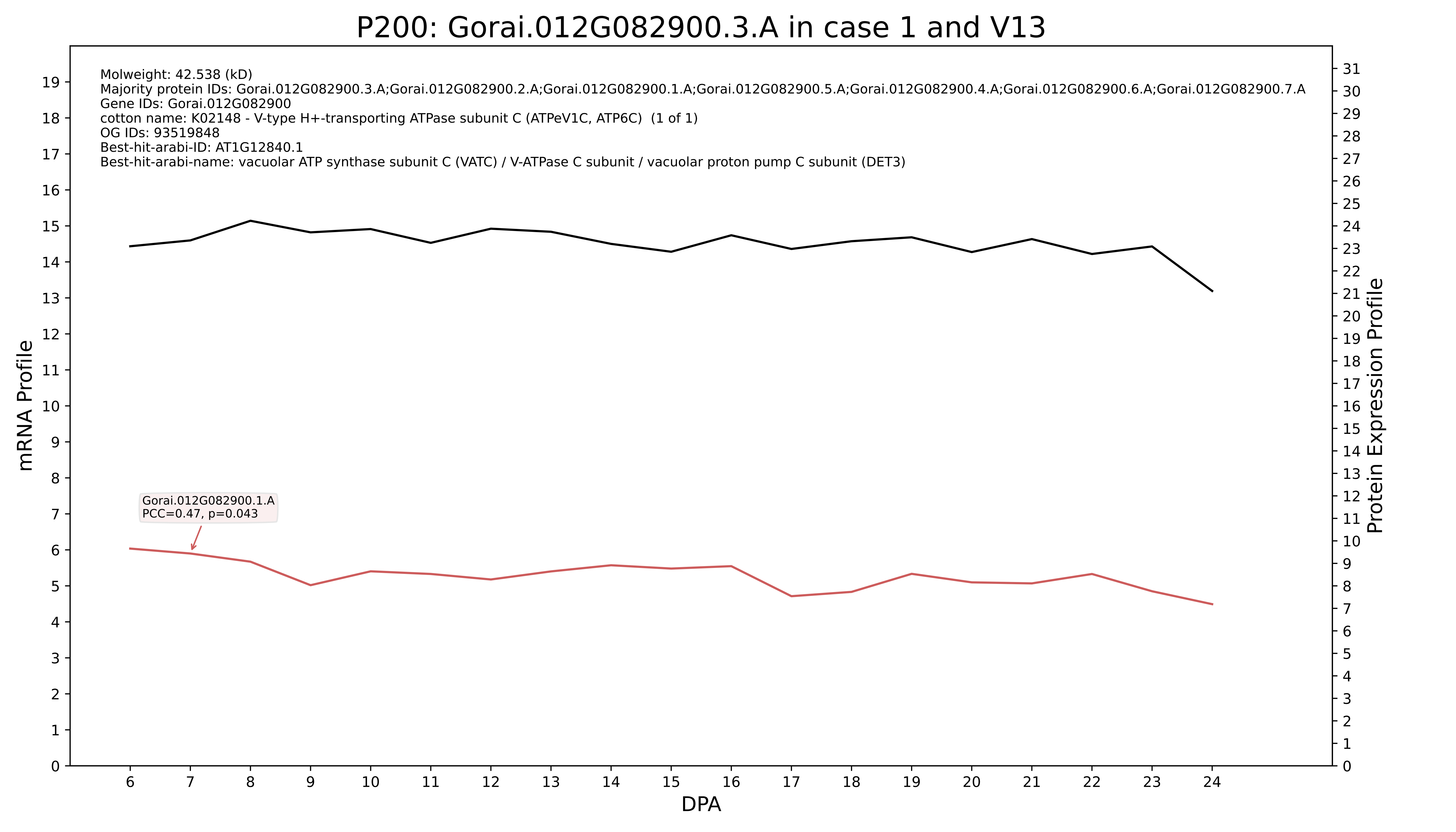
P200: Gorai.012G082900.3.A in case 1 and V13
Molweight: 42.538 (kD)
Majority protein IDs: Gorai.012G082900.3.A;Gorai.012G082900.2.A;Gorai.012G082900.1.A;Gorai.012G082900.5.A;Gorai.012G082900.4.A;Gorai.012G082900.6.A;Gorai.012G082900.7.A
Gene IDs: Gorai.012G082900
cotton name: K02148 - V-type H+-transporting ATPase subunit C (ATPeV1C, ATP6C) (1 of 1)
OG IDs: 93519848
Best-hit-arabi-ID: AT1G12840.1
Best-hit-arabi-name: vacuolar ATP synthase subunit C (VATC) / V-ATPase C subunit / vacuolar proton pump C subunit (DET3)
Gorai.012G082900.1.A
PCC=0.47, p=0.043
mRNA Profile
Protein Expression Profile
DPA

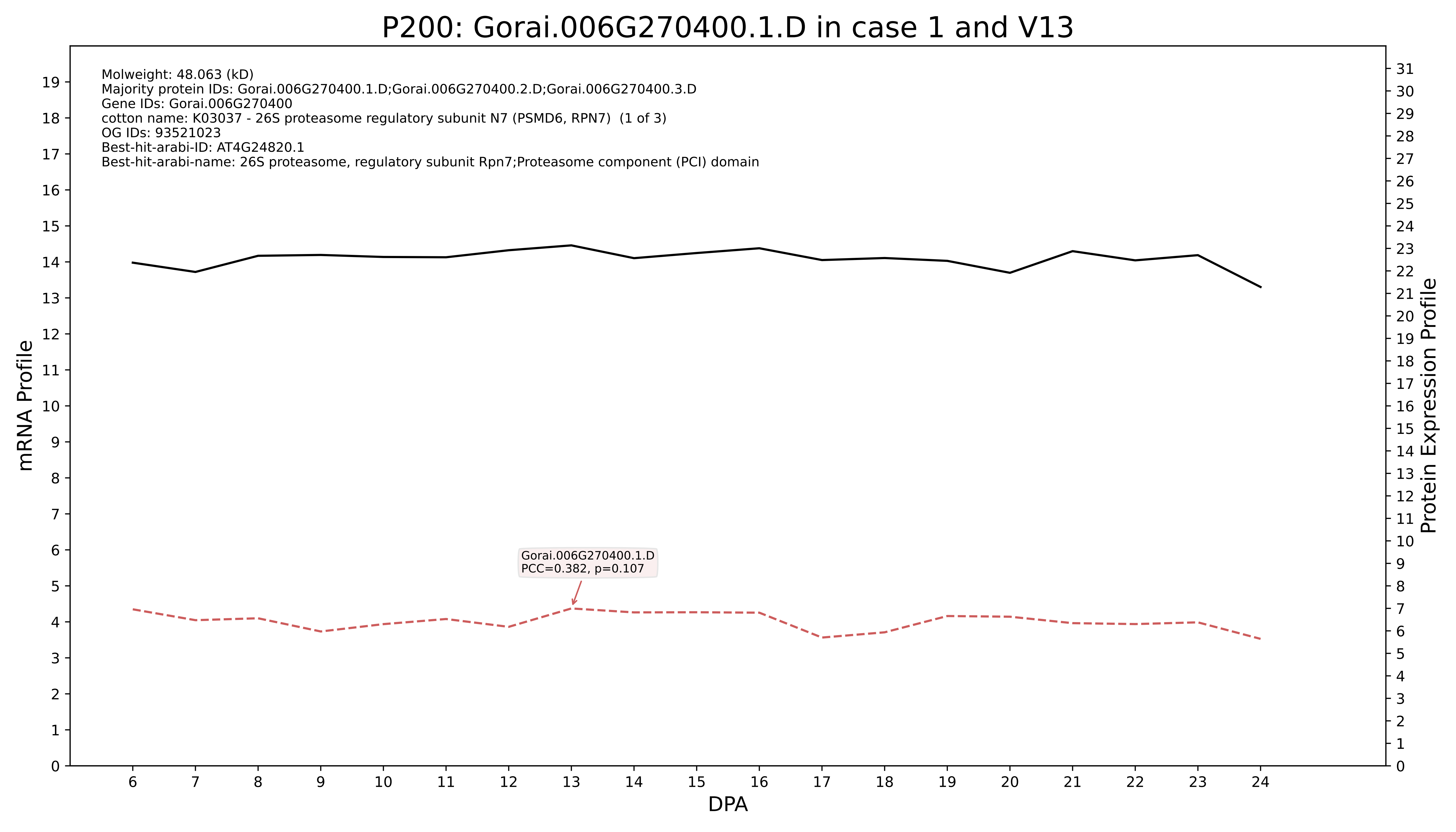
P200: Gorai.006G270400.1.D in case 1 and V13
Molweight: 48.063 (kD)
Majority protein IDs: Gorai.006G270400.1.D;Gorai.006G270400.2.D;Gorai.006G270400.3.D
Gene IDs: Gorai.006G270400
cotton name: K03037 - 26S proteasome regulatory subunit N7 (PSMD6, RPN7) (1 of 3)
OG IDs: 93521023
Best-hit-arabi-ID: AT4G24820.1
Best-hit-arabi-name: 26S proteasome, regulatory subunit Rpn7;Proteasome component (PCI) domain
Gorai.006G270400.1.D
PCC=0.382, p=0.107
mRNA Profile
Protein Expression Profile
DPA

### P200: Gorai.004G132600.1.A in case 1 and V13

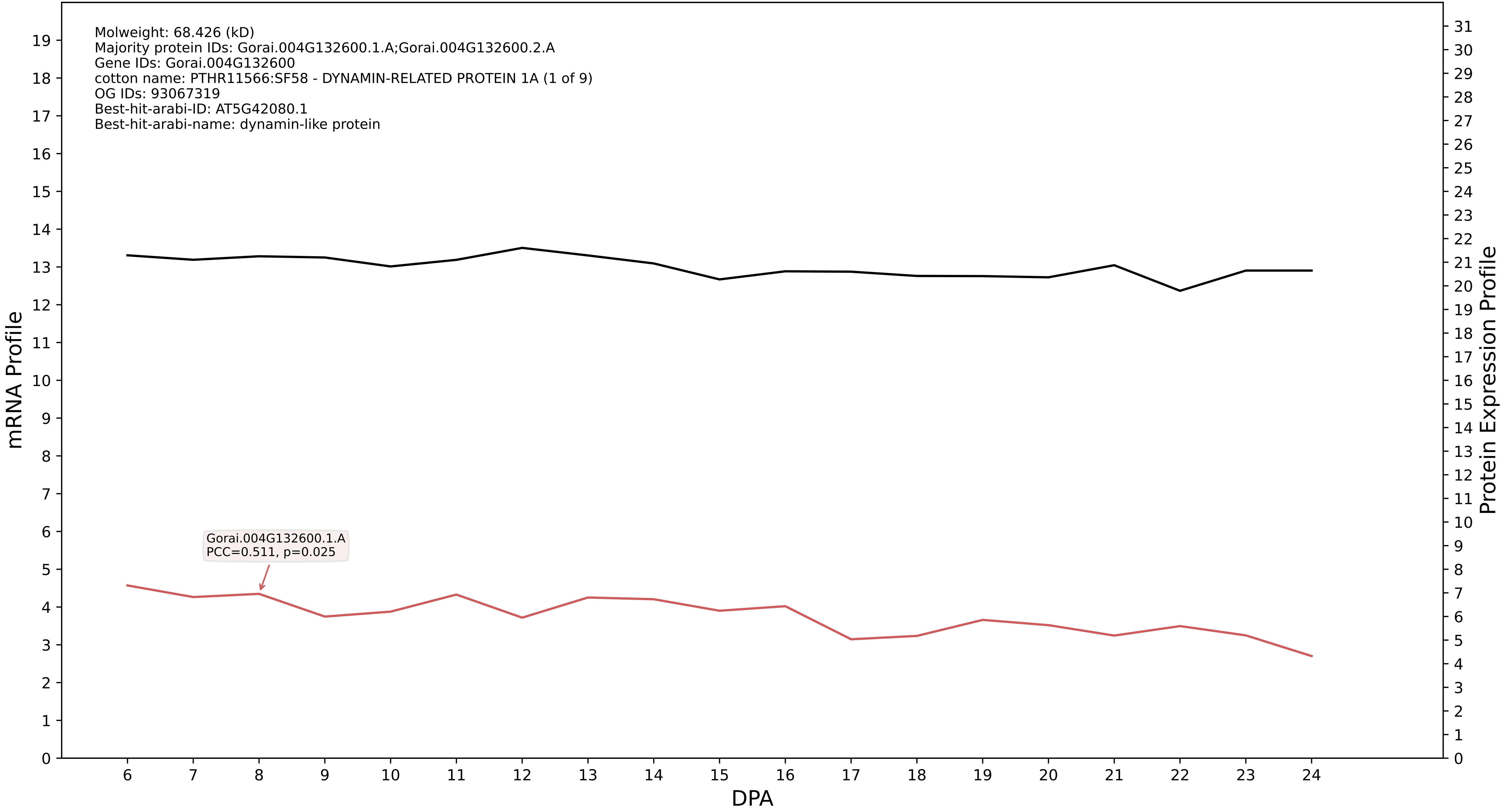

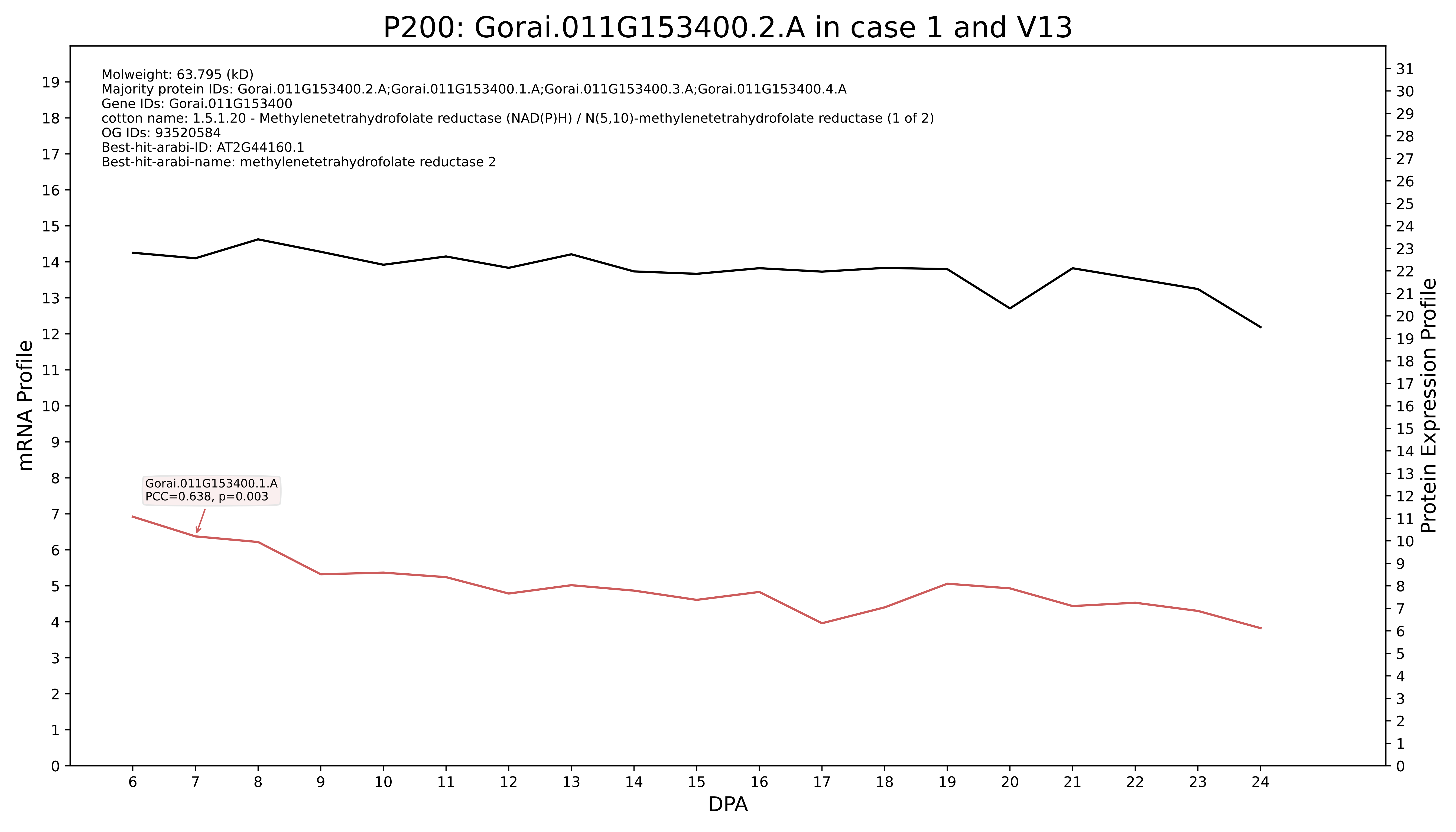
P200: Gorai.011G153400.2.A in case 1 and V13
Molweight: 63.795 (kD)
Majority protein IDs: Gorai.011G153400.2.A;Gorai.011G153400.1.A;Gorai.011G153400.3.A;Gorai.011G153400.4.A
Gene IDs: Gorai.011G153400
cotton name: 1.5.1.20 - Methylenetetrahydrofolate reductase (NAD(P)H) / N(5,10)-methylenetetrahydrofolate reductase (1 of 2)
OG IDs: 93520584
Best-hit-arabi-ID: AT2G44160.1
Best-hit-arabi-name: methylenetetrahydrofolate reductase 2
Gorai.011G153400.1.A
PCC=0.638, p=0.003
mRNA Profile
Protein Expression Profile
DPA

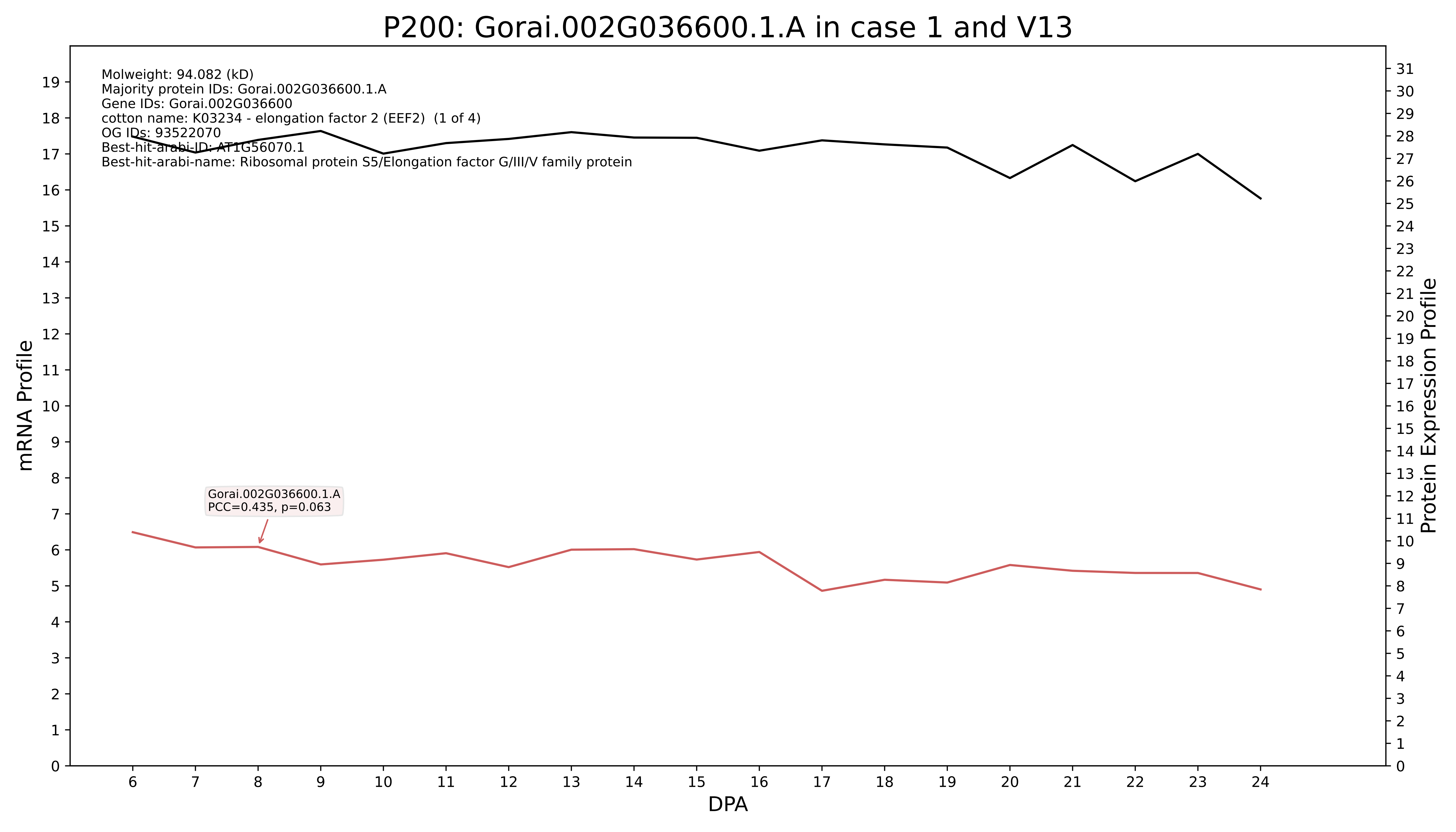

P200: Gorai.002G036600.1.A in case 1 and V13
Molweight: 94.082 (kD)
Majority protein IDs: Gorai.002G036600.1.A
Gene IDs: Gorai.002G036600
cotton name: K03234 - elongation factor 2 (EEF2) (1 of 4)
OG IDs: 93522070
Best-hit-arabi-ID: AT1G56070.1
Best-hit-arabi-name: Ribosomal protein S5/Elongation factor G/III/V family protein
Gorai.002G036600.1.A
PCC=0.435, p=0.063
mRNA Profile
Protein Expression Profile
DPA

P200: Gorai.010G035600.1.D in case 1 and V13

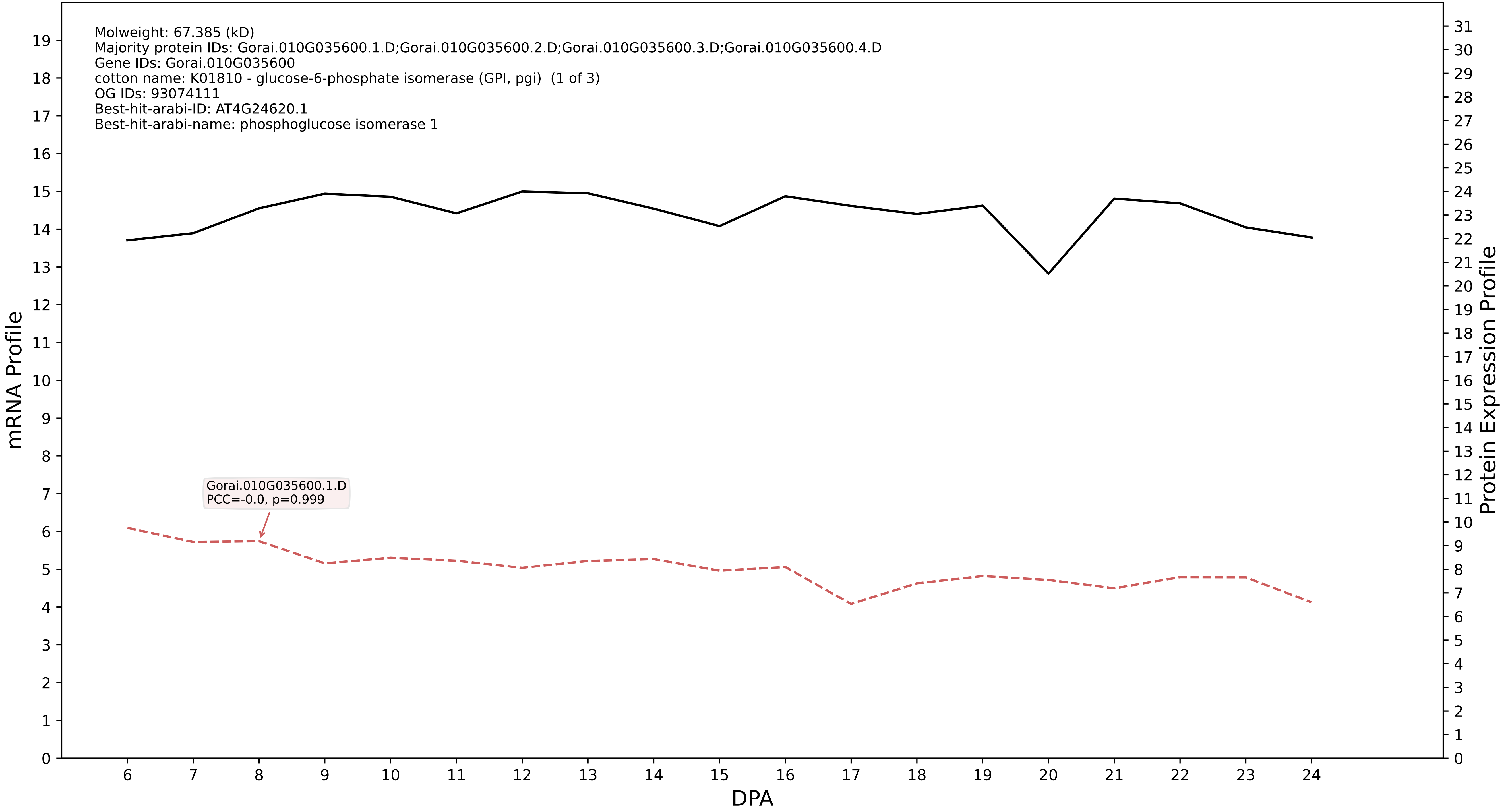

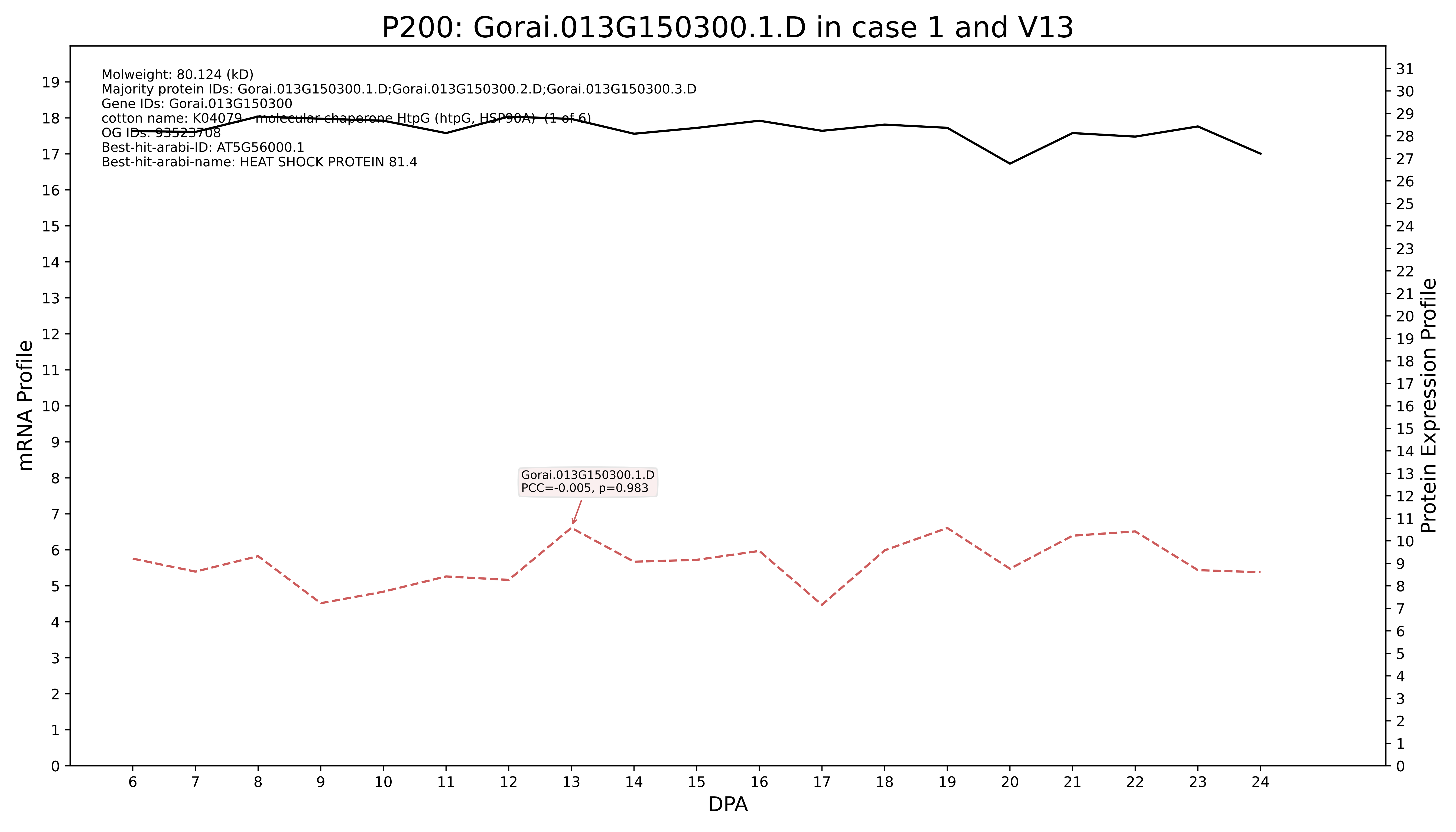
P200: Gorai.013G150300.1.D in case 1 and V13
Molweight: 80.124 (kD)
Majority protein IDs: Gorai.013G150300.1.D;Gorai.013G150300.2.D;Gorai.013G150300.3.D
Gene IDs: Gorai.013G150300
cotton name: K04079 molecular chaperone HtpG (htpG, HSP90A) (1 of 6)
OG IDs: 93523708
Best-hit-arabi-ID: AT5G56000.1
Best-hit-arabi-name: HEAT SHOCK PROTEIN 81.4
Gorai.013G150300.1.D
PCC=-0.005, p=0.983
mRNA Profile
Protein Expression Profile
DPA

### P200: Gorai.002G020200.1.D in case 1 and V13

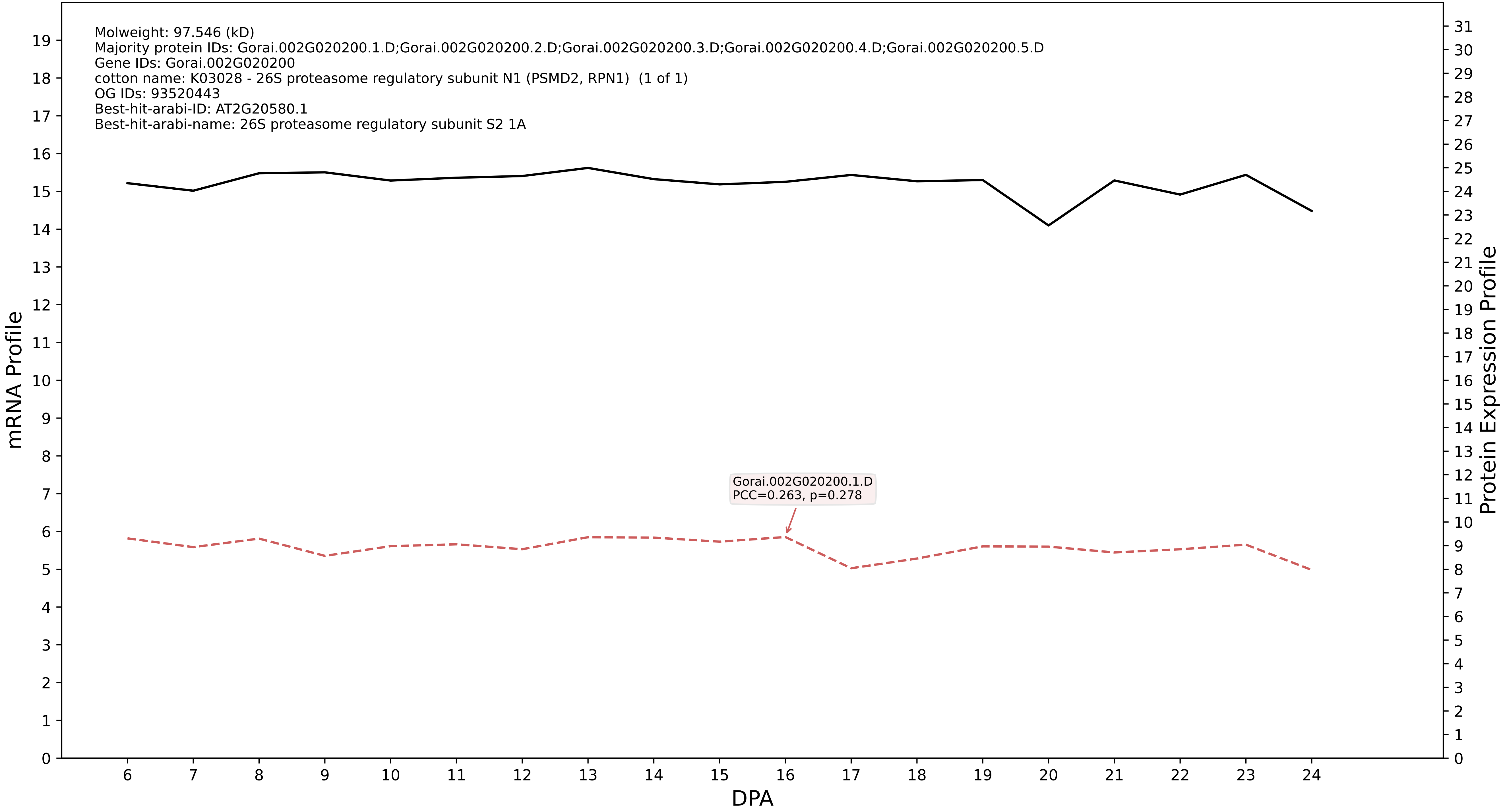

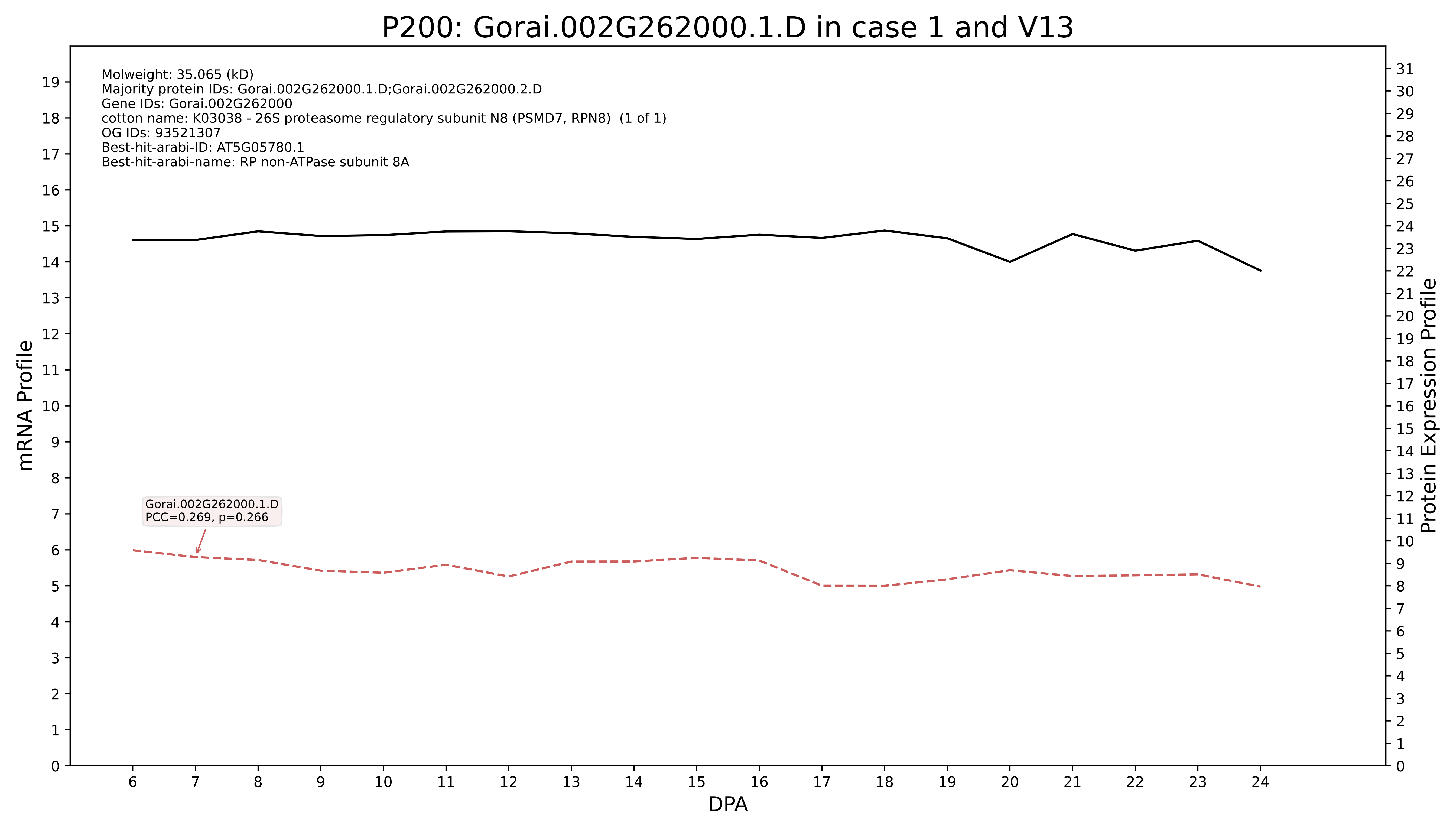

P200: Gorai.002G262000.1.D in case 1 and V13
Molweight: 35.065 (kD)
Majority protein IDs: Gorai.002G262000.1.D;Gorai.002G262000.2.D
Gene IDs: Gorai.002G262000
cotton name: K03038 - 26S proteasome regulatory subunit N8 (PSMD7, RPN8) (1 of 1)
OG IDs: 93521307
Best-hit-arabi-ID: AT5G05780.1
Best-hit-arabi-name: RP non-ATPase subunit 8A
Gorai.002G262000.1.D
PCC=0.269, p=0.266
mRNA Profile
Protein Expression Profile
DPA

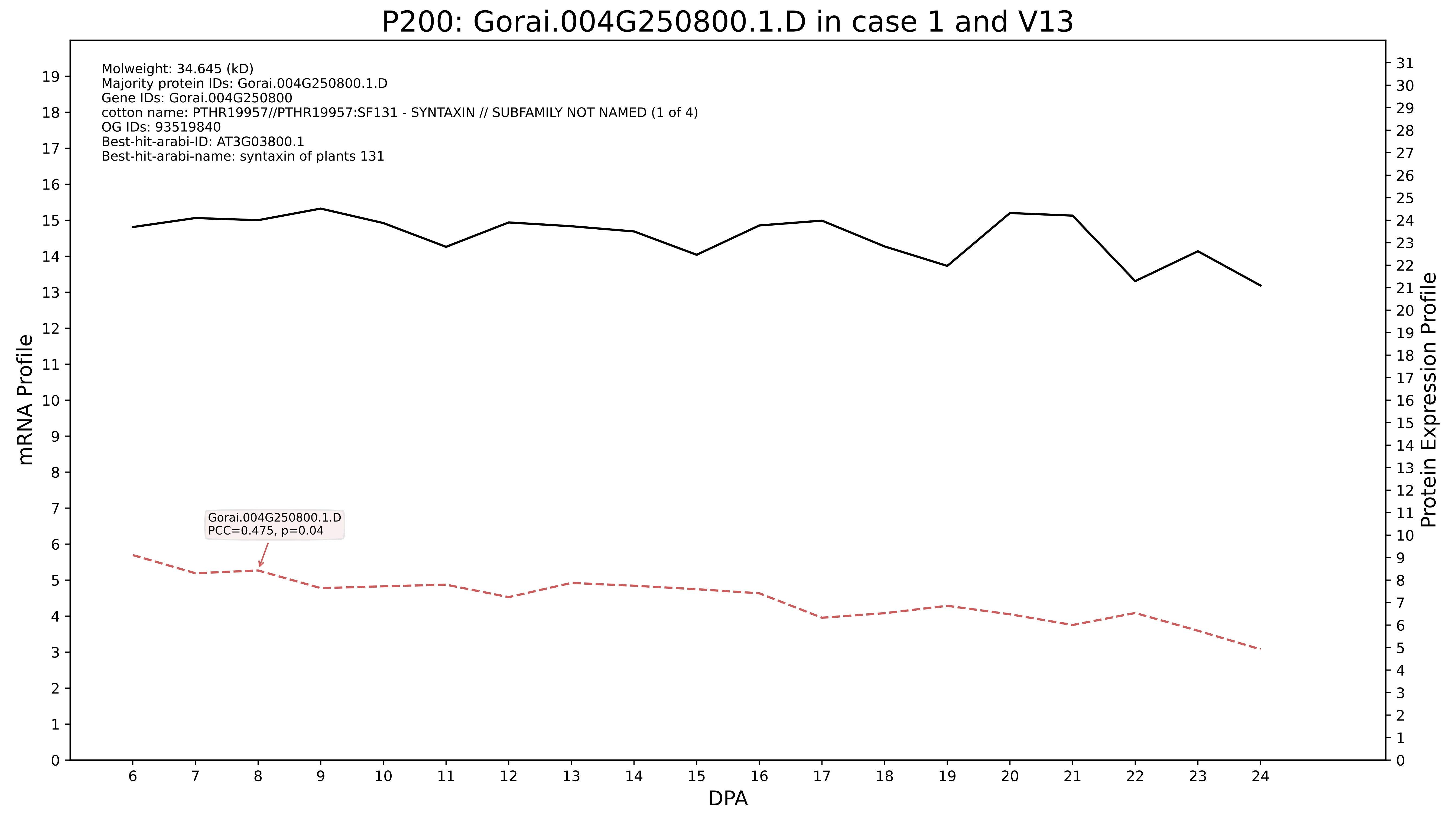
P200: Gorai.004G250800.1.D in case 1 and V13
Molweight: 34.645 (kD)
Majority protein IDs: Gorai.004G250800.1.D
Gene IDs: Gorai.004G250800
cotton name: PTHR19957//PTHR19957:SF131 - SYNTAXIN // SUBFAMILY NOT NAMED (1 of 4)
OG IDs: 93519840
Best-hit-arabi-ID: AT3G03800.1
Best-hit-arabi-name: syntaxin of plants 131
Gorai.004G250800.1.D
PCC=0.475, p=0.04
mRNA Profile
Protein Expression Profile
DPA

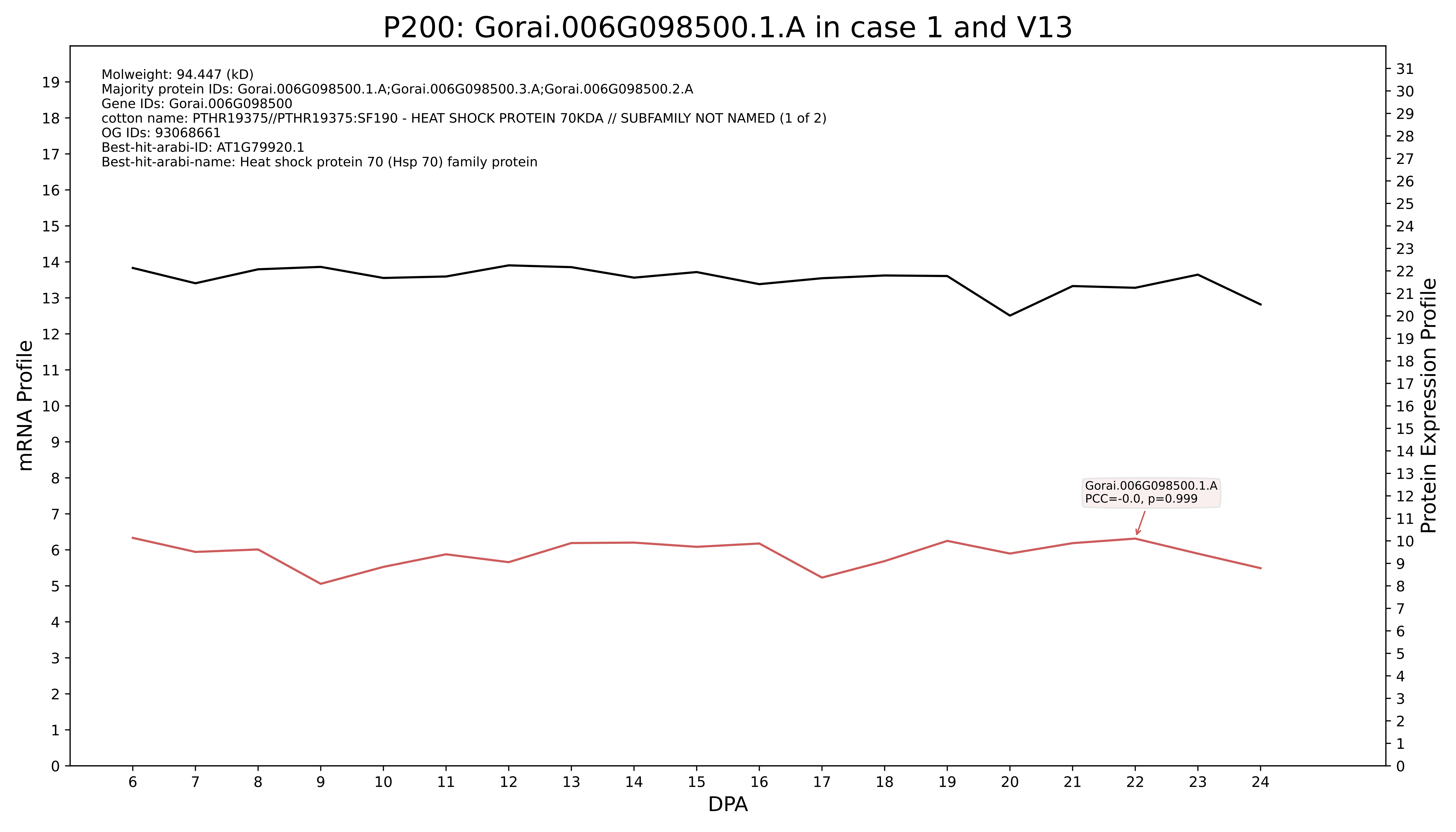
P200: Gorai.006G098500.1.A in case 1 and V13
Molweight: 94.447 (kD)
Majority protein IDs: Gorai.006G098500.1.A;Gorai.006G098500.3.A;Gorai.006G098500.2.A
Gene IDs: Gorai.006G098500
cotton name: PTHR19375//PTHR19375:SF190 - HEAT SHOCK PROTEIN 70KDA // SUBFAMILY NOT NAMED (1 of 2)
OG IDs: 93068661
Best-hit-arabi-ID: AT1G79920.1
Best-hit-arabi-name: Heat shock protein 70 (Hsp 70) family protein
Gorai.006G098500.1.A
PCC=-0.0, p=0.999
mRNA Profile
Protein Expression Profile
DPA

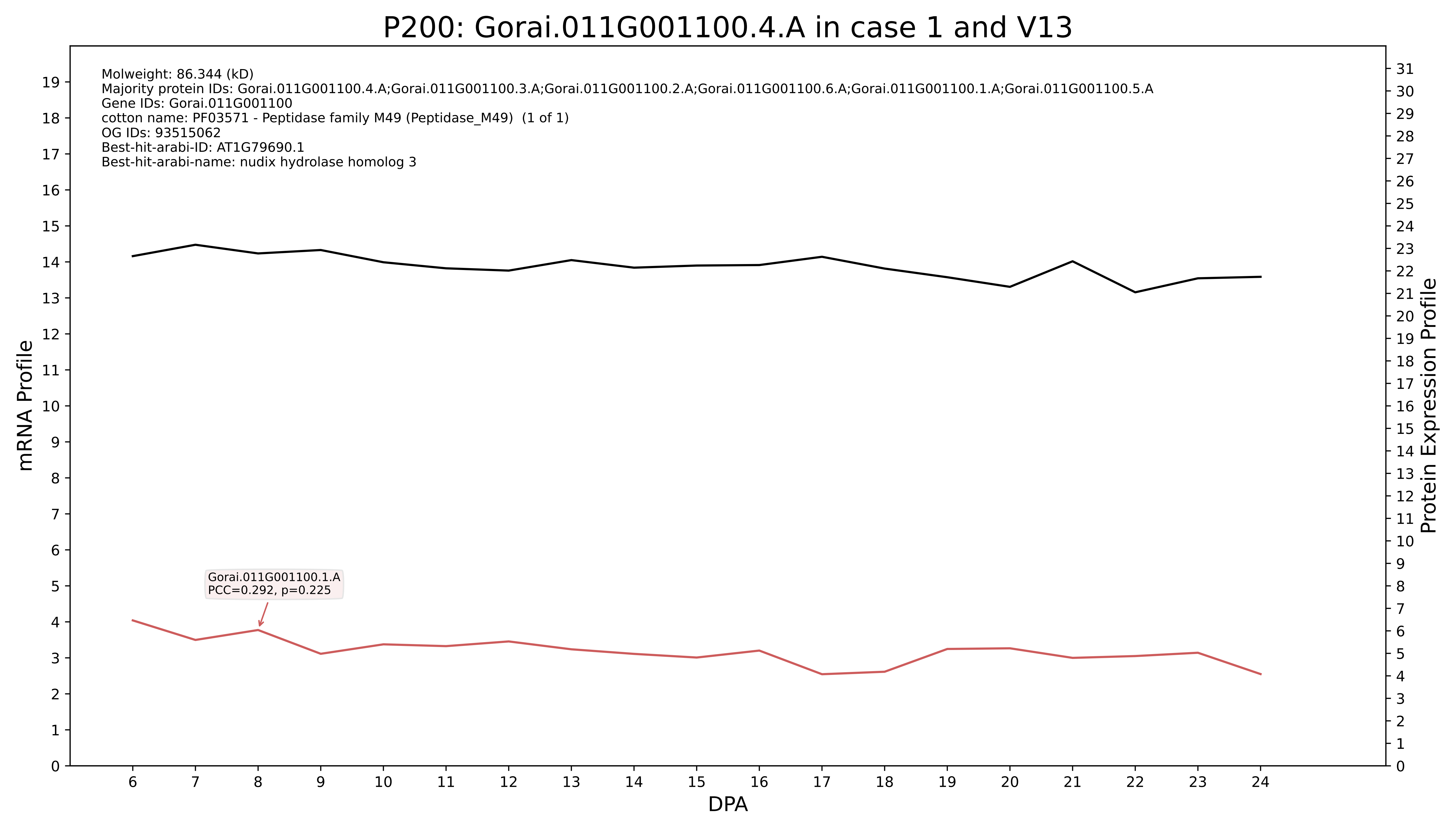
P200: Gorai.011G001100.4.A in case 1 and V13
Molweight: 86.344 (kD)
Majority protein IDs: Gorai.011G001100.4.A;Gorai.011G001100.3.A;Gorai.011G001100.2.A;Gorai.011G001100.6.A;Gorai.011G001100.1.A;Gorai.011G001100.5.A
Gene IDs: Gorai.011G001100
cotton name: PF03571 - Peptidase family M49 (Peptidase_M49) (1 of 1)
OG IDs: 93515062
Best-hit-arabi-ID: AT1G79690.1
Best-hit-arabi-name: nudix hydrolase homolog 3
Gorai.011G001100.1.A
PCC=0.292, p=0.225
mRNA Profile
Protein Expression Profile
DPA

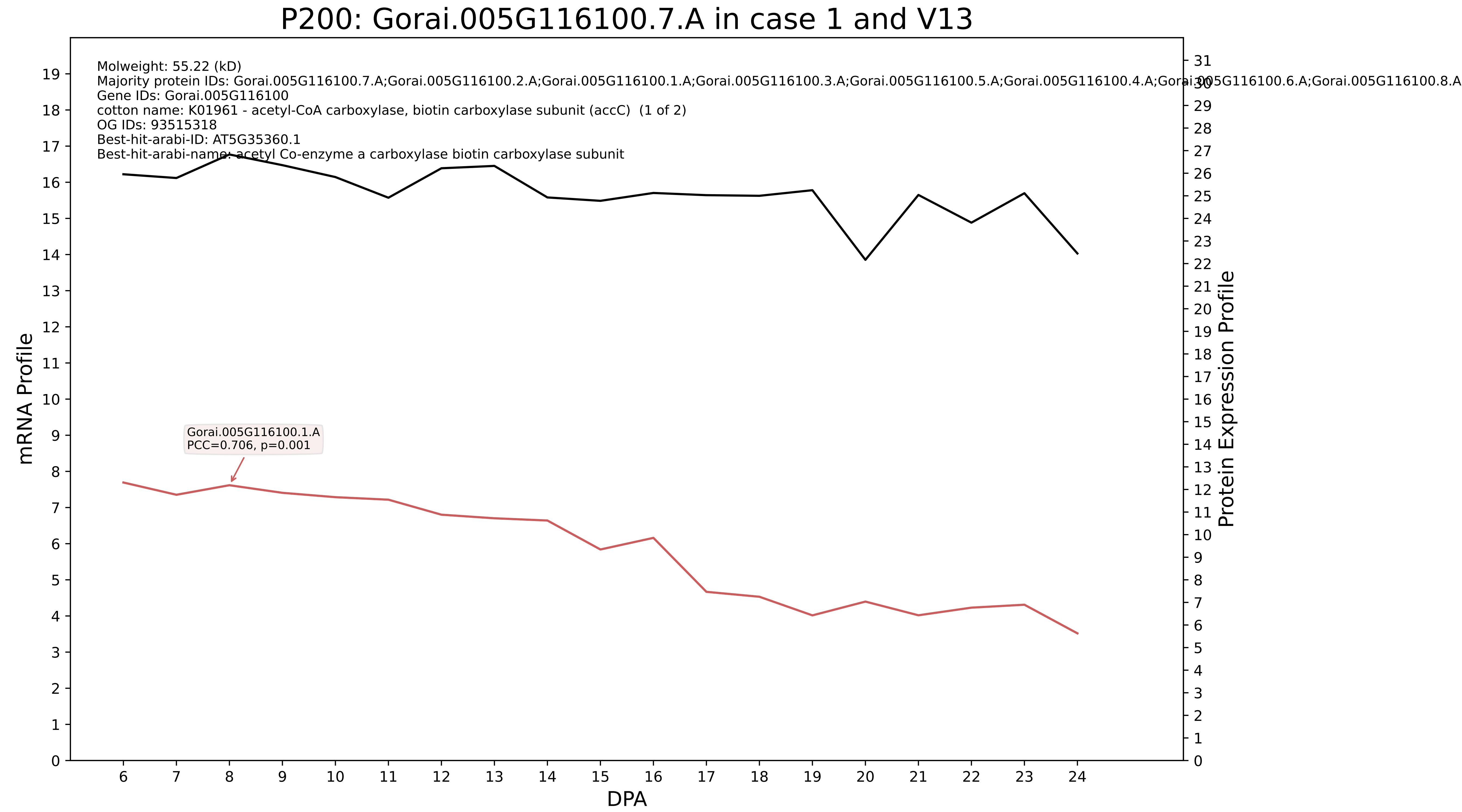
P200: Gorai.005G116100.7.A in case 1 and V13
Molweight: 55.22 (kD)
Majority protein IDs: Gorai.005G116100.7.A;Gorai.005G116100.2.A;Gorai.005G116100.1.A;Gorai.005G116100.3.A;Gorai.005G116100.5.A;Gorai.005G116100.4.A;Gorai.005G116100.6.A;Gorai.005G116100.8.A
Gene IDs: Gorai.005G116100
cotton name: K01961 - acetyl-CoA carboxylase, biotin carboxylase subunit (accC) (1 of 2)
OG IDs: 93515318
Best-hit-arabi-ID: AT5G35360.1
Best-hit-arabi-name: acetyl Co-enzyme a carboxylase biotin carboxylase subunit
Gorai.005G116100.1.A
PCC=0.706, p=0.001
mRNA Profile
Protein Expression Profile
DPA

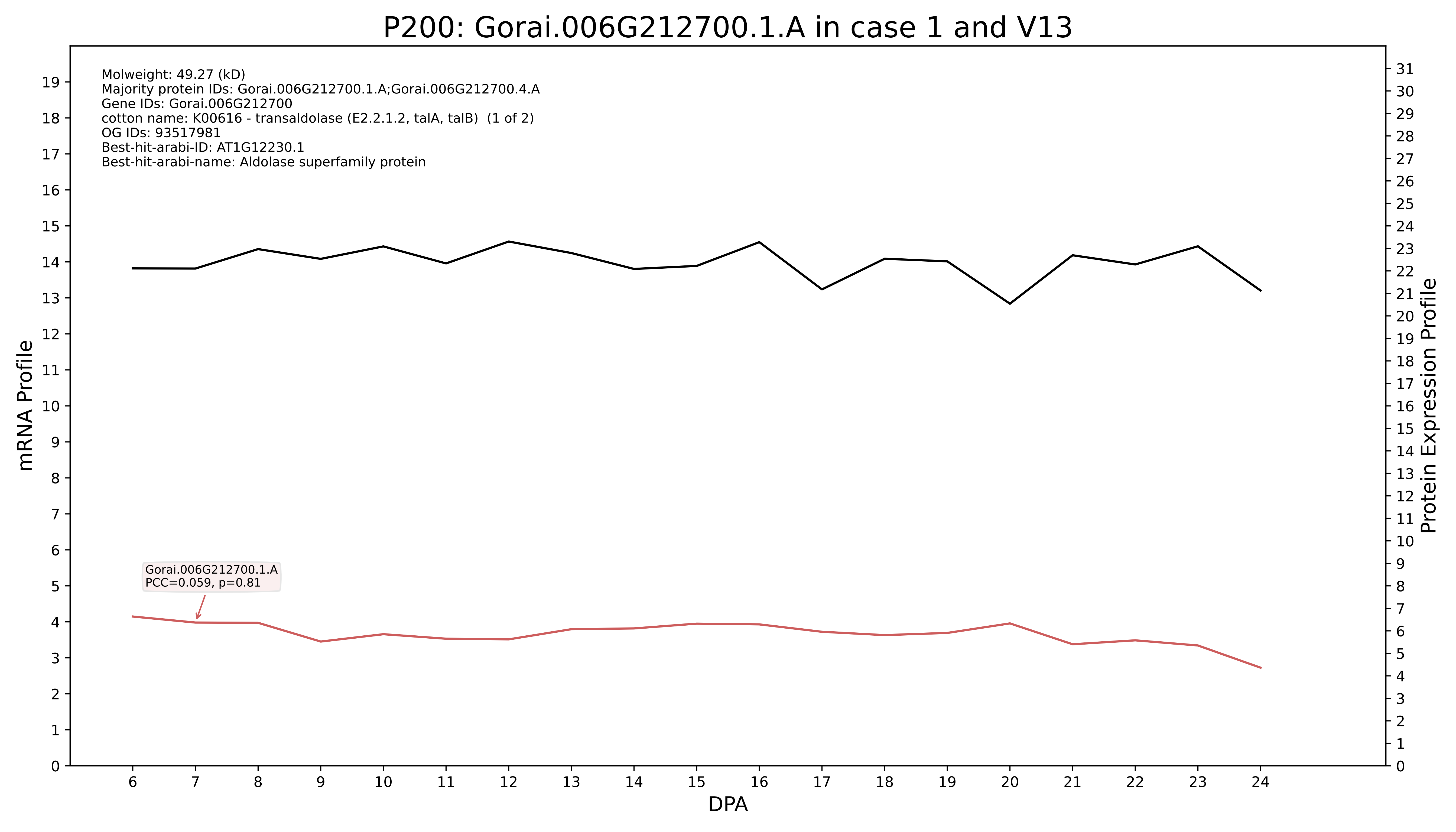
P200: Gorai.006G212700.1.A in case 1 and V13
Molweight: 49.27 (kD)
Majority protein IDs: Gorai.006G212700.1.A;Gorai.006G212700.4.A
Gene IDs: Gorai.006G212700
cotton name: K00616 - transaldolase (E2.2.1.2, talA, talB) (1 of 2)
OG IDs: 93517981
Best-hit-arabi-ID: AT1G12230.1
Best-hit-arabi-name: Aldolase superfamily protein
Gorai.006G212700.1.A
PCC=0.059, p=0.81
mRNA Profile
Protein Expression Profile
DPA
0 1 2 3 4 5 6 7 8 9 10 11 12 13 14 15 16 17 18 19
0 1 2 3 4 5 6 7 8 9 10 11 12 13 14 15 16 17 18 19 20 21 22 23 24 25 26 27 28 29 30 31
6 7 8 9 10 11 12 13 14 15 16 17 18 19 20 21 22 23 24

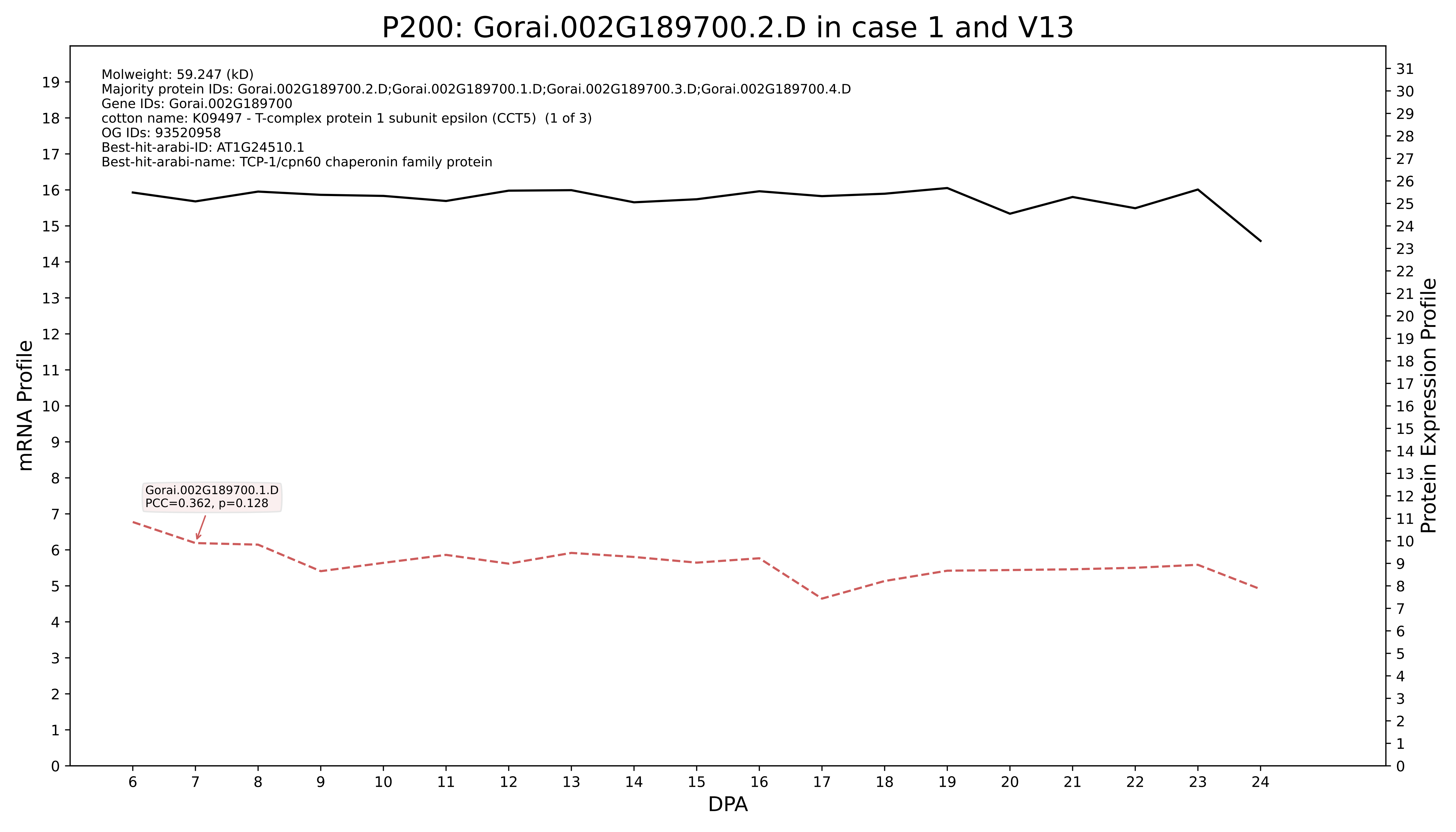
P200: Gorai.002G189700.2.D in case 1 and V13
Molweight: 59.247 (kD)
Majority protein IDs: Gorai.002G189700.2.D;Gorai.002G189700.1.D;Gorai.002G189700.3.D;Gorai.002G189700.4.D
Gene IDs: Gorai.002G189700
cotton name: K09497 - T-complex protein 1 subunit epsilon (CCT5) (1 of 3)
OG IDs: 93520958
Best-hit-arabi-ID: AT1G24510.1
Best-hit-arabi-name: TCP-1/cpn60 chaperonin family protein
Gorai.002G189700.1.D
PCC=0.362, p=0.128
mRNA Profile
Protein Expression Profile
DPA

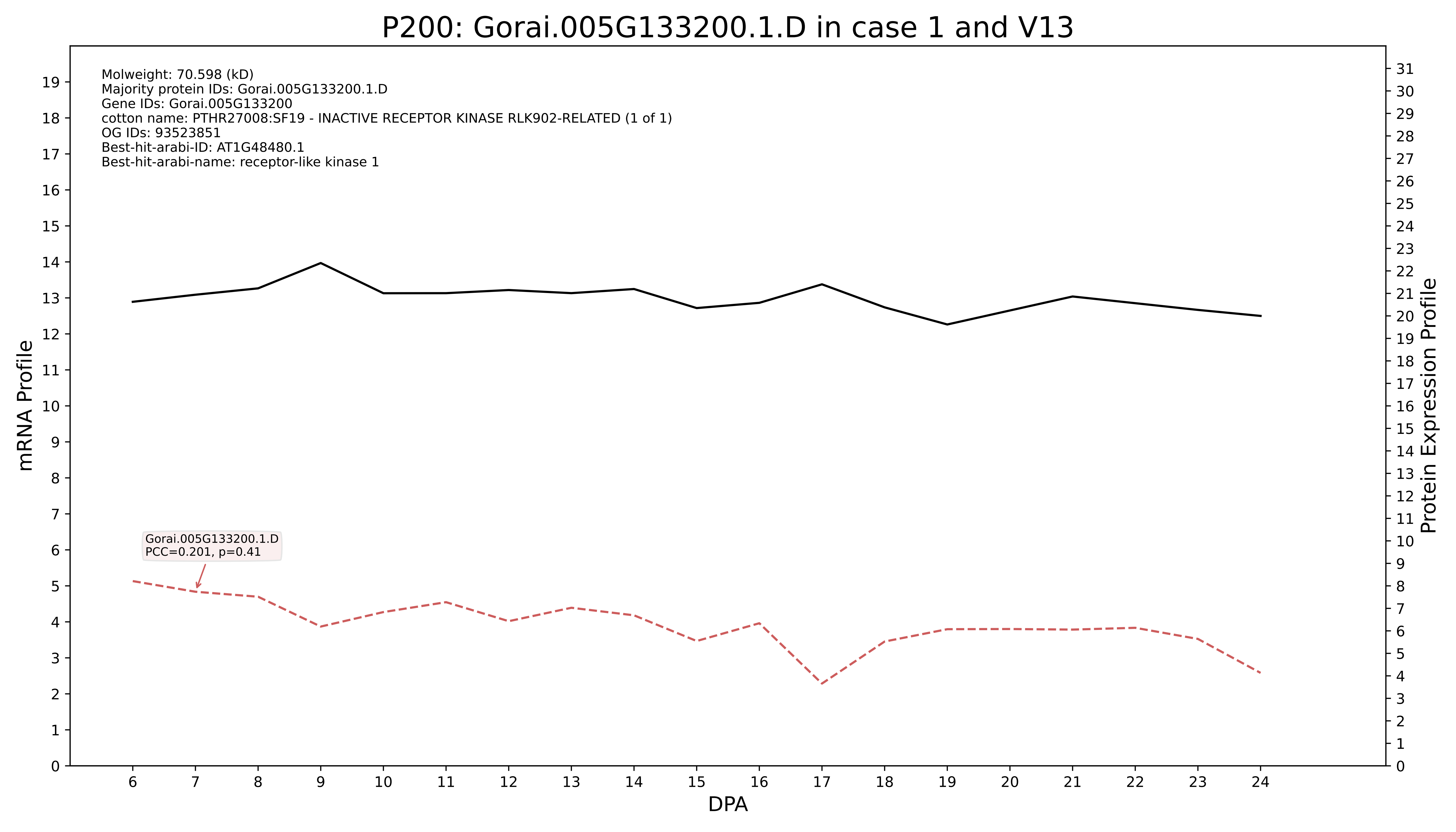

P200: Gorai.005G133200.1.D in case 1 and V13
Molweight: 70.598 (kD)
Majority protein IDs: Gorai.005G133200.1.D
Gene IDs: Gorai.005G133200
cotton name: PTHR27008:SF19 - INACTIVE RECEPTOR KINASE RLK902-RELATED (1 of 1)
OG IDs: 93523851
Best-hit-arabi-ID: AT1G48480.1
Best-hit-arabi-name: receptor-like kinase 1
Gorai.005G133200.1.D
PCC=0.201, p=0.41
mRNA Profile
Protein Expression Profile
DPA

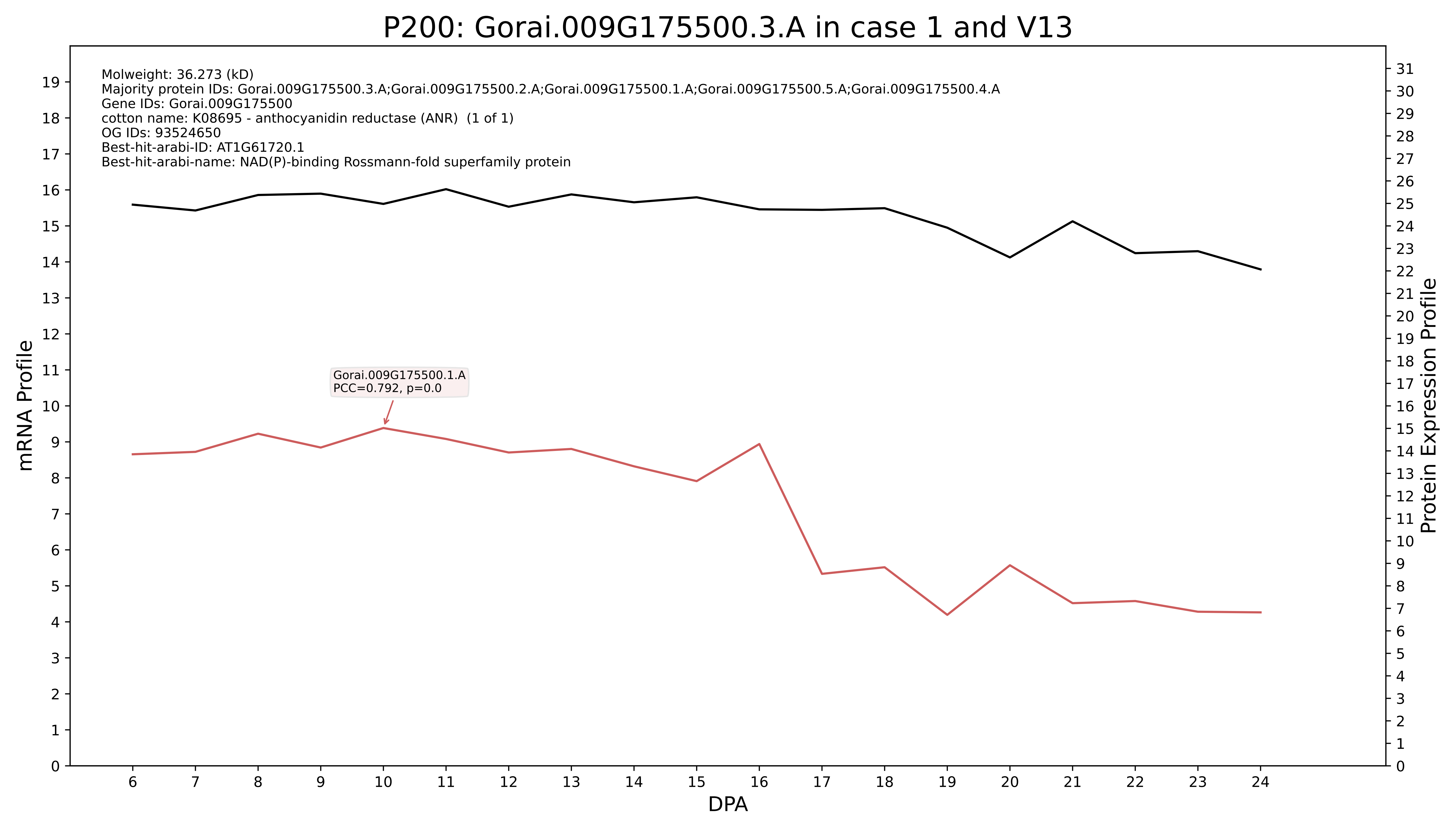

P200: Gorai.009G175500.3.A in case 1 and V13
Molweight: 36.273 (kD)
Majority protein IDs: Gorai.009G175500.3.A;Gorai.009G175500.2.A;Gorai.009G175500.1.A;Gorai.009G175500.5.A;Gorai.009G175500.4.A
Gene IDs: Gorai.009G175500
cotton name: K08695 - anthocyanidin reductase (ANR) (1 of 1)
OG IDs: 93524650
Best-hit-arabi-ID: AT1G61720.1
Best-hit-arabi-name: NAD(P)-binding Rossmann-fold superfamily protein
Gorai.009G175500.1.A
PCC=0.792, p=0.0
mRNA Profile
Protein Expression Profile
DPA

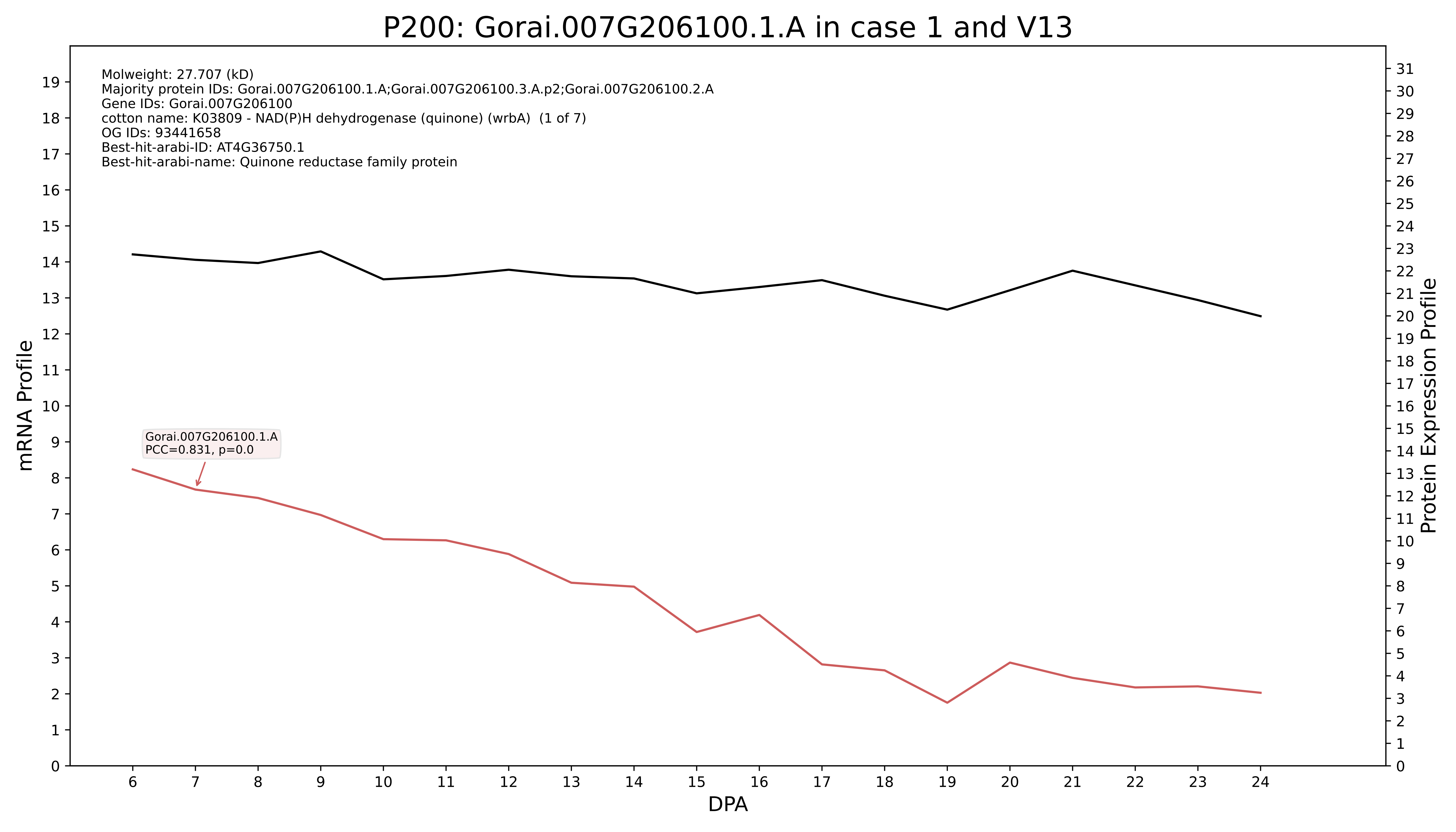
P200: Gorai.007G206100.1.A in case 1 and V13
Molweight: 27.707 (kD)
Majority protein IDs: Gorai.007G206100.1.A;Gorai.007G206100.3.A.p2;Gorai.007G206100.2.A
Gene IDs: Gorai.007G206100
cotton name: K03809 - NAD(P)H dehydrogenase (quinone) (wrbA) (1 of 7)
OG IDs: 93441658
Best-hit-arabi-ID: AT4G36750.1
Best-hit-arabi-name: Quinone reductase family protein
Gorai.007G206100.1.A
PCC=0.831, p=0.0
mRNA Profile
Protein Expression Profile
DPA

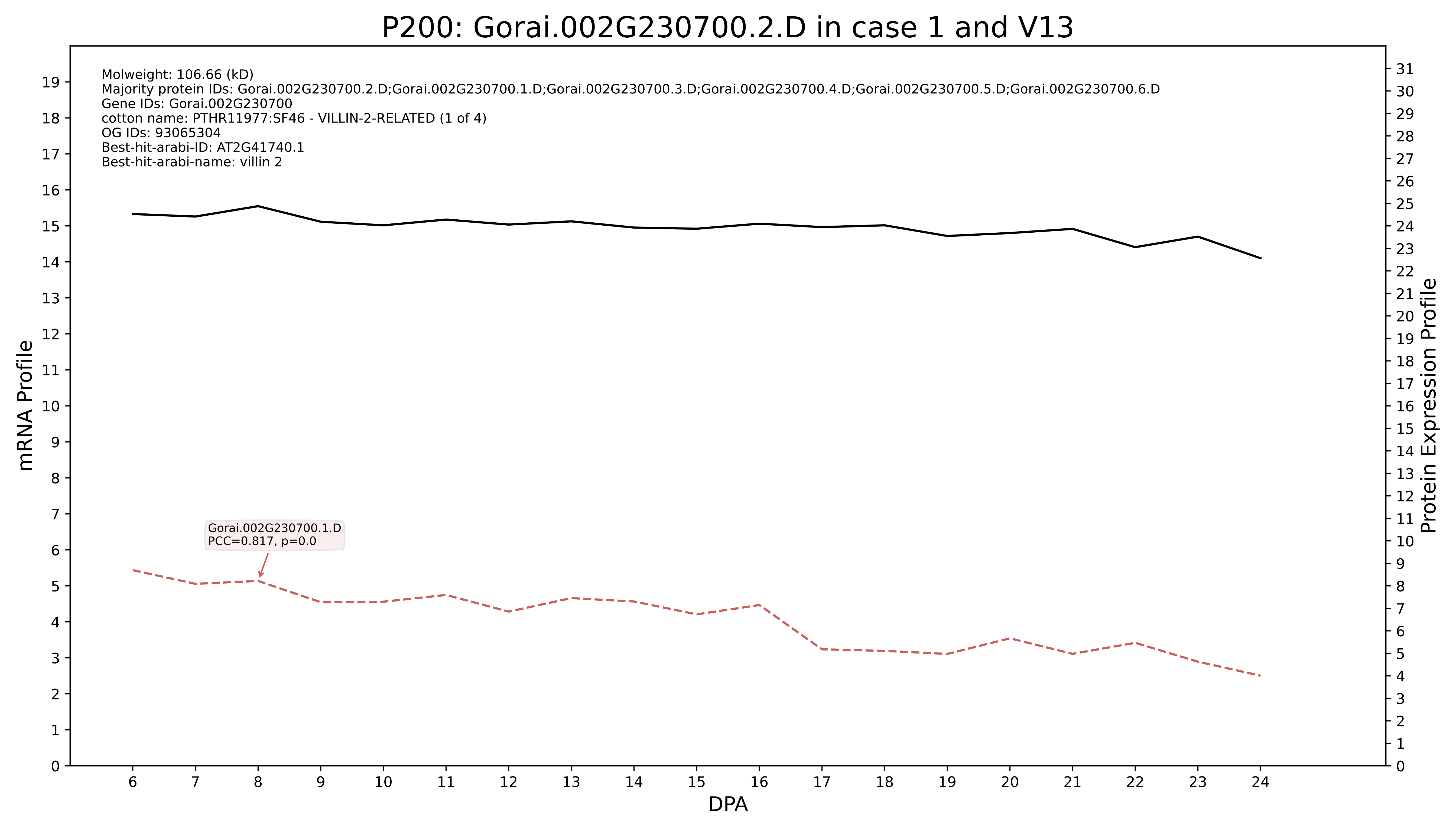

P200: Gorai.002G230700.2.D in case 1 and V13
Molweight: 106.66 (kD)
Majority protein IDs: Gorai.002G230700.2.D;Gorai.002G230700.1.D;Gorai.002G230700.3.D;Gorai.002G230700.4.D;Gorai.002G230700.5.D;Gorai.002G230700.6.D
Gene IDs: Gorai.002G230700
cotton name: PTHR11977:SF46 - VILLIN-2-RELATED (1 of 4)
OG IDs: 93065304
Best-hit-arabi-ID: AT2G41740.1
Best-hit-arabi-name: villin 2
Gorai.002G230700.1.D
PCC=0.817, p=0.0
mRNA Profile
Protein Expression Profile
DPA

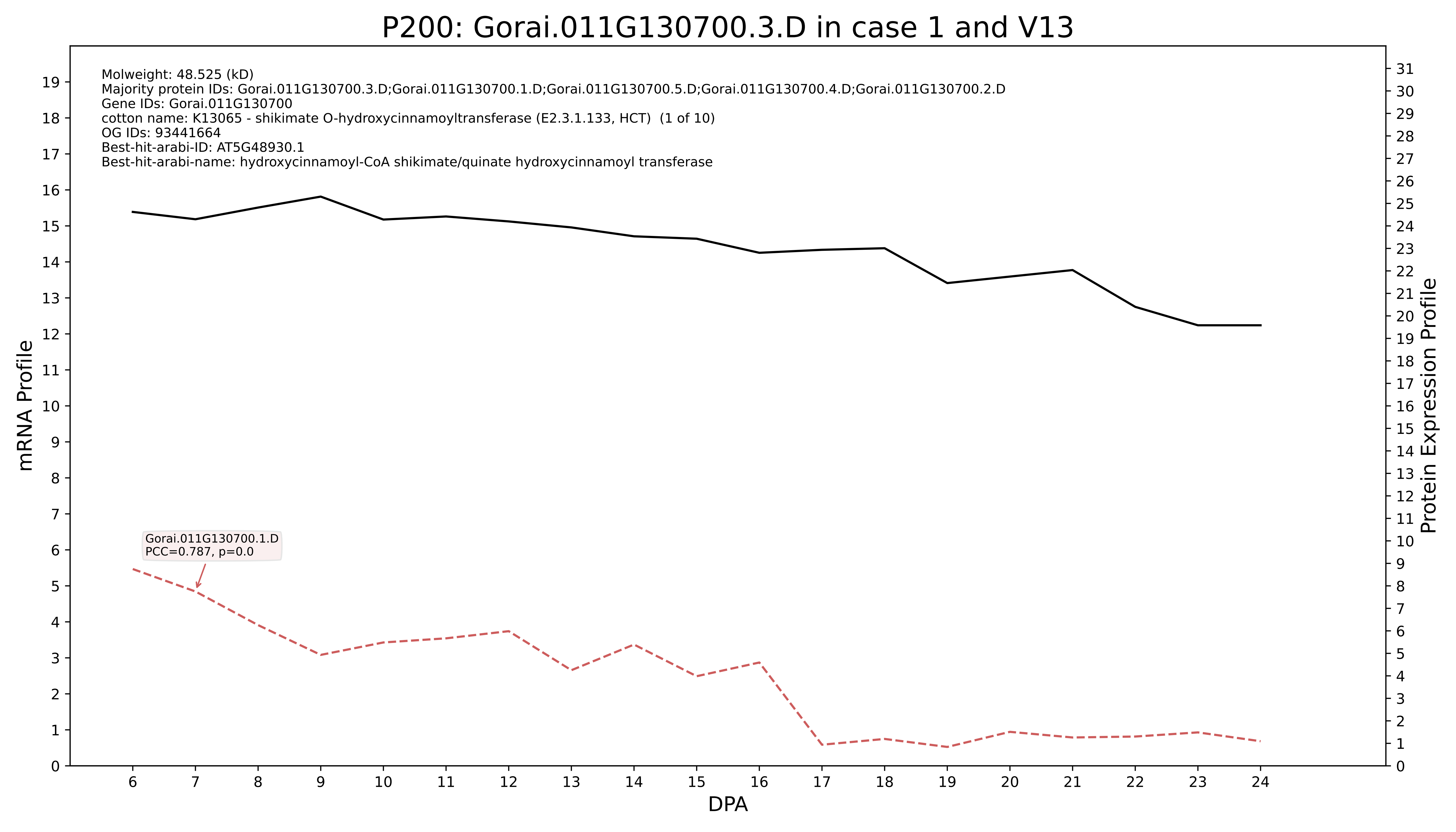

P200: Gorai.011G130700.3.D in case 1 and V13
Molweight: 48.525 (kD)
Majority protein IDs: Gorai.011G130700.3.D;Gorai.011G130700.1.D;Gorai.011G130700.5.D;Gorai.011G130700.4.D;Gorai.011G130700.2.D
Gene IDs: Gorai.011G130700
cotton name: K13065 - shikimate O-hydroxycinnamoyltransferase (E2.3.1.133, HCT) (1 of 10)
OG IDs: 93441664
Best-hit-arabi-ID: AT5G48930.1
Best-hit-arabi-name: hydroxycinnamoyl-CoA shikimate/quinate hydroxycinnamoyl transferase
Gorai.011G130700.1.D
PCC=0.787, p=0.0
mRNA Profile
Protein Expression Profile
DPA

### P200: Gorai.009G200600.2.D in case 1 and V13

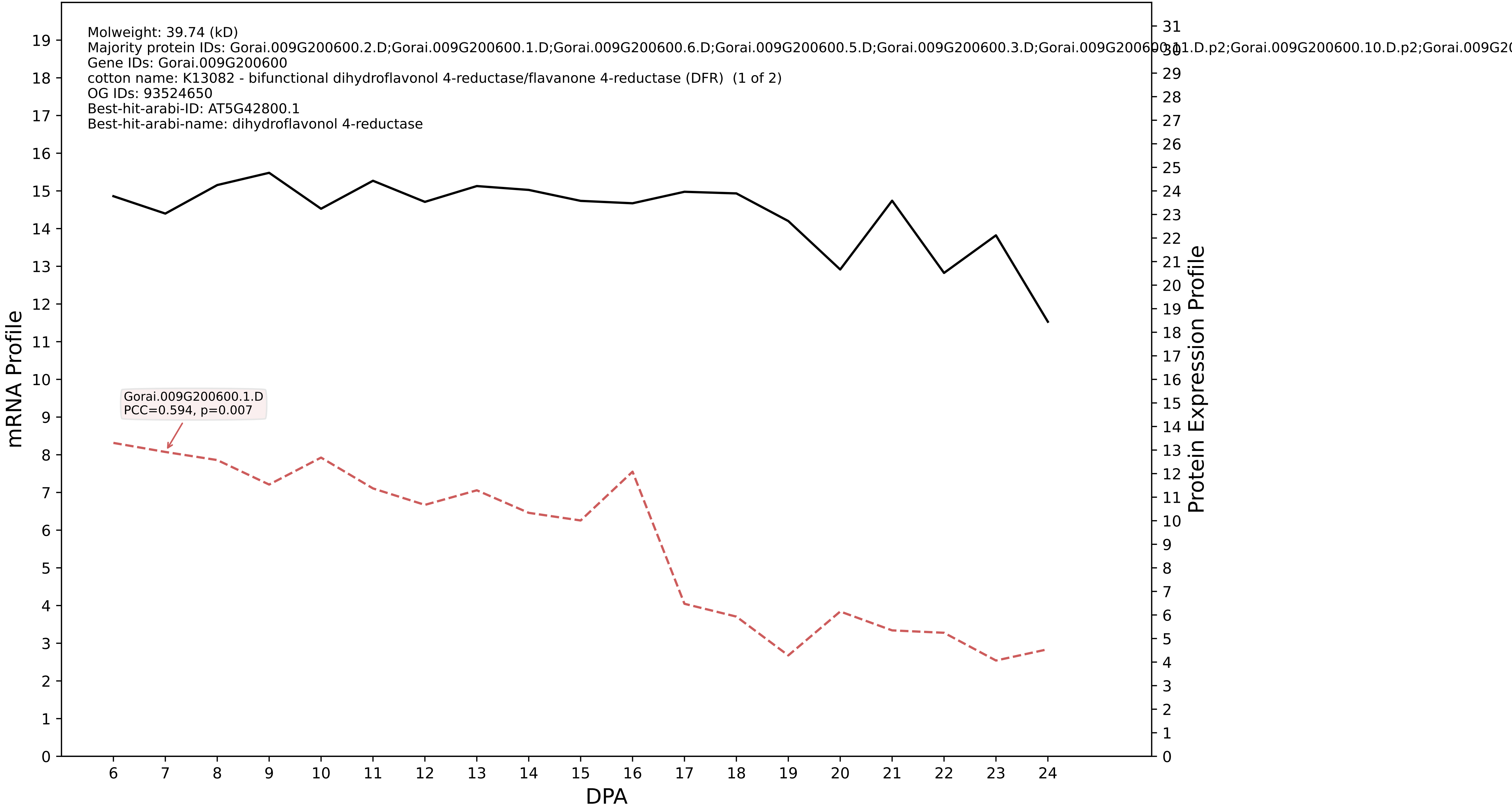

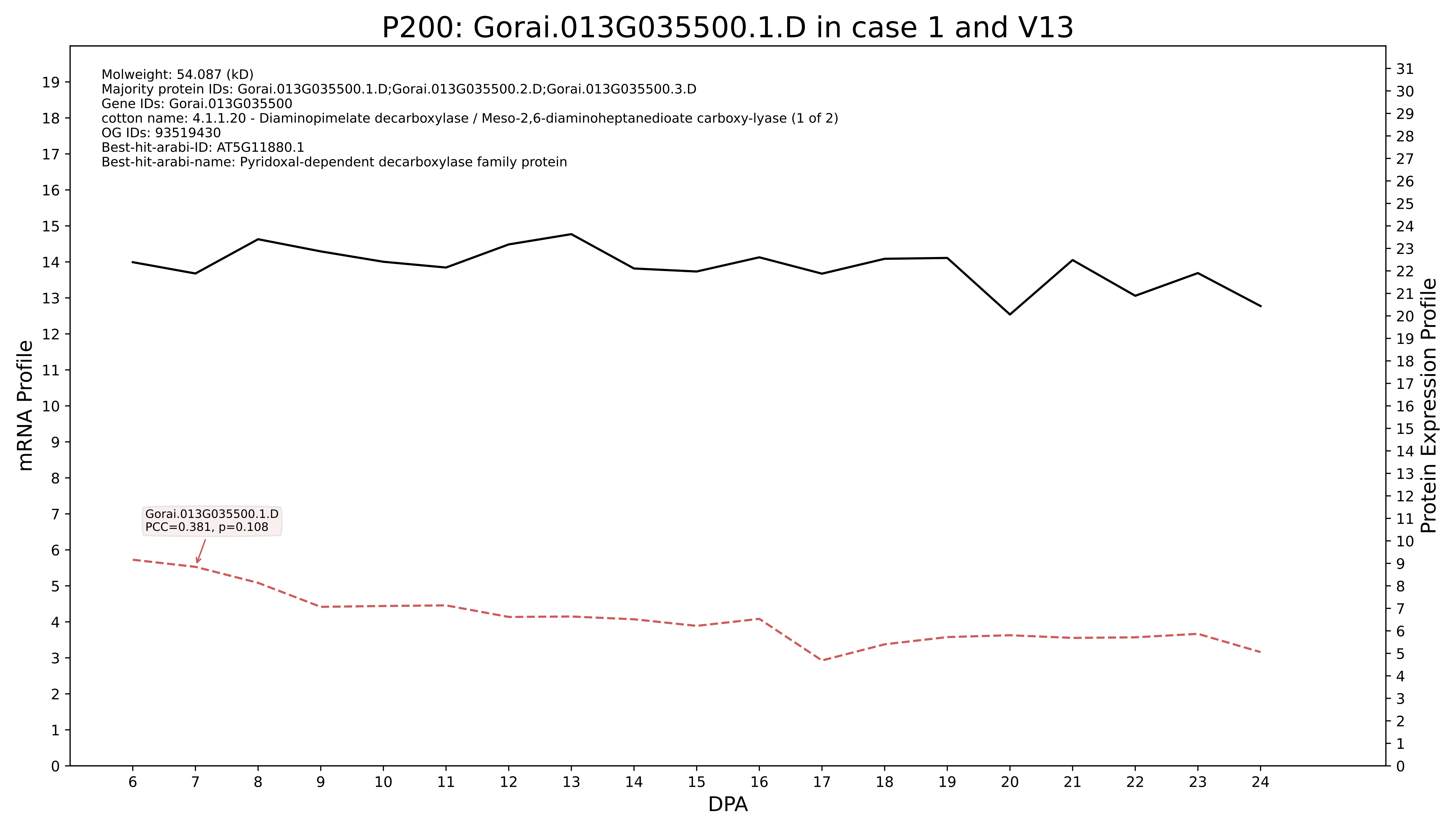
P200: Gorai.013G035500.1.D in case 1 and V13
Molweight: 54.087 (kD)
Majority protein IDs: Gorai.013G035500.1.D;Gorai.013G035500.2.D;Gorai.013G035500.3.D
Gene IDs: Gorai.013G035500
cotton name: 4.1.1.20 - Diaminopimelate decarboxylase / Meso-2,6-diaminoheptanedioate carboxy-lyase (1 of 2)
OG IDs: 93519430
Best-hit-arabi-ID: AT5G11880.1
Best-hit-arabi-name: Pyridoxal-dependent decarboxylase family protein
Gorai.013G035500.1.D
PCC=0.381, p=0.108
mRNA Profile
Protein Expression Profile
DPA

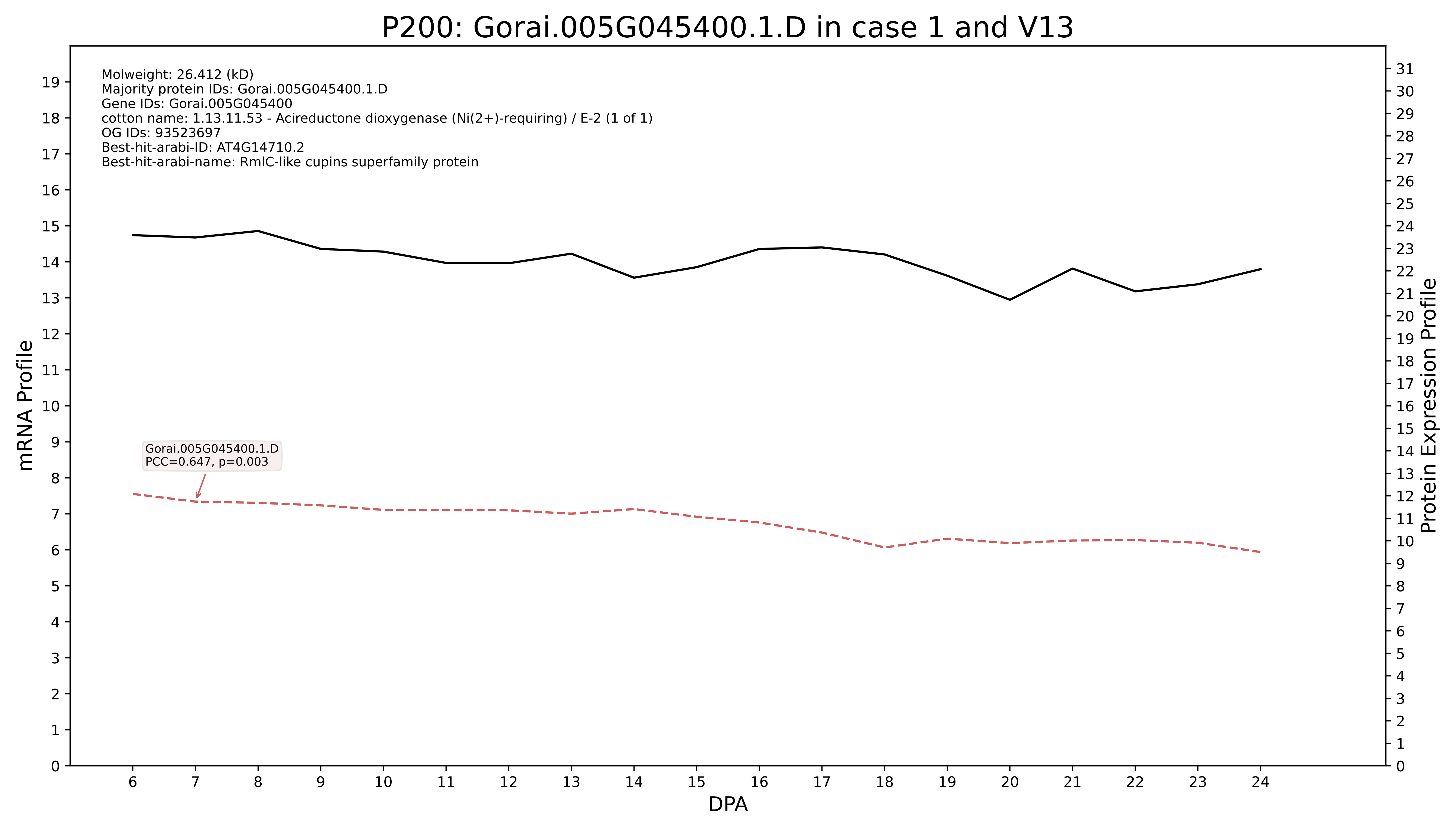
P200: Gorai.005G045400.1.D in case 1 and V13
Molweight: 26.412 (kD)
Majority protein IDs: Gorai.005G045400.1.D
Gene IDs: Gorai.005G045400
cotton name: 1.13.11.53 - Acireductone dioxygenase (Ni(2+)-requiring) / E-2 (1 of 1)
OG IDs: 93523697
Best-hit-arabi-ID: AT4G14710.2
Best-hit-arabi-name: RmlC-like cupins superfamily protein
Gorai.005G045400.1.D
PCC=0.647, p=0.003
mRNA Profile
Protein Expression Profile
DPA

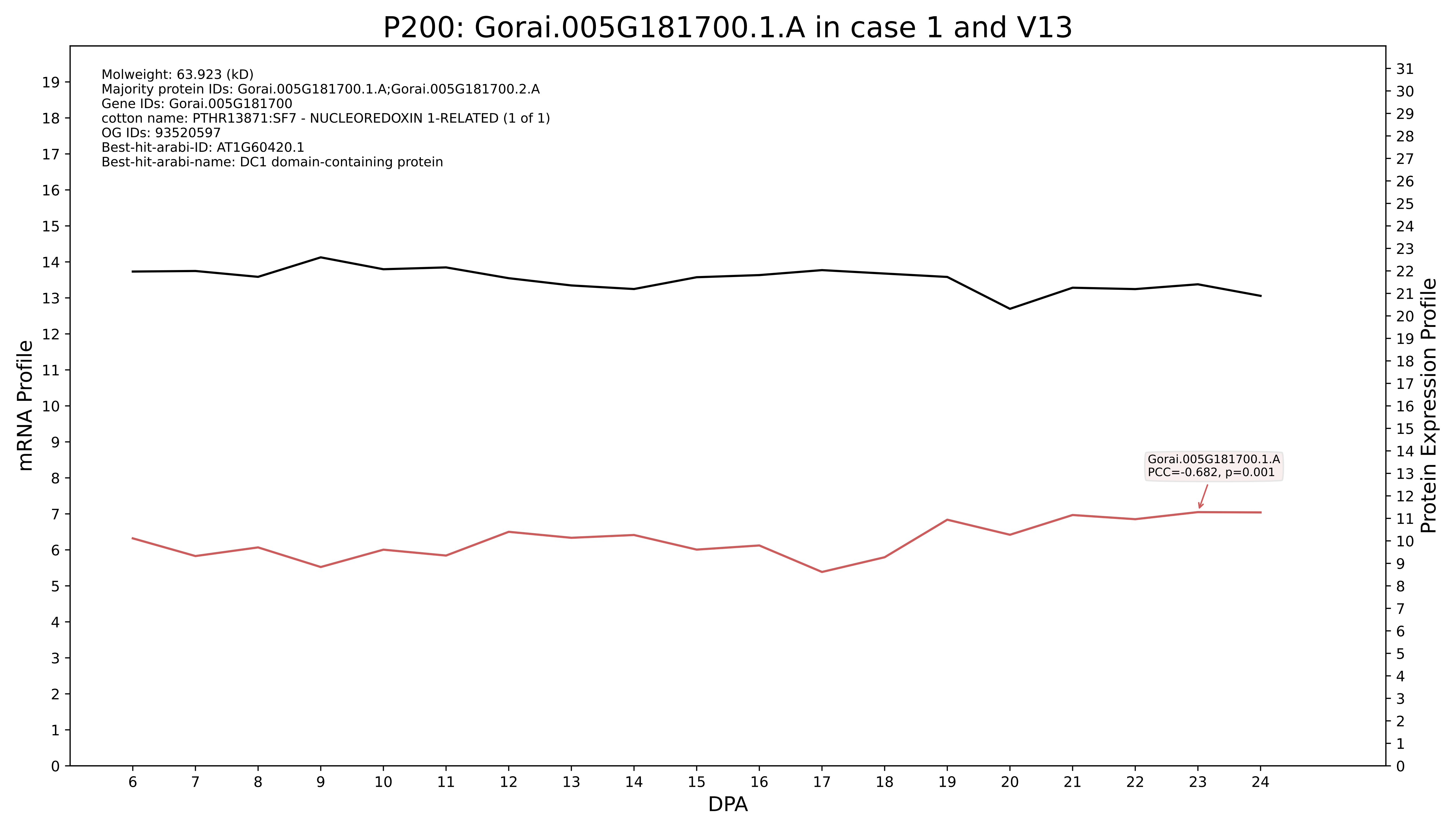

P200: Gorai.005G181700.1.A in case 1 and V13
Molweight: 63.923 (kD)
Majority protein IDs: Gorai.005G181700.1.A;Gorai.005G181700.2.A
Gene IDs: Gorai.005G181700
cotton name: PTHR13871:SF7 - NUCLEOREDOXIN 1-RELATED (1 of 1)
OG IDs: 93520597
Best-hit-arabi-ID: AT1G60420.1
Best-hit-arabi-name: DC1 domain-containing protein
Gorai.005G181700.1.A
PCC=-0.682, p=0.001
mRNA Profile
Protein Expression Profile
DPA

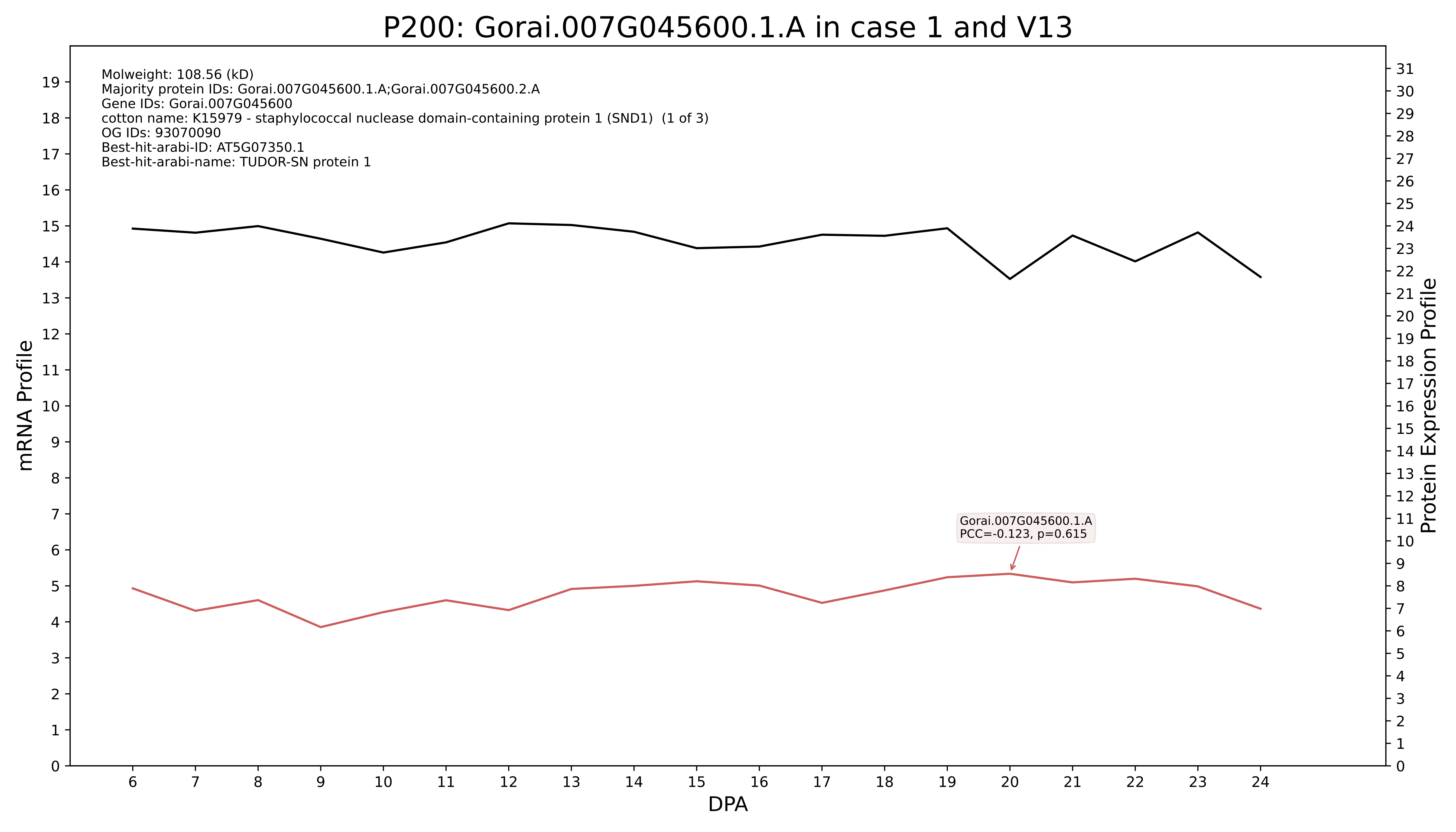
P200: Gorai.007G045600.1.A in case 1 and V13
Molweight: 108.56 (kD)
Majority protein IDs: Gorai.007G045600.1.A;Gorai.007G045600.2.A
Gene IDs: Gorai.007G045600
cotton name: K15979 - staphylococcal nuclease domain-containing protein 1 (SND1) (1 of 3)
OG IDs: 93070090
Best-hit-arabi-ID: AT5G07350.1
Best-hit-arabi-name: TUDOR-SN protein 1
Gorai.007G045600.1.A
PCC=-0.123, p=0.615
mRNA Profile
Protein Expression Profile
DPA

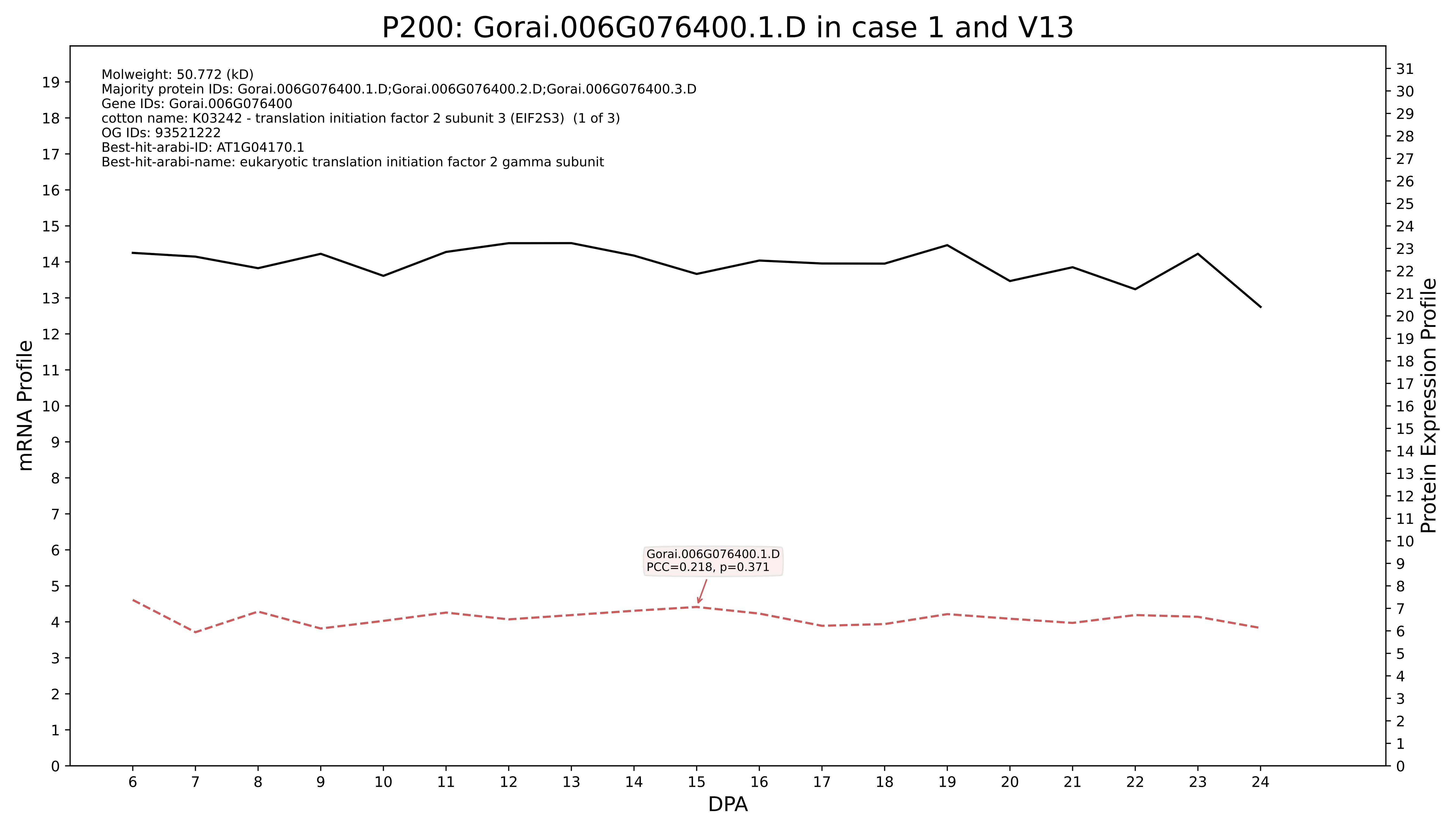

P200: Gorai.006G076400.1.D in case 1 and V13
Molweight: 50.772 (kD)
Majority protein IDs: Gorai.006G076400.1.D;Gorai.006G076400.2.D;Gorai.006G076400.3.D
Gene IDs: Gorai.006G076400
cotton name: K03242 - translation initiation factor 2 subunit 3 (EIF2S3) (1 of 3)
OG IDs: 93521222
Best-hit-arabi-ID: AT1G04170.1
Best-hit-arabi-name: eukaryotic translation initiation factor 2 gamma subunit
Gorai.006G076400.1.D
PCC=0.218, p=0.371
mRNA Profile
Protein Expression Profile
DPA

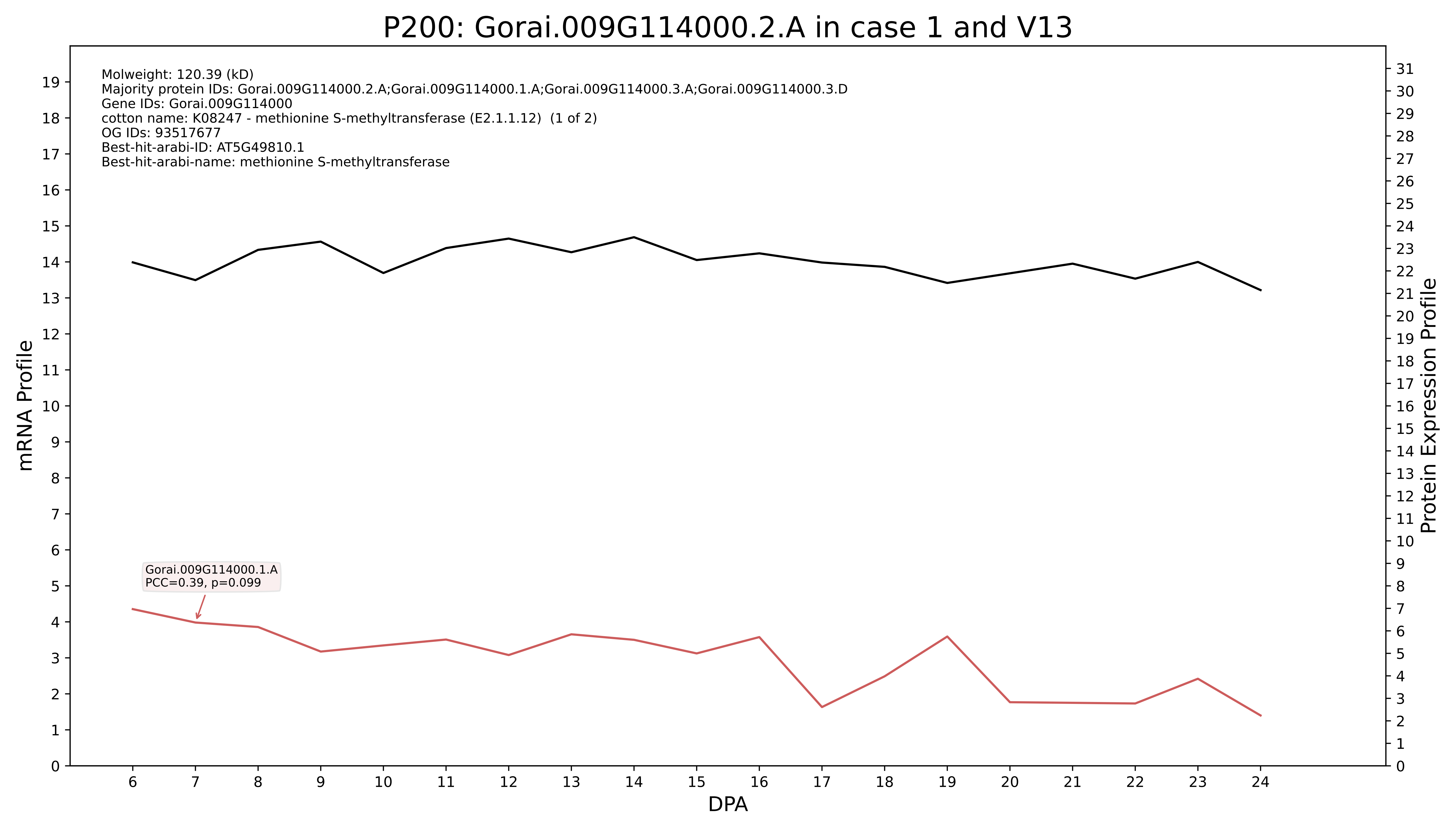
P200: Gorai.009G114000.2.A in case 1 and V13
Molweight: 120.39 (kD)
Majority protein IDs: Gorai.009G114000.2.A;Gorai.009G114000.1.A;Gorai.009G114000.3.A;Gorai.009G114000.3.D
Gene IDs: Gorai.009G114000
cotton name: K08247 - methionine S-methyltransferase (E2.1.1.12) (1 of 2)
OG IDs: 93517677
Best-hit-arabi-ID: AT5G49810.1
Best-hit-arabi-name: methionine S-methyltransferase
Gorai.009G114000.1.A
PCC=0.39, p=0.099
mRNA Profile
Protein Expression Profile
DPA

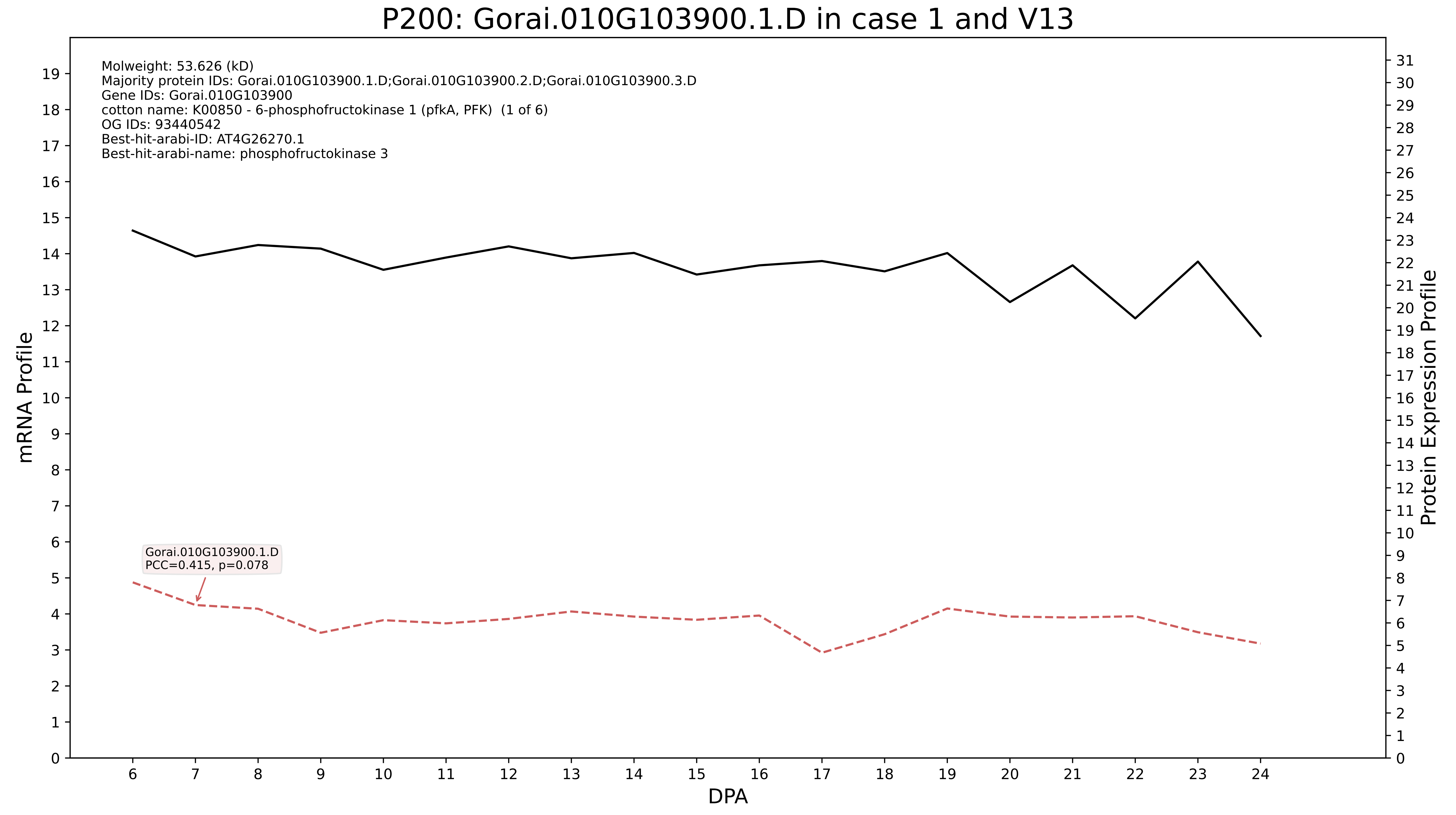
P200: Gorai.010G103900.1.D in case 1 and V13
Molweight: 53.626 (kD)
Majority protein IDs: Gorai.010G103900.1.D;Gorai.010G103900.2.D;Gorai.010G103900.3.D
Gene IDs: Gorai.010G103900
cotton name: K00850 - 6-phosphofructokinase 1 (pfkA, PFK) (1 of 6)
OG IDs: 93440542
Best-hit-arabi-ID: AT4G26270.1
Best-hit-arabi-name: phosphofructokinase 3
Gorai.010G103900.1.D
PCC=0.415, p=0.078
mRNA Profile
Protein Expression Profile
DPA

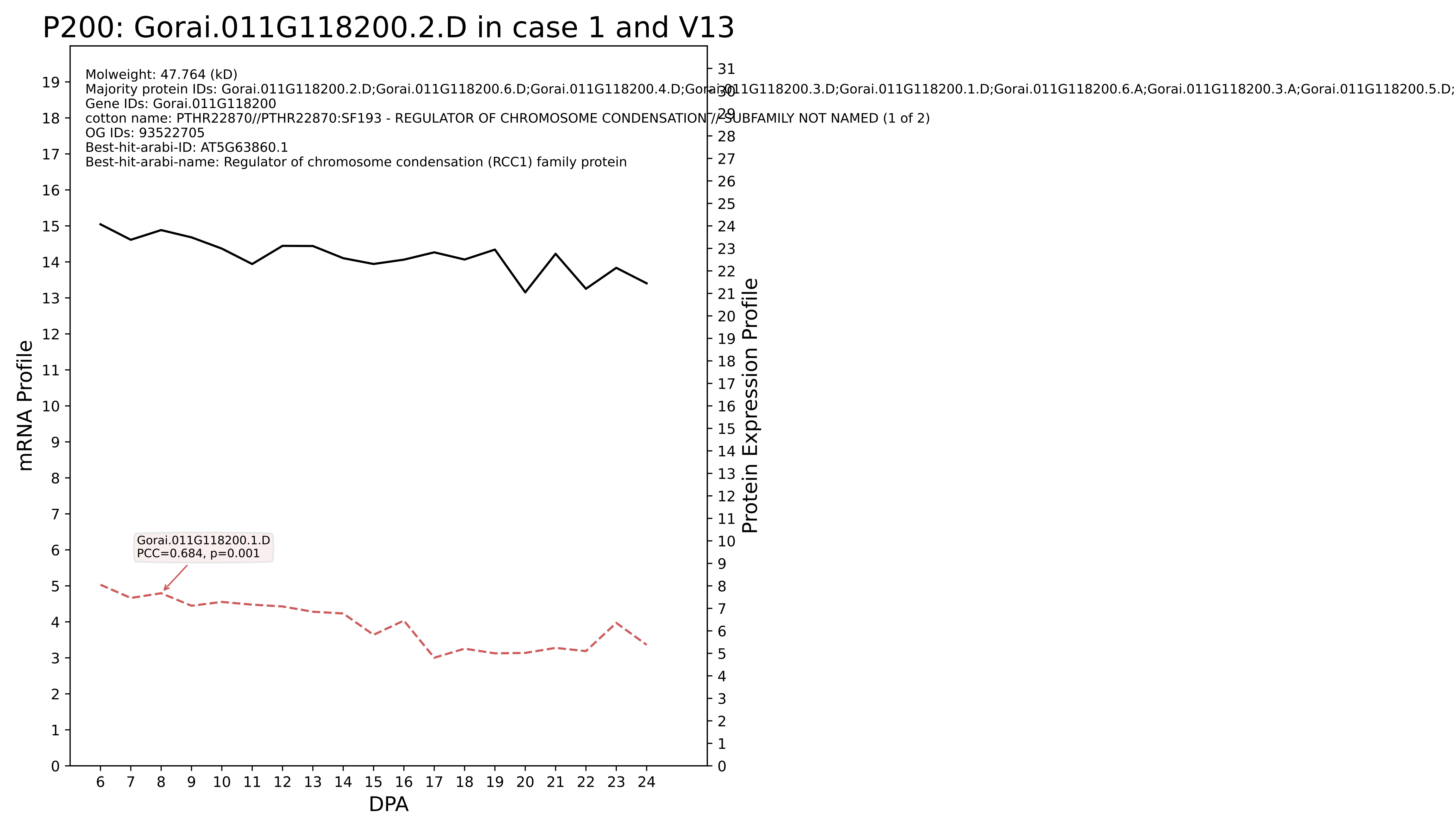

P200: Gorai.011G118200.2.D in case 1 and V13
Molweight: 47.764 (kD)
Majority protein IDs: Gorai.011G118200.2.D;Gorai.011G118200.6.D;Gorai.011G118200.4.D;Gorai.011G118200.3.D;Gorai.011G118200.1.D;Gorai.011G118200.6.A;Gorai.011G118200.3.A;Gorai.011G118200.5.D;
Gene IDs: Gorai.011G118200
cotton name: PTHR22870//PTHR22870:SF193 - REGULATOR OF CHROMOSOME CONDENSATION // SUBFAMILY NOT NAMED (1 of 2)
OG IDs: 93522705
Best-hit-arabi-ID: AT5G63860.1
Best-hit-arabi-name: Regulator of chromosome condensation (RCC1) family protein
Gorai.011G118200.1.D
PCC=0.684, p=0.001
mRNA Profile
Protein Expression Profile
DPA

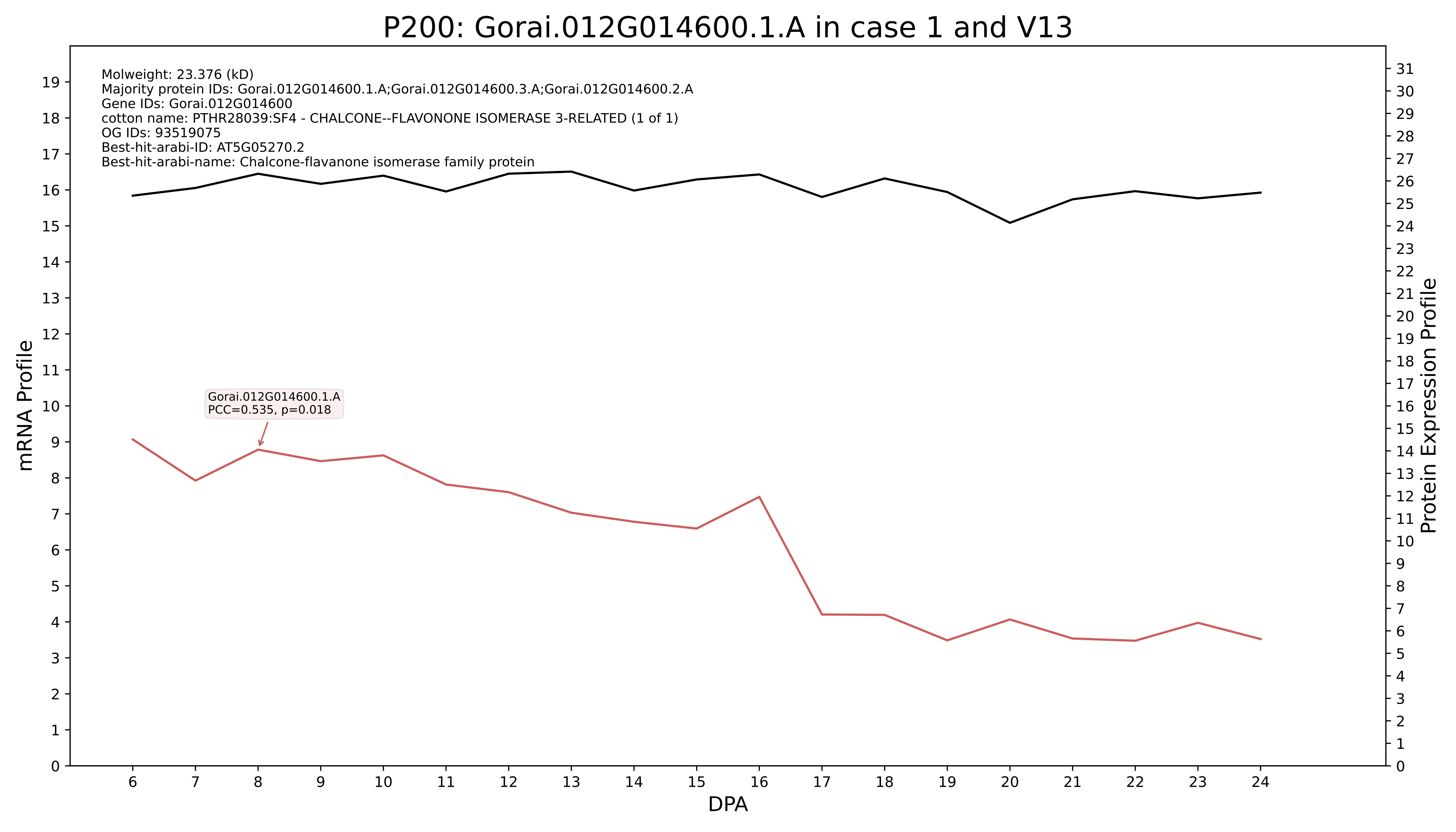

P200: Gorai.012G014600.1.A in case 1 and V13
Molweight: 23.376 (kD)
Majority protein IDs: Gorai.012G014600.1.A;Gorai.012G014600.3.A;Gorai.012G014600.2.A
Gene IDs: Gorai.012G014600
cotton name: PTHR28039:SF4 - CHALCONE--FLAVONONE ISOMERASE 3-RELATED (1 of 1)
OG IDs: 93519075
Best-hit-arabi-ID: AT5G05270.2
Best-hit-arabi-name: Chalcone-flavanone isomerase family protein
Gorai.012G014600.1.A
PCC=0.535, p=0.018
mRNA Profile
Protein Expression Profile
DPA

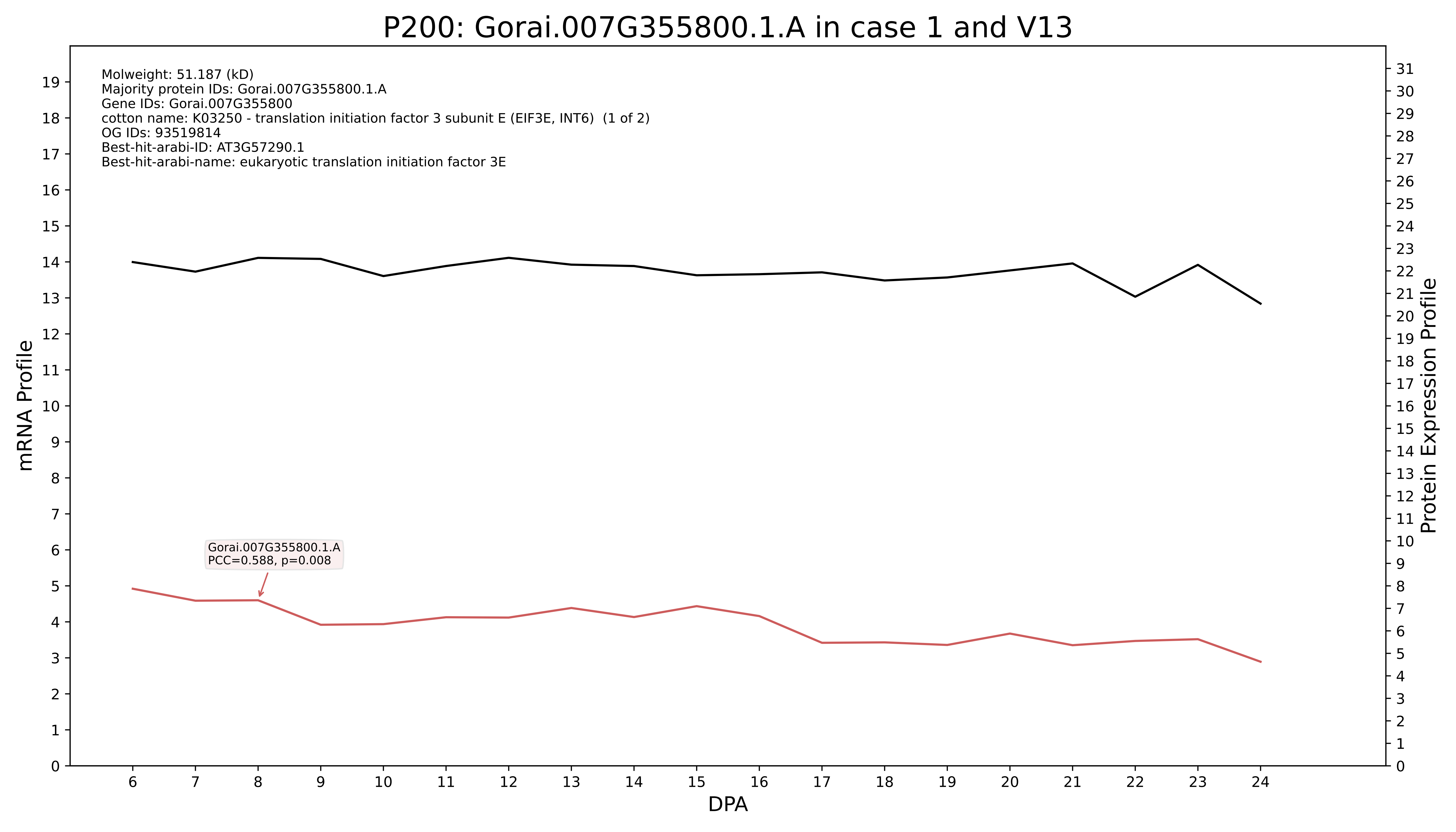

P200: Gorai.007G355800.1.A in case 1 and V13
Molweight: 51.187 (kD)
Majority protein IDs: Gorai.007G355800.1.A
Gene IDs: Gorai.007G355800
cotton name: K03250 - translation initiation factor 3 subunit E (EIF3E, INT6) (1 of 2)
OG IDs: 93519814
Best-hit-arabi-ID: AT3G57290.1
Best-hit-arabi-name: eukaryotic translation initiation factor 3E
Gorai.007G355800.1.A
PCC=0.588, p=0.008
mRNA Profile
Protein Expression Profile
DPA

### P200: Gorai.006G207300.2.D in case 1 and V13

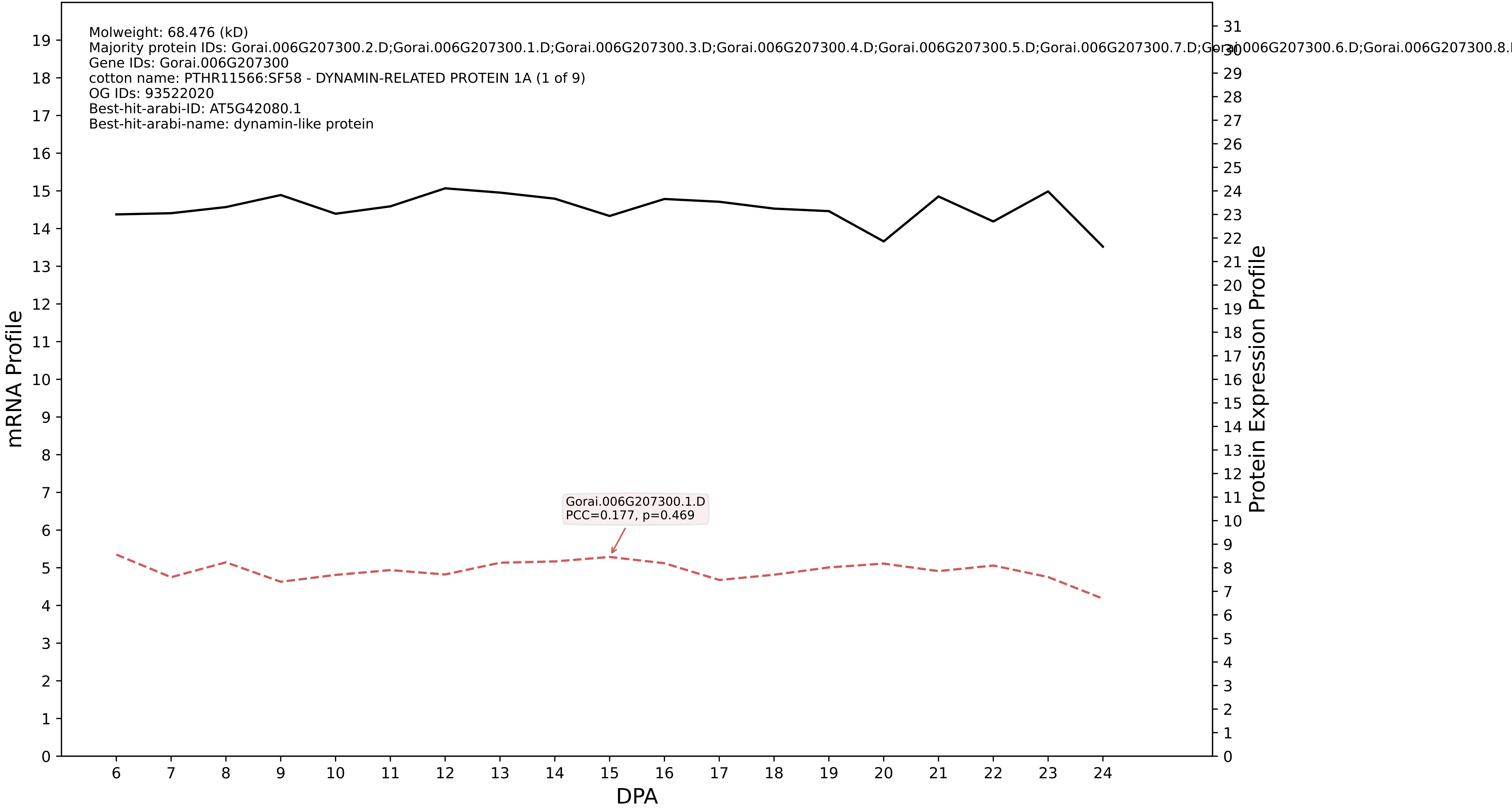

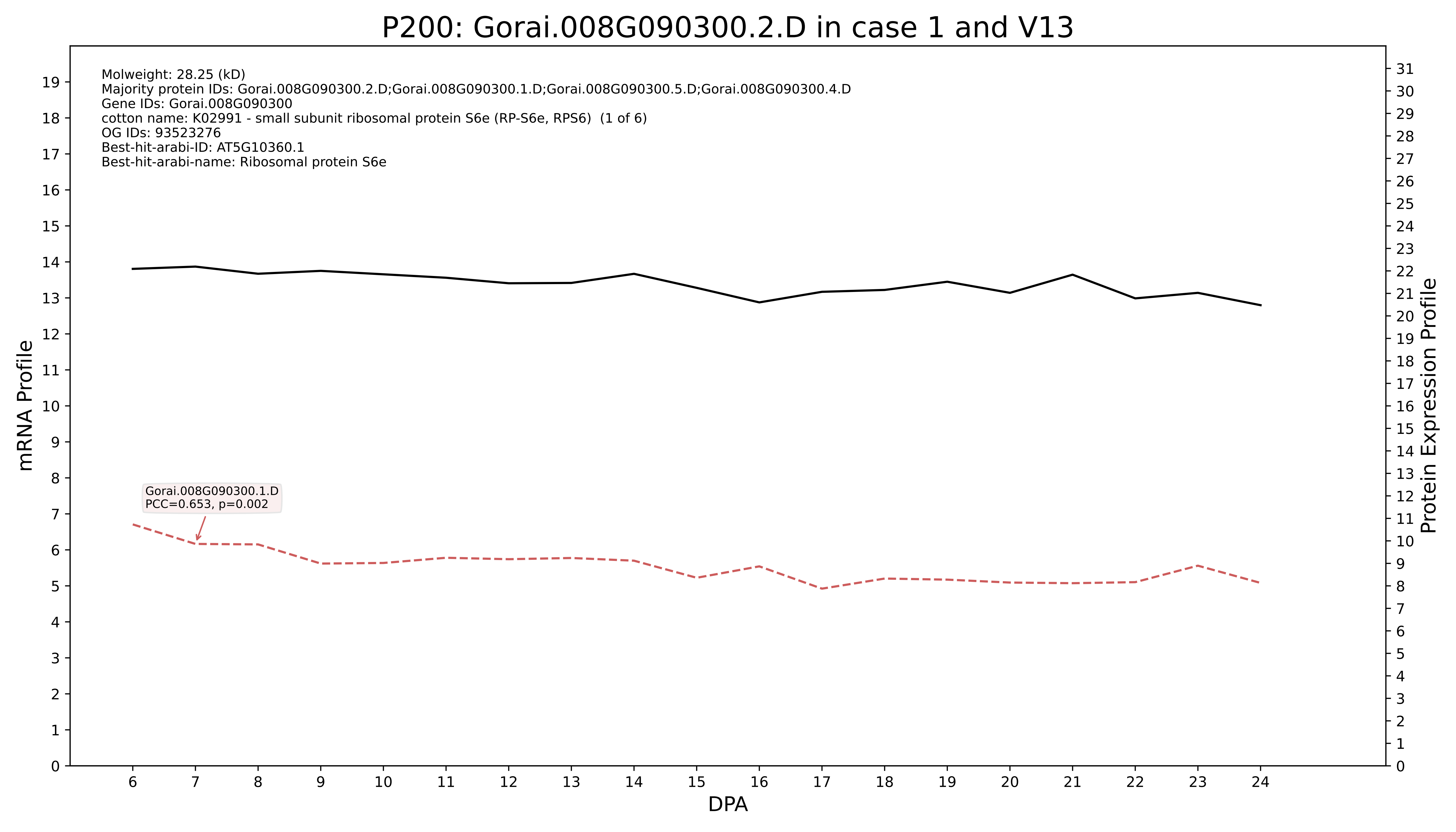
P200: Gorai.008G090300.2.D in case 1 and V13
Molweight: 28.25 (kD)
Majority protein IDs: Gorai.008G090300.2.D;Gorai.008G090300.1.D;Gorai.008G090300.5.D;Gorai.008G090300.4.D
Gene IDs: Gorai.008G090300
cotton name: K02991 - small subunit ribosomal protein S6e (RP-S6e, RPS6) (1 of 6)
OG IDs: 93523276
Best-hit-arabi-ID: AT5G10360.1
Best-hit-arabi-name: Ribosomal protein S6e
Gorai.008G090300.1.D
PCC=0.653, p=0.002
mRNA Profile
Protein Expression Profile
DPA

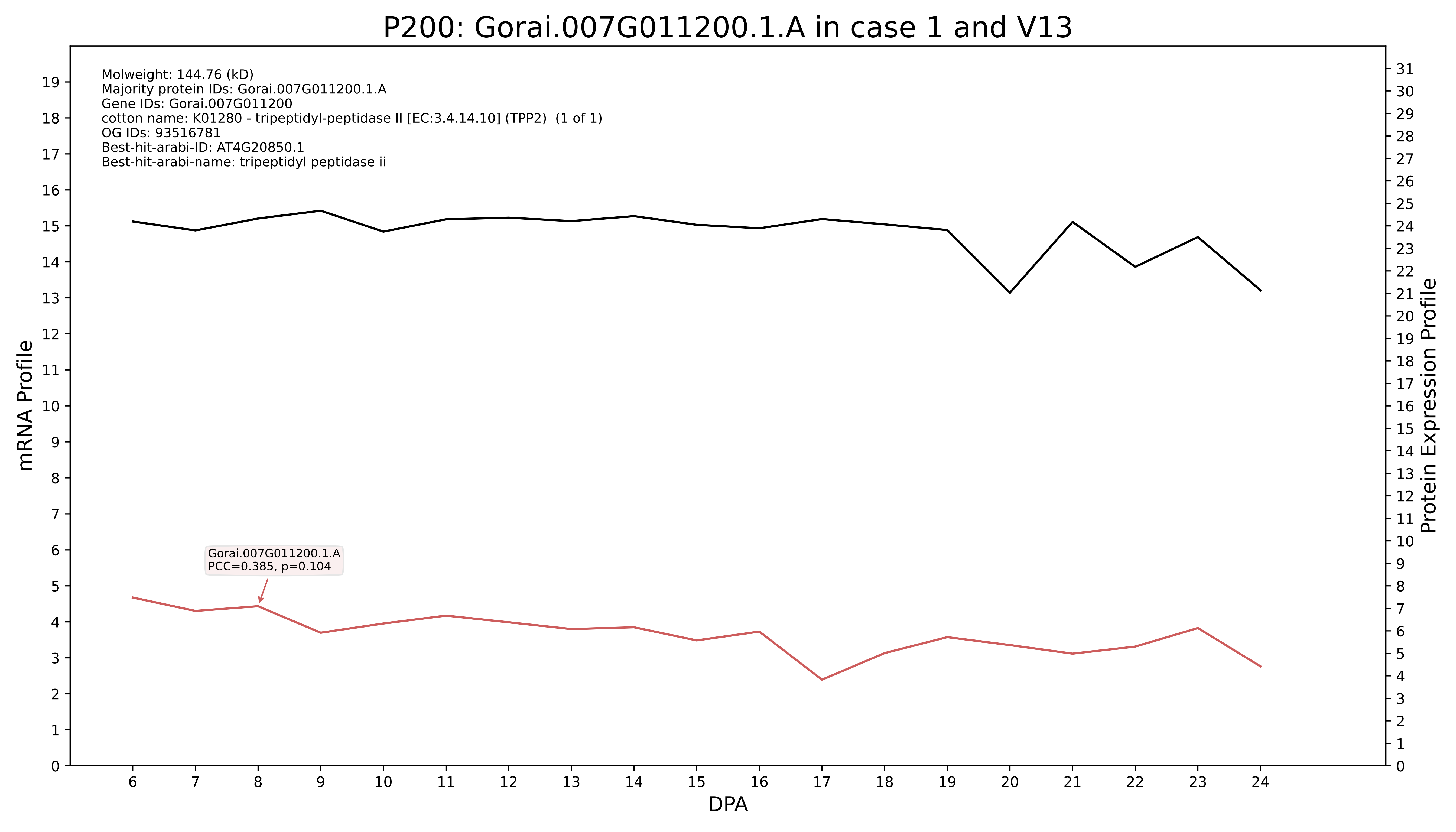

P200: Gorai.007G011200.1.A in case 1 and V13
Molweight: 144.76 (kD)
Majority protein IDs: Gorai.007G011200.1.A
Gene IDs: Gorai.007G011200
cotton name: K01280 - tripeptidyl-peptidase II [EC:3.4.14.10] (TPP2) (1 of 1)
OG IDs: 93516781
Best-hit-arabi-ID: AT4G20850.1
Best-hit-arabi-name: tripeptidyl peptidase ii
Gorai.007G011200.1.A
PCC=0.385, p=0.104
mRNA Profile
Protein Expression Profile
DPA

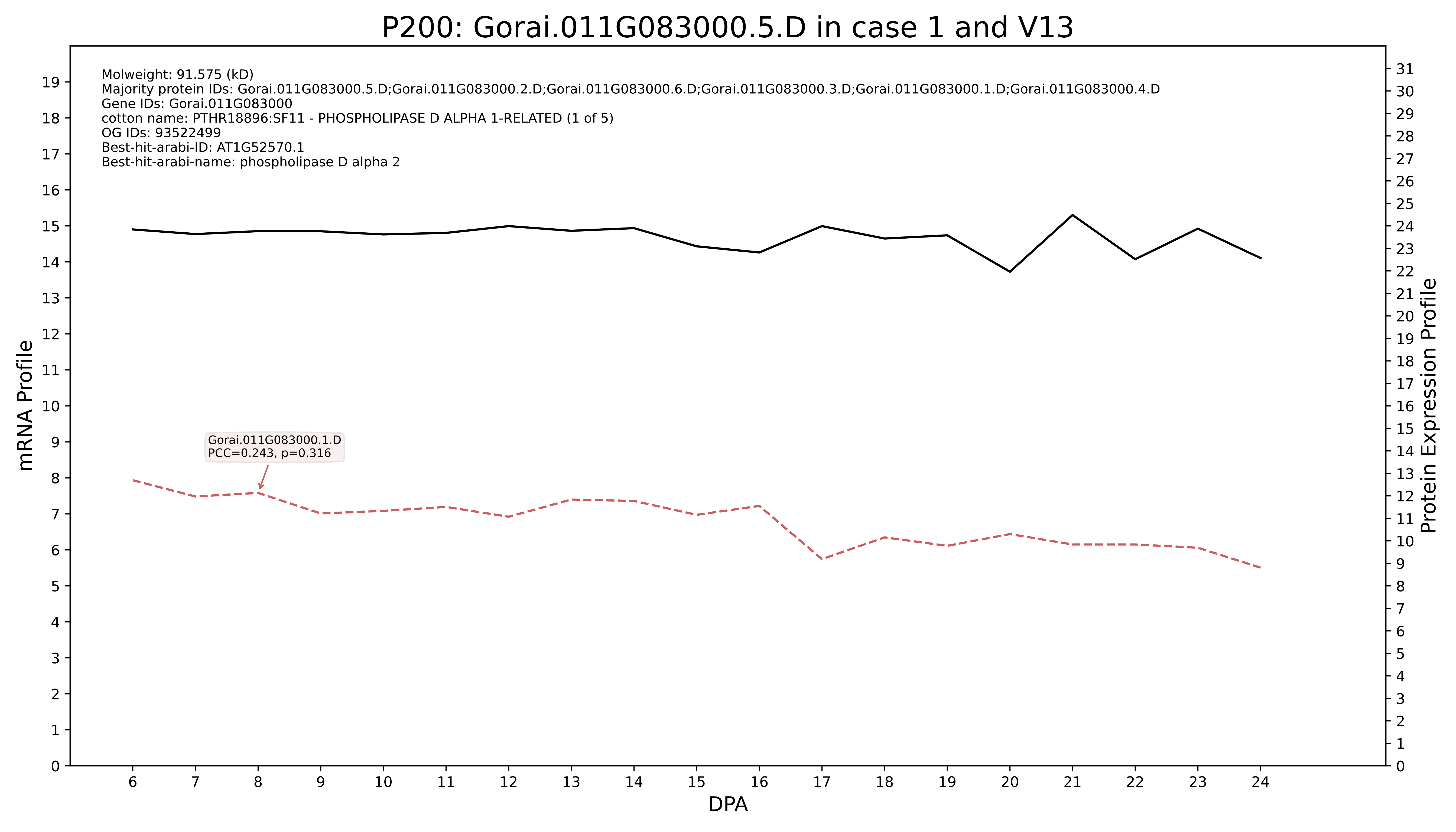
P200: Gorai.011G083000.5.D in case 1 and V13
Molweight: 91.575 (kD)
Majority protein IDs: Gorai.011G083000.5.D;Gorai.011G083000.2.D;Gorai.011G083000.6.D;Gorai.011G083000.3.D;Gorai.011G083000.1.D;Gorai.011G083000.4.D
Gene IDs: Gorai.011G083000
cotton name: PTHR18896:SF11 - PHOSPHOLIPASE D ALPHA 1-RELATED (1 of 5)
OG IDs: 93522499
Best-hit-arabi-ID: AT1G52570.1
Best-hit-arabi-name: phospholipase D alpha 2
Gorai.011G083000.1.D
PCC=0.243, p=0.316
mRNA Profile
Protein Expression Profile
DPA

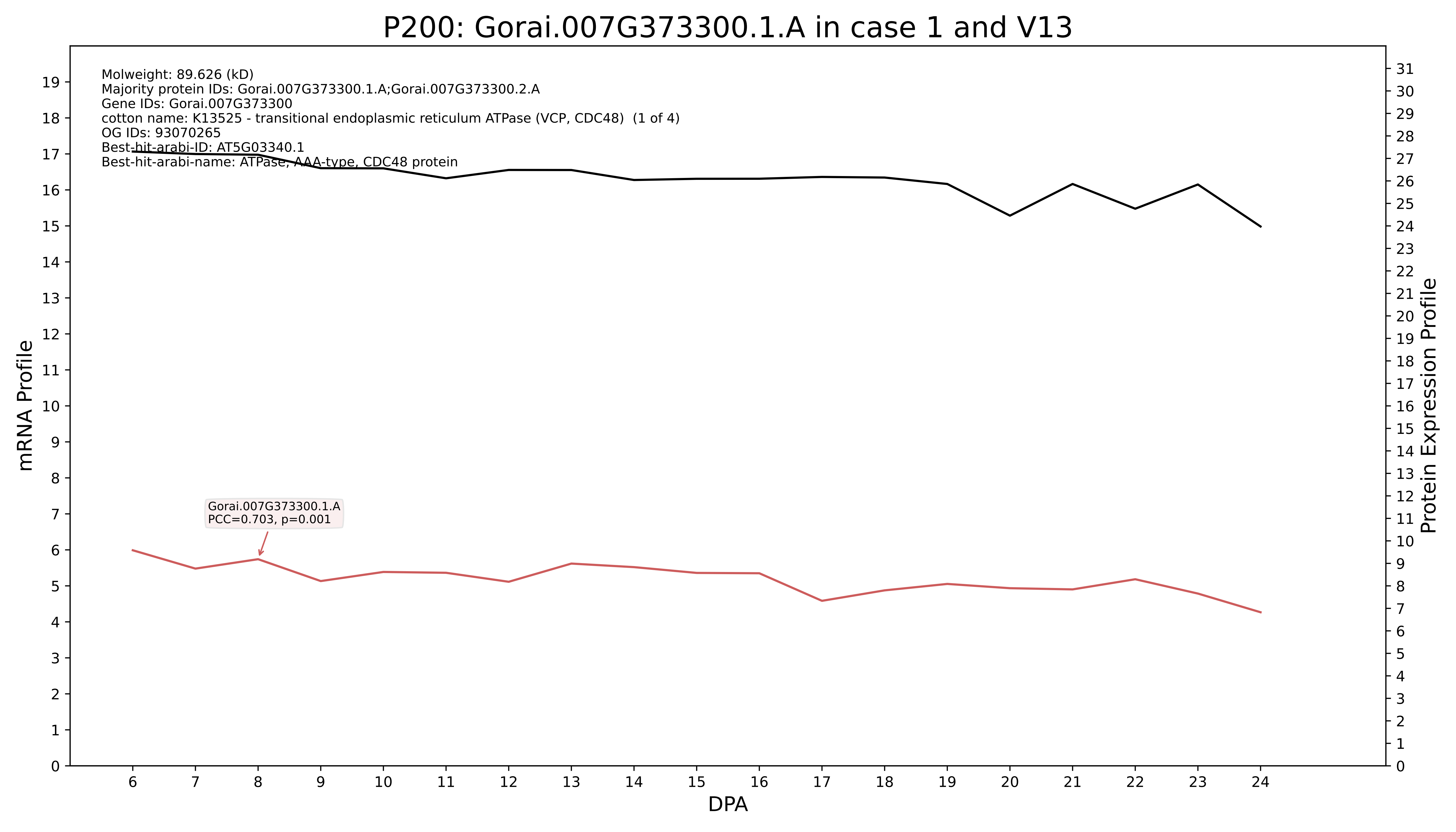
P200: Gorai.007G373300.1.A in case 1 and V13
Molweight: 89.626 (kD)
Majority protein IDs: Gorai.007G373300.1.A;Gorai.007G373300.2.A
Gene IDs: Gorai.007G373300
cotton name: K13525 - transitional endoplasmic reticulum ATPase (VCP, CDC48) (1 of 4)
OG IDs: 93070265
Best-hit-arabi-ID: AT5G03340.1
Best-hit-arabi-name: ATPase, AAA-type, CDC48 protein
Gorai.007G373300.1.A
PCC=0.703, p=0.001
mRNA Profile
Protein Expression Profile
DPA

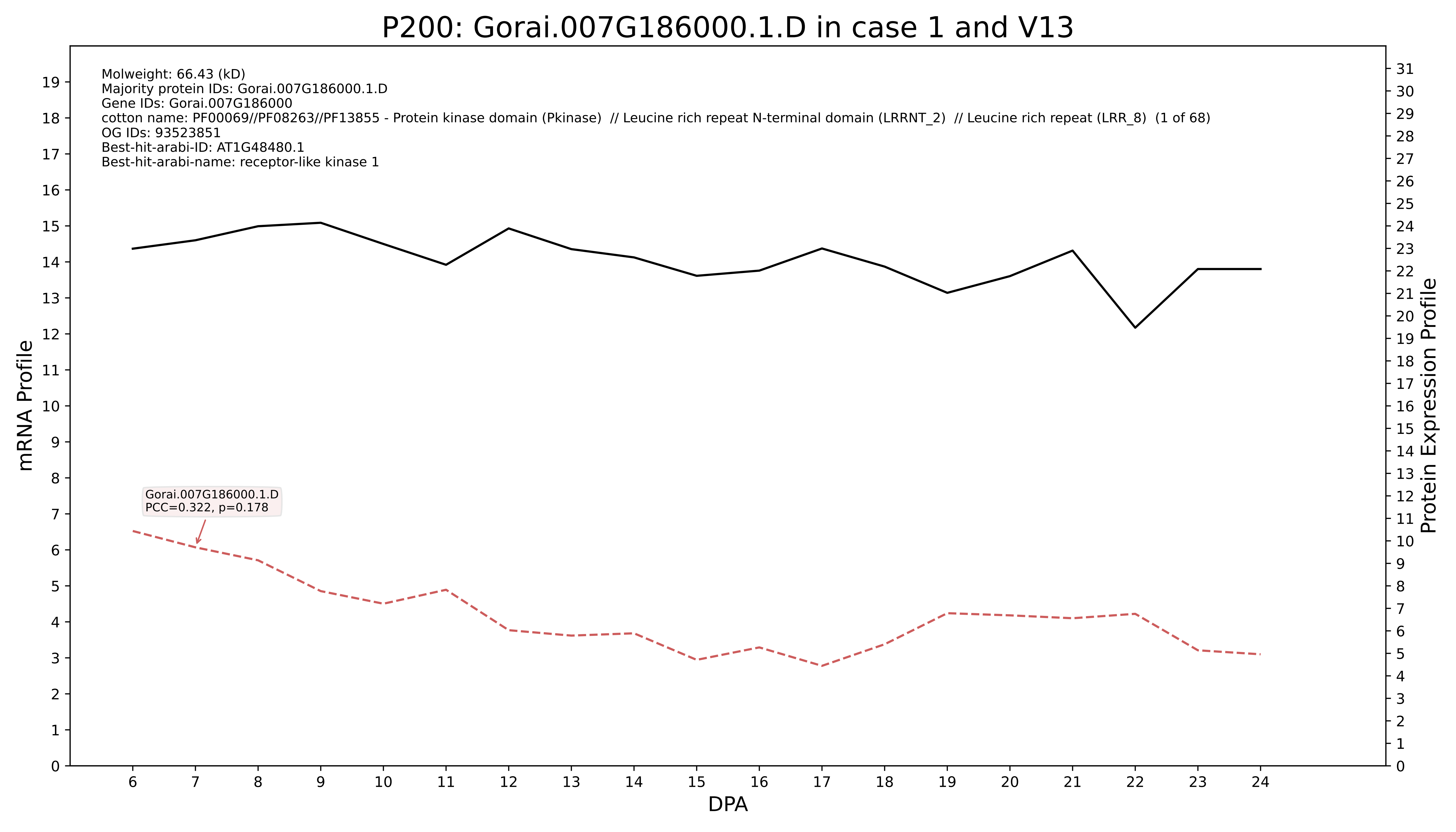

P200: Gorai.007G186000.1.D in case 1 and V13
Molweight: 66.43 (kD)
Majority protein IDs: Gorai.007G186000.1.D
Gene IDs: Gorai.007G186000
cotton name: PF00069//PF08263//PF13855 - Protein kinase domain (Pkinase) // Leucine rich repeat N-terminal domain (LRRNT_2) // Leucine rich repeat (LRR_8) (1 of 68)
OG IDs: 93523851
Best-hit-arabi-ID: AT1G48480.1
Best-hit-arabi-name: receptor-like kinase 1
Gorai.007G186000.1.D
PCC=0.322, p=0.178
mRNA Profile
Protein Expression Profile
DPA

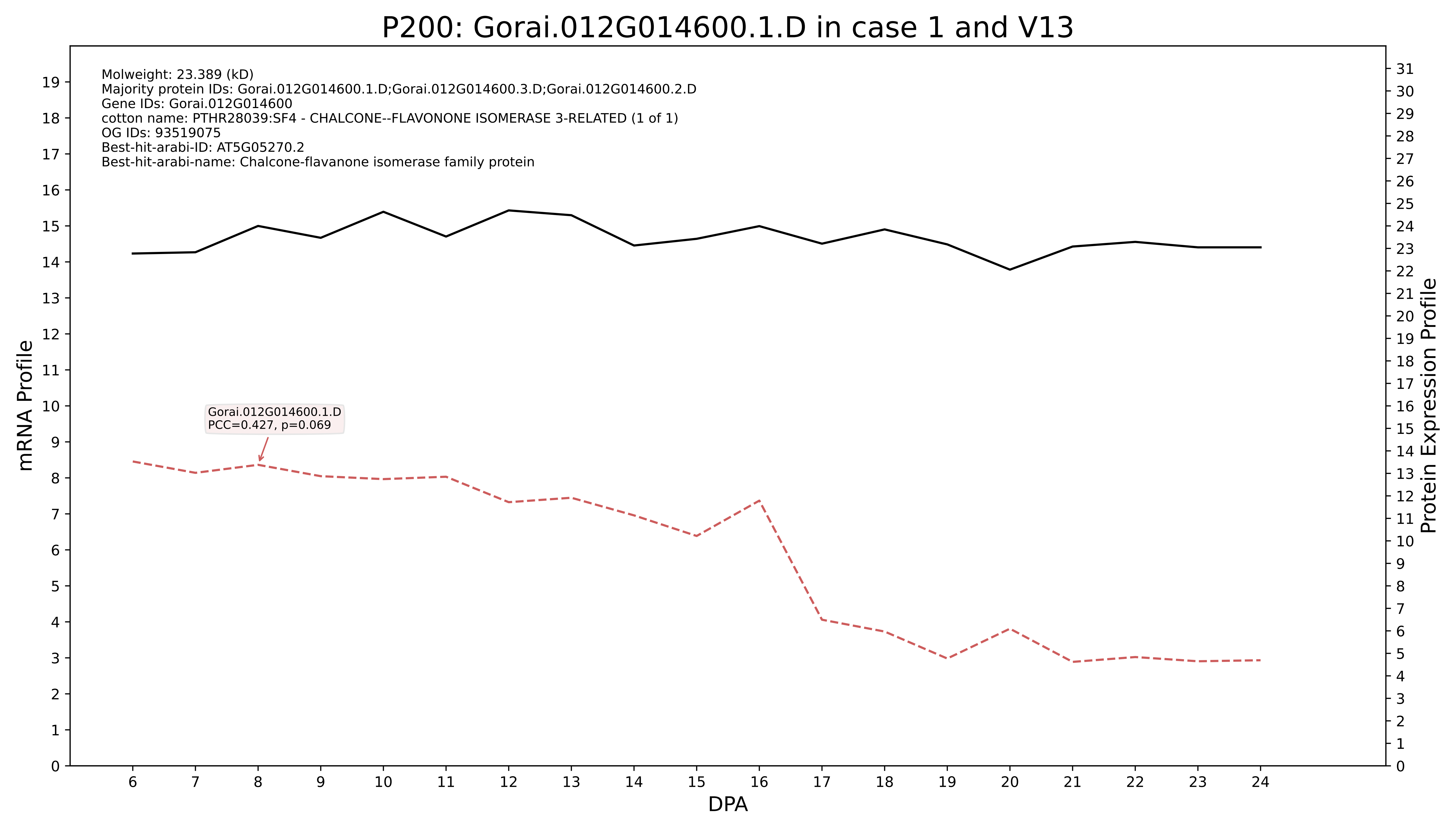
P200: Gorai.012G014600.1.D in case 1 and V13
Molweight: 23.389 (kD)
Majority protein IDs: Gorai.012G014600.1.D;Gorai.012G014600.3.D;Gorai.012G014600.2.D
Gene IDs: Gorai.012G014600
cotton name: PTHR28039:SF4 - CHALCONE--FLAVONONE ISOMERASE 3-RELATED (1 of 1)
OG IDs: 93519075
Best-hit-arabi-ID: AT5G05270.2
Best-hit-arabi-name: Chalcone-flavanone isomerase family protein
Gorai.012G014600.1.D
PCC=0.427, p=0.069
mRNA Profile
Protein Expression Profile
DPA

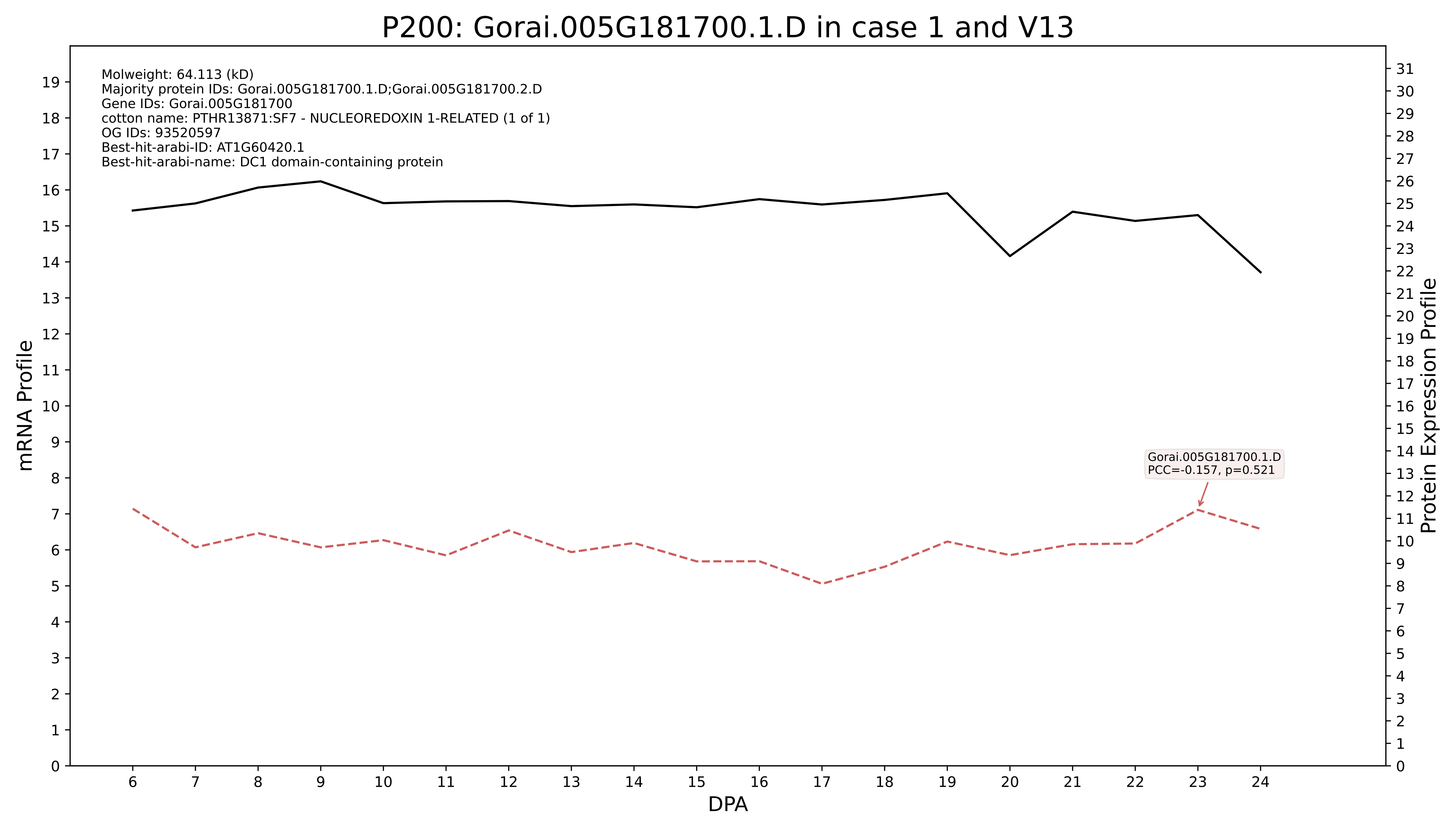
P200: Gorai.005G181700.1.D in case 1 and V13
Molweight: 64.113 (kD)
Majority protein IDs: Gorai.005G181700.1.D;Gorai.005G181700.2.D
Gene IDs: Gorai.005G181700
cotton name: PTHR13871:SF7 - NUCLEOREDOXIN 1-RELATED (1 of 1)
OG IDs: 93520597
Best-hit-arabi-ID: AT1G60420.1
Best-hit-arabi-name: DC1 domain-containing protein
Gorai.005G181700.1.D
PCC=-0.157, p=0.521
mRNA Profile
Protein Expression Profile
DPA

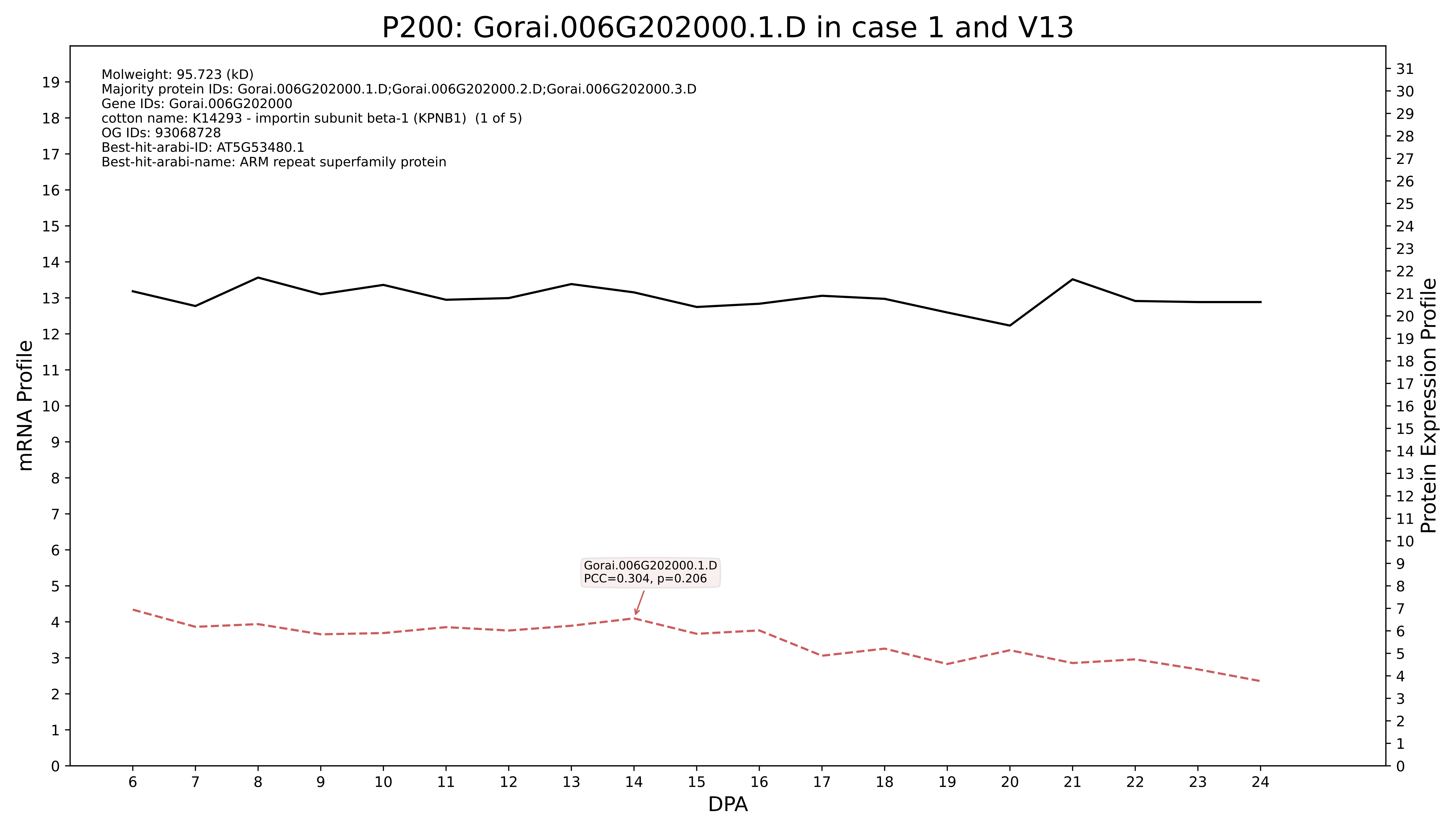
P200: Gorai.006G202000.1.D in case 1 and V13
Molweight: 95.723 (kD)
Majority protein IDs: Gorai.006G202000.1.D;Gorai.006G202000.2.D;Gorai.006G202000.3.D
Gene IDs: Gorai.006G202000
cotton name: K14293 - importin subunit beta-1 (KPNB1) (1 of 5)
OG IDs: 93068728
Best-hit-arabi-ID: AT5G53480.1
Best-hit-arabi-name: ARM repeat superfamily protein
Gorai.006G202000.1.D
PCC=0.304, p=0.206
mRNA Profile
Protein Expression Profile
DPA

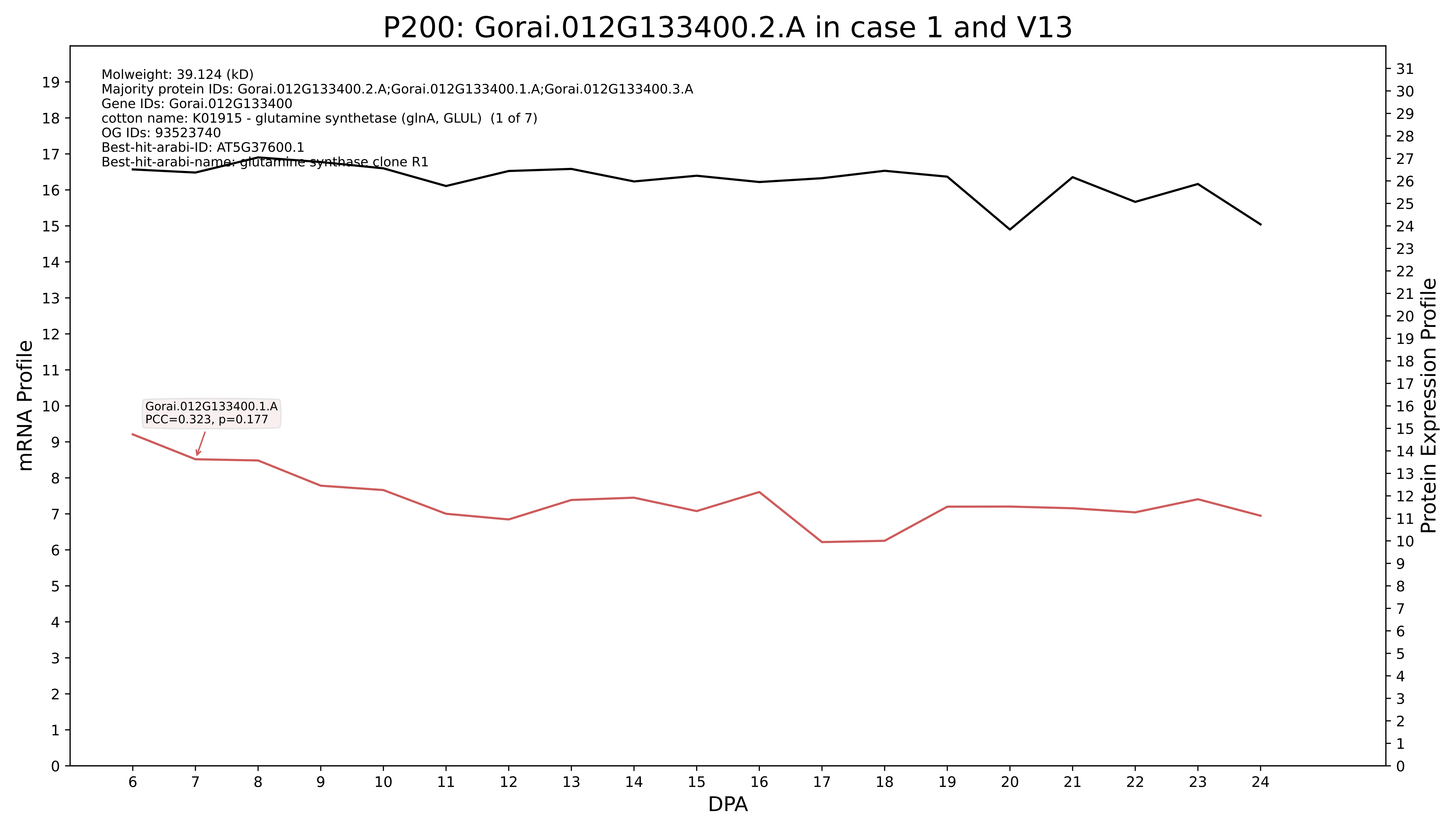
P200: Gorai.012G133400.2.A in case 1 and V13
Molweight: 39.124 (kD)
Majority protein IDs: Gorai.012G133400.2.A;Gorai.012G133400.1.A;Gorai.012G133400.3.A
Gene IDs: Gorai.012G133400
cotton name: K01915 - glutamine synthetase (glnA, GLUL) (1 of 7)
OG IDs: 93523740
Best-hit-arabi-ID: AT5G37600.1
Best-hit-arabi-name: glutamine synthase clone R1
Gorai.012G133400.1.A
PCC=0.323, p=0.177
mRNA Profile
Protein Expression Profile
DPA

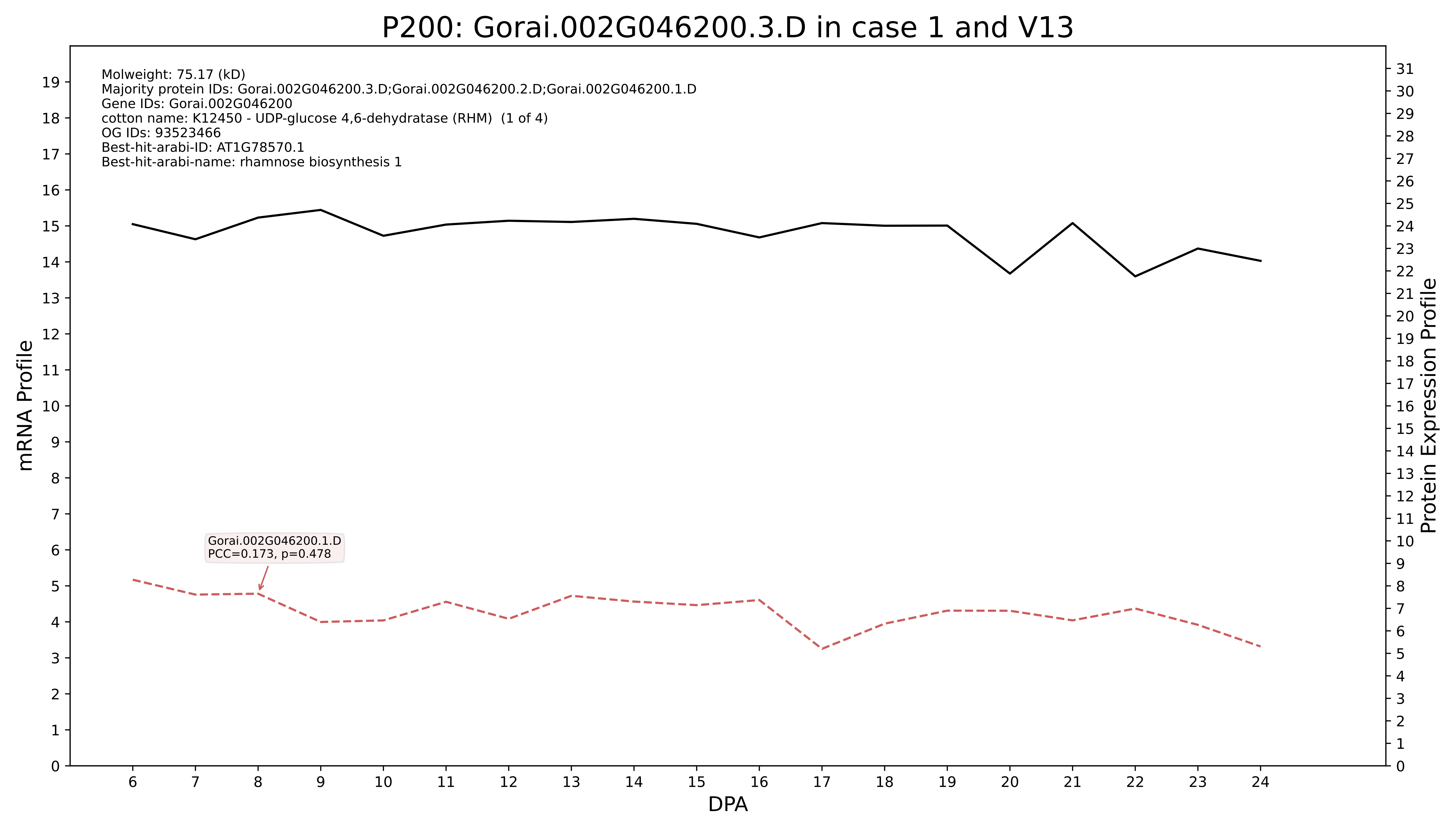
P200: Gorai.002G046200.3.D in case 1 and V13
Molweight: 75.17 (kD)
Majority protein IDs: Gorai.002G046200.3.D;Gorai.002G046200.2.D;Gorai.002G046200.1.D
Gene IDs: Gorai.002G046200
cotton name: K12450 - UDP-glucose 4,6-dehydratase (RHM) (1 of 4)
OG IDs: 93523466
Best-hit-arabi-ID: AT1G78570.1
Best-hit-arabi-name: rhamnose biosynthesis 1
Gorai.002G046200.1.D
PCC=0.173, p=0.478
mRNA Profile
Protein Expression Profile
DPA

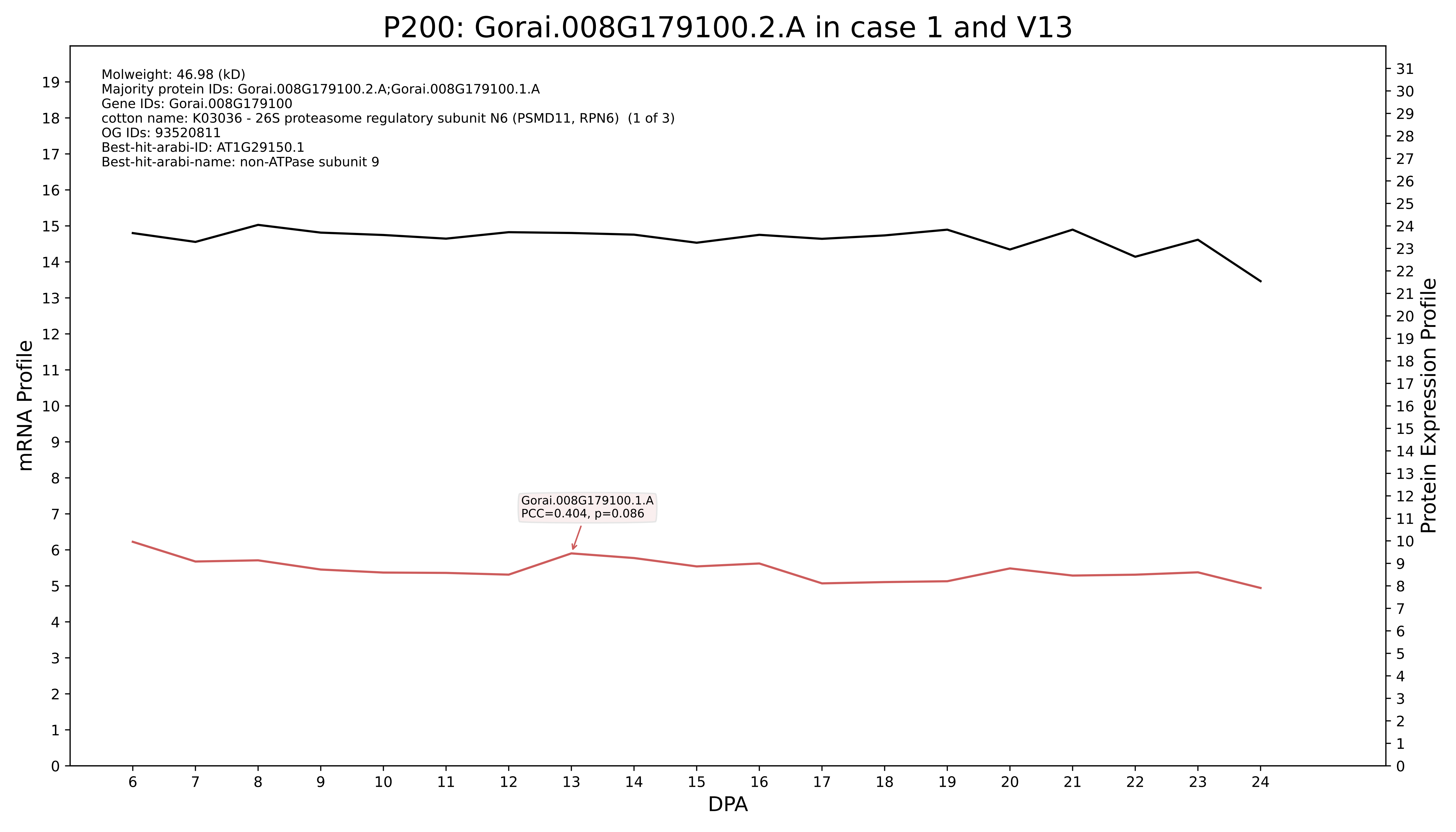

P200: Gorai.008G179100.2.A in case 1 and V13
Molweight: 46.98 (kD)
Majority protein IDs: Gorai.008G179100.2.A;Gorai.008G179100.1.A
Gene IDs: Gorai.008G179100
cotton name: K03036 - 26S proteasome regulatory subunit N6 (PSMD11, RPN6) (1 of 3)
OG IDs: 93520811
Best-hit-arabi-ID: AT1G29150.1
Best-hit-arabi-name: non-ATPase subunit 9
Gorai.008G179100.1.A
PCC=0.404, p=0.086
mRNA Profile
Protein Expression Profile
DPA

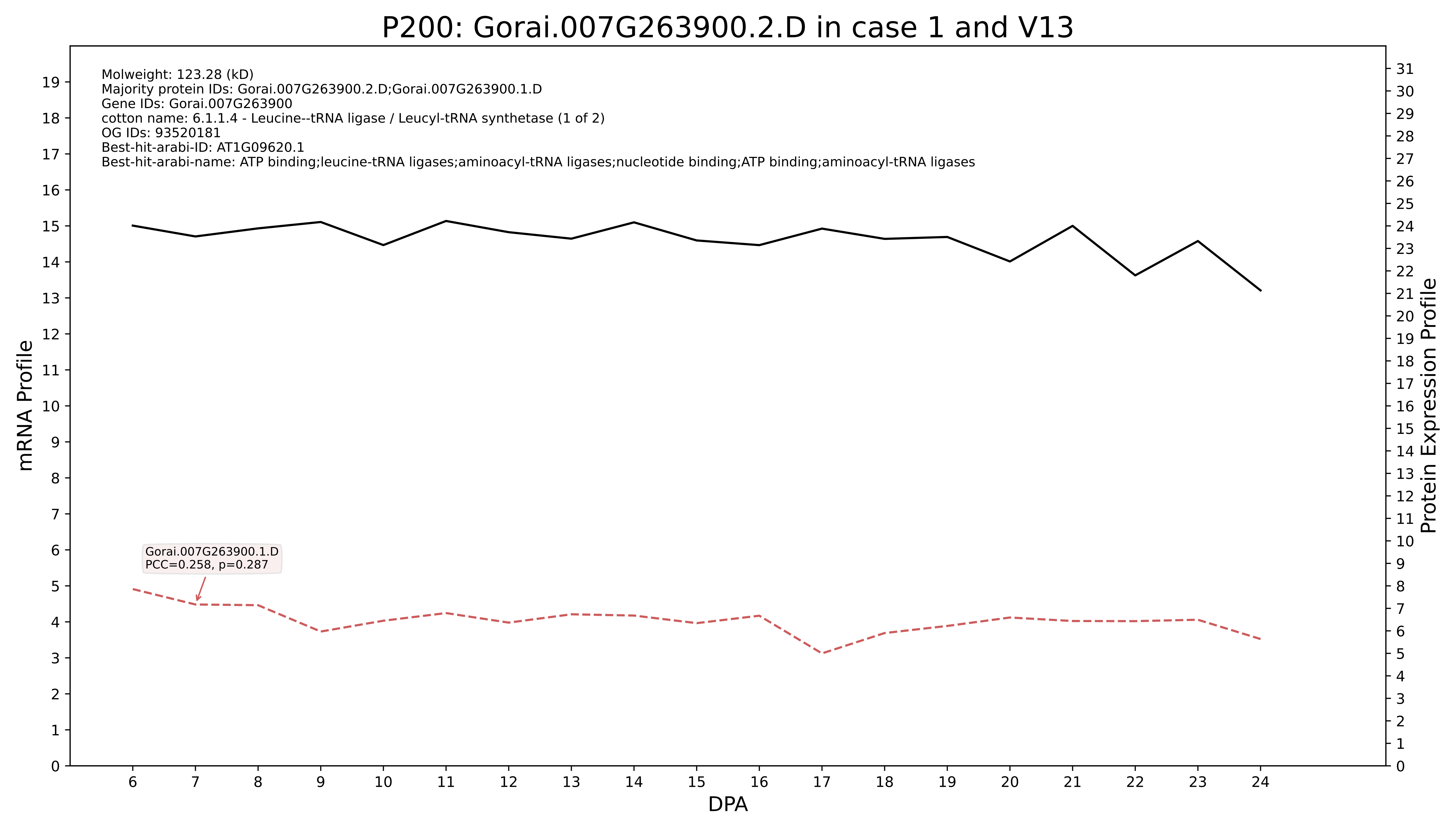
P200: Gorai.007G263900.2.D in case 1 and V13
Molweight: 123.28 (kD)
Majority protein IDs: Gorai.007G263900.2.D;Gorai.007G263900.1.D
Gene IDs: Gorai.007G263900
cotton name: 6.1.1.4 - Leucine--tRNA ligase / Leucyl-tRNA synthetase (1 of 2)
OG IDs: 93520181
Best-hit-arabi-ID: AT1G09620.1
Best-hit-arabi-name: ATP binding;leucine-tRNA ligases;aminoacyl-tRNA ligases;nucleotide binding;ATP binding;aminoacyl-tRNA ligases
Gorai.007G263900.1.D
PCC=0.258, p=0.287
mRNA Profile
Protein Expression Profile
DPA

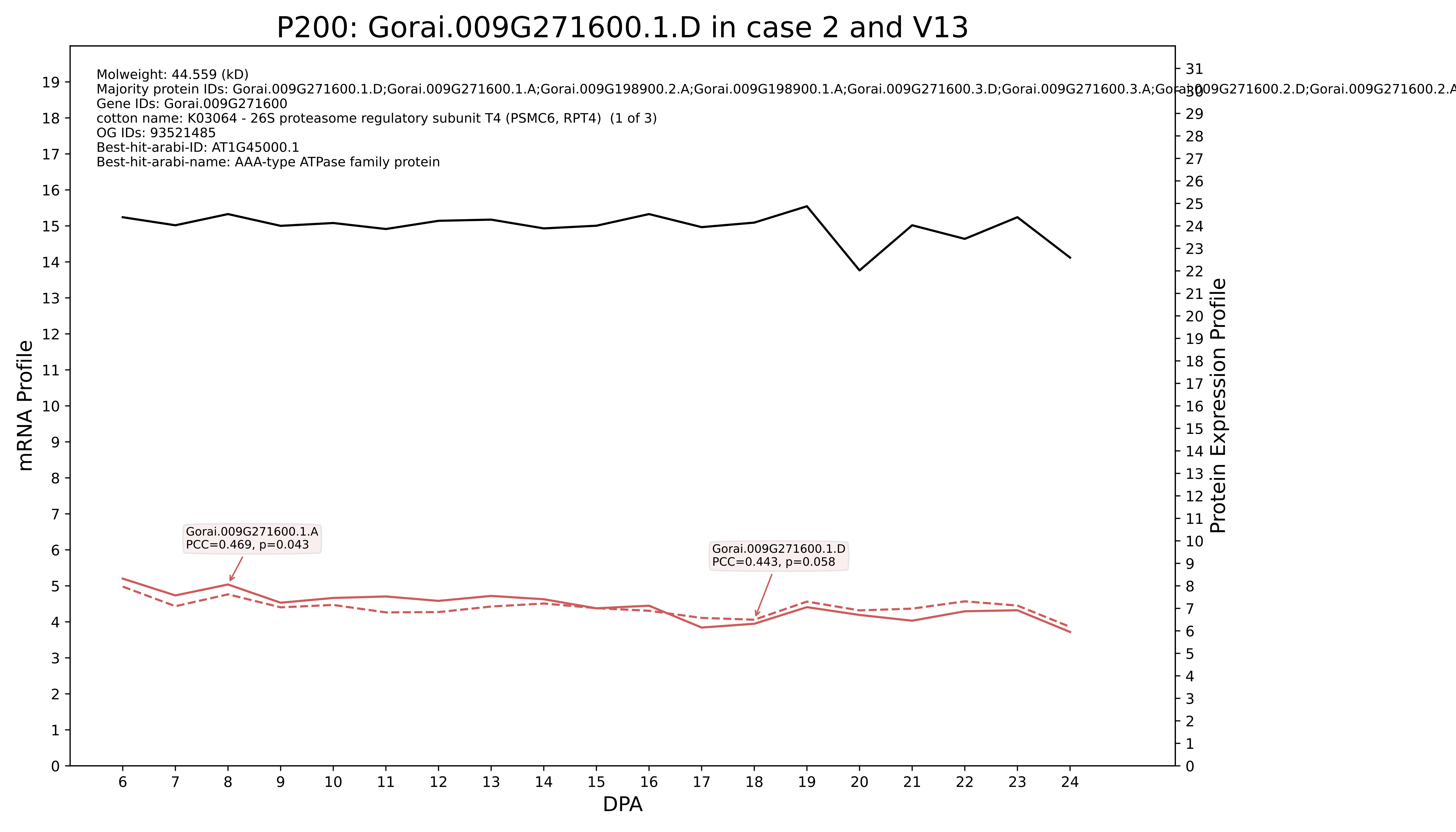
P200: Gorai.009G271600.1.D in case 2 and V13
Molweight: 44.559 (kD)
Majority protein IDs: Gorai.009G271600.1.D;Gorai.009G271600.1.A;Gorai.009G198900.2.A;Gorai.009G198900.1.A;Gorai.009G271600.3.D;Gorai.009G271600.3.A;Gorai.009G271600.2.D;Gorai.009G271600.2.A
Gene IDs: Gorai.009G271600
cotton name: K03064 - 26S proteasome regulatory subunit T4 (PSMC6, RPT4) (1 of 3)
OG IDs: 93521485
Best-hit-arabi-ID: AT1G45000.1
Best-hit-arabi-name: AAA-type ATPase family protein
Gorai.009G271600.1.A
PCC=0.469, p=0.043
Gorai.009G271600.1.D
PCC=0.443, p=0.058
mRNA Profile
Protein Expression Profile
DPA

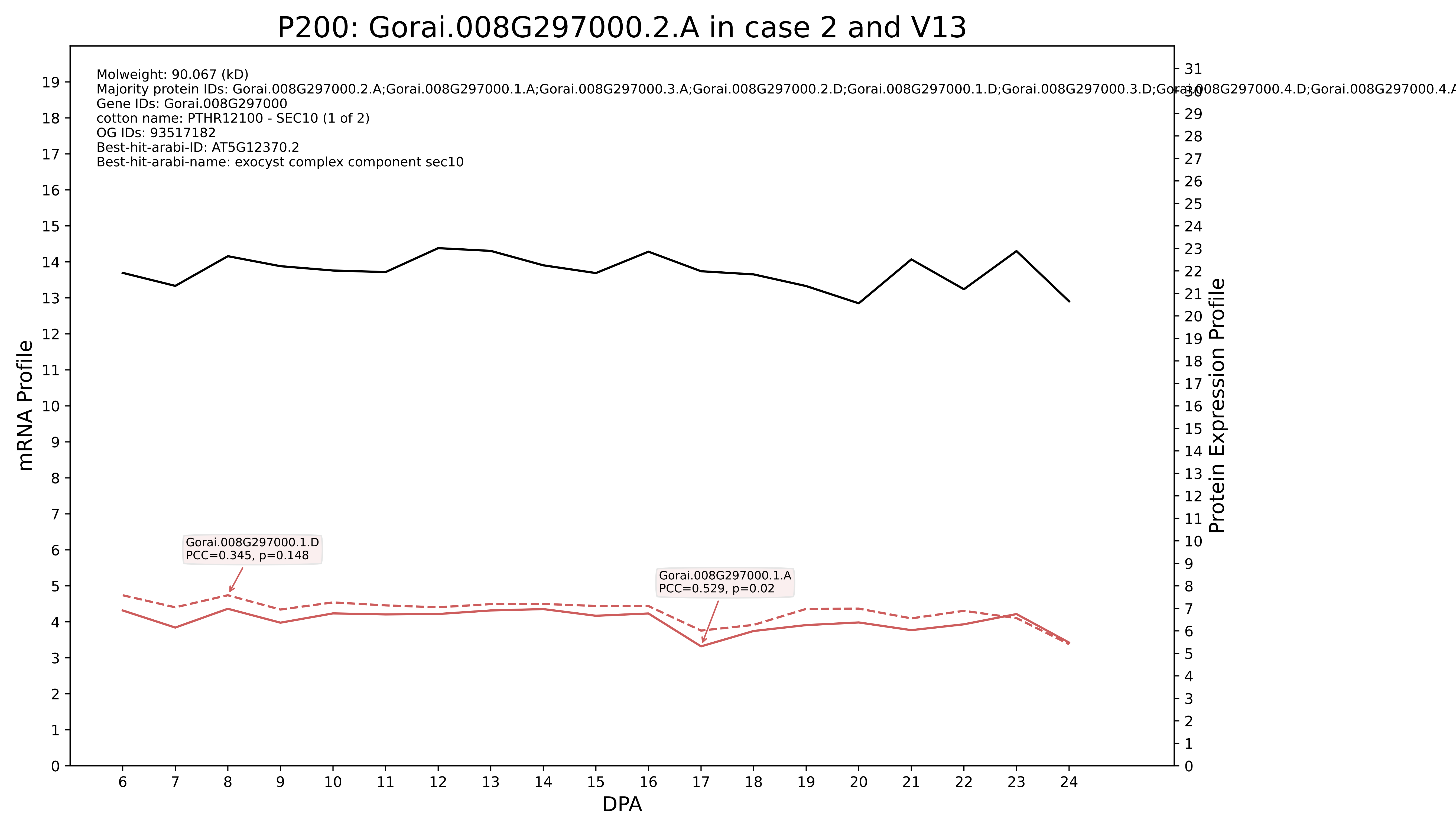
P200: Gorai.008G297000.2.A in case 2 and V13
Molweight: 90.067 (kD)
Majority protein IDs: Gorai.008G297000.2.A;Gorai.008G297000.1.A;Gorai.008G297000.3.A;Gorai.008G297000.2.D;Gorai.008G297000.1.D;Gorai.008G297000.3.D;Gorai.008G297000.4.D;Gorai.008G297000.4.A
Gene IDs: Gorai.008G297000
cotton name: PTHR12100 - SEC10 (1 of 2)
OG IDs: 93517182
Best-hit-arabi-ID: AT5G12370.2
Best-hit-arabi-name: exocyst complex component sec10
Gorai.008G297000.1.D
PCC=0.345, p=0.148
Gorai.008G297000.1.A
PCC=0.529, p=0.02
mRNA Profile
Protein Expression Profile
DPA

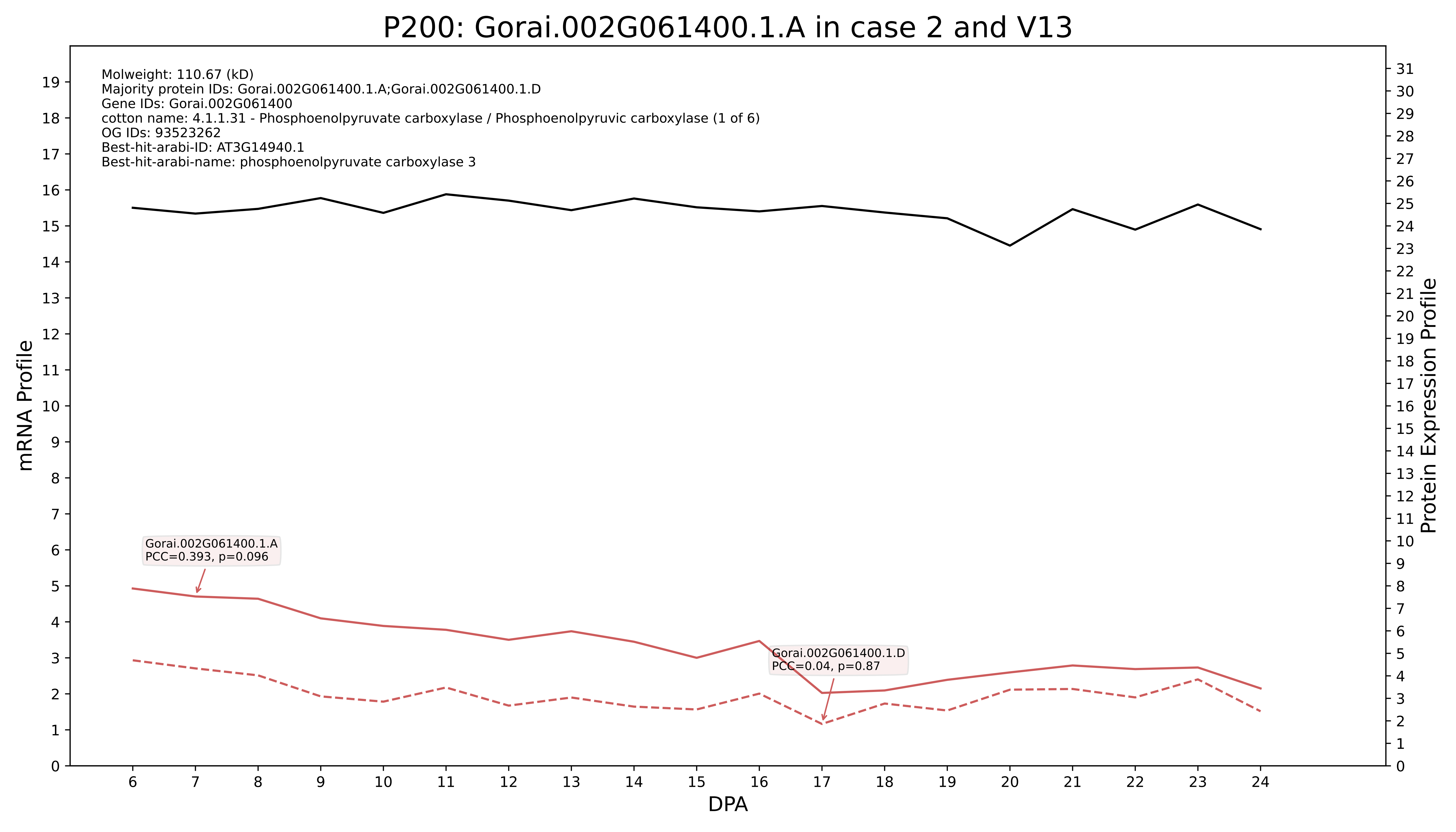
P200: Gorai.002G061400.1.A in case 2 and V13
Molweight: 110.67 (kD)
Majority protein IDs: Gorai.002G061400.1.A;Gorai.002G061400.1.D
Gene IDs: Gorai.002G061400
cotton name: 4.1.1.31 - Phosphoenolpyruvate carboxylase / Phosphoenolpyruvic carboxylase (1 of 6)
OG IDs: 93523262
Best-hit-arabi-ID: AT3G14940.1
Best-hit-arabi-name: phosphoenolpyruvate carboxylase 3
Gorai.002G061400.1.A
PCC=0.393, p=0.096
Gorai.002G061400.1.D
PCC=0.04, p=0.87
mRNA Profile
Protein Expression Profile
DPA

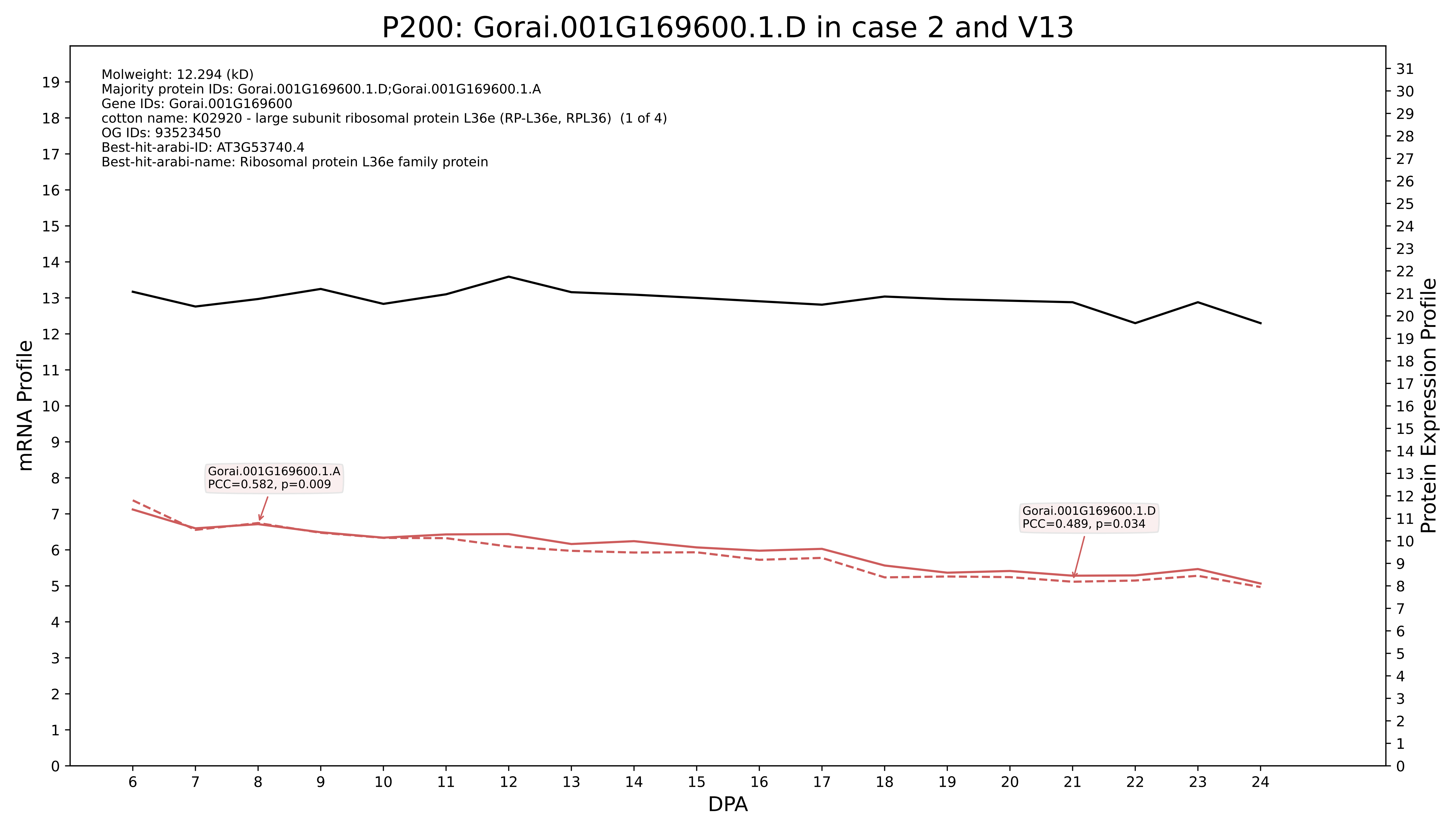

P200: Gorai.001G169600.1.D in case 2 and V13
Molweight: 12.294 (kD)
Majority protein IDs: Gorai.001G169600.1.D;Gorai.001G169600.1.A
Gene IDs: Gorai.001G169600
cotton name: K02920 - large subunit ribosomal protein L36e (RP-L36e, RPL36) (1 of 4)
OG IDs: 93523450
Best-hit-arabi-ID: AT3G53740.4
Best-hit-arabi-name: Ribosomal protein L36e family protein
Gorai.001G169600.1.A
PCC=0.582, p=0.009
Gorai.001G169600.1.D
PCC=0.489, p=0.034
mRNA Profile
Protein Expression Profile
DPA

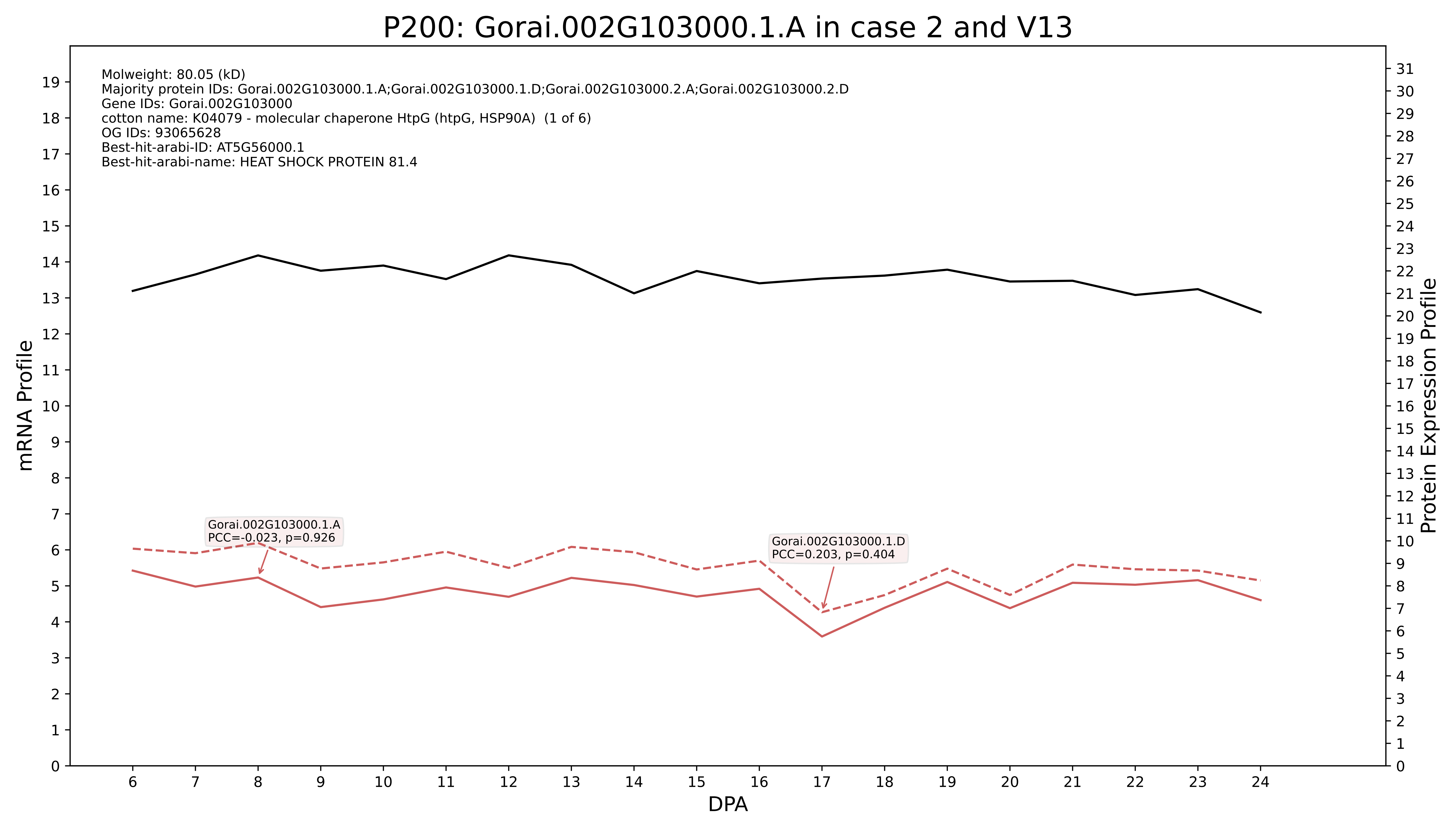

P200: Gorai.002G103000.1.A in case 2 and V13
Molweight: 80.05 (kD)
Majority protein IDs: Gorai.002G103000.1.A;Gorai.002G103000.1.D;Gorai.002G103000.2.A;Gorai.002G103000.2.D
Gene IDs: Gorai.002G103000
cotton name: K04079 - molecular chaperone HtpG (htpG, HSP90A) (1 of 6)
OG IDs: 93065628
Best-hit-arabi-ID: AT5G56000.1
Best-hit-arabi-name: HEAT SHOCK PROTEIN 81.4
Gorai.002G103000.1.A
PCC=-0.023, p=0.926
Gorai.002G103000.1.D
PCC=0.203, p=0.404
mRNA Profile
Protein Expression Profile
DPA

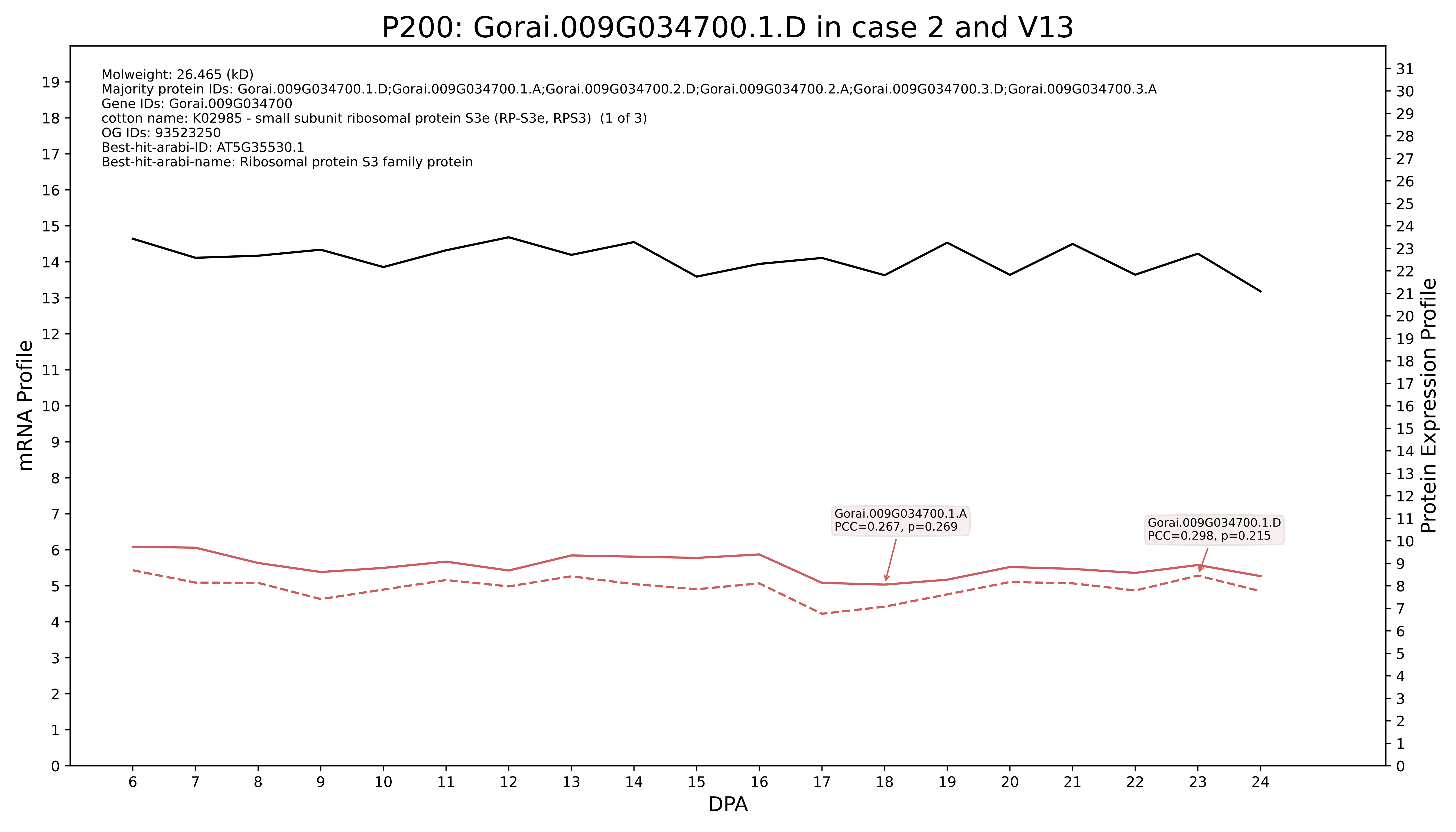
P200: Gorai.009G034700.1.D in case 2 and V13
Molweight: 26.465 (kD)
Majority protein IDs: Gorai.009G034700.1.D;Gorai.009G034700.1.A;Gorai.009G034700.2.D;Gorai.009G034700.2.A;Gorai.009G034700.3.D;Gorai.009G034700.3.A
Gene IDs: Gorai.009G034700
cotton name: K02985 - small subunit ribosomal protein S3e (RP-S3e, RPS3) (1 of 3)
OG IDs: 93523250
Best-hit-arabi-ID: AT5G35530.1
Best-hit-arabi-name: Ribosomal protein S3 family protein
Gorai.009G034700.1.A
PCC=0.267, p=0.269
Gorai.009G034700.1.D
PCC=0.298, p=0.215
mRNA Profile
Protein Expression Profile
DPA

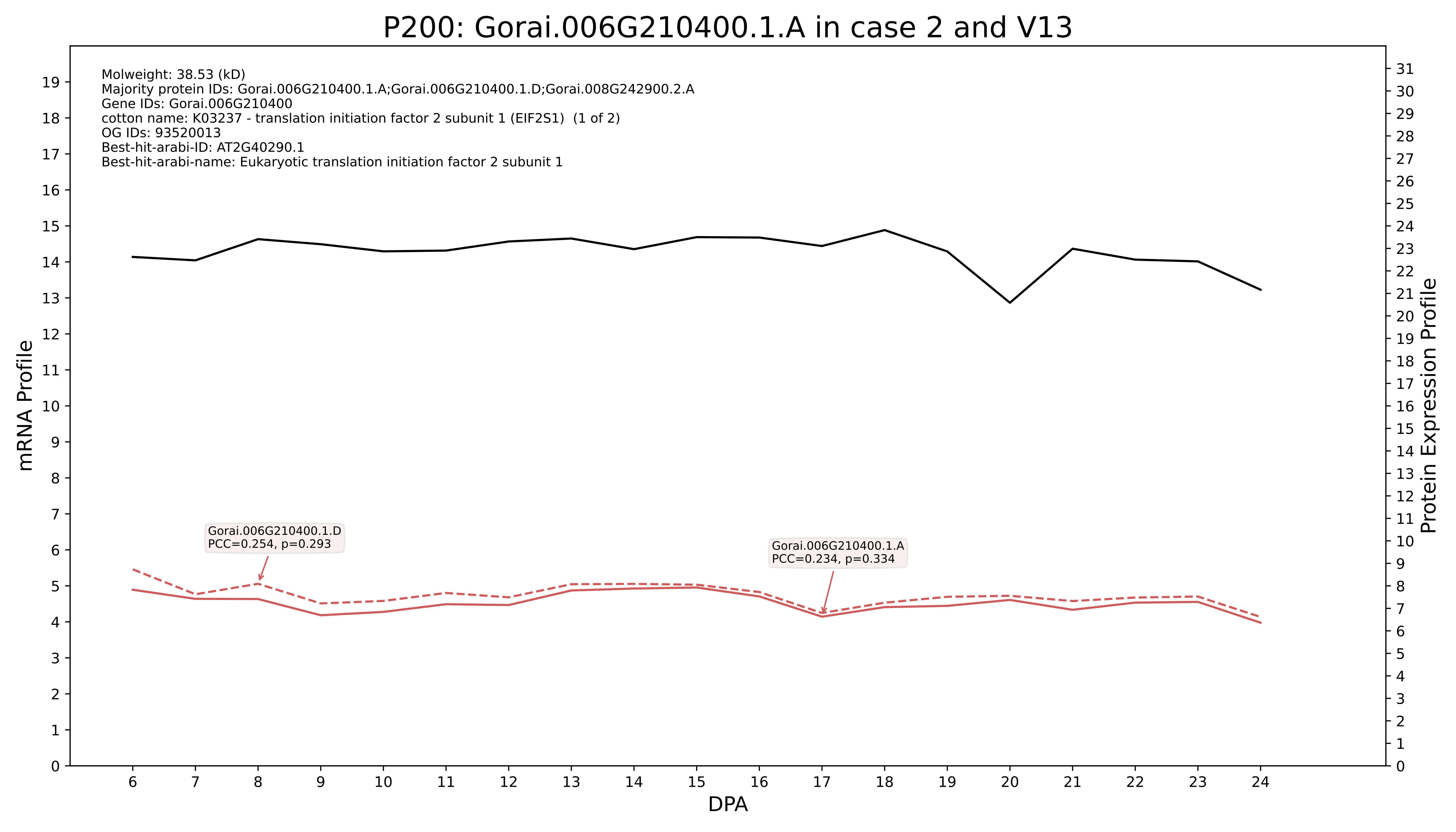
P200: Gorai.006G210400.1.A in case 2 and V13
Molweight: 38.53 (kD)
Majority protein IDs: Gorai.006G210400.1.A;Gorai.006G210400.1.D;Gorai.008G242900.2.A
Gene IDs: Gorai.006G210400
cotton name: K03237 - translation initiation factor 2 subunit 1 (EIF2S1) (1 of 2)
OG IDs: 93520013
Best-hit-arabi-ID: AT2G40290.1
Best-hit-arabi-name: Eukaryotic translation initiation factor 2 subunit 1
Gorai.006G210400.1.D
PCC=0.254, p=0.293
Gorai.006G210400.1.A
PCC=0.234, p=0.334
mRNA Profile
Protein Expression Profile
DPA

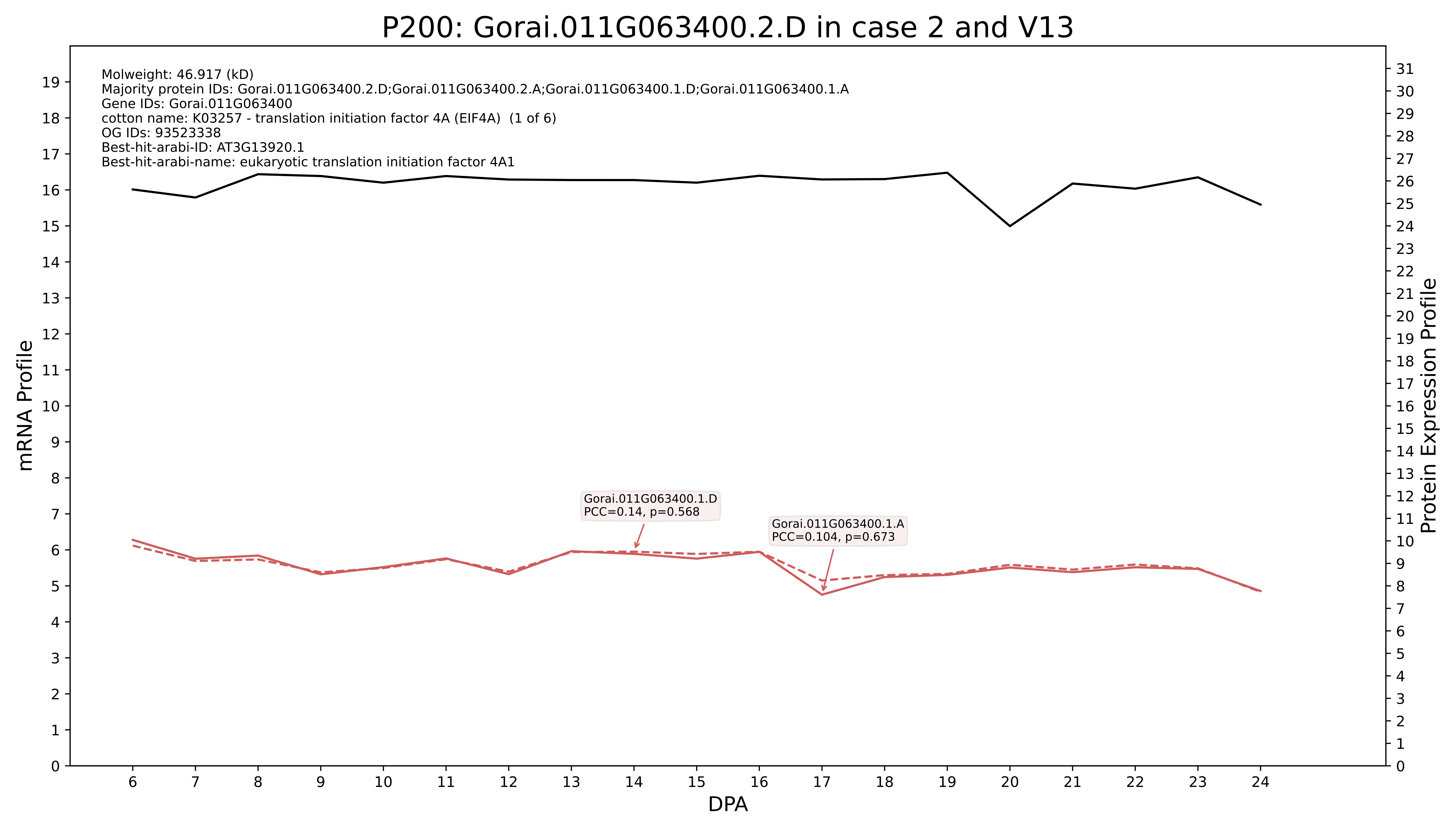
P200: Gorai.011G063400.2.D in case 2 and V13
Molweight: 46.917 (kD)
Majority protein IDs: Gorai.011G063400.2.D;Gorai.011G063400.2.A;Gorai.011G063400.1.D;Gorai.011G063400.1.A
Gene IDs: Gorai.011G063400
cotton name: K03257 - translation initiation factor 4A (EIF4A) (1 of 6)
OG IDs: 93523338
Best-hit-arabi-ID: AT3G13920.1
Best-hit-arabi-name: eukaryotic translation initiation factor 4A1
Gorai.011G063400.1.D
PCC=0.14, p=0.568
Gorai.011G063400.1.A
PCC=0.104, p=0.673
mRNA Profile
Protein Expression Profile
DPA

00: Gorai.005G045000.1.D in case 2 and V13

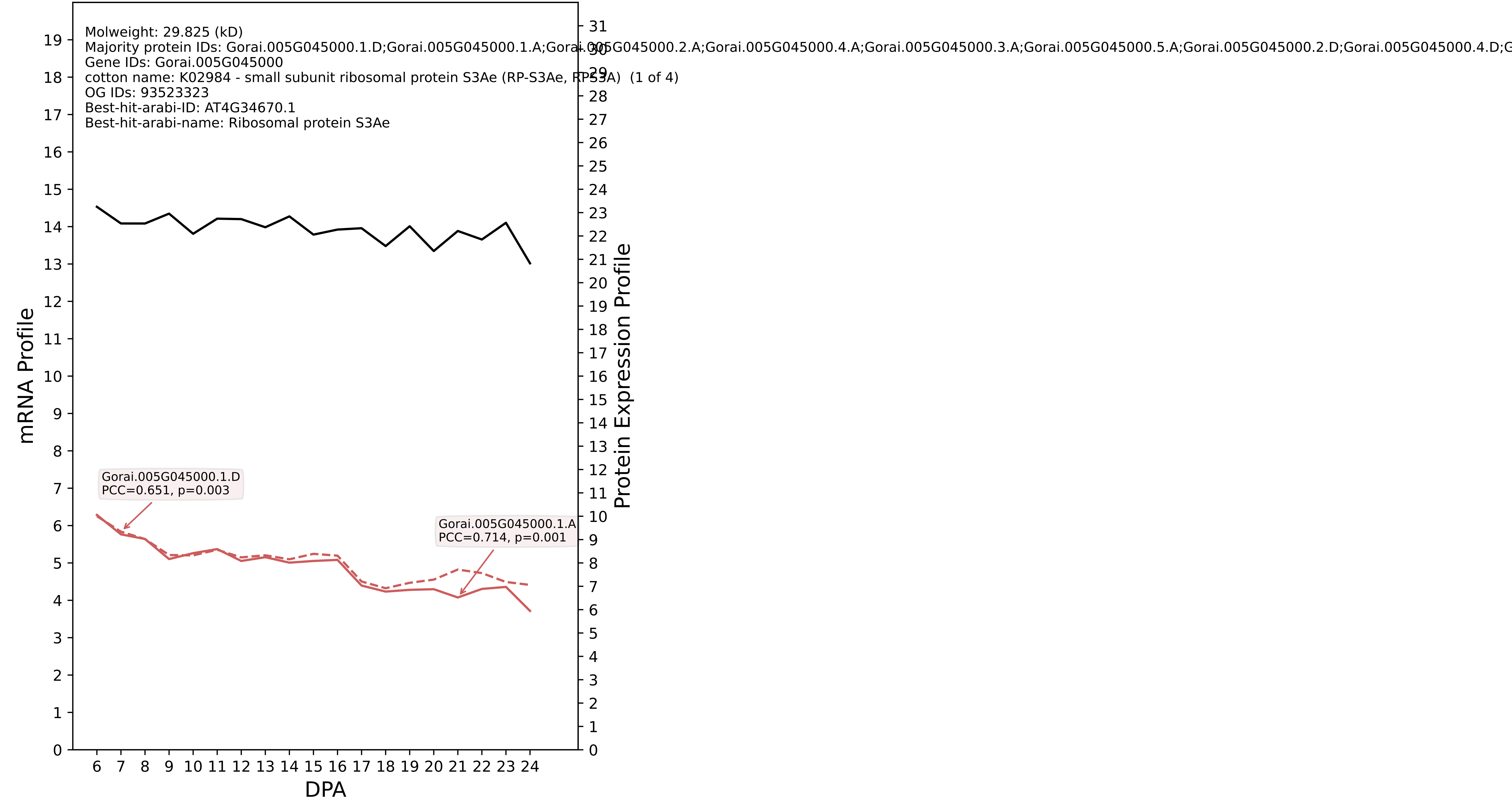

### 00: Gorai.012G070300.5.D in case 2 and V13

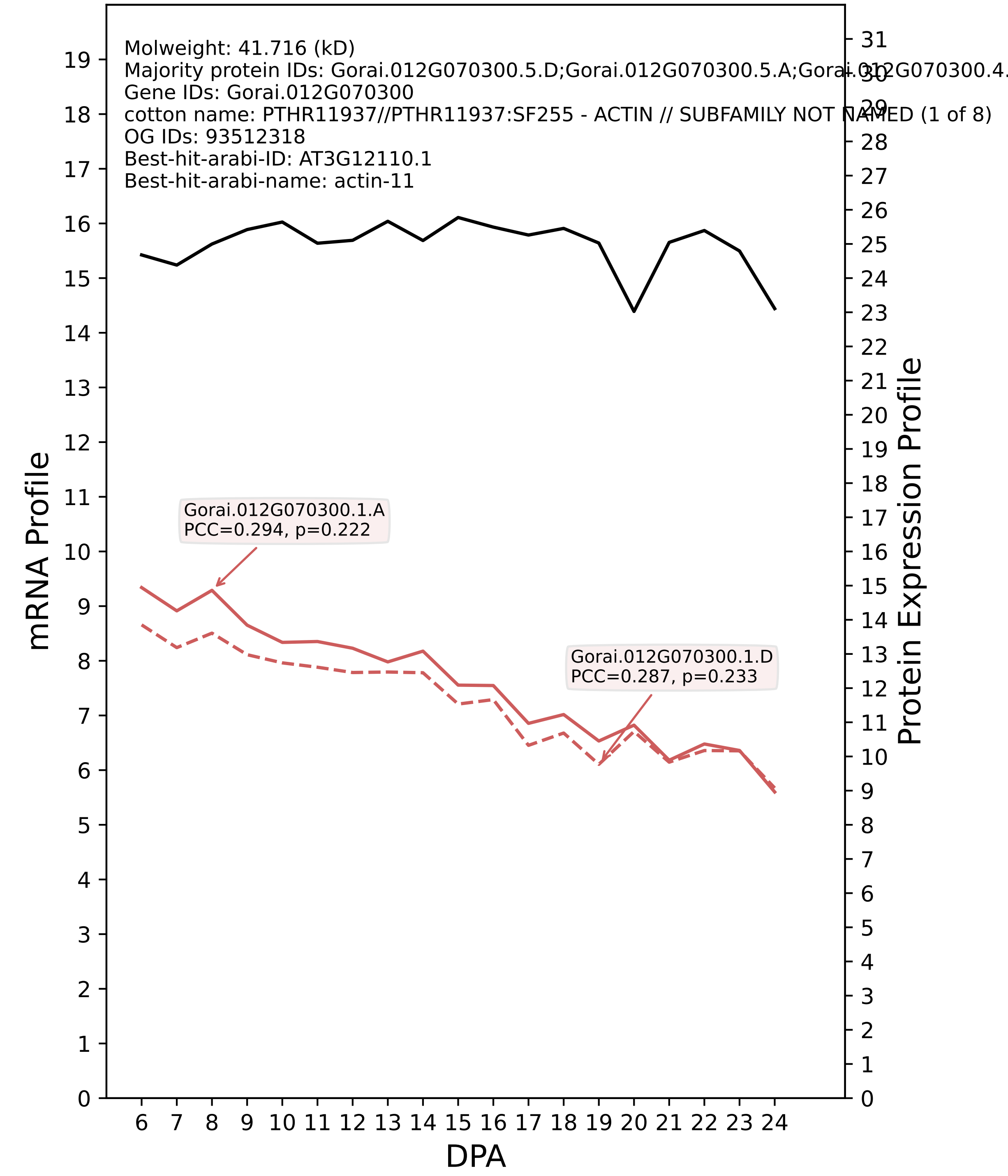

00: Gorai.011G000800.3.D in case 2 and V13

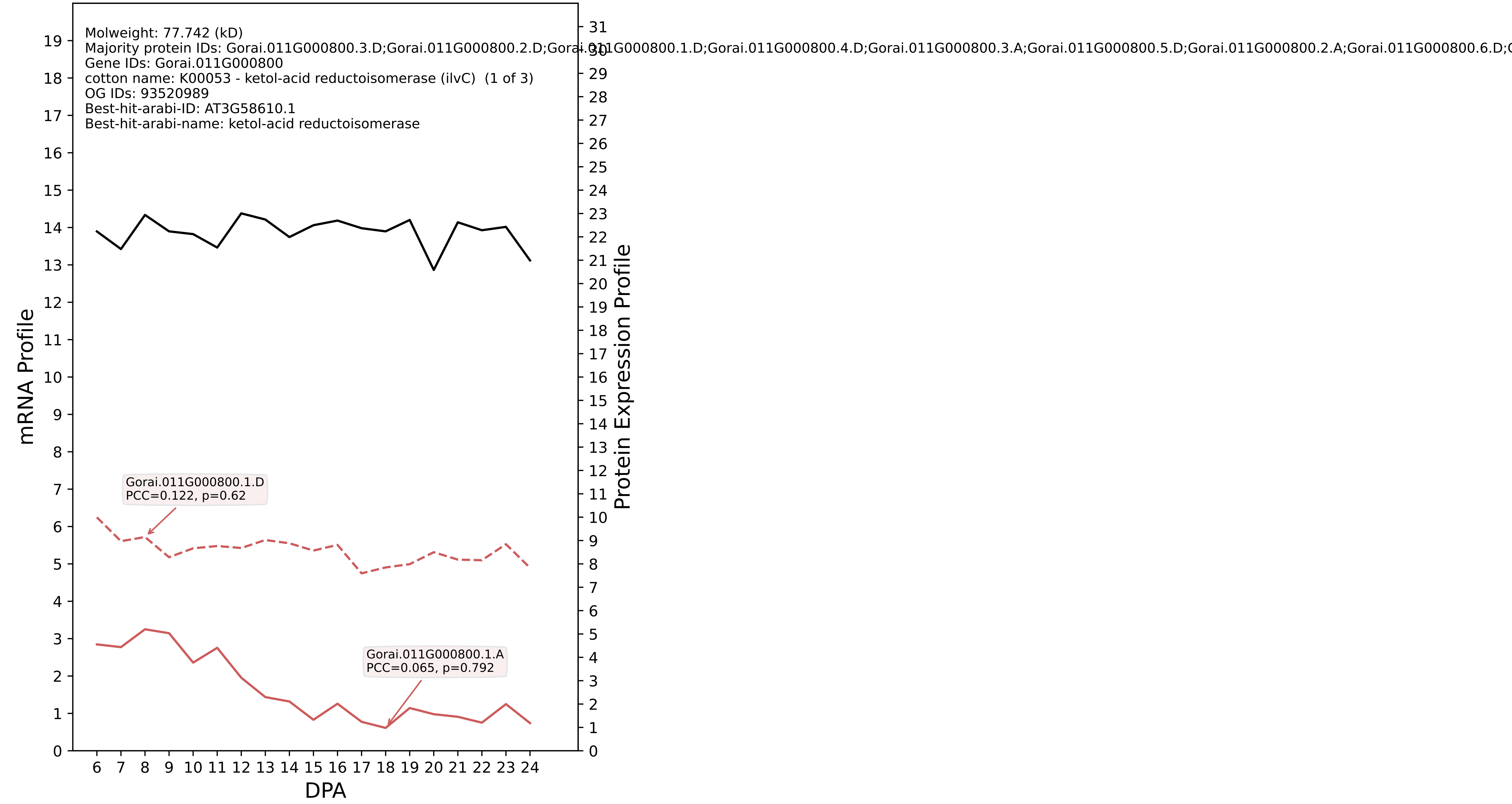

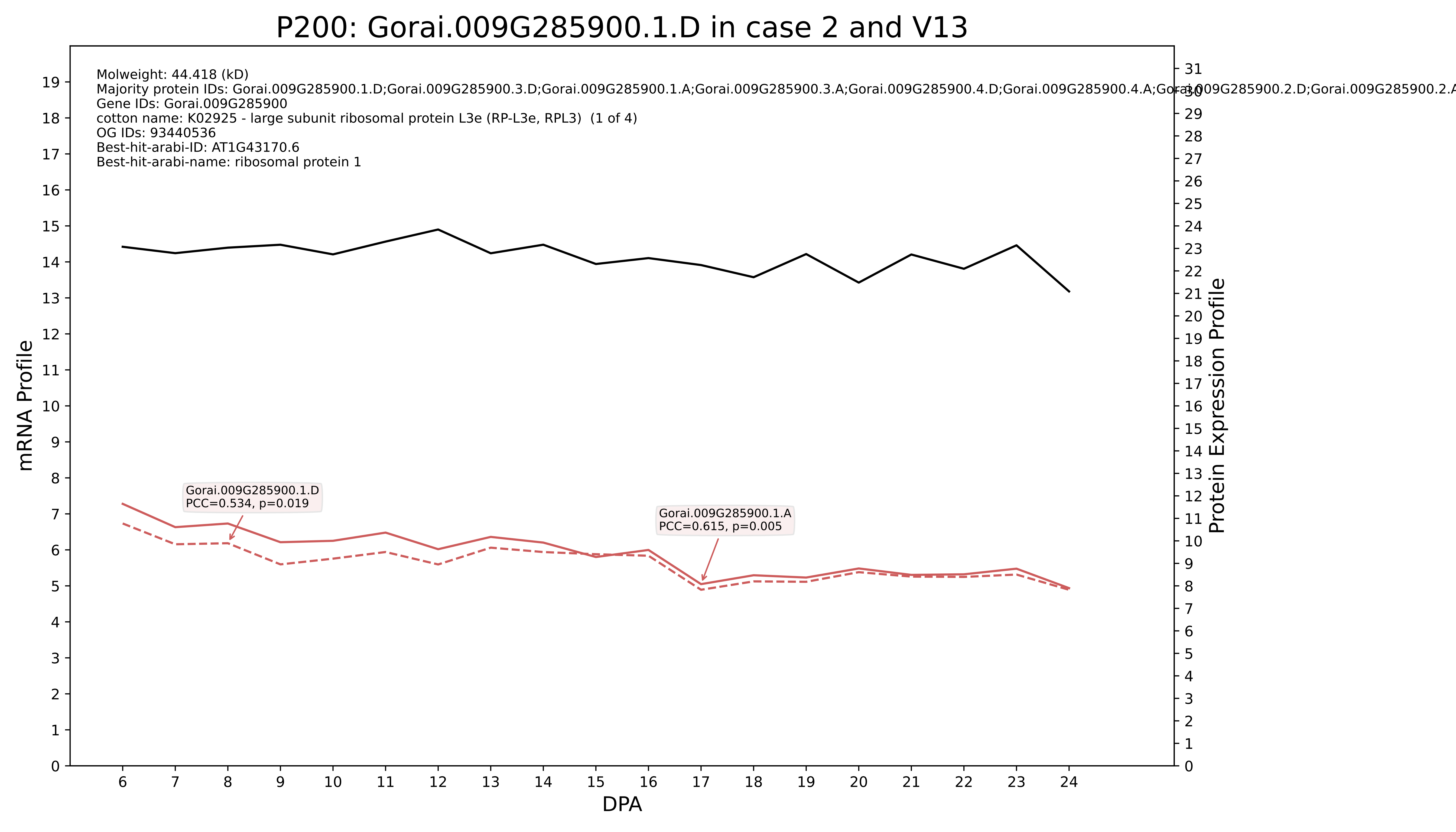
P200: Gorai.009G285900.1.D in case 2 and V13
Molweight: 44.418 (kD)
Majority protein IDs: Gorai.009G285900.1.D;Gorai.009G285900.3.D;Gorai.009G285900.1.A;Gorai.009G285900.3.A;Gorai.009G285900.4.D;Gorai.009G285900.4.A;Gorai.009G285900.2.D;Gorai.009G285900.2.A
Gene IDs: Gorai.009G285900
cotton name: K02925 - large subunit ribosomal protein L3e (RP-L3e, RPL3) (1 of 4)
OG IDs: 93440536
Best-hit-arabi-ID: AT1G43170.6
Best-hit-arabi-name: ribosomal protein 1
Gorai.009G285900.1.D
PCC=0.534, p=0.019
Gorai.009G285900.1.A
PCC=0.615, p=0.005
mRNA Profile
Protein Expression Profile
DPA

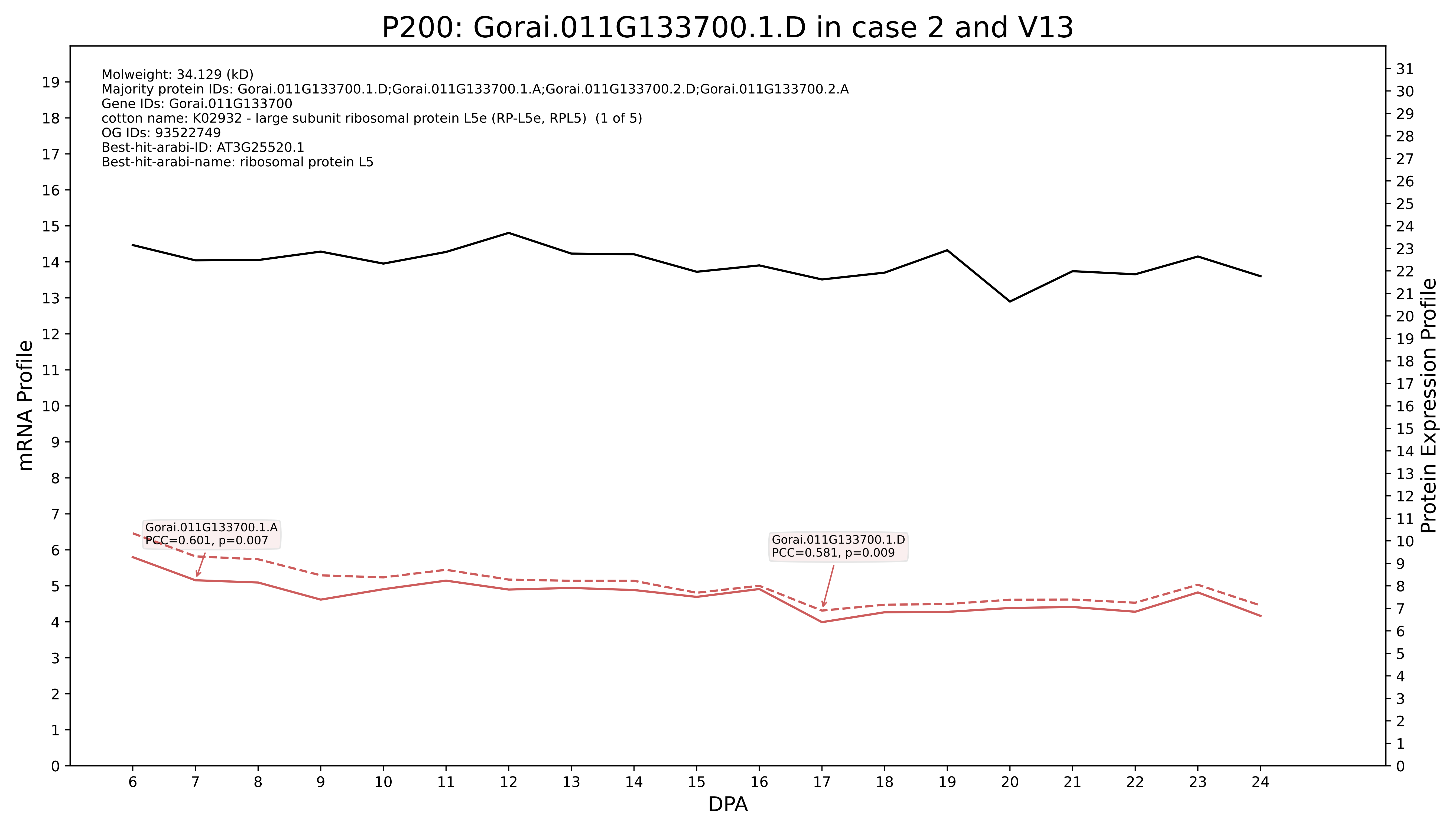
P200: Gorai.011G133700.1.D in case 2 and V13
Molweight: 34.129 (kD)
Majority protein IDs: Gorai.011G133700.1.D;Gorai.011G133700.1.A;Gorai.011G133700.2.D;Gorai.011G133700.2.A
Gene IDs: Gorai.011G133700
cotton name: K02932 - large subunit ribosomal protein L5e (RP-L5e, RPL5) (1 of 5)
OG IDs: 93522749
Best-hit-arabi-ID: AT3G25520.1
Best-hit-arabi-name: ribosomal protein L5
Gorai.011G133700.1.A
PCC=0.601, p=0.007
Gorai.011G133700.1.D
PCC=0.581, p=0.009
mRNA Profile
Protein Expression Profile
DPA

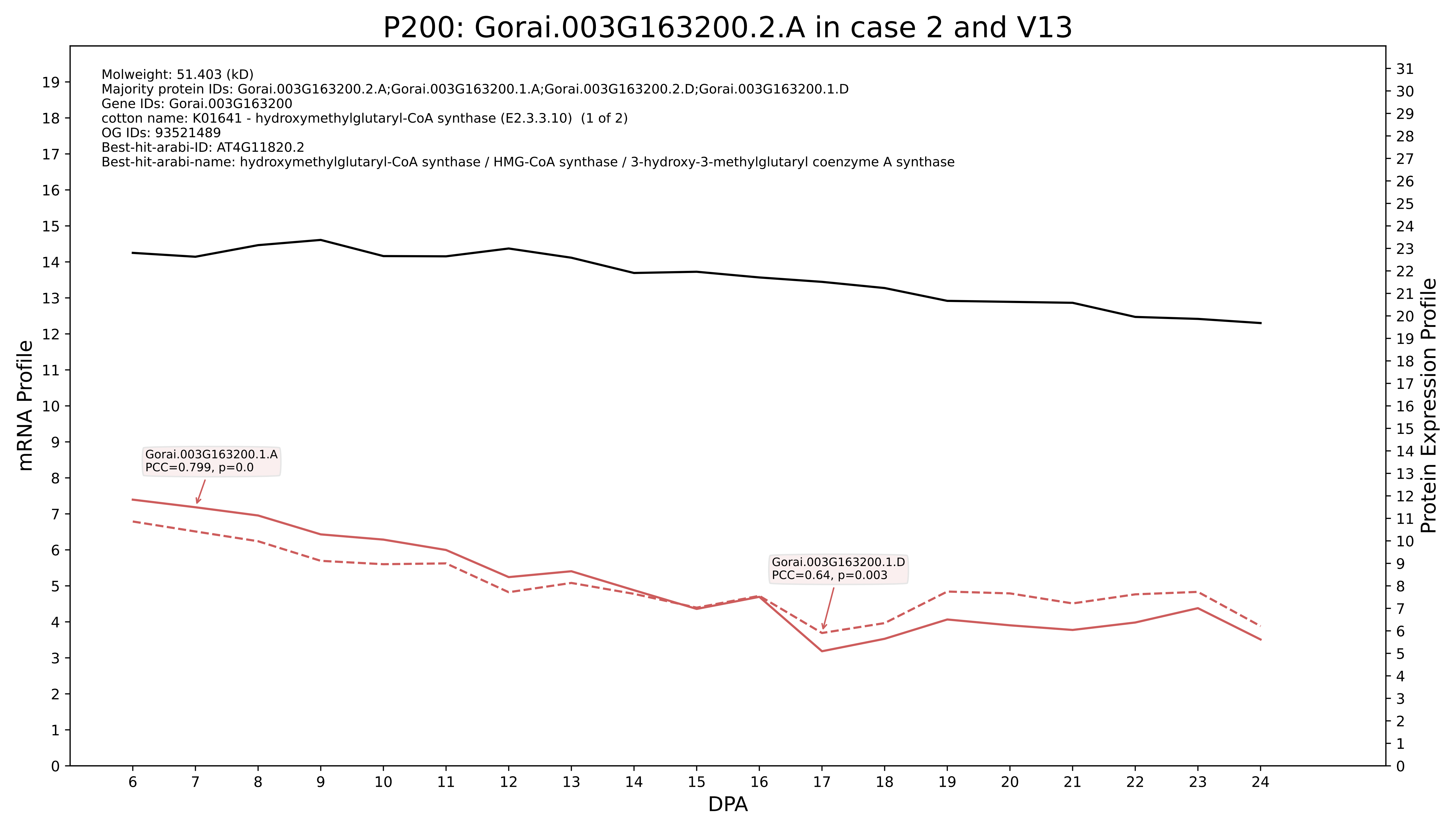

P200: Gorai.003G163200.2.A in case 2 and V13
Molweight: 51.403 (kD)
Majority protein IDs: Gorai.003G163200.2.A;Gorai.003G163200.1.A;Gorai.003G163200.2.D;Gorai.003G163200.1.D
Gene IDs: Gorai.003G163200
cotton name: K01641 - hydroxymethylglutaryl-CoA synthase (E2.3.3.10) (1 of 2)
OG IDs: 93521489
Best-hit-arabi-ID: AT4G11820.2
Best-hit-arabi-name: hydroxymethylglutaryl-CoA synthase / HMG-CoA synthase / 3-hydroxy-3-methylglutaryl coenzyme A synthase
Gorai.003G163200.1.A
PCC=0.799, p=0.0
Gorai.003G163200.1.D
PCC=0.64, p=0.003
mRNA Profile
Protein Expression Profile
DPA

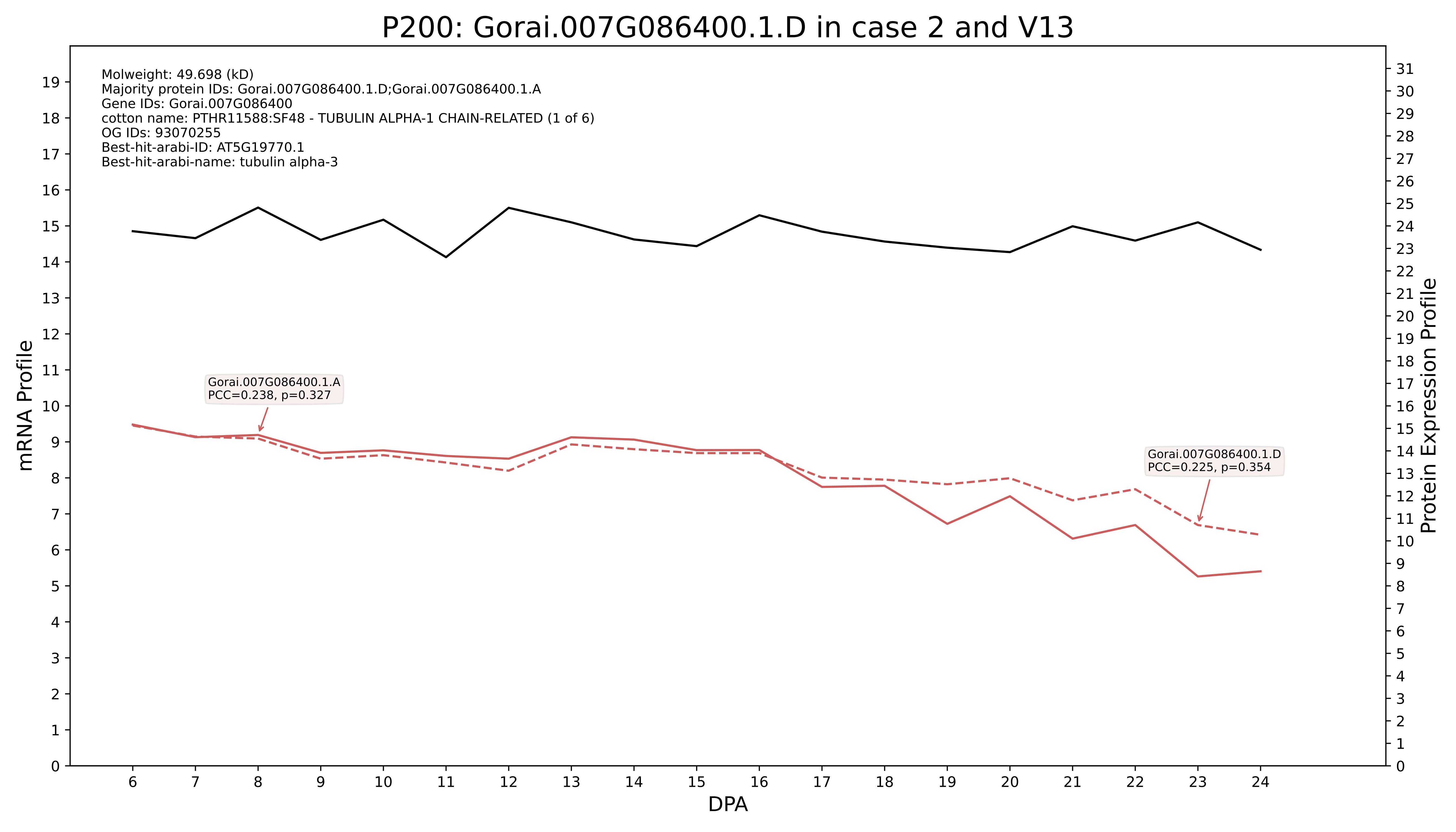

P200: Gorai.007G086400.1.D in case 2 and V13
Molweight: 49.698 (kD)
Majority protein IDs: Gorai.007G086400.1.D;Gorai.007G086400.1.A
Gene IDs: Gorai.007G086400
cotton name: PTHR11588:SF48 - TUBULIN ALPHA-1 CHAIN-RELATED (1 of 6)
OG IDs: 93070255
Best-hit-arabi-ID: AT5G19770.1
Best-hit-arabi-name: tubulin alpha-3
Gorai.007G086400.1.A
PCC=0.238, p=0.327
Gorai.007G086400.1.D
PCC=0.225, p=0.354
mRNA Profile
Protein Expression Profile
DPA

### P200: Gorai.009G420600.1.D in case 2 and V13

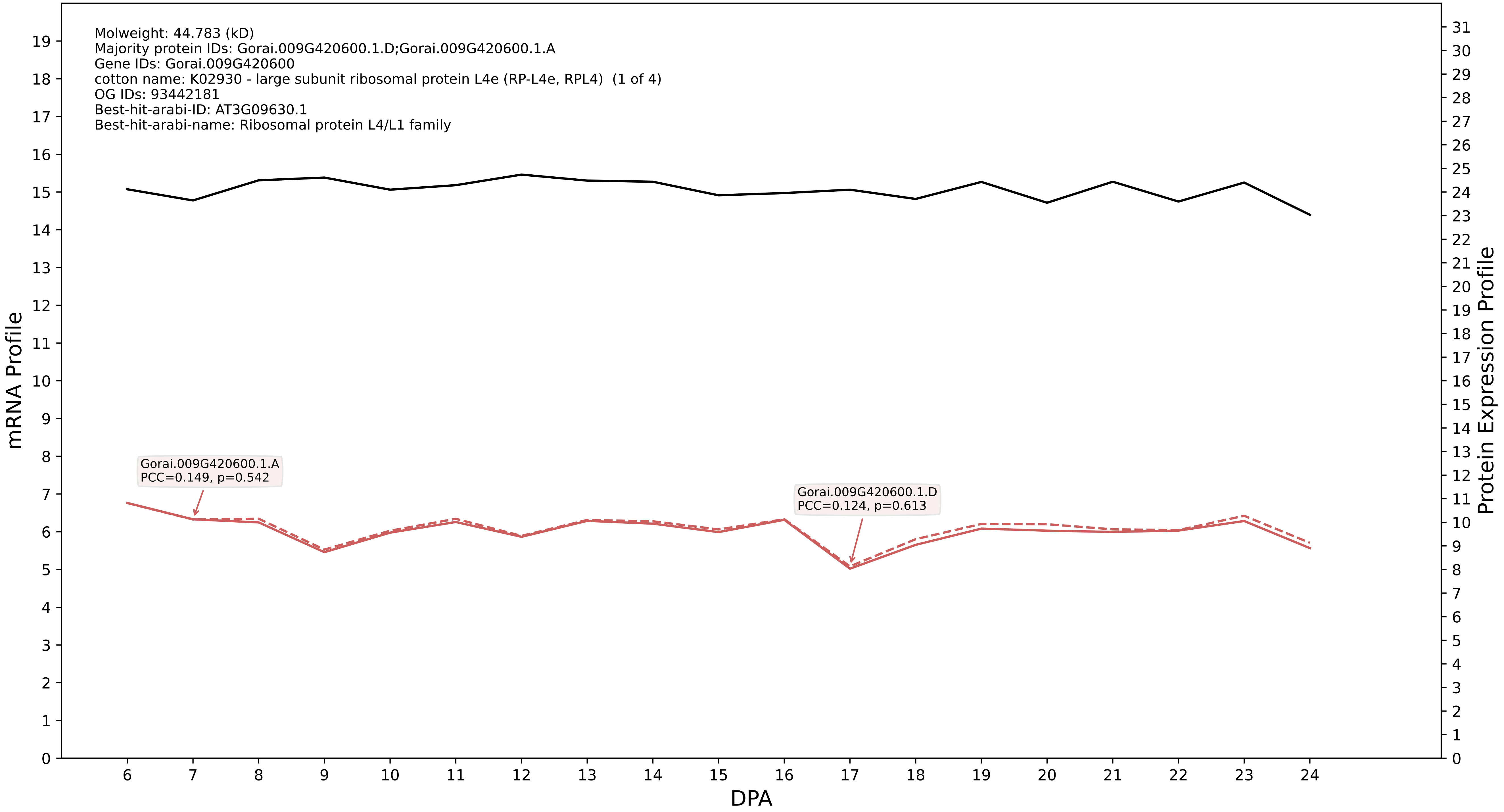

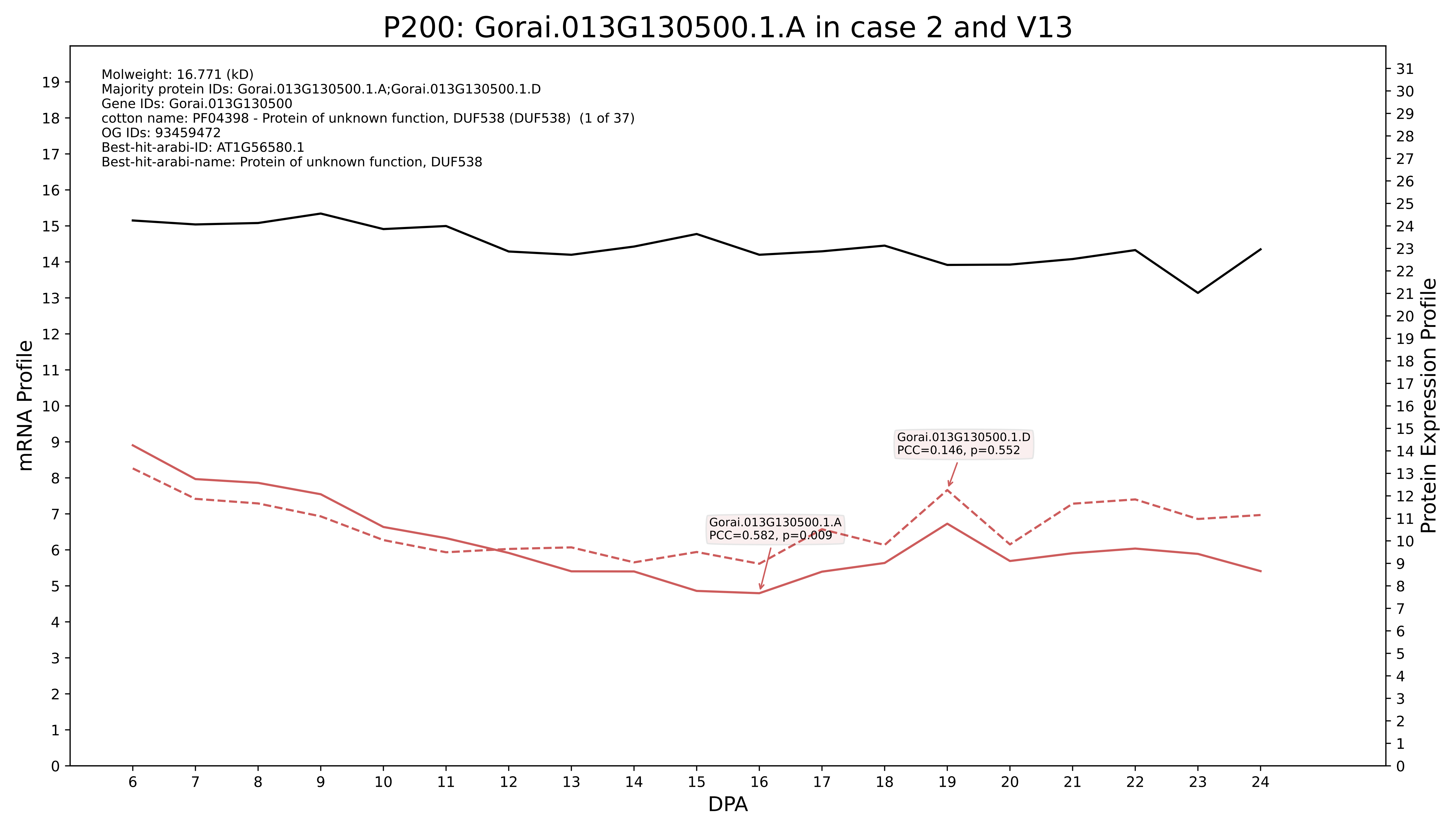
P200: Gorai.013G130500.1.A in case 2 and V13
Molweight: 16.771 (kD)
Majority protein IDs: Gorai.013G130500.1.A;Gorai.013G130500.1.D
Gene IDs: Gorai.013G130500
cotton name: PF04398 - Protein of unknown function, DUF538 (DUF538) (1 of 37)
OG IDs: 93459472
Best-hit-arabi-ID: AT1G56580.1
Best-hit-arabi-name: Protein of unknown function, DUF538
Gorai.013G130500.1.D
PCC=0.146, p=0.552
Gorai.013G130500.1.A
PCC=0.582, p=0.009
mRNA Profile
Protein Expression Profile
DPA

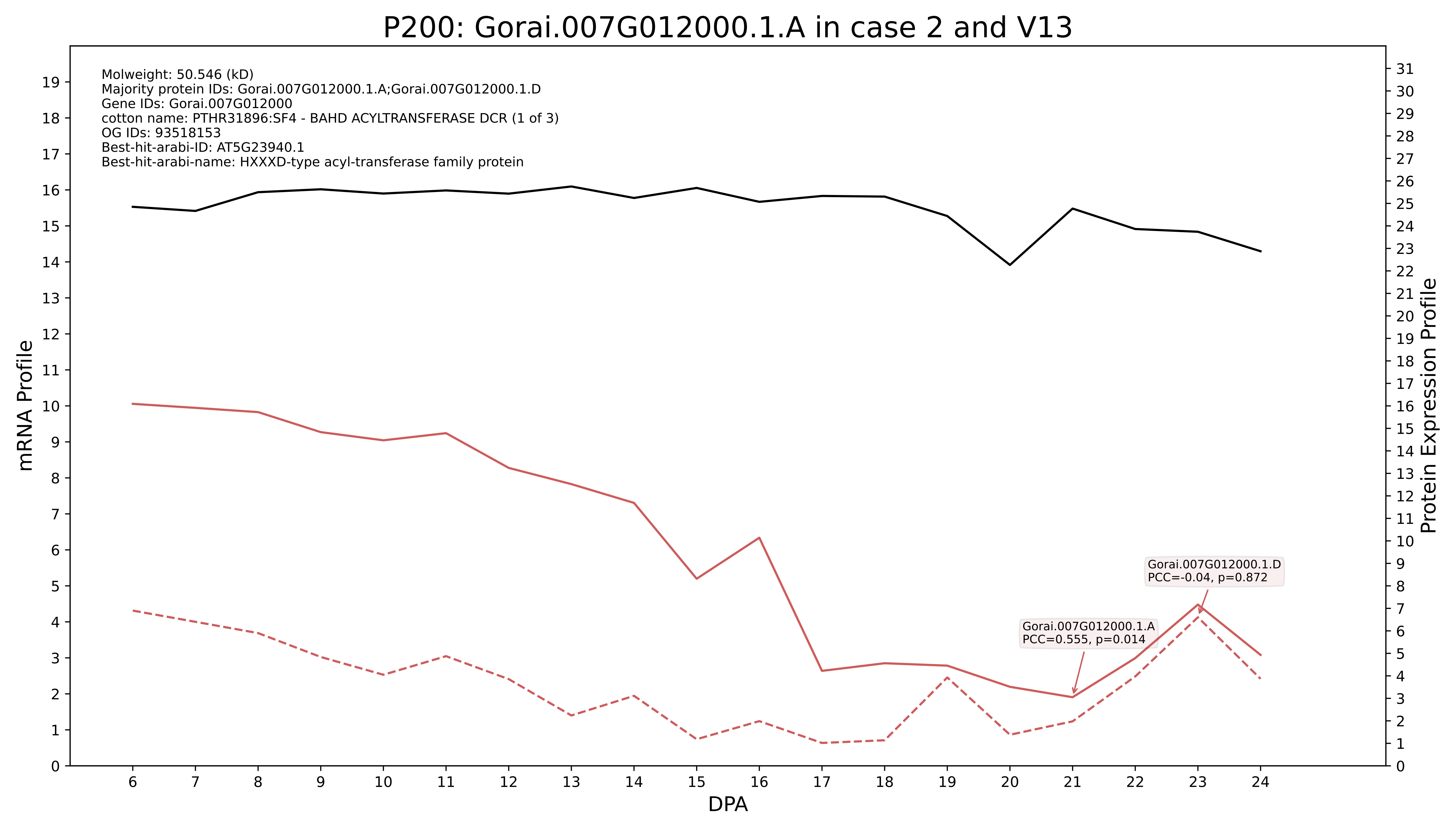

P200: Gorai.007G012000.1.A in case 2 and V13
Molweight: 50.546 (kD)
Majority protein IDs: Gorai.007G012000.1.A;Gorai.007G012000.1.D
Gene IDs: Gorai.007G012000
cotton name: PTHR31896:SF4 - BAHD ACYLTRANSFERASE DCR (1 of 3)
OG IDs: 93518153
Best-hit-arabi-ID: AT5G23940.1
Best-hit-arabi-name: HXXXD-type acyl-transferase family protein
Gorai.007G012000.1.D
PCC=-0.04, p=0.872
Gorai.007G012000.1.A
PCC=0.555, p=0.014
mRNA Profile
Protein Expression Profile
DPA

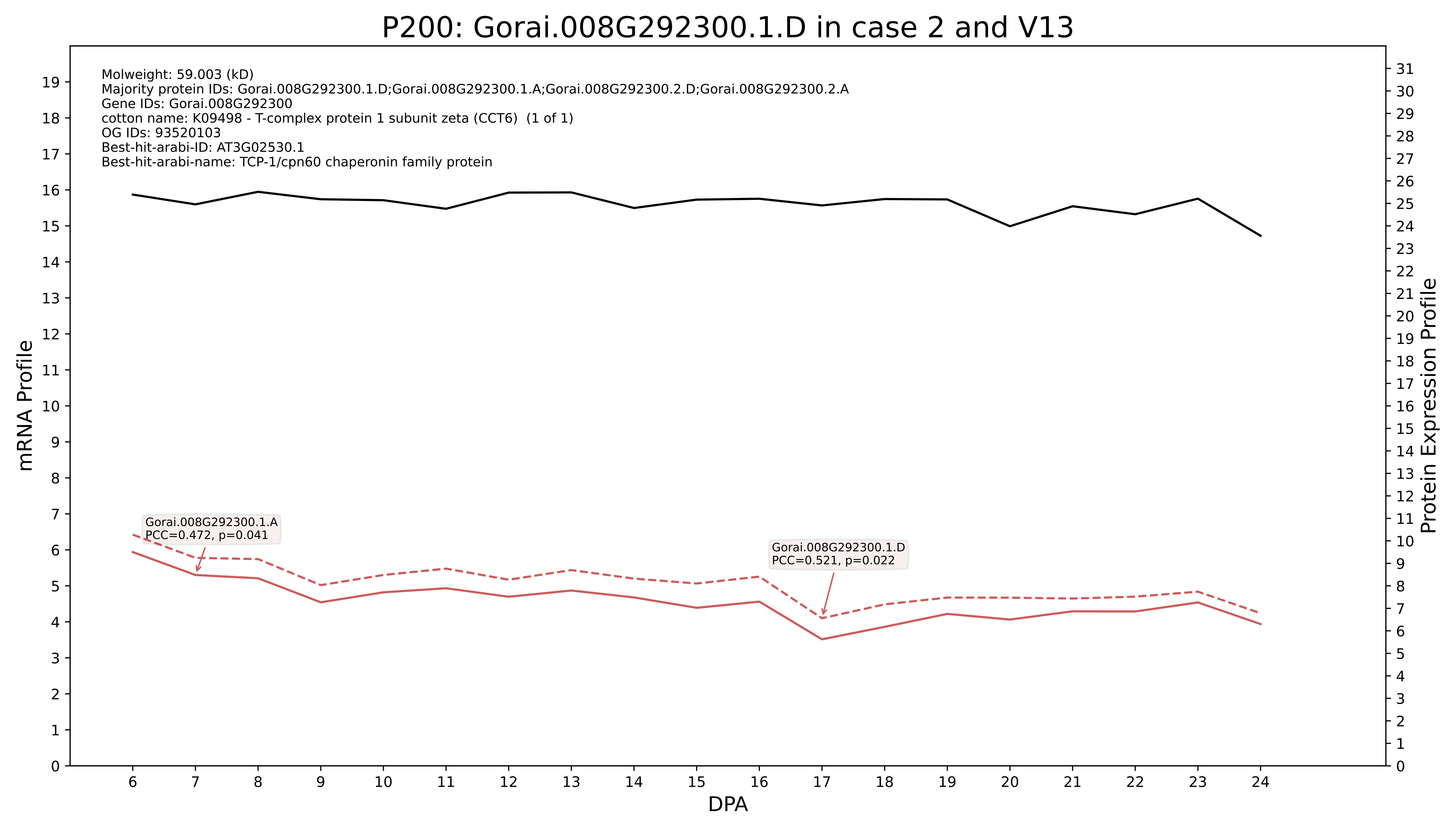

P200: Gorai.008G292300.1.D in case 2 and V13
Molweight: 59.003 (kD)
Majority protein IDs: Gorai.008G292300.1.D;Gorai.008G292300.1.A;Gorai.008G292300.2.D;Gorai.008G292300.2.A
Gene IDs: Gorai.008G292300
cotton name: K09498 - T-complex protein 1 subunit zeta (CCT6) (1 of 1)
OG IDs: 93520103
Best-hit-arabi-ID: AT3G02530.1
Best-hit-arabi-name: TCP-1/cpn60 chaperonin family protein
Gorai.008G292300.1.A
PCC=0.472, p=0.041
Gorai.008G292300.1.D
PCC=0.521, p=0.022
mRNA Profile
Protein Expression Profile
DPA

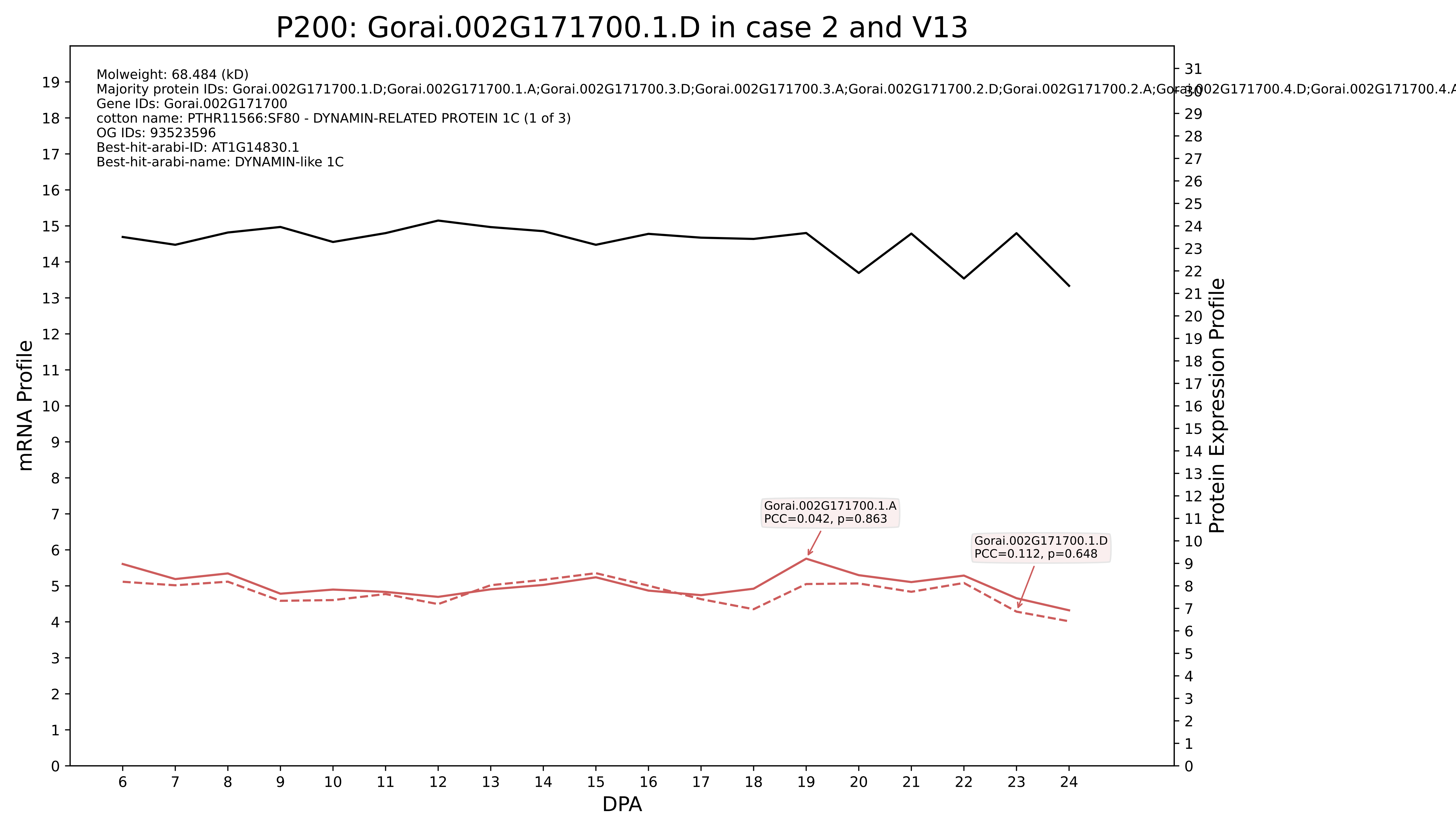
P200: Gorai.002G171700.1.D in case 2 and V13
Molweight: 68.484 (kD)
Majority protein IDs: Gorai.002G171700.1.D;Gorai.002G171700.1.A;Gorai.002G171700.3.D;Gorai.002G171700.3.A;Gorai.002G171700.2.D;Gorai.002G171700.2.A;Gorai.002G171700.4.D;Gorai.002G171700.4.A
Gene IDs: Gorai.002G171700
cotton name: PTHR11566:SF80 - DYNAMIN-RELATED PROTEIN 1C (1 of 3)
OG IDs: 93523596
Best-hit-arabi-ID: AT1G14830.1
Best-hit-arabi-name: DYNAMIN-like 1C
Gorai.002G171700.1.A
PCC=0.042, p=0.863
Gorai.002G171700.1.D
PCC=0.112, p=0.648
mRNA Profile
Protein Expression Profile
DPA

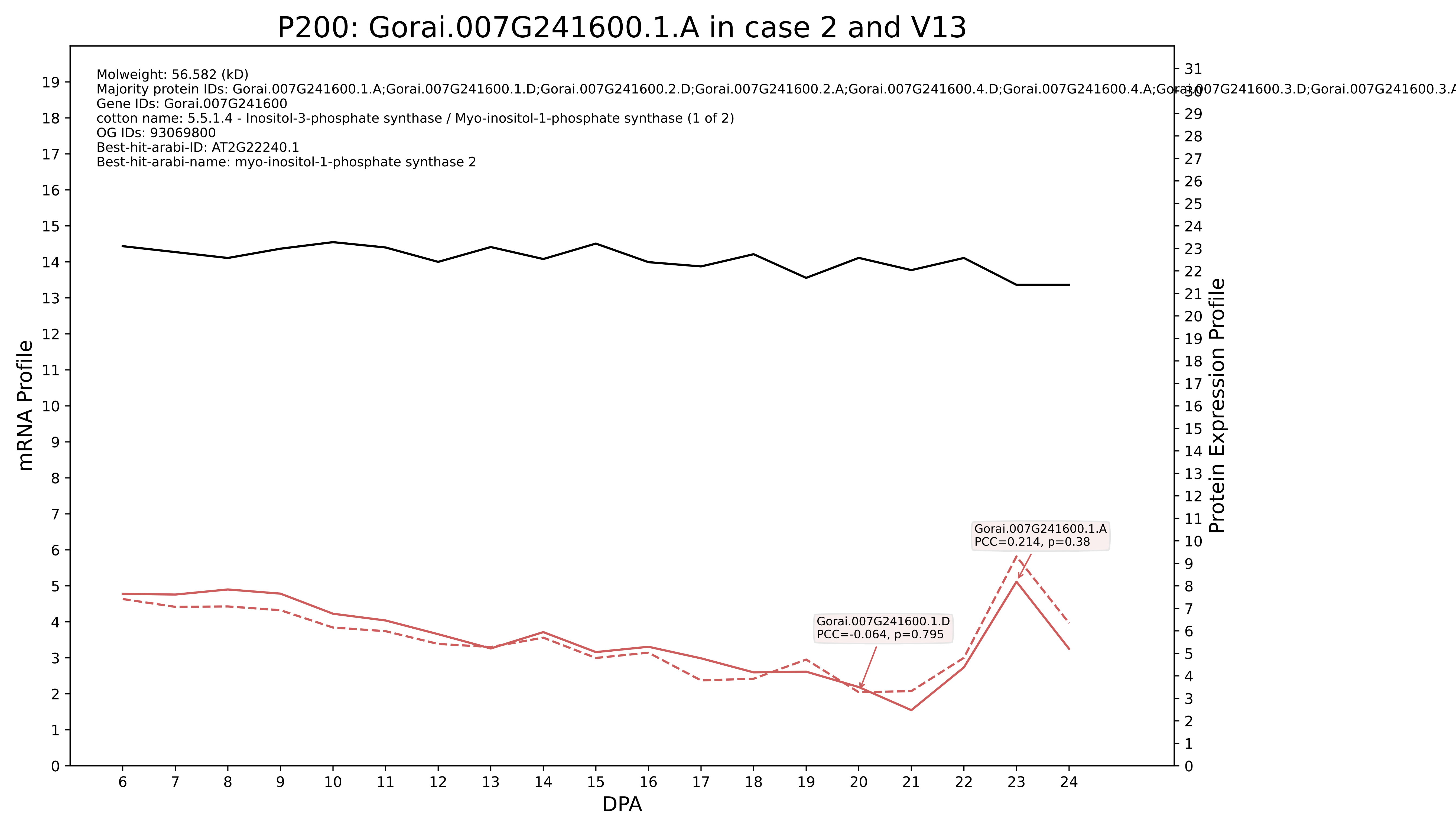
P200: Gorai.007G241600.1.A in case 2 and V13
Molweight: 56.582 (kD)
Majority protein IDs: Gorai.007G241600.1.A;Gorai.007G241600.1.D;Gorai.007G241600.2.D;Gorai.007G241600.2.A;Gorai.007G241600.4.D;Gorai.007G241600.4.A;Gorai.007G241600.3.D;Gorai.007G241600.3.A
Gene IDs: Gorai.007G241600
cotton name: 5.5.1.4 - Inositol-3-phosphate synthase / Myo-inositol-1-phosphate synthase (1 of 2)
OG IDs: 93069800
Best-hit-arabi-ID: AT2G22240.1
Best-hit-arabi-name: myo-inositol-1-phosphate synthase 2
Gorai.007G241600.1.A
PCC=0.214, p=0.38
Gorai.007G241600.1.D
PCC=-0.064, p=0.795
mRNA Profile
Protein Expression Profile
DPA

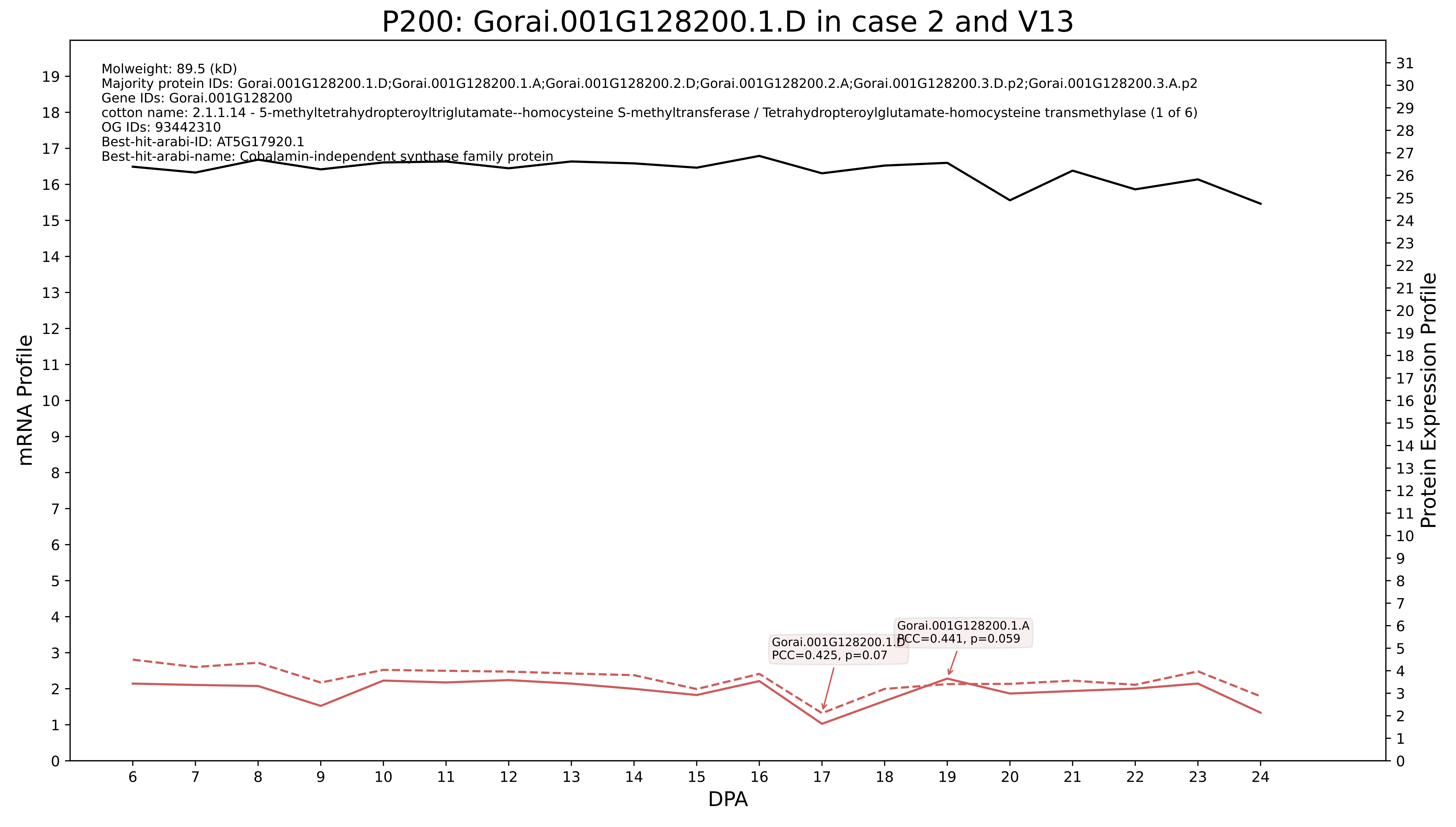
P200: Gorai.001G128200.1.D in case 2 and V13
Molweight: 89.5 (kD)
Majority protein IDs: Gorai.001G128200.1.D;Gorai.001G128200.1.A;Gorai.001G128200.2.D;Gorai.001G128200.2.A;Gorai.001G128200.3.D.p2;Gorai.001G128200.3.A.p2
Gene IDs: Gorai.001G128200
cotton name: 2.1.1.14 - 5-methyltetrahydropteroyltriglutamate--homocysteine S-methyltransferase / Tetrahydropteroylglutamate-homocysteine transmethylase (1 of 6)
OG IDs: 93442310
Best-hit-arabi-ID: AT5G17920.1
Best-hit-arabi-name: Cobalamin-independent synthase family protein
Gorai.001G128200.1.D
PCC=0.425, p=0.07
Gorai.001G128200.1.A
PCC=0.441, p=0.059
mRNA Profile
Protein Expression Profile
DPA

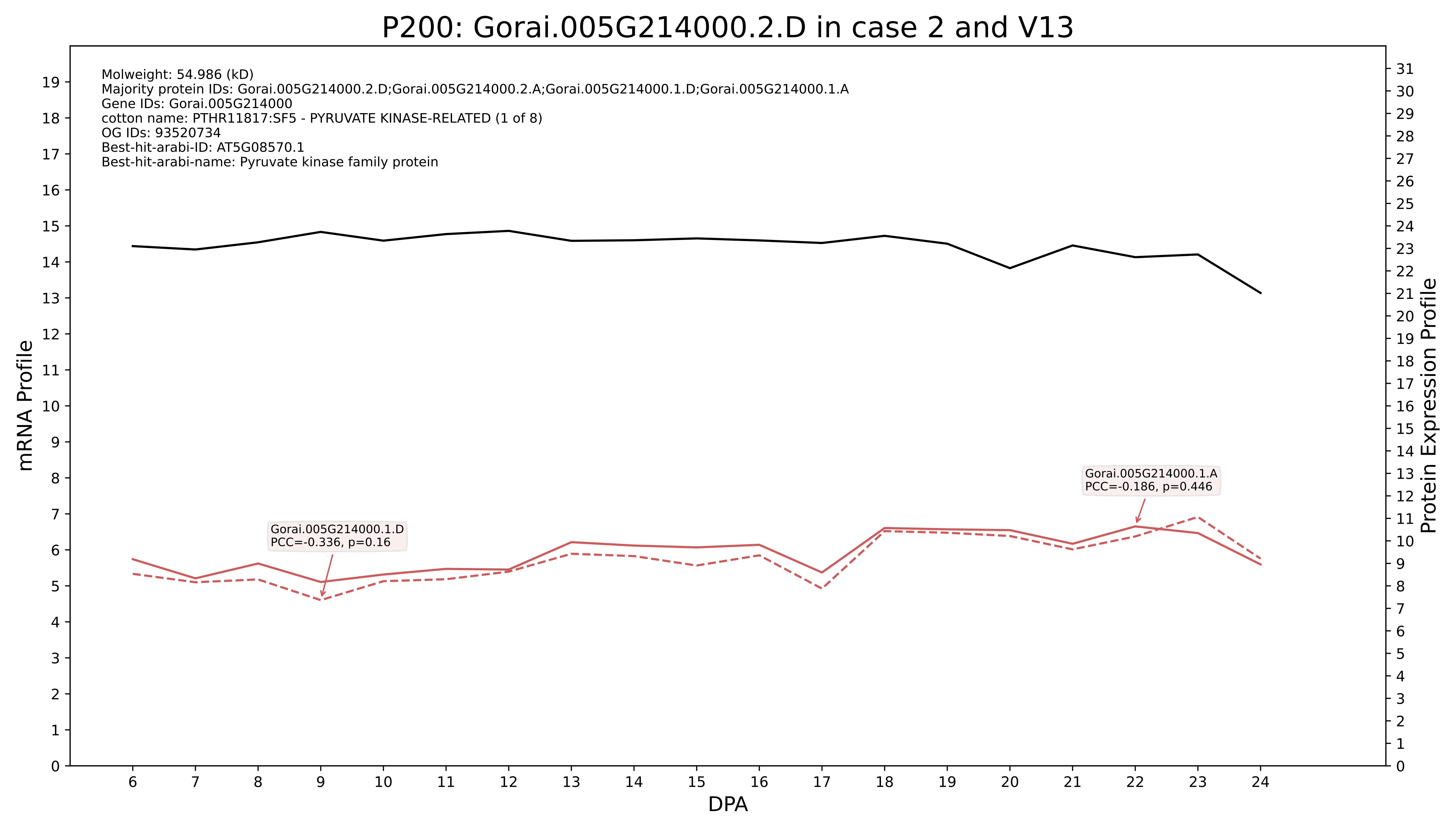
P200: Gorai.005G214000.2.D in case 2 and V13
Molweight: 54.986 (kD)
Majority protein IDs: Gorai.005G214000.2.D;Gorai.005G214000.2.A;Gorai.005G214000.1.D;Gorai.005G214000.1.A
Gene IDs: Gorai.005G214000
cotton name: PTHR11817:SF5 - PYRUVATE KINASE-RELATED (1 of 8)
OG IDs: 93520734
Best-hit-arabi-ID: AT5G08570.1
Best-hit-arabi-name: Pyruvate kinase family protein
Gorai.005G214000.1.D
PCC=-0.336, p=0.16
Gorai.005G214000.1.A
PCC=-0.186, p=0.446
mRNA Profile
Protein Expression Profile
DPA

012G055000.1.D in case 3 and V13

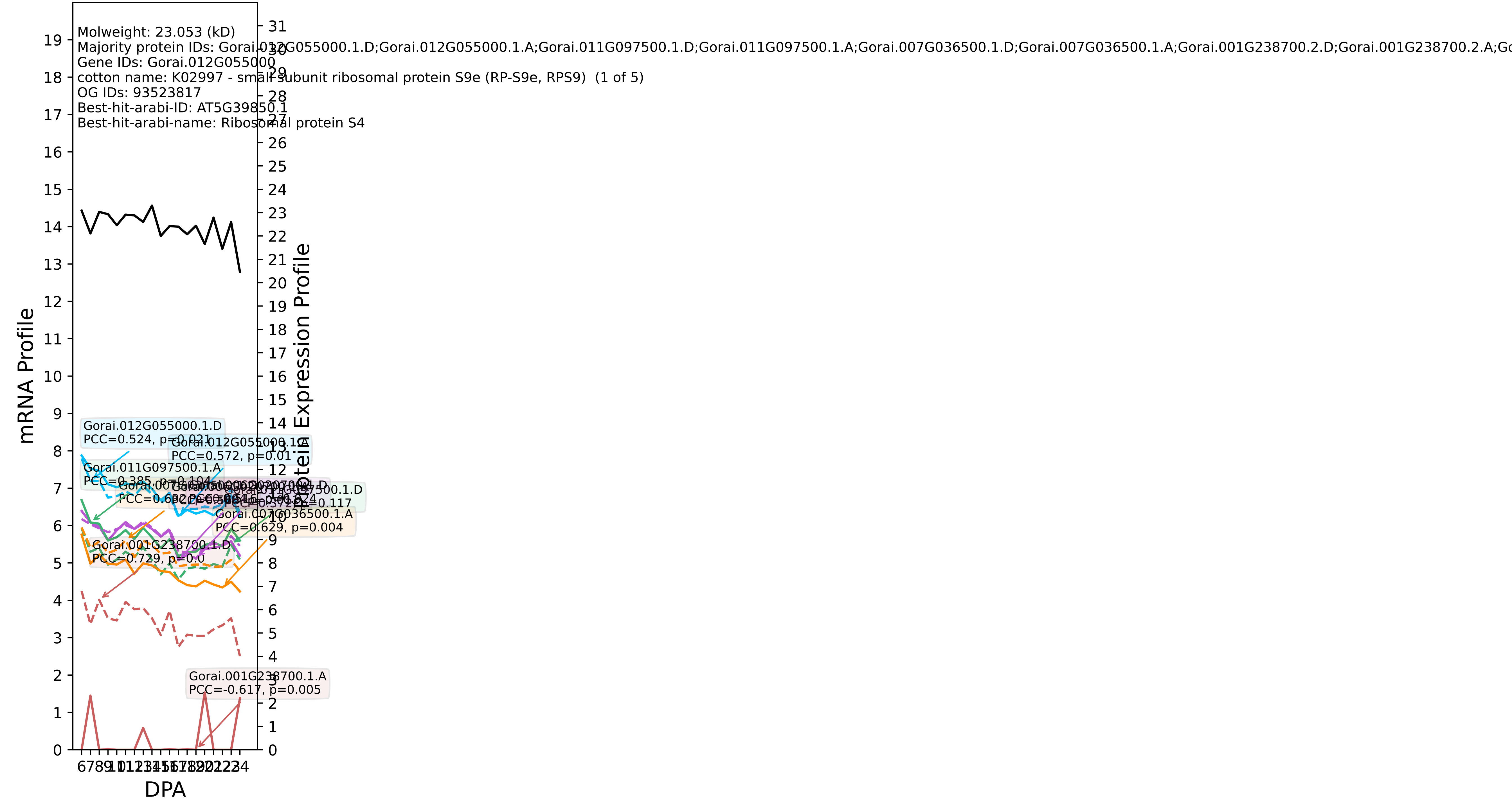

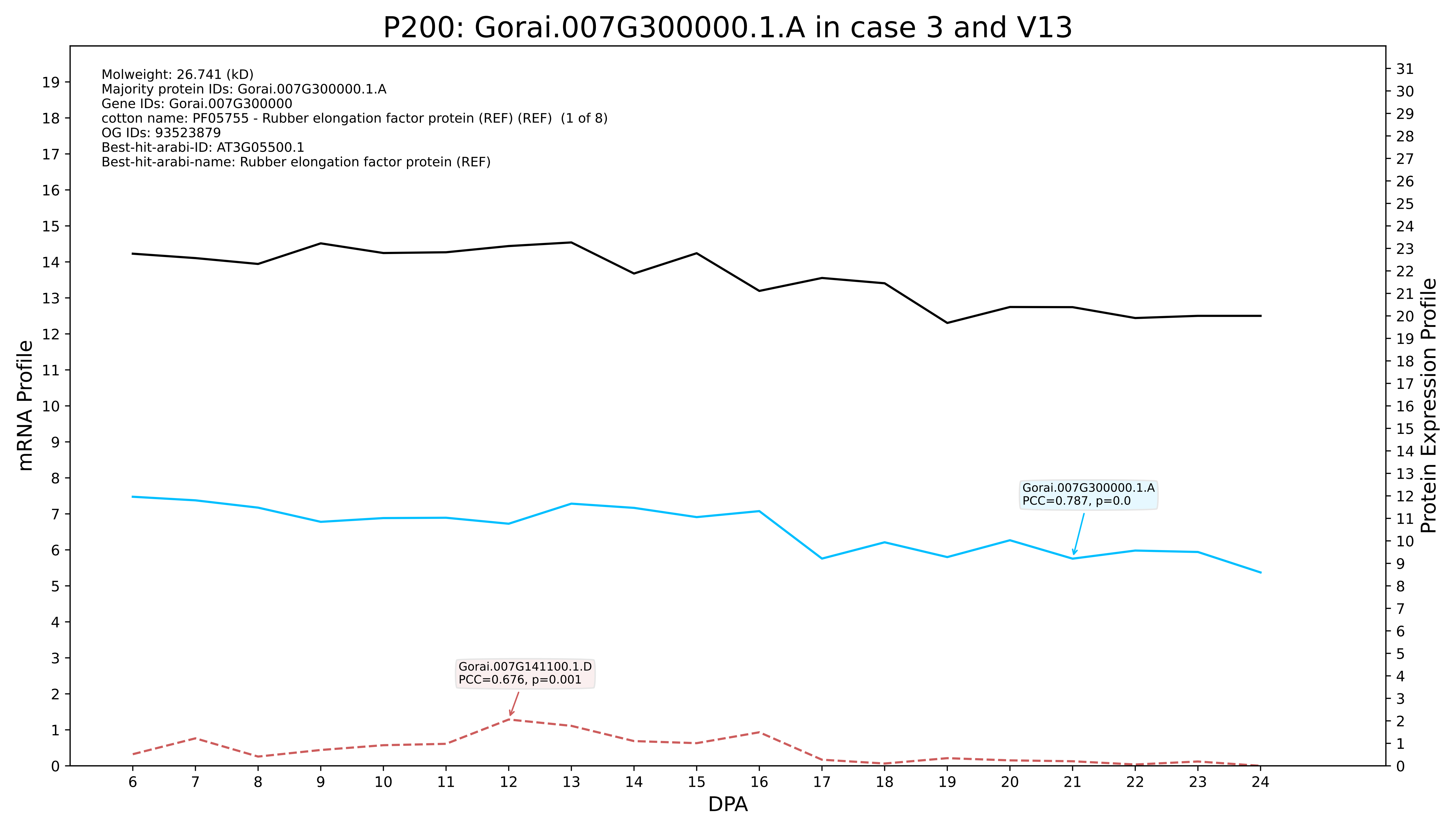
P200: Gorai.007G300000.1.A in case 3 and V13
Molweight: 26.741 (kD)
Majority protein IDs: Gorai.007G300000.1.A
Gene IDs: Gorai.007G300000
cotton name: PF05755 - Rubber elongation factor protein (REF) (REF) (1 of 8)
OG IDs: 93523879
Best-hit-arabi-ID: AT3G05500.1
Best-hit-arabi-name: Rubber elongation factor protein (REF)
Gorai.007G300000.1.A
PCC=0.787, p=0.0
Gorai.007G141100.1.D
PCC=0.676, p=0.001
mRNA Profile
Protein Expression Profile
DPA

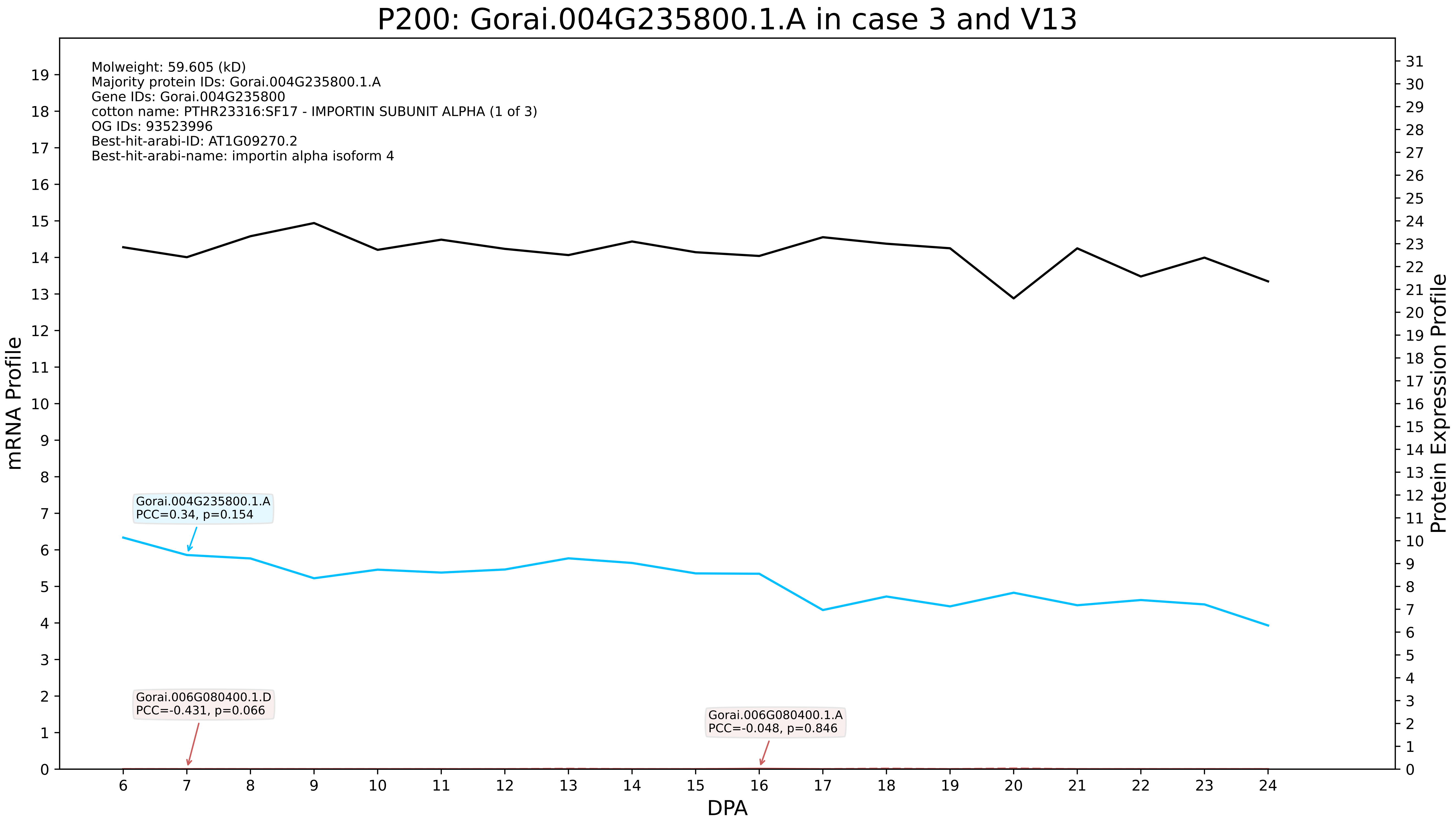
P200: Gorai.004G235800.1.A in case 3 and V13
Molweight: 59.605 (kD)
Majority protein IDs: Gorai.004G235800.1.A
Gene IDs: Gorai.004G235800
cotton name: PTHR23316:SF17 - IMPORTIN SUBUNIT ALPHA (1 of 3)
OG IDs: 93523996
Best-hit-arabi-ID: AT1G09270.2
Best-hit-arabi-name: importin alpha isoform 4
Gorai.004G235800.1.A
PCC=0.34, p=0.154
Gorai.006G080400.1.D
PCC=-0.431, p=0.066
Gorai.006G080400.1.A
PCC=-0.048, p=0.846
mRNA Profile
Protein Expression Profile
DPA

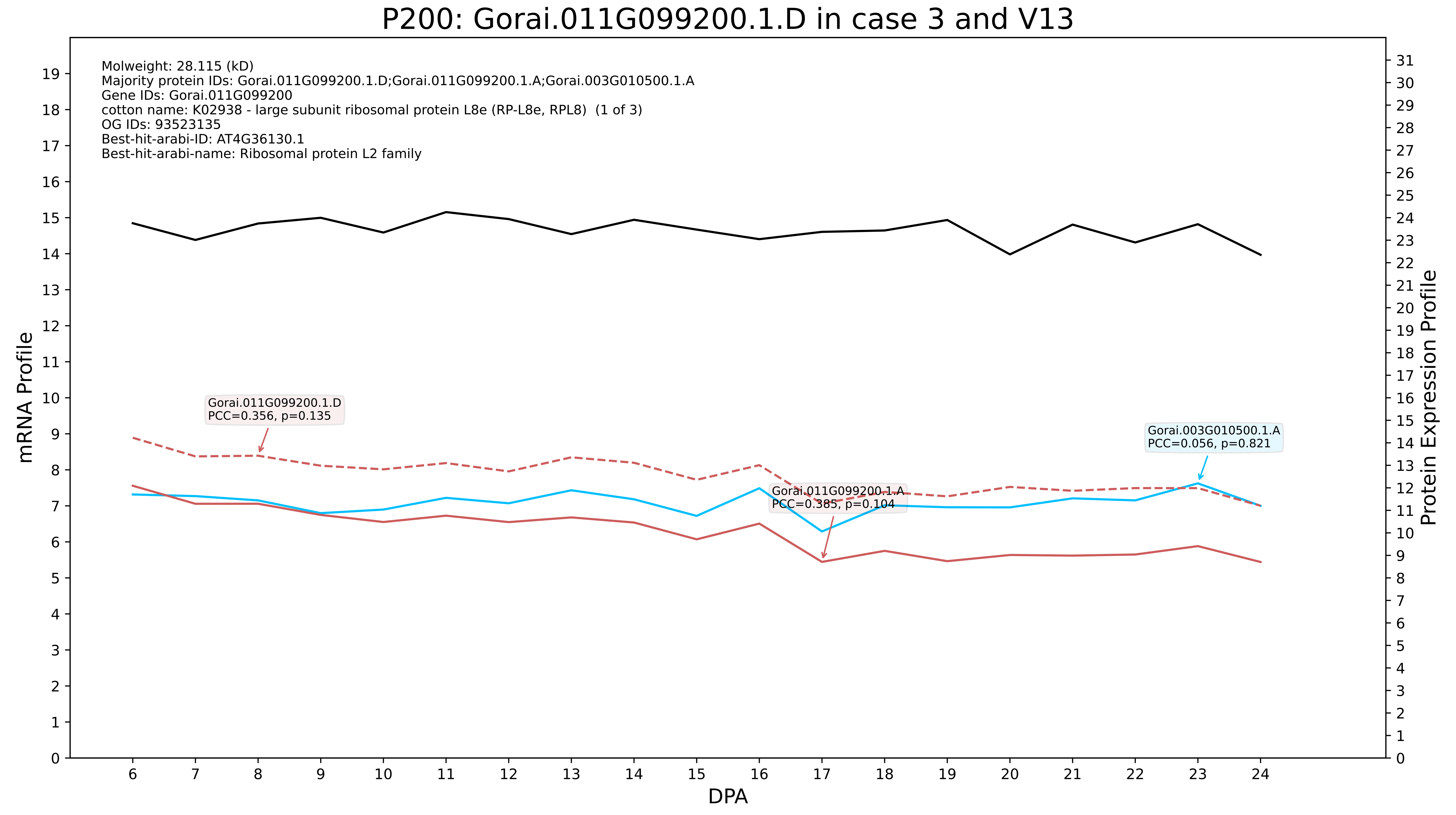

P200: Gorai.011G099200.1.D in case 3 and V13
Molweight: 28.115 (kD)
Majority protein IDs: Gorai.011G099200.1.D;Gorai.011G099200.1.A;Gorai.003G010500.1.A
Gene IDs: Gorai.011G099200
cotton name: K02938 - large subunit ribosomal protein L8e (RP-L8e, RPL8) (1 of 3)
OG IDs: 93523135
Best-hit-arabi-ID: AT4G36130.1
Best-hit-arabi-name: Ribosomal protein L2 family
Gorai.011G099200.1.D
PCC=0.356, p=0.135
Gorai.011G099200.1.A
PCC=0.385, p=0.104
Gorai.003G010500.1.A
PCC=0.056, p=0.821
mRNA Profile
Protein Expression Profile
DPA

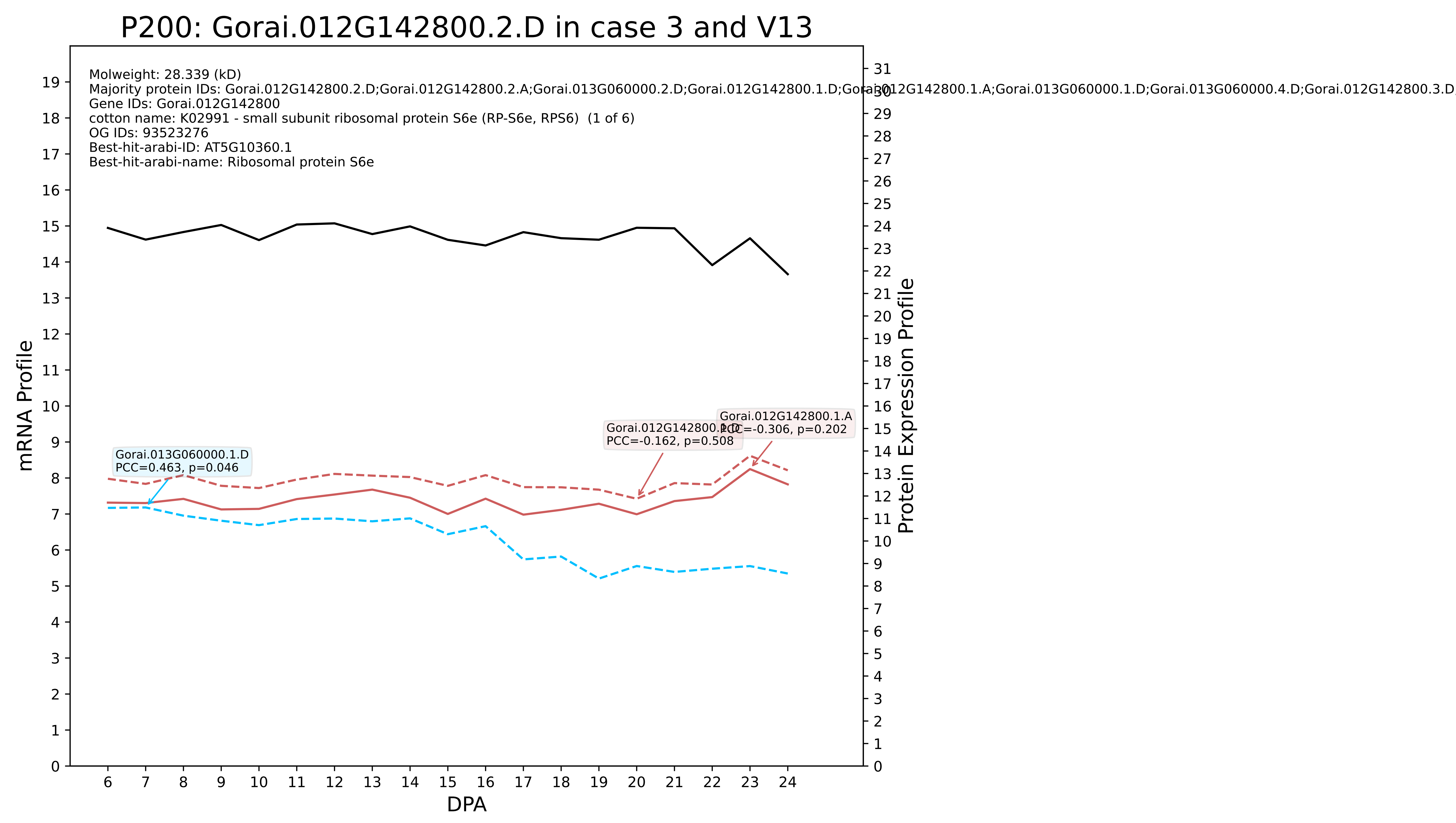
P200: Gorai.012G142800.2.D in case 3 and V13
Molweight: 28.339 (kD)
Majority protein IDs: Gorai.012G142800.2.D;Gorai.012G142800.2.A;Gorai.013G060000.2.D;Gorai.012G142800.1.D;Gorai.012G142800.1.A;Gorai.013G060000.1.D;Gorai.013G060000.4.D;Gorai.012G142800.3.D
Gene IDs: Gorai.012G142800
cotton name: K02991 - small subunit ribosomal protein S6e (RP-S6e, RPS6) (1 of 6)
OG IDs: 93523276
Best-hit-arabi-ID: AT5G10360.1
Best-hit-arabi-name: Ribosomal protein S6e
Gorai.013G060000.1.D
PCC=0.463, p=0.046
Gorai.012G142800.1.D
PCC=-0.162, p=0.508
Gorai.012G142800.1.A
PCC=-0.306, p=0.202
mRNA Profile
Protein Expression Profile
DPA

### P200: Gorai.005G259200.1.D in case 3 and V13

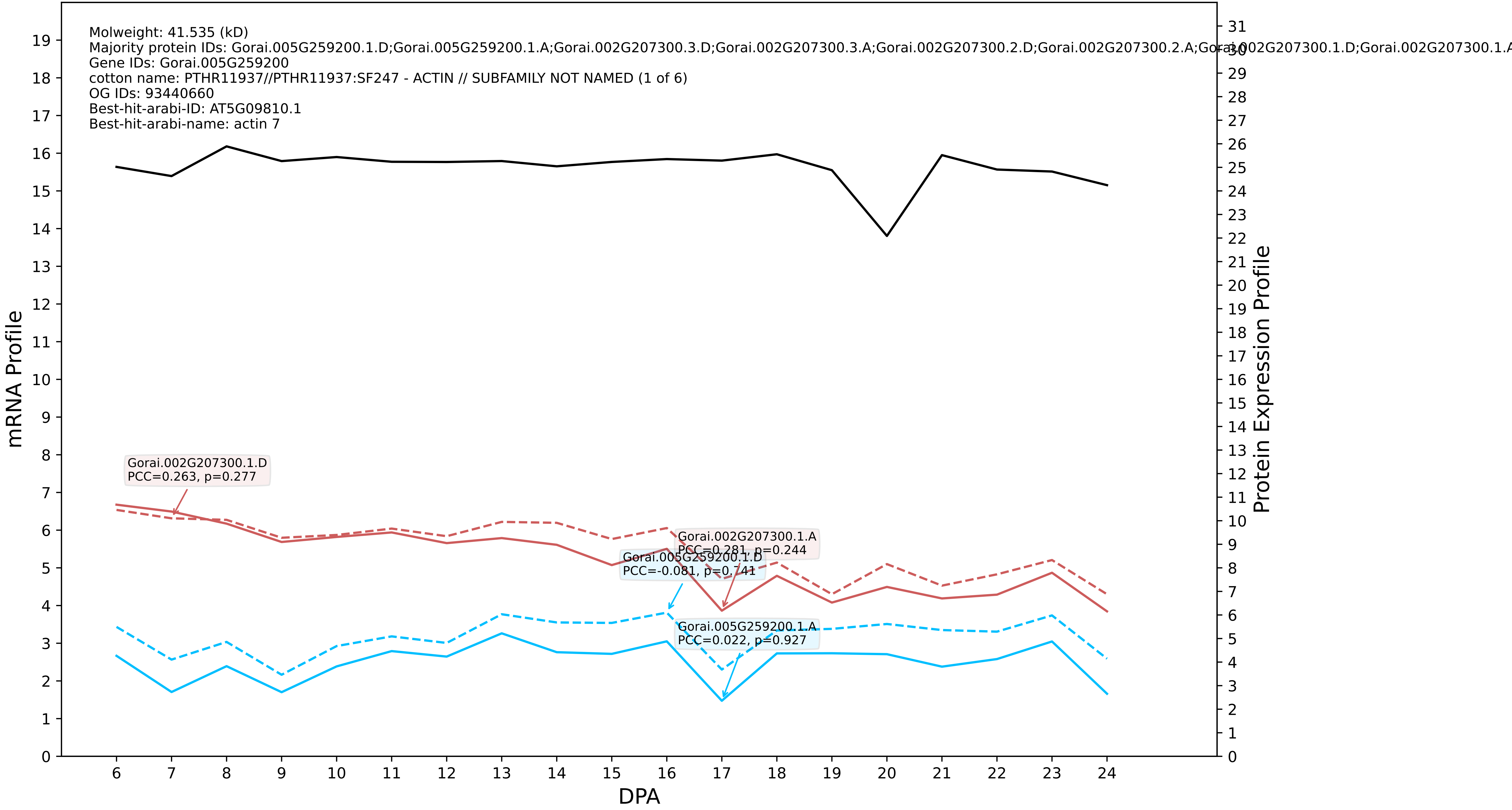

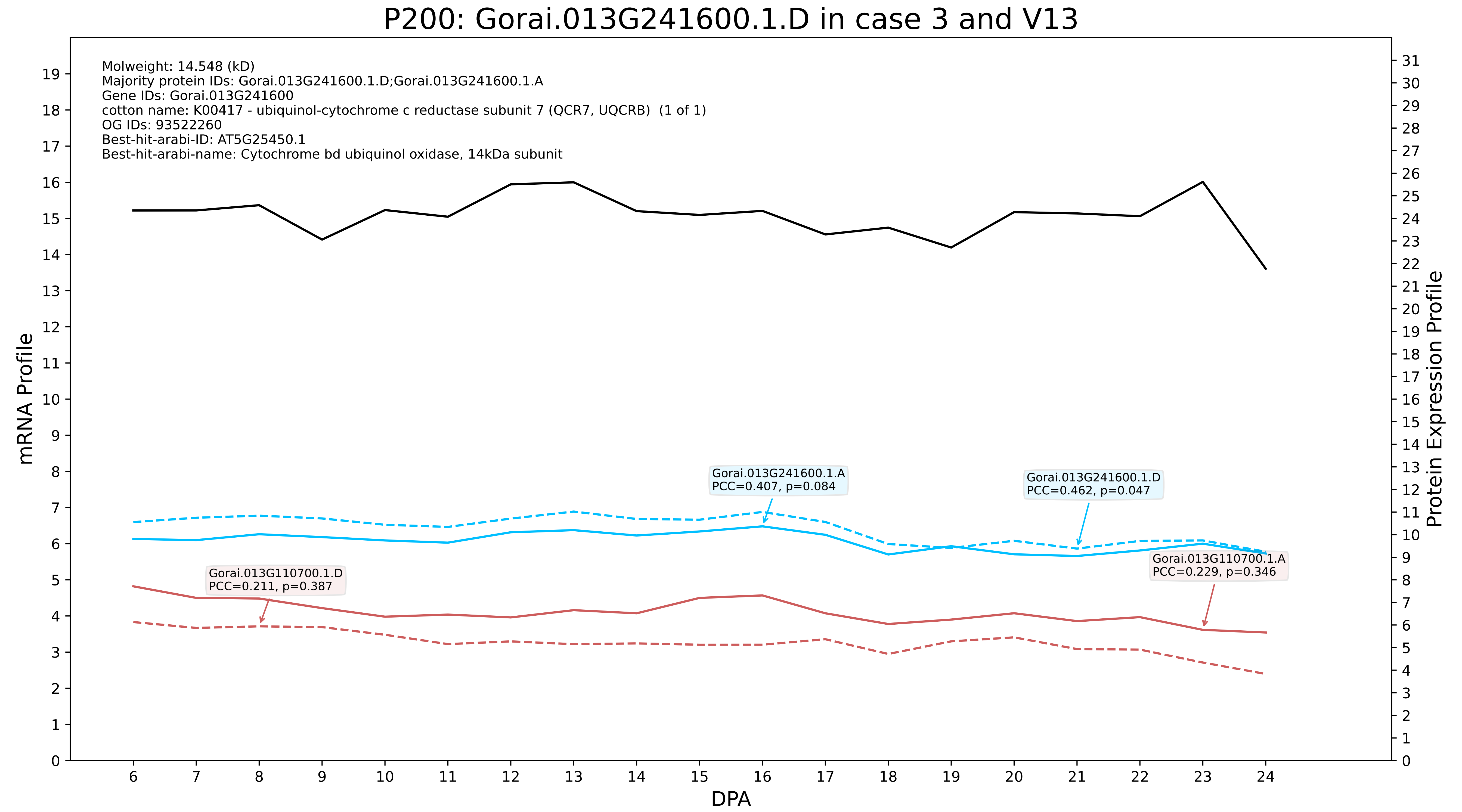
P200: Gorai.013G241600.1.D in case 3 and V13
Molweight: 14.548 (kD)
Majority protein IDs: Gorai.013G241600.1.D;Gorai.013G241600.1.A
Gene IDs: Gorai.013G241600
cotton name: K00417 - ubiquinol-cytochrome c reductase subunit 7 (QCR7, UQCRB) (1 of 1)
OG IDs: 93522260
Best-hit-arabi-ID: AT5G25450.1
Best-hit-arabi-name: Cytochrome bd ubiquinol oxidase, 14kDa subunit
Gorai.013G241600.1.A
PCC=0.407, p=0.084
Gorai.013G241600.1.D
PCC=0.462, p=0.047
Gorai.013G110700.1.D
PCC=0.211, p=0.387
Gorai.013G110700.1.A
PCC=0.229, p=0.346
mRNA Profile
Protein Expression Profile
DPA

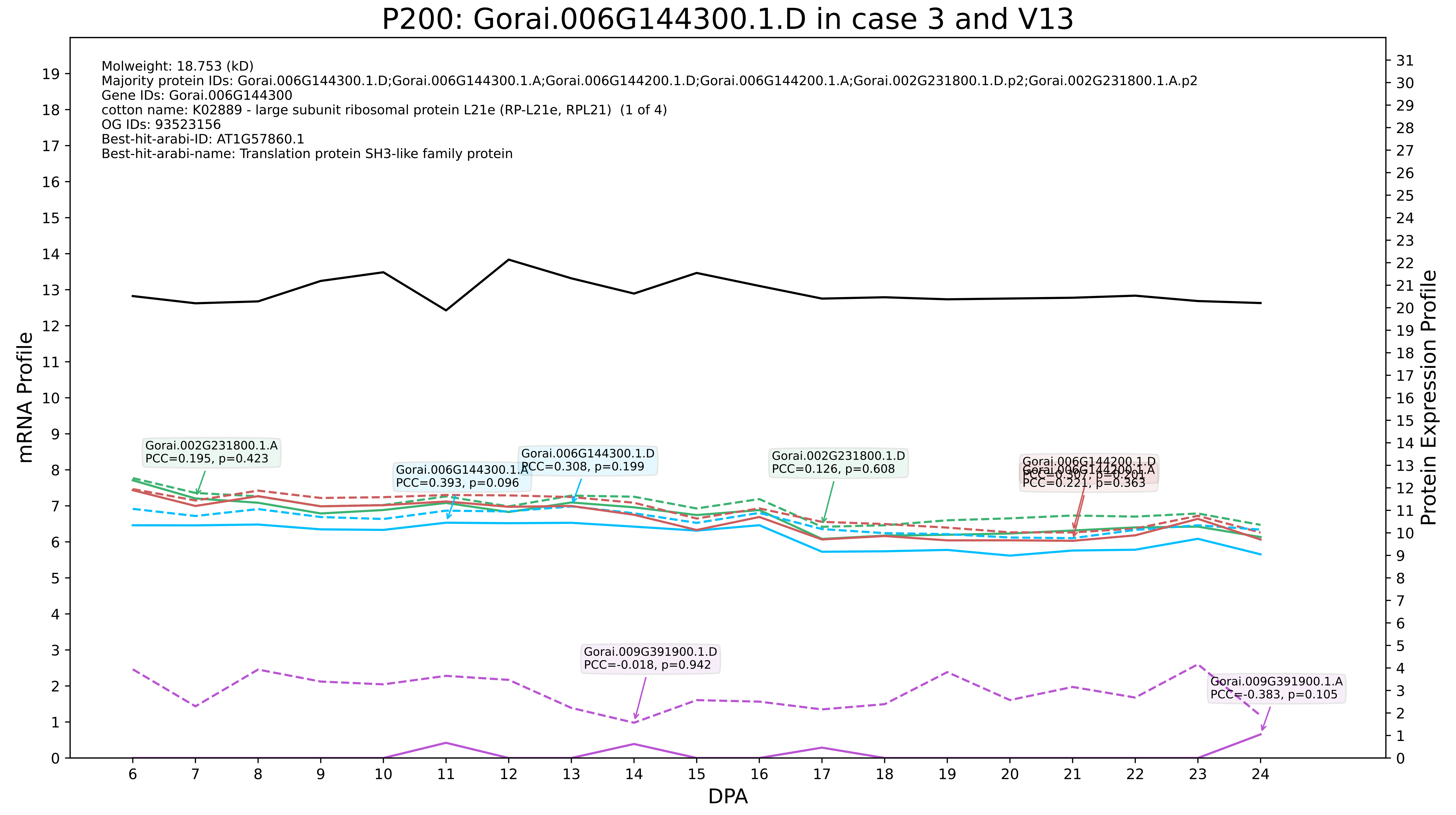
P200: Gorai.006G144300.1.D in case 3 and V13
Molweight: 18.753 (kD)
Majority protein IDs: Gorai.006G144300.1.D;Gorai.006G144300.1.A;Gorai.006G144200.1.D;Gorai.006G144200.1.A;Gorai.002G231800.1.D.p2;Gorai.002G231800.1.A.p2
Gene IDs: Gorai.006G144300
cotton name: K02889 - large subunit ribosomal protein L21e (RP-L21e, RPL21) (1 of 4)
OG IDs: 93523156
Best-hit-arabi-ID: AT1G57860.1
Best-hit-arabi-name: Translation protein SH3-like family protein
Gorai.002G231800.1.A
PCC=0.195, p=0.423
Gorai.006G144300.1.A
PCC=0.393, p=0.096
Gorai.006G144300.1.D
PCC=0.308, p=0.199
Gorai.002G231800.1.D
PCC=0.126, p=0.608
Gorai.006G144200.1.D
PCC=0.307, p=0.201
Gorai.006G144200.1.A
PCC=0.221, p=0.363
Gorai.009G391900.1.D
PCC=-0.018, p=0.942
Gorai.009G391900.1.A
PCC=-0.383, p=0.105
mRNA Profile
Protein Expression Profile
DPA

### P200: Gorai.010G236400.1.D in case 3 and V13

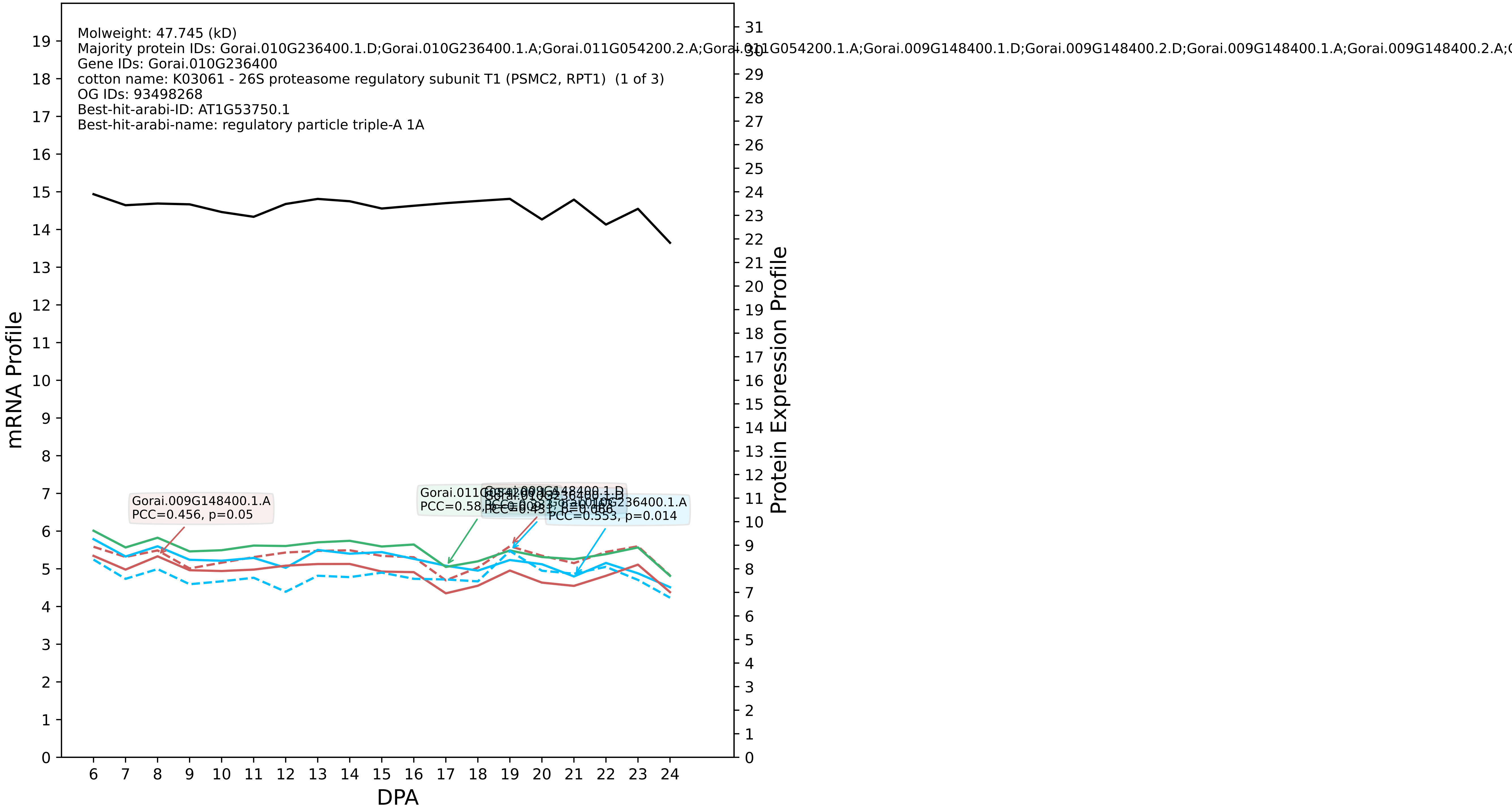

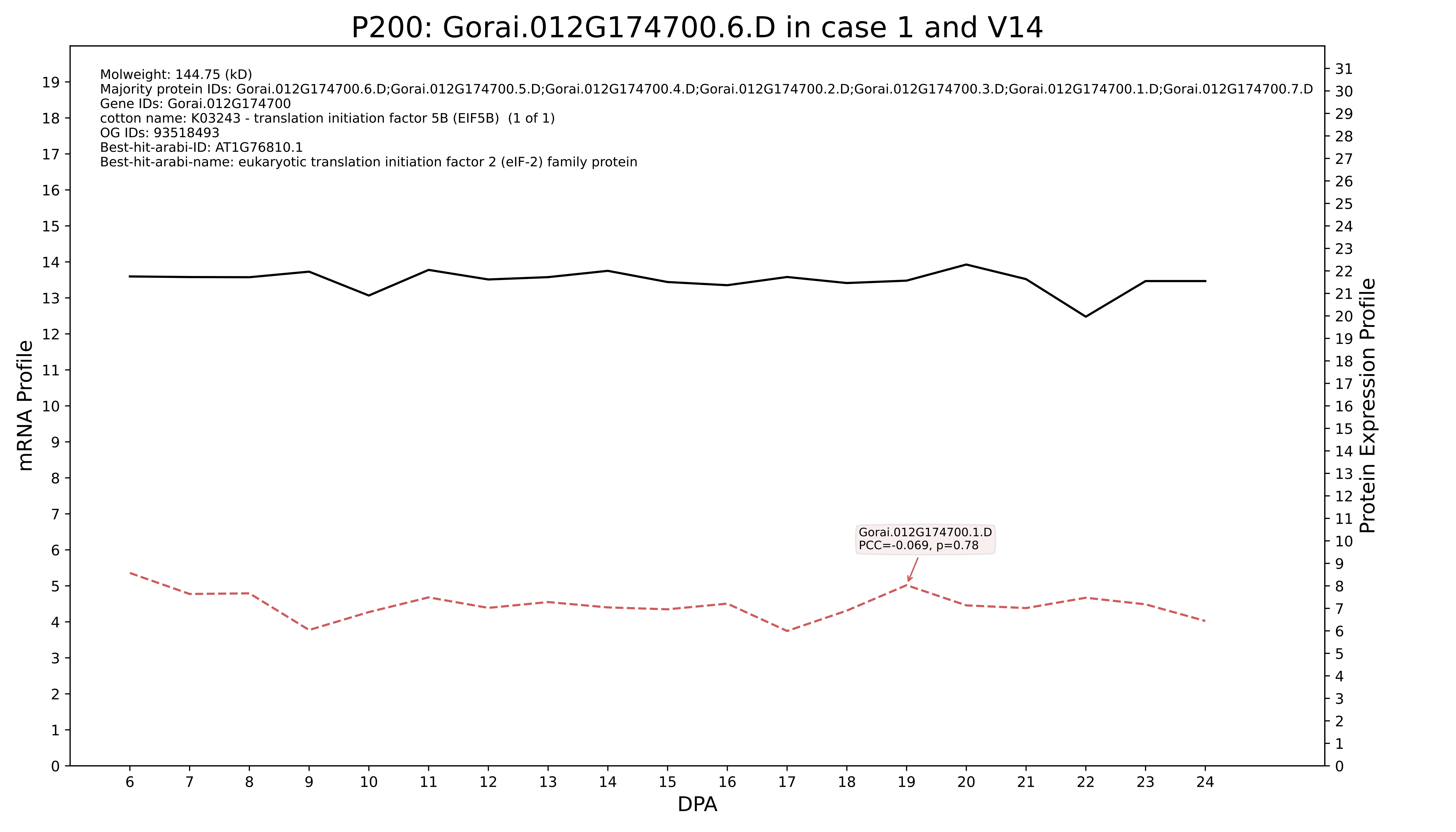
P200: Gorai.012G174700.6.D in case 1 and V14
Molweight: 144.75 (kD)
Majority protein IDs: Gorai.012G174700.6.D;Gorai.012G174700.5.D;Gorai.012G174700.4.D;Gorai.012G174700.2.D;Gorai.012G174700.3.D;Gorai.012G174700.1.D;Gorai.012G174700.7.D
Gene IDs: Gorai.012G174700
cotton name: K03243 - translation initiation factor 5B (EIF5B) (1 of 1)
OG IDs: 93518493
Best-hit-arabi-ID: AT1G76810.1
Best-hit-arabi-name: eukaryotic translation initiation factor 2 (eIF-2) family protein
Gorai.012G174700.1.D
PCC=-0.069, p=0.78
mRNA Profile
Protein Expression Profile
DPA

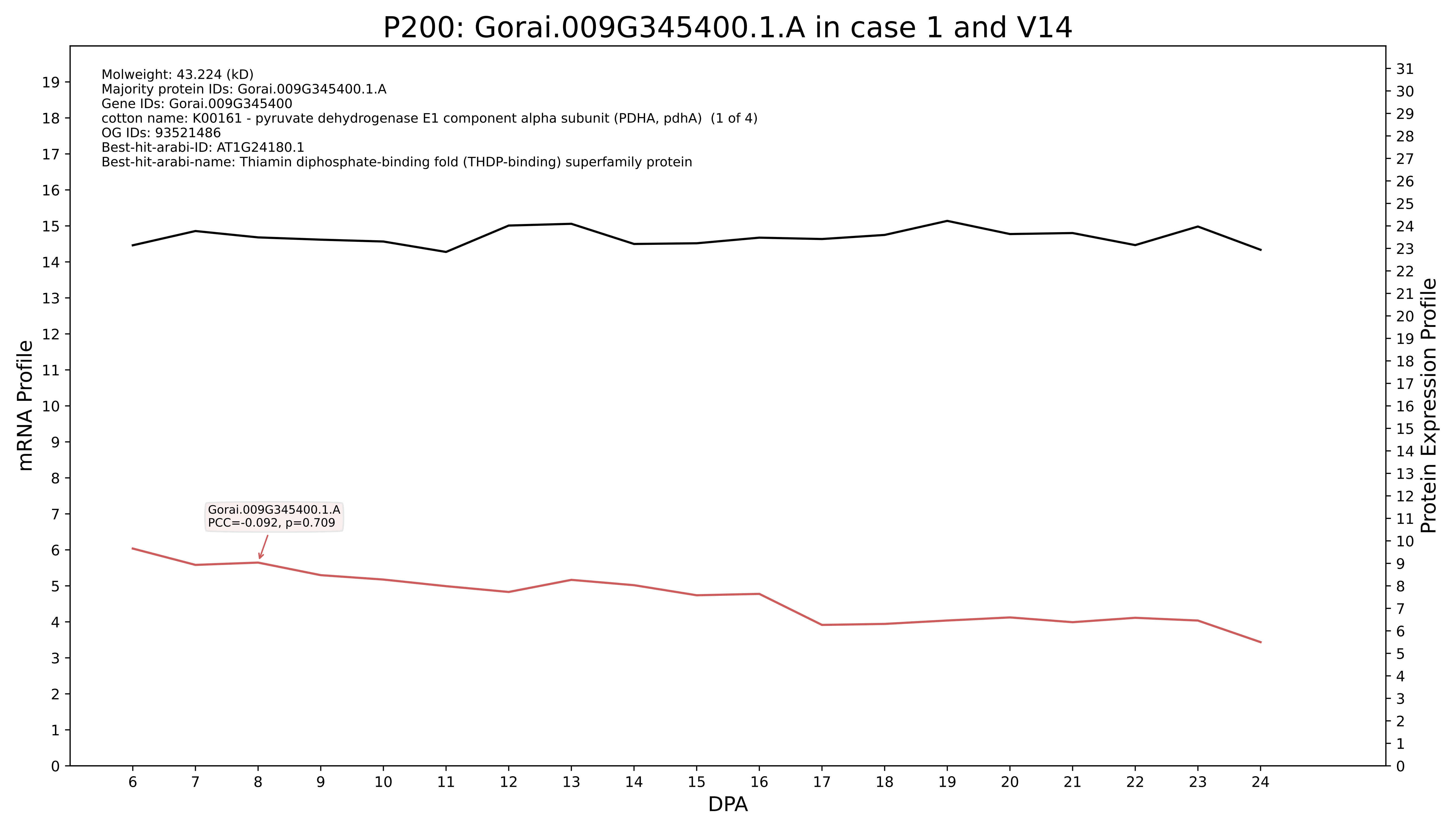
P200: Gorai.009G345400.1.A in case 1 and V14
Molweight: 43.224 (kD)
Majority protein IDs: Gorai.009G345400.1.A
Gene IDs: Gorai.009G345400
cotton name: K00161 - pyruvate dehydrogenase E1 component alpha subunit (PDHA, pdhA) (1 of 4)
OG IDs: 93521486
Best-hit-arabi-ID: AT1G24180.1
Best-hit-arabi-name: Thiamin diphosphate-binding fold (THDP-binding) superfamily protein
Gorai.009G345400.1.A
PCC=-0.092, p=0.709
mRNA Profile
Protein Expression Profile
DPA

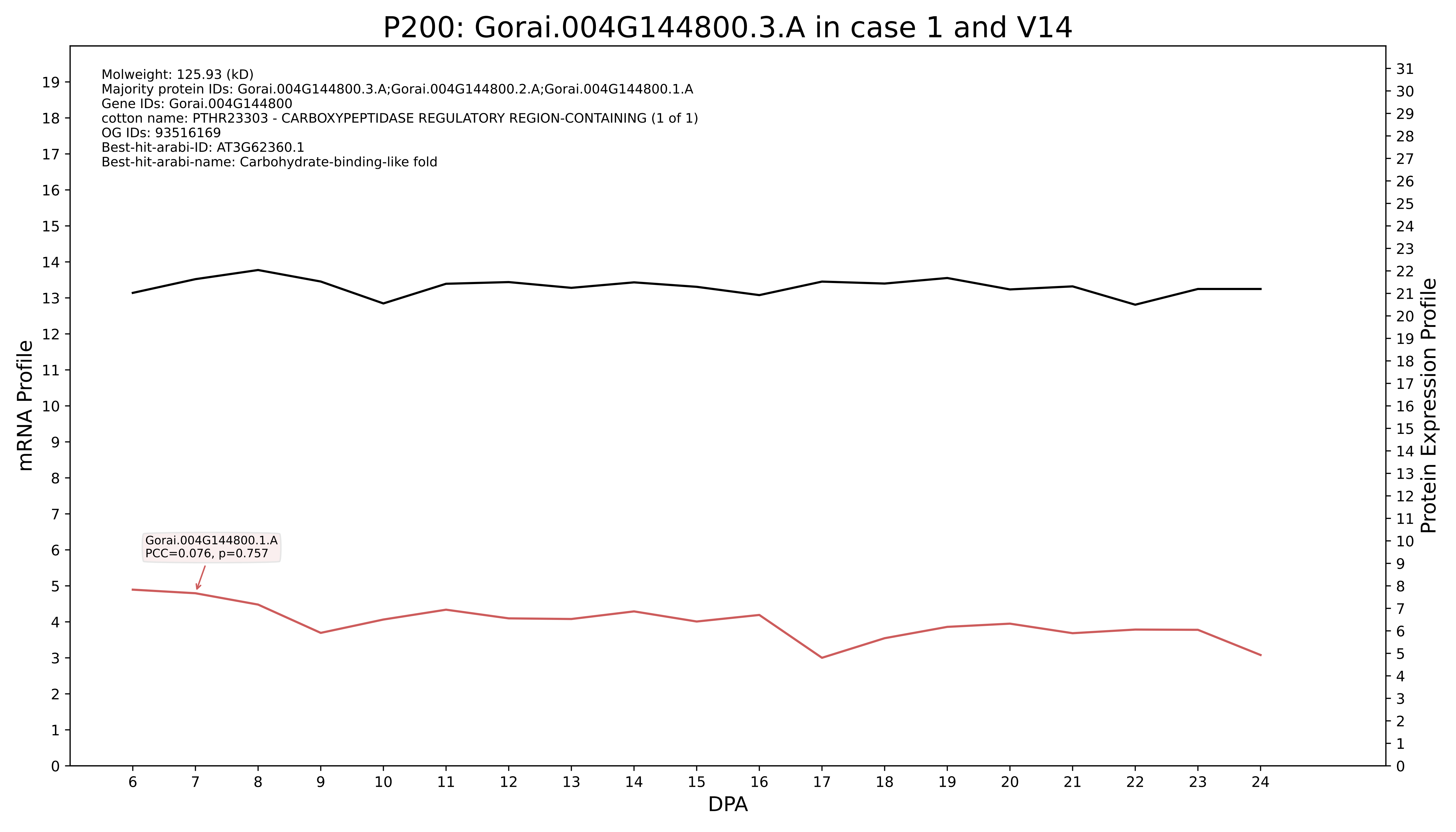

P200: Gorai.004G144800.3.A in case 1 and V14
Molweight: 125.93 (kD)
Majority protein IDs: Gorai.004G144800.3.A;Gorai.004G144800.2.A;Gorai.004G144800.1.A
Gene IDs: Gorai.004G144800
cotton name: PTHR23303 - CARBOXYPEPTIDASE REGULATORY REGION-CONTAINING (1 of 1)
OG IDs: 93516169
Best-hit-arabi-ID: AT3G62360.1
Best-hit-arabi-name: Carbohydrate-binding-like fold
Gorai.004G144800.1.A
PCC=0.076, p=0.757
mRNA Profile
Protein Expression Profile
DPA

### P200: Gorai.004G098000.1.D in case 1 and V14

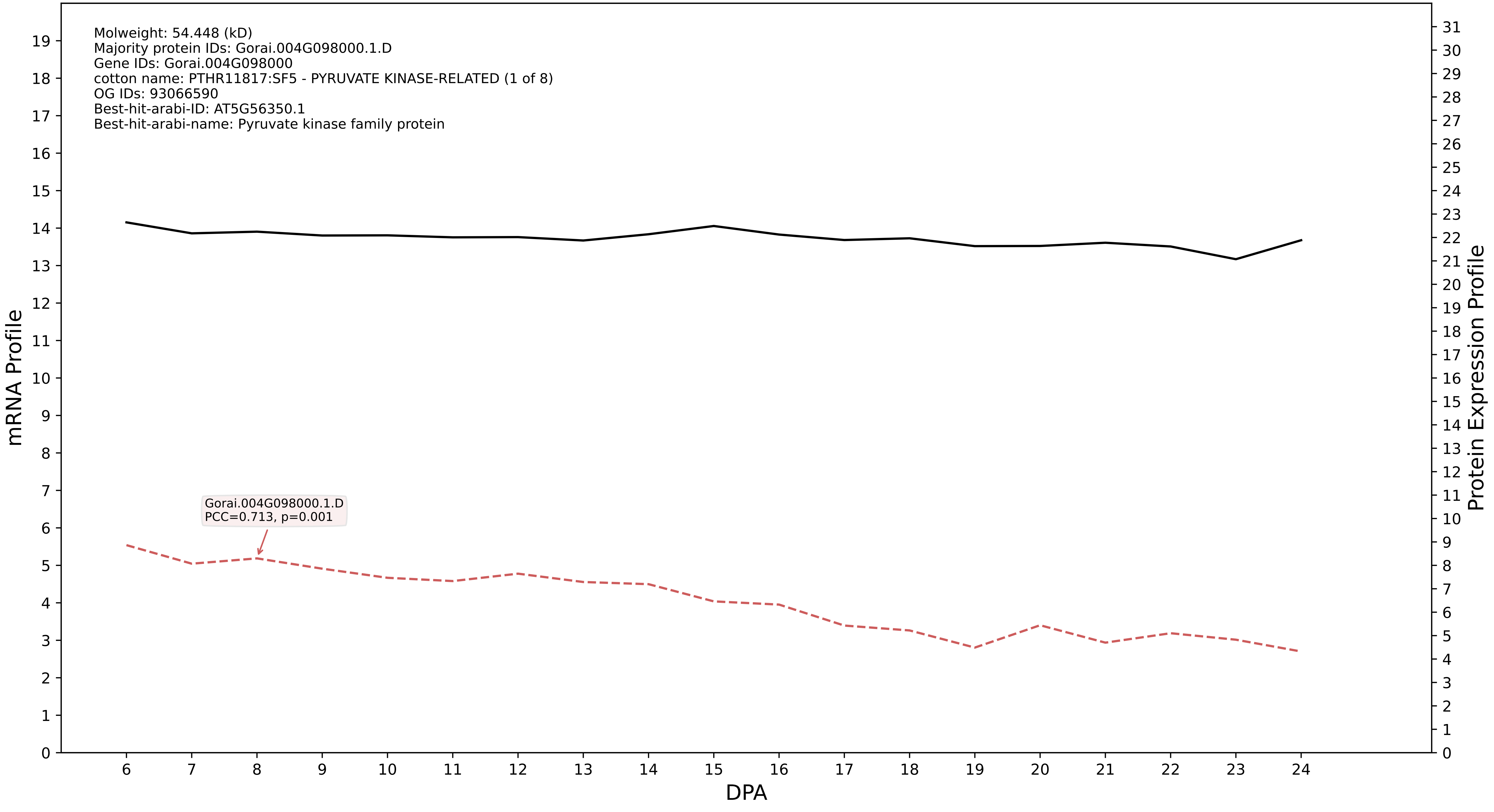

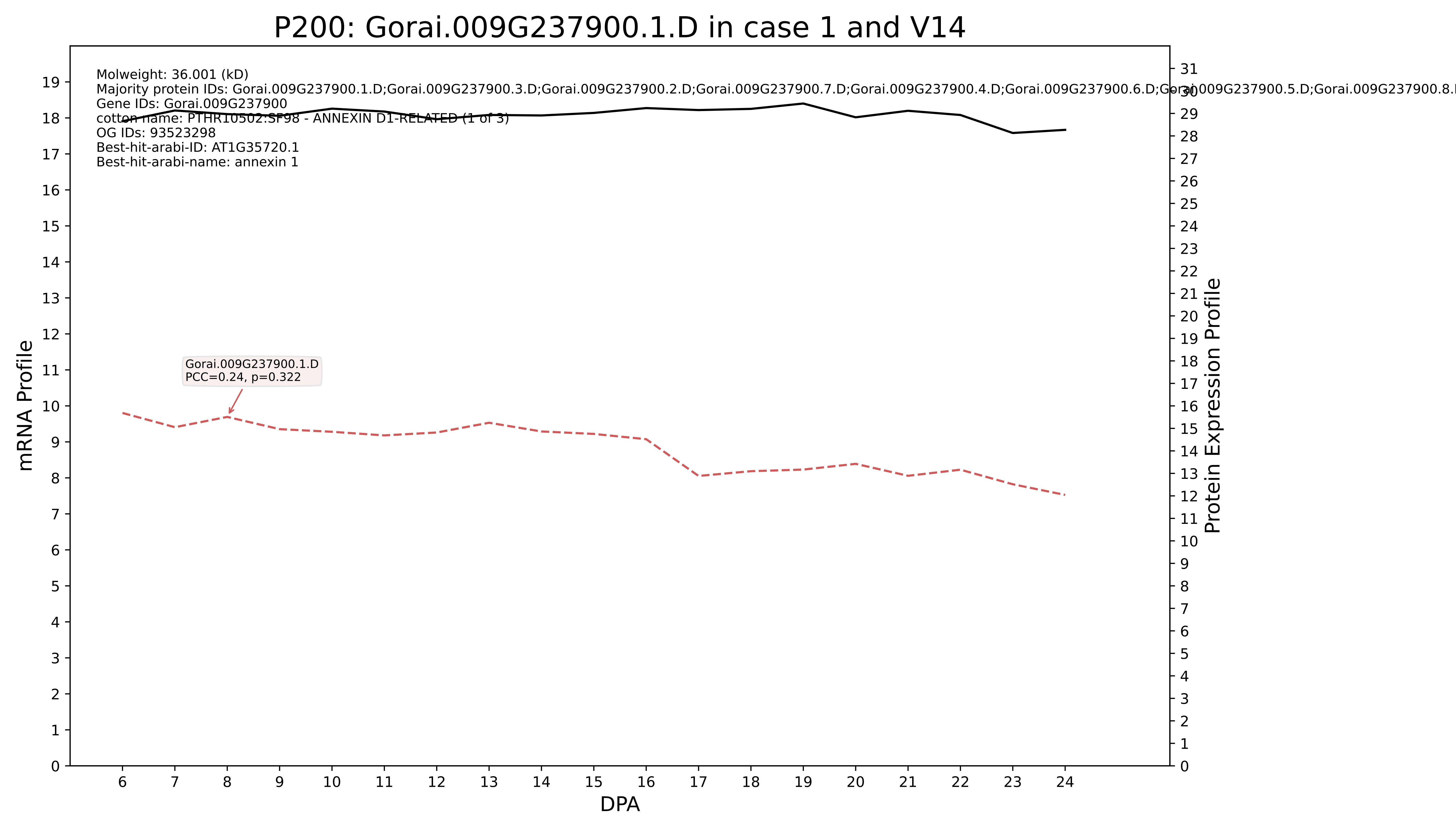
P200: Gorai.009G237900.1.D in case 1 and V14
Molweight: 36.001 (kD)
Majority protein IDs: Gorai.009G237900.1.D;Gorai.009G237900.3.D;Gorai.009G237900.2.D;Gorai.009G237900.7.D;Gorai.009G237900.4.D;Gorai.009G237900.6.D;Gorai.009G237900.5.D;Gorai.009G237900.8.D
Gene IDs: Gorai.009G237900
cotton name: PTHR10502:SF98 - ANNEXIN D1-RELATED (1 of 3)
OG IDs: 93523298
Best-hit-arabi-ID: AT1G35720.1
Best-hit-arabi-name: annexin 1
Gorai.009G237900.1.D
PCC=0.24, p=0.322
mRNA Profile
Protein Expression Profile
DPA

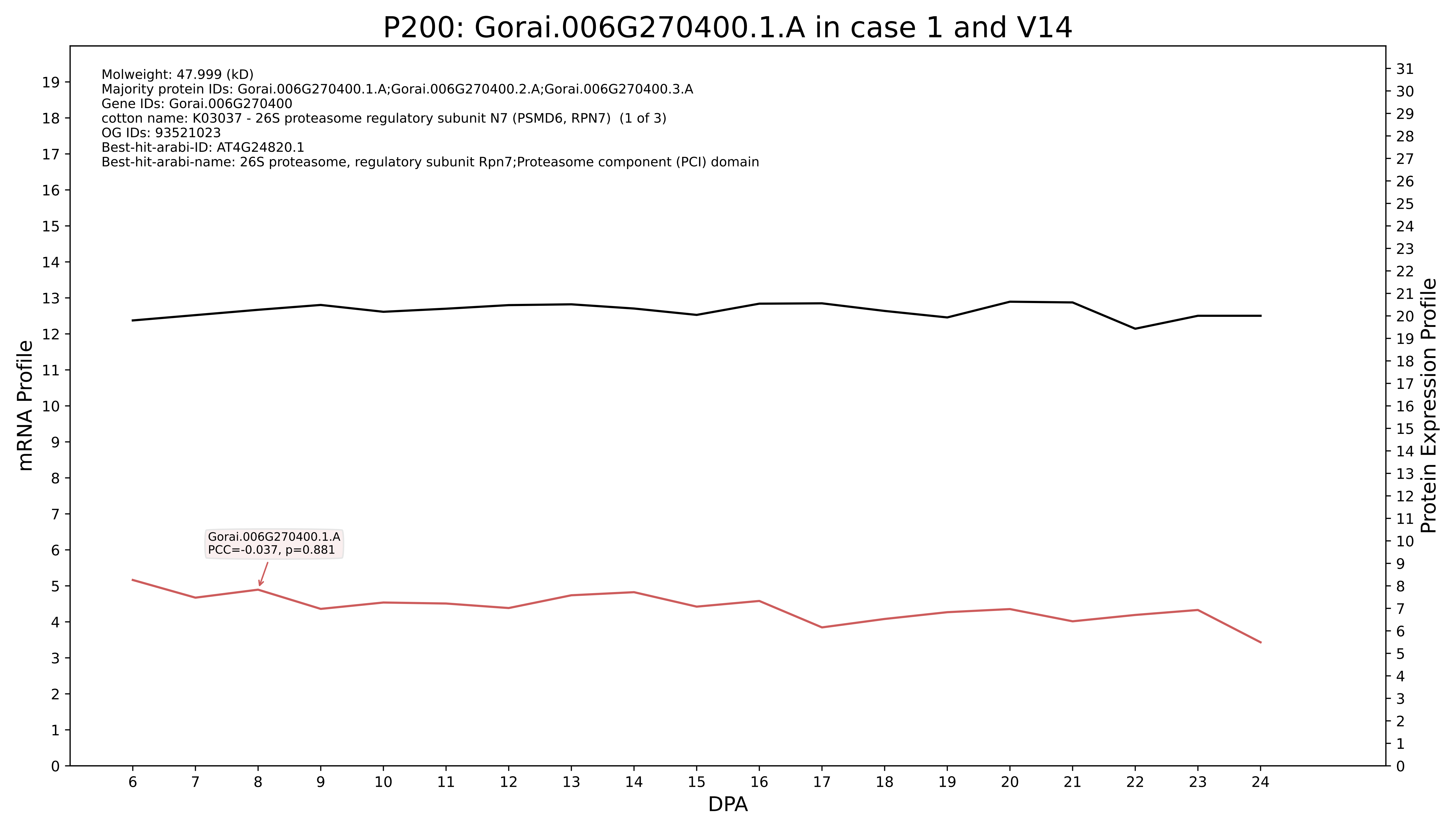
P200: Gorai.006G270400.1.A in case 1 and V14
Molweight: 47.999 (kD)
Majority protein IDs: Gorai.006G270400.1.A;Gorai.006G270400.2.A;Gorai.006G270400.3.A
Gene IDs: Gorai.006G270400
cotton name: K03037 - 26S proteasome regulatory subunit N7 (PSMD6, RPN7) (1 of 3)
OG IDs: 93521023
Best-hit-arabi-ID: AT4G24820.1
Best-hit-arabi-name: 26S proteasome, regulatory subunit Rpn7;Proteasome component (PCI) domain
Gorai.006G270400.1.A
PCC=-0.037, p=0.881
mRNA Profile
Protein Expression Profile
DPA

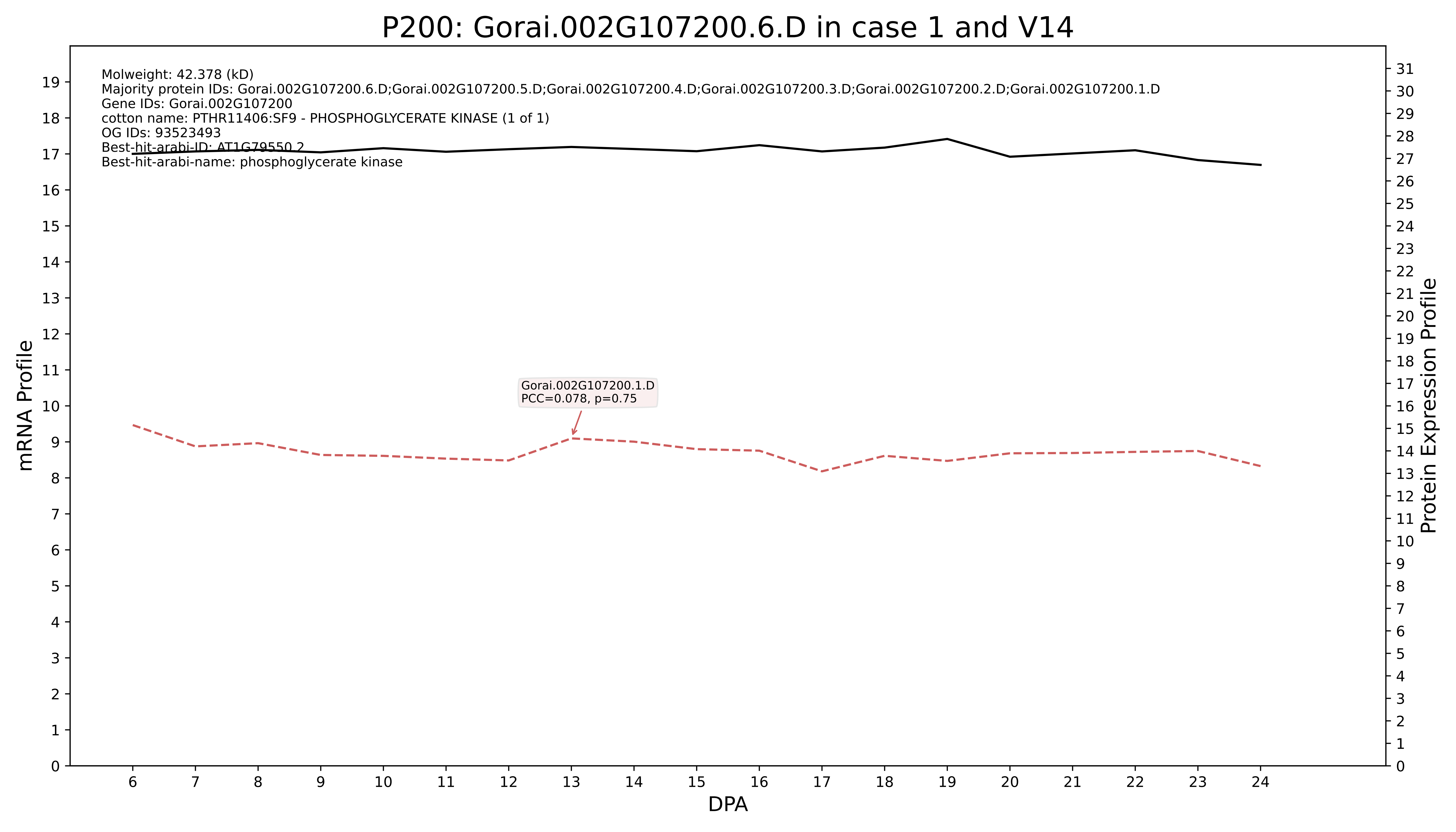
P200: Gorai.002G107200.6.D in case 1 and V14
Molweight: 42.378 (kD)
Majority protein IDs: Gorai.002G107200.6.D;Gorai.002G107200.5.D;Gorai.002G107200.4.D;Gorai.002G107200.3.D;Gorai.002G107200.2.D;Gorai.002G107200.1.D
Gene IDs: Gorai.002G107200
cotton name: PTHR11406:SF9 - PHOSPHOGLYCERATE KINASE (1 of 1)
OG IDs: 93523493
Best-hit-arabi-ID: AT1G79550.2
Best-hit-arabi-name: phosphoglycerate kinase
Gorai.002G107200.1.D
PCC=0.078, p=0.75
mRNA Profile
Protein Expression Profile
DPA

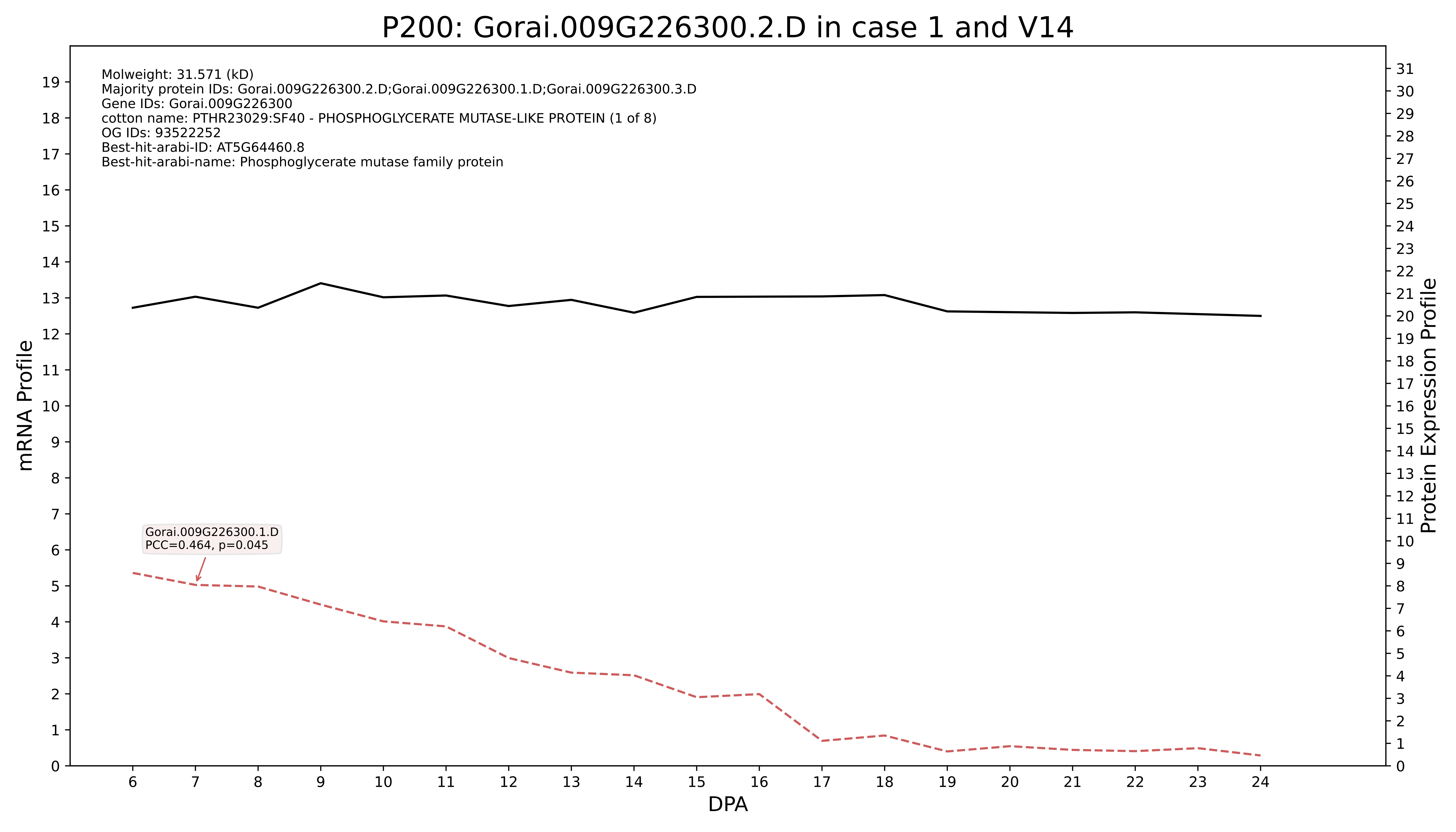

P200: Gorai.009G226300.2.D in case 1 and V14
Molweight: 31.571 (kD)
Majority protein IDs: Gorai.009G226300.2.D;Gorai.009G226300.1.D;Gorai.009G226300.3.D
Gene IDs: Gorai.009G226300
cotton name: PTHR23029:SF40 - PHOSPHOGLYCERATE MUTASE-LIKE PROTEIN (1 of 8)
OG IDs: 93522252
Best-hit-arabi-ID: AT5G64460.8
Best-hit-arabi-name: Phosphoglycerate mutase family protein
Gorai.009G226300.1.D
PCC=0.464, p=0.045
mRNA Profile
Protein Expression Profile
DPA

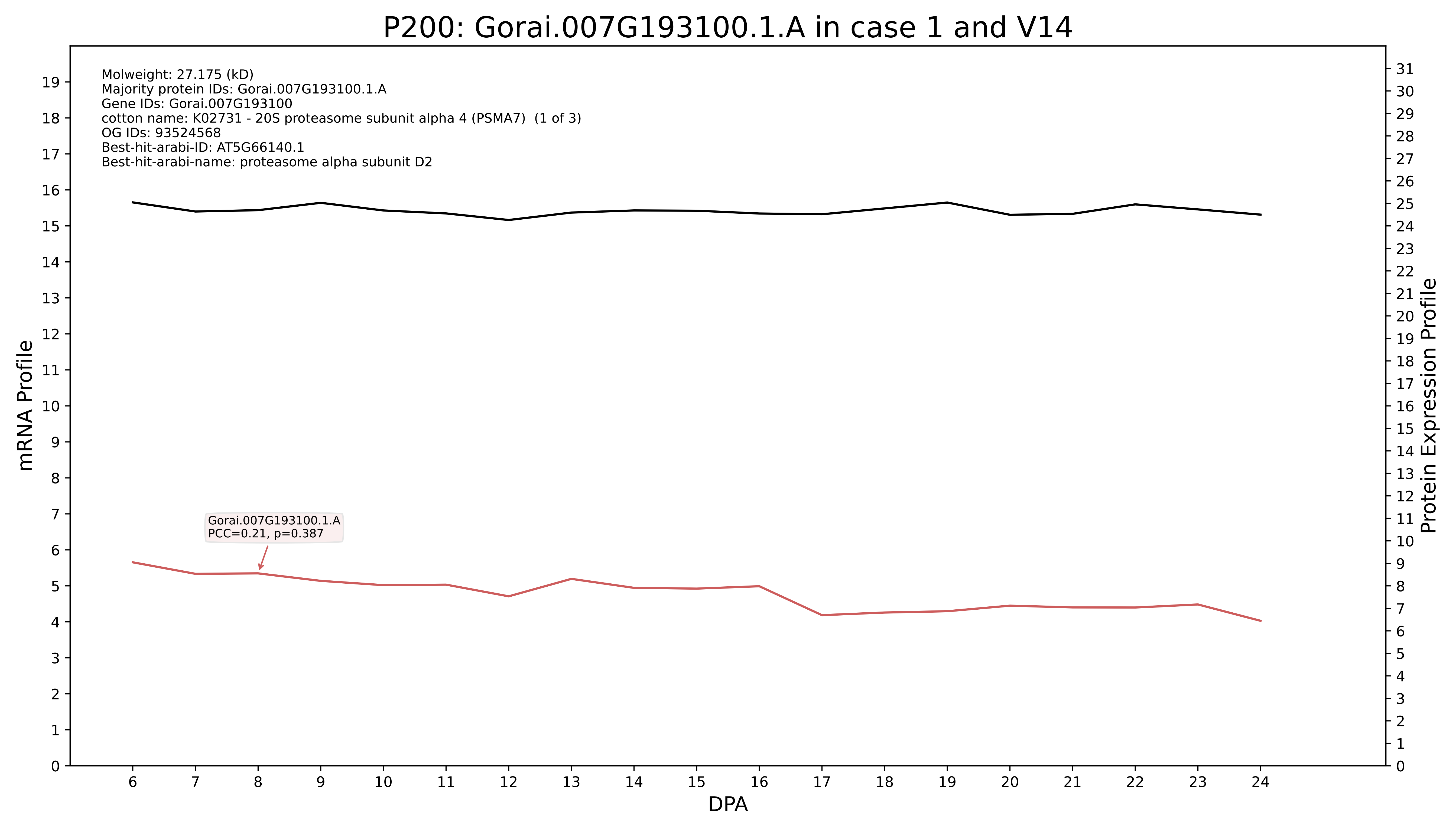
P200: Gorai.007G193100.1.A in case 1 and V14
Molweight: 27.175 (kD)
Majority protein IDs: Gorai.007G193100.1.A
Gene IDs: Gorai.007G193100
cotton name: K02731 - 20S proteasome subunit alpha 4 (PSMA7) (1 of 3)
OG IDs: 93524568
Best-hit-arabi-ID: AT5G66140.1
Best-hit-arabi-name: proteasome alpha subunit D2
Gorai.007G193100.1.A
PCC=0.21, p=0.387
mRNA Profile
Protein Expression Profile
DPA

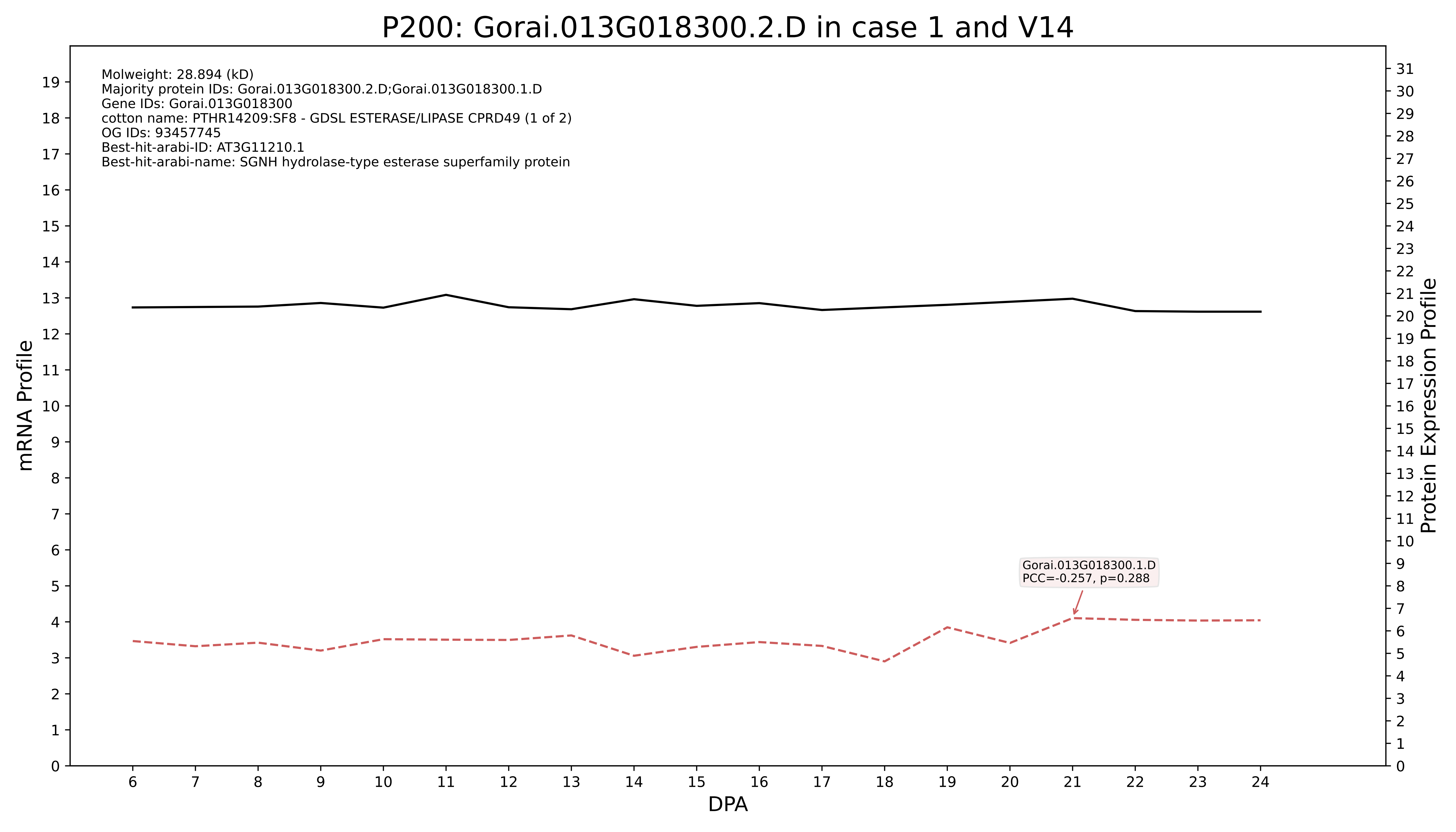
P200: Gorai.013G018300.2.D in case 1 and V14
Molweight: 28.894 (kD)
Majority protein IDs: Gorai.013G018300.2.D;Gorai.013G018300.1.D
Gene IDs: Gorai.013G018300
cotton name: PTHR14209:SF8 - GDSL ESTERASE/LIPASE CPRD49 (1 of 2)
OG IDs: 93457745
Best-hit-arabi-ID: AT3G11210.1
Best-hit-arabi-name: SGNH hydrolase-type esterase superfamily protein
Gorai.013G018300.1.D
PCC=-0.257, p=0.288
mRNA Profile
Protein Expression Profile
DPA

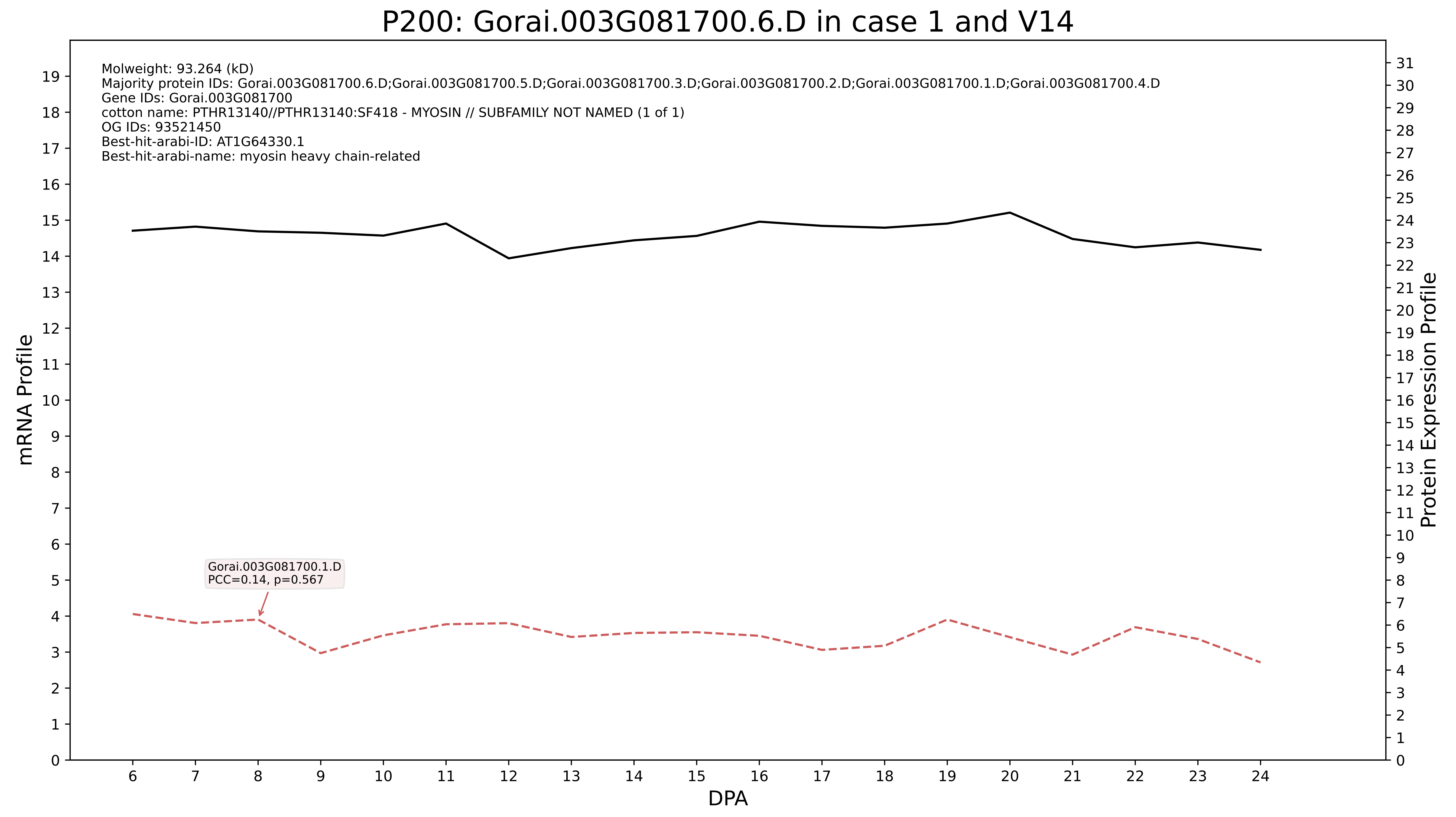

P200: Gorai.003G081700.6.D in case 1 and V14
Molweight: 93.264 (kD)
Majority protein IDs: Gorai.003G081700.6.D;Gorai.003G081700.5.D;Gorai.003G081700.3.D;Gorai.003G081700.2.D;Gorai.003G081700.1.D;Gorai.003G081700.4.D
Gene IDs: Gorai.003G081700
cotton name: PTHR13140//PTHR13140:SF418 - MYOSIN // SUBFAMILY NOT NAMED (1 of 1)
OG IDs: 93521450
Best-hit-arabi-ID: AT1G64330.1
Best-hit-arabi-name: myosin heavy chain-related
Gorai.003G081700.1.D
PCC=0.14, p=0.567
mRNA Profile
Protein Expression Profile
DPA

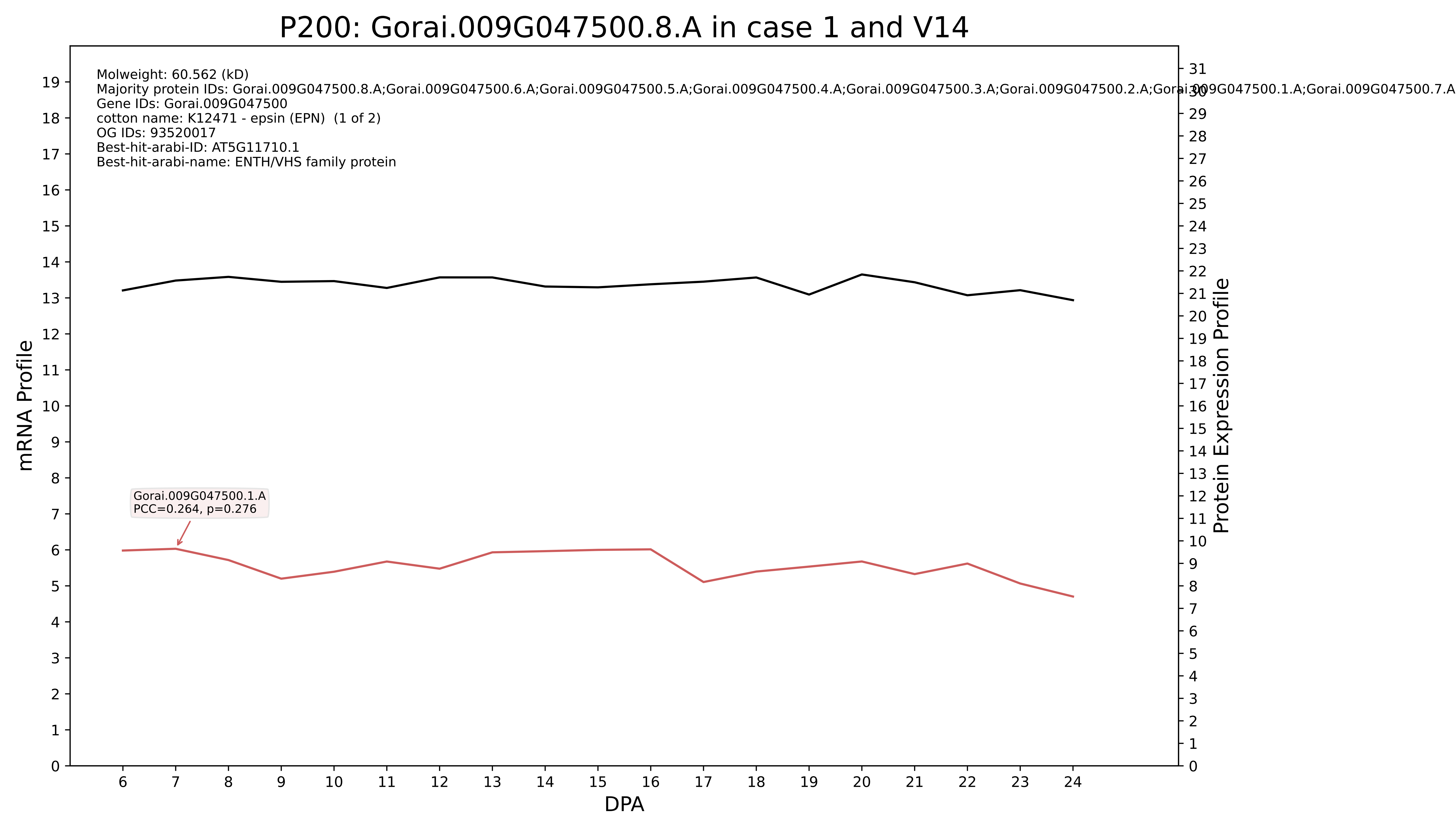
P200: Gorai.009G047500.8.A in case 1 and V14
Molweight: 60.562 (kD)
Majority protein IDs: Gorai.009G047500.8.A;Gorai.009G047500.6.A;Gorai.009G047500.5.A;Gorai.009G047500.4.A;Gorai.009G047500.3.A;Gorai.009G047500.2.A;Gorai.009G047500.1.A;Gorai.009G047500.7.A
Gene IDs: Gorai.009G047500
cotton name: K12471 - epsin (EPN) (1 of 2)
OG IDs: 93520017
Best-hit-arabi-ID: AT5G11710.1
Best-hit-arabi-name: ENTH/VHS family protein
Gorai.009G047500.1.A
PCC=0.264, p=0.276
mRNA Profile
Protein Expression Profile
DPA

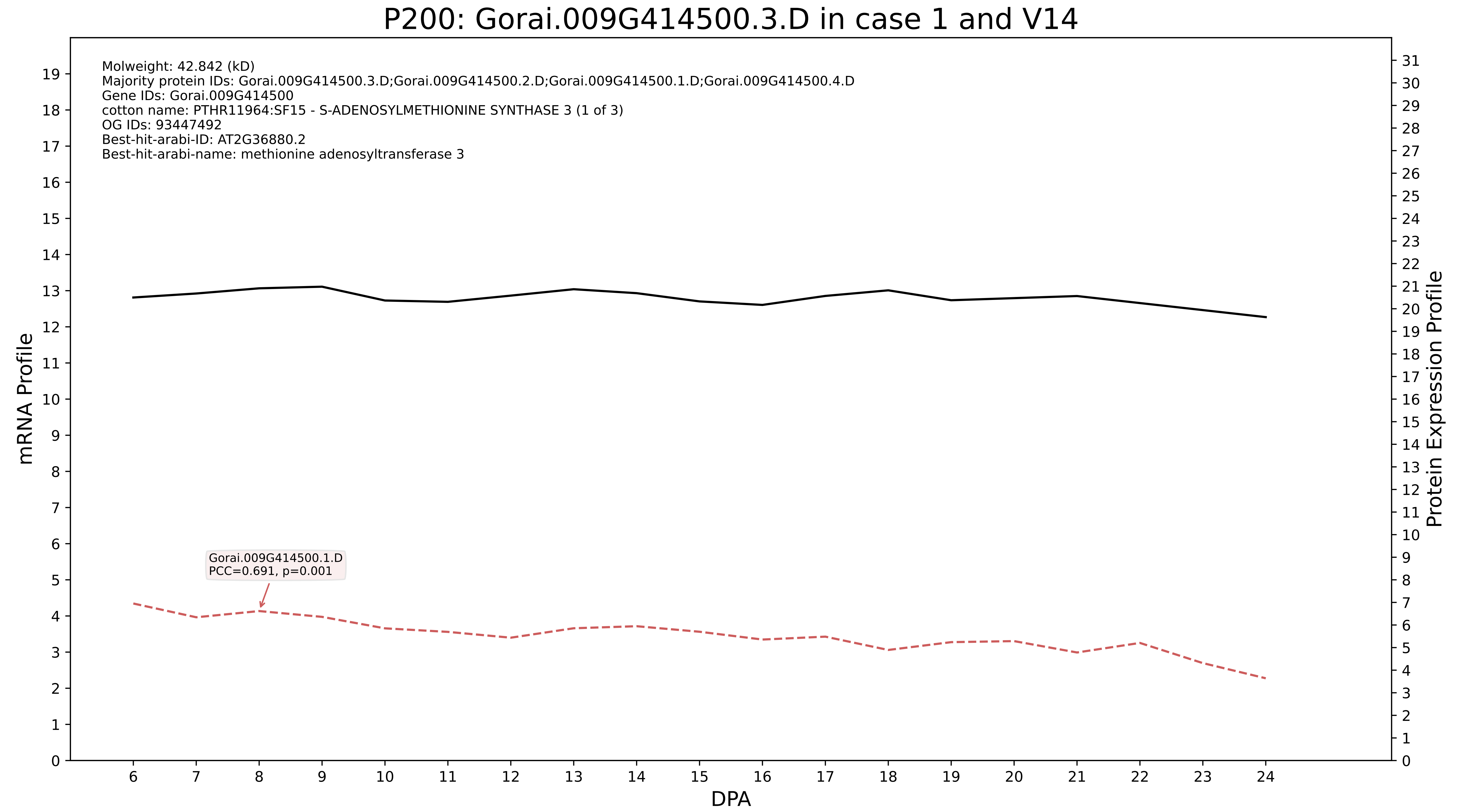
P200: Gorai.009G414500.3.D in case 1 and V14
Molweight: 42.842 (kD)
Majority protein IDs: Gorai.009G414500.3.D;Gorai.009G414500.2.D;Gorai.009G414500.1.D;Gorai.009G414500.4.D
Gene IDs: Gorai.009G414500
cotton name: PTHR11964:SF15 - S-ADENOSYLMETHIONINE SYNTHASE 3 (1 of 3)
OG IDs: 93447492
Best-hit-arabi-ID: AT2G36880.2
Best-hit-arabi-name: methionine adenosyltransferase 3
Gorai.009G414500.1.D
PCC=0.691, p=0.001
mRNA Profile
Protein Expression Profile
DPA

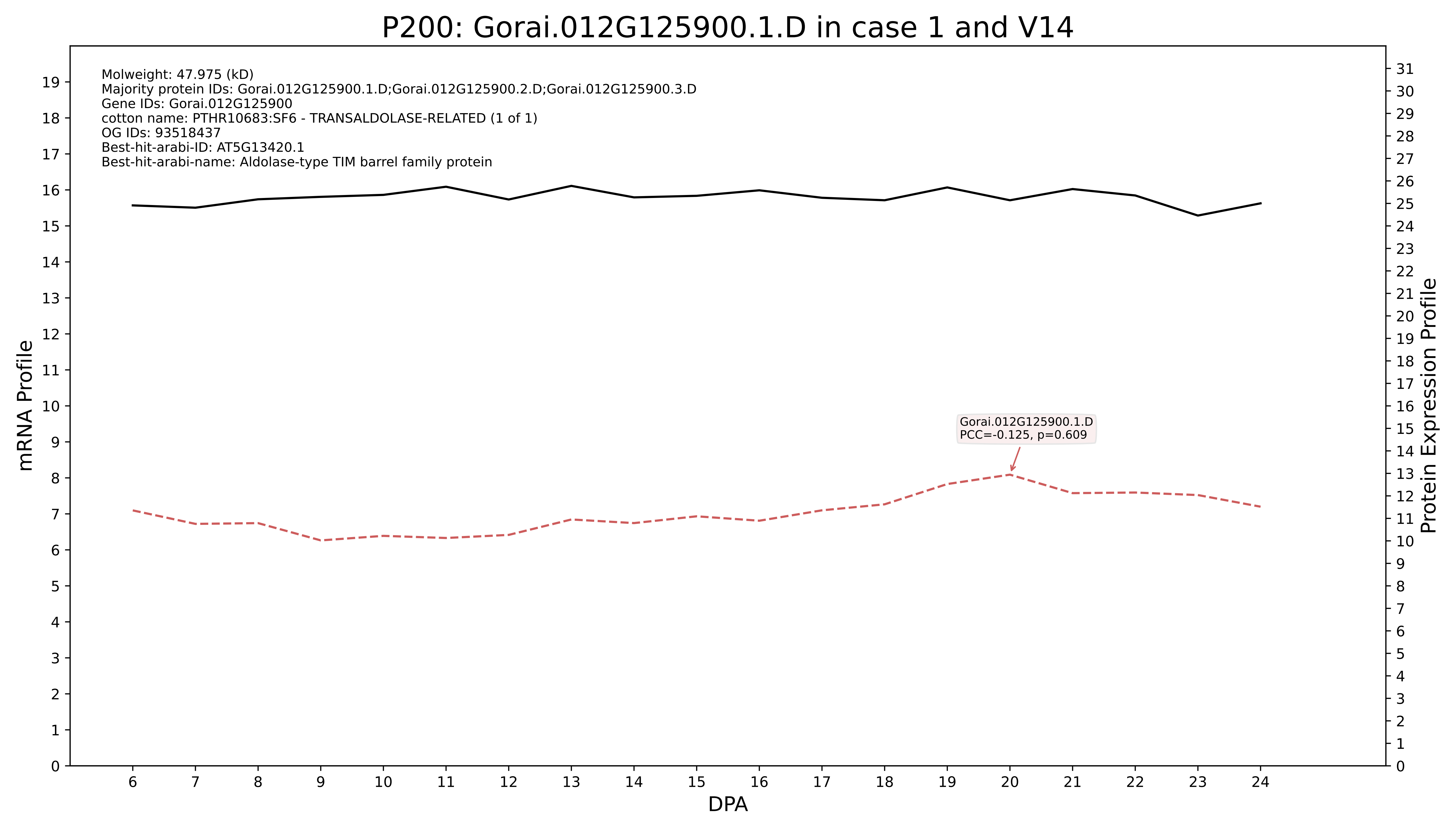
P200: Gorai.012G125900.1.D in case 1 and V14
Molweight: 47.975 (kD)
Majority protein IDs: Gorai.012G125900.1.D;Gorai.012G125900.2.D;Gorai.012G125900.3.D
Gene IDs: Gorai.012G125900
cotton name: PTHR10683:SF6 - TRANSALDOLASE-RELATED (1 of 1)
OG IDs: 93518437
Best-hit-arabi-ID: AT5G13420.1
Best-hit-arabi-name: Aldolase-type TIM barrel family protein
Gorai.012G125900.1.D
PCC=-0.125, p=0.609
mRNA Profile
Protein Expression Profile
DPA

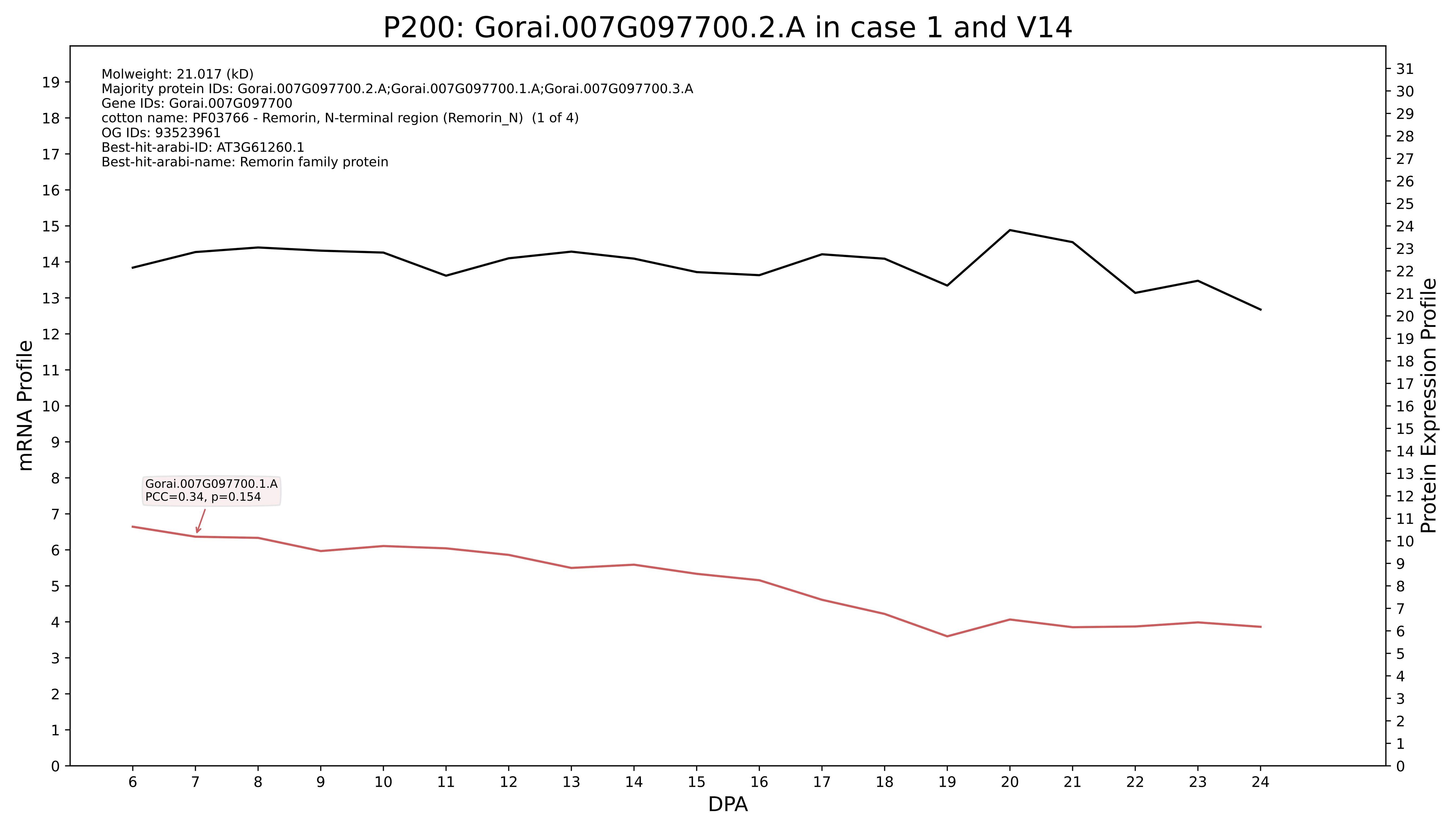

P200: Gorai.007G097700.2.A in case 1 and V14
Molweight: 21.017 (kD)
Majority protein IDs: Gorai.007G097700.2.A;Gorai.007G097700.1.A;Gorai.007G097700.3.A
Gene IDs: Gorai.007G097700
cotton name: PF03766 - Remorin, N-terminal region (Remorin_N) (1 of 4)
OG IDs: 93523961
Best-hit-arabi-ID: AT3G61260.1
Best-hit-arabi-name: Remorin family protein
Gorai.007G097700.1.A
PCC=0.34, p=0.154
mRNA Profile
Protein Expression Profile
DPA

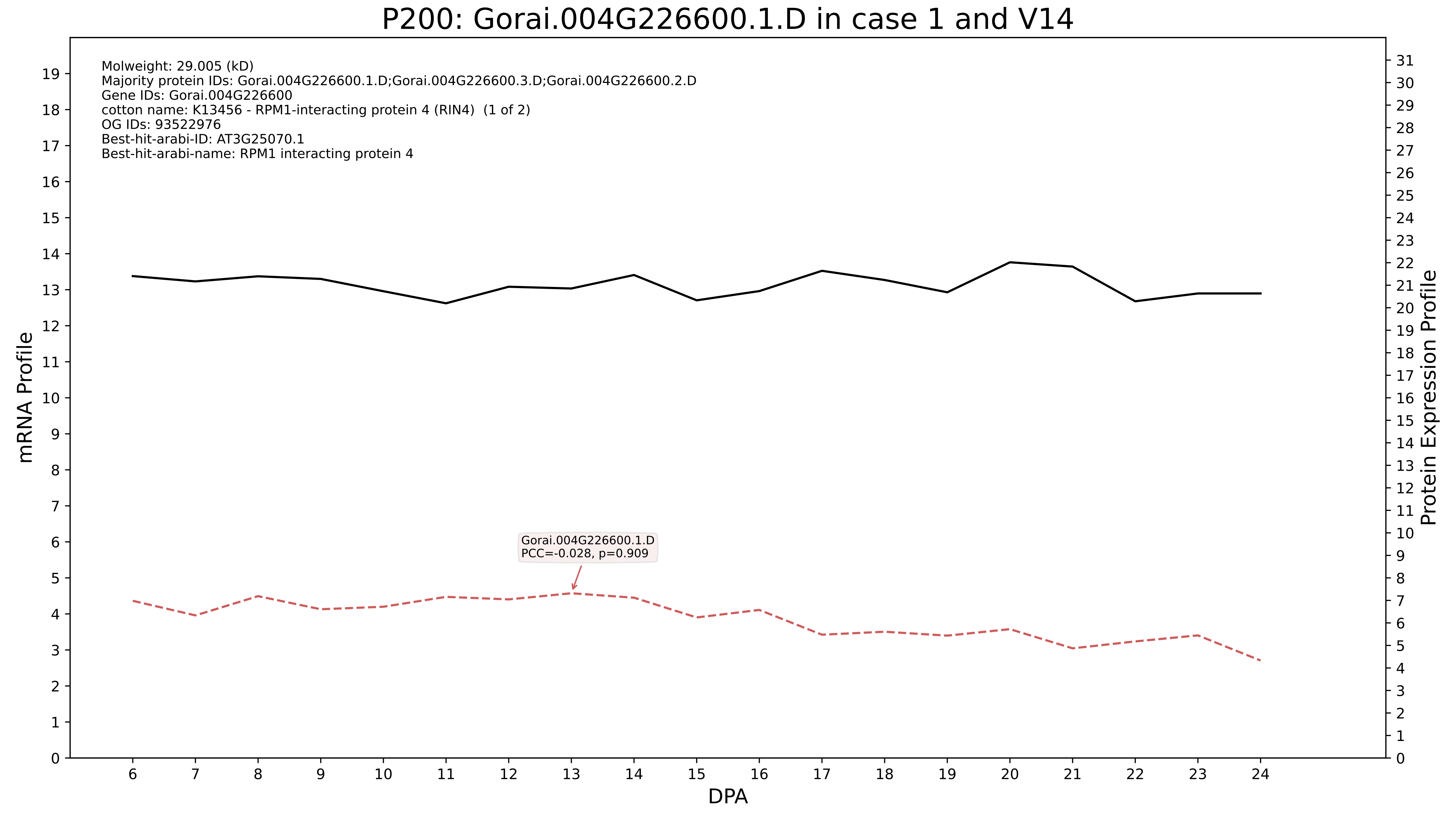
P200: Gorai.004G226600.1.D in case 1 and V14
Molweight: 29.005 (kD)
Majority protein IDs: Gorai.004G226600.1.D;Gorai.004G226600.3.D;Gorai.004G226600.2.D
Gene IDs: Gorai.004G226600
cotton name: K13456 - RPM1-interacting protein 4 (RIN4) (1 of 2)
OG IDs: 93522976
Best-hit-arabi-ID: AT3G25070.1
Best-hit-arabi-name: RPM1 interacting protein 4
Gorai.004G226600.1.D
PCC=-0.028, p=0.909
mRNA Profile
Protein Expression Profile
DPA

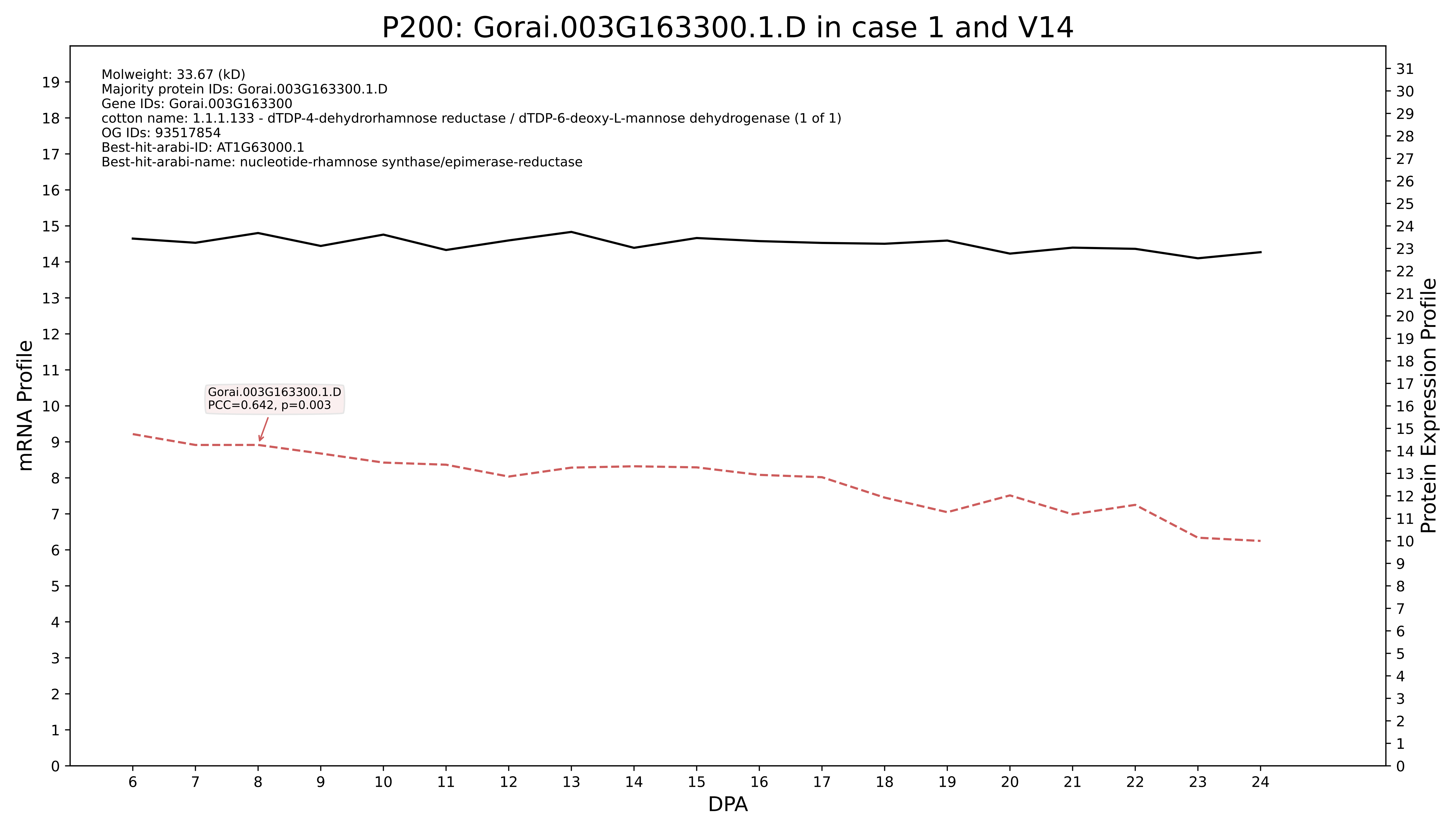

P200: Gorai.003G163300.1.D in case 1 and V14
Molweight: 33.67 (kD)
Majority protein IDs: Gorai.003G163300.1.D
Gene IDs: Gorai.003G163300
cotton name: 1.1.1.133 - dTDP-4-dehydrorhamnose reductase / dTDP-6-deoxy-L-mannose dehydrogenase (1 of 1)
OG IDs: 93517854
Best-hit-arabi-ID: AT1G63000.1
Best-hit-arabi-name: nucleotide-rhamnose synthase/epimerase-reductase
Gorai.003G163300.1.D
PCC=0.642, p=0.003
mRNA Profile
Protein Expression Profile
DPA

### P200: Gorai.005G172100.1.D in case 1 and V14

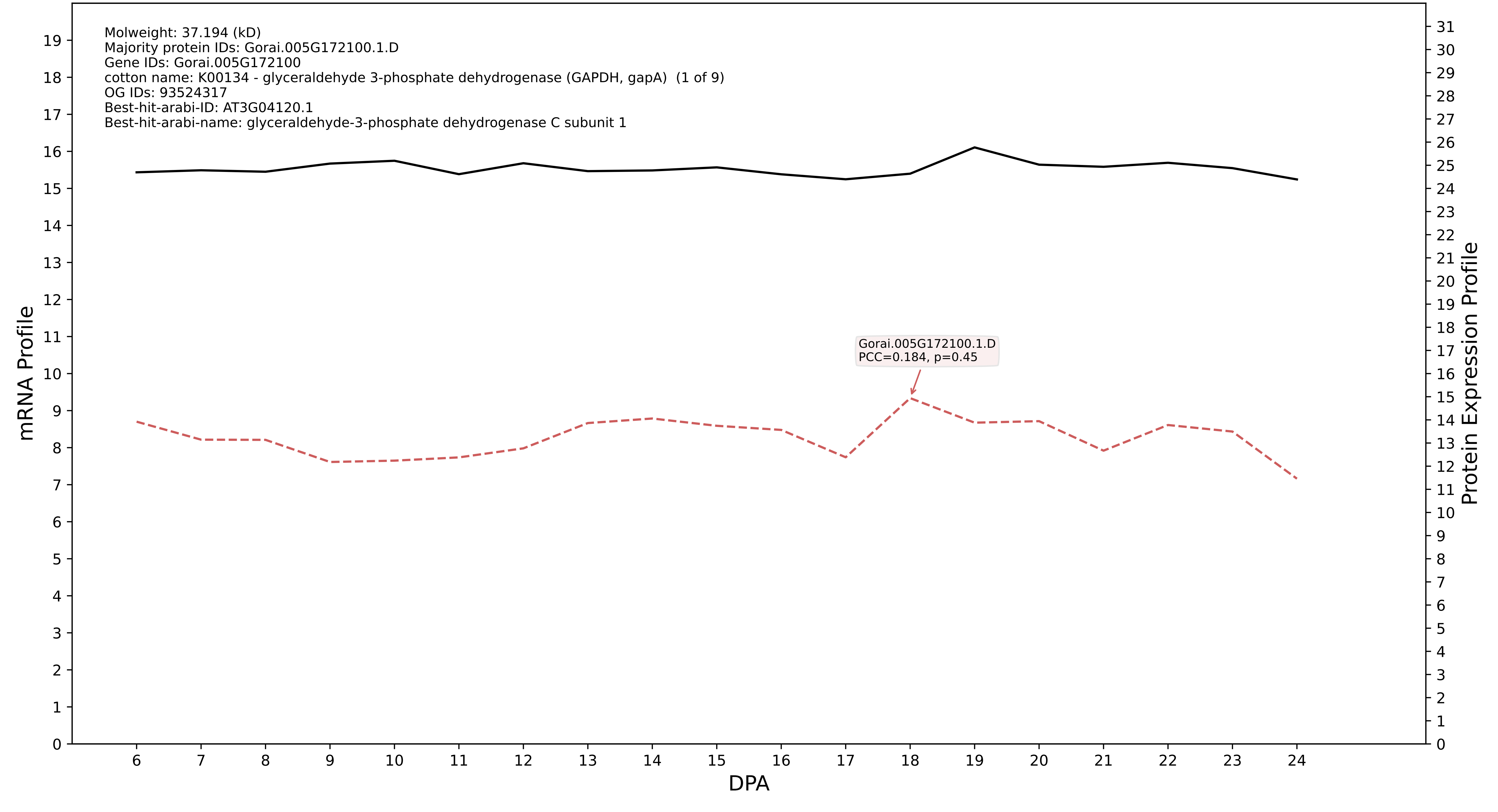

### P200: Gorai.005G115600.2.A in case 1 and V14

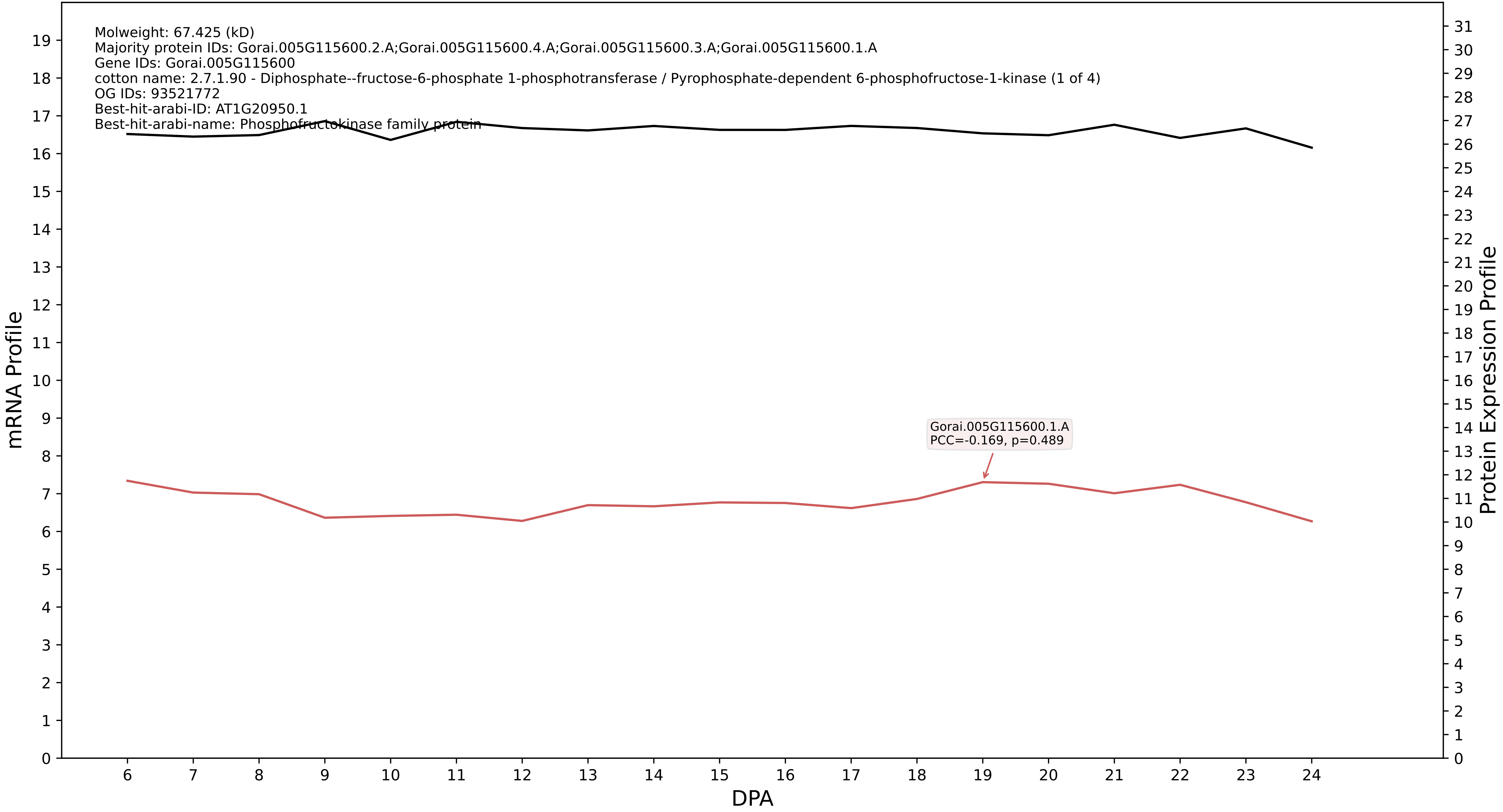

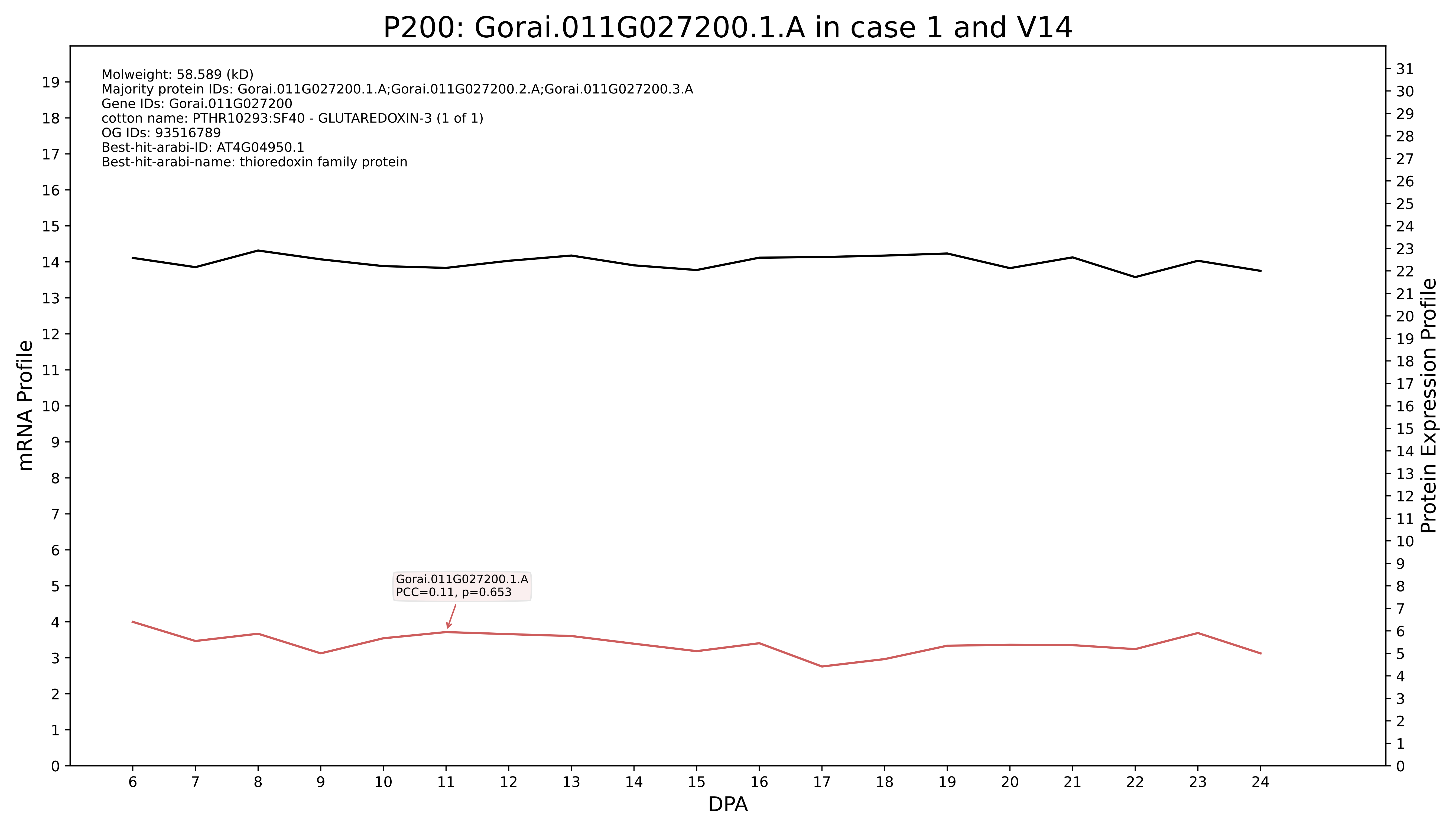
P200: Gorai.011G027200.1.A in case 1 and V14
Molweight: 58.589 (kD)
Majority protein IDs: Gorai.011G027200.1.A;Gorai.011G027200.2.A;Gorai.011G027200.3.A
Gene IDs: Gorai.011G027200
cotton name: PTHR10293:SF40 - GLUTAREDOXIN-3 (1 of 1)
OG IDs: 93516789
Best-hit-arabi-ID: AT4G04950.1
Best-hit-arabi-name: thioredoxin family protein
Gorai.011G027200.1.A
PCC=0.11, p=0.653
mRNA Profile
Protein Expression Profile
DPA

### P200: Gorai.010G080400.1.D in case 1 and V14

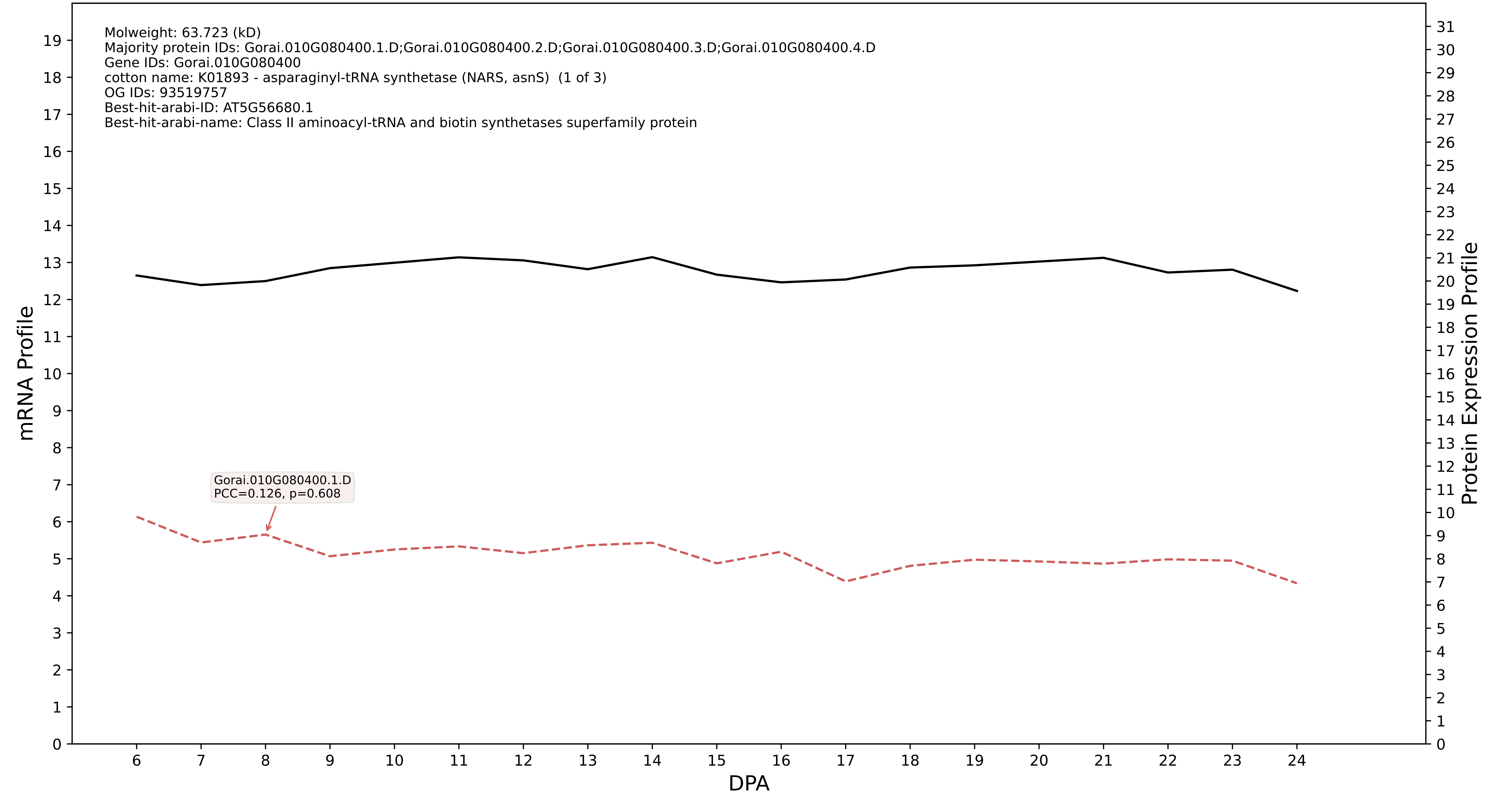

### P200: Gorai.009G102400.2.D in case 1 and V14

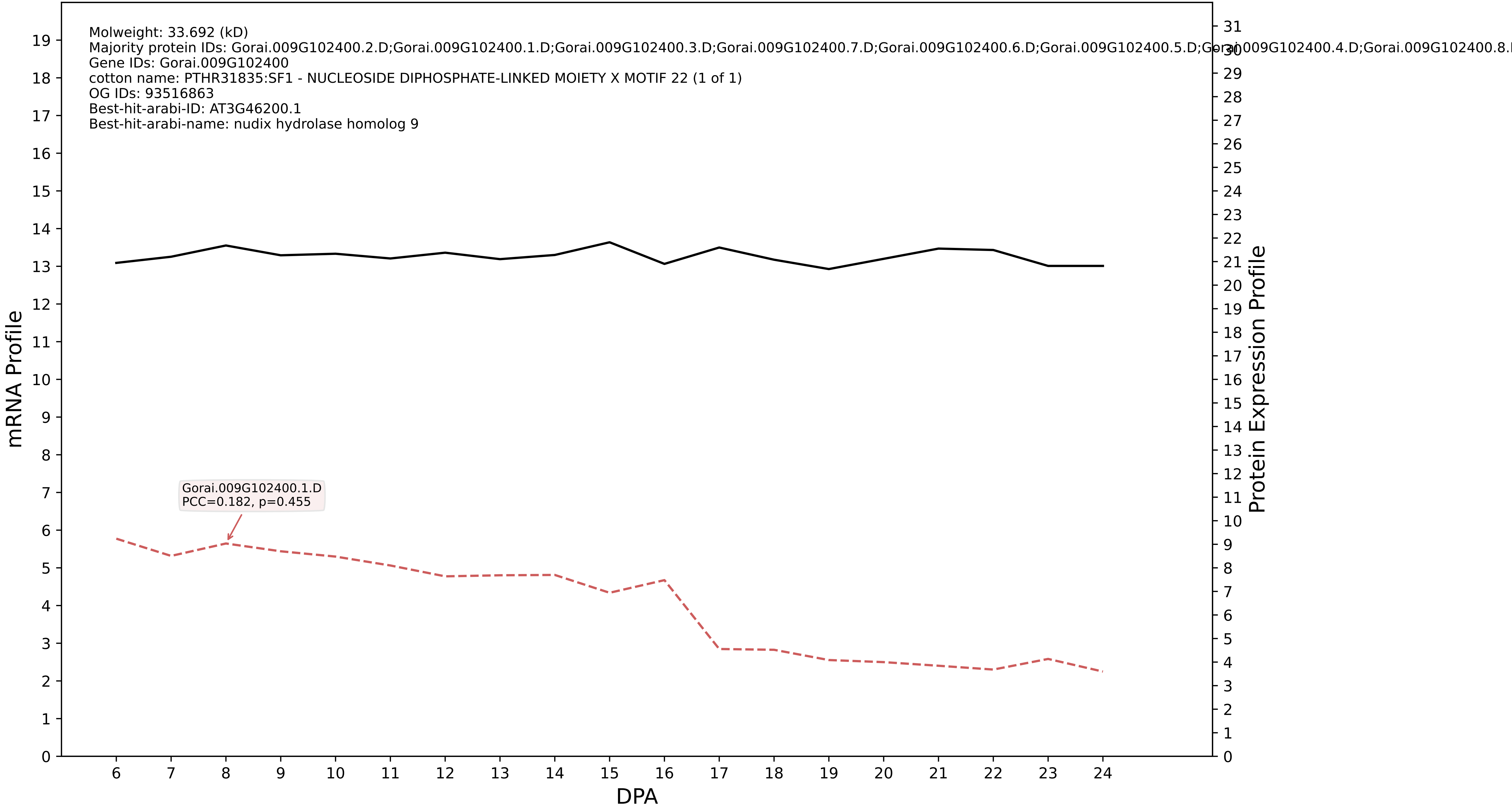

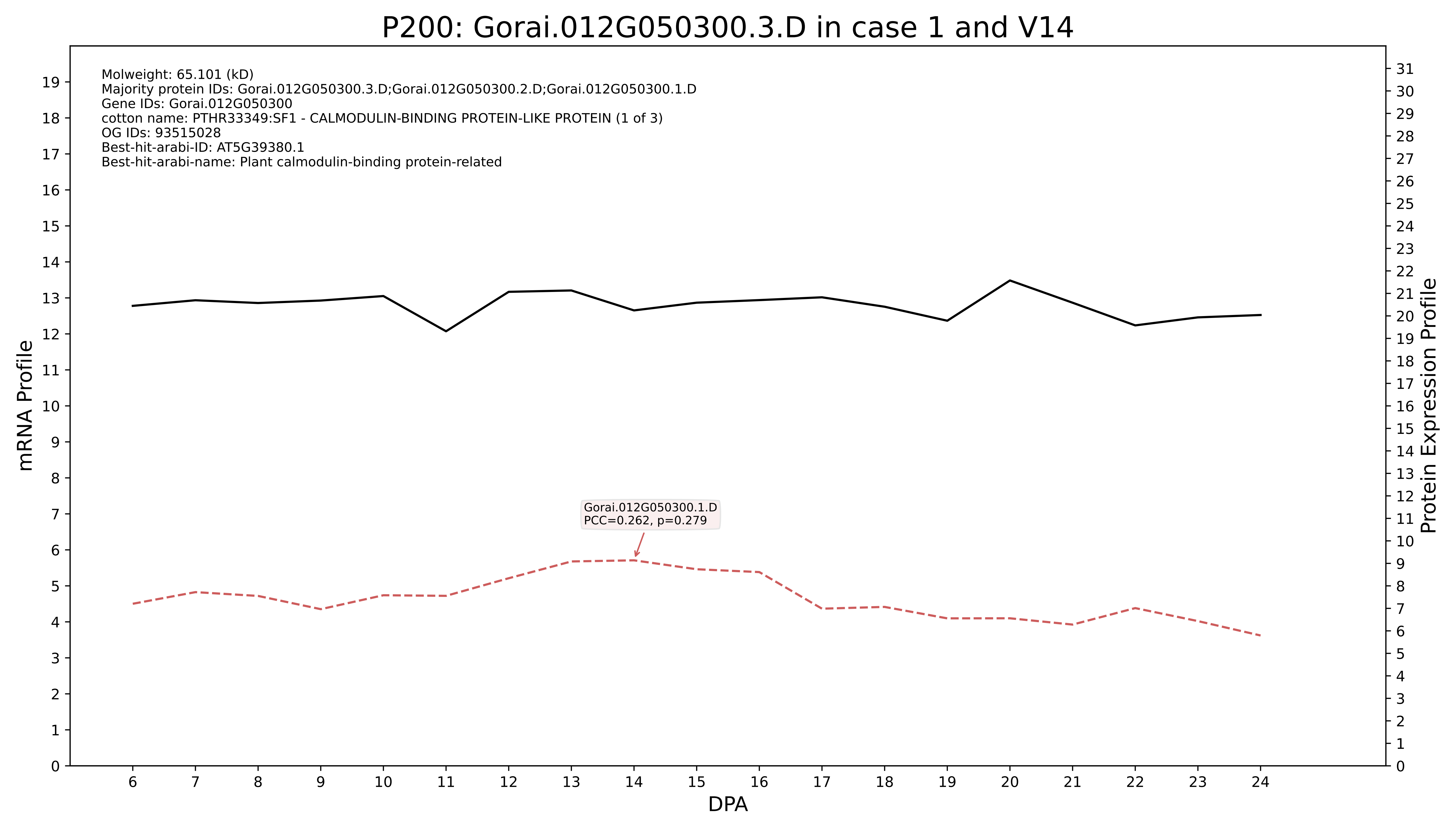
P200: Gorai.012G050300.3.D in case 1 and V14
Molweight: 65.101 (kD)
Majority protein IDs: Gorai.012G050300.3.D;Gorai.012G050300.2.D;Gorai.012G050300.1.D
Gene IDs: Gorai.012G050300
cotton name: PTHR33349:SF1 - CALMODULIN-BINDING PROTEIN-LIKE PROTEIN (1 of 3)
OG IDs: 93515028
Best-hit-arabi-ID: AT5G39380.1
Best-hit-arabi-name: Plant calmodulin-binding protein-related
Gorai.012G050300.1.D
PCC=0.262, p=0.279
mRNA Profile
Protein Expression Profile
DPA

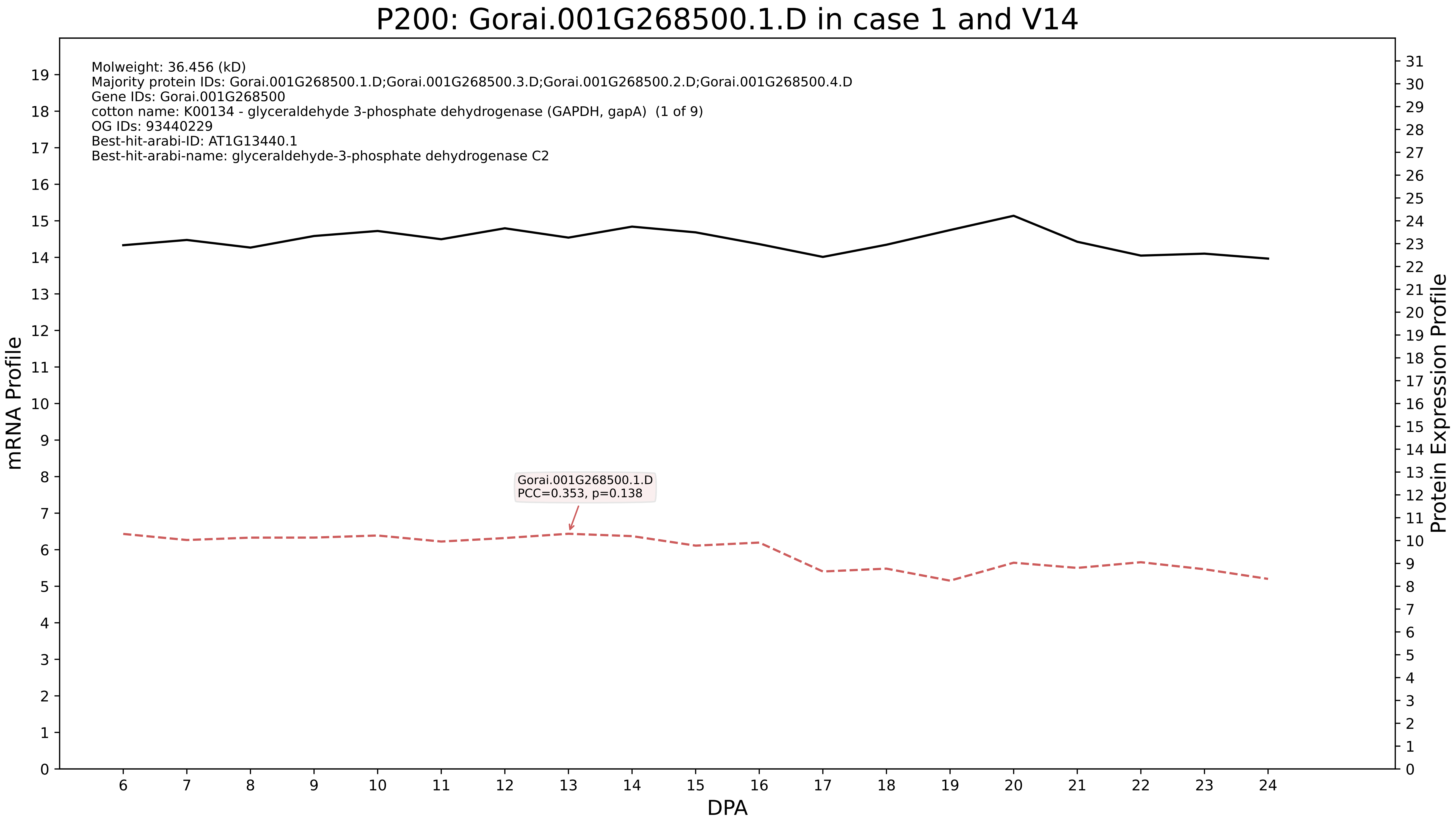
P200: Gorai.001G268500.1.D in case 1 and V14
Molweight: 36.456 (kD)
Majority protein IDs: Gorai.001G268500.1.D;Gorai.001G268500.3.D;Gorai.001G268500.2.D;Gorai.001G268500.4.D
Gene IDs: Gorai.001G268500
cotton name: K00134 - glyceraldehyde 3-phosphate dehydrogenase (GAPDH, gapA) (1 of 9)
OG IDs: 93440229
Best-hit-arabi-ID: AT1G13440.1
Best-hit-arabi-name: glyceraldehyde-3-phosphate dehydrogenase C2
Gorai.001G268500.1.D
PCC=0.353, p=0.138
mRNA Profile
Protein Expression Profile
DPA

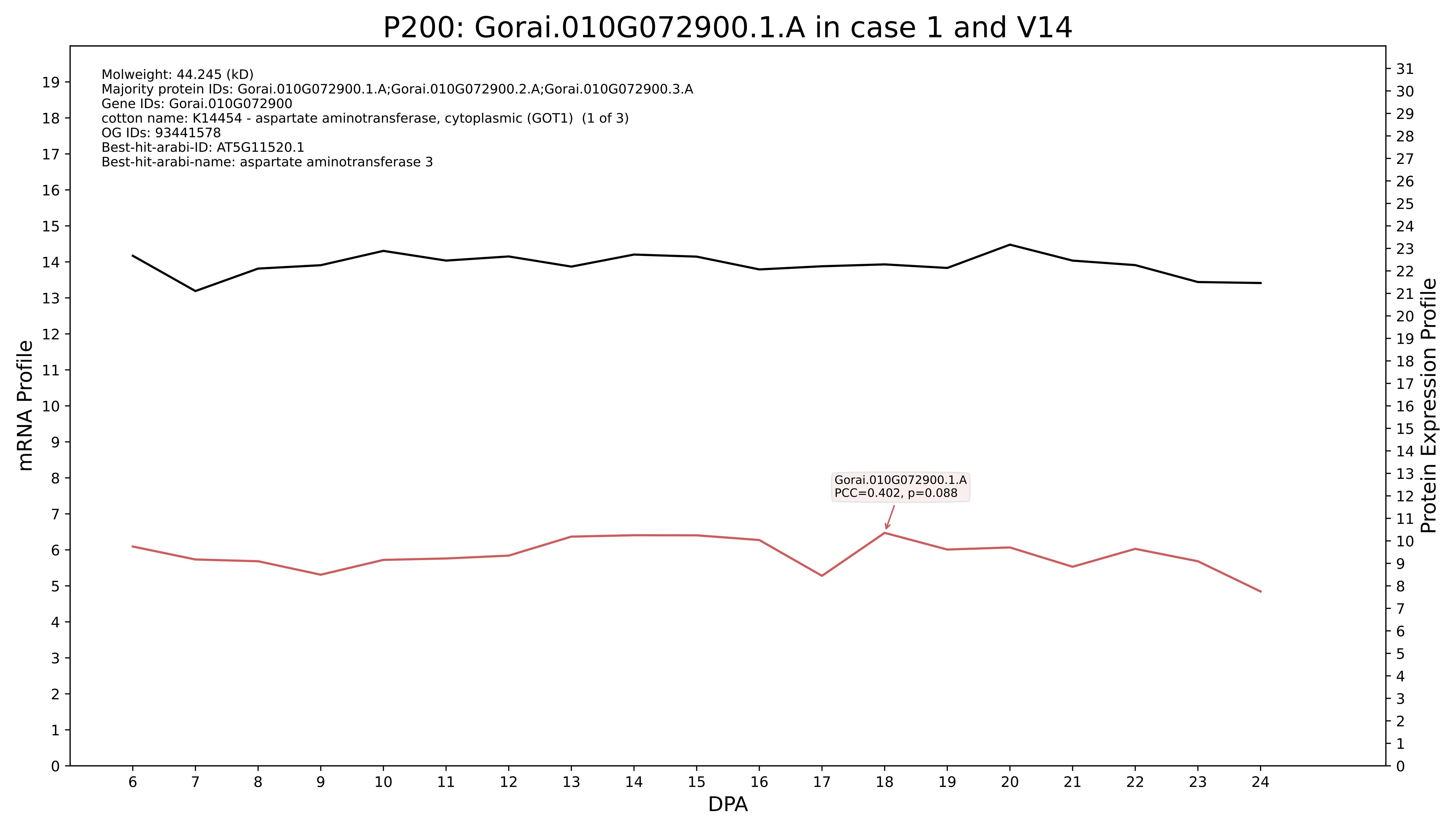
P200: Gorai.010G072900.1.A in case 1 and V14
Molweight: 44.245 (kD)
Majority protein IDs: Gorai.010G072900.1.A;Gorai.010G072900.2.A;Gorai.010G072900.3.A
Gene IDs: Gorai.010G072900
cotton name: K14454 - aspartate aminotransferase, cytoplasmic (GOT1) (1 of 3)
OG IDs: 93441578
Best-hit-arabi-ID: AT5G11520.1
Best-hit-arabi-name: aspartate aminotransferase 3
Gorai.010G072900.1.A
PCC=0.402, p=0.088
mRNA Profile
Protein Expression Profile
DPA

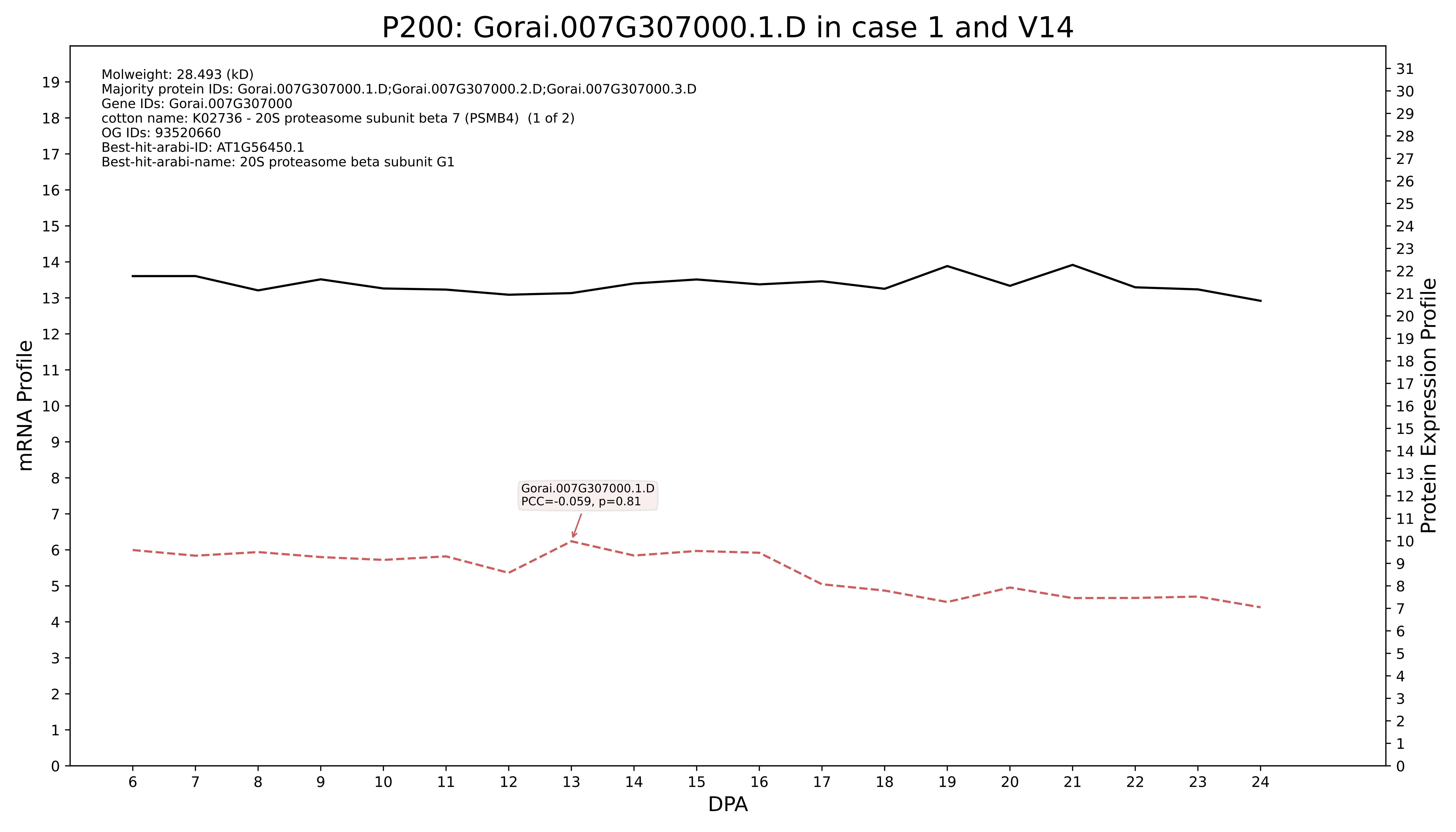
P200: Gorai.007G307000.1.D in case 1 and V14
Molweight: 28.493 (kD)
Majority protein IDs: Gorai.007G307000.1.D;Gorai.007G307000.2.D;Gorai.007G307000.3.D
Gene IDs: Gorai.007G307000
cotton name: K02736 - 20S proteasome subunit beta 7 (PSMB4) (1 of 2)
OG IDs: 93520660
Best-hit-arabi-ID: AT1G56450.1
Best-hit-arabi-name: 20S proteasome beta subunit G1
Gorai.007G307000.1.D
PCC=-0.059, p=0.81
mRNA Profile
Protein Expression Profile
DPA

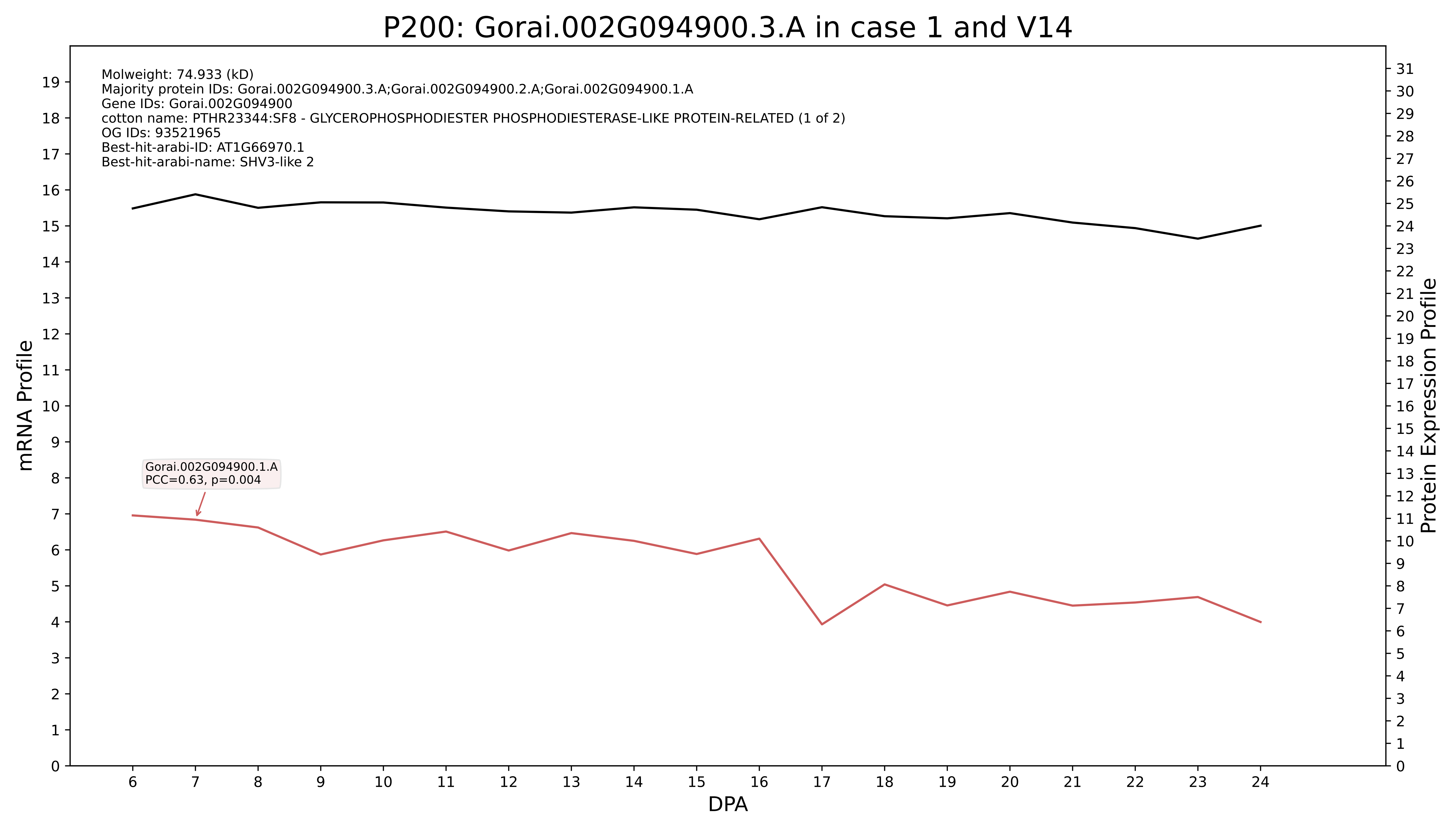
P200: Gorai.002G094900.3.A in case 1 and V14
Molweight: 74.933 (kD)
Majority protein IDs: Gorai.002G094900.3.A;Gorai.002G094900.2.A;Gorai.002G094900.1.A
Gene IDs: Gorai.002G094900
cotton name: PTHR23344:SF8 - GLYCEROPHOSPHODIESTER PHOSPHODIESTERASE-LIKE PROTEIN-RELATED (1 of 2)
OG IDs: 93521965
Best-hit-arabi-ID: AT1G66970.1
Best-hit-arabi-name: SHV3-like 2
Gorai.002G094900.1.A
PCC=0.63, p=0.004
mRNA Profile
Protein Expression Profile
DPA

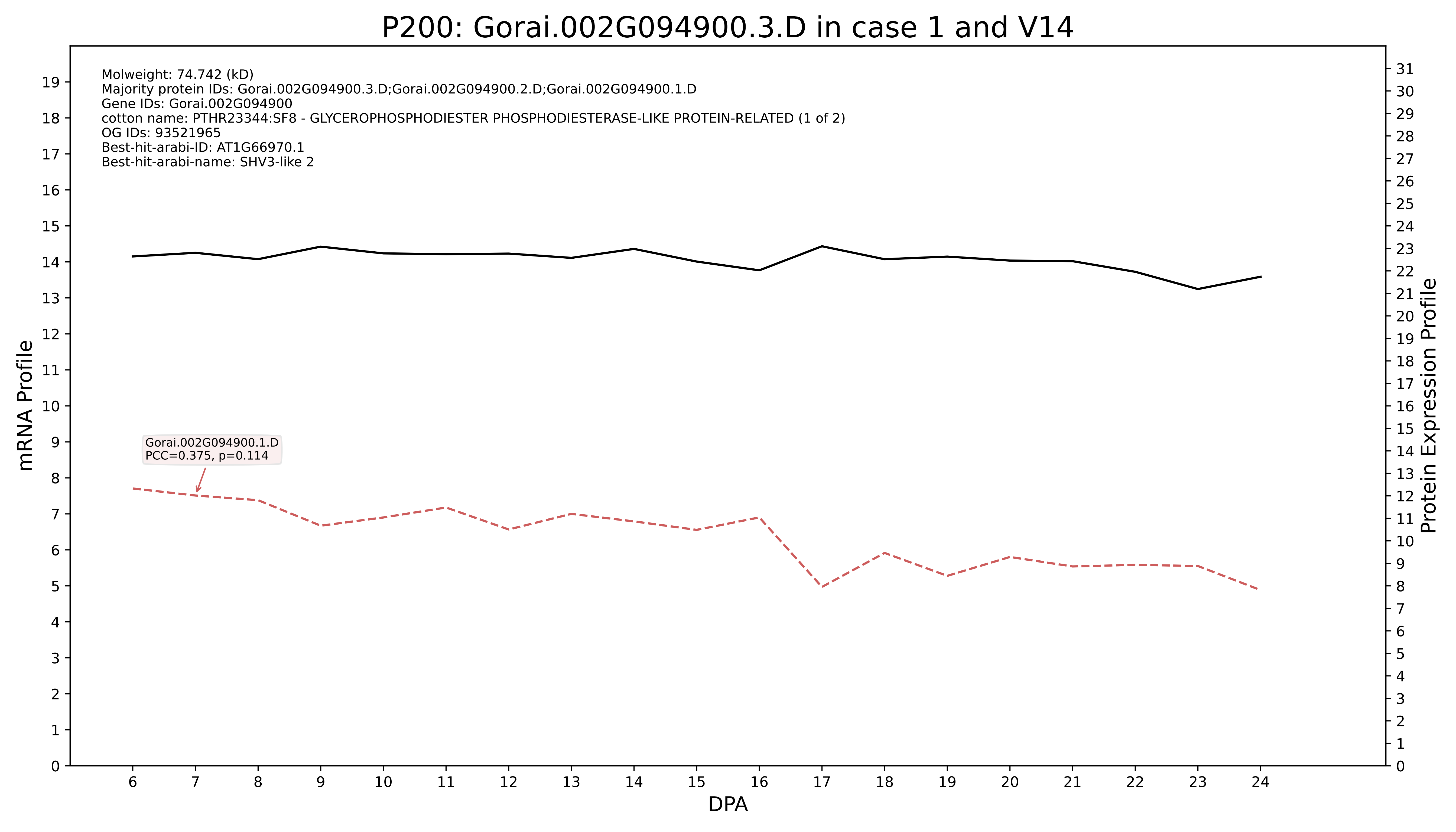
P200: Gorai.002G094900.3.D in case 1 and V14
Molweight: 74.742 (kD)
Majority protein IDs: Gorai.002G094900.3.D;Gorai.002G094900.2.D;Gorai.002G094900.1.D
Gene IDs: Gorai.002G094900
cotton name: PTHR23344:SF8 - GLYCEROPHOSPHODIESTER PHOSPHODIESTERASE-LIKE PROTEIN-RELATED (1 of 2)
OG IDs: 93521965
Best-hit-arabi-ID: AT1G66970.1
Best-hit-arabi-name: SHV3-like 2
Gorai.002G094900.1.D
PCC=0.375, p=0.114
mRNA Profile
Protein Expression Profile
DPA

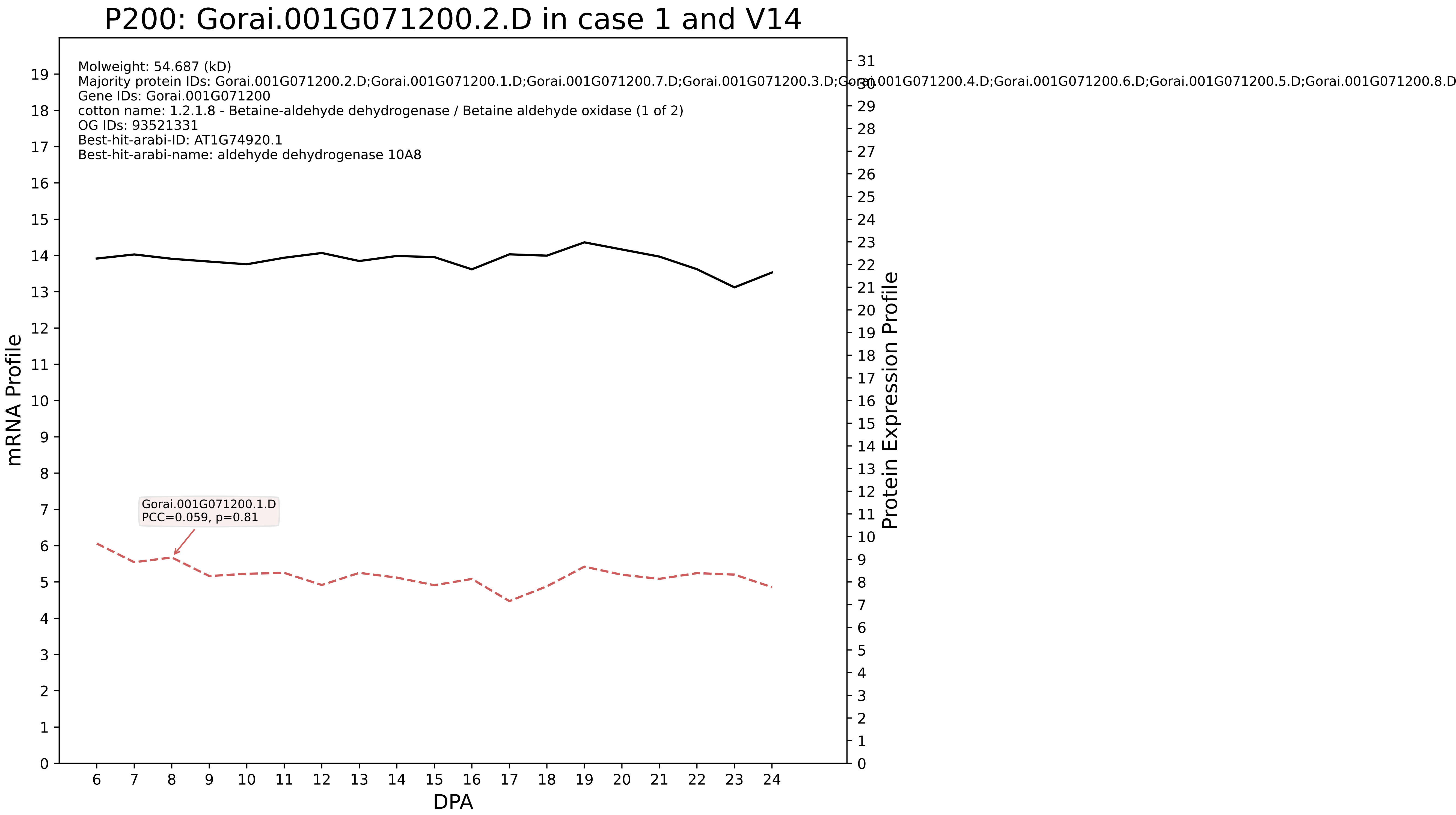
P200: Gorai.001G071200.2.D in case 1 and V14
Molweight: 54.687 (kD)
Majority protein IDs: Gorai.001G071200.2.D;Gorai.001G071200.1.D;Gorai.001G071200.7.D;Gorai.001G071200.3.D;Gorai.001G071200.4.D;Gorai.001G071200.6.D;Gorai.001G071200.5.D;Gorai.001G071200.8.D
Gene IDs: Gorai.001G071200
cotton name: 1.2.1.8 - Betaine-aldehyde dehydrogenase / Betaine aldehyde oxidase (1 of 2)
OG IDs: 93521331
Best-hit-arabi-ID: AT1G74920.1
Best-hit-arabi-name: aldehyde dehydrogenase 10A8
Gorai.001G071200.1.D
PCC=0.059, p=0.81
mRNA Profile
Protein Expression Profile
DPA

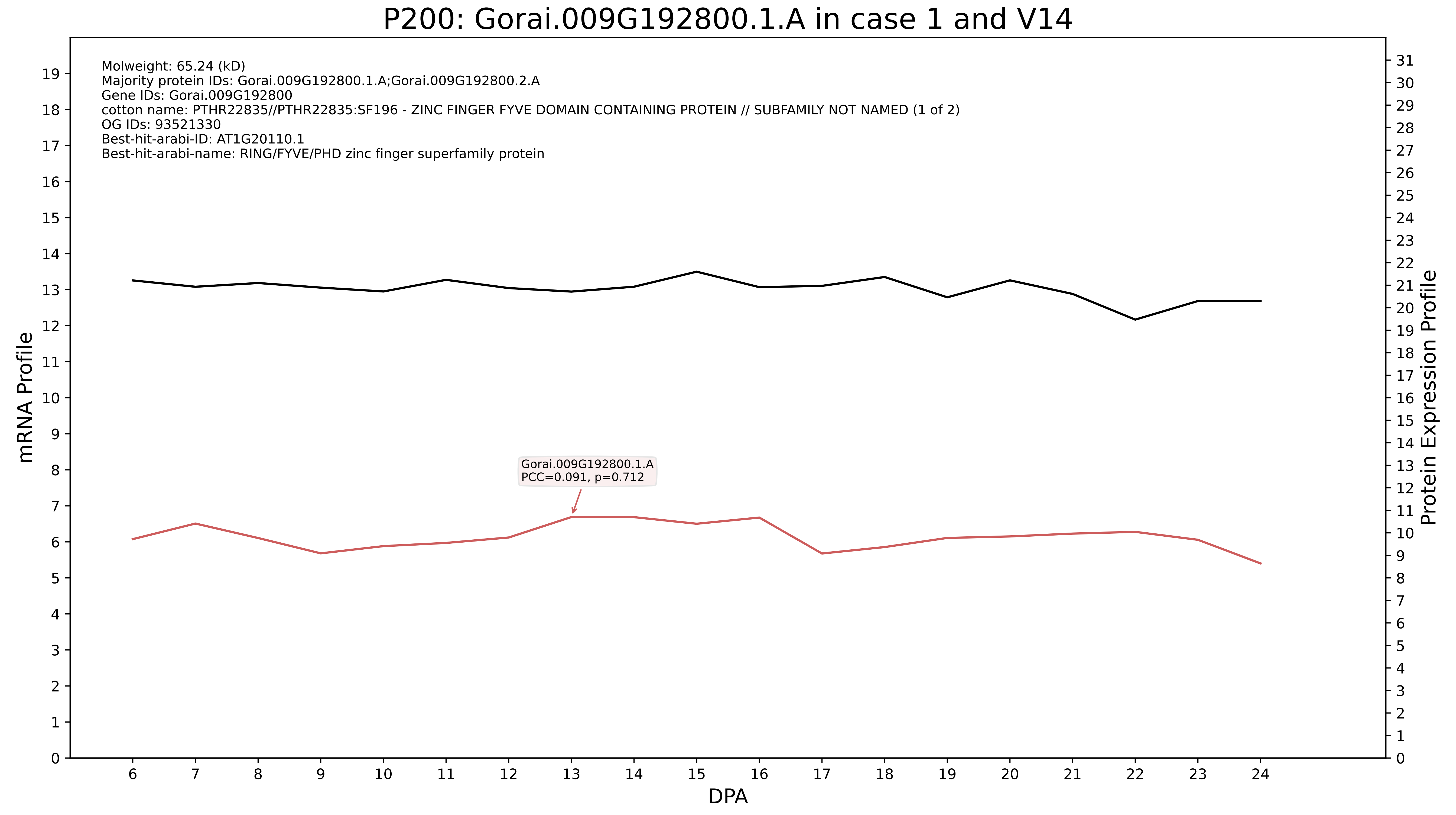

P200: Gorai.009G192800.1.A in case 1 and V14
Molweight: 65.24 (kD)
Majority protein IDs: Gorai.009G192800.1.A;Gorai.009G192800.2.A
Gene IDs: Gorai.009G192800
cotton name: PTHR22835//PTHR22835:SF196 - ZINC FINGER FYVE DOMAIN CONTAINING PROTEIN // SUBFAMILY NOT NAMED (1 of 2)
OG IDs: 93521330
Best-hit-arabi-ID: AT1G20110.1
Best-hit-arabi-name: RING/FYVE/PHD zinc finger superfamily protein
Gorai.009G192800.1.A
PCC=0.091, p=0.712
mRNA Profile
Protein Expression Profile
DPA

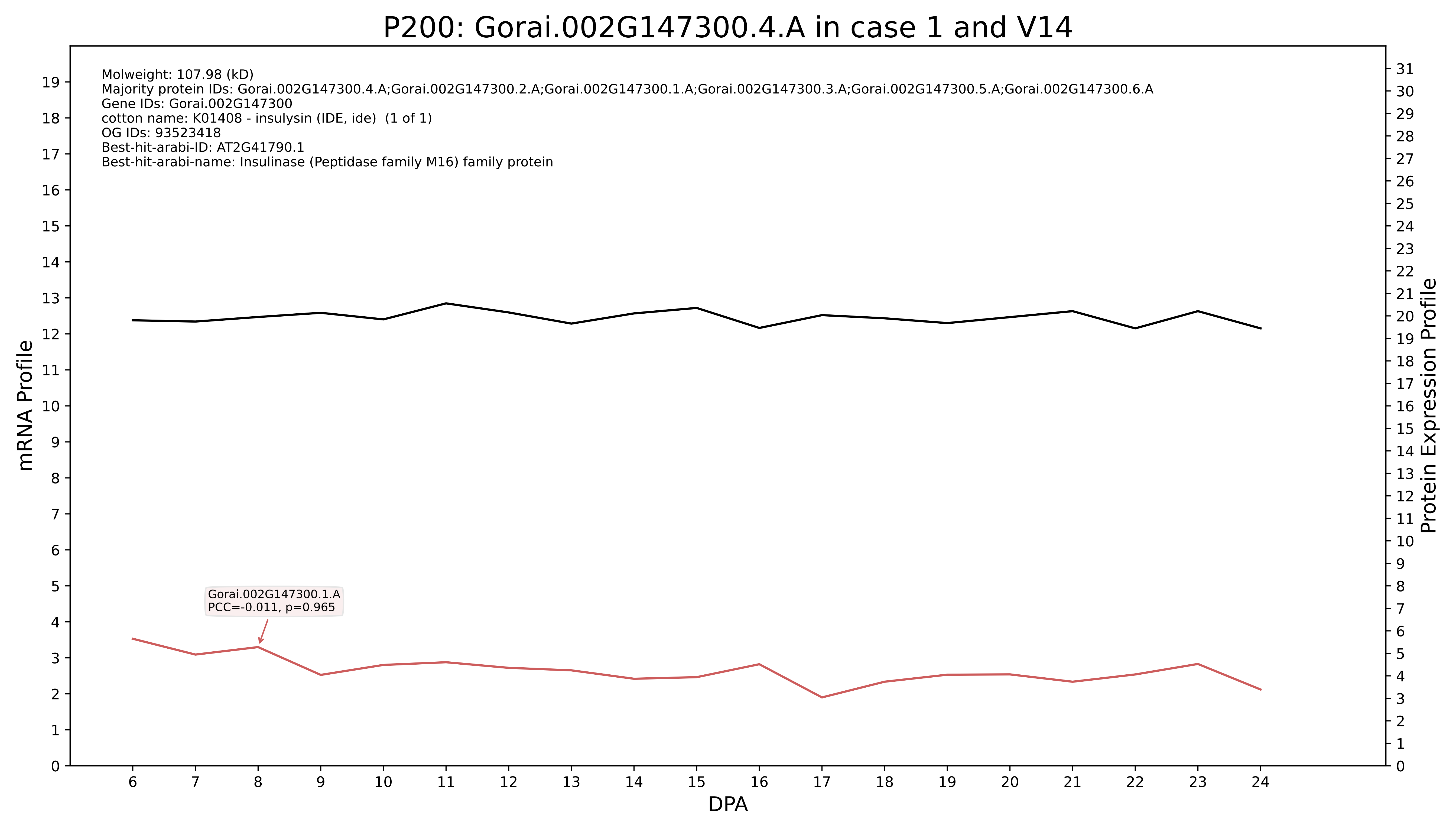
P200: Gorai.002G147300.4.A in case 1 and V14
Molweight: 107.98 (kD)
Majority protein IDs: Gorai.002G147300.4.A;Gorai.002G147300.2.A;Gorai.002G147300.1.A;Gorai.002G147300.3.A;Gorai.002G147300.5.A;Gorai.002G147300.6.A
Gene IDs: Gorai.002G147300
cotton name: K01408 - insulysin (IDE, ide) (1 of 1)
OG IDs: 93523418
Best-hit-arabi-ID: AT2G41790.1
Best-hit-arabi-name: Insulinase (Peptidase family M16) family protein
Gorai.002G147300.1.A
PCC=-0.011, p=0.965
mRNA Profile
Protein Expression Profile
DPA

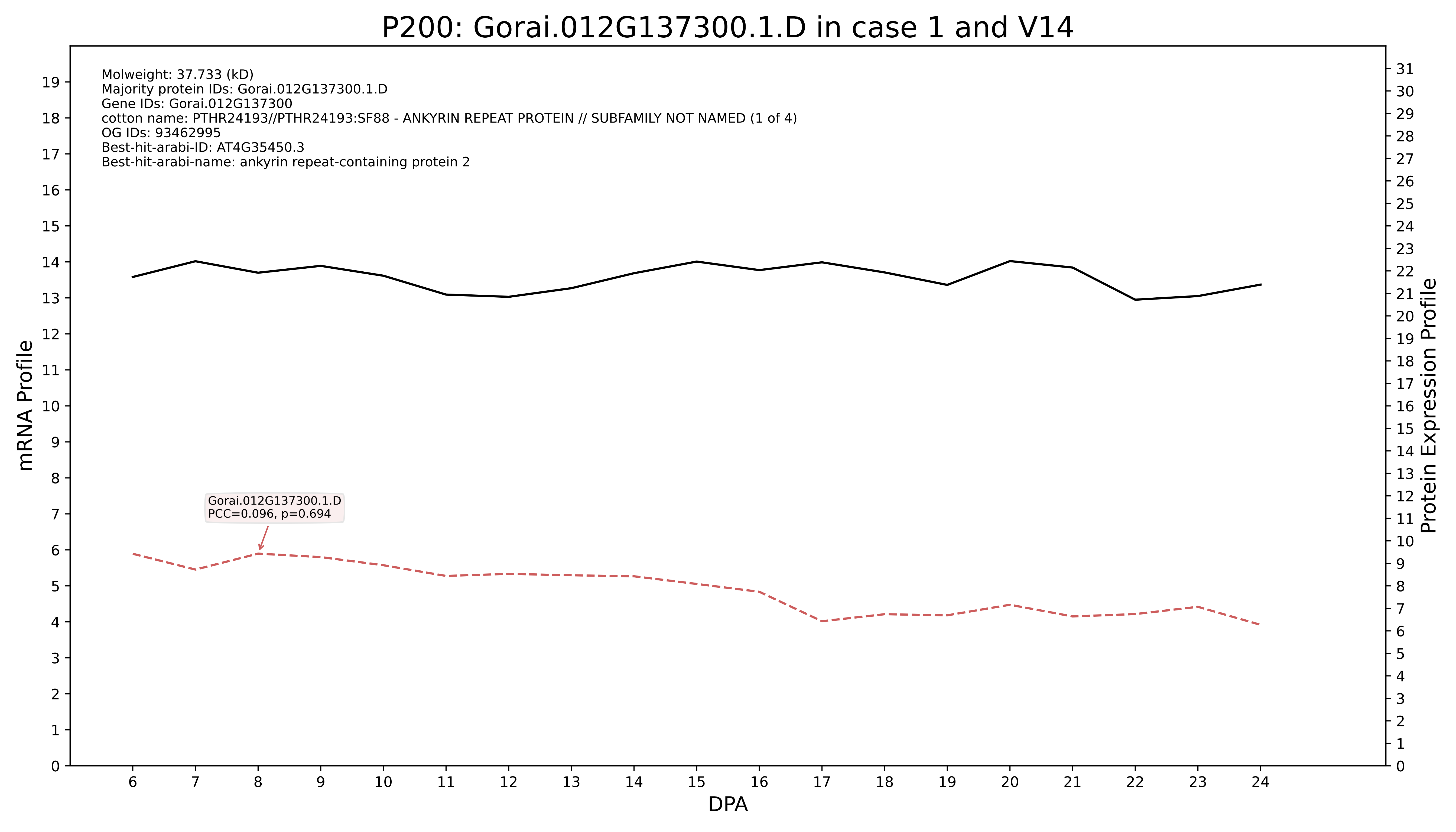
P200: Gorai.012G137300.1.D in case 1 and V14
Molweight: 37.733 (kD)
Majority protein IDs: Gorai.012G137300.1.D
Gene IDs: Gorai.012G137300
cotton name: PTHR24193//PTHR24193:SF88 - ANKYRIN REPEAT PROTEIN // SUBFAMILY NOT NAMED (1 of 4)
OG IDs: 93462995
Best-hit-arabi-ID: AT4G35450.3
Best-hit-arabi-name: ankyrin repeat-containing protein 2
Gorai.012G137300.1.D
PCC=0.096, p=0.694
mRNA Profile
Protein Expression Profile
DPA

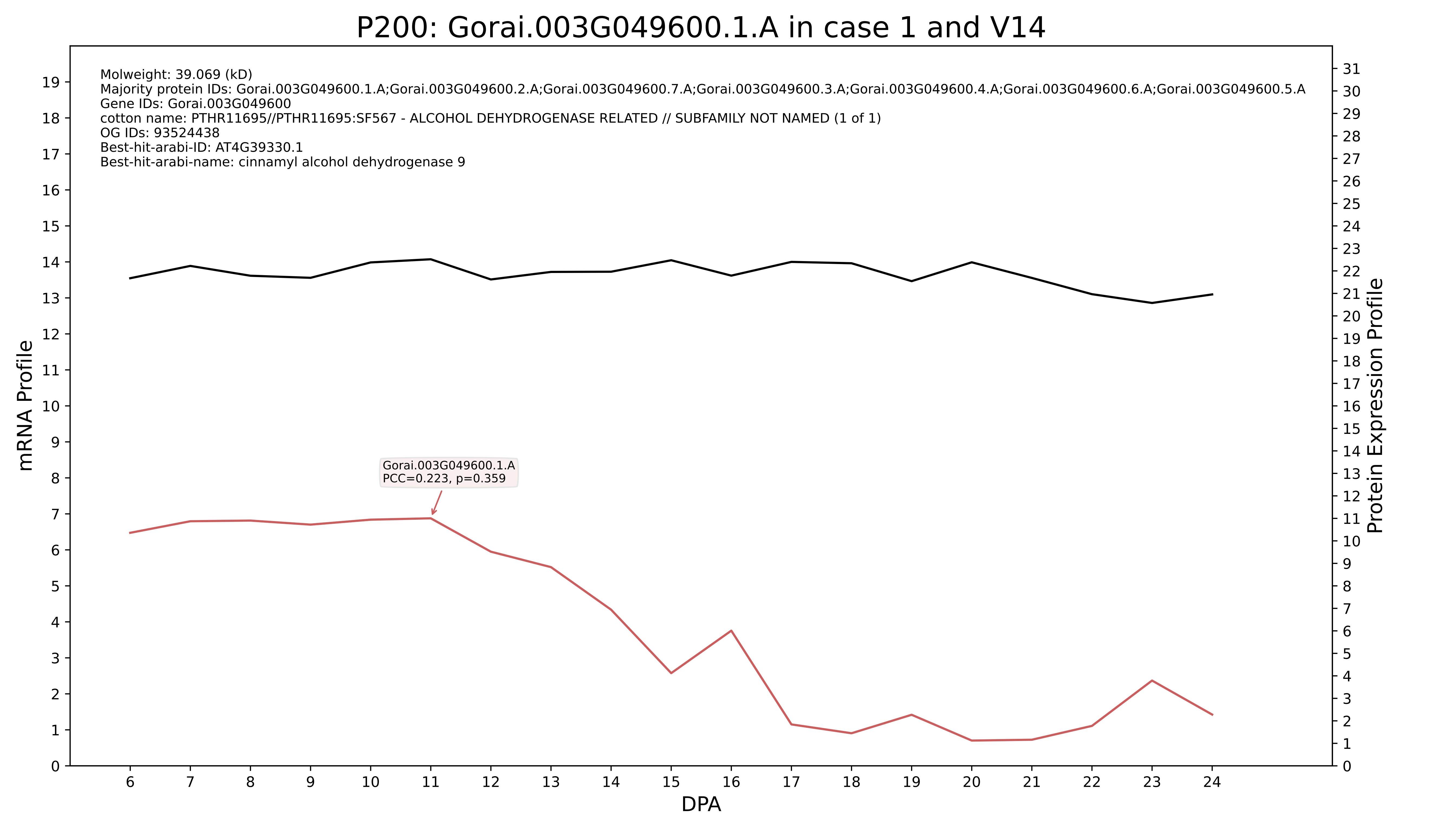
P200: Gorai.003G049600.1.A in case 1 and V14
Molweight: 39.069 (kD)
Majority protein IDs: Gorai.003G049600.1.A;Gorai.003G049600.2.A;Gorai.003G049600.7.A;Gorai.003G049600.3.A;Gorai.003G049600.4.A;Gorai.003G049600.6.A;Gorai.003G049600.5.A
Gene IDs: Gorai.003G049600
cotton name: PTHR11695//PTHR11695:SF567 - ALCOHOL DEHYDROGENASE RELATED // SUBFAMILY NOT NAMED (1 of 1)
OG IDs: 93524438
Best-hit-arabi-ID: AT4G39330.1
Best-hit-arabi-name: cinnamyl alcohol dehydrogenase 9
Gorai.003G049600.1.A
PCC=0.223, p=0.359
mRNA Profile
Protein Expression Profile
DPA

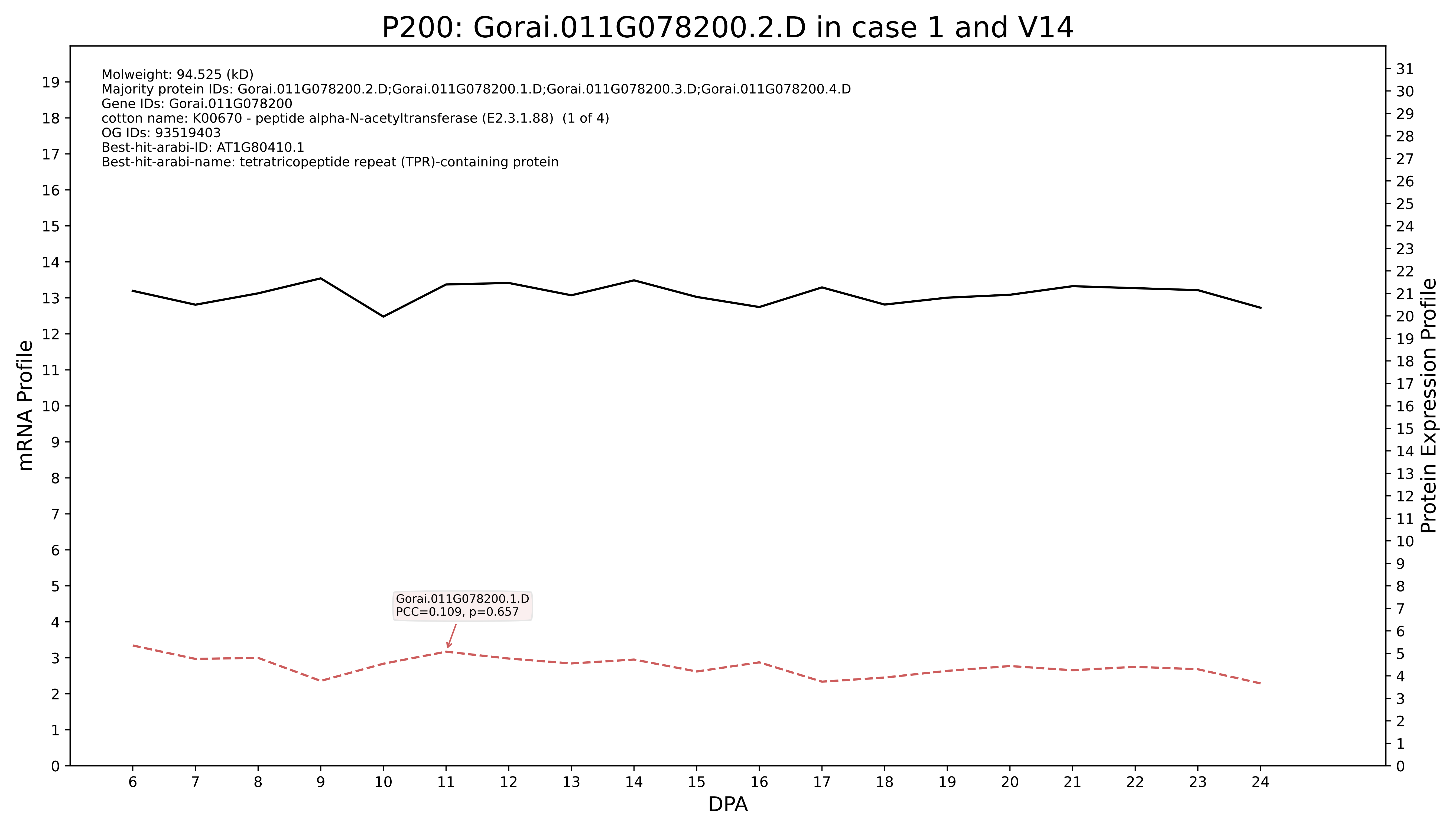
P200: Gorai.011G078200.2.D in case 1 and V14
Molweight: 94.525 (kD)
Majority protein IDs: Gorai.011G078200.2.D;Gorai.011G078200.1.D;Gorai.011G078200.3.D;Gorai.011G078200.4.D
Gene IDs: Gorai.011G078200
cotton name: K00670 - peptide alpha-N-acetyltransferase (E2.3.1.88) (1 of 4)
OG IDs: 93519403
Best-hit-arabi-ID: AT1G80410.1
Best-hit-arabi-name: tetratricopeptide repeat (TPR)-containing protein
Gorai.011G078200.1.D
PCC=0.109, p=0.657
mRNA Profile
Protein Expression Profile
DPA

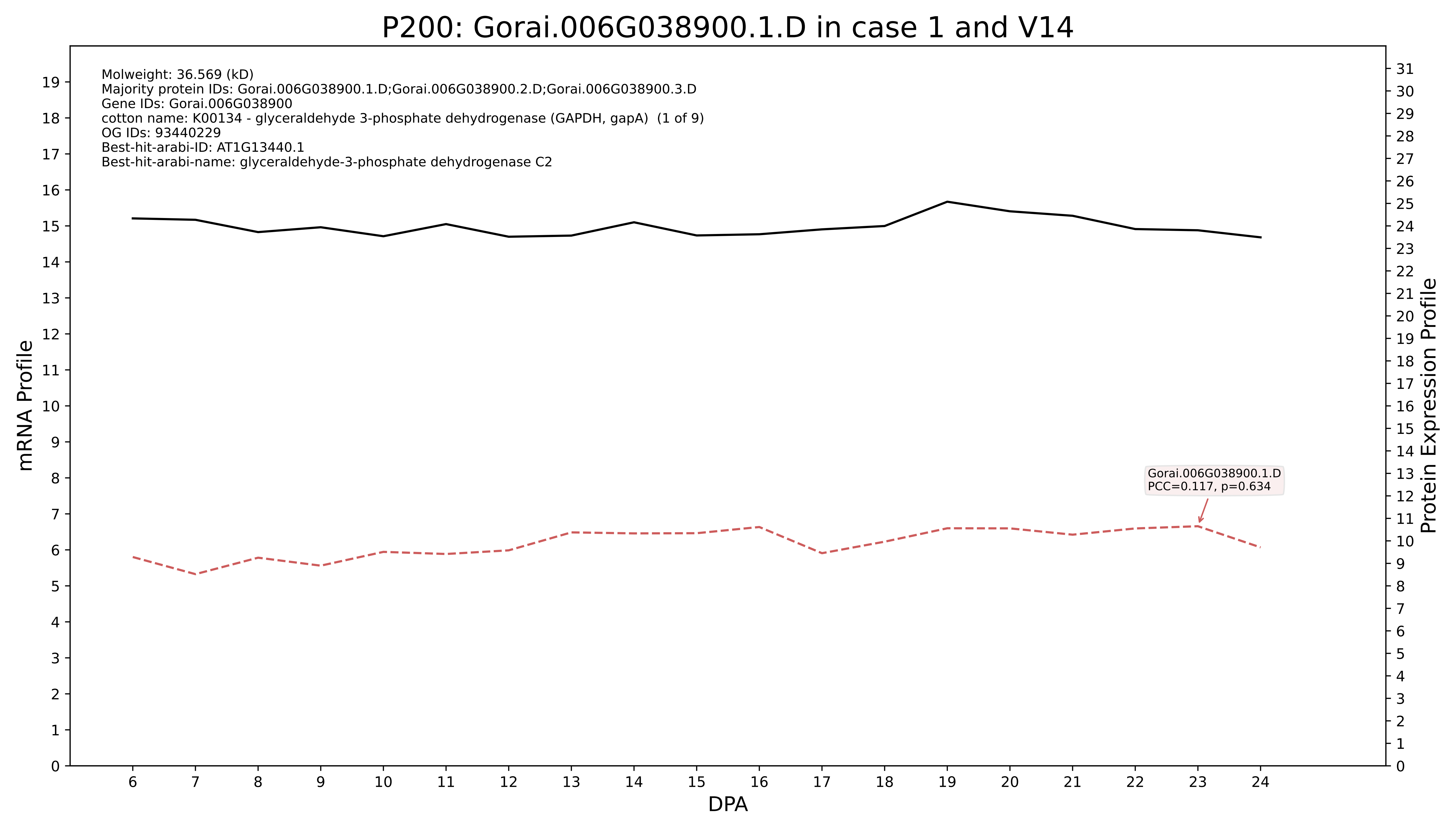
P200: Gorai.006G038900.1.D in case 1 and V14
Molweight: 36.569 (kD)
Majority protein IDs: Gorai.006G038900.1.D;Gorai.006G038900.2.D;Gorai.006G038900.3.D
Gene IDs: Gorai.006G038900
cotton name: K00134 - glyceraldehyde 3-phosphate dehydrogenase (GAPDH, gapA) (1 of 9)
OG IDs: 93440229
Best-hit-arabi-ID: AT1G13440.1
Best-hit-arabi-name: glyceraldehyde-3-phosphate dehydrogenase C2
Gorai.006G038900.1.D
PCC=0.117, p=0.634
mRNA Profile
Protein Expression Profile
DPA

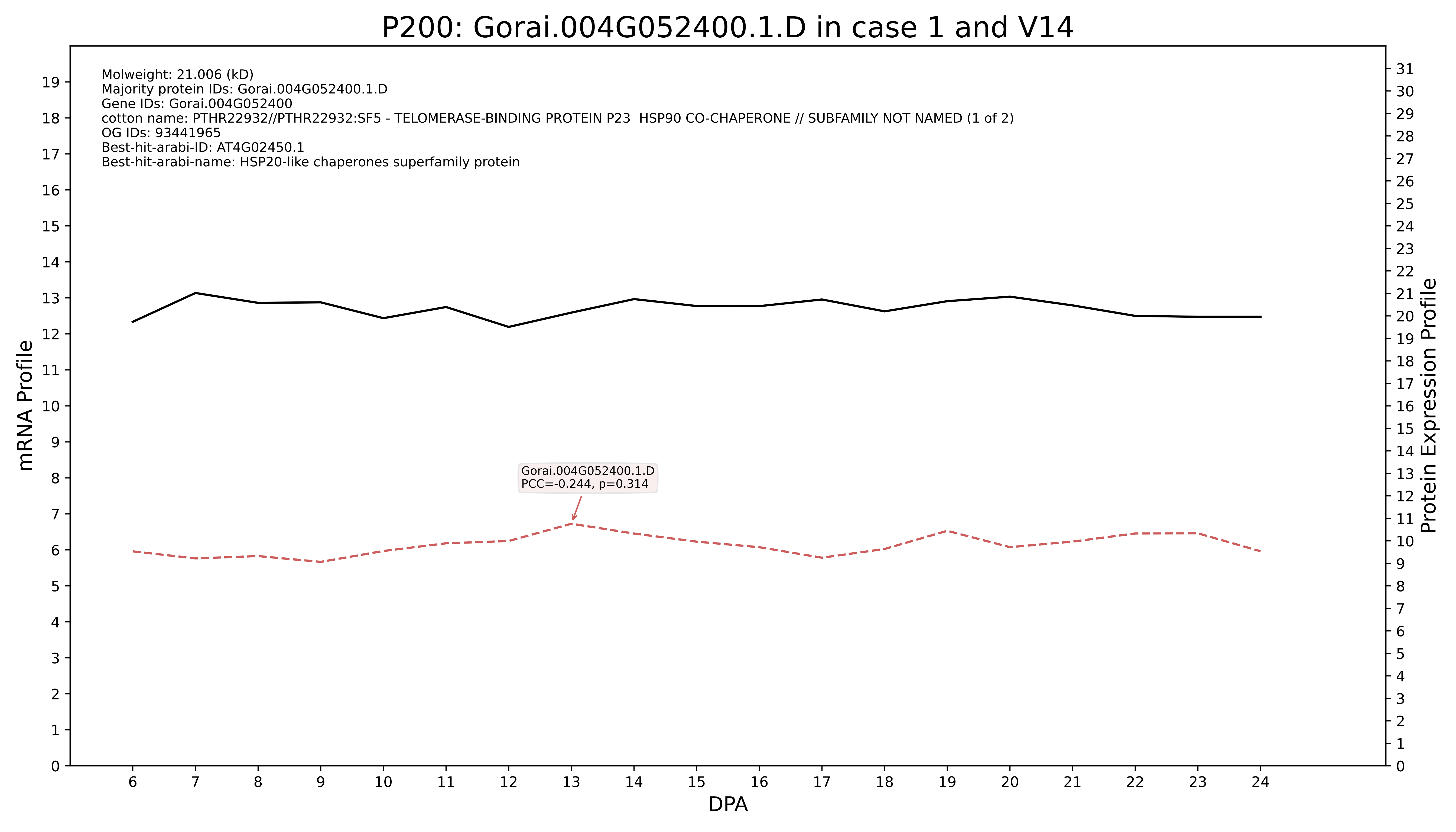
P200: Gorai.004G052400.1.D in case 1 and V14
Molweight: 21.006 (kD)
Majority protein IDs: Gorai.004G052400.1.D
Gene IDs: Gorai.004G052400
cotton name: PTHR22932//PTHR22932:SF5 - TELOMERASE-BINDING PROTEIN P23 HSP90 CO-CHAPERONE // SUBFAMILY NOT NAMED (1 of 2)
OG IDs: 93441965
Best-hit-arabi-ID: AT4G02450.1
Best-hit-arabi-name: HSP20-like chaperones superfamily protein
Gorai.004G052400.1.D
PCC=-0.244, p=0.314
mRNA Profile
Protein Expression Profile
DPA

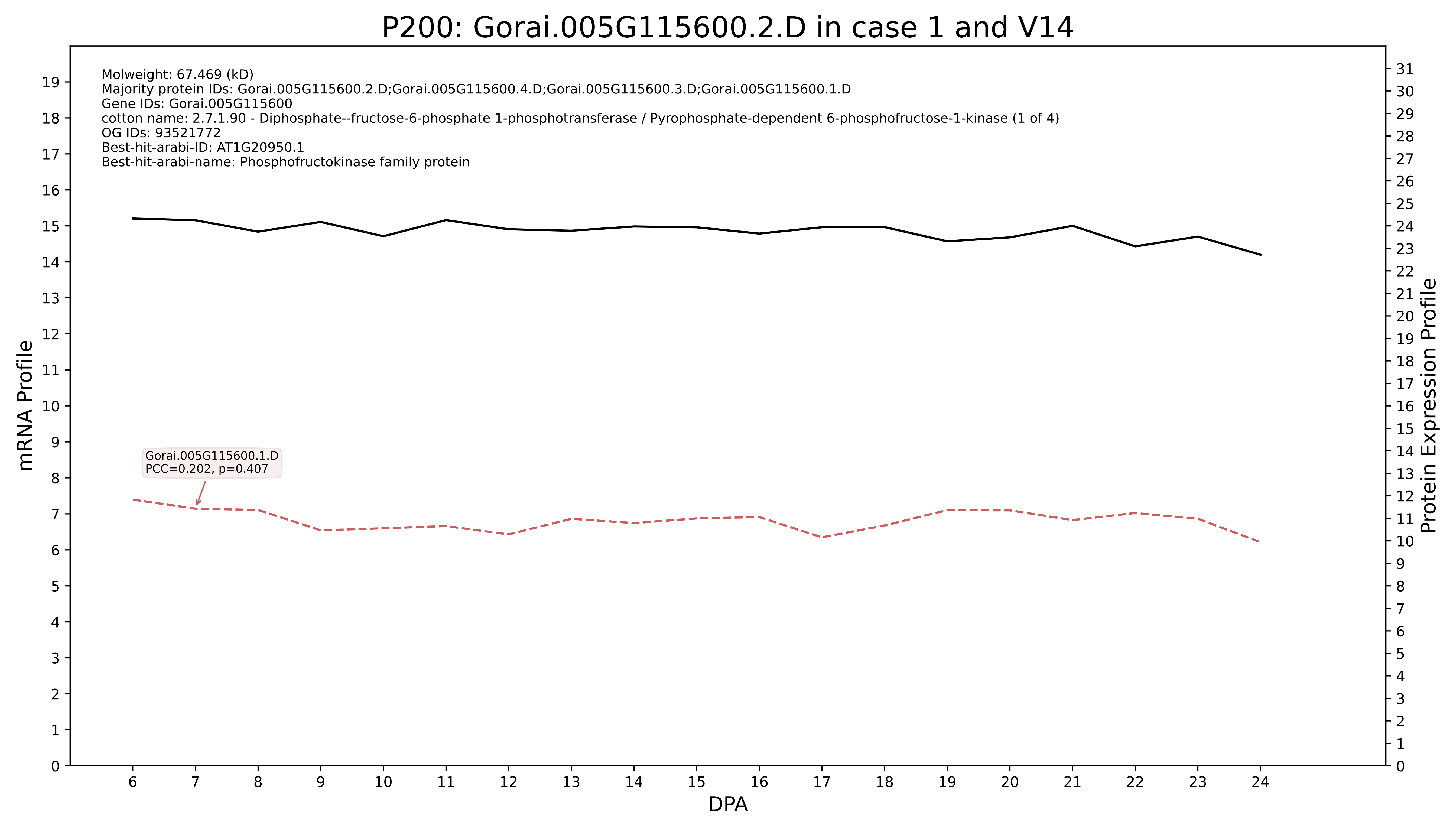
P200: Gorai.005G115600.2.D in case 1 and V14
Molweight: 67.469 (kD)
Majority protein IDs: Gorai.005G115600.2.D;Gorai.005G115600.4.D;Gorai.005G115600.3.D;Gorai.005G115600.1.D
Gene IDs: Gorai.005G115600
cotton name: 2.7.1.90 - Diphosphate--fructose-6-phosphate 1-phosphotransferase / Pyrophosphate-dependent 6-phosphofructose-1-kinase (1 of 4)
OG IDs: 93521772
Best-hit-arabi-ID: AT1G20950.1
Best-hit-arabi-name: Phosphofructokinase family protein
Gorai.005G115600.1.D
PCC=0.202, p=0.407
mRNA Profile
Protein Expression Profile
DPA

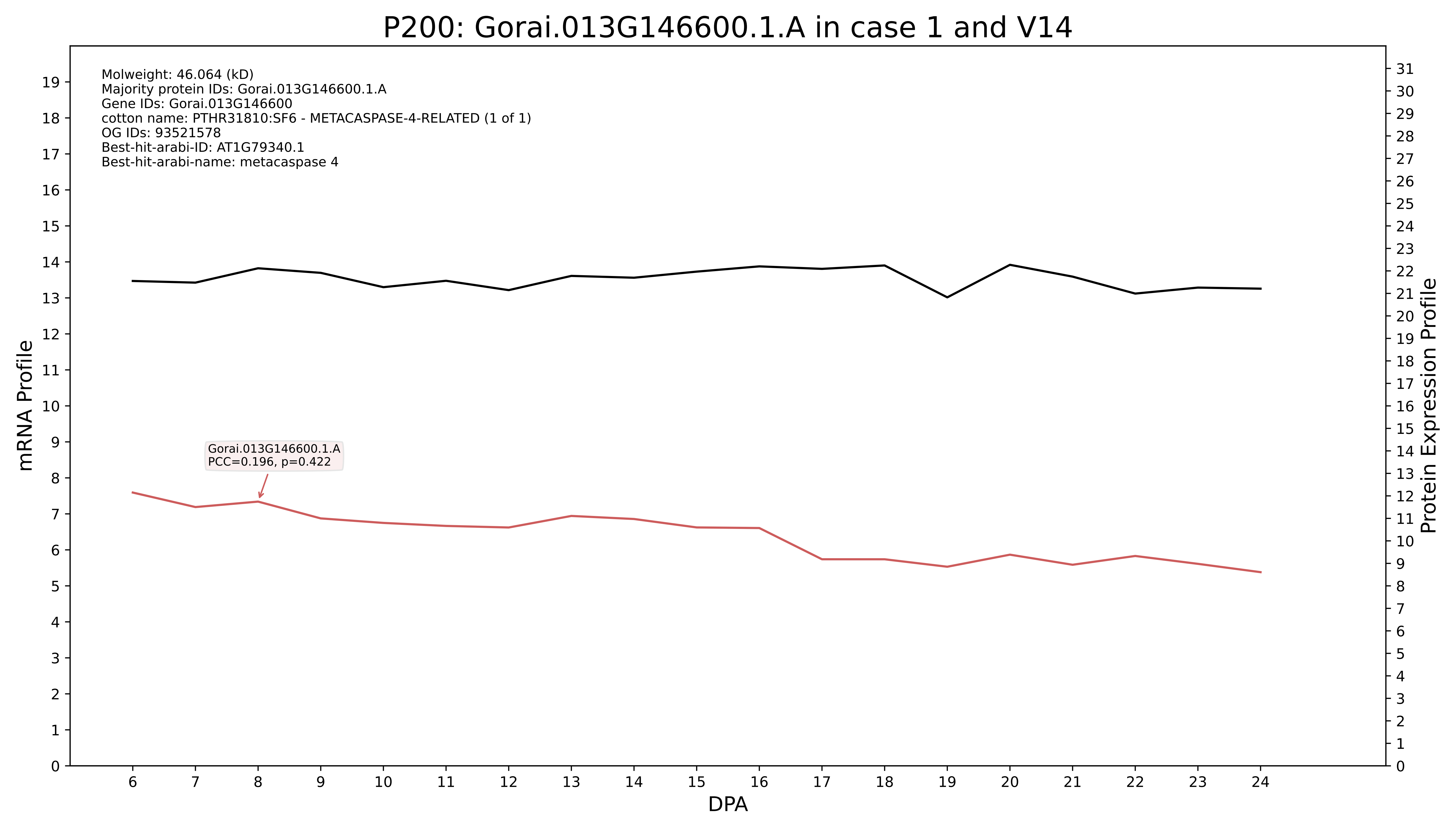
P200: Gorai.013G146600.1.A in case 1 and V14
Molweight: 46.064 (kD)
Majority protein IDs: Gorai.013G146600.1.A
Gene IDs: Gorai.013G146600
cotton name: PTHR31810:SF6 - METACASPASE-4-RELATED (1 of 1)
OG IDs: 93521578
Best-hit-arabi-ID: AT1G79340.1
Best-hit-arabi-name: metacaspase 4
Gorai.013G146600.1.A
PCC=0.196, p=0.422
mRNA Profile
Protein Expression Profile
DPA

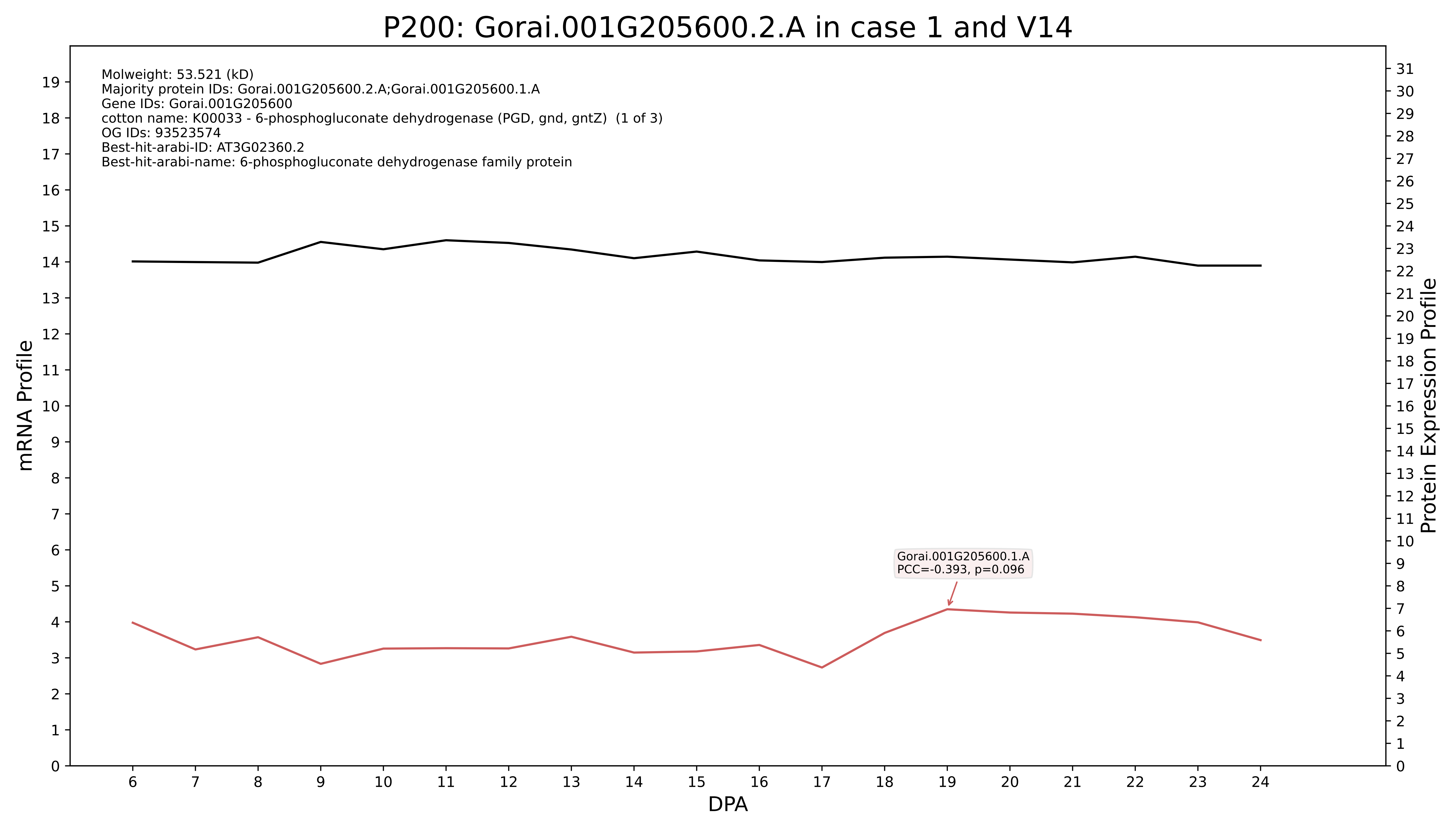
P200: Gorai.001G205600.2.A in case 1 and V14
Molweight: 53.521 (kD)
Majority protein IDs: Gorai.001G205600.2.A;Gorai.001G205600.1.A
Gene IDs: Gorai.001G205600
cotton name: K00033 - 6-phosphogluconate dehydrogenase (PGD, gnd, gntZ) (1 of 3)
OG IDs: 93523574
Best-hit-arabi-ID: AT3G02360.2
Best-hit-arabi-name: 6-phosphogluconate dehydrogenase family protein
Gorai.001G205600.1.A
PCC=-0.393, p=0.096
mRNA Profile
Protein Expression Profile
DPA

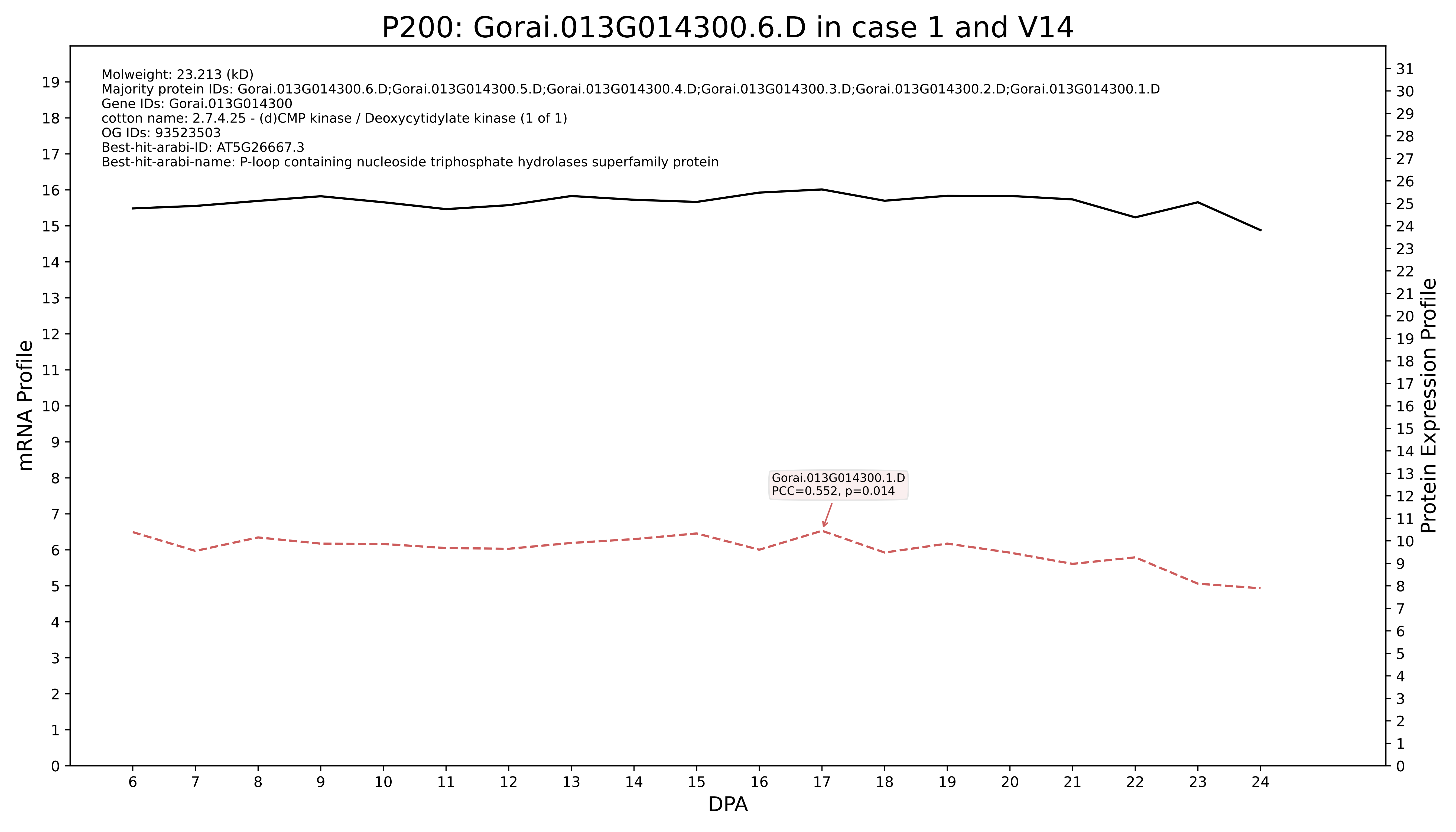

P200: Gorai.013G014300.6.D in case 1 and V14
Molweight: 23.213 (kD)
Majority protein IDs: Gorai.013G014300.6.D;Gorai.013G014300.5.D;Gorai.013G014300.4.D;Gorai.013G014300.3.D;Gorai.013G014300.2.D;Gorai.013G014300.1.D
Gene IDs: Gorai.013G014300
cotton name: 2.7.4.25 - (d)CMP kinase / Deoxycytidylate kinase (1 of 1)
OG IDs: 93523503
Best-hit-arabi-ID: AT5G26667.3
Best-hit-arabi-name: P-loop containing nucleoside triphosphate hydrolases superfamily protein
Gorai.013G014300.1.D
PCC=0.552, p=0.014
mRNA Profile
Protein Expression Profile
DPA

### P200: Gorai.007G011200.1.D in case 1 and V14

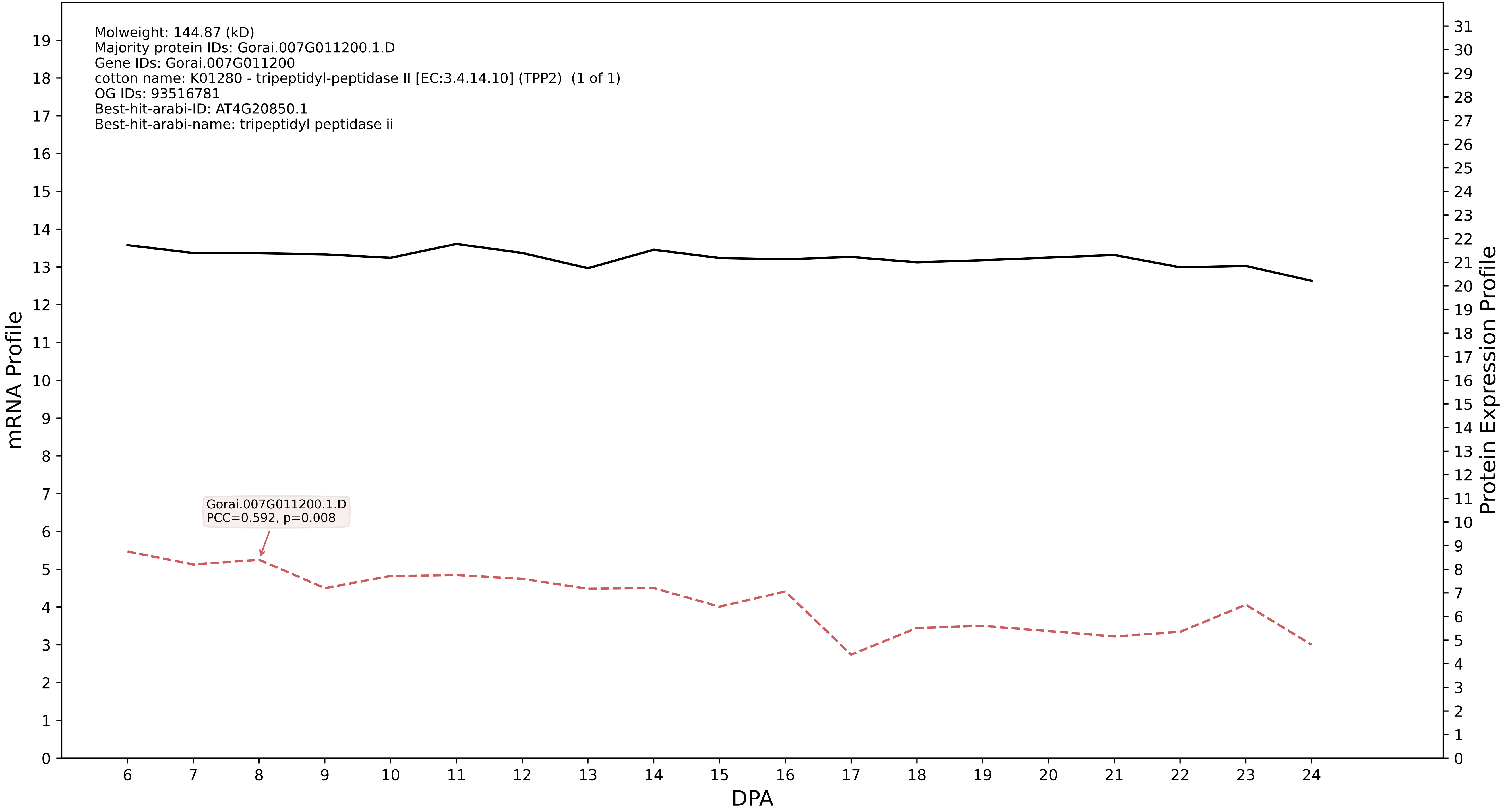

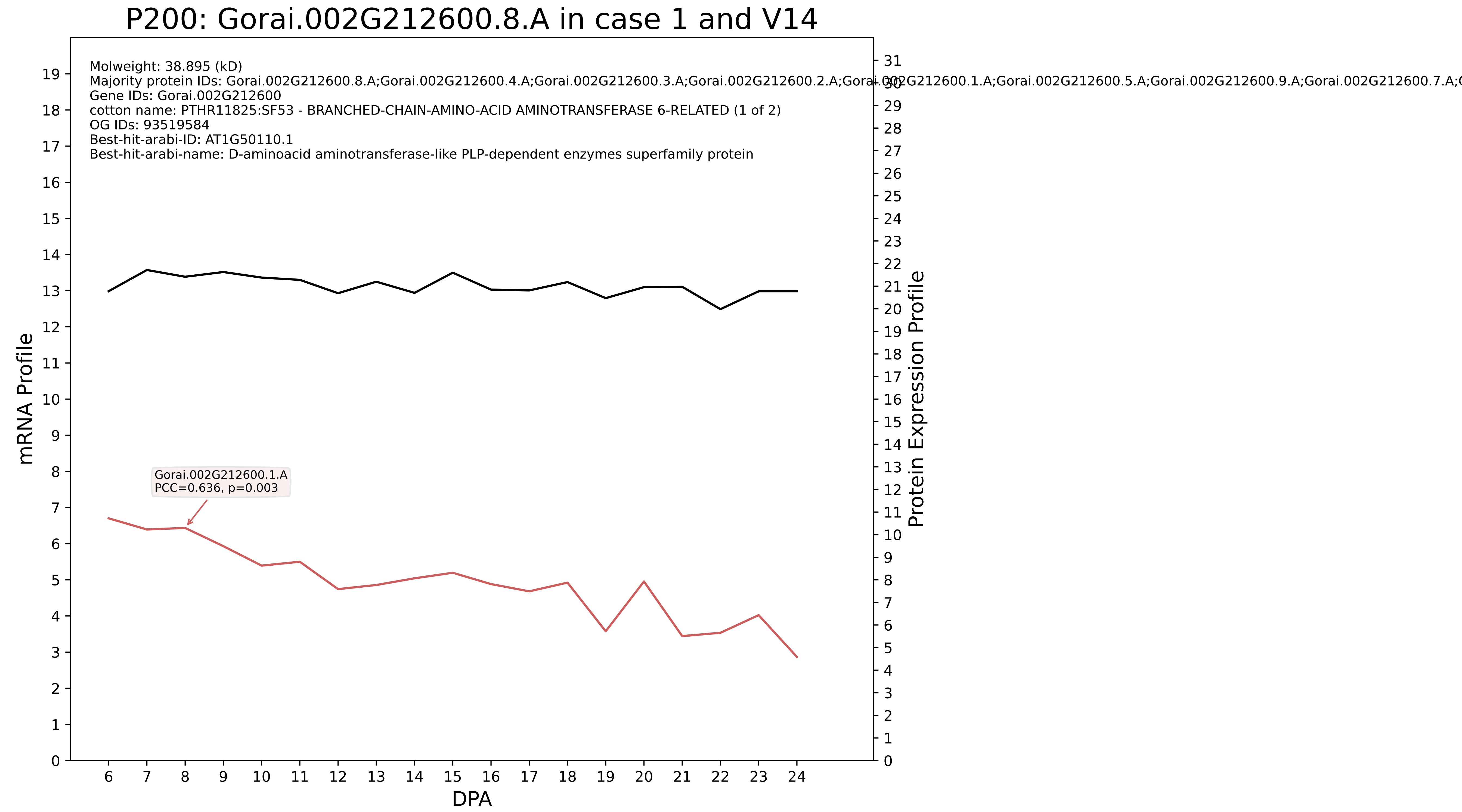
P200: Gorai.002G212600.8.A in case 1 and V14
Molweight: 38.895 (kD)
Majority protein IDs: Gorai.002G212600.8.A;Gorai.002G212600.4.A;Gorai.002G212600.3.A;Gorai.002G212600.2.A;Gorai.002G212600.1.A;Gorai.002G212600.5.A;Gorai.002G212600.9.A;Gorai.002G212600.7.A;G
Gene IDs: Gorai.002G212600
cotton name: PTHR11825:SF53 - BRANCHED-CHAIN-AMINO-ACID AMINOTRANSFERASE 6-RELATED (1 of 2)
OG IDs: 93519584
Best-hit-arabi-ID: AT1G50110.1
Best-hit-arabi-name: D-aminoacid aminotransferase-like PLP-dependent enzymes superfamily protein
Gorai.002G212600.1.A
PCC=0.636, p=0.003
mRNA Profile
Protein Expression Profile
DPA

### P200: Gorai.004G028000.1.D in case 1 and V14

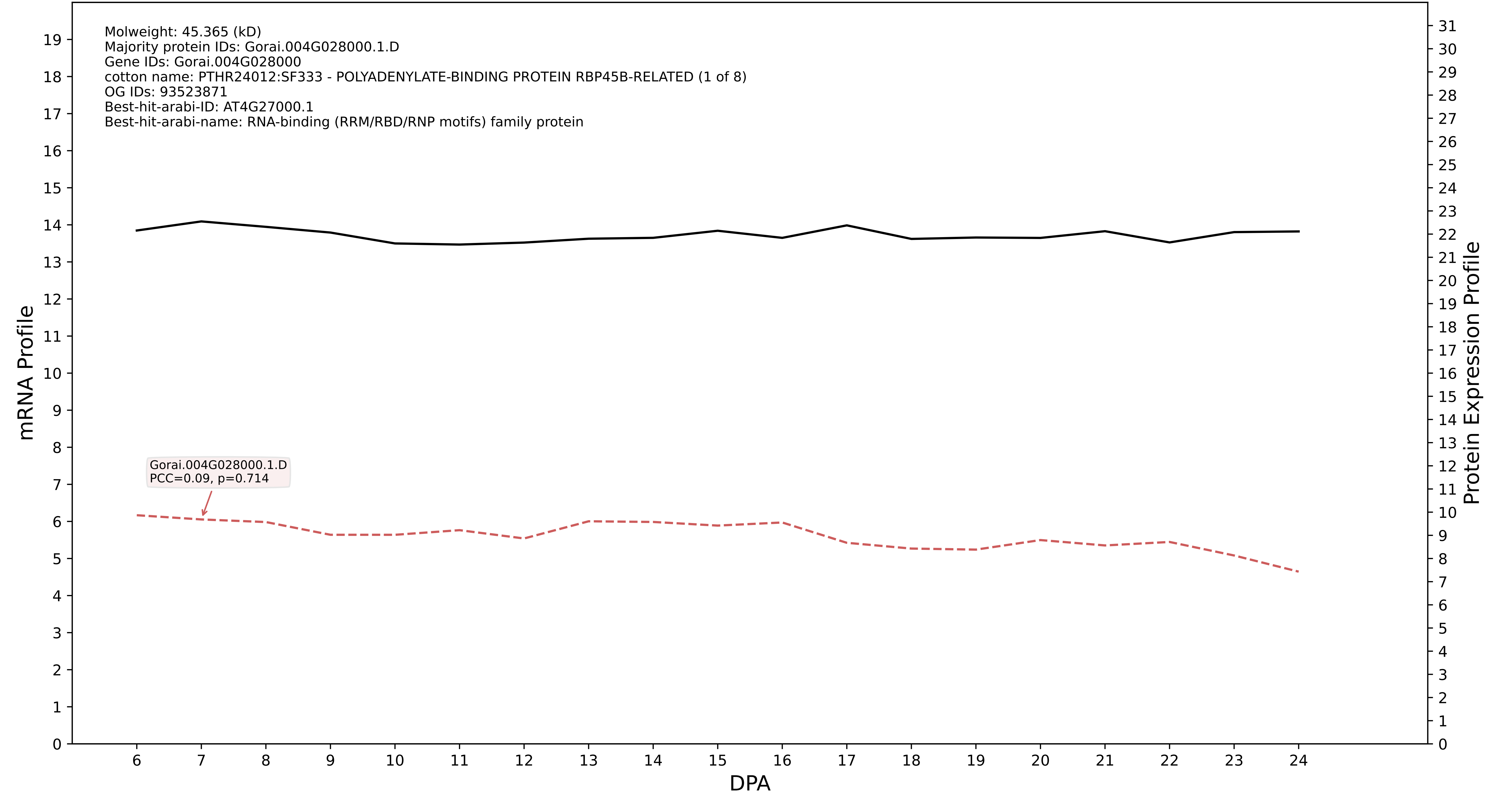

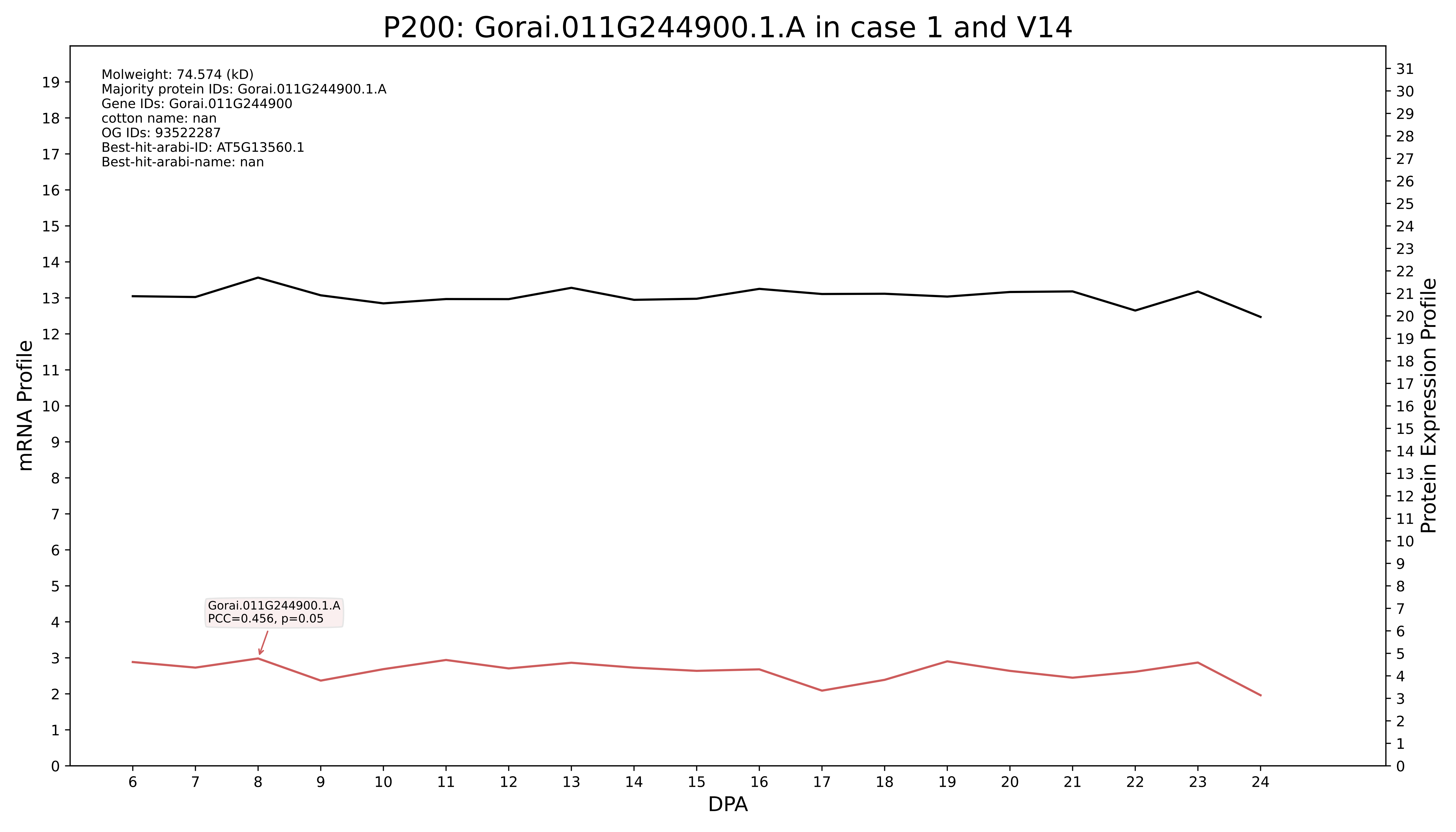
P200: Gorai.011G244900.1.A in case 1 and V14
Molweight: 74.574 (kD)
Majority protein IDs: Gorai.011G244900.1.A
Gene IDs: Gorai.011G244900
cotton name: nan
OG IDs: 93522287
Best-hit-arabi-ID: AT5G13560.1
Best-hit-arabi-name: nan
Gorai.011G244900.1.A
PCC=0.456, p=0.05
mRNA Profile
Protein Expression Profile
DPA

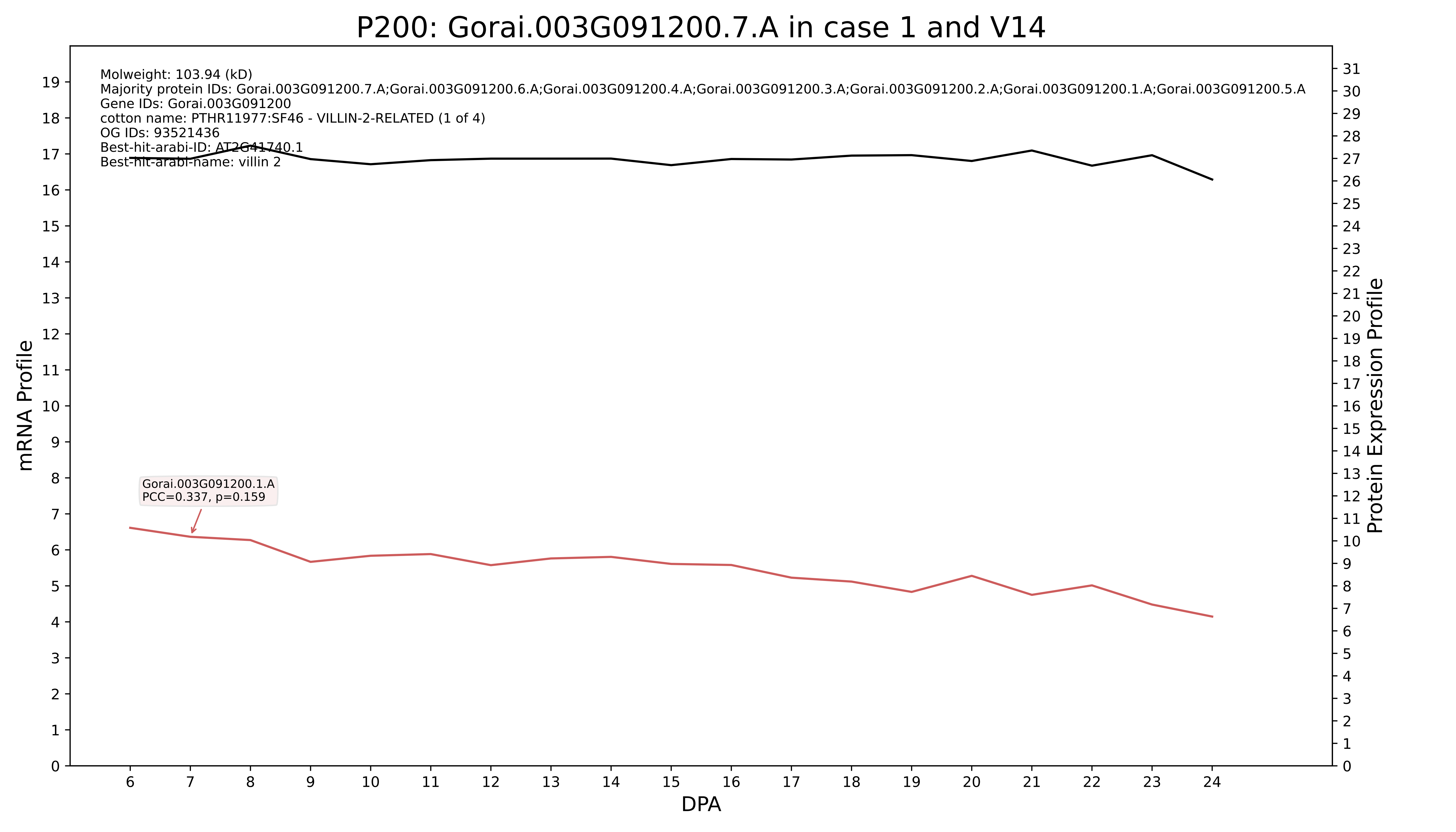
P200: Gorai.003G091200.7.A in case 1 and V14
Molweight: 103.94 (kD)
Majority protein IDs: Gorai.003G091200.7.A;Gorai.003G091200.6.A;Gorai.003G091200.4.A;Gorai.003G091200.3.A;Gorai.003G091200.2.A;Gorai.003G091200.1.A;Gorai.003G091200.5.A
Gene IDs: Gorai.003G091200
cotton name: PTHR11977:SF46 - VILLIN-2-RELATED (1 of 4)
OG IDs: 93521436
Best-hit-arabi-ID: AT2G41740.1
Best-hit-arabi-name: villin 2
Gorai.003G091200.1.A
PCC=0.337, p=0.159
mRNA Profile
Protein Expression Profile
DPA

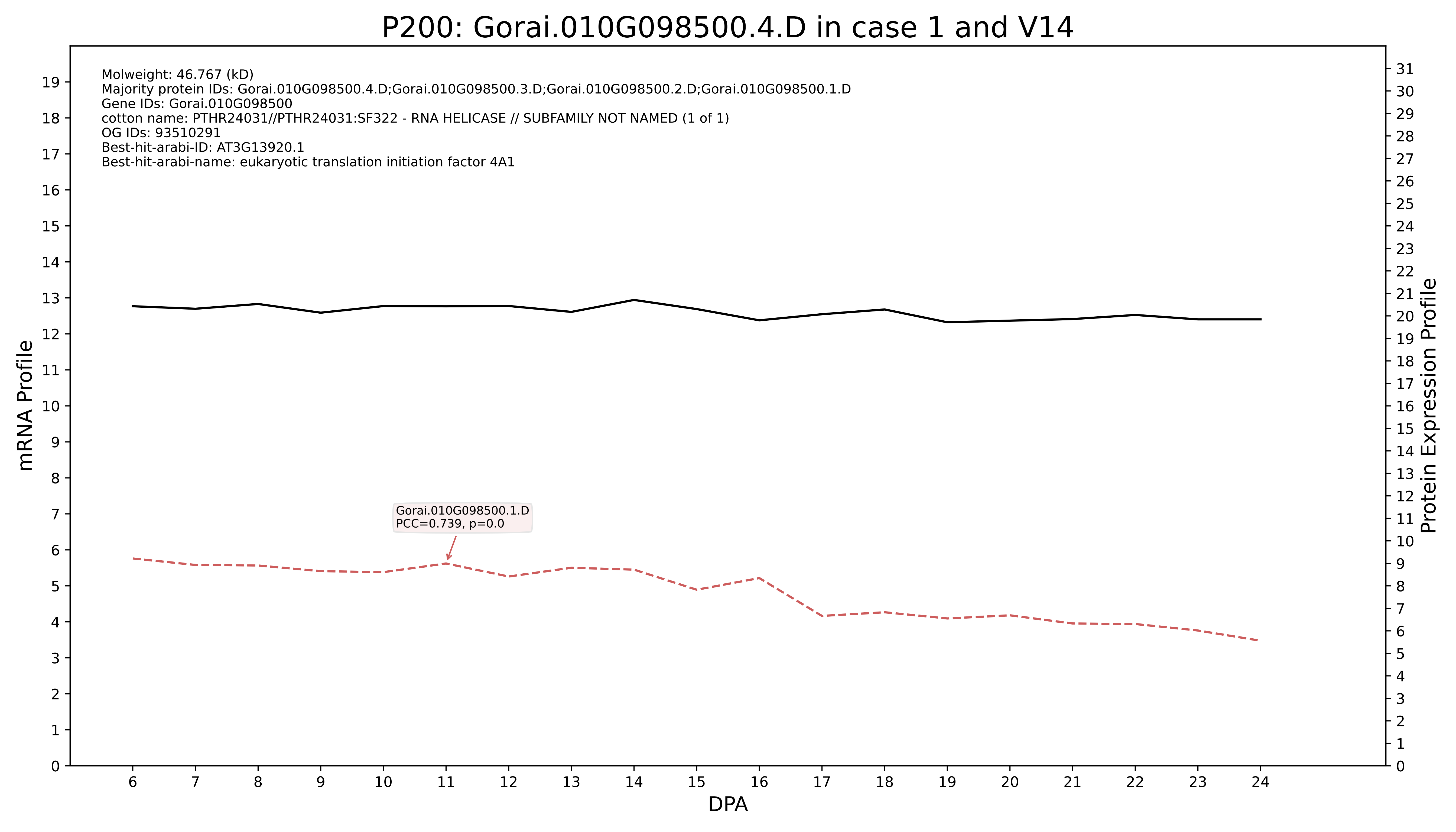
P200: Gorai.010G098500.4.D in case 1 and V14
Molweight: 46.767 (kD)
Majority protein IDs: Gorai.010G098500.4.D;Gorai.010G098500.3.D;Gorai.010G098500.2.D;Gorai.010G098500.1.D
Gene IDs: Gorai.010G098500
cotton name: PTHR24031//PTHR24031:SF322 - RNA HELICASE // SUBFAMILY NOT NAMED (1 of 1)
OG IDs: 93510291
Best-hit-arabi-ID: AT3G13920.1
Best-hit-arabi-name: eukaryotic translation initiation factor 4A1
Gorai.010G098500.1.D
PCC=0.739, p=0.0
mRNA Profile
Protein Expression Profile
DPA

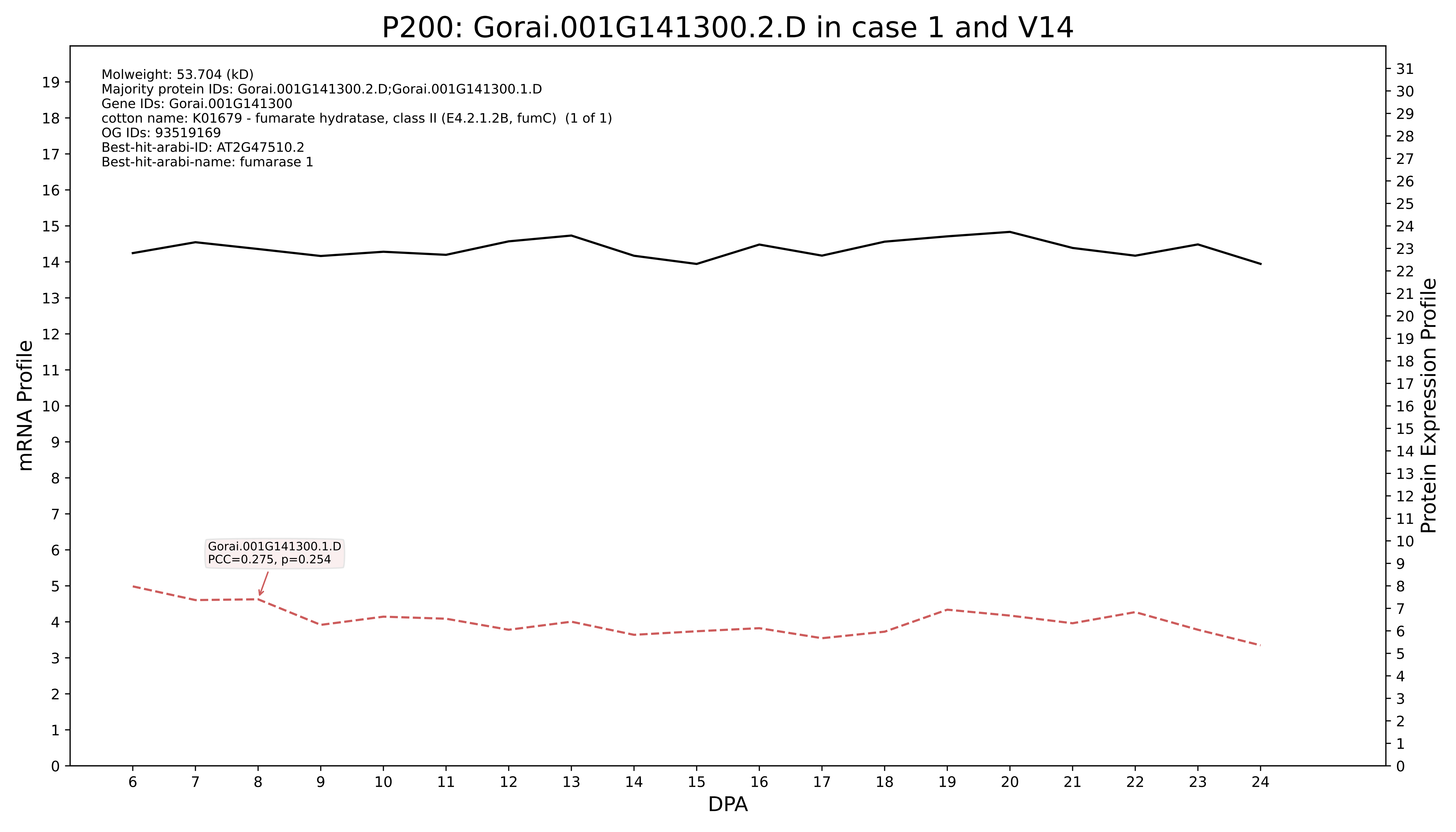
P200: Gorai.001G141300.2.D in case 1 and V14
Molweight: 53.704 (kD)
Majority protein IDs: Gorai.001G141300.2.D;Gorai.001G141300.1.D
Gene IDs: Gorai.001G141300
cotton name: K01679 - fumarate hydratase, class II (E4.2.1.2B, fumC) (1 of 1)
OG IDs: 93519169
Best-hit-arabi-ID: AT2G47510.2
Best-hit-arabi-name: fumarase 1
Gorai.001G141300.1.D
PCC=0.275, p=0.254
mRNA Profile
Protein Expression Profile
DPA

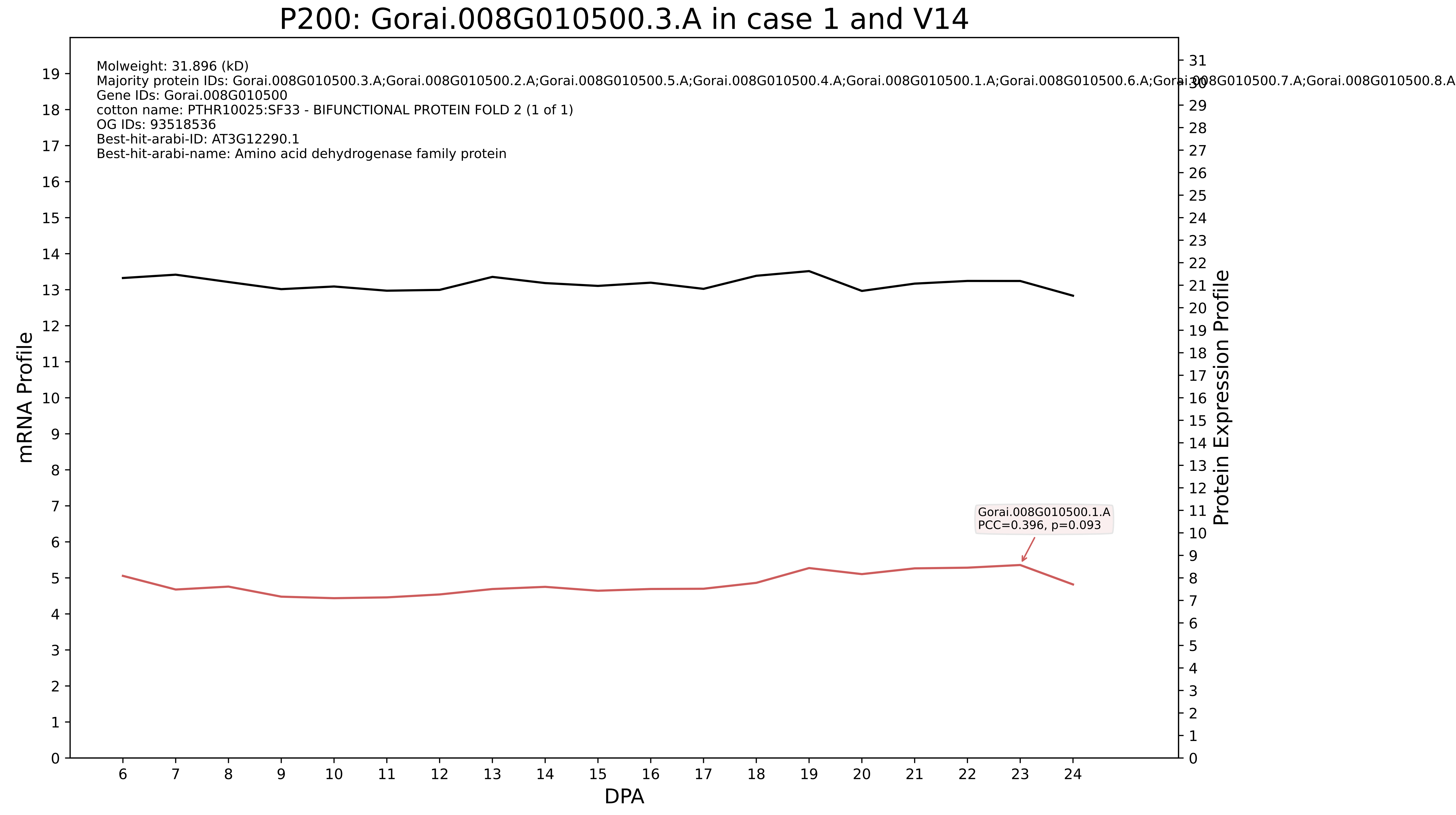

P200: Gorai.008G010500.3.A in case 1 and V14
Molweight: 31.896 (kD)
Majority protein IDs: Gorai.008G010500.3.A;Gorai.008G010500.2.A;Gorai.008G010500.5.A;Gorai.008G010500.4.A;Gorai.008G010500.1.A;Gorai.008G010500.6.A;Gorai.008G010500.7.A;Gorai.008G010500.8.A
Gene IDs: Gorai.008G010500
cotton name: PTHR10025:SF33 - BIFUNCTIONAL PROTEIN FOLD 2 (1 of 1)
OG IDs: 93518536
Best-hit-arabi-ID: AT3G12290.1
Best-hit-arabi-name: Amino acid dehydrogenase family protein
Gorai.008G010500.1.A
PCC=0.396, p=0.093
mRNA Profile
Protein Expression Profile
DPA

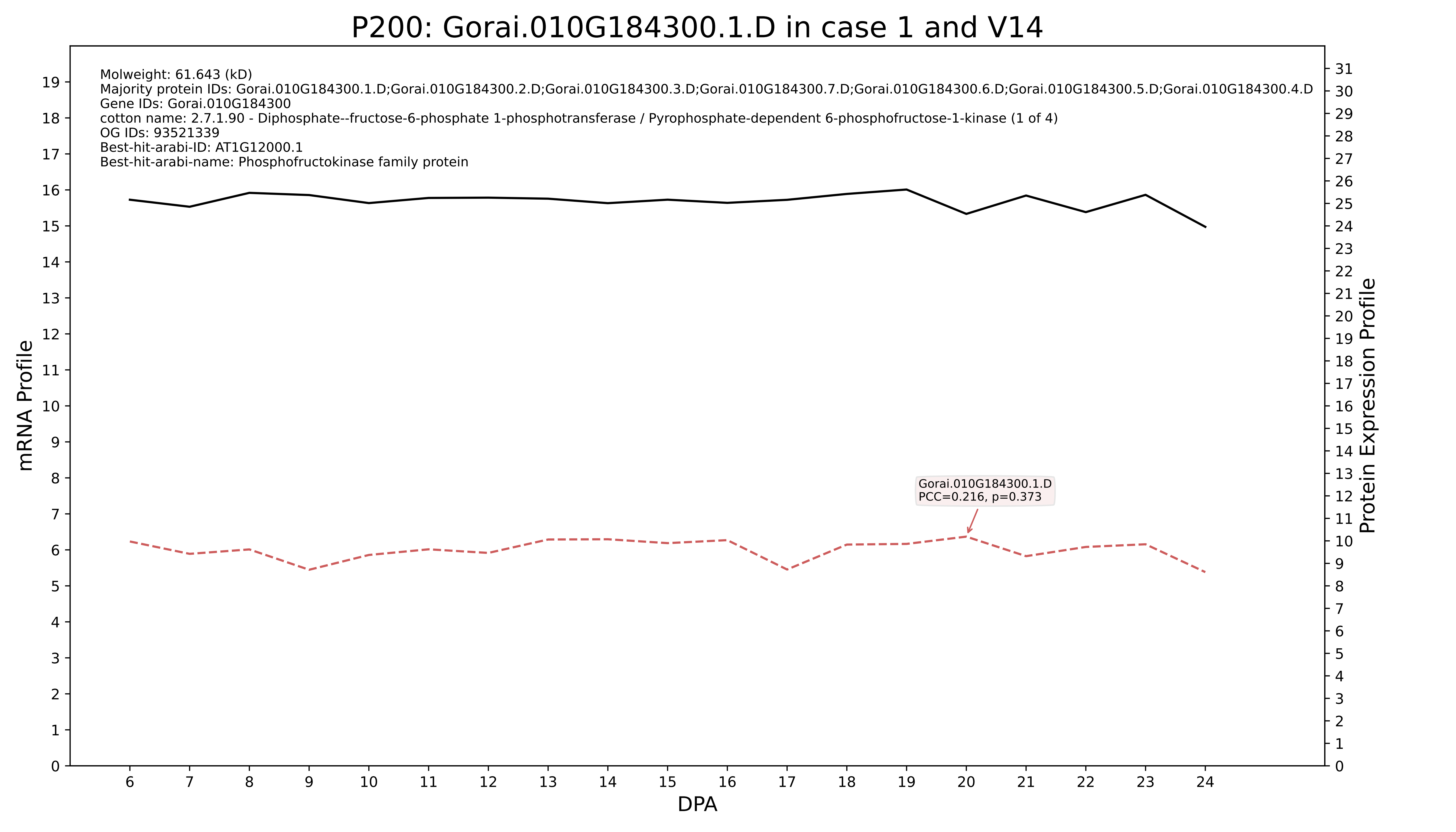

P200: Gorai.010G184300.1.D in case 1 and V14
Molweight: 61.643 (kD)
Majority protein IDs: Gorai.010G184300.1.D;Gorai.010G184300.2.D;Gorai.010G184300.3.D;Gorai.010G184300.7.D;Gorai.010G184300.6.D;Gorai.010G184300.5.D;Gorai.010G184300.4.D
Gene IDs: Gorai.010G184300
cotton name: 2.7.1.90 - Diphosphate--fructose-6-phosphate 1-phosphotransferase / Pyrophosphate-dependent 6-phosphofructose-1-kinase (1 of 4)
OG IDs: 93521339
Best-hit-arabi-ID: AT1G12000.1
Best-hit-arabi-name: Phosphofructokinase family protein
Gorai.010G184300.1.D
PCC=0.216, p=0.373
mRNA Profile
Protein Expression Profile
DPA

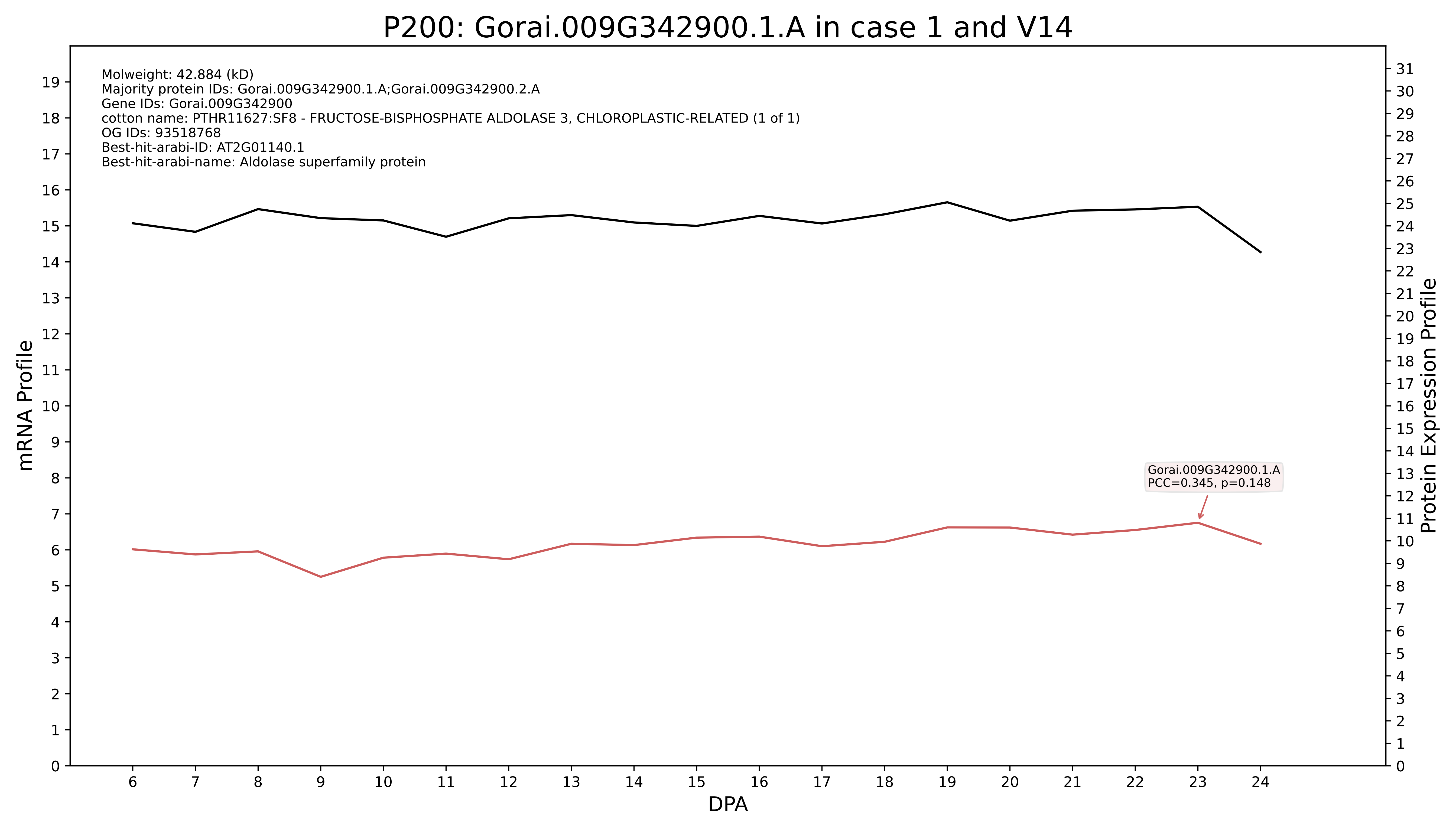
P200: Gorai.009G342900.1.A in case 1 and V14
Molweight: 42.884 (kD)
Majority protein IDs: Gorai.009G342900.1.A;Gorai.009G342900.2.A
Gene IDs: Gorai.009G342900
cotton name: PTHR11627:SF8 - FRUCTOSE-BISPHOSPHATE ALDOLASE 3, CHLOROPLASTIC-RELATED (1 of 1)
OG IDs: 93518768
Best-hit-arabi-ID: AT2G01140.1
Best-hit-arabi-name: Aldolase superfamily protein
Gorai.009G342900.1.A
PCC=0.345, p=0.148
mRNA Profile
Protein Expression Profile
DPA

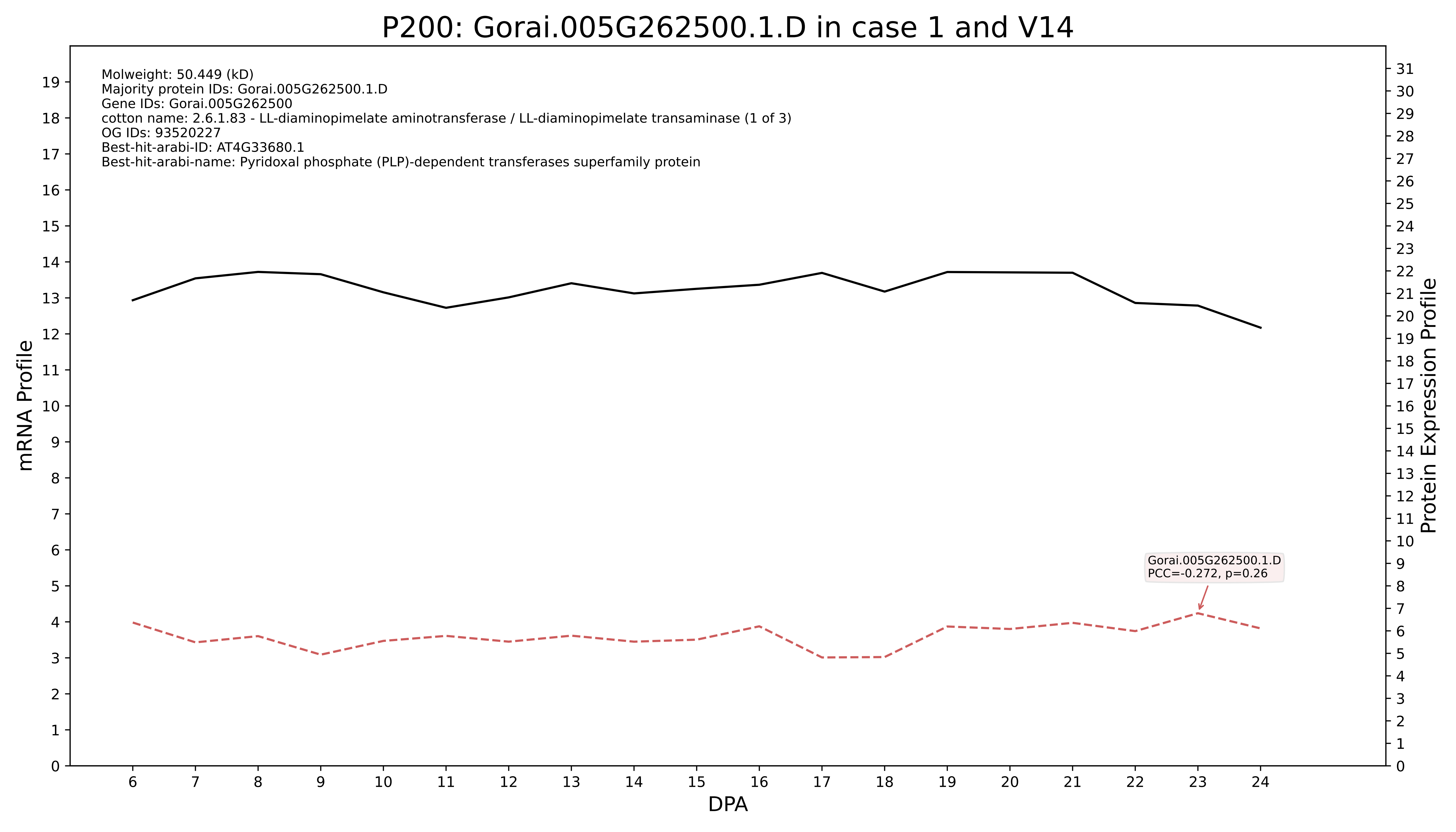
P200: Gorai.005G262500.1.D in case 1 and V14
Molweight: 50.449 (kD)
Majority protein IDs: Gorai.005G262500.1.D
Gene IDs: Gorai.005G262500
cotton name: 2.6.1.83 - LL-diaminopimelate aminotransferase / LL-diaminopimelate transaminase (1 of 3)
OG IDs: 93520227
Best-hit-arabi-ID: AT4G33680.1
Best-hit-arabi-name: Pyridoxal phosphate (PLP)-dependent transferases superfamily protein
Gorai.005G262500.1.D
PCC=-0.272, p=0.26
mRNA Profile
Protein Expression Profile
DPA

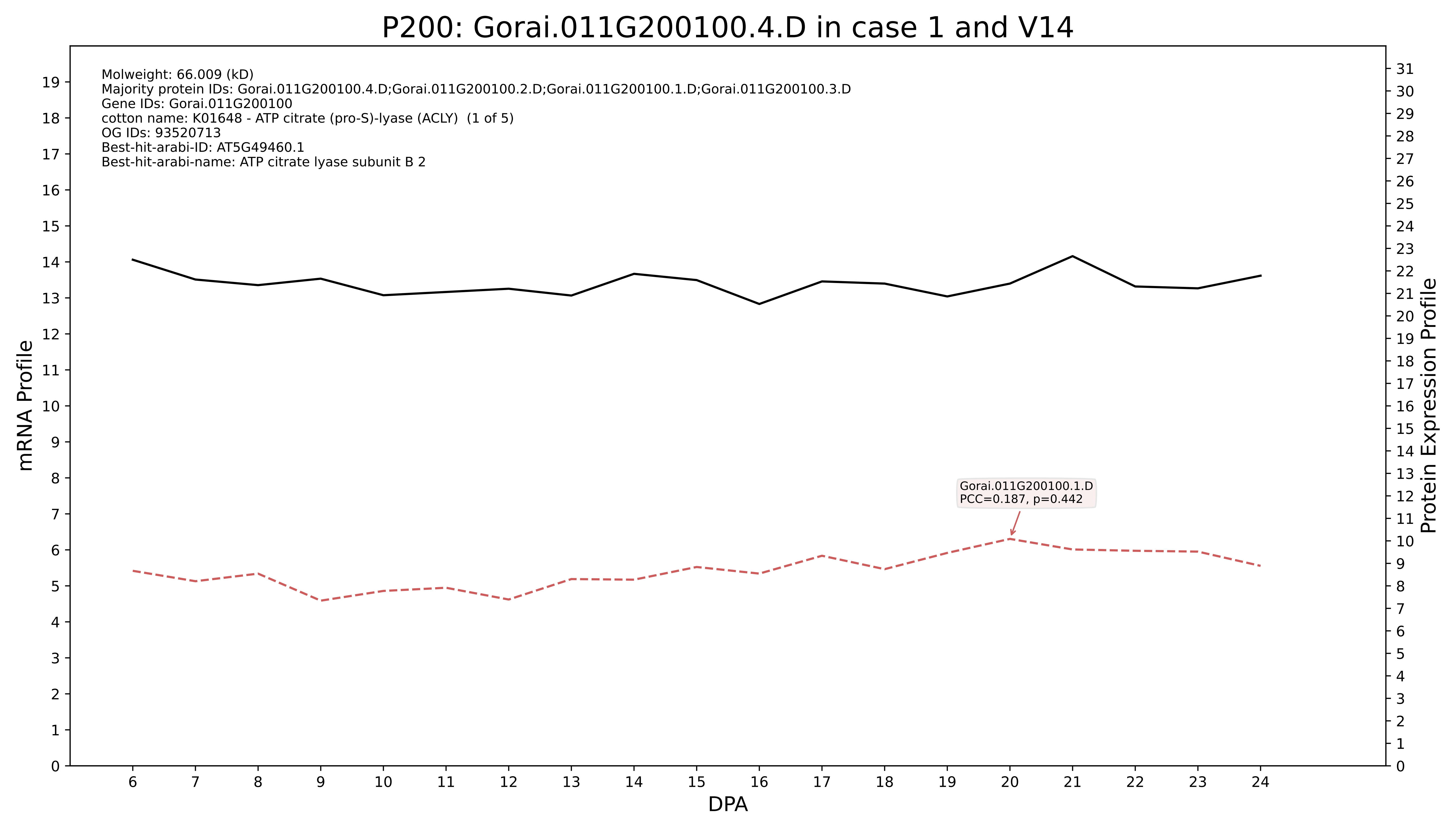
P200: Gorai.011G200100.4.D in case 1 and V14
Molweight: 66.009 (kD)
Majority protein IDs: Gorai.011G200100.4.D;Gorai.011G200100.2.D;Gorai.011G200100.1.D;Gorai.011G200100.3.D
Gene IDs: Gorai.011G200100
cotton name: K01648 - ATP citrate (pro-S)-lyase (ACLY) (1 of 5)
OG IDs: 93520713
Best-hit-arabi-ID: AT5G49460.1
Best-hit-arabi-name: ATP citrate lyase subunit B 2
Gorai.011G200100.1.D
PCC=0.187, p=0.442
mRNA Profile
Protein Expression Profile
DPA

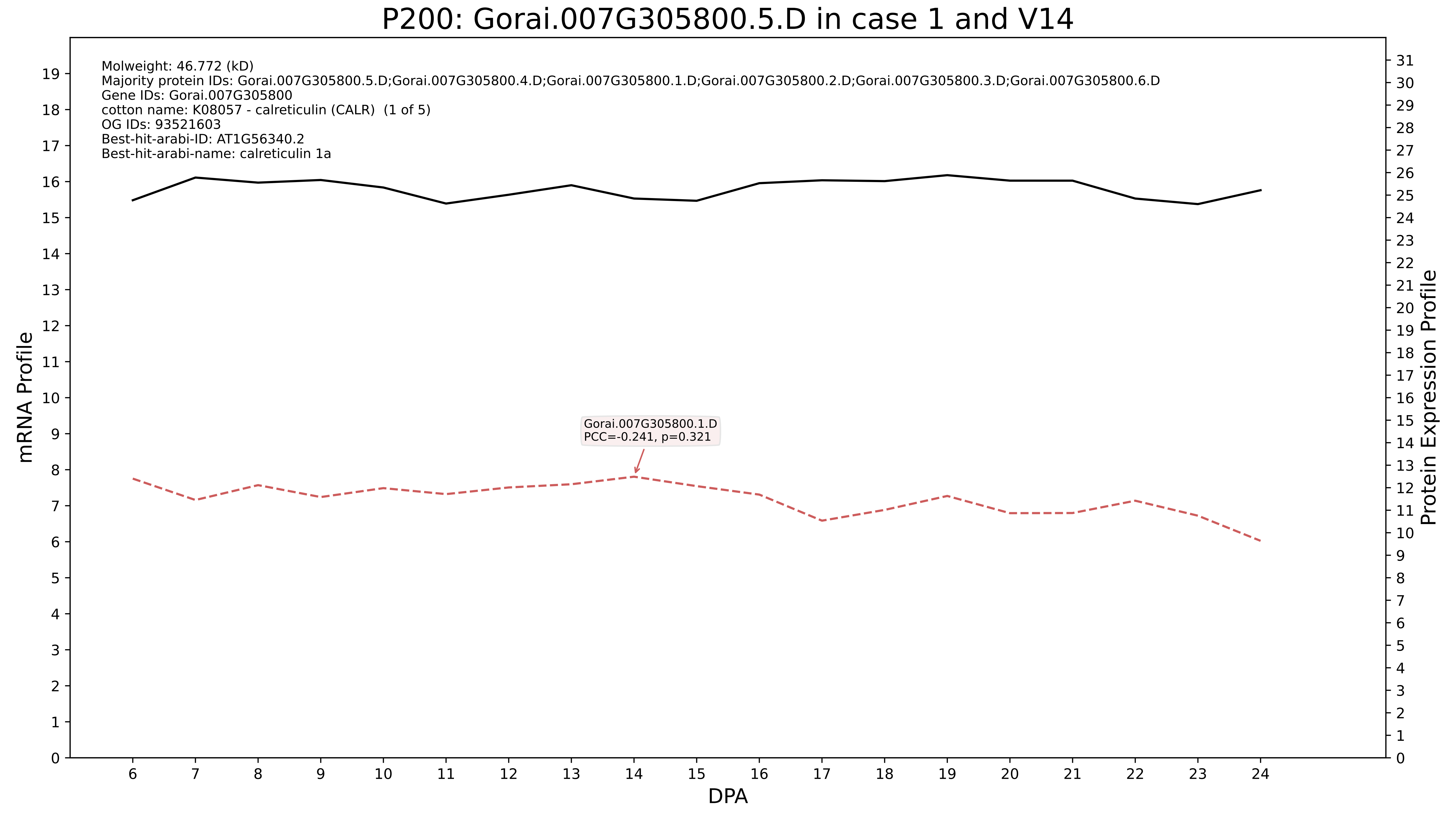
P200: Gorai.007G305800.5.D in case 1 and V14
Molweight: 46.772 (kD)
Majority protein IDs: Gorai.007G305800.5.D;Gorai.007G305800.4.D;Gorai.007G305800.1.D;Gorai.007G305800.2.D;Gorai.007G305800.3.D;Gorai.007G305800.6.D
Gene IDs: Gorai.007G305800
cotton name: K08057 - calreticulin (CALR) (1 of 5)
OG IDs: 93521603
Best-hit-arabi-ID: AT1G56340.2
Best-hit-arabi-name: calreticulin 1a
Gorai.007G305800.1.D
PCC=-0.241, p=0.321
mRNA Profile
Protein Expression Profile
DPA

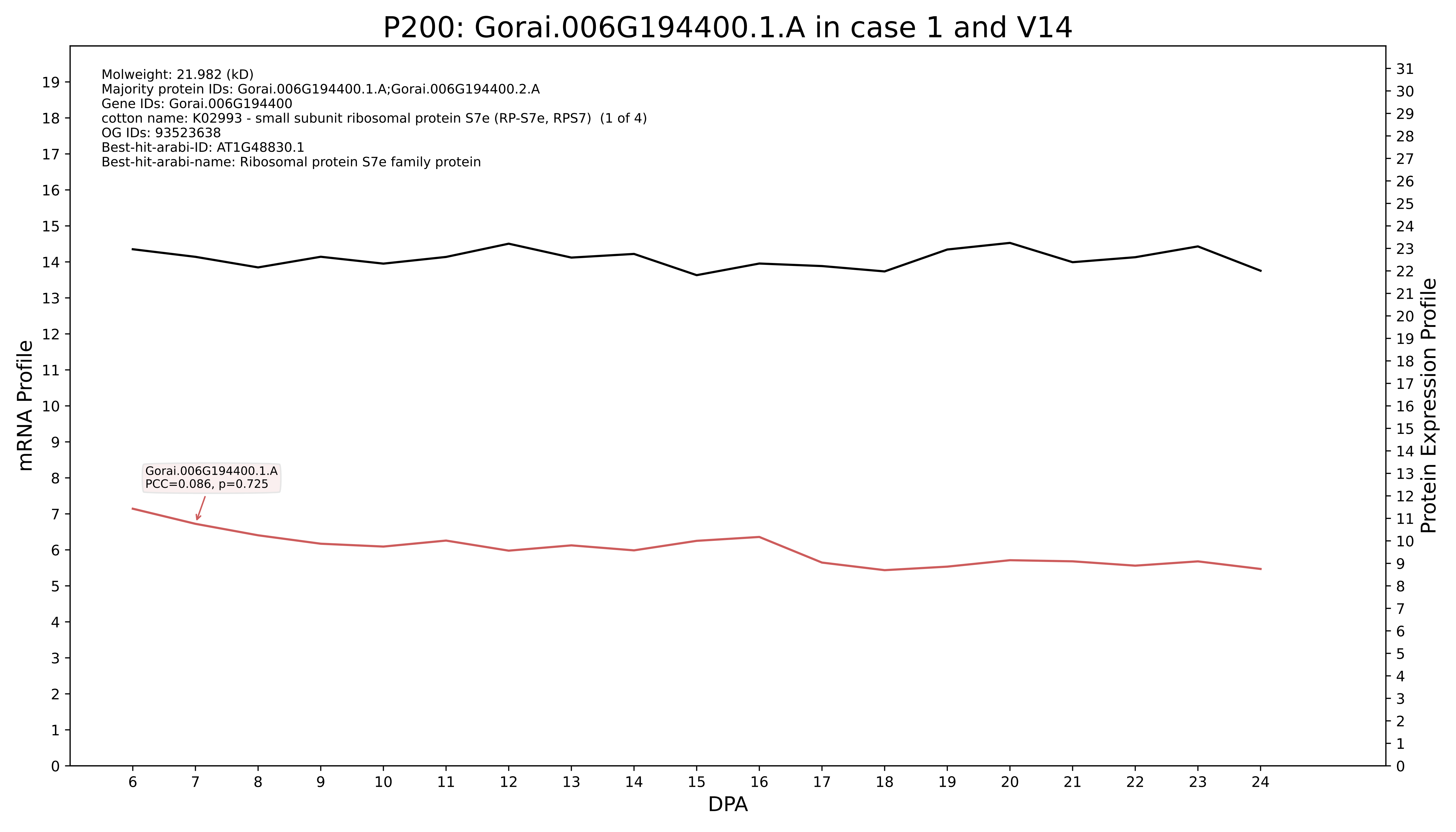
P200: Gorai.006G194400.1.A in case 1 and V14
Molweight: 21.982 (kD)
Majority protein IDs: Gorai.006G194400.1.A;Gorai.006G194400.2.A
Gene IDs: Gorai.006G194400
cotton name: K02993 - small subunit ribosomal protein S7e (RP-S7e, RPS7) (1 of 4)
OG IDs: 93523638
Best-hit-arabi-ID: AT1G48830.1
Best-hit-arabi-name: Ribosomal protein S7e family protein
Gorai.006G194400.1.A
PCC=0.086, p=0.725
mRNA Profile
Protein Expression Profile
DPA

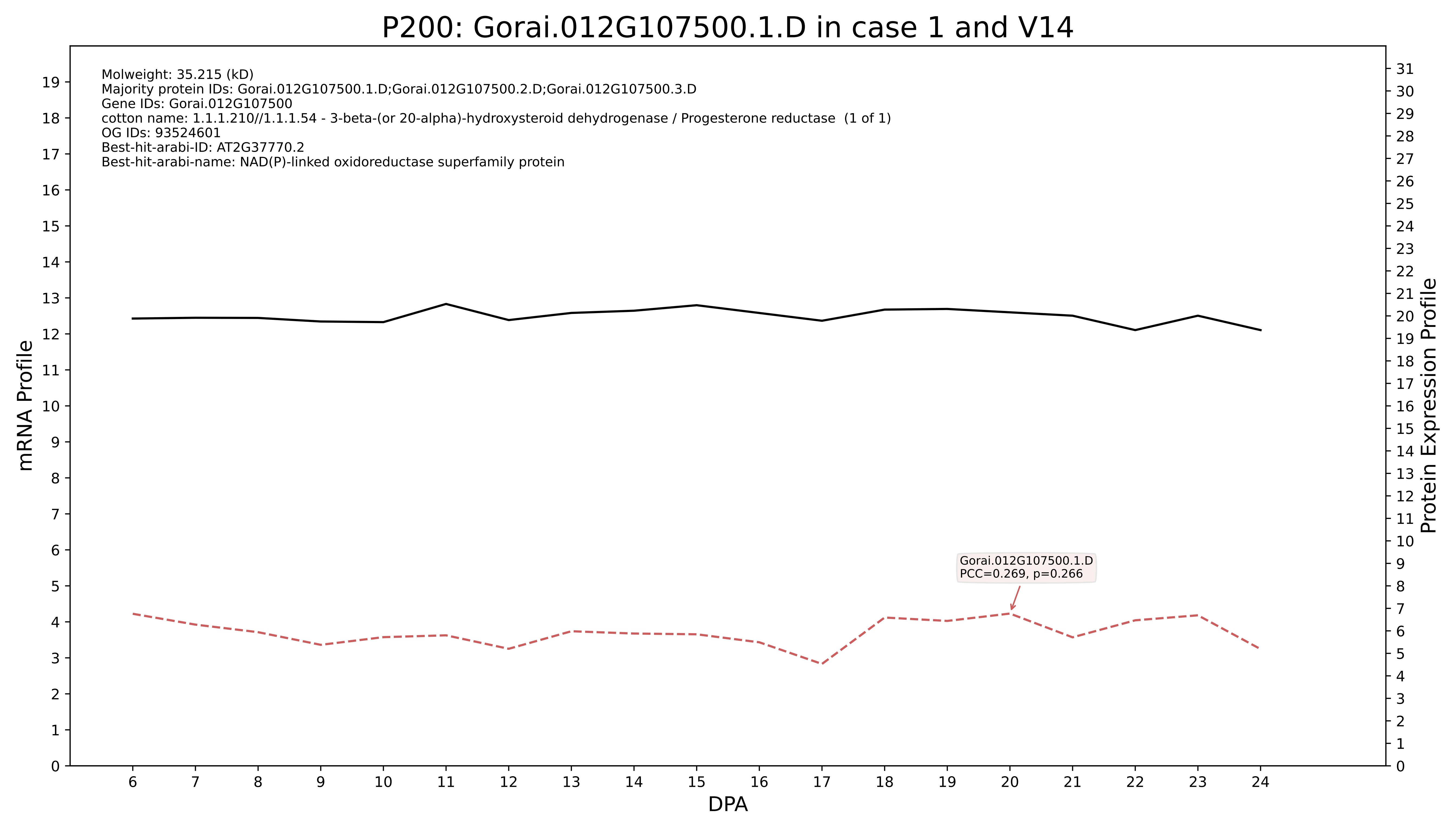

P200: Gorai.012G107500.1.D in case 1 and V14
Molweight: 35.215 (kD)
Majority protein IDs: Gorai.012G107500.1.D;Gorai.012G107500.2.D;Gorai.012G107500.3.D
Gene IDs: Gorai.012G107500
cotton name: 1.1.1.210//1.1.1.54 - 3-beta-(or 20-alpha)-hydroxysteroid dehydrogenase / Progesterone reductase (1 of 1)
OG IDs: 93524601
Best-hit-arabi-ID: AT2G37770.2
Best-hit-arabi-name: NAD(P)-linked oxidoreductase superfamily protein
Gorai.012G107500.1.D
PCC=0.269, p=0.266
mRNA Profile
Protein Expression Profile
DPA

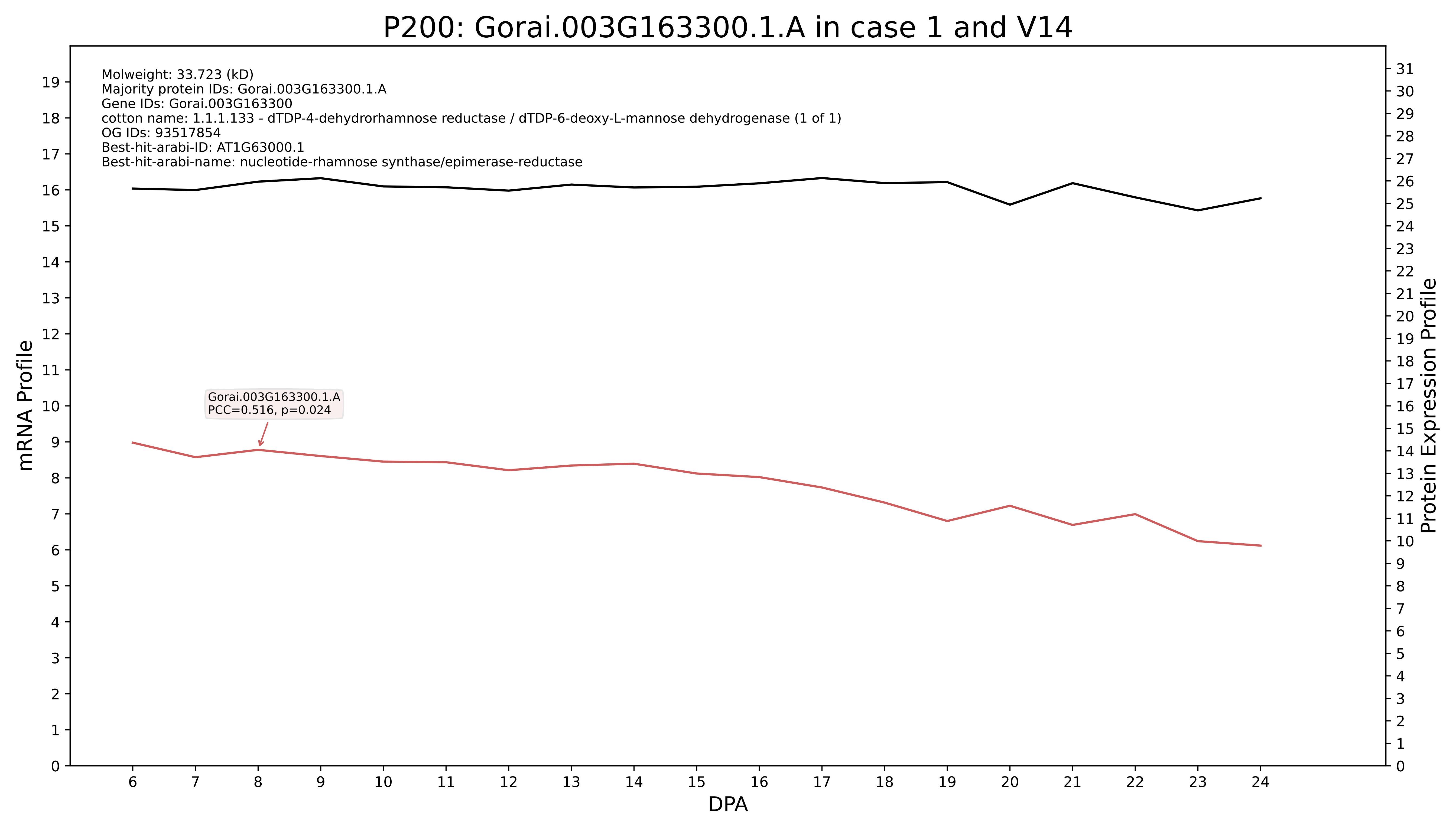
P200: Gorai.003G163300.1.A in case 1 and V14
Molweight: 33.723 (kD)
Majority protein IDs: Gorai.003G163300.1.A
Gene IDs: Gorai.003G163300
cotton name: 1.1.1.133 - dTDP-4-dehydrorhamnose reductase / dTDP-6-deoxy-L-mannose dehydrogenase (1 of 1)
OG IDs: 93517854
Best-hit-arabi-ID: AT1G63000.1
Best-hit-arabi-name: nucleotide-rhamnose synthase/epimerase-reductase
Gorai.003G163300.1.A
PCC=0.516, p=0.024
mRNA Profile
Protein Expression Profile
DPA

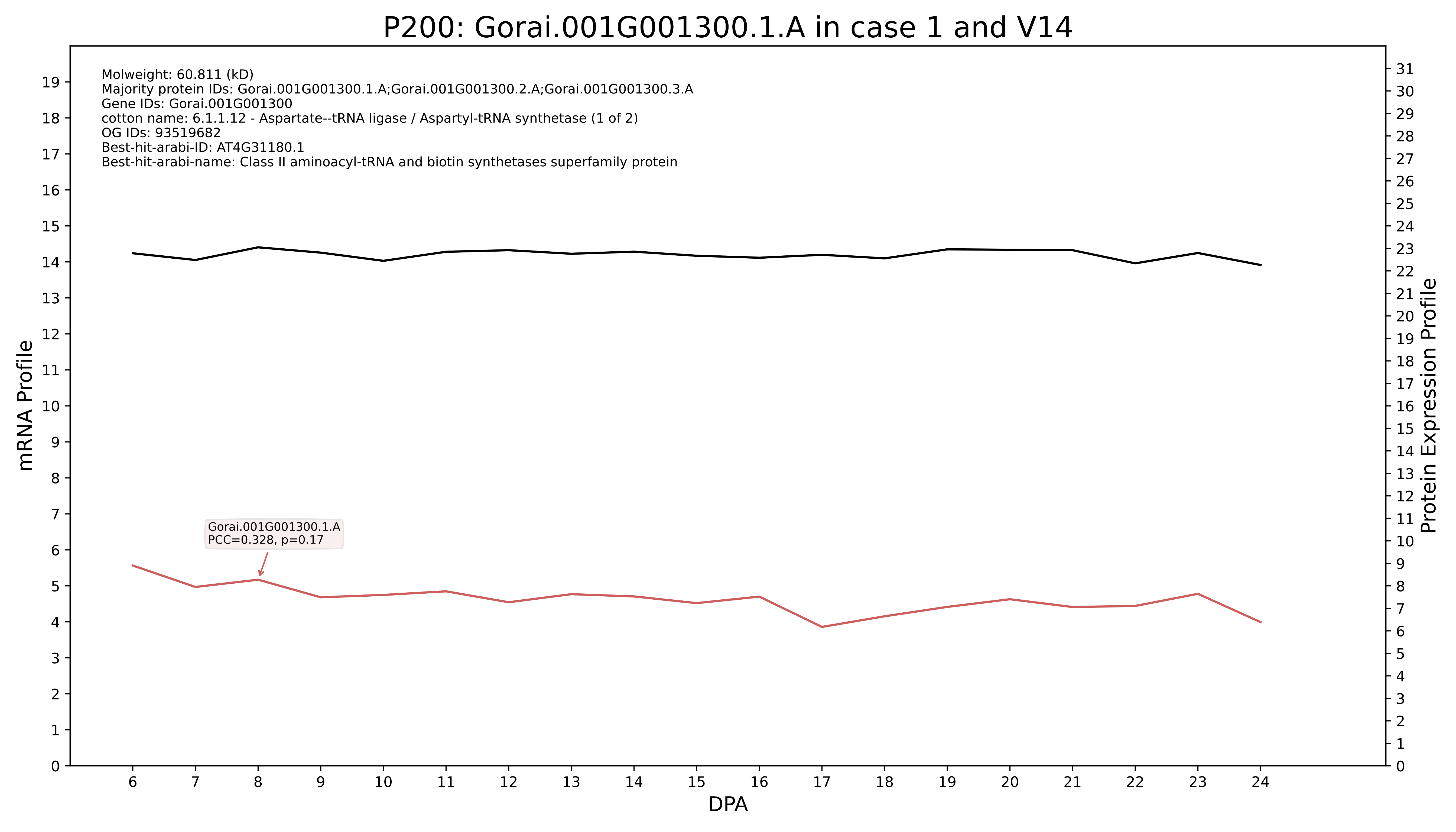
P200: Gorai.001G001300.1.A in case 1 and V14
Molweight: 60.811 (kD)
Majority protein IDs: Gorai.001G001300.1.A;Gorai.001G001300.2.A;Gorai.001G001300.3.A
Gene IDs: Gorai.001G001300
cotton name: 6.1.1.12 - Aspartate--tRNA ligase / Aspartyl-tRNA synthetase (1 of 2)
OG IDs: 93519682
Best-hit-arabi-ID: AT4G31180.1
Best-hit-arabi-name: Class II aminoacyl-tRNA and biotin synthetases superfamily protein
Gorai.001G001300.1.A
PCC=0.328, p=0.17
mRNA Profile
Protein Expression Profile
DPA

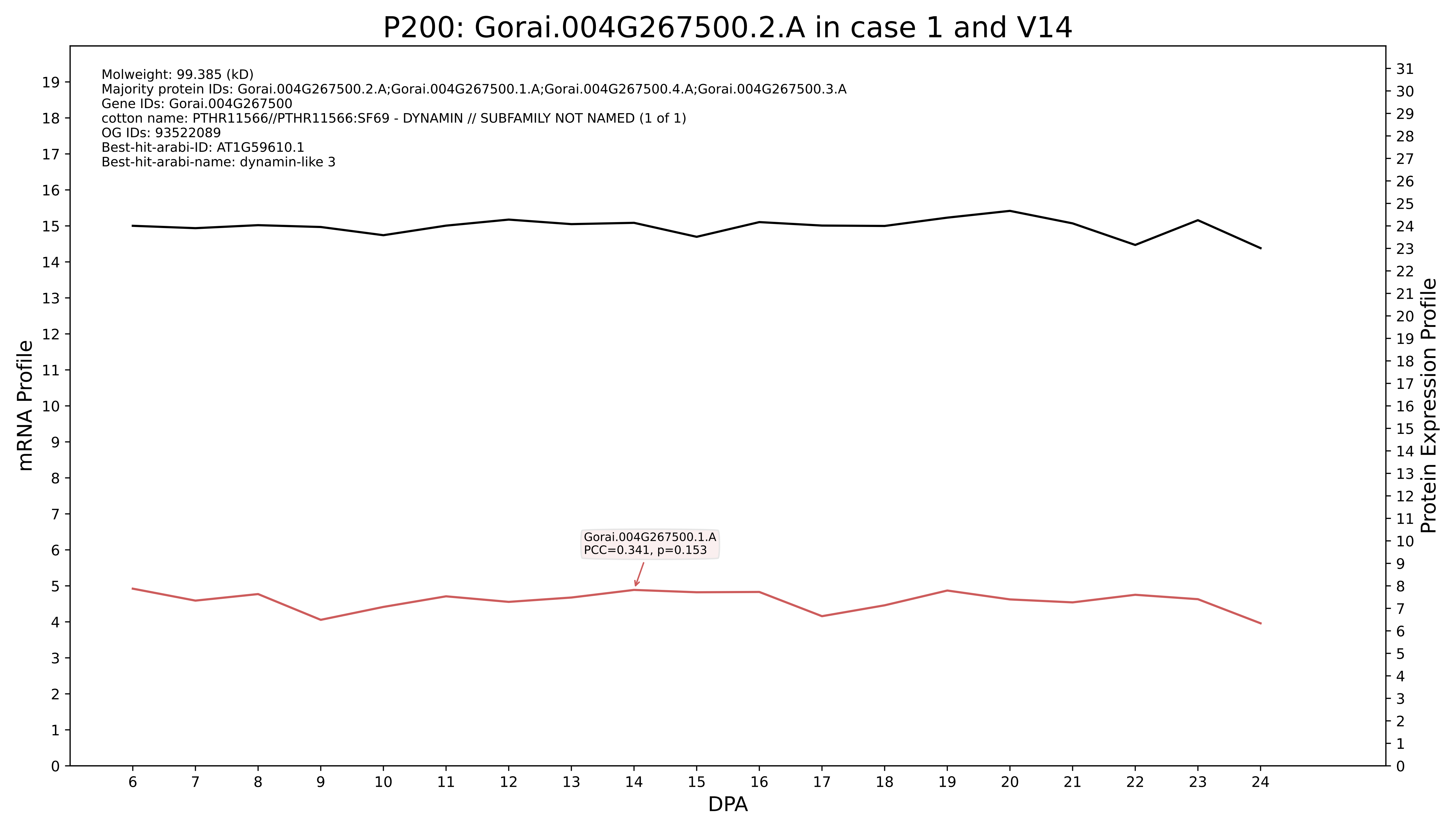
P200: Gorai.004G267500.2.A in case 1 and V14
Molweight: 99.385 (kD)
Majority protein IDs: Gorai.004G267500.2.A;Gorai.004G267500.1.A;Gorai.004G267500.4.A;Gorai.004G267500.3.A
Gene IDs: Gorai.004G267500
cotton name: PTHR11566//PTHR11566:SF69 - DYNAMIN // SUBFAMILY NOT NAMED (1 of 1)
OG IDs: 93522089
Best-hit-arabi-ID: AT1G59610.1
Best-hit-arabi-name: dynamin-like 3
Gorai.004G267500.1.A
PCC=0.341, p=0.153
mRNA Profile
Protein Expression Profile
DPA

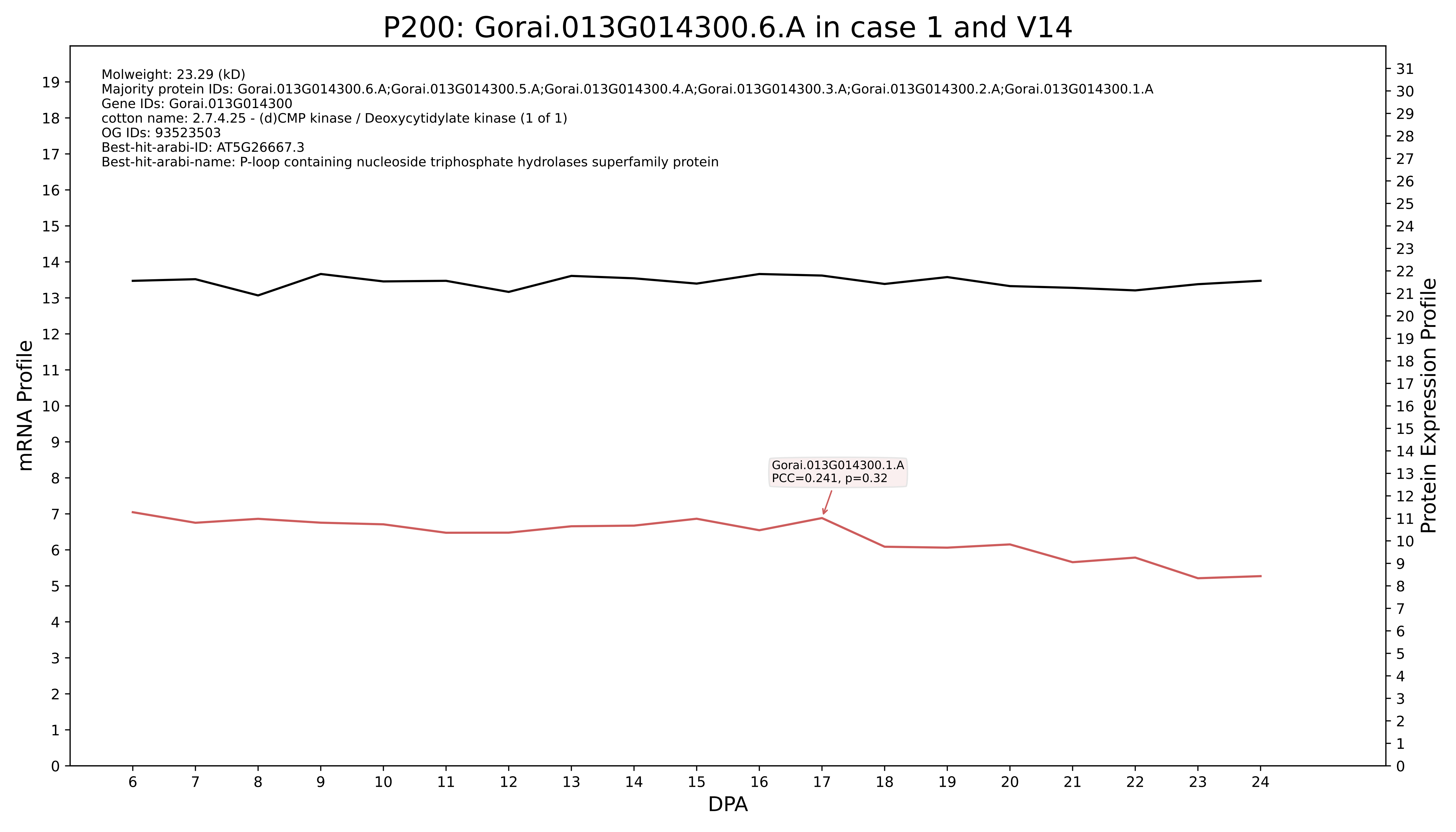
P200: Gorai.013G014300.6.A in case 1 and V14
Molweight: 23.29 (kD)
Majority protein IDs: Gorai.013G014300.6.A;Gorai.013G014300.5.A;Gorai.013G014300.4.A;Gorai.013G014300.3.A;Gorai.013G014300.2.A;Gorai.013G014300.1.A
Gene IDs: Gorai.013G014300
cotton name: 2.7.4.25 - (d)CMP kinase / Deoxycytidylate kinase (1 of 1)
OG IDs: 93523503
Best-hit-arabi-ID: AT5G26667.3
Best-hit-arabi-name: P-loop containing nucleoside triphosphate hydrolases superfamily protein
Gorai.013G014300.1.A
PCC=0.241, p=0.32
mRNA Profile
Protein Expression Profile
DPA

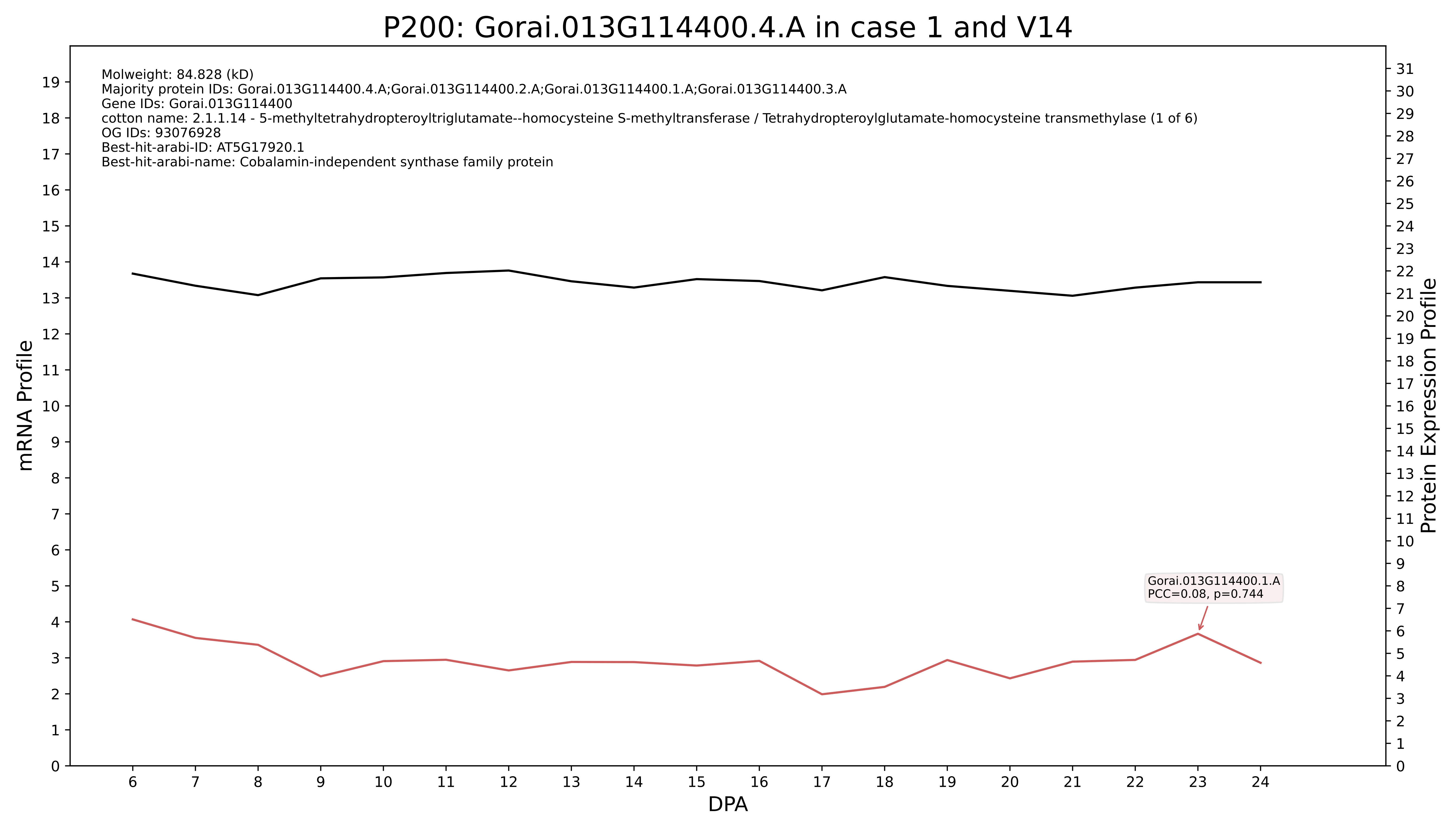
P200: Gorai.013G114400.4.A in case 1 and V14
Molweight: 84.828 (kD)
Majority protein IDs: Gorai.013G114400.4.A;Gorai.013G114400.2.A;Gorai.013G114400.1.A;Gorai.013G114400.3.A
Gene IDs: Gorai.013G114400
cotton name: 2.1.1.14 - 5-methyltetrahydropteroyltriglutamate--homocysteine S-methyltransferase / Tetrahydropteroylglutamate-homocysteine transmethylase (1 of 6)
OG IDs: 93076928
Best-hit-arabi-ID: AT5G17920.1
Best-hit-arabi-name: Cobalamin-independent synthase family protein
Gorai.013G114400.1.A
PCC=0.08, p=0.744
mRNA Profile
Protein Expression Profile
DPA

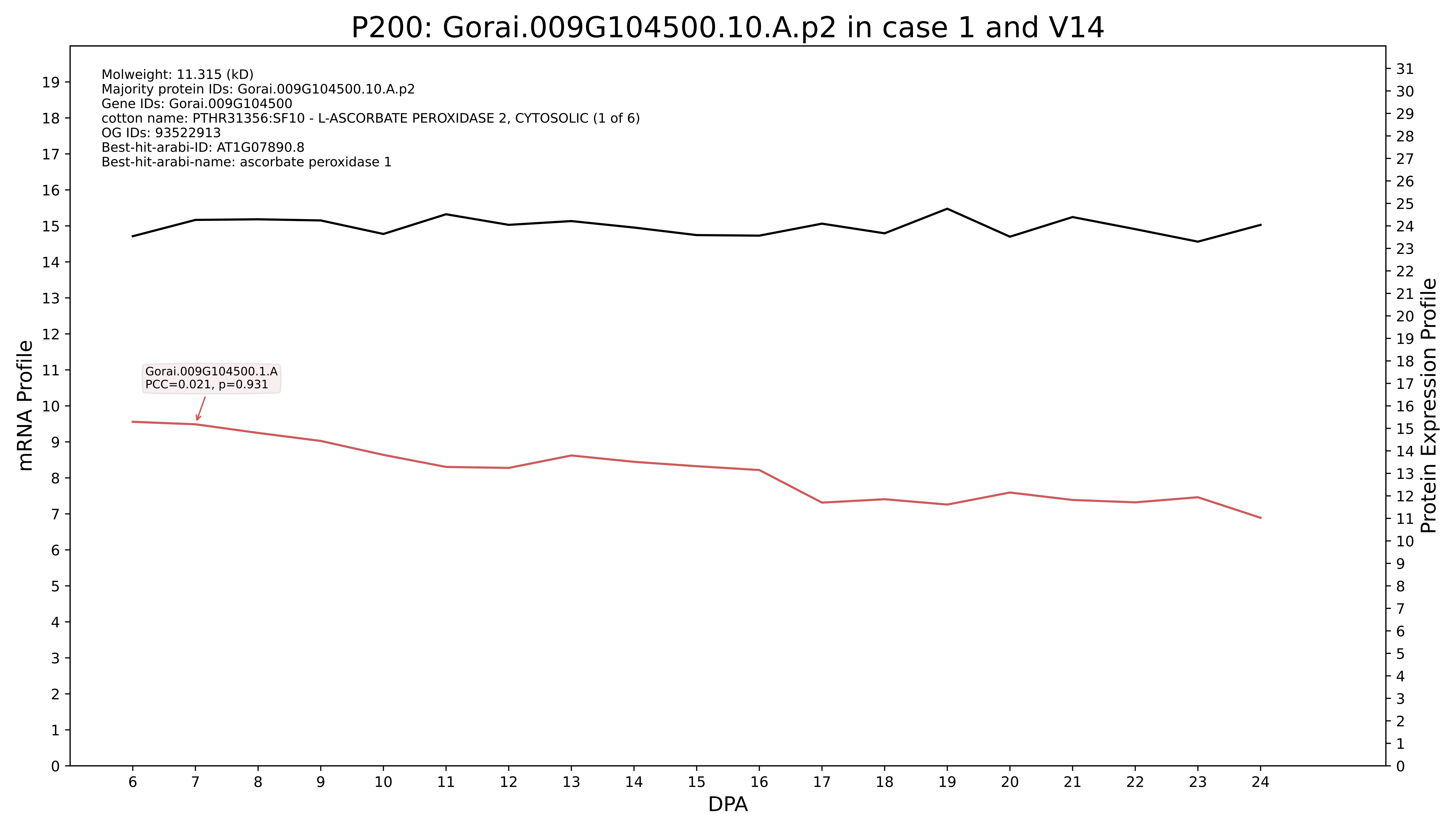

P200: Gorai.009G104500.10.A.p2 in case 1 and V14
Molweight: 11.315 (kD)
Majority protein IDs: Gorai.009G104500.10.A.p2
Gene IDs: Gorai.009G104500
cotton name: PTHR31356:SF10 - L-ASCORBATE PEROXIDASE 2, CYTOSOLIC (1 of 6)
OG IDs: 93522913
Best-hit-arabi-ID: AT1G07890.8
Best-hit-arabi-name: ascorbate peroxidase 1
Gorai.009G104500.1.A
PCC=0.021, p=0.931
mRNA Profile
Protein Expression Profile
DPA

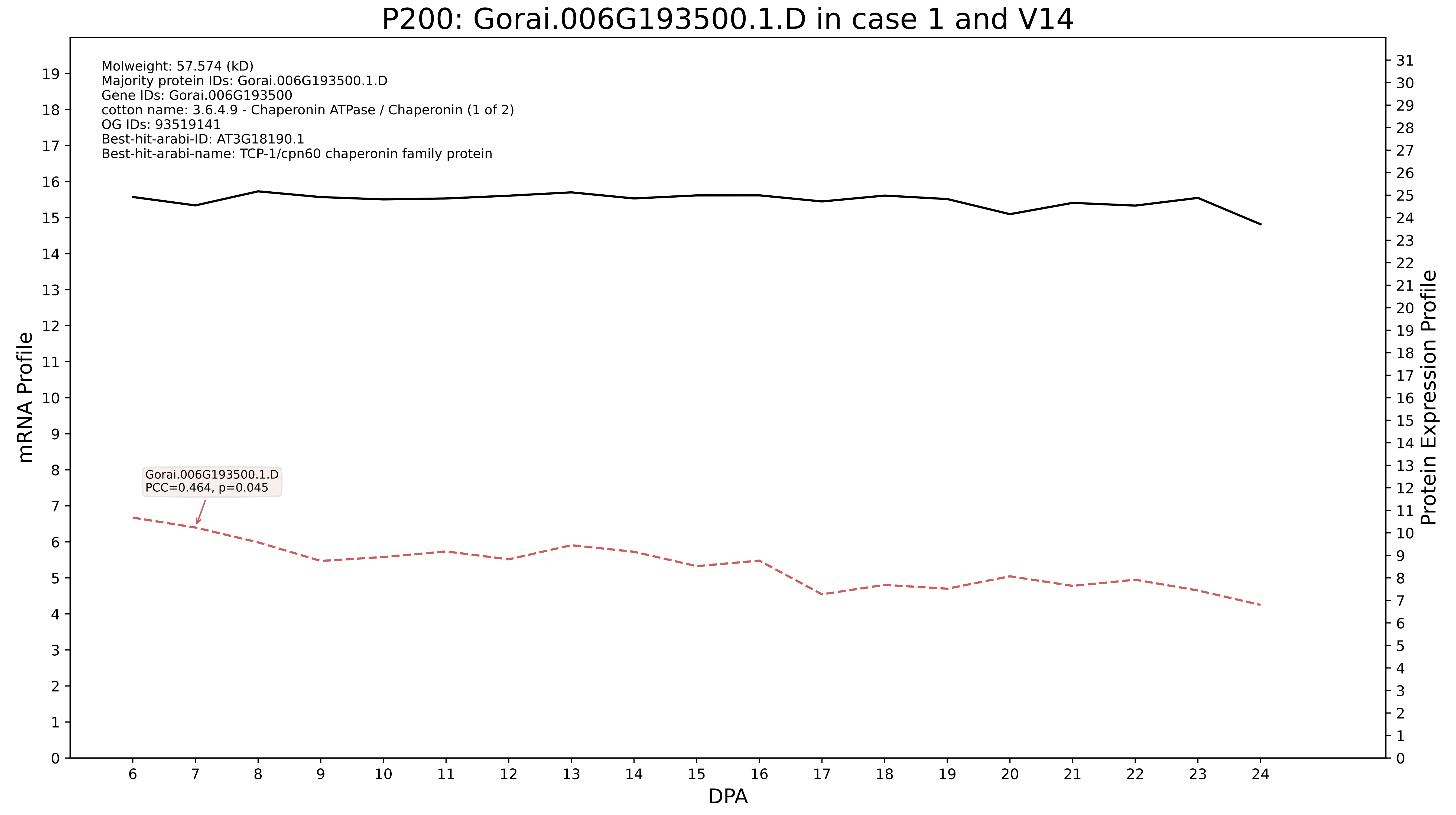
P200: Gorai.006G193500.1.D in case 1 and V14
Molweight: 57.574 (kD)
Majority protein IDs: Gorai.006G193500.1.D
Gene IDs: Gorai.006G193500
cotton name: 3.6.4.9 - Chaperonin ATPase / Chaperonin (1 of 2)
OG IDs: 93519141
Best-hit-arabi-ID: AT3G18190.1
Best-hit-arabi-name: TCP-1/cpn60 chaperonin family protein
Gorai.006G193500.1.D
PCC=0.464, p=0.045
mRNA Profile
Protein Expression Profile
DPA

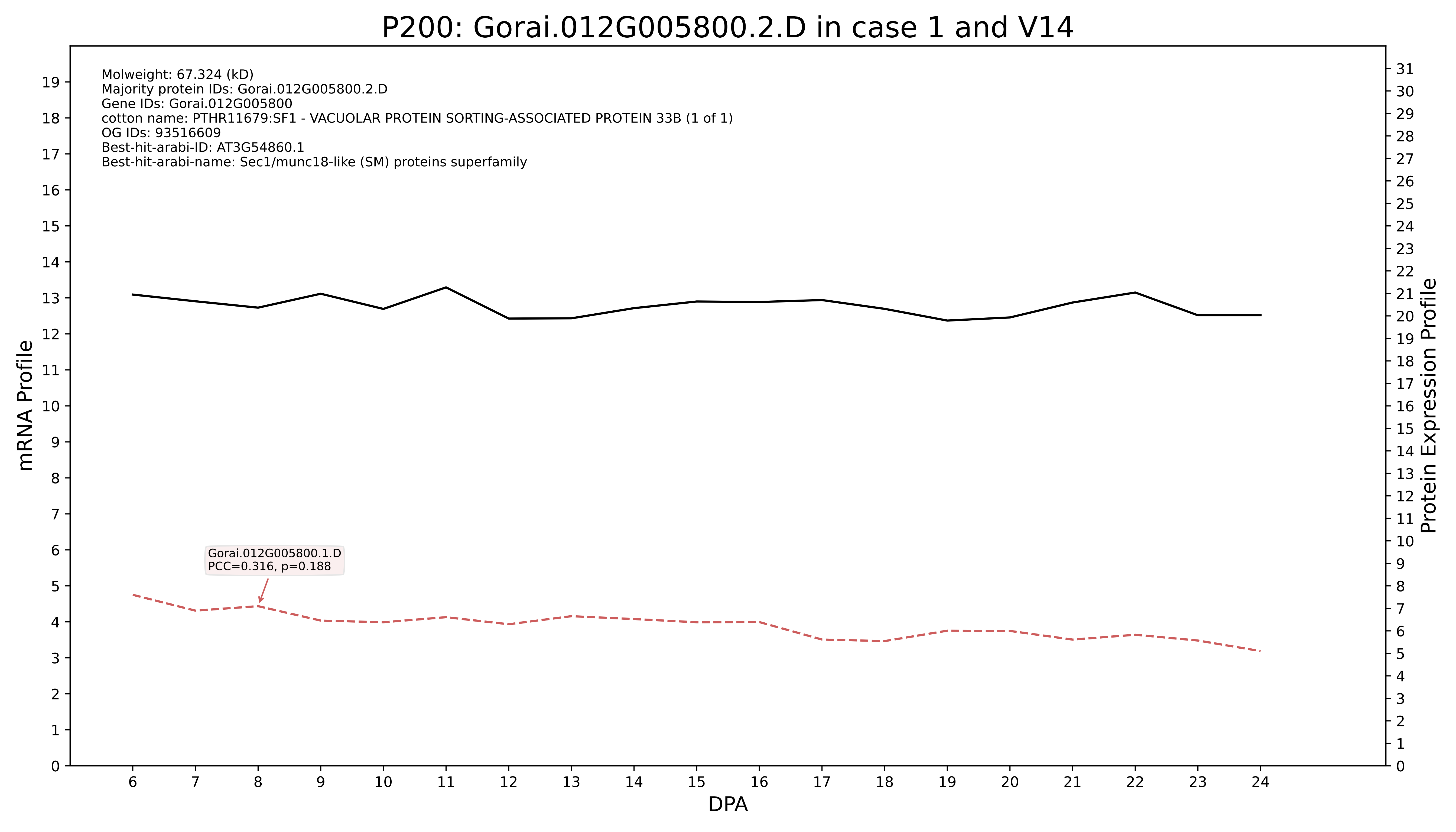
P200: Gorai.012G005800.2.D in case 1 and V14
Molweight: 67.324 (kD)
Majority protein IDs: Gorai.012G005800.2.D
Gene IDs: Gorai.012G005800
cotton name: PTHR11679:SF1 - VACUOLAR PROTEIN SORTING-ASSOCIATED PROTEIN 33B (1 of 1)
OG IDs: 93516609
Best-hit-arabi-ID: AT3G54860.1
Best-hit-arabi-name: Sec1/munc18-like (SM) proteins superfamily
Gorai.012G005800.1.D
PCC=0.316, p=0.188
mRNA Profile
Protein Expression Profile
DPA

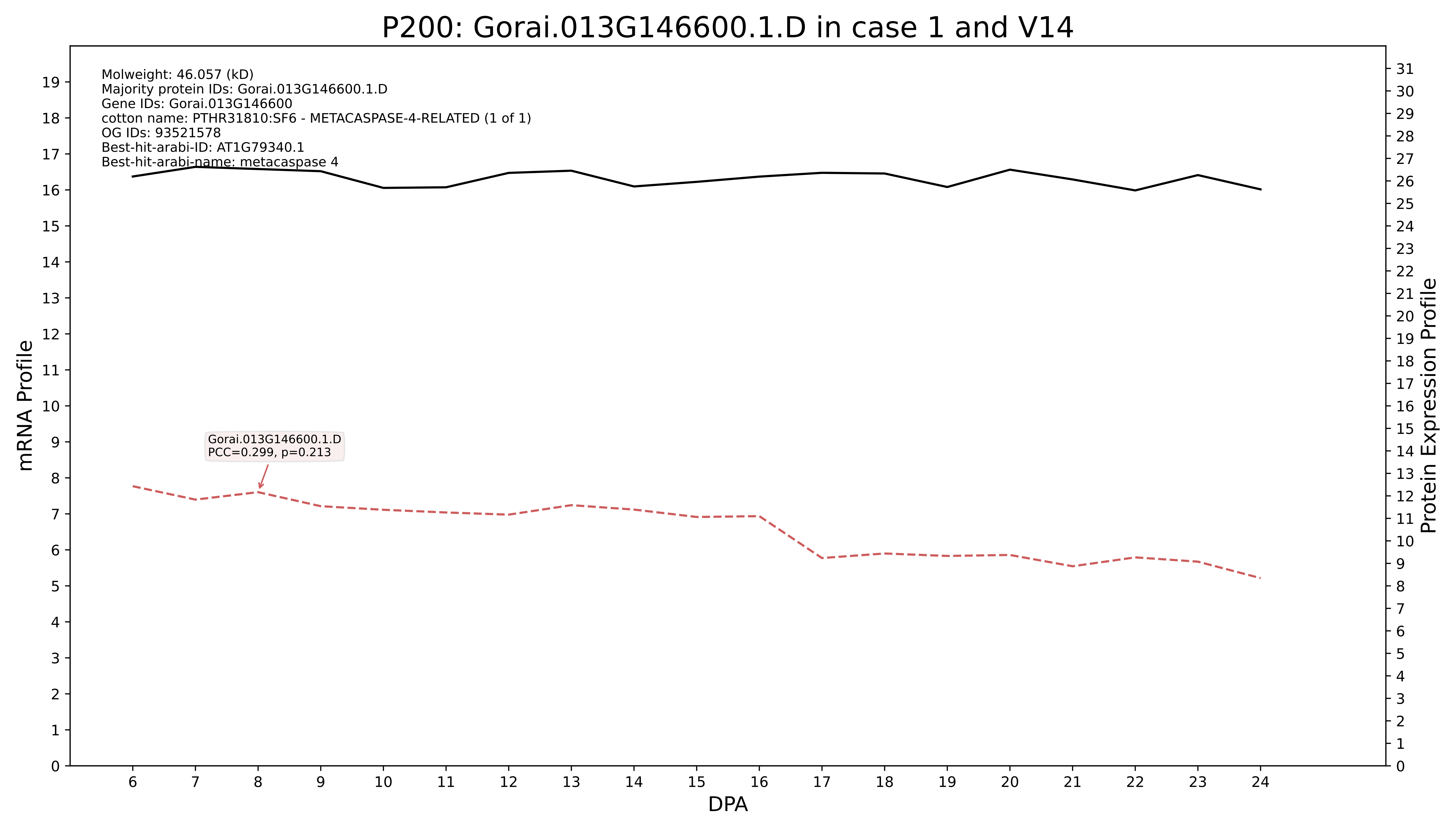

P200: Gorai.013G146600.1.D in case 1 and V14
Molweight: 46.057 (kD)
Majority protein IDs: Gorai.013G146600.1.D
Gene IDs: Gorai.013G146600
cotton name: PTHR31810:SF6 - METACASPASE-4-RELATED (1 of 1)
OG IDs: 93521578
Best-hit-arabi-ID: AT1G79340.1
Best-hit-arabi-name: metacaspase 4
Gorai.013G146600.1.D
PCC=0.299, p=0.213
mRNA Profile
Protein Expression Profile
DPA

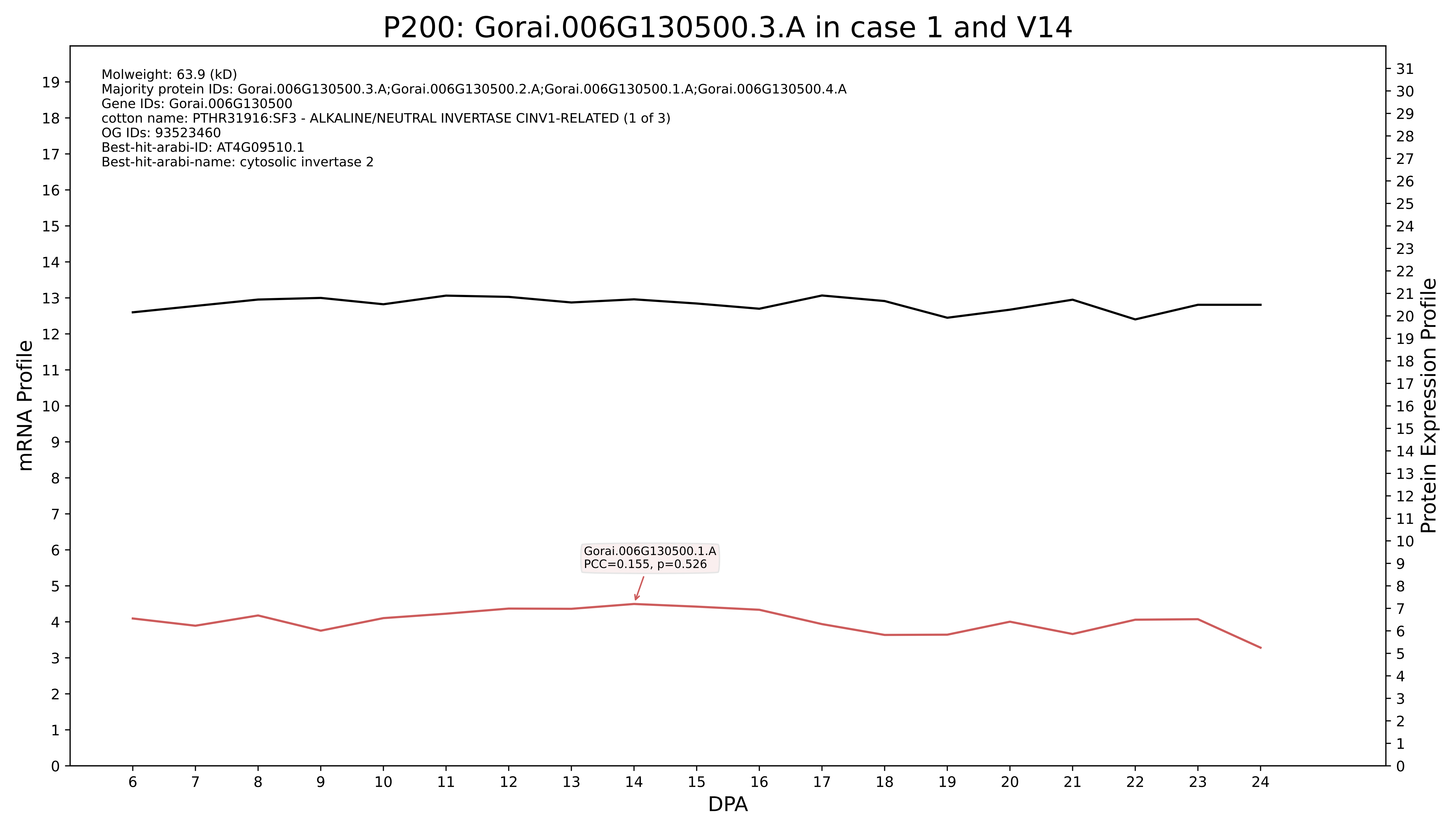
P200: Gorai.006G130500.3.A in case 1 and V14
Molweight: 63.9 (kD)
Majority protein IDs: Gorai.006G130500.3.A;Gorai.006G130500.2.A;Gorai.006G130500.1.A;Gorai.006G130500.4.A
Gene IDs: Gorai.006G130500
cotton name: PTHR31916:SF3 - ALKALINE/NEUTRAL INVERTASE CINV1-RELATED (1 of 3)
OG IDs: 93523460
Best-hit-arabi-ID: AT4G09510.1
Best-hit-arabi-name: cytosolic invertase 2
Gorai.006G130500.1.A
PCC=0.155, p=0.526
mRNA Profile
Protein Expression Profile
DPA

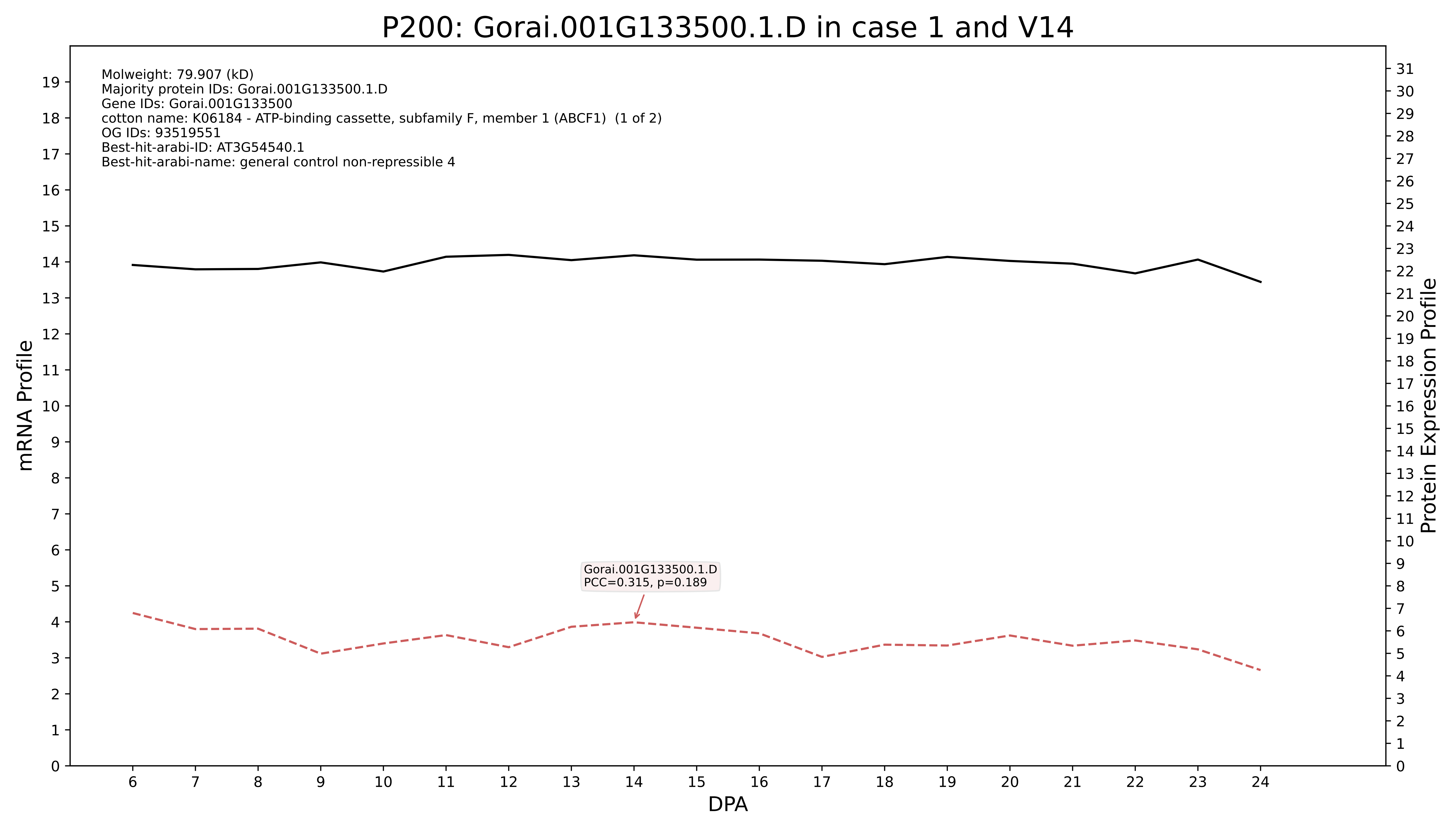
P200: Gorai.001G133500.1.D in case 1 and V14
Molweight: 79.907 (kD)
Majority protein IDs: Gorai.001G133500.1.D
Gene IDs: Gorai.001G133500
cotton name: K06184 - ATP-binding cassette, subfamily F, member 1 (ABCF1) (1 of 2)
OG IDs: 93519551
Best-hit-arabi-ID: AT3G54540.1
Best-hit-arabi-name: general control non-repressible 4
Gorai.001G133500.1.D
PCC=0.315, p=0.189
mRNA Profile
Protein Expression Profile
DPA

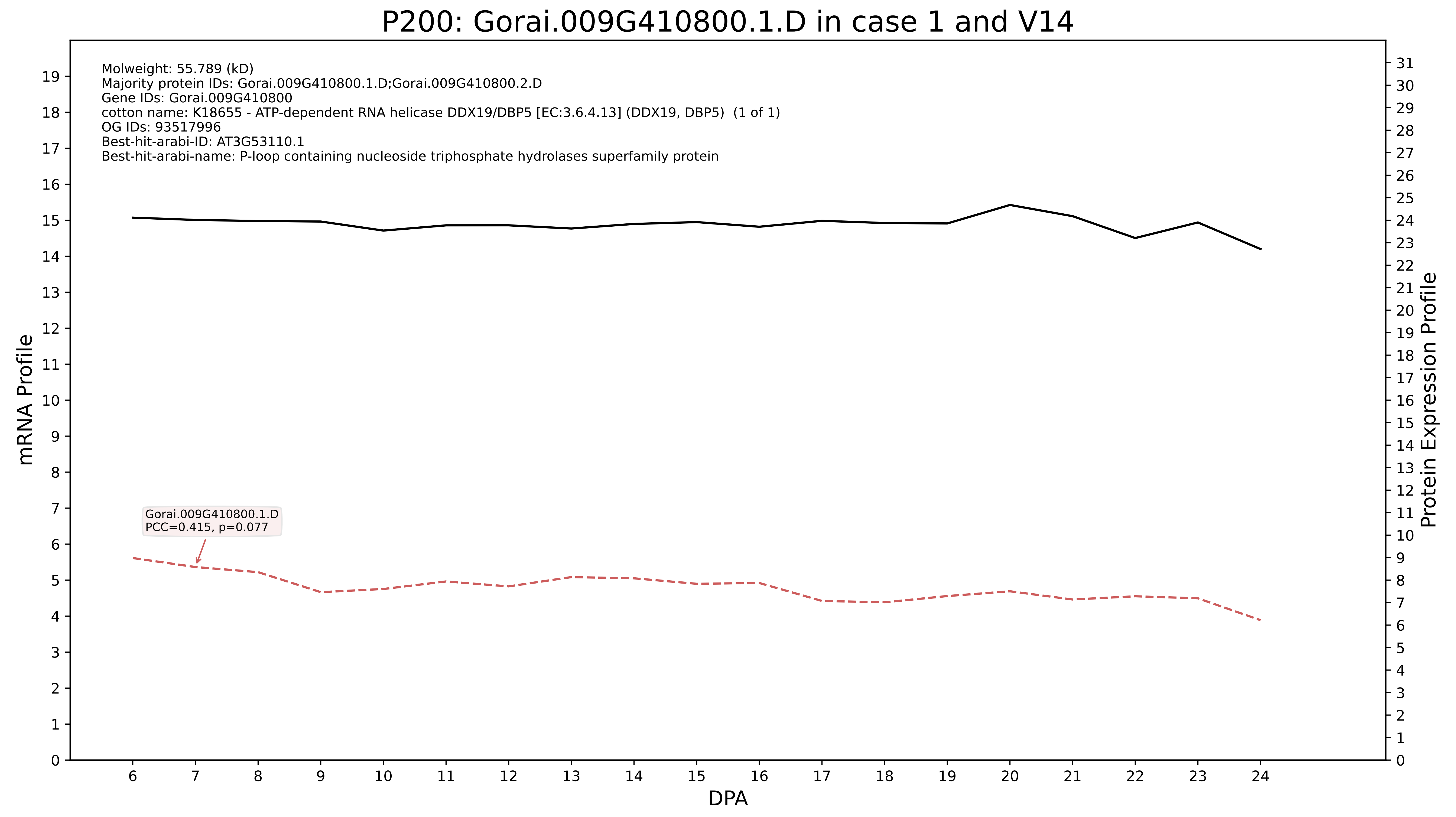

P200: Gorai.009G410800.1.D in case 1 and V14
Molweight: 55.789 (kD)
Majority protein IDs: Gorai.009G410800.1.D;Gorai.009G410800.2.D
Gene IDs: Gorai.009G410800
cotton name: K18655 - ATP-dependent RNA helicase DDX19/DBP5 [EC:3.6.4.13] (DDX19, DBP5) (1 of 1)
OG IDs: 93517996
Best-hit-arabi-ID: AT3G53110.1
Best-hit-arabi-name: P-loop containing nucleoside triphosphate hydrolases superfamily protein
Gorai.009G410800.1.D
PCC=0.415, p=0.077
mRNA Profile
Protein Expression Profile
DPA

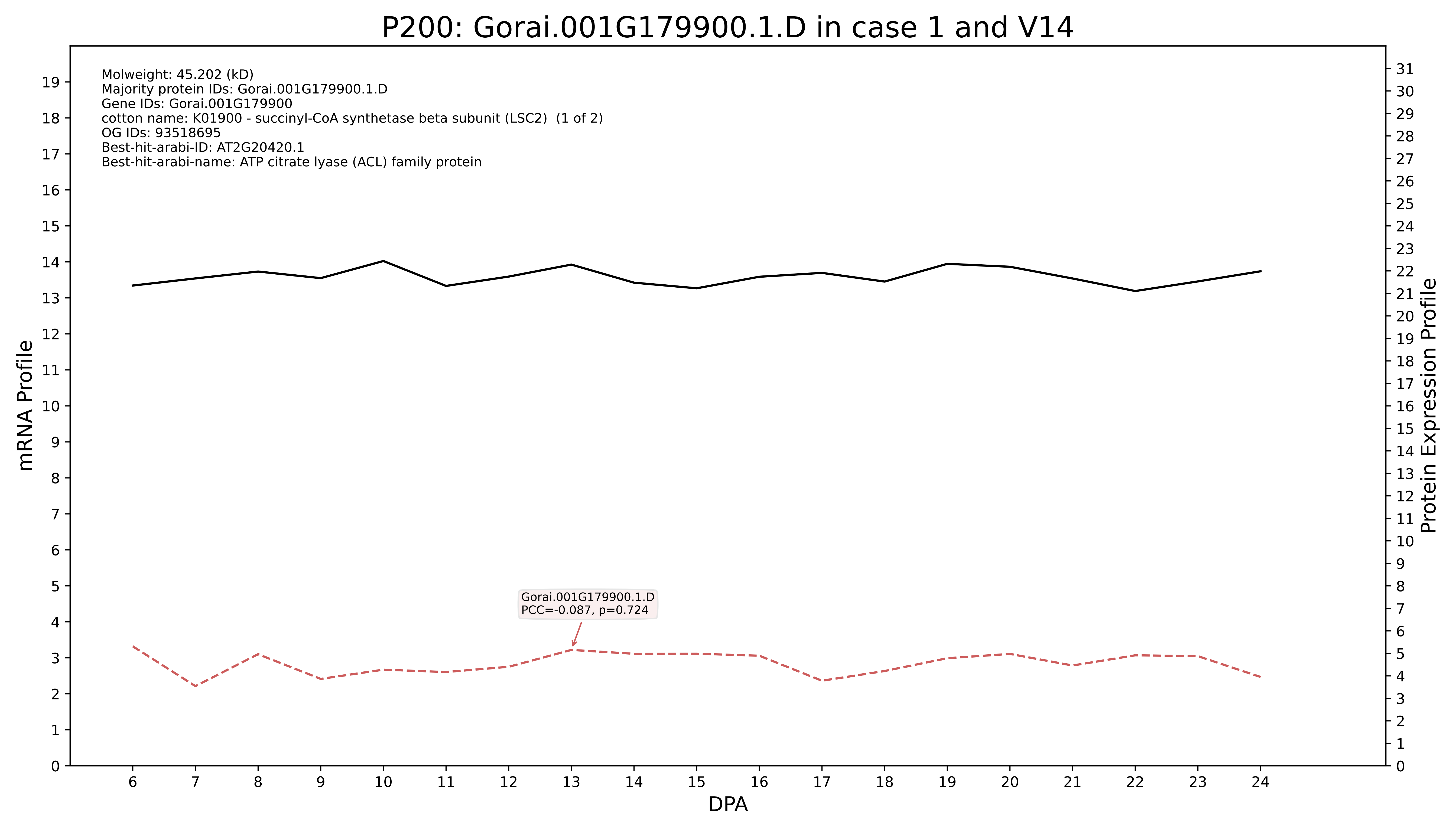
P200: Gorai.001G179900.1.D in case 1 and V14
Molweight: 45.202 (kD)
Majority protein IDs: Gorai.001G179900.1.D
Gene IDs: Gorai.001G179900
cotton name: K01900 - succinyl-CoA synthetase beta subunit (LSC2) (1 of 2)
OG IDs: 93518695
Best-hit-arabi-ID: AT2G20420.1
Best-hit-arabi-name: ATP citrate lyase (ACL) family protein
Gorai.001G179900.1.D
PCC=-0.087, p=0.724
mRNA Profile
Protein Expression Profile
DPA

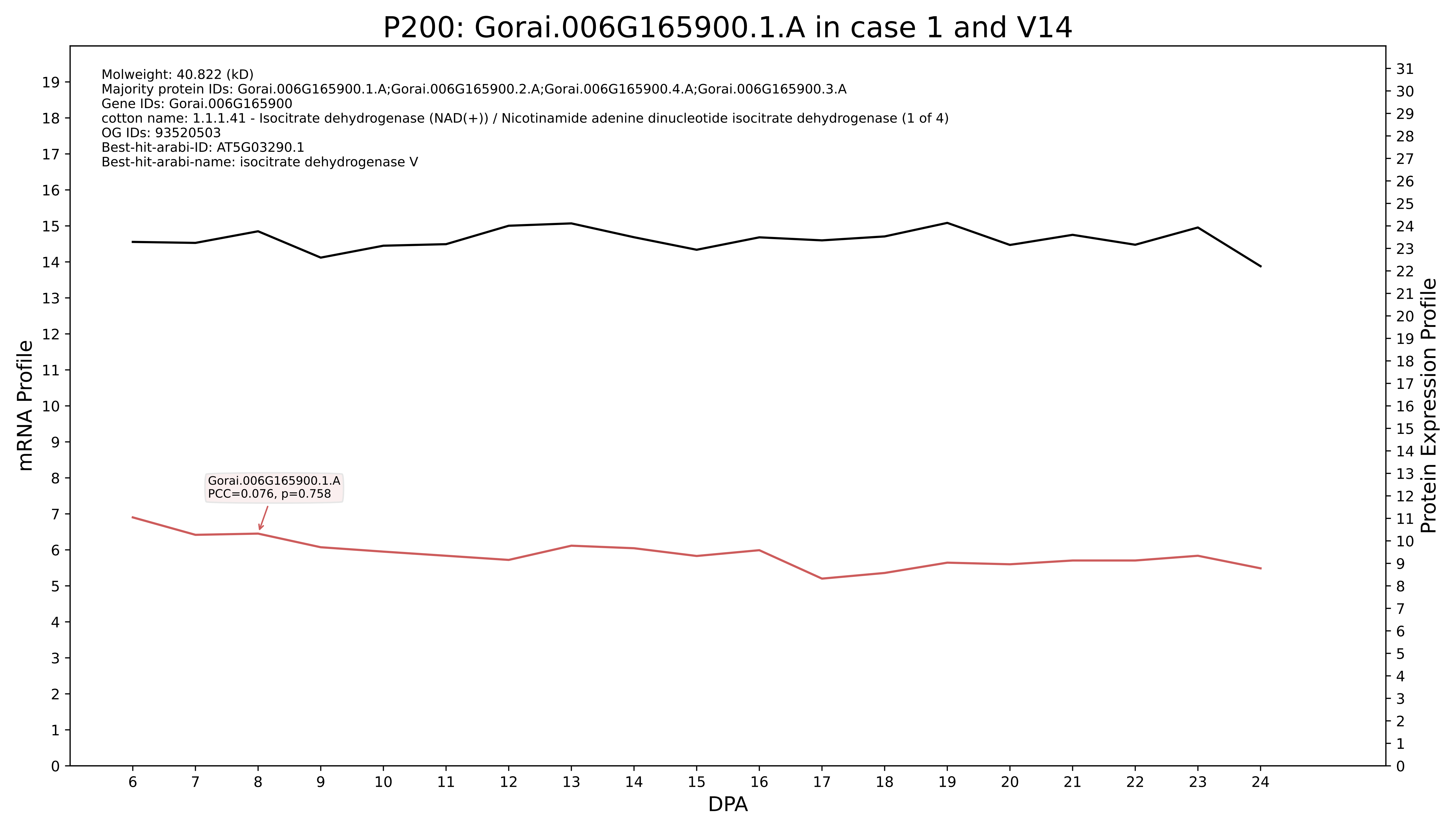
P200: Gorai.006G165900.1.A in case 1 and V14
Molweight: 40.822 (kD)
Majority protein IDs: Gorai.006G165900.1.A;Gorai.006G165900.2.A;Gorai.006G165900.4.A;Gorai.006G165900.3.A
Gene IDs: Gorai.006G165900
cotton name: 1.1.1.41 - Isocitrate dehydrogenase (NAD(+)) / Nicotinamide adenine dinucleotide isocitrate dehydrogenase (1 of 4)
OG IDs: 93520503
Best-hit-arabi-ID: AT5G03290.1
Best-hit-arabi-name: isocitrate dehydrogenase V
Gorai.006G165900.1.A
PCC=0.076, p=0.758
mRNA Profile
Protein Expression Profile
DPA

### P200: Gorai.001G050100.2.A in case 1 and V14

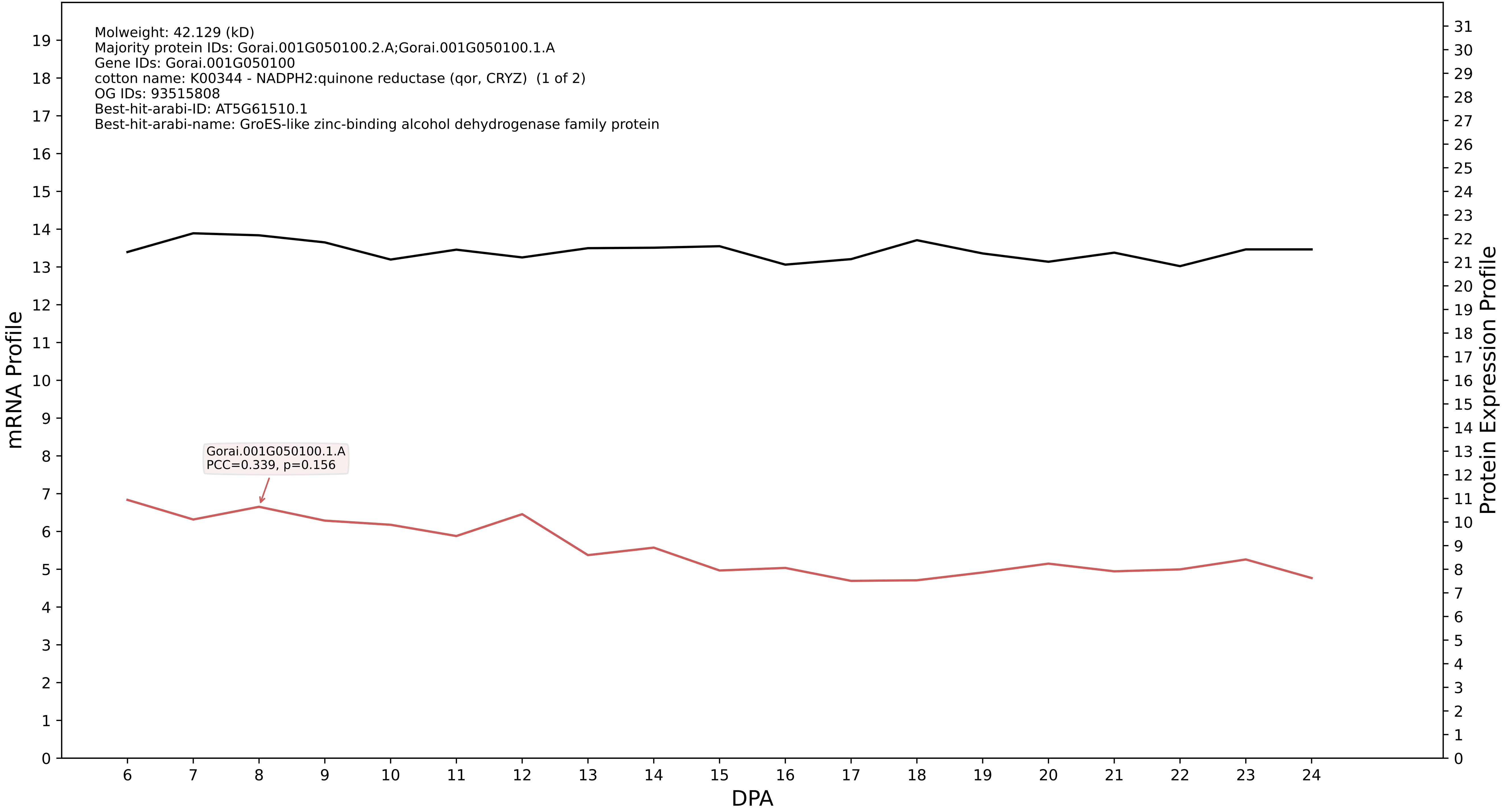

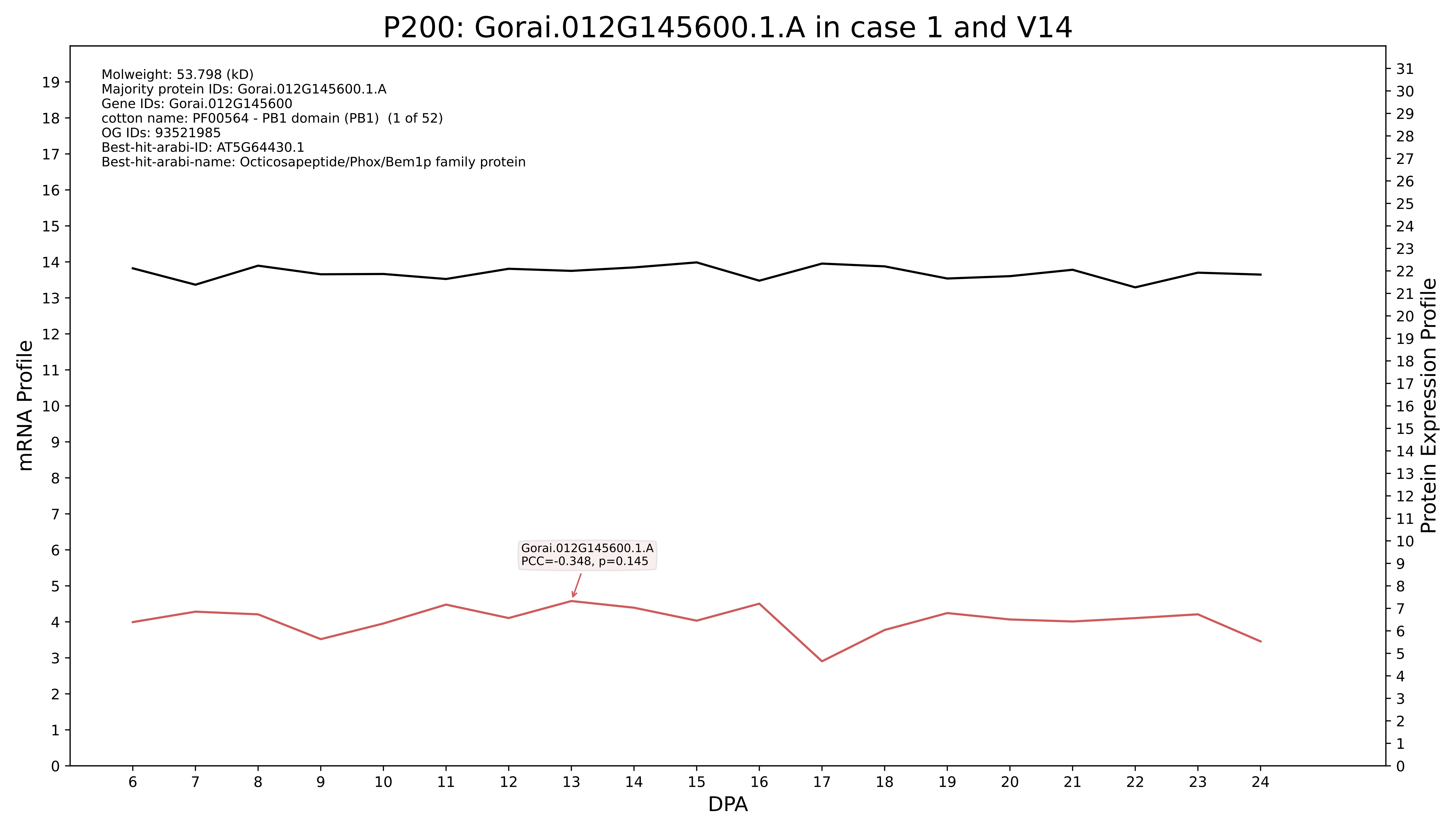

P200: Gorai.012G145600.1.A in case 1 and V14
Molweight: 53.798 (kD)
Majority protein IDs: Gorai.012G145600.1.A
Gene IDs: Gorai.012G145600
cotton name: PF00564 - PB1 domain (PB1) (1 of 52)
OG IDs: 93521985
Best-hit-arabi-ID: AT5G64430.1
Best-hit-arabi-name: Octicosapeptide/Phox/Bem1p family protein
Gorai.012G145600.1.A
PCC=-0.348, p=0.145
mRNA Profile
Protein Expression Profile
DPA

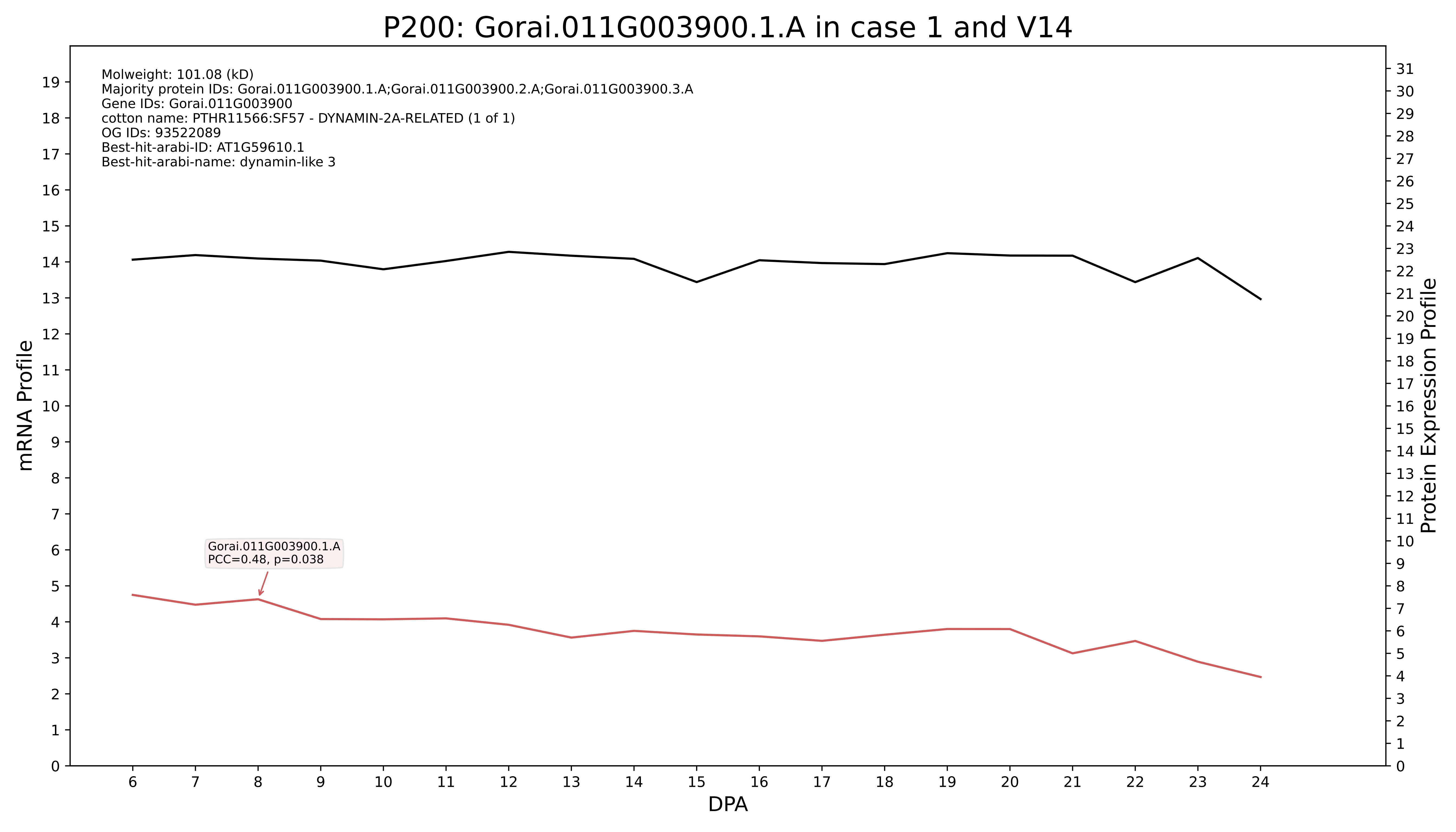

P200: Gorai.011G003900.1.A in case 1 and V14
Molweight: 101.08 (kD)
Majority protein IDs: Gorai.011G003900.1.A;Gorai.011G003900.2.A;Gorai.011G003900.3.A
Gene IDs: Gorai.011G003900
cotton name: PTHR11566:SF57 - DYNAMIN-2A-RELATED (1 of 1)
OG IDs: 93522089
Best-hit-arabi-ID: AT1G59610.1
Best-hit-arabi-name: dynamin-like 3
Gorai.011G003900.1.A
PCC=0.48, p=0.038
mRNA Profile
Protein Expression Profile
DPA

### P200: Gorai.011G112000.3.A in case 1 and V14

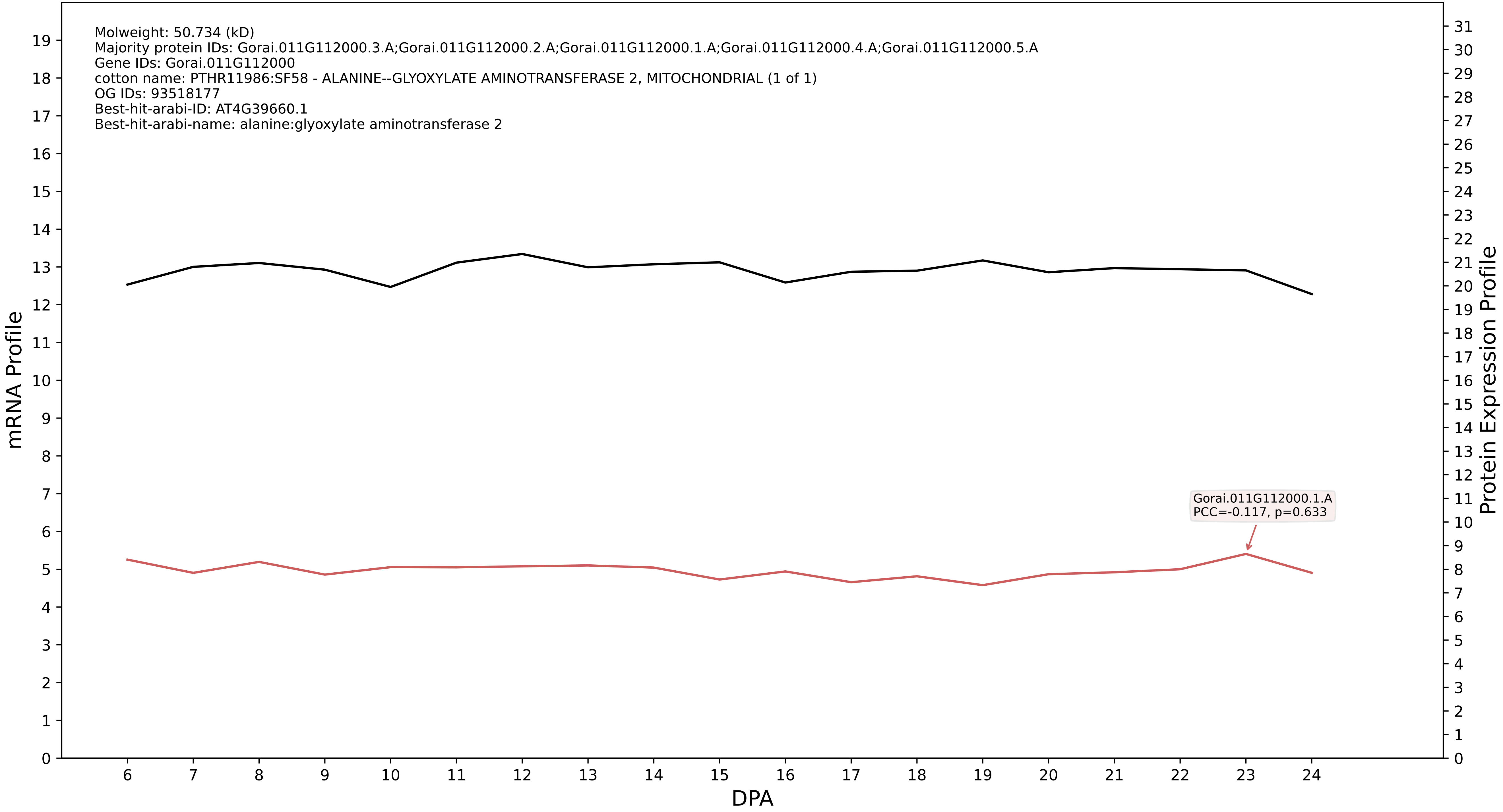

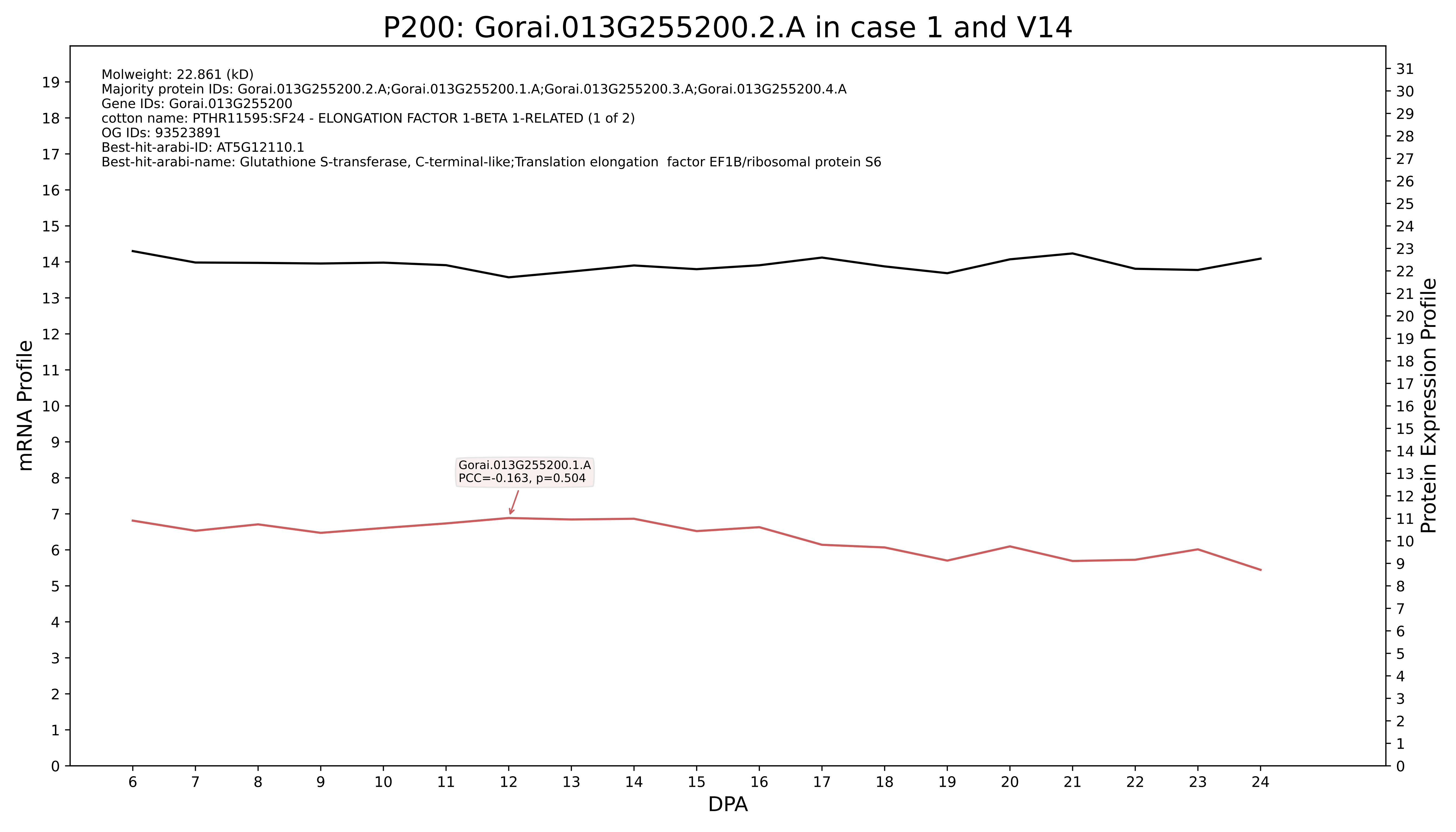

P200: Gorai.013G255200.2.A in case 1 and V14
Molweight: 22.861 (kD)
Majority protein IDs: Gorai.013G255200.2.A;Gorai.013G255200.1.A;Gorai.013G255200.3.A;Gorai.013G255200.4.A
Gene IDs: Gorai.013G255200
cotton name: PTHR11595:SF24 - ELONGATION FACTOR 1-BETA 1-RELATED (1 of 2)
OG IDs: 93523891
Best-hit-arabi-ID: AT5G12110.1
Best-hit-arabi-name: Glutathione S-transferase, C-terminal-like;Translation elongation factor EF1B/ribosomal protein S6
Gorai.013G255200.1.A
PCC=-0.163, p=0.504
mRNA Profile
Protein Expression Profile
DPA

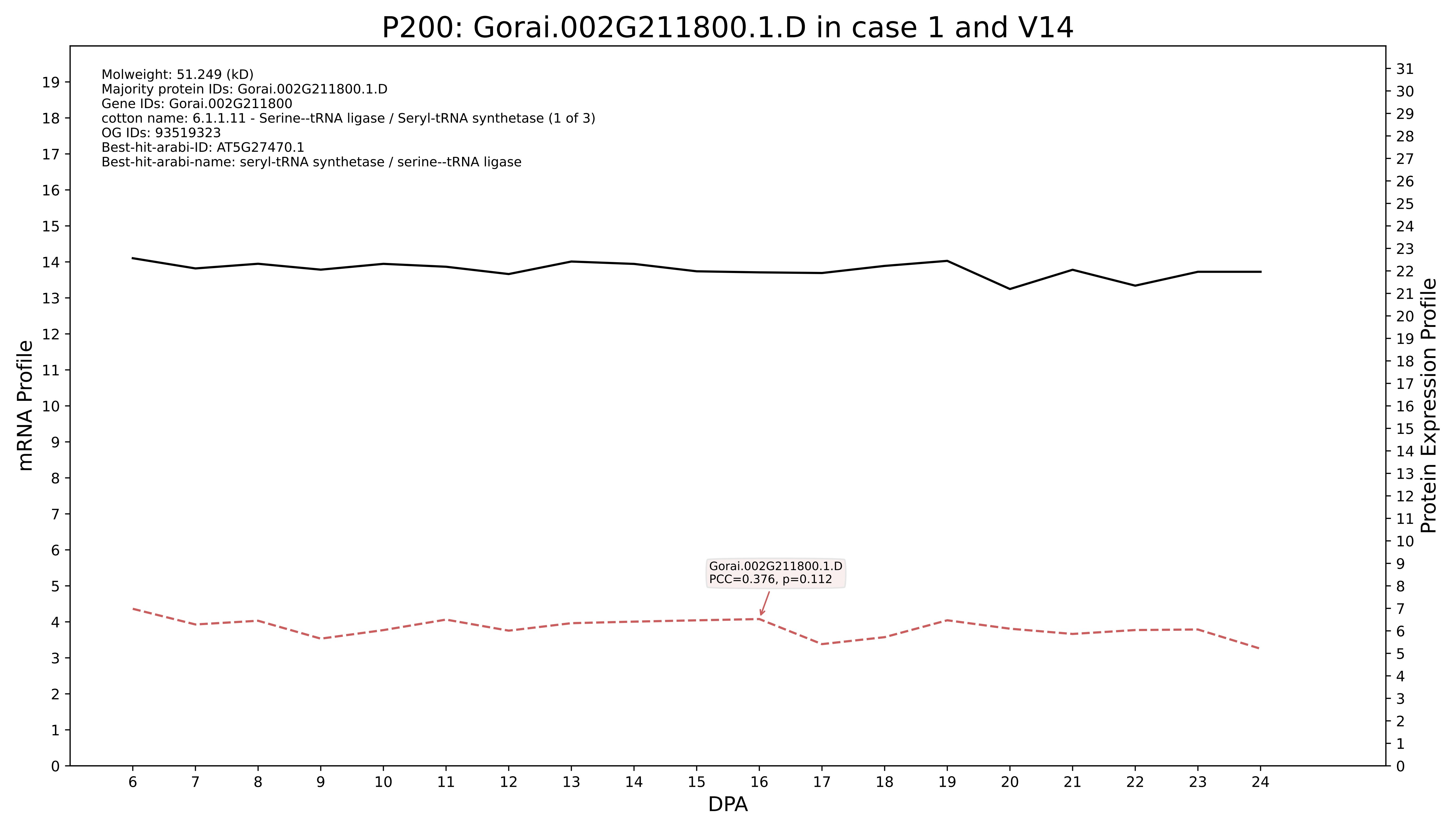

P200: Gorai.002G211800.1.D in case 1 and V14
Molweight: 51.249 (kD)
Majority protein IDs: Gorai.002G211800.1.D
Gene IDs: Gorai.002G211800
cotton name: 6.1.1.11 - Serine--tRNA ligase / Seryl-tRNA synthetase (1 of 3)
OG IDs: 93519323
Best-hit-arabi-ID: AT5G27470.1
Best-hit-arabi-name: seryl-tRNA synthetase / serine--tRNA ligase
Gorai.002G211800.1.D
PCC=0.376, p=0.112
mRNA Profile
Protein Expression Profile
DPA

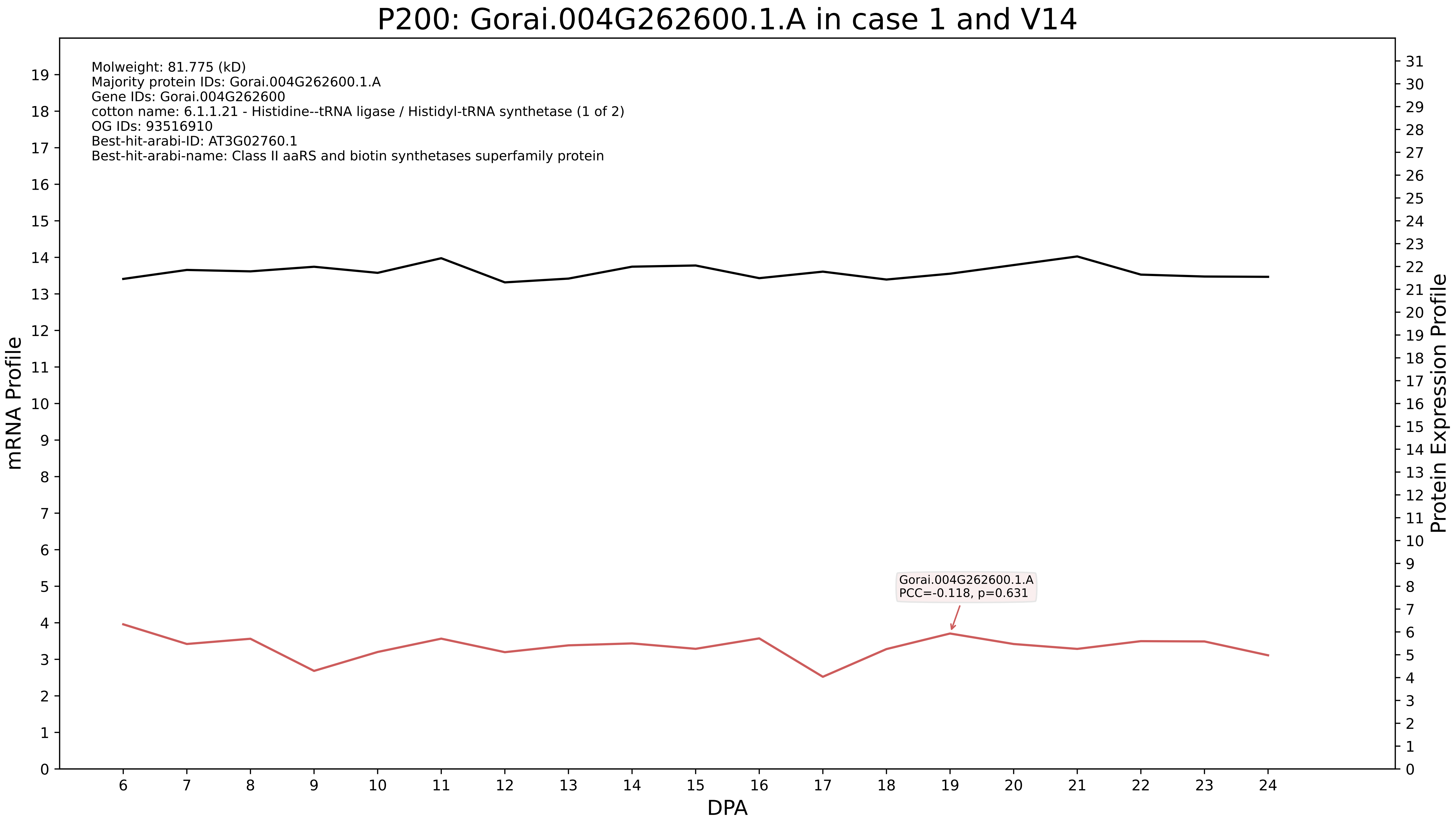
P200: Gorai.004G262600.1.A in case 1 and V14
Molweight: 81.775 (kD)
Majority protein IDs: Gorai.004G262600.1.A
Gene IDs: Gorai.004G262600
cotton name: 6.1.1.21 - Histidine--tRNA ligase / Histidyl-tRNA synthetase (1 of 2)
OG IDs: 93516910
Best-hit-arabi-ID: AT3G02760.1
Best-hit-arabi-name: Class II aaRS and biotin synthetases superfamily protein
Gorai.004G262600.1.A
PCC=-0.118, p=0.631
mRNA Profile
Protein Expression Profile
DPA

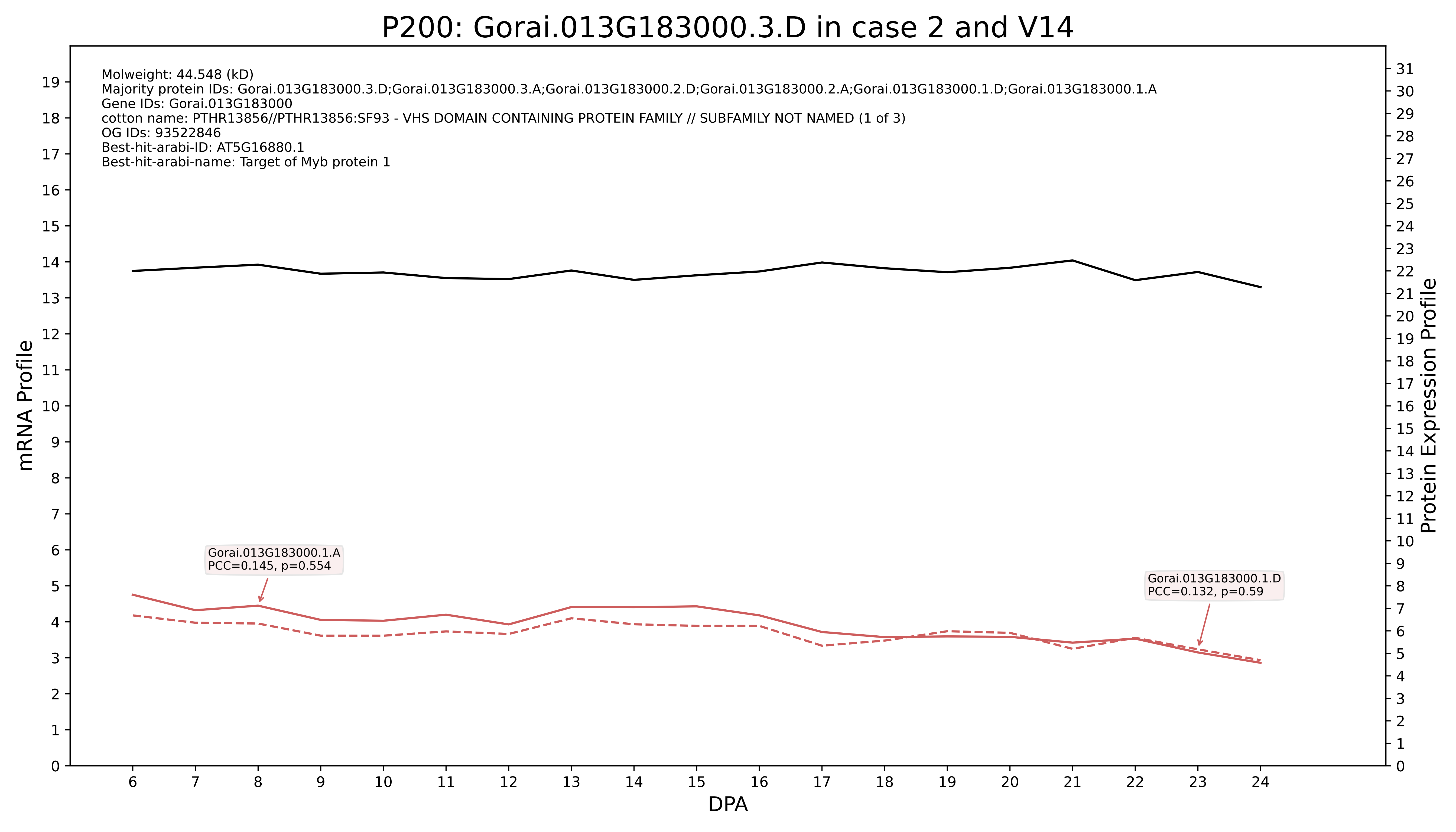
P200: Gorai.013G183000.3.D in case 2 and V14
Molweight: 44.548 (kD)
Majority protein IDs: Gorai.013G183000.3.D;Gorai.013G183000.3.A;Gorai.013G183000.2.D;Gorai.013G183000.2.A;Gorai.013G183000.1.D;Gorai.013G183000.1.A
Gene IDs: Gorai.013G183000
cotton name: PTHR13856//PTHR13856:SF93 - VHS DOMAIN CONTAINING PROTEIN FAMILY // SUBFAMILY NOT NAMED (1 of 3)
OG IDs: 93522846
Best-hit-arabi-ID: AT5G16880.1
Best-hit-arabi-name: Target of Myb protein 1
Gorai.013G183000.1.A
PCC=0.145, p=0.554
Gorai.013G183000.1.D
PCC=0.132, p=0.59
mRNA Profile
Protein Expression Profile
DPA

00: Gorai.001G121400.4.D in case 2 and V14

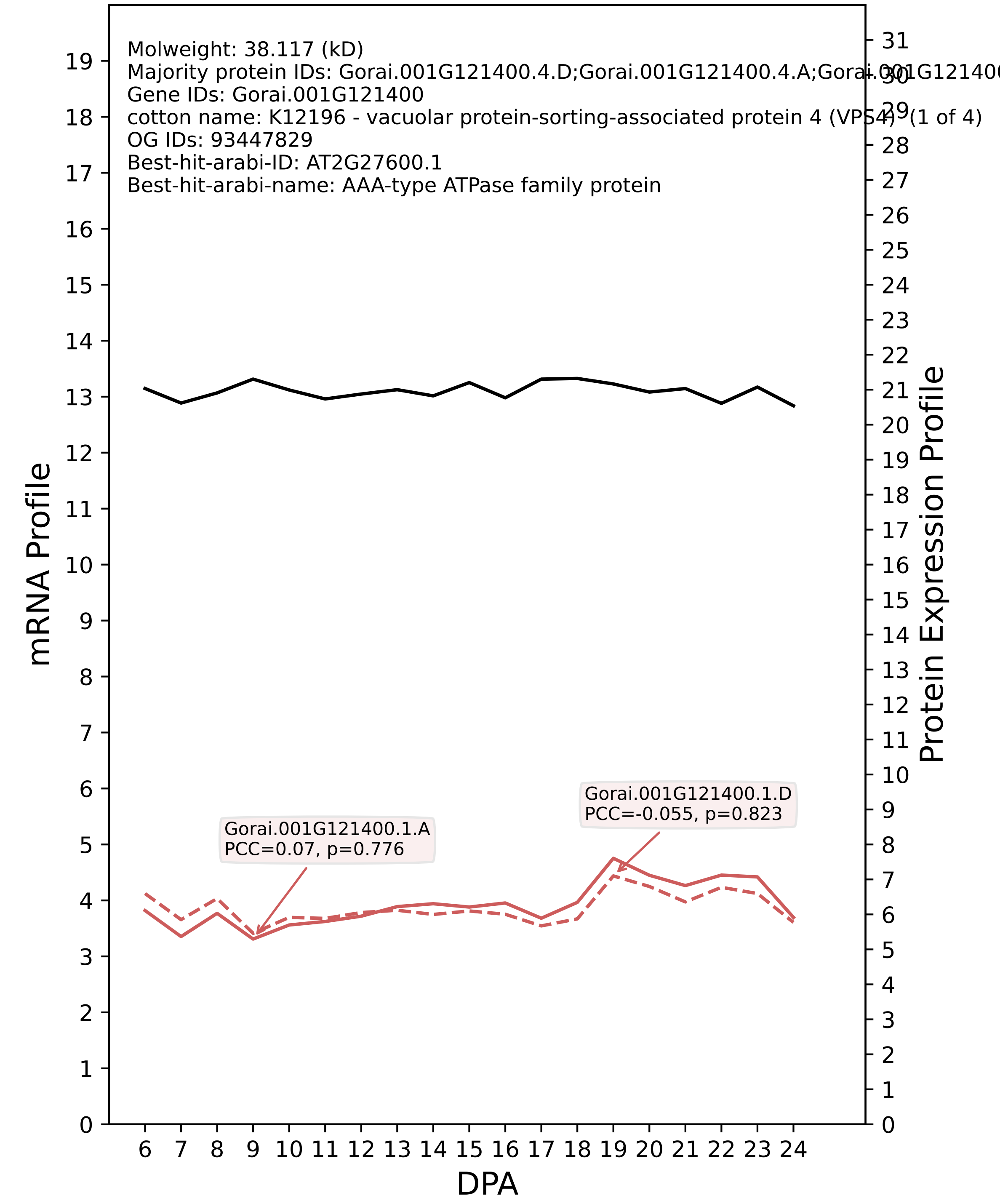

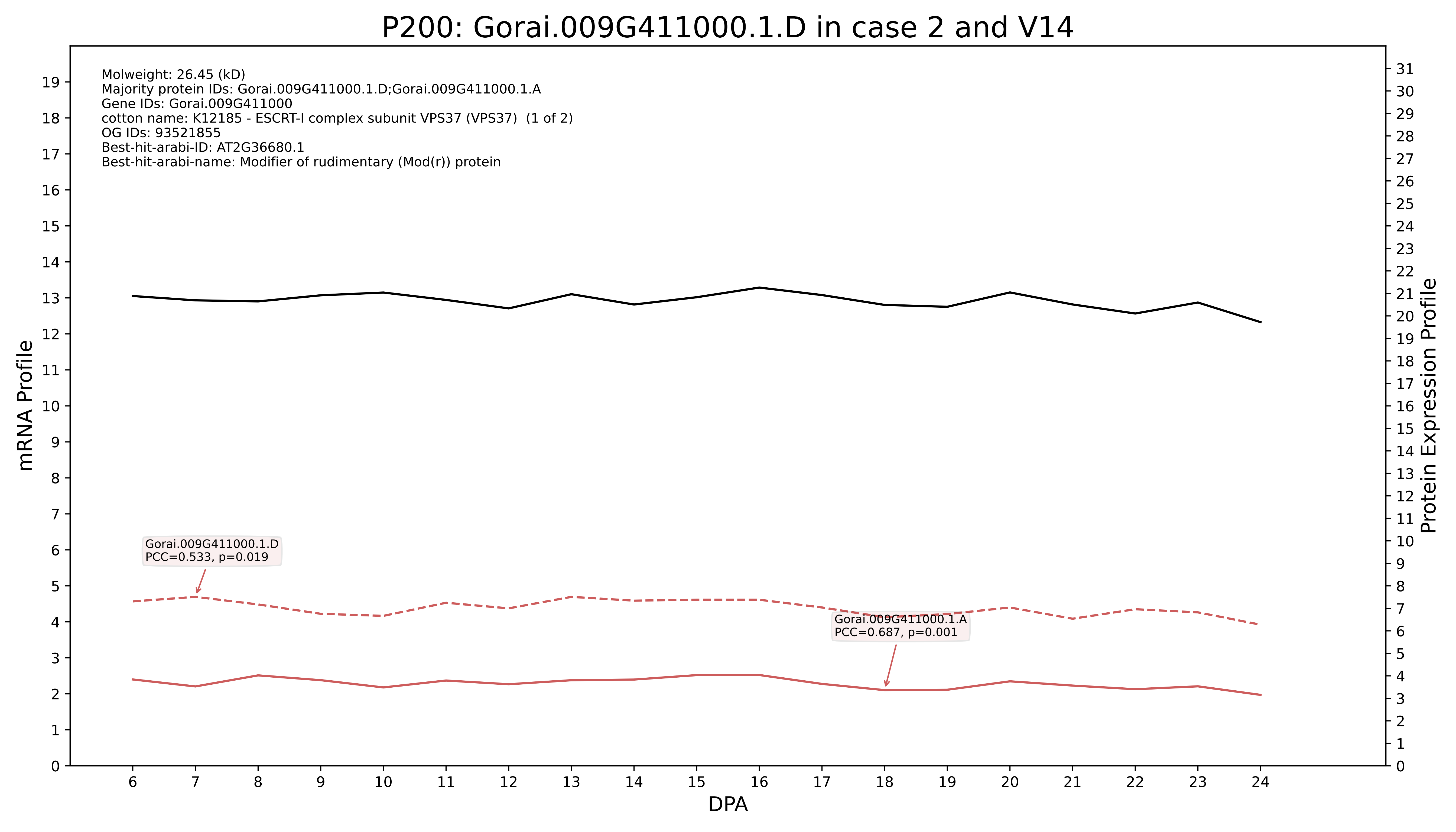
P200: Gorai.009G411000.1.D in case 2 and V14
Molweight: 26.45 (kD)
Majority protein IDs: Gorai.009G411000.1.D;Gorai.009G411000.1.A
Gene IDs: Gorai.009G411000
cotton name: K12185 - ESCRT-I complex subunit VPS37 (VPS37) (1 of 2)
OG IDs: 93521855
Best-hit-arabi-ID: AT2G36680.1
Best-hit-arabi-name: Modifier of rudimentary (Mod(r)) protein
Gorai.009G411000.1.D
PCC=0.533, p=0.019
Gorai.009G411000.1.A
PCC=0.687, p=0.001
mRNA Profile
Protein Expression Profile
DPA

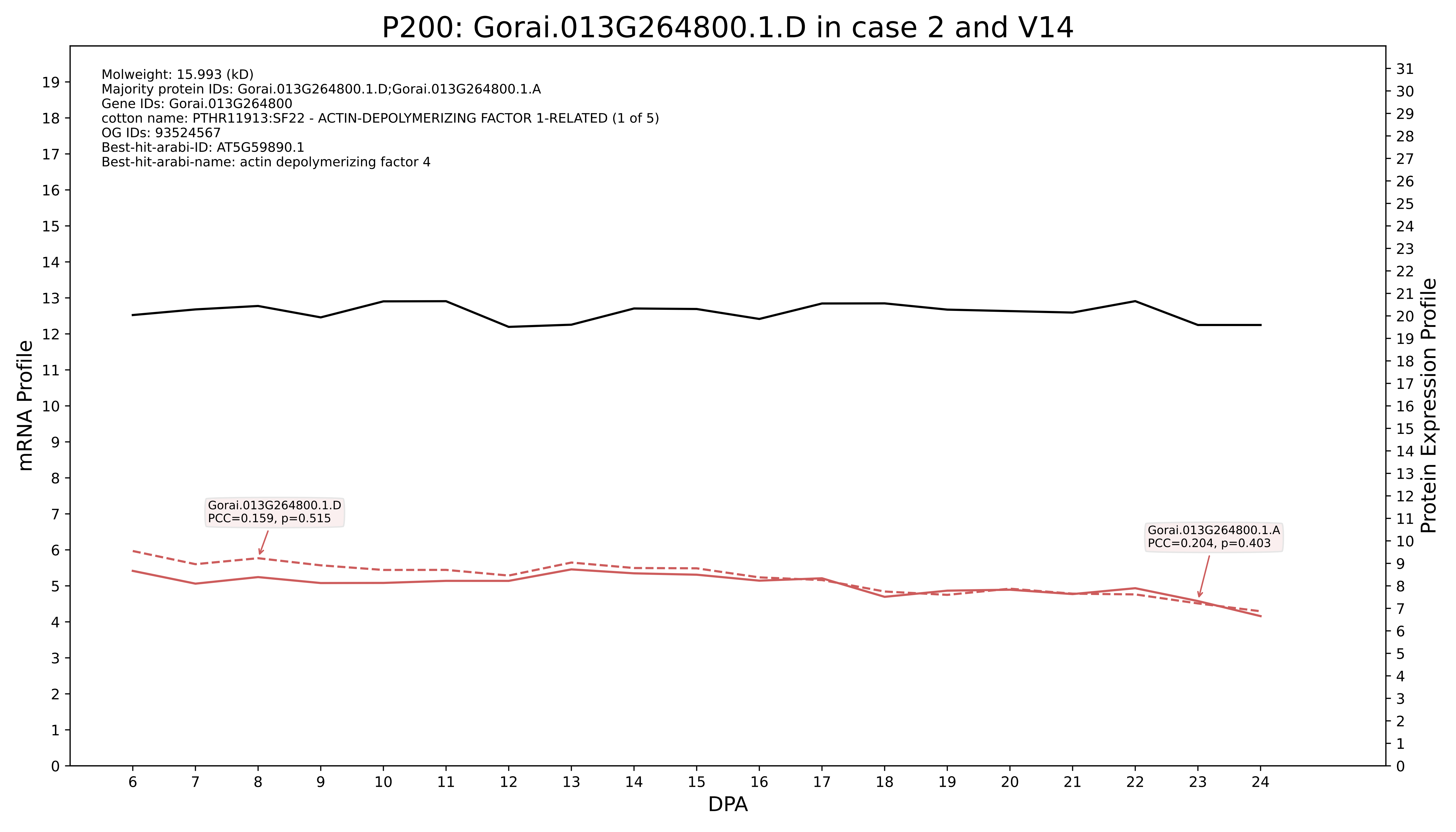
P200: Gorai.013G264800.1.D in case 2 and V14
Molweight: 15.993 (kD)
Majority protein IDs: Gorai.013G264800.1.D;Gorai.013G264800.1.A
Gene IDs: Gorai.013G264800
cotton name: PTHR11913:SF22 - ACTIN-DEPOLYMERIZING FACTOR 1-RELATED (1 of 5)
OG IDs: 93524567
Best-hit-arabi-ID: AT5G59890.1
Best-hit-arabi-name: actin depolymerizing factor 4
Gorai.013G264800.1.D
PCC=0.159, p=0.515
Gorai.013G264800.1.A
PCC=0.204, p=0.403
mRNA Profile
Protein Expression Profile
DPA

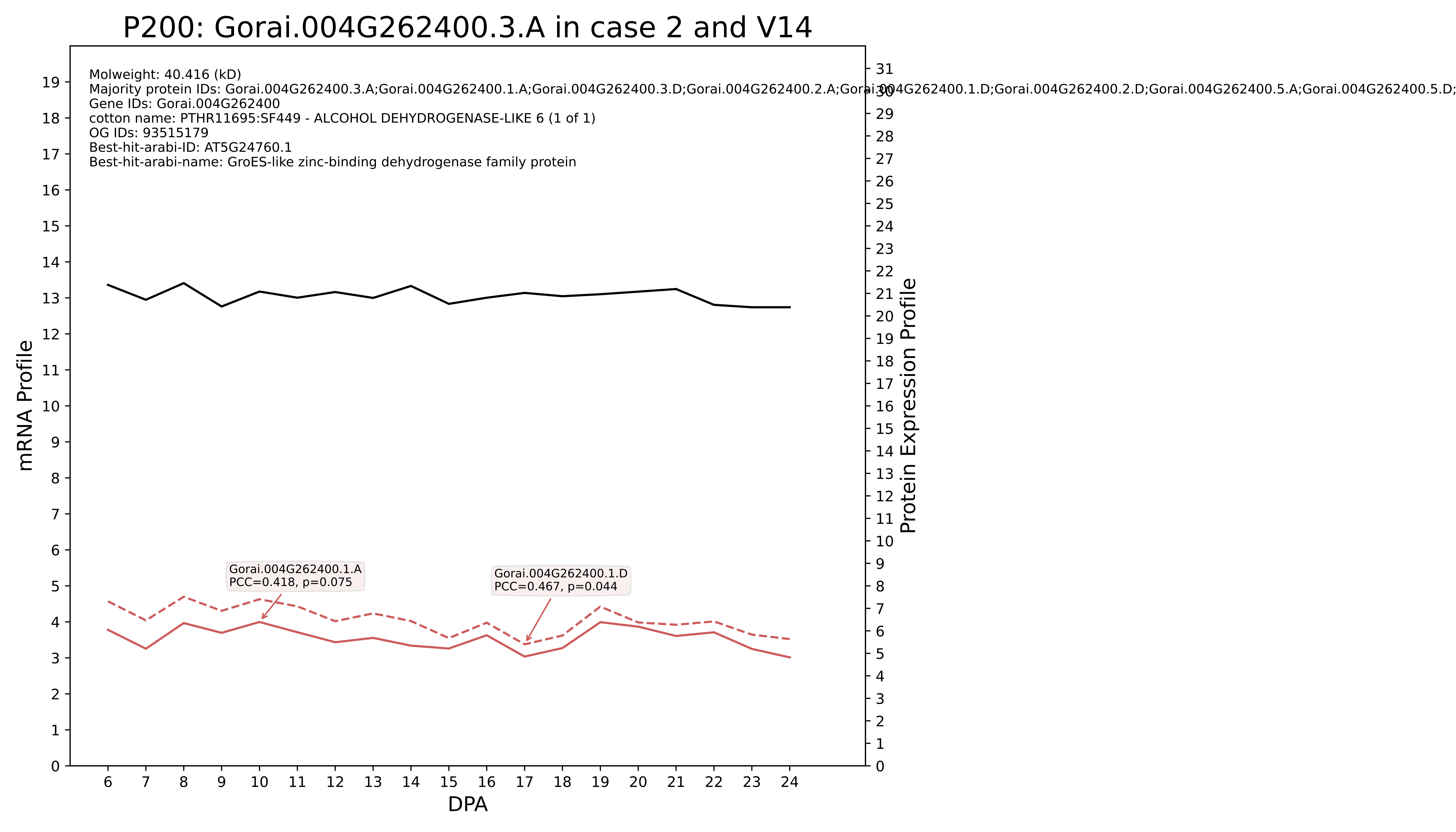
P200: Gorai.004G262400.3.A in case 2 and V14
Molweight: 40.416 (kD)
Majority protein IDs: Gorai.004G262400.3.A;Gorai.004G262400.1.A;Gorai.004G262400.3.D;Gorai.004G262400.2.A;Gorai.004G262400.1.D;Gorai.004G262400.2.D;Gorai.004G262400.5.A;Gorai.004G262400.5.D;
Gene IDs: Gorai.004G262400
cotton name: PTHR11695:SF449 - ALCOHOL DEHYDROGENASE-LIKE 6 (1 of 1)
OG IDs: 93515179
Best-hit-arabi-ID: AT5G24760.1
Best-hit-arabi-name: GroES-like zinc-binding dehydrogenase family protein
Gorai.004G262400.1.A
PCC=0.418, p=0.075
Gorai.004G262400.1.D
PCC=0.467, p=0.044
mRNA Profile
Protein Expression Profile
DPA

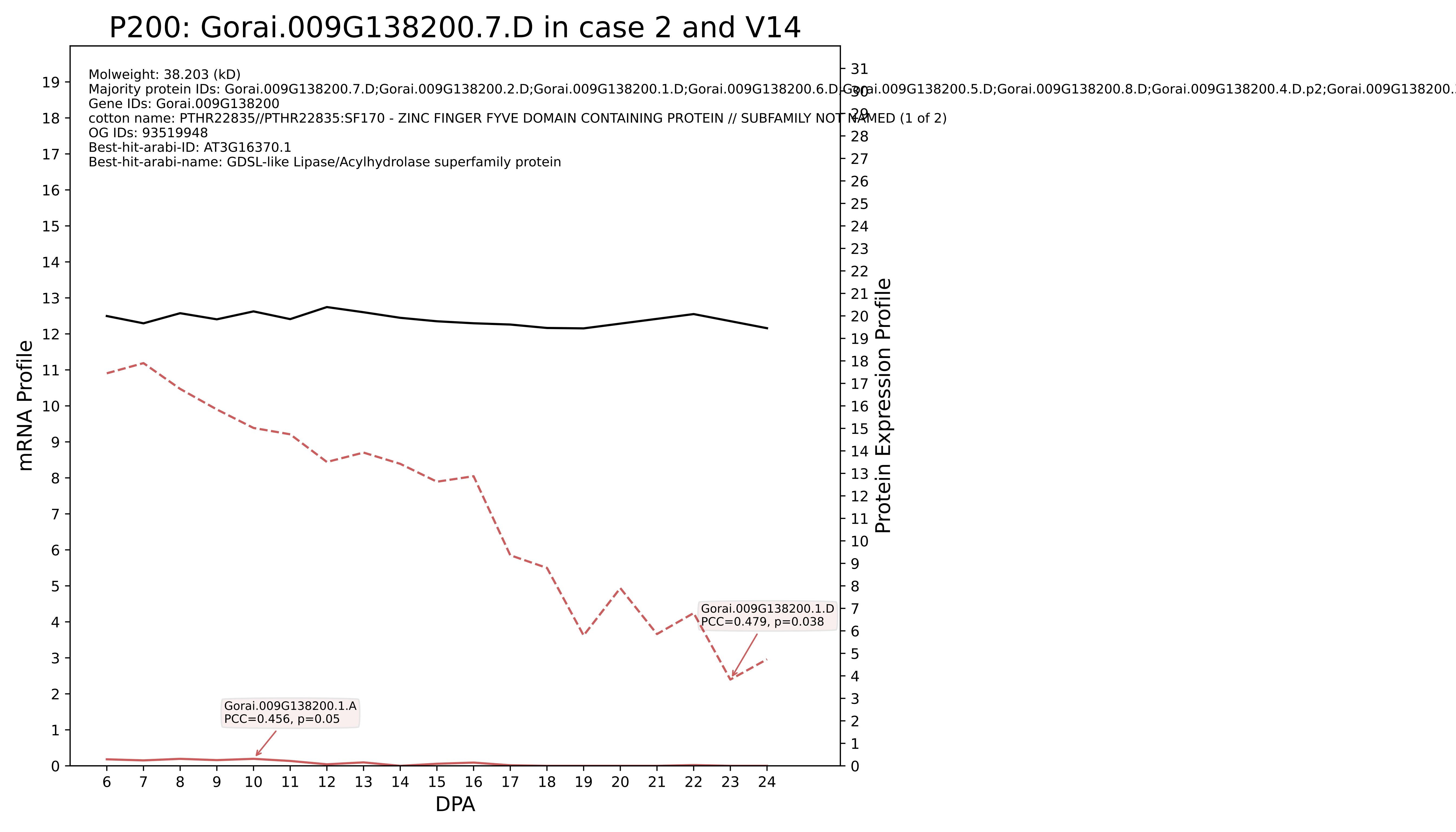
P200: Gorai.009G138200.7.D in case 2 and V14
Molweight: 38.203 (kD)
Majority protein IDs: Gorai.009G138200.7.D;Gorai.009G138200.2.D;Gorai.009G138200.1.D;Gorai.009G138200.6.D;Gorai.009G138200.5.D;Gorai.009G138200.8.D;Gorai.009G138200.4.D.p2;Gorai.009G138200.
Gene IDs: Gorai.009G138200
cotton name: PTHR22835//PTHR22835:SF170 - ZINC FINGER FYVE DOMAIN CONTAINING PROTEIN // SUBFAMILY NOT NAMED (1 of 2)
OG IDs: 93519948
Best-hit-arabi-ID: AT3G16370.1
Best-hit-arabi-name: GDSL-like Lipase/Acylhydrolase superfamily protein
Gorai.009G138200.1.D
PCC=0.479, p=0.038
Gorai.009G138200.1.A
PCC=0.456, p=0.05
mRNA Profile
Protein Expression Profile
DPA

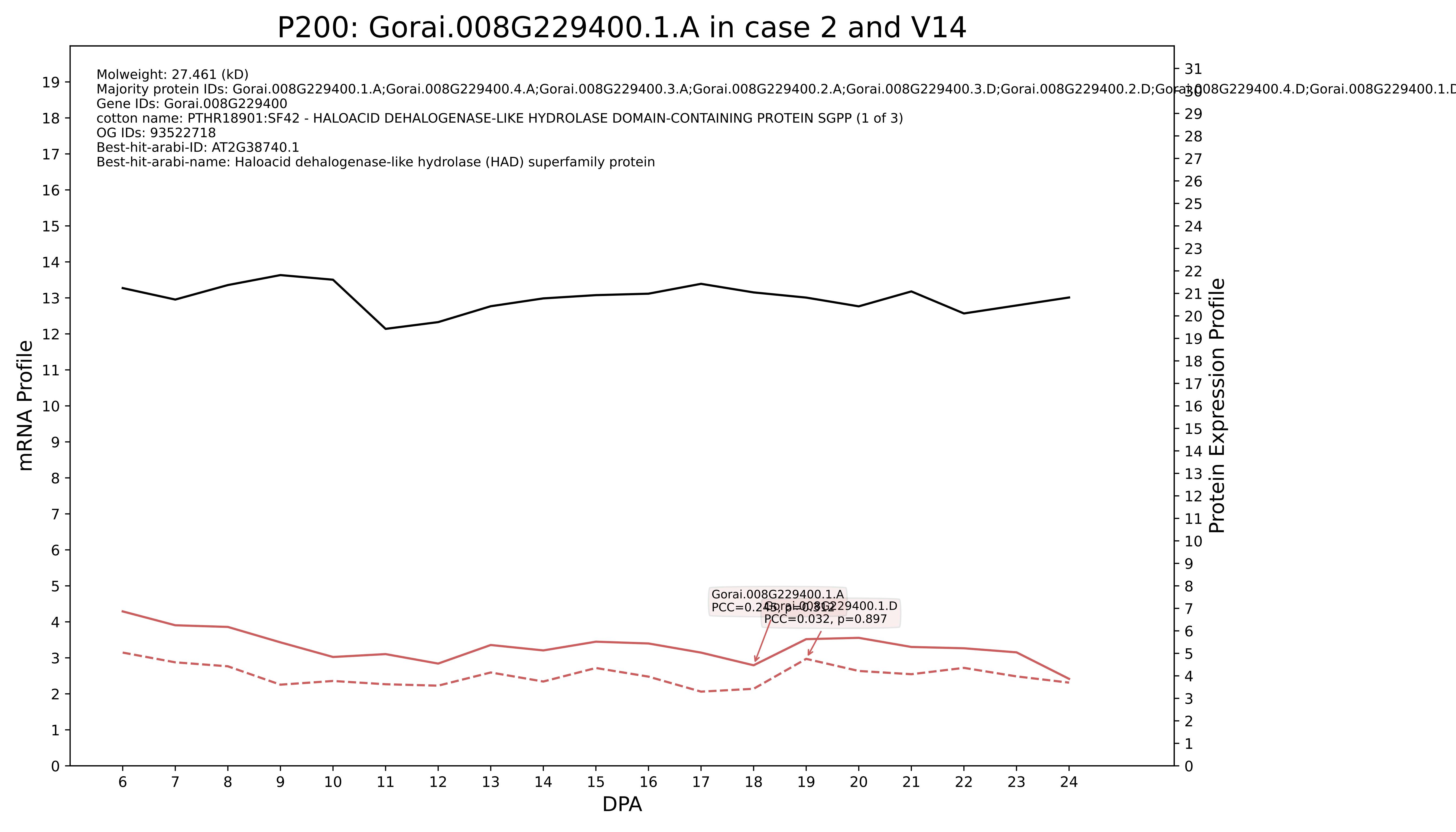
P200: Gorai.008G229400.1.A in case 2 and V14
Molweight: 27.461 (kD)
Majority protein IDs: Gorai.008G229400.1.A;Gorai.008G229400.4.A;Gorai.008G229400.3.A;Gorai.008G229400.2.A;Gorai.008G229400.3.D;Gorai.008G229400.2.D;Gorai.008G229400.4.D;Gorai.008G229400.1.D
Gene IDs: Gorai.008G229400
cotton name: PTHR18901:SF42 - HALOACID DEHALOGENASE-LIKE HYDROLASE DOMAIN-CONTAINING PROTEIN SGPP (1 of 3)
OG IDs: 93522718
Best-hit-arabi-ID: AT2G38740.1
Best-hit-arabi-name: Haloacid dehalogenase-like hydrolase (HAD) superfamily protein
Gorai.008G229400.1.A
Gorai.008G229400.1.D
PCC=0.032, p=0.897
mRNA Profile
Protein Expression Profile
DPA

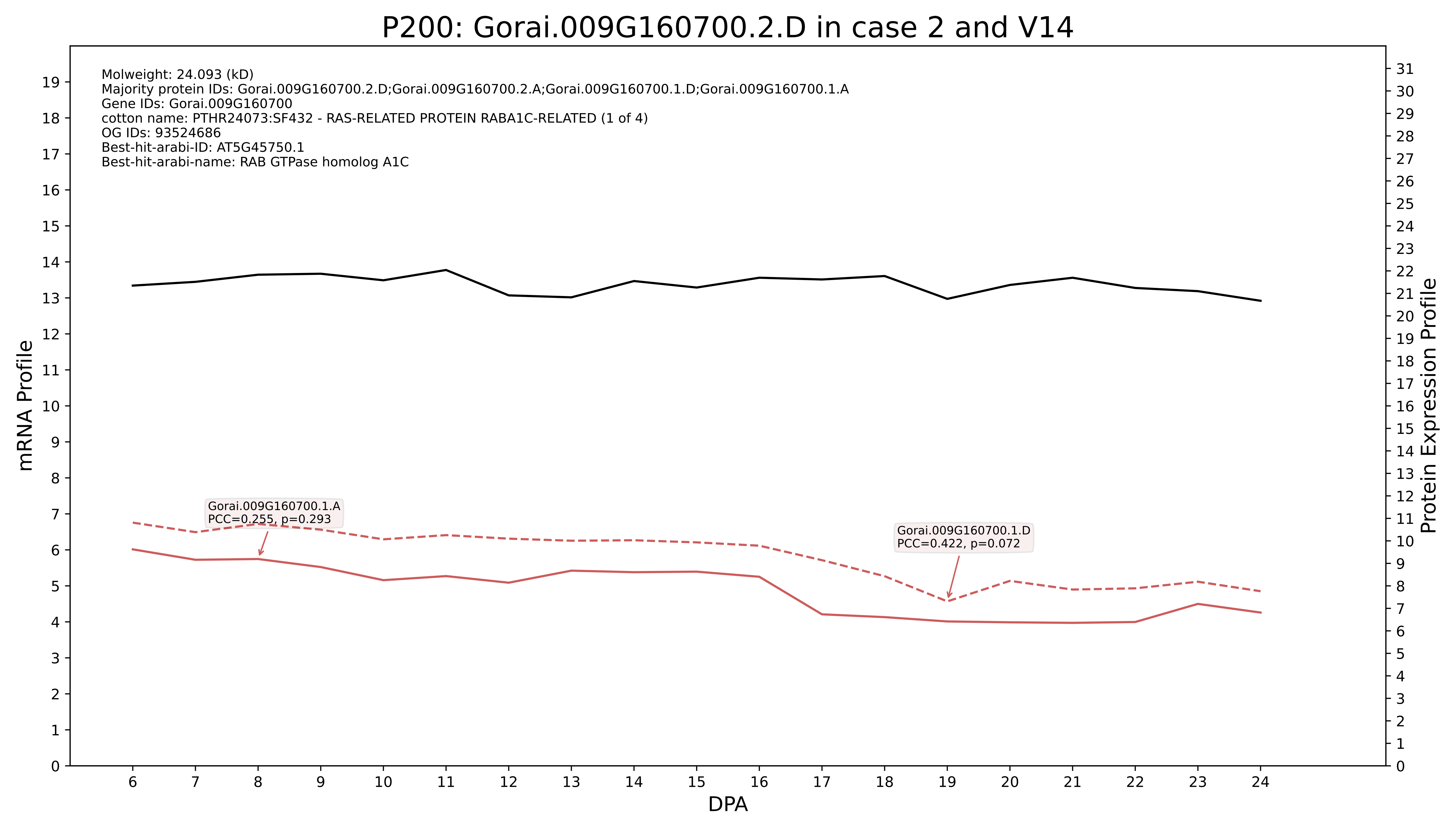

P200: Gorai.009G160700.2.D in case 2 and V14
Molweight: 24.093 (kD)
Majority protein IDs: Gorai.009G160700.2.D;Gorai.009G160700.2.A;Gorai.009G160700.1.D;Gorai.009G160700.1.A
Gene IDs: Gorai.009G160700
cotton name: PTHR24073:SF432 - RAS-RELATED PROTEIN RABA1C-RELATED (1 of 4)
OG IDs: 93524686
Best-hit-arabi-ID: AT5G45750.1
Best-hit-arabi-name: RAB GTPase homolog A1C
Gorai.009G160700.1.A
PCC=0.255, p=0.293
Gorai.009G160700.1.D
PCC=0.422, p=0.072
mRNA Profile
Protein Expression Profile
DPA

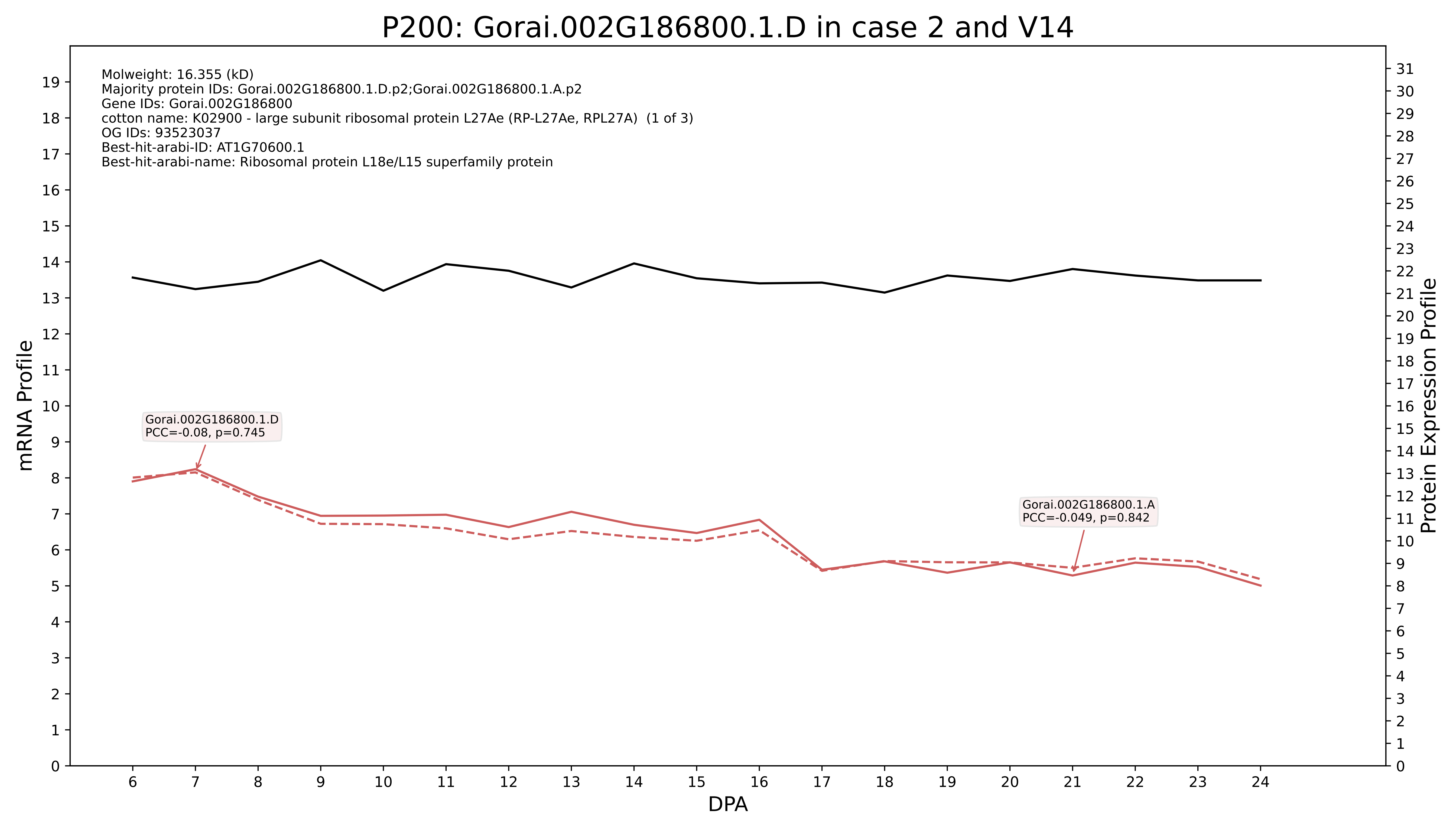

P200: Gorai.002G186800.1.D in case 2 and V14
Molweight: 16.355 (kD)
Majority protein IDs: Gorai.002G186800.1.D.p2;Gorai.002G186800.1.A.p2
Gene IDs: Gorai.002G186800
cotton name: K02900 - large subunit ribosomal protein L27Ae (RP-L27Ae, RPL27A) (1 of 3)
OG IDs: 93523037
Best-hit-arabi-ID: AT1G70600.1
Best-hit-arabi-name: Ribosomal protein L18e/L15 superfamily protein
Gorai.002G186800.1.D
PCC=-0.08, p=0.745
Gorai.002G186800.1.A
PCC=-0.049, p=0.842
mRNA Profile
Protein Expression Profile
DPA

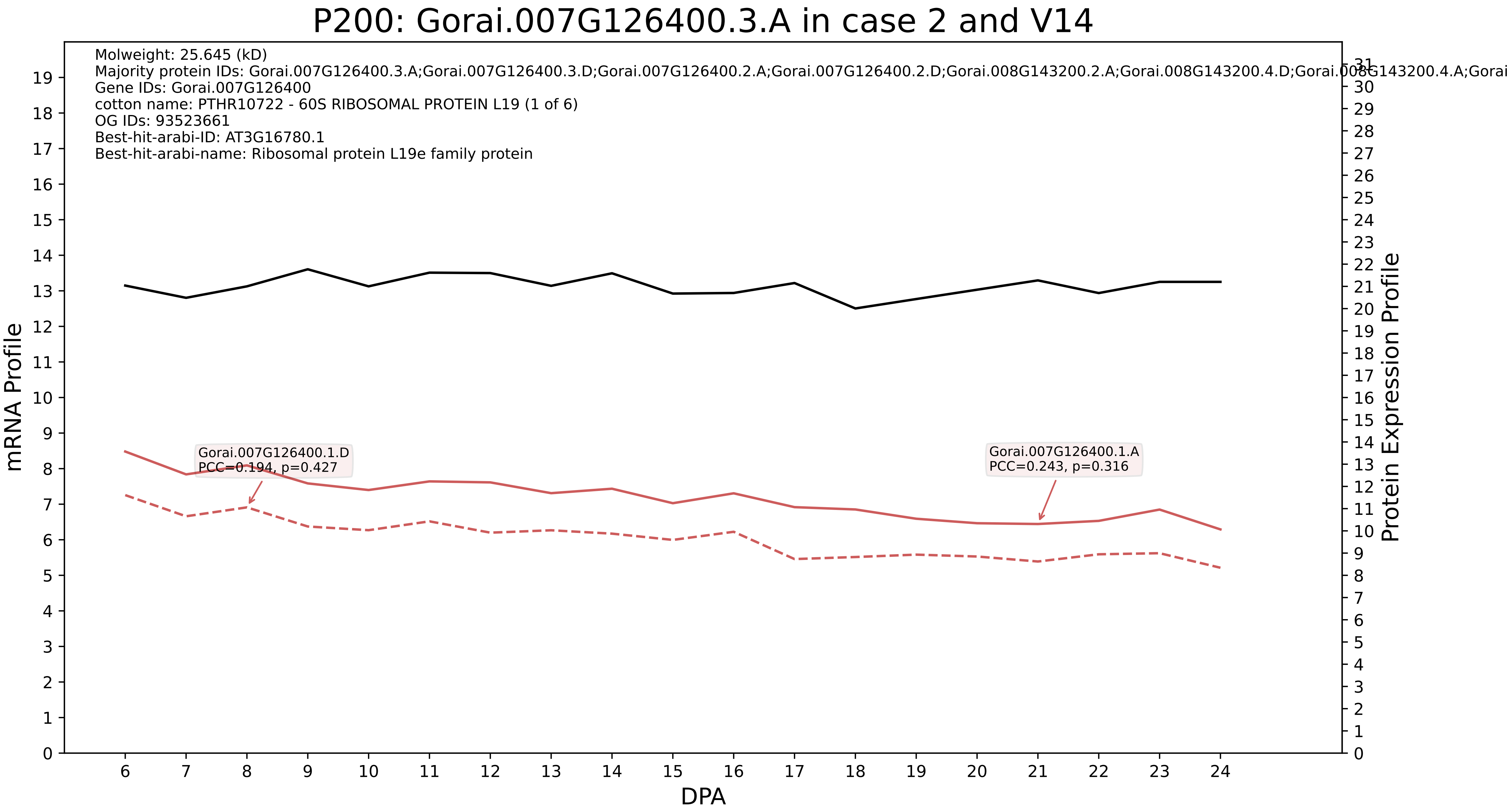

P200: Gorai.007G126400.3.A in case 2 and V14
Molweight: 25.645 (kD)
Majority protein IDs: Gorai.007G126400.3.A;Gorai.007G126400.3.D;Gorai.007G126400.2.A;Gorai.007G126400.2.D;Gorai.008G143200.2.A;Gorai.008G143200.4.D;Gorai.008G143200.4.A;Gorai
Gene IDs: Gorai.007G126400
cotton name: PTHR10722 - 60S RIBOSOMAL PROTEIN L19 (1 of 6)
OG IDs: 93523661
Best-hit-arabi-ID: AT3G16780.1
Best-hit-arabi-name: Ribosomal protein L19e family protein
Gorai.007G126400.1.D
PCC=0.194, p=0.427
Gorai.007G126400.1.A
PCC=0.243, p=0.316
mRNA Profile
Protein Expression Profile
DPA

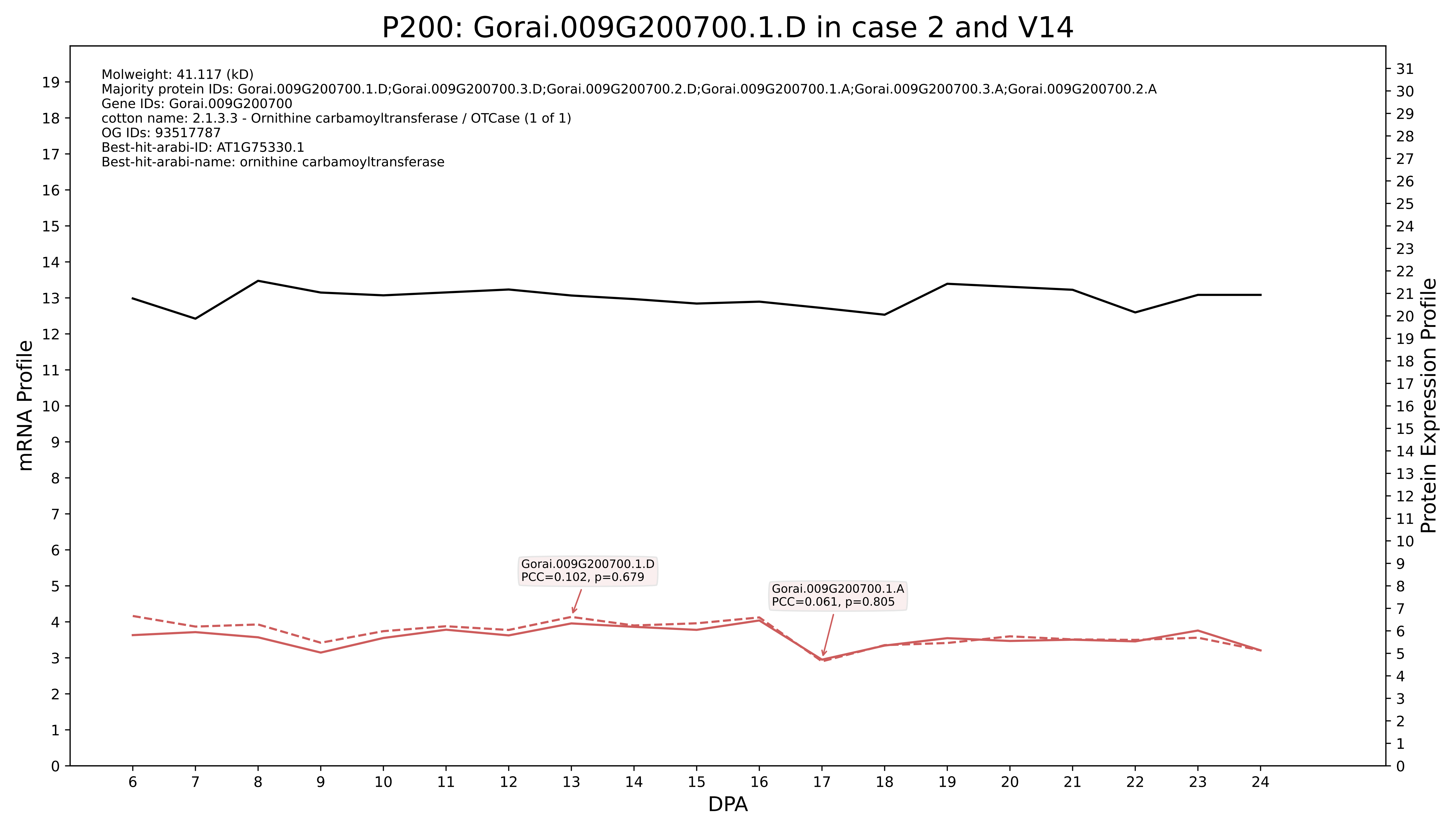
P200: Gorai.009G200700.1.D in case 2 and V14
Molweight: 41.117 (kD)
Majority protein IDs: Gorai.009G200700.1.D;Gorai.009G200700.3.D;Gorai.009G200700.2.D;Gorai.009G200700.1.A;Gorai.009G200700.3.A;Gorai.009G200700.2.A
Gene IDs: Gorai.009G200700
cotton name: 2.1.3.3 - Ornithine carbamoyltransferase / OTCase (1 of 1)
OG IDs: 93517787
Best-hit-arabi-ID: AT1G75330.1
Best-hit-arabi-name: ornithine carbamoyltransferase
Gorai.009G200700.1.D
PCC=0.102, p=0.679
Gorai.009G200700.1.A
PCC=0.061, p=0.805
mRNA Profile
Protein Expression Profile
DPA

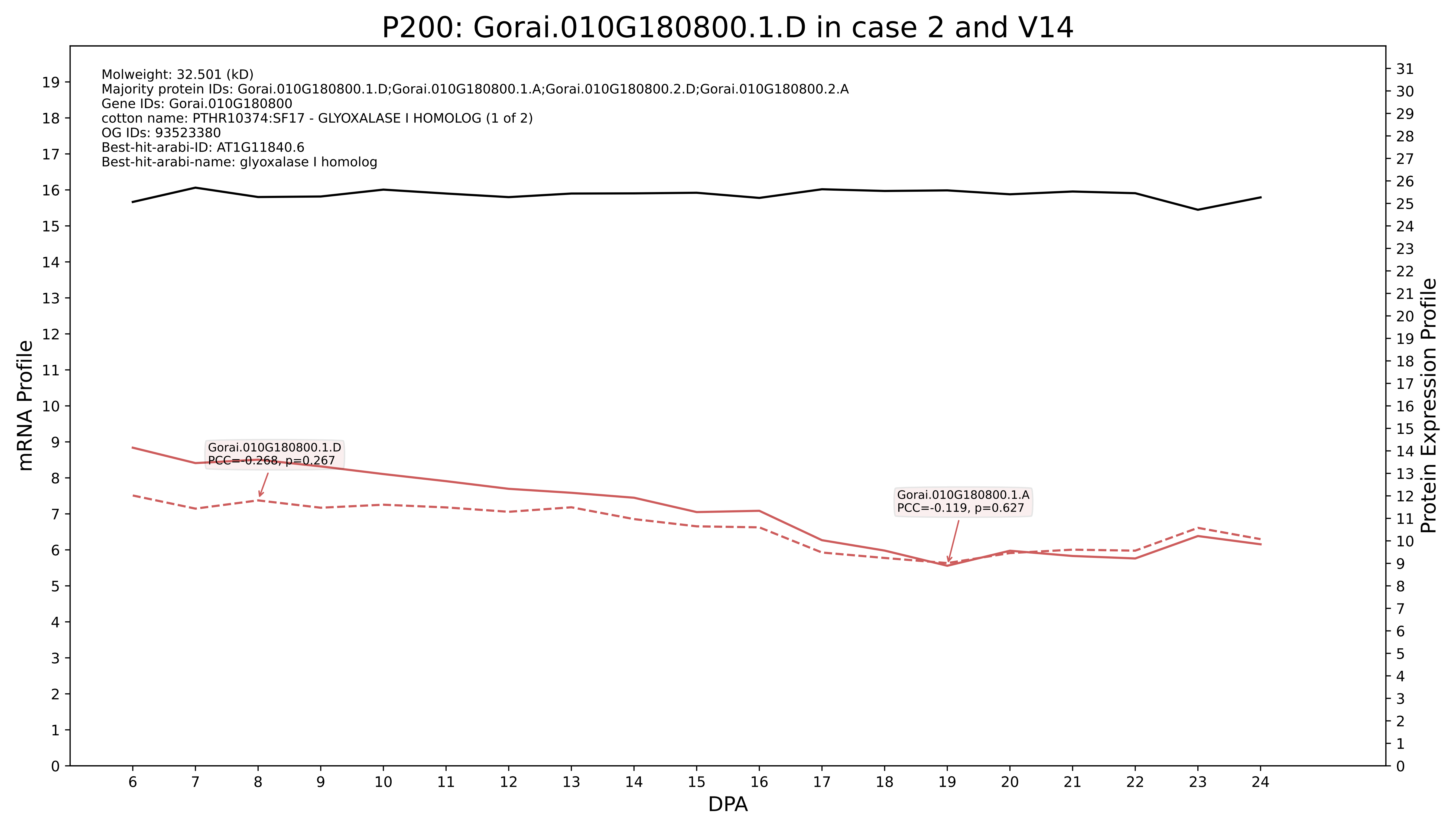

P200: Gorai.010G180800.1.D in case 2 and V14
Molweight: 32.501 (kD)
Majority protein IDs: Gorai.010G180800.1.D;Gorai.010G180800.1.A;Gorai.010G180800.2.D;Gorai.010G180800.2.A
Gene IDs: Gorai.010G180800
cotton name: PTHR10374:SF17 - GLYOXALASE I HOMOLOG (1 of 2)
OG IDs: 93523380
Best-hit-arabi-ID: AT1G11840.6
Best-hit-arabi-name: glyoxalase I homolog
Gorai.010G180800.1.D
PCC=-0.268, p=0.267
Gorai.010G180800.1.A
PCC=-0.119, p=0.627
mRNA Profile
Protein Expression Profile
DPA

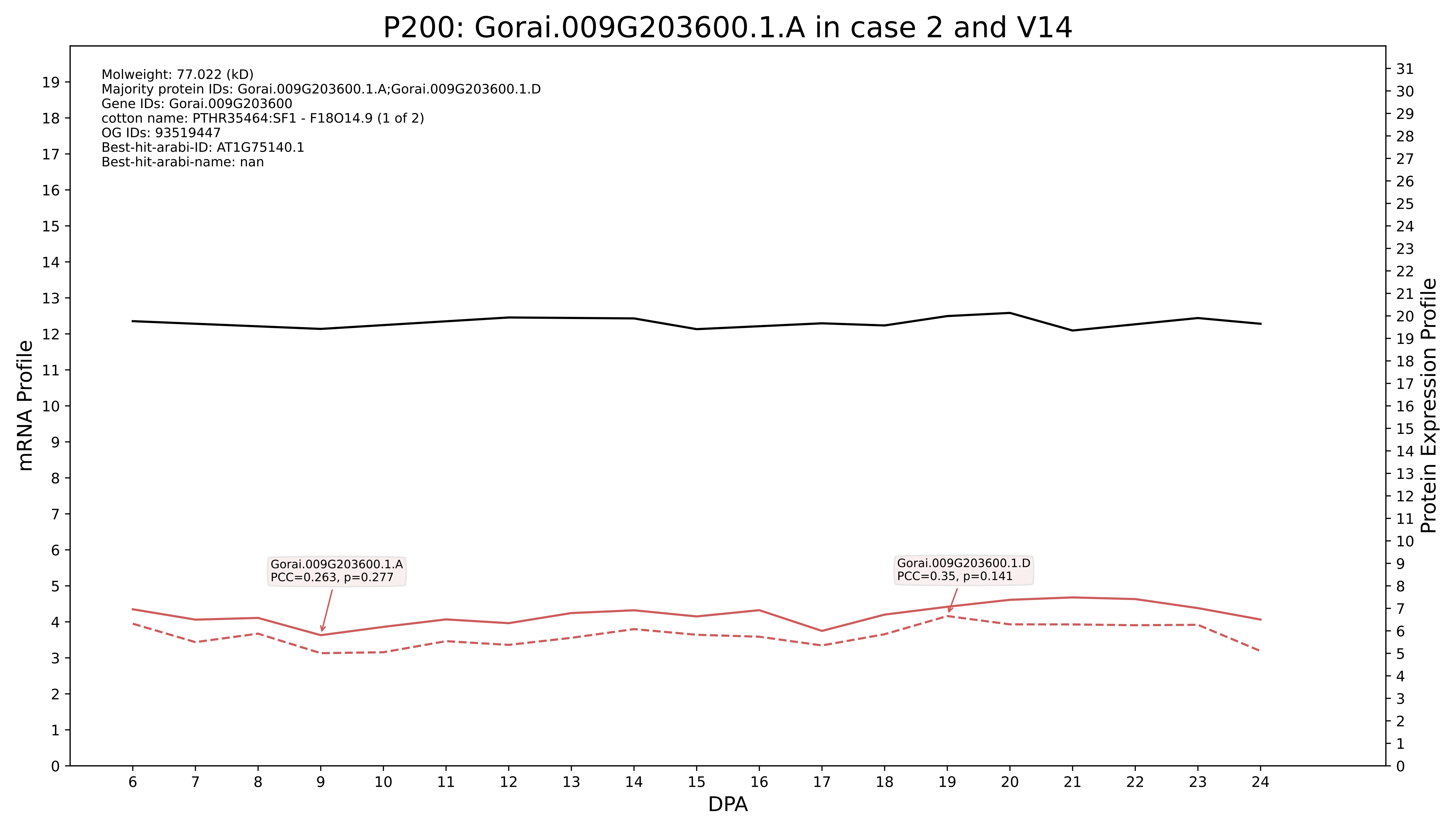
P200: Gorai.009G203600.1.A in case 2 and V14
Molweight: 77.022 (kD)
Majority protein IDs: Gorai.009G203600.1.A;Gorai.009G203600.1.D
Gene IDs: Gorai.009G203600
cotton name: PTHR35464:SF1 - F18O14.9 (1 of 2)
OG IDs: 93519447
Best-hit-arabi-ID: AT1G75140.1
Best-hit-arabi-name: nan
Gorai.009G203600.1.A
PCC=0.263, p=0.277
Gorai.009G203600.1.D
PCC=0.35, p=0.141
mRNA Profile
Protein Expression Profile
DPA

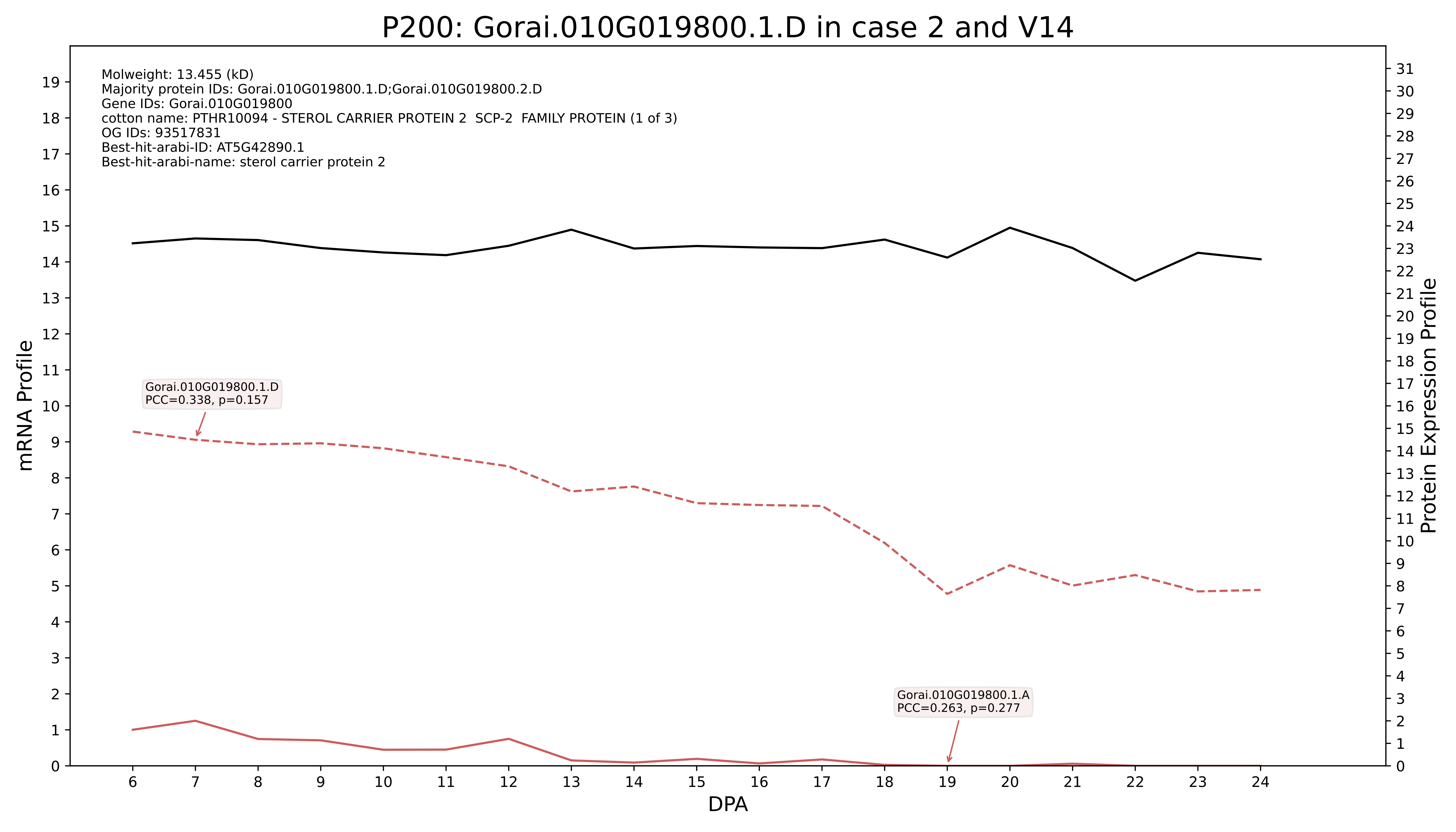

P200: Gorai.010G019800.1.D in case 2 and V14
Molweight: 13.455 (kD)
Majority protein IDs: Gorai.010G019800.1.D;Gorai.010G019800.2.D
Gene IDs: Gorai.010G019800
cotton name: PTHR10094 - STEROL CARRIER PROTEIN 2 SCP-2 FAMILY PROTEIN (1 of 3)
OG IDs: 93517831
Best-hit-arabi-ID: AT5G42890.1
Best-hit-arabi-name: sterol carrier protein 2
Gorai.010G019800.1.D
PCC=0.338, p=0.157
Gorai.010G019800.1.A
PCC=0.263, p=0.277
mRNA Profile
Protein Expression Profile
DPA

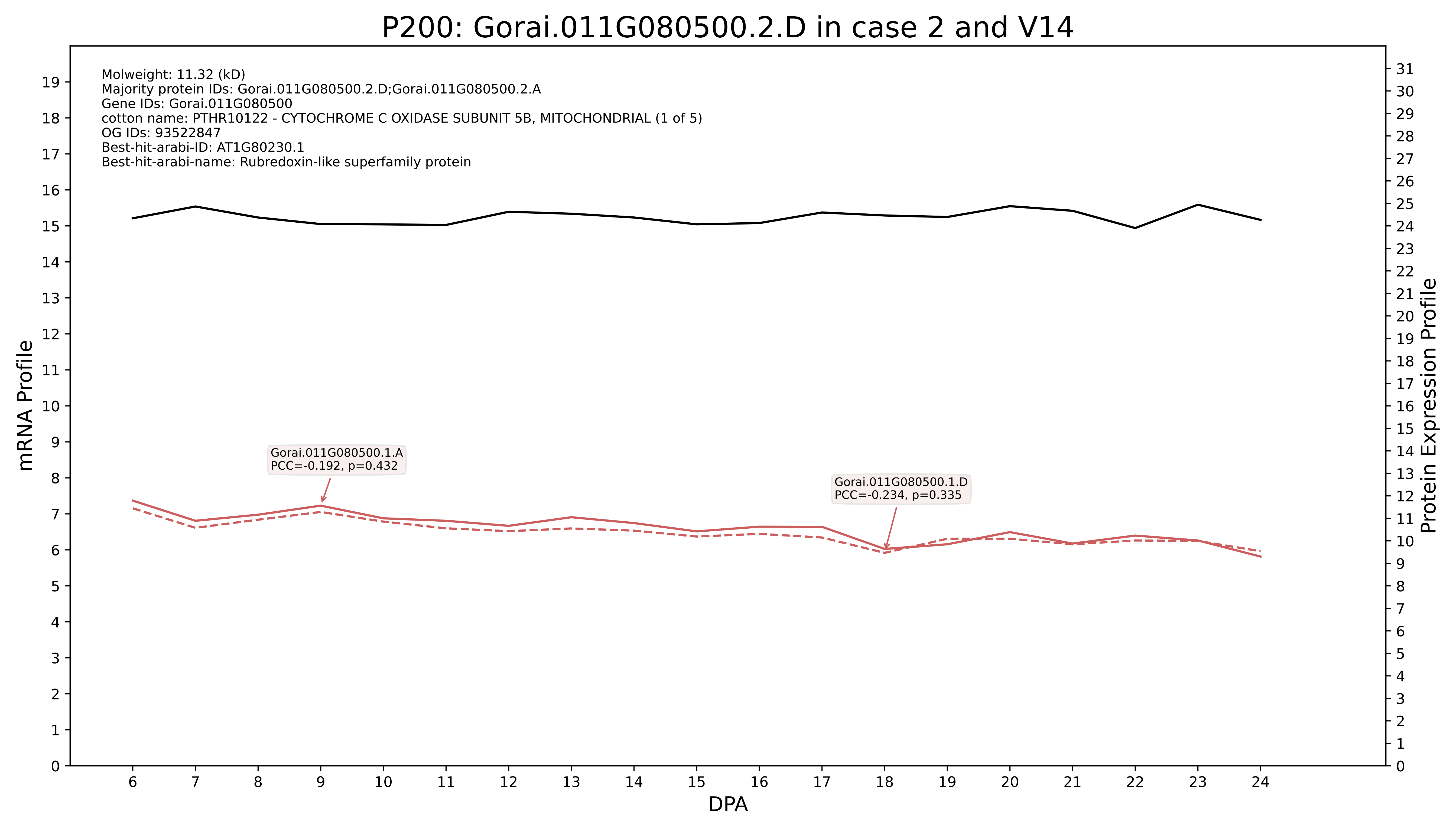

P200: Gorai.011G080500.2.D in case 2 and V14
Molweight: 11.32 (kD)
Majority protein IDs: Gorai.011G080500.2.D;Gorai.011G080500.2.A
Gene IDs: Gorai.011G080500
cotton name: PTHR10122 - CYTOCHROME C OXIDASE SUBUNIT 5B, MITOCHONDRIAL (1 of 5)
OG IDs: 93522847
Best-hit-arabi-ID: AT1G80230.1
Best-hit-arabi-name: Rubredoxin-like superfamily protein
Gorai.011G080500.1.A
PCC=-0.192, p=0.432
Gorai.011G080500.1.D
PCC=-0.234, p=0.335
mRNA Profile
Protein Expression Profile
DPA

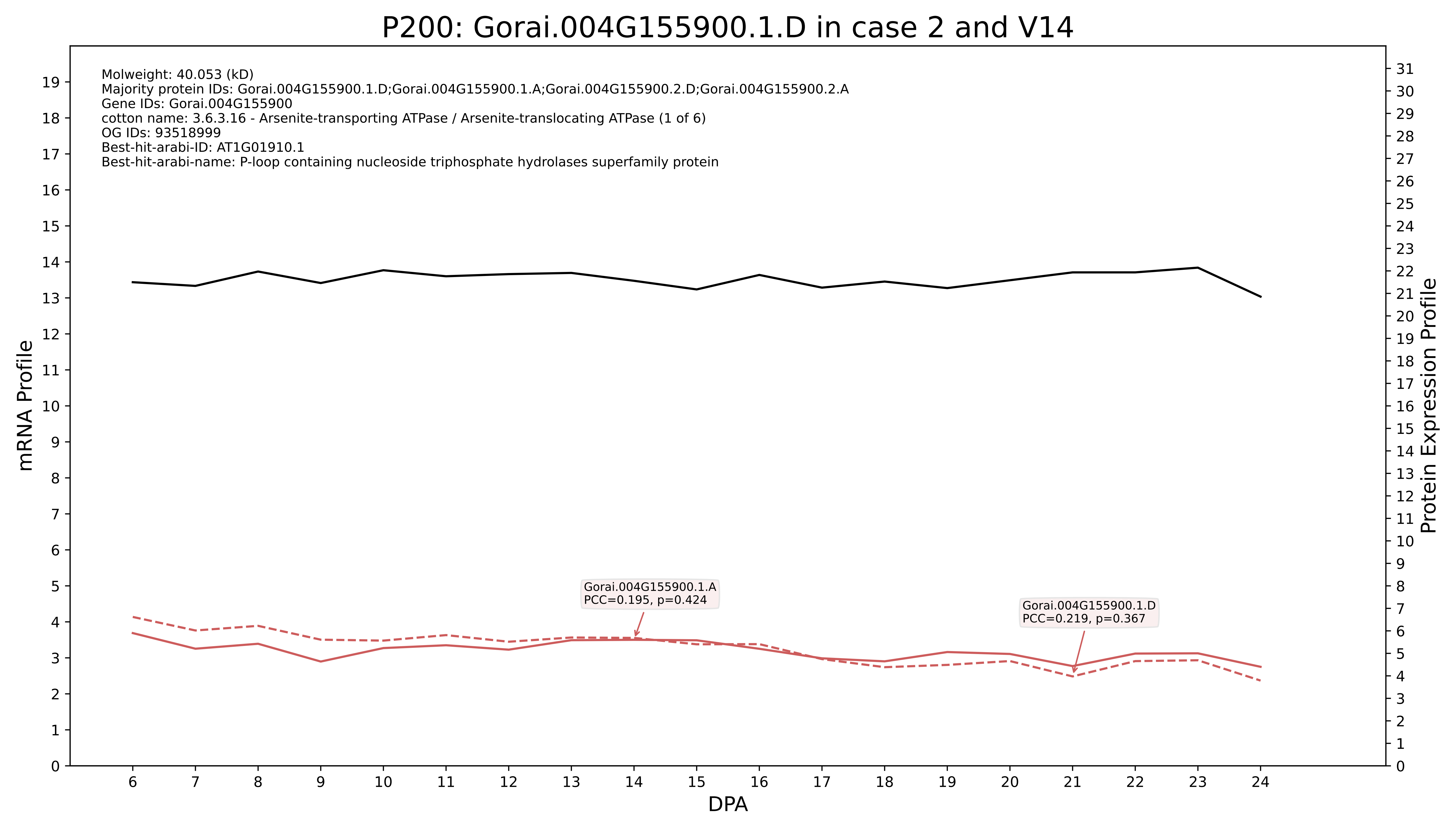

P200: Gorai.004G155900.1.D in case 2 and V14
Molweight: 40.053 (kD)
Majority protein IDs: Gorai.004G155900.1.D;Gorai.004G155900.1.A;Gorai.004G155900.2.D;Gorai.004G155900.2.A
Gene IDs: Gorai.004G155900
cotton name: 3.6.3.16 - Arsenite-transporting ATPase / Arsenite-translocating ATPase (1 of 6)
OG IDs: 93518999
Best-hit-arabi-ID: AT1G01910.1
Best-hit-arabi-name: P-loop containing nucleoside triphosphate hydrolases superfamily protein
Gorai.004G155900.1.A
PCC=0.195, p=0.424
Gorai.004G155900.1.D
PCC=0.219, p=0.367
mRNA Profile
Protein Expression Profile
DPA

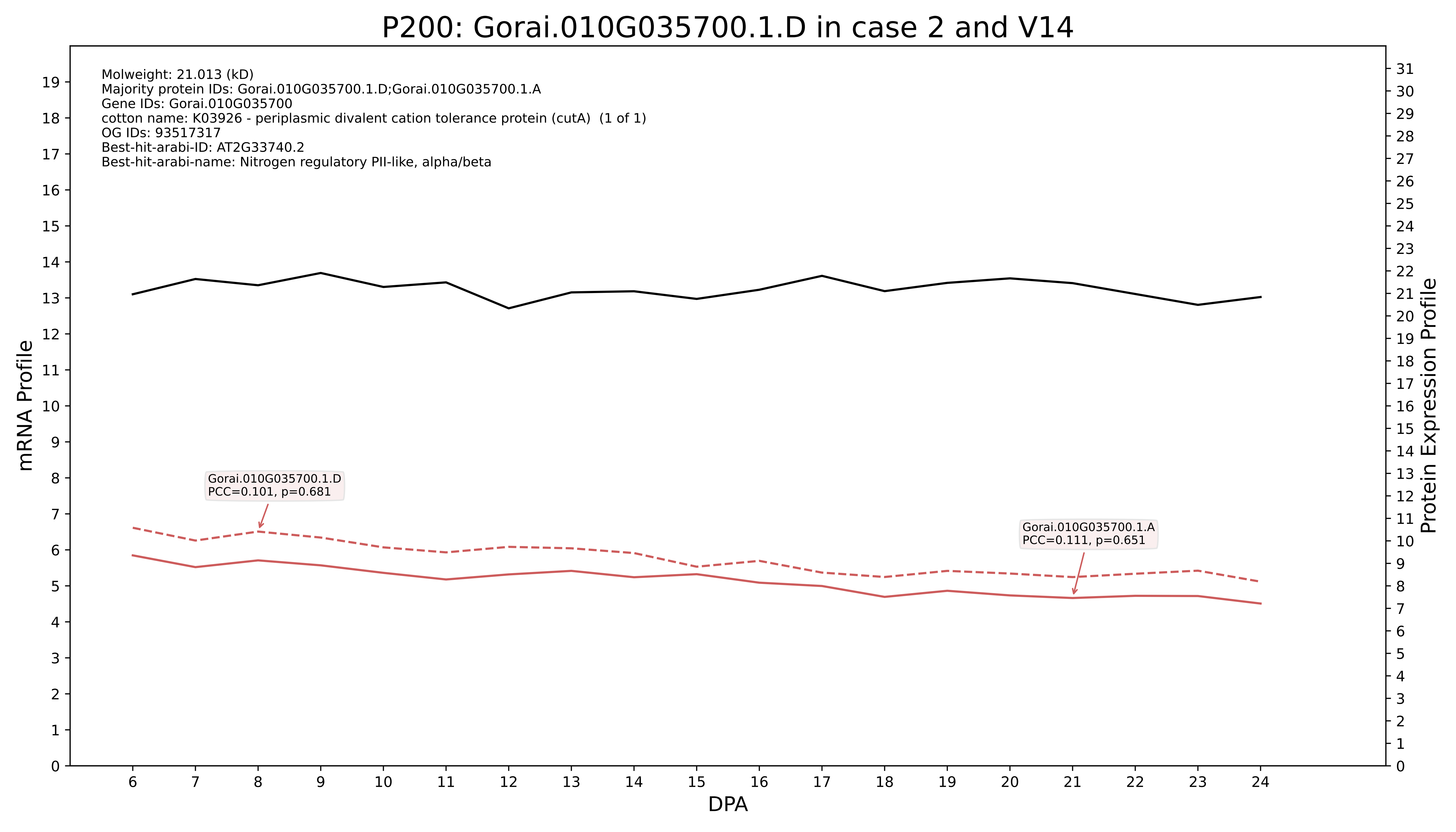
P200: Gorai.010G035700.1.D in case 2 and V14
Molweight: 21.013 (kD)
Majority protein IDs: Gorai.010G035700.1.D;Gorai.010G035700.1.A
Gene IDs: Gorai.010G035700
cotton name: K03926 - periplasmic divalent cation tolerance protein (cutA) (1 of 1)
OG IDs: 93517317
Best-hit-arabi-ID: AT2G33740.2
Best-hit-arabi-name: Nitrogen regulatory PII-like, alpha/beta
Gorai.010G035700.1.D
PCC=0.101, p=0.681
Gorai.010G035700.1.A
PCC=0.111, p=0.651
mRNA Profile
Protein Expression Profile
DPA

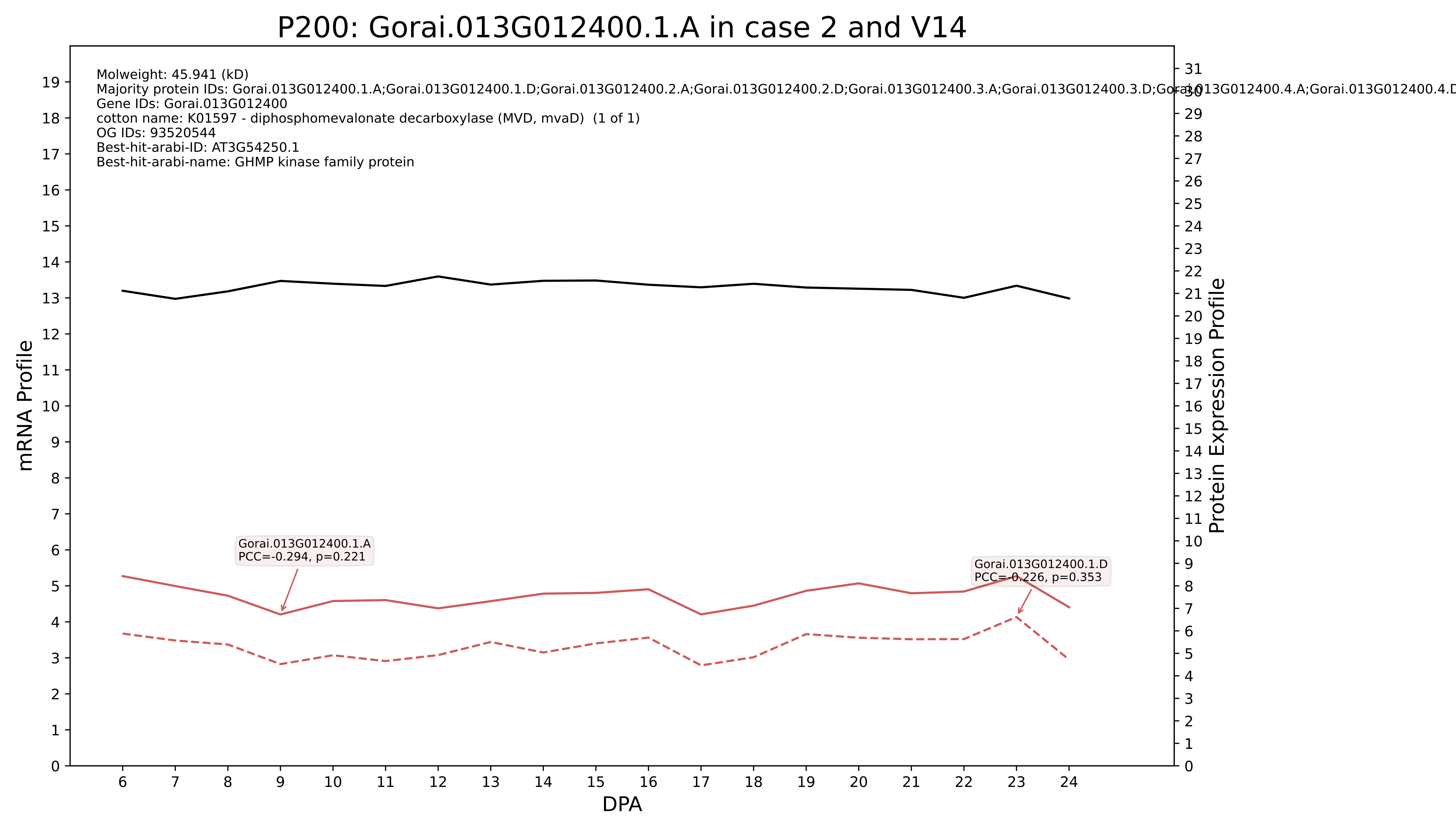

P200: Gorai.013G012400.1.A in case 2 and V14
Molweight: 45.941 (kD)
Majority protein IDs: Gorai.013G012400.1.A;Gorai.013G012400.1.D;Gorai.013G012400.2.A;Gorai.013G012400.2.D;Gorai.013G012400.3.A;Gorai.013G012400.3.D;Gorai.013G012400.4.A;Gorai.013G012400.4.D
Gene IDs: Gorai.013G012400
cotton name: K01597 - diphosphomevalonate decarboxylase (MVD, mvaD) (1 of 1)
OG IDs: 93520544
Best-hit-arabi-ID: AT3G54250.1
Best-hit-arabi-name: GHMP kinase family protein
Gorai.013G012400.1.A
PCC=-0.294, p=0.221
Gorai.013G012400.1.D
PCC=-0.226, p=0.353
mRNA Profile
Protein Expression Profile
DPA

### P200: Gorai.004G224000.1.D in case 2 and V14

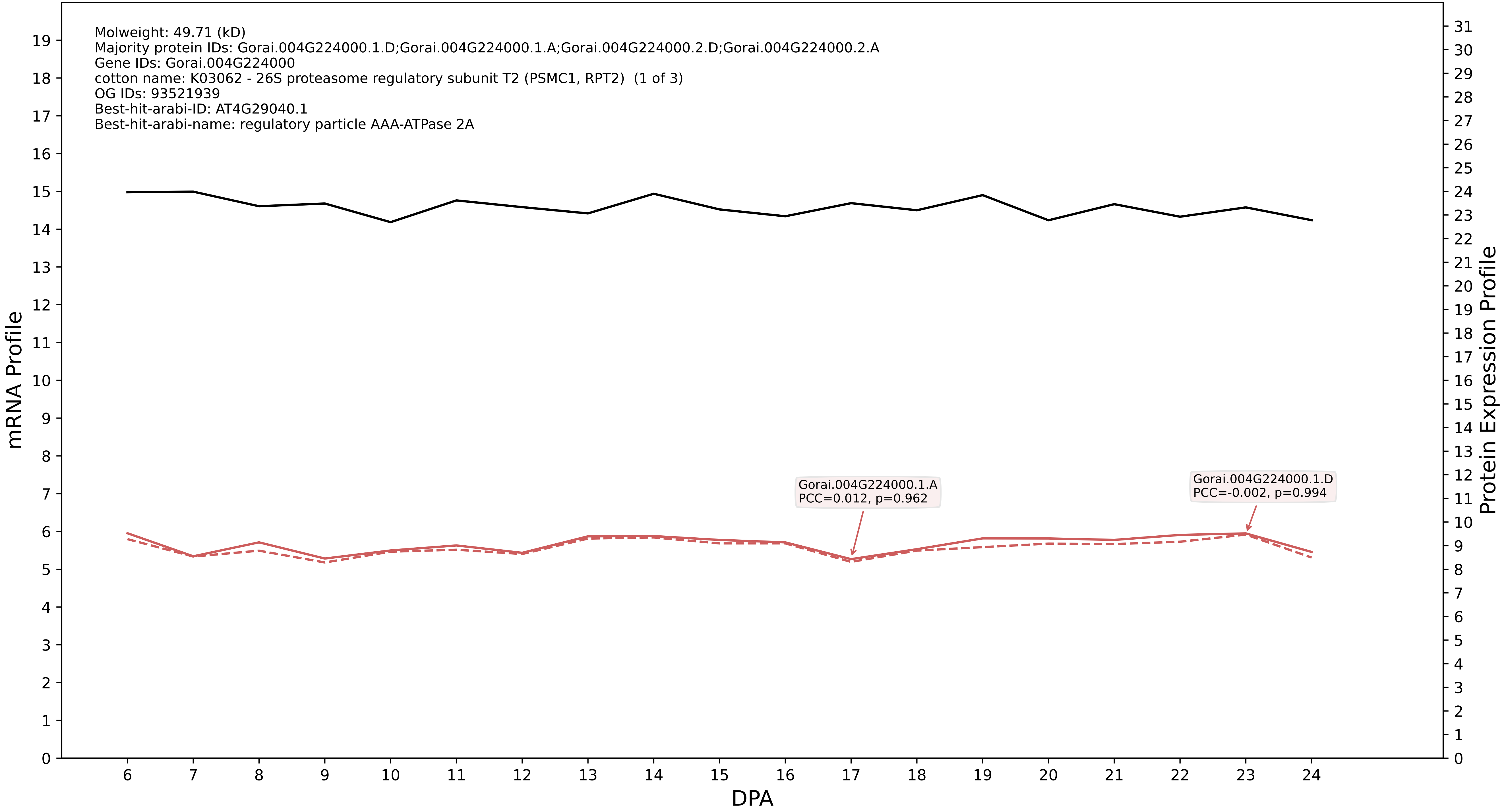

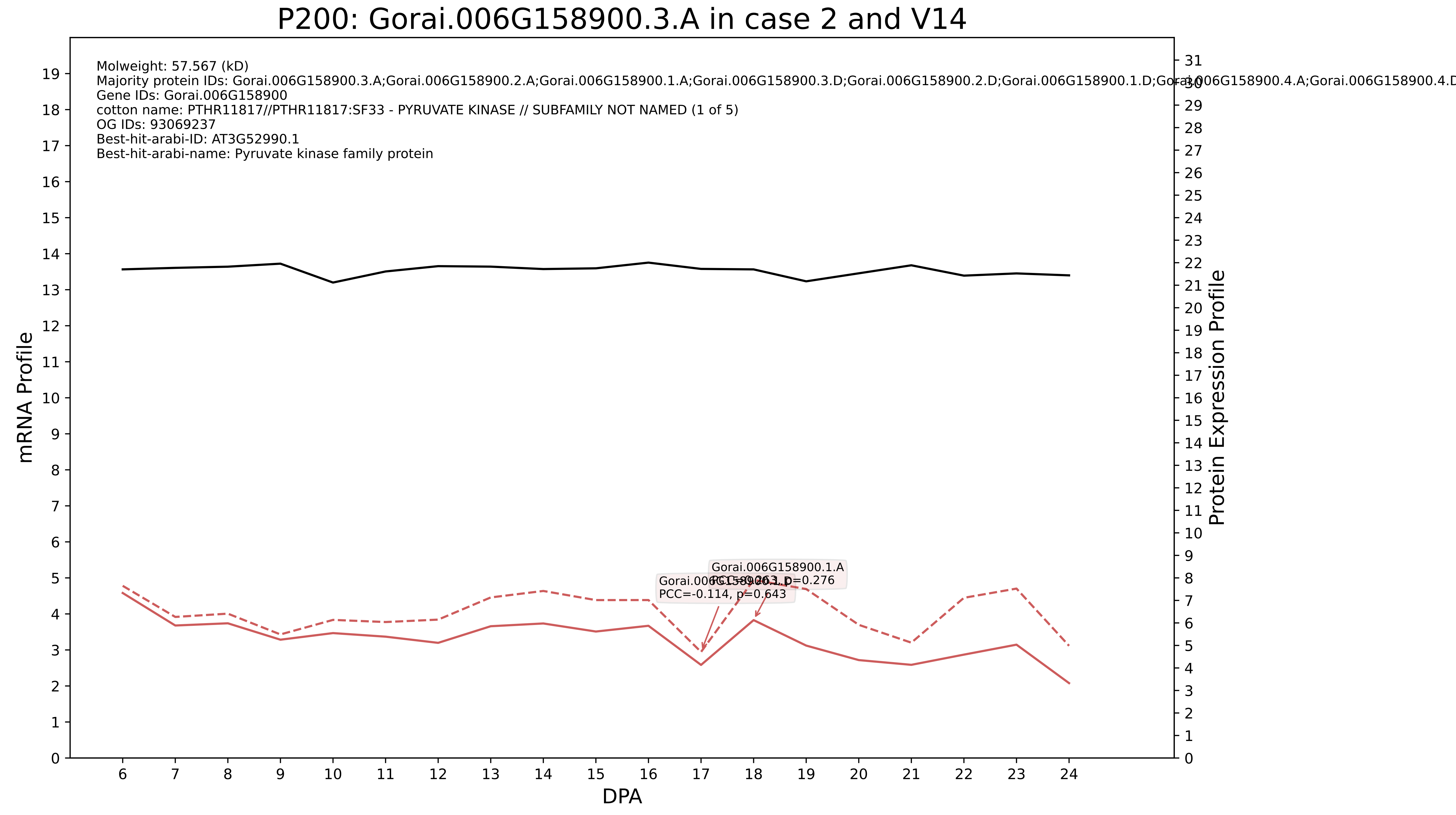
P200: Gorai.006G158900.3.A in case 2 and V14
Molweight: 57.567 (kD)
Majority protein IDs: Gorai.006G158900.3.A;Gorai.006G158900.2.A;Gorai.006G158900.1.A;Gorai.006G158900.3.D;Gorai.006G158900.2.D;Gorai.006G158900.1.D;Gorai.006G158900.4.A;Gorai.006G158900.4.D
Gene IDs: Gorai.006G158900
cotton name: PTHR11817//PTHR11817:SF33 - PYRUVATE KINASE // SUBFAMILY NOT NAMED (1 of 5)
OG IDs: 93069237
Best-hit-arabi-ID: AT3G52990.1
Best-hit-arabi-name: Pyruvate kinase family protein
Gorai.006G158900.1.A
PCC=0.263, p=0.276
PCC=-0.114, p=0.643
mRNA Profile
Protein Expression Profile
DPA

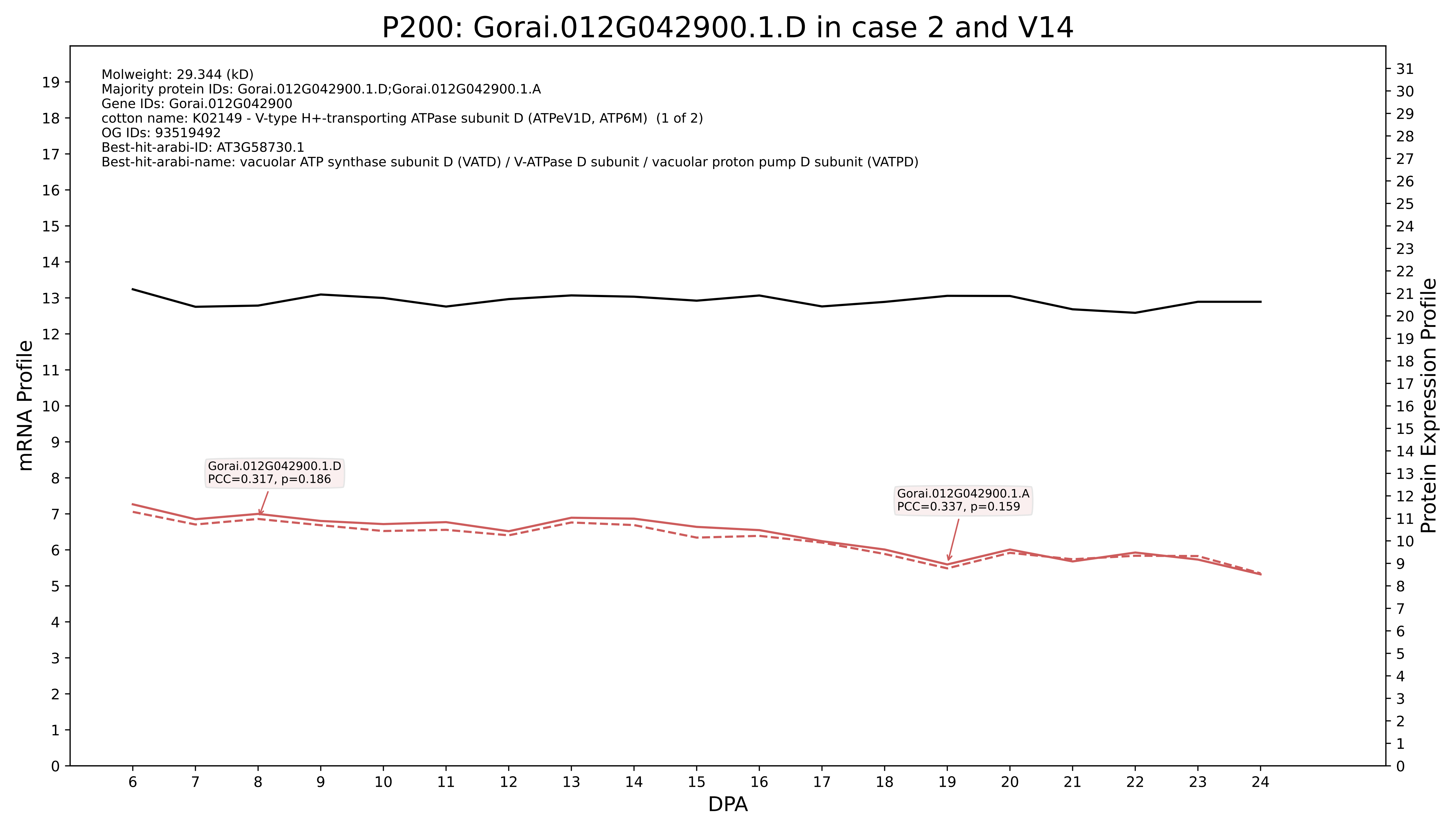
P200: Gorai.012G042900.1.D in case 2 and V14
Molweight: 29.344 (kD)
Majority protein IDs: Gorai.012G042900.1.D;Gorai.012G042900.1.A
Gene IDs: Gorai.012G042900
cotton name: K02149 - V-type H+-transporting ATPase subunit D (ATPeV1D, ATP6M) (1 of 2)
OG IDs: 93519492
Best-hit-arabi-ID: AT3G58730.1
Best-hit-arabi-name: vacuolar ATP synthase subunit D (VATD) / V-ATPase D subunit / vacuolar proton pump D subunit (VATPD)
Gorai.012G042900.1.D
PCC=0.317, p=0.186
Gorai.012G042900.1.A
PCC=0.337, p=0.159
mRNA Profile
Protein Expression Profile
DPA

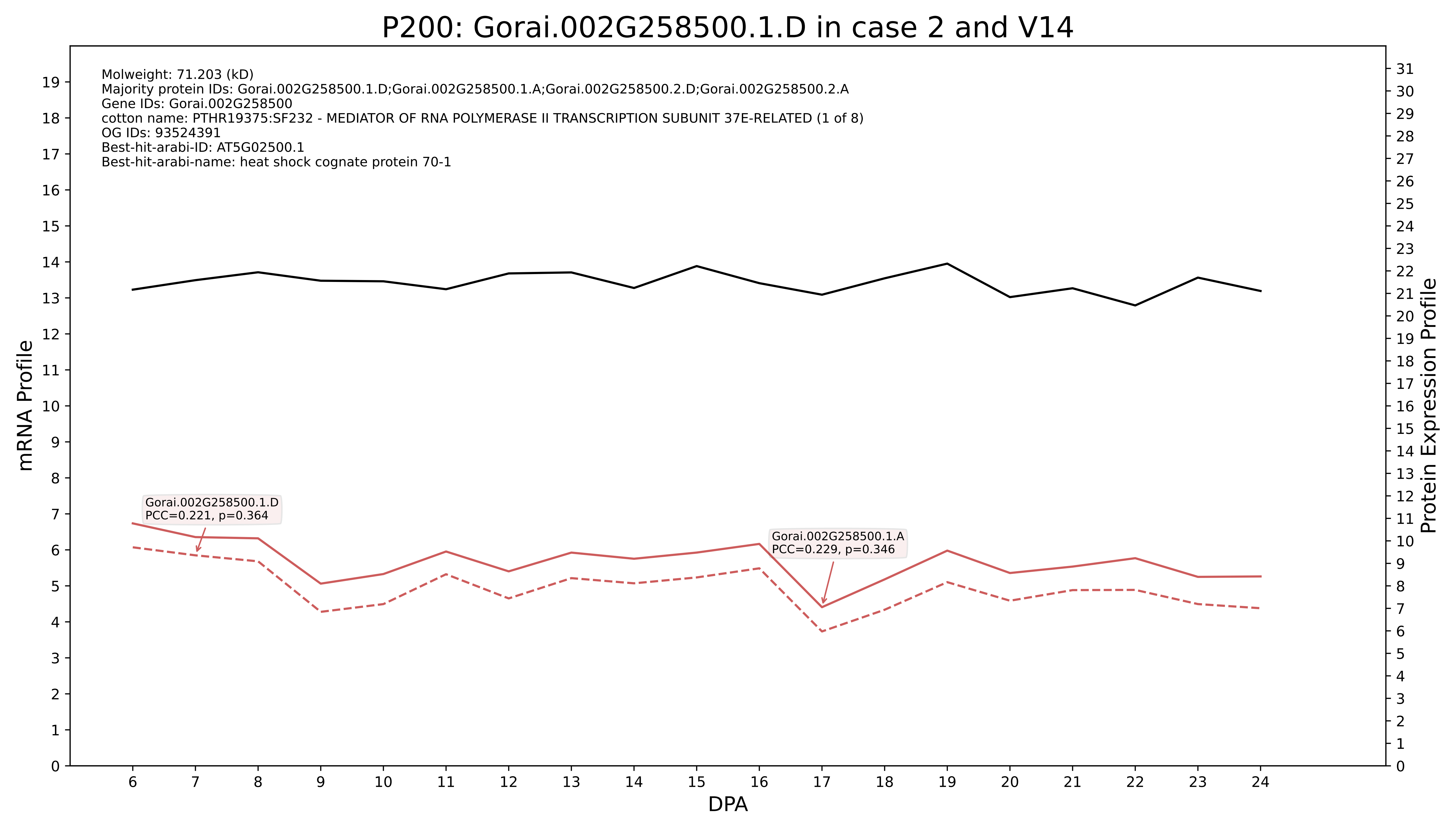
P200: Gorai.002G258500.1.D in case 2 and V14
Molweight: 71.203 (kD)
Majority protein IDs: Gorai.002G258500.1.D;Gorai.002G258500.1.A;Gorai.002G258500.2.D;Gorai.002G258500.2.A
Gene IDs: Gorai.002G258500
cotton name: PTHR19375:SF232 - MEDIATOR OF RNA POLYMERASE II TRANSCRIPTION SUBUNIT 37E-RELATED (1 of 8)
OG IDs: 93524391
Best-hit-arabi-ID: AT5G02500.1
Best-hit-arabi-name: heat shock cognate protein 70-1
Gorai.002G258500.1.D
PCC=0.221, p=0.364
Gorai.002G258500.1.A
PCC=0.229, p=0.346
mRNA Profile
Protein Expression Profile
DPA

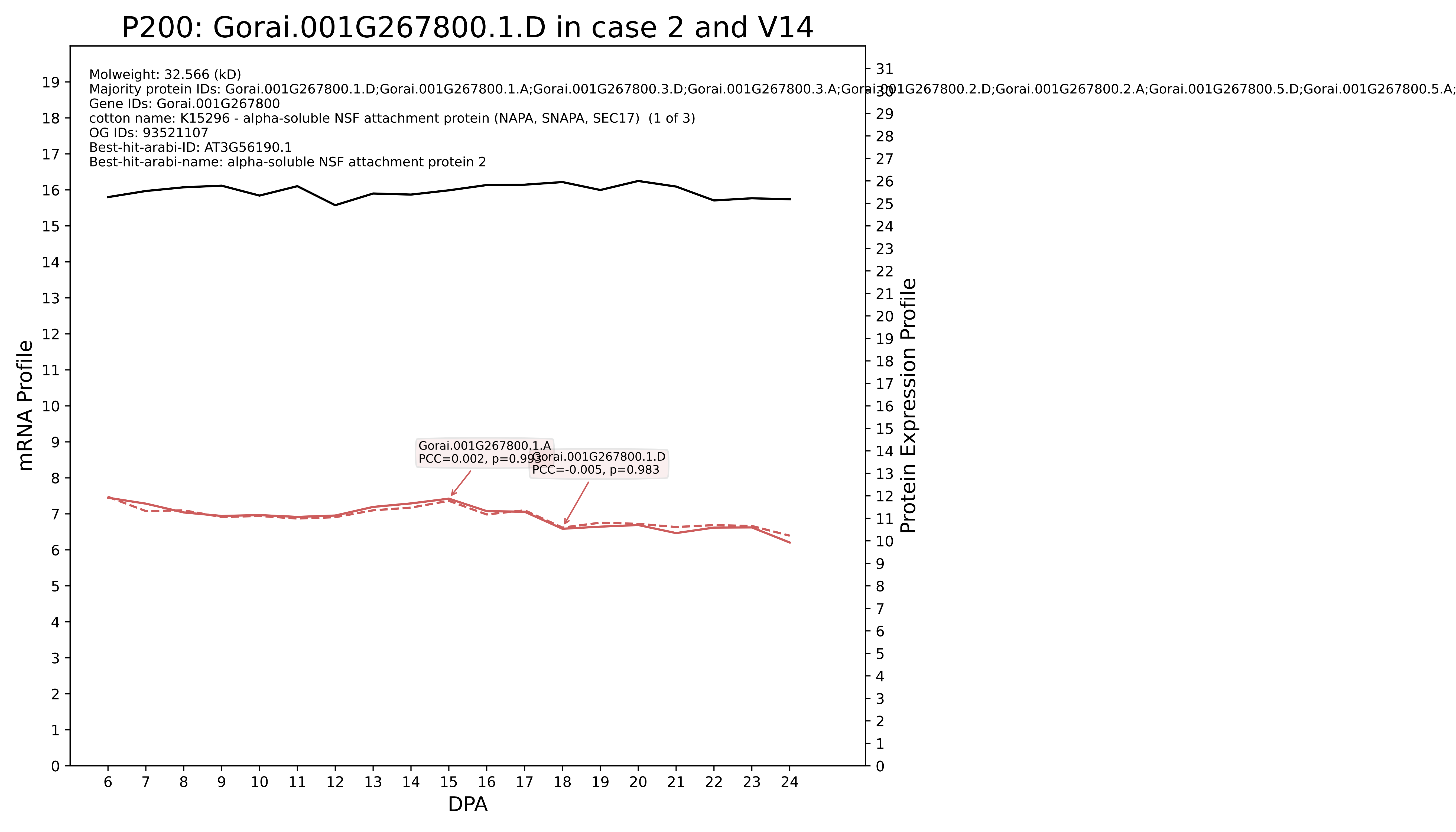

P200: Gorai.001G267800.1.D in case 2 and V14
Molweight: 32.566 (kD)
Majority protein IDs: Gorai.001G267800.1.D;Gorai.001G267800.1.A;Gorai.001G267800.3.D;Gorai.001G267800.3.A;Gorai.001G267800.2.D;Gorai.001G267800.2.A;Gorai.001G267800.5.D;Gorai.001G267800.5.A;
Gene IDs: Gorai.001G267800
cotton name: K15296 - alpha-soluble NSF attachment protein (NAPA, SNAPA, SEC17) (1 of 3)
OG IDs: 93521107
Best-hit-arabi-ID: AT3G56190.1
Best-hit-arabi-name: alpha-soluble NSF attachment protein 2
Gorai.001G267800.1.A
PCC=0.002, p=0.993
Gorai.001G267800.1.D
PCC=-0.005, p=0.983
mRNA Profile
Protein Expression Profile
DPA

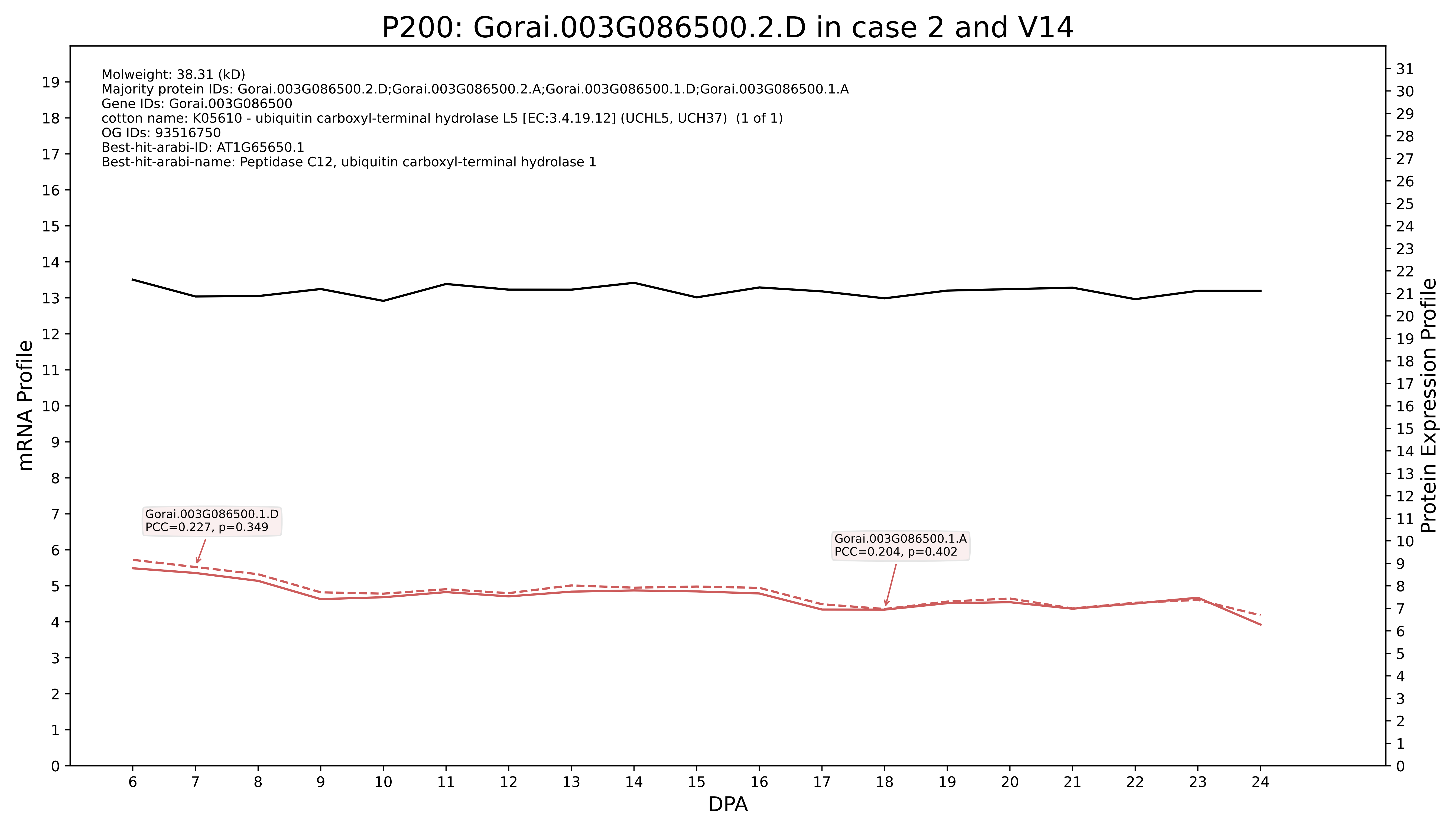
P200: Gorai.003G086500.2.D in case 2 and V14
Molweight: 38.31 (kD)
Majority protein IDs: Gorai.003G086500.2.D;Gorai.003G086500.2.A;Gorai.003G086500.1.D;Gorai.003G086500.1.A
Gene IDs: Gorai.003G086500
cotton name: K05610 - ubiquitin carboxyl-terminal hydrolase L5 [EC:3.4.19.12] (UCHL5, UCH37) (1 of 1)
OG IDs: 93516750
Best-hit-arabi-ID: AT1G65650.1
Best-hit-arabi-name: Peptidase C12, ubiquitin carboxyl-terminal hydrolase 1
Gorai.003G086500.1.D
PCC=0.227, p=0.349
Gorai.003G086500.1.A
PCC=0.204, p=0.402
mRNA Profile
Protein Expression Profile
DPA

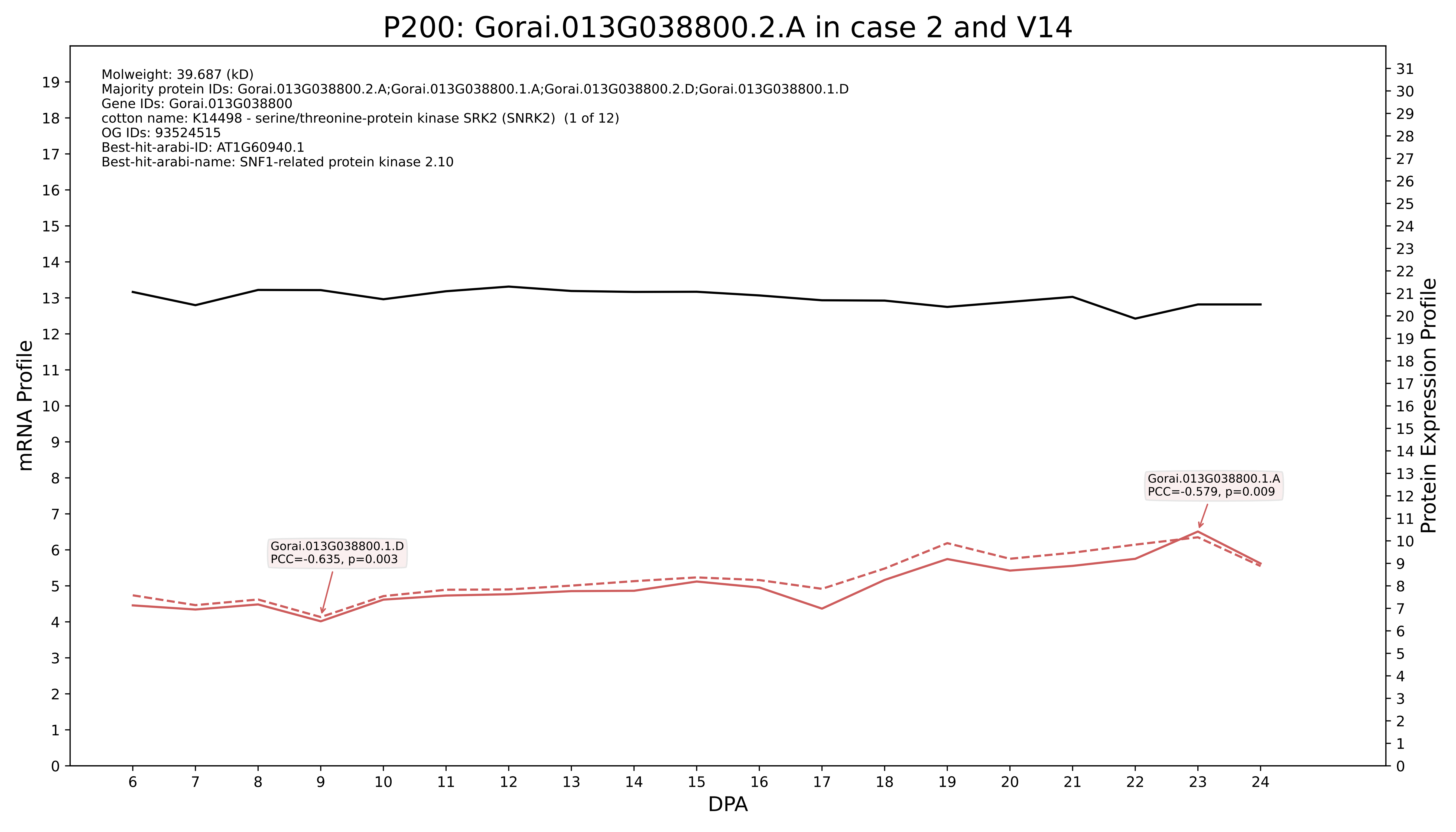

P200: Gorai.013G038800.2.A in case 2 and V14
Molweight: 39.687 (kD)
Majority protein IDs: Gorai.013G038800.2.A;Gorai.013G038800.1.A;Gorai.013G038800.2.D;Gorai.013G038800.1.D
Gene IDs: Gorai.013G038800
cotton name: K14498 - serine/threonine-protein kinase SRK2 (SNRK2) (1 of 12)
OG IDs: 93524515
Best-hit-arabi-ID: AT1G60940.1
Best-hit-arabi-name: SNF1-related protein kinase 2.10
Gorai.013G038800.1.A
PCC=-0.579, p=0.009
Gorai.013G038800.1.D
PCC=-0.635, p=0.003
mRNA Profile
Protein Expression Profile
DPA

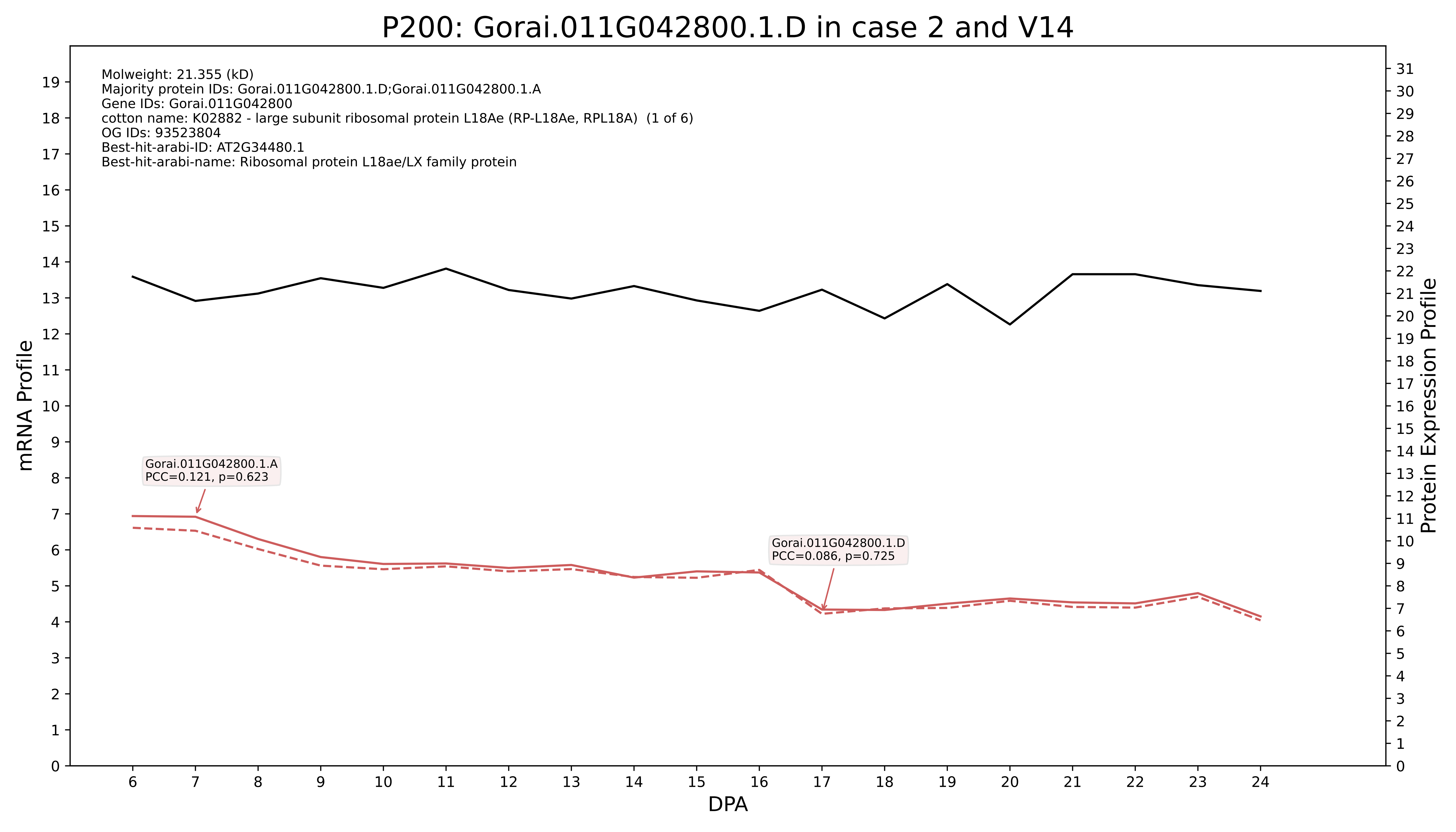
P200: Gorai.011G042800.1.D in case 2 and V14
Molweight: 21.355 (kD)
Majority protein IDs: Gorai.011G042800.1.D;Gorai.011G042800.1.A
Gene IDs: Gorai.011G042800
cotton name: K02882 - large subunit ribosomal protein L18Ae (RP-L18Ae, RPL18A) (1 of 6)
OG IDs: 93523804
Best-hit-arabi-ID: AT2G34480.1
Best-hit-arabi-name: Ribosomal protein L18ae/LX family protein
Gorai.011G042800.1.A
PCC=0.121, p=0.623
Gorai.011G042800.1.D
PCC=0.086, p=0.725
mRNA Profile
Protein Expression Profile
DPA

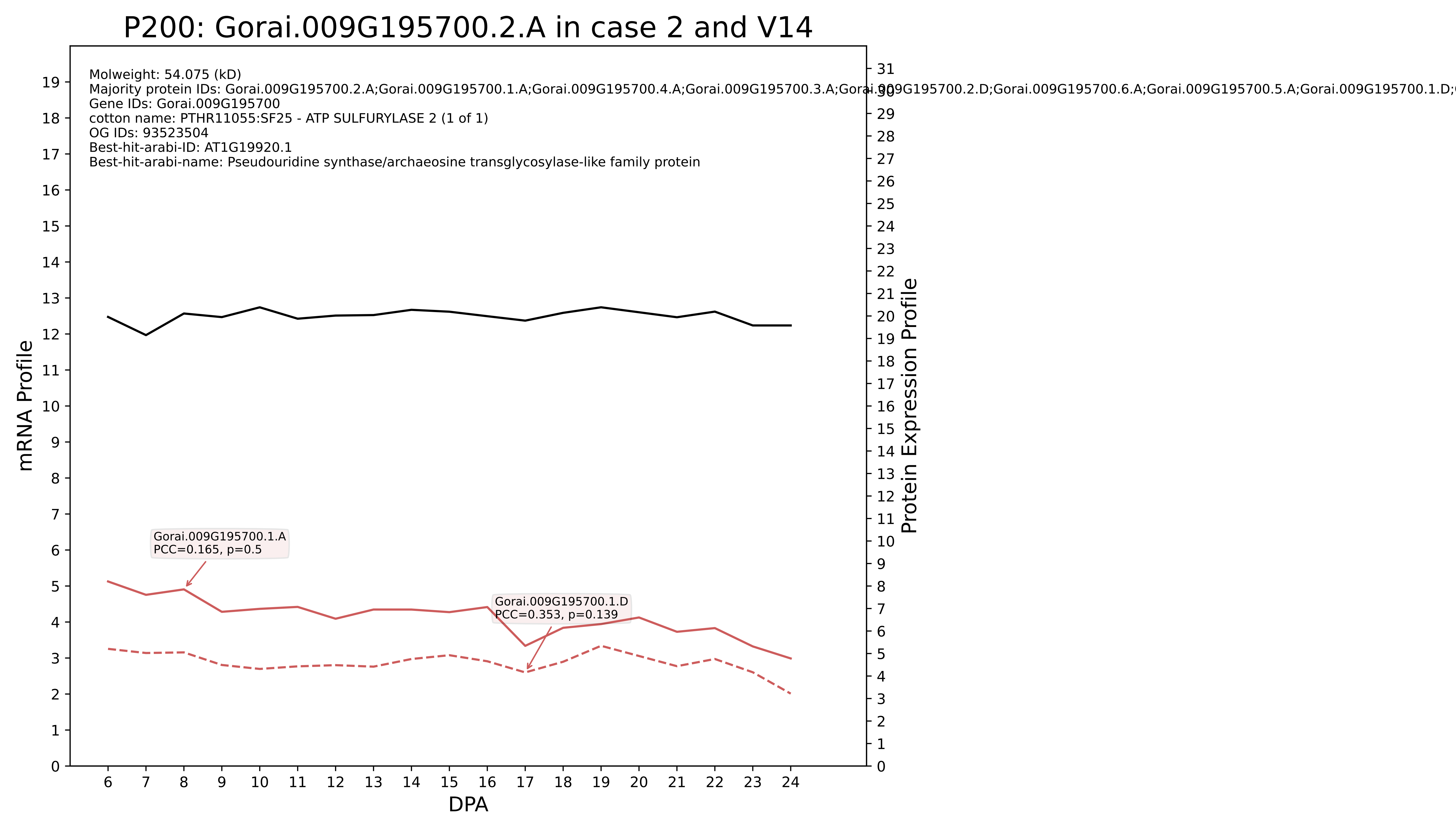
P200: Gorai.009G195700.2.A in case 2 and V14
Molweight: 54.075 (kD)
Majority protein IDs: Gorai.009G195700.2.A;Gorai.009G195700.1.A;Gorai.009G195700.4.A;Gorai.009G195700.3.A;Gorai.009G195700.2.D;Gorai.009G195700.6.A;Gorai.009G195700.5.A;Gorai.009G195700.1.D;
Gene IDs: Gorai.009G195700
cotton name: PTHR11055:SF25 - ATP SULFURYLASE 2 (1 of 1)
OG IDs: 93523504
Best-hit-arabi-ID: AT1G19920.1
Best-hit-arabi-name: Pseudouridine synthase/archaeosine transglycosylase-like family protein
Gorai.009G195700.1.A
PCC=0.165, p=0.5
Gorai.009G195700.1.D
PCC=0.353, p=0.139
mRNA Profile
Protein Expression Profile
DPA

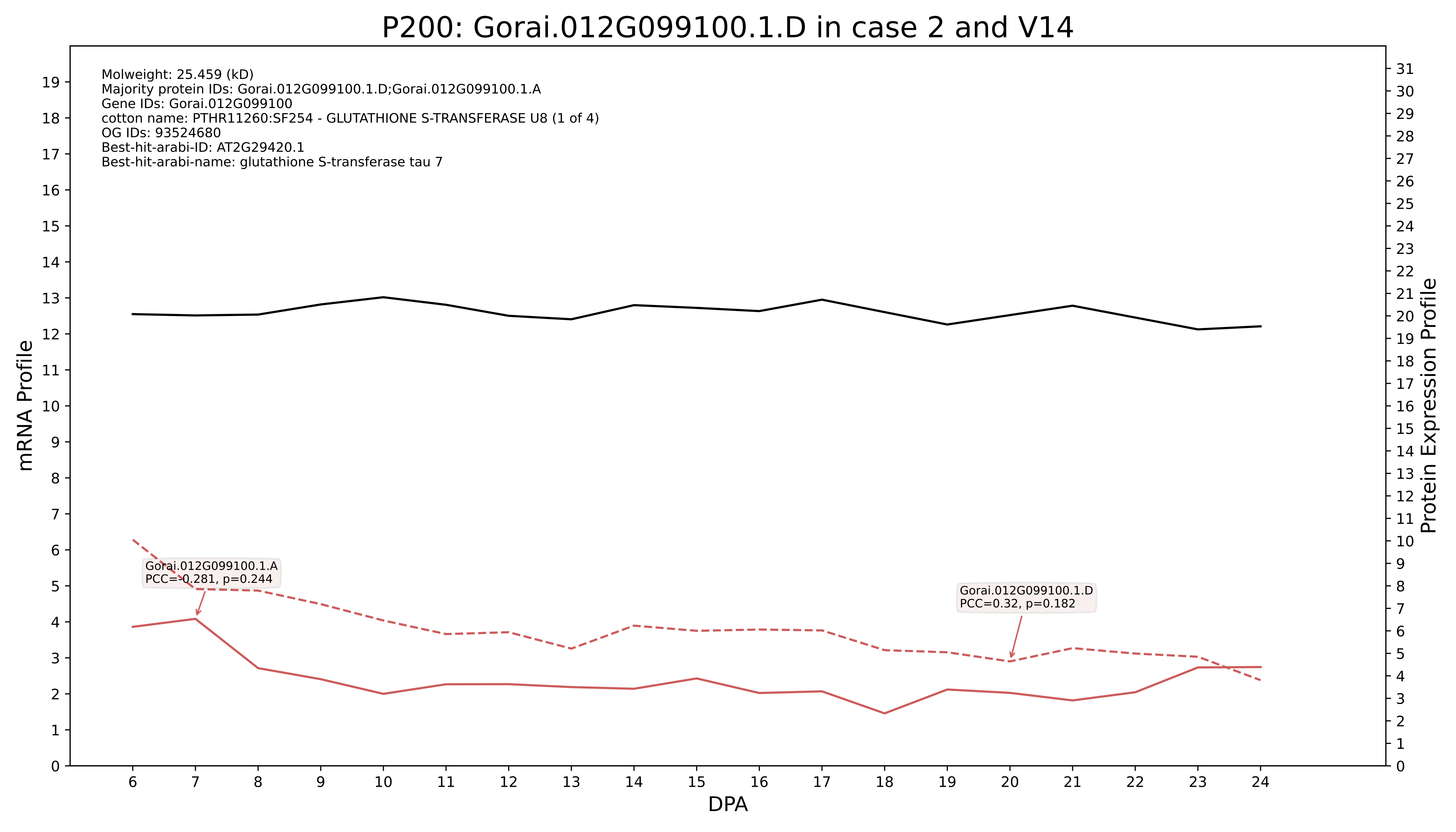

P200: Gorai.012G099100.1.D in case 2 and V14
Molweight: 25.459 (kD)
Majority protein IDs: Gorai.012G099100.1.D;Gorai.012G099100.1.A
Gene IDs: Gorai.012G099100
cotton name: PTHR11260:SF254 - GLUTATHIONE S-TRANSFERASE U8 (1 of 4)
OG IDs: 93524680
Best-hit-arabi-ID: AT2G29420.1
Best-hit-arabi-name: glutathione S-transferase tau 7
Gorai.012G099100.1.A
PCC=-0.281, p=0.244
Gorai.012G099100.1.D
PCC=0.32, p=0.182
mRNA Profile
Protein Expression Profile
DPA

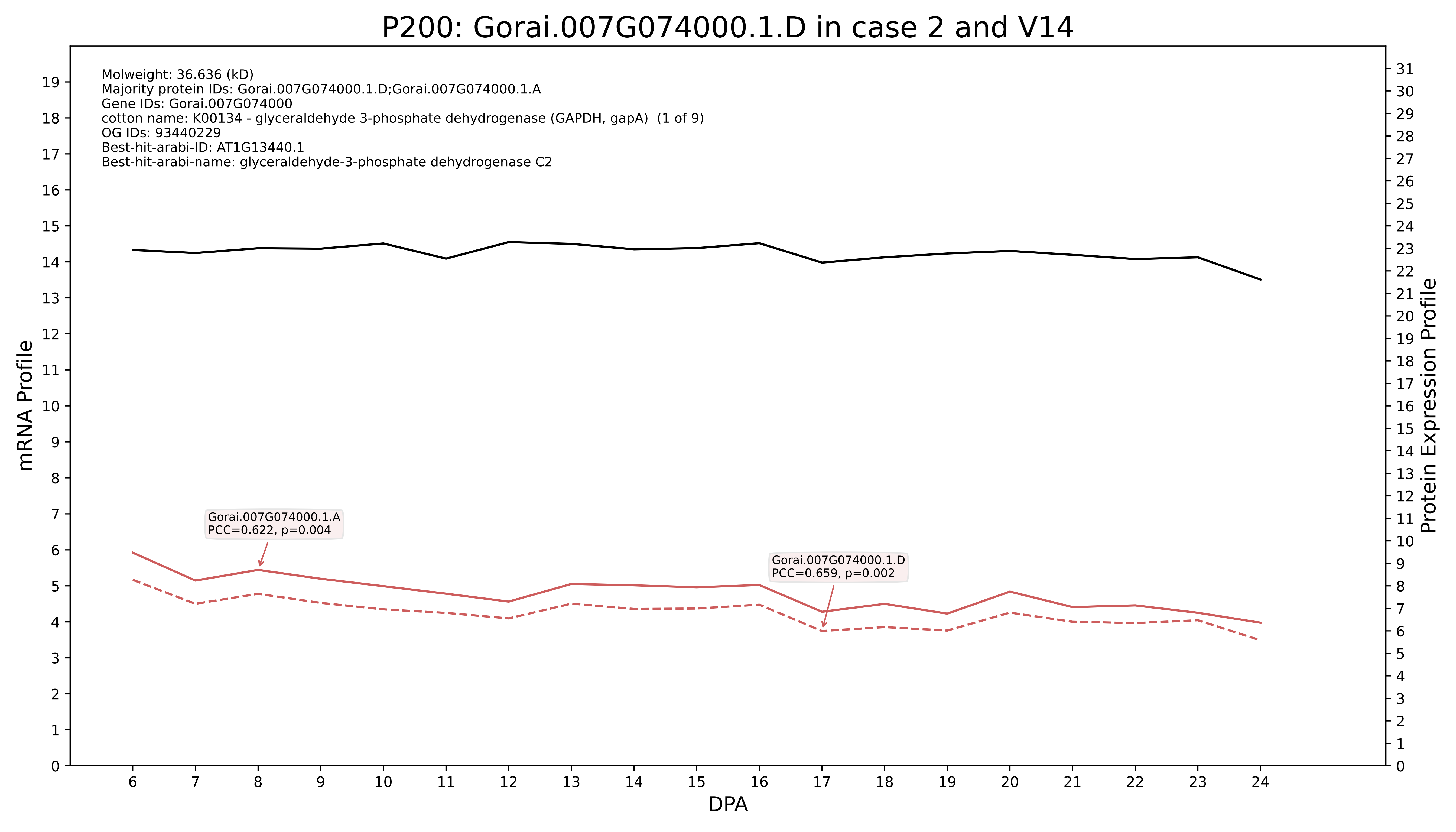
P200: Gorai.007G074000.1.D in case 2 and V14
Molweight: 36.636 (kD)
Majority protein IDs: Gorai.007G074000.1.D;Gorai.007G074000.1.A
Gene IDs: Gorai.007G074000
cotton name: K00134 - glyceraldehyde 3-phosphate dehydrogenase (GAPDH, gapA) (1 of 9)
OG IDs: 93440229
Best-hit-arabi-ID: AT1G13440.1
Best-hit-arabi-name: glyceraldehyde-3-phosphate dehydrogenase C2
Gorai.007G074000.1.A
PCC=0.622, p=0.004
Gorai.007G074000.1.D
PCC=0.659, p=0.002
mRNA Profile
Protein Expression Profile
DPA

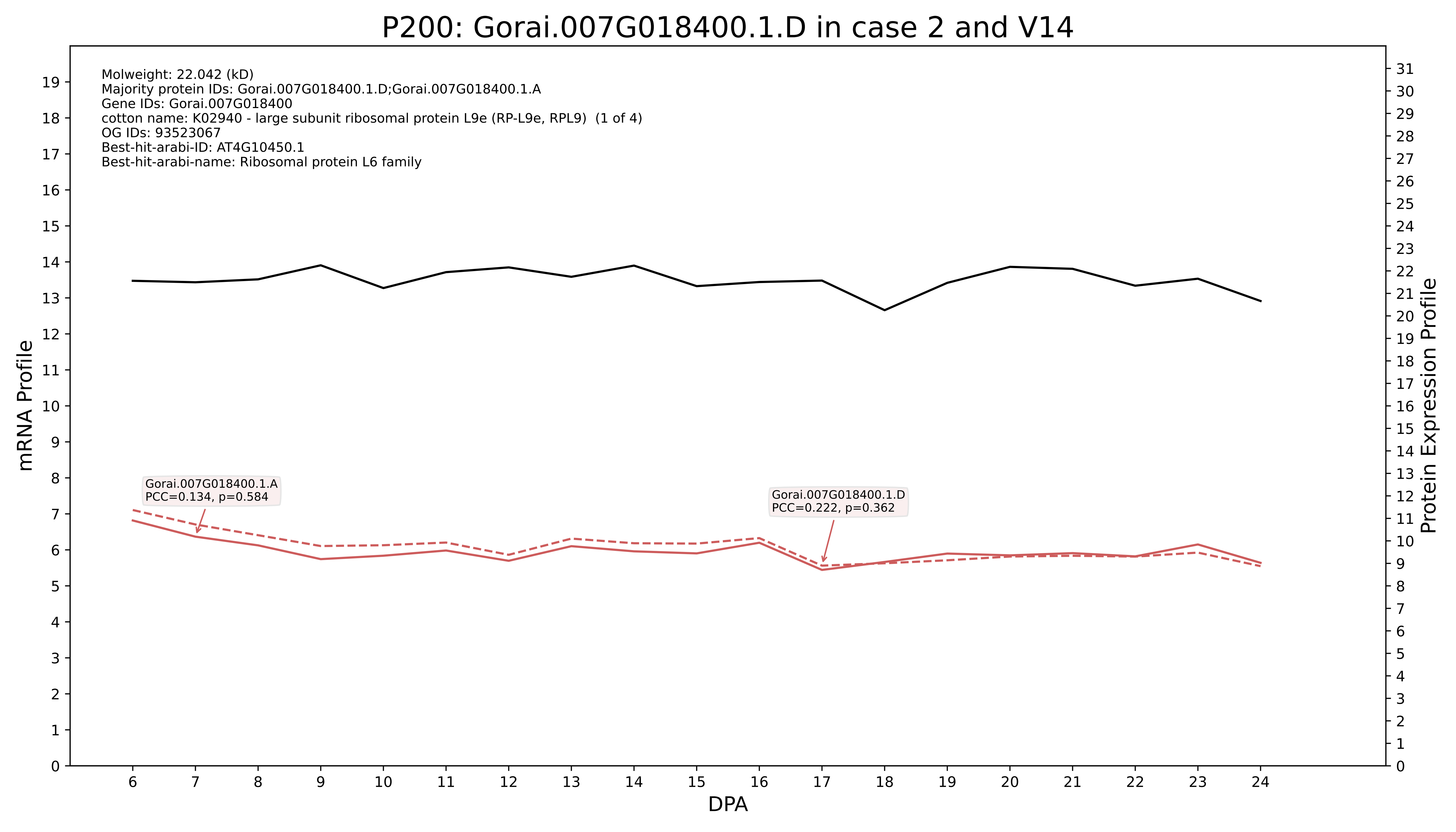
P200: Gorai.007G018400.1.D in case 2 and V14
Molweight: 22.042 (kD)
Majority protein IDs: Gorai.007G018400.1.D;Gorai.007G018400.1.A
Gene IDs: Gorai.007G018400
cotton name: K02940 - large subunit ribosomal protein L9e (RP-L9e, RPL9) (1 of 4)
OG IDs: 93523067
Best-hit-arabi-ID: AT4G10450.1
Best-hit-arabi-name: Ribosomal protein L6 family
Gorai.007G018400.1.A
PCC=0.134, p=0.584
Gorai.007G018400.1.D
PCC=0.222, p=0.362
mRNA Profile
Protein Expression Profile
DPA

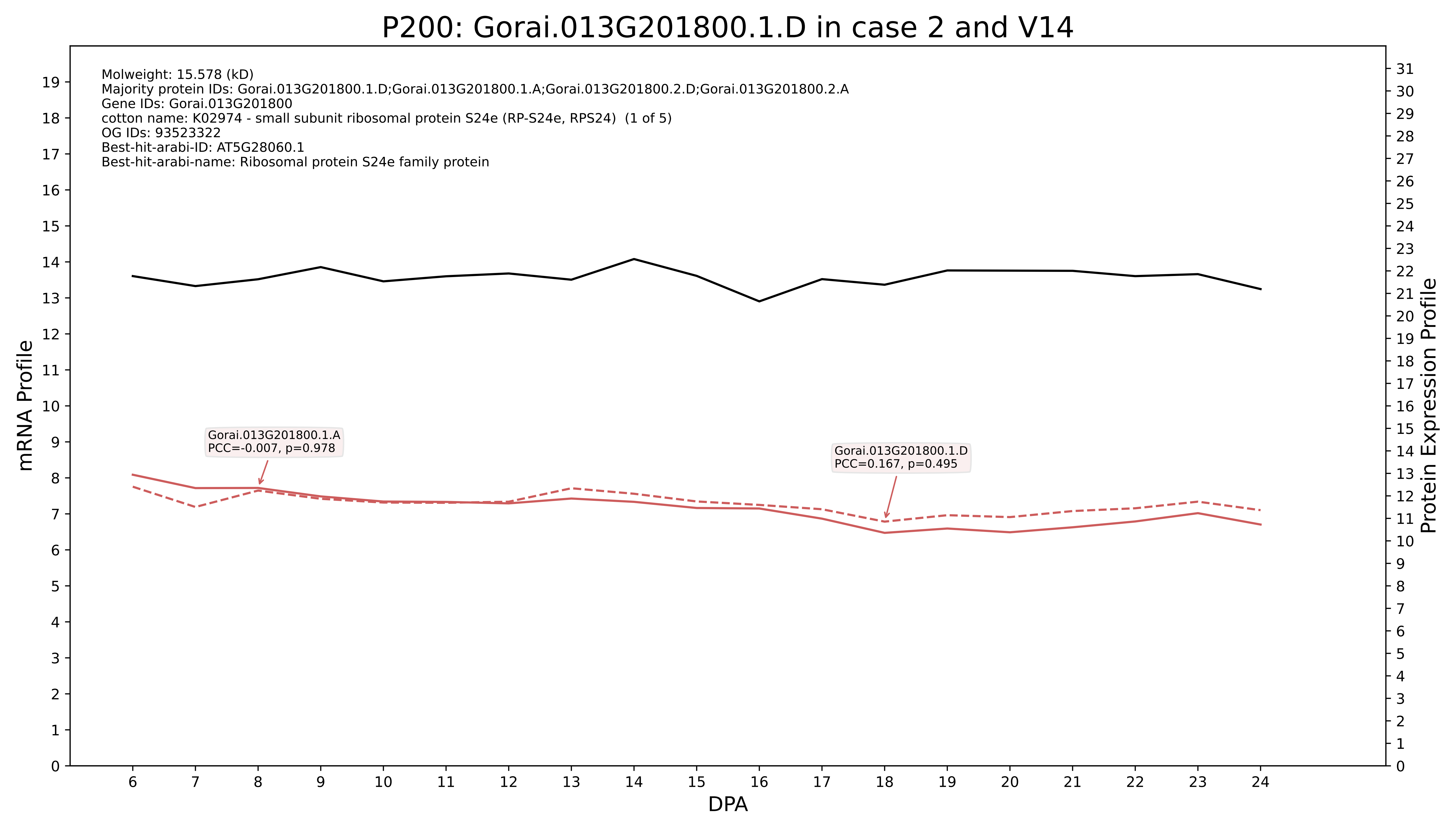
P200: Gorai.013G201800.1.D in case 2 and V14
Molweight: 15.578 (kD)
Majority protein IDs: Gorai.013G201800.1.D;Gorai.013G201800.1.A;Gorai.013G201800.2.D;Gorai.013G201800.2.A
Gene IDs: Gorai.013G201800
cotton name: K02974 - small subunit ribosomal protein S24e (RP-S24e, RPS24) (1 of 5)
OG IDs: 93523322
Best-hit-arabi-ID: AT5G28060.1
Best-hit-arabi-name: Ribosomal protein S24e family protein
Gorai.013G201800.1.A
PCC=-0.007, p=0.978
Gorai.013G201800.1.D
PCC=0.167, p=0.495
mRNA Profile
Protein Expression Profile
DPA

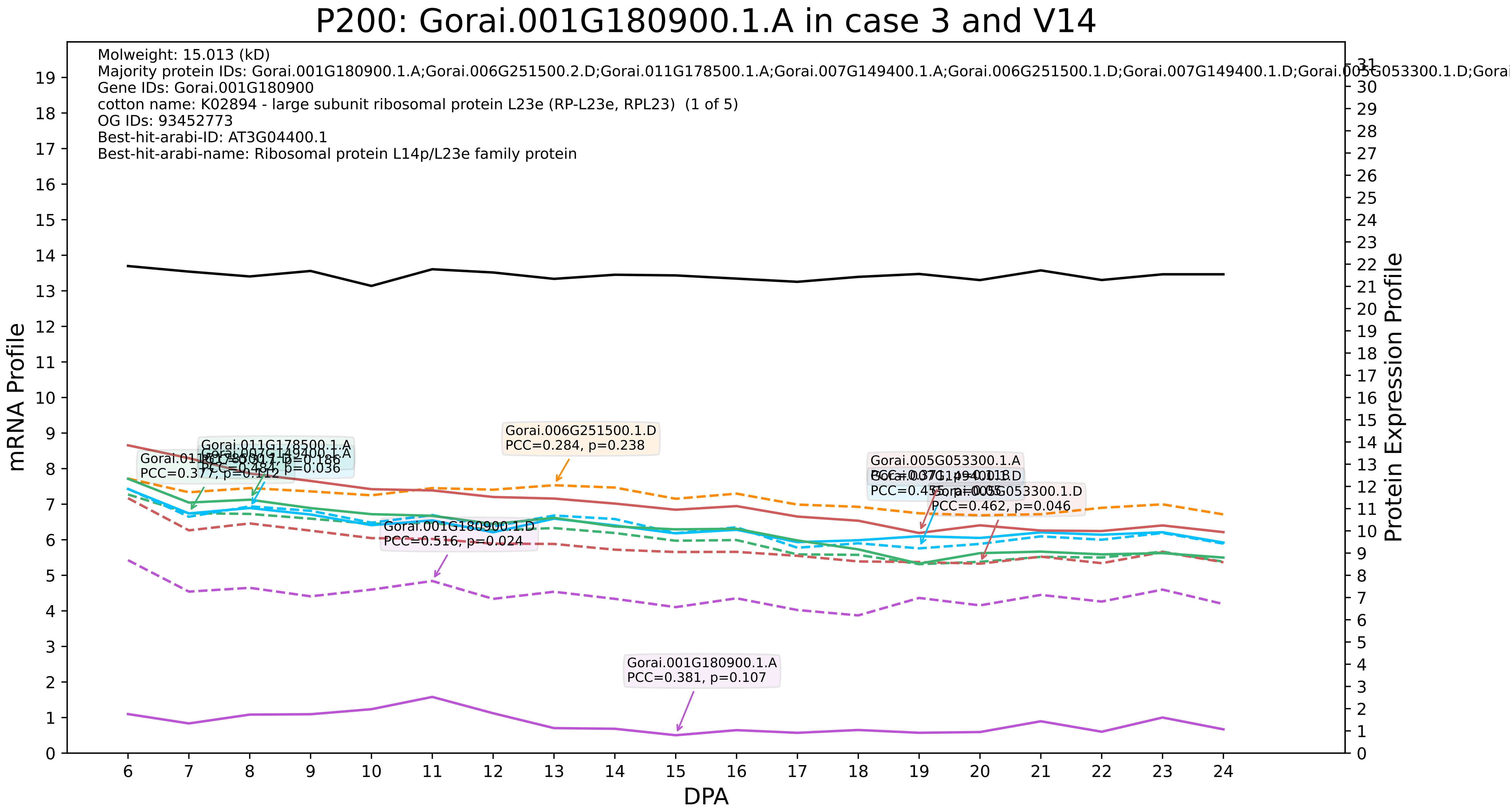
P200: Gorai.001G180900.1.A in case 3 and V14
Molweight: 15.013 (kD)
Majority protein IDs: Gorai.001G180900.1.A;Gorai.006G251500.2.D;Gorai.011G178500.1.A;Gorai.007G149400.1.A;Gorai.006G251500.1.D;Gorai.007G149400.1.D;Gorai.005G053300.1.D;Gorai
Gene IDs: Gorai.001G180900
cotton name: K02894 - large subunit ribosomal protein L23e (RP-L23e, RPL23) (1 of 5)
OG IDs: 93452773
Best-hit-arabi-ID: AT3G04400.1
Best-hit-arabi-name: Ribosomal protein L14p/L23e family protein
Gorai.006G251500.1.D
PCC=0.284, p=0.238
Gorai.011G178500.1.A
Gorai.001G180900.1.D
PCC=0.516, p=0.024
Gorai.005G053300.1.A
Gorai.005G053300.1.D
PCC=0.462, p=0.046
Gorai.001G180900.1.A
PCC=0.381, p=0.107
mRNA Profile
Protein Expression Profile
DPA

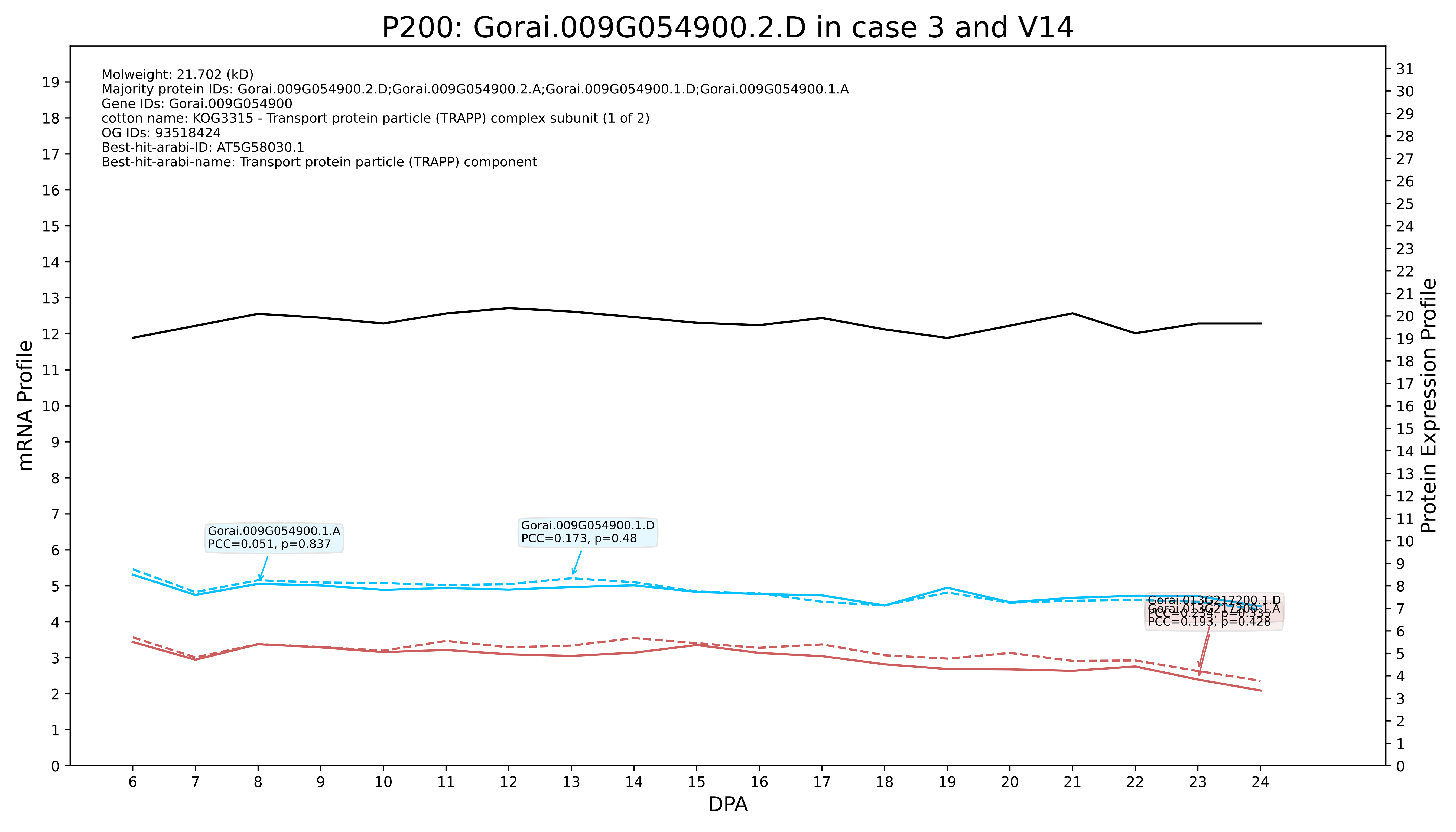
P200: Gorai.009G054900.2.D in case 3 and V14
Molweight: 21.702 (kD)
Majority protein IDs: Gorai.009G054900.2.D;Gorai.009G054900.2.A;Gorai.009G054900.1.D;Gorai.009G054900.1.A
Gene IDs: Gorai.009G054900
cotton name: KOG3315 - Transport protein particle (TRAPP) complex subunit (1 of 2)
OG IDs: 93518424
Best-hit-arabi-ID: AT5G58030.1
Best-hit-arabi-name: Transport protein particle (TRAPP) component
Gorai.009G054900.1.A
PCC=0.051, p=0.837
Gorai.009G054900.1.D
PCC=0.173, p=0.48
Gorai.013G217200.1.D
Gorai.013G217200.1.A
PCC=0.234, p=0.335
PCC=0.193, p=0.428
mRNA Profile
Protein Expression Profile
DPA

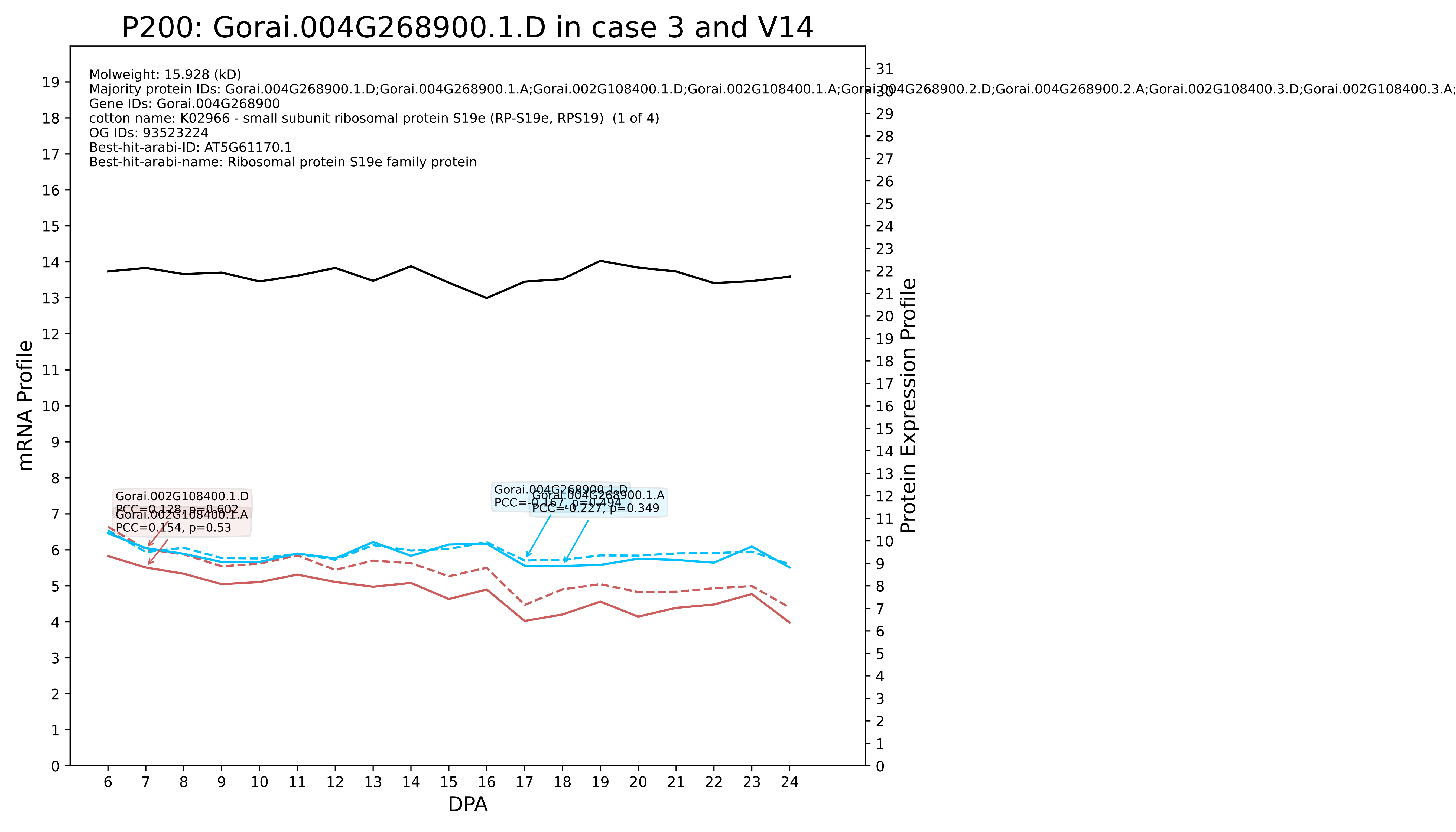
P200: Gorai.004G268900.1.D in case 3 and V14
Molweight: 15.928 (kD)
Majority protein IDs: Gorai.004G268900.1.D;Gorai.004G268900.1.A;Gorai.002G108400.1.D;Gorai.002G108400.1.A;Gorai.004G268900.2.D;Gorai.004G268900.2.A;Gorai.002G108400.3.D;Gorai.002G108400.3.A;
Gene IDs: Gorai.004G268900
cotton name: K02966 - small subunit ribosomal protein S19e (RP-S19e, RPS19) (1 of 4)
OG IDs: 93523224
Best-hit-arabi-ID: AT5G61170.1
Best-hit-arabi-name: Ribosomal protein S19e family protein
Gorai.002G108400.1.D
PCC=0.128, p=0.602
Gorai.002G108400.1.A
PCC=0.154, p=0.53
Gorai.004G268900.1.D
PCC=[illegible]
Gorai.004G268900.1.A
PCC=-0.227, p=0.349
mRNA Profile
Protein Expression Profile
DPA

### P200: Gorai.009G230200.2.D in case 3 and V14

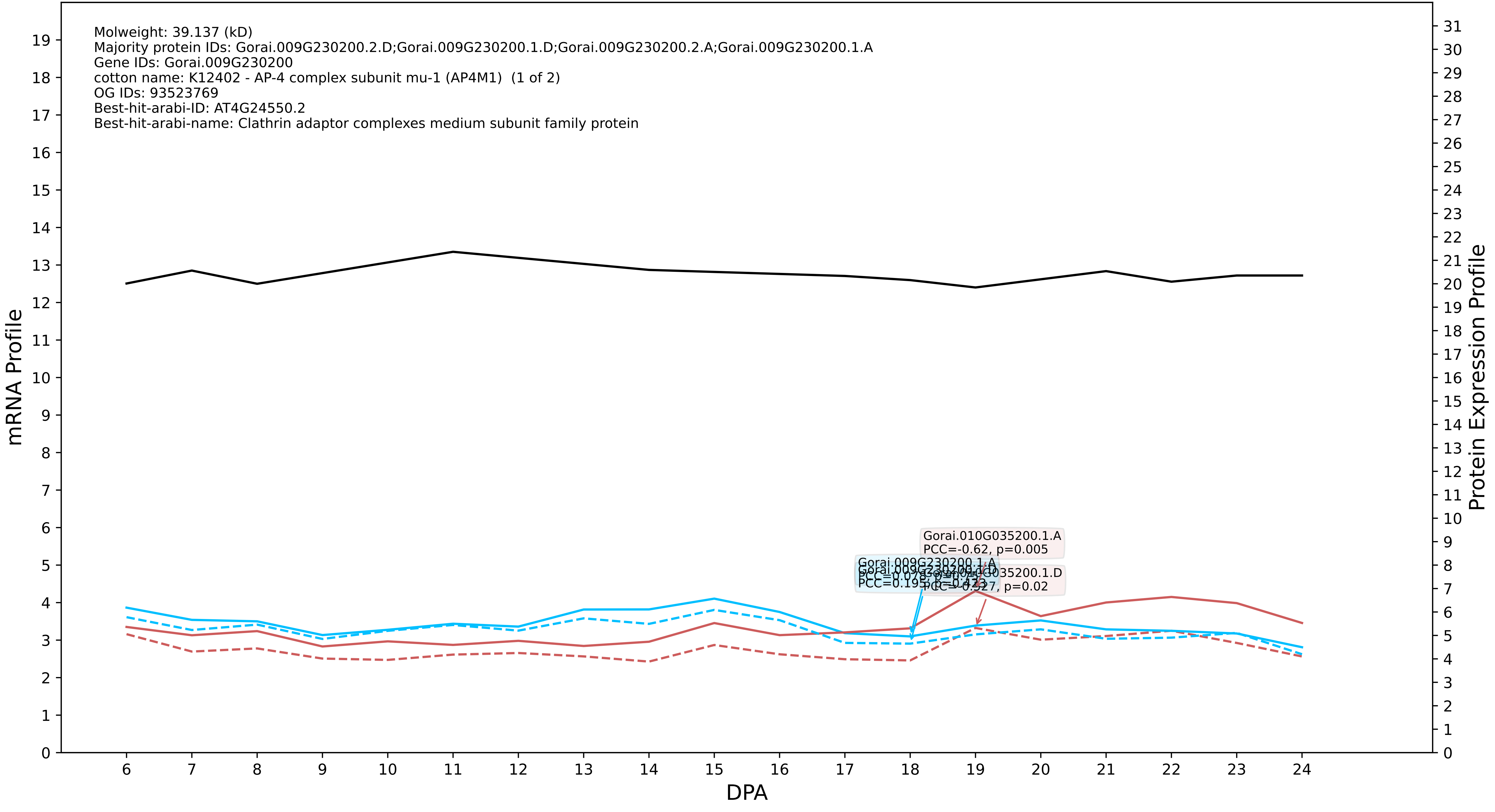

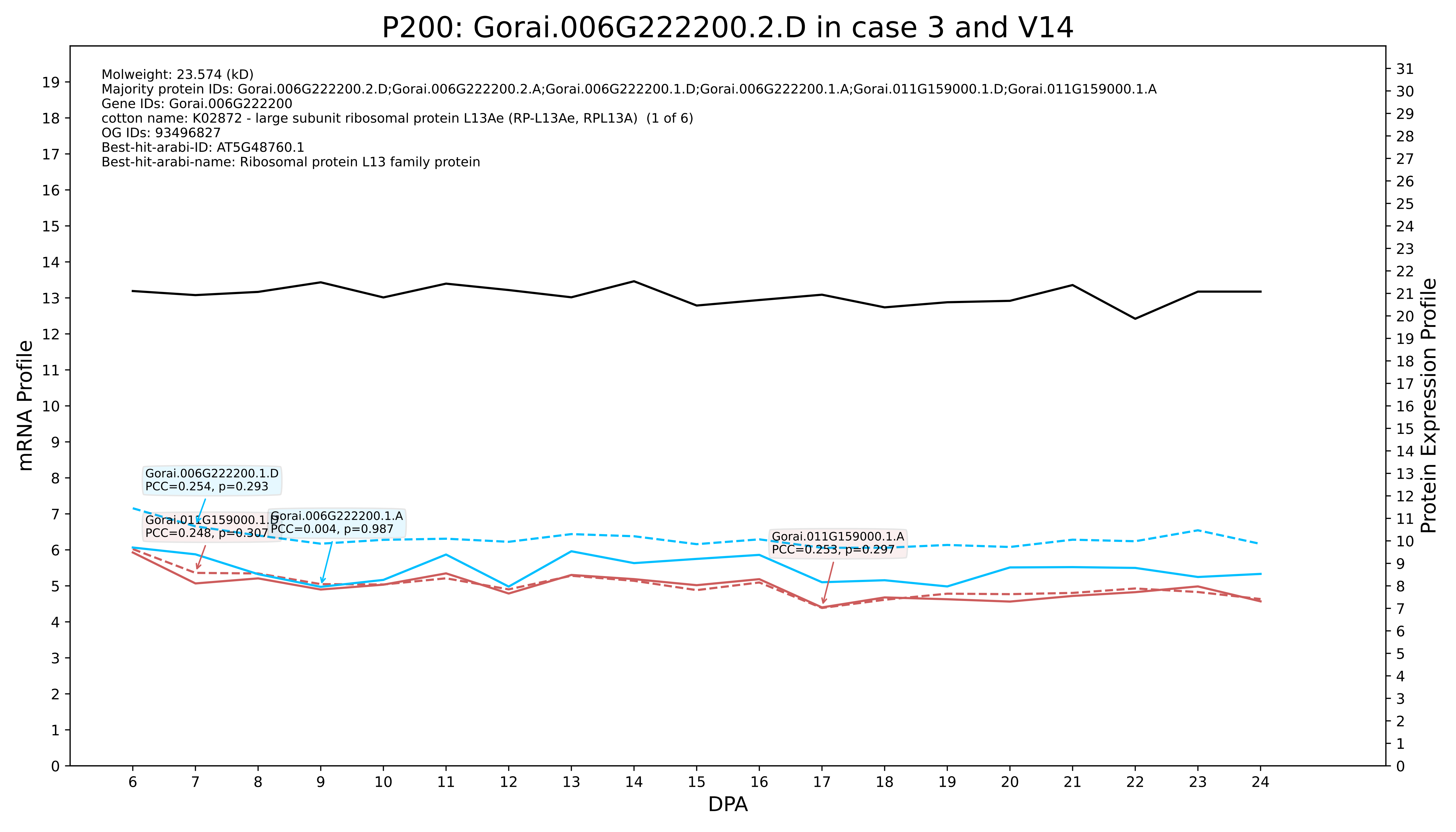
P200: Gorai.006G222200.2.D in case 3 and V14
Molweight: 23.574 (kD)
Majority protein IDs: Gorai.006G222200.2.D;Gorai.006G222200.2.A;Gorai.006G222200.1.D;Gorai.006G222200.1.A;Gorai.011G159000.1.D;Gorai.011G159000.1.A
Gene IDs: Gorai.006G222200
cotton name: K02872 - large subunit ribosomal protein L13Ae (RP-L13Ae, RPL13A) (1 of 6)
OG IDs: 93496827
Best-hit-arabi-ID: AT5G48760.1
Best-hit-arabi-name: Ribosomal protein L13 family protein
Gorai.006G222200.1.D
PCC=0.254, p=0.293
Gorai.011G159000.1.D
PCC=0.248, p=0.307
Gorai.006G222200.1.A
PCC=0.004, p=0.987
Gorai.011G159000.1.A
PCC=0.253, p=0.297
mRNA Profile
Protein Expression Profile
DPA

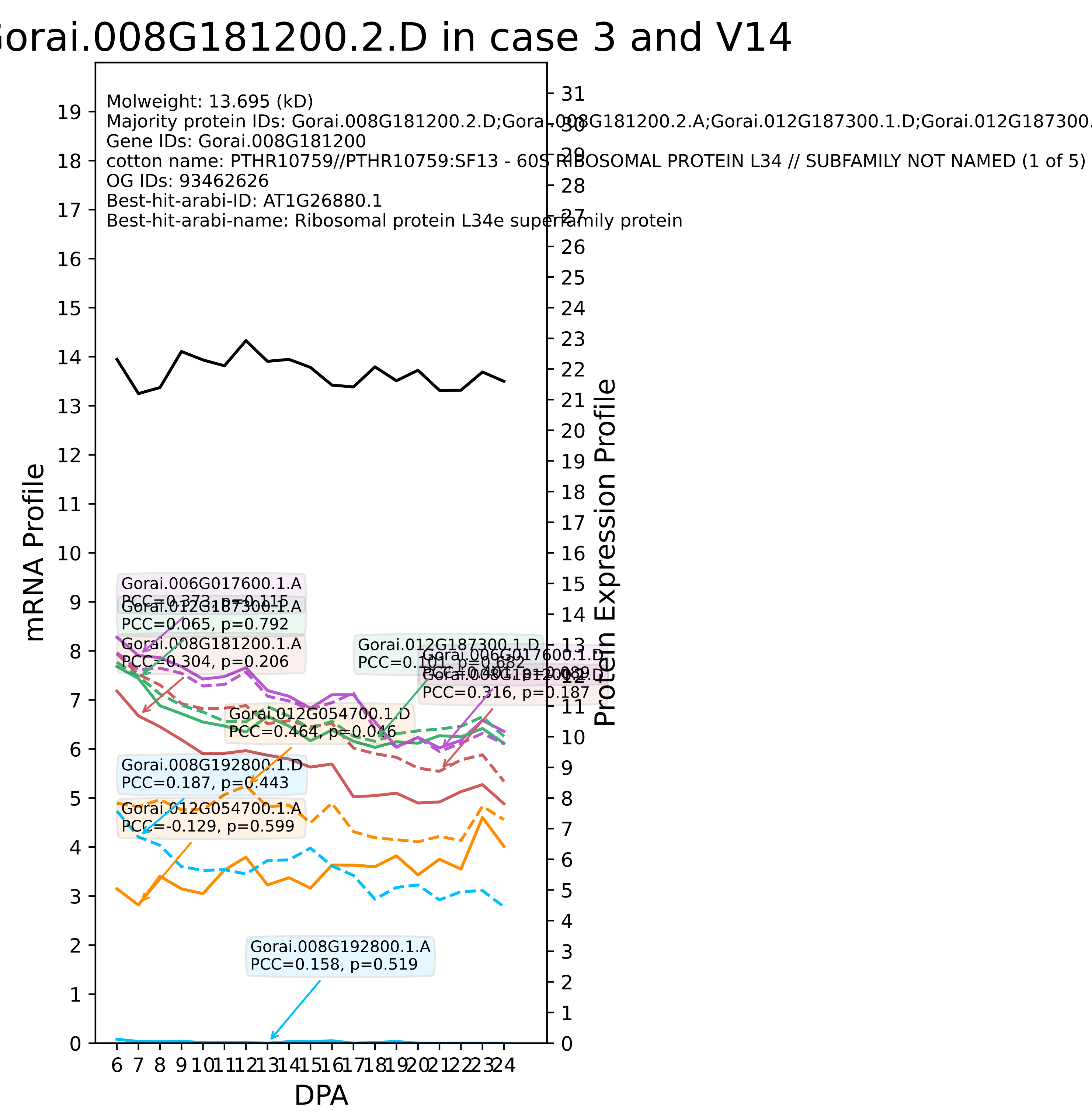
Gorai.008G181200.2.D in case 3 and V14
Molweight: 13.695 (kD)
Majority protein IDs: Gorai.008G181200.2.D;Gorai.008G181200.2.A;Gorai.012G187300.1.D;Gorai.012G187300.1.A;Gorai.008G192800.1.D;Gorai.006G017600.1.D;Gorai.008G181200.1.D;Gorai.008G181200.1.A;G
Gene IDs: Gorai.008G181200
cotton name: PTHR10759//PTHR10759:SF13 - 60S RIBOSOMAL PROTEIN L34 // SUBFAMILY NOT NAMED (1 of 5)
OG IDs: 93462626
Best-hit-arabi-ID: AT1G26880.1
Best-hit-arabi-name: Ribosomal protein L34e superfamily protein
Gorai.006G017600.1.A
PCC=0.373, p=0.115
Gorai.012G187300.1.A
PCC=0.065, p=0.792
Gorai.008G181200.1.A
PCC=0.304, p=0.206
Gorai.012G187300.1.D
PCC=0.101, p=0.682
Gorai.006G017600.1.D
Gorai.008G181200.1.D
PCC=0.316, p=0.187
Gorai.012G054700.1.D
PCC=0.464, p=0.046
Gorai.008G192800.1.D
PCC=0.187, p=0.443
Gorai.012G054700.1.A
PCC=-0.129, p=0.599
Gorai.008G192800.1.A
PCC=0.158, p=0.519
mRNA Profile
Protein Expression Profile
DPA

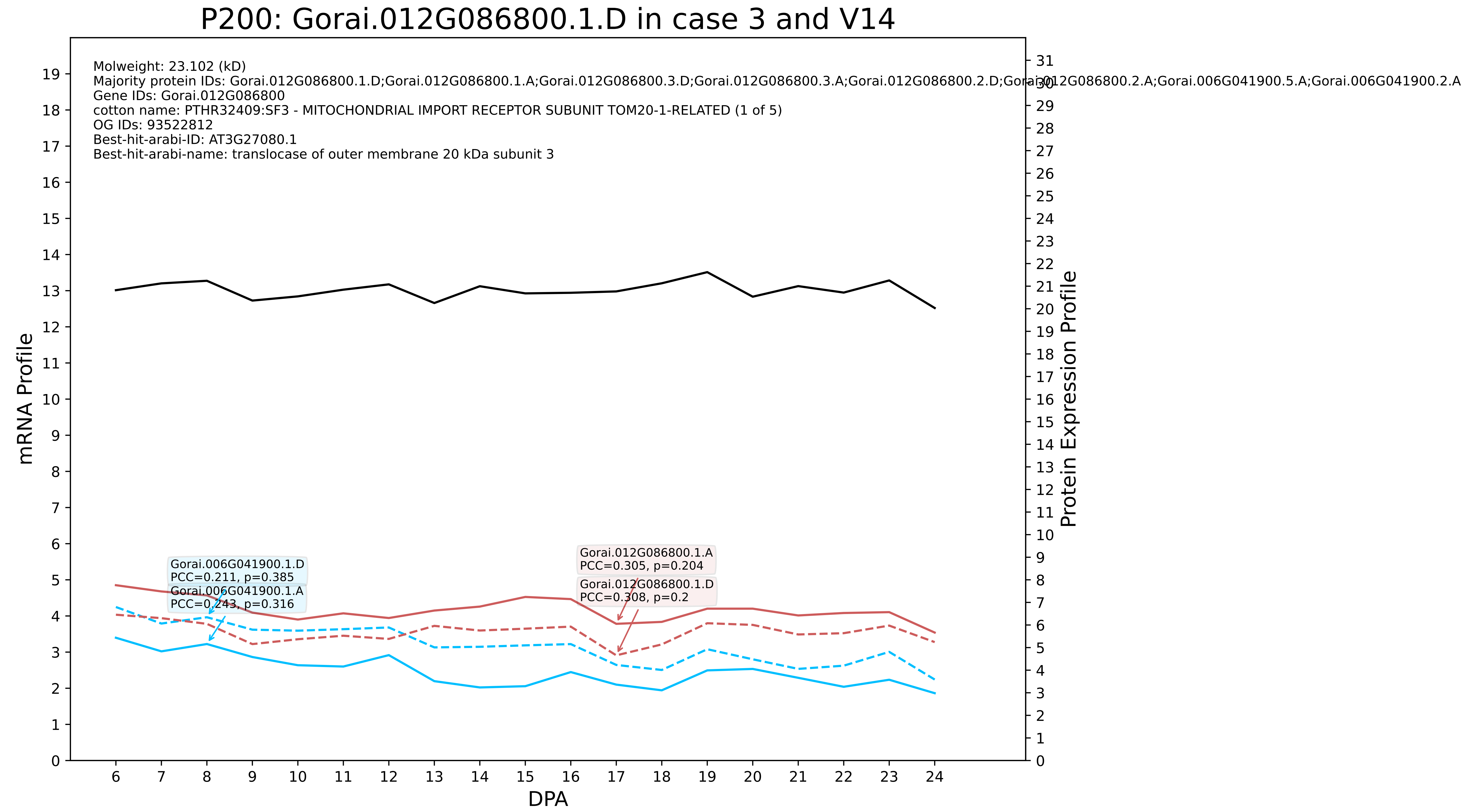
P200: Gorai.012G086800.1.D in case 3 and V14
Molweight: 23.102 (kD)
Majority protein IDs: Gorai.012G086800.1.D;Gorai.012G086800.1.A;Gorai.012G086800.3.D;Gorai.012G086800.3.A;Gorai.012G086800.2.D;Gorai.012G086800.2.A;Gorai.006G041900.5.A;Gorai.006G041900.2.A
Gene IDs: Gorai.012G086800
cotton name: PTHR32409:SF3 - MITOCHONDRIAL IMPORT RECEPTOR SUBUNIT TOM20-1-RELATED (1 of 5)
OG IDs: 93522812
Best-hit-arabi-ID: AT3G27080.1
Best-hit-arabi-name: translocase of outer membrane 20 kDa subunit 3
Gorai.006G041900.1.D
PCC=0.211, p=0.385
Gorai.006G041900.1.A
PCC=0.243, p=0.316
Gorai.012G086800.1.A
PCC=0.305, p=0.204
Gorai.012G086800.1.D
PCC=0.308, p=0.2
mRNA Profile
Protein Expression Profile
DPA

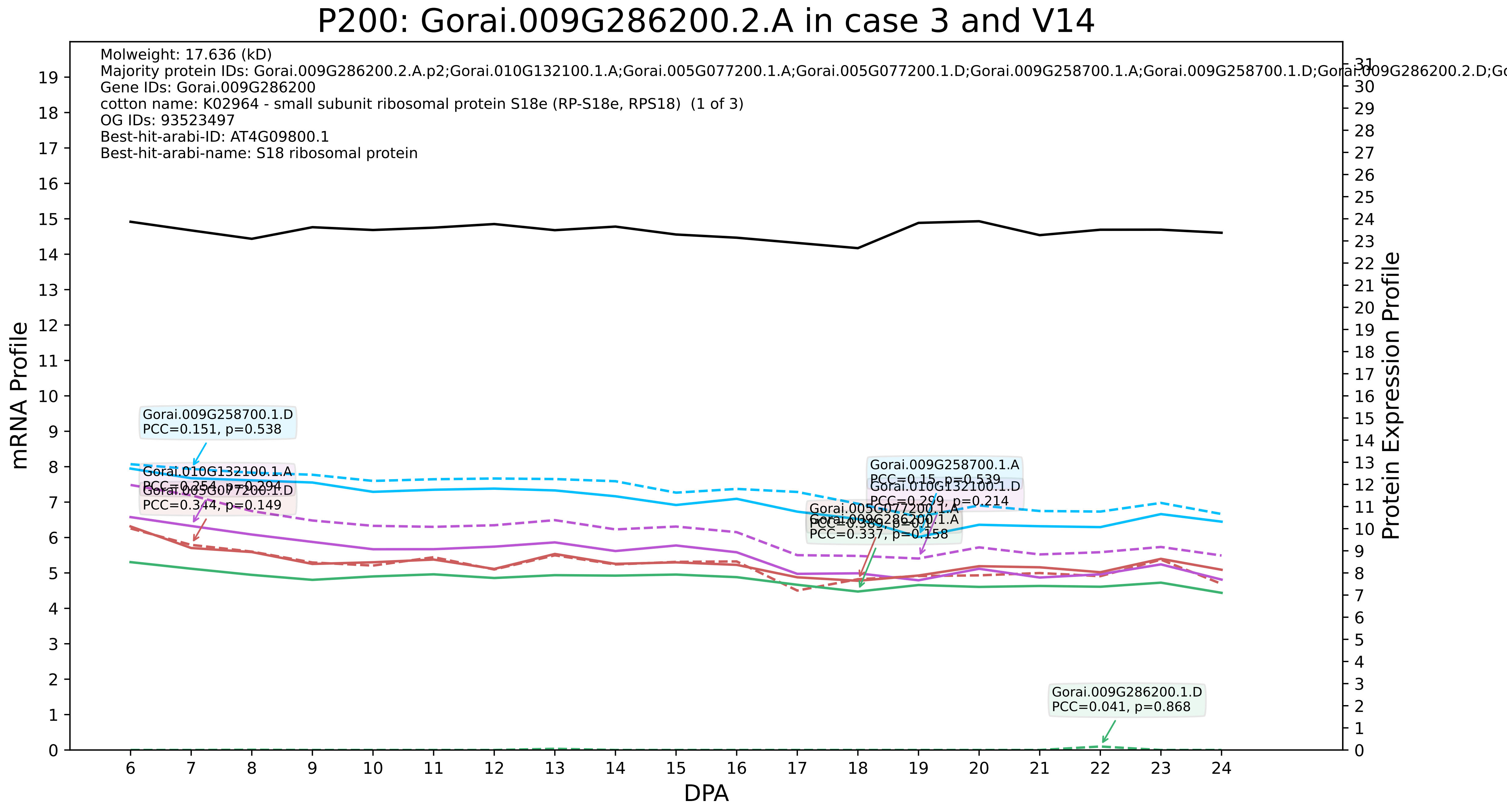

P200: Gorai.009G286200.2.A in case 3 and V14
Molweight: 17.636 (kD)
Majority protein IDs: Gorai.009G286200.2.A.p2;Gorai.010G132100.1.A;Gorai.005G077200.1.A;Gorai.005G077200.1.D;Gorai.009G258700.1.A;Gorai.009G258700.1.D;Gorai.009G286200.2.D;Go
Gene IDs: Gorai.009G286200
cotton name: K02964 - small subunit ribosomal protein S18e (RP-S18e, RPS18) (1 of 3)
OG IDs: 93523497
Best-hit-arabi-ID: AT4G09800.1
Best-hit-arabi-name: S18 ribosomal protein
mRNA Profile
Protein Expression Profile
DPA
Gorai.009G258700.1.D
PCC=0.151, p=0.538
Gorai.010G132100.1.A
PCC=0.254, p=0.294
Gorai.005G077200.1.D
PCC=0.344, p=0.149
Gorai.009G258700.1.A
PCC=0.15, p=0.539
Gorai.010G132100.1.D
PCC=0.299, p=0.214
Gorai.005G077200.1.A
Gorai.009G286200.1.A
PCC=0.337, p=0.158
Gorai.009G286200.1.D
PCC=0.041, p=0.868

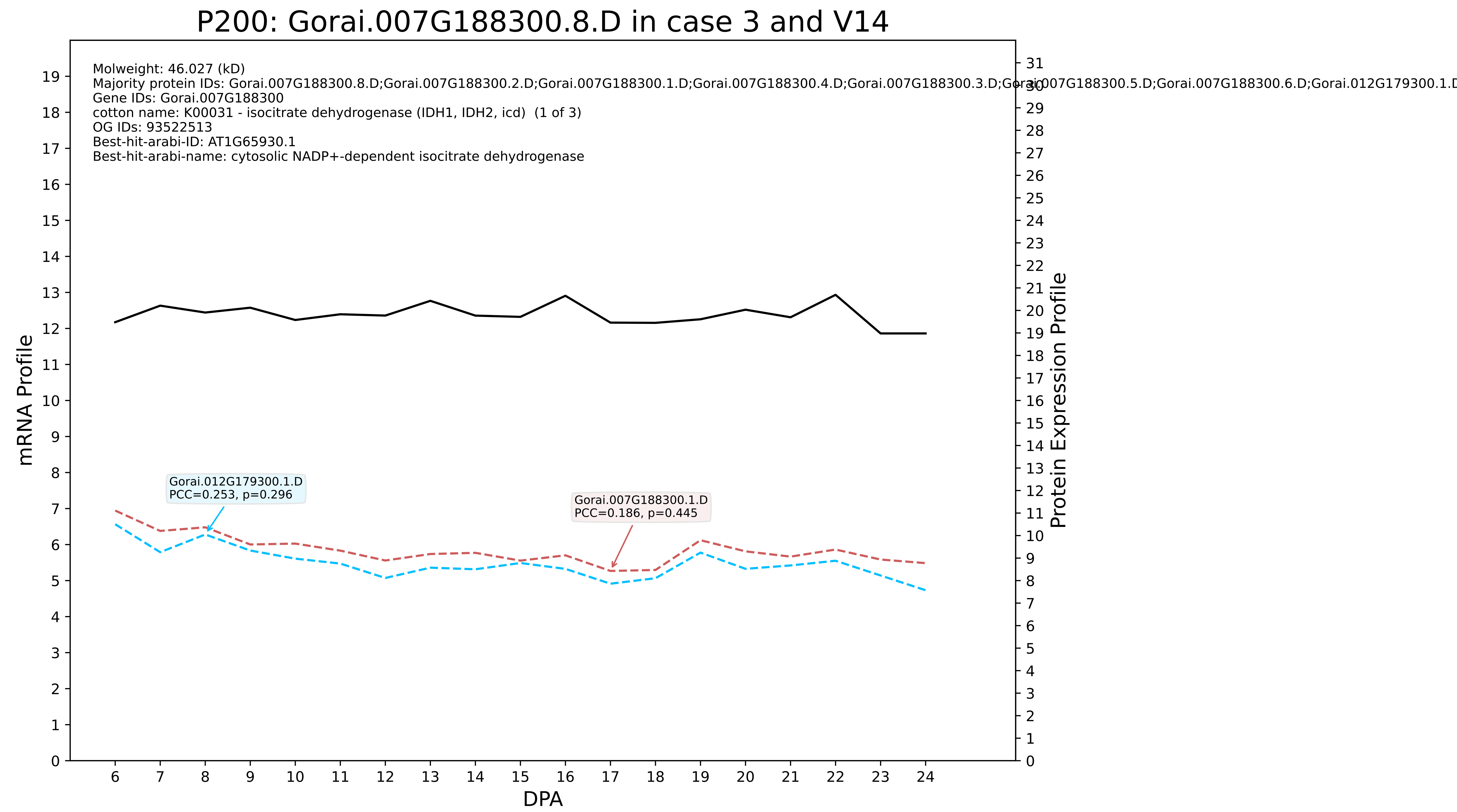
P200: Gorai.007G188300.8.D in case 3 and V14
Molweight: 46.027 (kD)
Majority protein IDs: Gorai.007G188300.8.D;Gorai.007G188300.2.D;Gorai.007G188300.1.D;Gorai.007G188300.4.D;Gorai.007G188300.3.D;Gorai.007G188300.5.D;Gorai.007G188300.6.D;Gorai.012G179300.1.D
Gene IDs: Gorai.007G188300
cotton name: K00031 - isocitrate dehydrogenase (IDH1, IDH2, icd) (1 of 3)
OG IDs: 93522513
Best-hit-arabi-ID: AT1G65930.1
Best-hit-arabi-name: cytosolic NADP+-dependent isocitrate dehydrogenase
Gorai.012G179300.1.D
PCC=0.253, p=0.296
Gorai.007G188300.1.D
PCC=0.186, p=0.445
mRNA Profile
Protein Expression Profile
DPA

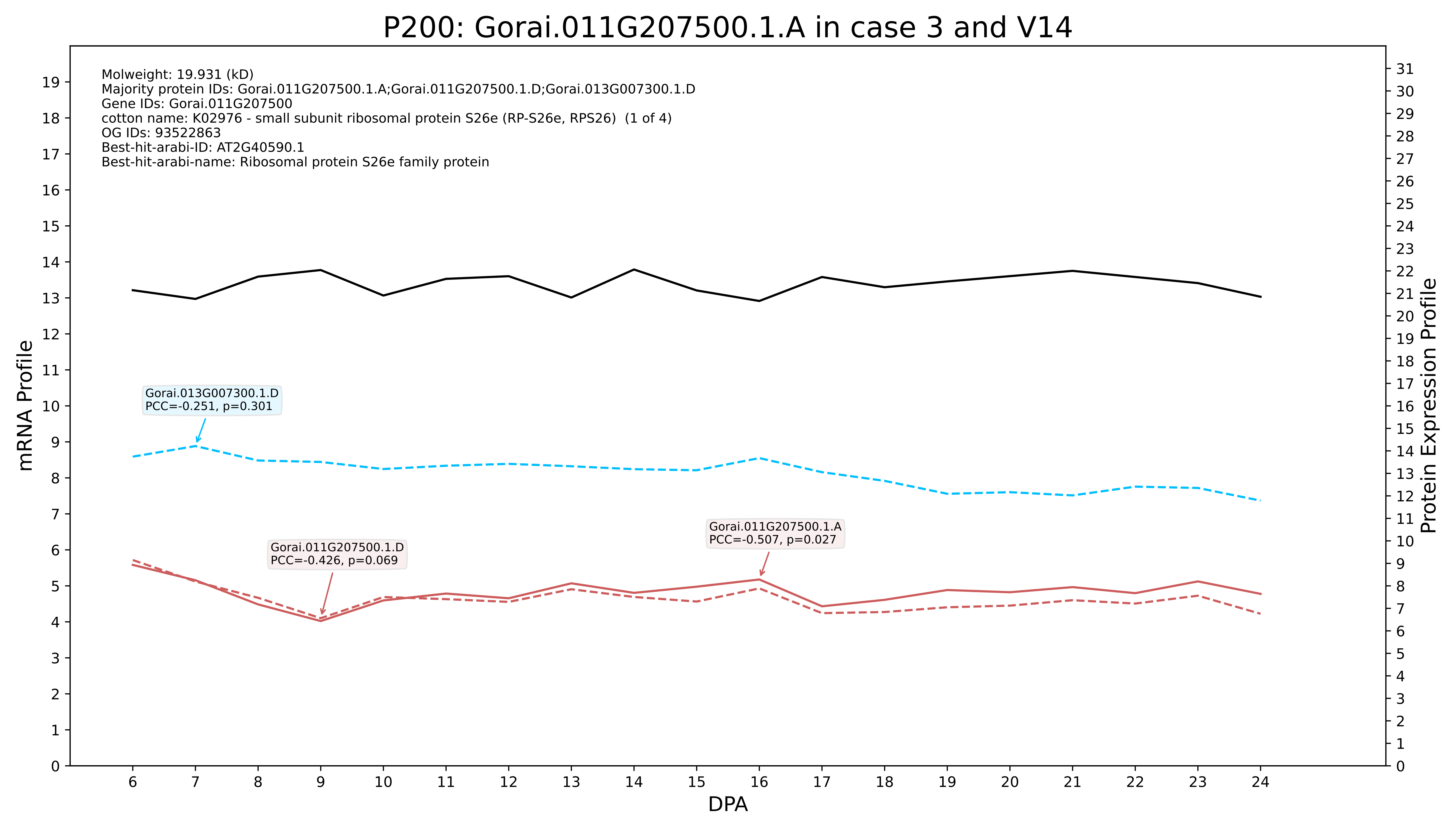
P200: Gorai.011G207500.1.A in case 3 and V14
Molweight: 19.931 (kD)
Majority protein IDs: Gorai.011G207500.1.A;Gorai.011G207500.1.D;Gorai.013G007300.1.D
Gene IDs: Gorai.011G207500
cotton name: K02976 - small subunit ribosomal protein S26e (RP-S26e, RPS26) (1 of 4)
OG IDs: 93522863
Best-hit-arabi-ID: AT2G40590.1
Best-hit-arabi-name: Ribosomal protein S26e family protein
Gorai.013G007300.1.D
PCC=-0.251, p=0.301
Gorai.011G207500.1.A
PCC=-0.507, p=0.027
Gorai.011G207500.1.D
PCC=-0.426, p=0.069
mRNA Profile
Protein Expression Profile
DPA

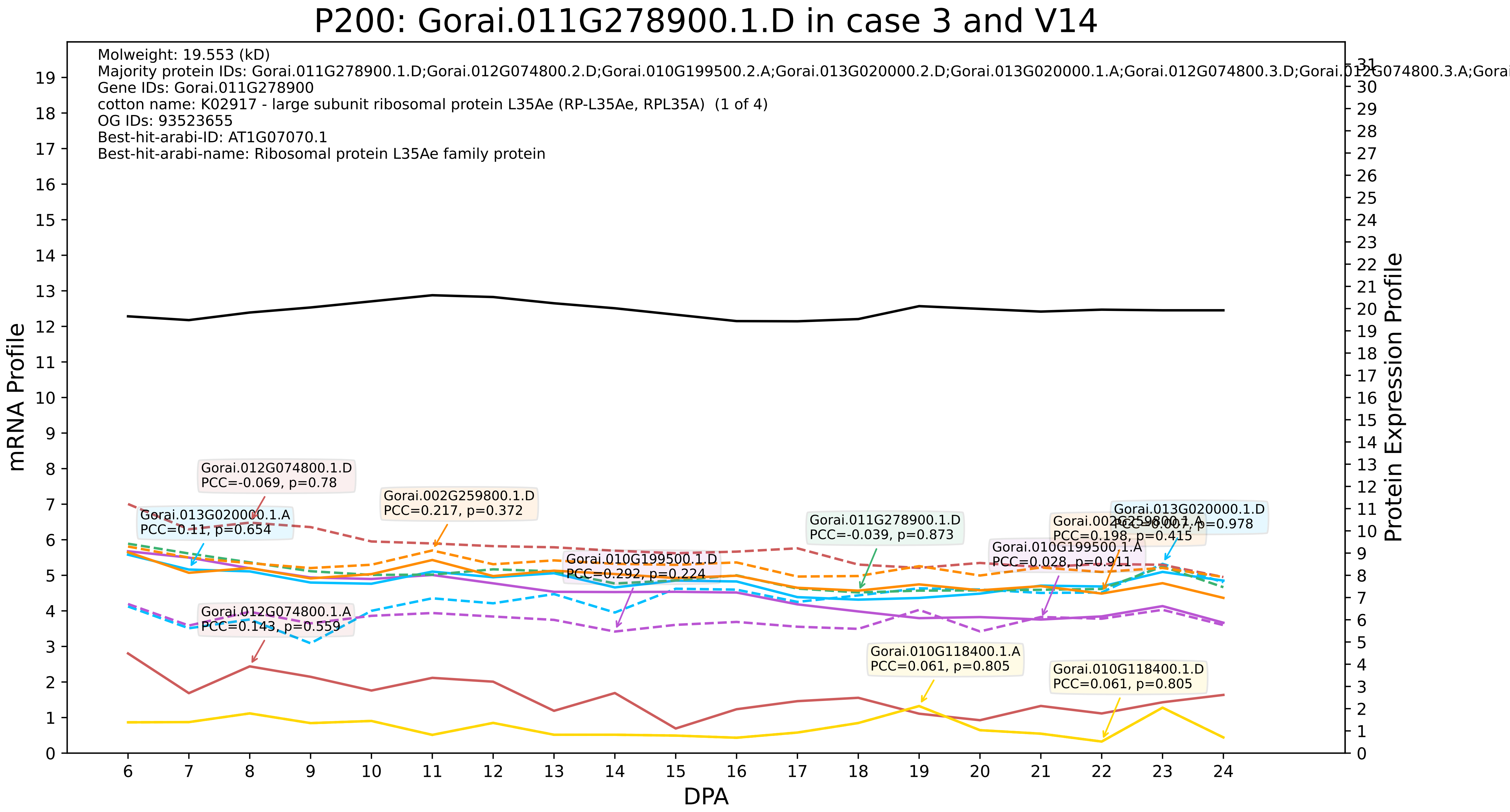
P200: Gorai.011G278900.1.D in case 3 and V14
Molweight: 19.553 (kD)
Majority protein IDs: Gorai.011G278900.1.D;Gorai.012G074800.2.D;Gorai.010G199500.2.A;Gorai.013G020000.2.D;Gorai.013G020000.1.A;Gorai.012G074800.3.D;Gorai.012G074800.3.A;Gorai
Gene IDs: Gorai.011G278900
cotton name: K02917 - large subunit ribosomal protein L35Ae (RP-L35Ae, RPL35A) (1 of 4)
OG IDs: 93523655
Best-hit-arabi-ID: AT1G07070.1
Best-hit-arabi-name: Ribosomal protein L35Ae family protein
Gorai.012G074800.1.D
PCC=-0.069, p=0.78
Gorai.013G020000.1.A
PCC=0.11, p=0.654
Gorai.002G259800.1.D
PCC=0.217, p=0.372
Gorai.011G278900.1.D
PCC=-0.039, p=0.873
Gorai.013G020000.1.D
PCC=0.007, p=0.978
Gorai.002G259800.1.A
PCC=0.198, p=0.415
Gorai.010G199500.1.A
PCC=0.028, p=0.911
Gorai.010G199500.1.D
PCC=0.292, p=0.224
Gorai.012G074800.1.A
PCC=0.143, p=0.559
Gorai.010G118400.1.A
PCC=0.061, p=0.805
Gorai.010G118400.1.D
PCC=0.061, p=0.805
mRNA Profile
Protein Expression Profile
DPA

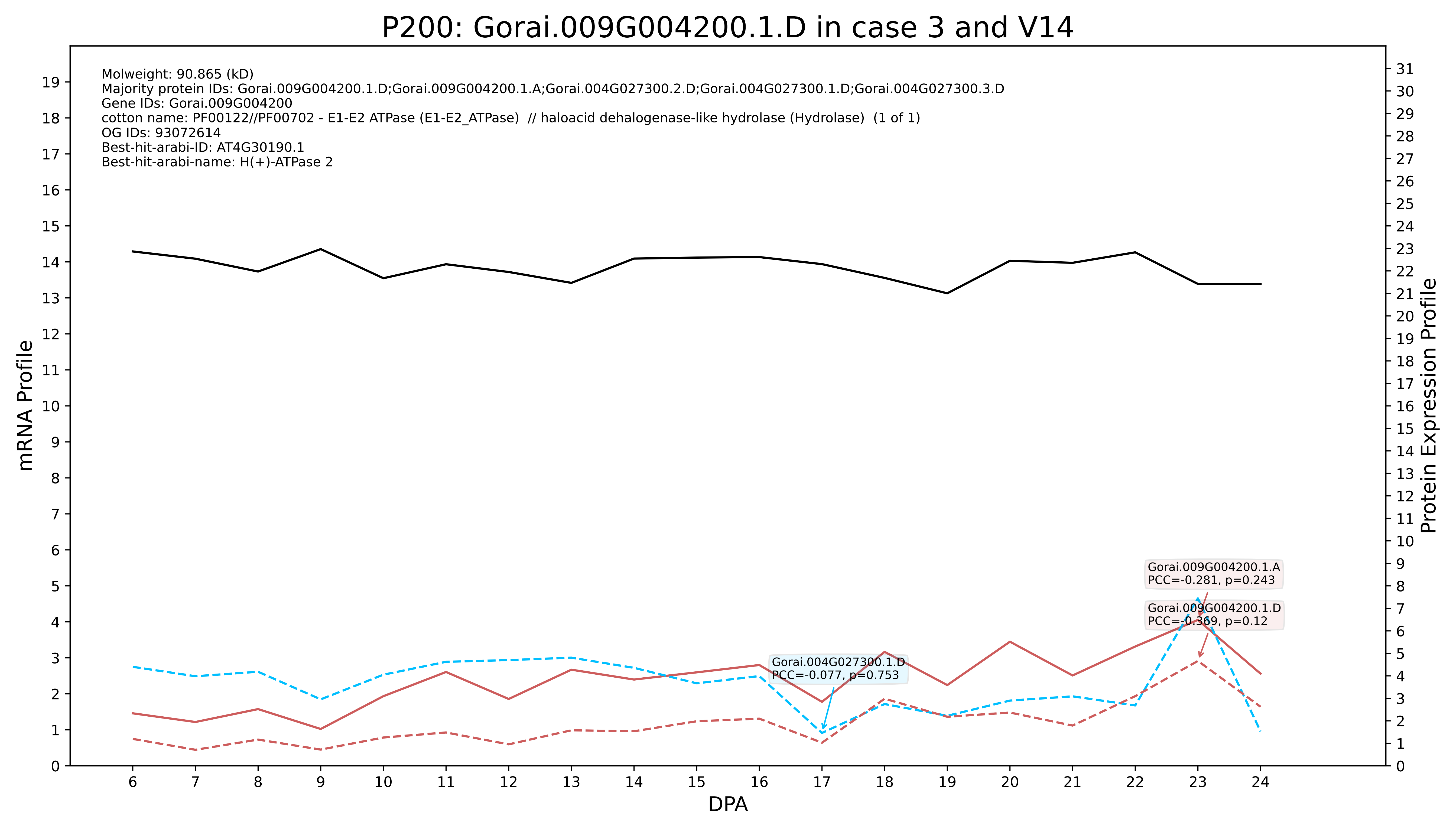
P200: Gorai.009G004200.1.D in case 3 and V14
Molweight: 90.865 (kD)
Majority protein IDs: Gorai.009G004200.1.D;Gorai.009G004200.1.A;Gorai.004G027300.2.D;Gorai.004G027300.1.D;Gorai.004G027300.3.D
Gene IDs: Gorai.009G004200
cotton name: PF00122//PF00702 - E1-E2 ATPase (E1-E2_ATPase) // haloacid dehalogenase-like hydrolase (Hydrolase) (1 of 1)
OG IDs: 93072614
Best-hit-arabi-ID: AT4G30190.1
Best-hit-arabi-name: H(+)-ATPase 2
Gorai.009G004200.1.A
PCC=-0.281, p=0.243
Gorai.009G004200.1.D
PCC=-0.369, p=0.12
Gorai.004G027300.1.D
PCC=-0.077, p=0.753
mRNA Profile
Protein Expression Profile
DPA

### P200: Gorai.007G122600.1.D in case 3 and V14

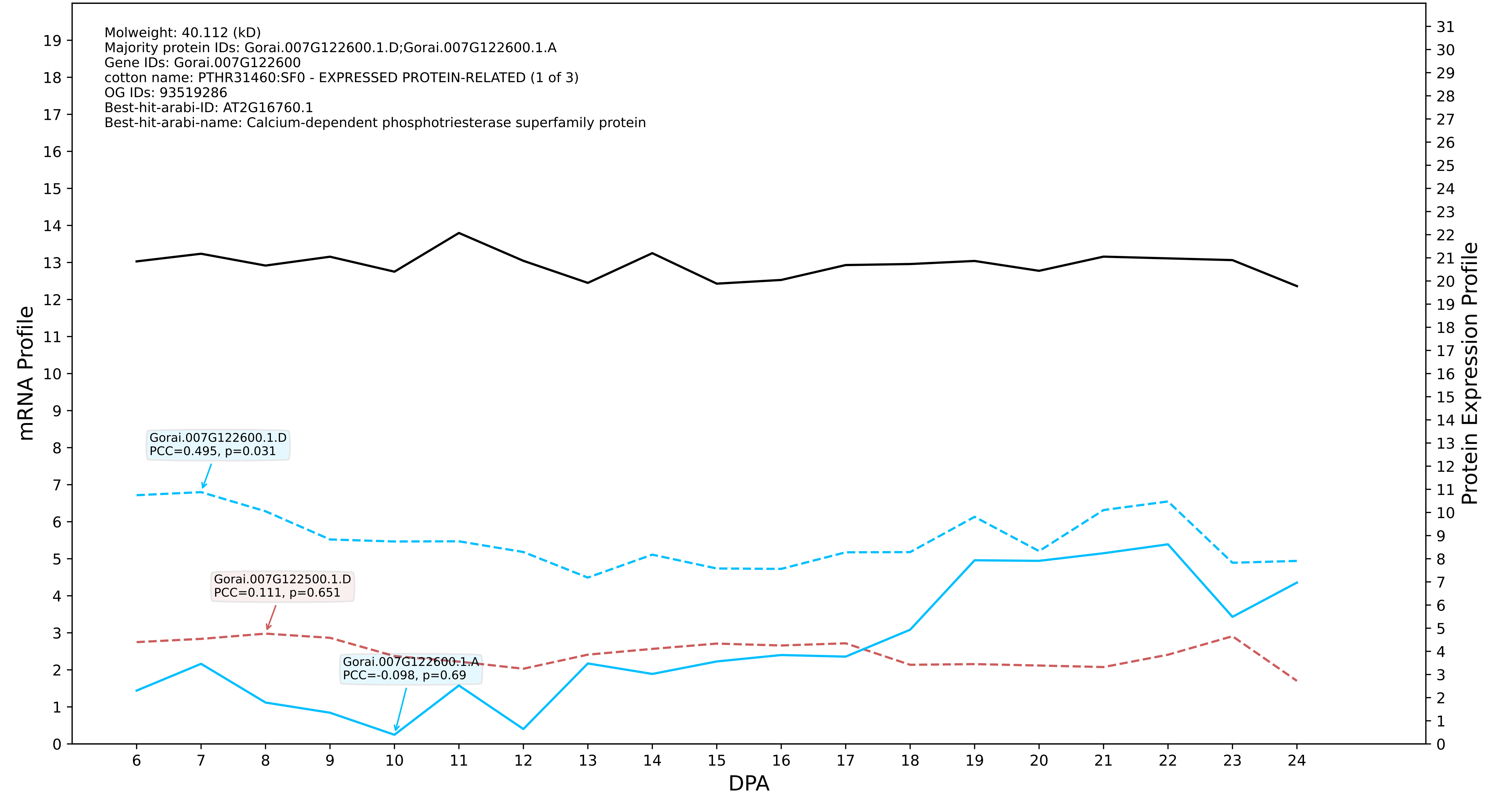

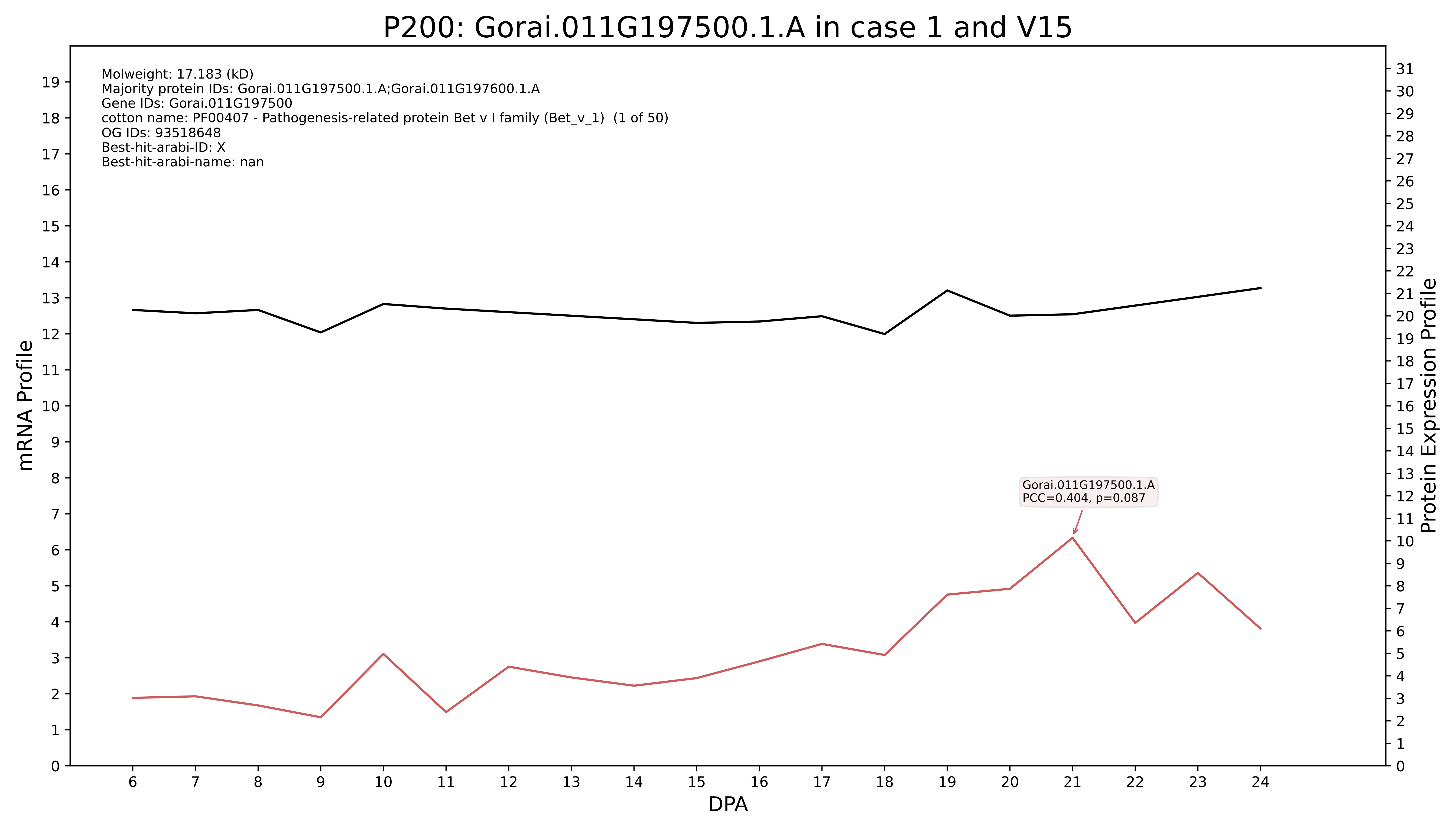
P200: Gorai.011G197500.1.A in case 1 and V15
Molweight: 17.183 (kD)
Majority protein IDs: Gorai.011G197500.1.A;Gorai.011G197600.1.A
Gene IDs: Gorai.011G197500
cotton name: PF00407 - Pathogenesis-related protein Bet v I family (Bet_v_1) (1 of 50)
OG IDs: 93518648
Best-hit-arabi-ID: X
Best-hit-arabi-name: nan
Gorai.011G197500.1.A
PCC=0.404, p=0.087
mRNA Profile
Protein Expression Profile
DPA

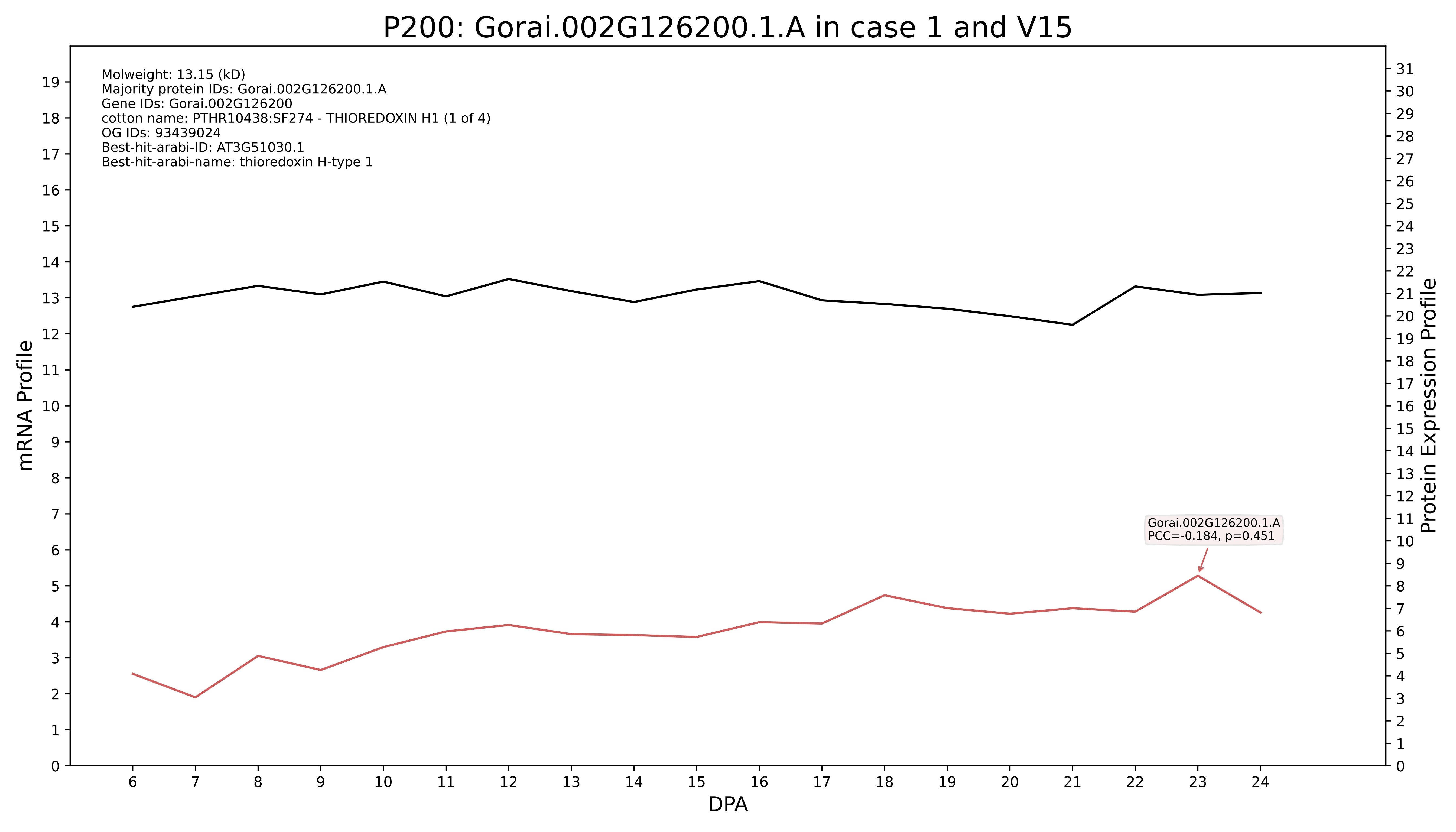
P200: Gorai.002G126200.1.A in case 1 and V15
Molweight: 13.15 (kD)
Majority protein IDs: Gorai.002G126200.1.A
Gene IDs: Gorai.002G126200
cotton name: PTHR10438:SF274 - THIOREDOXIN H1 (1 of 4)
OG IDs: 93439024
Best-hit-arabi-ID: AT3G51030.1
Best-hit-arabi-name: thioredoxin H-type 1
Gorai.002G126200.1.A
PCC=-0.184, p=0.451
mRNA Profile
Protein Expression Profile
DPA

### P200: Gorai.006G009900.4.A in case 1 and V15

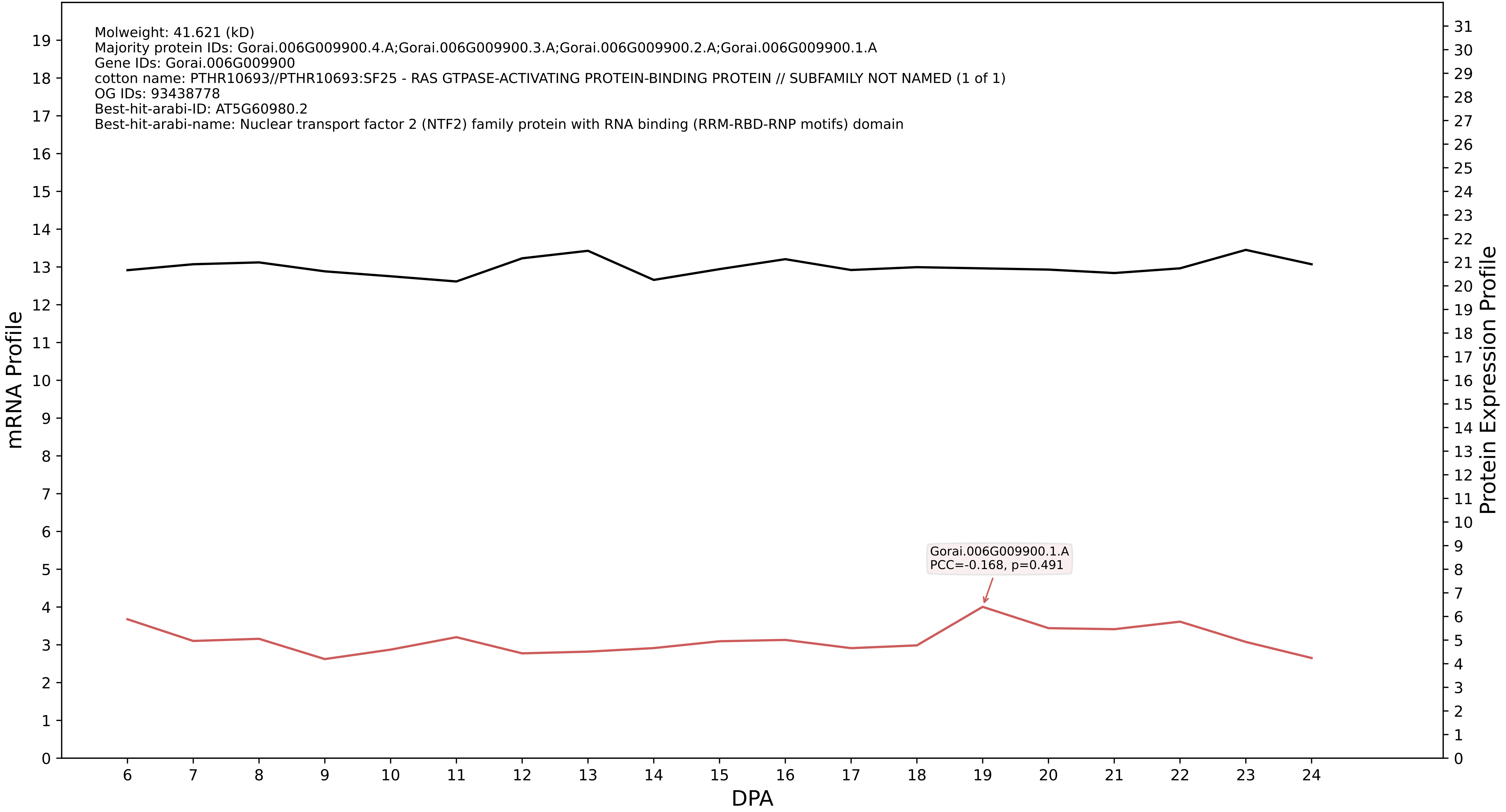

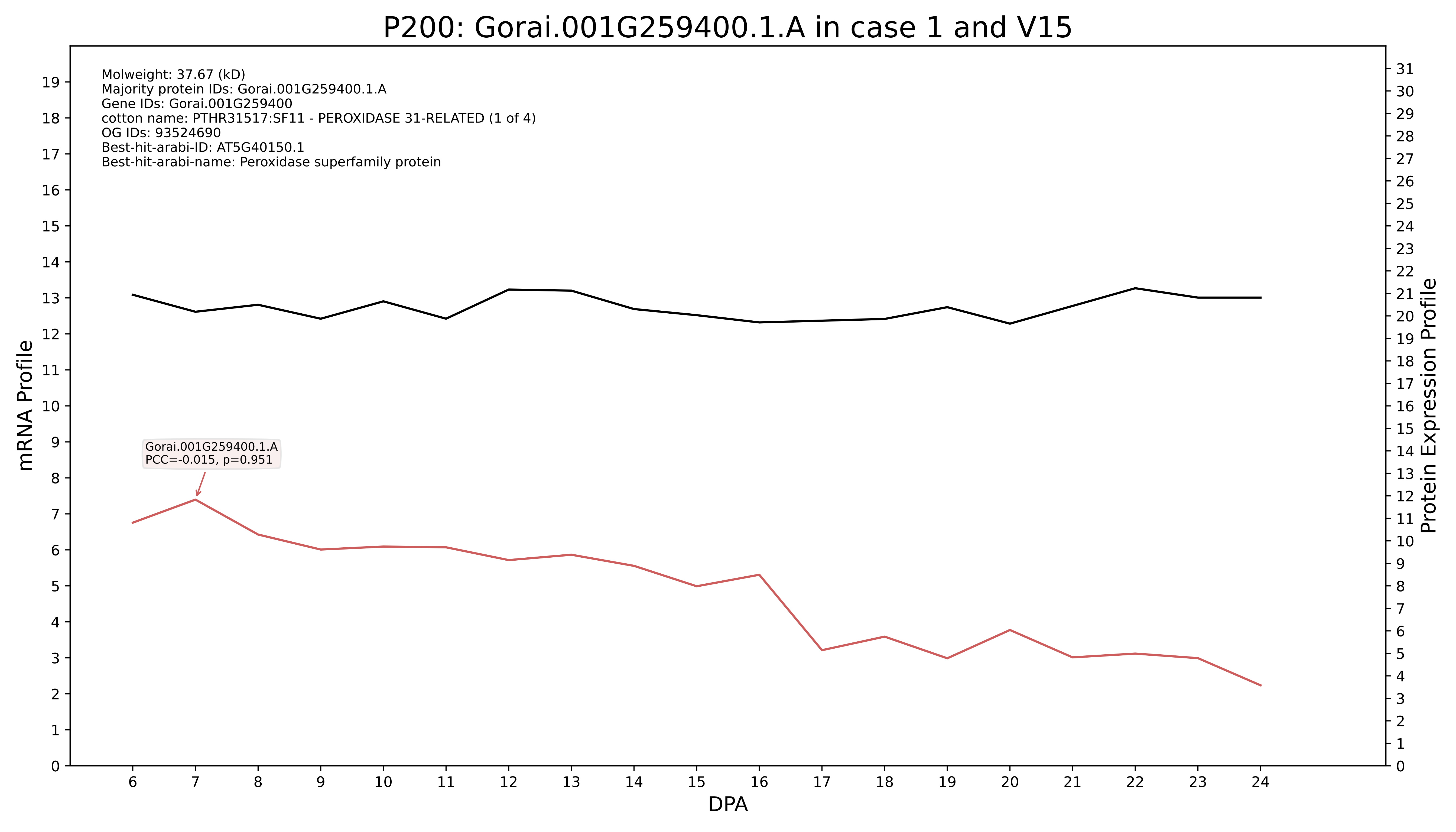
P200: Gorai.001G259400.1.A in case 1 and V15
Molweight: 37.67 (kD)
Majority protein IDs: Gorai.001G259400.1.A
Gene IDs: Gorai.001G259400
cotton name: PTHR31517:SF11 - PEROXIDASE 31-RELATED (1 of 4)
OG IDs: 93524690
Best-hit-arabi-ID: AT5G40150.1
Best-hit-arabi-name: Peroxidase superfamily protein
Gorai.001G259400.1.A
PCC=-0.015, p=0.951
mRNA Profile
Protein Expression Profile
DPA

### P200: Gorai.009G234600.3.D in case 1 and V15

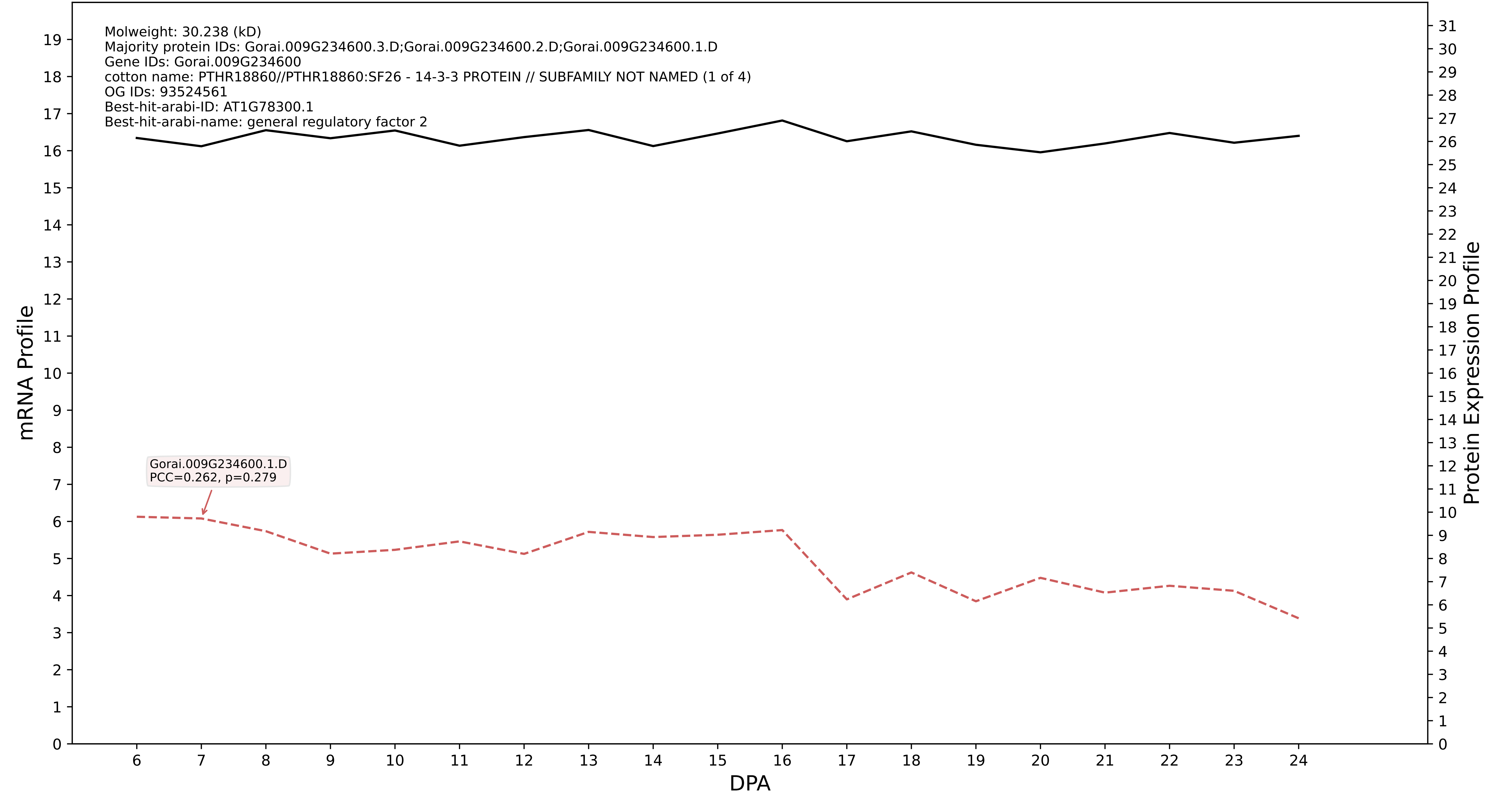

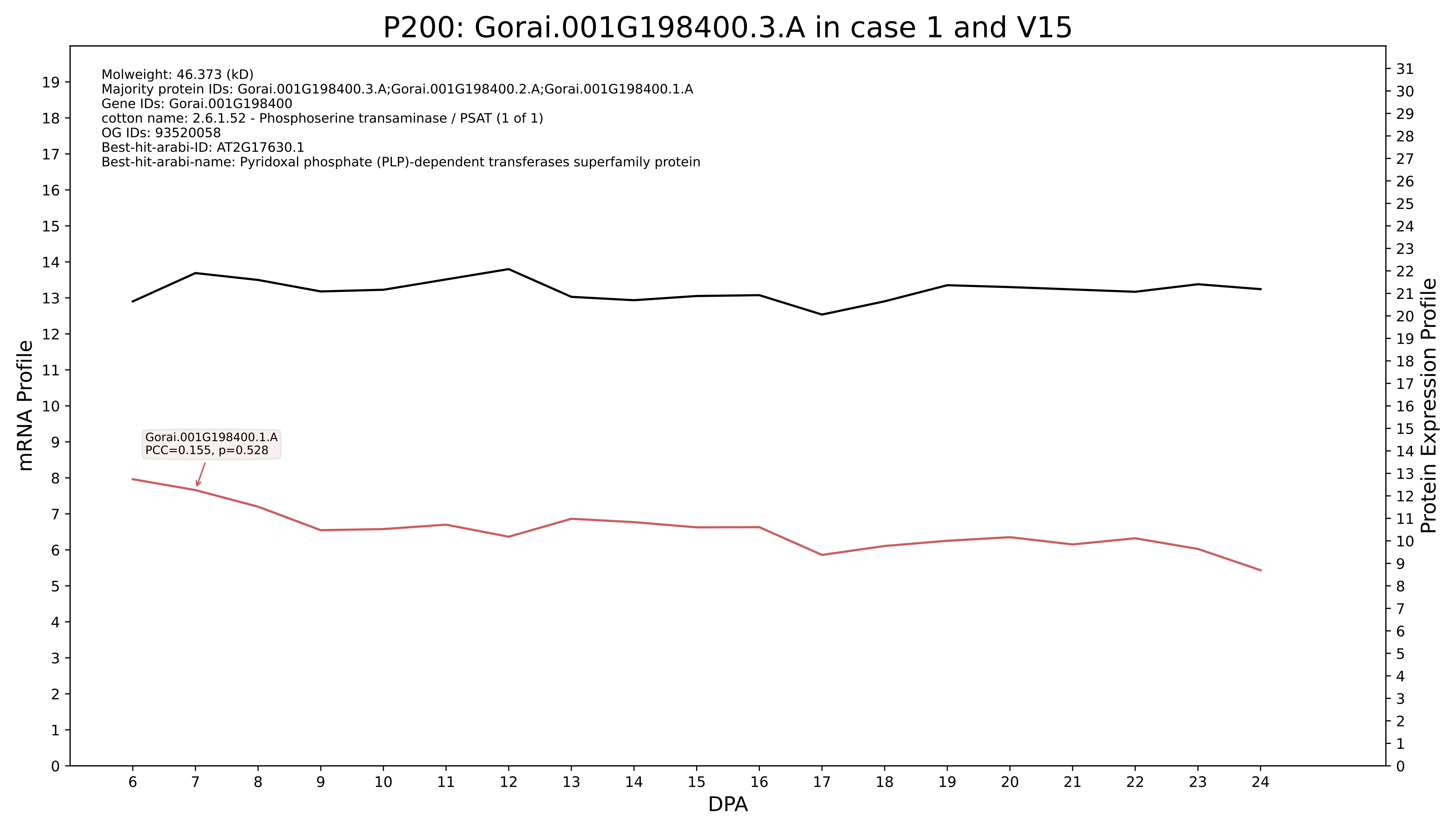
P200: Gorai.001G198400.3.A in case 1 and V15
Molweight: 46.373 (kD)
Majority protein IDs: Gorai.001G198400.3.A;Gorai.001G198400.2.A;Gorai.001G198400.1.A
Gene IDs: Gorai.001G198400
cotton name: 2.6.1.52 - Phosphoserine transaminase / PSAT (1 of 1)
OG IDs: 93520058
Best-hit-arabi-ID: AT2G17630.1
Best-hit-arabi-name: Pyridoxal phosphate (PLP)-dependent transferases superfamily protein
Gorai.001G198400.1.A
PCC=0.155, p=0.528
mRNA Profile
Protein Expression Profile
DPA

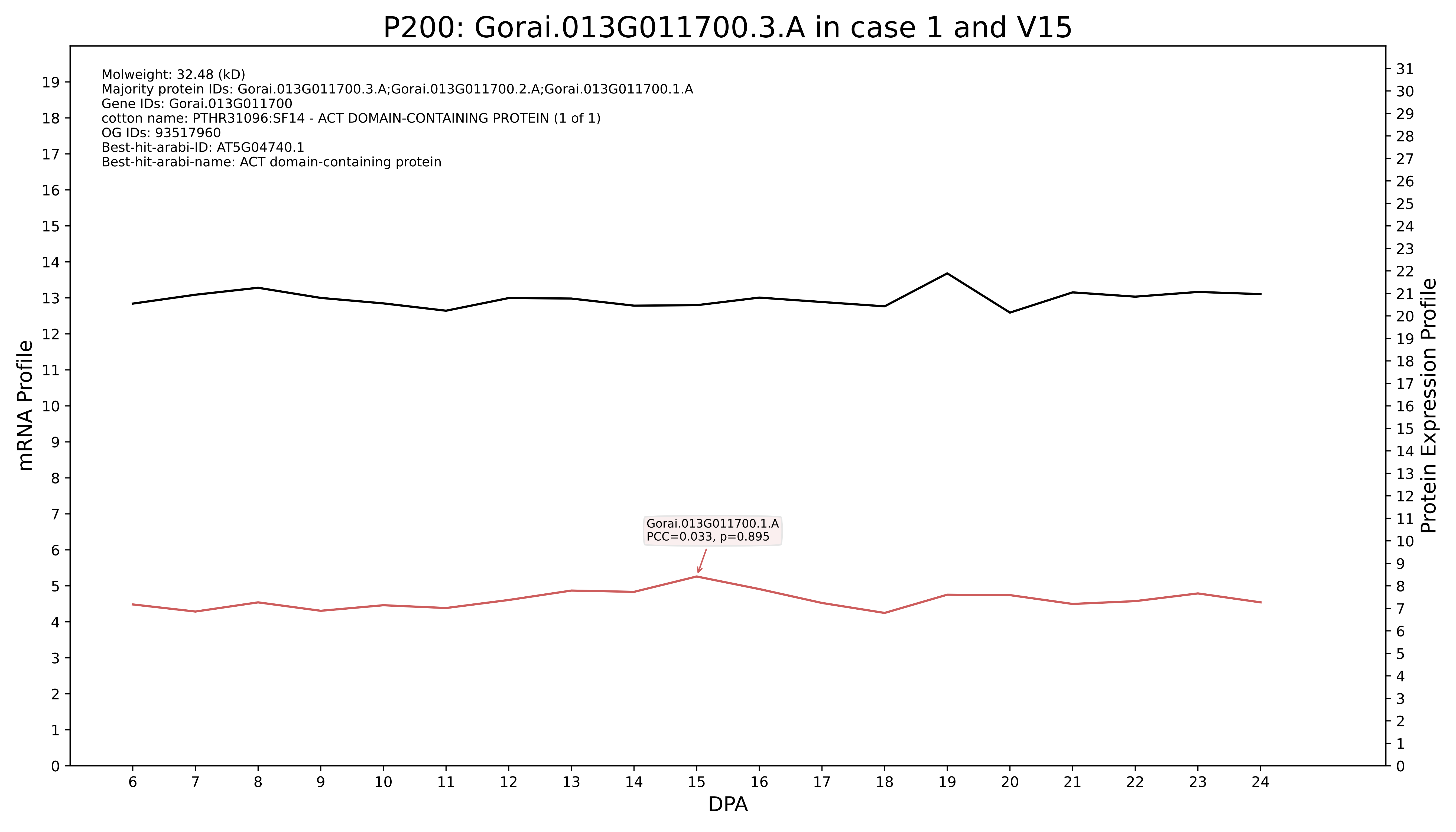
P200: Gorai.013G011700.3.A in case 1 and V15
Molweight: 32.48 (kD)
Majority protein IDs: Gorai.013G011700.3.A;Gorai.013G011700.2.A;Gorai.013G011700.1.A
Gene IDs: Gorai.013G011700
cotton name: PTHR31096:SF14 - ACT DOMAIN-CONTAINING PROTEIN (1 of 1)
OG IDs: 93517960
Best-hit-arabi-ID: AT5G04740.1
Best-hit-arabi-name: ACT domain-containing protein
Gorai.013G011700.1.A
PCC=0.033, p=0.895
mRNA Profile
Protein Expression Profile
DPA

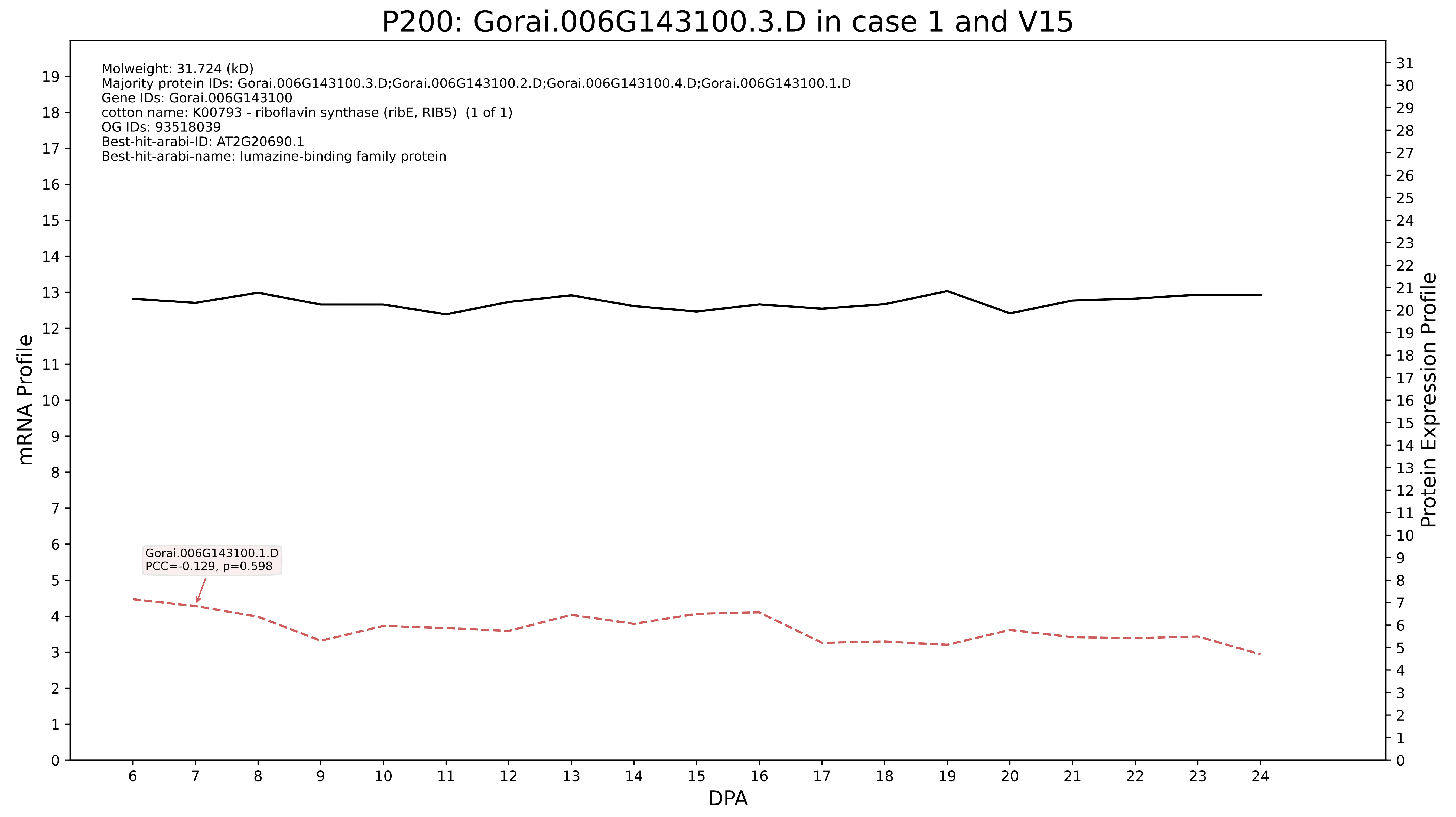
P200: Gorai.006G143100.3.D in case 1 and V15
Molweight: 31.724 (kD)
Majority protein IDs: Gorai.006G143100.3.D;Gorai.006G143100.2.D;Gorai.006G143100.4.D;Gorai.006G143100.1.D
Gene IDs: Gorai.006G143100
cotton name: K00793 - riboflavin synthase (ribE, RIB5) (1 of 1)
OG IDs: 93518039
Best-hit-arabi-ID: AT2G20690.1
Best-hit-arabi-name: lumazine-binding family protein
Gorai.006G143100.1.D
PCC=-0.129, p=0.598
mRNA Profile
Protein Expression Profile
DPA

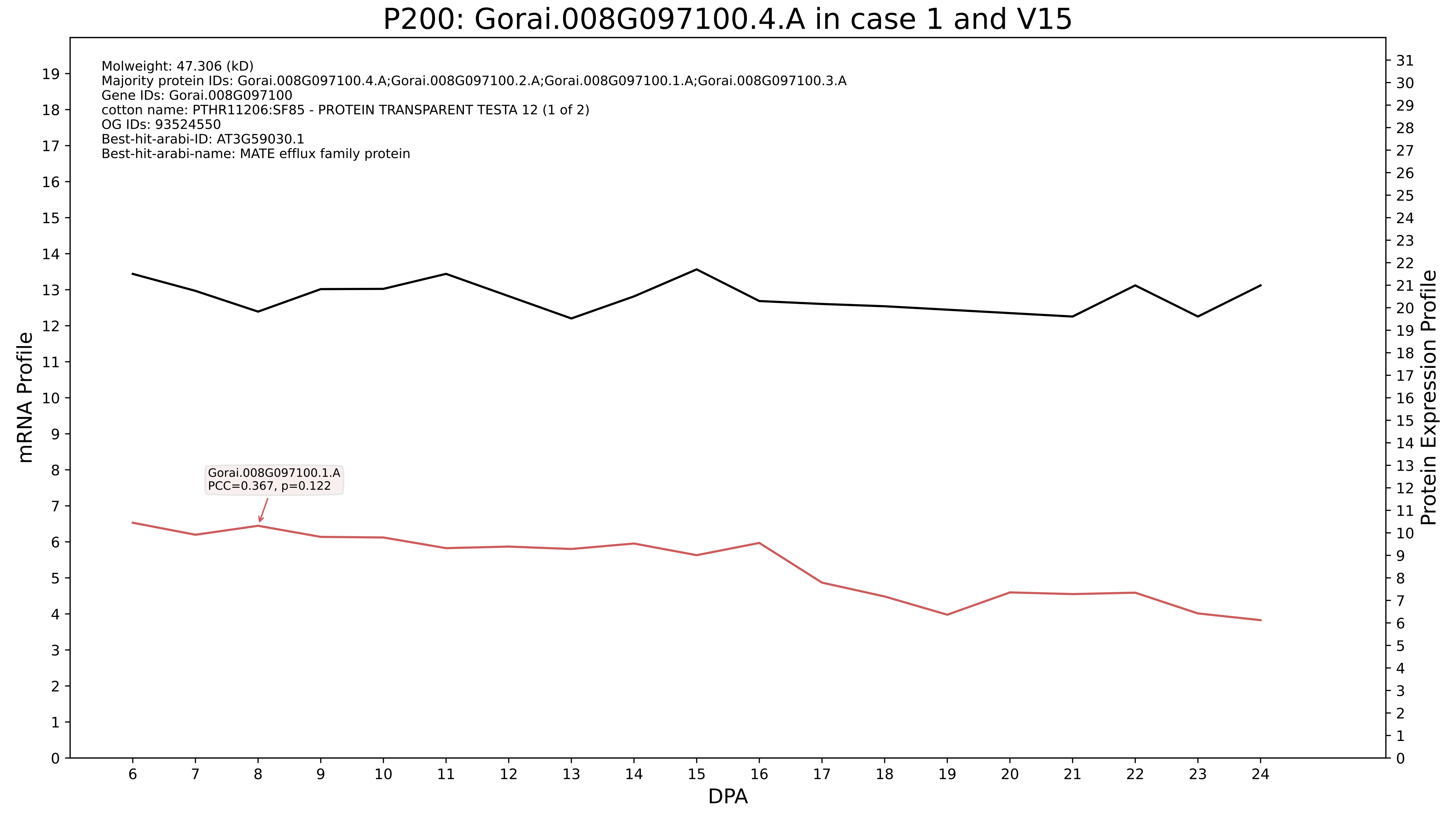

P200: Gorai.008G097100.4.A in case 1 and V15
Molweight: 47.306 (kD)
Majority protein IDs: Gorai.008G097100.4.A;Gorai.008G097100.2.A;Gorai.008G097100.1.A;Gorai.008G097100.3.A
Gene IDs: Gorai.008G097100
cotton name: PTHR11206:SF85 - PROTEIN TRANSPARENT TESTA 12 (1 of 2)
OG IDs: 93524550
Best-hit-arabi-ID: AT3G59030.1
Best-hit-arabi-name: MATE efflux family protein
Gorai.008G097100.1.A
PCC=0.367, p=0.122
mRNA Profile
Protein Expression Profile
DPA

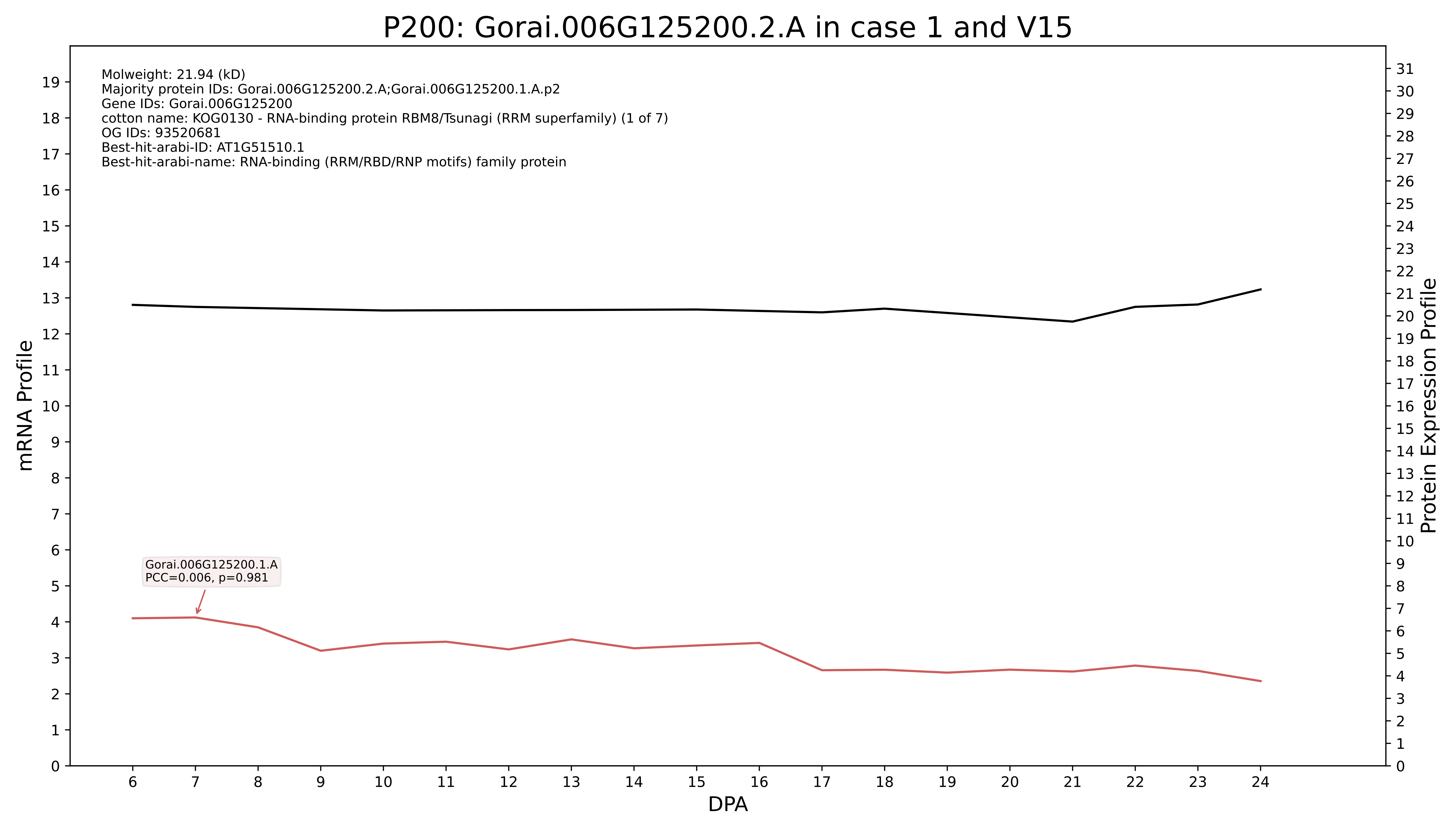
P200: Gorai.006G125200.2.A in case 1 and V15
Molweight: 21.94 (kD)
Majority protein IDs: Gorai.006G125200.2.A;Gorai.006G125200.1.A.p2
Gene IDs: Gorai.006G125200
cotton name: KOG0130 - RNA-binding protein RBM8/Tsunagi (RRM superfamily) (1 of 7)
OG IDs: 93520681
Best-hit-arabi-ID: AT1G51510.1
Best-hit-arabi-name: RNA-binding (RRM/RBD/RNP motifs) family protein
Gorai.006G125200.1.A
PCC=0.006, p=0.981
mRNA Profile
Protein Expression Profile
DPA

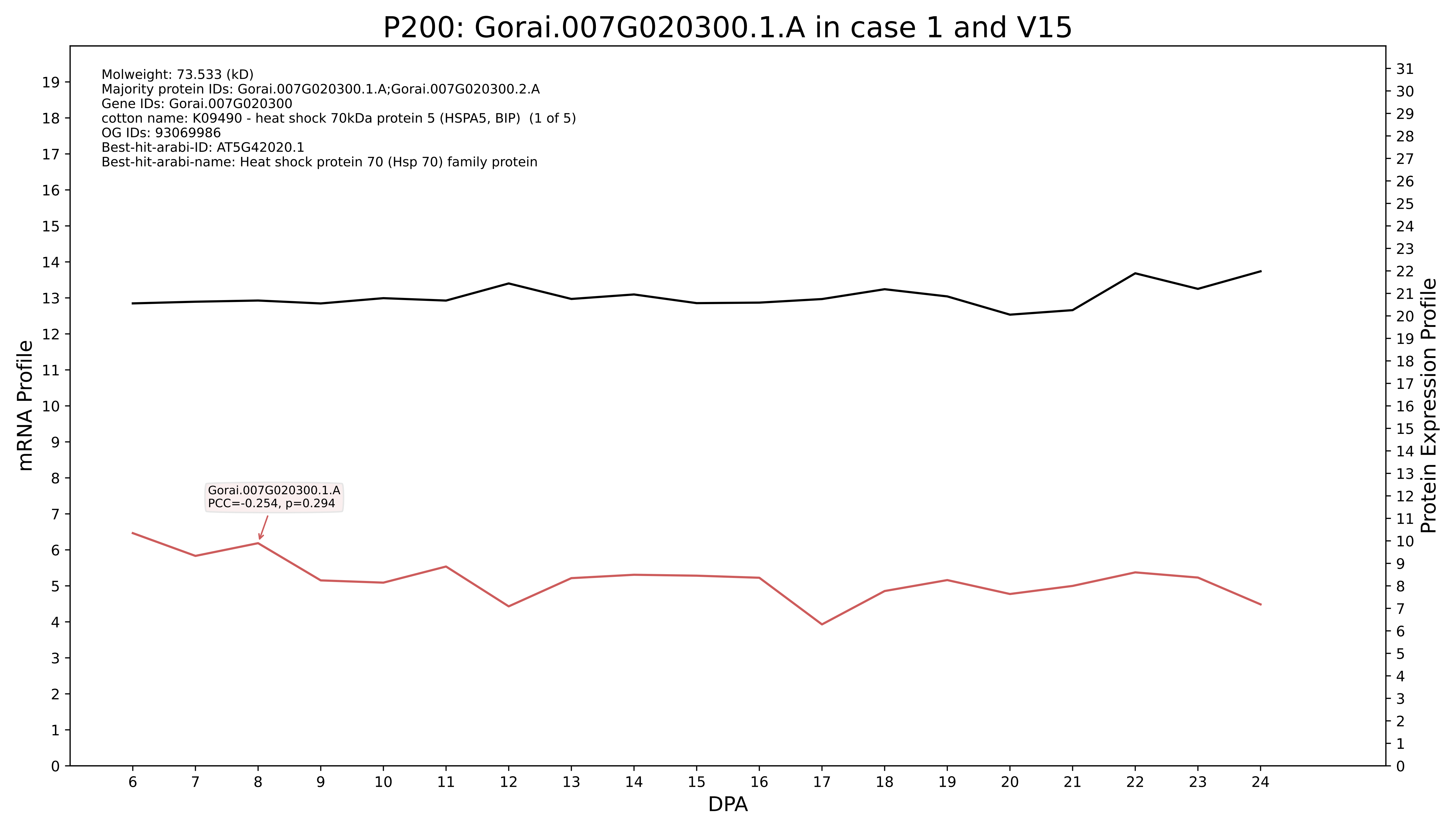
P200: Gorai.007G020300.1.A in case 1 and V15
Molweight: 73.533 (kD)
Majority protein IDs: Gorai.007G020300.1.A;Gorai.007G020300.2.A
Gene IDs: Gorai.007G020300
cotton name: K09490 - heat shock 70kDa protein 5 (HSPA5, BIP) (1 of 5)
OG IDs: 93069986
Best-hit-arabi-ID: AT5G42020.1
Best-hit-arabi-name: Heat shock protein 70 (Hsp 70) family protein
Gorai.007G020300.1.A
PCC=-0.254, p=0.294
mRNA Profile
Protein Expression Profile
DPA

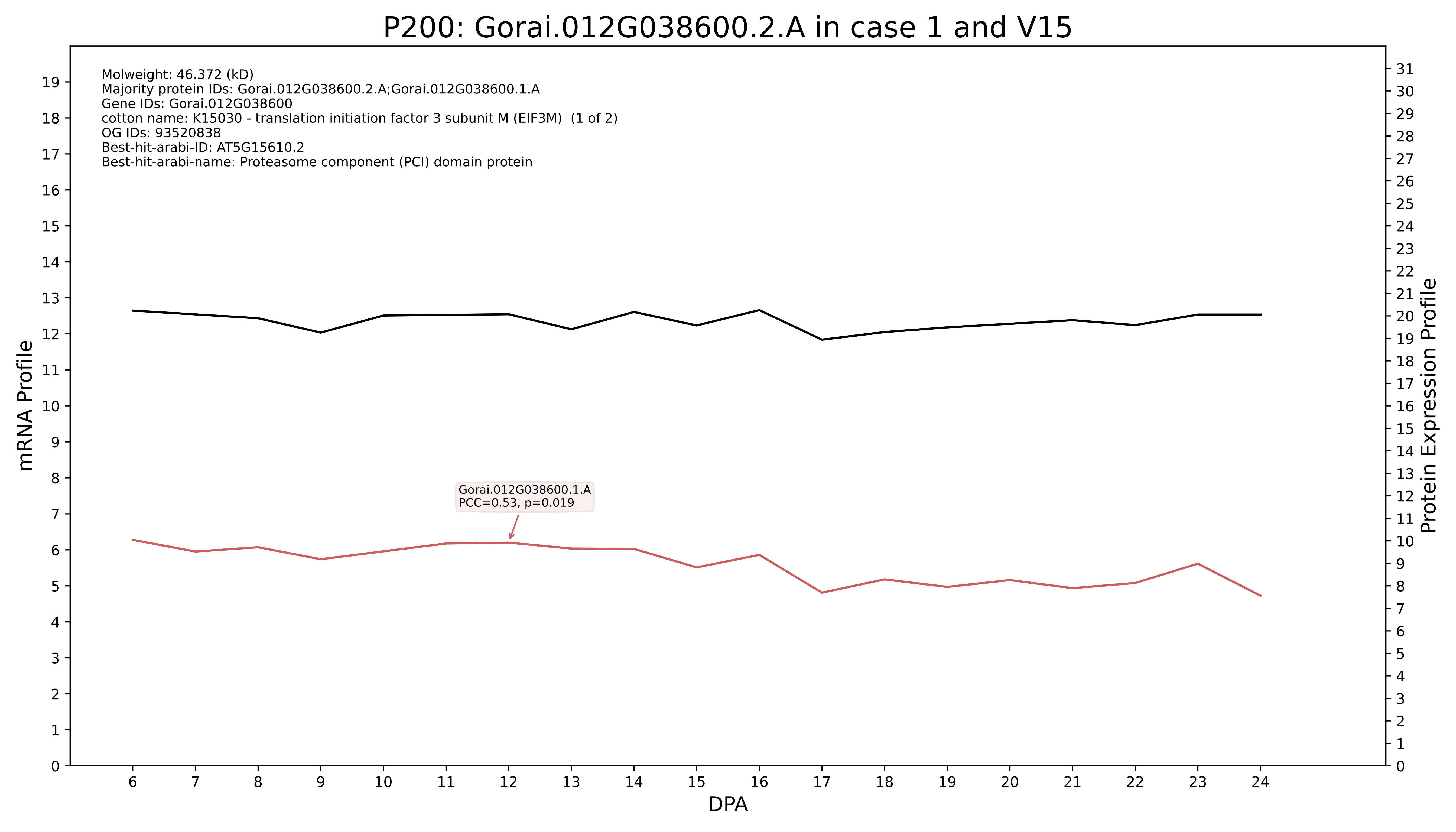
P200: Gorai.012G038600.2.A in case 1 and V15
Molweight: 46.372 (kD)
Majority protein IDs: Gorai.012G038600.2.A;Gorai.012G038600.1.A
Gene IDs: Gorai.012G038600
cotton name: K15030 - translation initiation factor 3 subunit M (EIF3M) (1 of 2)
OG IDs: 93520838
Best-hit-arabi-ID: AT5G15610.2
Best-hit-arabi-name: Proteasome component (PCI) domain protein
Gorai.012G038600.1.A
PCC=0.53, p=0.019
mRNA Profile
Protein Expression Profile
DPA

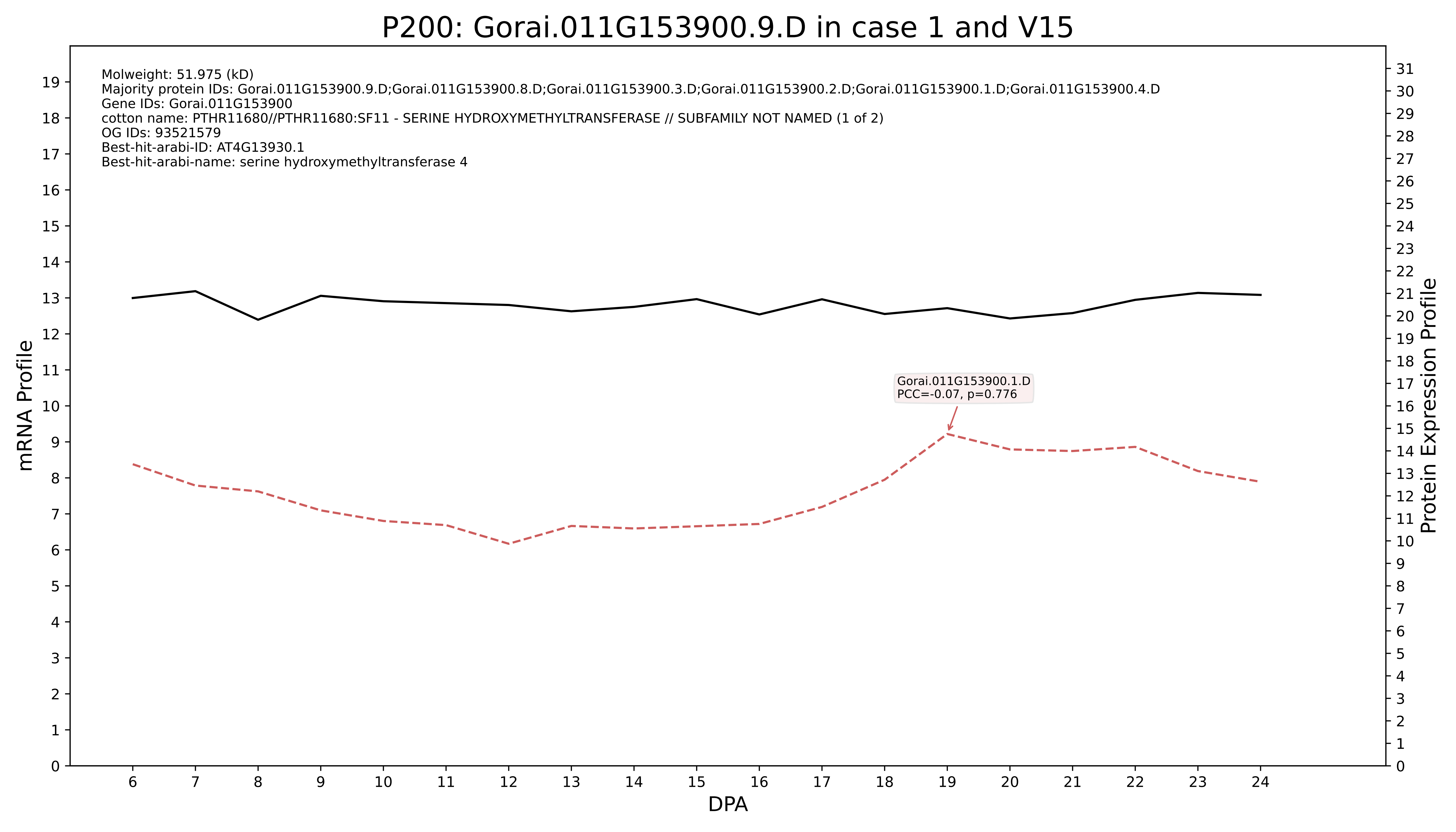
P200: Gorai.011G153900.9.D in case 1 and V15
Molweight: 51.975 (kD)
Majority protein IDs: Gorai.011G153900.9.D;Gorai.011G153900.8.D;Gorai.011G153900.3.D;Gorai.011G153900.2.D;Gorai.011G153900.1.D;Gorai.011G153900.4.D
Gene IDs: Gorai.011G153900
cotton name: PTHR11680//PTHR11680:SF11 - SERINE HYDROXYMETHYLTRANSFERASE // SUBFAMILY NOT NAMED (1 of 2)
OG IDs: 93521579
Best-hit-arabi-ID: AT4G13930.1
Best-hit-arabi-name: serine hydroxymethyltransferase 4
Gorai.011G153900.1.D
PCC=-0.07, p=0.776
mRNA Profile
Protein Expression Profile
DPA

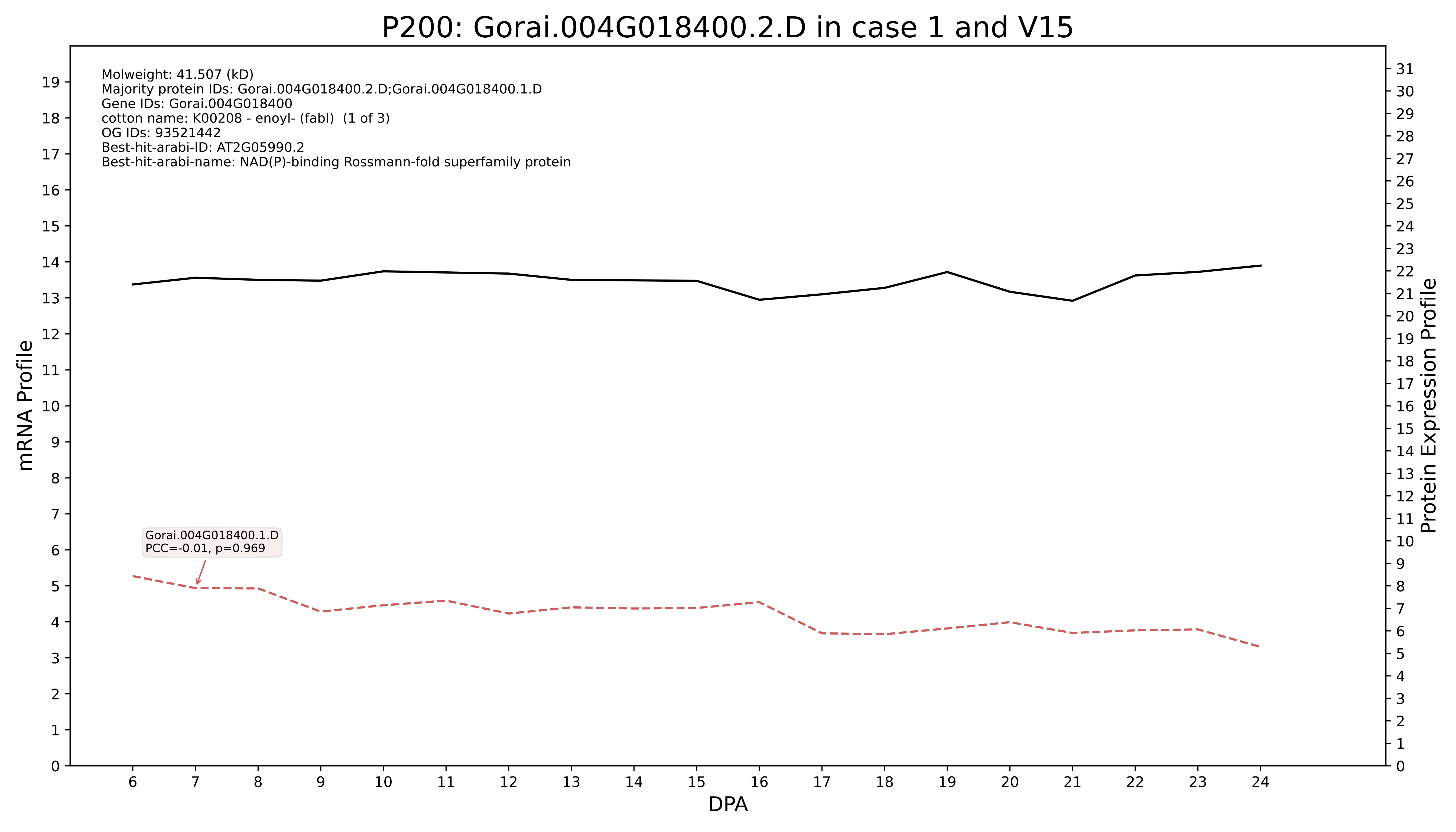

P200: Gorai.004G018400.2.D in case 1 and V15
Molweight: 41.507 (kD)
Majority protein IDs: Gorai.004G018400.2.D;Gorai.004G018400.1.D
Gene IDs: Gorai.004G018400
cotton name: K00208 - enoyl- (fabI) (1 of 3)
OG IDs: 93521442
Best-hit-arabi-ID: AT2G05990.2
Best-hit-arabi-name: NAD(P)-binding Rossmann-fold superfamily protein
Gorai.004G018400.1.D
PCC=-0.01, p=0.969
mRNA Profile
Protein Expression Profile
DPA

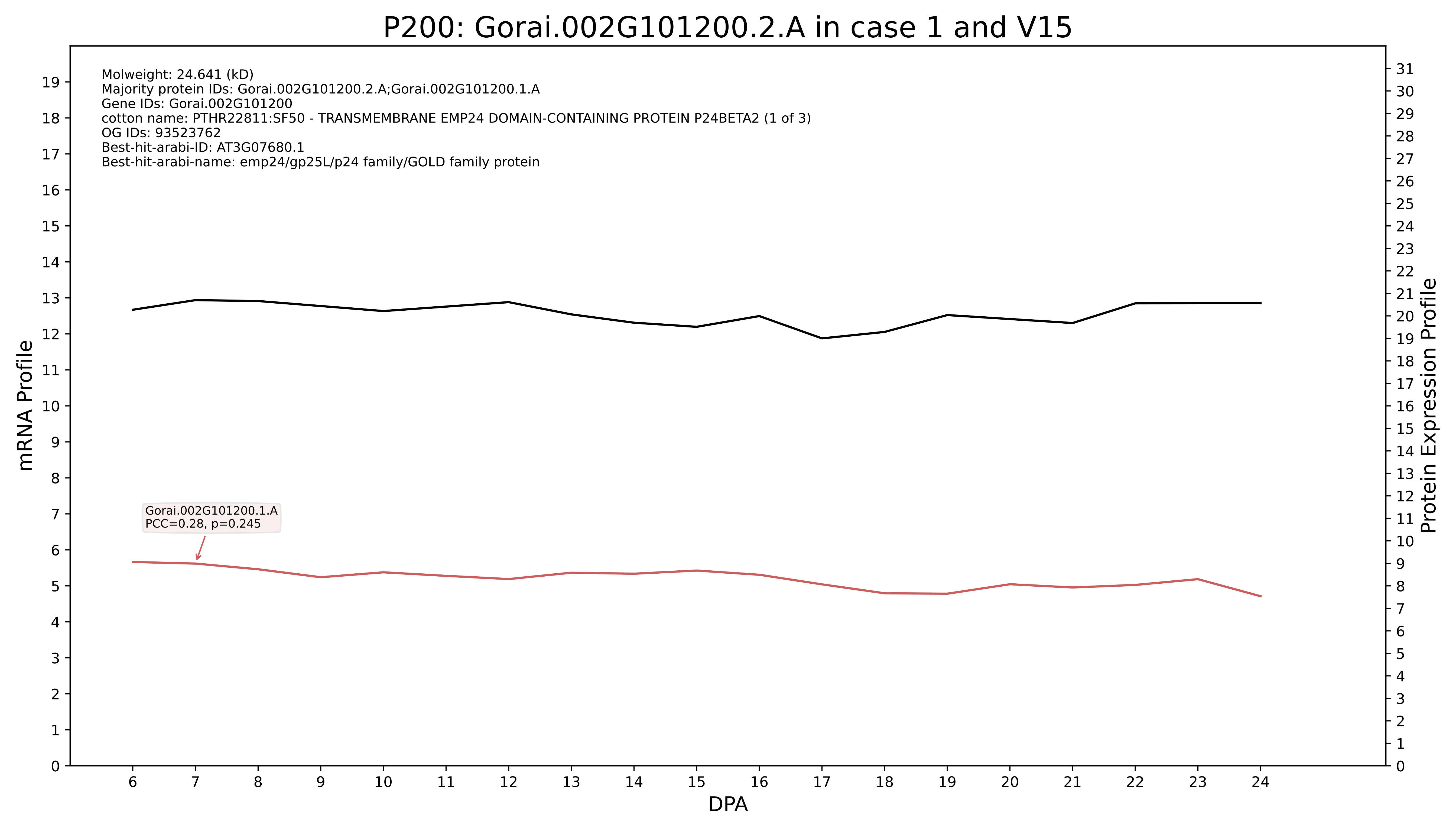
P200: Gorai.002G101200.2.A in case 1 and V15
Molweight: 24.641 (kD)
Majority protein IDs: Gorai.002G101200.2.A;Gorai.002G101200.1.A
Gene IDs: Gorai.002G101200
cotton name: PTHR22811:SF50 - TRANSMEMBRANE EMP24 DOMAIN-CONTAINING PROTEIN P24BETA2 (1 of 3)
OG IDs: 93523762
Best-hit-arabi-ID: AT3G07680.1
Best-hit-arabi-name: emp24/gp25L/p24 family/GOLD family protein
Gorai.002G101200.1.A
PCC=0.28, p=0.245
mRNA Profile
Protein Expression Profile
DPA

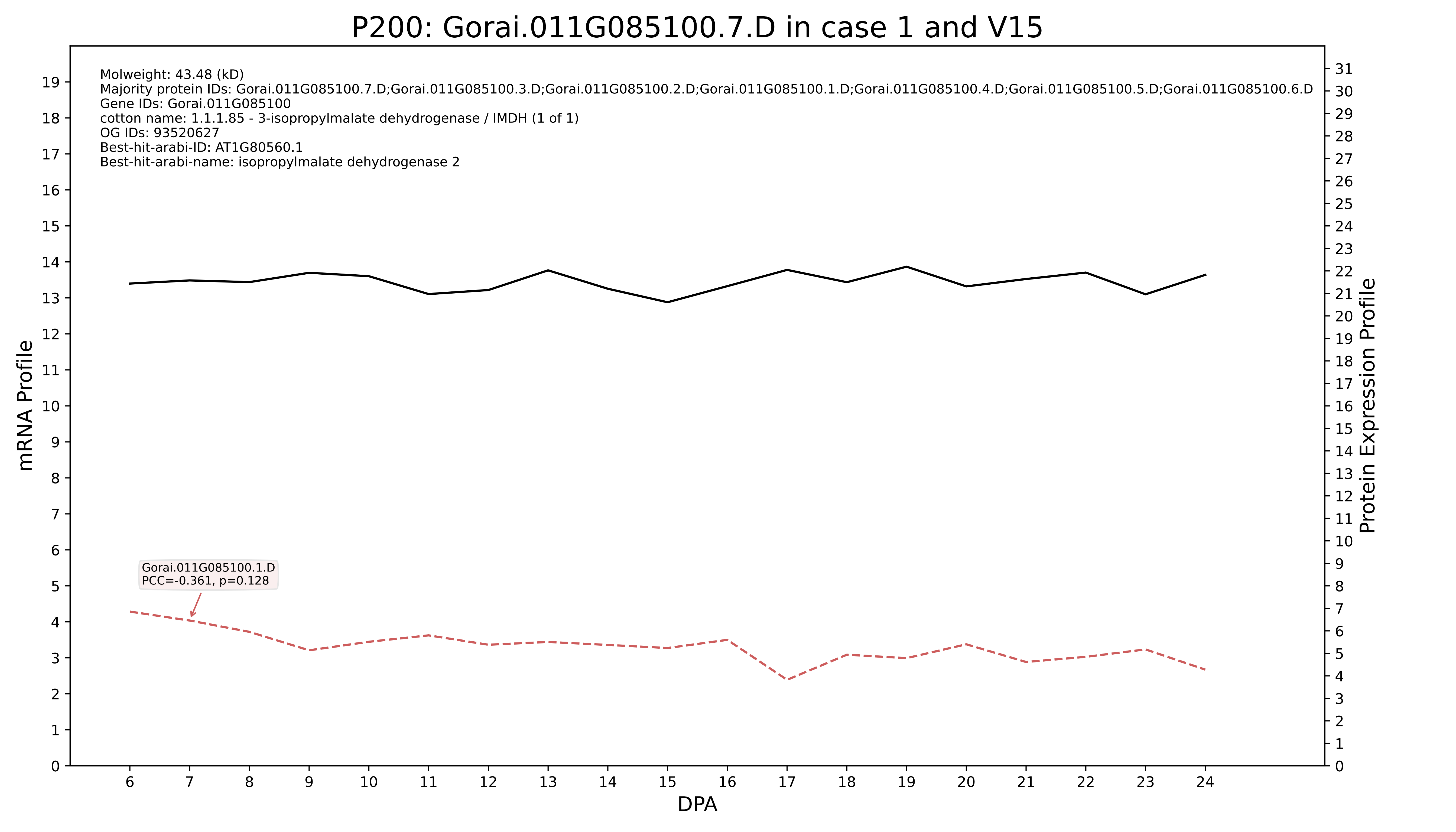
P200: Gorai.011G085100.7.D in case 1 and V15
Molweight: 43.48 (kD)
Majority protein IDs: Gorai.011G085100.7.D;Gorai.011G085100.3.D;Gorai.011G085100.2.D;Gorai.011G085100.1.D;Gorai.011G085100.4.D;Gorai.011G085100.5.D;Gorai.011G085100.6.D
Gene IDs: Gorai.011G085100
cotton name: 1.1.1.85 - 3-isopropylmalate dehydrogenase / IMDH (1 of 1)
OG IDs: 93520627
Best-hit-arabi-ID: AT1G80560.1
Best-hit-arabi-name: isopropylmalate dehydrogenase 2
Gorai.011G085100.1.D
PCC=-0.361, p=0.128
mRNA Profile
Protein Expression Profile
DPA

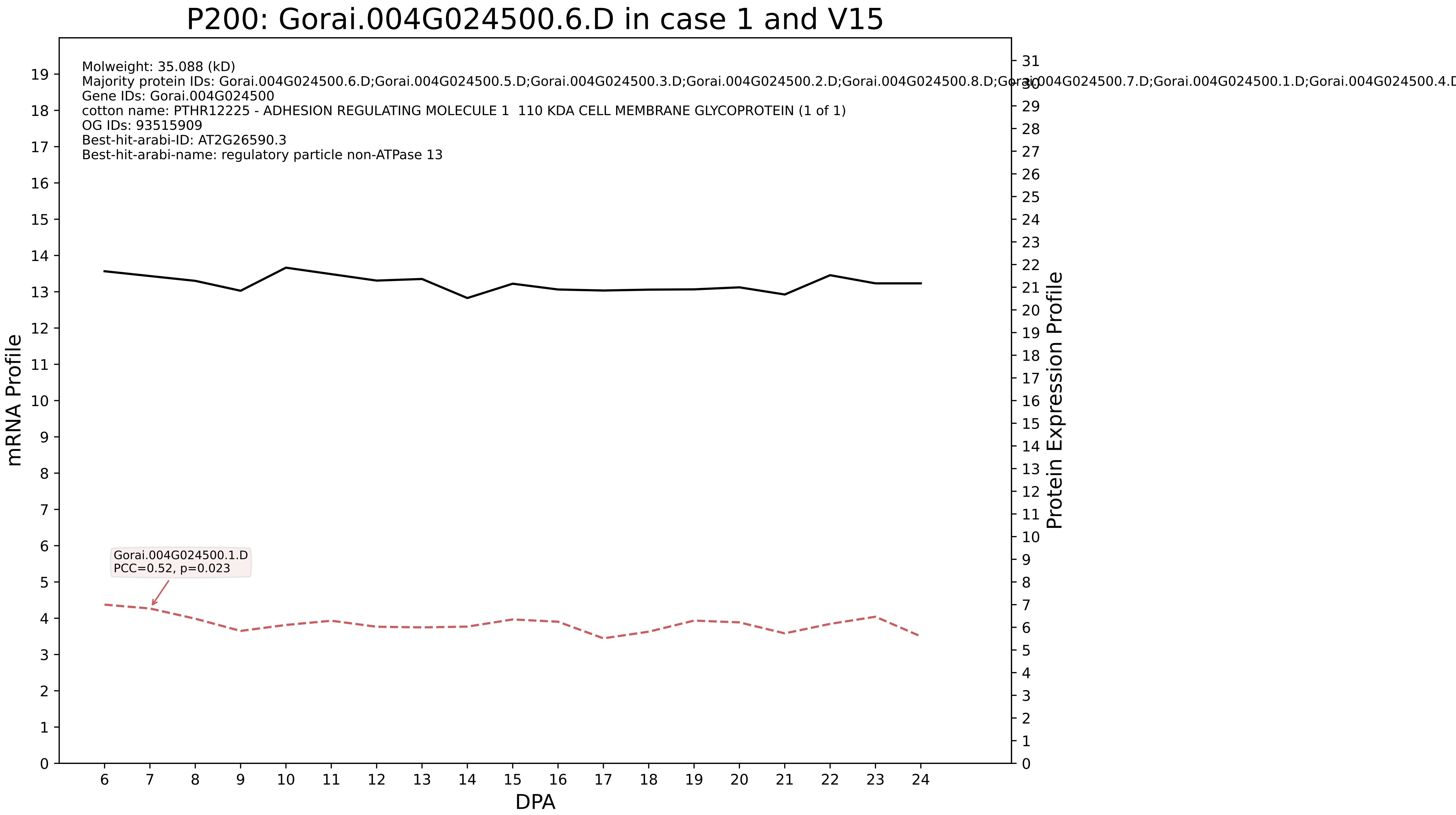
P200: Gorai.004G024500.6.D in case 1 and V15
Molweight: 35.088 (kD)
Majority protein IDs: Gorai.004G024500.6.D;Gorai.004G024500.5.D;Gorai.004G024500.3.D;Gorai.004G024500.2.D;Gorai.004G024500.8.D;Gorai.004G024500.7.D;Gorai.004G024500.1.D;Gorai.004G024500.4.D
Gene IDs: Gorai.004G024500
cotton name: PTHR12225 - ADHESION REGULATING MOLECULE 1 110 KDA CELL MEMBRANE GLYCOPROTEIN (1 of 1)
OG IDs: 93515909
Best-hit-arabi-ID: AT2G26590.3
Best-hit-arabi-name: regulatory particle non-ATPase 13
Gorai.004G024500.1.D
PCC=0.52, p=0.023
mRNA Profile
Protein Expression Profile
DPA

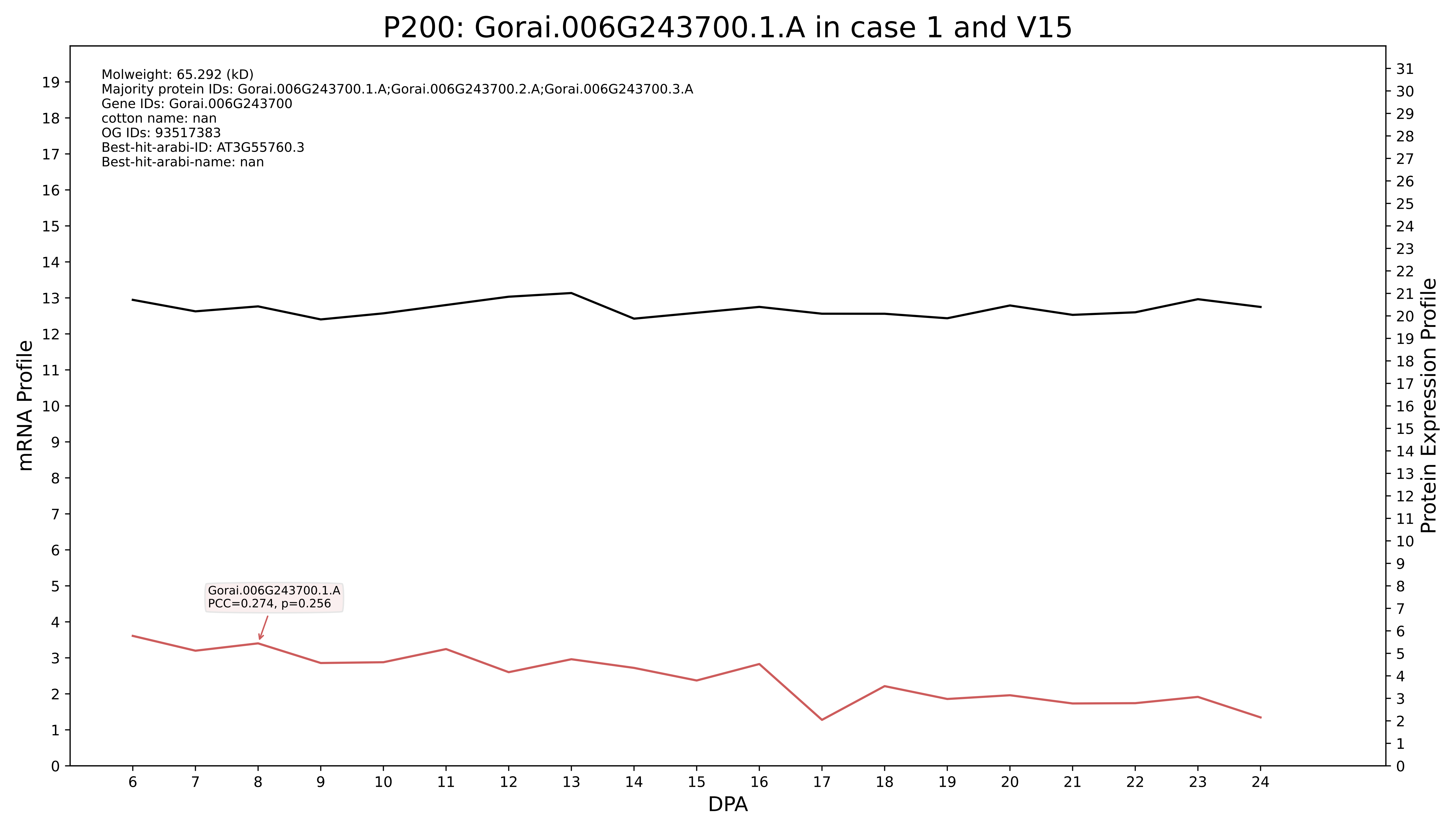
P200: Gorai.006G243700.1.A in case 1 and V15
Molweight: 65.292 (kD)
Majority protein IDs: Gorai.006G243700.1.A;Gorai.006G243700.2.A;Gorai.006G243700.3.A
Gene IDs: Gorai.006G243700
cotton name: nan
OG IDs: 93517383
Best-hit-arabi-ID: AT3G55760.3
Best-hit-arabi-name: nan
Gorai.006G243700.1.A
PCC=0.274, p=0.256
mRNA Profile
Protein Expression Profile
DPA

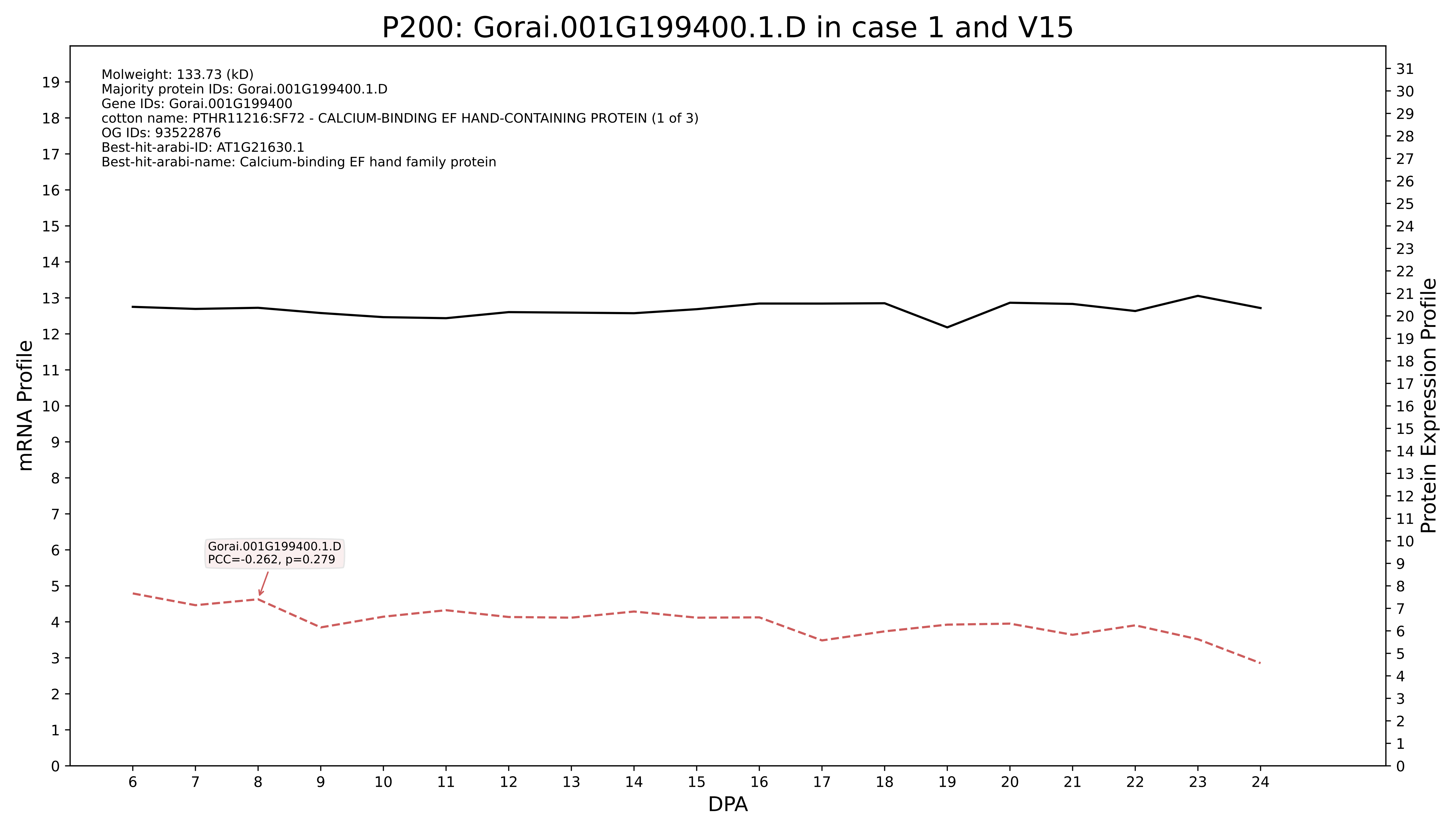
P200: Gorai.001G199400.1.D in case 1 and V15
Molweight: 133.73 (kD)
Majority protein IDs: Gorai.001G199400.1.D
Gene IDs: Gorai.001G199400
cotton name: PTHR11216:SF72 - CALCIUM-BINDING EF HAND-CONTAINING PROTEIN (1 of 3)
OG IDs: 93522876
Best-hit-arabi-ID: AT1G21630.1
Best-hit-arabi-name: Calcium-binding EF hand family protein
Gorai.001G199400.1.D
PCC=-0.262, p=0.279
mRNA Profile
Protein Expression Profile
DPA

### P200: Gorai.005G172100.1.A in case 1 and V15

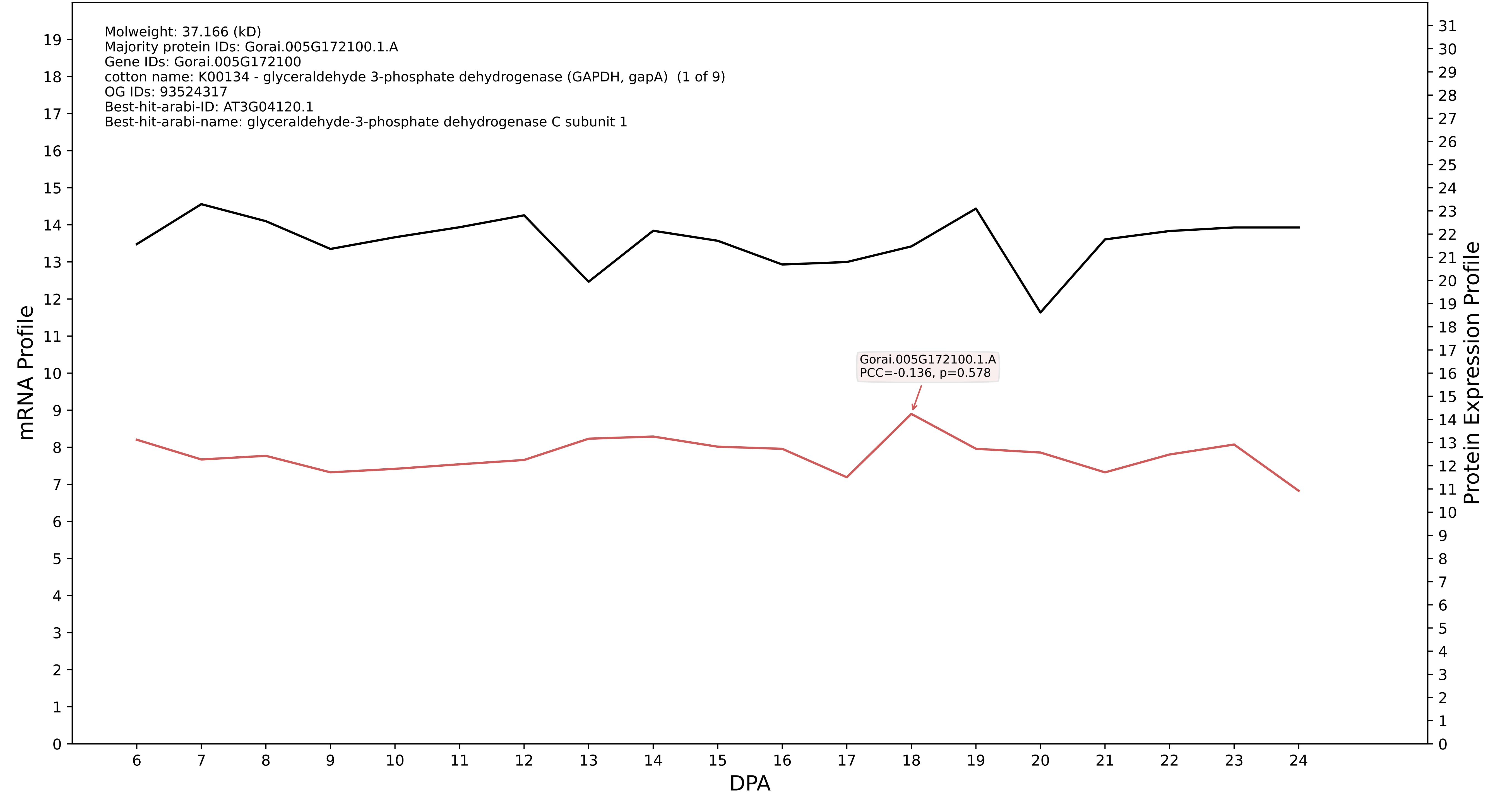

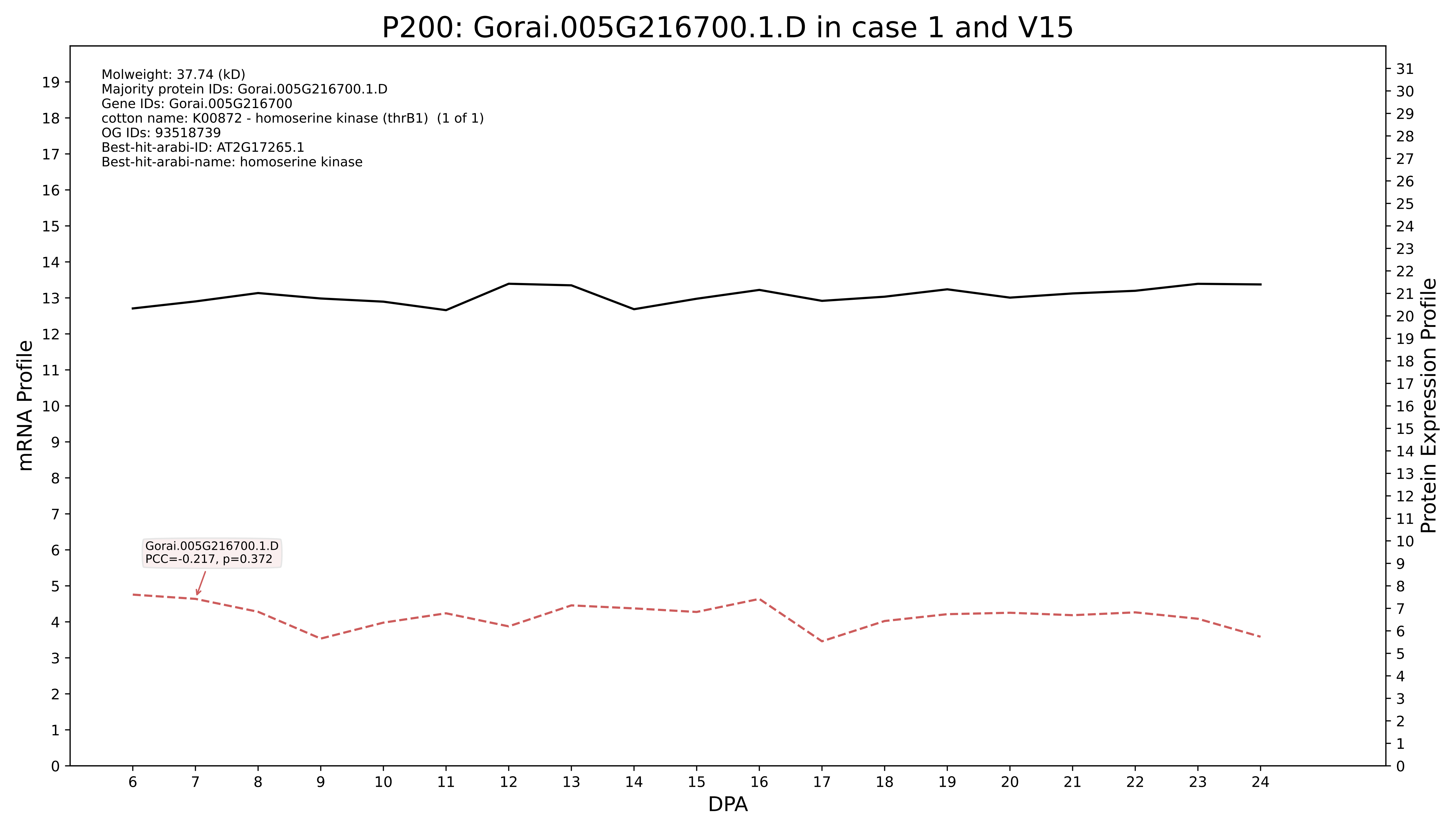
P200: Gorai.005G216700.1.D in case 1 and V15
Molweight: 37.74 (kD)
Majority protein IDs: Gorai.005G216700.1.D
Gene IDs: Gorai.005G216700
cotton name: K00872 - homoserine kinase (thrB1) (1 of 1)
OG IDs: 93518739
Best-hit-arabi-ID: AT2G17265.1
Best-hit-arabi-name: homoserine kinase
Gorai.005G216700.1.D
PCC=-0.217, p=0.372
mRNA Profile
Protein Expression Profile
DPA

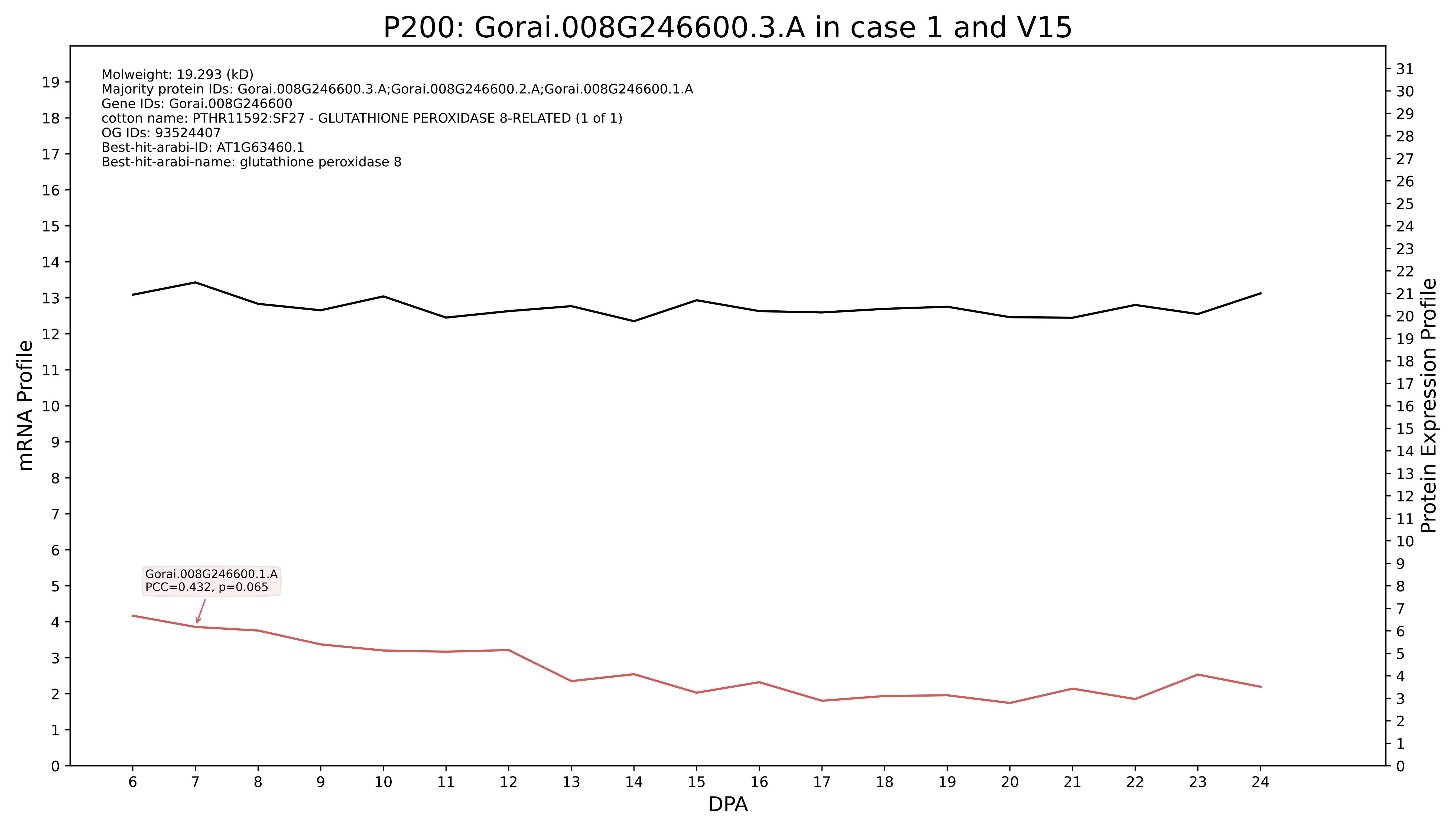
P200: Gorai.008G246600.3.A in case 1 and V15
Molweight: 19.293 (kD)
Majority protein IDs: Gorai.008G246600.3.A;Gorai.008G246600.2.A;Gorai.008G246600.1.A
Gene IDs: Gorai.008G246600
cotton name: PTHR11592:SF27 - GLUTATHIONE PEROXIDASE 8-RELATED (1 of 1)
OG IDs: 93524407
Best-hit-arabi-ID: AT1G63460.1
Best-hit-arabi-name: glutathione peroxidase 8
Gorai.008G246600.1.A
PCC=0.432, p=0.065
mRNA Profile
Protein Expression Profile
DPA

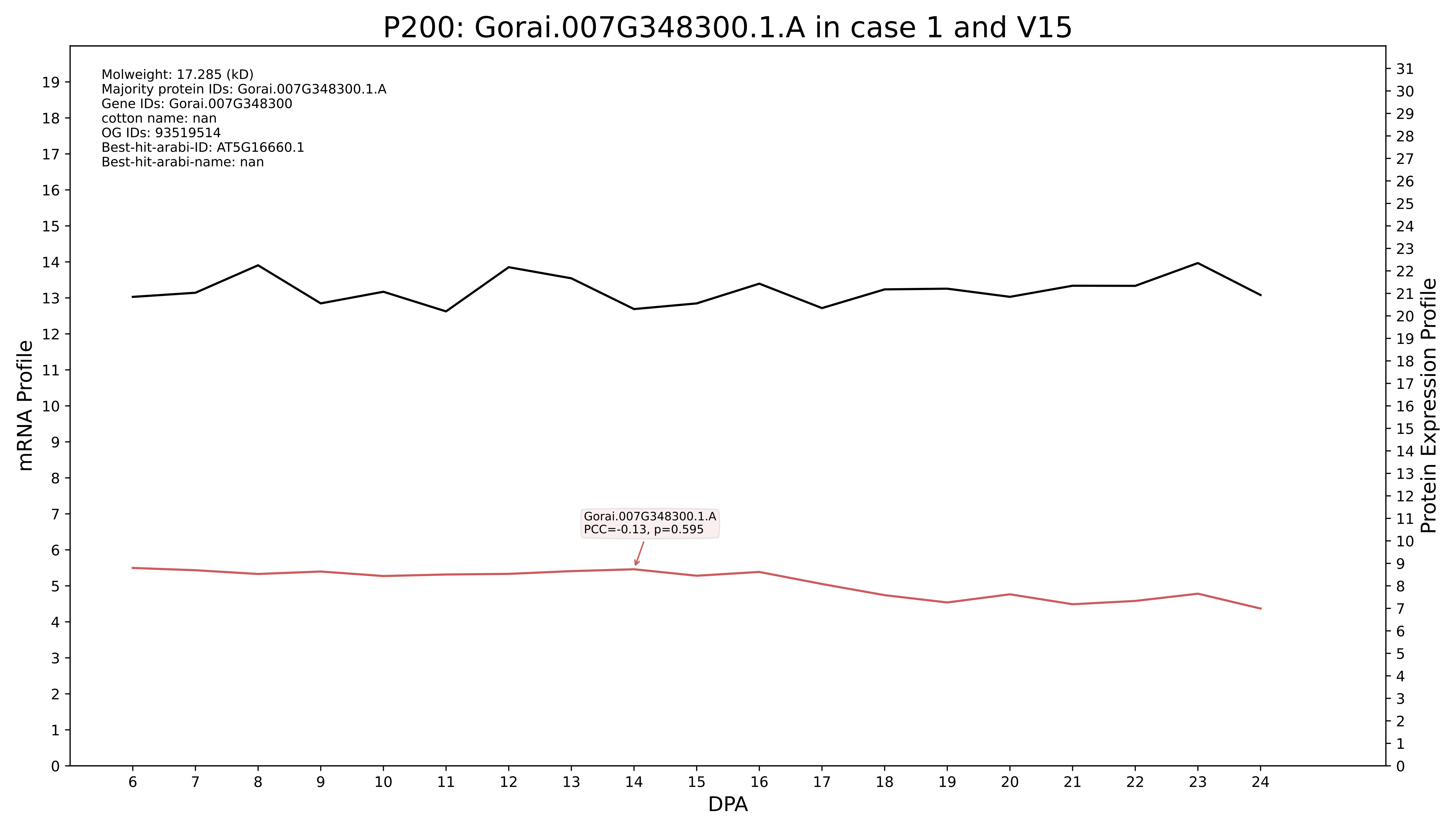
P200: Gorai.007G348300.1.A in case 1 and V15
Molweight: 17.285 (kD)
Majority protein IDs: Gorai.007G348300.1.A
Gene IDs: Gorai.007G348300
cotton name: nan
OG IDs: 93519514
Best-hit-arabi-ID: AT5G16660.1
Best-hit-arabi-name: nan
Gorai.007G348300.1.A
PCC=-0.13, p=0.595
mRNA Profile
Protein Expression Profile
DPA

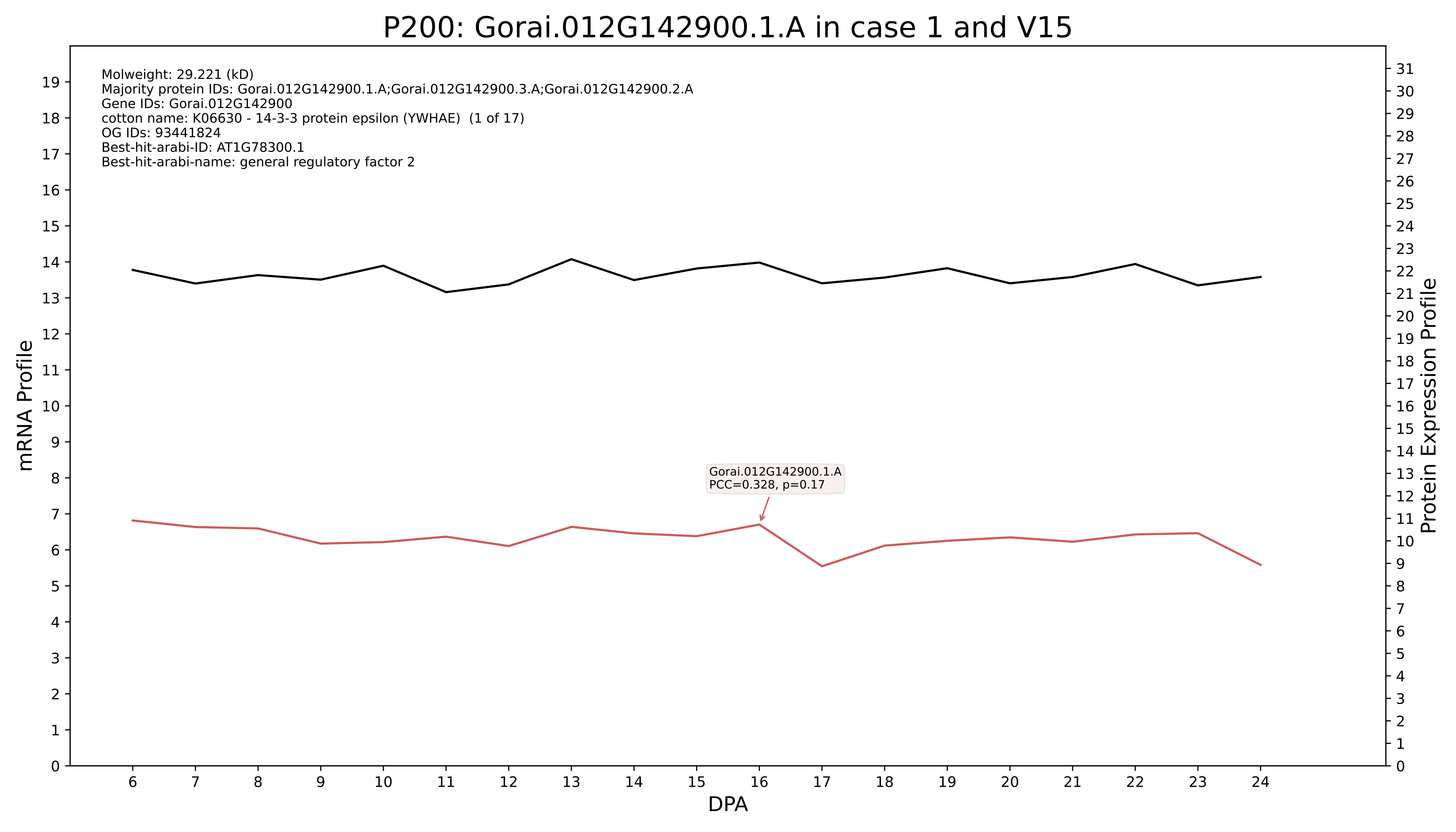

P200: Gorai.012G142900.1.A in case 1 and V15
Molweight: 29.221 (kD)
Majority protein IDs: Gorai.012G142900.1.A;Gorai.012G142900.3.A;Gorai.012G142900.2.A
Gene IDs: Gorai.012G142900
cotton name: K06630 - 14-3-3 protein epsilon (YWHAE) (1 of 17)
OG IDs: 93441824
Best-hit-arabi-ID: AT1G78300.1
Best-hit-arabi-name: general regulatory factor 2
Gorai.012G142900.1.A
PCC=0.328, p=0.17
mRNA Profile
Protein Expression Profile
DPA

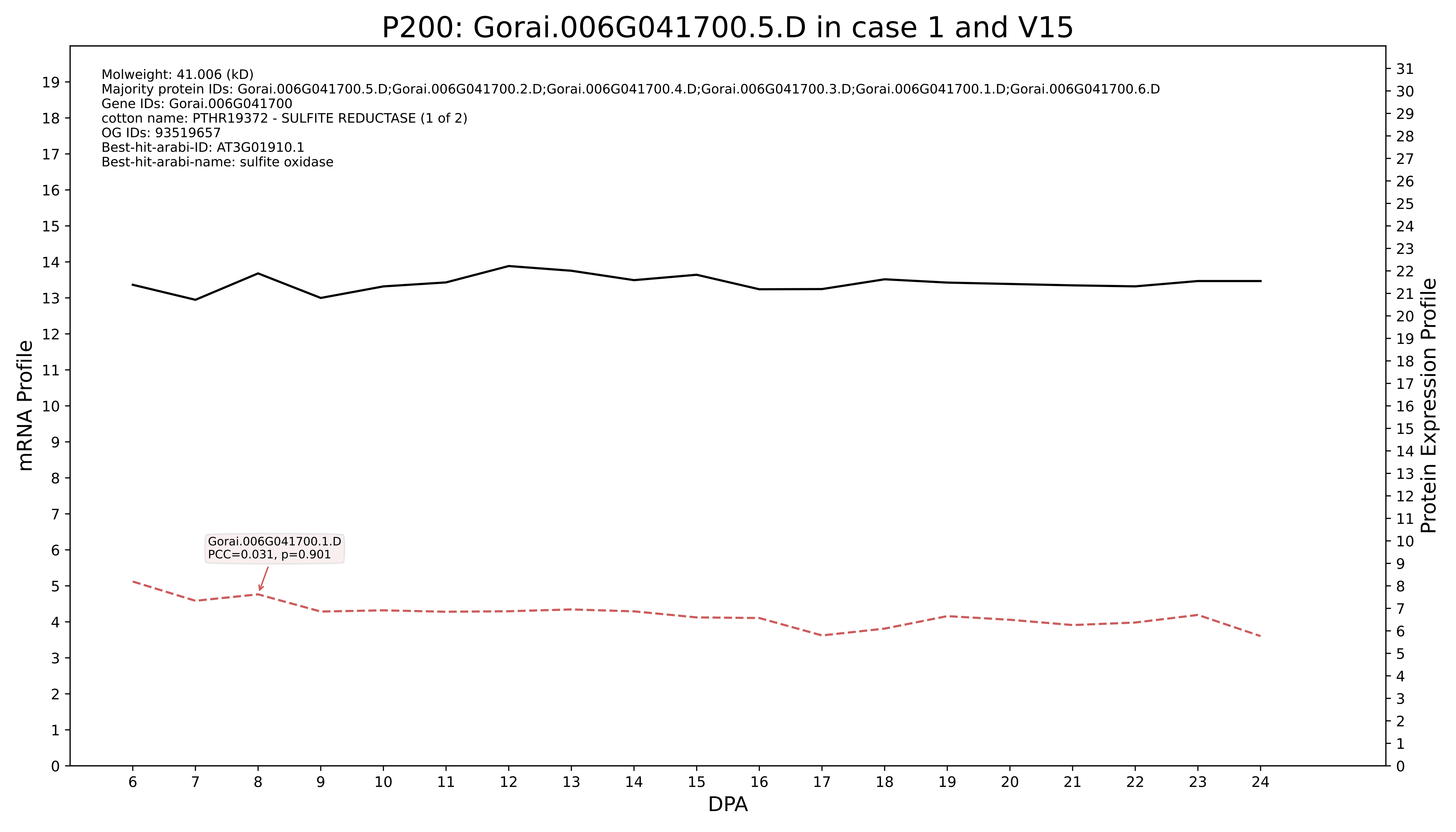
P200: Gorai.006G041700.5.D in case 1 and V15
Molweight: 41.006 (kD)
Majority protein IDs: Gorai.006G041700.5.D;Gorai.006G041700.2.D;Gorai.006G041700.4.D;Gorai.006G041700.3.D;Gorai.006G041700.1.D;Gorai.006G041700.6.D
Gene IDs: Gorai.006G041700
cotton name: PTHR19372 - SULFITE REDUCTASE (1 of 2)
OG IDs: 93519657
Best-hit-arabi-ID: AT3G01910.1
Best-hit-arabi-name: sulfite oxidase
Gorai.006G041700.1.D
PCC=0.031, p=0.901
mRNA Profile
Protein Expression Profile
DPA

### P200: Gorai.005G216700.1.A in case 1 and V15

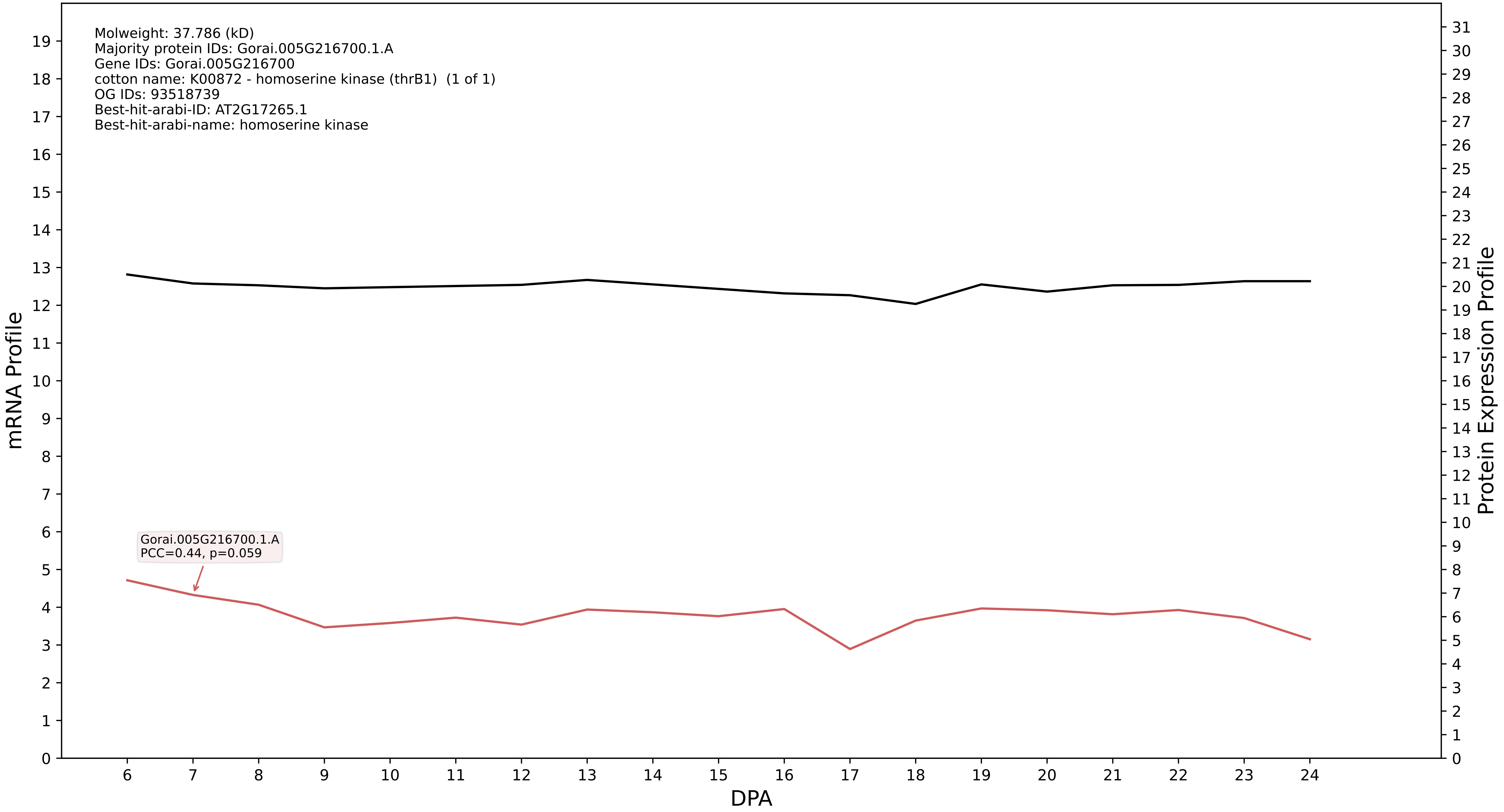

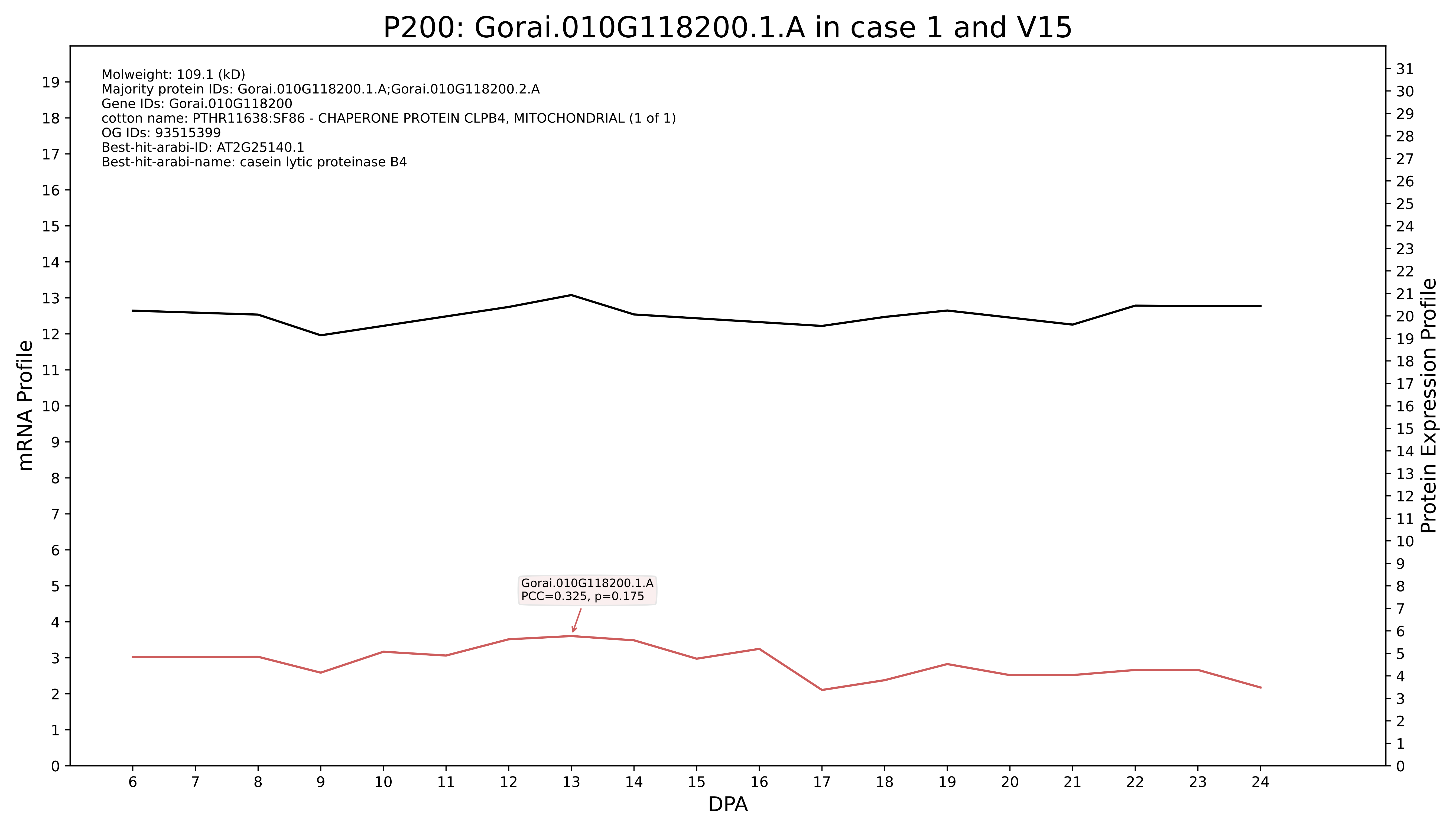

P200: Gorai.010G118200.1.A in case 1 and V15
Molweight: 109.1 (kD)
Majority protein IDs: Gorai.010G118200.1.A;Gorai.010G118200.2.A
Gene IDs: Gorai.010G118200
cotton name: PTHR11638:SF86 - CHAPERONE PROTEIN CLPB4, MITOCHONDRIAL (1 of 1)
OG IDs: 93515399
Best-hit-arabi-ID: AT2G25140.1
Best-hit-arabi-name: casein lytic proteinase B4
Gorai.010G118200.1.A
PCC=0.325, p=0.175
mRNA Profile
Protein Expression Profile
DPA

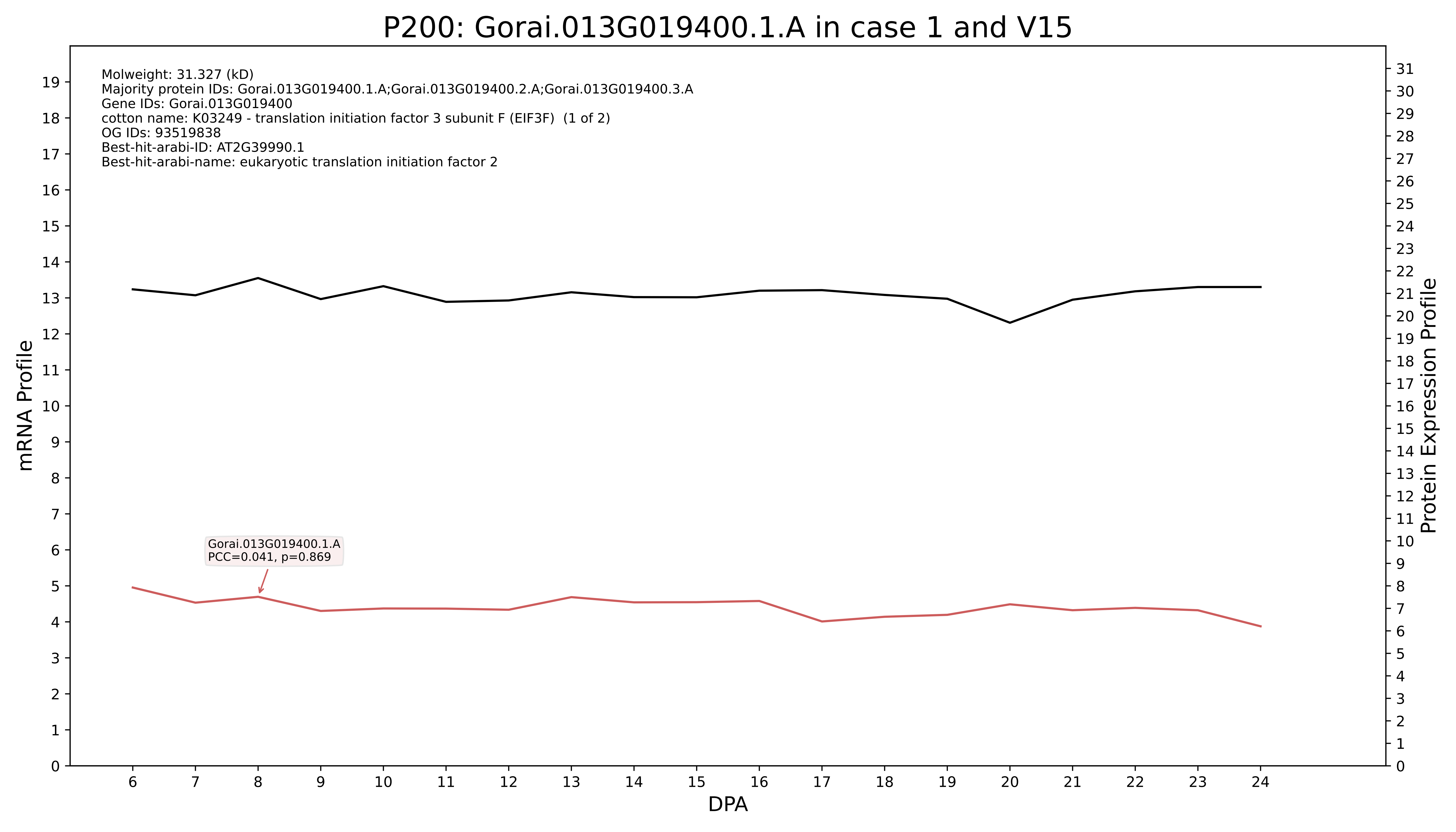
P200: Gorai.013G019400.1.A in case 1 and V15
Molweight: 31.327 (kD)
Majority protein IDs: Gorai.013G019400.1.A;Gorai.013G019400.2.A;Gorai.013G019400.3.A
Gene IDs: Gorai.013G019400
cotton name: K03249 - translation initiation factor 3 subunit F (EIF3F) (1 of 2)
OG IDs: 93519838
Best-hit-arabi-ID: AT2G39990.1
Best-hit-arabi-name: eukaryotic translation initiation factor 2
Gorai.013G019400.1.A
PCC=0.041, p=0.869
mRNA Profile
Protein Expression Profile
DPA

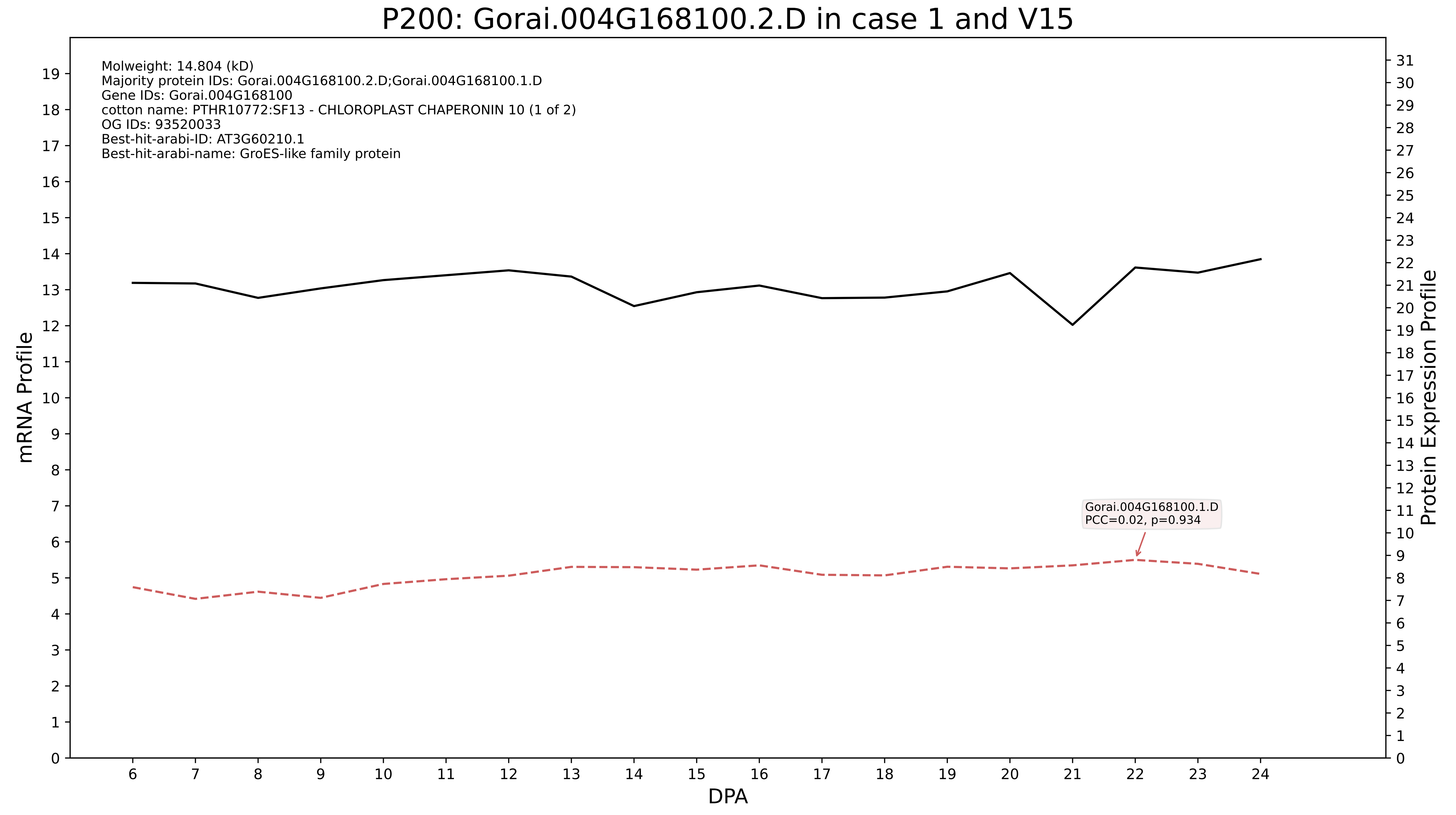
P200: Gorai.004G168100.2.D in case 1 and V15
Molweight: 14.804 (kD)
Majority protein IDs: Gorai.004G168100.2.D;Gorai.004G168100.1.D
Gene IDs: Gorai.004G168100
cotton name: PTHR10772:SF13 - CHLOROPLAST CHAPERONIN 10 (1 of 2)
OG IDs: 93520033
Best-hit-arabi-ID: AT3G60210.1
Best-hit-arabi-name: GroES-like family protein
Gorai.004G168100.1.D
PCC=0.02, p=0.934
mRNA Profile
Protein Expression Profile
DPA

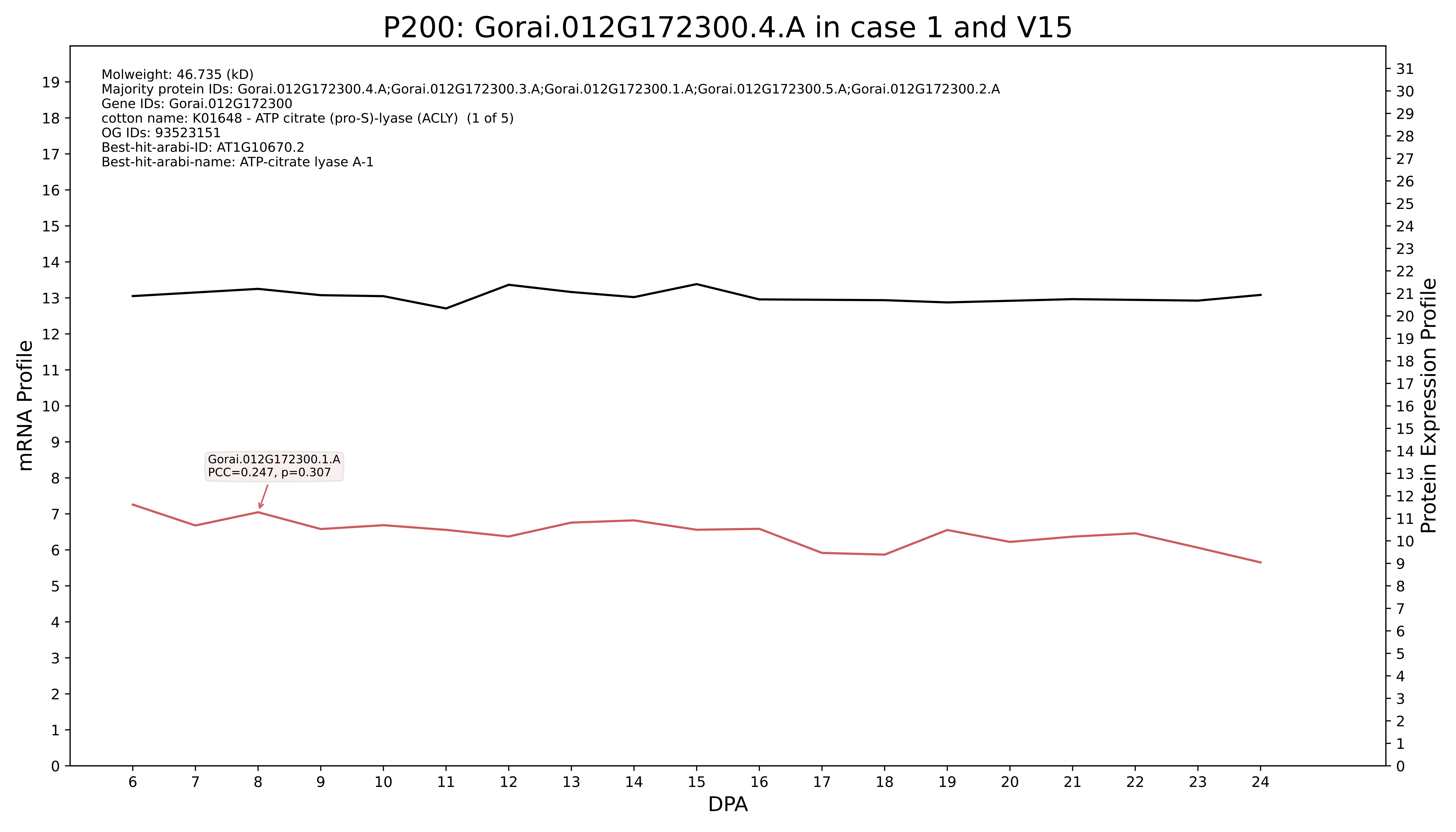
P200: Gorai.012G172300.4.A in case 1 and V15
Molweight: 46.735 (kD)
Majority protein IDs: Gorai.012G172300.4.A;Gorai.012G172300.3.A;Gorai.012G172300.1.A;Gorai.012G172300.5.A;Gorai.012G172300.2.A
Gene IDs: Gorai.012G172300
cotton name: K01648 - ATP citrate (pro-S)-lyase (ACLY) (1 of 5)
OG IDs: 93523151
Best-hit-arabi-ID: AT1G10670.2
Best-hit-arabi-name: ATP-citrate lyase A-1
Gorai.012G172300.1.A
PCC=0.247, p=0.307
mRNA Profile
Protein Expression Profile
DPA

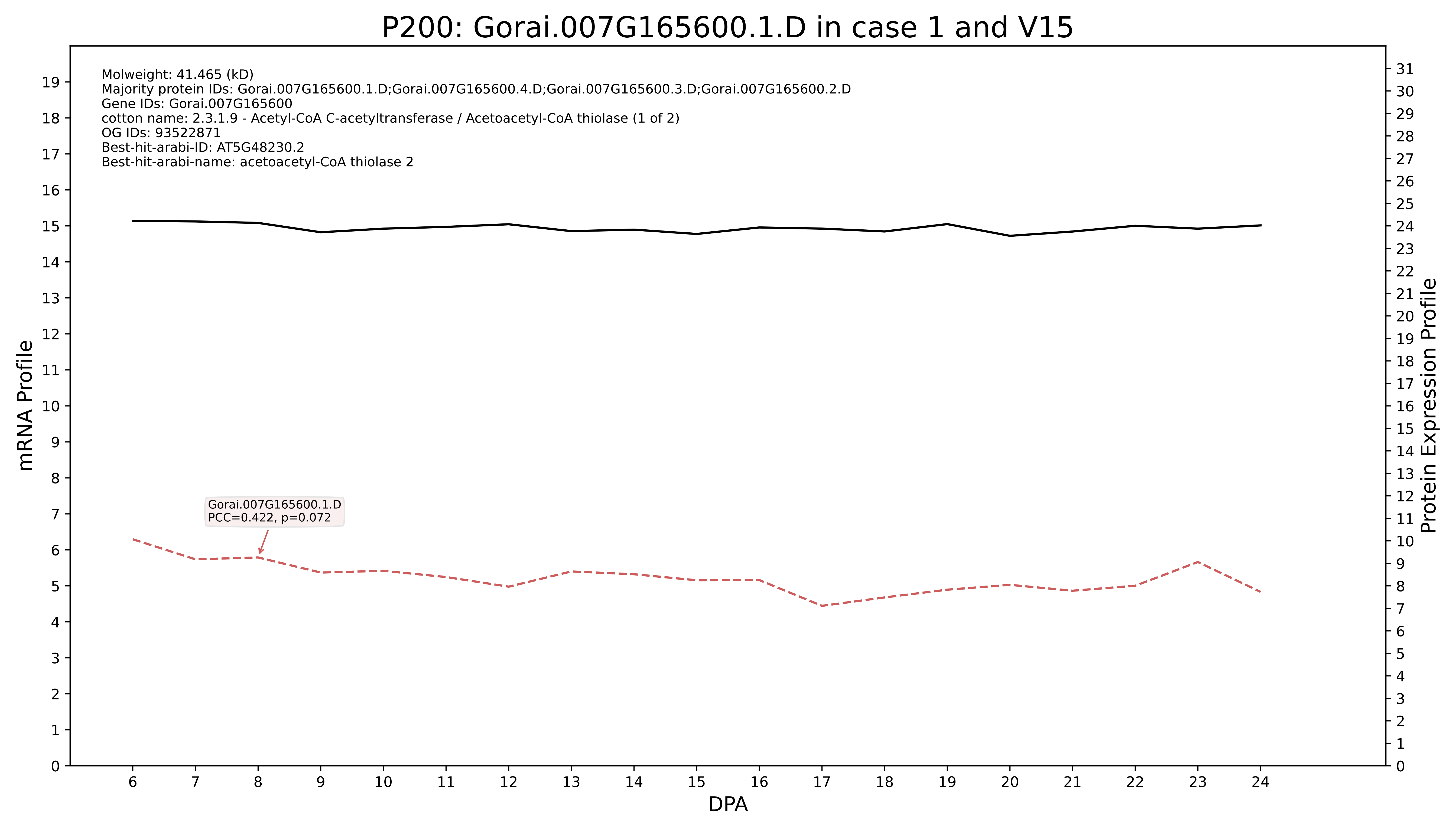

P200: Gorai.007G165600.1.D in case 1 and V15
Molweight: 41.465 (kD)
Majority protein IDs: Gorai.007G165600.1.D;Gorai.007G165600.4.D;Gorai.007G165600.3.D;Gorai.007G165600.2.D
Gene IDs: Gorai.007G165600
cotton name: 2.3.1.9 - Acetyl-CoA C-acetyltransferase / Acetoacetyl-CoA thiolase (1 of 2)
OG IDs: 93522871
Best-hit-arabi-ID: AT5G48230.2
Best-hit-arabi-name: acetoacetyl-CoA thiolase 2
Gorai.007G165600.1.D
PCC=0.422, p=0.072
mRNA Profile
Protein Expression Profile
DPA

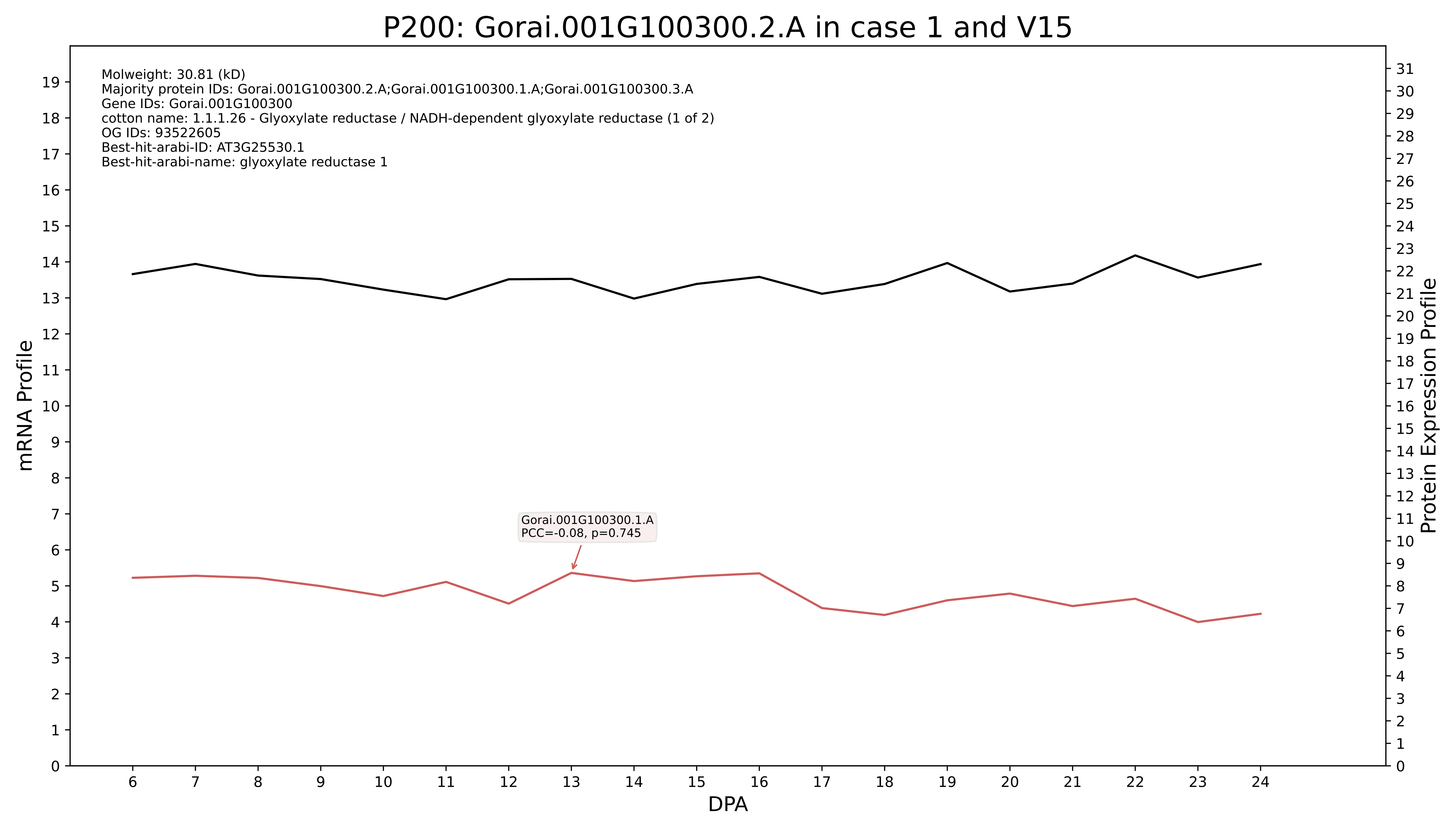

P200: Gorai.001G100300.2.A in case 1 and V15
Molweight: 30.81 (kD)
Majority protein IDs: Gorai.001G100300.2.A;Gorai.001G100300.1.A;Gorai.001G100300.3.A
Gene IDs: Gorai.001G100300
cotton name: 1.1.1.26 - Glyoxylate reductase / NADH-dependent glyoxylate reductase (1 of 2)
OG IDs: 93522605
Best-hit-arabi-ID: AT3G25530.1
Best-hit-arabi-name: glyoxylate reductase 1
Gorai.001G100300.1.A
PCC=-0.08, p=0.745
mRNA Profile
Protein Expression Profile
DPA

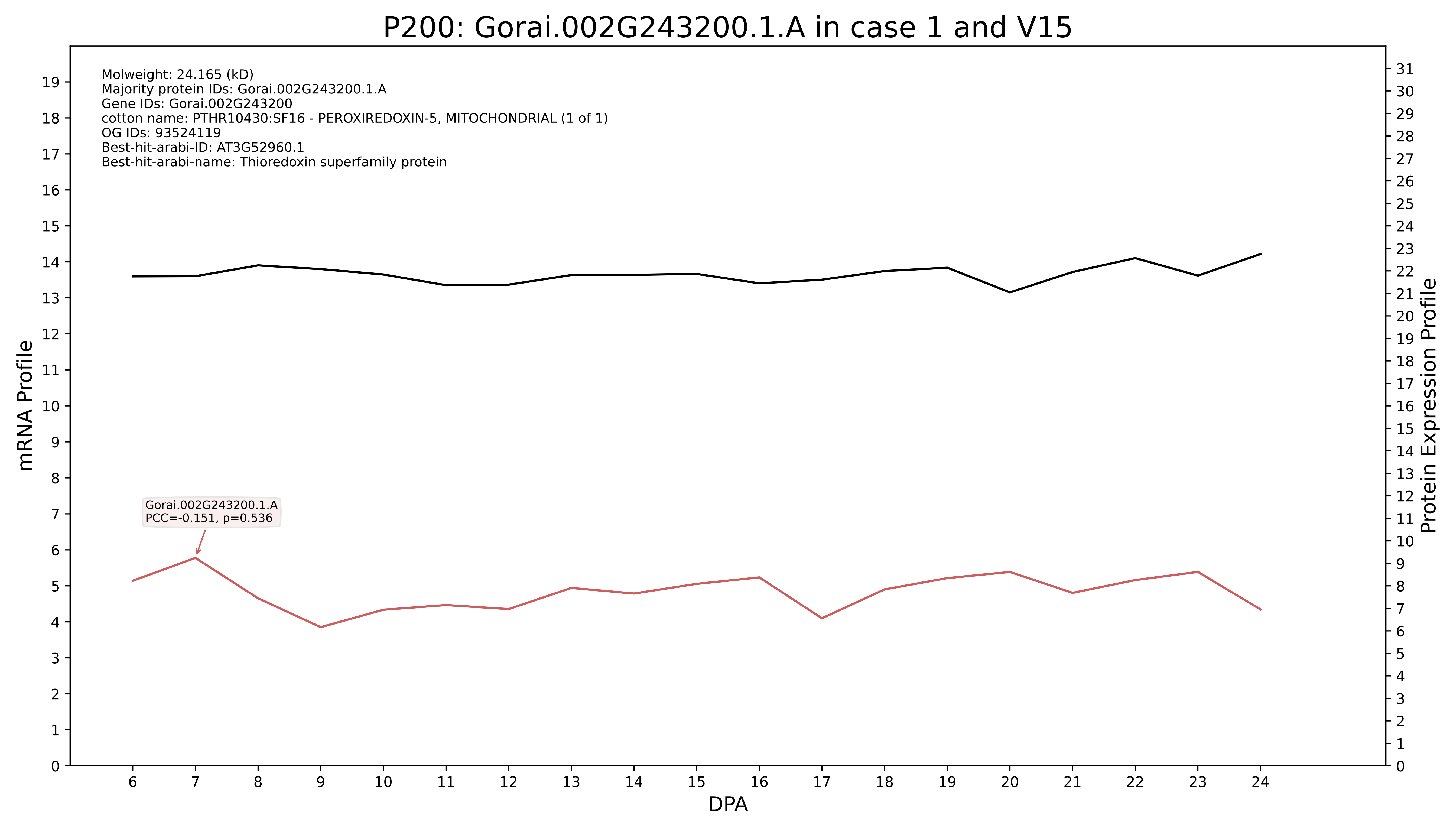
P200: Gorai.002G243200.1.A in case 1 and V15
Molweight: 24.165 (kD)
Majority protein IDs: Gorai.002G243200.1.A
Gene IDs: Gorai.002G243200
cotton name: PTHR10430:SF16 - PEROXIREDOXIN-5, MITOCHONDRIAL (1 of 1)
OG IDs: 93524119
Best-hit-arabi-ID: AT3G52960.1
Best-hit-arabi-name: Thioredoxin superfamily protein
Gorai.002G243200.1.A
PCC=-0.151, p=0.536
mRNA Profile
Protein Expression Profile
DPA

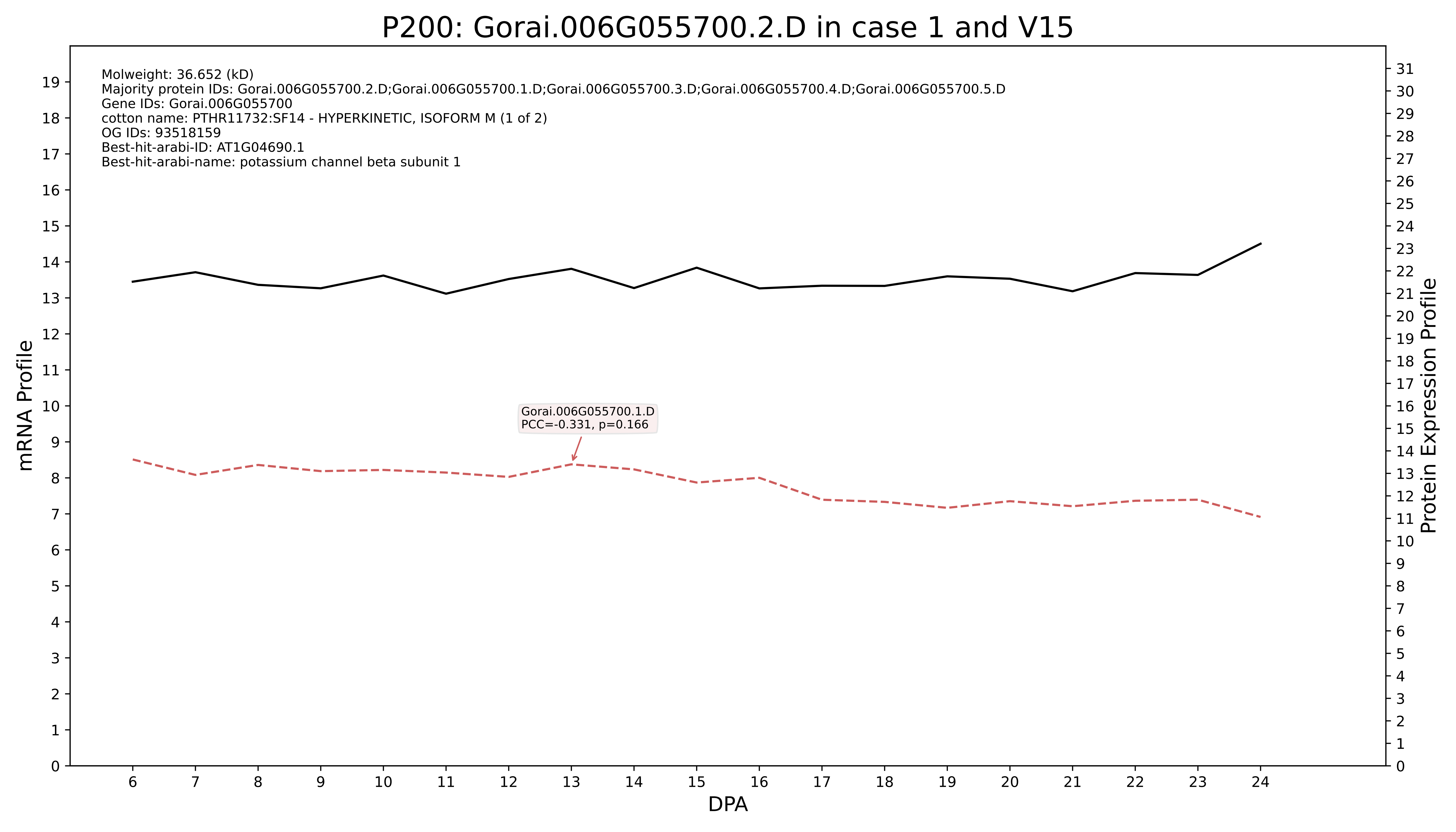
P200: Gorai.006G055700.2.D in case 1 and V15
Molweight: 36.652 (kD)
Majority protein IDs: Gorai.006G055700.2.D;Gorai.006G055700.1.D;Gorai.006G055700.3.D;Gorai.006G055700.4.D;Gorai.006G055700.5.D
Gene IDs: Gorai.006G055700
cotton name: PTHR11732:SF14 - HYPERKINETIC, ISOFORM M (1 of 2)
OG IDs: 93518159
Best-hit-arabi-ID: AT1G04690.1
Best-hit-arabi-name: potassium channel beta subunit 1
Gorai.006G055700.1.D
PCC=-0.331, p=0.166
mRNA Profile
Protein Expression Profile
DPA

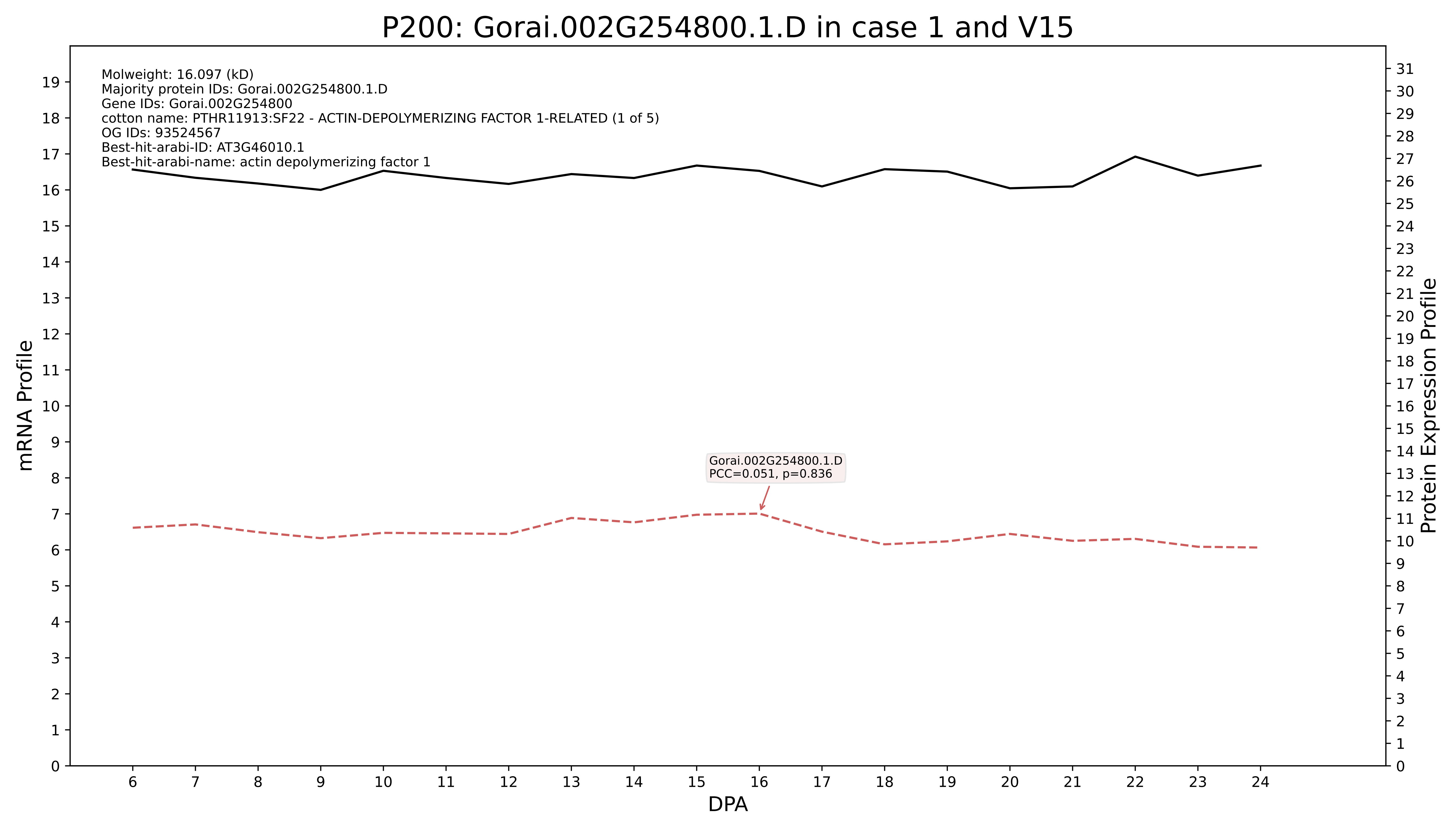
P200: Gorai.002G254800.1.D in case 1 and V15
Molweight: 16.097 (kD)
Majority protein IDs: Gorai.002G254800.1.D
Gene IDs: Gorai.002G254800
cotton name: PTHR11913:SF22 - ACTIN-DEPOLYMERIZING FACTOR 1-RELATED (1 of 5)
OG IDs: 93524567
Best-hit-arabi-ID: AT3G46010.1
Best-hit-arabi-name: actin depolymerizing factor 1
Gorai.002G254800.1.D
PCC=0.051, p=0.836
mRNA Profile
Protein Expression Profile
DPA

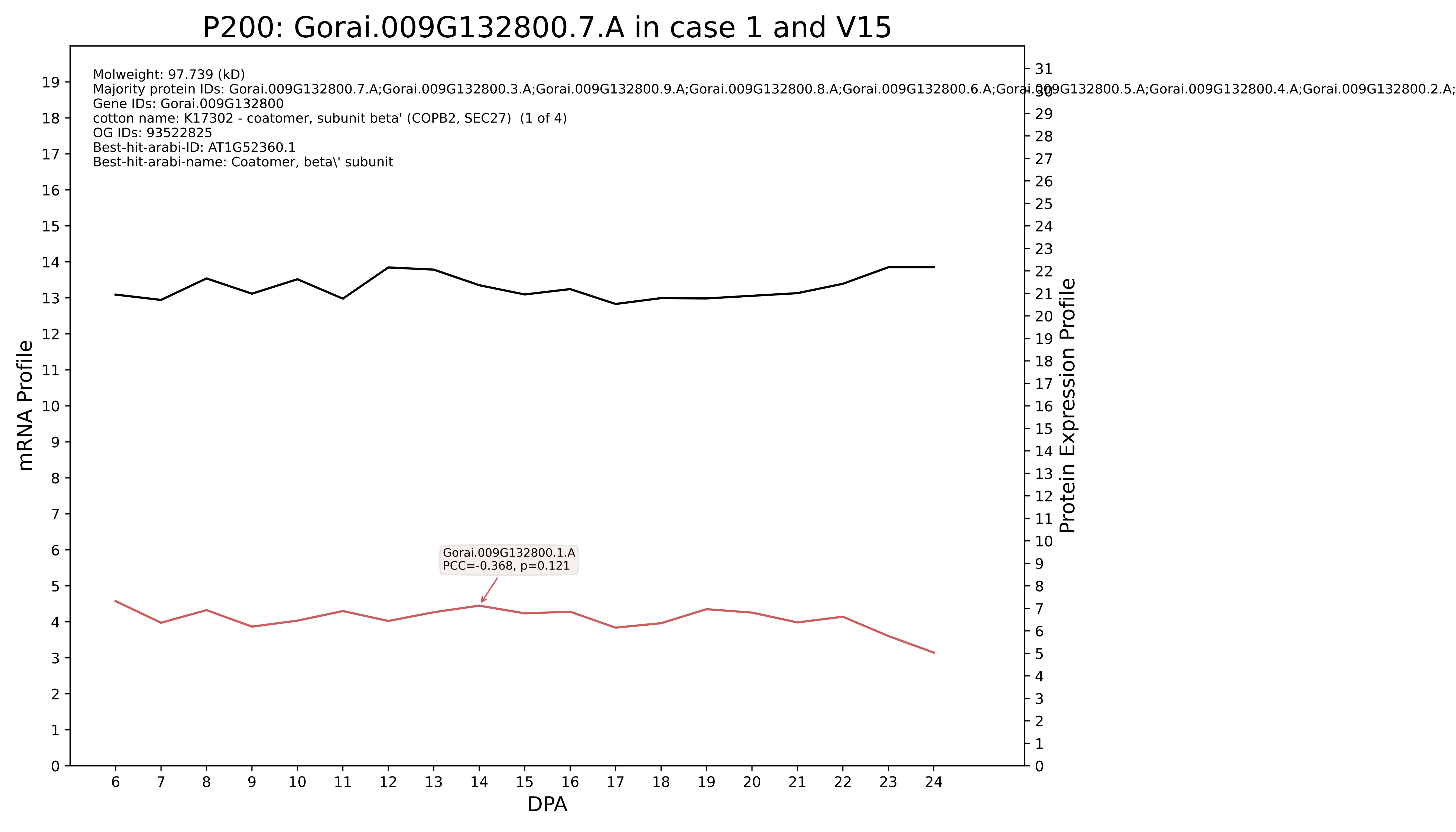
P200: Gorai.009G132800.7.A in case 1 and V15
Molweight: 97.739 (kD)
Majority protein IDs: Gorai.009G132800.7.A;Gorai.009G132800.3.A;Gorai.009G132800.9.A;Gorai.009G132800.8.A;Gorai.009G132800.6.A;Gorai.009G132800.5.A;Gorai.009G132800.4.A;Gorai.009G132800.2.A;
Gene IDs: Gorai.009G132800
cotton name: K17302 - coatomer, subunit beta' (COPB2, SEC27) (1 of 4)
OG IDs: 93522825
Best-hit-arabi-ID: AT1G52360.1
Best-hit-arabi-name: Coatomer, beta\' subunit
Gorai.009G132800.1.A
PCC=-0.368, p=0.121
mRNA Profile
Protein Expression Profile
DPA

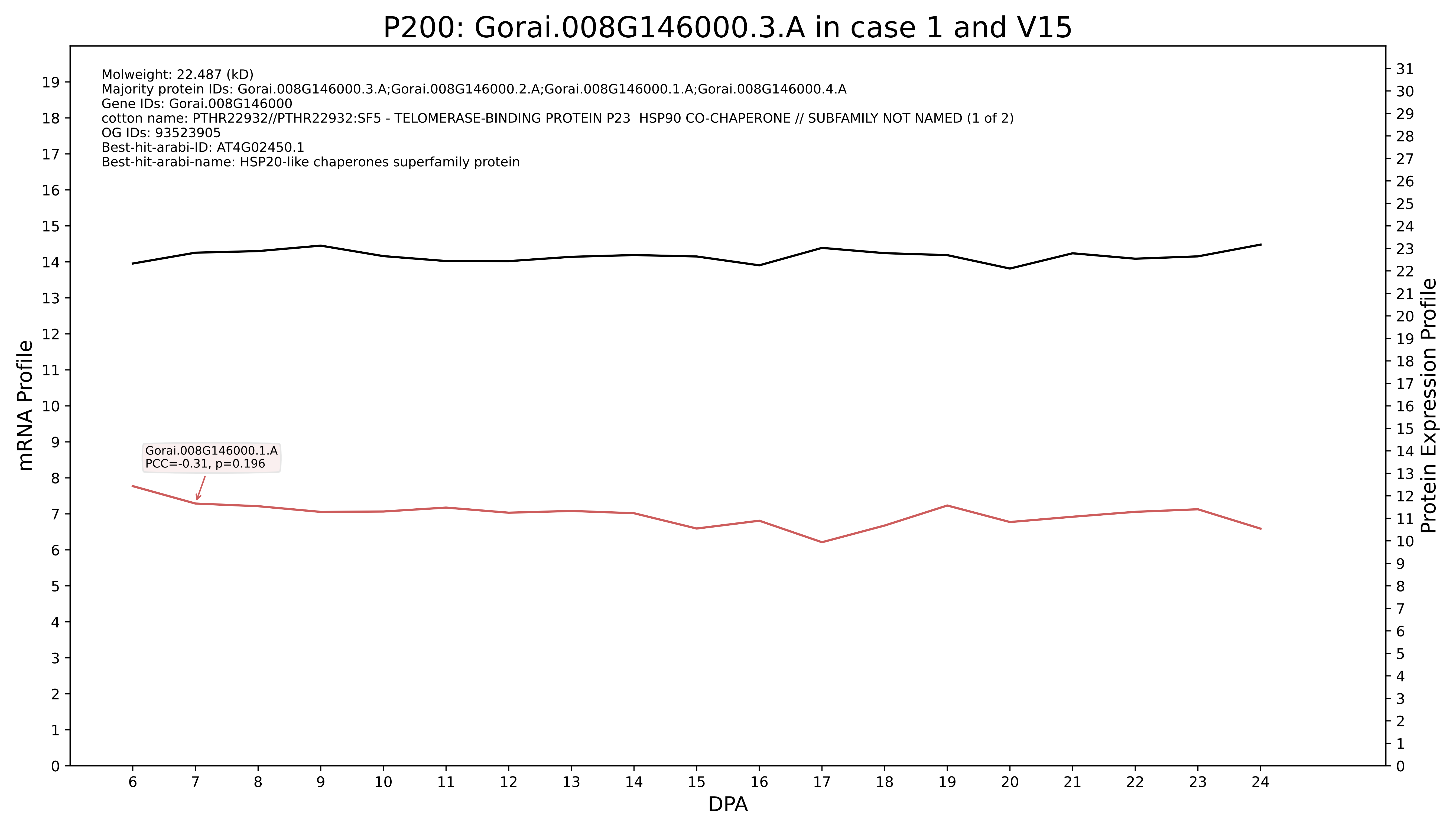
P200: Gorai.008G146000.3.A in case 1 and V15
Molweight: 22.487 (kD)
Majority protein IDs: Gorai.008G146000.3.A;Gorai.008G146000.2.A;Gorai.008G146000.1.A;Gorai.008G146000.4.A
Gene IDs: Gorai.008G146000
cotton name: PTHR22932//PTHR22932:SF5 - TELOMERASE-BINDING PROTEIN P23 HSP90 CO-CHAPERONE // SUBFAMILY NOT NAMED (1 of 2)
OG IDs: 93523905
Best-hit-arabi-ID: AT4G02450.1
Best-hit-arabi-name: HSP20-like chaperones superfamily protein
Gorai.008G146000.1.A
PCC=-0.31, p=0.196
mRNA Profile
Protein Expression Profile
DPA

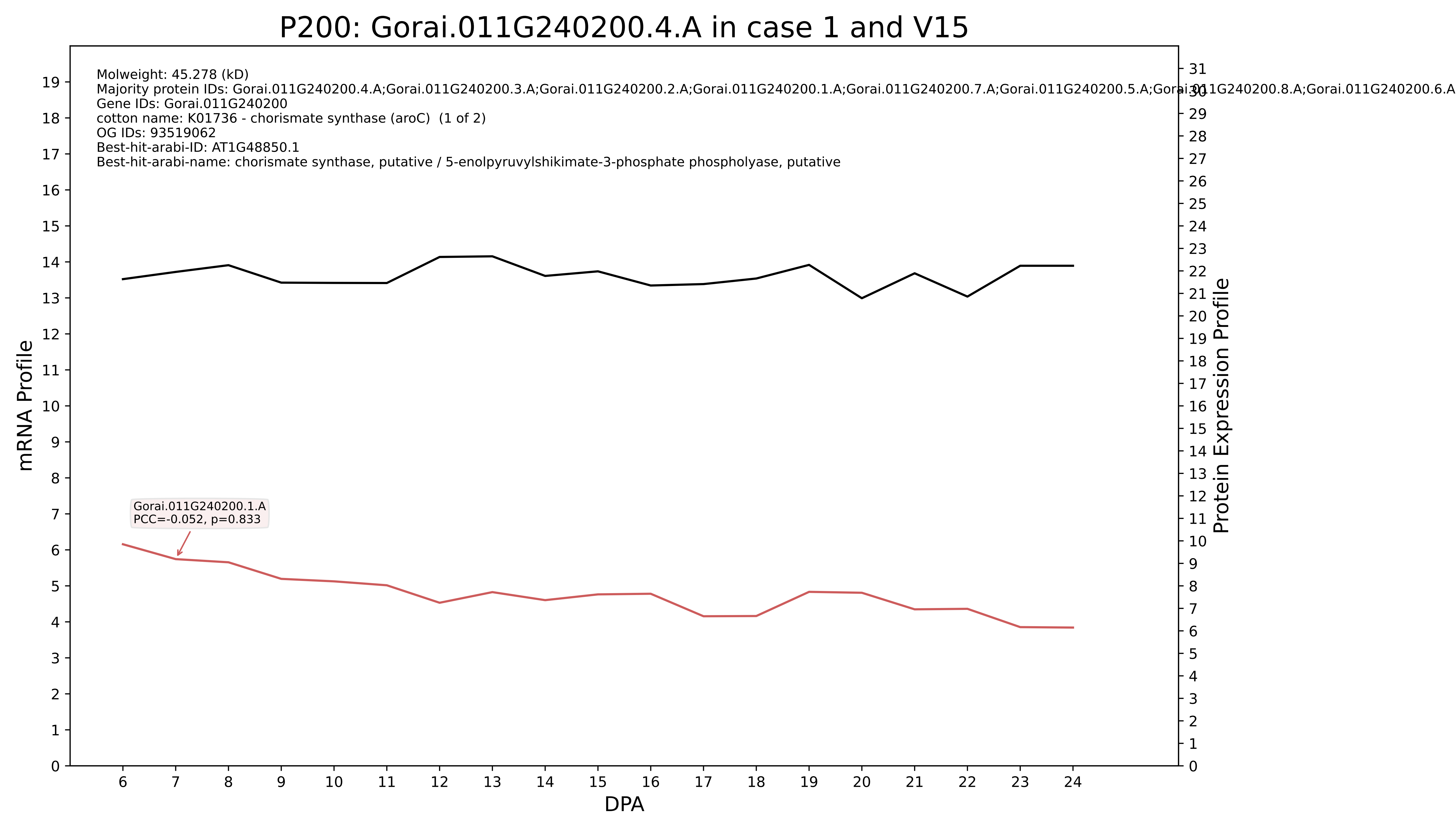
P200: Gorai.011G240200.4.A in case 1 and V15
Molweight: 45.278 (kD)
Majority protein IDs: Gorai.011G240200.4.A;Gorai.011G240200.3.A;Gorai.011G240200.2.A;Gorai.011G240200.1.A;Gorai.011G240200.7.A;Gorai.011G240200.5.A;Gorai.011G240200.8.A;Gorai.011G240200.6.A
Gene IDs: Gorai.011G240200
cotton name: K01736 - chorismate synthase (aroC) (1 of 2)
OG IDs: 93519062
Best-hit-arabi-ID: AT1G48850.1
Best-hit-arabi-name: chorismate synthase, putative / 5-enolpyruvylshikimate-3-phosphate phospholyase, putative
Gorai.011G240200.1.A
PCC=-0.052, p=0.833
mRNA Profile
Protein Expression Profile
DPA

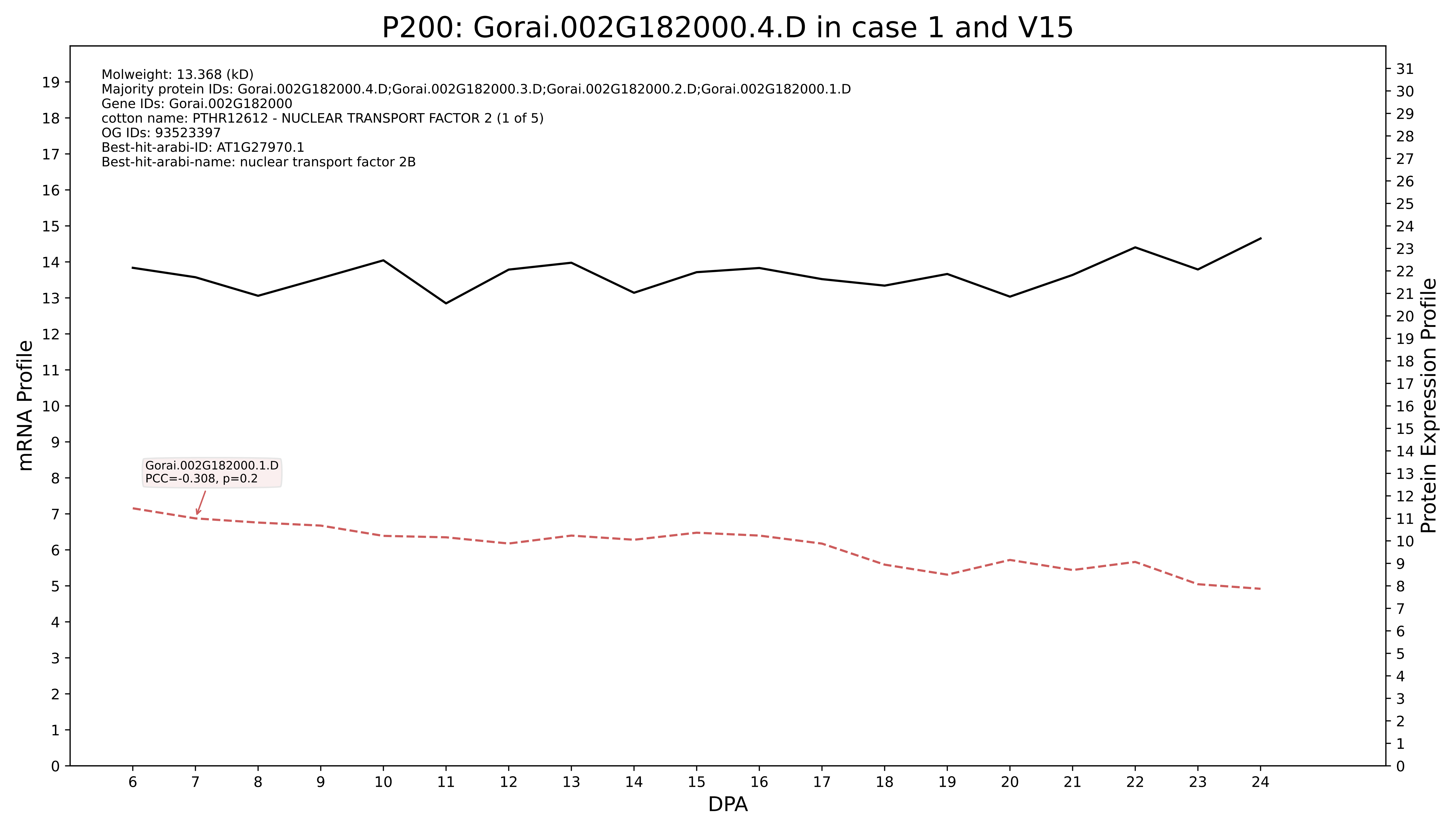

P200: Gorai.002G182000.4.D in case 1 and V15
Molweight: 13.368 (kD)
Majority protein IDs: Gorai.002G182000.4.D;Gorai.002G182000.3.D;Gorai.002G182000.2.D;Gorai.002G182000.1.D
Gene IDs: Gorai.002G182000
cotton name: PTHR12612 - NUCLEAR TRANSPORT FACTOR 2 (1 of 5)
OG IDs: 93523397
Best-hit-arabi-ID: AT1G27970.1
Best-hit-arabi-name: nuclear transport factor 2B
Gorai.002G182000.1.D
PCC=-0.308, p=0.2
mRNA Profile
Protein Expression Profile
DPA

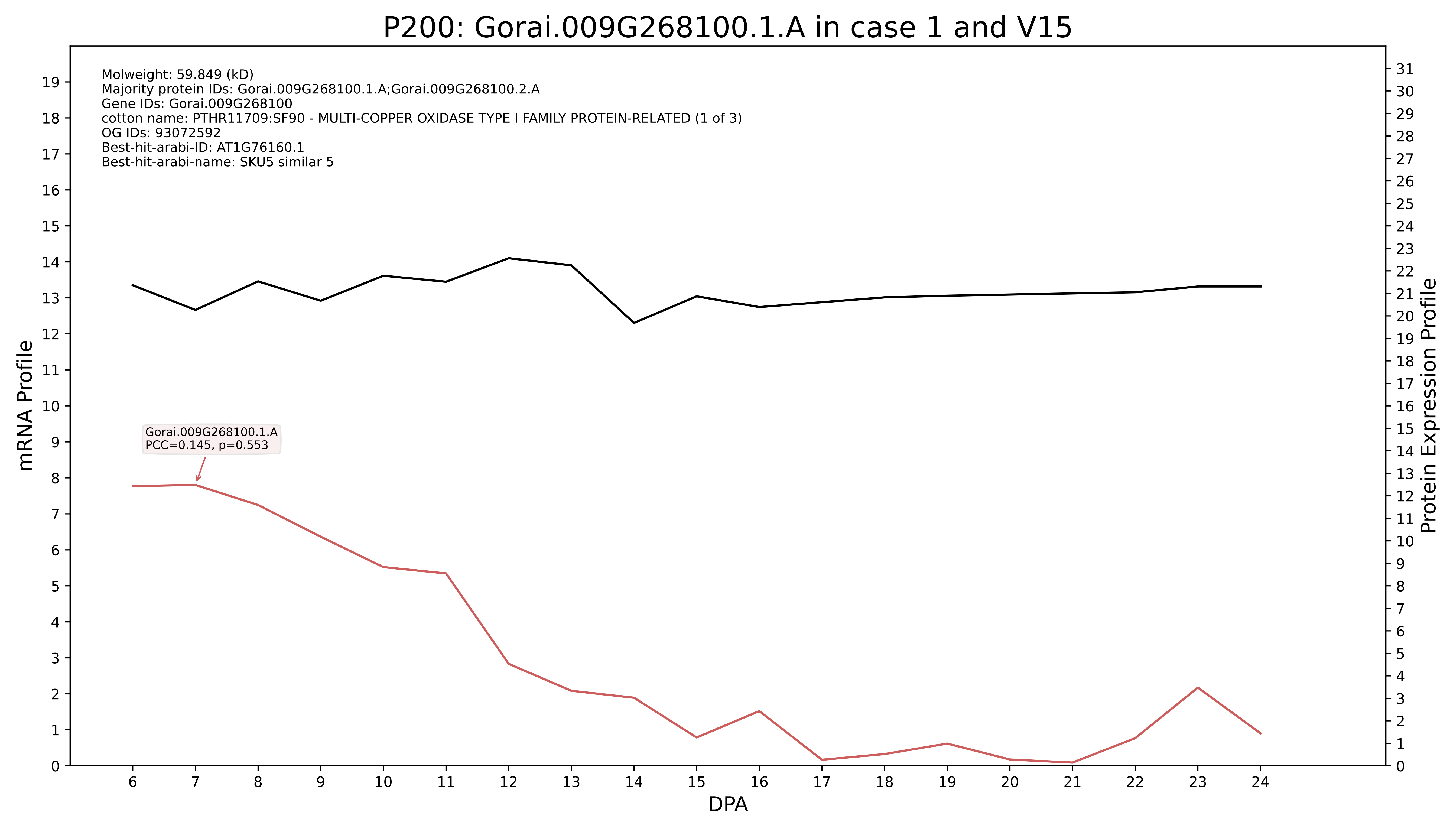
P200: Gorai.009G268100.1.A in case 1 and V15
Molweight: 59.849 (kD)
Majority protein IDs: Gorai.009G268100.1.A;Gorai.009G268100.2.A
Gene IDs: Gorai.009G268100
cotton name: PTHR11709:SF90 - MULTI-COPPER OXIDASE TYPE I FAMILY PROTEIN-RELATED (1 of 3)
OG IDs: 93072592
Best-hit-arabi-ID: AT1G76160.1
Best-hit-arabi-name: SKU5 similar 5
Gorai.009G268100.1.A
PCC=0.145, p=0.553
mRNA Profile
Protein Expression Profile
DPA

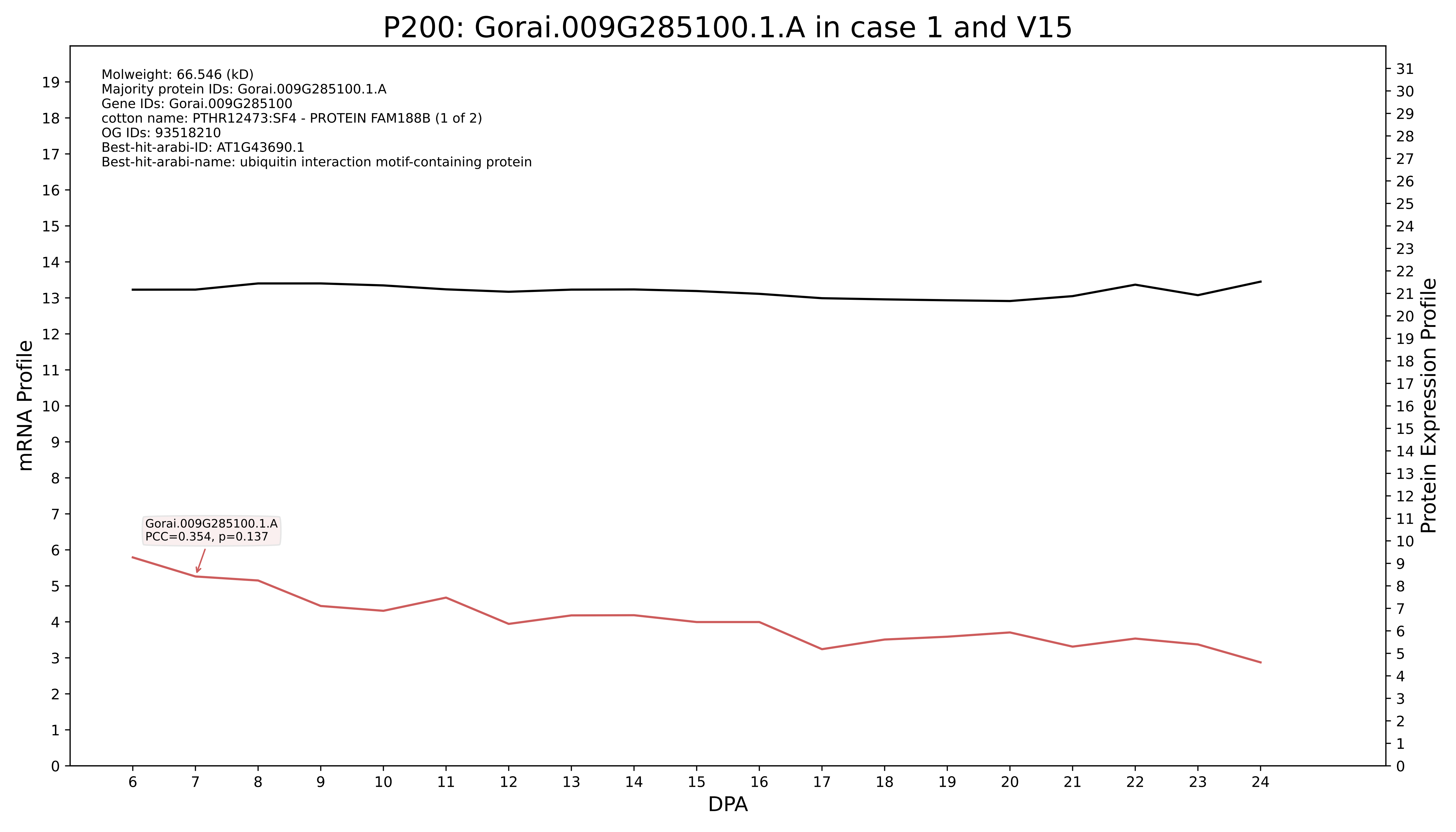
P200: Gorai.009G285100.1.A in case 1 and V15
Molweight: 66.546 (kD)
Majority protein IDs: Gorai.009G285100.1.A
Gene IDs: Gorai.009G285100
cotton name: PTHR12473:SF4 - PROTEIN FAM188B (1 of 2)
OG IDs: 93518210
Best-hit-arabi-ID: AT1G43690.1
Best-hit-arabi-name: ubiquitin interaction motif-containing protein
Gorai.009G285100.1.A
PCC=0.354, p=0.137
mRNA Profile
Protein Expression Profile
DPA

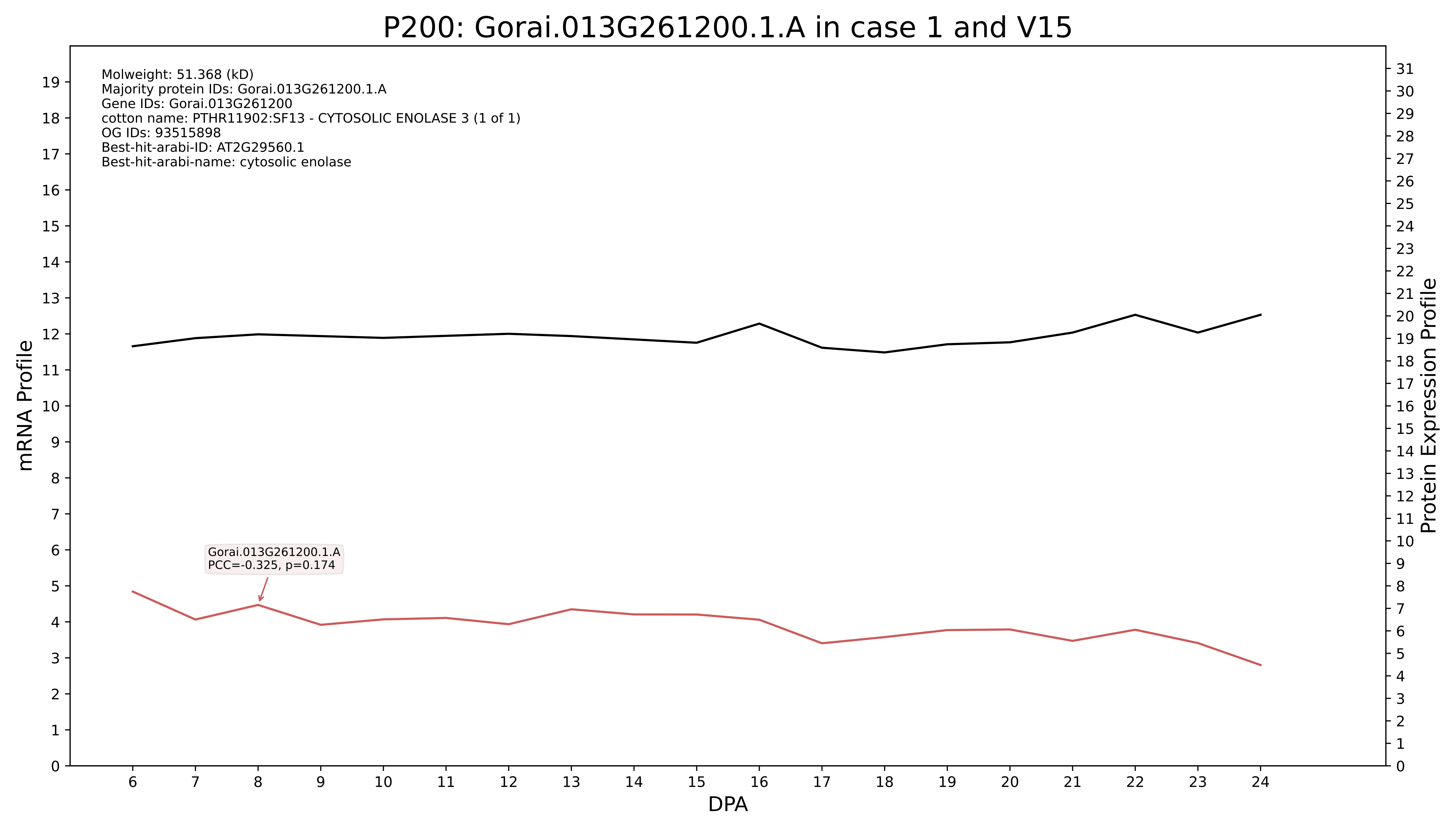
P200: Gorai.013G261200.1.A in case 1 and V15
Molweight: 51.368 (kD)
Majority protein IDs: Gorai.013G261200.1.A
Gene IDs: Gorai.013G261200
cotton name: PTHR11902:SF13 - CYTOSOLIC ENOLASE 3 (1 of 1)
OG IDs: 93515898
Best-hit-arabi-ID: AT2G29560.1
Best-hit-arabi-name: cytosolic enolase
Gorai.013G261200.1.A
PCC=-0.325, p=0.174
mRNA Profile
Protein Expression Profile
DPA

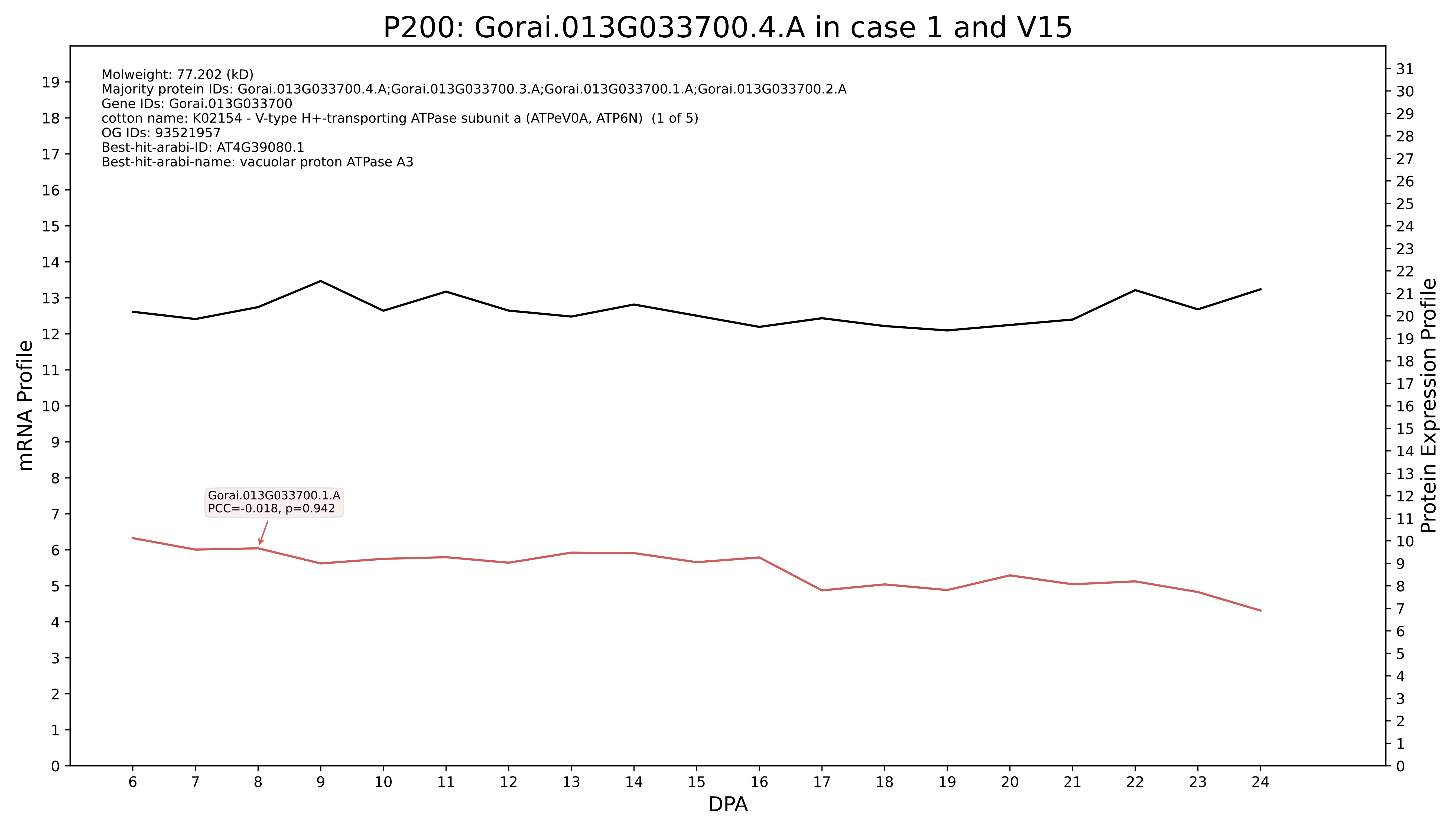

P200: Gorai.013G033700.4.A in case 1 and V15
Molweight: 77.202 (kD)
Majority protein IDs: Gorai.013G033700.4.A;Gorai.013G033700.3.A;Gorai.013G033700.1.A;Gorai.013G033700.2.A
Gene IDs: Gorai.013G033700
cotton name: K02154 - V-type H+-transporting ATPase subunit a (ATPeV0A, ATP6N) (1 of 5)
OG IDs: 93521957
Best-hit-arabi-ID: AT4G39080.1
Best-hit-arabi-name: vacuolar proton ATPase A3
Gorai.013G033700.1.A
PCC=-0.018, p=0.942
mRNA Profile
Protein Expression Profile
DPA

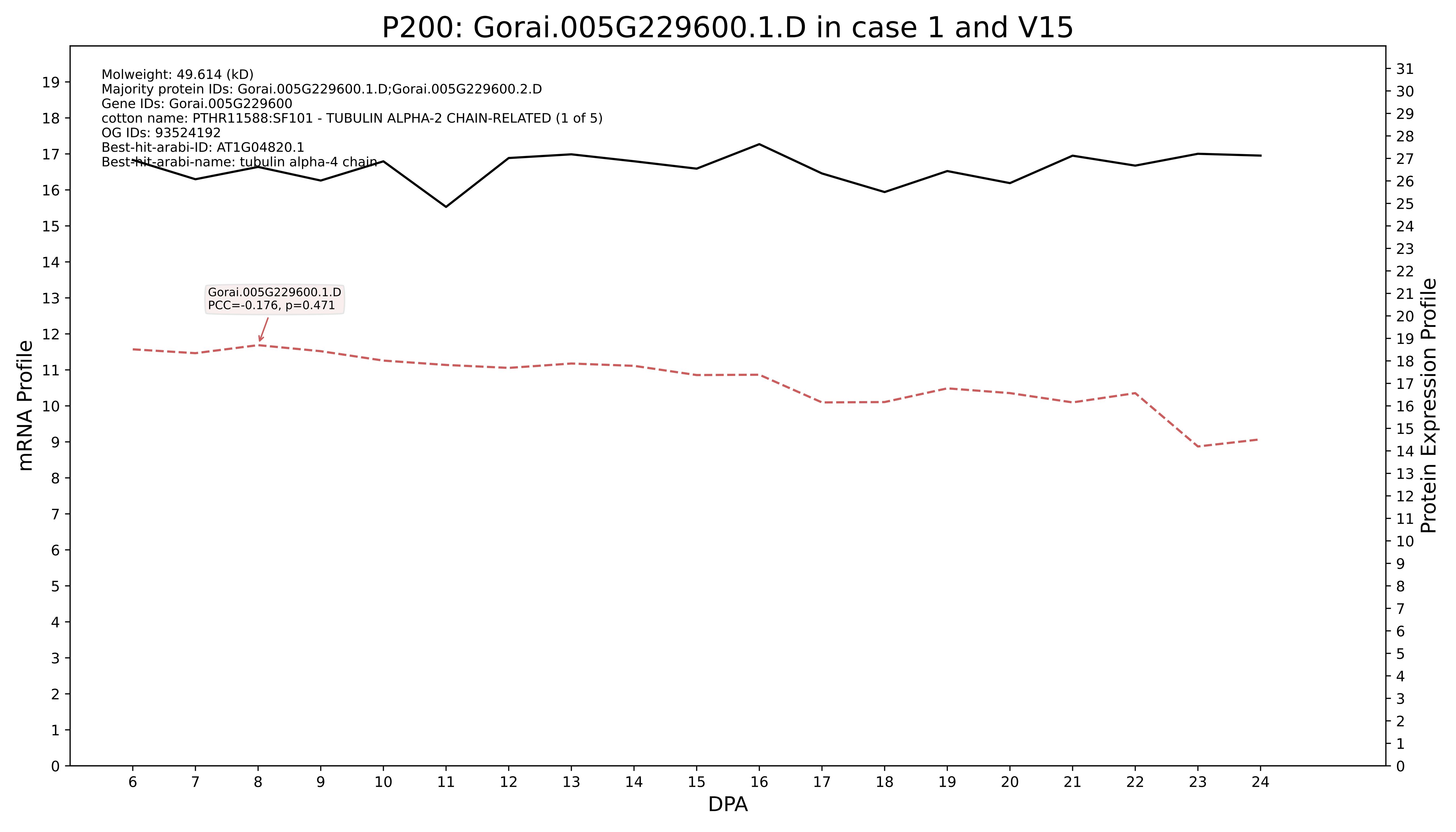
P200: Gorai.005G229600.1.D in case 1 and V15
Molweight: 49.614 (kD)
Majority protein IDs: Gorai.005G229600.1.D;Gorai.005G229600.2.D
Gene IDs: Gorai.005G229600
cotton name: PTHR11588:SF101 - TUBULIN ALPHA-2 CHAIN-RELATED (1 of 5)
OG IDs: 93524192
Best-hit-arabi-ID: AT1G04820.1
Best-hit-arabi-name: tubulin alpha-4 chain
Gorai.005G229600.1.D
PCC=-0.176, p=0.471
mRNA Profile
Protein Expression Profile
DPA

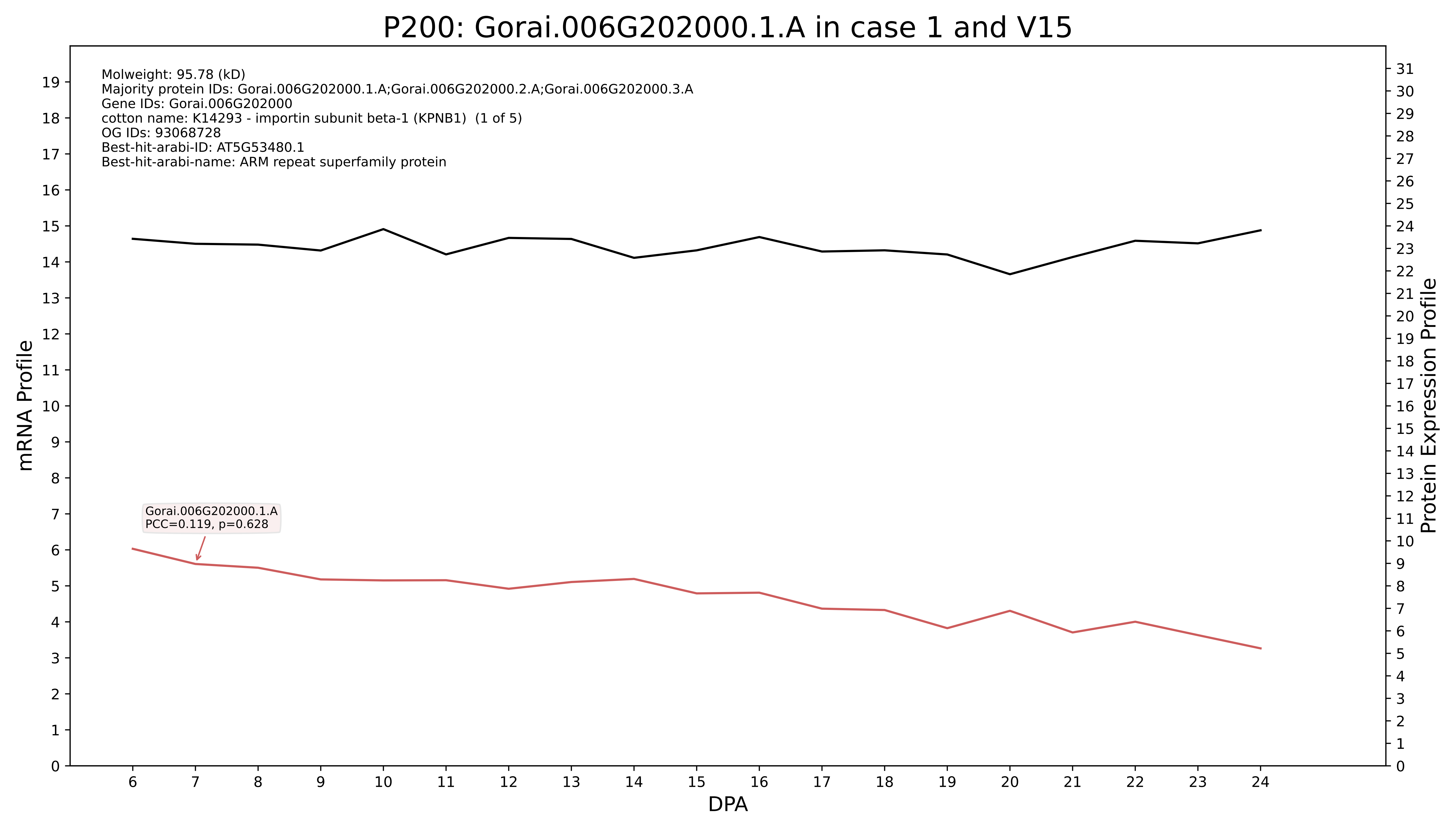
P200: Gorai.006G202000.1.A in case 1 and V15
Molweight: 95.78 (kD)
Majority protein IDs: Gorai.006G202000.1.A;Gorai.006G202000.2.A;Gorai.006G202000.3.A
Gene IDs: Gorai.006G202000
cotton name: K14293 - importin subunit beta-1 (KPNB1) (1 of 5)
OG IDs: 93068728
Best-hit-arabi-ID: AT5G53480.1
Best-hit-arabi-name: ARM repeat superfamily protein
Gorai.006G202000.1.A
PCC=0.119, p=0.628
mRNA Profile
Protein Expression Profile
DPA

### P200: Gorai.008G216000.1.D in case 1 and V15

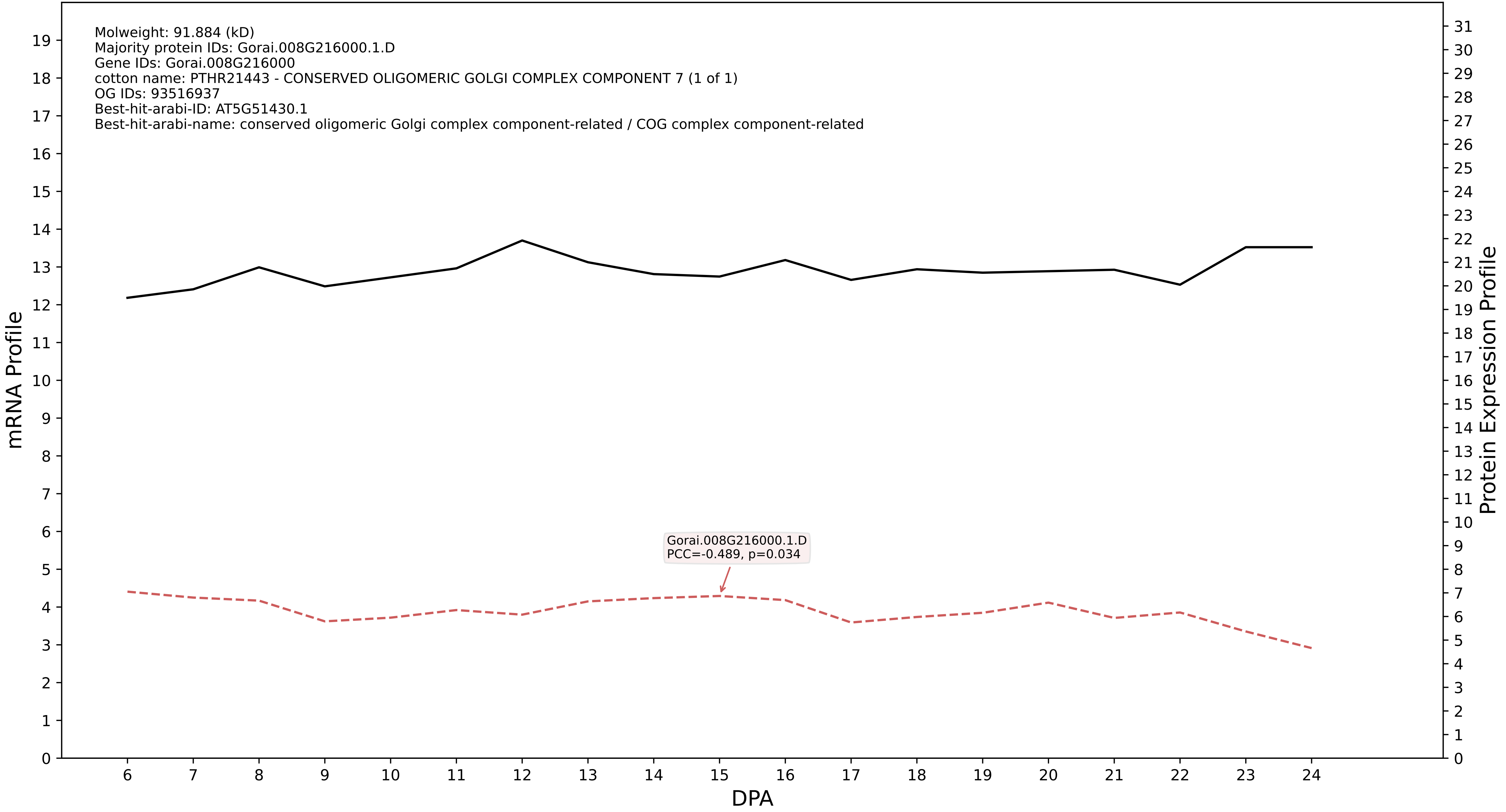

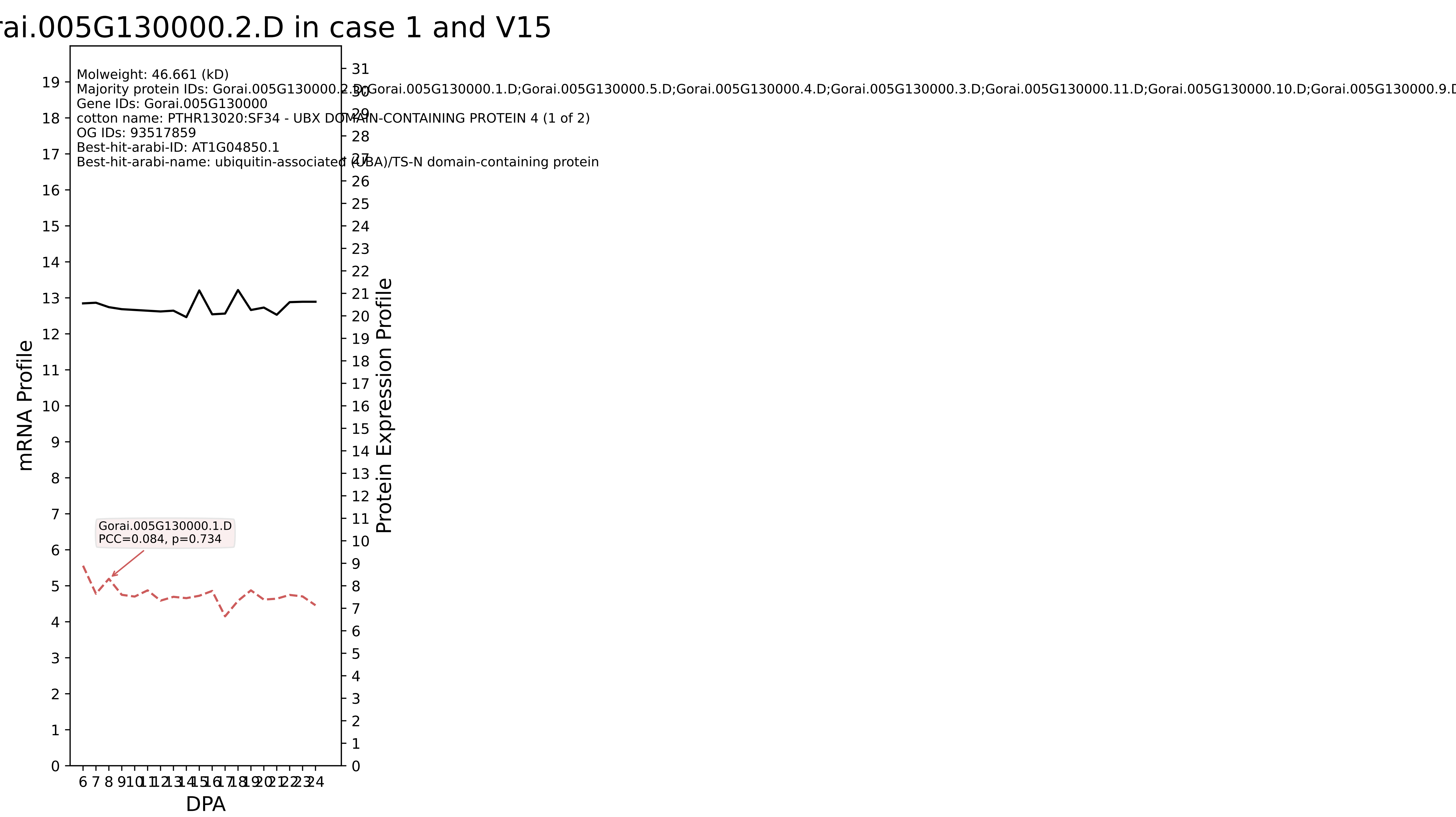
rai.005G130000.2.D in case 1 and V15
Molweight: 46.661 (kD)
Majority protein IDs: Gorai.005G130000.2.D;Gorai.005G130000.1.D;Gorai.005G130000.5.D;Gorai.005G130000.4.D;Gorai.005G130000.3.D;Gorai.005G130000.11.D;Gorai.005G130000.10.D;Gorai.005G130000.9.D
Gene IDs: Gorai.005G130000
cotton name: PTHR13020:SF34 - UBX DOMAIN-CONTAINING PROTEIN 4 (1 of 2)
OG IDs: 93517859
Best-hit-arabi-ID: AT1G04850.1
Best-hit-arabi-name: ubiquitin-associated (UBA)/TS-N domain-containing protein
Gorai.005G130000.1.D
PCC=0.084, p=0.734
mRNA Profile
Protein Expression Profile
DPA

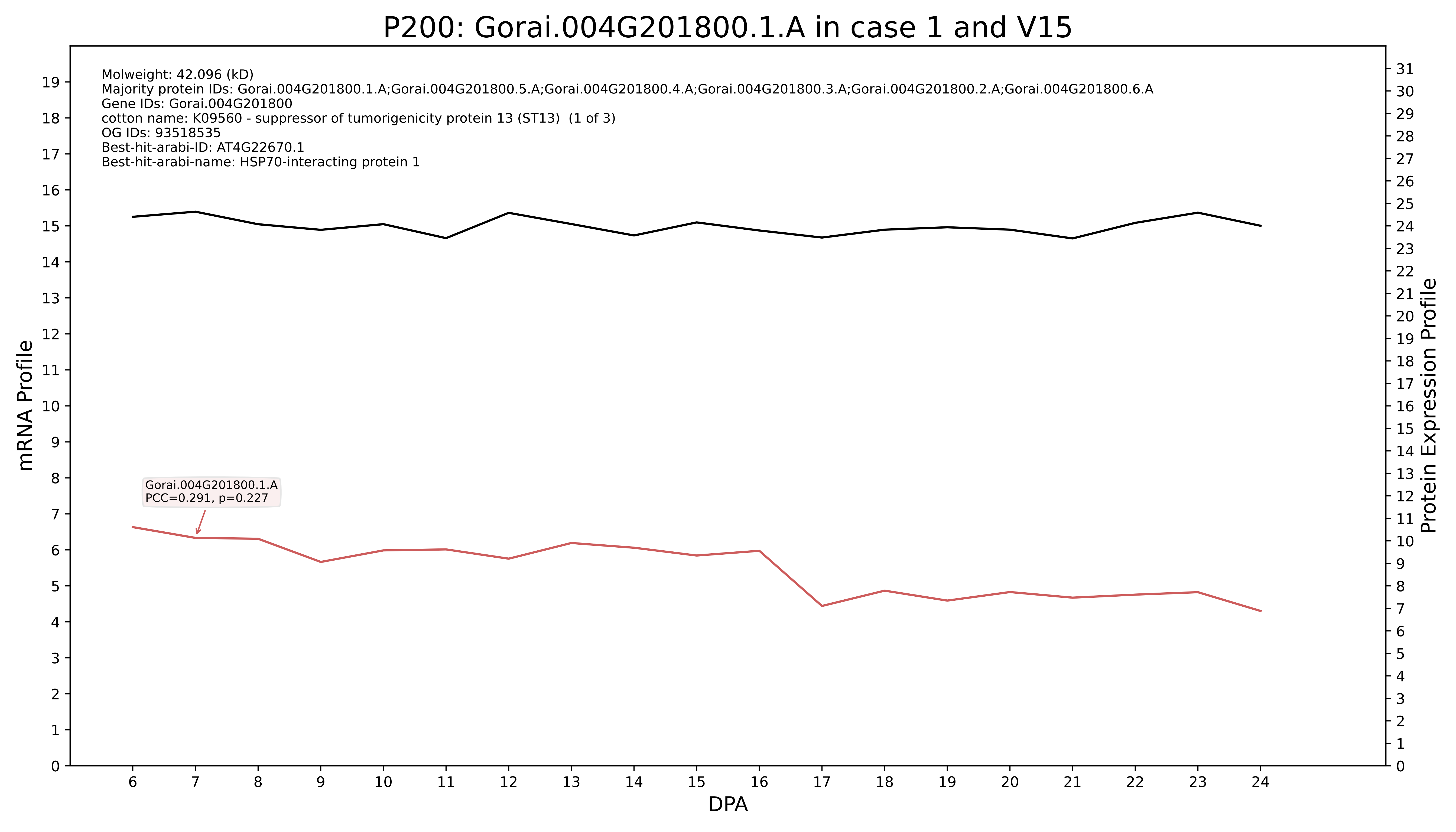
P200: Gorai.004G201800.1.A in case 1 and V15
Molweight: 42.096 (kD)
Majority protein IDs: Gorai.004G201800.1.A;Gorai.004G201800.5.A;Gorai.004G201800.4.A;Gorai.004G201800.3.A;Gorai.004G201800.2.A;Gorai.004G201800.6.A
Gene IDs: Gorai.004G201800
cotton name: K09560 - suppressor of tumorigenicity protein 13 (ST13) (1 of 3)
OG IDs: 93518535
Best-hit-arabi-ID: AT4G22670.1
Best-hit-arabi-name: HSP70-interacting protein 1
Gorai.004G201800.1.A
PCC=0.291, p=0.227
mRNA Profile
Protein Expression Profile
DPA

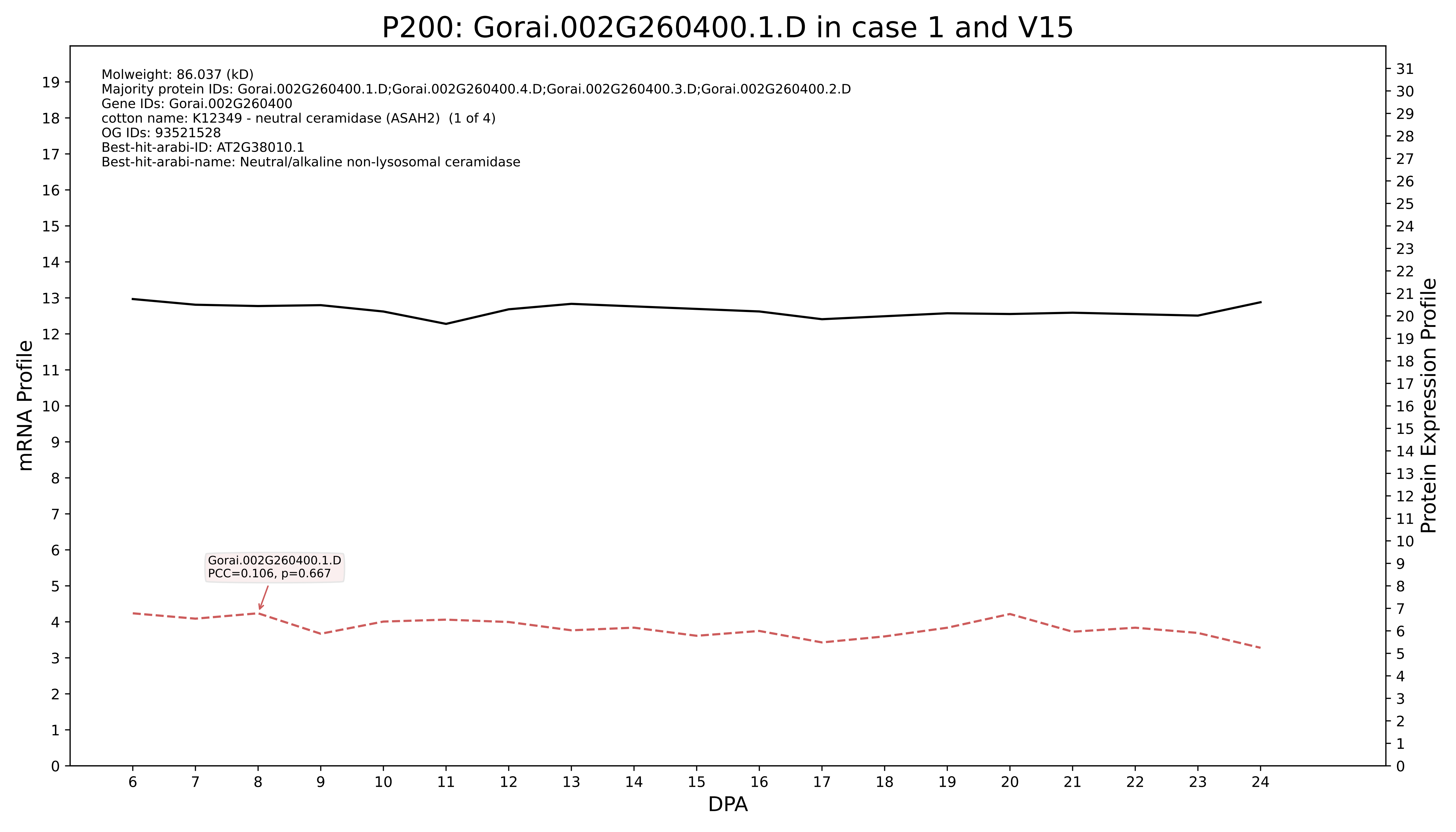

P200: Gorai.002G260400.1.D in case 1 and V15
Molweight: 86.037 (kD)
Majority protein IDs: Gorai.002G260400.1.D;Gorai.002G260400.4.D;Gorai.002G260400.3.D;Gorai.002G260400.2.D
Gene IDs: Gorai.002G260400
cotton name: K12349 - neutral ceramidase (ASAH2) (1 of 4)
OG IDs: 93521528
Best-hit-arabi-ID: AT2G38010.1
Best-hit-arabi-name: Neutral/alkaline non-lysosomal ceramidase
Gorai.002G260400.1.D
PCC=0.106, p=0.667
mRNA Profile
Protein Expression Profile
DPA

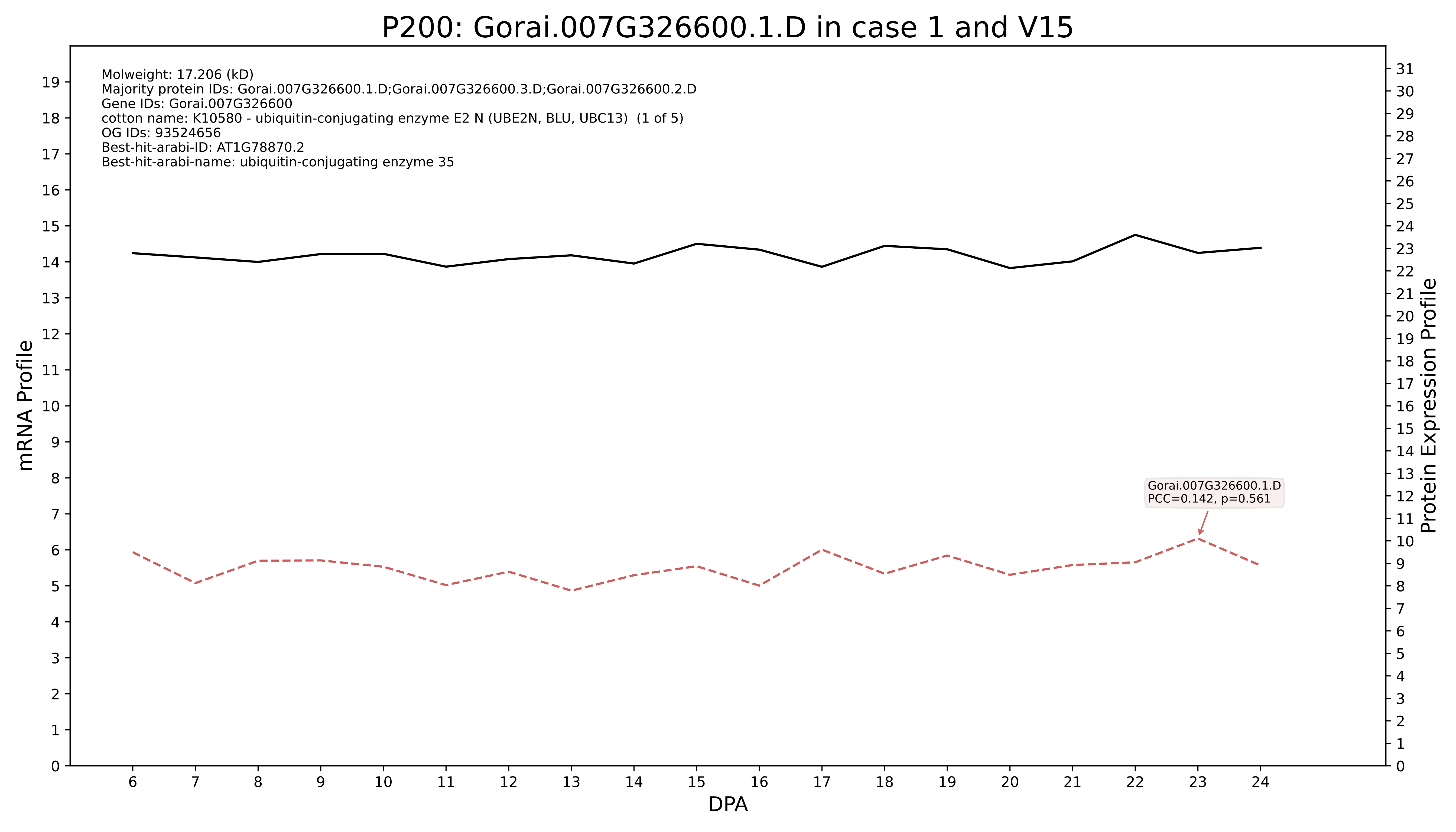
P200: Gorai.007G326600.1.D in case 1 and V15
Molweight: 17.206 (kD)
Majority protein IDs: Gorai.007G326600.1.D;Gorai.007G326600.3.D;Gorai.007G326600.2.D
Gene IDs: Gorai.007G326600
cotton name: K10580 - ubiquitin-conjugating enzyme E2 N (UBE2N, BLU, UBC13) (1 of 5)
OG IDs: 93524656
Best-hit-arabi-ID: AT1G78870.2
Best-hit-arabi-name: ubiquitin-conjugating enzyme 35
Gorai.007G326600.1.D
PCC=0.142, p=0.561
mRNA Profile
Protein Expression Profile
DPA

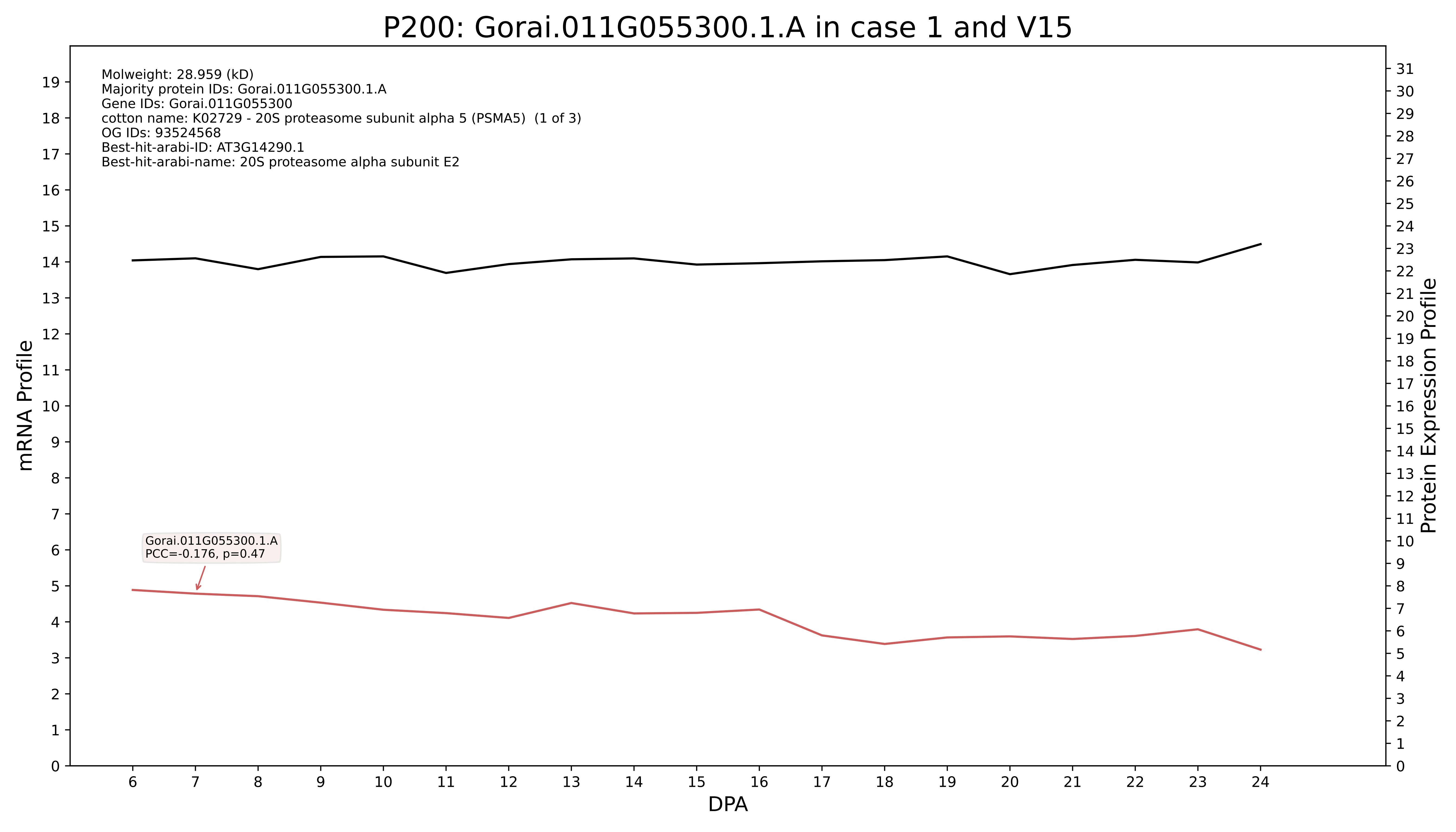
P200: Gorai.011G055300.1.A in case 1 and V15
Molweight: 28.959 (kD)
Majority protein IDs: Gorai.011G055300.1.A
Gene IDs: Gorai.011G055300
cotton name: K02729 - 20S proteasome subunit alpha 5 (PSMA5) (1 of 3)
OG IDs: 93524568
Best-hit-arabi-ID: AT3G14290.1
Best-hit-arabi-name: 20S proteasome alpha subunit E2
Gorai.011G055300.1.A
PCC=-0.176, p=0.47
mRNA Profile
Protein Expression Profile
DPA

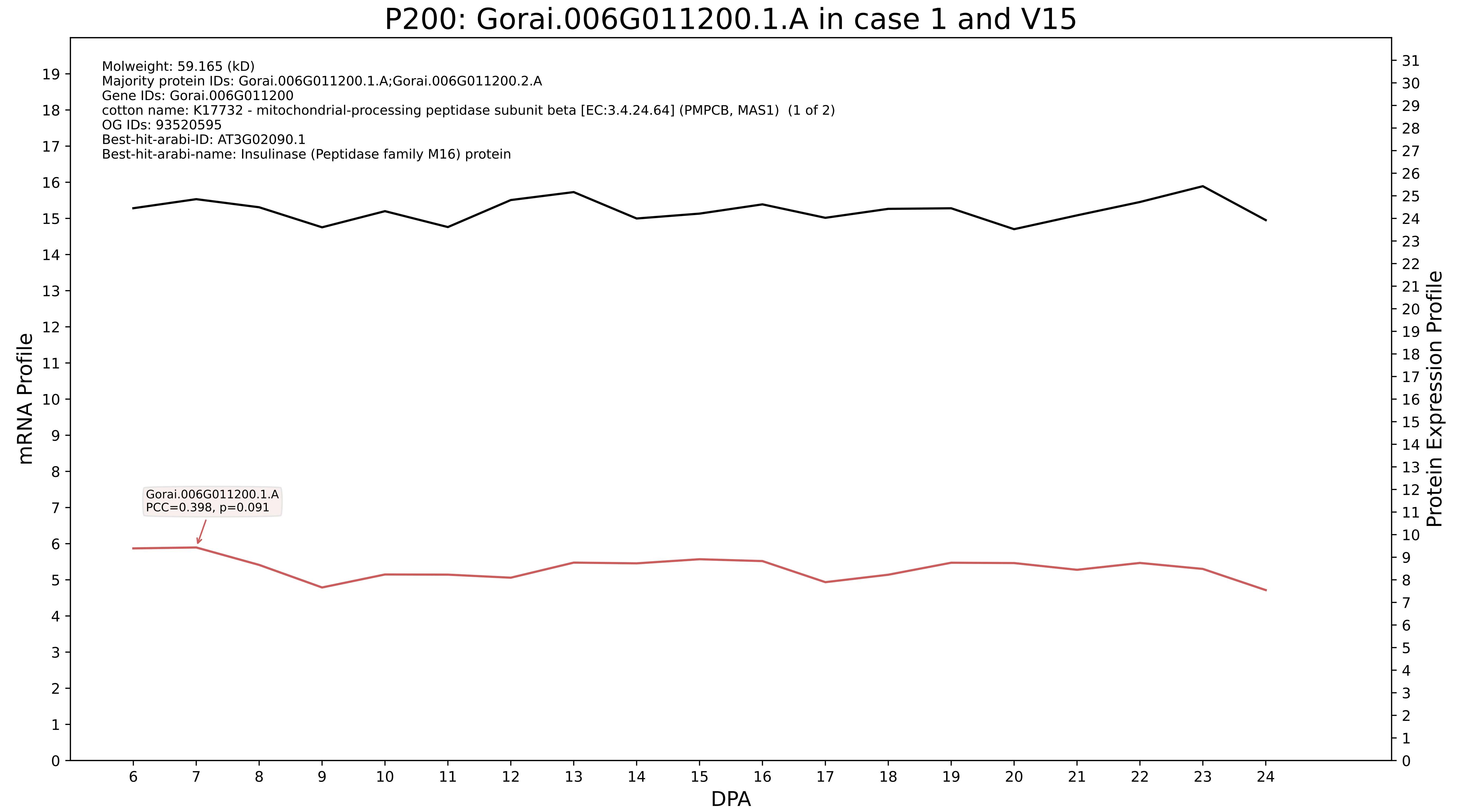
P200: Gorai.006G011200.1.A in case 1 and V15
Molweight: 59.165 (kD)
Majority protein IDs: Gorai.006G011200.1.A;Gorai.006G011200.2.A
Gene IDs: Gorai.006G011200
cotton name: K17732 - mitochondrial-processing peptidase subunit beta [EC:3.4.24.64] (PMPCB, MAS1) (1 of 2)
OG IDs: 93520595
Best-hit-arabi-ID: AT3G02090.1
Best-hit-arabi-name: Insulinase (Peptidase family M16) protein
Gorai.006G011200.1.A
PCC=0.398, p=0.091
mRNA Profile
Protein Expression Profile
DPA

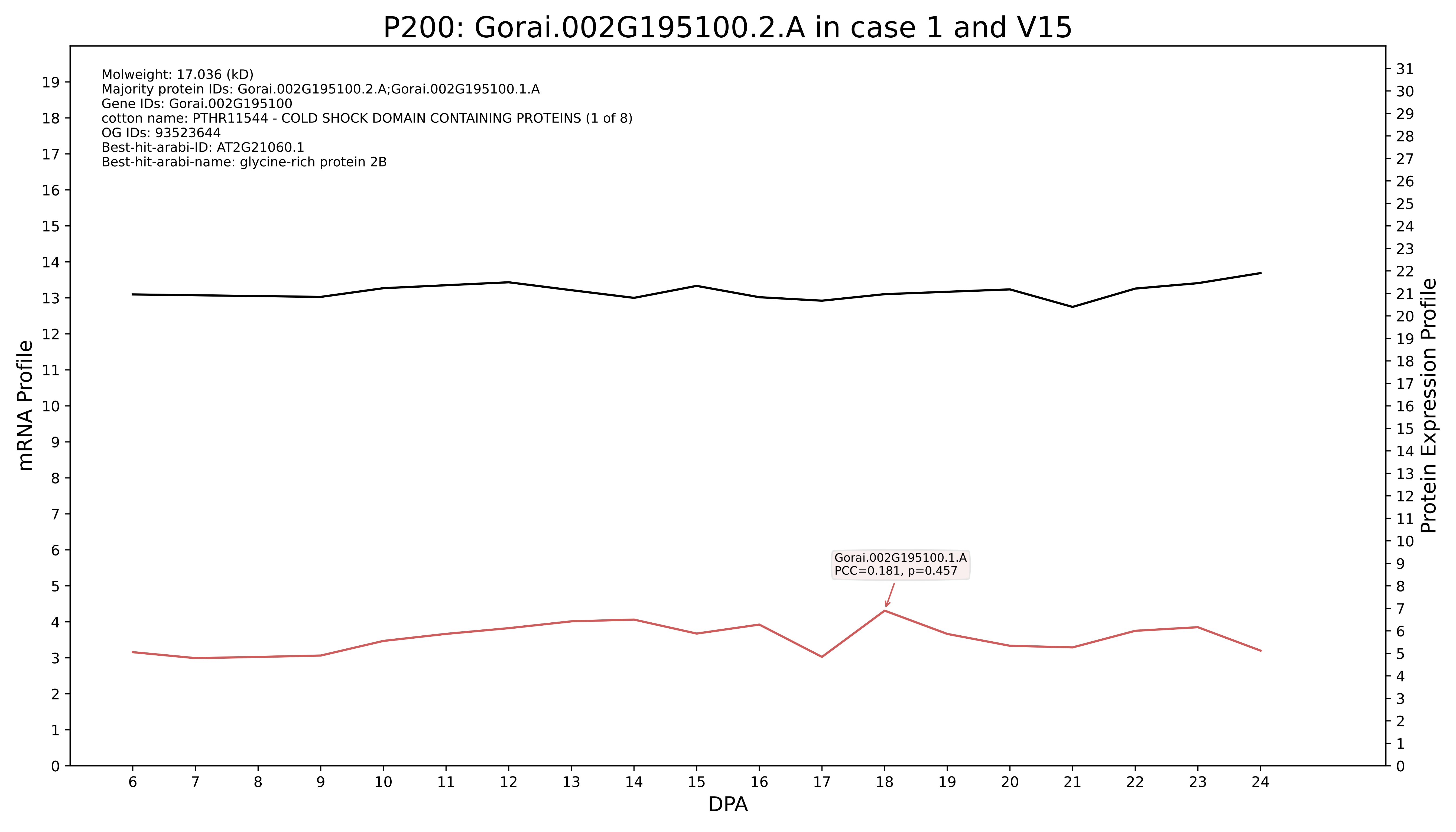

P200: Gorai.002G195100.2.A in case 1 and V15
Molweight: 17.036 (kD)
Majority protein IDs: Gorai.002G195100.2.A;Gorai.002G195100.1.A
Gene IDs: Gorai.002G195100
cotton name: PTHR11544 - COLD SHOCK DOMAIN CONTAINING PROTEINS (1 of 8)
OG IDs: 93523644
Best-hit-arabi-ID: AT2G21060.1
Best-hit-arabi-name: glycine-rich protein 2B
Gorai.002G195100.1.A
PCC=0.181, p=0.457
mRNA Profile
Protein Expression Profile
DPA

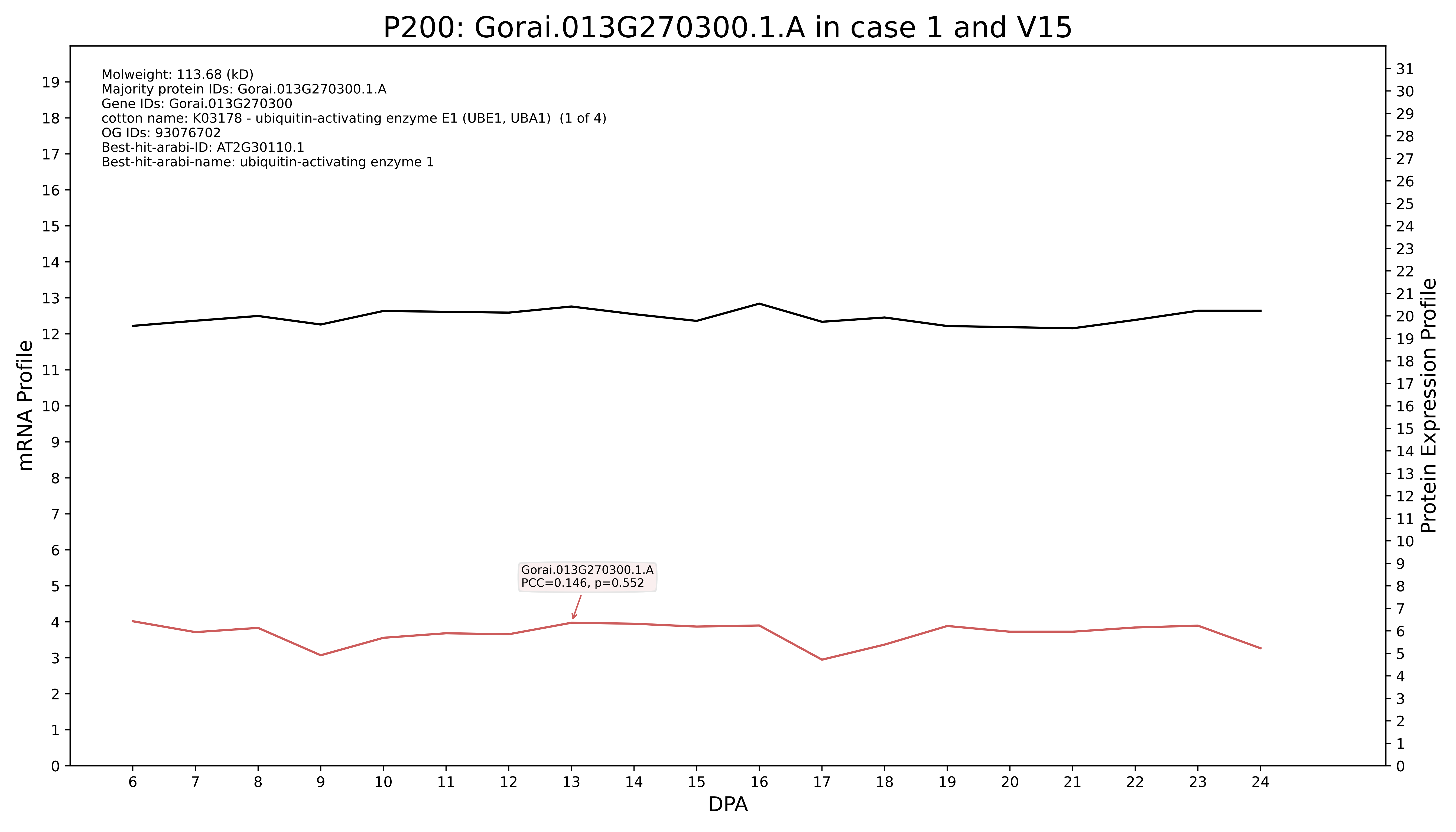
P200: Gorai.013G270300.1.A in case 1 and V15
Molweight: 113.68 (kD)
Majority protein IDs: Gorai.013G270300.1.A
Gene IDs: Gorai.013G270300
cotton name: K03178 - ubiquitin-activating enzyme E1 (UBE1, UBA1) (1 of 4)
OG IDs: 93076702
Best-hit-arabi-ID: AT2G30110.1
Best-hit-arabi-name: ubiquitin-activating enzyme 1
Gorai.013G270300.1.A
PCC=0.146, p=0.552
mRNA Profile
Protein Expression Profile
DPA

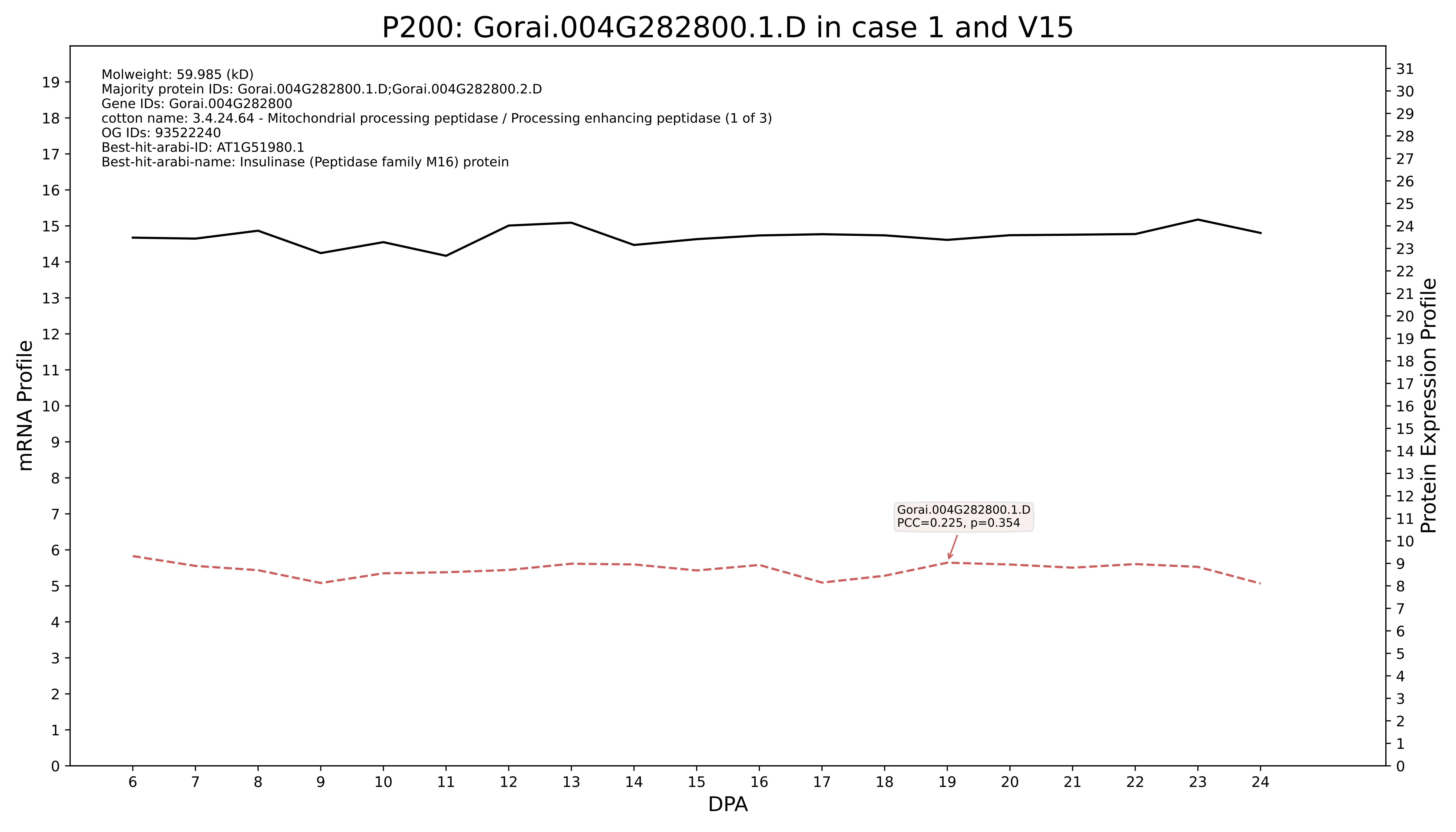

P200: Gorai.004G282800.1.D in case 1 and V15
Molweight: 59.985 (kD)
Majority protein IDs: Gorai.004G282800.1.D;Gorai.004G282800.2.D
Gene IDs: Gorai.004G282800
cotton name: 3.4.24.64 - Mitochondrial processing peptidase / Processing enhancing peptidase (1 of 3)
OG IDs: 93522240
Best-hit-arabi-ID: AT1G51980.1
Best-hit-arabi-name: Insulinase (Peptidase family M16) protein
Gorai.004G282800.1.D
PCC=0.225, p=0.354
mRNA Profile
Protein Expression Profile
DPA

P200: Gorai.009G194500.1.D in case 1 and V15

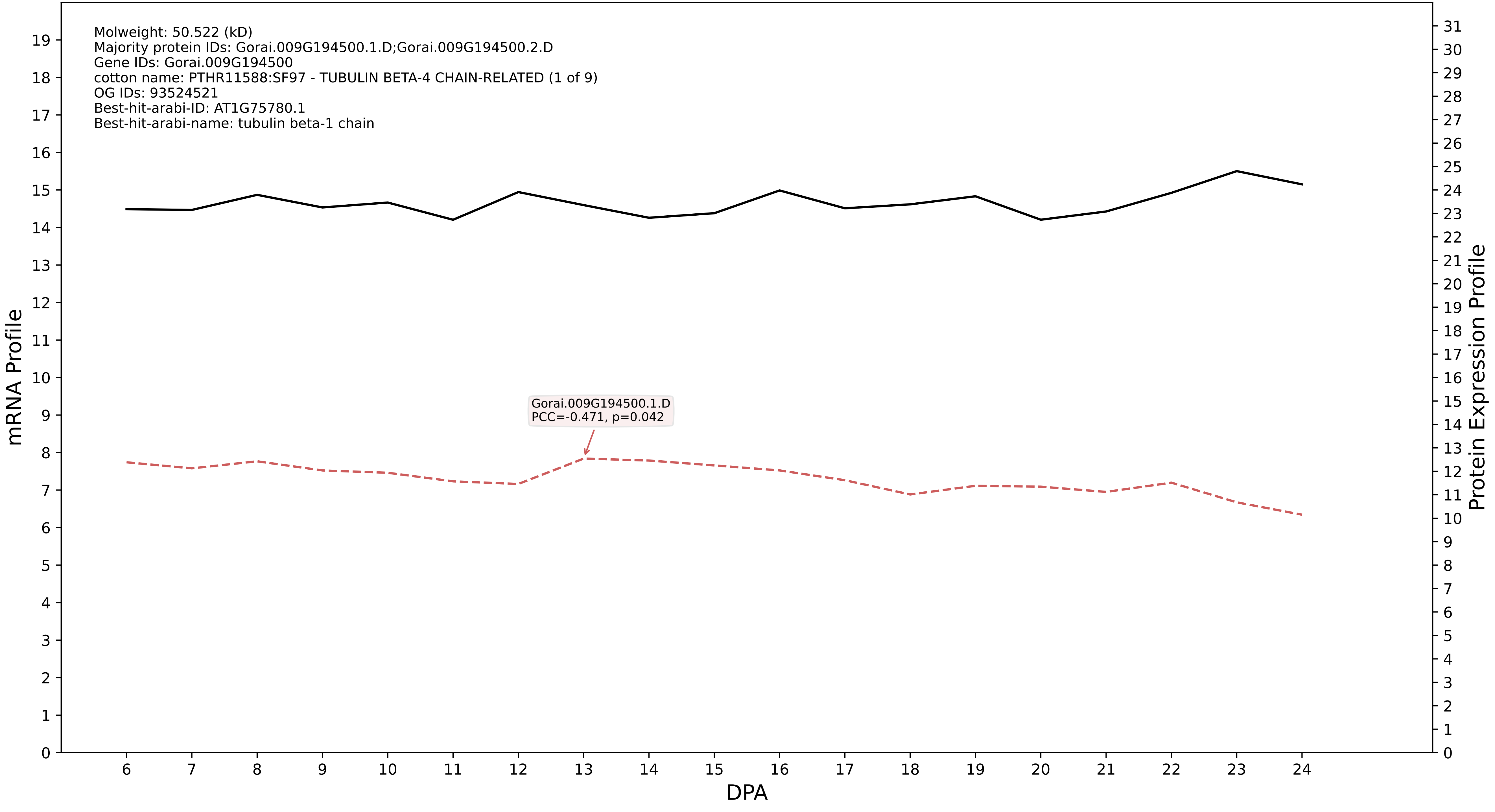

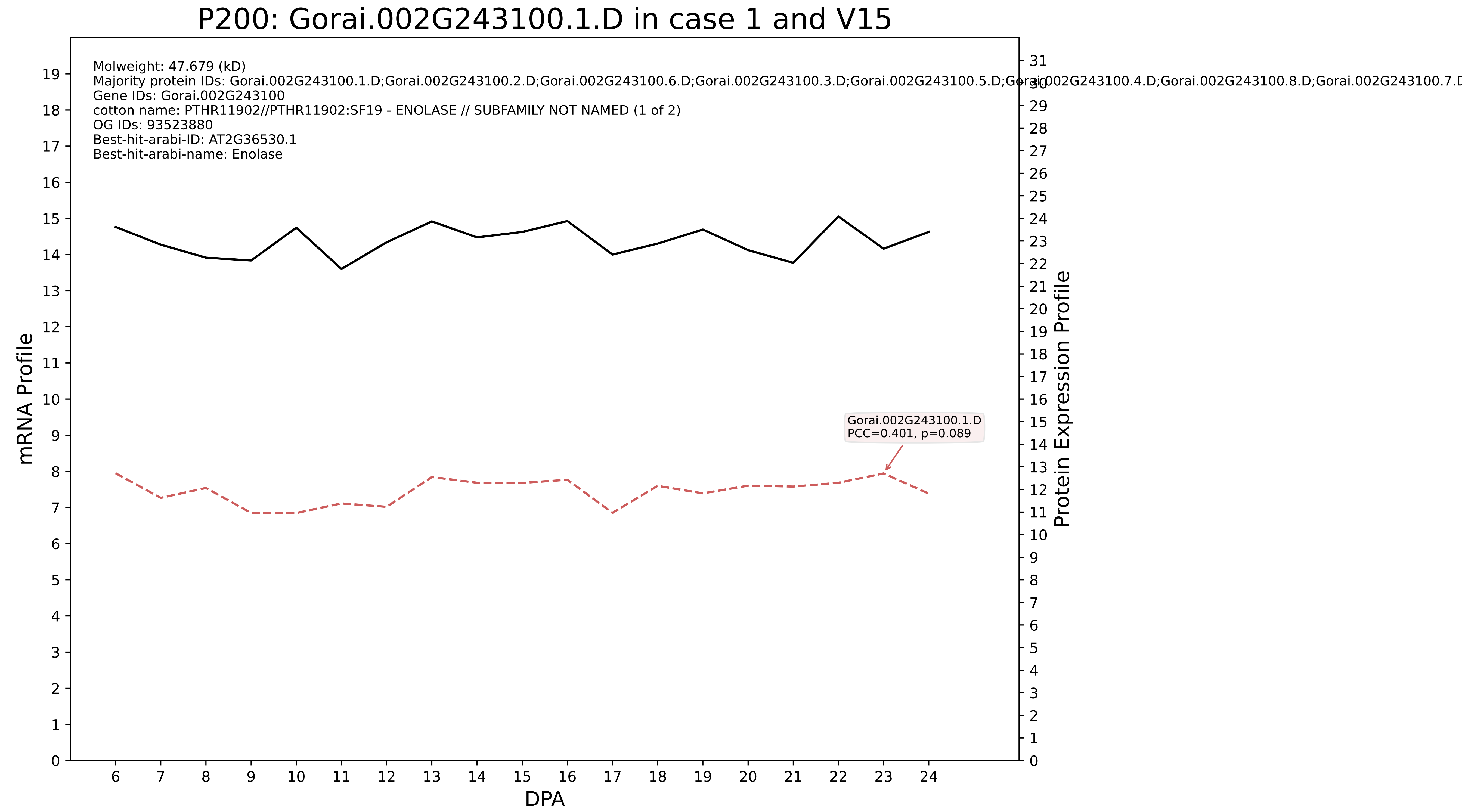

P200: Gorai.002G243100.1.D in case 1 and V15
Molweight: 47.679 (kD)
Majority protein IDs: Gorai.002G243100.1.D;Gorai.002G243100.2.D;Gorai.002G243100.6.D;Gorai.002G243100.3.D;Gorai.002G243100.5.D;Gorai.002G243100.4.D;Gorai.002G243100.8.D;Gorai.002G243100.7.D
Gene IDs: Gorai.002G243100
cotton name: PTHR11902//PTHR11902:SF19 - ENOLASE // SUBFAMILY NOT NAMED (1 of 2)
OG IDs: 93523880
Best-hit-arabi-ID: AT2G36530.1
Best-hit-arabi-name: Enolase
Gorai.002G243100.1.D
PCC=0.401, p=0.089
mRNA Profile
Protein Expression Profile
DPA

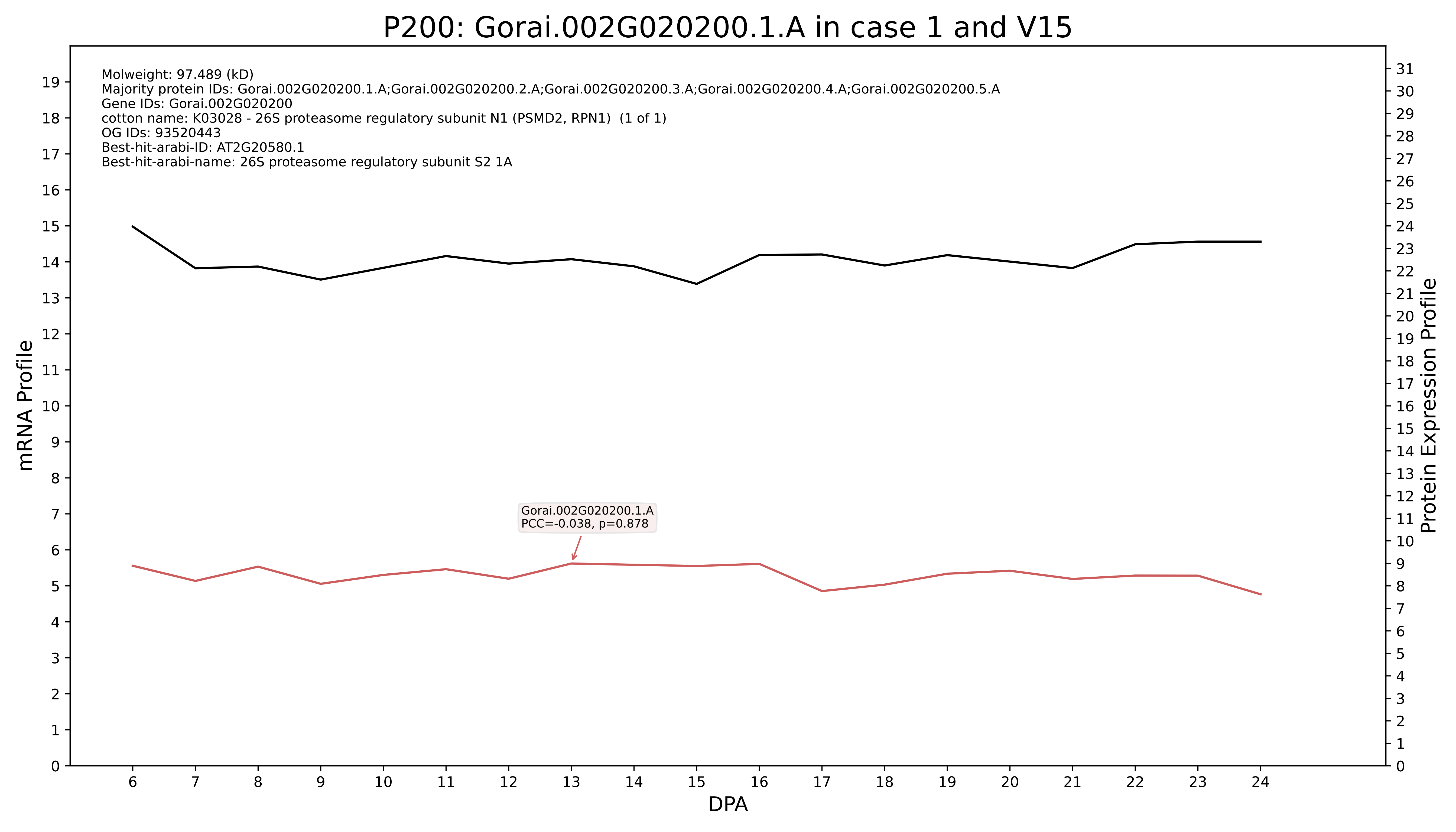
P200: Gorai.002G020200.1.A in case 1 and V15
Molweight: 97.489 (kD)
Majority protein IDs: Gorai.002G020200.1.A;Gorai.002G020200.2.A;Gorai.002G020200.3.A;Gorai.002G020200.4.A;Gorai.002G020200.5.A
Gene IDs: Gorai.002G020200
cotton name: K03028 - 26S proteasome regulatory subunit N1 (PSMD2, RPN1) (1 of 1)
OG IDs: 93520443
Best-hit-arabi-ID: AT2G20580.1
Best-hit-arabi-name: 26S proteasome regulatory subunit S2 1A
Gorai.002G020200.1.A
PCC=-0.038, p=0.878
mRNA Profile
Protein Expression Profile
DPA

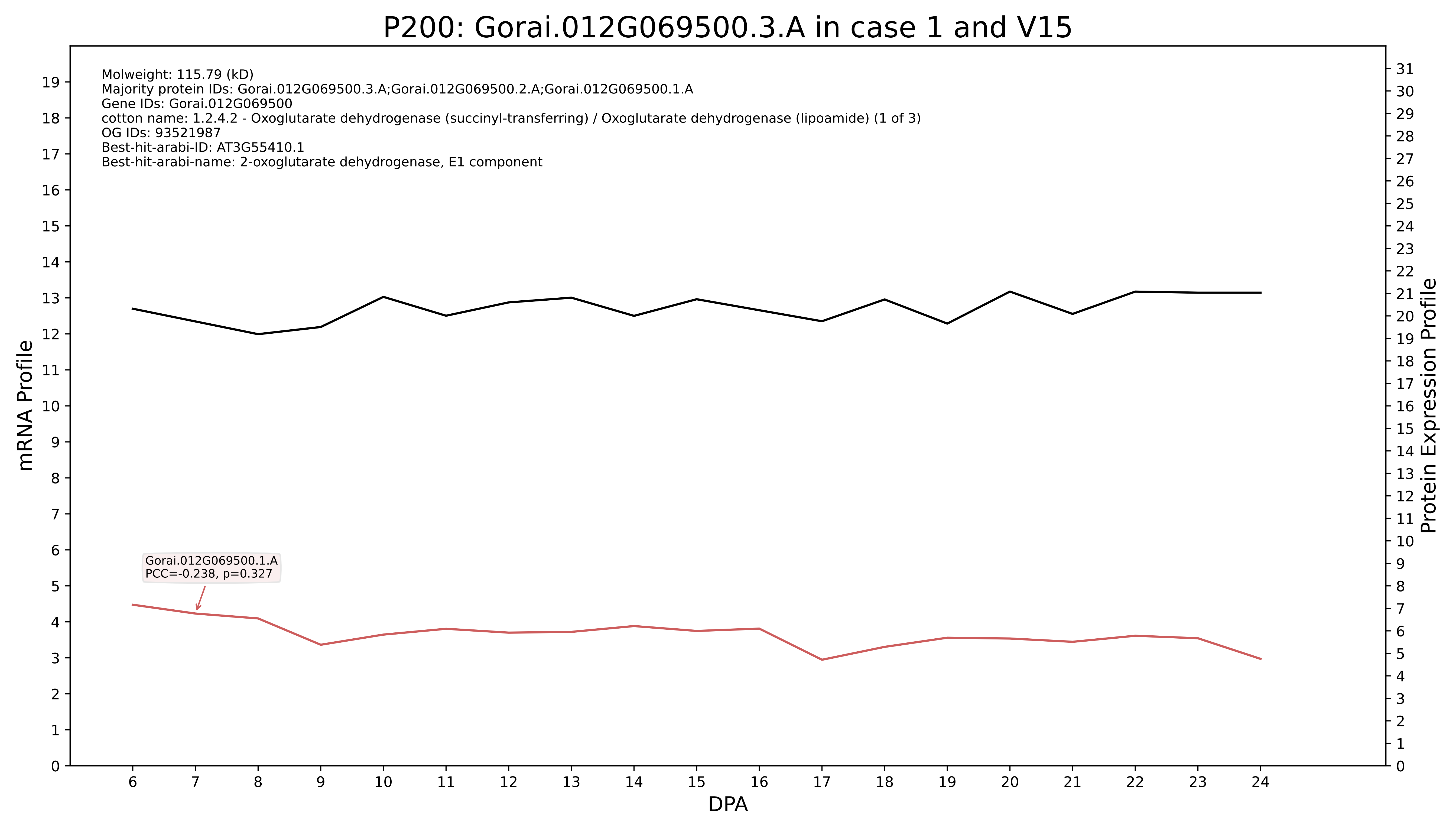
P200: Gorai.012G069500.3.A in case 1 and V15
Molweight: 115.79 (kD)
Majority protein IDs: Gorai.012G069500.3.A;Gorai.012G069500.2.A;Gorai.012G069500.1.A
Gene IDs: Gorai.012G069500
cotton name: 1.2.4.2 - Oxoglutarate dehydrogenase (succinyl-transferring) / Oxoglutarate dehydrogenase (lipoamide) (1 of 3)
OG IDs: 93521987
Best-hit-arabi-ID: AT3G55410.1
Best-hit-arabi-name: 2-oxoglutarate dehydrogenase, E1 component
Gorai.012G069500.1.A
PCC=-0.238, p=0.327
mRNA Profile
Protein Expression Profile
DPA

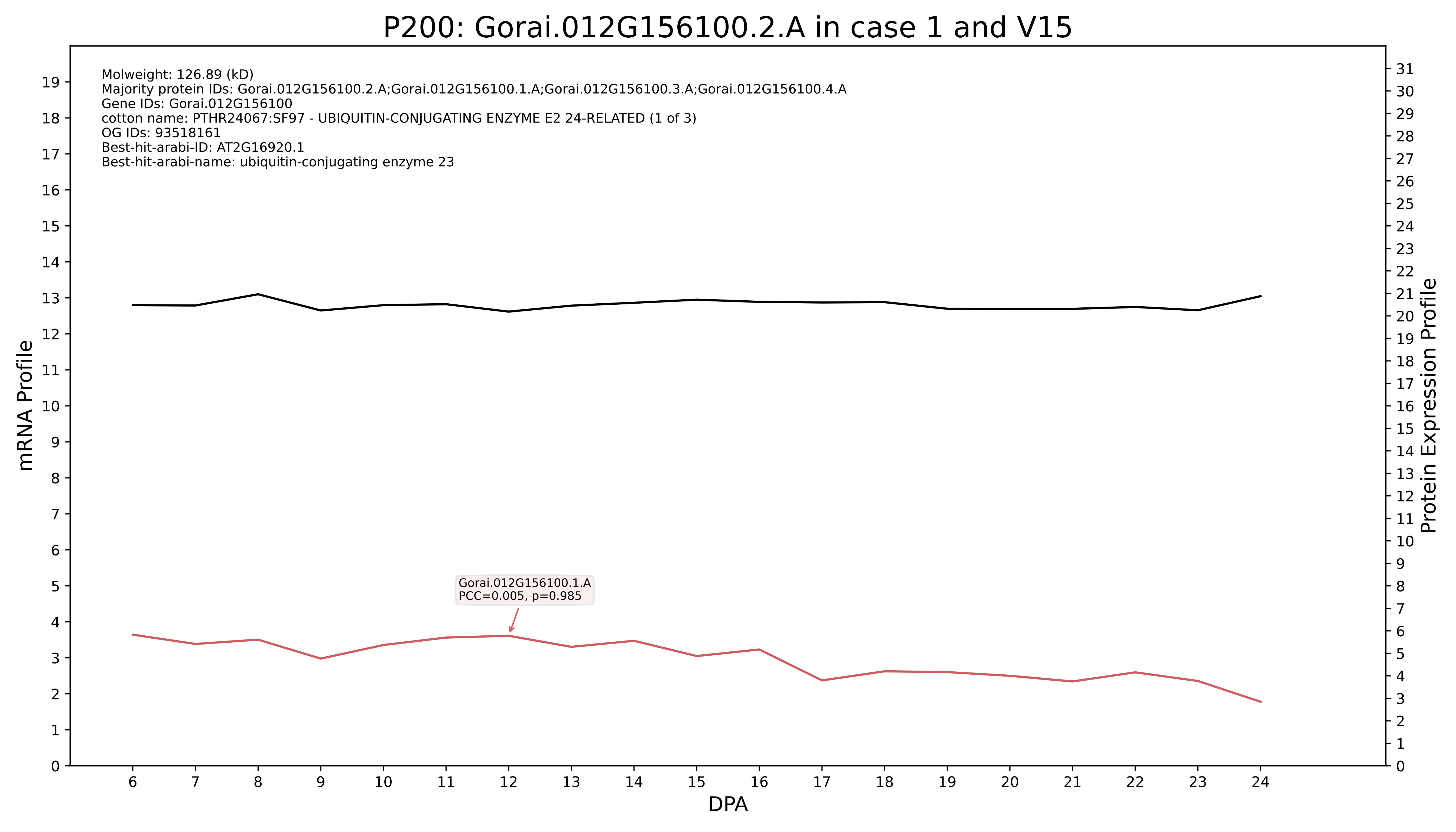
P200: Gorai.012G156100.2.A in case 1 and V15
Molweight: 126.89 (kD)
Majority protein IDs: Gorai.012G156100.2.A;Gorai.012G156100.1.A;Gorai.012G156100.3.A;Gorai.012G156100.4.A
Gene IDs: Gorai.012G156100
cotton name: PTHR24067:SF97 - UBIQUITIN-CONJUGATING ENZYME E2 24-RELATED (1 of 3)
OG IDs: 93518161
Best-hit-arabi-ID: AT2G16920.1
Best-hit-arabi-name: ubiquitin-conjugating enzyme 23
Gorai.012G156100.1.A
PCC=0.005, p=0.985
mRNA Profile
Protein Expression Profile
DPA

### P200: Gorai.001G197200.7.D in case 1 and V15

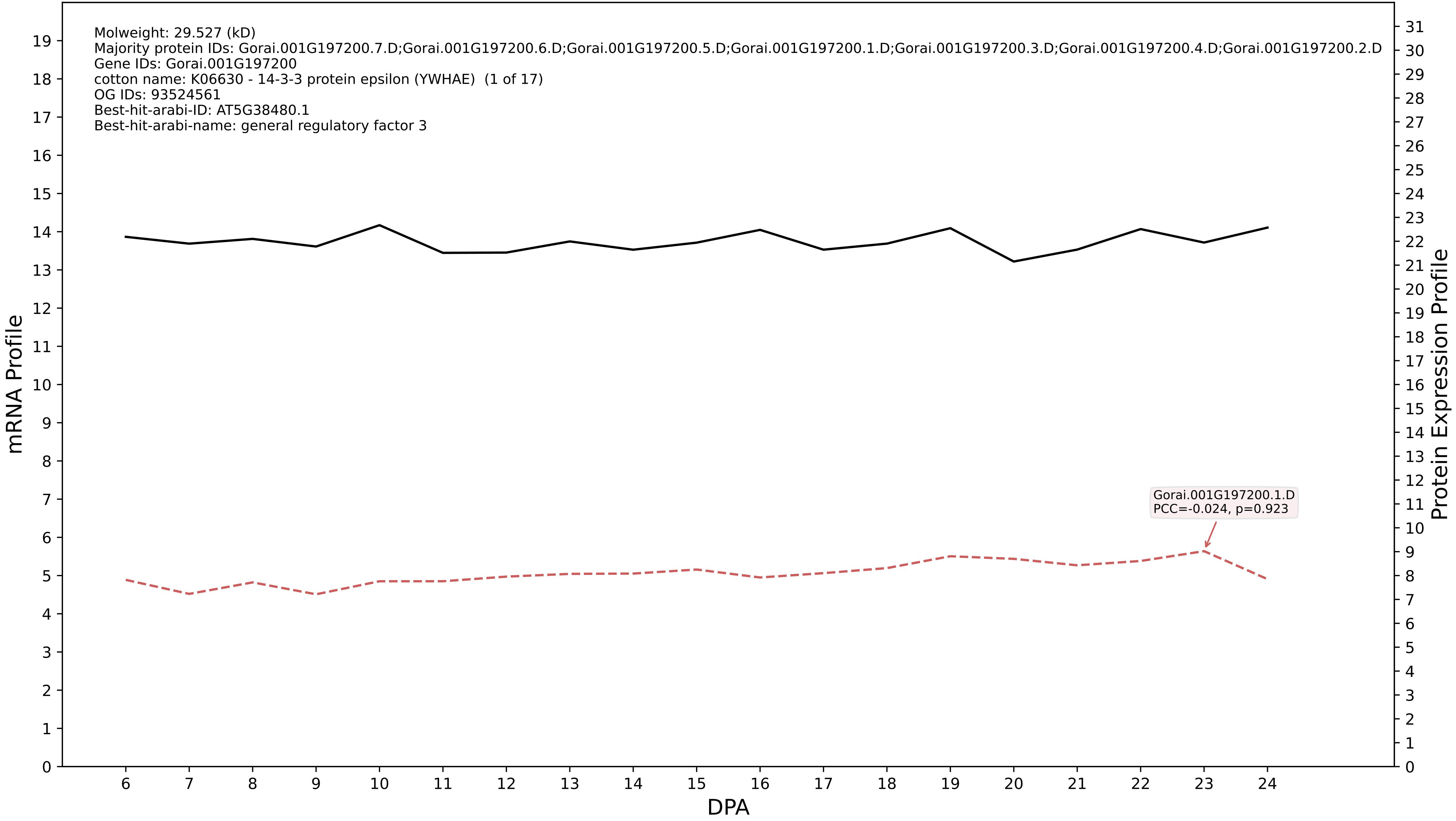

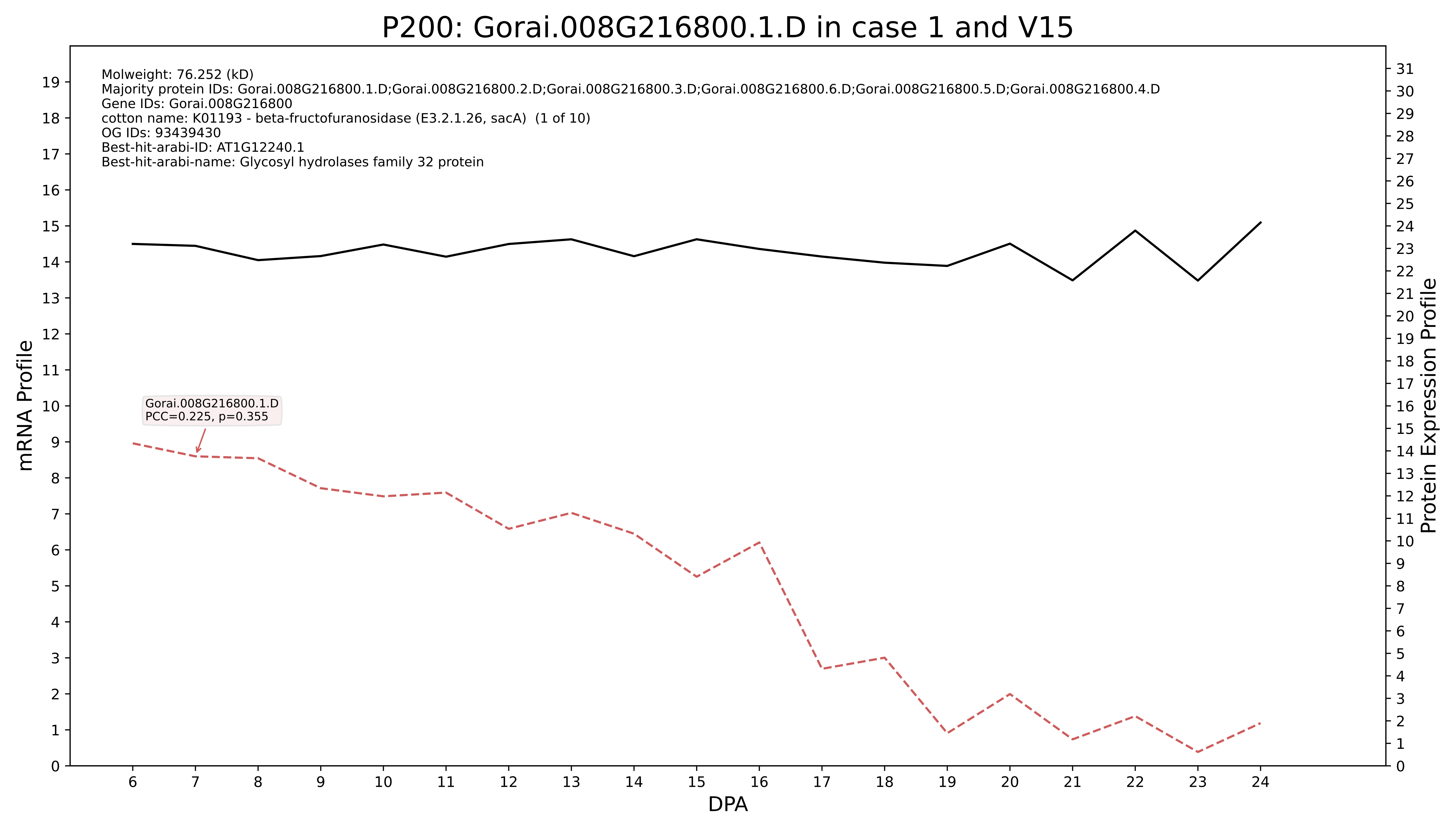

P200: Gorai.008G216800.1.D in case 1 and V15
Molweight: 76.252 (kD)
Majority protein IDs: Gorai.008G216800.1.D;Gorai.008G216800.2.D;Gorai.008G216800.3.D;Gorai.008G216800.6.D;Gorai.008G216800.5.D;Gorai.008G216800.4.D
Gene IDs: Gorai.008G216800
cotton name: K01193 - beta-fructofuranosidase (E3.2.1.26, sacA) (1 of 10)
OG IDs: 93439430
Best-hit-arabi-ID: AT1G12240.1
Best-hit-arabi-name: Glycosyl hydrolases family 32 protein
Gorai.008G216800.1.D
PCC=0.225, p=0.355
mRNA Profile
Protein Expression Profile
DPA

### P200: Gorai.011G080700.3.D in case 1 and V15

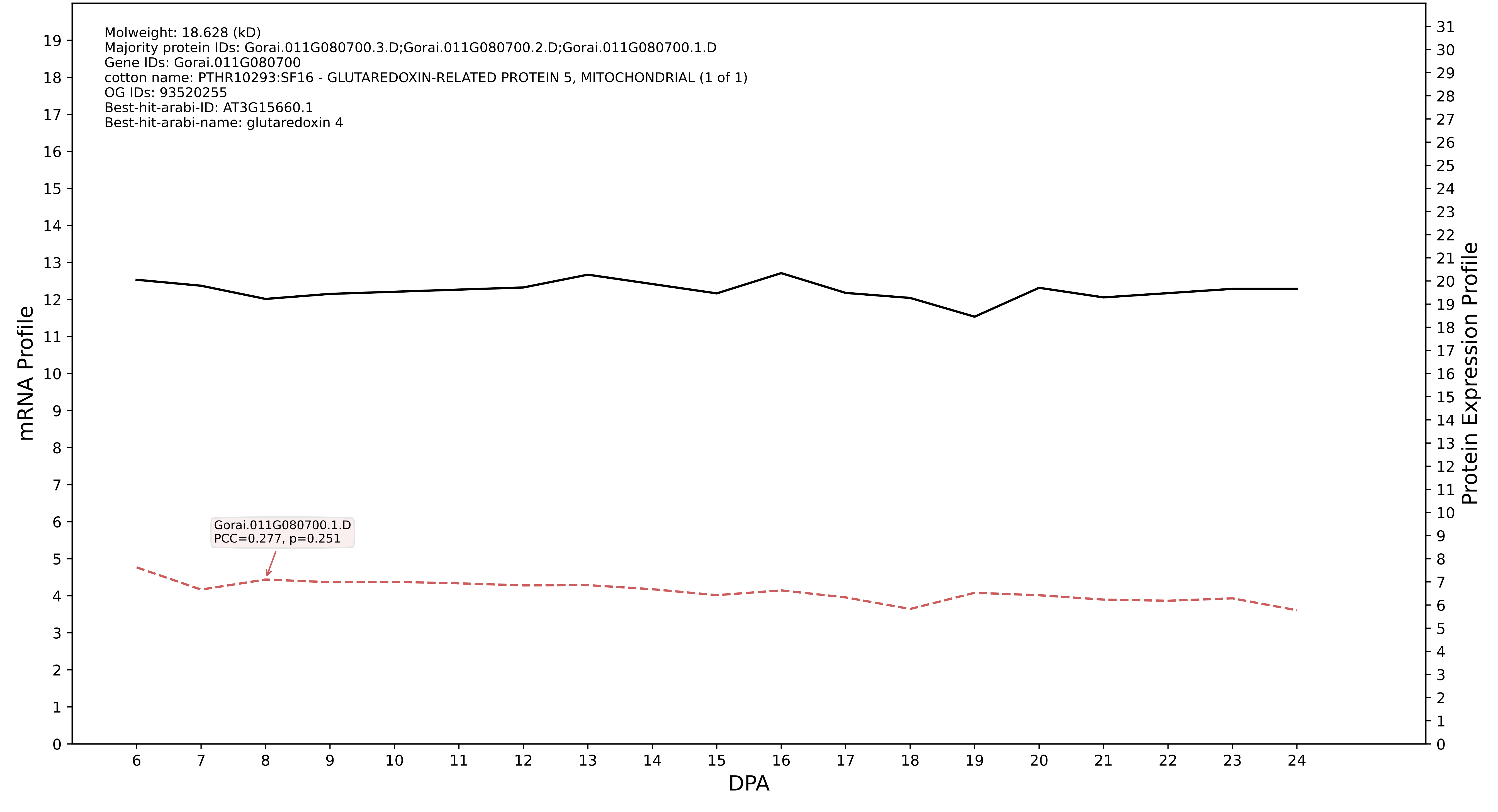

### P200: Gorai.009G000700.1.A in case 1 and V15

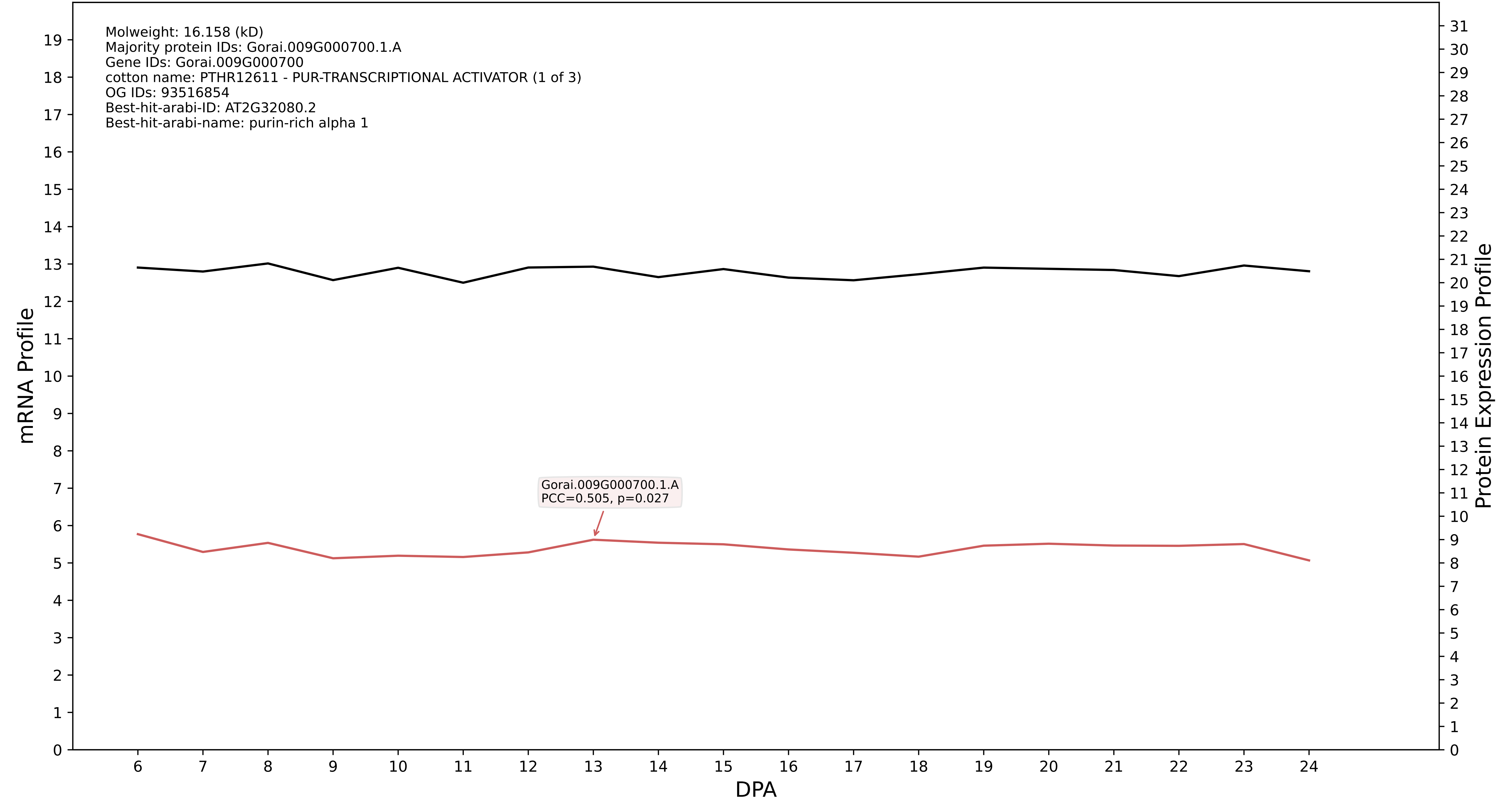

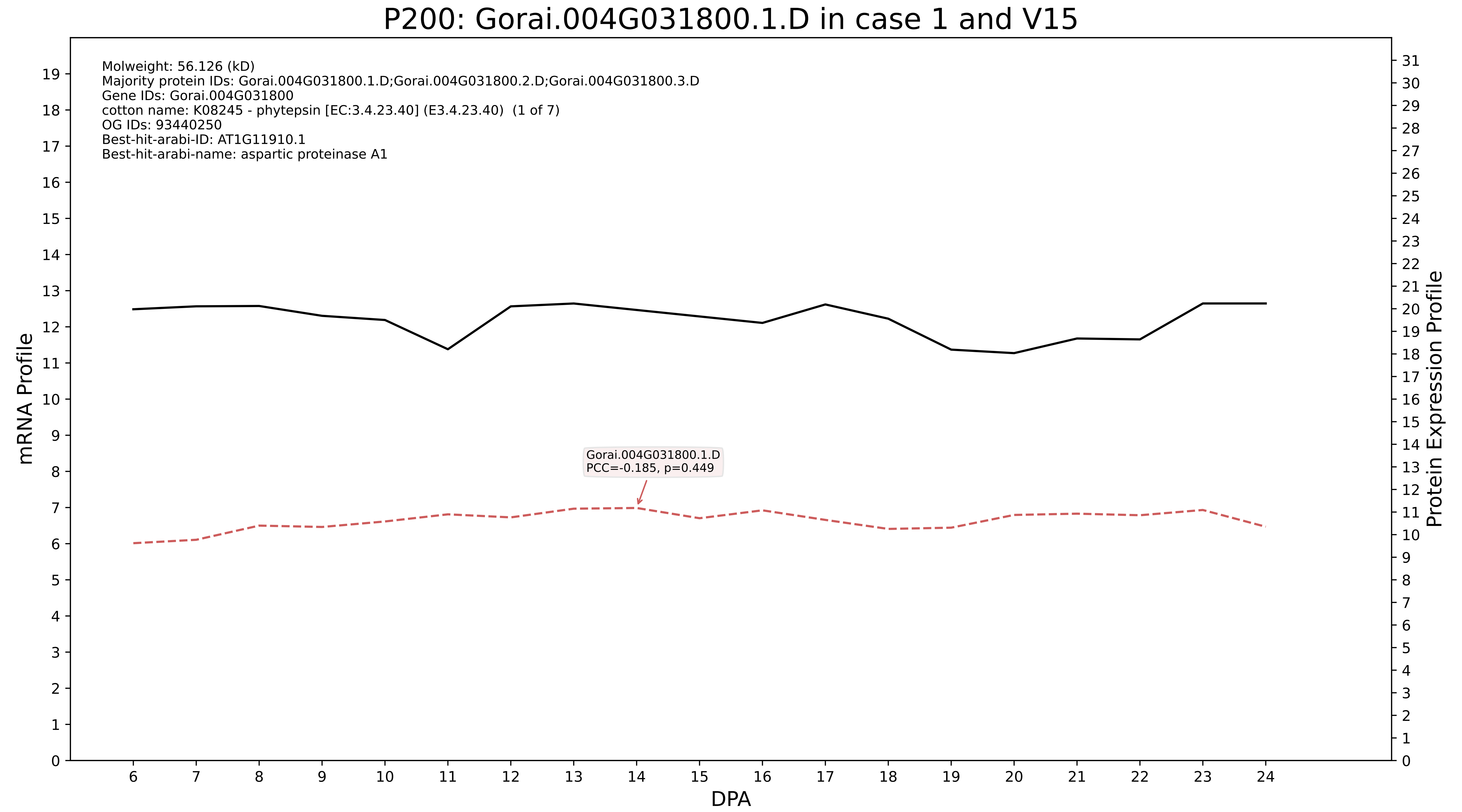
P200: Gorai.004G031800.1.D in case 1 and V15
Molweight: 56.126 (kD)
Majority protein IDs: Gorai.004G031800.1.D;Gorai.004G031800.2.D;Gorai.004G031800.3.D
Gene IDs: Gorai.004G031800
cotton name: K08245 - phytepsin [EC:3.4.23.40] (E3.4.23.40) (1 of 7)
OG IDs: 93440250
Best-hit-arabi-ID: AT1G11910.1
Best-hit-arabi-name: aspartic proteinase A1
Gorai.004G031800.1.D
PCC=-0.185, p=0.449
mRNA Profile
Protein Expression Profile
DPA

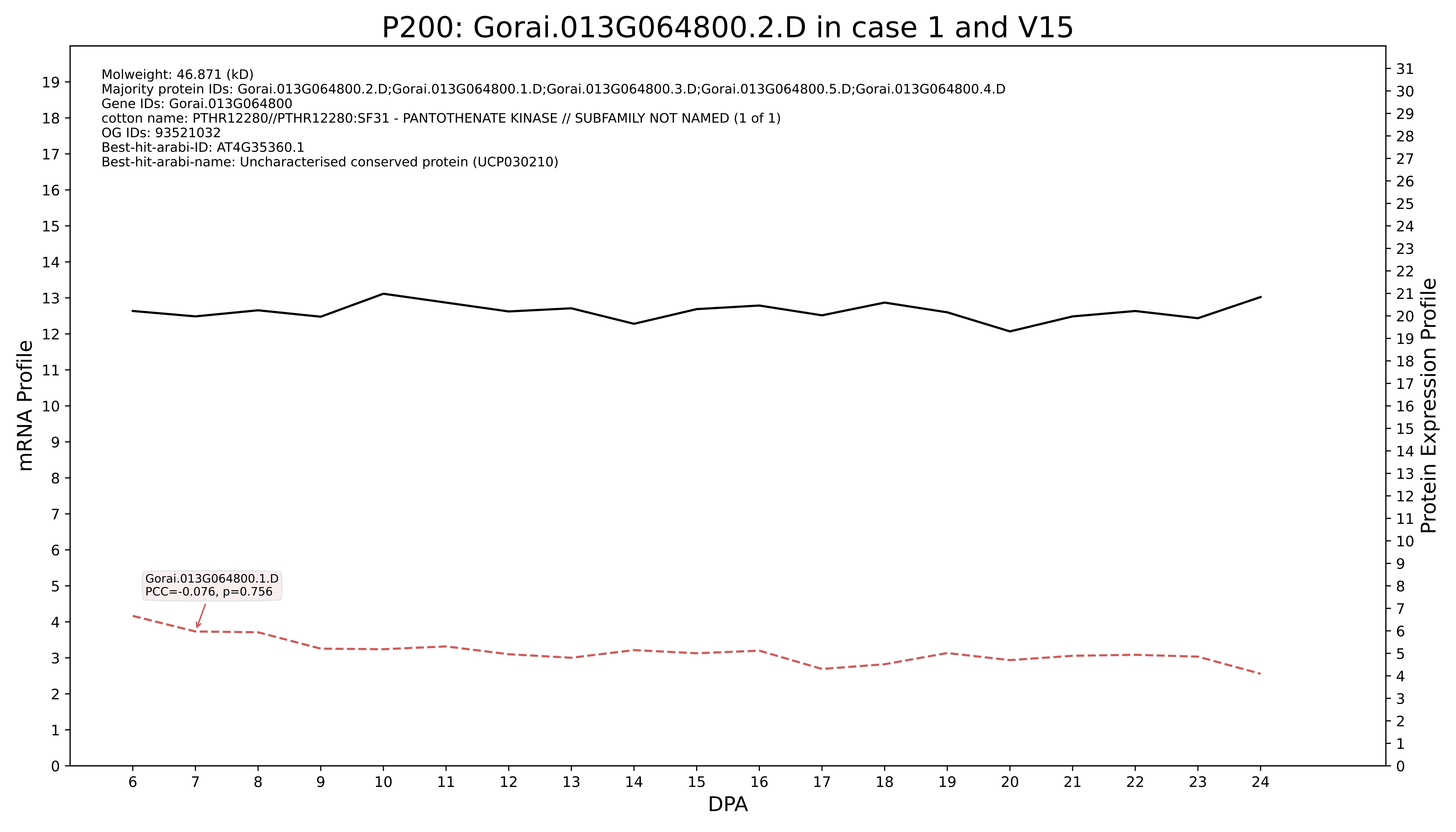
P200: Gorai.013G064800.2.D in case 1 and V15
Molweight: 46.871 (kD)
Majority protein IDs: Gorai.013G064800.2.D;Gorai.013G064800.1.D;Gorai.013G064800.3.D;Gorai.013G064800.5.D;Gorai.013G064800.4.D
Gene IDs: Gorai.013G064800
cotton name: PTHR12280//PTHR12280:SF31 - PANTOTHENATE KINASE // SUBFAMILY NOT NAMED (1 of 1)
OG IDs: 93521032
Best-hit-arabi-ID: AT4G35360.1
Best-hit-arabi-name: Uncharacterised conserved protein (UCP030210)
Gorai.013G064800.1.D
PCC=-0.076, p=0.756
mRNA Profile
Protein Expression Profile
DPA

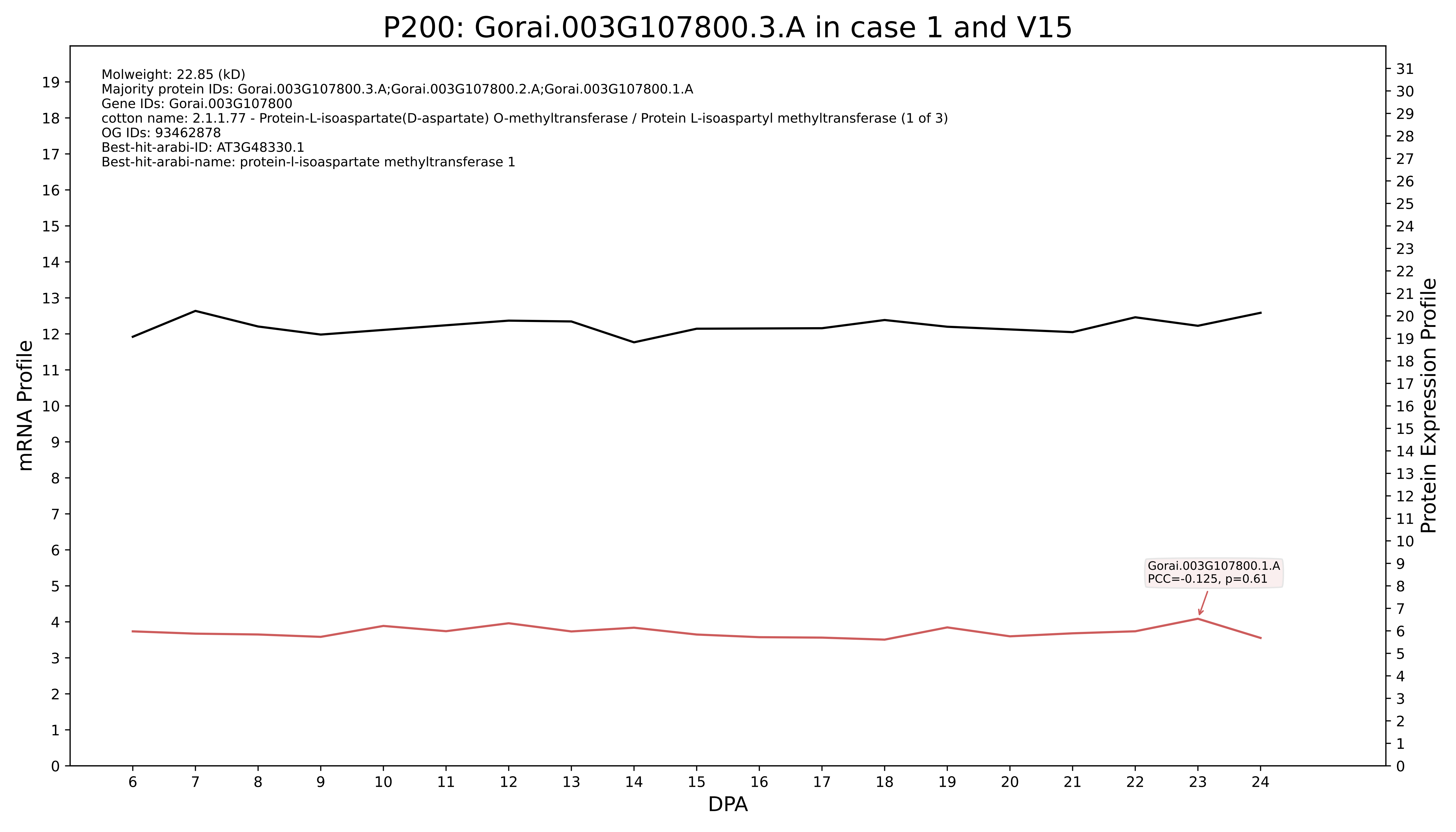
P200: Gorai.003G107800.3.A in case 1 and V15
Molweight: 22.85 (kD)
Majority protein IDs: Gorai.003G107800.3.A;Gorai.003G107800.2.A;Gorai.003G107800.1.A
Gene IDs: Gorai.003G107800
cotton name: 2.1.1.77 - Protein-L-isoaspartate(D-aspartate) O-methyltransferase / Protein L-isoaspartyl methyltransferase (1 of 3)
OG IDs: 93462878
Best-hit-arabi-ID: AT3G48330.1
Best-hit-arabi-name: protein-l-isoaspartate methyltransferase 1
Gorai.003G107800.1.A
PCC=-0.125, p=0.61
mRNA Profile
Protein Expression Profile
DPA

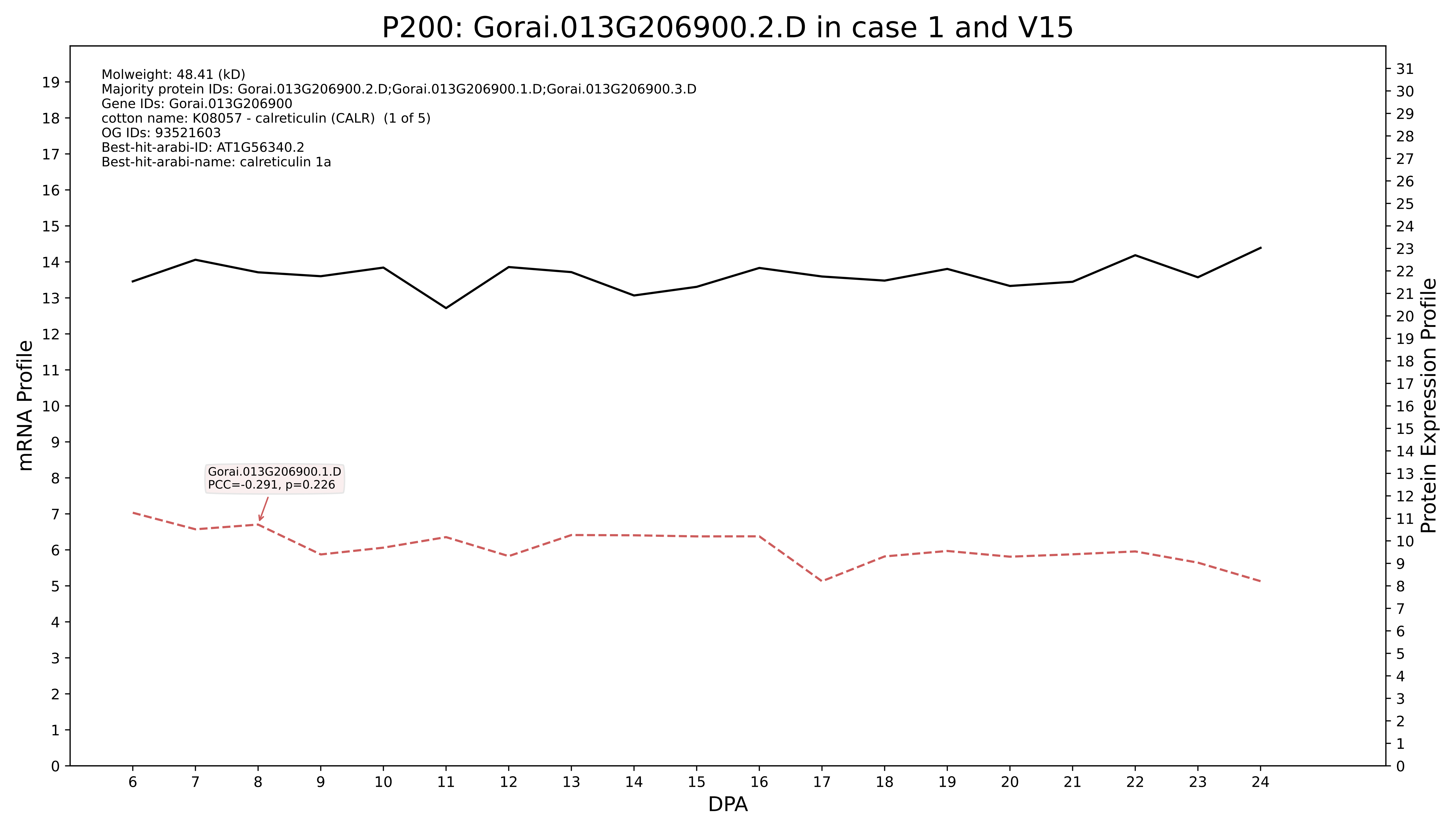
P200: Gorai.013G206900.2.D in case 1 and V15
Molweight: 48.41 (kD)
Majority protein IDs: Gorai.013G206900.2.D;Gorai.013G206900.1.D;Gorai.013G206900.3.D
Gene IDs: Gorai.013G206900
cotton name: K08057 - calreticulin (CALR) (1 of 5)
OG IDs: 93521603
Best-hit-arabi-ID: AT1G56340.2
Best-hit-arabi-name: calreticulin 1a
Gorai.013G206900.1.D
PCC=-0.291, p=0.226
mRNA Profile
Protein Expression Profile
DPA

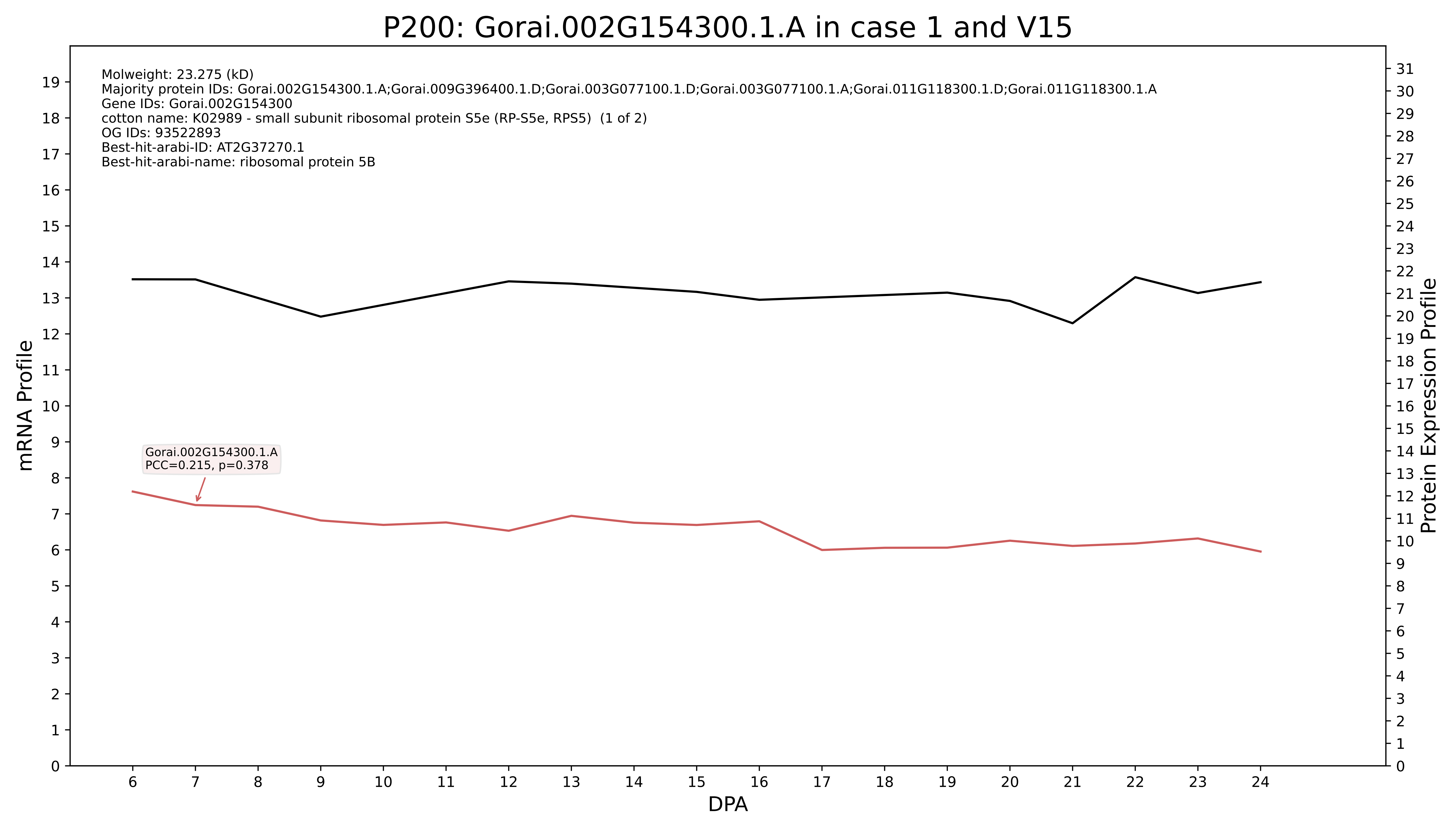

P200: Gorai.002G154300.1.A in case 1 and V15
Molweight: 23.275 (kD)
Majority protein IDs: Gorai.002G154300.1.A;Gorai.009G396400.1.D;Gorai.003G077100.1.D;Gorai.003G077100.1.A;Gorai.011G118300.1.D;Gorai.011G118300.1.A
Gene IDs: Gorai.002G154300
cotton name: K02989 - small subunit ribosomal protein S5e (RP-S5e, RPS5) (1 of 2)
OG IDs: 93522893
Best-hit-arabi-ID: AT2G37270.1
Best-hit-arabi-name: ribosomal protein 5B
Gorai.002G154300.1.A
PCC=0.215, p=0.378
mRNA Profile
Protein Expression Profile
DPA

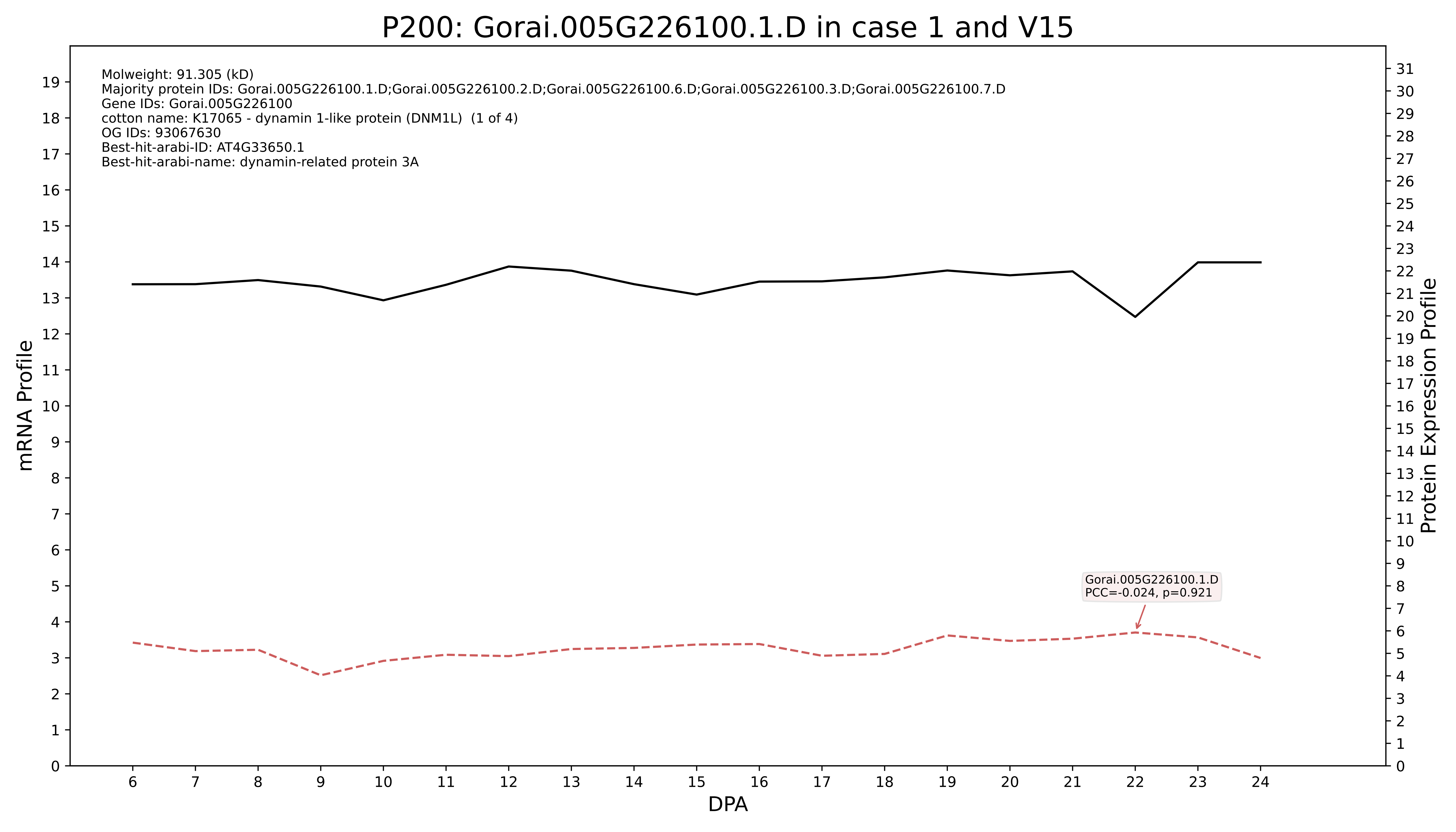
P200: Gorai.005G226100.1.D in case 1 and V15
Molweight: 91.305 (kD)
Majority protein IDs: Gorai.005G226100.1.D;Gorai.005G226100.2.D;Gorai.005G226100.6.D;Gorai.005G226100.3.D;Gorai.005G226100.7.D
Gene IDs: Gorai.005G226100
cotton name: K17065 - dynamin 1-like protein (DNM1L) (1 of 4)
OG IDs: 93067630
Best-hit-arabi-ID: AT4G33650.1
Best-hit-arabi-name: dynamin-related protein 3A
Gorai.005G226100.1.D
PCC=-0.024, p=0.921
mRNA Profile
Protein Expression Profile
DPA

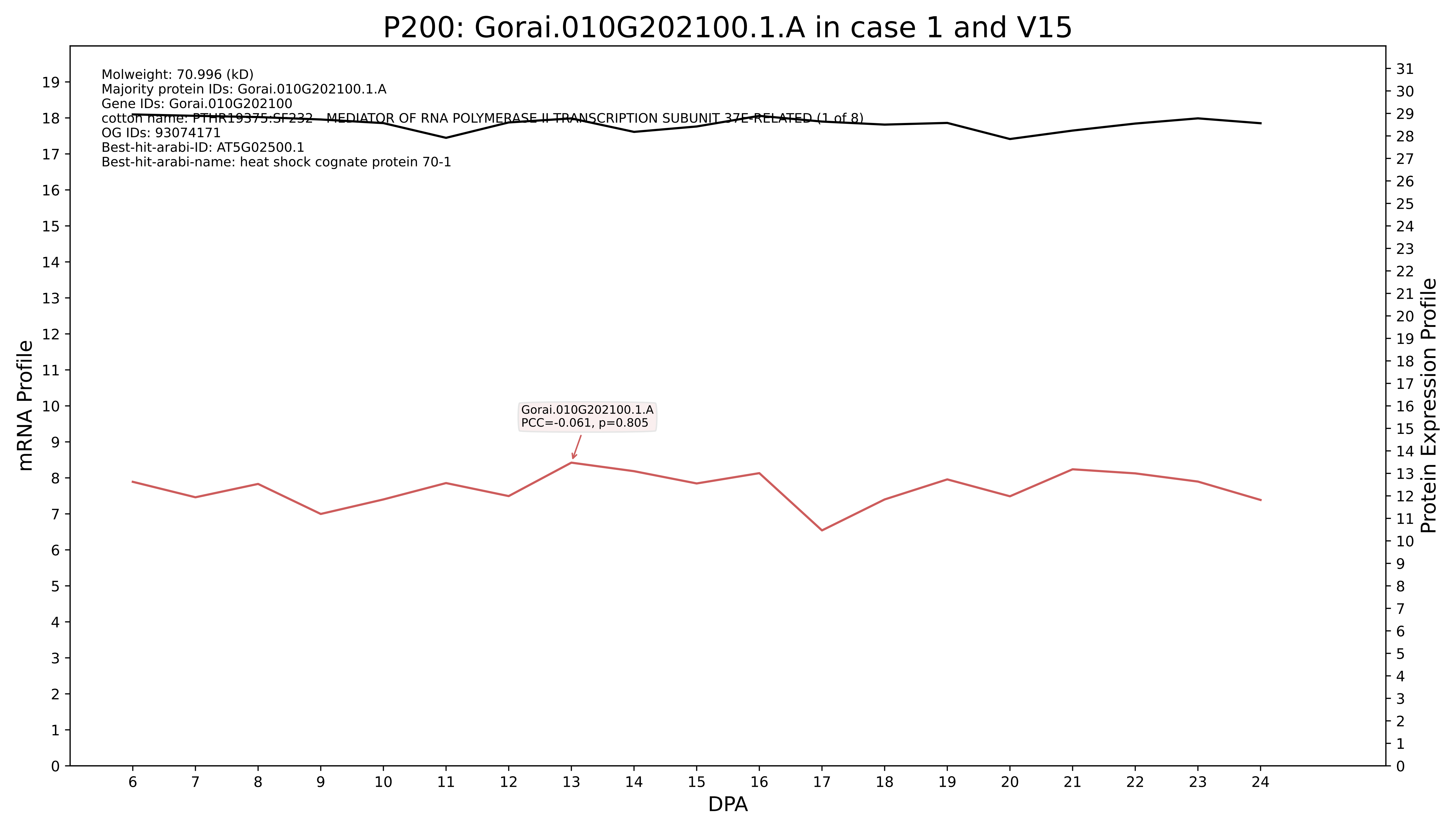
P200: Gorai.010G202100.1.A in case 1 and V15
Molweight: 70.996 (kD)
Majority protein IDs: Gorai.010G202100.1.A
Gene IDs: Gorai.010G202100
cotton name: PTHR19375:SF232 - MEDIATOR OF RNA POLYMERASE II TRANSCRIPTION SUBUNIT 37E-RELATED (1 of 8)
OG IDs: 93074171
Best-hit-arabi-ID: AT5G02500.1
Best-hit-arabi-name: heat shock cognate protein 70-1
Gorai.010G202100.1.A
PCC=-0.061, p=0.805
mRNA Profile
Protein Expression Profile
DPA

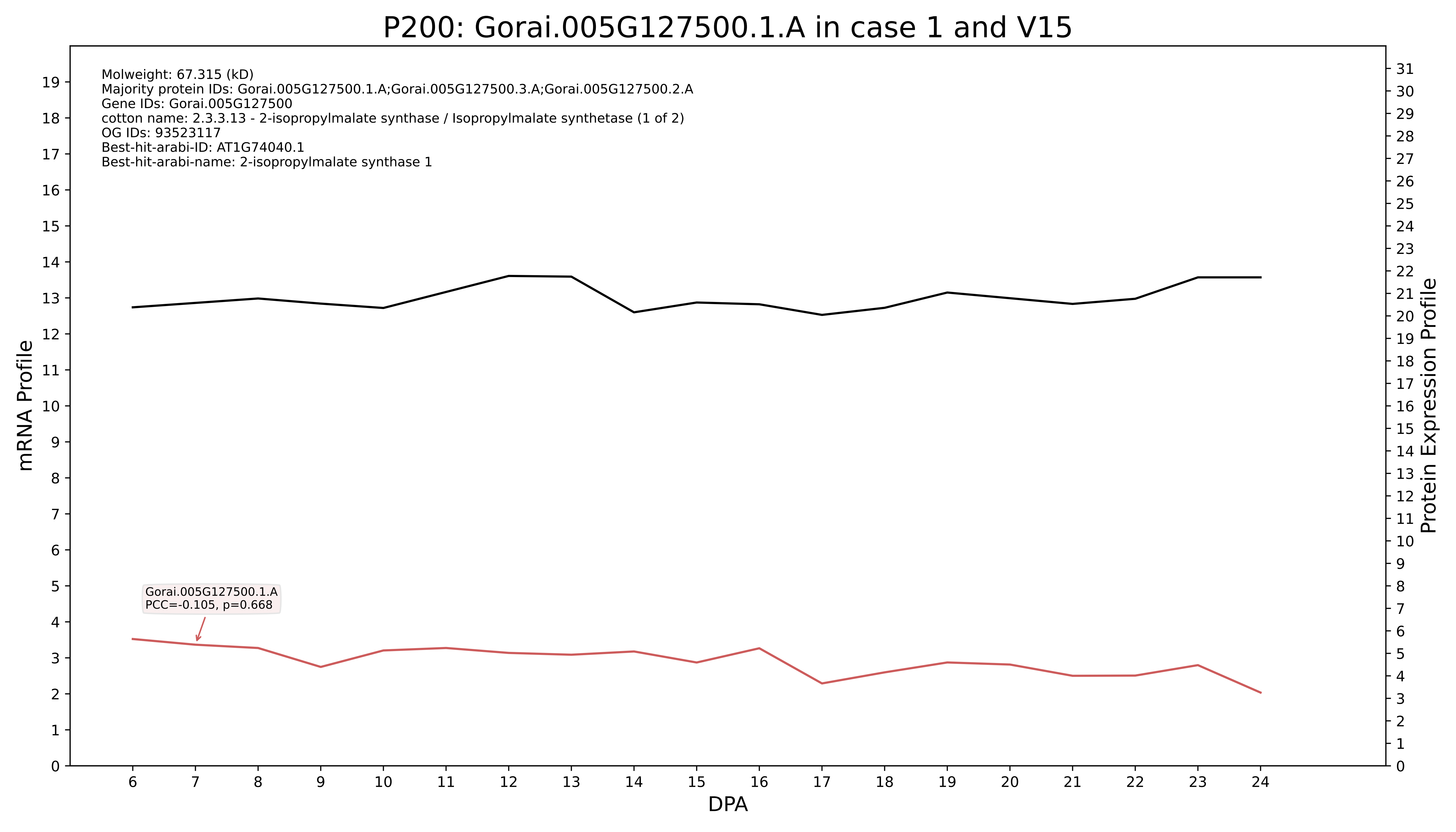
P200: Gorai.005G127500.1.A in case 1 and V15
Molweight: 67.315 (kD)
Majority protein IDs: Gorai.005G127500.1.A;Gorai.005G127500.3.A;Gorai.005G127500.2.A
Gene IDs: Gorai.005G127500
cotton name: 2.3.3.13 - 2-isopropylmalate synthase / Isopropylmalate synthetase (1 of 2)
OG IDs: 93523117
Best-hit-arabi-ID: AT1G74040.1
Best-hit-arabi-name: 2-isopropylmalate synthase 1
Gorai.005G127500.1.A
PCC=-0.105, p=0.668
mRNA Profile
Protein Expression Profile
DPA

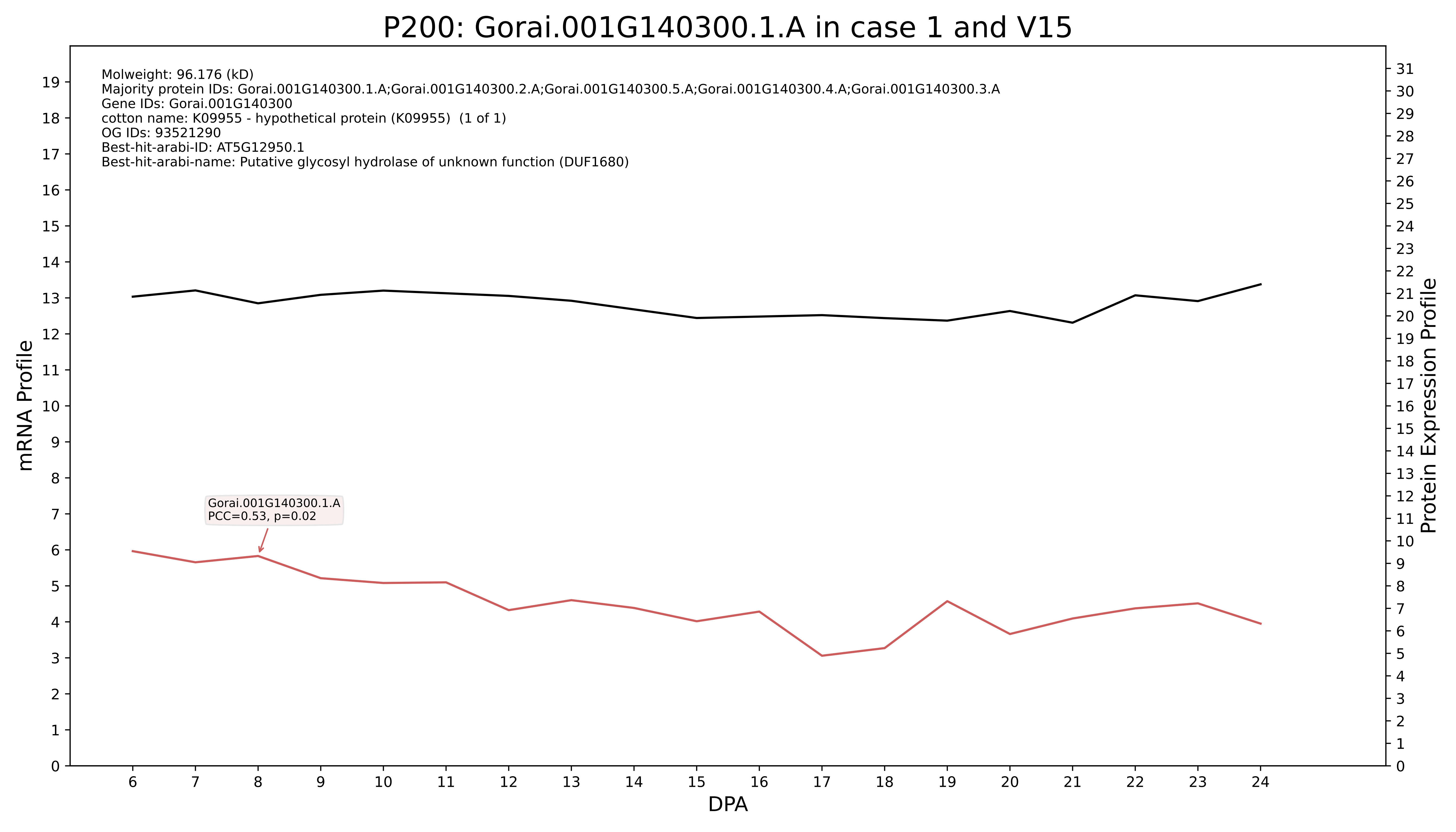
P200: Gorai.001G140300.1.A in case 1 and V15
Molweight: 96.176 (kD)
Majority protein IDs: Gorai.001G140300.1.A;Gorai.001G140300.2.A;Gorai.001G140300.5.A;Gorai.001G140300.4.A;Gorai.001G140300.3.A
Gene IDs: Gorai.001G140300
cotton name: K09955 - hypothetical protein (K09955) (1 of 1)
OG IDs: 93521290
Best-hit-arabi-ID: AT5G12950.1
Best-hit-arabi-name: Putative glycosyl hydrolase of unknown function (DUF1680)
Gorai.001G140300.1.A
PCC=0.53, p=0.02
mRNA Profile
Protein Expression Profile
DPA

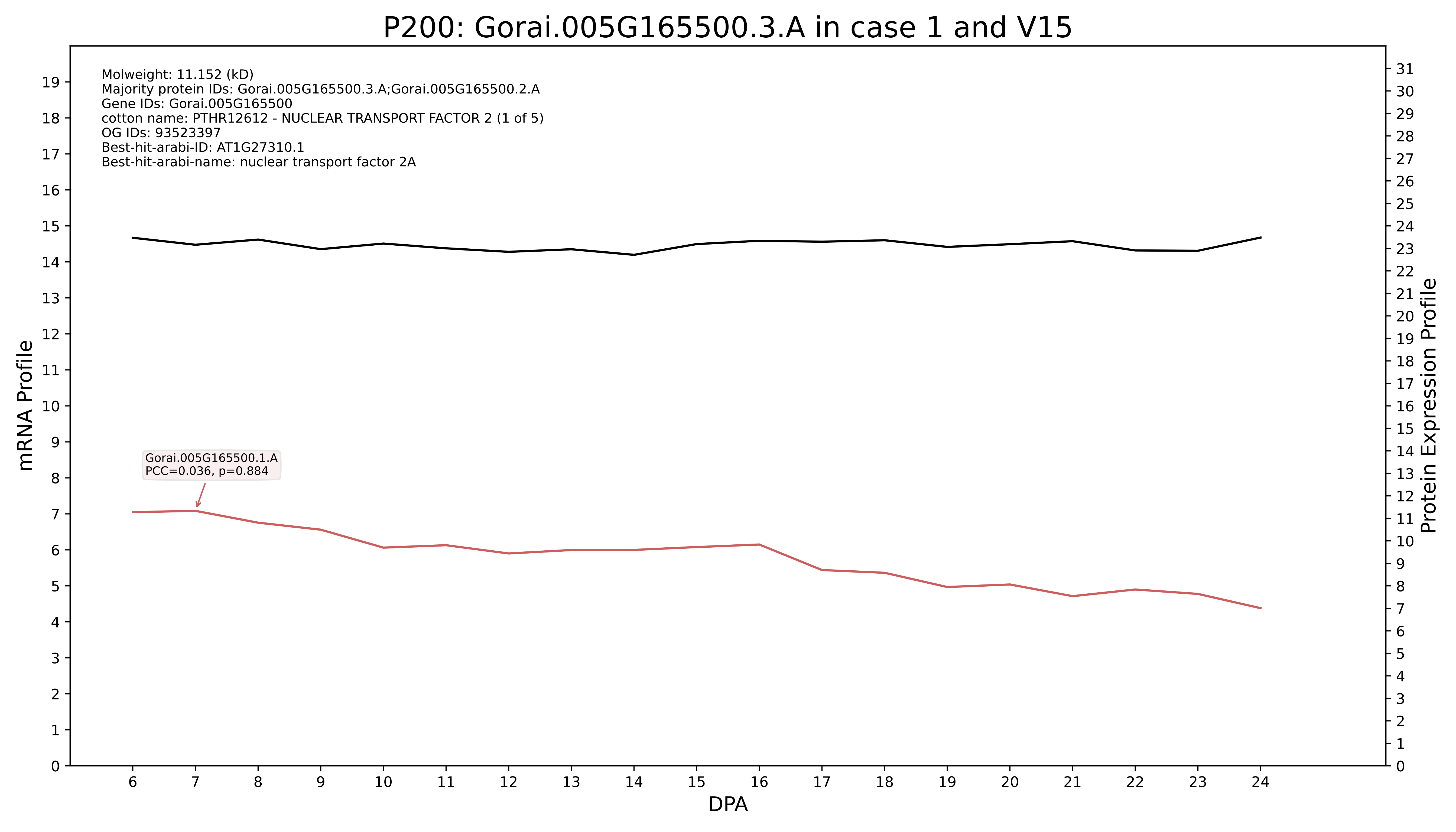
P200: Gorai.005G165500.3.A in case 1 and V15
Molweight: 11.152 (kD)
Majority protein IDs: Gorai.005G165500.3.A;Gorai.005G165500.2.A
Gene IDs: Gorai.005G165500
cotton name: PTHR12612 - NUCLEAR TRANSPORT FACTOR 2 (1 of 5)
OG IDs: 93523397
Best-hit-arabi-ID: AT1G27310.1
Best-hit-arabi-name: nuclear transport factor 2A
Gorai.005G165500.1.A
PCC=0.036, p=0.884
mRNA Profile
Protein Expression Profile
DPA

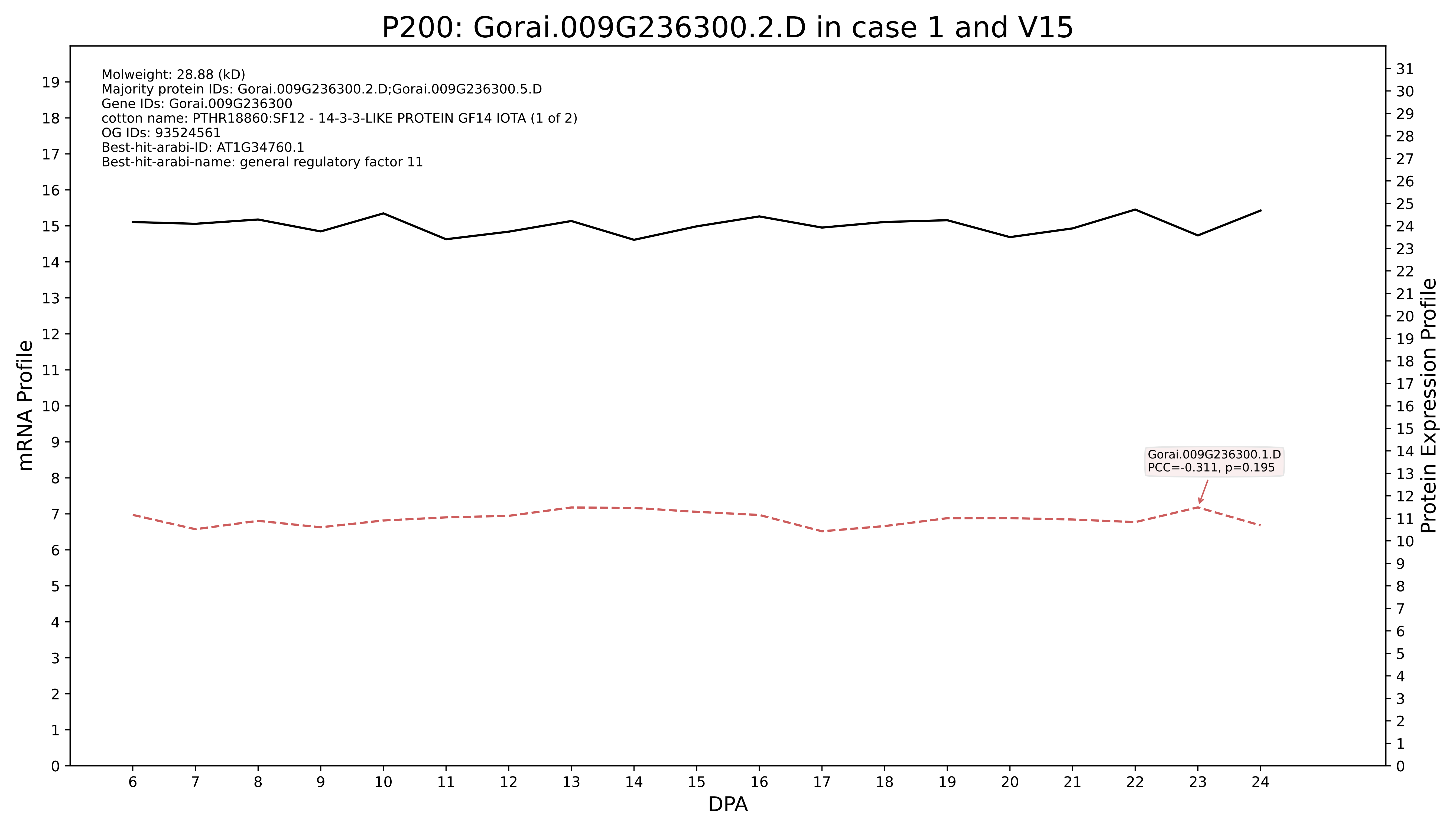
P200: Gorai.009G236300.2.D in case 1 and V15
Molweight: 28.88 (kD)
Majority protein IDs: Gorai.009G236300.2.D;Gorai.009G236300.5.D
Gene IDs: Gorai.009G236300
cotton name: PTHR18860:SF12 - 14-3-3-LIKE PROTEIN GF14 IOTA (1 of 2)
OG IDs: 93524561
Best-hit-arabi-ID: AT1G34760.1
Best-hit-arabi-name: general regulatory factor 11
Gorai.009G236300.1.D
PCC=-0.311, p=0.195
mRNA Profile
Protein Expression Profile
DPA

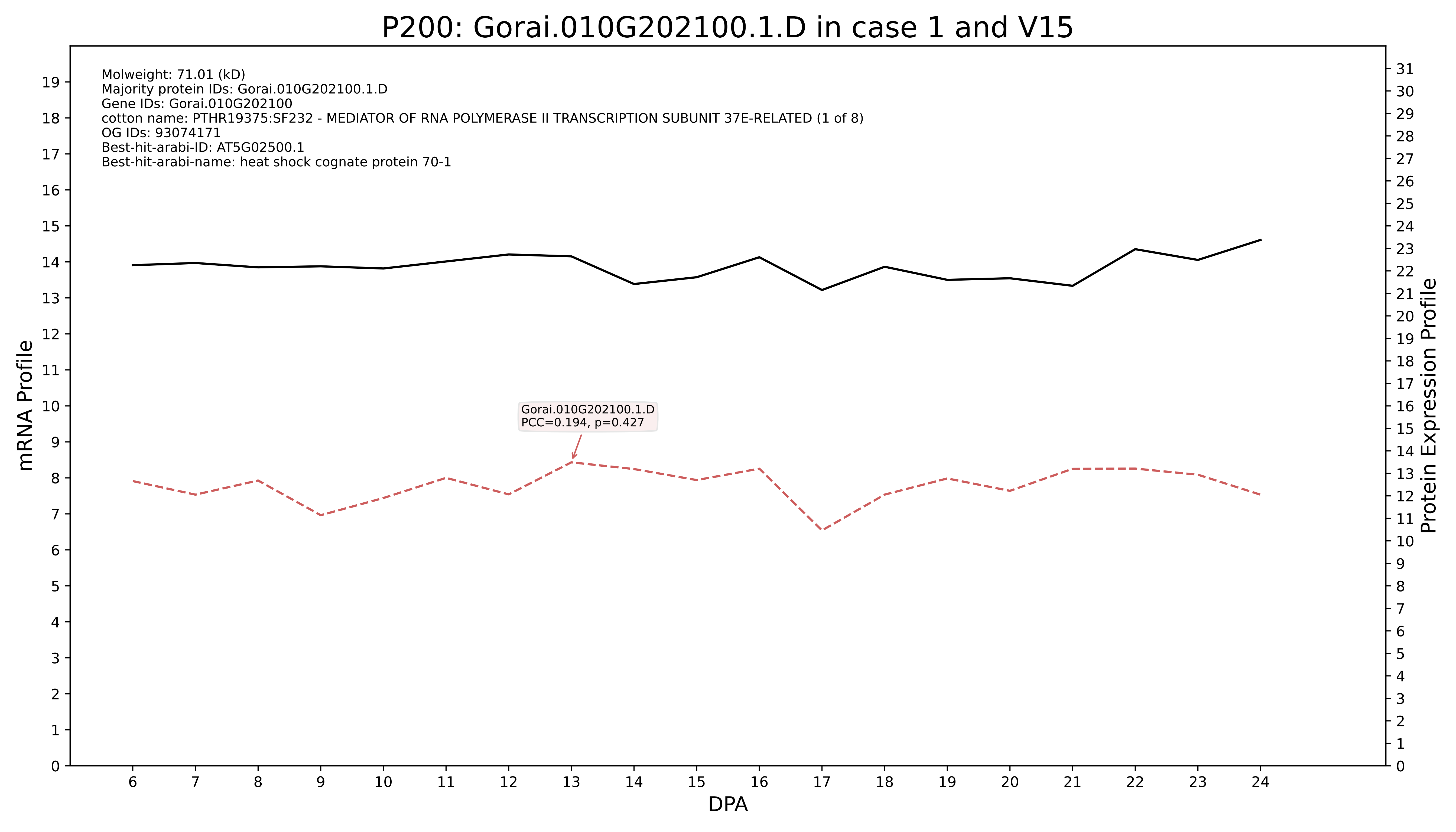
P200: Gorai.010G202100.1.D in case 1 and V15
Molweight: 71.01 (kD)
Majority protein IDs: Gorai.010G202100.1.D
Gene IDs: Gorai.010G202100
cotton name: PTHR19375:SF232 - MEDIATOR OF RNA POLYMERASE II TRANSCRIPTION SUBUNIT 37E-RELATED (1 of 8)
OG IDs: 93074171
Best-hit-arabi-ID: AT5G02500.1
Best-hit-arabi-name: heat shock cognate protein 70-1
Gorai.010G202100.1.D
PCC=0.194, p=0.427
mRNA Profile
Protein Expression Profile
DPA

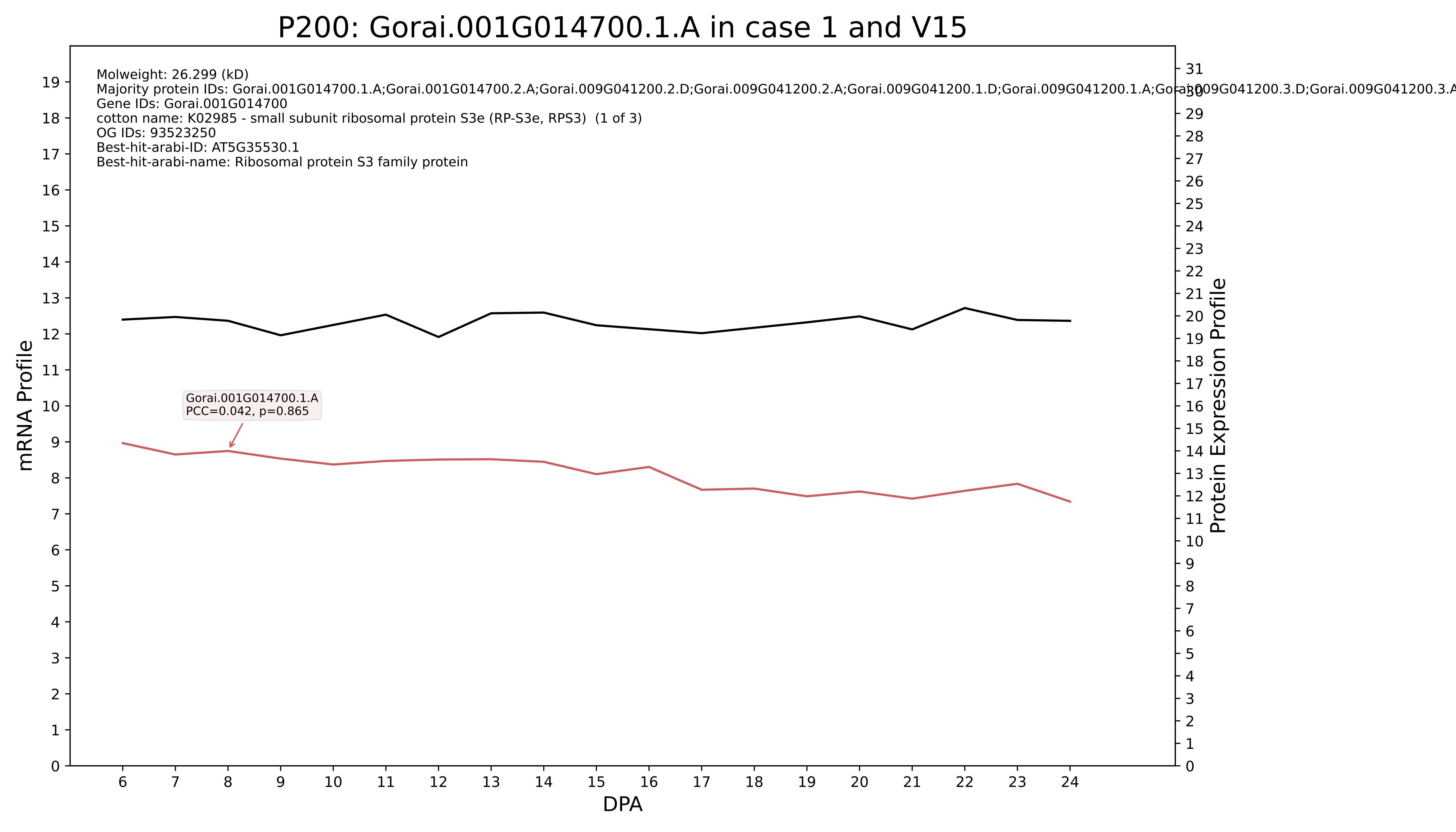
P200: Gorai.001G014700.1.A in case 1 and V15
Molweight: 26.299 (kD)
Majority protein IDs: Gorai.001G014700.1.A;Gorai.001G014700.2.A;Gorai.009G041200.2.D;Gorai.009G041200.2.A;Gorai.009G041200.1.D;Gorai.009G041200.1.A;Gorai.009G041200.3.D;Gorai.009G041200.3.A
Gene IDs: Gorai.001G014700
cotton name: K02985 - small subunit ribosomal protein S3e (RP-S3e, RPS3) (1 of 3)
OG IDs: 93523250
Best-hit-arabi-ID: AT5G35530.1
Best-hit-arabi-name: Ribosomal protein S3 family protein
Gorai.001G014700.1.A
PCC=0.042, p=0.865
mRNA Profile
Protein Expression Profile
DPA

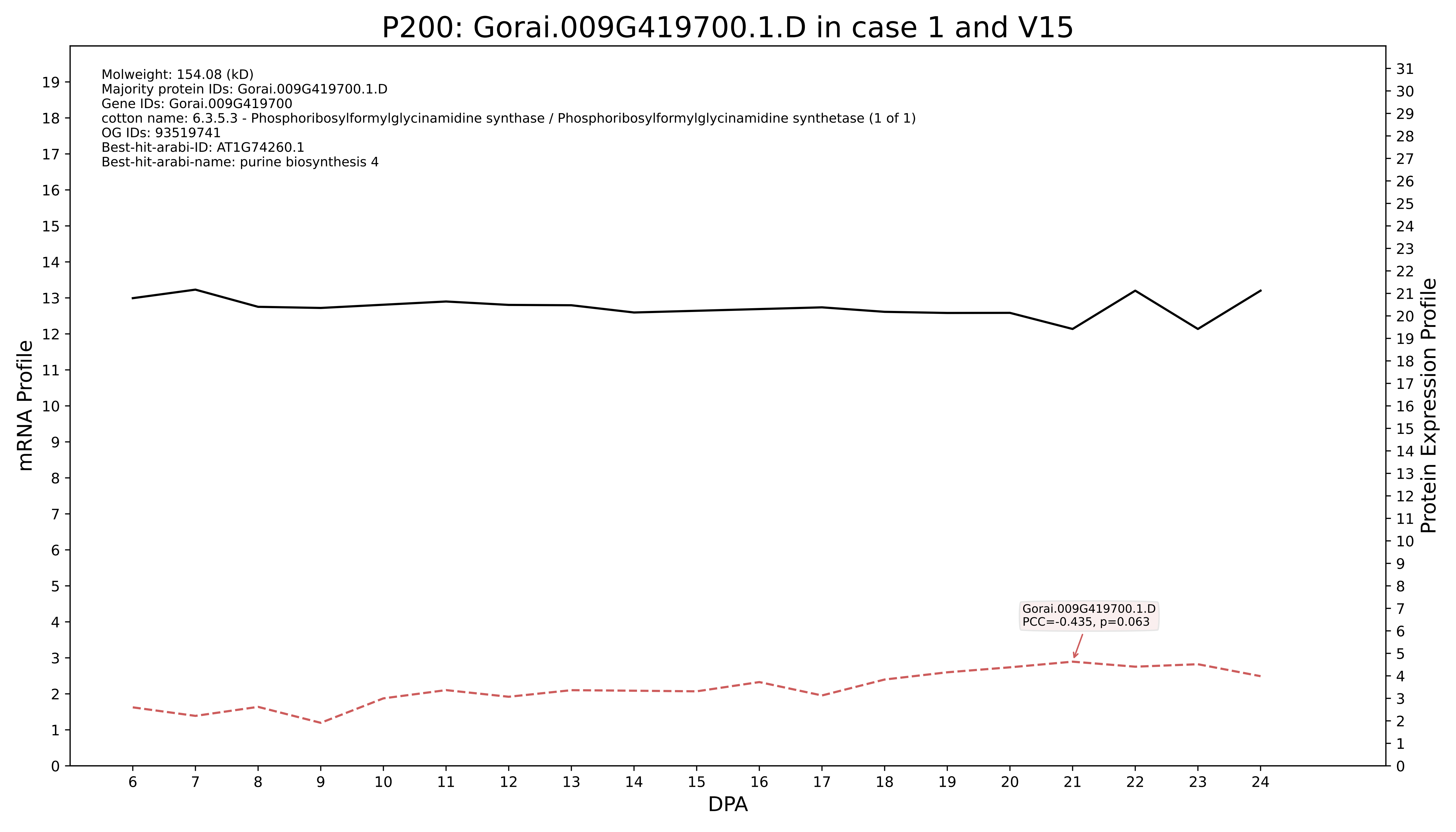
P200: Gorai.009G419700.1.D in case 1 and V15
Molweight: 154.08 (kD)
Majority protein IDs: Gorai.009G419700.1.D
Gene IDs: Gorai.009G419700
cotton name: 6.3.5.3 - Phosphoribosylformylglycinamidine synthase / Phosphoribosylformylglycinamidine synthetase (1 of 1)
OG IDs: 93519741
Best-hit-arabi-ID: AT1G74260.1
Best-hit-arabi-name: purine biosynthesis 4
Gorai.009G419700.1.D
PCC=-0.435, p=0.063
mRNA Profile
Protein Expression Profile
DPA

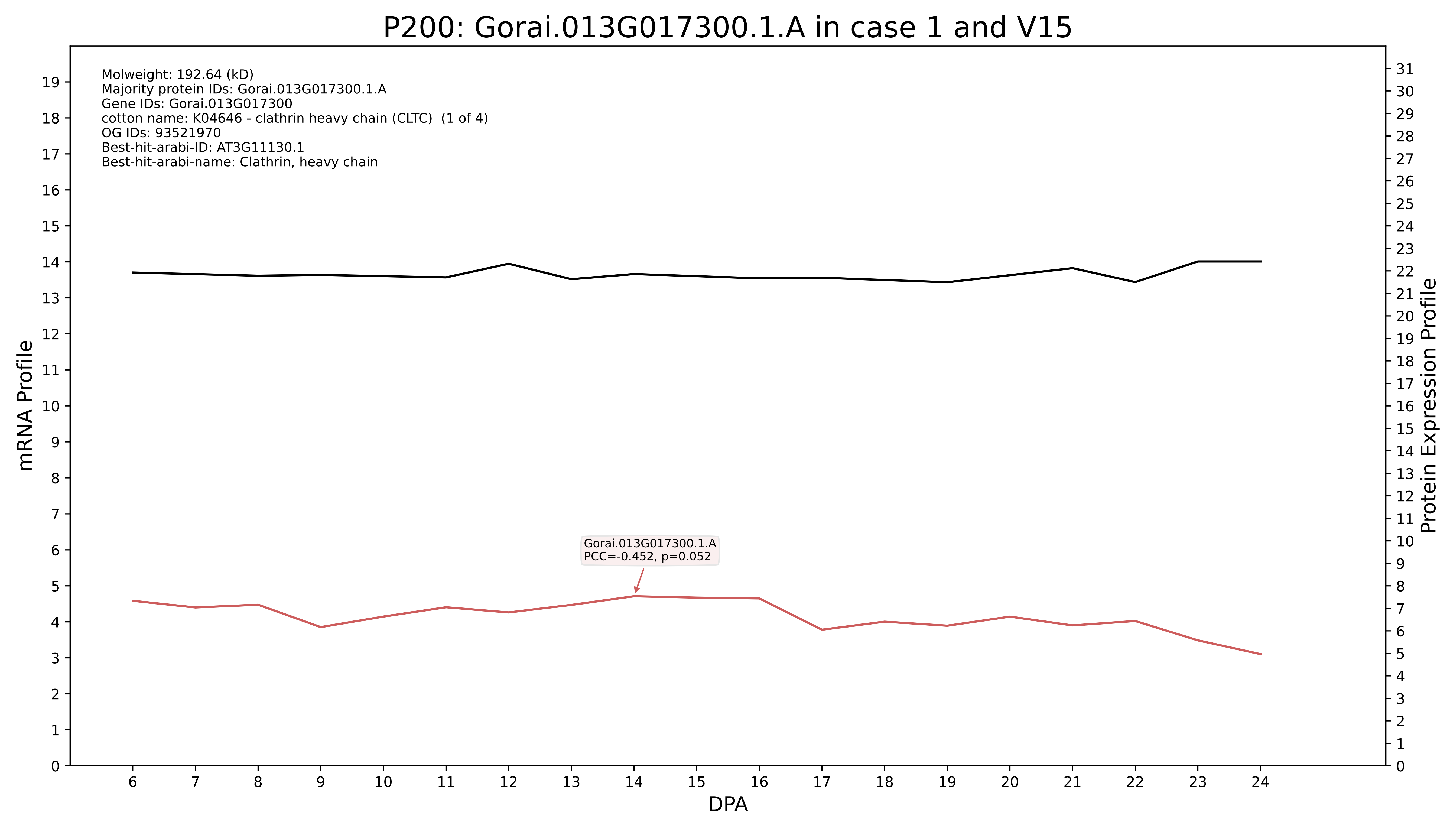
P200: Gorai.013G017300.1.A in case 1 and V15
Molweight: 192.64 (kD)
Majority protein IDs: Gorai.013G017300.1.A
Gene IDs: Gorai.013G017300
cotton name: K04646 - clathrin heavy chain (CLTC) (1 of 4)
OG IDs: 93521970
Best-hit-arabi-ID: AT3G11130.1
Best-hit-arabi-name: Clathrin, heavy chain
Gorai.013G017300.1.A
PCC=-0.452, p=0.052
mRNA Profile
Protein Expression Profile
DPA

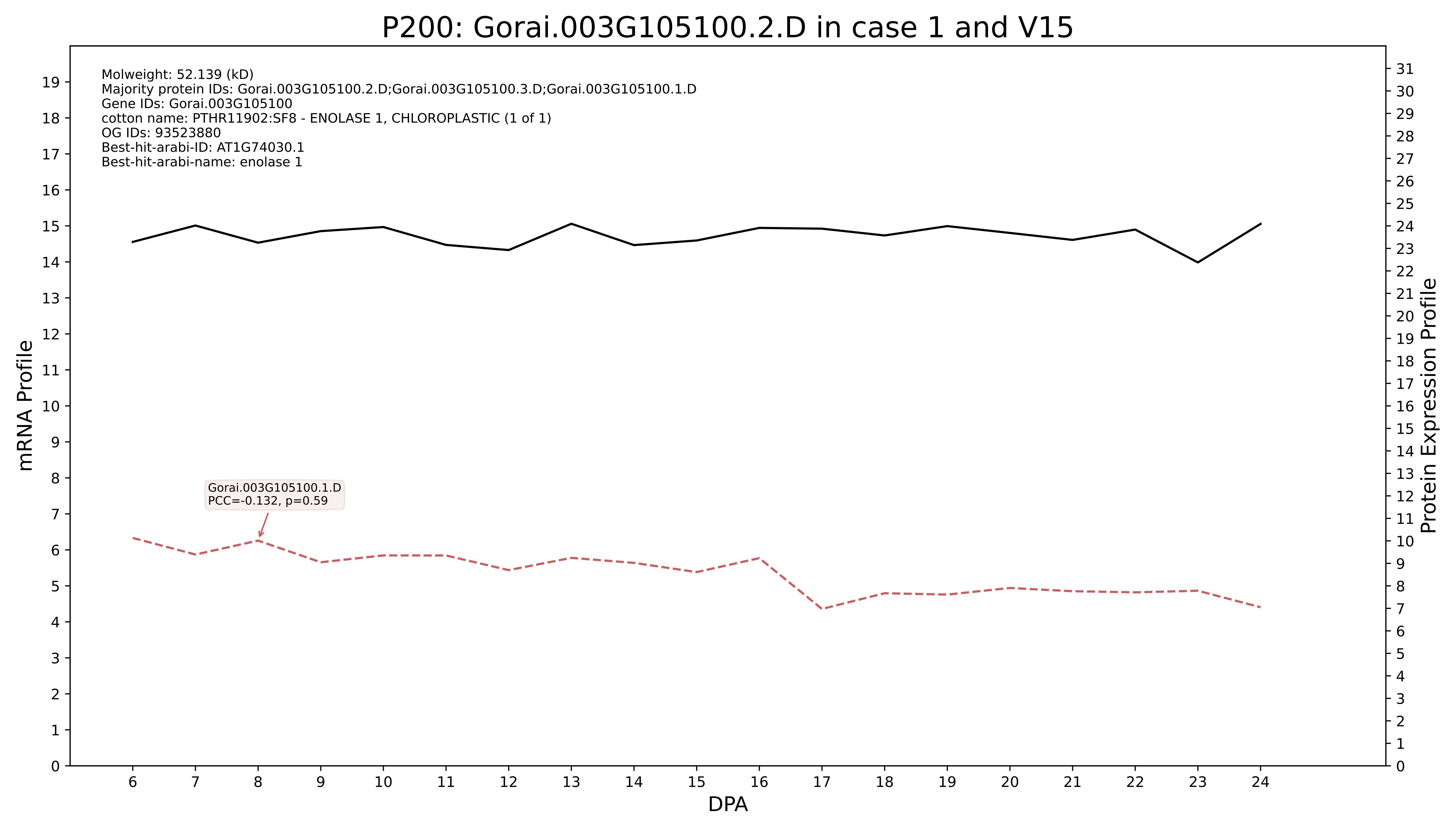

P200: Gorai.003G105100.2.D in case 1 and V15
Molweight: 52.139 (kD)
Majority protein IDs: Gorai.003G105100.2.D;Gorai.003G105100.3.D;Gorai.003G105100.1.D
Gene IDs: Gorai.003G105100
cotton name: PTHR11902:SF8 - ENOLASE 1, CHLOROPLASTIC (1 of 1)
OG IDs: 93523880
Best-hit-arabi-ID: AT1G74030.1
Best-hit-arabi-name: enolase 1
Gorai.003G105100.1.D
PCC=-0.132, p=0.59
mRNA Profile
Protein Expression Profile
DPA

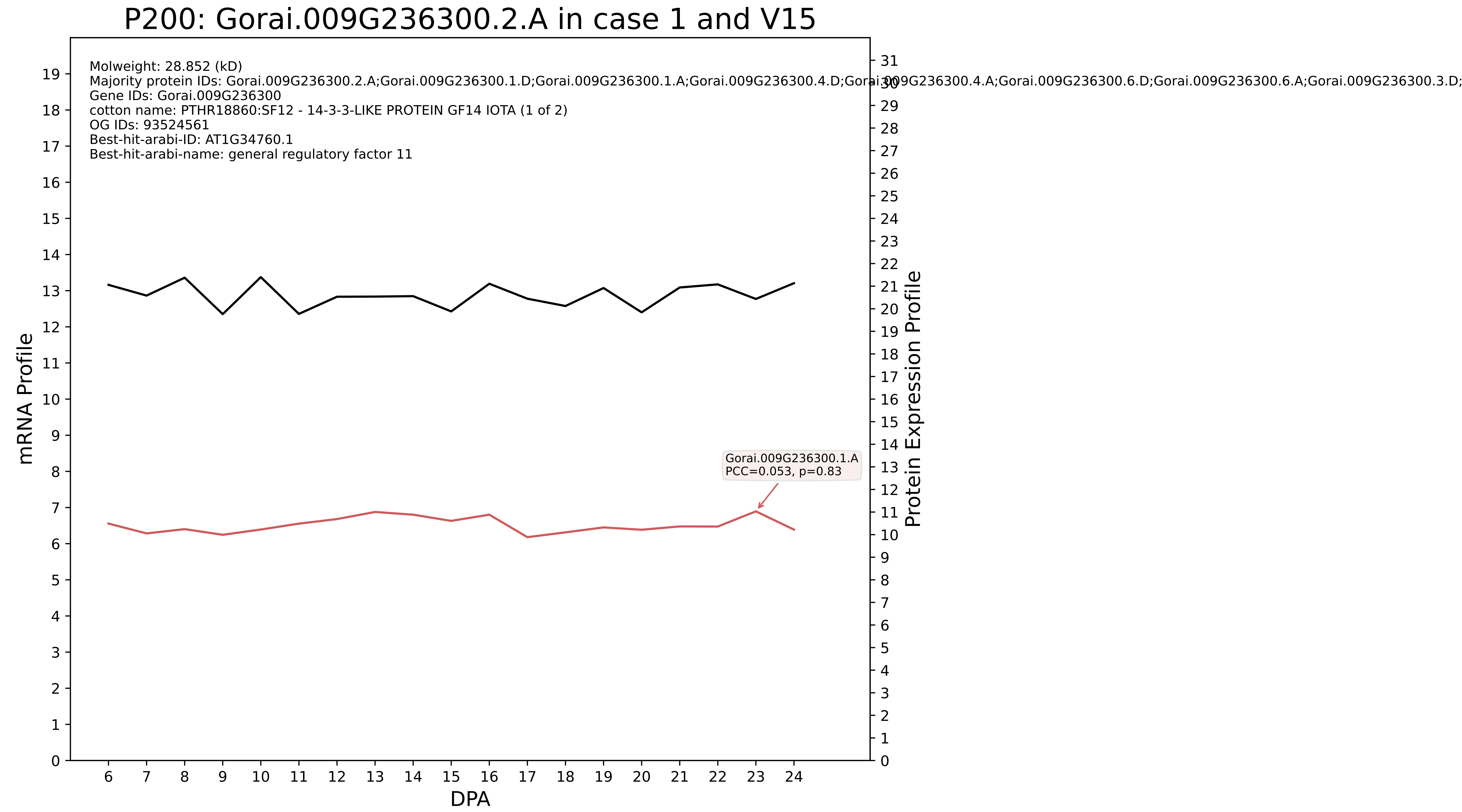
P200: Gorai.009G236300.2.A in case 1 and V15
Molweight: 28.852 (kD)
Majority protein IDs: Gorai.009G236300.2.A;Gorai.009G236300.1.D;Gorai.009G236300.1.A;Gorai.009G236300.4.D;Gorai.009G236300.4.A;Gorai.009G236300.6.D;Gorai.009G236300.6.A;Gorai.009G236300.3.D;
Gene IDs: Gorai.009G236300
cotton name: PTHR18860:SF12 - 14-3-3-LIKE PROTEIN GF14 IOTA (1 of 2)
OG IDs: 93524561
Best-hit-arabi-ID: AT1G34760.1
Best-hit-arabi-name: general regulatory factor 11
Gorai.009G236300.1.A
PCC=0.053, p=0.83
mRNA Profile
Protein Expression Profile
DPA

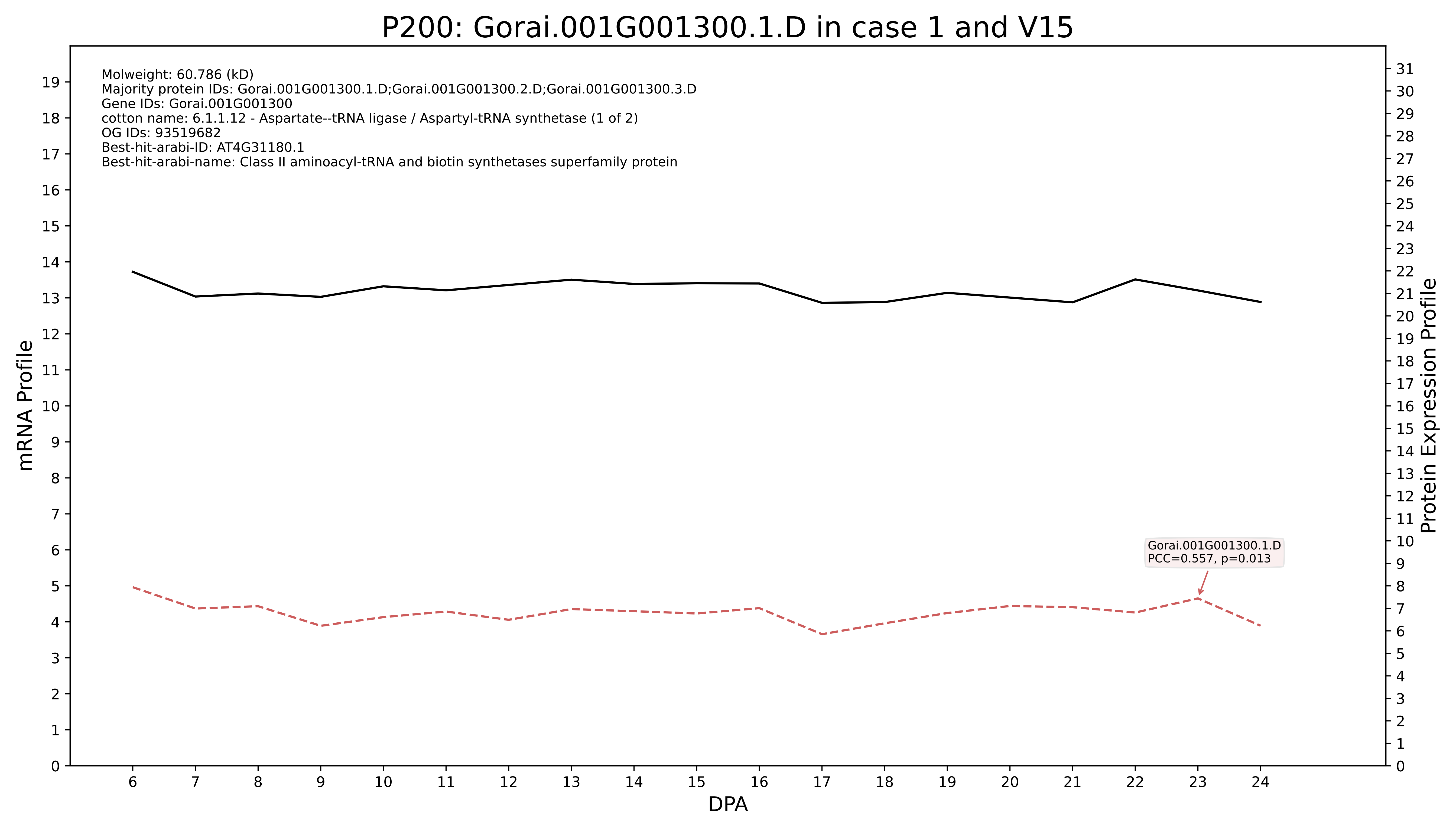
P200: Gorai.001G001300.1.D in case 1 and V15
Molweight: 60.786 (kD)
Majority protein IDs: Gorai.001G001300.1.D;Gorai.001G001300.2.D;Gorai.001G001300.3.D
Gene IDs: Gorai.001G001300
cotton name: 6.1.1.12 - Aspartate--tRNA ligase / Aspartyl-tRNA synthetase (1 of 2)
OG IDs: 93519682
Best-hit-arabi-ID: AT4G31180.1
Best-hit-arabi-name: Class II aminoacyl-tRNA and biotin synthetases superfamily protein
Gorai.001G001300.1.D
PCC=0.557, p=0.013
mRNA Profile
Protein Expression Profile
DPA

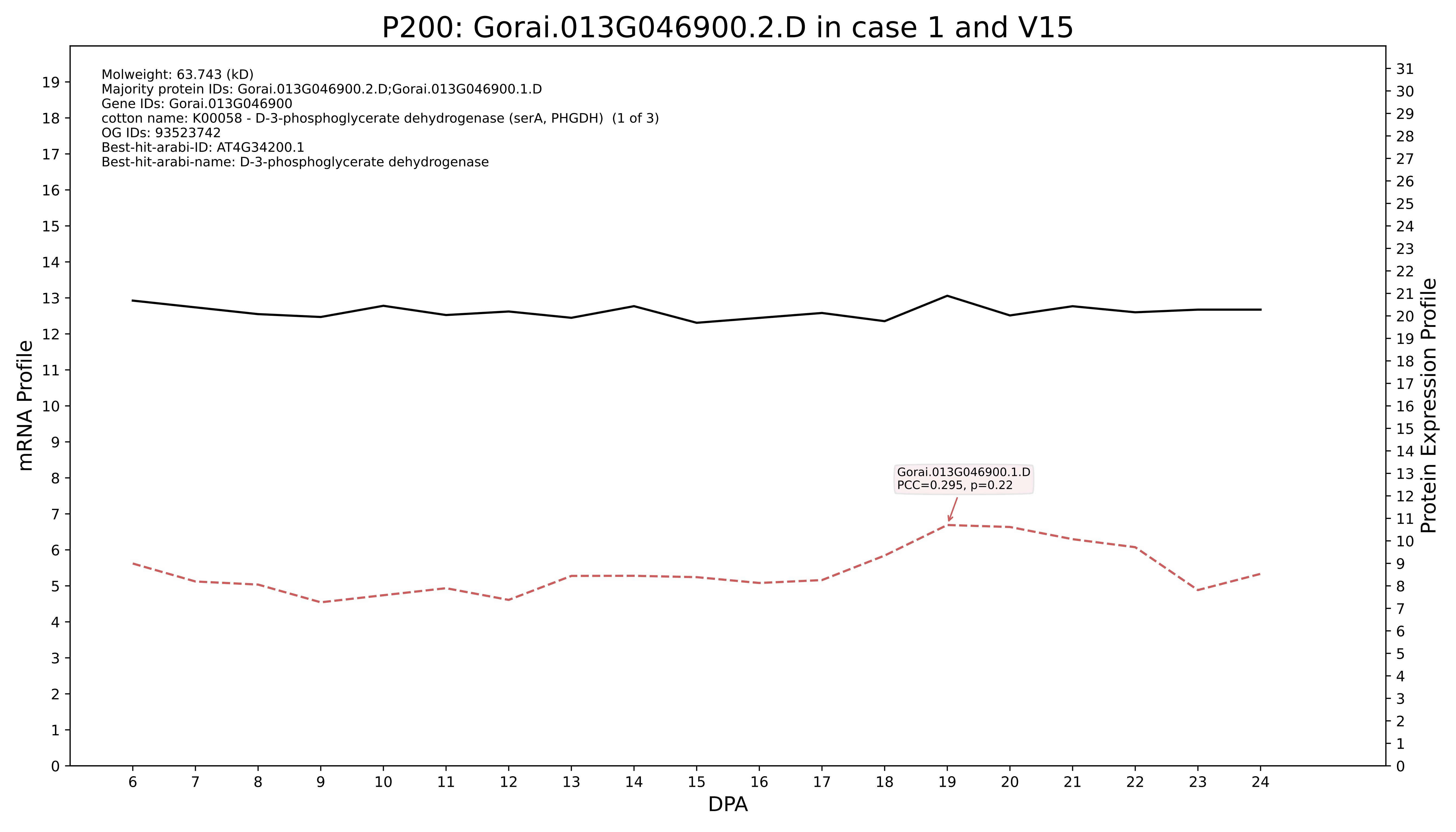

P200: Gorai.013G046900.2.D in case 1 and V15
Molweight: 63.743 (kD)
Majority protein IDs: Gorai.013G046900.2.D;Gorai.013G046900.1.D
Gene IDs: Gorai.013G046900
cotton name: K00058 - D-3-phosphoglycerate dehydrogenase (serA, PHGDH) (1 of 3)
OG IDs: 93523742
Best-hit-arabi-ID: AT4G34200.1
Best-hit-arabi-name: D-3-phosphoglycerate dehydrogenase
Gorai.013G046900.1.D
PCC=0.295, p=0.22
mRNA Profile
Protein Expression Profile
DPA

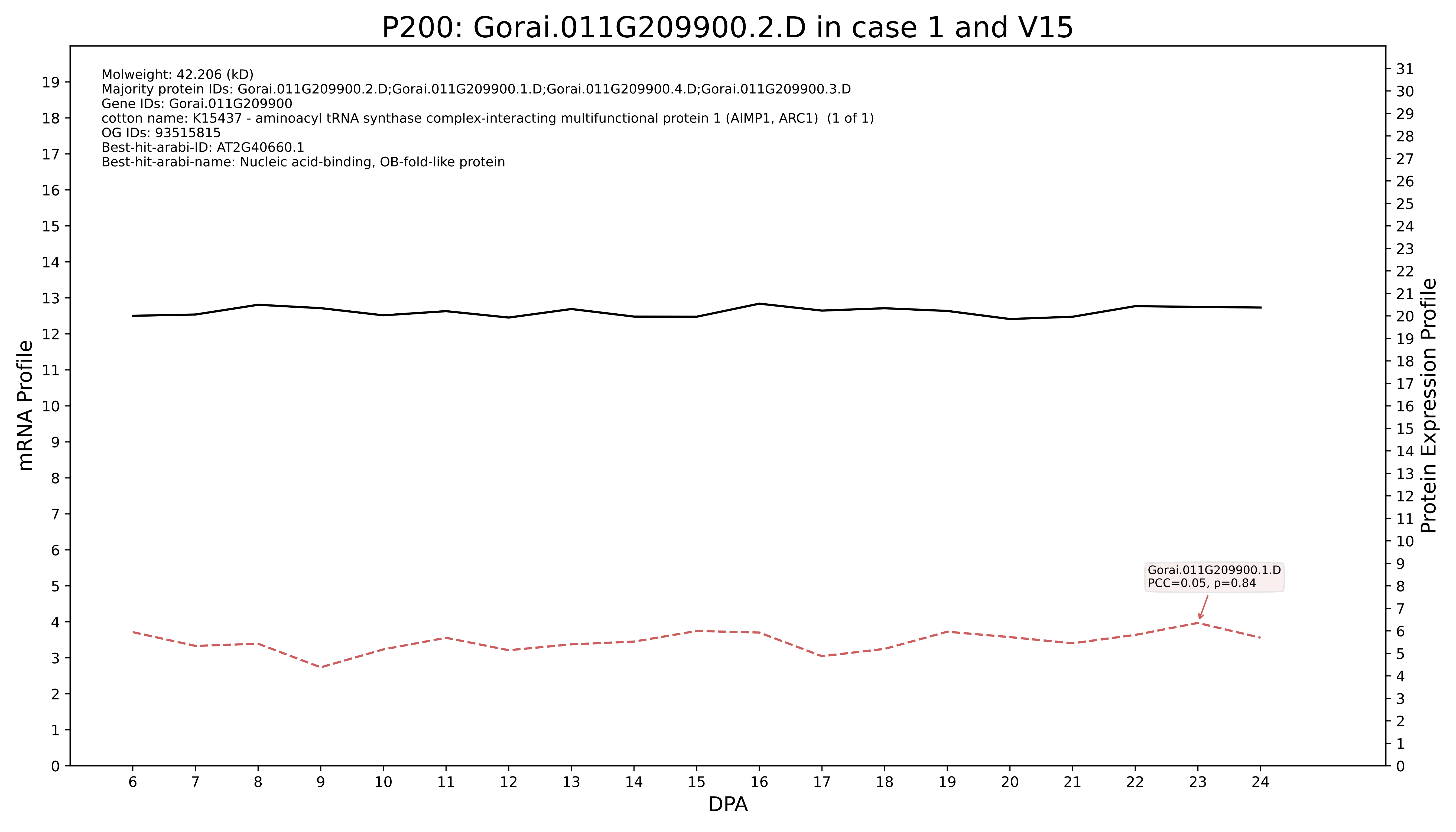

P200: Gorai.011G209900.2.D in case 1 and V15
Molweight: 42.206 (kD)
Majority protein IDs: Gorai.011G209900.2.D;Gorai.011G209900.1.D;Gorai.011G209900.4.D;Gorai.011G209900.3.D
Gene IDs: Gorai.011G209900
cotton name: K15437 - aminoacyl tRNA synthase complex-interacting multifunctional protein 1 (AIMP1, ARC1) (1 of 1)
OG IDs: 93515815
Best-hit-arabi-ID: AT2G40660.1
Best-hit-arabi-name: Nucleic acid-binding, OB-fold-like protein
Gorai.011G209900.1.D
PCC=0.05, p=0.84
mRNA Profile
Protein Expression Profile
DPA

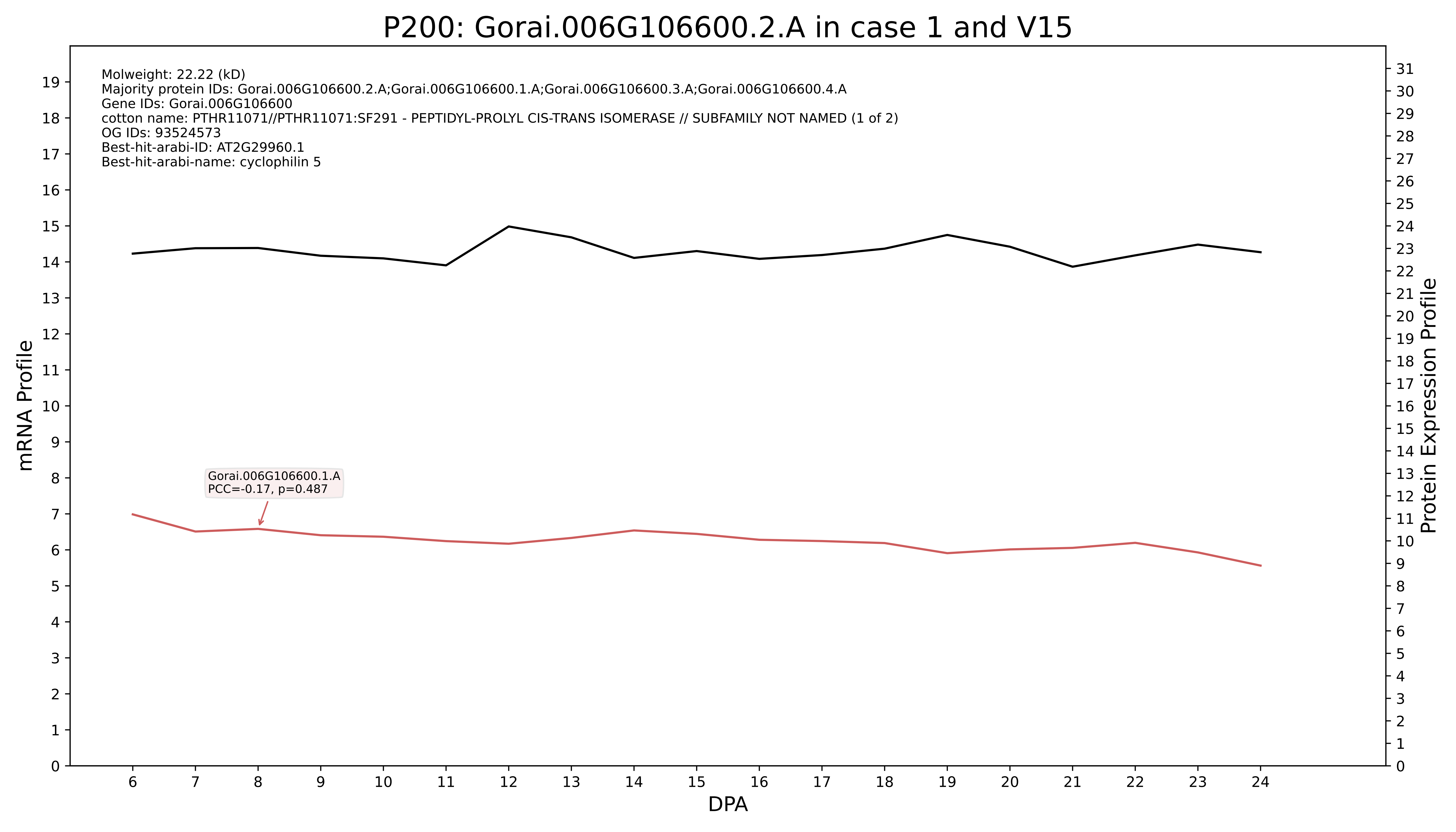

P200: Gorai.006G106600.2.A in case 1 and V15
Molweight: 22.22 (kD)
Majority protein IDs: Gorai.006G106600.2.A;Gorai.006G106600.1.A;Gorai.006G106600.3.A;Gorai.006G106600.4.A
Gene IDs: Gorai.006G106600
cotton name: PTHR11071//PTHR11071:SF291 - PEPTIDYL-PROLYL CIS-TRANS ISOMERASE // SUBFAMILY NOT NAMED (1 of 2)
OG IDs: 93524573
Best-hit-arabi-ID: AT2G29960.1
Best-hit-arabi-name: cyclophilin 5
Gorai.006G106600.1.A
PCC=-0.17, p=0.487
mRNA Profile
Protein Expression Profile
DPA

### P200: Gorai.011G283100.1.D in case 1 and V15

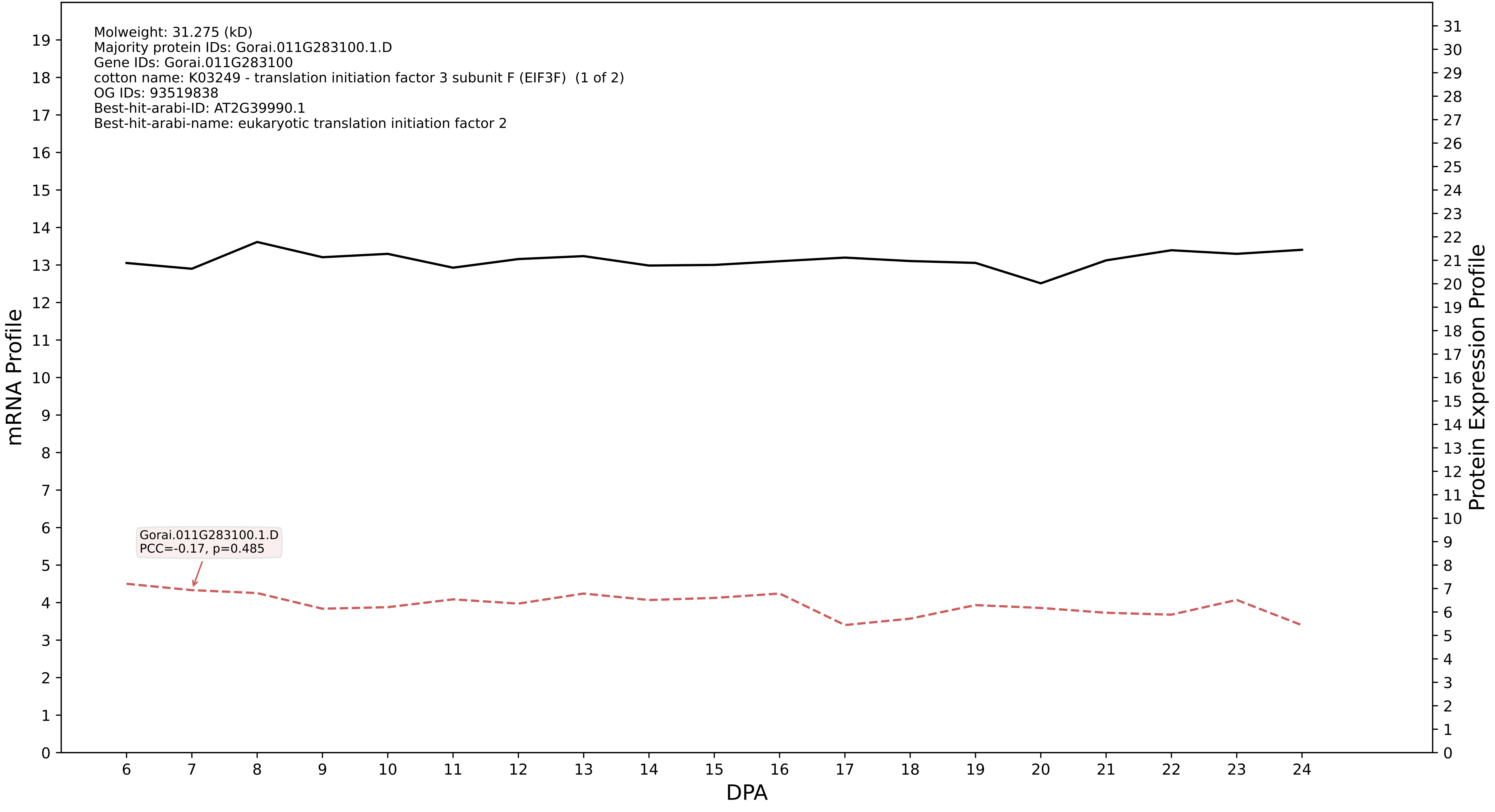

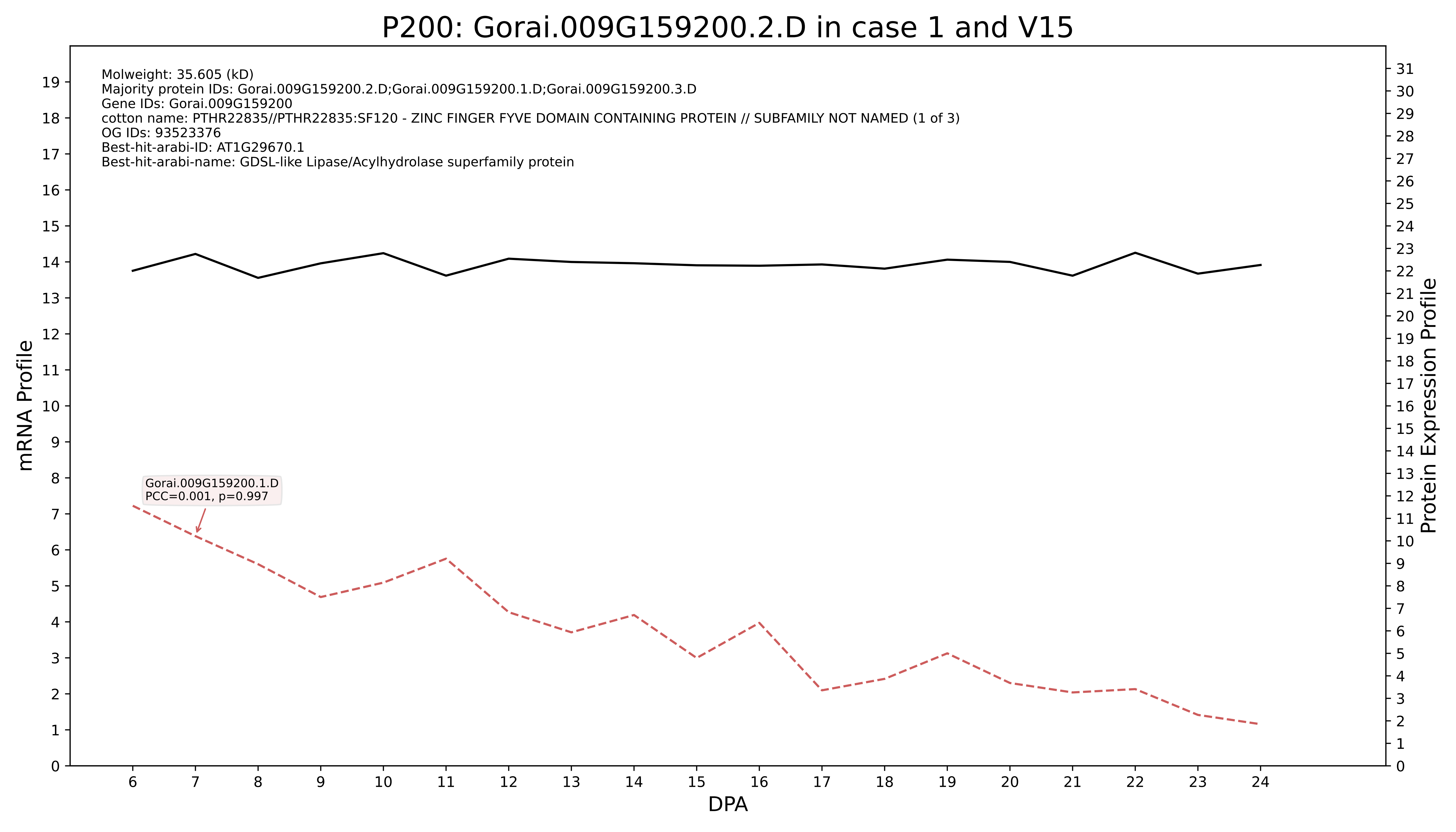
P200: Gorai.009G159200.2.D in case 1 and V15
Molweight: 35.605 (kD)
Majority protein IDs: Gorai.009G159200.2.D;Gorai.009G159200.1.D;Gorai.009G159200.3.D
Gene IDs: Gorai.009G159200
cotton name: PTHR22835//PTHR22835:SF120 - ZINC FINGER FYVE DOMAIN CONTAINING PROTEIN // SUBFAMILY NOT NAMED (1 of 3)
OG IDs: 93523376
Best-hit-arabi-ID: AT1G29670.1
Best-hit-arabi-name: GDSL-like Lipase/Acylhydrolase superfamily protein
Gorai.009G159200.1.D
PCC=0.001, p=0.997
mRNA Profile
Protein Expression Profile
DPA

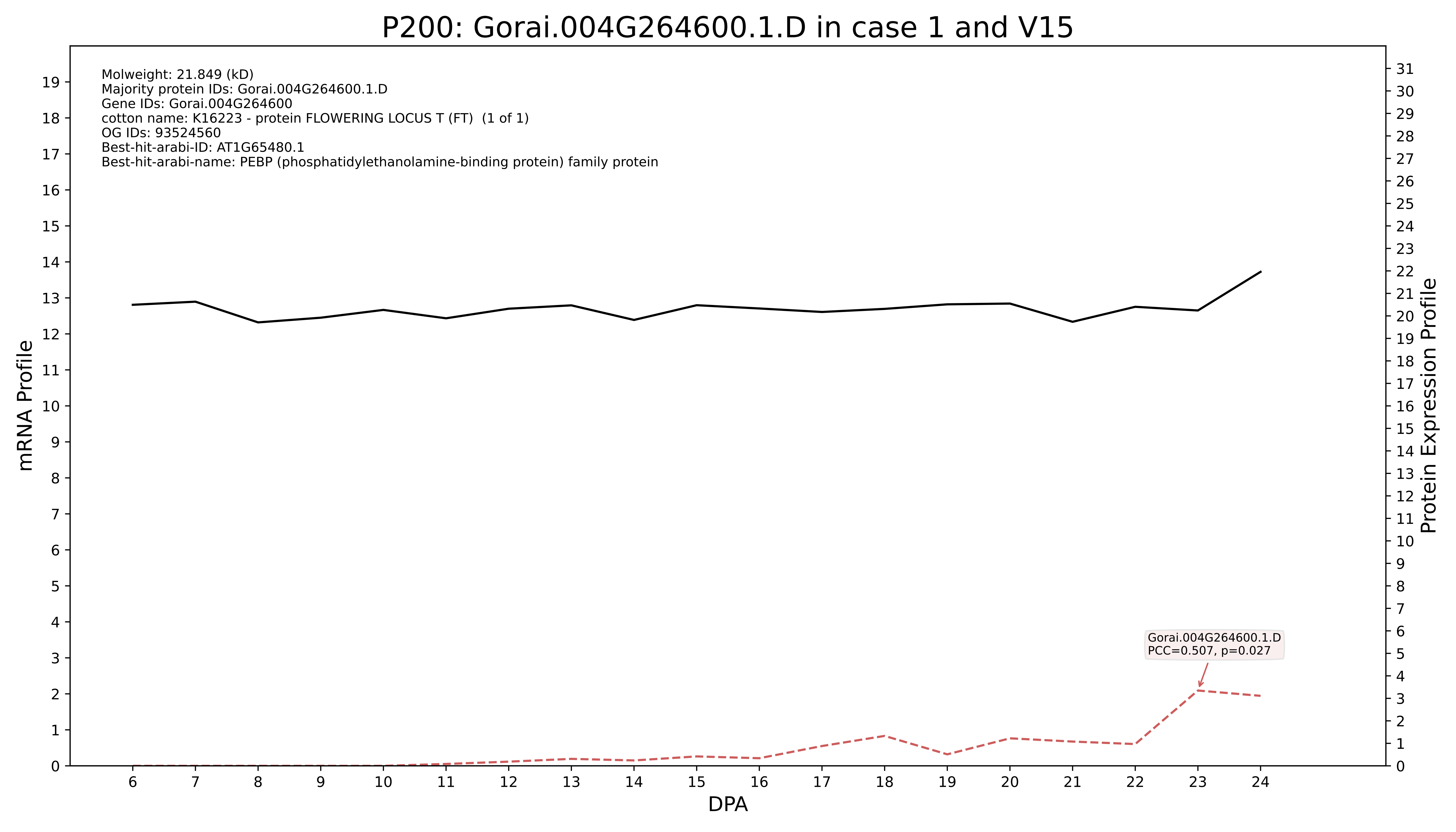

P200: Gorai.004G264600.1.D in case 1 and V15
Molweight: 21.849 (kD)
Majority protein IDs: Gorai.004G264600.1.D
Gene IDs: Gorai.004G264600
cotton name: K16223 - protein FLOWERING LOCUS T (FT) (1 of 1)
OG IDs: 93524560
Best-hit-arabi-ID: AT1G65480.1
Best-hit-arabi-name: PEBP (phosphatidylethanolamine-binding protein) family protein
Gorai.004G264600.1.D
PCC=0.507, p=0.027
mRNA Profile
Protein Expression Profile
DPA

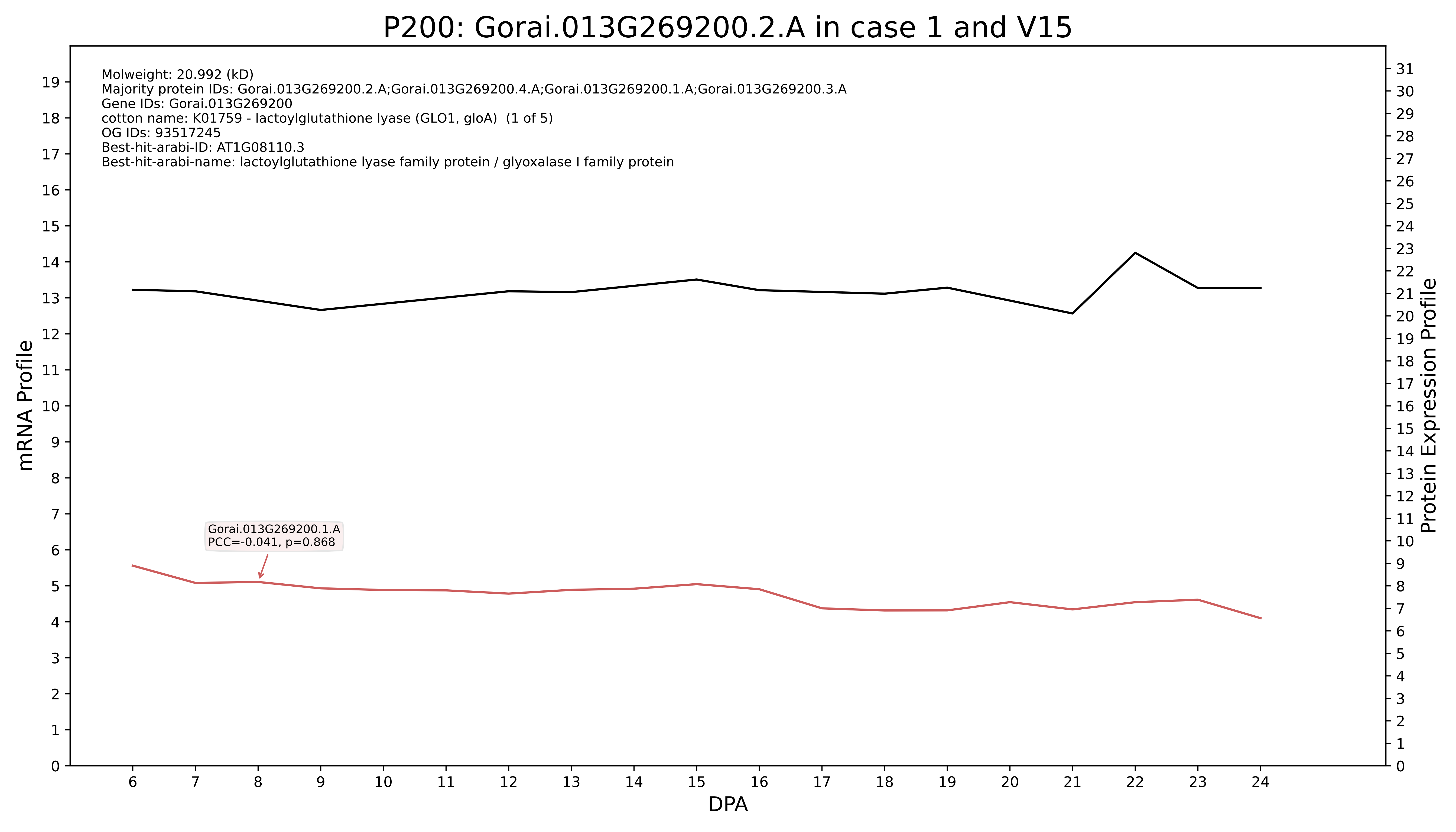
P200: Gorai.013G269200.2.A in case 1 and V15
Molweight: 20.992 (kD)
Majority protein IDs: Gorai.013G269200.2.A;Gorai.013G269200.4.A;Gorai.013G269200.1.A;Gorai.013G269200.3.A
Gene IDs: Gorai.013G269200
cotton name: K01759 - lactoylglutathione lyase (GLO1, gloA) (1 of 5)
OG IDs: 93517245
Best-hit-arabi-ID: AT1G08110.3
Best-hit-arabi-name: lactoylglutathione lyase family protein / glyoxalase I family protein
Gorai.013G269200.1.A
PCC=-0.041, p=0.868
mRNA Profile
Protein Expression Profile
DPA

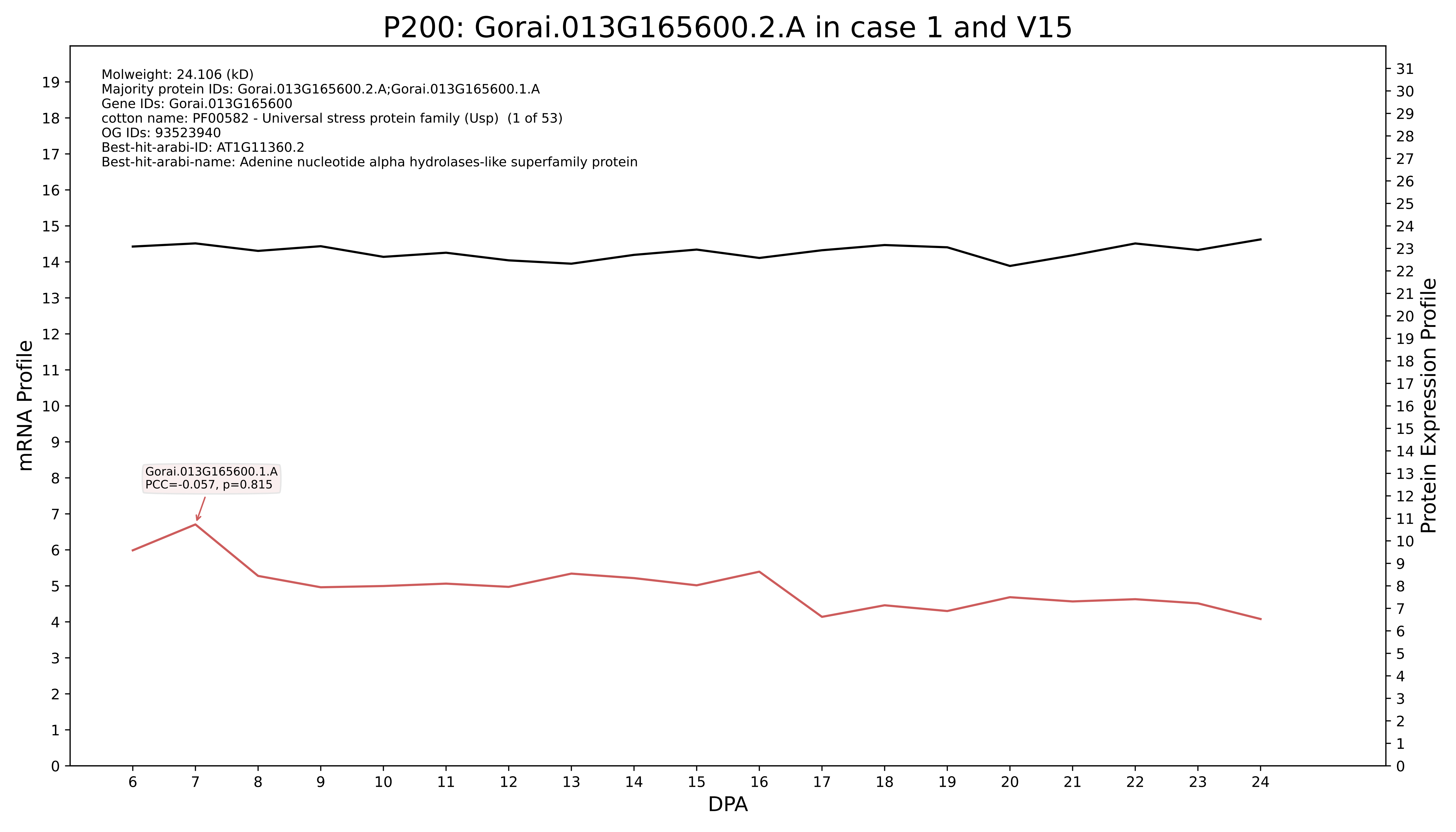

P200: Gorai.013G165600.2.A in case 1 and V15
Molweight: 24.106 (kD)
Majority protein IDs: Gorai.013G165600.2.A;Gorai.013G165600.1.A
Gene IDs: Gorai.013G165600
cotton name: PF00582 - Universal stress protein family (Usp) (1 of 53)
OG IDs: 93523940
Best-hit-arabi-ID: AT1G11360.2
Best-hit-arabi-name: Adenine nucleotide alpha hydrolases-like superfamily protein
Gorai.013G165600.1.A
PCC=-0.057, p=0.815
mRNA Profile
Protein Expression Profile
DPA

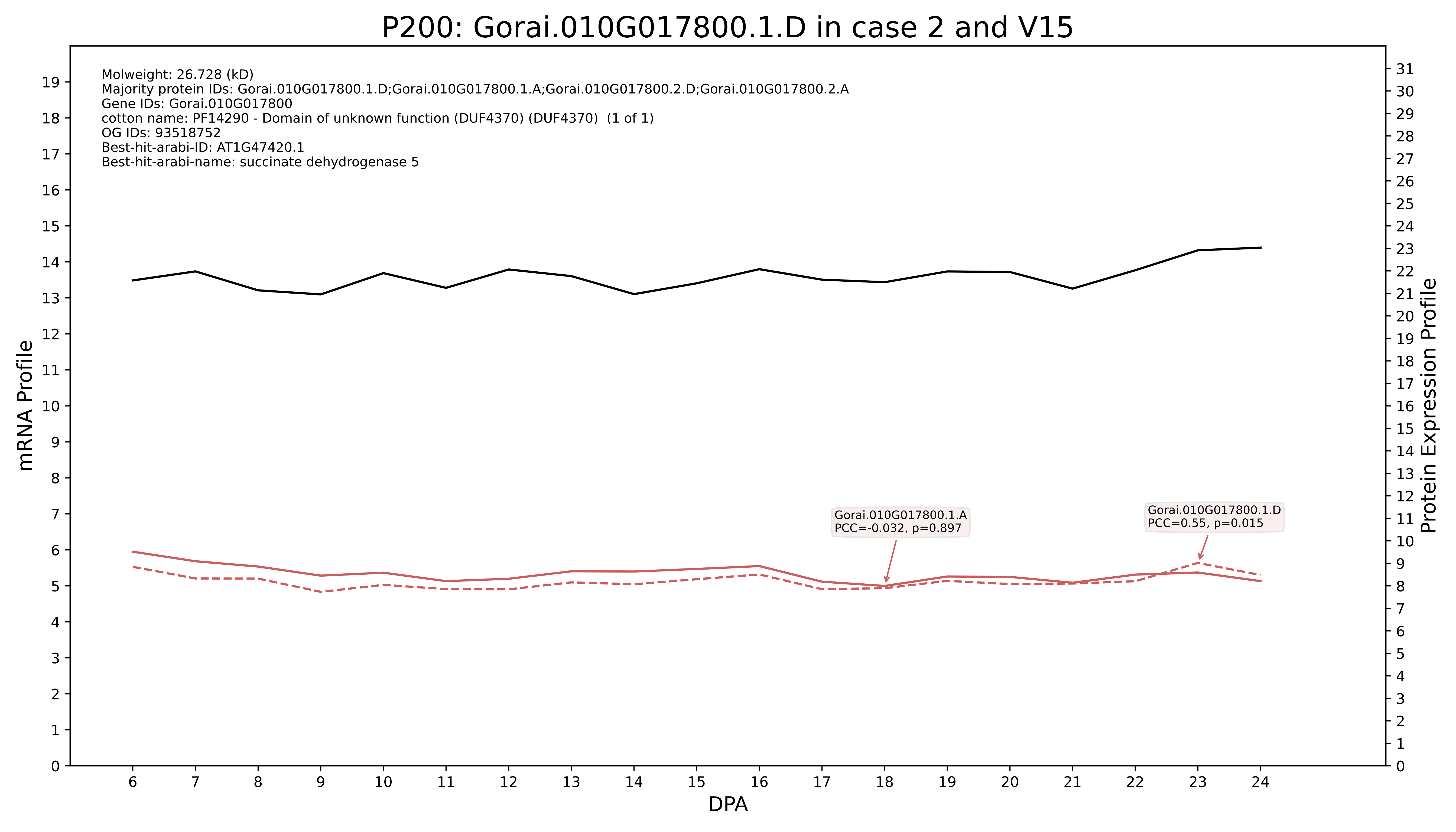
P200: Gorai.010G017800.1.D in case 2 and V15
Molweight: 26.728 (kD)
Majority protein IDs: Gorai.010G017800.1.D;Gorai.010G017800.1.A;Gorai.010G017800.2.D;Gorai.010G017800.2.A
Gene IDs: Gorai.010G017800
cotton name: PF14290 - Domain of unknown function (DUF4370) (DUF4370) (1 of 1)
OG IDs: 93518752
Best-hit-arabi-ID: AT1G47420.1
Best-hit-arabi-name: succinate dehydrogenase 5
Gorai.010G017800.1.A
PCC=-0.032, p=0.897
Gorai.010G017800.1.D
PCC=0.55, p=0.015
mRNA Profile
Protein Expression Profile
DPA

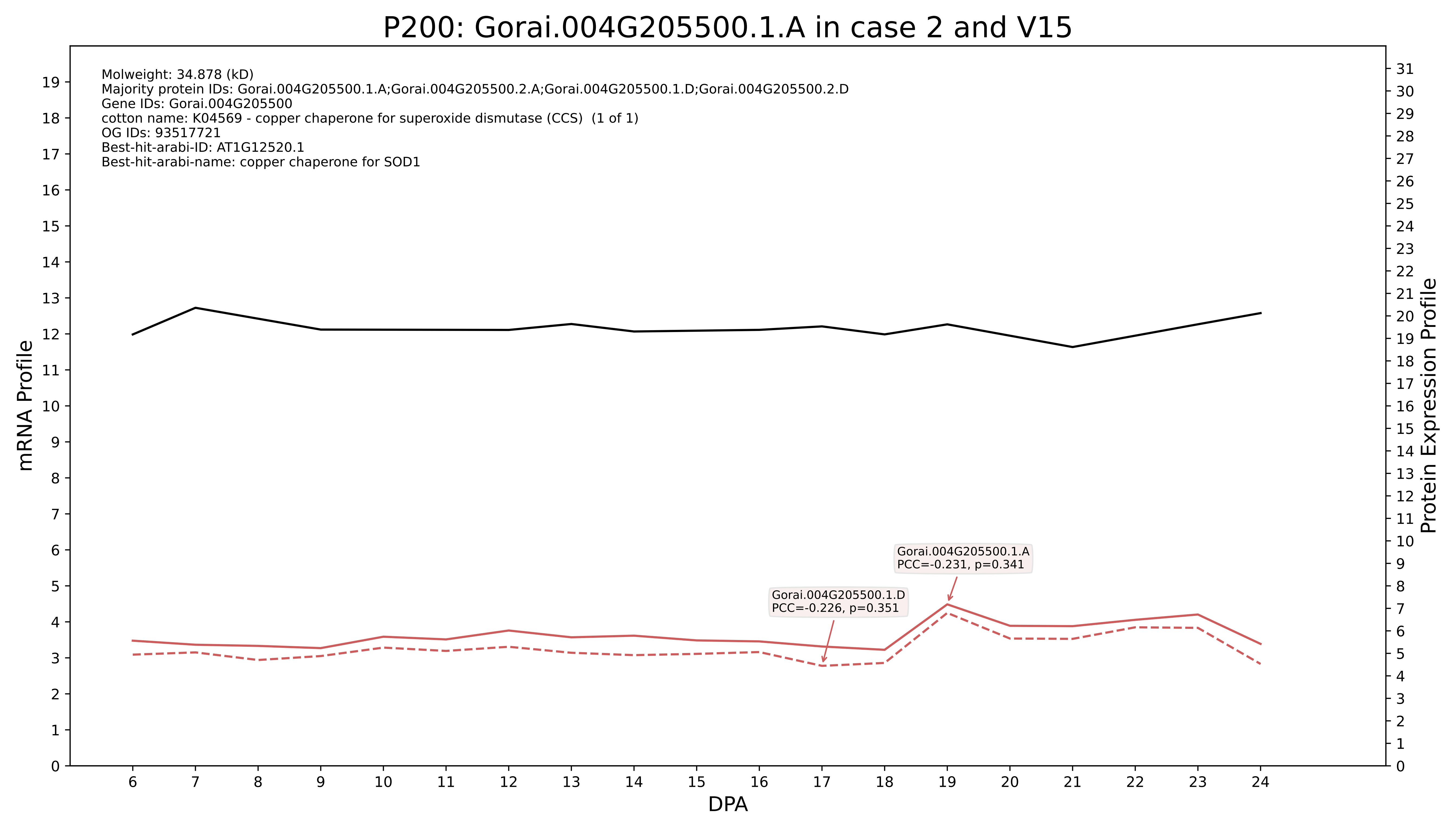
P200: Gorai.004G205500.1.A in case 2 and V15
Molweight: 34.878 (kD)
Majority protein IDs: Gorai.004G205500.1.A;Gorai.004G205500.2.A;Gorai.004G205500.1.D;Gorai.004G205500.2.D
Gene IDs: Gorai.004G205500
cotton name: K04569 - copper chaperone for superoxide dismutase (CCS) (1 of 1)
OG IDs: 93517721
Best-hit-arabi-ID: AT1G12520.1
Best-hit-arabi-name: copper chaperone for SOD1
Gorai.004G205500.1.A
PCC=-0.231, p=0.341
Gorai.004G205500.1.D
PCC=-0.226, p=0.351
mRNA Profile
Protein Expression Profile
DPA

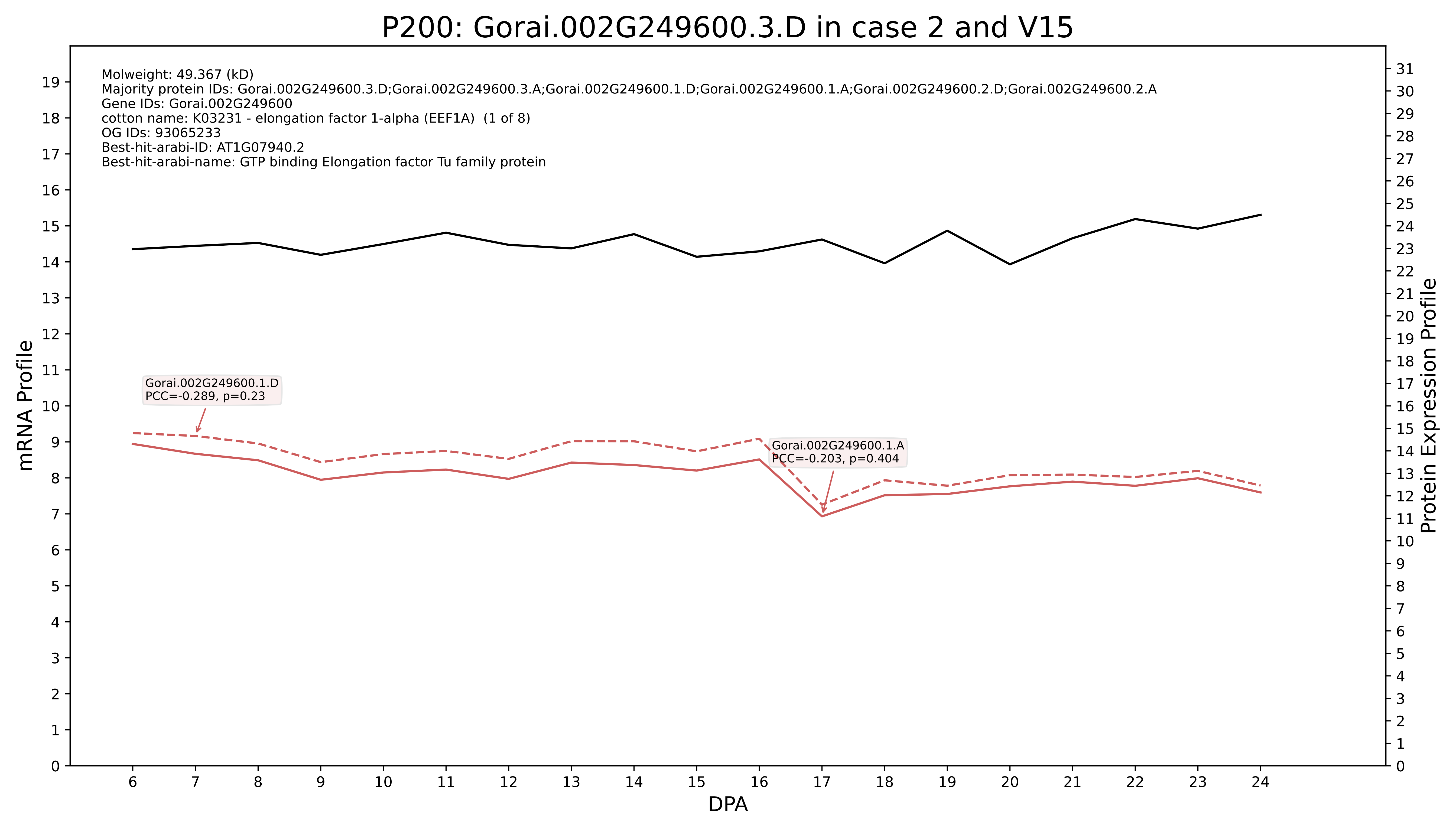
P200: Gorai.002G249600.3.D in case 2 and V15
Molweight: 49.367 (kD)
Majority protein IDs: Gorai.002G249600.3.D;Gorai.002G249600.3.A;Gorai.002G249600.1.D;Gorai.002G249600.1.A;Gorai.002G249600.2.D;Gorai.002G249600.2.A
Gene IDs: Gorai.002G249600
cotton name: K03231 - elongation factor 1-alpha (EEF1A) (1 of 8)
OG IDs: 93065233
Best-hit-arabi-ID: AT1G07940.2
Best-hit-arabi-name: GTP binding Elongation factor Tu family protein
Gorai.002G249600.1.D
PCC=-0.289, p=0.23
Gorai.002G249600.1.A
PCC=-0.203, p=0.404
mRNA Profile
Protein Expression Profile
DPA

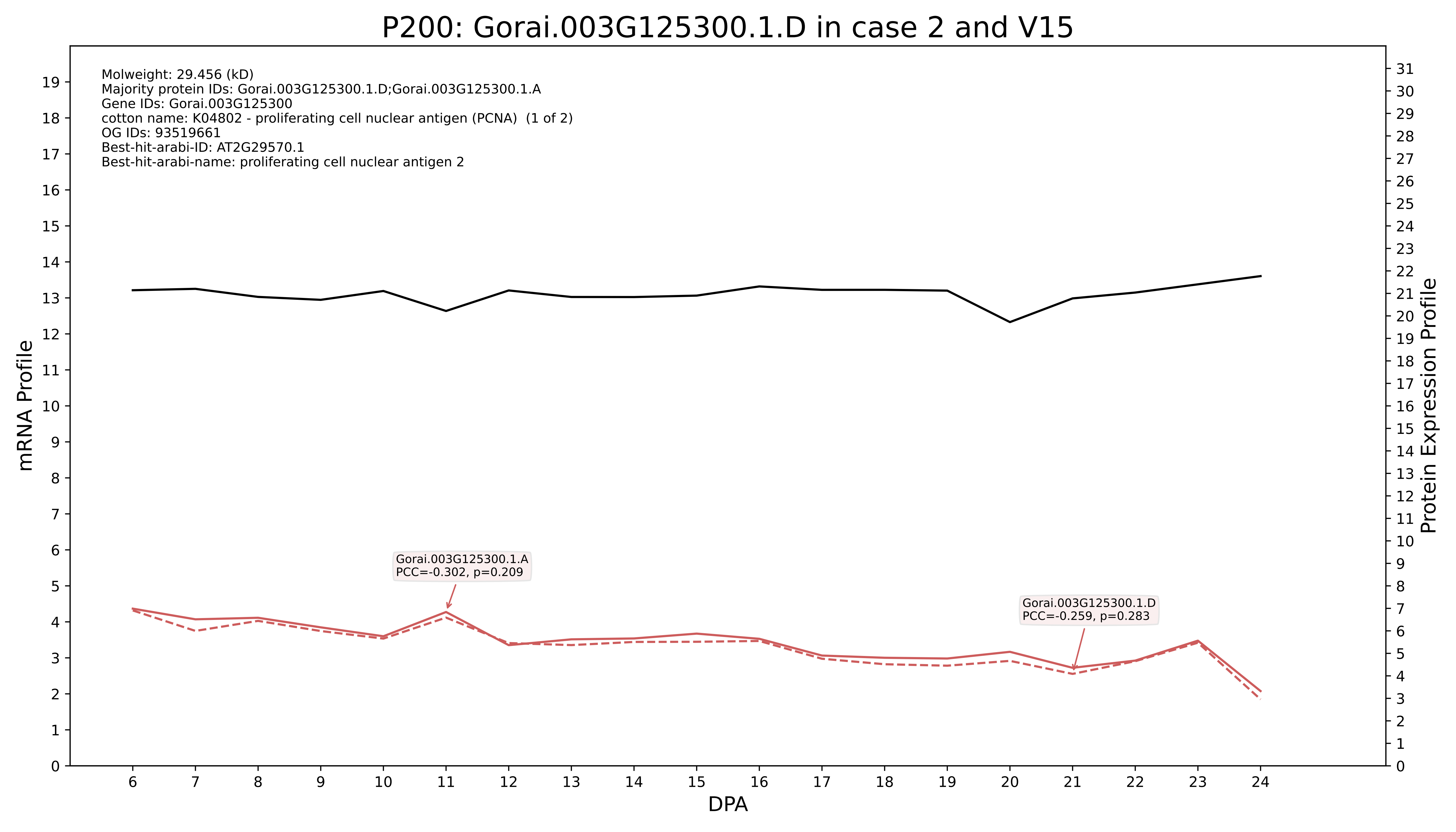

P200: Gorai.003G125300.1.D in case 2 and V15
Molweight: 29.456 (kD)
Majority protein IDs: Gorai.003G125300.1.D;Gorai.003G125300.1.A
Gene IDs: Gorai.003G125300
cotton name: K04802 - proliferating cell nuclear antigen (PCNA) (1 of 2)
OG IDs: 93519661
Best-hit-arabi-ID: AT2G29570.1
Best-hit-arabi-name: proliferating cell nuclear antigen 2
Gorai.003G125300.1.A
PCC=-0.302, p=0.209
Gorai.003G125300.1.D
PCC=-0.259, p=0.283
mRNA Profile
Protein Expression Profile
DPA

00: Gorai.001G213900.5.D in case 2 and V15

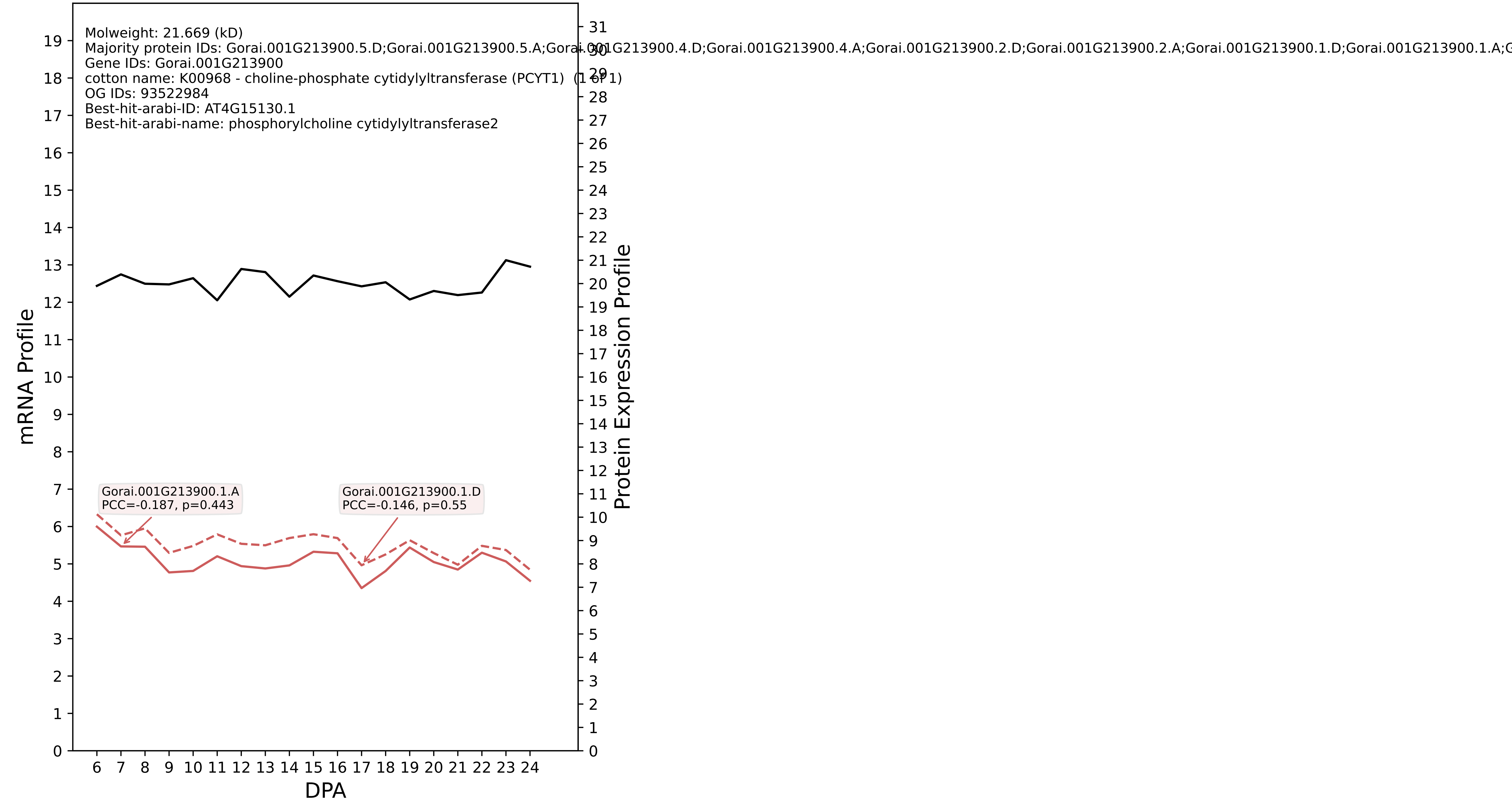

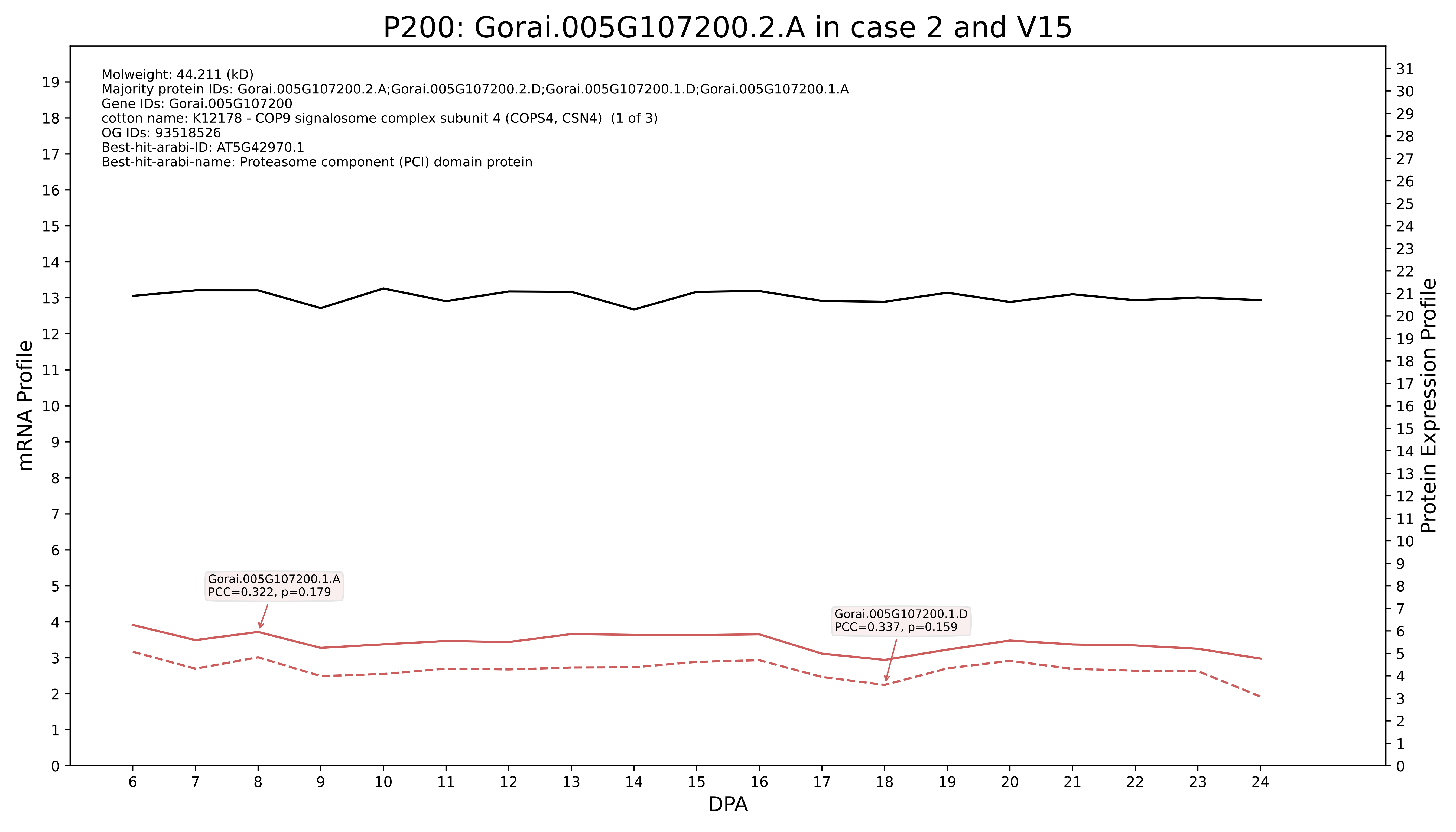
P200: Gorai.005G107200.2.A in case 2 and V15
Molweight: 44.211 (kD)
Majority protein IDs: Gorai.005G107200.2.A;Gorai.005G107200.2.D;Gorai.005G107200.1.D;Gorai.005G107200.1.A
Gene IDs: Gorai.005G107200
cotton name: K12178 - COP9 signalosome complex subunit 4 (COPS4, CSN4) (1 of 3)
OG IDs: 93518526
Best-hit-arabi-ID: AT5G42970.1
Best-hit-arabi-name: Proteasome component (PCI) domain protein
Gorai.005G107200.1.A
PCC=0.322, p=0.179
Gorai.005G107200.1.D
PCC=0.337, p=0.159
mRNA Profile
Protein Expression Profile
DPA

007G150200.6.A in case 2 and V15

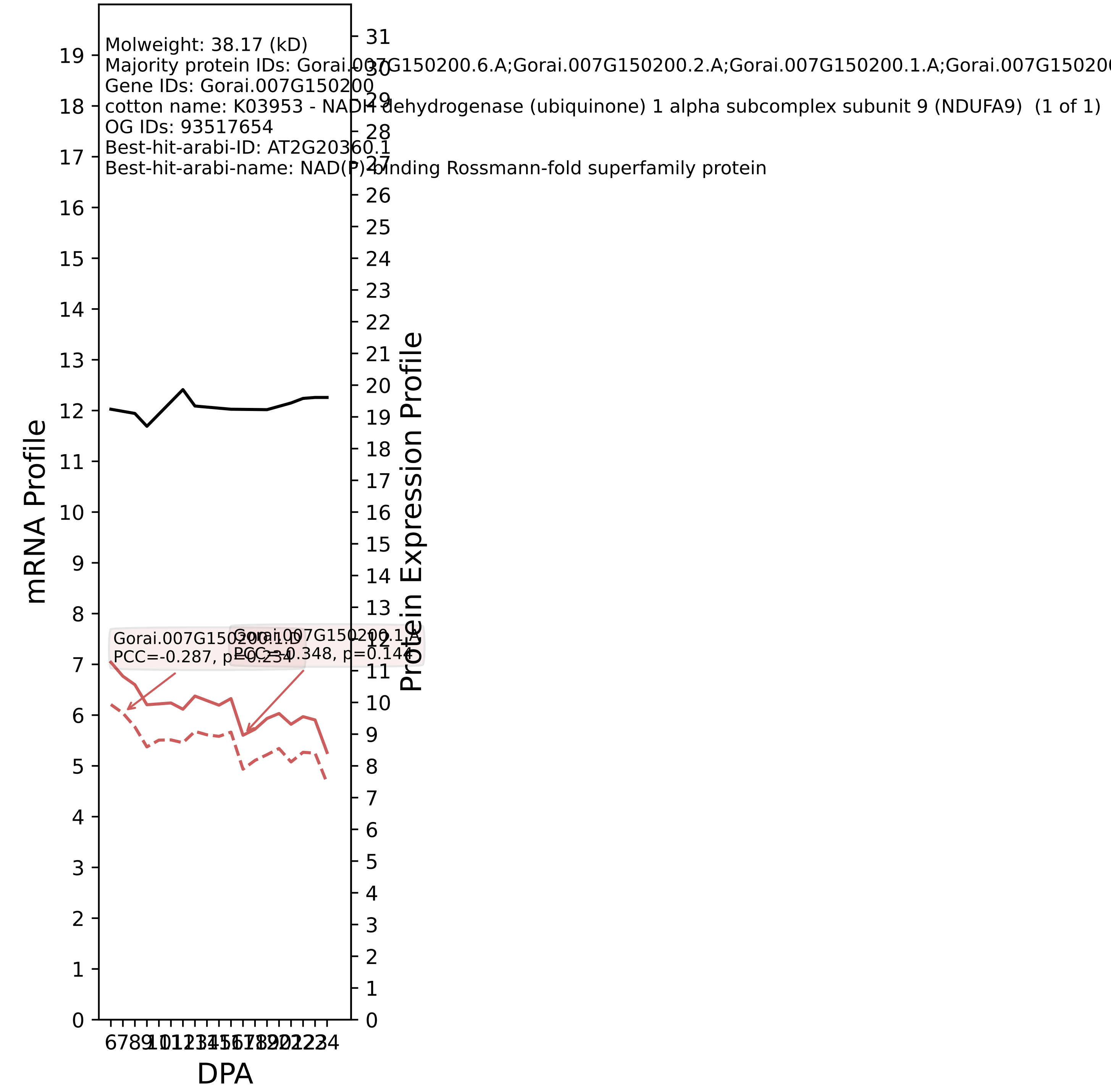

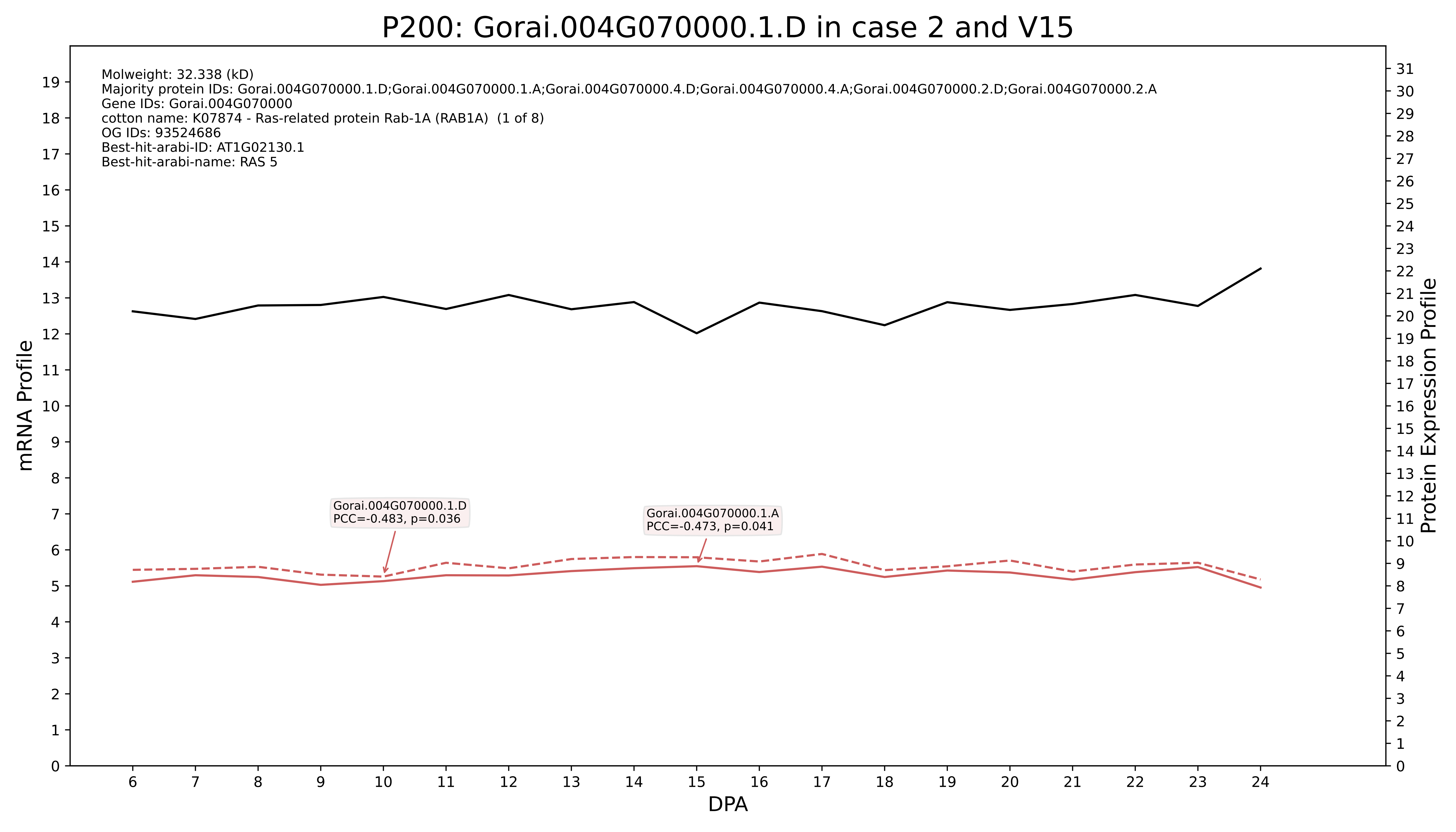
P200: Gorai.004G070000.1.D in case 2 and V15
Molweight: 32.338 (kD)
Majority protein IDs: Gorai.004G070000.1.D;Gorai.004G070000.1.A;Gorai.004G070000.4.D;Gorai.004G070000.4.A;Gorai.004G070000.2.D;Gorai.004G070000.2.A
Gene IDs: Gorai.004G070000
cotton name: K07874 - Ras-related protein Rab-1A (RAB1A) (1 of 8)
OG IDs: 93524686
Best-hit-arabi-ID: AT1G02130.1
Best-hit-arabi-name: RAS 5
Gorai.004G070000.1.D
PCC=-0.483, p=0.036
Gorai.004G070000.1.A
PCC=-0.473, p=0.041
mRNA Profile
Protein Expression Profile
DPA

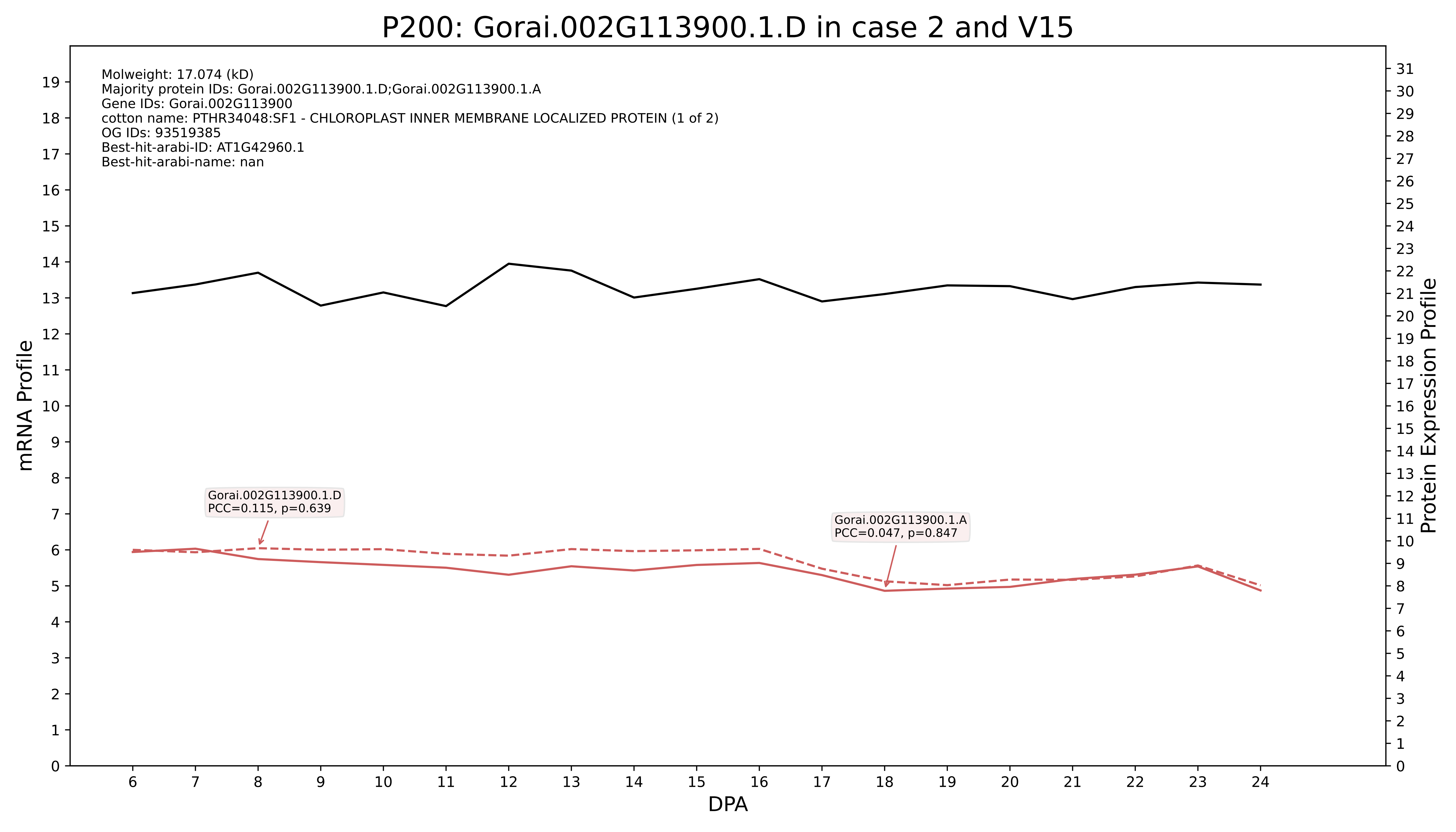

P200: Gorai.002G113900.1.D in case 2 and V15
Molweight: 17.074 (kD)
Majority protein IDs: Gorai.002G113900.1.D;Gorai.002G113900.1.A
Gene IDs: Gorai.002G113900
cotton name: PTHR34048:SF1 - CHLOROPLAST INNER MEMBRANE LOCALIZED PROTEIN (1 of 2)
OG IDs: 93519385
Best-hit-arabi-ID: AT1G42960.1
Best-hit-arabi-name: nan
Gorai.002G113900.1.D
PCC=0.115, p=0.639
Gorai.002G113900.1.A
PCC=0.047, p=0.847
mRNA Profile
Protein Expression Profile
DPA

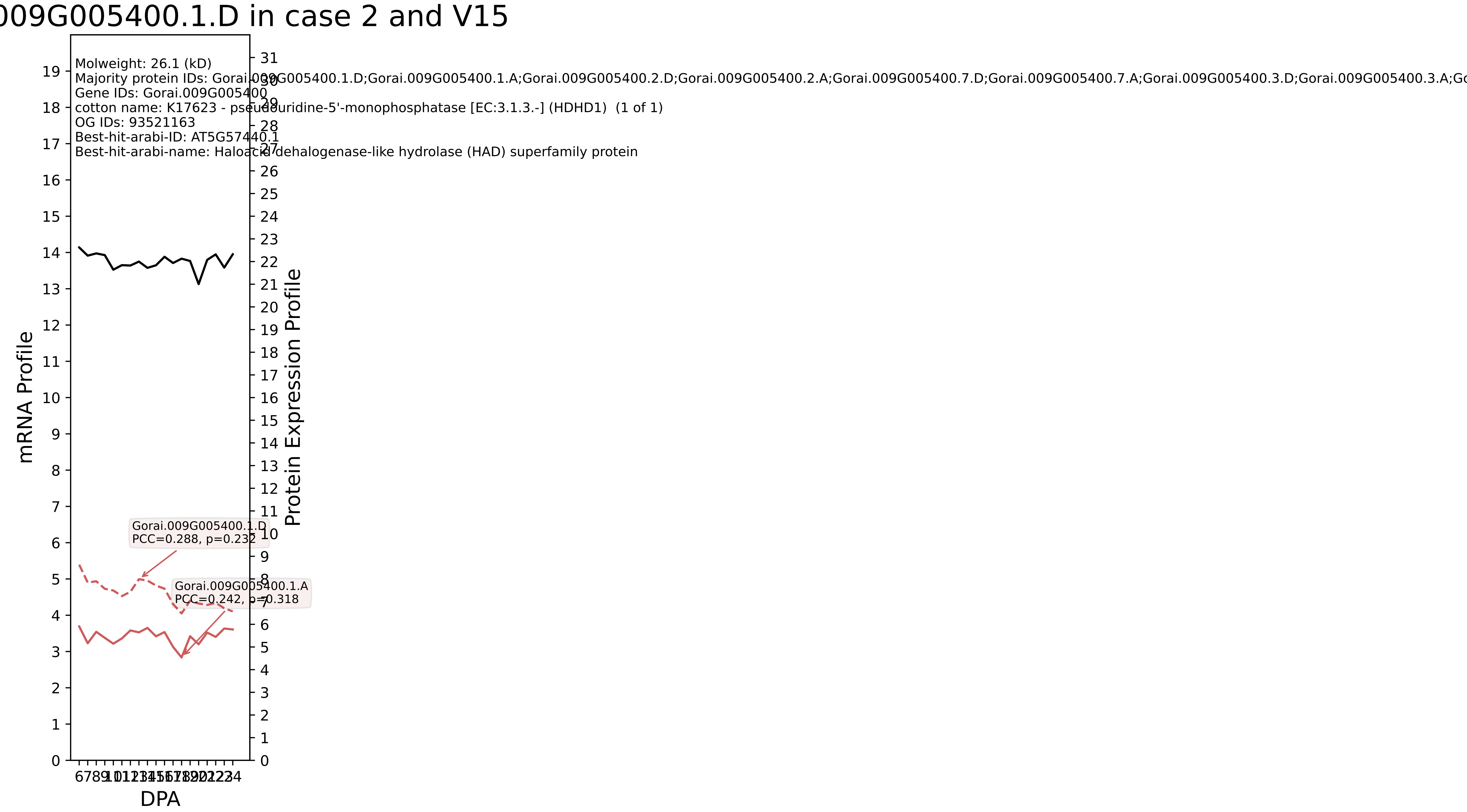

009G005400.1.D in case 2 and V15
Molweight: 26.1 (kD)
Majority protein IDs: Gorai.009G005400.1.D;Gorai.009G005400.1.A;Gorai.009G005400.2.D;Gorai.009G005400.2.A;Gorai.009G005400.7.D;Gorai.009G005400.7.A;Gorai.009G005400.3.D;Gorai.009G005400.3.A;Go
Gene IDs: Gorai.009G005400
cotton name: K17623 - pseudouridine-5'-monophosphatase [EC:3.1.3.-] (HDHD1) (1 of 1)
OG IDs: 93521163
Best-hit-arabi-ID: AT5G57440.1
Best-hit-arabi-name: Haloacid dehalogenase-like hydrolase (HAD) superfamily protein
Gorai.009G005400.1.D
PCC=0.288, p=0.232
Gorai.009G005400.1.A
PCC=0.242, p=0.318
mRNA Profile
Protein Expression Profile
DPA

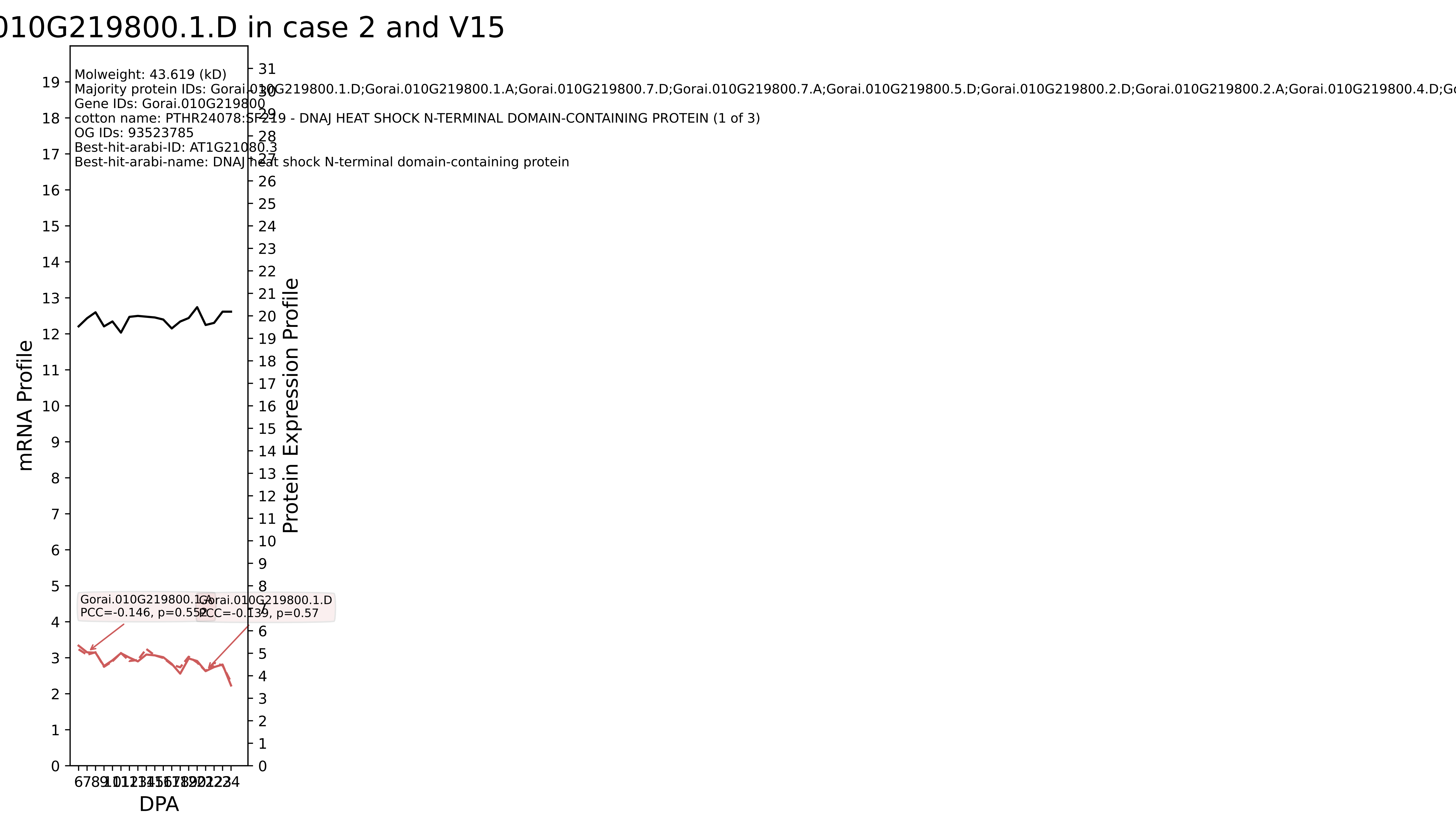
010G219800.1.D in case 2 and V15
Molweight: 43.619 (kD)
Majority protein IDs: Gorai.010G219800.1.D;Gorai.010G219800.1.A;Gorai.010G219800.7.D;Gorai.010G219800.7.A;Gorai.010G219800.5.D;Gorai.010G219800.2.D;Gorai.010G219800.2.A;Gorai.010G219800.4.D;Go
Gene IDs: Gorai.010G219800
cotton name: PTHR24078:SF219 - DNAJ HEAT SHOCK N-TERMINAL DOMAIN-CONTAINING PROTEIN (1 of 3)
OG IDs: 93523785
Best-hit-arabi-ID: AT1G21080.3
Best-hit-arabi-name: DNAJ heat shock N-terminal domain-containing protein
Gorai.010G219800.1.A
PCC=-0.146, p=0.552
Gorai.010G219800.1.D
PCC=-0.139, p=0.57
mRNA Profile
Protein Expression Profile
DPA

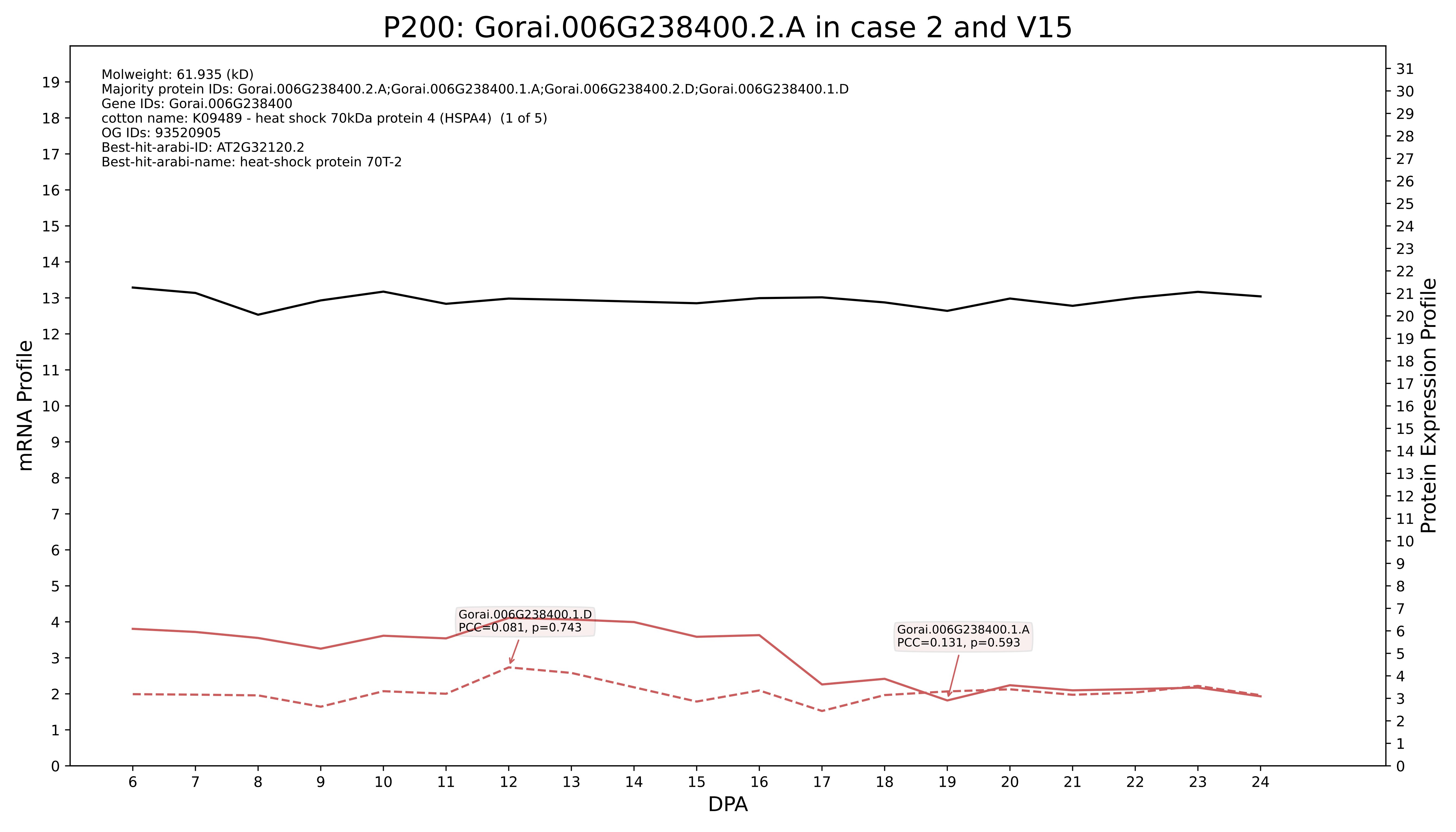
P200: Gorai.006G238400.2.A in case 2 and V15
Molweight: 61.935 (kD)
Majority protein IDs: Gorai.006G238400.2.A;Gorai.006G238400.1.A;Gorai.006G238400.2.D;Gorai.006G238400.1.D
Gene IDs: Gorai.006G238400
cotton name: K09489 - heat shock 70kDa protein 4 (HSPA4) (1 of 5)
OG IDs: 93520905
Best-hit-arabi-ID: AT2G32120.2
Best-hit-arabi-name: heat-shock protein 70T-2
Gorai.006G238400.1.D
PCC=0.081, p=0.743
Gorai.006G238400.1.A
PCC=0.131, p=0.593
mRNA Profile
Protein Expression Profile
DPA

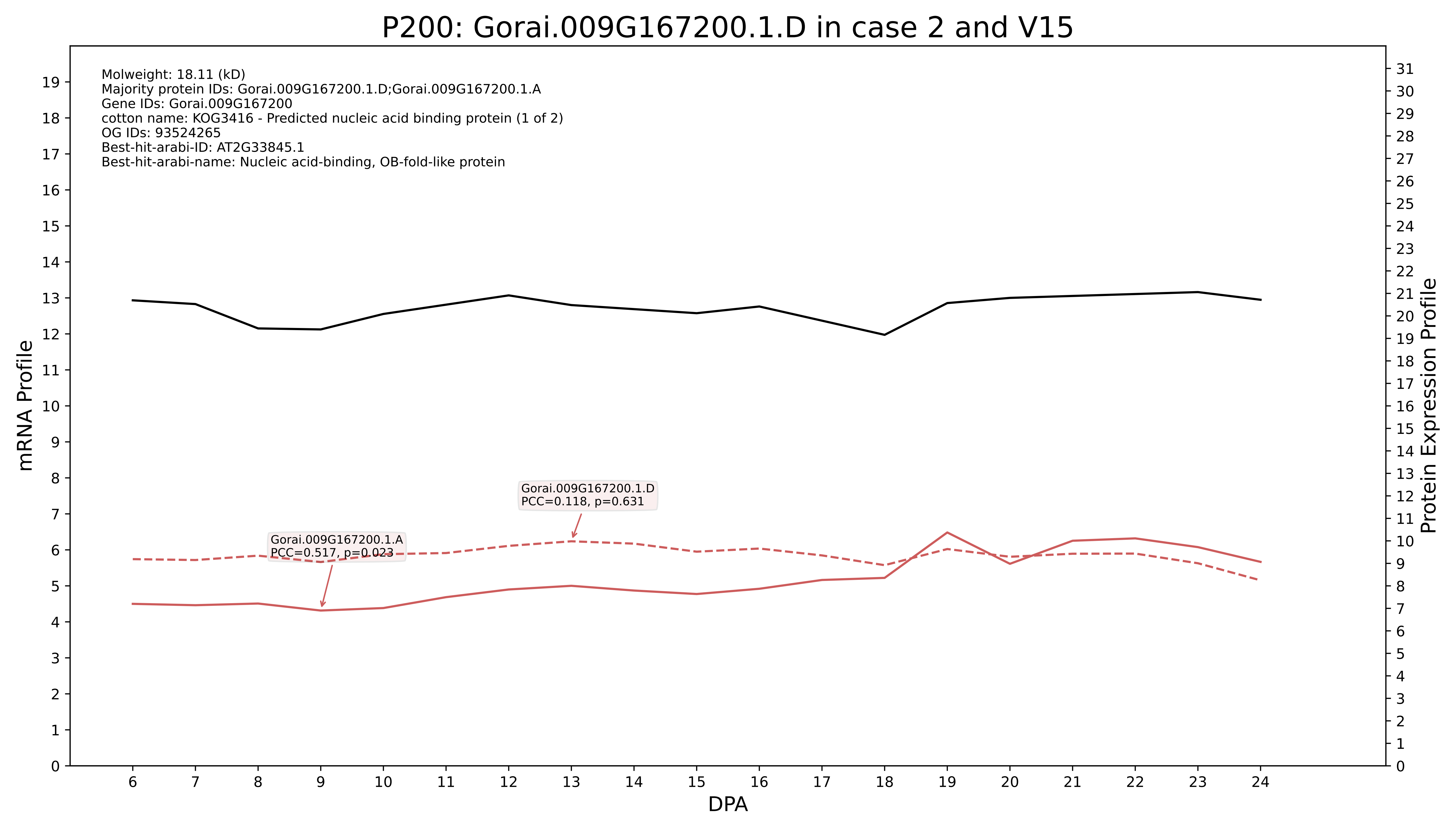
P200: Gorai.009G167200.1.D in case 2 and V15
Molweight: 18.11 (kD)
Majority protein IDs: Gorai.009G167200.1.D;Gorai.009G167200.1.A
Gene IDs: Gorai.009G167200
cotton name: KOG3416 - Predicted nucleic acid binding protein (1 of 2)
OG IDs: 93524265
Best-hit-arabi-ID: AT2G33845.1
Best-hit-arabi-name: Nucleic acid-binding, OB-fold-like protein
Gorai.009G167200.1.D
PCC=0.118, p=0.631
Gorai.009G167200.1.A
PCC=0.517, p=0.023
mRNA Profile
Protein Expression Profile
DPA

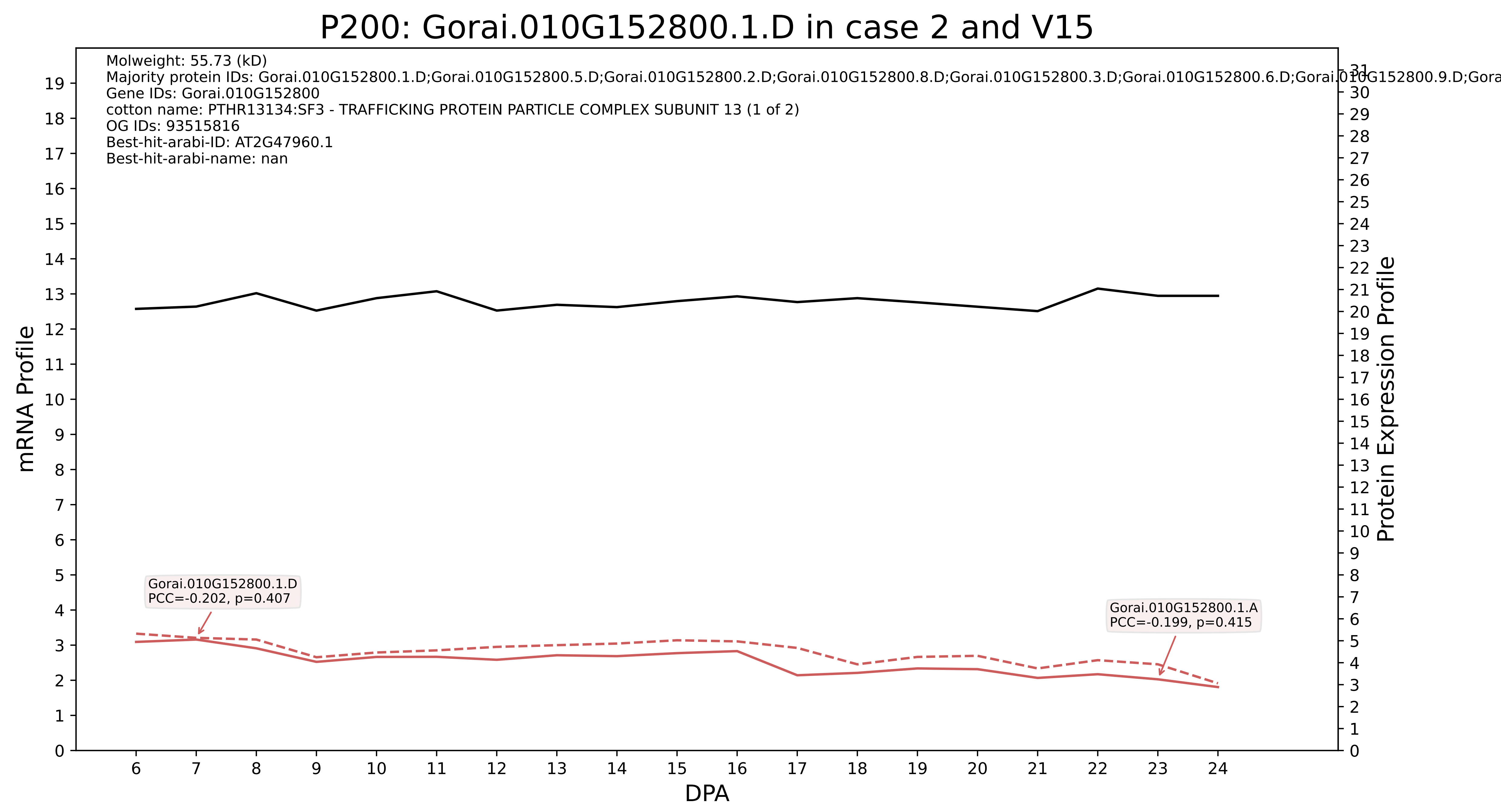

P200: Gorai.010G152800.1.D in case 2 and V15
Molweight: 55.73 (kD)
Majority protein IDs: Gorai.010G152800.1.D;Gorai.010G152800.5.D;Gorai.010G152800.2.D;Gorai.010G152800.8.D;Gorai.010G152800.3.D;Gorai.010G152800.6.D;Gorai.010G152800.9.D;Gora
Gene IDs: Gorai.010G152800
cotton name: PTHR13134:SF3 - TRAFFICKING PROTEIN PARTICLE COMPLEX SUBUNIT 13 (1 of 2)
OG IDs: 93515816
Best-hit-arabi-ID: AT2G47960.1
Best-hit-arabi-name: nan
Gorai.010G152800.1.D
PCC=-0.202, p=0.407
Gorai.010G152800.1.A
PCC=-0.199, p=0.415
mRNA Profile
Protein Expression Profile
DPA

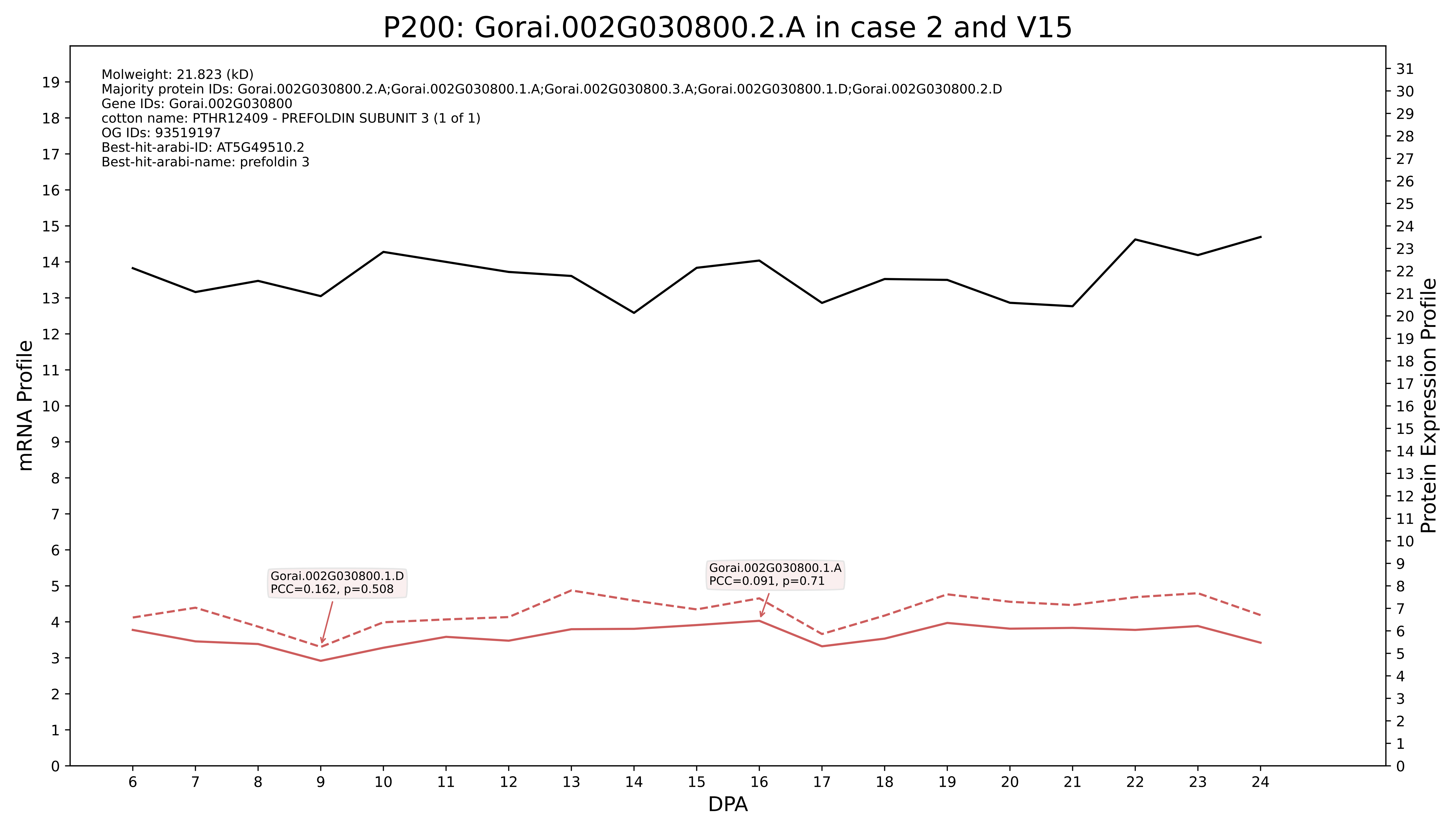
P200: Gorai.002G030800.2.A in case 2 and V15
Molweight: 21.823 (kD)
Majority protein IDs: Gorai.002G030800.2.A;Gorai.002G030800.1.A;Gorai.002G030800.3.A;Gorai.002G030800.1.D;Gorai.002G030800.2.D
Gene IDs: Gorai.002G030800
cotton name: PTHR12409 - PREFOLDIN SUBUNIT 3 (1 of 1)
OG IDs: 93519197
Best-hit-arabi-ID: AT5G49510.2
Best-hit-arabi-name: prefoldin 3
Gorai.002G030800.1.D
PCC=0.162, p=0.508
Gorai.002G030800.1.A
PCC=0.091, p=0.71
mRNA Profile
Protein Expression Profile
DPA

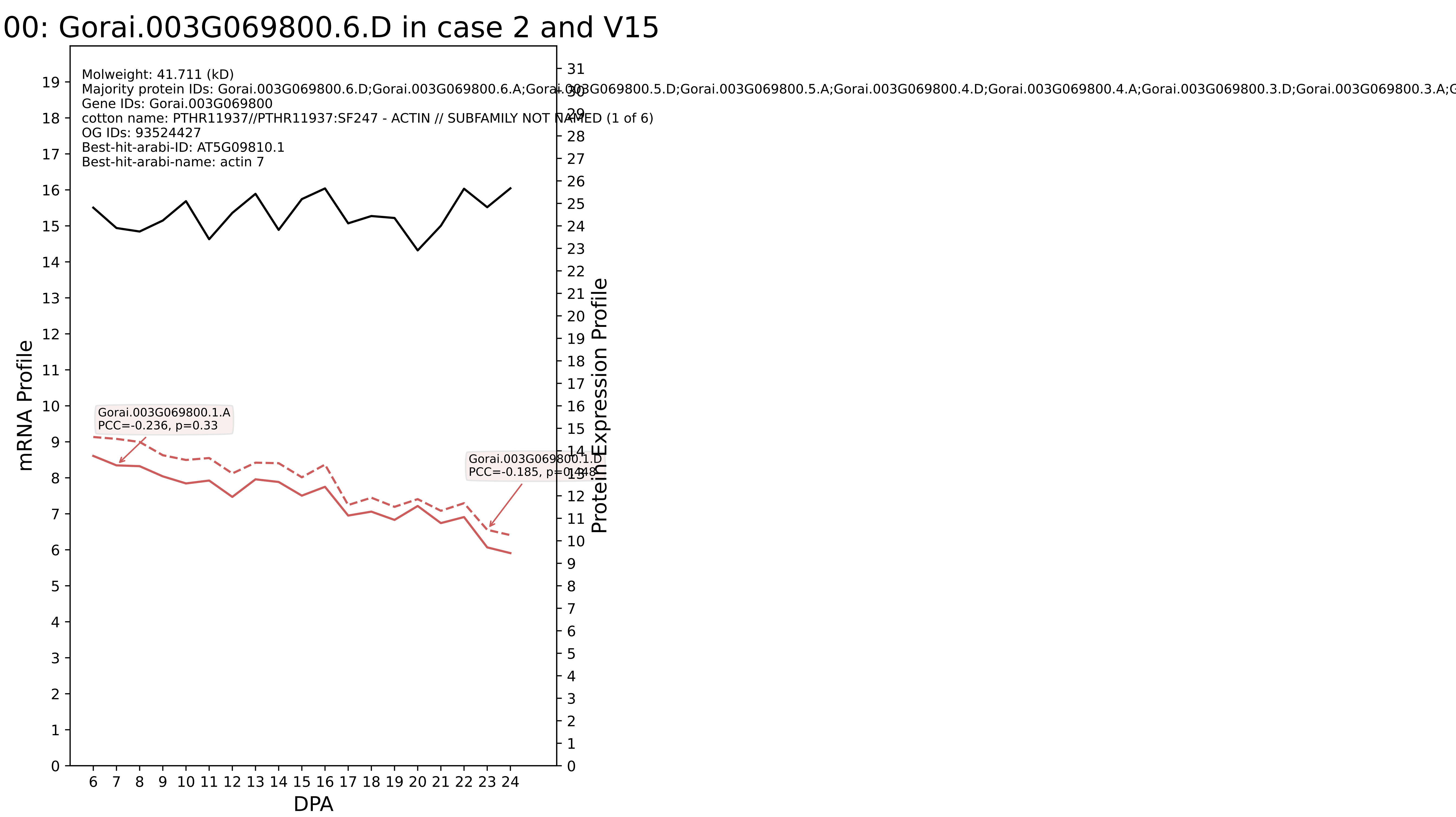
00: Gorai.003G069800.6.D in case 2 and V15
Molweight: 41.711 (kD)
Majority protein IDs: Gorai.003G069800.6.D;Gorai.003G069800.6.A;Gorai.003G069800.5.D;Gorai.003G069800.5.A;Gorai.003G069800.4.D;Gorai.003G069800.4.A;Gorai.003G069800.3.D;Gorai.003G069800.3.A;G
Gene IDs: Gorai.003G069800
cotton name: PTHR11937//PTHR11937:SF247 - ACTIN // SUBFAMILY NOT NAMED (1 of 6)
OG IDs: 93524427
Best-hit-arabi-ID: AT5G09810.1
Best-hit-arabi-name: actin 7
Gorai.003G069800.1.A
PCC=-0.236, p=0.33
Gorai.003G069800.1.D
PCC=-0.185, p=0.448
mRNA Profile
Protein Expression Profile
DPA

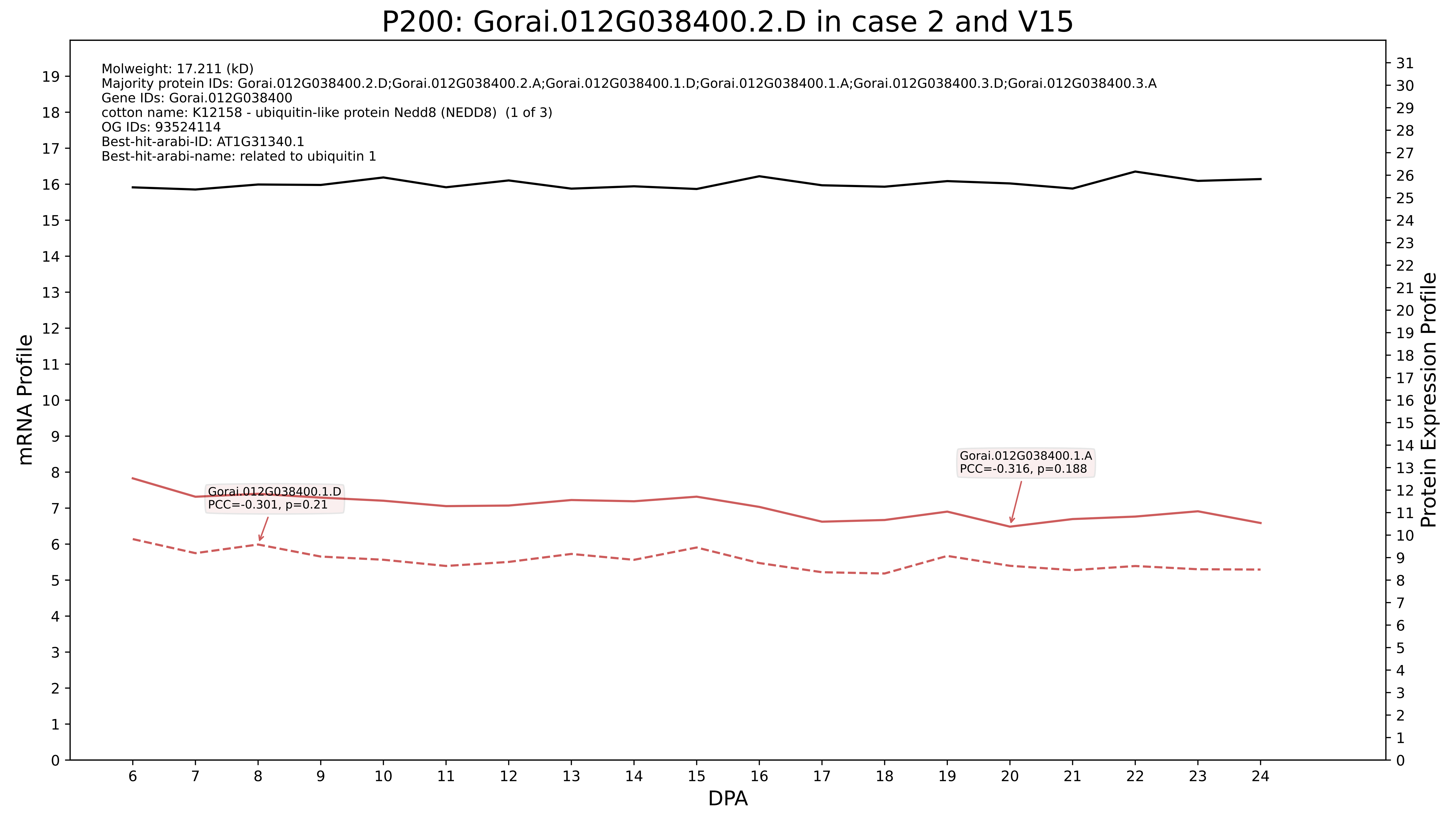
P200: Gorai.012G038400.2.D in case 2 and V15
Molweight: 17.211 (kD)
Majority protein IDs: Gorai.012G038400.2.D;Gorai.012G038400.2.A;Gorai.012G038400.1.D;Gorai.012G038400.1.A;Gorai.012G038400.3.D;Gorai.012G038400.3.A
Gene IDs: Gorai.012G038400
cotton name: K12158 - ubiquitin-like protein Nedd8 (NEDD8) (1 of 3)
OG IDs: 93524114
Best-hit-arabi-ID: AT1G31340.1
Best-hit-arabi-name: related to ubiquitin 1
Gorai.012G038400.1.D
PCC=-0.301, p=0.21
Gorai.012G038400.1.A
PCC=-0.316, p=0.188
mRNA Profile
Protein Expression Profile
DPA

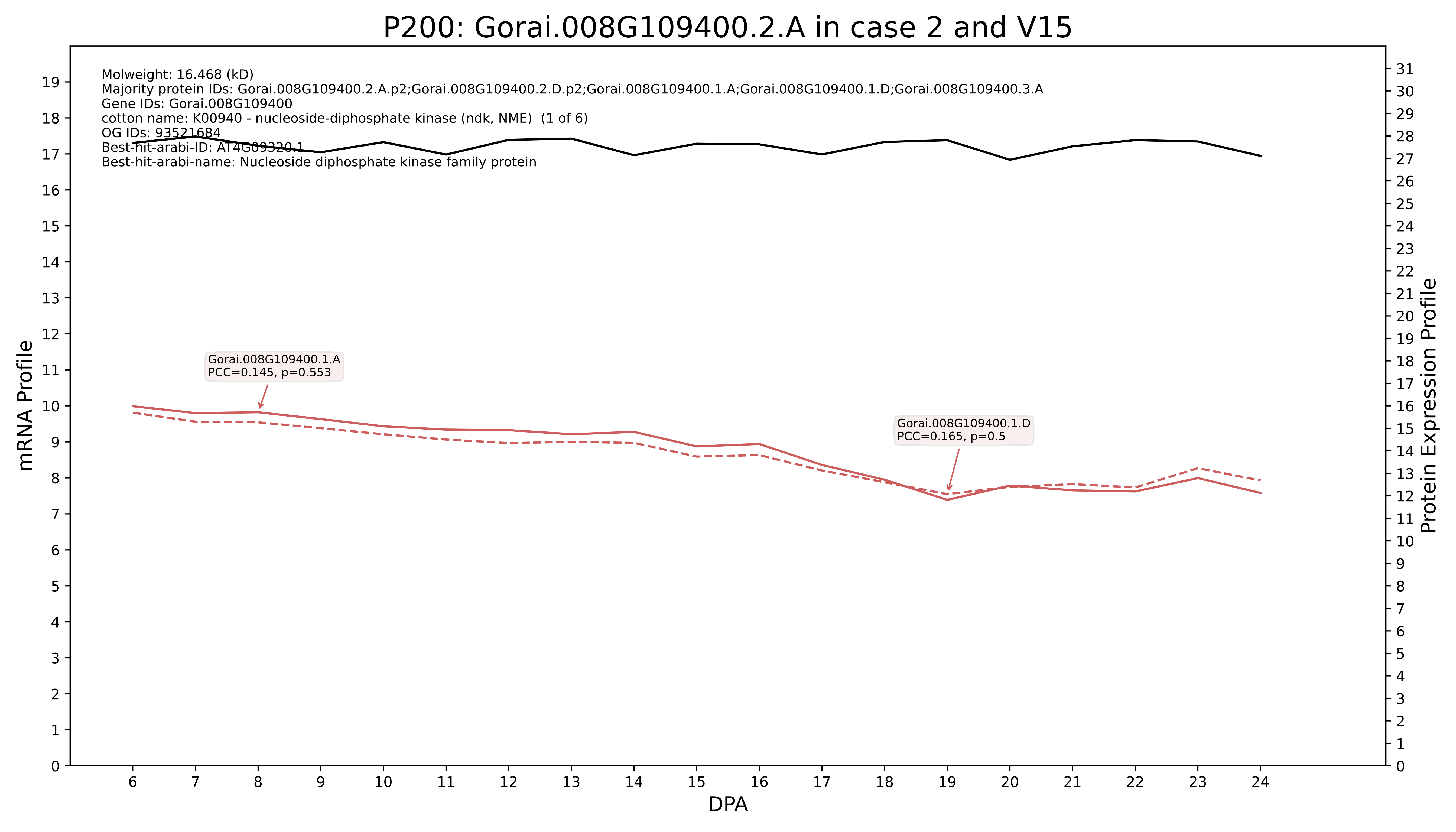
P200: Gorai.008G109400.2.A in case 2 and V15
Molweight: 16.468 (kD)
Majority protein IDs: Gorai.008G109400.2.A.p2;Gorai.008G109400.2.D.p2;Gorai.008G109400.1.A;Gorai.008G109400.1.D;Gorai.008G109400.3.A
Gene IDs: Gorai.008G109400
cotton name: K00940 - nucleoside-diphosphate kinase (ndk, NME) (1 of 6)
OG IDs: 93521684
Best-hit-arabi-ID: AT4G09320.1
Best-hit-arabi-name: Nucleoside diphosphate kinase family protein
Gorai.008G109400.1.A
PCC=0.145, p=0.553
Gorai.008G109400.1.D
PCC=0.165, p=0.5
mRNA Profile
Protein Expression Profile
DPA

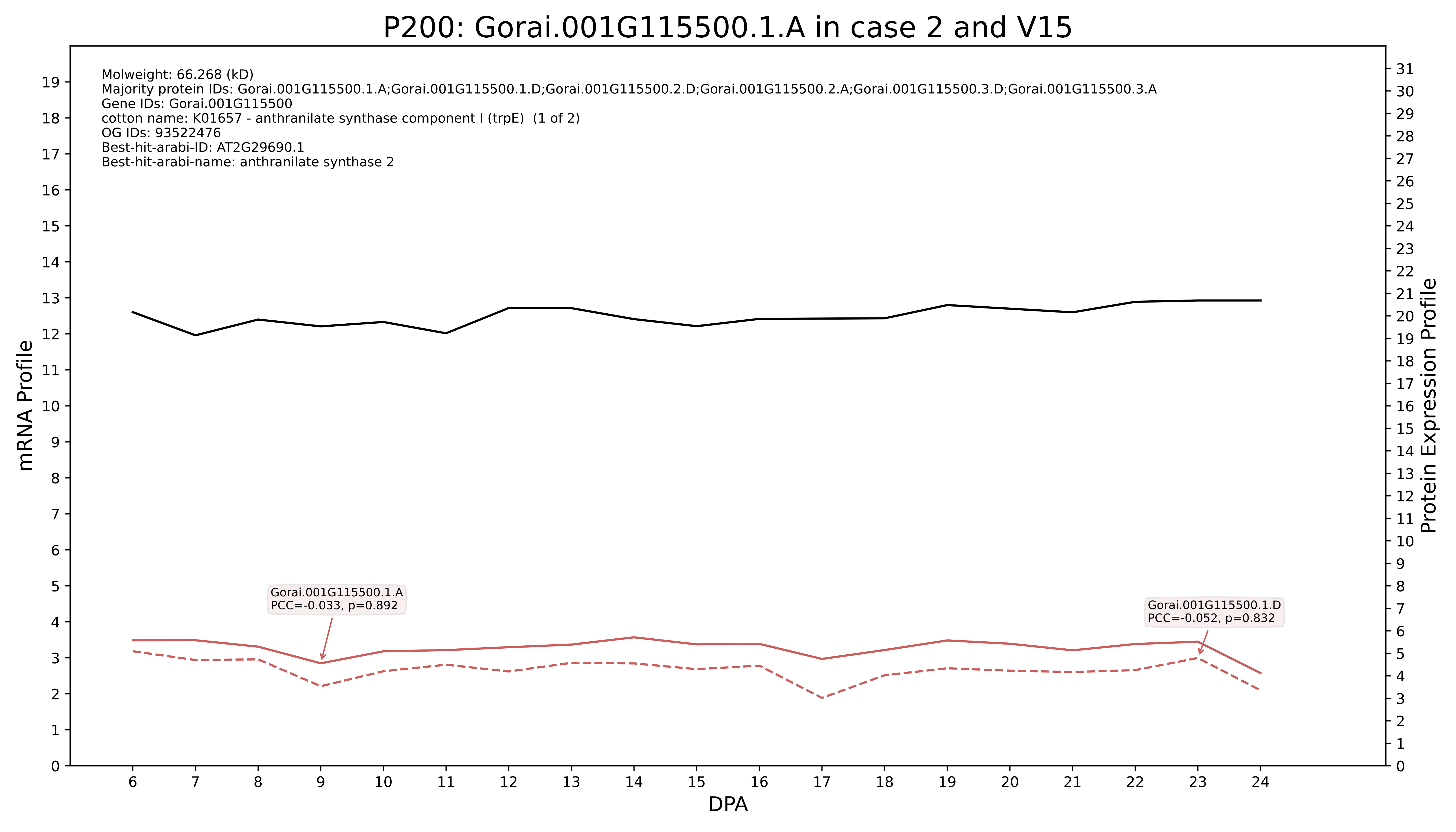
P200: Gorai.001G115500.1.A in case 2 and V15
Molweight: 66.268 (kD)
Majority protein IDs: Gorai.001G115500.1.A;Gorai.001G115500.1.D;Gorai.001G115500.2.D;Gorai.001G115500.2.A;Gorai.001G115500.3.D;Gorai.001G115500.3.A
Gene IDs: Gorai.001G115500
cotton name: K01657 - anthranilate synthase component I (trpE) (1 of 2)
OG IDs: 93522476
Best-hit-arabi-ID: AT2G29690.1
Best-hit-arabi-name: anthranilate synthase 2
Gorai.001G115500.1.A
PCC=-0.033, p=0.892
Gorai.001G115500.1.D
PCC=-0.052, p=0.832
mRNA Profile
Protein Expression Profile
DPA

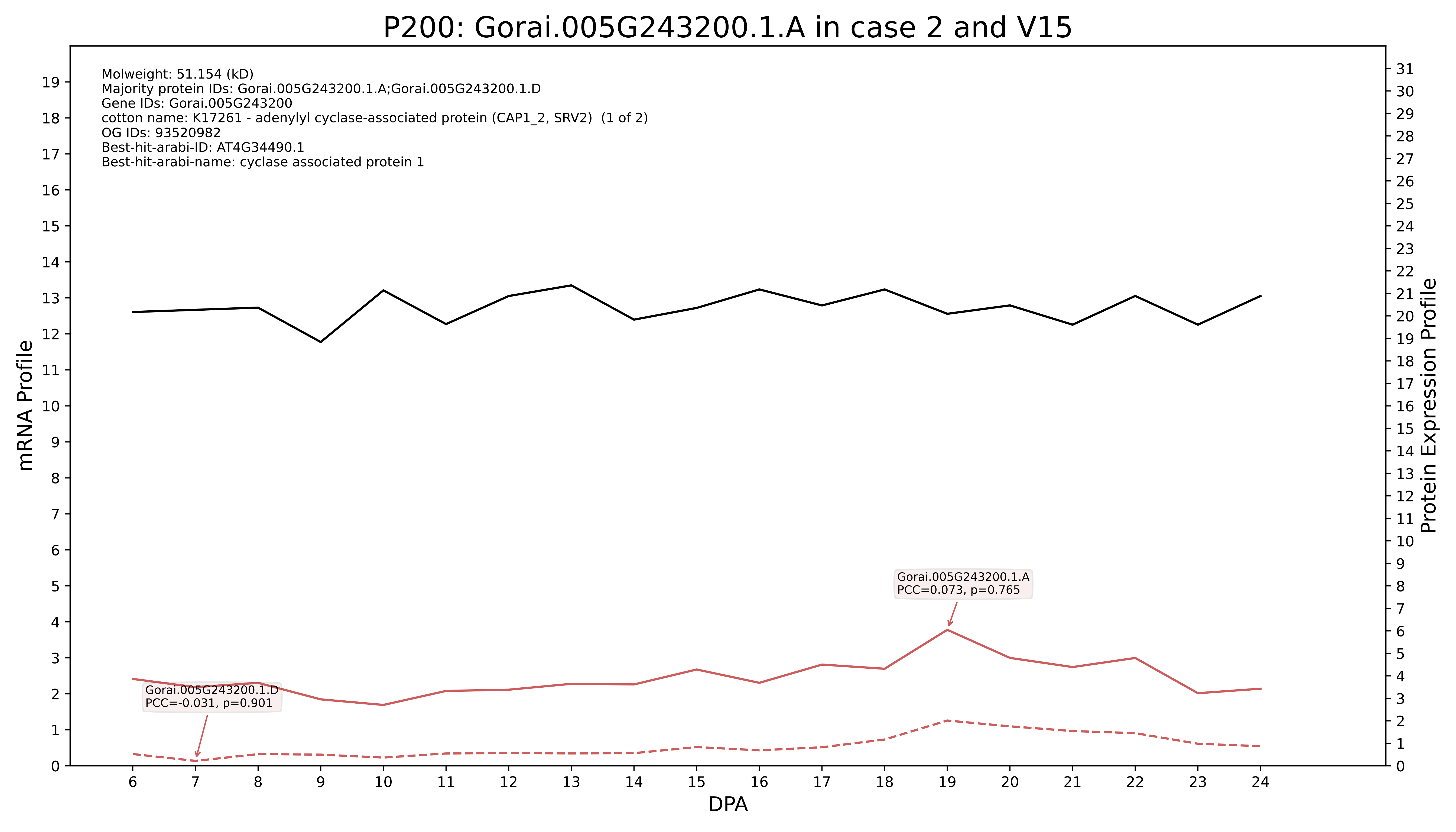
P200: Gorai.005G243200.1.A in case 2 and V15
Molweight: 51.154 (kD)
Majority protein IDs: Gorai.005G243200.1.A;Gorai.005G243200.1.D
Gene IDs: Gorai.005G243200
cotton name: K17261 - adenylyl cyclase-associated protein (CAP1_2, SRV2) (1 of 2)
OG IDs: 93520982
Best-hit-arabi-ID: AT4G34490.1
Best-hit-arabi-name: cyclase associated protein 1
Gorai.005G243200.1.A
PCC=0.073, p=0.765
Gorai.005G243200.1.D
PCC=-0.031, p=0.901
mRNA Profile
Protein Expression Profile
DPA

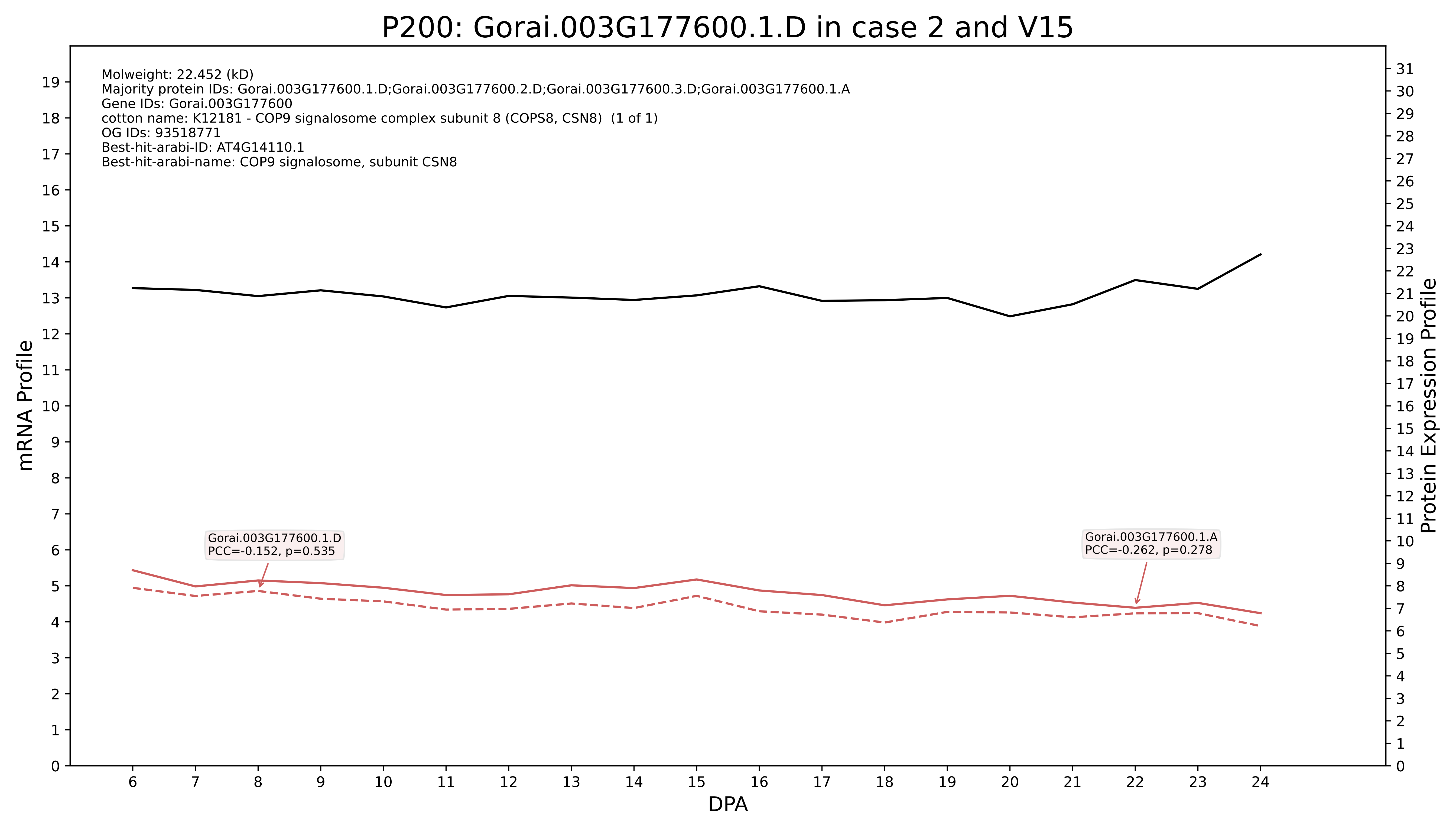
P200: Gorai.003G177600.1.D in case 2 and V15
Molweight: 22.452 (kD)
Majority protein IDs: Gorai.003G177600.1.D;Gorai.003G177600.2.D;Gorai.003G177600.3.D;Gorai.003G177600.1.A
Gene IDs: Gorai.003G177600
cotton name: K12181 - COP9 signalosome complex subunit 8 (COPS8, CSN8) (1 of 1)
OG IDs: 93518771
Best-hit-arabi-ID: AT4G14110.1
Best-hit-arabi-name: COP9 signalosome, subunit CSN8
Gorai.003G177600.1.D
PCC=-0.152, p=0.535
Gorai.003G177600.1.A
PCC=-0.262, p=0.278
mRNA Profile
Protein Expression Profile
DPA

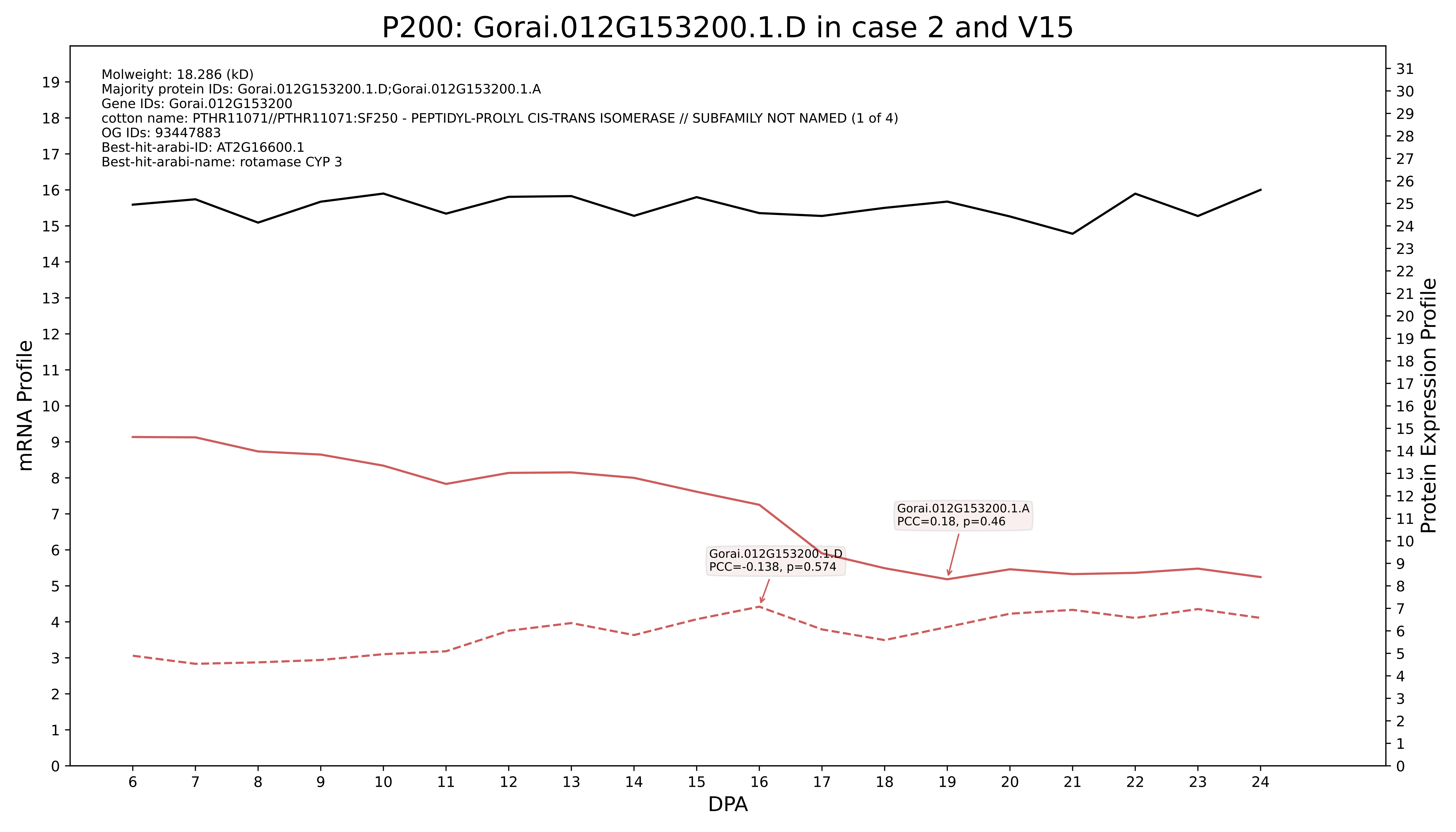
P200: Gorai.012G153200.1.D in case 2 and V15
Molweight: 18.286 (kD)
Majority protein IDs: Gorai.012G153200.1.D;Gorai.012G153200.1.A
Gene IDs: Gorai.012G153200
cotton name: PTHR11071//PTHR11071:SF250 - PEPTIDYL-PROLYL CIS-TRANS ISOMERASE // SUBFAMILY NOT NAMED (1 of 4)
OG IDs: 93447883
Best-hit-arabi-ID: AT2G16600.1
Best-hit-arabi-name: rotamase CYP 3
Gorai.012G153200.1.A
PCC=0.18, p=0.46
Gorai.012G153200.1.D
PCC=-0.138, p=0.574
mRNA Profile
Protein Expression Profile
DPA

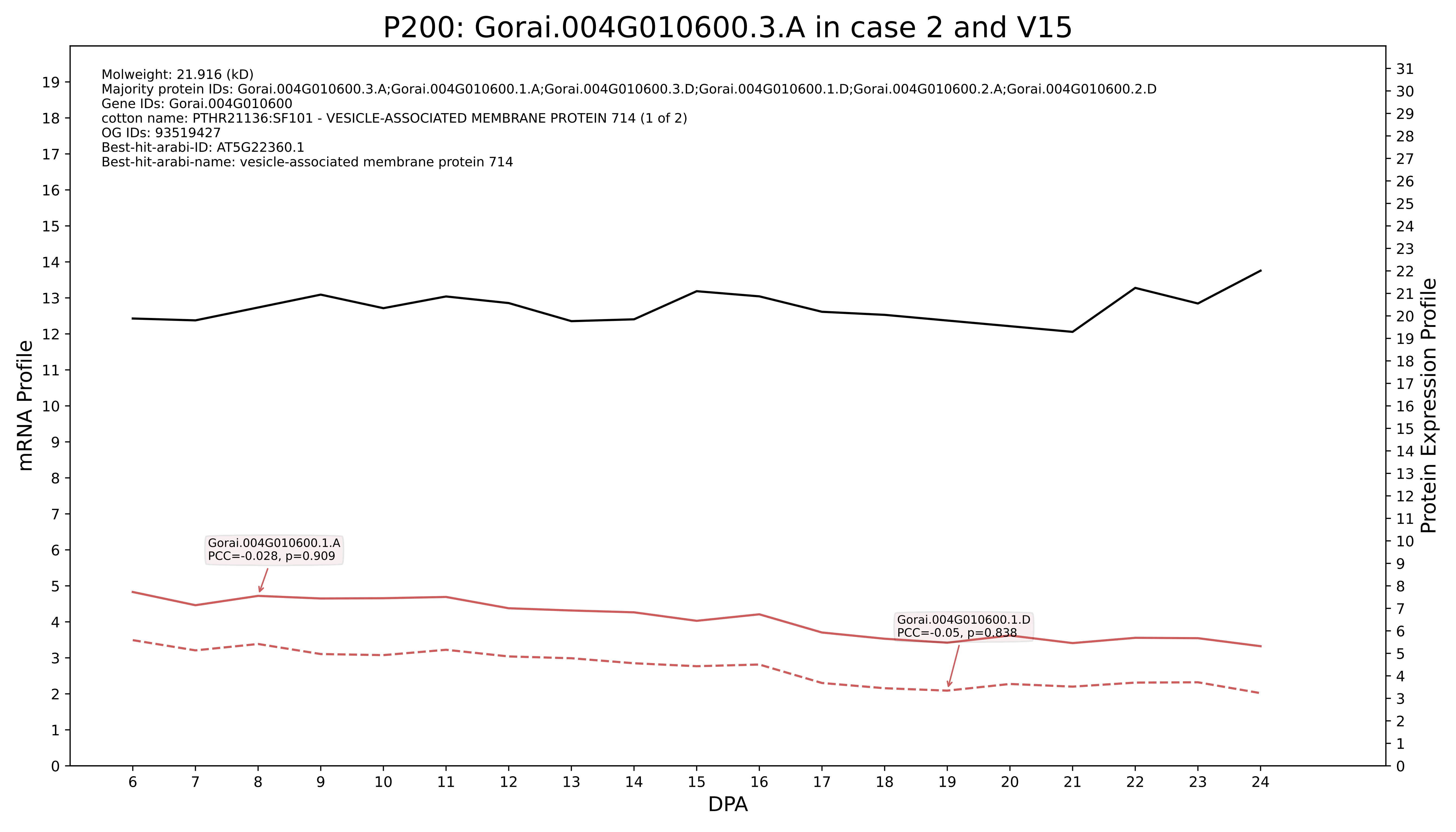

P200: Gorai.004G010600.3.A in case 2 and V15
Molweight: 21.916 (kD)
Majority protein IDs: Gorai.004G010600.3.A;Gorai.004G010600.1.A;Gorai.004G010600.3.D;Gorai.004G010600.1.D;Gorai.004G010600.2.A;Gorai.004G010600.2.D
Gene IDs: Gorai.004G010600
cotton name: PTHR21136:SF101 - VESICLE-ASSOCIATED MEMBRANE PROTEIN 714 (1 of 2)
OG IDs: 93519427
Best-hit-arabi-ID: AT5G22360.1
Best-hit-arabi-name: vesicle-associated membrane protein 714
Gorai.004G010600.1.A
PCC=-0.028, p=0.909
Gorai.004G010600.1.D
PCC=-0.05, p=0.838
mRNA Profile
Protein Expression Profile
DPA

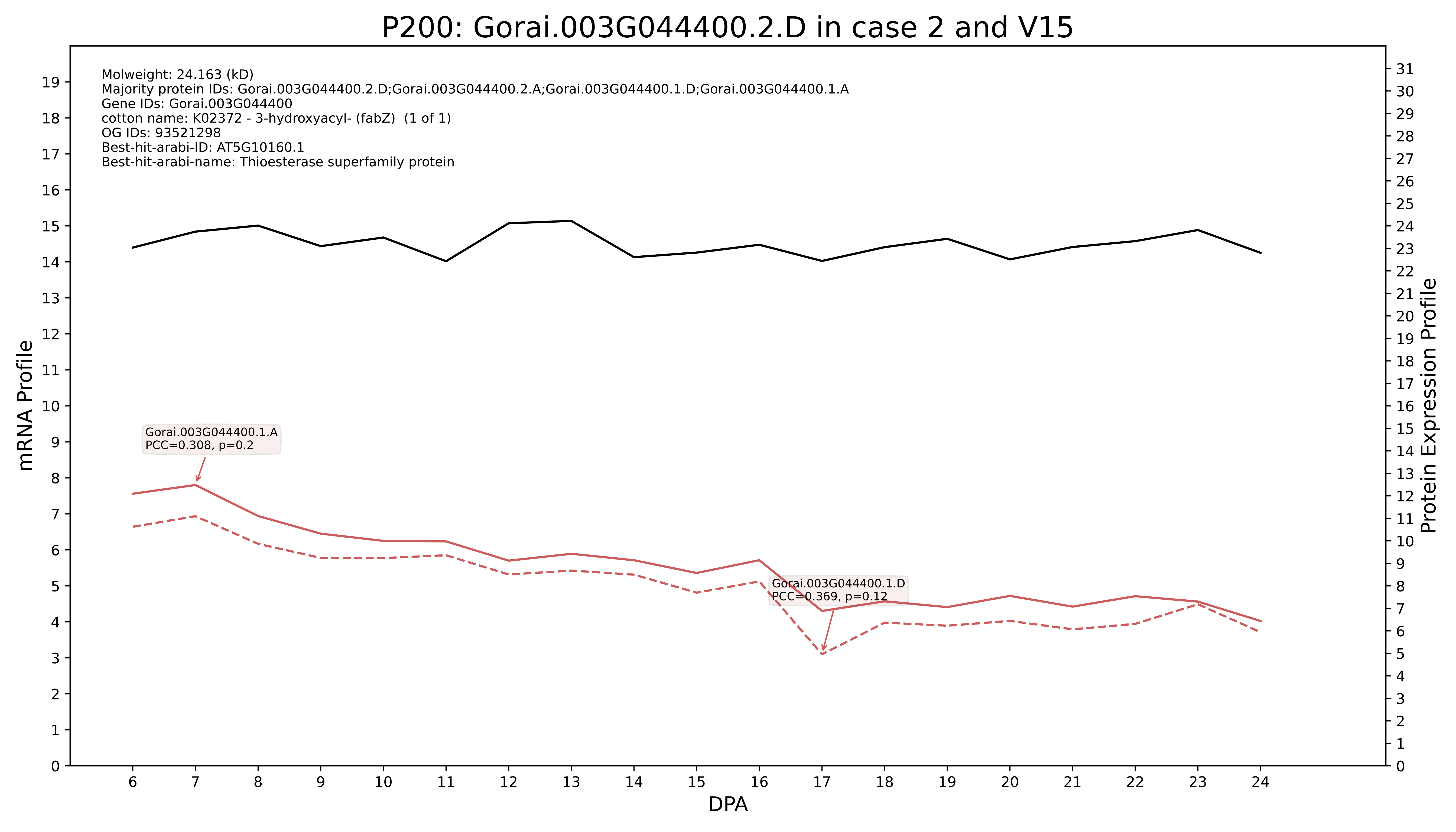

P200: Gorai.003G044400.2.D in case 2 and V15
Molweight: 24.163 (kD)
Majority protein IDs: Gorai.003G044400.2.D;Gorai.003G044400.2.A;Gorai.003G044400.1.D;Gorai.003G044400.1.A
Gene IDs: Gorai.003G044400
cotton name: K02372 - 3-hydroxyacyl- (fabZ) (1 of 1)
OG IDs: 93521298
Best-hit-arabi-ID: AT5G10160.1
Best-hit-arabi-name: Thioesterase superfamily protein
Gorai.003G044400.1.A
PCC=0.308, p=0.2
Gorai.003G044400.1.D
PCC=0.369, p=0.12
mRNA Profile
Protein Expression Profile
DPA

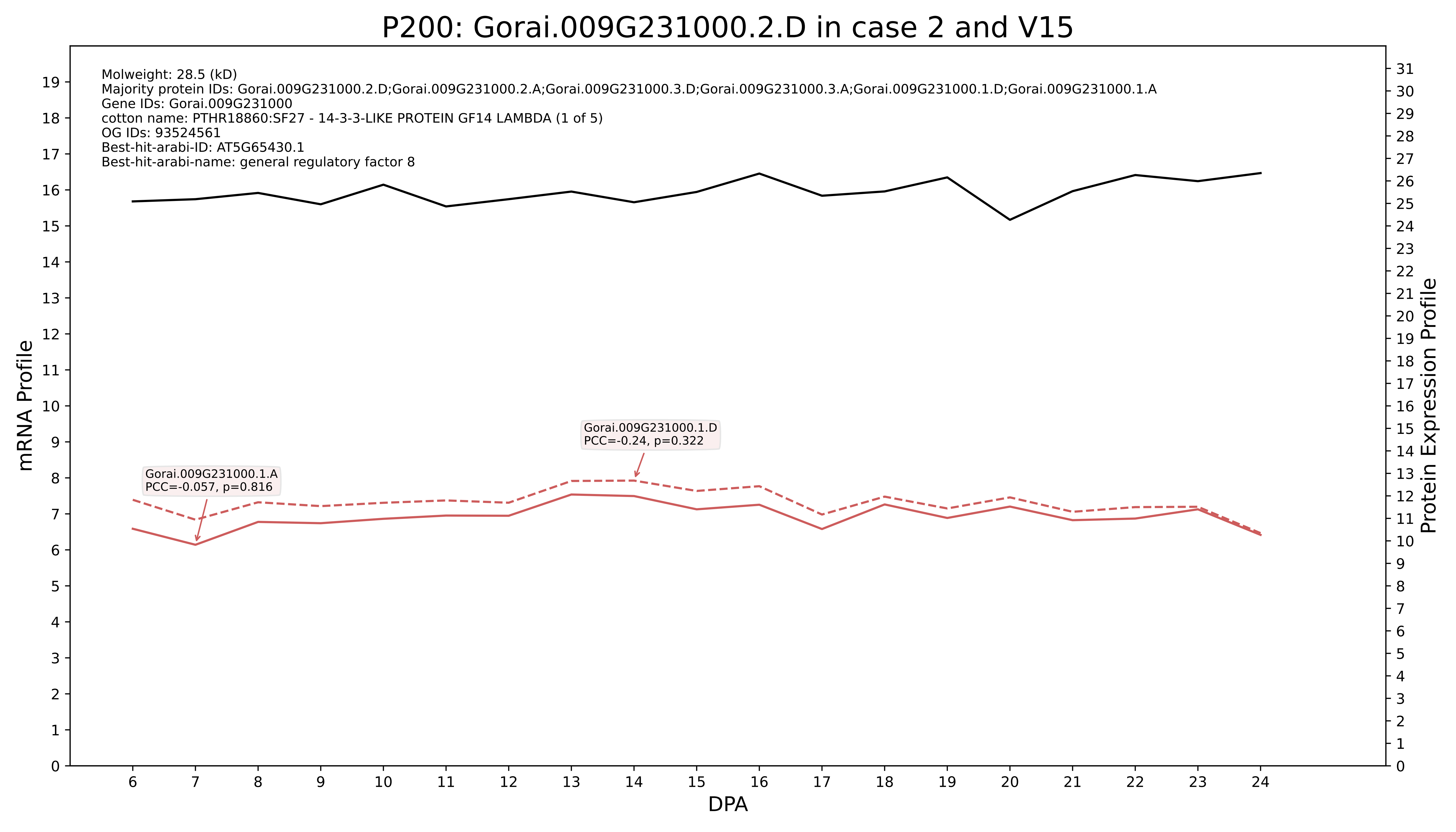

P200: Gorai.009G231000.2.D in case 2 and V15
Molweight: 28.5 (kD)
Majority protein IDs: Gorai.009G231000.2.D;Gorai.009G231000.2.A;Gorai.009G231000.3.D;Gorai.009G231000.3.A;Gorai.009G231000.1.D;Gorai.009G231000.1.A
Gene IDs: Gorai.009G231000
cotton name: PTHR18860:SF27 - 14-3-3-LIKE PROTEIN GF14 LAMBDA (1 of 5)
OG IDs: 93524561
Best-hit-arabi-ID: AT5G65430.1
Best-hit-arabi-name: general regulatory factor 8
Gorai.009G231000.1.D
PCC=-0.24, p=0.322
Gorai.009G231000.1.A
PCC=-0.057, p=0.816
mRNA Profile
Protein Expression Profile
DPA

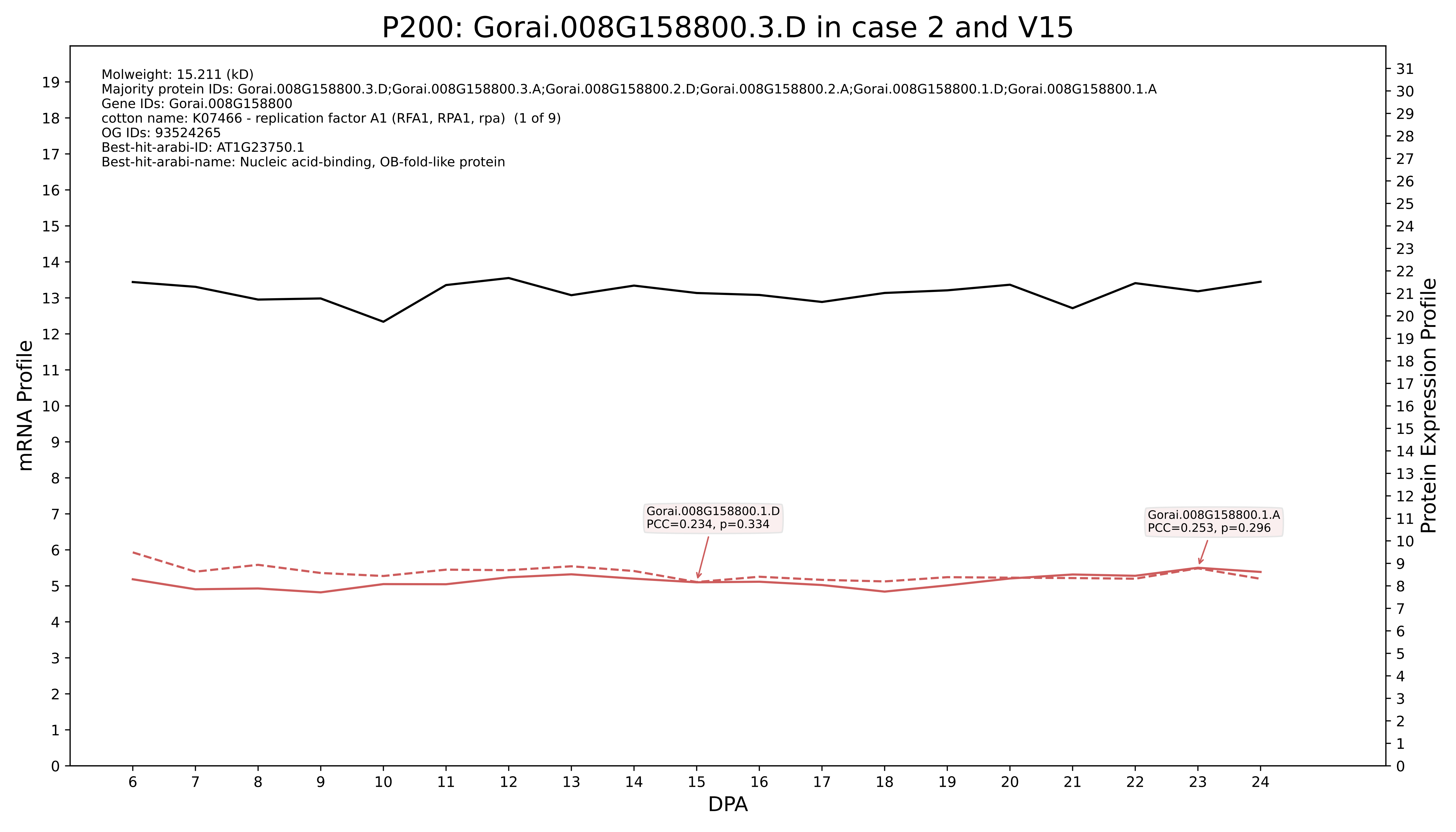

P200: Gorai.008G158800.3.D in case 2 and V15
Molweight: 15.211 (kD)
Majority protein IDs: Gorai.008G158800.3.D;Gorai.008G158800.3.A;Gorai.008G158800.2.D;Gorai.008G158800.2.A;Gorai.008G158800.1.D;Gorai.008G158800.1.A
Gene IDs: Gorai.008G158800
cotton name: K07466 - replication factor A1 (RFA1, RPA1, rpa) (1 of 9)
OG IDs: 93524265
Best-hit-arabi-ID: AT1G23750.1
Best-hit-arabi-name: Nucleic acid-binding, OB-fold-like protein
Gorai.008G158800.1.D
PCC=0.234, p=0.334
Gorai.008G158800.1.A
PCC=0.253, p=0.296
mRNA Profile
Protein Expression Profile
DPA

00: Gorai.005G242800.2.D in case 2 and V15

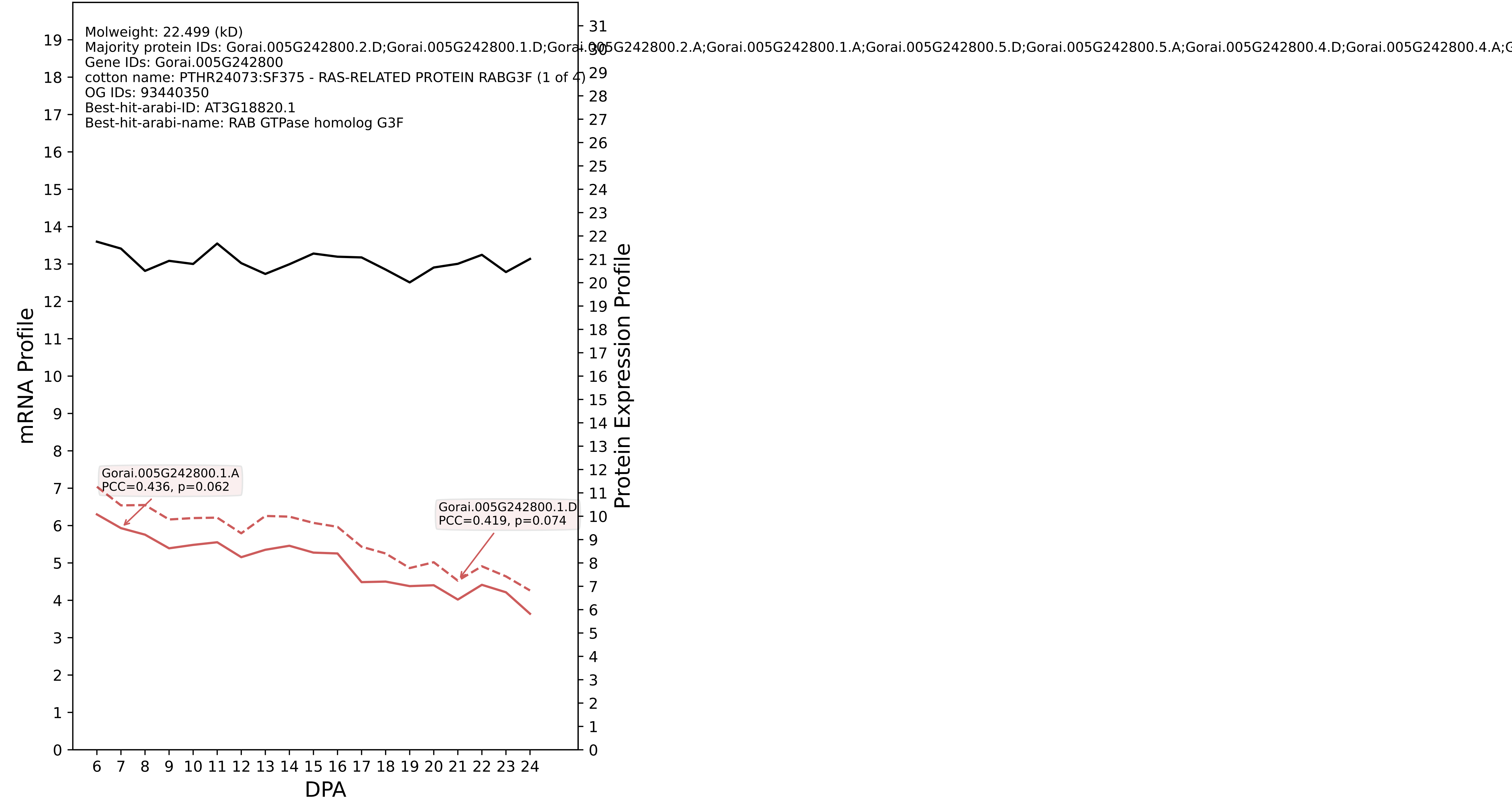

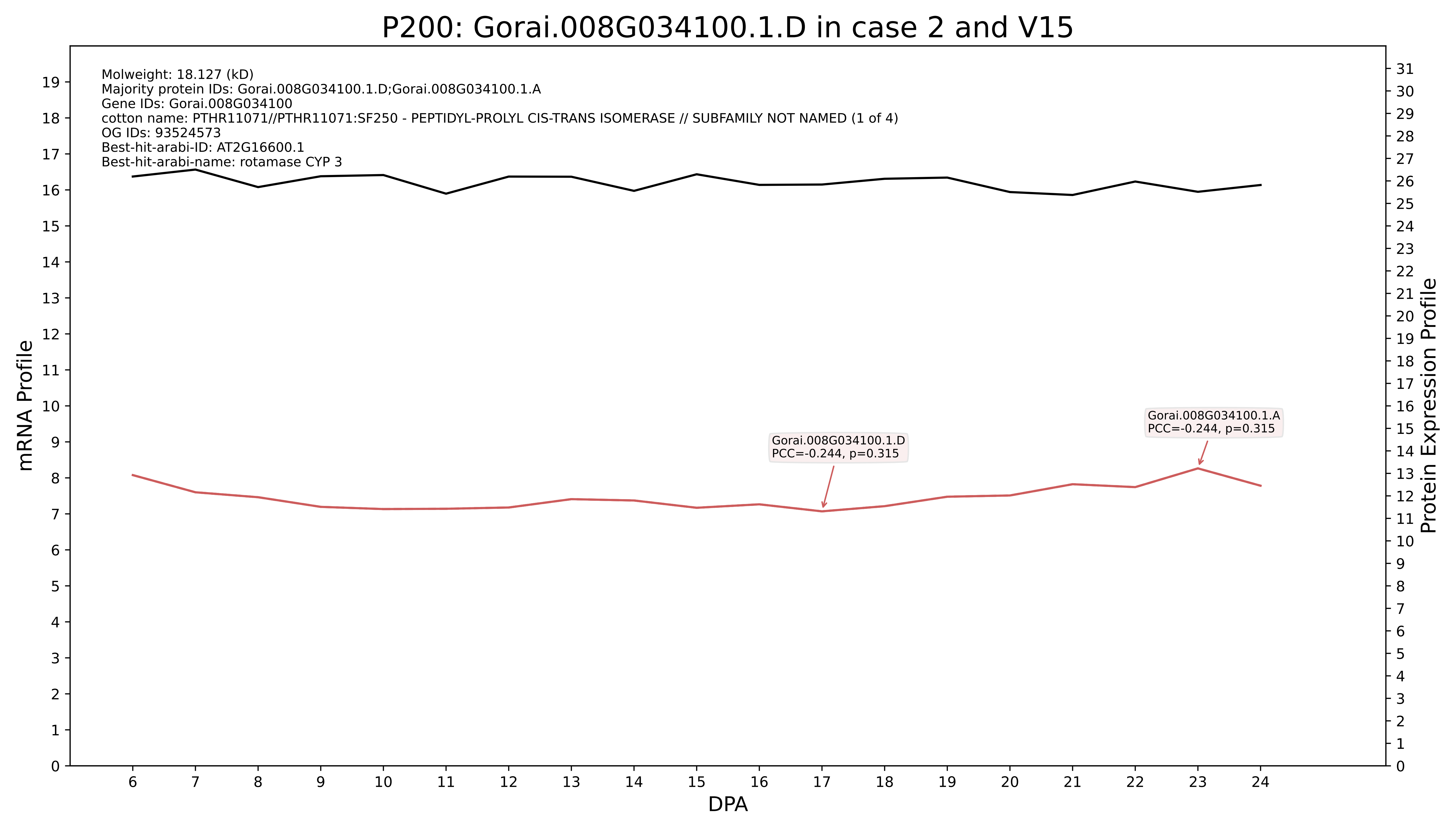
P200: Gorai.008G034100.1.D in case 2 and V15
Molweight: 18.127 (kD)
Majority protein IDs: Gorai.008G034100.1.D;Gorai.008G034100.1.A
Gene IDs: Gorai.008G034100
cotton name: PTHR11071//PTHR11071:SF250 - PEPTIDYL-PROLYL CIS-TRANS ISOMERASE // SUBFAMILY NOT NAMED (1 of 4)
OG IDs: 93524573
Best-hit-arabi-ID: AT2G16600.1
Best-hit-arabi-name: rotamase CYP 3
Gorai.008G034100.1.D
PCC=-0.244, p=0.315
Gorai.008G034100.1.A
PCC=-0.244, p=0.315
mRNA Profile
Protein Expression Profile
DPA

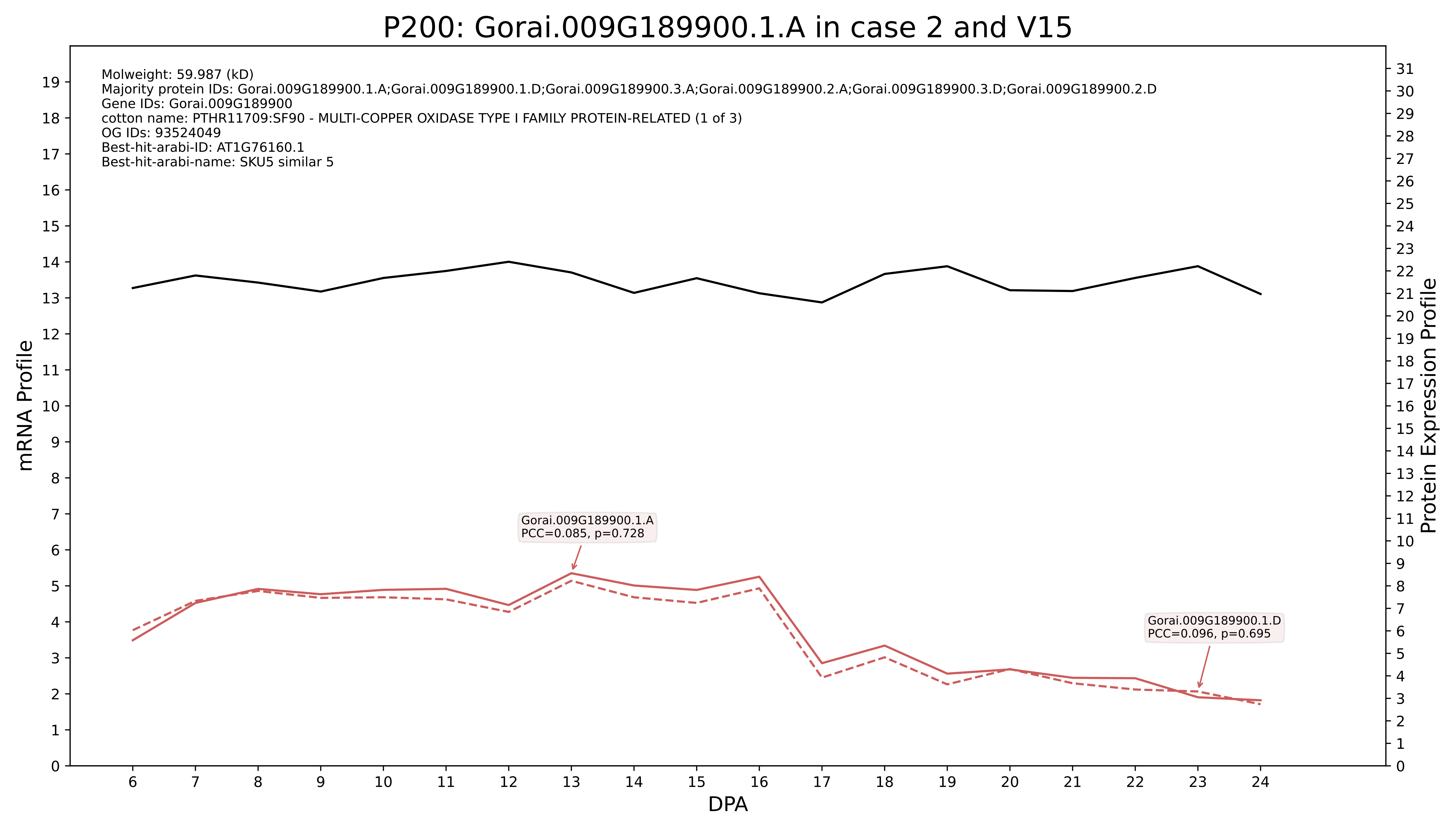
P200: Gorai.009G189900.1.A in case 2 and V15
Molweight: 59.987 (kD)
Majority protein IDs: Gorai.009G189900.1.A;Gorai.009G189900.1.D;Gorai.009G189900.3.A;Gorai.009G189900.2.A;Gorai.009G189900.3.D;Gorai.009G189900.2.D
Gene IDs: Gorai.009G189900
cotton name: PTHR11709:SF90 - MULTI-COPPER OXIDASE TYPE I FAMILY PROTEIN-RELATED (1 of 3)
OG IDs: 93524049
Best-hit-arabi-ID: AT1G76160.1
Best-hit-arabi-name: SKU5 similar 5
Gorai.009G189900.1.A
PCC=0.085, p=0.728
Gorai.009G189900.1.D
PCC=0.096, p=0.695
mRNA Profile
Protein Expression Profile
DPA

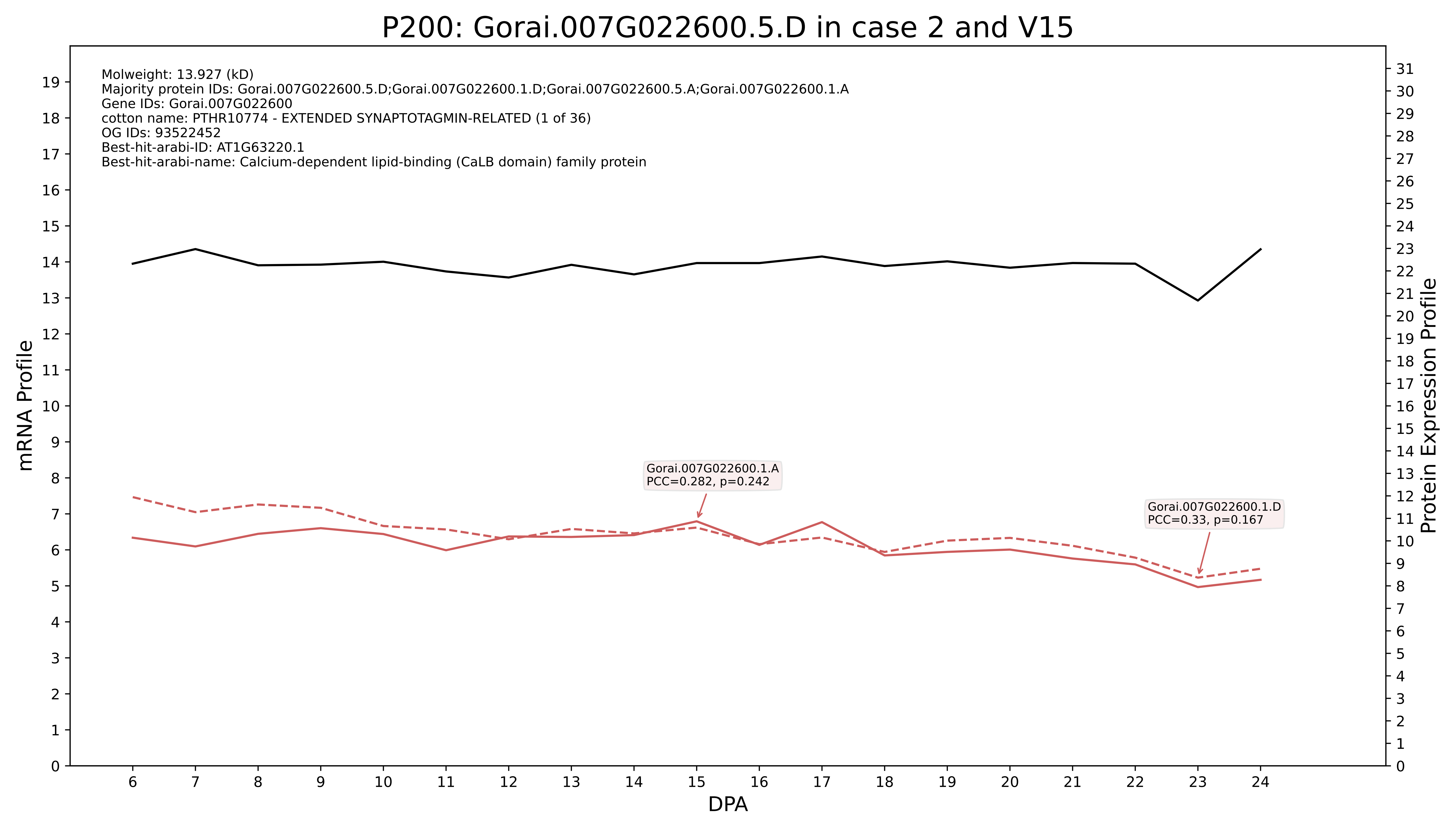
P200: Gorai.007G022600.5.D in case 2 and V15
Molweight: 13.927 (kD)
Majority protein IDs: Gorai.007G022600.5.D;Gorai.007G022600.1.D;Gorai.007G022600.5.A;Gorai.007G022600.1.A
Gene IDs: Gorai.007G022600
cotton name: PTHR10774 - EXTENDED SYNAPTOTAGMIN-RELATED (1 of 36)
OG IDs: 93522452
Best-hit-arabi-ID: AT1G63220.1
Best-hit-arabi-name: Calcium-dependent lipid-binding (CaLB domain) family protein
Gorai.007G022600.1.A
PCC=0.282, p=0.242
Gorai.007G022600.1.D
PCC=0.33, p=0.167
mRNA Profile
Protein Expression Profile
DPA

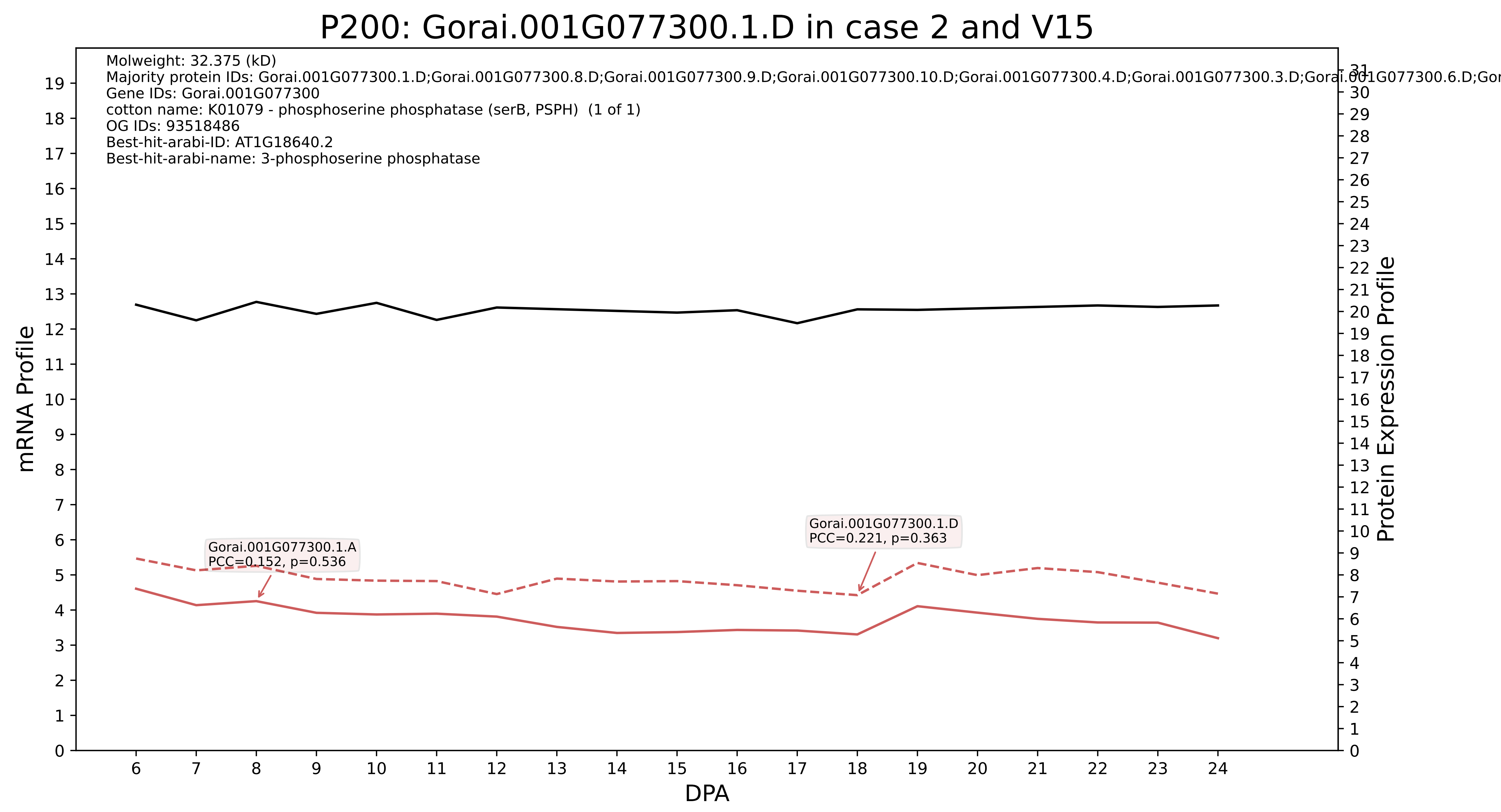

P200: Gorai.001G077300.1.D in case 2 and V15
Molweight: 32.375 (kD)
Majority protein IDs: Gorai.001G077300.1.D;Gorai.001G077300.8.D;Gorai.001G077300.9.D;Gorai.001G077300.10.D;Gorai.001G077300.4.D;Gorai.001G077300.3.D;Gorai.001G077300.6.D;Gor
Gene IDs: Gorai.001G077300
cotton name: K01079 - phosphoserine phosphatase (serB, PSPH) (1 of 1)
OG IDs: 93518486
Best-hit-arabi-ID: AT1G18640.2
Best-hit-arabi-name: 3-phosphoserine phosphatase
Gorai.001G077300.1.A
PCC=0.152, p=0.536
Gorai.001G077300.1.D
PCC=0.221, p=0.363
mRNA Profile
Protein Expression Profile
DPA

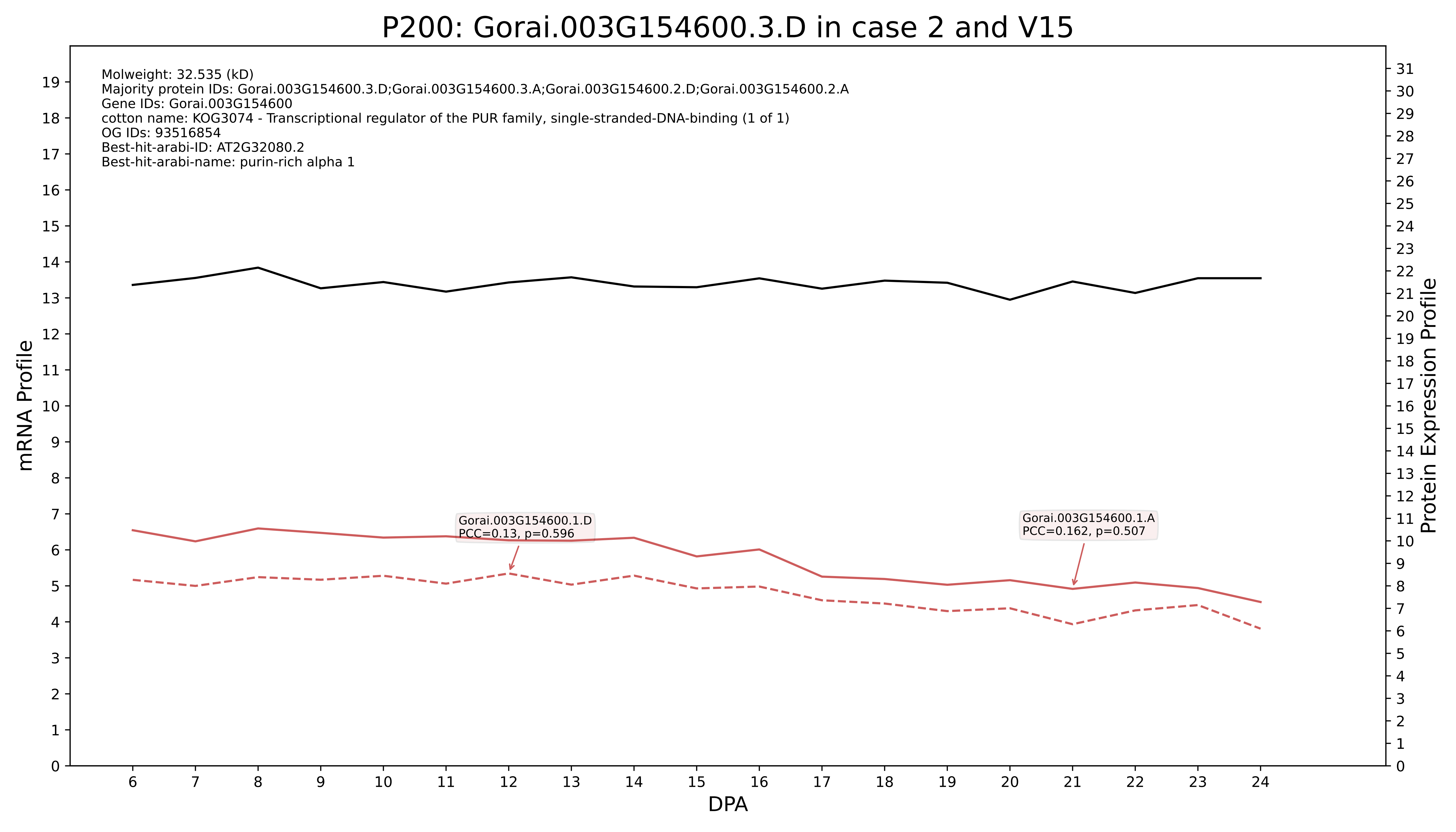

P200: Gorai.003G154600.3.D in case 2 and V15
Molweight: 32.535 (kD)
Majority protein IDs: Gorai.003G154600.3.D;Gorai.003G154600.3.A;Gorai.003G154600.2.D;Gorai.003G154600.2.A
Gene IDs: Gorai.003G154600
cotton name: KOG3074 - Transcriptional regulator of the PUR family, single-stranded-DNA-binding (1 of 1)
OG IDs: 93516854
Best-hit-arabi-ID: AT2G32080.2
Best-hit-arabi-name: purin-rich alpha 1
Gorai.003G154600.1.D
PCC=0.13, p=0.596
Gorai.003G154600.1.A
PCC=0.162, p=0.507
mRNA Profile
Protein Expression Profile
DPA

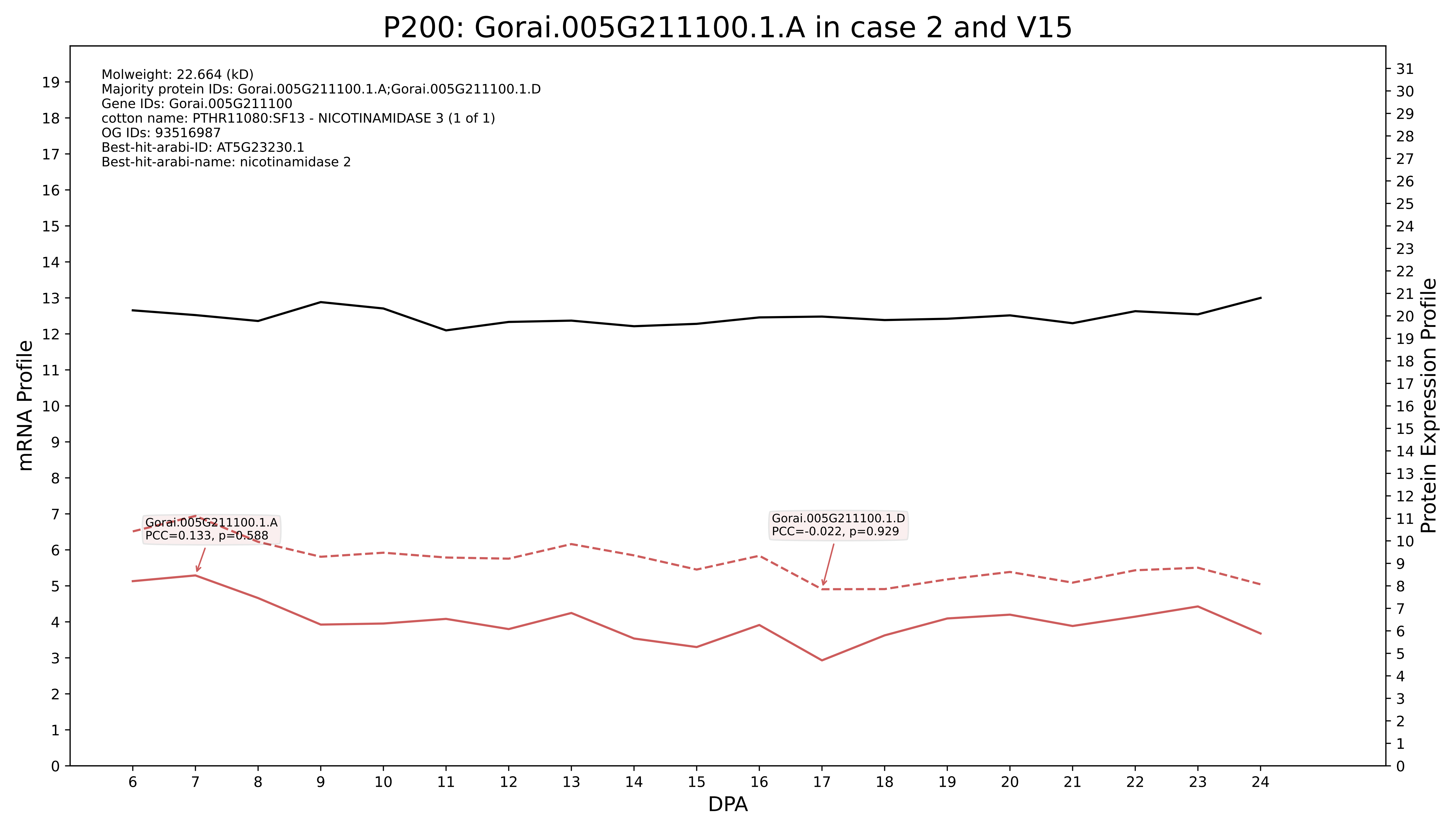
P200: Gorai.005G211100.1.A in case 2 and V15
Molweight: 22.664 (kD)
Majority protein IDs: Gorai.005G211100.1.A;Gorai.005G211100.1.D
Gene IDs: Gorai.005G211100
cotton name: PTHR11080:SF13 - NICOTINAMIDASE 3 (1 of 1)
OG IDs: 93516987
Best-hit-arabi-ID: AT5G23230.1
Best-hit-arabi-name: nicotinamidase 2
Gorai.005G211100.1.A
PCC=0.133, p=0.588
Gorai.005G211100.1.D
PCC=-0.022, p=0.929
mRNA Profile
Protein Expression Profile
DPA

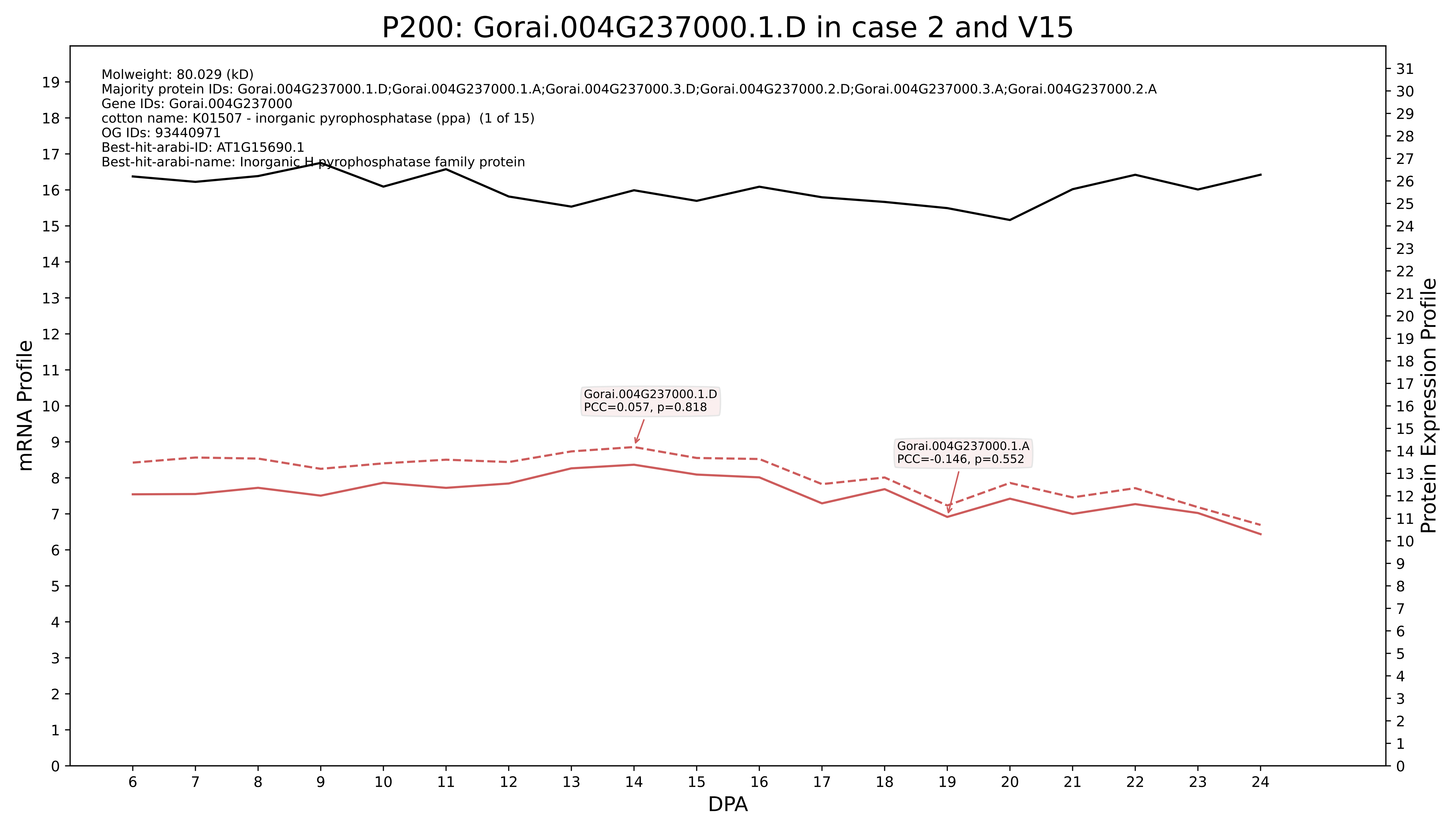

P200: Gorai.004G237000.1.D in case 2 and V15
Molweight: 80.029 (kD)
Majority protein IDs: Gorai.004G237000.1.D;Gorai.004G237000.1.A;Gorai.004G237000.3.D;Gorai.004G237000.2.D;Gorai.004G237000.3.A;Gorai.004G237000.2.A
Gene IDs: Gorai.004G237000
cotton name: K01507 - inorganic pyrophosphatase (ppa) (1 of 15)
OG IDs: 93440971
Best-hit-arabi-ID: AT1G15690.1
Best-hit-arabi-name: Inorganic H pyrophosphatase family protein
Gorai.004G237000.1.D
PCC=0.057, p=0.818
Gorai.004G237000.1.A
PCC=-0.146, p=0.552
mRNA Profile
Protein Expression Profile
DPA

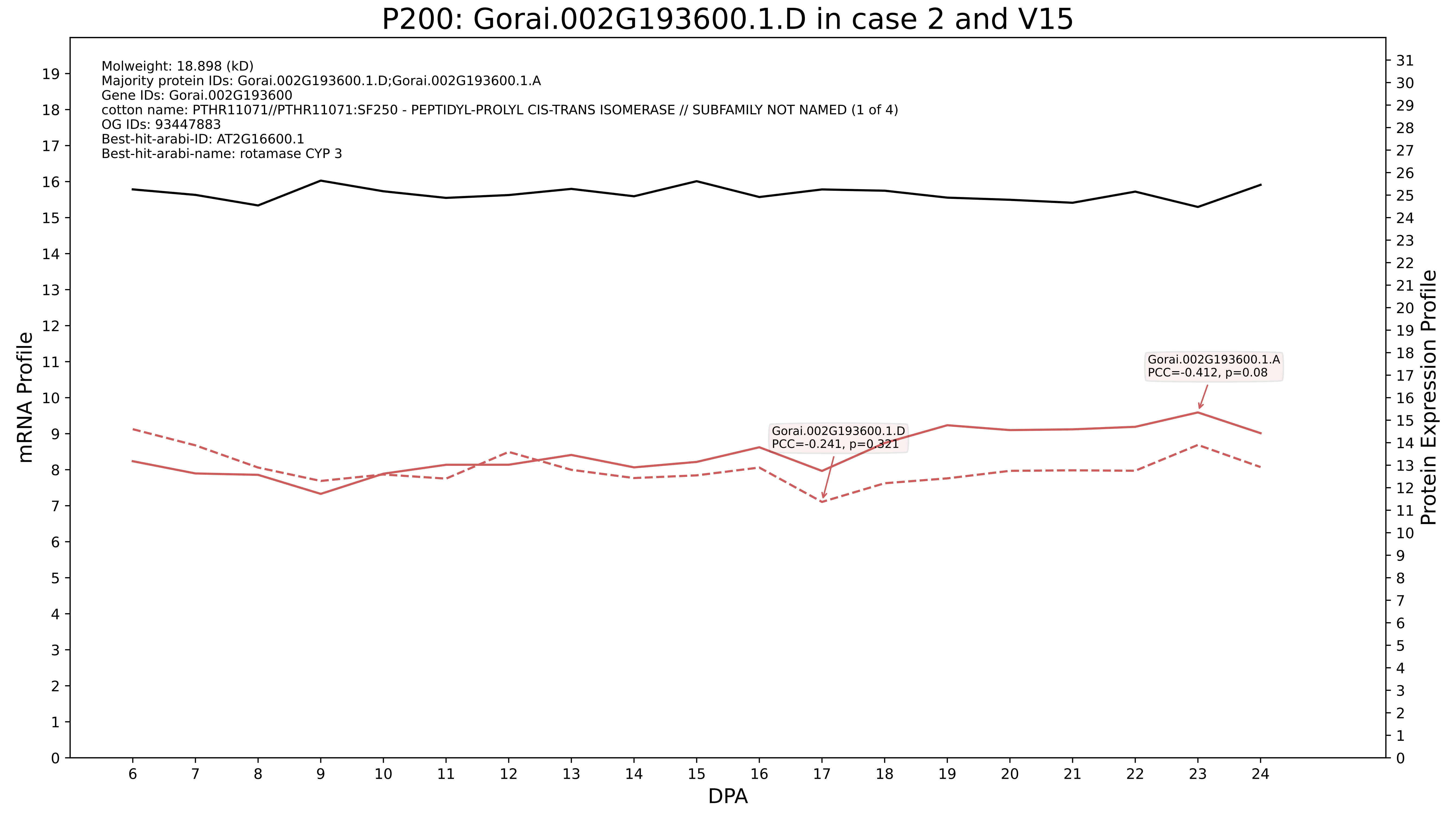

P200: Gorai.002G193600.1.D in case 2 and V15
Molweight: 18.898 (kD)
Majority protein IDs: Gorai.002G193600.1.D;Gorai.002G193600.1.A
Gene IDs: Gorai.002G193600
cotton name: PTHR11071//PTHR11071:SF250 - PEPTIDYL-PROLYL CIS-TRANS ISOMERASE // SUBFAMILY NOT NAMED (1 of 4)
OG IDs: 93447883
Best-hit-arabi-ID: AT2G16600.1
Best-hit-arabi-name: rotamase CYP 3
Gorai.002G193600.1.A
PCC=-0.412, p=0.08
Gorai.002G193600.1.D
PCC=-0.241, p=0.321
mRNA Profile
Protein Expression Profile
DPA

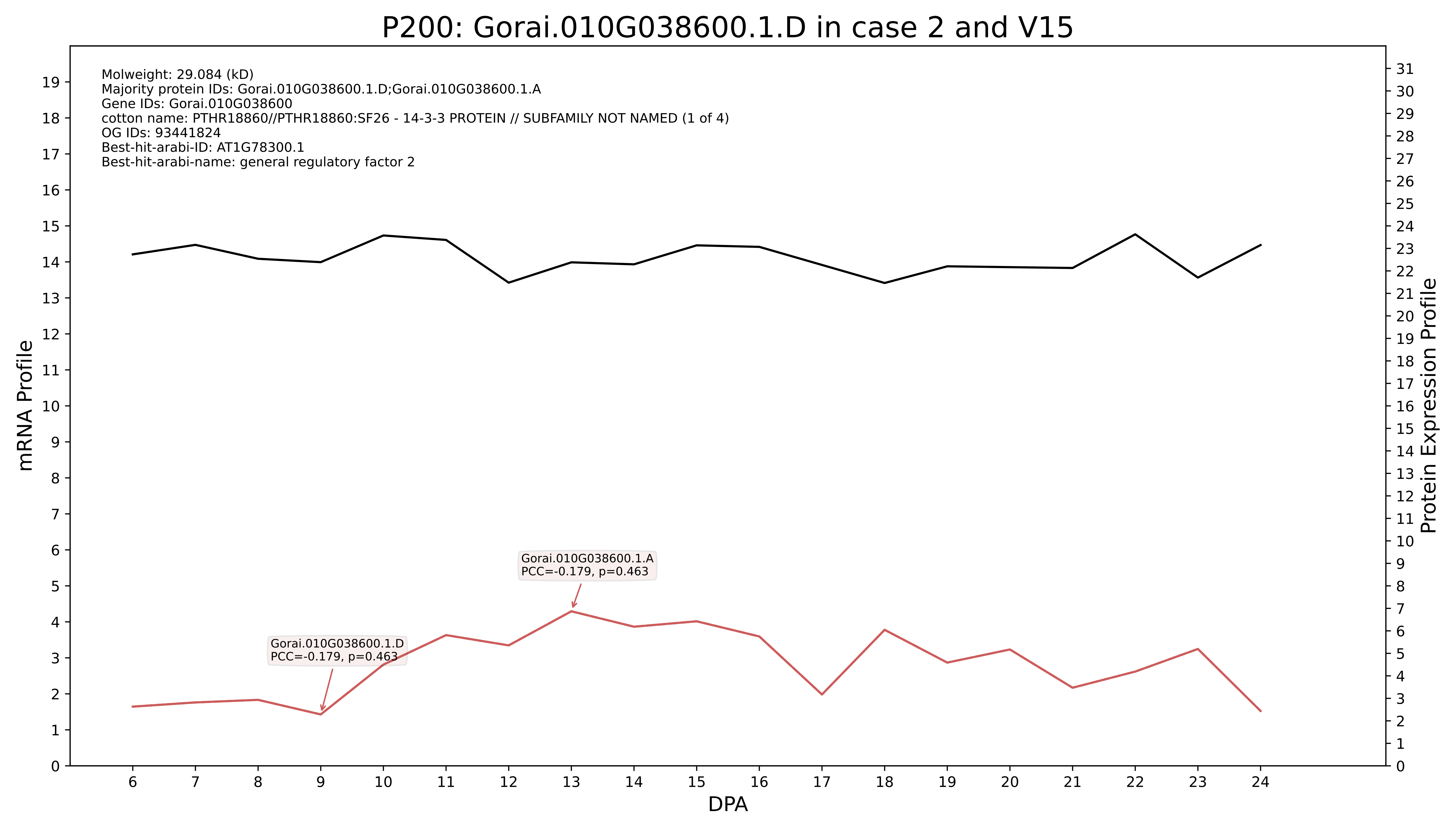

P200: Gorai.010G038600.1.D in case 2 and V15
Molweight: 29.084 (kD)
Majority protein IDs: Gorai.010G038600.1.D;Gorai.010G038600.1.A
Gene IDs: Gorai.010G038600
cotton name: PTHR18860//PTHR18860:SF26 - 14-3-3 PROTEIN // SUBFAMILY NOT NAMED (1 of 4)
OG IDs: 93441824
Best-hit-arabi-ID: AT1G78300.1
Best-hit-arabi-name: general regulatory factor 2
Gorai.010G038600.1.A
PCC=-0.179, p=0.463
Gorai.010G038600.1.D
PCC=-0.179, p=0.463
mRNA Profile
Protein Expression Profile
DPA

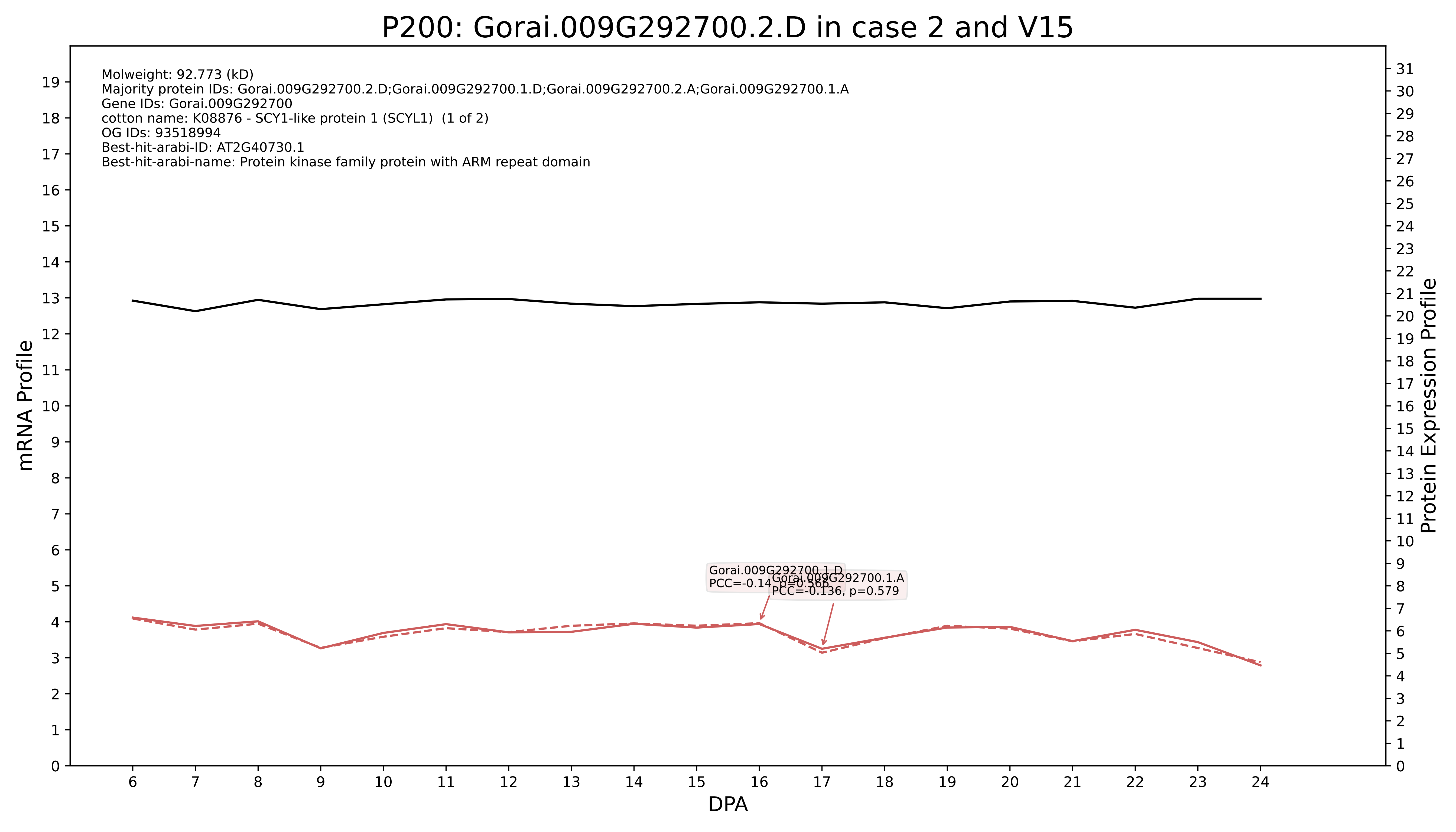
P200: Gorai.009G292700.2.D in case 2 and V15
Molweight: 92.773 (kD)
Majority protein IDs: Gorai.009G292700.2.D;Gorai.009G292700.1.D;Gorai.009G292700.2.A;Gorai.009G292700.1.A
Gene IDs: Gorai.009G292700
cotton name: K08876 - SCY1-like protein 1 (SCYL1) (1 of 2)
OG IDs: 93518994
Best-hit-arabi-ID: AT2G40730.1
Best-hit-arabi-name: Protein kinase family protein with ARM repeat domain
Gorai.009G292700.1.D
Gorai.009G292700.1.A
PCC=-0.136, p=0.579
mRNA Profile
Protein Expression Profile
DPA

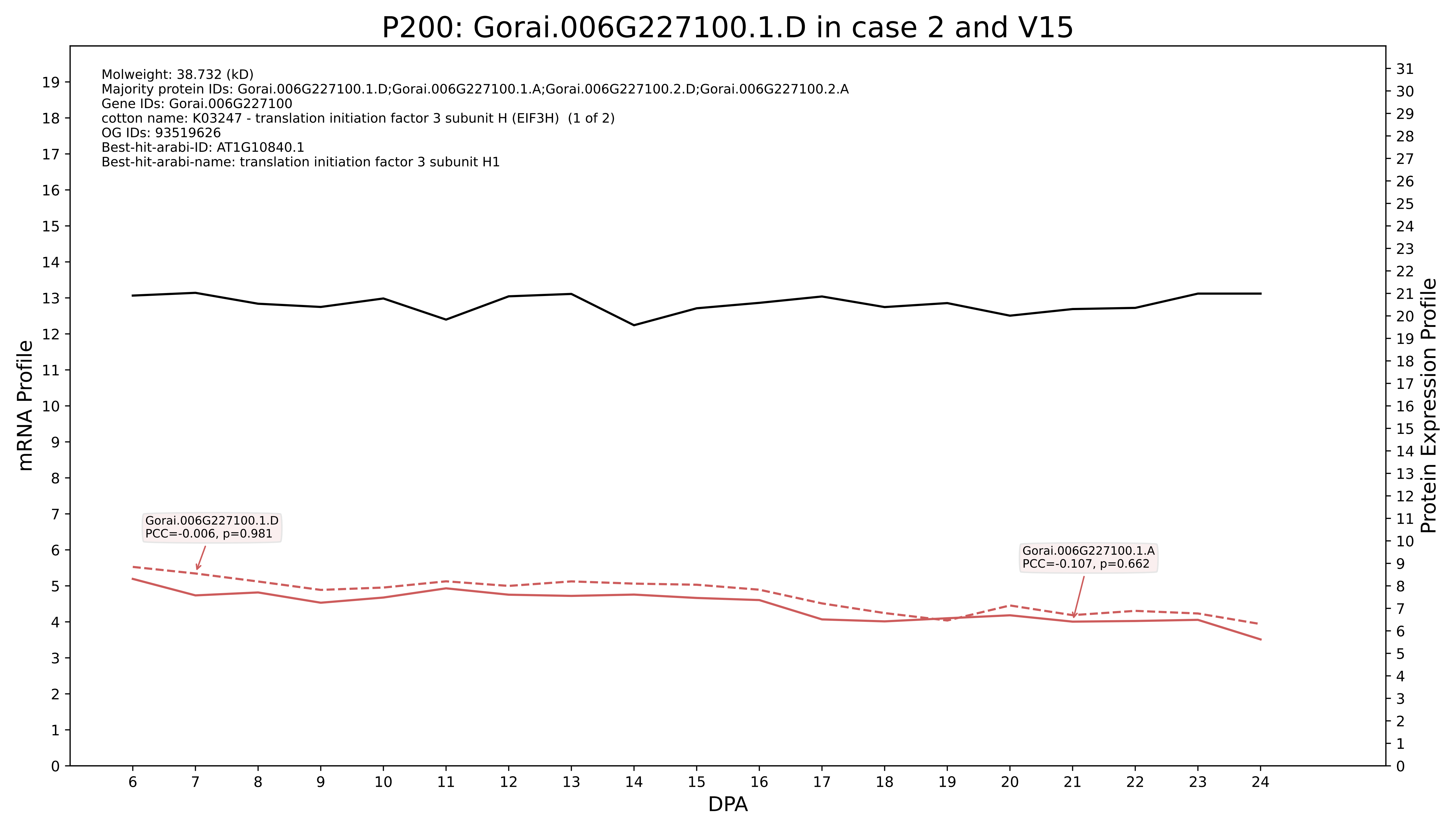
P200: Gorai.006G227100.1.D in case 2 and V15
Molweight: 38.732 (kD)
Majority protein IDs: Gorai.006G227100.1.D;Gorai.006G227100.1.A;Gorai.006G227100.2.D;Gorai.006G227100.2.A
Gene IDs: Gorai.006G227100
cotton name: K03247 - translation initiation factor 3 subunit H (EIF3H) (1 of 2)
OG IDs: 93519626
Best-hit-arabi-ID: AT1G10840.1
Best-hit-arabi-name: translation initiation factor 3 subunit H1
Gorai.006G227100.1.D
PCC=-0.006, p=0.981
Gorai.006G227100.1.A
PCC=-0.107, p=0.662
mRNA Profile
Protein Expression Profile
DPA

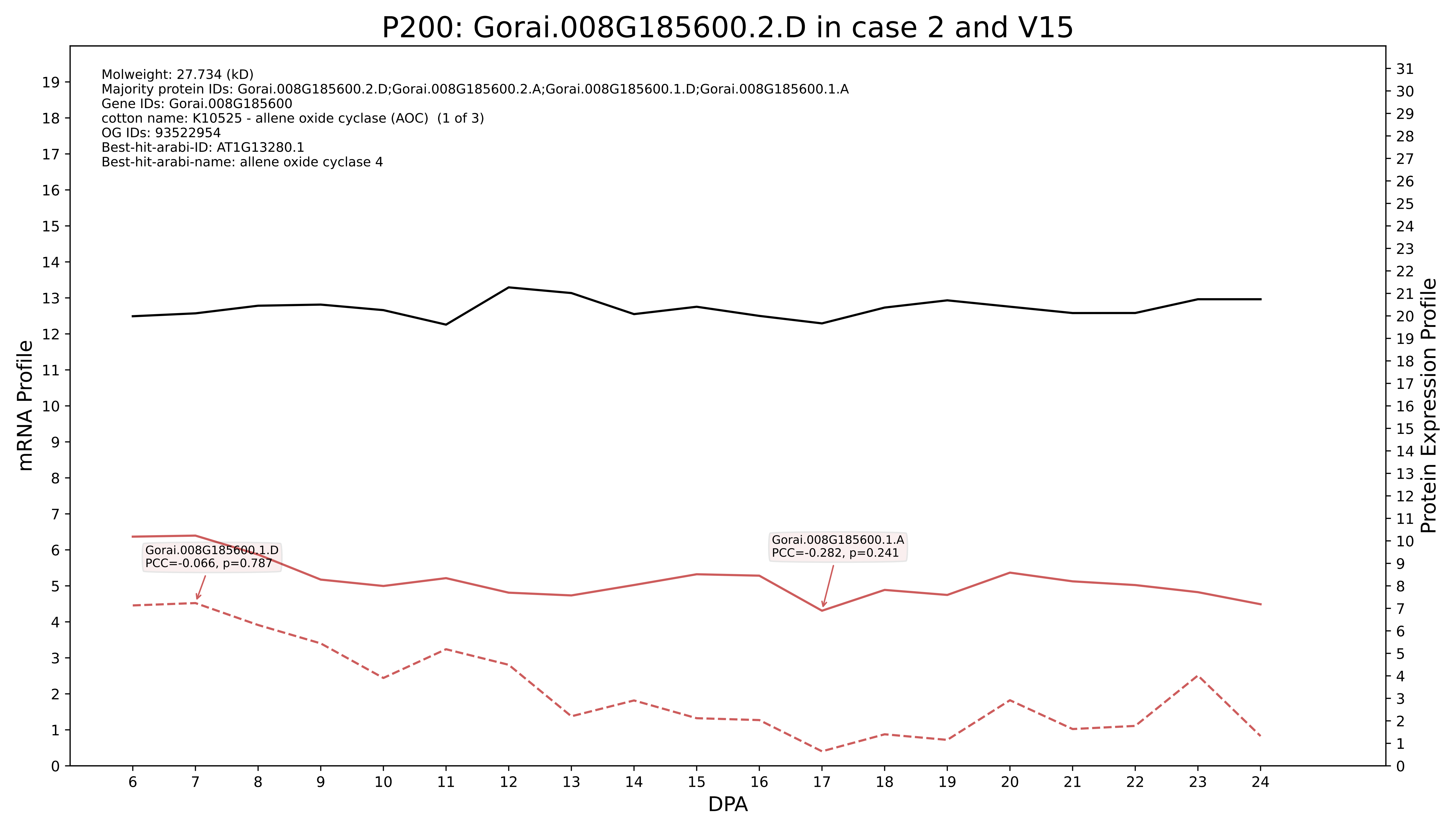
P200: Gorai.008G185600.2.D in case 2 and V15
Molweight: 27.734 (kD)
Majority protein IDs: Gorai.008G185600.2.D;Gorai.008G185600.2.A;Gorai.008G185600.1.D;Gorai.008G185600.1.A
Gene IDs: Gorai.008G185600
cotton name: K10525 - allene oxide cyclase (AOC) (1 of 3)
OG IDs: 93522954
Best-hit-arabi-ID: AT1G13280.1
Best-hit-arabi-name: allene oxide cyclase 4
Gorai.008G185600.1.D
PCC=-0.066, p=0.787
Gorai.008G185600.1.A
PCC=-0.282, p=0.241
mRNA Profile
Protein Expression Profile
DPA

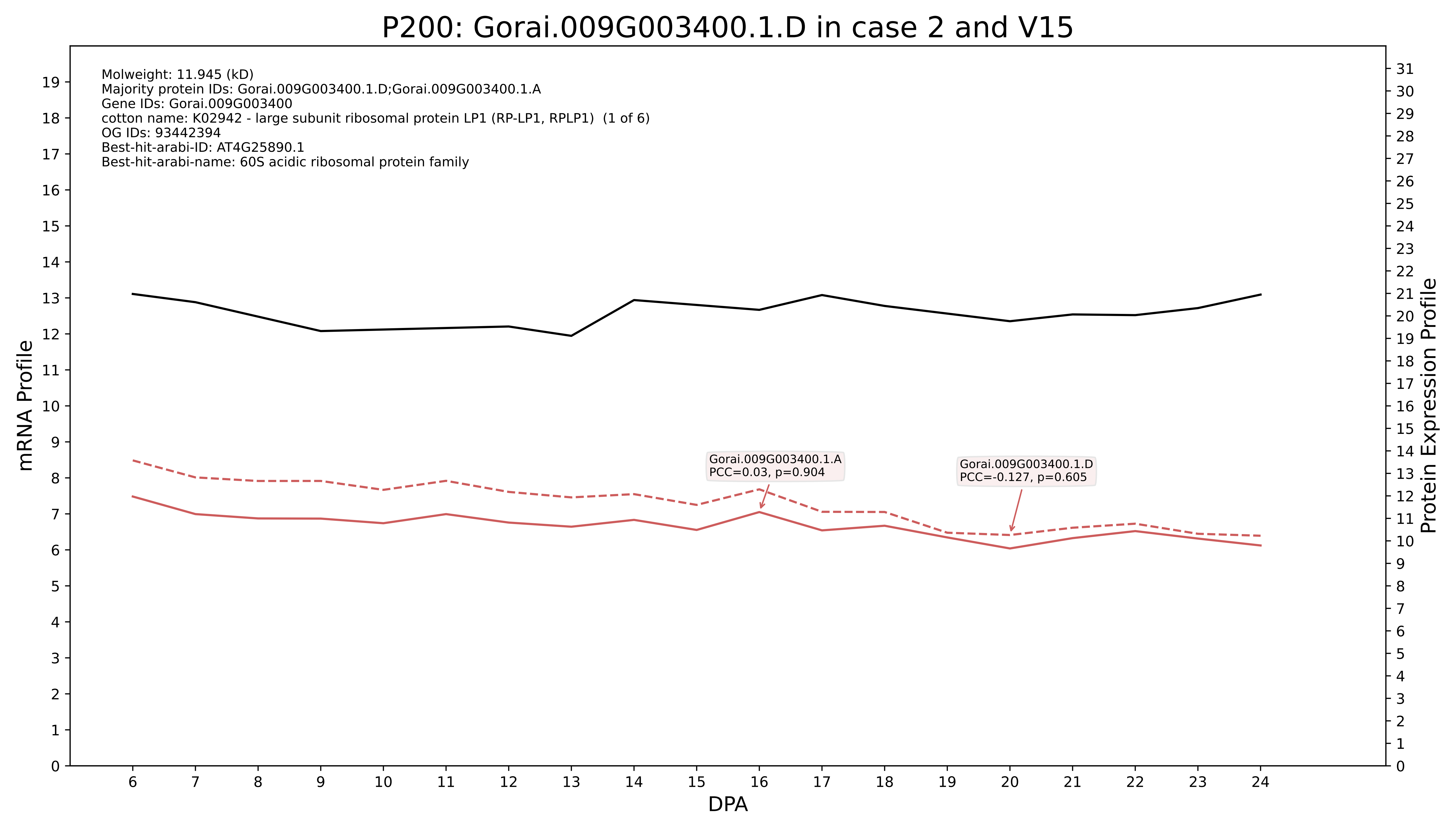
P200: Gorai.009G003400.1.D in case 2 and V15
Molweight: 11.945 (kD)
Majority protein IDs: Gorai.009G003400.1.D;Gorai.009G003400.1.A
Gene IDs: Gorai.009G003400
cotton name: K02942 - large subunit ribosomal protein LP1 (RP-LP1, RPLP1) (1 of 6)
OG IDs: 93442394
Best-hit-arabi-ID: AT4G25890.1
Best-hit-arabi-name: 60S acidic ribosomal protein family
Gorai.009G003400.1.A
PCC=0.03, p=0.904
Gorai.009G003400.1.D
PCC=-0.127, p=0.605
mRNA Profile
Protein Expression Profile
DPA

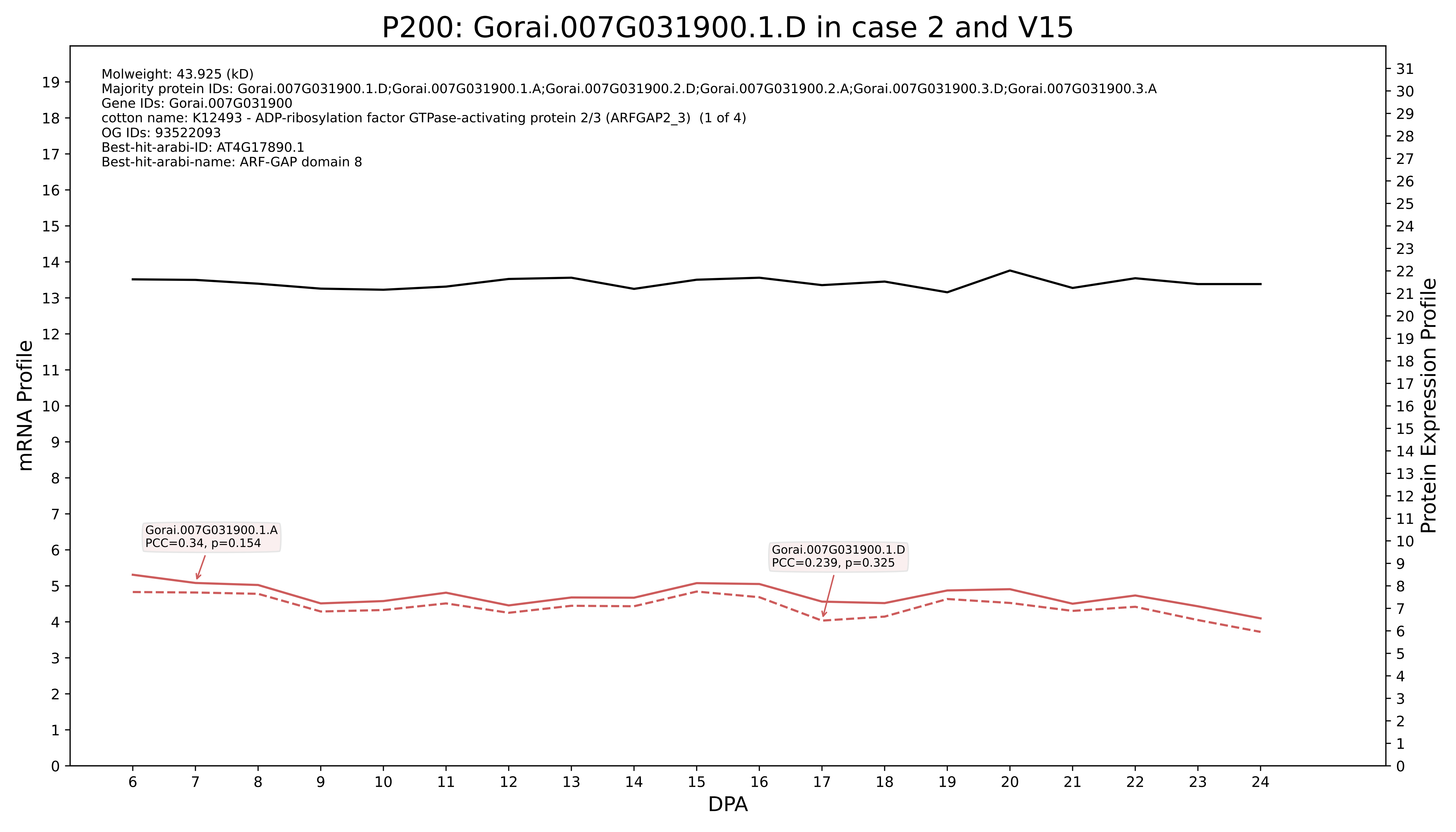
P200: Gorai.007G031900.1.D in case 2 and V15
Molweight: 43.925 (kD)
Majority protein IDs: Gorai.007G031900.1.D;Gorai.007G031900.1.A;Gorai.007G031900.2.D;Gorai.007G031900.2.A;Gorai.007G031900.3.D;Gorai.007G031900.3.A
Gene IDs: Gorai.007G031900
cotton name: K12493 - ADP-ribosylation factor GTPase-activating protein 2/3 (ARFGAP2_3) (1 of 4)
OG IDs: 93522093
Best-hit-arabi-ID: AT4G17890.1
Best-hit-arabi-name: ARF-GAP domain 8
Gorai.007G031900.1.A
PCC=0.34, p=0.154
Gorai.007G031900.1.D
PCC=0.239, p=0.325
mRNA Profile
Protein Expression Profile
DPA

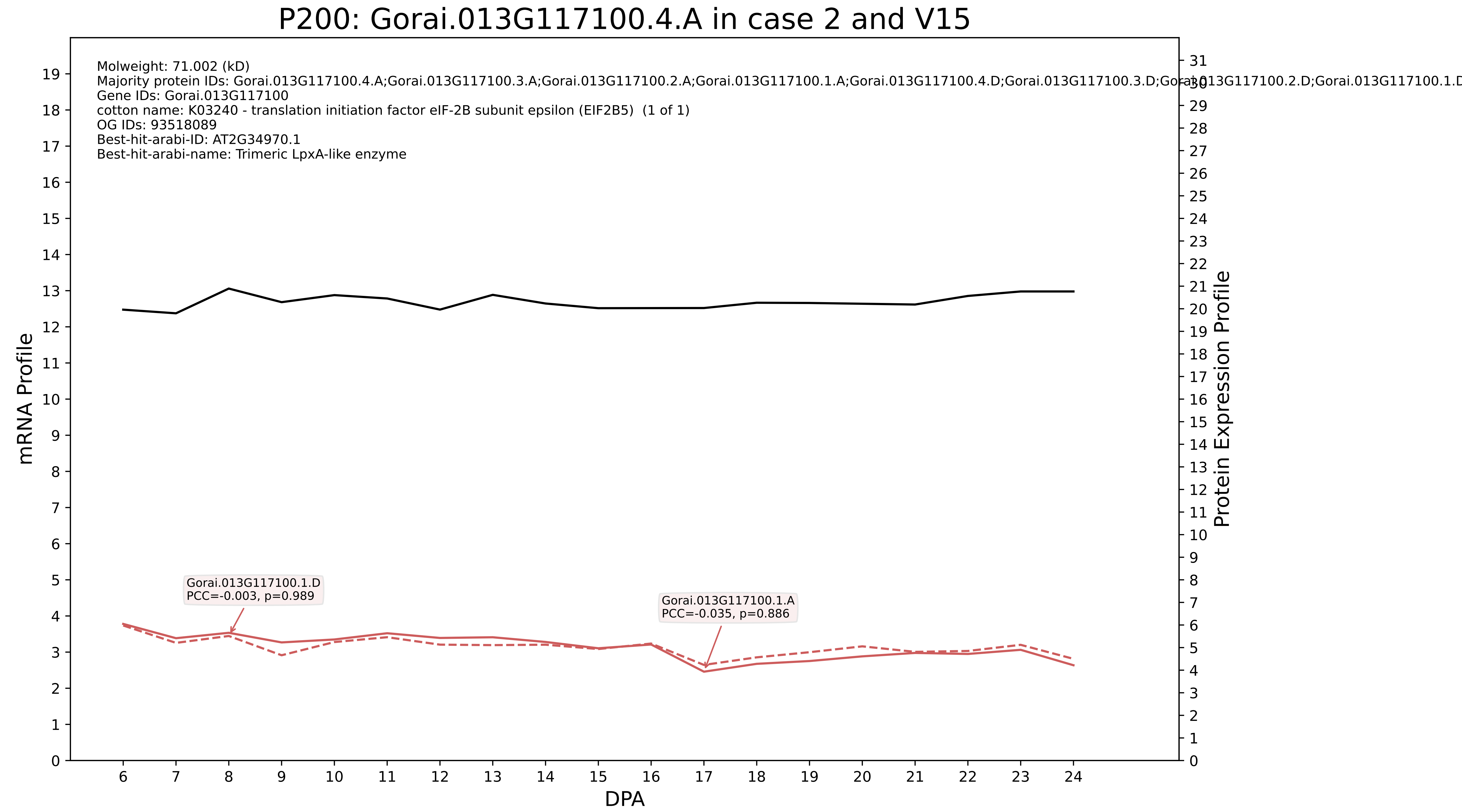
P200: Gorai.013G117100.4.A in case 2 and V15
Molweight: 71.002 (kD)
Majority protein IDs: Gorai.013G117100.4.A;Gorai.013G117100.3.A;Gorai.013G117100.2.A;Gorai.013G117100.1.A;Gorai.013G117100.4.D;Gorai.013G117100.3.D;Gorai.013G117100.2.D;Gorai.013G117100.1.D
Gene IDs: Gorai.013G117100
cotton name: K03240 - translation initiation factor eIF-2B subunit epsilon (EIF2B5) (1 of 1)
OG IDs: 93518089
Best-hit-arabi-ID: AT2G34970.1
Best-hit-arabi-name: Trimeric LpxA-like enzyme
Gorai.013G117100.1.D
PCC=-0.003, p=0.989
Gorai.013G117100.1.A
PCC=-0.035, p=0.886
mRNA Profile
Protein Expression Profile
DPA

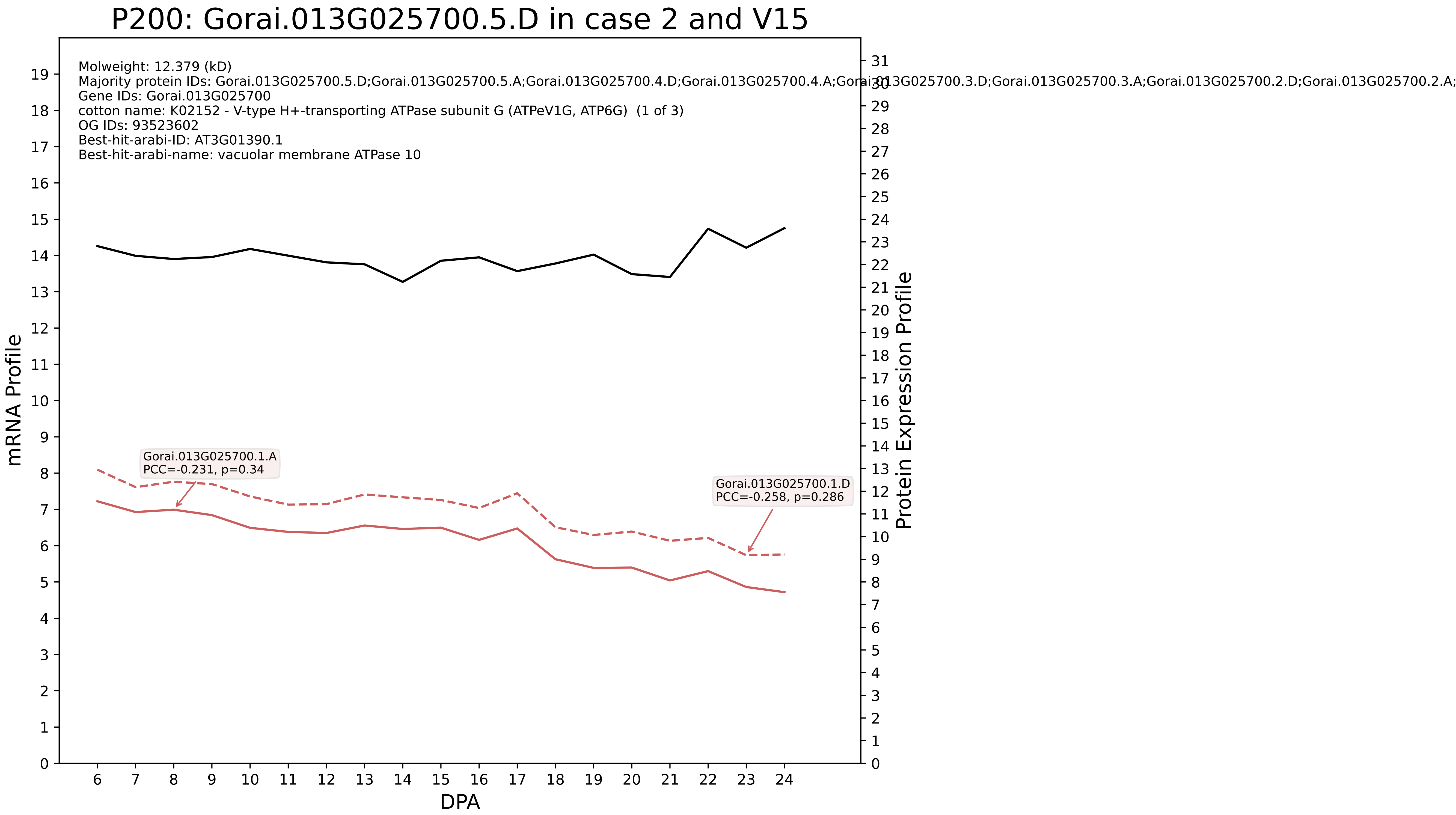
P200: Gorai.013G025700.5.D in case 2 and V15
Molweight: 12.379 (kD)
Majority protein IDs: Gorai.013G025700.5.D;Gorai.013G025700.5.A;Gorai.013G025700.4.D;Gorai.013G025700.4.A;Gorai.013G025700.3.D;Gorai.013G025700.3.A;Gorai.013G025700.2.D;Gorai.013G025700.2.A;
Gene IDs: Gorai.013G025700
cotton name: K02152 - V-type H+-transporting ATPase subunit G (ATPeV1G, ATP6G) (1 of 3)
OG IDs: 93523602
Best-hit-arabi-ID: AT3G01390.1
Best-hit-arabi-name: vacuolar membrane ATPase 10
Gorai.013G025700.1.A
PCC=-0.231, p=0.34
Gorai.013G025700.1.D
PCC=-0.258, p=0.286
mRNA Profile
Protein Expression Profile
DPA

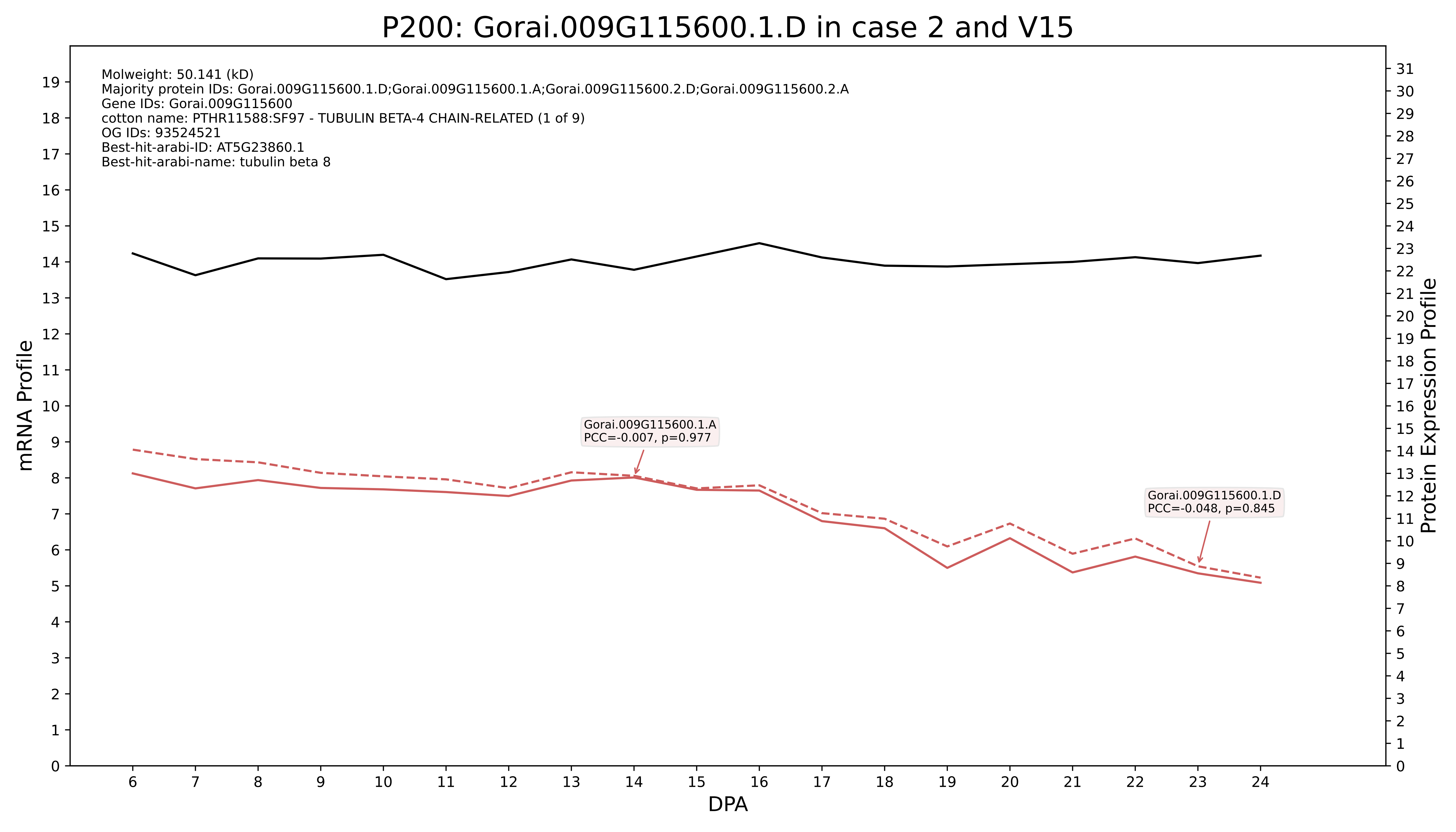

P200: Gorai.009G115600.1.D in case 2 and V15
Molweight: 50.141 (kD)
Majority protein IDs: Gorai.009G115600.1.D;Gorai.009G115600.1.A;Gorai.009G115600.2.D;Gorai.009G115600.2.A
Gene IDs: Gorai.009G115600
cotton name: PTHR11588:SF97 - TUBULIN BETA-4 CHAIN-RELATED (1 of 9)
OG IDs: 93524521
Best-hit-arabi-ID: AT5G23860.1
Best-hit-arabi-name: tubulin beta 8
Gorai.009G115600.1.A
PCC=-0.007, p=0.977
Gorai.009G115600.1.D
PCC=-0.048, p=0.845
mRNA Profile
Protein Expression Profile
DPA

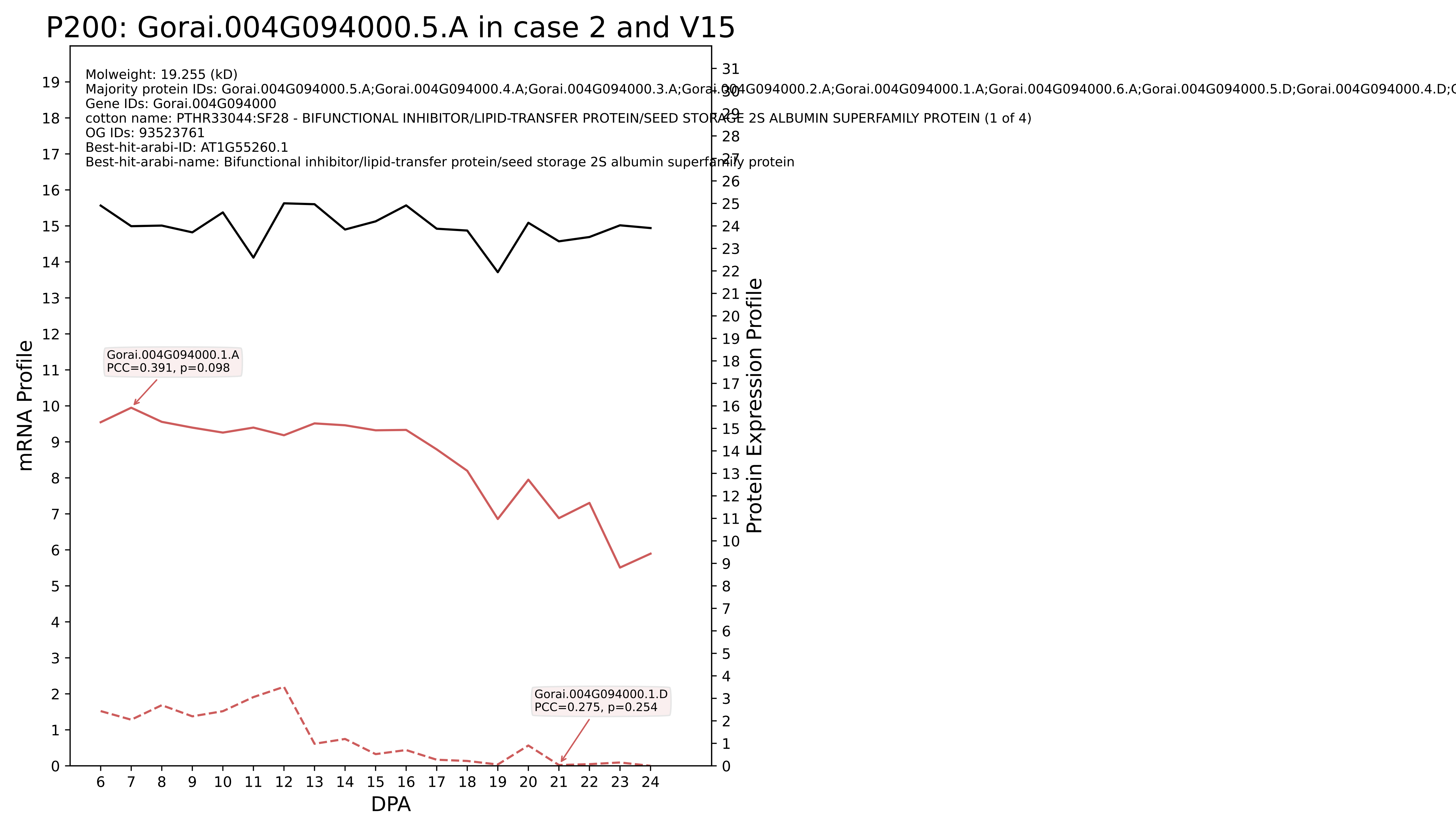
P200: Gorai.004G094000.5.A in case 2 and V15
Molweight: 19.255 (kD)
Majority protein IDs: Gorai.004G094000.5.A;Gorai.004G094000.4.A;Gorai.004G094000.3.A;Gorai.004G094000.2.A;Gorai.004G094000.1.A;Gorai.004G094000.6.A;Gorai.004G094000.5.D;Gorai.004G094000.4.D;G
Gene IDs: Gorai.004G094000
cotton name: PTHR33044:SF28 - BIFUNCTIONAL INHIBITOR/LIPID-TRANSFER PROTEIN/SEED STORAGE 2S ALBUMIN SUPERFAMILY PROTEIN (1 of 4)
OG IDs: 93523761
Best-hit-arabi-ID: AT1G55260.1
Best-hit-arabi-name: Bifunctional inhibitor/lipid-transfer protein/seed storage 2S albumin superfamily protein
Gorai.004G094000.1.A
PCC=0.391, p=0.098
Gorai.004G094000.1.D
PCC=0.275, p=0.254
mRNA Profile
Protein Expression Profile
DPA

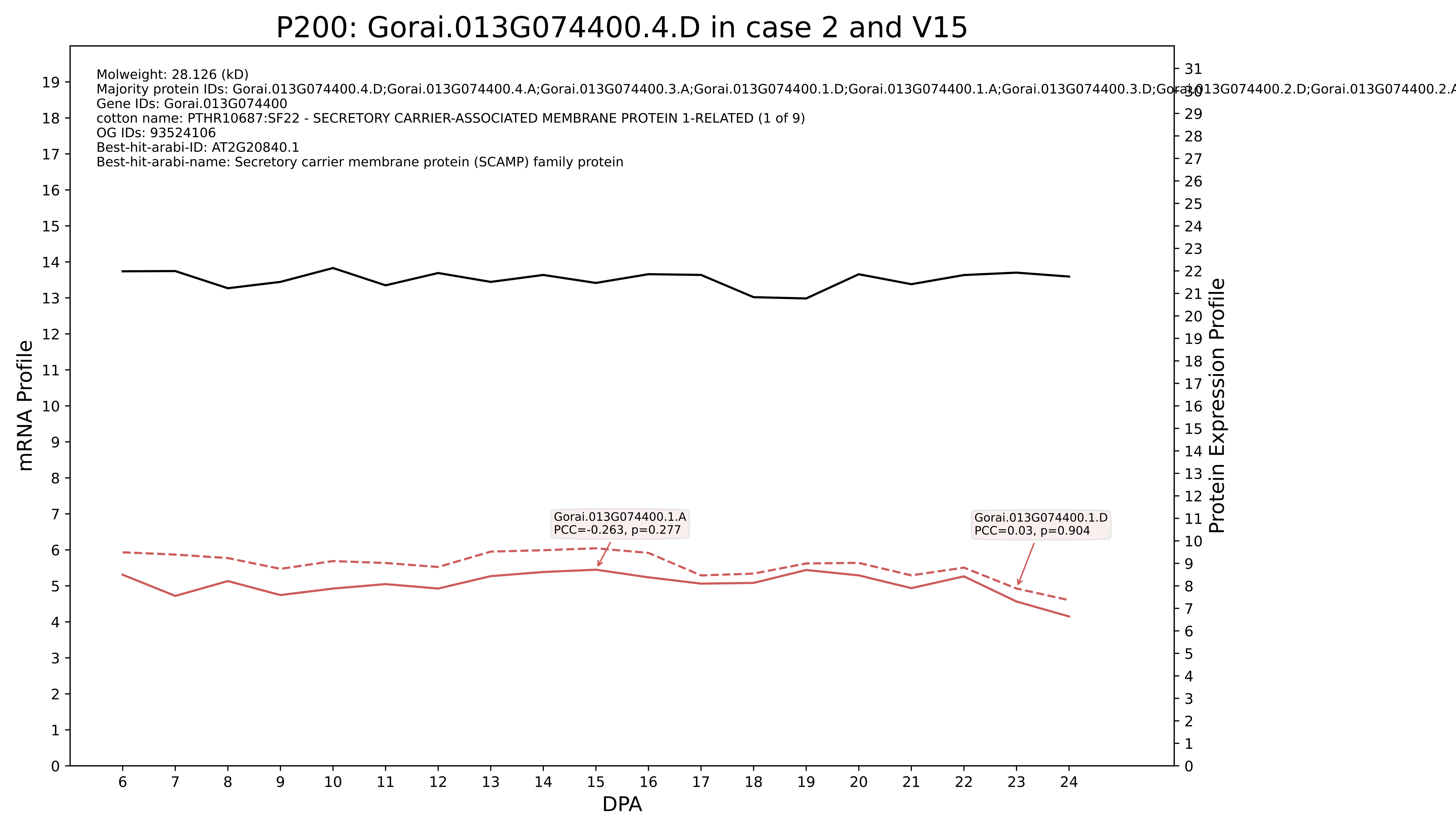

P200: Gorai.013G074400.4.D in case 2 and V15
Molweight: 28.126 (kD)
Majority protein IDs: Gorai.013G074400.4.D;Gorai.013G074400.4.A;Gorai.013G074400.3.A;Gorai.013G074400.1.D;Gorai.013G074400.1.A;Gorai.013G074400.3.D;Gorai.013G074400.2.D;Gorai.013G074400.2.A
Gene IDs: Gorai.013G074400
cotton name: PTHR10687:SF22 - SECRETORY CARRIER-ASSOCIATED MEMBRANE PROTEIN 1-RELATED (1 of 9)
OG IDs: 93524106
Best-hit-arabi-ID: AT2G20840.1
Best-hit-arabi-name: Secretory carrier membrane protein (SCAMP) family protein
Gorai.013G074400.1.A
PCC=-0.263, p=0.277
Gorai.013G074400.1.D
PCC=0.03, p=0.904
mRNA Profile
Protein Expression Profile
DPA

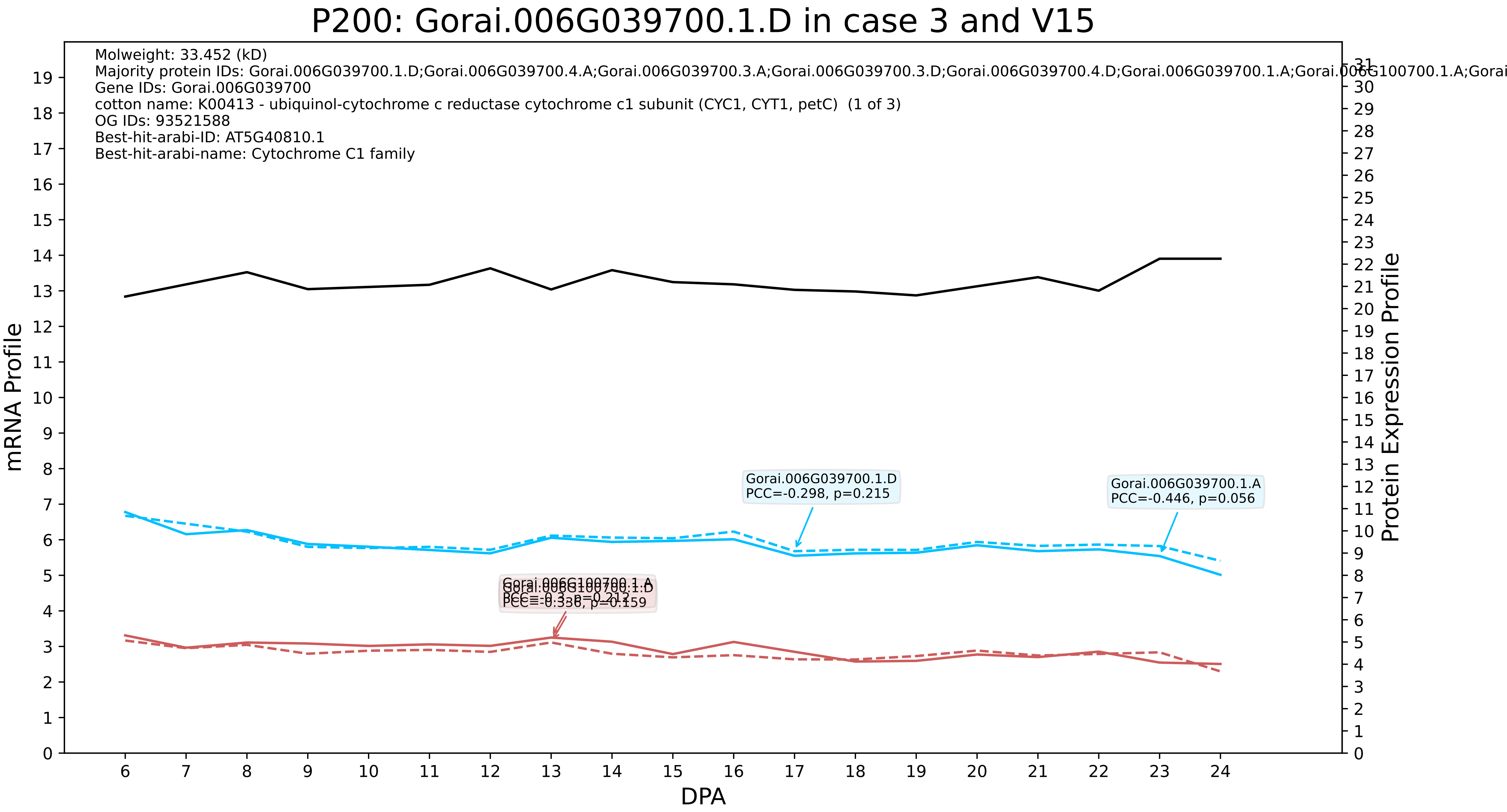

P200: Gorai.006G039700.1.D in case 3 and V15
Molweight: 33.452 (kD)
Majority protein IDs: Gorai.006G039700.1.D;Gorai.006G039700.4.A;Gorai.006G039700.3.A;Gorai.006G039700.3.D;Gorai.006G039700.4.D;Gorai.006G039700.1.A;Gorai.006G100700.1.A;Gorai
Gene IDs: Gorai.006G039700
cotton name: K00413 - ubiquinol-cytochrome c reductase cytochrome c1 subunit (CYC1, CYT1, petC) (1 of 3)
OG IDs: 93521588
Best-hit-arabi-ID: AT5G40810.1
Best-hit-arabi-name: Cytochrome C1 family
Gorai.006G039700.1.D
PCC=-0.298, p=0.215
Gorai.006G039700.1.A
PCC=-0.446, p=0.056
mRNA Profile
Protein Expression Profile
DPA

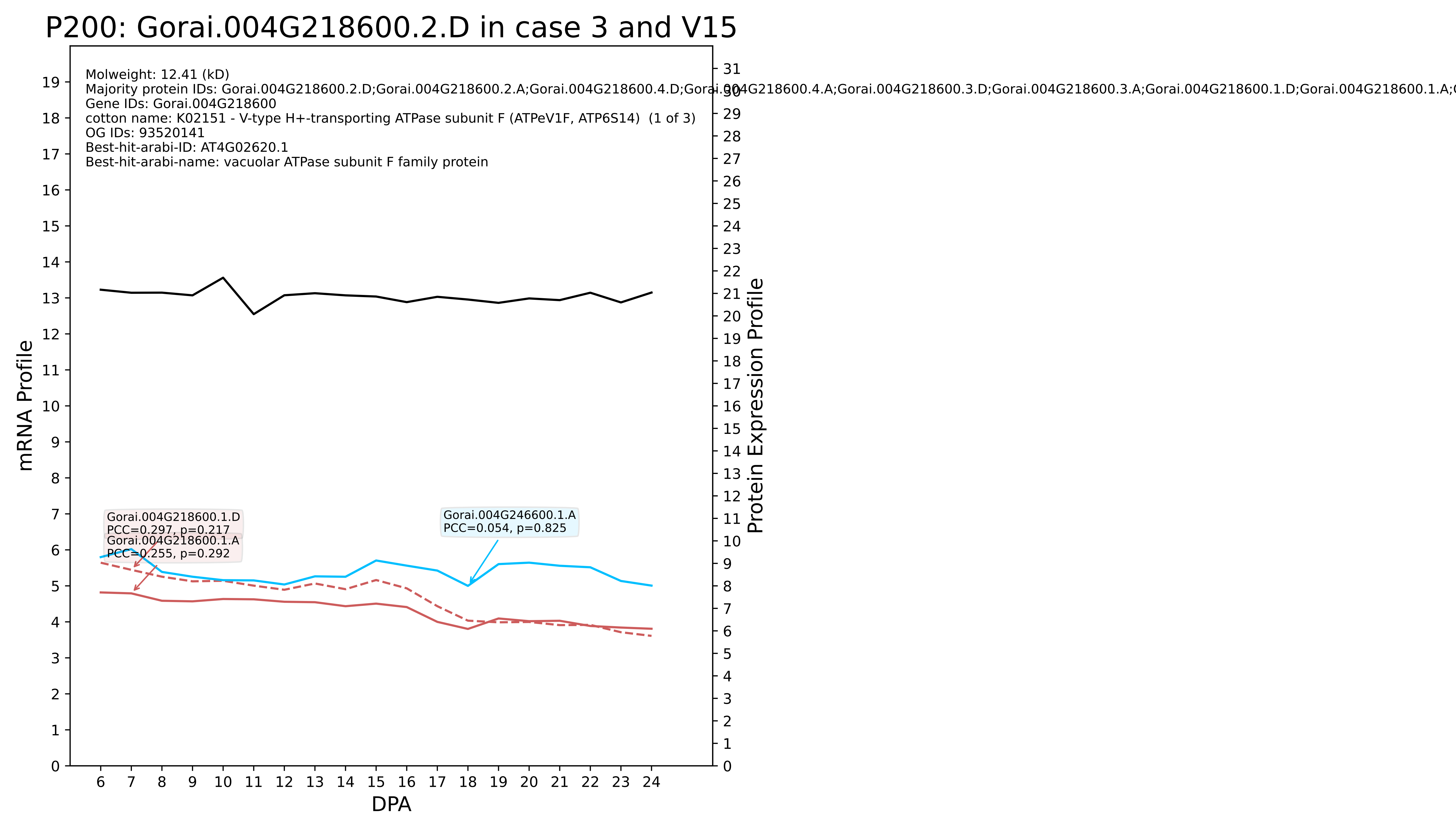
P200: Gorai.004G218600.2.D in case 3 and V15
Molweight: 12.41 (kD)
Majority protein IDs: Gorai.004G218600.2.D;Gorai.004G218600.2.A;Gorai.004G218600.4.D;Gorai.004G218600.4.A;Gorai.004G218600.3.D;Gorai.004G218600.3.A;Gorai.004G218600.1.D;Gorai.004G218600.1.A;
Gene IDs: Gorai.004G218600
cotton name: K02151 - V-type H+-transporting ATPase subunit F (ATPeV1F, ATP6S14) (1 of 3)
OG IDs: 93520141
Best-hit-arabi-ID: AT4G02620.1
Best-hit-arabi-name: vacuolar ATPase subunit F family protein
Gorai.004G218600.1.D
PCC=0.297, p=0.217
Gorai.004G218600.1.A
PCC=0.255, p=0.292
Gorai.004G246600.1.A
PCC=0.054, p=0.825
mRNA Profile
Protein Expression Profile
DPA

### P200: Gorai.013G213000.1.A in case 3 and V15

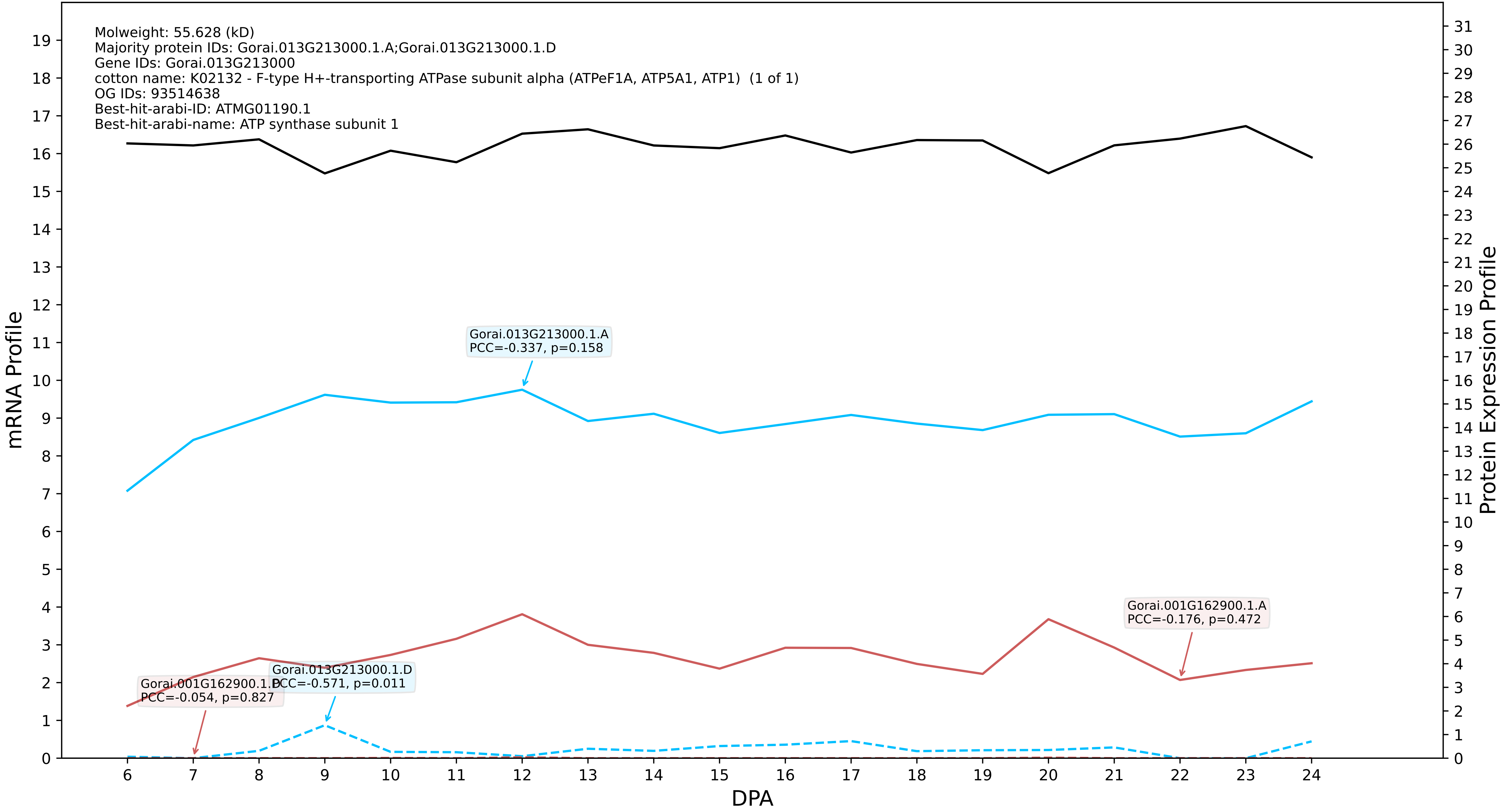

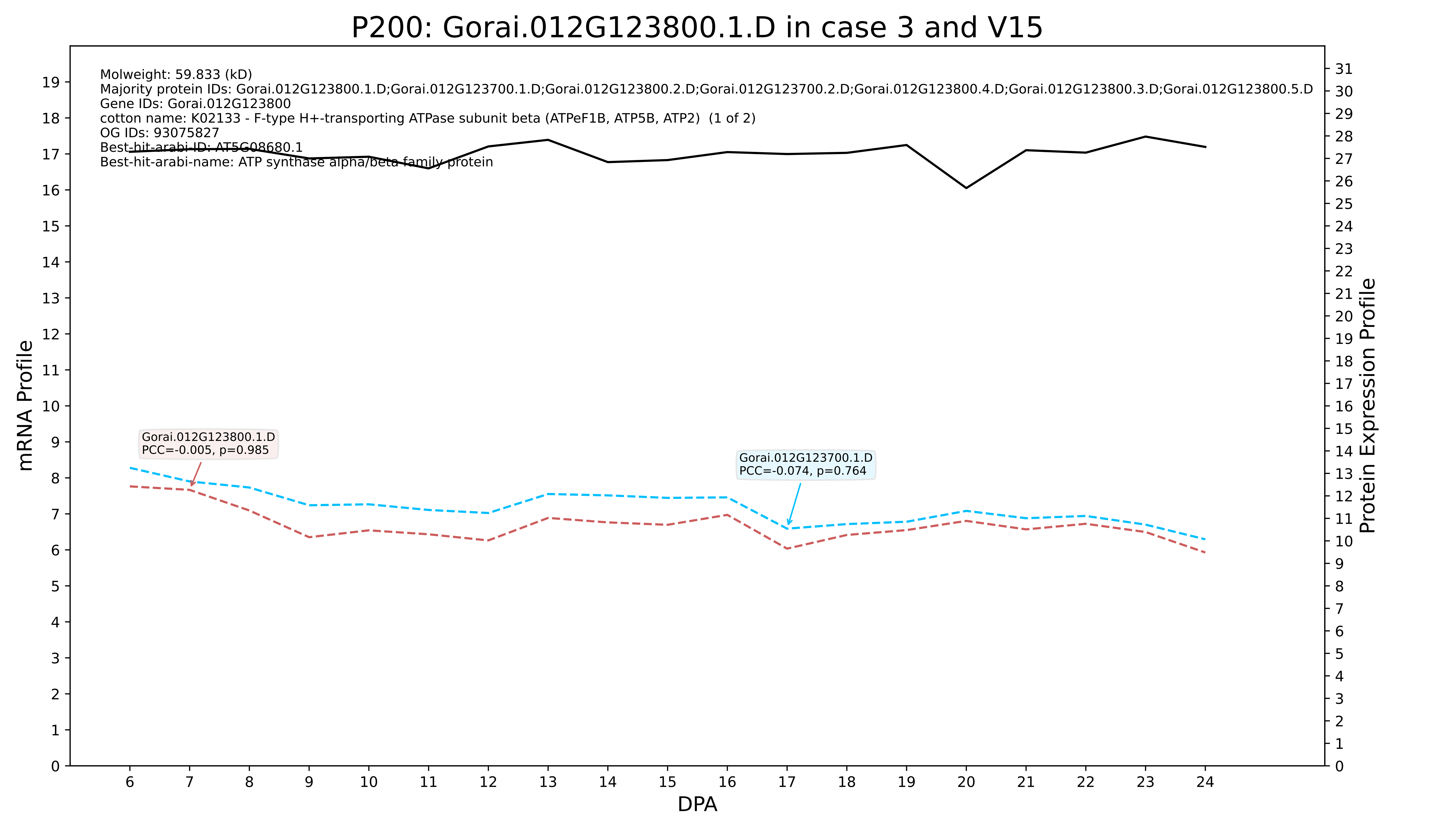

P200: Gorai.012G123800.1.D in case 3 and V15
Molweight: 59.833 (kD)
Majority protein IDs: Gorai.012G123800.1.D;Gorai.012G123700.1.D;Gorai.012G123800.2.D;Gorai.012G123700.2.D;Gorai.012G123800.4.D;Gorai.012G123800.3.D;Gorai.012G123800.5.D
Gene IDs: Gorai.012G123800
cotton name: K02133 - F-type H+-transporting ATPase subunit beta (ATPeF1B, ATP5B, ATP2) (1 of 2)
OG IDs: 93075827
Best-hit-arabi-ID: AT5G08680.1
Best-hit-arabi-name: ATP synthase alpha/beta family protein
Gorai.012G123800.1.D
PCC=-0.005, p=0.985
Gorai.012G123700.1.D
PCC=-0.074, p=0.764
mRNA Profile
Protein Expression Profile
DPA

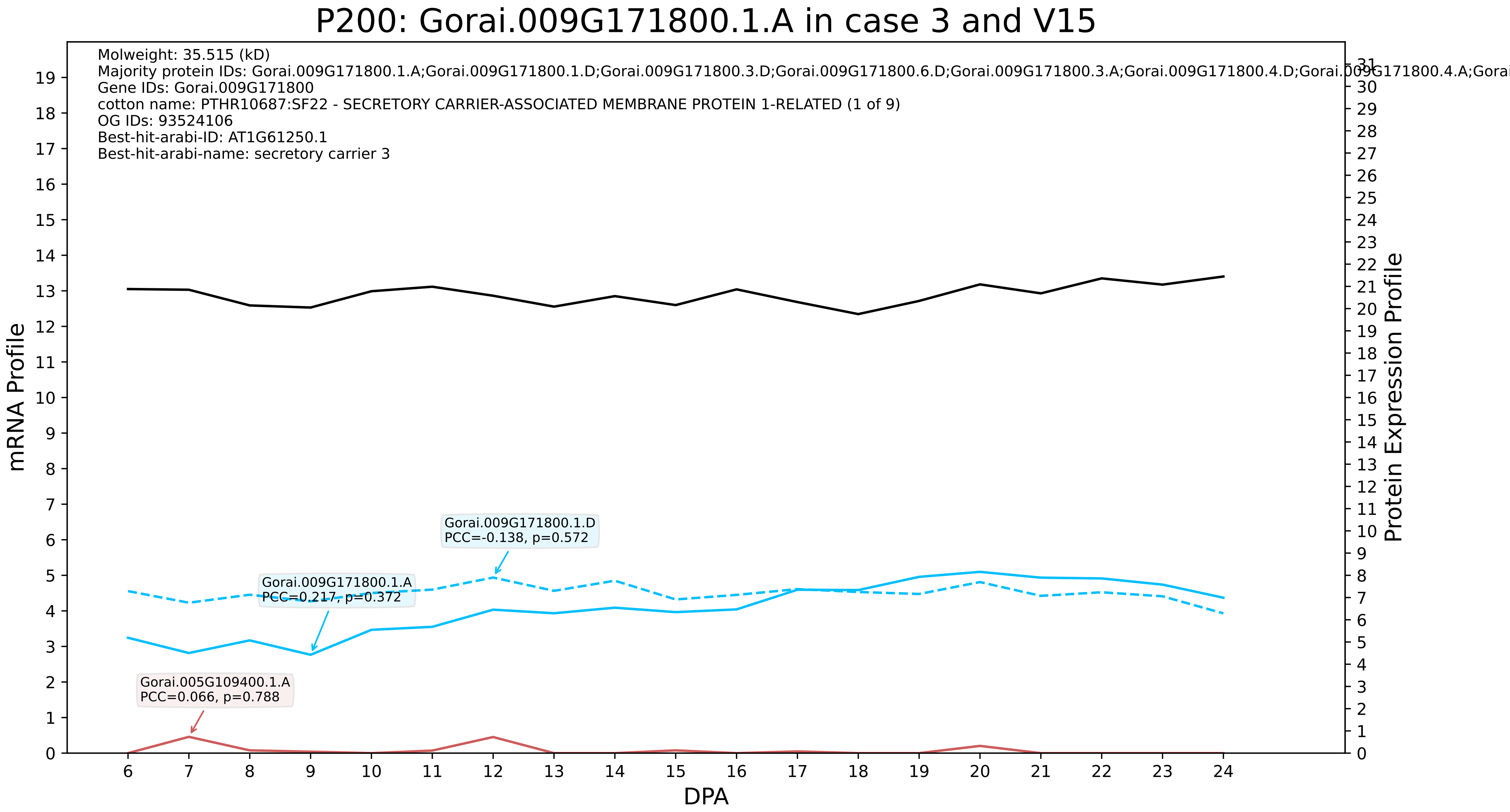
P200: Gorai.009G171800.1.A in case 3 and V15
Molweight: 35.515 (kD)
Majority protein IDs: Gorai.009G171800.1.A;Gorai.009G171800.1.D;Gorai.009G171800.3.D;Gorai.009G171800.6.D;Gorai.009G171800.3.A;Gorai.009G171800.4.D;Gorai.009G171800.4.A;Gorai
Gene IDs: Gorai.009G171800
cotton name: PTHR10687:SF22 - SECRETORY CARRIER-ASSOCIATED MEMBRANE PROTEIN 1-RELATED (1 of 9)
OG IDs: 93524106
Best-hit-arabi-ID: AT1G61250.1
Best-hit-arabi-name: secretory carrier 3
Gorai.009G171800.1.D
PCC=-0.138, p=0.572
Gorai.009G171800.1.A
PCC=0.217, p=0.372
Gorai.005G109400.1.A
PCC=0.066, p=0.788
mRNA Profile
Protein Expression Profile
DPA

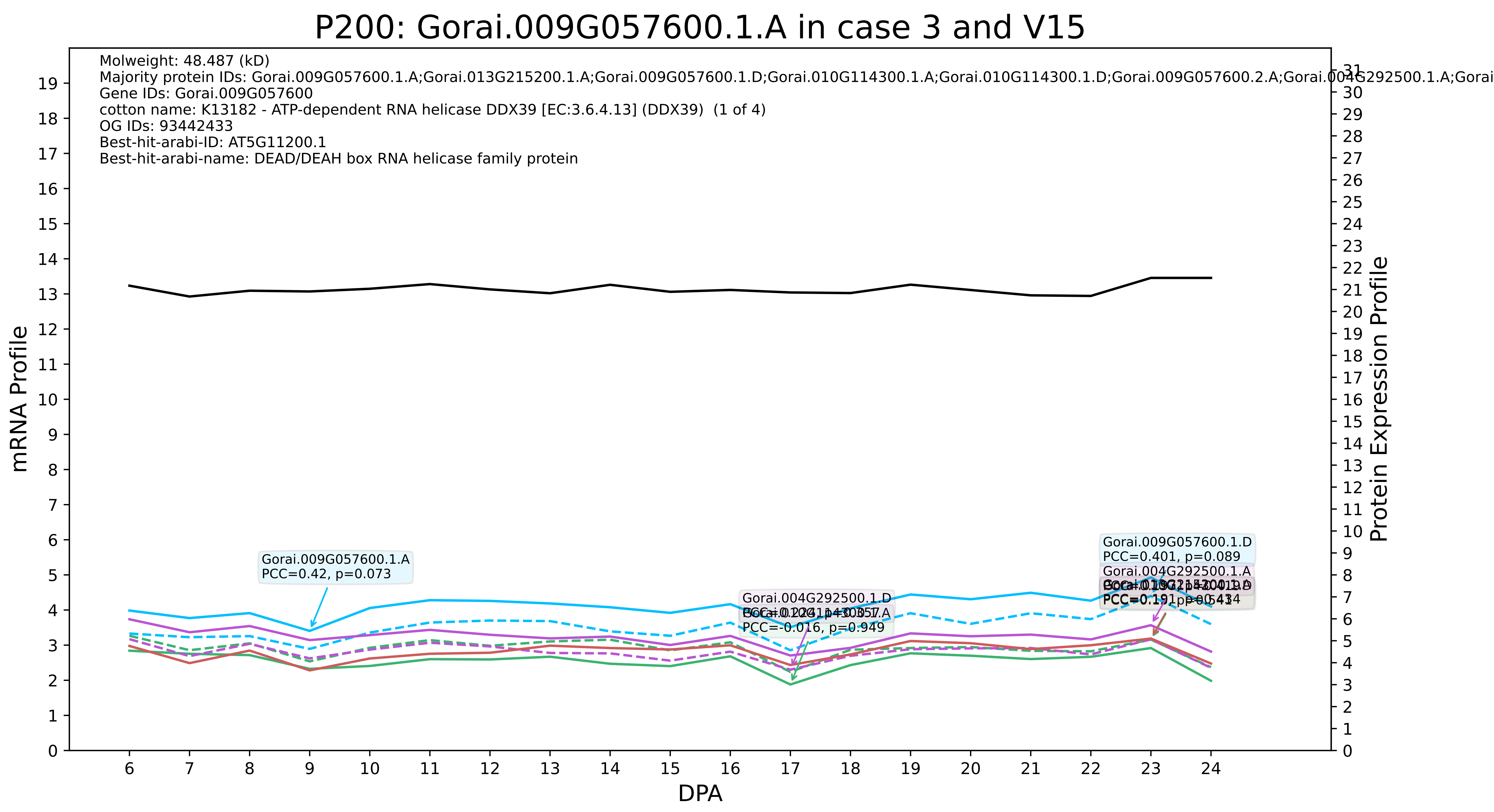

P200: Gorai.009G057600.1.A in case 3 and V15
Molweight: 48.487 (kD)
Majority protein IDs: Gorai.009G057600.1.A;Gorai.013G215200.1.A;Gorai.009G057600.1.D;Gorai.010G114300.1.A;Gorai.010G114300.1.D;Gorai.009G057600.2.A;Gorai.004G292500.1.A;Gorai
Gene IDs: Gorai.009G057600
cotton name: K13182 - ATP-dependent RNA helicase DDX39 [EC:3.6.4.13] (DDX39) (1 of 4)
OG IDs: 93442433
Best-hit-arabi-ID: AT5G11200.1
Best-hit-arabi-name: DEAD/DEAH box RNA helicase family protein
Gorai.009G057600.1.A
PCC=0.42, p=0.073
Gorai.004G292500.1.D
PCC=-0.016, p=0.949
Gorai.009G057600.1.D
PCC=0.401, p=0.089
Gorai.004G292500.1.A
mRNA Profile
Protein Expression Profile
DPA

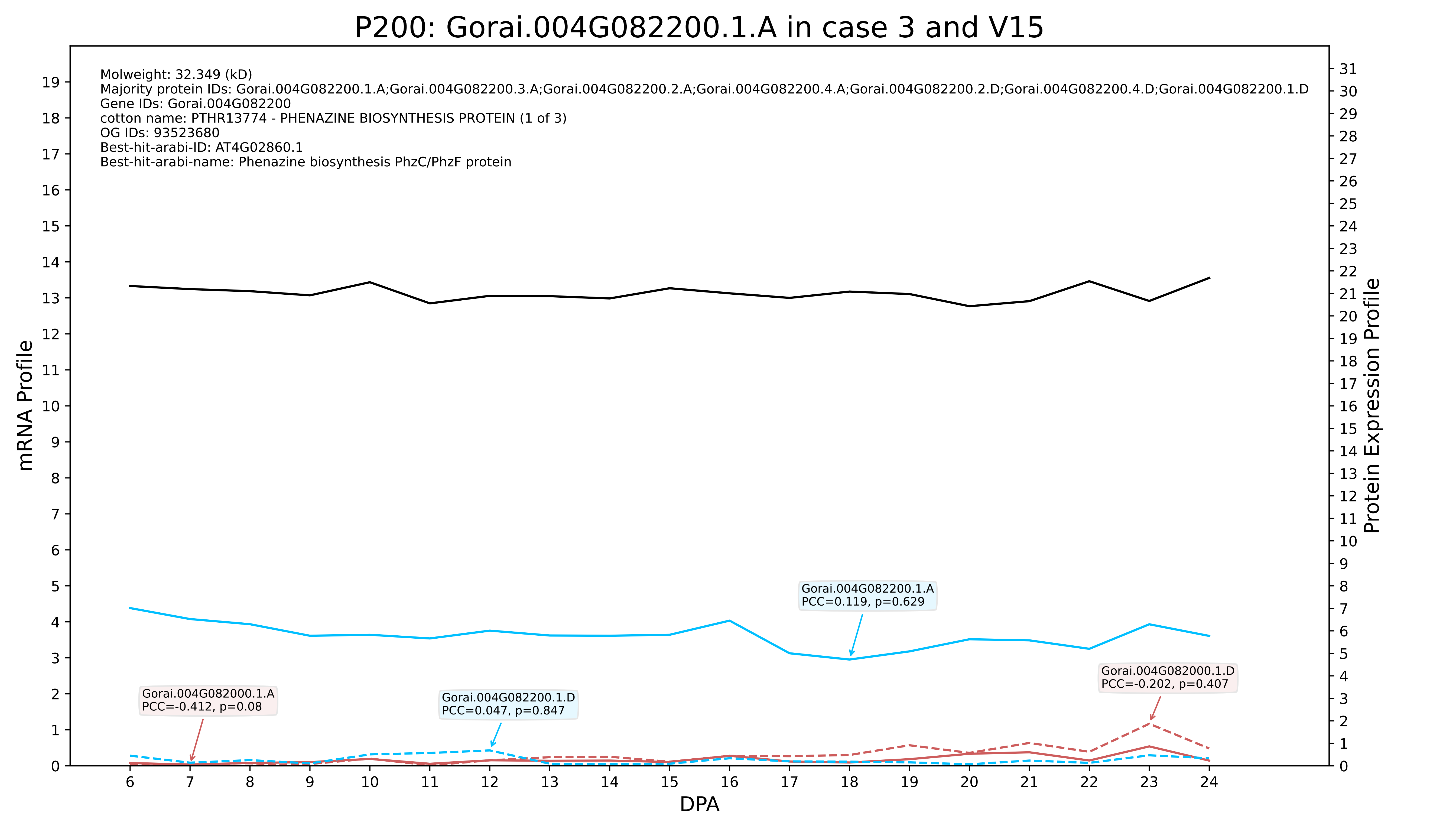

P200: Gorai.004G082200.1.A in case 3 and V15
Molweight: 32.349 (kD)
Majority protein IDs: Gorai.004G082200.1.A;Gorai.004G082200.3.A;Gorai.004G082200.2.A;Gorai.004G082200.4.A;Gorai.004G082200.2.D;Gorai.004G082200.4.D;Gorai.004G082200.1.D
Gene IDs: Gorai.004G082200
cotton name: PTHR13774 - PHENAZINE BIOSYNTHESIS PROTEIN (1 of 3)
OG IDs: 93523680
Best-hit-arabi-ID: AT4G02860.1
Best-hit-arabi-name: Phenazine biosynthesis PhzC/PhzF protein
Gorai.004G082200.1.A
PCC=0.119, p=0.629
Gorai.004G082000.1.D
PCC=-0.202, p=0.407
Gorai.004G082000.1.A
PCC=-0.412, p=0.08
Gorai.004G082200.1.D
PCC=0.047, p=0.847
mRNA Profile
Protein Expression Profile
DPA

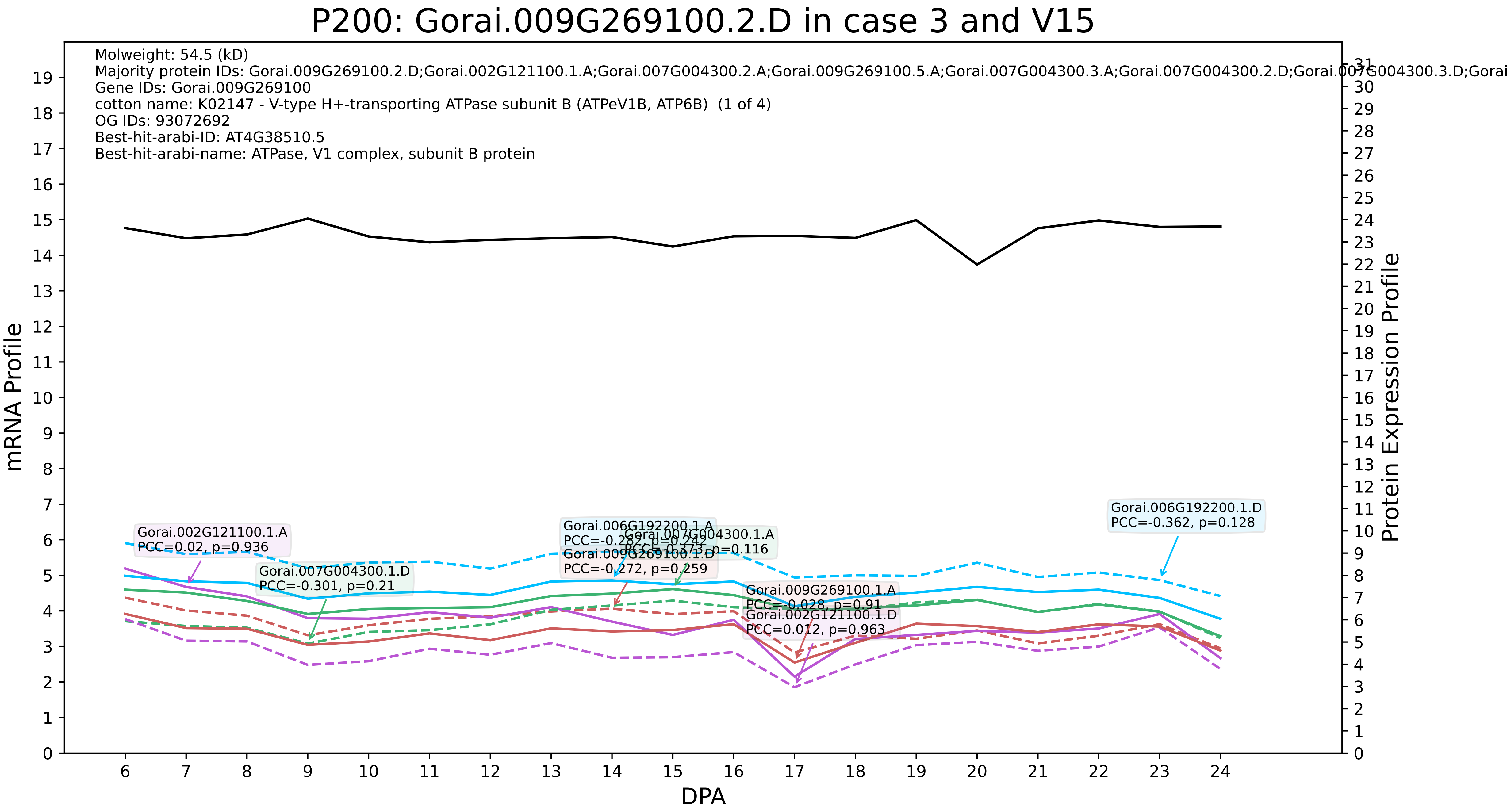

P200: Gorai.009G269100.2.D in case 3 and V15
Molweight: 54.5 (kD)
Majority protein IDs: Gorai.009G269100.2.D;Gorai.002G121100.1.A;Gorai.007G004300.2.A;Gorai.009G269100.5.A;Gorai.007G004300.3.A;Gorai.007G004300.2.D;Gorai.007G004300.3.D;Gorai
Gene IDs: Gorai.009G269100
cotton name: K02147 - V-type H+-transporting ATPase subunit B (ATPeV1B, ATP6B) (1 of 4)
OG IDs: 93072692
Best-hit-arabi-ID: AT4G38510.5
Best-hit-arabi-name: ATPase, V1 complex, subunit B protein
Gorai.002G121100.1.A
PCC=0.02, p=0.936
Gorai.007G004300.1.D
PCC=-0.301, p=0.21
Gorai.006G192200.1.A
PCC=-0.282, p=0.242
Gorai.007G004300.1.A
PCC=-0.373, p=0.116
Gorai.009G269100.1.D
PCC=-0.272, p=0.259
Gorai.009G269100.1.A
PCC=-0.028, p=0.91
Gorai.002G121100.1.D
PCC=0.012, p=0.963
Gorai.006G192200.1.D
PCC=-0.362, p=0.128
mRNA Profile
Protein Expression Profile
DPA

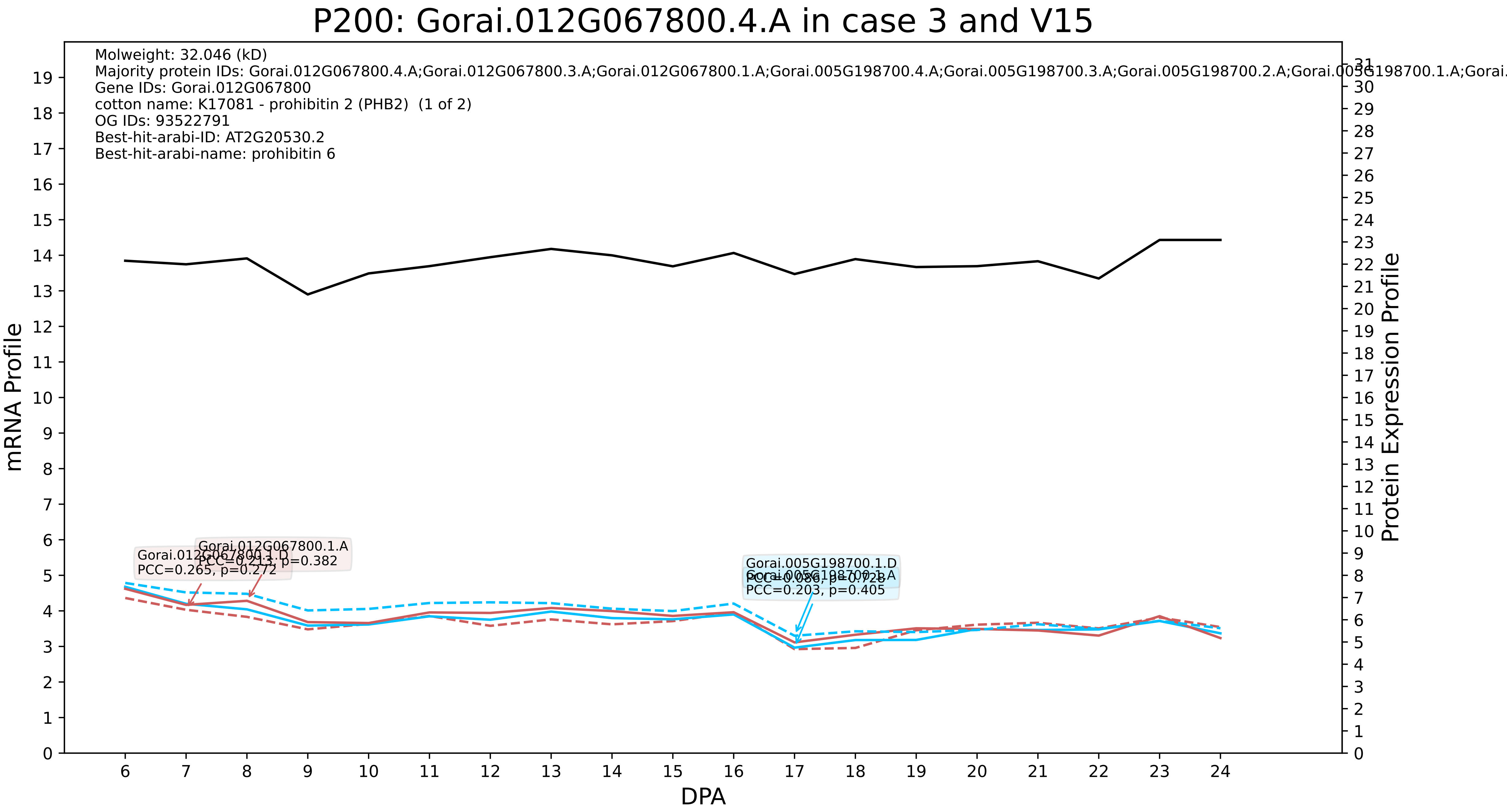

P200: Gorai.012G067800.4.A in case 3 and V15
Molweight: 32.046 (kD)
Majority protein IDs: Gorai.012G067800.4.A;Gorai.012G067800.3.A;Gorai.012G067800.1.A;Gorai.005G198700.4.A;Gorai.005G198700.3.A;Gorai.005G198700.2.A;Gorai.005G198700.1.A;Gorai.
Gene IDs: Gorai.012G067800
cotton name: K17081 - prohibitin 2 (PHB2) (1 of 2)
OG IDs: 93522791
Best-hit-arabi-ID: AT2G20530.2
Best-hit-arabi-name: prohibitin 6
Gorai.012G067800.1.D
PCC=0.265, p=0.272
Gorai.012G067800.1.A
PCC=0.213, p=0.382
Gorai.005G198700.1.D
PCC=0.086, p=0.726
Gorai.005G198700.1.A
PCC=0.203, p=0.405
mRNA Profile
Protein Expression Profile
DPA

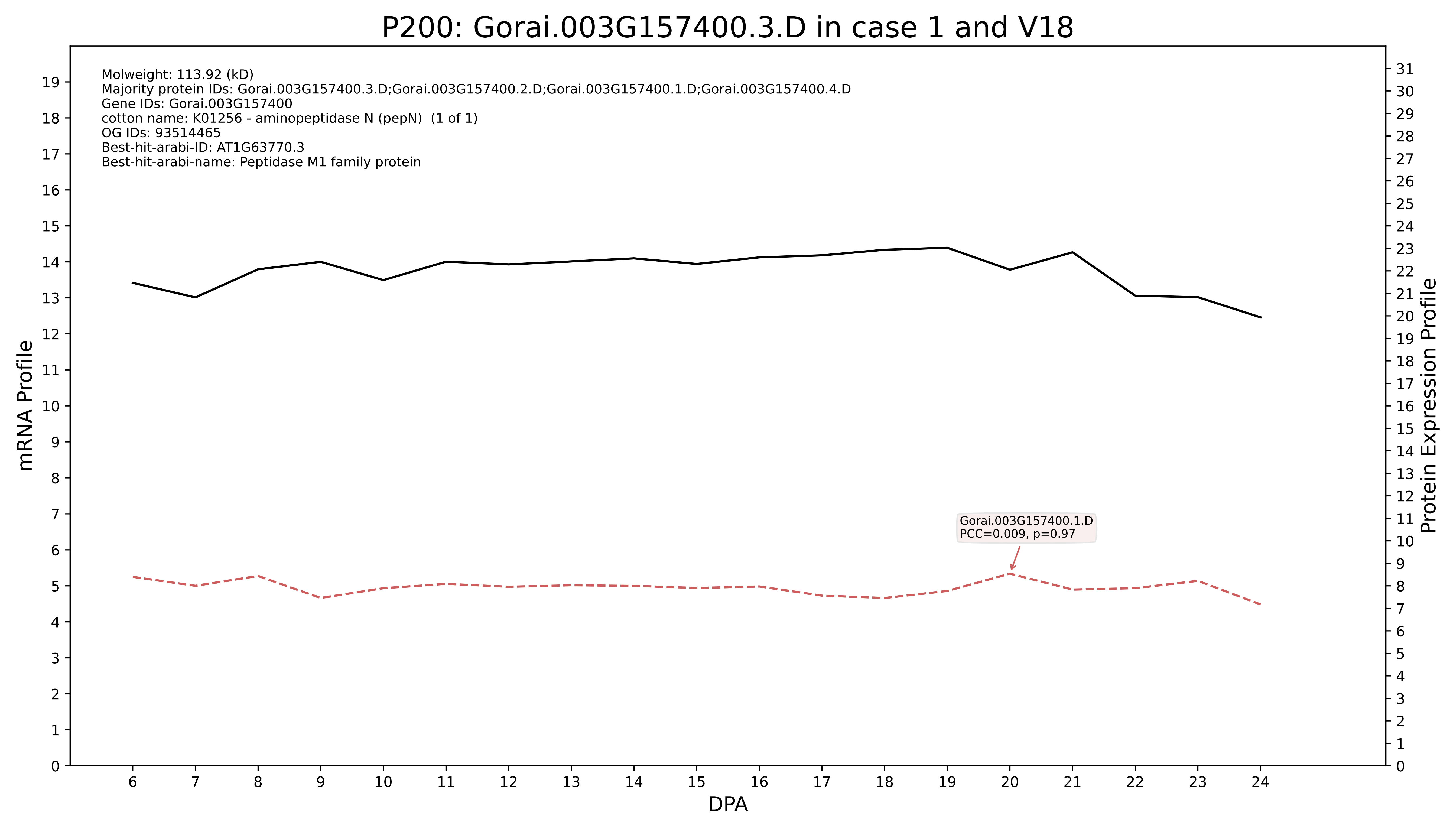
P200: Gorai.003G157400.3.D in case 1 and V18
Molweight: 113.92 (kD)
Majority protein IDs: Gorai.003G157400.3.D;Gorai.003G157400.2.D;Gorai.003G157400.1.D;Gorai.003G157400.4.D
Gene IDs: Gorai.003G157400
cotton name: K01256 - aminopeptidase N (pepN) (1 of 1)
OG IDs: 93514465
Best-hit-arabi-ID: AT1G63770.3
Best-hit-arabi-name: Peptidase M1 family protein
Gorai.003G157400.1.D
PCC=0.009, p=0.97
mRNA Profile
Protein Expression Profile
DPA

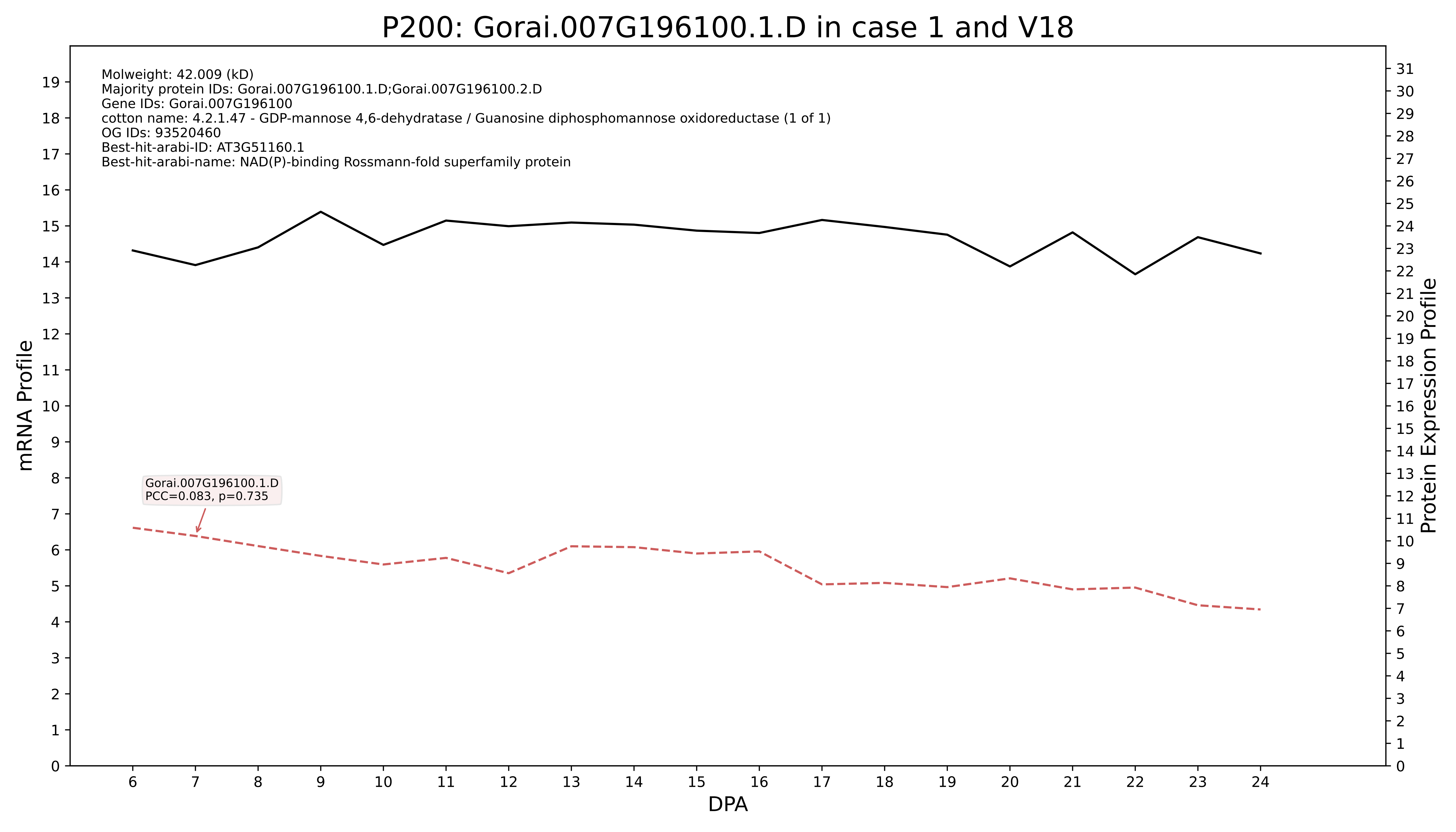
P200: Gorai.007G196100.1.D in case 1 and V18
Molweight: 42.009 (kD)
Majority protein IDs: Gorai.007G196100.1.D;Gorai.007G196100.2.D
Gene IDs: Gorai.007G196100
cotton name: 4.2.1.47 - GDP-mannose 4,6-dehydratase / Guanosine diphosphomannose oxidoreductase (1 of 1)
OG IDs: 93520460
Best-hit-arabi-ID: AT3G51160.1
Best-hit-arabi-name: NAD(P)-binding Rossmann-fold superfamily protein
Gorai.007G196100.1.D
PCC=0.083, p=0.735
mRNA Profile
Protein Expression Profile
DPA

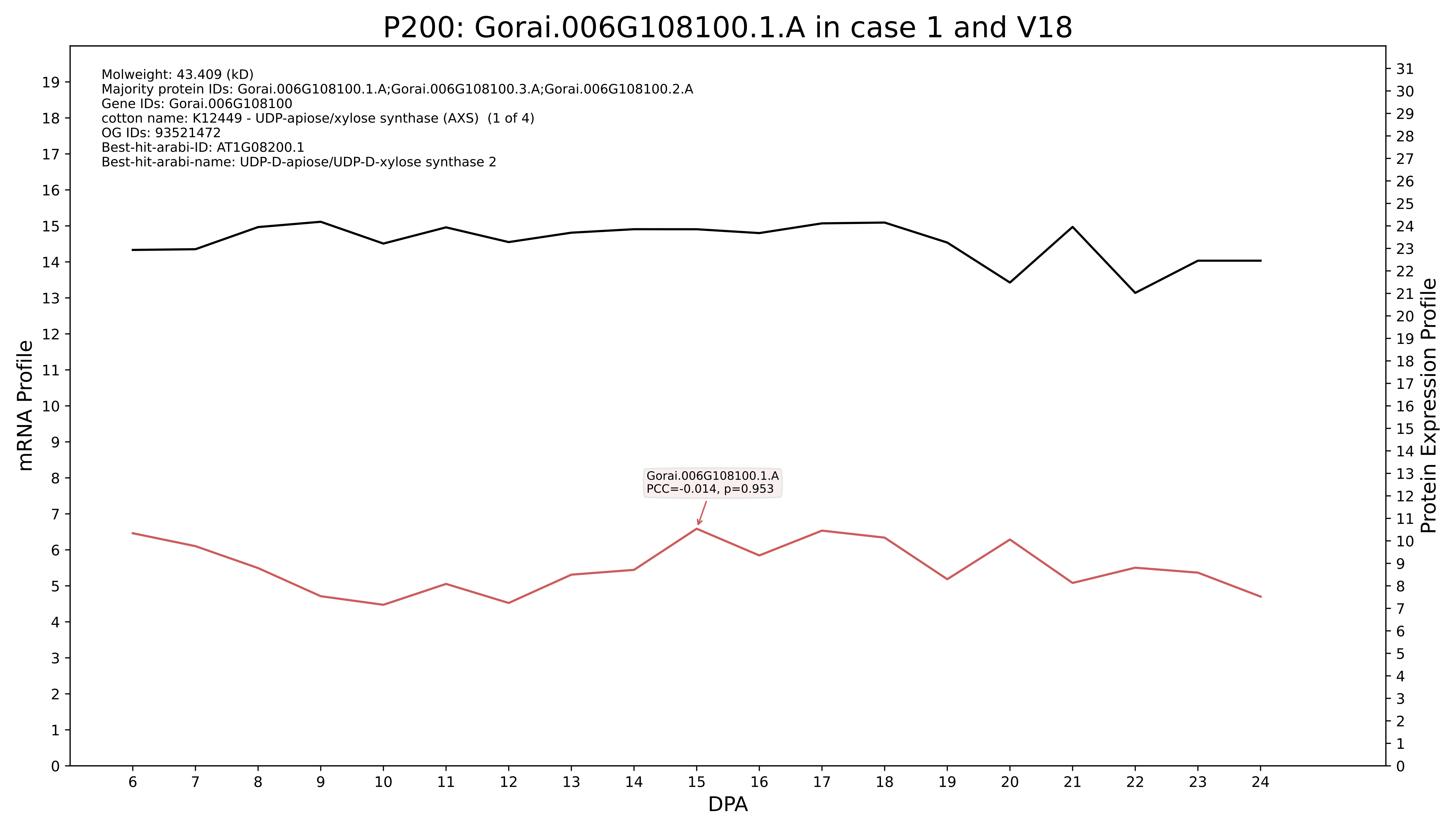
P200: Gorai.006G108100.1.A in case 1 and V18
Molweight: 43.409 (kD)
Majority protein IDs: Gorai.006G108100.1.A;Gorai.006G108100.3.A;Gorai.006G108100.2.A
Gene IDs: Gorai.006G108100
cotton name: K12449 - UDP-apiose/xylose synthase (AXS) (1 of 4)
OG IDs: 93521472
Best-hit-arabi-ID: AT1G08200.1
Best-hit-arabi-name: UDP-D-apiose/UDP-D-xylose synthase 2
Gorai.006G108100.1.A
PCC=-0.014, p=0.953
mRNA Profile
Protein Expression Profile
DPA

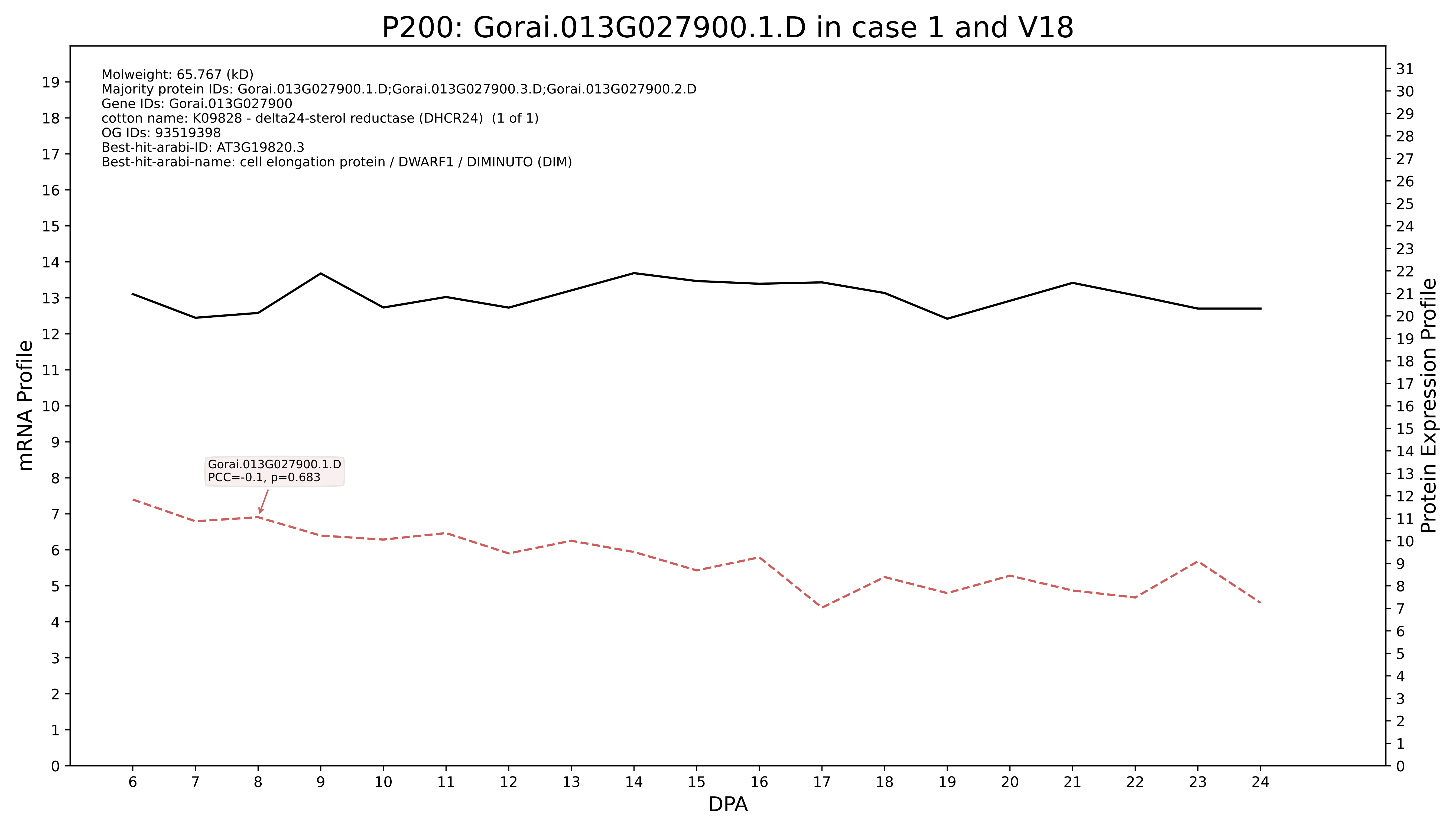

P200: Gorai.013G027900.1.D in case 1 and V18
Molweight: 65.767 (kD)
Majority protein IDs: Gorai.013G027900.1.D;Gorai.013G027900.3.D;Gorai.013G027900.2.D
Gene IDs: Gorai.013G027900
cotton name: K09828 - delta24-sterol reductase (DHCR24) (1 of 1)
OG IDs: 93519398
Best-hit-arabi-ID: AT3G19820.3
Best-hit-arabi-name: cell elongation protein / DWARF1 / DIMINUTO (DIM)
Gorai.013G027900.1.D
PCC=-0.1, p=0.683
mRNA Profile
Protein Expression Profile
DPA

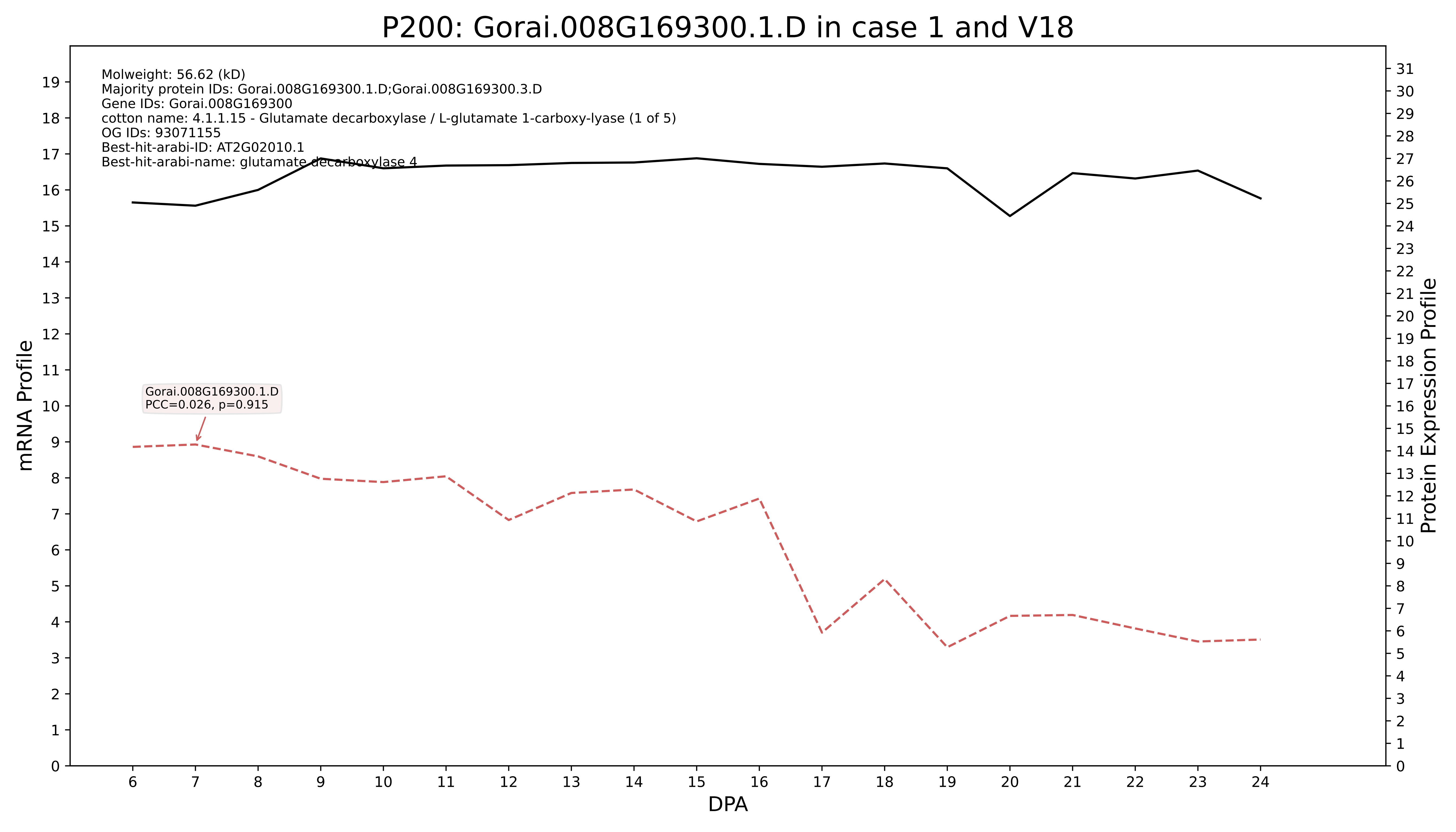

P200: Gorai.008G169300.1.D in case 1 and V18
Molweight: 56.62 (kD)
Majority protein IDs: Gorai.008G169300.1.D;Gorai.008G169300.3.D
Gene IDs: Gorai.008G169300
cotton name: 4.1.1.15 - Glutamate decarboxylase / L-glutamate 1-carboxy-lyase (1 of 5)
OG IDs: 93071155
Best-hit-arabi-ID: AT2G02010.1
Best-hit-arabi-name: glutamate decarboxylase 4
Gorai.008G169300.1.D
PCC=0.026, p=0.915
mRNA Profile
Protein Expression Profile
DPA

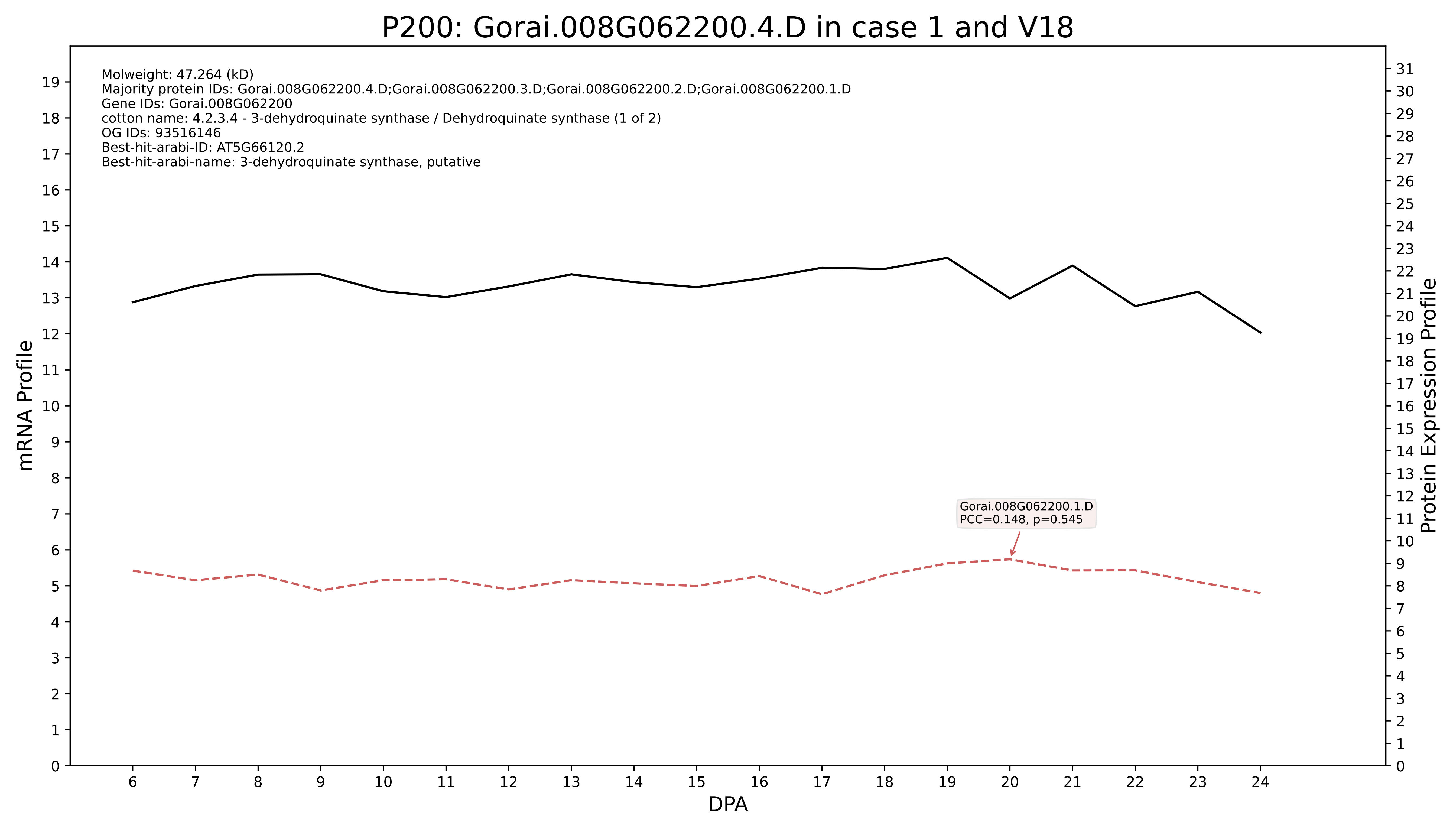
P200: Gorai.008G062200.4.D in case 1 and V18
Molweight: 47.264 (kD)
Majority protein IDs: Gorai.008G062200.4.D;Gorai.008G062200.3.D;Gorai.008G062200.2.D;Gorai.008G062200.1.D
Gene IDs: Gorai.008G062200
cotton name: 4.2.3.4 - 3-dehydroquinate synthase / Dehydroquinate synthase (1 of 2)
OG IDs: 93516146
Best-hit-arabi-ID: AT5G66120.2
Best-hit-arabi-name: 3-dehydroquinate synthase, putative
Gorai.008G062200.1.D
PCC=0.148, p=0.545
mRNA Profile
Protein Expression Profile
DPA

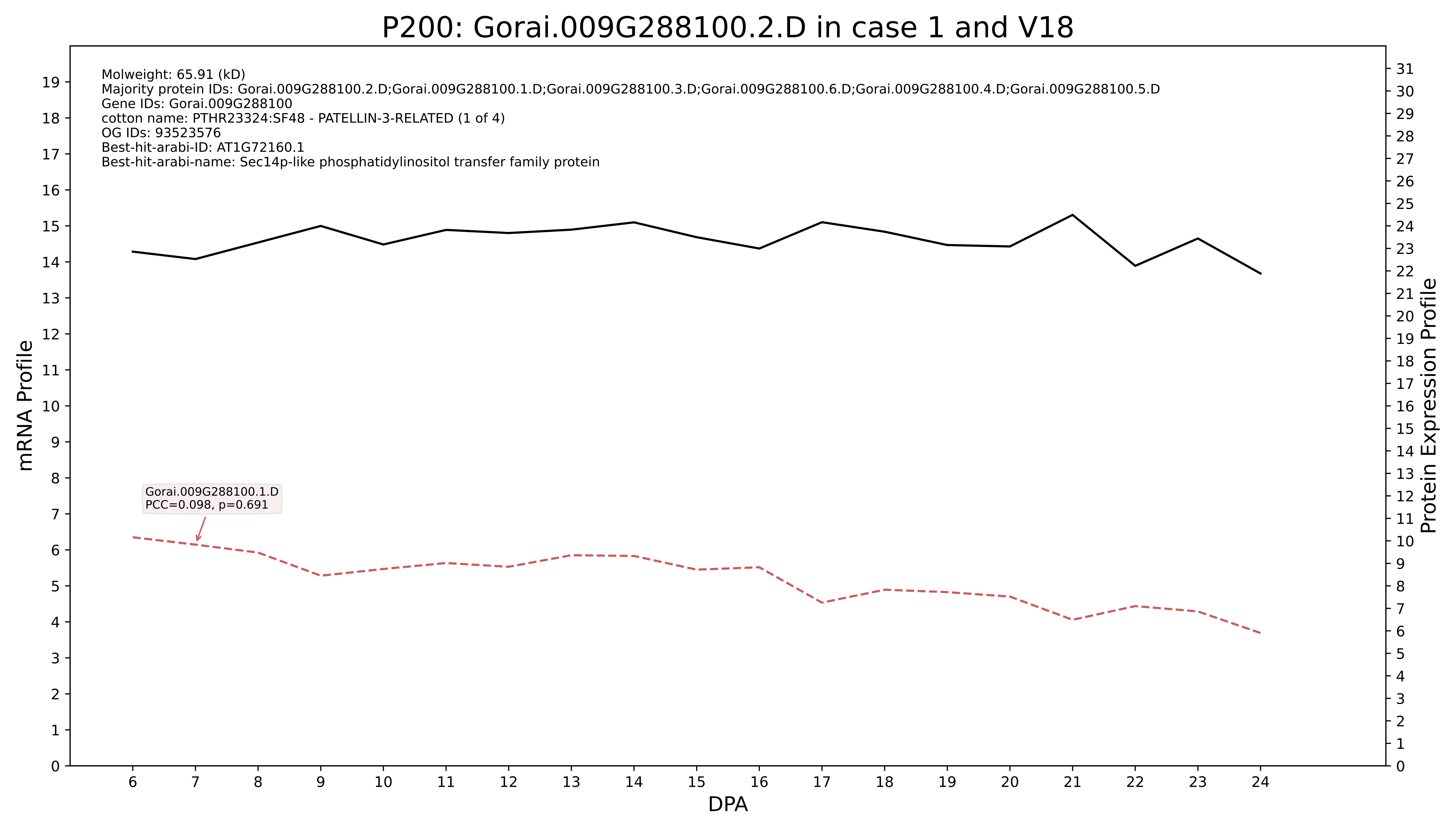
P200: Gorai.009G288100.2.D in case 1 and V18
Molweight: 65.91 (kD)
Majority protein IDs: Gorai.009G288100.2.D;Gorai.009G288100.1.D;Gorai.009G288100.3.D;Gorai.009G288100.6.D;Gorai.009G288100.4.D;Gorai.009G288100.5.D
Gene IDs: Gorai.009G288100
cotton name: PTHR23324:SF48 - PATELLIN-3-RELATED (1 of 4)
OG IDs: 93523576
Best-hit-arabi-ID: AT1G72160.1
Best-hit-arabi-name: Sec14p-like phosphatidylinositol transfer family protein
Gorai.009G288100.1.D
PCC=0.098, p=0.691
mRNA Profile
Protein Expression Profile
DPA

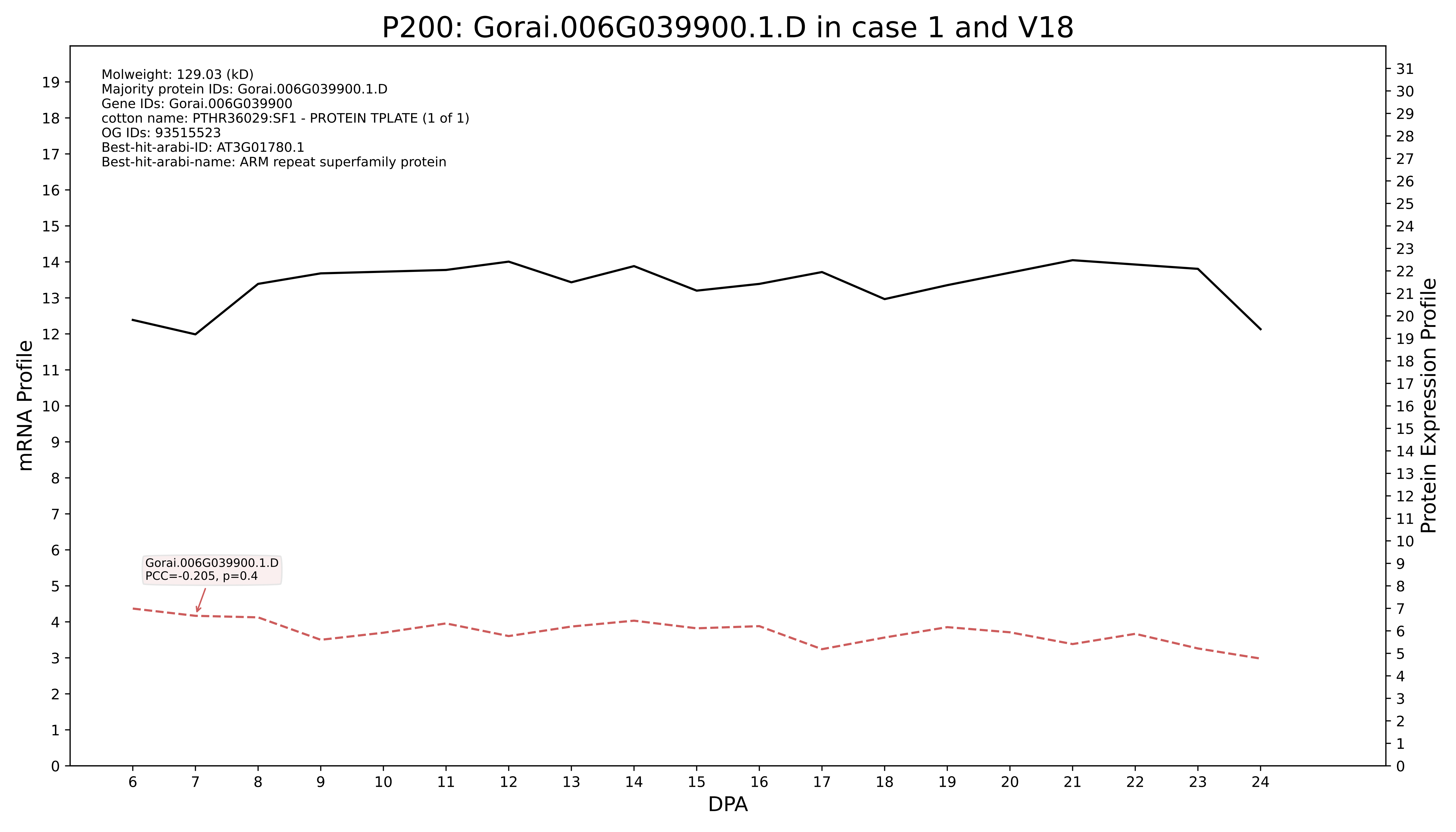
P200: Gorai.006G039900.1.D in case 1 and V18
Molweight: 129.03 (kD)
Majority protein IDs: Gorai.006G039900.1.D
Gene IDs: Gorai.006G039900
cotton name: PTHR36029:SF1 - PROTEIN TPLATE (1 of 1)
OG IDs: 93515523
Best-hit-arabi-ID: AT3G01780.1
Best-hit-arabi-name: ARM repeat superfamily protein
Gorai.006G039900.1.D
PCC=-0.205, p=0.4
mRNA Profile
Protein Expression Profile
DPA

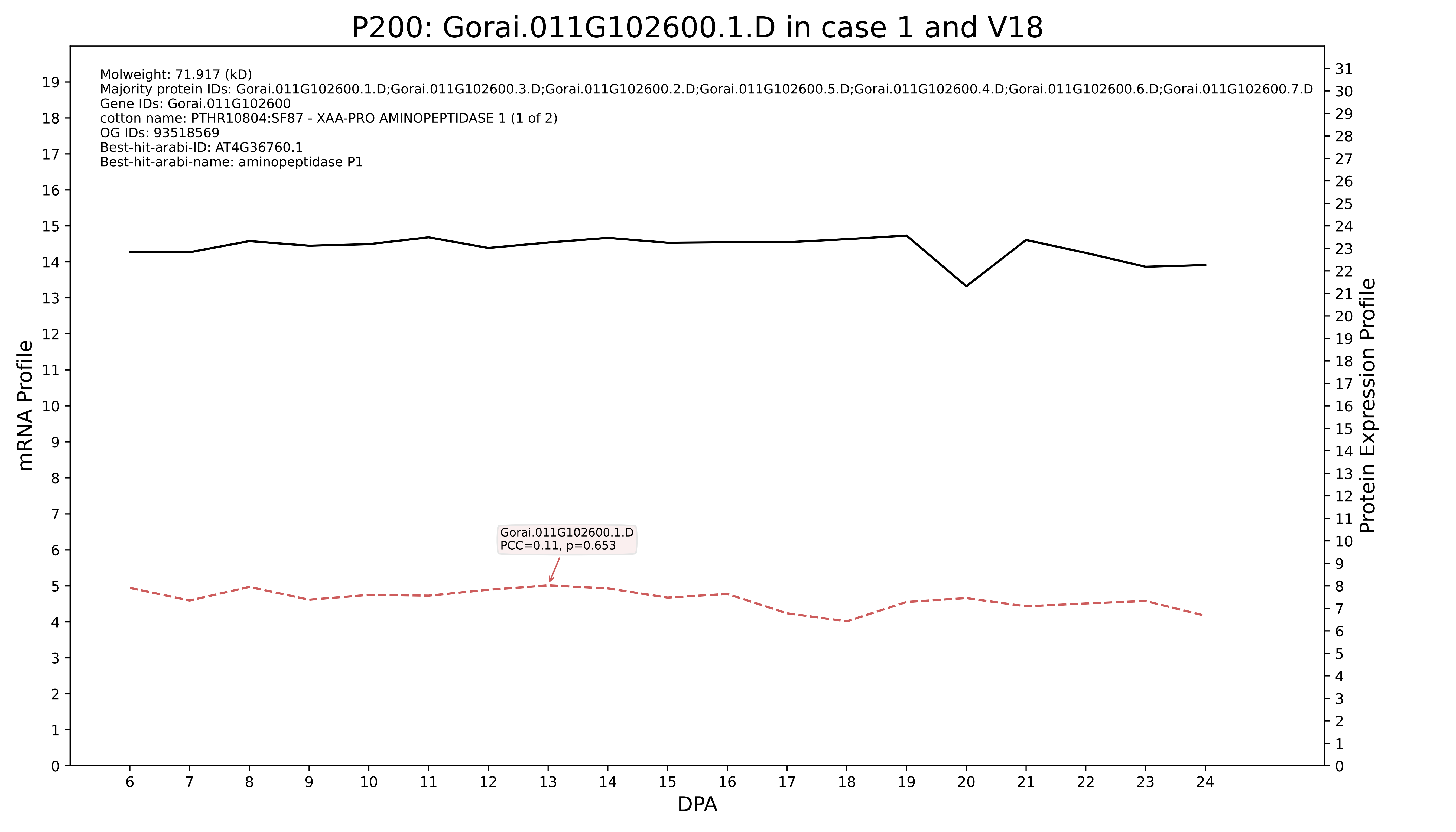

P200: Gorai.011G102600.1.D in case 1 and V18
Molweight: 71.917 (kD)
Majority protein IDs: Gorai.011G102600.1.D;Gorai.011G102600.3.D;Gorai.011G102600.2.D;Gorai.011G102600.5.D;Gorai.011G102600.4.D;Gorai.011G102600.6.D;Gorai.011G102600.7.D
Gene IDs: Gorai.011G102600
cotton name: PTHR10804:SF87 - XAA-PRO AMINOPEPTIDASE 1 (1 of 2)
OG IDs: 93518569
Best-hit-arabi-ID: AT4G36760.1
Best-hit-arabi-name: aminopeptidase P1
Gorai.011G102600.1.D
PCC=0.11, p=0.653
mRNA Profile
Protein Expression Profile
DPA

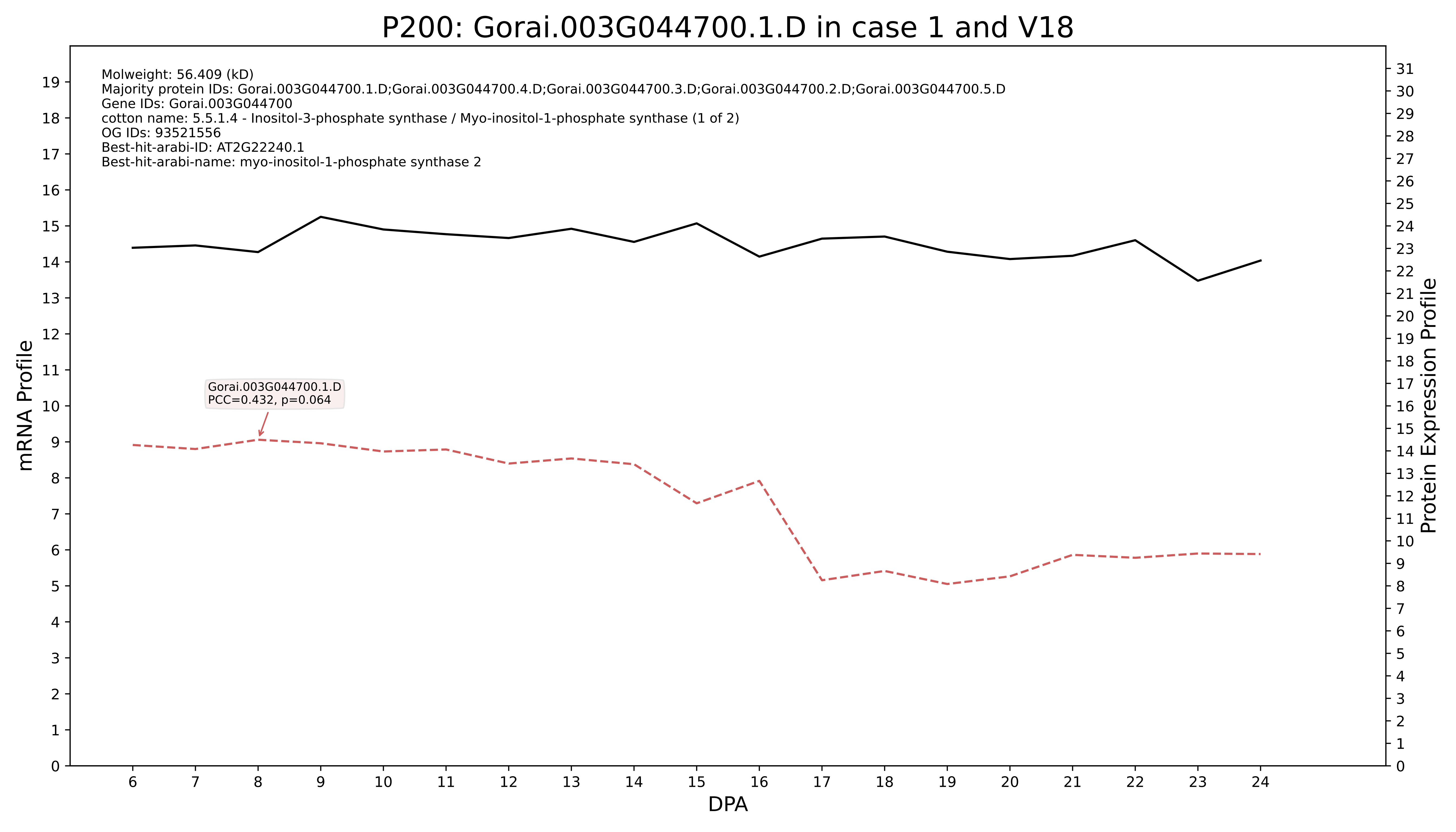
P200: Gorai.003G044700.1.D in case 1 and V18
Molweight: 56.409 (kD)
Majority protein IDs: Gorai.003G044700.1.D;Gorai.003G044700.4.D;Gorai.003G044700.3.D;Gorai.003G044700.2.D;Gorai.003G044700.5.D
Gene IDs: Gorai.003G044700
cotton name: 5.5.1.4 - Inositol-3-phosphate synthase / Myo-inositol-1-phosphate synthase (1 of 2)
OG IDs: 93521556
Best-hit-arabi-ID: AT2G22240.1
Best-hit-arabi-name: myo-inositol-1-phosphate synthase 2
Gorai.003G044700.1.D
PCC=0.432, p=0.064
mRNA Profile
Protein Expression Profile
DPA

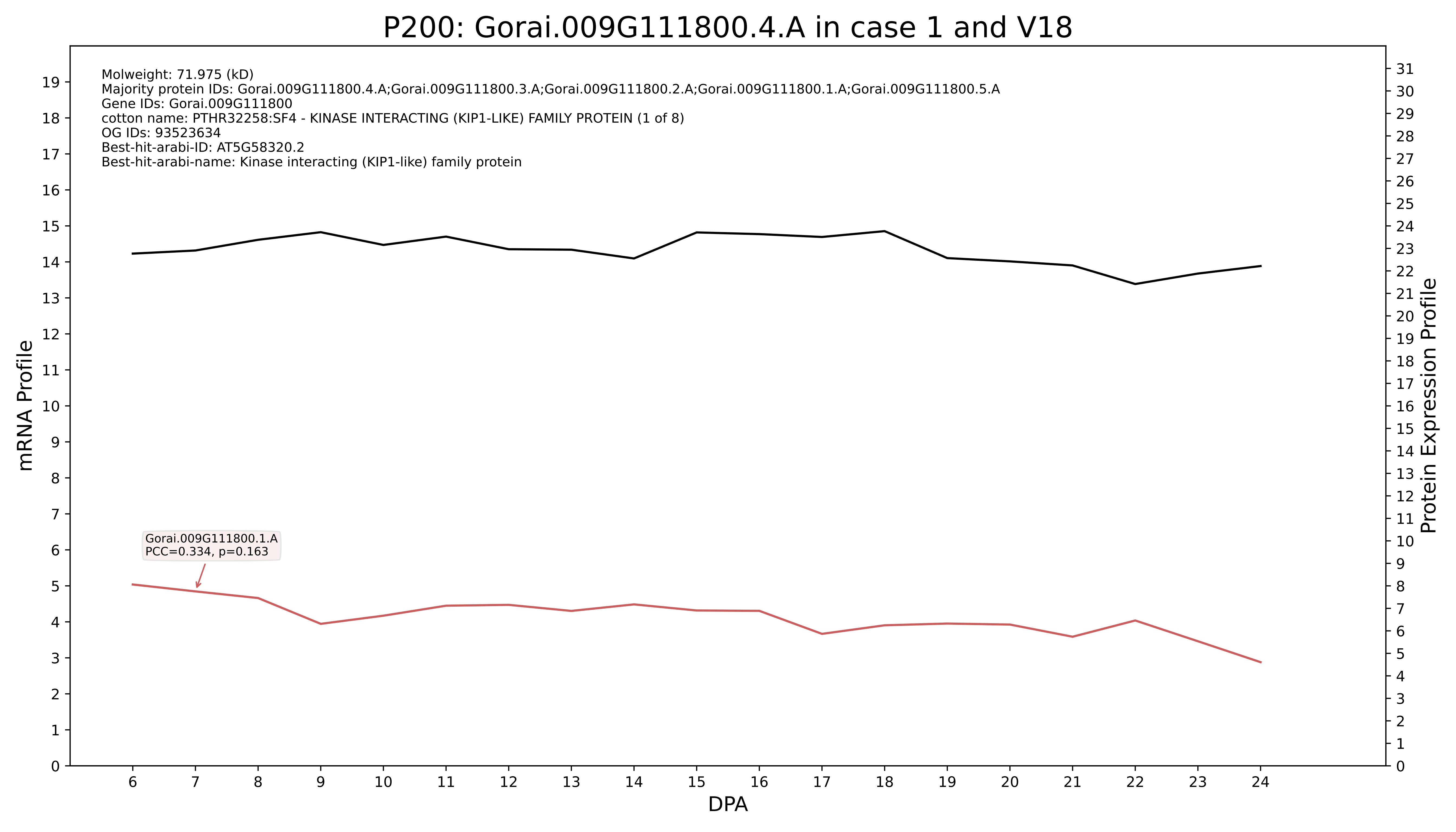
P200: Gorai.009G111800.4.A in case 1 and V18
Molweight: 71.975 (kD)
Majority protein IDs: Gorai.009G111800.4.A;Gorai.009G111800.3.A;Gorai.009G111800.2.A;Gorai.009G111800.1.A;Gorai.009G111800.5.A
Gene IDs: Gorai.009G111800
cotton name: PTHR32258:SF4 - KINASE INTERACTING (KIP1-LIKE) FAMILY PROTEIN (1 of 8)
OG IDs: 93523634
Best-hit-arabi-ID: AT5G58320.2
Best-hit-arabi-name: Kinase interacting (KIP1-like) family protein
Gorai.009G111800.1.A
PCC=0.334, p=0.163
mRNA Profile
Protein Expression Profile
DPA

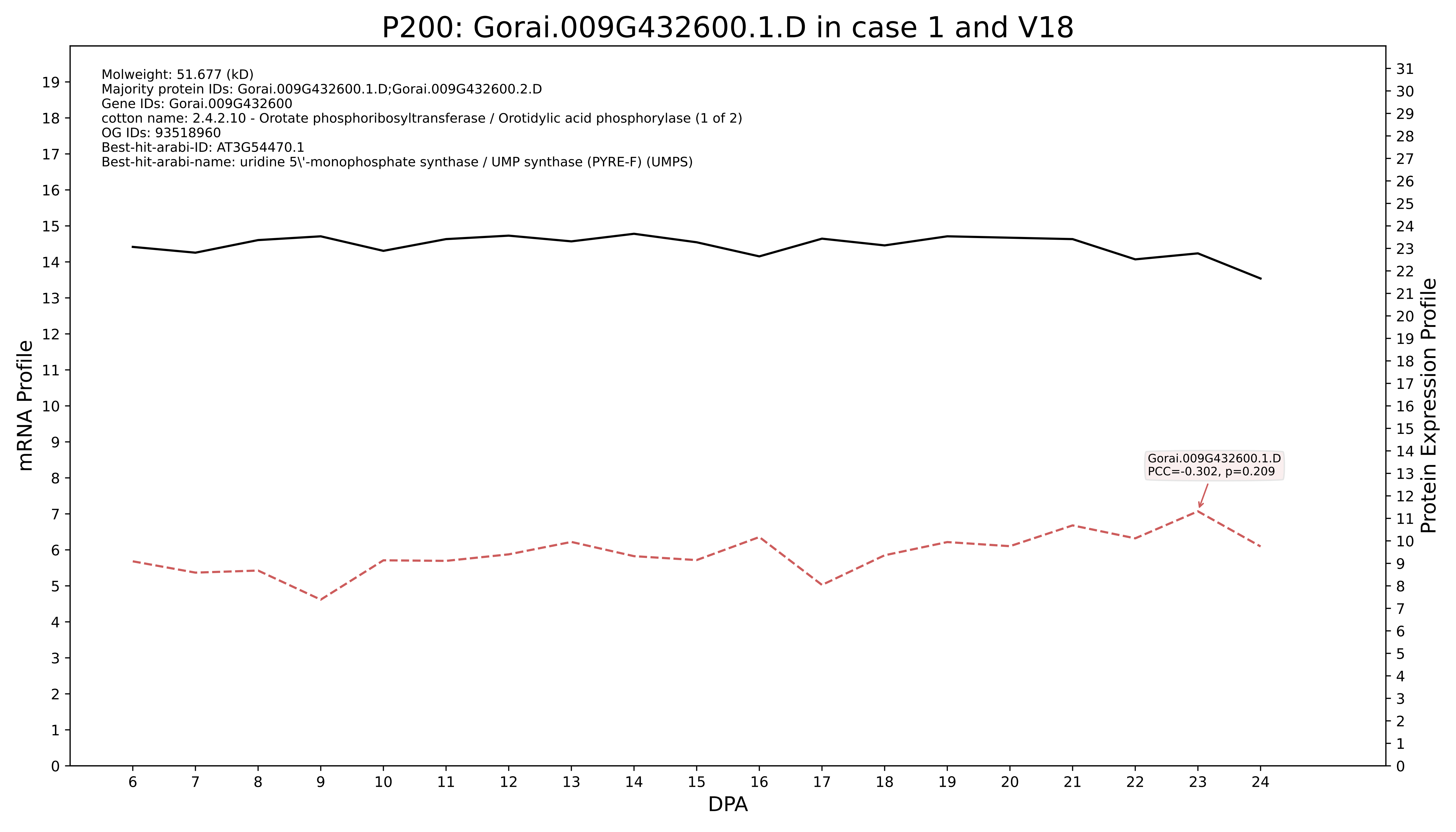

P200: Gorai.009G432600.1.D in case 1 and V18
Molweight: 51.677 (kD)
Majority protein IDs: Gorai.009G432600.1.D;Gorai.009G432600.2.D
Gene IDs: Gorai.009G432600
cotton name: 2.4.2.10 - Orotate phosphoribosyltransferase / Orotidylic acid phosphorylase (1 of 2)
OG IDs: 93518960
Best-hit-arabi-ID: AT3G54470.1
Best-hit-arabi-name: uridine 5\'-monophosphate synthase / UMP synthase (PYRE-F) (UMPS)
Gorai.009G432600.1.D
PCC=-0.302, p=0.209
mRNA Profile
Protein Expression Profile
DPA

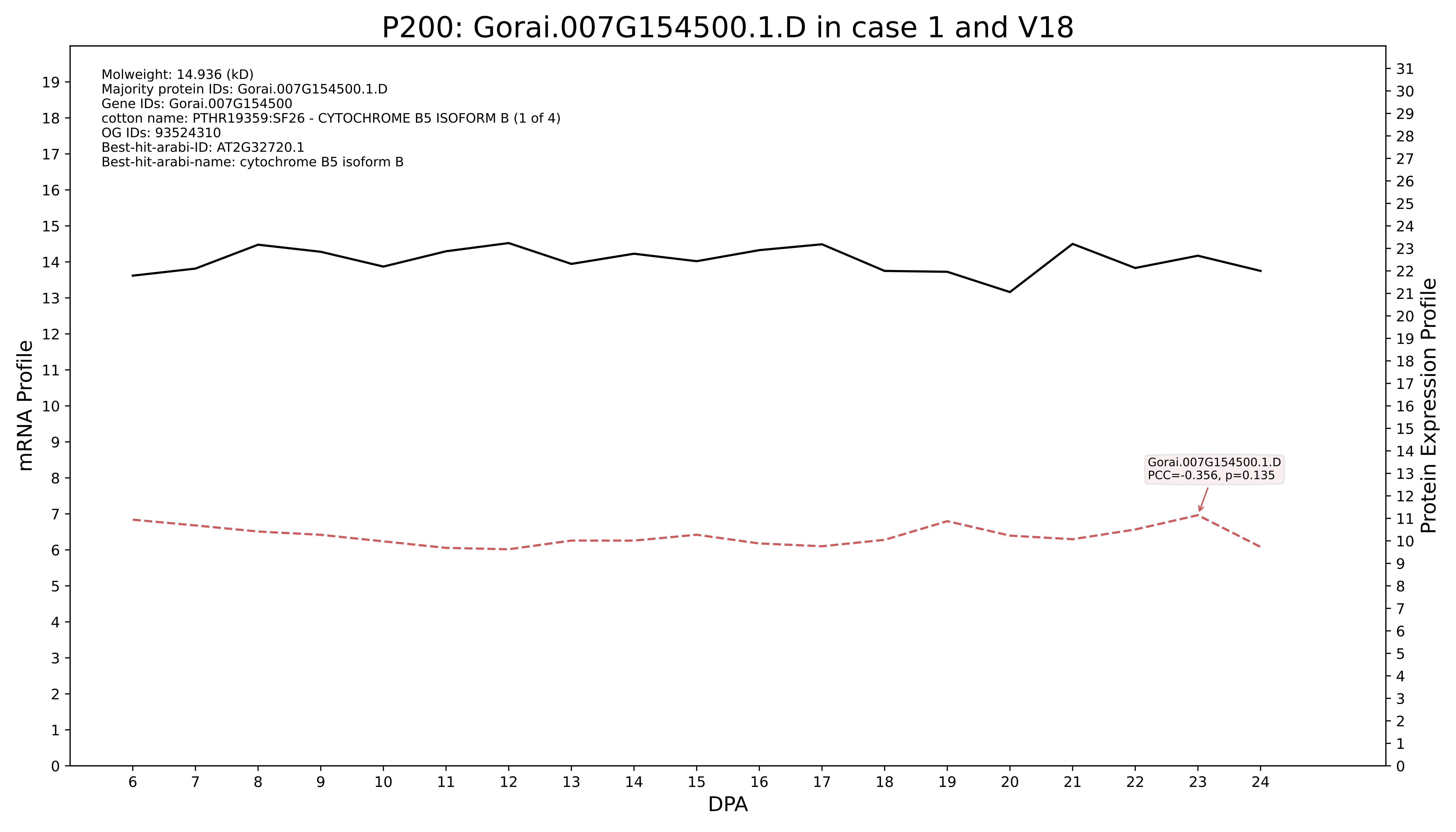

P200: Gorai.007G154500.1.D in case 1 and V18
Molweight: 14.936 (kD)
Majority protein IDs: Gorai.007G154500.1.D
Gene IDs: Gorai.007G154500
cotton name: PTHR19359:SF26 - CYTOCHROME B5 ISOFORM B (1 of 4)
OG IDs: 93524310
Best-hit-arabi-ID: AT2G32720.1
Best-hit-arabi-name: cytochrome B5 isoform B
Gorai.007G154500.1.D
PCC=-0.356, p=0.135
mRNA Profile
Protein Expression Profile
DPA

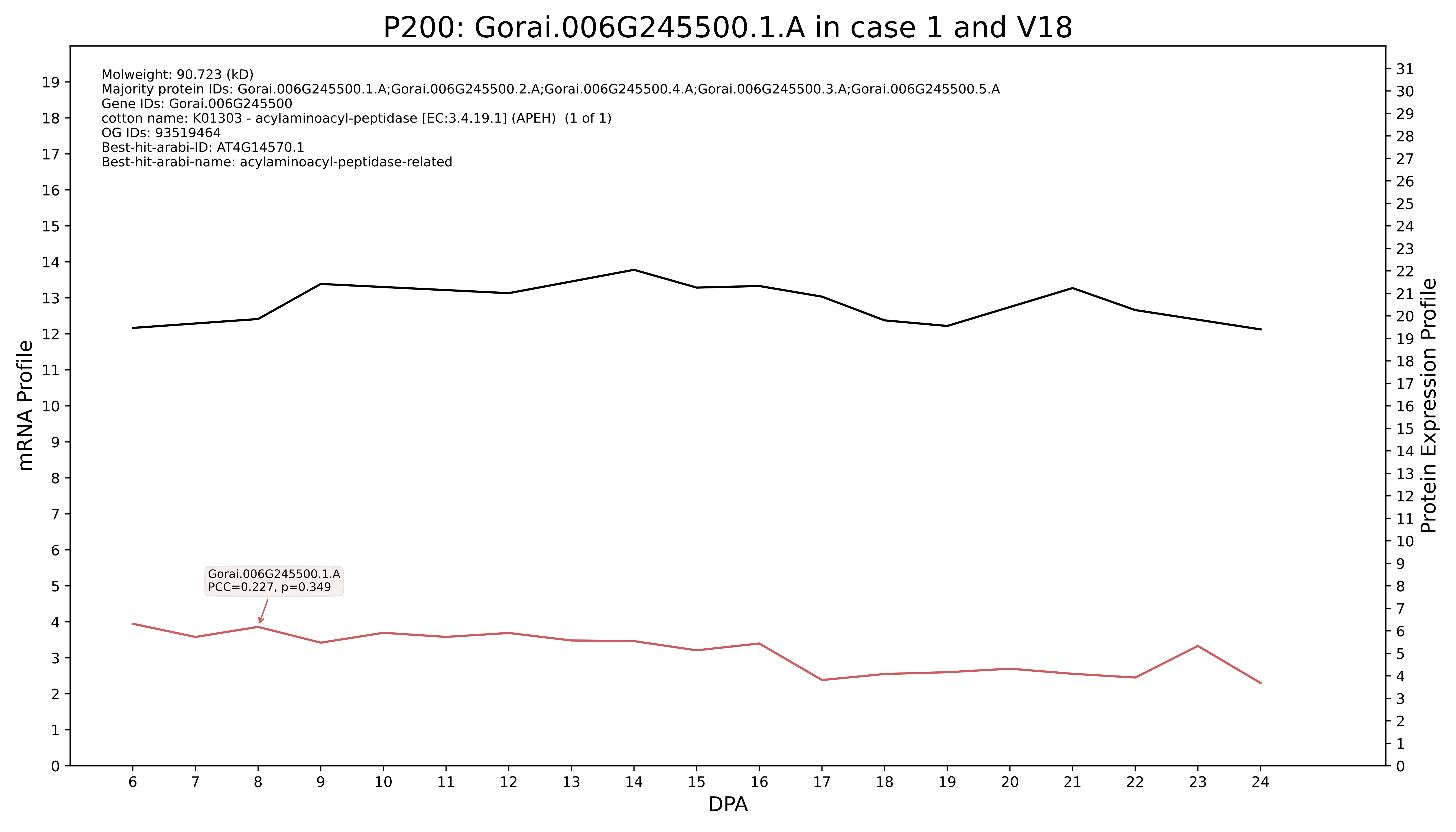
P200: Gorai.006G245500.1.A in case 1 and V18
Molweight: 90.723 (kD)
Majority protein IDs: Gorai.006G245500.1.A;Gorai.006G245500.2.A;Gorai.006G245500.4.A;Gorai.006G245500.3.A;Gorai.006G245500.5.A
Gene IDs: Gorai.006G245500
cotton name: K01303 - acylaminoacyl-peptidase [EC:3.4.19.1] (APEH) (1 of 1)
OG IDs: 93519464
Best-hit-arabi-ID: AT4G14570.1
Best-hit-arabi-name: acylaminoacyl-peptidase-related
Gorai.006G245500.1.A
PCC=0.227, p=0.349
mRNA Profile
Protein Expression Profile
DPA

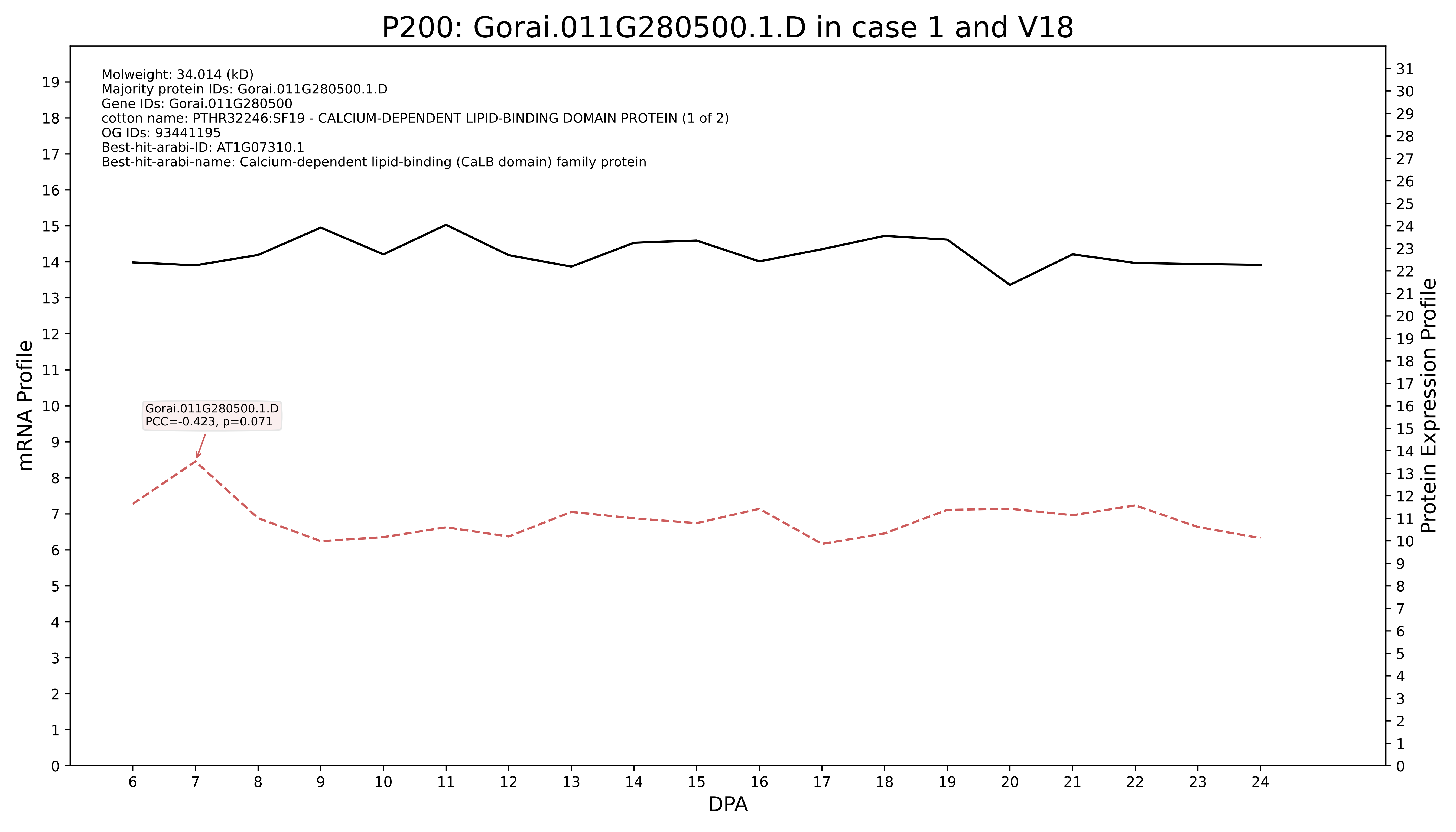

P200: Gorai.011G280500.1.D in case 1 and V18
Molweight: 34.014 (kD)
Majority protein IDs: Gorai.011G280500.1.D
Gene IDs: Gorai.011G280500
cotton name: PTHR32246:SF19 - CALCIUM-DEPENDENT LIPID-BINDING DOMAIN PROTEIN (1 of 2)
OG IDs: 93441195
Best-hit-arabi-ID: AT1G07310.1
Best-hit-arabi-name: Calcium-dependent lipid-binding (CaLB domain) family protein
Gorai.011G280500.1.D
PCC=-0.423, p=0.071
mRNA Profile
Protein Expression Profile
DPA

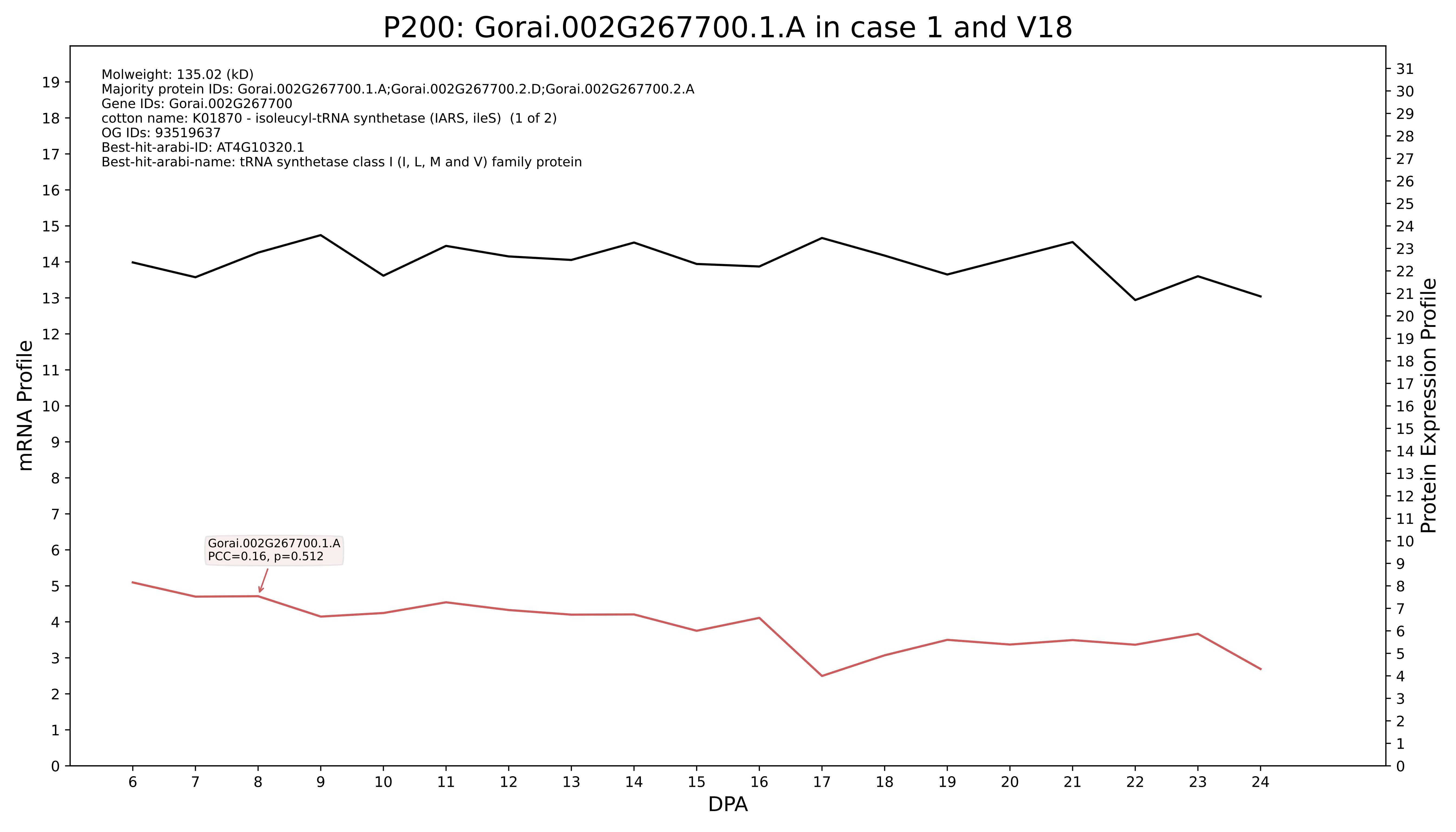

P200: Gorai.002G267700.1.A in case 1 and V18
Molweight: 135.02 (kD)
Majority protein IDs: Gorai.002G267700.1.A;Gorai.002G267700.2.D;Gorai.002G267700.2.A
Gene IDs: Gorai.002G267700
cotton name: K01870 - isoleucyl-tRNA synthetase (IARS, ileS) (1 of 2)
OG IDs: 93519637
Best-hit-arabi-ID: AT4G10320.1
Best-hit-arabi-name: tRNA synthetase class I (I, L, M and V) family protein
Gorai.002G267700.1.A
PCC=0.16, p=0.512
mRNA Profile
Protein Expression Profile
DPA

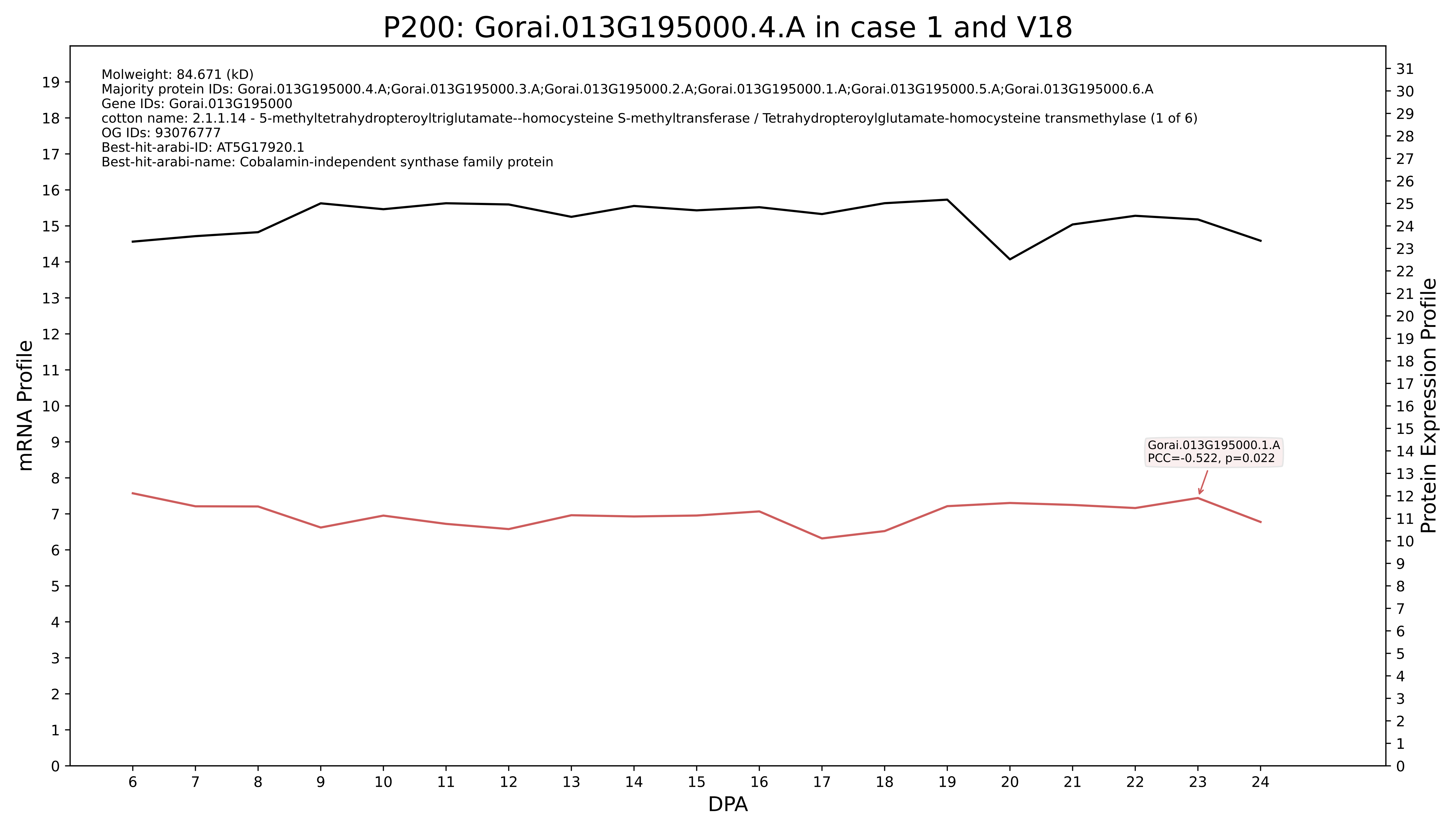
P200: Gorai.013G195000.4.A in case 1 and V18
Molweight: 84.671 (kD)
Majority protein IDs: Gorai.013G195000.4.A;Gorai.013G195000.3.A;Gorai.013G195000.2.A;Gorai.013G195000.1.A;Gorai.013G195000.5.A;Gorai.013G195000.6.A
Gene IDs: Gorai.013G195000
cotton name: 2.1.1.14 - 5-methyltetrahydropteroyltriglutamate--homocysteine S-methyltransferase / Tetrahydropteroylglutamate-homocysteine transmethylase (1 of 6)
OG IDs: 93076777
Best-hit-arabi-ID: AT5G17920.1
Best-hit-arabi-name: Cobalamin-independent synthase family protein
Gorai.013G195000.1.A
PCC=-0.522, p=0.022
mRNA Profile
Protein Expression Profile
DPA

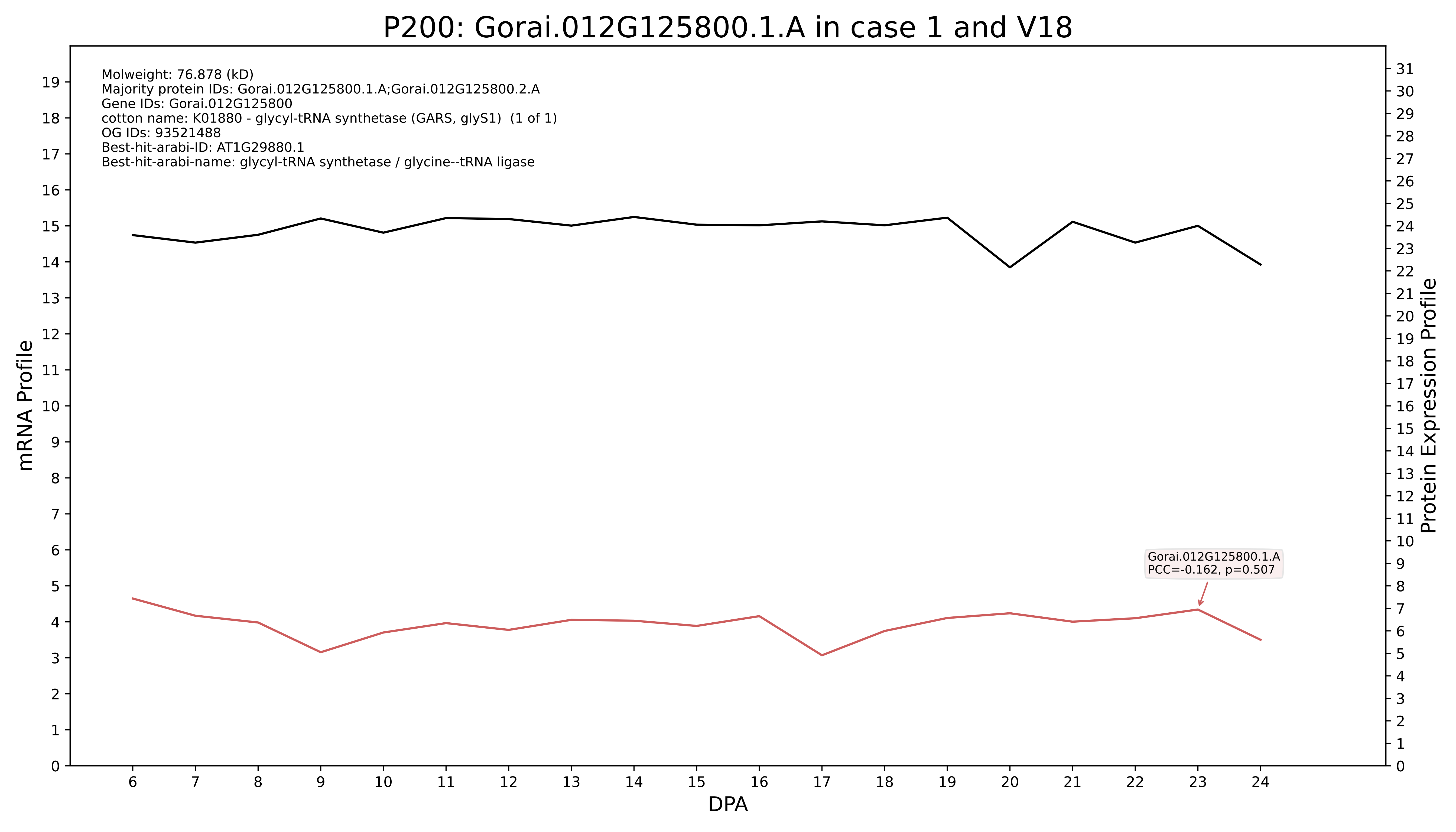
P200: Gorai.012G125800.1.A in case 1 and V18
Molweight: 76.878 (kD)
Majority protein IDs: Gorai.012G125800.1.A;Gorai.012G125800.2.A
Gene IDs: Gorai.012G125800
cotton name: K01880 - glycyl-tRNA synthetase (GARS, glyS1) (1 of 1)
OG IDs: 93521488
Best-hit-arabi-ID: AT1G29880.1
Best-hit-arabi-name: glycyl-tRNA synthetase / glycine--tRNA ligase
Gorai.012G125800.1.A
PCC=-0.162, p=0.507
mRNA Profile
Protein Expression Profile
DPA

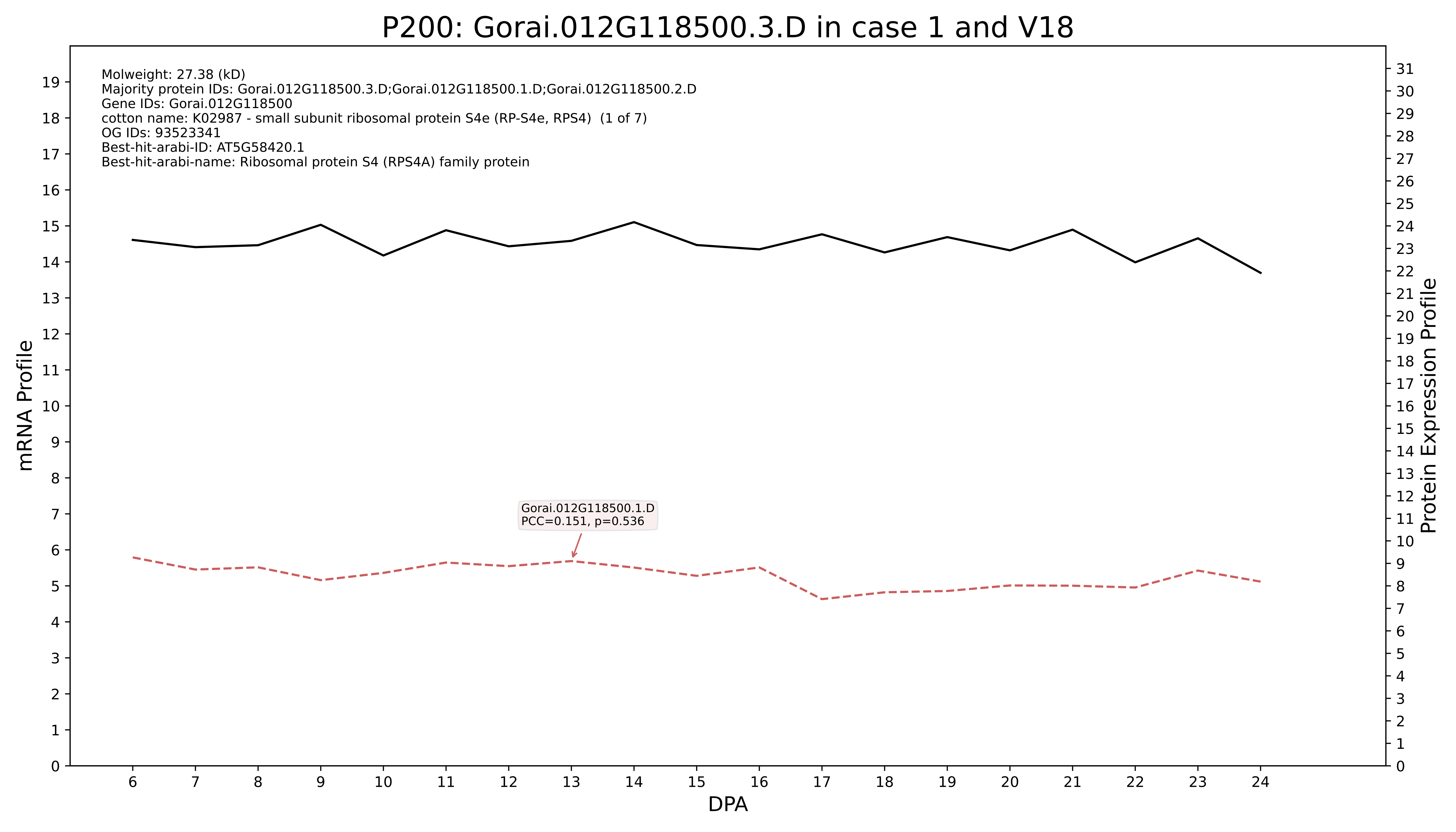
P200: Gorai.012G118500.3.D in case 1 and V18
Molweight: 27.38 (kD)
Majority protein IDs: Gorai.012G118500.3.D;Gorai.012G118500.1.D;Gorai.012G118500.2.D
Gene IDs: Gorai.012G118500
cotton name: K02987 - small subunit ribosomal protein S4e (RP-S4e, RPS4) (1 of 7)
OG IDs: 93523341
Best-hit-arabi-ID: AT5G58420.1
Best-hit-arabi-name: Ribosomal protein S4 (RPS4A) family protein
Gorai.012G118500.1.D
PCC=0.151, p=0.536
mRNA Profile
Protein Expression Profile
DPA

### P200: Gorai.012G146500.1.D in case 1 and V18

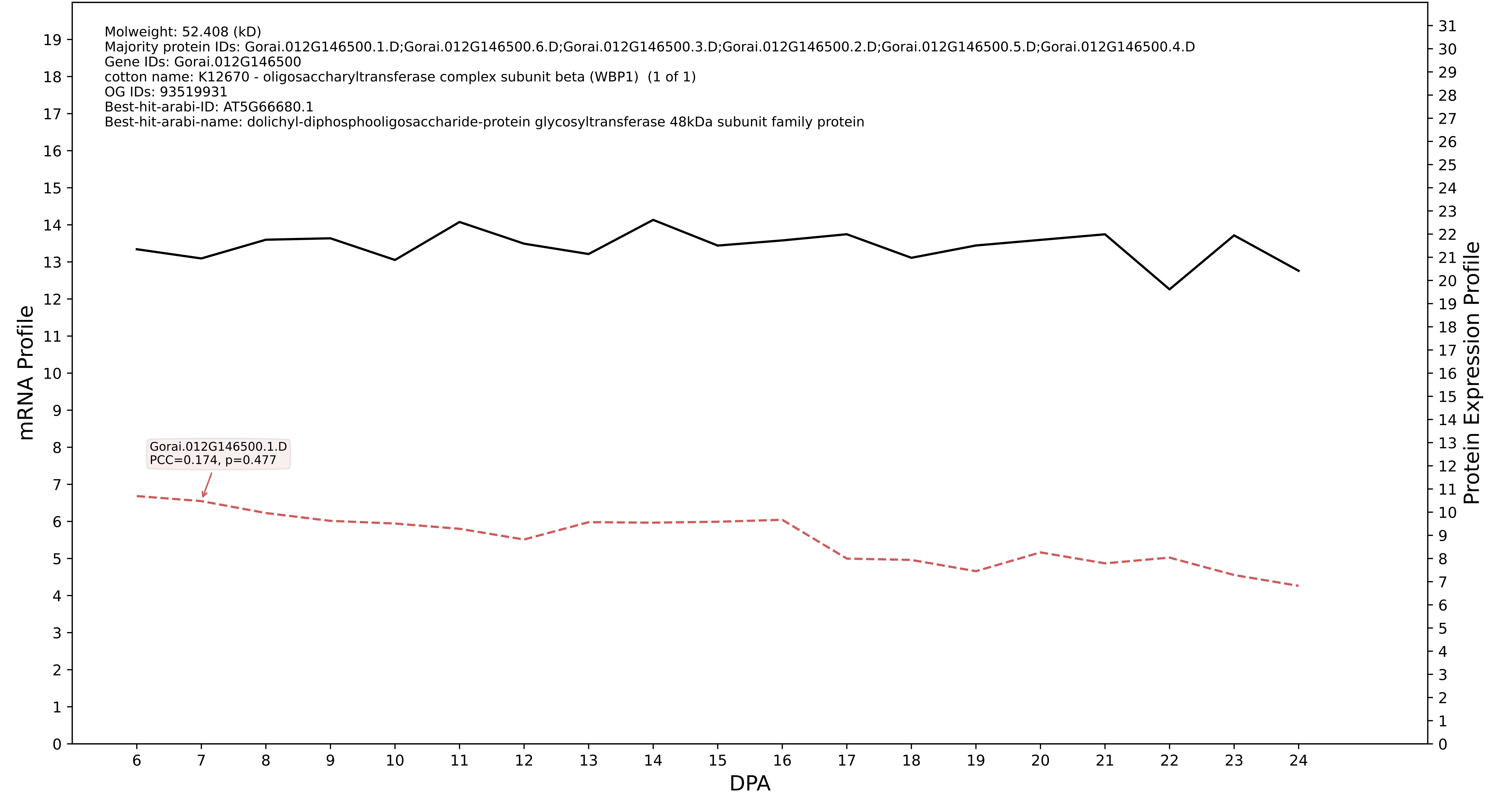

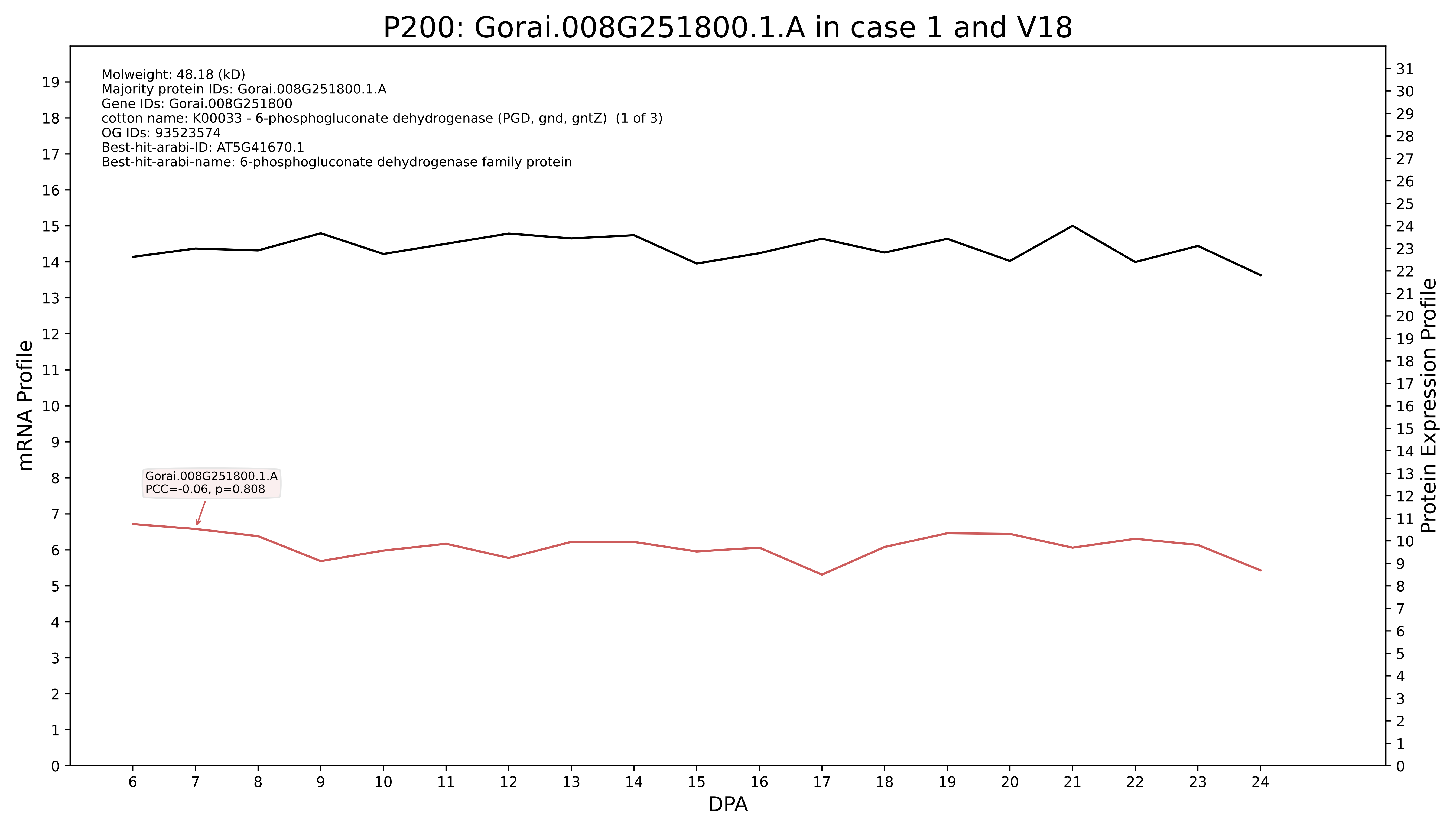

P200: Gorai.008G251800.1.A in case 1 and V18
Molweight: 48.18 (kD)
Majority protein IDs: Gorai.008G251800.1.A
Gene IDs: Gorai.008G251800
cotton name: K00033 - 6-phosphogluconate dehydrogenase (PGD, gnd, gntZ) (1 of 3)
OG IDs: 93523574
Best-hit-arabi-ID: AT5G41670.1
Best-hit-arabi-name: 6-phosphogluconate dehydrogenase family protein
Gorai.008G251800.1.A
PCC=-0.06, p=0.808
mRNA Profile
Protein Expression Profile
DPA

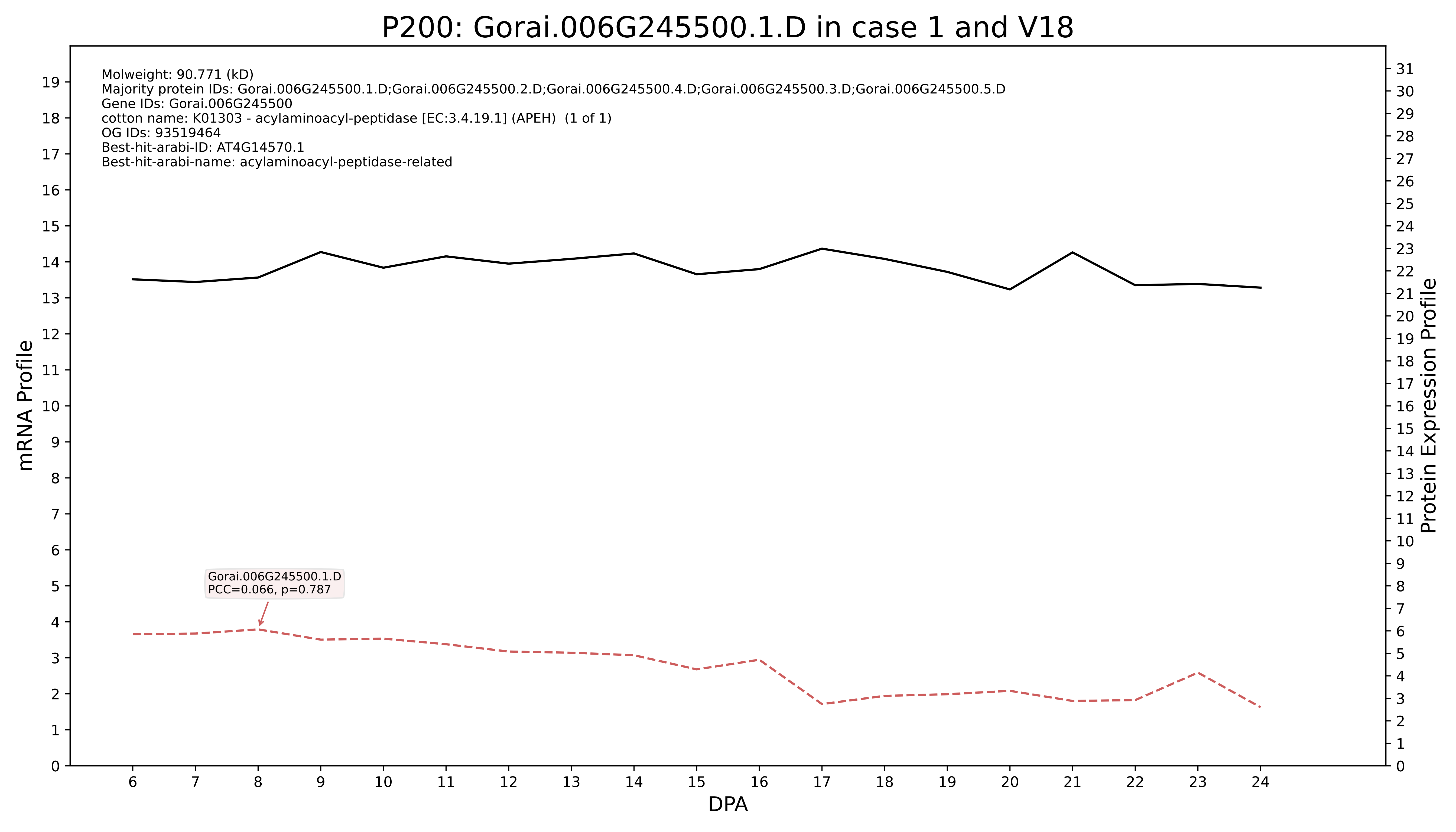
P200: Gorai.006G245500.1.D in case 1 and V18
Molweight: 90.771 (kD)
Majority protein IDs: Gorai.006G245500.1.D;Gorai.006G245500.2.D;Gorai.006G245500.4.D;Gorai.006G245500.3.D;Gorai.006G245500.5.D
Gene IDs: Gorai.006G245500
cotton name: K01303 - acylaminoacyl-peptidase [EC:3.4.19.1] (APEH) (1 of 1)
OG IDs: 93519464
Best-hit-arabi-ID: AT4G14570.1
Best-hit-arabi-name: acylaminoacyl-peptidase-related
Gorai.006G245500.1.D
PCC=0.066, p=0.787
mRNA Profile
Protein Expression Profile
DPA

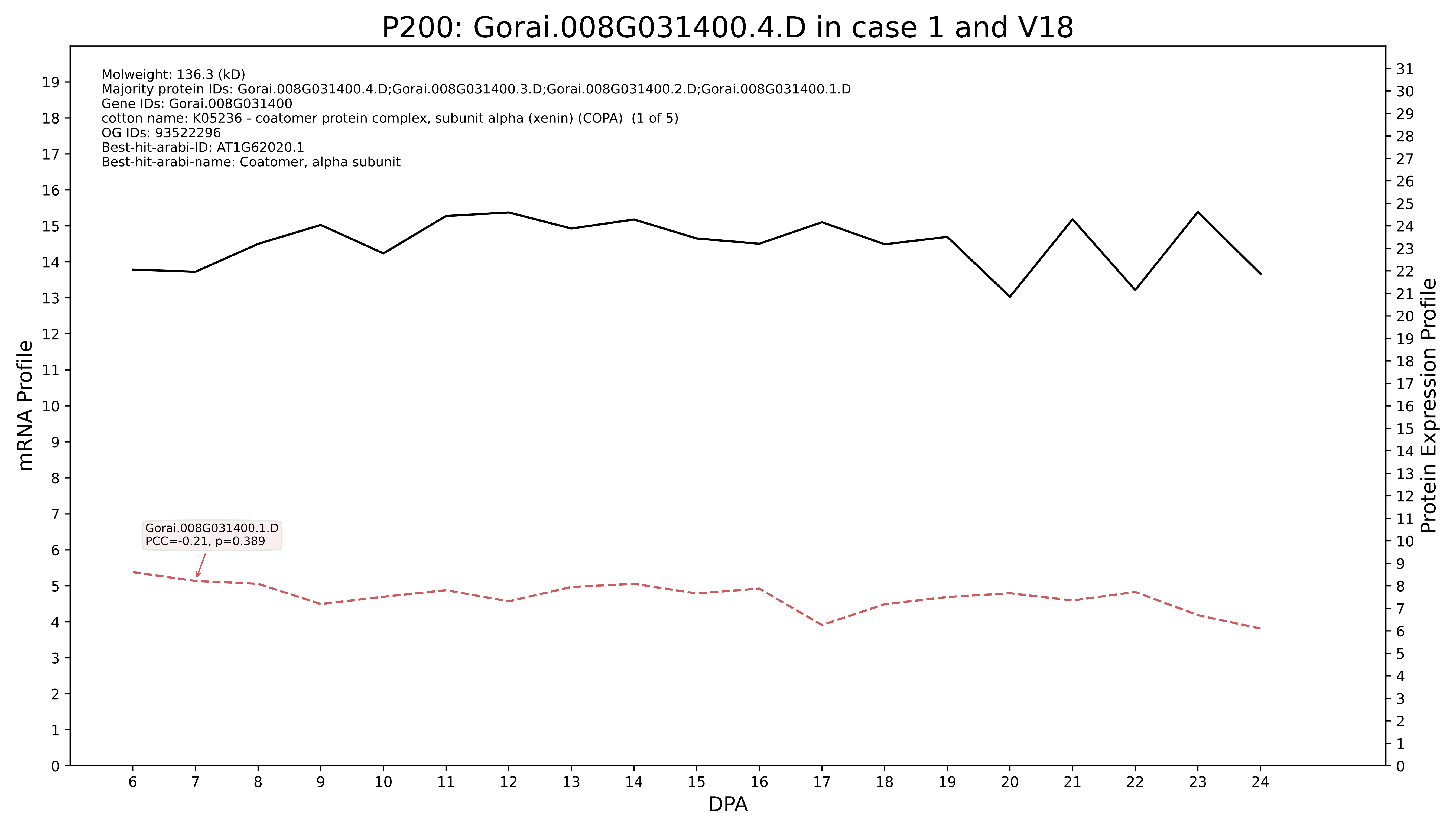

P200: Gorai.008G031400.4.D in case 1 and V18
Molweight: 136.3 (kD)
Majority protein IDs: Gorai.008G031400.4.D;Gorai.008G031400.3.D;Gorai.008G031400.2.D;Gorai.008G031400.1.D
Gene IDs: Gorai.008G031400
cotton name: K05236 - coatomer protein complex, subunit alpha (xenin) (COPA) (1 of 5)
OG IDs: 93522296
Best-hit-arabi-ID: AT1G62020.1
Best-hit-arabi-name: Coatomer, alpha subunit
Gorai.008G031400.1.D
PCC=-0.21, p=0.389
mRNA Profile
Protein Expression Profile
DPA

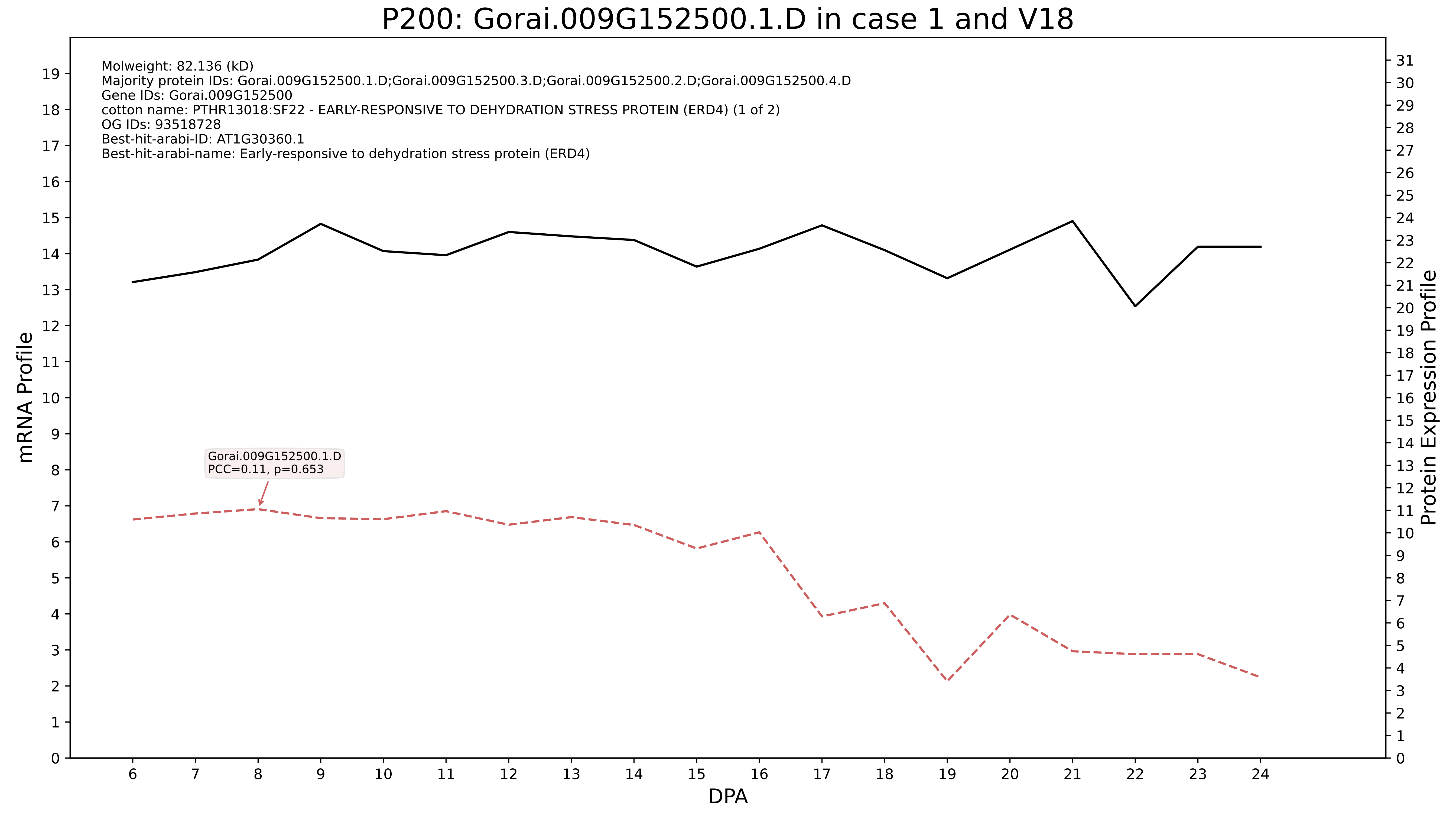
P200: Gorai.009G152500.1.D in case 1 and V18
Molweight: 82.136 (kD)
Majority protein IDs: Gorai.009G152500.1.D;Gorai.009G152500.3.D;Gorai.009G152500.2.D;Gorai.009G152500.4.D
Gene IDs: Gorai.009G152500
cotton name: PTHR13018:SF22 - EARLY-RESPONSIVE TO DEHYDRATION STRESS PROTEIN (ERD4) (1 of 2)
OG IDs: 93518728
Best-hit-arabi-ID: AT1G30360.1
Best-hit-arabi-name: Early-responsive to dehydration stress protein (ERD4)
Gorai.009G152500.1.D
PCC=0.11, p=0.653
mRNA Profile
Protein Expression Profile
DPA

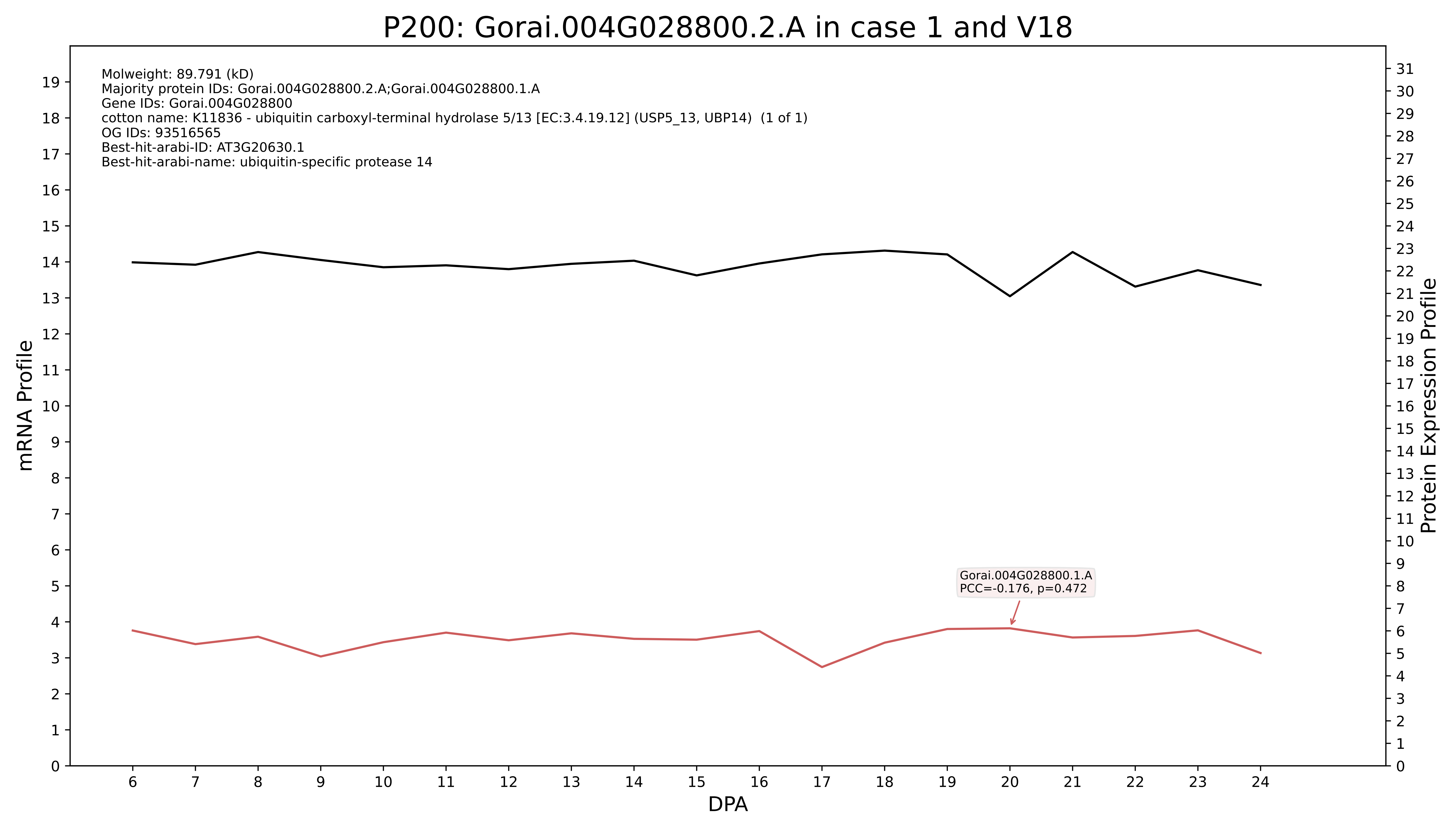
P200: Gorai.004G028800.2.A in case 1 and V18
Molweight: 89.791 (kD)
Majority protein IDs: Gorai.004G028800.2.A;Gorai.004G028800.1.A
Gene IDs: Gorai.004G028800
cotton name: K11836 - ubiquitin carboxyl-terminal hydrolase 5/13 [EC:3.4.19.12] (USP5_13, UBP14) (1 of 1)
OG IDs: 93516565
Best-hit-arabi-ID: AT3G20630.1
Best-hit-arabi-name: ubiquitin-specific protease 14
Gorai.004G028800.1.A
PCC=-0.176, p=0.472
mRNA Profile
Protein Expression Profile
DPA

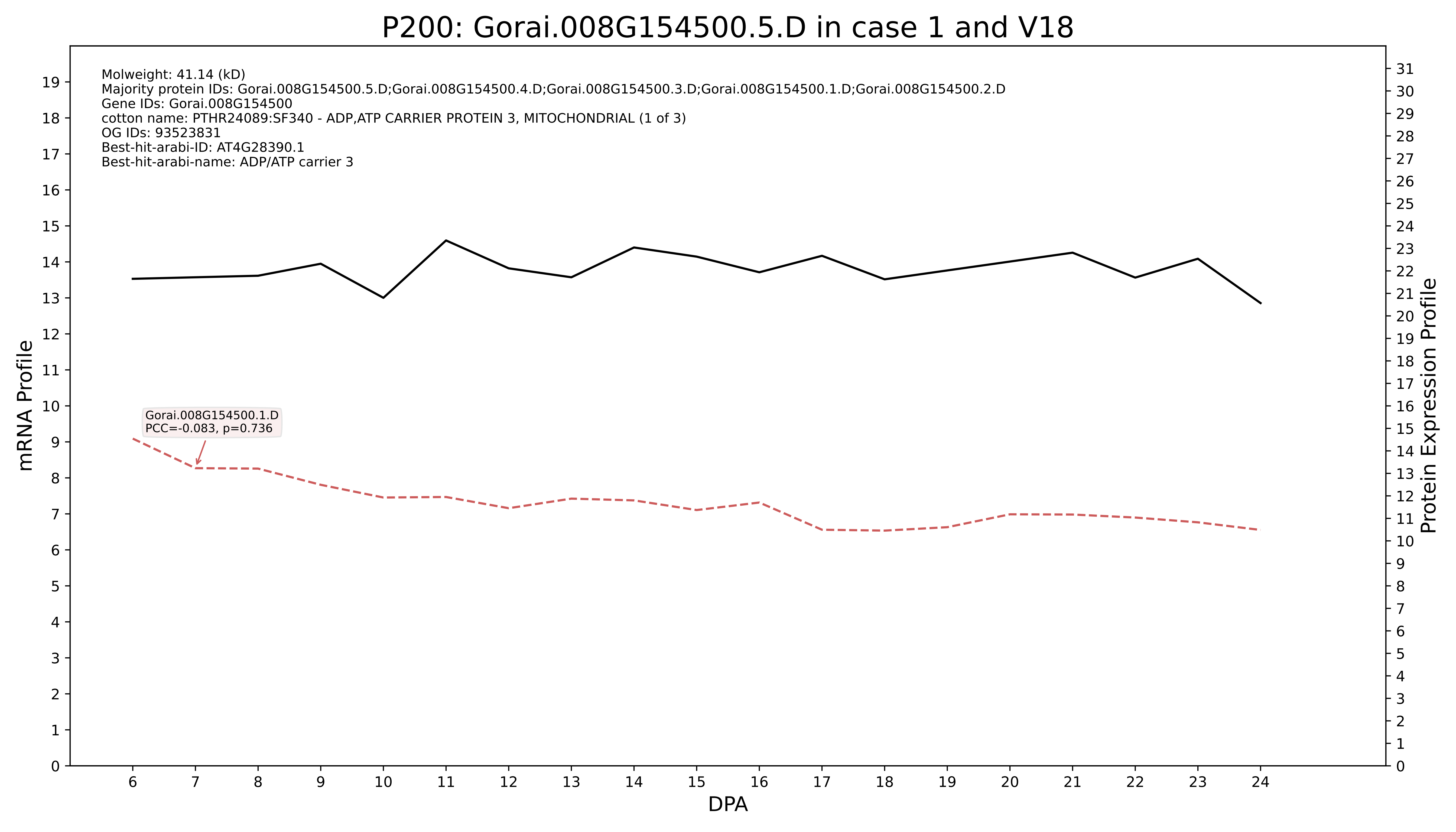
P200: Gorai.008G154500.5.D in case 1 and V18
Molweight: 41.14 (kD)
Majority protein IDs: Gorai.008G154500.5.D;Gorai.008G154500.4.D;Gorai.008G154500.3.D;Gorai.008G154500.1.D;Gorai.008G154500.2.D
Gene IDs: Gorai.008G154500
cotton name: PTHR24089:SF340 - ADP,ATP CARRIER PROTEIN 3, MITOCHONDRIAL (1 of 3)
OG IDs: 93523831
Best-hit-arabi-ID: AT4G28390.1
Best-hit-arabi-name: ADP/ATP carrier 3
Gorai.008G154500.1.D
PCC=-0.083, p=0.736
mRNA Profile
Protein Expression Profile
DPA

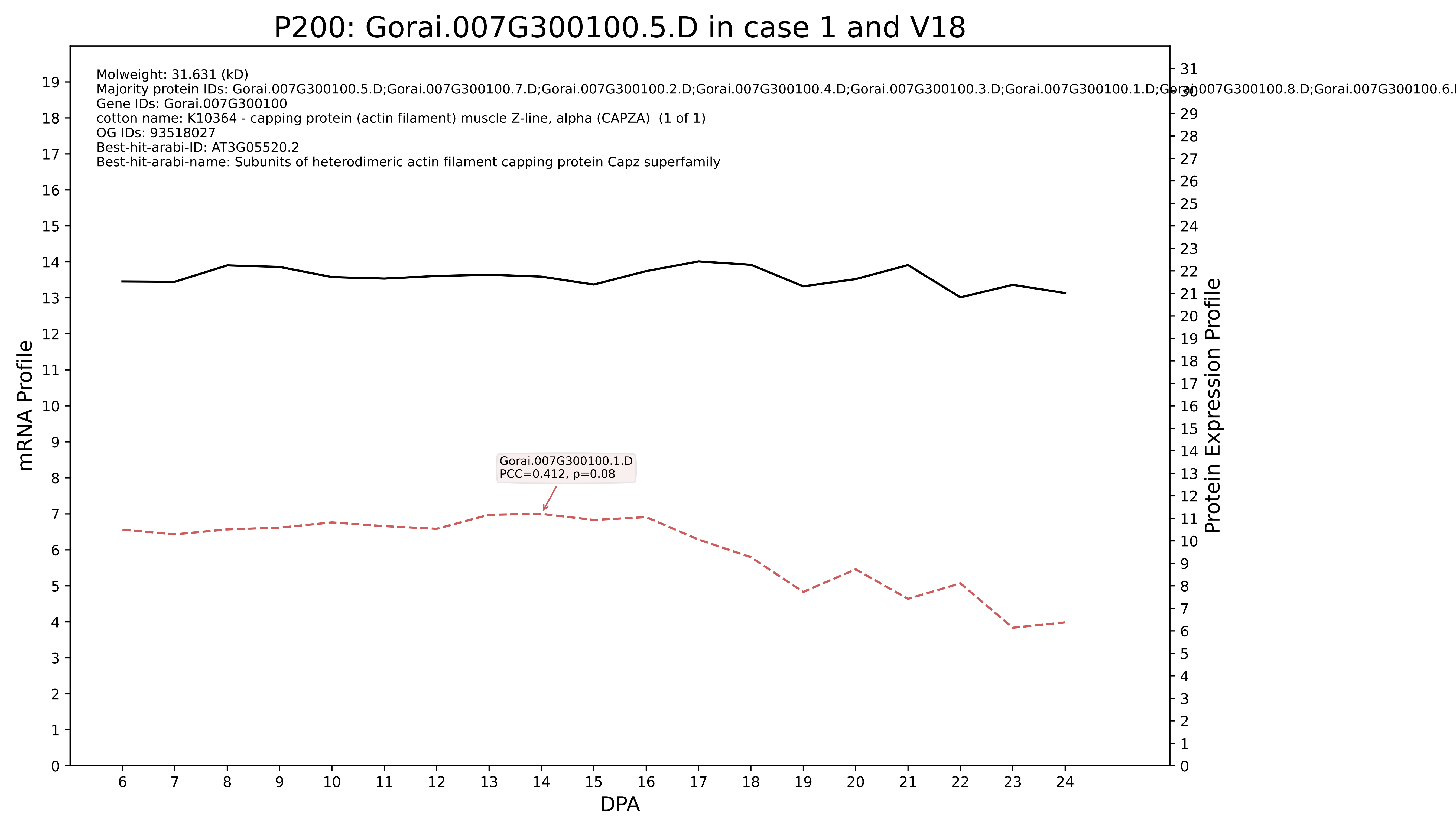
P200: Gorai.007G300100.5.D in case 1 and V18
Molweight: 31.631 (kD)
Majority protein IDs: Gorai.007G300100.5.D;Gorai.007G300100.7.D;Gorai.007G300100.2.D;Gorai.007G300100.4.D;Gorai.007G300100.3.D;Gorai.007G300100.1.D;Gorai.007G300100.8.D;Gorai.007G300100.6.D
Gene IDs: Gorai.007G300100
cotton name: K10364 - capping protein (actin filament) muscle Z-line, alpha (CAPZA) (1 of 1)
OG IDs: 93518027
Best-hit-arabi-ID: AT3G05520.2
Best-hit-arabi-name: Subunits of heterodimeric actin filament capping protein Capz superfamily
Gorai.007G300100.1.D
PCC=0.412, p=0.08
mRNA Profile
Protein Expression Profile
DPA

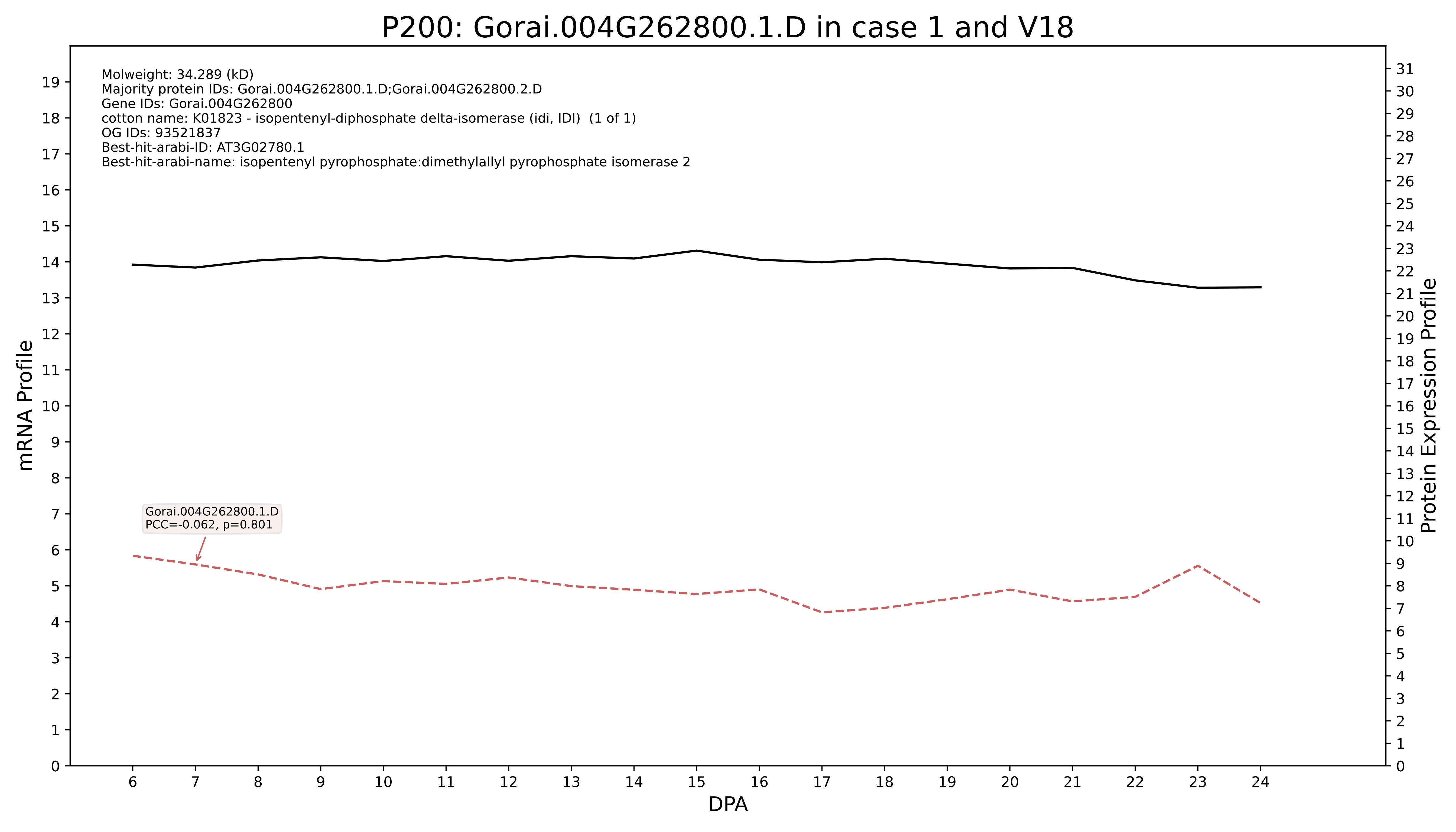

P200: Gorai.004G262800.1.D in case 1 and V18
Molweight: 34.289 (kD)
Majority protein IDs: Gorai.004G262800.1.D;Gorai.004G262800.2.D
Gene IDs: Gorai.004G262800
cotton name: K01823 - isopentenyl-diphosphate delta-isomerase (idi, IDI) (1 of 1)
OG IDs: 93521837
Best-hit-arabi-ID: AT3G02780.1
Best-hit-arabi-name: isopentenyl pyrophosphate:dimethylallyl pyrophosphate isomerase 2
Gorai.004G262800.1.D
PCC=-0.062, p=0.801
mRNA Profile
Protein Expression Profile
DPA

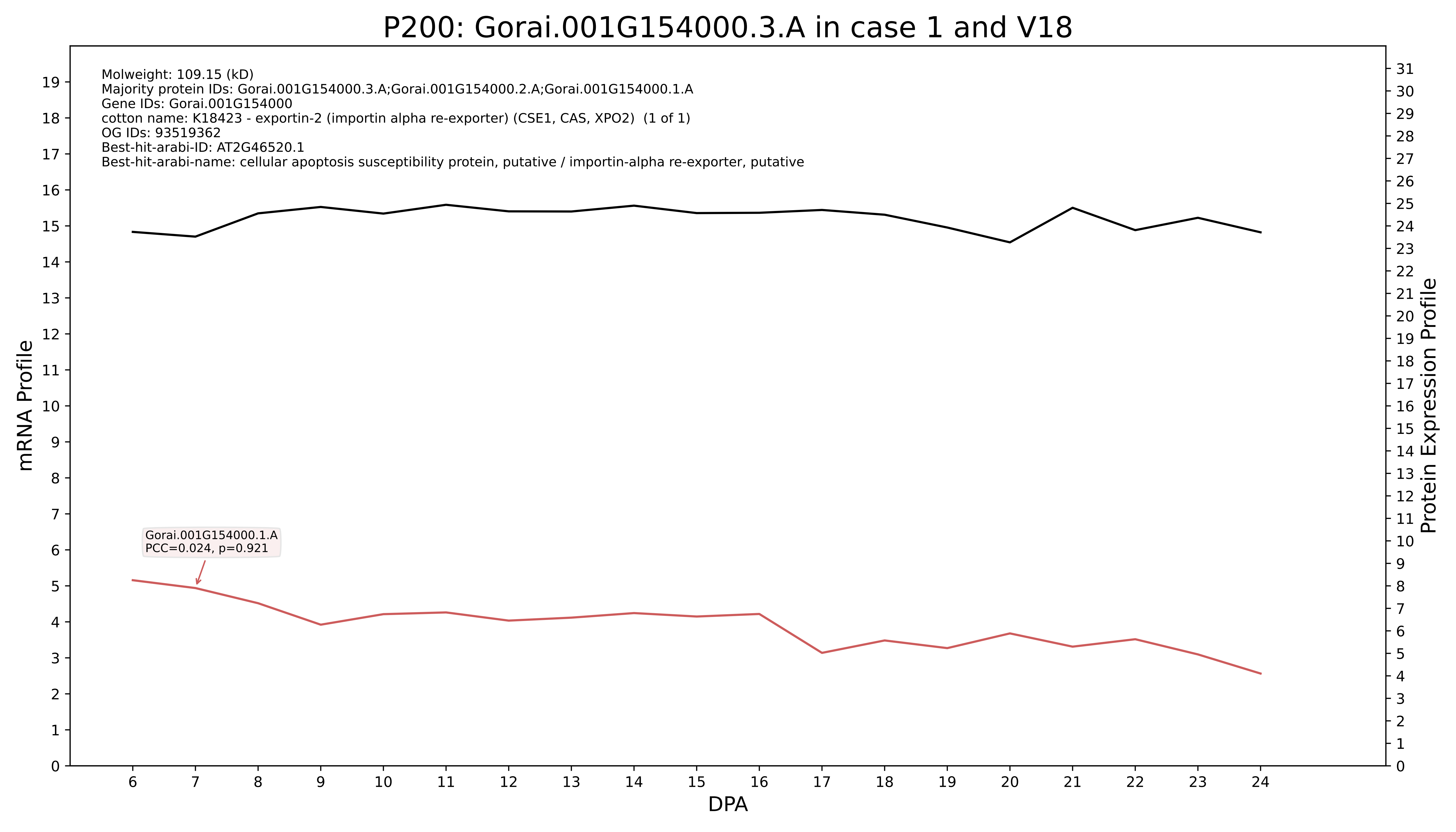
P200: Gorai.001G154000.3.A in case 1 and V18
Molweight: 109.15 (kD)
Majority protein IDs: Gorai.001G154000.3.A;Gorai.001G154000.2.A;Gorai.001G154000.1.A
Gene IDs: Gorai.001G154000
cotton name: K18423 - exportin-2 (importin alpha re-exporter) (CSE1, CAS, XPO2) (1 of 1)
OG IDs: 93519362
Best-hit-arabi-ID: AT2G46520.1
Best-hit-arabi-name: cellular apoptosis susceptibility protein, putative / importin-alpha re-exporter, putative
Gorai.001G154000.1.A
PCC=0.024, p=0.921
mRNA Profile
Protein Expression Profile
DPA

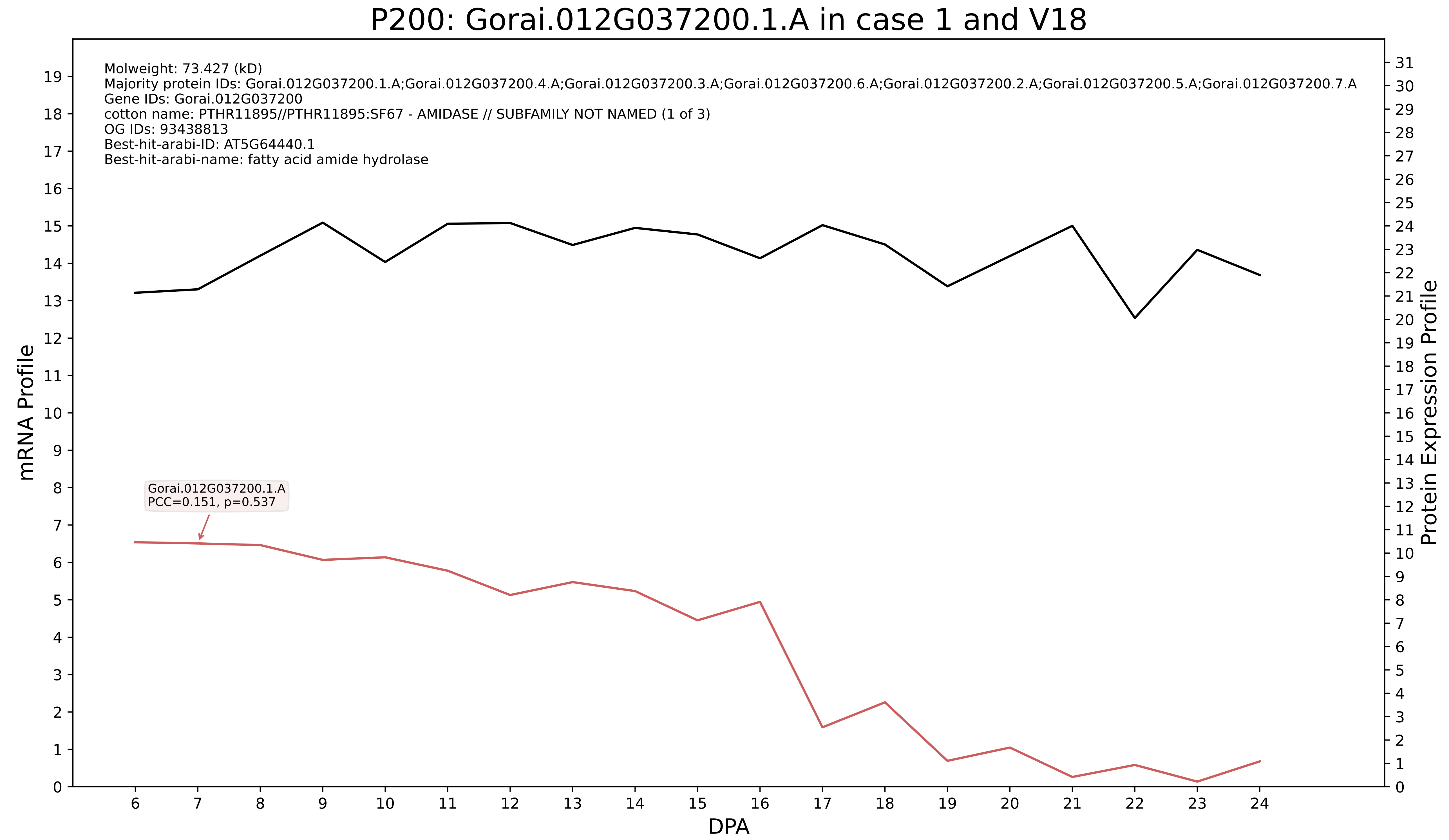

P200: Gorai.012G037200.1.A in case 1 and V18
Molweight: 73.427 (kD)
Majority protein IDs: Gorai.012G037200.1.A;Gorai.012G037200.4.A;Gorai.012G037200.3.A;Gorai.012G037200.6.A;Gorai.012G037200.2.A;Gorai.012G037200.5.A;Gorai.012G037200.7.A
Gene IDs: Gorai.012G037200
cotton name: PTHR11895//PTHR11895:SF67 - AMIDASE // SUBFAMILY NOT NAMED (1 of 3)
OG IDs: 93438813
Best-hit-arabi-ID: AT5G64440.1
Best-hit-arabi-name: fatty acid amide hydrolase
Gorai.012G037200.1.A
PCC=0.151, p=0.537
mRNA Profile
Protein Expression Profile
DPA

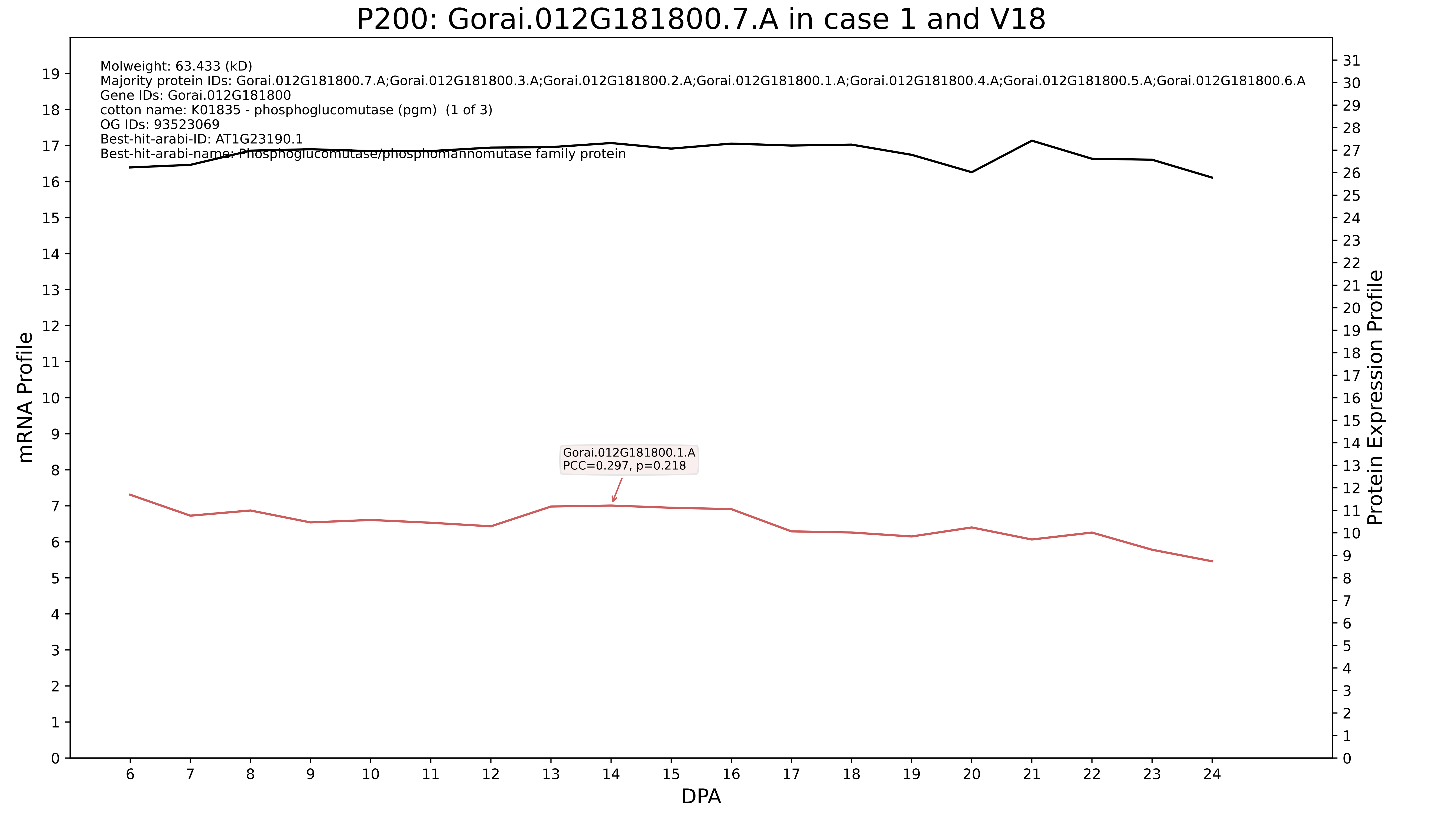

P200: Gorai.012G181800.7.A in case 1 and V18
Molweight: 63.433 (kD)
Majority protein IDs: Gorai.012G181800.7.A;Gorai.012G181800.3.A;Gorai.012G181800.2.A;Gorai.012G181800.1.A;Gorai.012G181800.4.A;Gorai.012G181800.5.A;Gorai.012G181800.6.A
Gene IDs: Gorai.012G181800
cotton name: K01835 - phosphoglucomutase (pgm) (1 of 3)
OG IDs: 93523069
Best-hit-arabi-ID: AT1G23190.1
Best-hit-arabi-name: Phosphoglucomutase/phosphomannomutase family protein
Gorai.012G181800.1.A
PCC=0.297, p=0.218
mRNA Profile
Protein Expression Profile
DPA

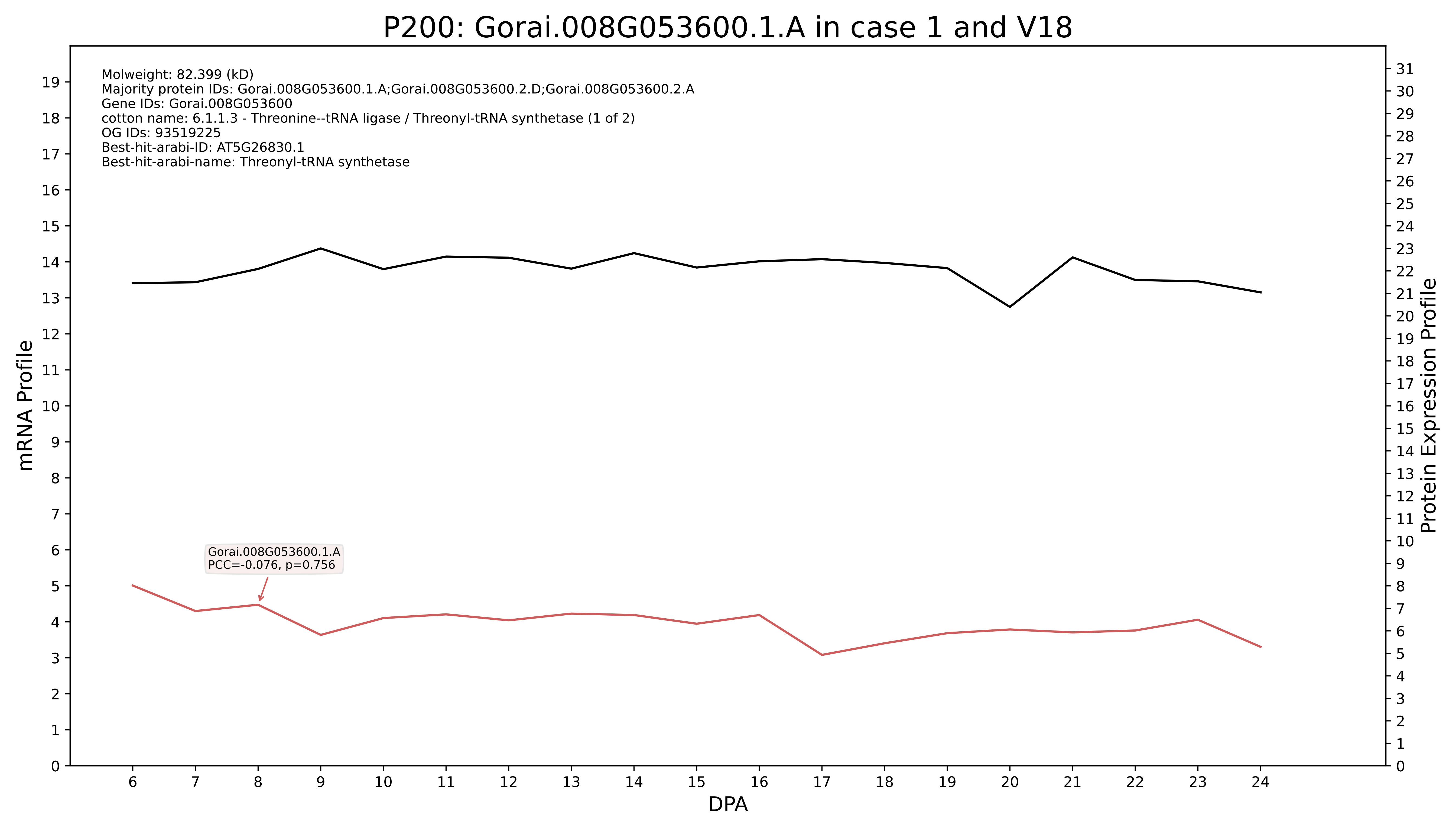
P200: Gorai.008G053600.1.A in case 1 and V18
Molweight: 82.399 (kD)
Majority protein IDs: Gorai.008G053600.1.A;Gorai.008G053600.2.D;Gorai.008G053600.2.A
Gene IDs: Gorai.008G053600
cotton name: 6.1.1.3 - Threonine--tRNA ligase / Threonyl-tRNA synthetase (1 of 2)
OG IDs: 93519225
Best-hit-arabi-ID: AT5G26830.1
Best-hit-arabi-name: Threonyl-tRNA synthetase
Gorai.008G053600.1.A
PCC=-0.076, p=0.756
mRNA Profile
Protein Expression Profile
DPA

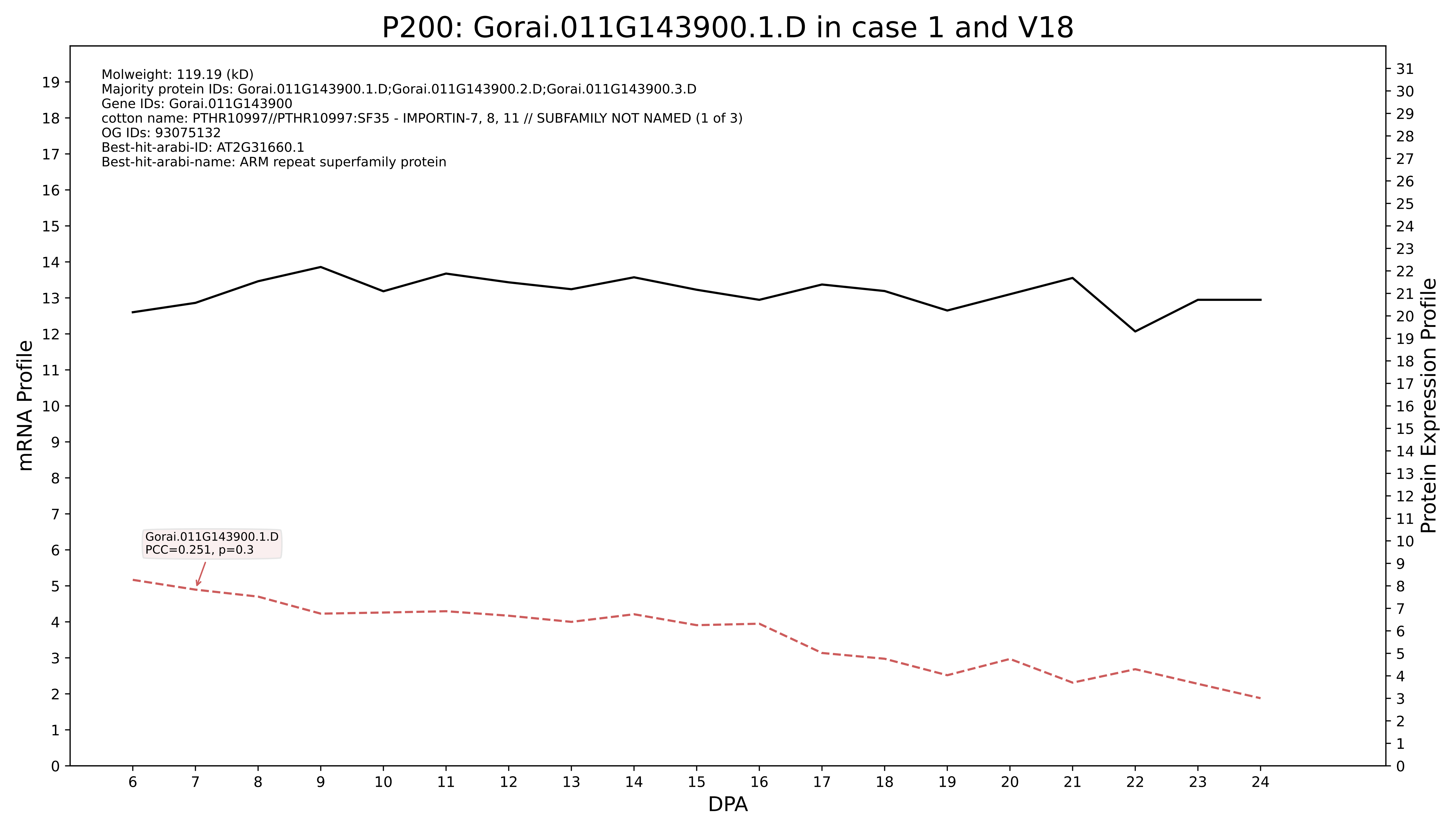

P200: Gorai.011G143900.1.D in case 1 and V18
Molweight: 119.19 (kD)
Majority protein IDs: Gorai.011G143900.1.D;Gorai.011G143900.2.D;Gorai.011G143900.3.D
Gene IDs: Gorai.011G143900
cotton name: PTHR10997//PTHR10997:SF35 - IMPORTIN-7, 8, 11 // SUBFAMILY NOT NAMED (1 of 3)
OG IDs: 93075132
Best-hit-arabi-ID: AT2G31660.1
Best-hit-arabi-name: ARM repeat superfamily protein
Gorai.011G143900.1.D
PCC=0.251, p=0.3
mRNA Profile
Protein Expression Profile
DPA

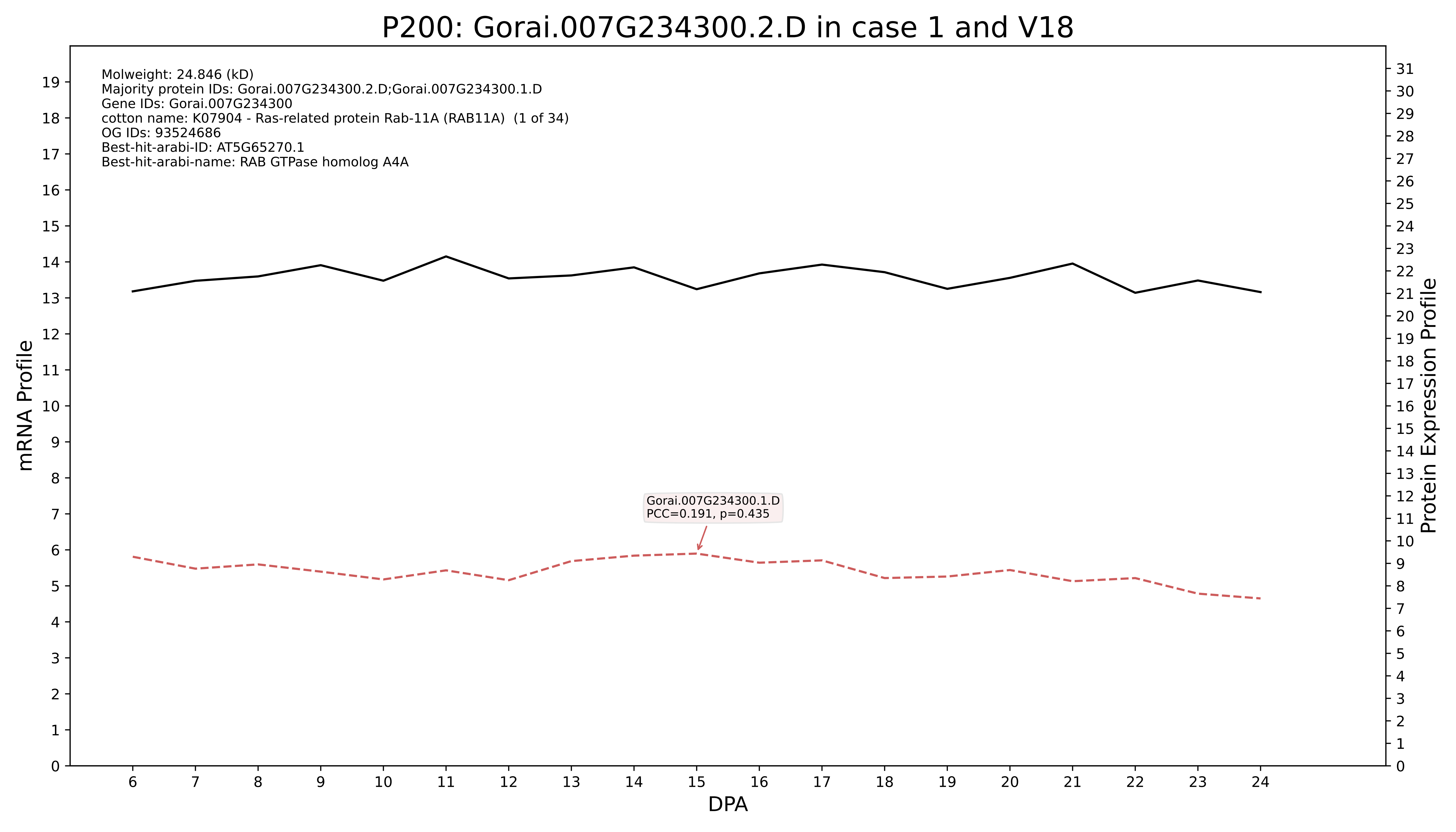

P200: Gorai.007G234300.2.D in case 1 and V18
Molweight: 24.846 (kD)
Majority protein IDs: Gorai.007G234300.2.D;Gorai.007G234300.1.D
Gene IDs: Gorai.007G234300
cotton name: K07904 - Ras-related protein Rab-11A (RAB11A) (1 of 34)
OG IDs: 93524686
Best-hit-arabi-ID: AT5G65270.1
Best-hit-arabi-name: RAB GTPase homolog A4A
Gorai.007G234300.1.D
PCC=0.191, p=0.435
mRNA Profile
Protein Expression Profile
DPA

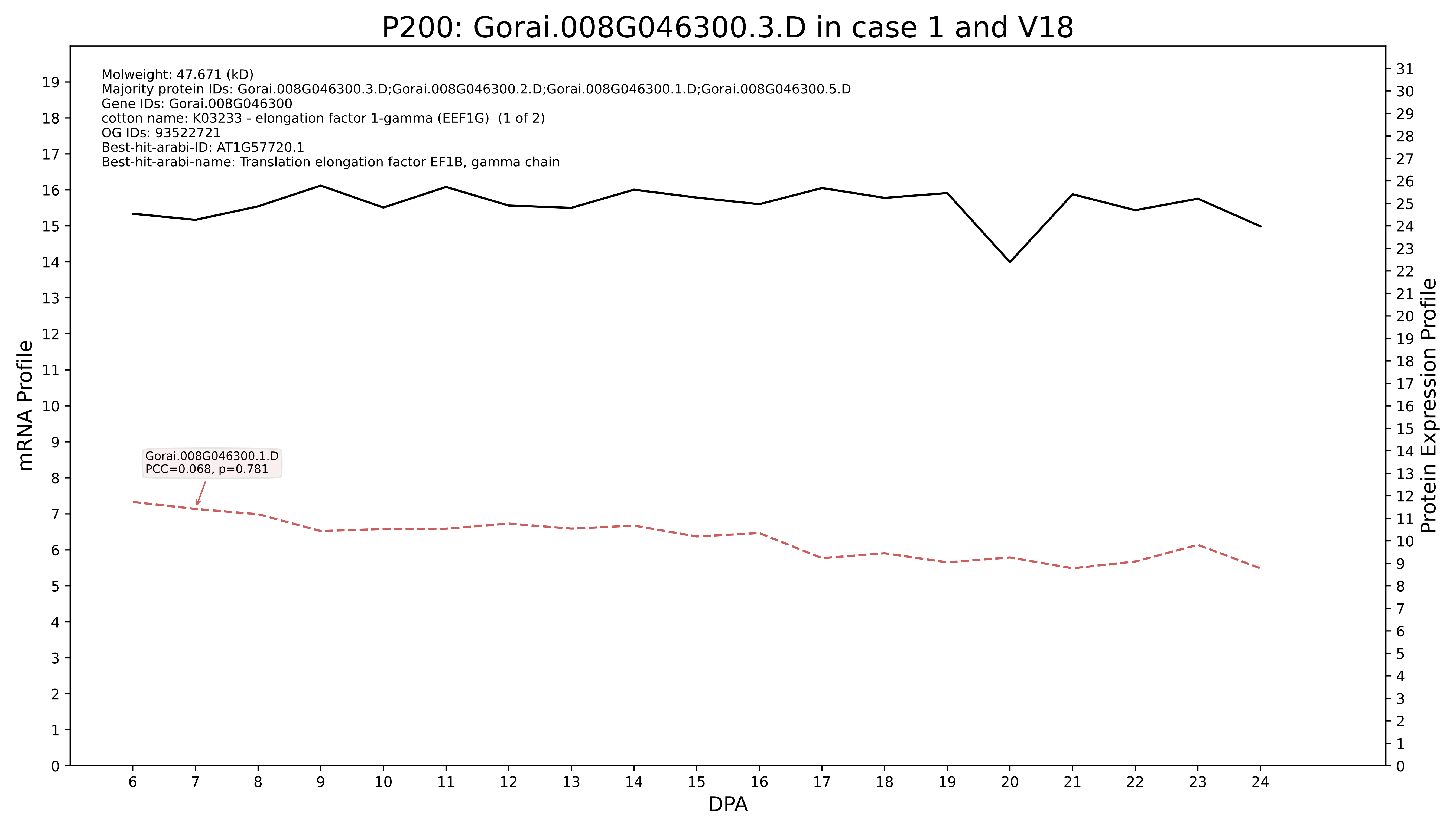
P200: Gorai.008G046300.3.D in case 1 and V18
Molweight: 47.671 (kD)
Majority protein IDs: Gorai.008G046300.3.D;Gorai.008G046300.2.D;Gorai.008G046300.1.D;Gorai.008G046300.5.D
Gene IDs: Gorai.008G046300
cotton name: K03233 - elongation factor 1-gamma (EEF1G) (1 of 2)
OG IDs: 93522721
Best-hit-arabi-ID: AT1G57720.1
Best-hit-arabi-name: Translation elongation factor EF1B, gamma chain
Gorai.008G046300.1.D
PCC=0.068, p=0.781
mRNA Profile
Protein Expression Profile
DPA

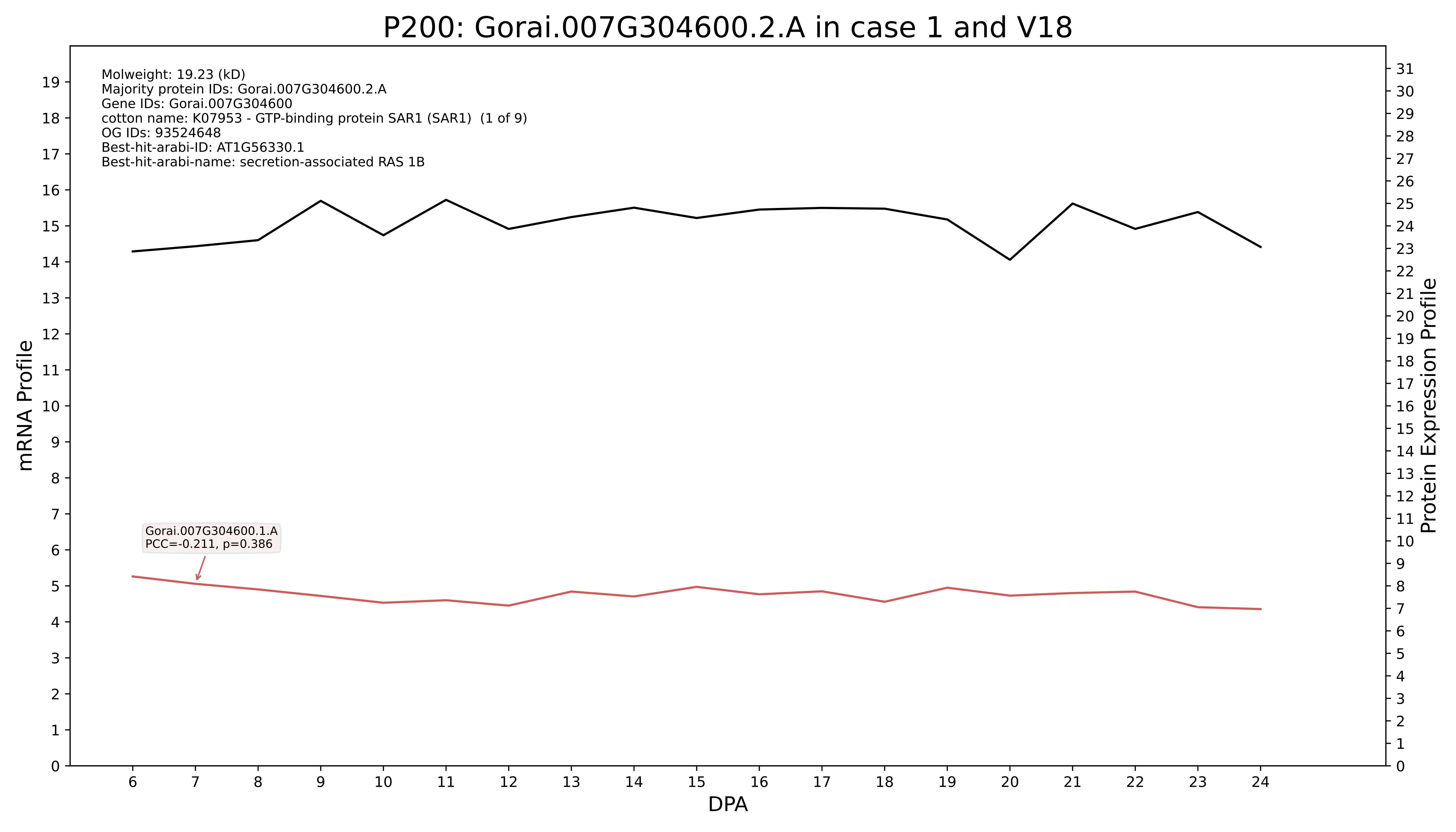
P200: Gorai.007G304600.2.A in case 1 and V18
Molweight: 19.23 (kD)
Majority protein IDs: Gorai.007G304600.2.A
Gene IDs: Gorai.007G304600
cotton name: K07953 - GTP-binding protein SAR1 (SAR1) (1 of 9)
OG IDs: 93524648
Best-hit-arabi-ID: AT1G56330.1
Best-hit-arabi-name: secretion-associated RAS 1B
Gorai.007G304600.1.A
PCC=-0.211, p=0.386
mRNA Profile
Protein Expression Profile
DPA

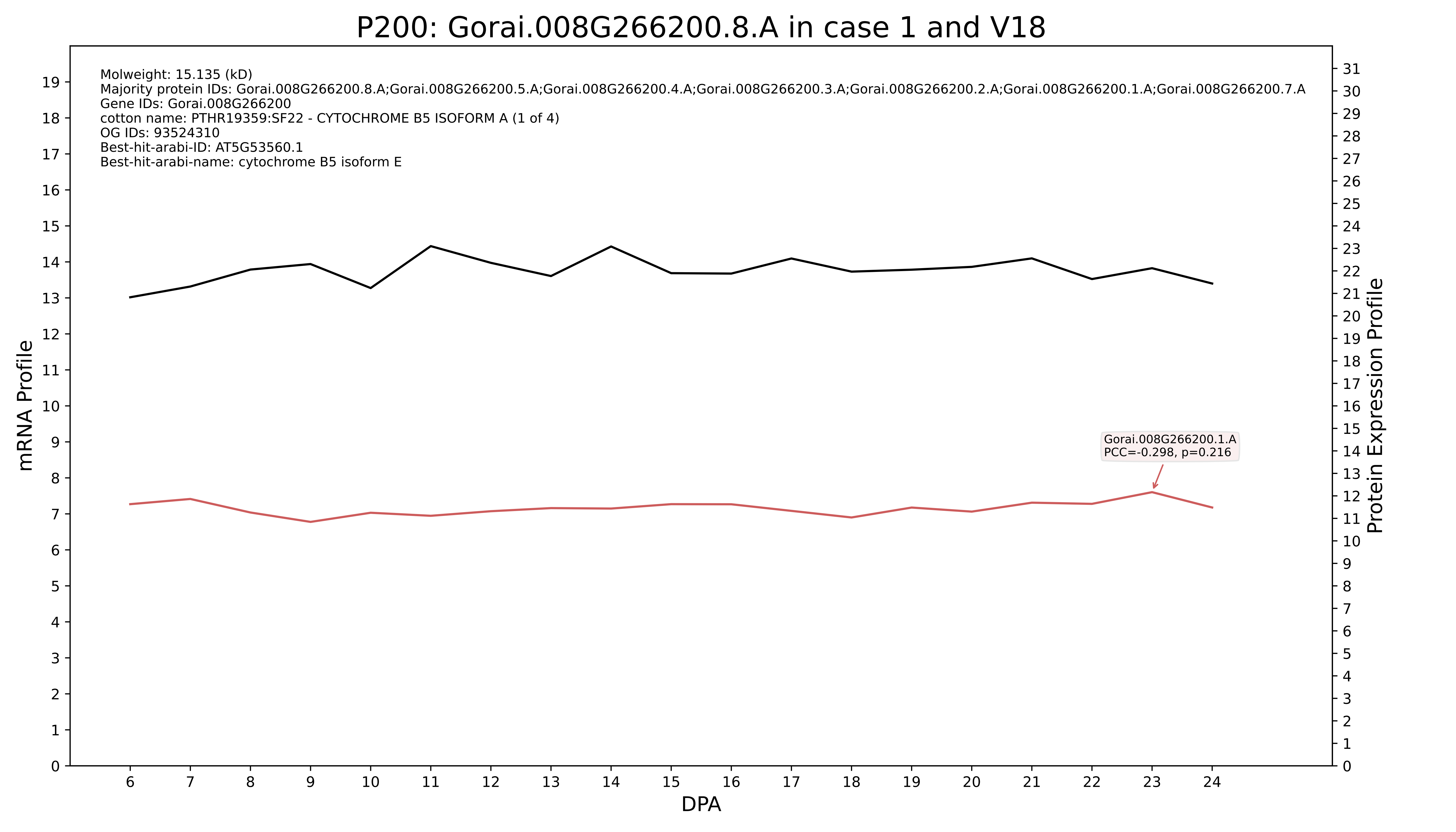
P200: Gorai.008G266200.8.A in case 1 and V18
Molweight: 15.135 (kD)
Majority protein IDs: Gorai.008G266200.8.A;Gorai.008G266200.5.A;Gorai.008G266200.4.A;Gorai.008G266200.3.A;Gorai.008G266200.2.A;Gorai.008G266200.1.A;Gorai.008G266200.7.A
Gene IDs: Gorai.008G266200
cotton name: PTHR19359:SF22 - CYTOCHROME B5 ISOFORM A (1 of 4)
OG IDs: 93524310
Best-hit-arabi-ID: AT5G53560.1
Best-hit-arabi-name: cytochrome B5 isoform E
Gorai.008G266200.1.A
PCC=-0.298, p=0.216
mRNA Profile
Protein Expression Profile
DPA

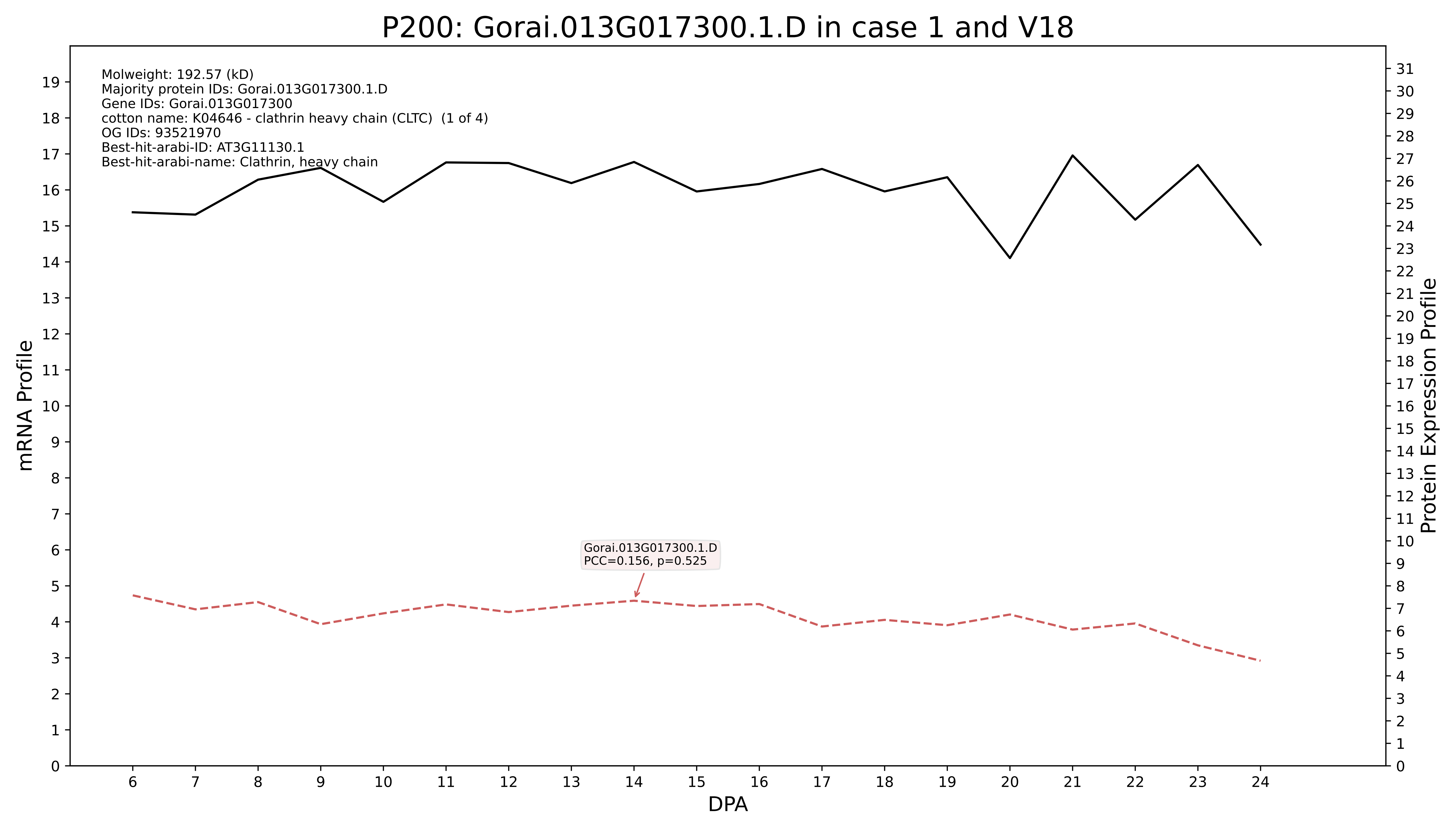
P200: Gorai.013G017300.1.D in case 1 and V18
Molweight: 192.57 (kD)
Majority protein IDs: Gorai.013G017300.1.D
Gene IDs: Gorai.013G017300
cotton name: K04646 - clathrin heavy chain (CLTC) (1 of 4)
OG IDs: 93521970
Best-hit-arabi-ID: AT3G11130.1
Best-hit-arabi-name: Clathrin, heavy chain
Gorai.013G017300.1.D
PCC=0.156, p=0.525
mRNA Profile
Protein Expression Profile
DPA

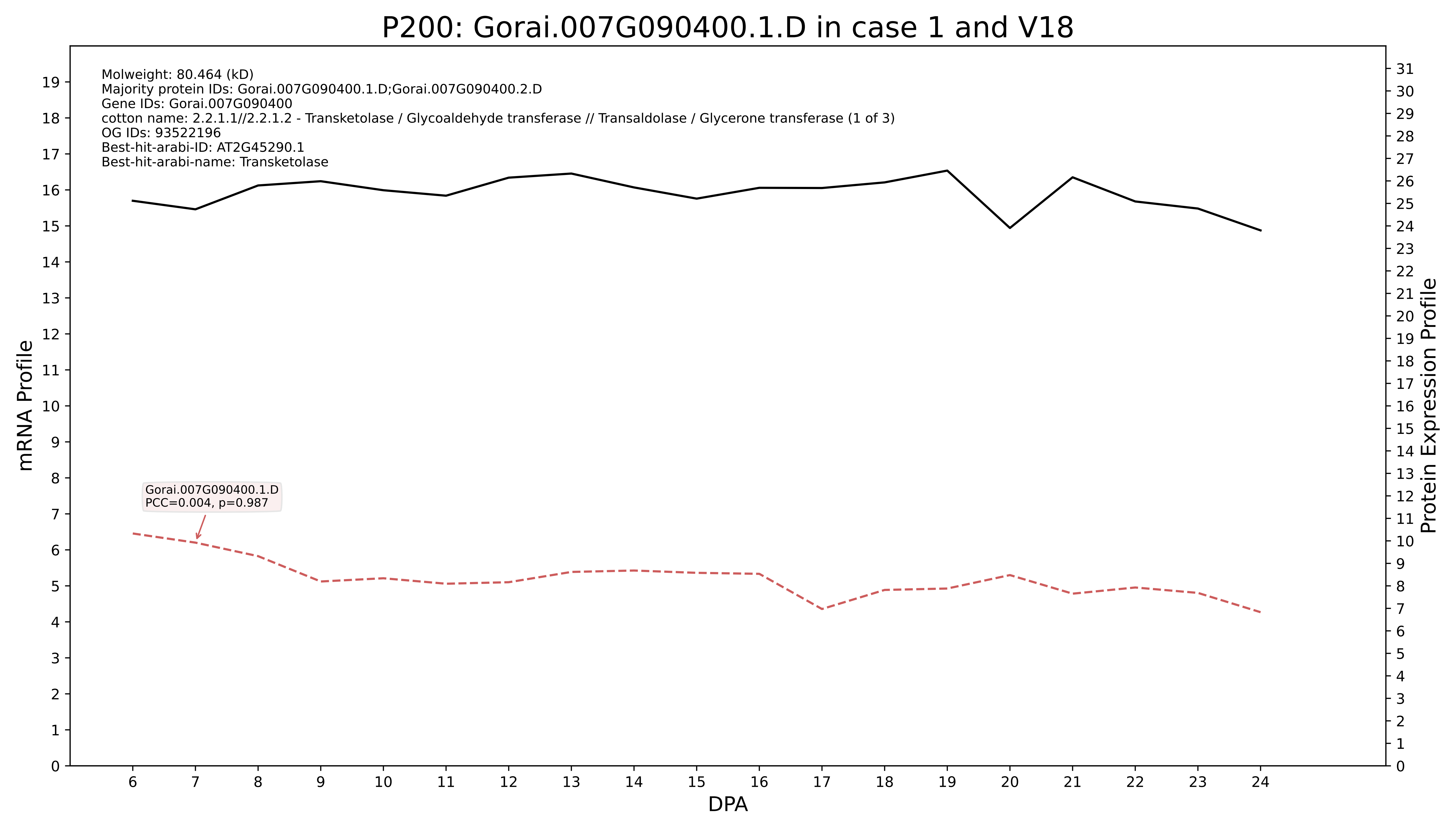

P200: Gorai.007G090400.1.D in case 1 and V18
Molweight: 80.464 (kD)
Majority protein IDs: Gorai.007G090400.1.D;Gorai.007G090400.2.D
Gene IDs: Gorai.007G090400
cotton name: 2.2.1.1//2.2.1.2 - Transketolase / Glycoaldehyde transferase // Transaldolase / Glycerone transferase (1 of 3)
OG IDs: 93522196
Best-hit-arabi-ID: AT2G45290.1
Best-hit-arabi-name: Transketolase
Gorai.007G090400.1.D
PCC=0.004, p=0.987
mRNA Profile
Protein Expression Profile
DPA

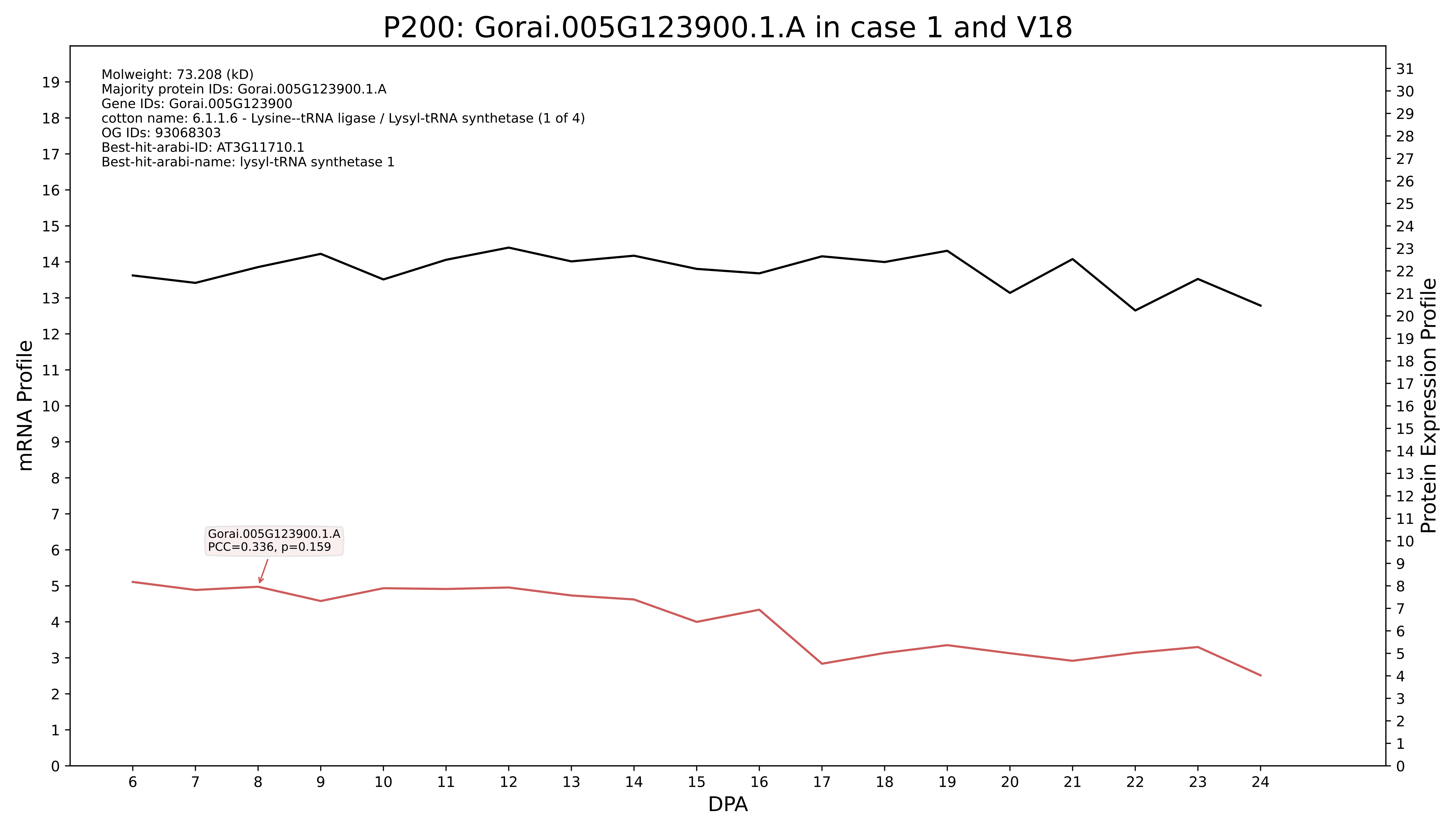

P200: Gorai.005G123900.1.A in case 1 and V18
Molweight: 73.208 (kD)
Majority protein IDs: Gorai.005G123900.1.A
Gene IDs: Gorai.005G123900
cotton name: 6.1.1.6 - Lysine--tRNA ligase / Lysyl-tRNA synthetase (1 of 4)
OG IDs: 93068303
Best-hit-arabi-ID: AT3G11710.1
Best-hit-arabi-name: lysyl-tRNA synthetase 1
Gorai.005G123900.1.A
PCC=0.336, p=0.159
mRNA Profile
Protein Expression Profile
DPA

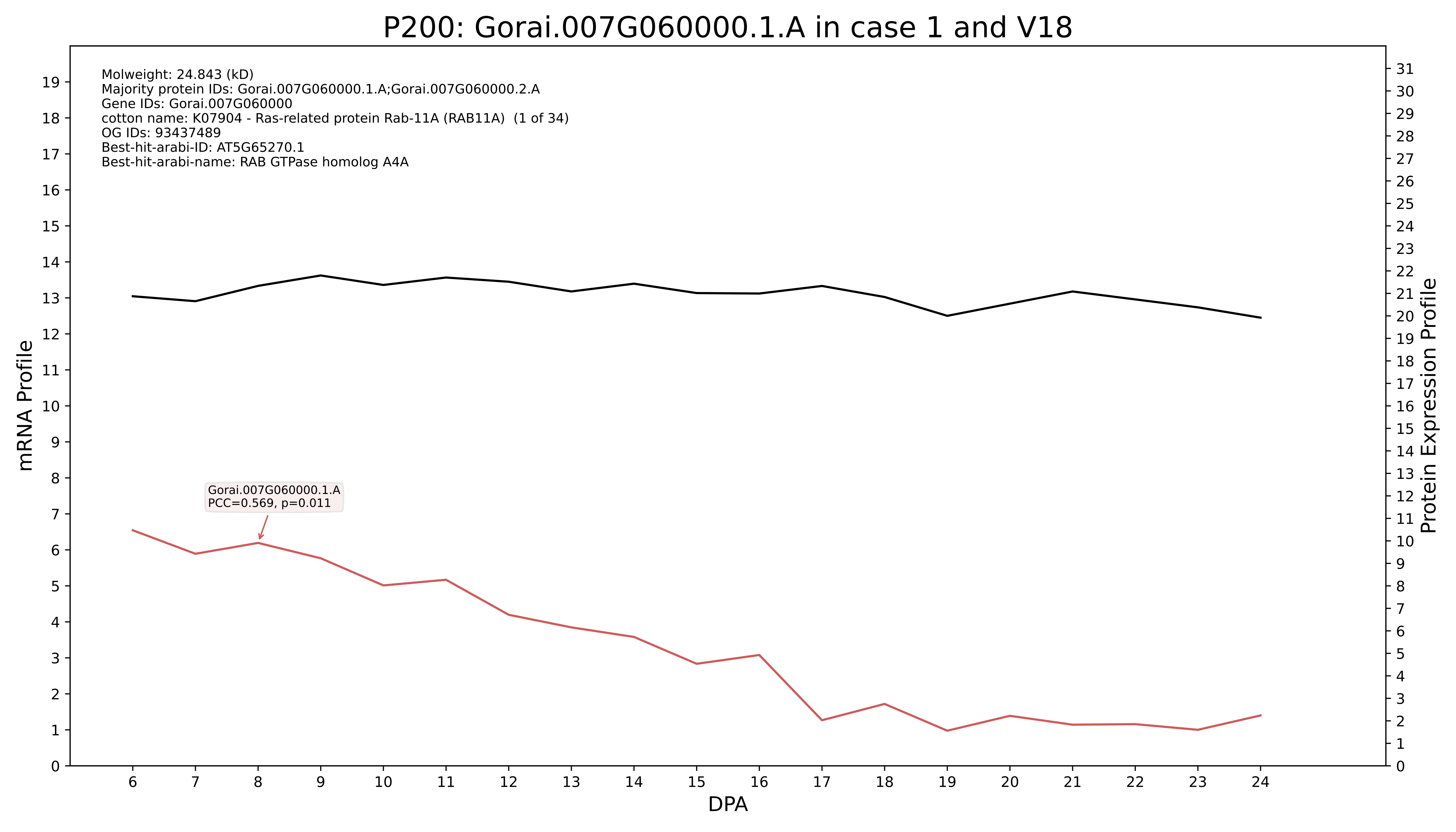
P200: Gorai.007G060000.1.A in case 1 and V18
Molweight: 24.843 (kD)
Majority protein IDs: Gorai.007G060000.1.A;Gorai.007G060000.2.A
Gene IDs: Gorai.007G060000
cotton name: K07904 - Ras-related protein Rab-11A (RAB11A) (1 of 34)
OG IDs: 93437489
Best-hit-arabi-ID: AT5G65270.1
Best-hit-arabi-name: RAB GTPase homolog A4A
Gorai.007G060000.1.A
PCC=0.569, p=0.011
mRNA Profile
Protein Expression Profile
DPA

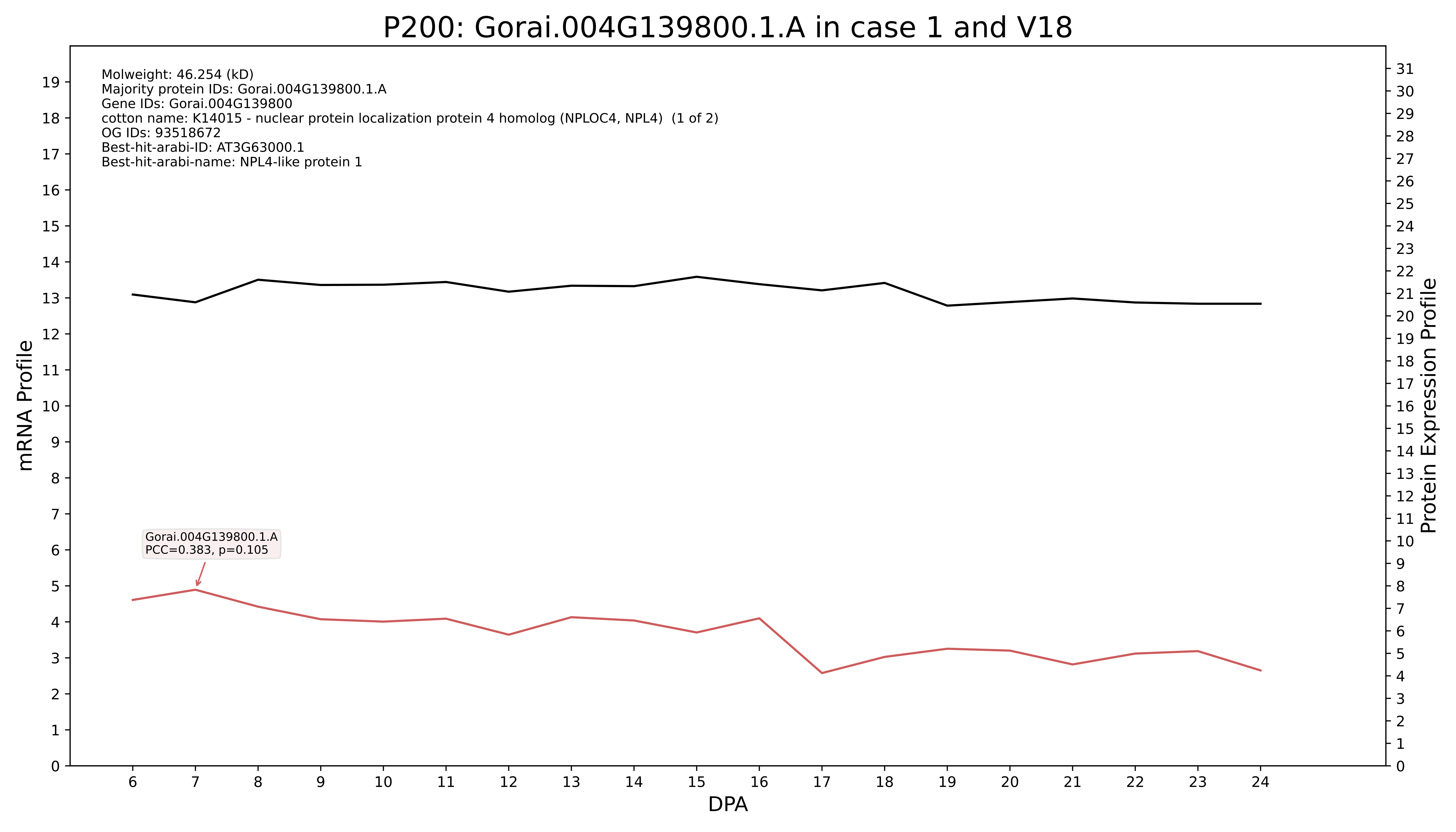

P200: Gorai.004G139800.1.A in case 1 and V18
Molweight: 46.254 (kD)
Majority protein IDs: Gorai.004G139800.1.A
Gene IDs: Gorai.004G139800
cotton name: K14015 - nuclear protein localization protein 4 homolog (NPLOC4, NPL4) (1 of 2)
OG IDs: 93518672
Best-hit-arabi-ID: AT3G63000.1
Best-hit-arabi-name: NPL4-like protein 1
Gorai.004G139800.1.A
PCC=0.383, p=0.105
mRNA Profile
Protein Expression Profile
DPA

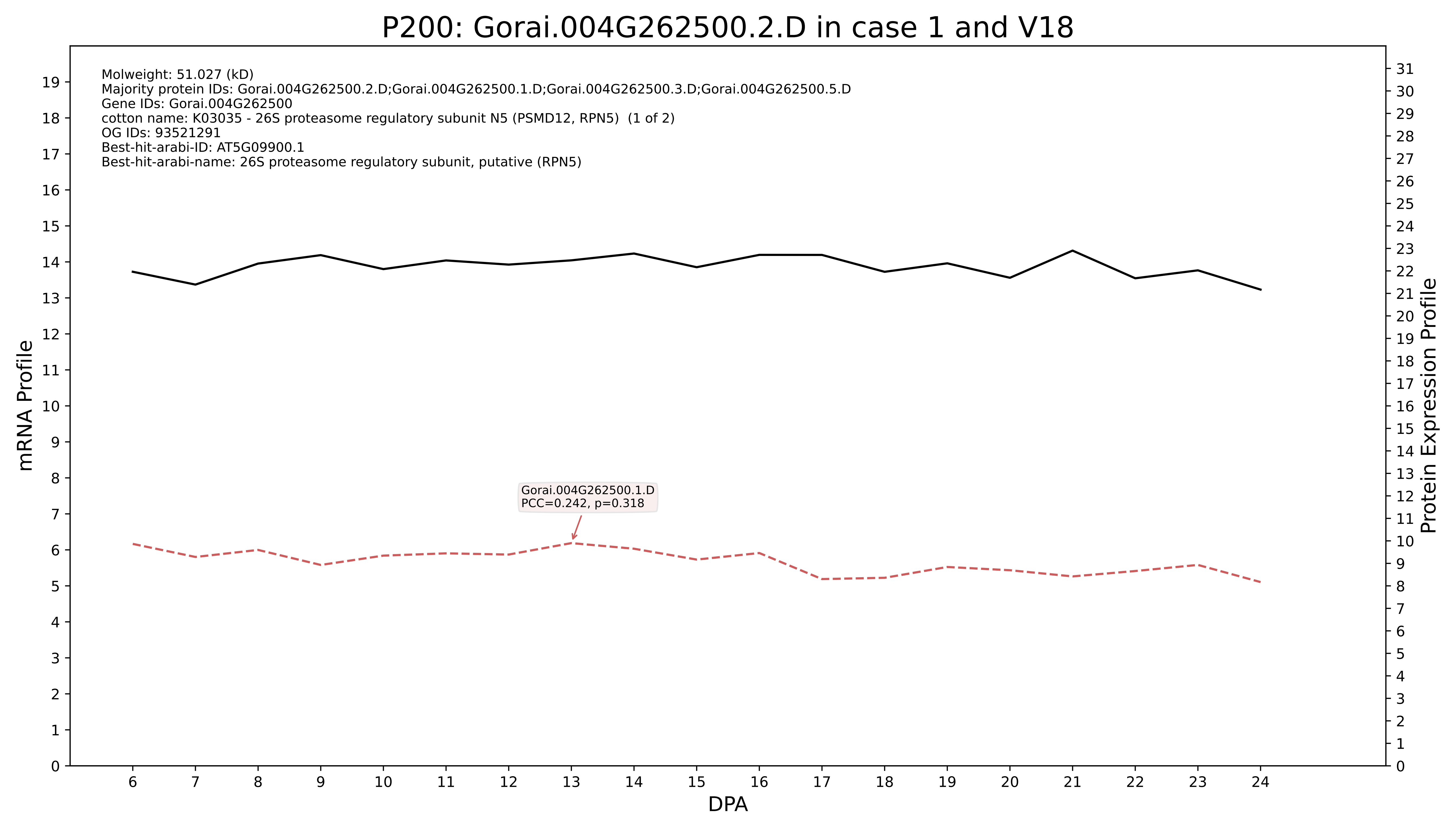
P200: Gorai.004G262500.2.D in case 1 and V18
Molweight: 51.027 (kD)
Majority protein IDs: Gorai.004G262500.2.D;Gorai.004G262500.1.D;Gorai.004G262500.3.D;Gorai.004G262500.5.D
Gene IDs: Gorai.004G262500
cotton name: K03035 - 26S proteasome regulatory subunit N5 (PSMD12, RPN5) (1 of 2)
OG IDs: 93521291
Best-hit-arabi-ID: AT5G09900.1
Best-hit-arabi-name: 26S proteasome regulatory subunit, putative (RPN5)
Gorai.004G262500.1.D
PCC=0.242, p=0.318
mRNA Profile
Protein Expression Profile
DPA

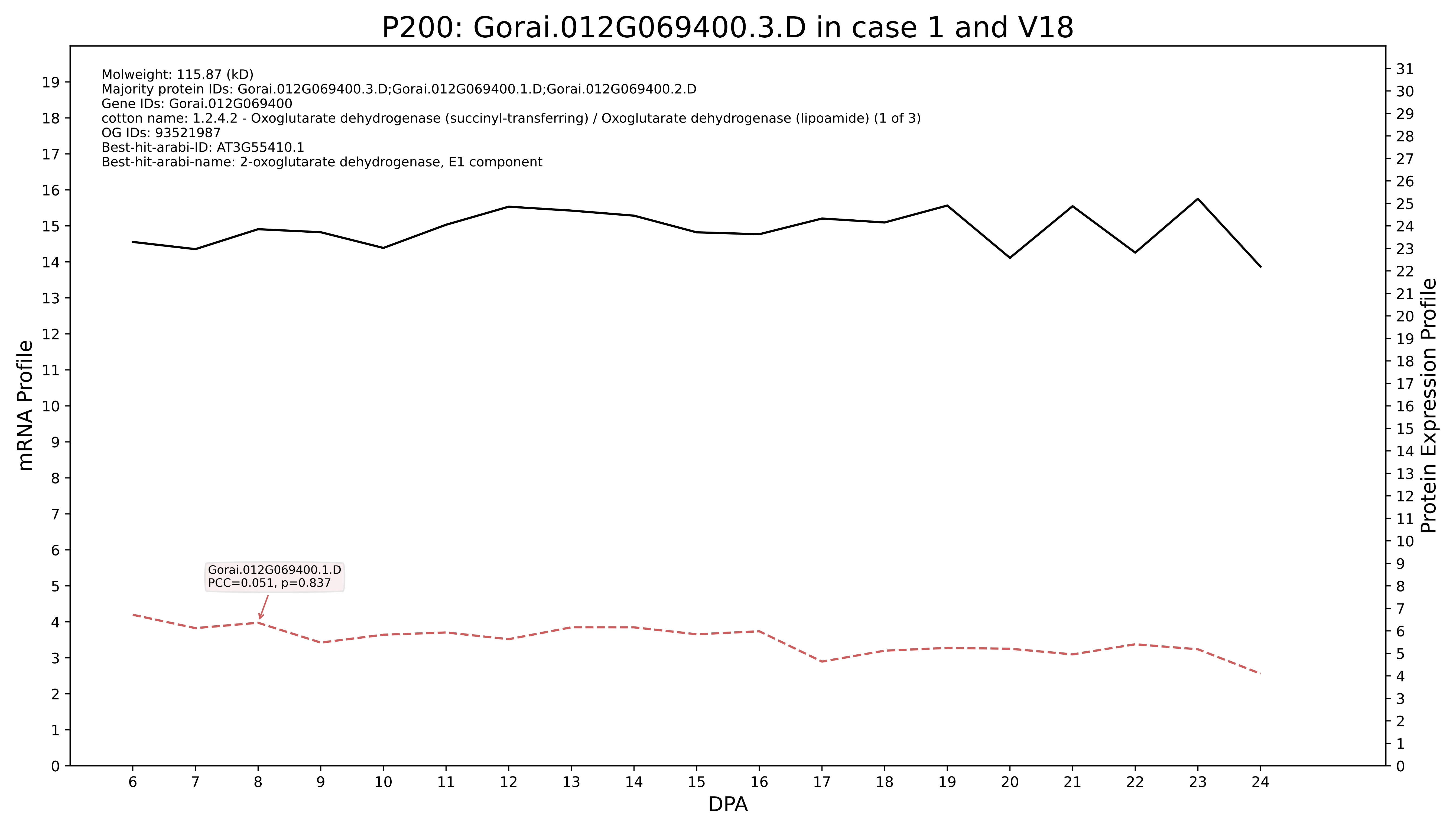
P200: Gorai.012G069400.3.D in case 1 and V18
Molweight: 115.87 (kD)
Majority protein IDs: Gorai.012G069400.3.D;Gorai.012G069400.1.D;Gorai.012G069400.2.D
Gene IDs: Gorai.012G069400
cotton name: 1.2.4.2 - Oxoglutarate dehydrogenase (succinyl-transferring) / Oxoglutarate dehydrogenase (lipoamide) (1 of 3)
OG IDs: 93521987
Best-hit-arabi-ID: AT3G55410.1
Best-hit-arabi-name: 2-oxoglutarate dehydrogenase, E1 component
Gorai.012G069400.1.D
PCC=0.051, p=0.837
mRNA Profile
Protein Expression Profile
DPA

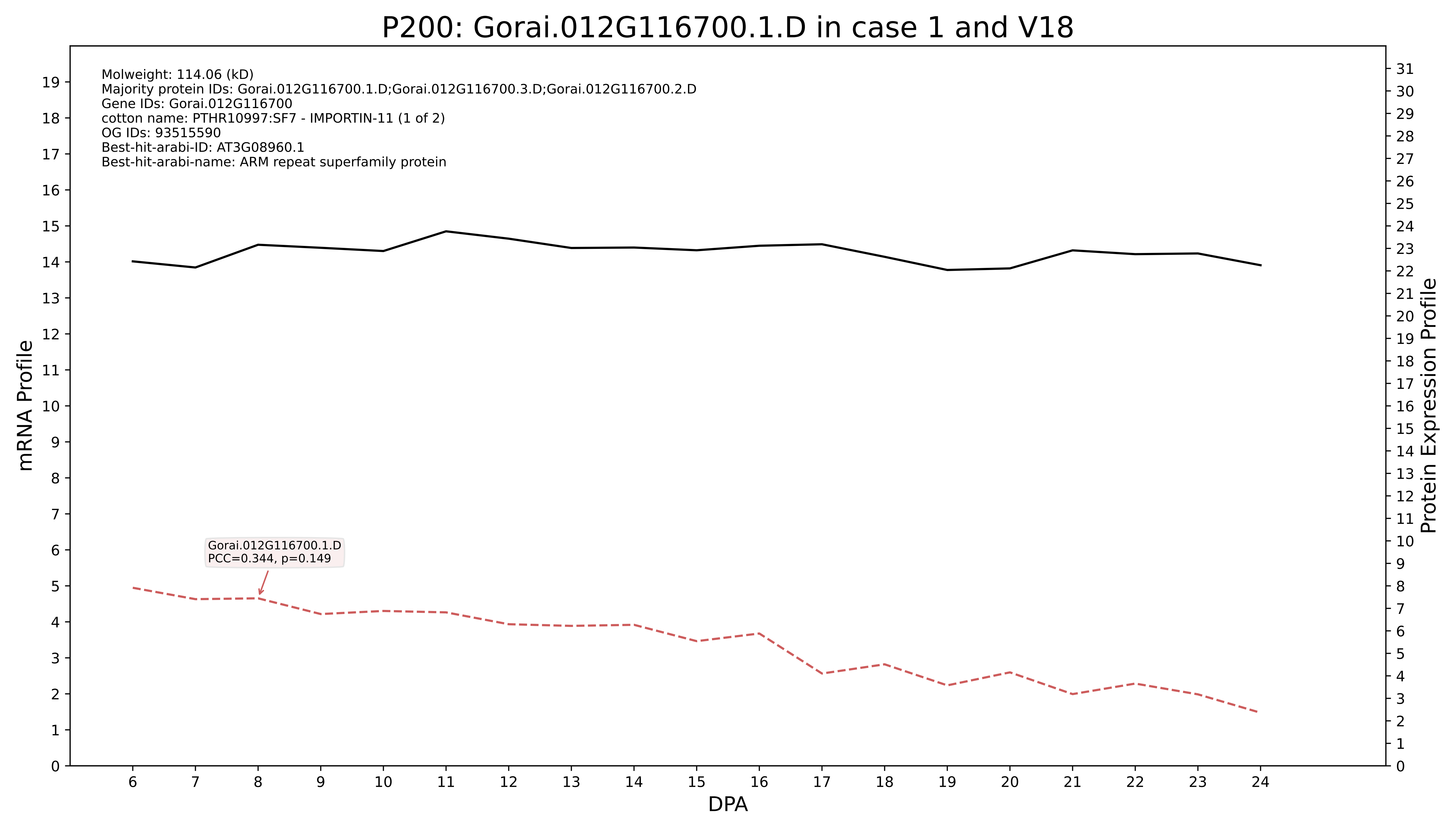
P200: Gorai.012G116700.1.D in case 1 and V18
Molweight: 114.06 (kD)
Majority protein IDs: Gorai.012G116700.1.D;Gorai.012G116700.3.D;Gorai.012G116700.2.D
Gene IDs: Gorai.012G116700
cotton name: PTHR10997:SF7 - IMPORTIN-11 (1 of 2)
OG IDs: 93515590
Best-hit-arabi-ID: AT3G08960.1
Best-hit-arabi-name: ARM repeat superfamily protein
Gorai.012G116700.1.D
PCC=0.344, p=0.149
mRNA Profile
Protein Expression Profile
DPA

### P200: Gorai.001G014100.1.A in case 1 and V18

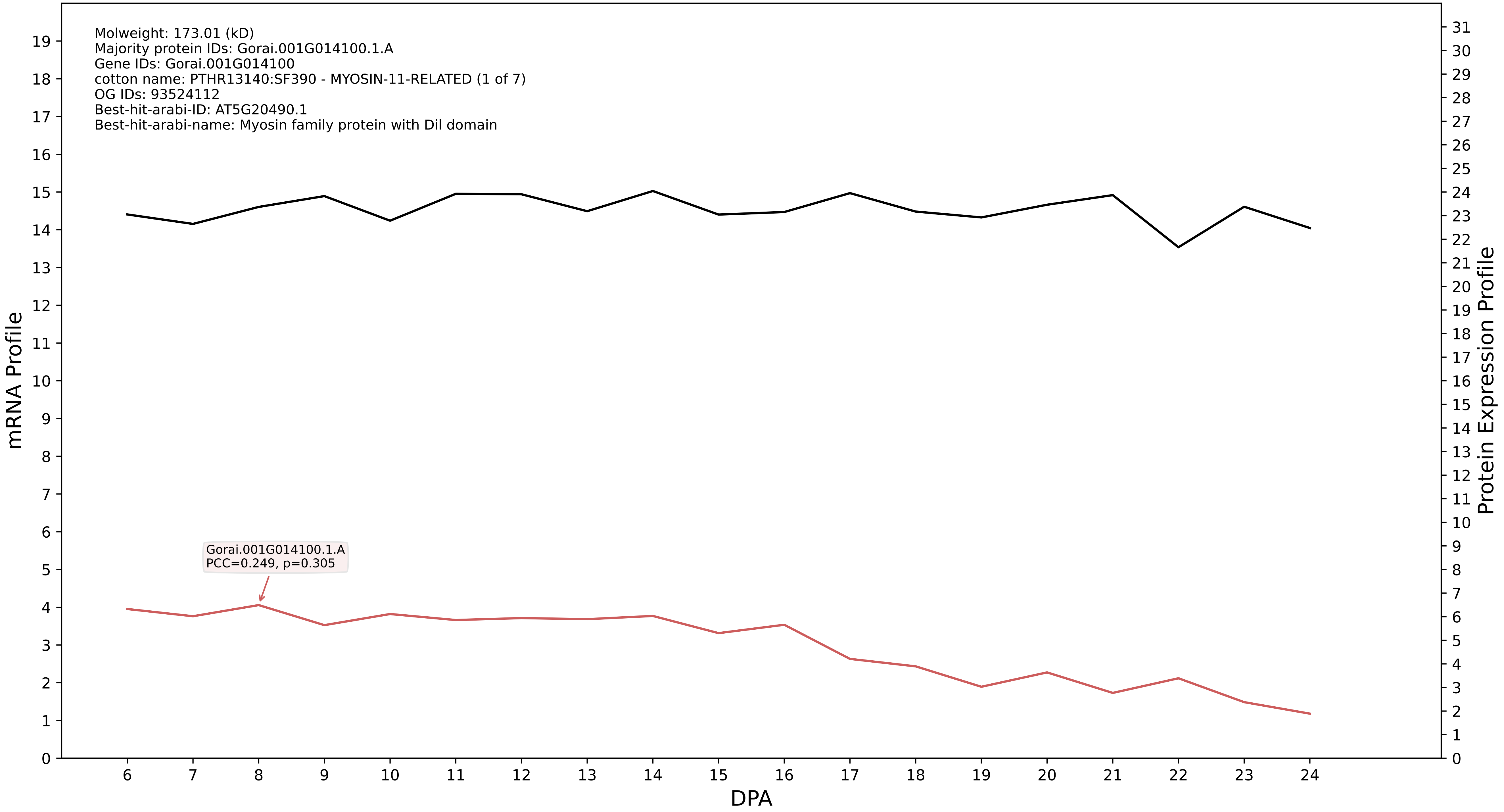

### P200: Gorai.008G046300.3.A in case 1 and V18

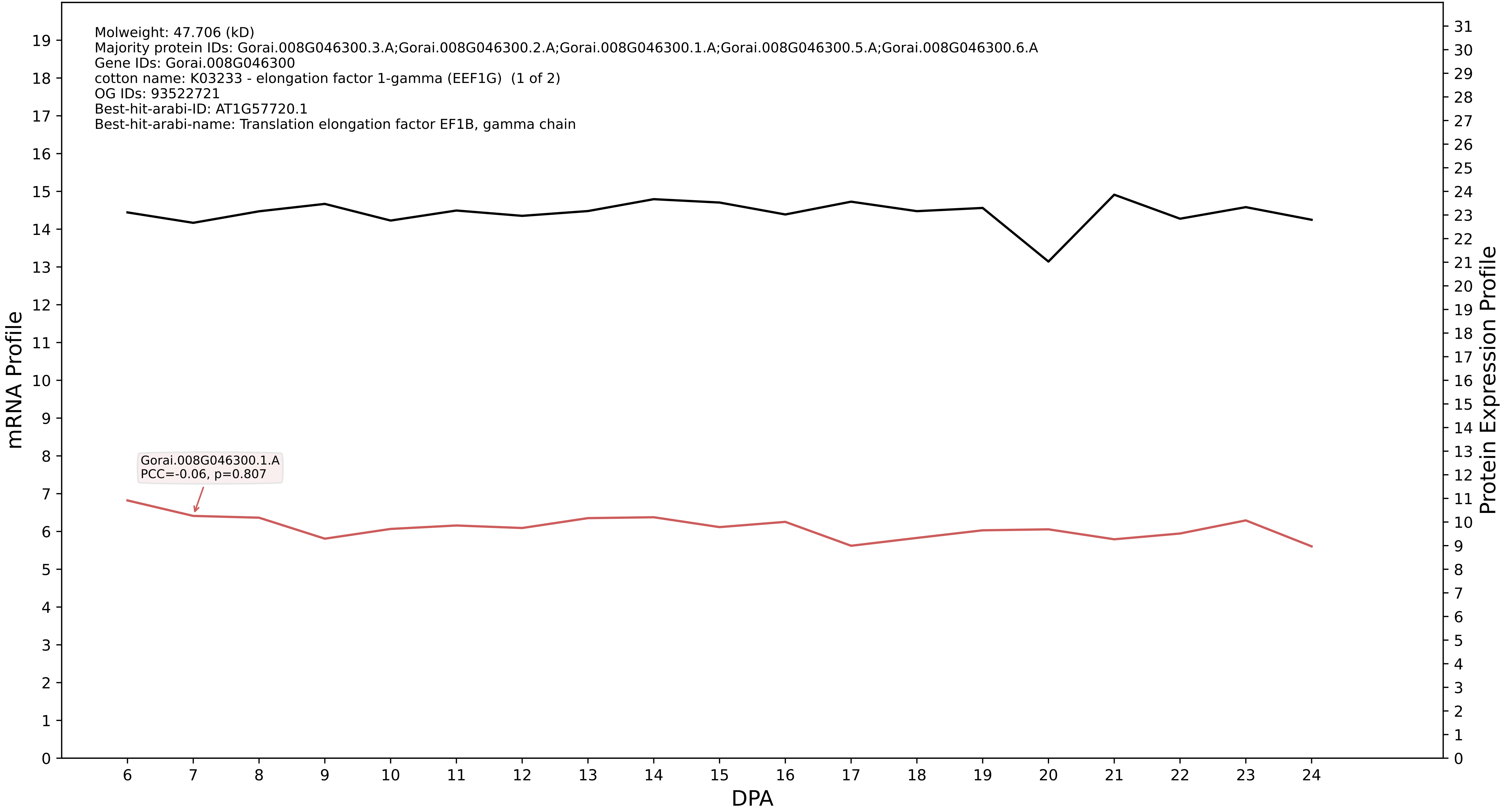

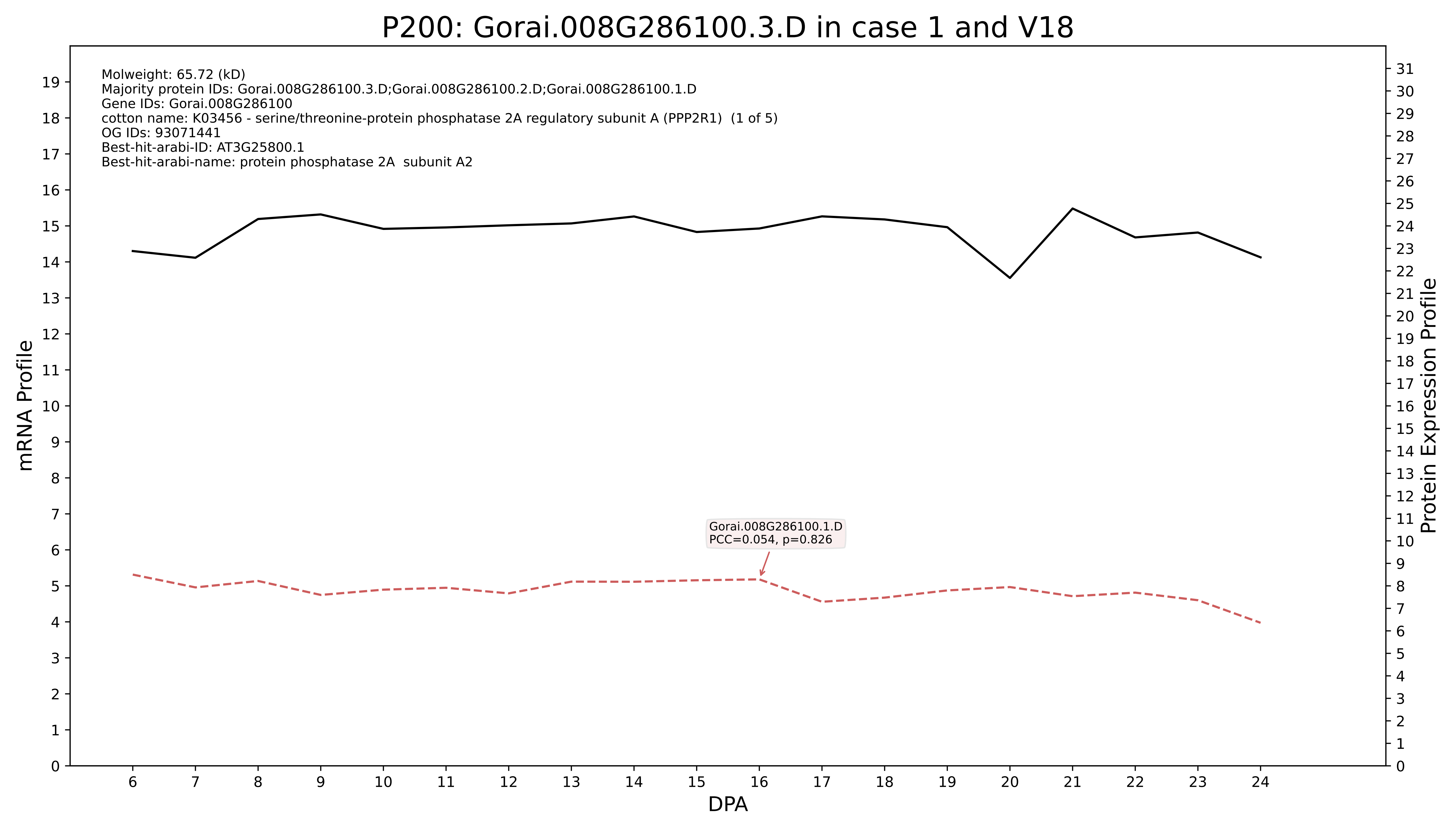
P200: Gorai.008G286100.3.D in case 1 and V18
Molweight: 65.72 (kD)
Majority protein IDs: Gorai.008G286100.3.D;Gorai.008G286100.2.D;Gorai.008G286100.1.D
Gene IDs: Gorai.008G286100
cotton name: K03456 - serine/threonine-protein phosphatase 2A regulatory subunit A (PPP2R1) (1 of 5)
OG IDs: 93071441
Best-hit-arabi-ID: AT3G25800.1
Best-hit-arabi-name: protein phosphatase 2A subunit A2
Gorai.008G286100.1.D
PCC=0.054, p=0.826
mRNA Profile
Protein Expression Profile
DPA

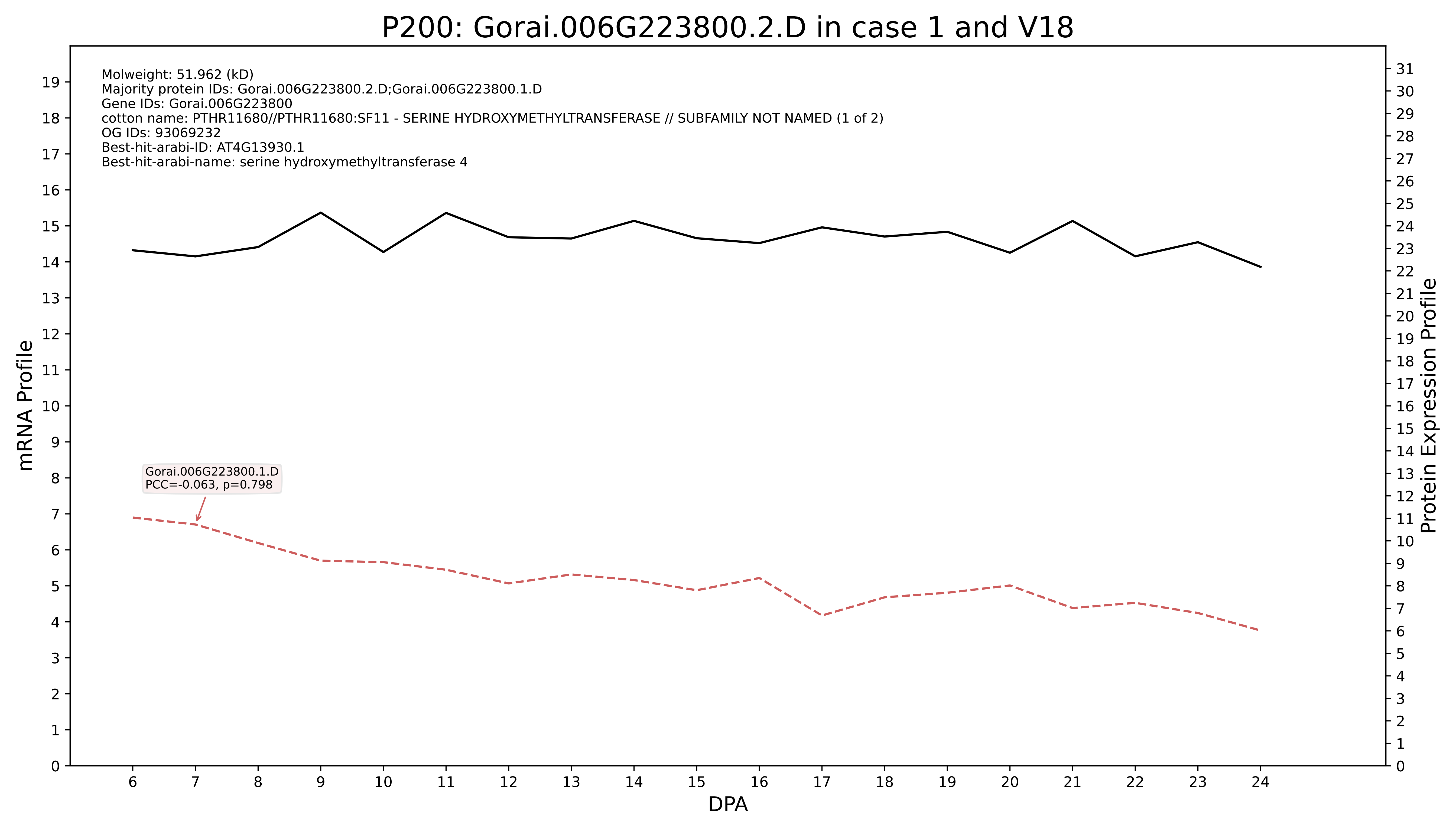

P200: Gorai.006G223800.2.D in case 1 and V18
Molweight: 51.962 (kD)
Majority protein IDs: Gorai.006G223800.2.D;Gorai.006G223800.1.D
Gene IDs: Gorai.006G223800
cotton name: PTHR11680//PTHR11680:SF11 - SERINE HYDROXYMETHYLTRANSFERASE // SUBFAMILY NOT NAMED (1 of 2)
OG IDs: 93069232
Best-hit-arabi-ID: AT4G13930.1
Best-hit-arabi-name: serine hydroxymethyltransferase 4
Gorai.006G223800.1.D
PCC=-0.063, p=0.798
mRNA Profile
Protein Expression Profile
DPA

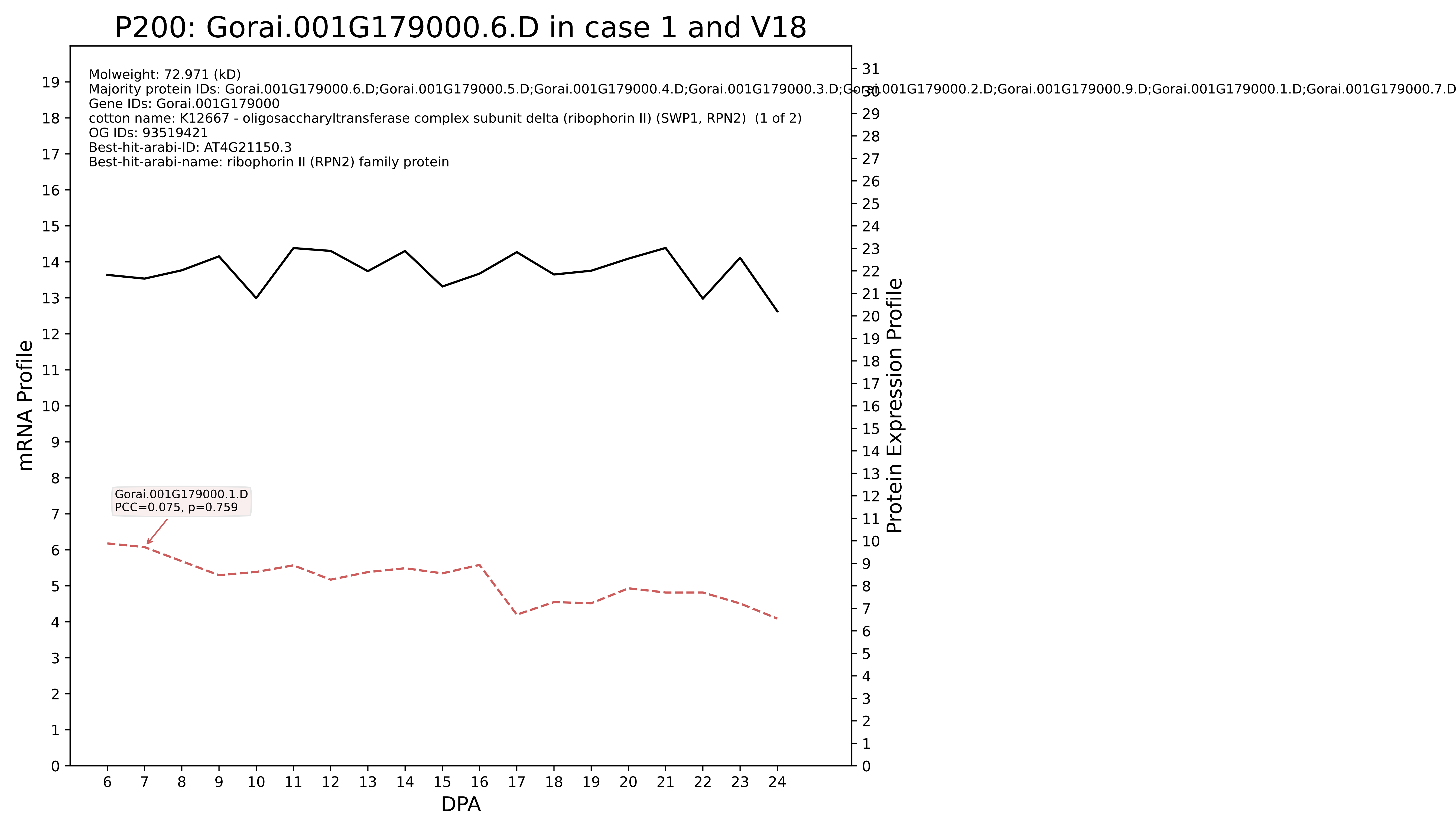
P200: Gorai.001G179000.6.D in case 1 and V18
Molweight: 72.971 (kD)
Majority protein IDs: Gorai.001G179000.6.D;Gorai.001G179000.5.D;Gorai.001G179000.4.D;Gorai.001G179000.3.D;Gorai.001G179000.2.D;Gorai.001G179000.9.D;Gorai.001G179000.1.D;Gorai.001G179000.7.D
Gene IDs: Gorai.001G179000
cotton name: K12667 - oligosaccharyltransferase complex subunit delta (ribophorin II) (SWP1, RPN2) (1 of 2)
OG IDs: 93519421
Best-hit-arabi-ID: AT4G21150.3
Best-hit-arabi-name: ribophorin II (RPN2) family protein
Gorai.001G179000.1.D
PCC=0.075, p=0.759
mRNA Profile
Protein Expression Profile
DPA

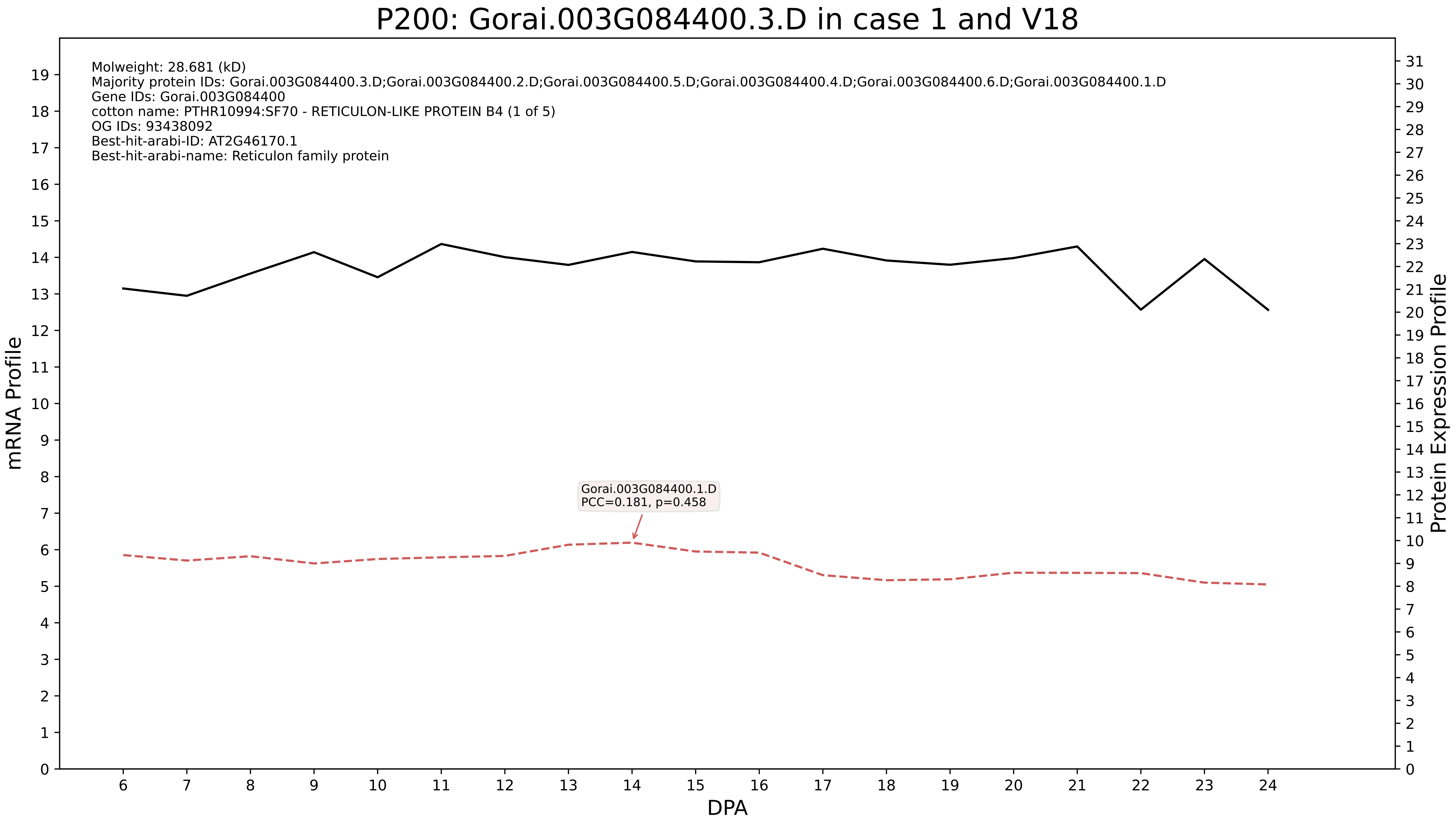
P200: Gorai.003G084400.3.D in case 1 and V18
Molweight: 28.681 (kD)
Majority protein IDs: Gorai.003G084400.3.D;Gorai.003G084400.2.D;Gorai.003G084400.5.D;Gorai.003G084400.4.D;Gorai.003G084400.6.D;Gorai.003G084400.1.D
Gene IDs: Gorai.003G084400
cotton name: PTHR10994:SF70 - RETICULON-LIKE PROTEIN B4 (1 of 5)
OG IDs: 93438092
Best-hit-arabi-ID: AT2G46170.1
Best-hit-arabi-name: Reticulon family protein
Gorai.003G084400.1.D
PCC=0.181, p=0.458
mRNA Profile
Protein Expression Profile
DPA

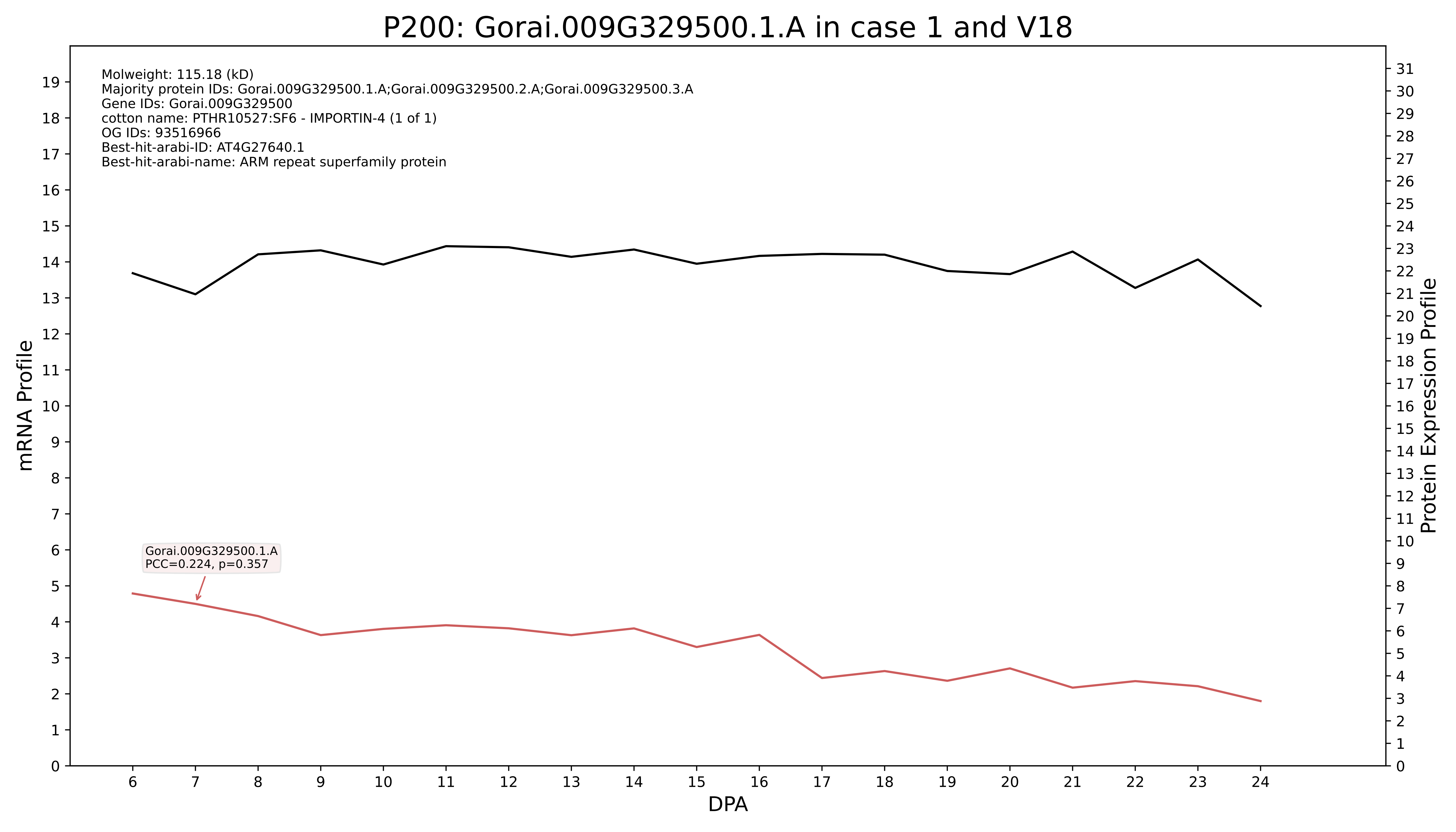
P200: Gorai.009G329500.1.A in case 1 and V18
Molweight: 115.18 (kD)
Majority protein IDs: Gorai.009G329500.1.A;Gorai.009G329500.2.A;Gorai.009G329500.3.A
Gene IDs: Gorai.009G329500
cotton name: PTHR10527:SF6 - IMPORTIN-4 (1 of 1)
OG IDs: 93516966
Best-hit-arabi-ID: AT4G27640.1
Best-hit-arabi-name: ARM repeat superfamily protein
Gorai.009G329500.1.A
PCC=0.224, p=0.357
mRNA Profile
Protein Expression Profile
DPA

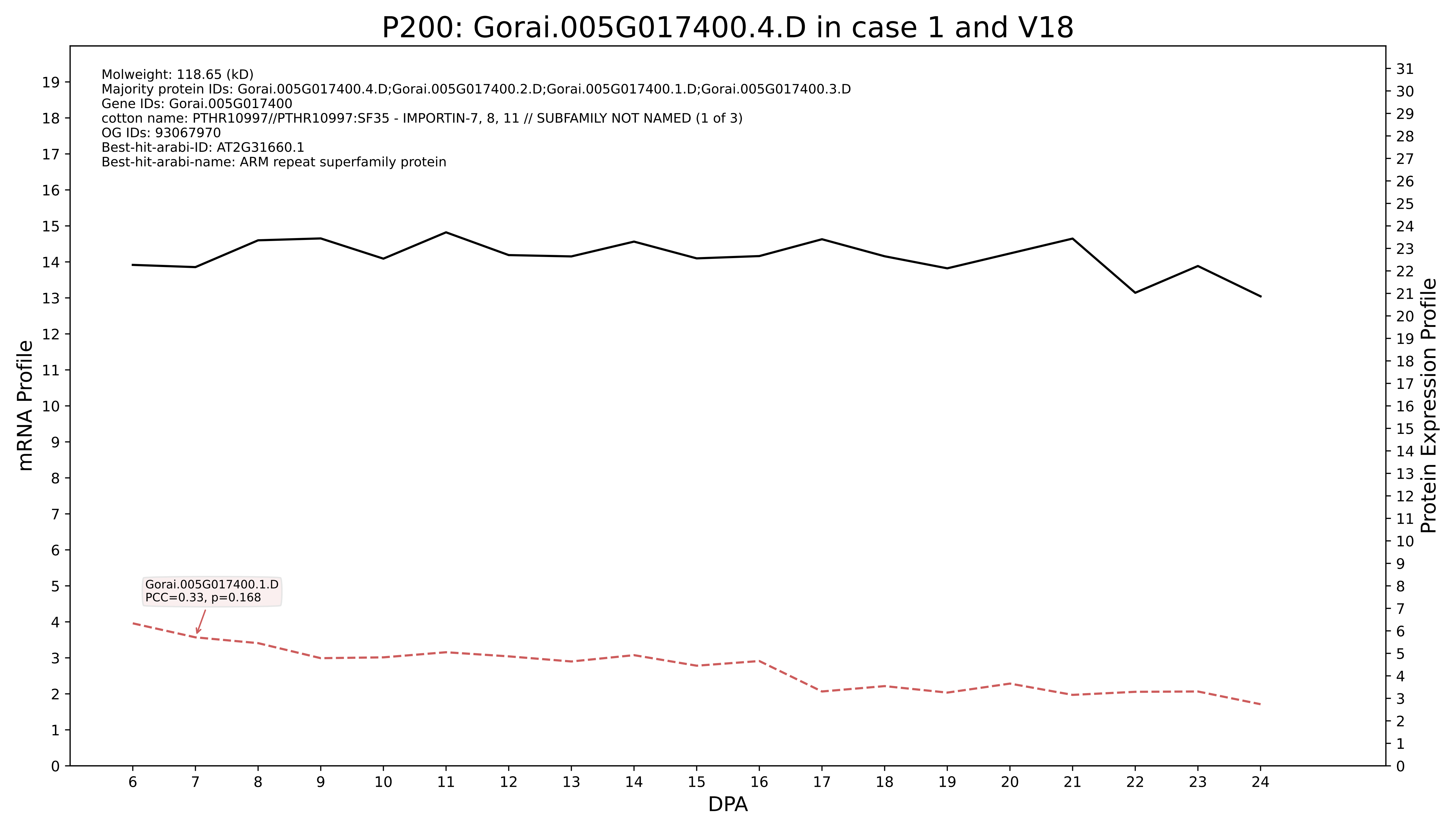

P200: Gorai.005G017400.4.D in case 1 and V18
Molweight: 118.65 (kD)
Majority protein IDs: Gorai.005G017400.4.D;Gorai.005G017400.2.D;Gorai.005G017400.1.D;Gorai.005G017400.3.D
Gene IDs: Gorai.005G017400
cotton name: PTHR10997//PTHR10997:SF35 - IMPORTIN-7, 8, 11 // SUBFAMILY NOT NAMED (1 of 3)
OG IDs: 93067970
Best-hit-arabi-ID: AT2G31660.1
Best-hit-arabi-name: ARM repeat superfamily protein
Gorai.005G017400.1.D
PCC=0.33, p=0.168
mRNA Profile
Protein Expression Profile
DPA

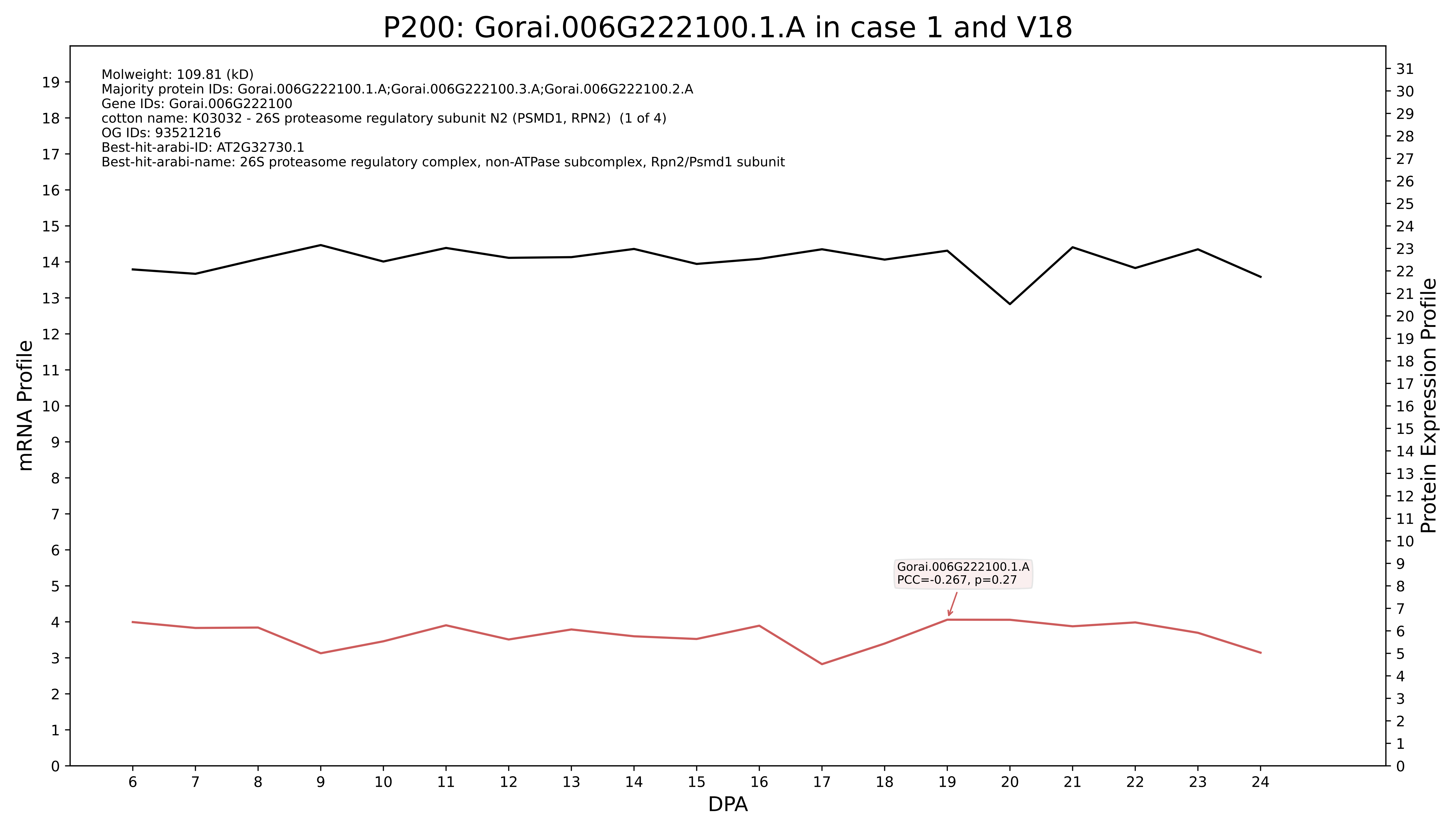
P200: Gorai.006G222100.1.A in case 1 and V18
Molweight: 109.81 (kD)
Majority protein IDs: Gorai.006G222100.1.A;Gorai.006G222100.3.A;Gorai.006G222100.2.A
Gene IDs: Gorai.006G222100
cotton name: K03032 - 26S proteasome regulatory subunit N2 (PSMD1, RPN2) (1 of 4)
OG IDs: 93521216
Best-hit-arabi-ID: AT2G32730.1
Best-hit-arabi-name: 26S proteasome regulatory complex, non-ATPase subcomplex, Rpn2/Psmd1 subunit
Gorai.006G222100.1.A
PCC=-0.267, p=0.27
mRNA Profile
Protein Expression Profile
DPA

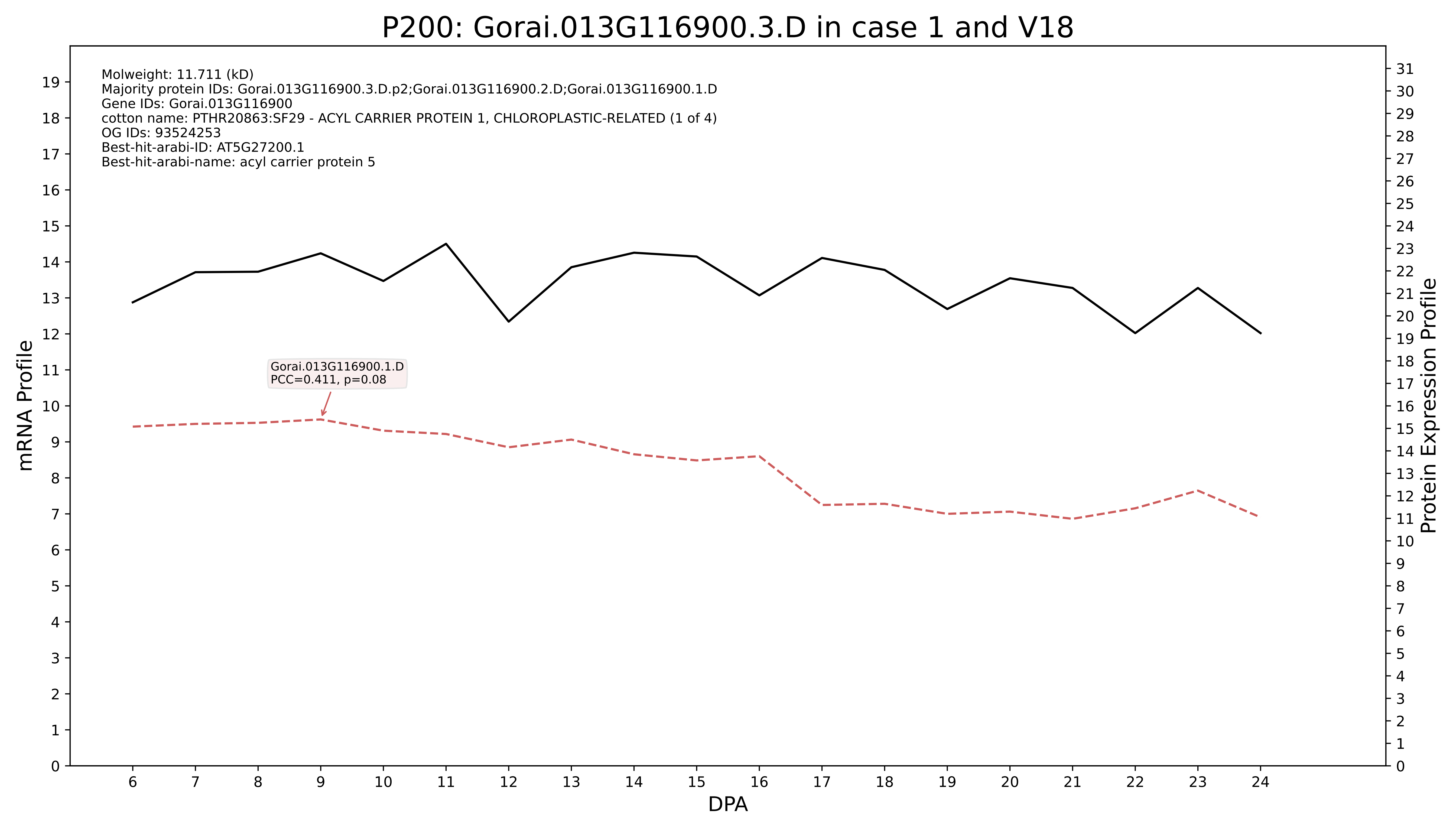

P200: Gorai.013G116900.3.D in case 1 and V18
Molweight: 11.711 (kD)
Majority protein IDs: Gorai.013G116900.3.D.p2;Gorai.013G116900.2.D;Gorai.013G116900.1.D
Gene IDs: Gorai.013G116900
cotton name: PTHR20863:SF29 - ACYL CARRIER PROTEIN 1, CHLOROPLASTIC-RELATED (1 of 4)
OG IDs: 93524253
Best-hit-arabi-ID: AT5G27200.1
Best-hit-arabi-name: acyl carrier protein 5
Gorai.013G116900.1.D
PCC=0.411, p=0.08
mRNA Profile
Protein Expression Profile
DPA

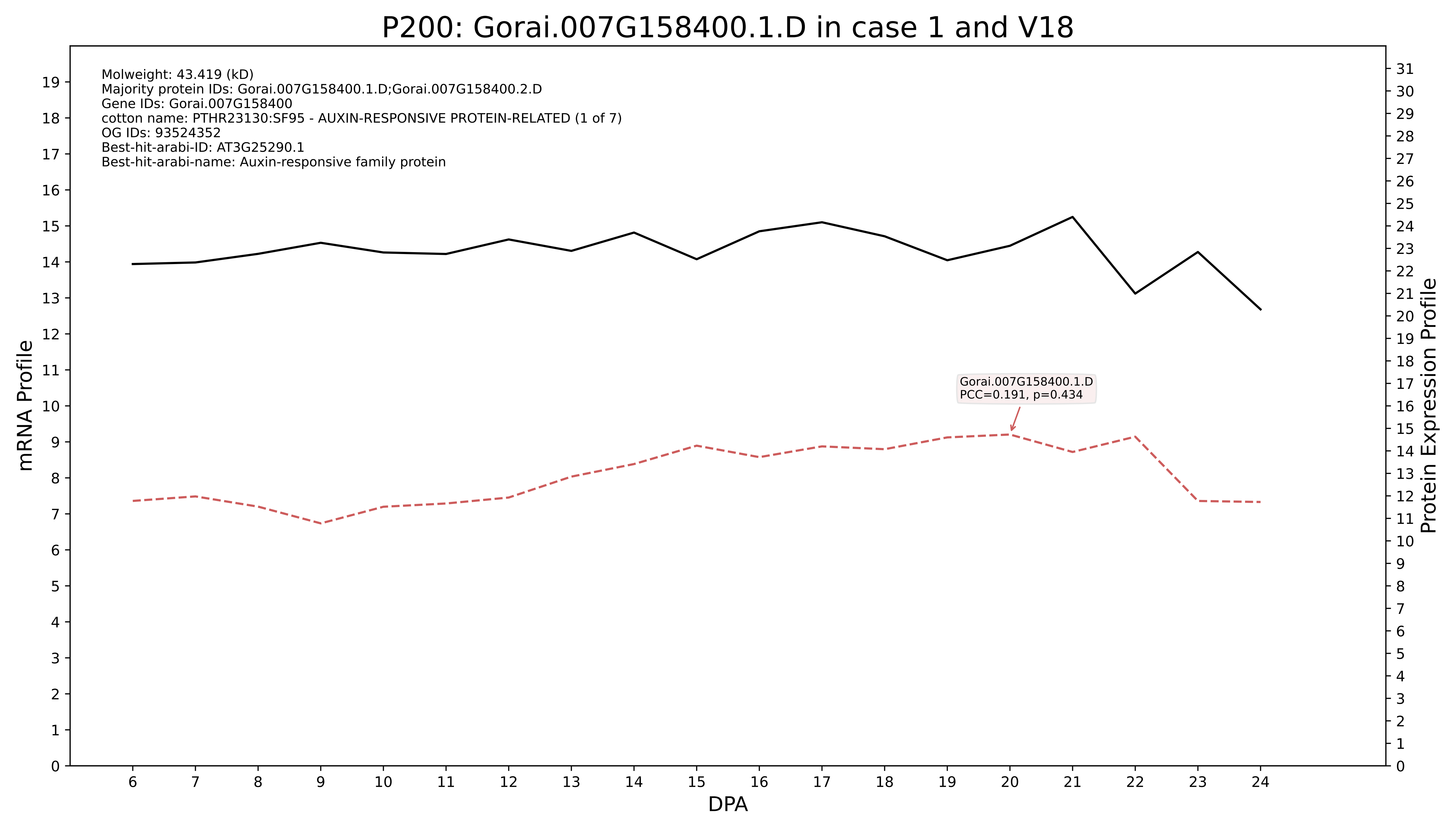

P200: Gorai.007G158400.1.D in case 1 and V18
Molweight: 43.419 (kD)
Majority protein IDs: Gorai.007G158400.1.D;Gorai.007G158400.2.D
Gene IDs: Gorai.007G158400
cotton name: PTHR23130:SF95 - AUXIN-RESPONSIVE PROTEIN-RELATED (1 of 7)
OG IDs: 93524352
Best-hit-arabi-ID: AT3G25290.1
Best-hit-arabi-name: Auxin-responsive family protein
Gorai.007G158400.1.D
PCC=0.191, p=0.434
mRNA Profile
Protein Expression Profile
DPA

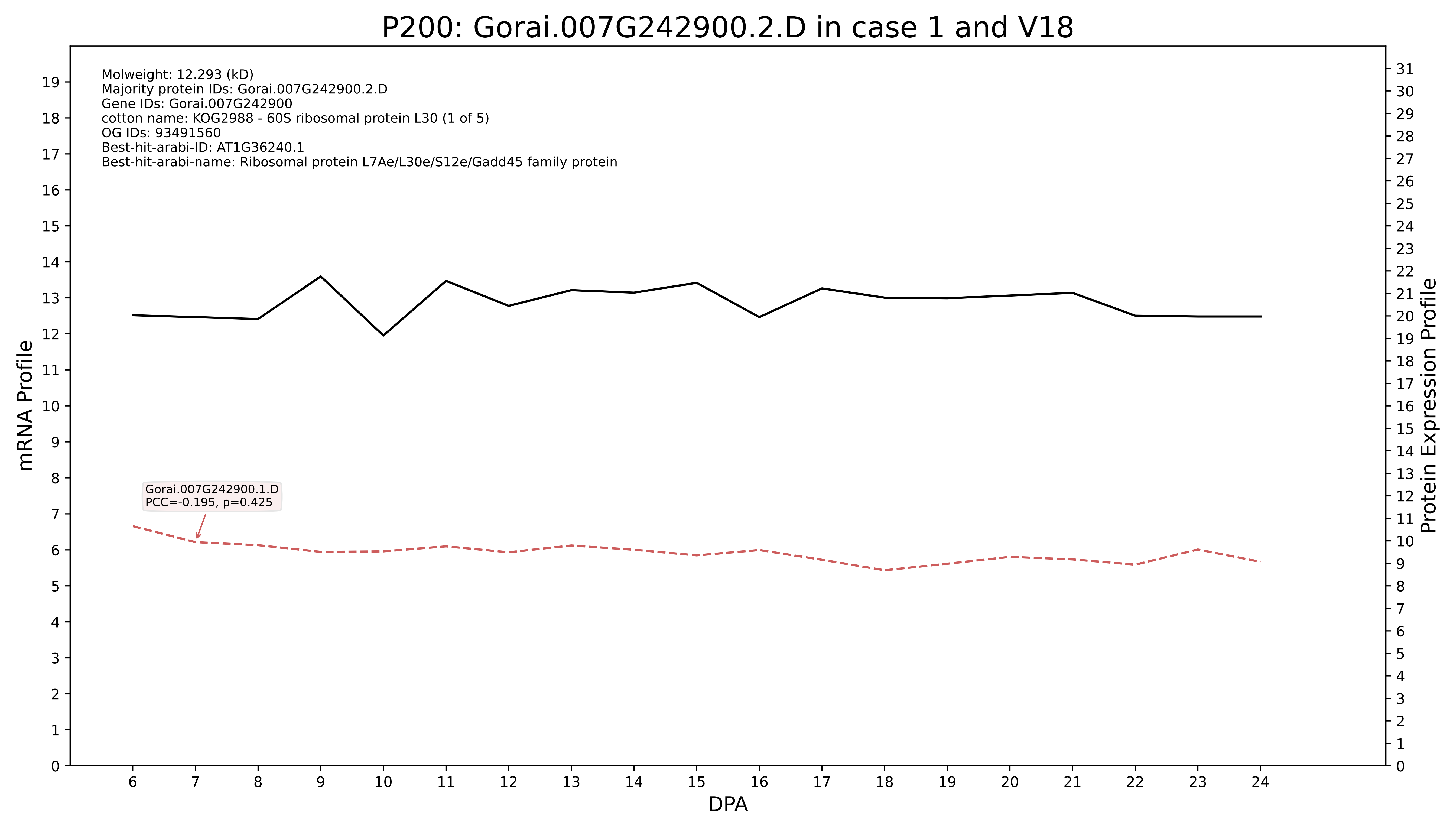
P200: Gorai.007G242900.2.D in case 1 and V18
Molweight: 12.293 (kD)
Majority protein IDs: Gorai.007G242900.2.D
Gene IDs: Gorai.007G242900
cotton name: KOG2988 - 60S ribosomal protein L30 (1 of 5)
OG IDs: 93491560
Best-hit-arabi-ID: AT1G36240.1
Best-hit-arabi-name: Ribosomal protein L7Ae/L30e/S12e/Gadd45 family protein
Gorai.007G242900.1.D
PCC=-0.195, p=0.425
mRNA Profile
Protein Expression Profile
DPA

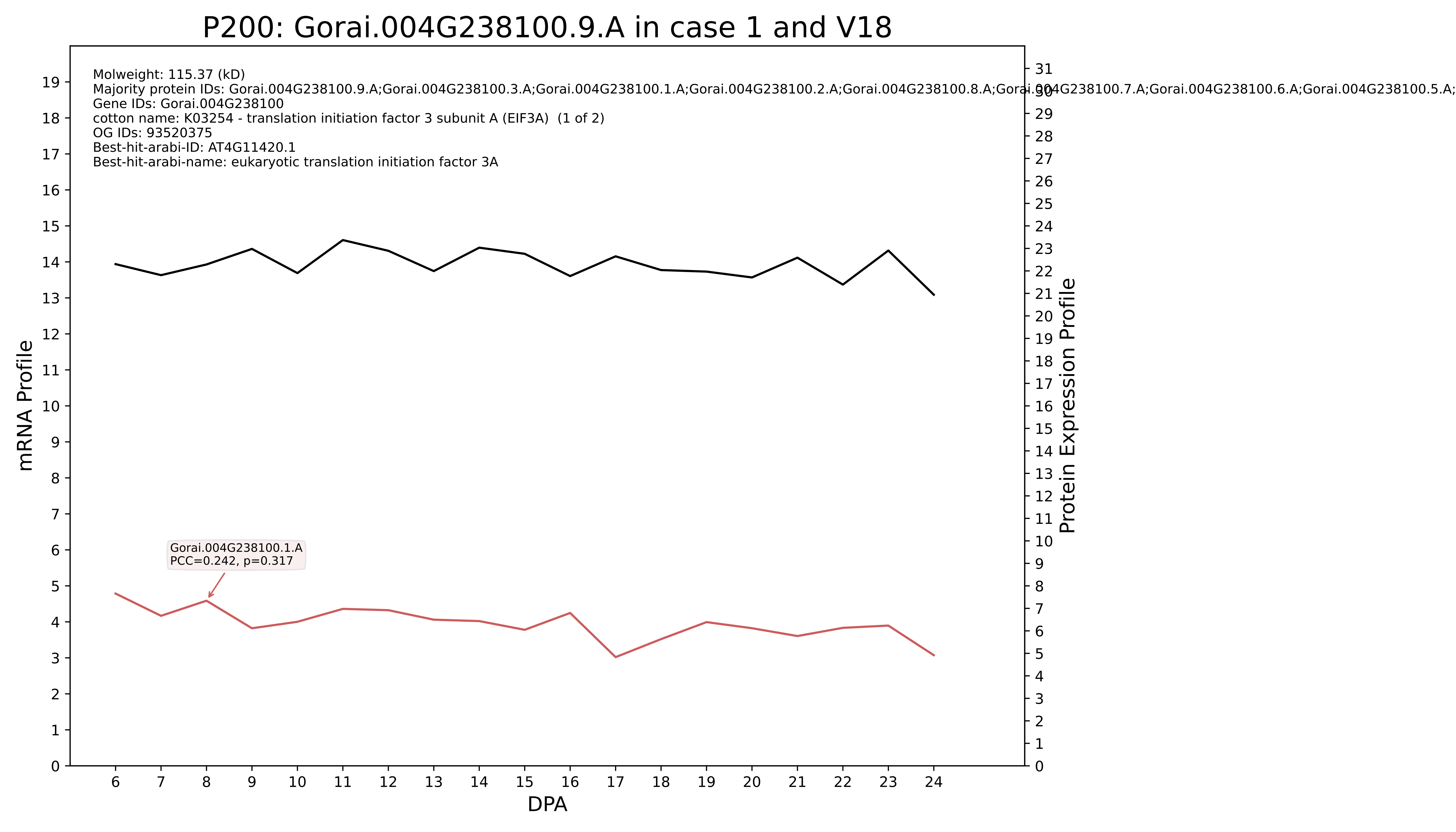
P200: Gorai.004G238100.9.A in case 1 and V18
Molweight: 115.37 (kD)
Majority protein IDs: Gorai.004G238100.9.A;Gorai.004G238100.3.A;Gorai.004G238100.1.A;Gorai.004G238100.2.A;Gorai.004G238100.8.A;Gorai.004G238100.7.A;Gorai.004G238100.6.A;Gorai.004G238100.5.A;
Gene IDs: Gorai.004G238100
cotton name: K03254 - translation initiation factor 3 subunit A (EIF3A) (1 of 2)
OG IDs: 93520375
Best-hit-arabi-ID: AT4G11420.1
Best-hit-arabi-name: eukaryotic translation initiation factor 3A
Gorai.004G238100.1.A
PCC=0.242, p=0.317
mRNA Profile
Protein Expression Profile
DPA

#### P200: Gorai.012G017900.1.D in case 1 and V18

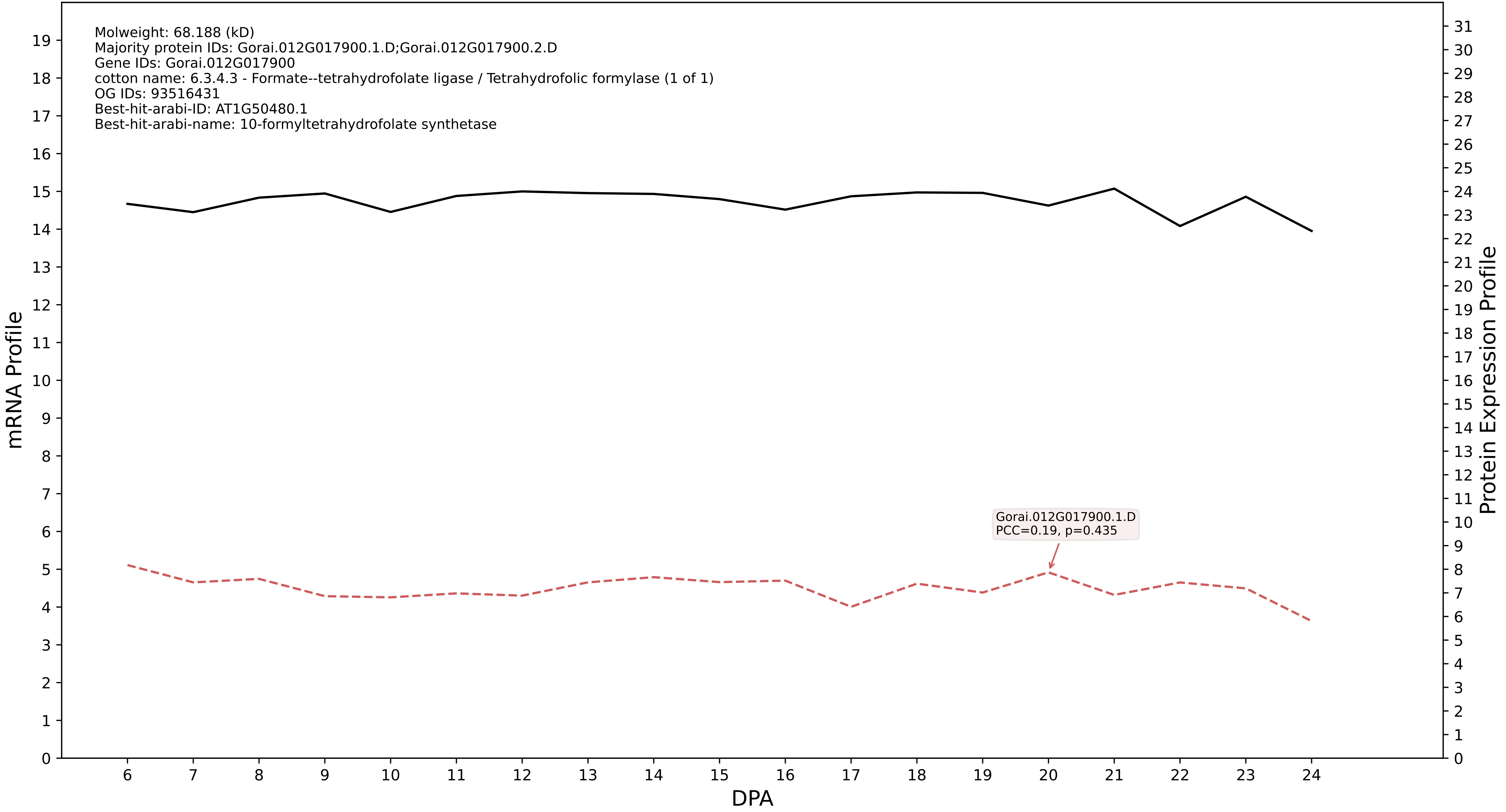

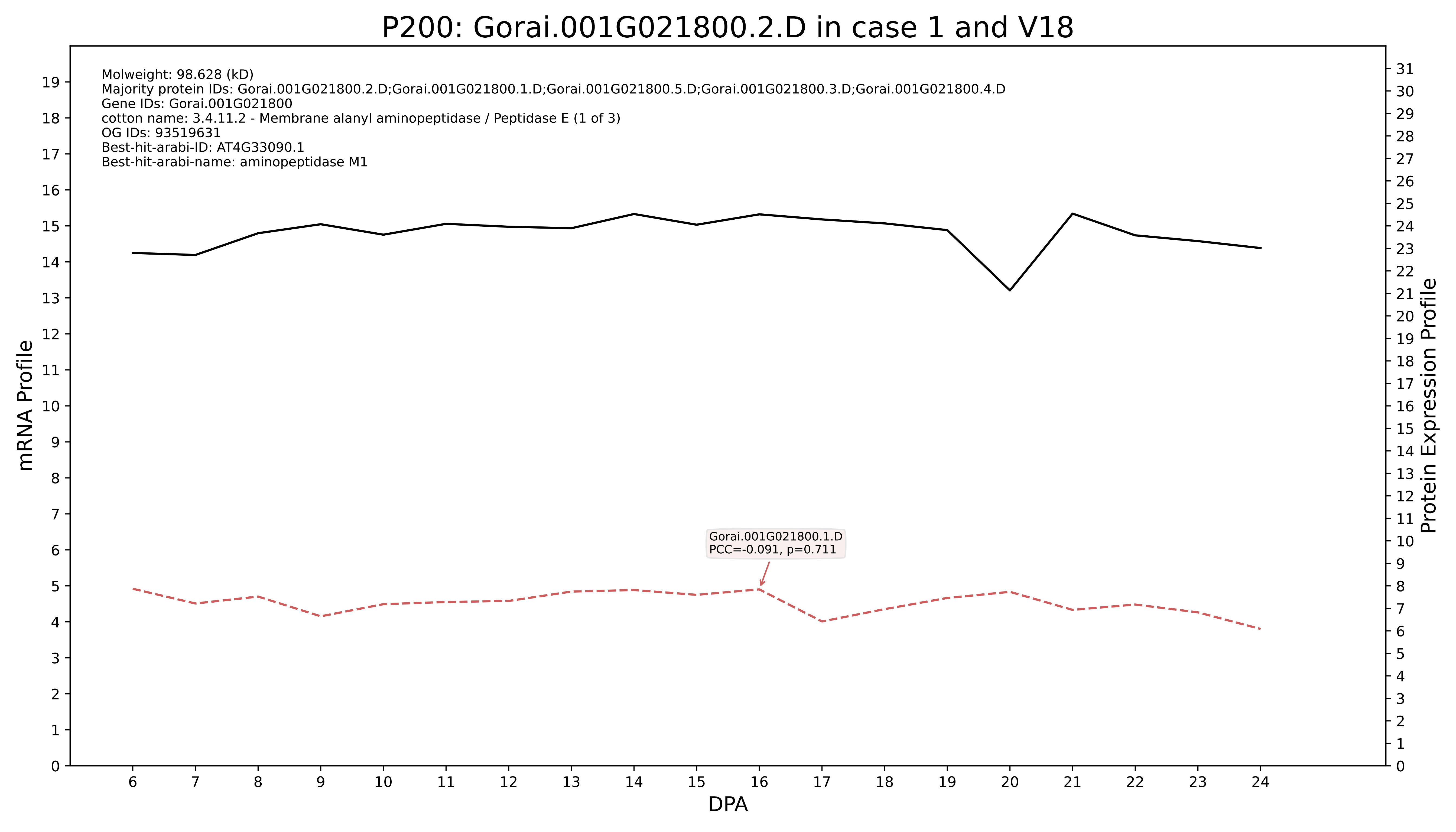
P200: Gorai.001G021800.2.D in case 1 and V18
Molweight: 98.628 (kD)
Majority protein IDs: Gorai.001G021800.2.D;Gorai.001G021800.1.D;Gorai.001G021800.5.D;Gorai.001G021800.3.D;Gorai.001G021800.4.D
Gene IDs: Gorai.001G021800
cotton name: 3.4.11.2 - Membrane alanyl aminopeptidase / Peptidase E (1 of 3)
OG IDs: 93519631
Best-hit-arabi-ID: AT4G33090.1
Best-hit-arabi-name: aminopeptidase M1
Gorai.001G021800.1.D
PCC=-0.091, p=0.711
mRNA Profile
Protein Expression Profile
DPA

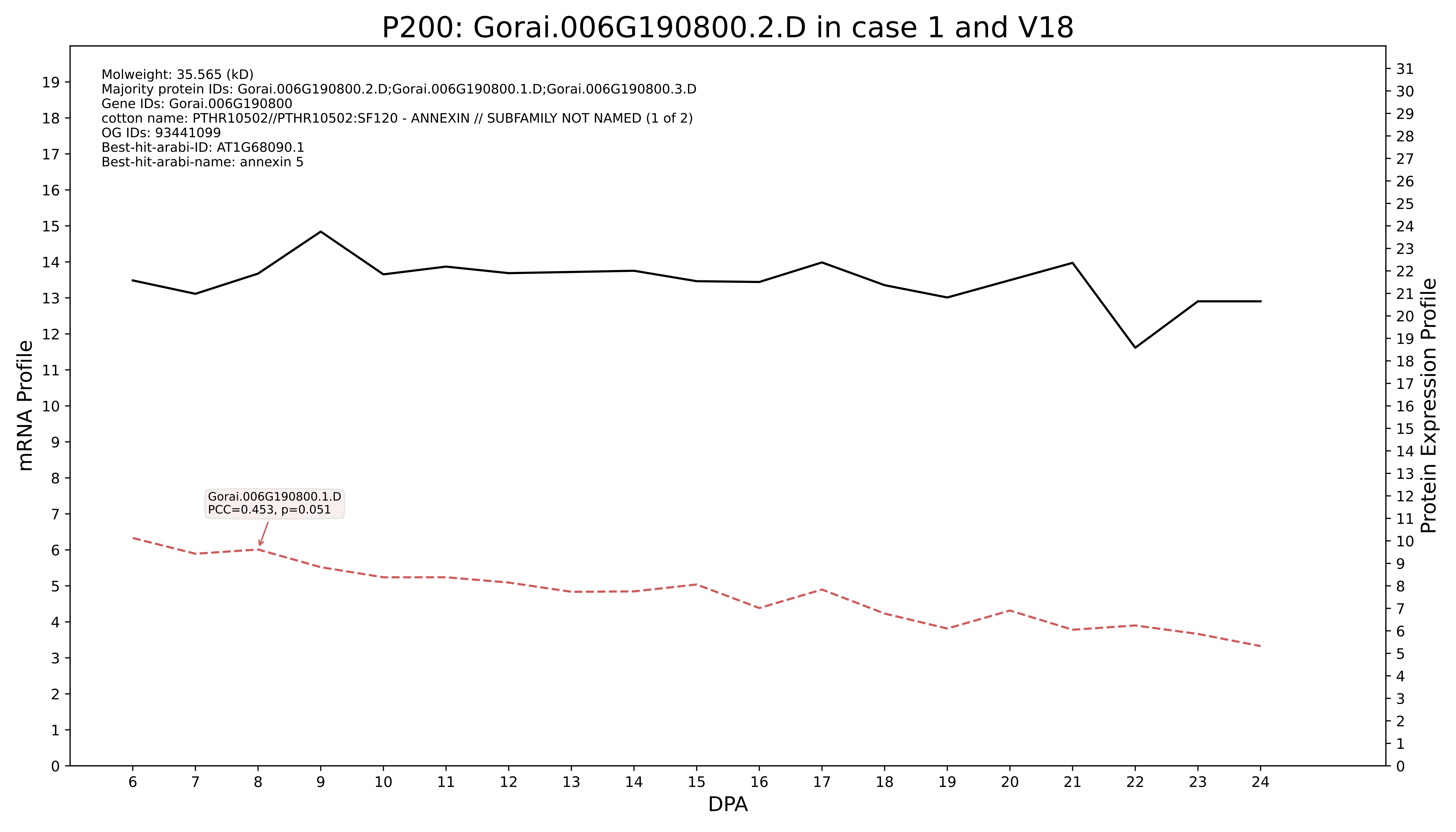
P200: Gorai.006G190800.2.D in case 1 and V18
Molweight: 35.565 (kD)
Majority protein IDs: Gorai.006G190800.2.D;Gorai.006G190800.1.D;Gorai.006G190800.3.D
Gene IDs: Gorai.006G190800
cotton name: PTHR10502//PTHR10502:SF120 - ANNEXIN // SUBFAMILY NOT NAMED (1 of 2)
OG IDs: 93441099
Best-hit-arabi-ID: AT1G68090.1
Best-hit-arabi-name: annexin 5
Gorai.006G190800.1.D
PCC=0.453, p=0.051
mRNA Profile
Protein Expression Profile
DPA

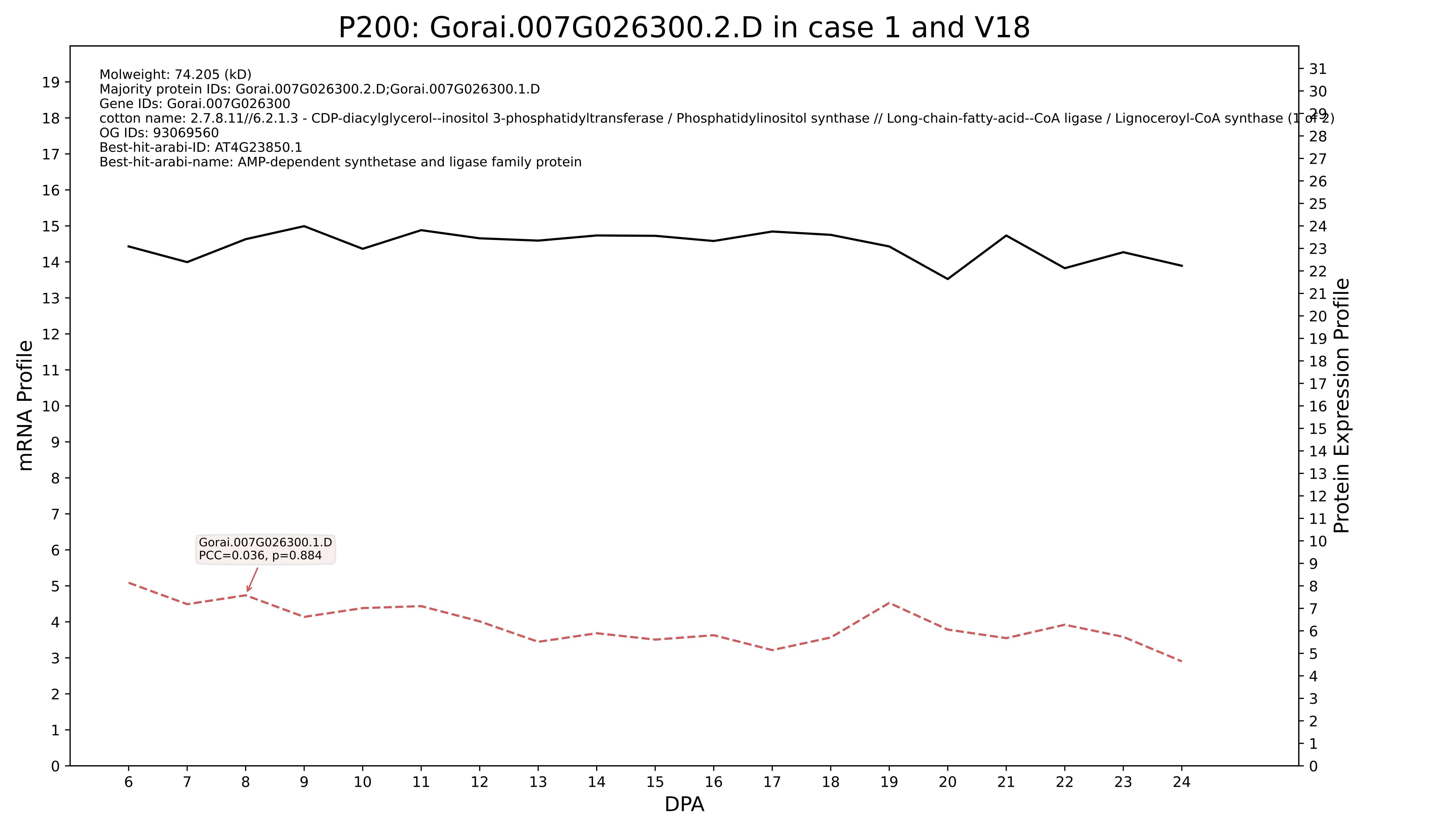

P200: Gorai.007G026300.2.D in case 1 and V18
Molweight: 74.205 (kD)
Majority protein IDs: Gorai.007G026300.2.D;Gorai.007G026300.1.D
Gene IDs: Gorai.007G026300
cotton name: 2.7.8.11//6.2.1.3 - CDP-diacylglycerol--inositol 3-phosphatidyltransferase / Phosphatidylinositol synthase // Long-chain-fatty-acid--CoA ligase / Lignoceroyl-CoA synthase (1 of 2)
OG IDs: 93069560
Best-hit-arabi-ID: AT4G23850.1
Best-hit-arabi-name: AMP-dependent synthetase and ligase family protein
Gorai.007G026300.1.D
PCC=0.036, p=0.884
mRNA Profile
Protein Expression Profile
DPA

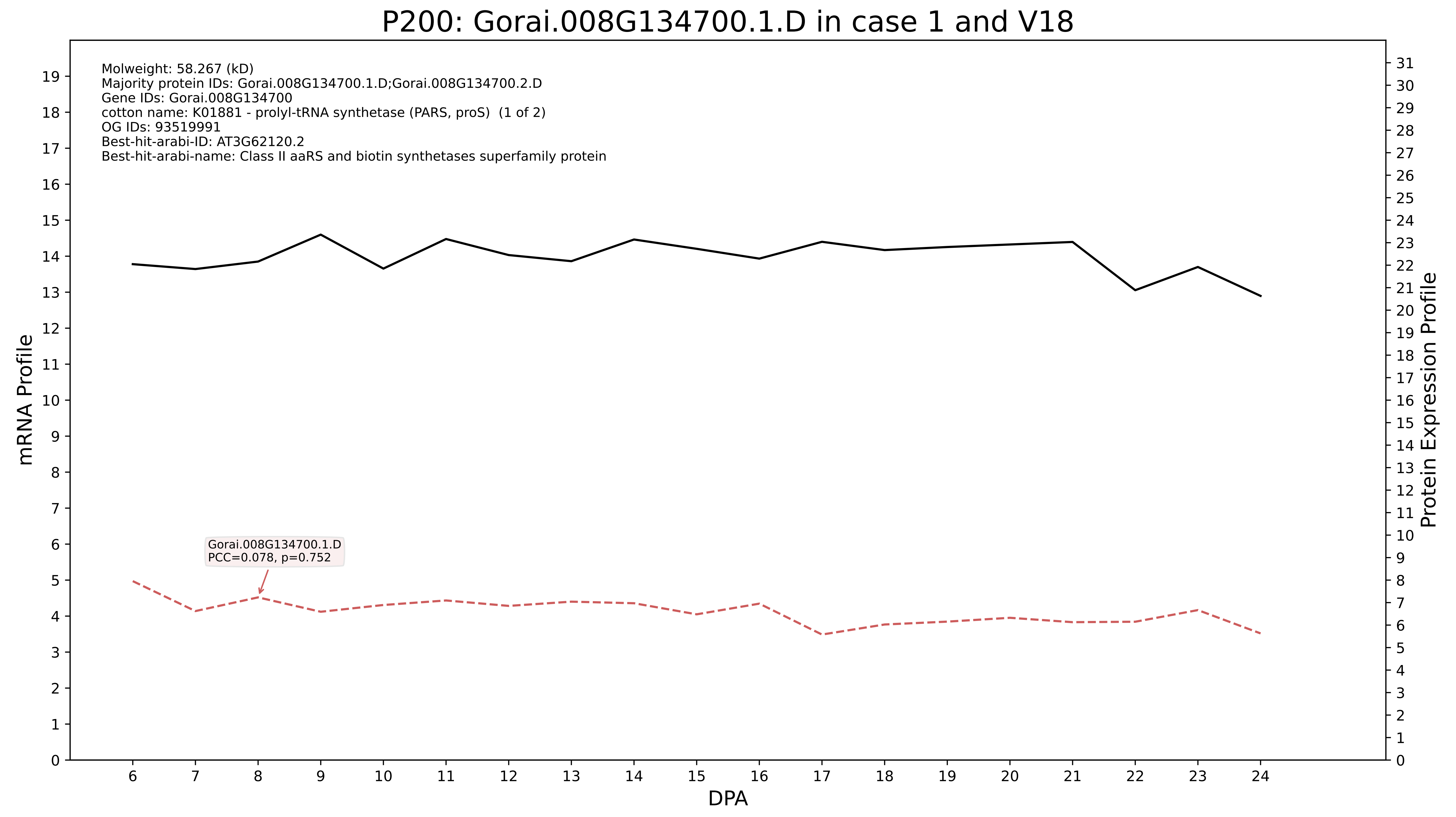

P200: Gorai.008G134700.1.D in case 1 and V18
Molweight: 58.267 (kD)
Majority protein IDs: Gorai.008G134700.1.D;Gorai.008G134700.2.D
Gene IDs: Gorai.008G134700
cotton name: K01881 - prolyl-tRNA synthetase (PARS, proS) (1 of 2)
OG IDs: 93519991
Best-hit-arabi-ID: AT3G62120.2
Best-hit-arabi-name: Class II aaRS and biotin synthetases superfamily protein
Gorai.008G134700.1.D
PCC=0.078, p=0.752
mRNA Profile
Protein Expression Profile
DPA

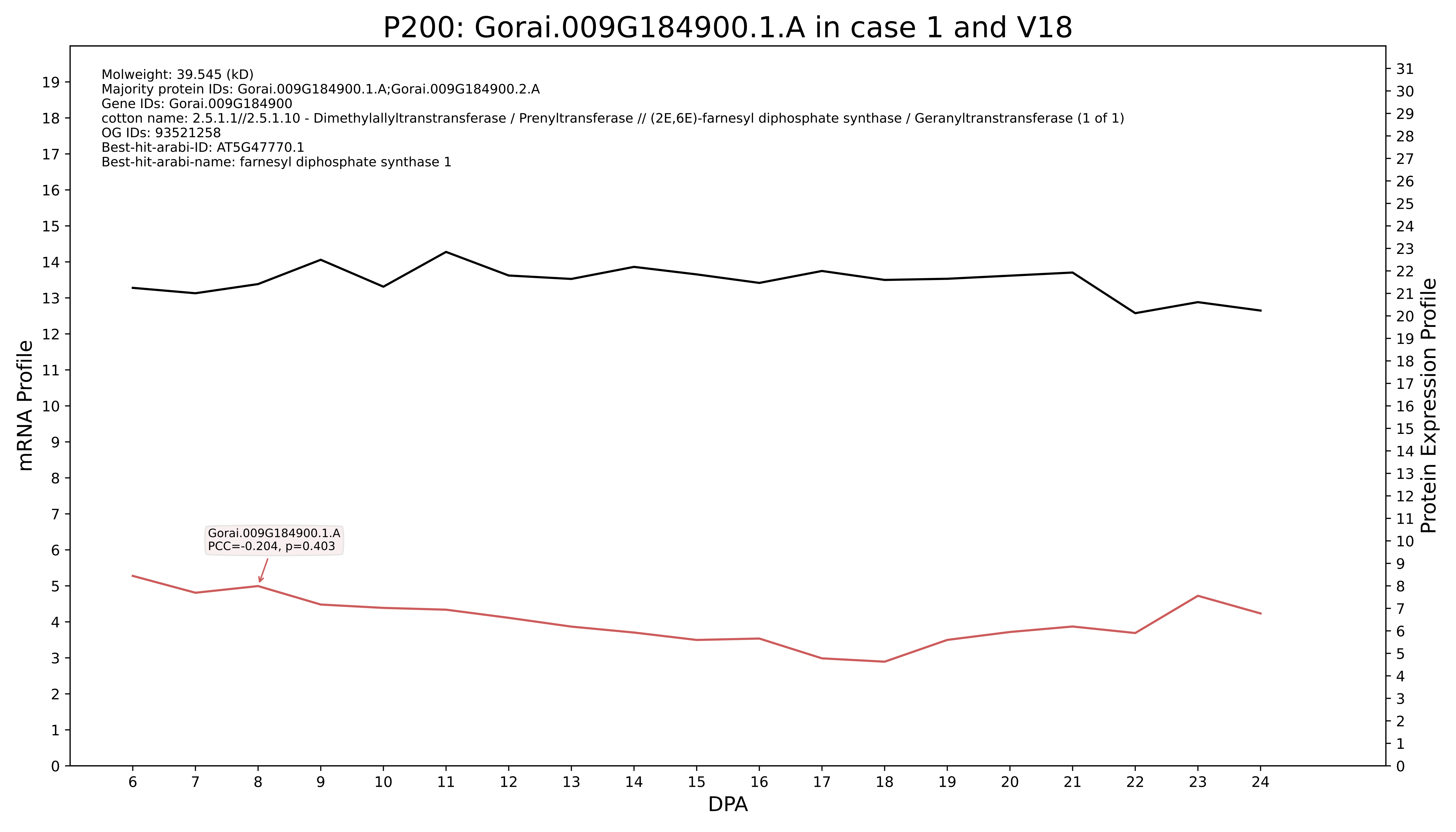
P200: Gorai.009G184900.1.A in case 1 and V18
Molweight: 39.545 (kD)
Majority protein IDs: Gorai.009G184900.1.A;Gorai.009G184900.2.A
Gene IDs: Gorai.009G184900
cotton name: 2.5.1.1//2.5.1.10 - Dimethylallyltranstransferase / Prenyltransferase // (2E,6E)-farnesyl diphosphate synthase / Geranyltranstransferase (1 of 1)
OG IDs: 93521258
Best-hit-arabi-ID: AT5G47770.1
Best-hit-arabi-name: farnesyl diphosphate synthase 1
Gorai.009G184900.1.A
PCC=-0.204, p=0.403
mRNA Profile
Protein Expression Profile
DPA

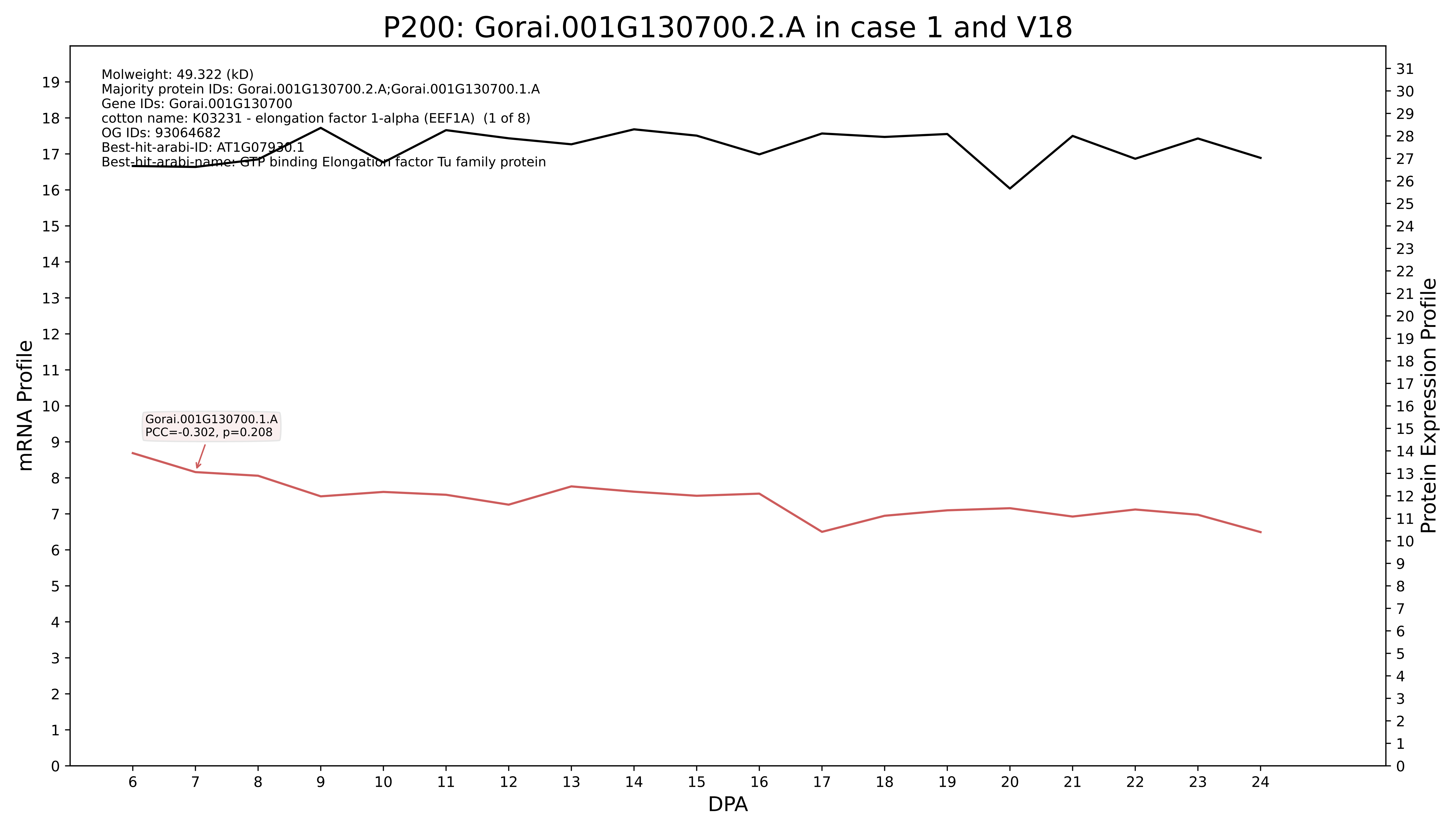
P200: Gorai.001G130700.2.A in case 1 and V18
Molweight: 49.322 (kD)
Majority protein IDs: Gorai.001G130700.2.A;Gorai.001G130700.1.A
Gene IDs: Gorai.001G130700
cotton name: K03231 - elongation factor 1-alpha (EEF1A) (1 of 8)
OG IDs: 93064682
Best-hit-arabi-ID: AT1G07930.1
Best-hit-arabi-name: GTP binding Elongation factor Tu family protein
Gorai.001G130700.1.A
PCC=-0.302, p=0.208
mRNA Profile
Protein Expression Profile
DPA

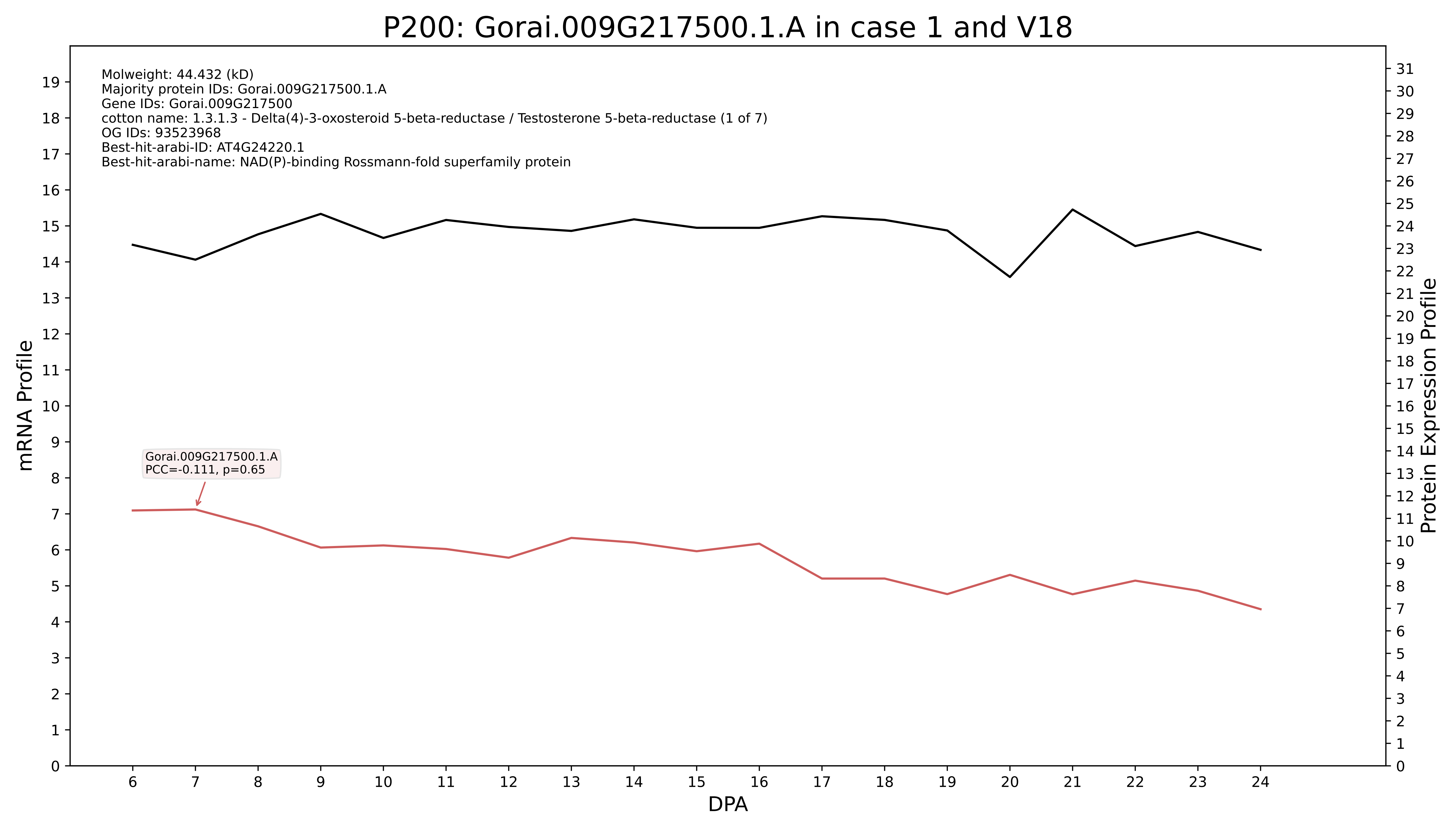
P200: Gorai.009G217500.1.A in case 1 and V18
Molweight: 44.432 (kD)
Majority protein IDs: Gorai.009G217500.1.A
Gene IDs: Gorai.009G217500
cotton name: 1.3.1.3 - Delta(4)-3-oxosteroid 5-beta-reductase / Testosterone 5-beta-reductase (1 of 7)
OG IDs: 93523968
Best-hit-arabi-ID: AT4G24220.1
Best-hit-arabi-name: NAD(P)-binding Rossmann-fold superfamily protein
Gorai.009G217500.1.A
PCC=-0.111, p=0.65
mRNA Profile
Protein Expression Profile
DPA

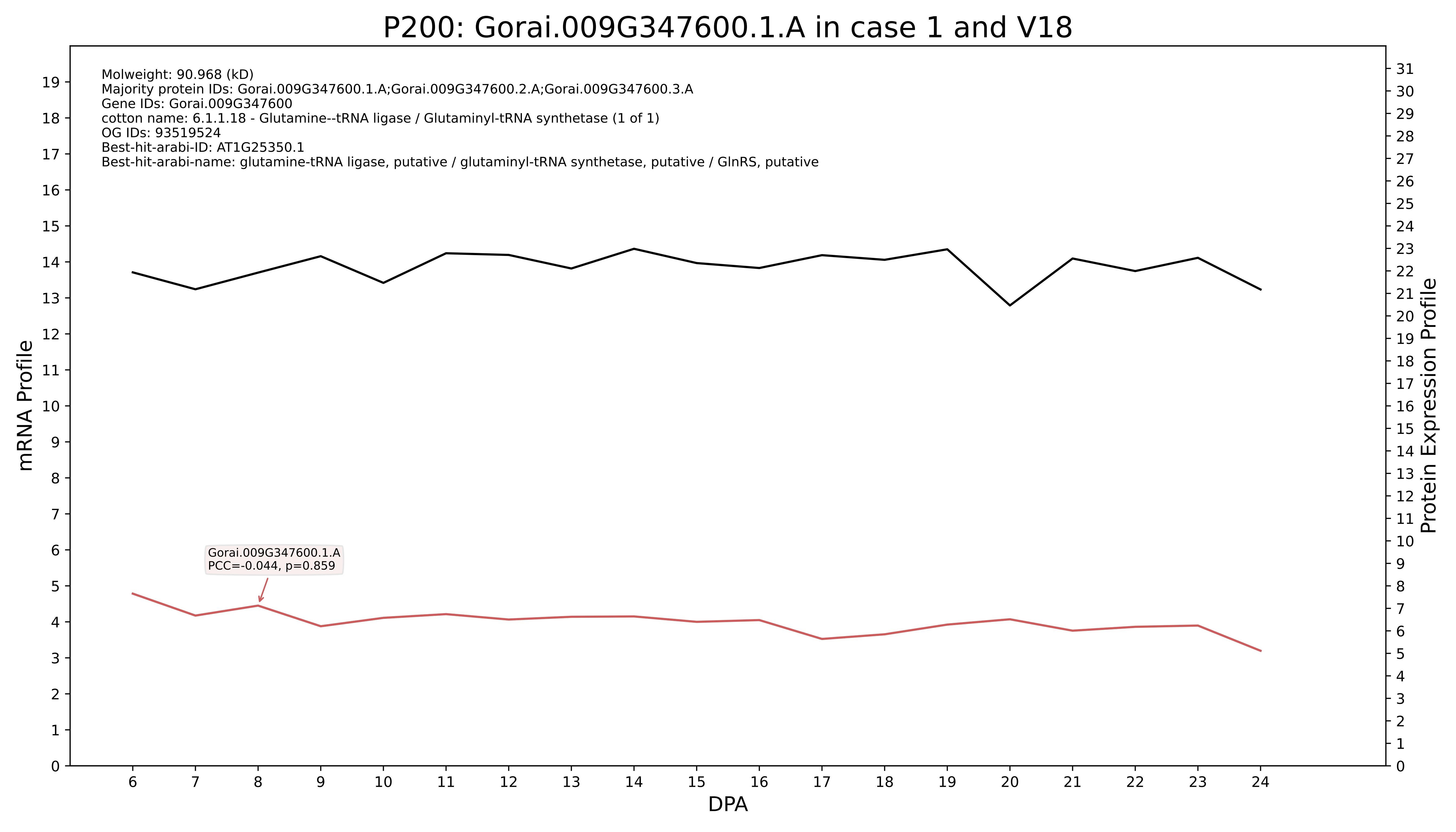
P200: Gorai.009G347600.1.A in case 1 and V18
Molweight: 90.968 (kD)
Majority protein IDs: Gorai.009G347600.1.A;Gorai.009G347600.2.A;Gorai.009G347600.3.A
Gene IDs: Gorai.009G347600
cotton name: 6.1.1.18 - Glutamine--tRNA ligase / Glutaminyl-tRNA synthetase (1 of 1)
OG IDs: 93519524
Best-hit-arabi-ID: AT1G25350.1
Best-hit-arabi-name: glutamine-tRNA ligase, putative / glutaminyl-tRNA synthetase, putative / GlnRS, putative
Gorai.009G347600.1.A
PCC=-0.044, p=0.859
mRNA Profile
Protein Expression Profile
DPA
0 1 2 3 4 5 6 7 8 9 10 11 12 13 14 15 16 17 18 19
0 1 2 3 4 5 6 7 8 9 10 11 12 13 14 15 16 17 18 19 20 21 22 23 24 25 26 27 28 29 30 31
6 7 8 9 10 11 12 13 14 15 16 17 18 19 20 21 22 23 24

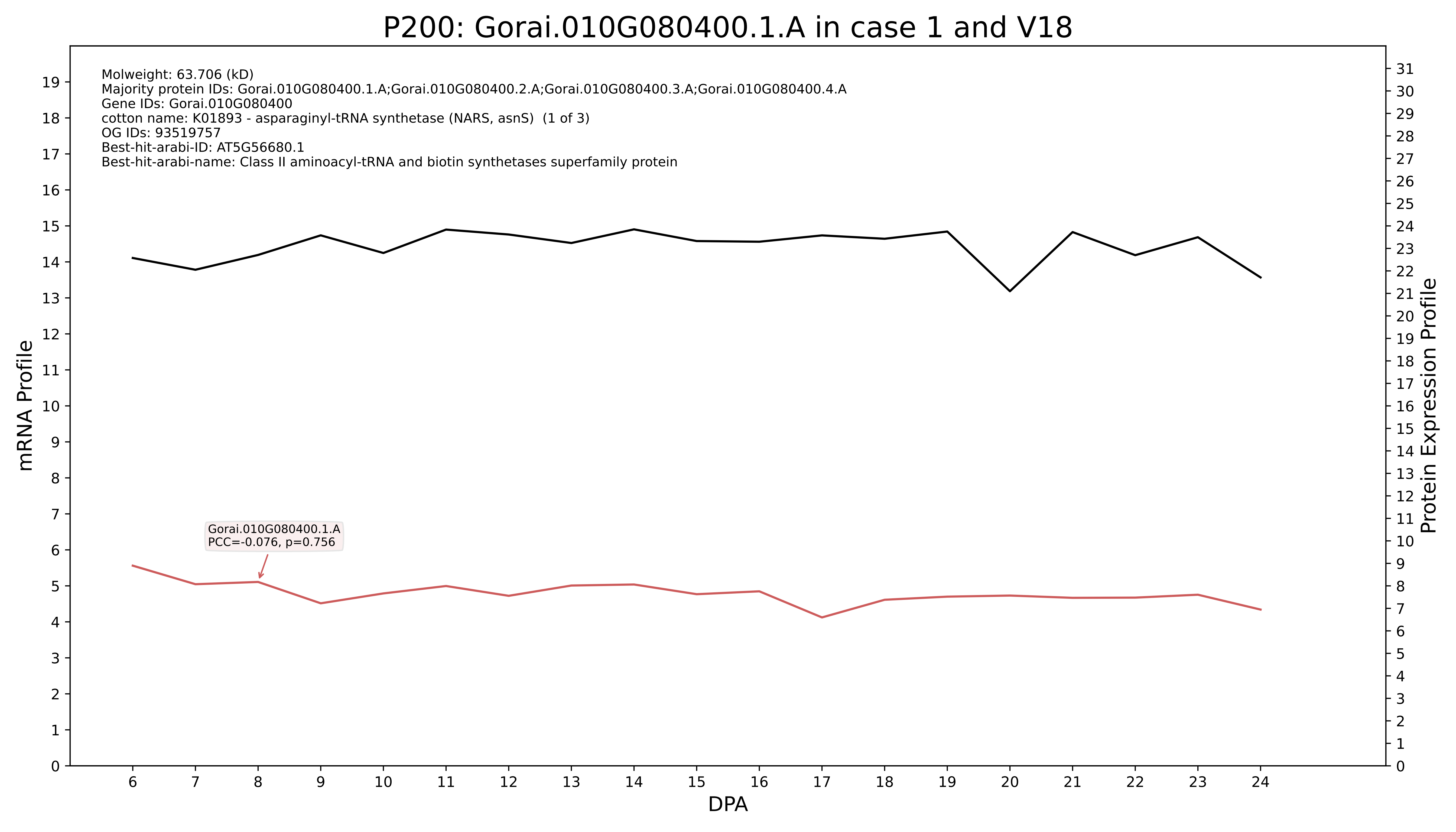
P200: Gorai.010G080400.1.A in case 1 and V18
Molweight: 63.706 (kD)
Majority protein IDs: Gorai.010G080400.1.A;Gorai.010G080400.2.A;Gorai.010G080400.3.A;Gorai.010G080400.4.A
Gene IDs: Gorai.010G080400
cotton name: K01893 - asparaginyl-tRNA synthetase (NARS, asnS) (1 of 3)
OG IDs: 93519757
Best-hit-arabi-ID: AT5G56680.1
Best-hit-arabi-name: Class II aminoacyl-tRNA and biotin synthetases superfamily protein
Gorai.010G080400.1.A
PCC=-0.076, p=0.756
mRNA Profile
Protein Expression Profile
DPA

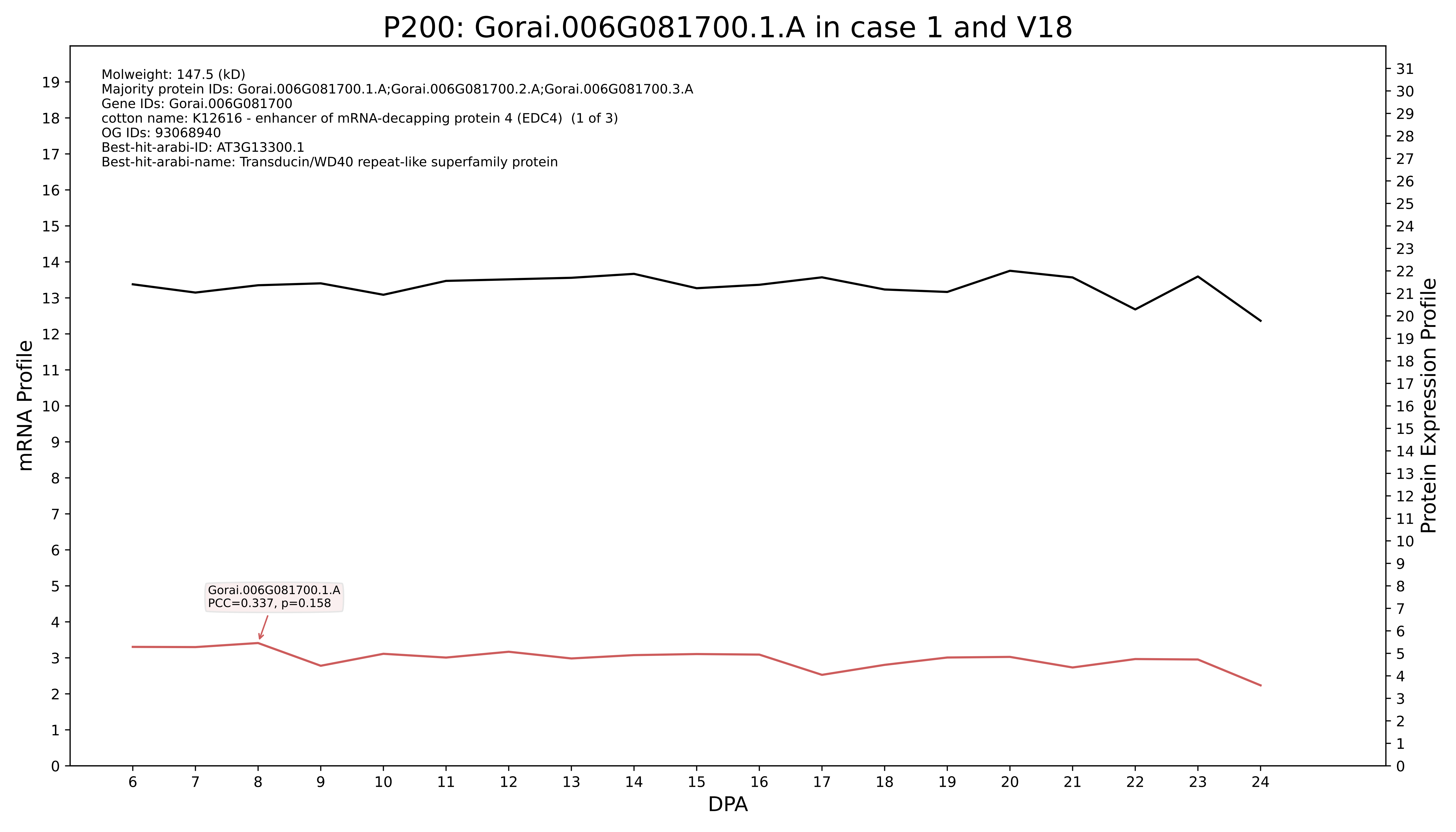
P200: Gorai.006G081700.1.A in case 1 and V18
Molweight: 147.5 (kD)
Majority protein IDs: Gorai.006G081700.1.A;Gorai.006G081700.2.A;Gorai.006G081700.3.A
Gene IDs: Gorai.006G081700
cotton name: K12616 - enhancer of mRNA-decapping protein 4 (EDC4) (1 of 3)
OG IDs: 93068940
Best-hit-arabi-ID: AT3G13300.1
Best-hit-arabi-name: Transducin/WD40 repeat-like superfamily protein
Gorai.006G081700.1.A
PCC=0.337, p=0.158
mRNA Profile
Protein Expression Profile
DPA

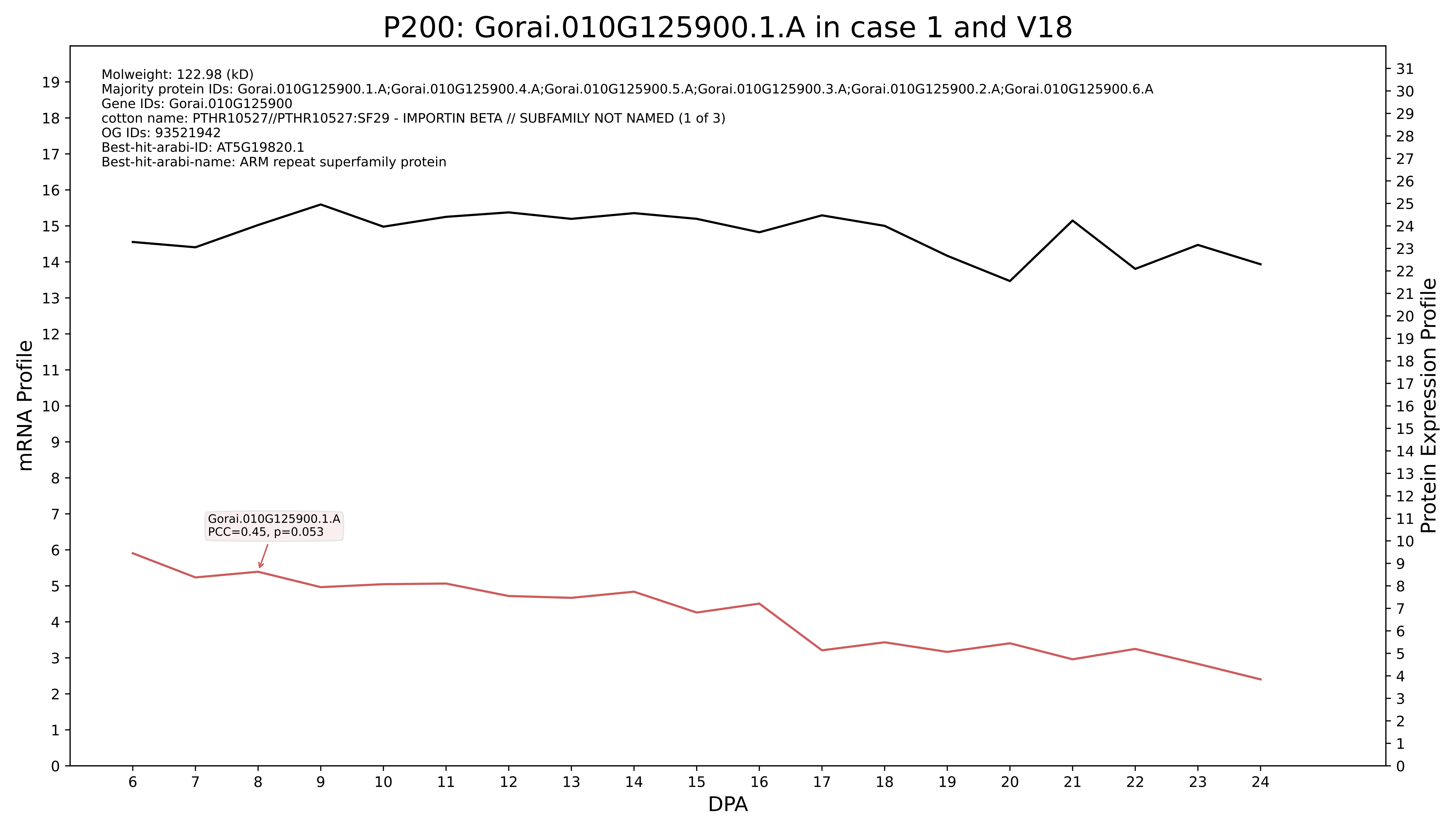

P200: Gorai.010G125900.1.A in case 1 and V18
Molweight: 122.98 (kD)
Majority protein IDs: Gorai.010G125900.1.A;Gorai.010G125900.4.A;Gorai.010G125900.5.A;Gorai.010G125900.3.A;Gorai.010G125900.2.A;Gorai.010G125900.6.A
Gene IDs: Gorai.010G125900
cotton name: PTHR10527//PTHR10527:SF29 - IMPORTIN BETA // SUBFAMILY NOT NAMED (1 of 3)
OG IDs: 93521942
Best-hit-arabi-ID: AT5G19820.1
Best-hit-arabi-name: ARM repeat superfamily protein
Gorai.010G125900.1.A
PCC=0.45, p=0.053
mRNA Profile
Protein Expression Profile
DPA

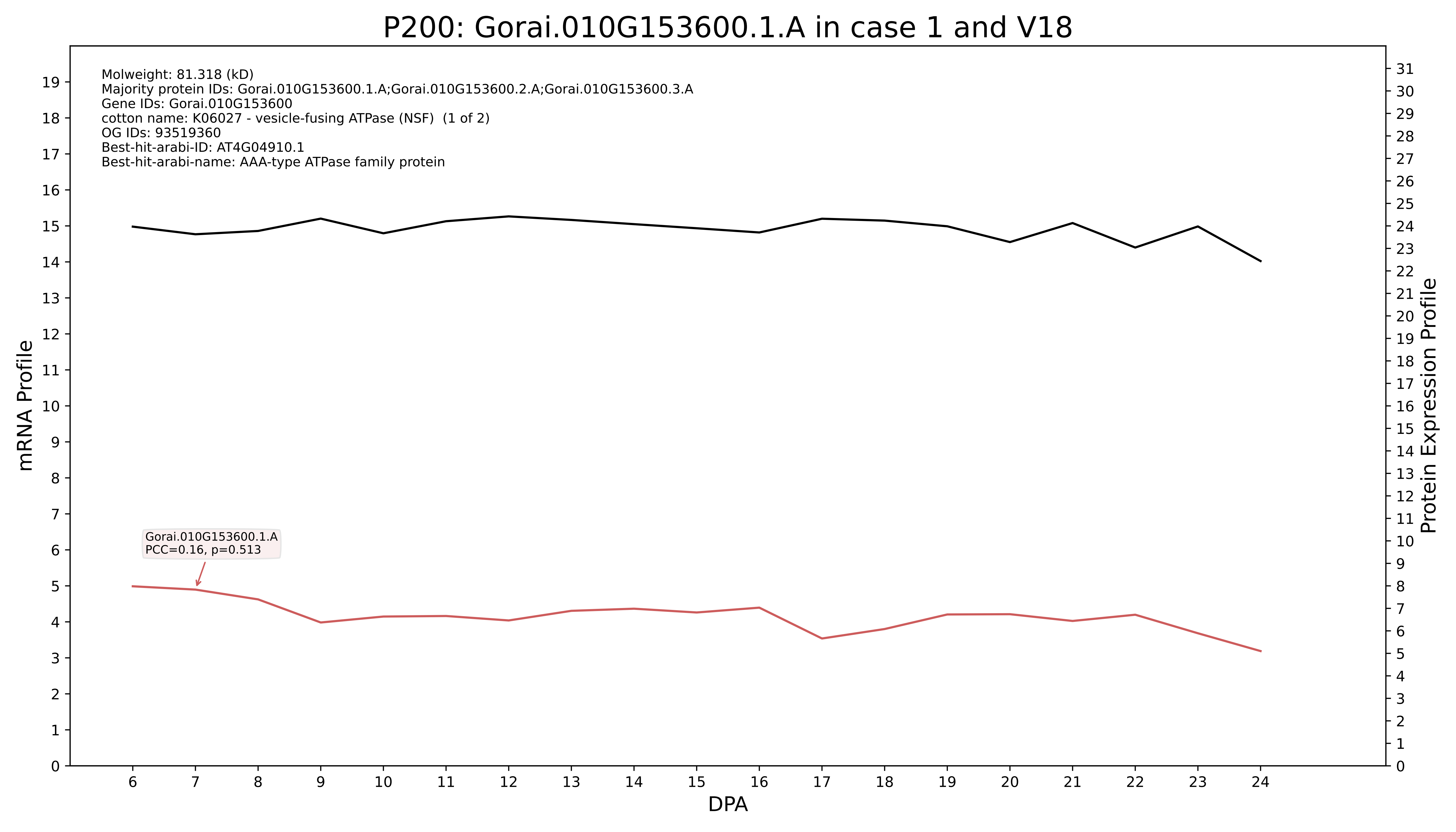

P200: Gorai.010G153600.1.A in case 1 and V18
Molweight: 81.318 (kD)
Majority protein IDs: Gorai.010G153600.1.A;Gorai.010G153600.2.A;Gorai.010G153600.3.A
Gene IDs: Gorai.010G153600
cotton name: K06027 - vesicle-fusing ATPase (NSF) (1 of 2)
OG IDs: 93519360
Best-hit-arabi-ID: AT4G04910.1
Best-hit-arabi-name: AAA-type ATPase family protein
Gorai.010G153600.1.A
PCC=0.16, p=0.513
mRNA Profile
Protein Expression Profile
DPA

### P200: Gorai.008G177900.1.D in case 1 and V18

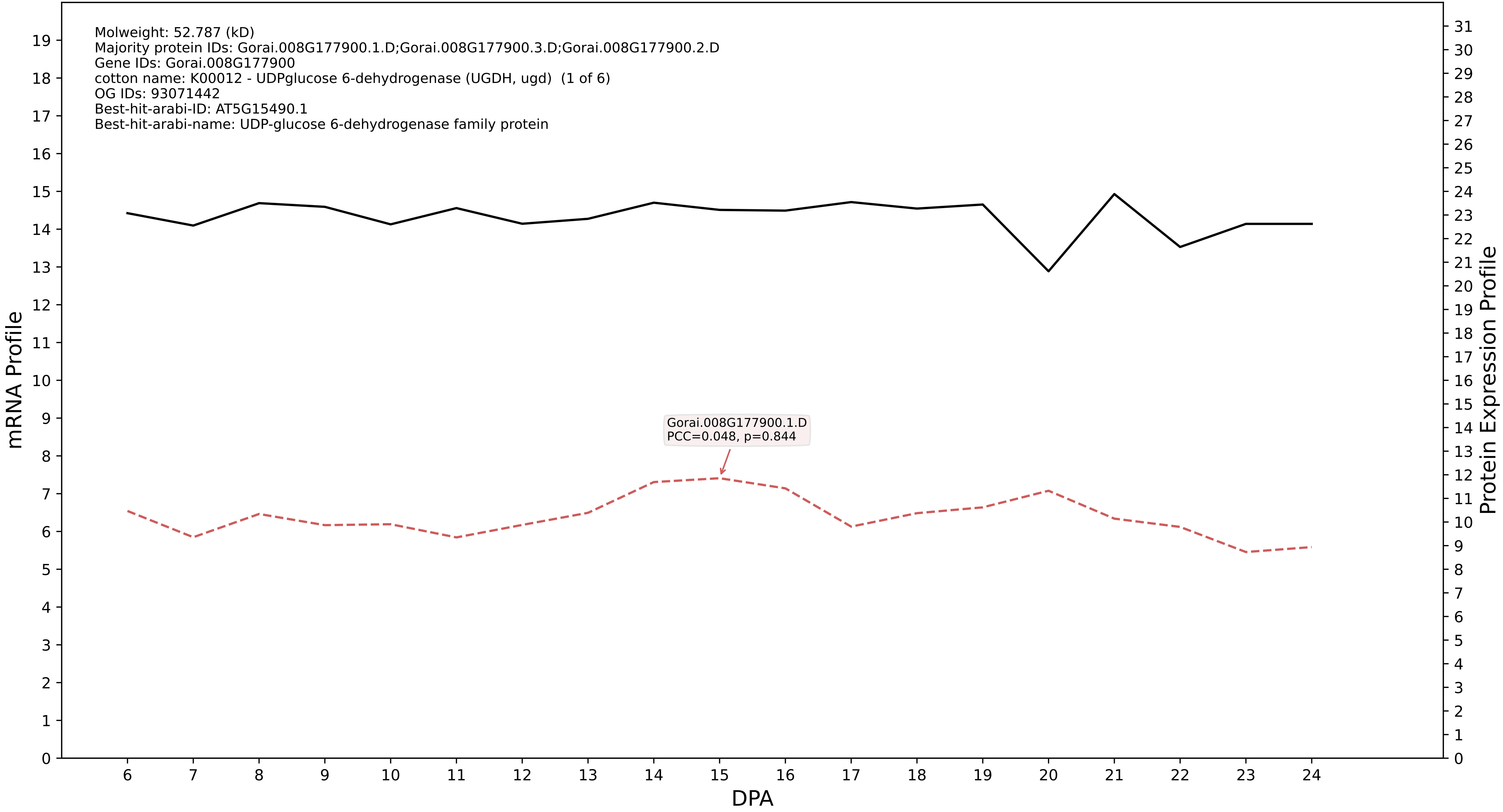

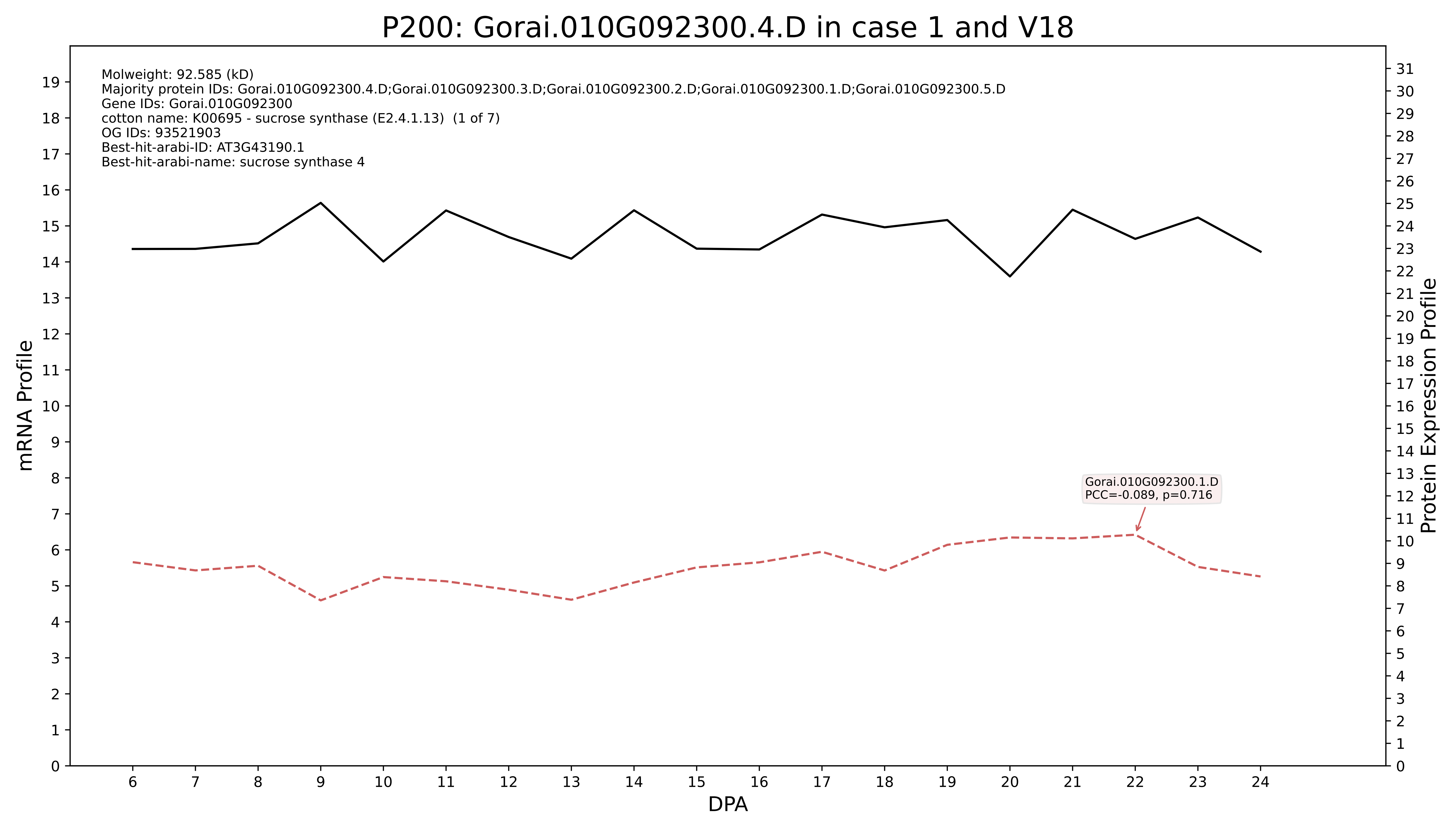
P200: Gorai.010G092300.4.D in case 1 and V18
Molweight: 92.585 (kD)
Majority protein IDs: Gorai.010G092300.4.D;Gorai.010G092300.3.D;Gorai.010G092300.2.D;Gorai.010G092300.1.D;Gorai.010G092300.5.D
Gene IDs: Gorai.010G092300
cotton name: K00695 - sucrose synthase (E2.4.1.13) (1 of 7)
OG IDs: 93521903
Best-hit-arabi-ID: AT3G43190.1
Best-hit-arabi-name: sucrose synthase 4
Gorai.010G092300.1.D
PCC=-0.089, p=0.716
mRNA Profile
Protein Expression Profile
DPA

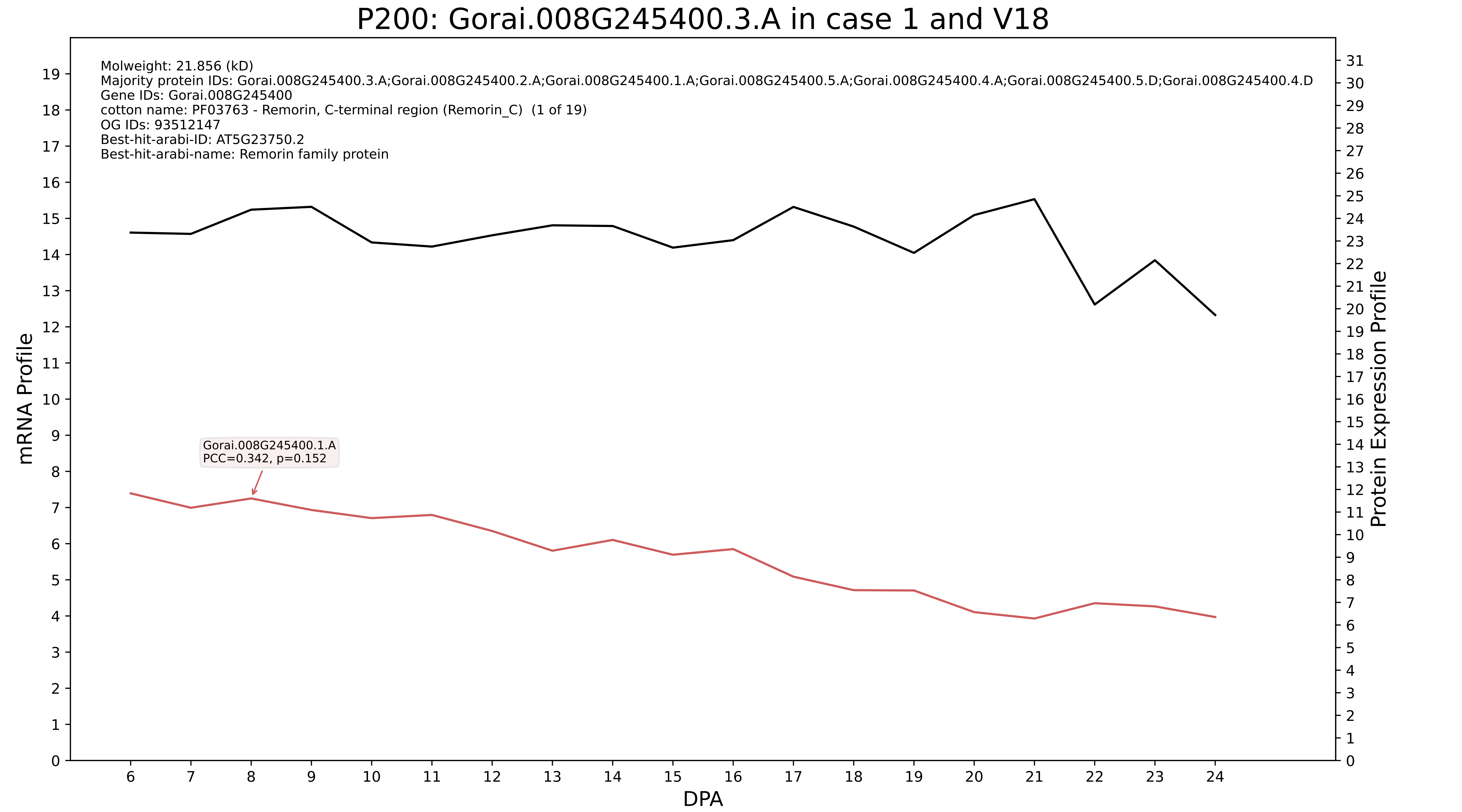

P200: Gorai.008G245400.3.A in case 1 and V18
Molweight: 21.856 (kD)
Majority protein IDs: Gorai.008G245400.3.A;Gorai.008G245400.2.A;Gorai.008G245400.1.A;Gorai.008G245400.5.A;Gorai.008G245400.4.A;Gorai.008G245400.5.D;Gorai.008G245400.4.D
Gene IDs: Gorai.008G245400
cotton name: PF03763 - Remorin, C-terminal region (Remorin_C) (1 of 19)
OG IDs: 93512147
Best-hit-arabi-ID: AT5G23750.2
Best-hit-arabi-name: Remorin family protein
Gorai.008G245400.1.A
PCC=0.342, p=0.152
mRNA Profile
Protein Expression Profile
DPA

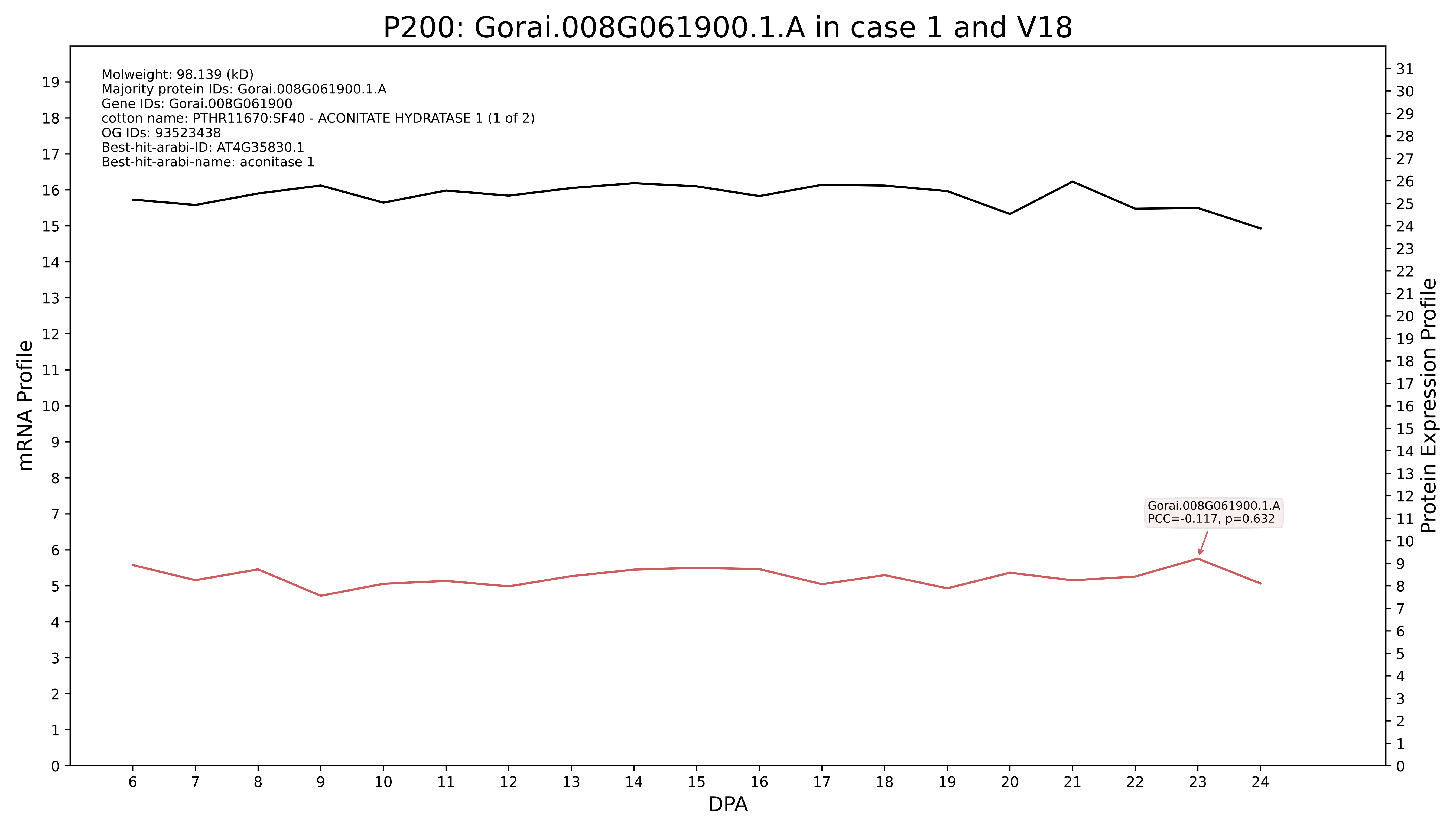
P200: Gorai.008G061900.1.A in case 1 and V18
Molweight: 98.139 (kD)
Majority protein IDs: Gorai.008G061900.1.A
Gene IDs: Gorai.008G061900
cotton name: PTHR11670:SF40 - ACONITATE HYDRATASE 1 (1 of 2)
OG IDs: 93523438
Best-hit-arabi-ID: AT4G35830.1
Best-hit-arabi-name: aconitase 1
Gorai.008G061900.1.A
PCC=-0.117, p=0.632
mRNA Profile
Protein Expression Profile
DPA

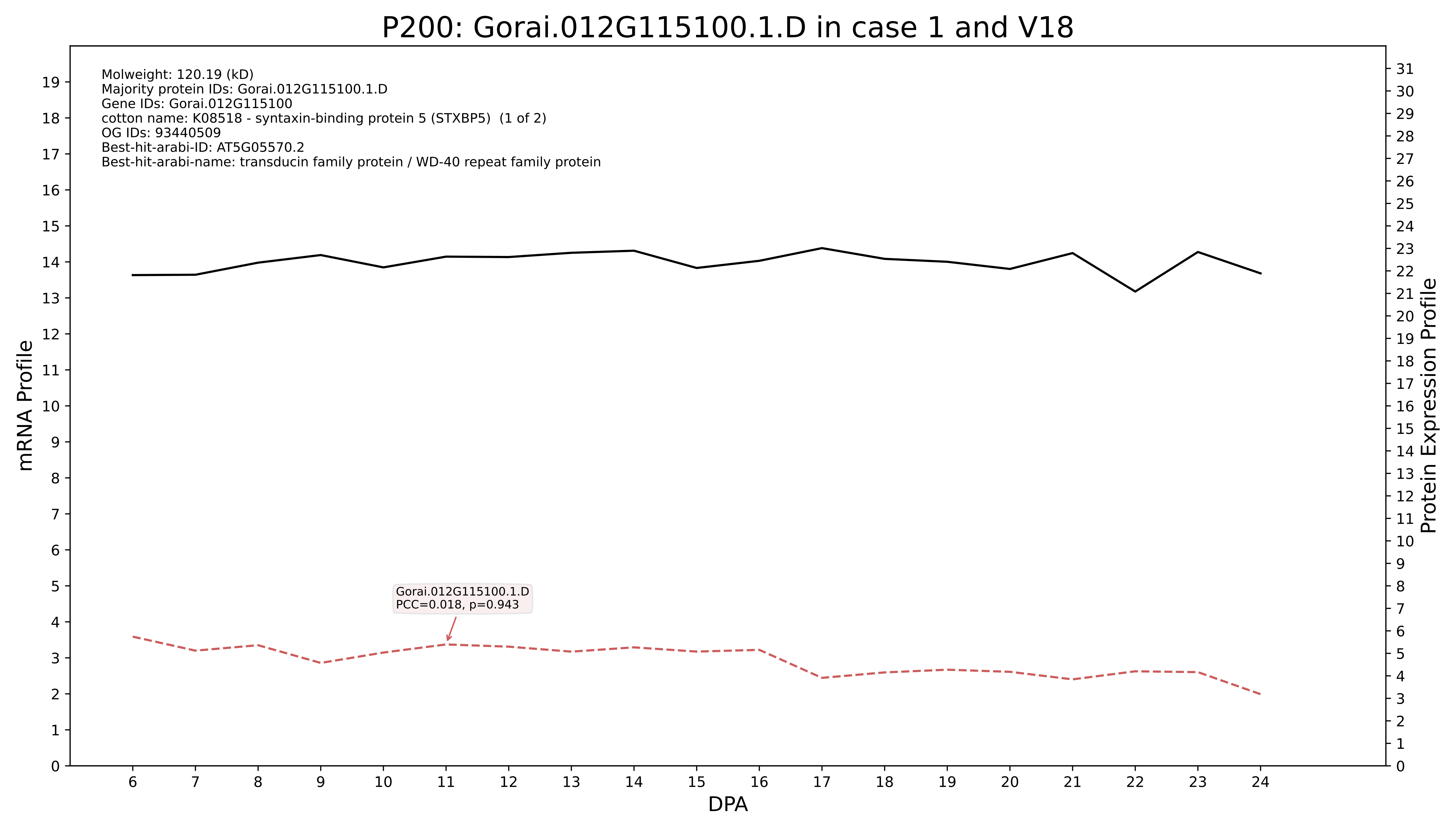

P200: Gorai.012G115100.1.D in case 1 and V18
Molweight: 120.19 (kD)
Majority protein IDs: Gorai.012G115100.1.D
Gene IDs: Gorai.012G115100
cotton name: K08518 - syntaxin-binding protein 5 (STXBP5) (1 of 2)
OG IDs: 93440509
Best-hit-arabi-ID: AT5G05570.2
Best-hit-arabi-name: transducin family protein / WD-40 repeat family protein
Gorai.012G115100.1.D
PCC=0.018, p=0.943
mRNA Profile
Protein Expression Profile
DPA

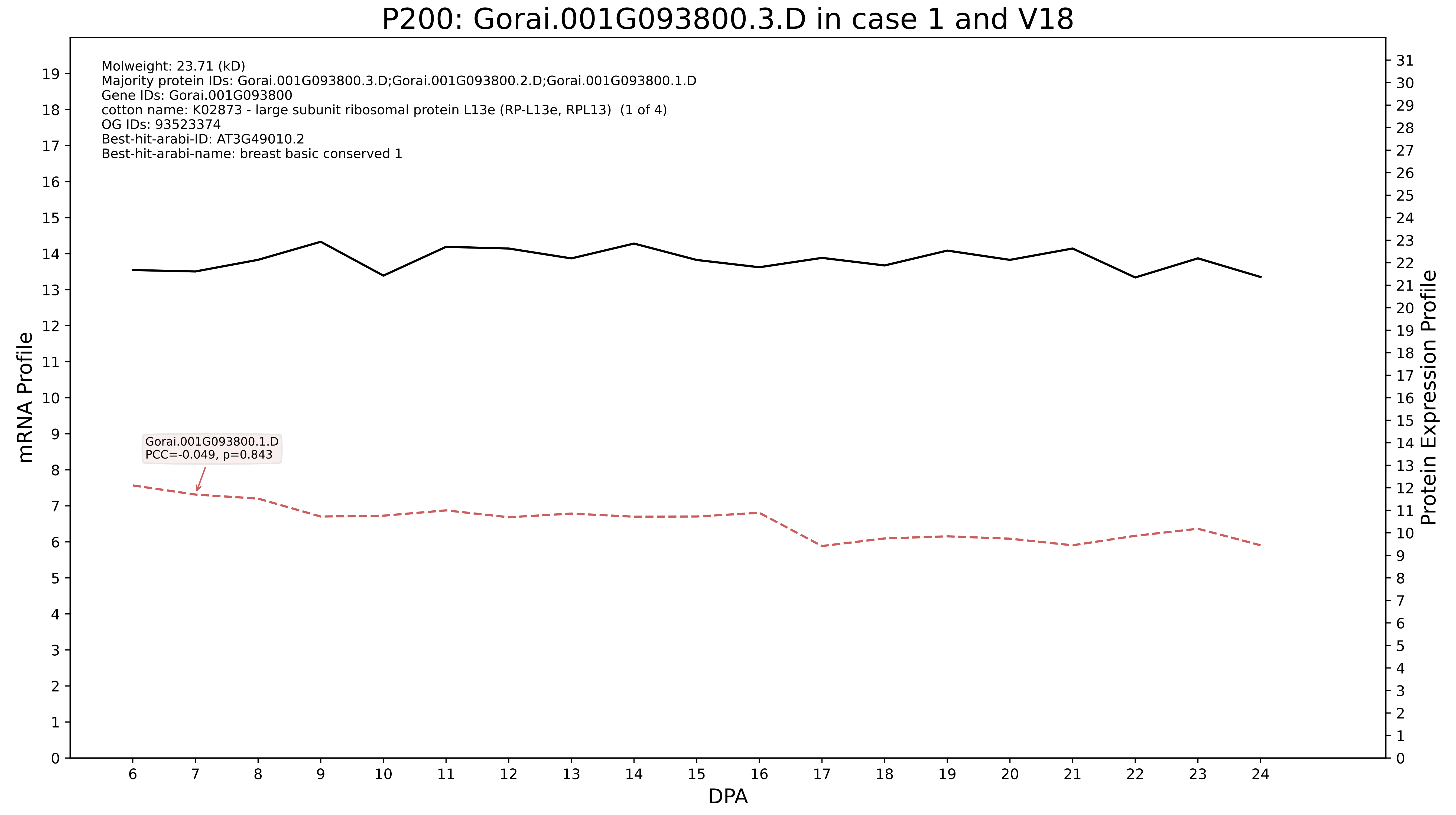
P200: Gorai.001G093800.3.D in case 1 and V18
Molweight: 23.71 (kD)
Majority protein IDs: Gorai.001G093800.3.D;Gorai.001G093800.2.D;Gorai.001G093800.1.D
Gene IDs: Gorai.001G093800
cotton name: K02873 - large subunit ribosomal protein L13e (RP-L13e, RPL13) (1 of 4)
OG IDs: 93523374
Best-hit-arabi-ID: AT3G49010.2
Best-hit-arabi-name: breast basic conserved 1
Gorai.001G093800.1.D
PCC=-0.049, p=0.843
mRNA Profile
Protein Expression Profile
DPA

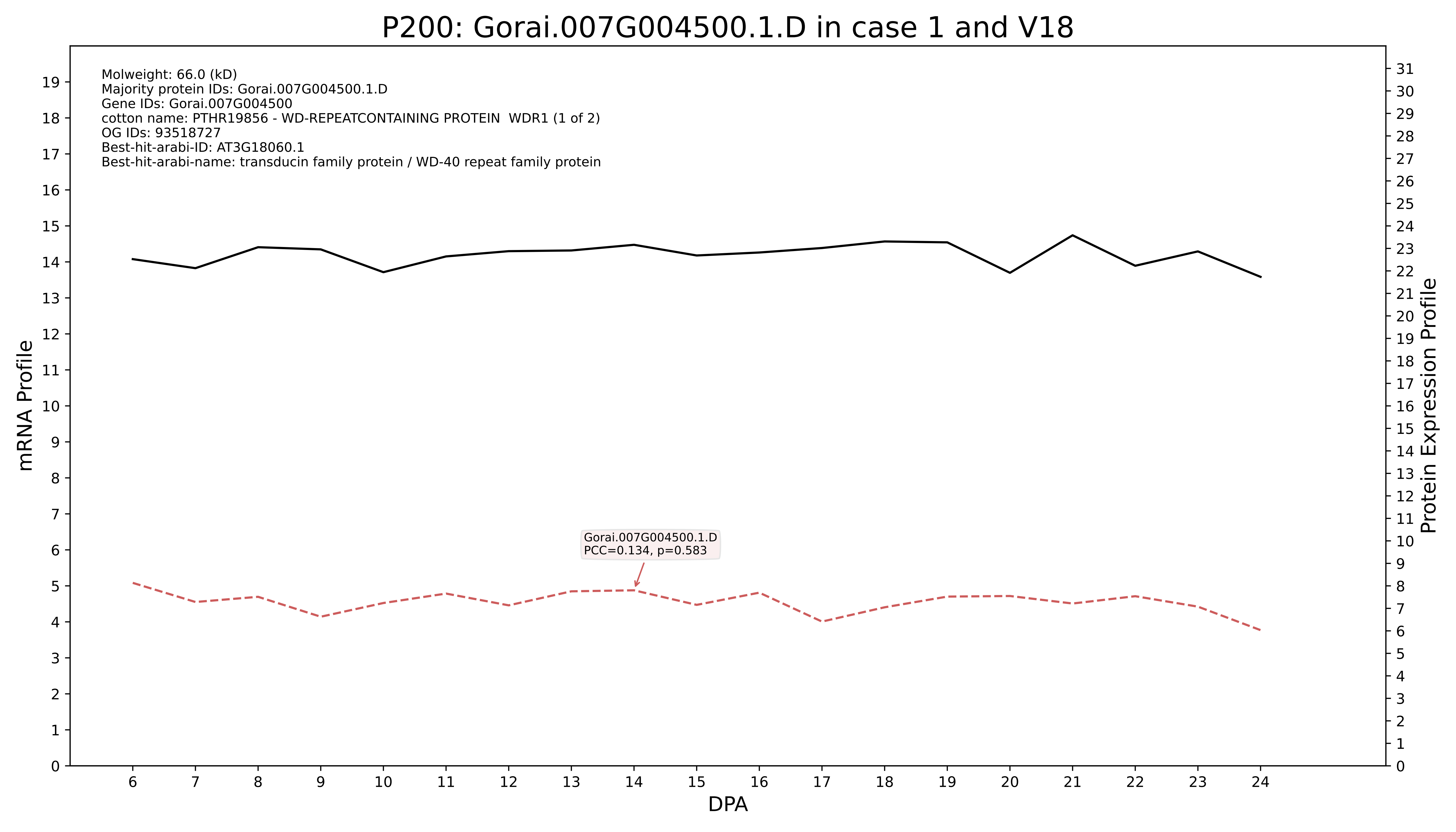
P200: Gorai.007G004500.1.D in case 1 and V18
Molweight: 66.0 (kD)
Majority protein IDs: Gorai.007G004500.1.D
Gene IDs: Gorai.007G004500
cotton name: PTHR19856 - WD-REPEATCONTAINING PROTEIN WDR1 (1 of 2)
OG IDs: 93518727
Best-hit-arabi-ID: AT3G18060.1
Best-hit-arabi-name: transducin family protein / WD-40 repeat family protein
Gorai.007G004500.1.D
PCC=0.134, p=0.583
mRNA Profile
Protein Expression Profile
DPA

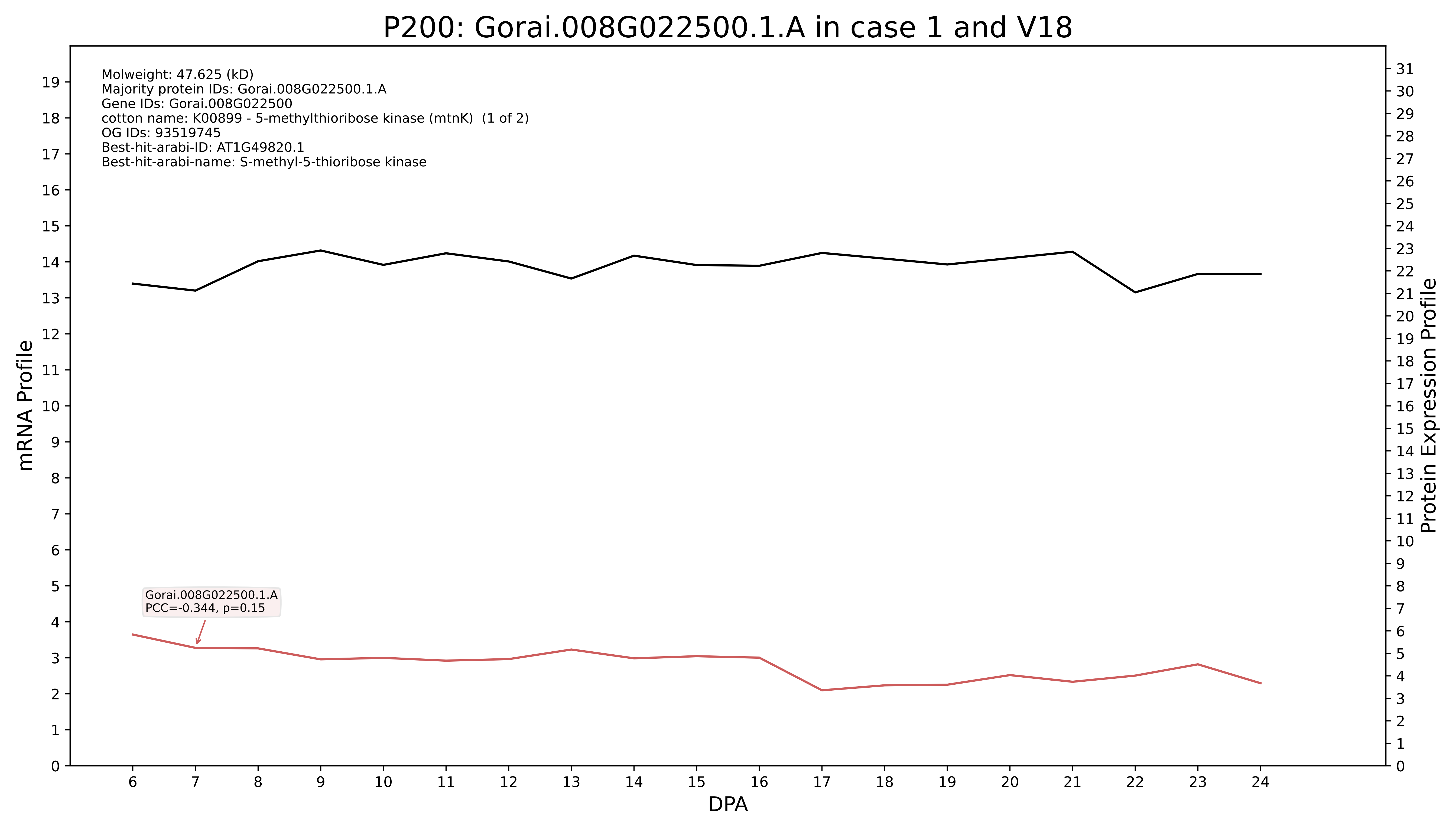
P200: Gorai.008G022500.1.A in case 1 and V18
Molweight: 47.625 (kD)
Majority protein IDs: Gorai.008G022500.1.A
Gene IDs: Gorai.008G022500
cotton name: K00899 - 5-methylthioribose kinase (mtnK) (1 of 2)
OG IDs: 93519745
Best-hit-arabi-ID: AT1G49820.1
Best-hit-arabi-name: S-methyl-5-thioribose kinase
Gorai.008G022500.1.A
PCC=-0.344, p=0.15
mRNA Profile
Protein Expression Profile
DPA

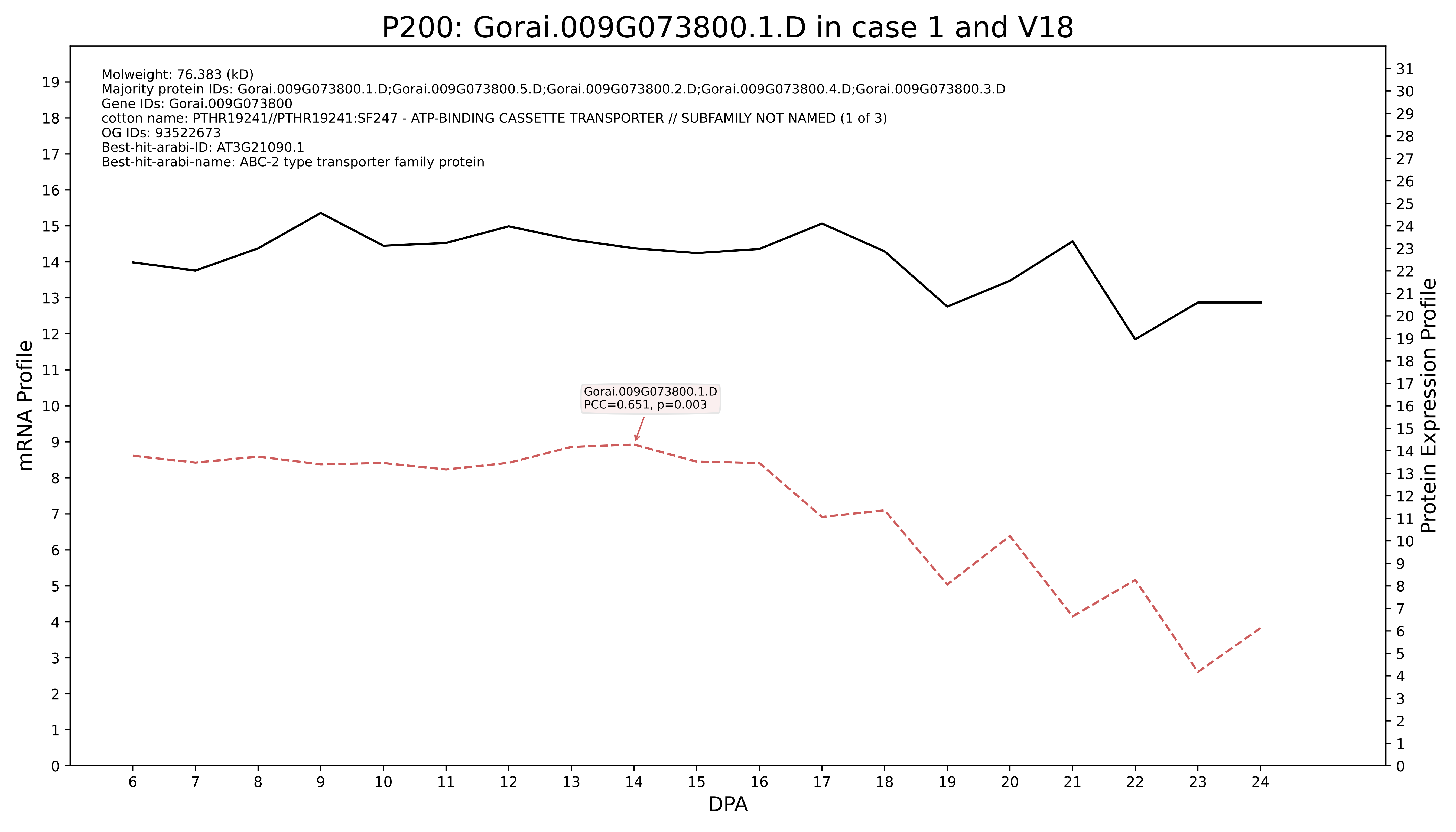
P200: Gorai.009G073800.1.D in case 1 and V18
Molweight: 76.383 (kD)
Majority protein IDs: Gorai.009G073800.1.D;Gorai.009G073800.5.D;Gorai.009G073800.2.D;Gorai.009G073800.4.D;Gorai.009G073800.3.D
Gene IDs: Gorai.009G073800
cotton name: PTHR19241//PTHR19241:SF247 - ATP-BINDING CASSETTE TRANSPORTER // SUBFAMILY NOT NAMED (1 of 3)
OG IDs: 93522673
Best-hit-arabi-ID: AT3G21090.1
Best-hit-arabi-name: ABC-2 type transporter family protein
Gorai.009G073800.1.D
PCC=0.651, p=0.003
mRNA Profile
Protein Expression Profile
DPA

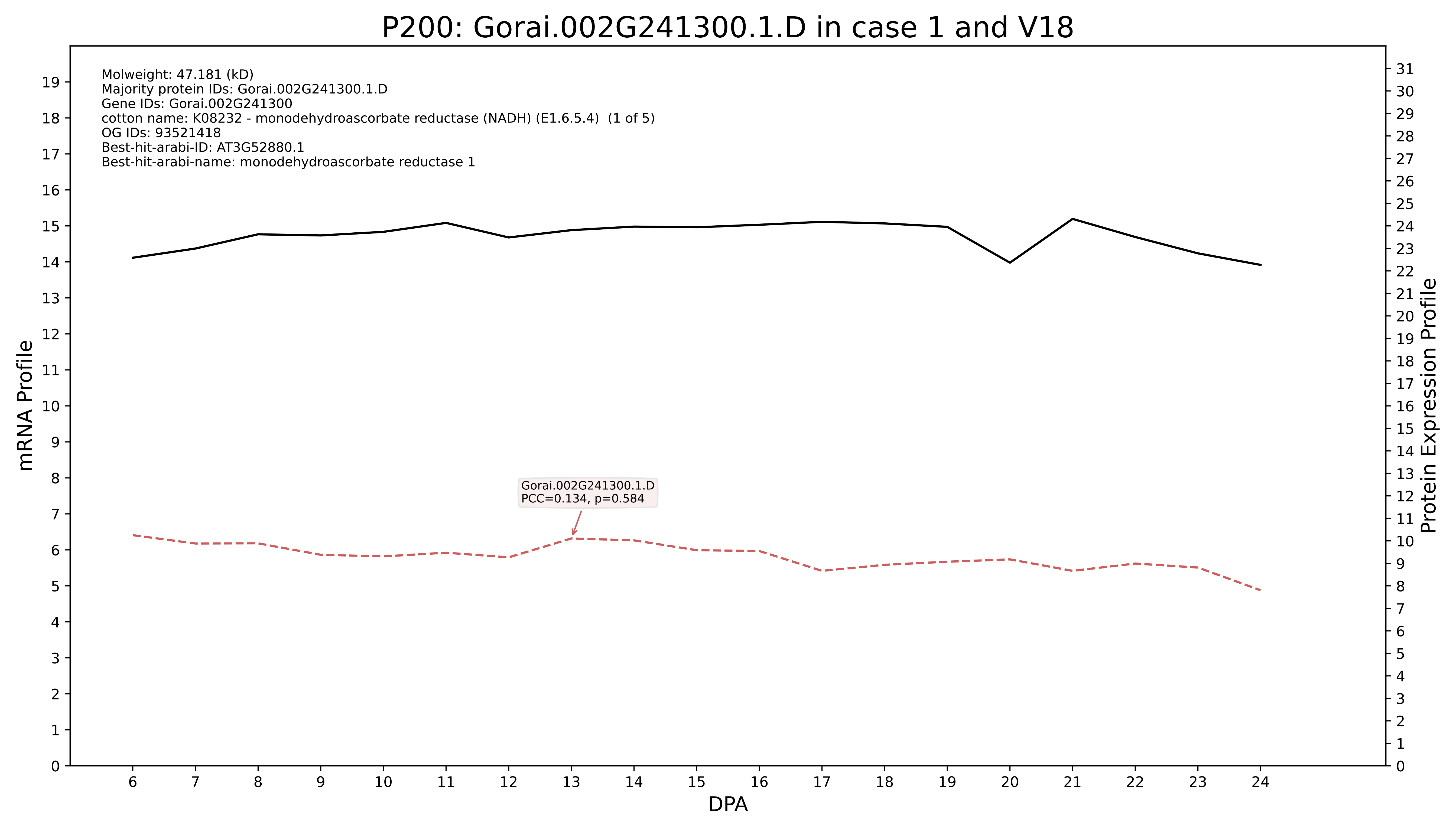

P200: Gorai.002G241300.1.D in case 1 and V18
Molweight: 47.181 (kD)
Majority protein IDs: Gorai.002G241300.1.D
Gene IDs: Gorai.002G241300
cotton name: K08232 - monodehydroascorbate reductase (NADH) (E1.6.5.4) (1 of 5)
OG IDs: 93521418
Best-hit-arabi-ID: AT3G52880.1
Best-hit-arabi-name: monodehydroascorbate reductase 1
Gorai.002G241300.1.D
PCC=0.134, p=0.584
mRNA Profile
Protein Expression Profile
DPA

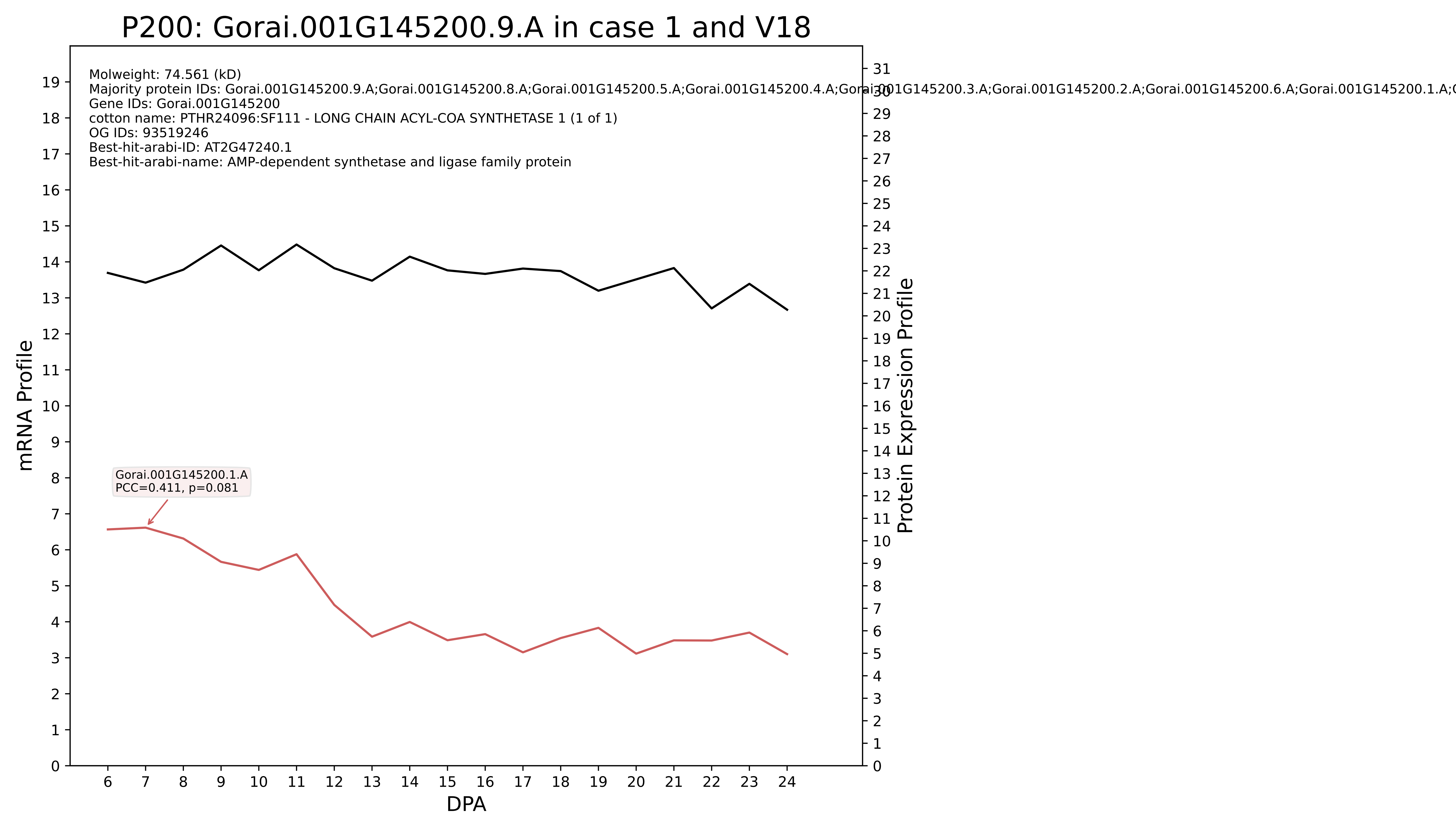

P200: Gorai.001G145200.9.A in case 1 and V18
Molweight: 74.561 (kD)
Majority protein IDs: Gorai.001G145200.9.A;Gorai.001G145200.8.A;Gorai.001G145200.5.A;Gorai.001G145200.4.A;Gorai.001G145200.3.A;Gorai.001G145200.2.A;Gorai.001G145200.6.A;Gorai.001G145200.1.A;G
Gene IDs: Gorai.001G145200
cotton name: PTHR24096:SF111 - LONG CHAIN ACYL-COA SYNTHETASE 1 (1 of 1)
OG IDs: 93519246
Best-hit-arabi-ID: AT2G47240.1
Best-hit-arabi-name: AMP-dependent synthetase and ligase family protein
Gorai.001G145200.1.A
PCC=0.411, p=0.081
mRNA Profile
Protein Expression Profile
DPA

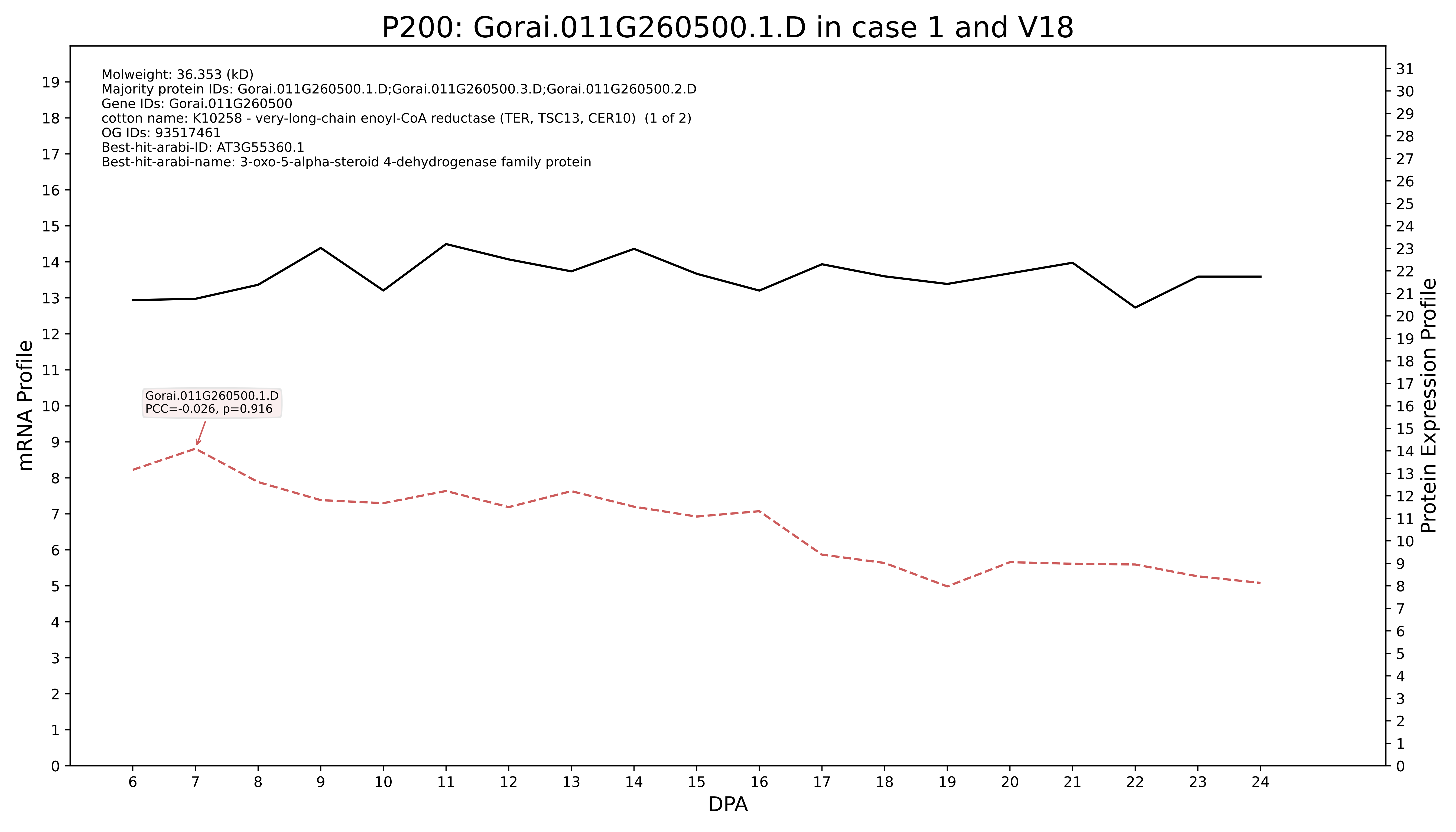
P200: Gorai.011G260500.1.D in case 1 and V18
Molweight: 36.353 (kD)
Majority protein IDs: Gorai.011G260500.1.D;Gorai.011G260500.3.D;Gorai.011G260500.2.D
Gene IDs: Gorai.011G260500
cotton name: K10258 - very-long-chain enoyl-CoA reductase (TER, TSC13, CER10) (1 of 2)
OG IDs: 93517461
Best-hit-arabi-ID: AT3G55360.1
Best-hit-arabi-name: 3-oxo-5-alpha-steroid 4-dehydrogenase family protein
Gorai.011G260500.1.D
PCC=-0.026, p=0.916
mRNA Profile
Protein Expression Profile
DPA

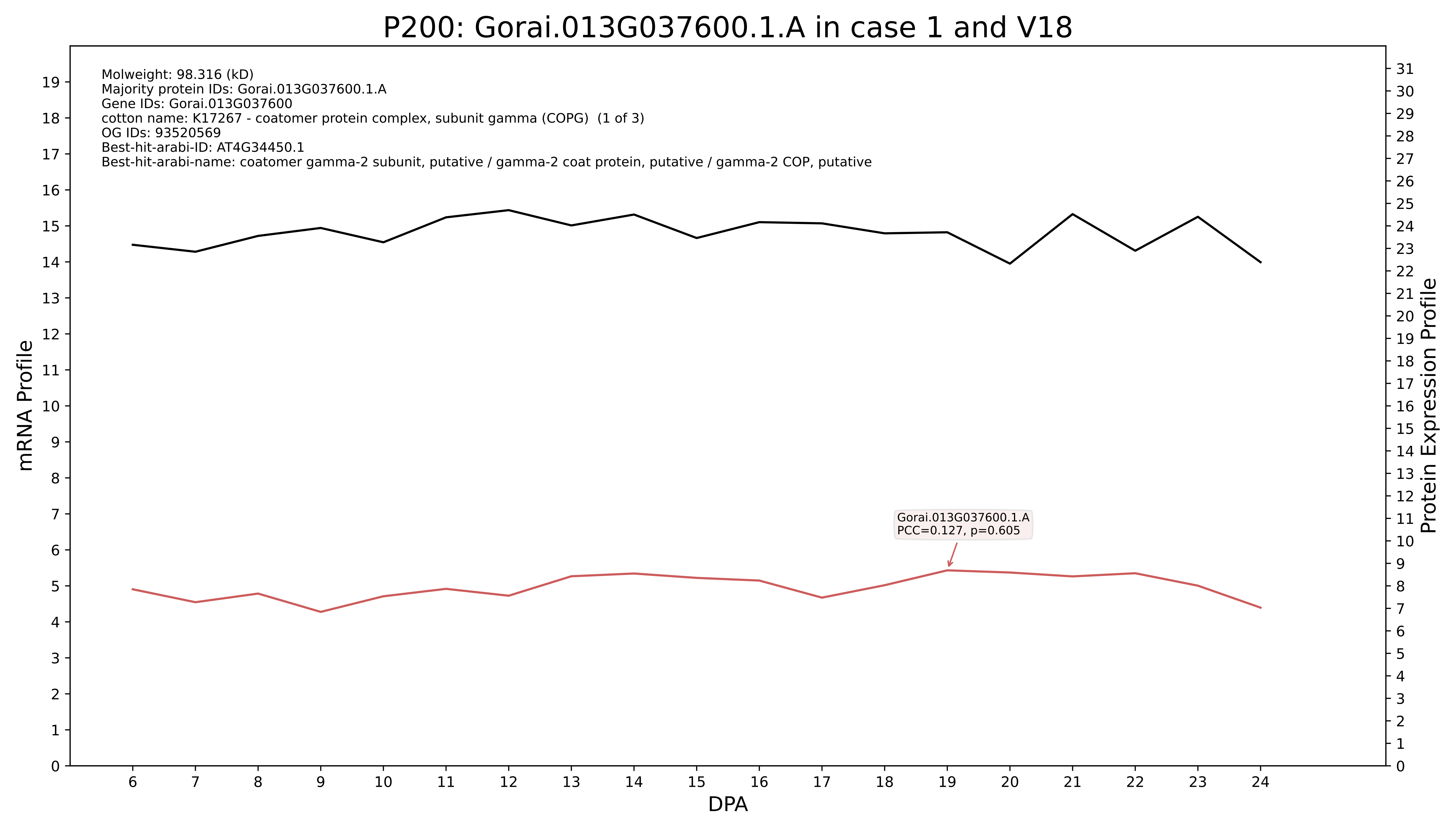
P200: Gorai.013G037600.1.A in case 1 and V18
Molweight: 98.316 (kD)
Majority protein IDs: Gorai.013G037600.1.A
Gene IDs: Gorai.013G037600
cotton name: K17267 - coatomer protein complex, subunit gamma (COPG) (1 of 3)
OG IDs: 93520569
Best-hit-arabi-ID: AT4G34450.1
Best-hit-arabi-name: coatomer gamma-2 subunit, putative / gamma-2 coat protein, putative / gamma-2 COP, putative
Gorai.013G037600.1.A
PCC=0.127, p=0.605
mRNA Profile
Protein Expression Profile
DPA

P200: Gorai.006G073400.1.A in case 1 and V18

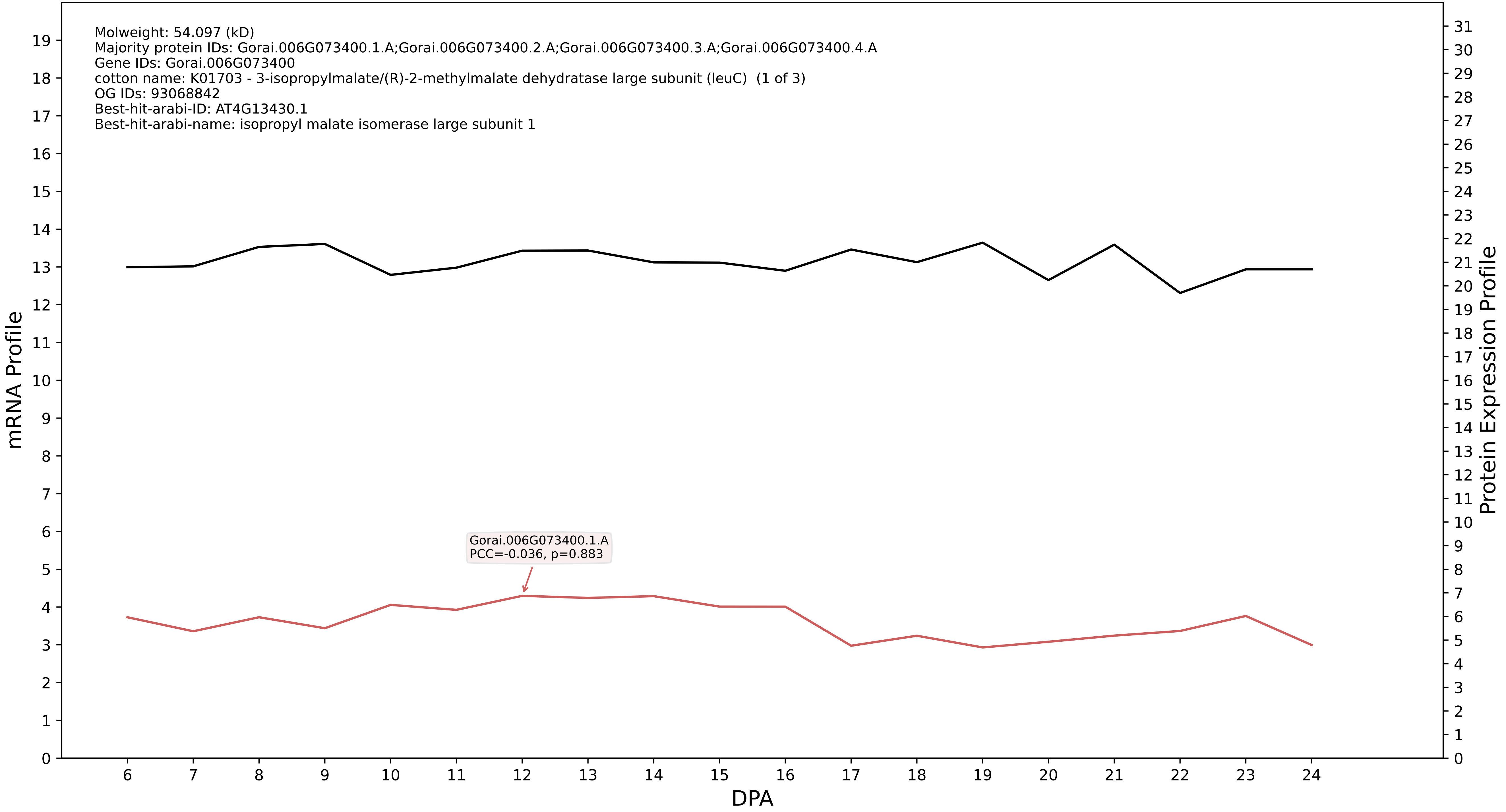

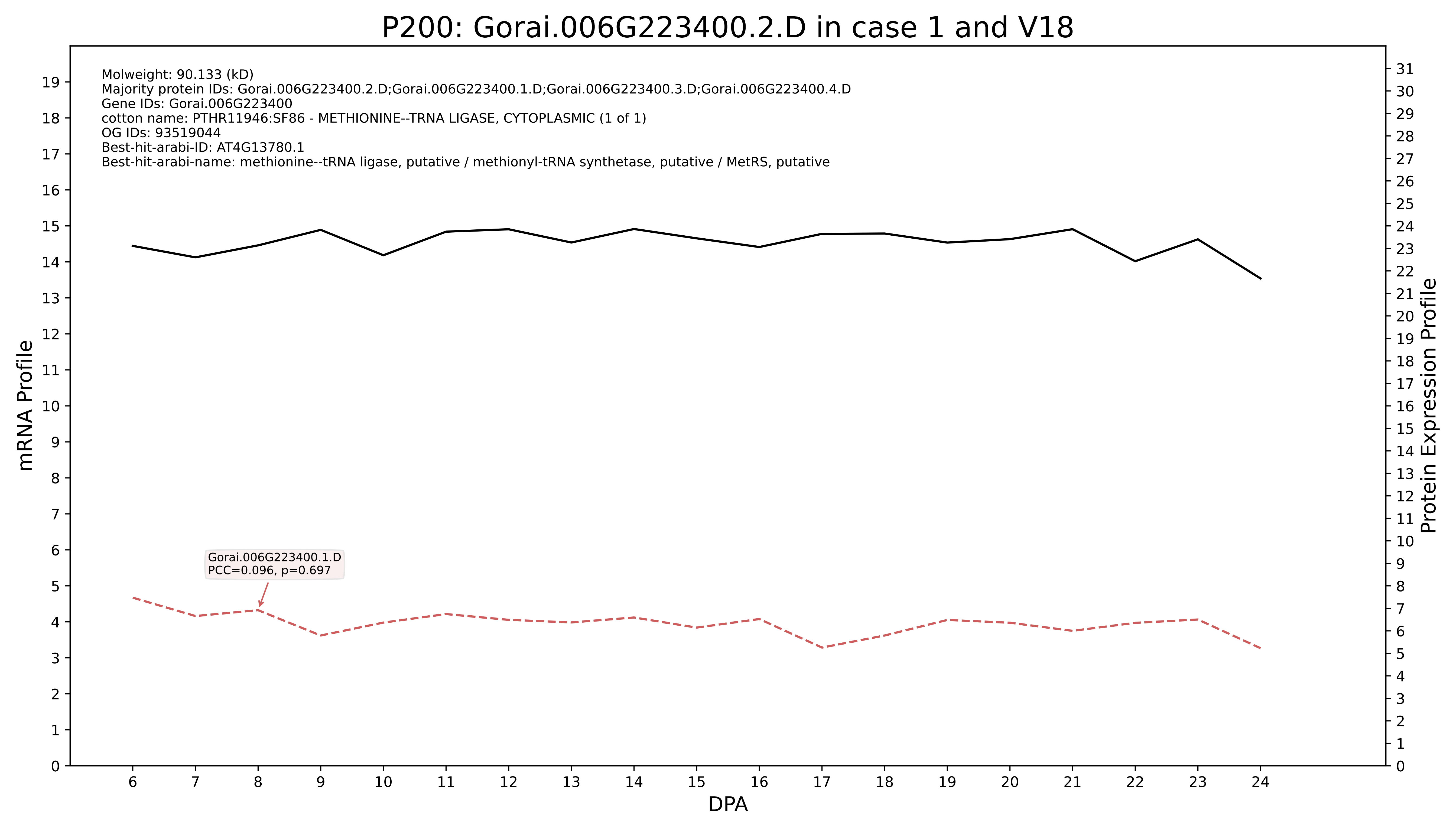
P200: Gorai.006G223400.2.D in case 1 and V18
Molweight: 90.133 (kD)
Majority protein IDs: Gorai.006G223400.2.D;Gorai.006G223400.1.D;Gorai.006G223400.3.D;Gorai.006G223400.4.D
Gene IDs: Gorai.006G223400
cotton name: PTHR11946:SF86 - METHIONINE--TRNA LIGASE, CYTOPLASMIC (1 of 1)
OG IDs: 93519044
Best-hit-arabi-ID: AT4G13780.1
Best-hit-arabi-name: methionine--tRNA ligase, putative / methionyl-tRNA synthetase, putative / MetRS, putative
Gorai.006G223400.1.D
PCC=0.096, p=0.697
mRNA Profile
Protein Expression Profile
DPA

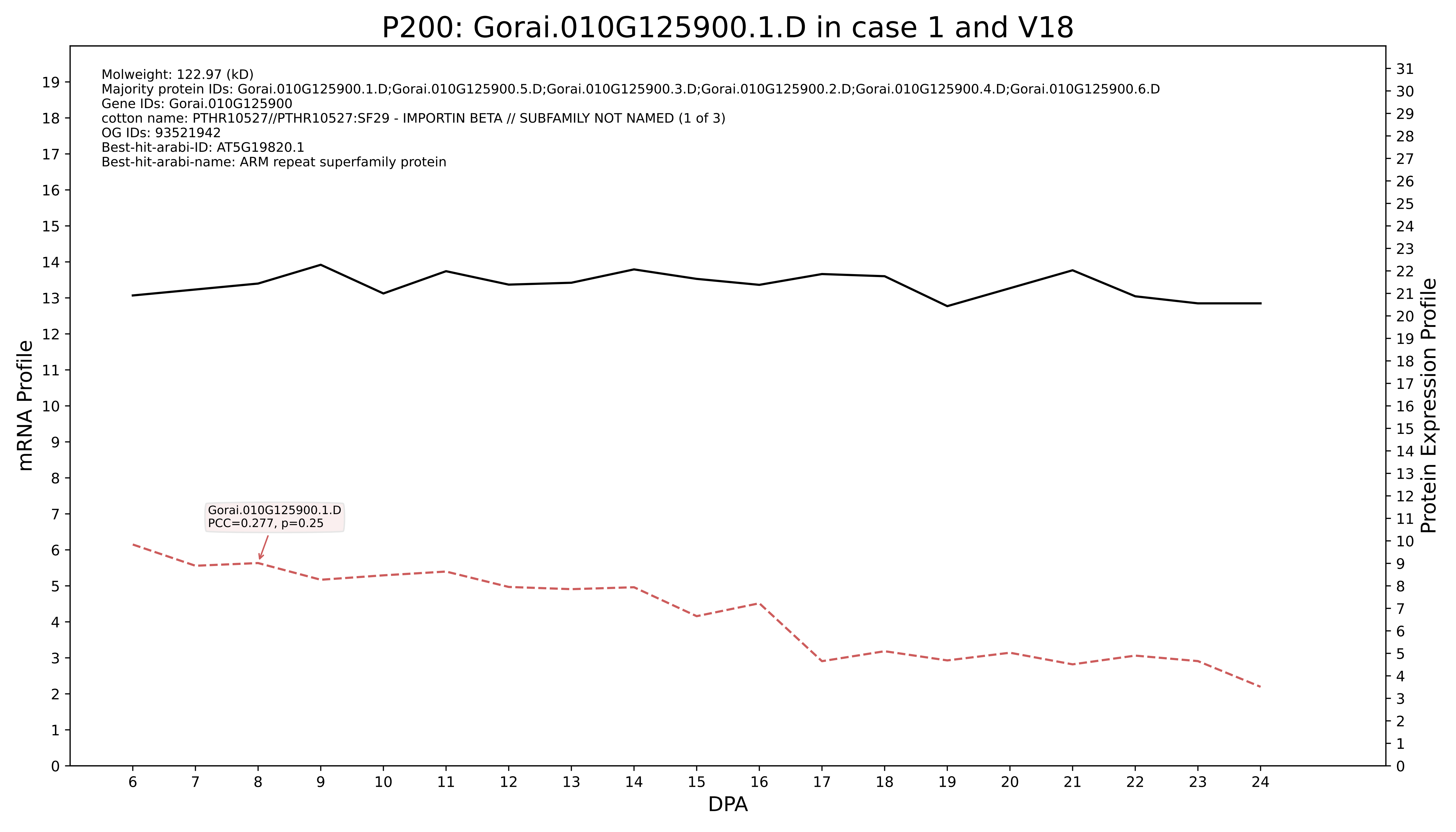
P200: Gorai.010G125900.1.D in case 1 and V18
Molweight: 122.97 (kD)
Majority protein IDs: Gorai.010G125900.1.D;Gorai.010G125900.5.D;Gorai.010G125900.3.D;Gorai.010G125900.2.D;Gorai.010G125900.4.D;Gorai.010G125900.6.D
Gene IDs: Gorai.010G125900
cotton name: PTHR10527//PTHR10527:SF29 - IMPORTIN BETA // SUBFAMILY NOT NAMED (1 of 3)
OG IDs: 93521942
Best-hit-arabi-ID: AT5G19820.1
Best-hit-arabi-name: ARM repeat superfamily protein
Gorai.010G125900.1.D
PCC=0.277, p=0.25
mRNA Profile
Protein Expression Profile
DPA

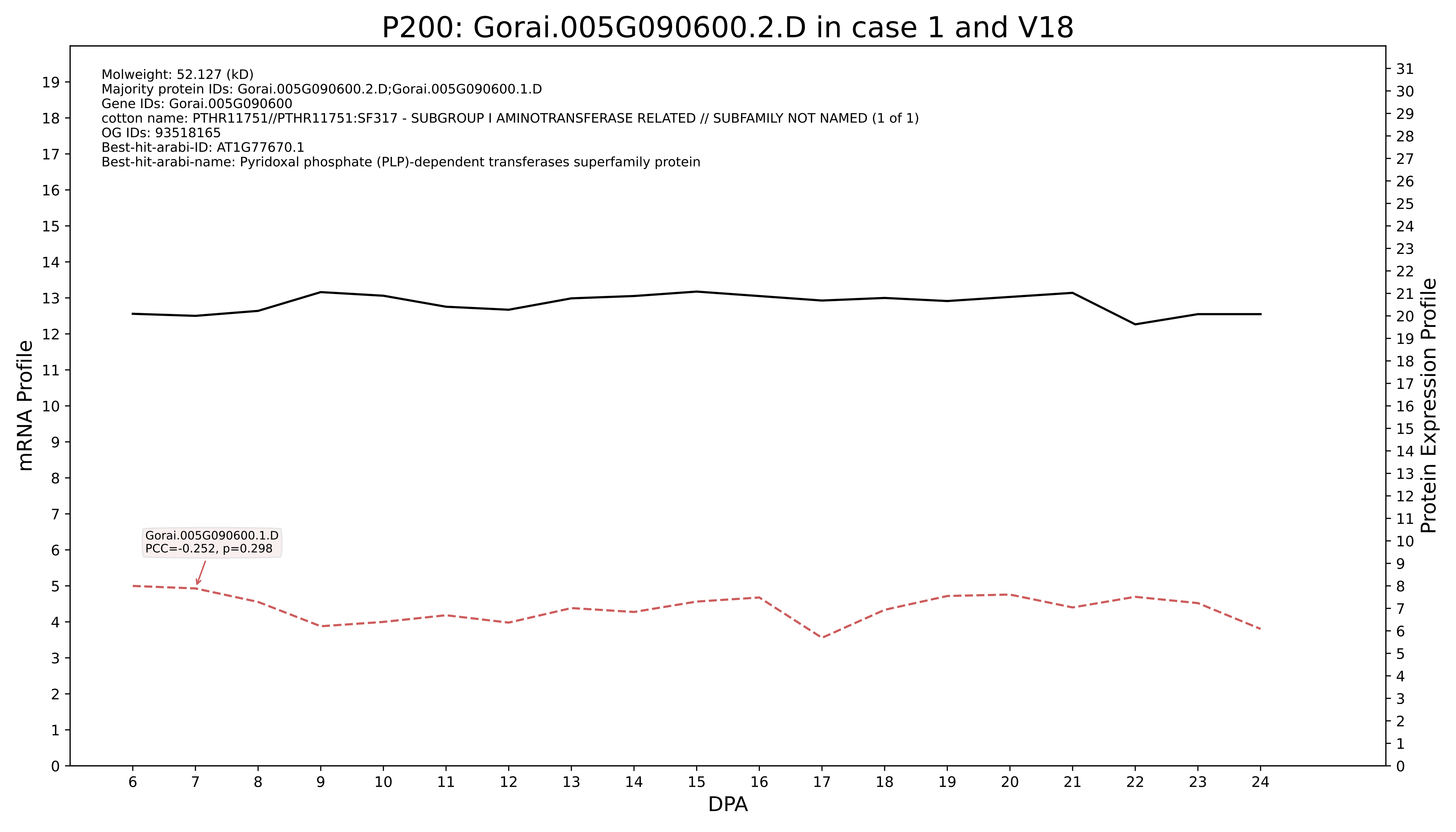
P200: Gorai.005G090600.2.D in case 1 and V18
Molweight: 52.127 (kD)
Majority protein IDs: Gorai.005G090600.2.D;Gorai.005G090600.1.D
Gene IDs: Gorai.005G090600
cotton name: PTHR11751//PTHR11751:SF317 - SUBGROUP I AMINOTRANSFERASE RELATED // SUBFAMILY NOT NAMED (1 of 1)
OG IDs: 93518165
Best-hit-arabi-ID: AT1G77670.1
Best-hit-arabi-name: Pyridoxal phosphate (PLP)-dependent transferases superfamily protein
Gorai.005G090600.1.D
PCC=-0.252, p=0.298
mRNA Profile
Protein Expression Profile
DPA

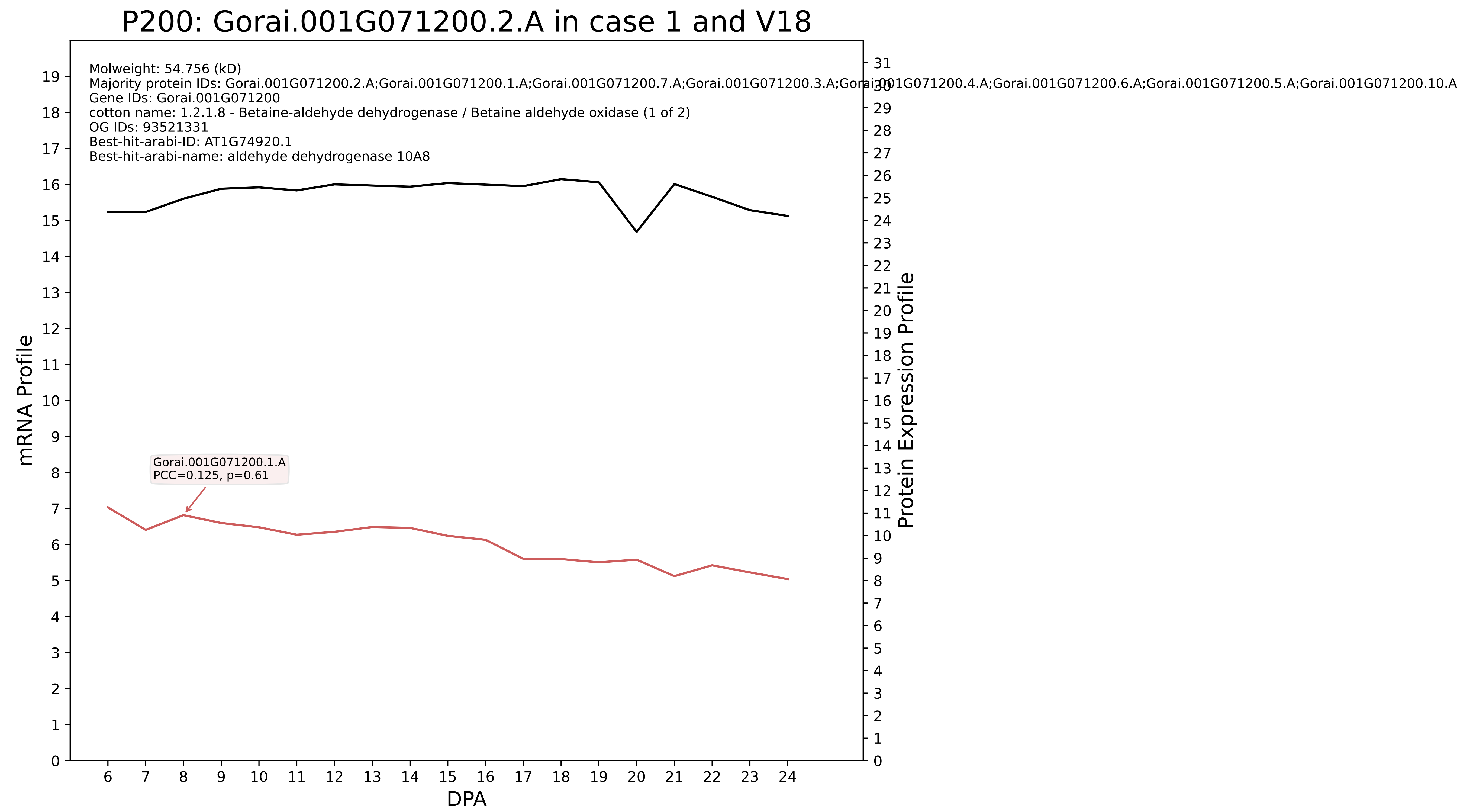

P200: Gorai.001G071200.2.A in case 1 and V18
Molweight: 54.756 (kD)
Majority protein IDs: Gorai.001G071200.2.A;Gorai.001G071200.1.A;Gorai.001G071200.7.A;Gorai.001G071200.3.A;Gorai.001G071200.4.A;Gorai.001G071200.6.A;Gorai.001G071200.5.A;Gorai.001G071200.10.A
Gene IDs: Gorai.001G071200
cotton name: 1.2.1.8 - Betaine-aldehyde dehydrogenase / Betaine aldehyde oxidase (1 of 2)
OG IDs: 93521331
Best-hit-arabi-ID: AT1G74920.1
Best-hit-arabi-name: aldehyde dehydrogenase 10A8
Gorai.001G071200.1.A
PCC=0.125, p=0.61
mRNA Profile
Protein Expression Profile
DPA

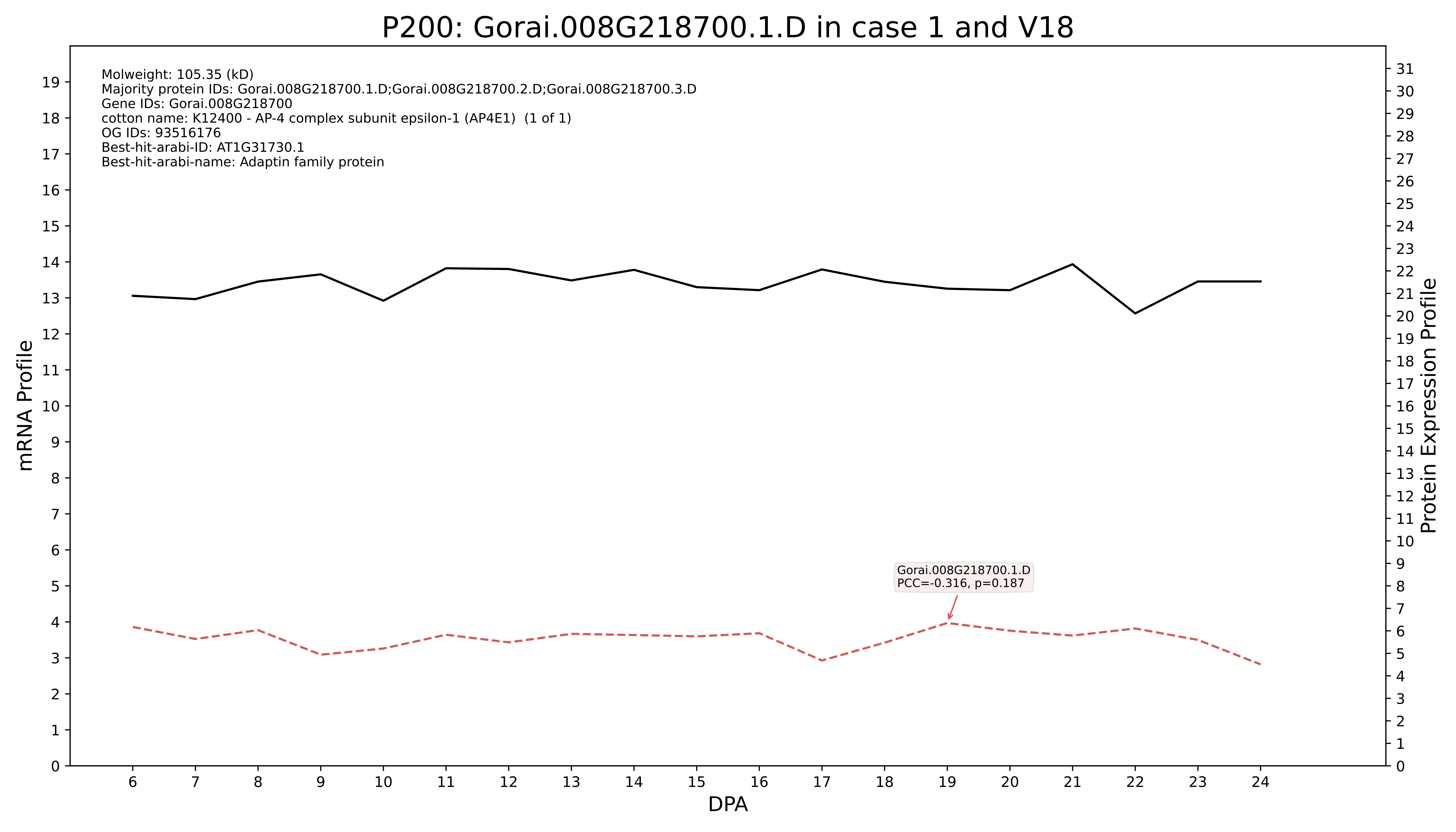
P200: Gorai.008G218700.1.D in case 1 and V18
Molweight: 105.35 (kD)
Majority protein IDs: Gorai.008G218700.1.D;Gorai.008G218700.2.D;Gorai.008G218700.3.D
Gene IDs: Gorai.008G218700
cotton name: K12400 - AP-4 complex subunit epsilon-1 (AP4E1) (1 of 1)
OG IDs: 93516176
Best-hit-arabi-ID: AT1G31730.1
Best-hit-arabi-name: Adaptin family protein
Gorai.008G218700.1.D
PCC=-0.316, p=0.187
mRNA Profile
Protein Expression Profile
DPA

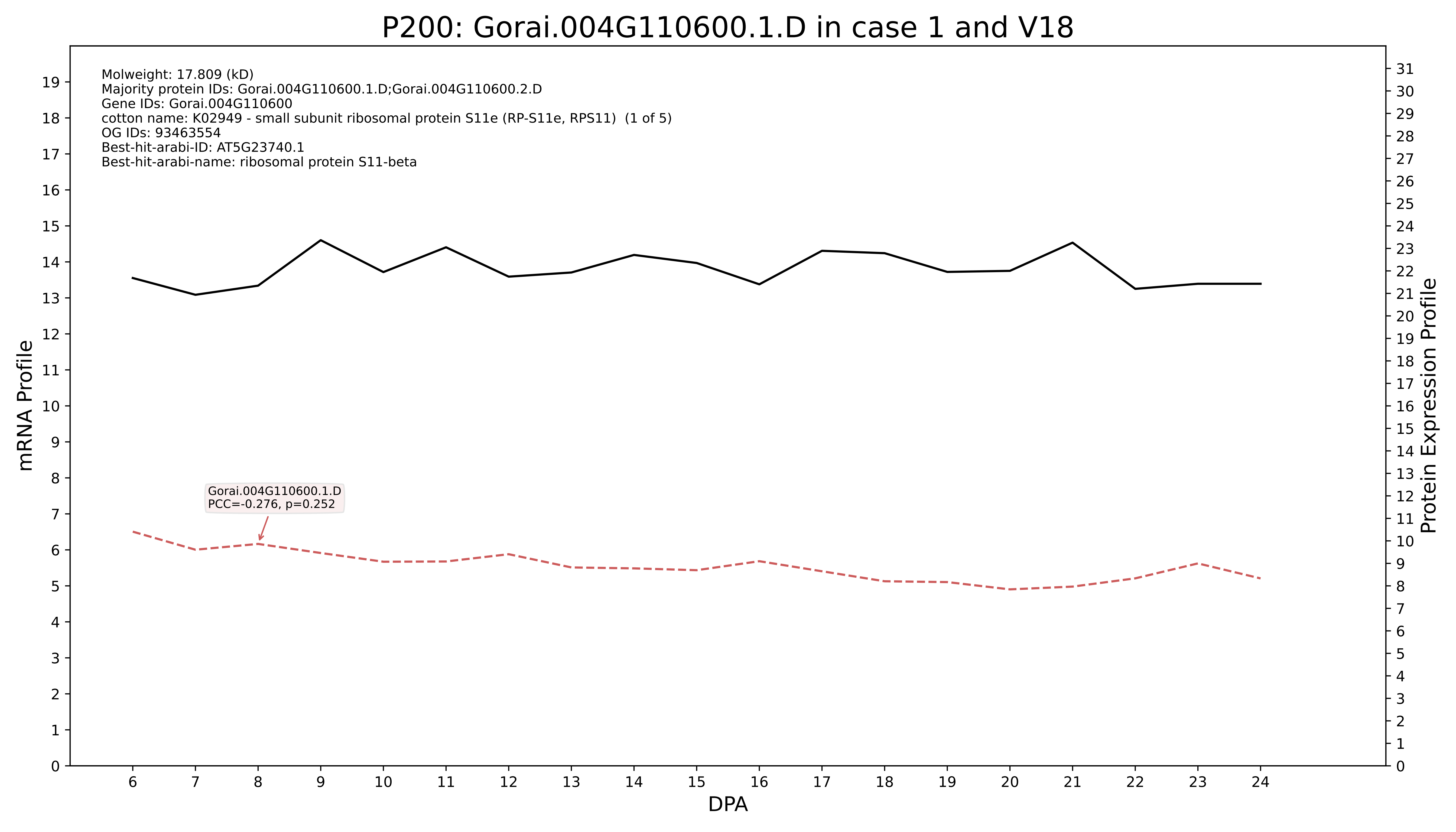

P200: Gorai.004G110600.1.D in case 1 and V18
Molweight: 17.809 (kD)
Majority protein IDs: Gorai.004G110600.1.D;Gorai.004G110600.2.D
Gene IDs: Gorai.004G110600
cotton name: K02949 - small subunit ribosomal protein S11e (RP-S11e, RPS11) (1 of 5)
OG IDs: 93463554
Best-hit-arabi-ID: AT5G23740.1
Best-hit-arabi-name: ribosomal protein S11-beta
Gorai.004G110600.1.D
PCC=-0.276, p=0.252
mRNA Profile
Protein Expression Profile
DPA

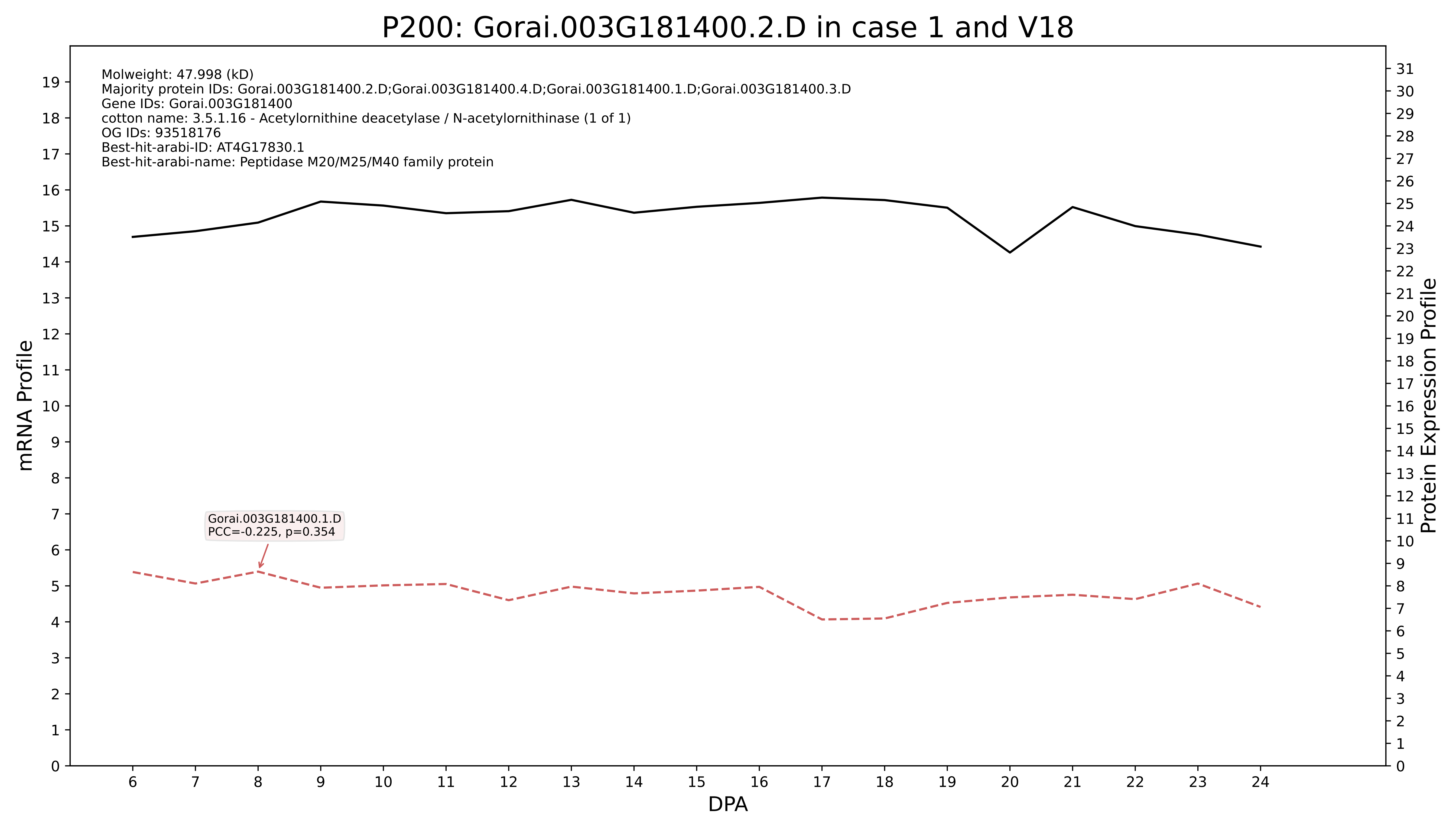

P200: Gorai.003G181400.2.D in case 1 and V18
Molweight: 47.998 (kD)
Majority protein IDs: Gorai.003G181400.2.D;Gorai.003G181400.4.D;Gorai.003G181400.1.D;Gorai.003G181400.3.D
Gene IDs: Gorai.003G181400
cotton name: 3.5.1.16 - Acetylornithine deacetylase / N-acetylornithinase (1 of 1)
OG IDs: 93518176
Best-hit-arabi-ID: AT4G17830.1
Best-hit-arabi-name: Peptidase M20/M25/M40 family protein
Gorai.003G181400.1.D
PCC=-0.225, p=0.354
mRNA Profile
Protein Expression Profile
DPA

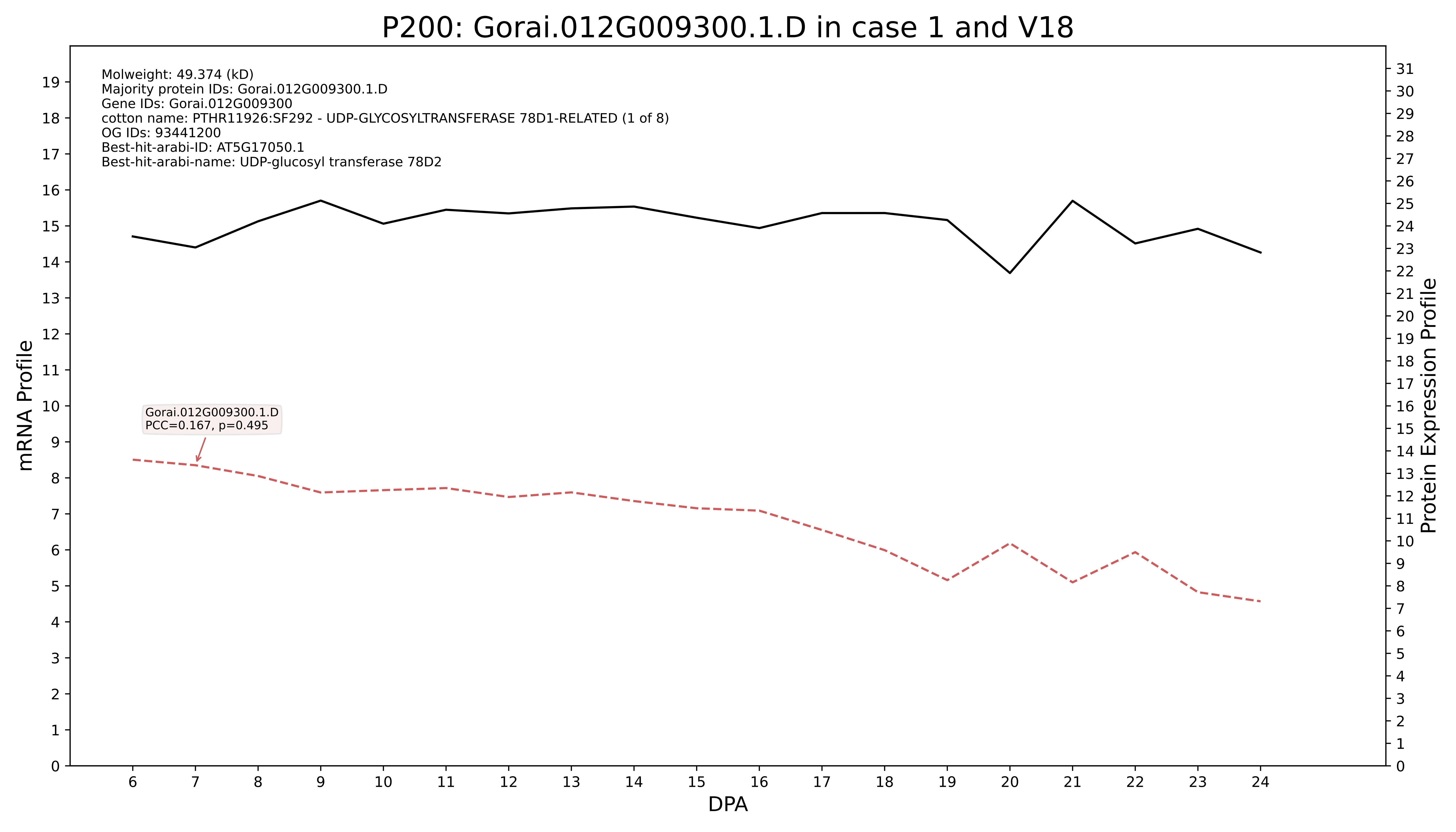
P200: Gorai.012G009300.1.D in case 1 and V18
Molweight: 49.374 (kD)
Majority protein IDs: Gorai.012G009300.1.D
Gene IDs: Gorai.012G009300
cotton name: PTHR11926:SF292 - UDP-GLYCOSYLTRANSFERASE 78D1-RELATED (1 of 8)
OG IDs: 93441200
Best-hit-arabi-ID: AT5G17050.1
Best-hit-arabi-name: UDP-glucosyl transferase 78D2
Gorai.012G009300.1.D
PCC=0.167, p=0.495
mRNA Profile
Protein Expression Profile
DPA

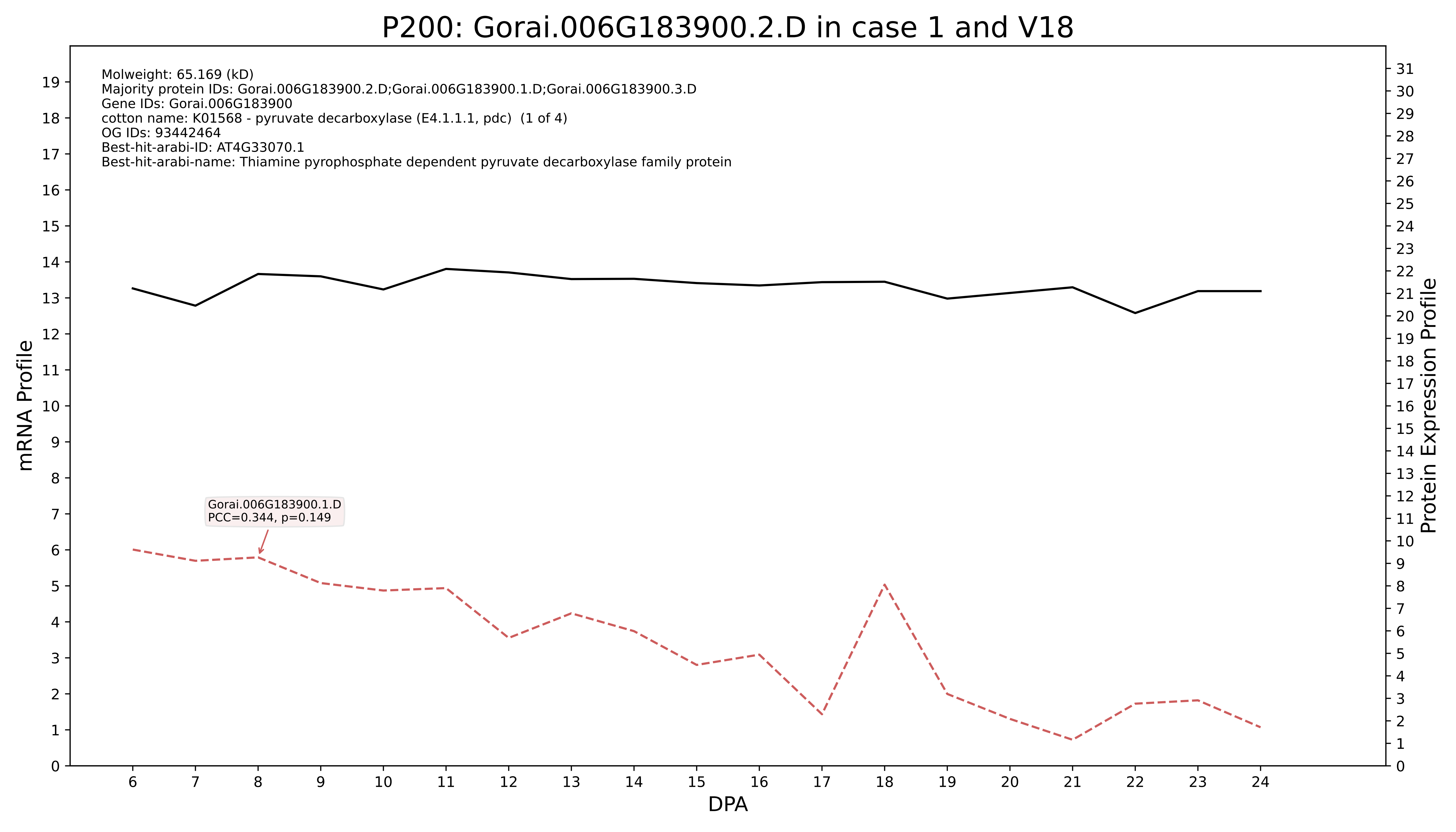
P200: Gorai.006G183900.2.D in case 1 and V18
Molweight: 65.169 (kD)
Majority protein IDs: Gorai.006G183900.2.D;Gorai.006G183900.1.D;Gorai.006G183900.3.D
Gene IDs: Gorai.006G183900
cotton name: K01568 - pyruvate decarboxylase (E4.1.1.1, pdc) (1 of 4)
OG IDs: 93442464
Best-hit-arabi-ID: AT4G33070.1
Best-hit-arabi-name: Thiamine pyrophosphate dependent pyruvate decarboxylase family protein
Gorai.006G183900.1.D
PCC=0.344, p=0.149
mRNA Profile
Protein Expression Profile
DPA

### P200: Gorai.007G301700.2.D in case 1 and V18

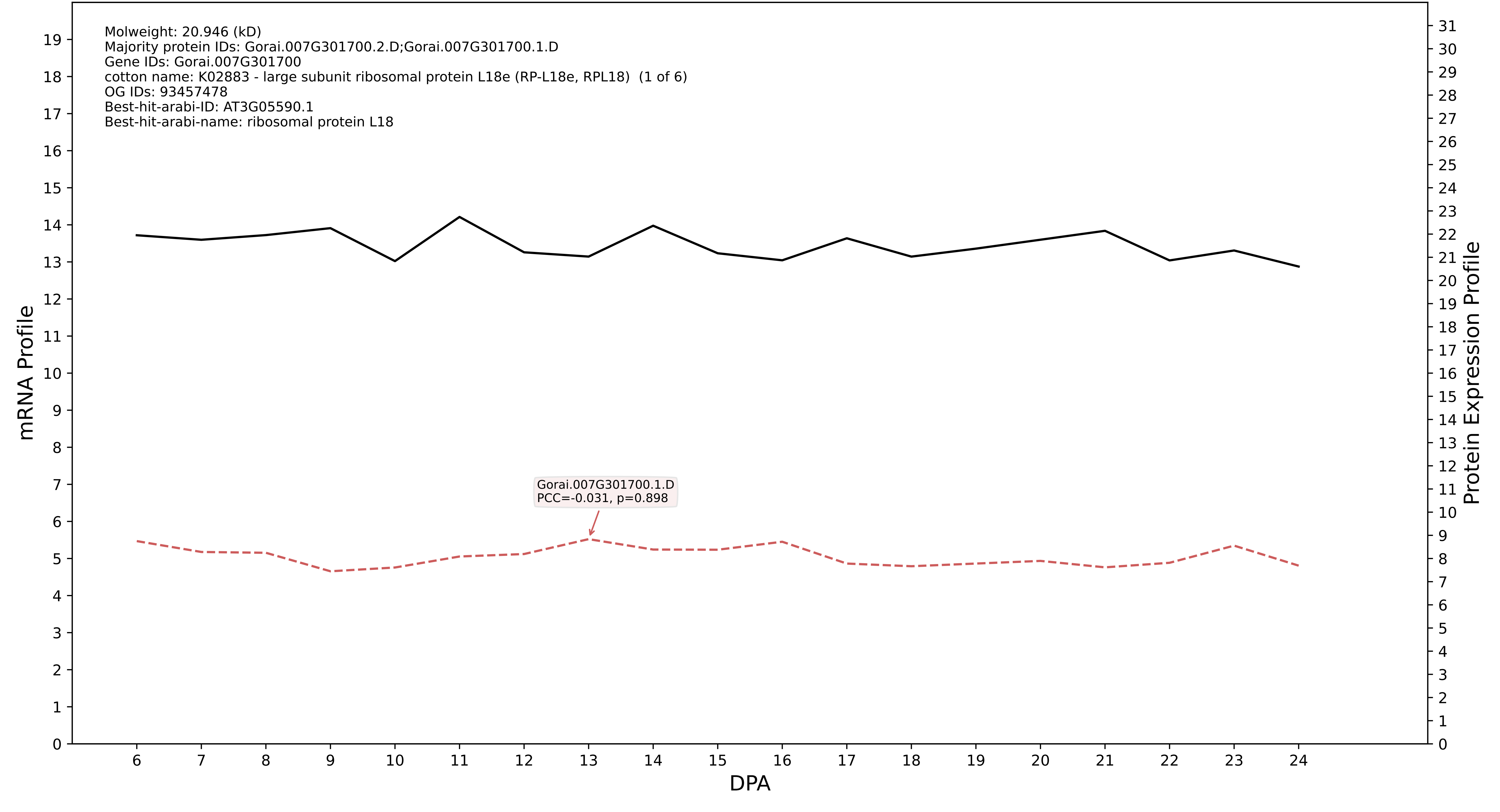

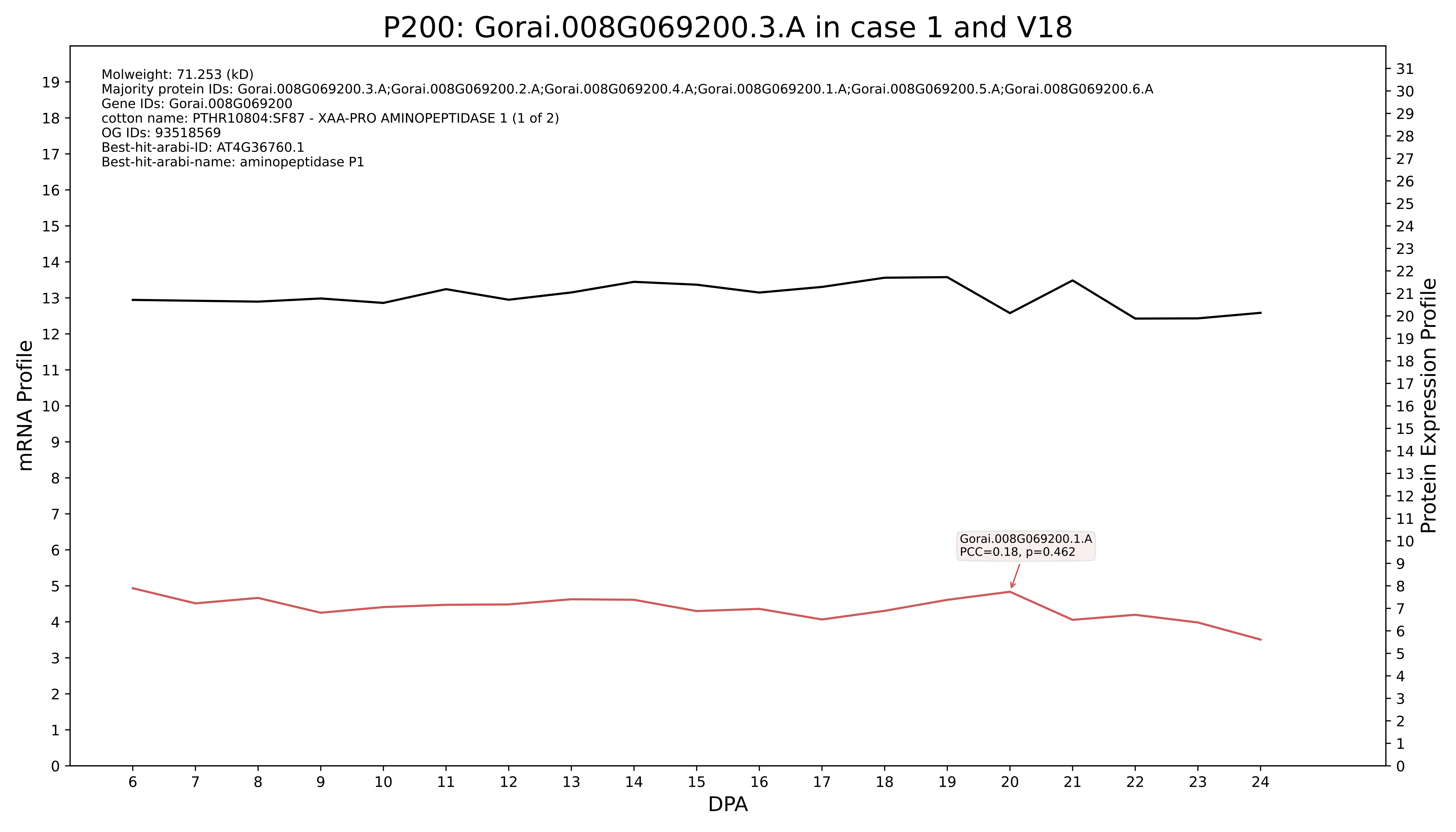
P200: Gorai.008G069200.3.A in case 1 and V18
Molweight: 71.253 (kD)
Majority protein IDs: Gorai.008G069200.3.A;Gorai.008G069200.2.A;Gorai.008G069200.4.A;Gorai.008G069200.1.A;Gorai.008G069200.5.A;Gorai.008G069200.6.A
Gene IDs: Gorai.008G069200
cotton name: PTHR10804:SF87 - XAA-PRO AMINOPEPTIDASE 1 (1 of 2)
OG IDs: 93518569
Best-hit-arabi-ID: AT4G36760.1
Best-hit-arabi-name: aminopeptidase P1
Gorai.008G069200.1.A
PCC=0.18, p=0.462
mRNA Profile
Protein Expression Profile
DPA

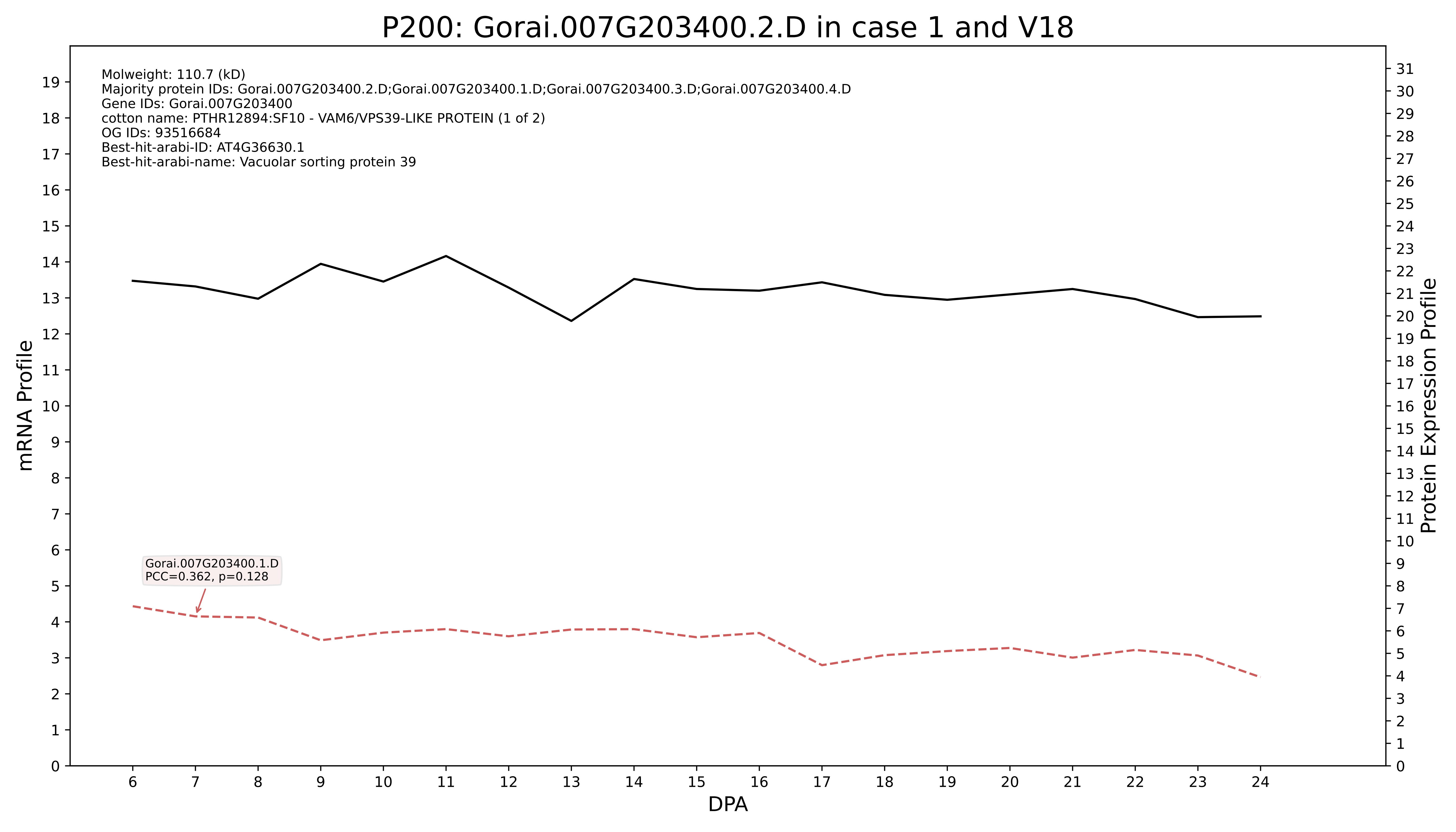
P200: Gorai.007G203400.2.D in case 1 and V18
Molweight: 110.7 (kD)
Majority protein IDs: Gorai.007G203400.2.D;Gorai.007G203400.1.D;Gorai.007G203400.3.D;Gorai.007G203400.4.D
Gene IDs: Gorai.007G203400
cotton name: PTHR12894:SF10 - VAM6/VPS39-LIKE PROTEIN (1 of 2)
OG IDs: 93516684
Best-hit-arabi-ID: AT4G36630.1
Best-hit-arabi-name: Vacuolar sorting protein 39
Gorai.007G203400.1.D
PCC=0.362, p=0.128
mRNA Profile
Protein Expression Profile
DPA

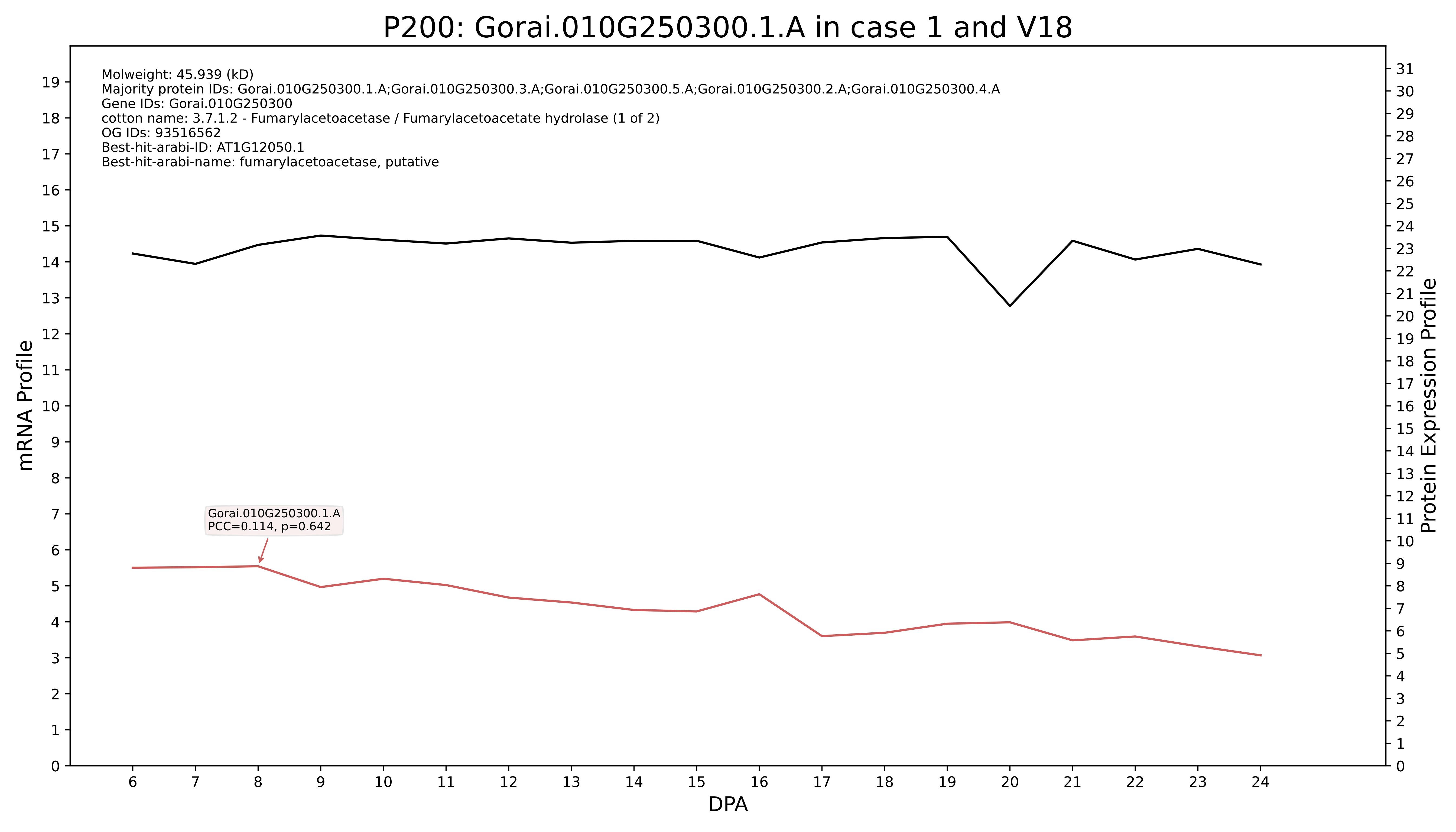
P200: Gorai.010G250300.1.A in case 1 and V18
Molweight: 45.939 (kD)
Majority protein IDs: Gorai.010G250300.1.A;Gorai.010G250300.3.A;Gorai.010G250300.5.A;Gorai.010G250300.2.A;Gorai.010G250300.4.A
Gene IDs: Gorai.010G250300
cotton name: 3.7.1.2 - Fumarylacetoacetase / Fumarylacetoacetate hydrolase (1 of 2)
OG IDs: 93516562
Best-hit-arabi-ID: AT1G12050.1
Best-hit-arabi-name: fumarylacetoacetase, putative
Gorai.010G250300.1.A
PCC=0.114, p=0.642
mRNA Profile
Protein Expression Profile
DPA

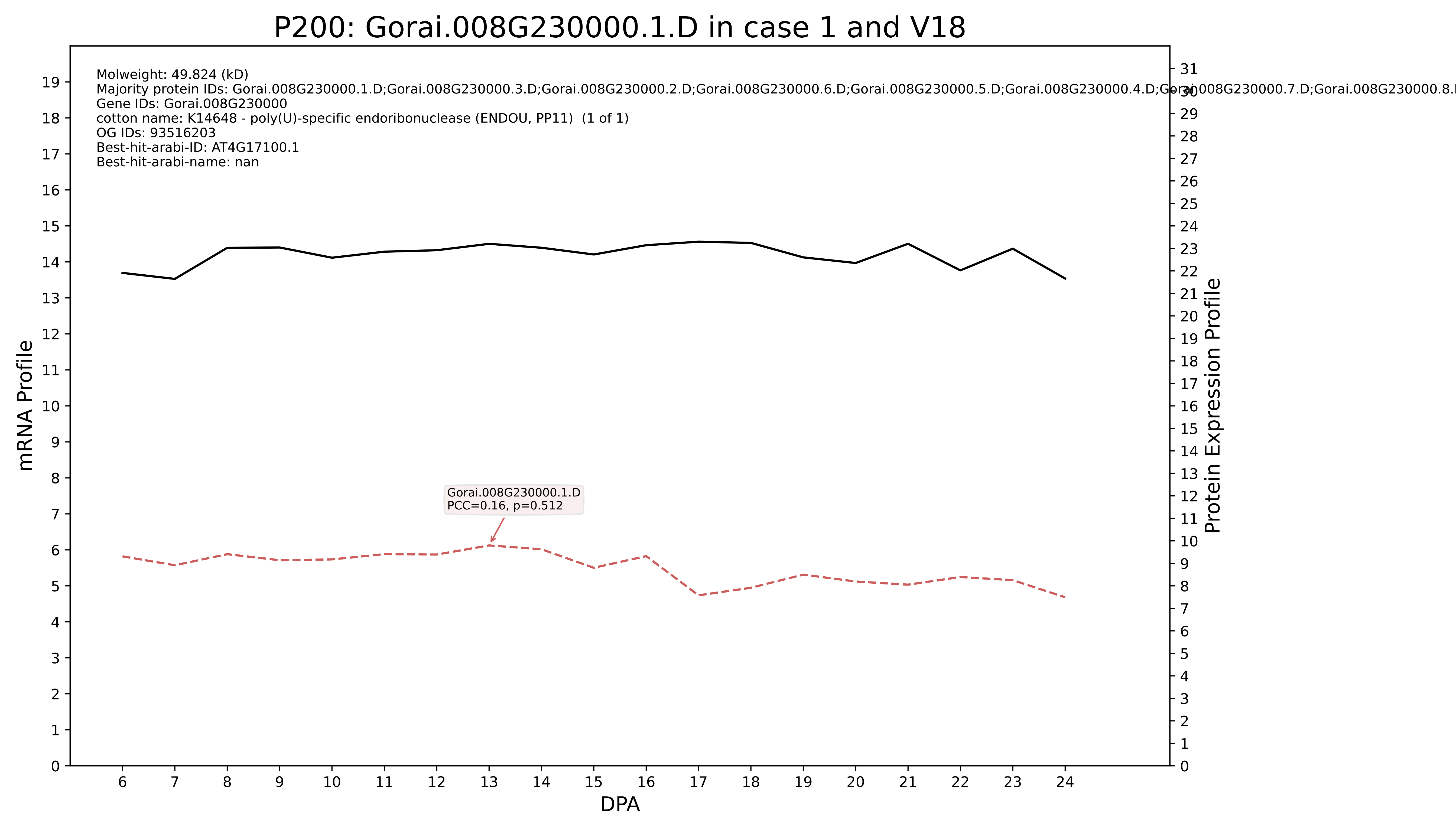
P200: Gorai.008G230000.1.D in case 1 and V18
Molweight: 49.824 (kD)
Majority protein IDs: Gorai.008G230000.1.D;Gorai.008G230000.3.D;Gorai.008G230000.2.D;Gorai.008G230000.6.D;Gorai.008G230000.5.D;Gorai.008G230000.4.D;Gorai.008G230000.7.D;Gorai.008G230000.8.
Gene IDs: Gorai.008G230000
cotton name: K14648 - poly(U)-specific endoribonuclease (ENDOU, PP11) (1 of 1)
OG IDs: 93516203
Best-hit-arabi-ID: AT4G17100.1
Best-hit-arabi-name: nan
Gorai.008G230000.1.D
PCC=0.16, p=0.512
mRNA Profile
Protein Expression Profile
DPA

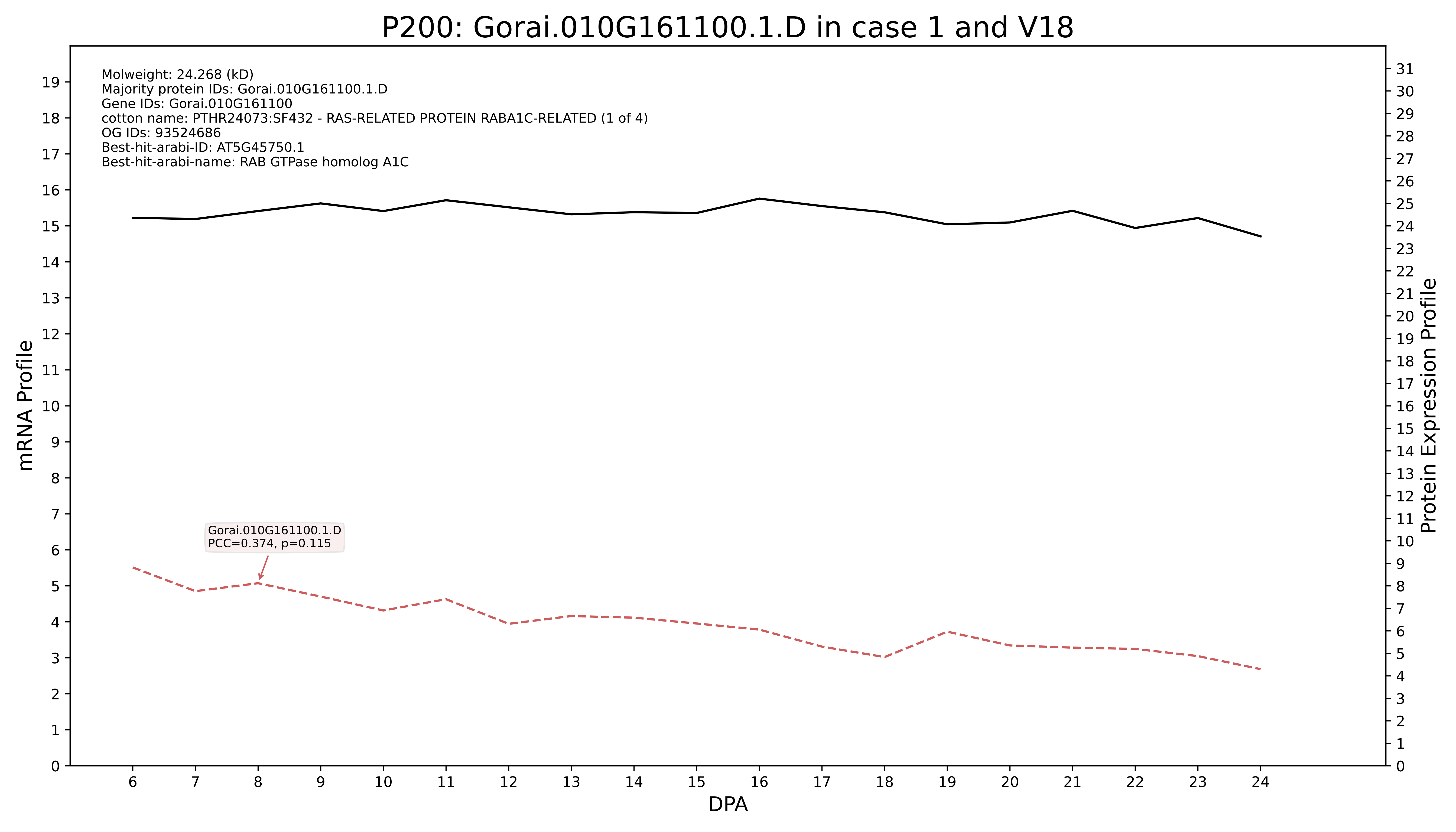
P200: Gorai.010G161100.1.D in case 1 and V18
Molweight: 24.268 (kD)
Majority protein IDs: Gorai.010G161100.1.D
Gene IDs: Gorai.010G161100
cotton name: PTHR24073:SF432 - RAS-RELATED PROTEIN RABA1C-RELATED (1 of 4)
OG IDs: 93524686
Best-hit-arabi-ID: AT5G45750.1
Best-hit-arabi-name: RAB GTPase homolog A1C
Gorai.010G161100.1.D
PCC=0.374, p=0.115
mRNA Profile
Protein Expression Profile
DPA

### P200: Gorai.006G103500.2.D in case 1 and V18

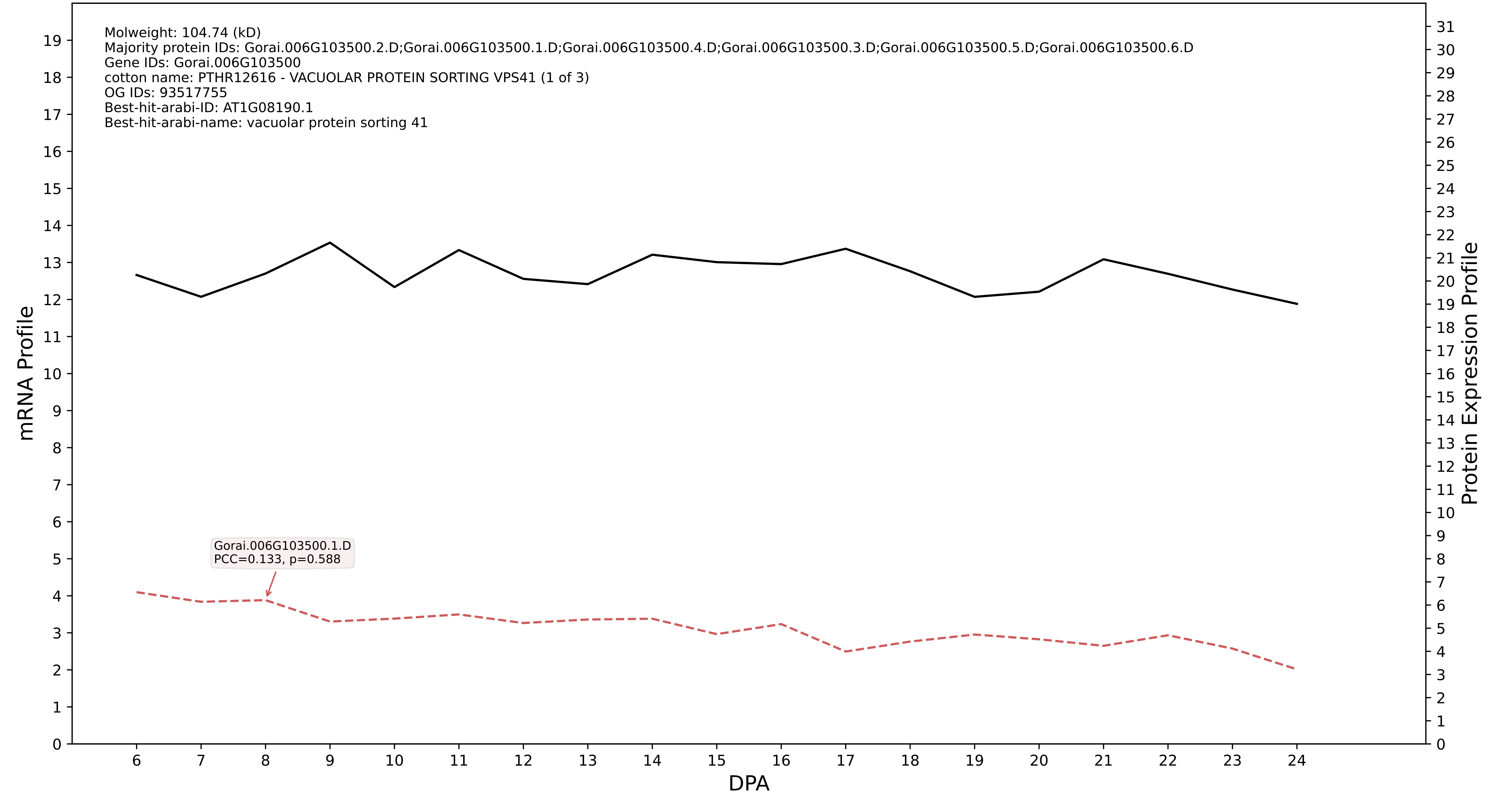

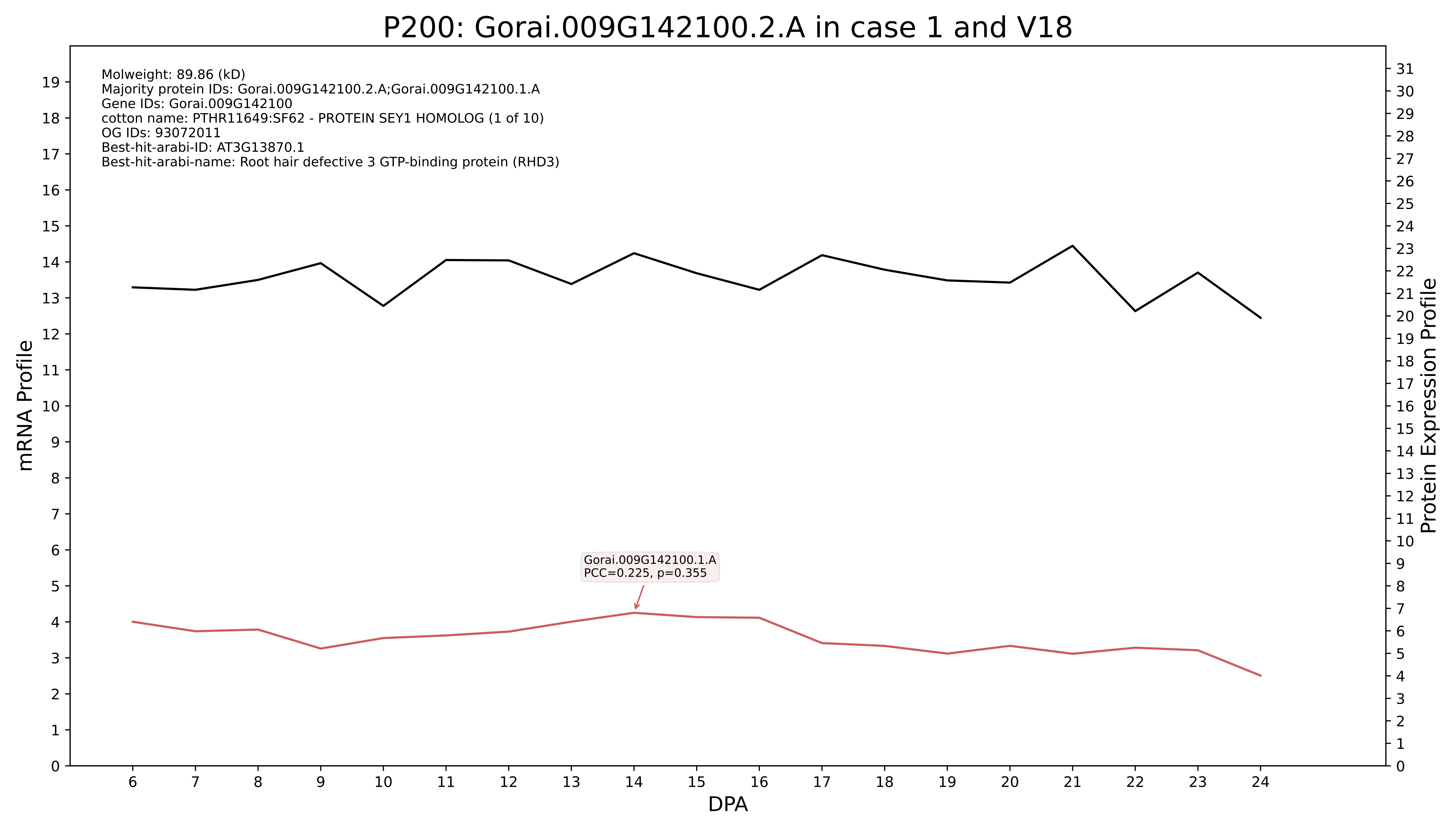

P200: Gorai.009G142100.2.A in case 1 and V18
Molweight: 89.86 (kD)
Majority protein IDs: Gorai.009G142100.2.A;Gorai.009G142100.1.A
Gene IDs: Gorai.009G142100
cotton name: PTHR11649:SF62 - PROTEIN SEY1 HOMOLOG (1 of 10)
OG IDs: 93072011
Best-hit-arabi-ID: AT3G13870.1
Best-hit-arabi-name: Root hair defective 3 GTP-binding protein (RHD3)
Gorai.009G142100.1.A
PCC=0.225, p=0.355
mRNA Profile
Protein Expression Profile
DPA

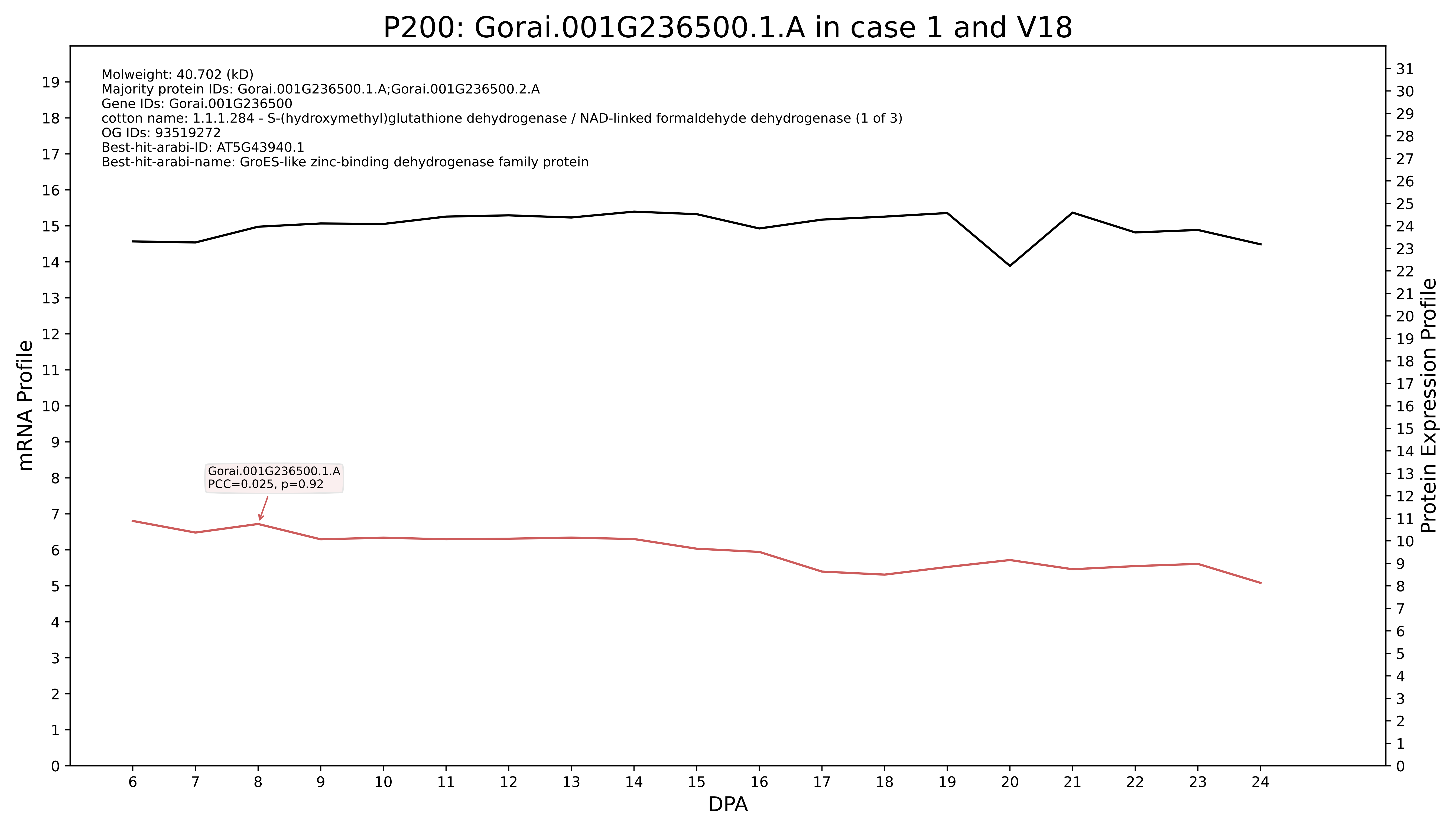
P200: Gorai.001G236500.1.A in case 1 and V18
Molweight: 40.702 (kD)
Majority protein IDs: Gorai.001G236500.1.A;Gorai.001G236500.2.A
Gene IDs: Gorai.001G236500
cotton name: 1.1.1.284 - S-(hydroxymethyl)glutathione dehydrogenase / NAD-linked formaldehyde dehydrogenase (1 of 3)
OG IDs: 93519272
Best-hit-arabi-ID: AT5G43940.1
Best-hit-arabi-name: GroES-like zinc-binding dehydrogenase family protein
Gorai.001G236500.1.A
PCC=0.025, p=0.92
mRNA Profile
Protein Expression Profile
DPA

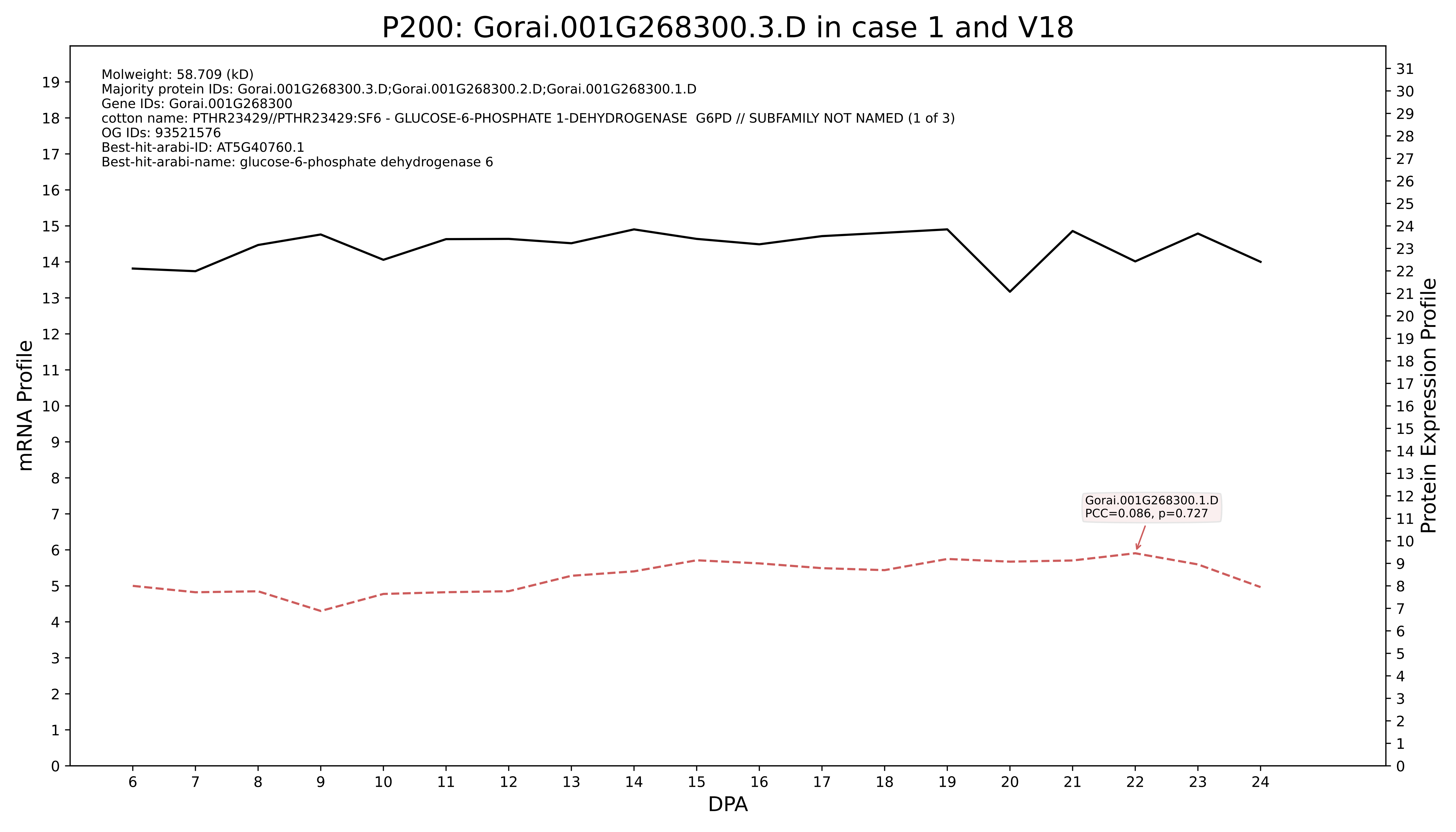
P200: Gorai.001G268300.3.D in case 1 and V18
Molweight: 58.709 (kD)
Majority protein IDs: Gorai.001G268300.3.D;Gorai.001G268300.2.D;Gorai.001G268300.1.D
Gene IDs: Gorai.001G268300
cotton name: PTHR23429//PTHR23429:SF6 - GLUCOSE-6-PHOSPHATE 1-DEHYDROGENASE G6PD // SUBFAMILY NOT NAMED (1 of 3)
OG IDs: 93521576
Best-hit-arabi-ID: AT5G40760.1
Best-hit-arabi-name: glucose-6-phosphate dehydrogenase 6
Gorai.001G268300.1.D
PCC=0.086, p=0.727
mRNA Profile
Protein Expression Profile
DPA

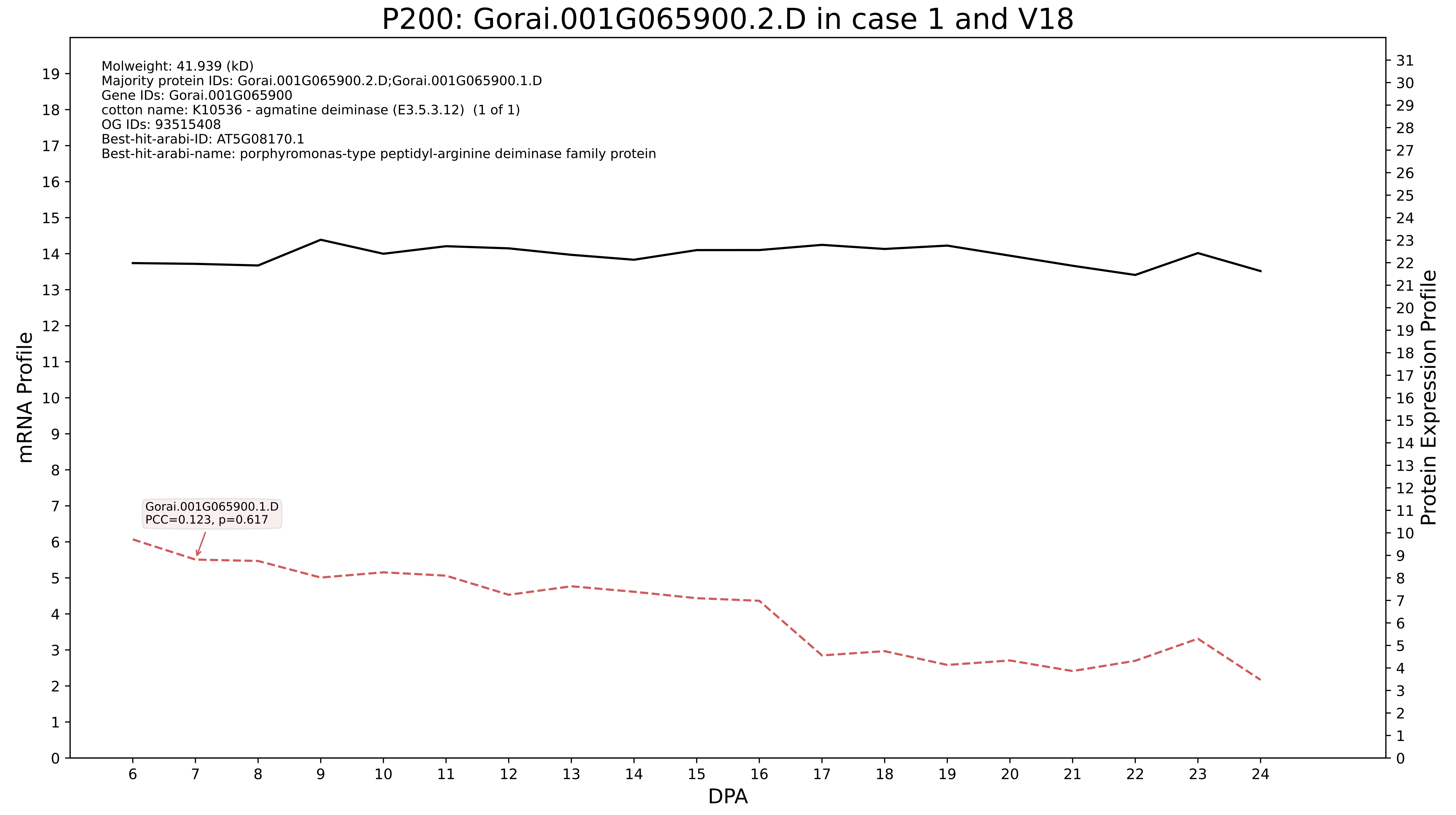

P200: Gorai.001G065900.2.D in case 1 and V18
Molweight: 41.939 (kD)
Majority protein IDs: Gorai.001G065900.2.D;Gorai.001G065900.1.D
Gene IDs: Gorai.001G065900
cotton name: K10536 - agmatine deiminase (E3.5.3.12) (1 of 1)
OG IDs: 93515408
Best-hit-arabi-ID: AT5G08170.1
Best-hit-arabi-name: porphyromonas-type peptidyl-arginine deiminase family protein
Gorai.001G065900.1.D
PCC=0.123, p=0.617
mRNA Profile
Protein Expression Profile
DPA

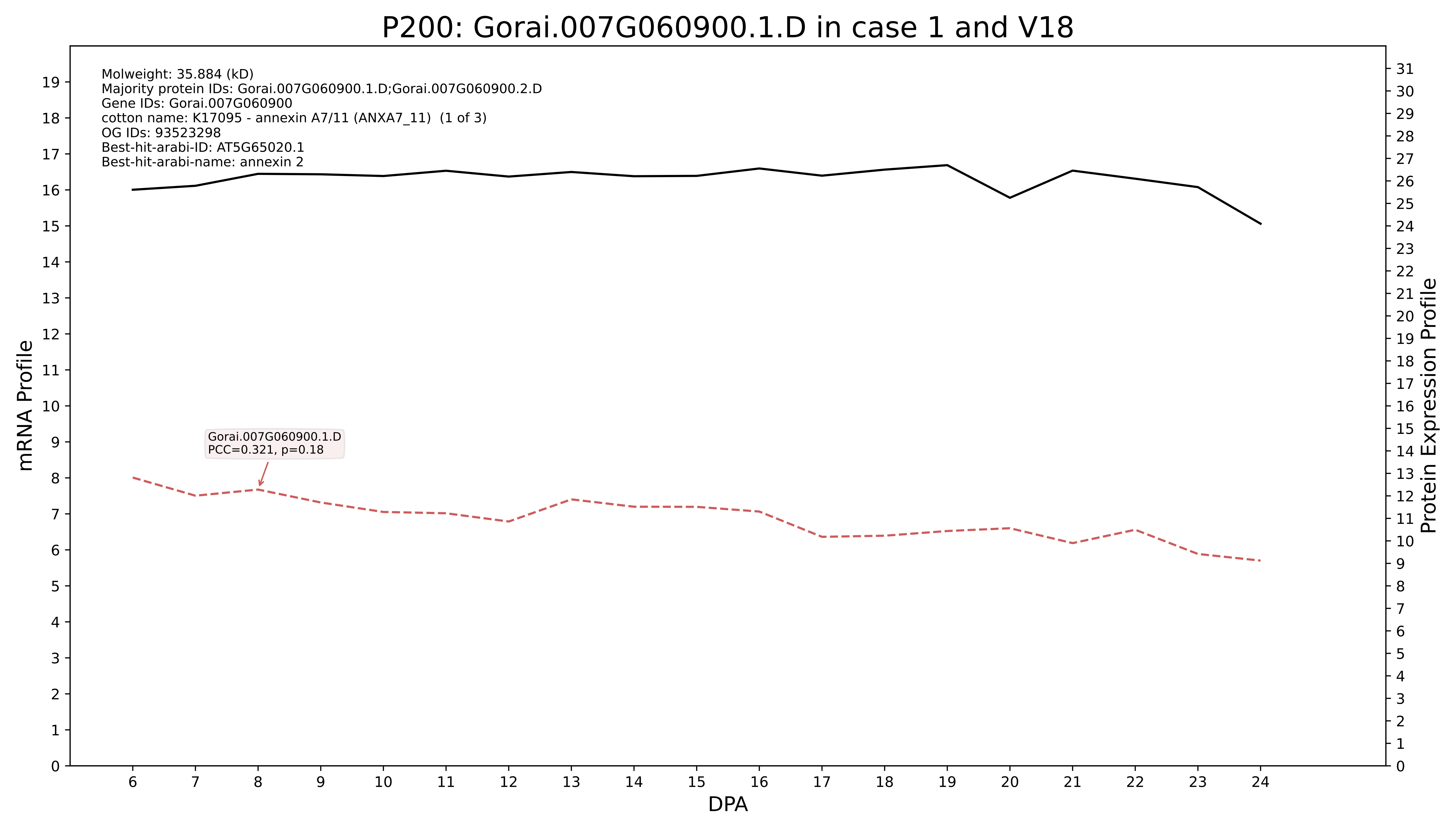
P200: Gorai.007G060900.1.D in case 1 and V18
Molweight: 35.884 (kD)
Majority protein IDs: Gorai.007G060900.1.D;Gorai.007G060900.2.D
Gene IDs: Gorai.007G060900
cotton name: K17095 - annexin A7/11 (ANXA7_11) (1 of 3)
OG IDs: 93523298
Best-hit-arabi-ID: AT5G65020.1
Best-hit-arabi-name: annexin 2
Gorai.007G060900.1.D
PCC=0.321, p=0.18
mRNA Profile
Protein Expression Profile
DPA

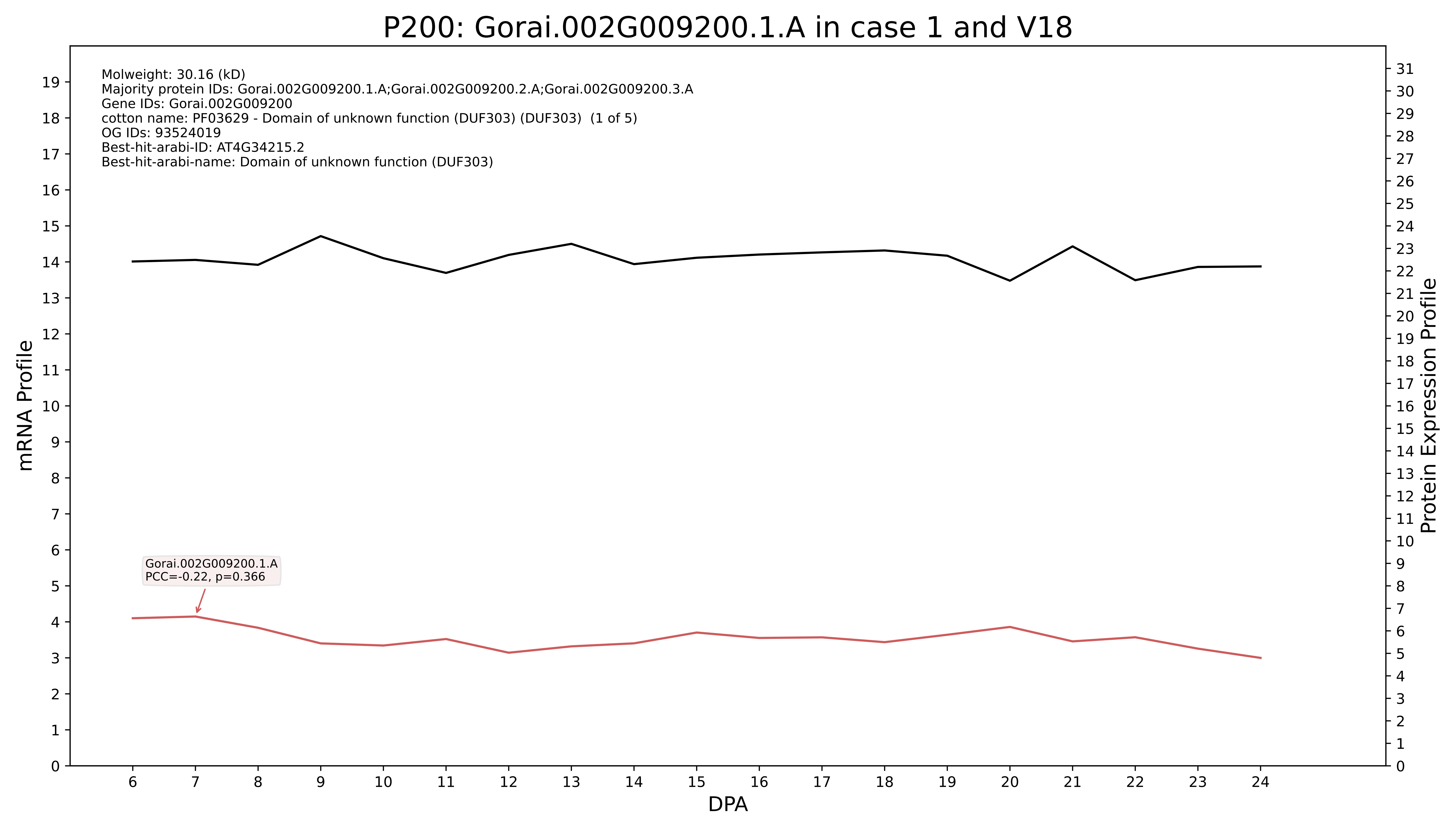
P200: Gorai.002G009200.1.A in case 1 and V18
Molweight: 30.16 (kD)
Majority protein IDs: Gorai.002G009200.1.A;Gorai.002G009200.2.A;Gorai.002G009200.3.A
Gene IDs: Gorai.002G009200
cotton name: PF03629 - Domain of unknown function (DUF303) (DUF303) (1 of 5)
OG IDs: 93524019
Best-hit-arabi-ID: AT4G34215.2
Best-hit-arabi-name: Domain of unknown function (DUF303)
Gorai.002G009200.1.A
PCC=-0.22, p=0.366
mRNA Profile
Protein Expression Profile
DPA

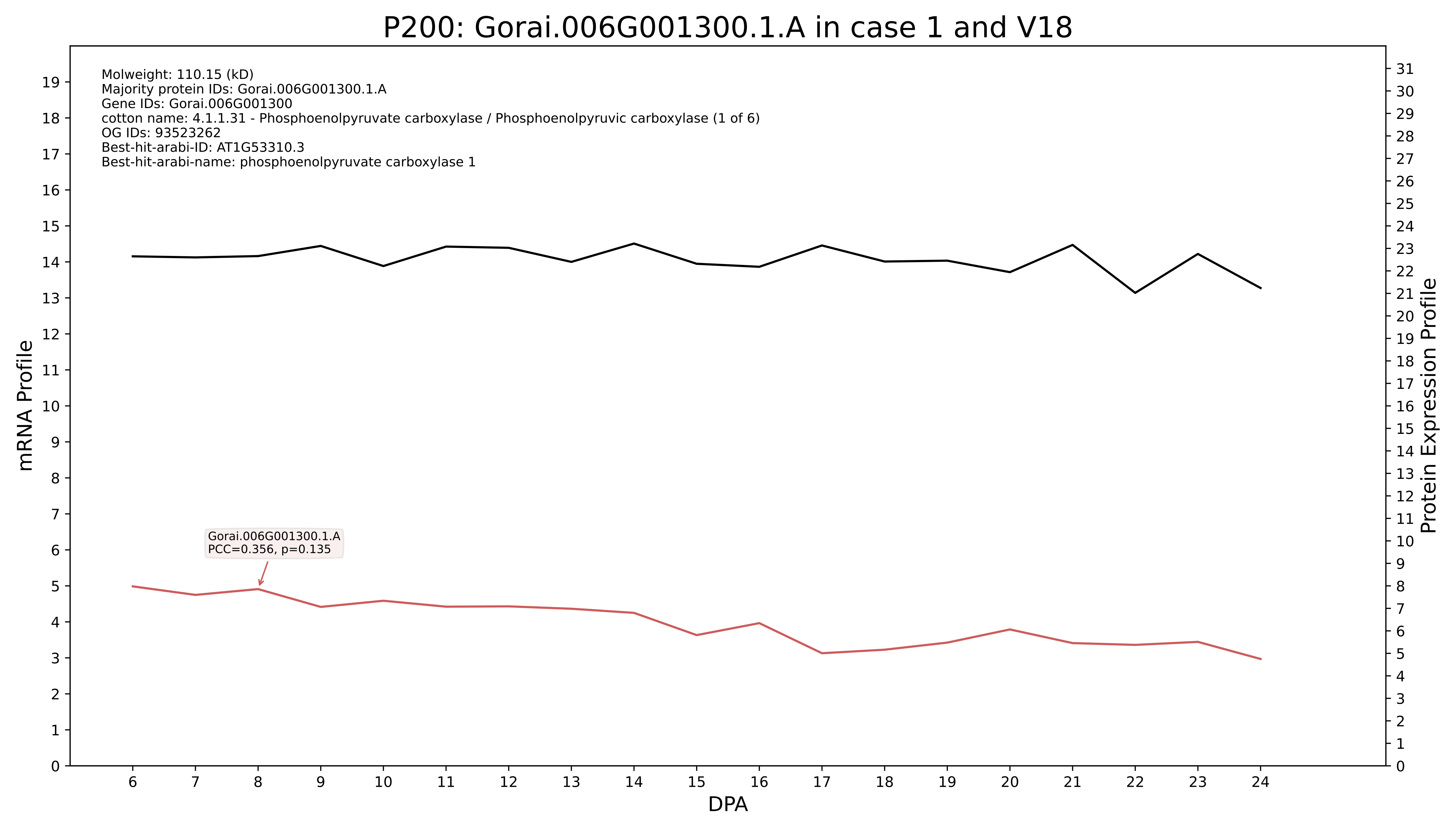
P200: Gorai.006G001300.1.A in case 1 and V18
Molweight: 110.15 (kD)
Majority protein IDs: Gorai.006G001300.1.A
Gene IDs: Gorai.006G001300
cotton name: 4.1.1.31 - Phosphoenolpyruvate carboxylase / Phosphoenolpyruvic carboxylase (1 of 6)
OG IDs: 93523262
Best-hit-arabi-ID: AT1G53310.3
Best-hit-arabi-name: phosphoenolpyruvate carboxylase 1
Gorai.006G001300.1.A
PCC=0.356, p=0.135
mRNA Profile
Protein Expression Profile
DPA

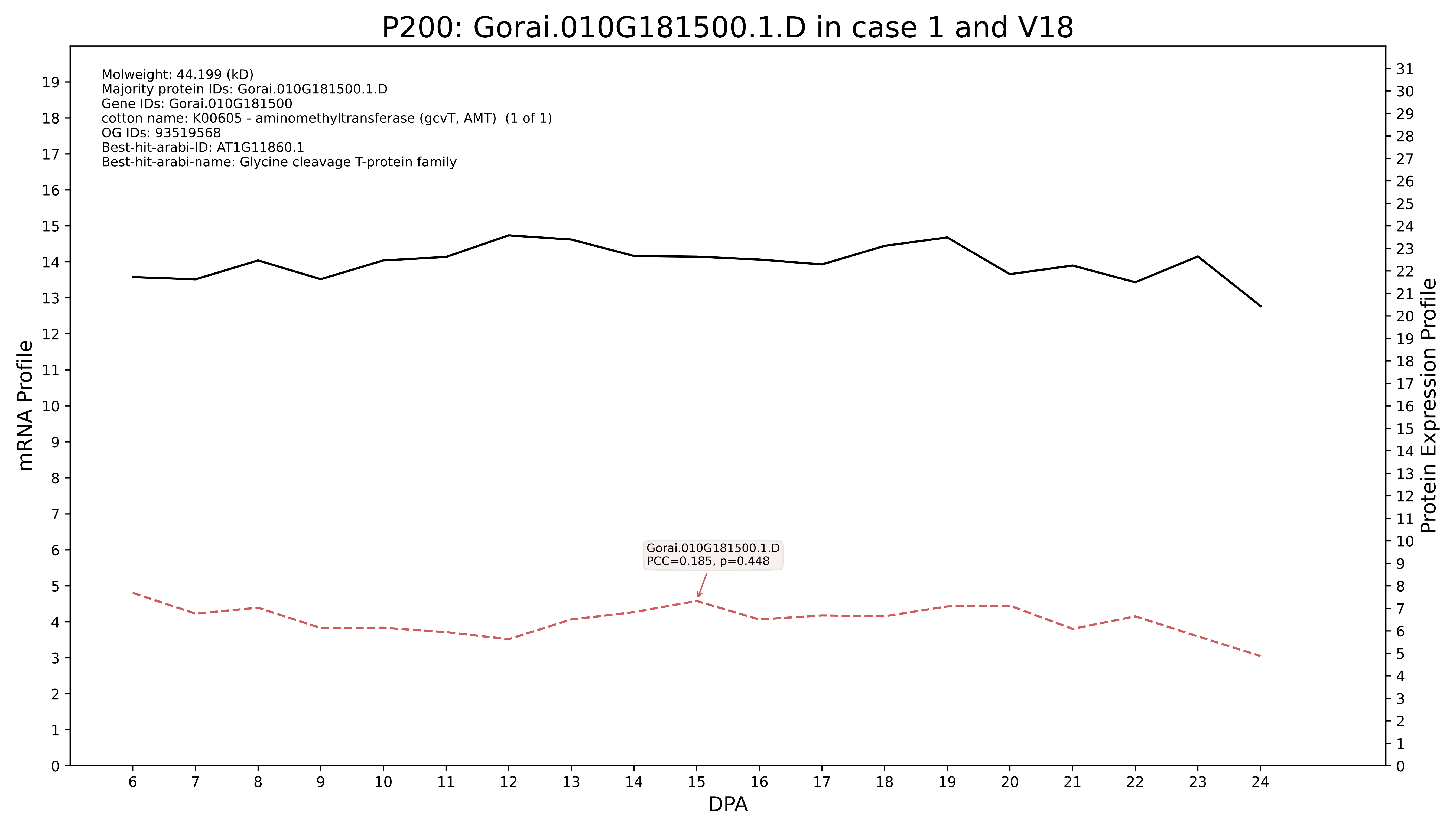
P200: Gorai.010G181500.1.D in case 1 and V18
Molweight: 44.199 (kD)
Majority protein IDs: Gorai.010G181500.1.D
Gene IDs: Gorai.010G181500
cotton name: K00605 - aminomethyltransferase (gcvT, AMT) (1 of 1)
OG IDs: 93519568
Best-hit-arabi-ID: AT1G11860.1
Best-hit-arabi-name: Glycine cleavage T-protein family
Gorai.010G181500.1.D
PCC=0.185, p=0.448
mRNA Profile
Protein Expression Profile
DPA

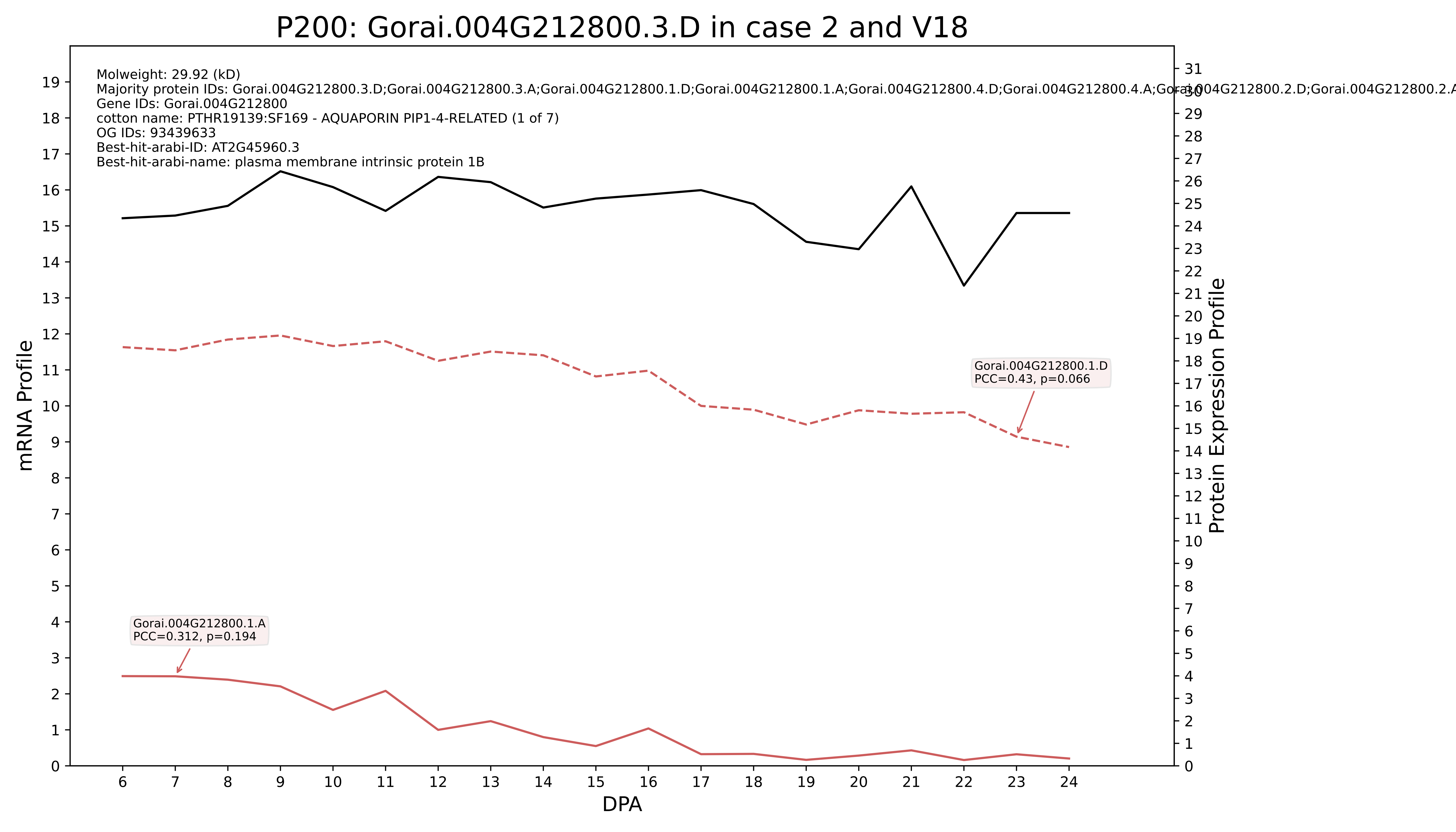
P200: Gorai.004G212800.3.D in case 2 and V18
Molweight: 29.92 (kD)
Majority protein IDs: Gorai.004G212800.3.D;Gorai.004G212800.3.A;Gorai.004G212800.1.D;Gorai.004G212800.1.A;Gorai.004G212800.4.D;Gorai.004G212800.4.A;Gorai.004G212800.2.D;Gorai.004G212800.2.A
Gene IDs: Gorai.004G212800
cotton name: PTHR19139:SF169 - AQUAPORIN PIP1-4-RELATED (1 of 7)
OG IDs: 93439633
Best-hit-arabi-ID: AT2G45960.3
Best-hit-arabi-name: plasma membrane intrinsic protein 1B
Gorai.004G212800.1.D
PCC=0.43, p=0.066
Gorai.004G212800.1.A
PCC=0.312, p=0.194
mRNA Profile
Protein Expression Profile
DPA

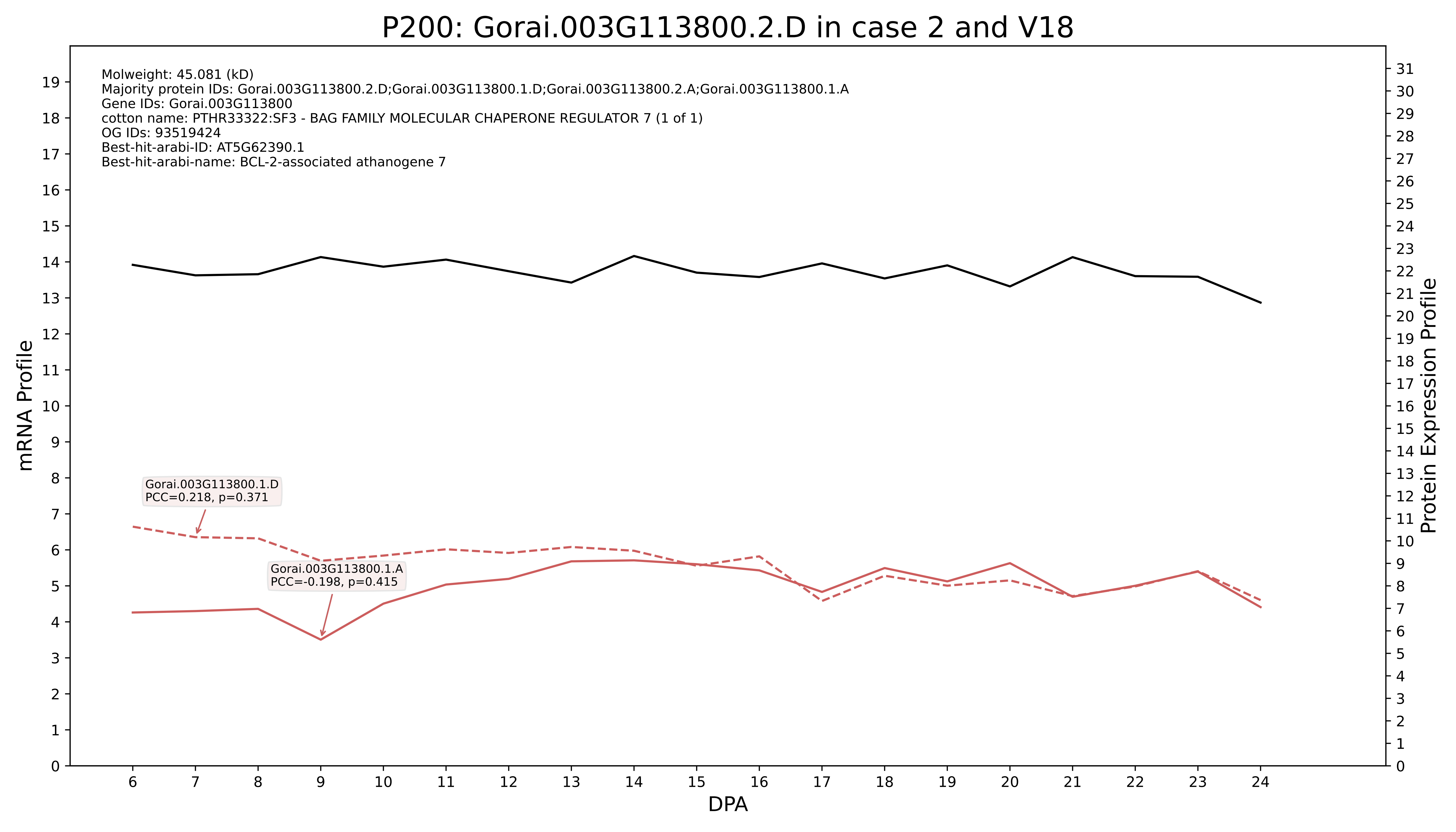
P200: Gorai.003G113800.2.D in case 2 and V18
Molweight: 45.081 (kD)
Majority protein IDs: Gorai.003G113800.2.D;Gorai.003G113800.1.D;Gorai.003G113800.2.A;Gorai.003G113800.1.A
Gene IDs: Gorai.003G113800
cotton name: PTHR33322:SF3 - BAG FAMILY MOLECULAR CHAPERONE REGULATOR 7 (1 of 1)
OG IDs: 93519424
Best-hit-arabi-ID: AT5G62390.1
Best-hit-arabi-name: BCL-2-associated athanogene 7
Gorai.003G113800.1.D
PCC=0.218, p=0.371
Gorai.003G113800.1.A
PCC=-0.198, p=0.415
mRNA Profile
Protein Expression Profile
DPA

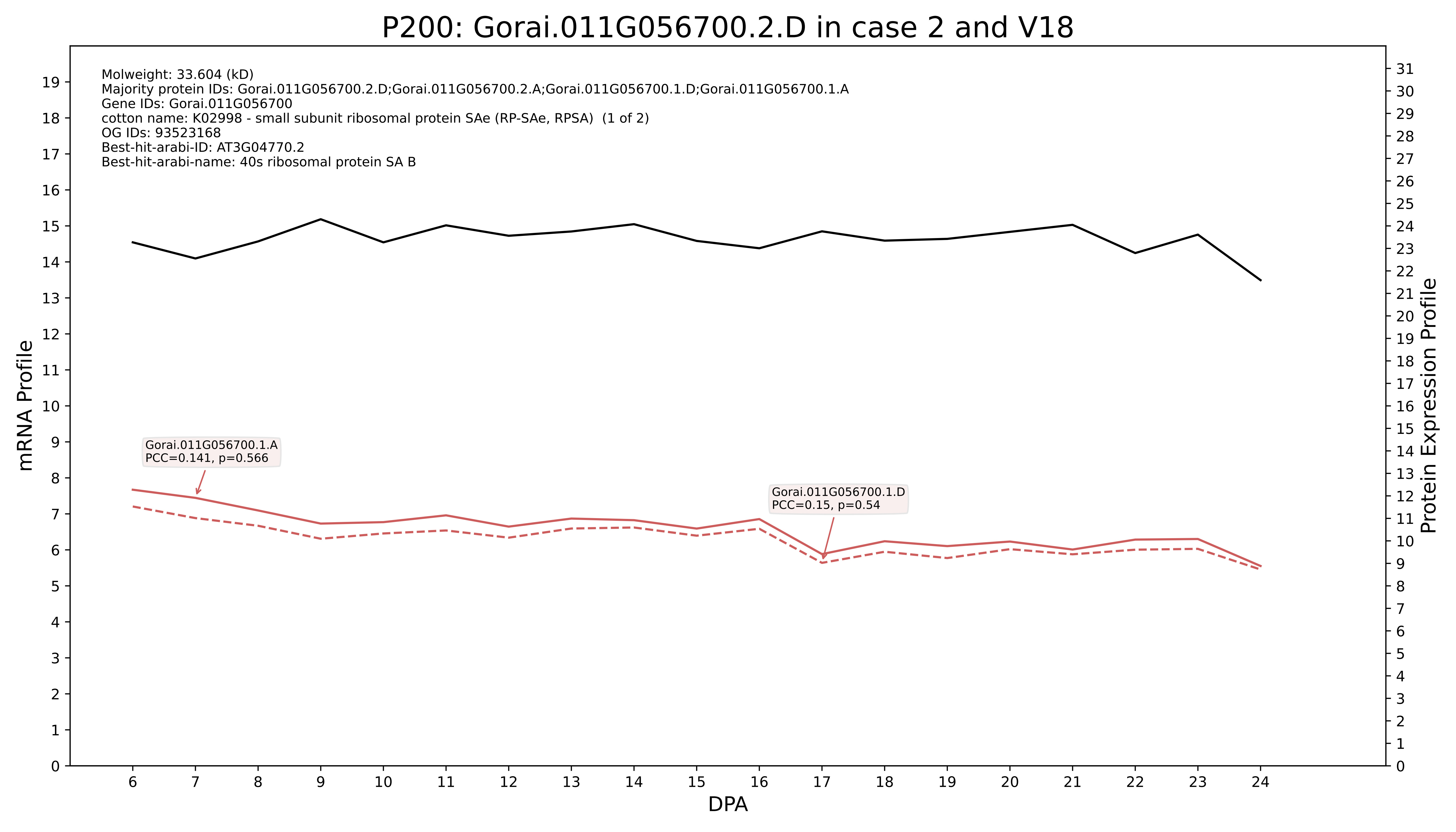

P200: Gorai.011G056700.2.D in case 2 and V18
Molweight: 33.604 (kD)
Majority protein IDs: Gorai.011G056700.2.D;Gorai.011G056700.2.A;Gorai.011G056700.1.D;Gorai.011G056700.1.A
Gene IDs: Gorai.011G056700
cotton name: K02998 - small subunit ribosomal protein SAe (RP-SAe, RPSA) (1 of 2)
OG IDs: 93523168
Best-hit-arabi-ID: AT3G04770.2
Best-hit-arabi-name: 40s ribosomal protein SA B
Gorai.011G056700.1.A
PCC=0.141, p=0.566
Gorai.011G056700.1.D
PCC=0.15, p=0.54
mRNA Profile
Protein Expression Profile
DPA

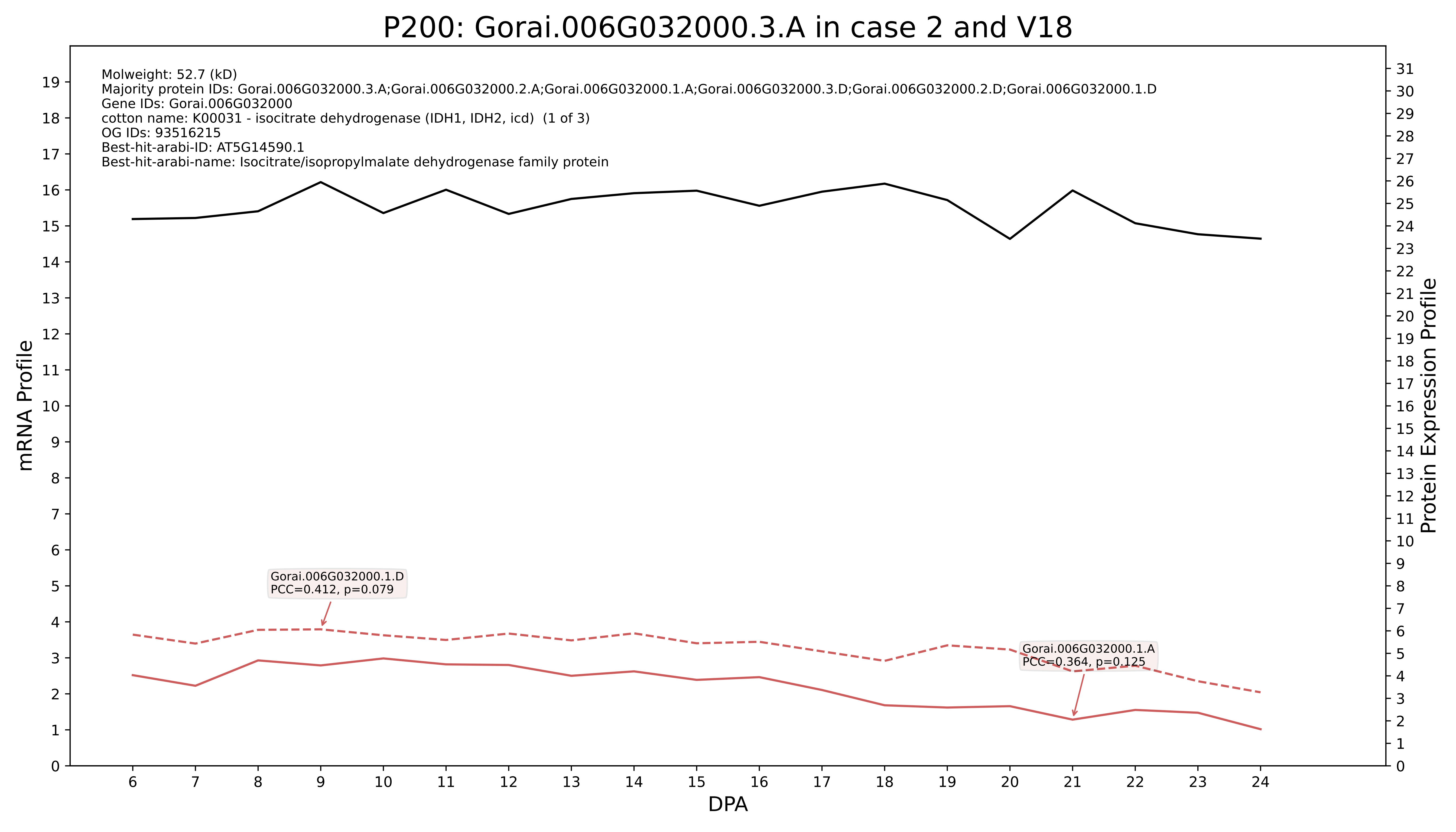
P200: Gorai.006G032000.3.A in case 2 and V18
Molweight: 52.7 (kD)
Majority protein IDs: Gorai.006G032000.3.A;Gorai.006G032000.2.A;Gorai.006G032000.1.A;Gorai.006G032000.3.D;Gorai.006G032000.2.D;Gorai.006G032000.1.D
Gene IDs: Gorai.006G032000
cotton name: K00031 - isocitrate dehydrogenase (IDH1, IDH2, icd) (1 of 3)
OG IDs: 93516215
Best-hit-arabi-ID: AT5G14590.1
Best-hit-arabi-name: Isocitrate/isopropylmalate dehydrogenase family protein
Gorai.006G032000.1.D
PCC=0.412, p=0.079
Gorai.006G032000.1.A
PCC=0.364, p=0.125
mRNA Profile
Protein Expression Profile
DPA

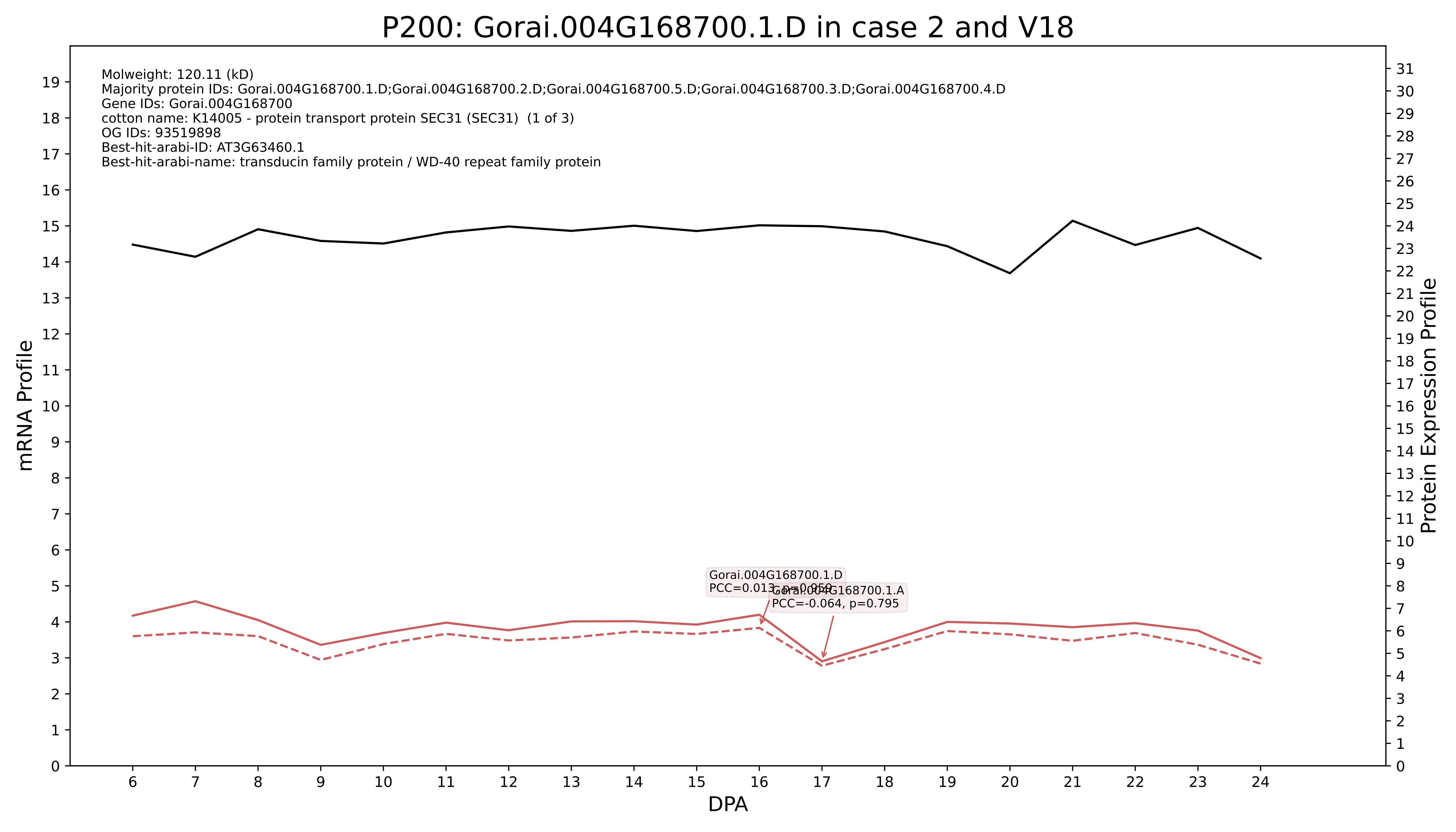
P200: Gorai.004G168700.1.D in case 2 and V18
Molweight: 120.11 (kD)
Majority protein IDs: Gorai.004G168700.1.D;Gorai.004G168700.2.D;Gorai.004G168700.5.D;Gorai.004G168700.3.D;Gorai.004G168700.4.D
Gene IDs: Gorai.004G168700
cotton name: K14005 - protein transport protein SEC31 (SEC31) (1 of 3)
OG IDs: 93519898
Best-hit-arabi-ID: AT3G63460.1
Best-hit-arabi-name: transducin family protein / WD-40 repeat family protein
Gorai.004G168700.1.A
PCC=-0.064, p=0.795
mRNA Profile
Protein Expression Profile
DPA

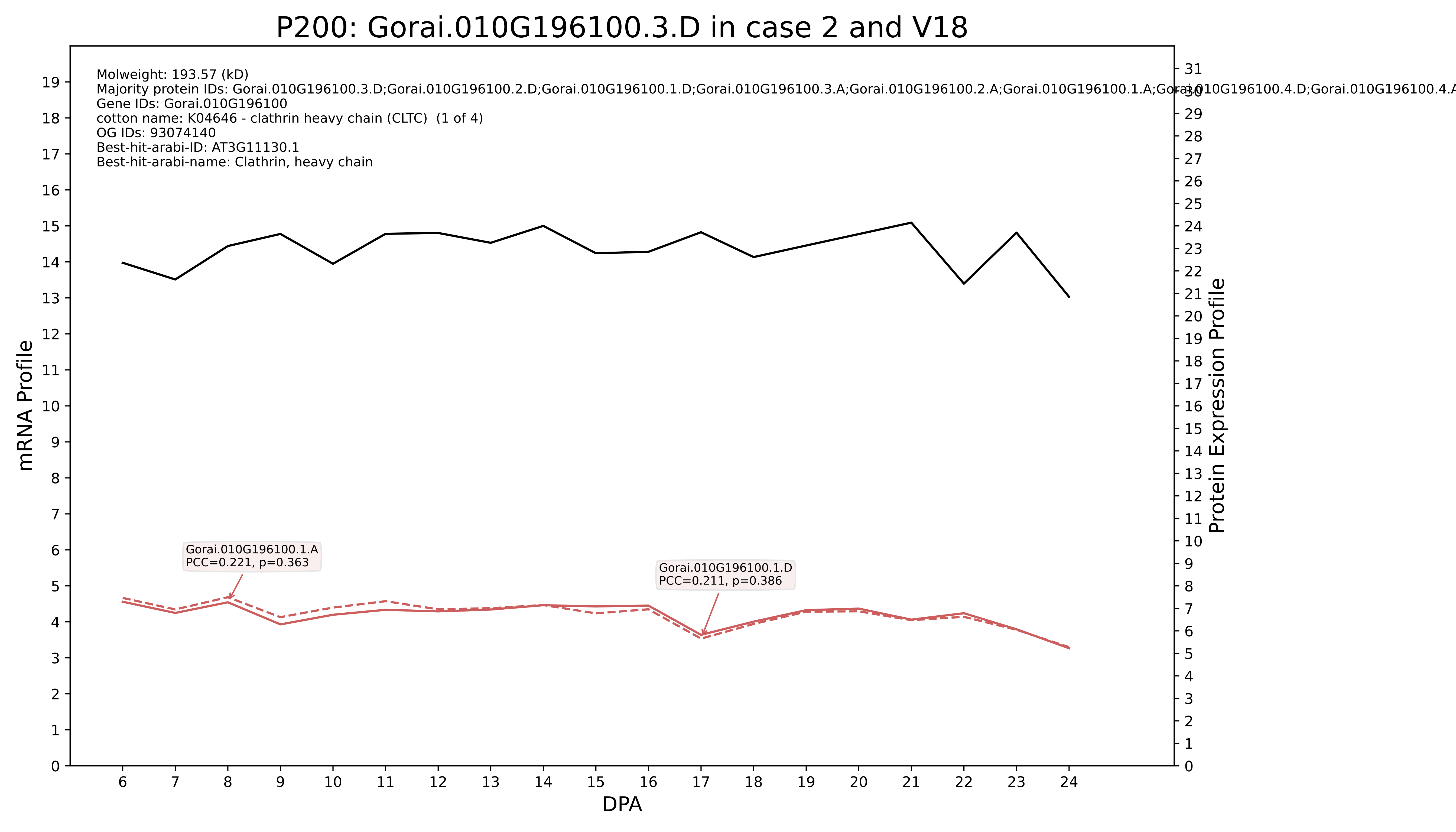
P200: Gorai.010G196100.3.D in case 2 and V18
Molweight: 193.57 (kD)
Majority protein IDs: Gorai.010G196100.3.D;Gorai.010G196100.2.D;Gorai.010G196100.1.D;Gorai.010G196100.3.A;Gorai.010G196100.2.A;Gorai.010G196100.1.A;Gorai.010G196100.4.D;Gorai.010G196100.4.A
Gene IDs: Gorai.010G196100
cotton name: K04646 - clathrin heavy chain (CLTC) (1 of 4)
OG IDs: 93074140
Best-hit-arabi-ID: AT3G11130.1
Best-hit-arabi-name: Clathrin, heavy chain
Gorai.010G196100.1.A
PCC=0.221, p=0.363
Gorai.010G196100.1.D
PCC=0.211, p=0.386
mRNA Profile
Protein Expression Profile
DPA

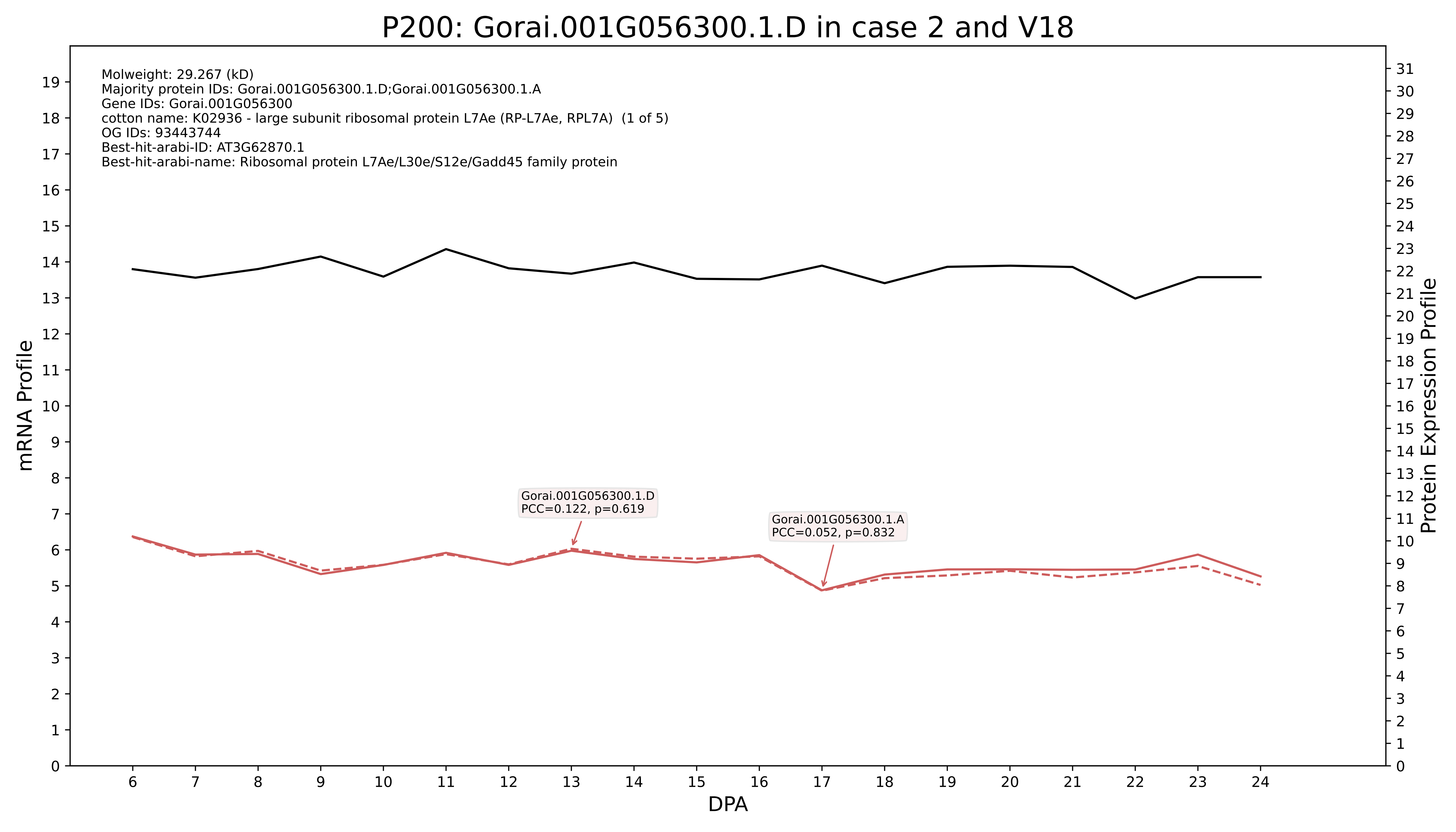
P200: Gorai.001G056300.1.D in case 2 and V18
Molweight: 29.267 (kD)
Majority protein IDs: Gorai.001G056300.1.D;Gorai.001G056300.1.A
Gene IDs: Gorai.001G056300
cotton name: K02936 - large subunit ribosomal protein L7Ae (RP-L7Ae, RPL7A) (1 of 5)
OG IDs: 93443744
Best-hit-arabi-ID: AT3G62870.1
Best-hit-arabi-name: Ribosomal protein L7Ae/L30e/S12e/Gadd45 family protein
Gorai.001G056300.1.D
PCC=0.122, p=0.619
Gorai.001G056300.1.A
PCC=0.052, p=0.832
mRNA Profile
Protein Expression Profile
DPA

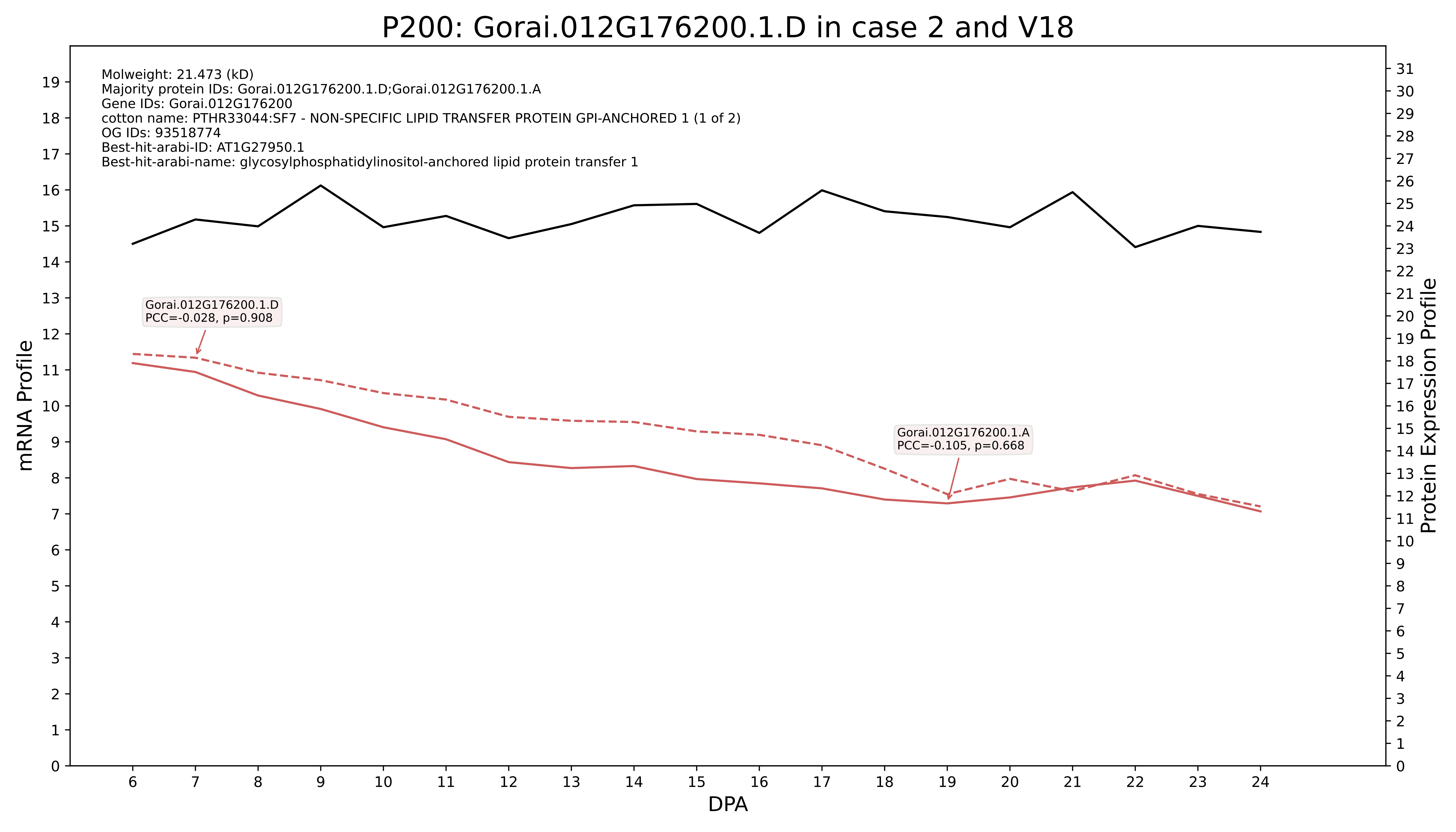
P200: Gorai.012G176200.1.D in case 2 and V18
Molweight: 21.473 (kD)
Majority protein IDs: Gorai.012G176200.1.D;Gorai.012G176200.1.A
Gene IDs: Gorai.012G176200
cotton name: PTHR33044:SF7 - NON-SPECIFIC LIPID TRANSFER PROTEIN GPI-ANCHORED 1 (1 of 2)
OG IDs: 93518774
Best-hit-arabi-ID: AT1G27950.1
Best-hit-arabi-name: glycosylphosphatidylinositol-anchored lipid protein transfer 1
Gorai.012G176200.1.D
PCC=-0.028, p=0.908
Gorai.012G176200.1.A
PCC=-0.105, p=0.668
mRNA Profile
Protein Expression Profile
DPA

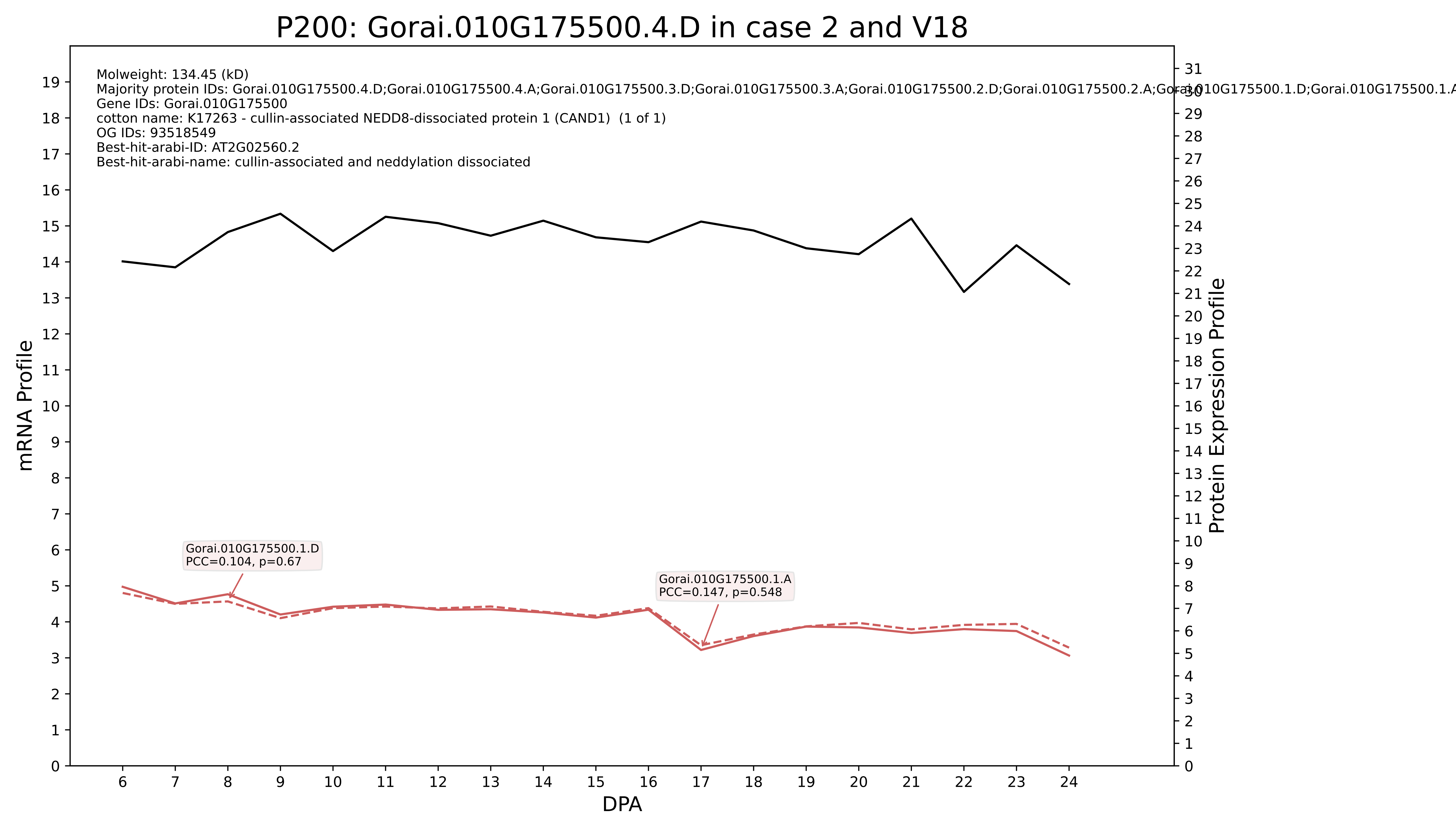

P200: Gorai.010G175500.4.D in case 2 and V18
Molweight: 134.45 (kD)
Majority protein IDs: Gorai.010G175500.4.D;Gorai.010G175500.4.A;Gorai.010G175500.3.D;Gorai.010G175500.3.A;Gorai.010G175500.2.D;Gorai.010G175500.2.A;Gorai.010G175500.1.D;Gorai.010G175500.1.A
Gene IDs: Gorai.010G175500
cotton name: K17263 - cullin-associated NEDD8-dissociated protein 1 (CAND1) (1 of 1)
OG IDs: 93518549
Best-hit-arabi-ID: AT2G02560.2
Best-hit-arabi-name: cullin-associated and neddylation dissociated
Gorai.010G175500.1.D
PCC=0.104, p=0.67
Gorai.010G175500.1.A
PCC=0.147, p=0.548
mRNA Profile
Protein Expression Profile
DPA

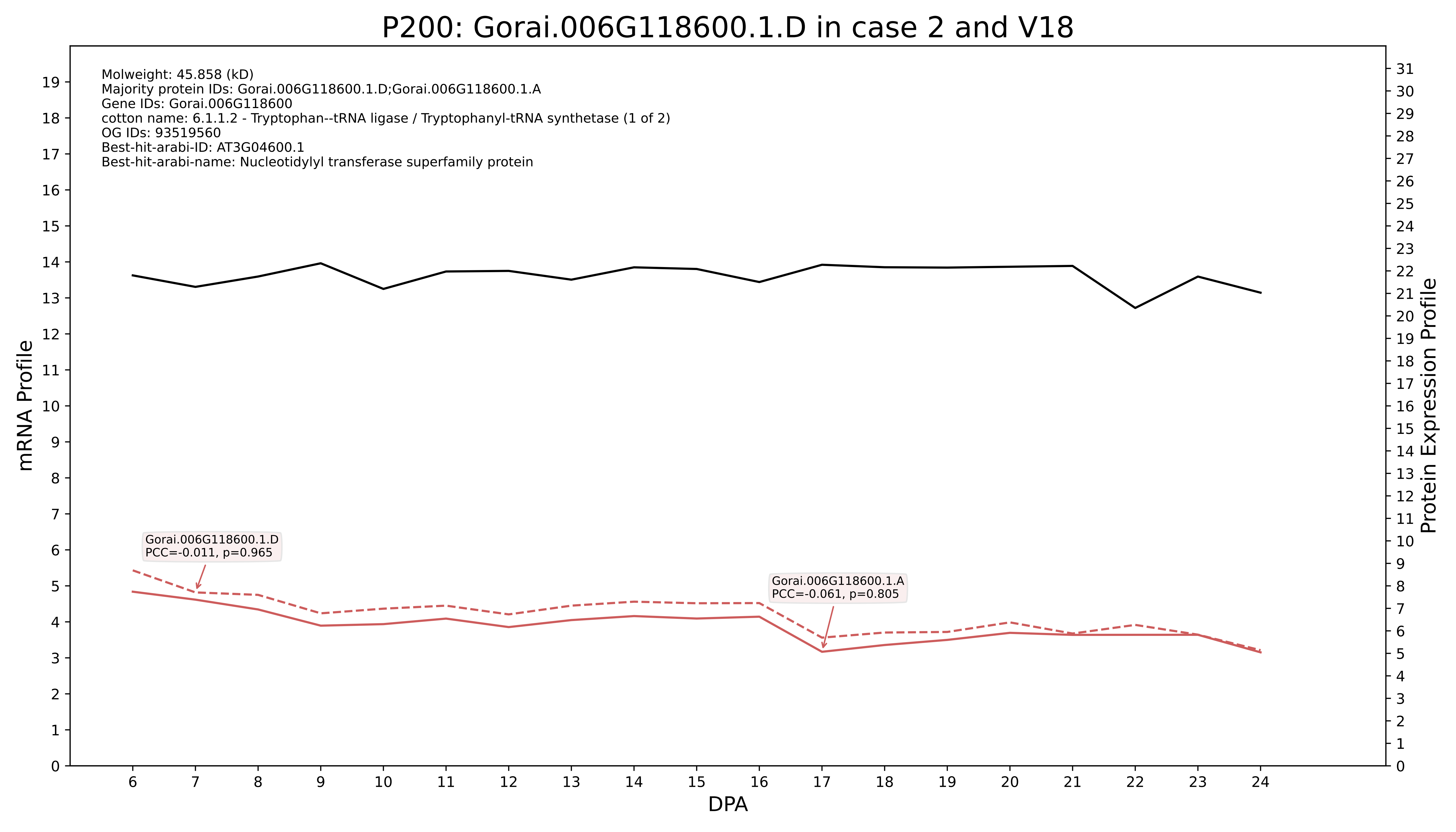
P200: Gorai.006G118600.1.D in case 2 and V18
Molweight: 45.858 (kD)
Majority protein IDs: Gorai.006G118600.1.D;Gorai.006G118600.1.A
Gene IDs: Gorai.006G118600
cotton name: 6.1.1.2 - Tryptophan--tRNA ligase / Tryptophanyl-tRNA synthetase (1 of 2)
OG IDs: 93519560
Best-hit-arabi-ID: AT3G04600.1
Best-hit-arabi-name: Nucleotidylyl transferase superfamily protein
Gorai.006G118600.1.D
PCC=-0.011, p=0.965
Gorai.006G118600.1.A
PCC=-0.061, p=0.805
mRNA Profile
Protein Expression Profile
DPA

200: Gorai.001G147400.6.A in case 2 and V18

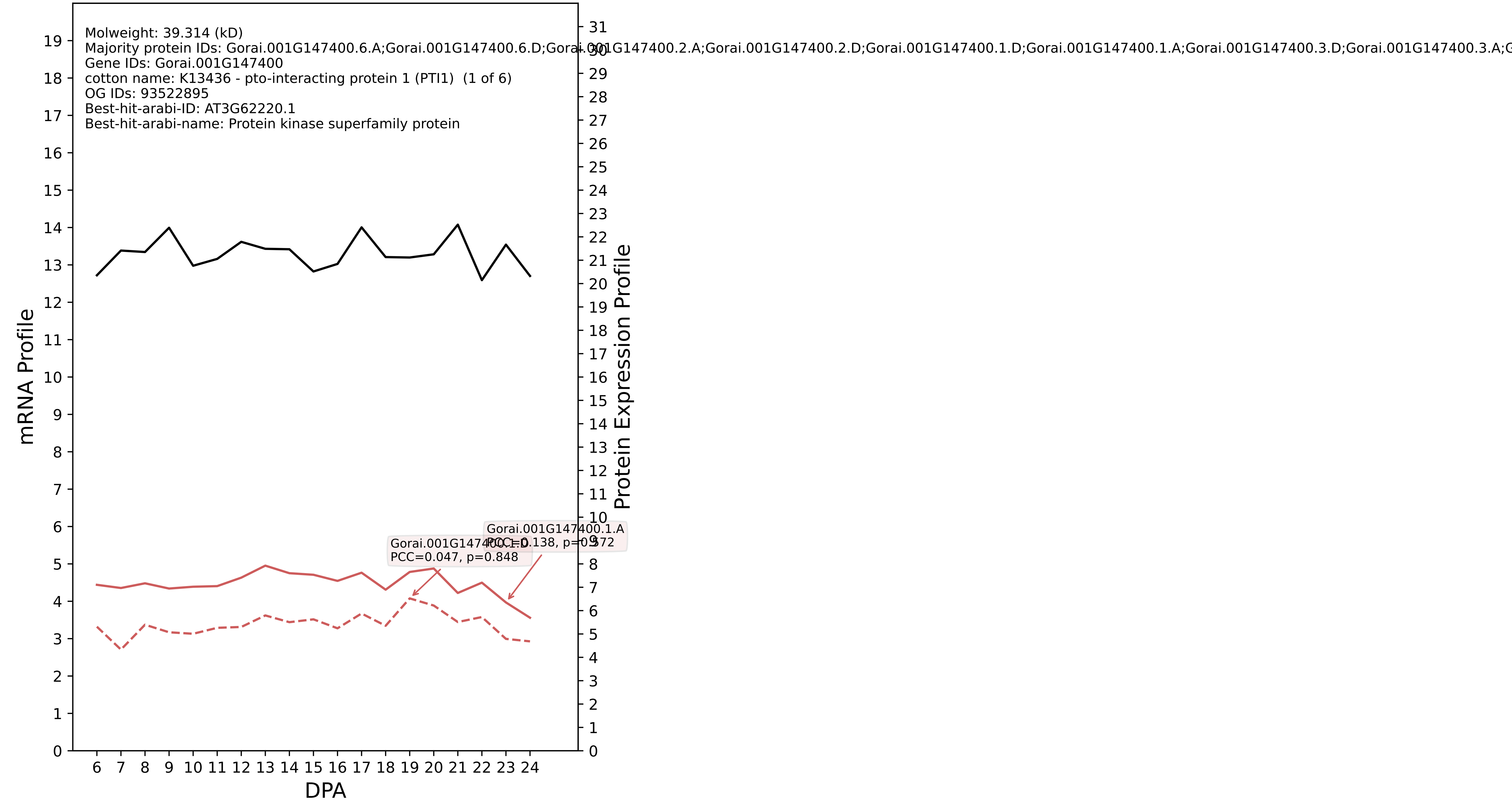

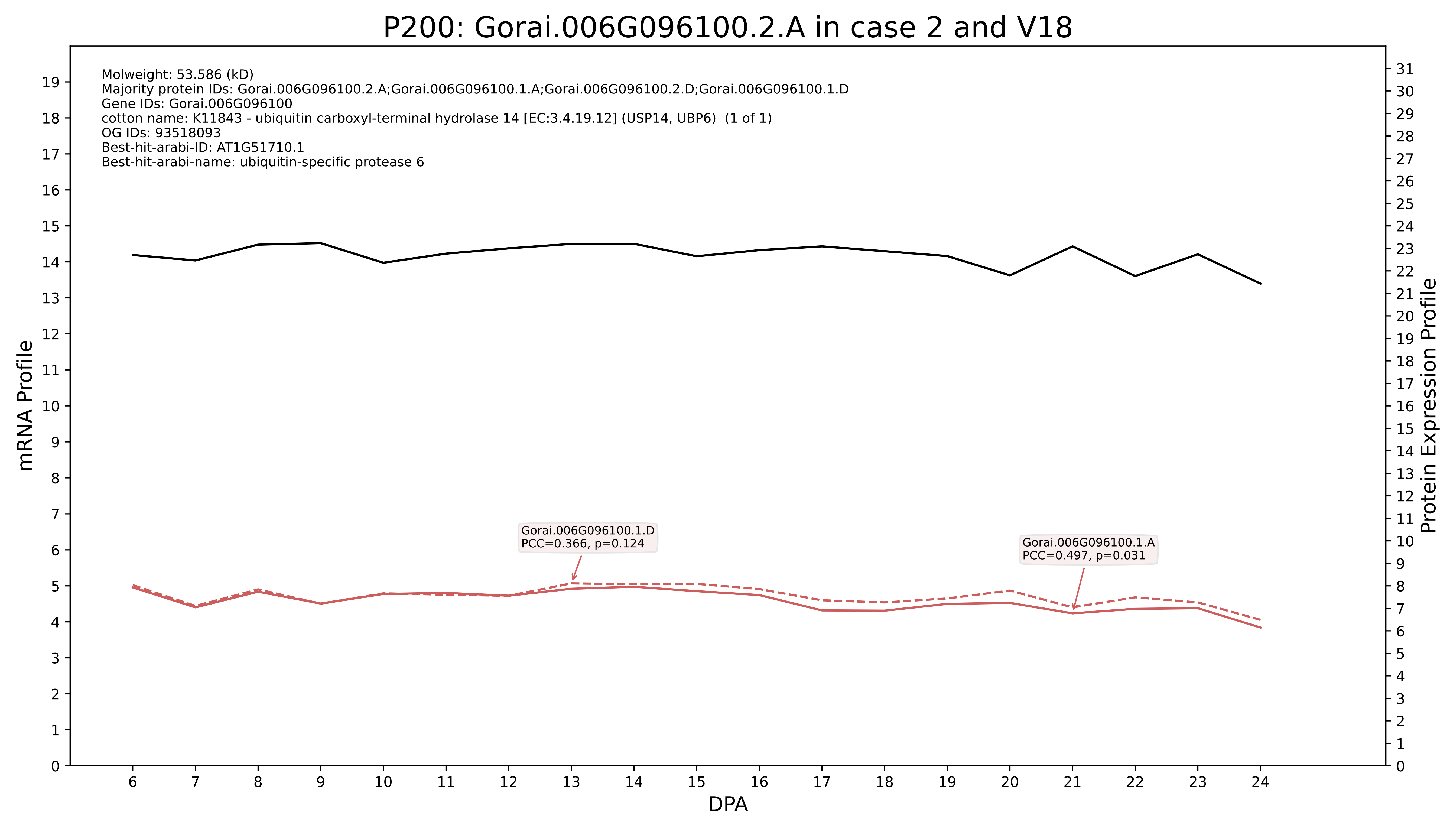

P200: Gorai.006G096100.2.A in case 2 and V18
Molweight: 53.586 (kD)
Majority protein IDs: Gorai.006G096100.2.A;Gorai.006G096100.1.A;Gorai.006G096100.2.D;Gorai.006G096100.1.D
Gene IDs: Gorai.006G096100
cotton name: K11843 - ubiquitin carboxyl-terminal hydrolase 14 [EC:3.4.19.12] (USP14, UBP6) (1 of 1)
OG IDs: 93518093
Best-hit-arabi-ID: AT1G51710.1
Best-hit-arabi-name: ubiquitin-specific protease 6
Gorai.006G096100.1.D
PCC=0.366, p=0.124
Gorai.006G096100.1.A
PCC=0.497, p=0.031
mRNA Profile
Protein Expression Profile
DPA

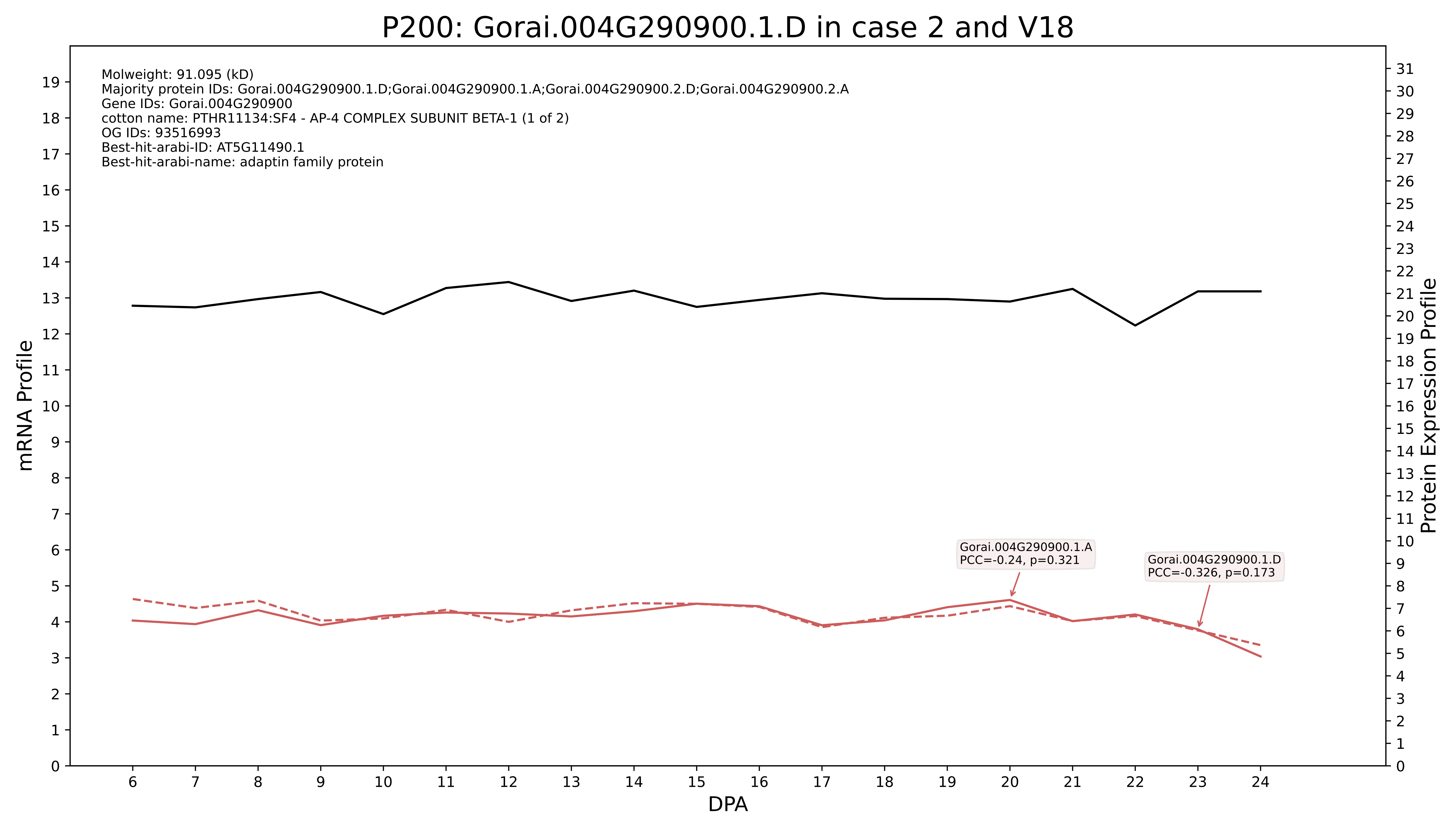
P200: Gorai.004G290900.1.D in case 2 and V18
Molweight: 91.095 (kD)
Majority protein IDs: Gorai.004G290900.1.D;Gorai.004G290900.1.A;Gorai.004G290900.2.D;Gorai.004G290900.2.A
Gene IDs: Gorai.004G290900
cotton name: PTHR11134:SF4 - AP-4 COMPLEX SUBUNIT BETA-1 (1 of 2)
OG IDs: 93516993
Best-hit-arabi-ID: AT5G11490.1
Best-hit-arabi-name: adaptin family protein
Gorai.004G290900.1.A
PCC=-0.24, p=0.321
Gorai.004G290900.1.D
PCC=-0.326, p=0.173
mRNA Profile
Protein Expression Profile
DPA

### P200: Gorai.004G139500.2.D in case 2 and V18

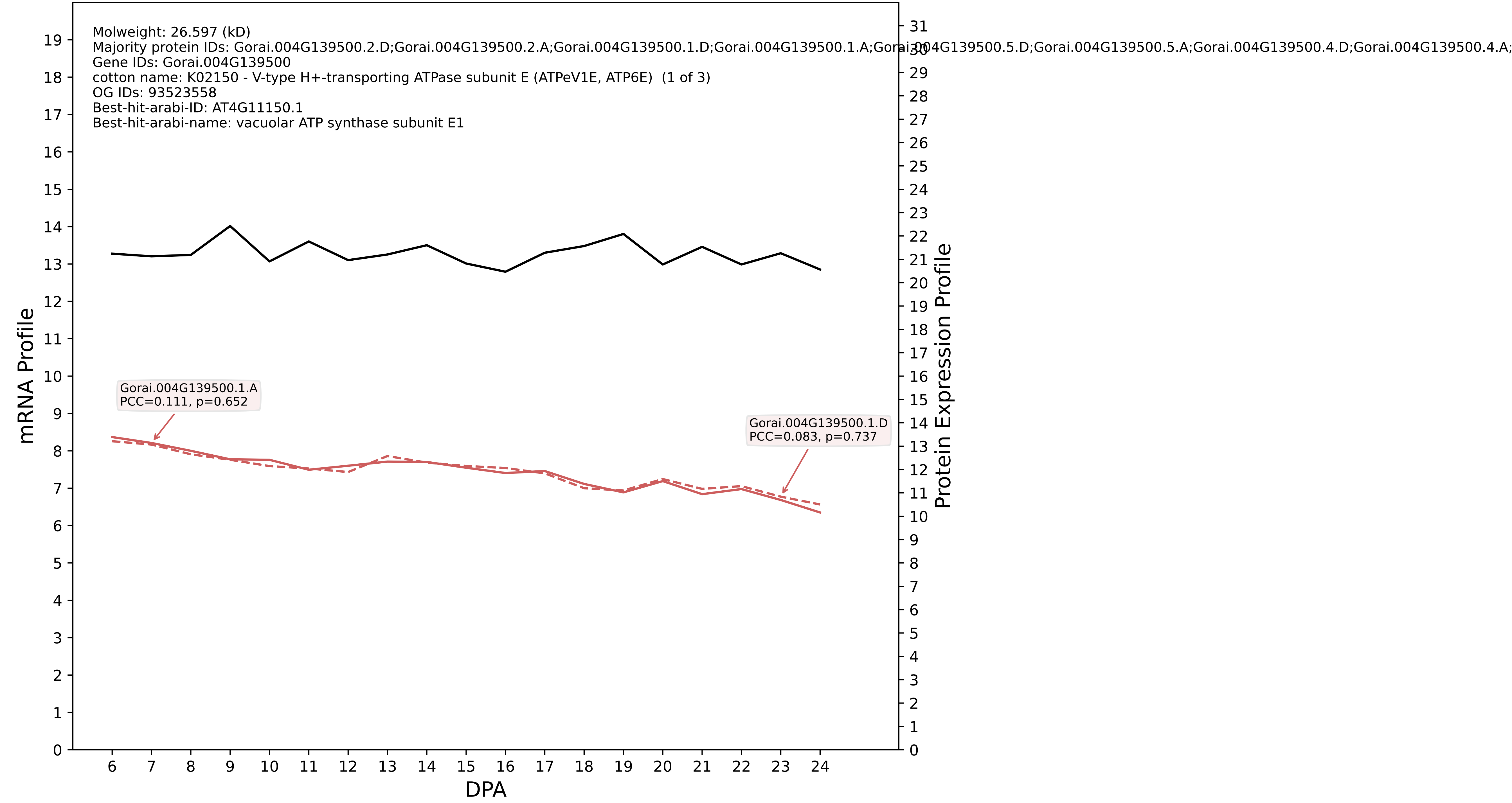

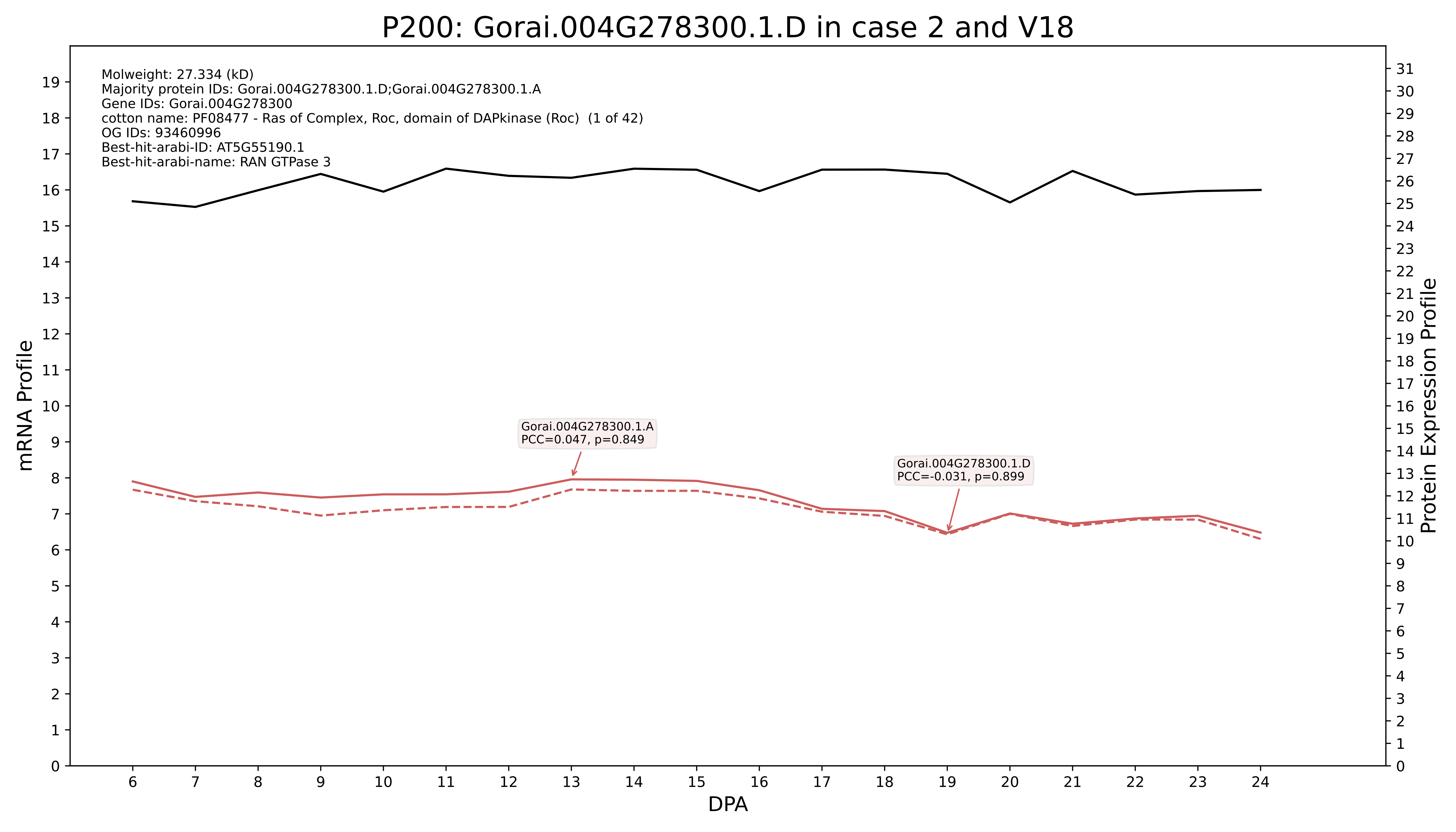
P200: Gorai.004G278300.1.D in case 2 and V18
Molweight: 27.334 (kD)
Majority protein IDs: Gorai.004G278300.1.D;Gorai.004G278300.1.A
Gene IDs: Gorai.004G278300
cotton name: PF08477 - Ras of Complex, Roc, domain of DAPkinase (Roc) (1 of 42)
OG IDs: 93460996
Best-hit-arabi-ID: AT5G55190.1
Best-hit-arabi-name: RAN GTPase 3
Gorai.004G278300.1.A
PCC=0.047, p=0.849
Gorai.004G278300.1.D
PCC=-0.031, p=0.899
mRNA Profile
Protein Expression Profile
DPA

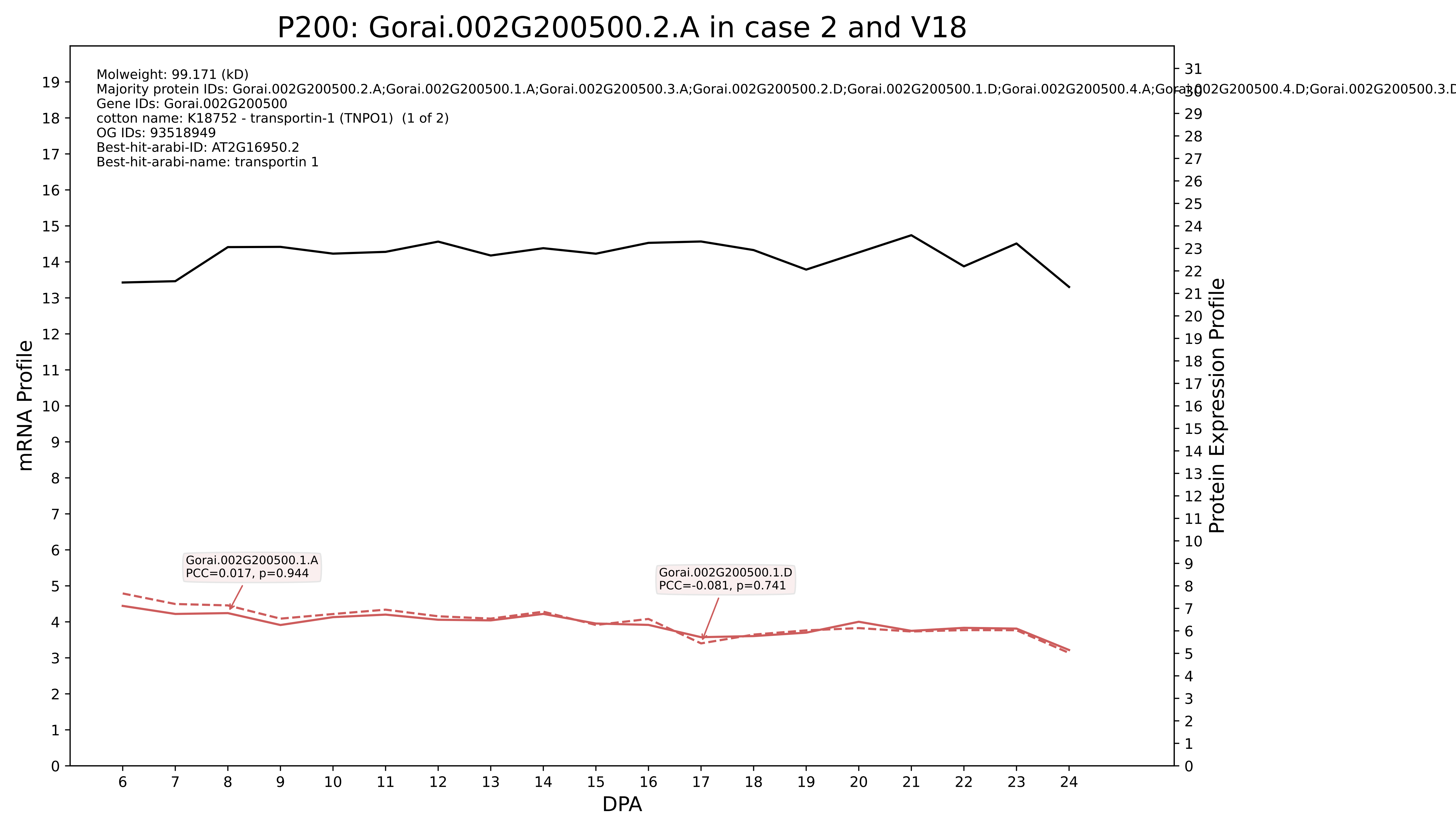
P200: Gorai.002G200500.2.A in case 2 and V18
Molweight: 99.171 (kD)
Majority protein IDs: Gorai.002G200500.2.A;Gorai.002G200500.1.A;Gorai.002G200500.3.A;Gorai.002G200500.2.D;Gorai.002G200500.1.D;Gorai.002G200500.4.A;Gorai.002G200500.4.D;Gorai.002G200500.3.D
Gene IDs: Gorai.002G200500
cotton name: K18752 - transportin-1 (TNPO1) (1 of 2)
OG IDs: 93518949
Best-hit-arabi-ID: AT2G16950.2
Best-hit-arabi-name: transportin 1
Gorai.002G200500.1.A
PCC=0.017, p=0.944
Gorai.002G200500.1.D
PCC=-0.081, p=0.741
mRNA Profile
Protein Expression Profile
DPA

### 00: Gorai.005G216500.1.D in case 2 and V18

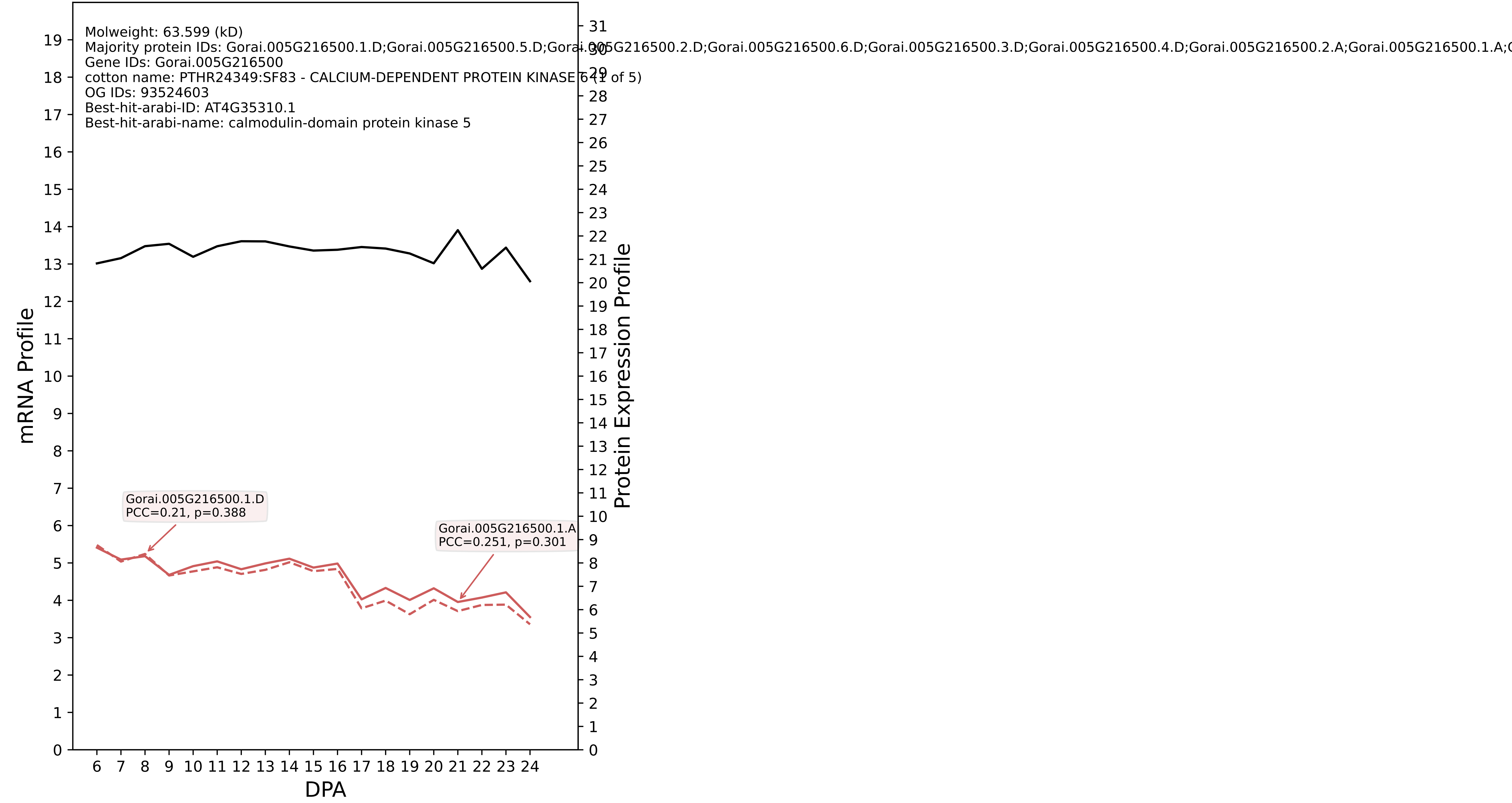

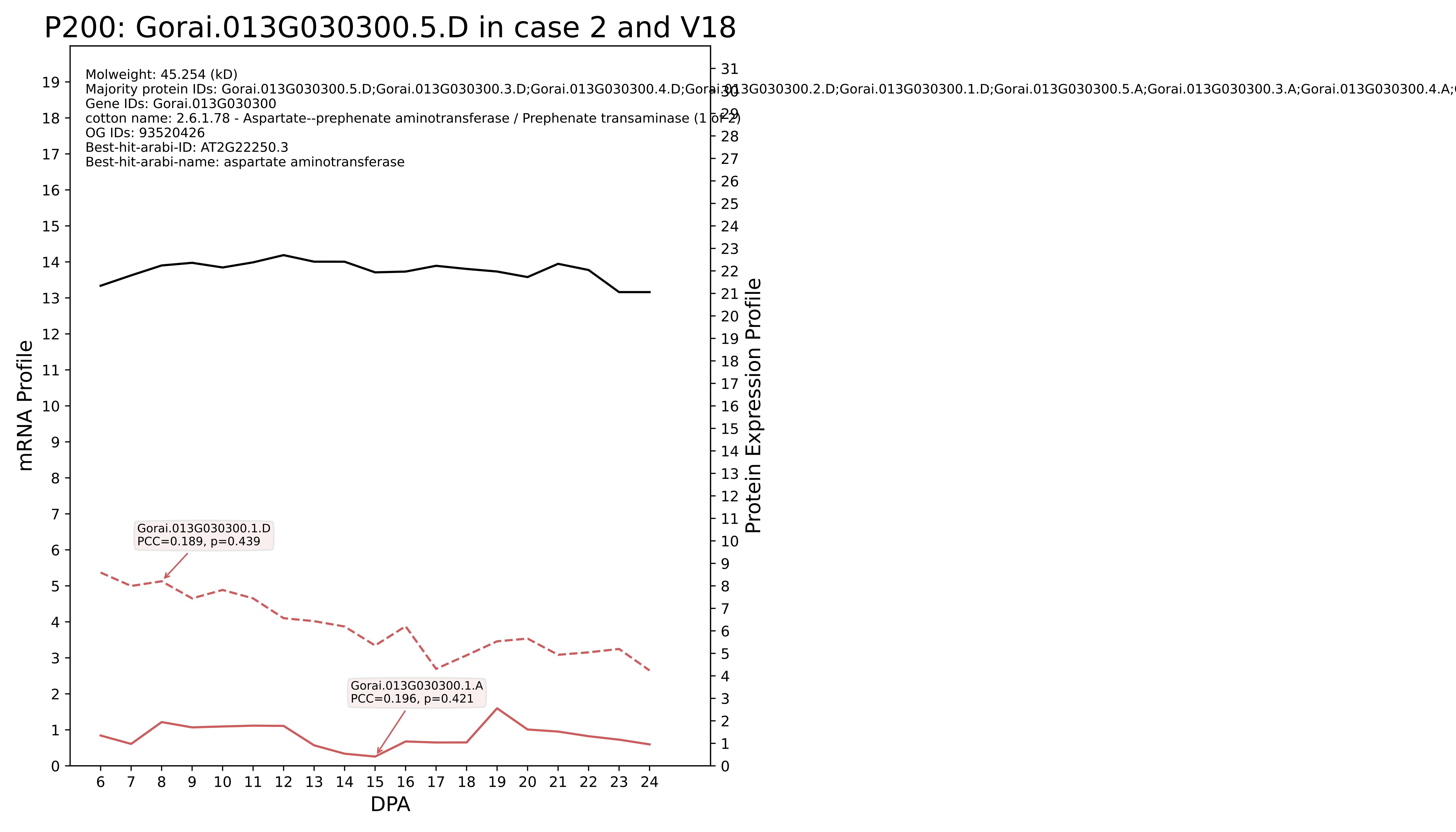
P200: Gorai.013G030300.5.D in case 2 and V18
Molweight: 45.254 (kD)
Majority protein IDs: Gorai.013G030300.5.D;Gorai.013G030300.3.D;Gorai.013G030300.4.D;Gorai.013G030300.2.D;Gorai.013G030300.1.D;Gorai.013G030300.5.A;Gorai.013G030300.3.A;Gorai.013G030300.4.A;
Gene IDs: Gorai.013G030300
cotton name: 2.6.1.78 - Aspartate--prephenate aminotransferase / Prephenate transaminase (1 of 2)
OG IDs: 93520426
Best-hit-arabi-ID: AT2G22250.3
Best-hit-arabi-name: aspartate aminotransferase
Gorai.013G030300.1.D
PCC=0.189, p=0.439
Gorai.013G030300.1.A
PCC=0.196, p=0.421
mRNA Profile
Protein Expression Profile
DPA

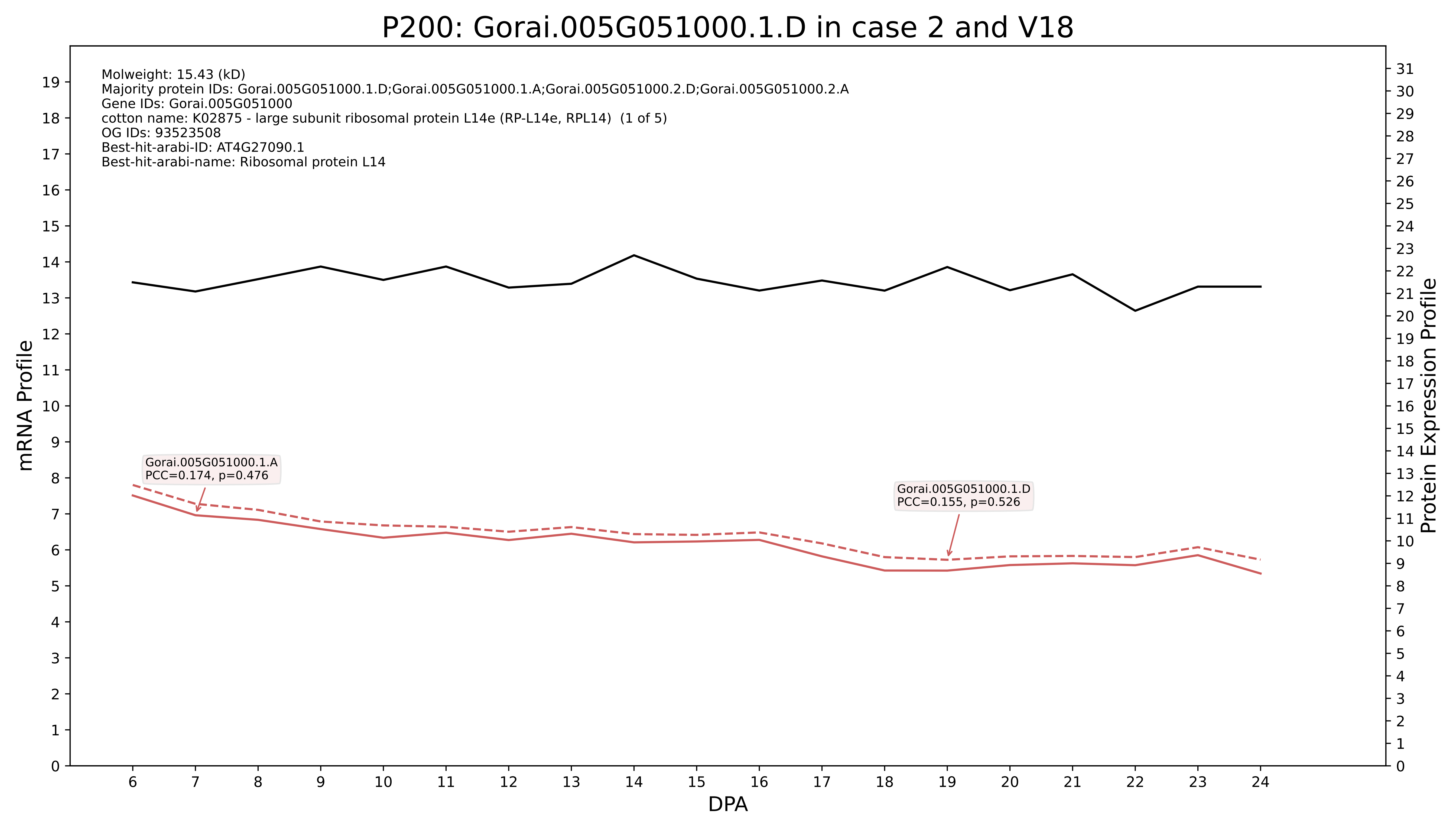
P200: Gorai.005G051000.1.D in case 2 and V18
Molweight: 15.43 (kD)
Majority protein IDs: Gorai.005G051000.1.D;Gorai.005G051000.1.A;Gorai.005G051000.2.D;Gorai.005G051000.2.A
Gene IDs: Gorai.005G051000
cotton name: K02875 - large subunit ribosomal protein L14e (RP-L14e, RPL14) (1 of 5)
OG IDs: 93523508
Best-hit-arabi-ID: AT4G27090.1
Best-hit-arabi-name: Ribosomal protein L14
Gorai.005G051000.1.A
PCC=0.174, p=0.476
Gorai.005G051000.1.D
PCC=0.155, p=0.526
mRNA Profile
Protein Expression Profile
DPA

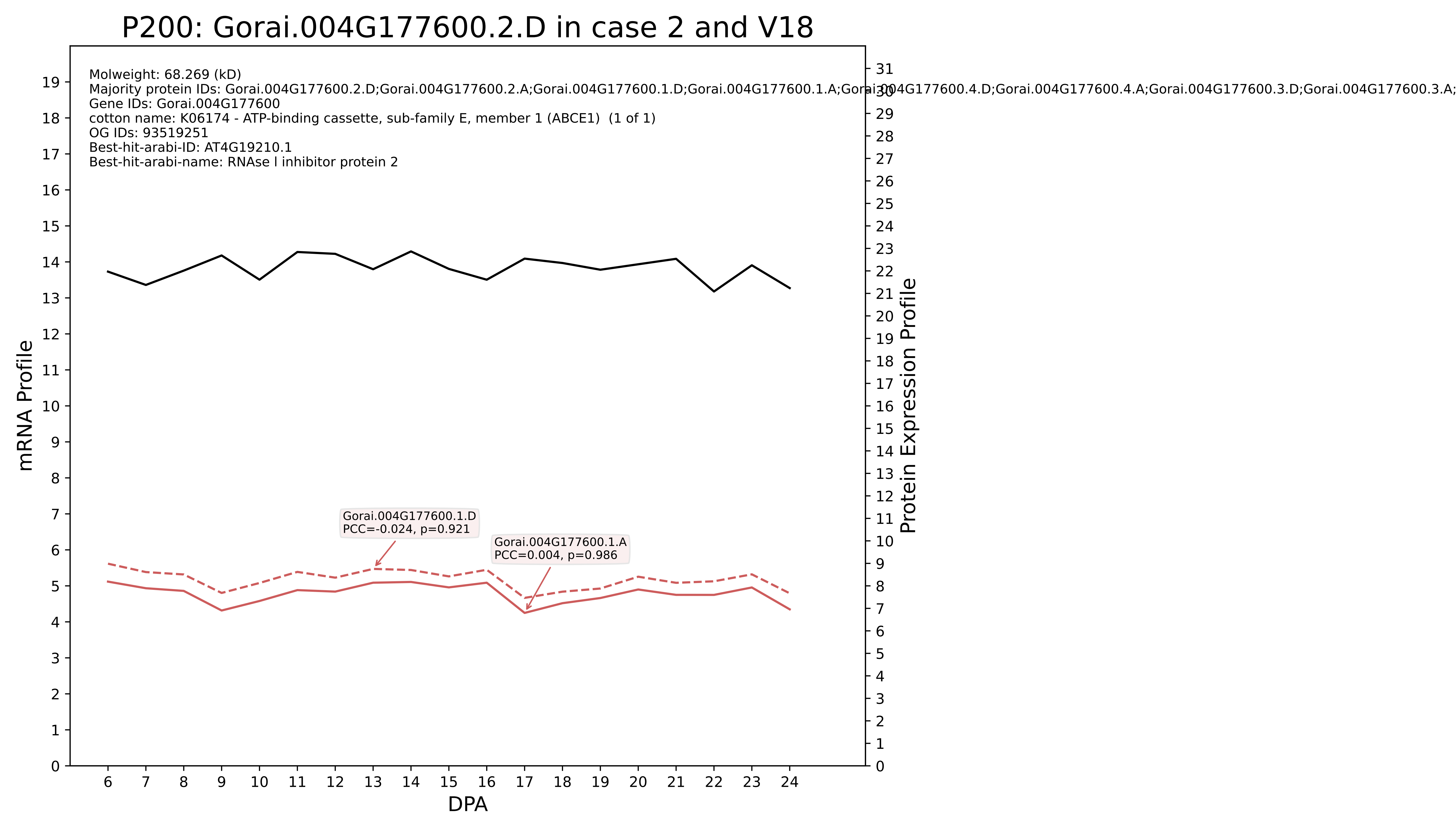

P200: Gorai.004G177600.2.D in case 2 and V18
Molweight: 68.269 (kD)
Majority protein IDs: Gorai.004G177600.2.D;Gorai.004G177600.2.A;Gorai.004G177600.1.D;Gorai.004G177600.1.A;Gorai.004G177600.4.D;Gorai.004G177600.4.A;Gorai.004G177600.3.D;Gorai.004G177600.3.A;
Gene IDs: Gorai.004G177600
cotton name: K06174 - ATP-binding cassette, sub-family E, member 1 (ABCE1) (1 of 1)
OG IDs: 93519251
Best-hit-arabi-ID: AT4G19210.1
Best-hit-arabi-name: RNAse l inhibitor protein 2
Gorai.004G177600.1.D
PCC=-0.024, p=0.921
Gorai.004G177600.1.A
PCC=0.004, p=0.986
mRNA Profile
Protein Expression Profile
DPA

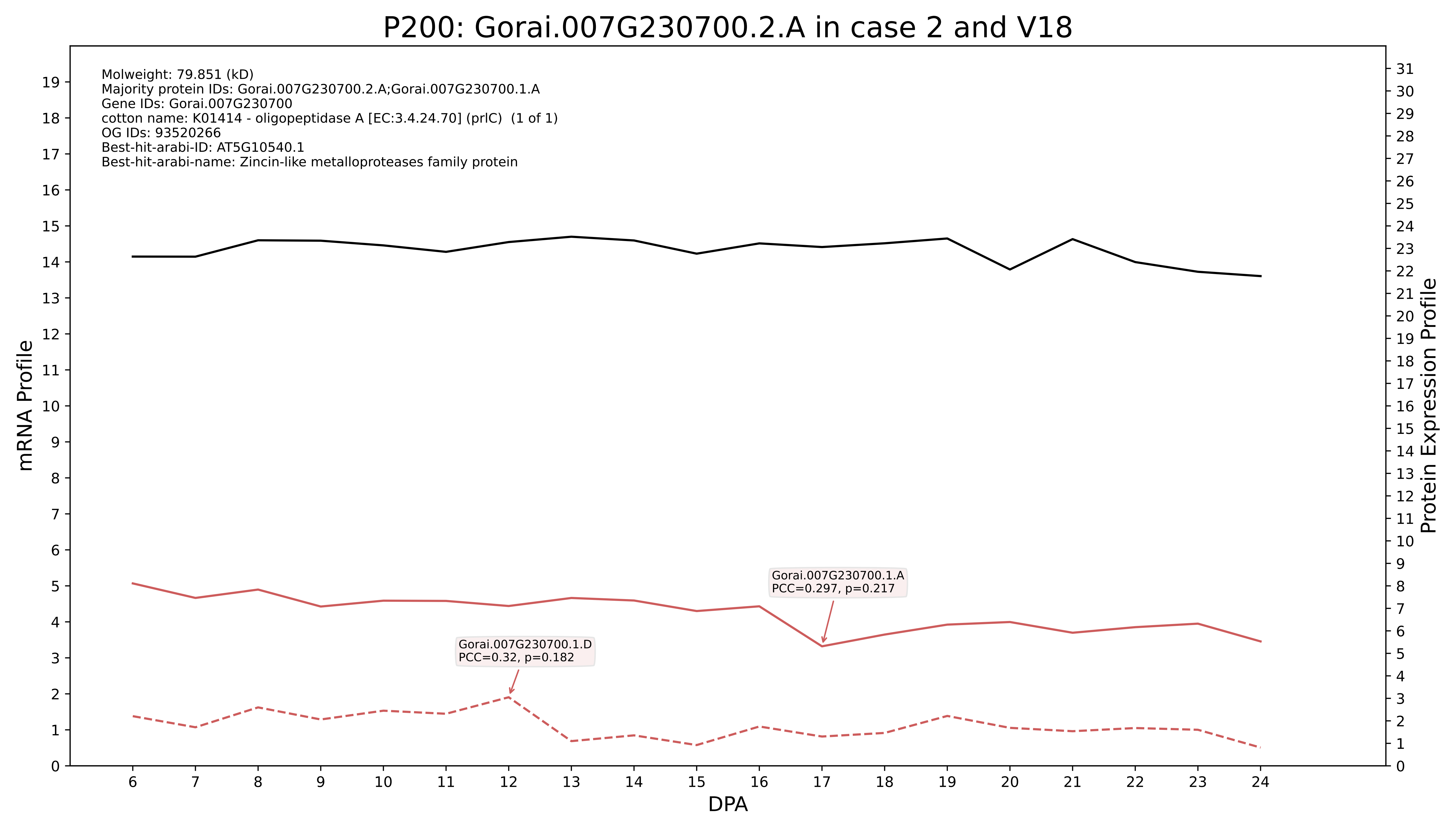
P200: Gorai.007G230700.2.A in case 2 and V18
Molweight: 79.851 (kD)
Majority protein IDs: Gorai.007G230700.2.A;Gorai.007G230700.1.A
Gene IDs: Gorai.007G230700
cotton name: K01414 - oligopeptidase A [EC:3.4.24.70] (prlC) (1 of 1)
OG IDs: 93520266
Best-hit-arabi-ID: AT5G10540.1
Best-hit-arabi-name: Zincin-like metalloproteases family protein
Gorai.007G230700.1.A
PCC=0.297, p=0.217
Gorai.007G230700.1.D
PCC=0.32, p=0.182
mRNA Profile
Protein Expression Profile
DPA

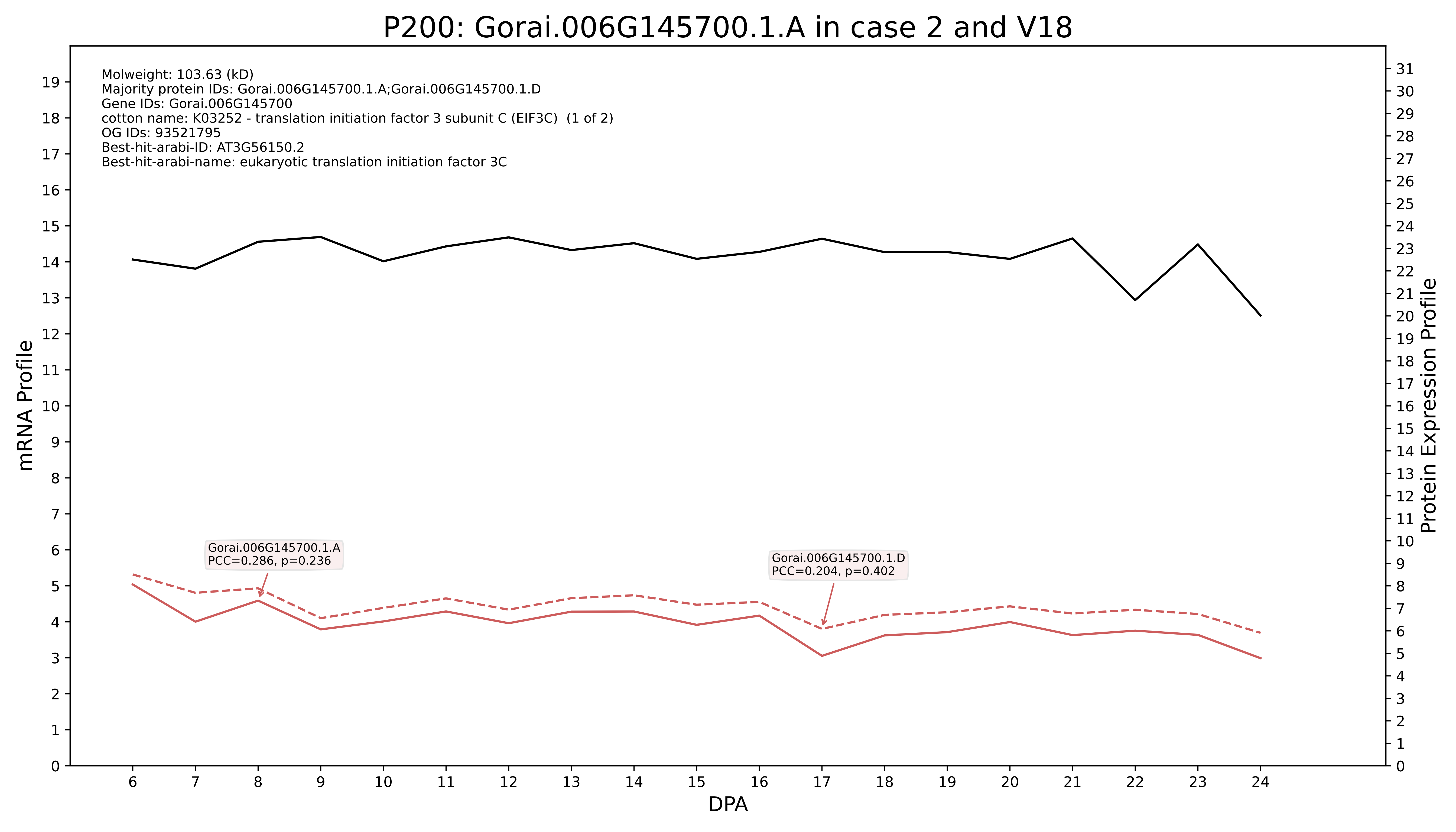
P200: Gorai.006G145700.1.A in case 2 and V18
Molweight: 103.63 (kD)
Majority protein IDs: Gorai.006G145700.1.A;Gorai.006G145700.1.D
Gene IDs: Gorai.006G145700
cotton name: K03252 - translation initiation factor 3 subunit C (EIF3C) (1 of 2)
OG IDs: 93521795
Best-hit-arabi-ID: AT3G56150.2
Best-hit-arabi-name: eukaryotic translation initiation factor 3C
Gorai.006G145700.1.A
PCC=0.286, p=0.236
Gorai.006G145700.1.D
PCC=0.204, p=0.402
mRNA Profile
Protein Expression Profile
DPA

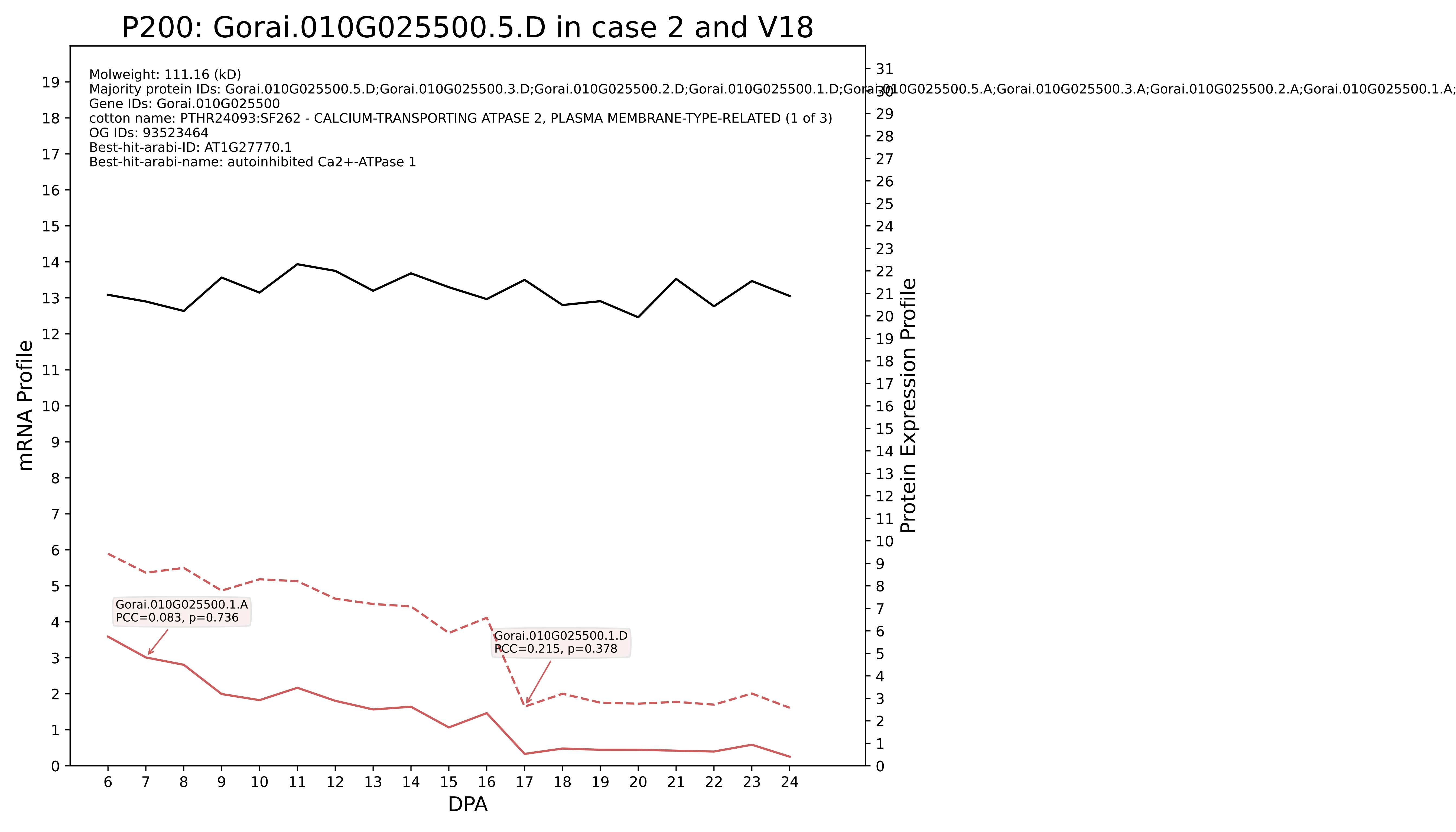
P200: Gorai.010G025500.5.D in case 2 and V18
Molweight: 111.16 (kD)
Majority protein IDs: Gorai.010G025500.5.D;Gorai.010G025500.3.D;Gorai.010G025500.2.D;Gorai.010G025500.1.D;Gorai.010G025500.5.A;Gorai.010G025500.3.A;Gorai.010G025500.2.A;Gorai.010G025500.1.A;
Gene IDs: Gorai.010G025500
cotton name: PTHR24093:SF262 - CALCIUM-TRANSPORTING ATPASE 2, PLASMA MEMBRANE-TYPE-RELATED (1 of 3)
OG IDs: 93523464
Best-hit-arabi-ID: AT1G27770.1
Best-hit-arabi-name: autoinhibited Ca2+-ATPase 1
Gorai.010G025500.1.A
PCC=0.083, p=0.736
Gorai.010G025500.1.D
PCC=0.215, p=0.378
mRNA Profile
Protein Expression Profile
DPA

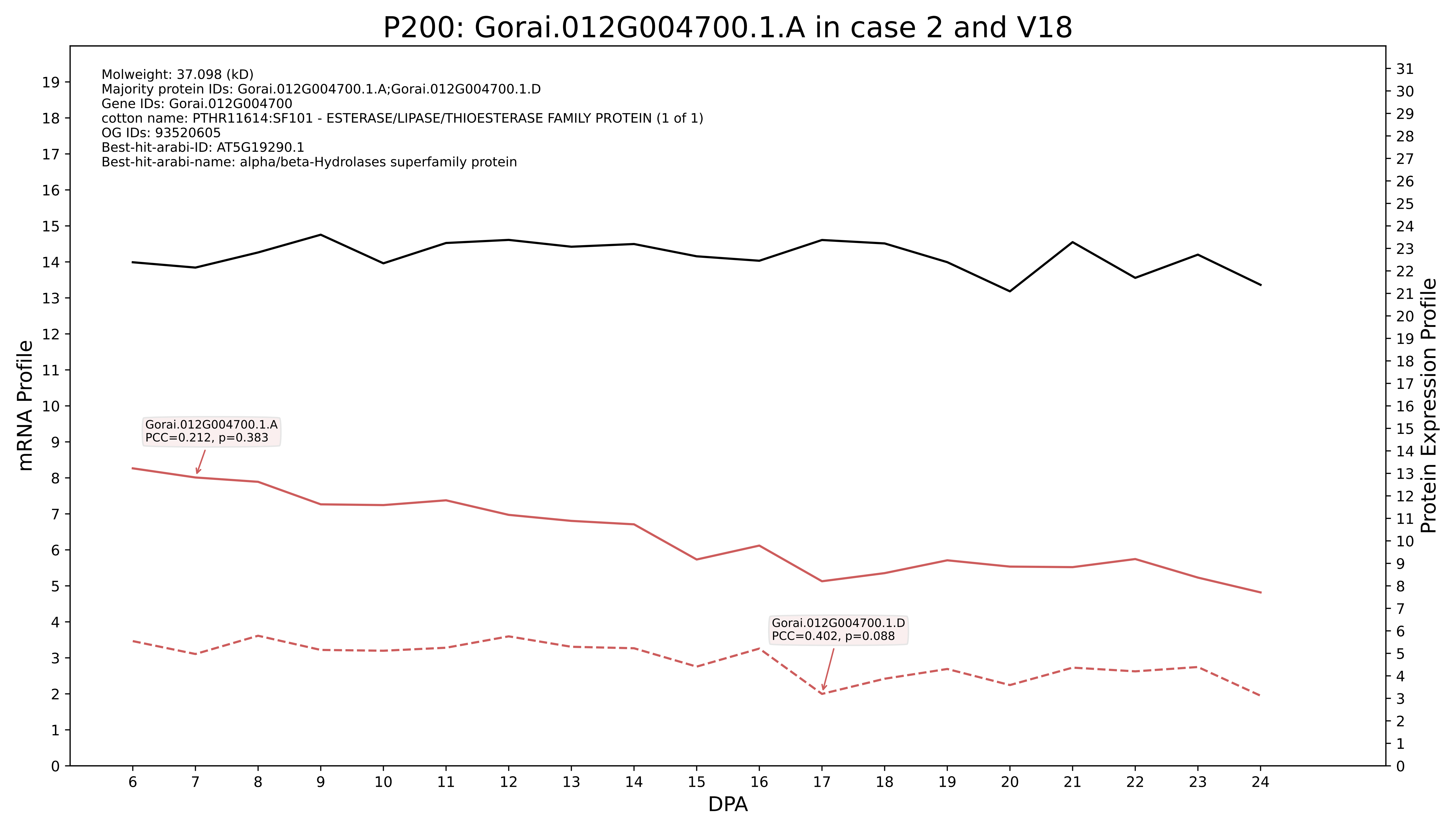

P200: Gorai.012G004700.1.A in case 2 and V18
Molweight: 37.098 (kD)
Majority protein IDs: Gorai.012G004700.1.A;Gorai.012G004700.1.D
Gene IDs: Gorai.012G004700
cotton name: PTHR11614:SF101 - ESTERASE/LIPASE/THIOESTERASE FAMILY PROTEIN (1 of 1)
OG IDs: 93520605
Best-hit-arabi-ID: AT5G19290.1
Best-hit-arabi-name: alpha/beta-Hydrolases superfamily protein
Gorai.012G004700.1.A
PCC=0.212, p=0.383
Gorai.012G004700.1.D
PCC=0.402, p=0.088
mRNA Profile
Protein Expression Profile
DPA

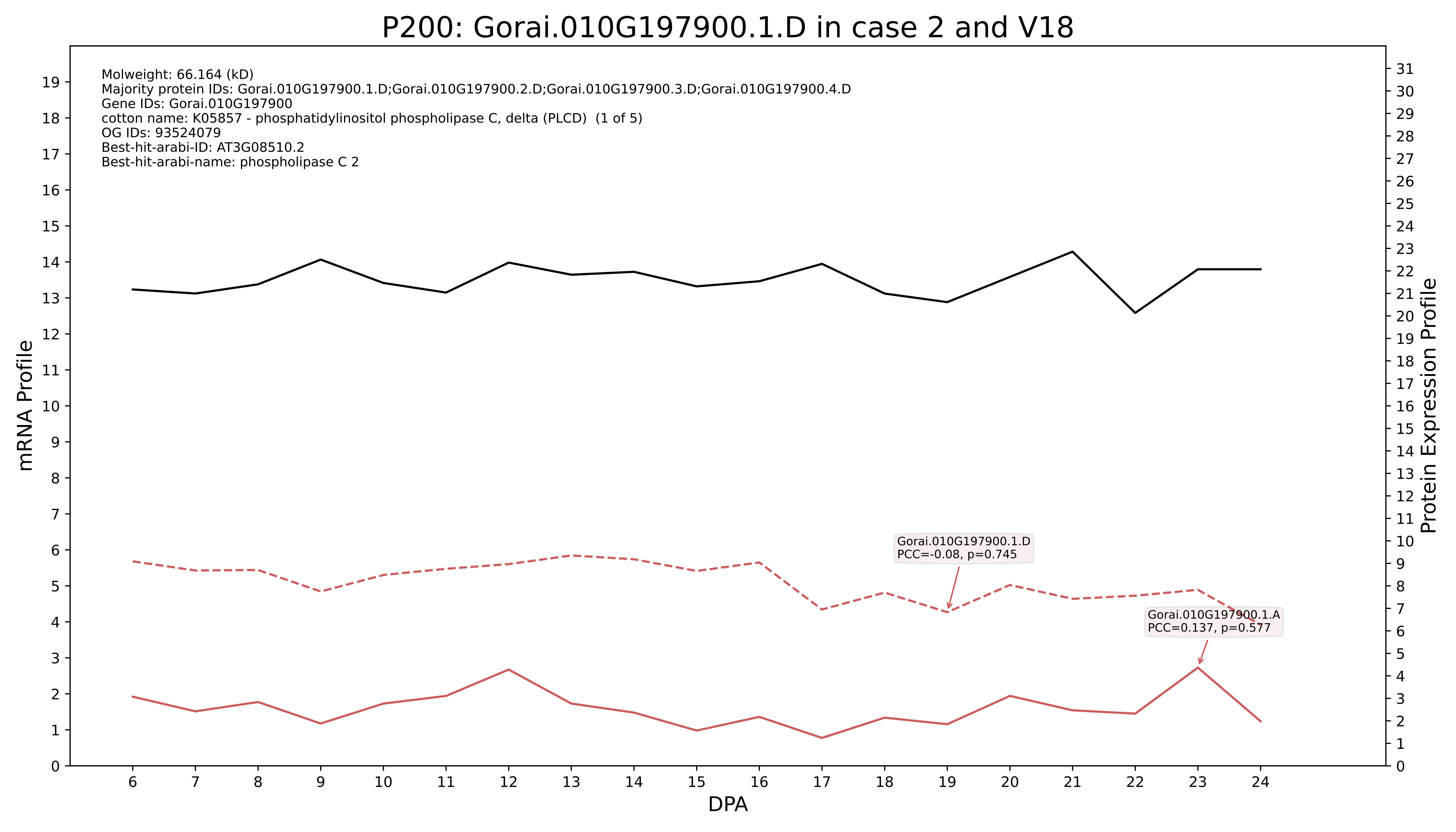
P200: Gorai.010G197900.1.D in case 2 and V18
Molweight: 66.164 (kD)
Majority protein IDs: Gorai.010G197900.1.D;Gorai.010G197900.2.D;Gorai.010G197900.3.D;Gorai.010G197900.4.D
Gene IDs: Gorai.010G197900
cotton name: K05857 - phosphatidylinositol phospholipase C, delta (PLCD) (1 of 5)
OG IDs: 93524079
Best-hit-arabi-ID: AT3G08510.2
Best-hit-arabi-name: phospholipase C 2
Gorai.010G197900.1.D
PCC=-0.08, p=0.745
Gorai.010G197900.1.A
PCC=0.137, p=0.577
mRNA Profile
Protein Expression Profile
DPA

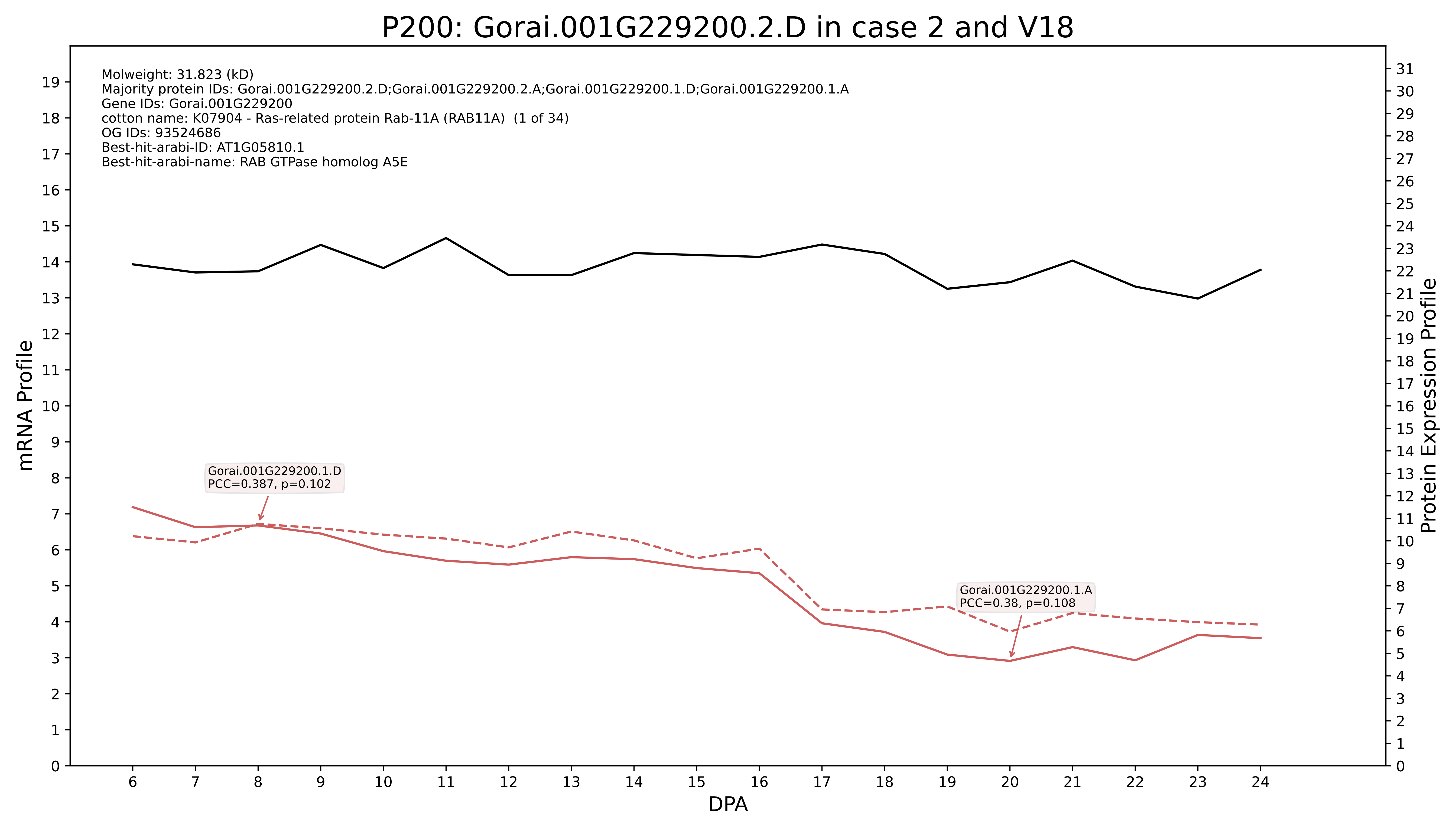
P200: Gorai.001G229200.2.D in case 2 and V18
Molweight: 31.823 (kD)
Majority protein IDs: Gorai.001G229200.2.D;Gorai.001G229200.2.A;Gorai.001G229200.1.D;Gorai.001G229200.1.A
Gene IDs: Gorai.001G229200
cotton name: K07904 - Ras-related protein Rab-11A (RAB11A) (1 of 34)
OG IDs: 93524686
Best-hit-arabi-ID: AT1G05810.1
Best-hit-arabi-name: RAB GTPase homolog A5E
Gorai.001G229200.1.D
PCC=0.387, p=0.102
Gorai.001G229200.1.A
PCC=0.38, p=0.108
mRNA Profile
Protein Expression Profile
DPA

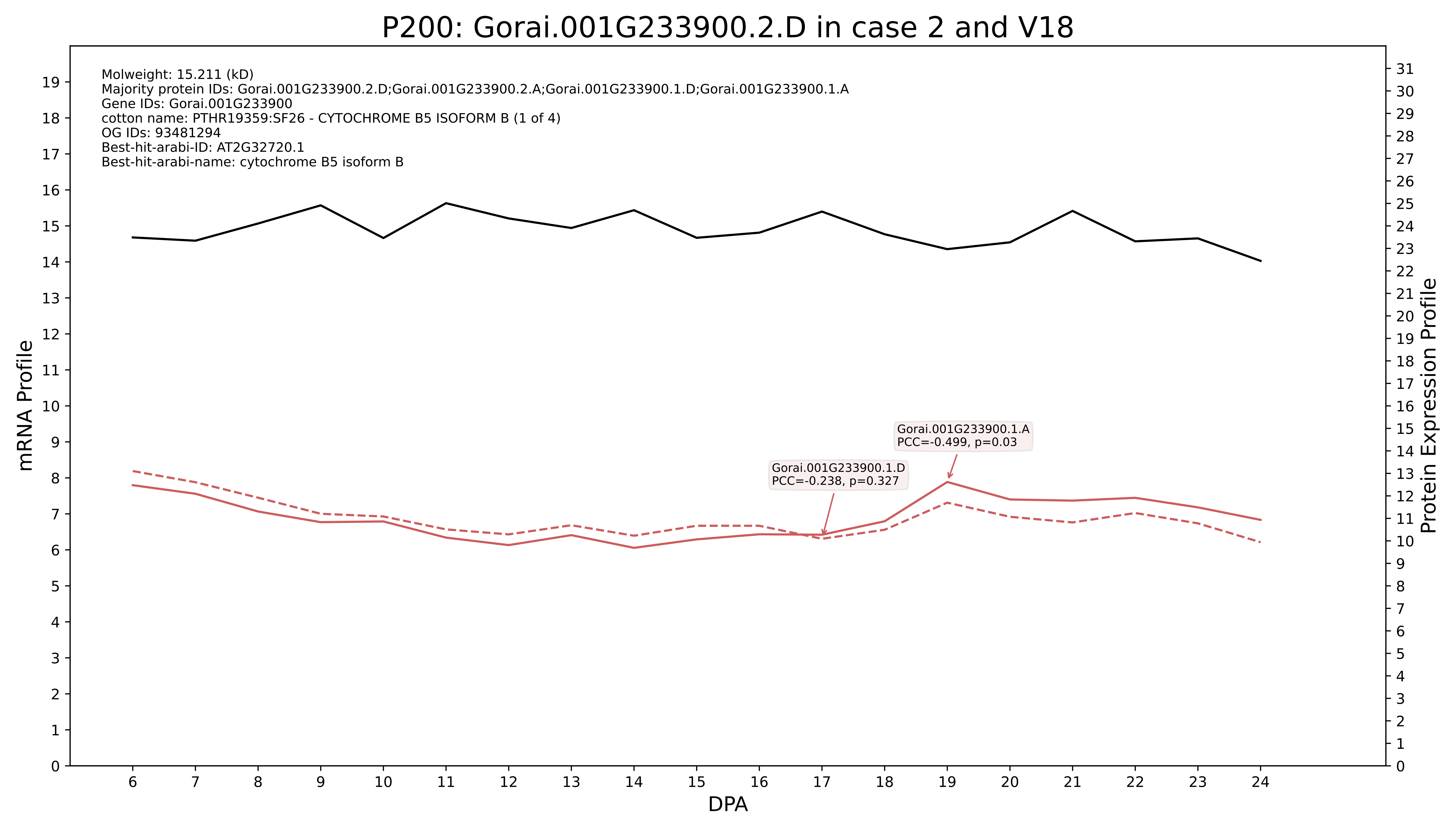
P200: Gorai.001G233900.2.D in case 2 and V18
Molweight: 15.211 (kD)
Majority protein IDs: Gorai.001G233900.2.D;Gorai.001G233900.2.A;Gorai.001G233900.1.D;Gorai.001G233900.1.A
Gene IDs: Gorai.001G233900
cotton name: PTHR19359:SF26 - CYTOCHROME B5 ISOFORM B (1 of 4)
OG IDs: 93481294
Best-hit-arabi-ID: AT2G32720.1
Best-hit-arabi-name: cytochrome B5 isoform B
Gorai.001G233900.1.A
PCC=-0.499, p=0.03
Gorai.001G233900.1.D
PCC=-0.238, p=0.327
mRNA Profile
Protein Expression Profile
DPA

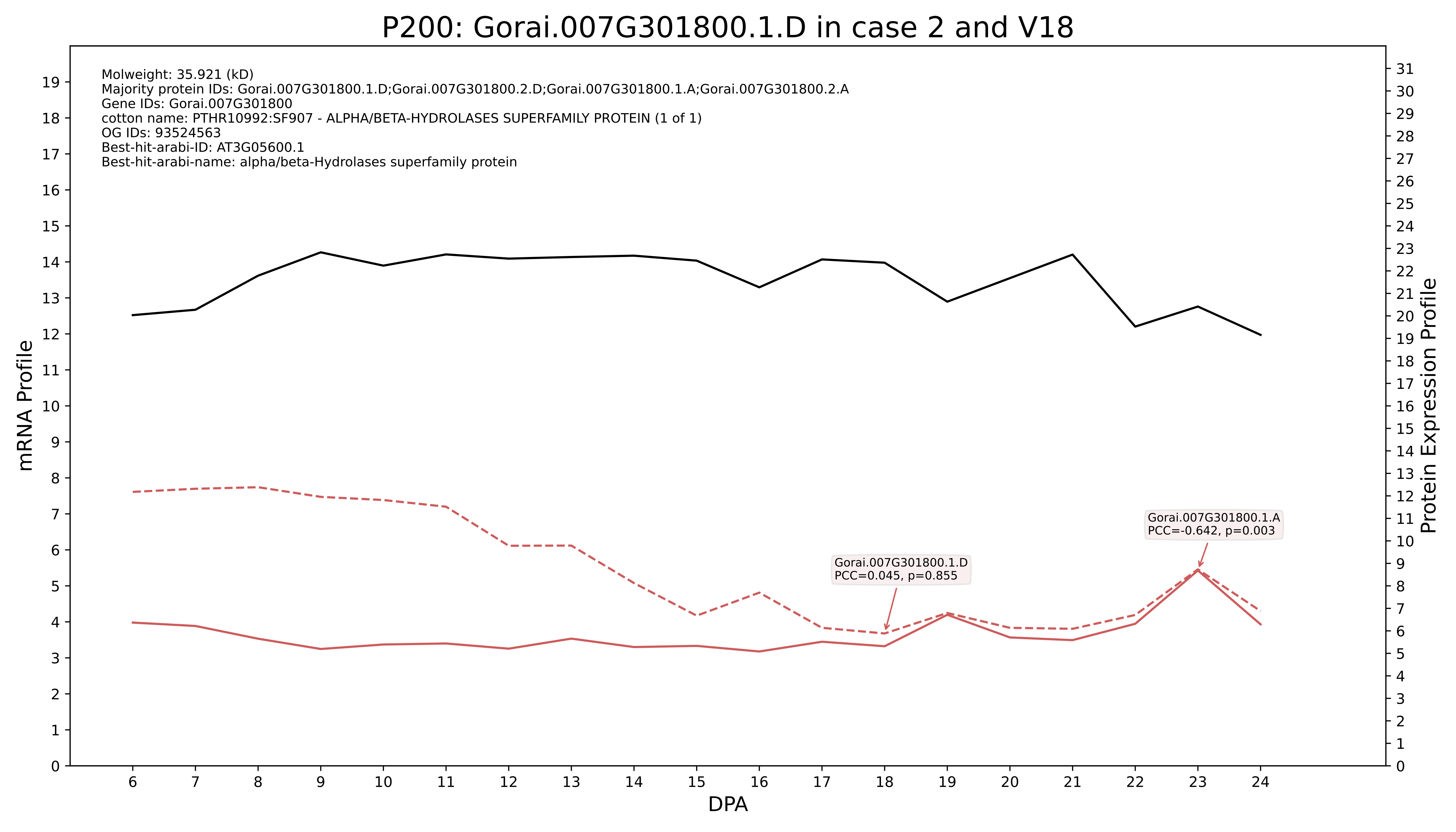
P200: Gorai.007G301800.1.D in case 2 and V18
Molweight: 35.921 (kD)
Majority protein IDs: Gorai.007G301800.1.D;Gorai.007G301800.2.D;Gorai.007G301800.1.A;Gorai.007G301800.2.A
Gene IDs: Gorai.007G301800
cotton name: PTHR10992:SF907 - ALPHA/BETA-HYDROLASES SUPERFAMILY PROTEIN (1 of 1)
OG IDs: 93524563
Best-hit-arabi-ID: AT3G05600.1
Best-hit-arabi-name: alpha/beta-Hydrolases superfamily protein
Gorai.007G301800.1.D
PCC=0.045, p=0.855
Gorai.007G301800.1.A
PCC=-0.642, p=0.003
mRNA Profile
Protein Expression Profile
DPA

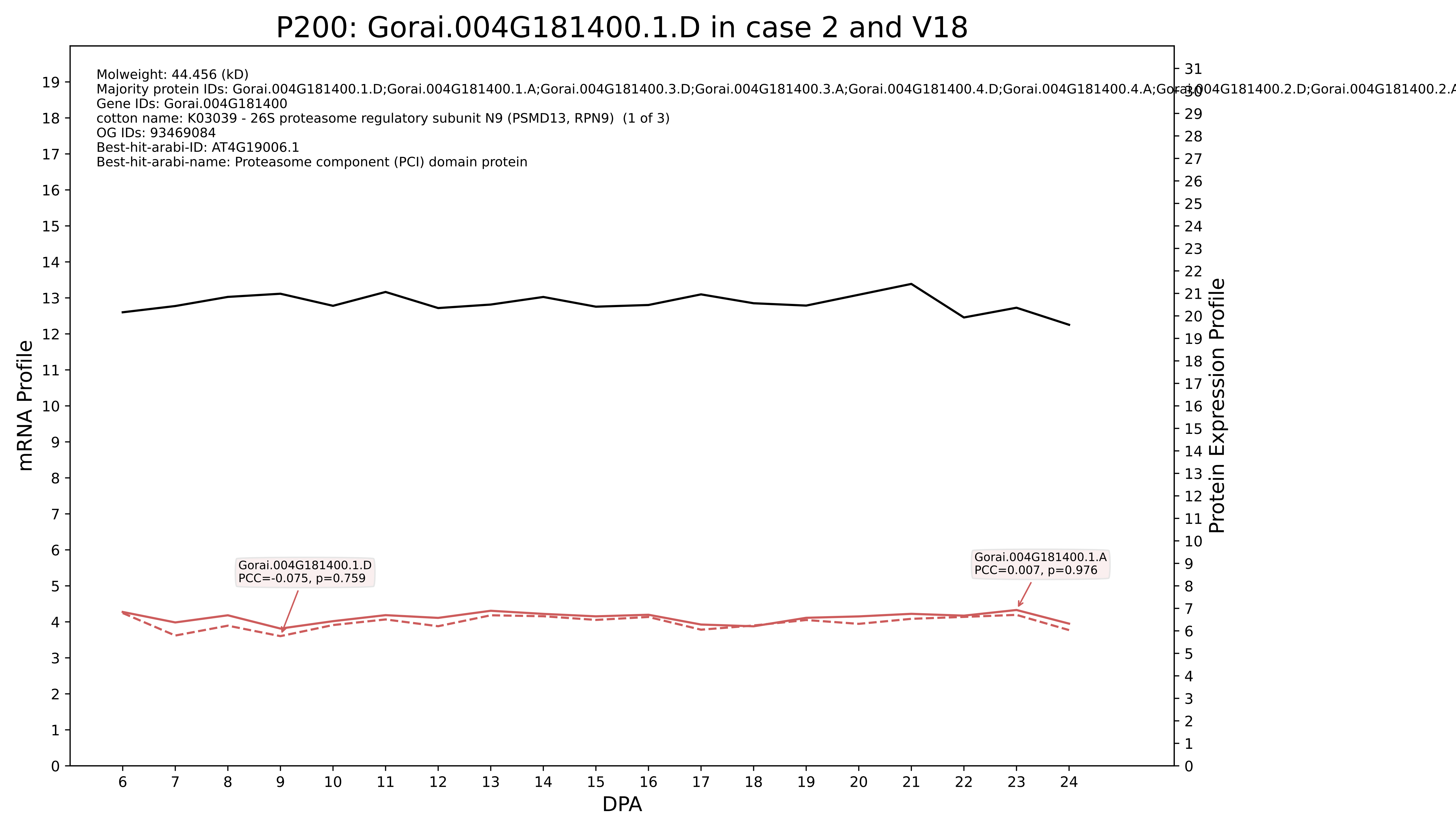
P200: Gorai.004G181400.1.D in case 2 and V18
Molweight: 44.456 (kD)
Majority protein IDs: Gorai.004G181400.1.D;Gorai.004G181400.1.A;Gorai.004G181400.3.D;Gorai.004G181400.3.A;Gorai.004G181400.4.D;Gorai.004G181400.4.A;Gorai.004G181400.2.D;Gorai.004G181400.2.A
Gene IDs: Gorai.004G181400
cotton name: K03039 - 26S proteasome regulatory subunit N9 (PSMD13, RPN9) (1 of 3)
OG IDs: 93469084
Best-hit-arabi-ID: AT4G19006.1
Best-hit-arabi-name: Proteasome component (PCI) domain protein
Gorai.004G181400.1.D
PCC=-0.075, p=0.759
Gorai.004G181400.1.A
PCC=0.007, p=0.976
mRNA Profile
Protein Expression Profile
DPA

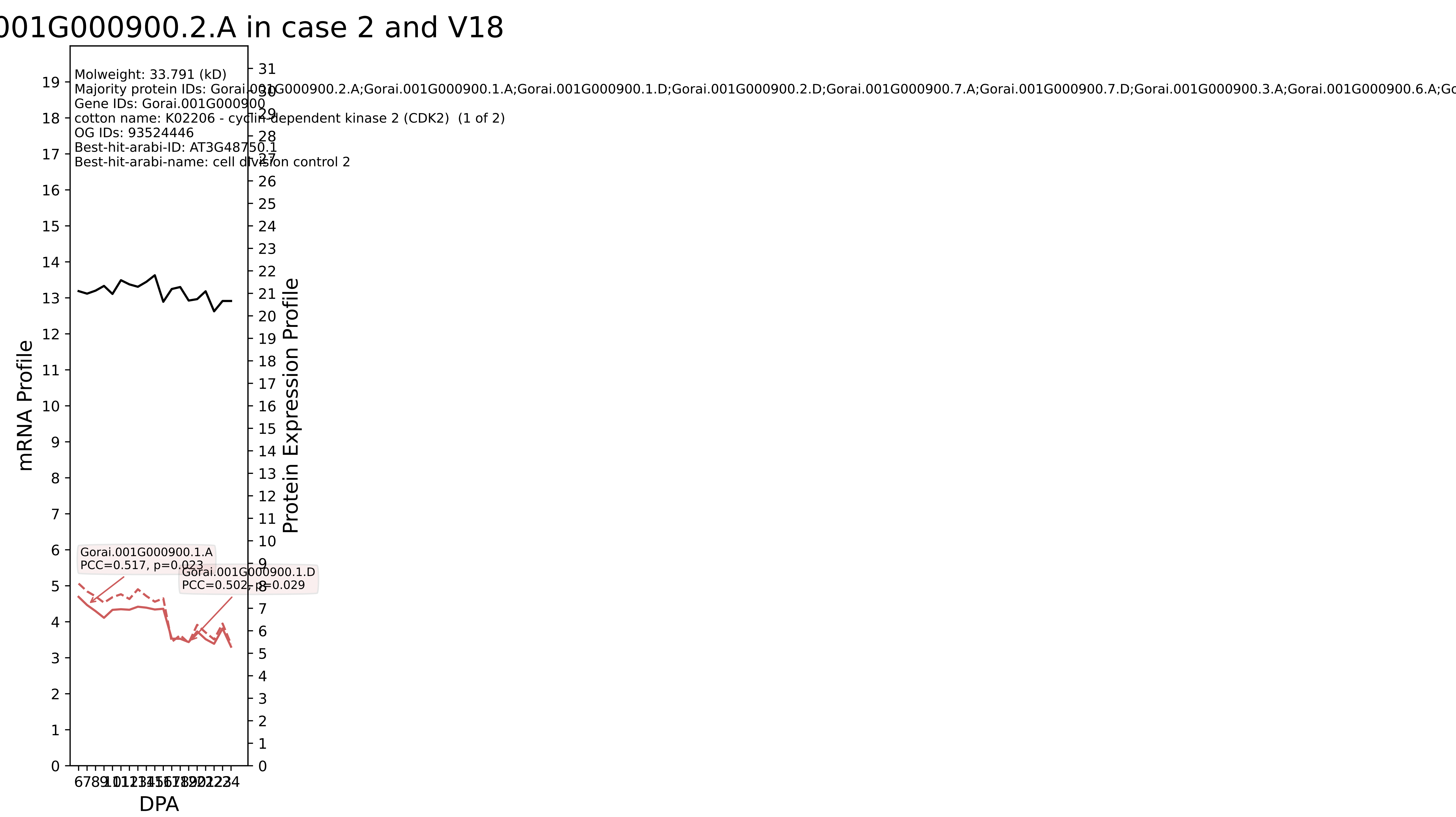

001G000900.2.A in case 2 and V18
Molweight: 33.791 (kD)
Majority protein IDs: Gorai.001G000900.2.A;Gorai.001G000900.1.A;Gorai.001G000900.1.D;Gorai.001G000900.2.D;Gorai.001G000900.7.A;Gorai.001G000900.7.D;Gorai.001G000900.3.A;Gorai.001G000900.6.A;Go
Gene IDs: Gorai.001G000900
cotton name: K02206 - cyclin-dependent kinase 2 (CDK2) (1 of 2)
OG IDs: 93524446
Best-hit-arabi-ID: AT3G48750.1
Best-hit-arabi-name: cell division control 2
Gorai.001G000900.1.A
PCC=0.517, p=0.023
Gorai.001G000900.1.D
PCC=0.502, p=0.029
mRNA Profile
Protein Expression Profile
DPA

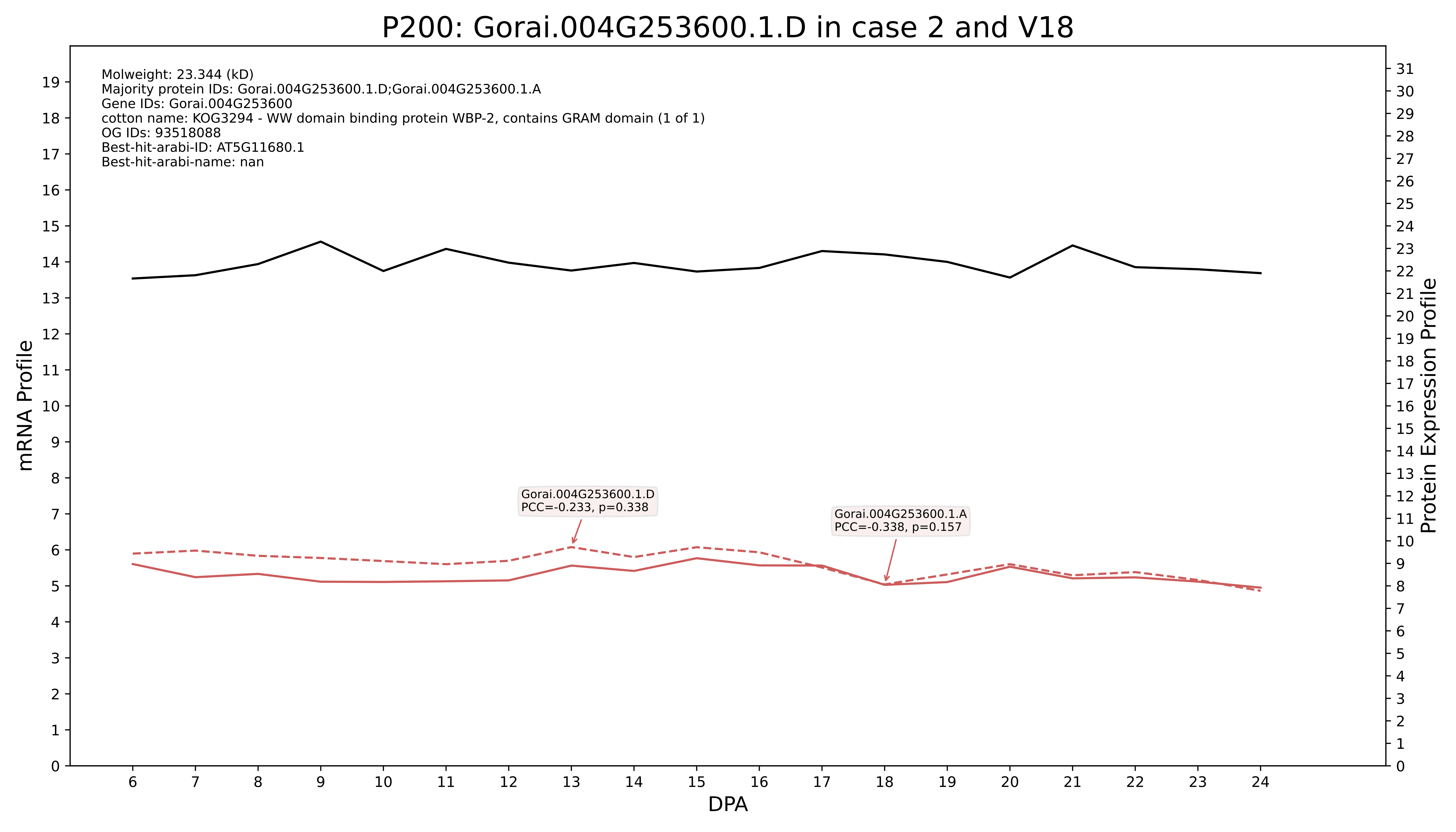
P200: Gorai.004G253600.1.D in case 2 and V18
Molweight: 23.344 (kD)
Majority protein IDs: Gorai.004G253600.1.D;Gorai.004G253600.1.A
Gene IDs: Gorai.004G253600
cotton name: KOG3294 - WW domain binding protein WBP-2, contains GRAM domain (1 of 1)
OG IDs: 93518088
Best-hit-arabi-ID: AT5G11680.1
Best-hit-arabi-name: nan
Gorai.004G253600.1.D
PCC=-0.233, p=0.338
Gorai.004G253600.1.A
PCC=-0.338, p=0.157
mRNA Profile
Protein Expression Profile
DPA

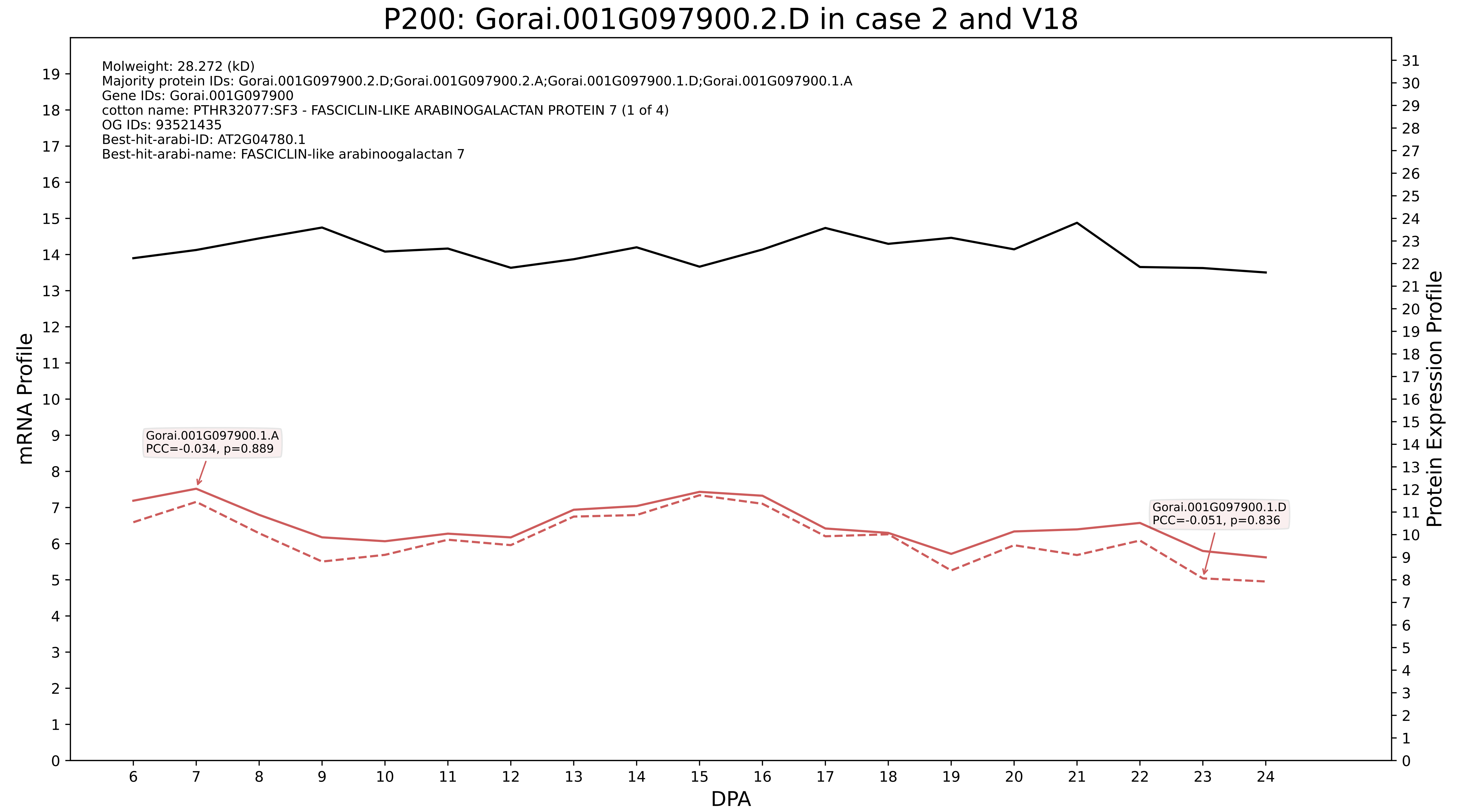

P200: Gorai.001G097900.2.D in case 2 and V18
Molweight: 28.272 (kD)
Majority protein IDs: Gorai.001G097900.2.D;Gorai.001G097900.2.A;Gorai.001G097900.1.D;Gorai.001G097900.1.A
Gene IDs: Gorai.001G097900
cotton name: PTHR32077:SF3 - FASCICLIN-LIKE ARABINOGALACTAN PROTEIN 7 (1 of 4)
OG IDs: 93521435
Best-hit-arabi-ID: AT2G04780.1
Best-hit-arabi-name: FASCICLIN-like arabinoogalactan 7
Gorai.001G097900.1.A
PCC=-0.034, p=0.889
Gorai.001G097900.1.D
PCC=-0.051, p=0.836
mRNA Profile
Protein Expression Profile
DPA

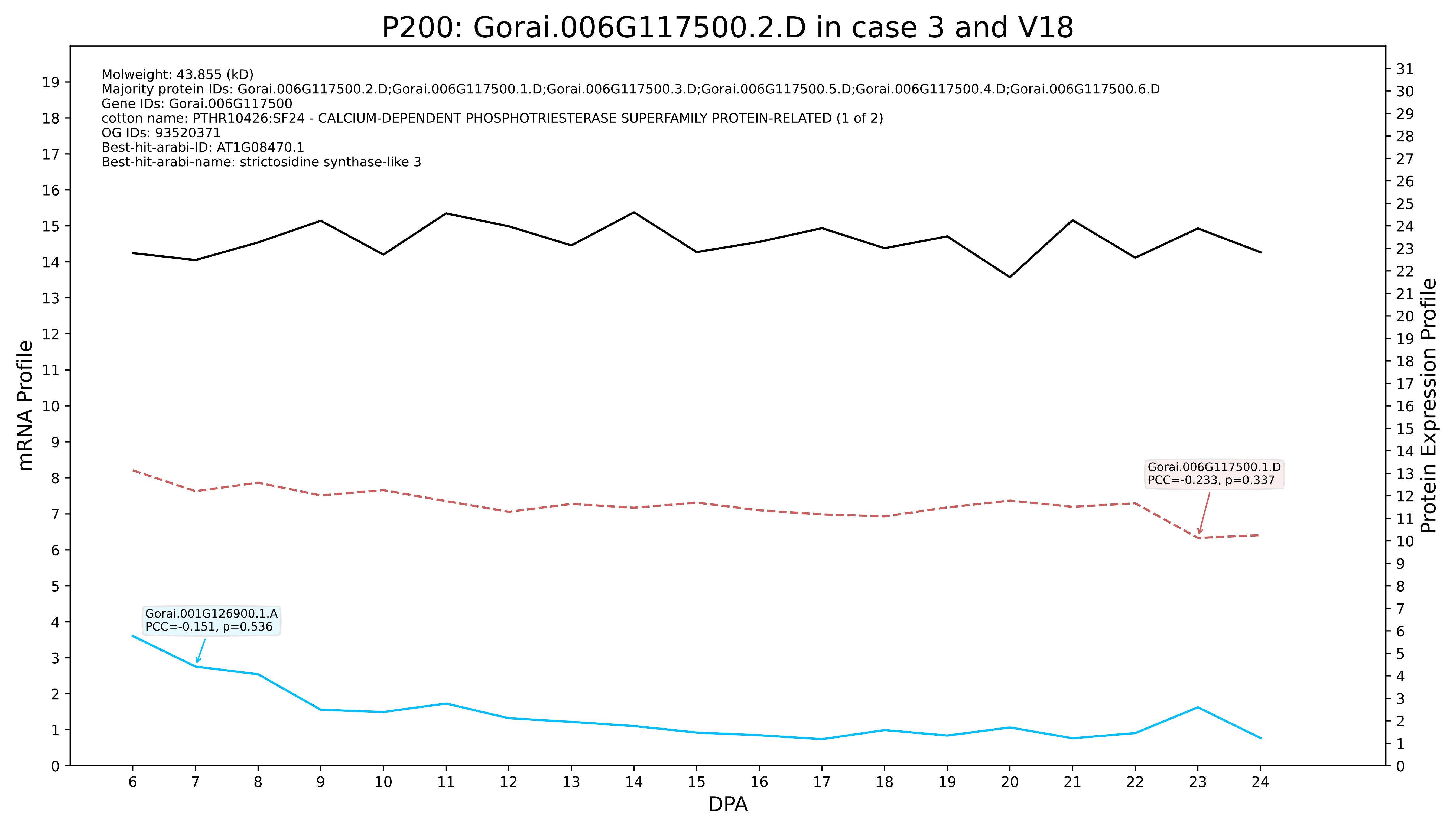

P200: Gorai.006G117500.2.D in case 3 and V18
Molweight: 43.855 (kD)
Majority protein IDs: Gorai.006G117500.2.D;Gorai.006G117500.1.D;Gorai.006G117500.3.D;Gorai.006G117500.5.D;Gorai.006G117500.4.D;Gorai.006G117500.6.D
Gene IDs: Gorai.006G117500
cotton name: PTHR10426:SF24 - CALCIUM-DEPENDENT PHOSPHOTRIESTERASE SUPERFAMILY PROTEIN-RELATED (1 of 2)
OG IDs: 93520371
Best-hit-arabi-ID: AT1G08470.1
Best-hit-arabi-name: strictosidine synthase-like 3
Gorai.006G117500.1.D
PCC=-0.233, p=0.337
Gorai.001G126900.1.A
PCC=-0.151, p=0.536
mRNA Profile
Protein Expression Profile
DPA

### P200: Gorai.011G038200.2.D in case 3 and V18

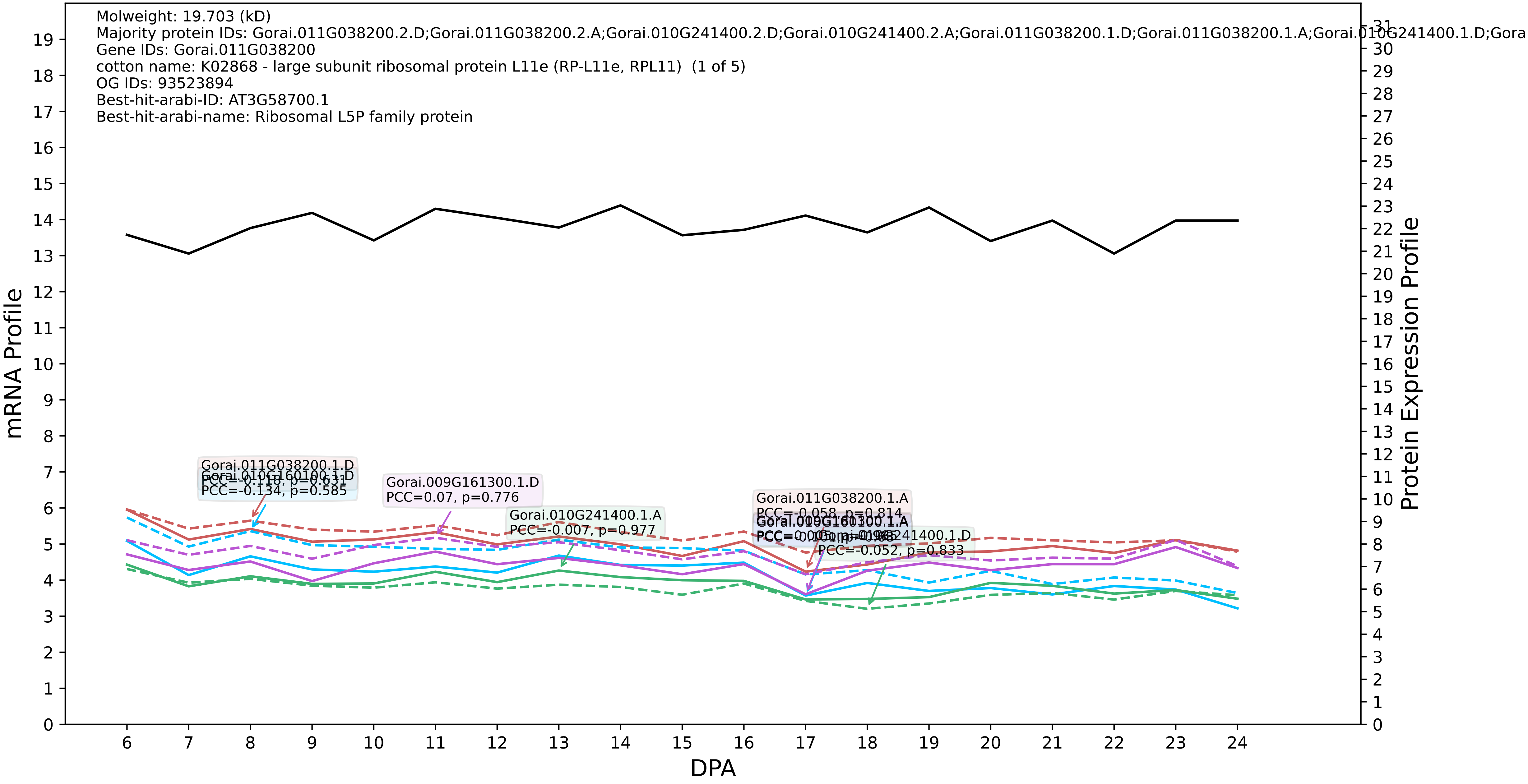

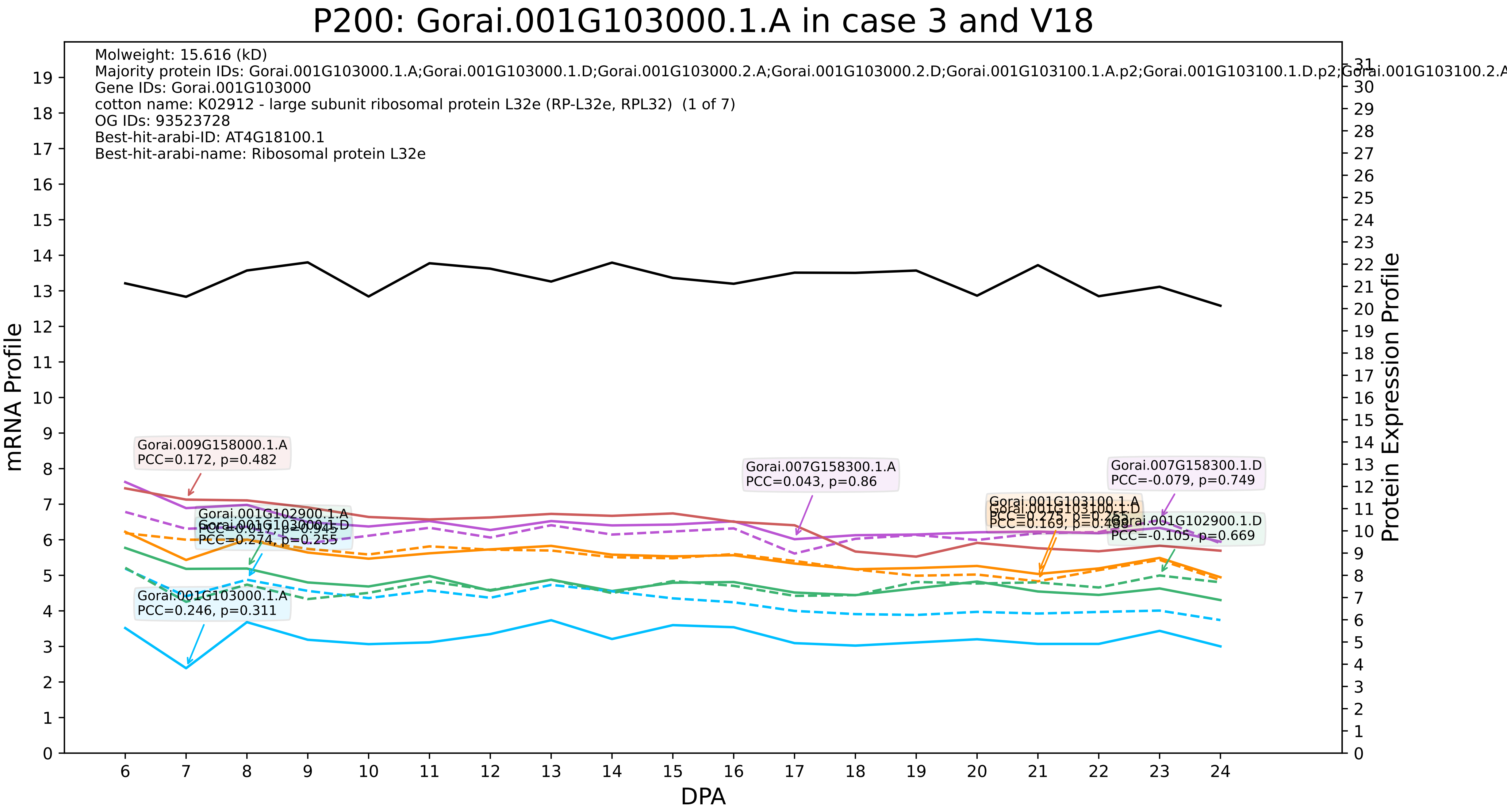

P200: Gorai.001G103000.1.A in case 3 and V18
Molweight: 15.616 (kD)
Majority protein IDs: Gorai.001G103000.1.A;Gorai.001G103000.1.D;Gorai.001G103000.2.A;Gorai.001G103000.2.D;Gorai.001G103100.1.A.p2;Gorai.001G103100.1.D.p2;Gorai.001G103100.2.A
Gene IDs: Gorai.001G103000
cotton name: K02912 - large subunit ribosomal protein L32e (RP-L32e, RPL32) (1 of 7)
OG IDs: 93523728
Best-hit-arabi-ID: AT4G18100.1
Best-hit-arabi-name: Ribosomal protein L32e
Gorai.009G158000.1.A
PCC=0.172, p=0.482
Gorai.007G158300.1.A
PCC=0.043, p=0.86
Gorai.007G158300.1.D
PCC=-0.079, p=0.749
Gorai.001G102900.1.A
PCC=0.274, p=0.255
Gorai.001G102900.1.D
PCC=-0.105, p=0.669
Gorai.001G103000.1.A
PCC=0.246, p=0.311
mRNA Profile
Protein Expression Profile
DPA

### P200: Gorai.003G136100.4.D in case 3 and V18

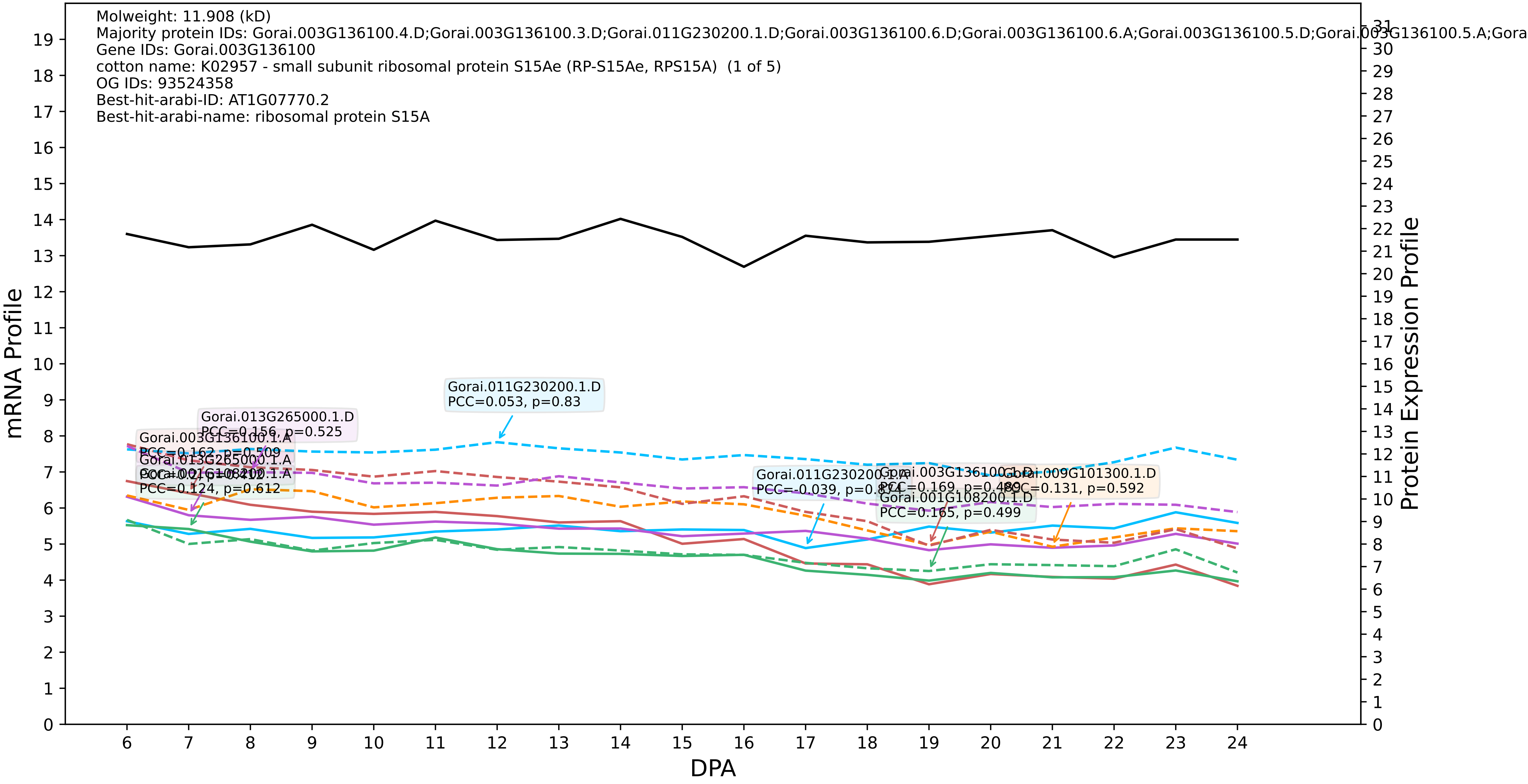

### P200: Gorai.006G112000.4.D in case 3 and V18

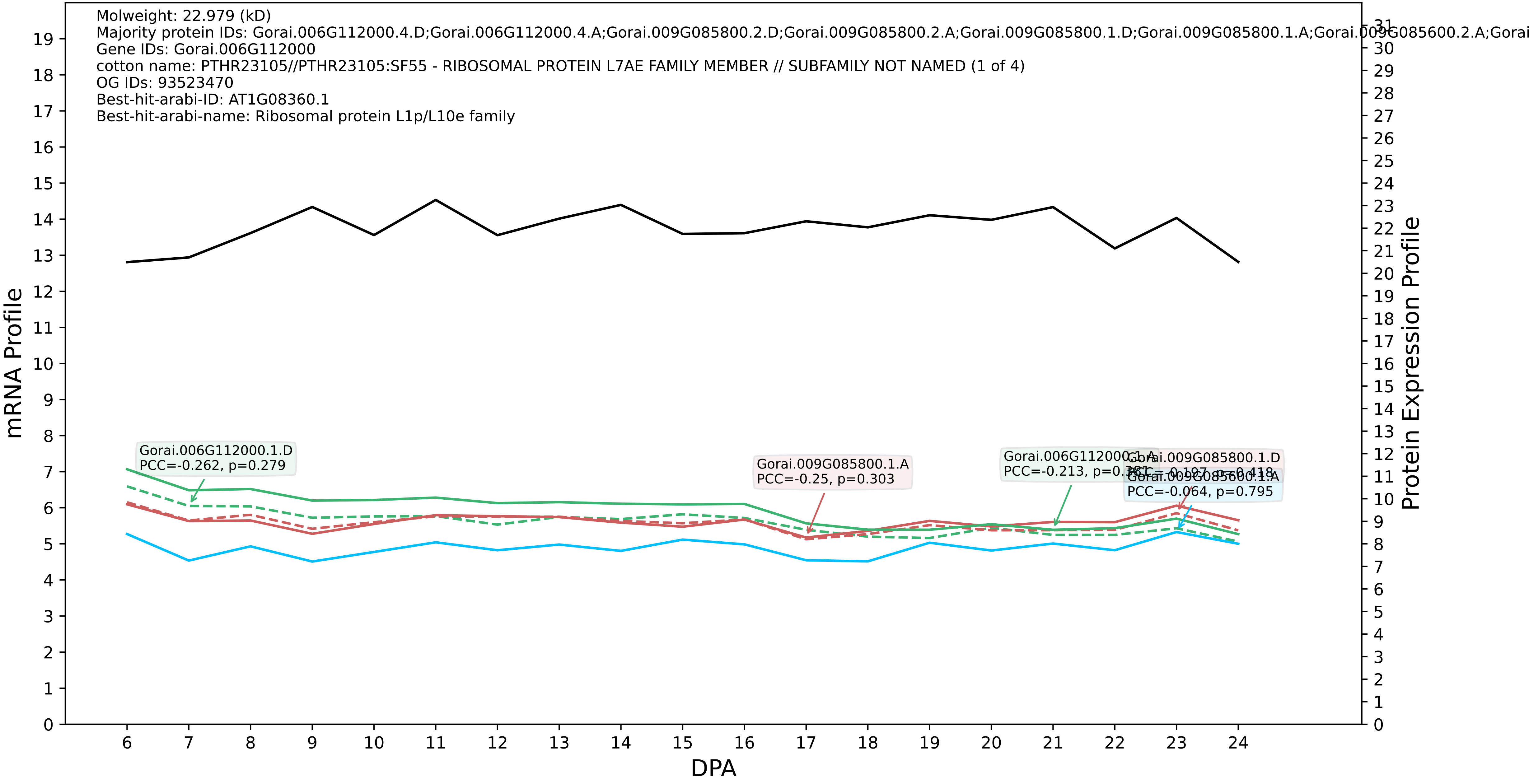

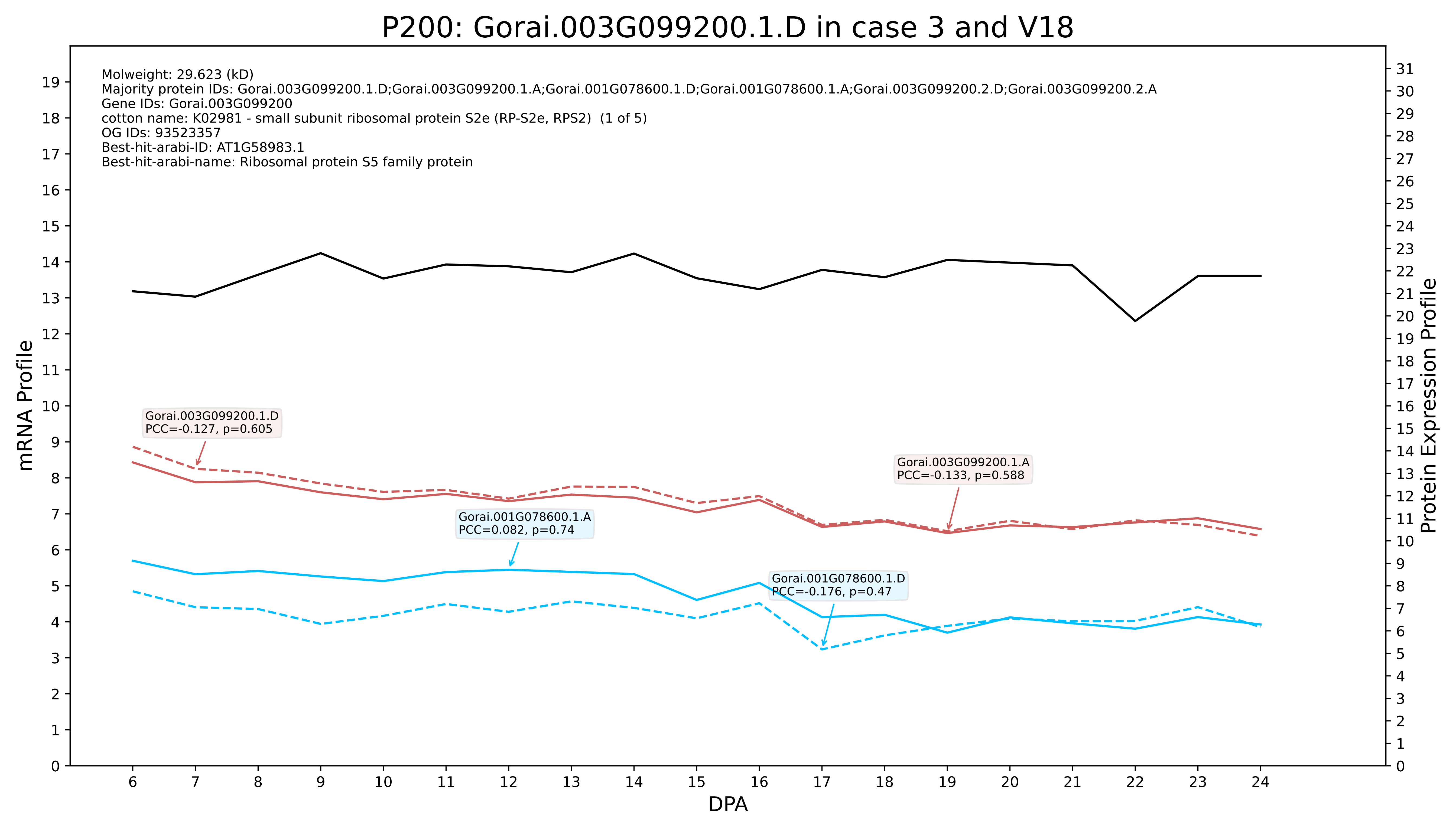
P200: Gorai.003G099200.1.D in case 3 and V18
Molweight: 29.623 (kD)
Majority protein IDs: Gorai.003G099200.1.D;Gorai.003G099200.1.A;Gorai.001G078600.1.D;Gorai.001G078600.1.A;Gorai.003G099200.2.D;Gorai.003G099200.2.A
Gene IDs: Gorai.003G099200
cotton name: K02981 - small subunit ribosomal protein S2e (RP-S2e, RPS2) (1 of 5)
OG IDs: 93523357
Best-hit-arabi-ID: AT1G58983.1
Best-hit-arabi-name: Ribosomal protein S5 family protein
Gorai.003G099200.1.D
PCC=-0.127, p=0.605
Gorai.003G099200.1.A
PCC=-0.133, p=0.588
Gorai.001G078600.1.A
PCC=0.082, p=0.74
Gorai.001G078600.1.D
PCC=-0.176, p=0.47
mRNA Profile
Protein Expression Profile
DPA

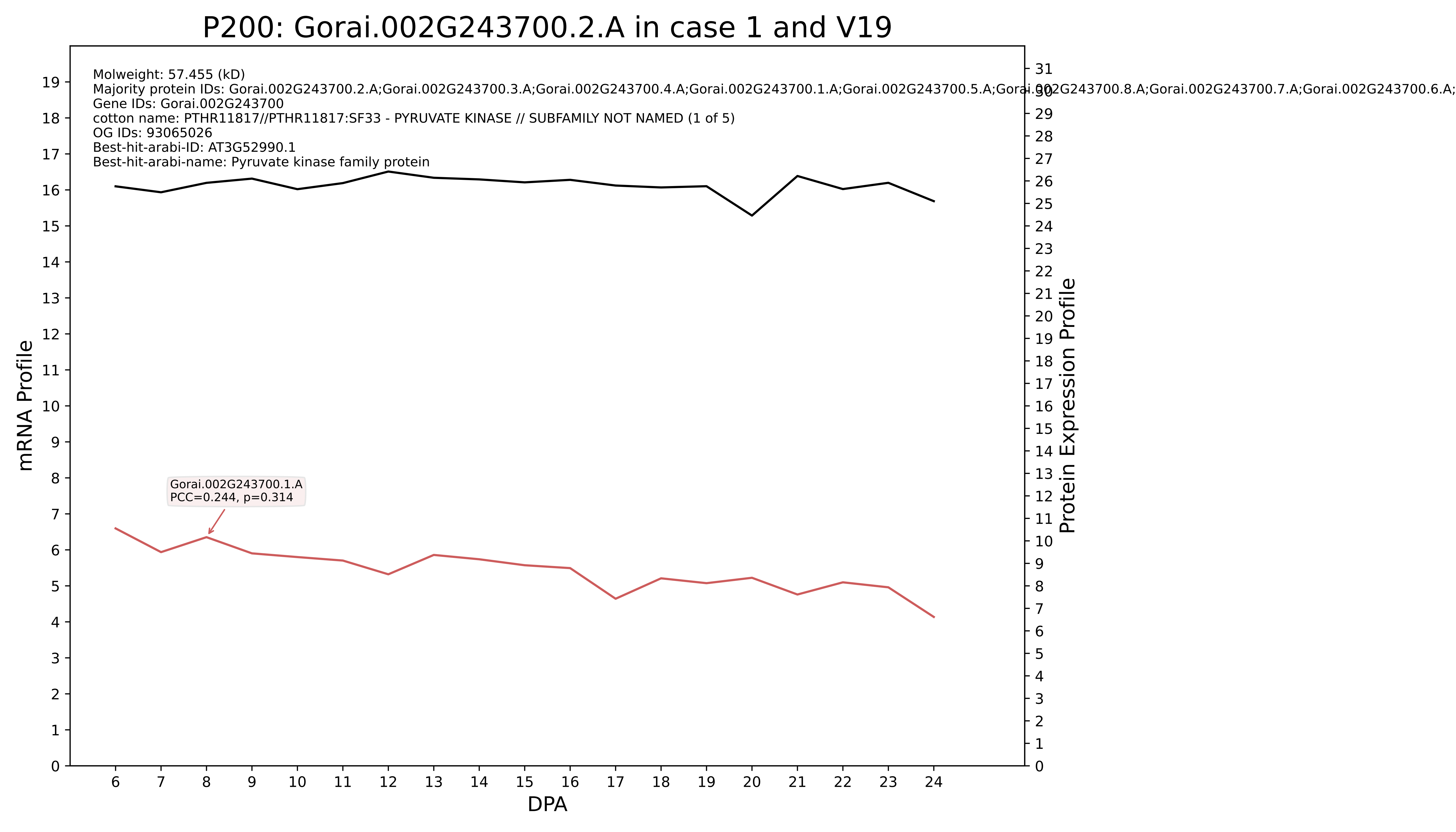
P200: Gorai.002G243700.2.A in case 1 and V19
Molweight: 57.455 (kD)
Majority protein IDs: Gorai.002G243700.2.A;Gorai.002G243700.3.A;Gorai.002G243700.4.A;Gorai.002G243700.1.A;Gorai.002G243700.5.A;Gorai.002G243700.8.A;Gorai.002G243700.7.A;Gorai.002G243700.6.A;
Gene IDs: Gorai.002G243700
cotton name: PTHR11817//PTHR11817:SF33 - PYRUVATE KINASE // SUBFAMILY NOT NAMED (1 of 5)
OG IDs: 93065026
Best-hit-arabi-ID: AT3G52990.1
Best-hit-arabi-name: Pyruvate kinase family protein
Gorai.002G243700.1.A
PCC=0.244, p=0.314
mRNA Profile
Protein Expression Profile
DPA

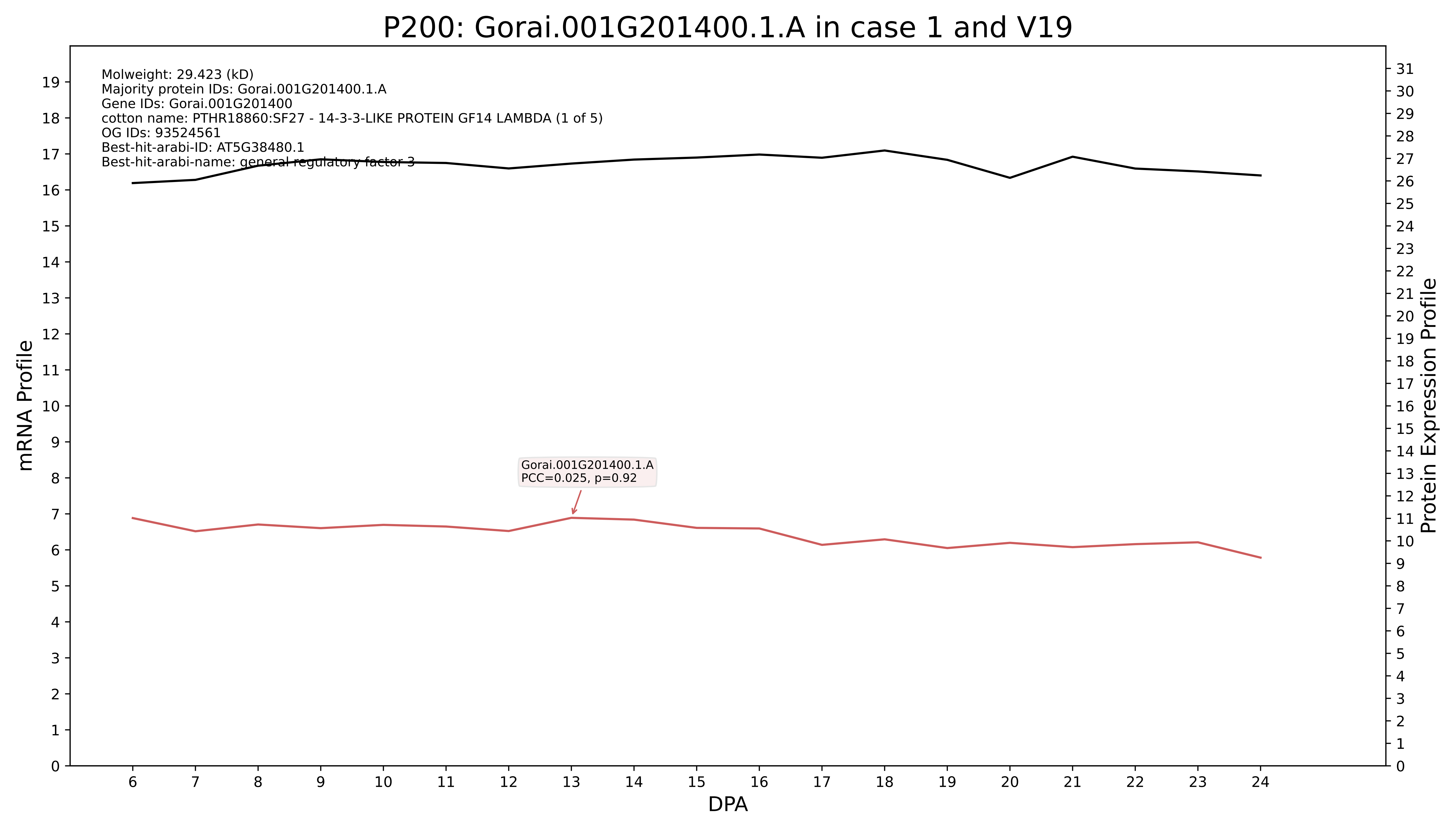

P200: Gorai.001G201400.1.A in case 1 and V19
Molweight: 29.423 (kD)
Majority protein IDs: Gorai.001G201400.1.A
Gene IDs: Gorai.001G201400
cotton name: PTHR18860:SF27 - 14-3-3-LIKE PROTEIN GF14 LAMBDA (1 of 5)
OG IDs: 93524561
Best-hit-arabi-ID: AT5G38480.1
Best-hit-arabi-name: general regulatory factor 3
Gorai.001G201400.1.A
PCC=0.025, p=0.92
mRNA Profile
Protein Expression Profile
DPA

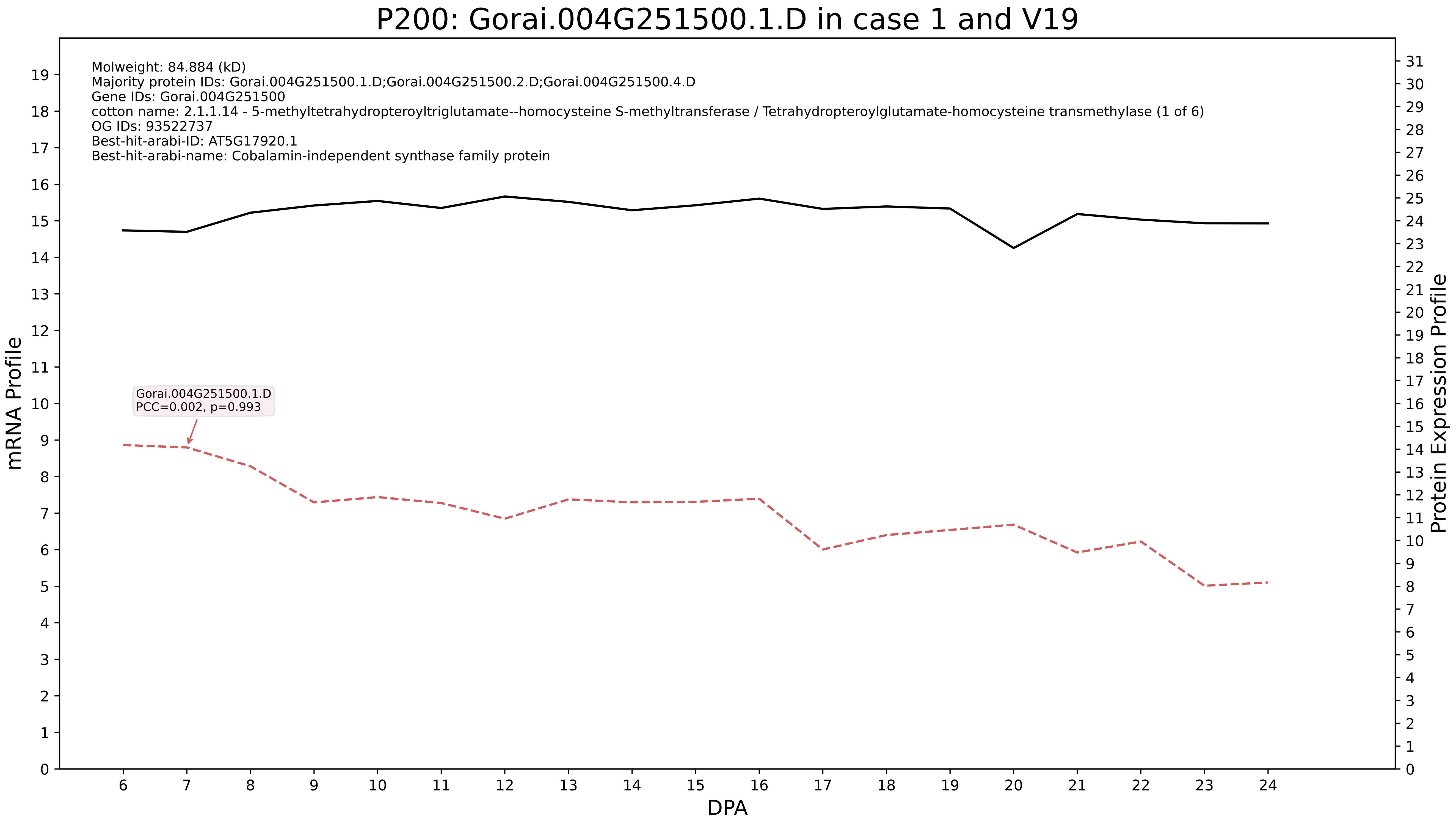
P200: Gorai.004G251500.1.D in case 1 and V19
Molweight: 84.884 (kD)
Majority protein IDs: Gorai.004G251500.1.D;Gorai.004G251500.2.D;Gorai.004G251500.4.D
Gene IDs: Gorai.004G251500
cotton name: 2.1.1.14 - 5-methyltetrahydropteroyltriglutamate--homocysteine S-methyltransferase / Tetrahydropteroylglutamate-homocysteine transmethylase (1 of 6)
OG IDs: 93522737
Best-hit-arabi-ID: AT5G17920.1
Best-hit-arabi-name: Cobalamin-independent synthase family protein
Gorai.004G251500.1.D
PCC=0.002, p=0.993
mRNA Profile
Protein Expression Profile
DPA

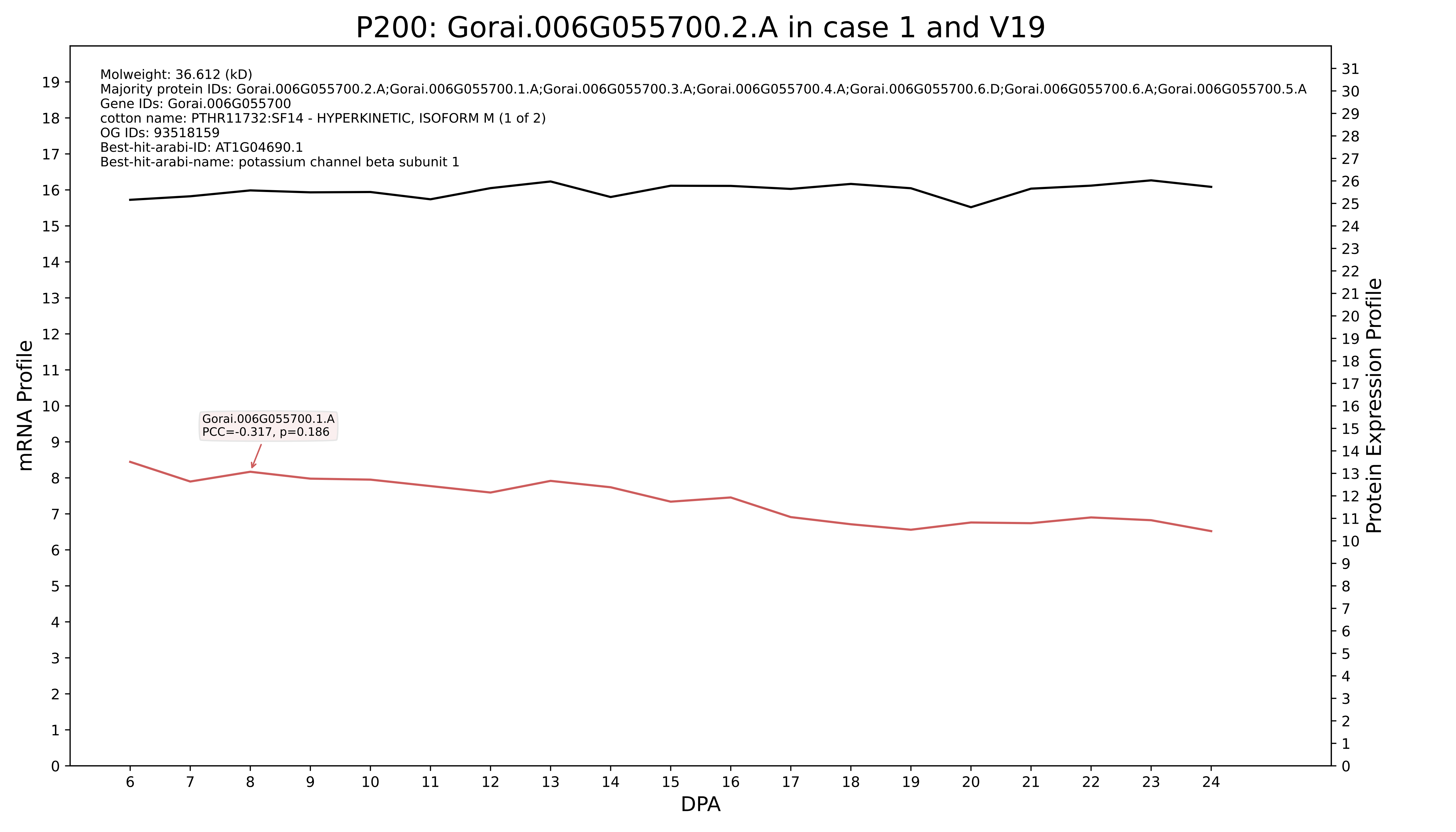
P200: Gorai.006G055700.2.A in case 1 and V19
Molweight: 36.612 (kD)
Majority protein IDs: Gorai.006G055700.2.A;Gorai.006G055700.1.A;Gorai.006G055700.3.A;Gorai.006G055700.4.A;Gorai.006G055700.6.D;Gorai.006G055700.6.A;Gorai.006G055700.5.A
Gene IDs: Gorai.006G055700
cotton name: PTHR11732:SF14 - HYPERKINETIC, ISOFORM M (1 of 2)
OG IDs: 93518159
Best-hit-arabi-ID: AT1G04690.1
Best-hit-arabi-name: potassium channel beta subunit 1
Gorai.006G055700.1.A
PCC=-0.317, p=0.186
mRNA Profile
Protein Expression Profile
DPA

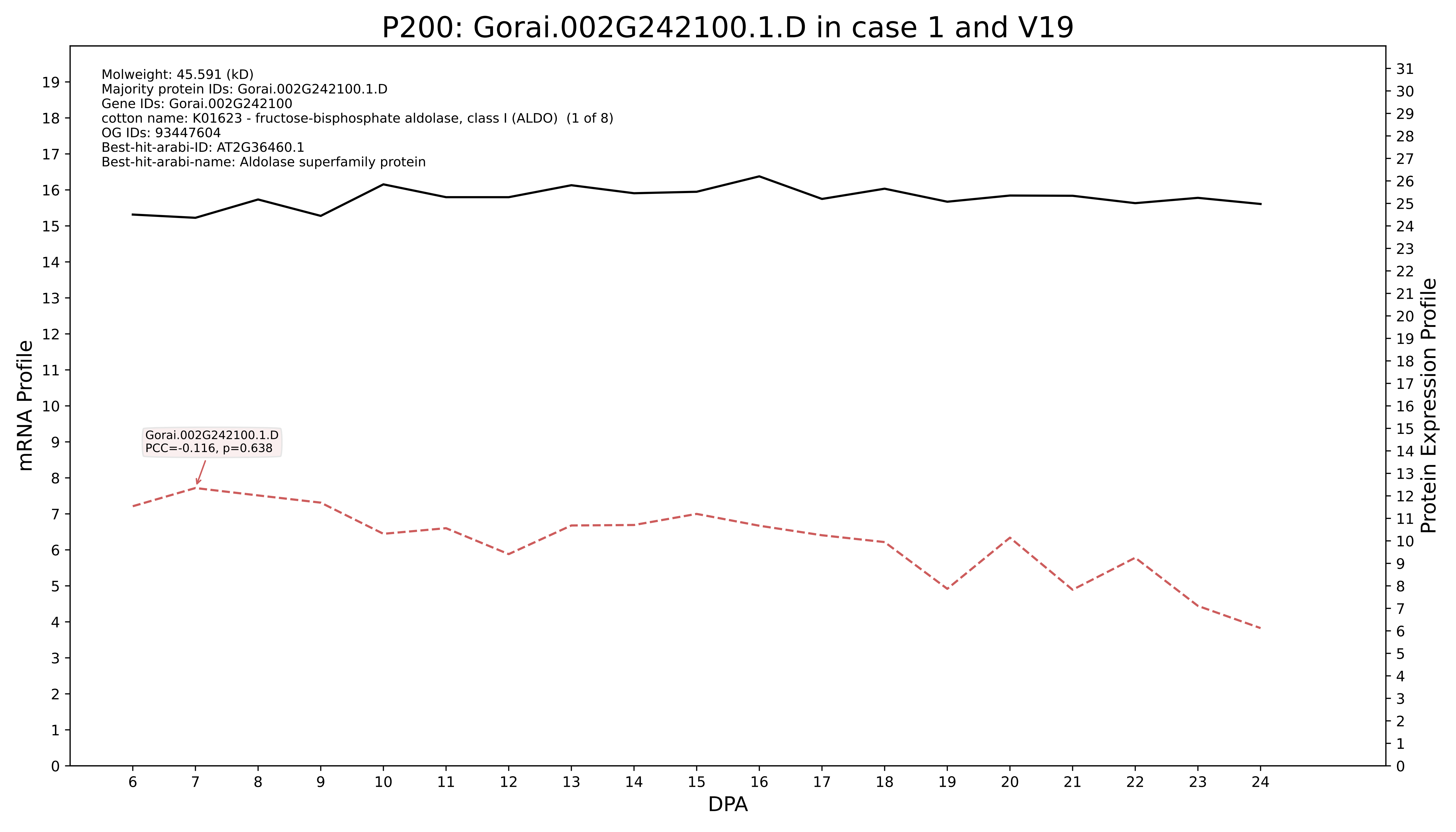

P200: Gorai.002G242100.1.D in case 1 and V19
Molweight: 45.591 (kD)
Majority protein IDs: Gorai.002G242100.1.D
Gene IDs: Gorai.002G242100
cotton name: K01623 - fructose-bisphosphate aldolase, class I (ALDO) (1 of 8)
OG IDs: 93447604
Best-hit-arabi-ID: AT2G36460.1
Best-hit-arabi-name: Aldolase superfamily protein
Gorai.002G242100.1.D
PCC=-0.116, p=0.638
mRNA Profile
Protein Expression Profile
DPA

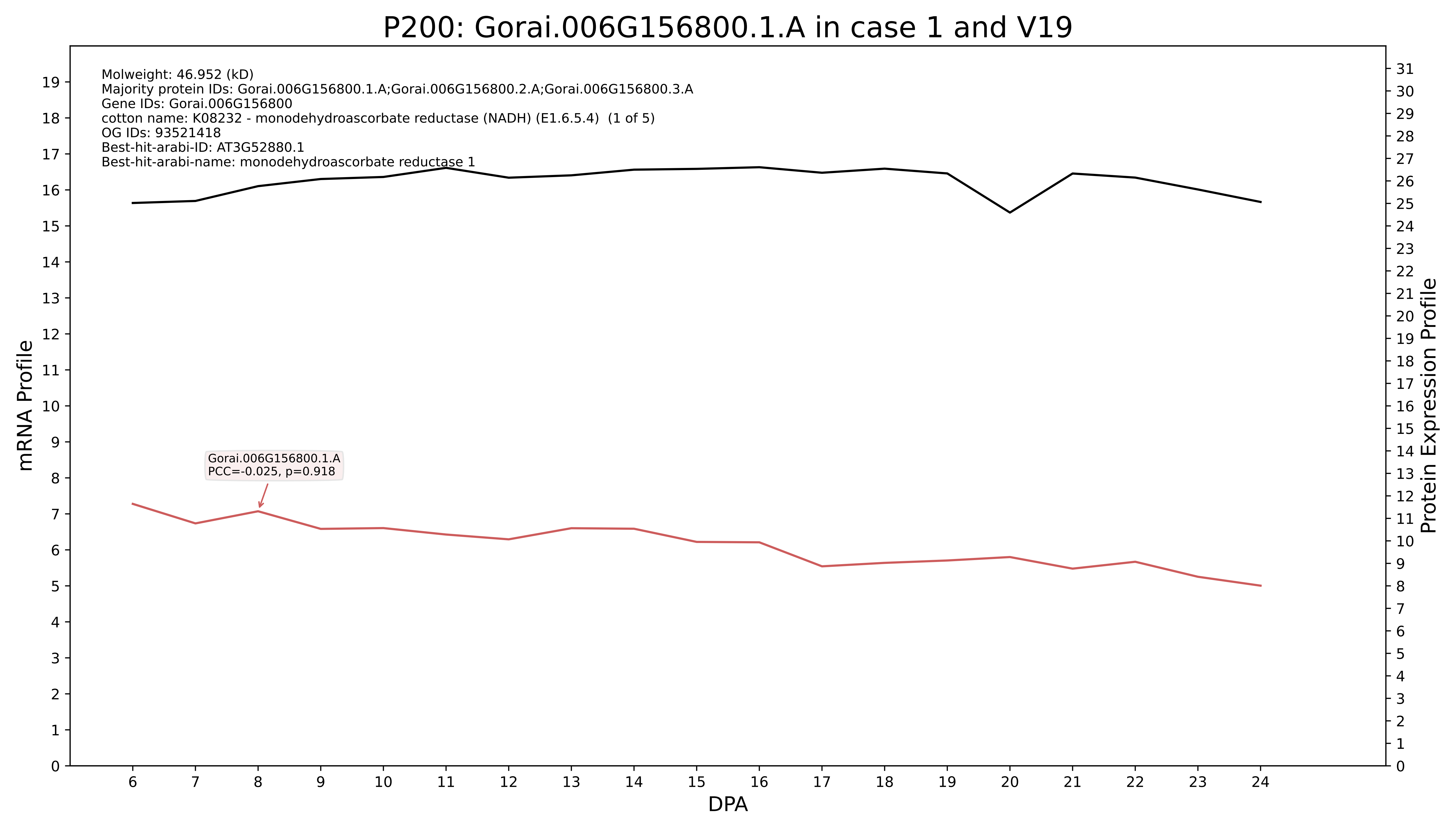

P200: Gorai.006G156800.1.A in case 1 and V19
Molweight: 46.952 (kD)
Majority protein IDs: Gorai.006G156800.1.A;Gorai.006G156800.2.A;Gorai.006G156800.3.A
Gene IDs: Gorai.006G156800
cotton name: K08232 - monodehydroascorbate reductase (NADH) (E1.6.5.4) (1 of 5)
OG IDs: 93521418
Best-hit-arabi-ID: AT3G52880.1
Best-hit-arabi-name: monodehydroascorbate reductase 1
Gorai.006G156800.1.A
PCC=-0.025, p=0.918
mRNA Profile
Protein Expression Profile
DPA

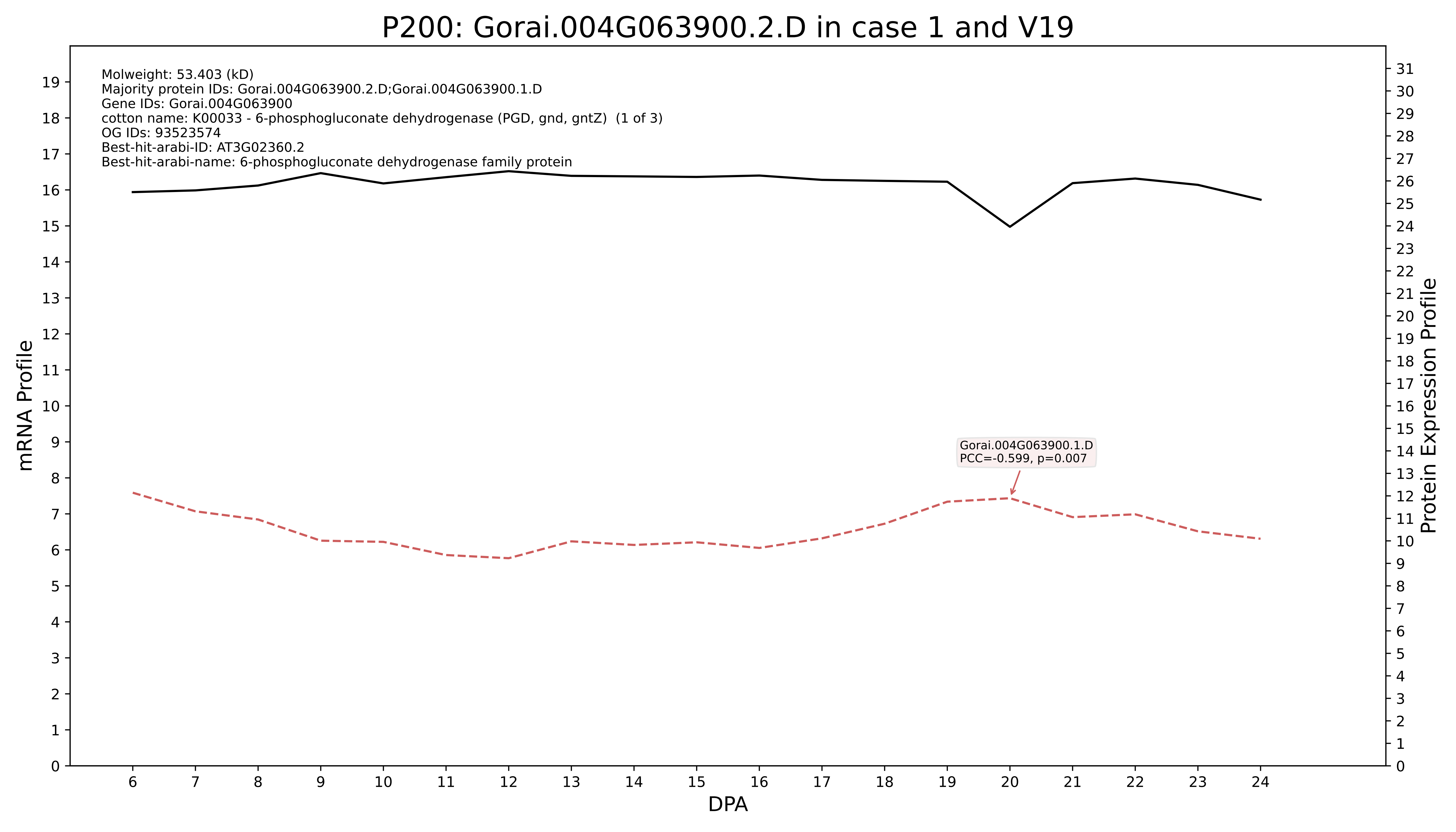
P200: Gorai.004G063900.2.D in case 1 and V19
Molweight: 53.403 (kD)
Majority protein IDs: Gorai.004G063900.2.D;Gorai.004G063900.1.D
Gene IDs: Gorai.004G063900
cotton name: K00033 - 6-phosphogluconate dehydrogenase (PGD, gnd, gntZ) (1 of 3)
OG IDs: 93523574
Best-hit-arabi-ID: AT3G02360.2
Best-hit-arabi-name: 6-phosphogluconate dehydrogenase family protein
Gorai.004G063900.1.D
PCC=-0.599, p=0.007
mRNA Profile
Protein Expression Profile
DPA

### P200: Gorai.007G375600.2.D in case 1 and V19

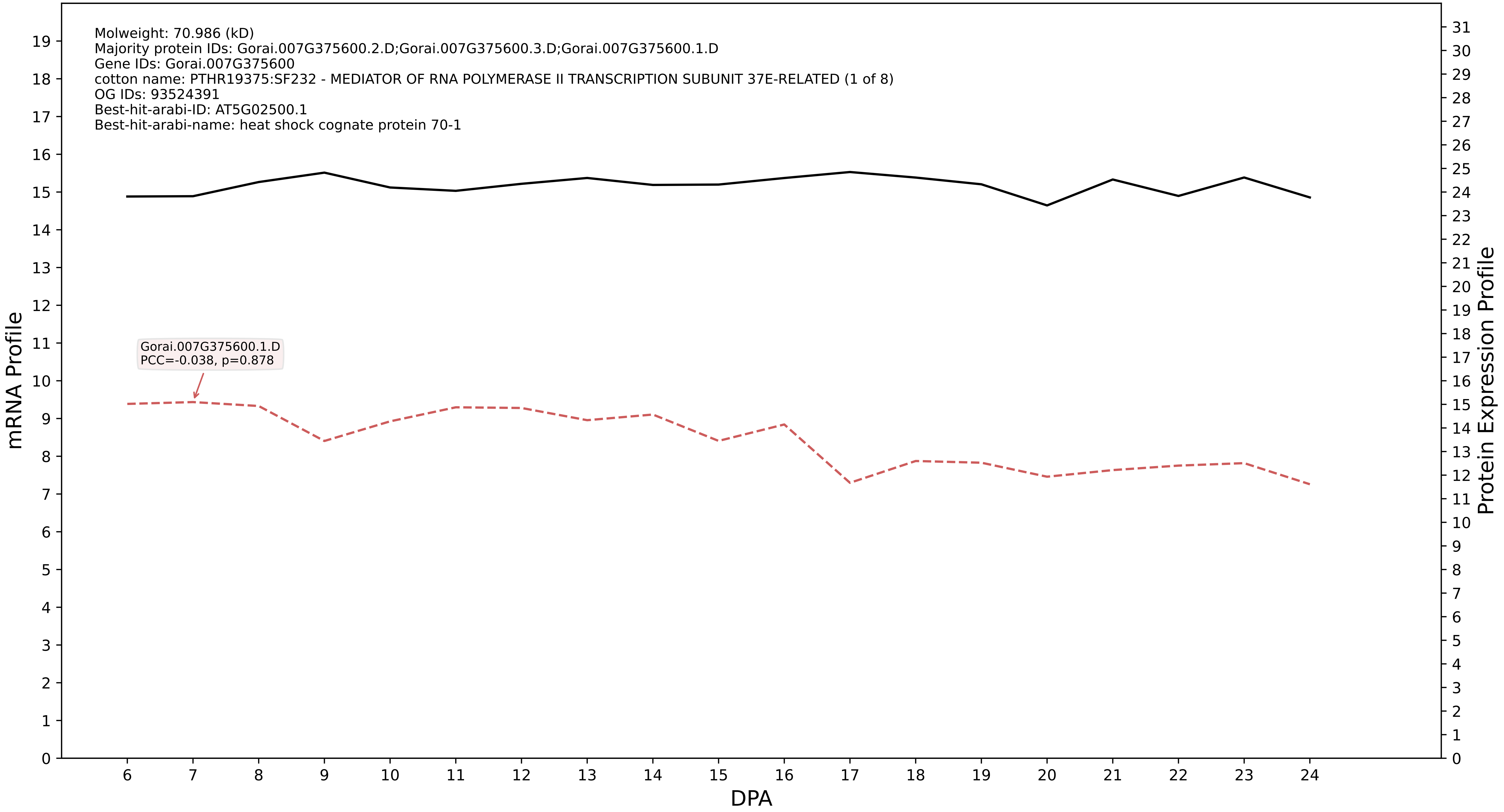

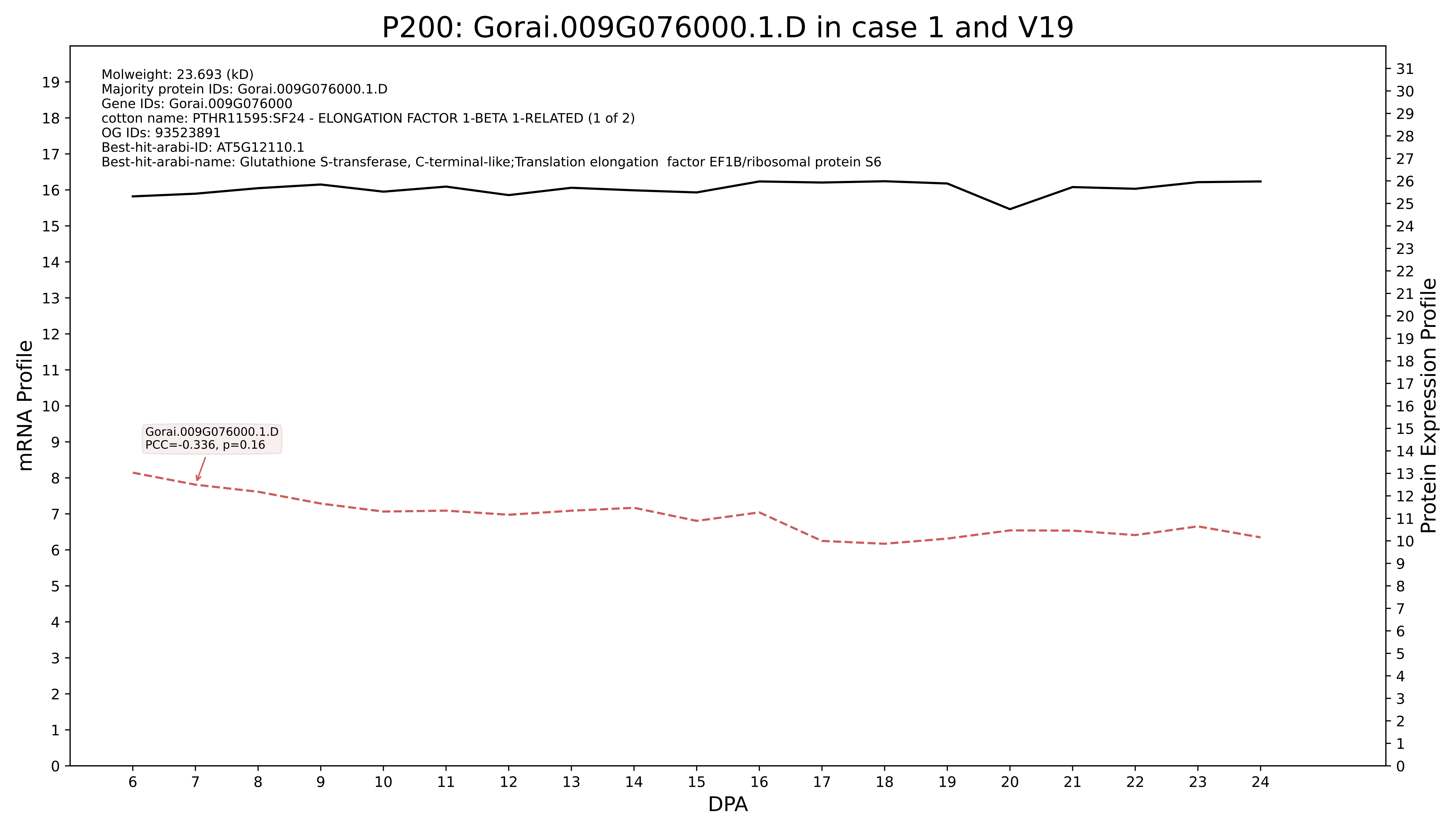
P200: Gorai.009G076000.1.D in case 1 and V19
Molweight: 23.693 (kD)
Majority protein IDs: Gorai.009G076000.1.D
Gene IDs: Gorai.009G076000
cotton name: PTHR11595:SF24 - ELONGATION FACTOR 1-BETA 1-RELATED (1 of 2)
OG IDs: 93523891
Best-hit-arabi-ID: AT5G12110.1
Best-hit-arabi-name: Glutathione S-transferase, C-terminal-like;Translation elongation factor EF1B/ribosomal protein S6
Gorai.009G076000.1.D
PCC=-0.336, p=0.16
mRNA Profile
Protein Expression Profile
DPA

### P200: Gorai.001G062000.3.D in case 1 and V19

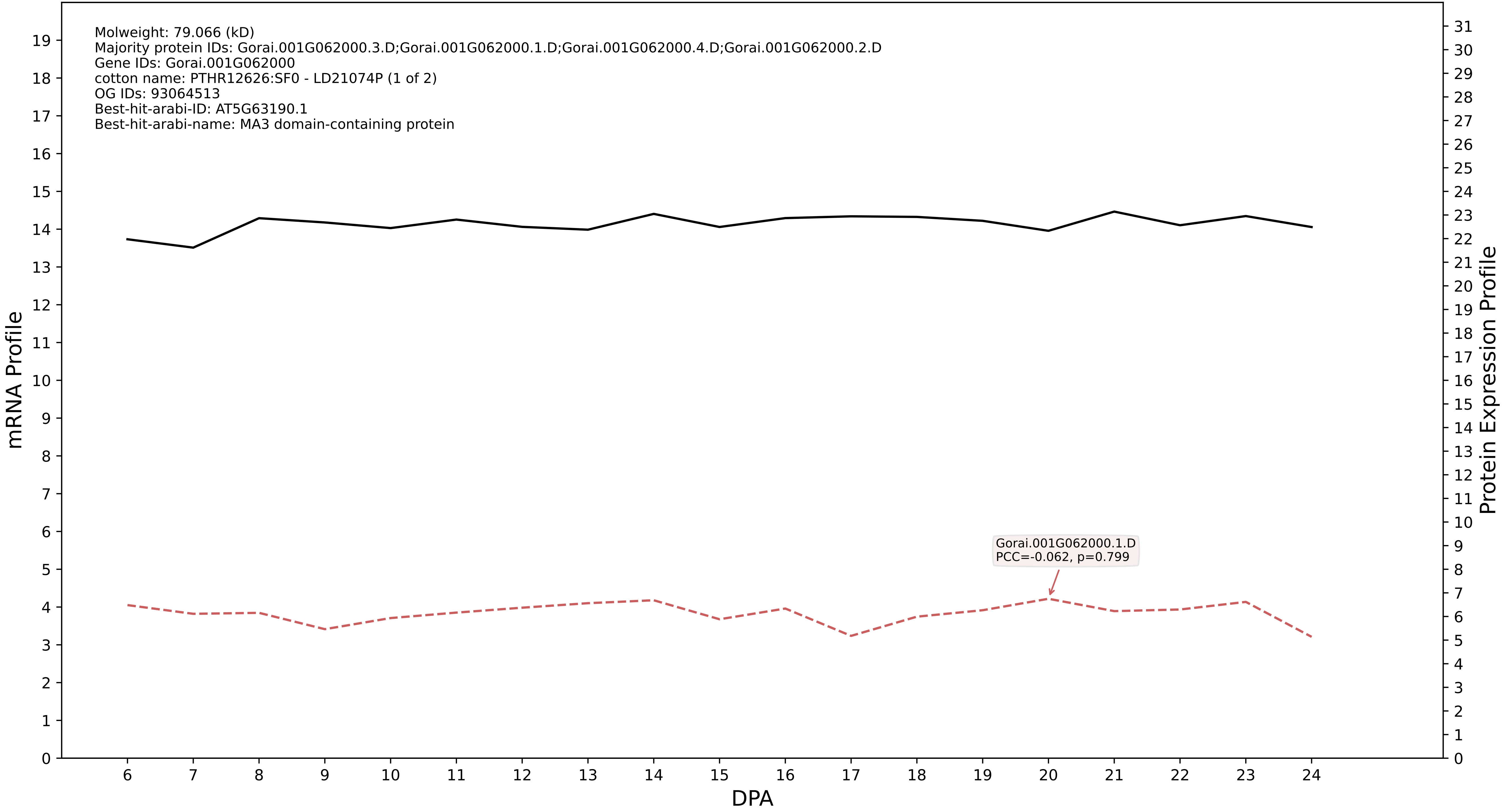

### P200: Gorai.007G314500.2.A in case 1 and V19

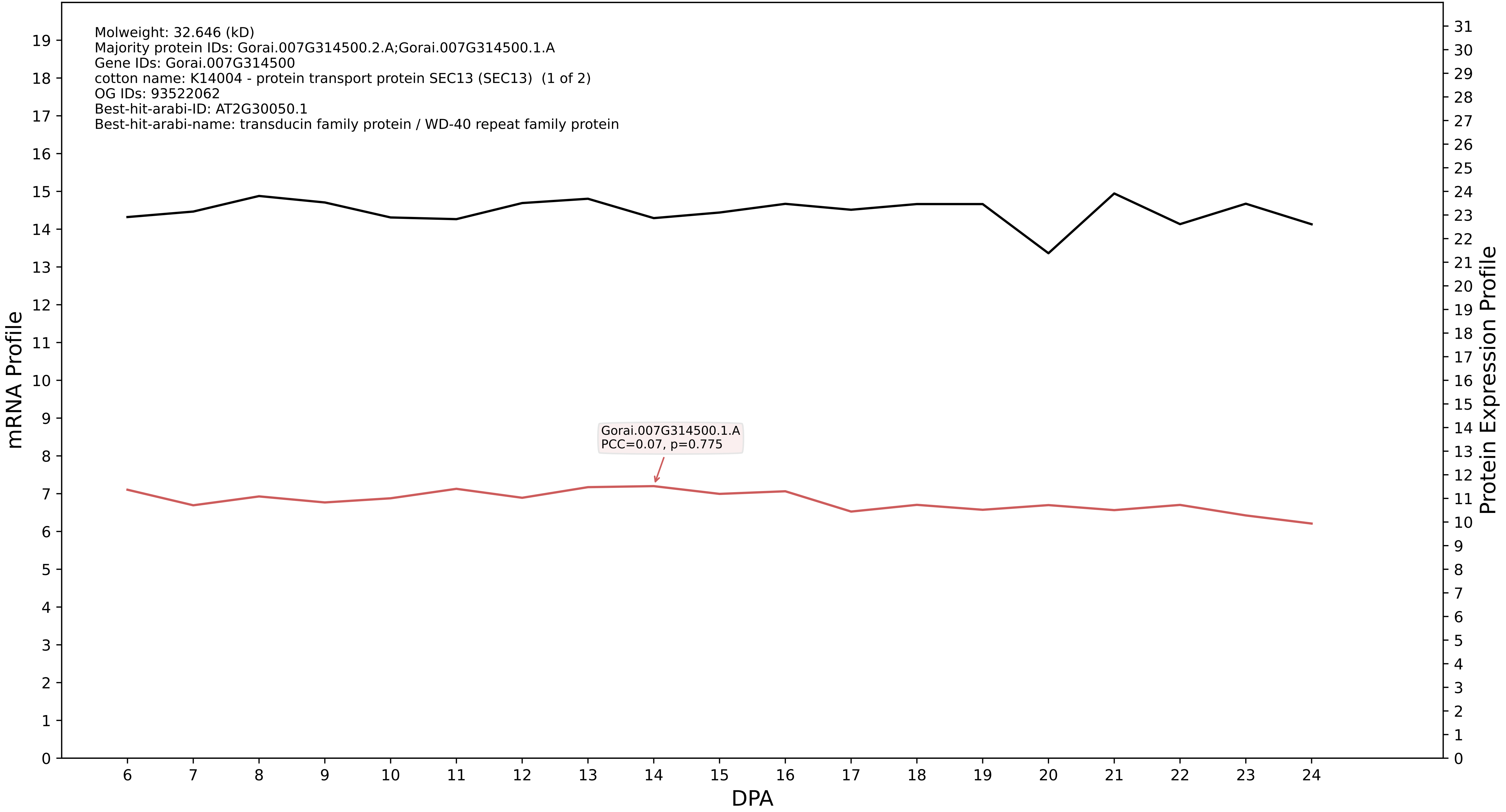

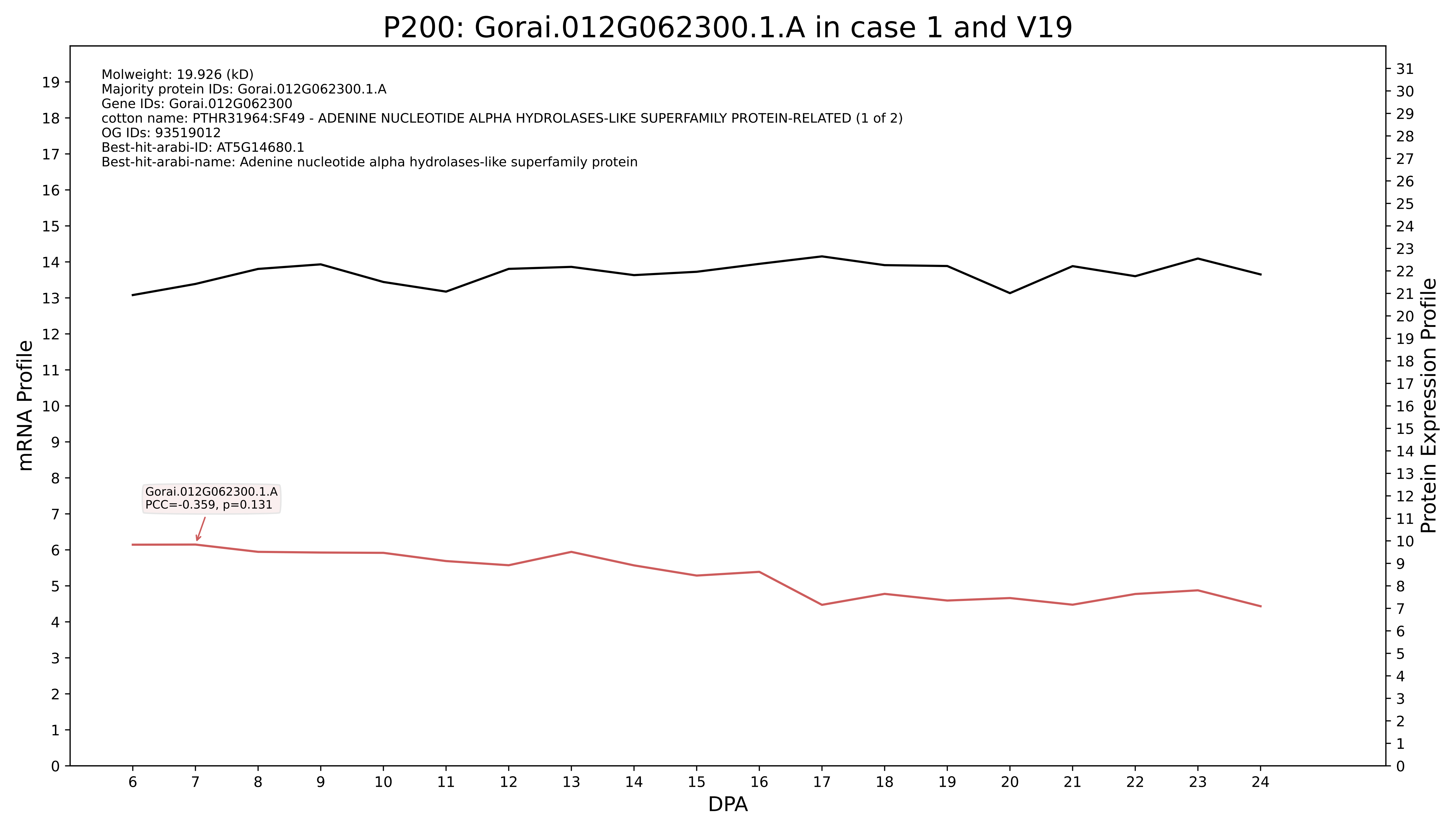
P200: Gorai.012G062300.1.A in case 1 and V19
Molweight: 19.926 (kD)
Majority protein IDs: Gorai.012G062300.1.A
Gene IDs: Gorai.012G062300
cotton name: PTHR31964:SF49 - ADENINE NUCLEOTIDE ALPHA HYDROLASES-LIKE SUPERFAMILY PROTEIN-RELATED (1 of 2)
OG IDs: 93519012
Best-hit-arabi-ID: AT5G14680.1
Best-hit-arabi-name: Adenine nucleotide alpha hydrolases-like superfamily protein
Gorai.012G062300.1.A
PCC=-0.359, p=0.131
mRNA Profile
Protein Expression Profile
DPA

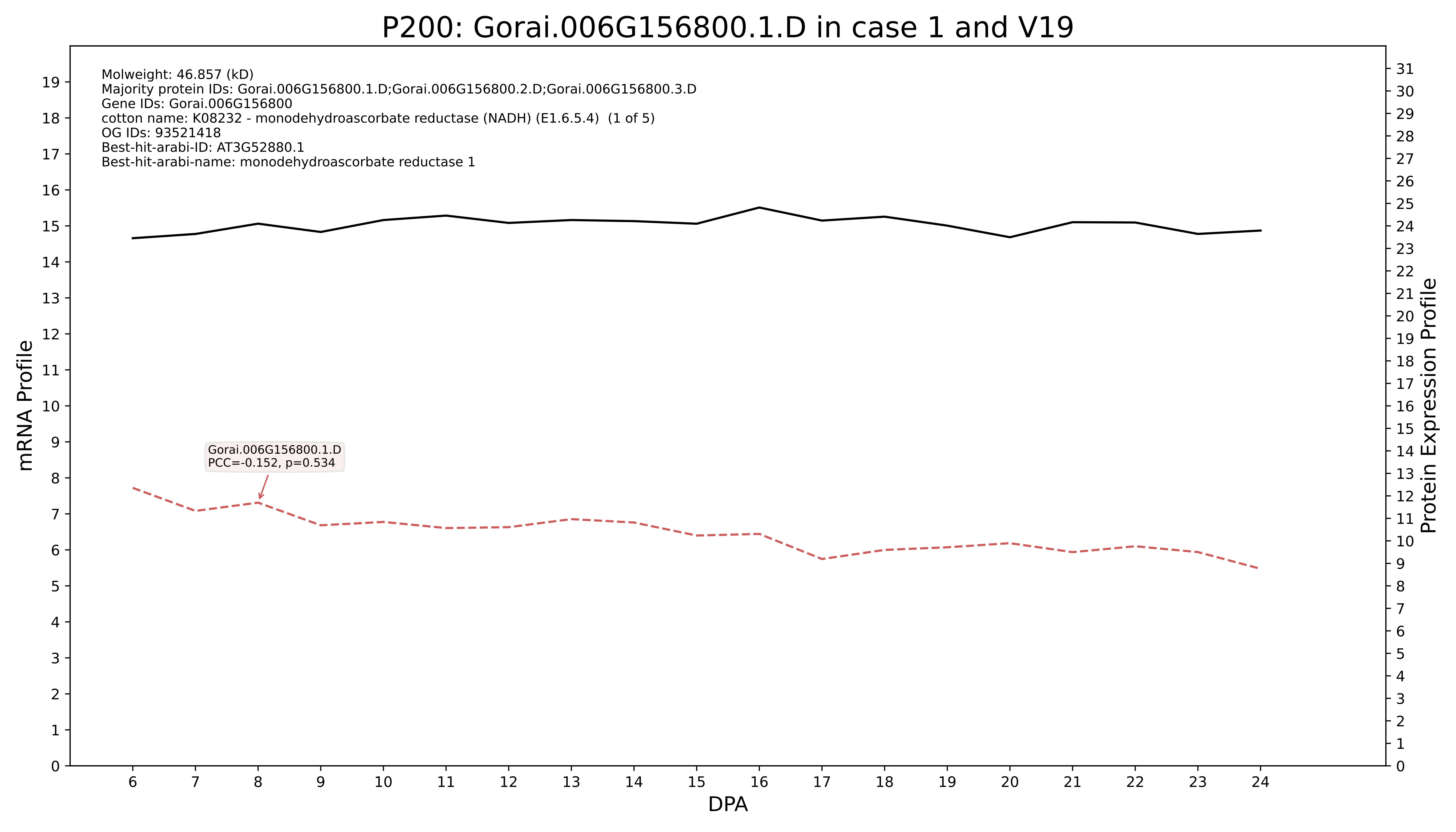
P200: Gorai.006G156800.1.D in case 1 and V19
Molweight: 46.857 (kD)
Majority protein IDs: Gorai.006G156800.1.D;Gorai.006G156800.2.D;Gorai.006G156800.3.D
Gene IDs: Gorai.006G156800
cotton name: K08232 - monodehydroascorbate reductase (NADH) (E1.6.5.4) (1 of 5)
OG IDs: 93521418
Best-hit-arabi-ID: AT3G52880.1
Best-hit-arabi-name: monodehydroascorbate reductase 1
Gorai.006G156800.1.D
PCC=-0.152, p=0.534
mRNA Profile
Protein Expression Profile
DPA

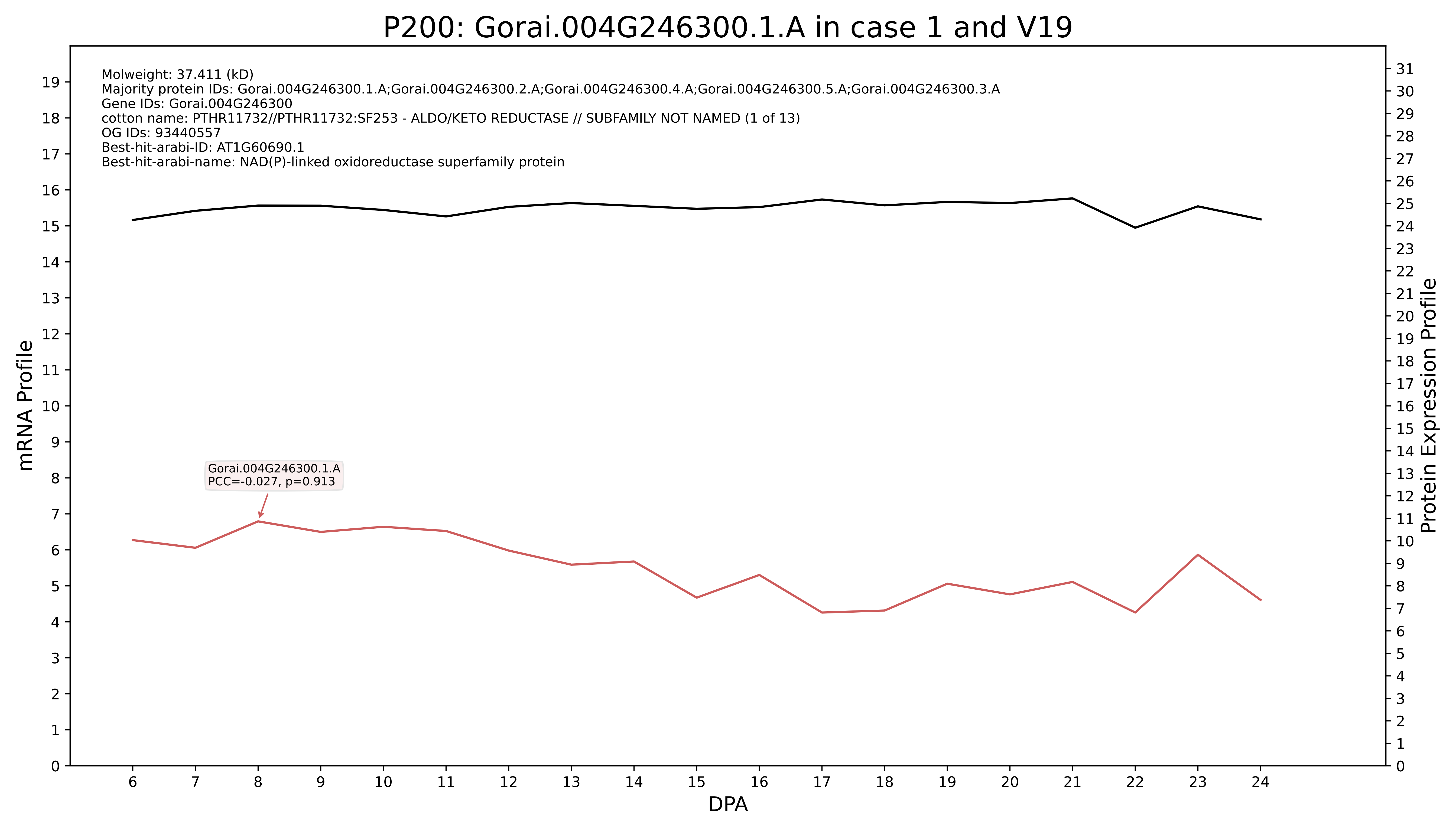
P200: Gorai.004G246300.1.A in case 1 and V19
Molweight: 37.411 (kD)
Majority protein IDs: Gorai.004G246300.1.A;Gorai.004G246300.2.A;Gorai.004G246300.4.A;Gorai.004G246300.5.A;Gorai.004G246300.3.A
Gene IDs: Gorai.004G246300
cotton name: PTHR11732//PTHR11732:SF253 - ALDO/KETO REDUCTASE // SUBFAMILY NOT NAMED (1 of 13)
OG IDs: 93440557
Best-hit-arabi-ID: AT1G60690.1
Best-hit-arabi-name: NAD(P)-linked oxidoreductase superfamily protein
Gorai.004G246300.1.A
PCC=-0.027, p=0.913
mRNA Profile
Protein Expression Profile
DPA

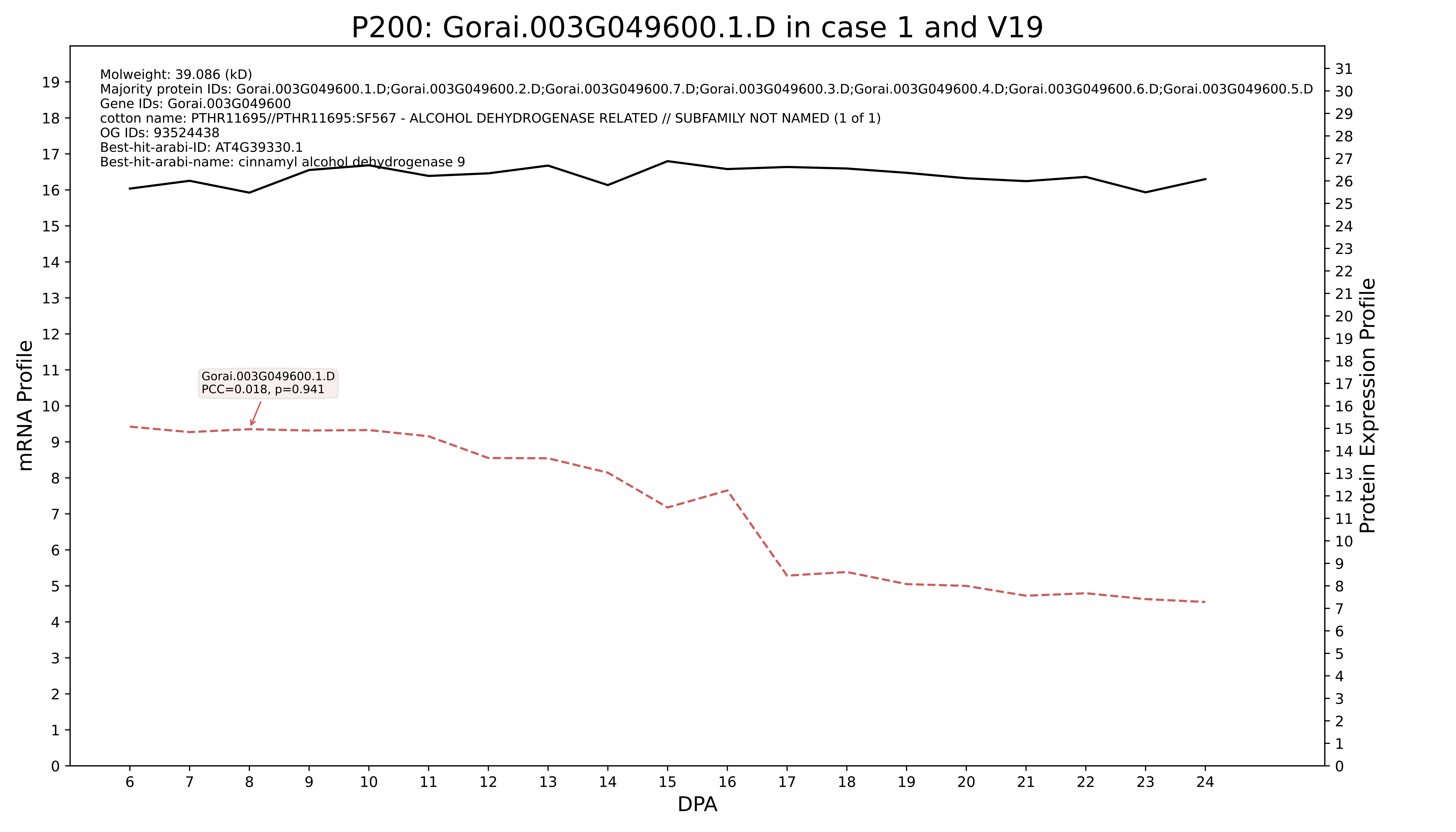
P200: Gorai.003G049600.1.D in case 1 and V19
Molweight: 39.086 (kD)
Majority protein IDs: Gorai.003G049600.1.D;Gorai.003G049600.2.D;Gorai.003G049600.7.D;Gorai.003G049600.3.D;Gorai.003G049600.4.D;Gorai.003G049600.6.D;Gorai.003G049600.5.D
Gene IDs: Gorai.003G049600
cotton name: PTHR11695//PTHR11695:SF567 - ALCOHOL DEHYDROGENASE RELATED // SUBFAMILY NOT NAMED (1 of 1)
OG IDs: 93524438
Best-hit-arabi-ID: AT4G39330.1
Best-hit-arabi-name: cinnamyl alcohol dehydrogenase 9
Gorai.003G049600.1.D
PCC=0.018, p=0.941
mRNA Profile
Protein Expression Profile
DPA

### P200: Gorai.012G125800.1.D in case 1 and V19

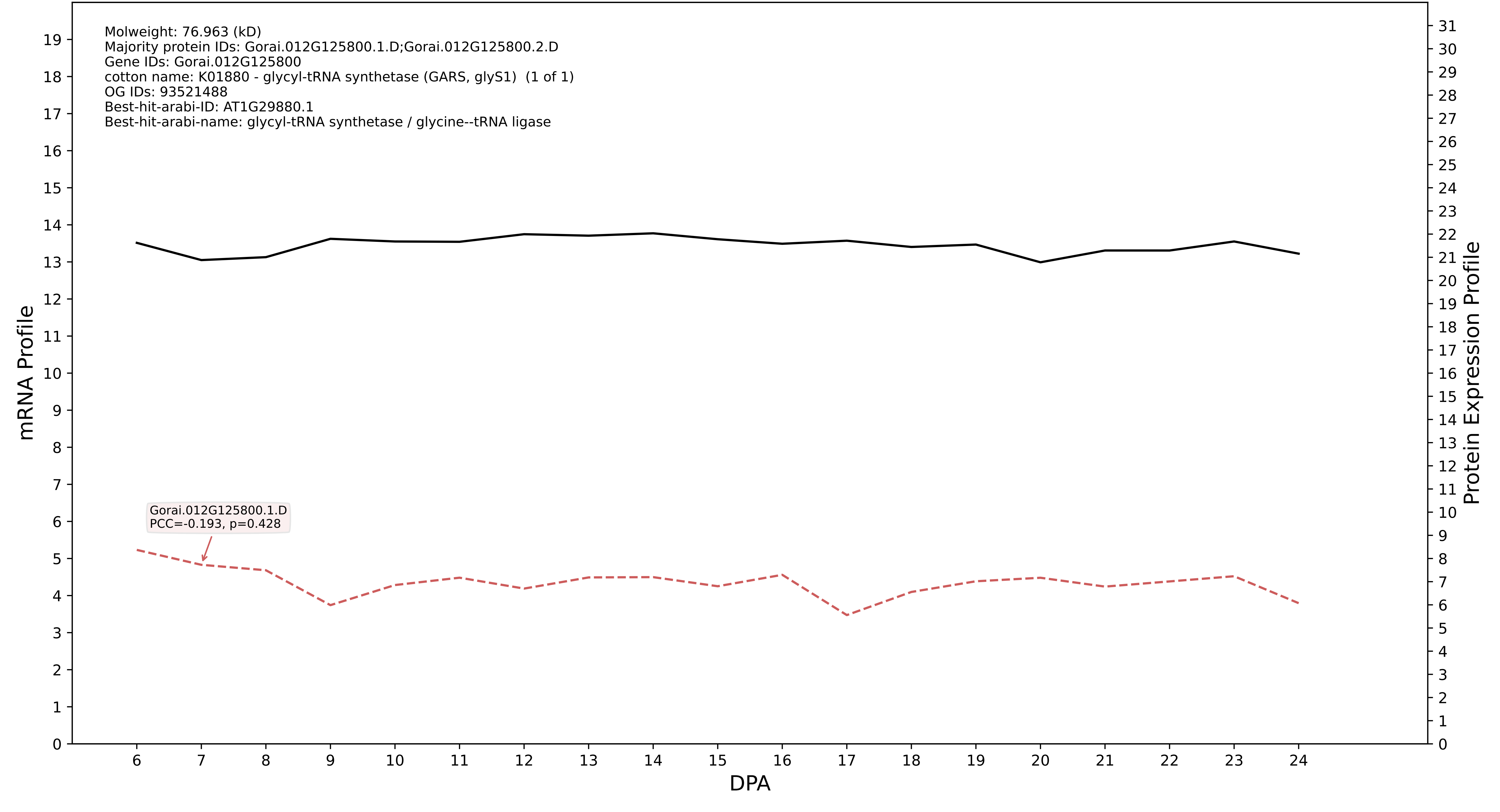

### P200: Gorai.005G007500.1.A in case 1 and V19

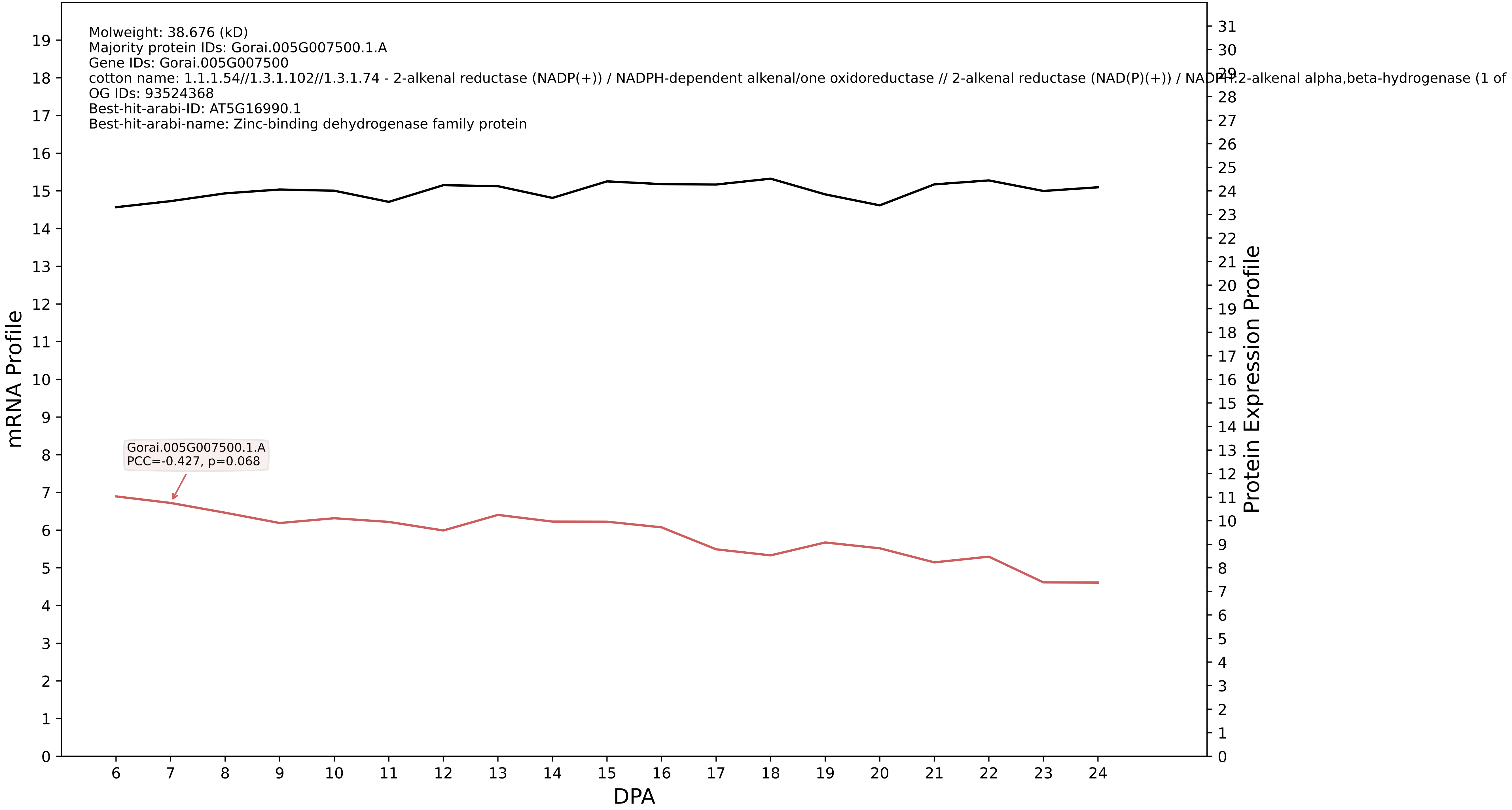

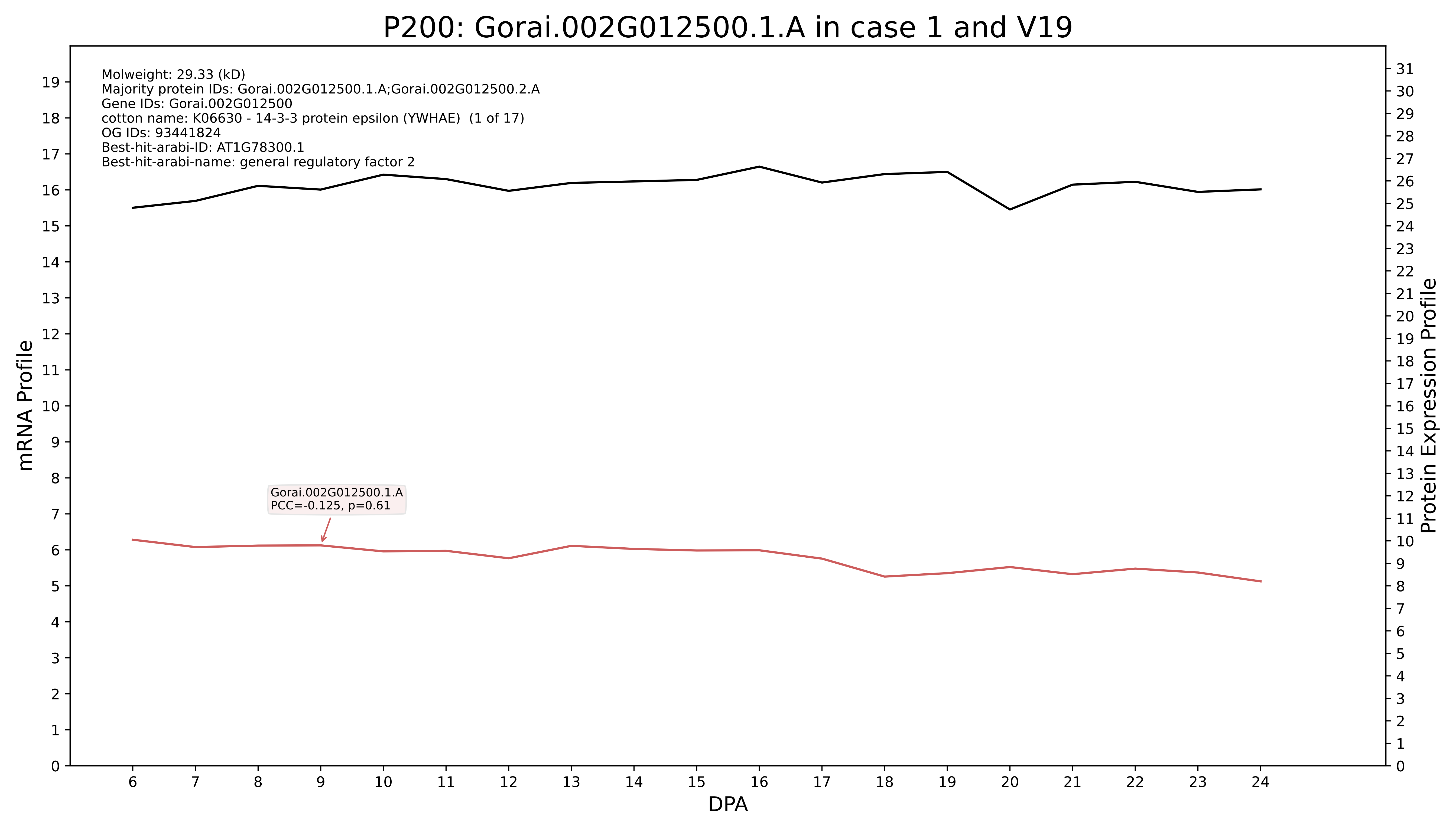
P200: Gorai.002G012500.1.A in case 1 and V19
Molweight: 29.33 (kD)
Majority protein IDs: Gorai.002G012500.1.A;Gorai.002G012500.2.A
Gene IDs: Gorai.002G012500
cotton name: K06630 - 14-3-3 protein epsilon (YWHAE) (1 of 17)
OG IDs: 93441824
Best-hit-arabi-ID: AT1G78300.1
Best-hit-arabi-name: general regulatory factor 2
Gorai.002G012500.1.A
PCC=-0.125, p=0.61
mRNA Profile
Protein Expression Profile
DPA

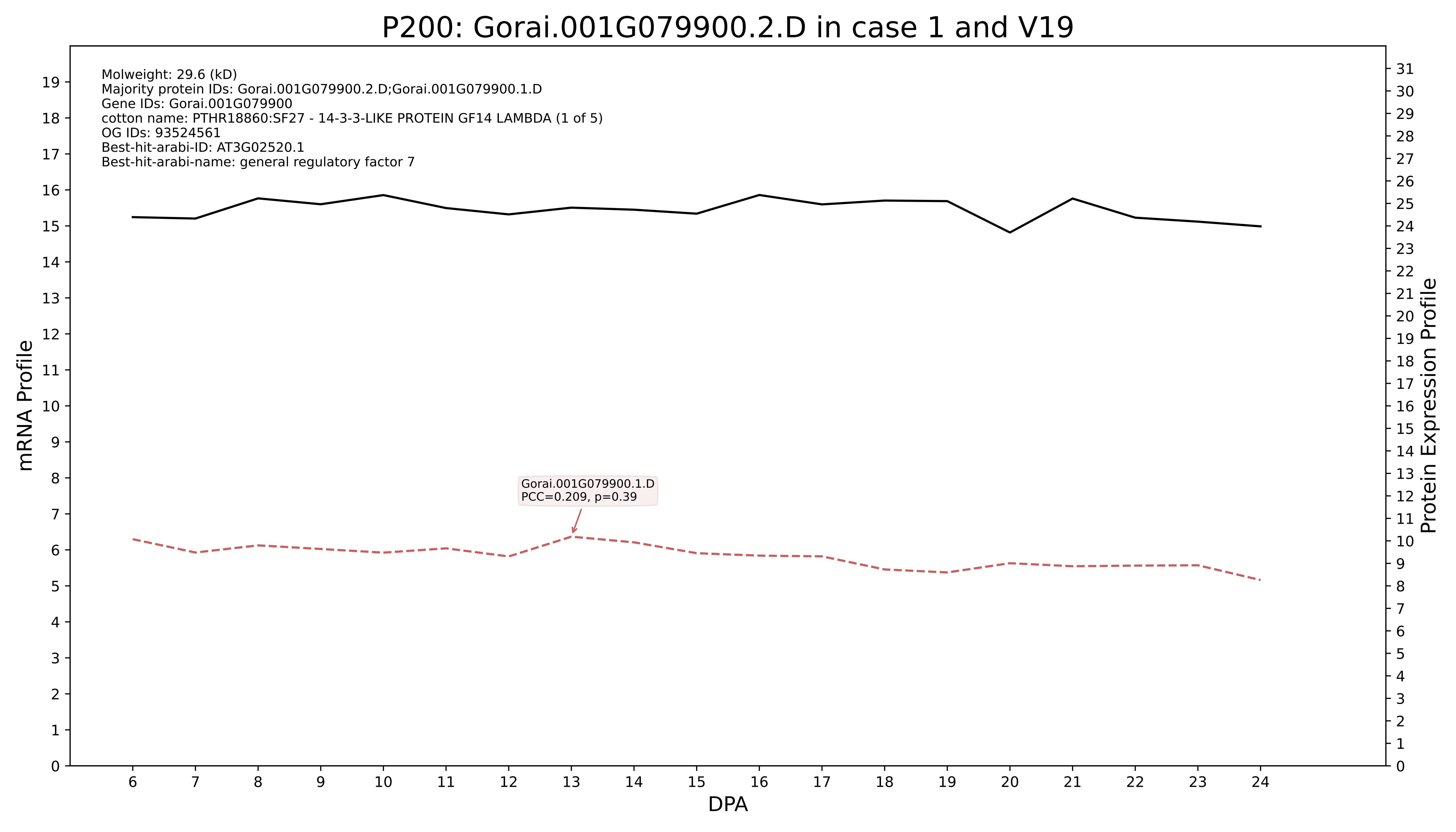
P200: Gorai.001G079900.2.D in case 1 and V19
Molweight: 29.6 (kD)
Majority protein IDs: Gorai.001G079900.2.D;Gorai.001G079900.1.D
Gene IDs: Gorai.001G079900
cotton name: PTHR18860:SF27 - 14-3-3-LIKE PROTEIN GF14 LAMBDA (1 of 5)
OG IDs: 93524561
Best-hit-arabi-ID: AT3G02520.1
Best-hit-arabi-name: general regulatory factor 7
Gorai.001G079900.1.D
PCC=0.209, p=0.39
mRNA Profile
Protein Expression Profile
DPA

### P200: Gorai.006G153000.4.A in case 1 and V19

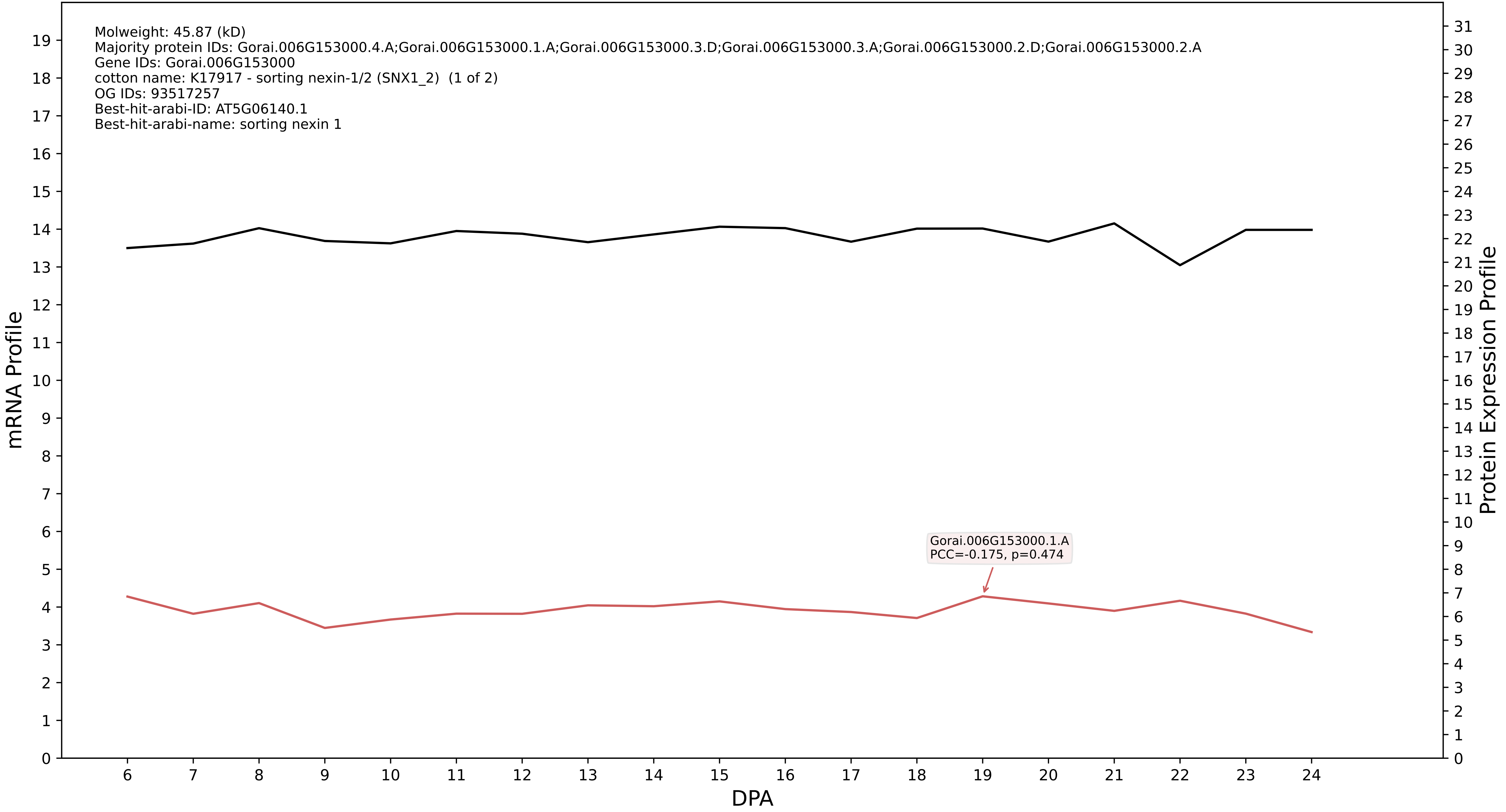

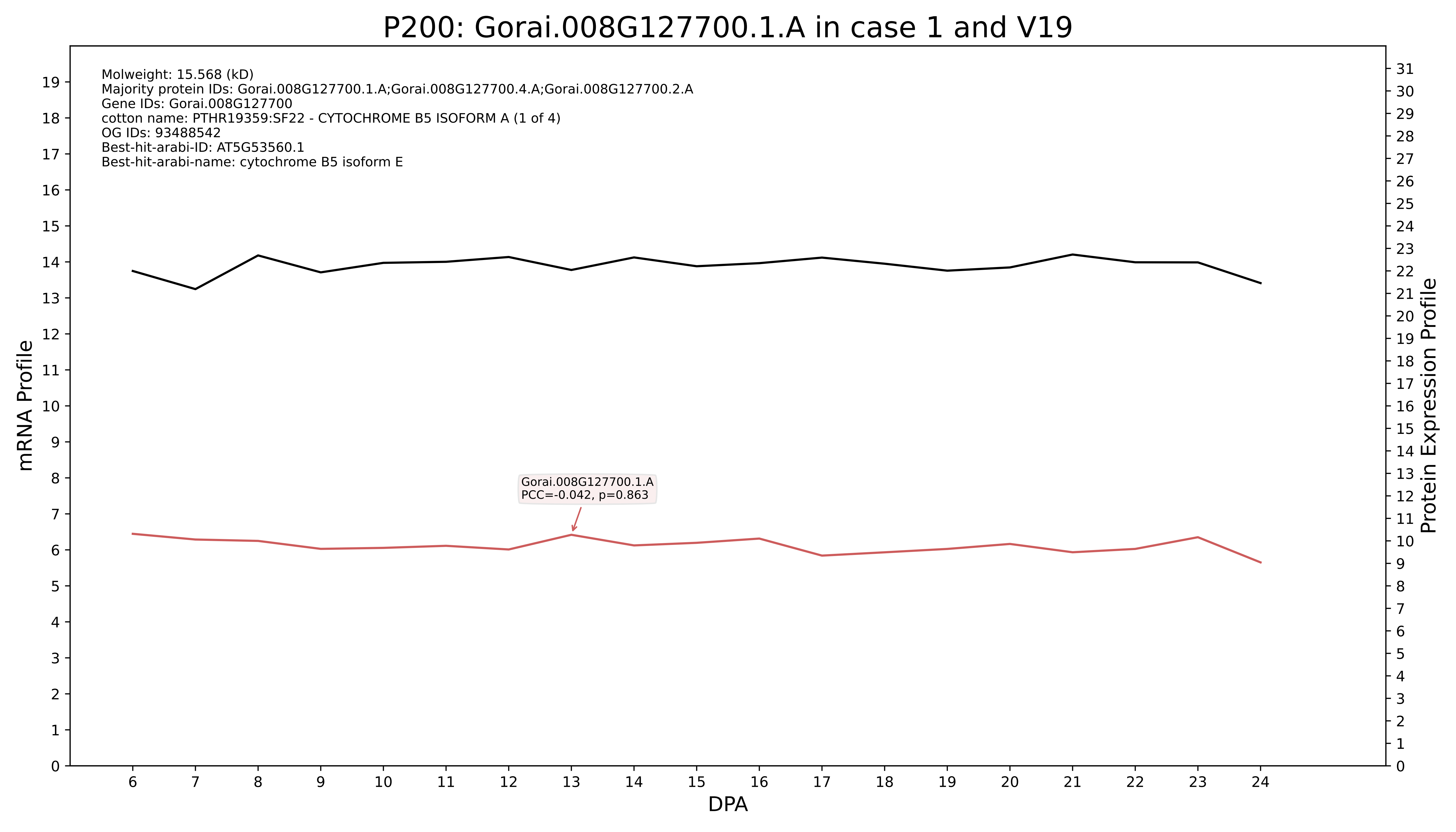

P200: Gorai.008G127700.1.A in case 1 and V19
Molweight: 15.568 (kD)
Majority protein IDs: Gorai.008G127700.1.A;Gorai.008G127700.4.A;Gorai.008G127700.2.A
Gene IDs: Gorai.008G127700
cotton name: PTHR19359:SF22 - CYTOCHROME B5 ISOFORM A (1 of 4)
OG IDs: 93488542
Best-hit-arabi-ID: AT5G53560.1
Best-hit-arabi-name: cytochrome B5 isoform E
Gorai.008G127700.1.A
PCC=-0.042, p=0.863
mRNA Profile
Protein Expression Profile
DPA

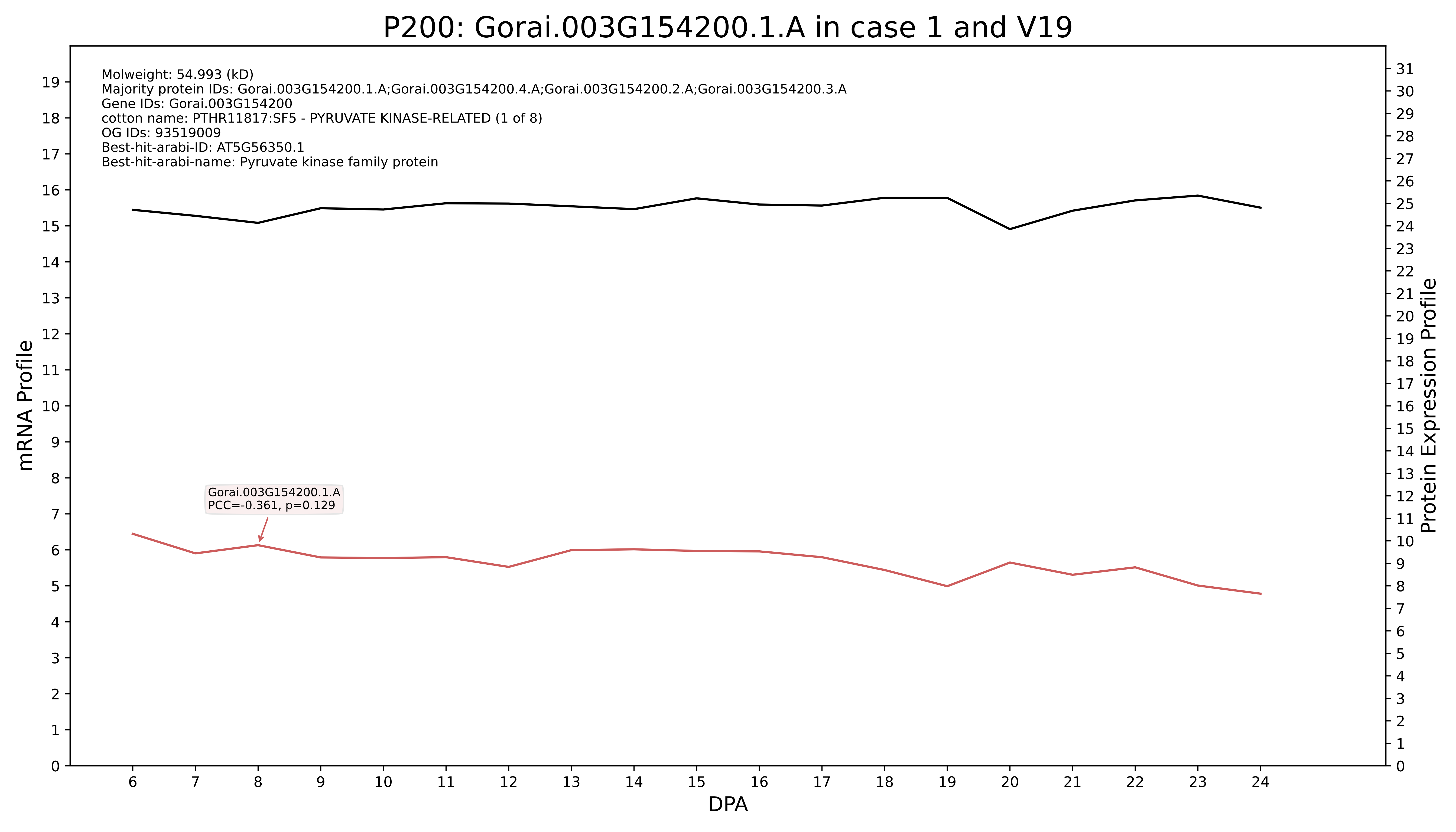

P200: Gorai.003G154200.1.A in case 1 and V19
Molweight: 54.993 (kD)
Majority protein IDs: Gorai.003G154200.1.A;Gorai.003G154200.4.A;Gorai.003G154200.2.A;Gorai.003G154200.3.A
Gene IDs: Gorai.003G154200
cotton name: PTHR11817:SF5 - PYRUVATE KINASE-RELATED (1 of 8)
OG IDs: 93519009
Best-hit-arabi-ID: AT5G56350.1
Best-hit-arabi-name: Pyruvate kinase family protein
Gorai.003G154200.1.A
PCC=-0.361, p=0.129
mRNA Profile
Protein Expression Profile
DPA

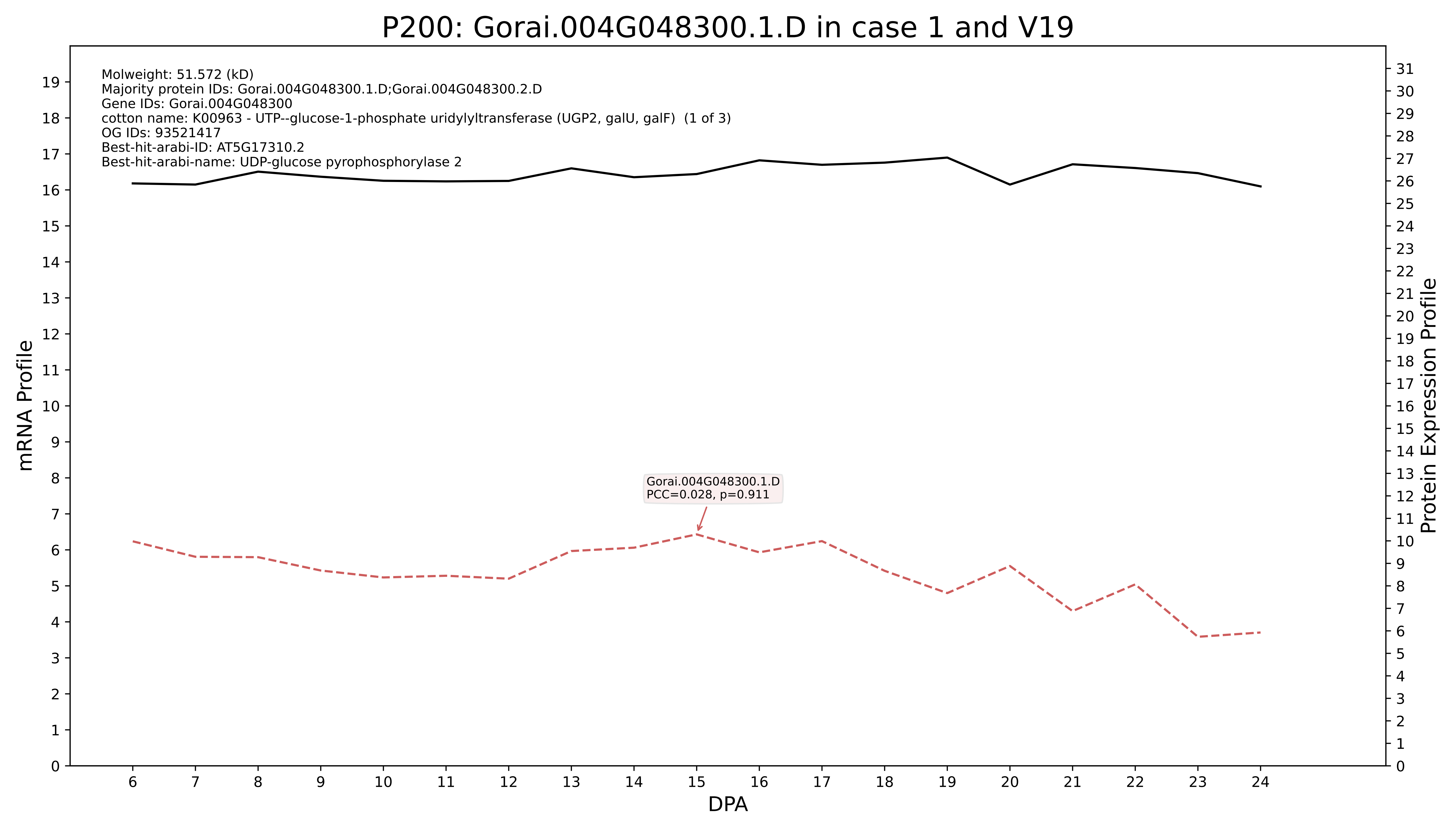
P200: Gorai.004G048300.1.D in case 1 and V19
Molweight: 51.572 (kD)
Majority protein IDs: Gorai.004G048300.1.D;Gorai.004G048300.2.D
Gene IDs: Gorai.004G048300
cotton name: K00963 - UTP--glucose-1-phosphate uridylyltransferase (UGP2, galU, galF) (1 of 3)
OG IDs: 93521417
Best-hit-arabi-ID: AT5G17310.2
Best-hit-arabi-name: UDP-glucose pyrophosphorylase 2
Gorai.004G048300.1.D
PCC=0.028, p=0.911
mRNA Profile
Protein Expression Profile
DPA

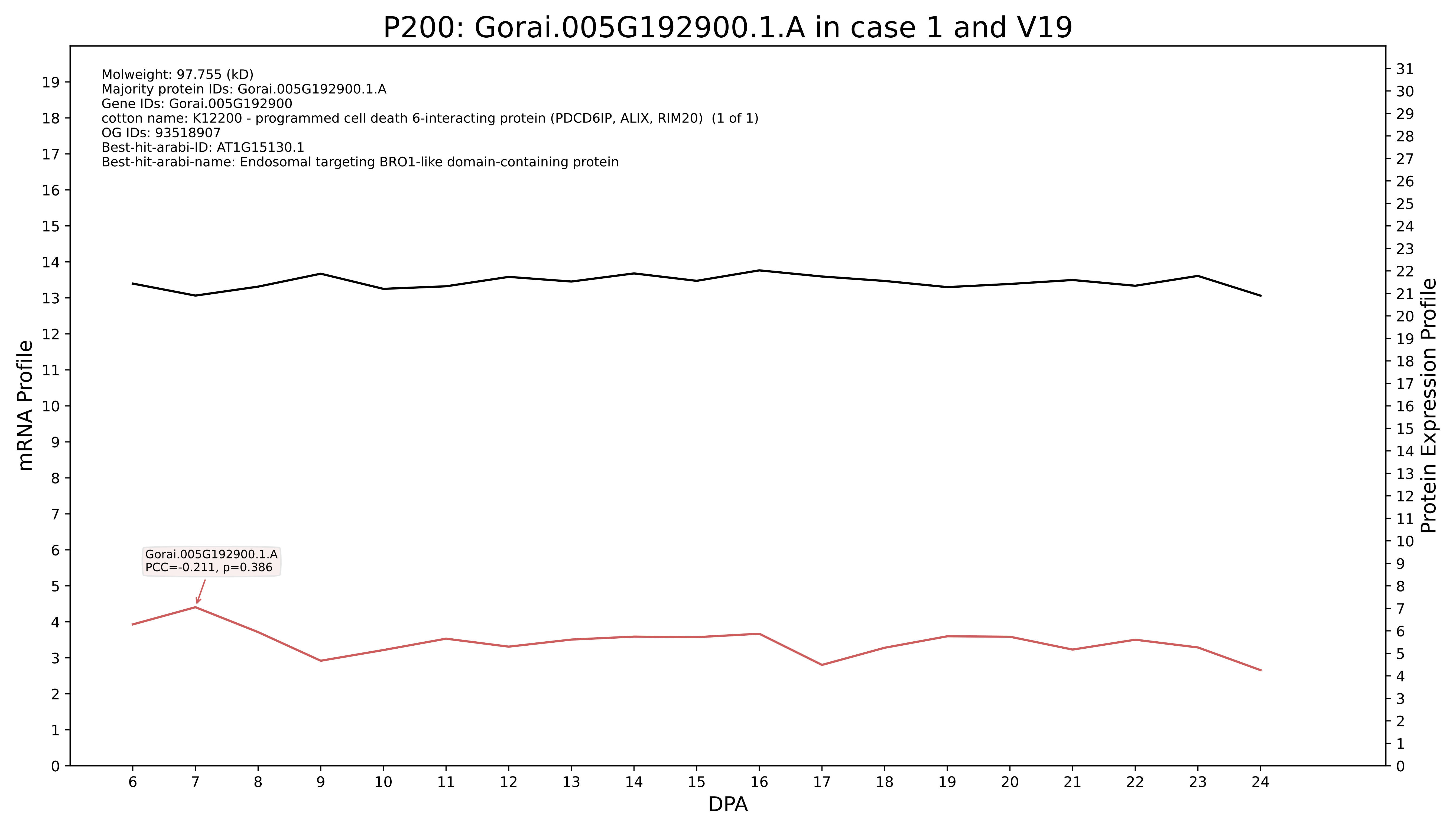
P200: Gorai.005G192900.1.A in case 1 and V19
Molweight: 97.755 (kD)
Majority protein IDs: Gorai.005G192900.1.A
Gene IDs: Gorai.005G192900
cotton name: K12200 - programmed cell death 6-interacting protein (PDCD6IP, ALIX, RIM20) (1 of 1)
OG IDs: 93518907
Best-hit-arabi-ID: AT1G15130.1
Best-hit-arabi-name: Endosomal targeting BRO1-like domain-containing protein
Gorai.005G192900.1.A
PCC=-0.211, p=0.386
mRNA Profile
Protein Expression Profile
DPA

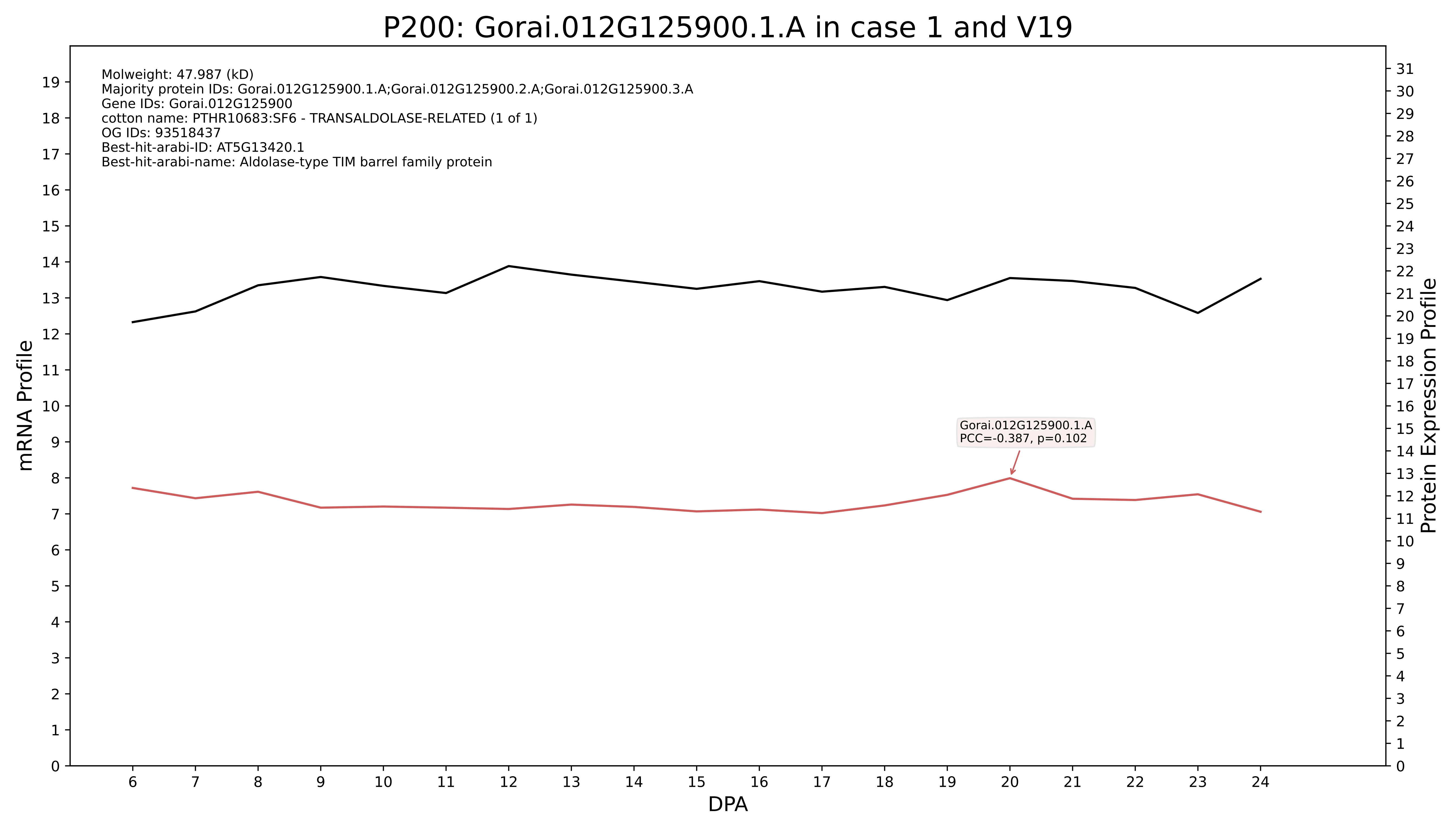
P200: Gorai.012G125900.1.A in case 1 and V19
Molweight: 47.987 (kD)
Majority protein IDs: Gorai.012G125900.1.A;Gorai.012G125900.2.A;Gorai.012G125900.3.A
Gene IDs: Gorai.012G125900
cotton name: PTHR10683:SF6 - TRANSALDOLASE-RELATED (1 of 1)
OG IDs: 93518437
Best-hit-arabi-ID: AT5G13420.1
Best-hit-arabi-name: Aldolase-type TIM barrel family protein
Gorai.012G125900.1.A
PCC=-0.387, p=0.102
mRNA Profile
Protein Expression Profile
DPA

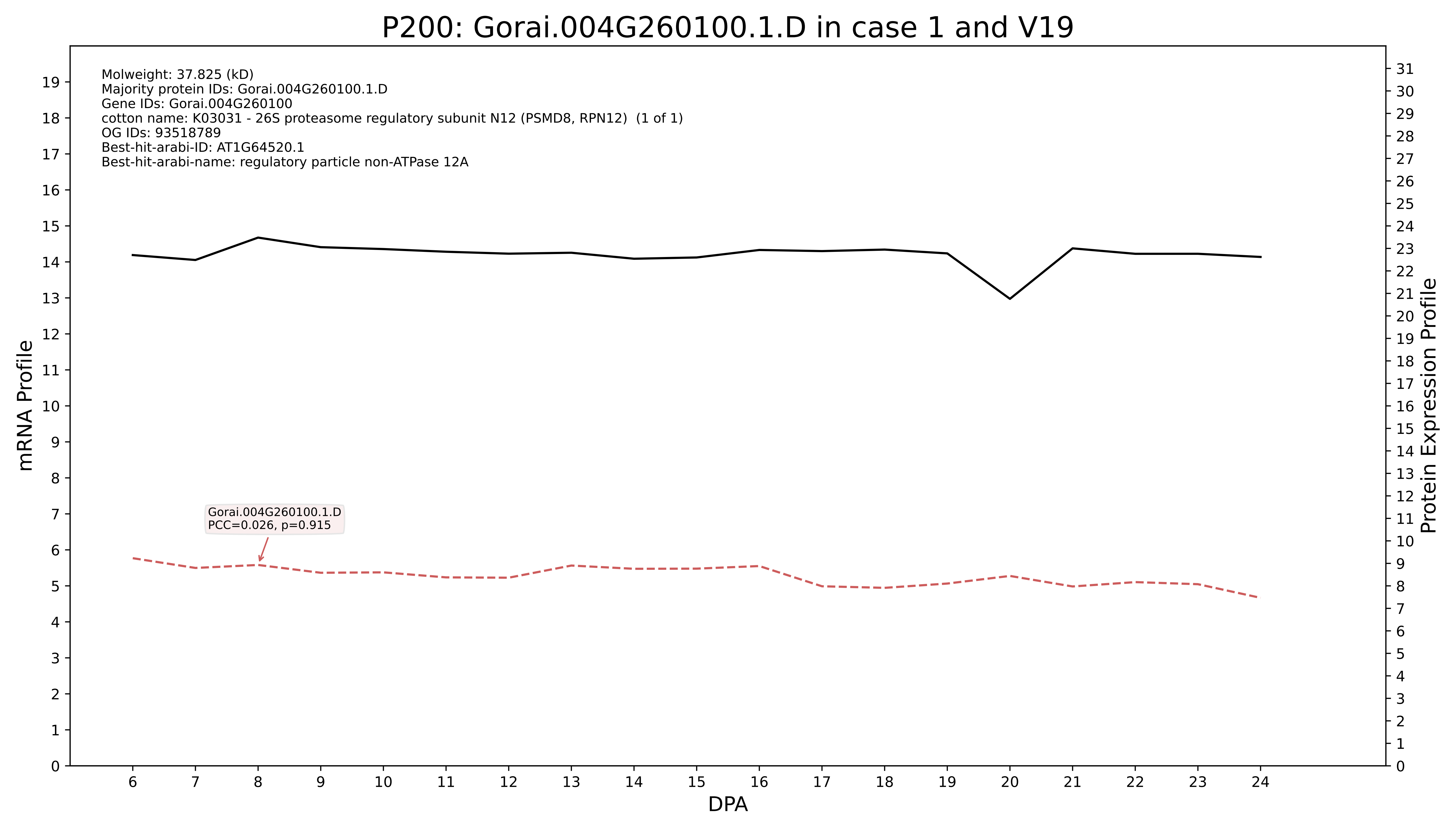
P200: Gorai.004G260100.1.D in case 1 and V19
Molweight: 37.825 (kD)
Majority protein IDs: Gorai.004G260100.1.D
Gene IDs: Gorai.004G260100
cotton name: K03031 - 26S proteasome regulatory subunit N12 (PSMD8, RPN12) (1 of 1)
OG IDs: 93518789
Best-hit-arabi-ID: AT1G64520.1
Best-hit-arabi-name: regulatory particle non-ATPase 12A
Gorai.004G260100.1.D
PCC=0.026, p=0.915
mRNA Profile
Protein Expression Profile
DPA

### P200: Gorai.006G148600.1.A in case 1 and V19

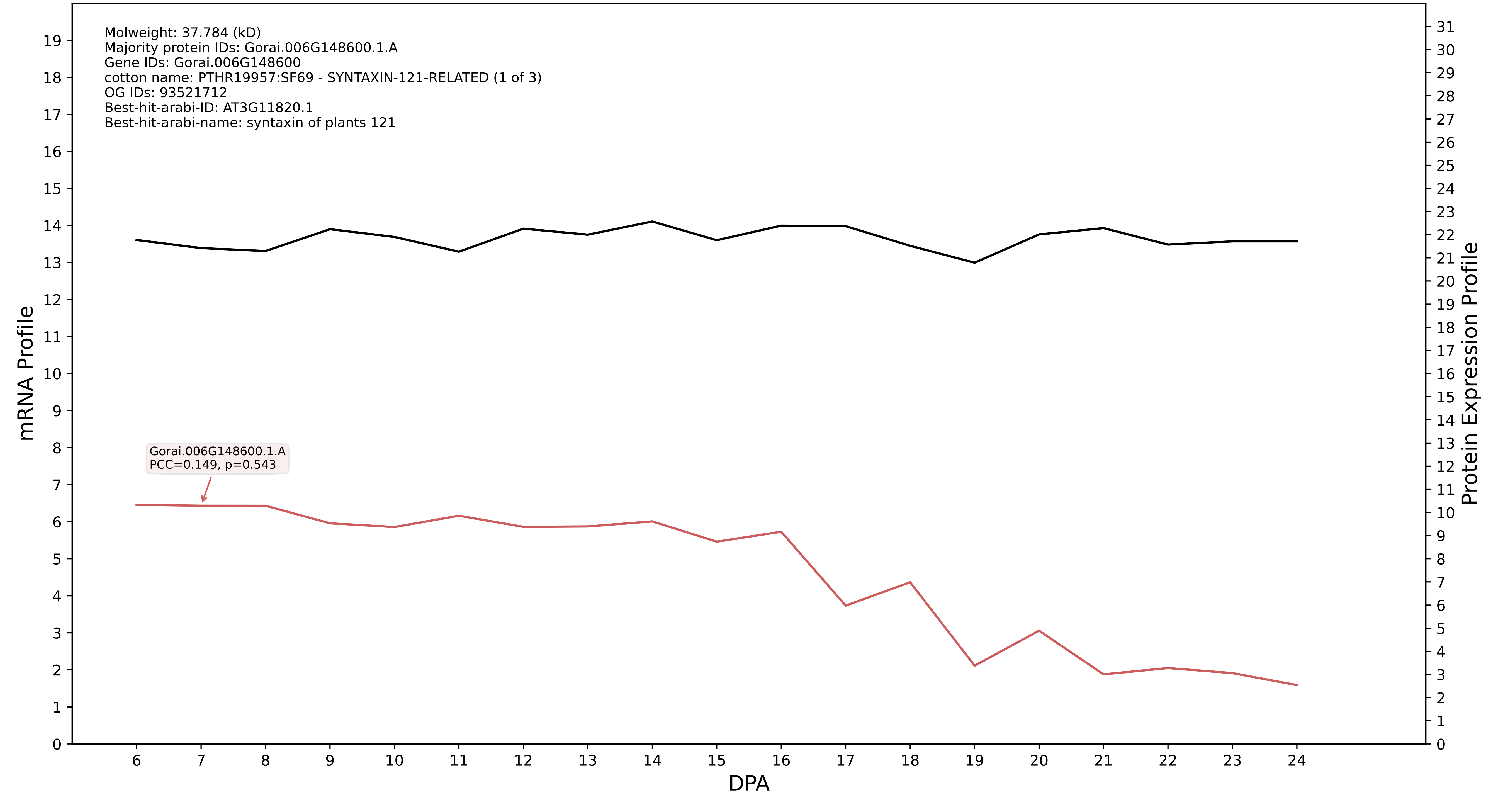

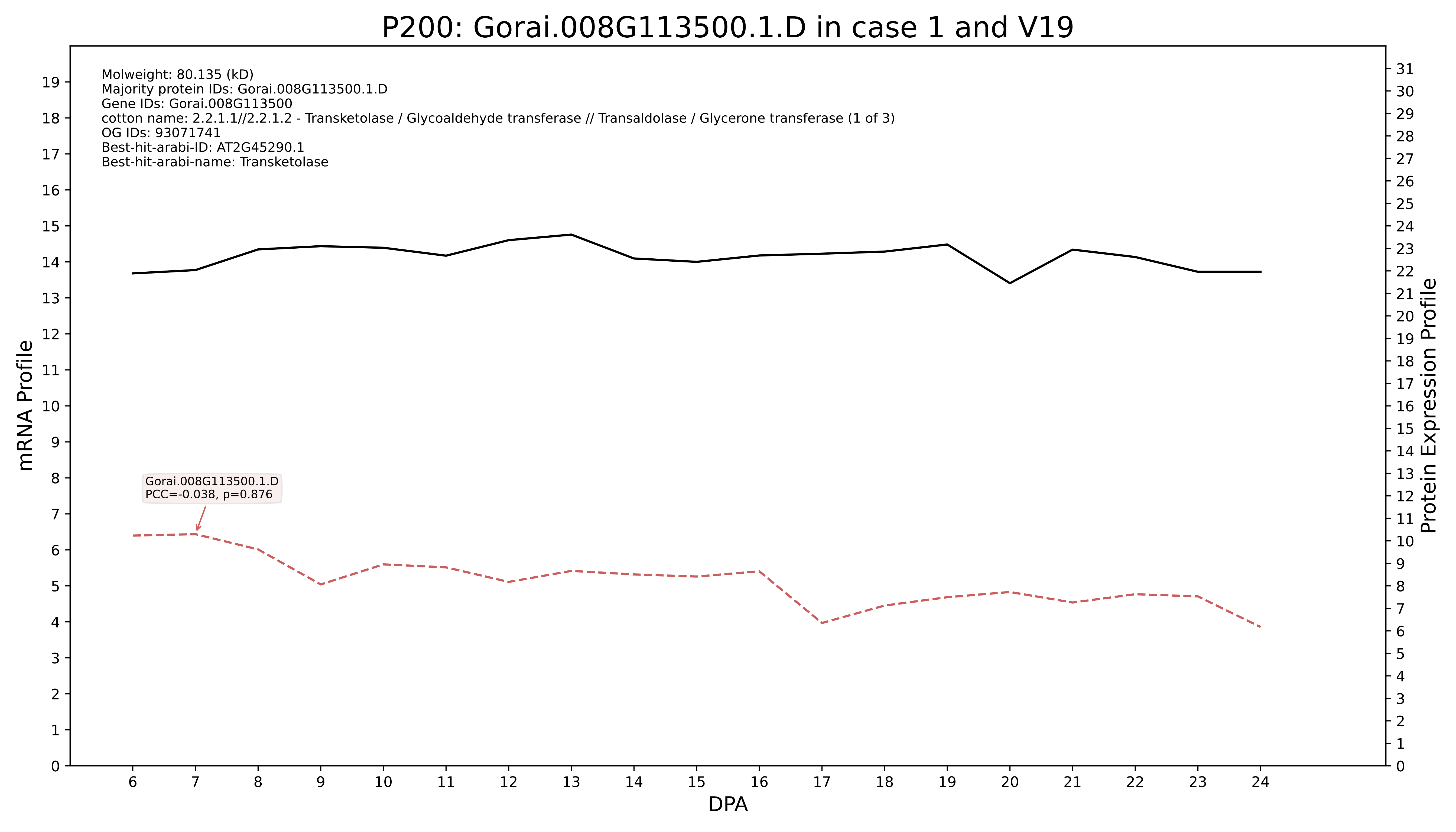
P200: Gorai.008G113500.1.D in case 1 and V19
Molweight: 80.135 (kD)
Majority protein IDs: Gorai.008G113500.1.D
Gene IDs: Gorai.008G113500
cotton name: 2.2.1.1//2.2.1.2 - Transketolase / Glycoaldehyde transferase // Transaldolase / Glycerone transferase (1 of 3)
OG IDs: 93071741
Best-hit-arabi-ID: AT2G45290.1
Best-hit-arabi-name: Transketolase
Gorai.008G113500.1.D
PCC=-0.038, p=0.876
mRNA Profile
Protein Expression Profile
DPA

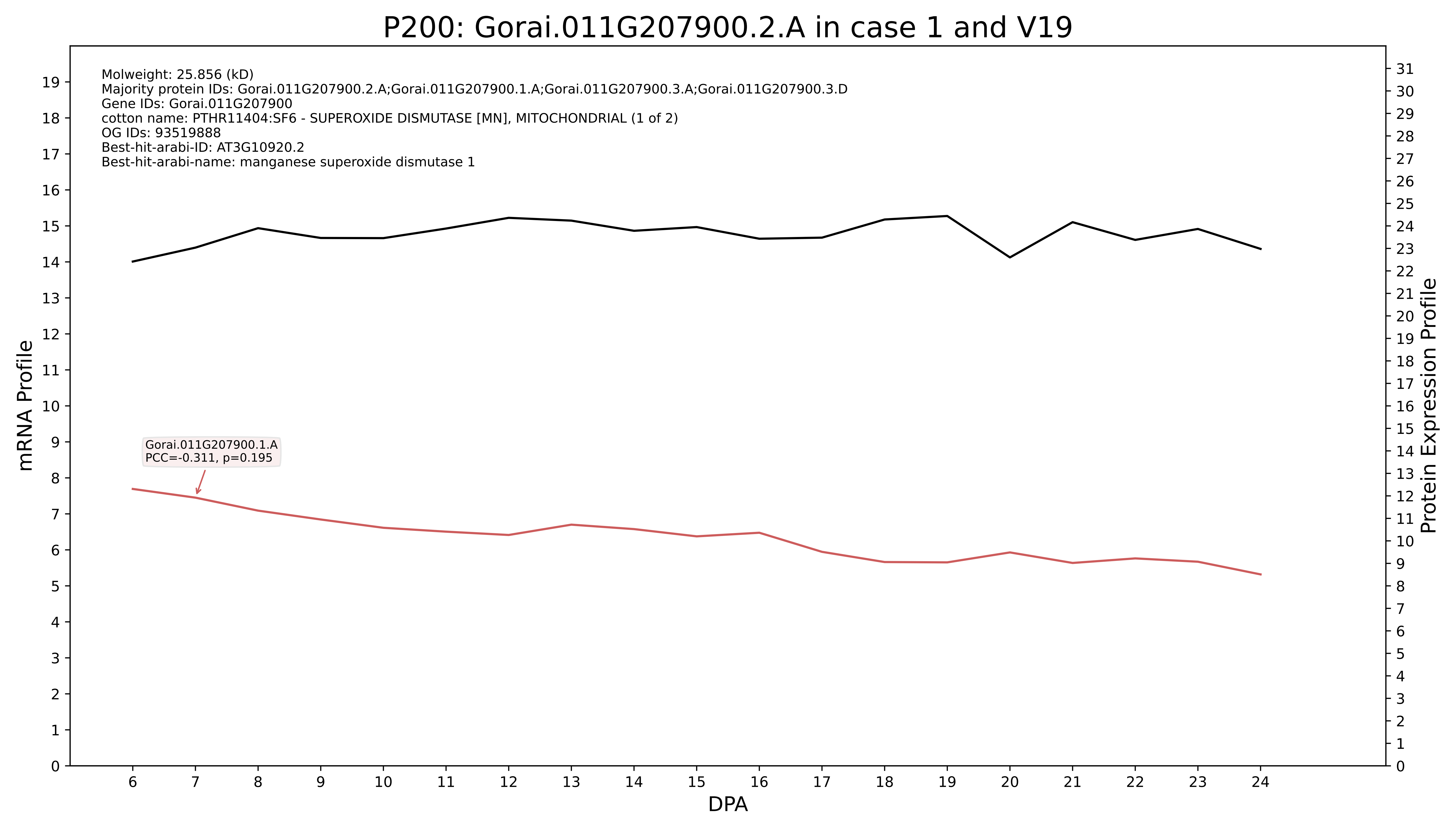
P200: Gorai.011G207900.2.A in case 1 and V19
Molweight: 25.856 (kD)
Majority protein IDs: Gorai.011G207900.2.A;Gorai.011G207900.1.A;Gorai.011G207900.3.A;Gorai.011G207900.3.D
Gene IDs: Gorai.011G207900
cotton name: PTHR11404:SF6 - SUPEROXIDE DISMUTASE [MN], MITOCHONDRIAL (1 of 2)
OG IDs: 93519888
Best-hit-arabi-ID: AT3G10920.2
Best-hit-arabi-name: manganese superoxide dismutase 1
Gorai.011G207900.1.A
PCC=-0.311, p=0.195
mRNA Profile
Protein Expression Profile
DPA

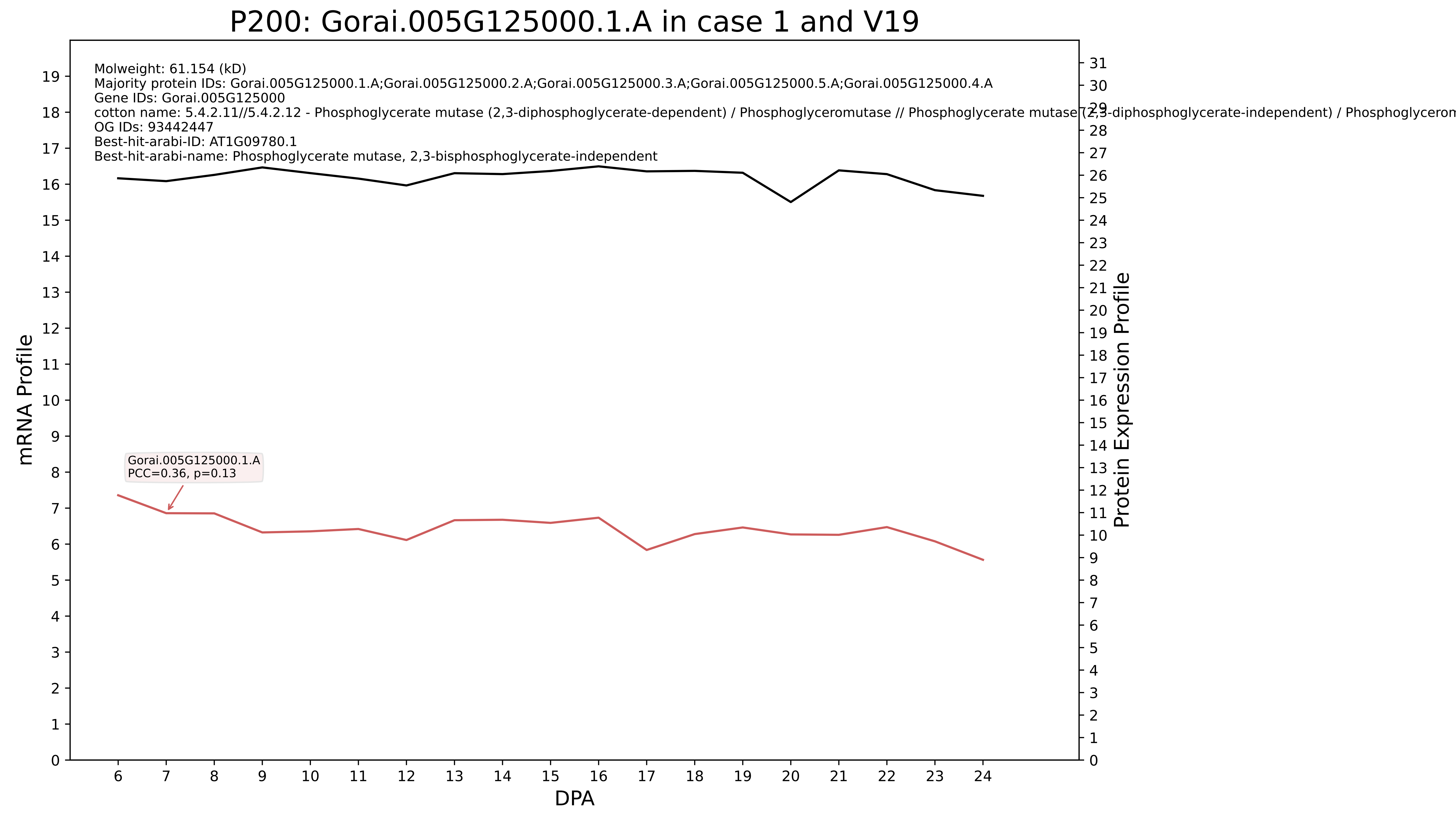
P200: Gorai.005G125000.1.A in case 1 and V19
Molweight: 61.154 (kD)
Majority protein IDs: Gorai.005G125000.1.A;Gorai.005G125000.2.A;Gorai.005G125000.3.A;Gorai.005G125000.5.A;Gorai.005G125000.4.A
Gene IDs: Gorai.005G125000
cotton name: 5.4.2.11//5.4.2.12 - Phosphoglycerate mutase (2,3-diphosphoglycerate-dependent) / Phosphoglyceromutase // Phosphoglycerate mutase (2,3-diphosphoglycerate-independent) / Phosphoglyceron
OG IDs: 93442447
Best-hit-arabi-ID: AT1G09780.1
Best-hit-arabi-name: Phosphoglycerate mutase, 2,3-bisphosphoglycerate-independent
Gorai.005G125000.1.A
PCC=0.36, p=0.13
mRNA Profile
Protein Expression Profile
DPA

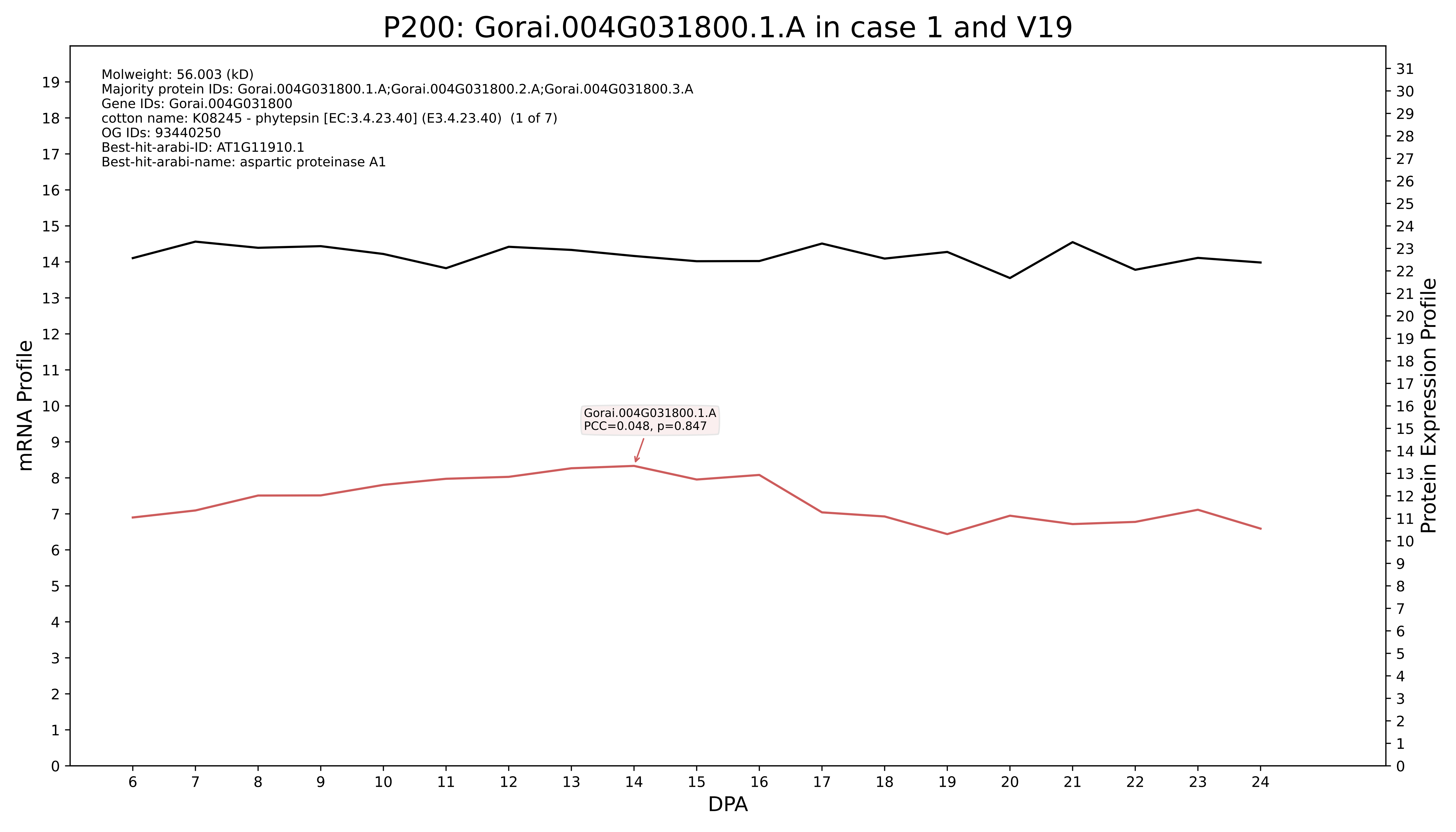
P200: Gorai.004G031800.1.A in case 1 and V19
Molweight: 56.003 (kD)
Majority protein IDs: Gorai.004G031800.1.A;Gorai.004G031800.2.A;Gorai.004G031800.3.A
Gene IDs: Gorai.004G031800
cotton name: K08245 - phytepsin [EC:3.4.23.40] (E3.4.23.40) (1 of 7)
OG IDs: 93440250
Best-hit-arabi-ID: AT1G11910.1
Best-hit-arabi-name: aspartic proteinase A1
Gorai.004G031800.1.A
PCC=0.048, p=0.847
mRNA Profile
Protein Expression Profile
DPA

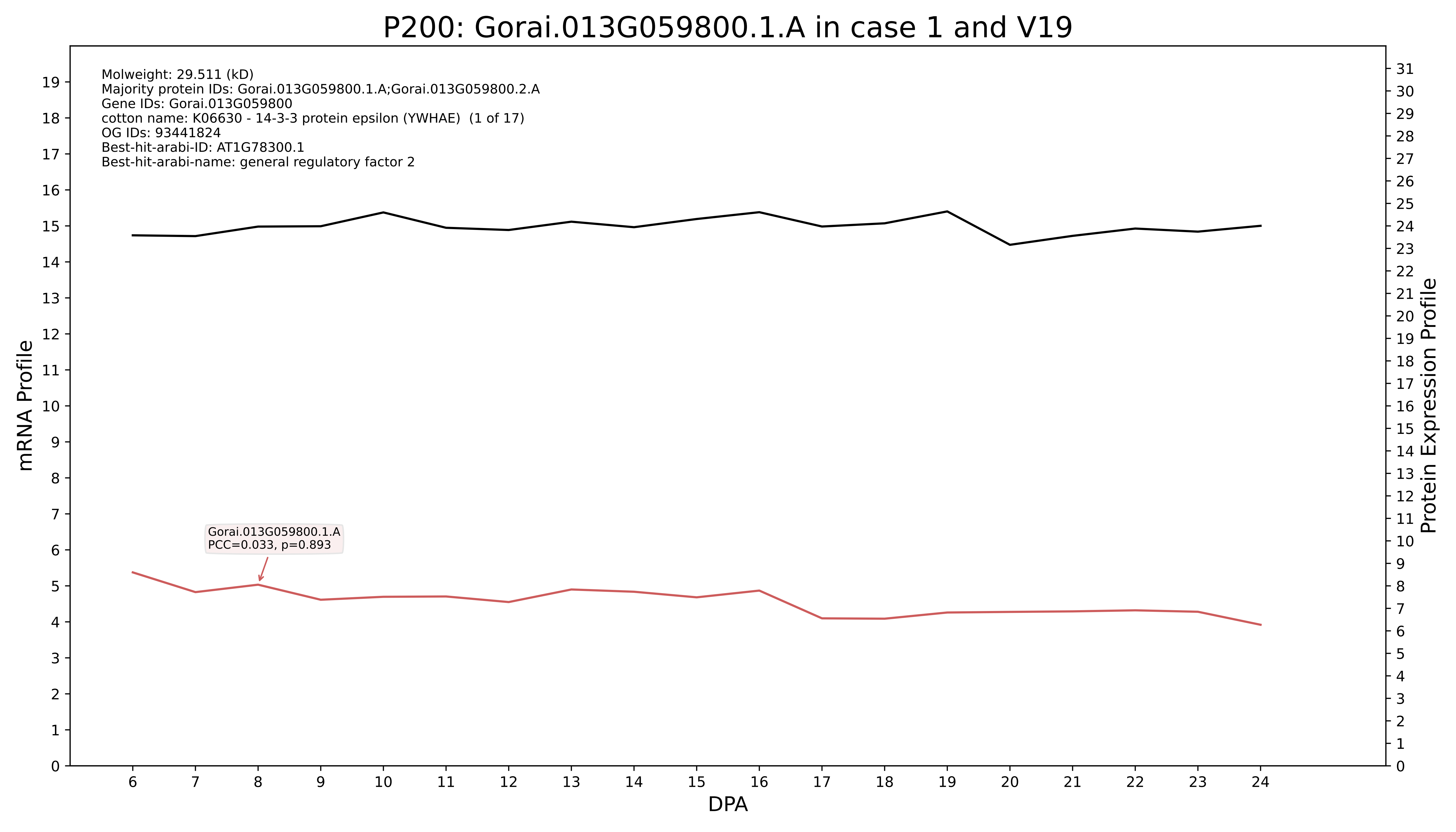

P200: Gorai.013G059800.1.A in case 1 and V19
Molweight: 29.511 (kD)
Majority protein IDs: Gorai.013G059800.1.A;Gorai.013G059800.2.A
Gene IDs: Gorai.013G059800
cotton name: K06630 - 14-3-3 protein epsilon (YWHAE) (1 of 17)
OG IDs: 93441824
Best-hit-arabi-ID: AT1G78300.1
Best-hit-arabi-name: general regulatory factor 2
Gorai.013G059800.1.A
PCC=0.033, p=0.893
mRNA Profile
Protein Expression Profile
DPA

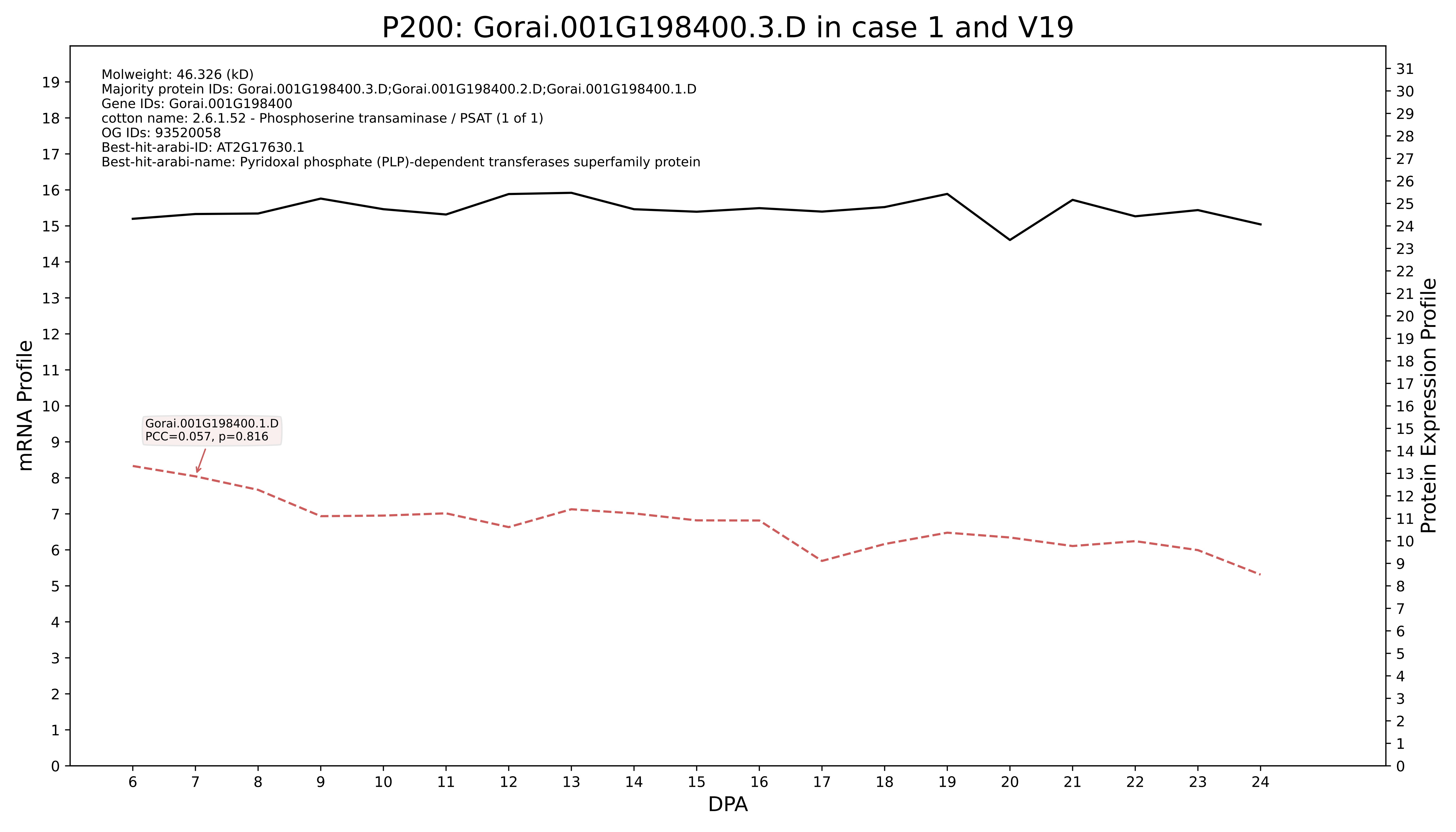
P200: Gorai.001G198400.3.D in case 1 and V19
Molweight: 46.326 (kD)
Majority protein IDs: Gorai.001G198400.3.D;Gorai.001G198400.2.D;Gorai.001G198400.1.D
Gene IDs: Gorai.001G198400
cotton name: 2.6.1.52 - Phosphoserine transaminase / PSAT (1 of 1)
OG IDs: 93520058
Best-hit-arabi-ID: AT2G17630.1
Best-hit-arabi-name: Pyridoxal phosphate (PLP)-dependent transferases superfamily protein
Gorai.001G198400.1.D
PCC=0.057, p=0.816
mRNA Profile
Protein Expression Profile
DPA

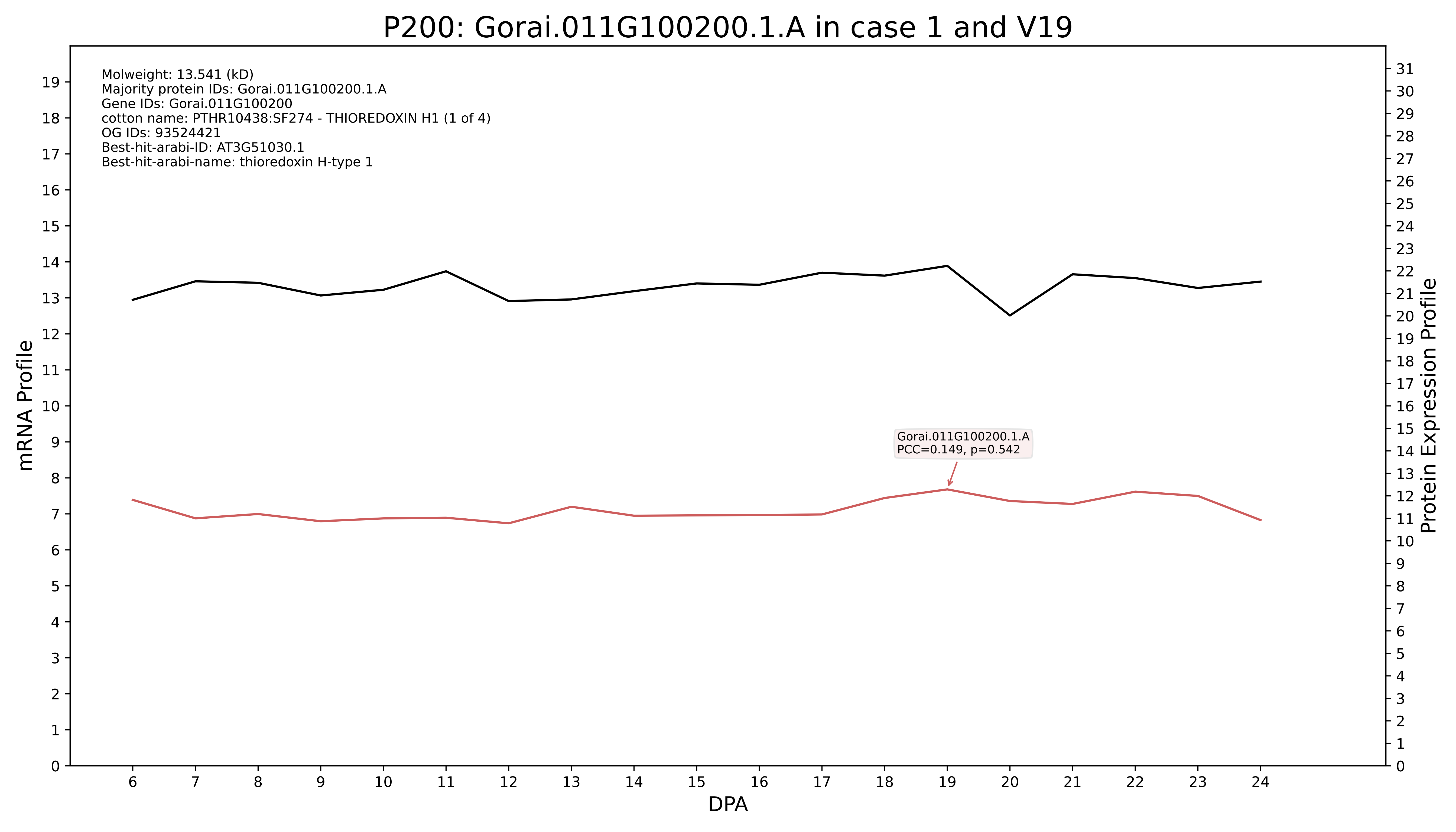
P200: Gorai.011G100200.1.A in case 1 and V19
Molweight: 13.541 (kD)
Majority protein IDs: Gorai.011G100200.1.A
Gene IDs: Gorai.011G100200
cotton name: PTHR10438:SF274 - THIOREDOXIN H1 (1 of 4)
OG IDs: 93524421
Best-hit-arabi-ID: AT3G51030.1
Best-hit-arabi-name: thioredoxin H-type 1
Gorai.011G100200.1.A
PCC=0.149, p=0.542
mRNA Profile
Protein Expression Profile
DPA

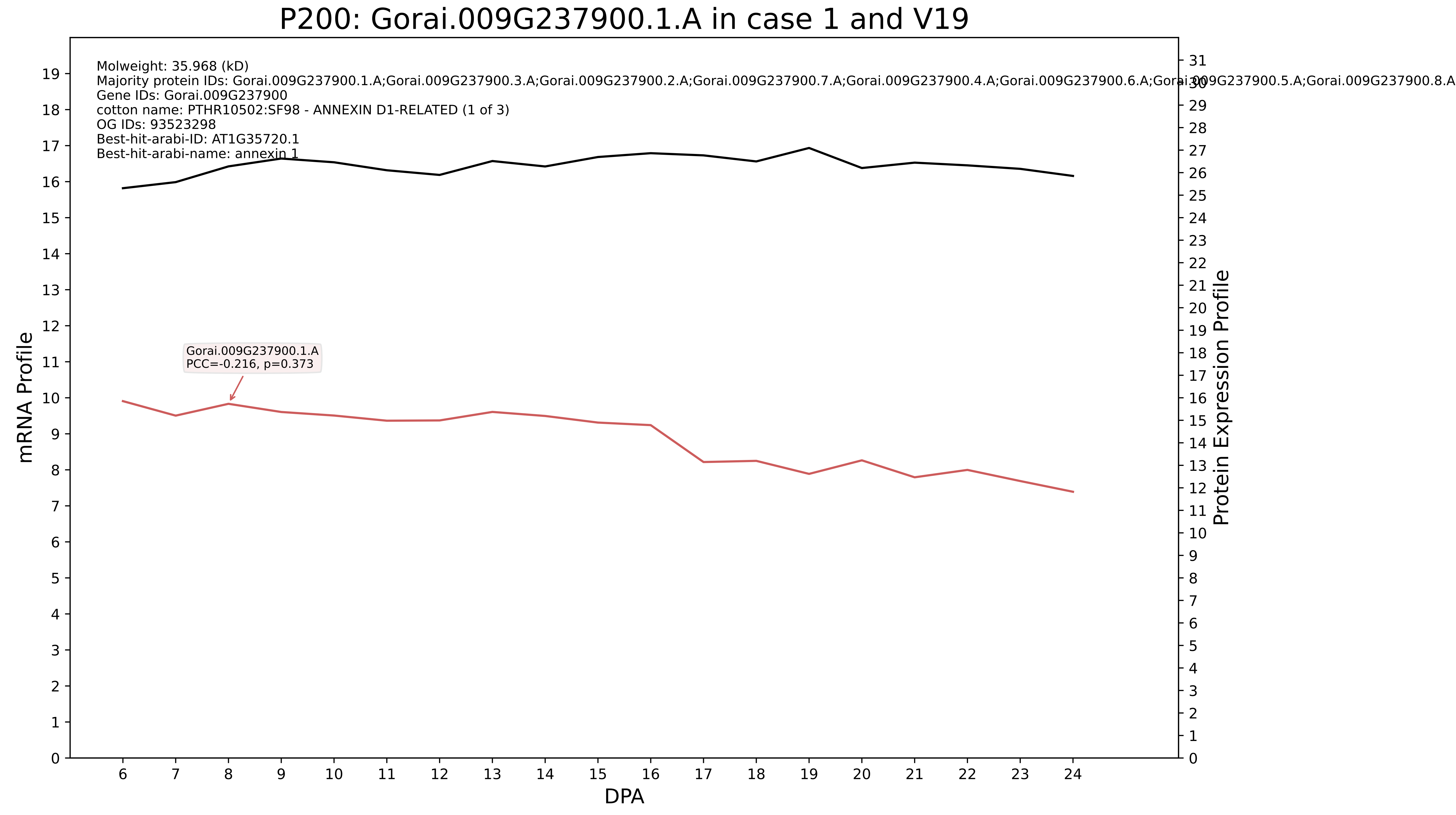
P200: Gorai.009G237900.1.A in case 1 and V19
Molweight: 35.968 (kD)
Majority protein IDs: Gorai.009G237900.1.A;Gorai.009G237900.3.A;Gorai.009G237900.2.A;Gorai.009G237900.7.A;Gorai.009G237900.4.A;Gorai.009G237900.6.A;Gorai.009G237900.5.A;Gorai.009G237900.8.A
Gene IDs: Gorai.009G237900
cotton name: PTHR10502:SF98 - ANNEXIN D1-RELATED (1 of 3)
OG IDs: 93523298
Best-hit-arabi-ID: AT1G35720.1
Best-hit-arabi-name: annexin 1
Gorai.009G237900.1.A
PCC=-0.216, p=0.373
mRNA Profile
Protein Expression Profile
DPA

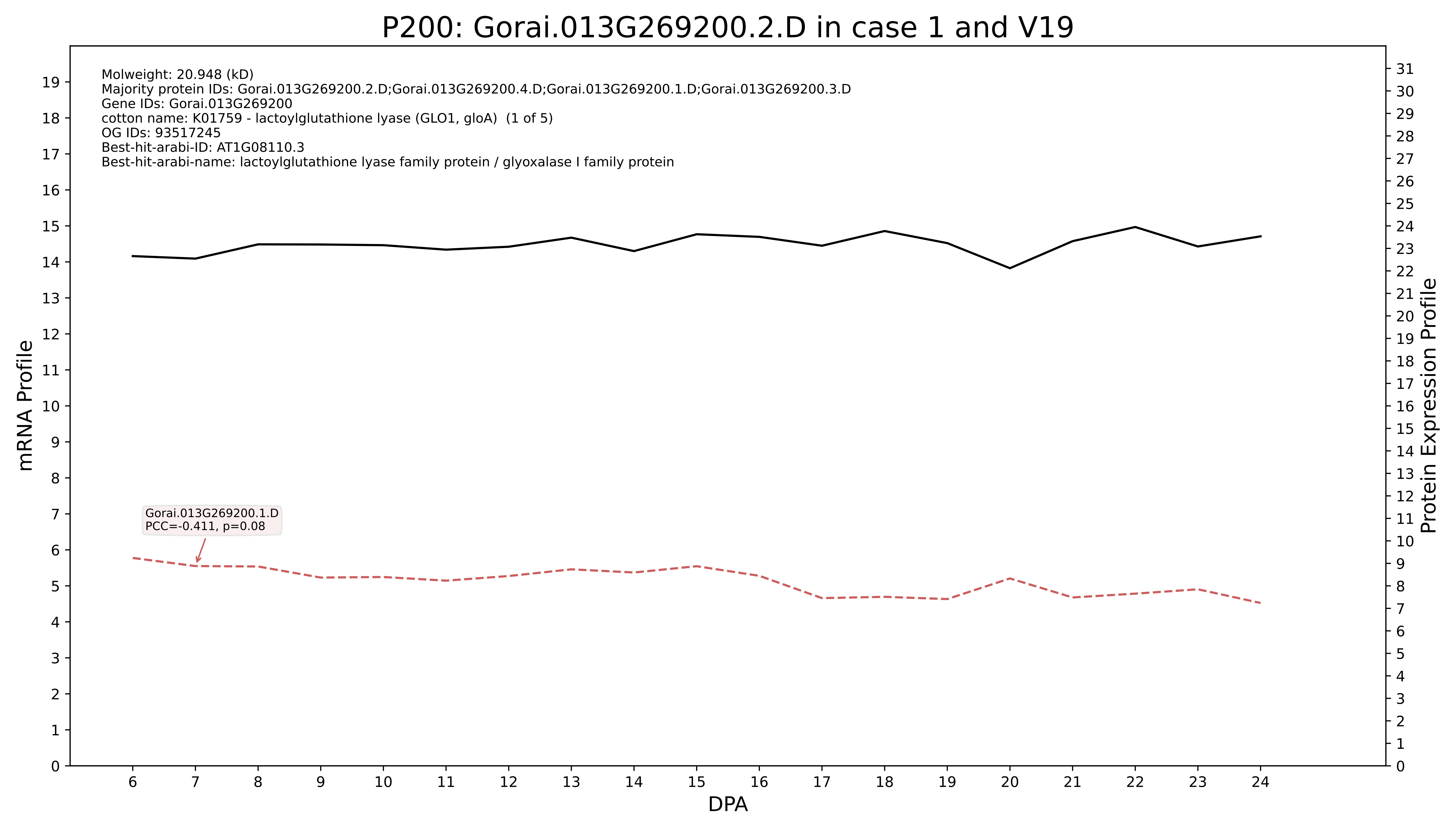
P200: Gorai.013G269200.2.D in case 1 and V19
Molweight: 20.948 (kD)
Majority protein IDs: Gorai.013G269200.2.D;Gorai.013G269200.4.D;Gorai.013G269200.1.D;Gorai.013G269200.3.D
Gene IDs: Gorai.013G269200
cotton name: K01759 - lactoylglutathione lyase (GLO1, gloA) (1 of 5)
OG IDs: 93517245
Best-hit-arabi-ID: AT1G08110.3
Best-hit-arabi-name: lactoylglutathione lyase family protein / glyoxalase I family protein
Gorai.013G269200.1.D
PCC=-0.411, p=0.08
mRNA Profile
Protein Expression Profile
DPA

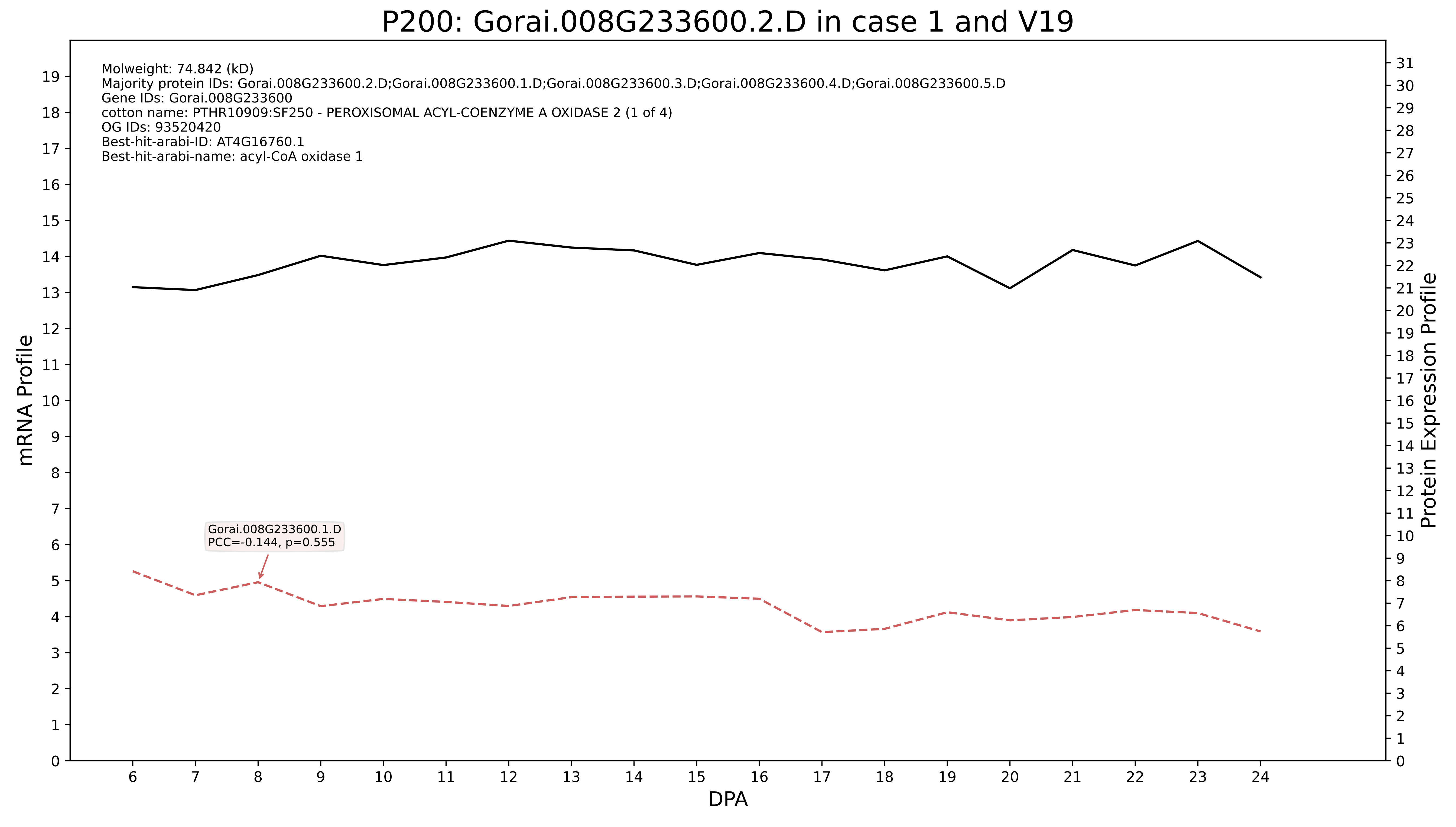
P200: Gorai.008G233600.2.D in case 1 and V19
Molweight: 74.842 (kD)
Majority protein IDs: Gorai.008G233600.2.D;Gorai.008G233600.1.D;Gorai.008G233600.3.D;Gorai.008G233600.4.D;Gorai.008G233600.5.D
Gene IDs: Gorai.008G233600
cotton name: PTHR10909:SF250 - PEROXISOMAL ACYL-COENZYME A OXIDASE 2 (1 of 4)
OG IDs: 93520420
Best-hit-arabi-ID: AT4G16760.1
Best-hit-arabi-name: acyl-CoA oxidase 1
Gorai.008G233600.1.D
PCC=-0.144, p=0.555
mRNA Profile
Protein Expression Profile
DPA

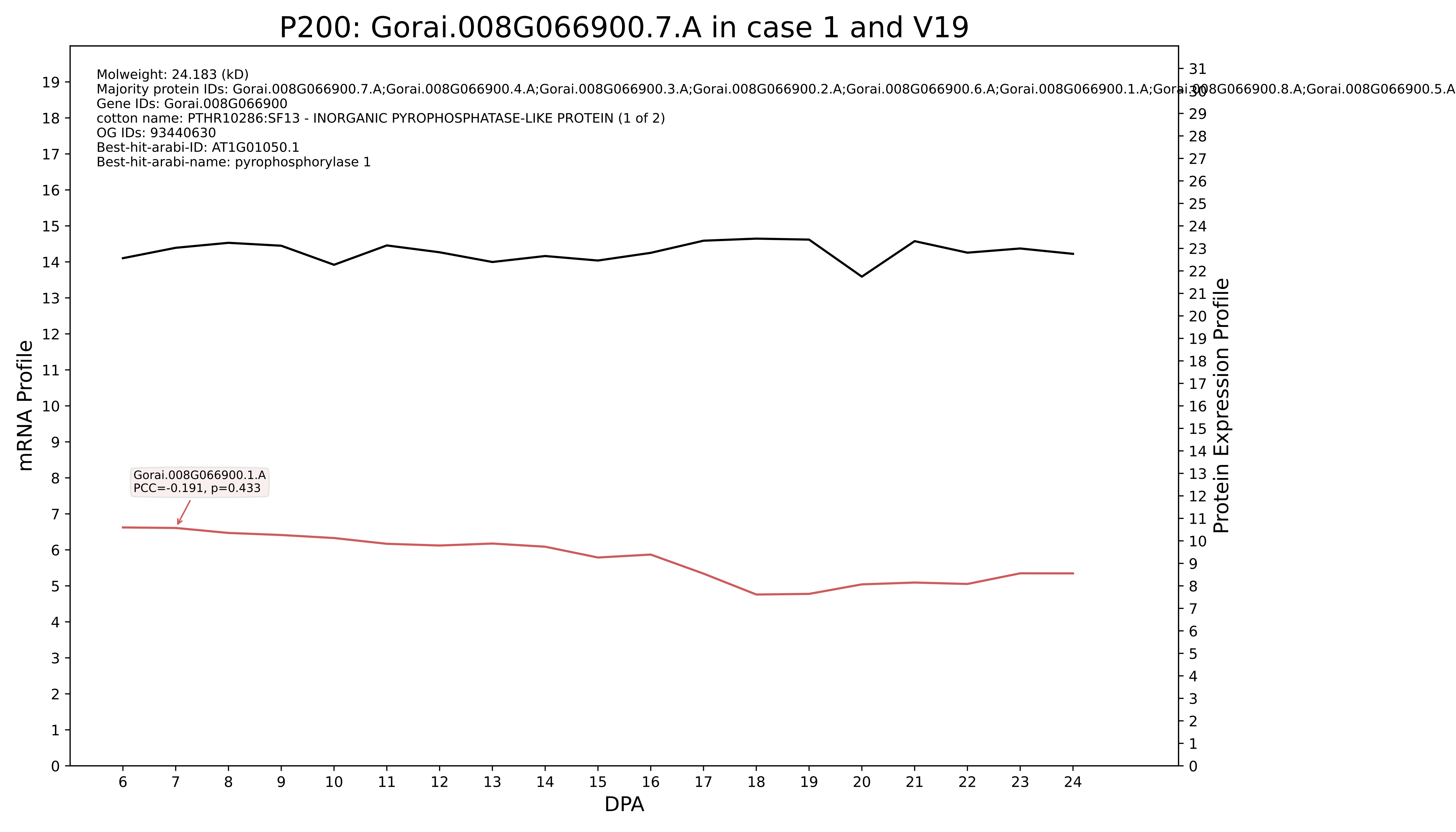

P200: Gorai.008G066900.7.A in case 1 and V19
Molweight: 24.183 (kD)
Majority protein IDs: Gorai.008G066900.7.A;Gorai.008G066900.4.A;Gorai.008G066900.3.A;Gorai.008G066900.2.A;Gorai.008G066900.6.A;Gorai.008G066900.1.A;Gorai.008G066900.8.A;Gorai.008G066900.5.A
Gene IDs: Gorai.008G066900
cotton name: PTHR10286:SF13 - INORGANIC PYROPHOSPHATASE-LIKE PROTEIN (1 of 2)
OG IDs: 93440630
Best-hit-arabi-ID: AT1G01050.1
Best-hit-arabi-name: pyrophosphorylase 1
Gorai.008G066900.1.A
PCC=-0.191, p=0.433
mRNA Profile
Protein Expression Profile
DPA

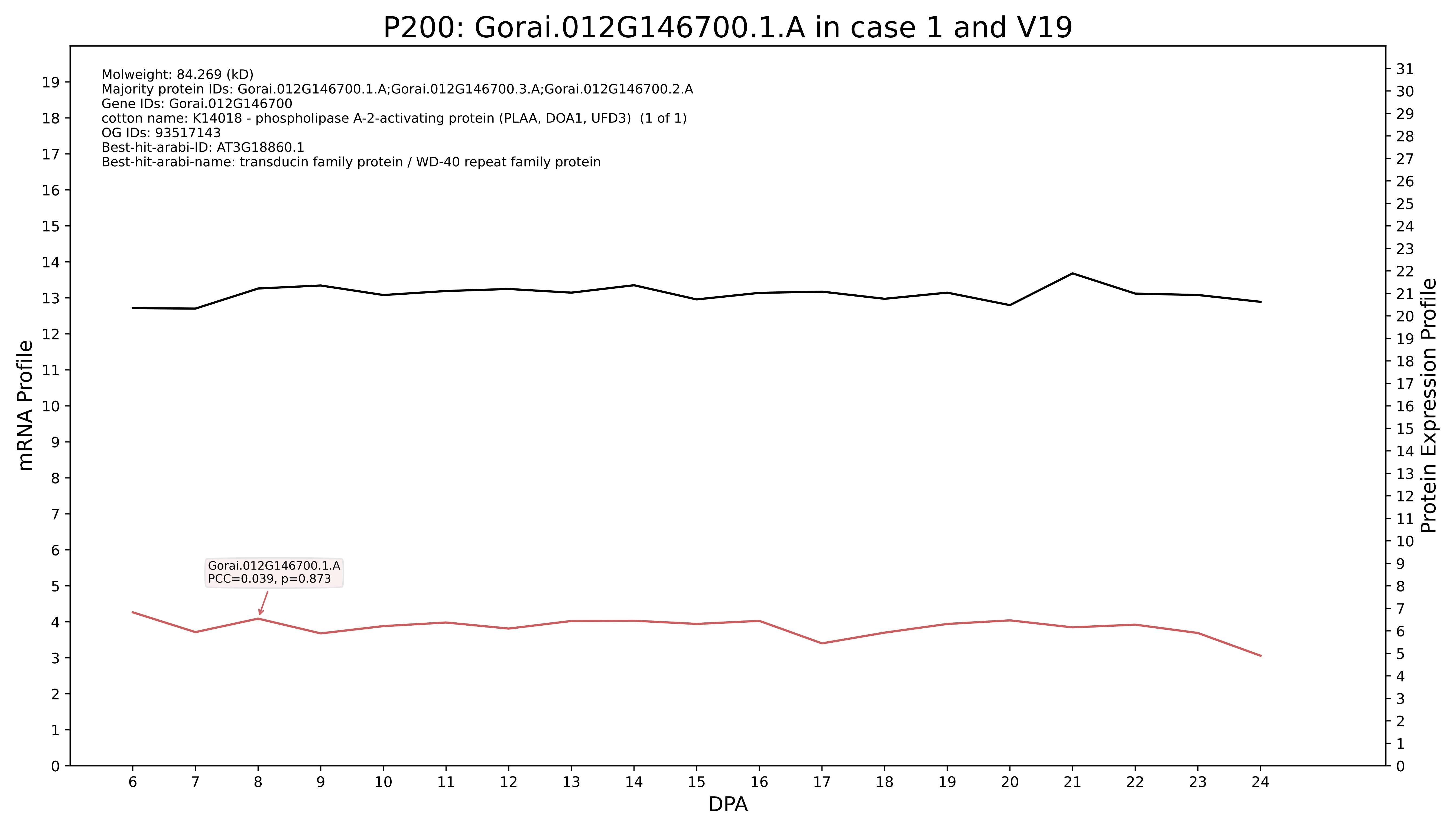

P200: Gorai.012G146700.1.A in case 1 and V19
Molweight: 84.269 (kD)
Majority protein IDs: Gorai.012G146700.1.A;Gorai.012G146700.3.A;Gorai.012G146700.2.A
Gene IDs: Gorai.012G146700
cotton name: K14018 - phospholipase A-2-activating protein (PLAA, DOA1, UFD3) (1 of 1)
OG IDs: 93517143
Best-hit-arabi-ID: AT3G18860.1
Best-hit-arabi-name: transducin family protein / WD-40 repeat family protein
Gorai.012G146700.1.A
PCC=0.039, p=0.873
mRNA Profile
Protein Expression Profile
DPA

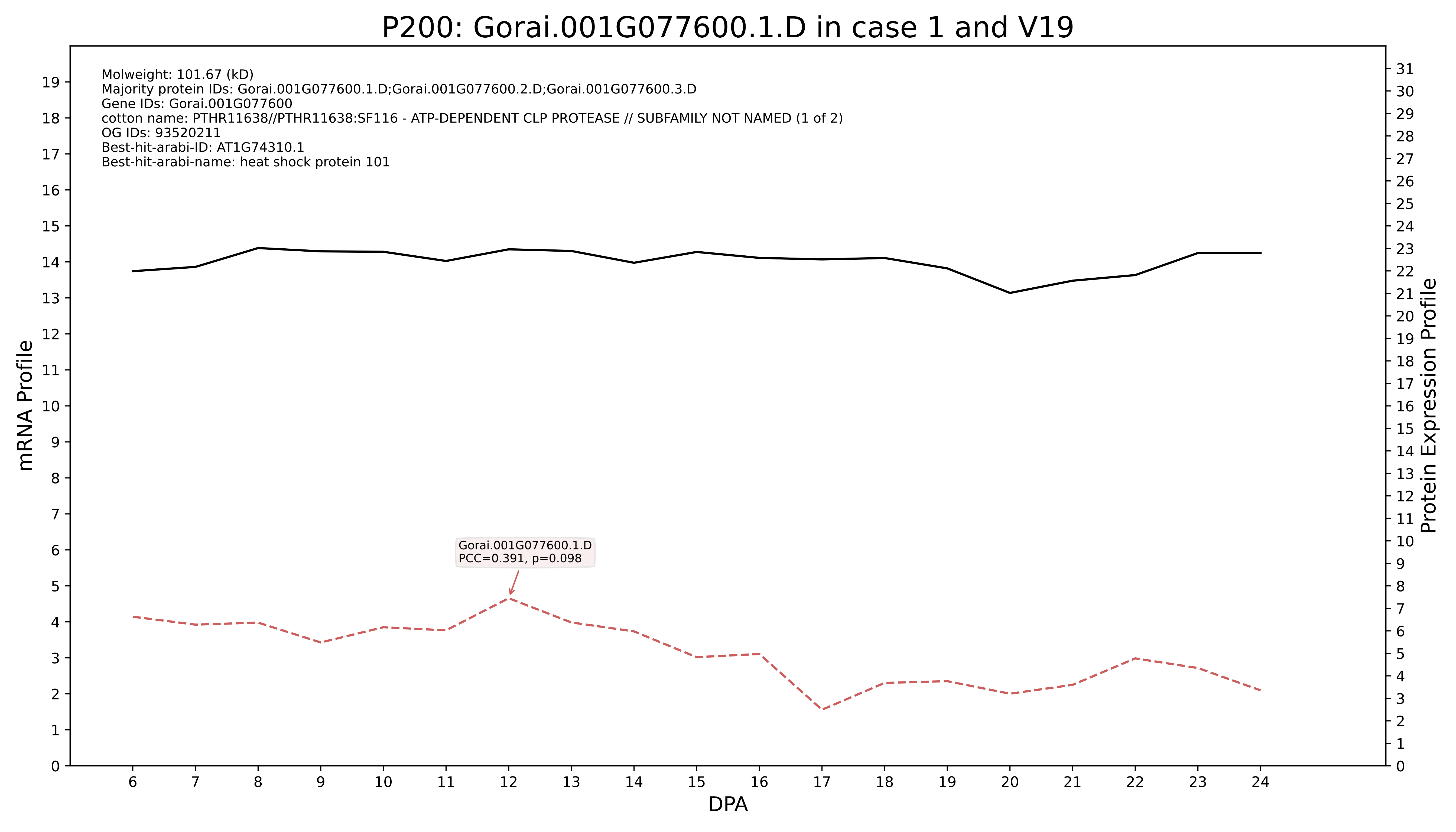
P200: Gorai.001G077600.1.D in case 1 and V19
Molweight: 101.67 (kD)
Majority protein IDs: Gorai.001G077600.1.D;Gorai.001G077600.2.D;Gorai.001G077600.3.D
Gene IDs: Gorai.001G077600
cotton name: PTHR11638//PTHR11638:SF116 - ATP-DEPENDENT CLP PROTEASE // SUBFAMILY NOT NAMED (1 of 2)
OG IDs: 93520211
Best-hit-arabi-ID: AT1G74310.1
Best-hit-arabi-name: heat shock protein 101
Gorai.001G077600.1.D
PCC=0.391, p=0.098
mRNA Profile
Protein Expression Profile
DPA

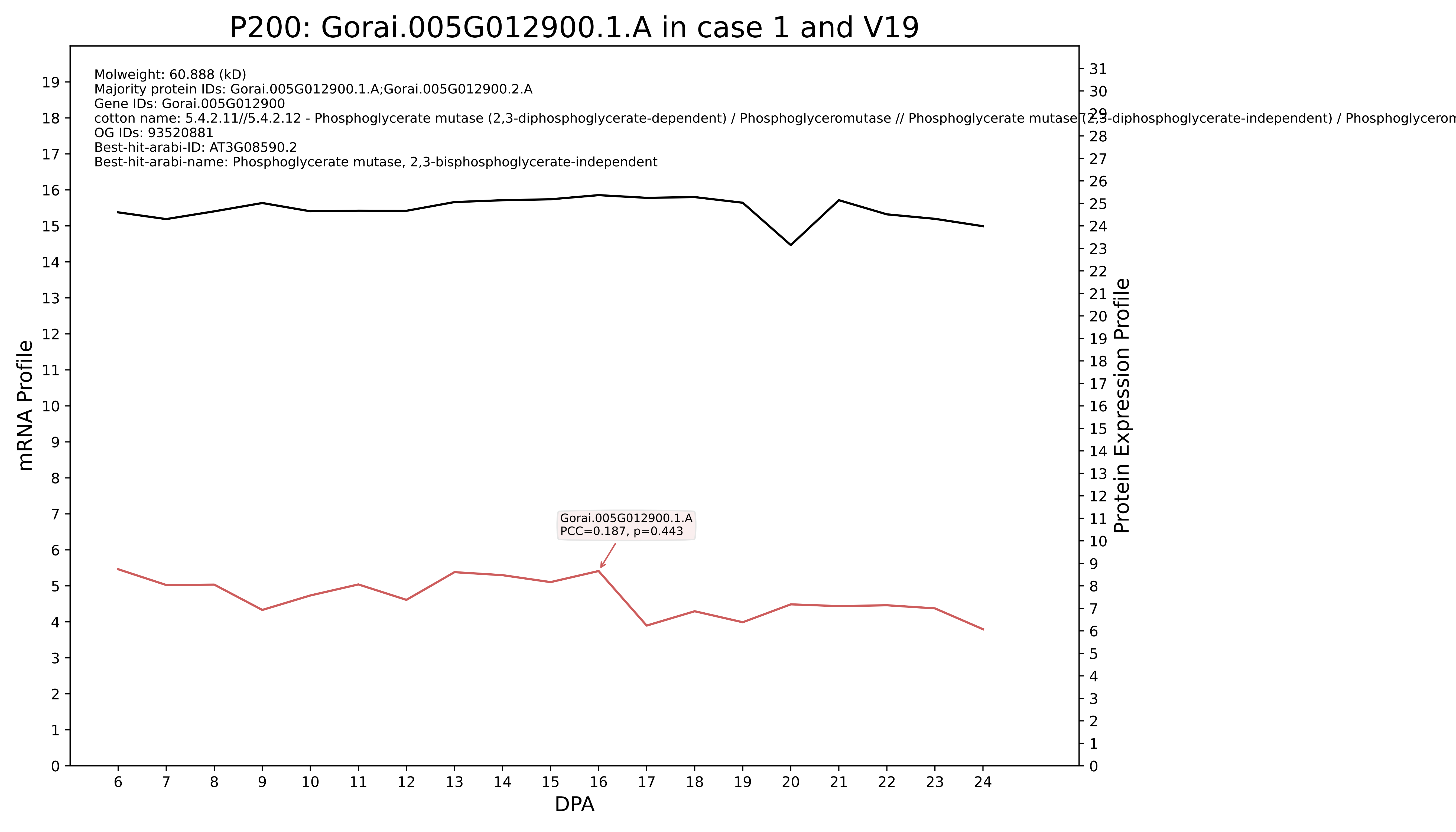
P200: Gorai.005G012900.1.A in case 1 and V19
Molweight: 60.888 (kD)
Majority protein IDs: Gorai.005G012900.1.A;Gorai.005G012900.2.A
Gene IDs: Gorai.005G012900
cotton name: 5.4.2.11//5.4.2.12 - Phosphoglycerate mutase (2,3-diphosphoglycerate-dependent) / Phosphoglyceromutase // Phosphoglycerate mutase (2,3-diphosphoglycerate-independent) / Phosphoglyceron
OG IDs: 93520881
Best-hit-arabi-ID: AT3G08590.2
Best-hit-arabi-name: Phosphoglycerate mutase, 2,3-bisphosphoglycerate-independent
Gorai.005G012900.1.A
PCC=0.187, p=0.443
mRNA Profile
Protein Expression Profile
DPA

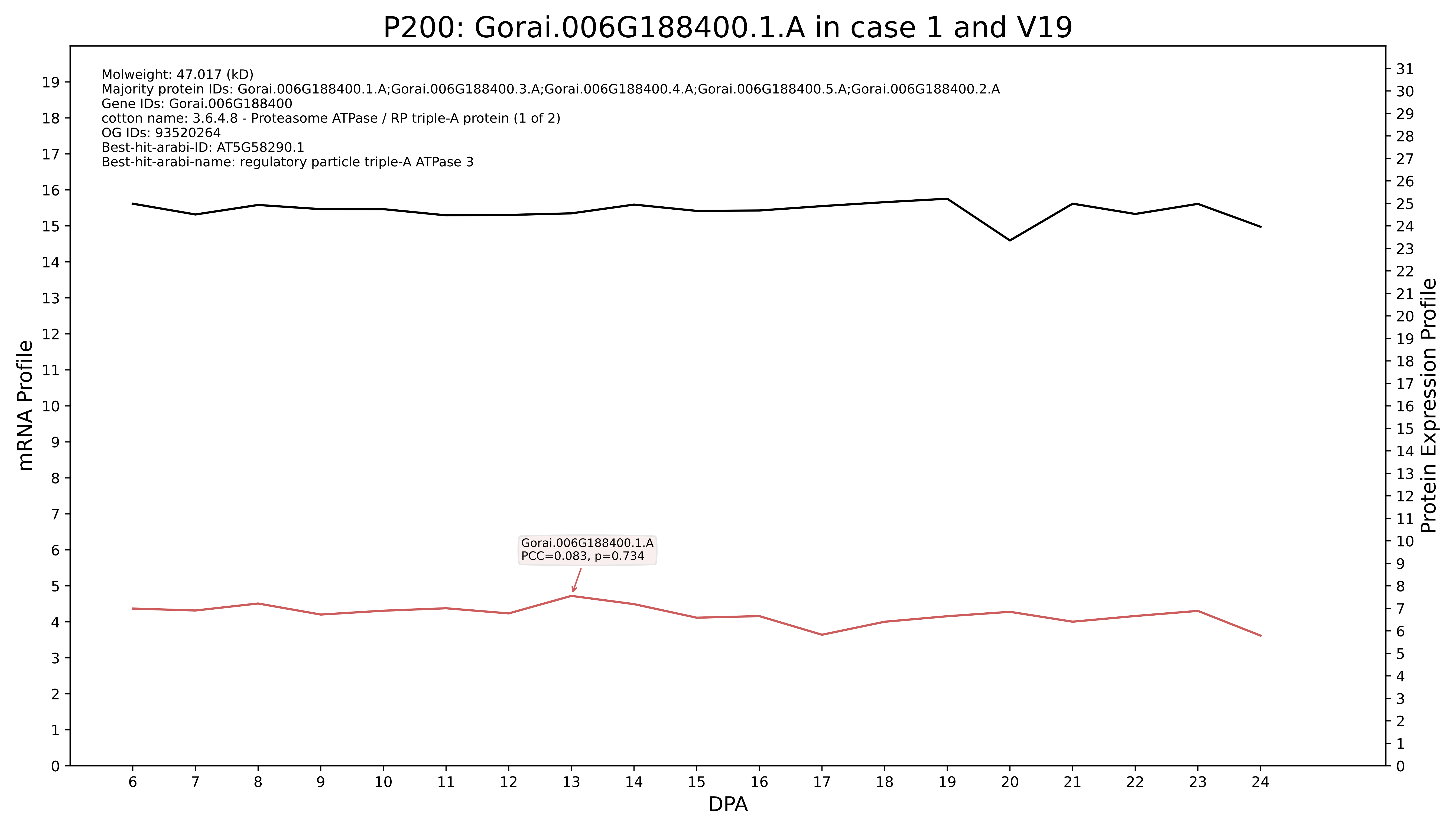
P200: Gorai.006G188400.1.A in case 1 and V19
Molweight: 47.017 (kD)
Majority protein IDs: Gorai.006G188400.1.A;Gorai.006G188400.3.A;Gorai.006G188400.4.A;Gorai.006G188400.5.A;Gorai.006G188400.2.A
Gene IDs: Gorai.006G188400
cotton name: 3.6.4.8 - Proteasome ATPase / RP triple-A protein (1 of 2)
OG IDs: 93520264
Best-hit-arabi-ID: AT5G58290.1
Best-hit-arabi-name: regulatory particle triple-A ATPase 3
Gorai.006G188400.1.A
PCC=0.083, p=0.734
mRNA Profile
Protein Expression Profile
DPA

#### P200: Gorai.007G057400.2.D in case 1 and V19

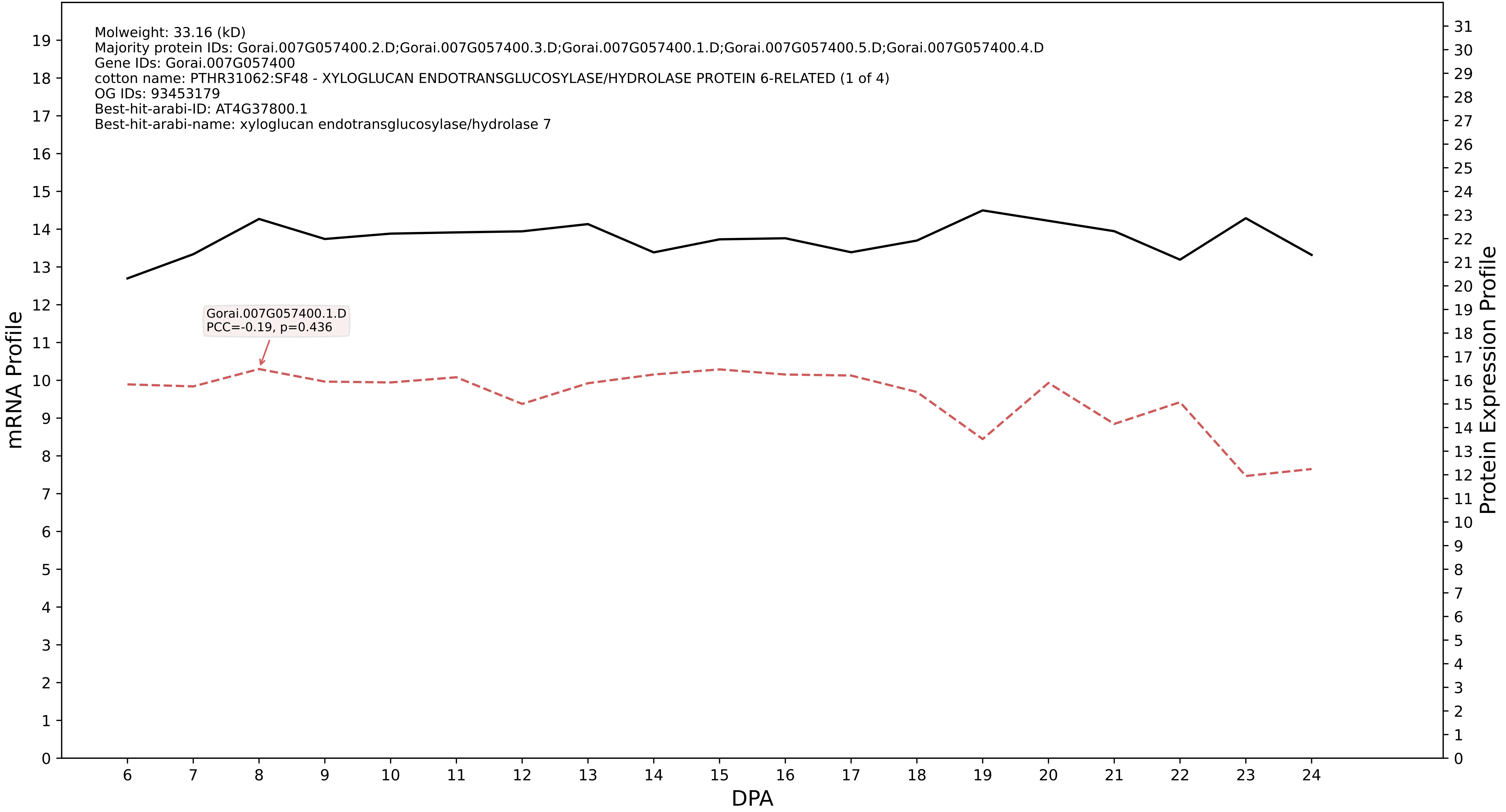

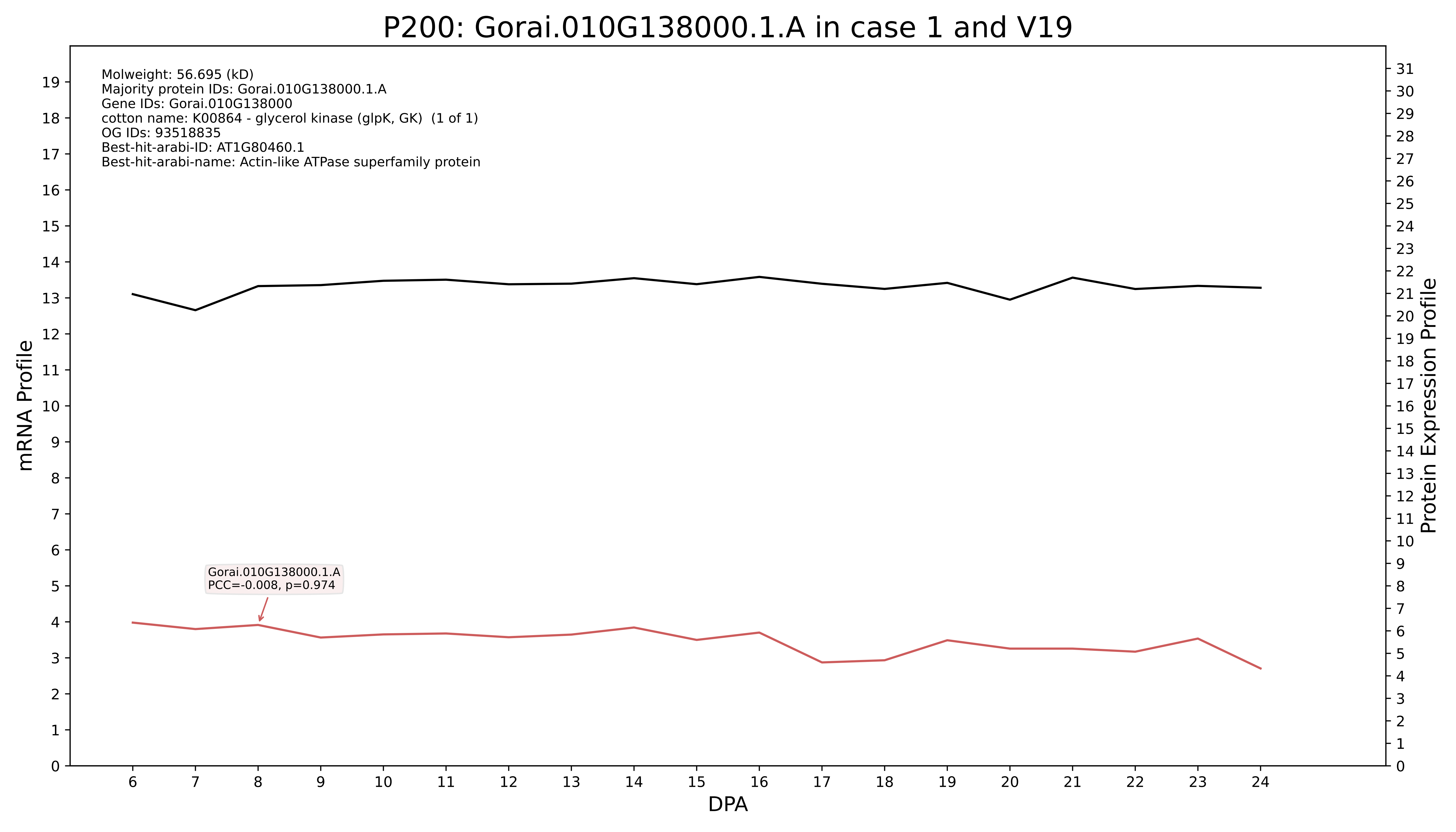
P200: Gorai.010G138000.1.A in case 1 and V19
Molweight: 56.695 (kD)
Majority protein IDs: Gorai.010G138000.1.A
Gene IDs: Gorai.010G138000
cotton name: K00864 - glycerol kinase (glpK, GK) (1 of 1)
OG IDs: 93518835
Best-hit-arabi-ID: AT1G80460.1
Best-hit-arabi-name: Actin-like ATPase superfamily protein
Gorai.010G138000.1.A
PCC=-0.008, p=0.974
mRNA Profile
Protein Expression Profile
DPA

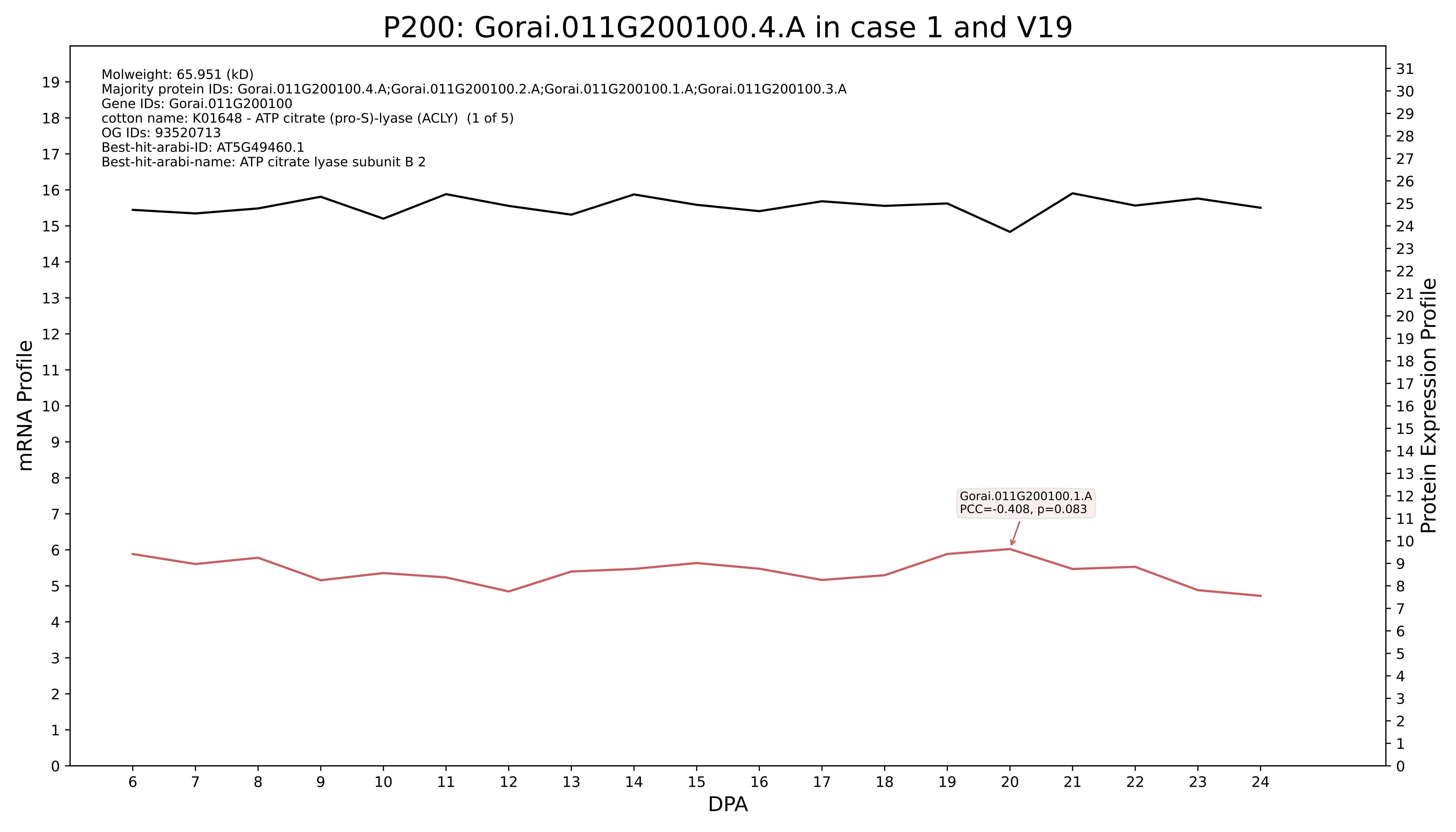

P200: Gorai.011G200100.4.A in case 1 and V19
Molweight: 65.951 (kD)
Majority protein IDs: Gorai.011G200100.4.A;Gorai.011G200100.2.A;Gorai.011G200100.1.A;Gorai.011G200100.3.A
Gene IDs: Gorai.011G200100
cotton name: K01648 - ATP citrate (pro-S)-lyase (ACLY) (1 of 5)
OG IDs: 93520713
Best-hit-arabi-ID: AT5G49460.1
Best-hit-arabi-name: ATP citrate lyase subunit B 2
Gorai.011G200100.1.A
PCC=-0.408, p=0.083
mRNA Profile
Protein Expression Profile
DPA

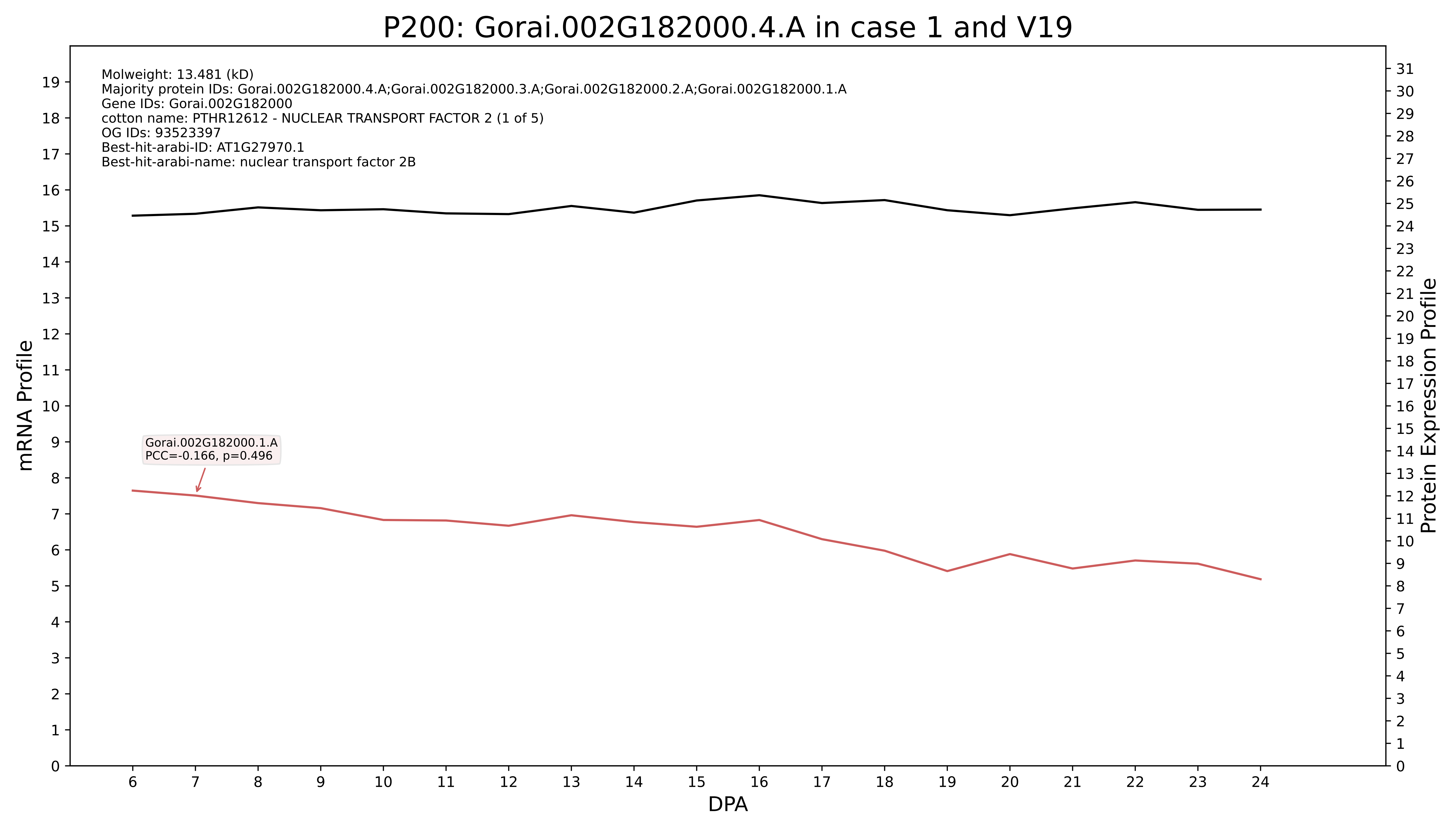

P200: Gorai.002G182000.4.A in case 1 and V19
Molweight: 13.481 (kD)
Majority protein IDs: Gorai.002G182000.4.A;Gorai.002G182000.3.A;Gorai.002G182000.2.A;Gorai.002G182000.1.A
Gene IDs: Gorai.002G182000
cotton name: PTHR12612 - NUCLEAR TRANSPORT FACTOR 2 (1 of 5)
OG IDs: 93523397
Best-hit-arabi-ID: AT1G27970.1
Best-hit-arabi-name: nuclear transport factor 2B
Gorai.002G182000.1.A
PCC=-0.166, p=0.496
mRNA Profile
Protein Expression Profile
DPA

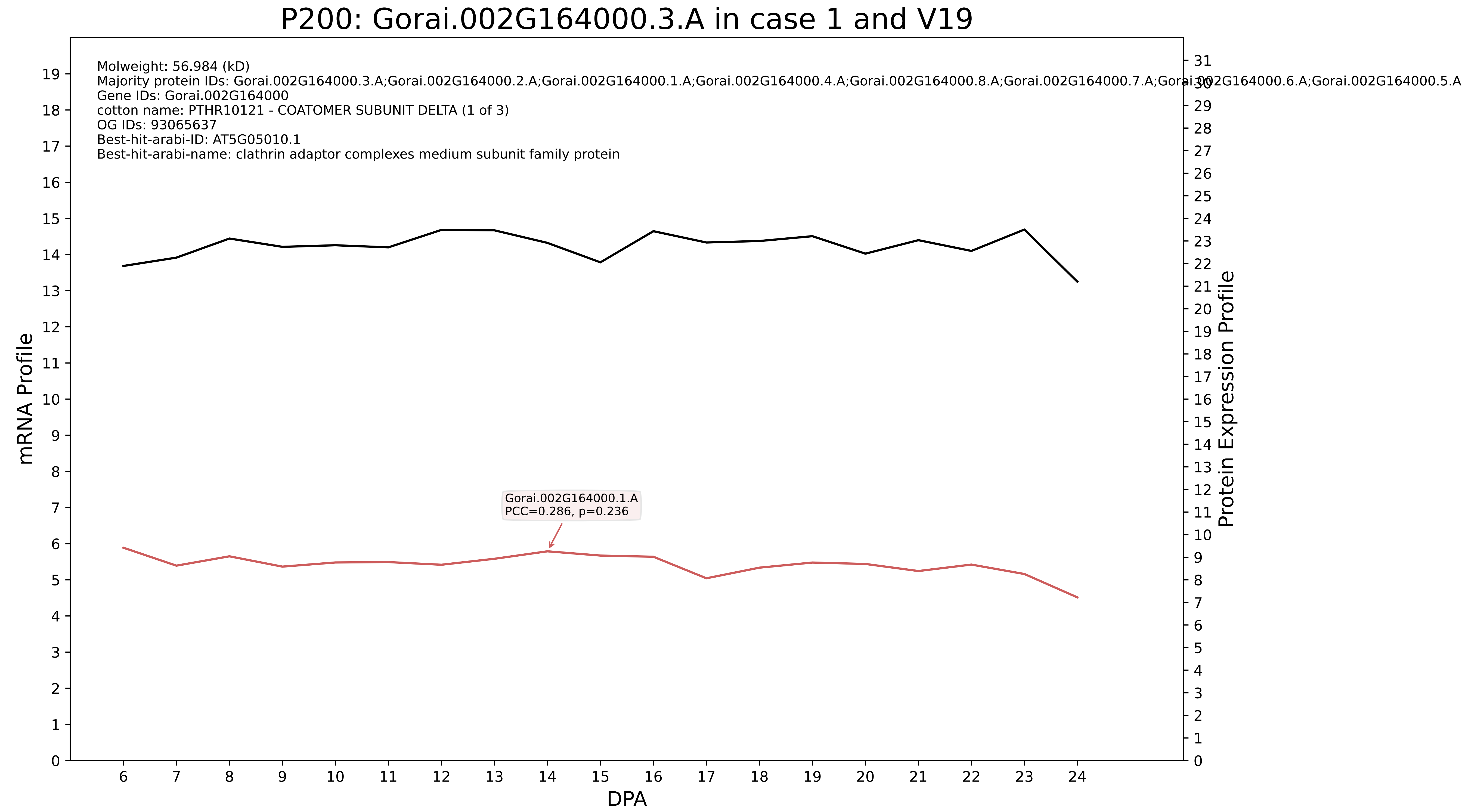
P200: Gorai.002G164000.3.A in case 1 and V19
Molweight: 56.984 (kD)
Majority protein IDs: Gorai.002G164000.3.A;Gorai.002G164000.2.A;Gorai.002G164000.1.A;Gorai.002G164000.4.A;Gorai.002G164000.8.A;Gorai.002G164000.7.A;Gorai.002G164000.6.A;Gorai.002G164000.5.A
Gene IDs: Gorai.002G164000
cotton name: PTHR10121 - COATOMER SUBUNIT DELTA (1 of 3)
OG IDs: 93065637
Best-hit-arabi-ID: AT5G05010.1
Best-hit-arabi-name: clathrin adaptor complexes medium subunit family protein
Gorai.002G164000.1.A
PCC=0.286, p=0.236
mRNA Profile
Protein Expression Profile
DPA

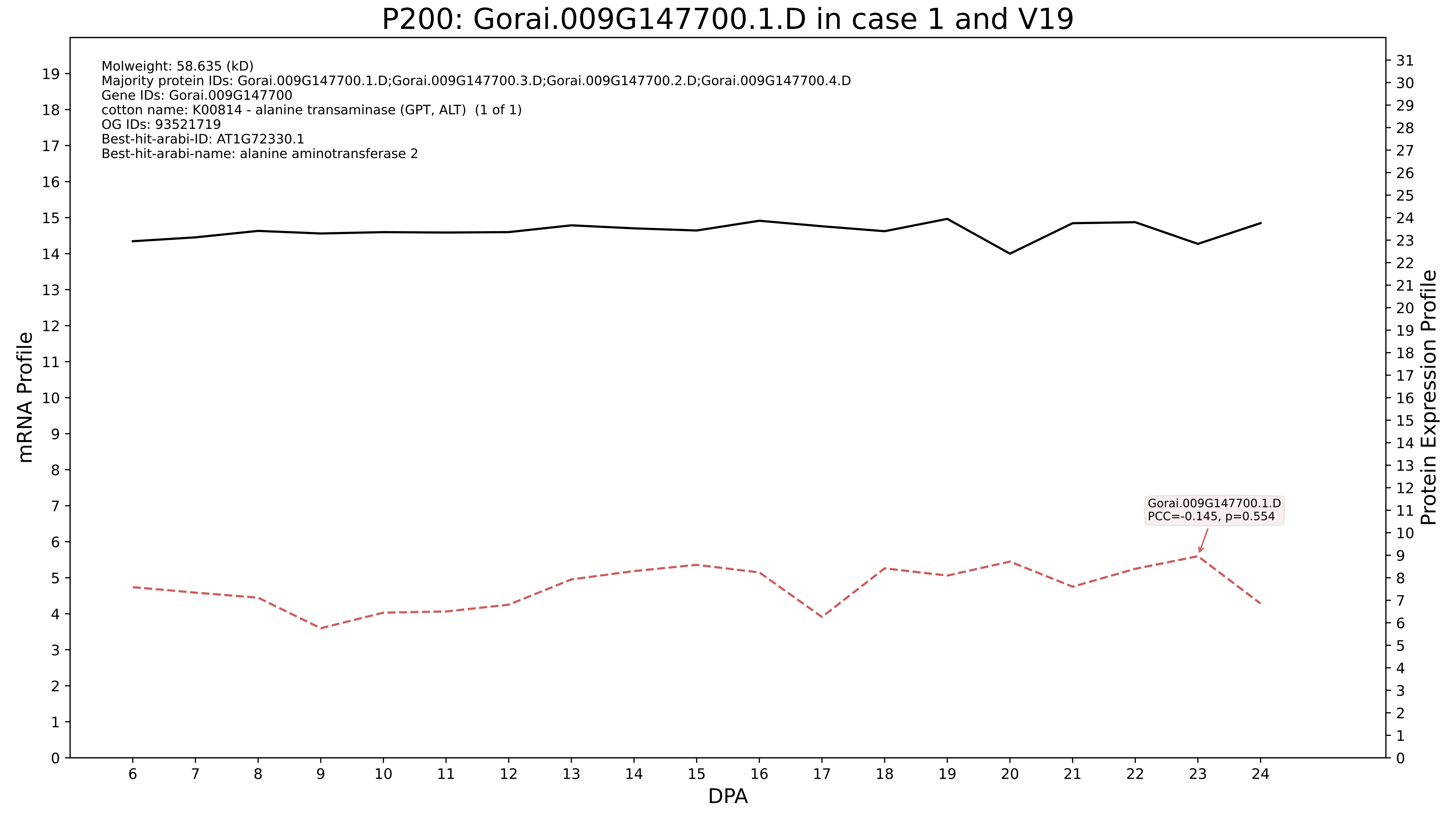

P200: Gorai.009G147700.1.D in case 1 and V19
Molweight: 58.635 (kD)
Majority protein IDs: Gorai.009G147700.1.D;Gorai.009G147700.3.D;Gorai.009G147700.2.D;Gorai.009G147700.4.D
Gene IDs: Gorai.009G147700
cotton name: K00814 - alanine transaminase (GPT, ALT) (1 of 1)
OG IDs: 93521719
Best-hit-arabi-ID: AT1G72330.1
Best-hit-arabi-name: alanine aminotransferase 2
Gorai.009G147700.1.D
PCC=-0.145, p=0.554
mRNA Profile
Protein Expression Profile
DPA

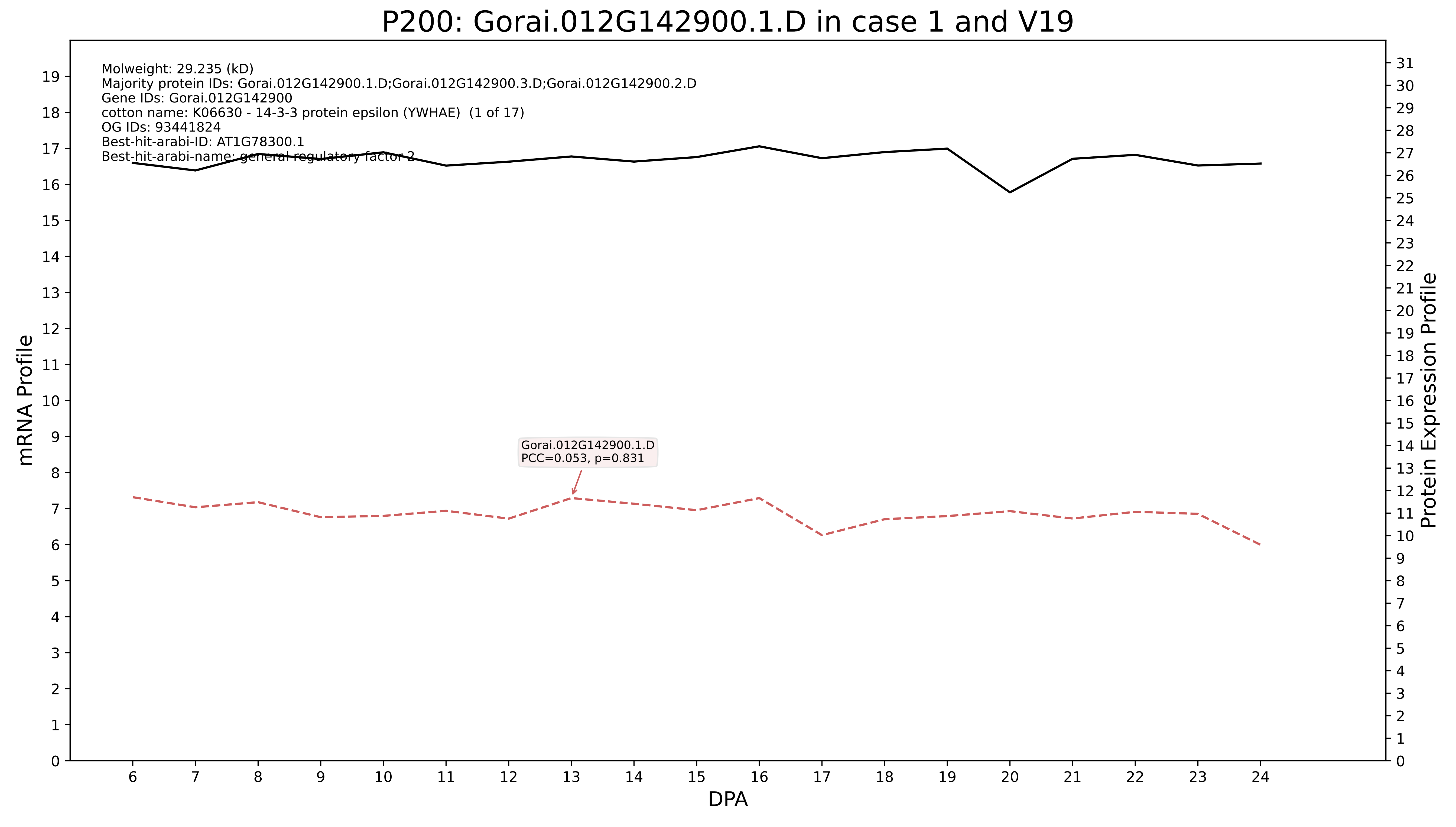
P200: Gorai.012G142900.1.D in case 1 and V19
Molweight: 29.235 (kD)
Majority protein IDs: Gorai.012G142900.1.D;Gorai.012G142900.3.D;Gorai.012G142900.2.D
Gene IDs: Gorai.012G142900
cotton name: K06630 - 14-3-3 protein epsilon (YWHAE) (1 of 17)
OG IDs: 93441824
Best-hit-arabi-ID: AT1G78300.1
Best-hit-arabi-name: general regulatory factor 2
Gorai.012G142900.1.D
PCC=0.053, p=0.831
mRNA Profile
Protein Expression Profile
DPA
6 7 8 9 10 11 12 13 14 15 16 17 18 19 20 21 22 23 24
0 1 2 3 4 5 6 7 8 9 10 11 12 13 14 15 16 17 18 19
0 1 2 3 4 5 6 7 8 9 10 11 12 13 14 15 16 17 18 19 20 21 22 23 24 25 26 27 28 29 30 31

### P200: Gorai.002G141000.5.D in case 1 and V19

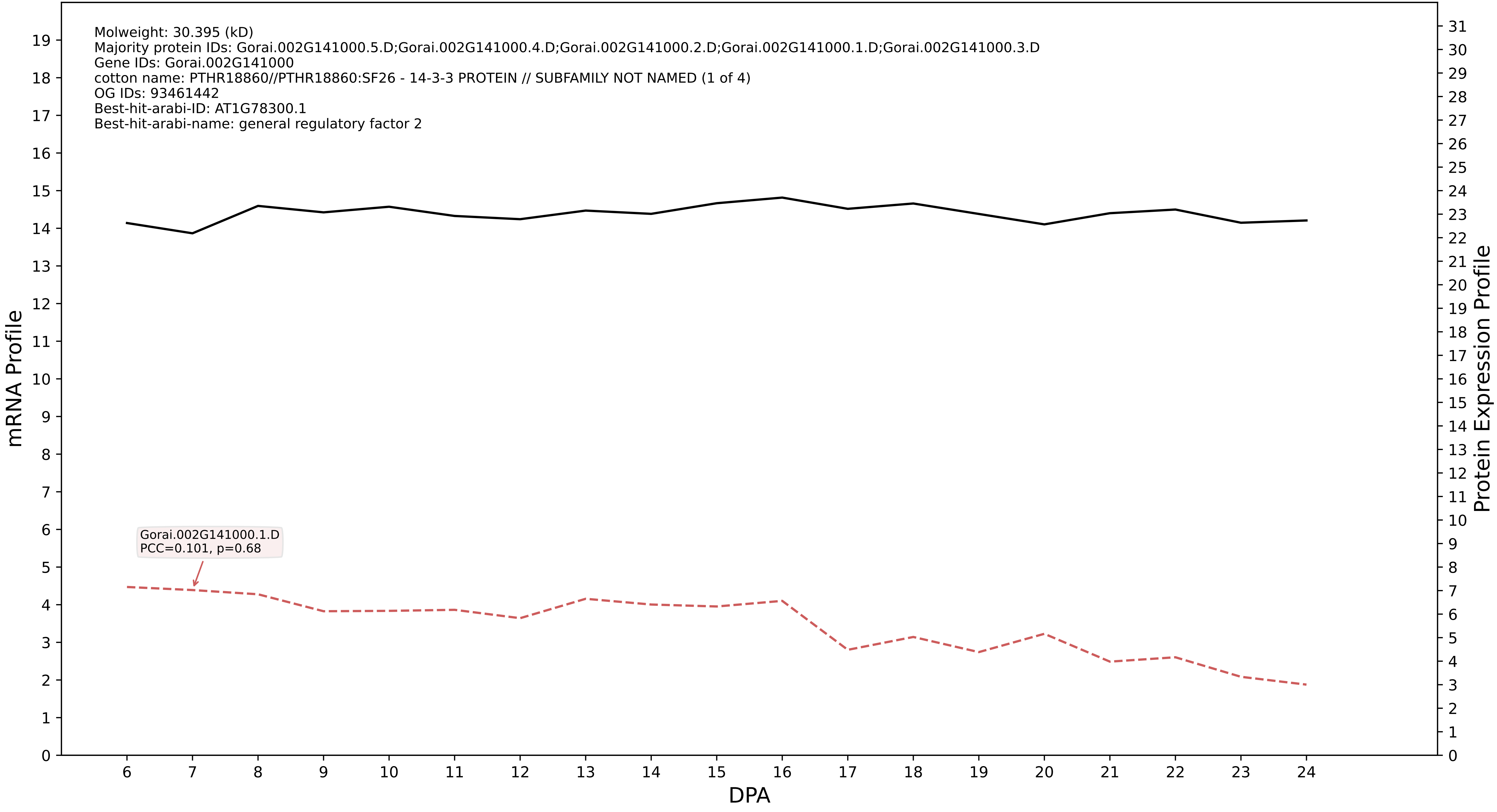

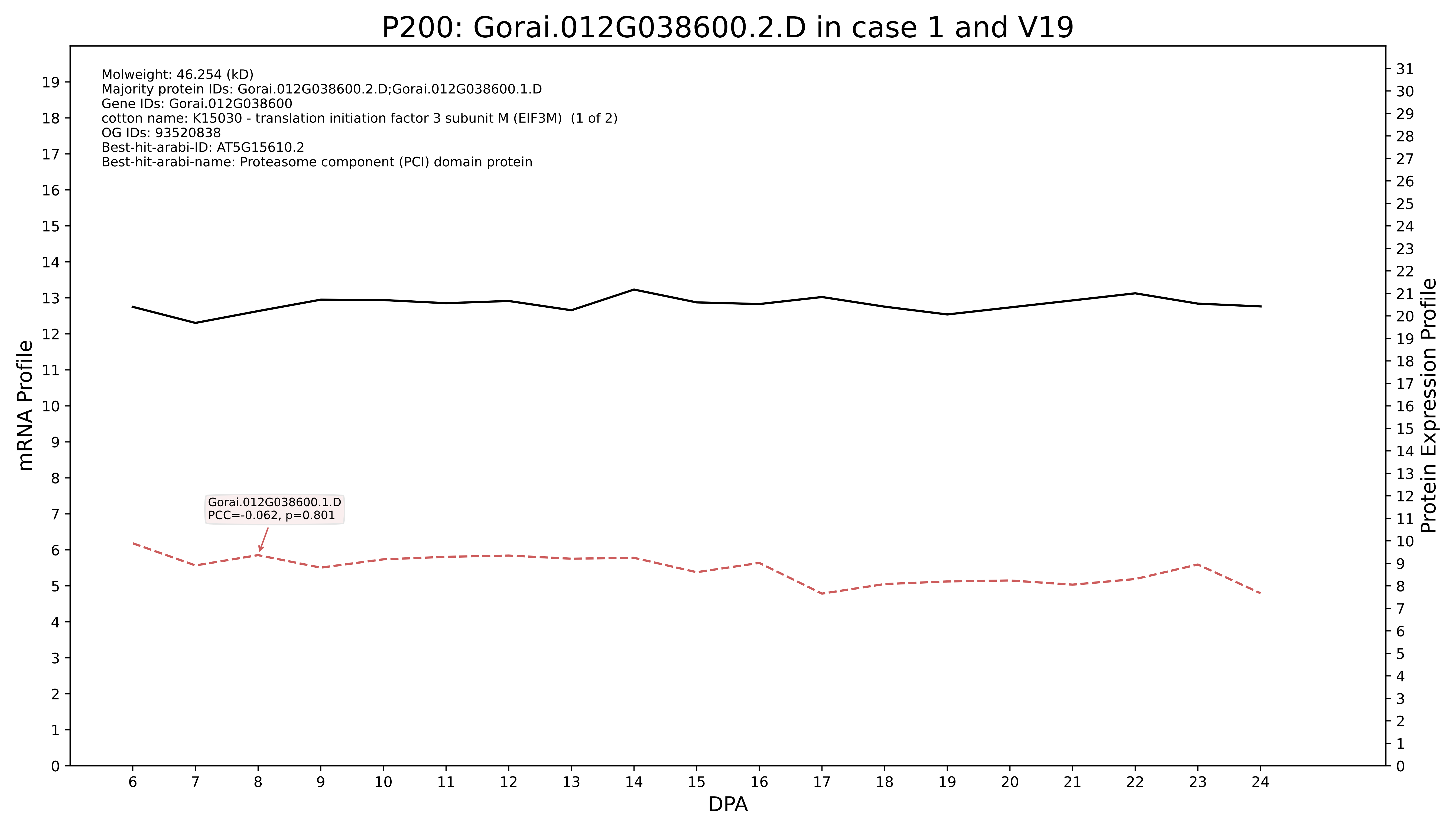
P200: Gorai.012G038600.2.D in case 1 and V19
Molweight: 46.254 (kD)
Majority protein IDs: Gorai.012G038600.2.D;Gorai.012G038600.1.D
Gene IDs: Gorai.012G038600
cotton name: K15030 - translation initiation factor 3 subunit M (EIF3M) (1 of 2)
OG IDs: 93520838
Best-hit-arabi-ID: AT5G15610.2
Best-hit-arabi-name: Proteasome component (PCI) domain protein
Gorai.012G038600.1.D
PCC=-0.062, p=0.801
mRNA Profile
Protein Expression Profile
DPA

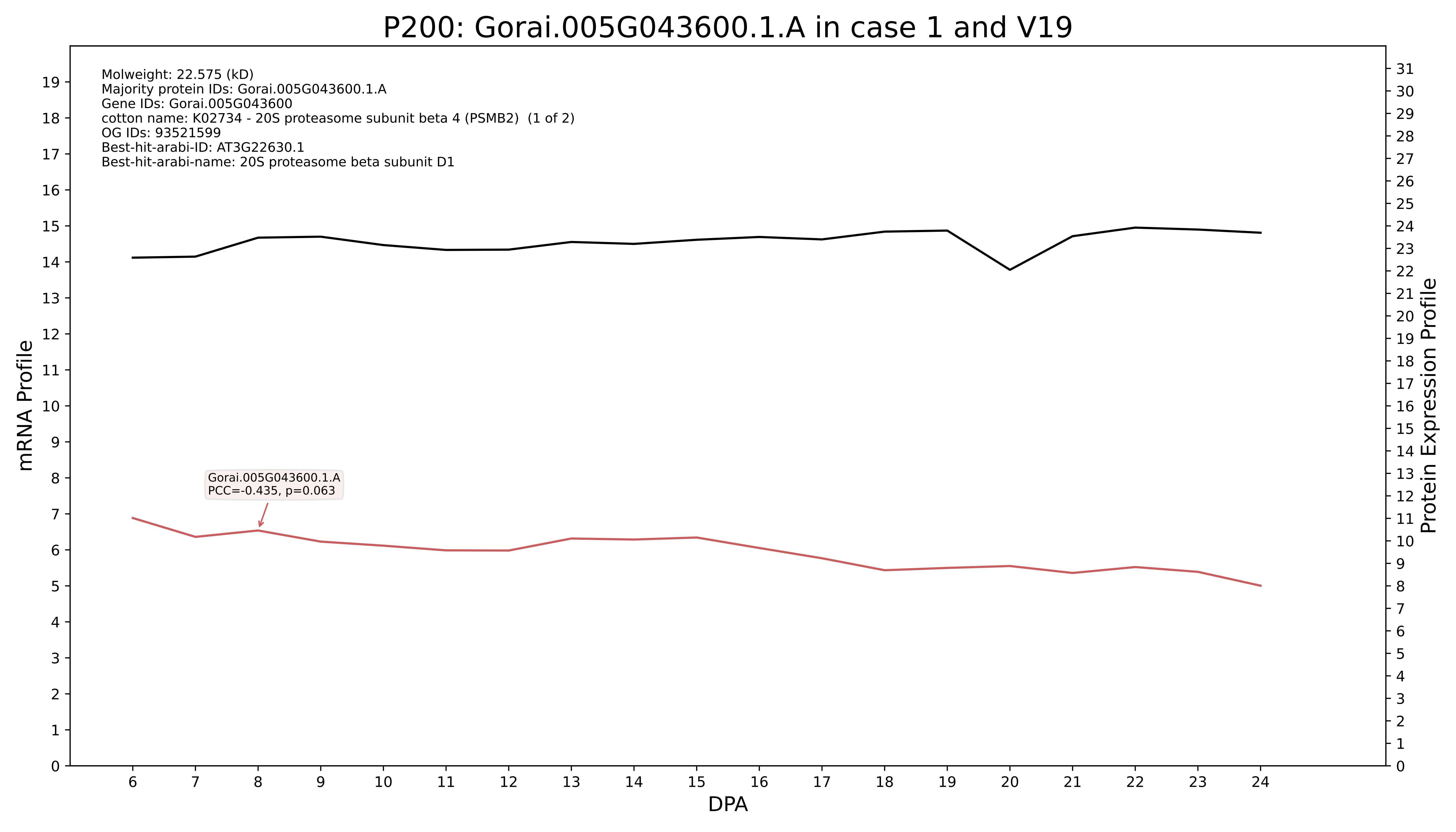
P200: Gorai.005G043600.1.A in case 1 and V19
Molweight: 22.575 (kD)
Majority protein IDs: Gorai.005G043600.1.A
Gene IDs: Gorai.005G043600
cotton name: K02734 - 20S proteasome subunit beta 4 (PSMB2) (1 of 2)
OG IDs: 93521599
Best-hit-arabi-ID: AT3G22630.1
Best-hit-arabi-name: 20S proteasome beta subunit D1
Gorai.005G043600.1.A
PCC=-0.435, p=0.063
mRNA Profile
Protein Expression Profile
DPA

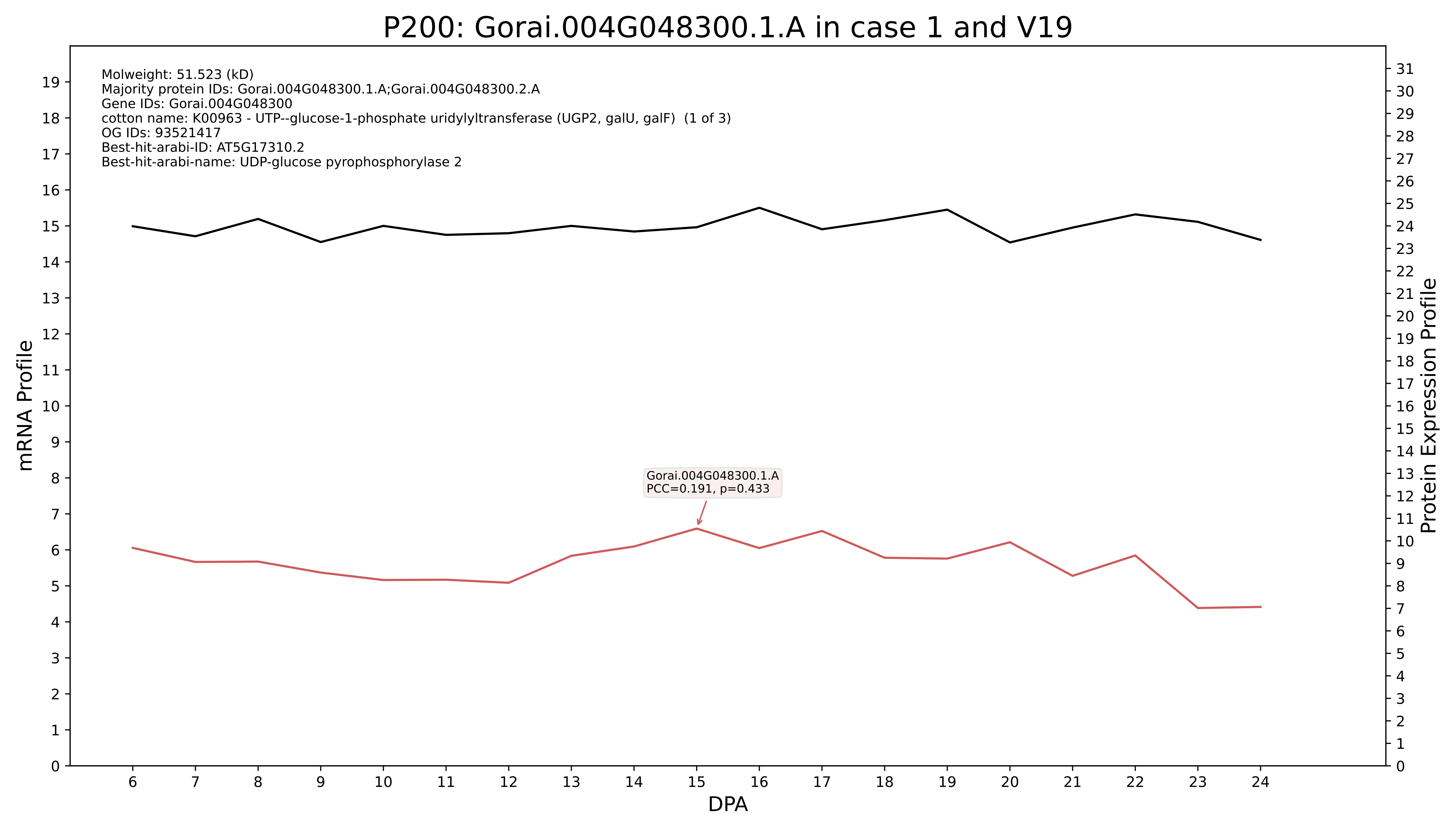

P200: Gorai.004G048300.1.A in case 1 and V19
Molweight: 51.523 (kD)
Majority protein IDs: Gorai.004G048300.1.A;Gorai.004G048300.2.A
Gene IDs: Gorai.004G048300
cotton name: K00963 - UTP--glucose-1-phosphate uridylyltransferase (UGP2, galU, galF) (1 of 3)
OG IDs: 93521417
Best-hit-arabi-ID: AT5G17310.2
Best-hit-arabi-name: UDP-glucose pyrophosphorylase 2
Gorai.004G048300.1.A
PCC=0.191, p=0.433
mRNA Profile
Protein Expression Profile
DPA

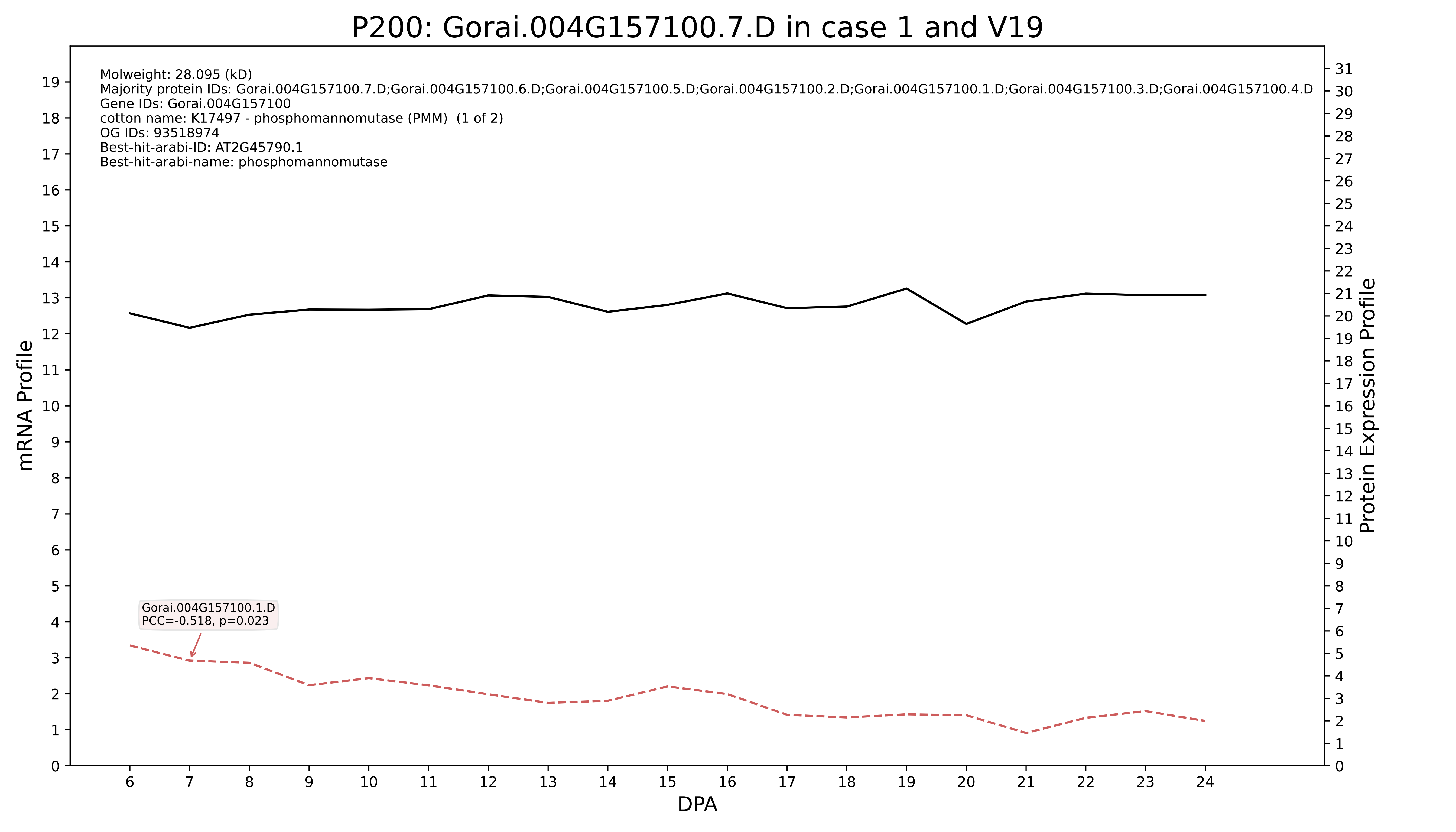
P200: Gorai.004G157100.7.D in case 1 and V19
Molweight: 28.095 (kD)
Majority protein IDs: Gorai.004G157100.7.D;Gorai.004G157100.6.D;Gorai.004G157100.5.D;Gorai.004G157100.2.D;Gorai.004G157100.1.D;Gorai.004G157100.3.D;Gorai.004G157100.4.D
Gene IDs: Gorai.004G157100
cotton name: K17497 - phosphomannomutase (PMM) (1 of 2)
OG IDs: 93518974
Best-hit-arabi-ID: AT2G45790.1
Best-hit-arabi-name: phosphomannomutase
Gorai.004G157100.1.D
PCC=-0.518, p=0.023
mRNA Profile
Protein Expression Profile
DPA

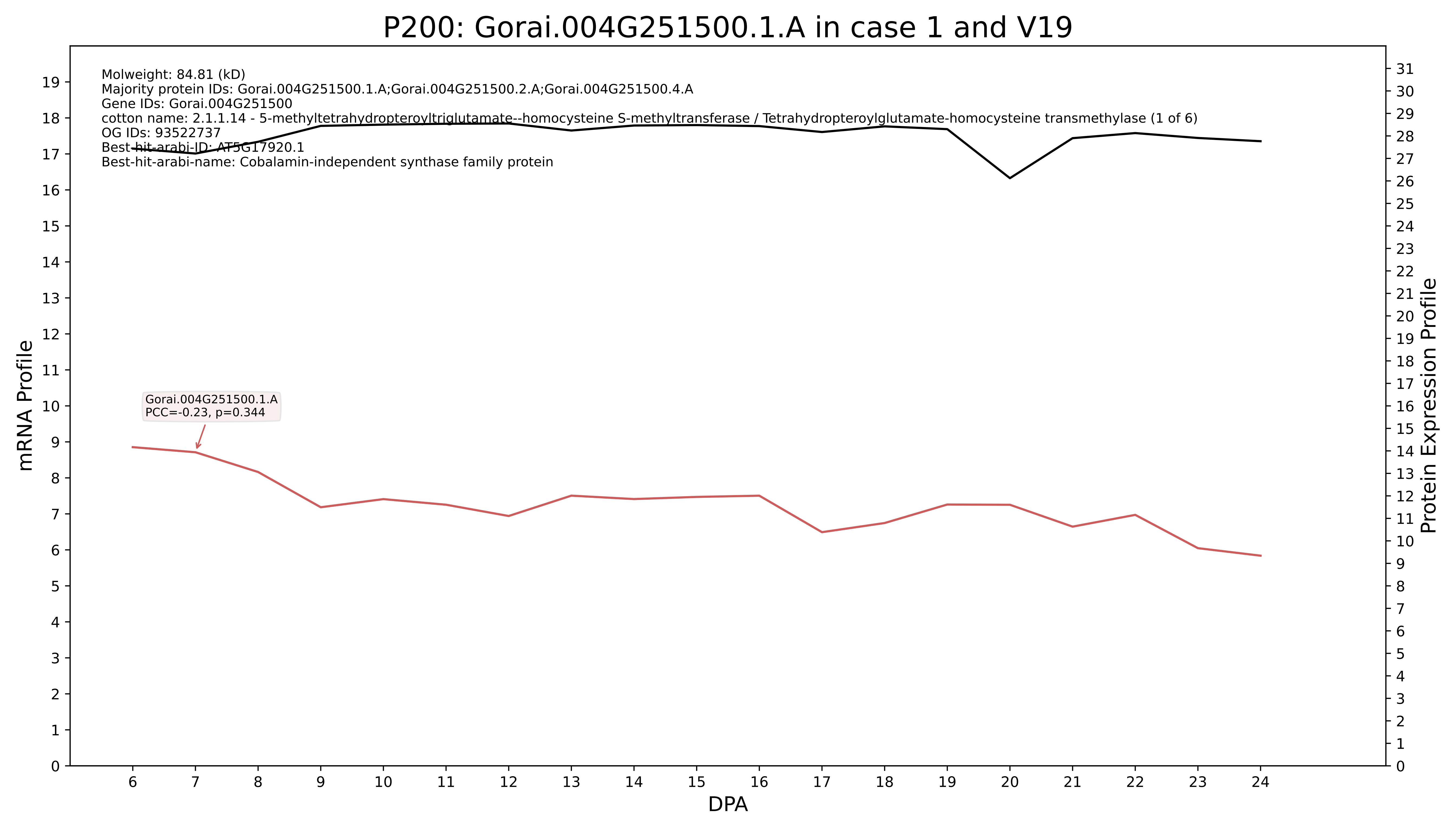

P200: Gorai.004G251500.1.A in case 1 and V19
Molweight: 84.81 (kD)
Majority protein IDs: Gorai.004G251500.1.A;Gorai.004G251500.2.A;Gorai.004G251500.4.A
Gene IDs: Gorai.004G251500
cotton name: 2.1.1.14 - 5-methyltetrahydropteroyltriglutamate--homocysteine S-methyltransferase / Tetrahydropteroylglutamate-homocysteine transmethylase (1 of 6)
OG IDs: 93522737
Best-hit-arabi-ID: AT5G17920.1
Best-hit-arabi-name: Cobalamin-independent synthase family protein
Gorai.004G251500.1.A
PCC=-0.23, p=0.344
mRNA Profile
Protein Expression Profile
DPA

P200: Gorai.002G024600.1.D in case 1 and V19

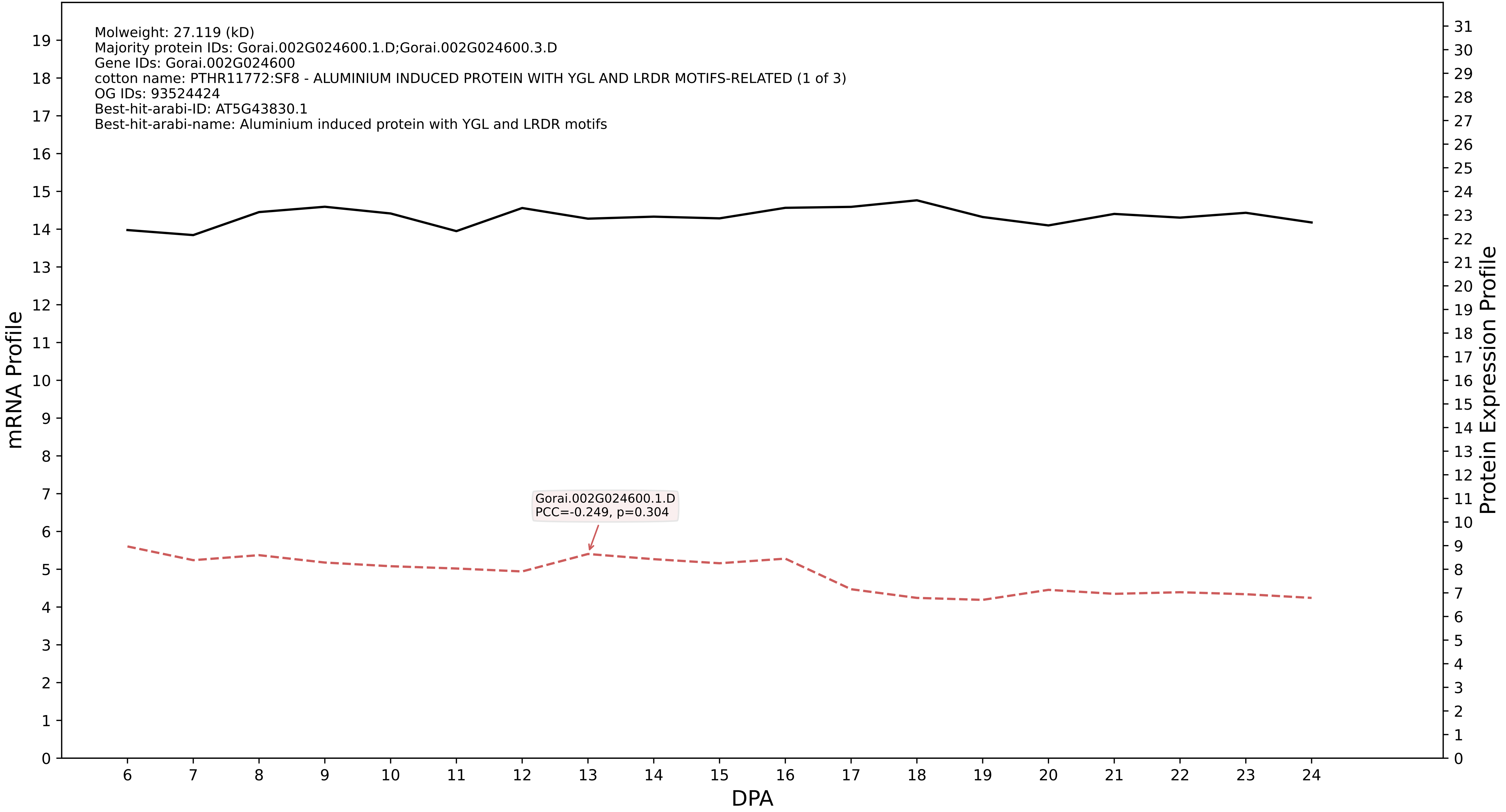

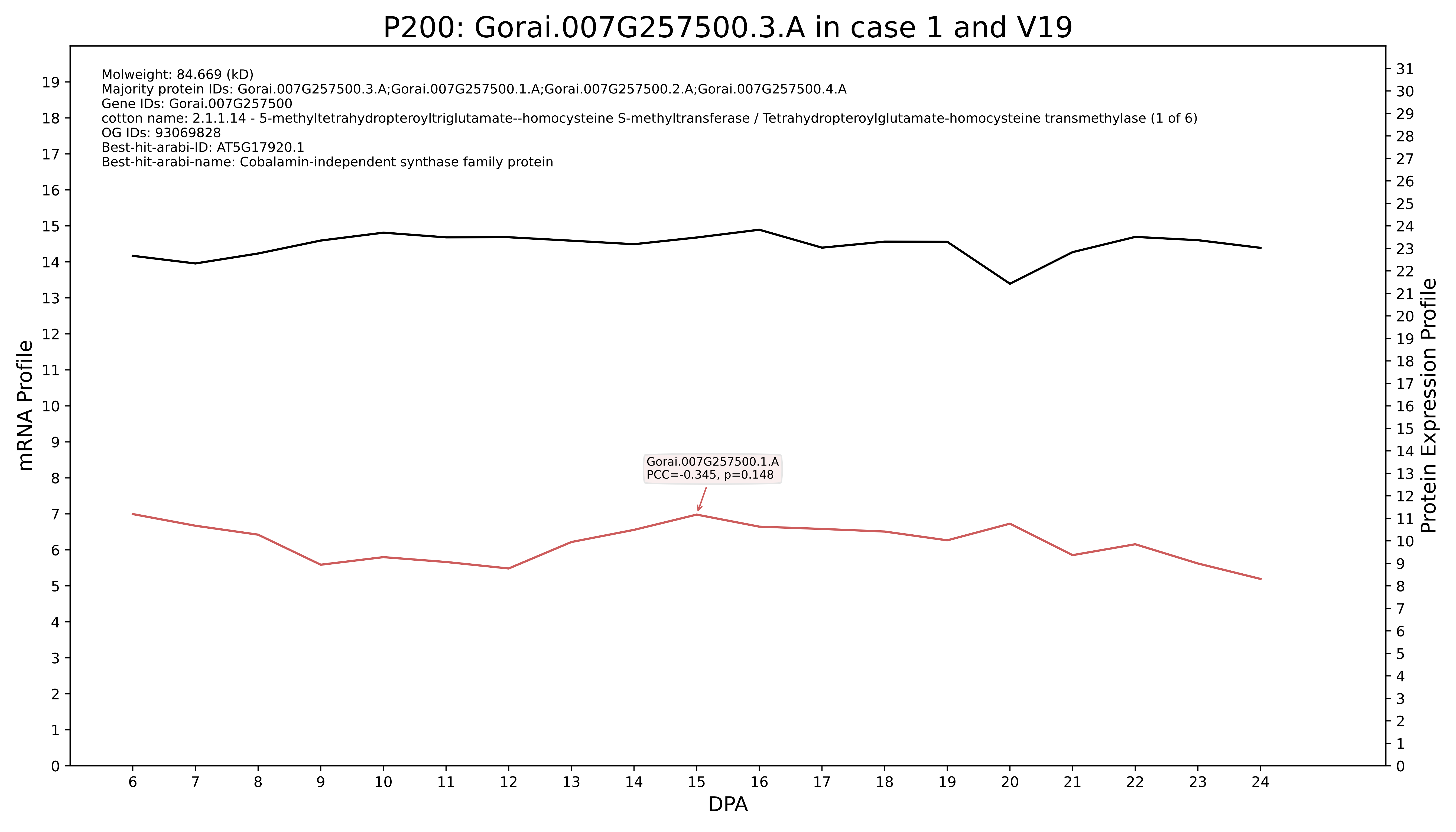
P200: Gorai.007G257500.3.A in case 1 and V19
Molweight: 84.669 (kD)
Majority protein IDs: Gorai.007G257500.3.A;Gorai.007G257500.1.A;Gorai.007G257500.2.A;Gorai.007G257500.4.A
Gene IDs: Gorai.007G257500
cotton name: 2.1.1.14 - 5-methyltetrahydropteroyltriglutamate--homocysteine S-methyltransferase / Tetrahydropteroylglutamate-homocysteine transmethylase (1 of 6)
OG IDs: 93069828
Best-hit-arabi-ID: AT5G17920.1
Best-hit-arabi-name: Cobalamin-independent synthase family protein
Gorai.007G257500.1.A
PCC=-0.345, p=0.148
mRNA Profile
Protein Expression Profile
DPA

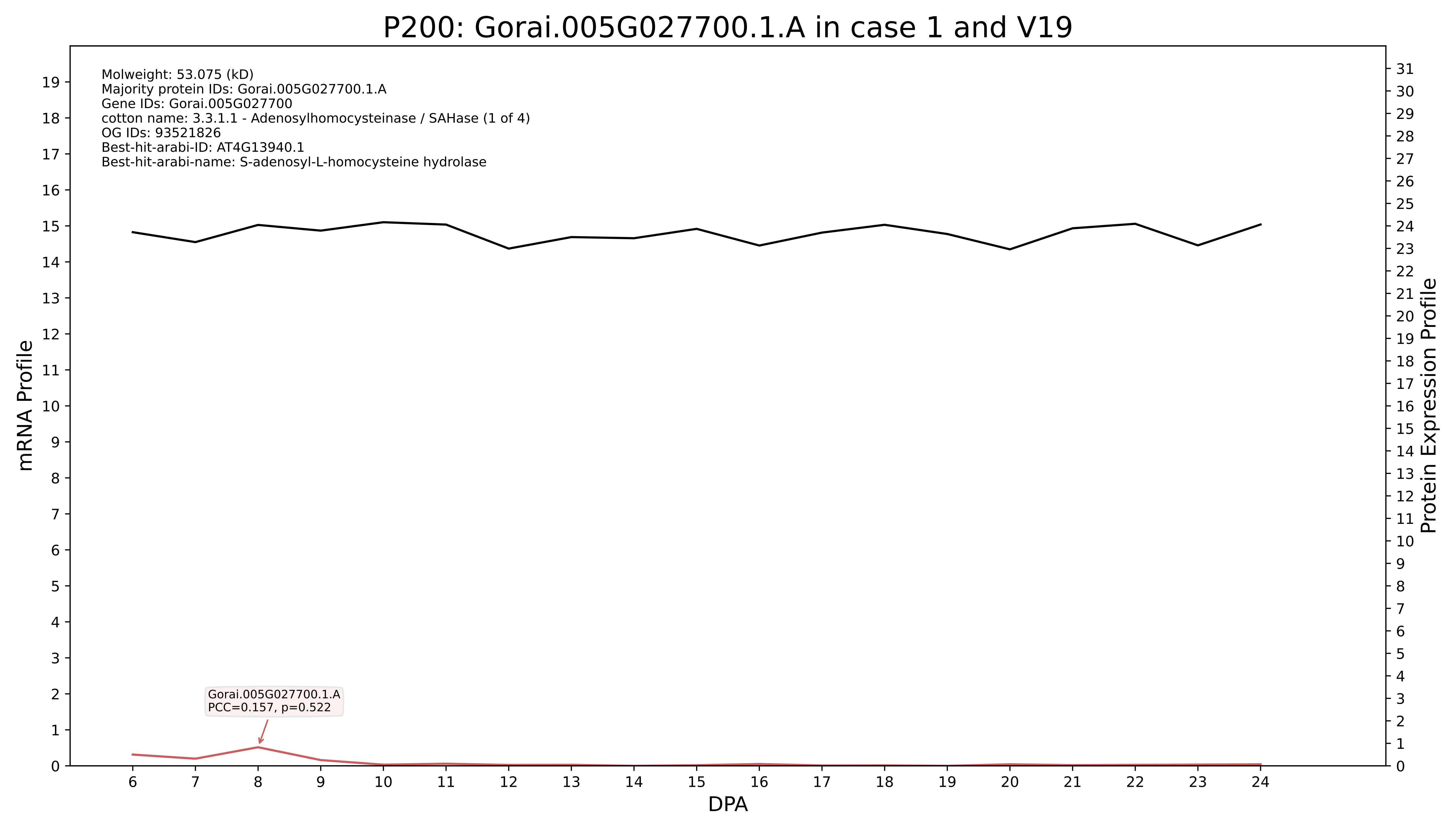
P200: Gorai.005G027700.1.A in case 1 and V19
Molweight: 53.075 (kD)
Majority protein IDs: Gorai.005G027700.1.A
Gene IDs: Gorai.005G027700
cotton name: 3.3.1.1 - Adenosylhomocysteinase / SAHase (1 of 4)
OG IDs: 93521826
Best-hit-arabi-ID: AT4G13940.1
Best-hit-arabi-name: S-adenosyl-L-homocysteine hydrolase
Gorai.005G027700.1.A
PCC=0.157, p=0.522
mRNA Profile
Protein Expression Profile
DPA

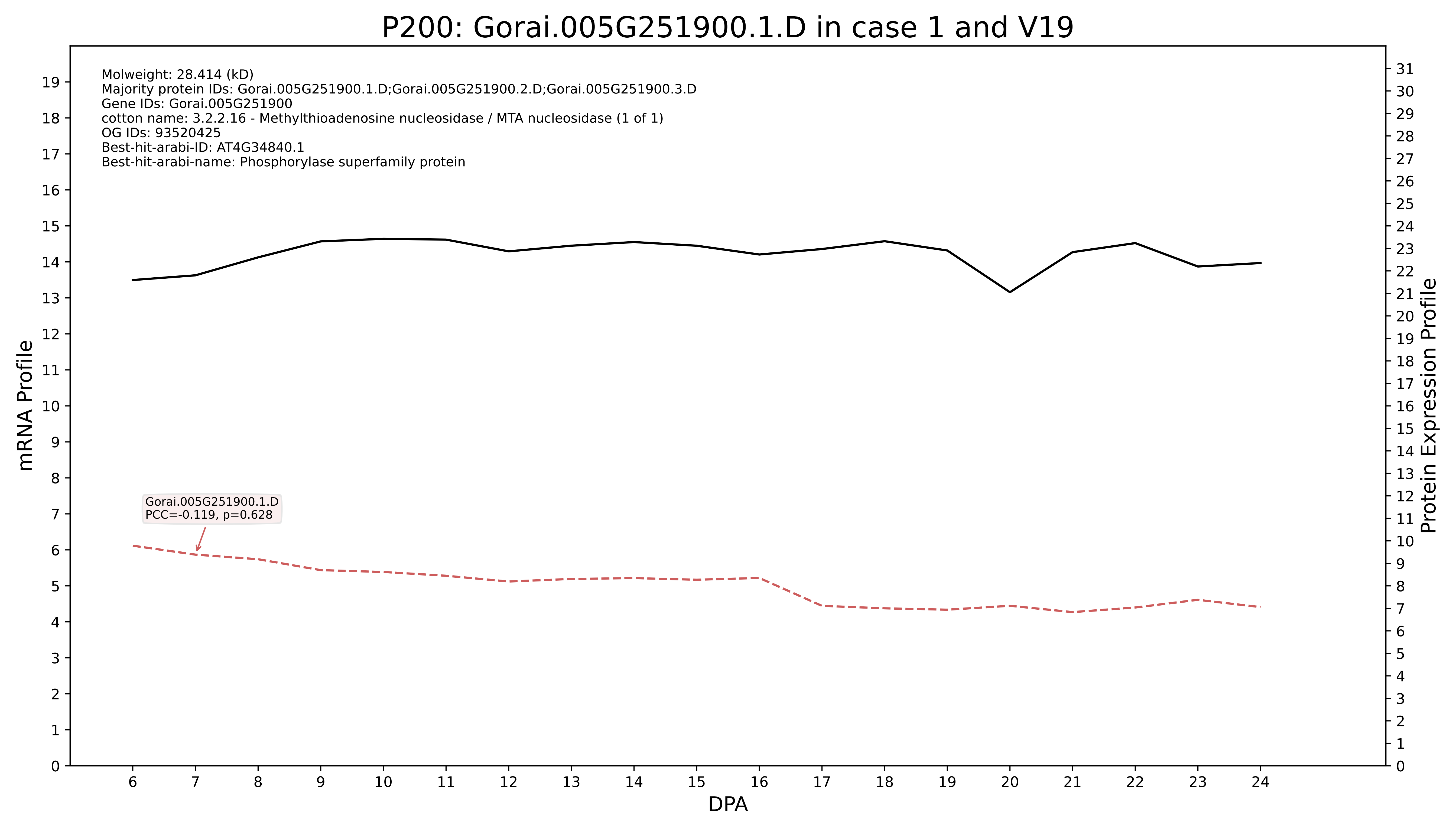
P200: Gorai.005G251900.1.D in case 1 and V19
Molweight: 28.414 (kD)
Majority protein IDs: Gorai.005G251900.1.D;Gorai.005G251900.2.D;Gorai.005G251900.3.D
Gene IDs: Gorai.005G251900
cotton name: 3.2.2.16 - Methylthioadenosine nucleosidase / MTA nucleosidase (1 of 1)
OG IDs: 93520425
Best-hit-arabi-ID: AT4G34840.1
Best-hit-arabi-name: Phosphorylase superfamily protein
Gorai.005G251900.1.D
PCC=-0.119, p=0.628
mRNA Profile
Protein Expression Profile
DPA

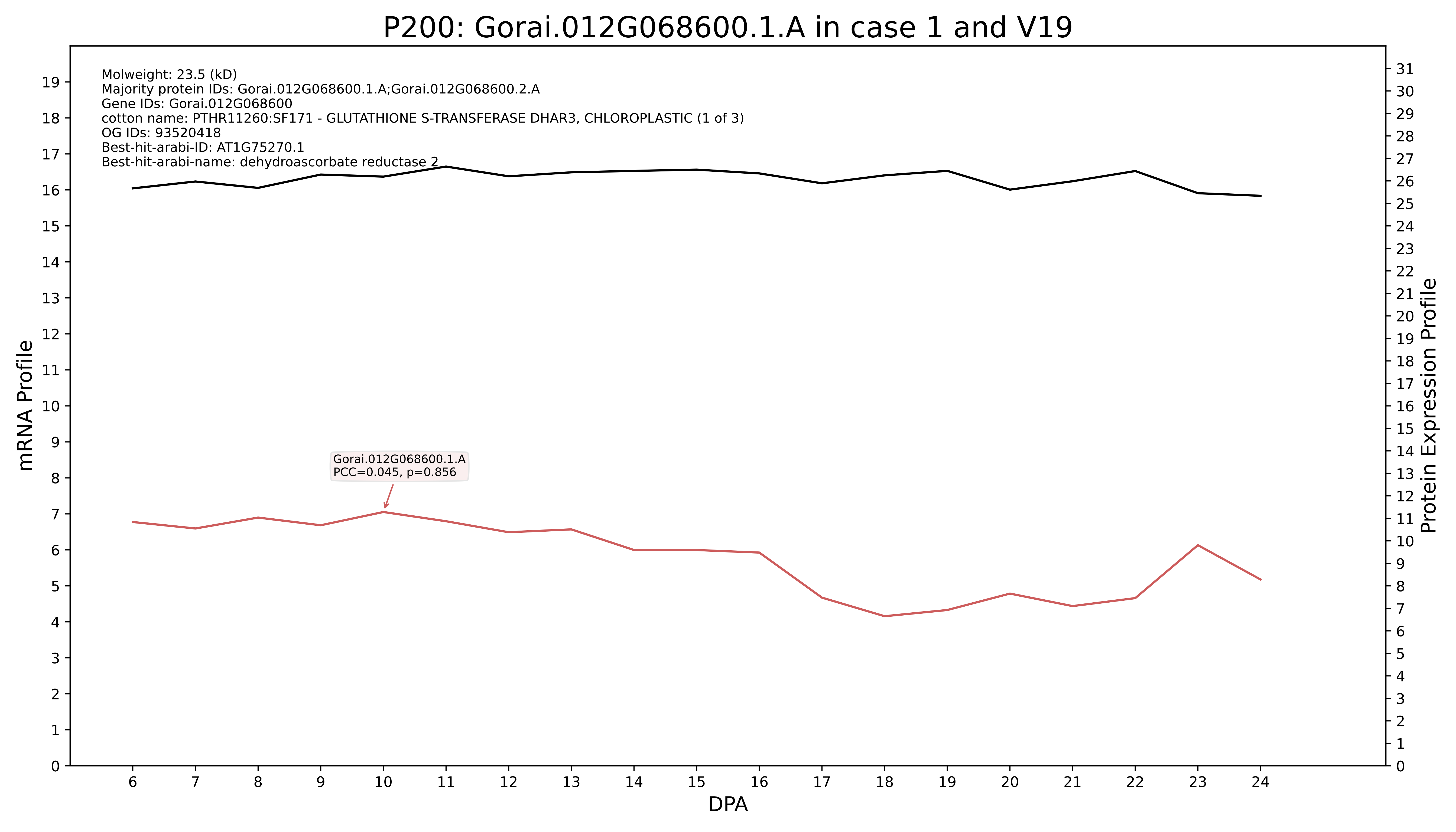
P200: Gorai.012G068600.1.A in case 1 and V19
Molweight: 23.5 (kD)
Majority protein IDs: Gorai.012G068600.1.A;Gorai.012G068600.2.A
Gene IDs: Gorai.012G068600
cotton name: PTHR11260:SF171 - GLUTATHIONE S-TRANSFERASE DHAR3, CHLOROPLASTIC (1 of 3)
OG IDs: 93520418
Best-hit-arabi-ID: AT1G75270.1
Best-hit-arabi-name: dehydroascorbate reductase 2
Gorai.012G068600.1.A
PCC=0.045, p=0.856
mRNA Profile
Protein Expression Profile
DPA

### P200: Gorai.013G172400.1.D in case 1 and V19

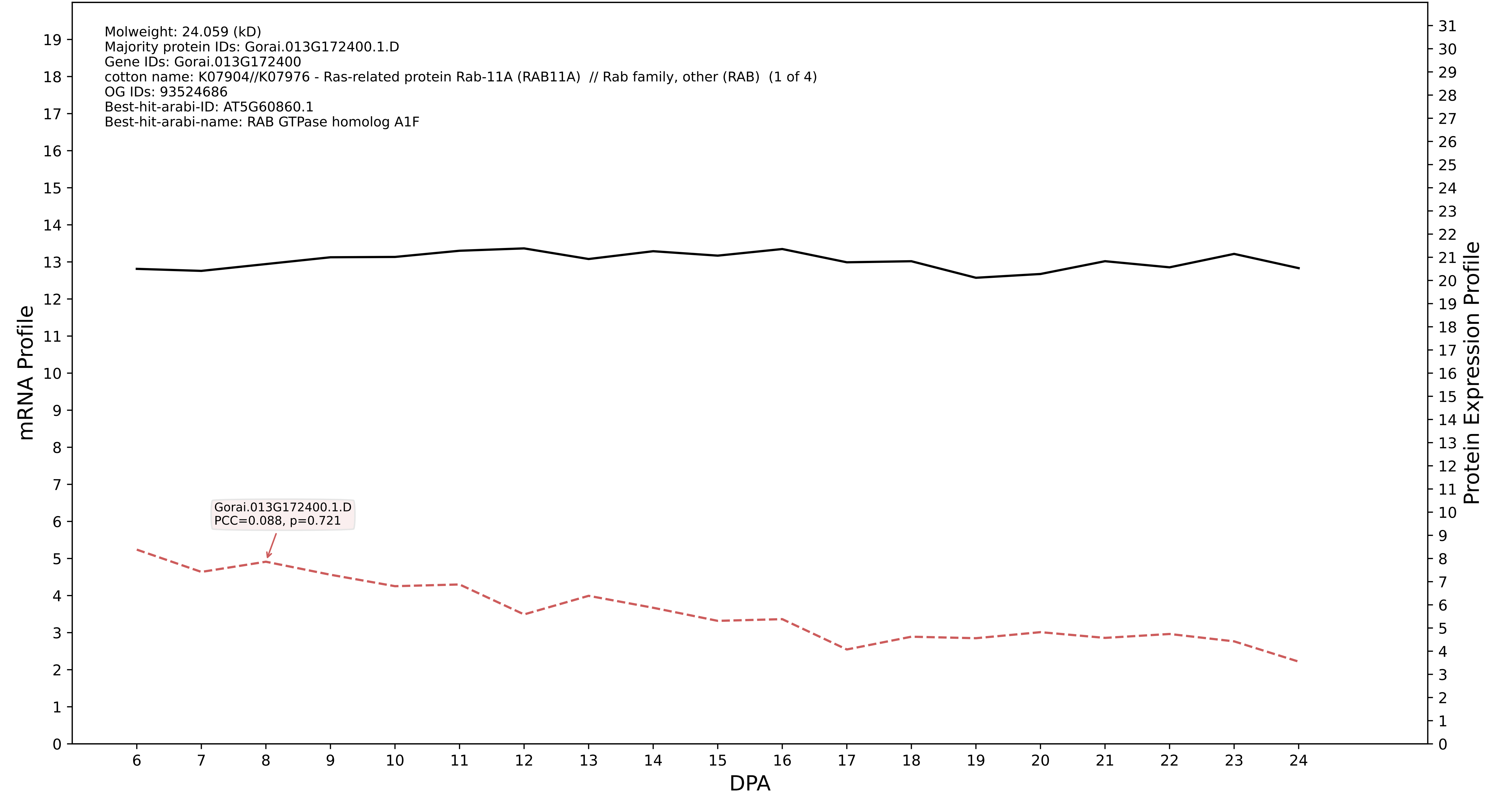

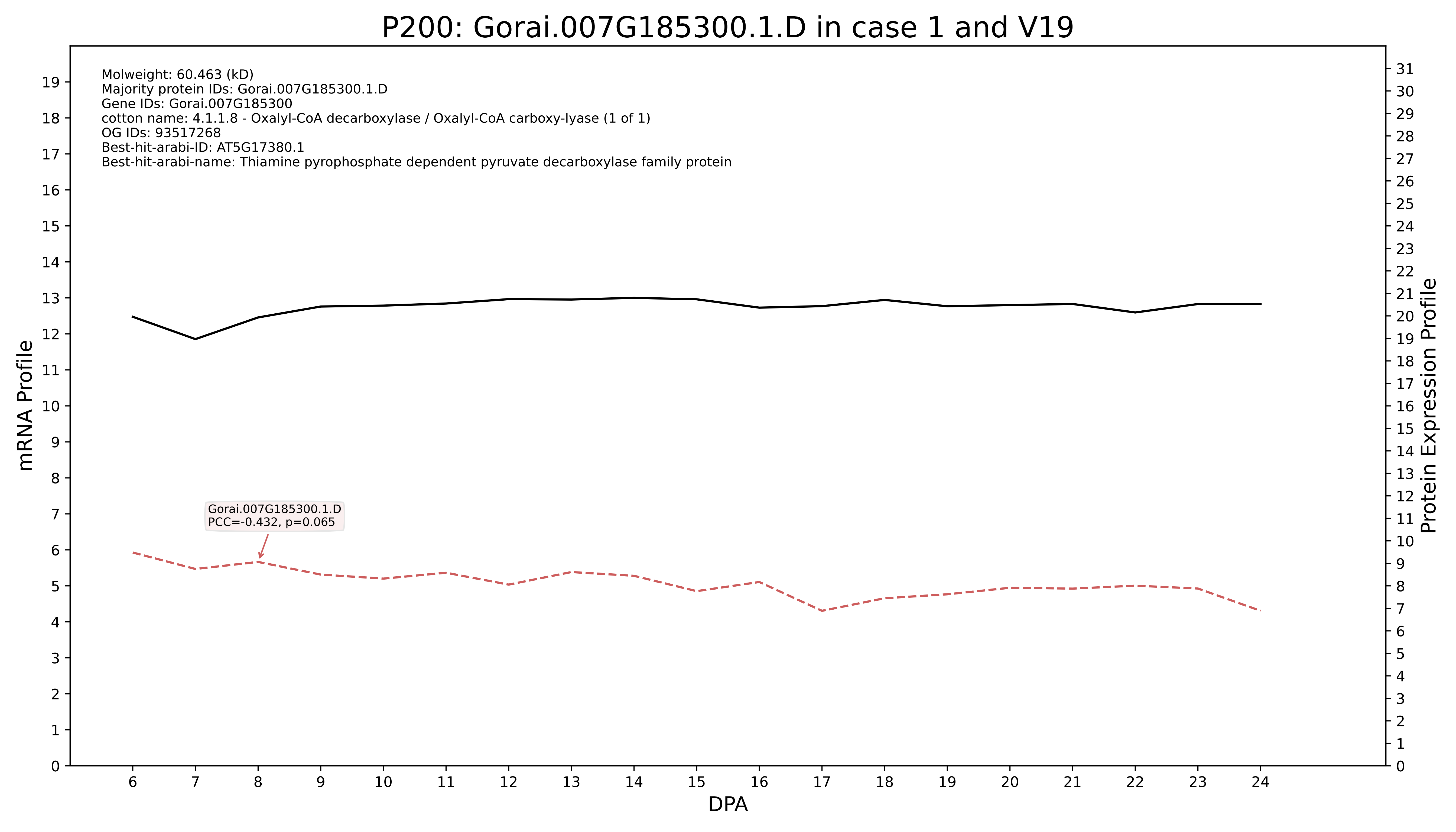
P200: Gorai.007G185300.1.D in case 1 and V19
Molweight: 60.463 (kD)
Majority protein IDs: Gorai.007G185300.1.D
Gene IDs: Gorai.007G185300
cotton name: 4.1.1.8 - Oxalyl-CoA decarboxylase / Oxalyl-CoA carboxy-lyase (1 of 1)
OG IDs: 93517268
Best-hit-arabi-ID: AT5G17380.1
Best-hit-arabi-name: Thiamine pyrophosphate dependent pyruvate decarboxylase family protein
Gorai.007G185300.1.D
PCC=-0.432, p=0.065
mRNA Profile
Protein Expression Profile
DPA

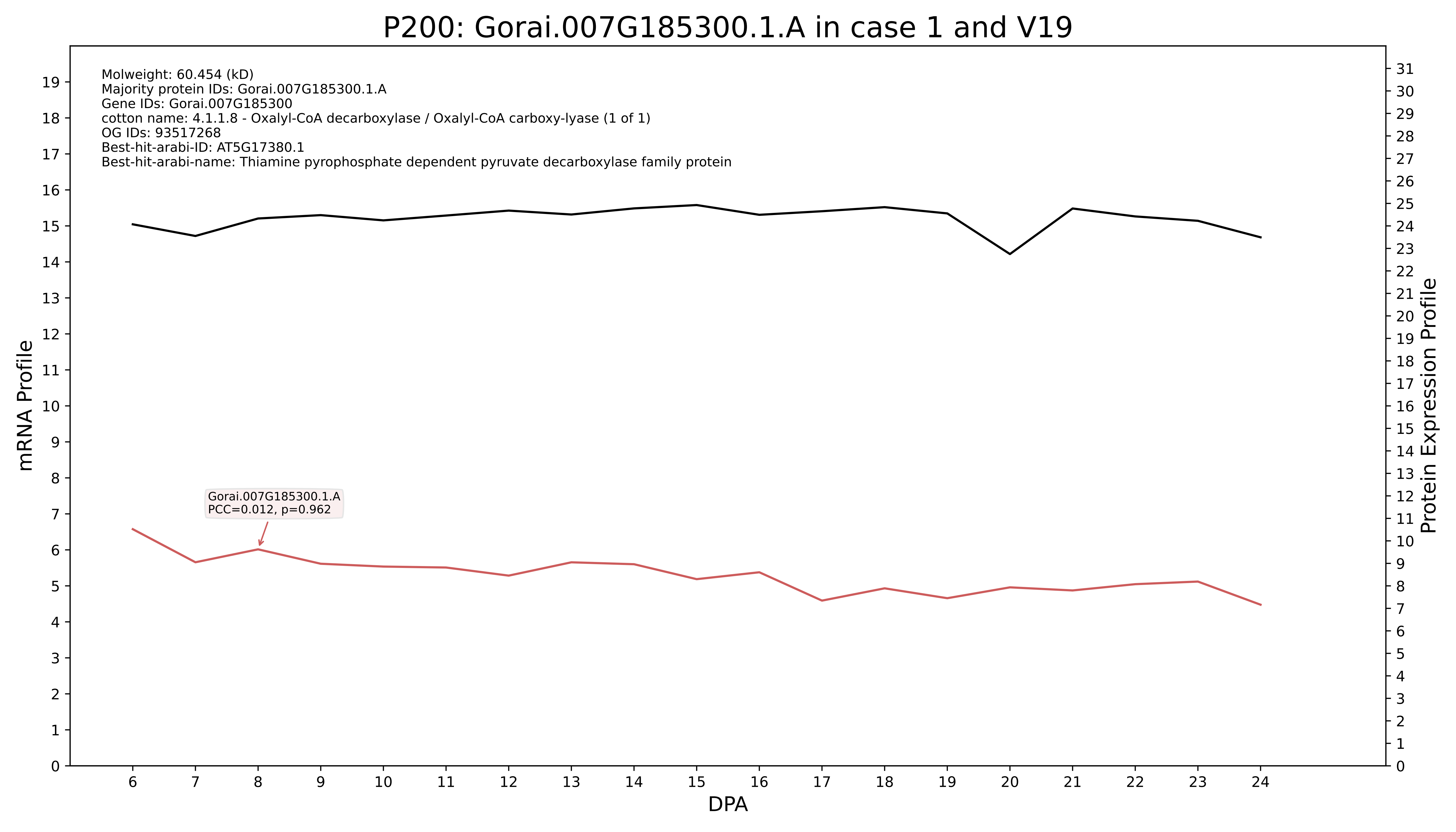
P200: Gorai.007G185300.1.A in case 1 and V19
Molweight: 60.454 (kD)
Majority protein IDs: Gorai.007G185300.1.A
Gene IDs: Gorai.007G185300
cotton name: 4.1.1.8 - Oxalyl-CoA decarboxylase / Oxalyl-CoA carboxy-lyase (1 of 1)
OG IDs: 93517268
Best-hit-arabi-ID: AT5G17380.1
Best-hit-arabi-name: Thiamine pyrophosphate dependent pyruvate decarboxylase family protein
Gorai.007G185300.1.A
PCC=0.012, p=0.962
mRNA Profile
Protein Expression Profile
DPA

### P200: Gorai.005G119100.1.A in case 1 and V19

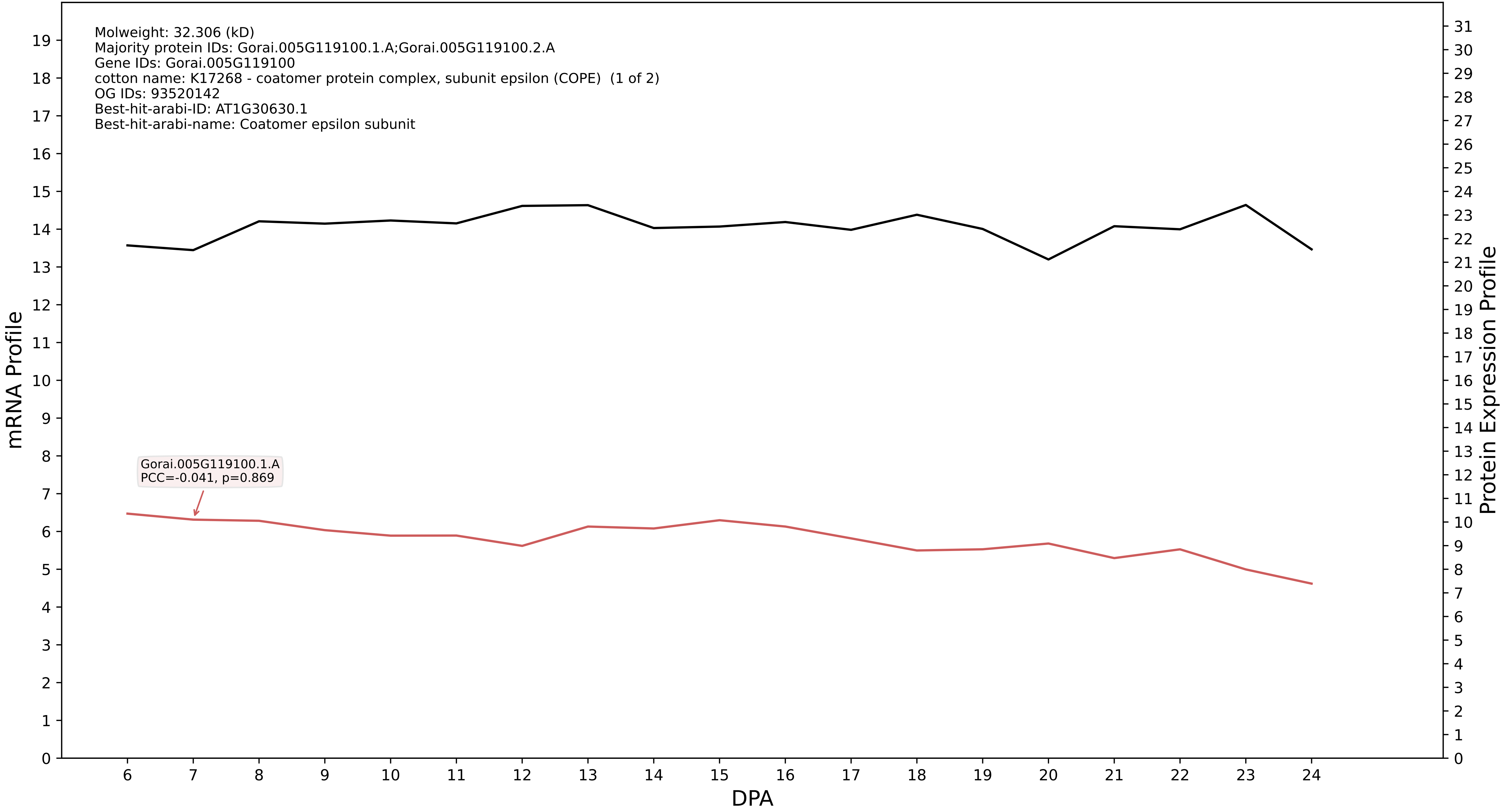

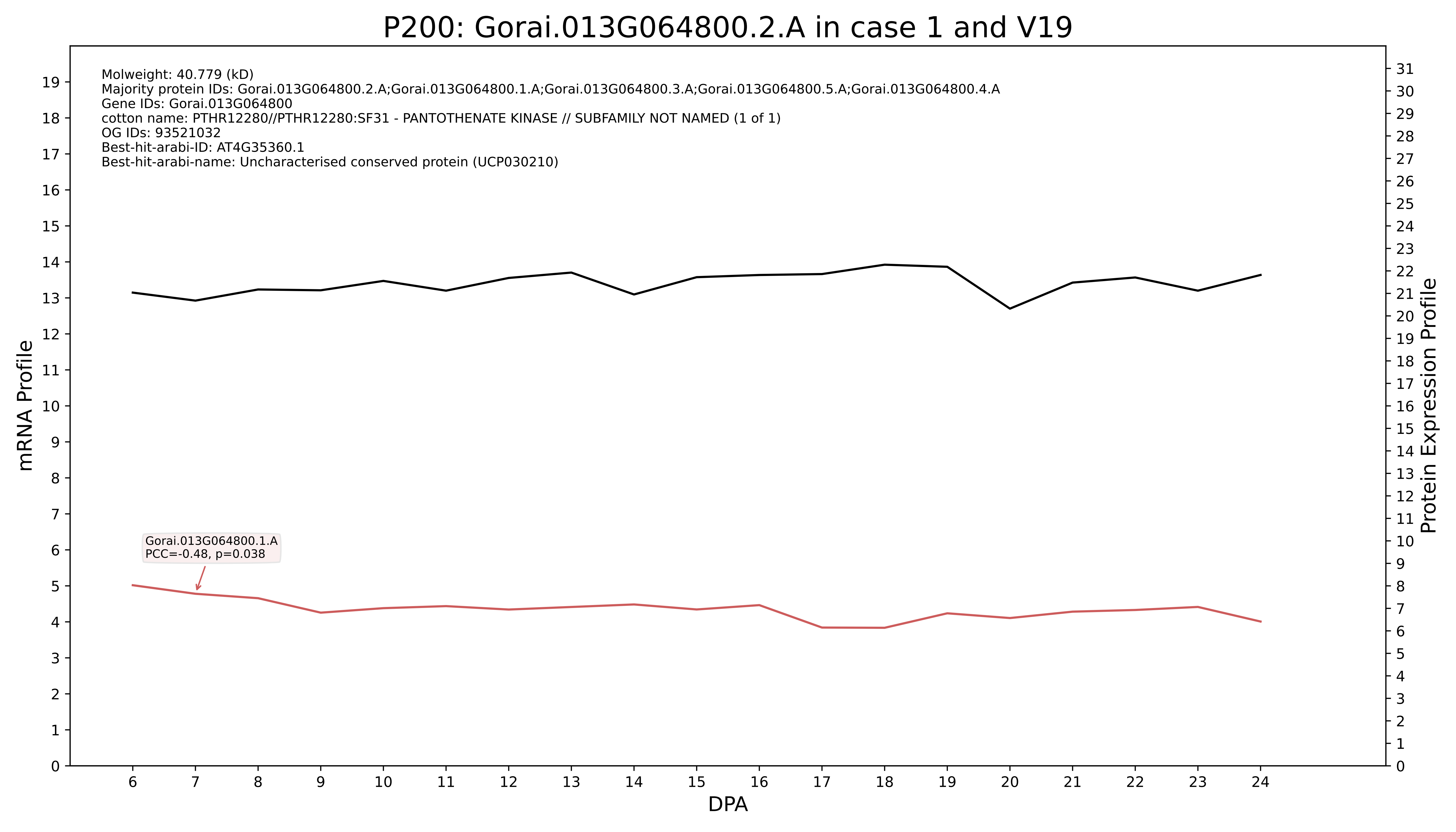

P200: Gorai.013G064800.2.A in case 1 and V19
Molweight: 40.779 (kD)
Majority protein IDs: Gorai.013G064800.2.A;Gorai.013G064800.1.A;Gorai.013G064800.3.A;Gorai.013G064800.5.A;Gorai.013G064800.4.A
Gene IDs: Gorai.013G064800
cotton name: PTHR12280//PTHR12280:SF31 - PANTOTHENATE KINASE // SUBFAMILY NOT NAMED (1 of 1)
OG IDs: 93521032
Best-hit-arabi-ID: AT4G35360.1
Best-hit-arabi-name: Uncharacterised conserved protein (UCP030210)
Gorai.013G064800.1.A
PCC=-0.48, p=0.038
mRNA Profile
Protein Expression Profile
DPA

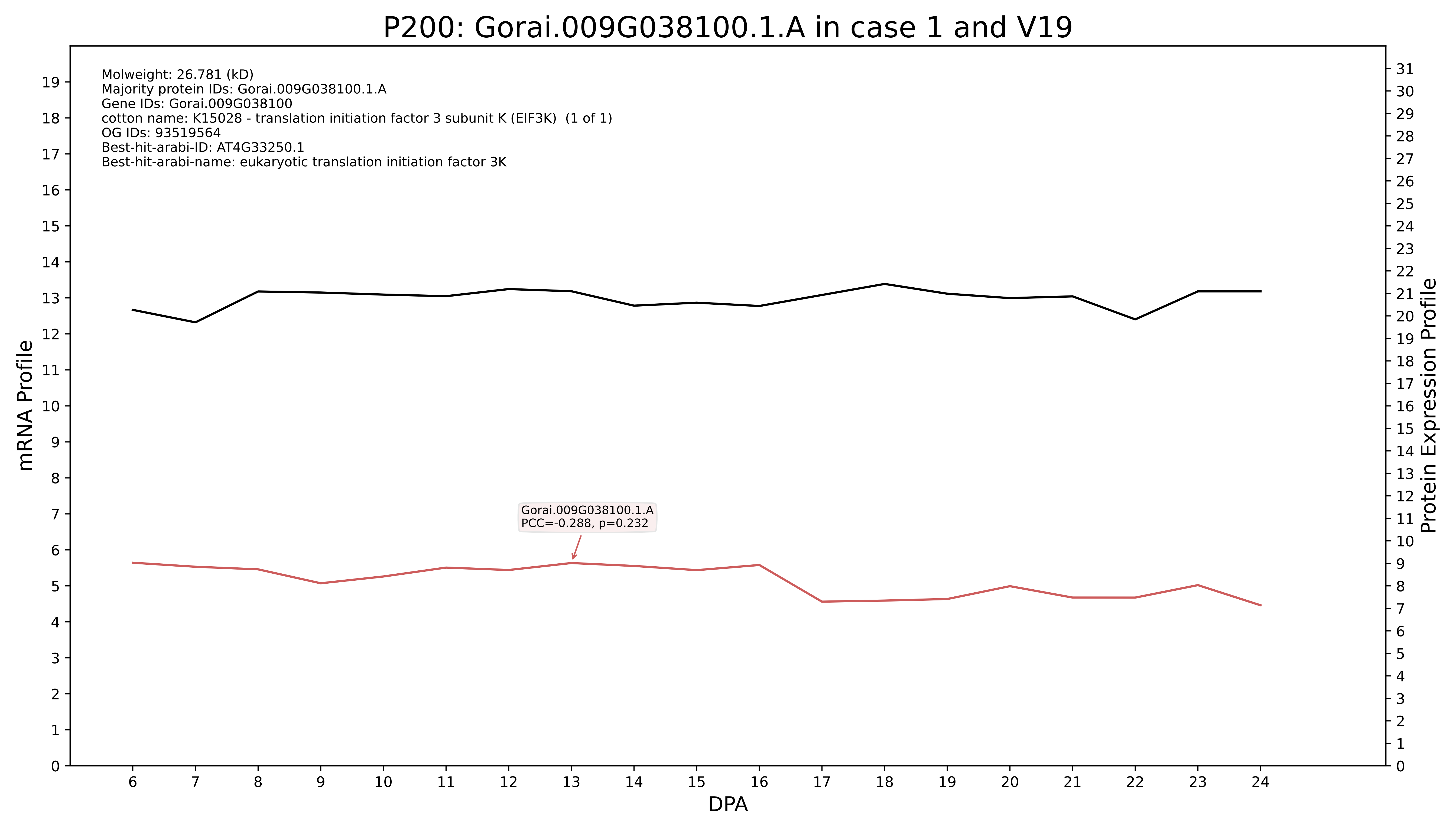
P200: Gorai.009G038100.1.A in case 1 and V19
Molweight: 26.781 (kD)
Majority protein IDs: Gorai.009G038100.1.A
Gene IDs: Gorai.009G038100
cotton name: K15028 - translation initiation factor 3 subunit K (EIF3K) (1 of 1)
OG IDs: 93519564
Best-hit-arabi-ID: AT4G33250.1
Best-hit-arabi-name: eukaryotic translation initiation factor 3K
Gorai.009G038100.1.A
PCC=-0.288, p=0.232
mRNA Profile
Protein Expression Profile
DPA

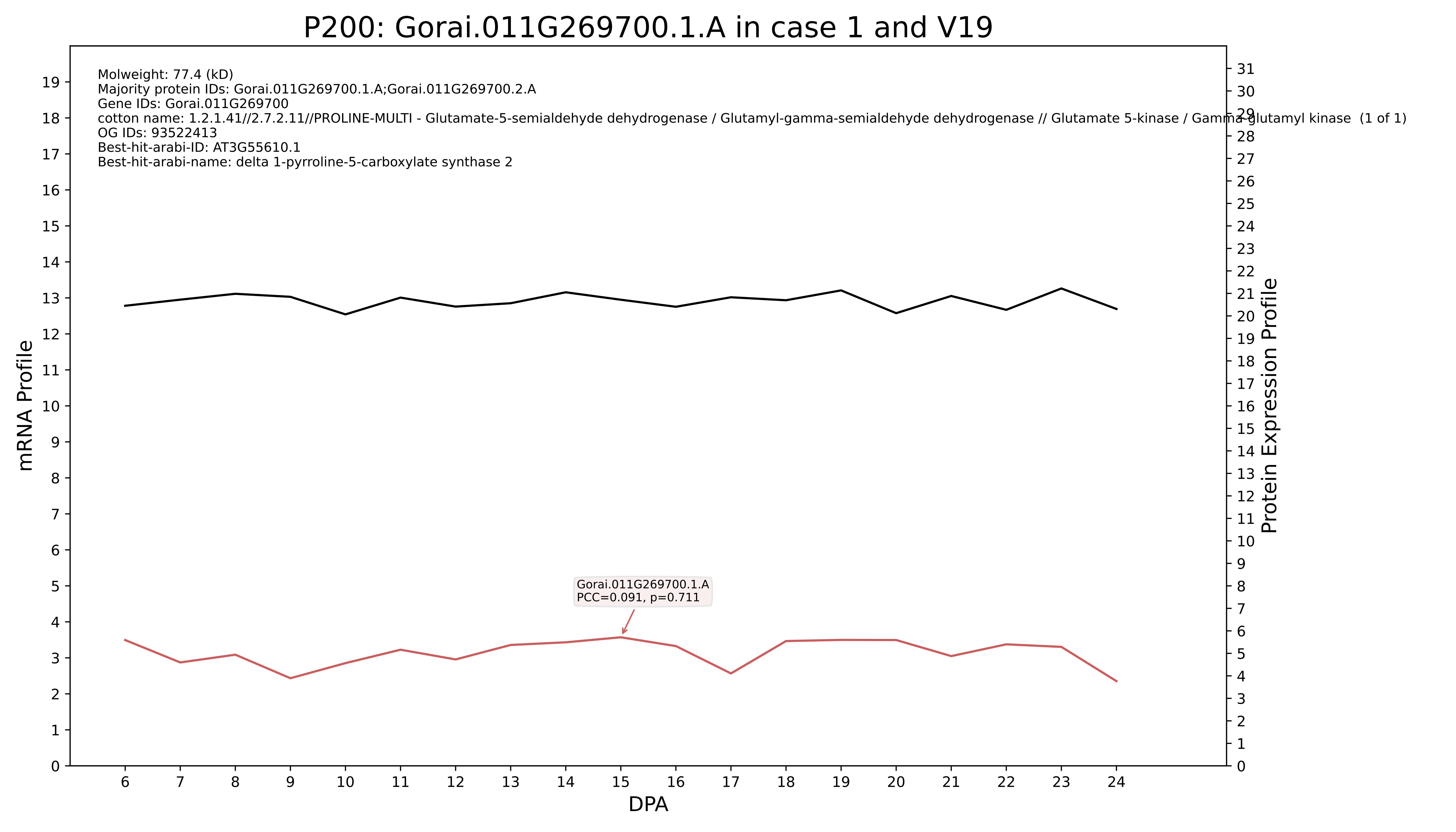
P200: Gorai.011G269700.1.A in case 1 and V19
Molweight: 77.4 (kD)
Majority protein IDs: Gorai.011G269700.1.A;Gorai.011G269700.2.A
Gene IDs: Gorai.011G269700
cotton name: 1.2.1.41//2.7.2.11//PROLINE-MULTI - Glutamate-5-semialdehyde dehydrogenase / Glutamyl-gamma-semialdehyde dehydrogenase // Glutamate 5-kinase / Gamma-glutamyl kinase (1 of 1)
OG IDs: 93522413
Best-hit-arabi-ID: AT3G55610.1
Best-hit-arabi-name: delta 1-pyrroline-5-carboxylate synthase 2
Gorai.011G269700.1.A
PCC=0.091, p=0.711
mRNA Profile
Protein Expression Profile
DPA

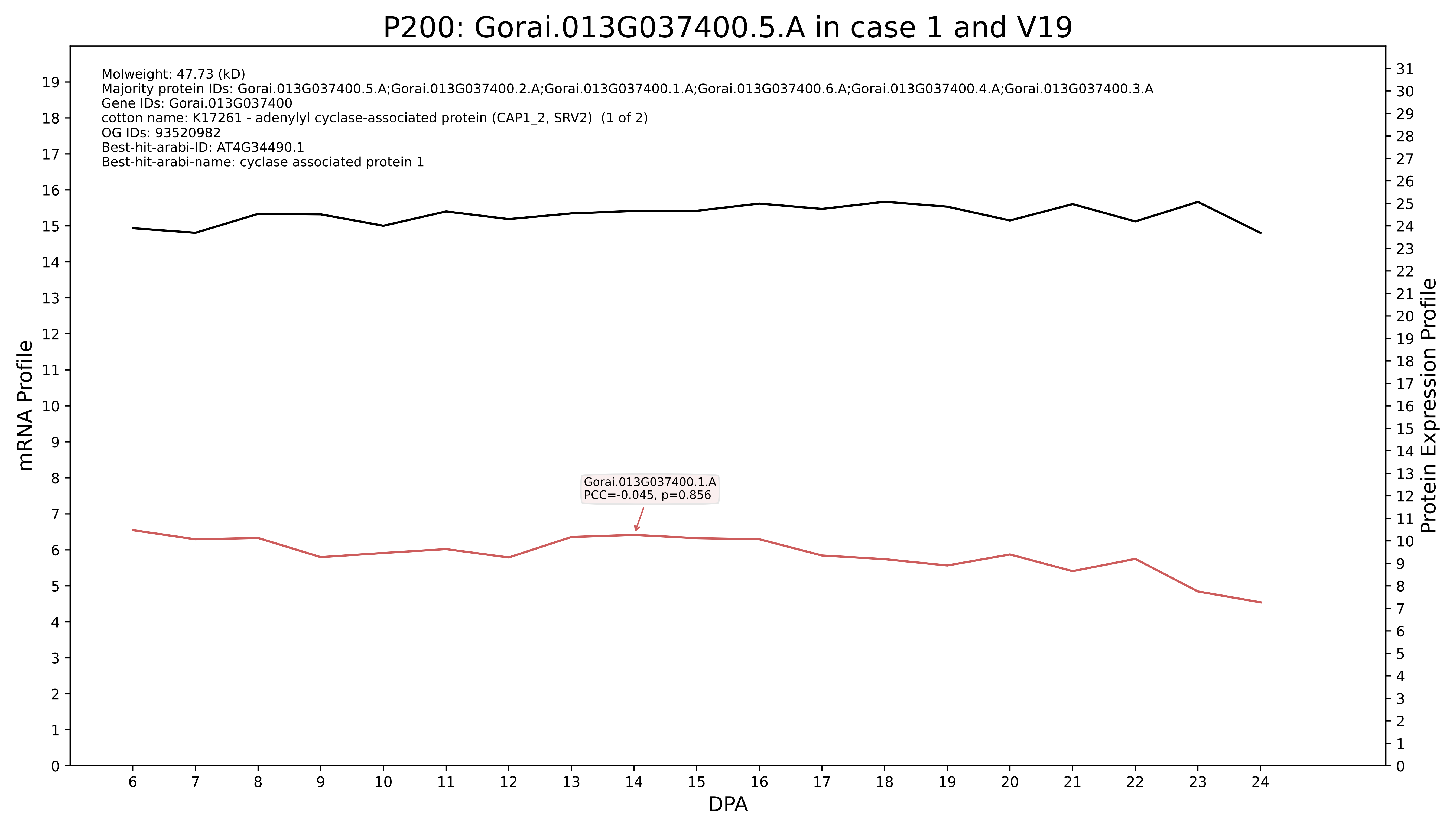
P200: Gorai.013G037400.5.A in case 1 and V19
Molweight: 47.73 (kD)
Majority protein IDs: Gorai.013G037400.5.A;Gorai.013G037400.2.A;Gorai.013G037400.1.A;Gorai.013G037400.6.A;Gorai.013G037400.4.A;Gorai.013G037400.3.A
Gene IDs: Gorai.013G037400
cotton name: K17261 - adenylyl cyclase-associated protein (CAP1_2, SRV2) (1 of 2)
OG IDs: 93520982
Best-hit-arabi-ID: AT4G34490.1
Best-hit-arabi-name: cyclase associated protein 1
Gorai.013G037400.1.A
PCC=-0.045, p=0.856
mRNA Profile
Protein Expression Profile
DPA

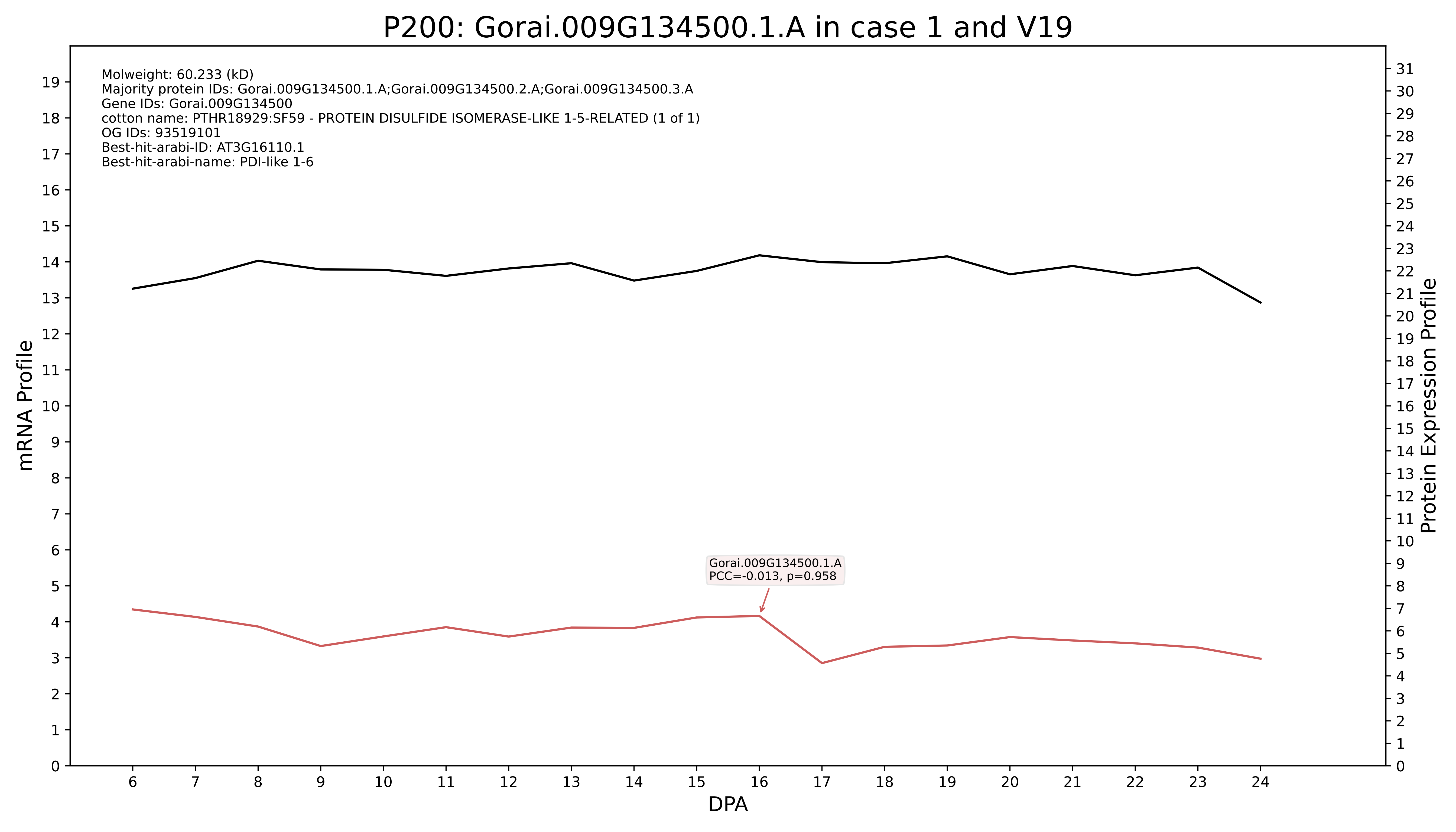
P200: Gorai.009G134500.1.A in case 1 and V19
Molweight: 60.233 (kD)
Majority protein IDs: Gorai.009G134500.1.A;Gorai.009G134500.2.A;Gorai.009G134500.3.A
Gene IDs: Gorai.009G134500
cotton name: PTHR18929:SF59 - PROTEIN DISULFIDE ISOMERASE-LIKE 1-5-RELATED (1 of 1)
OG IDs: 93519101
Best-hit-arabi-ID: AT3G16110.1
Best-hit-arabi-name: PDI-like 1-6
Gorai.009G134500.1.A
PCC=-0.013, p=0.958
mRNA Profile
Protein Expression Profile
DPA

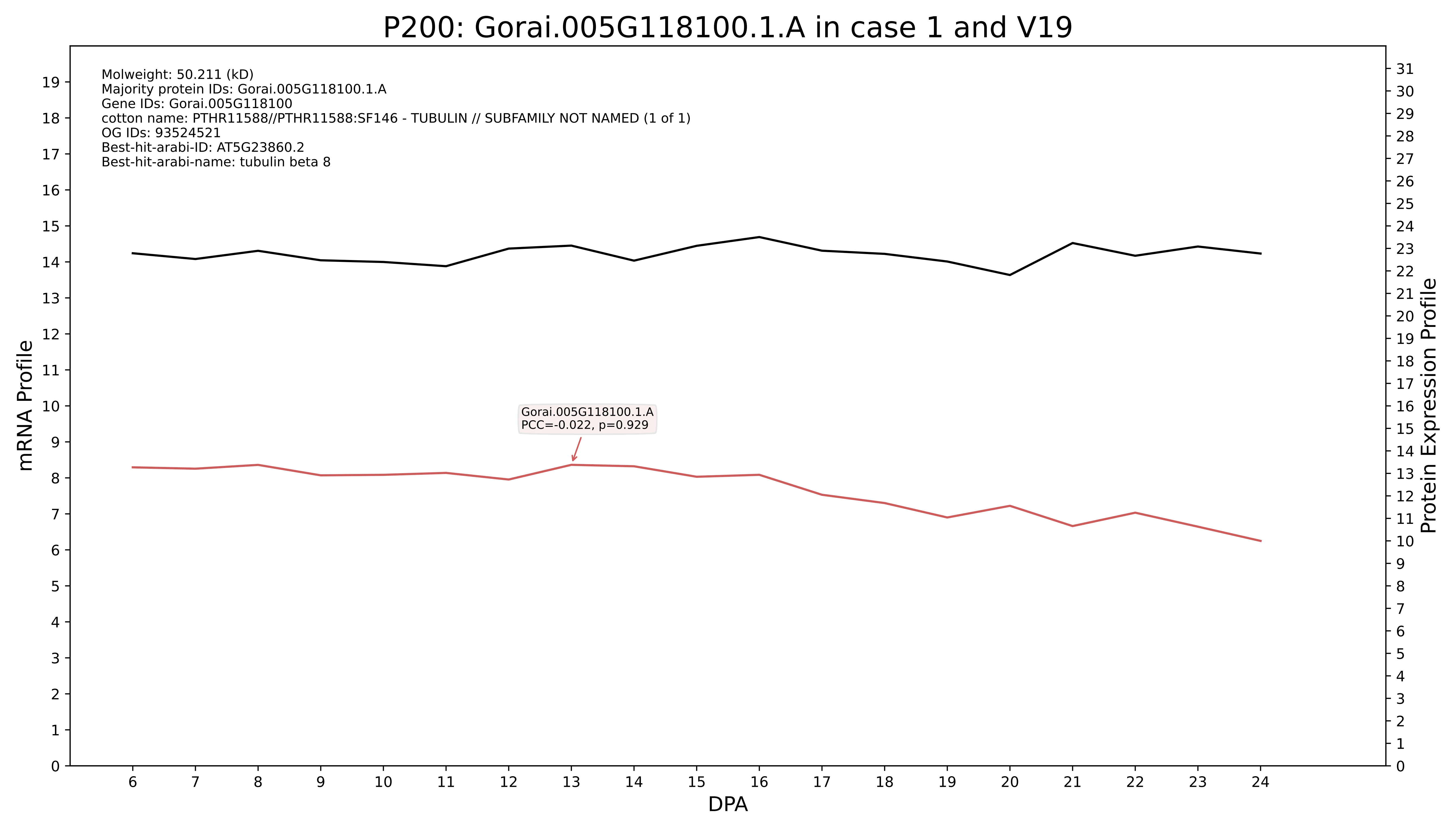
P200: Gorai.005G118100.1.A in case 1 and V19
Molweight: 50.211 (kD)
Majority protein IDs: Gorai.005G118100.1.A
Gene IDs: Gorai.005G118100
cotton name: PTHR11588//PTHR11588:SF146 - TUBULIN // SUBFAMILY NOT NAMED (1 of 1)
OG IDs: 93524521
Best-hit-arabi-ID: AT5G23860.2
Best-hit-arabi-name: tubulin beta 8
Gorai.005G118100.1.A
PCC=-0.022, p=0.929
mRNA Profile
Protein Expression Profile
DPA

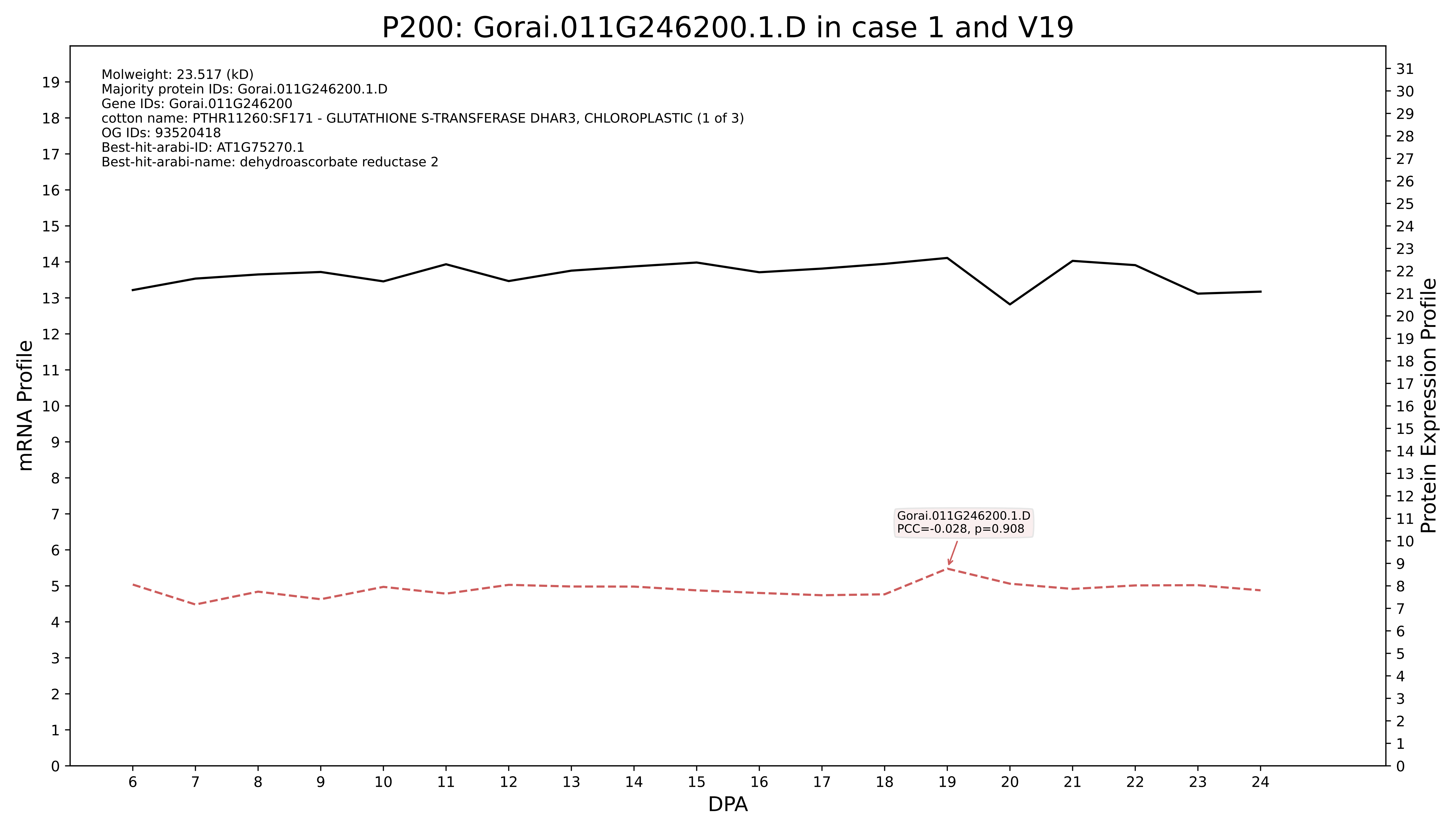
P200: Gorai.011G246200.1.D in case 1 and V19
Molweight: 23.517 (kD)
Majority protein IDs: Gorai.011G246200.1.D
Gene IDs: Gorai.011G246200
cotton name: PTHR11260:SF171 - GLUTATHIONE S-TRANSFERASE DHAR3, CHLOROPLASTIC (1 of 3)
OG IDs: 93520418
Best-hit-arabi-ID: AT1G75270.1
Best-hit-arabi-name: dehydroascorbate reductase 2
Gorai.011G246200.1.D
PCC=-0.028, p=0.908
mRNA Profile
Protein Expression Profile
DPA

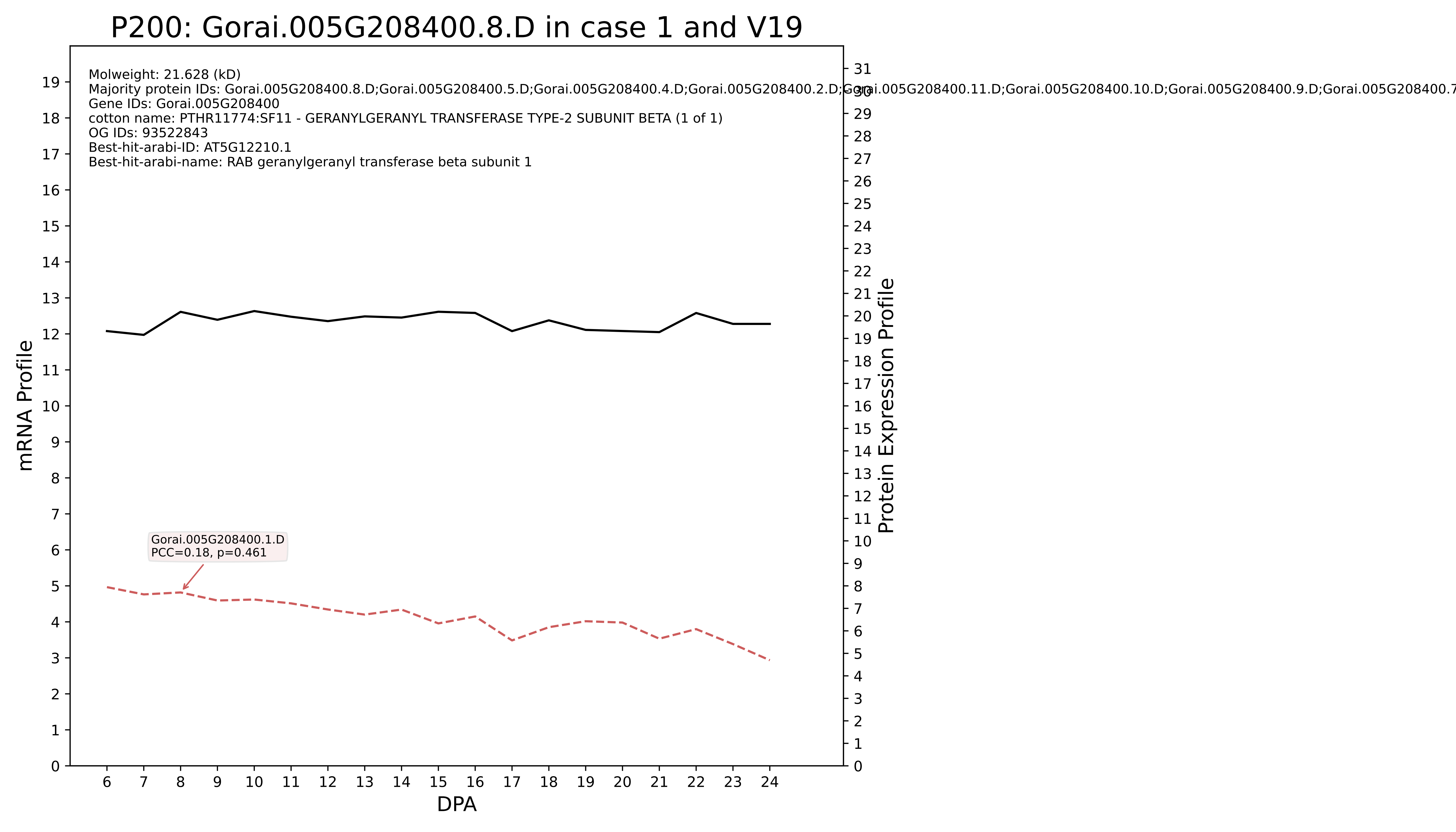
P200: Gorai.005G208400.8.D in case 1 and V19
Molweight: 21.628 (kD)
Majority protein IDs: Gorai.005G208400.8.D;Gorai.005G208400.5.D;Gorai.005G208400.4.D;Gorai.005G208400.2.D;Gorai.005G208400.11.D;Gorai.005G208400.10.D;Gorai.005G208400.9.D;Gorai.005G208400.7
Gene IDs: Gorai.005G208400
cotton name: PTHR11774:SF11 - GERANYLGERANYL TRANSFERASE TYPE-2 SUBUNIT BETA (1 of 1)
OG IDs: 93522843
Best-hit-arabi-ID: AT5G12210.1
Best-hit-arabi-name: RAB geranylgeranyl transferase beta subunit 1
Gorai.005G208400.1.D
PCC=0.18, p=0.461
mRNA Profile
Protein Expression Profile
DPA

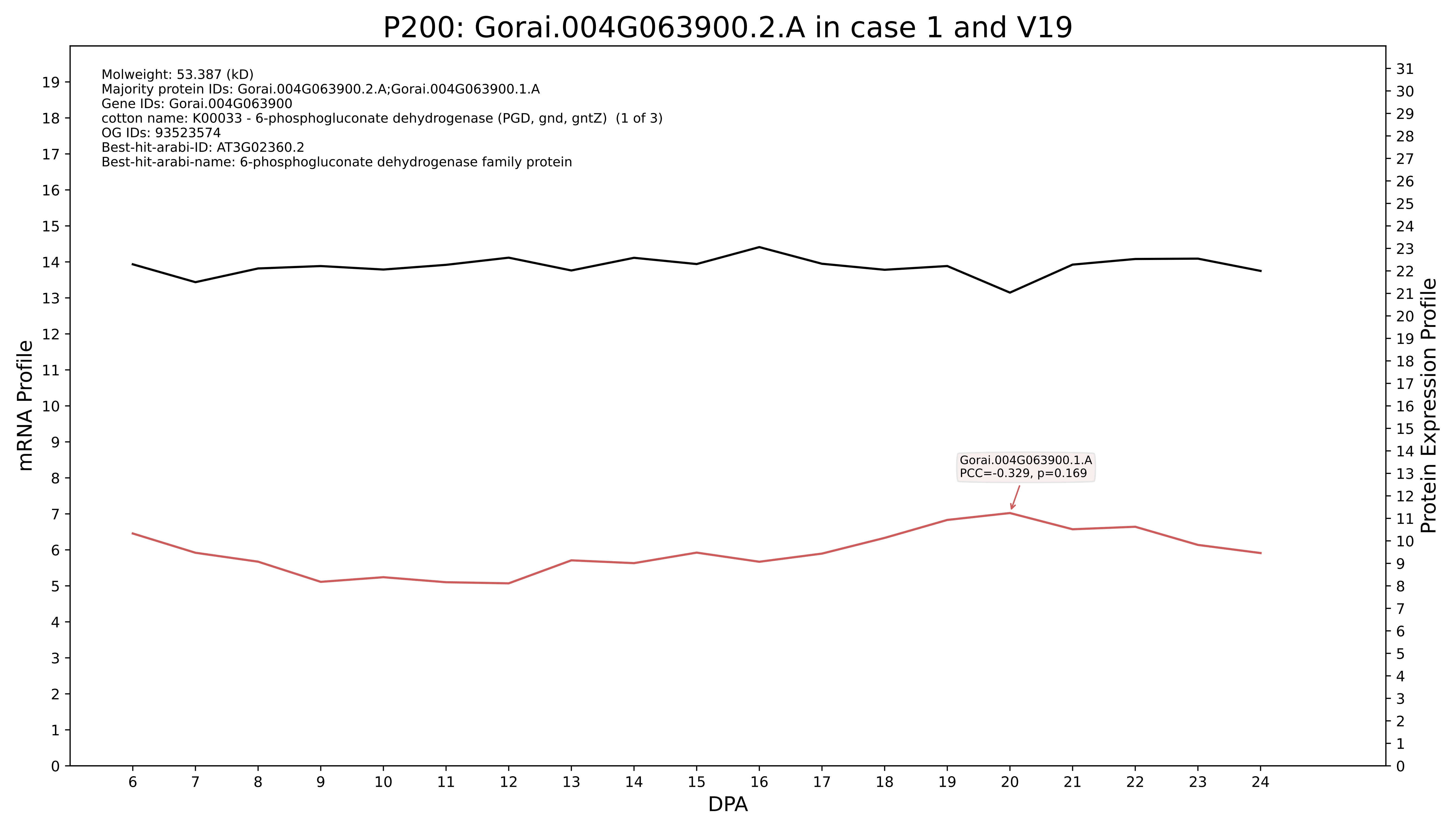
P200: Gorai.004G063900.2.A in case 1 and V19
Molweight: 53.387 (kD)
Majority protein IDs: Gorai.004G063900.2.A;Gorai.004G063900.1.A
Gene IDs: Gorai.004G063900
cotton name: K00033 - 6-phosphogluconate dehydrogenase (PGD, gnd, gntZ) (1 of 3)
OG IDs: 93523574
Best-hit-arabi-ID: AT3G02360.2
Best-hit-arabi-name: 6-phosphogluconate dehydrogenase family protein
Gorai.004G063900.1.A
PCC=-0.329, p=0.169
mRNA Profile
Protein Expression Profile
DPA

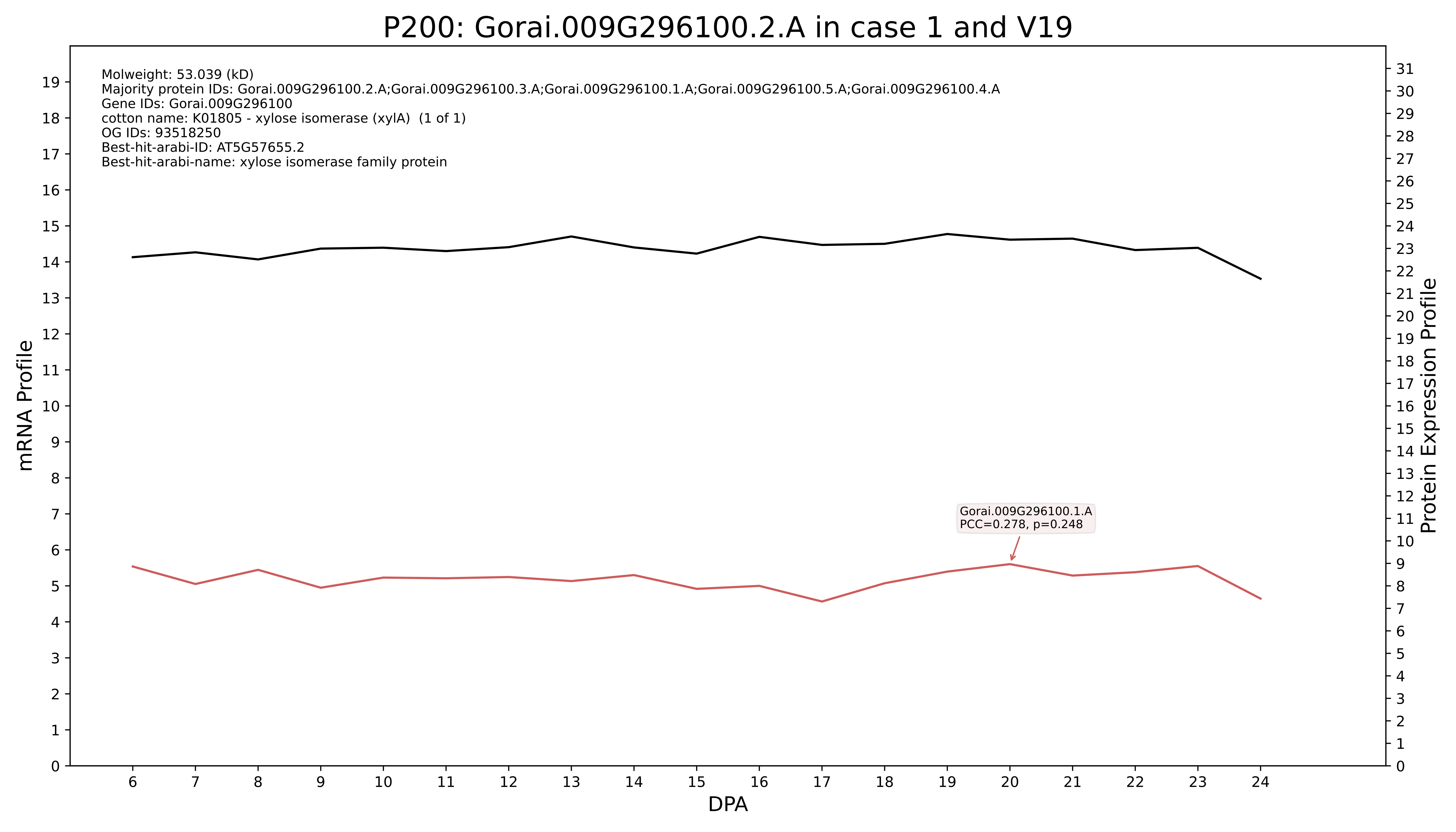

P200: Gorai.009G296100.2.A in case 1 and V19
Molweight: 53.039 (kD)
Majority protein IDs: Gorai.009G296100.2.A;Gorai.009G296100.3.A;Gorai.009G296100.1.A;Gorai.009G296100.5.A;Gorai.009G296100.4.A
Gene IDs: Gorai.009G296100
cotton name: K01805 - xylose isomerase (xylA) (1 of 1)
OG IDs: 93518250
Best-hit-arabi-ID: AT5G57655.2
Best-hit-arabi-name: xylose isomerase family protein
Gorai.009G296100.1.A
PCC=0.278, p=0.248
mRNA Profile
Protein Expression Profile
DPA

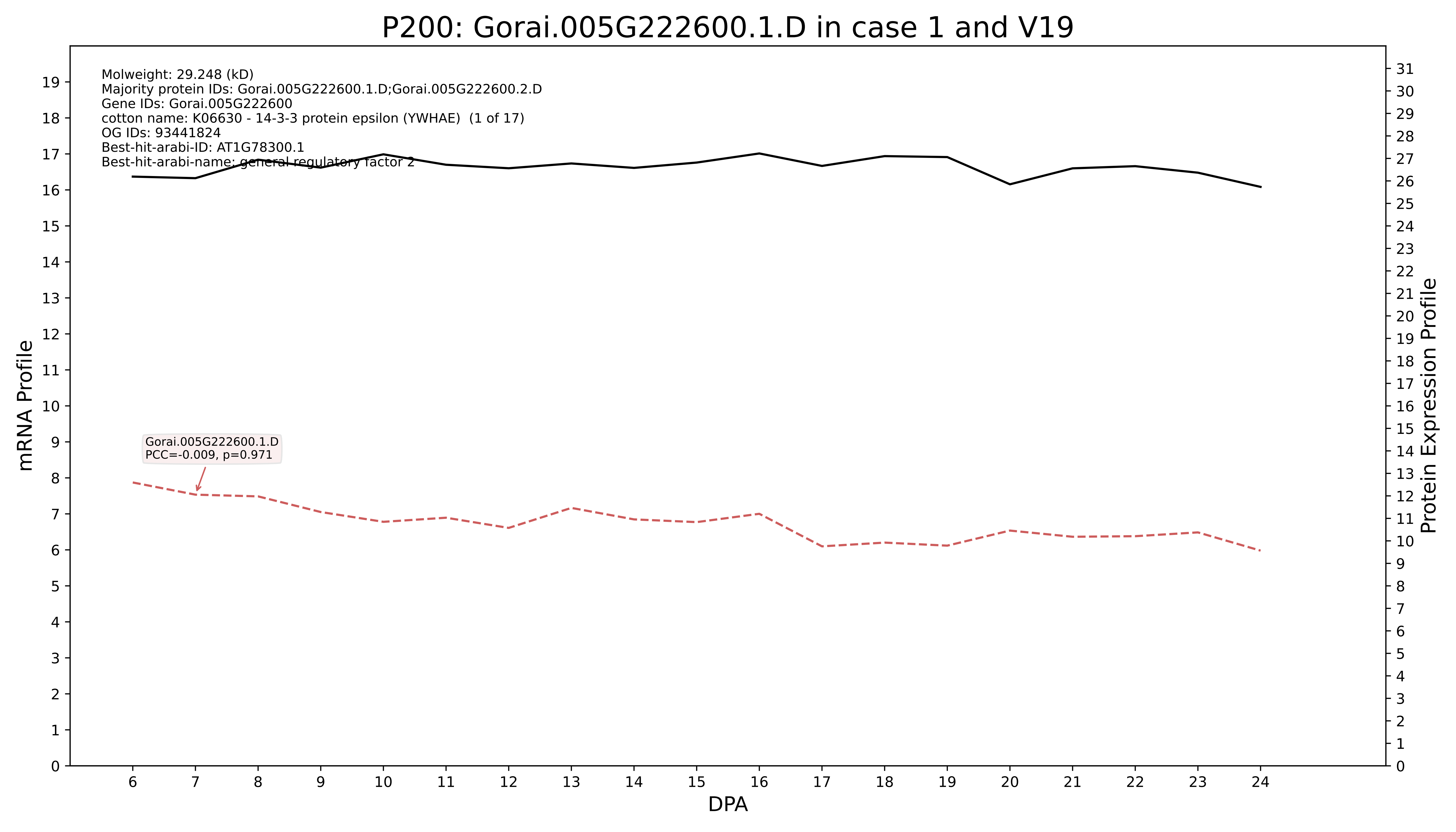
P200: Gorai.005G222600.1.D in case 1 and V19
Molweight: 29.248 (kD)
Majority protein IDs: Gorai.005G222600.1.D;Gorai.005G222600.2.D
Gene IDs: Gorai.005G222600
cotton name: K06630 - 14-3-3 protein epsilon (YWHAE) (1 of 17)
OG IDs: 93441824
Best-hit-arabi-ID: AT1G78300.1
Best-hit-arabi-name: general regulatory factor 2
Gorai.005G222600.1.D
PCC=-0.009, p=0.971
mRNA Profile
Protein Expression Profile
DPA

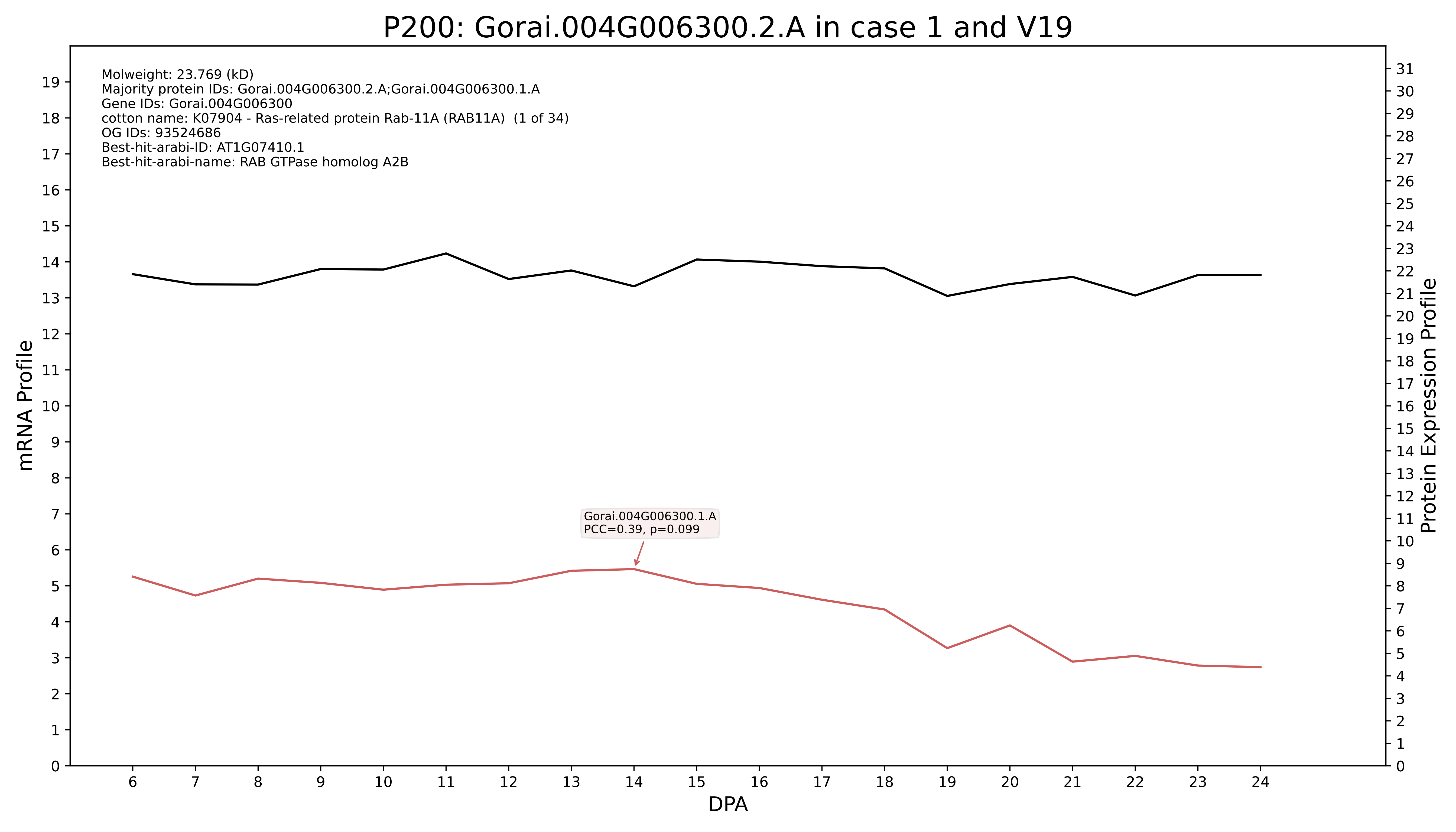
P200: Gorai.004G006300.2.A in case 1 and V19
Molweight: 23.769 (kD)
Majority protein IDs: Gorai.004G006300.2.A;Gorai.004G006300.1.A
Gene IDs: Gorai.004G006300
cotton name: K07904 - Ras-related protein Rab-11A (RAB11A) (1 of 34)
OG IDs: 93524686
Best-hit-arabi-ID: AT1G07410.1
Best-hit-arabi-name: RAB GTPase homolog A2B
Gorai.004G006300.1.A
PCC=0.39, p=0.099
mRNA Profile
Protein Expression Profile
DPA

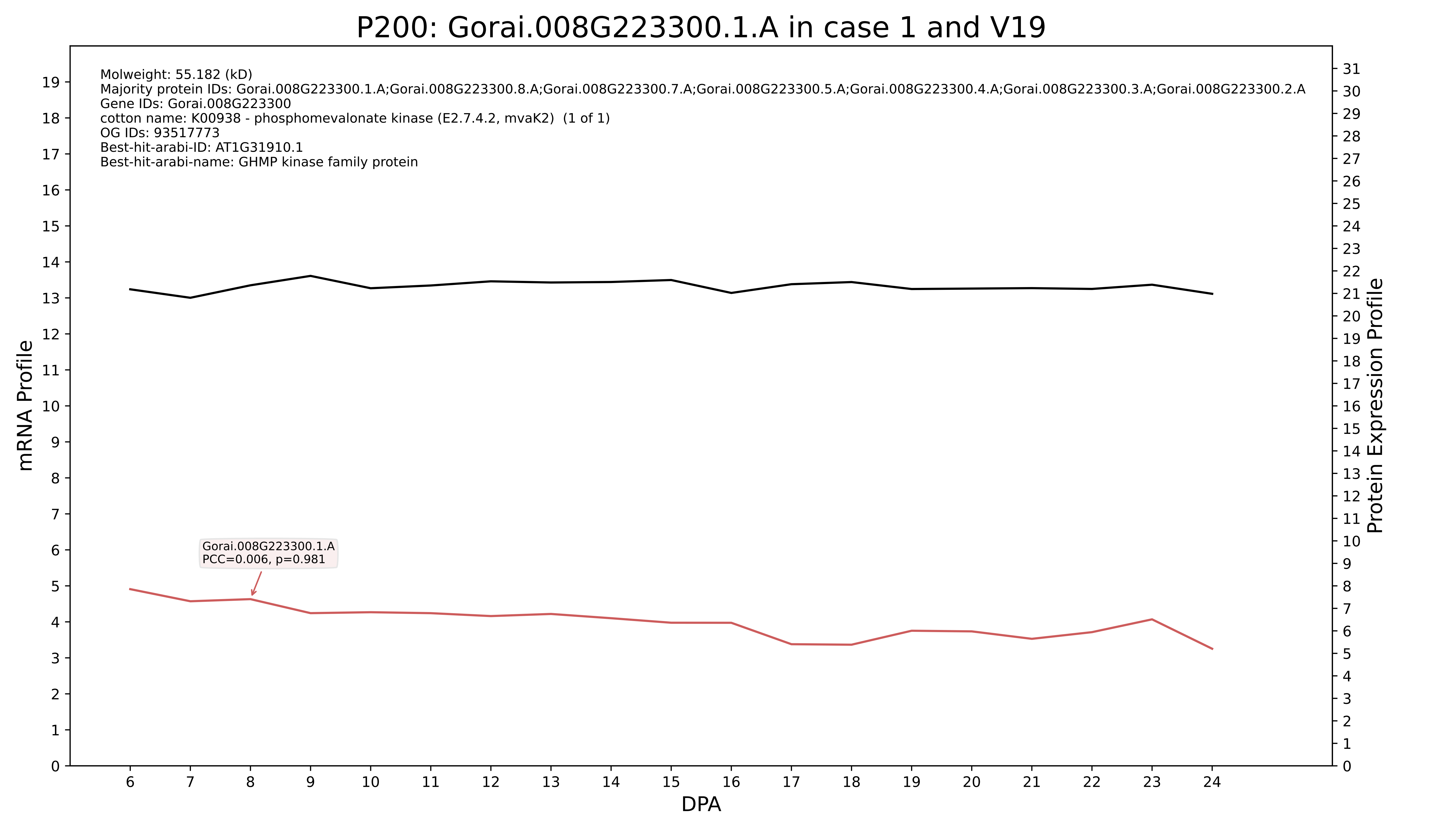

P200: Gorai.008G223300.1.A in case 1 and V19
Molweight: 55.182 (kD)
Majority protein IDs: Gorai.008G223300.1.A;Gorai.008G223300.8.A;Gorai.008G223300.7.A;Gorai.008G223300.5.A;Gorai.008G223300.4.A;Gorai.008G223300.3.A;Gorai.008G223300.2.A
Gene IDs: Gorai.008G223300
cotton name: K00938 - phosphomevalonate kinase (E2.7.4.2, mvaK2) (1 of 1)
OG IDs: 93517773
Best-hit-arabi-ID: AT1G31910.1
Best-hit-arabi-name: GHMP kinase family protein
Gorai.008G223300.1.A
PCC=0.006, p=0.981
mRNA Profile
Protein Expression Profile
DPA

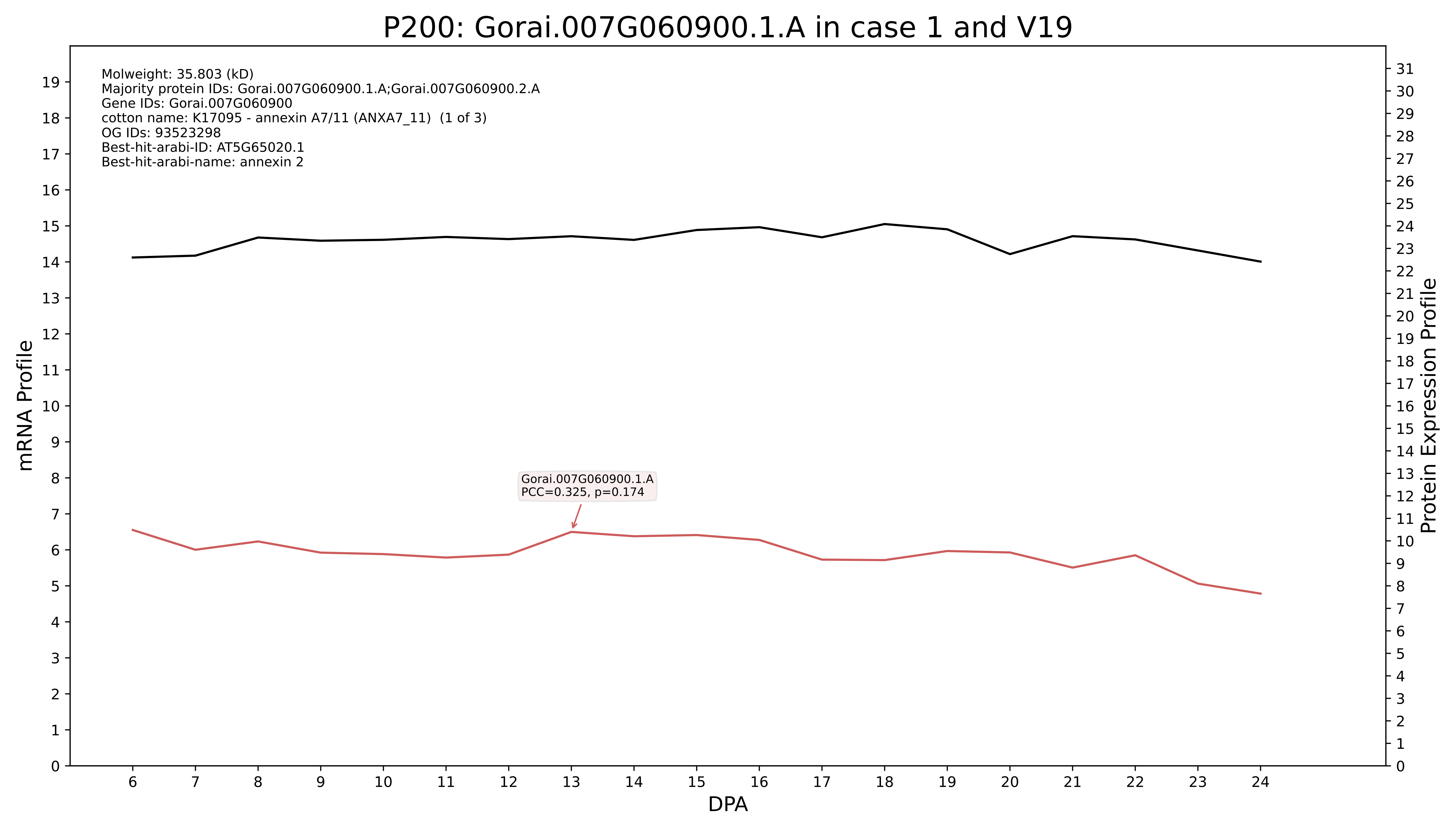
P200: Gorai.007G060900.1.A in case 1 and V19
Molweight: 35.803 (kD)
Majority protein IDs: Gorai.007G060900.1.A;Gorai.007G060900.2.A
Gene IDs: Gorai.007G060900
cotton name: K17095 - annexin A7/11 (ANXA7_11) (1 of 3)
OG IDs: 93523298
Best-hit-arabi-ID: AT5G65020.1
Best-hit-arabi-name: annexin 2
Gorai.007G060900.1.A
PCC=0.325, p=0.174
mRNA Profile
Protein Expression Profile
DPA

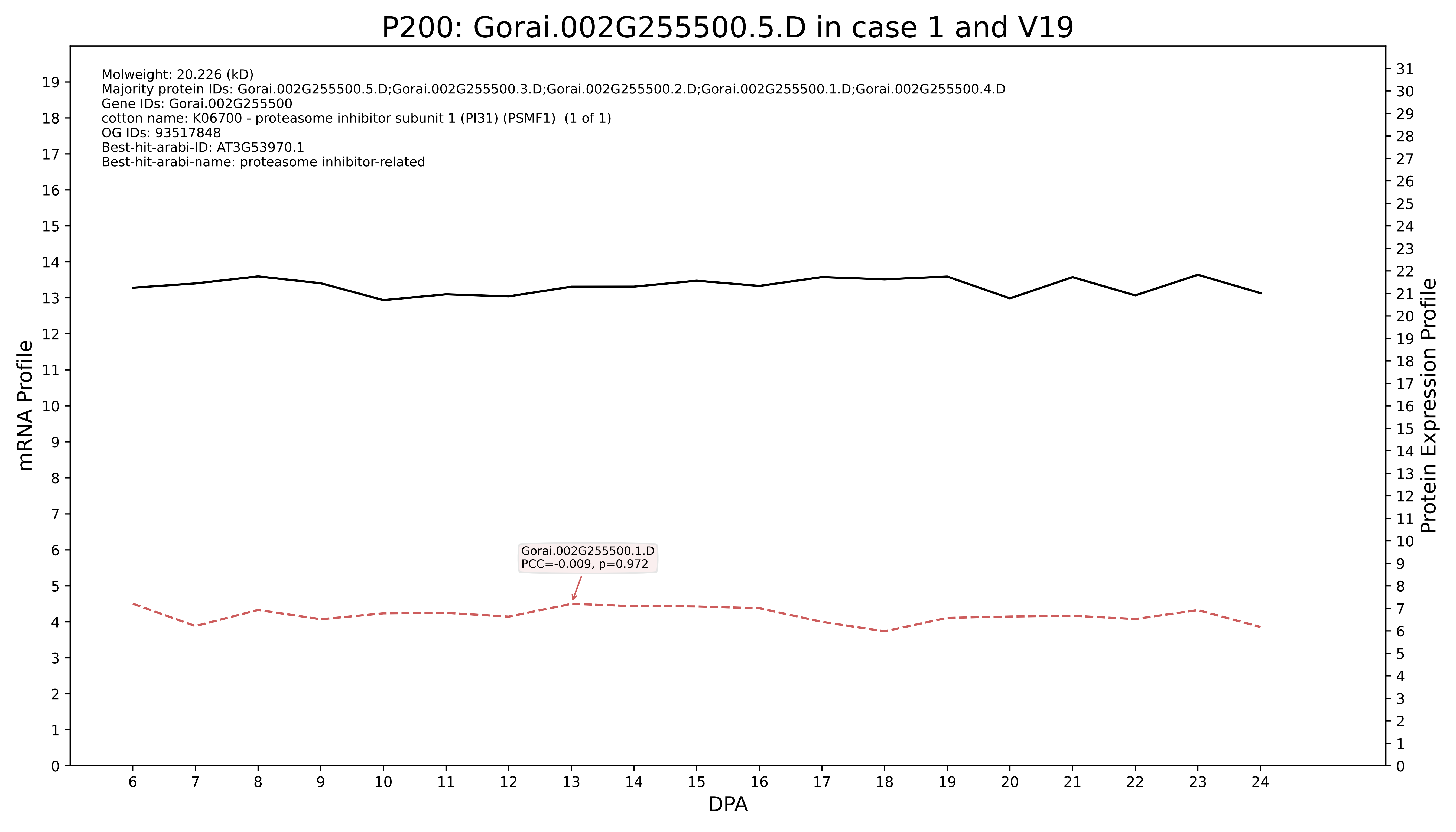
P200: Gorai.002G255500.5.D in case 1 and V19
Molweight: 20.226 (kD)
Majority protein IDs: Gorai.002G255500.5.D;Gorai.002G255500.3.D;Gorai.002G255500.2.D;Gorai.002G255500.1.D;Gorai.002G255500.4.D
Gene IDs: Gorai.002G255500
cotton name: K06700 - proteasome inhibitor subunit 1 (PI31) (PSMF1) (1 of 1)
OG IDs: 93517848
Best-hit-arabi-ID: AT3G53970.1
Best-hit-arabi-name: proteasome inhibitor-related
Gorai.002G255500.1.D
PCC=-0.009, p=0.972
mRNA Profile
Protein Expression Profile
DPA

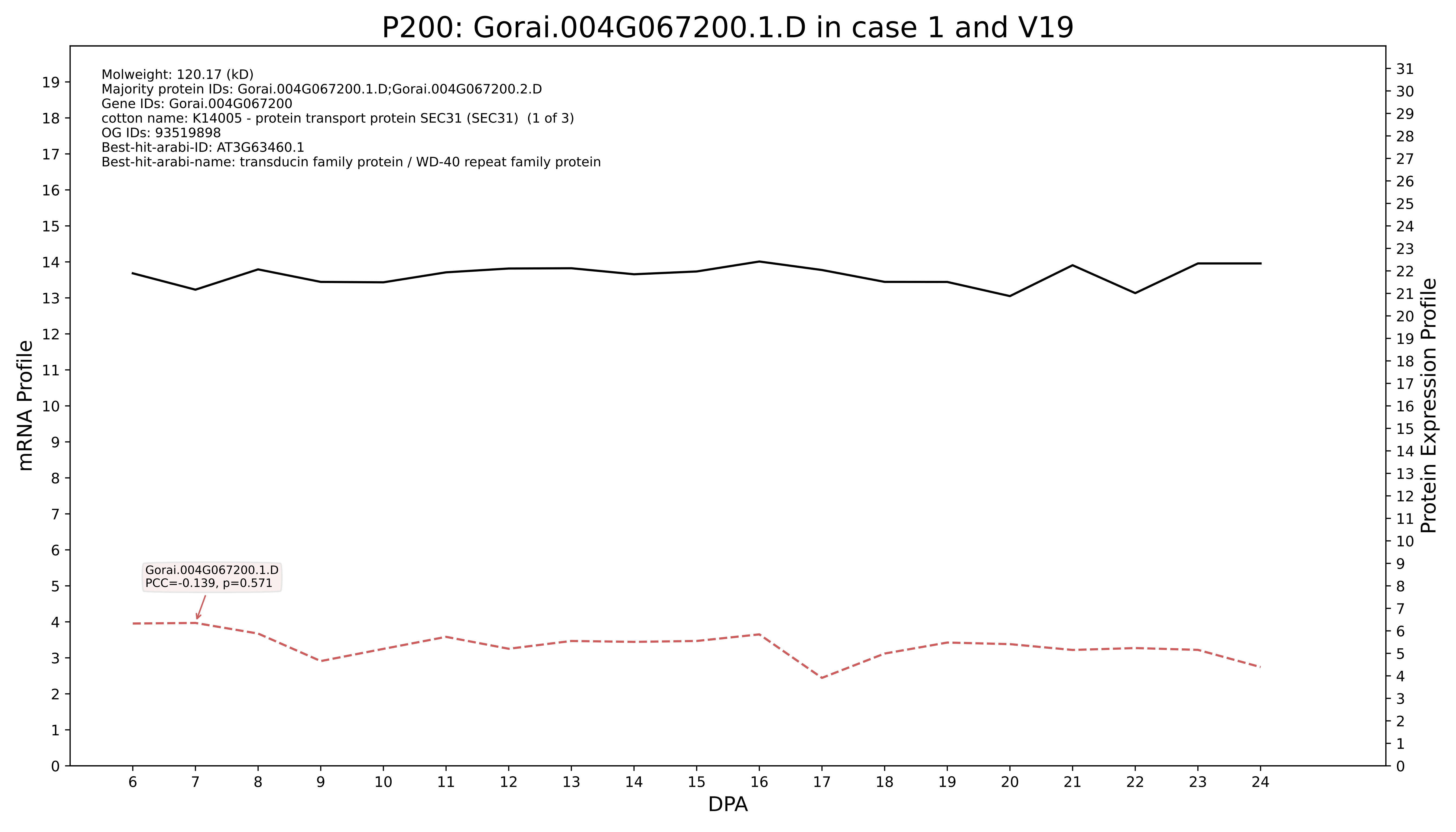
P200: Gorai.004G067200.1.D in case 1 and V19
Molweight: 120.17 (kD)
Majority protein IDs: Gorai.004G067200.1.D;Gorai.004G067200.2.D
Gene IDs: Gorai.004G067200
cotton name: K14005 - protein transport protein SEC31 (SEC31) (1 of 3)
OG IDs: 93519898
Best-hit-arabi-ID: AT3G63460.1
Best-hit-arabi-name: transducin family protein / WD-40 repeat family protein
Gorai.004G067200.1.D
PCC=-0.139, p=0.571
mRNA Profile
Protein Expression Profile
DPA

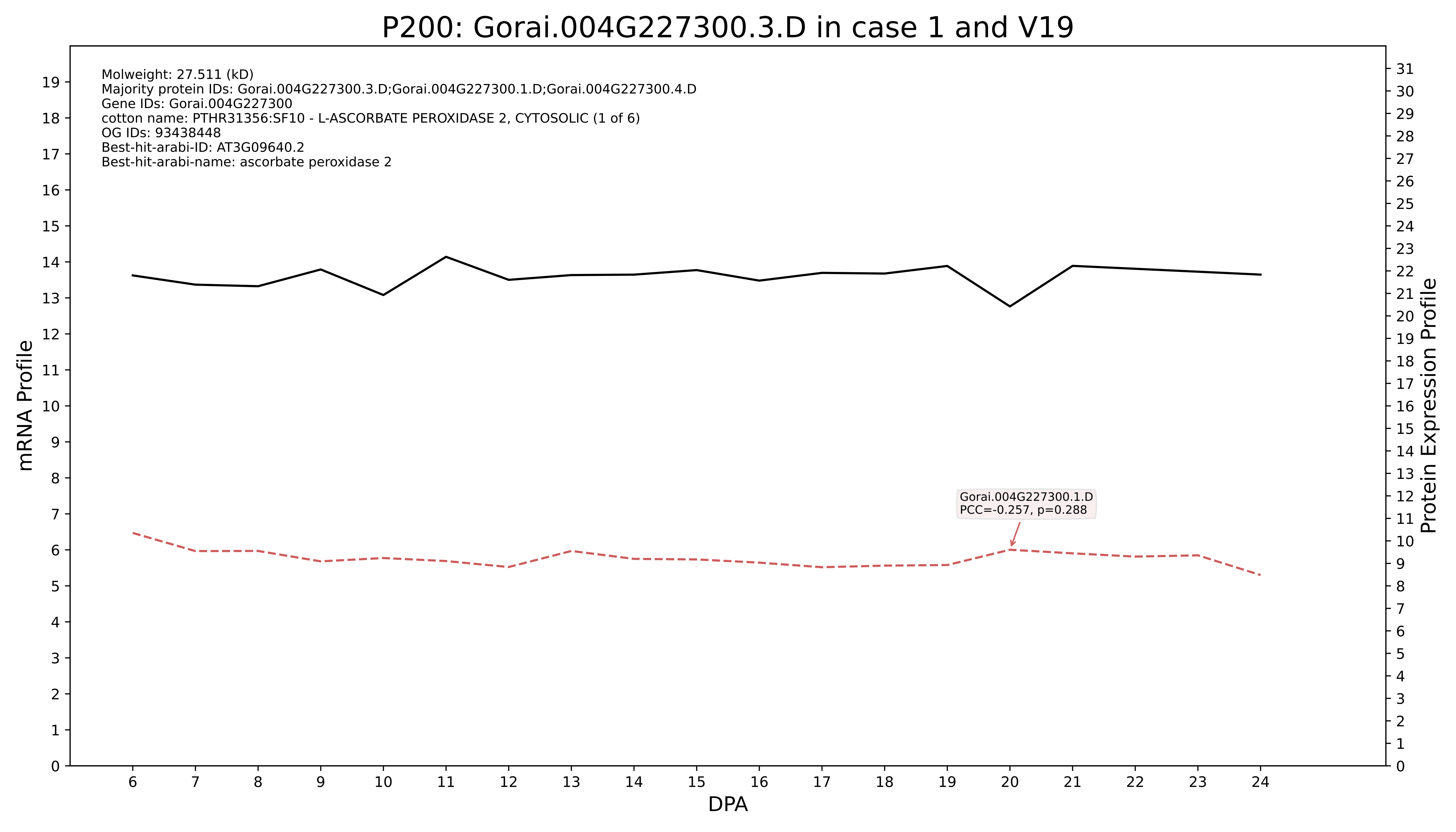
P200: Gorai.004G227300.3.D in case 1 and V19
Molweight: 27.511 (kD)
Majority protein IDs: Gorai.004G227300.3.D;Gorai.004G227300.1.D;Gorai.004G227300.4.D
Gene IDs: Gorai.004G227300
cotton name: PTHR31356:SF10 - L-ASCORBATE PEROXIDASE 2, CYTOSOLIC (1 of 6)
OG IDs: 93438448
Best-hit-arabi-ID: AT3G09640.2
Best-hit-arabi-name: ascorbate peroxidase 2
Gorai.004G227300.1.D
PCC=-0.257, p=0.288
mRNA Profile
Protein Expression Profile
DPA

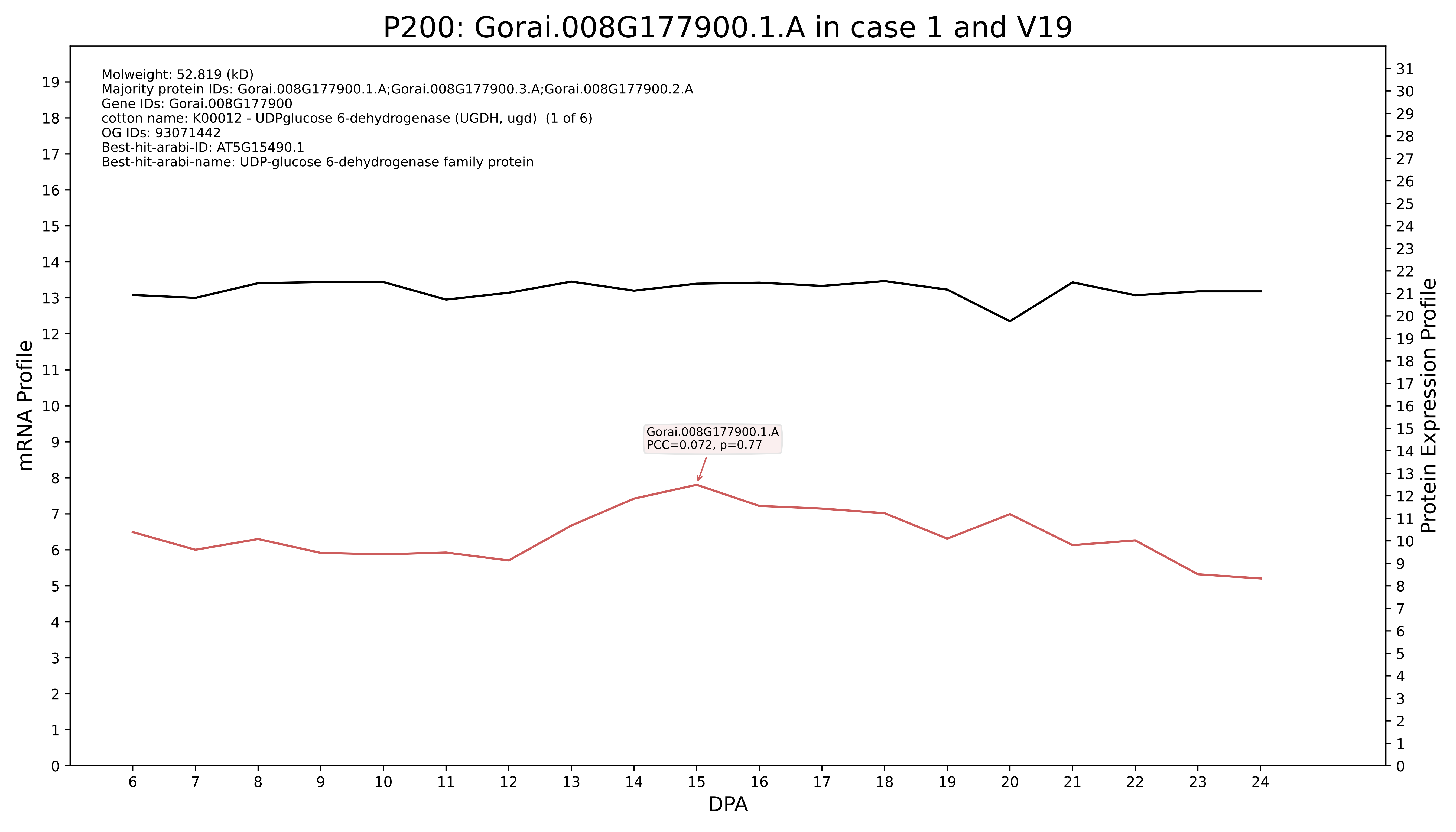
P200: Gorai.008G177900.1.A in case 1 and V19
Molweight: 52.819 (kD)
Majority protein IDs: Gorai.008G177900.1.A;Gorai.008G177900.3.A;Gorai.008G177900.2.A
Gene IDs: Gorai.008G177900
cotton name: K00012 - UDPglucose 6-dehydrogenase (UGDH, ugd) (1 of 6)
OG IDs: 93071442
Best-hit-arabi-ID: AT5G15490.1
Best-hit-arabi-name: UDP-glucose 6-dehydrogenase family protein
Gorai.008G177900.1.A
PCC=0.072, p=0.77
mRNA Profile
Protein Expression Profile
DPA

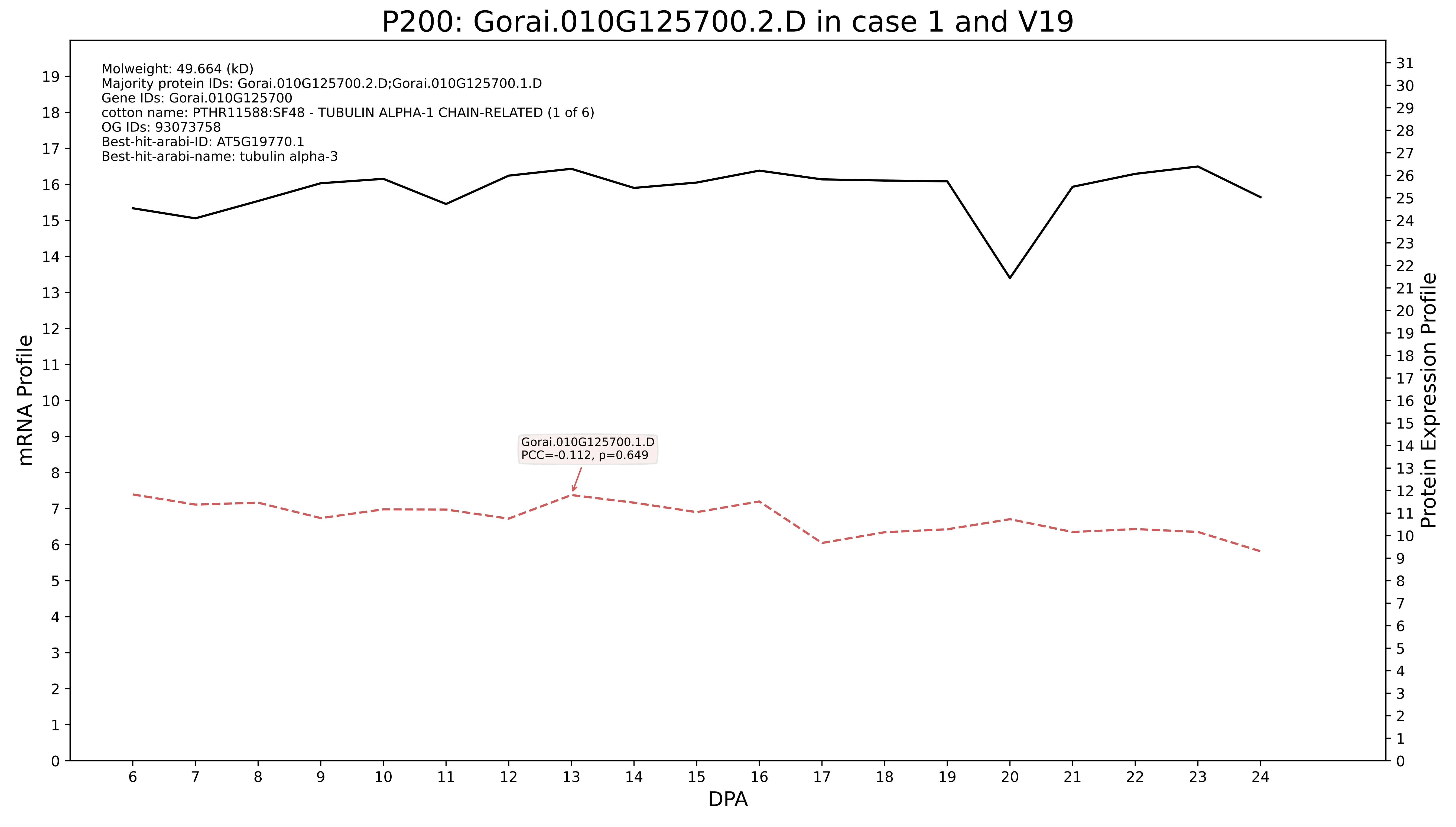
P200: Gorai.010G125700.2.D in case 1 and V19
Molweight: 49.664 (kD)
Majority protein IDs: Gorai.010G125700.2.D;Gorai.010G125700.1.D
Gene IDs: Gorai.010G125700
cotton name: PTHR11588:SF48 - TUBULIN ALPHA-1 CHAIN-RELATED (1 of 6)
OG IDs: 93073758
Best-hit-arabi-ID: AT5G19770.1
Best-hit-arabi-name: tubulin alpha-3
Gorai.010G125700.1.D
PCC=-0.112, p=0.649
mRNA Profile
Protein Expression Profile
DPA

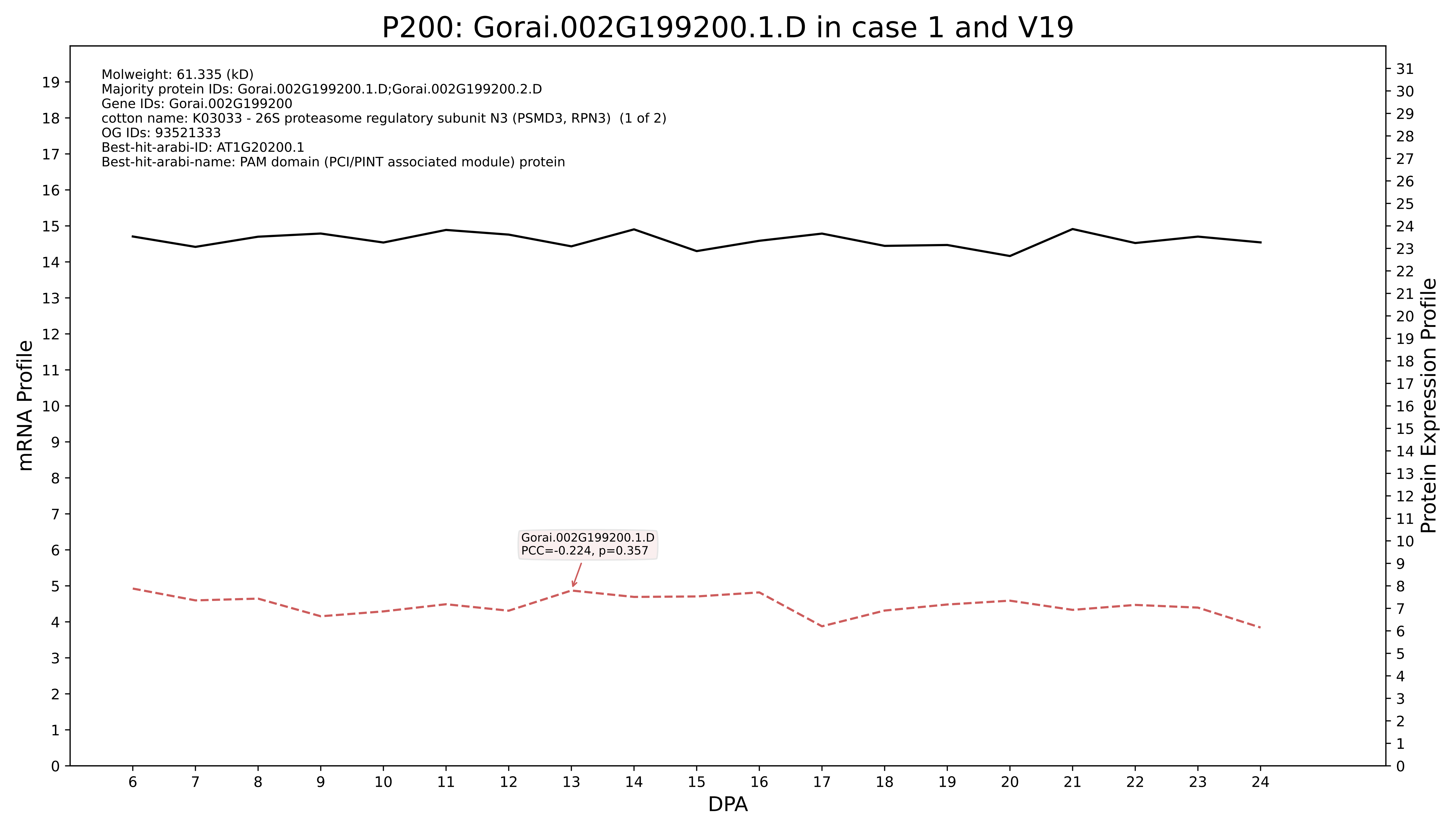
P200: Gorai.002G199200.1.D in case 1 and V19
Molweight: 61.335 (kD)
Majority protein IDs: Gorai.002G199200.1.D;Gorai.002G199200.2.D
Gene IDs: Gorai.002G199200
cotton name: K03033 - 26S proteasome regulatory subunit N3 (PSMD3, RPN3) (1 of 2)
OG IDs: 93521333
Best-hit-arabi-ID: AT1G20200.1
Best-hit-arabi-name: PAM domain (PCI/PINT associated module) protein
Gorai.002G199200.1.D
PCC=-0.224, p=0.357
mRNA Profile
Protein Expression Profile
DPA

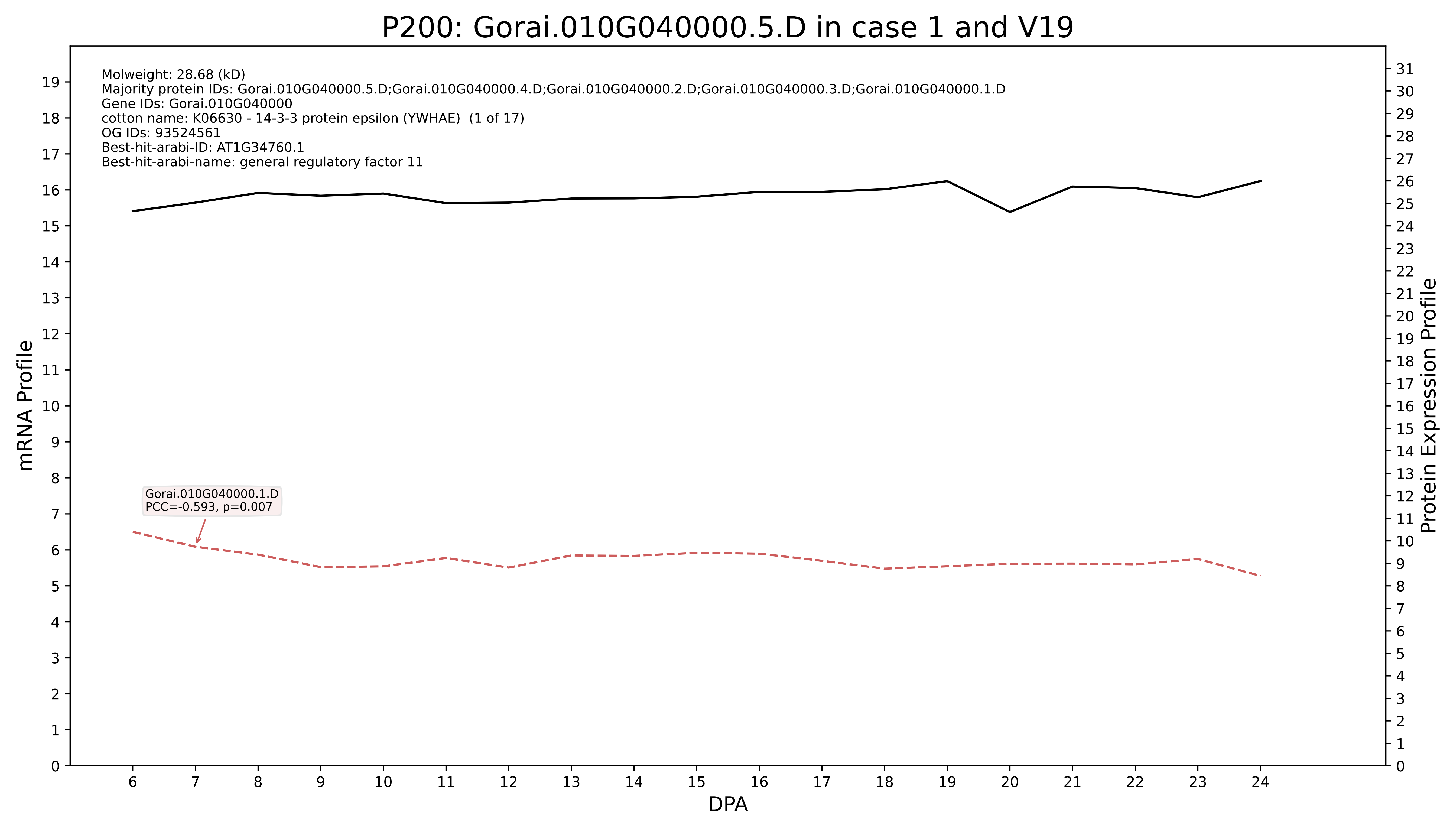
P200: Gorai.010G040000.5.D in case 1 and V19
Molweight: 28.68 (kD)
Majority protein IDs: Gorai.010G040000.5.D;Gorai.010G040000.4.D;Gorai.010G040000.2.D;Gorai.010G040000.3.D;Gorai.010G040000.1.D
Gene IDs: Gorai.010G040000
cotton name: K06630 - 14-3-3 protein epsilon (YWHAE) (1 of 17)
OG IDs: 93524561
Best-hit-arabi-ID: AT1G34760.1
Best-hit-arabi-name: general regulatory factor 11
Gorai.010G040000.1.D
PCC=-0.593, p=0.007
mRNA Profile
Protein Expression Profile
DPA

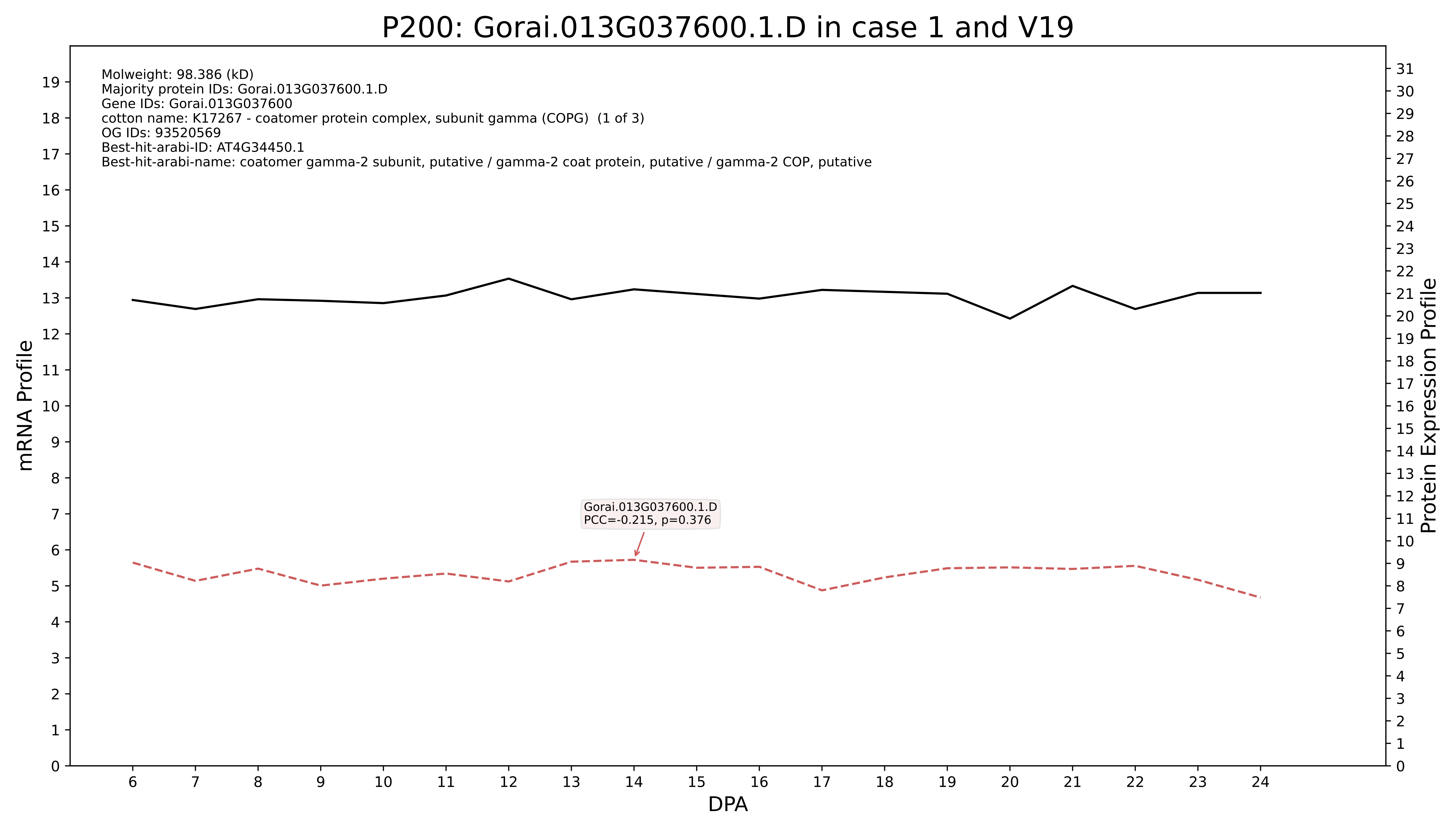

P200: Gorai.013G037600.1.D in case 1 and V19
Molweight: 98.386 (kD)
Majority protein IDs: Gorai.013G037600.1.D
Gene IDs: Gorai.013G037600
cotton name: K17267 - coatomer protein complex, subunit gamma (COPG) (1 of 3)
OG IDs: 93520569
Best-hit-arabi-ID: AT4G34450.1
Best-hit-arabi-name: coatomer gamma-2 subunit, putative / gamma-2 coat protein, putative / gamma-2 COP, putative
Gorai.013G037600.1.D
PCC=-0.215, p=0.376
mRNA Profile
Protein Expression Profile
DPA

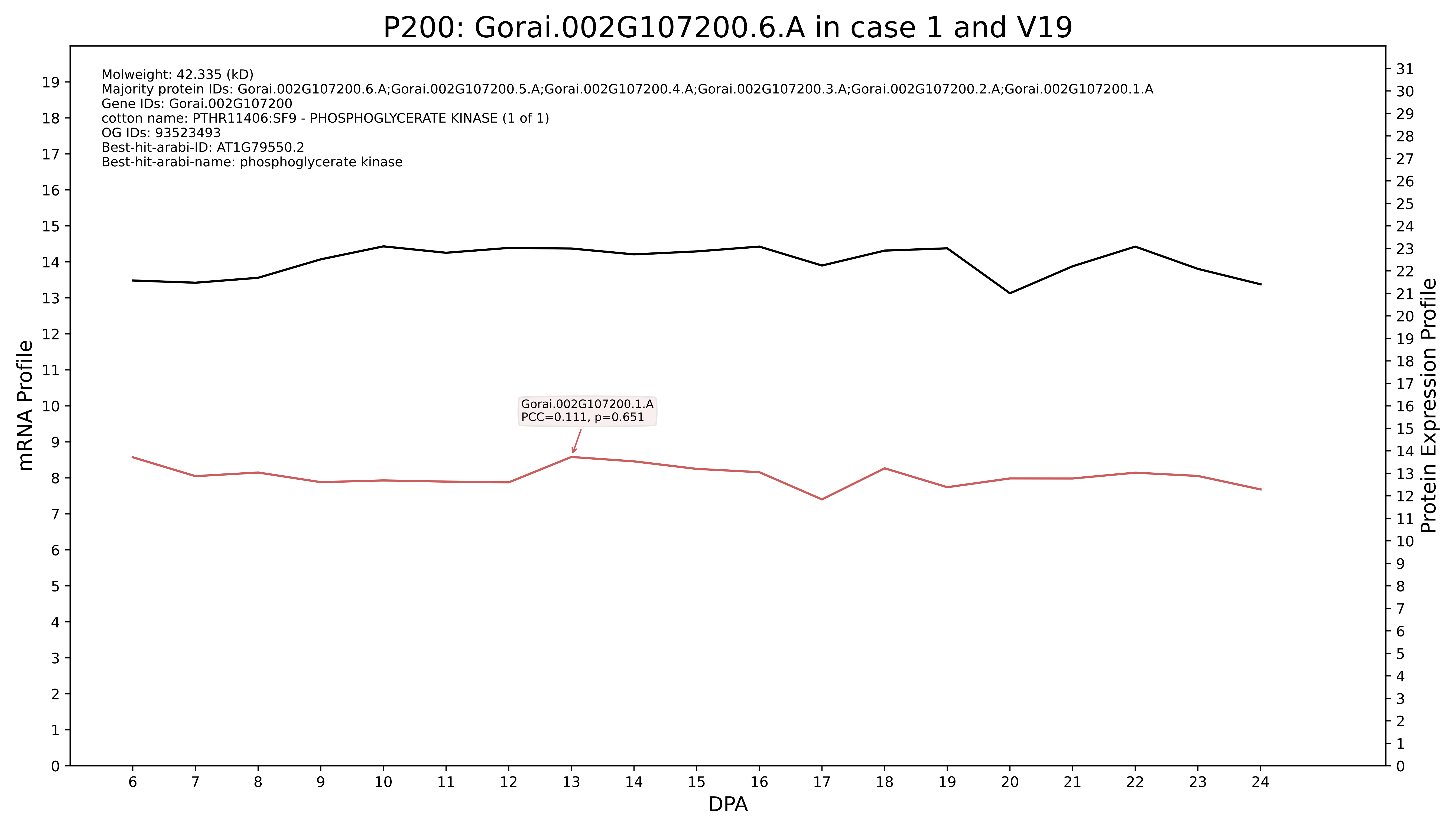
P200: Gorai.002G107200.6.A in case 1 and V19
Molweight: 42.335 (kD)
Majority protein IDs: Gorai.002G107200.6.A;Gorai.002G107200.5.A;Gorai.002G107200.4.A;Gorai.002G107200.3.A;Gorai.002G107200.2.A;Gorai.002G107200.1.A
Gene IDs: Gorai.002G107200
cotton name: PTHR11406:SF9 - PHOSPHOGLYCERATE KINASE (1 of 1)
OG IDs: 93523493
Best-hit-arabi-ID: AT1G79550.2
Best-hit-arabi-name: phosphoglycerate kinase
Gorai.002G107200.1.A
PCC=0.111, p=0.651
mRNA Profile
Protein Expression Profile
DPA

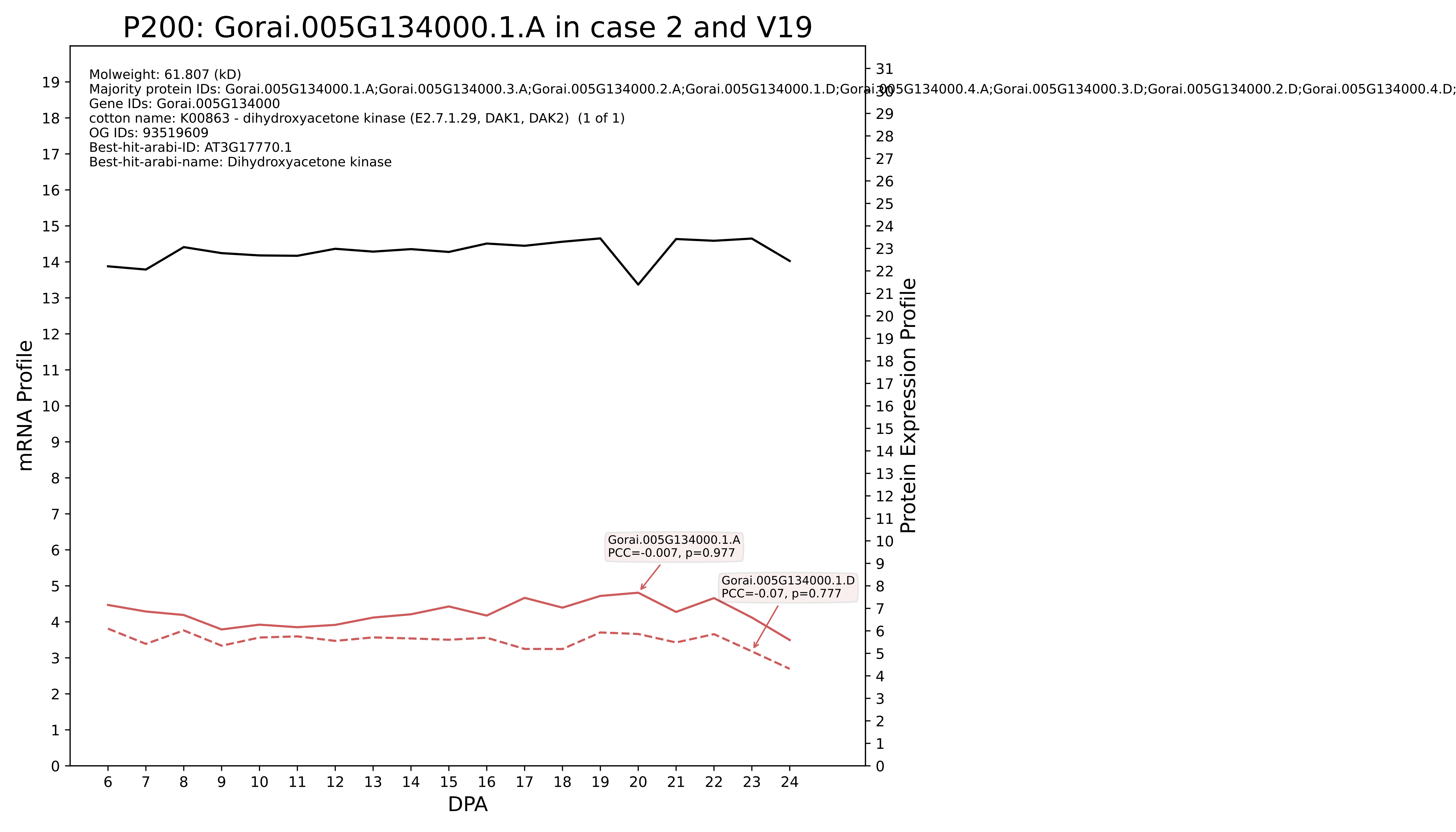
P200: Gorai.005G134000.1.A in case 2 and V19
Molweight: 61.807 (kD)
Majority protein IDs: Gorai.005G134000.1.A;Gorai.005G134000.3.A;Gorai.005G134000.2.A;Gorai.005G134000.1.D;Gorai.005G134000.4.A;Gorai.005G134000.3.D;Gorai.005G134000.2.D;Gorai.005G134000.4.D;
Gene IDs: Gorai.005G134000
cotton name: K00863 - dihydroxyacetone kinase (E2.7.1.29, DAK1, DAK2) (1 of 1)
OG IDs: 93519609
Best-hit-arabi-ID: AT3G17770.1
Best-hit-arabi-name: Dihydroxyacetone kinase
Gorai.005G134000.1.A
PCC=-0.007, p=0.977
Gorai.005G134000.1.D
PCC=-0.07, p=0.777
mRNA Profile
Protein Expression Profile
DPA

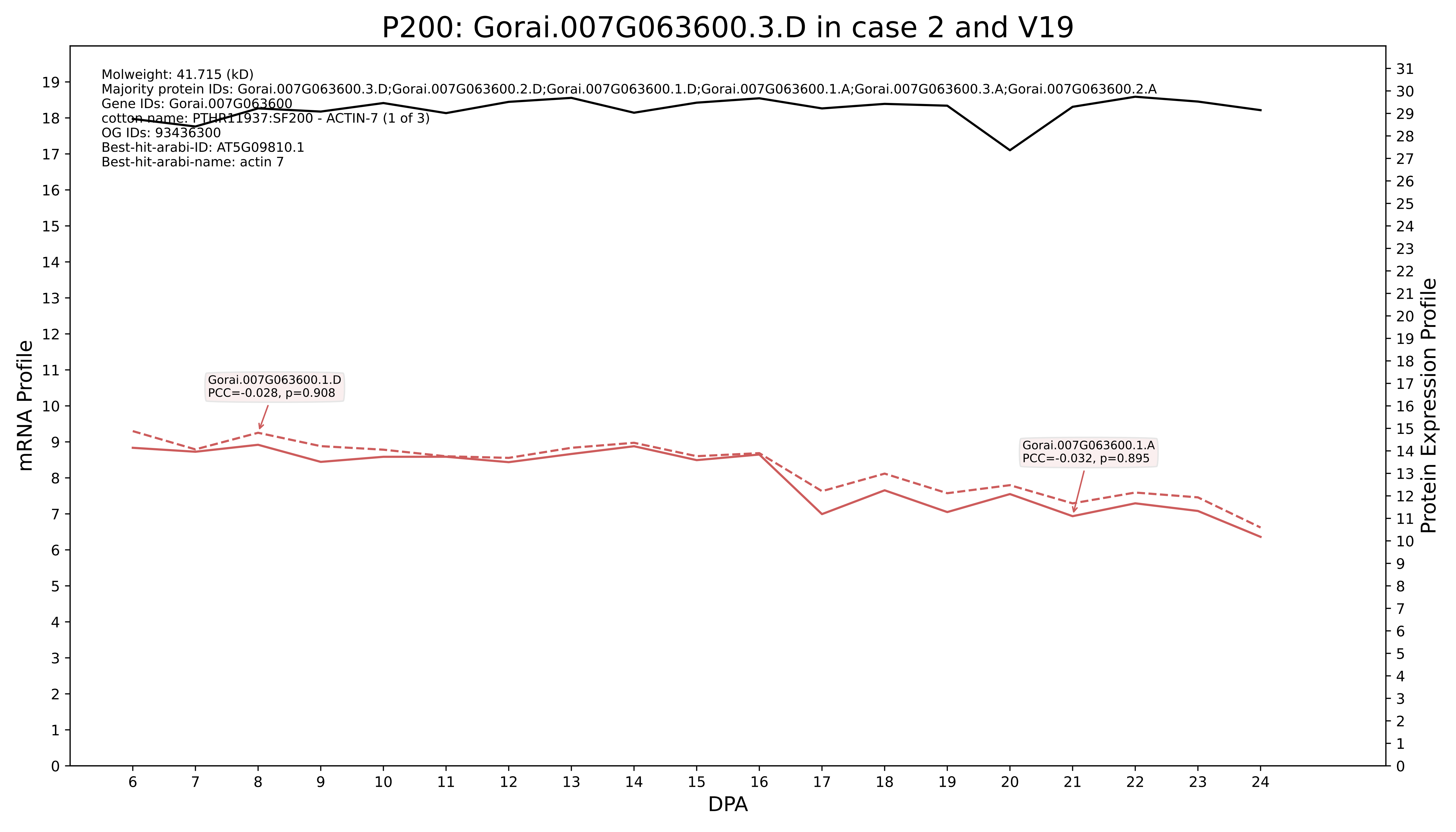
P200: Gorai.007G063600.3.D in case 2 and V19
Molweight: 41.715 (kD)
Majority protein IDs: Gorai.007G063600.3.D;Gorai.007G063600.2.D;Gorai.007G063600.1.D;Gorai.007G063600.1.A;Gorai.007G063600.3.A;Gorai.007G063600.2.A
Gene IDs: Gorai.007G063600
cotton-name: PTHR11937:SF200 - ACTIN-7 (1 of 3)
OG IDs: 93436300
Best-hit-arabi-ID: AT5G09810.1
Best-hit-arabi-name: actin 7
Gorai.007G063600.1.D
PCC=-0.028, p=0.908
Gorai.007G063600.1.A
PCC=-0.032, p=0.895
mRNA Profile
Protein Expression Profile
DPA

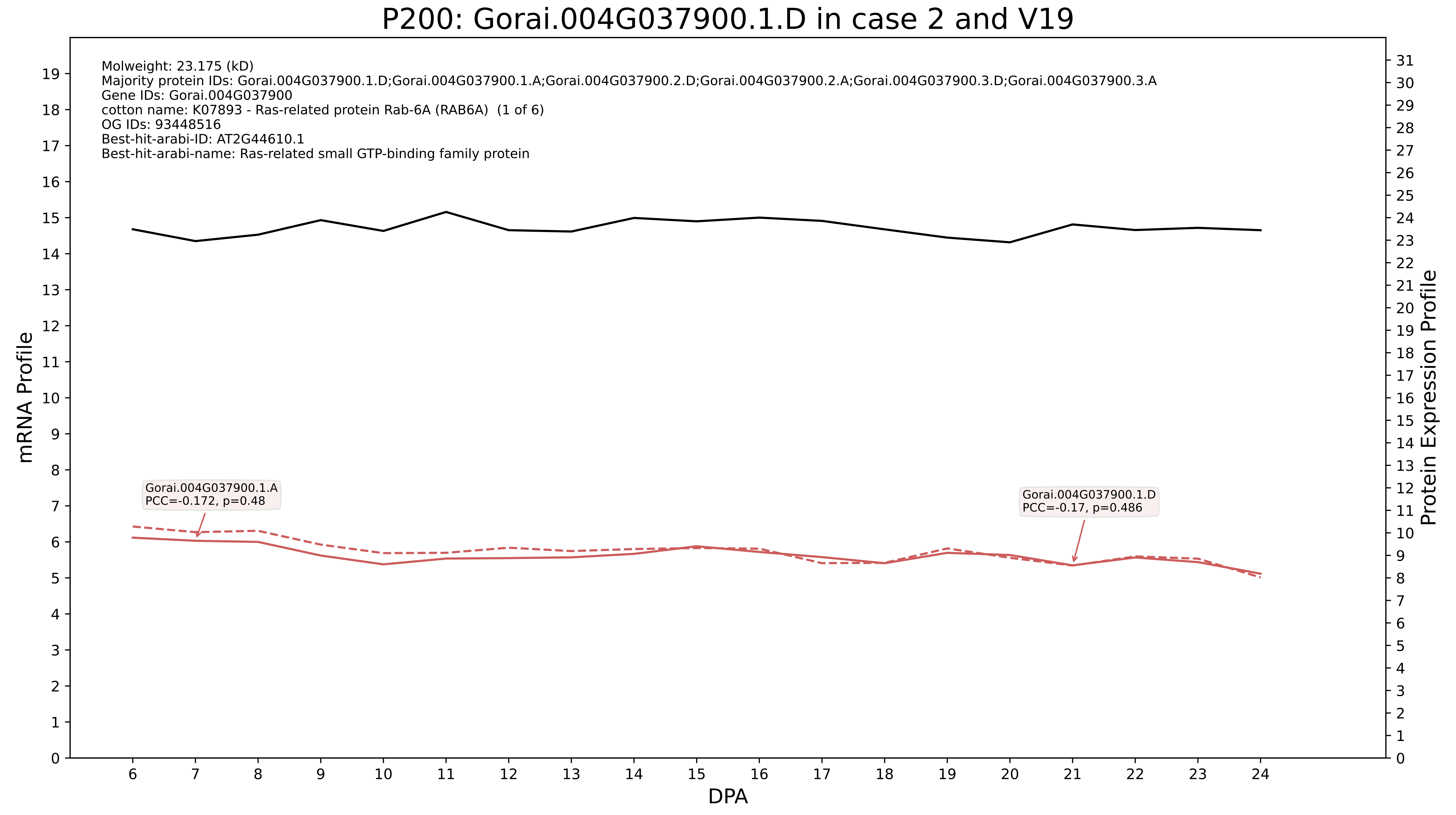
P200: Gorai.004G037900.1.D in case 2 and V19
Molweight: 23.175 (kD)
Majority protein IDs: Gorai.004G037900.1.D;Gorai.004G037900.1.A;Gorai.004G037900.2.D;Gorai.004G037900.2.A;Gorai.004G037900.3.D;Gorai.004G037900.3.A
Gene IDs: Gorai.004G037900
cotton name: K07893 - Ras-related protein Rab-6A (RAB6A) (1 of 6)
OG IDs: 93448516
Best-hit-arabi-ID: AT2G44610.1
Best-hit-arabi-name: Ras-related small GTP-binding family protein
Gorai.004G037900.1.A
PCC=-0.172, p=0.48
Gorai.004G037900.1.D
PCC=-0.17, p=0.486
mRNA Profile
Protein Expression Profile
DPA

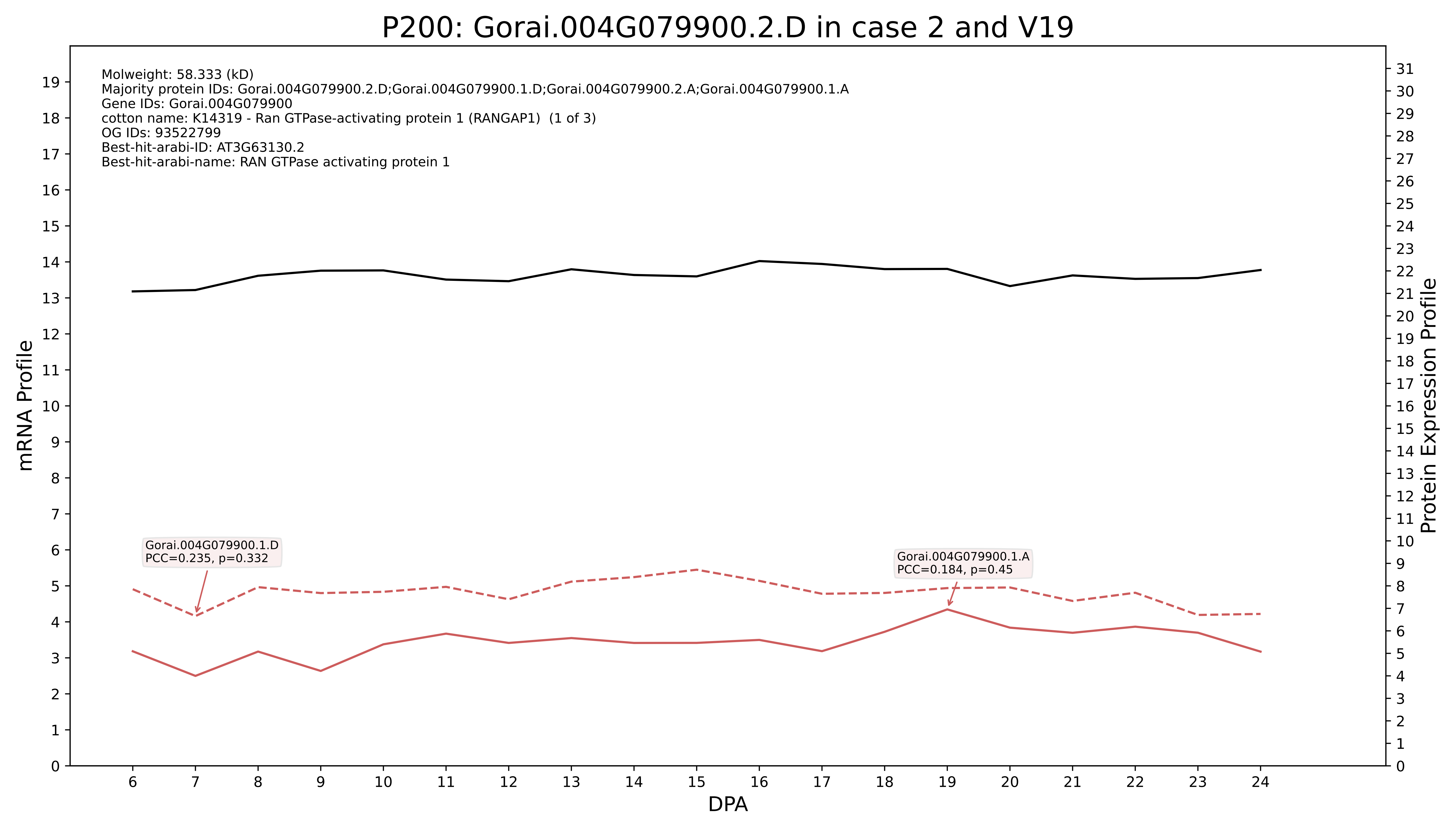

P200: Gorai.004G079900.2.D in case 2 and V19
Molweight: 58.333 (kD)
Majority protein IDs: Gorai.004G079900.2.D;Gorai.004G079900.1.D;Gorai.004G079900.2.A;Gorai.004G079900.1.A
Gene IDs: Gorai.004G079900
cotton name: K14319 - Ran GTPase-activating protein 1 (RANGAP1) (1 of 3)
OG IDs: 93522799
Best-hit-arabi-ID: AT3G63130.2
Best-hit-arabi-name: RAN GTPase activating protein 1
Gorai.004G079900.1.D
PCC=0.235, p=0.332
Gorai.004G079900.1.A
PCC=0.184, p=0.45
mRNA Profile
Protein Expression Profile
DPA

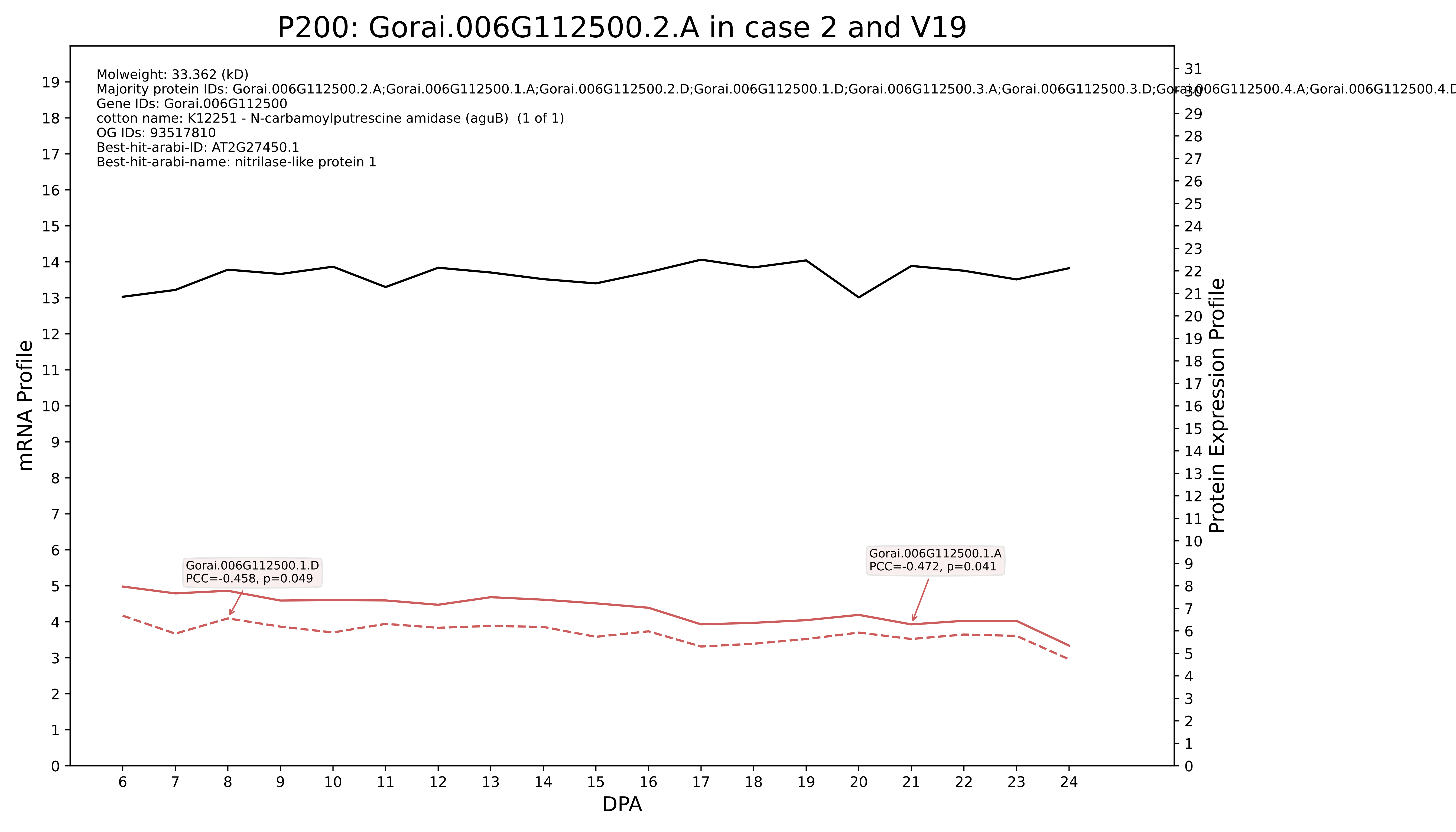
P200: Gorai.006G112500.2.A in case 2 and V19
Molweight: 33.362 (kD)
Majority protein IDs: Gorai.006G112500.2.A;Gorai.006G112500.1.A;Gorai.006G112500.2.D;Gorai.006G112500.1.D;Gorai.006G112500.3.A;Gorai.006G112500.3.D;Gorai.006G112500.4.A;Gorai.006G112500.4.D
Gene IDs: Gorai.006G112500
cotton name: K12251 - N-carbamoylputrescine amidase (aguB) (1 of 1)
OG IDs: 93517810
Best-hit-arabi-ID: AT2G27450.1
Best-hit-arabi-name: nitrilase-like protein 1
Gorai.006G112500.1.D
PCC=-0.458, p=0.049
Gorai.006G112500.1.A
PCC=-0.472, p=0.041
mRNA Profile
Protein Expression Profile
DPA

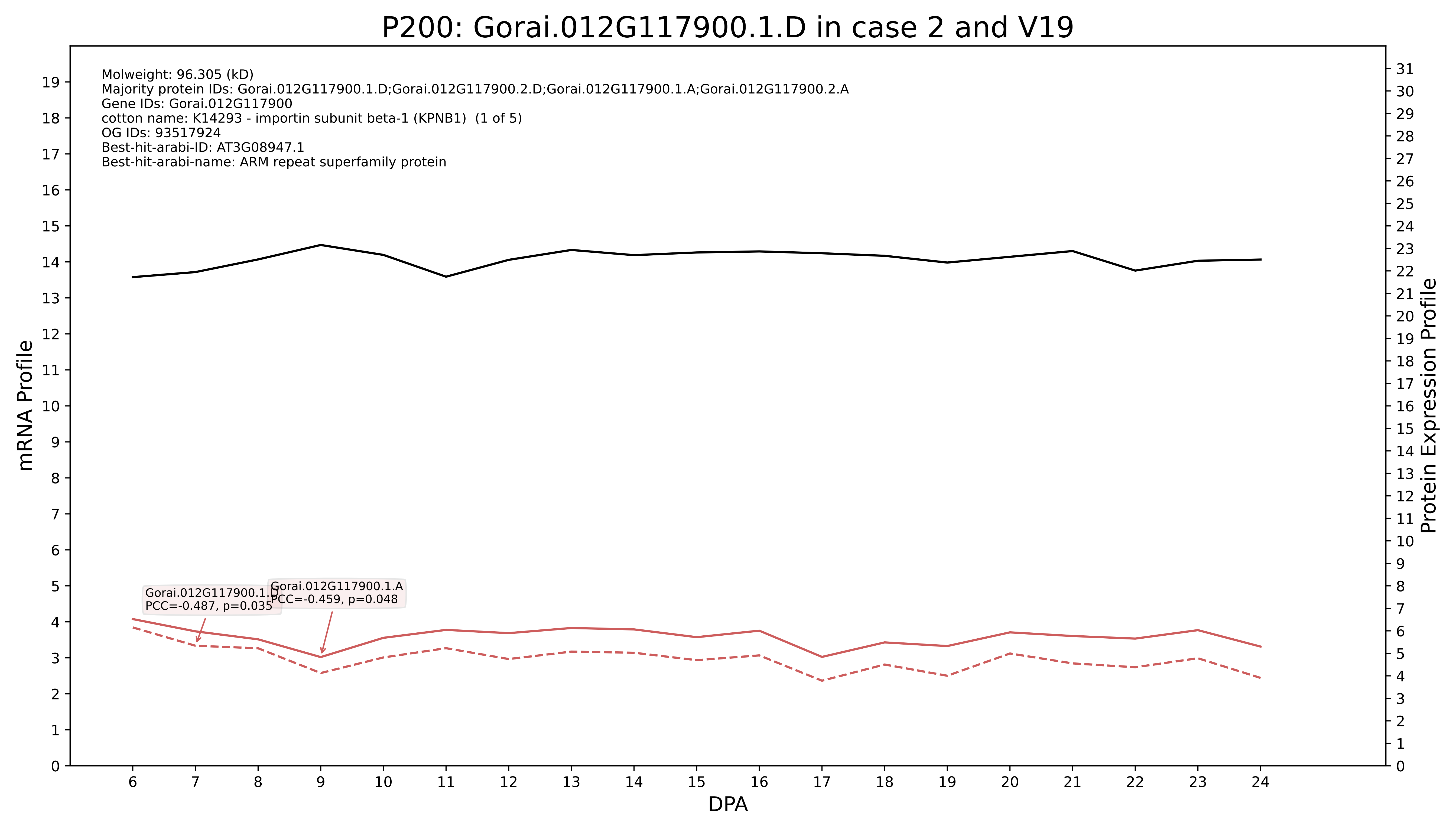
P200: Gorai.012G117900.1.D in case 2 and V19
Molweight: 96.305 (kD)
Majority protein IDs: Gorai.012G117900.1.D;Gorai.012G117900.2.D;Gorai.012G117900.1.A;Gorai.012G117900.2.A
Gene IDs: Gorai.012G117900
cotton name: K14293 - importin subunit beta-1 (KPNB1) (1 of 5)
OG IDs: 93517924
Best-hit-arabi-ID: AT3G08947.1
Best-hit-arabi-name: ARM repeat superfamily protein
Gorai.012G117900.1.D
PCC=-0.487, p=0.035
Gorai.012G117900.1.A
PCC=-0.459, p=0.048
mRNA Profile
Protein Expression Profile
DPA

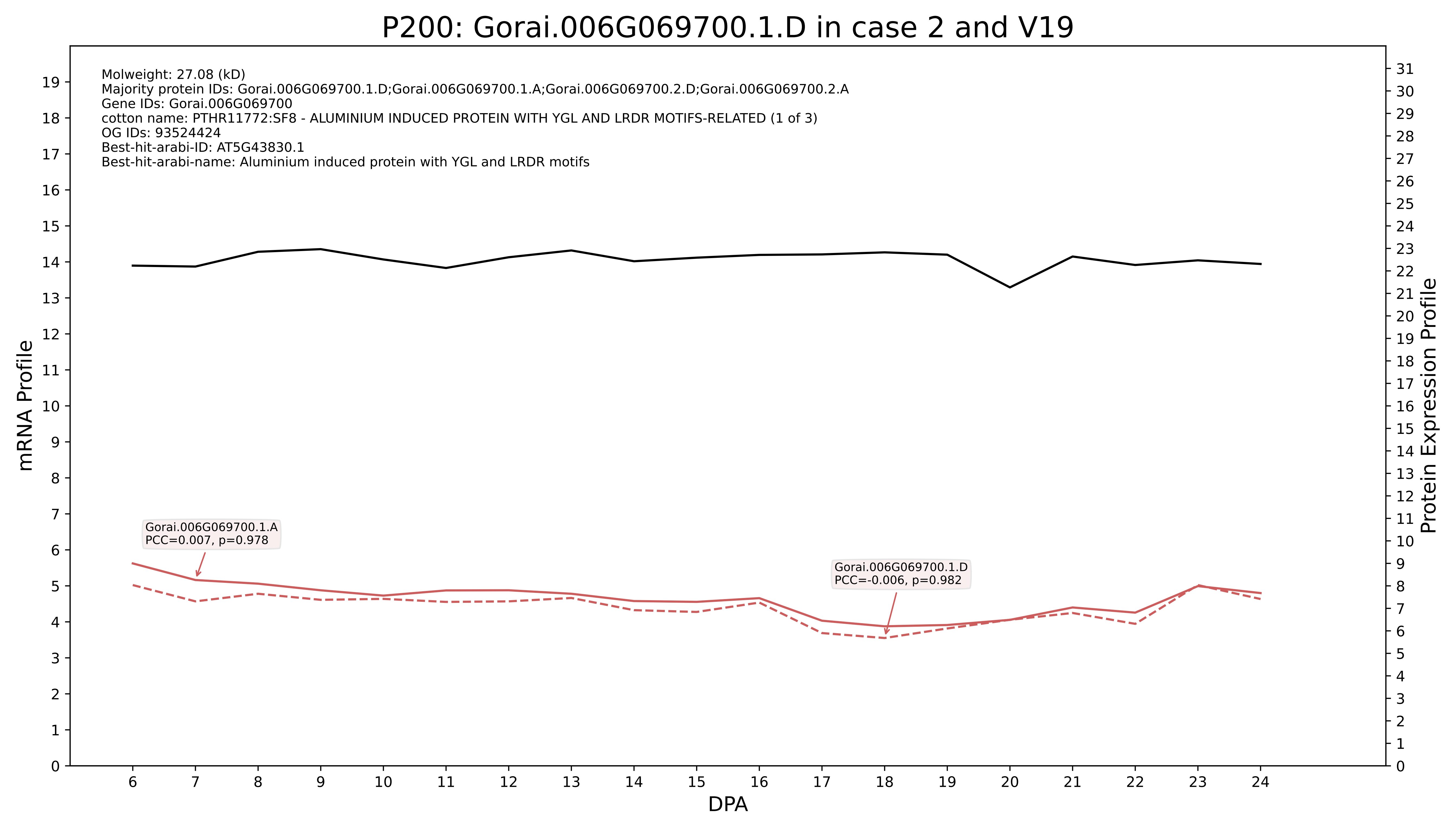

P200: Gorai.006G069700.1.D in case 2 and V19
Molweight: 27.08 (kD)
Majority protein IDs: Gorai.006G069700.1.D;Gorai.006G069700.1.A;Gorai.006G069700.2.D;Gorai.006G069700.2.A
Gene IDs: Gorai.006G069700
cotton name: PTHR11772:SF8 - ALUMINIUM INDUCED PROTEIN WITH YGL AND LRDR MOTIFS-RELATED (1 of 3)
OG IDs: 93524424
Best-hit-arabi-ID: AT5G43830.1
Best-hit-arabi-name: Aluminium induced protein with YGL and LRDR motifs
Gorai.006G069700.1.A
PCC=0.007, p=0.978
Gorai.006G069700.1.D
PCC=-0.006, p=0.982
mRNA Profile
Protein Expression Profile
DPA

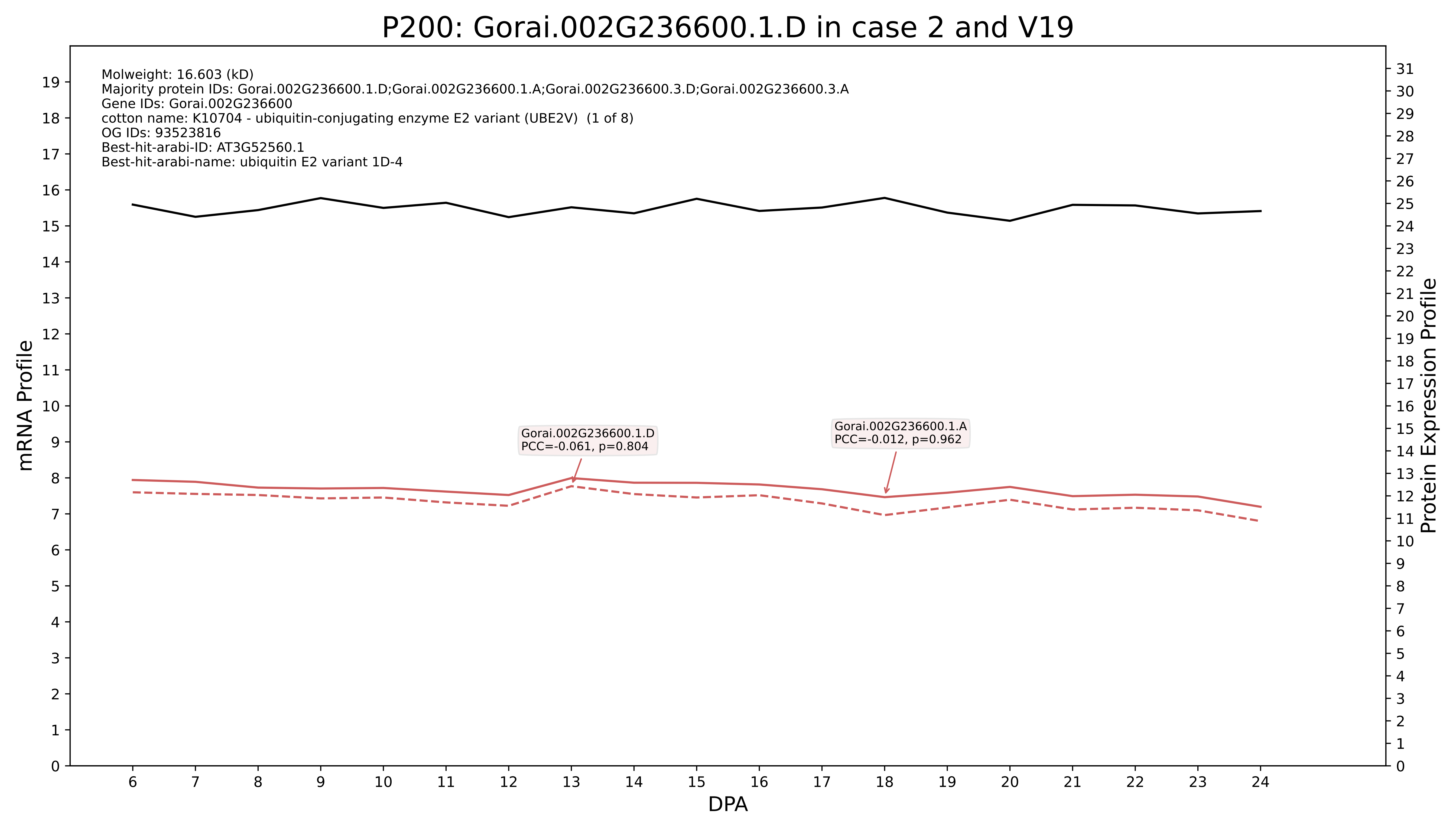

P200: Gorai.002G236600.1.D in case 2 and V19
Molweight: 16.603 (kD)
Majority protein IDs: Gorai.002G236600.1.D;Gorai.002G236600.1.A;Gorai.002G236600.3.D;Gorai.002G236600.3.A
Gene IDs: Gorai.002G236600
cotton name: K10704 - ubiquitin-conjugating enzyme E2 variant (UBE2V) (1 of 8)
OG IDs: 93523816
Best-hit-arabi-ID: AT3G52560.1
Best-hit-arabi-name: ubiquitin E2 variant 1D-4
Gorai.002G236600.1.D
PCC=-0.061, p=0.804
Gorai.002G236600.1.A
PCC=-0.012, p=0.962
mRNA Profile
Protein Expression Profile
DPA

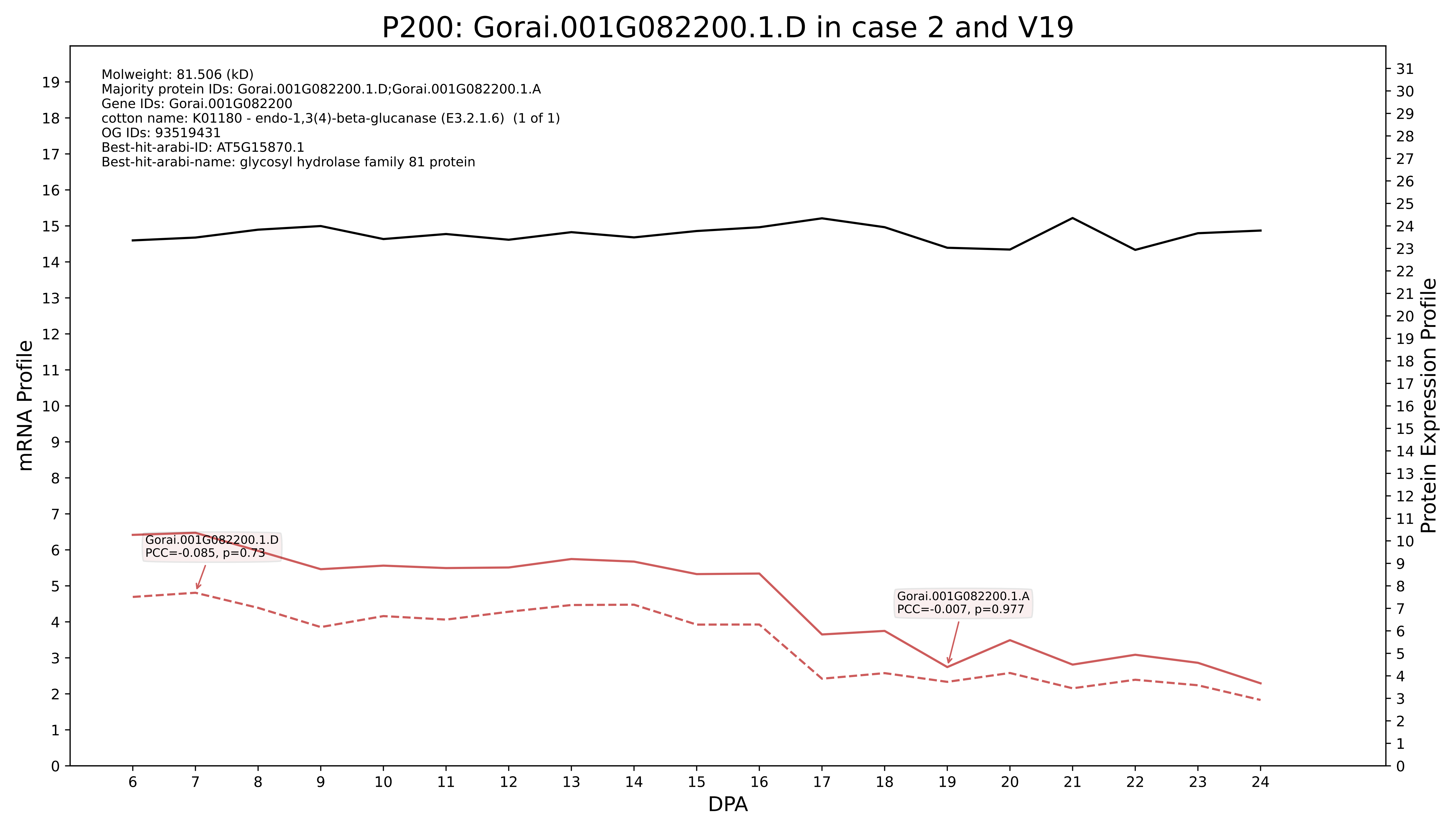
P200: Gorai.001G082200.1.D in case 2 and V19
Molweight: 81.506 (kD)
Majority protein IDs: Gorai.001G082200.1.D;Gorai.001G082200.1.A
Gene IDs: Gorai.001G082200
cotton name: K01180 - endo-1,3(4)-beta-glucanase (E3.2.1.6) (1 of 1)
OG IDs: 93519431
Best-hit-arabi-ID: AT5G15870.1
Best-hit-arabi-name: glycosyl hydrolase family 81 protein
Gorai.001G082200.1.D
PCC=-0.085, p=0.73
Gorai.001G082200.1.A
PCC=-0.007, p=0.977
mRNA Profile
Protein Expression Profile
DPA

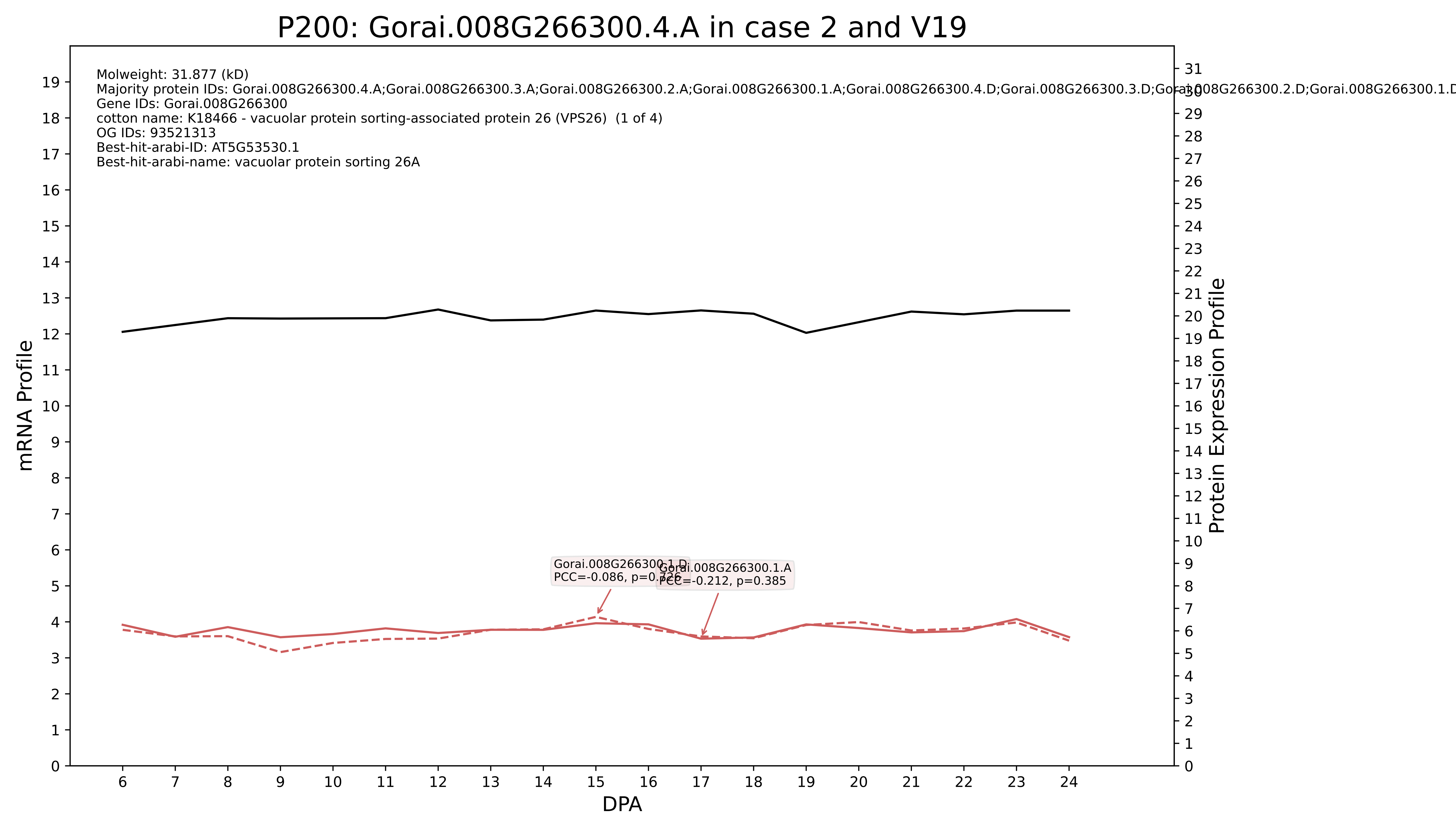

P200: Gorai.008G266300.4.A in case 2 and V19
Molweight: 31.877 (kD)
Majority protein IDs: Gorai.008G266300.4.A;Gorai.008G266300.3.A;Gorai.008G266300.2.A;Gorai.008G266300.1.A;Gorai.008G266300.4.D;Gorai.008G266300.3.D;Gorai.008G266300.2.D;Gorai.008G266300.1.D
Gene IDs: Gorai.008G266300
cotton name: K18466 - vacuolar protein sorting-associated protein 26 (VPS26) (1 of 4)
OG IDs: 93521313
Best-hit-arabi-ID: AT5G53530.1
Best-hit-arabi-name: vacuolar protein sorting 26A
Gorai.008G266300.1.D
PCC=-0.086, p=0.726
Gorai.008G266300.1.A
PCC=-0.212, p=0.385
mRNA Profile
Protein Expression Profile
DPA

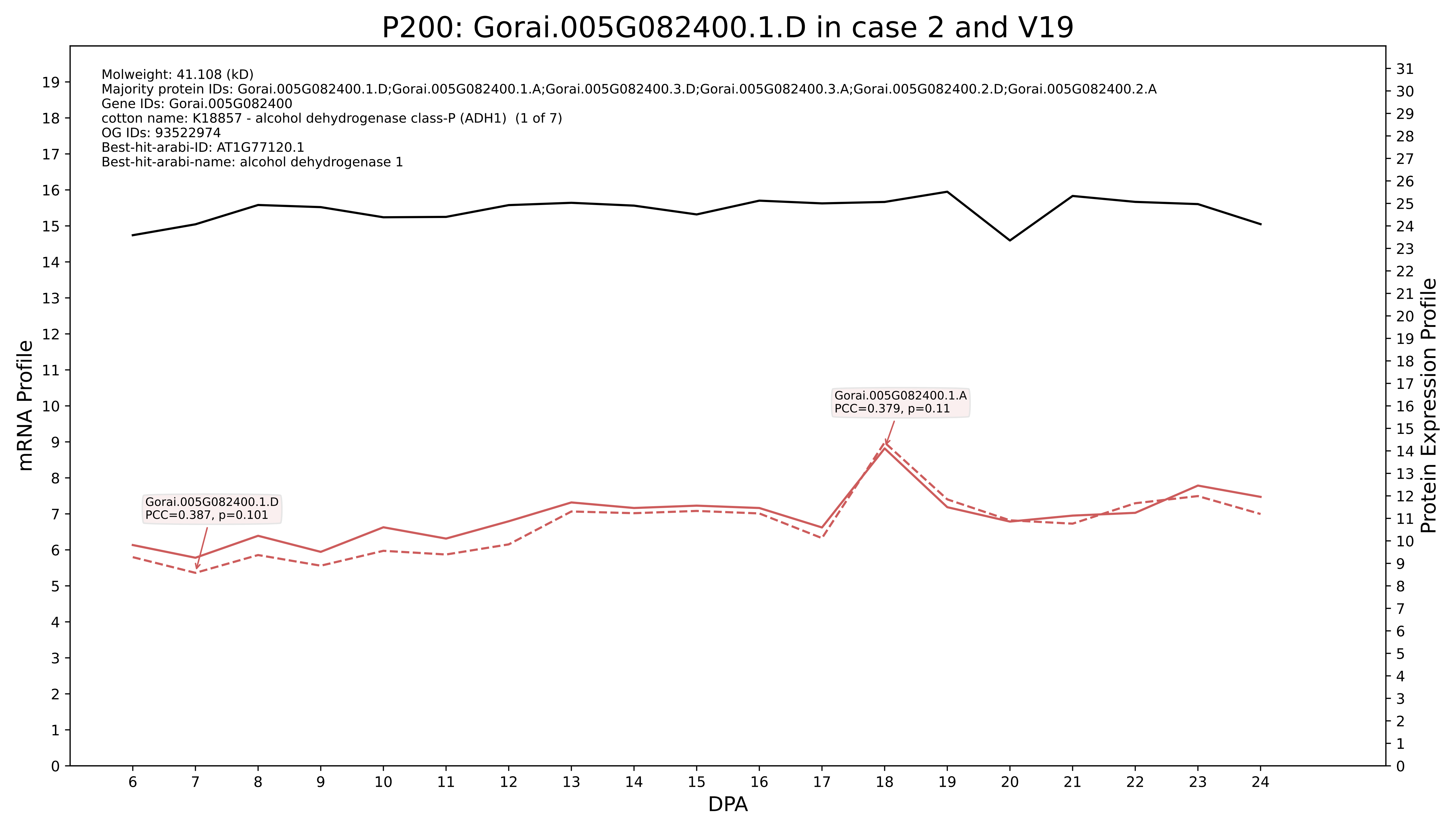
P200: Gorai.005G082400.1.D in case 2 and V19
Molweight: 41.108 (kD)
Majority protein IDs: Gorai.005G082400.1.D;Gorai.005G082400.1.A;Gorai.005G082400.3.D;Gorai.005G082400.3.A;Gorai.005G082400.2.D;Gorai.005G082400.2.A
Gene IDs: Gorai.005G082400
cotton name: K18857 - alcohol dehydrogenase class-P (ADH1) (1 of 7)
OG IDs: 93522974
Best-hit-arabi-ID: AT1G77120.1
Best-hit-arabi-name: alcohol dehydrogenase 1
Gorai.005G082400.1.A
PCC=0.379, p=0.11
Gorai.005G082400.1.D
PCC=0.387, p=0.101
mRNA Profile
Protein Expression Profile
DPA

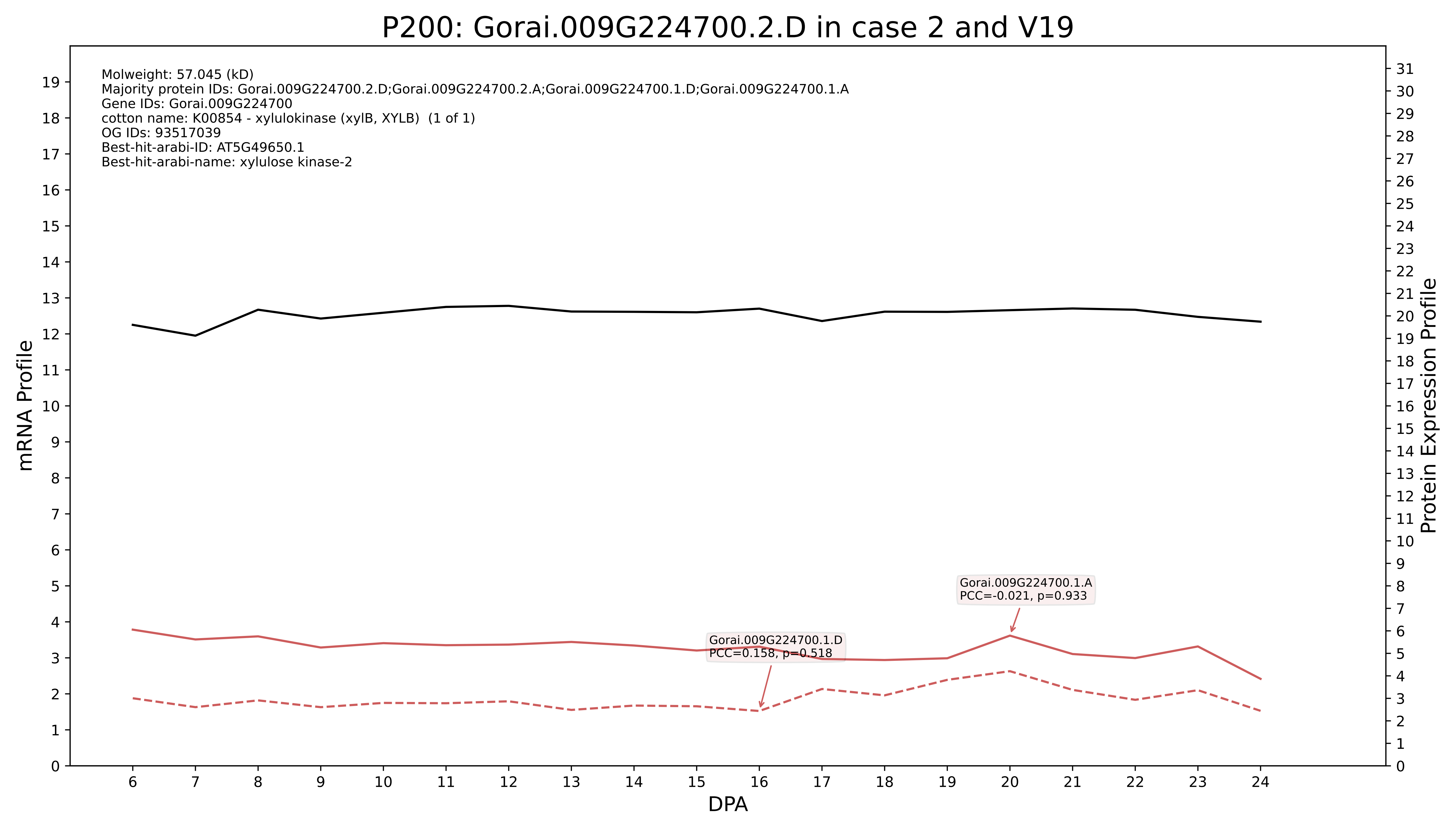
P200: Gorai.009G224700.2.D in case 2 and V19
Molweight: 57.045 (kD)
Majority protein IDs: Gorai.009G224700.2.D;Gorai.009G224700.2.A;Gorai.009G224700.1.D;Gorai.009G224700.1.A
Gene IDs: Gorai.009G224700
cotton name: K00854 - xylulokinase (xylB, XYLB) (1 of 1)
OG IDs: 93517039
Best-hit-arabi-ID: AT5G49650.1
Best-hit-arabi-name: xylulose kinase-2
Gorai.009G224700.1.A
PCC=-0.021, p=0.933
Gorai.009G224700.1.D
PCC=0.158, p=0.518
mRNA Profile
Protein Expression Profile
DPA

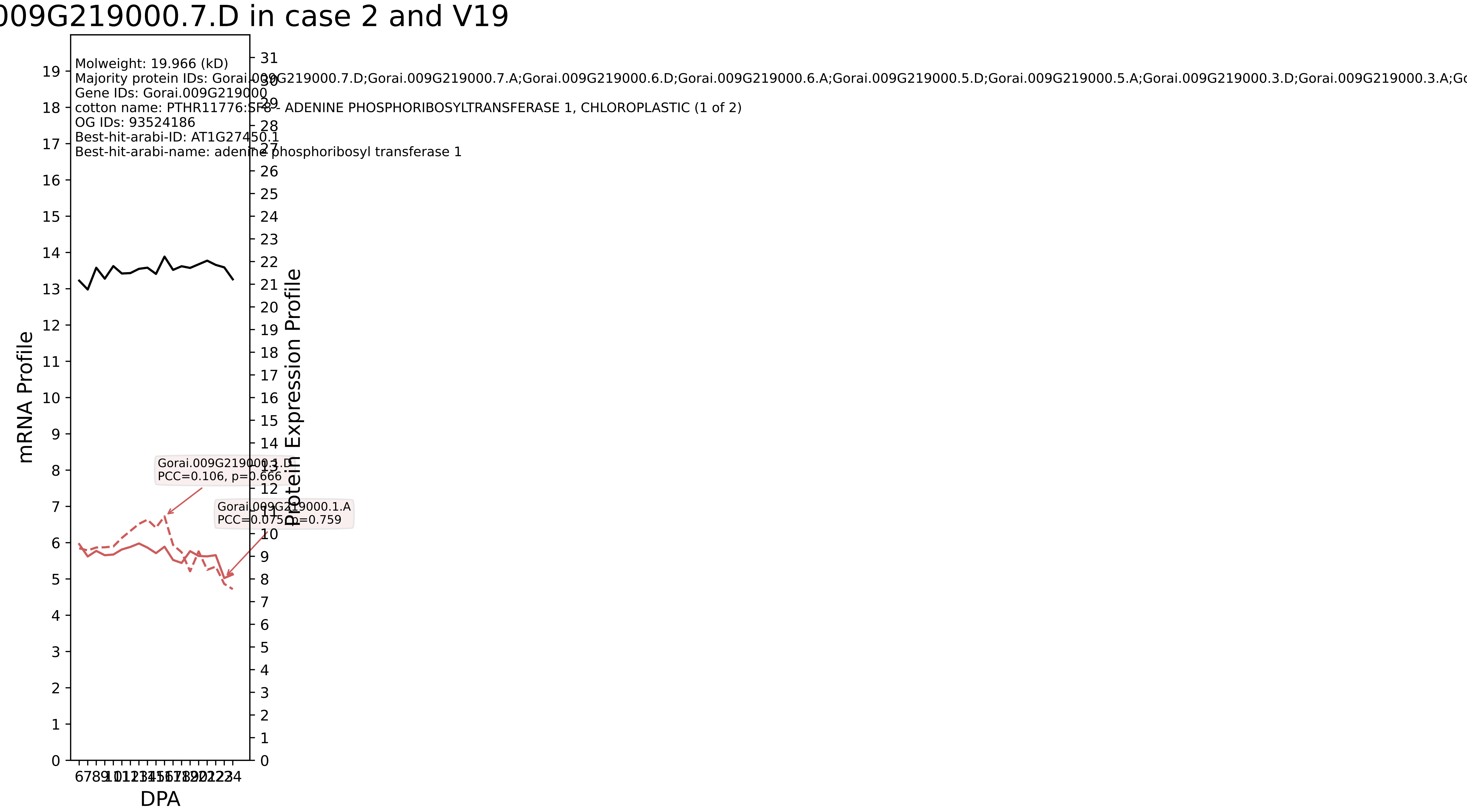
009G219000.7.D in case 2 and V19
Molweight: 19.966 (kD)
Majority protein IDs: Gorai.009G219000.7.D;Gorai.009G219000.7.A;Gorai.009G219000.6.D;Gorai.009G219000.6.A;Gorai.009G219000.5.D;Gorai.009G219000.5.A;Gorai.009G219000.3.D;Gorai.009G219000.3.A;Go
Gene IDs: Gorai.009G219000
cotton name: PTHR11776:SF8 - ADENINE PHOSPHORIBOSYLTRANSFERASE 1, CHLOROPLASTIC (1 of 2)
OG IDs: 93524186
Best-hit-arabi-ID: AT1G27450.1
Best-hit-arabi-name: adenine phosphoribosyl transferase 1
Gorai.009G219000.1.D
PCC=0.106, p=0.666
Gorai.009G219000.1.A
PCC=0.075, p=0.759
mRNA Profile
Protein Expression Profile
DPA

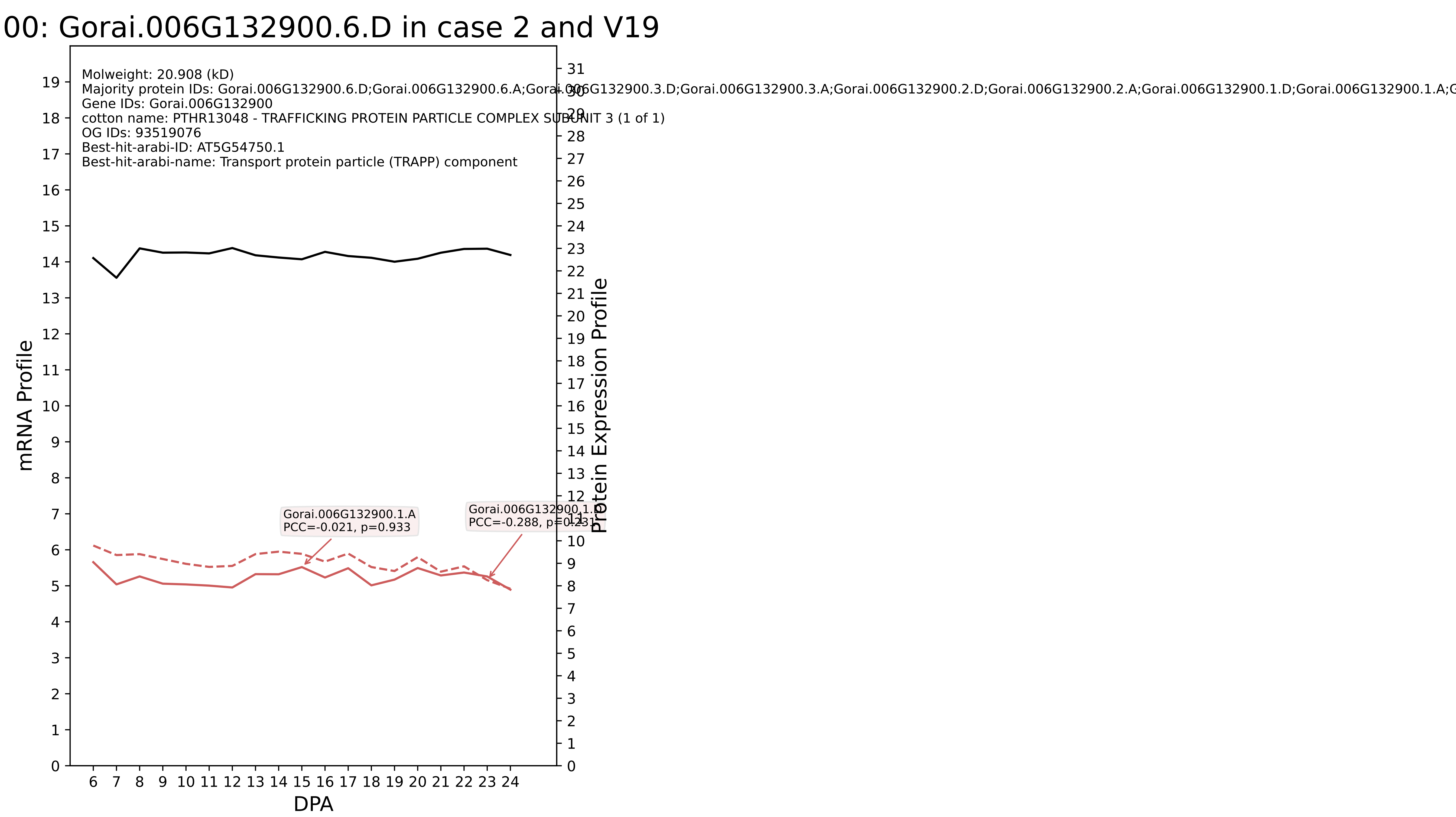
00: Gorai.006G132900.6.D in case 2 and V19
Molweight: 20.908 (kD)
Majority protein IDs: Gorai.006G132900.6.D;Gorai.006G132900.6.A;Gorai.006G132900.3.D;Gorai.006G132900.3.A;Gorai.006G132900.2.D;Gorai.006G132900.2.A;Gorai.006G132900.1.D;Gorai.006G132900.1.A;G
Gene IDs: Gorai.006G132900
cotton name: PTHR13048 - TRAFFICKING PROTEIN PARTICLE COMPLEX SUBUNIT 3 (1 of 1)
OG IDs: 93519076
Best-hit-arabi-ID: AT5G54750.1
Best-hit-arabi-name: Transport protein particle (TRAPP) component
Gorai.006G132900.1.A
PCC=-0.021, p=0.933
PCC=-0.288, p=0.231
mRNA Profile
Protein Expression Profile
DPA

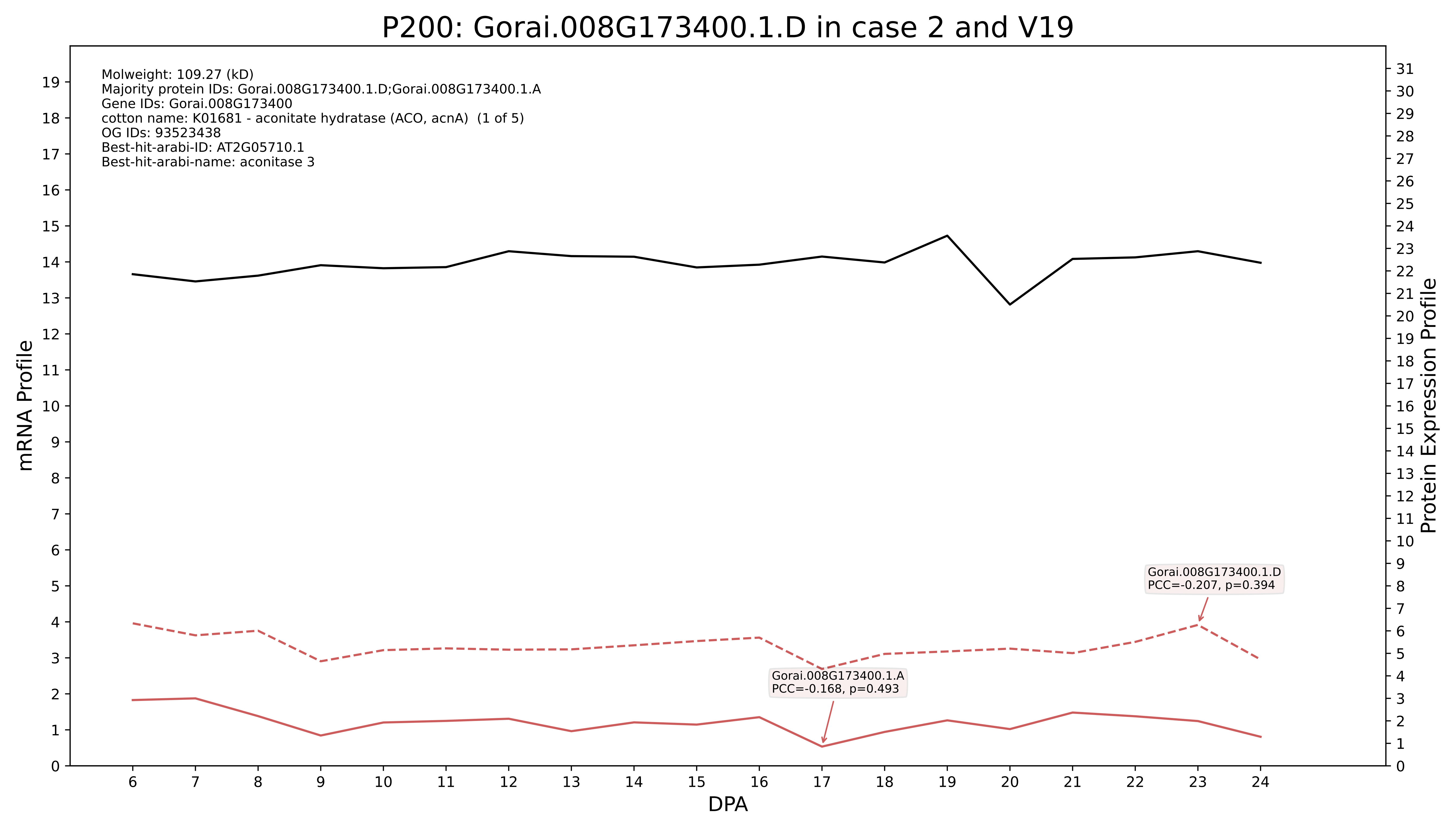
P200: Gorai.008G173400.1.D in case 2 and V19
Molweight: 109.27 (kD)
Majority protein IDs: Gorai.008G173400.1.D;Gorai.008G173400.1.A
Gene IDs: Gorai.008G173400
cotton name: K01681 - aconitate hydratase (ACO, acnA) (1 of 5)
OG IDs: 93523438
Best-hit-arabi-ID: AT2G05710.1
Best-hit-arabi-name: aconitase 3
Gorai.008G173400.1.D
PCC=-0.207, p=0.394
Gorai.008G173400.1.A
PCC=-0.168, p=0.493
mRNA Profile
Protein Expression Profile
DPA

### P200: Gorai.002G268600.3.D in case 2 and V19

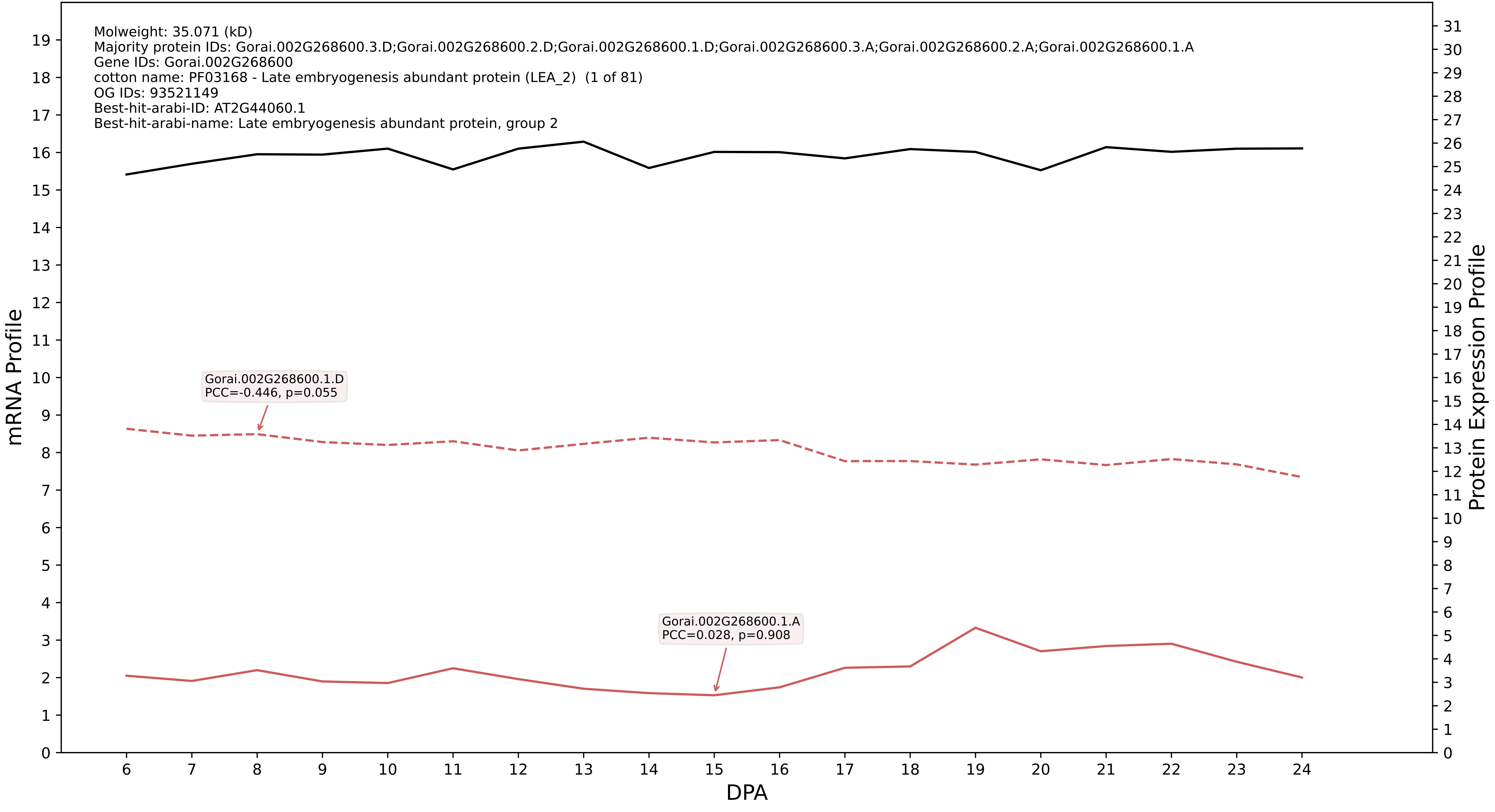

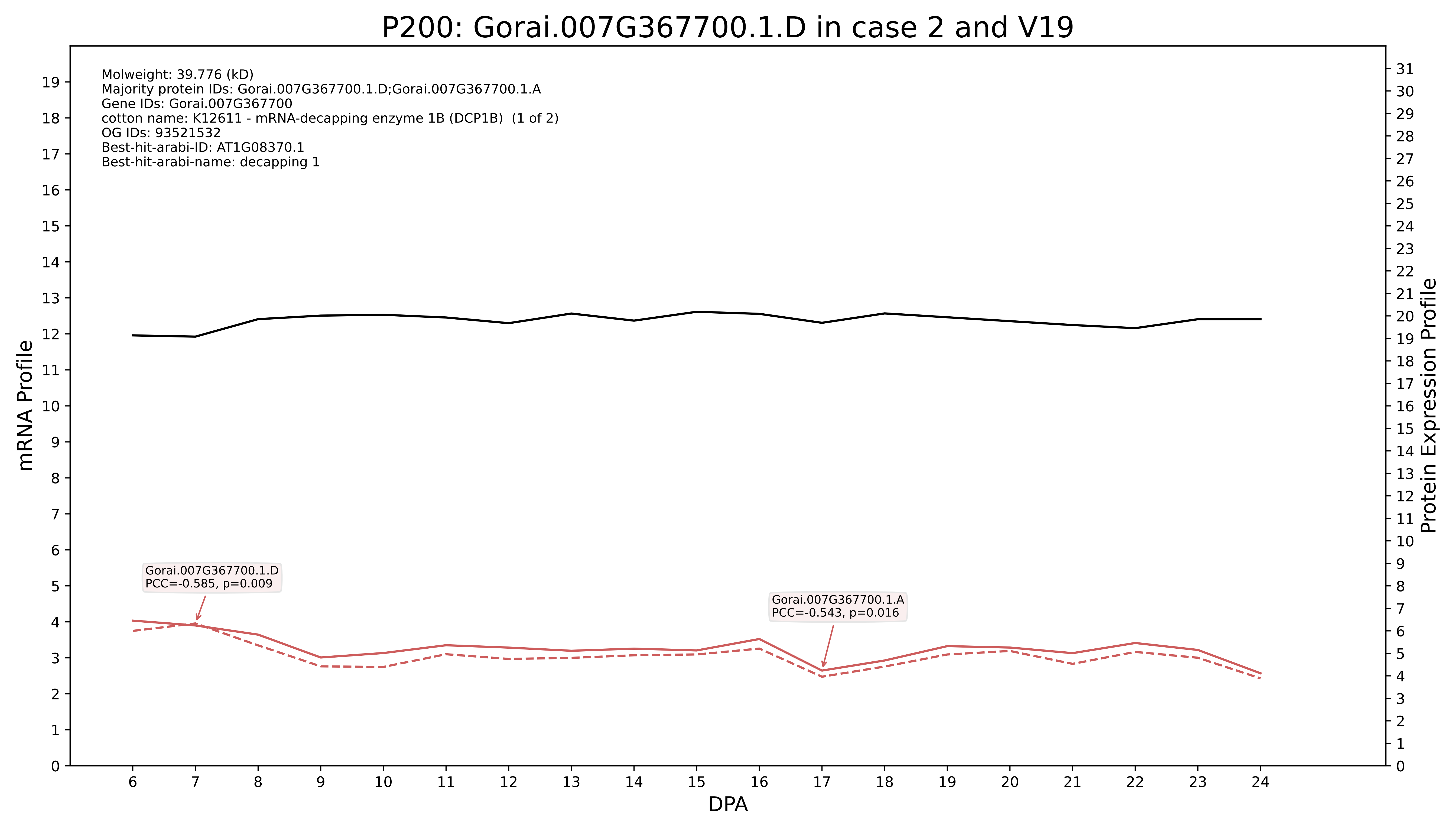
P200: Gorai.007G367700.1.D in case 2 and V19
Molweight: 39.776 (kD)
Majority protein IDs: Gorai.007G367700.1.D;Gorai.007G367700.1.A
Gene IDs: Gorai.007G367700
cotton name: K12611 - mRNA-decapping enzyme 1B (DCP1B) (1 of 2)
OG IDs: 93521532
Best-hit-arabi-ID: AT1G08370.1
Best-hit-arabi-name: decapping 1
Gorai.007G367700.1.D
PCC=-0.585, p=0.009
Gorai.007G367700.1.A
PCC=-0.543, p=0.016
mRNA Profile
Protein Expression Profile
DPA

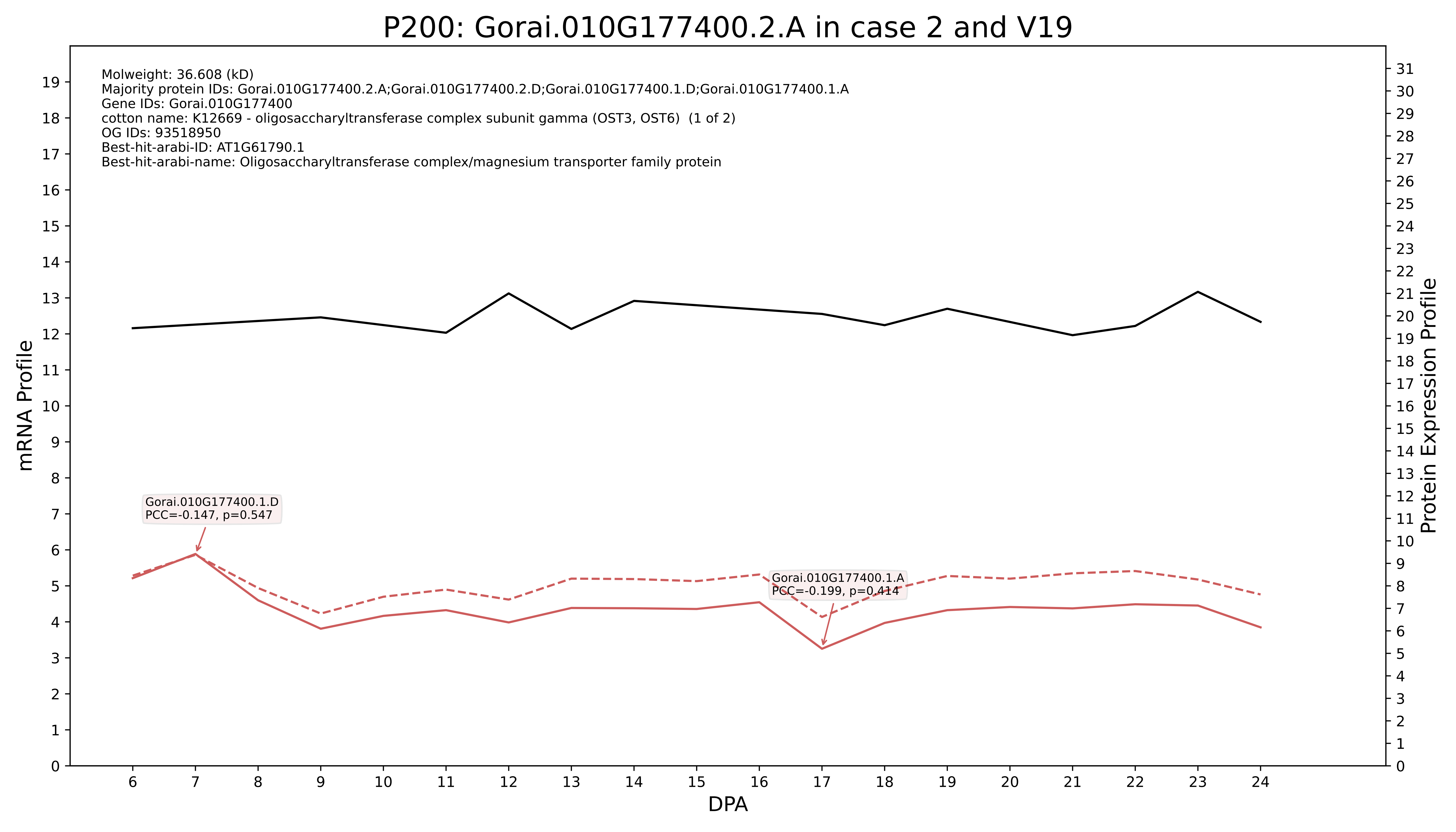
P200: Gorai.010G177400.2.A in case 2 and V19
Molweight: 36.608 (kD)
Majority protein IDs: Gorai.010G177400.2.A;Gorai.010G177400.2.D;Gorai.010G177400.1.D;Gorai.010G177400.1.A
Gene IDs: Gorai.010G177400
cotton name: K12669 - oligosaccharyltransferase complex subunit gamma (OST3, OST6) (1 of 2)
OG IDs: 93518950
Best-hit-arabi-ID: AT1G61790.1
Best-hit-arabi-name: Oligosaccharyltransferase complex/magnesium transporter family protein
Gorai.010G177400.1.D
PCC=-0.147, p=0.547
Gorai.010G177400.1.A
PCC=-0.199, p=0.414
mRNA Profile
Protein Expression Profile
DPA

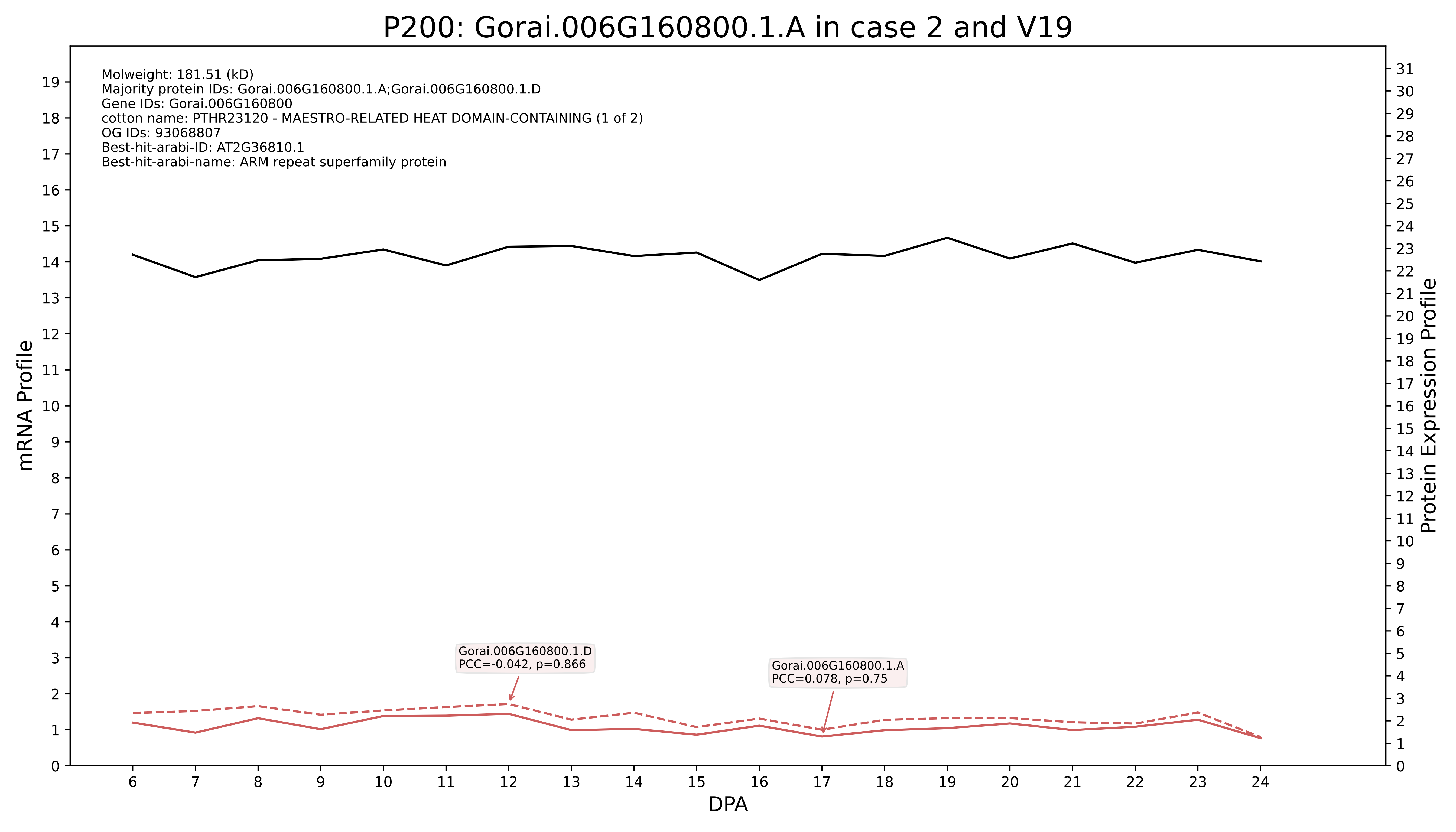
P200: Gorai.006G160800.1.A in case 2 and V19
Molweight: 181.51 (kD)
Majority protein IDs: Gorai.006G160800.1.A;Gorai.006G160800.1.D
Gene IDs: Gorai.006G160800
cotton name: PTHR23120 - MAESTRO-RELATED HEAT DOMAIN-CONTAINING (1 of 2)
OG IDs: 93068807
Best-hit-arabi-ID: AT2G36810.1
Best-hit-arabi-name: ARM repeat superfamily protein
Gorai.006G160800.1.D
PCC=-0.042, p=0.866
Gorai.006G160800.1.A
PCC=0.078, p=0.75
mRNA Profile
Protein Expression Profile
DPA

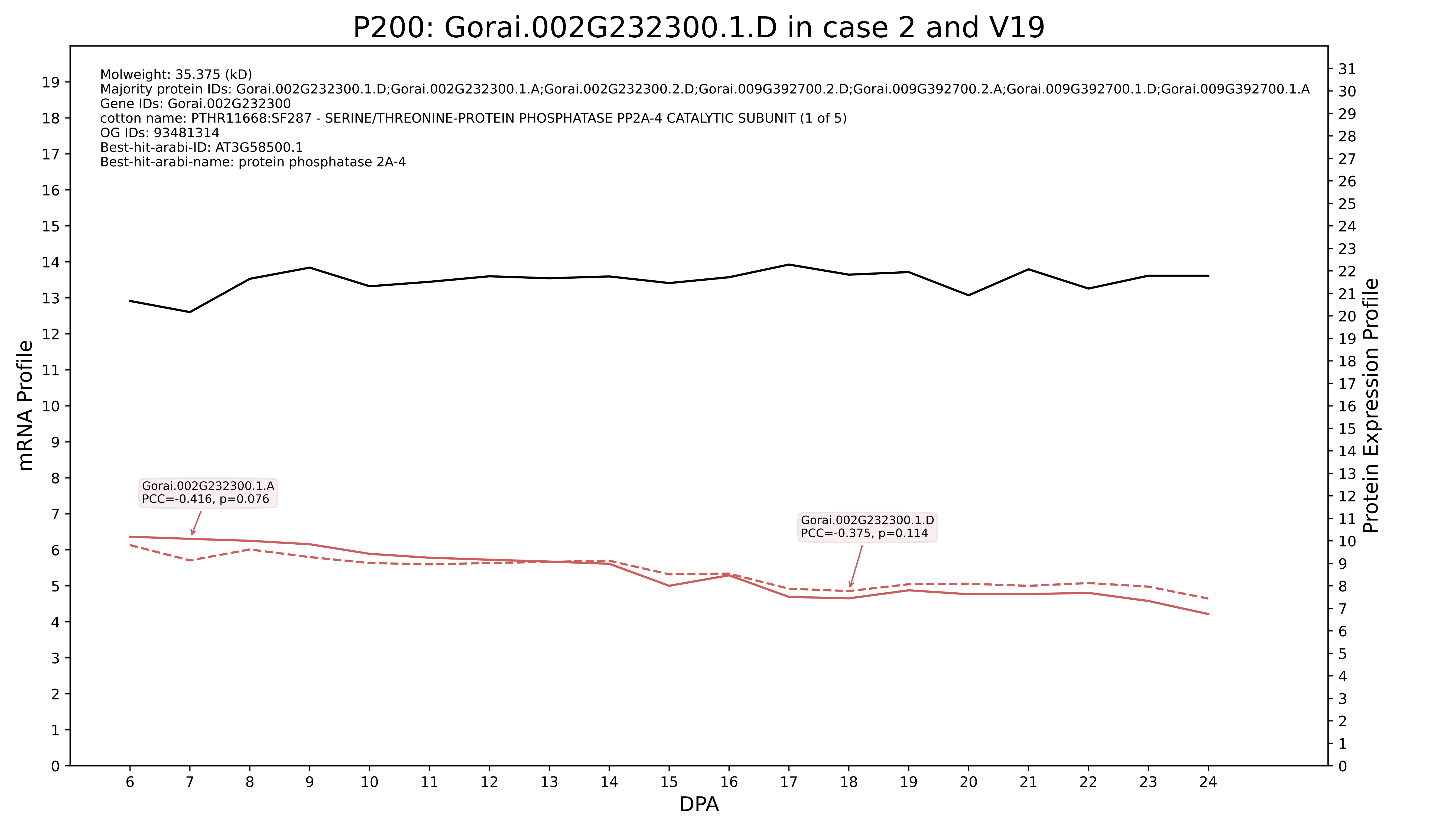
P200: Gorai.002G232300.1.D in case 2 and V19
Molweight: 35.375 (kD)
Majority protein IDs: Gorai.002G232300.1.D;Gorai.002G232300.1.A;Gorai.002G232300.2.D;Gorai.009G392700.2.D;Gorai.009G392700.2.A;Gorai.009G392700.1.D;Gorai.009G392700.1.A
Gene IDs: Gorai.002G232300
cotton name: PTHR11668:SF287 - SERINE/THREONINE-PROTEIN PHOSPHATASE PP2A-4 CATALYTIC SUBUNIT (1 of 5)
OG IDs: 93481314
Best-hit-arabi-ID: AT3G58500.1
Best-hit-arabi-name: protein phosphatase 2A-4
Gorai.002G232300.1.A
PCC=-0.416, p=0.076
Gorai.002G232300.1.D
PCC=-0.375, p=0.114
mRNA Profile
Protein Expression Profile
DPA

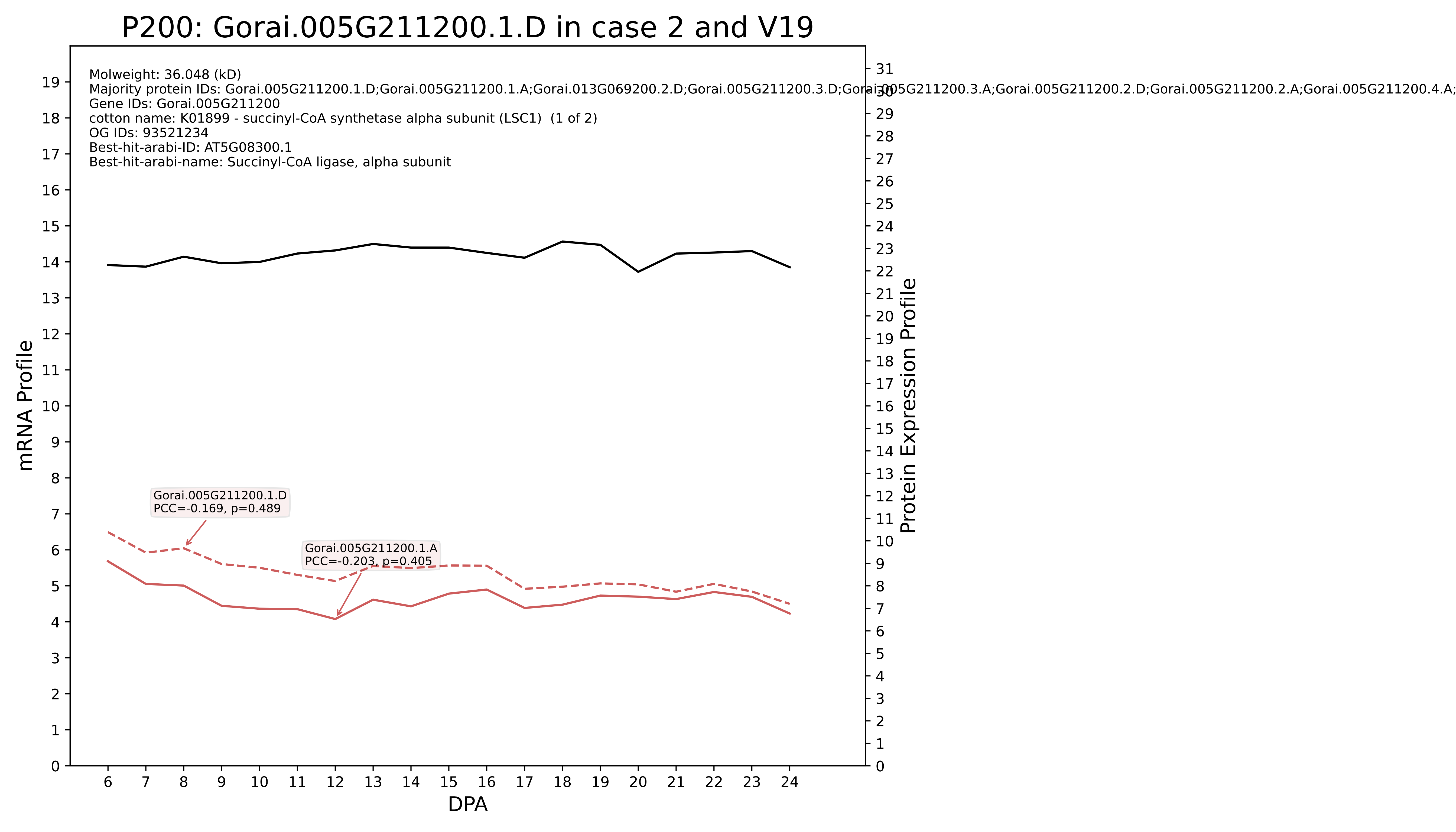
P200: Gorai.005G211200.1.D in case 2 and V19
Molweight: 36.048 (kD)
Majority protein IDs: Gorai.005G211200.1.D;Gorai.005G211200.1.A;Gorai.013G069200.2.D;Gorai.005G211200.3.D;Gorai.005G211200.3.A;Gorai.005G211200.2.D;Gorai.005G211200.2.A;Gorai.005G211200.4.A;
Gene IDs: Gorai.005G211200
cotton name: K01899 - succinyl-CoA synthetase alpha subunit (LSC1) (1 of 2)
OG IDs: 93521234
Best-hit-arabi-ID: AT5G08300.1
Best-hit-arabi-name: Succinyl-CoA ligase, alpha subunit
Gorai.005G211200.1.D
PCC=-0.169, p=0.489
Gorai.005G211200.1.A
PCC=-0.203, p=0.405
mRNA Profile
Protein Expression Profile
DPA

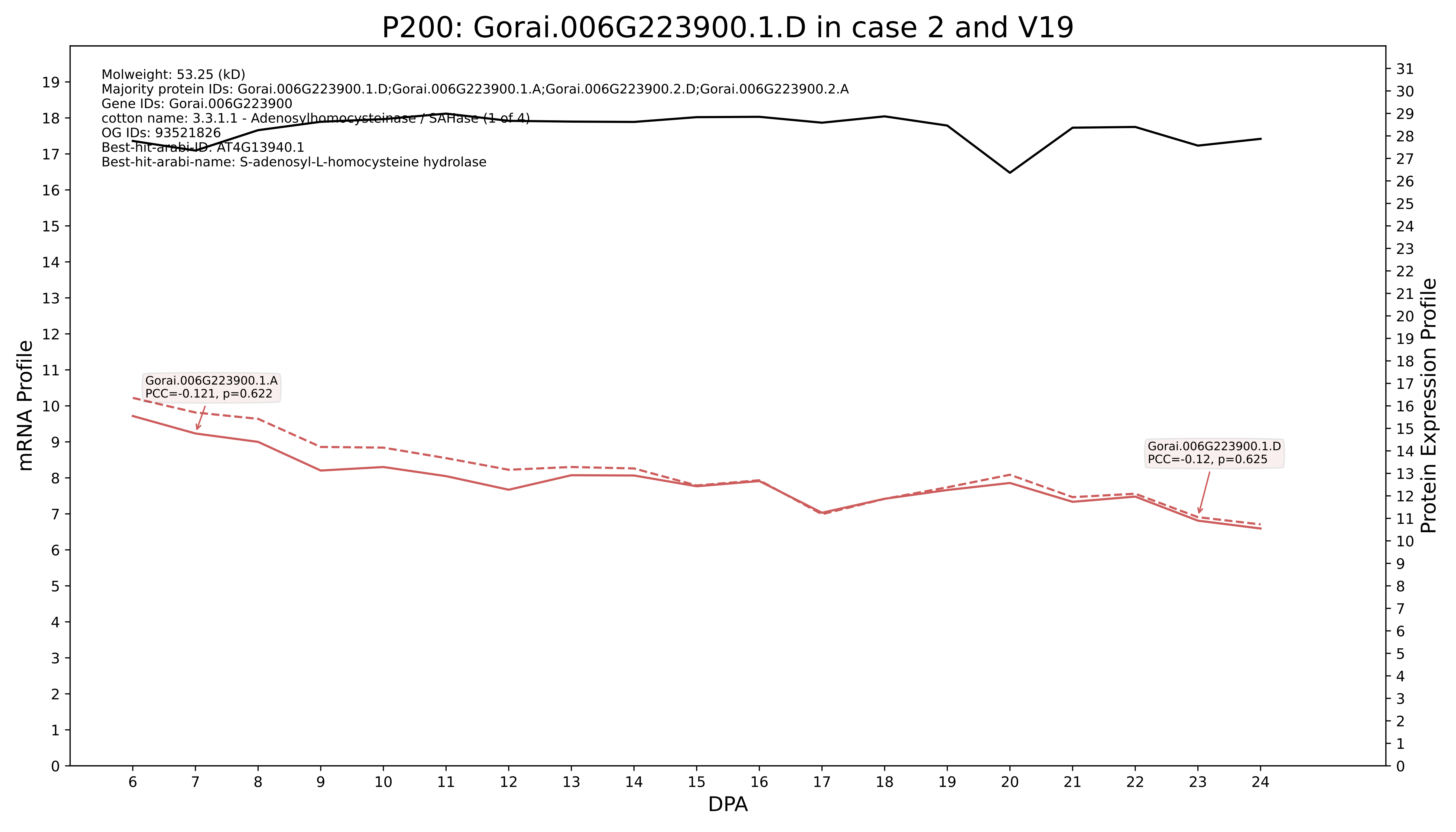
P200: Gorai.006G223900.1.D in case 2 and V19
Molweight: 53.25 (kD)
Majority protein IDs: Gorai.006G223900.1.D;Gorai.006G223900.1.A;Gorai.006G223900.2.D;Gorai.006G223900.2.A
Gene IDs: Gorai.006G223900
cotton name: 3.3.1.1 - Adenosylhomocysteinase / SAHase (1 of 4)
OG IDs: 93521826
Best-hit-arabi-ID: AT4G13940.1
Best-hit-arabi-name: S-adenosyl-L-homocysteine hydrolase
Gorai.006G223900.1.A
PCC=-0.121, p=0.622
Gorai.006G223900.1.D
PCC=-0.12, p=0.625
mRNA Profile
Protein Expression Profile
DPA

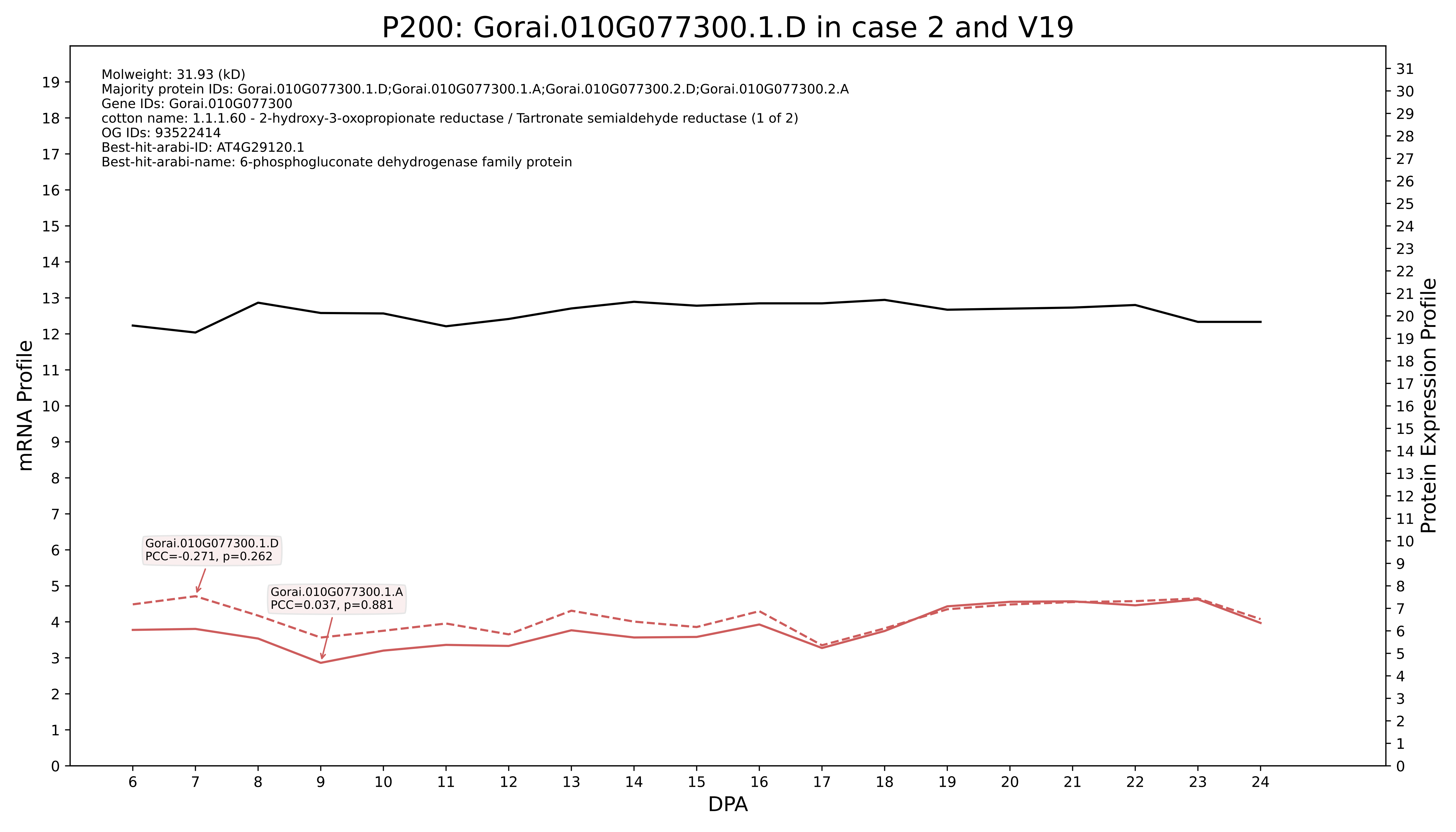
P200: Gorai.010G077300.1.D in case 2 and V19
Molweight: 31.93 (kD)
Majority protein IDs: Gorai.010G077300.1.D;Gorai.010G077300.1.A;Gorai.010G077300.2.D;Gorai.010G077300.2.A
Gene IDs: Gorai.010G077300
cotton name: 1.1.1.60 - 2-hydroxy-3-oxopropionate reductase / Tartronate semialdehyde reductase (1 of 2)
OG IDs: 93522414
Best-hit-arabi-ID: AT4G29120.1
Best-hit-arabi-name: 6-phosphogluconate dehydrogenase family protein
Gorai.010G077300.1.D
PCC=-0.271, p=0.262
Gorai.010G077300.1.A
PCC=0.037, p=0.881
mRNA Profile
Protein Expression Profile
DPA

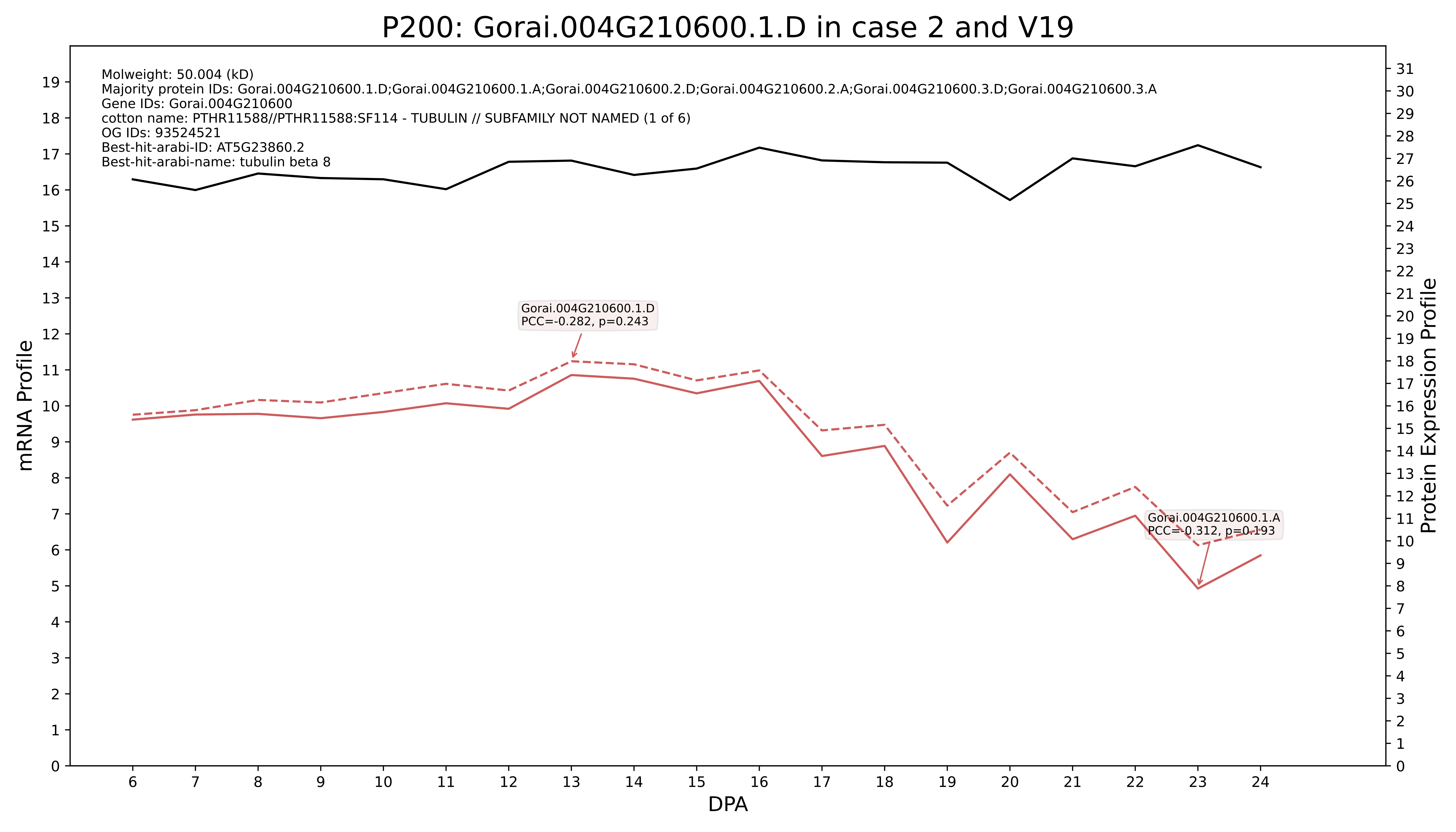
P200: Gorai.004G210600.1.D in case 2 and V19
Molweight: 50.004 (kD)
Majority protein IDs: Gorai.004G210600.1.D;Gorai.004G210600.1.A;Gorai.004G210600.2.D;Gorai.004G210600.2.A;Gorai.004G210600.3.D;Gorai.004G210600.3.A
Gene IDs: Gorai.004G210600
cotton name: PTHR11588//PTHR11588:SF114 - TUBULIN // SUBFAMILY NOT NAMED (1 of 6)
OG IDs: 93524521
Best-hit-arabi-ID: AT5G23860.2
Best-hit-arabi-name: tubulin beta 8
Gorai.004G210600.1.D
PCC=-0.282, p=0.243
Gorai.004G210600.1.A
PCC=-0.312, p=0.193
mRNA Profile
Protein Expression Profile
DPA

### P200: Gorai.007G049400.1.A in case 2 and V19

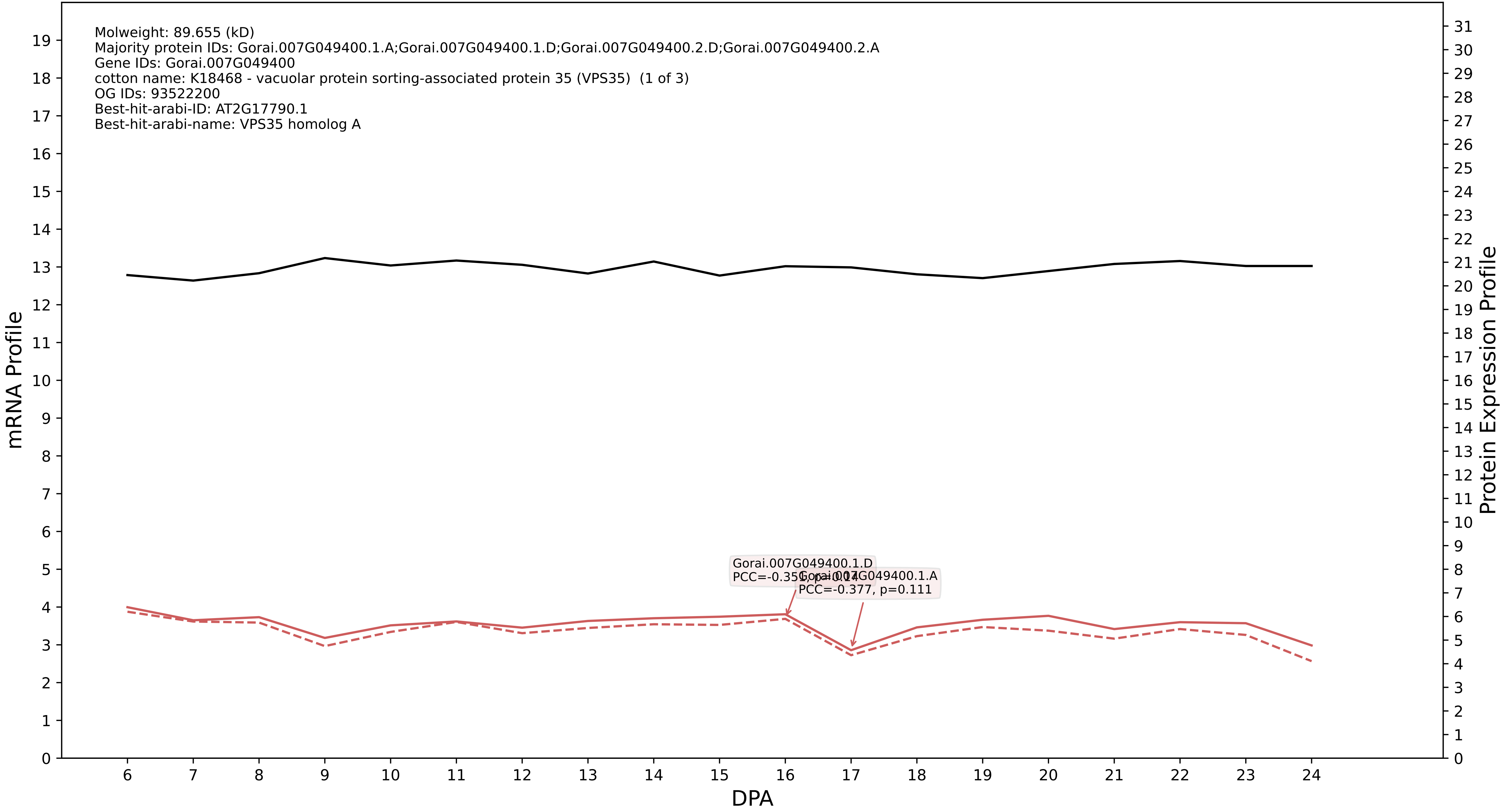

00: Gorai.003G094300.2.D in case 2 and V19

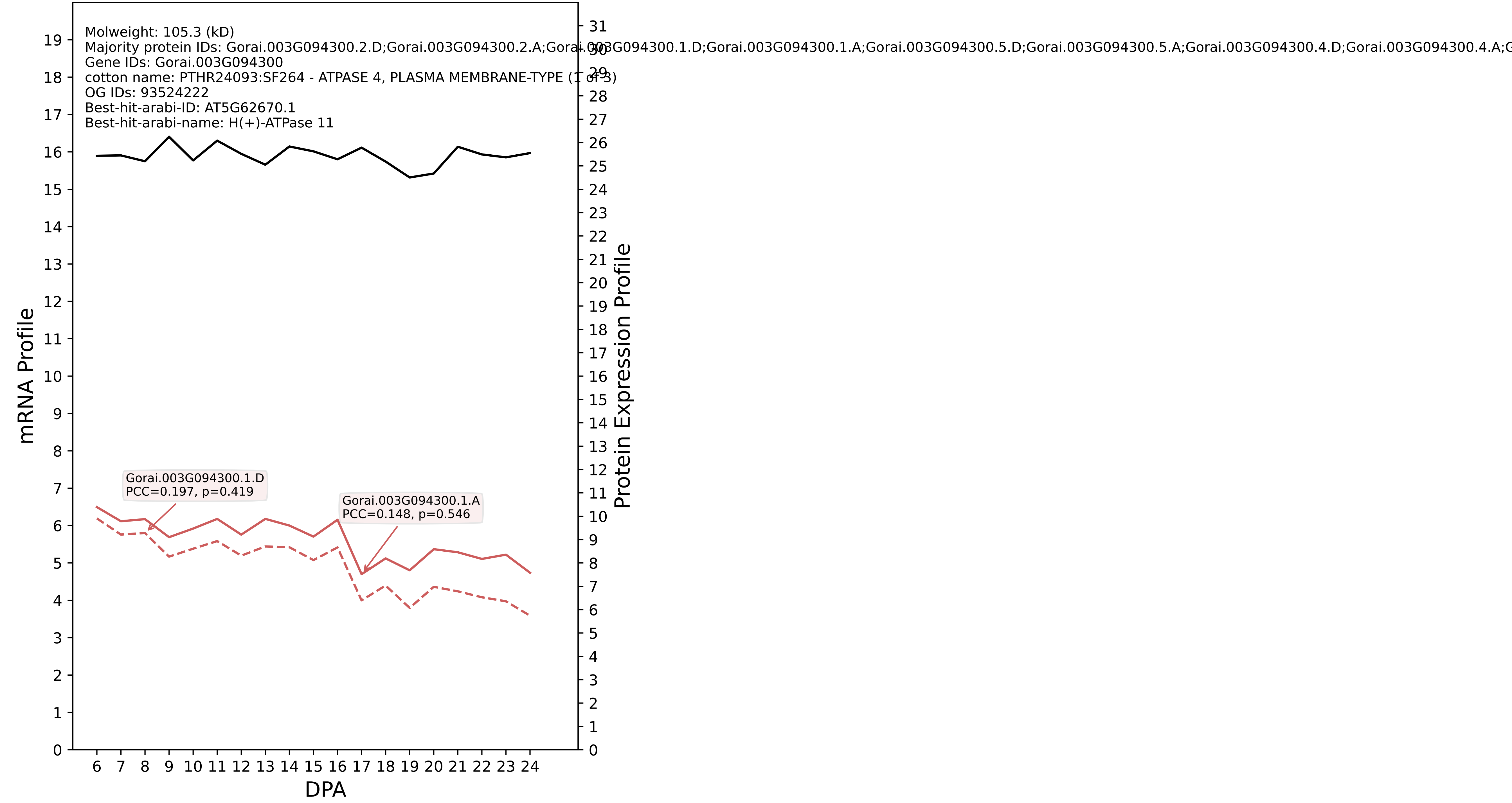

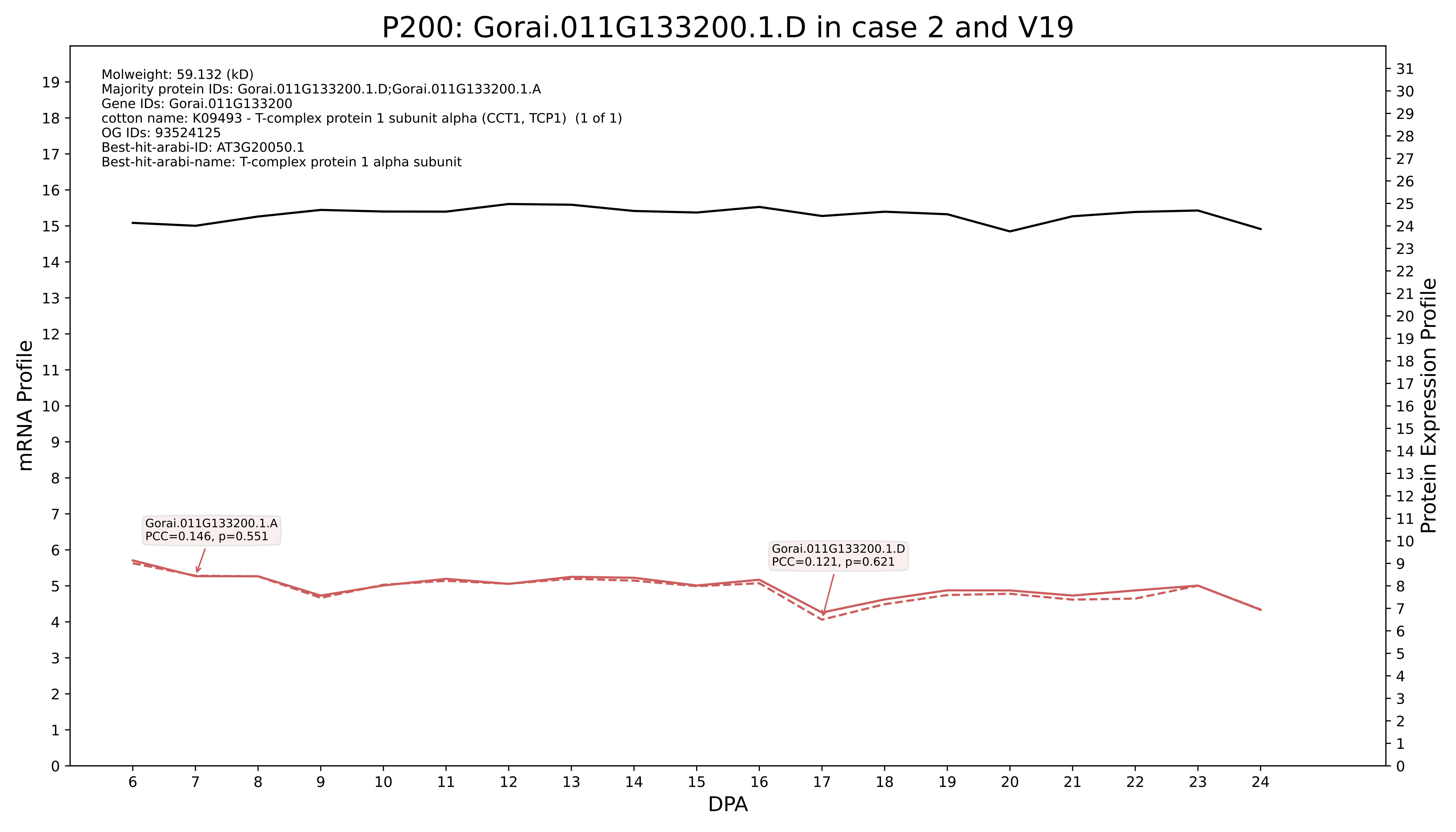
P200: Gorai.011G133200.1.D in case 2 and V19
Molweight: 59.132 (kD)
Majority protein IDs: Gorai.011G133200.1.D;Gorai.011G133200.1.A
Gene IDs: Gorai.011G133200
cotton name: K09493 - T-complex protein 1 subunit alpha (CCT1, TCP1) (1 of 1)
OG IDs: 93524125
Best-hit-arabi-ID: AT3G20050.1
Best-hit-arabi-name: T-complex protein 1 alpha subunit
Gorai.011G133200.1.A
PCC=0.146, p=0.551
Gorai.011G133200.1.D
PCC=0.121, p=0.621
mRNA Profile
Protein Expression Profile
DPA

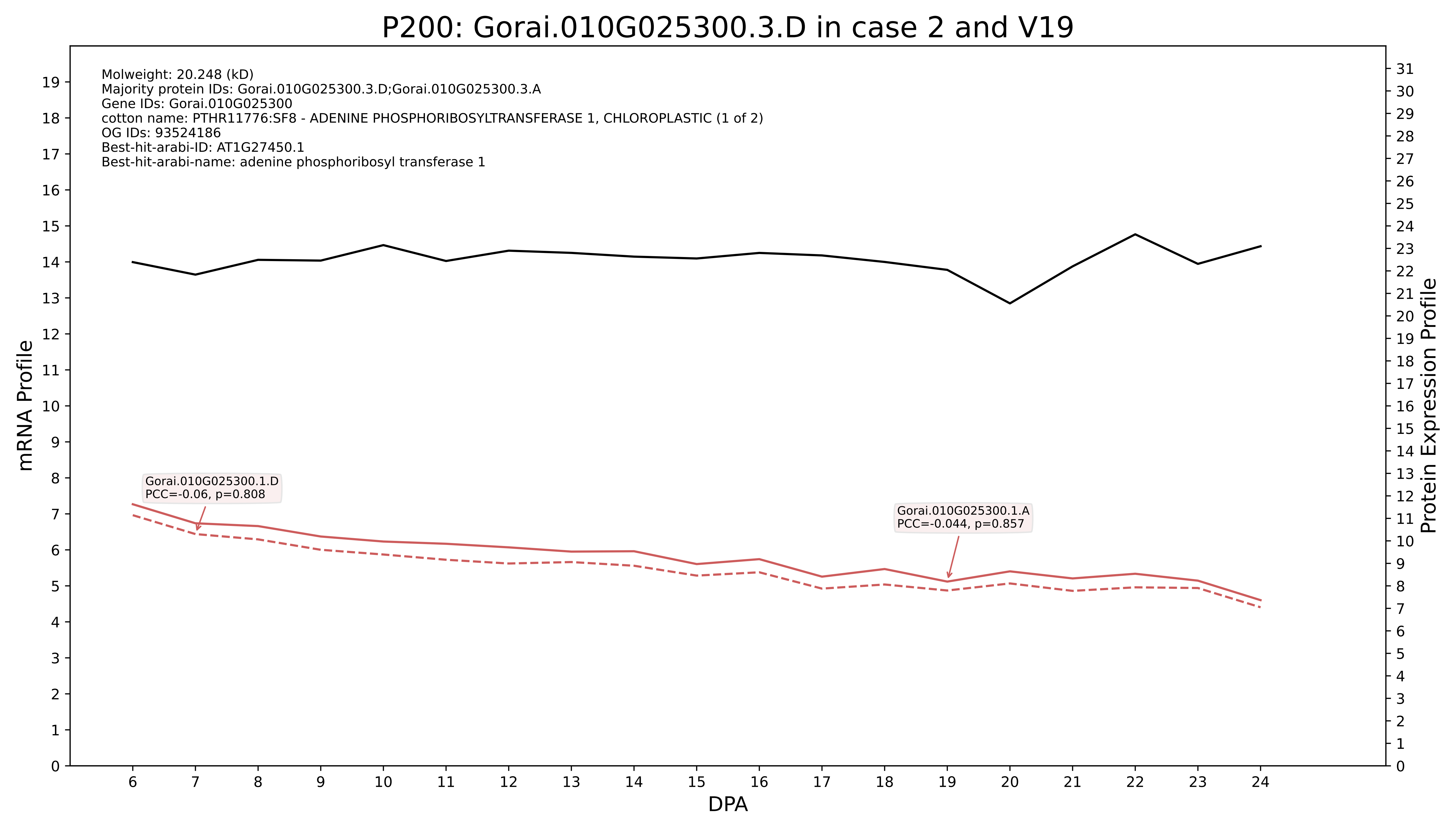
P200: Gorai.010G025300.3.D in case 2 and V19
Molweight: 20.248 (kD)
Majority protein IDs: Gorai.010G025300.3.D;Gorai.010G025300.3.A
Gene IDs: Gorai.010G025300
cotton name: PTHR11776:SF8 - ADENINE PHOSPHORIBOSYLTRANSFERASE 1, CHLOROPLASTIC (1 of 2)
OG IDs: 93524186
Best-hit-arabi-ID: AT1G27450.1
Best-hit-arabi-name: adenine phosphoribosyl transferase 1
Gorai.010G025300.1.D
PCC=-0.06, p=0.808
Gorai.010G025300.1.A
PCC=-0.044, p=0.857
mRNA Profile
Protein Expression Profile
DPA

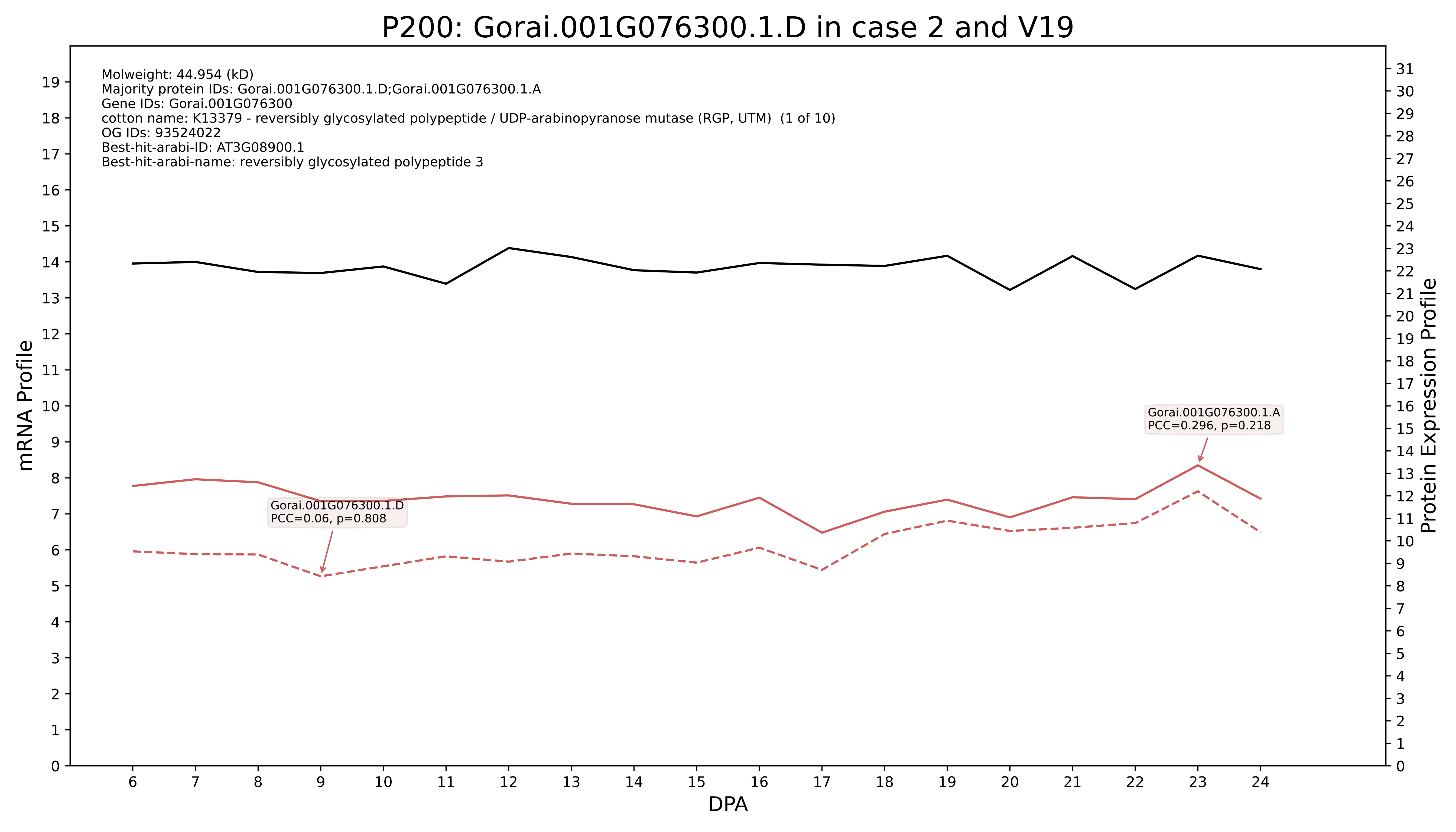
P200: Gorai.001G076300.1.D in case 2 and V19
Molweight: 44.954 (kD)
Majority protein IDs: Gorai.001G076300.1.D;Gorai.001G076300.1.A
Gene IDs: Gorai.001G076300
cotton name: K13379 - reversibly glycosylated polypeptide / UDP-arabinopyranose mutase (RGP, UTM) (1 of 10)
OG IDs: 93524022
Best-hit-arabi-ID: AT3G08900.1
Best-hit-arabi-name: reversibly glycosylated polypeptide 3
Gorai.001G076300.1.A
PCC=0.296, p=0.218
Gorai.001G076300.1.D
PCC=0.06, p=0.808
mRNA Profile
Protein Expression Profile
DPA

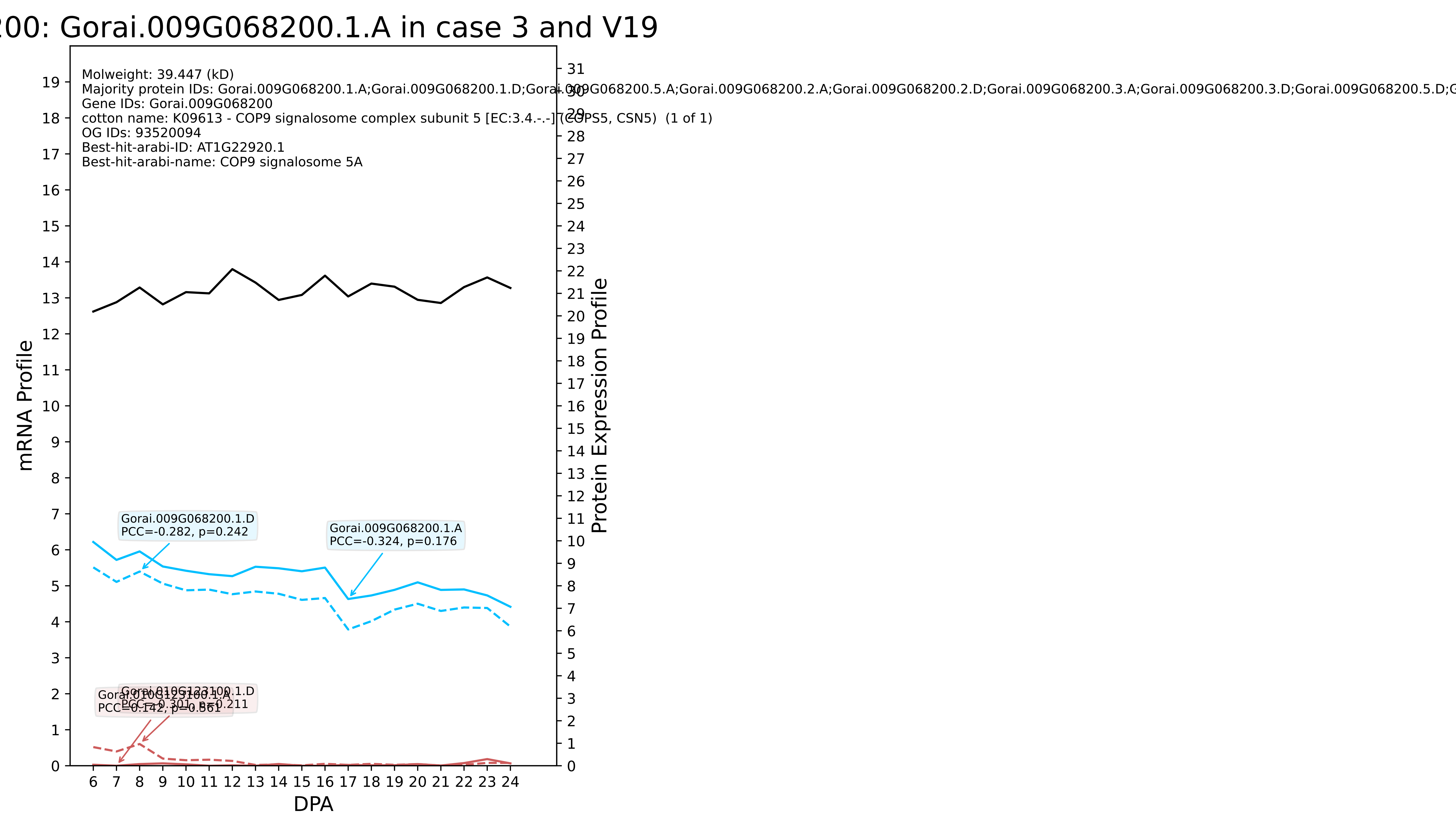

200: Gorai.009G068200.1.A in case 3 and V19
Molweight: 39.447 (kD)
Majority protein IDs: Gorai.009G068200.1.A;Gorai.009G068200.1.D;Gorai.009G068200.5.A;Gorai.009G068200.2.A;Gorai.009G068200.2.D;Gorai.009G068200.3.A;Gorai.009G068200.3.D;Gorai.009G068200.5.D;G
Gene IDs: Gorai.009G068200
cotton name: K09613 - COP9 signalosome complex subunit 5 [EC:3.4.-.-] (COPS5, CSN5) (1 of 1)
OG IDs: 93520094
Best-hit-arabi-ID: AT1G22920.1
Best-hit-arabi-name: COP9 signalosome 5A
Gorai.009G068200.1.D
PCC=-0.282, p=0.242
Gorai.009G068200.1.A
PCC=-0.324, p=0.176
Gorai.010G123100.1.D
PCC=-0.301, p=0.211
Gorai.010G123100.1.A
PCC=-0.142, p=0.561
mRNA Profile
Protein Expression Profile
DPA

### P200: Gorai.013G131000.1.D in case 3 and V19

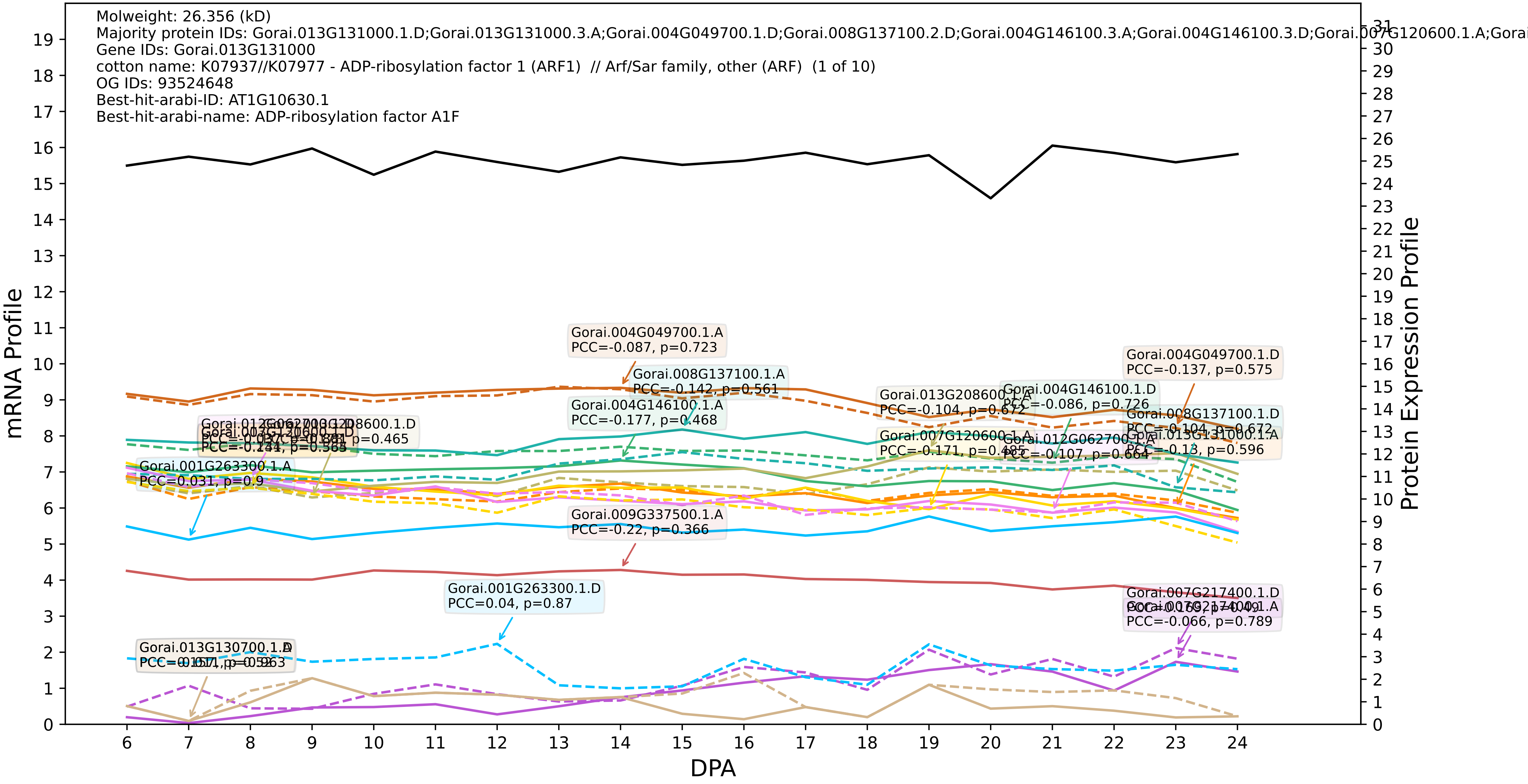

P200: Gorai.008G149500.1.D in case 3 and V19

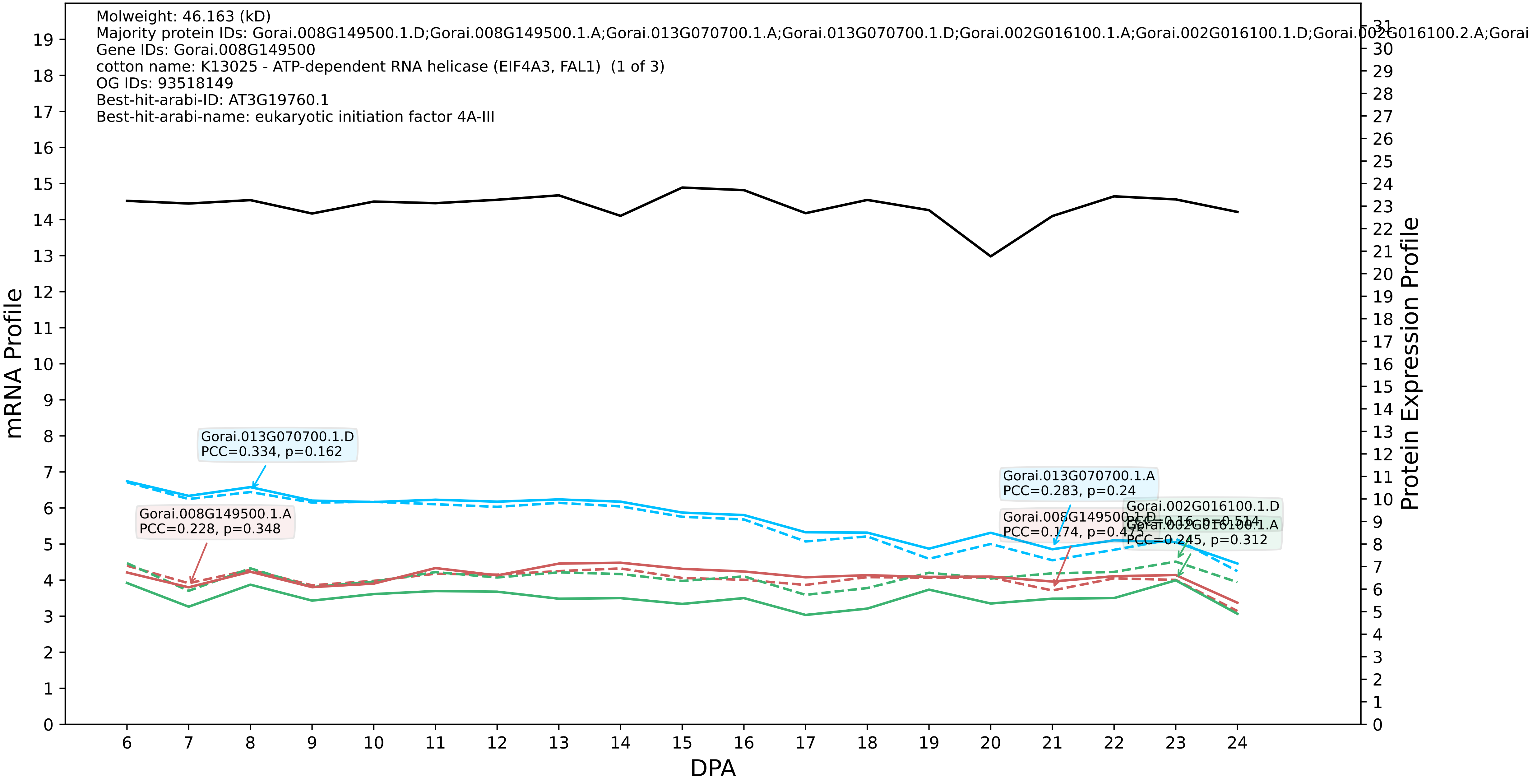

### 006G025300.1.D in case 3 and V19

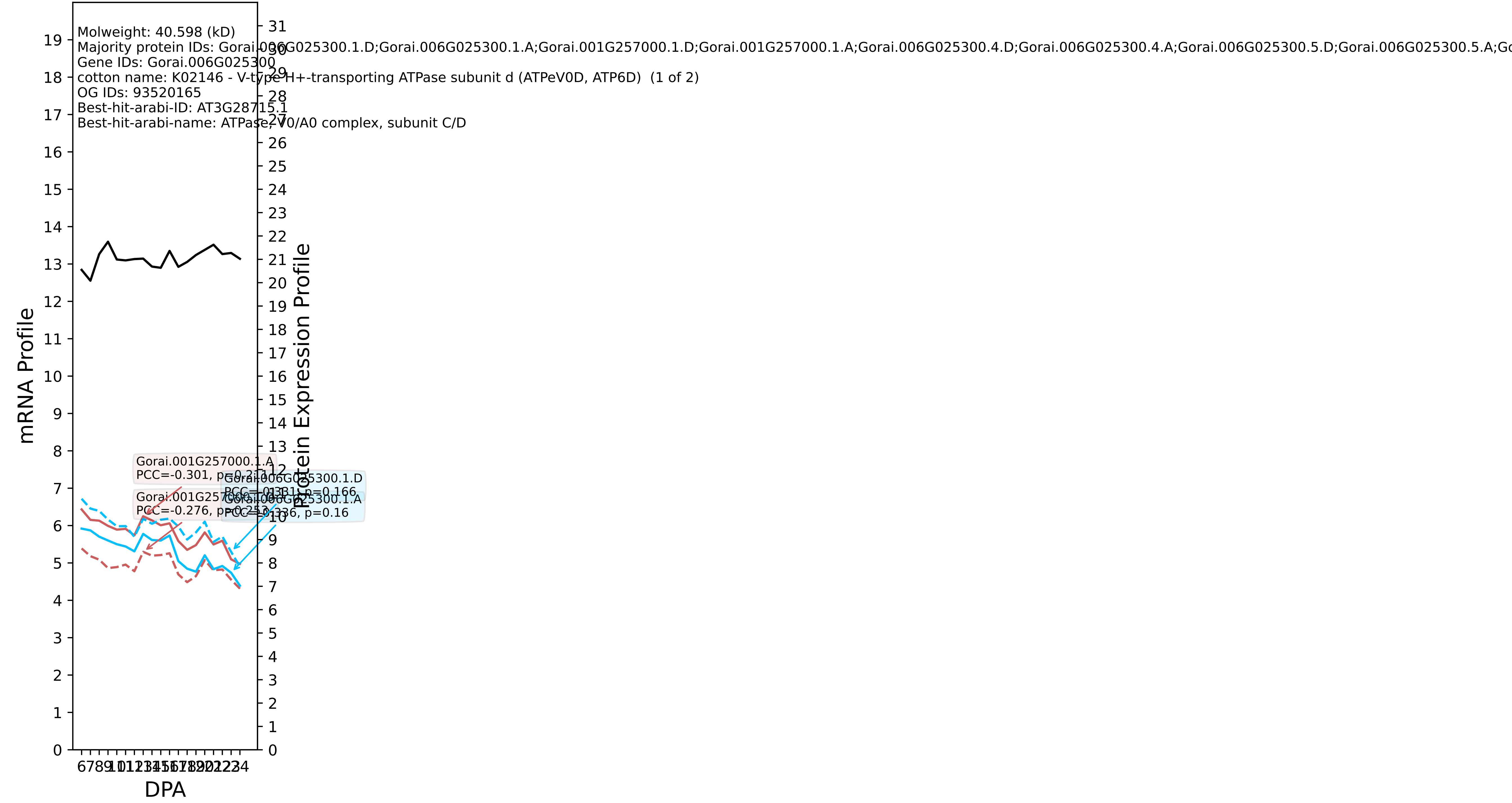

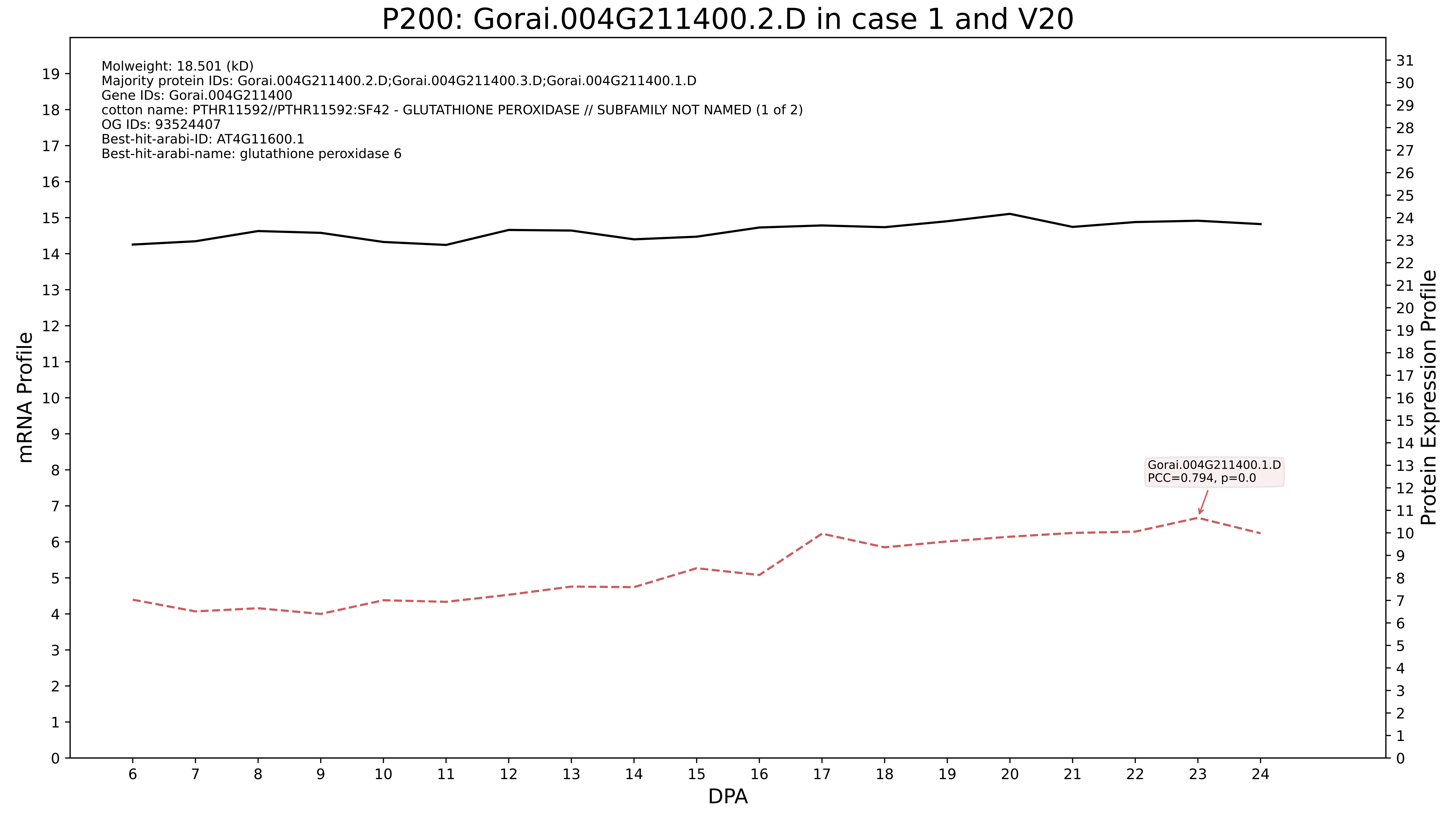
P200: Gorai.004G211400.2.D in case 1 and V20
Molweight: 18.501 (kD)
Majority protein IDs: Gorai.004G211400.2.D;Gorai.004G211400.3.D;Gorai.004G211400.1.D
Gene IDs: Gorai.004G211400
cotton name: PTHR11592//PTHR11592:SF42 - GLUTATHIONE PEROXIDASE // SUBFAMILY NOT NAMED (1 of 2)
OG IDs: 93524407
Best-hit-arabi-ID: AT4G11600.1
Best-hit-arabi-name: glutathione peroxidase 6
Gorai.004G211400.1.D
PCC=0.794, p=0.0
mRNA Profile
Protein Expression Profile
DPA

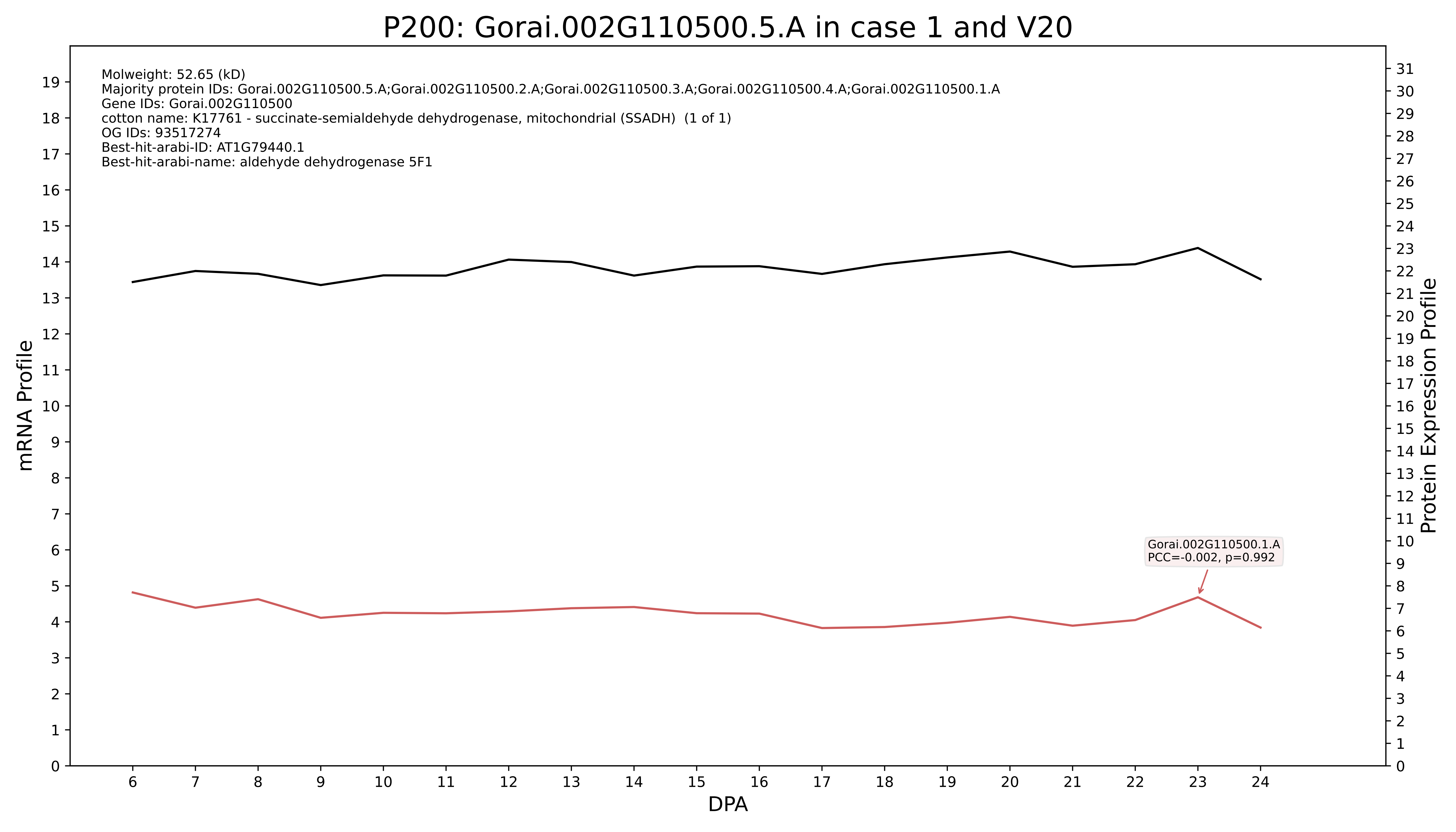

P200: Gorai.002G110500.5.A in case 1 and V20
Molweight: 52.65 (kD)
Majority protein IDs: Gorai.002G110500.5.A;Gorai.002G110500.2.A;Gorai.002G110500.3.A;Gorai.002G110500.4.A;Gorai.002G110500.1.A
Gene IDs: Gorai.002G110500
cotton name: K17761 - succinate-semialdehyde dehydrogenase, mitochondrial (SSADH) (1 of 1)
OG IDs: 93517274
Best-hit-arabi-ID: AT1G79440.1
Best-hit-arabi-name: aldehyde dehydrogenase 5F1
Gorai.002G110500.1.A
PCC=-0.002, p=0.992
mRNA Profile
Protein Expression Profile
DPA

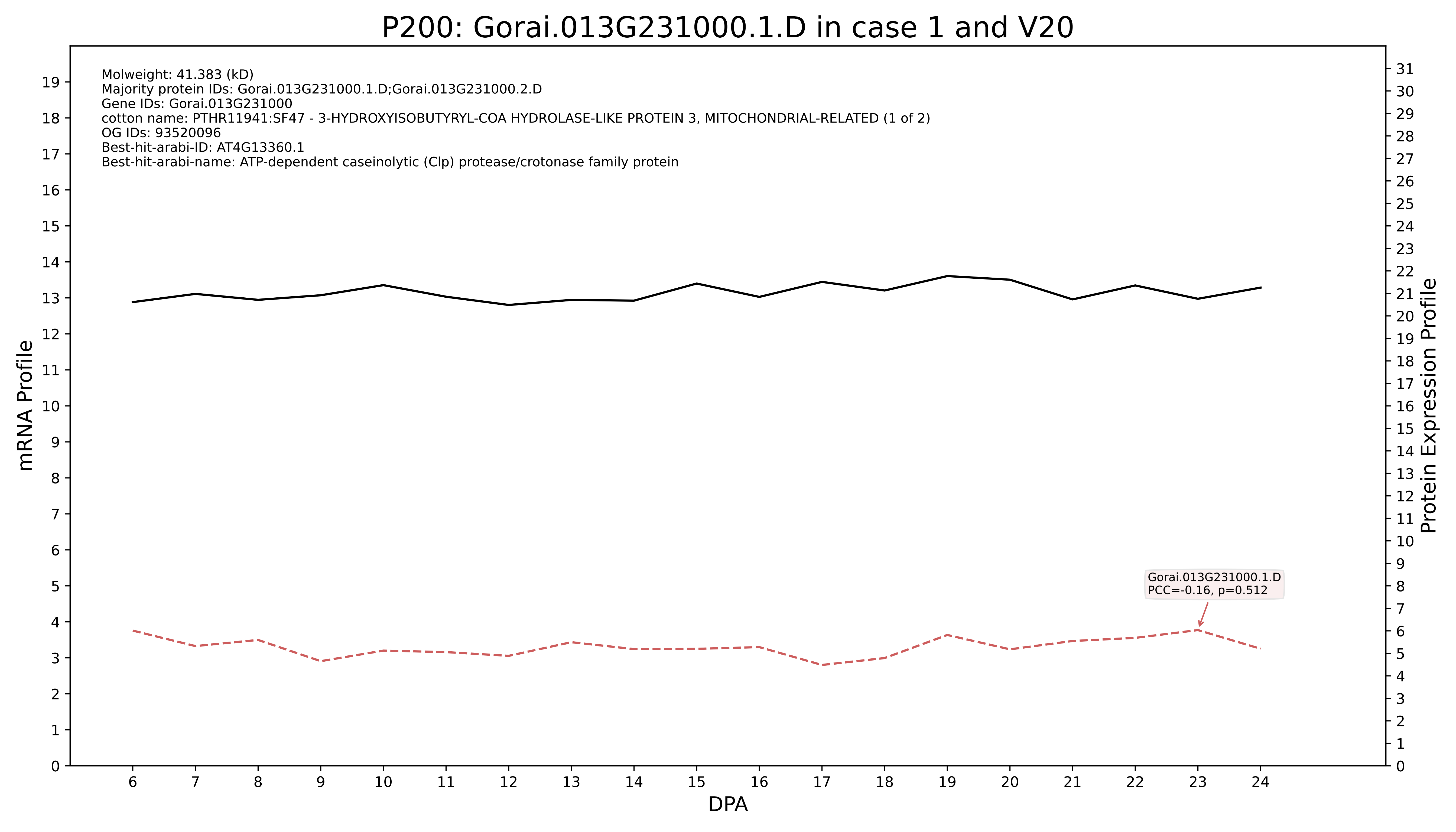
P200: Gorai.013G231000.1.D in case 1 and V20
Molweight: 41.383 (kD)
Majority protein IDs: Gorai.013G231000.1.D;Gorai.013G231000.2.D
Gene IDs: Gorai.013G231000
cotton name: PTHR11941:SF47 - 3-HYDROXYISOBUTYRYL-COA HYDROLASE-LIKE PROTEIN 3, MITOCHONDRIAL-RELATED (1 of 2)
OG IDs: 93520096
Best-hit-arabi-ID: AT4G13360.1
Best-hit-arabi-name: ATP-dependent caseinolytic (Clp) protease/crotonase family protein
Gorai.013G231000.1.D
PCC=-0.16, p=0.512
mRNA Profile
Protein Expression Profile
DPA

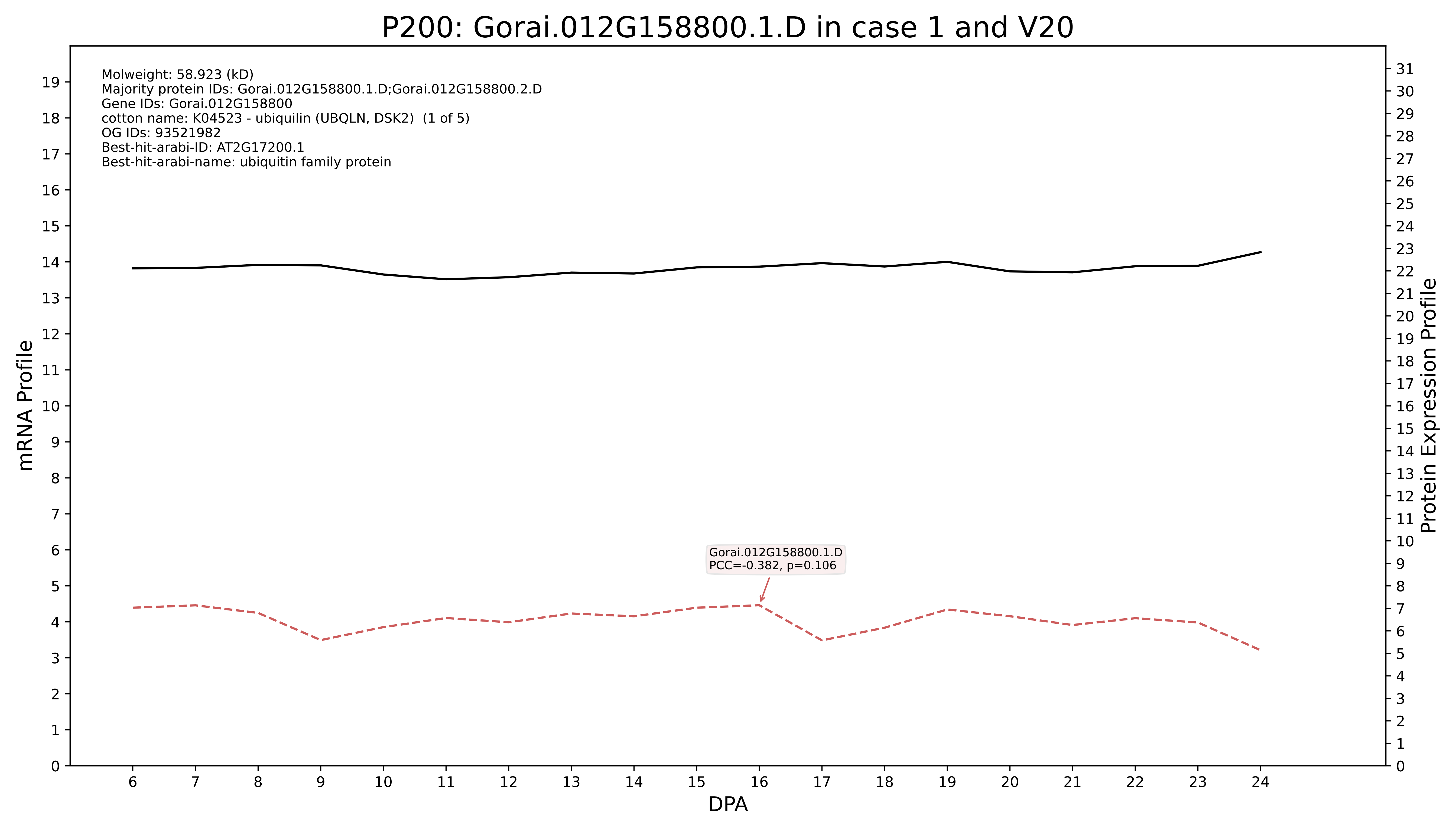

P200: Gorai.012G158800.1.D in case 1 and V20
Molweight: 58.923 (kD)
Majority protein IDs: Gorai.012G158800.1.D;Gorai.012G158800.2.D
Gene IDs: Gorai.012G158800
cotton name: K04523 - ubiquilin (UBQLN, DSK2) (1 of 5)
OG IDs: 93521982
Best-hit-arabi-ID: AT2G17200.1
Best-hit-arabi-name: ubiquitin family protein
Gorai.012G158800.1.D
PCC=-0.382, p=0.106
mRNA Profile
Protein Expression Profile
DPA

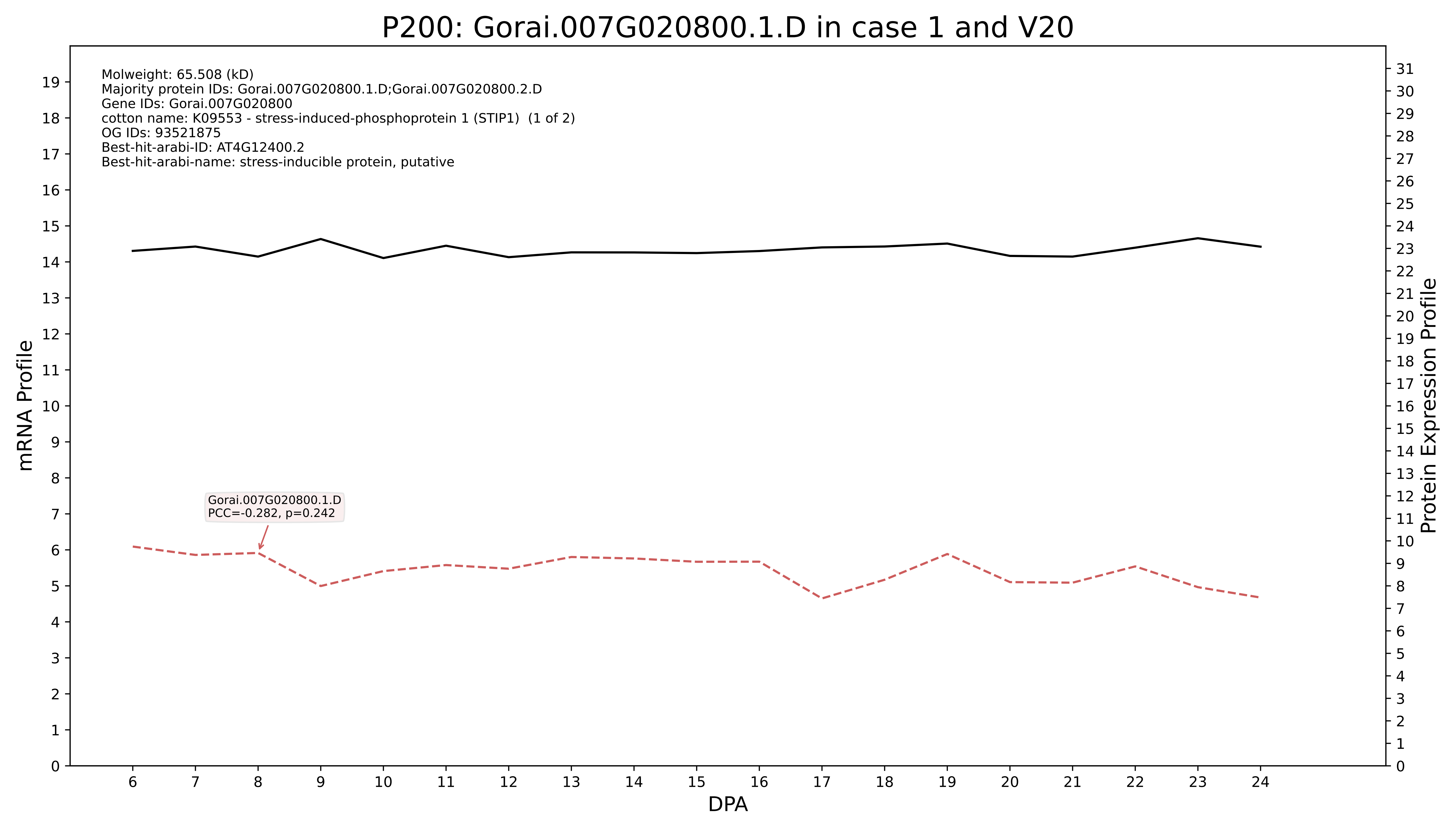
P200: Gorai.007G020800.1.D in case 1 and V20
Molweight: 65.508 (kD)
Majority protein IDs: Gorai.007G020800.1.D;Gorai.007G020800.2.D
Gene IDs: Gorai.007G020800
cotton name: K09553 - stress-induced-phosphoprotein 1 (STIP1) (1 of 2)
OG IDs: 93521875
Best-hit-arabi-ID: AT4G12400.2
Best-hit-arabi-name: stress-inducible protein, putative
Gorai.007G020800.1.D
PCC=-0.282, p=0.242
mRNA Profile
Protein Expression Profile
DPA

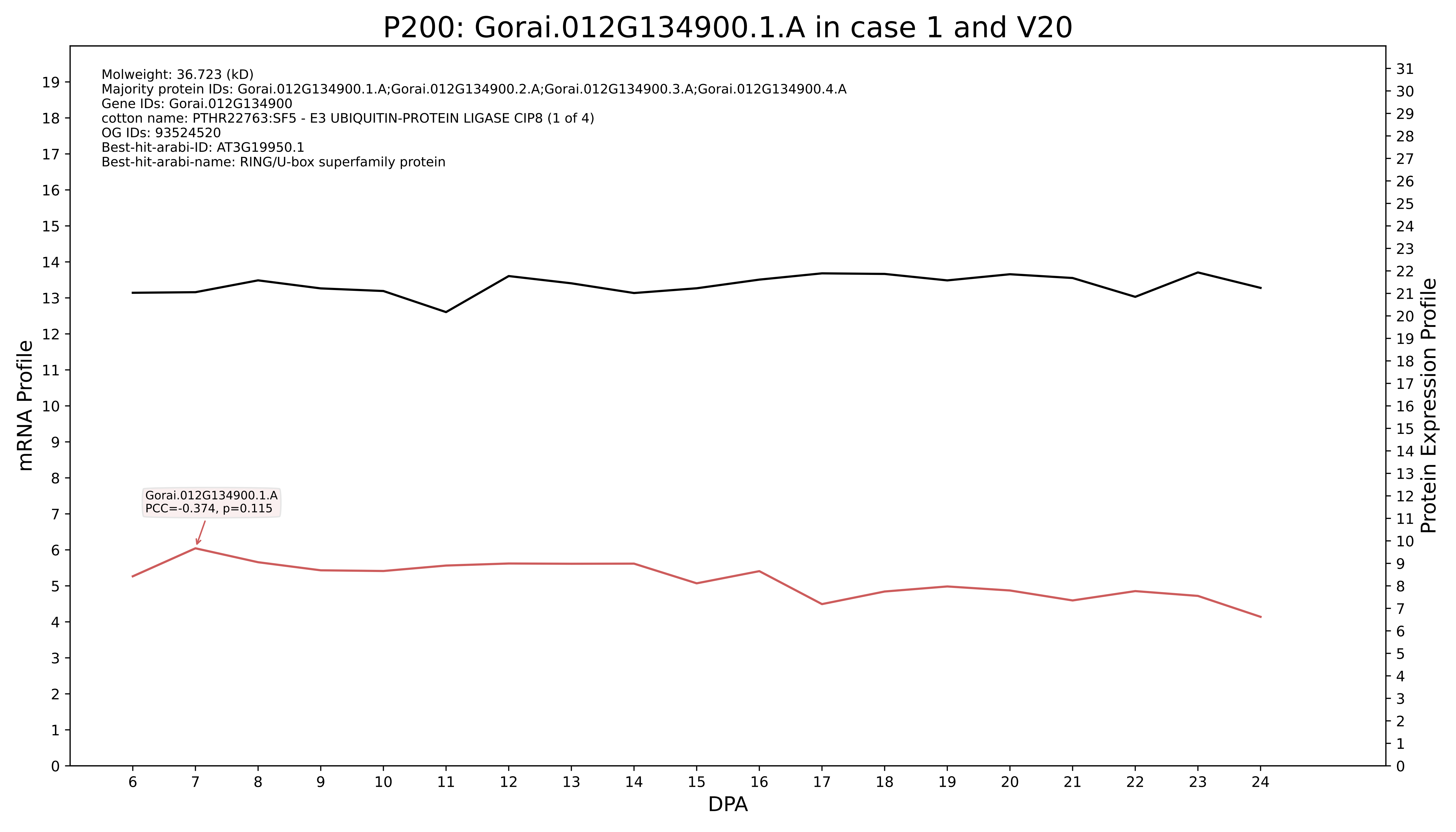

P200: Gorai.012G134900.1.A in case 1 and V20
Molweight: 36.723 (kD)
Majority protein IDs: Gorai.012G134900.1.A;Gorai.012G134900.2.A;Gorai.012G134900.3.A;Gorai.012G134900.4.A
Gene IDs: Gorai.012G134900
cotton name: PTHR22763:SF5 - E3 UBIQUITIN-PROTEIN LIGASE CIP8 (1 of 4)
OG IDs: 93524520
Best-hit-arabi-ID: AT3G19950.1
Best-hit-arabi-name: RING/U-box superfamily protein
Gorai.012G134900.1.A
PCC=-0.374, p=0.115
mRNA Profile
Protein Expression Profile
DPA

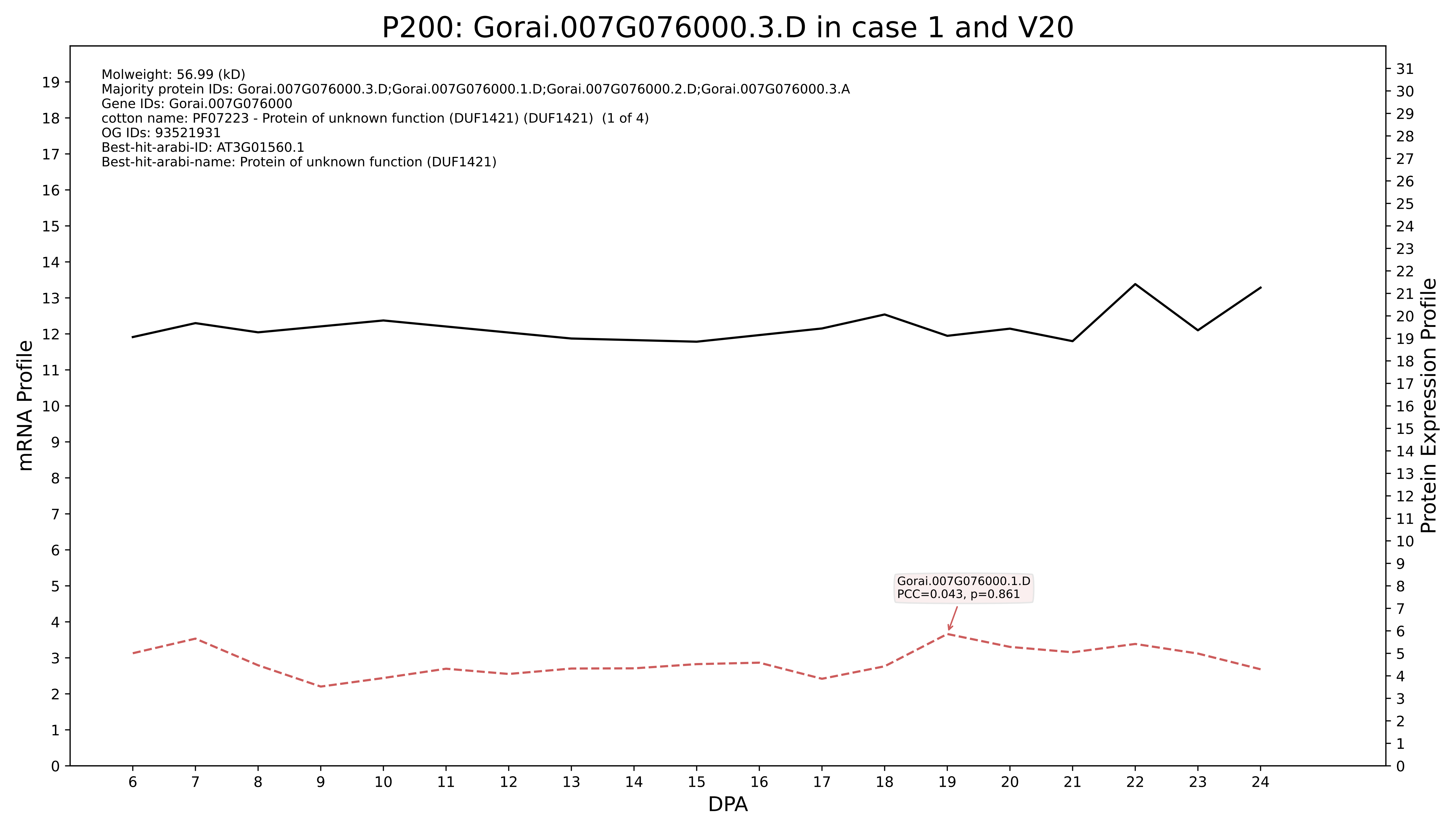
P200: Gorai.007G076000.3.D in case 1 and V20
Molweight: 56.99 (kD)
Majority protein IDs: Gorai.007G076000.3.D;Gorai.007G076000.1.D;Gorai.007G076000.2.D;Gorai.007G076000.3.A
Gene IDs: Gorai.007G076000
cotton name: PF07223 - Protein of unknown function (DUF1421) (DUF1421) (1 of 4)
OG IDs: 93521931
Best-hit-arabi-ID: AT3G01560.1
Best-hit-arabi-name: Protein of unknown function (DUF1421)
Gorai.007G076000.1.D
PCC=0.043, p=0.861
mRNA Profile
Protein Expression Profile
DPA

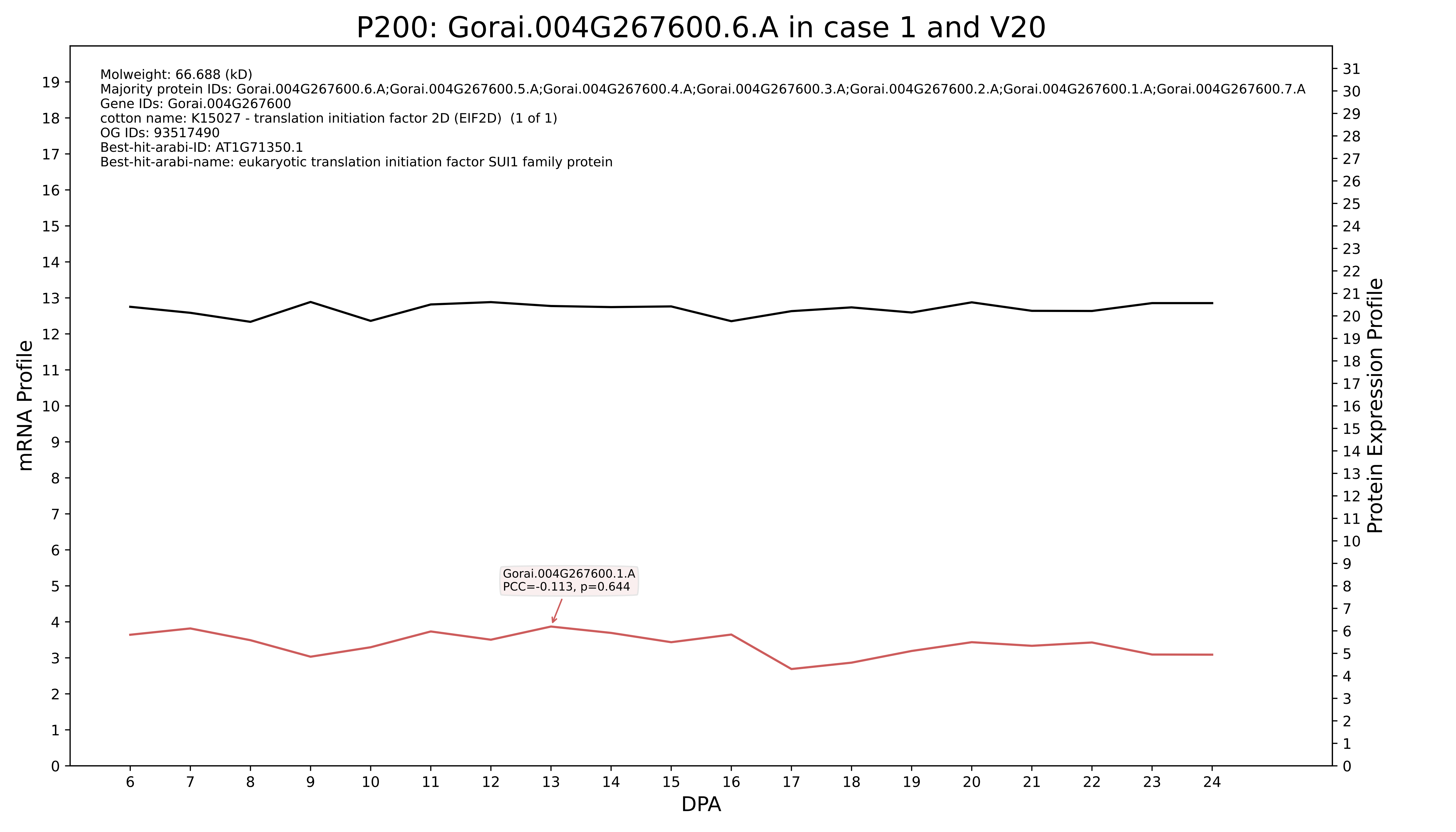
P200: Gorai.004G267600.6.A in case 1 and V20
Molweight: 66.688 (kD)
Majority protein IDs: Gorai.004G267600.6.A;Gorai.004G267600.5.A;Gorai.004G267600.4.A;Gorai.004G267600.3.A;Gorai.004G267600.2.A;Gorai.004G267600.1.A;Gorai.004G267600.7.A
Gene IDs: Gorai.004G267600
cotton name: K15027 - translation initiation factor 2D (EIF2D) (1 of 1)
OG IDs: 93517490
Best-hit-arabi-ID: AT1G71350.1
Best-hit-arabi-name: eukaryotic translation initiation factor SUI1 family protein
Gorai.004G267600.1.A
PCC=-0.113, p=0.644
mRNA Profile
Protein Expression Profile
DPA

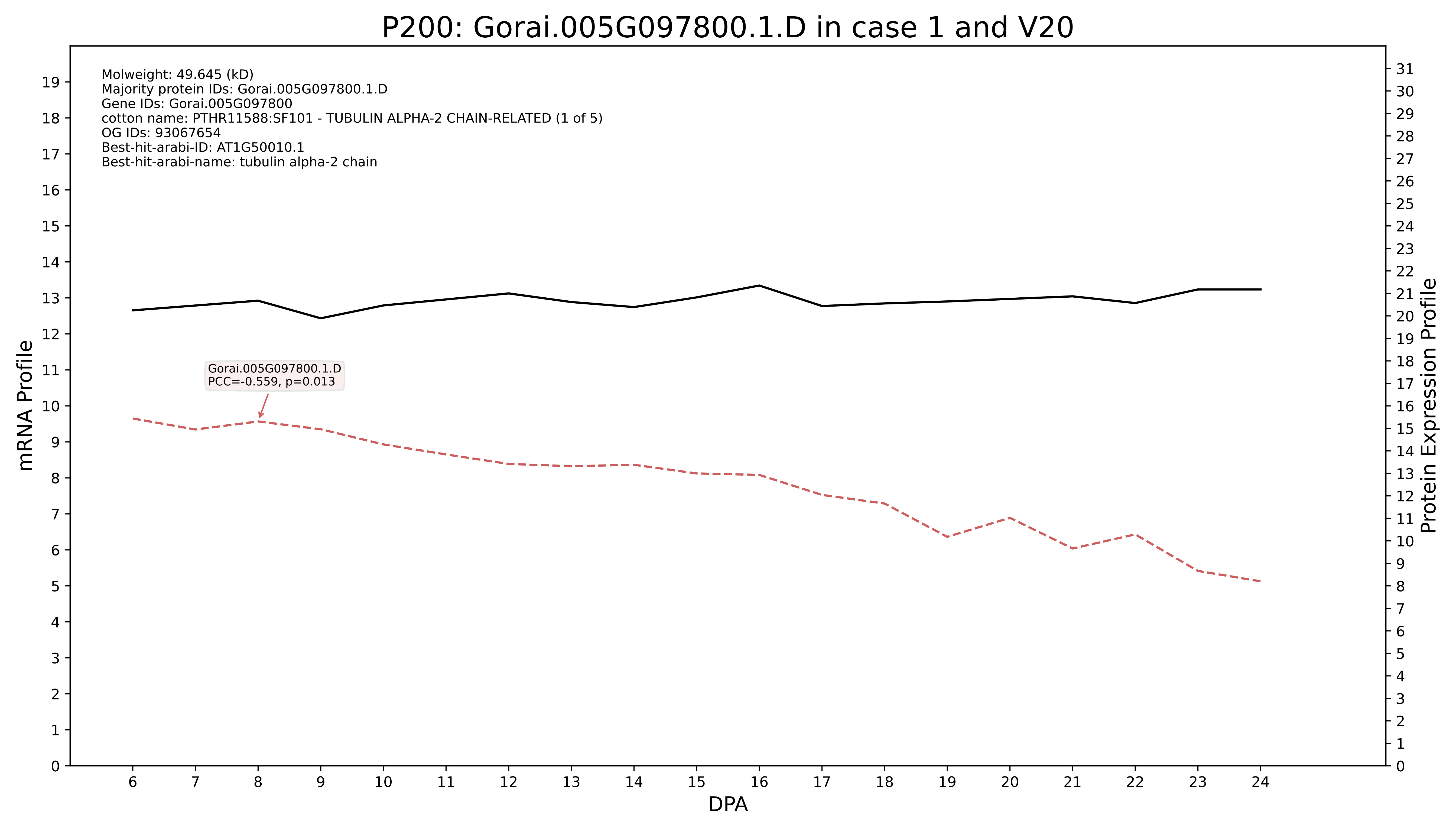
P200: Gorai.005G097800.1.D in case 1 and V20
Molweight: 49.645 (kD)
Majority protein IDs: Gorai.005G097800.1.D
Gene IDs: Gorai.005G097800
cotton name: PTHR11588:SF101 - TUBULIN ALPHA-2 CHAIN-RELATED (1 of 5)
OG IDs: 93067654
Best-hit-arabi-ID: AT1G50010.1
Best-hit-arabi-name: tubulin alpha-2 chain
Gorai.005G097800.1.D
PCC=-0.559, p=0.013
mRNA Profile
Protein Expression Profile
DPA

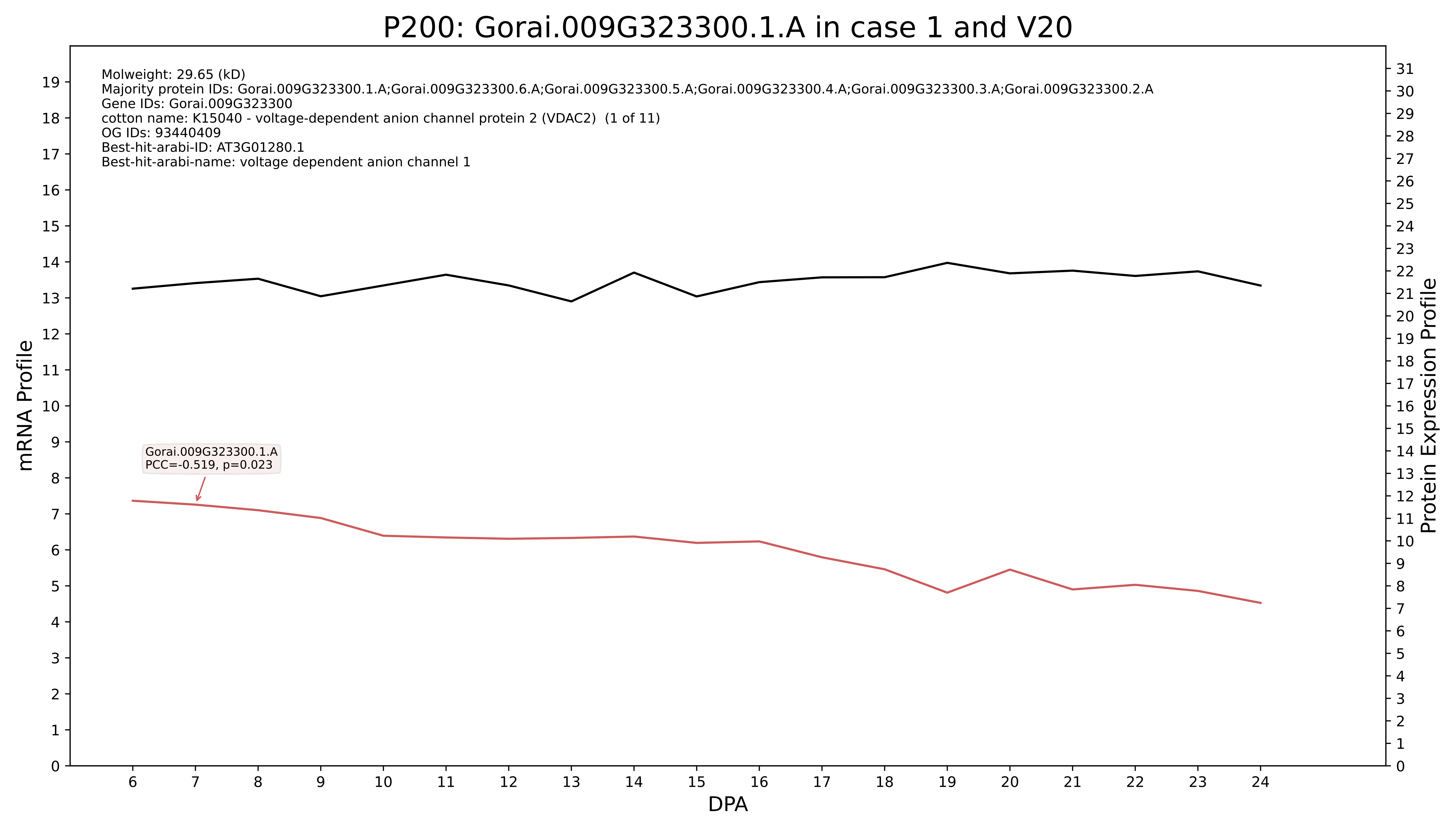
P200: Gorai.009G323300.1.A in case 1 and V20
Molweight: 29.65 (kD)
Majority protein IDs: Gorai.009G323300.1.A;Gorai.009G323300.6.A;Gorai.009G323300.5.A;Gorai.009G323300.4.A;Gorai.009G323300.3.A;Gorai.009G323300.2.A
Gene IDs: Gorai.009G323300
cotton name: K15040 - voltage-dependent anion channel protein 2 (VDAC2) (1 of 11)
OG IDs: 93440409
Best-hit-arabi-ID: AT3G01280.1
Best-hit-arabi-name: voltage dependent anion channel 1
Gorai.009G323300.1.A
PCC=-0.519, p=0.023
mRNA Profile
Protein Expression Profile
DPA

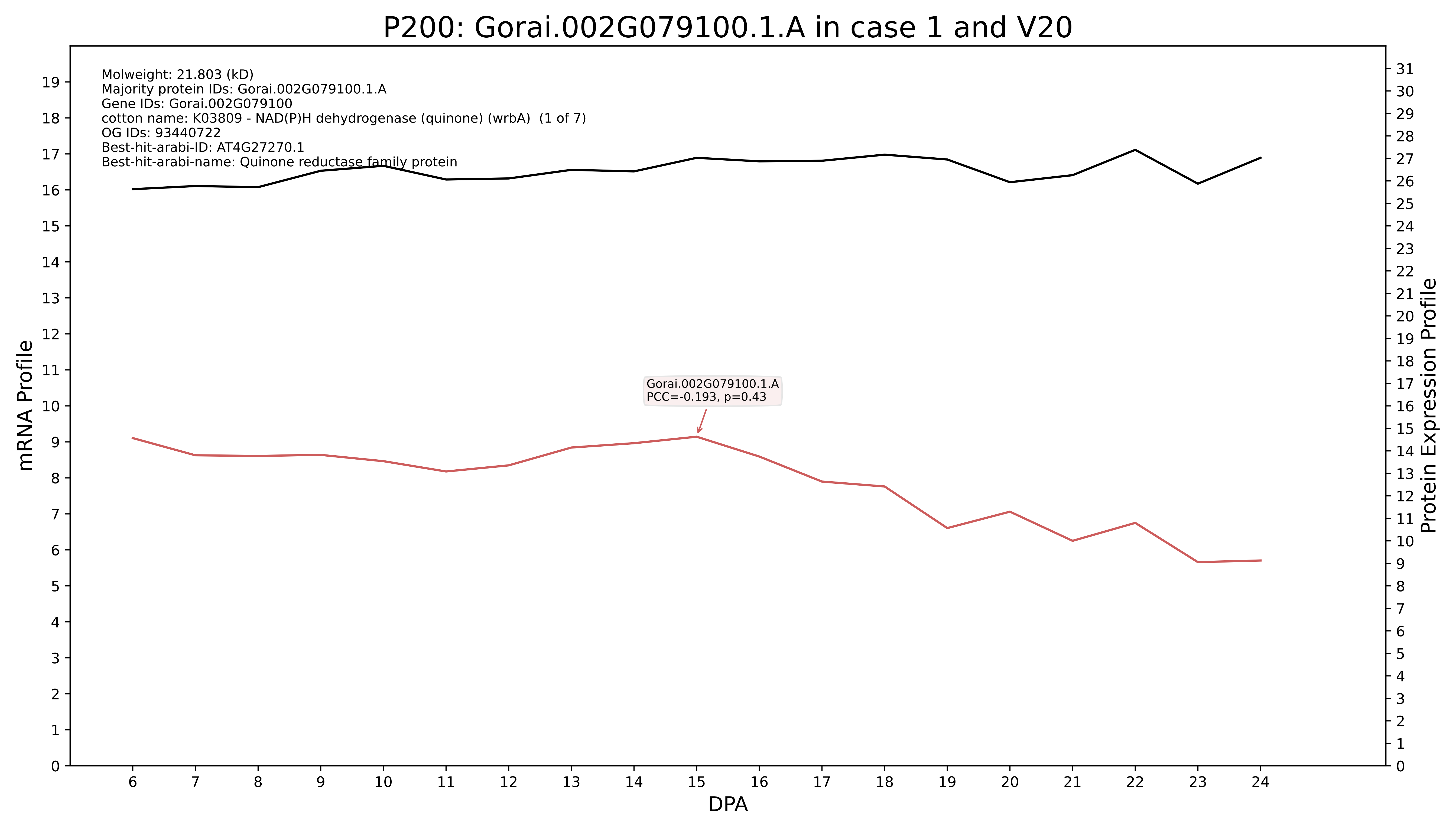
P200: Gorai.002G079100.1.A in case 1 and V20
Molweight: 21.803 (kD)
Majority protein IDs: Gorai.002G079100.1.A
Gene IDs: Gorai.002G079100
cotton name: K03809 - NAD(P)H dehydrogenase (quinone) (wrbA) (1 of 7)
OG IDs: 93440722
Best-hit-arabi-ID: AT4G27270.1
Best-hit-arabi-name: Quinone reductase family protein
Gorai.002G079100.1.A
PCC=-0.193, p=0.43
mRNA Profile
Protein Expression Profile
DPA

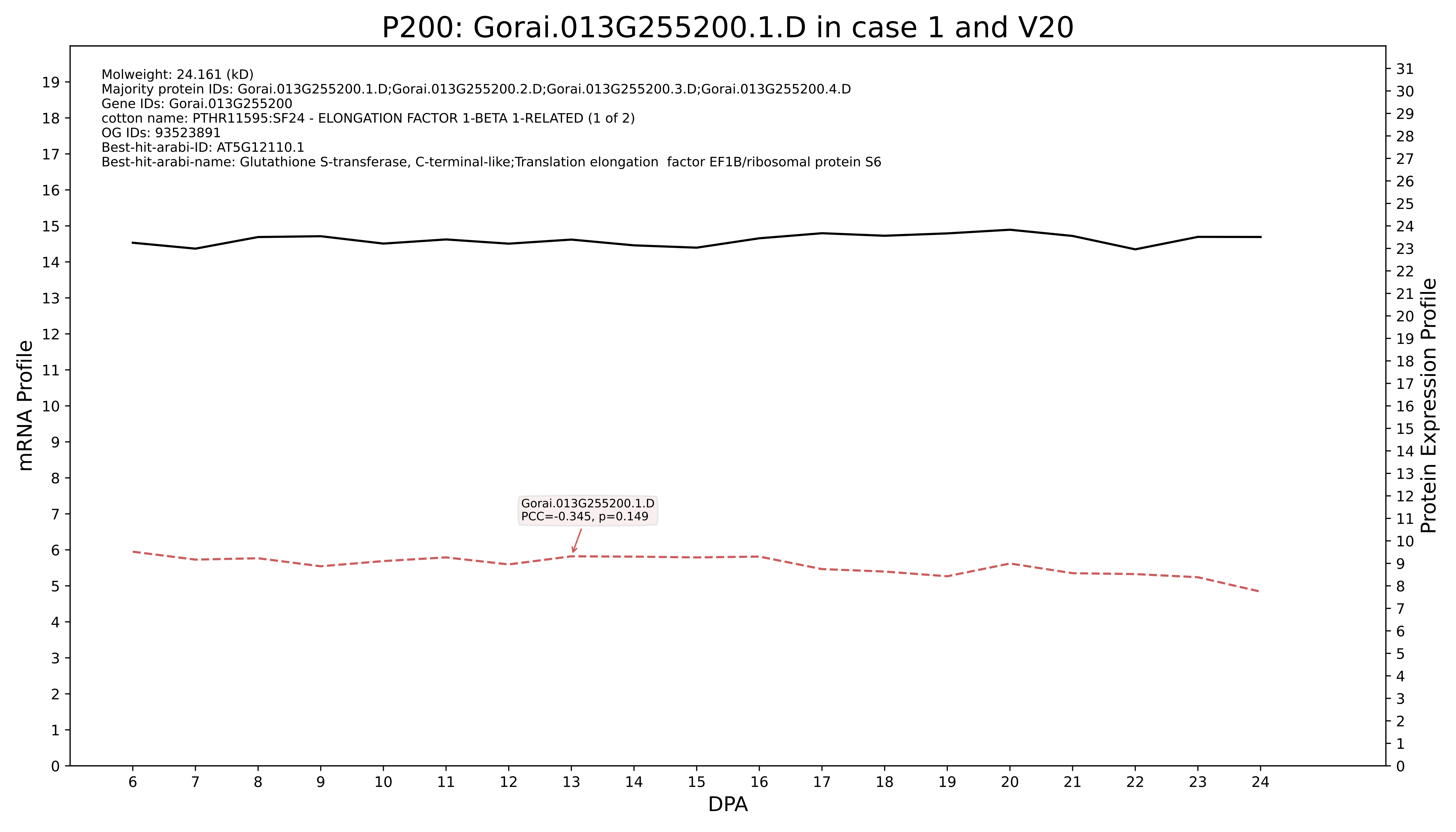

P200: Gorai.013G255200.1.D in case 1 and V20
Molweight: 24.161 (kD)
Majority protein IDs: Gorai.013G255200.1.D;Gorai.013G255200.2.D;Gorai.013G255200.3.D;Gorai.013G255200.4.D
Gene IDs: Gorai.013G255200
cotton name: PTHR11595:SF24 - ELONGATION FACTOR 1-BETA 1-RELATED (1 of 2)
OG IDs: 93523891
Best-hit-arabi-ID: AT5G12110.1
Best-hit-arabi-name: Glutathione S-transferase, C-terminal-like;Translation elongation factor EF1B/ribosomal protein S6
Gorai.013G255200.1.D
PCC=-0.345, p=0.149
mRNA Profile
Protein Expression Profile
DPA

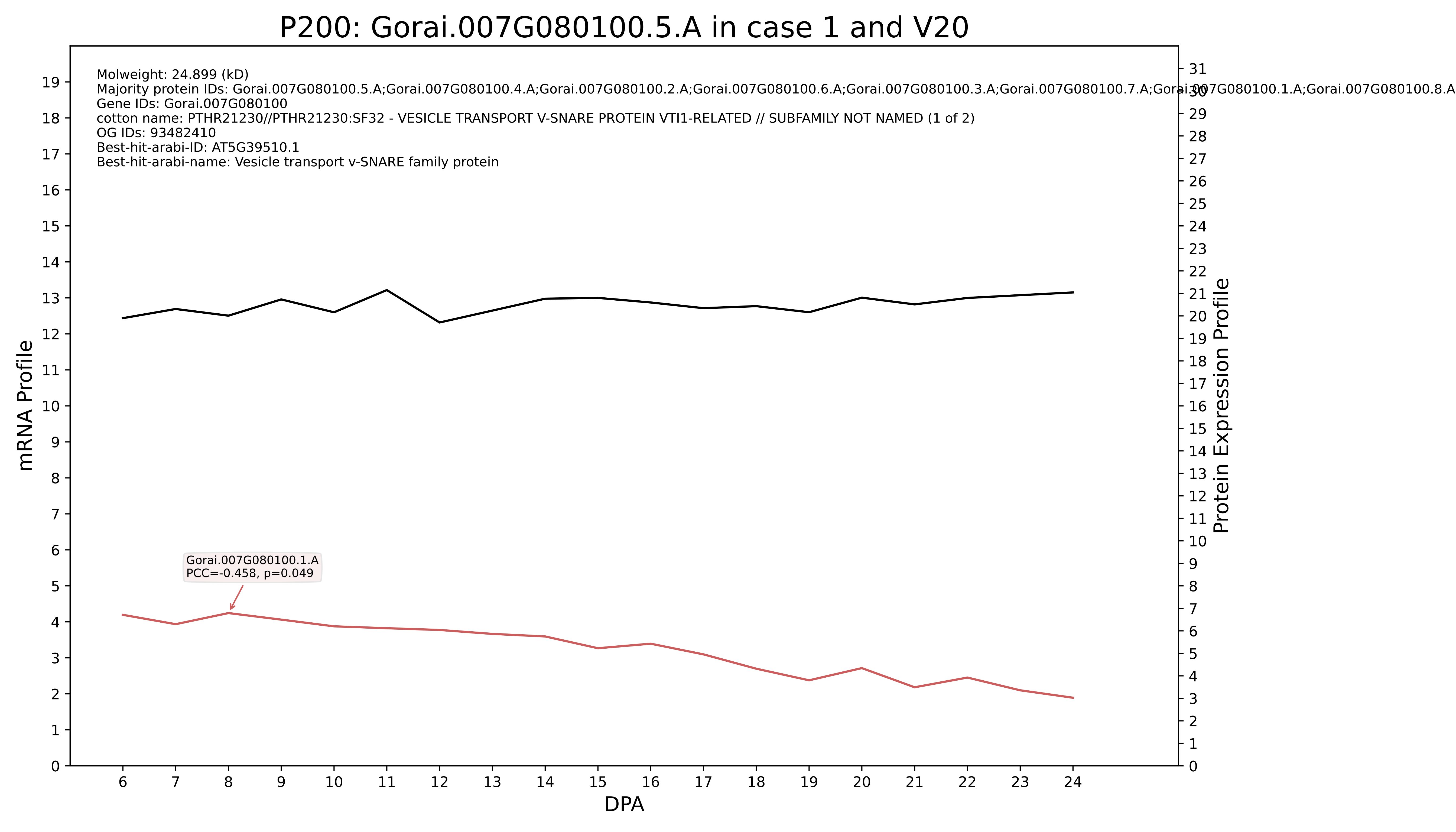

P200: Gorai.007G080100.5.A in case 1 and V20
Molweight: 24.899 (kD)
Majority protein IDs: Gorai.007G080100.5.A;Gorai.007G080100.4.A;Gorai.007G080100.2.A;Gorai.007G080100.6.A;Gorai.007G080100.3.A;Gorai.007G080100.7.A;Gorai.007G080100.1.A;Gorai.007G080100.8.A
Gene IDs: Gorai.007G080100
cotton name: PTHR21230//PTHR21230:SF32 - VESICLE TRANSPORT V-SNARE PROTEIN VTI1-RELATED // SUBFAMILY NOT NAMED (1 of 2)
OG IDs: 93482410
Best-hit-arabi-ID: AT5G39510.1
Best-hit-arabi-name: Vesicle transport v-SNARE family protein
Gorai.007G080100.1.A
PCC=-0.458, p=0.049
mRNA Profile
Protein Expression Profile
DPA

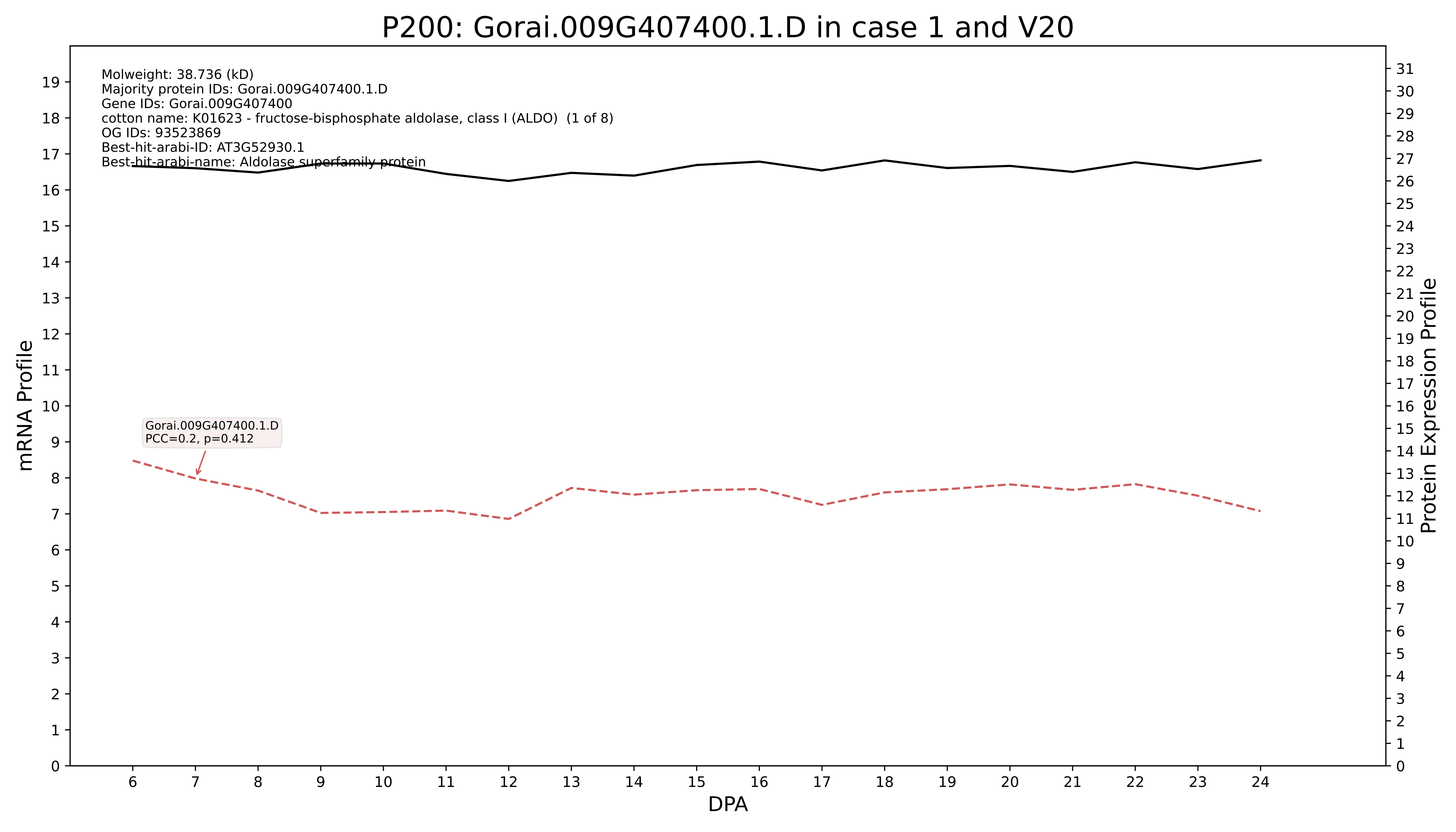

P200: Gorai.009G407400.1.D in case 1 and V20
Molweight: 38.736 (kD)
Majority protein IDs: Gorai.009G407400.1.D
Gene IDs: Gorai.009G407400
cotton name: K01623 - fructose-bisphosphate aldolase, class I (ALDO) (1 of 8)
OG IDs: 93523869
Best-hit-arabi-ID: AT3G52930.1
Best-hit-arabi-name: Aldolase superfamily protein
Gorai.009G407400.1.D
PCC=0.2, p=0.412
mRNA Profile
Protein Expression Profile
DPA

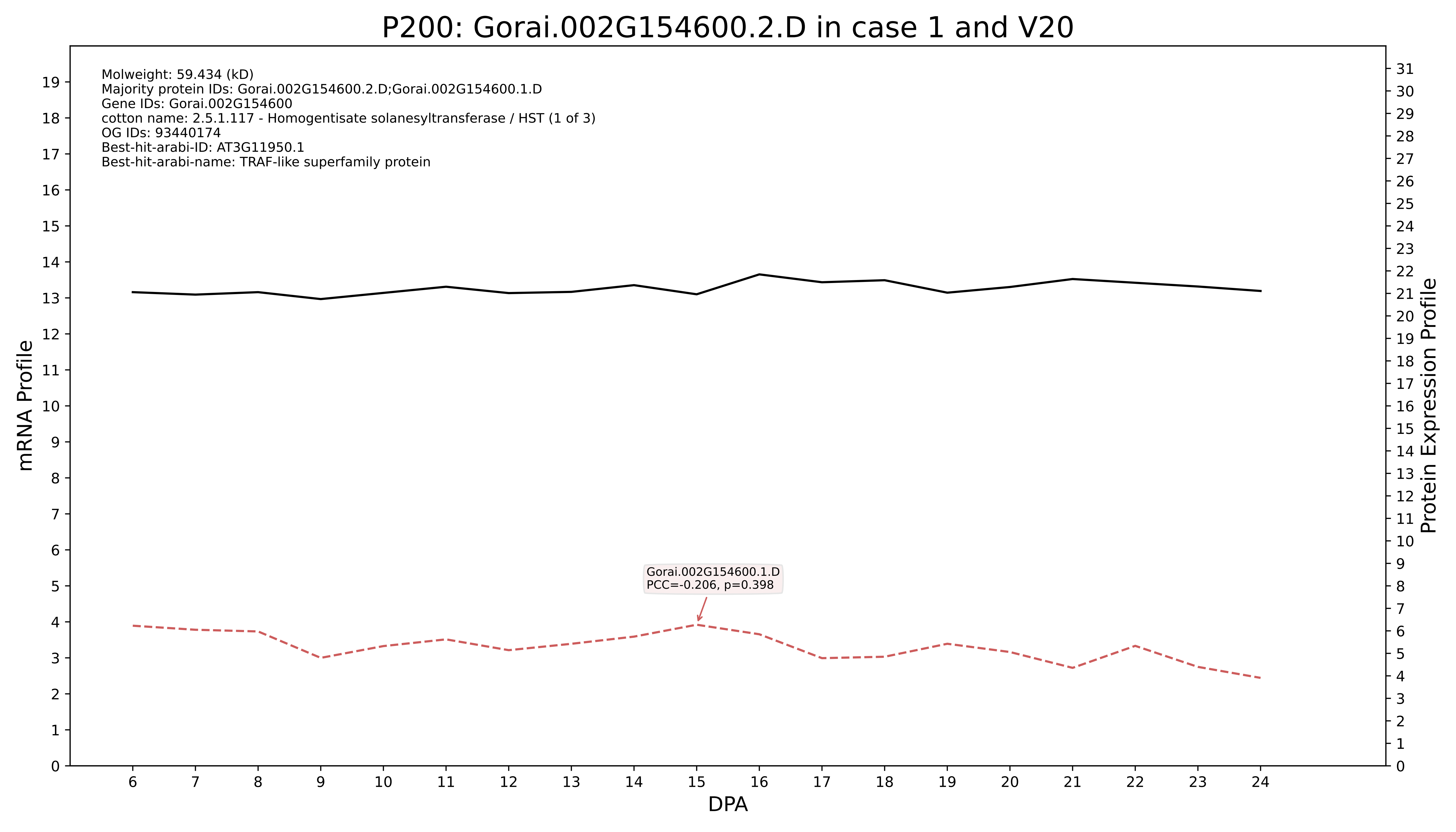
P200: Gorai.002G154600.2.D in case 1 and V20
Molweight: 59.434 (kD)
Majority protein IDs: Gorai.002G154600.2.D;Gorai.002G154600.1.D
Gene IDs: Gorai.002G154600
cotton name: 2.5.1.117 - Homogentisate solanesyltransferase / HST (1 of 3)
OG IDs: 93440174
Best-hit-arabi-ID: AT3G11950.1
Best-hit-arabi-name: TRAF-like superfamily protein
Gorai.002G154600.1.D
PCC=-0.206, p=0.398
mRNA Profile
Protein Expression Profile
DPA

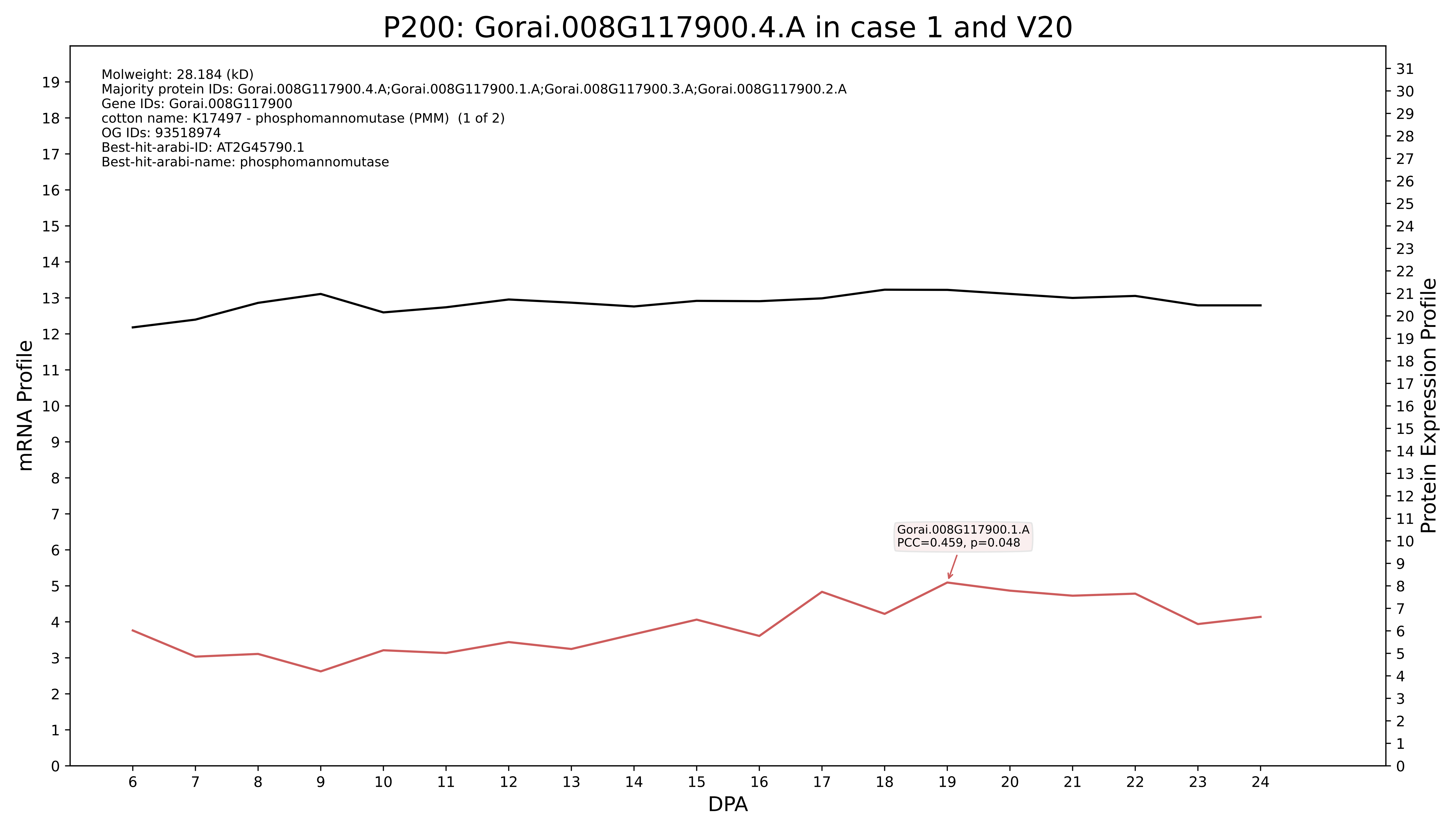
P200: Gorai.008G117900.4.A in case 1 and V20
Molweight: 28.184 (kD)
Majority protein IDs: Gorai.008G117900.4.A;Gorai.008G117900.1.A;Gorai.008G117900.3.A;Gorai.008G117900.2.A
Gene IDs: Gorai.008G117900
cotton name: K17497 - phosphomannomutase (PMM) (1 of 2)
OG IDs: 93518974
Best-hit-arabi-ID: AT2G45790.1
Best-hit-arabi-name: phosphomannomutase
Gorai.008G117900.1.A
PCC=0.459, p=0.048
mRNA Profile
Protein Expression Profile
DPA

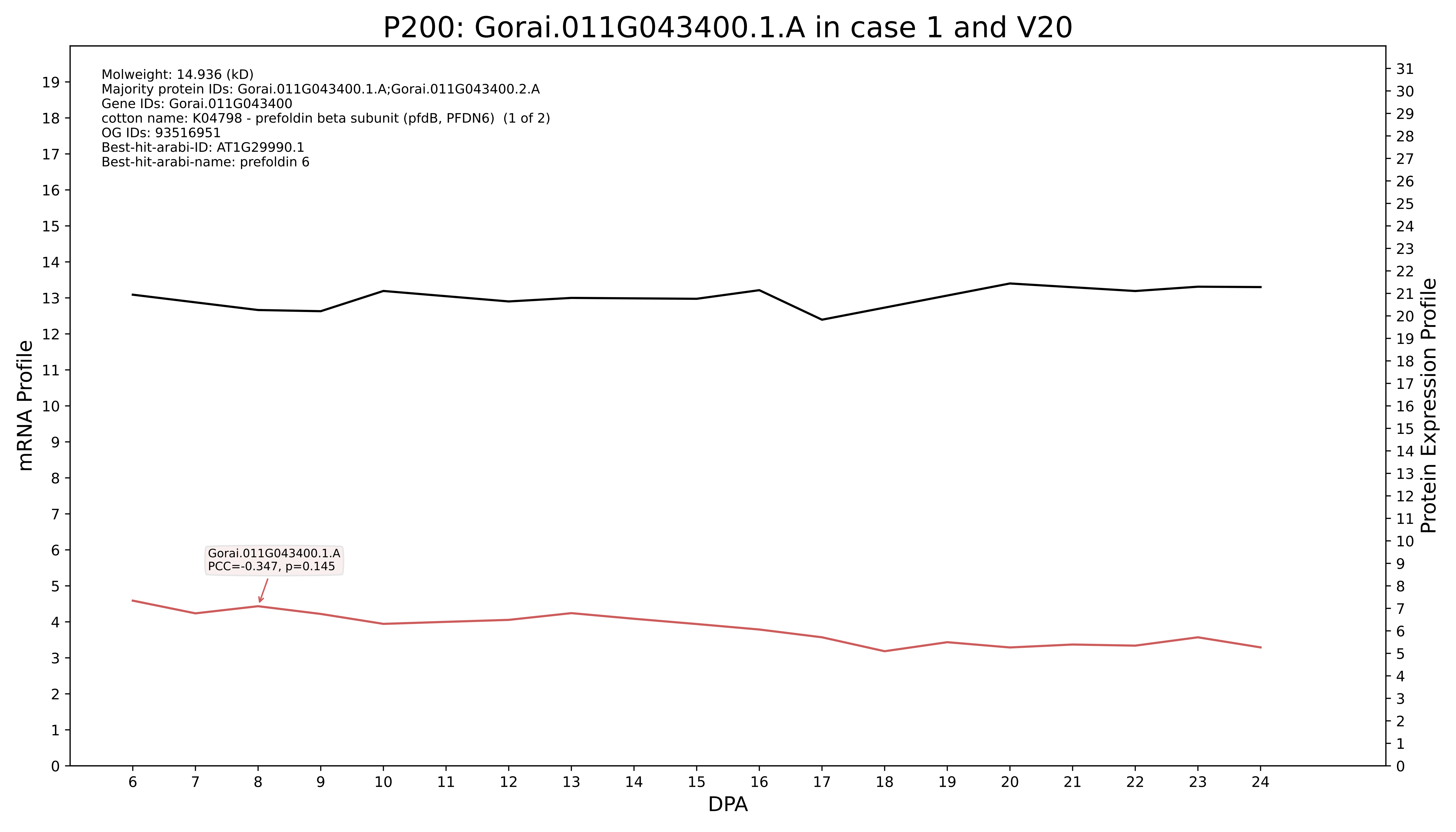

P200: Gorai.011G043400.1.A in case 1 and V20
Molweight: 14.936 (kD)
Majority protein IDs: Gorai.011G043400.1.A;Gorai.011G043400.2.A
Gene IDs: Gorai.011G043400
cotton name: K04798 - prefoldin beta subunit (pfdB, PFDN6) (1 of 2)
OG IDs: 93516951
Best-hit-arabi-ID: AT1G29990.1
Best-hit-arabi-name: prefoldin 6
Gorai.011G043400.1.A
PCC=-0.347, p=0.145
mRNA Profile
Protein Expression Profile
DPA

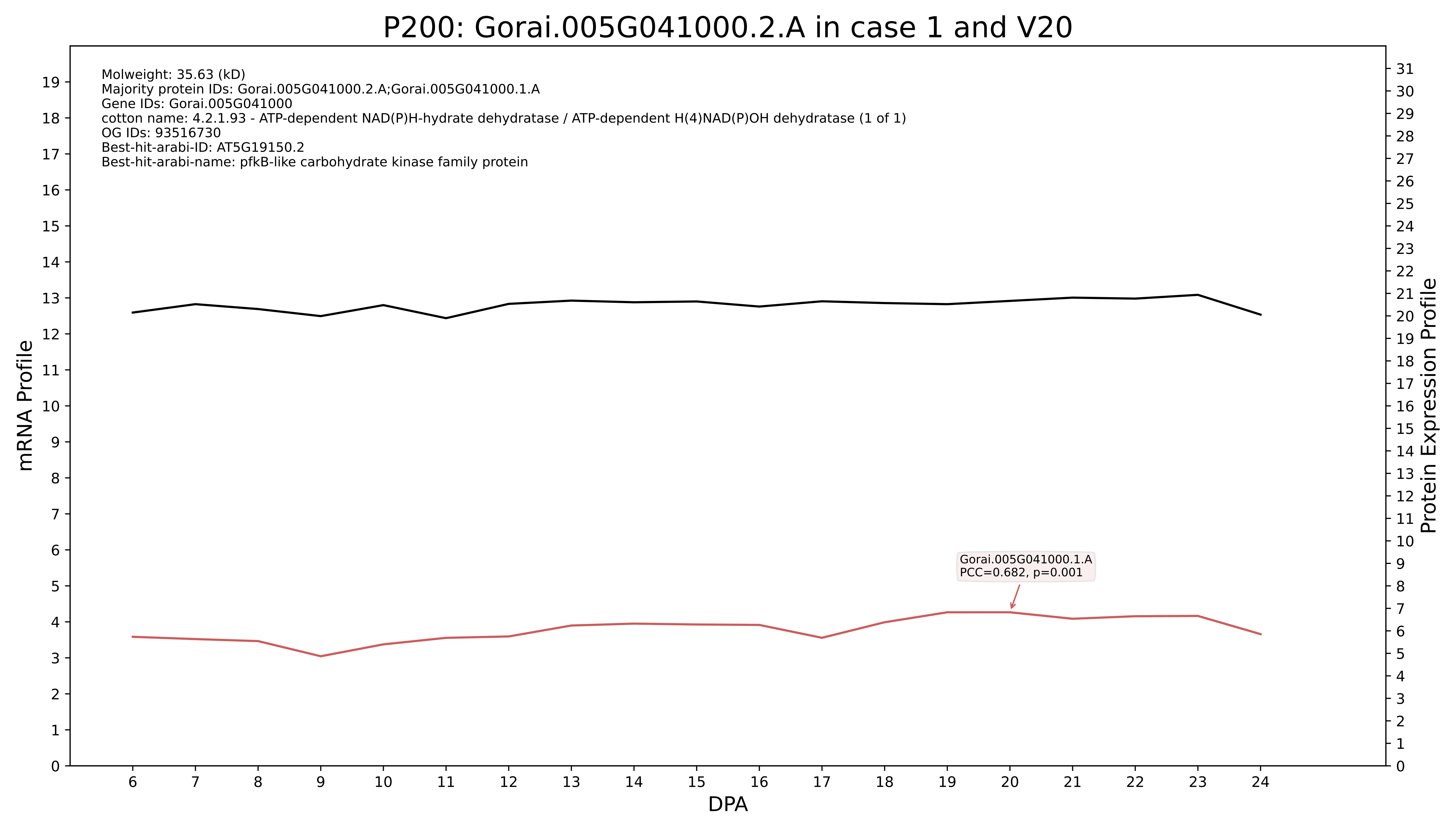
P200: Gorai.005G041000.2.A in case 1 and V20
Molweight: 35.63 (kD)
Majority protein IDs: Gorai.005G041000.2.A;Gorai.005G041000.1.A
Gene IDs: Gorai.005G041000
cotton name: 4.2.1.93 - ATP-dependent NAD(P)H-hydrate dehydratase / ATP-dependent H(4)NAD(P)OH dehydratase (1 of 1)
OG IDs: 93516730
Best-hit-arabi-ID: AT5G19150.2
Best-hit-arabi-name: pfkB-like carbohydrate kinase family protein
Gorai.005G041000.1.A
PCC=0.682, p=0.001
mRNA Profile
Protein Expression Profile
DPA

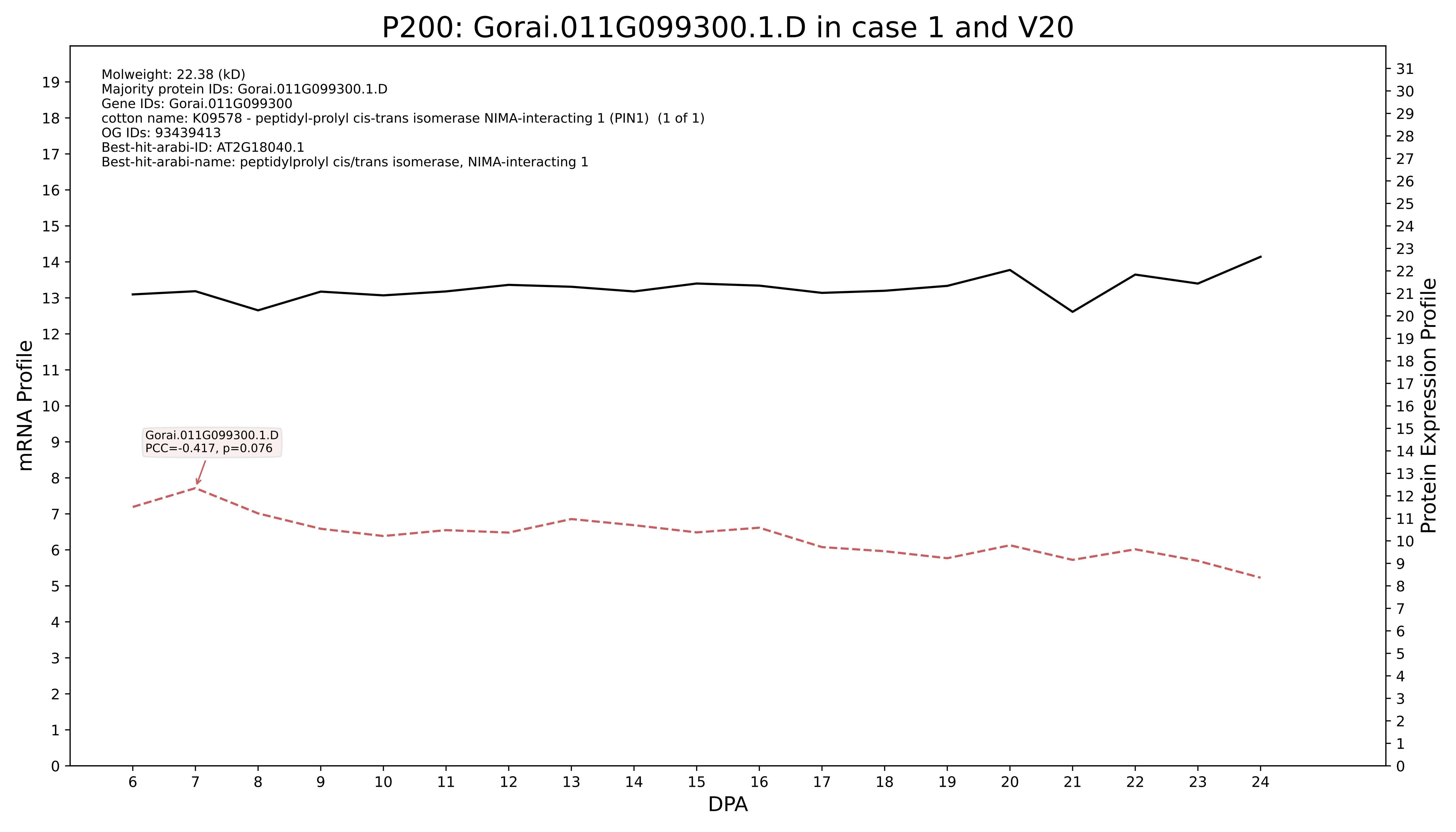
P200: Gorai.011G099300.1.D in case 1 and V20
Molweight: 22.38 (kD)
Majority protein IDs: Gorai.011G099300.1.D
Gene IDs: Gorai.011G099300
cotton name: K09578 - peptidyl-prolyl cis-trans isomerase NIMA-interacting 1 (PIN1) (1 of 1)
OG IDs: 93439413
Best-hit-arabi-ID: AT2G18040.1
Best-hit-arabi-name: peptidylprolyl cis/trans isomerase, NIMA-interacting 1
Gorai.011G099300.1.D
PCC=-0.417, p=0.076
mRNA Profile
Protein Expression Profile
DPA

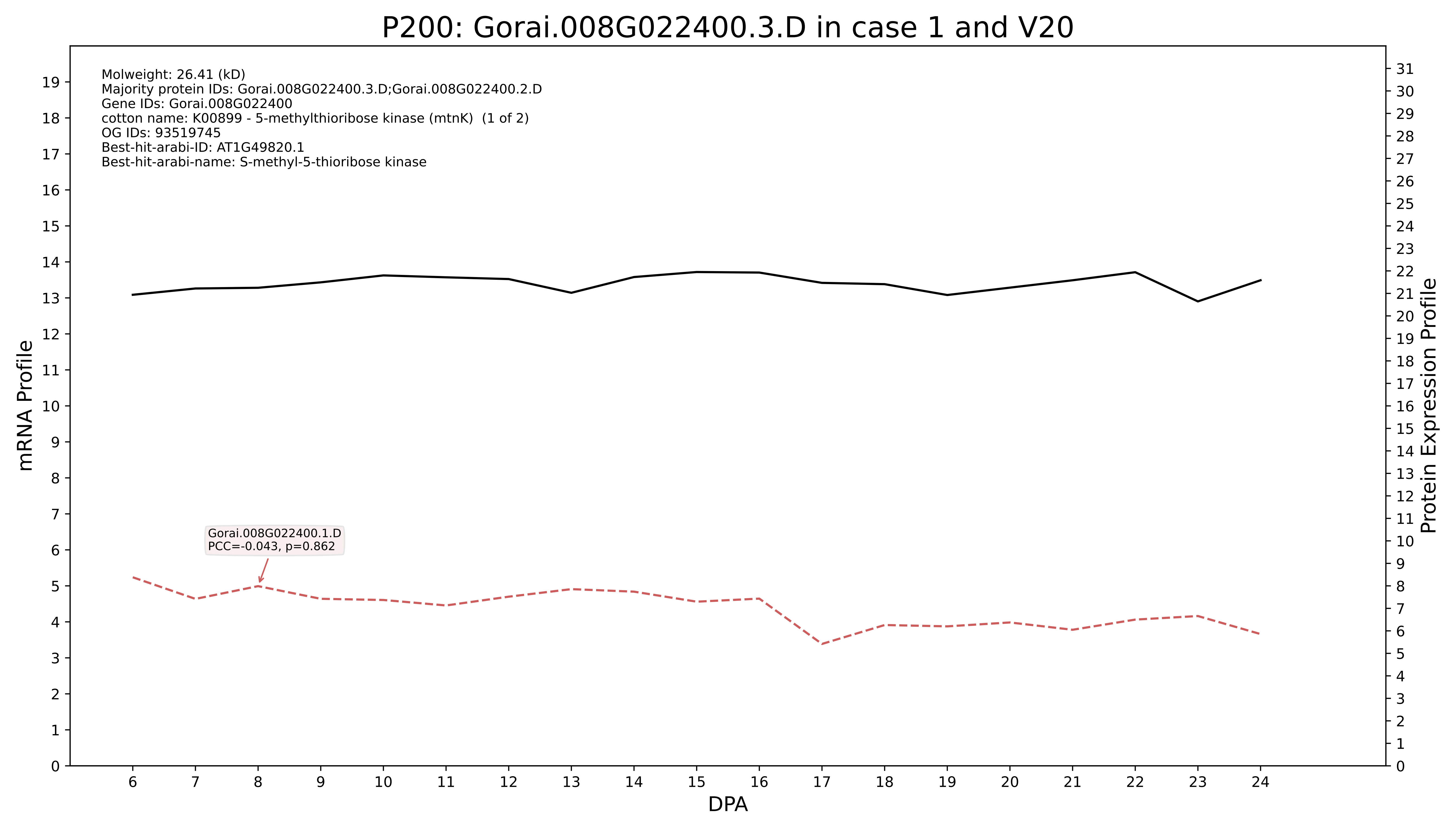

P200: Gorai.008G022400.3.D in case 1 and V20
Molweight: 26.41 (kD)
Majority protein IDs: Gorai.008G022400.3.D;Gorai.008G022400.2.D
Gene IDs: Gorai.008G022400
cotton name: K00899 - 5-methylthioribose kinase (mtnK) (1 of 2)
OG IDs: 93519745
Best-hit-arabi-ID: AT1G49820.1
Best-hit-arabi-name: S-methyl-5-thioribose kinase
Gorai.008G022400.1.D
PCC=-0.043, p=0.862
mRNA Profile
Protein Expression Profile
DPA

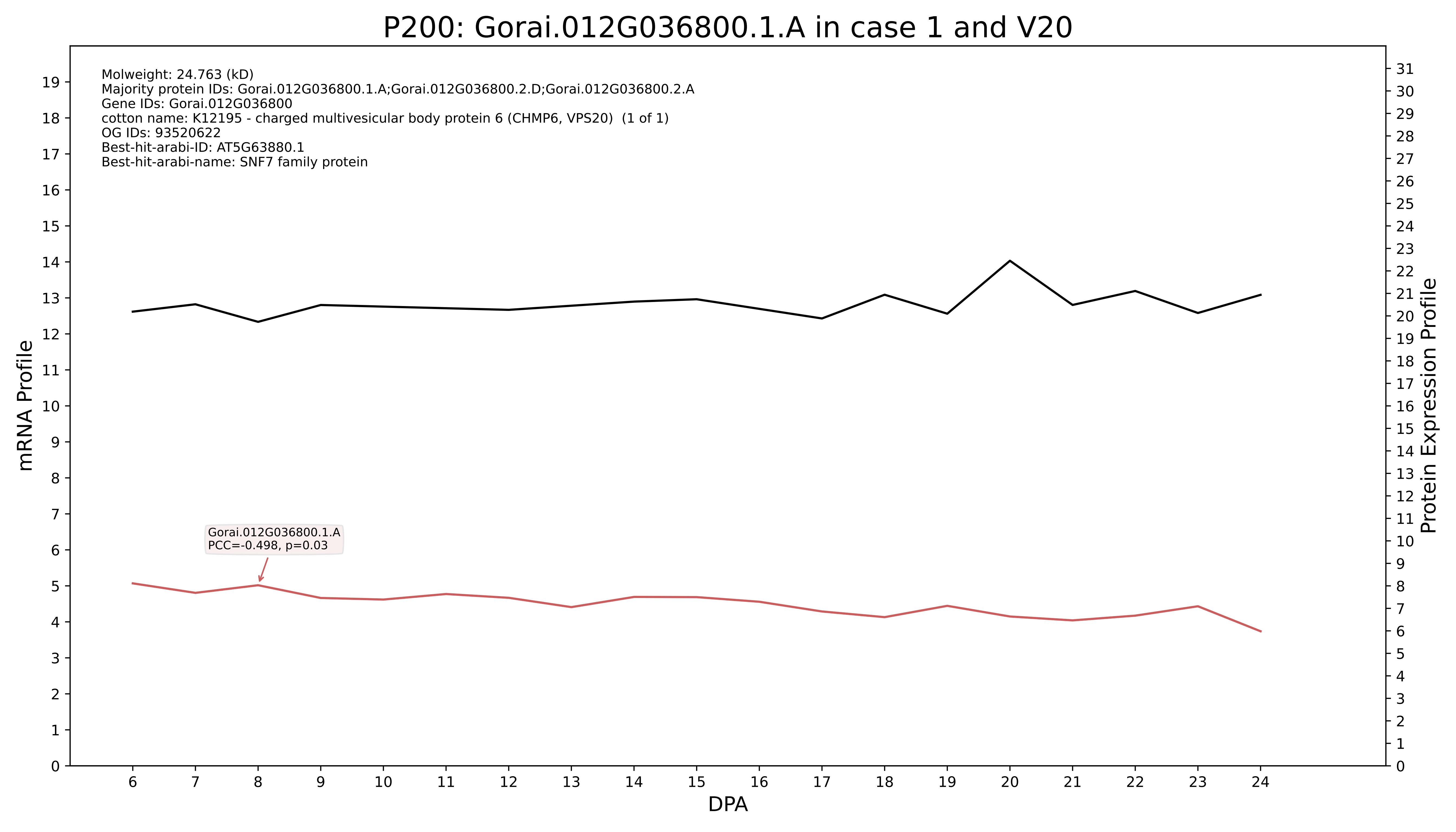
P200: Gorai.012G036800.1.A in case 1 and V20
Molweight: 24.763 (kD)
Majority protein IDs: Gorai.012G036800.1.A;Gorai.012G036800.2.D;Gorai.012G036800.2.A
Gene IDs: Gorai.012G036800
cotton name: K12195 - charged multivesicular body protein 6 (CHMP6, VPS20) (1 of 1)
OG IDs: 93520622
Best-hit-arabi-ID: AT5G63880.1
Best-hit-arabi-name: SNF7 family protein
Gorai.012G036800.1.A
PCC=-0.498, p=0.03
mRNA Profile
Protein Expression Profile
DPA

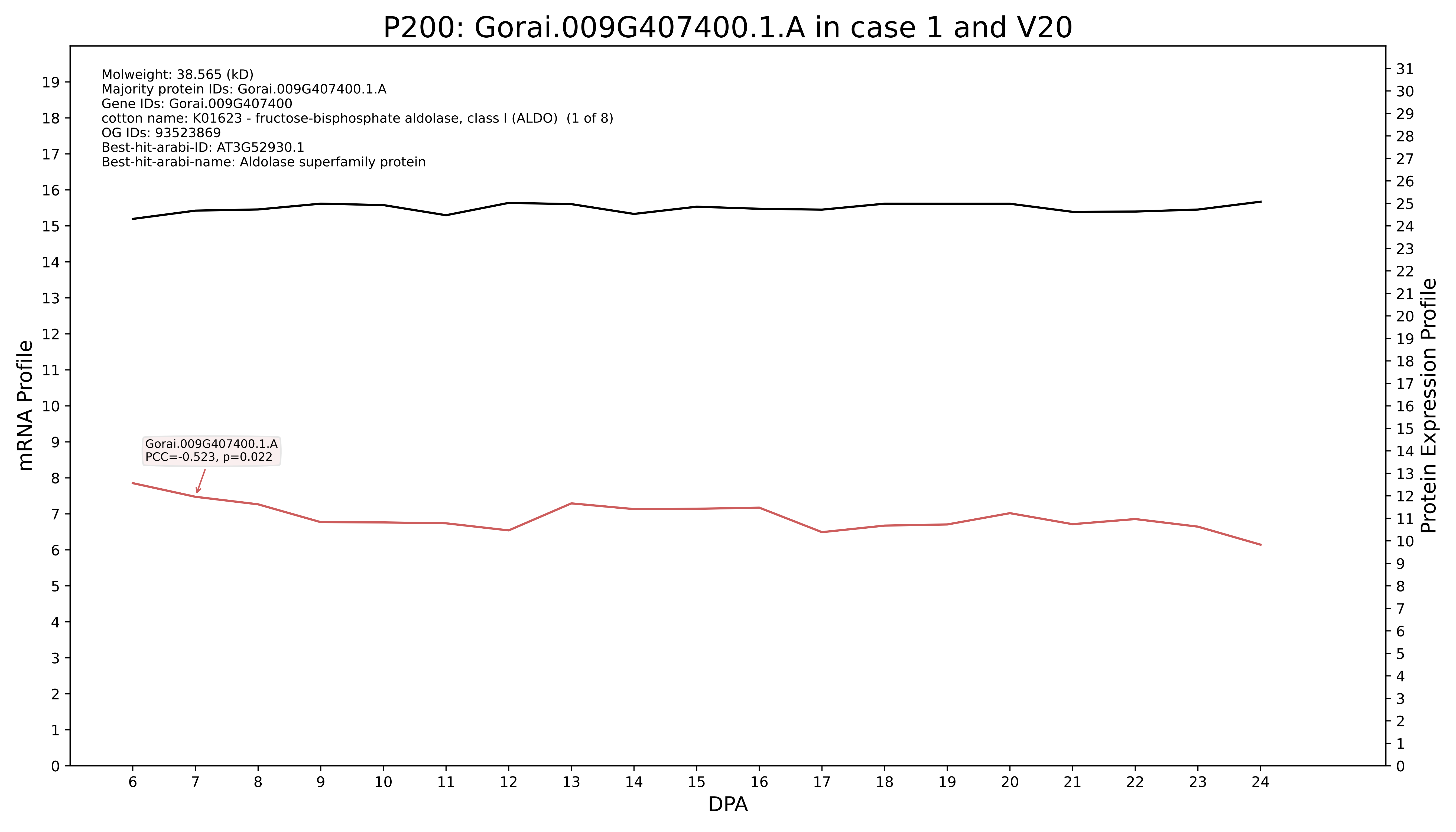
P200: Gorai.009G407400.1.A in case 1 and V20
Molweight: 38.565 (kD)
Majority protein IDs: Gorai.009G407400.1.A
Gene IDs: Gorai.009G407400
cotton name: K01623 - fructose-bisphosphate aldolase, class I (ALDO) (1 of 8)
OG IDs: 93523869
Best-hit-arabi-ID: AT3G52930.1
Best-hit-arabi-name: Aldolase superfamily protein
Gorai.009G407400.1.A
PCC=-0.523, p=0.022
mRNA Profile
Protein Expression Profile
DPA

### P200: Gorai.002G094300.8.D in case 1 and V20

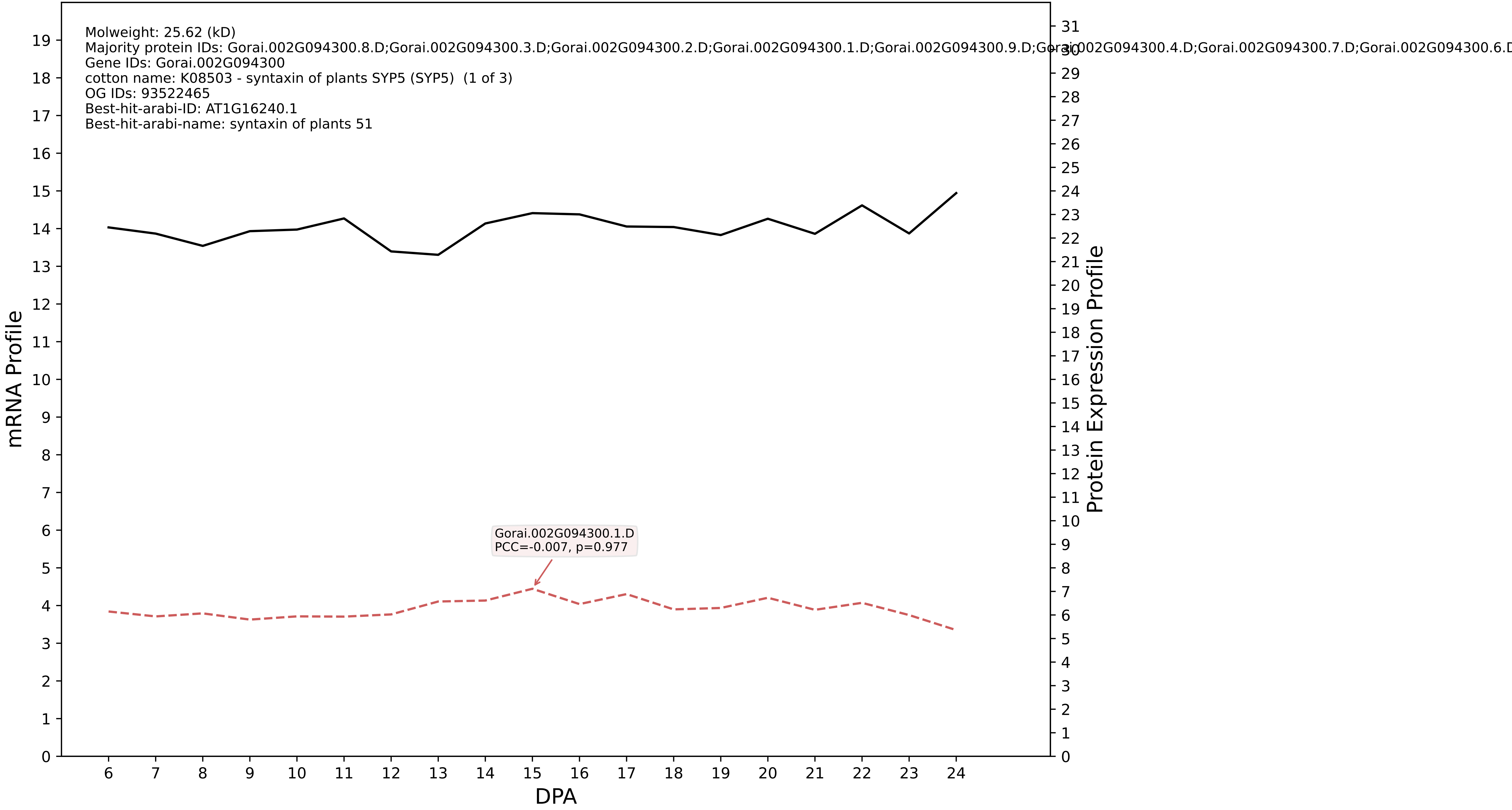

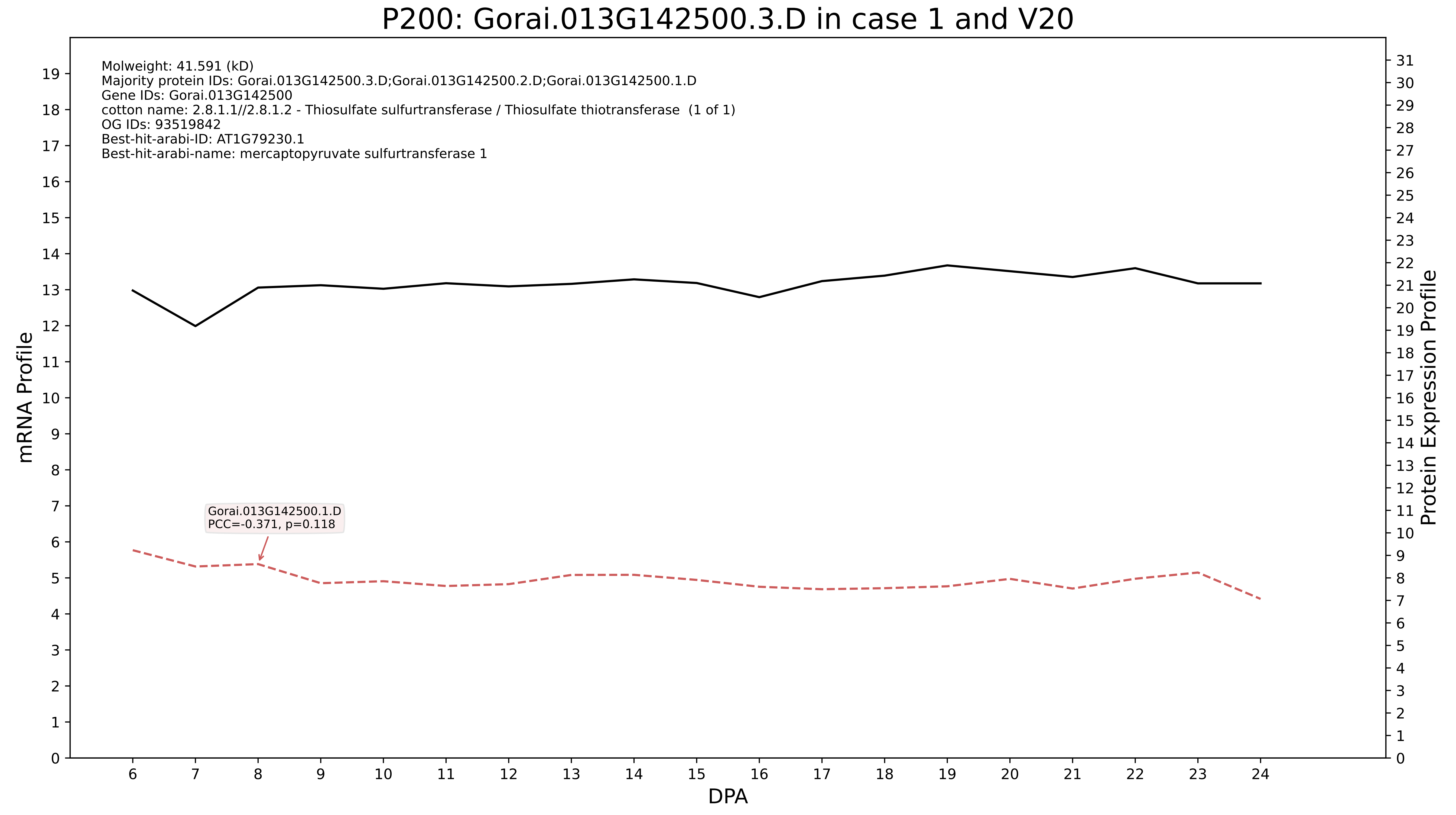
P200: Gorai.013G142500.3.D in case 1 and V20
Molweight: 41.591 (kD)
Majority protein IDs: Gorai.013G142500.3.D;Gorai.013G142500.2.D;Gorai.013G142500.1.D
Gene IDs: Gorai.013G142500
cotton name: 2.8.1.1//2.8.1.2 - Thiosulfate sulfurtransferase / Thiosulfate thiotransferase (1 of 1)
OG IDs: 93519842
Best-hit-arabi-ID: AT1G79230.1
Best-hit-arabi-name: mercaptopyruvate sulfurtransferase 1
Gorai.013G142500.1.D
PCC=-0.371, p=0.118
mRNA Profile
Protein Expression Profile
DPA

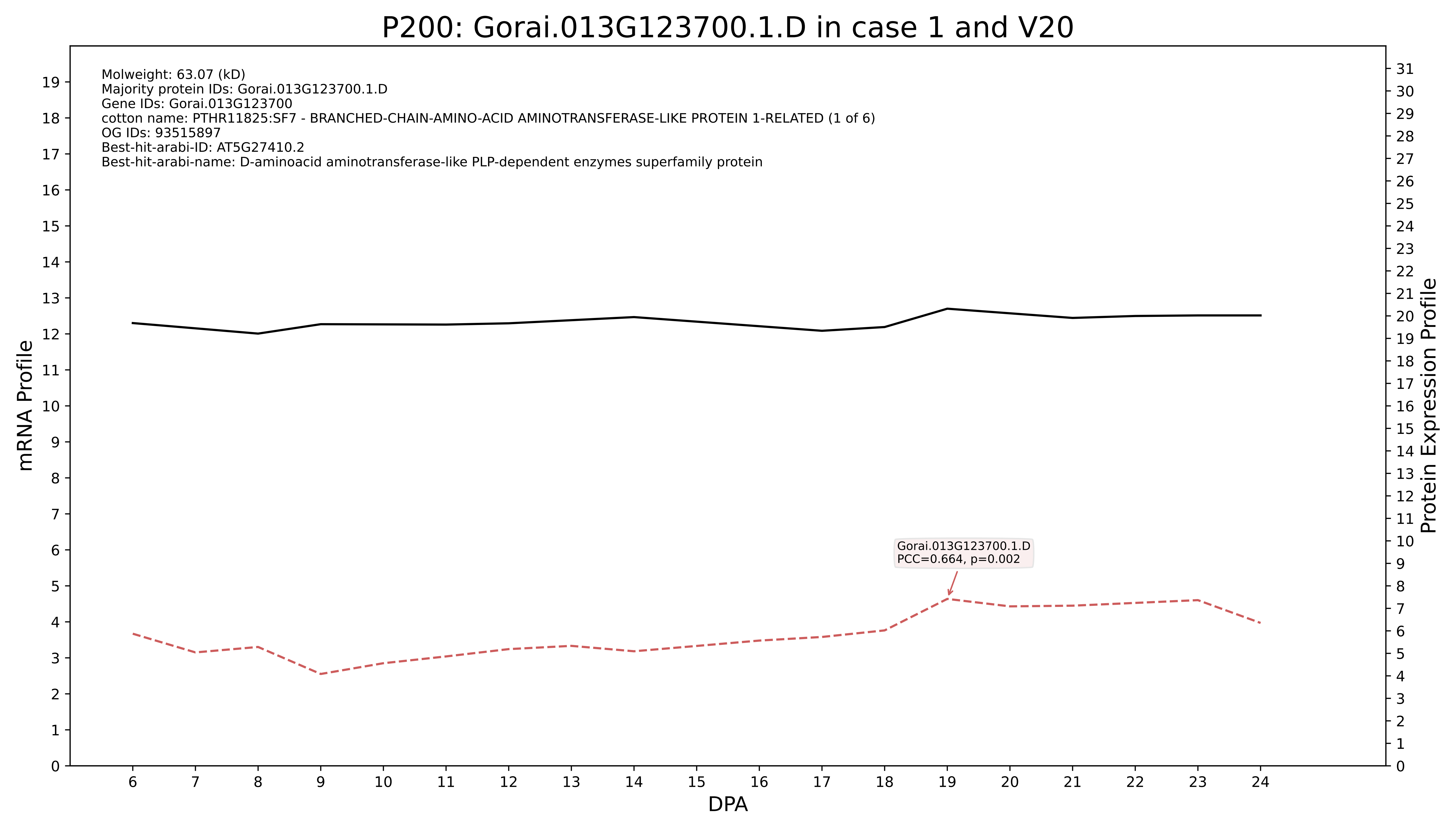
P200: Gorai.013G123700.1.D in case 1 and V20
Molweight: 63.07 (kD)
Majority protein IDs: Gorai.013G123700.1.D
Gene IDs: Gorai.013G123700
cotton name: PTHR11825:SF7 - BRANCHED-CHAIN-AMINO-ACID AMINOTRANSFERASE-LIKE PROTEIN 1-RELATED (1 of 6)
OG IDs: 93515897
Best-hit-arabi-ID: AT5G27410.2
Best-hit-arabi-name: D-aminoacid aminotransferase-like PLP-dependent enzymes superfamily protein
Gorai.013G123700.1.D
PCC=0.664, p=0.002
mRNA Profile
Protein Expression Profile
DPA

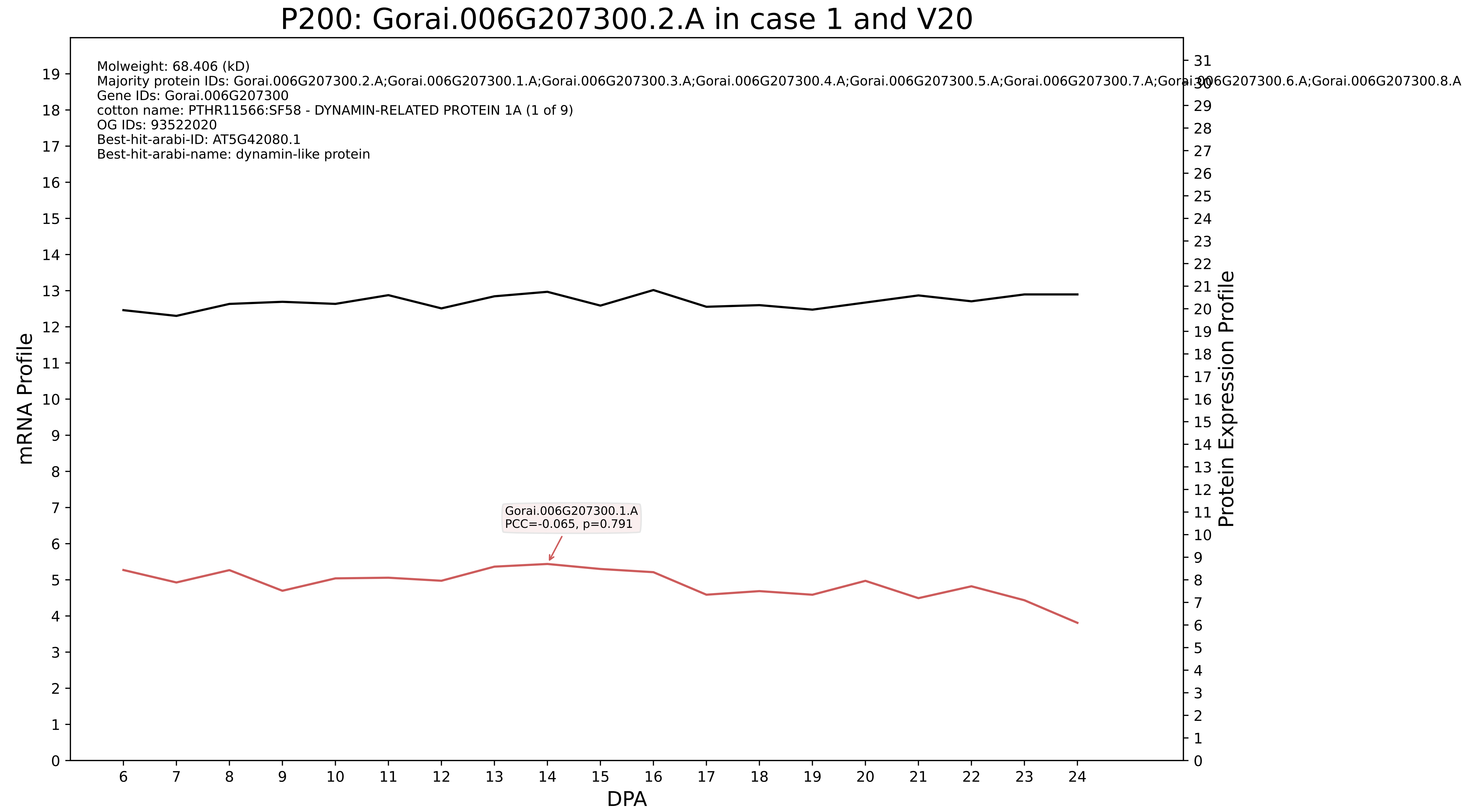
P200: Gorai.006G207300.2.A in case 1 and V20
Molweight: 68.406 (kD)
Majority protein IDs: Gorai.006G207300.2.A;Gorai.006G207300.1.A;Gorai.006G207300.3.A;Gorai.006G207300.4.A;Gorai.006G207300.5.A;Gorai.006G207300.7.A;Gorai.006G207300.6.A;Gorai.006G207300.8.A
Gene IDs: Gorai.006G207300
cotton name: PTHR11566:SF58 - DYNAMIN-RELATED PROTEIN 1A (1 of 9)
OG IDs: 93522020
Best-hit-arabi-ID: AT5G42080.1
Best-hit-arabi-name: dynamin-like protein
Gorai.006G207300.1.A
PCC=-0.065, p=0.791
mRNA Profile
Protein Expression Profile
DPA

### P200: Gorai.009G407500.5.A in case 1 and V20

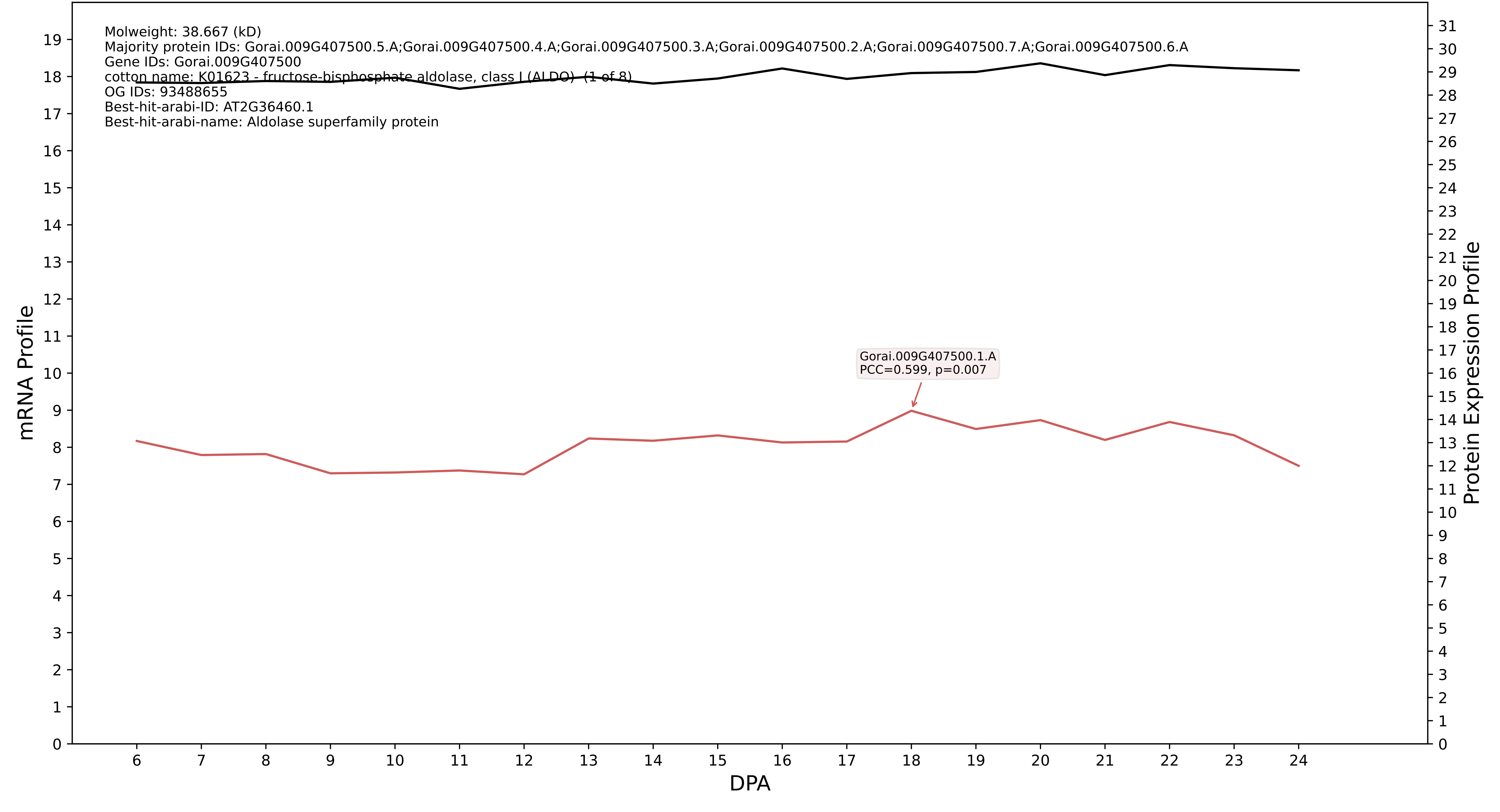

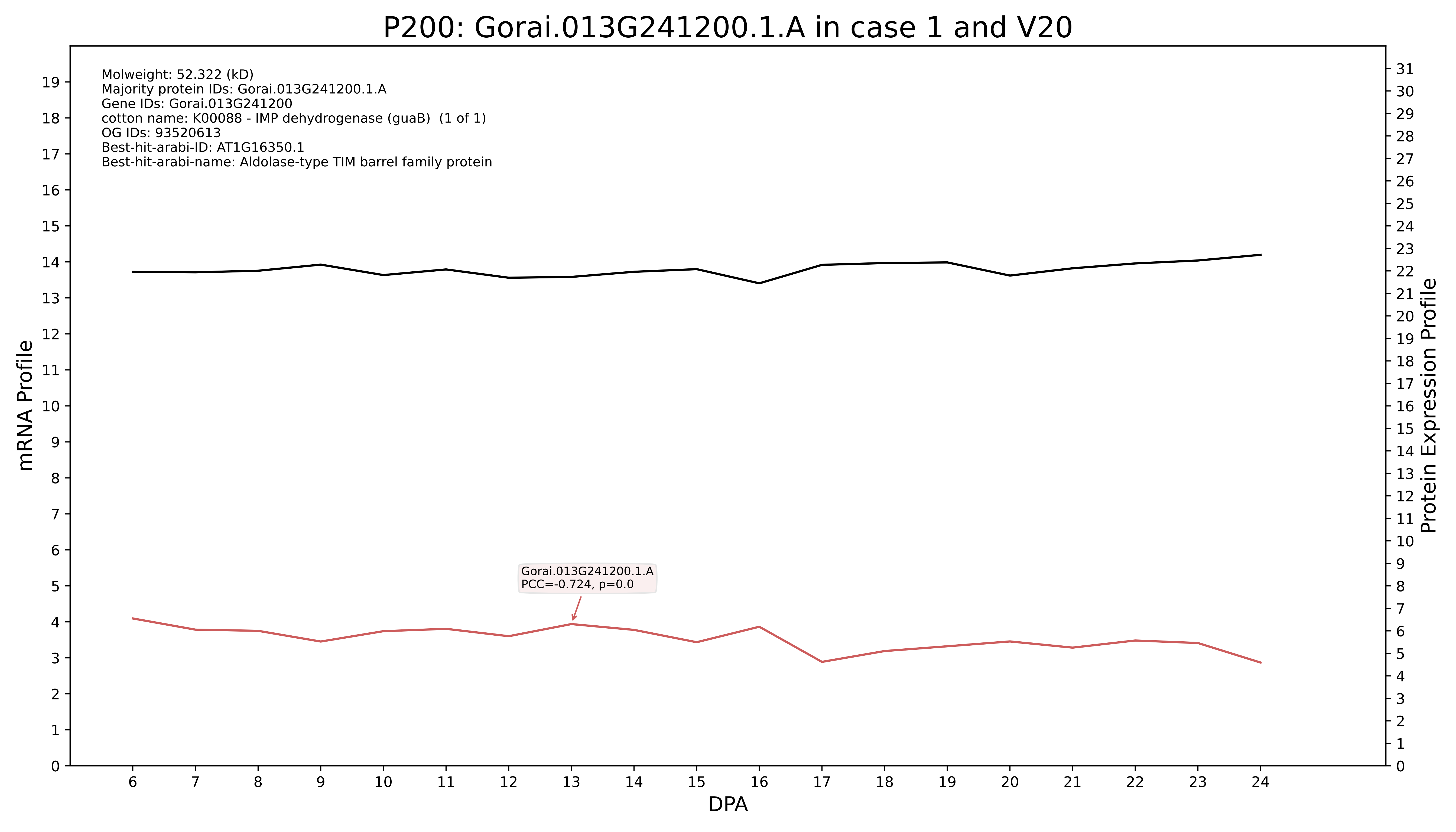
P200: Gorai.013G241200.1.A in case 1 and V20
Molweight: 52.322 (kD)
Majority protein IDs: Gorai.013G241200.1.A
Gene IDs: Gorai.013G241200
cotton name: K00088 - IMP dehydrogenase (guaB) (1 of 1)
OG IDs: 93520613
Best-hit-arabi-ID: AT1G16350.1
Best-hit-arabi-name: Aldolase-type TIM barrel family protein
Gorai.013G241200.1.A
PCC=-0.724, p=0.0
mRNA Profile
Protein Expression Profile
DPA

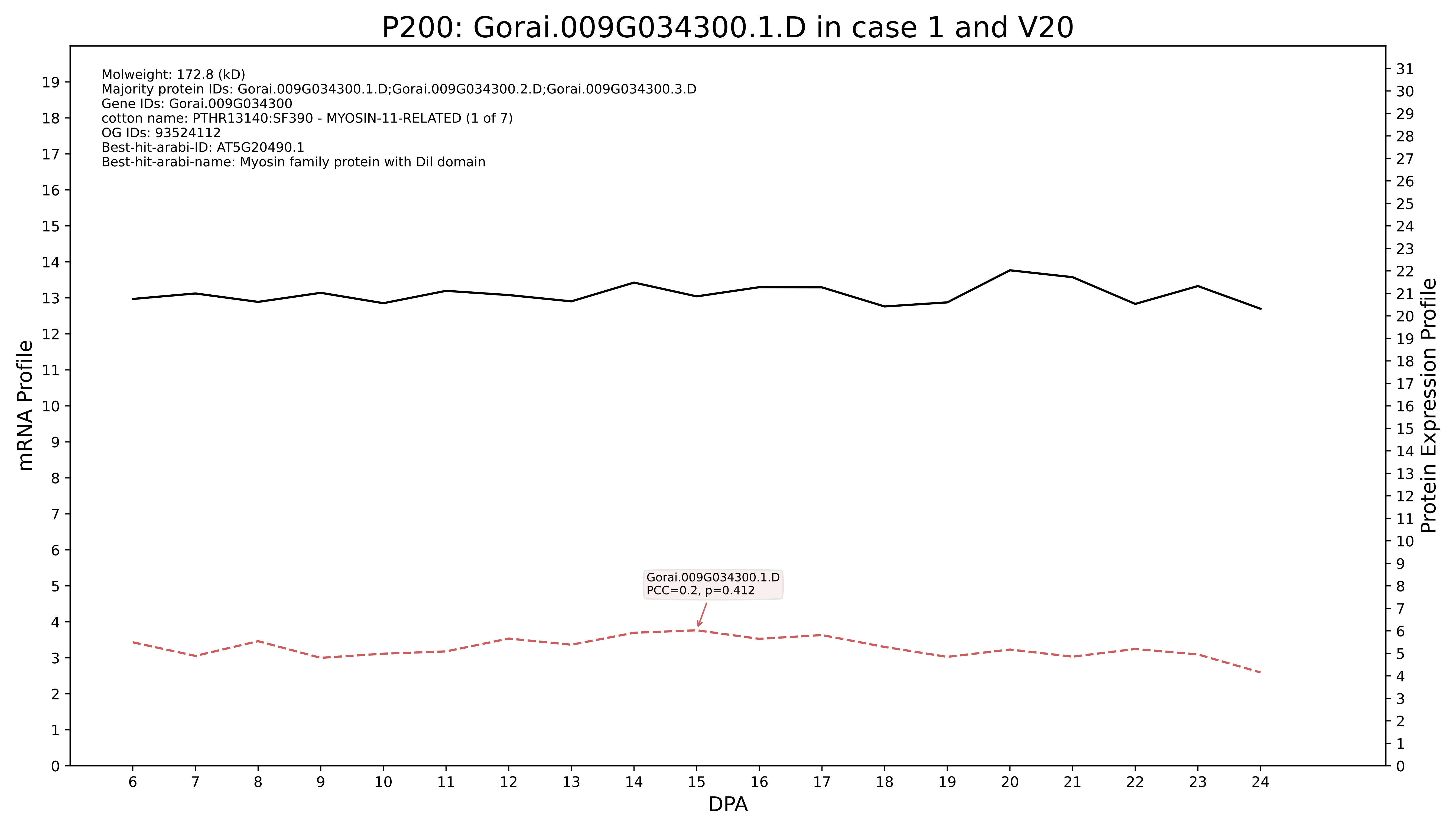
P200: Gorai.009G034300.1.D in case 1 and V20
Molweight: 172.8 (kD)
Majority protein IDs: Gorai.009G034300.1.D;Gorai.009G034300.2.D;Gorai.009G034300.3.D
Gene IDs: Gorai.009G034300
cotton name: PTHR13140:SF390 - MYOSIN-11-RELATED (1 of 7)
OG IDs: 93524112
Best-hit-arabi-ID: AT5G20490.1
Best-hit-arabi-name: Myosin family protein with Dil domain
Gorai.009G034300.1.D
PCC=0.2, p=0.412
mRNA Profile
Protein Expression Profile
DPA

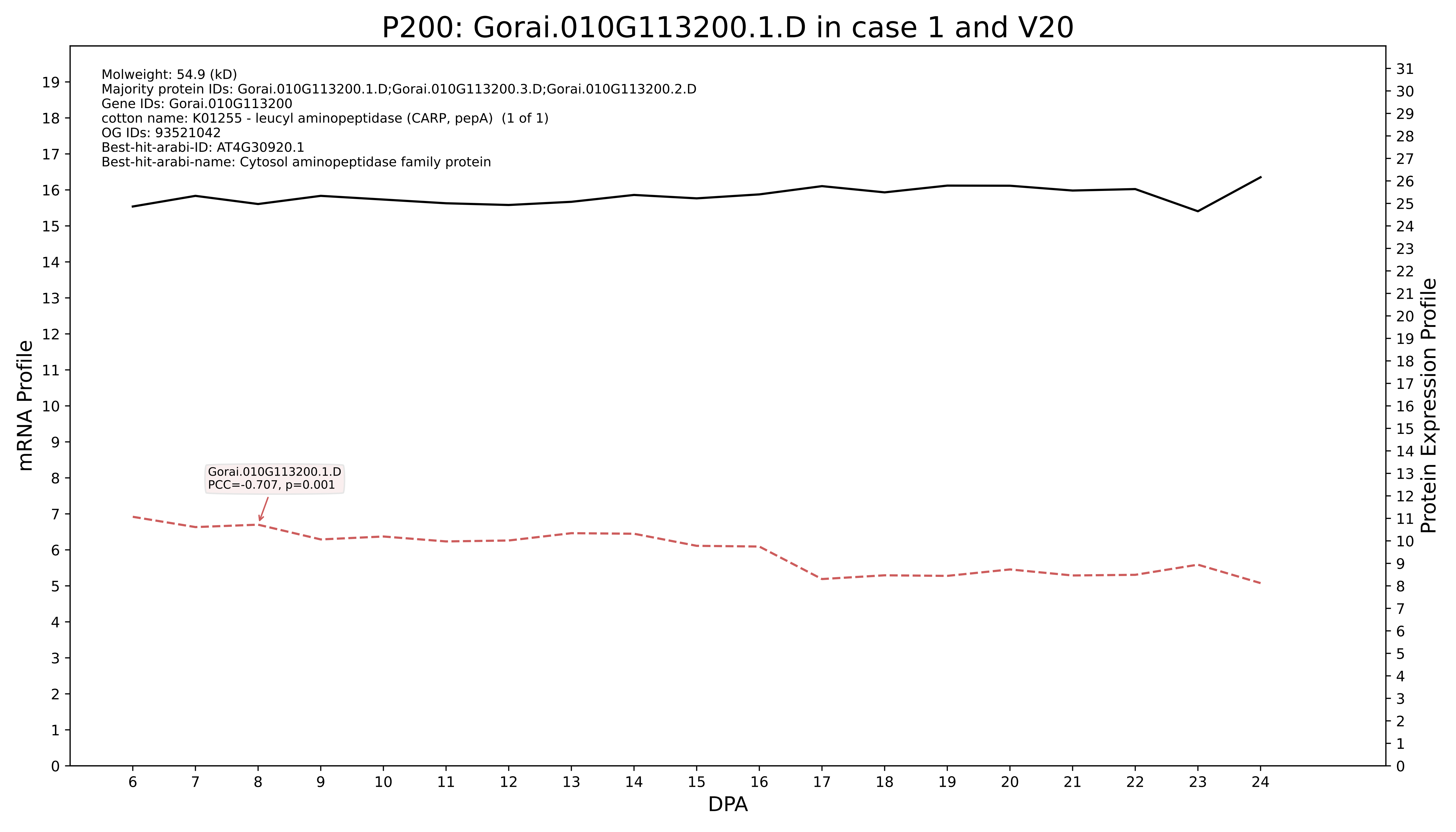
P200: Gorai.010G113200.1.D in case 1 and V20
Molweight: 54.9 (kD)
Majority protein IDs: Gorai.010G113200.1.D;Gorai.010G113200.3.D;Gorai.010G113200.2.D
Gene IDs: Gorai.010G113200
cotton name: K01255 - leucyl aminopeptidase (CARP, pepA) (1 of 1)
OG IDs: 93521042
Best-hit-arabi-ID: AT4G30920.1
Best-hit-arabi-name: Cytosol aminopeptidase family protein
Gorai.010G113200.1.D
PCC=-0.707, p=0.001
mRNA Profile
Protein Expression Profile
DPA

### P200: Gorai.012G063600.1.A in case 1 and V20

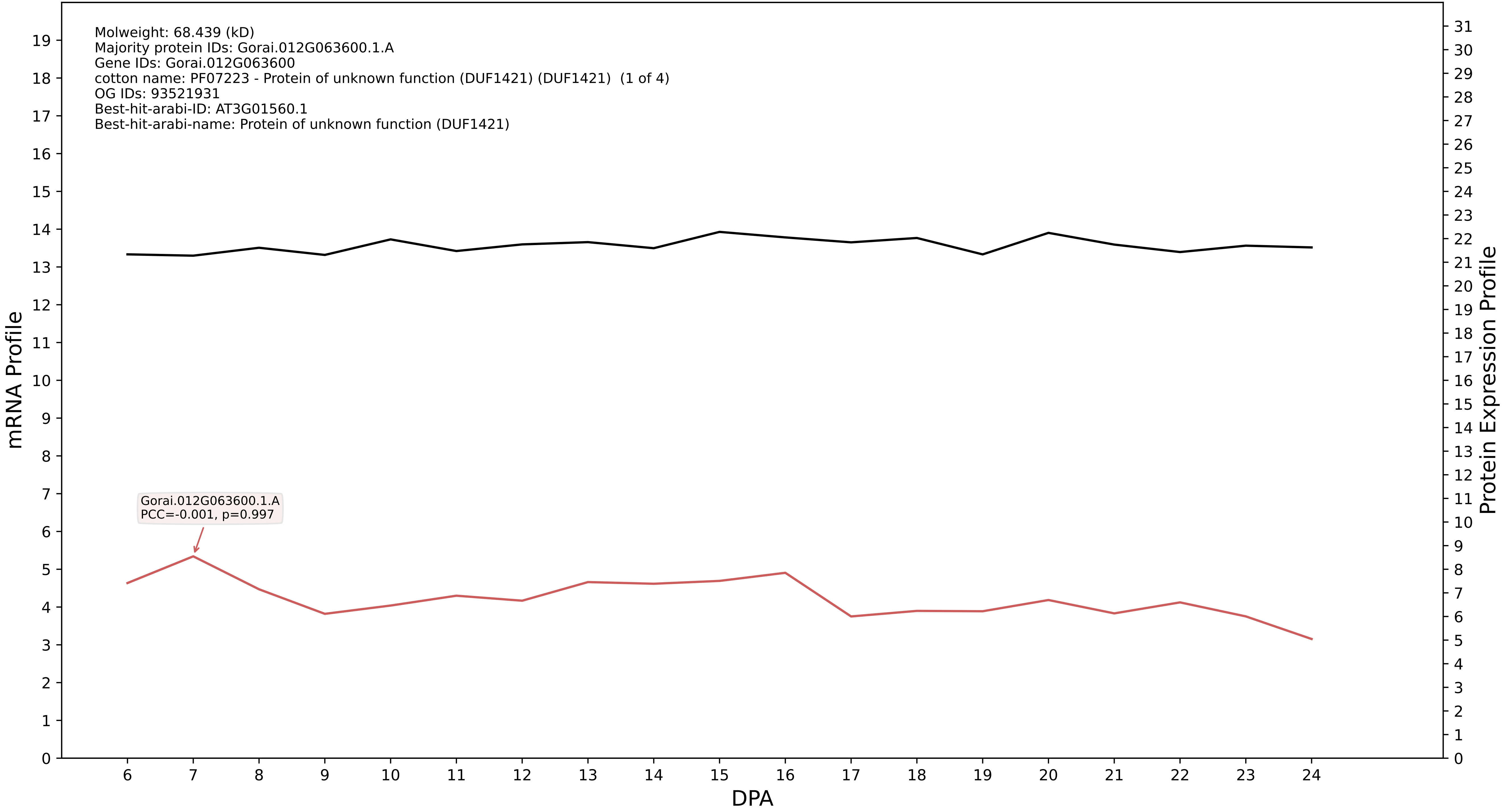

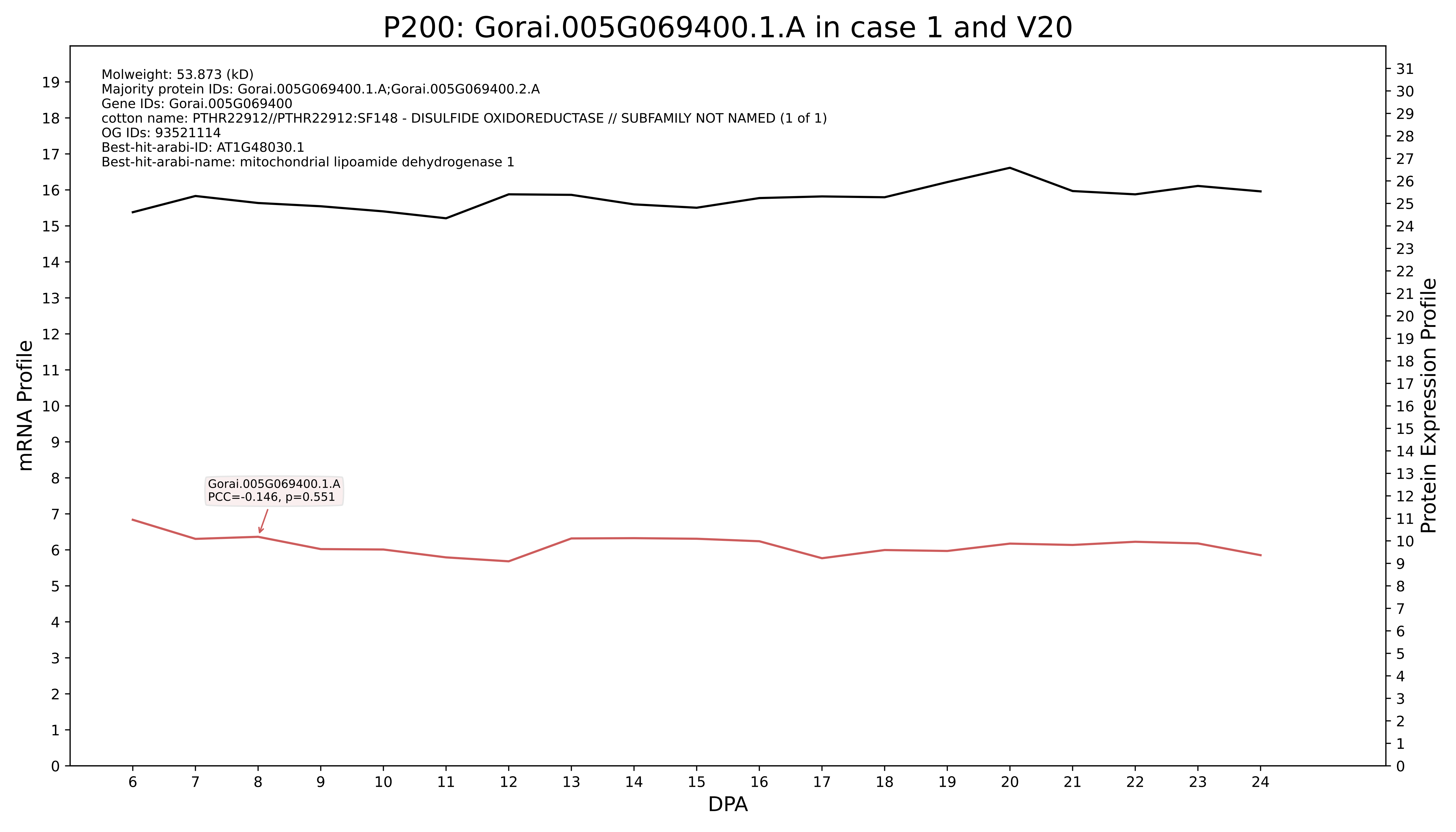
P200: Gorai.005G069400.1.A in case 1 and V20
Molweight: 53.873 (kD)
Majority protein IDs: Gorai.005G069400.1.A;Gorai.005G069400.2.A
Gene IDs: Gorai.005G069400
cotton name: PTHR22912//PTHR22912:SF148 - DISULFIDE OXIDOREDUCTASE // SUBFAMILY NOT NAMED (1 of 1)
OG IDs: 93521114
Best-hit-arabi-ID: AT1G48030.1
Best-hit-arabi-name: mitochondrial lipoamide dehydrogenase 1
Gorai.005G069400.1.A
PCC=-0.146, p=0.551
mRNA Profile
Protein Expression Profile
DPA

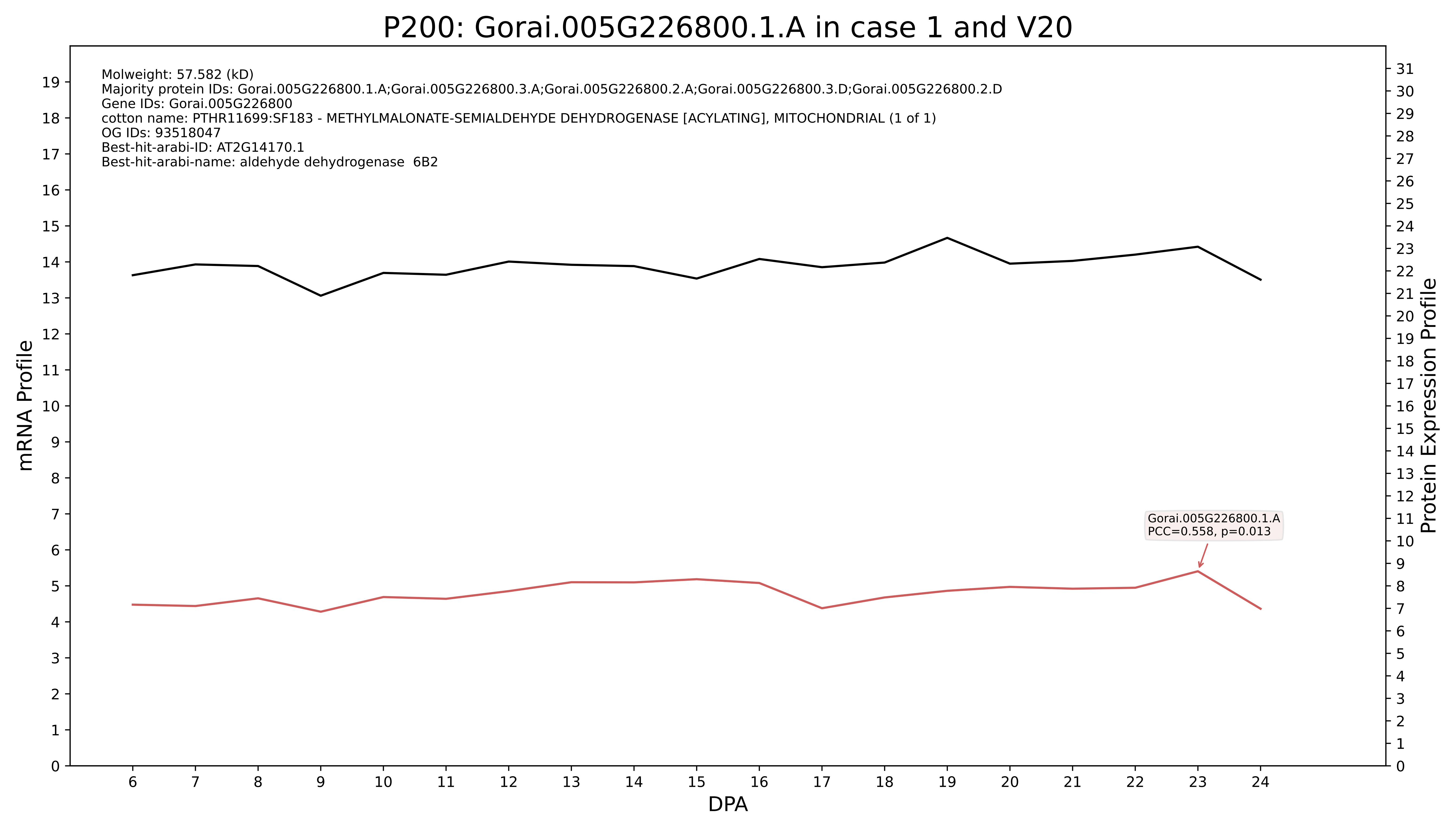
P200: Gorai.005G226800.1.A in case 1 and V20
Molweight: 57.582 (kD)
Majority protein IDs: Gorai.005G226800.1.A;Gorai.005G226800.3.A;Gorai.005G226800.2.A;Gorai.005G226800.3.D;Gorai.005G226800.2.D
Gene IDs: Gorai.005G226800
cotton name: PTHR11699:SF183 - METHYLMALONATE-SEMIALDEHYDE DEHYDROGENASE [ACYLATING], MITOCHONDRIAL (1 of 1)
OG IDs: 93518047
Best-hit-arabi-ID: AT2G14170.1
Best-hit-arabi-name: aldehyde dehydrogenase 6B2
Gorai.005G226800.1.A
PCC=0.558, p=0.013
mRNA Profile
Protein Expression Profile
DPA

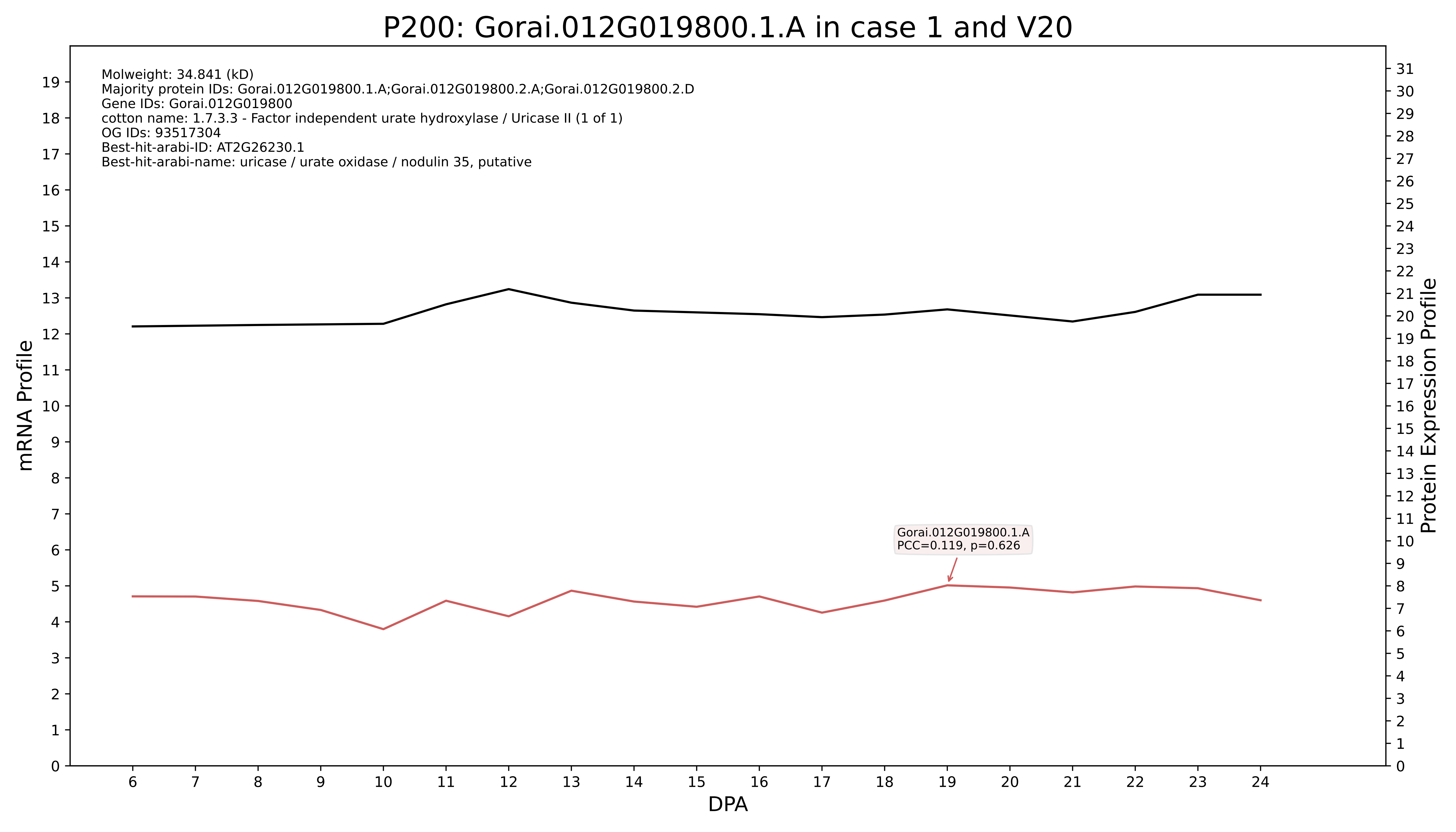
P200: Gorai.012G019800.1.A in case 1 and V20
Molweight: 34.841 (kD)
Majority protein IDs: Gorai.012G019800.1.A;Gorai.012G019800.2.A;Gorai.012G019800.2.D
Gene IDs: Gorai.012G019800
cotton name: 1.7.3.3 - Factor independent urate hydroxylase / Uricase II (1 of 1)
OG IDs: 93517304
Best-hit-arabi-ID: AT2G26230.1
Best-hit-arabi-name: uricase / urate oxidase / nodulin 35, putative
Gorai.012G019800.1.A
PCC=0.119, p=0.626
mRNA Profile
Protein Expression Profile
DPA

### P200: Gorai.013G159800.2.D in case 1 and V20

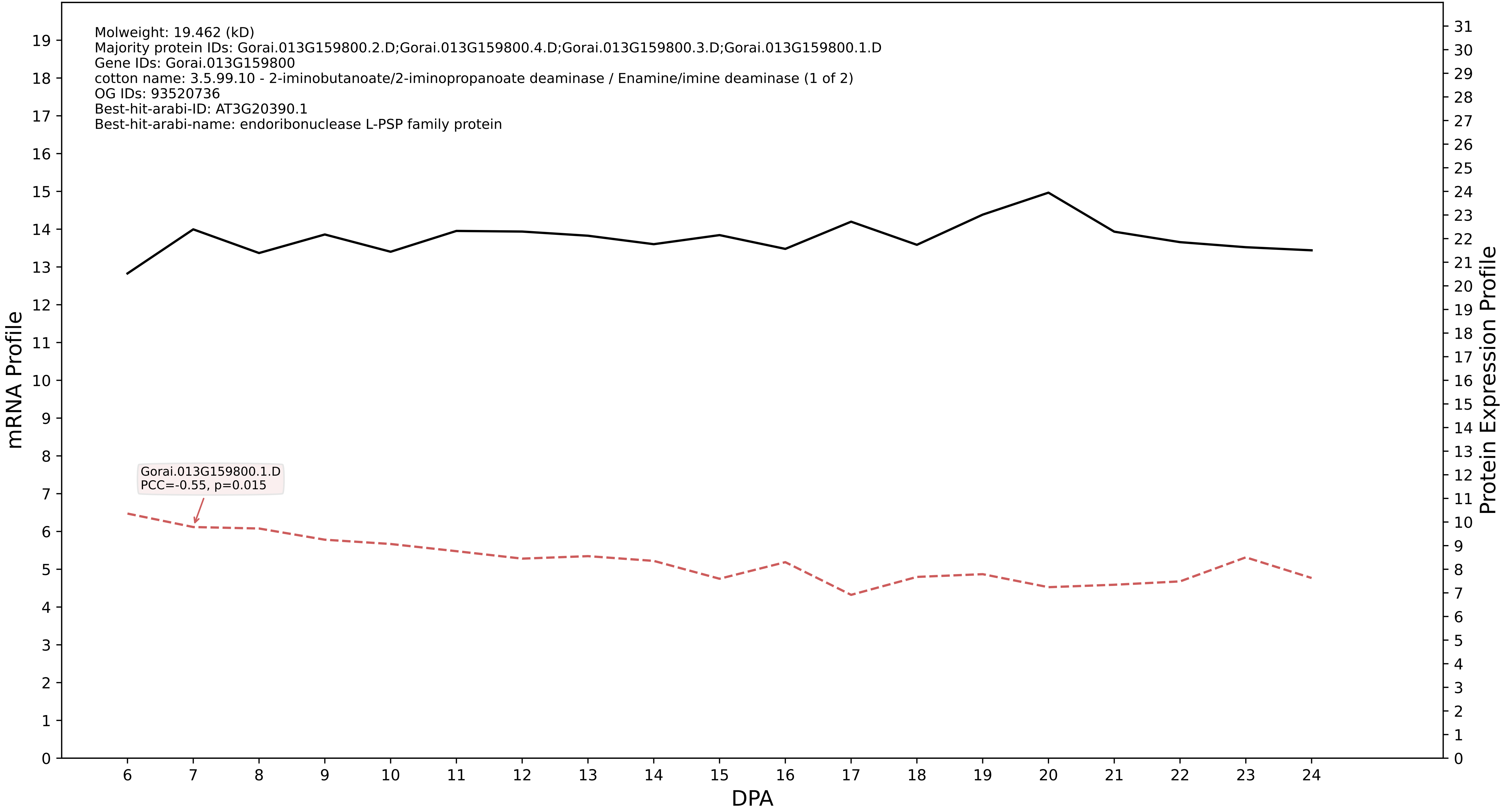

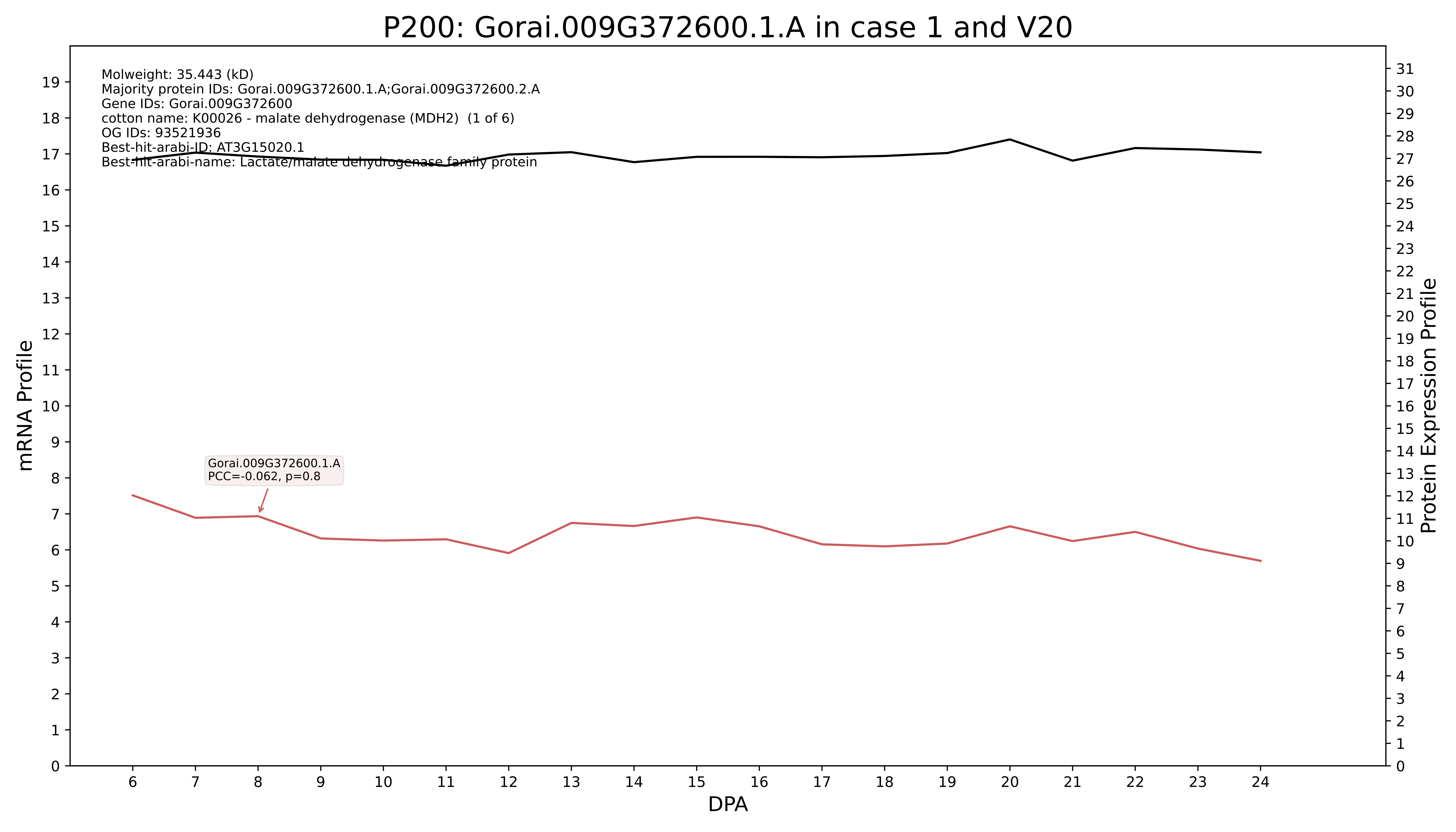
P200: Gorai.009G372600.1.A in case 1 and V20
Molweight: 35.443 (kD)
Majority protein IDs: Gorai.009G372600.1.A;Gorai.009G372600.2.A
Gene IDs: Gorai.009G372600
cotton name: K00026 - malate dehydrogenase (MDH2) (1 of 6)
OG IDs: 93521936
Best-hit-arabi-ID: AT3G15020.1
Best-hit-arabi-name: Lactate/malate dehydrogenase family protein
Gorai.009G372600.1.A
PCC=-0.062, p=0.8
mRNA Profile
Protein Expression Profile
DPA

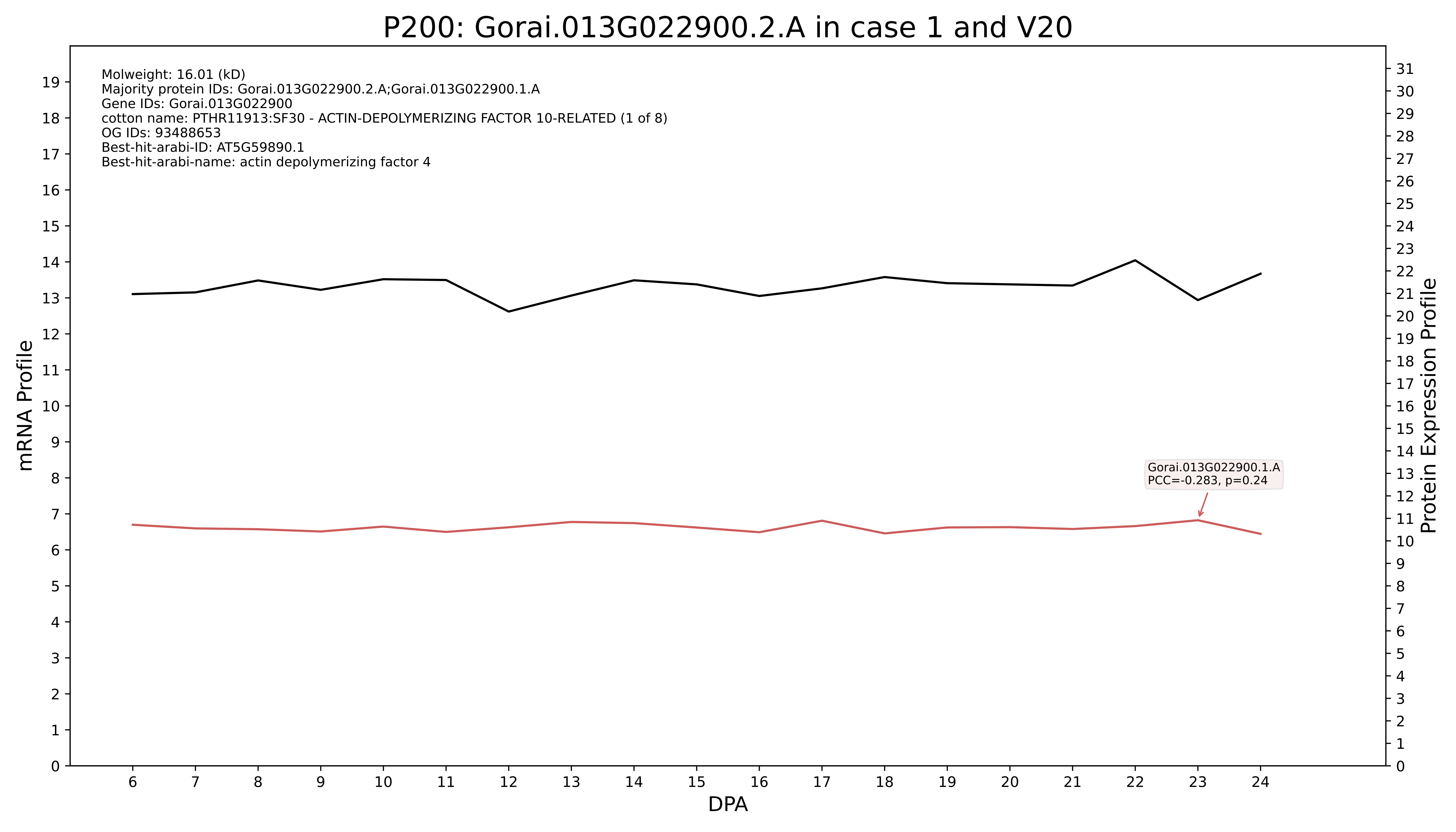
P200: Gorai.013G022900.2.A in case 1 and V20
Molweight: 16.01 (kD)
Majority protein IDs: Gorai.013G022900.2.A;Gorai.013G022900.1.A
Gene IDs: Gorai.013G022900
cotton name: PTHR11913:SF30 - ACTIN-DEPOLYMERIZING FACTOR 10-RELATED (1 of 8)
OG IDs: 93488653
Best-hit-arabi-ID: AT5G59890.1
Best-hit-arabi-name: actin depolymerizing factor 4
Gorai.013G022900.1.A
PCC=-0.283, p=0.24
mRNA Profile
Protein Expression Profile
DPA

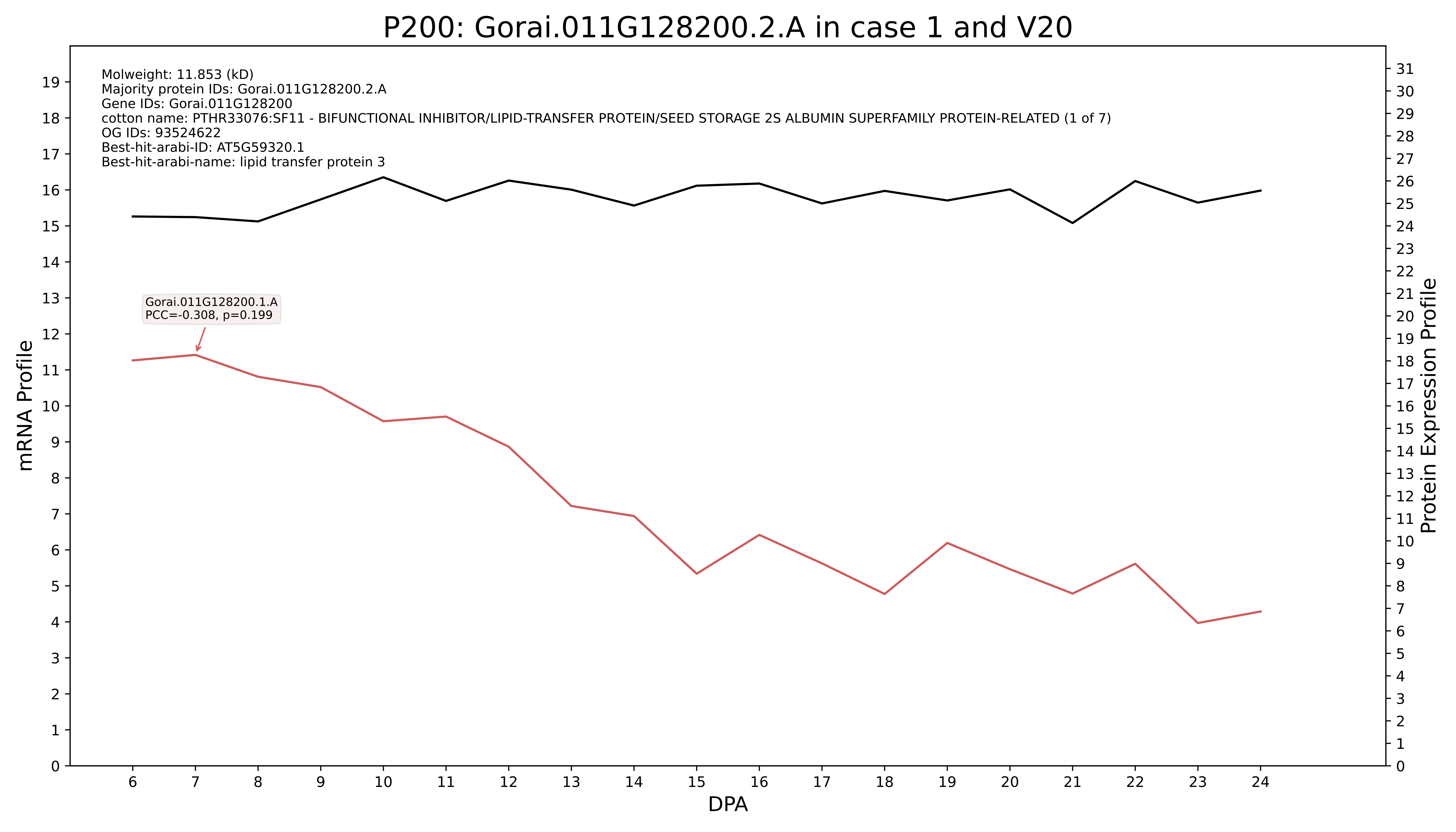

P200: Gorai.011G128200.2.A in case 1 and V20
Molweight: 11.853 (kD)
Majority protein IDs: Gorai.011G128200.2.A
Gene IDs: Gorai.011G128200
cotton name: PTHR33076:SF11 - BIFUNCTIONAL INHIBITOR/LIPID-TRANSFER PROTEIN/SEED STORAGE 2S ALBUMIN SUPERFAMILY PROTEIN-RELATED (1 of 7)
OG IDs: 93524622
Best-hit-arabi-ID: AT5G59320.1
Best-hit-arabi-name: lipid transfer protein 3
Gorai.011G128200.1.A
PCC=-0.308, p=0.199
mRNA Profile
Protein Expression Profile
DPA

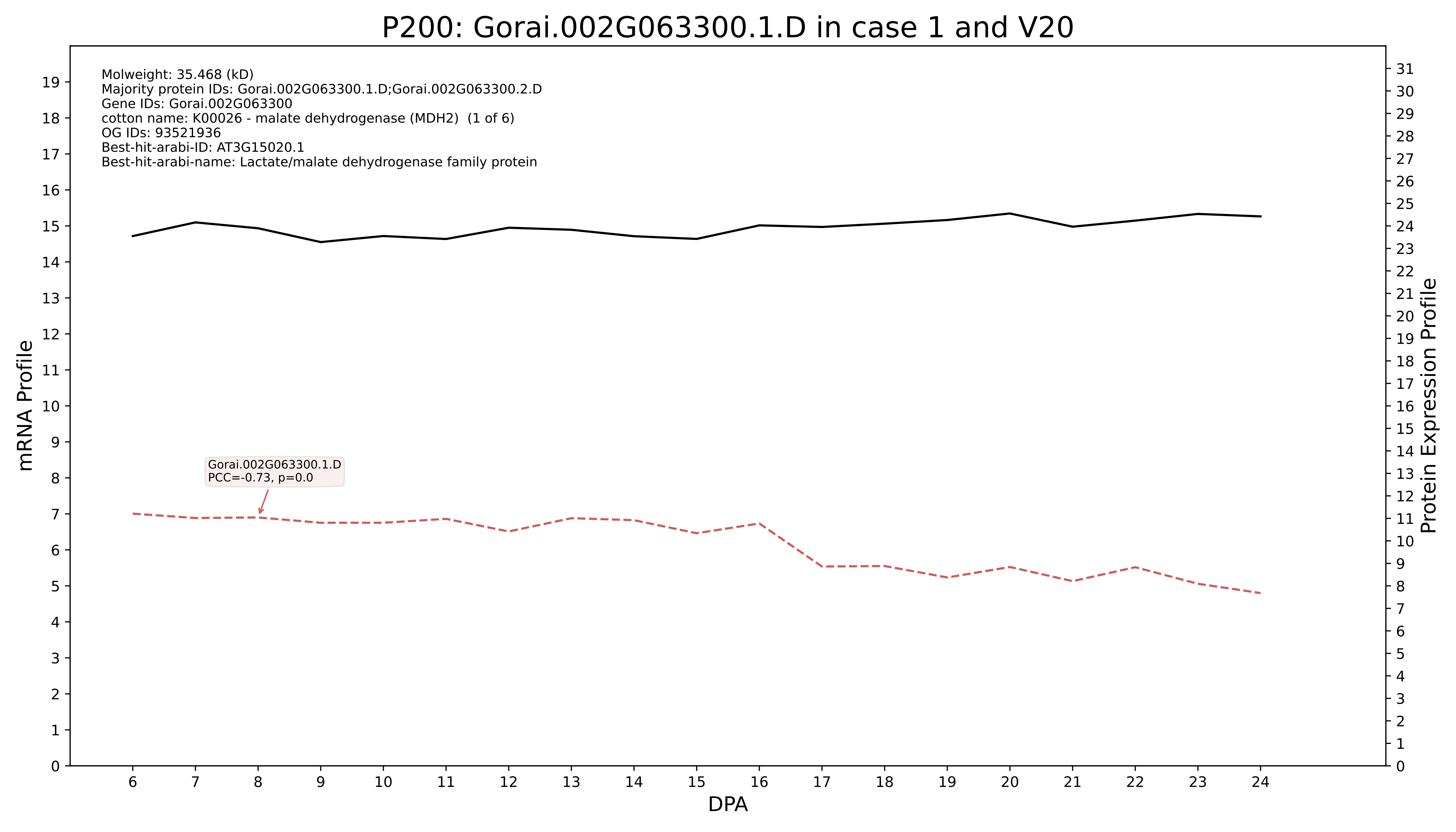

P200: Gorai.002G063300.1.D in case 1 and V20
Molweight: 35.468 (kD)
Majority protein IDs: Gorai.002G063300.1.D;Gorai.002G063300.2.D
Gene IDs: Gorai.002G063300
cotton name: K00026 - malate dehydrogenase (MDH2) (1 of 6)
OG IDs: 93521936
Best-hit-arabi-ID: AT3G15020.1
Best-hit-arabi-name: Lactate/malate dehydrogenase family protein
Gorai.002G063300.1.D
PCC=-0.73, p=0.0
mRNA Profile
Protein Expression Profile
DPA

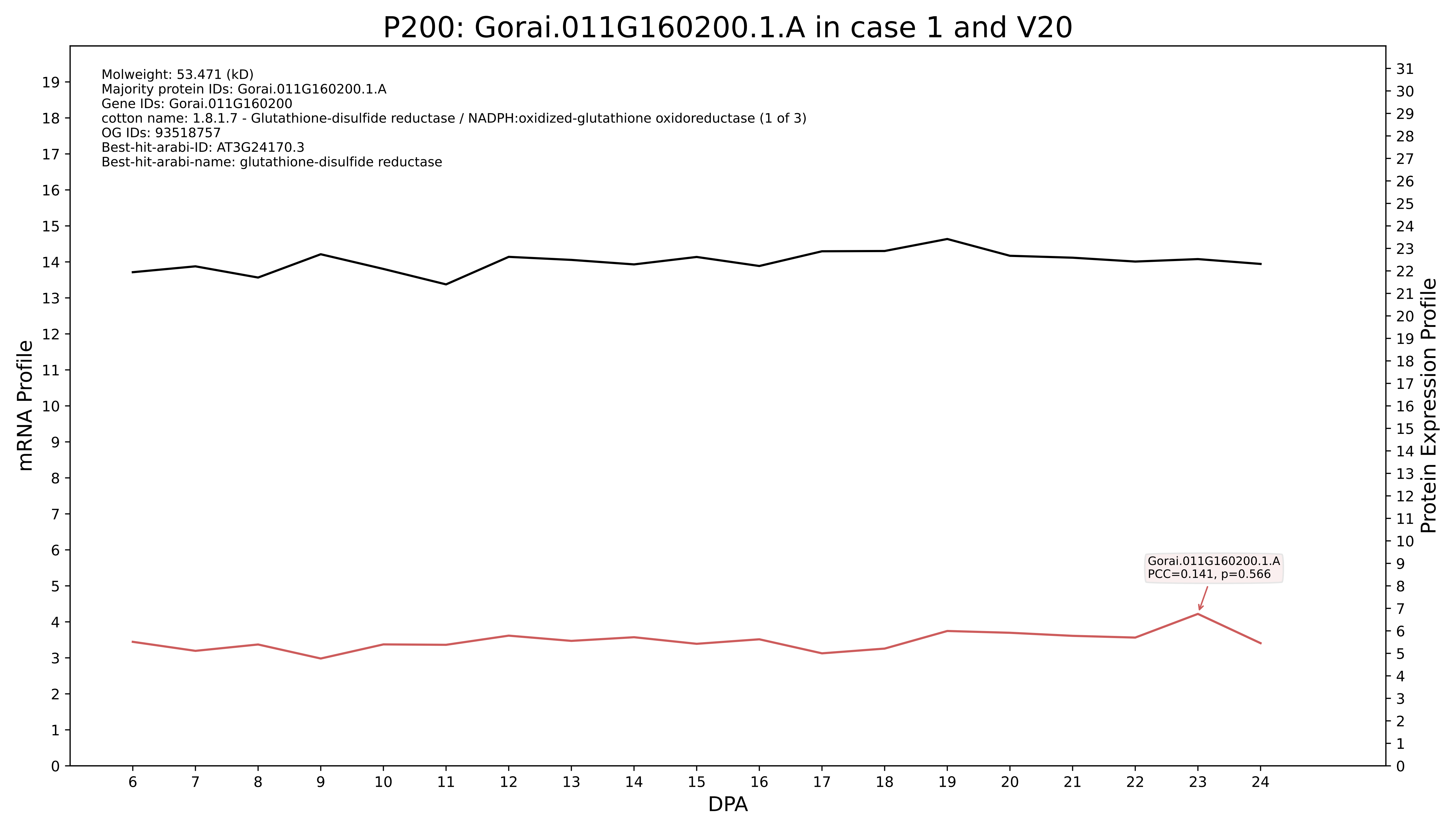

P200: Gorai.011G160200.1.A in case 1 and V20
Molweight: 53.471 (kD)
Majority protein IDs: Gorai.011G160200.1.A
Gene IDs: Gorai.011G160200
cotton name: 1.8.1.7 - Glutathione-disulfide reductase / NADPH:oxidized-glutathione oxidoreductase (1 of 3)
OG IDs: 93518757
Best-hit-arabi-ID: AT3G24170.3
Best-hit-arabi-name: glutathione-disulfide reductase
Gorai.011G160200.1.A
PCC=0.141, p=0.566
mRNA Profile
Protein Expression Profile
DPA

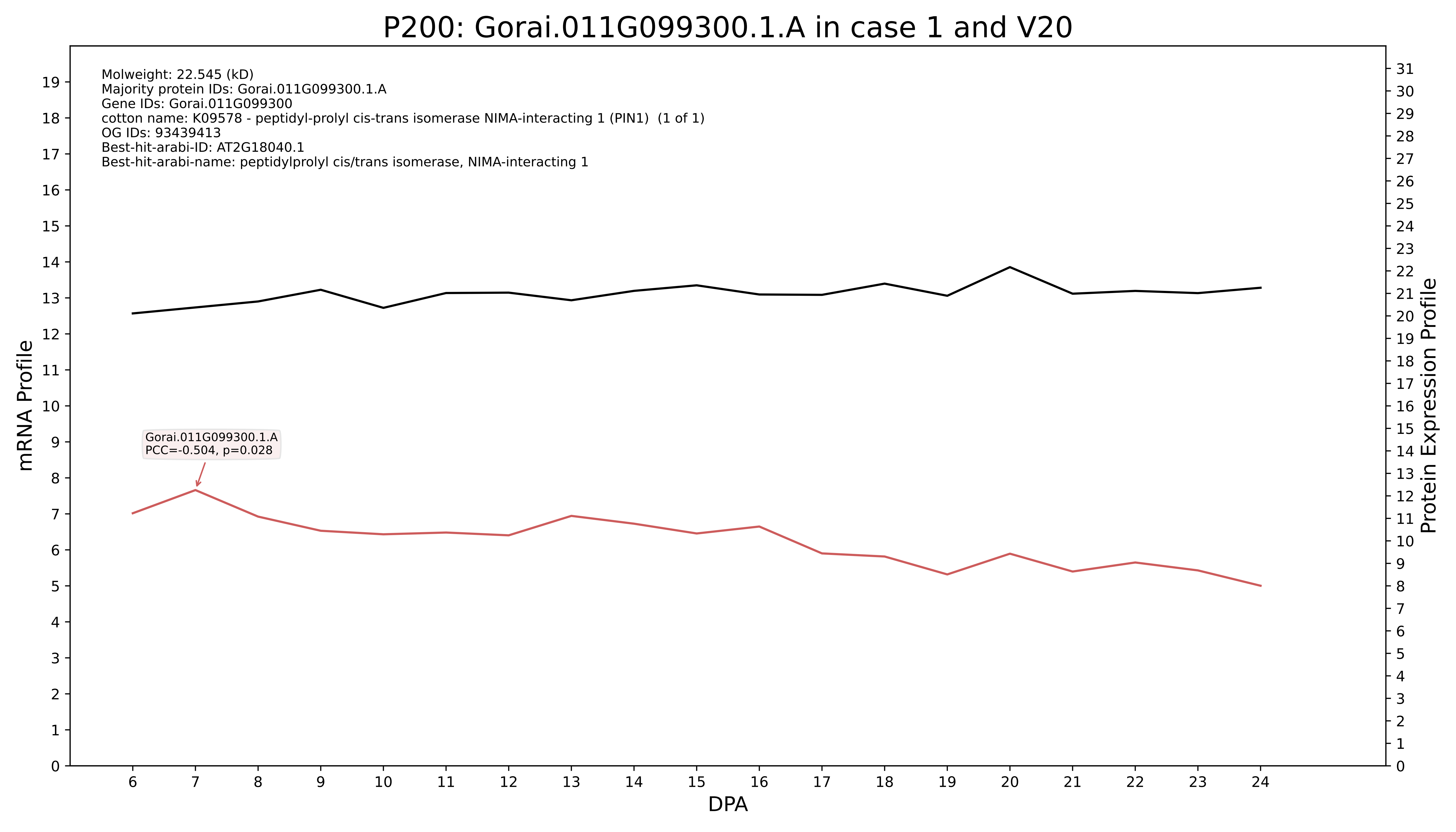

P200: Gorai.011G099300.1.A in case 1 and V20
Molweight: 22.545 (kD)
Majority protein IDs: Gorai.011G099300.1.A
Gene IDs: Gorai.011G099300
cotton name: K09578 - peptidyl-prolyl cis-trans isomerase NIMA-interacting 1 (PIN1) (1 of 1)
OG IDs: 93439413
Best-hit-arabi-ID: AT2G18040.1
Best-hit-arabi-name: peptidylprolyl cis/trans isomerase, NIMA-interacting 1
Gorai.011G099300.1.A
PCC=-0.504, p=0.028
mRNA Profile
Protein Expression Profile
DPA

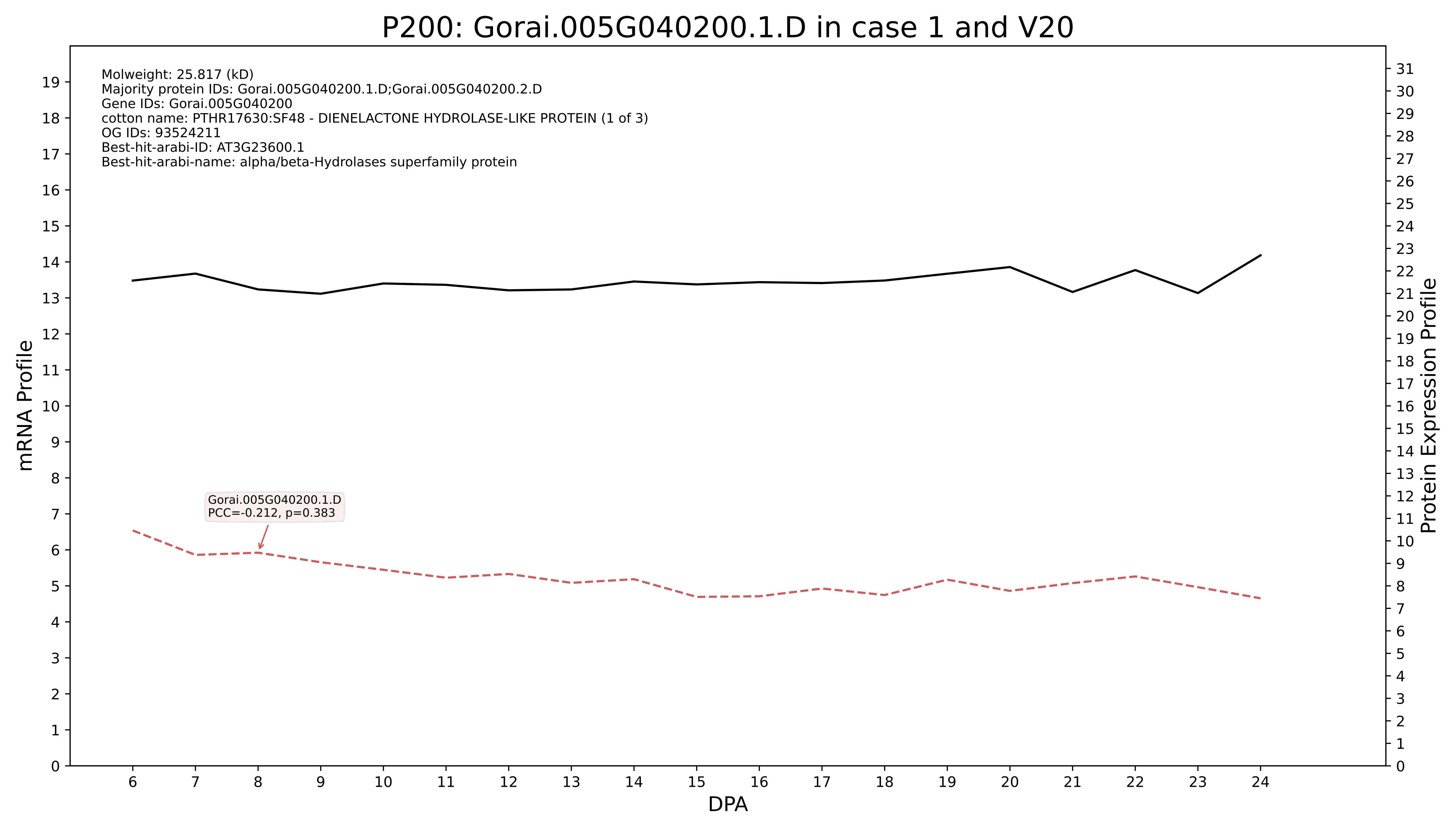
P200: Gorai.005G040200.1.D in case 1 and V20
Molweight: 25.817 (kD)
Majority protein IDs: Gorai.005G040200.1.D;Gorai.005G040200.2.D
Gene IDs: Gorai.005G040200
cotton name: PTHR17630:SF48 - DIENELACTONE HYDROLASE-LIKE PROTEIN (1 of 3)
OG IDs: 93524211
Best-hit-arabi-ID: AT3G23600.1
Best-hit-arabi-name: alpha/beta-Hydrolases superfamily protein
Gorai.005G040200.1.D
PCC=-0.212, p=0.383
mRNA Profile
Protein Expression Profile
DPA

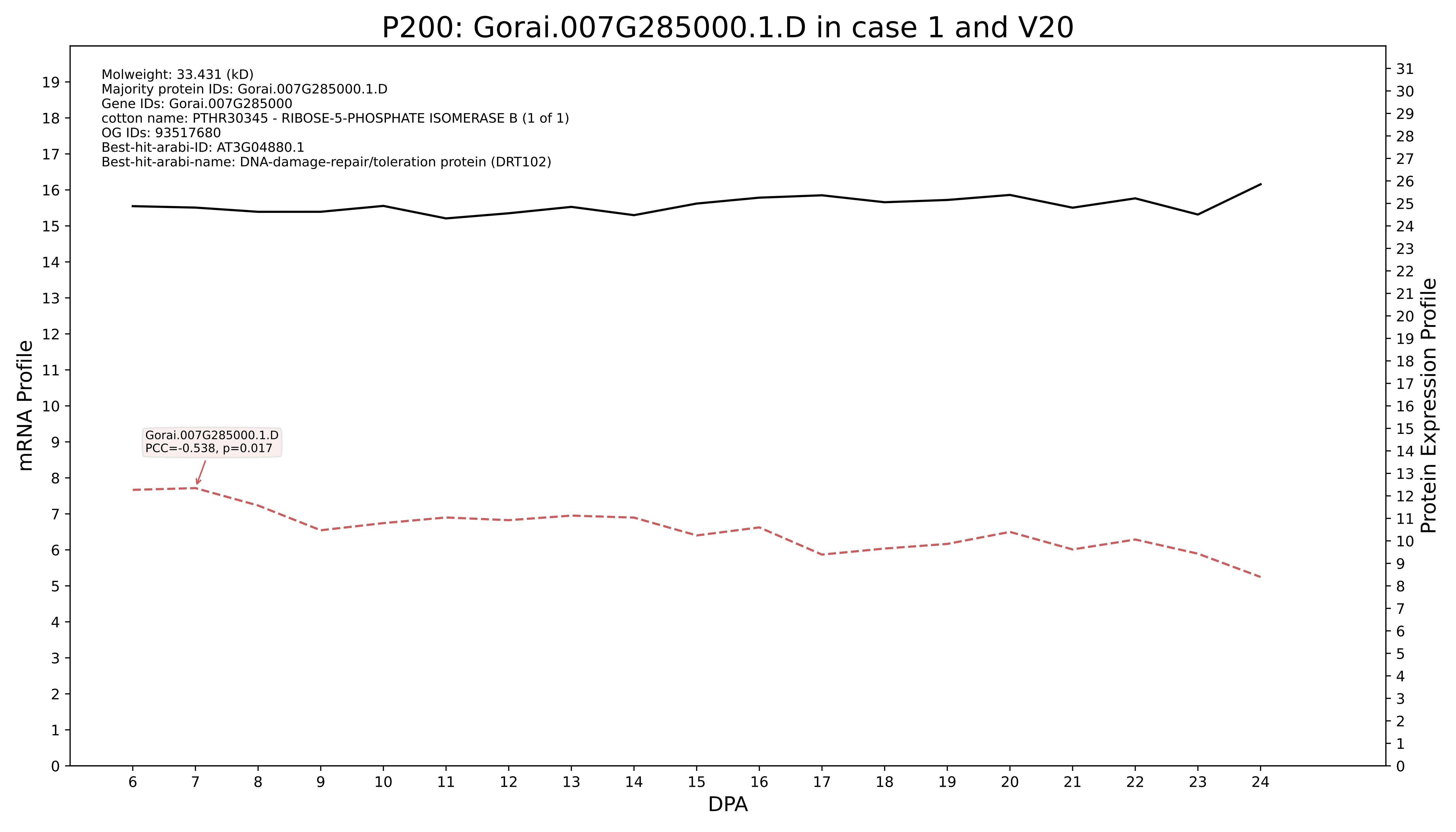
P200: Gorai.007G285000.1.D in case 1 and V20
Molweight: 33.431 (kD)
Majority protein IDs: Gorai.007G285000.1.D
Gene IDs: Gorai.007G285000
cotton name: PTHR30345 - RIBOSE-5-PHOSPHATE ISOMERASE B (1 of 1)
OG IDs: 93517680
Best-hit-arabi-ID: AT3G04880.1
Best-hit-arabi-name: DNA-damage-repair/toleration protein (DRT102)
Gorai.007G285000.1.D
PCC=-0.538, p=0.017
mRNA Profile
Protein Expression Profile
DPA

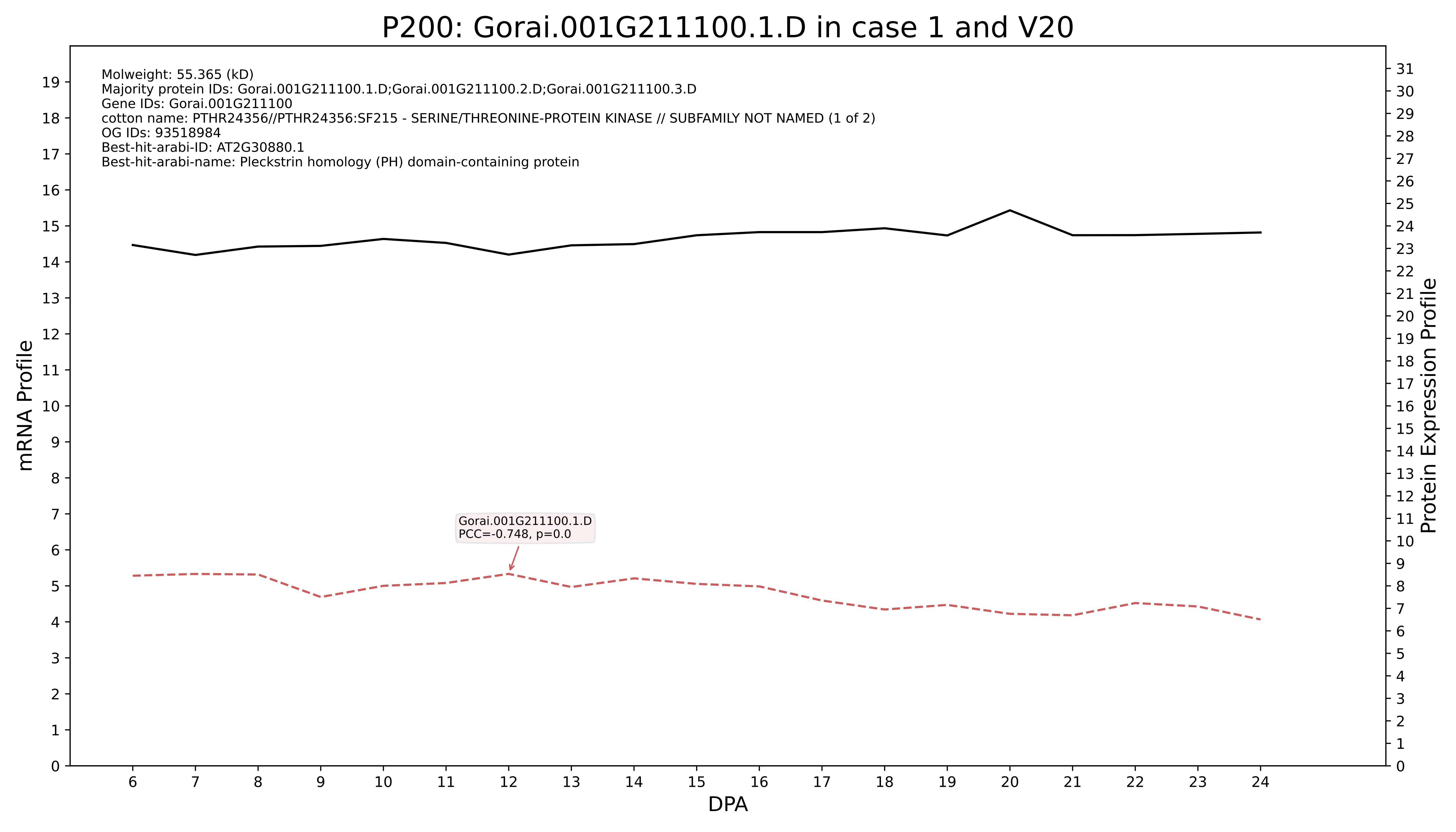

P200: Gorai.001G211100.1.D in case 1 and V20
Molweight: 55.365 (kD)
Majority protein IDs: Gorai.001G211100.1.D;Gorai.001G211100.2.D;Gorai.001G211100.3.D
Gene IDs: Gorai.001G211100
cotton name: PTHR24356//PTHR24356:SF215 - SERINE/THREONINE-PROTEIN KINASE // SUBFAMILY NOT NAMED (1 of 2)
OG IDs: 93518984
Best-hit-arabi-ID: AT2G30880.1
Best-hit-arabi-name: Pleckstrin homology (PH) domain-containing protein
Gorai.001G211100.1.D
PCC=-0.748, p=0.0
mRNA Profile
Protein Expression Profile
DPA

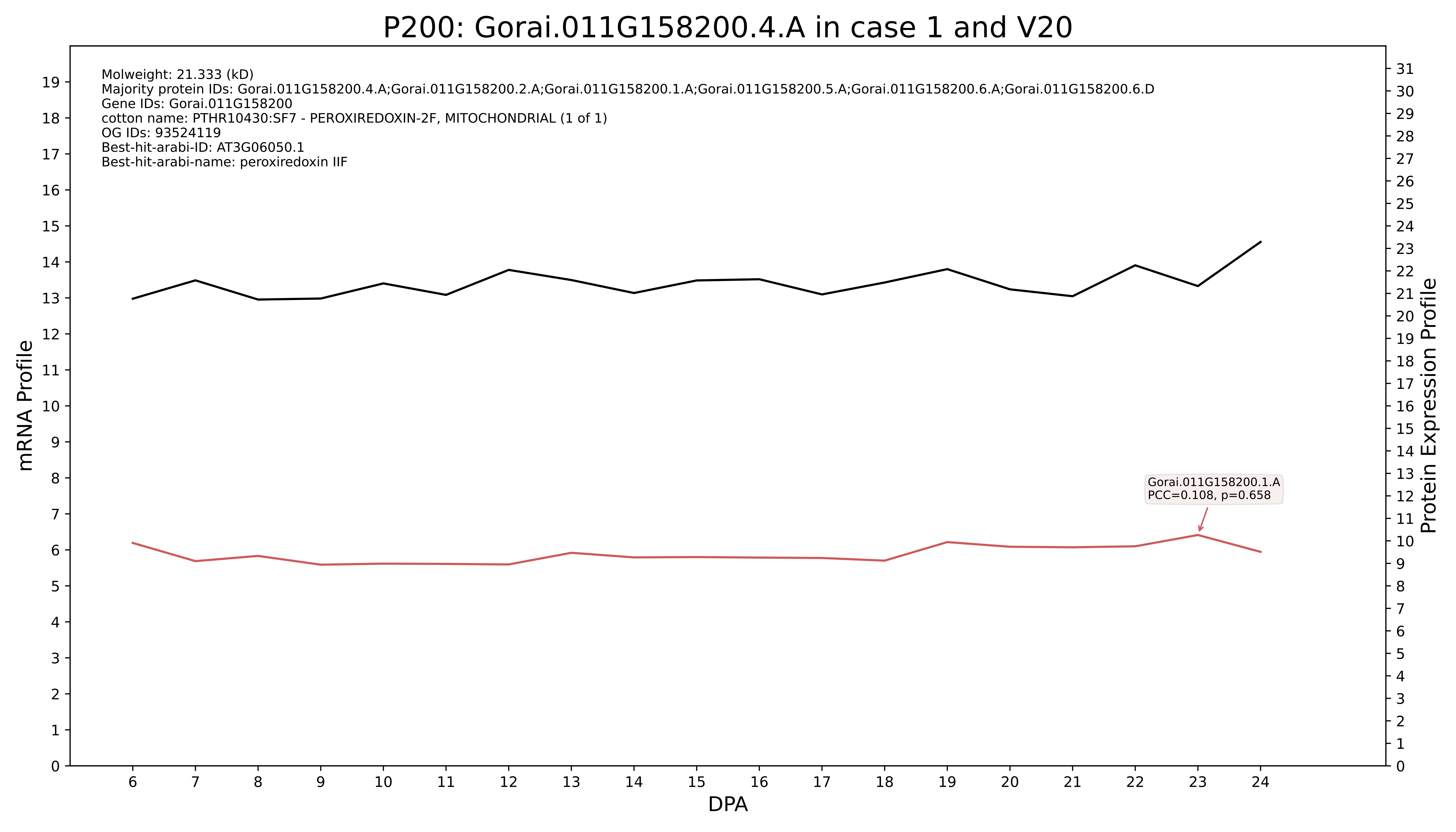
P200: Gorai.011G158200.4.A in case 1 and V20
Molweight: 21.333 (kD)
Majority protein IDs: Gorai.011G158200.4.A;Gorai.011G158200.2.A;Gorai.011G158200.1.A;Gorai.011G158200.5.A;Gorai.011G158200.6.A;Gorai.011G158200.6.D
Gene IDs: Gorai.011G158200
cotton name: PTHR10430:SF7 - PEROXIREDOXIN-2F, MITOCHONDRIAL (1 of 1)
OG IDs: 93524119
Best-hit-arabi-ID: AT3G06050.1
Best-hit-arabi-name: peroxiredoxin IIF
Gorai.011G158200.1.A
PCC=0.108, p=0.658
mRNA Profile
Protein Expression Profile
DPA

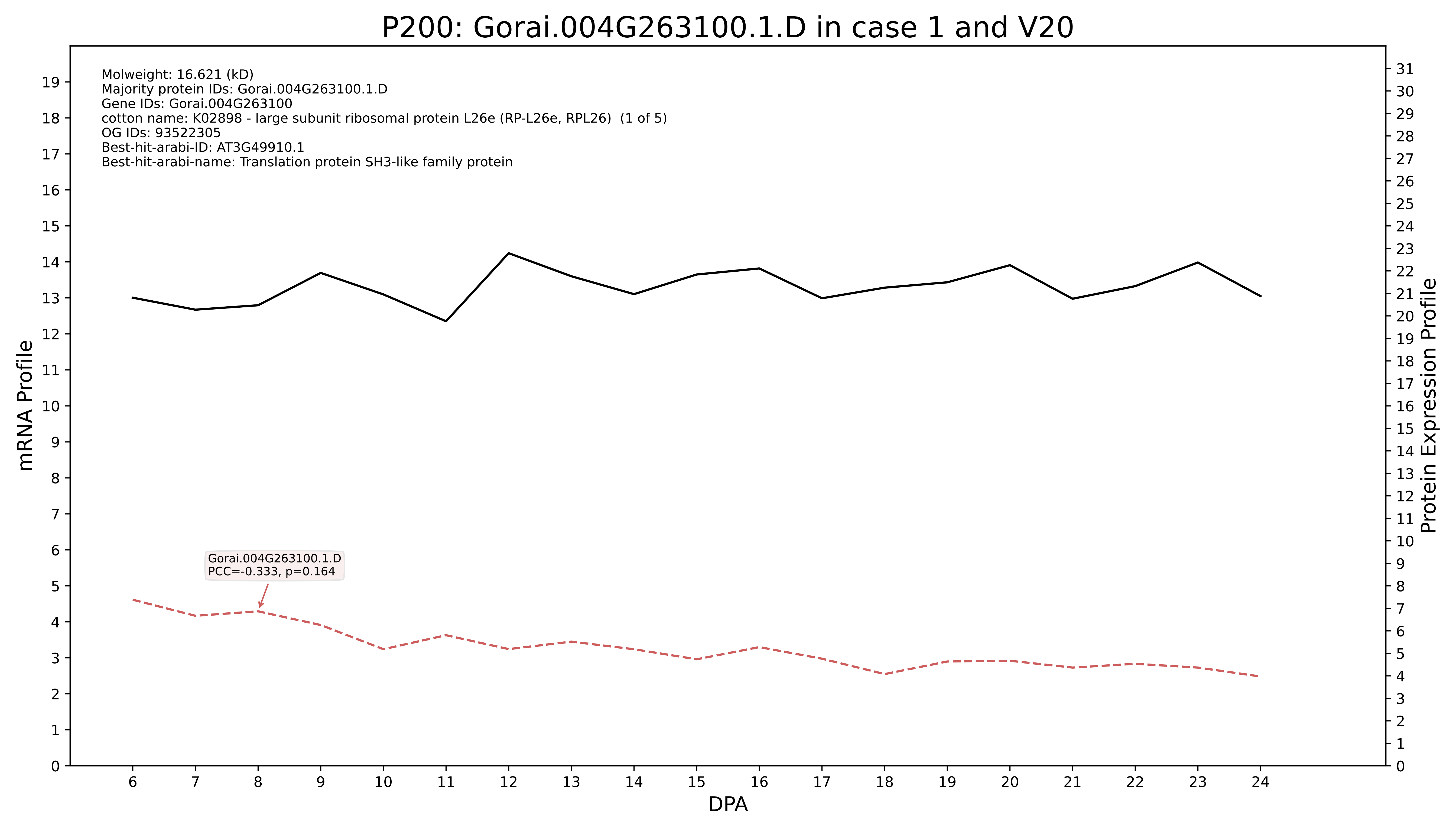
P200: Gorai.004G263100.1.D in case 1 and V20
Molweight: 16.621 (kD)
Majority protein IDs: Gorai.004G263100.1.D
Gene IDs: Gorai.004G263100
cotton name: K02898 - large subunit ribosomal protein L26e (RP-L26e, RPL26) (1 of 5)
OG IDs: 93522305
Best-hit-arabi-ID: AT3G49910.1
Best-hit-arabi-name: Translation protein SH3-like family protein
Gorai.004G263100.1.D
PCC=-0.333, p=0.164
mRNA Profile
Protein Expression Profile
DPA

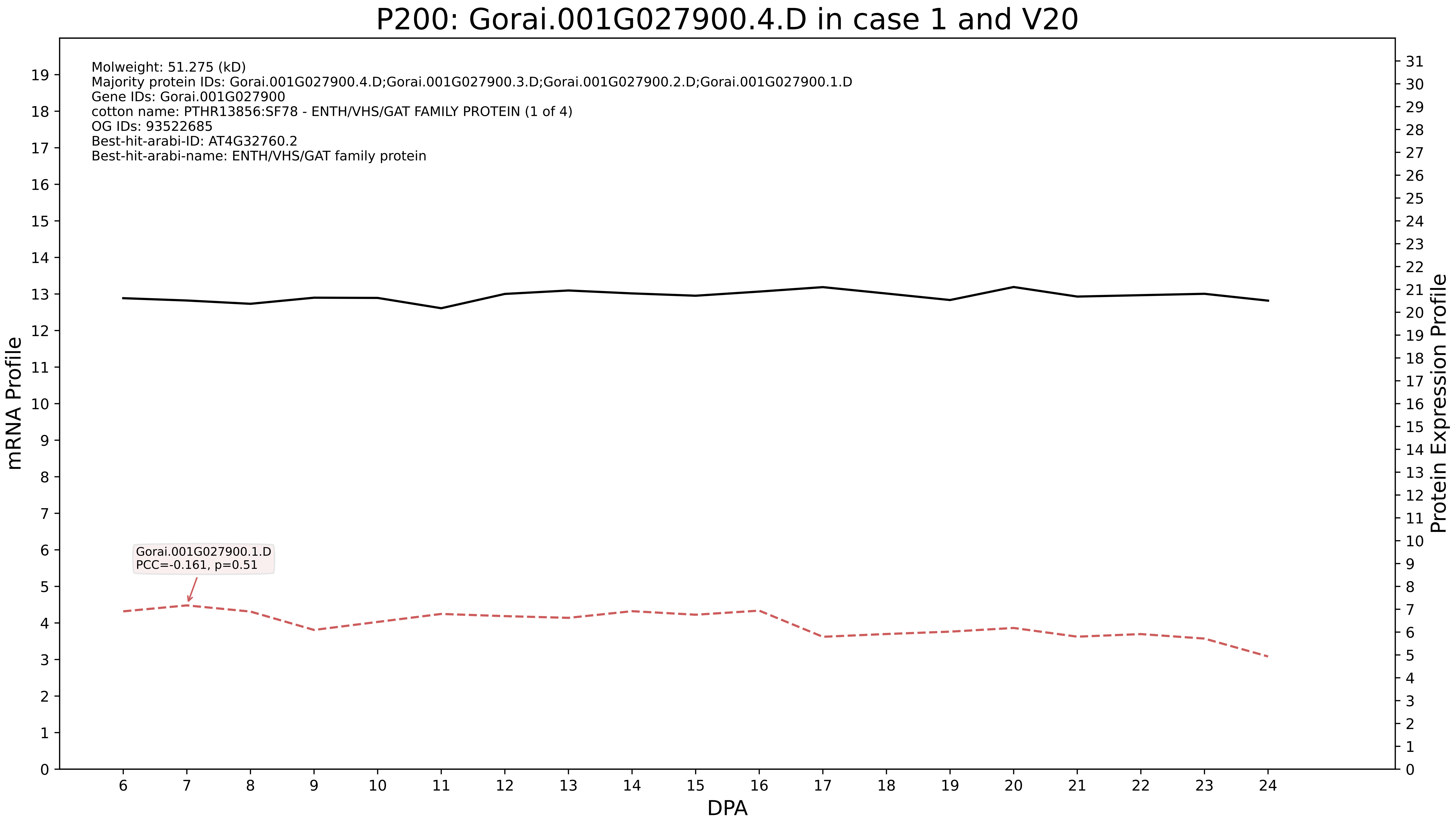
P200: Gorai.001G027900.4.D in case 1 and V20
Molweight: 51.275 (kD)
Majority protein IDs: Gorai.001G027900.4.D;Gorai.001G027900.3.D;Gorai.001G027900.2.D;Gorai.001G027900.1.D
Gene IDs: Gorai.001G027900
cotton name: PTHR13856:SF78 - ENTH/VHS/GAT FAMILY PROTEIN (1 of 4)
OG IDs: 93522685
Best-hit-arabi-ID: AT4G32760.2
Best-hit-arabi-name: ENTH/VHS/GAT family protein
Gorai.001G027900.1.D
PCC=-0.161, p=0.51
mRNA Profile
Protein Expression Profile
DPA

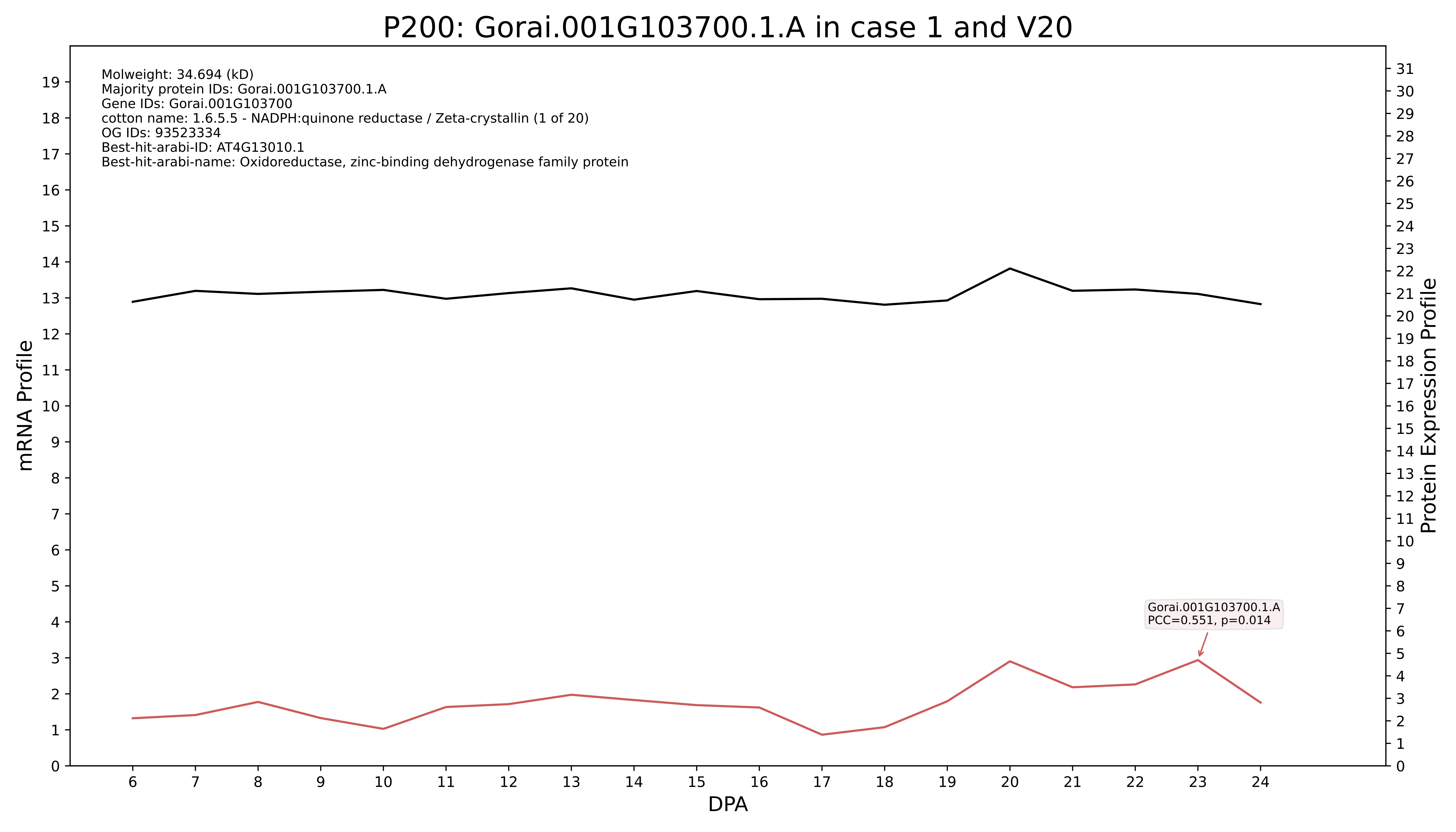
P200: Gorai.001G103700.1.A in case 1 and V20
Molweight: 34.694 (kD)
Majority protein IDs: Gorai.001G103700.1.A
Gene IDs: Gorai.001G103700
cotton name: 1.6.5.5 - NADPH:quinone reductase / Zeta-crystallin (1 of 20)
OG IDs: 93523334
Best-hit-arabi-ID: AT4G13010.1
Best-hit-arabi-name: Oxidoreductase, zinc-binding dehydrogenase family protein
Gorai.001G103700.1.A
PCC=0.551, p=0.014
mRNA Profile
Protein Expression Profile
DPA

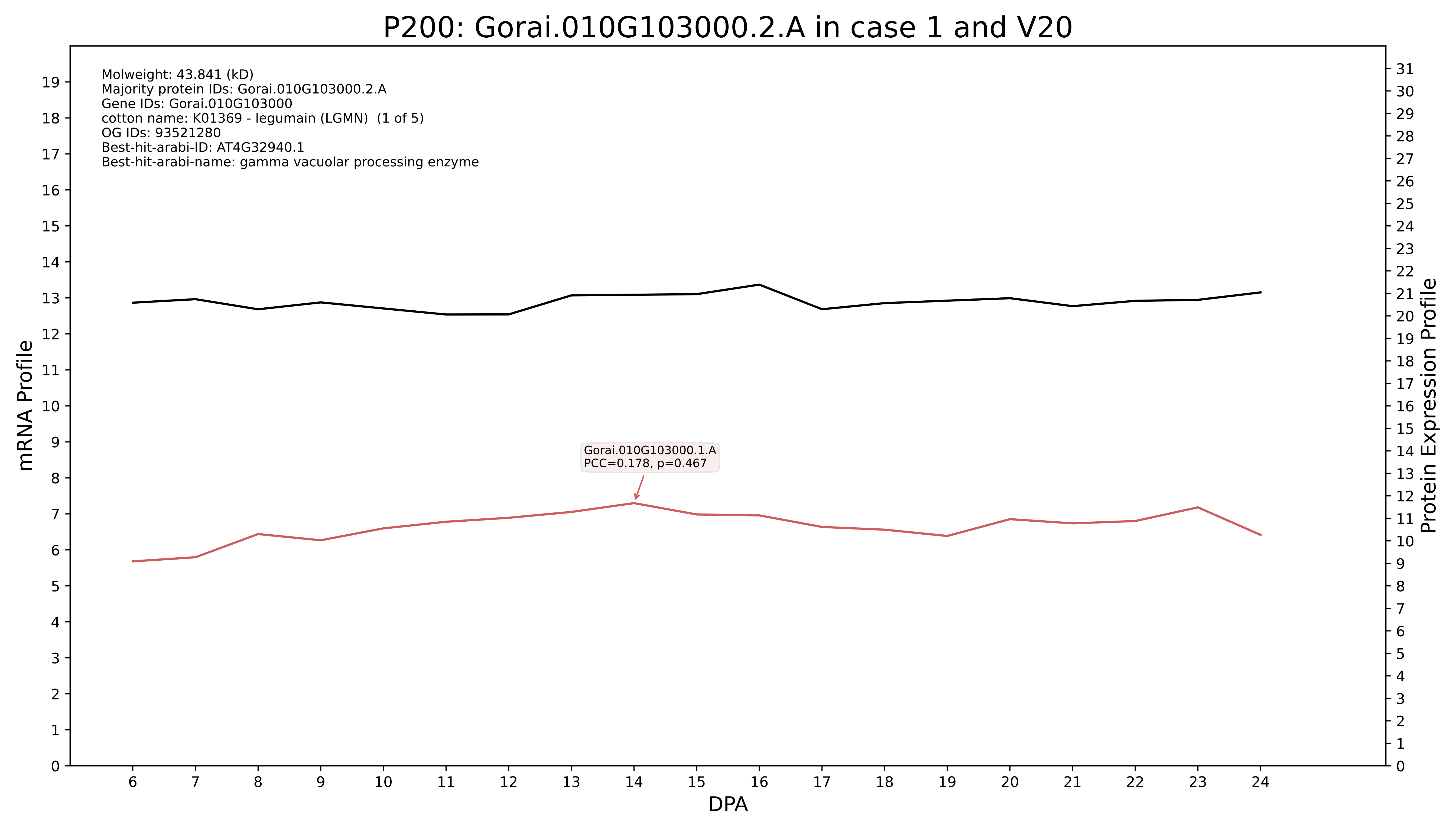
P200: Gorai.010G103000.2.A in case 1 and V20
Molweight: 43.841 (kD)
Majority protein IDs: Gorai.010G103000.2.A
Gene IDs: Gorai.010G103000
cotton name: K01369 - legumain (LGMN) (1 of 5)
OG IDs: 93521280
Best-hit-arabi-ID: AT4G32940.1
Best-hit-arabi-name: gamma vacuolar processing enzyme
Gorai.010G103000.1.A
PCC=0.178, p=0.467
mRNA Profile
Protein Expression Profile
DPA

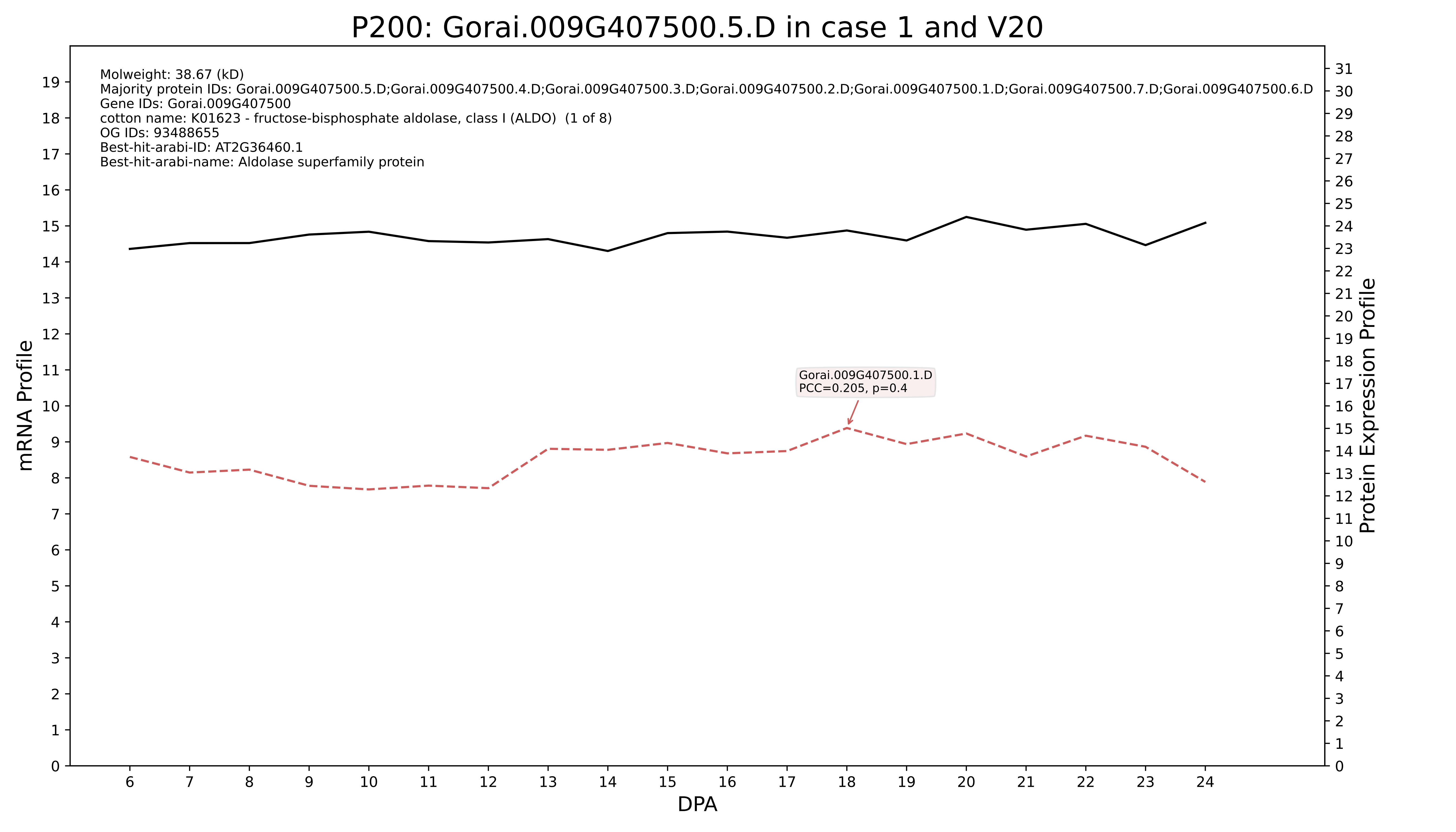
P200: Gorai.009G407500.5.D in case 1 and V20
Molweight: 38.67 (kD)
Majority protein IDs: Gorai.009G407500.5.D;Gorai.009G407500.4.D;Gorai.009G407500.3.D;Gorai.009G407500.2.D;Gorai.009G407500.1.D;Gorai.009G407500.7.D;Gorai.009G407500.6.D
Gene IDs: Gorai.009G407500
cotton name: K01623 - fructose-bisphosphate aldolase, class I (ALDO) (1 of 8)
OG IDs: 93488655
Best-hit-arabi-ID: AT2G36460.1
Best-hit-arabi-name: Aldolase superfamily protein
Gorai.009G407500.1.D
PCC=0.205, p=0.4
mRNA Profile
Protein Expression Profile
DPA

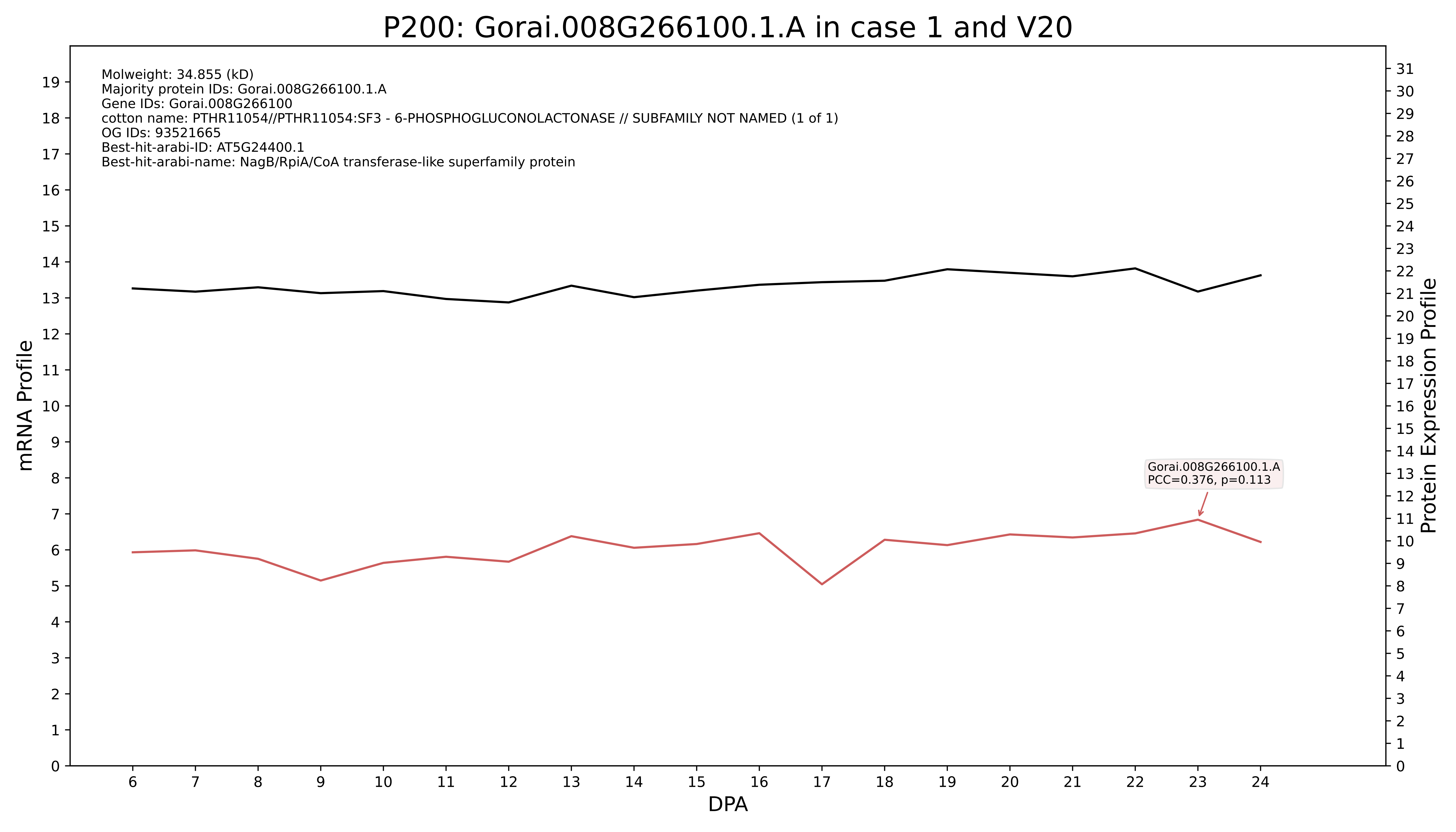
P200: Gorai.008G266100.1.A in case 1 and V20
Molweight: 34.855 (kD)
Majority protein IDs: Gorai.008G266100.1.A
Gene IDs: Gorai.008G266100
cotton name: PTHR11054//PTHR11054:SF3 - 6-PHOSPHOGLUCONOLACTONASE // SUBFAMILY NOT NAMED (1 of 1)
OG IDs: 93521665
Best-hit-arabi-ID: AT5G24400.1
Best-hit-arabi-name: NagB/RpiA/CoA transferase-like superfamily protein
Gorai.008G266100.1.A
PCC=0.376, p=0.113
mRNA Profile
Protein Expression Profile
DPA

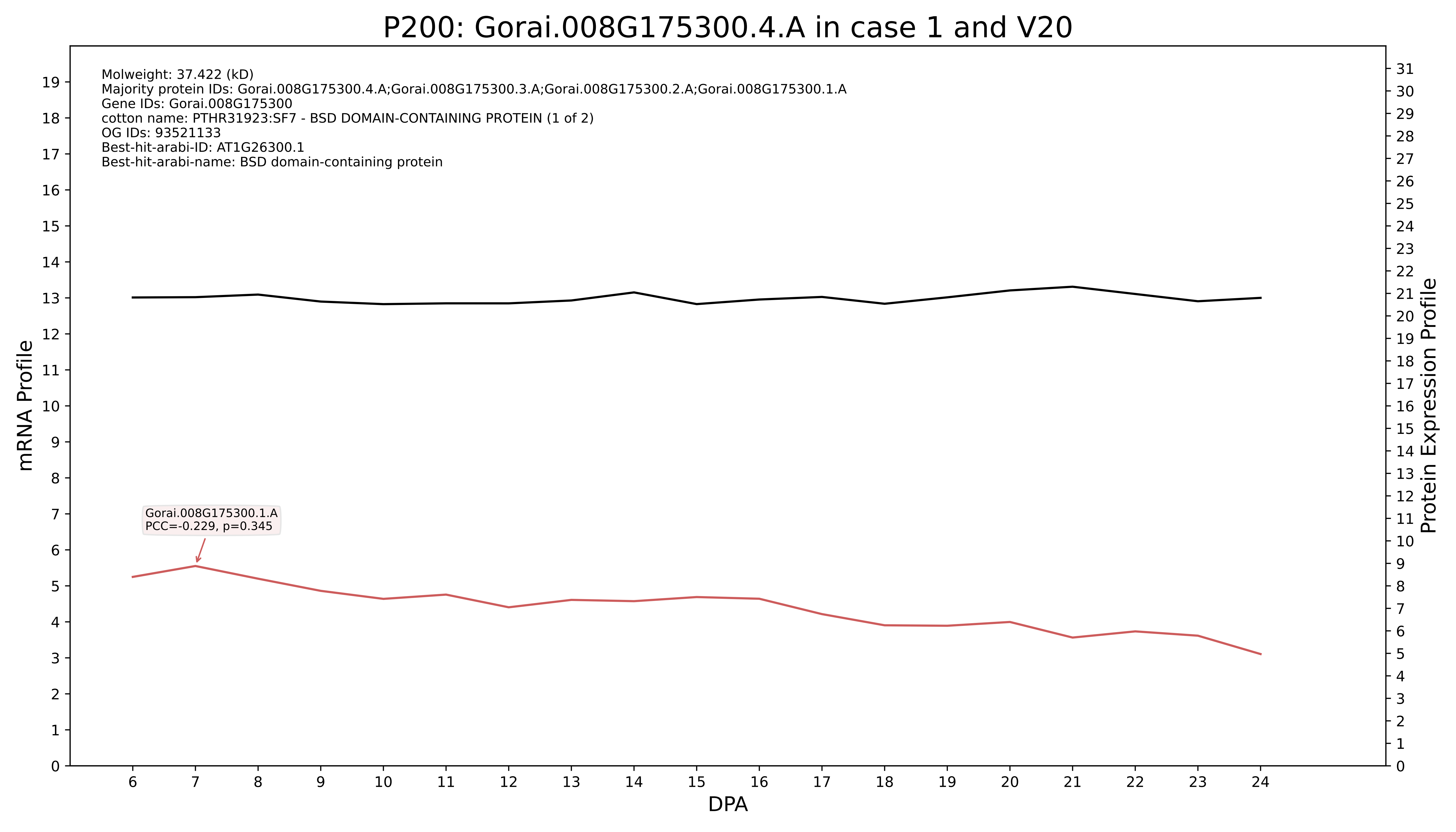
P200: Gorai.008G175300.4.A in case 1 and V20
Molweight: 37.422 (kD)
Majority protein IDs: Gorai.008G175300.4.A;Gorai.008G175300.3.A;Gorai.008G175300.2.A;Gorai.008G175300.1.A
Gene IDs: Gorai.008G175300
cotton name: PTHR31923:SF7 - BSD DOMAIN-CONTAINING PROTEIN (1 of 2)
OG IDs: 93521133
Best-hit-arabi-ID: AT1G26300.1
Best-hit-arabi-name: BSD domain-containing protein
Gorai.008G175300.1.A
PCC=-0.229, p=0.345
mRNA Profile
Protein Expression Profile
DPA

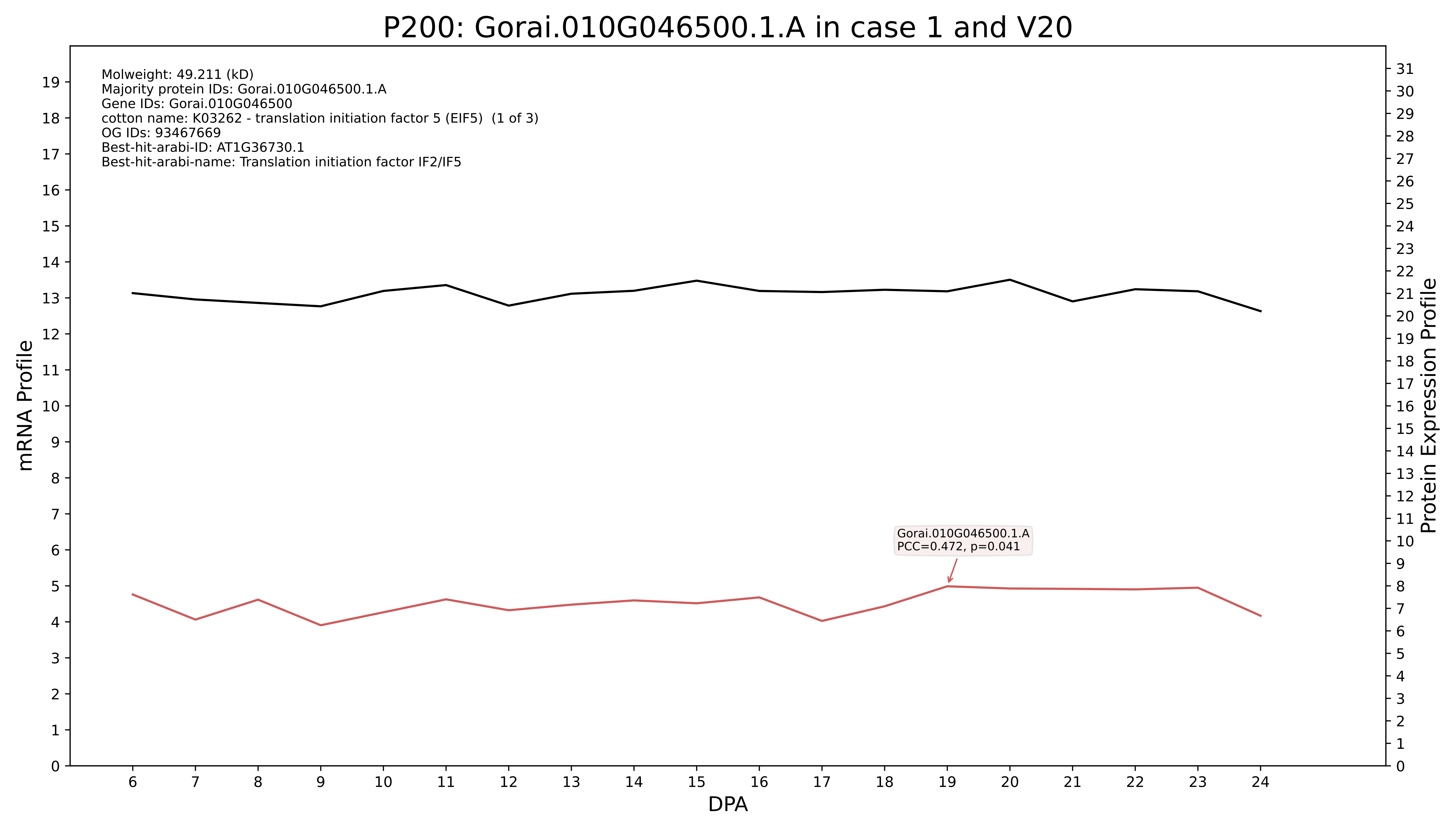
P200: Gorai.010G046500.1.A in case 1 and V20
Molweight: 49.211 (kD)
Majority protein IDs: Gorai.010G046500.1.A
Gene IDs: Gorai.010G046500
cotton name: K03262 - translation initiation factor 5 (EIF5) (1 of 3)
OG IDs: 93467669
Best-hit-arabi-ID: AT1G36730.1
Best-hit-arabi-name: Translation initiation factor IF2/IF5
Gorai.010G046500.1.A
PCC=0.472, p=0.041
mRNA Profile
Protein Expression Profile
DPA

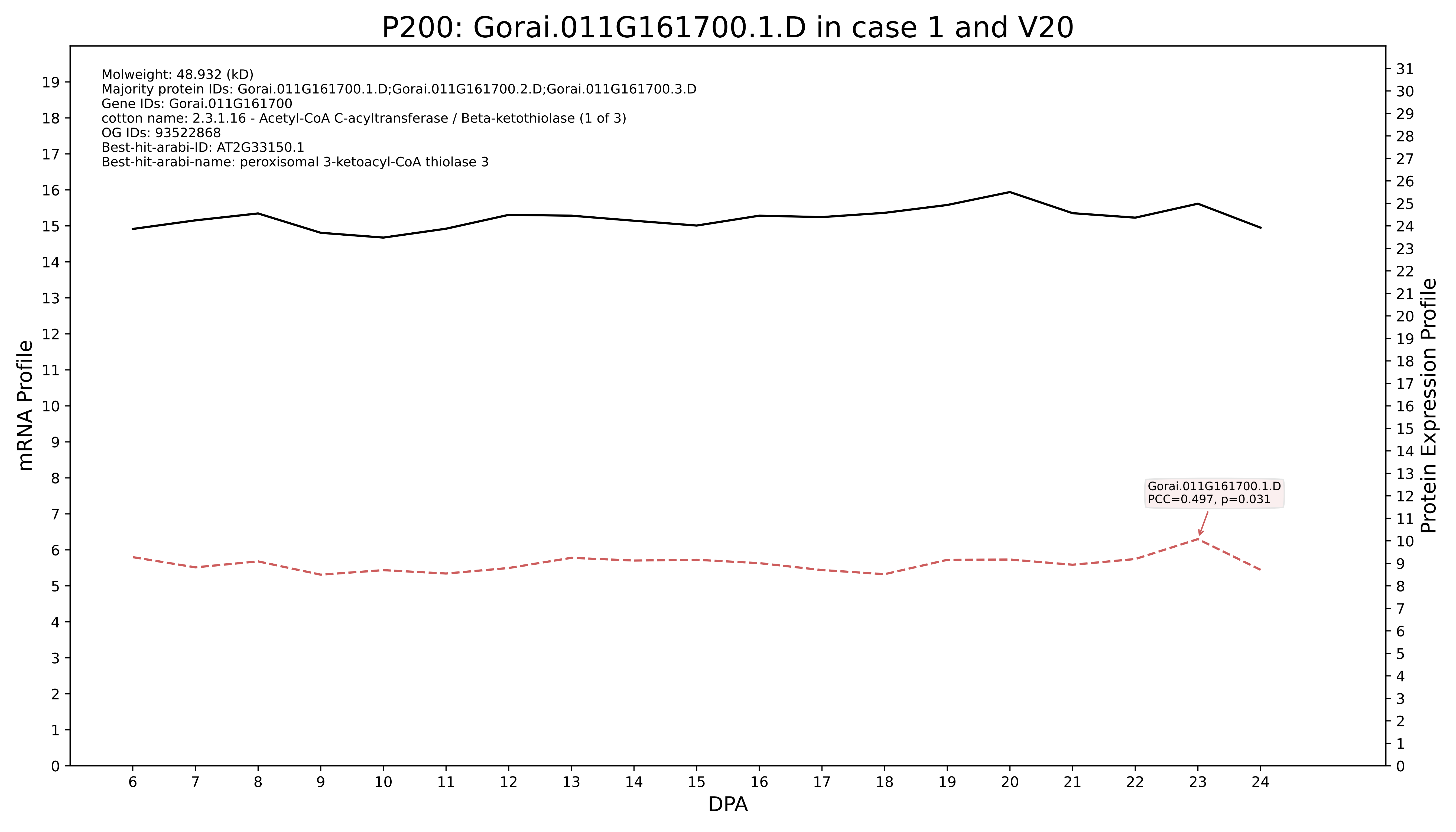
P200: Gorai.011G161700.1.D in case 1 and V20
Molweight: 48.932 (kD)
Majority protein IDs: Gorai.011G161700.1.D;Gorai.011G161700.2.D;Gorai.011G161700.3.D
Gene IDs: Gorai.011G161700
cotton name: 2.3.1.16 - Acetyl-CoA C-acyltransferase / Beta-ketothiolase (1 of 3)
OG IDs: 93522868
Best-hit-arabi-ID: AT2G33150.1
Best-hit-arabi-name: peroxisomal 3-ketoacyl-CoA thiolase 3
Gorai.011G161700.1.D
PCC=0.497, p=0.031
mRNA Profile
Protein Expression Profile
DPA

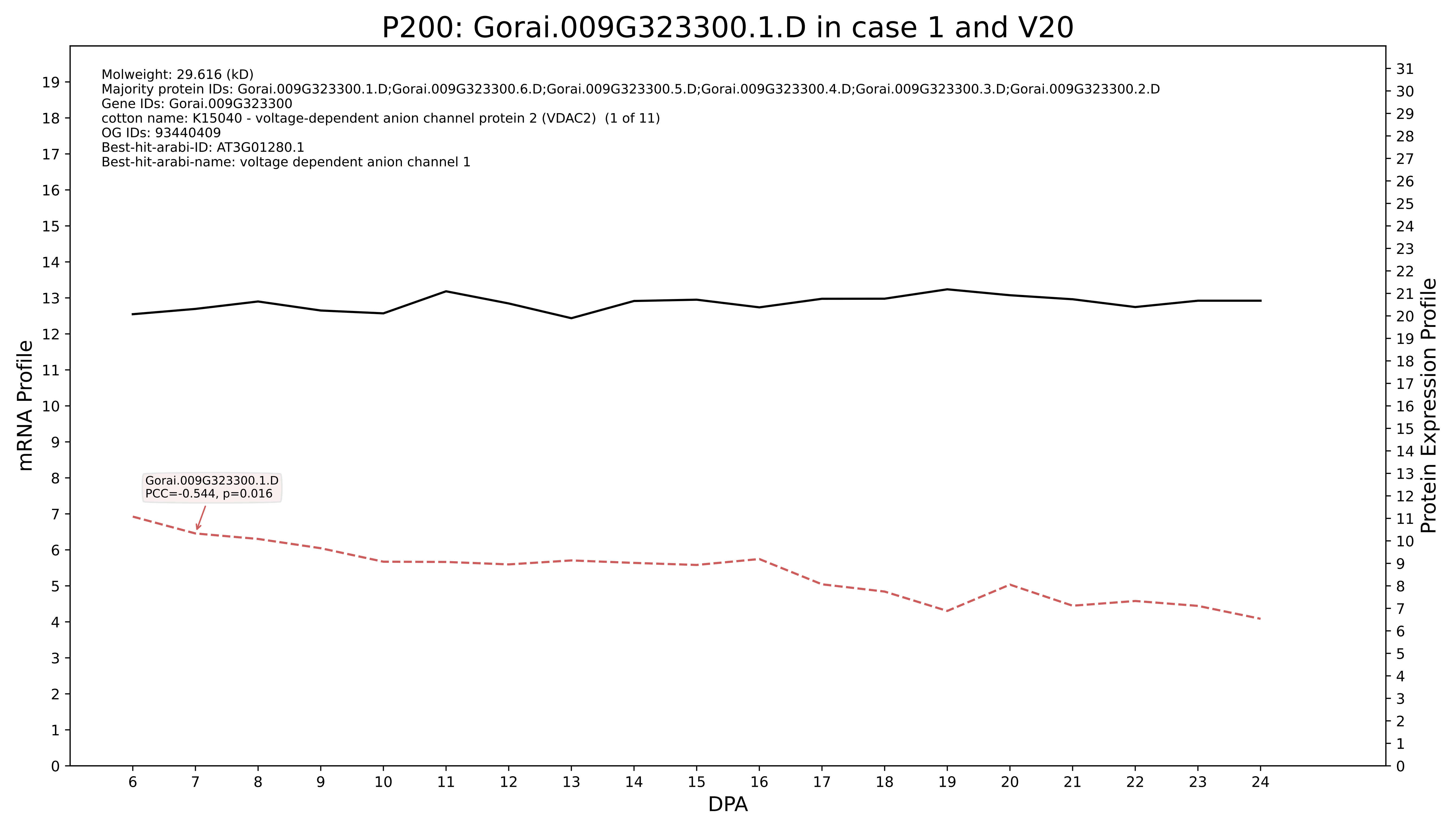

P200: Gorai.009G323300.1.D in case 1 and V20
Molweight: 29.616 (kD)
Majority protein IDs: Gorai.009G323300.1.D;Gorai.009G323300.6.D;Gorai.009G323300.5.D;Gorai.009G323300.4.D;Gorai.009G323300.3.D;Gorai.009G323300.2.D
Gene IDs: Gorai.009G323300
cotton name: K15040 - voltage-dependent anion channel protein 2 (VDAC2) (1 of 11)
OG IDs: 93440409
Best-hit-arabi-ID: AT3G01280.1
Best-hit-arabi-name: voltage dependent anion channel 1
Gorai.009G323300.1.D
PCC=-0.544, p=0.016
mRNA Profile
Protein Expression Profile
DPA

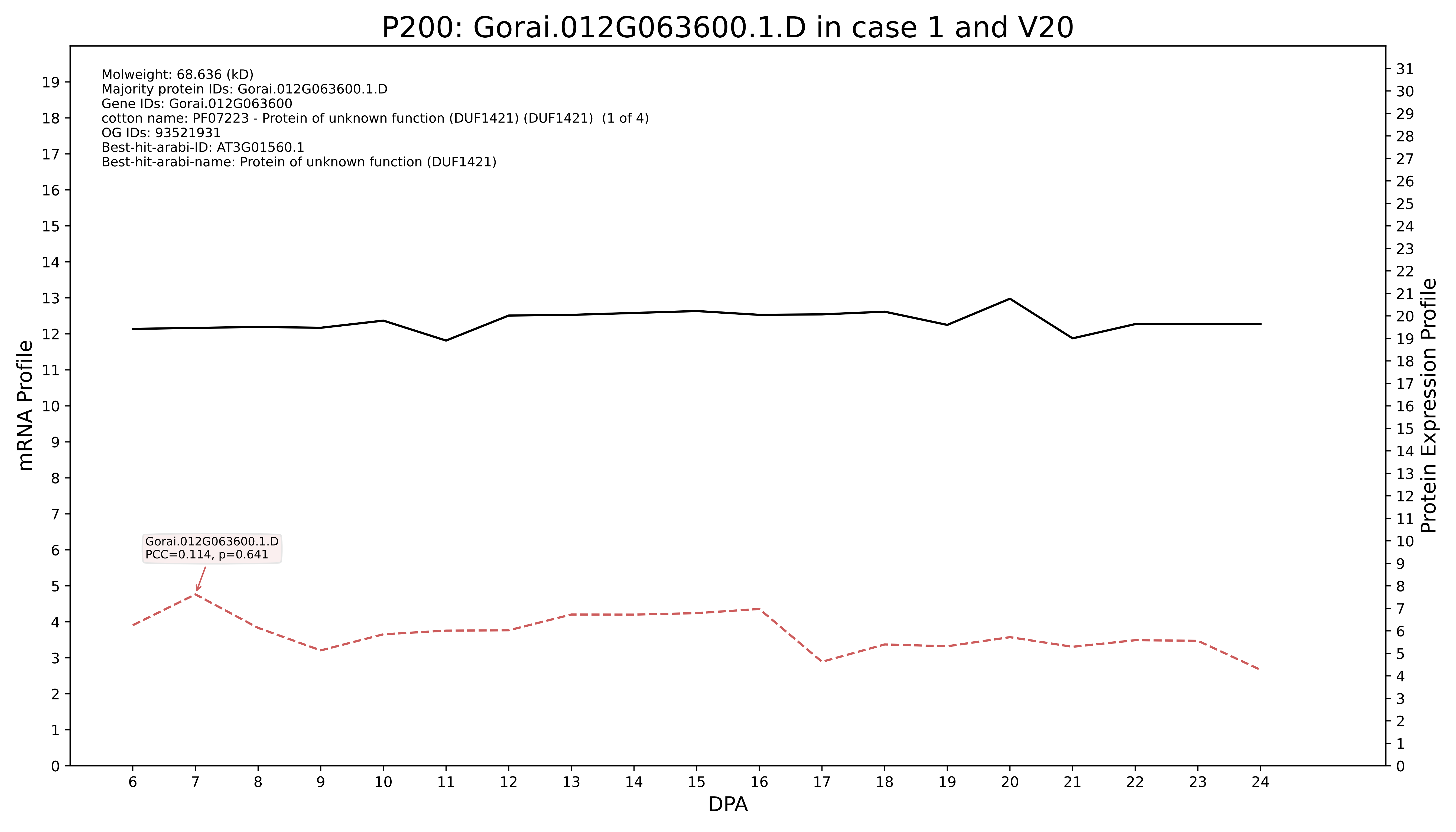

P200: Gorai.012G063600.1.D in case 1 and V20
Molweight: 68.636 (kD)
Majority protein IDs: Gorai.012G063600.1.D
Gene IDs: Gorai.012G063600
cotton name: PF07223 - Protein of unknown function (DUF1421) (DUF1421) (1 of 4)
OG IDs: 93521931
Best-hit-arabi-ID: AT3G01560.1
Best-hit-arabi-name: Protein of unknown function (DUF1421)
Gorai.012G063600.1.D
PCC=0.114, p=0.641
mRNA Profile
Protein Expression Profile
DPA

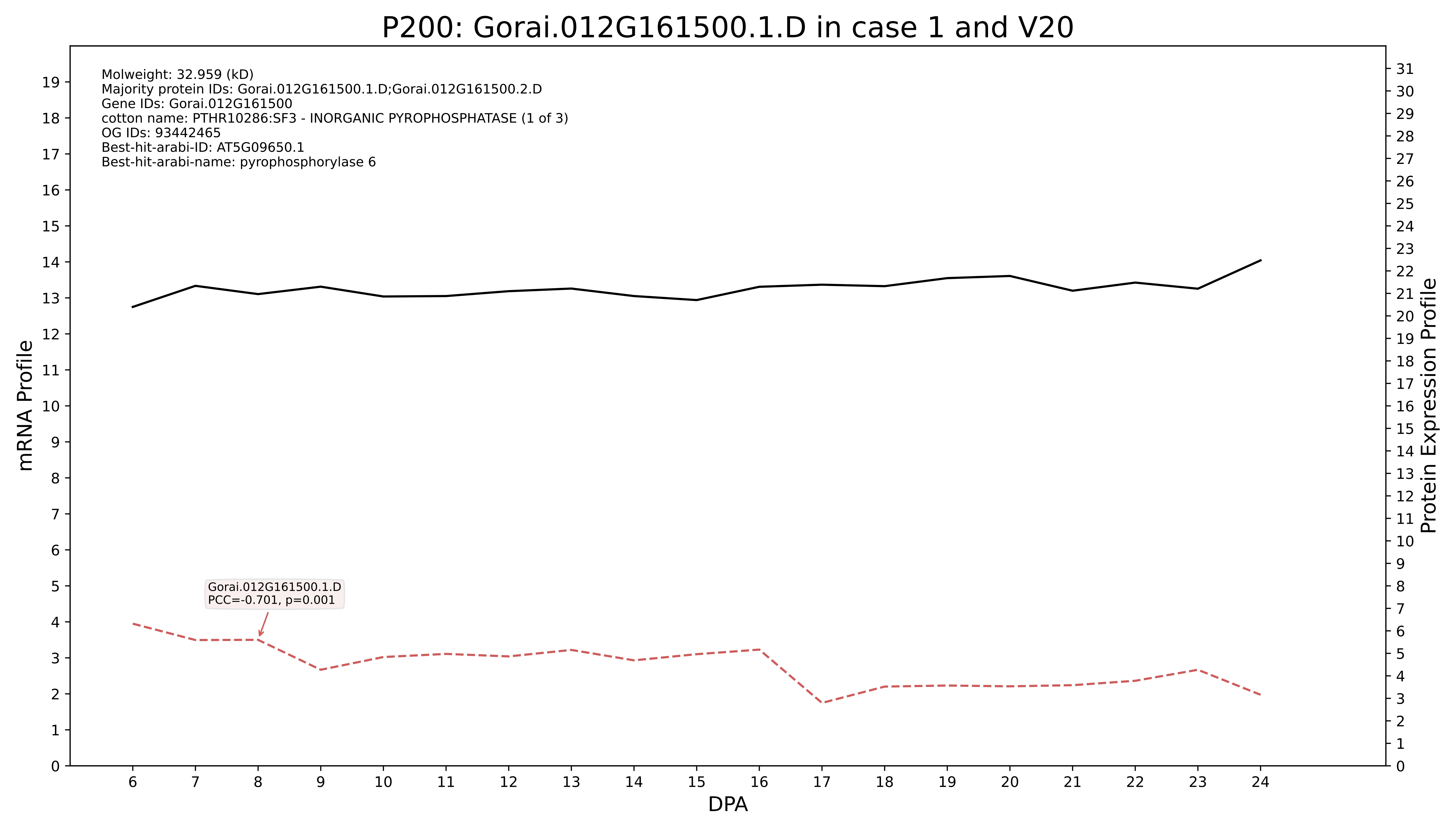
P200: Gorai.012G161500.1.D in case 1 and V20
Molweight: 32.959 (kD)
Majority protein IDs: Gorai.012G161500.1.D;Gorai.012G161500.2.D
Gene IDs: Gorai.012G161500
cotton name: PTHR10286:SF3 - INORGANIC PYROPHOSPHATASE (1 of 3)
OG IDs: 93442465
Best-hit-arabi-ID: AT5G09650.1
Best-hit-arabi-name: pyrophosphorylase 6
Gorai.012G161500.1.D
PCC=-0.701, p=0.001
mRNA Profile
Protein Expression Profile
DPA

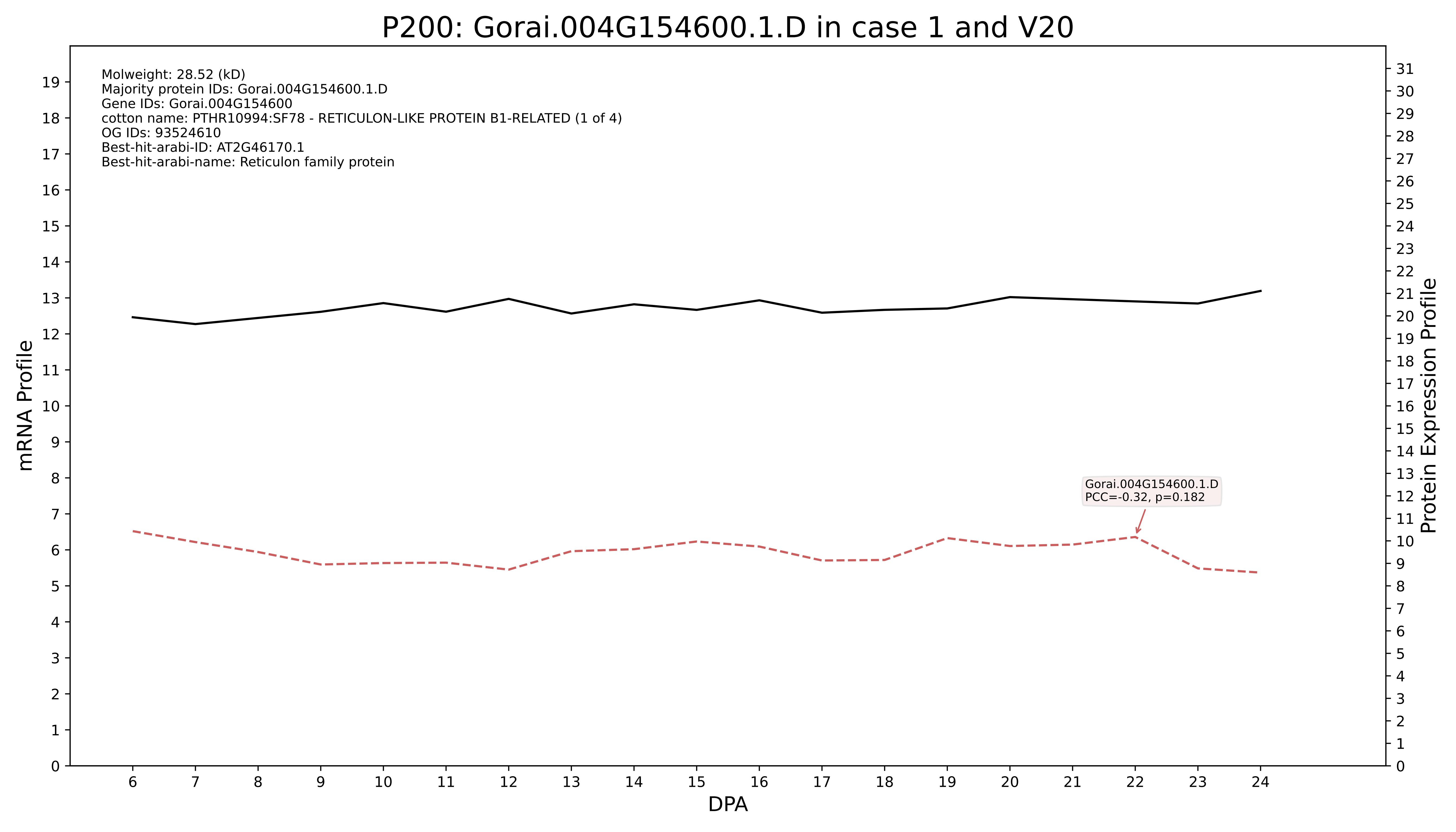

P200: Gorai.004G154600.1.D in case 1 and V20
Molweight: 28.52 (kD)
Majority protein IDs: Gorai.004G154600.1.D
Gene IDs: Gorai.004G154600
cotton name: PTHR10994:SF78 - RETICULON-LIKE PROTEIN B1-RELATED (1 of 4)
OG IDs: 93524610
Best-hit-arabi-ID: AT2G46170.1
Best-hit-arabi-name: Reticulon family protein
Gorai.004G154600.1.D
PCC=-0.32, p=0.182
mRNA Profile
Protein Expression Profile
DPA

### P200: Gorai.008G026600.1.A in case 1 and V20

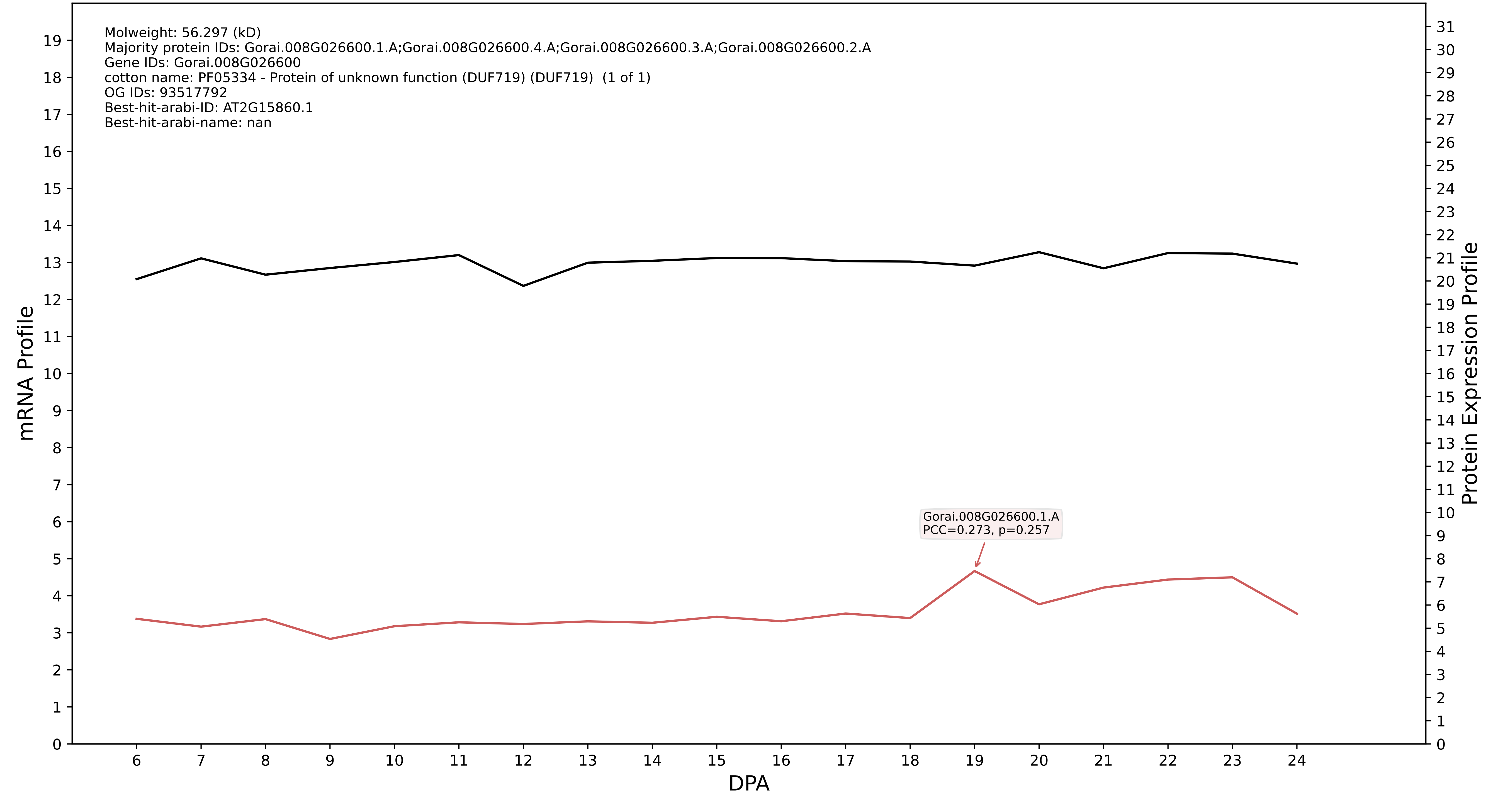

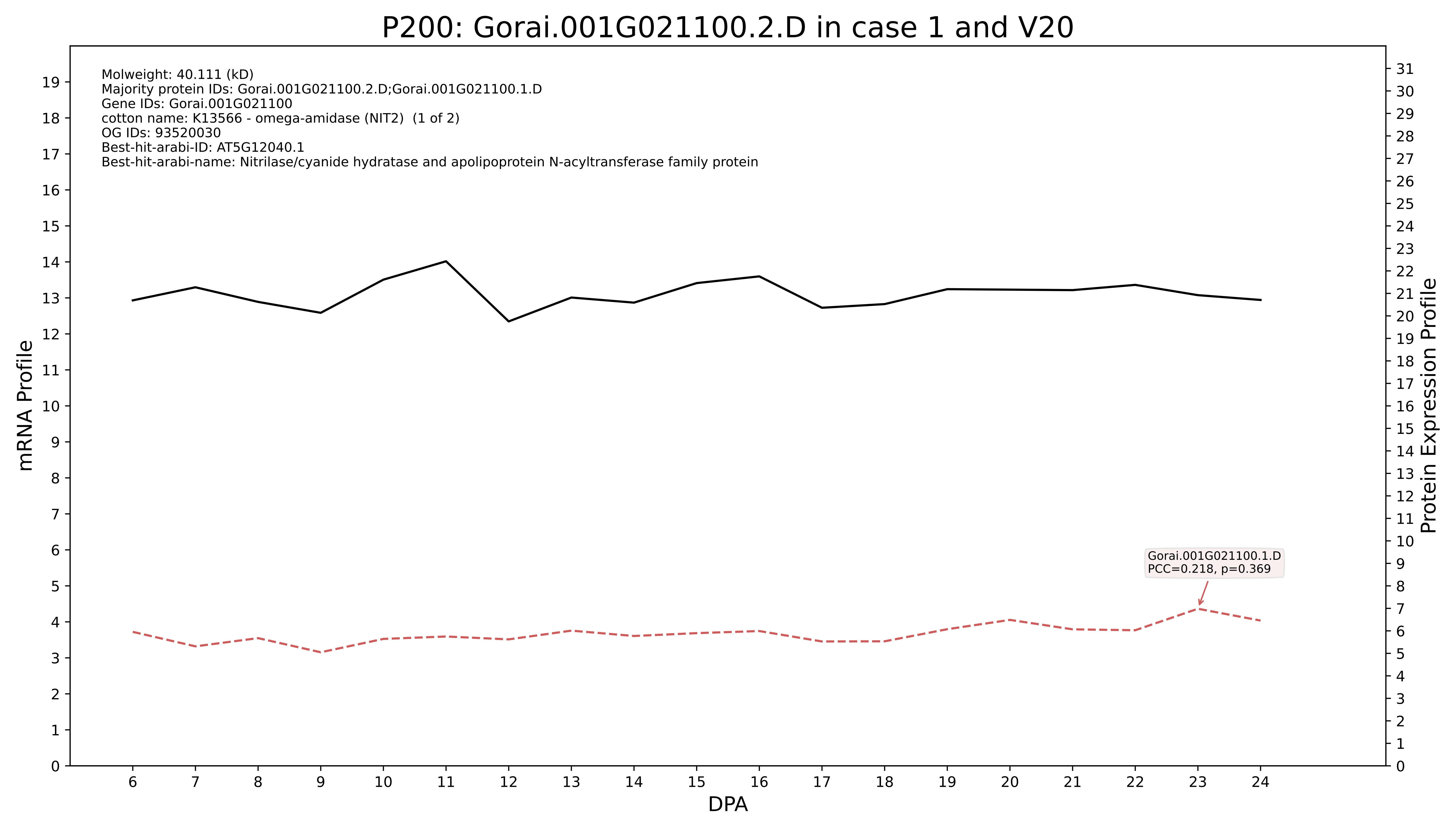

P200: Gorai.001G021100.2.D in case 1 and V20
Molweight: 40.111 (kD)
Majority protein IDs: Gorai.001G021100.2.D;Gorai.001G021100.1.D
Gene IDs: Gorai.001G021100
cotton name: K13566 - omega-amidase (NIT2) (1 of 2)
OG IDs: 93520030
Best-hit-arabi-ID: AT5G12040.1
Best-hit-arabi-name: Nitrilase/cyanide hydratase and apolipoprotein N-acyltransferase family protein
Gorai.001G021100.1.D
PCC=0.218, p=0.369
mRNA Profile
Protein Expression Profile
DPA

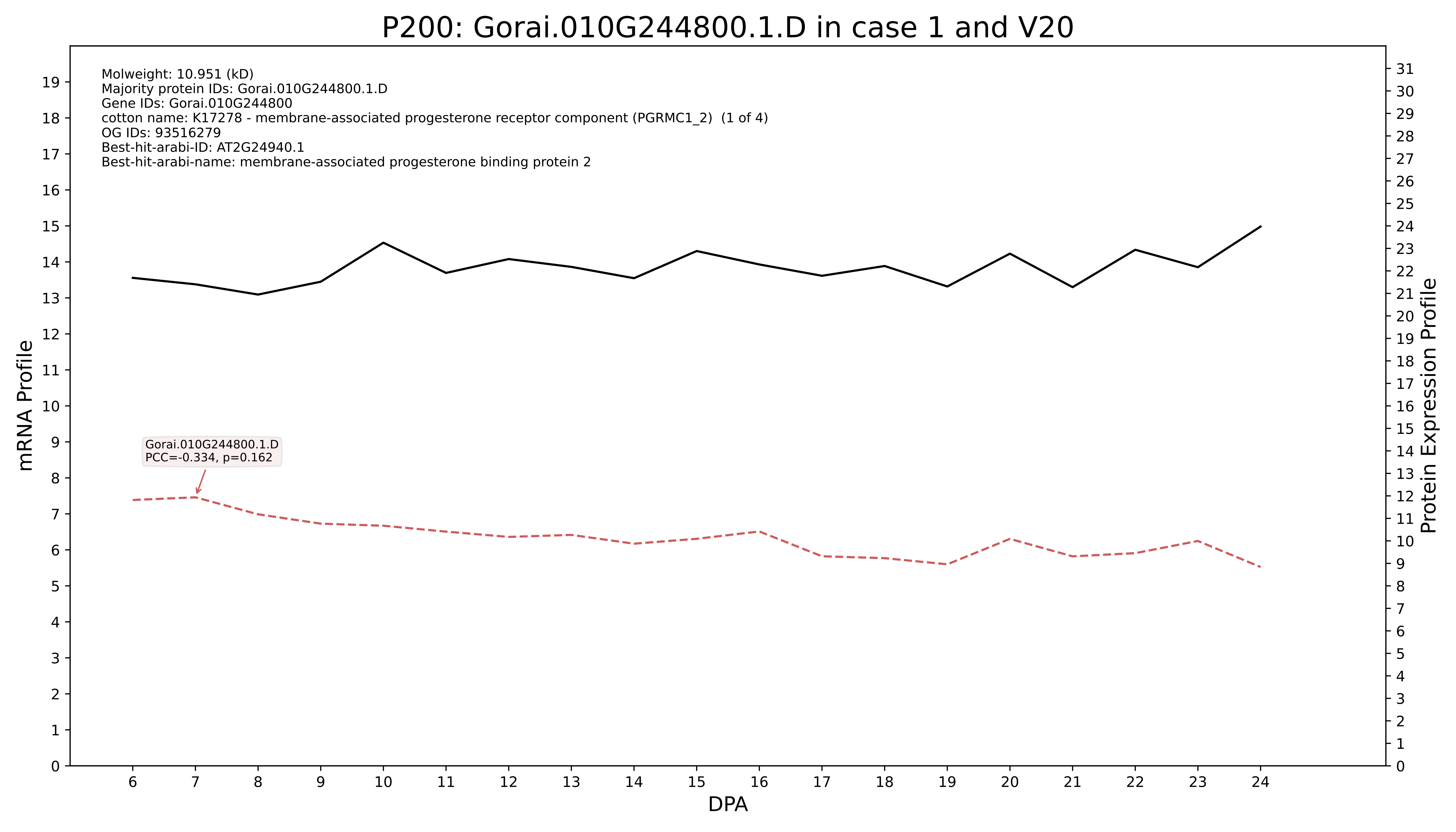
P200: Gorai.010G244800.1.D in case 1 and V20
Molweight: 10.951 (kD)
Majority protein IDs: Gorai.010G244800.1.D
Gene IDs: Gorai.010G244800
cotton name: K17278 - membrane-associated progesterone receptor component (PGRMC1_2) (1 of 4)
OG IDs: 93516279
Best-hit-arabi-ID: AT2G24940.1
Best-hit-arabi-name: membrane-associated progesterone binding protein 2
Gorai.010G244800.1.D
PCC=-0.334, p=0.162
mRNA Profile
Protein Expression Profile
DPA

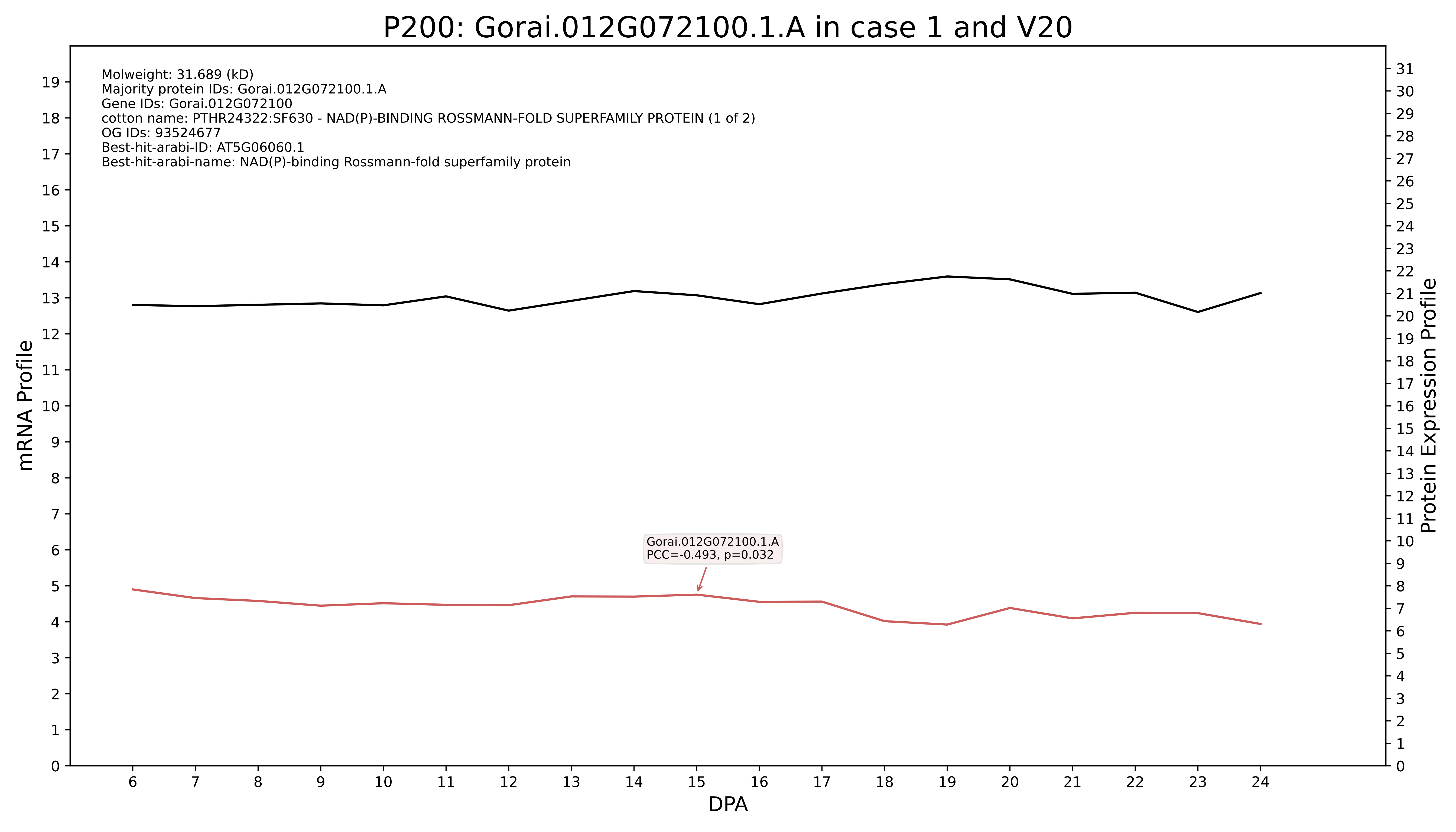

P200: Gorai.012G072100.1.A in case 1 and V20
Molweight: 31.689 (kD)
Majority protein IDs: Gorai.012G072100.1.A
Gene IDs: Gorai.012G072100
cotton name: PTHR24322:SF630 - NAD(P)-BINDING ROSSMANN-FOLD SUPERFAMILY PROTEIN (1 of 2)
OG IDs: 93524677
Best-hit-arabi-ID: AT5G06060.1
Best-hit-arabi-name: NAD(P)-binding Rossmann-fold superfamily protein
Gorai.012G072100.1.A
PCC=-0.493, p=0.032
mRNA Profile
Protein Expression Profile
DPA

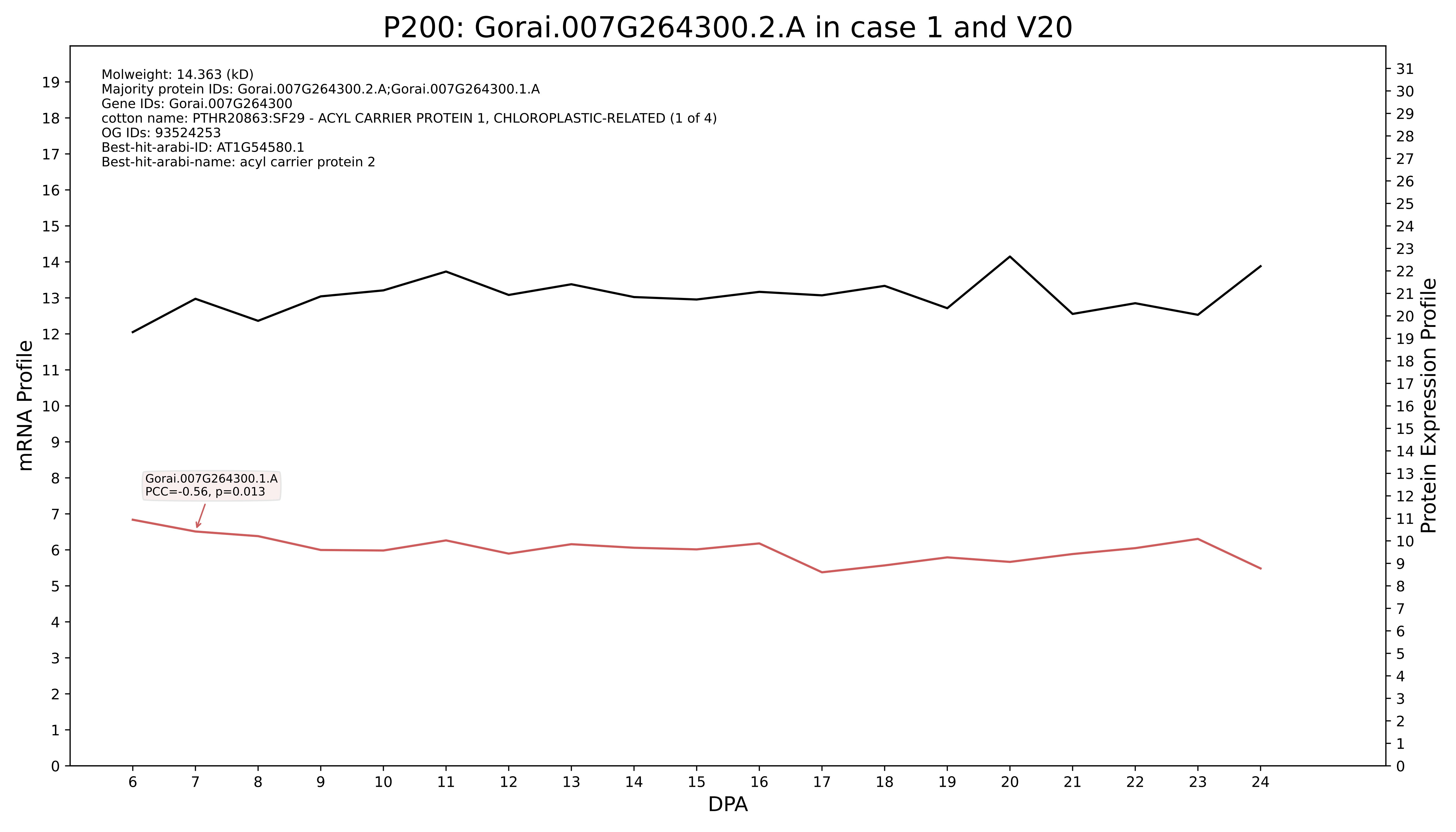

P200: Gorai.007G264300.2.A in case 1 and V20
Molweight: 14.363 (kD)
Majority protein IDs: Gorai.007G264300.2.A;Gorai.007G264300.1.A
Gene IDs: Gorai.007G264300
cotton name: PTHR20863:SF29 - ACYL CARRIER PROTEIN 1, CHLOROPLASTIC-RELATED (1 of 4)
OG IDs: 93524253
Best-hit-arabi-ID: AT1G54580.1
Best-hit-arabi-name: acyl carrier protein 2
Gorai.007G264300.1.A
PCC=-0.56, p=0.013
mRNA Profile
Protein Expression Profile
DPA

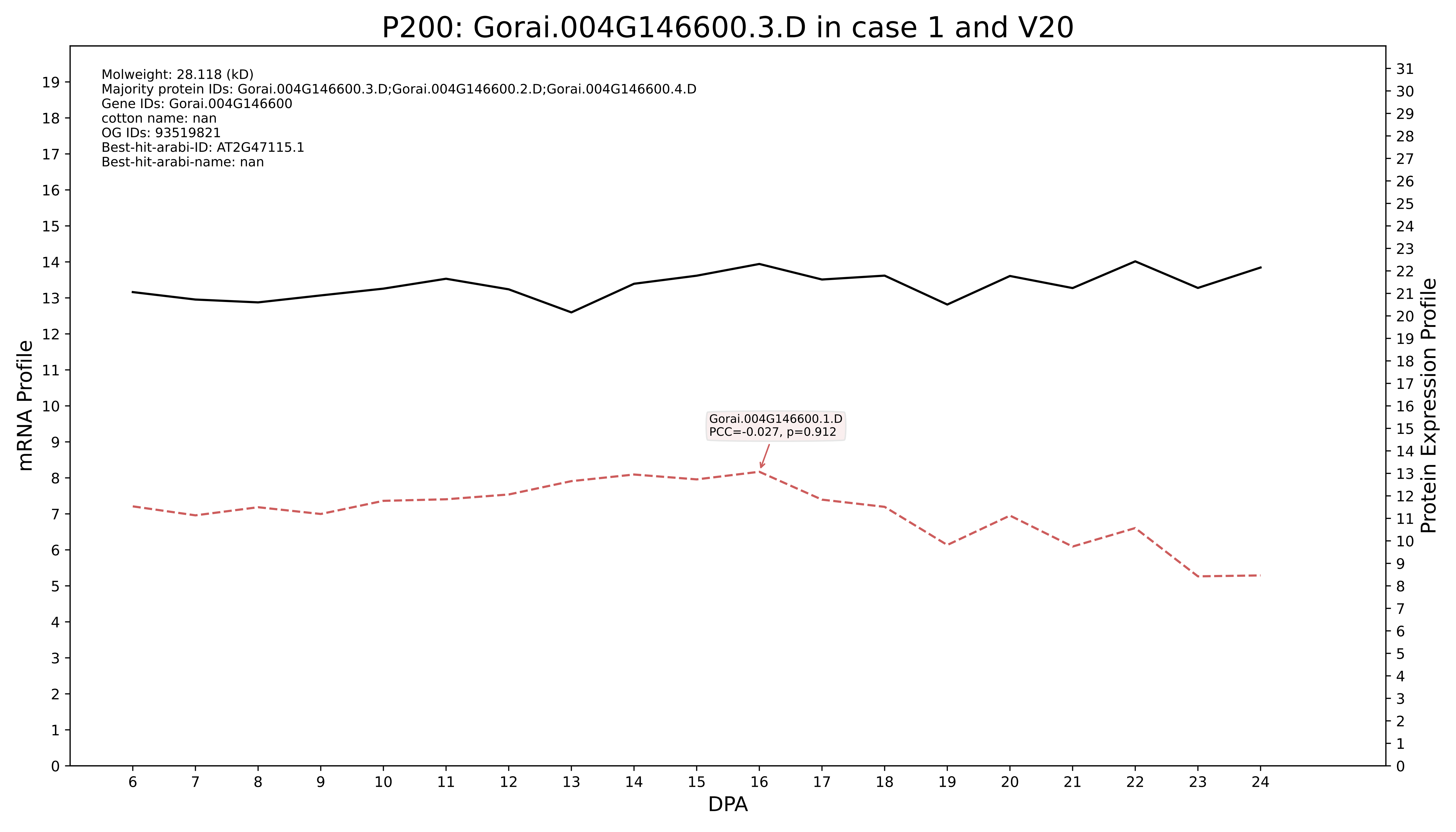
P200: Gorai.004G146600.3.D in case 1 and V20
Molweight: 28.118 (kD)
Majority protein IDs: Gorai.004G146600.3.D;Gorai.004G146600.2.D;Gorai.004G146600.4.D
Gene IDs: Gorai.004G146600
cotton name: nan
OG IDs: 93519821
Best-hit-arabi-ID: AT2G47115.1
Best-hit-arabi-name: nan
Gorai.004G146600.1.D
PCC=-0.027, p=0.912
mRNA Profile
Protein Expression Profile
DPA

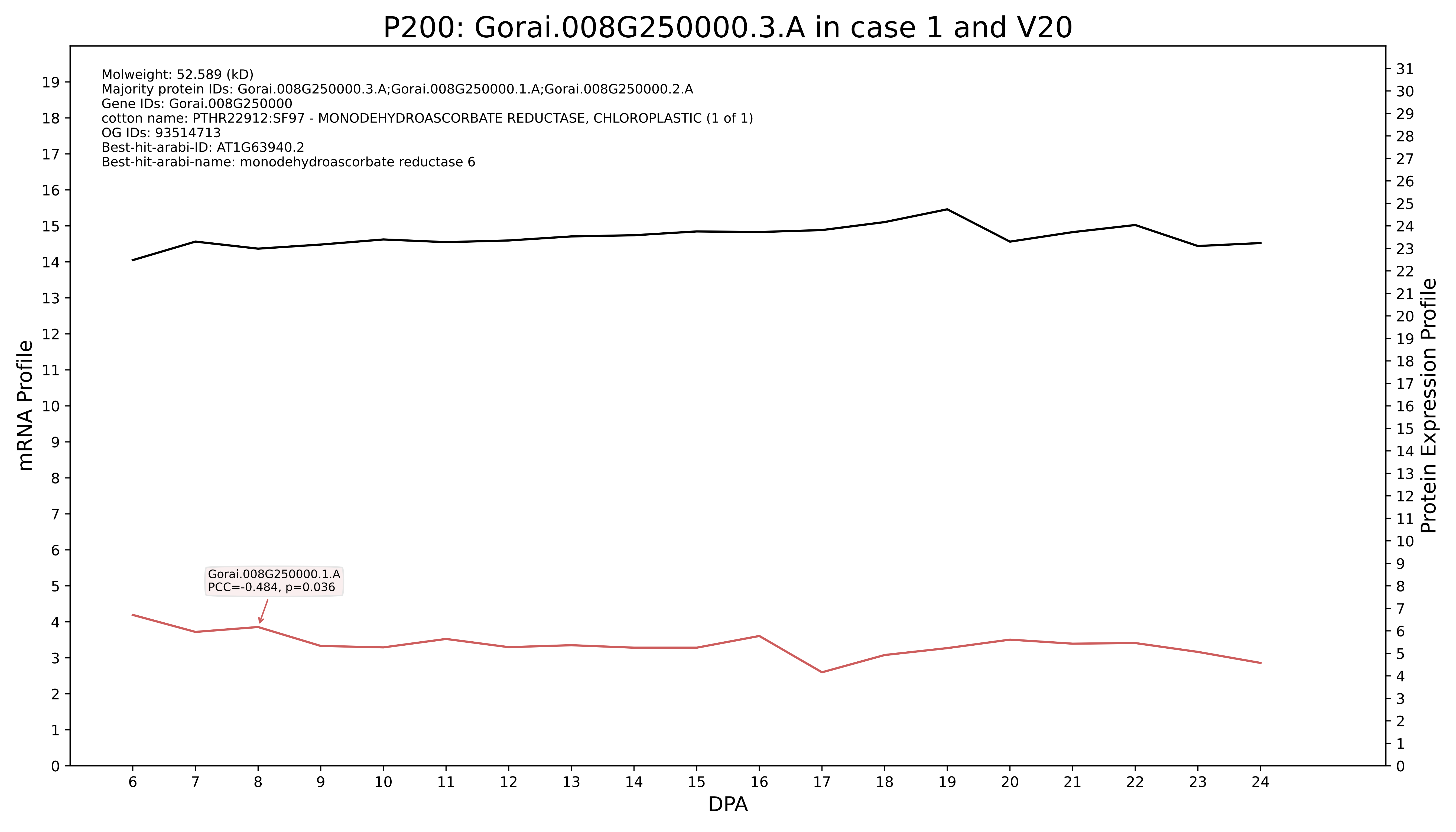

P200: Gorai.008G250000.3.A in case 1 and V20
Molweight: 52.589 (kD)
Majority protein IDs: Gorai.008G250000.3.A;Gorai.008G250000.1.A;Gorai.008G250000.2.A
Gene IDs: Gorai.008G250000
cotton name: PTHR22912:SF97 - MONODEHYDROASCORBATE REDUCTASE, CHLOROPLASTIC (1 of 1)
OG IDs: 93514713
Best-hit-arabi-ID: AT1G63940.2
Best-hit-arabi-name: monodehydroascorbate reductase 6
Gorai.008G250000.1.A
PCC=-0.484, p=0.036
mRNA Profile
Protein Expression Profile
DPA

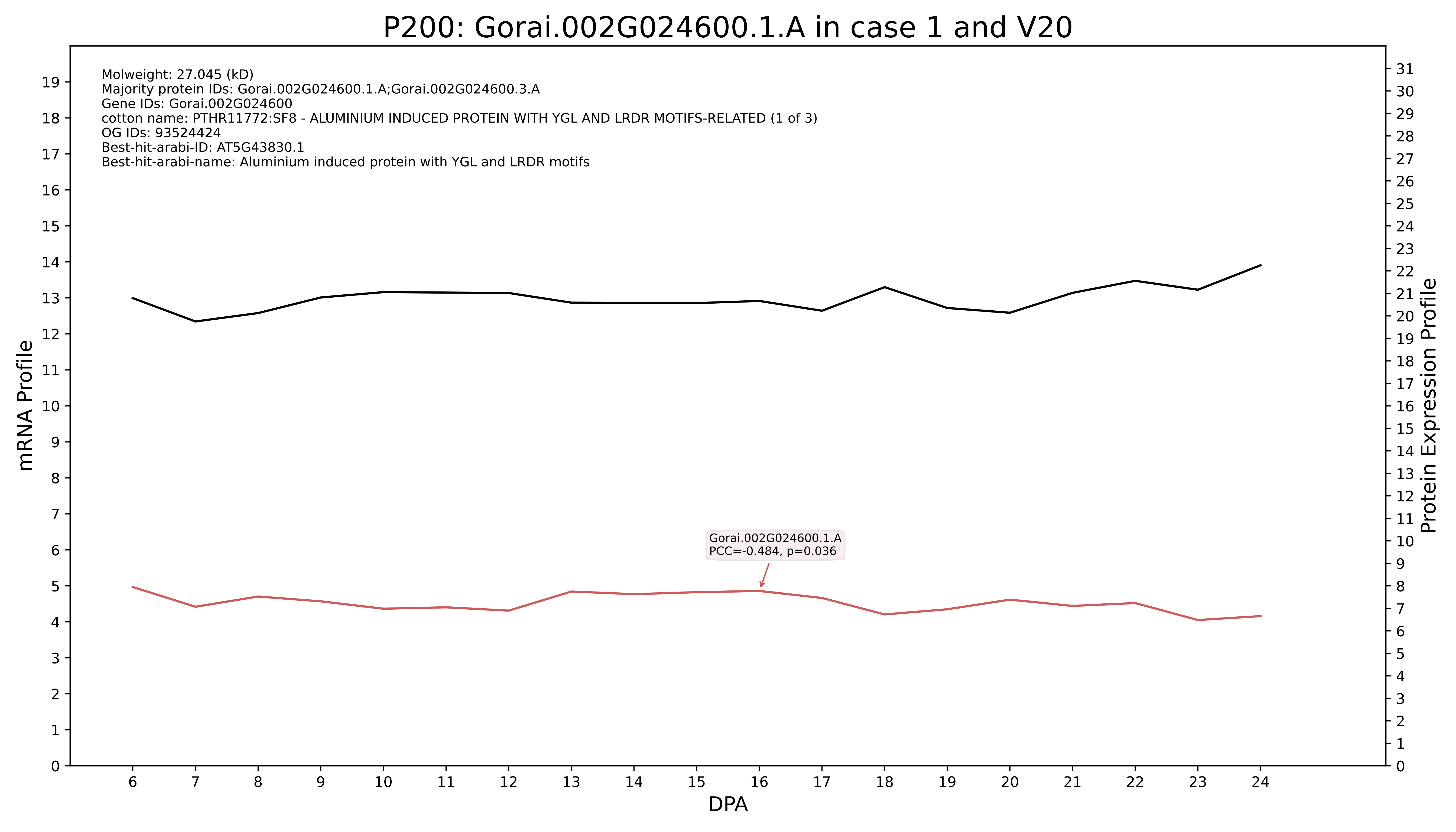
P200: Gorai.002G024600.1.A in case 1 and V20
Molweight: 27.045 (kD)
Majority protein IDs: Gorai.002G024600.1.A;Gorai.002G024600.3.A
Gene IDs: Gorai.002G024600
cotton name: PTHR11772:SF8 - ALUMINIUM INDUCED PROTEIN WITH YGL AND LRDR MOTIFS-RELATED (1 of 3)
OG IDs: 93524424
Best-hit-arabi-ID: AT5G43830.1
Best-hit-arabi-name: Aluminium induced protein with YGL and LRDR motifs
Gorai.002G024600.1.A
PCC=-0.484, p=0.036
mRNA Profile
Protein Expression Profile
DPA

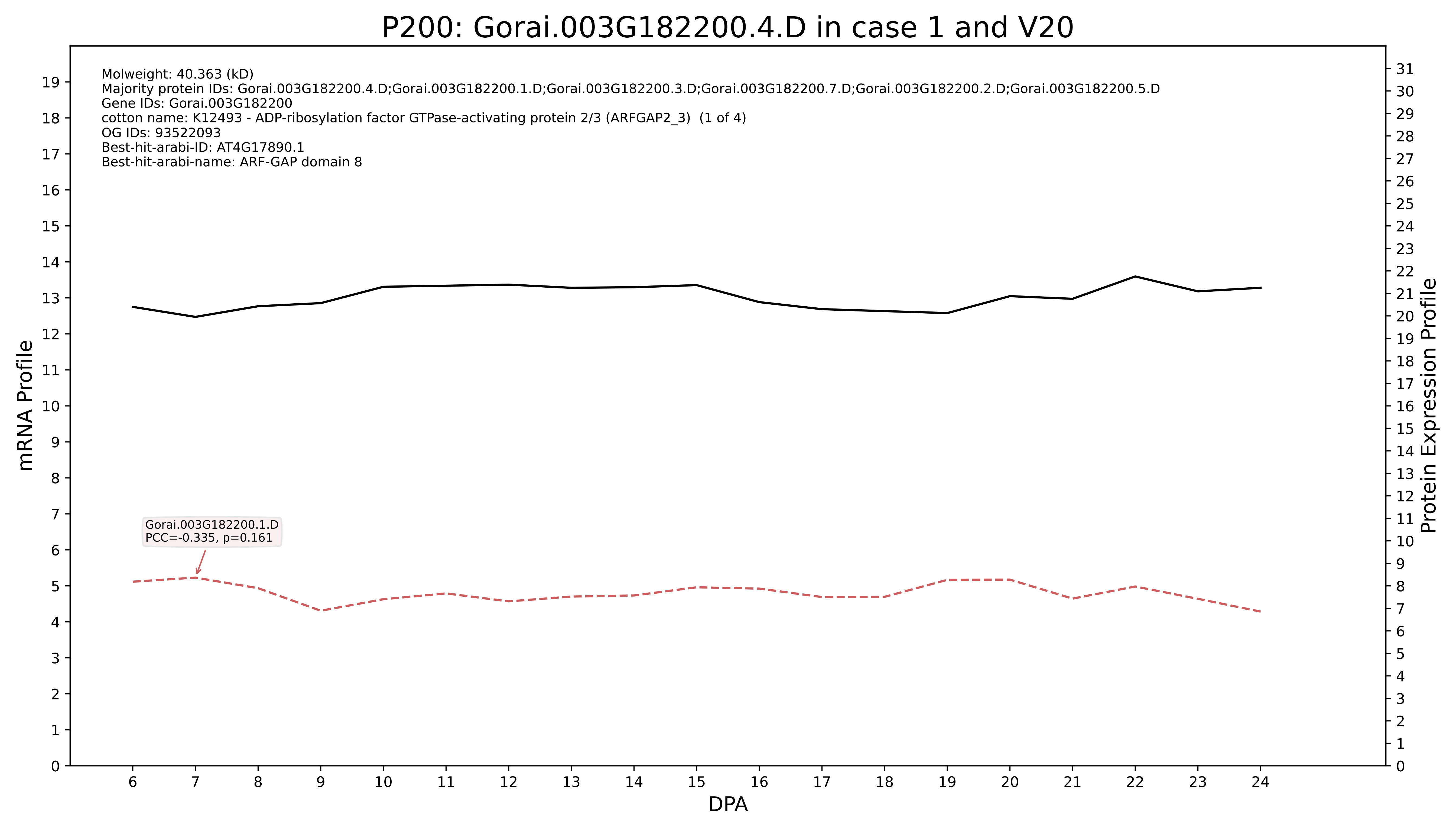

P200: Gorai.003G182200.4.D in case 1 and V20
Molweight: 40.363 (kD)
Majority protein IDs: Gorai.003G182200.4.D;Gorai.003G182200.1.D;Gorai.003G182200.3.D;Gorai.003G182200.7.D;Gorai.003G182200.2.D;Gorai.003G182200.5.D
Gene IDs: Gorai.003G182200
cotton name: K12493 - ADP-ribosylation factor GTPase-activating protein 2/3 (ARFGAP2_3) (1 of 4)
OG IDs: 93522093
Best-hit-arabi-ID: AT4G17890.1
Best-hit-arabi-name: ARF-GAP domain 8
Gorai.003G182200.1.D
PCC=-0.335, p=0.161
mRNA Profile
Protein Expression Profile
DPA

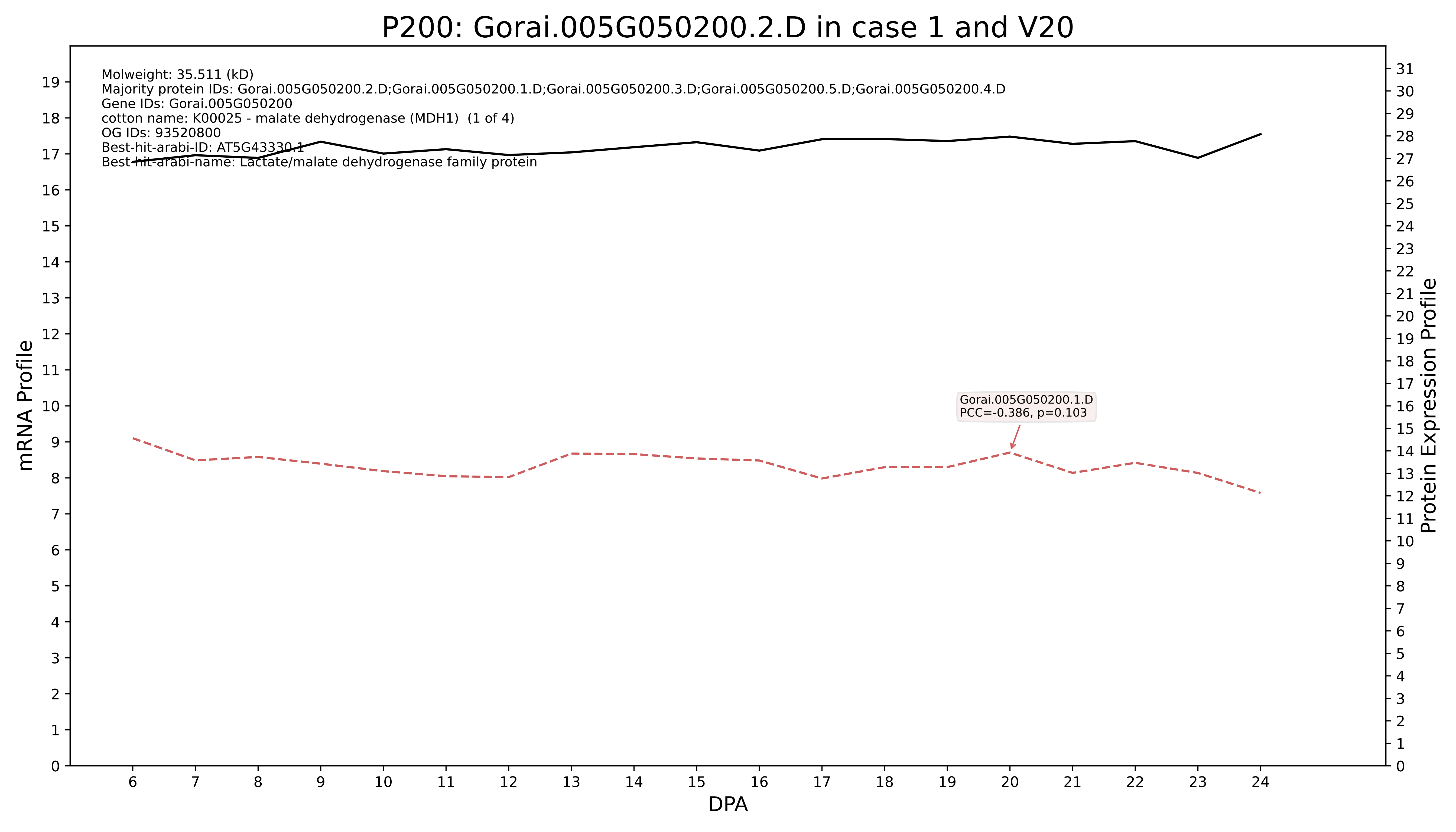
P200: Gorai.005G050200.2.D in case 1 and V20
Molweight: 35.511 (kD)
Majority protein IDs: Gorai.005G050200.2.D;Gorai.005G050200.1.D;Gorai.005G050200.3.D;Gorai.005G050200.5.D;Gorai.005G050200.4.D
Gene IDs: Gorai.005G050200
cotton name: K00025 - malate dehydrogenase (MDH1) (1 of 4)
OG IDs: 93520800
Best-hit-arabi-ID: AT5G43330.1
Best-hit-arabi-name: Lactate/malate dehydrogenase family protein
Gorai.005G050200.1.D
PCC=-0.386, p=0.103
mRNA Profile
Protein Expression Profile
DPA

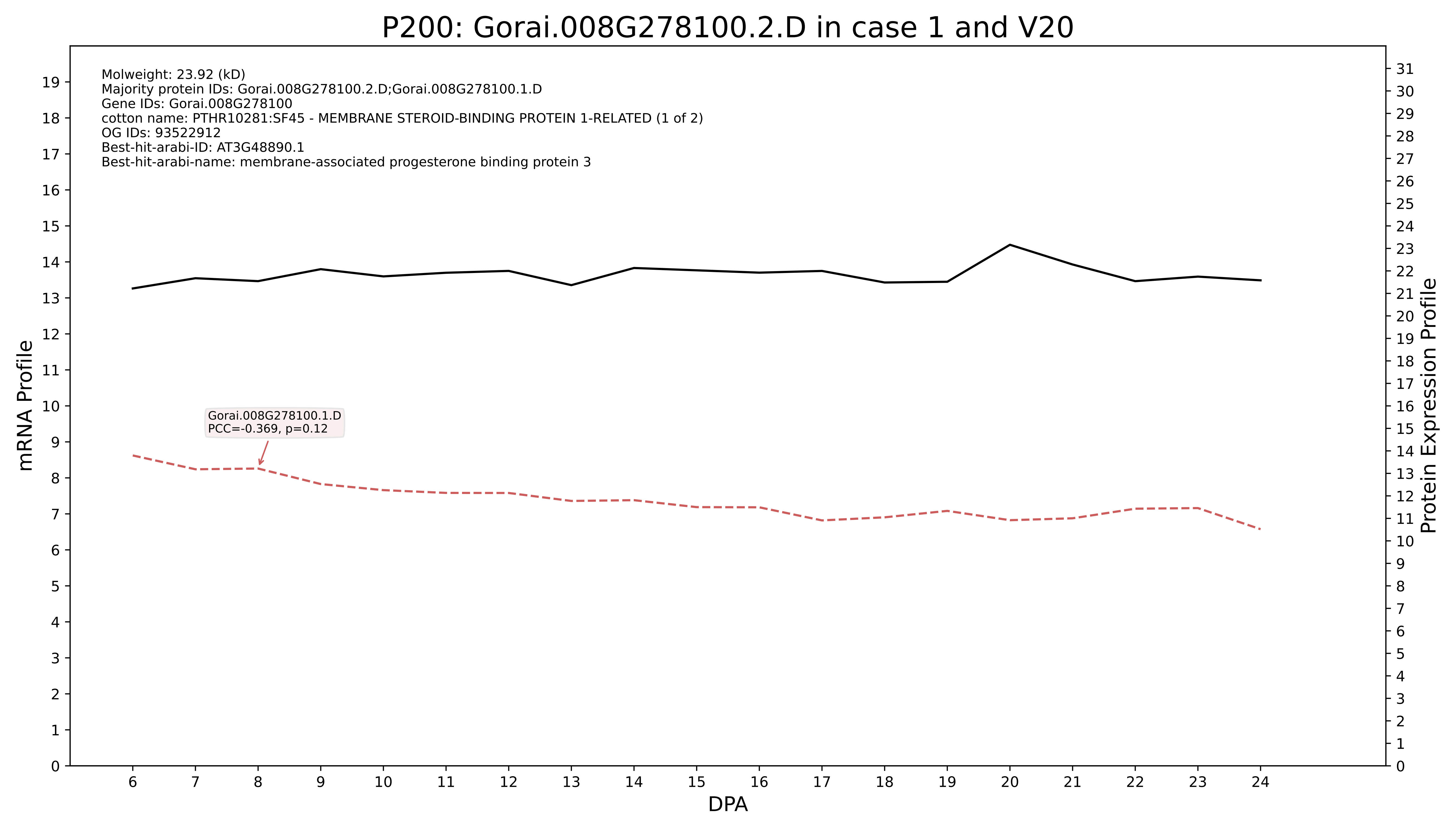
P200: Gorai.008G278100.2.D in case 1 and V20
Molweight: 23.92 (kD)
Majority protein IDs: Gorai.008G278100.2.D;Gorai.008G278100.1.D
Gene IDs: Gorai.008G278100
cotton name: PTHR10281:SF45 - MEMBRANE STEROID-BINDING PROTEIN 1-RELATED (1 of 2)
OG IDs: 93522912
Best-hit-arabi-ID: AT3G48890.1
Best-hit-arabi-name: membrane-associated progesterone binding protein 3
Gorai.008G278100.1.D
PCC=-0.369, p=0.12
mRNA Profile
Protein Expression Profile
DPA

### P200: Gorai.005G266800.1.D in case 1 and V20

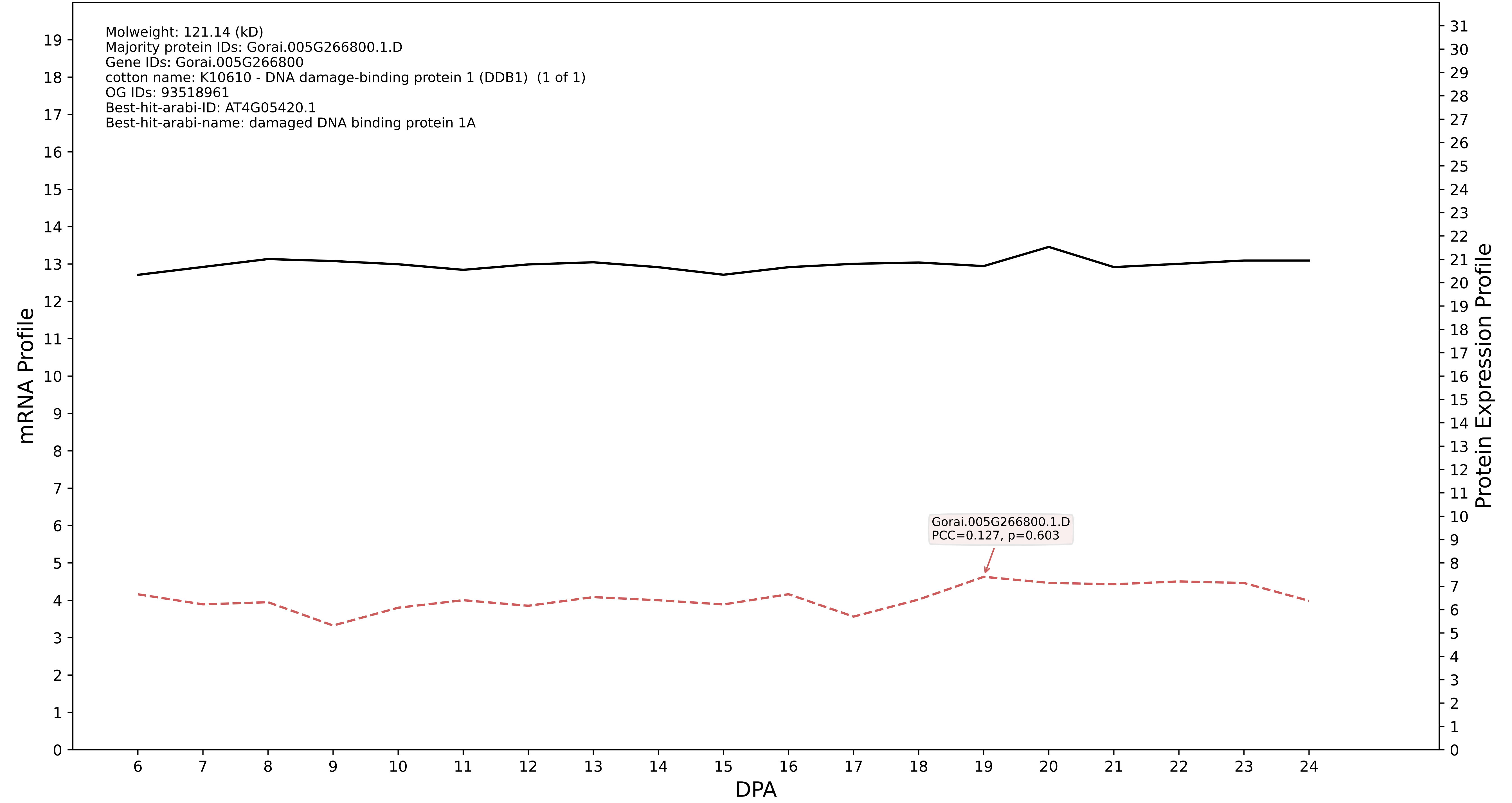

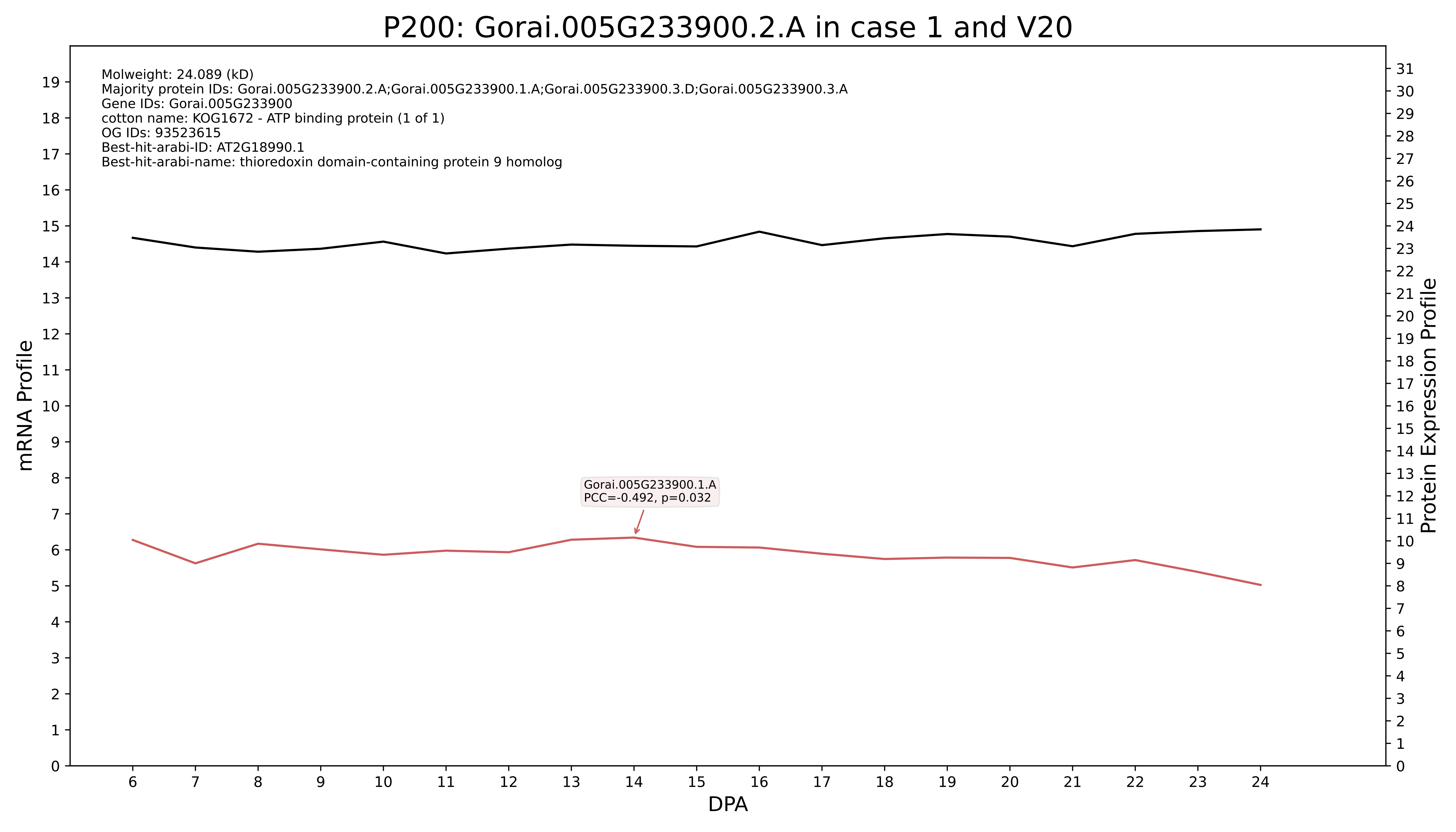

P200: Gorai.005G233900.2.A in case 1 and V20
Molweight: 24.089 (kD)
Majority protein IDs: Gorai.005G233900.2.A;Gorai.005G233900.1.A;Gorai.005G233900.3.D;Gorai.005G233900.3.A
Gene IDs: Gorai.005G233900
cotton name: KOG1672 - ATP binding protein (1 of 1)
OG IDs: 93523615
Best-hit-arabi-ID: AT2G18990.1
Best-hit-arabi-name: thioredoxin domain-containing protein 9 homolog
Gorai.005G233900.1.A
PCC=-0.492, p=0.032
mRNA Profile
Protein Expression Profile
DPA

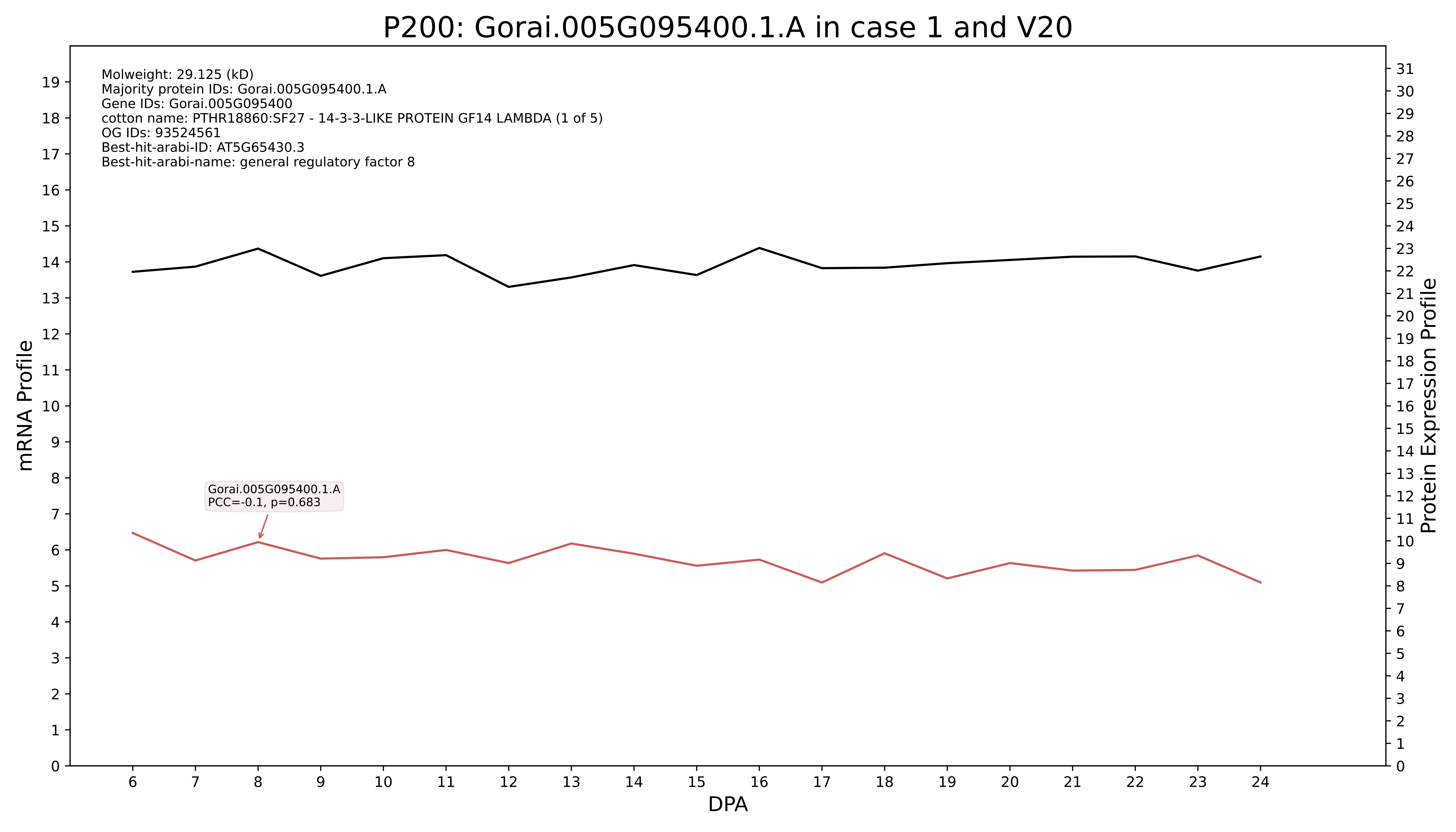
P200: Gorai.005G095400.1.A in case 1 and V20
Molweight: 29.125 (kD)
Majority protein IDs: Gorai.005G095400.1.A
Gene IDs: Gorai.005G095400
cotton name: PTHR18860:SF27 - 14-3-3-LIKE PROTEIN GF14 LAMBDA (1 of 5)
OG IDs: 93524561
Best-hit-arabi-ID: AT5G65430.3
Best-hit-arabi-name: general regulatory factor 8
Gorai.005G095400.1.A
PCC=-0.1, p=0.683
mRNA Profile
Protein Expression Profile
DPA

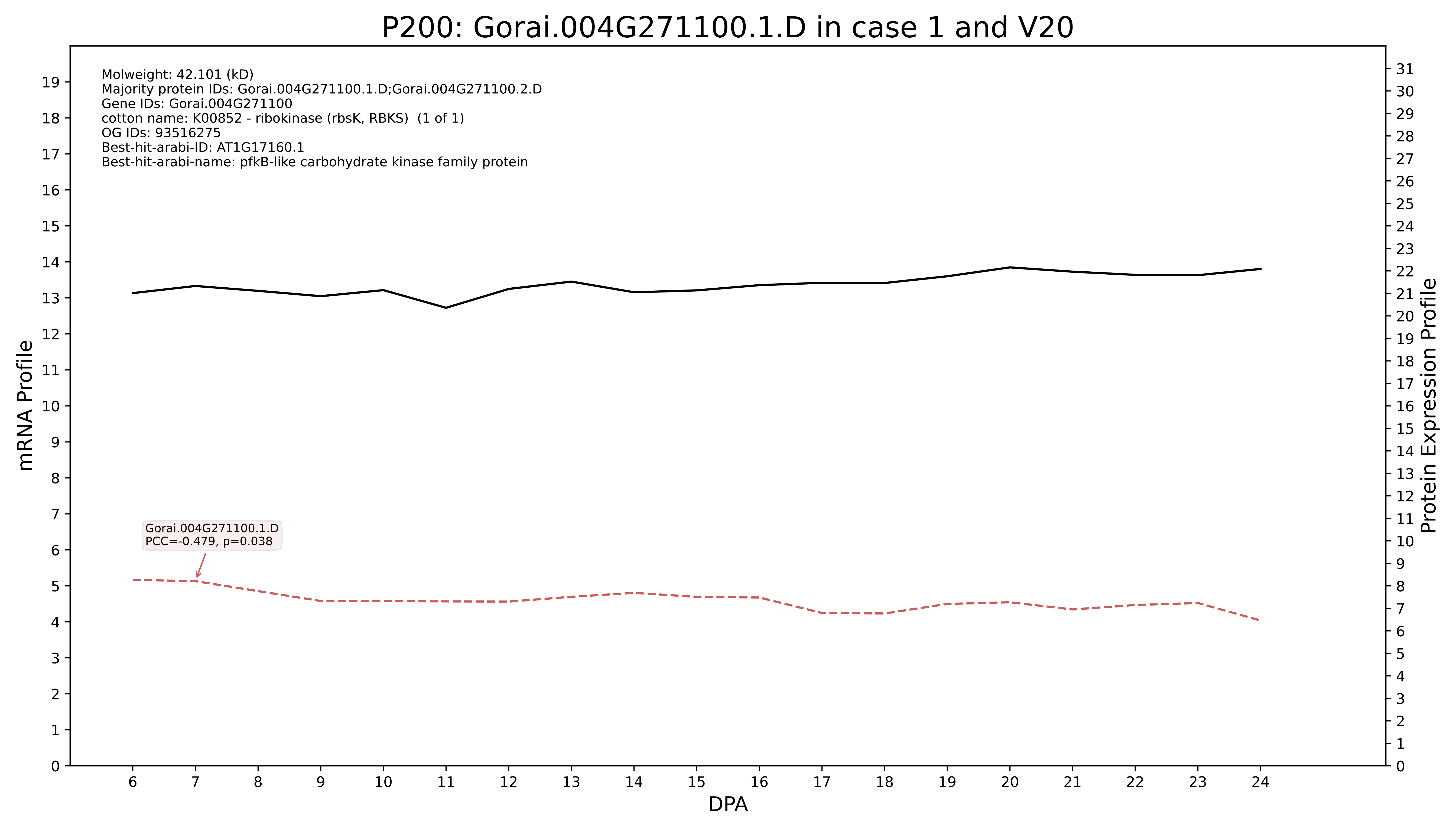
P200: Gorai.004G271100.1.D in case 1 and V20
Molweight: 42.101 (kD)
Majority protein IDs: Gorai.004G271100.1.D;Gorai.004G271100.2.D
Gene IDs: Gorai.004G271100
cotton name: K00852 - ribokinase (rbsK, RBKS) (1 of 1)
OG IDs: 93516275
Best-hit-arabi-ID: AT1G17160.1
Best-hit-arabi-name: pfkB-like carbohydrate kinase family protein
Gorai.004G271100.1.D
PCC=-0.479, p=0.038
mRNA Profile
Protein Expression Profile
DPA

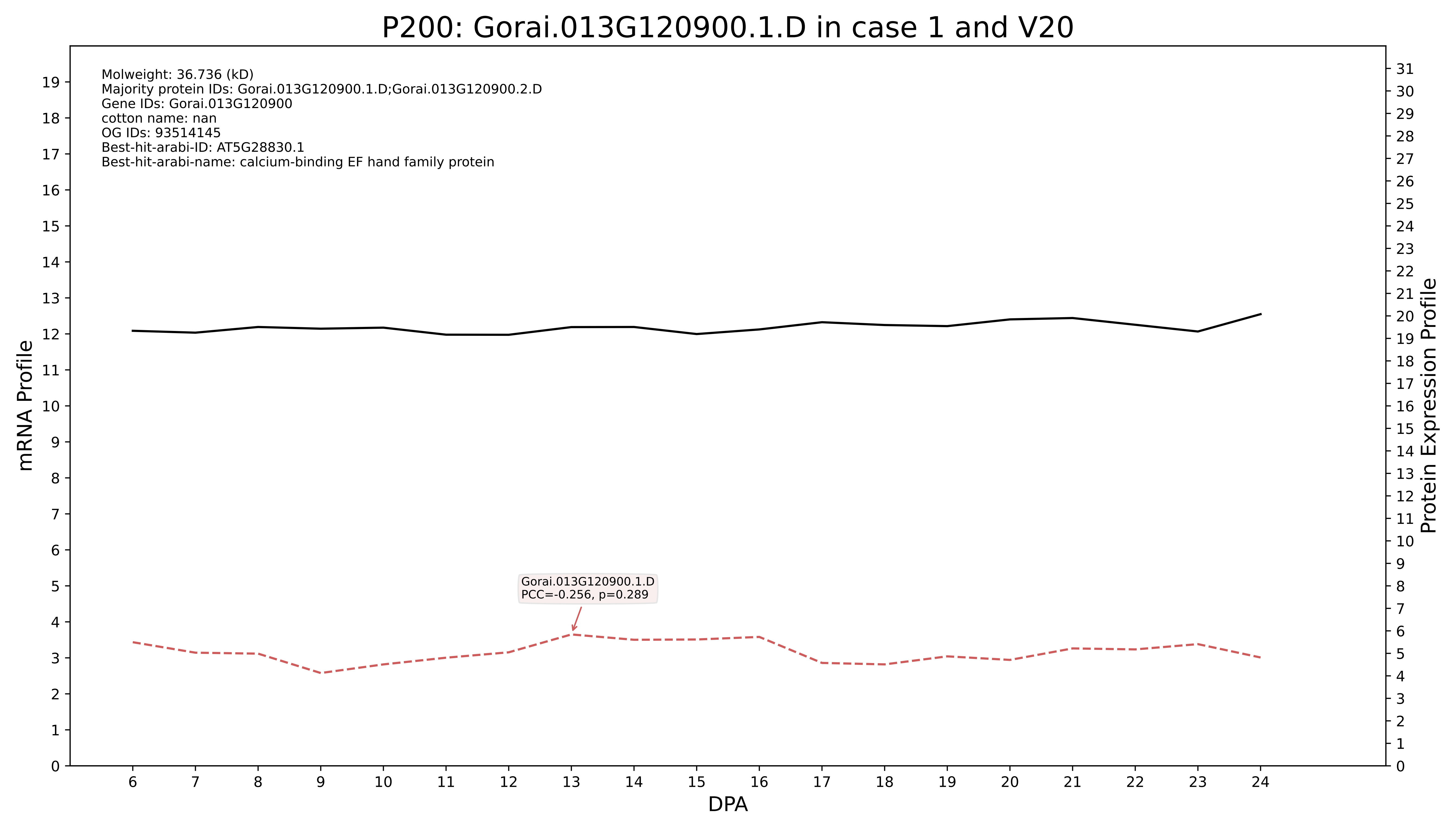
P200: Gorai.013G120900.1.D in case 1 and V20
Molweight: 36.736 (kD)
Majority protein IDs: Gorai.013G120900.1.D;Gorai.013G120900.2.D
Gene IDs: Gorai.013G120900
cotton name: nan
OG IDs: 93514145
Best-hit-arabi-ID: AT5G28830.1
Best-hit-arabi-name: calcium-binding EF hand family protein
Gorai.013G120900.1.D
PCC=-0.256, p=0.289
mRNA Profile
Protein Expression Profile
DPA

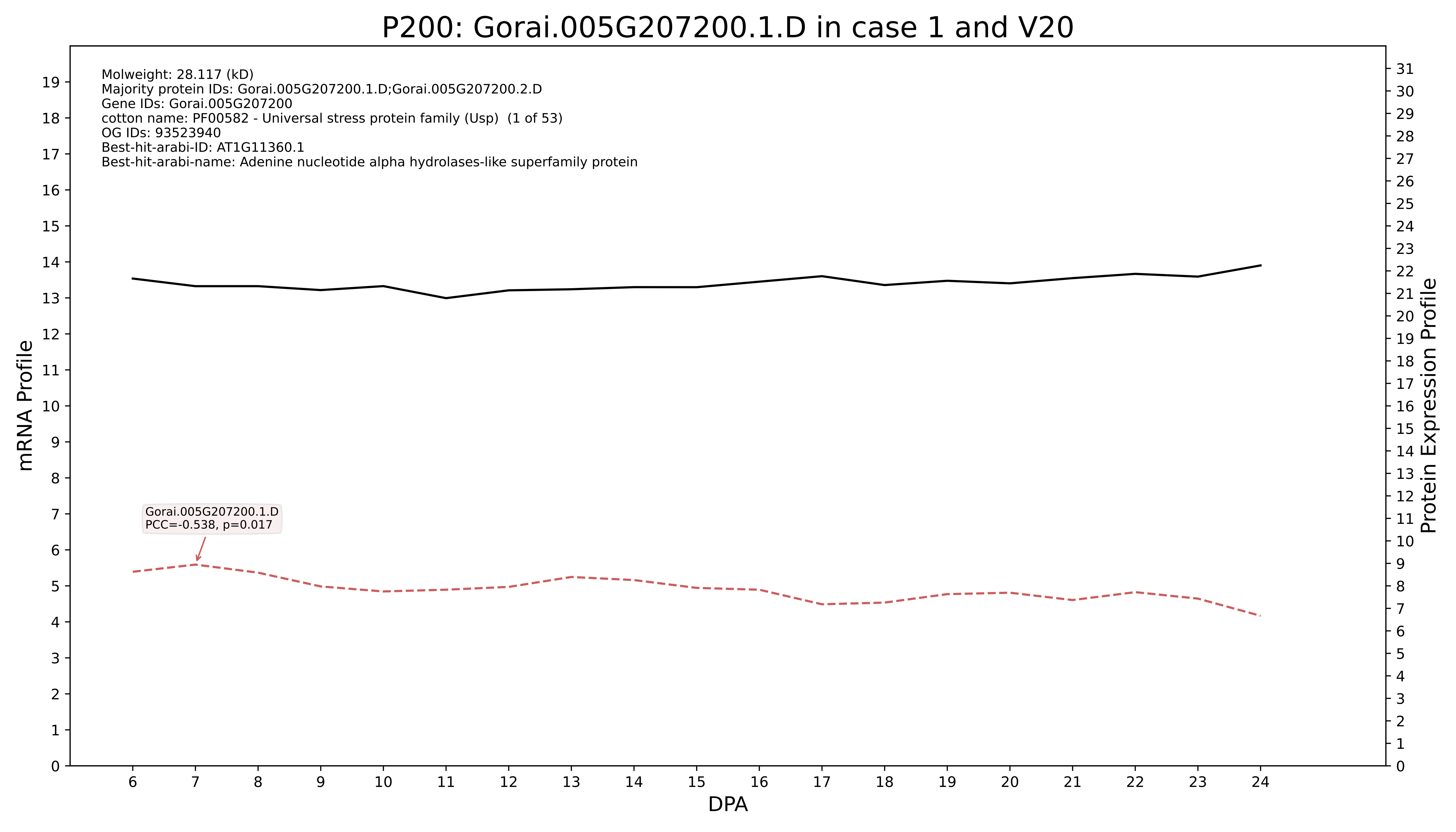
P200: Gorai.005G207200.1.D in case 1 and V20
Molweight: 28.117 (kD)
Majority protein IDs: Gorai.005G207200.1.D;Gorai.005G207200.2.D
Gene IDs: Gorai.005G207200
cotton name: PF00582 - Universal stress protein family (Usp) (1 of 53)
OG IDs: 93523940
Best-hit-arabi-ID: AT1G11360.1
Best-hit-arabi-name: Adenine nucleotide alpha hydrolases-like superfamily protein
Gorai.005G207200.1.D
PCC=-0.538, p=0.017
mRNA Profile
Protein Expression Profile
DPA

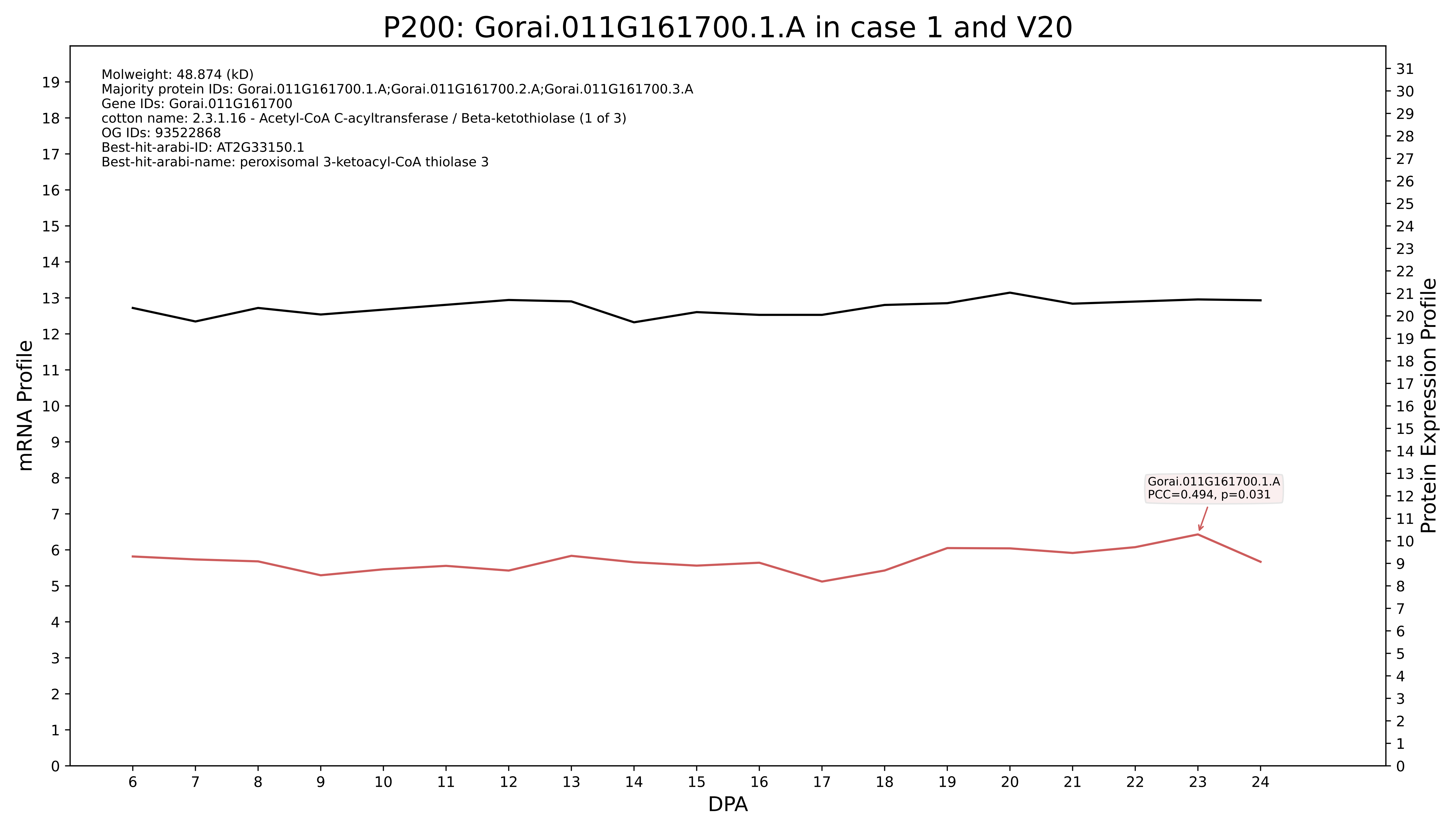

P200: Gorai.011G161700.1.A in case 1 and V20
Molweight: 48.874 (kD)
Majority protein IDs: Gorai.011G161700.1.A;Gorai.011G161700.2.A;Gorai.011G161700.3.A
Gene IDs: Gorai.011G161700
cotton name: 2.3.1.16 - Acetyl-CoA C-acyltransferase / Beta-ketothiolase (1 of 3)
OG IDs: 93522868
Best-hit-arabi-ID: AT2G33150.1
Best-hit-arabi-name: peroxisomal 3-ketoacyl-CoA thiolase 3
Gorai.011G161700.1.A
PCC=0.494, p=0.031
mRNA Profile
Protein Expression Profile
DPA

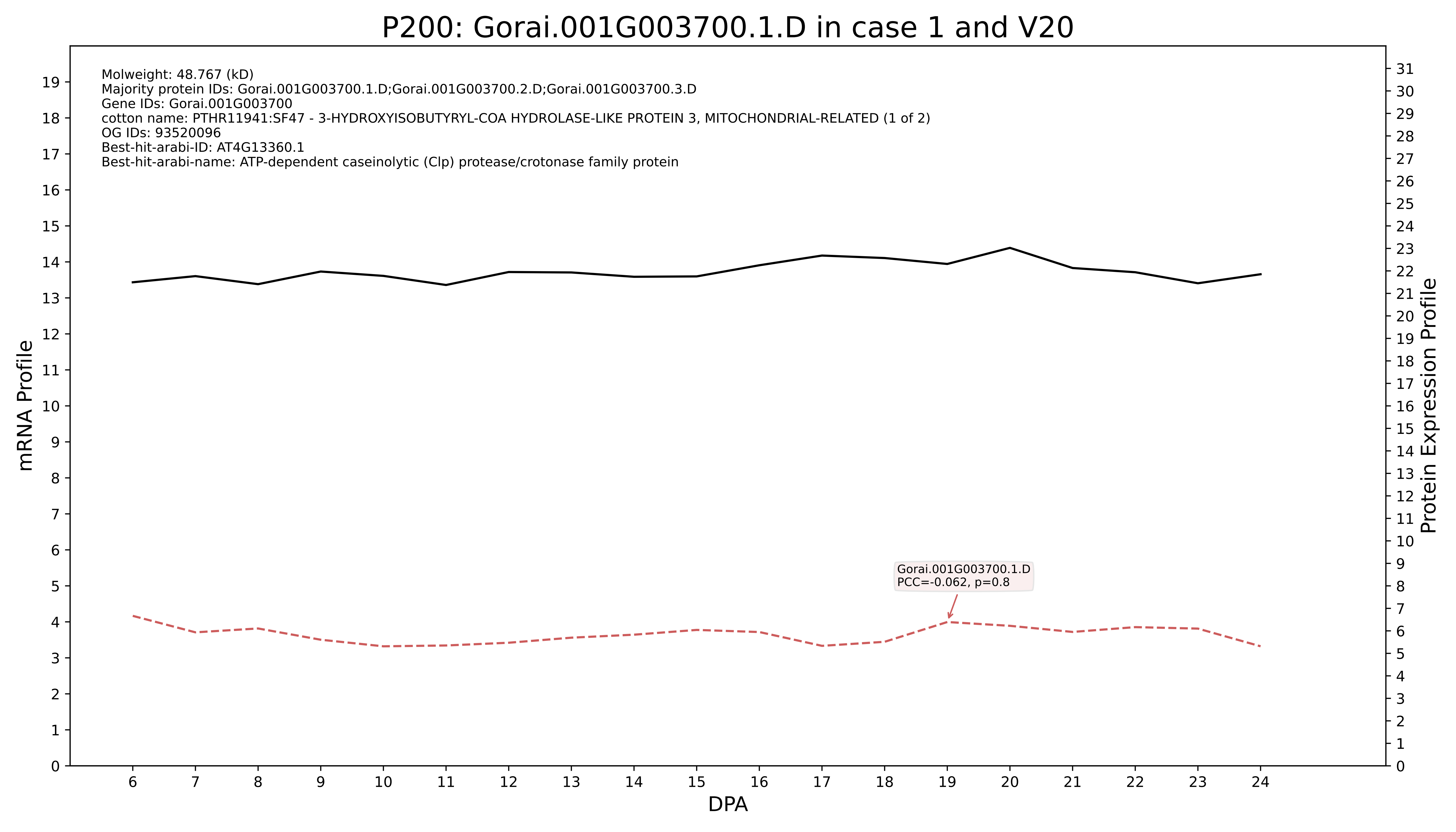
P200: Gorai.001G003700.1.D in case 1 and V20
Molweight: 48.767 (kD)
Majority protein IDs: Gorai.001G003700.1.D;Gorai.001G003700.2.D;Gorai.001G003700.3.D
Gene IDs: Gorai.001G003700
cotton name: PTHR11941:SF47 - 3-HYDROXYISOBUTYRYL-COA HYDROLASE-LIKE PROTEIN 3, MITOCHONDRIAL-RELATED (1 of 2)
OG IDs: 93520096
Best-hit-arabi-ID: AT4G13360.1
Best-hit-arabi-name: ATP-dependent caseinolytic (Clp) protease/crotonase family protein
Gorai.001G003700.1.D
PCC=-0.062, p=0.8
mRNA Profile
Protein Expression Profile
DPA

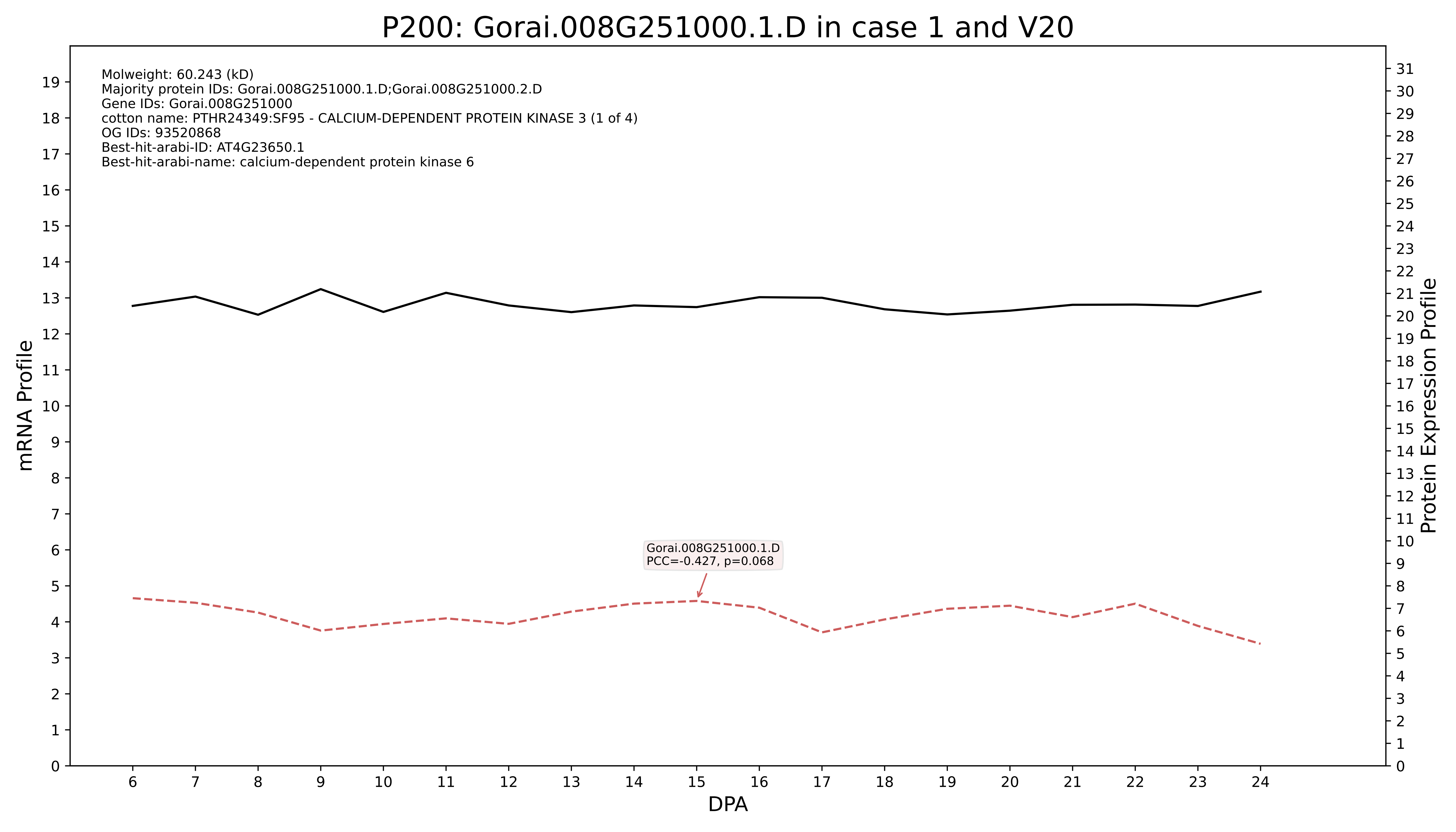

P200: Gorai.008G251000.1.D in case 1 and V20
Molweight: 60.243 (kD)
Majority protein IDs: Gorai.008G251000.1.D;Gorai.008G251000.2.D
Gene IDs: Gorai.008G251000
cotton name: PTHR24349:SF95 - CALCIUM-DEPENDENT PROTEIN KINASE 3 (1 of 4)
OG IDs: 93520868
Best-hit-arabi-ID: AT4G23650.1
Best-hit-arabi-name: calcium-dependent protein kinase 6
Gorai.008G251000.1.D
PCC=-0.427, p=0.068
mRNA Profile
Protein Expression Profile
DPA

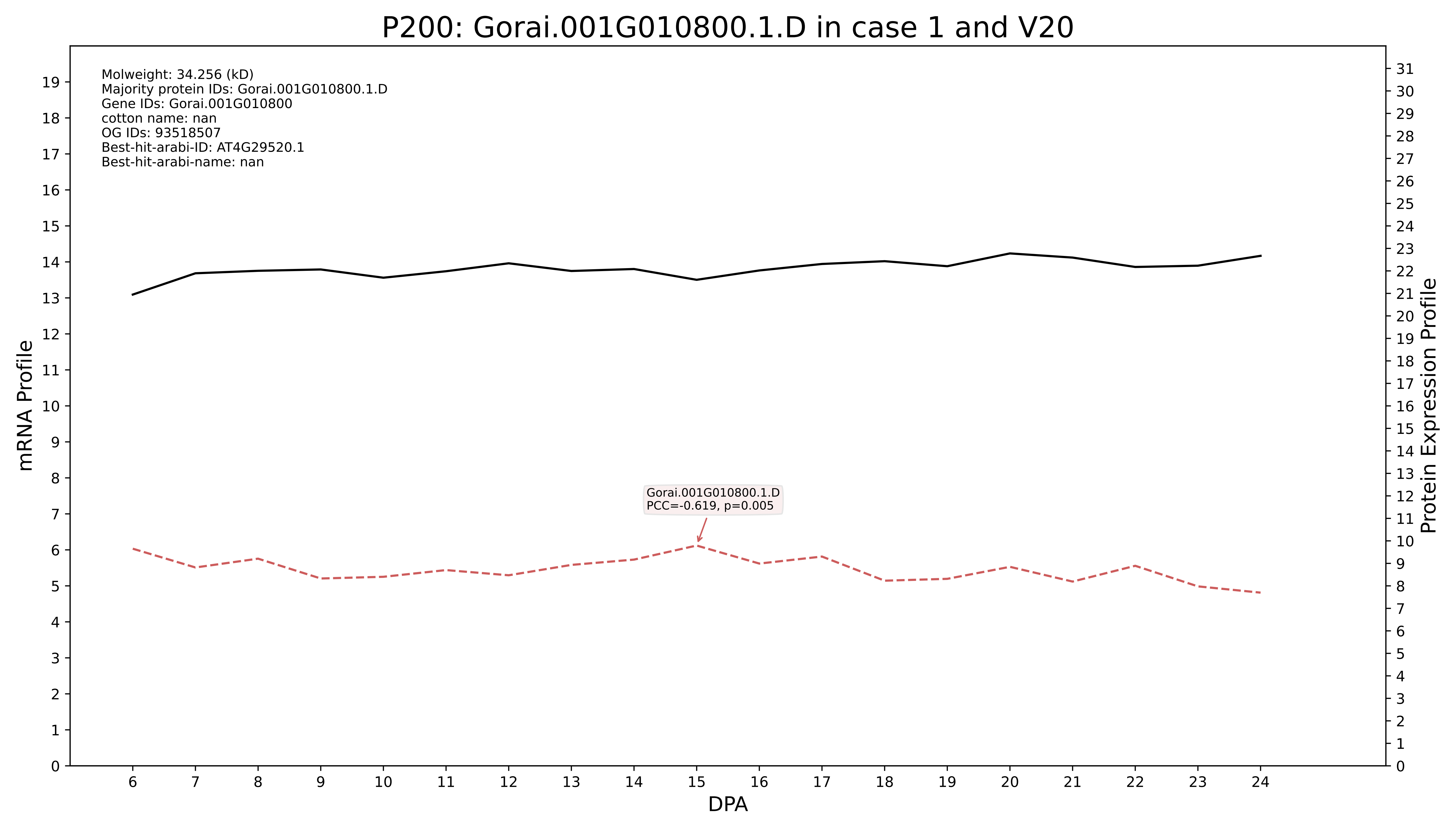

P200: Gorai.001G010800.1.D in case 1 and V20
Molweight: 34.256 (kD)
Majority protein IDs: Gorai.001G010800.1.D
Gene IDs: Gorai.001G010800
cotton name: nan
OG IDs: 93518507
Best-hit-arabi-ID: AT4G29520.1
Best-hit-arabi-name: nan
Gorai.001G010800.1.D
PCC=-0.619, p=0.005
mRNA Profile
Protein Expression Profile
DPA

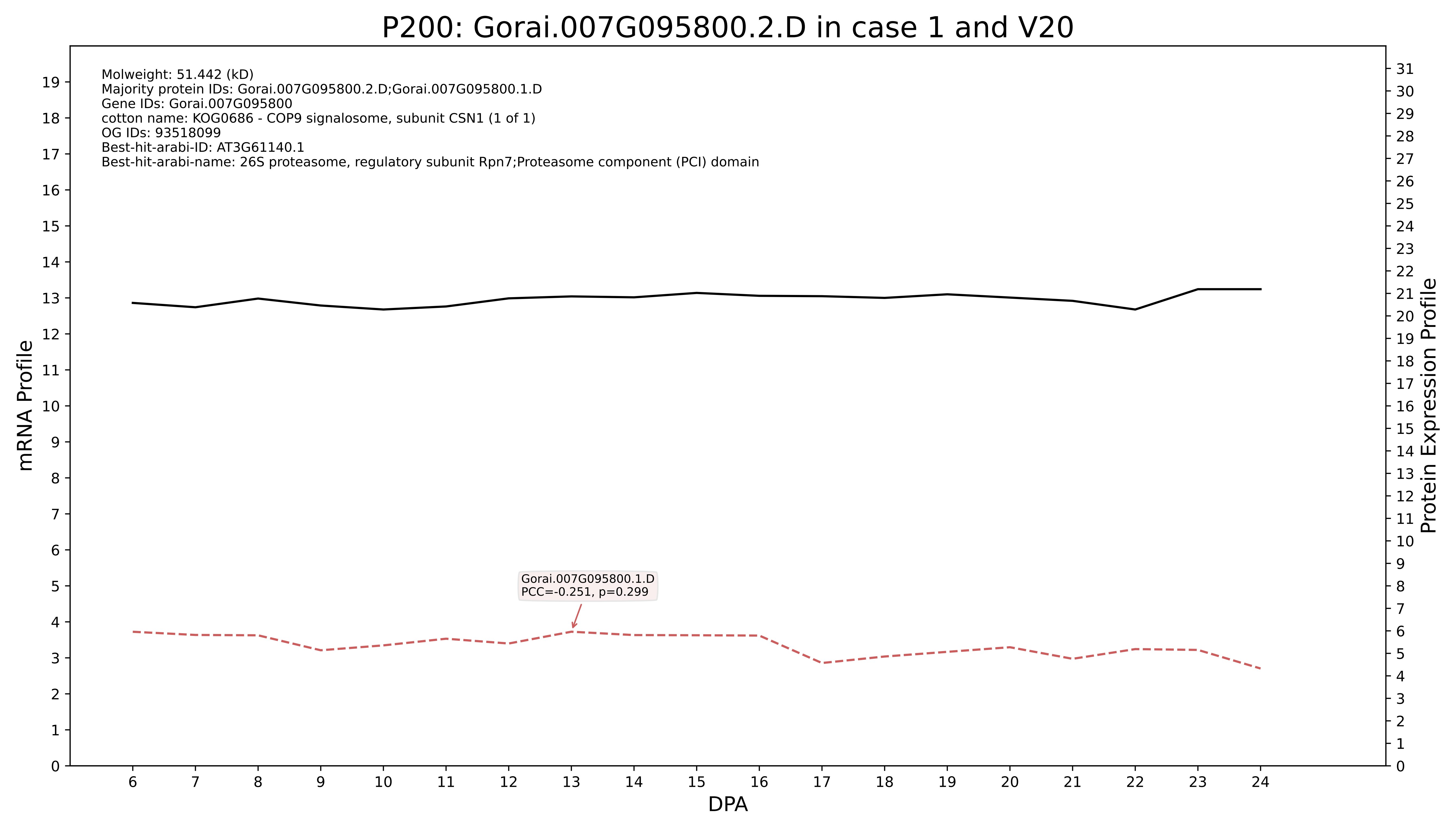
P200: Gorai.007G095800.2.D in case 1 and V20
Molweight: 51.442 (kD)
Majority protein IDs: Gorai.007G095800.2.D;Gorai.007G095800.1.D
Gene IDs: Gorai.007G095800
cotton name: KOG0686 - COP9 signalosome, subunit CSN1 (1 of 1)
OG IDs: 93518099
Best-hit-arabi-ID: AT3G61140.1
Best-hit-arabi-name: 26S proteasome, regulatory subunit Rpn7;Proteasome component (PCI) domain
Gorai.007G095800.1.D
PCC=-0.251, p=0.299
mRNA Profile
Protein Expression Profile
DPA

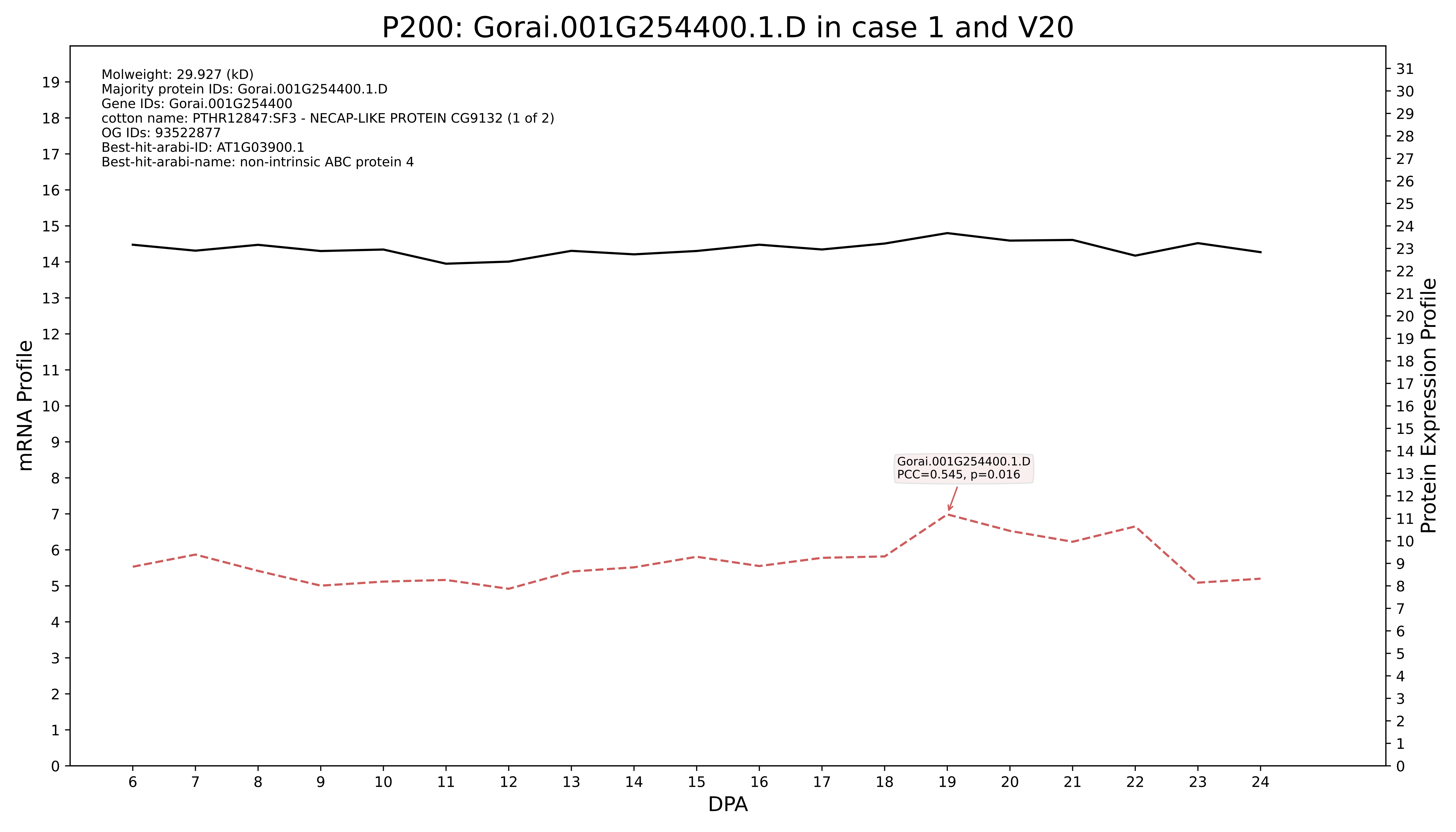
P200: Gorai.001G254400.1.D in case 1 and V20
Molweight: 29.927 (kD)
Majority protein IDs: Gorai.001G254400.1.D
Gene IDs: Gorai.001G254400
cotton name: PTHR12847:SF3 - NECAP-LIKE PROTEIN CG9132 (1 of 2)
OG IDs: 93522877
Best-hit-arabi-ID: AT1G03900.1
Best-hit-arabi-name: non-intrinsic ABC protein 4
Gorai.001G254400.1.D
PCC=0.545, p=0.016
mRNA Profile
Protein Expression Profile
DPA

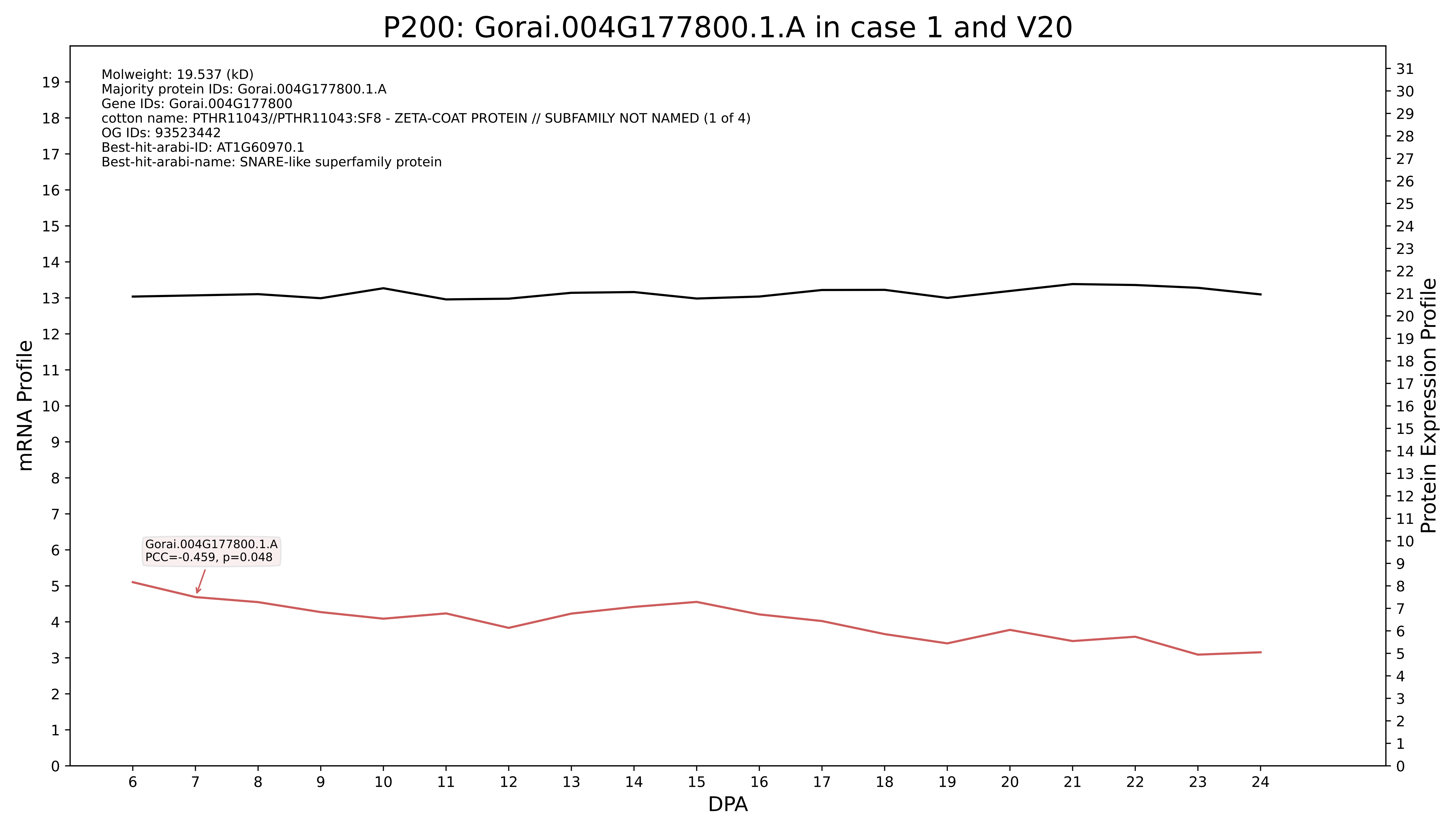

P200: Gorai.004G177800.1.A in case 1 and V20
Molweight: 19.537 (kD)
Majority protein IDs: Gorai.004G177800.1.A
Gene IDs: Gorai.004G177800
cotton name: PTHR11043//PTHR11043:SF8 - ZETA-COAT PROTEIN // SUBFAMILY NOT NAMED (1 of 4)
OG IDs: 93523442
Best-hit-arabi-ID: AT1G60970.1
Best-hit-arabi-name: SNARE-like superfamily protein
Gorai.004G177800.1.A
PCC=-0.459, p=0.048
mRNA Profile
Protein Expression Profile
DPA

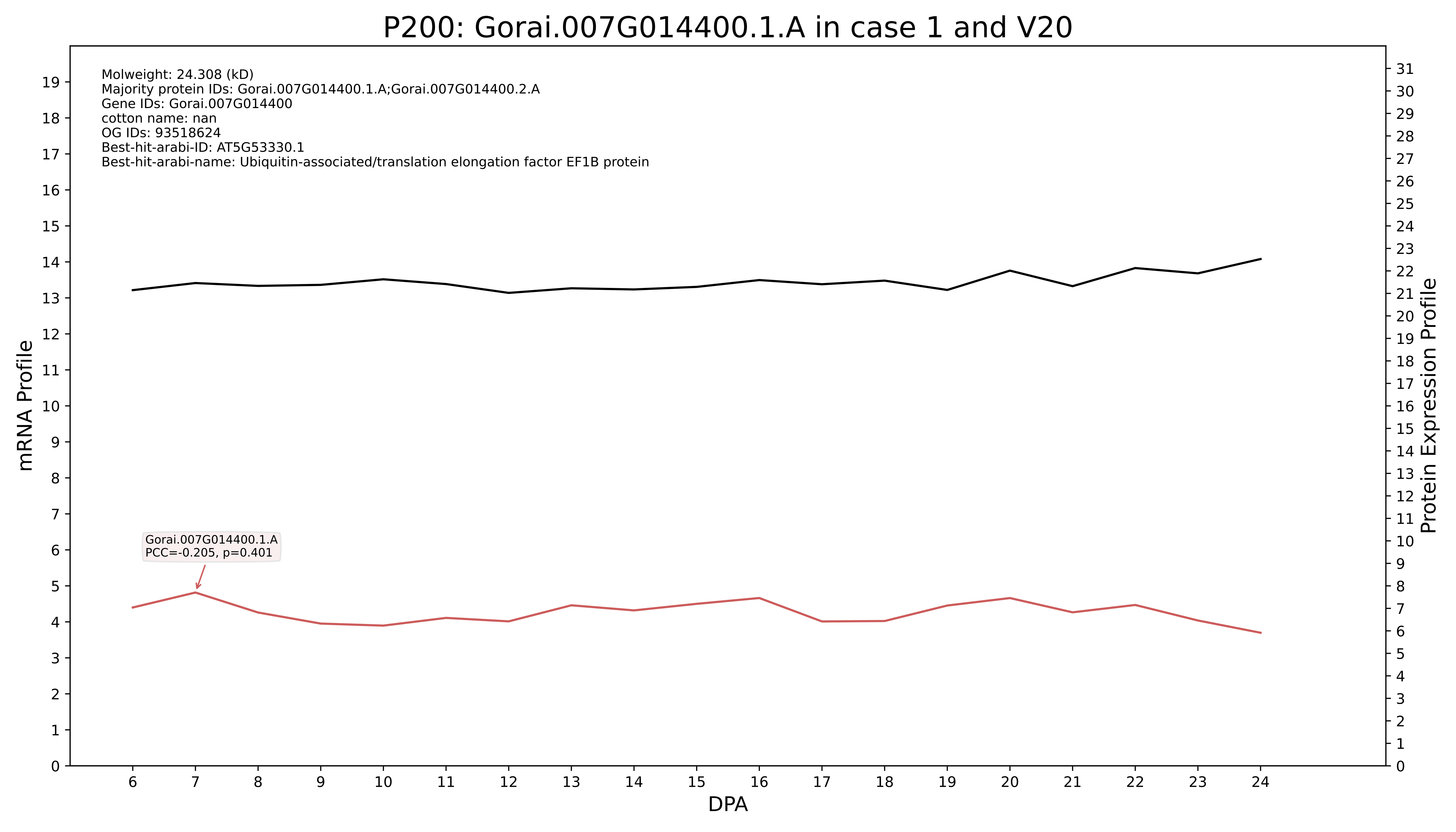

P200: Gorai.007G014400.1.A in case 1 and V20
Molweight: 24.308 (kD)
Majority protein IDs: Gorai.007G014400.1.A;Gorai.007G014400.2.A
Gene IDs: Gorai.007G014400
cotton name: nan
OG IDs: 93518624
Best-hit-arabi-ID: AT5G53330.1
Best-hit-arabi-name: Ubiquitin-associated/translation elongation factor EF1B protein
Gorai.007G014400.1.A
PCC=-0.205, p=0.401
mRNA Profile
Protein Expression Profile
DPA

### P200: Gorai.011G234100.3.A in case 1 and V20

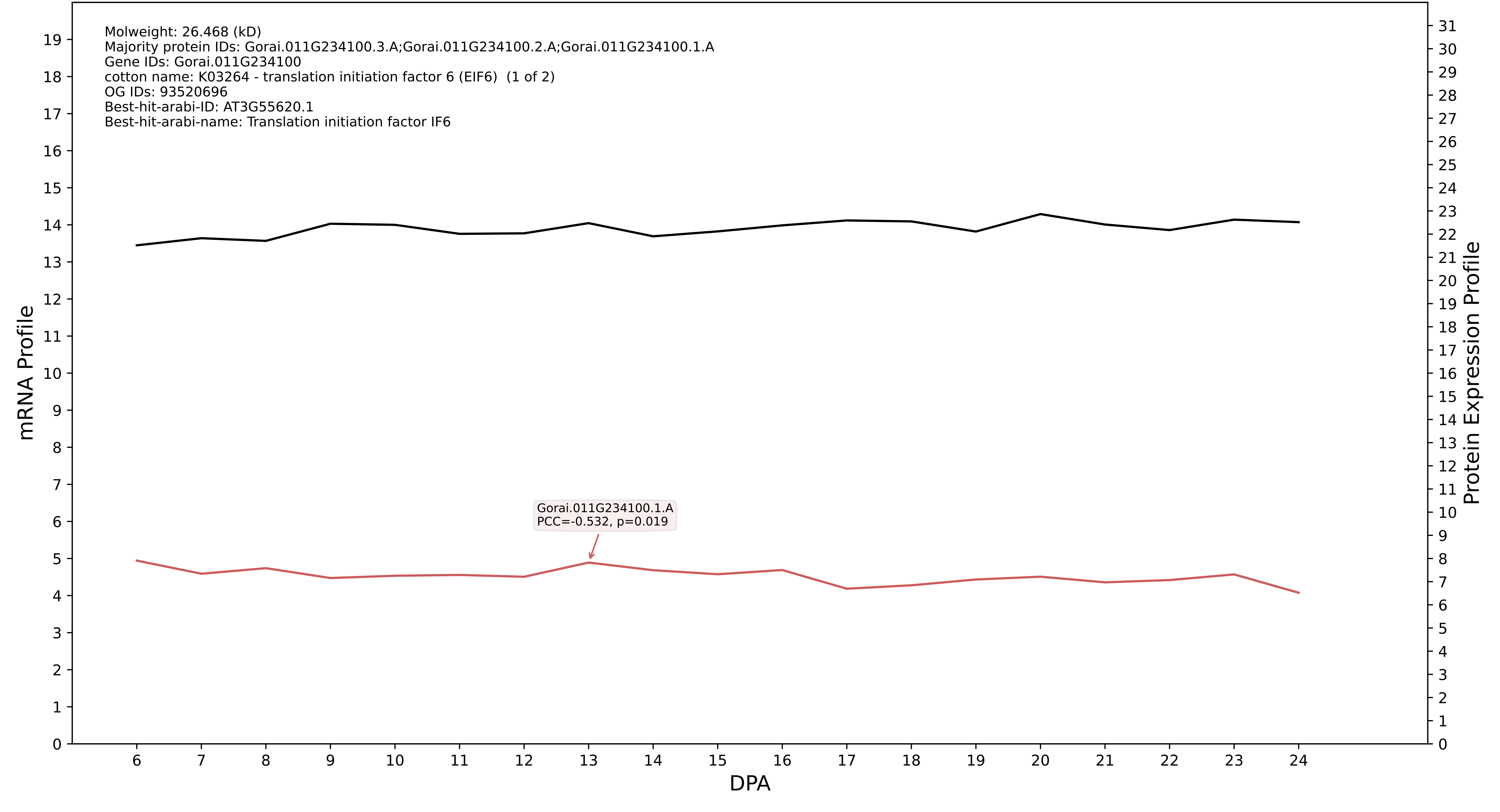

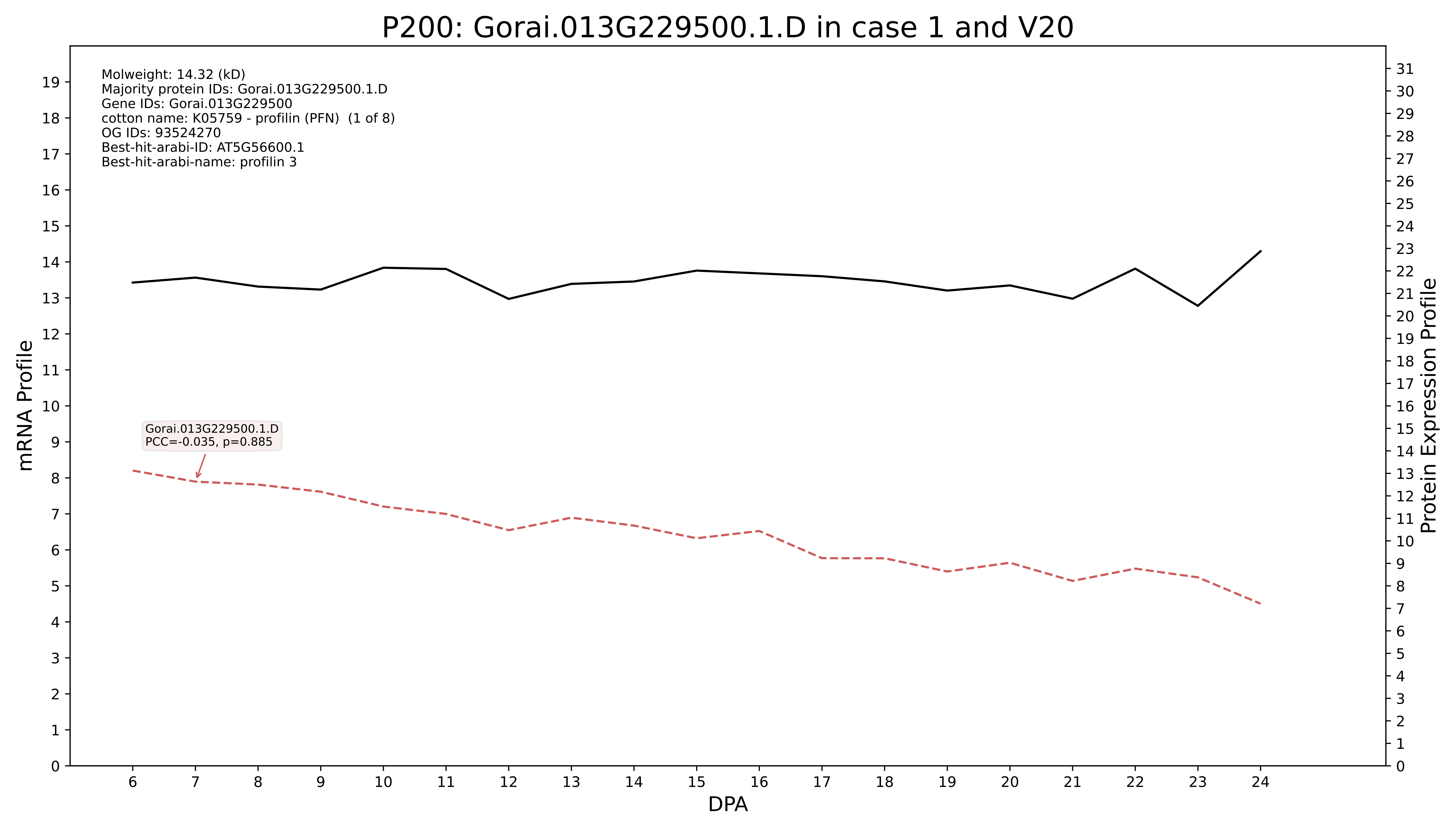

P200: Gorai.013G229500.1.D in case 1 and V20
Molweight: 14.32 (kD)
Majority protein IDs: Gorai.013G229500.1.D
Gene IDs: Gorai.013G229500
cotton name: K05759 - profilin (PFN) (1 of 8)
OG IDs: 93524270
Best-hit-arabi-ID: AT5G56600.1
Best-hit-arabi-name: profilin 3
Gorai.013G229500.1.D
PCC=-0.035, p=0.885
mRNA Profile
Protein Expression Profile
DPA

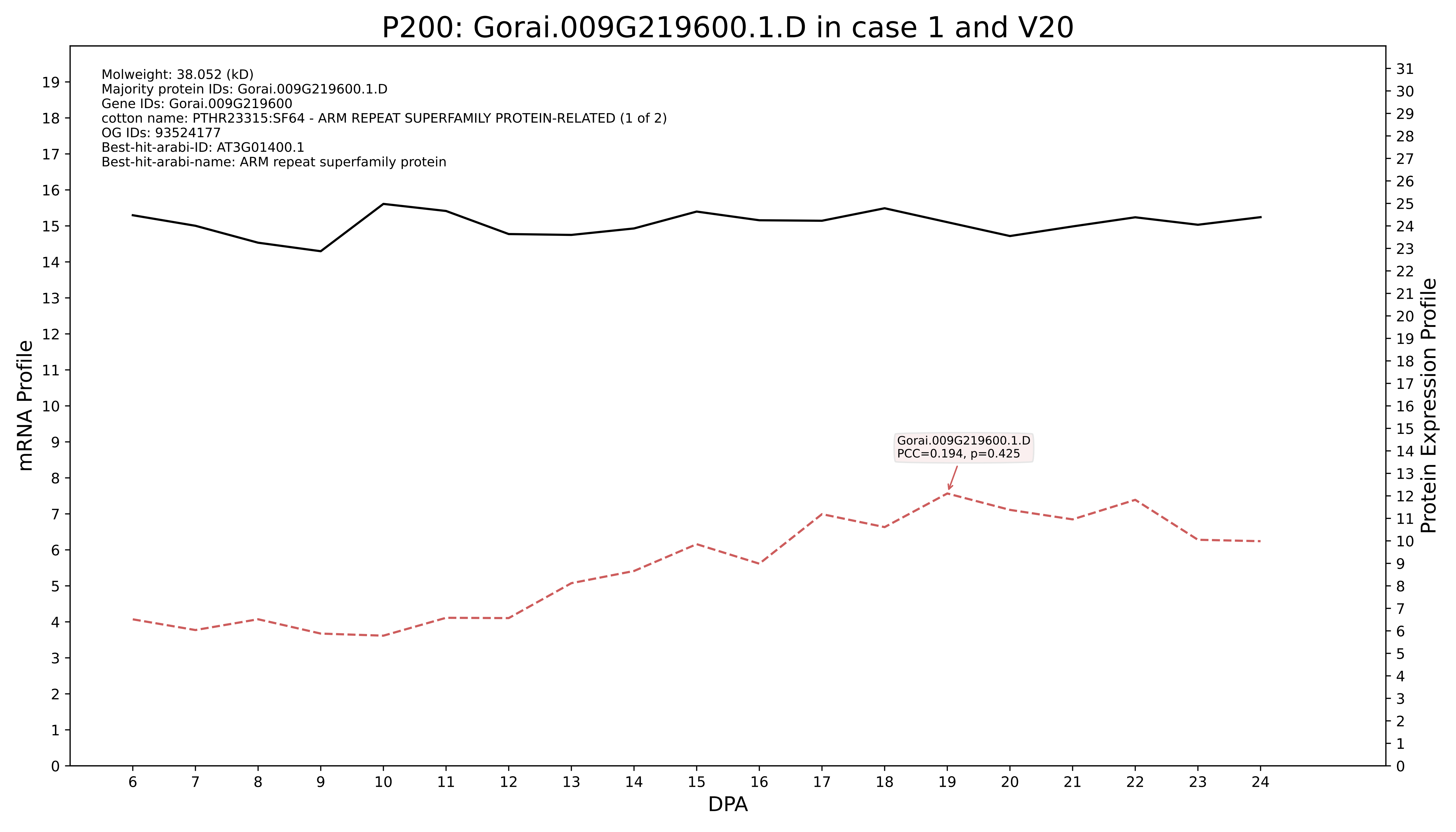

P200: Gorai.009G219600.1.D in case 1 and V20
Molweight: 38.052 (kD)
Majority protein IDs: Gorai.009G219600.1.D
Gene IDs: Gorai.009G219600
cotton name: PTHR23315:SF64 - ARM REPEAT SUPERFAMILY PROTEIN-RELATED (1 of 2)
OG IDs: 93524177
Best-hit-arabi-ID: AT3G01400.1
Best-hit-arabi-name: ARM repeat superfamily protein
Gorai.009G219600.1.D
PCC=0.194, p=0.425
mRNA Profile
Protein Expression Profile
DPA

### P200: Gorai.009G296100.2.D in case 1 and V20

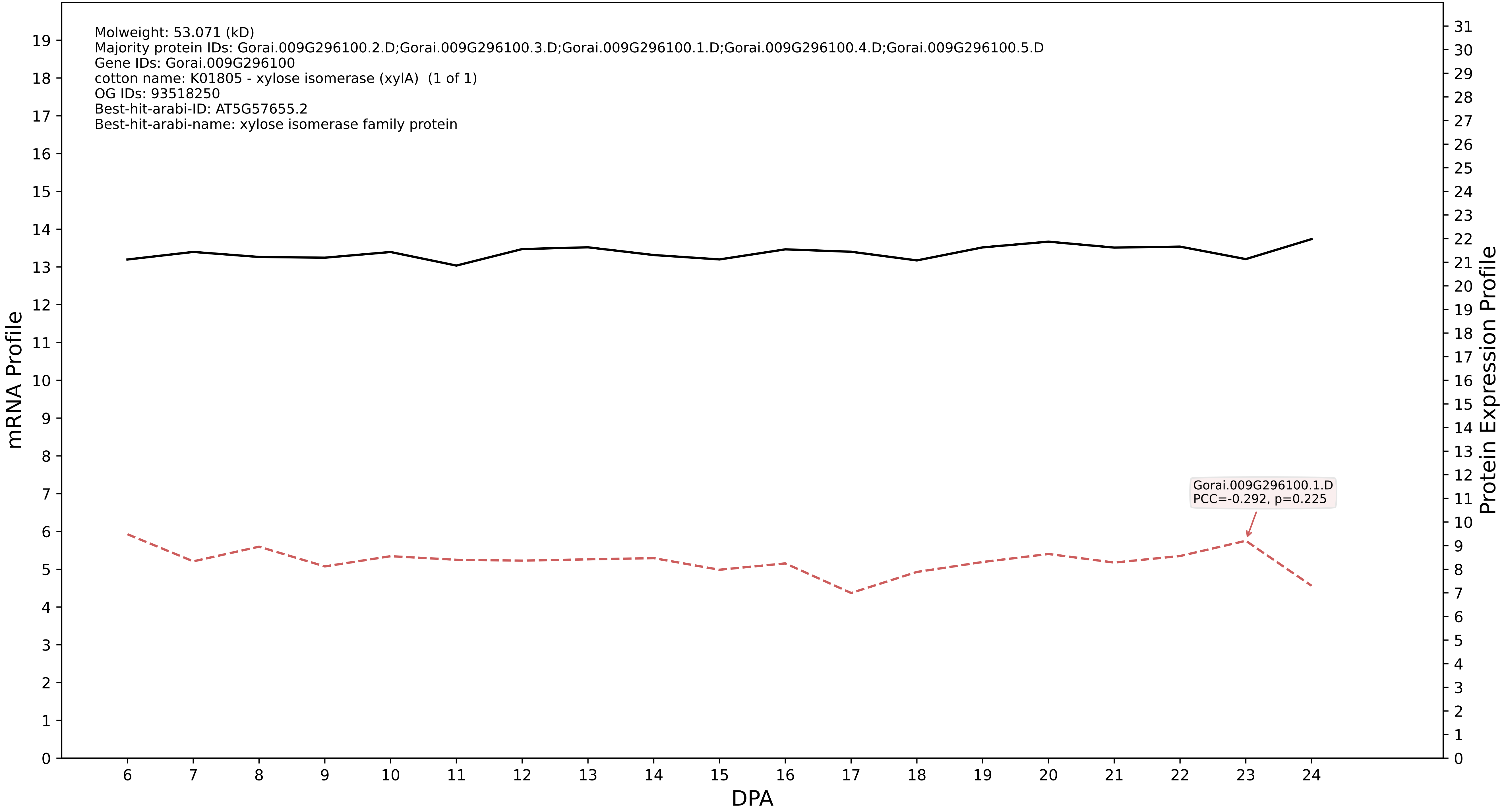

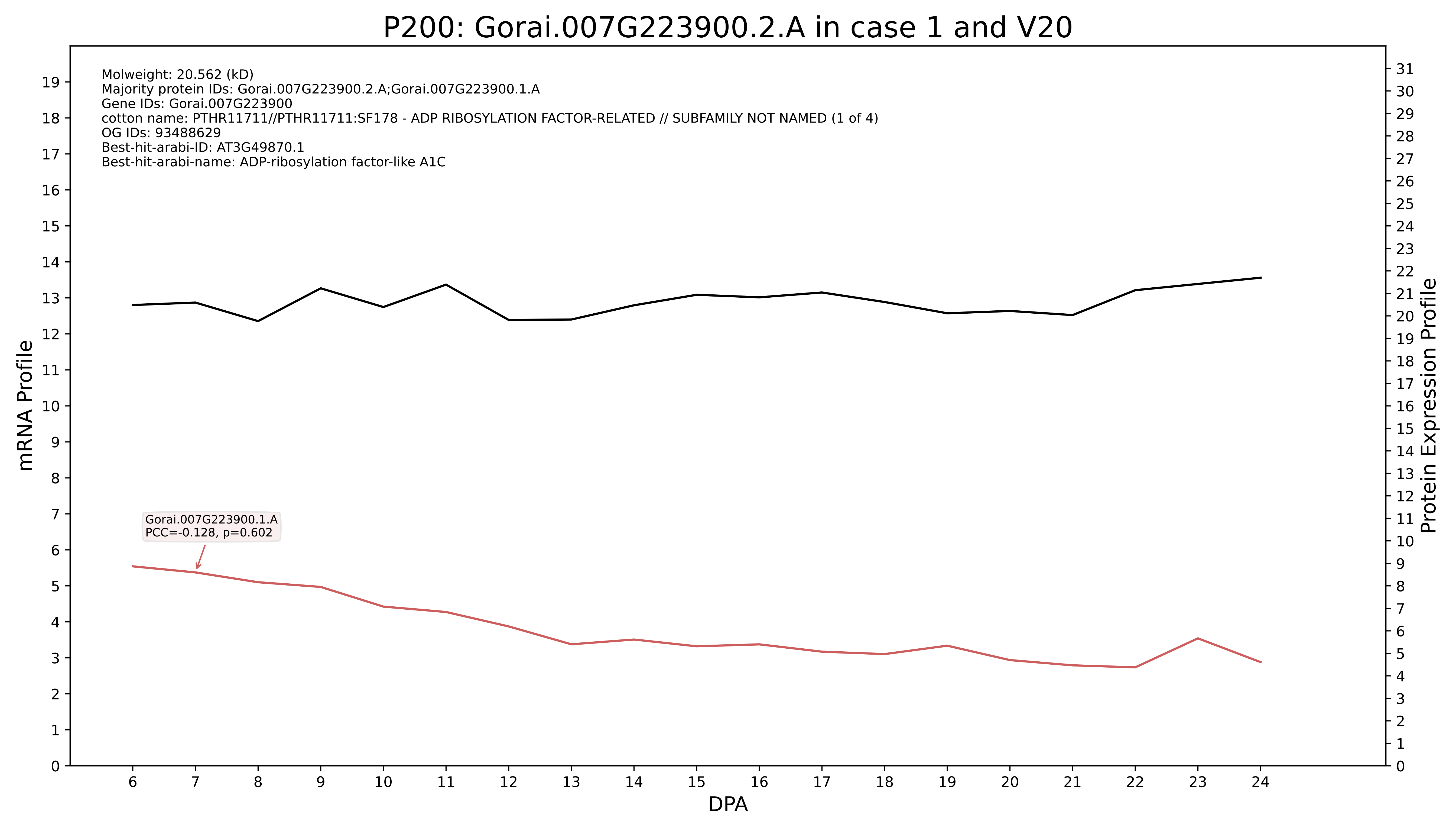
P200: Gorai.007G223900.2.A in case 1 and V20
Molweight: 20.562 (kD)
Majority protein IDs: Gorai.007G223900.2.A;Gorai.007G223900.1.A
Gene IDs: Gorai.007G223900
cotton name: PTHR11711//PTHR11711:SF178 - ADP RIBOSYLATION FACTOR-RELATED // SUBFAMILY NOT NAMED (1 of 4)
OG IDs: 93488629
Best-hit-arabi-ID: AT3G49870.1
Best-hit-arabi-name: ADP-ribosylation factor-like A1C
Gorai.007G223900.1.A
PCC=-0.128, p=0.602
mRNA Profile
Protein Expression Profile
DPA

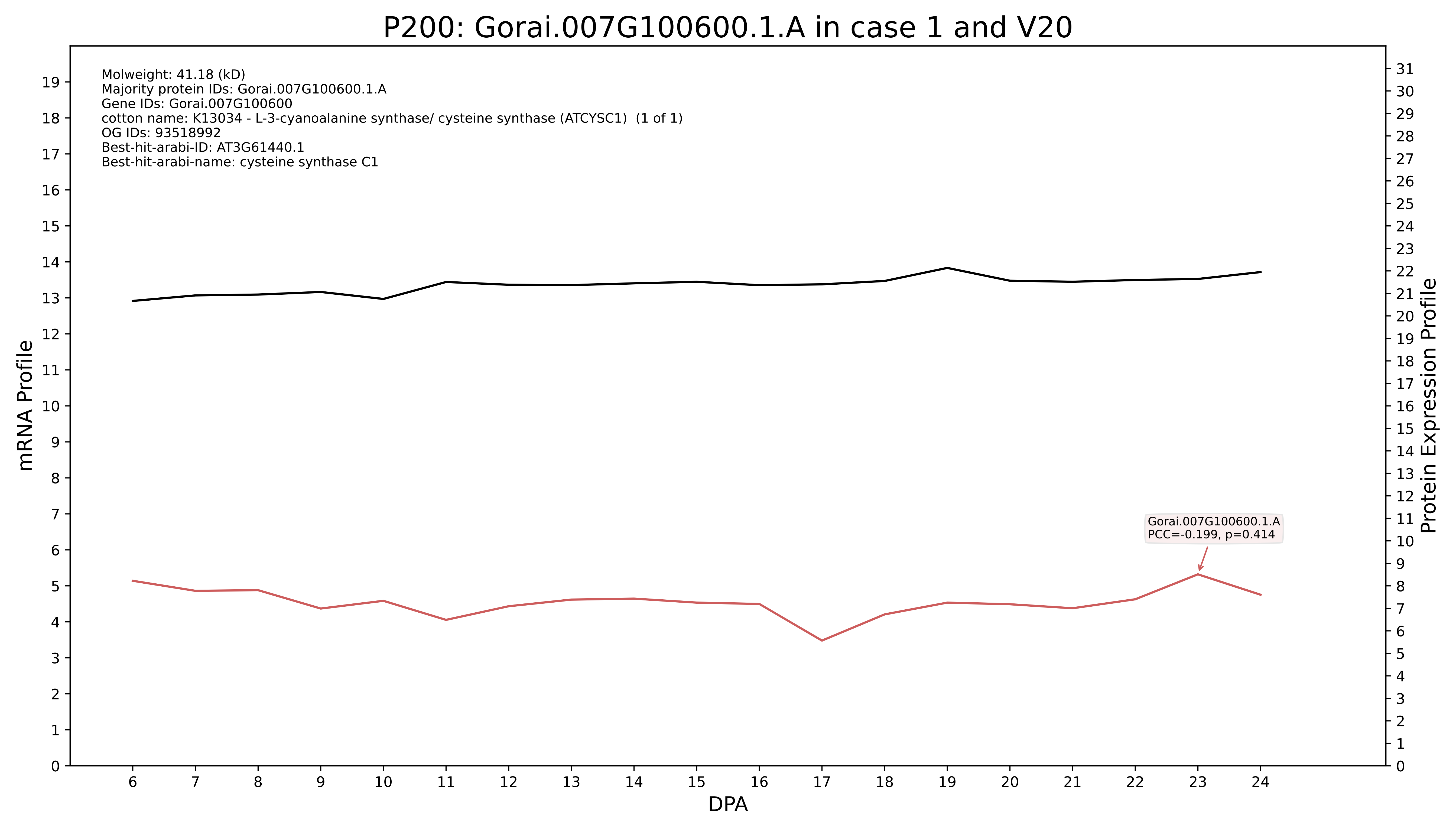
P200: Gorai.007G100600.1.A in case 1 and V20
Molweight: 41.18 (kD)
Majority protein IDs: Gorai.007G100600.1.A
Gene IDs: Gorai.007G100600
cotton name: K13034 - L-3-cyanoalanine synthase/ cysteine synthase (ATCYSC1) (1 of 1)
OG IDs: 93518992
Best-hit-arabi-ID: AT3G61440.1
Best-hit-arabi-name: cysteine synthase C1
Gorai.007G100600.1.A
PCC=-0.199, p=0.414
mRNA Profile
Protein Expression Profile
DPA

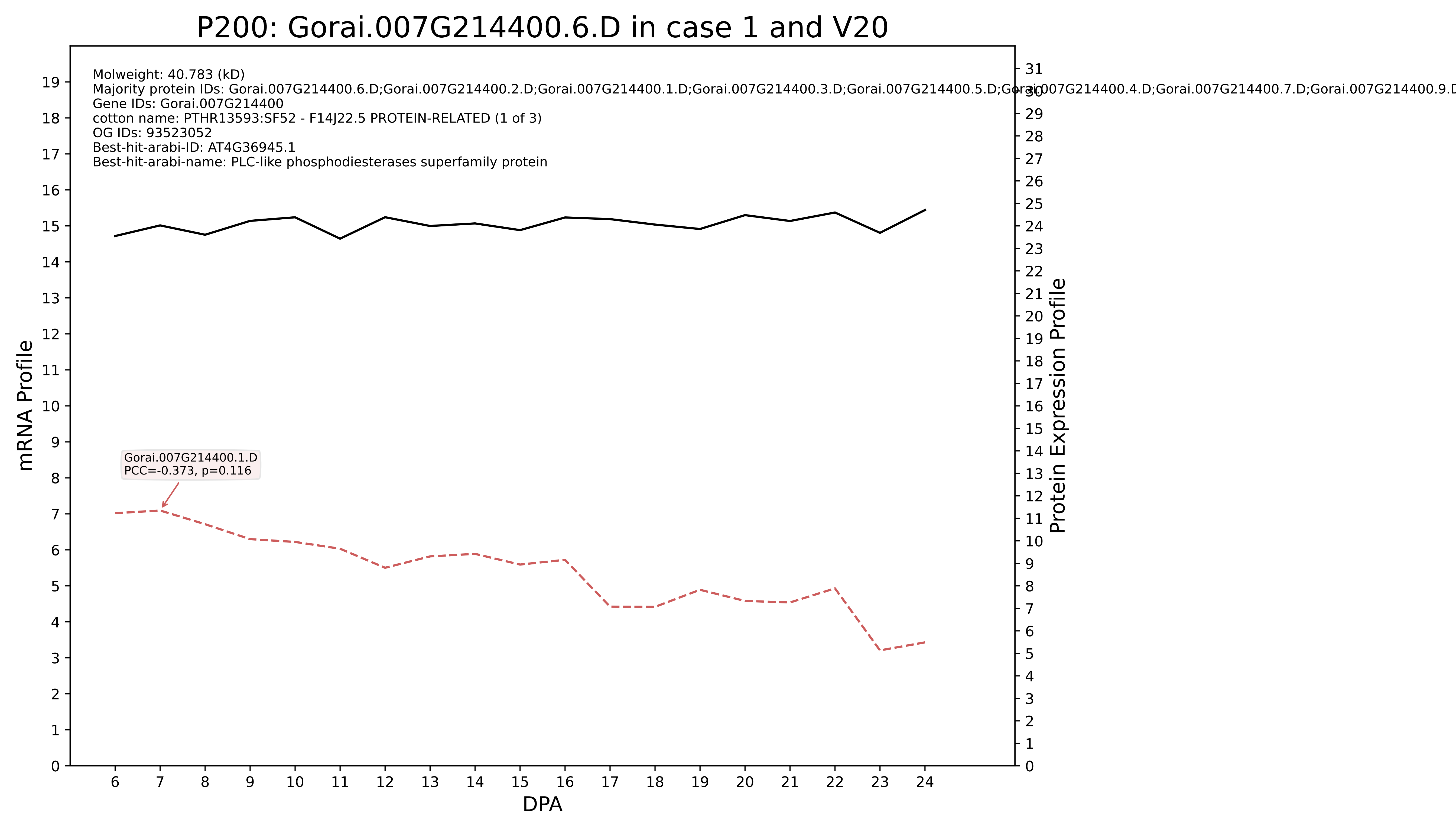
P200: Gorai.007G214400.6.D in case 1 and V20
Molweight: 40.783 (kD)
Majority protein IDs: Gorai.007G214400.6.D;Gorai.007G214400.2.D;Gorai.007G214400.1.D;Gorai.007G214400.3.D;Gorai.007G214400.5.D;Gorai.007G214400.4.D;Gorai.007G214400.7.D;Gorai.007G214400.9.D
Gene IDs: Gorai.007G214400
cotton name: PTHR13593:SF52 - F14J22.5 PROTEIN-RELATED (1 of 3)
OG IDs: 93523052
Best-hit-arabi-ID: AT4G36945.1
Best-hit-arabi-name: PLC-like phosphodiesterases superfamily protein
Gorai.007G214400.1.D
PCC=-0.373, p=0.116
mRNA Profile
Protein Expression Profile
DPA

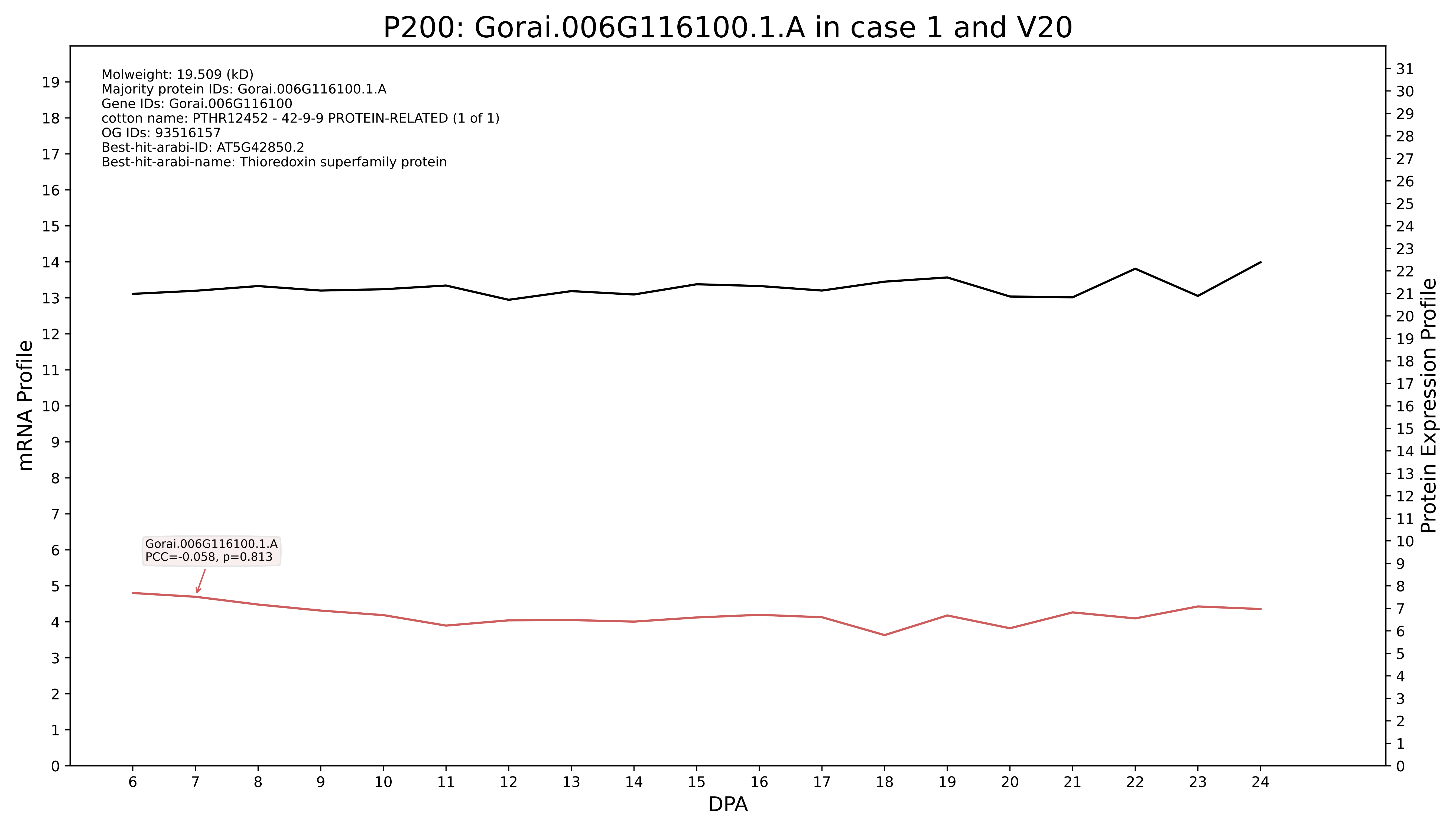

P200: Gorai.006G116100.1.A in case 1 and V20
Molweight: 19.509 (kD)
Majority protein IDs: Gorai.006G116100.1.A
Gene IDs: Gorai.006G116100
cotton name: PTHR12452 - 42-9-9 PROTEIN-RELATED (1 of 1)
OG IDs: 93516157
Best-hit-arabi-ID: AT5G42850.2
Best-hit-arabi-name: Thioredoxin superfamily protein
Gorai.006G116100.1.A
PCC=-0.058, p=0.813
mRNA Profile
Protein Expression Profile
DPA

### P200: Gorai.008G010200.1.D in case 1 and V20

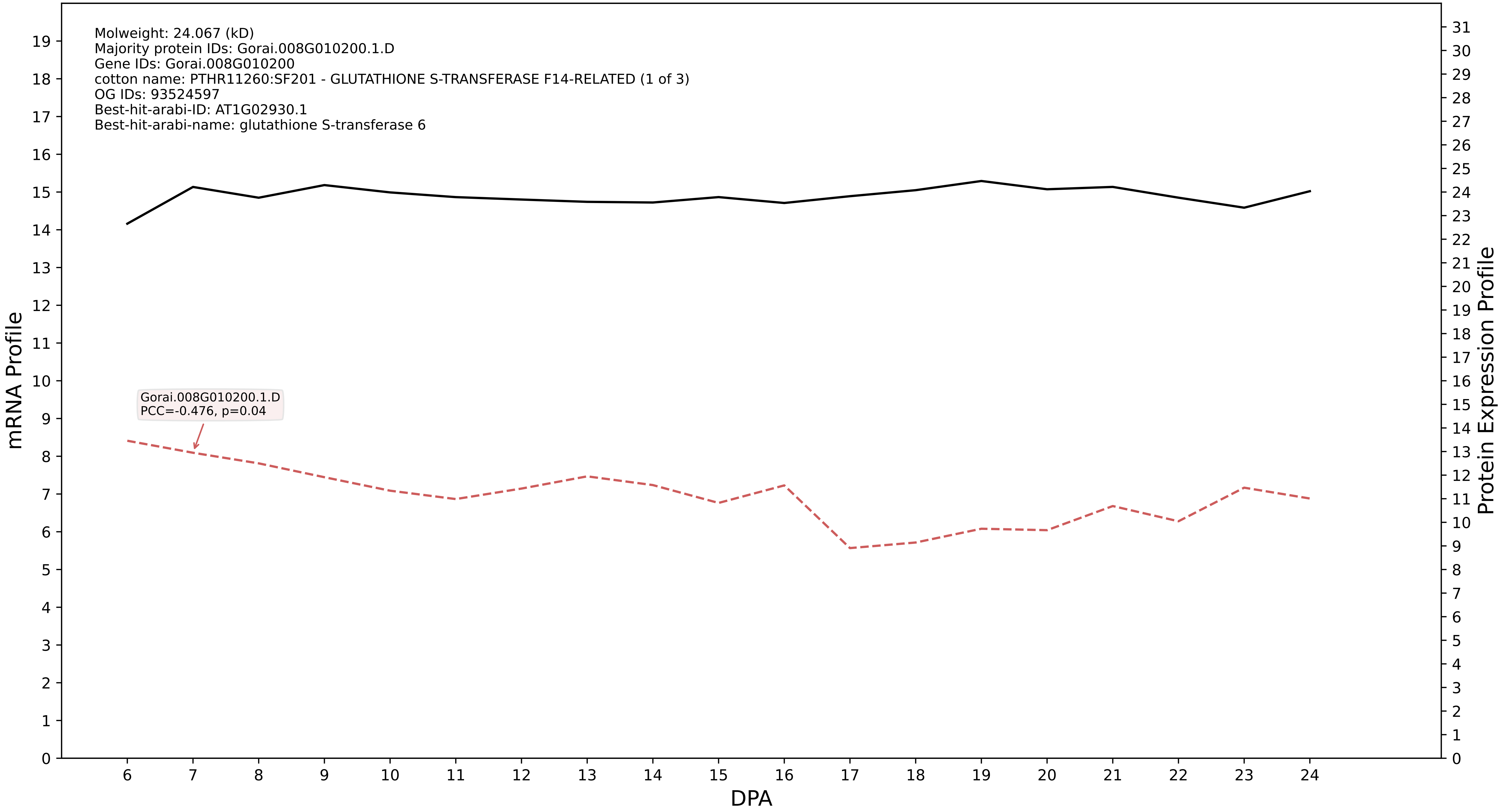

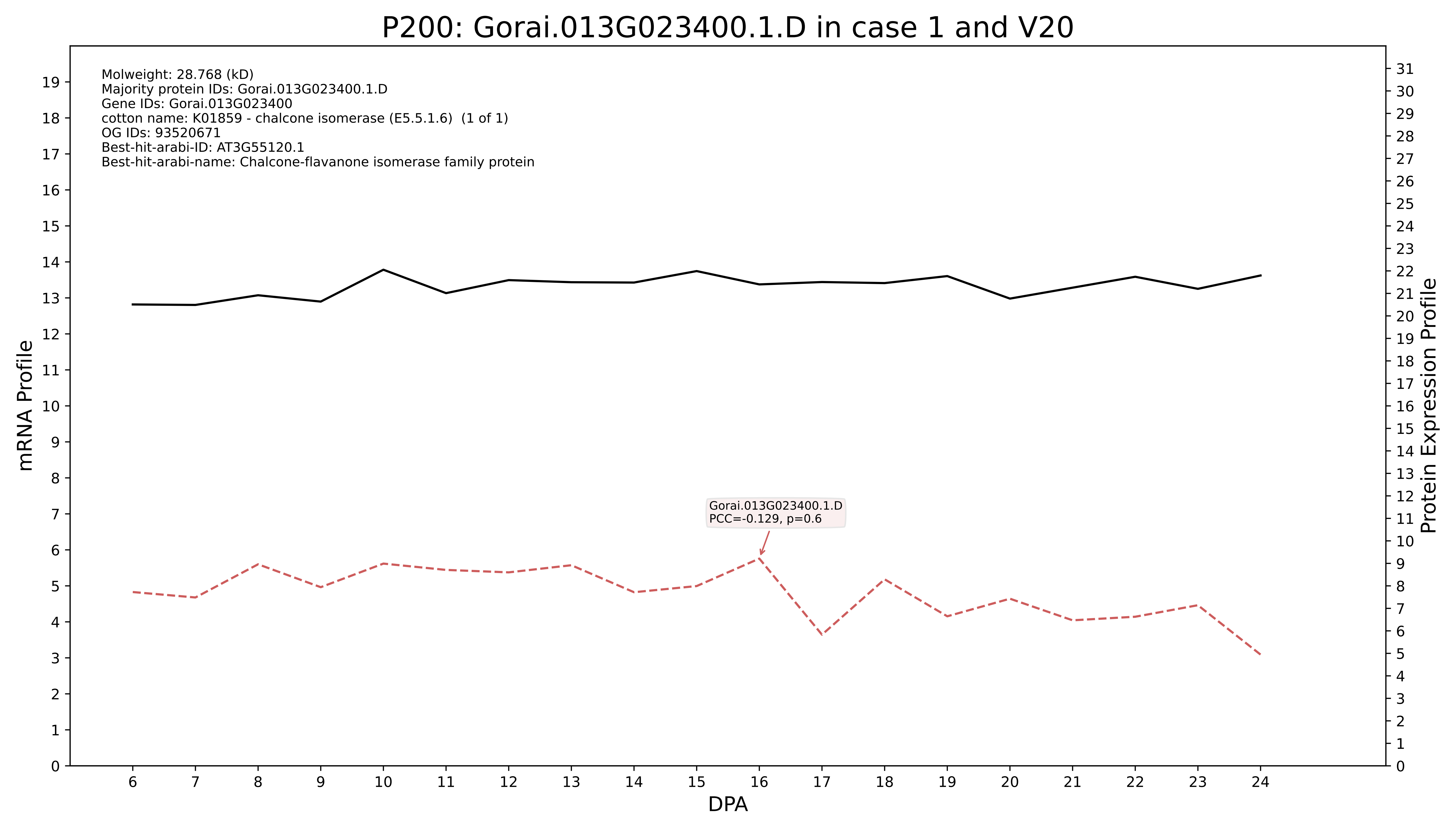
P200: Gorai.013G023400.1.D in case 1 and V20
Molweight: 28.768 (kD)
Majority protein IDs: Gorai.013G023400.1.D
Gene IDs: Gorai.013G023400
cotton name: K01859 - chalcone isomerase (E5.5.1.6) (1 of 1)
OG IDs: 93520671
Best-hit-arabi-ID: AT3G55120.1
Best-hit-arabi-name: Chalcone-flavanone isomerase family protein
Gorai.013G023400.1.D
PCC=-0.129, p=0.6
mRNA Profile
Protein Expression Profile
DPA

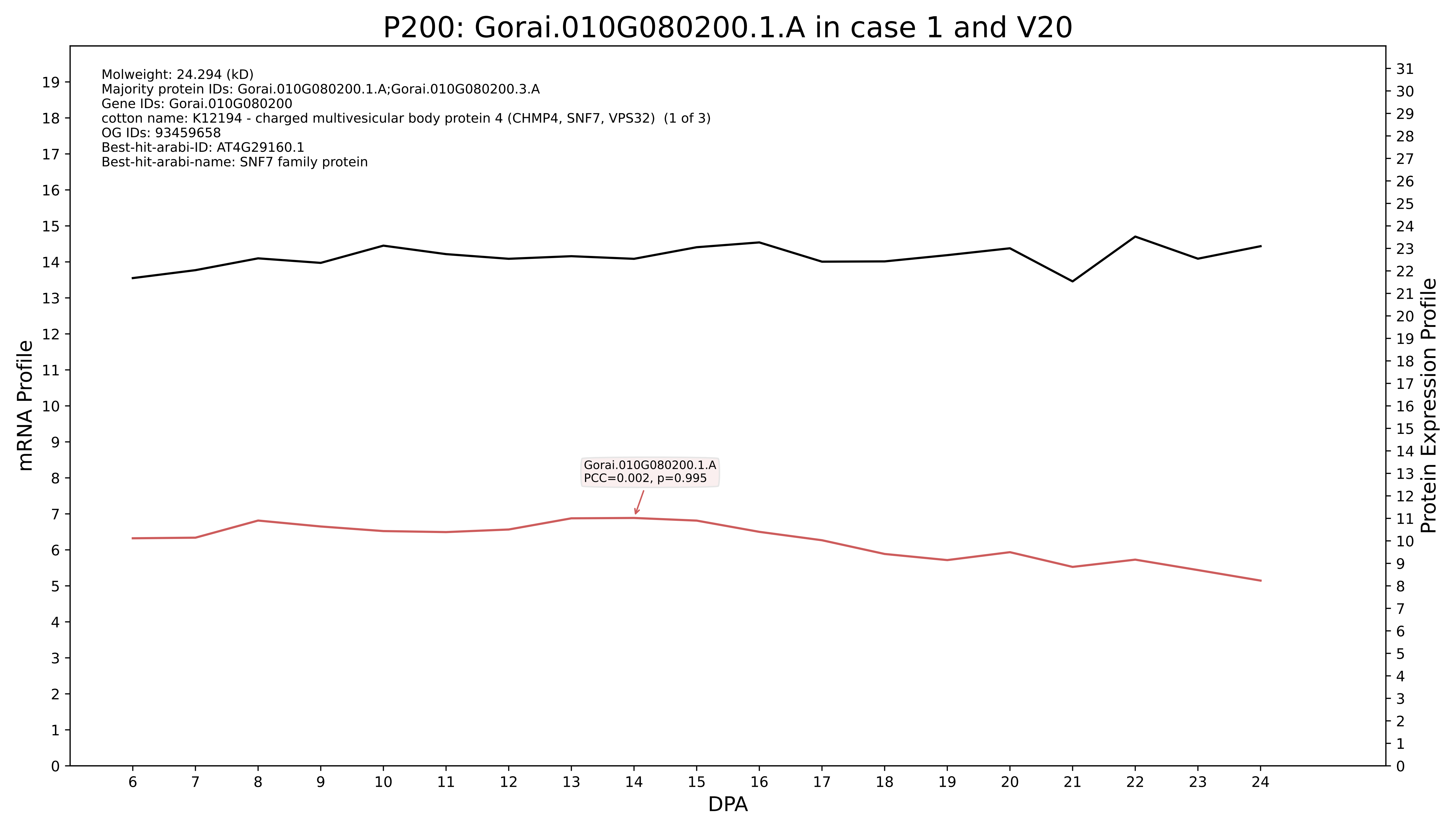
P200: Gorai.010G080200.1.A in case 1 and V20
Molweight: 24.294 (kD)
Majority protein IDs: Gorai.010G080200.1.A;Gorai.010G080200.3.A
Gene IDs: Gorai.010G080200
cotton name: K12194 - charged multivesicular body protein 4 (CHMP4, SNF7, VPS32) (1 of 3)
OG IDs: 93459658
Best-hit-arabi-ID: AT4G29160.1
Best-hit-arabi-name: SNF7 family protein
Gorai.010G080200.1.A
PCC=0.002, p=0.995
mRNA Profile
Protein Expression Profile
DPA

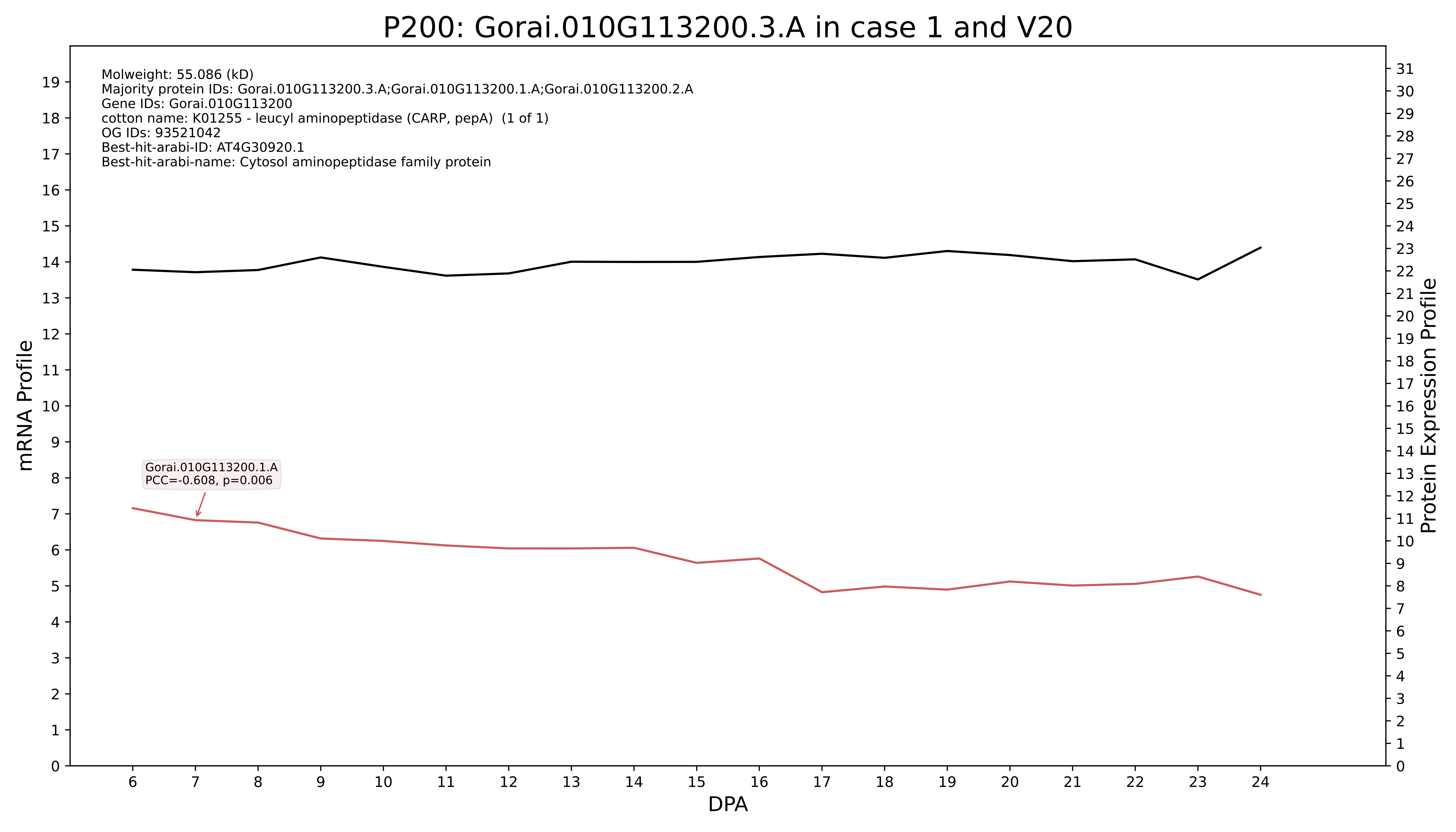

P200: Gorai.010G113200.3.A in case 1 and V20
Molweight: 55.086 (kD)
Majority protein IDs: Gorai.010G113200.3.A;Gorai.010G113200.1.A;Gorai.010G113200.2.A
Gene IDs: Gorai.010G113200
cotton name: K01255 - leucyl aminopeptidase (CARP, pepA) (1 of 1)
OG IDs: 93521042
Best-hit-arabi-ID: AT4G30920.1
Best-hit-arabi-name: Cytosol aminopeptidase family protein
Gorai.010G113200.1.A
PCC=-0.608, p=0.006
mRNA Profile
Protein Expression Profile
DPA

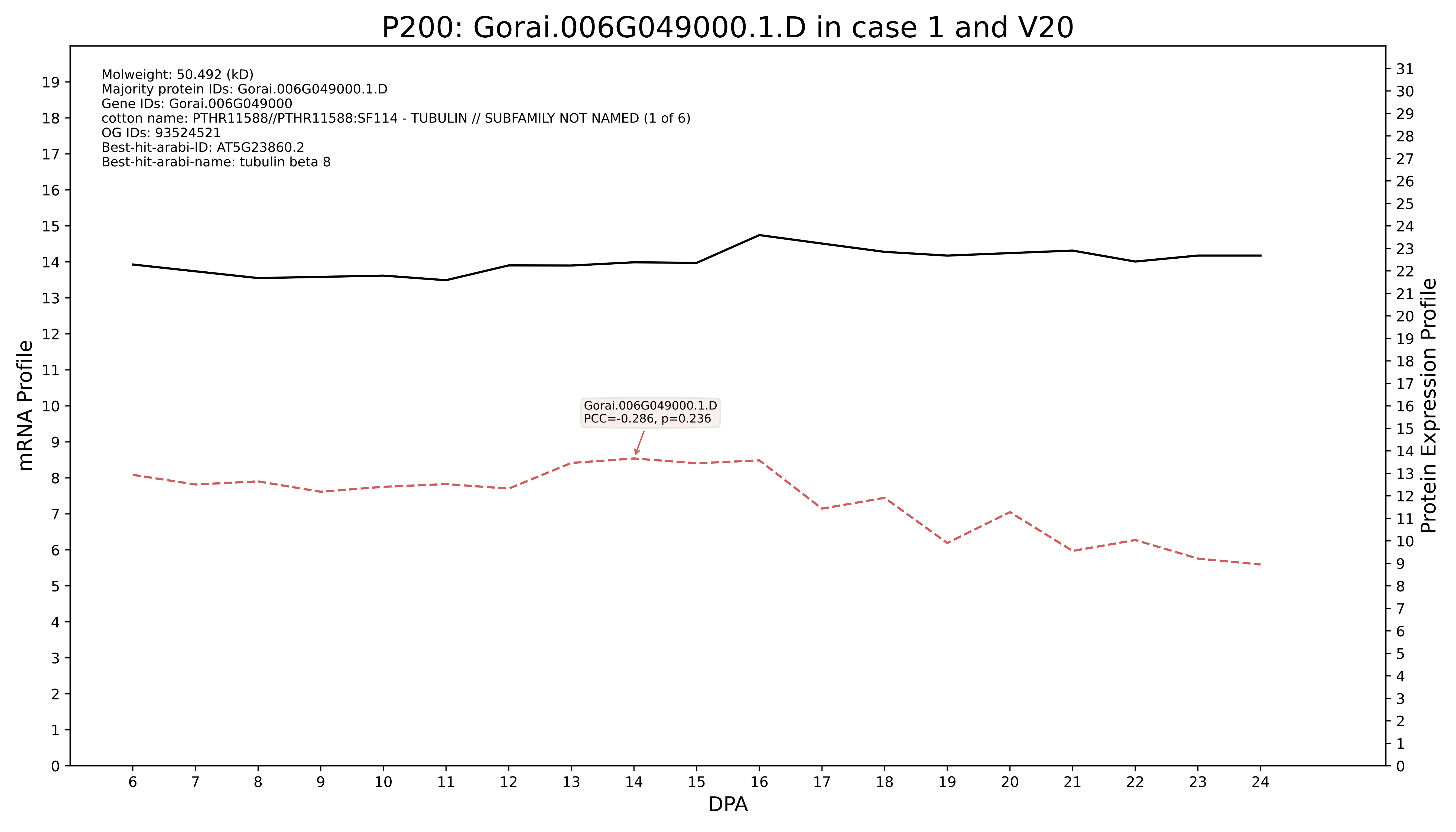

P200: Gorai.006G049000.1.D in case 1 and V20
Molweight: 50.492 (kD)
Majority protein IDs: Gorai.006G049000.1.D
Gene IDs: Gorai.006G049000
cotton name: PTHR11588//PTHR11588:SF114 - TUBULIN // SUBFAMILY NOT NAMED (1 of 6)
OG IDs: 93524521
Best-hit-arabi-ID: AT5G23860.2
Best-hit-arabi-name: tubulin beta 8
Gorai.006G049000.1.D
PCC=-0.286, p=0.236
mRNA Profile
Protein Expression Profile
DPA

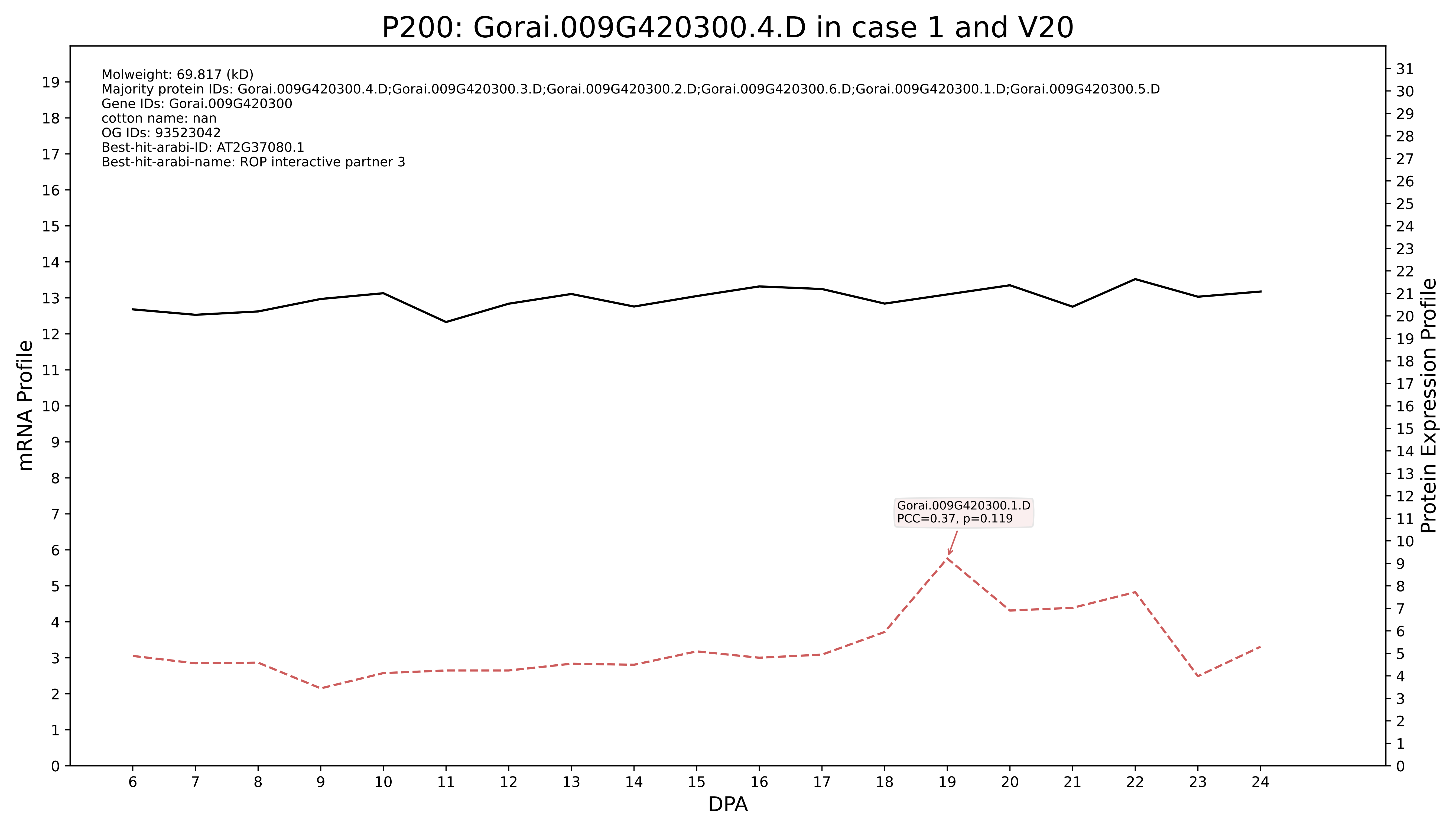
P200: Gorai.009G420300.4.D in case 1 and V20
Molweight: 69.817 (kD)
Majority protein IDs: Gorai.009G420300.4.D;Gorai.009G420300.3.D;Gorai.009G420300.2.D;Gorai.009G420300.6.D;Gorai.009G420300.1.D;Gorai.009G420300.5.D
Gene IDs: Gorai.009G420300
cotton name: nan
OG IDs: 93523042
Best-hit-arabi-ID: AT2G37080.1
Best-hit-arabi-name: ROP interactive partner 3
Gorai.009G420300.1.D
PCC=0.37, p=0.119
mRNA Profile
Protein Expression Profile
DPA

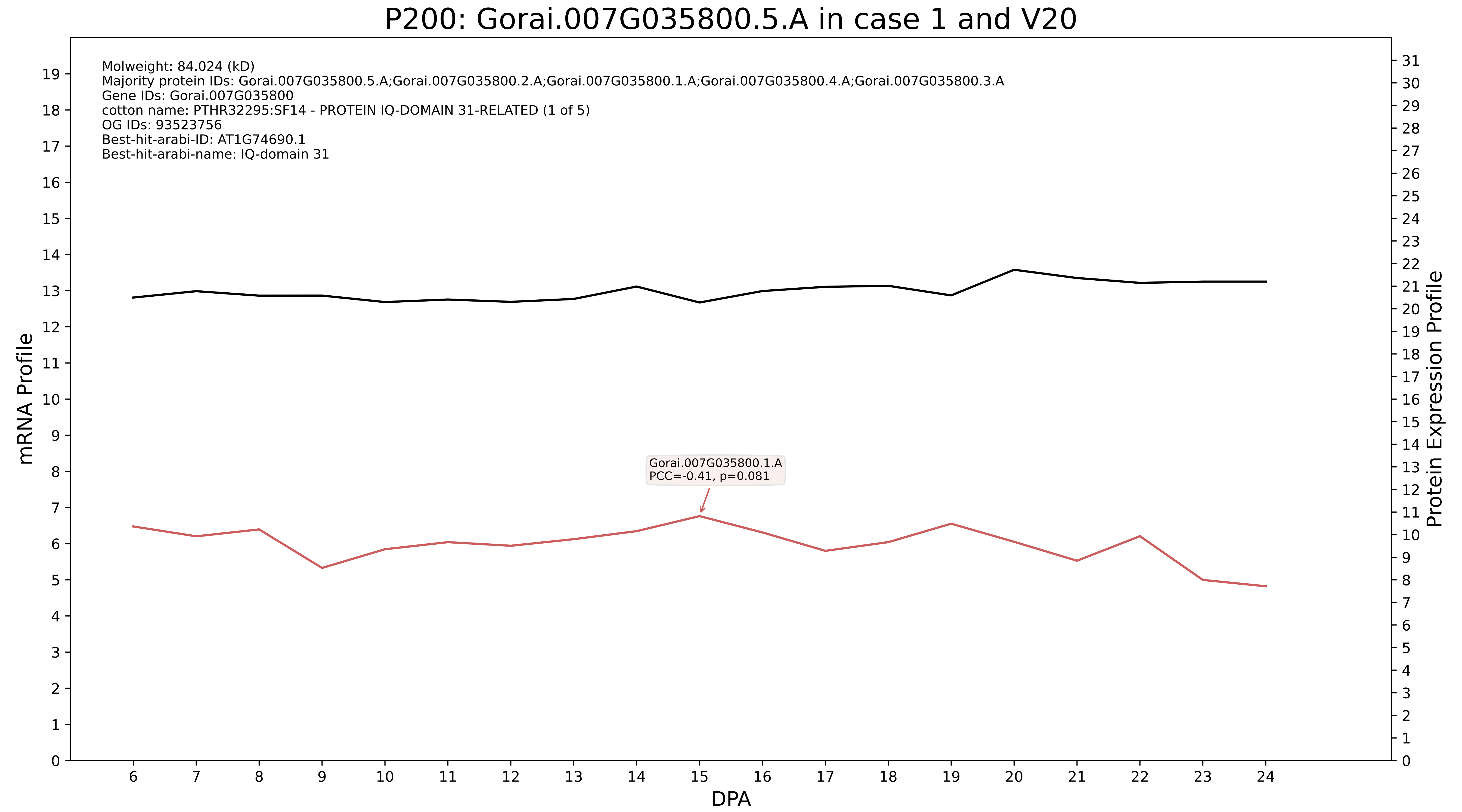

P200: Gorai.007G035800.5.A in case 1 and V20
Molweight: 84.024 (kD)
Majority protein IDs: Gorai.007G035800.5.A;Gorai.007G035800.2.A;Gorai.007G035800.1.A;Gorai.007G035800.4.A;Gorai.007G035800.3.A
Gene IDs: Gorai.007G035800
cotton name: PTHR32295:SF14 - PROTEIN IQ-DOMAIN 31-RELATED (1 of 5)
OG IDs: 93523756
Best-hit-arabi-ID: AT1G74690.1
Best-hit-arabi-name: IQ-domain 31
Gorai.007G035800.1.A
PCC=-0.41, p=0.081
mRNA Profile
Protein Expression Profile
DPA

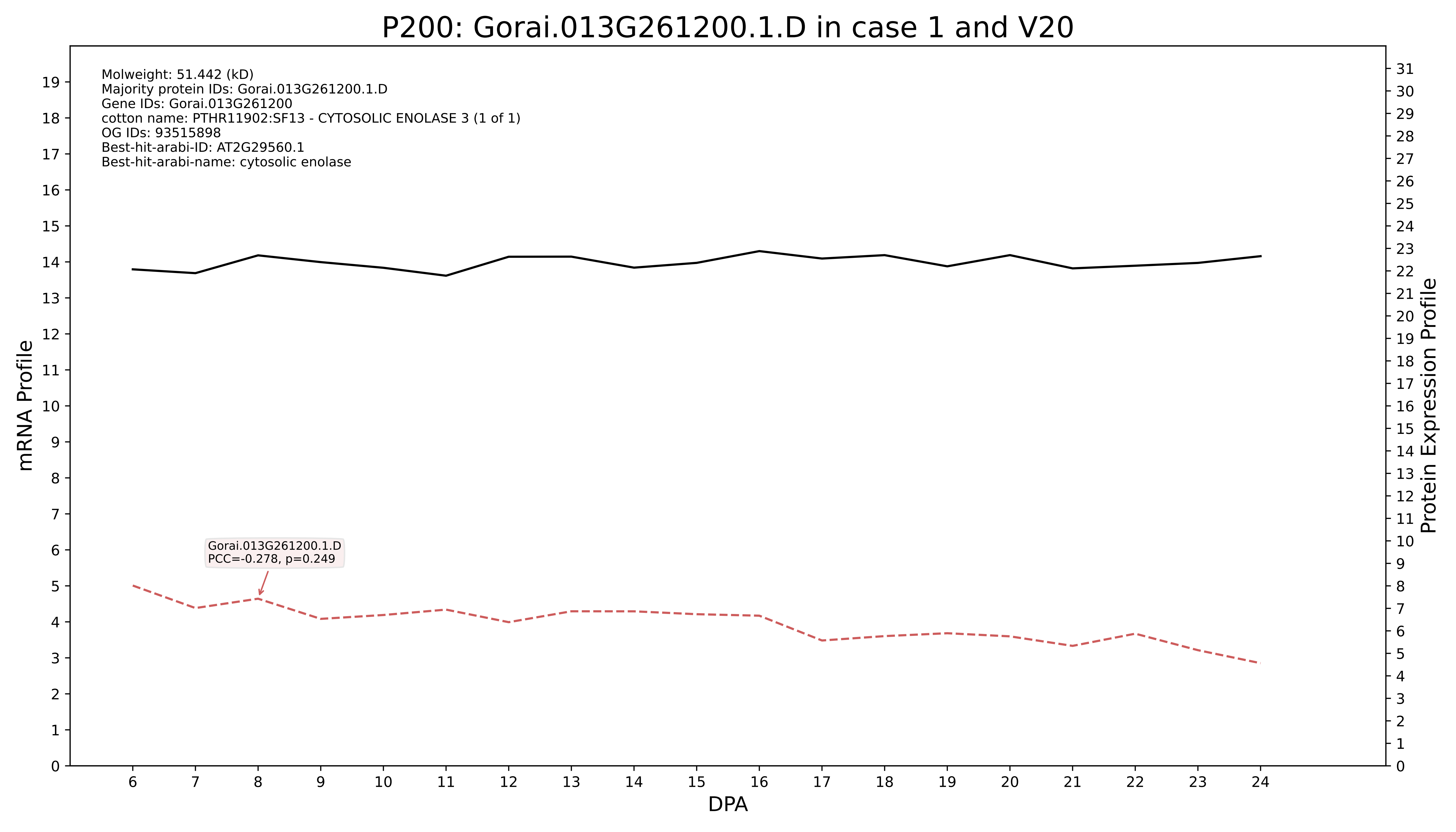
P200: Gorai.013G261200.1.D in case 1 and V20
Molweight: 51.442 (kD)
Majority protein IDs: Gorai.013G261200.1.D
Gene IDs: Gorai.013G261200
cotton name: PTHR11902:SF13 - CYTOSOLIC ENOLASE 3 (1 of 1)
OG IDs: 93515898
Best-hit-arabi-ID: AT2G29560.1
Best-hit-arabi-name: cytosolic enolase
Gorai.013G261200.1.D
PCC=-0.278, p=0.249
mRNA Profile
Protein Expression Profile
DPA

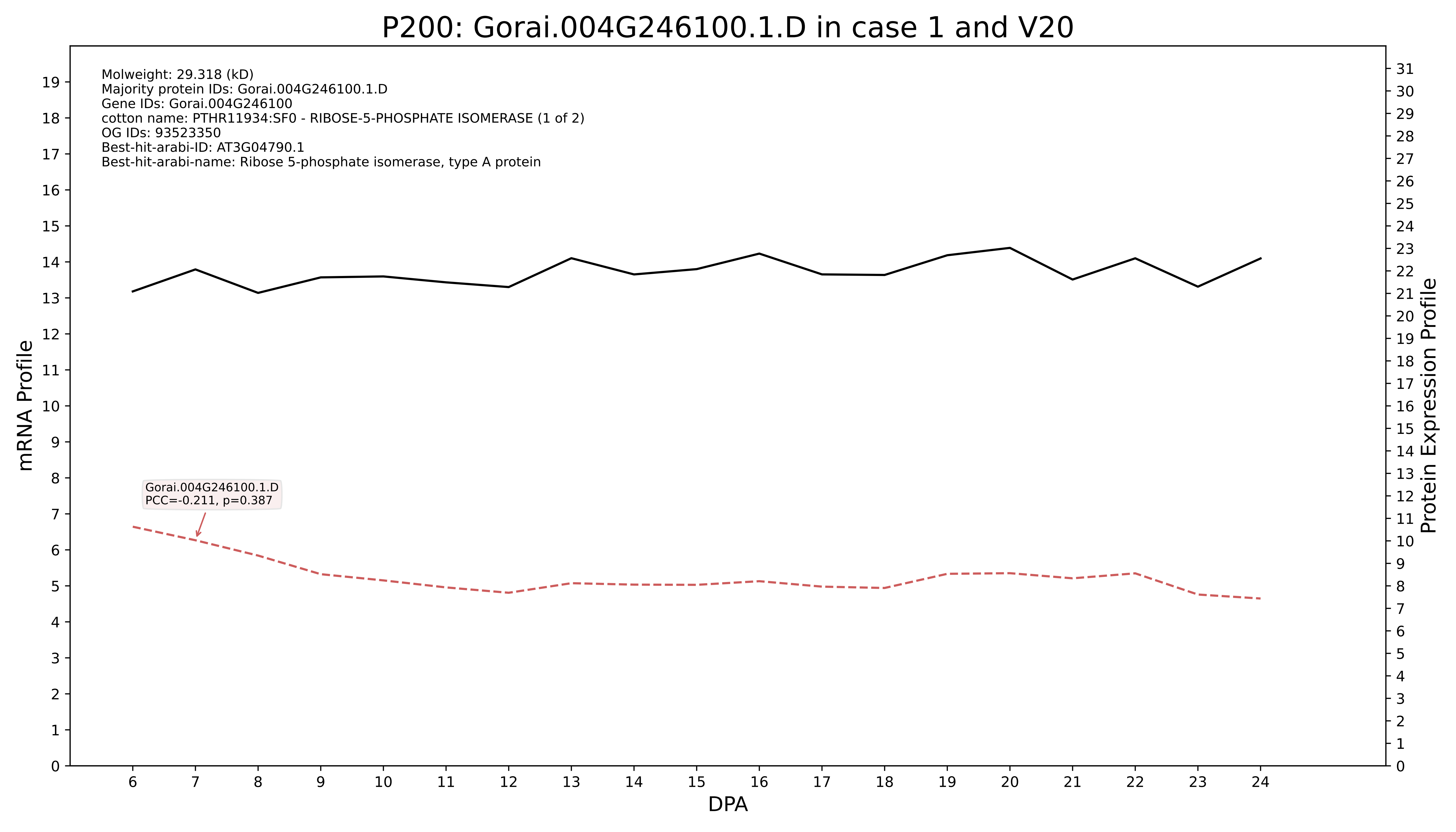
P200: Gorai.004G246100.1.D in case 1 and V20
Molweight: 29.318 (kD)
Majority protein IDs: Gorai.004G246100.1.D
Gene IDs: Gorai.004G246100
cotton name: PTHR11934:SF0 - RIBOSE-5-PHOSPHATE ISOMERASE (1 of 2)
OG IDs: 93523350
Best-hit-arabi-ID: AT3G04790.1
Best-hit-arabi-name: Ribose 5-phosphate isomerase, type A protein
Gorai.004G246100.1.D
PCC=-0.211, p=0.387
mRNA Profile
Protein Expression Profile
DPA

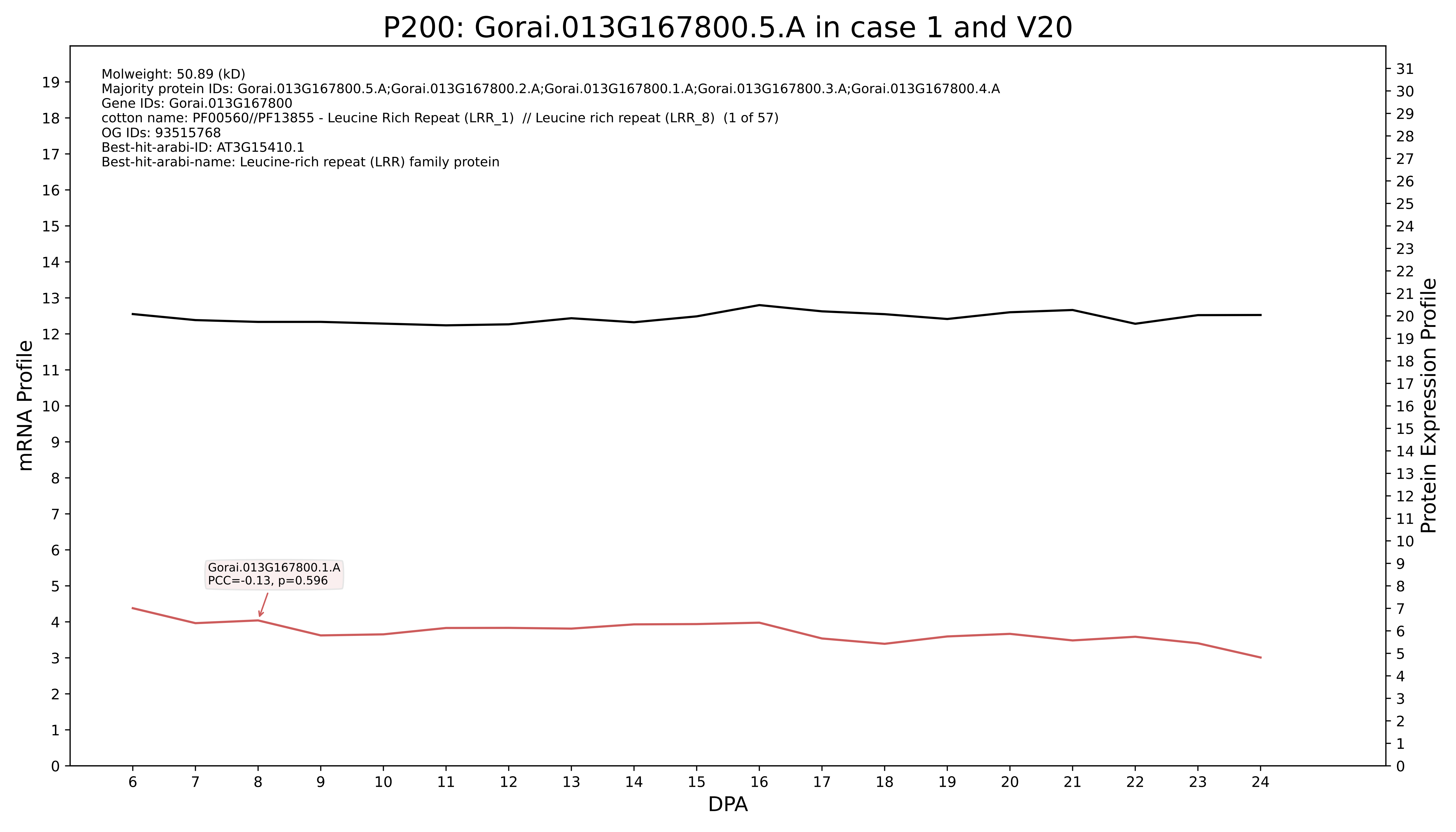
P200: Gorai.013G167800.5.A in case 1 and V20
Molweight: 50.89 (kD)
Majority protein IDs: Gorai.013G167800.5.A;Gorai.013G167800.2.A;Gorai.013G167800.1.A;Gorai.013G167800.3.A;Gorai.013G167800.4.A
Gene IDs: Gorai.013G167800
cotton name: PF00560//PF13855 - Leucine Rich Repeat (LRR_1) // Leucine rich repeat (LRR_8) (1 of 57)
OG IDs: 93515768
Best-hit-arabi-ID: AT3G15410.1
Best-hit-arabi-name: Leucine-rich repeat (LRR) family protein
Gorai.013G167800.1.A
PCC=-0.13, p=0.596
mRNA Profile
Protein Expression Profile
DPA

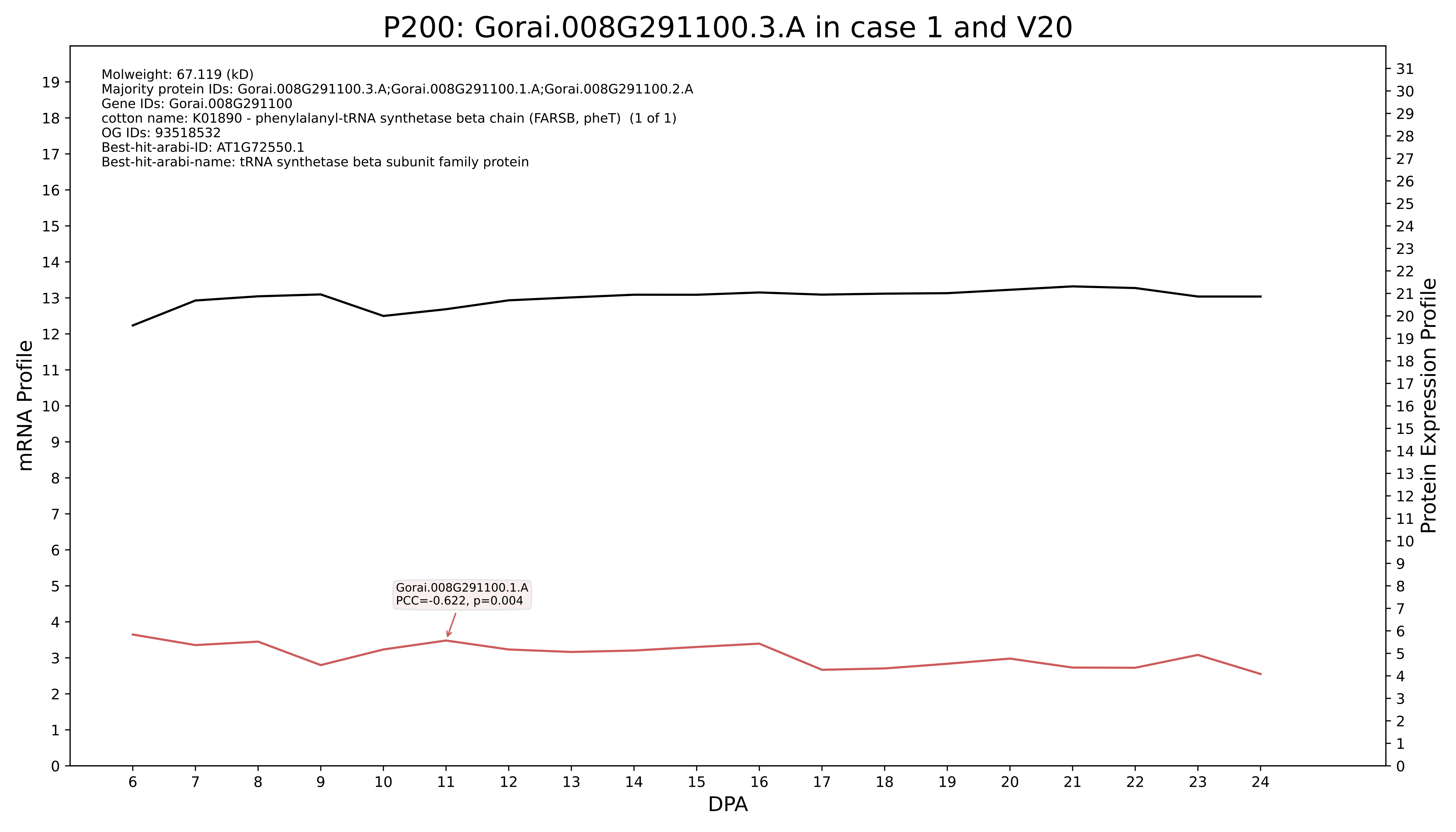

P200: Gorai.008G291100.3.A in case 1 and V20
Molweight: 67.119 (kD)
Majority protein IDs: Gorai.008G291100.3.A;Gorai.008G291100.1.A;Gorai.008G291100.2.A
Gene IDs: Gorai.008G291100
cotton name: K01890 - phenylalanyl-tRNA synthetase beta chain (FARSB, pheT) (1 of 1)
OG IDs: 93518532
Best-hit-arabi-ID: AT1G72550.1
Best-hit-arabi-name: tRNA synthetase beta subunit family protein
Gorai.008G291100.1.A
PCC=-0.622, p=0.004
mRNA Profile
Protein Expression Profile
DPA

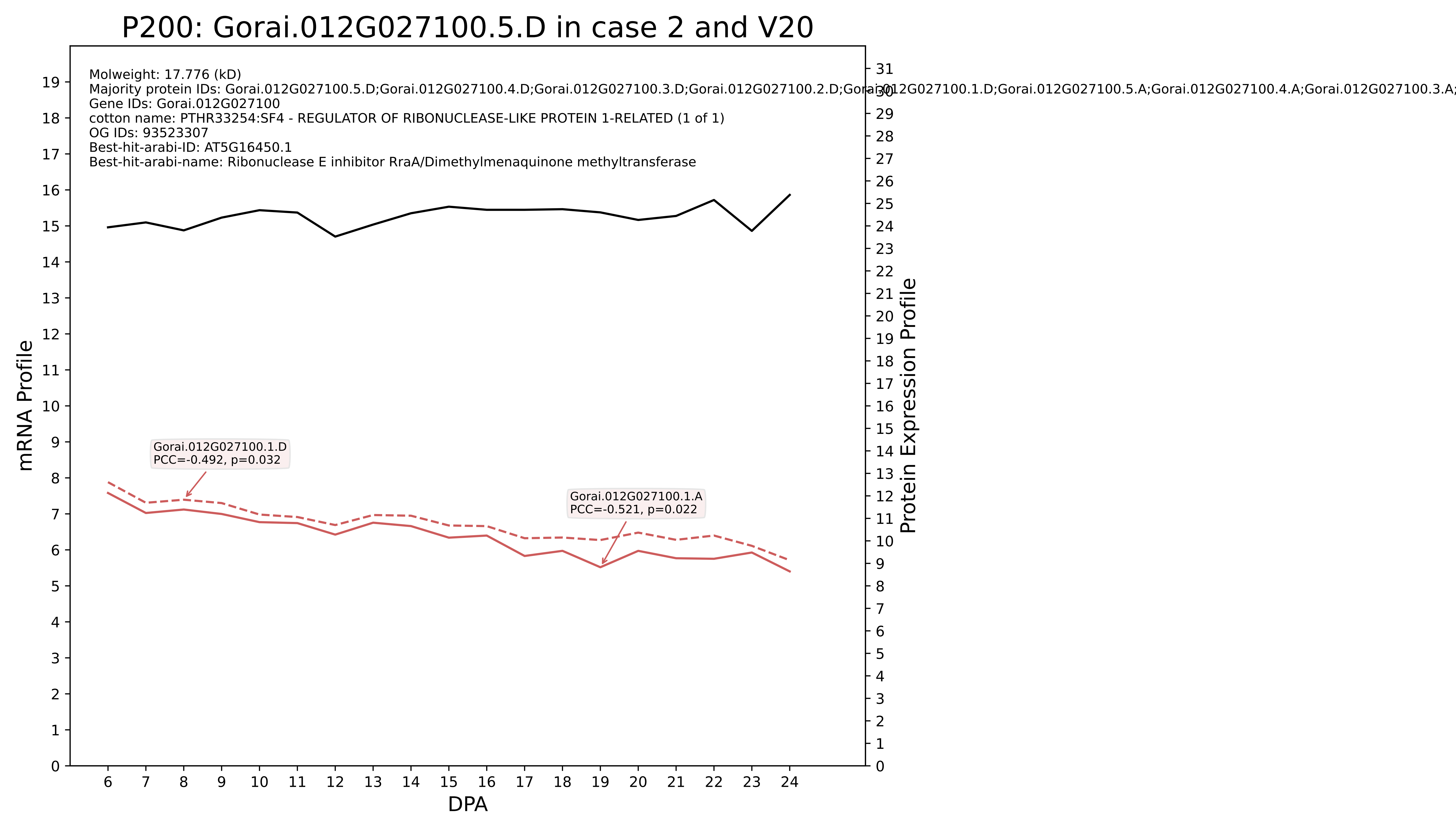
P200: Gorai.012G027100.5.D in case 2 and V20
Molweight: 17.776 (kD)
Majority protein IDs: Gorai.012G027100.5.D;Gorai.012G027100.4.D;Gorai.012G027100.3.D;Gorai.012G027100.2.D;Gorai.012G027100.1.D;Gorai.012G027100.5.A;Gorai.012G027100.4.A;Gorai.012G027100.3.A;
Gene IDs: Gorai.012G027100
cotton name: PTHR33254:SF4 - REGULATOR OF RIBONUCLEASE-LIKE PROTEIN 1-RELATED (1 of 1)
OG IDs: 93523307
Best-hit-arabi-ID: AT5G16450.1
Best-hit-arabi-name: Ribonuclease E inhibitor RraA/Dimethylmenaquinone methyltransferase
Gorai.012G027100.1.D
PCC=-0.492, p=0.032
Gorai.012G027100.1.A
PCC=-0.521, p=0.022
mRNA Profile
Protein Expression Profile
DPA

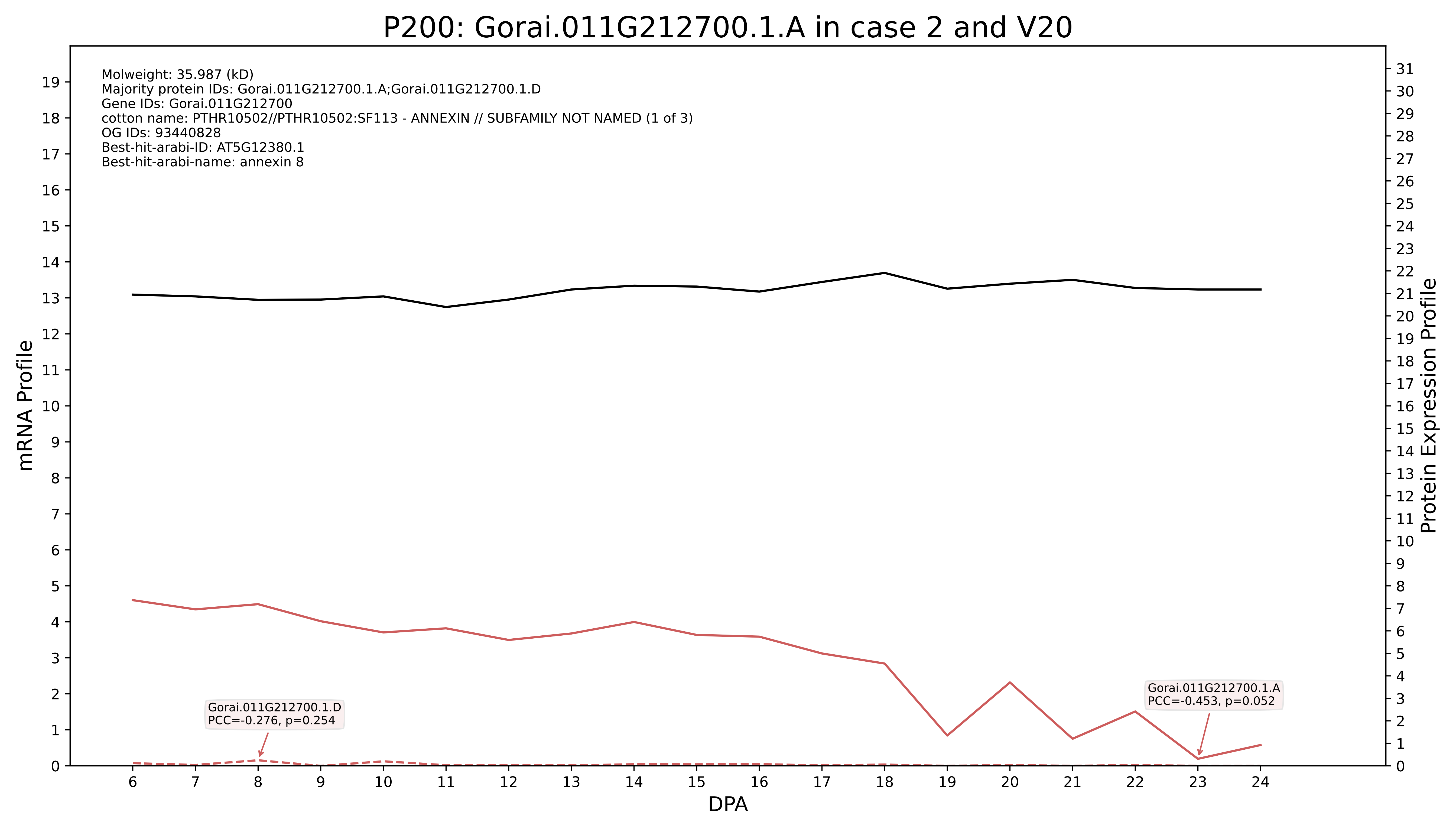
P200: Gorai.011G212700.1.A in case 2 and V20
Molweight: 35.987 (kD)
Majority protein IDs: Gorai.011G212700.1.A;Gorai.011G212700.1.D
Gene IDs: Gorai.011G212700
cotton name: PTHR10502//PTHR10502:SF113 - ANNEXIN // SUBFAMILY NOT NAMED (1 of 3)
OG IDs: 93440828
Best-hit-arabi-ID: AT5G12380.1
Best-hit-arabi-name: annexin 8
Gorai.011G212700.1.D
PCC=-0.276, p=0.254
Gorai.011G212700.1.A
PCC=-0.453, p=0.052
mRNA Profile
Protein Expression Profile
DPA

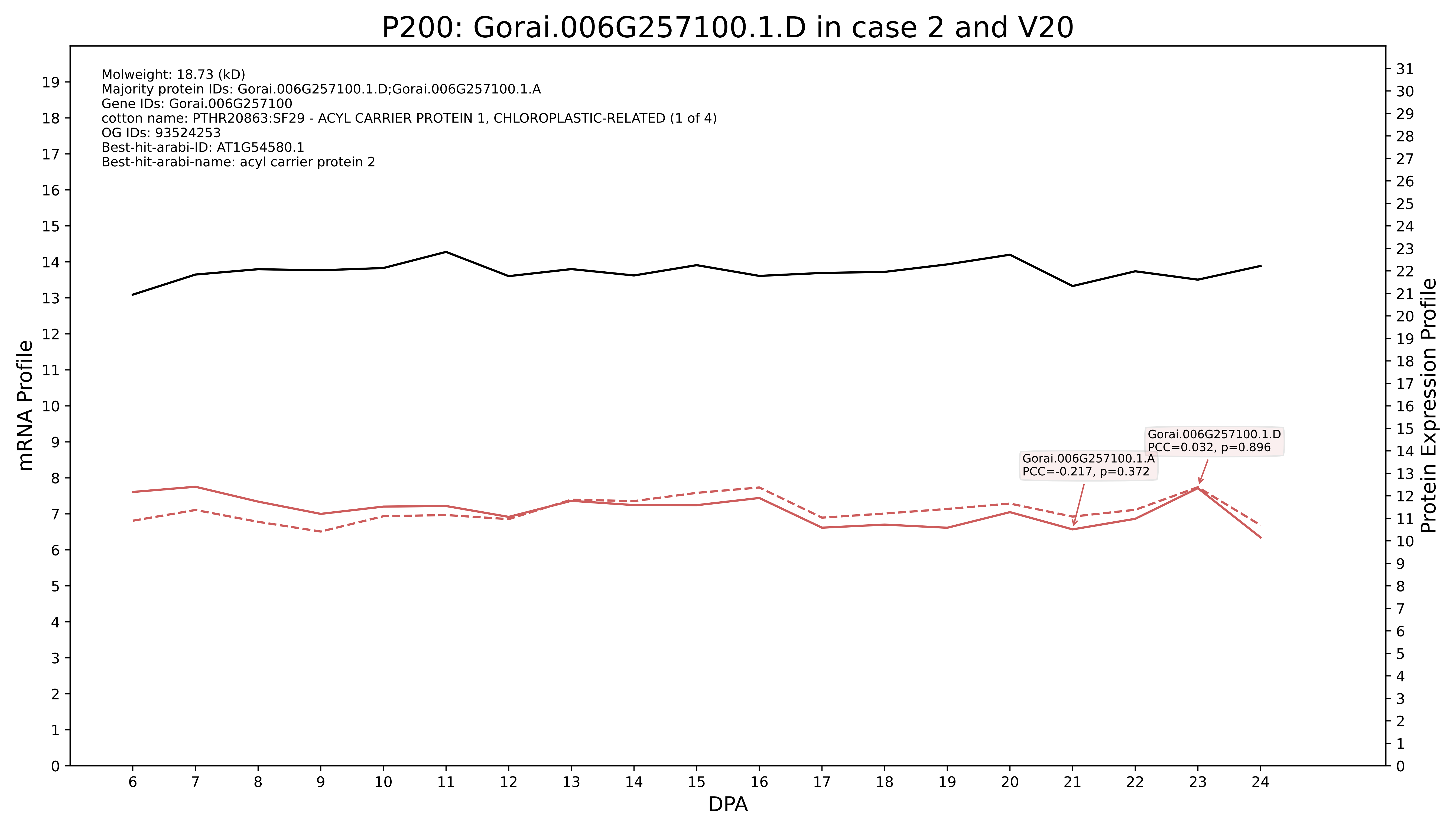

P200: Gorai.006G257100.1.D in case 2 and V20
Molweight: 18.73 (kD)
Majority protein IDs: Gorai.006G257100.1.D;Gorai.006G257100.1.A
Gene IDs: Gorai.006G257100
cotton name: PTHR20863:SF29 - ACYL CARRIER PROTEIN 1, CHLOROPLASTIC-RELATED (1 of 4)
OG IDs: 93524253
Best-hit-arabi-ID: AT1G54580.1
Best-hit-arabi-name: acyl carrier protein 2
Gorai.006G257100.1.A
PCC=-0.217, p=0.372
Gorai.006G257100.1.D
PCC=0.032, p=0.896
mRNA Profile
Protein Expression Profile
DPA

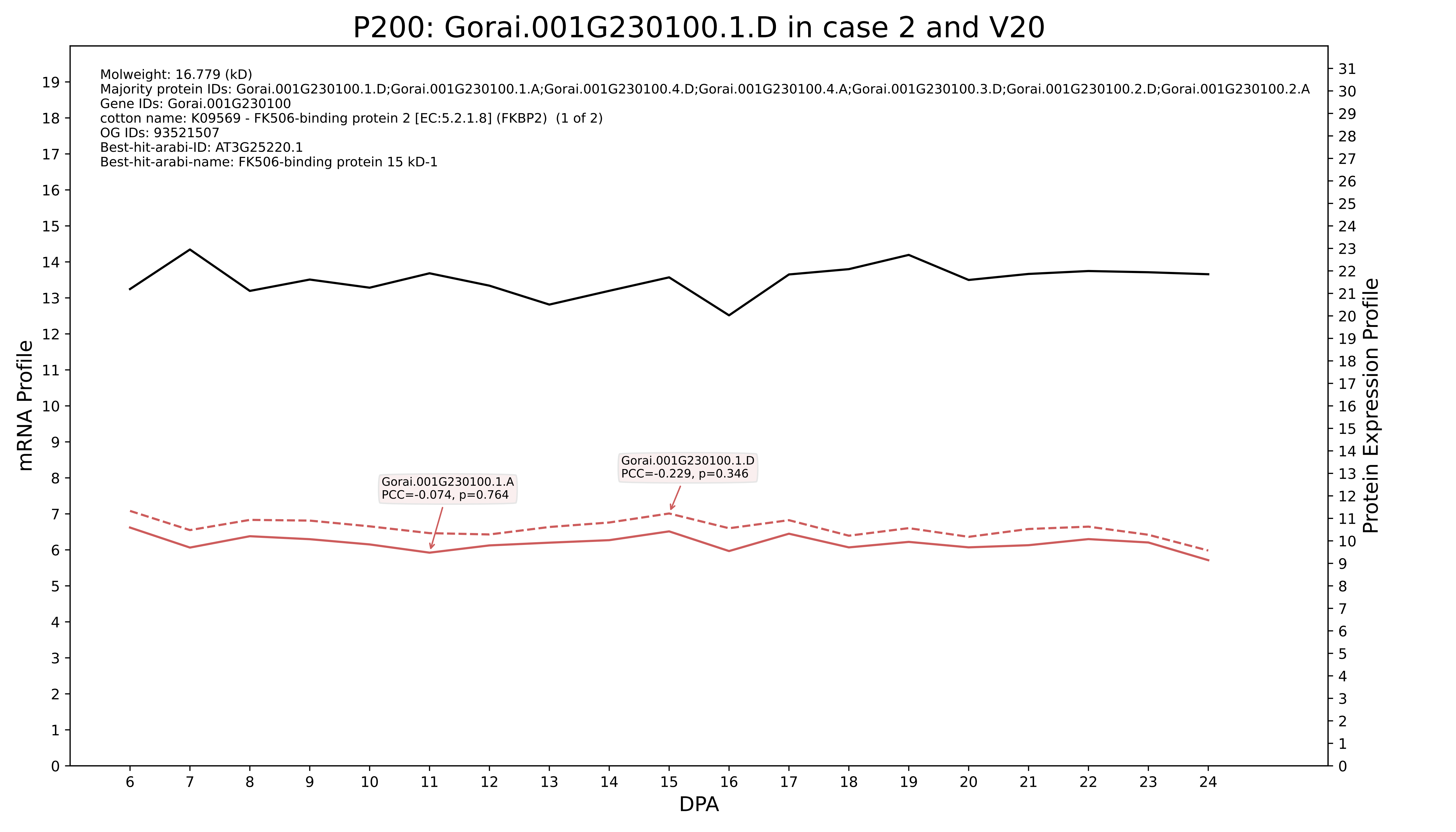

P200: Gorai.001G230100.1.D in case 2 and V20
Molweight: 16.779 (kD)
Majority protein IDs: Gorai.001G230100.1.D;Gorai.001G230100.1.A;Gorai.001G230100.4.D;Gorai.001G230100.4.A;Gorai.001G230100.3.D;Gorai.001G230100.2.D;Gorai.001G230100.2.A
Gene IDs: Gorai.001G230100
cotton name: K09569 - FK506-binding protein 2 [EC:5.2.1.8] (FKBP2) (1 of 2)
OG IDs: 93521507
Best-hit-arabi-ID: AT3G25220.1
Best-hit-arabi-name: FK506-binding protein 15 kD-1
Gorai.001G230100.1.A
PCC=-0.074, p=0.764
Gorai.001G230100.1.D
PCC=-0.229, p=0.346
mRNA Profile
Protein Expression Profile
DPA

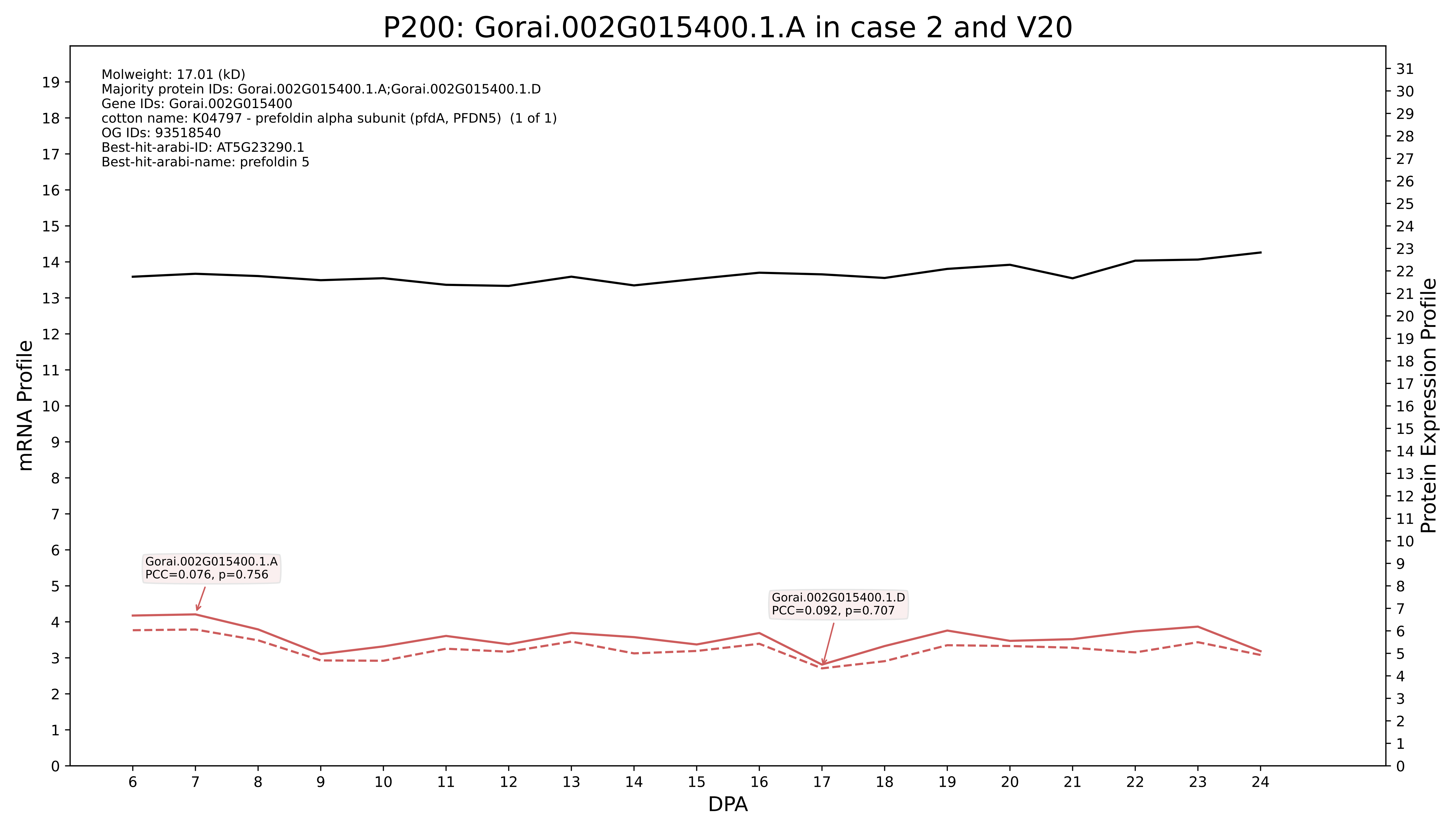
P200: Gorai.002G015400.1.A in case 2 and V20
Molweight: 17.01 (kD)
Majority protein IDs: Gorai.002G015400.1.A;Gorai.002G015400.1.D
Gene IDs: Gorai.002G015400
cotton name: K04797 - prefoldin alpha subunit (pfdA, PFDN5) (1 of 1)
OG IDs: 93518540
Best-hit-arabi-ID: AT5G23290.1
Best-hit-arabi-name: prefoldin 5
Gorai.002G015400.1.A
PCC=0.076, p=0.756
Gorai.002G015400.1.D
PCC=0.092, p=0.707
mRNA Profile
Protein Expression Profile
DPA

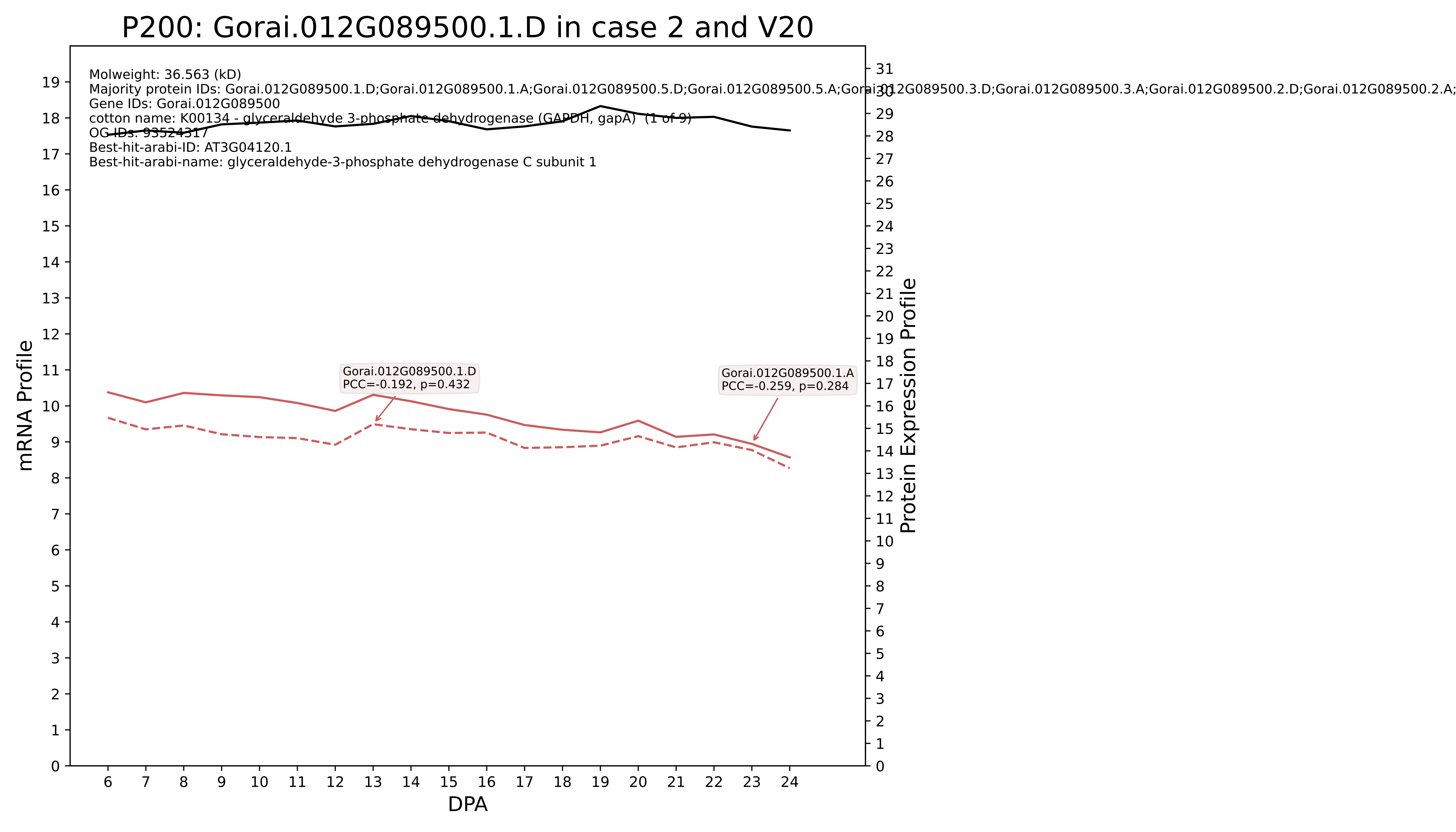
P200: Gorai.012G089500.1.D in case 2 and V20
Molweight: 36.563 (kD)
Majority protein IDs: Gorai.012G089500.1.D;Gorai.012G089500.1.A;Gorai.012G089500.5.D;Gorai.012G089500.5.A;Gorai.012G089500.3.D;Gorai.012G089500.3.A;Gorai.012G089500.2.D;Gorai.012G089500.2.A;
Gene IDs: Gorai.012G089500
cotton name: K00134 - glyceraldehyde 3-phosphate dehydrogenase (GAPDH, gapA) (1 of 9)
OG-IDs: 93524317
Best-hit-arabi-ID: AT3G04120.1
Best-hit-arabi-name: glyceraldehyde-3-phosphate dehydrogenase C subunit 1
Gorai.012G089500.1.D
PCC=-0.192, p=0.432
Gorai.012G089500.1.A
PCC=-0.259, p=0.284
mRNA Profile
Protein Expression Profile
DPA

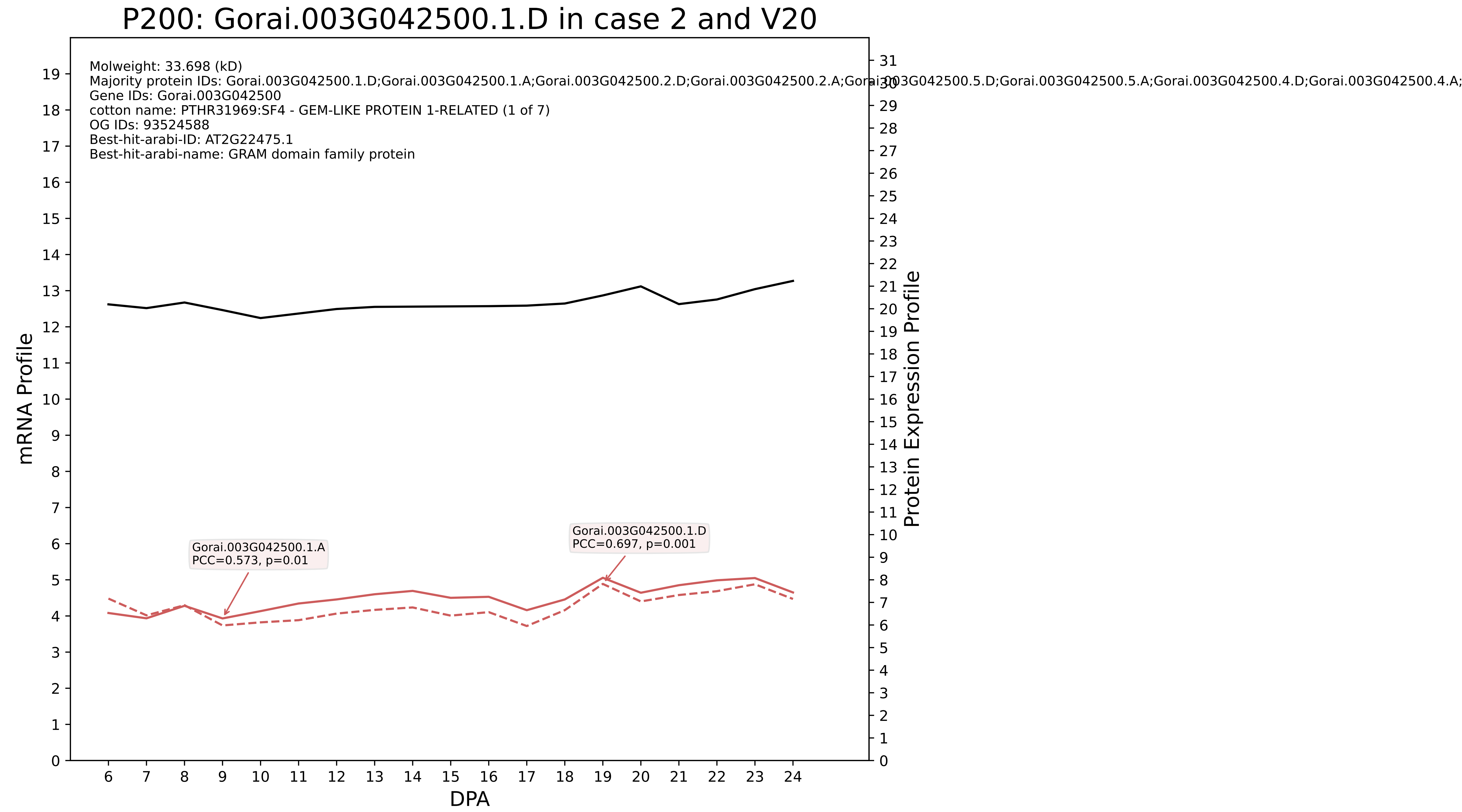
P200: Gorai.003G042500.1.D in case 2 and V20
Molweight: 33.698 (kD)
Majority protein IDs: Gorai.003G042500.1.D;Gorai.003G042500.1.A;Gorai.003G042500.2.D;Gorai.003G042500.2.A;Gorai.003G042500.5.D;Gorai.003G042500.5.A;Gorai.003G042500.4.D;Gorai.003G042500.4.A;
Gene IDs: Gorai.003G042500
cotton name: PTHR31969:SF4 - GEM-LIKE PROTEIN 1-RELATED (1 of 7)
OG IDs: 93524588
Best-hit-arabi-ID: AT2G22475.1
Best-hit-arabi-name: GRAM domain family protein
Gorai.003G042500.1.A
PCC=0.573, p=0.01
Gorai.003G042500.1.D
PCC=0.697, p=0.001
mRNA Profile
Protein Expression Profile
DPA

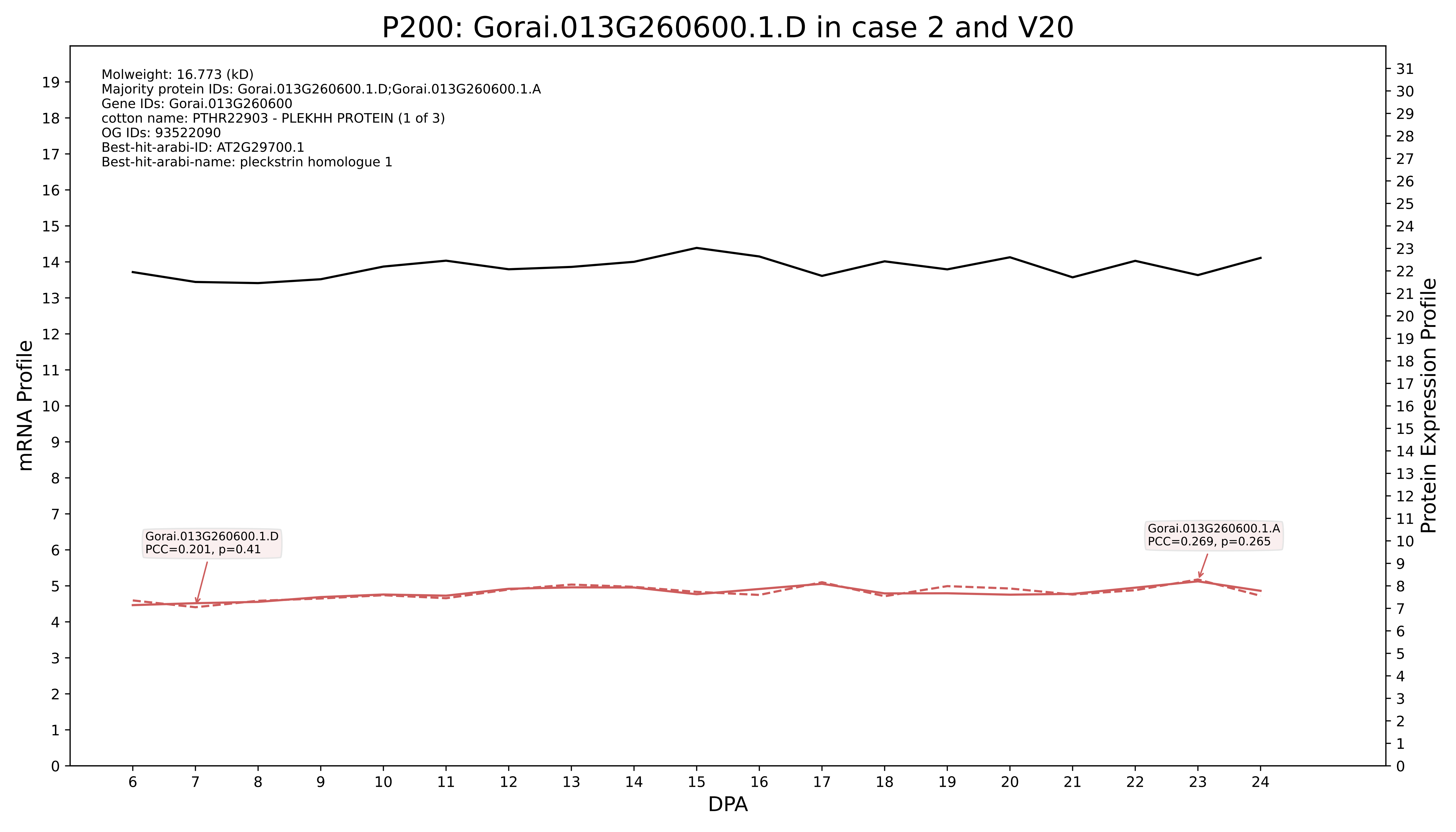
P200: Gorai.013G260600.1.D in case 2 and V20
Molweight: 16.773 (kD)
Majority protein IDs: Gorai.013G260600.1.D;Gorai.013G260600.1.A
Gene IDs: Gorai.013G260600
cotton name: PTHR22903 - PLEKHH PROTEIN (1 of 3)
OG IDs: 93522090
Best-hit-arabi-ID: AT2G29700.1
Best-hit-arabi-name: pleckstrin homologue 1
Gorai.013G260600.1.D
PCC=0.201, p=0.41
Gorai.013G260600.1.A
PCC=0.269, p=0.265
mRNA Profile
Protein Expression Profile
DPA

### P200: Gorai.011G178800.1.D in case 2 and V20

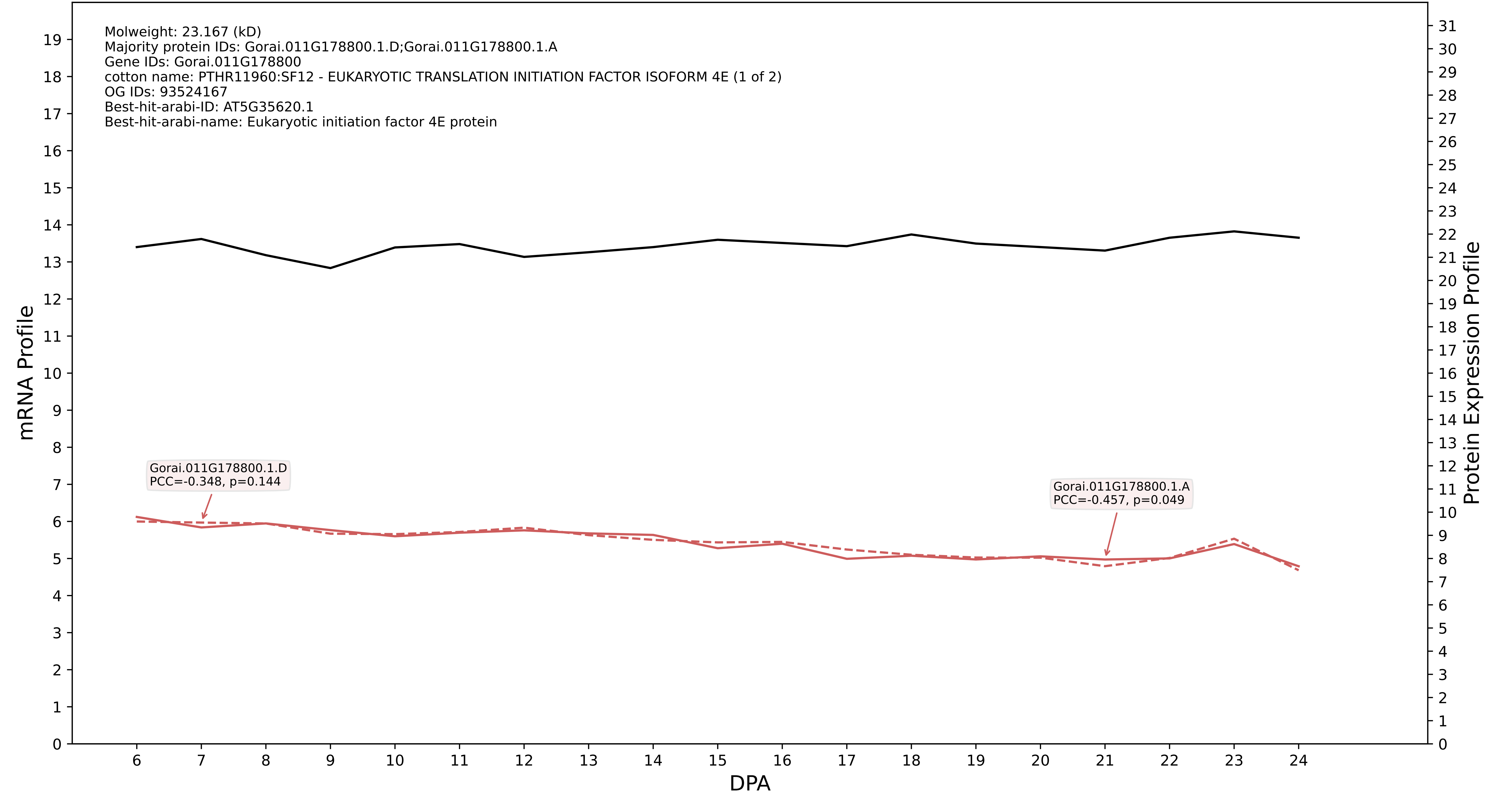

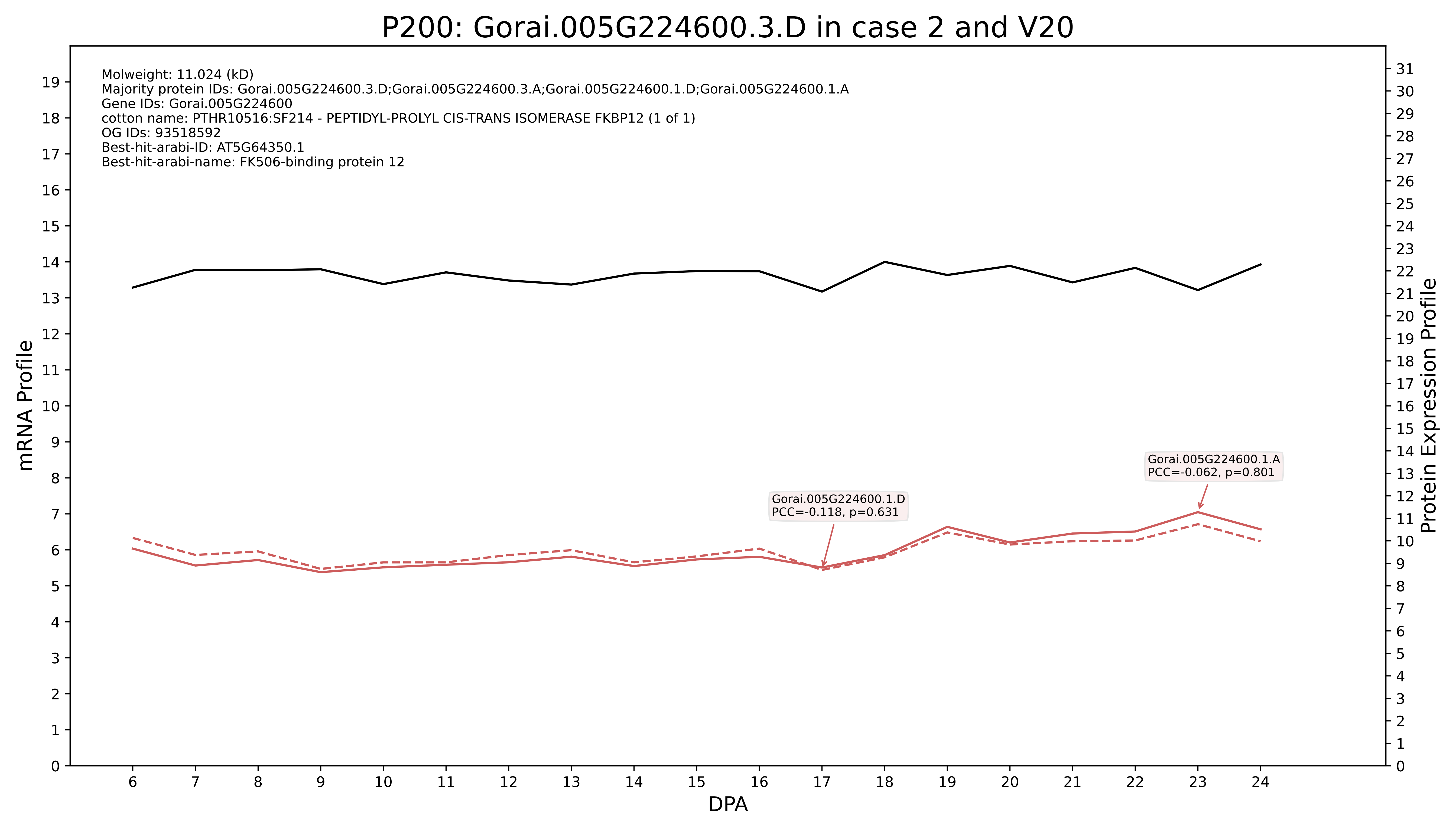
P200: Gorai.005G224600.3.D in case 2 and V20
Molweight: 11.024 (kD)
Majority protein IDs: Gorai.005G224600.3.D;Gorai.005G224600.3.A;Gorai.005G224600.1.D;Gorai.005G224600.1.A
Gene IDs: Gorai.005G224600
cotton name: PTHR10516:SF214 - PEPTIDYL-PROLYL CIS-TRANS ISOMERASE FKBP12 (1 of 1)
OG IDs: 93518592
Best-hit-arabi-ID: AT5G64350.1
Best-hit-arabi-name: FK506-binding protein 12
Gorai.005G224600.1.D
PCC=-0.118, p=0.631
Gorai.005G224600.1.A
PCC=-0.062, p=0.801
mRNA Profile
Protein Expression Profile
DPA

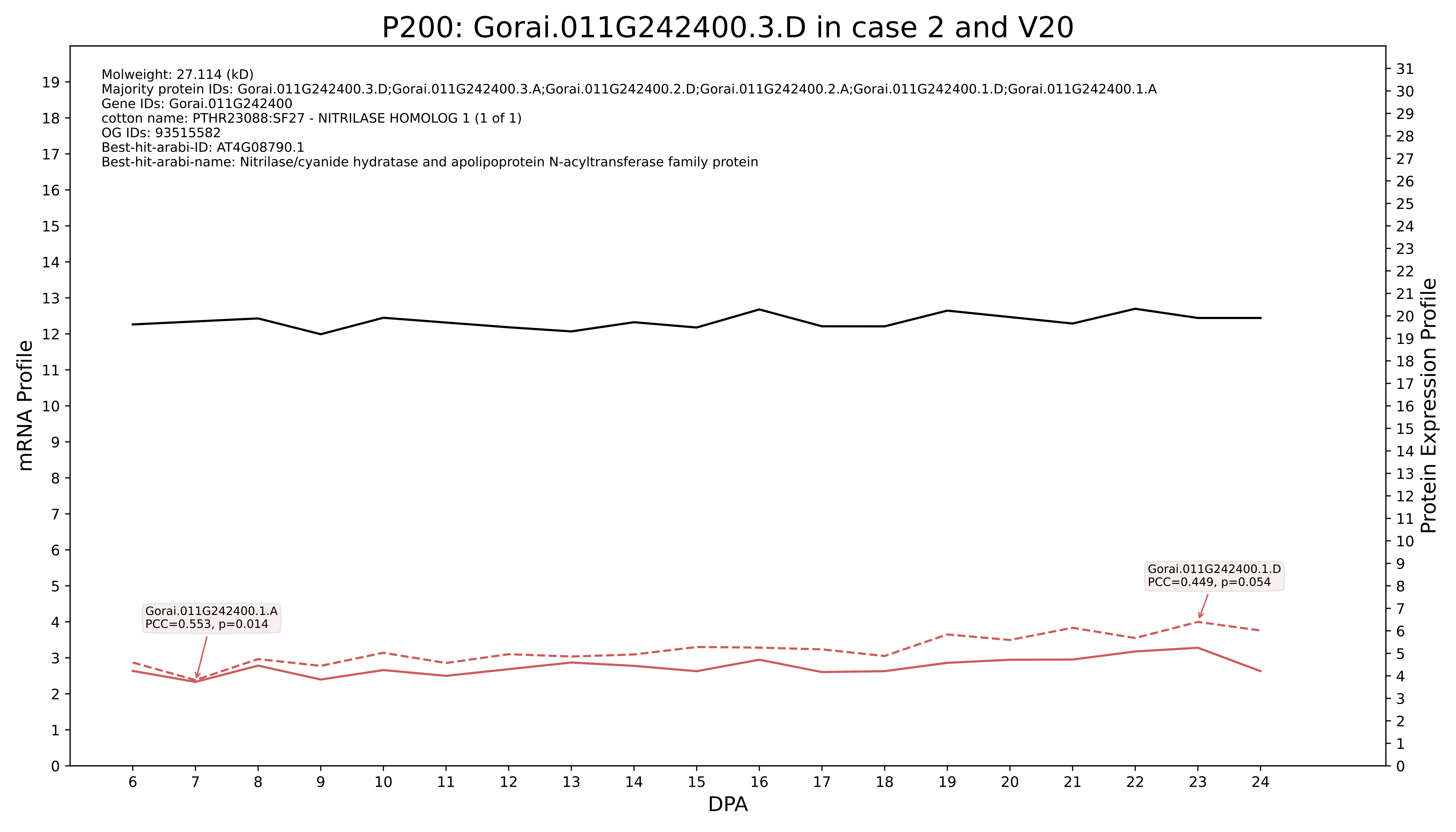
P200: Gorai.011G242400.3.D in case 2 and V20
Molweight: 27.114 (kD)
Majority protein IDs: Gorai.011G242400.3.D;Gorai.011G242400.3.A;Gorai.011G242400.2.D;Gorai.011G242400.2.A;Gorai.011G242400.1.D;Gorai.011G242400.1.A
Gene IDs: Gorai.011G242400
cotton name: PTHR23088:SF27 - NITRILASE HOMOLOG 1 (1 of 1)
OG IDs: 93515582
Best-hit-arabi-ID: AT4G08790.1
Best-hit-arabi-name: Nitrilase/cyanide hydratase and apolipoprotein N-acyltransferase family protein
Gorai.011G242400.1.D
PCC=0.449, p=0.054
Gorai.011G242400.1.A
PCC=0.553, p=0.014
mRNA Profile
Protein Expression Profile
DPA
6 7 8 9 10 11 12 13 14 15 16 17 18 19 20 21 22 23 24

### P200: Gorai.009G034900.3.D in case 2 and V20

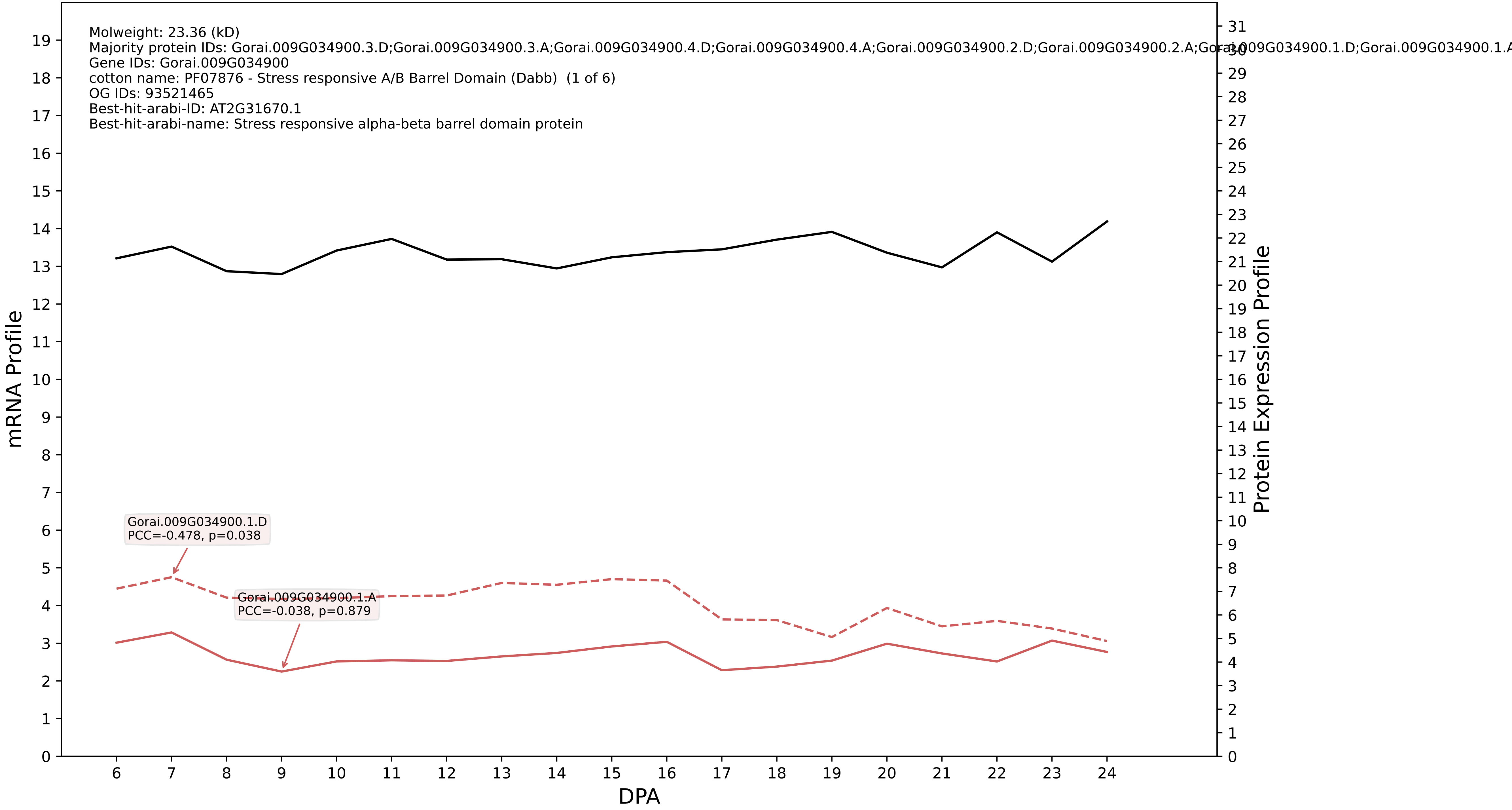

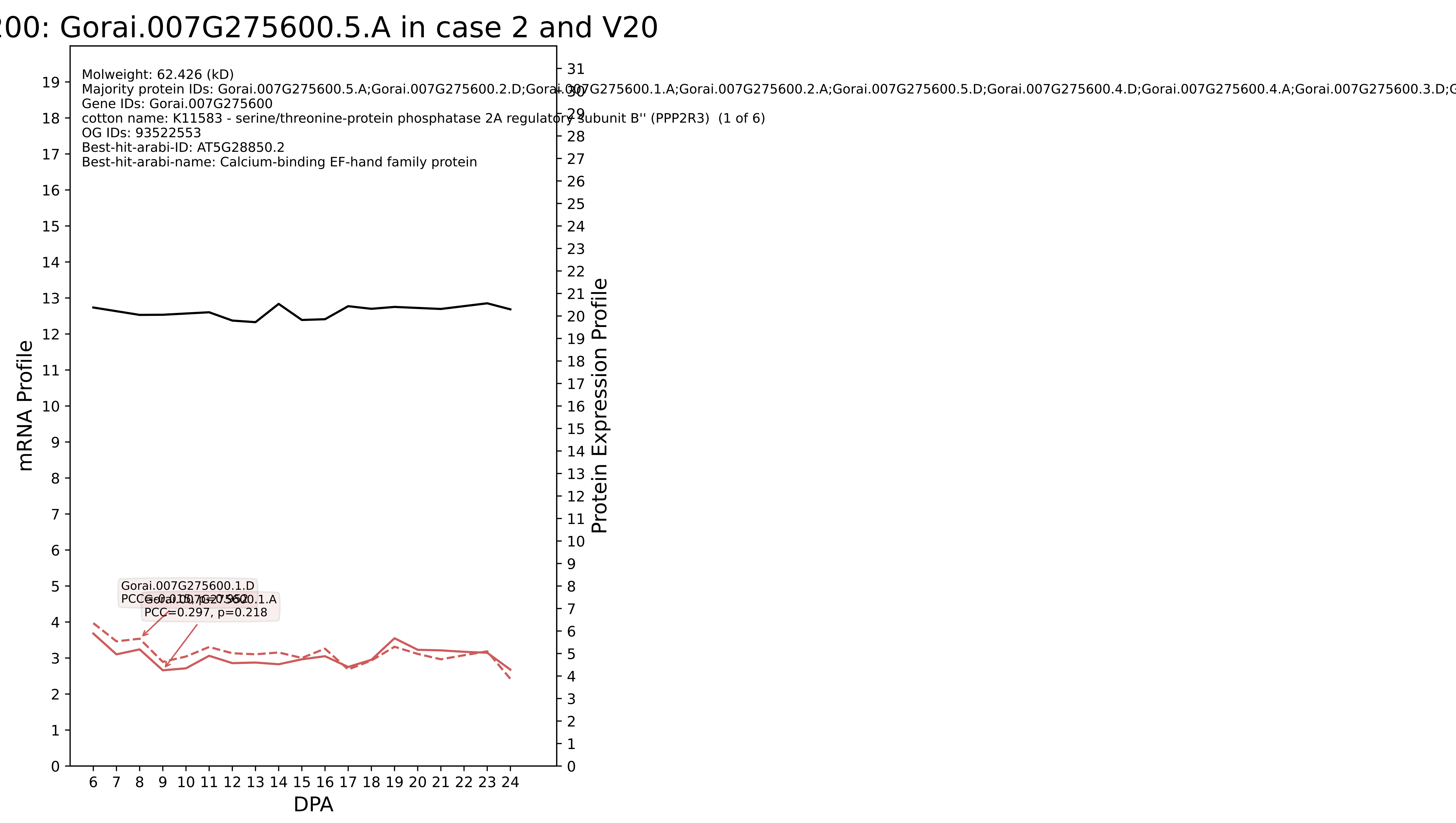
200: Gorai.007G275600.5.A in case 2 and V20
Molweight: 62.426 (kD)
Majority protein IDs: Gorai.007G275600.5.A;Gorai.007G275600.2.D;Gorai.007G275600.1.A;Gorai.007G275600.2.A;Gorai.007G275600.5.D;Gorai.007G275600.4.D;Gorai.007G275600.4.A;Gorai.007G275600.3.D;G
Gene IDs: Gorai.007G275600
cotton name: K11583 - serine/threonine-protein phosphatase 2A regulatory subunit B'' (PPP2R3) (1 of 6)
OG IDs: 93522553
Best-hit-arabi-ID: AT5G28850.2
Best-hit-arabi-name: Calcium-binding EF-hand family protein
Gorai.007G275600.1.D
PCC=0.015, p=0.950
Gorai.007G275600.1.A
PCC=0.297, p=0.218
mRNA Profile
Protein Expression Profile
DPA

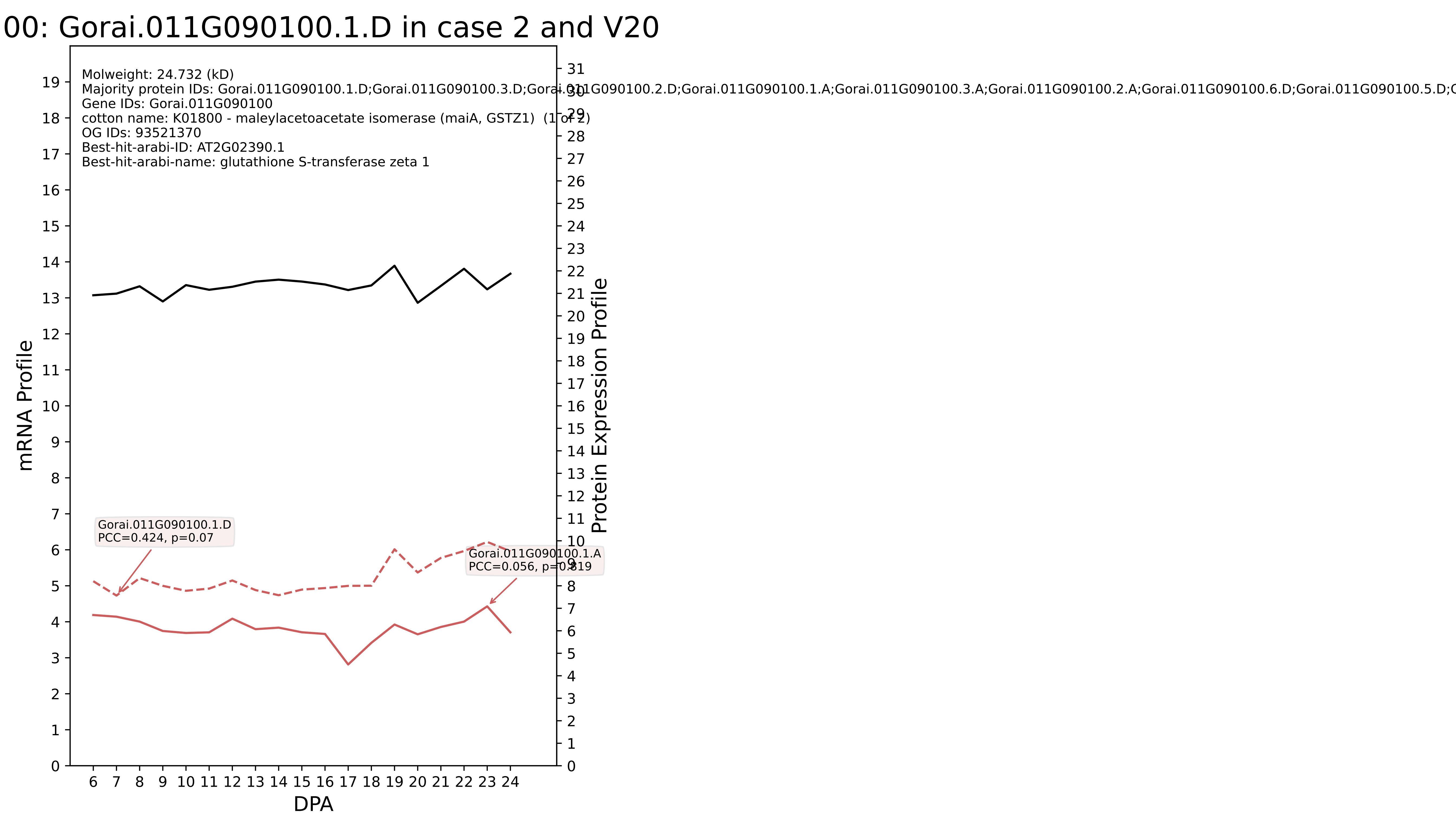
00: Gorai.011G090100.1.D in case 2 and V20
Molweight: 24.732 (kD)
Majority protein IDs: Gorai.011G090100.1.D;Gorai.011G090100.3.D;Gorai.011G090100.2.D;Gorai.011G090100.1.A;Gorai.011G090100.3.A;Gorai.011G090100.2.A;Gorai.011G090100.6.D;Gorai.011G090100.5.D;G
Gene IDs: Gorai.011G090100
cotton name: K01800 - maleylacetoacetate isomerase (maiA, GSTZ1) (1 of 2)
OG IDs: 93521370
Best-hit-arabi-ID: AT2G02390.1
Best-hit-arabi-name: glutathione S-transferase zeta 1
Gorai.011G090100.1.D
PCC=0.424, p=0.07
Gorai.011G090100.1.A
PCC=0.056, p=0.819
mRNA Profile
Protein Expression Profile
DPA

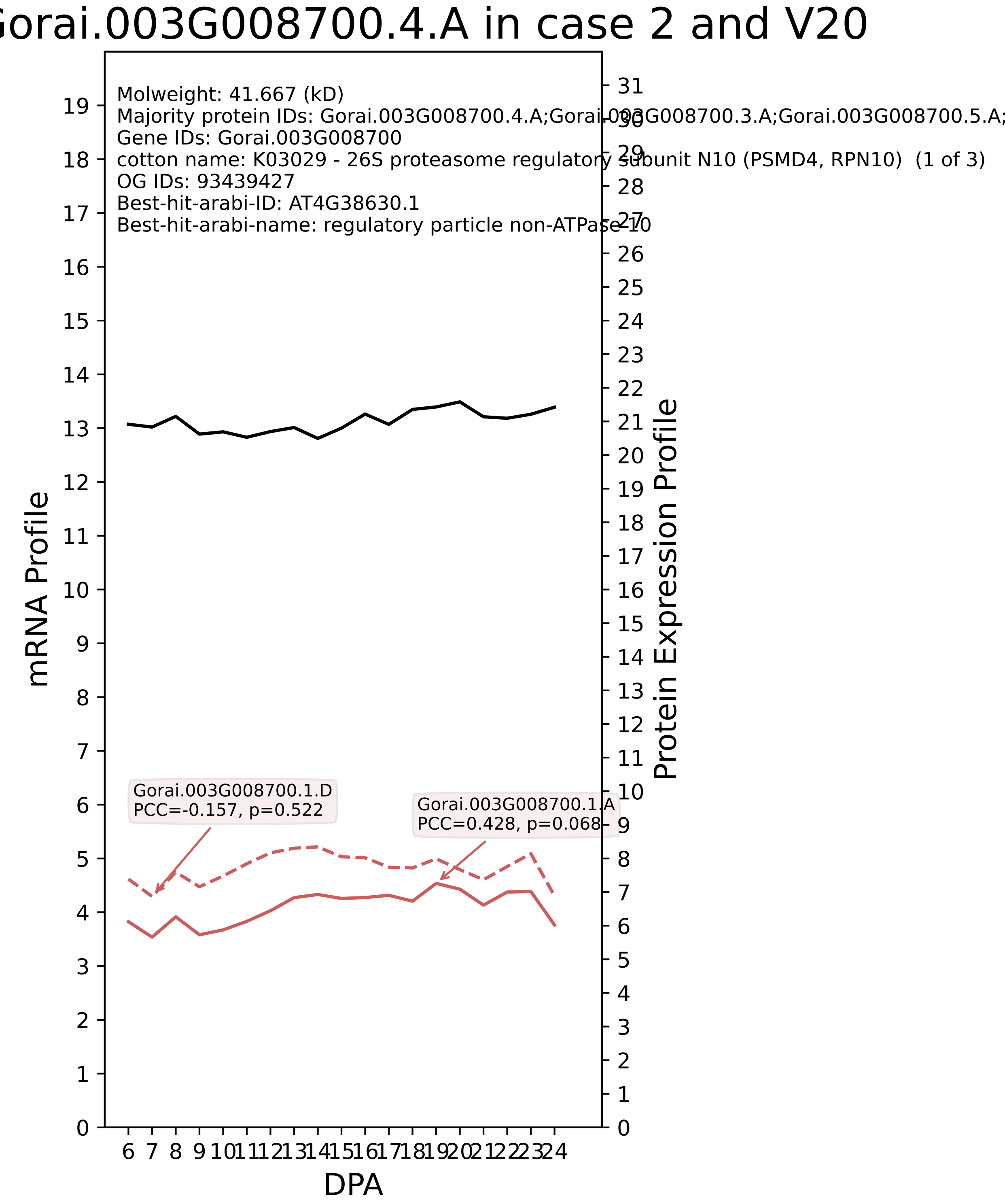
Gorai.003G008700.4.A in case 2 and V20
Molweight: 41.667 (kD)
Majority protein IDs: Gorai.003G008700.4.A;Gorai.003G008700.3.A;Gorai.003G008700.5.A;Gorai.003G008700.2.A;Gorai.003G008700.1.A;Gorai.003G008700.4.D;Gorai.003G008700.3.D;Gorai.003G008700.5.D;G
Gene IDs: Gorai.003G008700
cotton name: K03029 - 26S proteasome regulatory subunit N10 (PSMD4, RPN10) (1 of 3)
OG IDs: 93439427
Best-hit-arabi-ID: AT4G38630.1
Best-hit-arabi-name: regulatory particle non-ATPase 10
Gorai.003G008700.1.D
PCC=-0.157, p=0.522
Gorai.003G008700.1.A
PCC=0.428, p=0.068
mRNA Profile
Protein Expression Profile
DPA

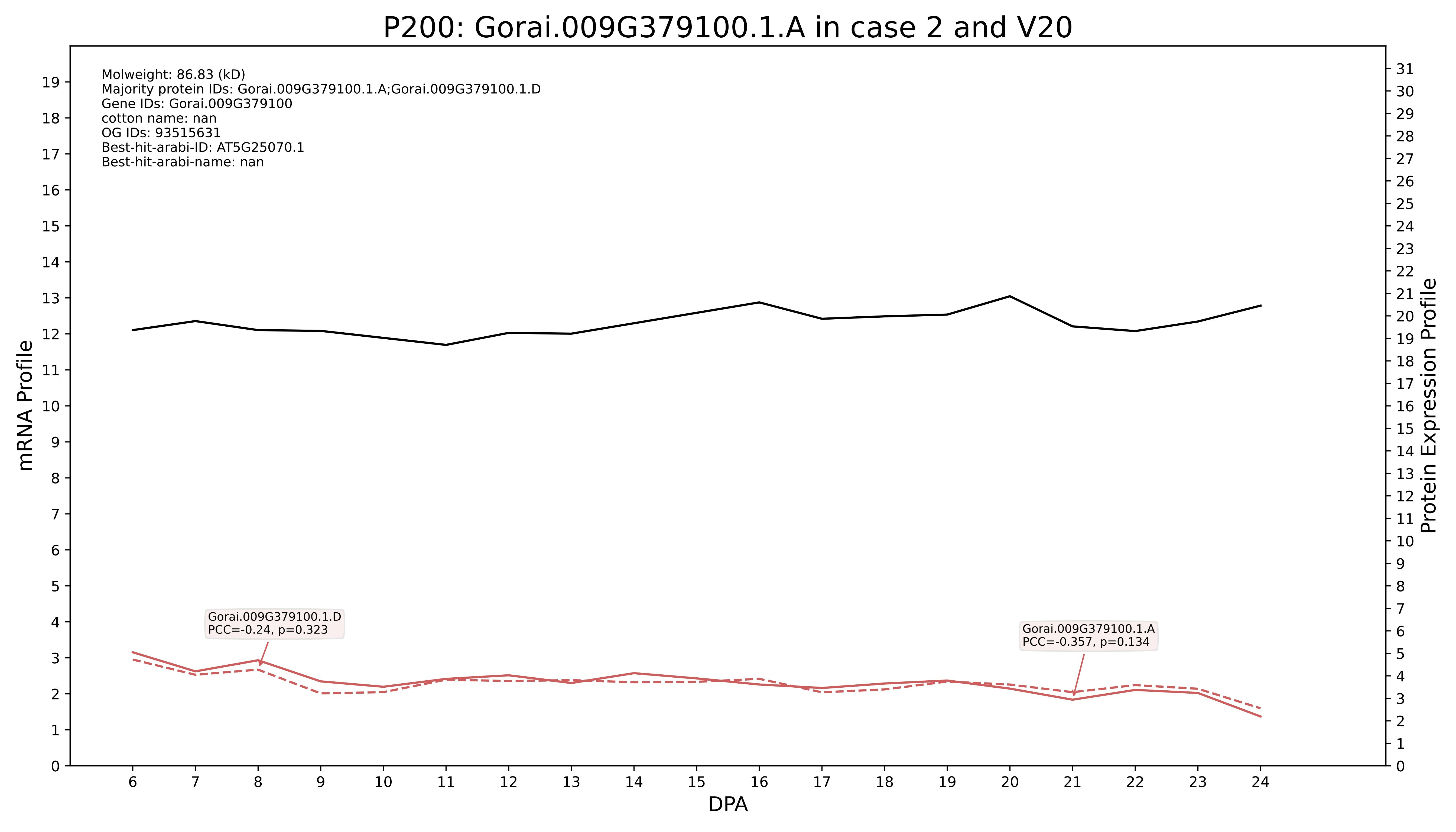
P200: Gorai.009G379100.1.A in case 2 and V20
Molweight: 86.83 (kD)
Majority protein IDs: Gorai.009G379100.1.A;Gorai.009G379100.1.D
Gene IDs: Gorai.009G379100
cotton name: nan
OG IDs: 93515631
Best-hit-arabi-ID: AT5G25070.1
Best-hit-arabi-name: nan
Gorai.009G379100.1.D
PCC=-0.24, p=0.323
Gorai.009G379100.1.A
PCC=-0.357, p=0.134
mRNA Profile
Protein Expression Profile
DPA

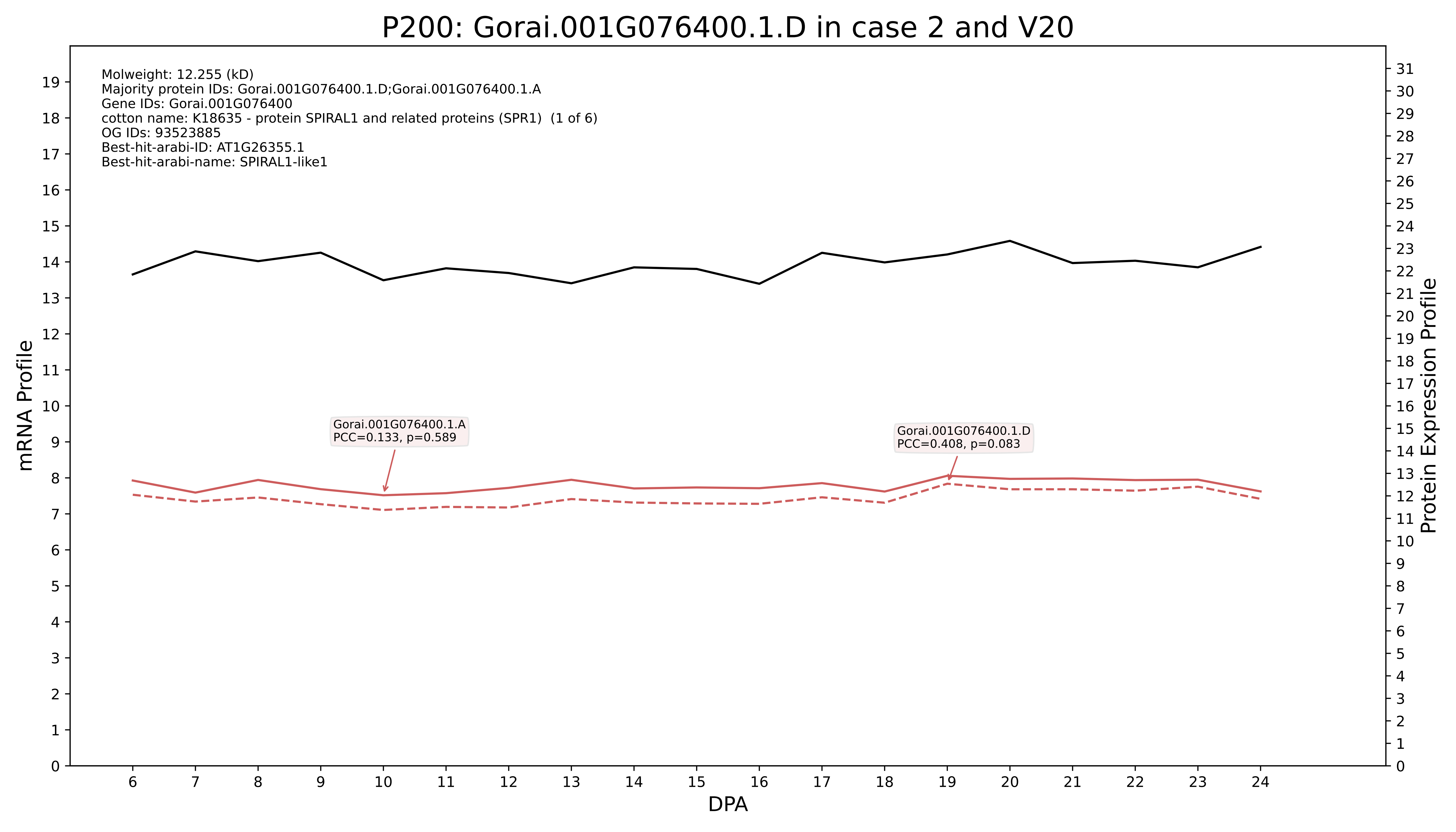
P200: Gorai.001G076400.1.D in case 2 and V20
Molweight: 12.255 (kD)
Majority protein IDs: Gorai.001G076400.1.D;Gorai.001G076400.1.A
Gene IDs: Gorai.001G076400
cotton name: K18635 - protein SPIRAL1 and related proteins (SPR1) (1 of 6)
OG IDs: 93523885
Best-hit-arabi-ID: AT1G26355.1
Best-hit-arabi-name: SPIRAL1-like1
Gorai.001G076400.1.A
PCC=0.133, p=0.589
Gorai.001G076400.1.D
PCC=0.408, p=0.083
mRNA Profile
Protein Expression Profile
DPA

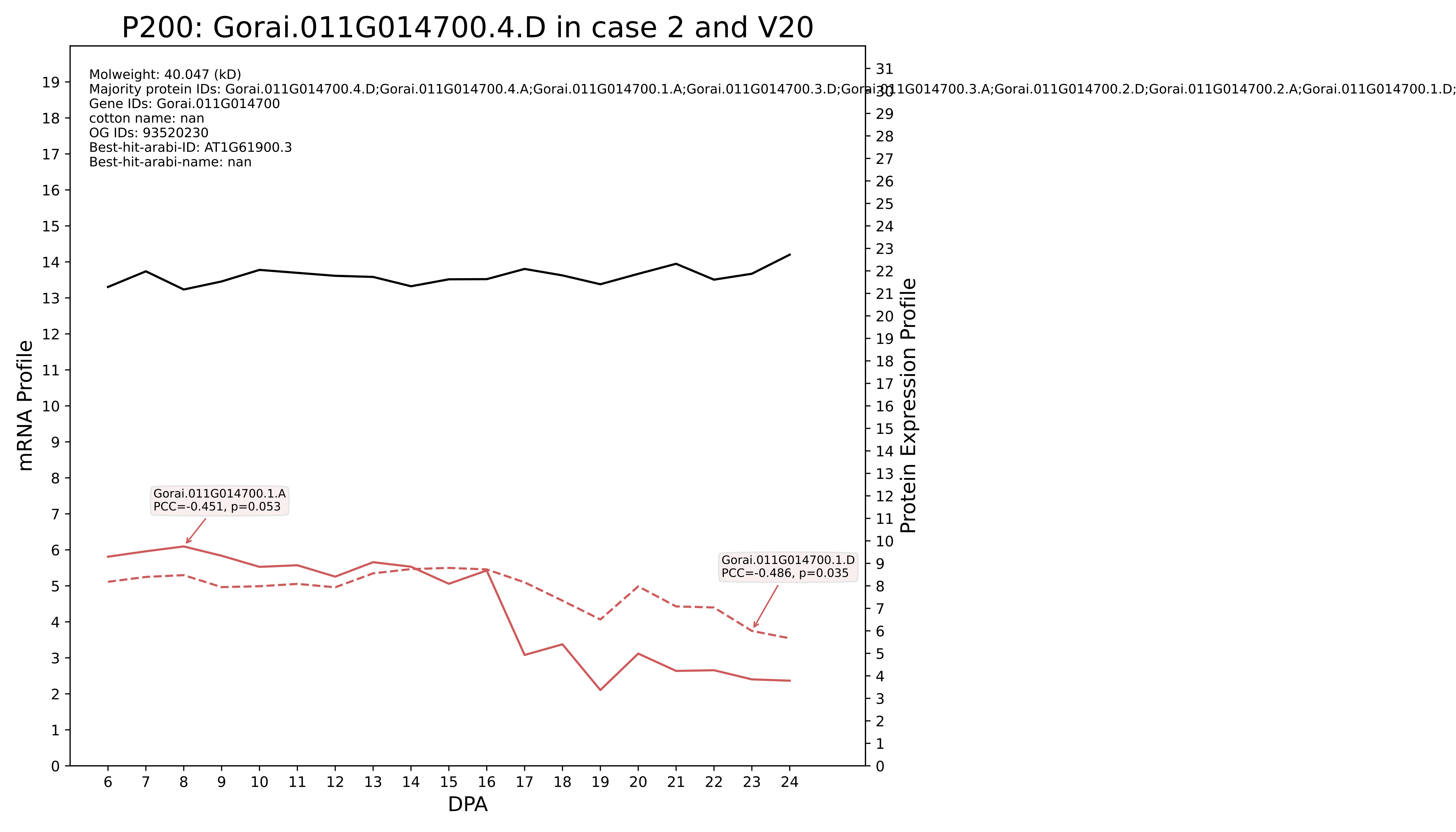
P200: Gorai.011G014700.4.D in case 2 and V20
Molweight: 40.047 (kD)
Majority protein IDs: Gorai.011G014700.4.D;Gorai.011G014700.4.A;Gorai.011G014700.1.A;Gorai.011G014700.3.D;Gorai.011G014700.3.A;Gorai.011G014700.2.D;Gorai.011G014700.2.A;Gorai.011G014700.1.D;
Gene IDs: Gorai.011G014700
cotton name: nan
OG IDs: 93520230
Best-hit-arabi-ID: AT1G61900.3
Best-hit-arabi-name: nan
Gorai.011G014700.1.A
PCC=-0.451, p=0.053
Gorai.011G014700.1.D
PCC=-0.486, p=0.035
mRNA Profile
Protein Expression Profile
DPA

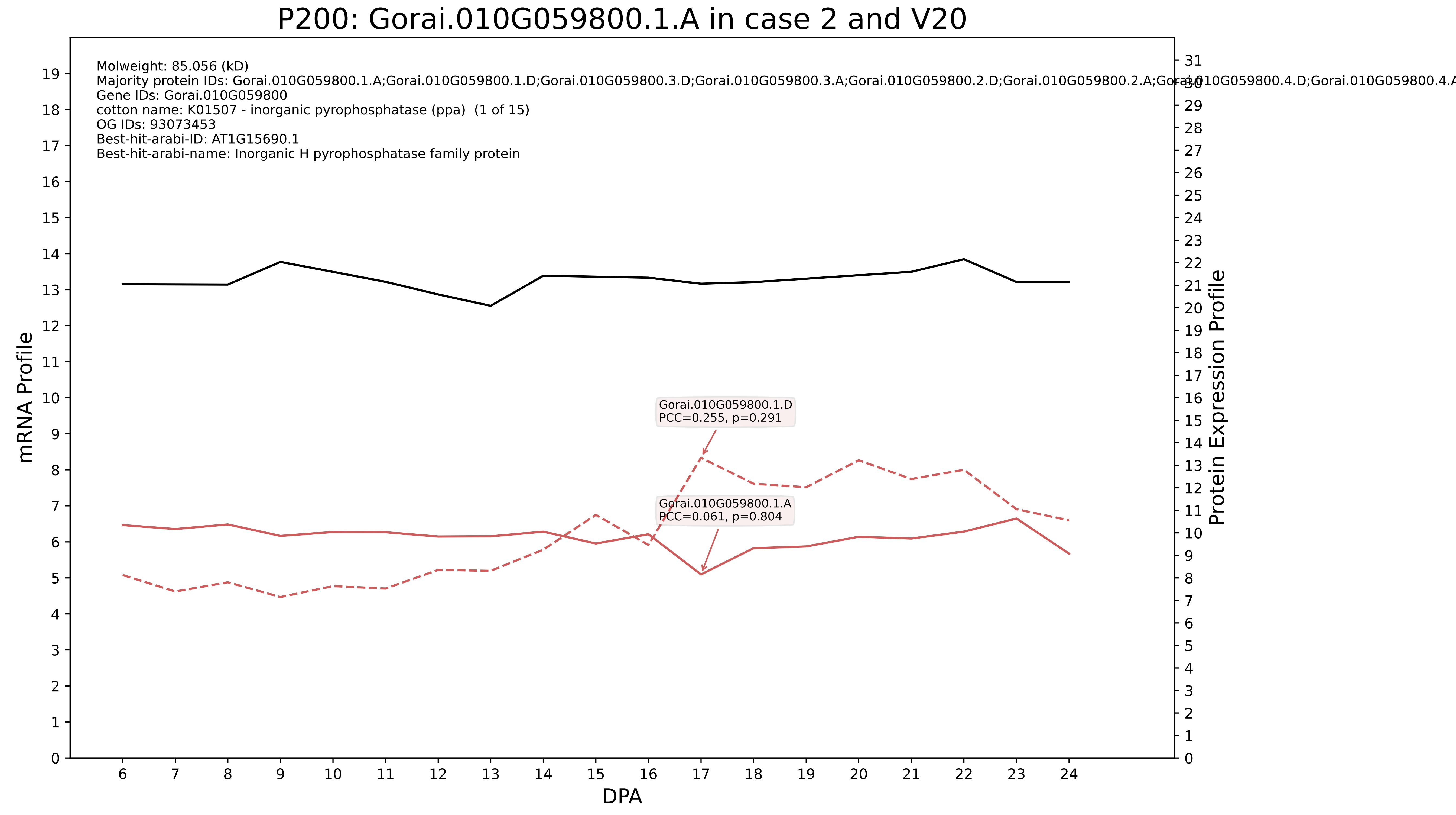
P200: Gorai.010G059800.1.A in case 2 and V20
Molweight: 85.056 (kD)
Majority protein IDs: Gorai.010G059800.1.A;Gorai.010G059800.1.D;Gorai.010G059800.3.D;Gorai.010G059800.3.A;Gorai.010G059800.2.D;Gorai.010G059800.2.A;Gorai.010G059800.4.D;Gorai.010G059800.4.A
Gene IDs: Gorai.010G059800
cotton name: K01507 - inorganic pyrophosphatase (ppa) (1 of 15)
OG IDs: 93073453
Best-hit-arabi-ID: AT1G15690.1
Best-hit-arabi-name: Inorganic H pyrophosphatase family protein
Gorai.010G059800.1.D
PCC=0.255, p=0.291
Gorai.010G059800.1.A
PCC=0.061, p=0.804
mRNA Profile
Protein Expression Profile
DPA

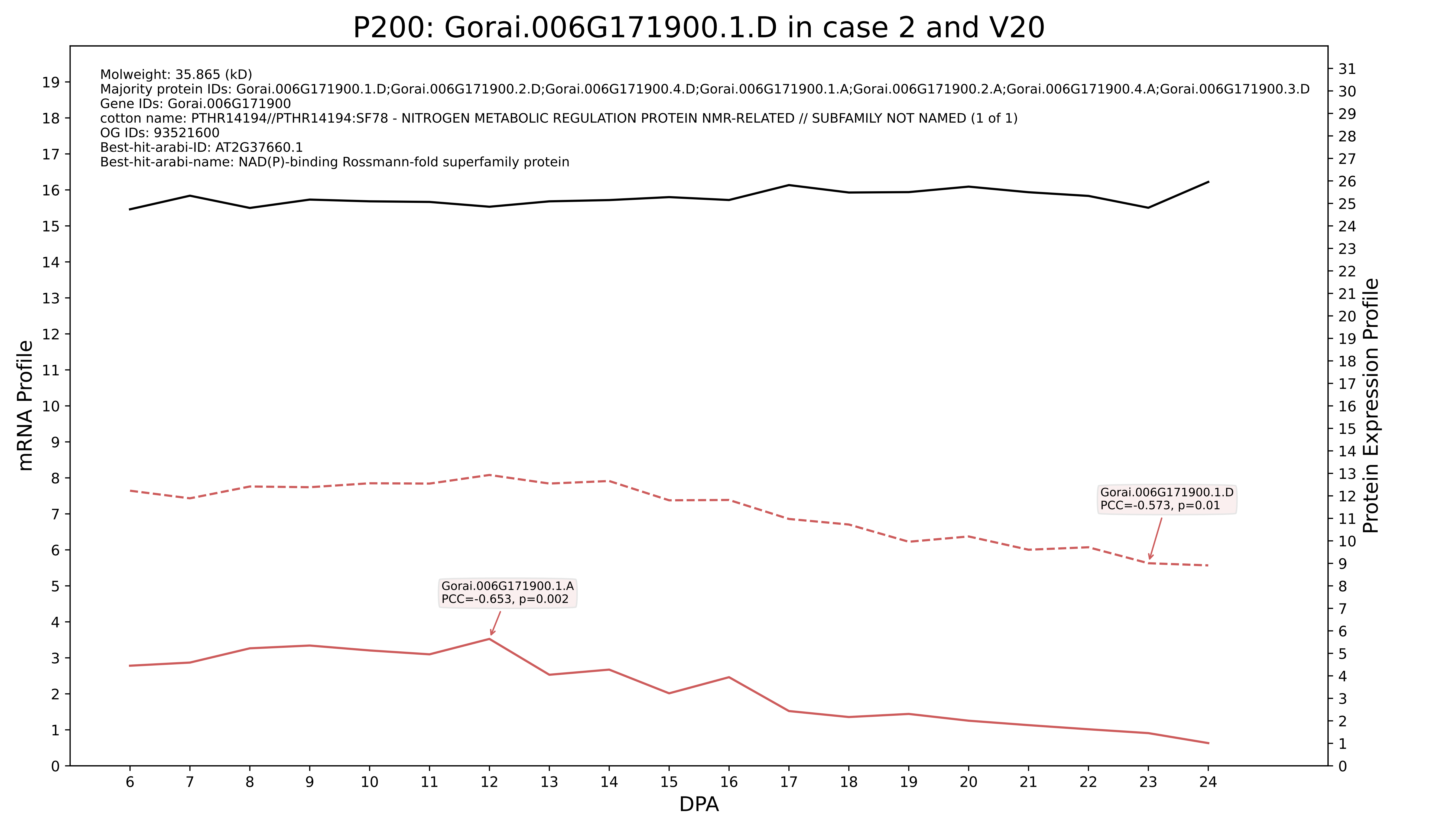
P200: Gorai.006G171900.1.D in case 2 and V20
Molweight: 35.865 (kD)
Majority protein IDs: Gorai.006G171900.1.D;Gorai.006G171900.2.D;Gorai.006G171900.4.D;Gorai.006G171900.1.A;Gorai.006G171900.2.A;Gorai.006G171900.4.A;Gorai.006G171900.3.D
Gene IDs: Gorai.006G171900
cotton name: PTHR14194//PTHR14194:SF78 - NITROGEN METABOLIC REGULATION PROTEIN NMR-RELATED // SUBFAMILY NOT NAMED (1 of 1)
OG IDs: 93521600
Best-hit-arabi-ID: AT2G37660.1
Best-hit-arabi-name: NAD(P)-binding Rossmann-fold superfamily protein
Gorai.006G171900.1.D
PCC=-0.573, p=0.01
Gorai.006G171900.1.A
PCC=-0.653, p=0.002
mRNA Profile
Protein Expression Profile
DPA

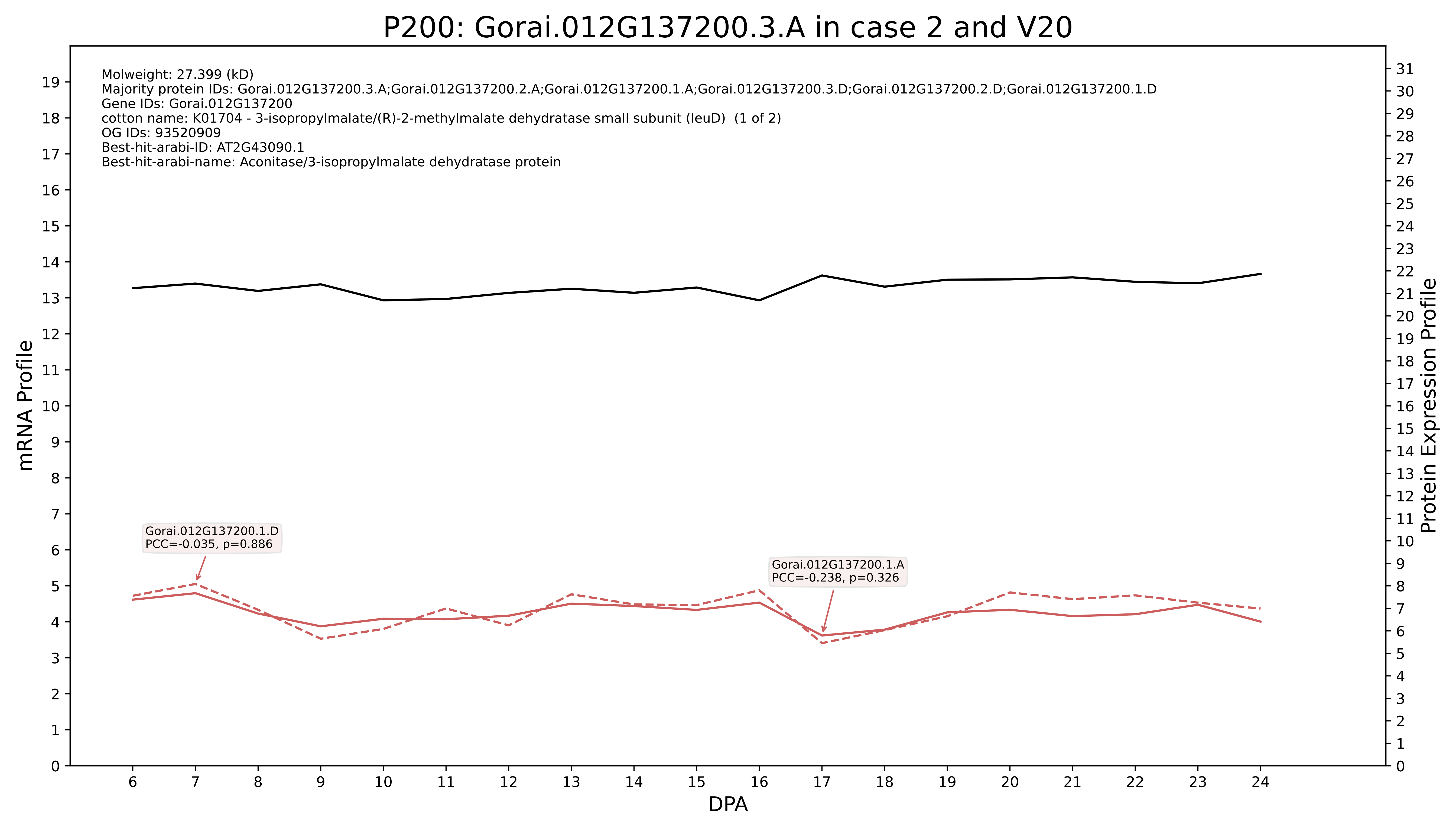
P200: Gorai.012G137200.3.A in case 2 and V20
Molweight: 27.399 (kD)
Majority protein IDs: Gorai.012G137200.3.A;Gorai.012G137200.2.A;Gorai.012G137200.1.A;Gorai.012G137200.3.D;Gorai.012G137200.2.D;Gorai.012G137200.1.D
Gene IDs: Gorai.012G137200
cotton name: K01704 - 3-isopropylmalate/(R)-2-methylmalate dehydratase small subunit (leuD) (1 of 2)
OG IDs: 93520909
Best-hit-arabi-ID: AT2G43090.1
Best-hit-arabi-name: Aconitase/3-isopropylmalate dehydratase protein
Gorai.012G137200.1.D
PCC=-0.035, p=0.886
Gorai.012G137200.1.A
PCC=-0.238, p=0.326
mRNA Profile
Protein Expression Profile
DPA

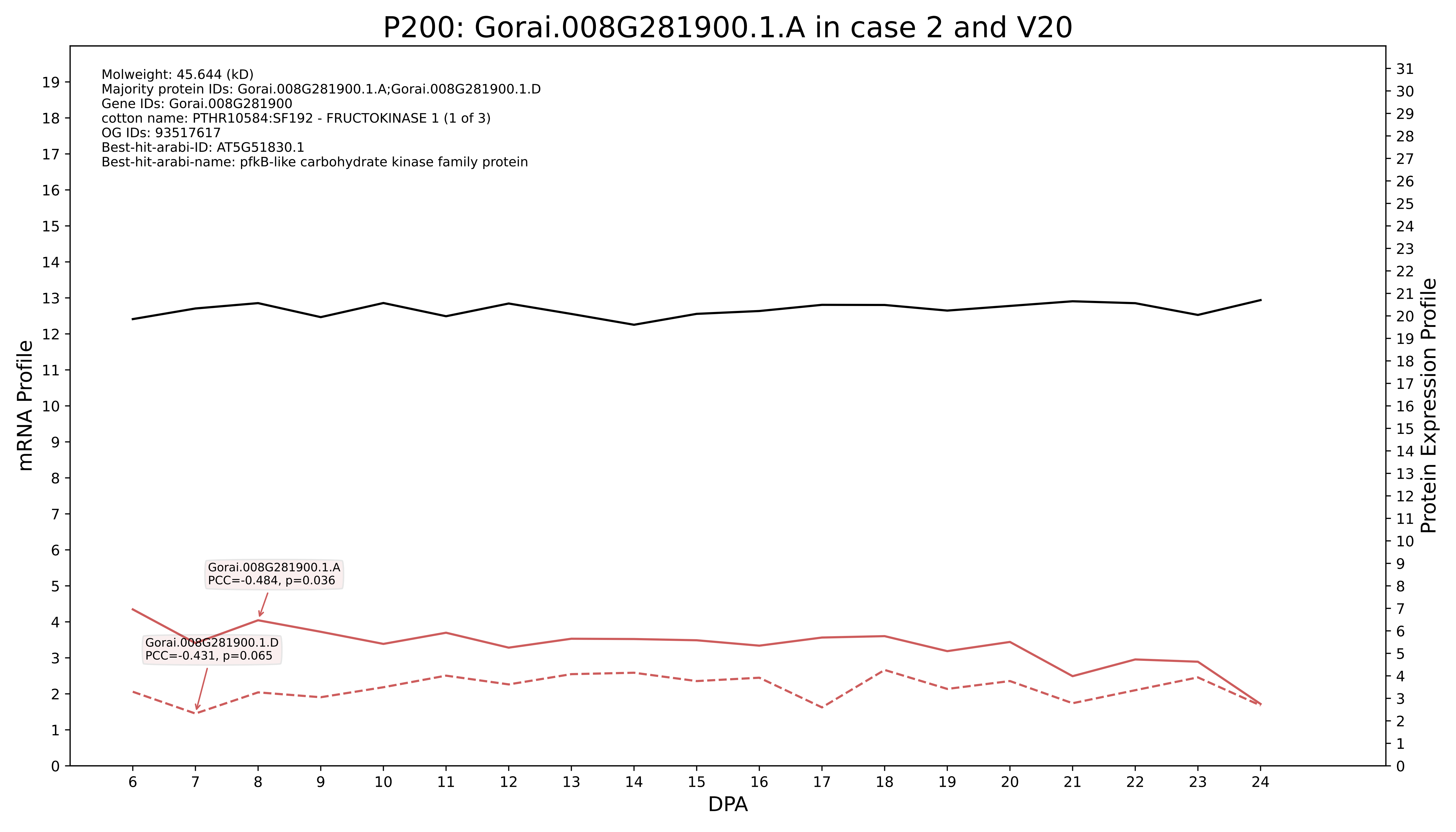
P200: Gorai.008G281900.1.A in case 2 and V20
Molweight: 45.644 (kD)
Majority protein IDs: Gorai.008G281900.1.A;Gorai.008G281900.1.D
Gene IDs: Gorai.008G281900
cotton name: PTHR10584:SF192 - FRUCTOKINASE 1 (1 of 3)
OG IDs: 93517617
Best-hit-arabi-ID: AT5G51830.1
Best-hit-arabi-name: pfkB-like carbohydrate kinase family protein
Gorai.008G281900.1.A
PCC=-0.484, p=0.036
Gorai.008G281900.1.D
PCC=-0.431, p=0.065
mRNA Profile
Protein Expression Profile
DPA

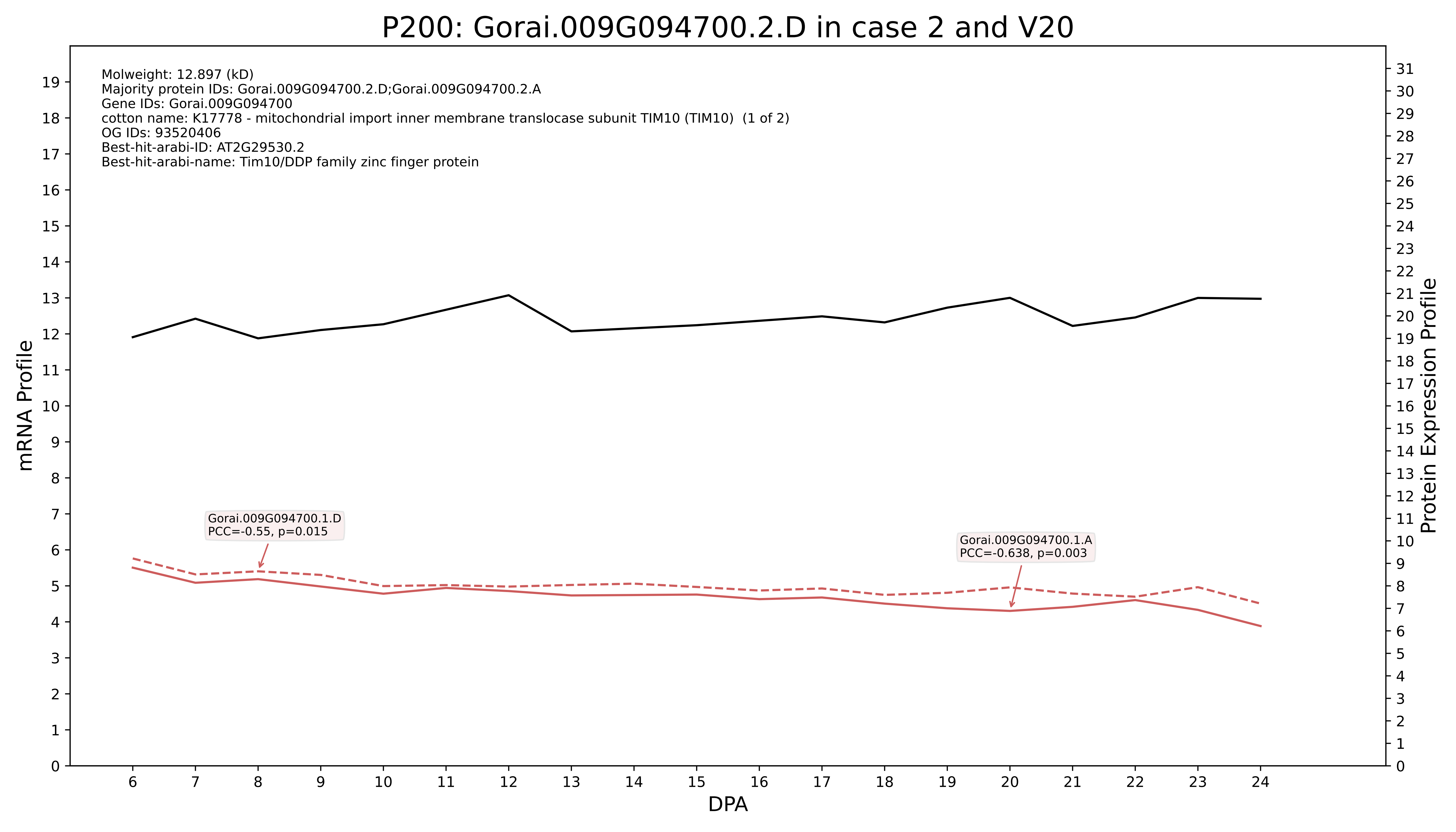
P200: Gorai.009G094700.2.D in case 2 and V20
Molweight: 12.897 (kD)
Majority protein IDs: Gorai.009G094700.2.D;Gorai.009G094700.2.A
Gene IDs: Gorai.009G094700
cotton name: K17778 - mitochondrial import inner membrane translocase subunit TIM10 (TIM10) (1 of 2)
OG IDs: 93520406
Best-hit-arabi-ID: AT2G29530.2
Best-hit-arabi-name: Tim10/DDP family zinc finger protein
Gorai.009G094700.1.D
PCC=-0.55, p=0.015
Gorai.009G094700.1.A
PCC=-0.638, p=0.003
mRNA Profile
Protein Expression Profile
DPA

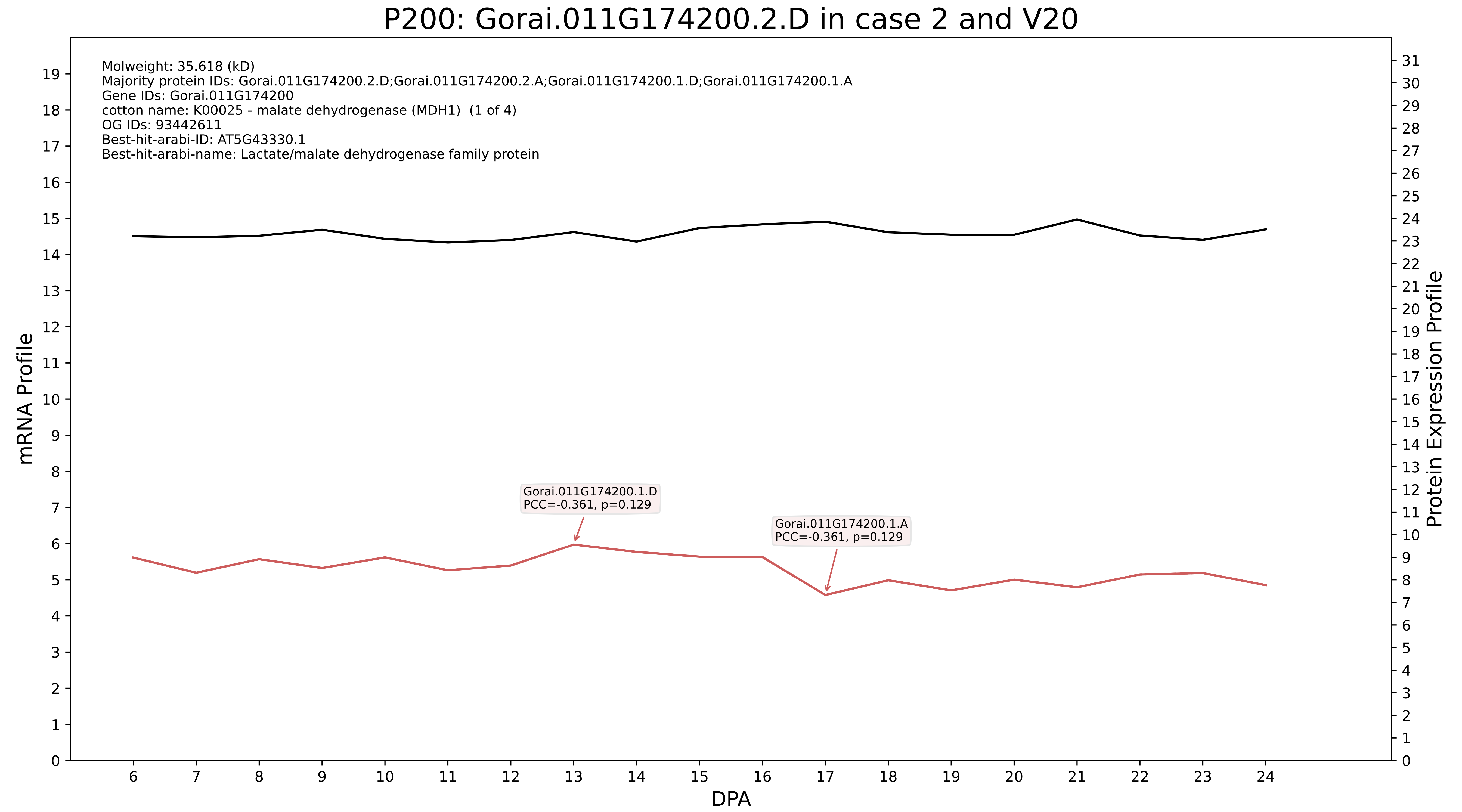
P200: Gorai.011G174200.2.D in case 2 and V20
Molweight: 35.618 (kD)
Majority protein IDs: Gorai.011G174200.2.D;Gorai.011G174200.2.A;Gorai.011G174200.1.D;Gorai.011G174200.1.A
Gene IDs: Gorai.011G174200
cotton name: K00025 - malate dehydrogenase (MDH1) (1 of 4)
OG IDs: 93442611
Best-hit-arabi-ID: AT5G43330.1
Best-hit-arabi-name: Lactate/malate dehydrogenase family protein
Gorai.011G174200.1.D
PCC=-0.361, p=0.129
Gorai.011G174200.1.A
PCC=-0.361, p=0.129
mRNA Profile
Protein Expression Profile
DPA

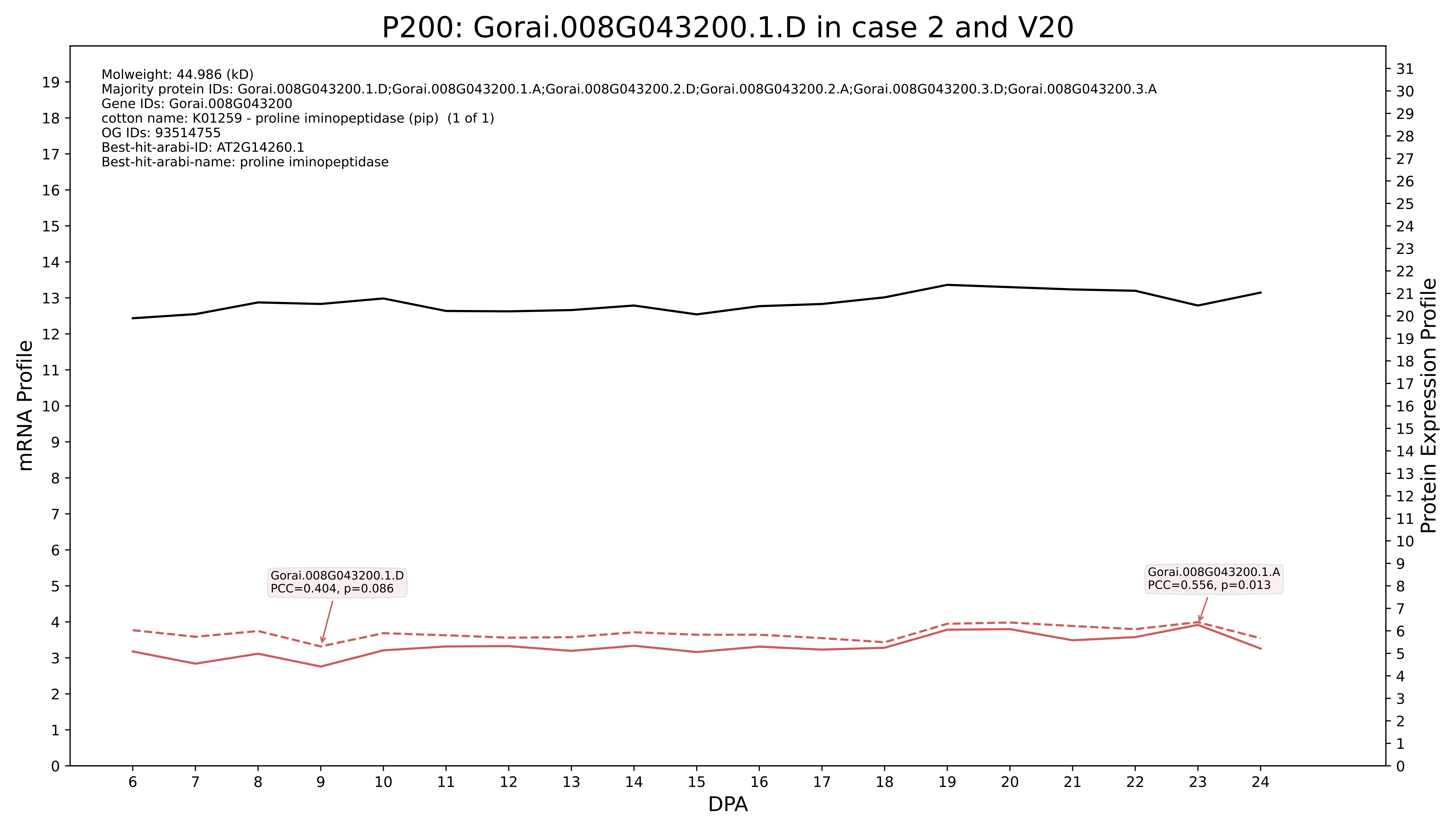

P200: Gorai.008G043200.1.D in case 2 and V20
Molweight: 44.986 (kD)
Majority protein IDs: Gorai.008G043200.1.D;Gorai.008G043200.1.A;Gorai.008G043200.2.D;Gorai.008G043200.2.A;Gorai.008G043200.3.D;Gorai.008G043200.3.A
Gene IDs: Gorai.008G043200
cotton name: K01259 - proline iminopeptidase (pip) (1 of 1)
OG IDs: 93514755
Best-hit-arabi-ID: AT2G14260.1
Best-hit-arabi-name: proline iminopeptidase
Gorai.008G043200.1.D
PCC=0.404, p=0.086
Gorai.008G043200.1.A
PCC=0.556, p=0.013
mRNA Profile
Protein Expression Profile
DPA

### P200: Gorai.005G093700.2.D in case 2 and V20

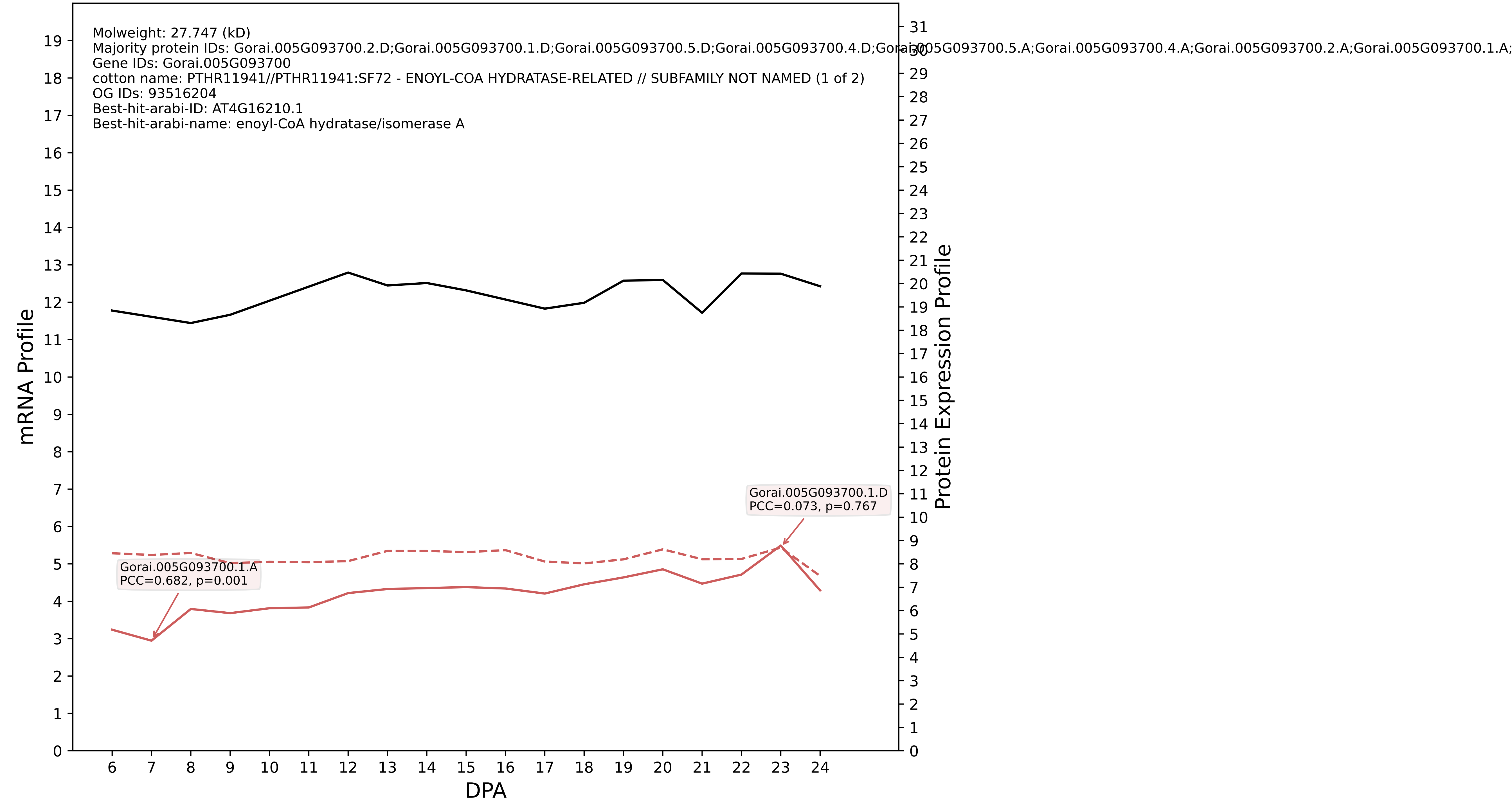

### P200: Gorai.004G147800.1.A in case 2 and V20

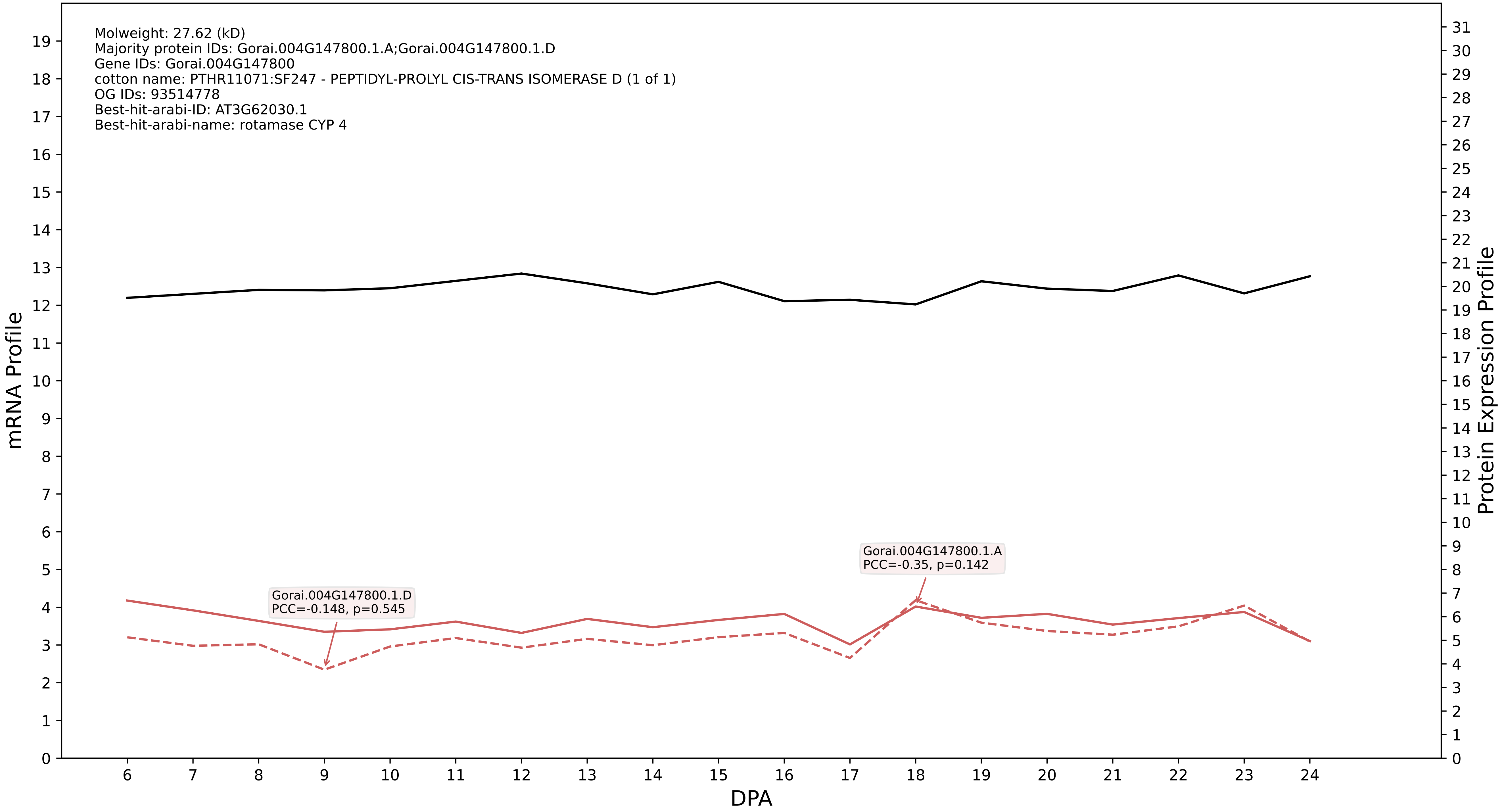

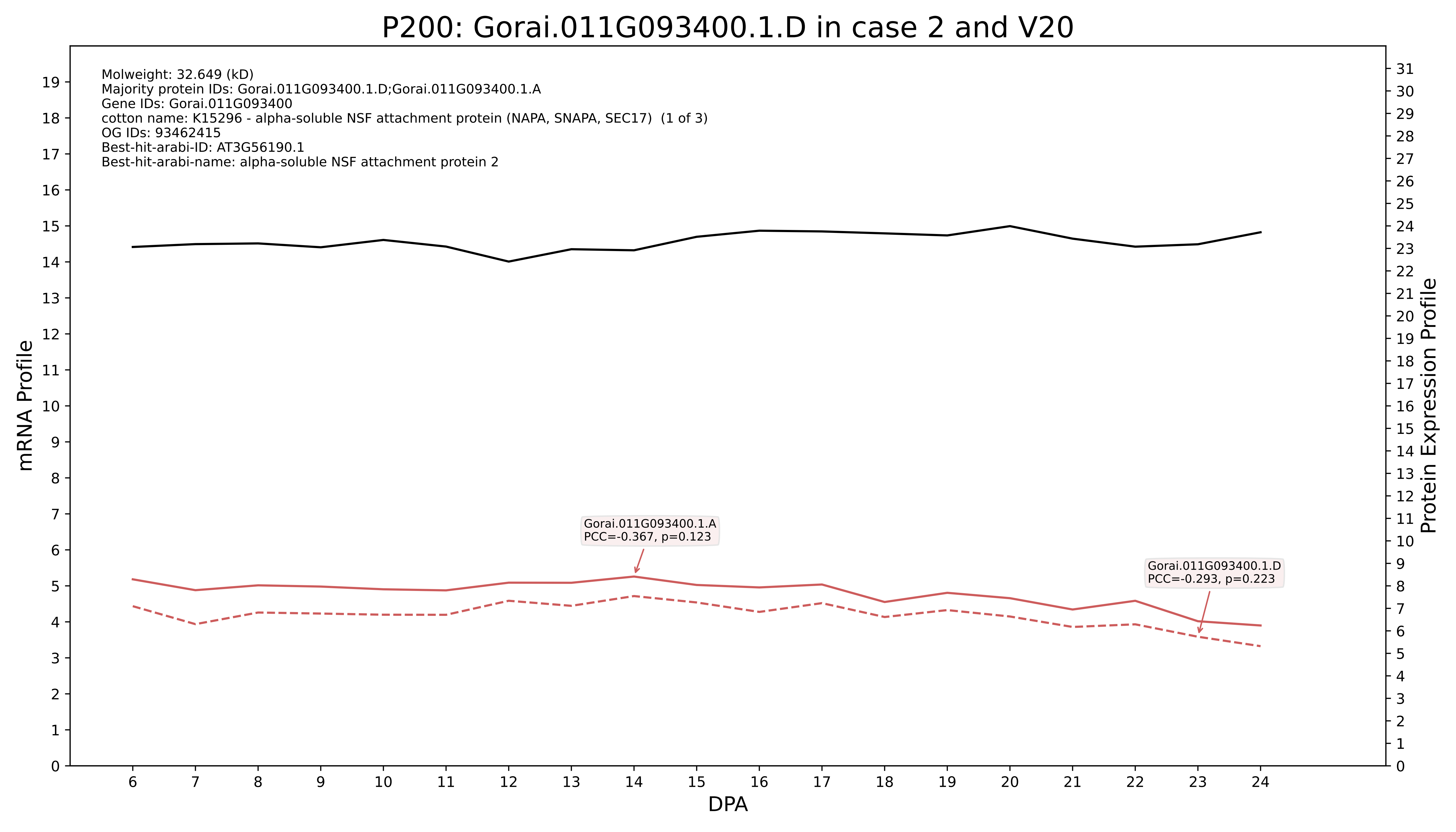
P200: Gorai.011G093400.1.D in case 2 and V20
Molweight: 32.649 (kD)
Majority protein IDs: Gorai.011G093400.1.D;Gorai.011G093400.1.A
Gene IDs: Gorai.011G093400
cotton name: K15296 - alpha-soluble NSF attachment protein (NAPA, SNAPA, SEC17) (1 of 3)
OG IDs: 93462415
Best-hit-arabi-ID: AT3G56190.1
Best-hit-arabi-name: alpha-soluble NSF attachment protein 2
Gorai.011G093400.1.A
PCC=-0.367, p=0.123
Gorai.011G093400.1.D
PCC=-0.293, p=0.223
mRNA Profile
Protein Expression Profile
DPA

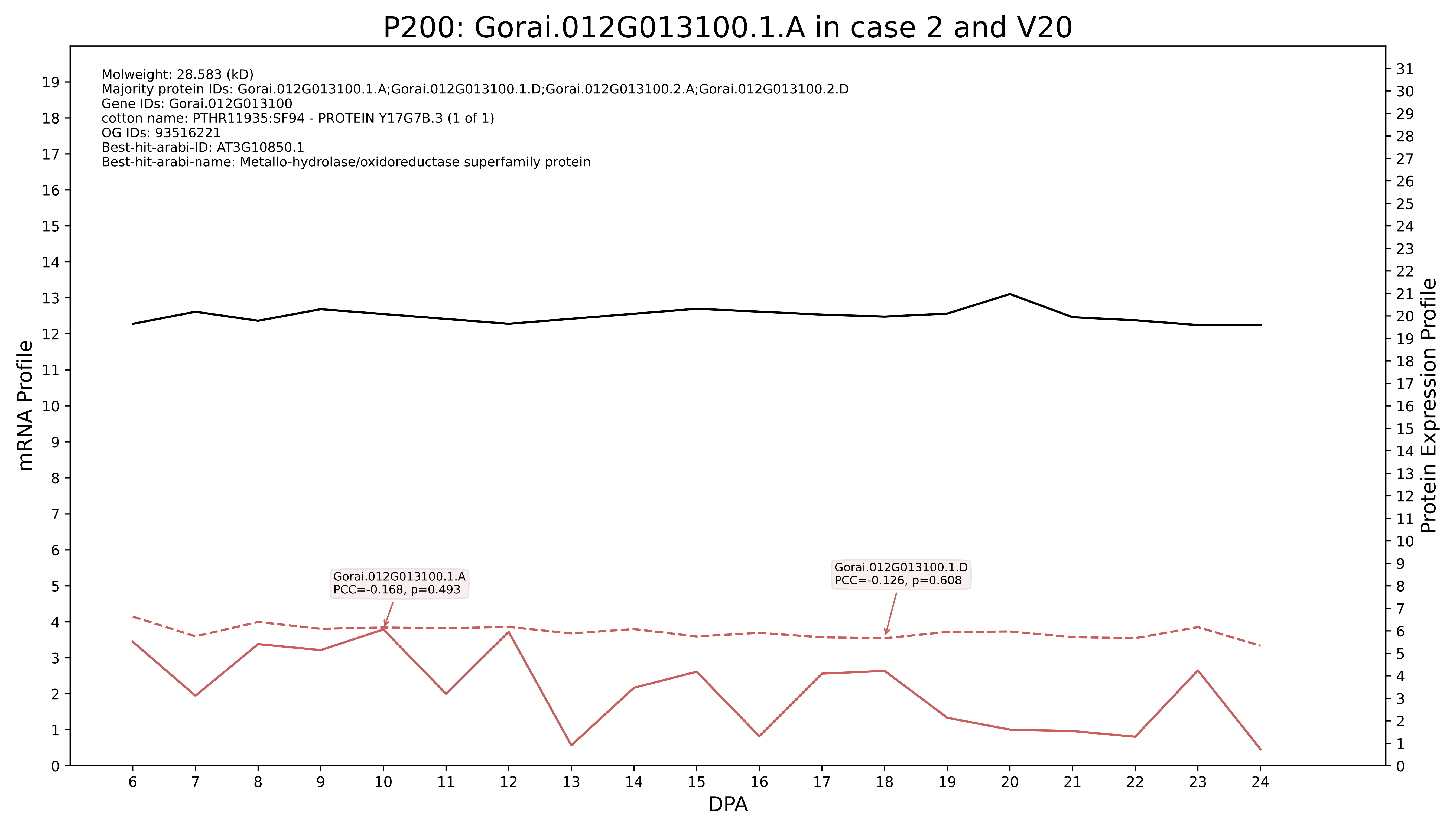
P200: Gorai.012G013100.1.A in case 2 and V20
Molweight: 28.583 (kD)
Majority protein IDs: Gorai.012G013100.1.A;Gorai.012G013100.1.D;Gorai.012G013100.2.A;Gorai.012G013100.2.D
Gene IDs: Gorai.012G013100
cotton name: PTHR11935:SF94 - PROTEIN Y17G7B.3 (1 of 1)
OG IDs: 93516221
Best-hit-arabi-ID: AT3G10850.1
Best-hit-arabi-name: Metallo-hydrolase/oxidoreductase superfamily protein
Gorai.012G013100.1.A
PCC=-0.168, p=0.493
Gorai.012G013100.1.D
PCC=-0.126, p=0.608
mRNA Profile
Protein Expression Profile
DPA

005G146400.7.D in case 2 and V20

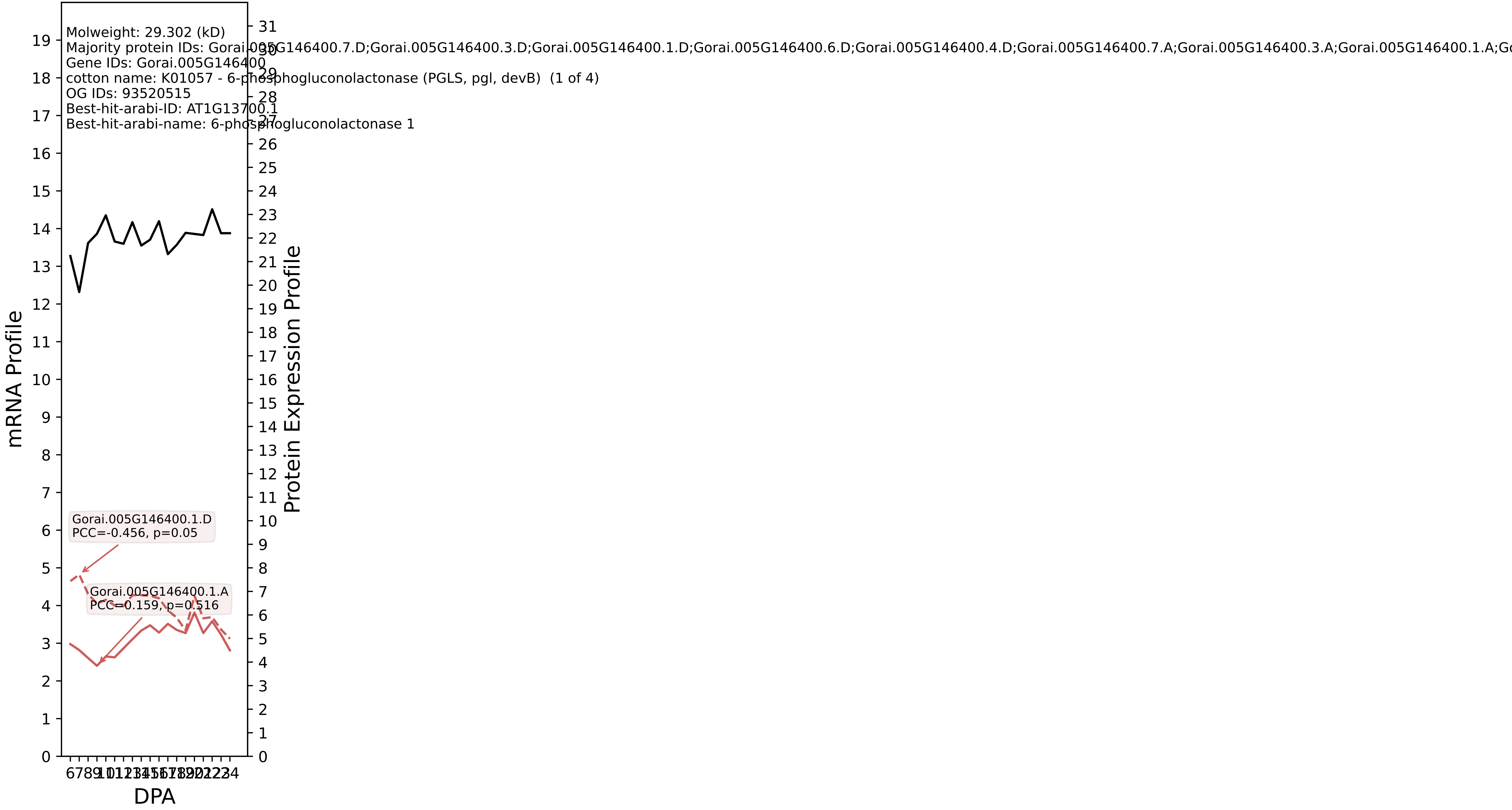

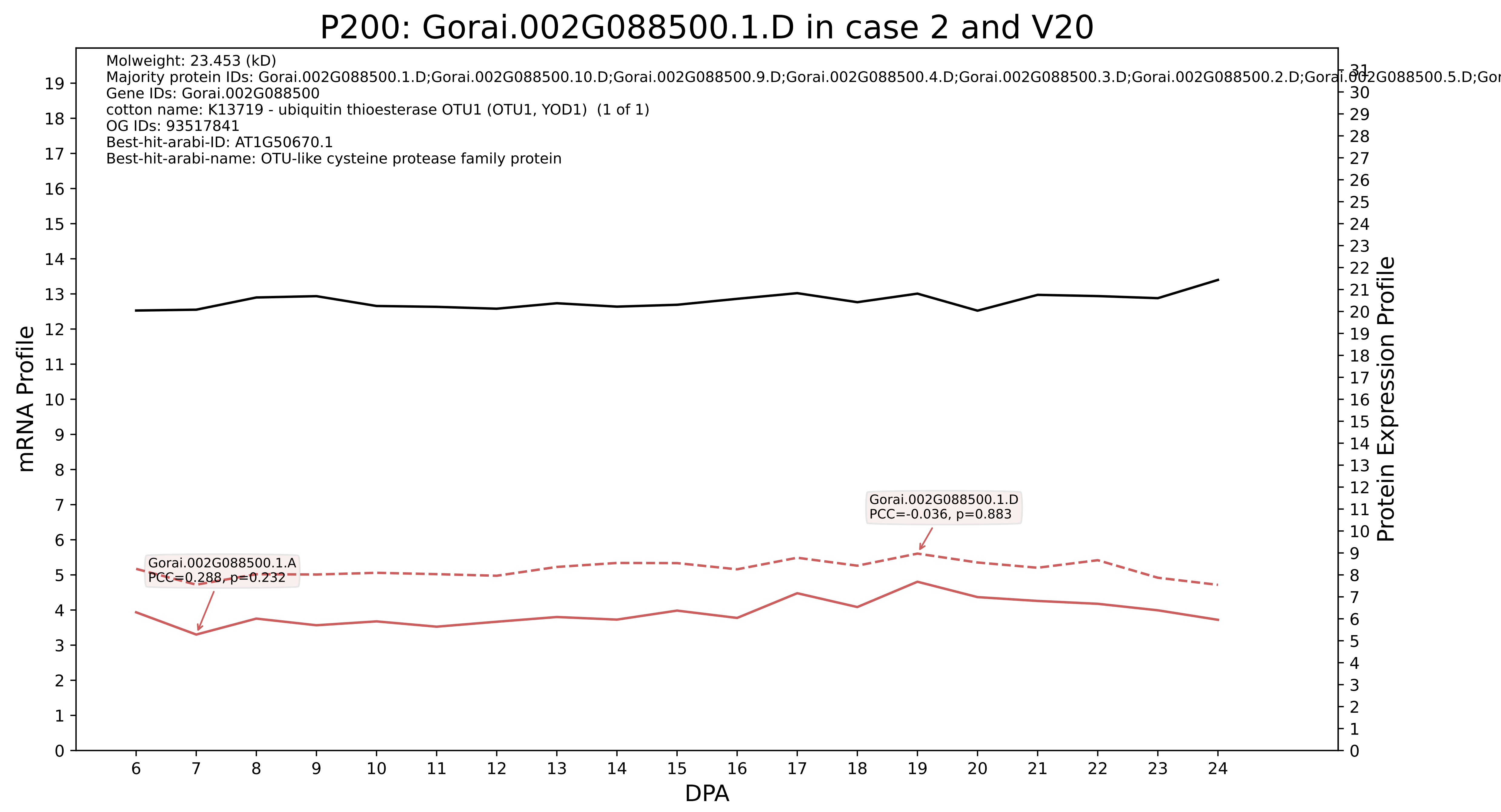

P200: Gorai.002G088500.1.D in case 2 and V20
Molweight: 23.453 (kD)
Majority protein IDs: Gorai.002G088500.1.D;Gorai.002G088500.10.D;Gorai.002G088500.9.D;Gorai.002G088500.4.D;Gorai.002G088500.3.D;Gorai.002G088500.2.D;Gorai.002G088500.5.D;Gor
Gene IDs: Gorai.002G088500
cotton name: K13719 - ubiquitin thioesterase OTU1 (OTU1, YOD1) (1 of 1)
OG IDs: 93517841
Best-hit-arabi-ID: AT1G50670.1
Best-hit-arabi-name: OTU-like cysteine protease family protein
Gorai.002G088500.1.D
PCC=-0.036, p=0.883
Gorai.002G088500.1.A
PCC=0.288, p=0.232
mRNA Profile
Protein Expression Profile
DPA

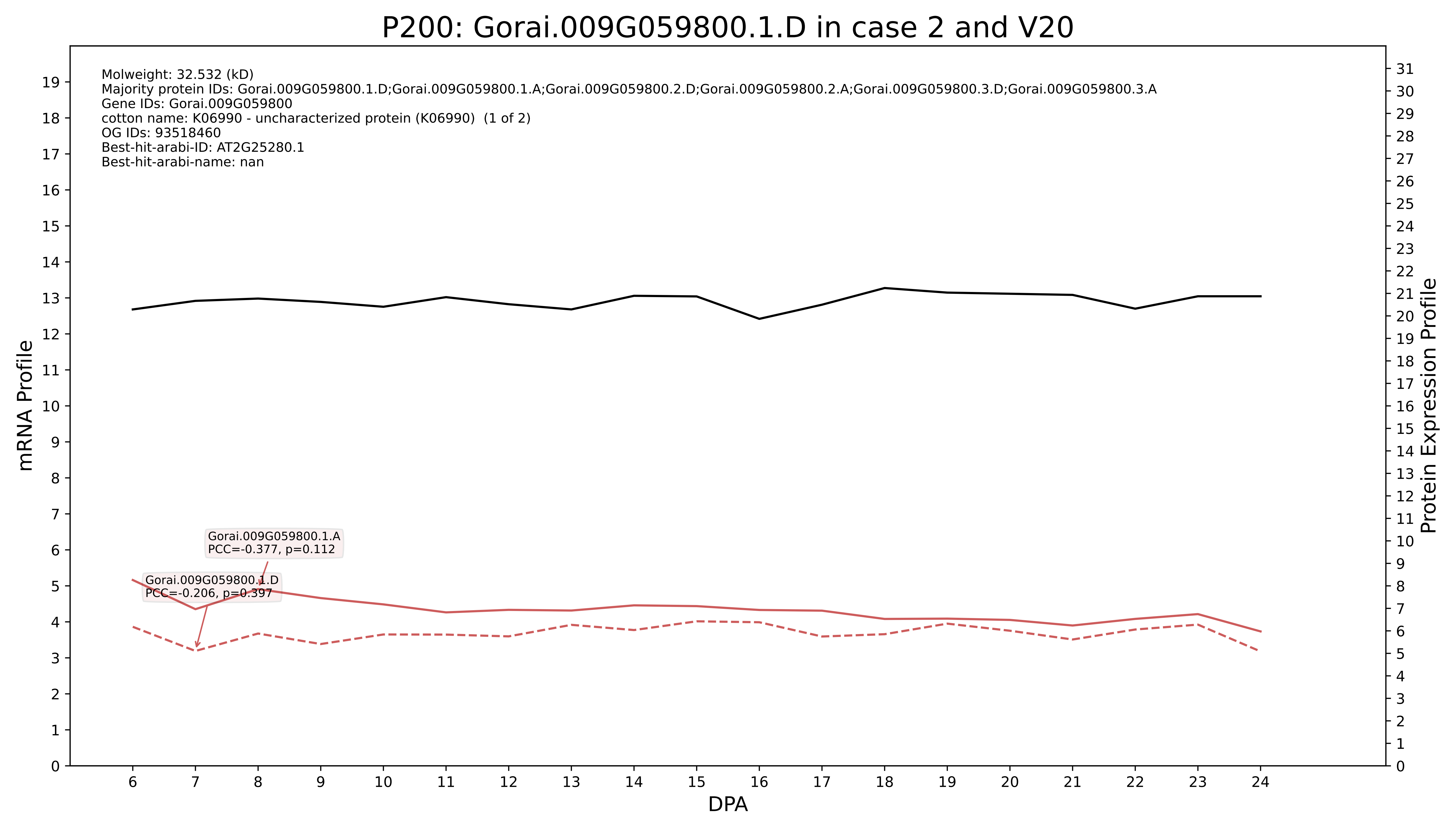

P200: Gorai.009G059800.1.D in case 2 and V20
Molweight: 32.532 (kD)
Majority protein IDs: Gorai.009G059800.1.D;Gorai.009G059800.1.A;Gorai.009G059800.2.D;Gorai.009G059800.2.A;Gorai.009G059800.3.D;Gorai.009G059800.3.A
Gene IDs: Gorai.009G059800
cotton name: K06990 - uncharacterized protein (K06990) (1 of 2)
OG IDs: 93518460
Best-hit-arabi-ID: AT2G25280.1
Best-hit-arabi-name: nan
Gorai.009G059800.1.A
PCC=-0.377, p=0.112
Gorai.009G059800.1.D
PCC=-0.206, p=0.397
mRNA Profile
Protein Expression Profile
DPA

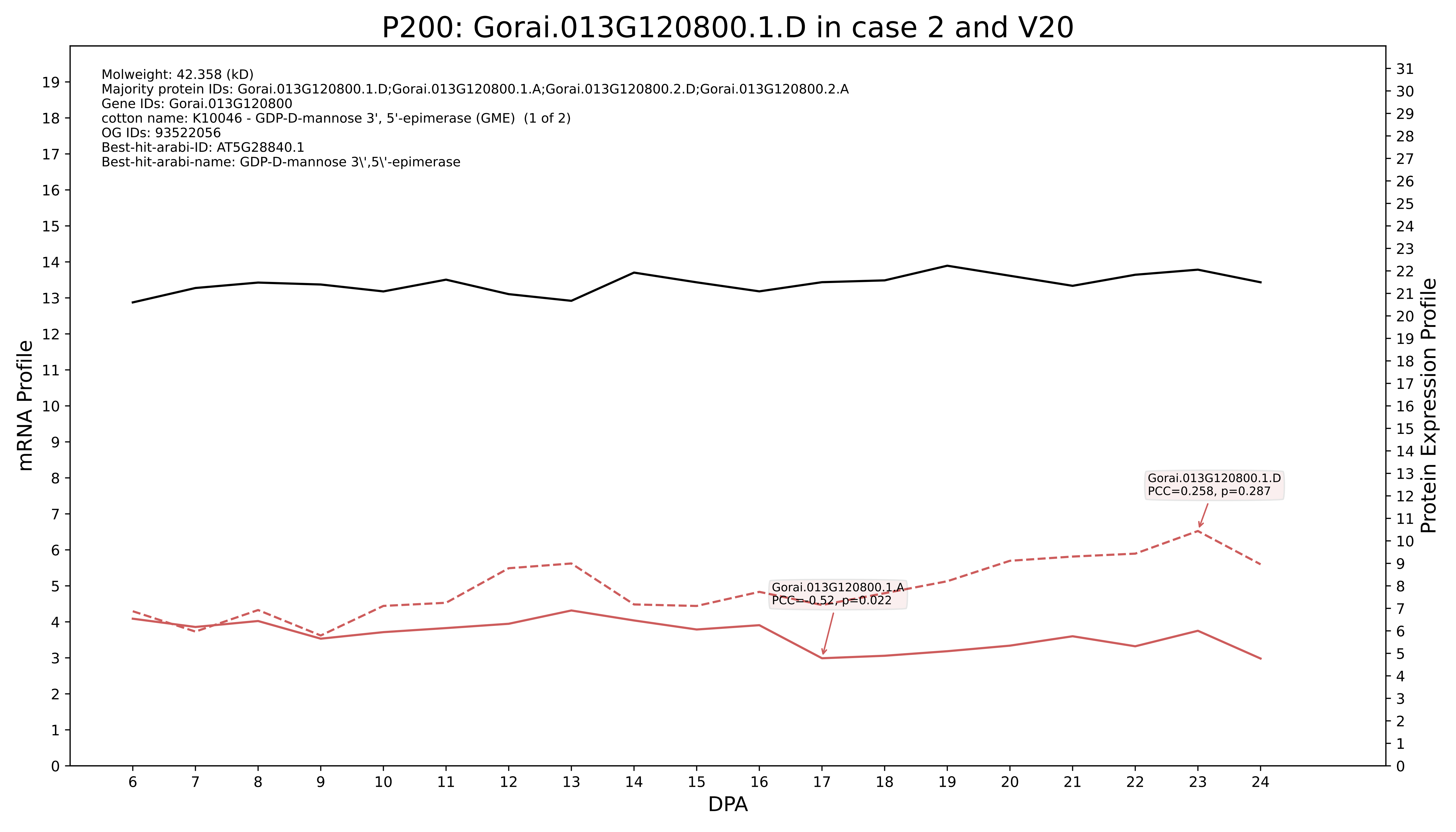
P200: Gorai.013G120800.1.D in case 2 and V20
Molweight: 42.358 (kD)
Majority protein IDs: Gorai.013G120800.1.D;Gorai.013G120800.1.A;Gorai.013G120800.2.D;Gorai.013G120800.2.A
Gene IDs: Gorai.013G120800
cotton name: K10046 - GDP-D-mannose 3', 5'-epimerase (GME) (1 of 2)
OG IDs: 93522056
Best-hit-arabi-ID: AT5G28840.1
Best-hit-arabi-name: GDP-D-mannose 3\',5\'-epimerase
Gorai.013G120800.1.D
PCC=0.258, p=0.287
Gorai.013G120800.1.A
PCC=-0.52, p=0.022
mRNA Profile
Protein Expression Profile
DPA

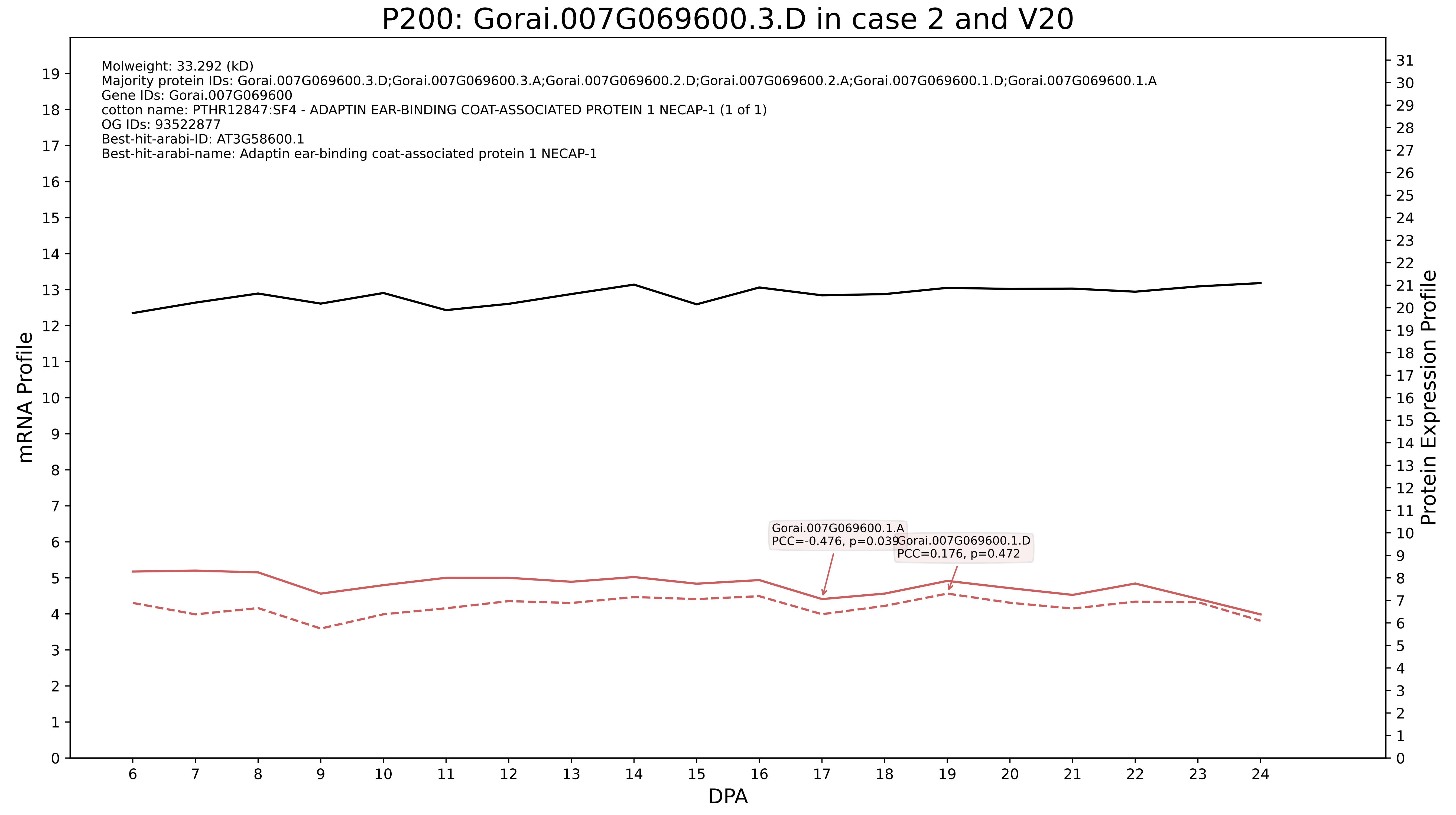
P200: Gorai.007G069600.3.D in case 2 and V20
Molweight: 33.292 (kD)
Majority protein IDs: Gorai.007G069600.3.D;Gorai.007G069600.3.A;Gorai.007G069600.2.D;Gorai.007G069600.2.A;Gorai.007G069600.1.D;Gorai.007G069600.1.A
Gene IDs: Gorai.007G069600
cotton name: PTHR12847:SF4 - ADAPTIN EAR-BINDING COAT-ASSOCIATED PROTEIN 1 NECAP-1 (1 of 1)
OG IDs: 93522877
Best-hit-arabi-ID: AT3G58600.1
Best-hit-arabi-name: Adaptin ear-binding coat-associated protein 1 NECAP-1
Gorai.007G069600.1.A
PCC=-0.476, p=0.039
Gorai.007G069600.1.D
PCC=0.176, p=0.472
mRNA Profile
Protein Expression Profile
DPA

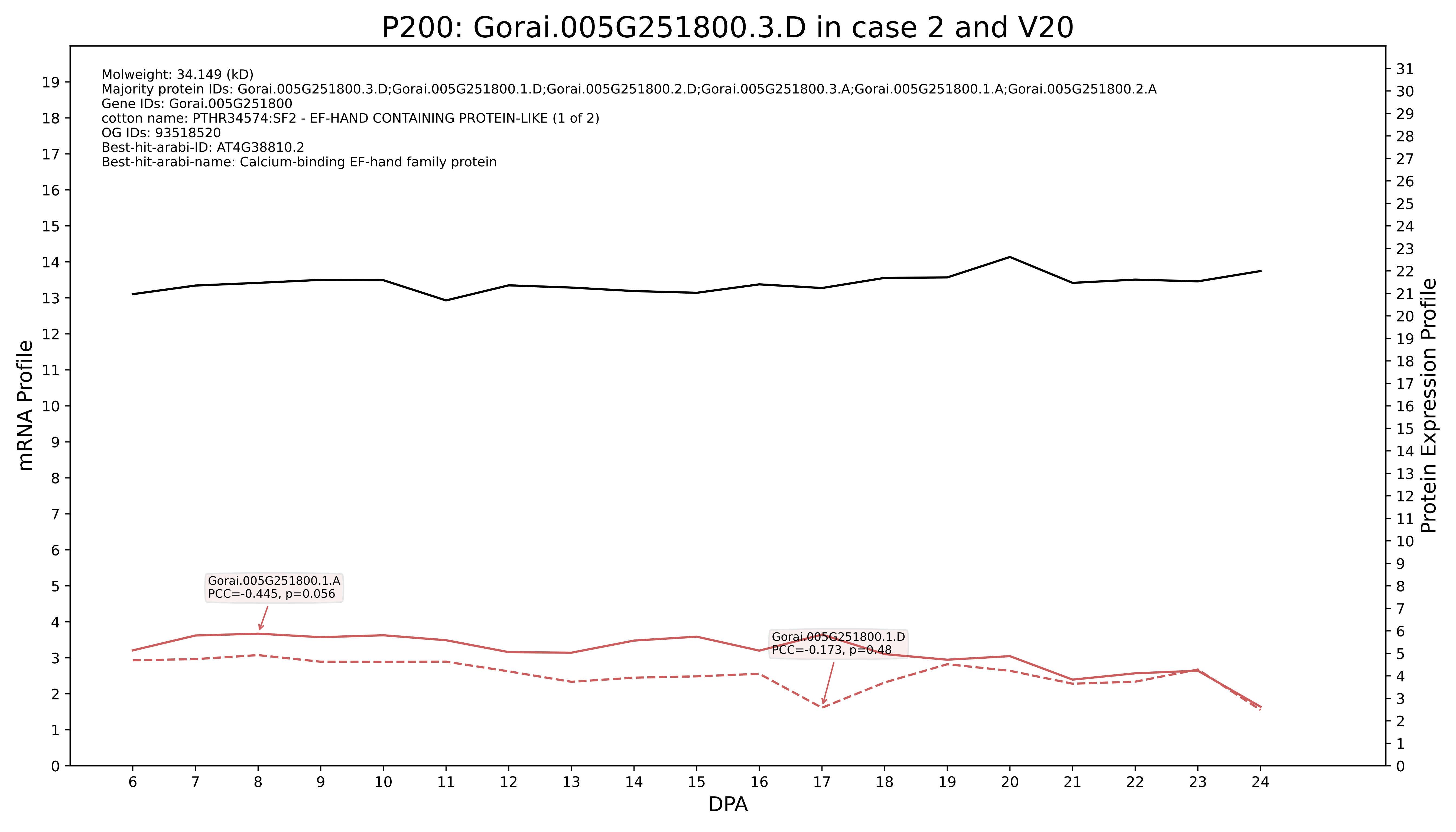
P200: Gorai.005G251800.3.D in case 2 and V20
Molweight: 34.149 (kD)
Majority protein IDs: Gorai.005G251800.3.D;Gorai.005G251800.1.D;Gorai.005G251800.2.D;Gorai.005G251800.3.A;Gorai.005G251800.1.A;Gorai.005G251800.2.A
Gene IDs: Gorai.005G251800
cotton name: PTHR34574:SF2 - EF-HAND CONTAINING PROTEIN-LIKE (1 of 2)
OG IDs: 93518520
Best-hit-arabi-ID: AT4G38810.2
Best-hit-arabi-name: Calcium-binding EF-hand family protein
Gorai.005G251800.1.A
PCC=-0.445, p=0.056
Gorai.005G251800.1.D
PCC=-0.173, p=0.48
mRNA Profile
Protein Expression Profile
DPA

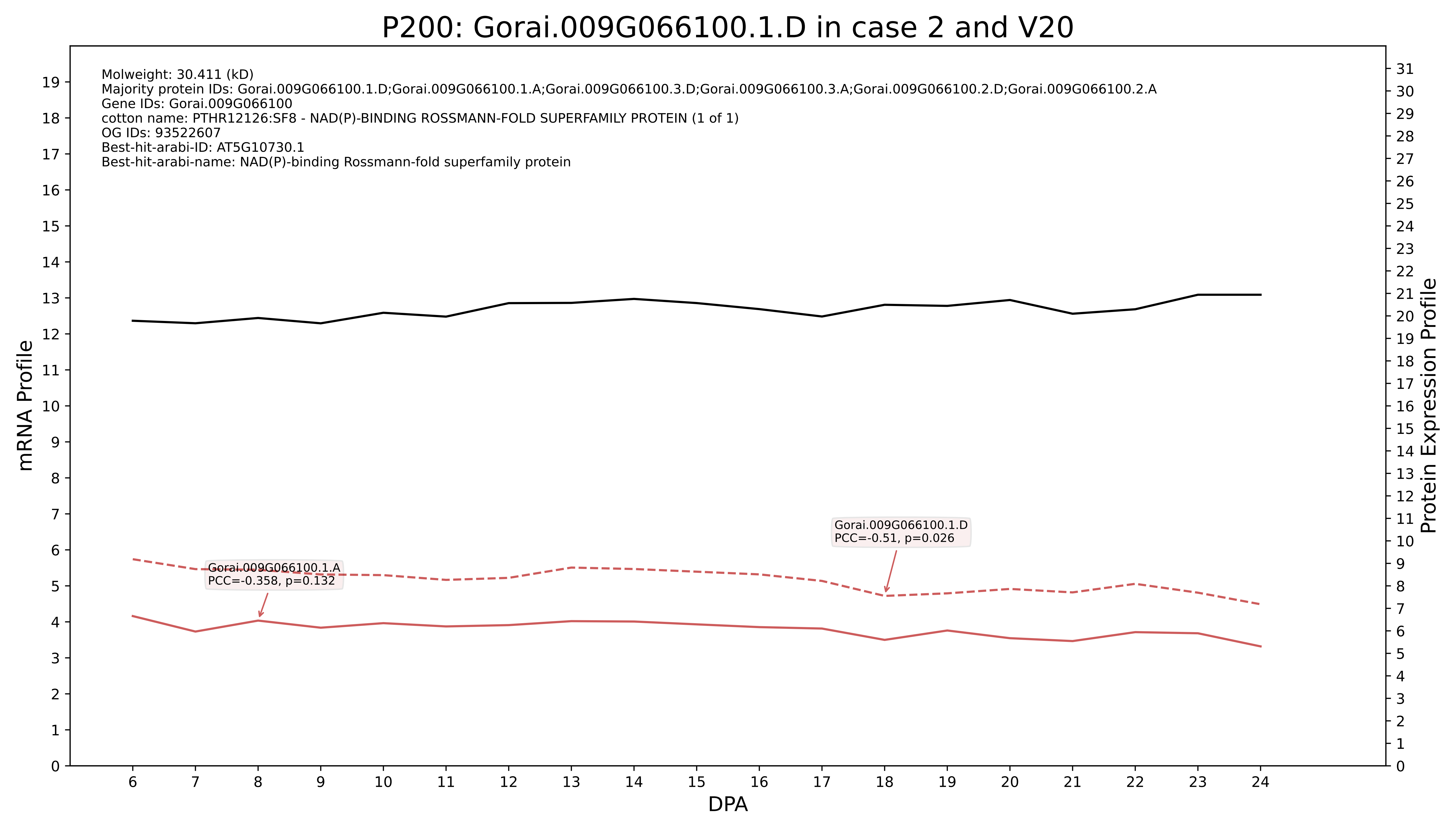
P200: Gorai.009G066100.1.D in case 2 and V20
Molweight: 30.411 (kD)
Majority protein IDs: Gorai.009G066100.1.D;Gorai.009G066100.1.A;Gorai.009G066100.3.D;Gorai.009G066100.3.A;Gorai.009G066100.2.D;Gorai.009G066100.2.A
Gene IDs: Gorai.009G066100
cotton name: PTHR12126:SF8 - NAD(P)-BINDING ROSSMANN-FOLD SUPERFAMILY PROTEIN (1 of 1)
OG IDs: 93522607
Best-hit-arabi-ID: AT5G10730.1
Best-hit-arabi-name: NAD(P)-binding Rossmann-fold superfamily protein
Gorai.009G066100.1.D
PCC=-0.51, p=0.026
Gorai.009G066100.1.A
PCC=-0.358, p=0.132
mRNA Profile
Protein Expression Profile
DPA

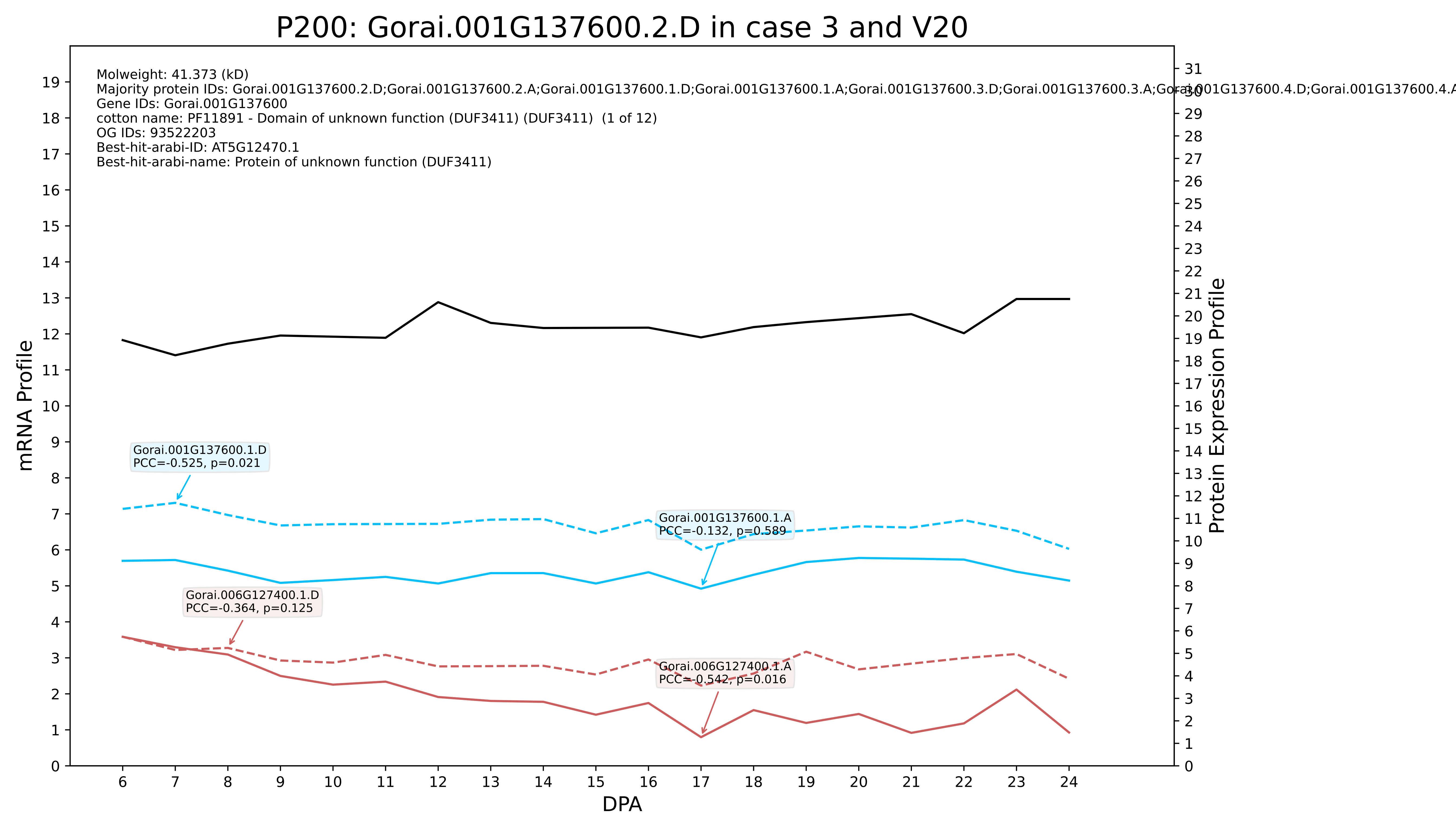

P200: Gorai.001G137600.2.D in case 3 and V20
Molweight: 41.373 (kD)
Majority protein IDs: Gorai.001G137600.2.D;Gorai.001G137600.2.A;Gorai.001G137600.1.D;Gorai.001G137600.1.A;Gorai.001G137600.3.D;Gorai.001G137600.3.A;Gorai.001G137600.4.D;Gorai.001G137600.4.A
Gene IDs: Gorai.001G137600
cotton name: PF11891 - Domain of unknown function (DUF3411) (DUF3411) (1 of 12)
OG IDs: 93522203
Best-hit-arabi-ID: AT5G12470.1
Best-hit-arabi-name: Protein of unknown function (DUF3411)
Gorai.001G137600.1.D
PCC=-0.525, p=0.021
Gorai.001G137600.1.A
PCC=-0.132, p=0.589
Gorai.006G127400.1.D
PCC=-0.364, p=0.125
Gorai.006G127400.1.A
PCC=-0.542, p=0.016
mRNA Profile
Protein Expression Profile
DPA

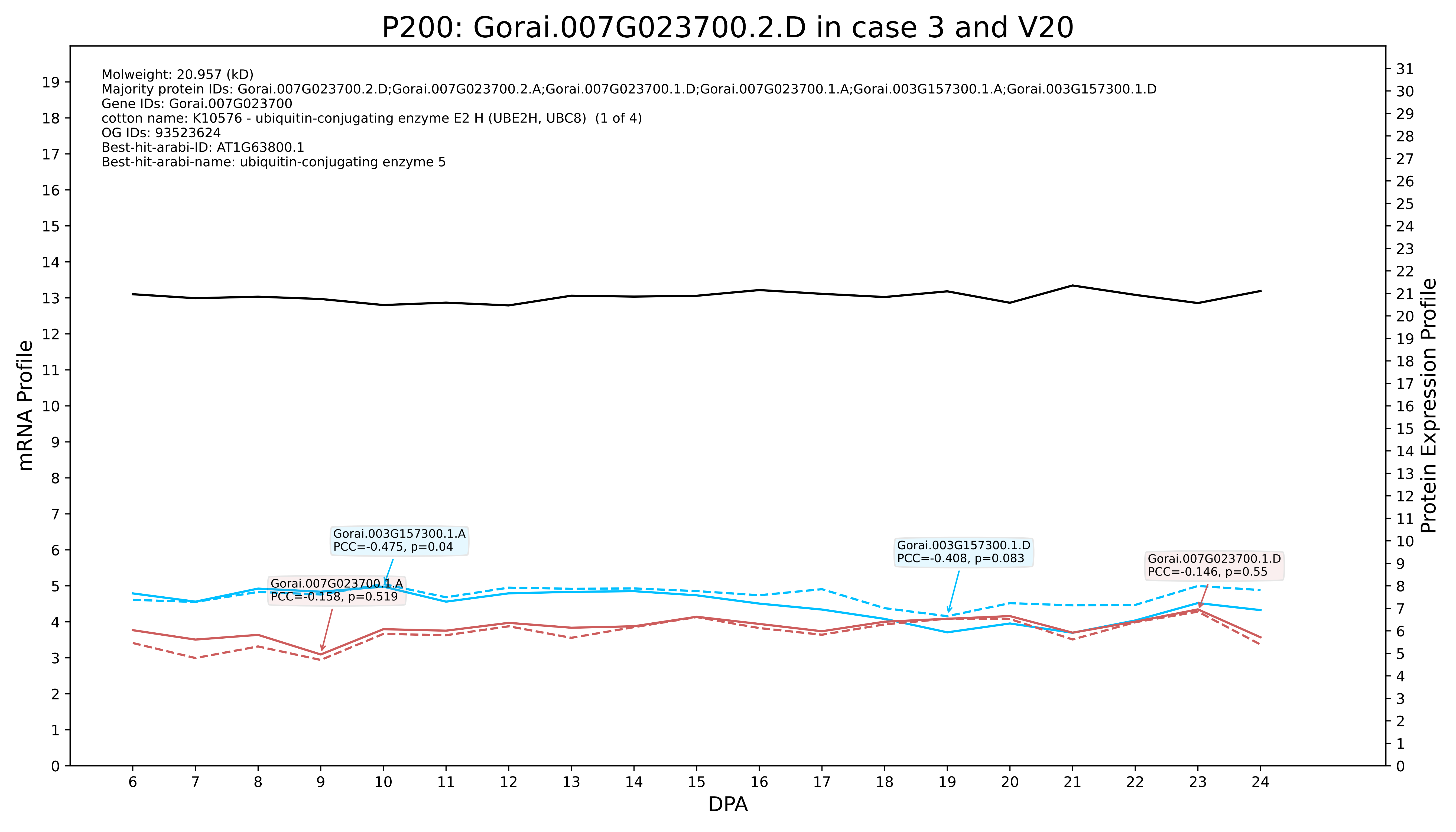

P200: Gorai.007G023700.2.D in case 3 and V20
Molweight: 20.957 (kD)
Majority protein IDs: Gorai.007G023700.2.D;Gorai.007G023700.2.A;Gorai.007G023700.1.D;Gorai.007G023700.1.A;Gorai.003G157300.1.A;Gorai.003G157300.1.D
Gene IDs: Gorai.007G023700
cotton name: K10576 - ubiquitin-conjugating enzyme E2 H (UBE2H, UBC8) (1 of 4)
OG IDs: 93523624
Best-hit-arabi-ID: AT1G63800.1
Best-hit-arabi-name: ubiquitin-conjugating enzyme 5
Gorai.003G157300.1.A
PCC=-0.475, p=0.04
Gorai.007G023700.1.A
PCC=-0.158, p=0.519
Gorai.003G157300.1.D
PCC=-0.408, p=0.083
Gorai.007G023700.1.D
PCC=-0.146, p=0.55
mRNA Profile
Protein Expression Profile
DPA

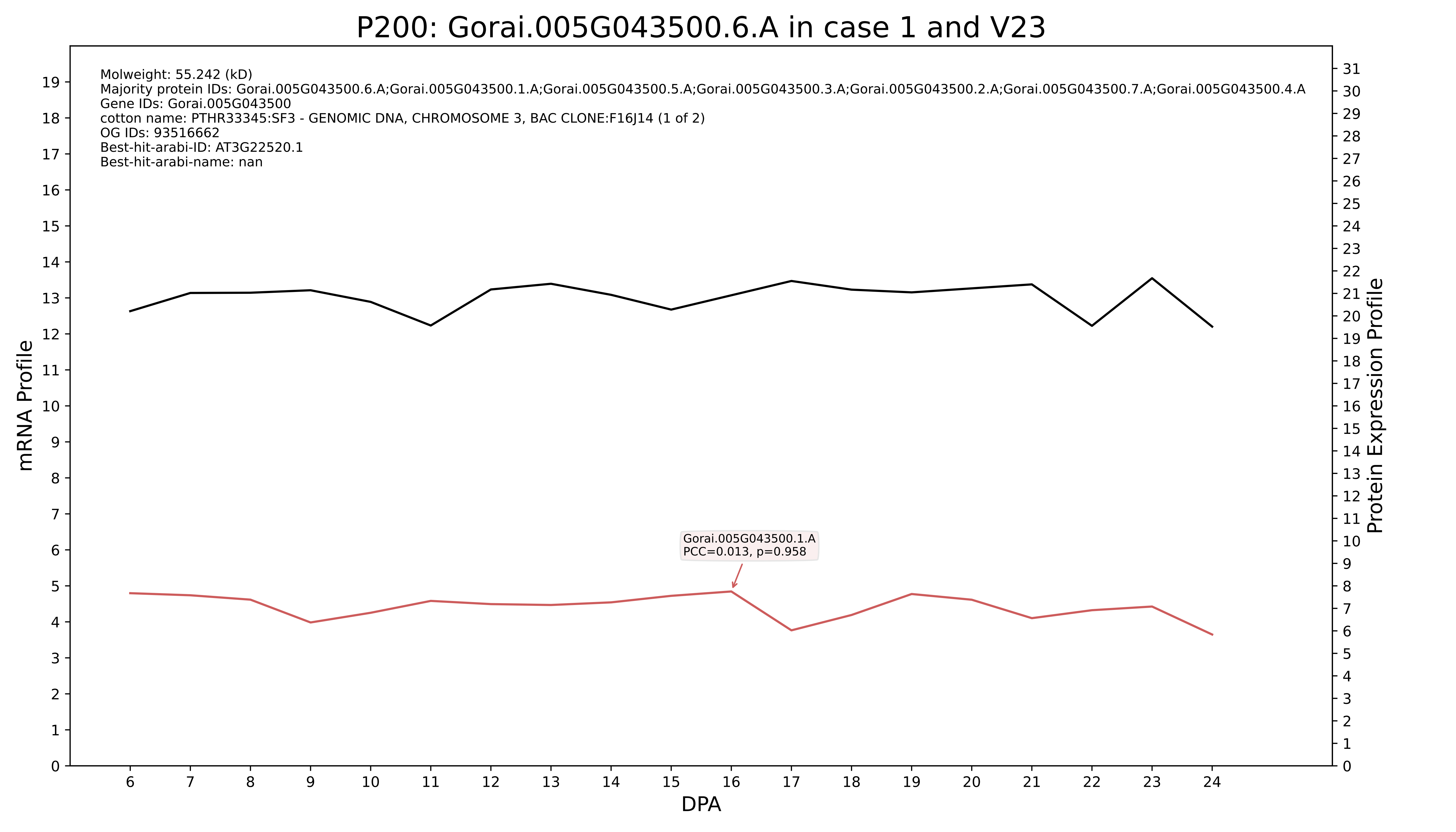
P200: Gorai.005G043500.6.A in case 1 and V23
Molweight: 55.242 (kD)
Majority protein IDs: Gorai.005G043500.6.A;Gorai.005G043500.1.A;Gorai.005G043500.5.A;Gorai.005G043500.3.A;Gorai.005G043500.2.A;Gorai.005G043500.7.A;Gorai.005G043500.4.A
Gene IDs: Gorai.005G043500
cotton name: PTHR33345:SF3 - GENOMIC DNA, CHROMOSOME 3, BAC CLONE:F16J14 (1 of 2)
OG IDs: 93516662
Best-hit-arabi-ID: AT3G22520.1
Best-hit-arabi-name: nan
Gorai.005G043500.1.A
PCC=0.013, p=0.958
mRNA Profile
Protein Expression Profile
DPA

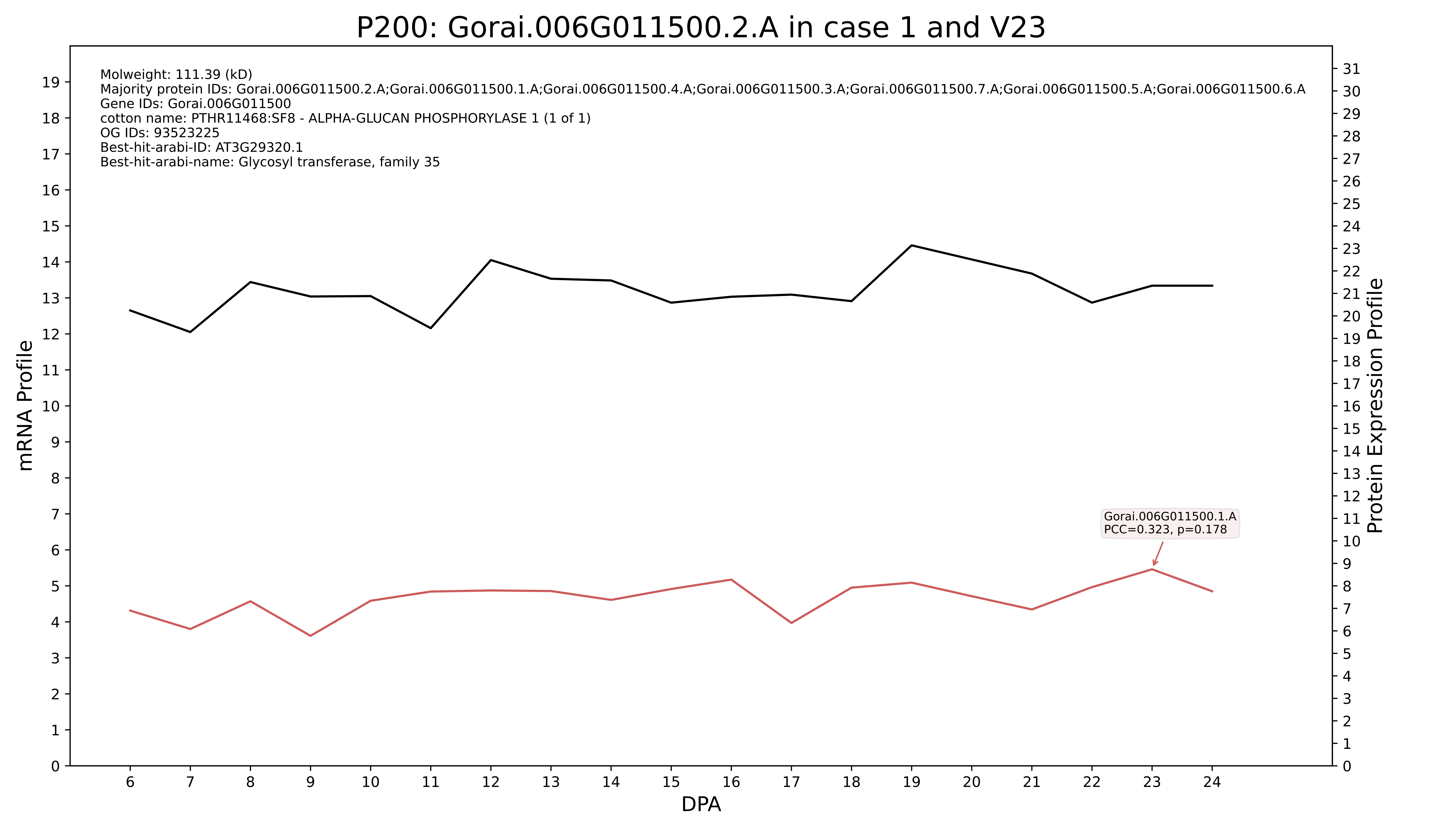
P200: Gorai.006G011500.2.A in case 1 and V23
Molweight: 111.39 (kD)
Majority protein IDs: Gorai.006G011500.2.A;Gorai.006G011500.1.A;Gorai.006G011500.4.A;Gorai.006G011500.3.A;Gorai.006G011500.7.A;Gorai.006G011500.5.A;Gorai.006G011500.6.A
Gene IDs: Gorai.006G011500
cotton name: PTHR11468:SF8 - ALPHA-GLUCAN PHOSPHORYLASE 1 (1 of 1)
OG IDs: 93523225
Best-hit-arabi-ID: AT3G29320.1
Best-hit-arabi-name: Glycosyl transferase, family 35
Gorai.006G011500.1.A
PCC=0.323, p=0.178
mRNA Profile
Protein Expression Profile
DPA

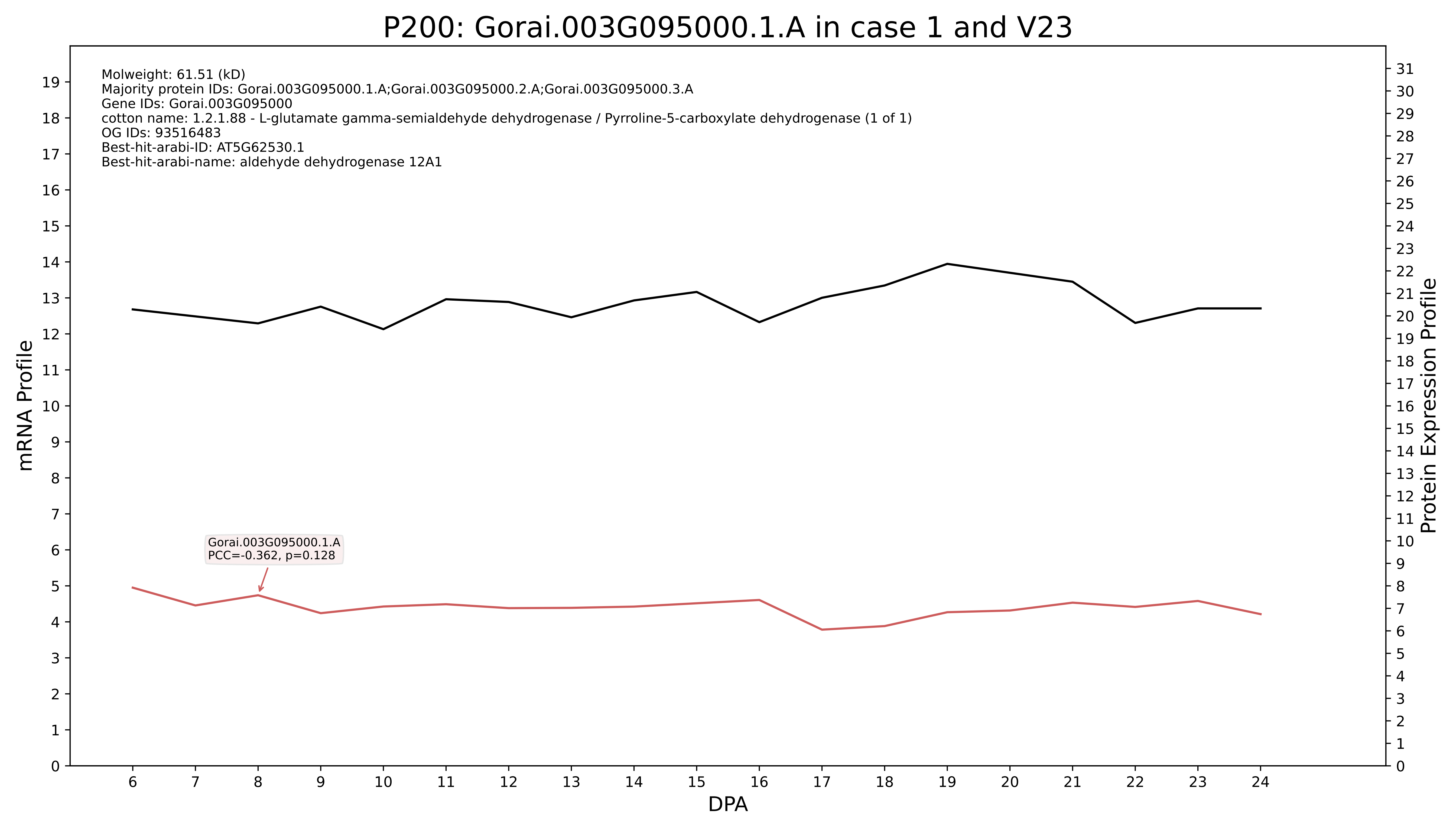
P200: Gorai.003G095000.1.A in case 1 and V23
Molweight: 61.51 (kD)
Majority protein IDs: Gorai.003G095000.1.A;Gorai.003G095000.2.A;Gorai.003G095000.3.A
Gene IDs: Gorai.003G095000
cotton name: 1.2.1.88 - L-glutamate gamma-semialdehyde dehydrogenase / Pyrroline-5-carboxylate dehydrogenase (1 of 1)
OG IDs: 93516483
Best-hit-arabi-ID: AT5G62530.1
Best-hit-arabi-name: aldehyde dehydrogenase 12A1
Gorai.003G095000.1.A
PCC=-0.362, p=0.128
mRNA Profile
Protein Expression Profile
DPA

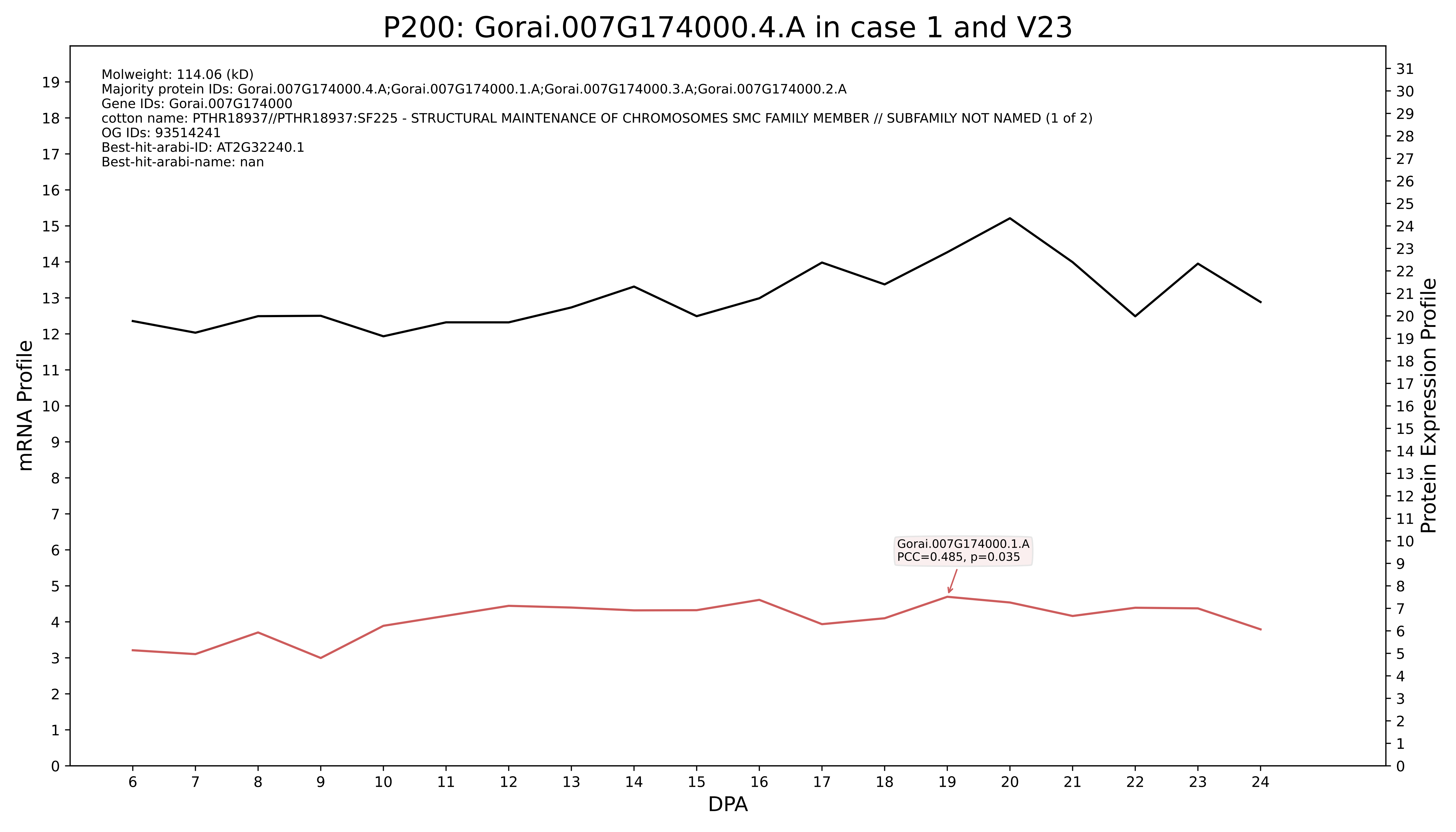
P200: Gorai.007G174000.4.A in case 1 and V23
Molweight: 114.06 (kD)
Majority protein IDs: Gorai.007G174000.4.A;Gorai.007G174000.1.A;Gorai.007G174000.3.A;Gorai.007G174000.2.A
Gene IDs: Gorai.007G174000
cotton name: PTHR18937//PTHR18937:SF225 - STRUCTURAL MAINTENANCE OF CHROMOSOMES SMC FAMILY MEMBER // SUBFAMILY NOT NAMED (1 of 2)
OG IDs: 93514241
Best-hit-arabi-ID: AT2G32240.1
Best-hit-arabi-name: nan
Gorai.007G174000.1.A
PCC=0.485, p=0.035
mRNA Profile
Protein Expression Profile
DPA

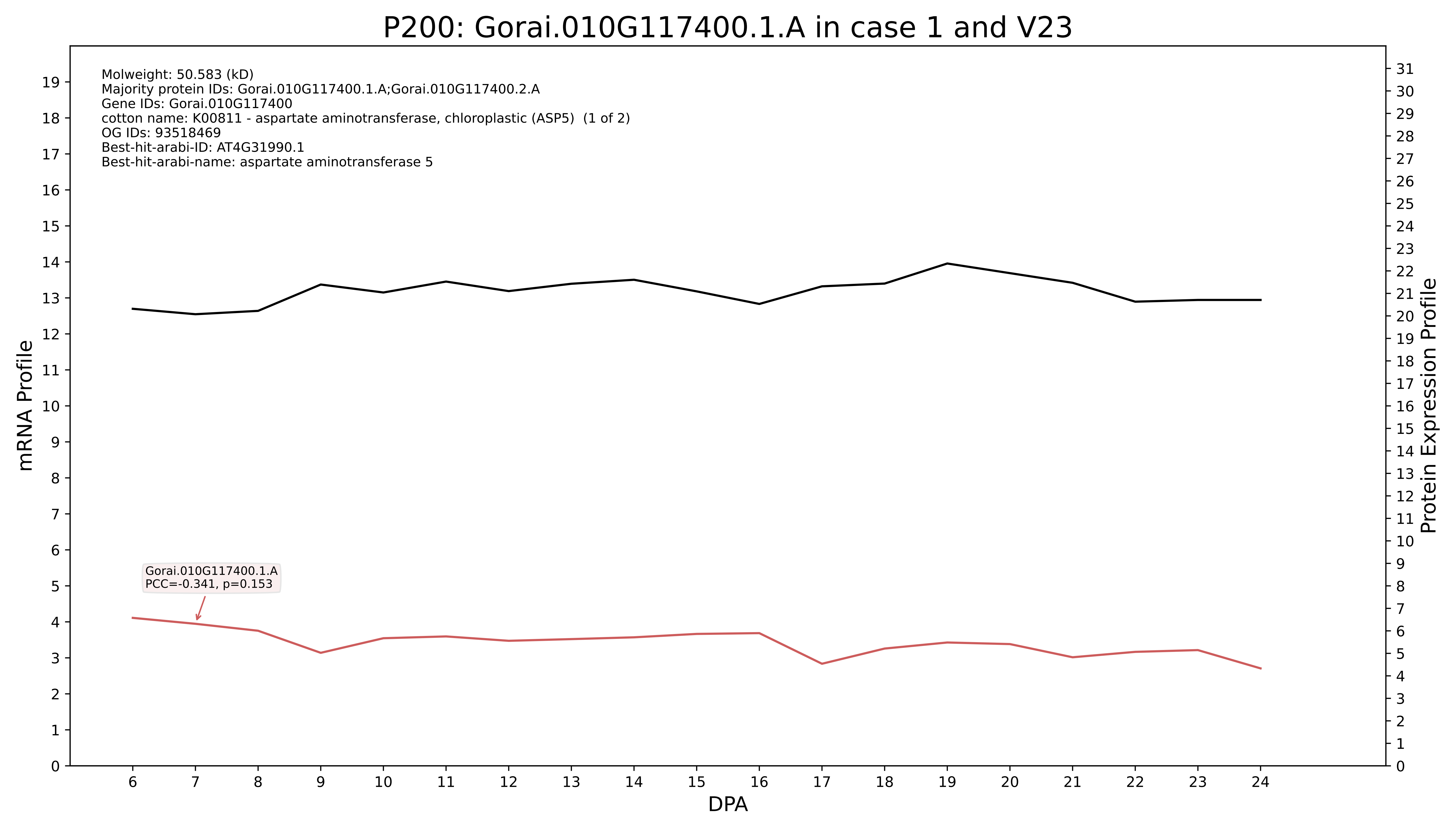
P200: Gorai.010G117400.1.A in case 1 and V23
Molweight: 50.583 (kD)
Majority protein IDs: Gorai.010G117400.1.A;Gorai.010G117400.2.A
Gene IDs: Gorai.010G117400
cotton name: K00811 - aspartate aminotransferase, chloroplastic (ASP5) (1 of 2)
OG IDs: 93518469
Best-hit-arabi-ID: AT4G31990.1
Best-hit-arabi-name: aspartate aminotransferase 5
Gorai.010G117400.1.A
PCC=-0.341, p=0.153
mRNA Profile
Protein Expression Profile
DPA

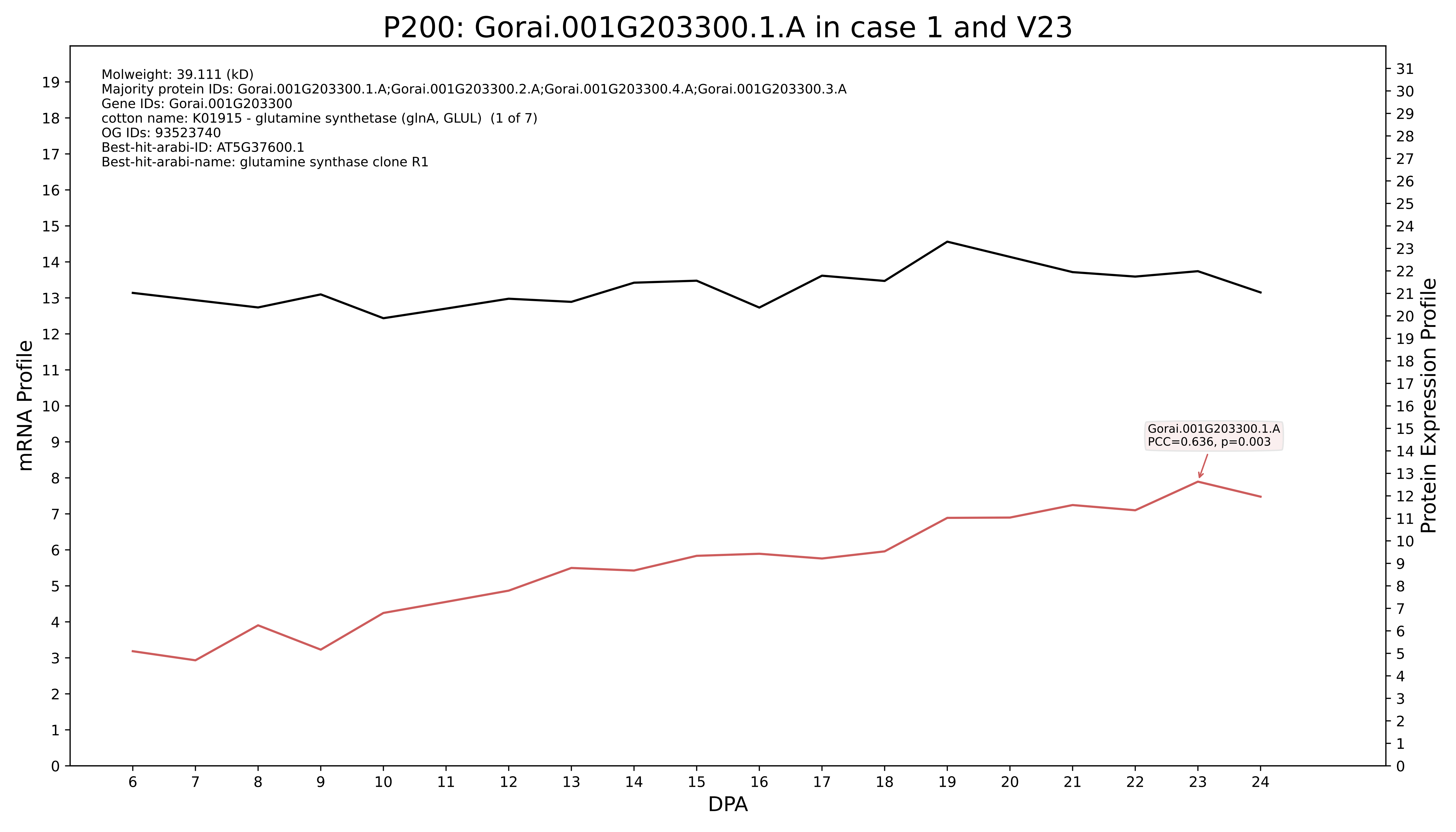
P200: Gorai.001G203300.1.A in case 1 and V23
Molweight: 39.111 (kD)
Majority protein IDs: Gorai.001G203300.1.A;Gorai.001G203300.2.A;Gorai.001G203300.4.A;Gorai.001G203300.3.A
Gene IDs: Gorai.001G203300
cotton name: K01915 - glutamine synthetase (glnA, GLUL) (1 of 7)
OG IDs: 93523740
Best-hit-arabi-ID: AT5G37600.1
Best-hit-arabi-name: glutamine synthase clone R1
Gorai.001G203300.1.A
PCC=0.636, p=0.003
mRNA Profile
Protein Expression Profile
DPA

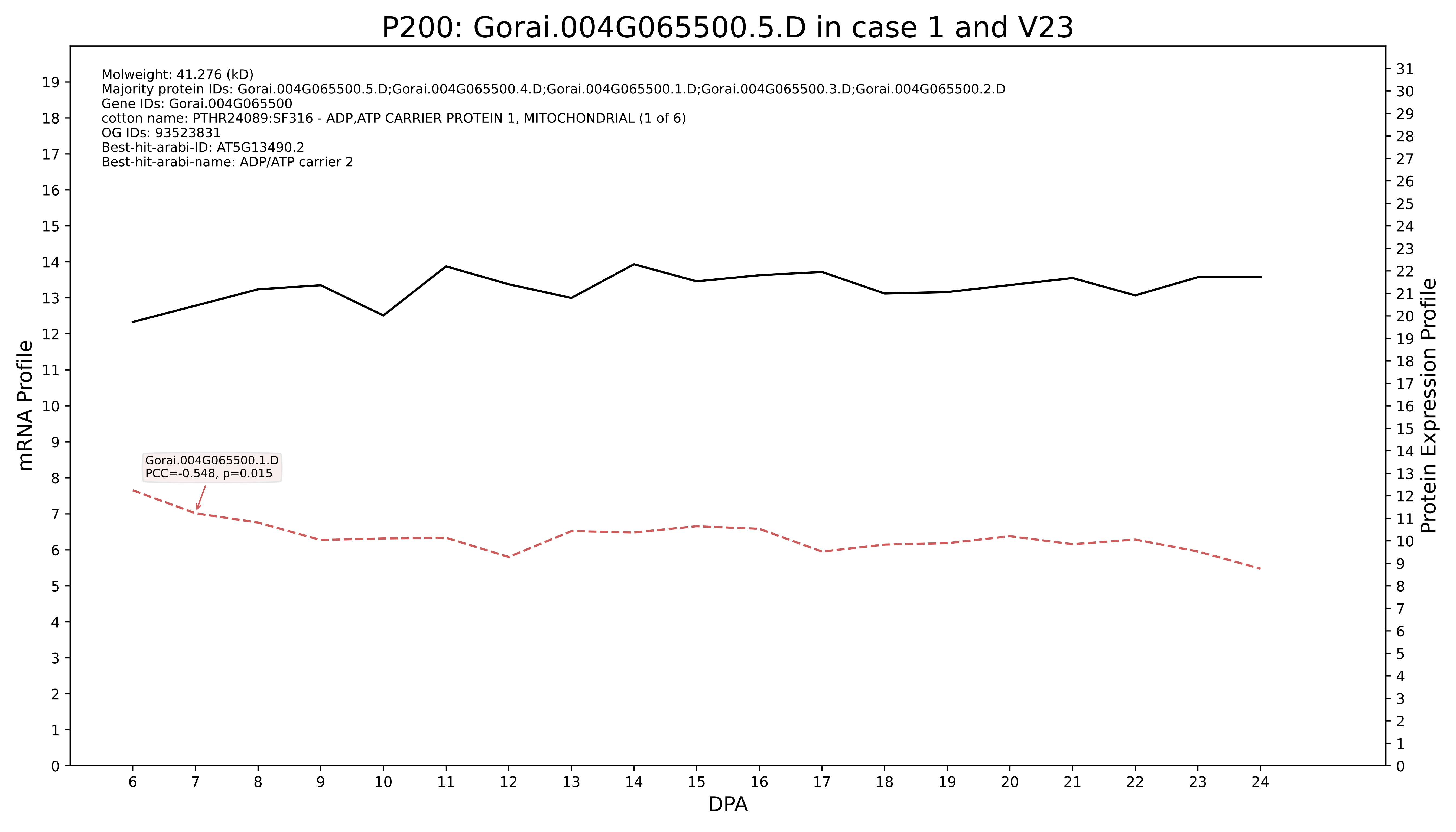
P200: Gorai.004G065500.5.D in case 1 and V23
Molweight: 41.276 (kD)
Majority protein IDs: Gorai.004G065500.5.D;Gorai.004G065500.4.D;Gorai.004G065500.1.D;Gorai.004G065500.3.D;Gorai.004G065500.2.D
Gene IDs: Gorai.004G065500
cotton name: PTHR24089:SF316 - ADP,ATP CARRIER PROTEIN 1, MITOCHONDRIAL (1 of 6)
OG IDs: 93523831
Best-hit-arabi-ID: AT5G13490.2
Best-hit-arabi-name: ADP/ATP carrier 2
Gorai.004G065500.1.D
PCC=-0.548, p=0.015
mRNA Profile
Protein Expression Profile
DPA

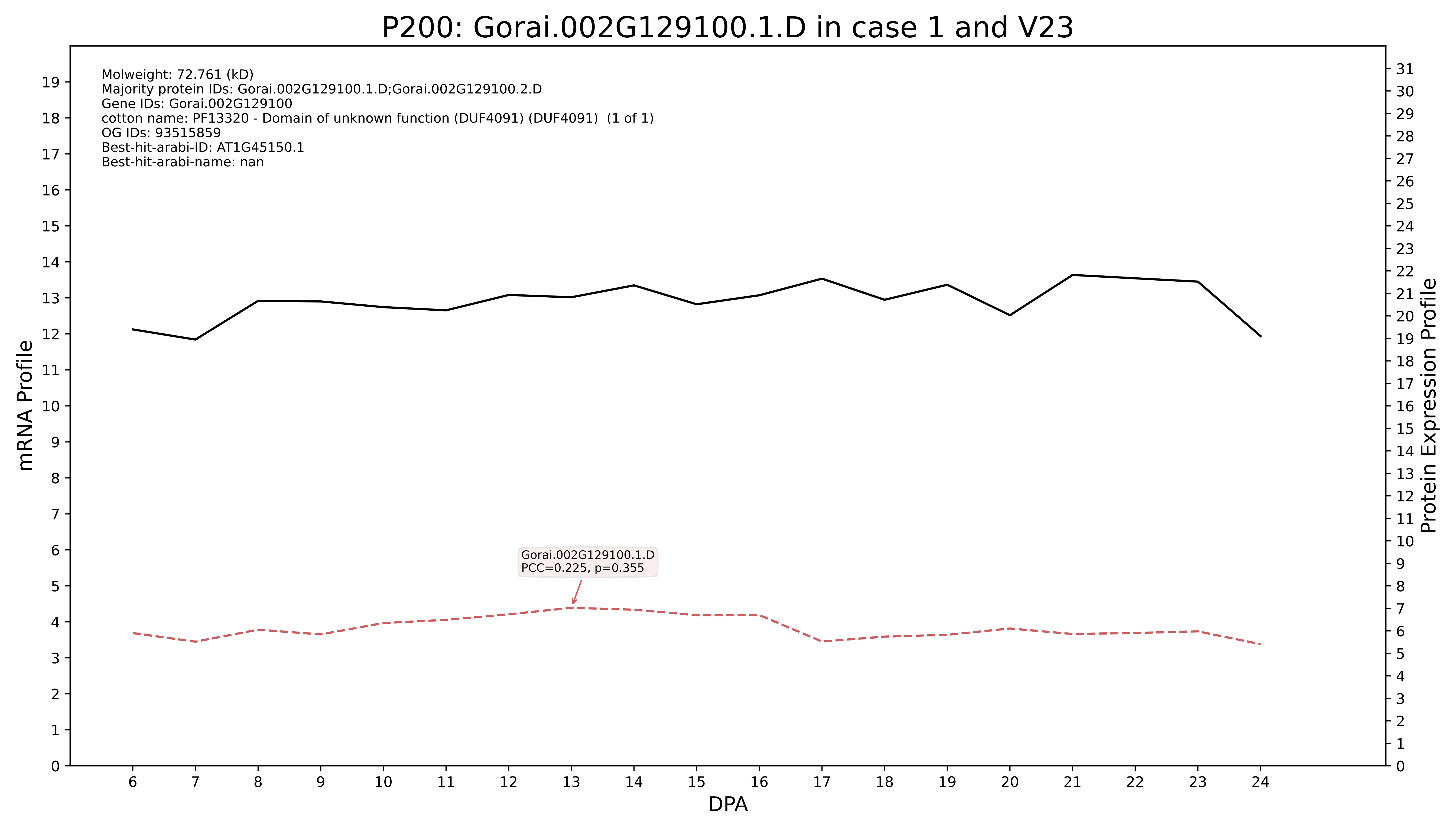

P200: Gorai.002G129100.1.D in case 1 and V23
Molweight: 72.761 (kD)
Majority protein IDs: Gorai.002G129100.1.D;Gorai.002G129100.2.D
Gene IDs: Gorai.002G129100
cotton name: PF13320 - Domain of unknown function (DUF4091) (DUF4091) (1 of 1)
OG IDs: 93515859
Best-hit-arabi-ID: AT1G45150.1
Best-hit-arabi-name: nan
Gorai.002G129100.1.D
PCC=0.225, p=0.355
mRNA Profile
Protein Expression Profile
DPA

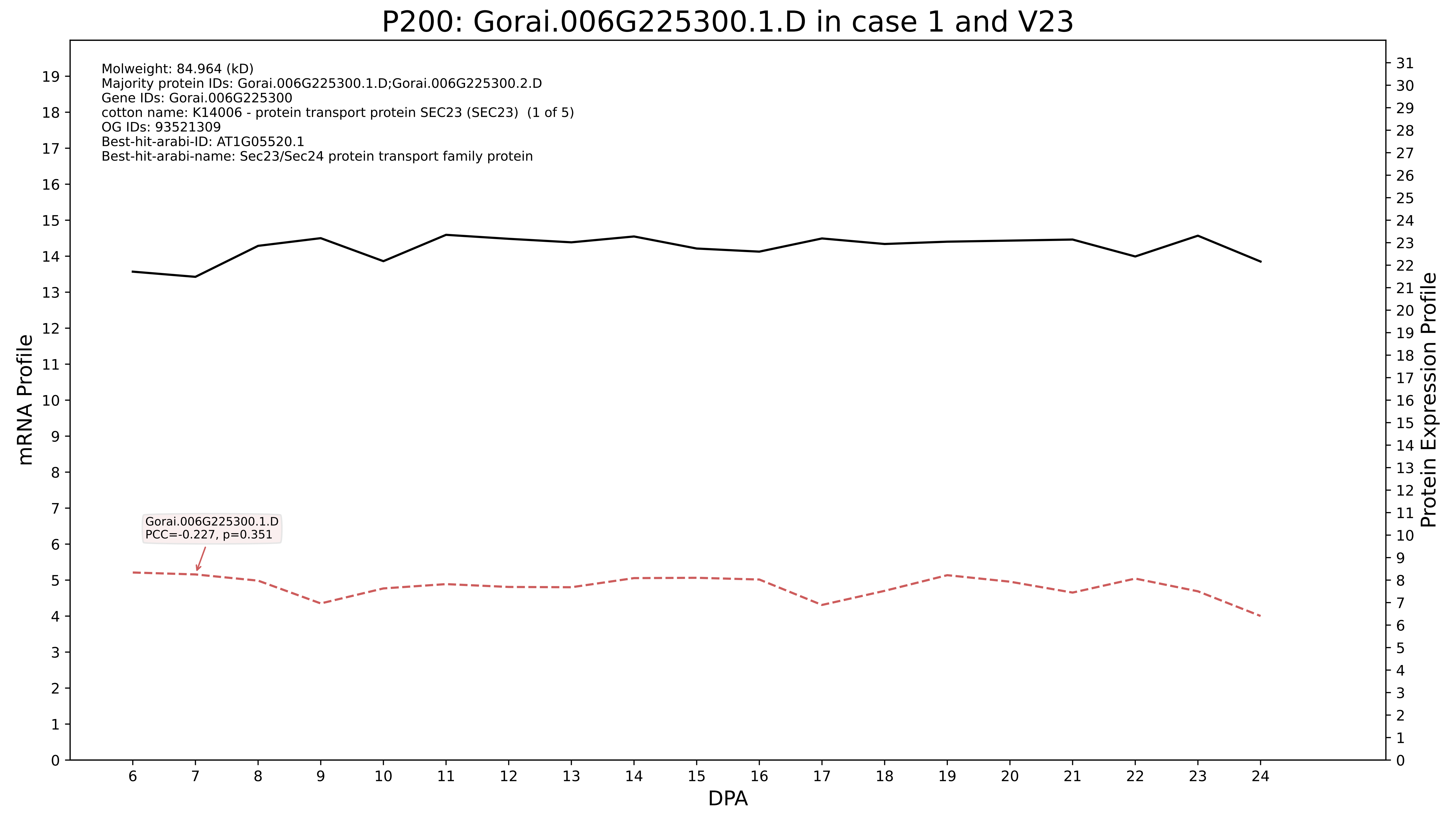
P200: Gorai.006G225300.1.D in case 1 and V23
Molweight: 84.964 (kD)
Majority protein IDs: Gorai.006G225300.1.D;Gorai.006G225300.2.D
Gene IDs: Gorai.006G225300
cotton name: K14006 - protein transport protein SEC23 (SEC23) (1 of 5)
OG IDs: 93521309
Best-hit-arabi-ID: AT1G05520.1
Best-hit-arabi-name: Sec23/Sec24 protein transport family protein
Gorai.006G225300.1.D
PCC=-0.227, p=0.351
mRNA Profile
Protein Expression Profile
DPA

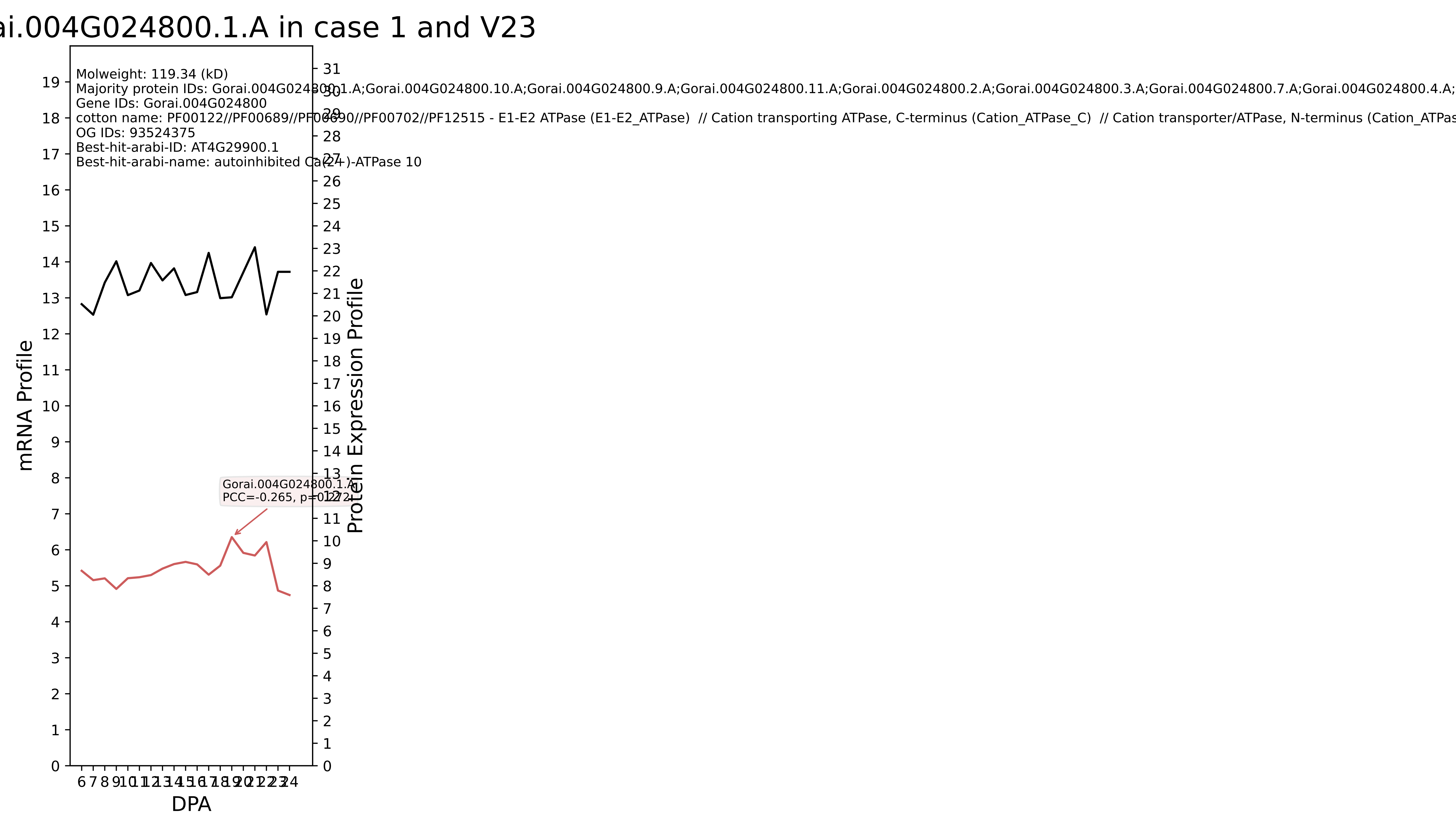
ai.004G024800.1.A in case 1 and V23
Molweight: 119.34 (kD)
Majority protein IDs: Gorai.004G024800.1.A;Gorai.004G024800.10.A;Gorai.004G024800.9.A;Gorai.004G024800.11.A;Gorai.004G024800.2.A;Gorai.004G024800.3.A;Gorai.004G024800.7.A;Gorai.004G024800.4.A;
Gene IDs: Gorai.004G024800
cotton name: PF00122//PF00689//PF00690//PF00702//PF12515 - E1-E2 ATPase (E1-E2_ATPase) // Cation transporting ATPase, C-terminus (Cation_ATPase_C) // Cation transporter/ATPase, N-terminus (Cation_ATPas
OG IDs: 93524375
Best-hit-arabi-ID: AT4G29900.1
Best-hit-arabi-name: autoinhibited Ca(2+)-ATPase 10
Gorai.004G024800.1.A
PCC=-0.265, p=0.272
mRNA Profile
Protein Expression Profile
DPA

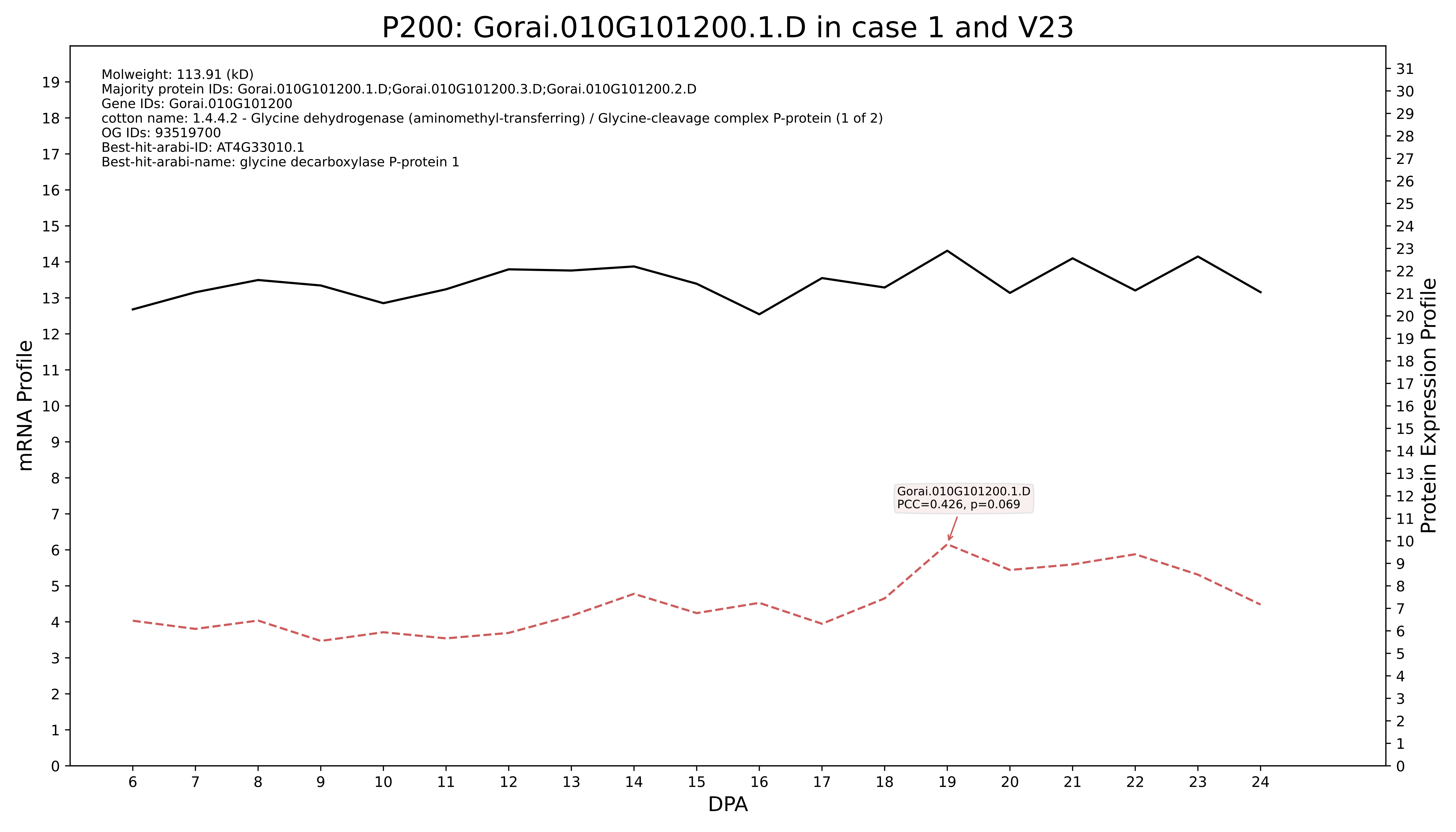
P200: Gorai.010G101200.1.D in case 1 and V23
Molweight: 113.91 (kD)
Majority protein IDs: Gorai.010G101200.1.D;Gorai.010G101200.3.D;Gorai.010G101200.2.D
Gene IDs: Gorai.010G101200
cotton name: 1.4.4.2 - Glycine dehydrogenase (aminomethyl-transferring) / Glycine-cleavage complex P-protein (1 of 2)
OG IDs: 93519700
Best-hit-arabi-ID: AT4G33010.1
Best-hit-arabi-name: glycine decarboxylase P-protein 1
Gorai.010G101200.1.D
PCC=0.426, p=0.069
mRNA Profile
Protein Expression Profile
DPA

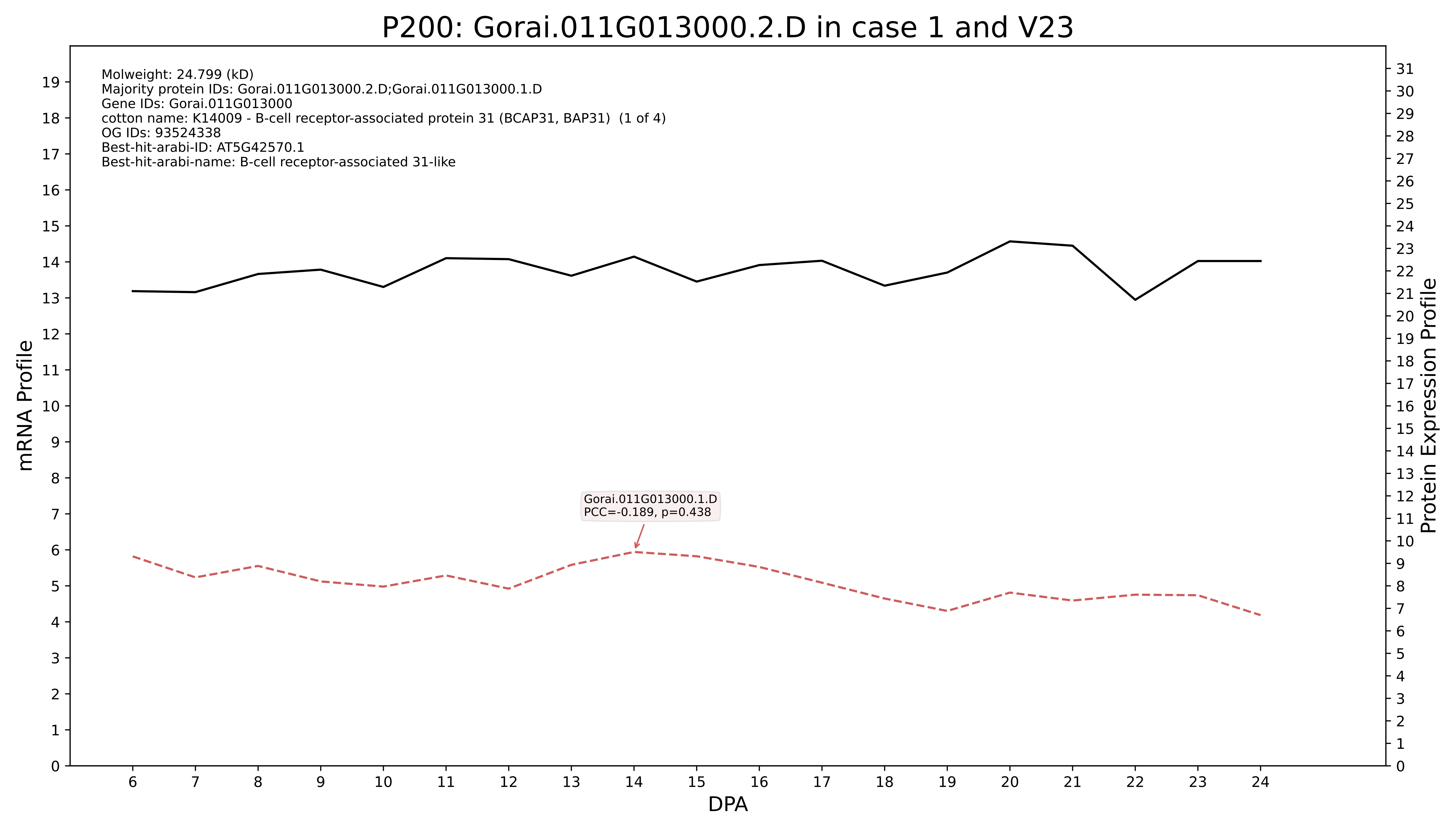
P200: Gorai.011G013000.2.D in case 1 and V23
Molweight: 24.799 (kD)
Majority protein IDs: Gorai.011G013000.2.D;Gorai.011G013000.1.D
Gene IDs: Gorai.011G013000
cotton name: K14009 - B-cell receptor-associated protein 31 (BCAP31, BAP31) (1 of 4)
OG IDs: 93524338
Best-hit-arabi-ID: AT5G42570.1
Best-hit-arabi-name: B-cell receptor-associated 31-like
Gorai.011G013000.1.D
PCC=-0.189, p=0.438
mRNA Profile
Protein Expression Profile
DPA

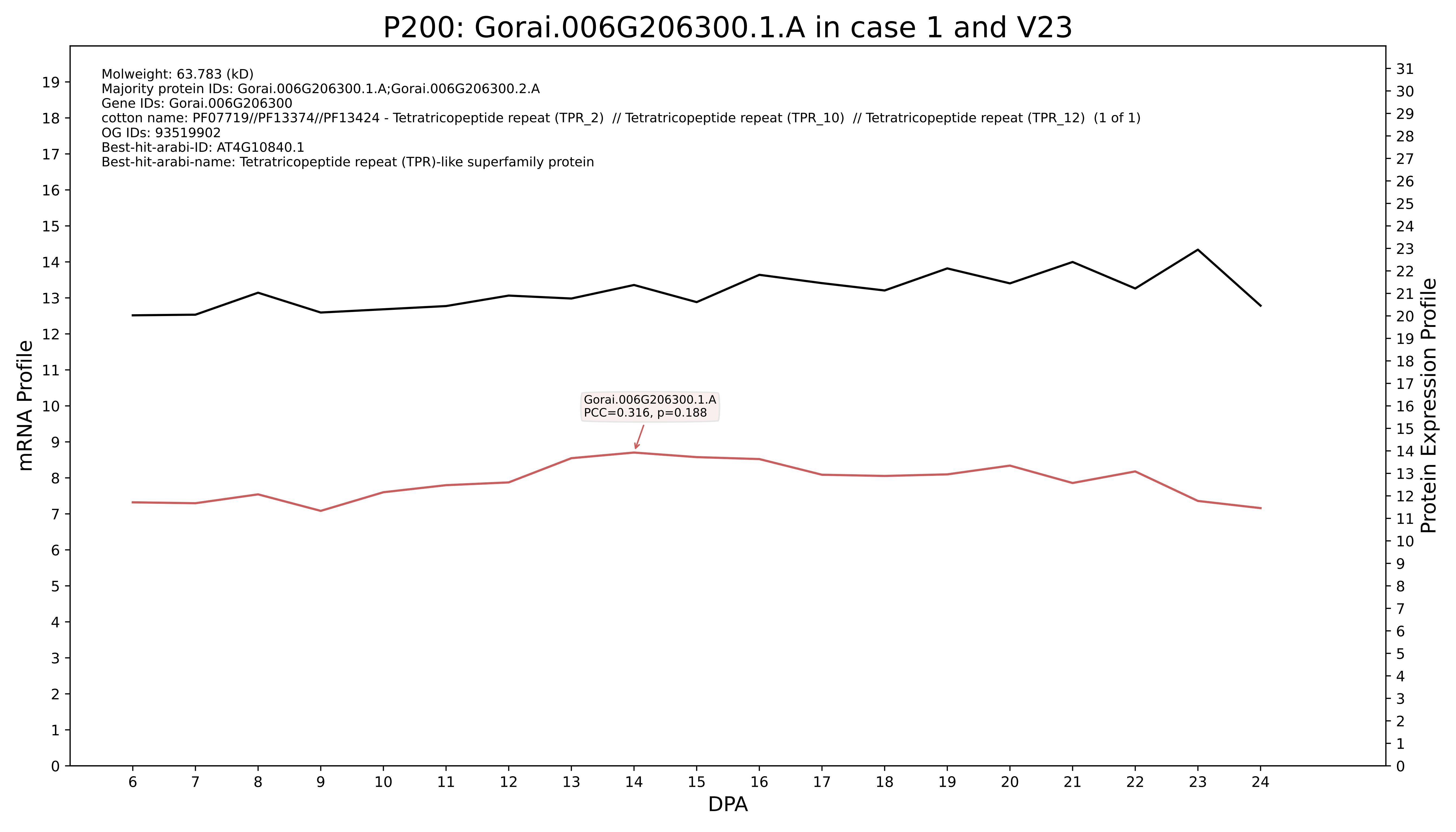
P200: Gorai.006G206300.1.A in case 1 and V23
Molweight: 63.783 (kD)
Majority protein IDs: Gorai.006G206300.1.A;Gorai.006G206300.2.A
Gene IDs: Gorai.006G206300
cotton name: PF07719//PF13374//PF13424 - Tetratricopeptide repeat (TPR_2) // Tetratricopeptide repeat (TPR_10) // Tetratricopeptide repeat (TPR_12) (1 of 1)
OG IDs: 93519902
Best-hit-arabi-ID: AT4G10840.1
Best-hit-arabi-name: Tetratricopeptide repeat (TPR)-like superfamily protein
Gorai.006G206300.1.A
PCC=0.316, p=0.188
mRNA Profile
Protein Expression Profile
DPA

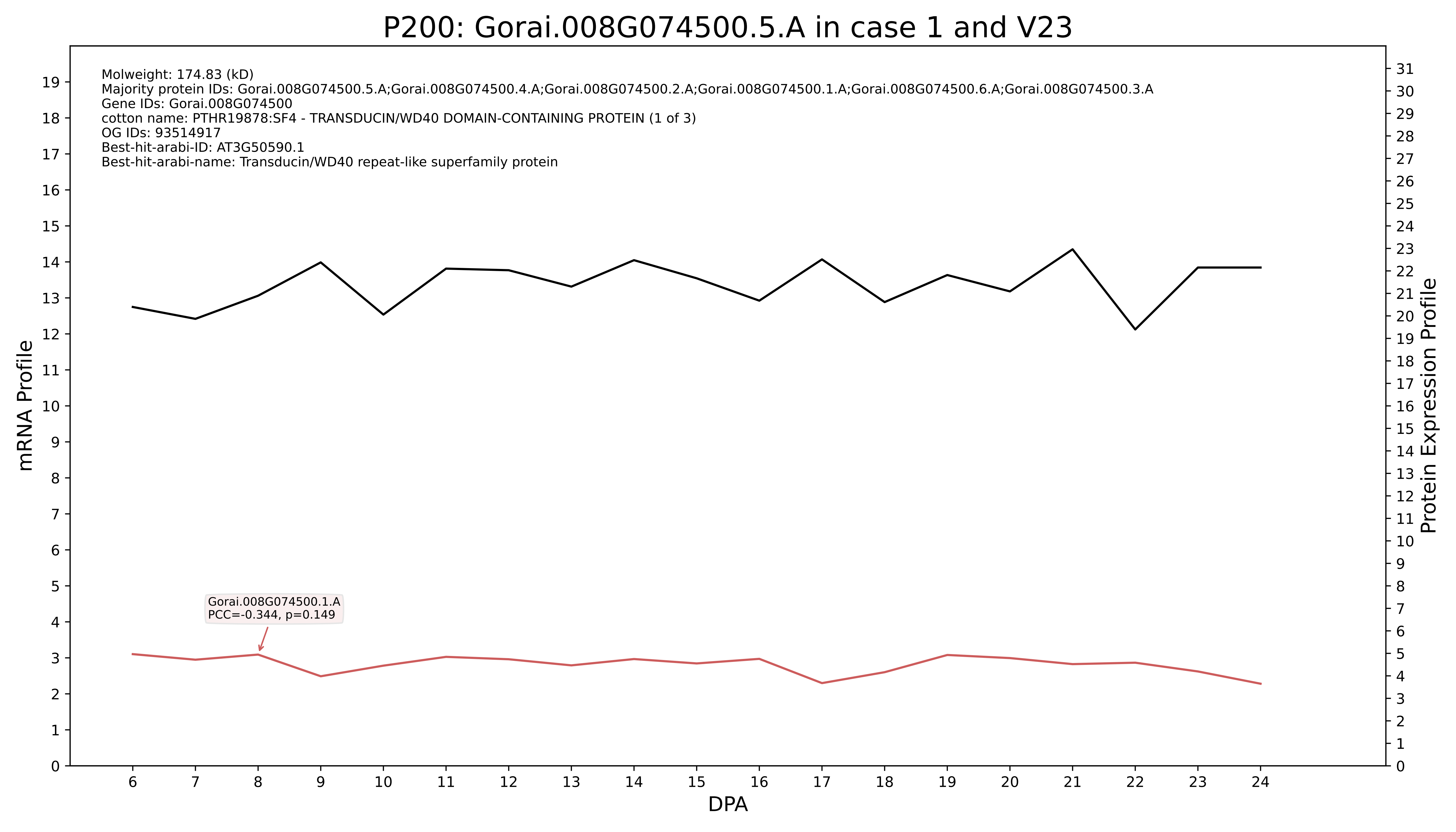
P200: Gorai.008G074500.5.A in case 1 and V23
Molweight: 174.83 (kD)
Majority protein IDs: Gorai.008G074500.5.A;Gorai.008G074500.4.A;Gorai.008G074500.2.A;Gorai.008G074500.1.A;Gorai.008G074500.6.A;Gorai.008G074500.3.A
Gene IDs: Gorai.008G074500
cotton name: PTHR19878:SF4 - TRANSDUCIN/WD40 DOMAIN-CONTAINING PROTEIN (1 of 3)
OG IDs: 93514917
Best-hit-arabi-ID: AT3G50590.1
Best-hit-arabi-name: Transducin/WD40 repeat-like superfamily protein
Gorai.008G074500.1.A
PCC=-0.344, p=0.149
mRNA Profile
Protein Expression Profile
DPA

### P200: Gorai.008G187900.1.A in case 1 and V23

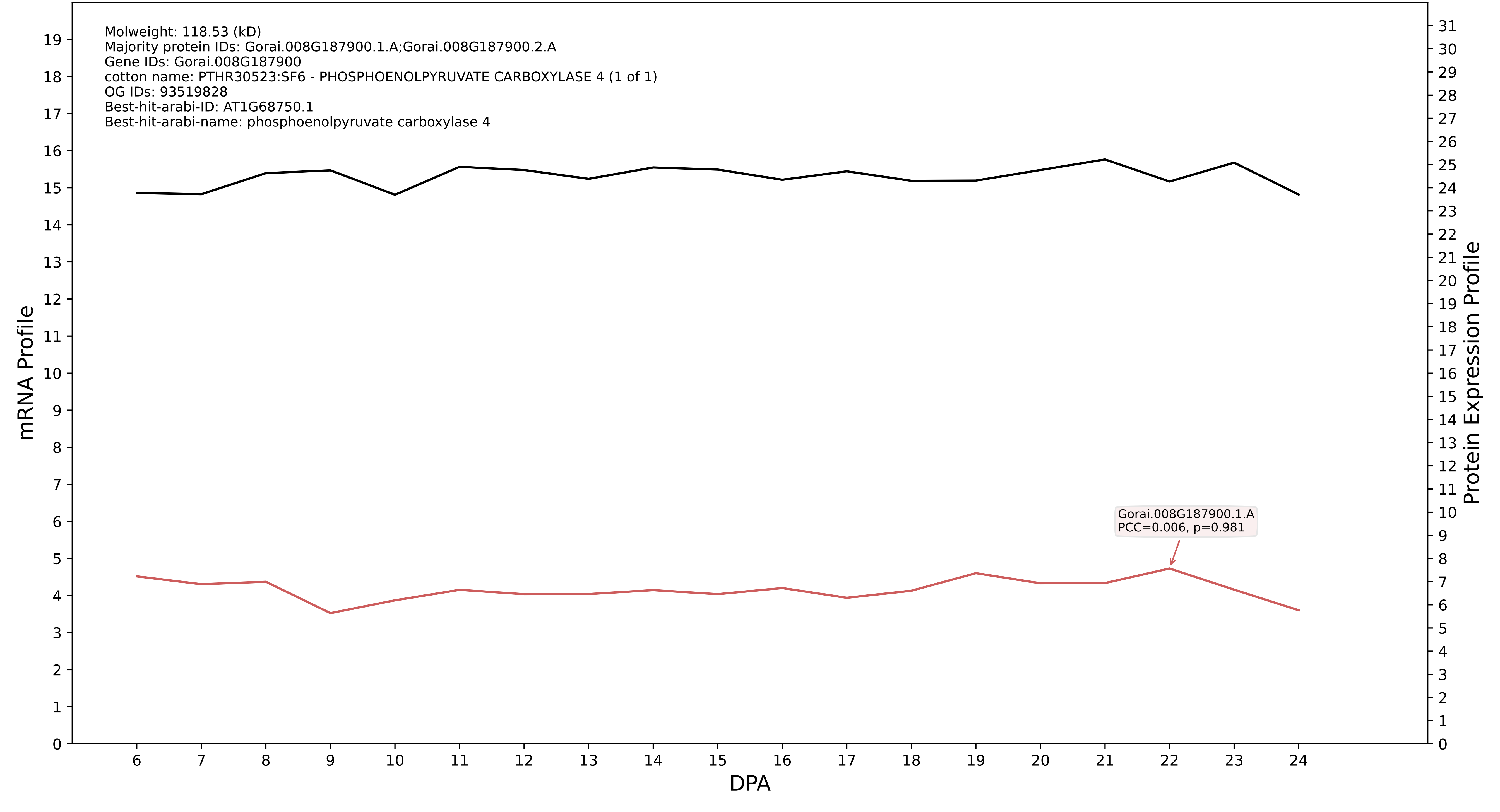

### P200: Gorai.001G261400.1.A in case 1 and V23

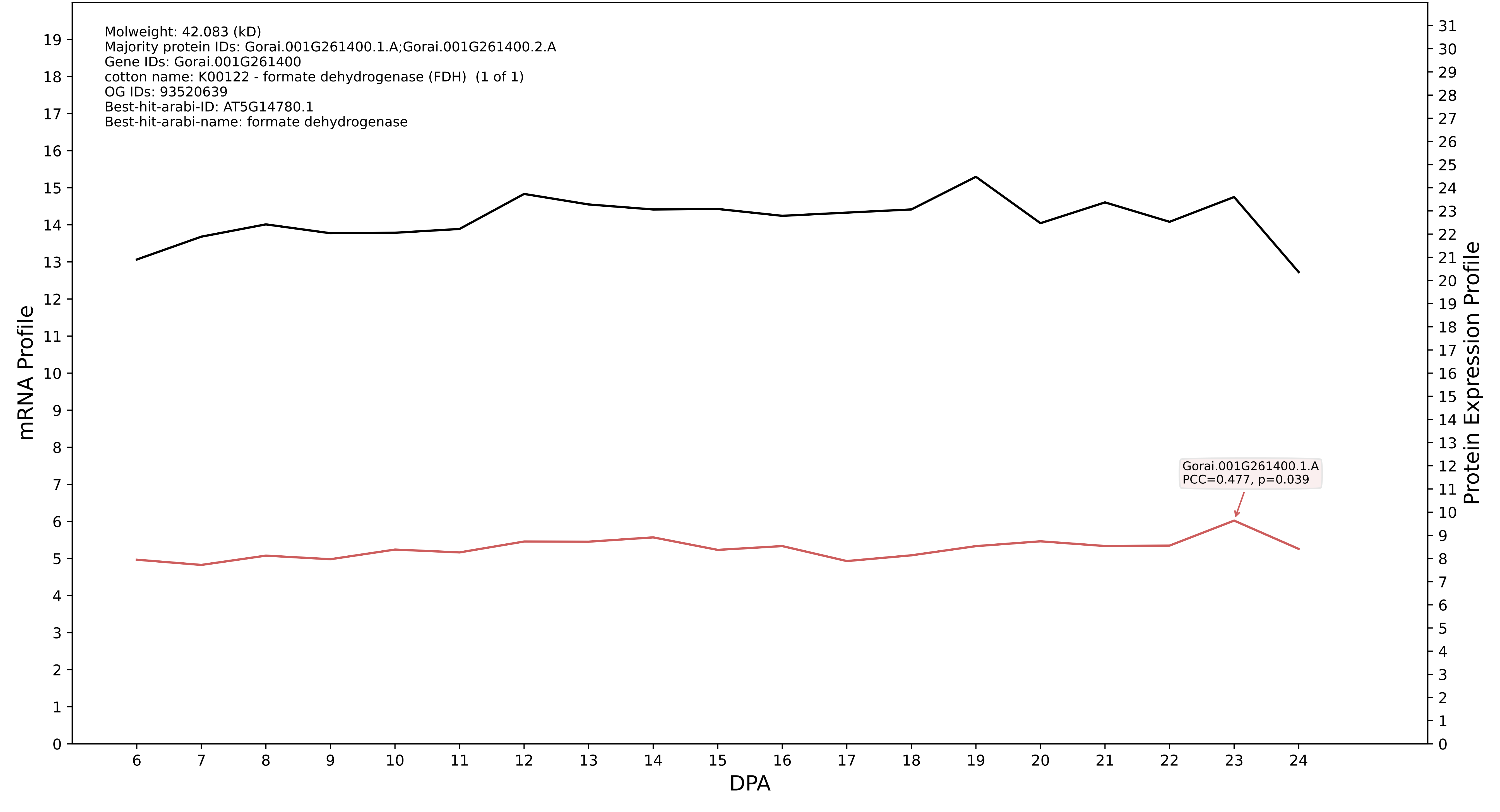

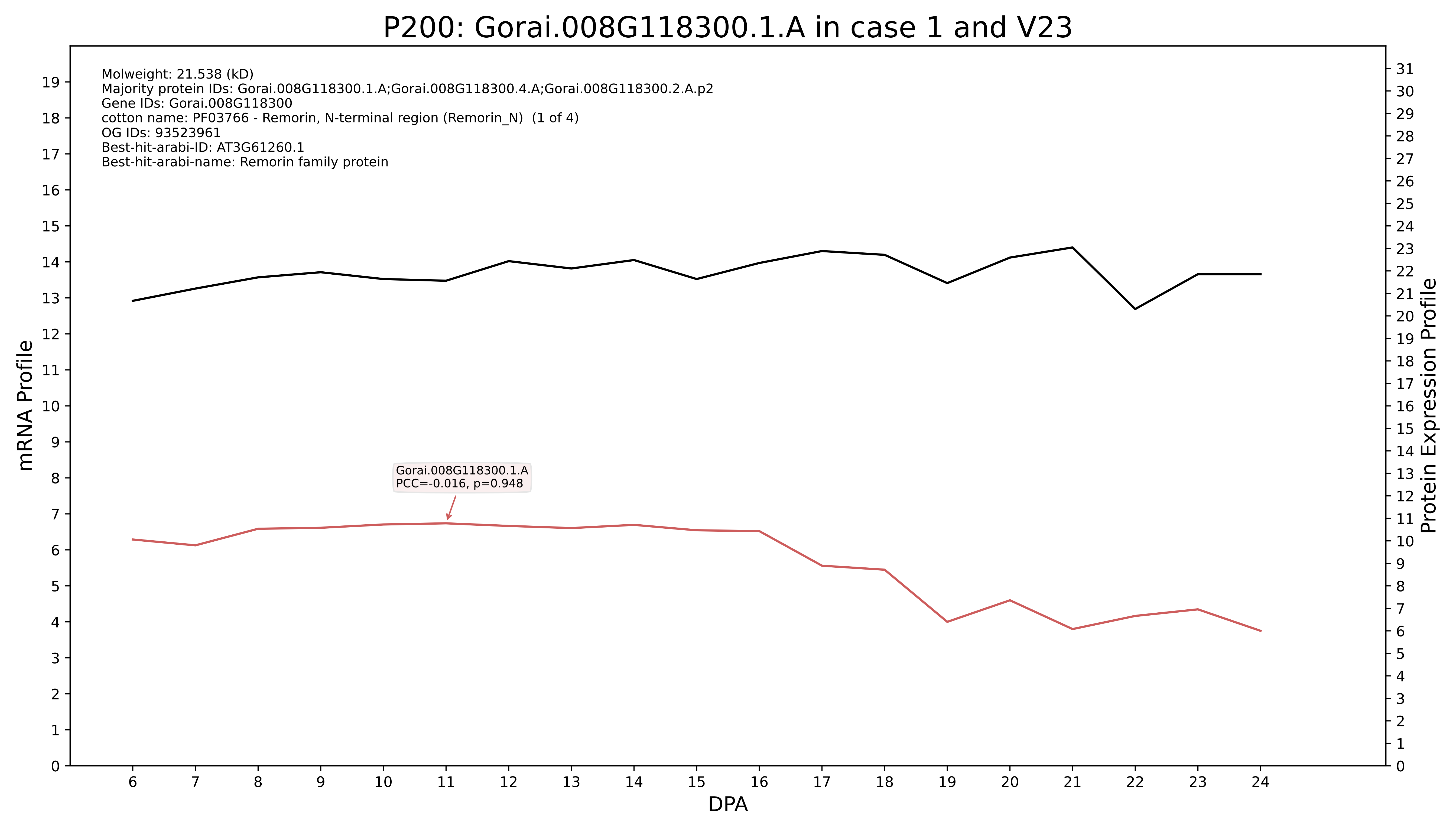
P200: Gorai.008G118300.1.A in case 1 and V23
Molweight: 21.538 (kD)
Majority protein IDs: Gorai.008G118300.1.A;Gorai.008G118300.4.A;Gorai.008G118300.2.A.p2
Gene IDs: Gorai.008G118300
cotton name: PF03766 - Remorin, N-terminal region (Remorin_N) (1 of 4)
OG IDs: 93523961
Best-hit-arabi-ID: AT3G61260.1
Best-hit-arabi-name: Remorin family protein
Gorai.008G118300.1.A
PCC=-0.016, p=0.948
mRNA Profile
Protein Expression Profile
DPA

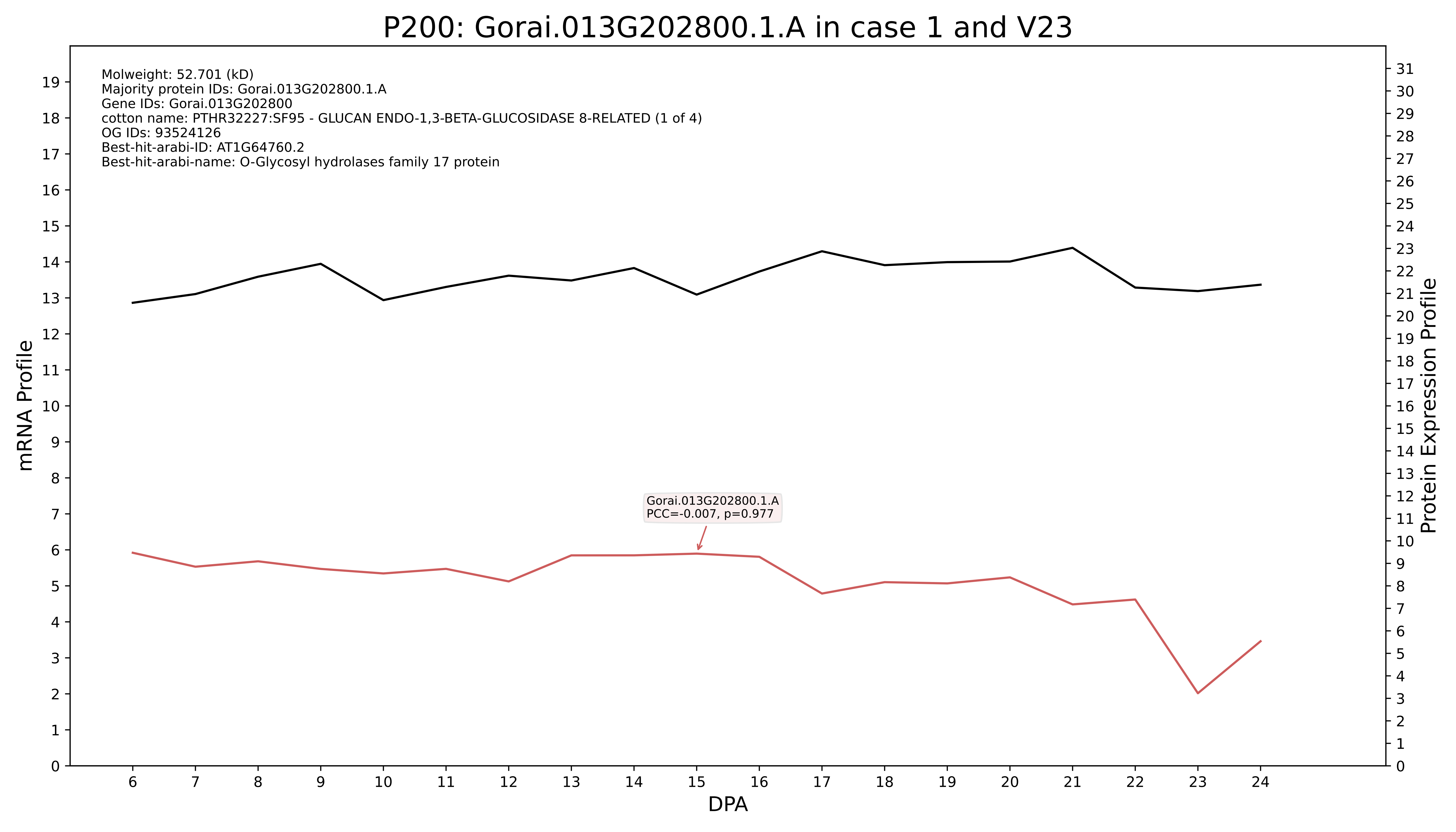
P200: Gorai.013G202800.1.A in case 1 and V23
Molweight: 52.701 (kD)
Majority protein IDs: Gorai.013G202800.1.A
Gene IDs: Gorai.013G202800
cotton name: PTHR32227:SF95 - GLUCAN ENDO-1,3-BETA-GLUCOSIDASE 8-RELATED (1 of 4)
OG IDs: 93524126
Best-hit-arabi-ID: AT1G64760.2
Best-hit-arabi-name: O-Glycosyl hydrolases family 17 protein
Gorai.013G202800.1.A
PCC=-0.007, p=0.977
mRNA Profile
Protein Expression Profile
DPA

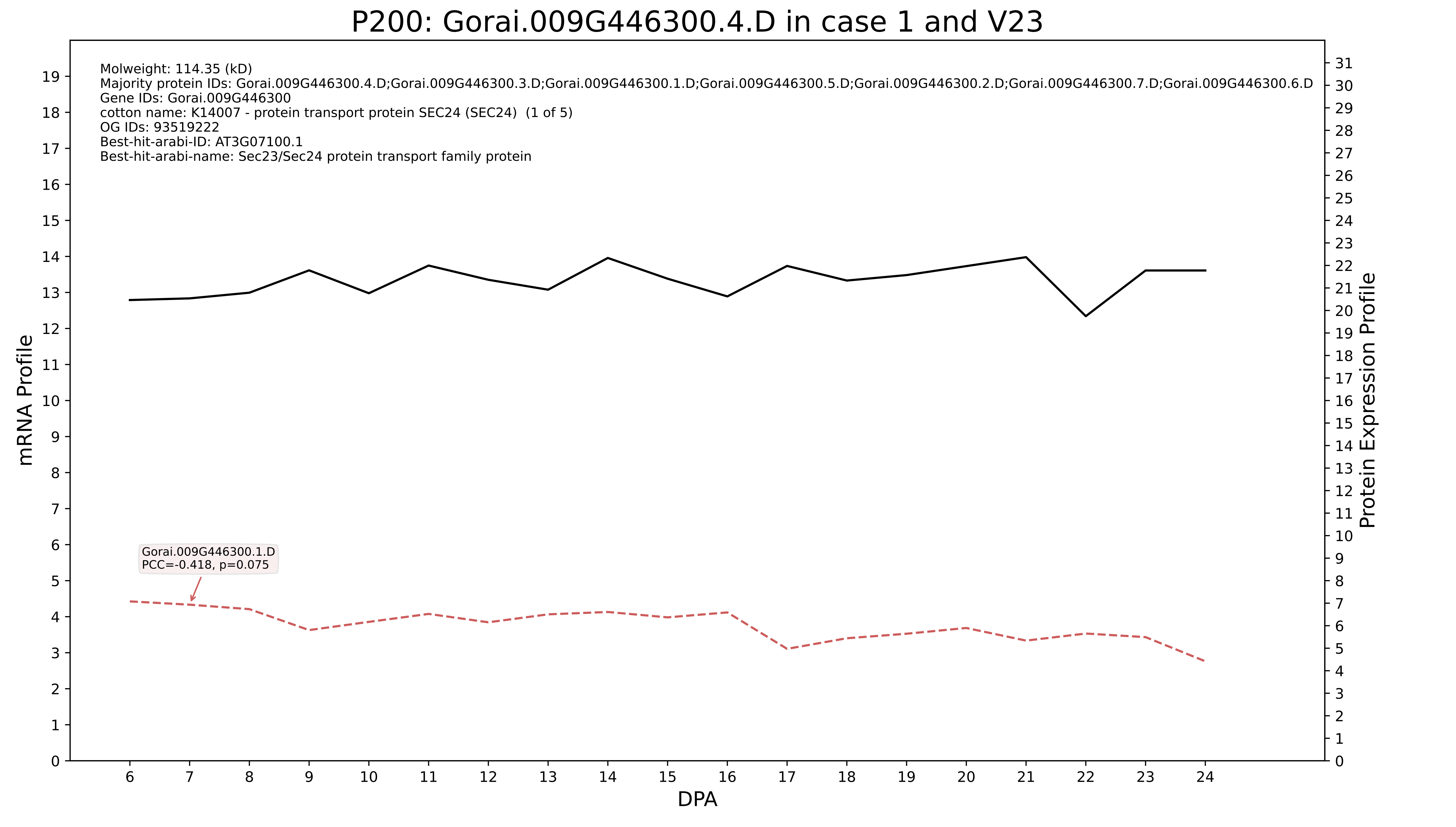
P200: Gorai.009G446300.4.D in case 1 and V23
Molweight: 114.35 (kD)
Majority protein IDs: Gorai.009G446300.4.D;Gorai.009G446300.3.D;Gorai.009G446300.1.D;Gorai.009G446300.5.D;Gorai.009G446300.2.D;Gorai.009G446300.7.D;Gorai.009G446300.6.D
Gene IDs: Gorai.009G446300
cotton name: K14007 - protein transport protein SEC24 (SEC24) (1 of 5)
OG IDs: 93519222
Best-hit-arabi-ID: AT3G07100.1
Best-hit-arabi-name: Sec23/Sec24 protein transport family protein
Gorai.009G446300.1.D
PCC=-0.418, p=0.075
mRNA Profile
Protein Expression Profile
DPA
0 1 2 3 4 5 6 7 8 9 10 11 12 13 14 15 16 17 18 19
0 1 2 3 4 5 6 7 8 9 10 11 12 13 14 15 16 17 18 19 20 21 22 23 24 25 26 27 28 29 30 31
6 7 8 9 10 11 12 13 14 15 16 17 18 19 20 21 22 23 24

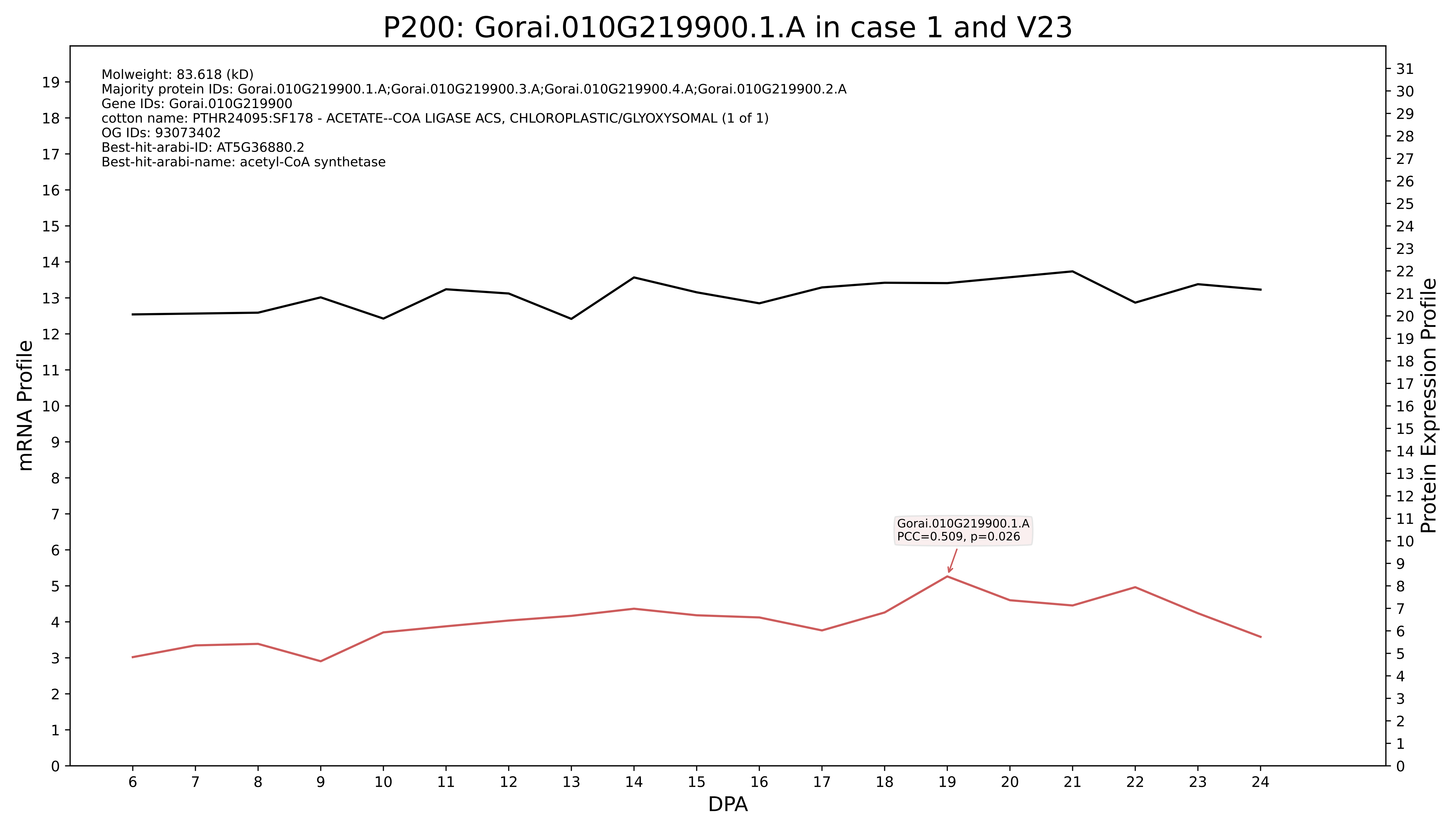
P200: Gorai.010G219900.1.A in case 1 and V23
Molweight: 83.618 (kD)
Majority protein IDs: Gorai.010G219900.1.A;Gorai.010G219900.3.A;Gorai.010G219900.4.A;Gorai.010G219900.2.A
Gene IDs: Gorai.010G219900
cotton name: PTHR24095:SF178 - ACETATE--COA LIGASE ACS, CHLOROPLASTIC/GLYOXYSOMAL (1 of 1)
OG IDs: 93073402
Best-hit-arabi-ID: AT5G36880.2
Best-hit-arabi-name: acetyl-CoA synthetase
Gorai.010G219900.1.A
PCC=0.509, p=0.026
mRNA Profile
Protein Expression Profile
DPA

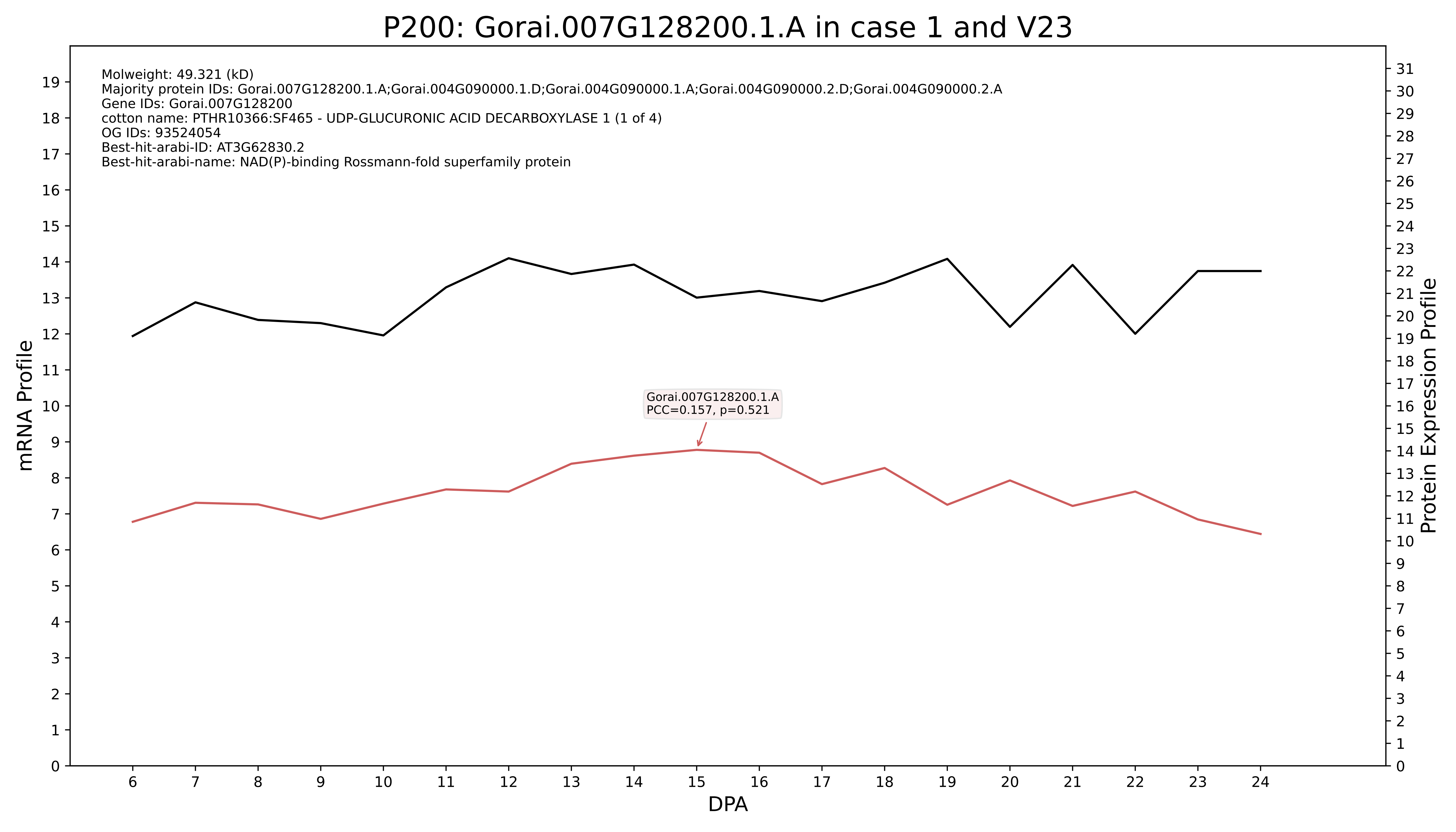
P200: Gorai.007G128200.1.A in case 1 and V23
Molweight: 49.321 (kD)
Majority protein IDs: Gorai.007G128200.1.A;Gorai.004G090000.1.D;Gorai.004G090000.1.A;Gorai.004G090000.2.D;Gorai.004G090000.2.A
Gene IDs: Gorai.007G128200
cotton name: PTHR10366:SF465 - UDP-GLUCURONIC ACID DECARBOXYLASE 1 (1 of 4)
OG IDs: 93524054
Best-hit-arabi-ID: AT3G62830.2
Best-hit-arabi-name: NAD(P)-binding Rossmann-fold superfamily protein
Gorai.007G128200.1.A
PCC=0.157, p=0.521
mRNA Profile
Protein Expression Profile
DPA

### P200: Gorai.003G154700.3.A in case 1 and V23

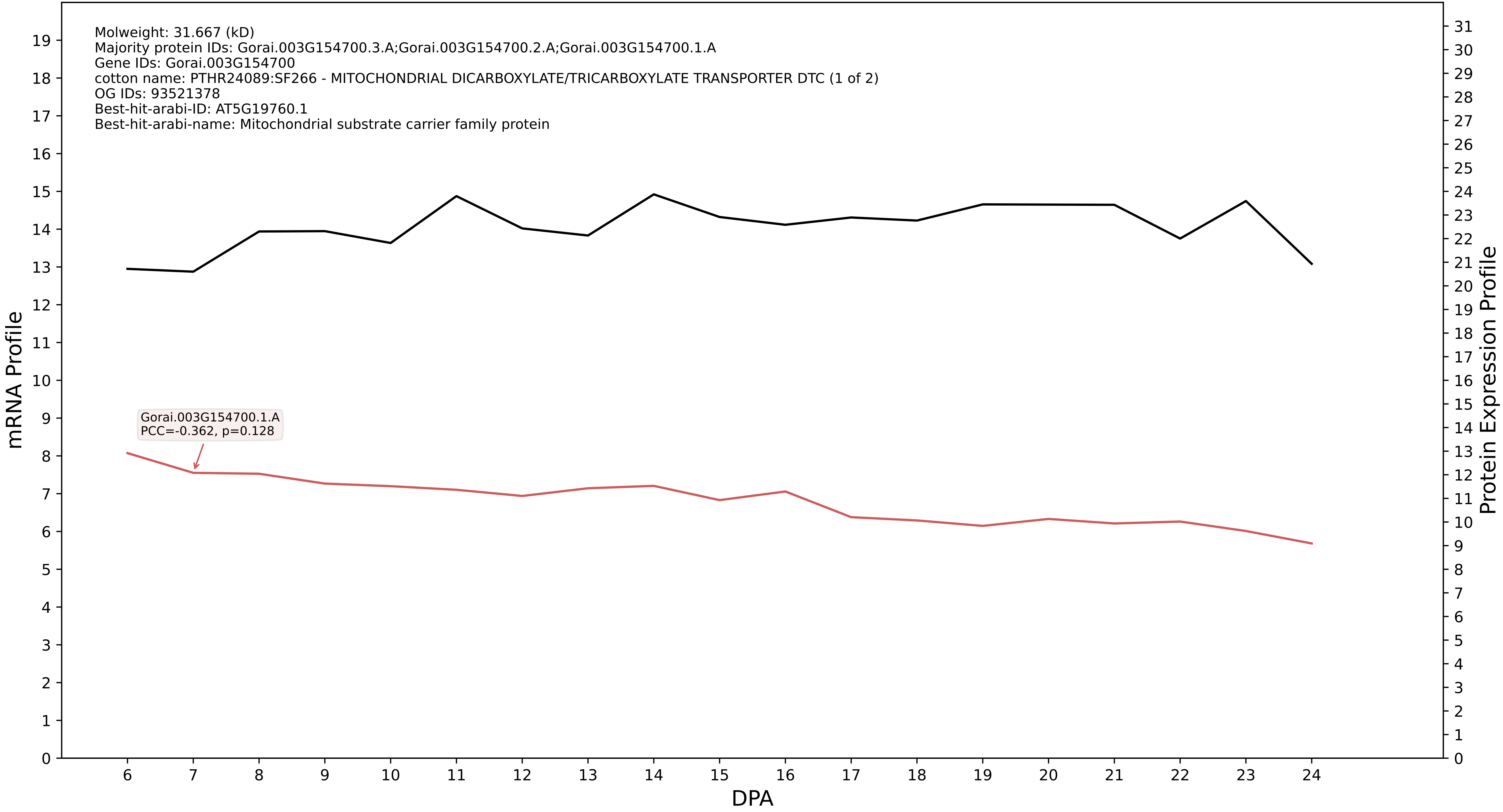

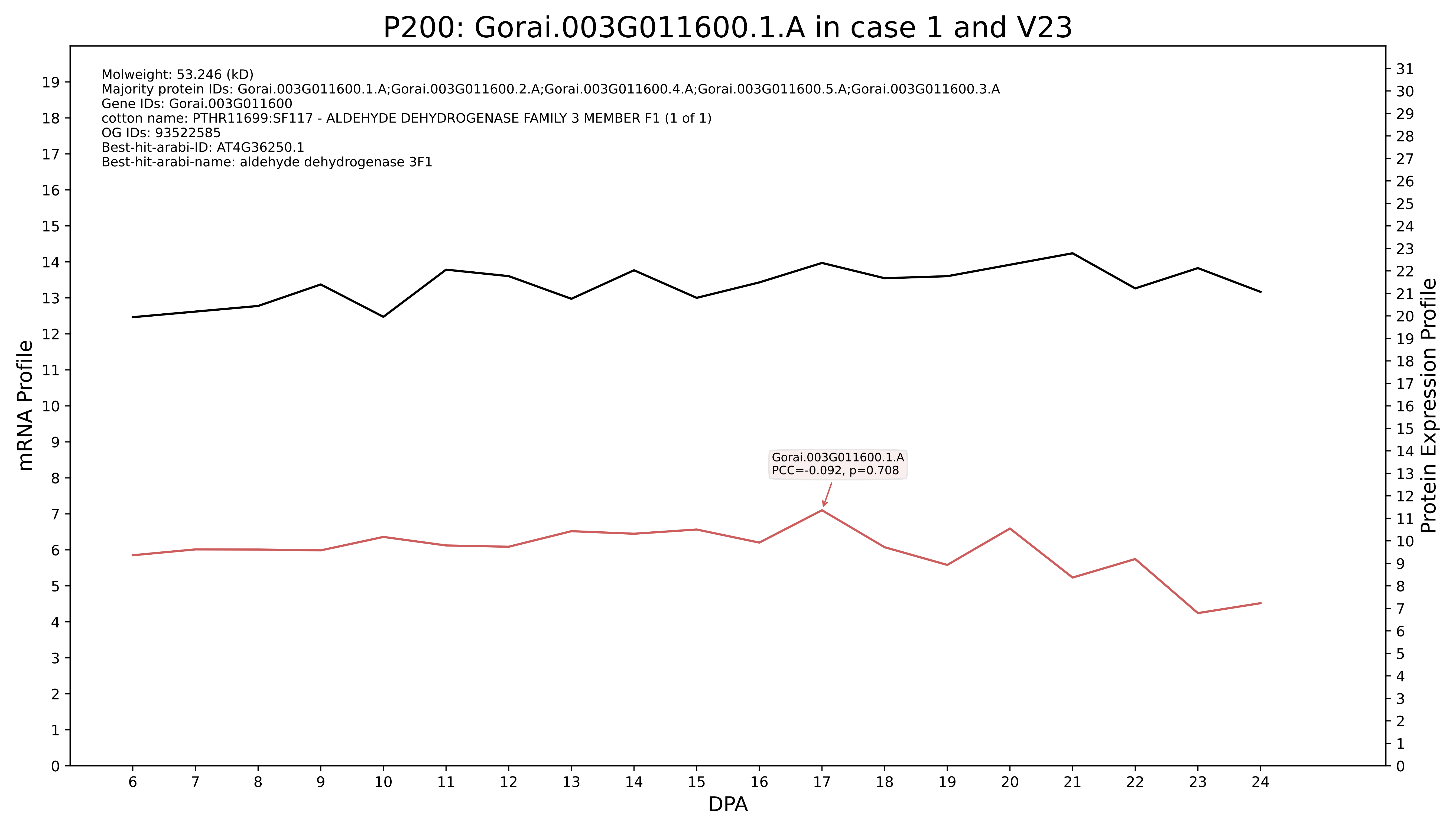
P200: Gorai.003G011600.1.A in case 1 and V23
Molweight: 53.246 (kD)
Majority protein IDs: Gorai.003G011600.1.A;Gorai.003G011600.2.A;Gorai.003G011600.4.A;Gorai.003G011600.5.A;Gorai.003G011600.3.A
Gene IDs: Gorai.003G011600
cotton name: PTHR11699:SF117 - ALDEHYDE DEHYDROGENASE FAMILY 3 MEMBER F1 (1 of 1)
OG IDs: 93522585
Best-hit-arabi-ID: AT4G36250.1
Best-hit-arabi-name: aldehyde dehydrogenase 3F1
Gorai.003G011600.1.A
PCC=-0.092, p=0.708
mRNA Profile
Protein Expression Profile
DPA

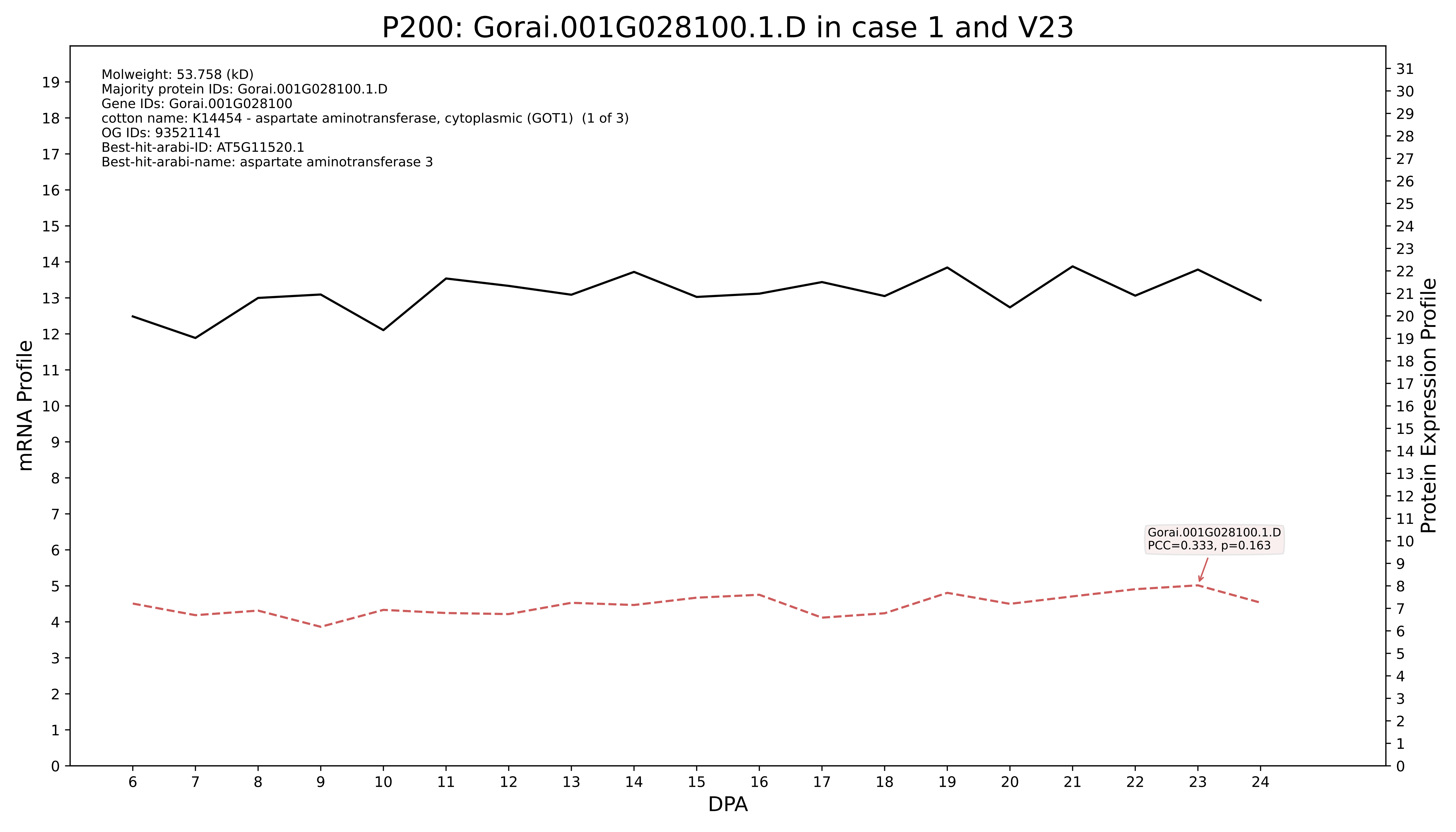
P200: Gorai.001G028100.1.D in case 1 and V23
Molweight: 53.758 (kD)
Majority protein IDs: Gorai.001G028100.1.D
Gene IDs: Gorai.001G028100
cotton name: K14454 - aspartate aminotransferase, cytoplasmic (GOT1) (1 of 3)
OG IDs: 93521141
Best-hit-arabi-ID: AT5G11520.1
Best-hit-arabi-name: aspartate aminotransferase 3
Gorai.001G028100.1.D
PCC=0.333, p=0.163
mRNA Profile
Protein Expression Profile
DPA

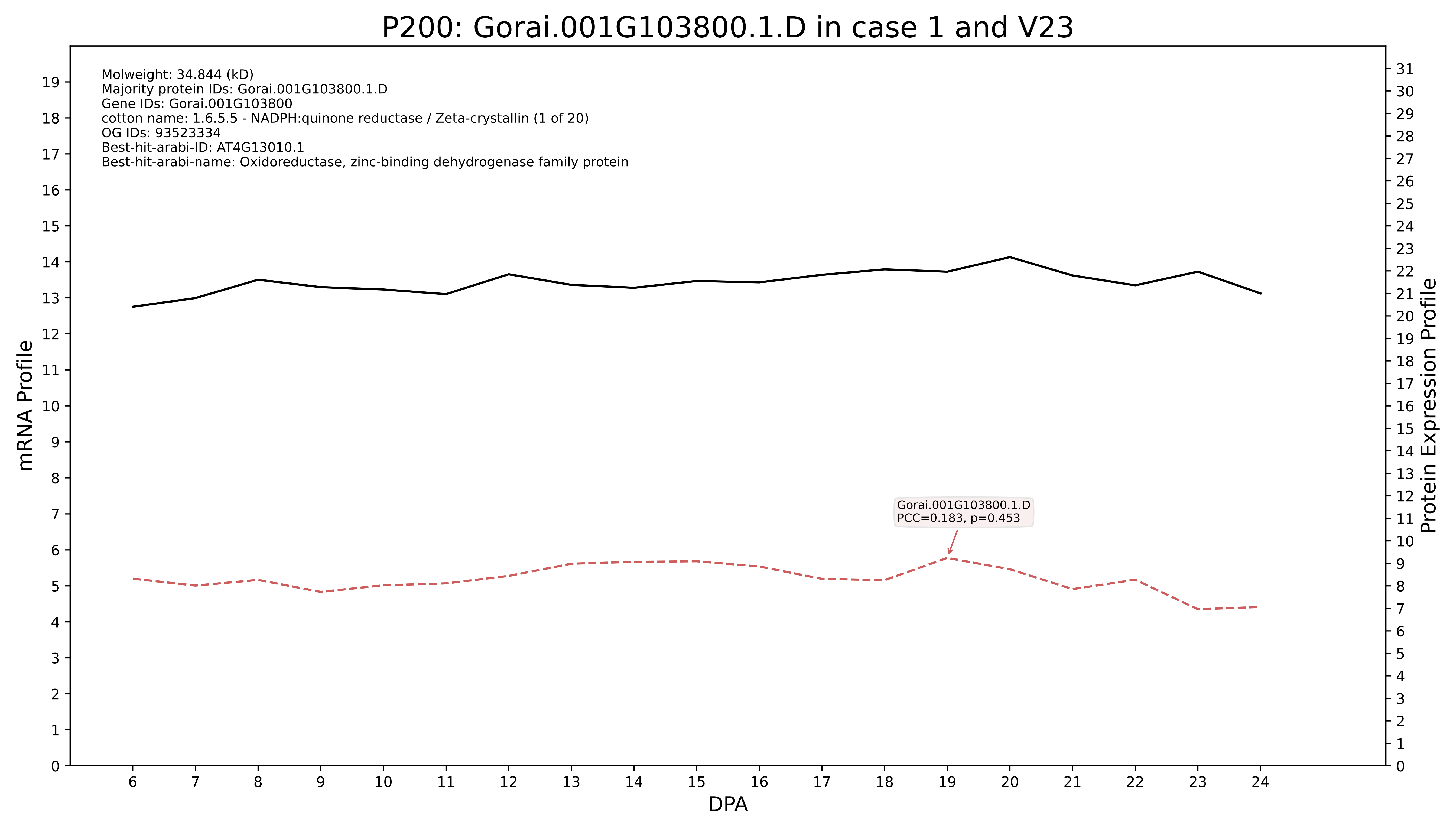
P200: Gorai.001G103800.1.D in case 1 and V23
Molweight: 34.844 (kD)
Majority protein IDs: Gorai.001G103800.1.D
Gene IDs: Gorai.001G103800
cotton name: 1.6.5.5 - NADPH:quinone reductase / Zeta-crystallin (1 of 20)
OG IDs: 93523334
Best-hit-arabi-ID: AT4G13010.1
Best-hit-arabi-name: Oxidoreductase, zinc-binding dehydrogenase family protein
Gorai.001G103800.1.D
PCC=0.183, p=0.453
mRNA Profile
Protein Expression Profile
DPA

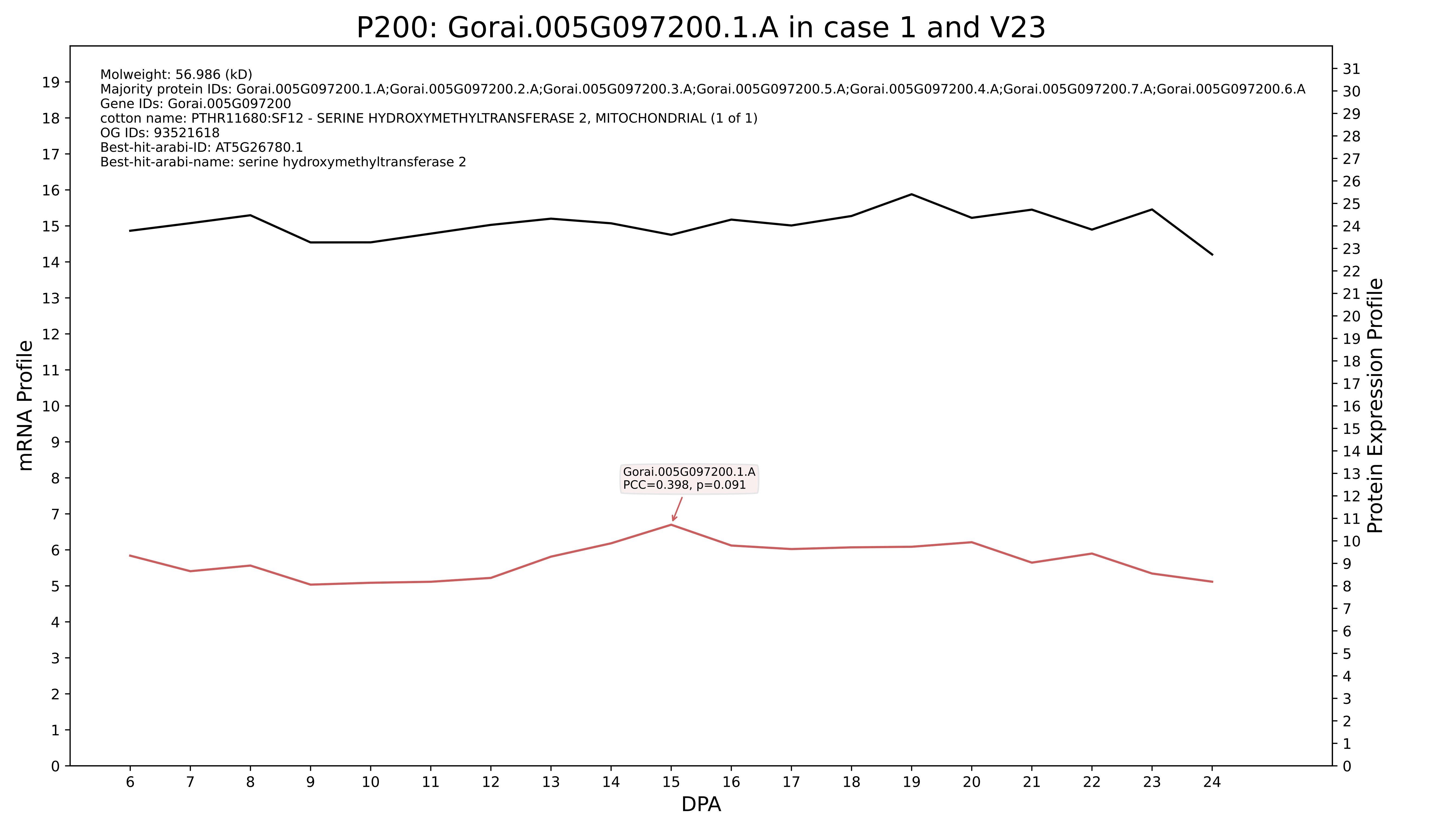

P200: Gorai.005G097200.1.A in case 1 and V23
Molweight: 56.986 (kD)
Majority protein IDs: Gorai.005G097200.1.A;Gorai.005G097200.2.A;Gorai.005G097200.3.A;Gorai.005G097200.5.A;Gorai.005G097200.4.A;Gorai.005G097200.7.A;Gorai.005G097200.6.A
Gene IDs: Gorai.005G097200
cotton name: PTHR11680:SF12 - SERINE HYDROXYMETHYLTRANSFERASE 2, MITOCHONDRIAL (1 of 1)
OG IDs: 93521618
Best-hit-arabi-ID: AT5G26780.1
Best-hit-arabi-name: serine hydroxymethyltransferase 2
Gorai.005G097200.1.A
PCC=0.398, p=0.091
mRNA Profile
Protein Expression Profile
DPA

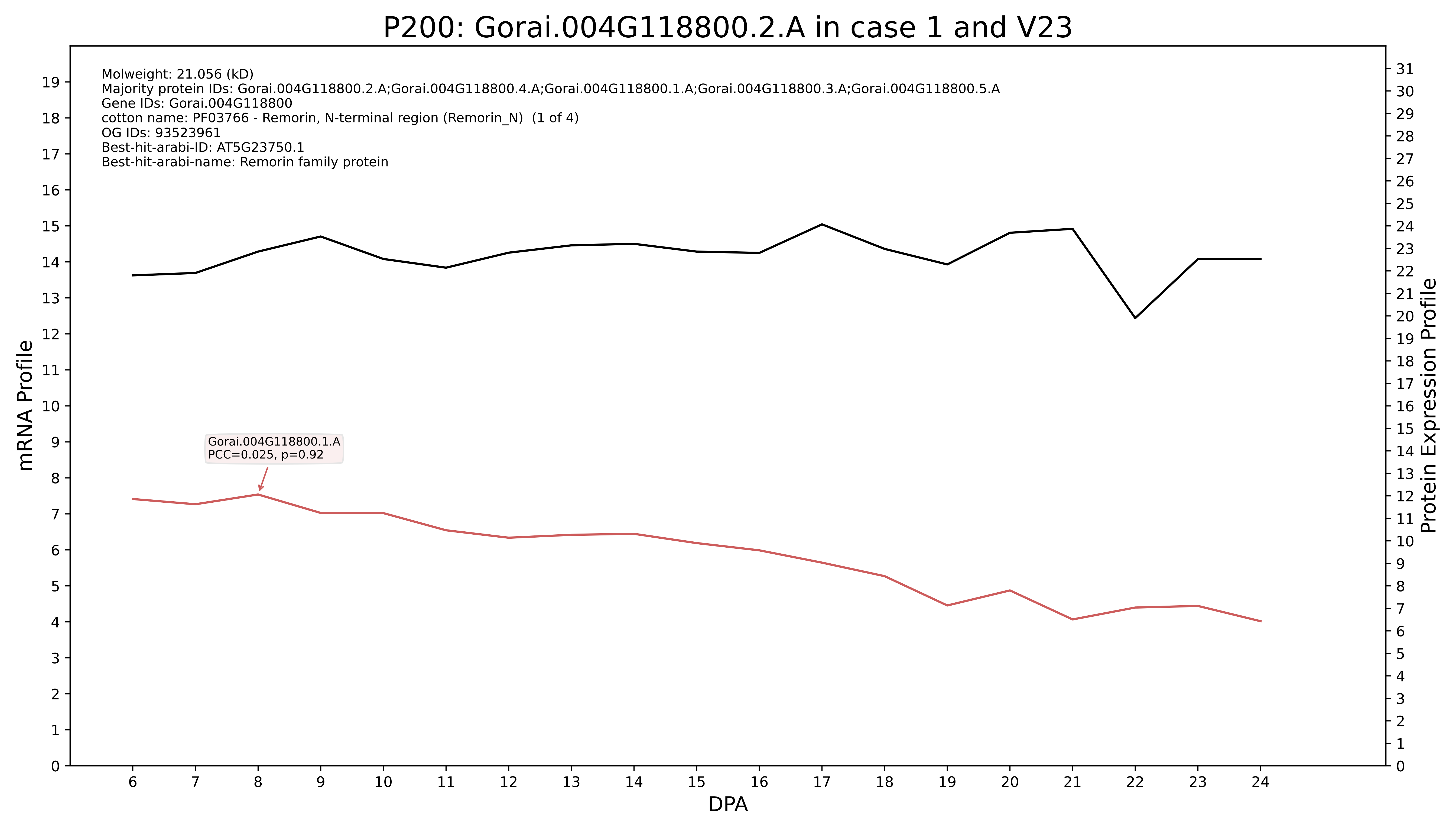
P200: Gorai.004G118800.2.A in case 1 and V23
Molweight: 21.056 (kD)
Majority protein IDs: Gorai.004G118800.2.A;Gorai.004G118800.4.A;Gorai.004G118800.1.A;Gorai.004G118800.3.A;Gorai.004G118800.5.A
Gene IDs: Gorai.004G118800
cotton name: PF03766 - Remorin, N-terminal region (Remorin_N) (1 of 4)
OG IDs: 93523961
Best-hit-arabi-ID: AT5G23750.1
Best-hit-arabi-name: Remorin family protein
Gorai.004G118800.1.A
PCC=0.025, p=0.92
mRNA Profile
Protein Expression Profile
DPA

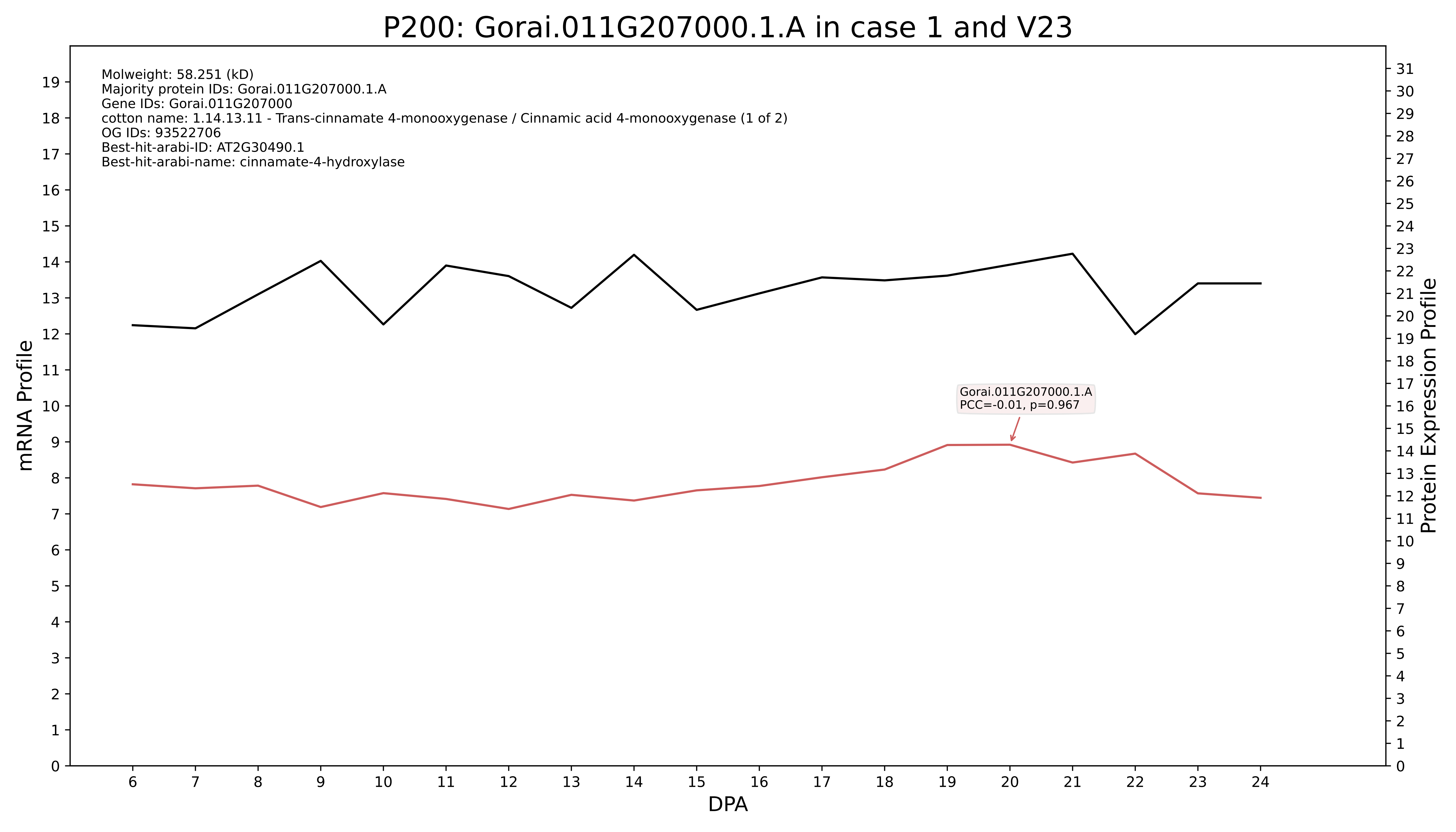

P200: Gorai.011G207000.1.A in case 1 and V23
Molweight: 58.251 (kD)
Majority protein IDs: Gorai.011G207000.1.A
Gene IDs: Gorai.011G207000
cotton name: 1.14.13.11 - Trans-cinnamate 4-monooxygenase / Cinnamic acid 4-monooxygenase (1 of 2)
OG IDs: 93522706
Best-hit-arabi-ID: AT2G30490.1
Best-hit-arabi-name: cinnamate-4-hydroxylase
Gorai.011G207000.1.A
PCC=-0.01, p=0.967
mRNA Profile
Protein Expression Profile
DPA

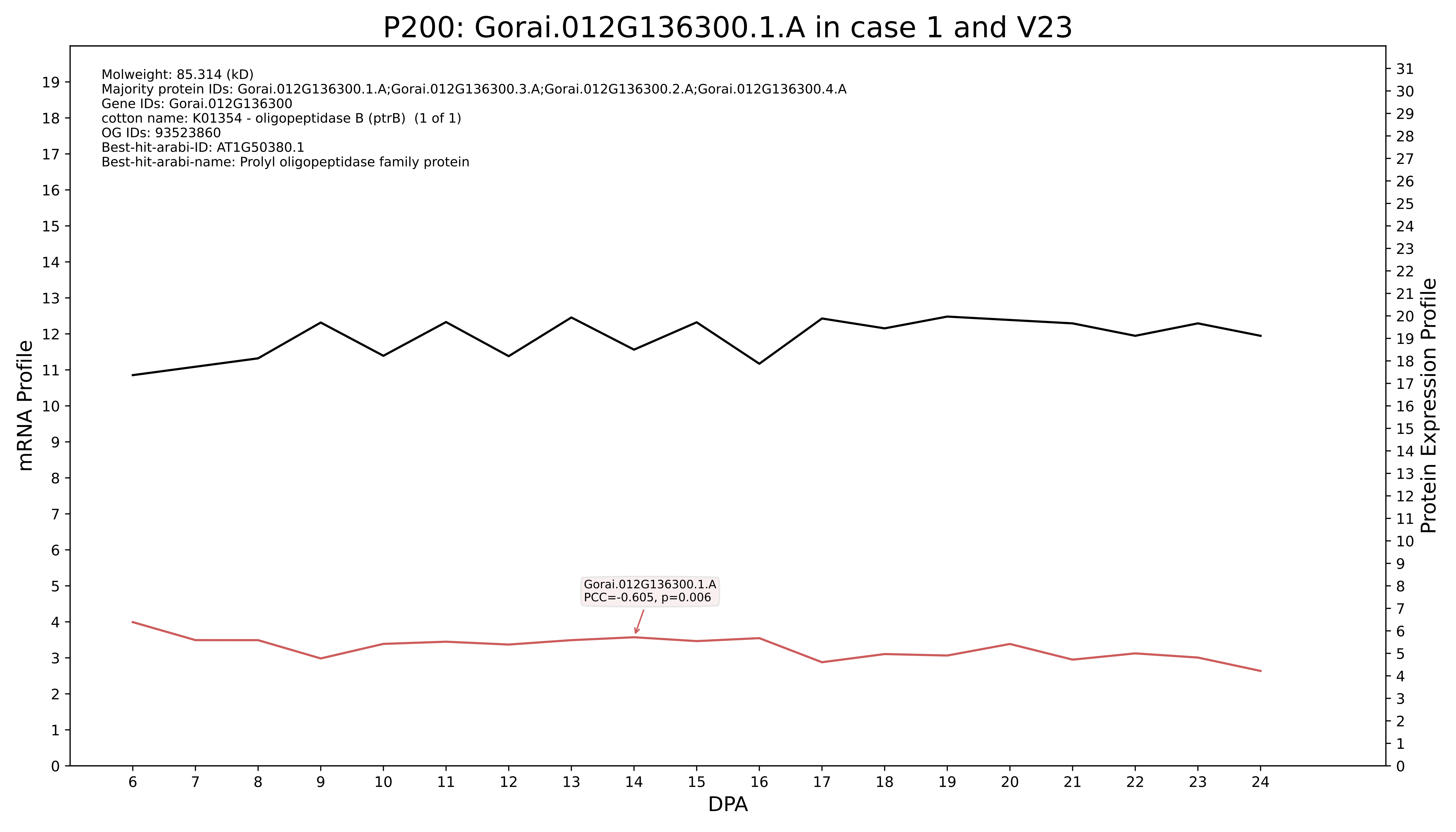
P200: Gorai.012G136300.1.A in case 1 and V23
Molweight: 85.314 (kD)
Majority protein IDs: Gorai.012G136300.1.A;Gorai.012G136300.3.A;Gorai.012G136300.2.A;Gorai.012G136300.4.A
Gene IDs: Gorai.012G136300
cotton name: K01354 - oligopeptidase B (ptrB) (1 of 1)
OG IDs: 93523860
Best-hit-arabi-ID: AT1G50380.1
Best-hit-arabi-name: Prolyl oligopeptidase family protein
Gorai.012G136300.1.A
PCC=-0.605, p=0.006
mRNA Profile
Protein Expression Profile
DPA

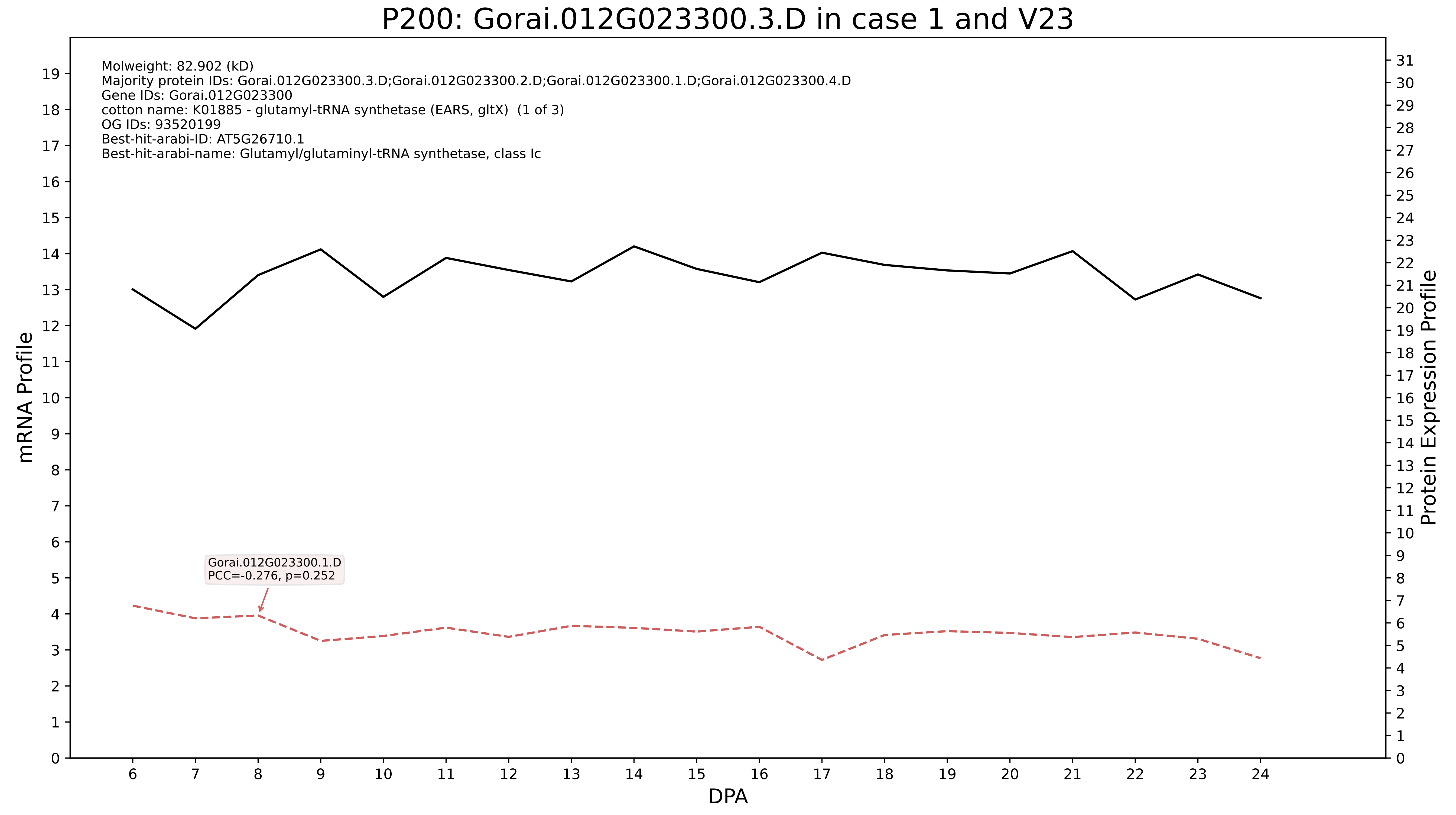
P200: Gorai.012G023300.3.D in case 1 and V23
Molweight: 82.902 (kD)
Majority protein IDs: Gorai.012G023300.3.D;Gorai.012G023300.2.D;Gorai.012G023300.1.D;Gorai.012G023300.4.D
Gene IDs: Gorai.012G023300
cotton name: K01885 - glutamyl-tRNA synthetase (EARS, gltX) (1 of 3)
OG IDs: 93520199
Best-hit-arabi-ID: AT5G26710.1
Best-hit-arabi-name: Glutamyl/glutaminyl-tRNA synthetase, class Ic
Gorai.012G023300.1.D
PCC=-0.276, p=0.252
mRNA Profile
Protein Expression Profile
DPA

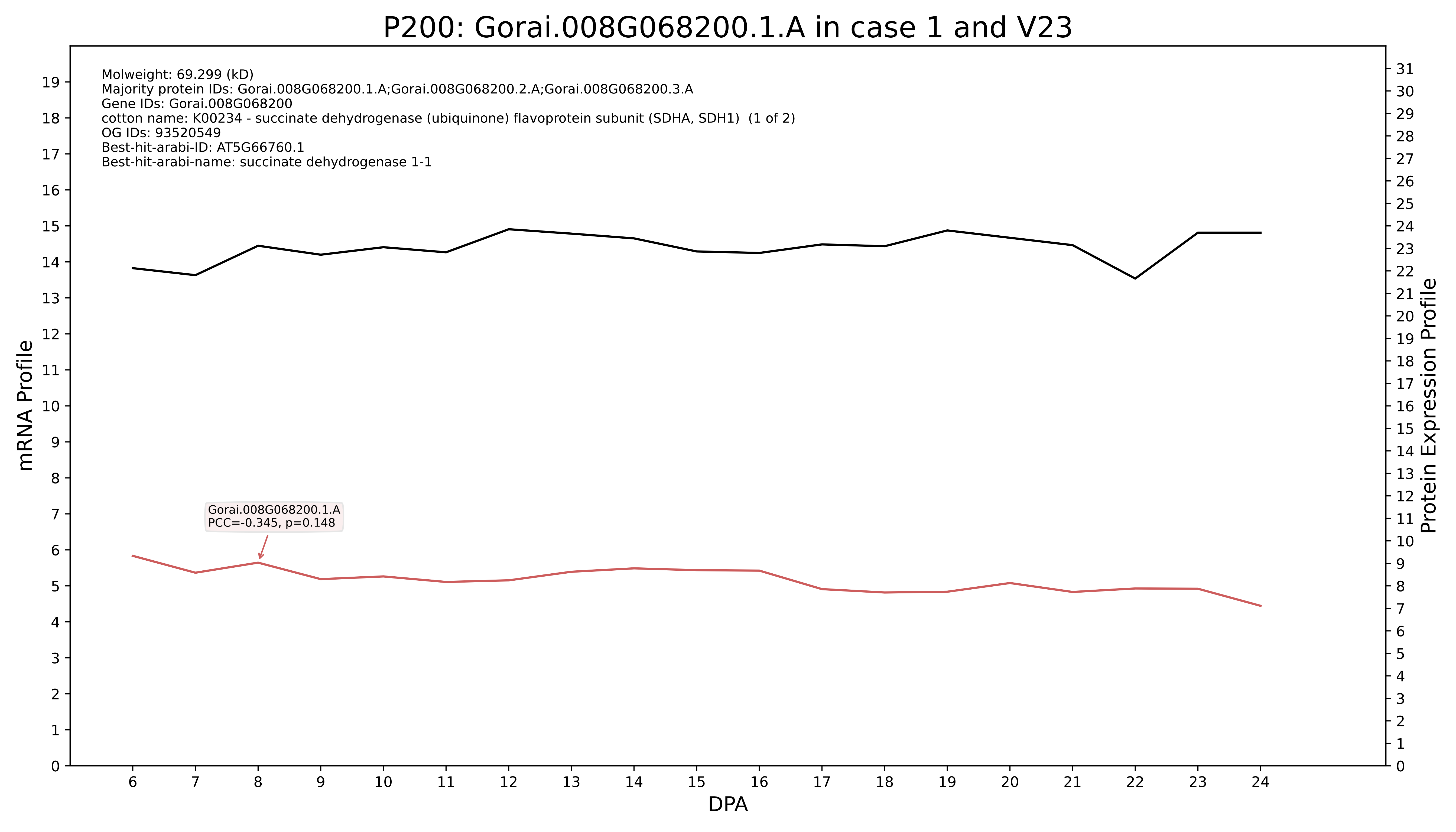

P200: Gorai.008G068200.1.A in case 1 and V23
Molweight: 69.299 (kD)
Majority protein IDs: Gorai.008G068200.1.A;Gorai.008G068200.2.A;Gorai.008G068200.3.A
Gene IDs: Gorai.008G068200
cotton name: K00234 - succinate dehydrogenase (ubiquinone) flavoprotein subunit (SDHA, SDH1) (1 of 2)
OG IDs: 93520549
Best-hit-arabi-ID: AT5G66760.1
Best-hit-arabi-name: succinate dehydrogenase 1-1
Gorai.008G068200.1.A
PCC=-0.345, p=0.148
mRNA Profile
Protein Expression Profile
DPA

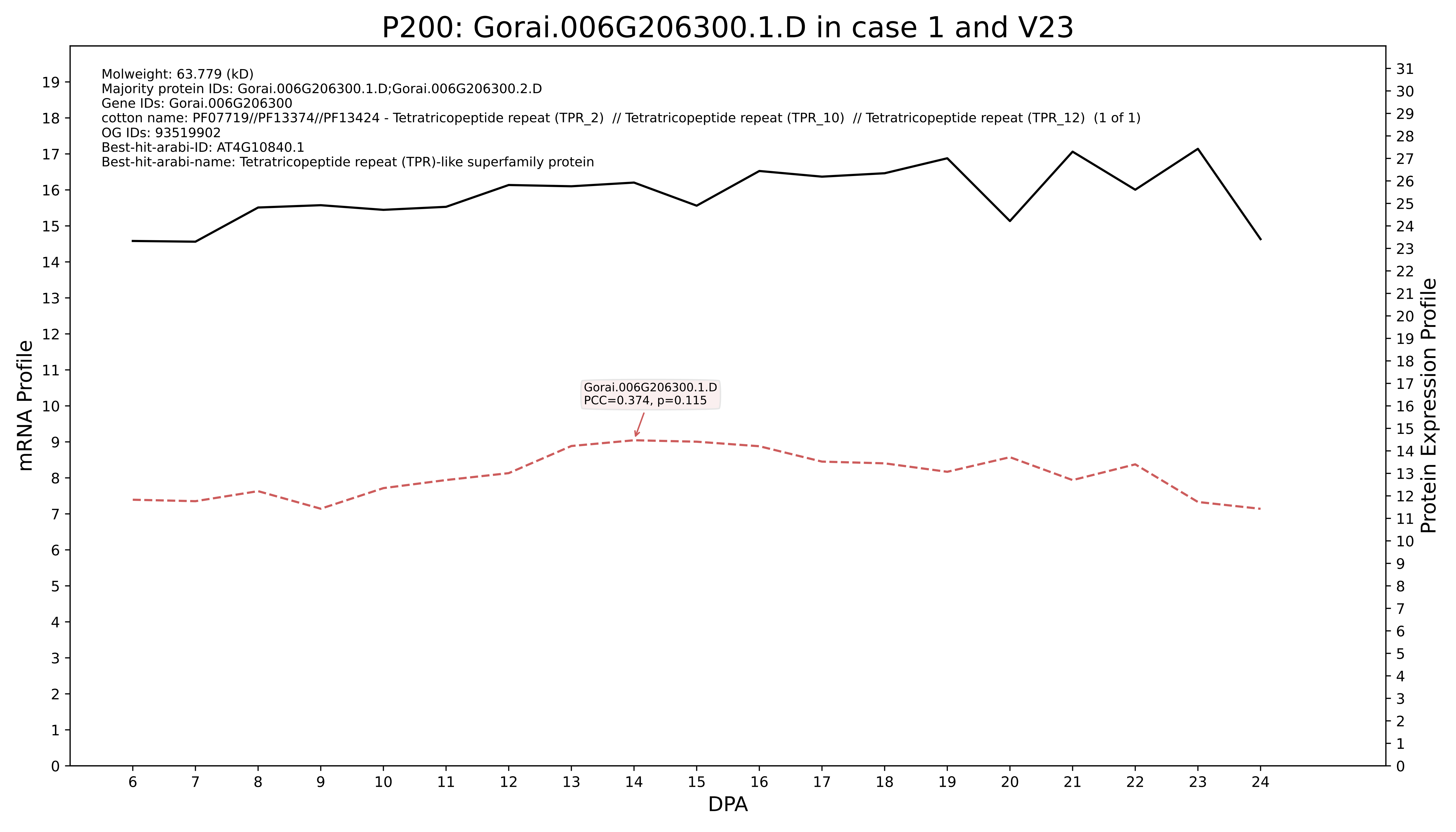

P200: Gorai.006G206300.1.D in case 1 and V23
Molweight: 63.779 (kD)
Majority protein IDs: Gorai.006G206300.1.D;Gorai.006G206300.2.D
Gene IDs: Gorai.006G206300
cotton name: PF07719//PF13374//PF13424 - Tetratricopeptide repeat (TPR_2) // Tetratricopeptide repeat (TPR_10) // Tetratricopeptide repeat (TPR_12) (1 of 1)
OG IDs: 93519902
Best-hit-arabi-ID: AT4G10840.1
Best-hit-arabi-name: Tetratricopeptide repeat (TPR)-like superfamily protein
Gorai.006G206300.1.D
PCC=0.374, p=0.115
mRNA Profile
Protein Expression Profile
DPA

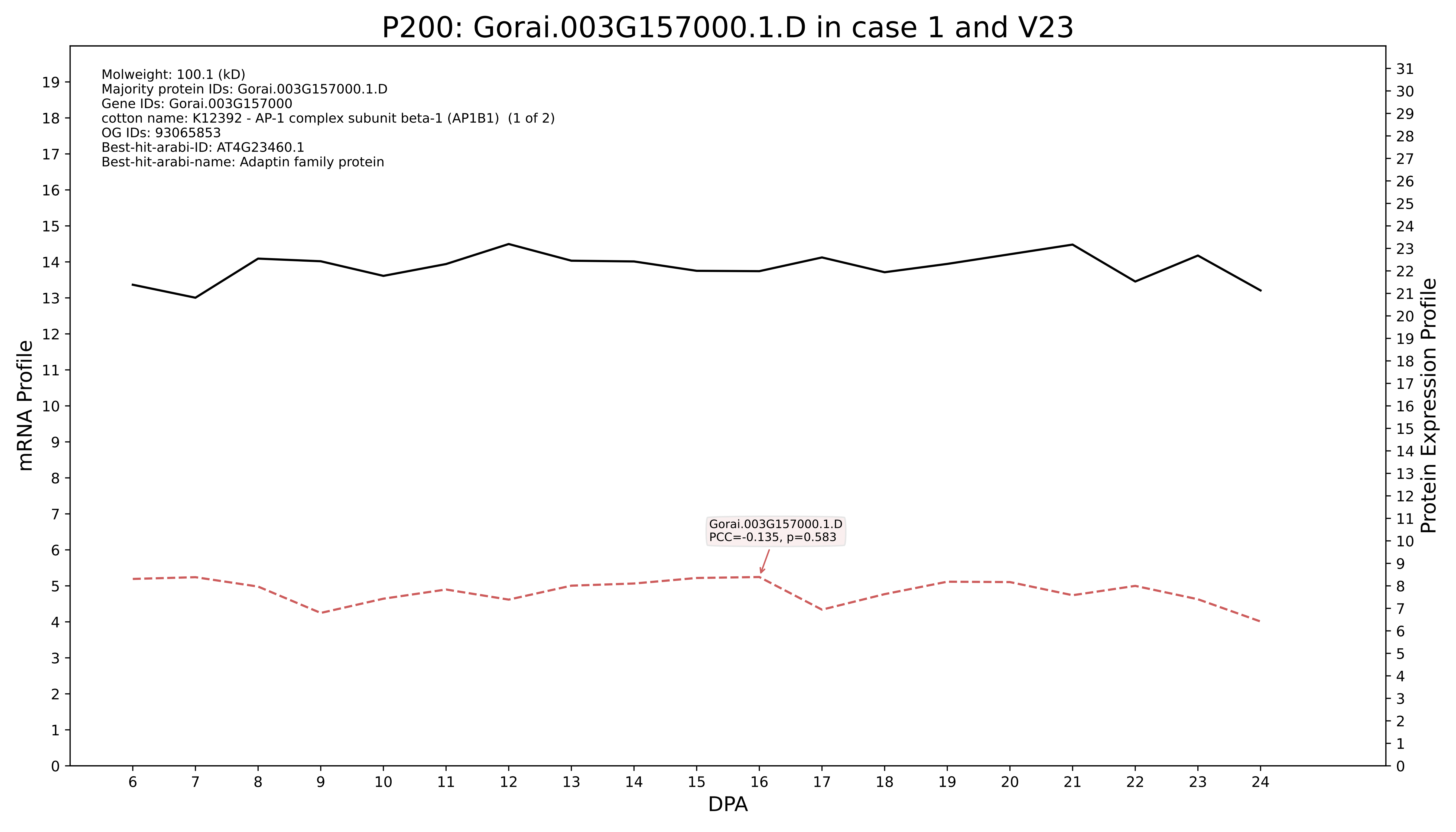
P200: Gorai.003G157000.1.D in case 1 and V23
Molweight: 100.1 (kD)
Majority protein IDs: Gorai.003G157000.1.D
Gene IDs: Gorai.003G157000
cotton name: K12392 - AP-1 complex subunit beta-1 (AP1B1) (1 of 2)
OG IDs: 93065853
Best-hit-arabi-ID: AT4G23460.1
Best-hit-arabi-name: Adaptin family protein
Gorai.003G157000.1.D
PCC=-0.135, p=0.583
mRNA Profile
Protein Expression Profile
DPA

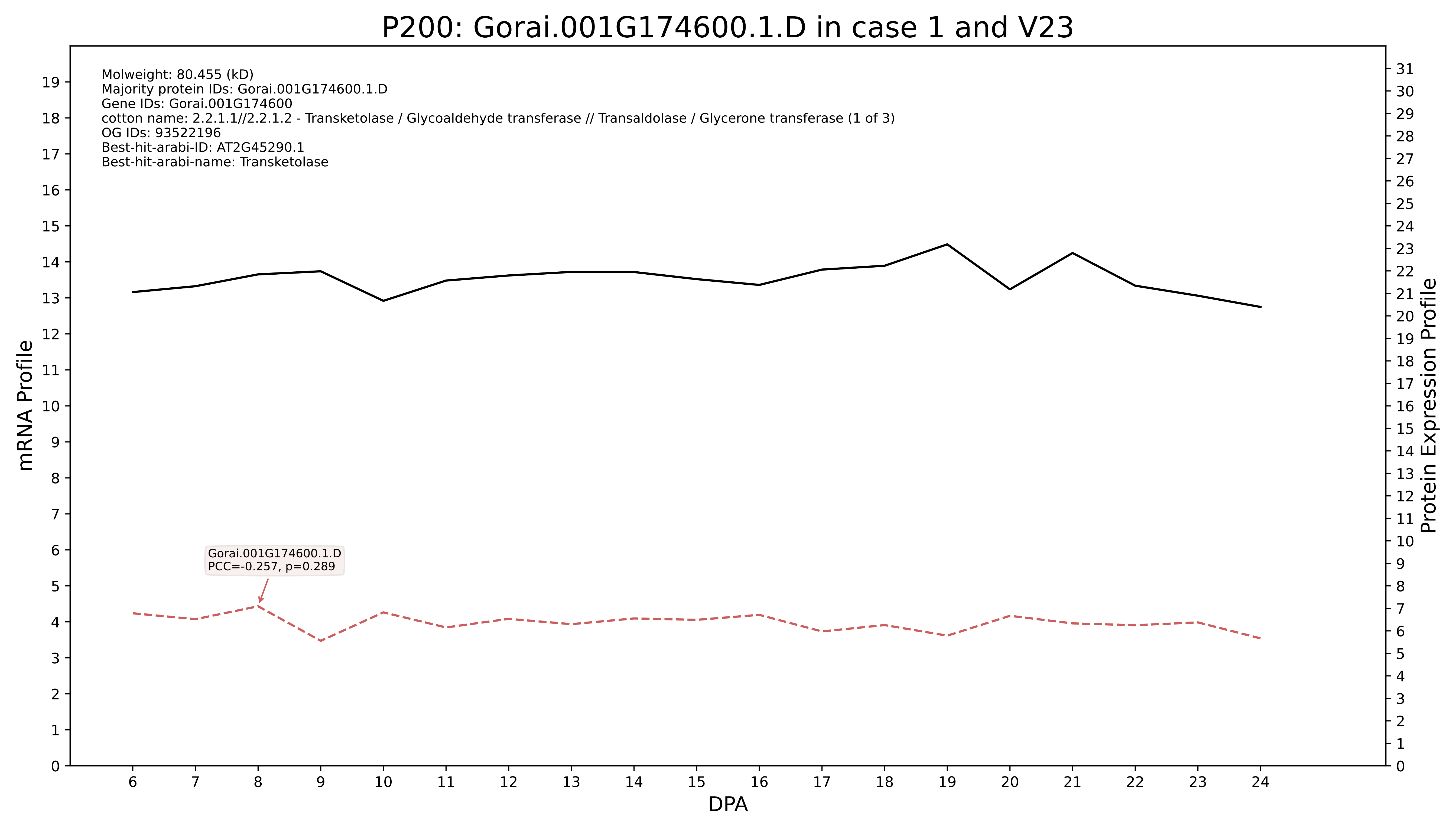

P200: Gorai.001G174600.1.D in case 1 and V23
Molweight: 80.455 (kD)
Majority protein IDs: Gorai.001G174600.1.D
Gene IDs: Gorai.001G174600
cotton name: 2.2.1.1//2.2.1.2 - Transketolase / Glycoaldehyde transferase // Transaldolase / Glycerone transferase (1 of 3)
OG IDs: 93522196
Best-hit-arabi-ID: AT2G45290.1
Best-hit-arabi-name: Transketolase
Gorai.001G174600.1.D
PCC=-0.257, p=0.289
mRNA Profile
Protein Expression Profile
DPA

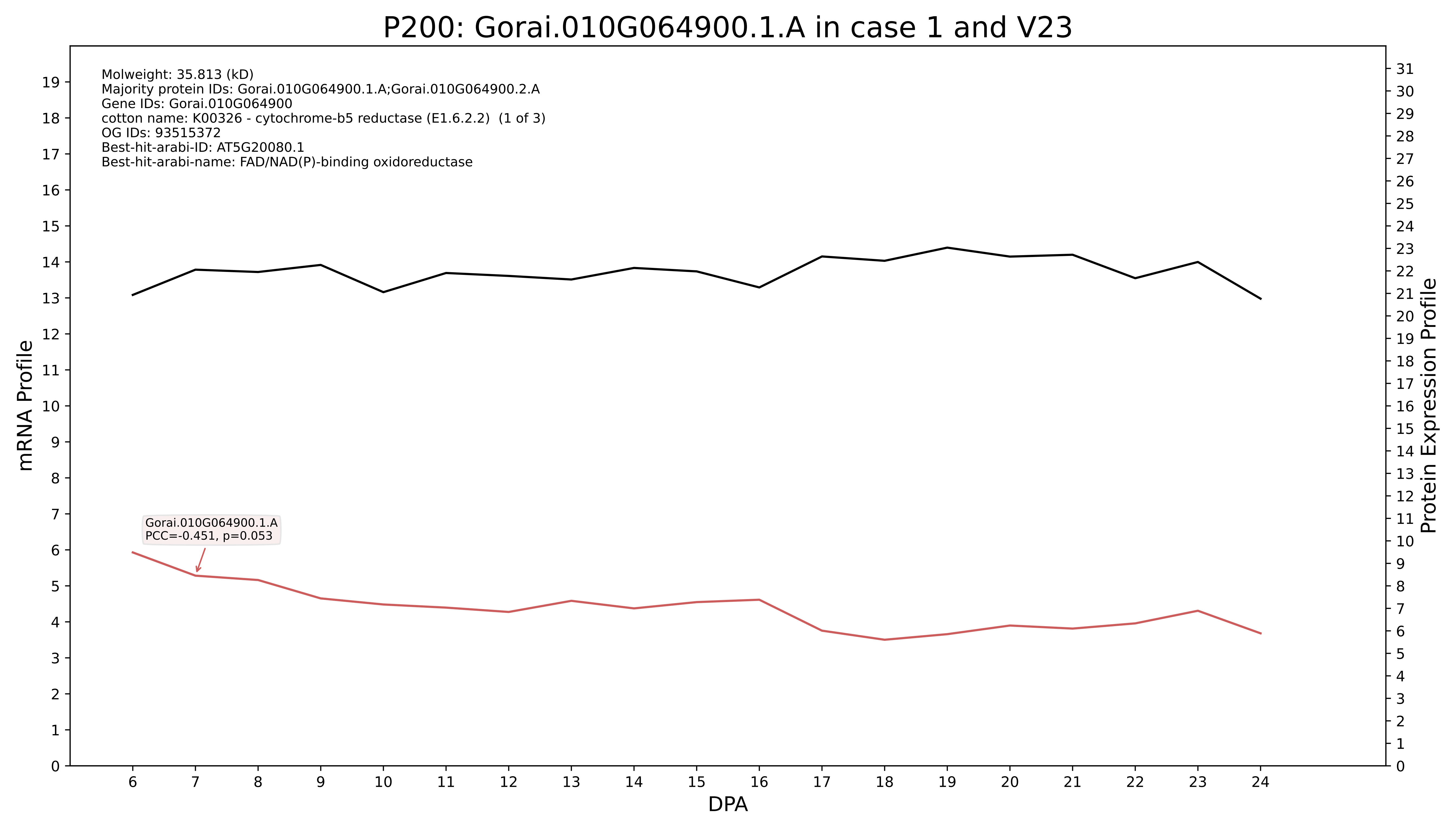
P200: Gorai.010G064900.1.A in case 1 and V23
Molweight: 35.813 (kD)
Majority protein IDs: Gorai.010G064900.1.A;Gorai.010G064900.2.A
Gene IDs: Gorai.010G064900
cotton name: K00326 - cytochrome-b5 reductase (E1.6.2.2) (1 of 3)
OG IDs: 93515372
Best-hit-arabi-ID: AT5G20080.1
Best-hit-arabi-name: FAD/NAD(P)-binding oxidoreductase
Gorai.010G064900.1.A
PCC=-0.451, p=0.053
mRNA Profile
Protein Expression Profile
DPA

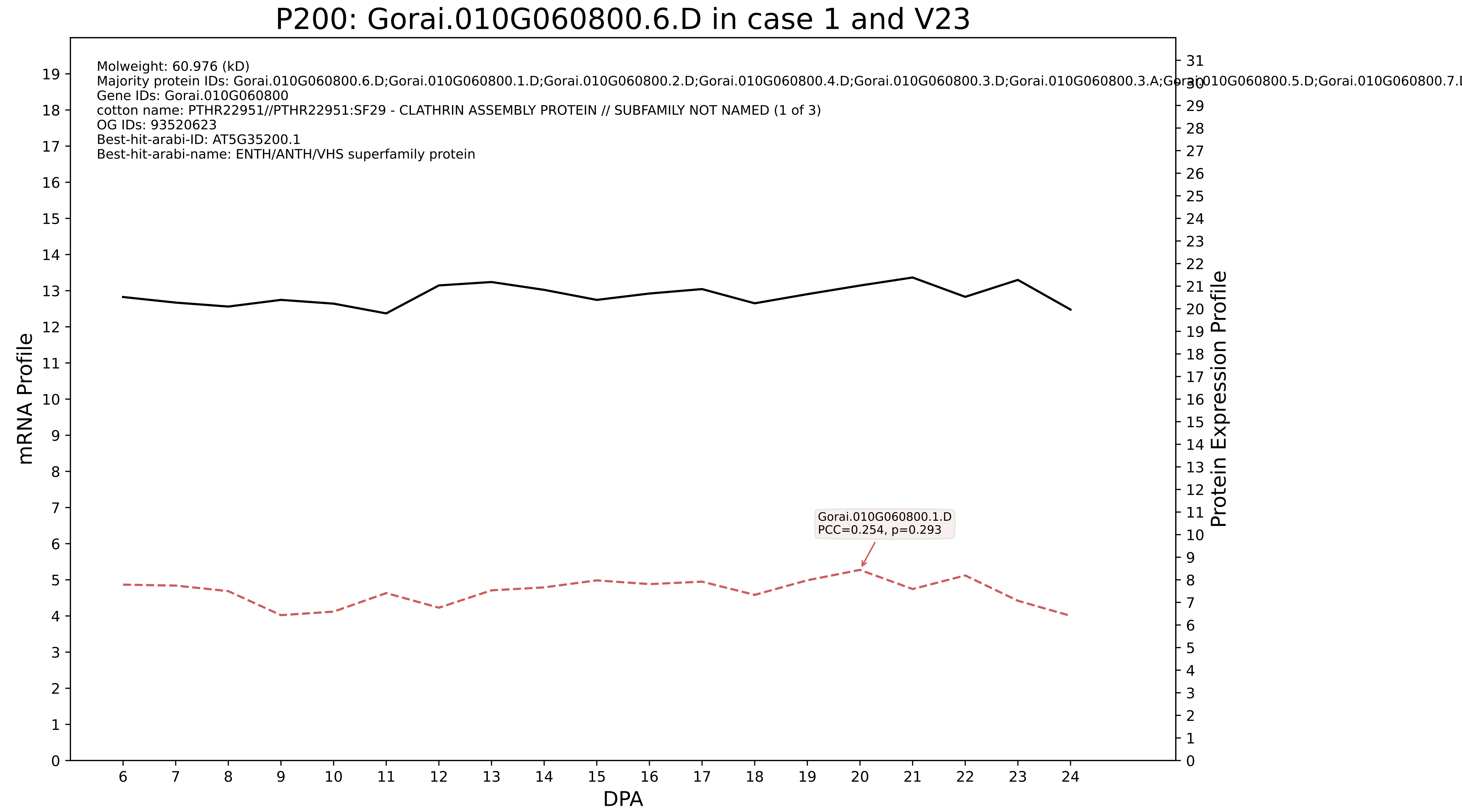
P200: Gorai.010G060800.6.D in case 1 and V23
Molweight: 60.976 (kD)
Majority protein IDs: Gorai.010G060800.6.D;Gorai.010G060800.1.D;Gorai.010G060800.2.D;Gorai.010G060800.4.D;Gorai.010G060800.3.D;Gorai.010G060800.3.A;Gorai.010G060800.5.D;Gorai.010G060800.7.D
Gene IDs: Gorai.010G060800
cotton name: PTHR22951//PTHR22951:SF29 - CLATHRIN ASSEMBLY PROTEIN // SUBFAMILY NOT NAMED (1 of 3)
OG IDs: 93520623
Best-hit-arabi-ID: AT5G35200.1
Best-hit-arabi-name: ENTH/ANTH/VHS superfamily protein
Gorai.010G060800.1.D
PCC=0.254, p=0.293
mRNA Profile
Protein Expression Profile
DPA

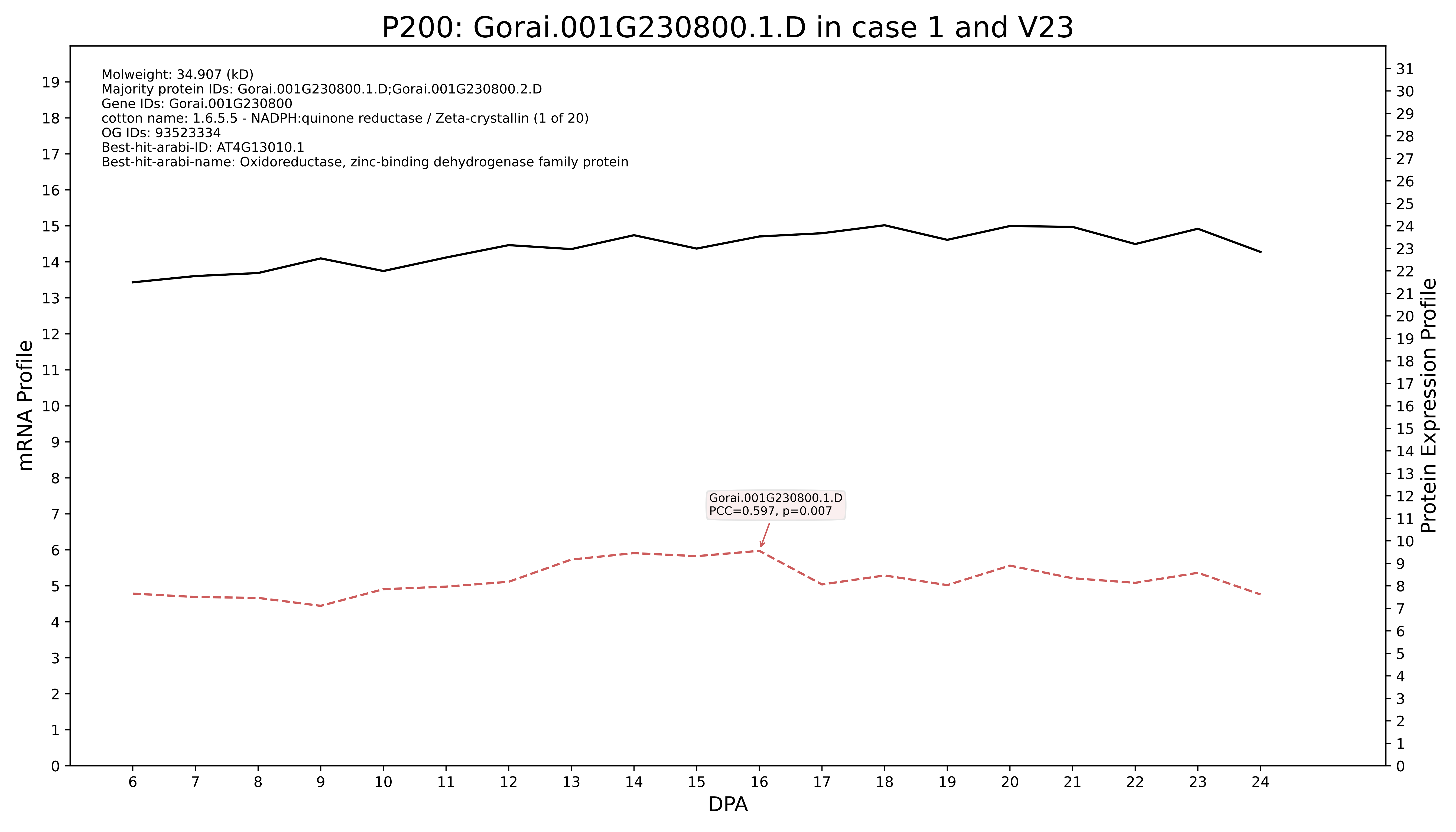

P200: Gorai.001G230800.1.D in case 1 and V23
Molweight: 34.907 (kD)
Majority protein IDs: Gorai.001G230800.1.D;Gorai.001G230800.2.D
Gene IDs: Gorai.001G230800
cotton name: 1.6.5.5 - NADPH:quinone reductase / Zeta-crystallin (1 of 20)
OG IDs: 93523334
Best-hit-arabi-ID: AT4G13010.1
Best-hit-arabi-name: Oxidoreductase, zinc-binding dehydrogenase family protein
Gorai.001G230800.1.D
PCC=0.597, p=0.007
mRNA Profile
Protein Expression Profile
DPA

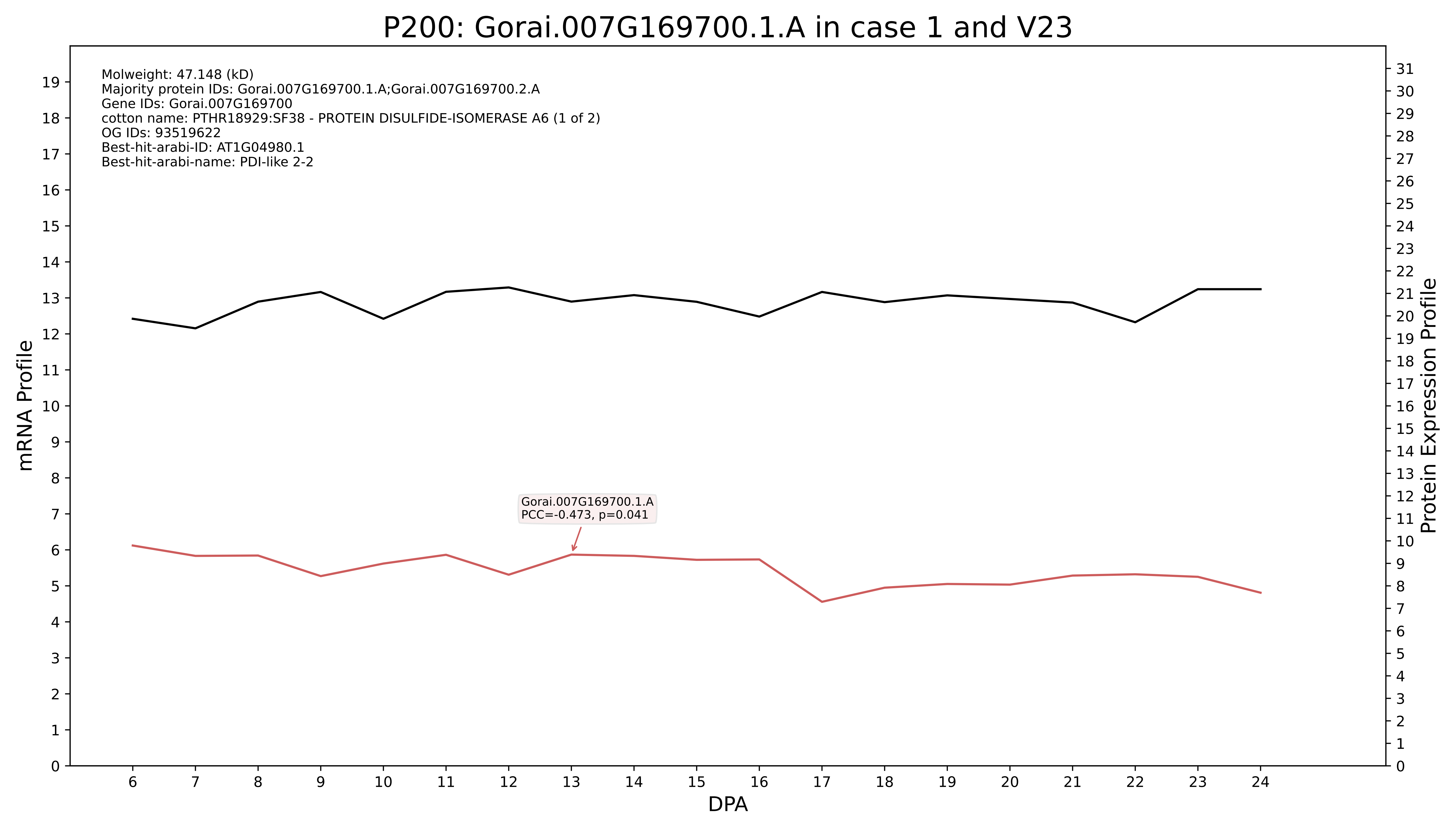

P200: Gorai.007G169700.1.A in case 1 and V23
Molweight: 47.148 (kD)
Majority protein IDs: Gorai.007G169700.1.A;Gorai.007G169700.2.A
Gene IDs: Gorai.007G169700
cotton name: PTHR18929:SF38 - PROTEIN DISULFIDE-ISOMERASE A6 (1 of 2)
OG IDs: 93519622
Best-hit-arabi-ID: AT1G04980.1
Best-hit-arabi-name: PDI-like 2-2
Gorai.007G169700.1.A
PCC=-0.473, p=0.041
mRNA Profile
Protein Expression Profile
DPA

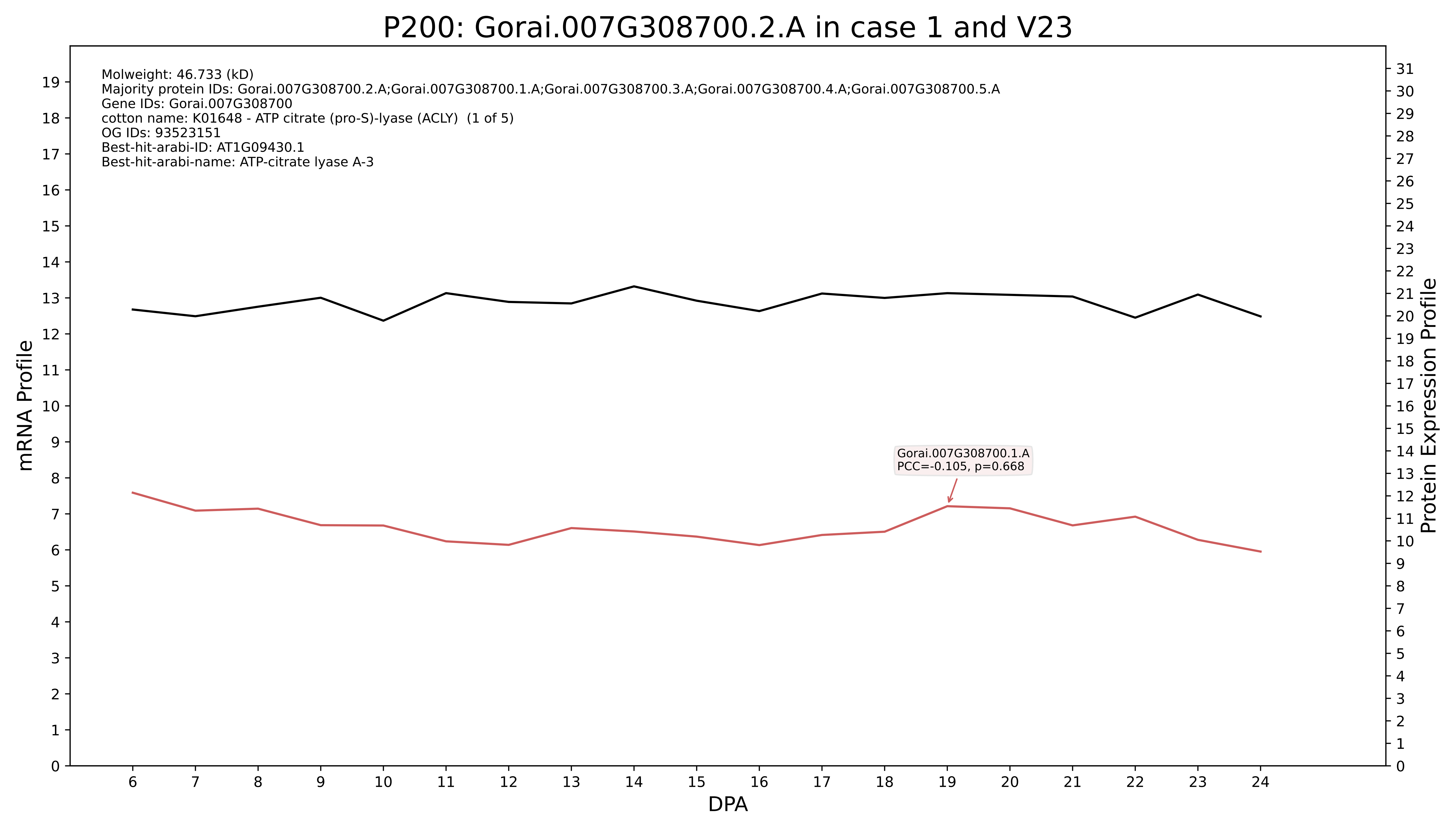
P200: Gorai.007G308700.2.A in case 1 and V23
Molweight: 46.733 (kD)
Majority protein IDs: Gorai.007G308700.2.A;Gorai.007G308700.1.A;Gorai.007G308700.3.A;Gorai.007G308700.4.A;Gorai.007G308700.5.A
Gene IDs: Gorai.007G308700
cotton name: K01648 - ATP citrate (pro-S)-lyase (ACLY) (1 of 5)
OG IDs: 93523151
Best-hit-arabi-ID: AT1G09430.1
Best-hit-arabi-name: ATP-citrate lyase A-3
Gorai.007G308700.1.A
PCC=-0.105, p=0.668
mRNA Profile
Protein Expression Profile
DPA

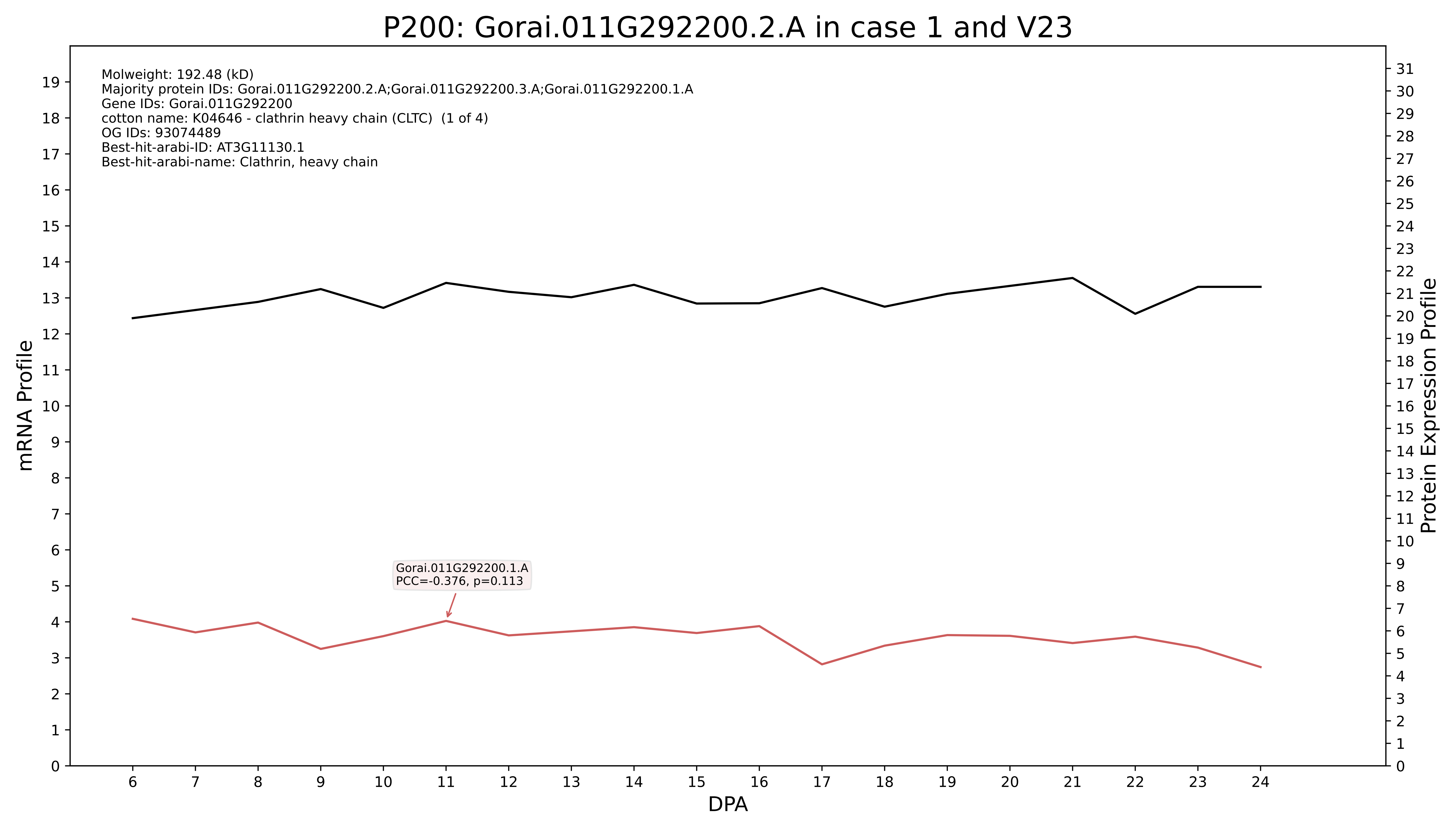
P200: Gorai.011G292200.2.A in case 1 and V23
Molweight: 192.48 (kD)
Majority protein IDs: Gorai.011G292200.2.A;Gorai.011G292200.3.A;Gorai.011G292200.1.A
Gene IDs: Gorai.011G292200
cotton name: K04646 - clathrin heavy chain (CLTC) (1 of 4)
OG IDs: 93074489
Best-hit-arabi-ID: AT3G11130.1
Best-hit-arabi-name: Clathrin, heavy chain
Gorai.011G292200.1.A
PCC=-0.376, p=0.113
mRNA Profile
Protein Expression Profile
DPA

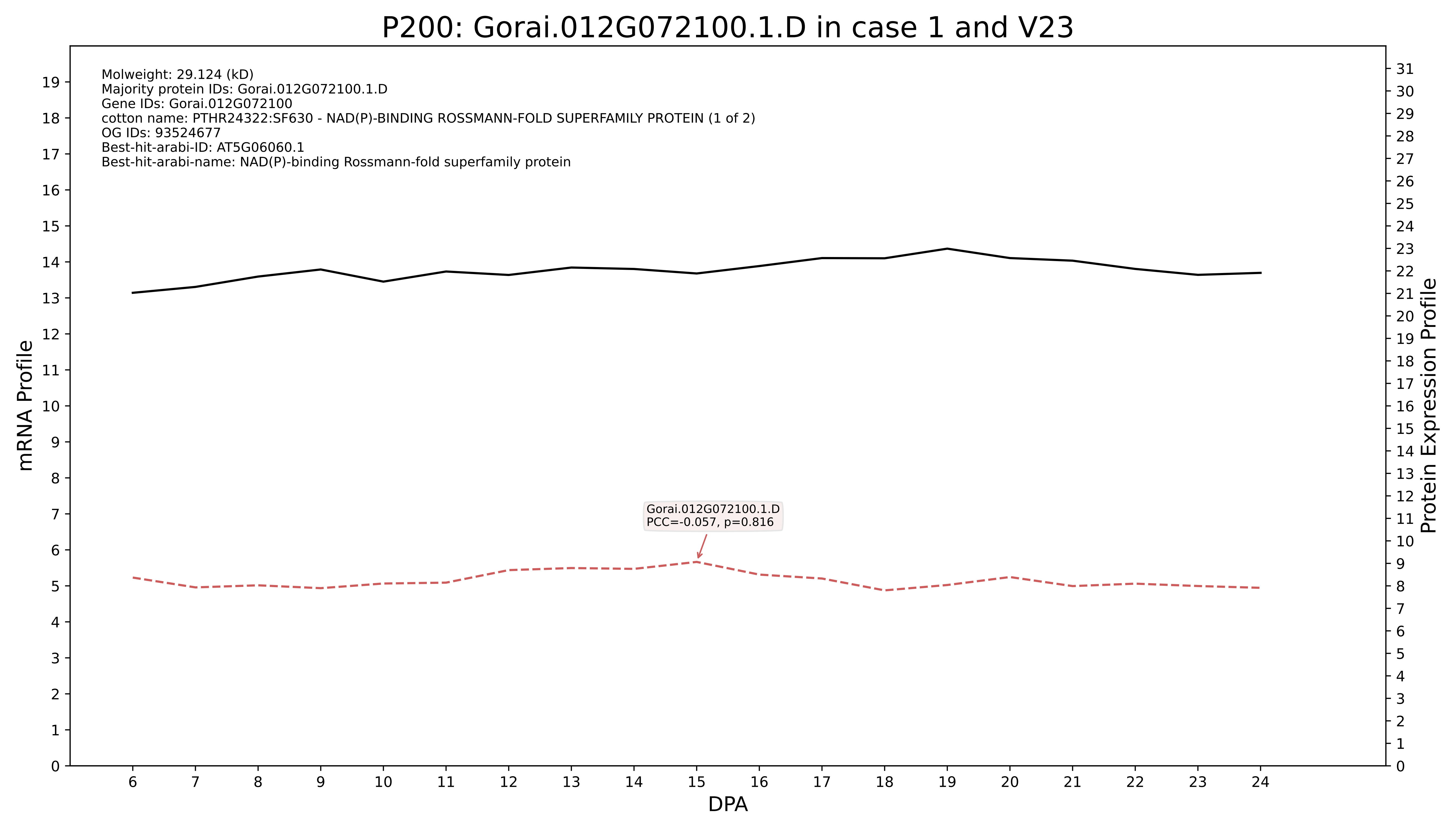
P200: Gorai.012G072100.1.D in case 1 and V23
Molweight: 29.124 (kD)
Majority protein IDs: Gorai.012G072100.1.D
Gene IDs: Gorai.012G072100
cotton name: PTHR24322:SF630 - NAD(P)-BINDING ROSSMANN-FOLD SUPERFAMILY PROTEIN (1 of 2)
OG IDs: 93524677
Best-hit-arabi-ID: AT5G06060.1
Best-hit-arabi-name: NAD(P)-binding Rossmann-fold superfamily protein
Gorai.012G072100.1.D
PCC=-0.057, p=0.816
mRNA Profile
Protein Expression Profile
DPA

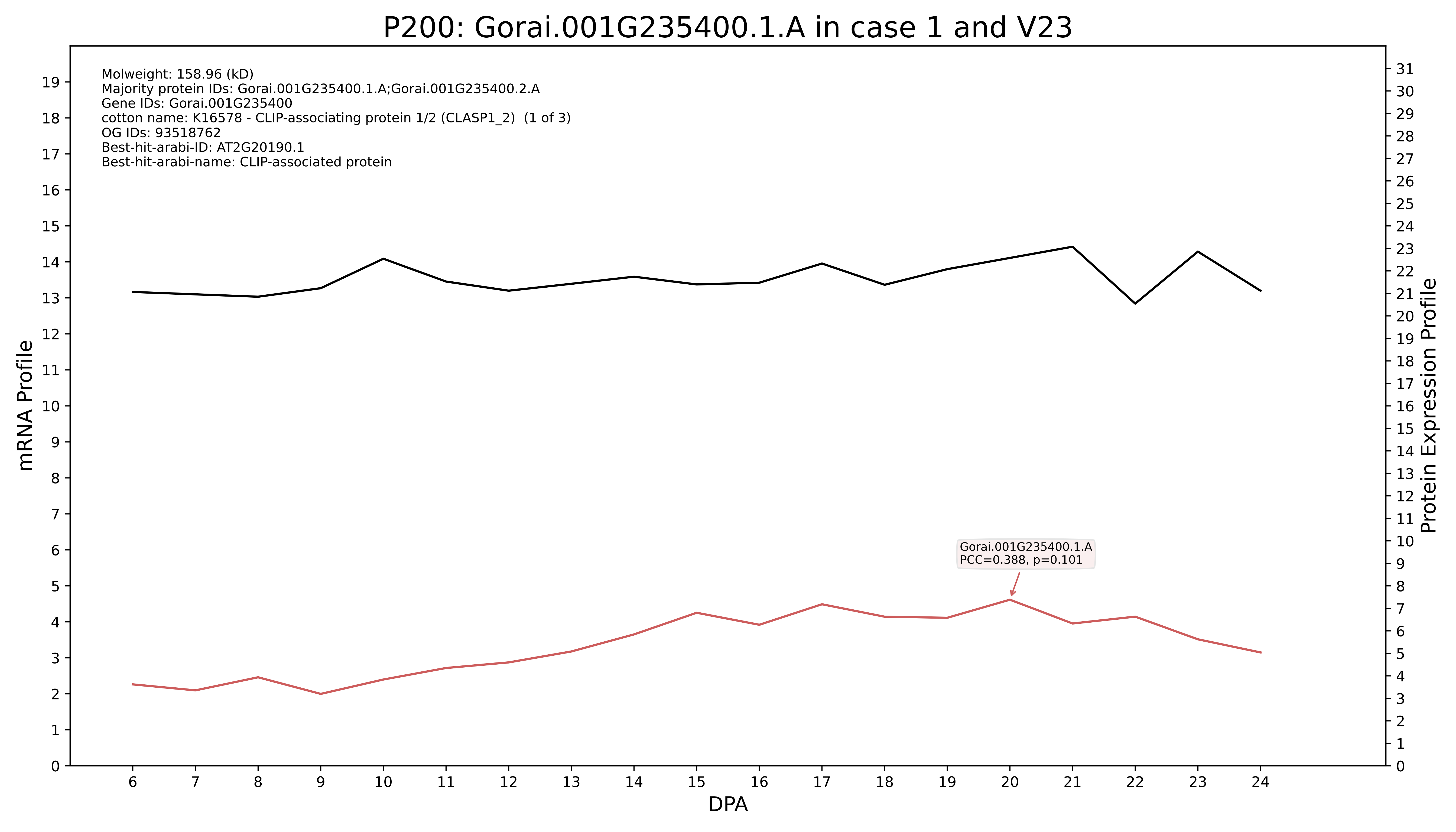

P200: Gorai.001G235400.1.A in case 1 and V23
Molweight: 158.96 (kD)
Majority protein IDs: Gorai.001G235400.1.A;Gorai.001G235400.2.A
Gene IDs: Gorai.001G235400
cotton name: K16578 - CLIP-associating protein 1/2 (CLASP1_2) (1 of 3)
OG IDs: 93518762
Best-hit-arabi-ID: AT2G20190.1
Best-hit-arabi-name: CLIP-associated protein
Gorai.001G235400.1.A
PCC=0.388, p=0.101
mRNA Profile
Protein Expression Profile
DPA

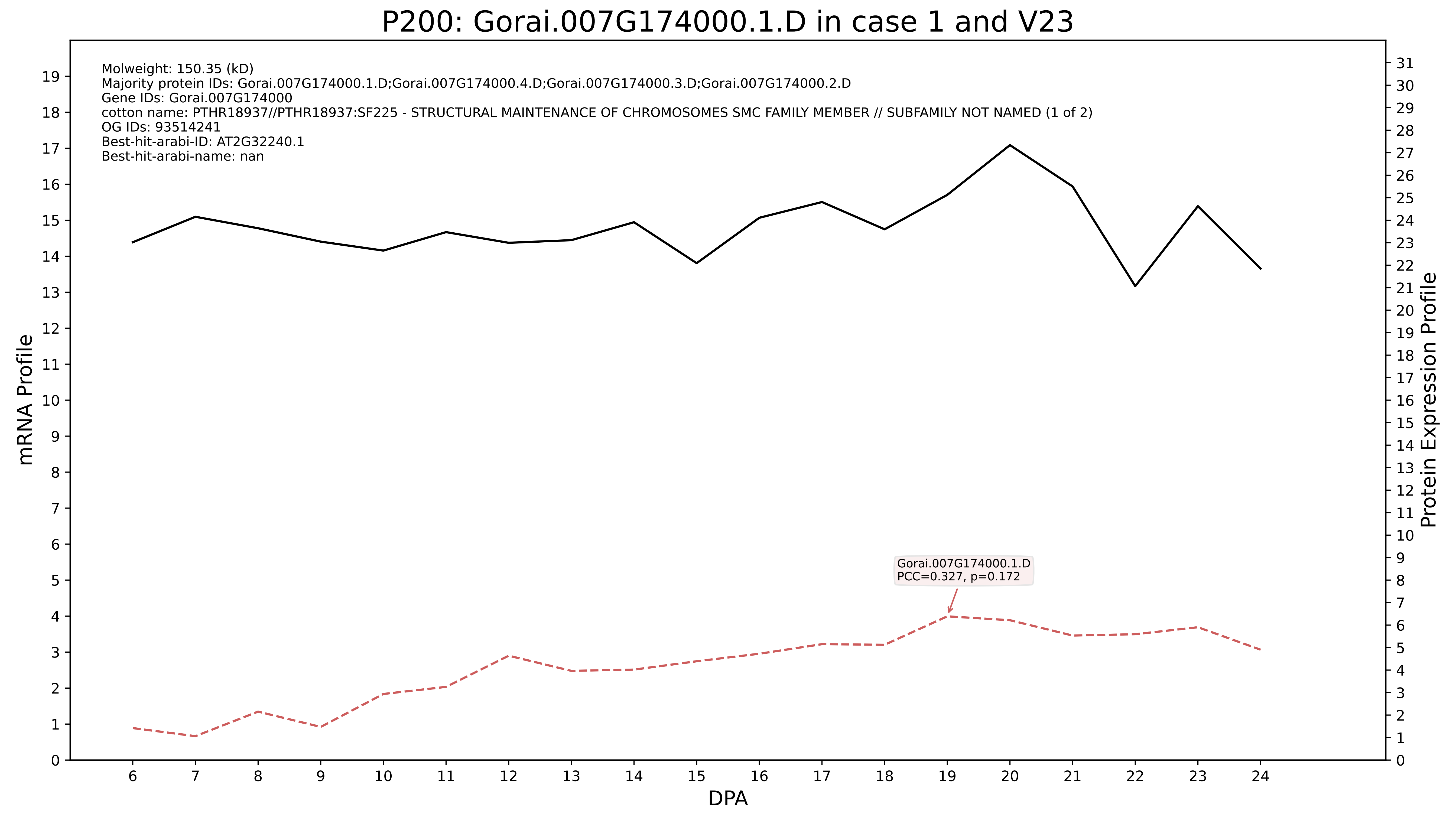
P200: Gorai.007G174000.1.D in case 1 and V23
Molweight: 150.35 (kD)
Majority protein IDs: Gorai.007G174000.1.D;Gorai.007G174000.4.D;Gorai.007G174000.3.D;Gorai.007G174000.2.D
Gene IDs: Gorai.007G174000
cotton name: PTHR18937//PTHR18937:SF225 - STRUCTURAL MAINTENANCE OF CHROMOSOMES SMC FAMILY MEMBER // SUBFAMILY NOT NAMED (1 of 2)
OG IDs: 93514241
Best-hit-arabi-ID: AT2G32240.1
Best-hit-arabi-name: nan
Gorai.007G174000.1.D
PCC=0.327, p=0.172
mRNA Profile
Protein Expression Profile
DPA

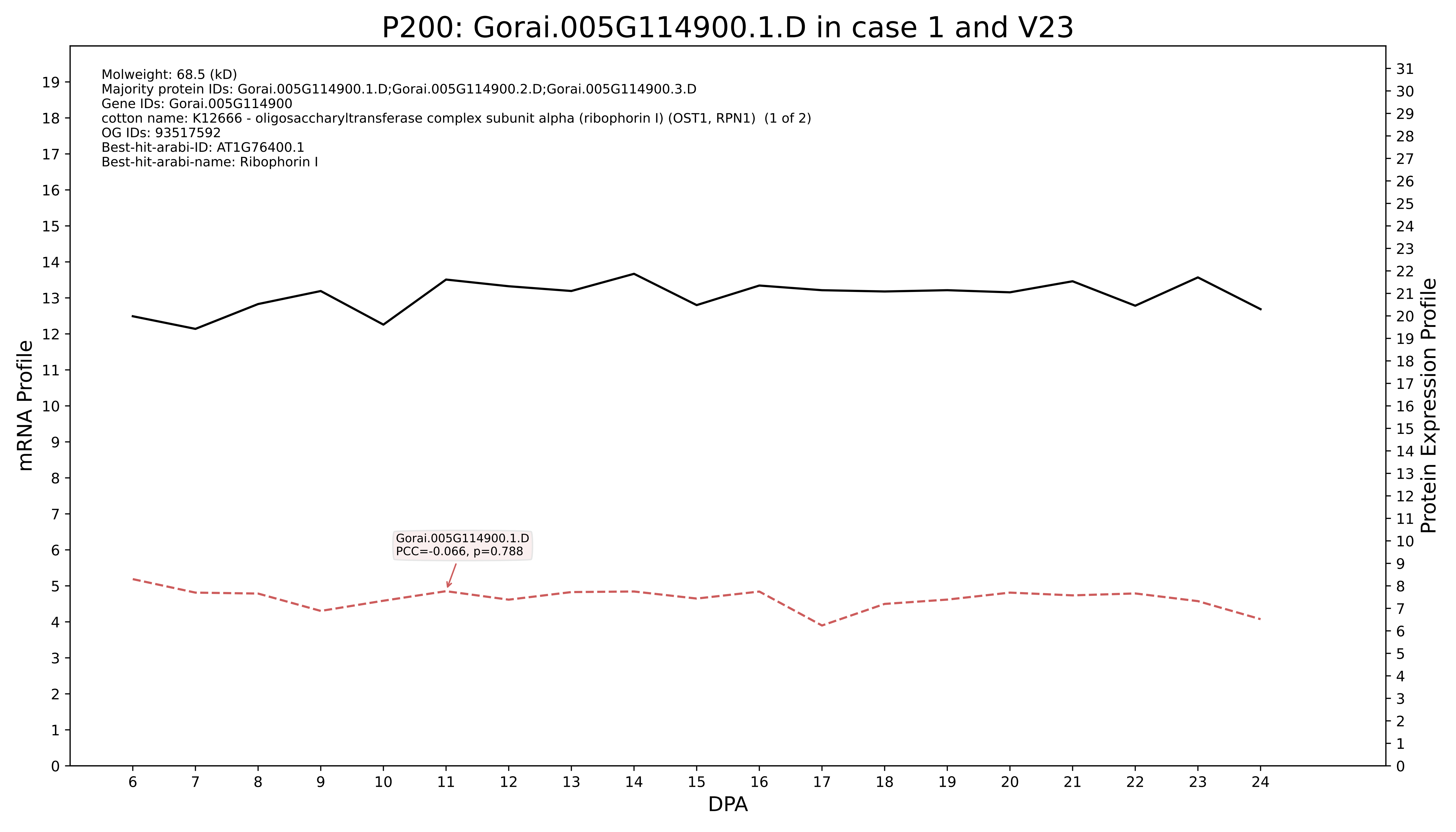

P200: Gorai.005G114900.1.D in case 1 and V23
Molweight: 68.5 (kD)
Majority protein IDs: Gorai.005G114900.1.D;Gorai.005G114900.2.D;Gorai.005G114900.3.D
Gene IDs: Gorai.005G114900
cotton name: K12666 - oligosaccharyltransferase complex subunit alpha (ribophorin I) (OST1, RPN1) (1 of 2)
OG IDs: 93517592
Best-hit-arabi-ID: AT1G76400.1
Best-hit-arabi-name: Ribophorin I
Gorai.005G114900.1.D
PCC=-0.066, p=0.788
mRNA Profile
Protein Expression Profile
DPA

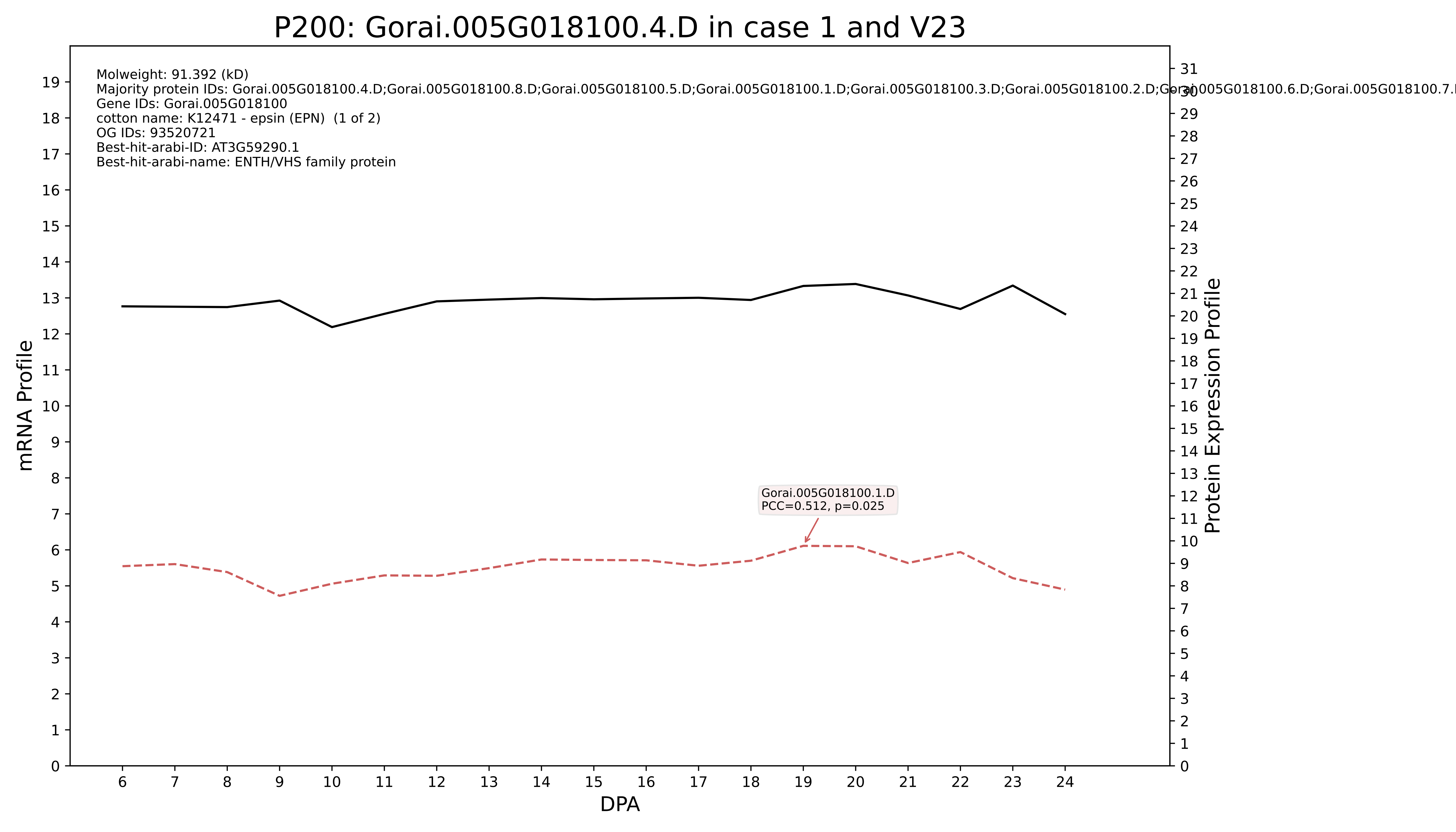
P200: Gorai.005G018100.4.D in case 1 and V23
Molweight: 91.392 (kD)
Majority protein IDs: Gorai.005G018100.4.D;Gorai.005G018100.8.D;Gorai.005G018100.5.D;Gorai.005G018100.1.D;Gorai.005G018100.3.D;Gorai.005G018100.2.D;Gorai.005G018100.6.D;Gorai.005G018100.7.D
Gene IDs: Gorai.005G018100
cotton name: K12471 - epsin (EPN) (1 of 2)
OG IDs: 93520721
Best-hit-arabi-ID: AT3G59290.1
Best-hit-arabi-name: ENTH/VHS family protein
Gorai.005G018100.1.D
PCC=0.512, p=0.025
mRNA Profile
Protein Expression Profile
DPA

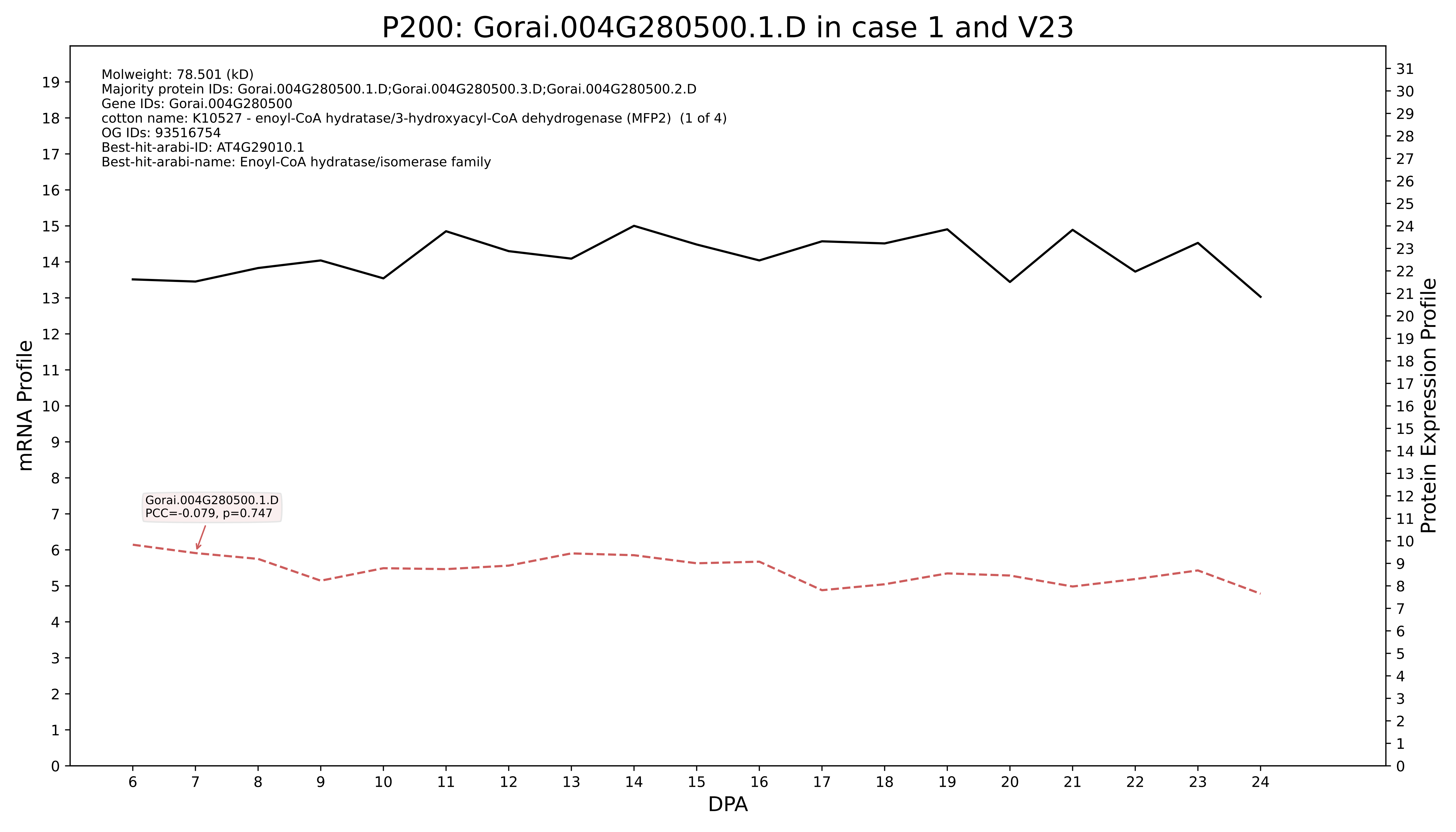
P200: Gorai.004G280500.1.D in case 1 and V23
Molweight: 78.501 (kD)
Majority protein IDs: Gorai.004G280500.1.D;Gorai.004G280500.3.D;Gorai.004G280500.2.D
Gene IDs: Gorai.004G280500
cotton name: K10527 - enoyl-CoA hydratase/3-hydroxyacyl-CoA dehydrogenase (MFP2) (1 of 4)
OG IDs: 93516754
Best-hit-arabi-ID: AT4G29010.1
Best-hit-arabi-name: Enoyl-CoA hydratase/isomerase family
Gorai.004G280500.1.D
PCC=-0.079, p=0.747
mRNA Profile
Protein Expression Profile
DPA

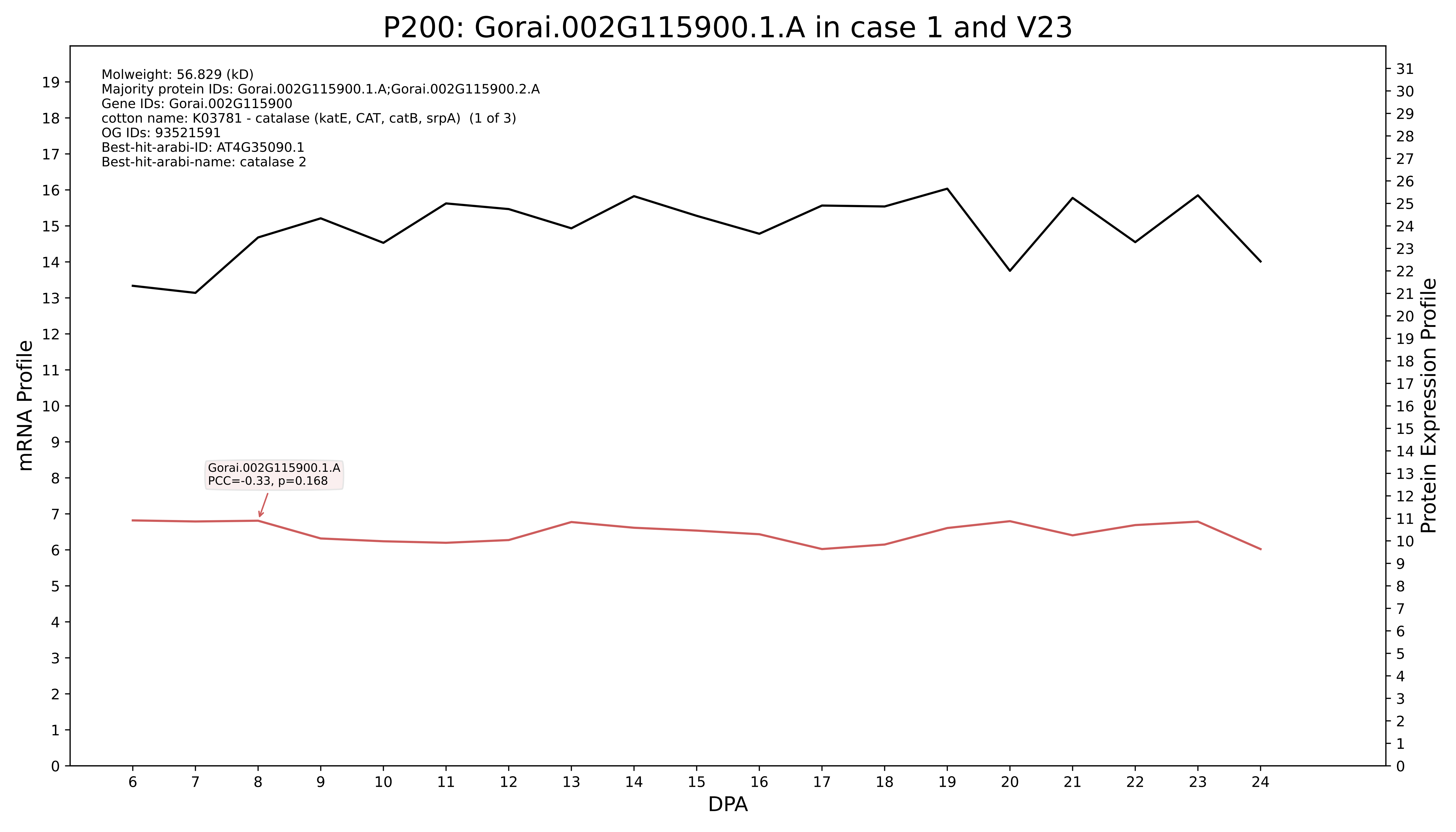

P200: Gorai.002G115900.1.A in case 1 and V23
Molweight: 56.829 (kD)
Majority protein IDs: Gorai.002G115900.1.A;Gorai.002G115900.2.A
Gene IDs: Gorai.002G115900
cotton name: K03781 - catalase (katE, CAT, catB, srpA) (1 of 3)
OG IDs: 93521591
Best-hit-arabi-ID: AT4G35090.1
Best-hit-arabi-name: catalase 2
Gorai.002G115900.1.A
PCC=-0.33, p=0.168
mRNA Profile
Protein Expression Profile
DPA

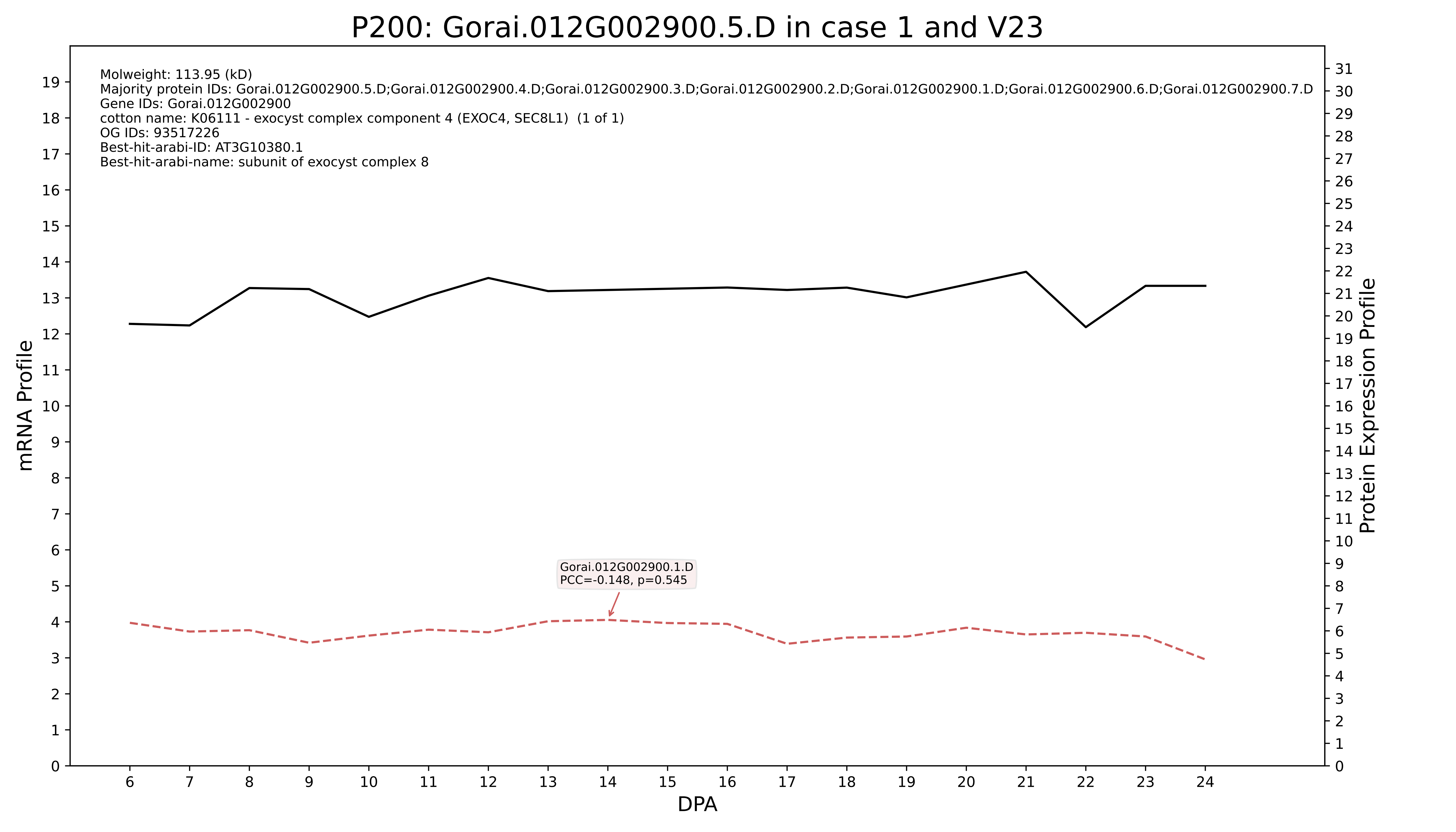
P200: Gorai.012G002900.5.D in case 1 and V23
Molweight: 113.95 (kD)
Majority protein IDs: Gorai.012G002900.5.D;Gorai.012G002900.4.D;Gorai.012G002900.3.D;Gorai.012G002900.2.D;Gorai.012G002900.1.D;Gorai.012G002900.6.D;Gorai.012G002900.7.D
Gene IDs: Gorai.012G002900
cotton name: K06111 - exocyst complex component 4 (EXOC4, SEC8L1) (1 of 1)
OG IDs: 93517226
Best-hit-arabi-ID: AT3G10380.1
Best-hit-arabi-name: subunit of exocyst complex 8
Gorai.012G002900.1.D
PCC=-0.148, p=0.545
mRNA Profile
Protein Expression Profile
DPA

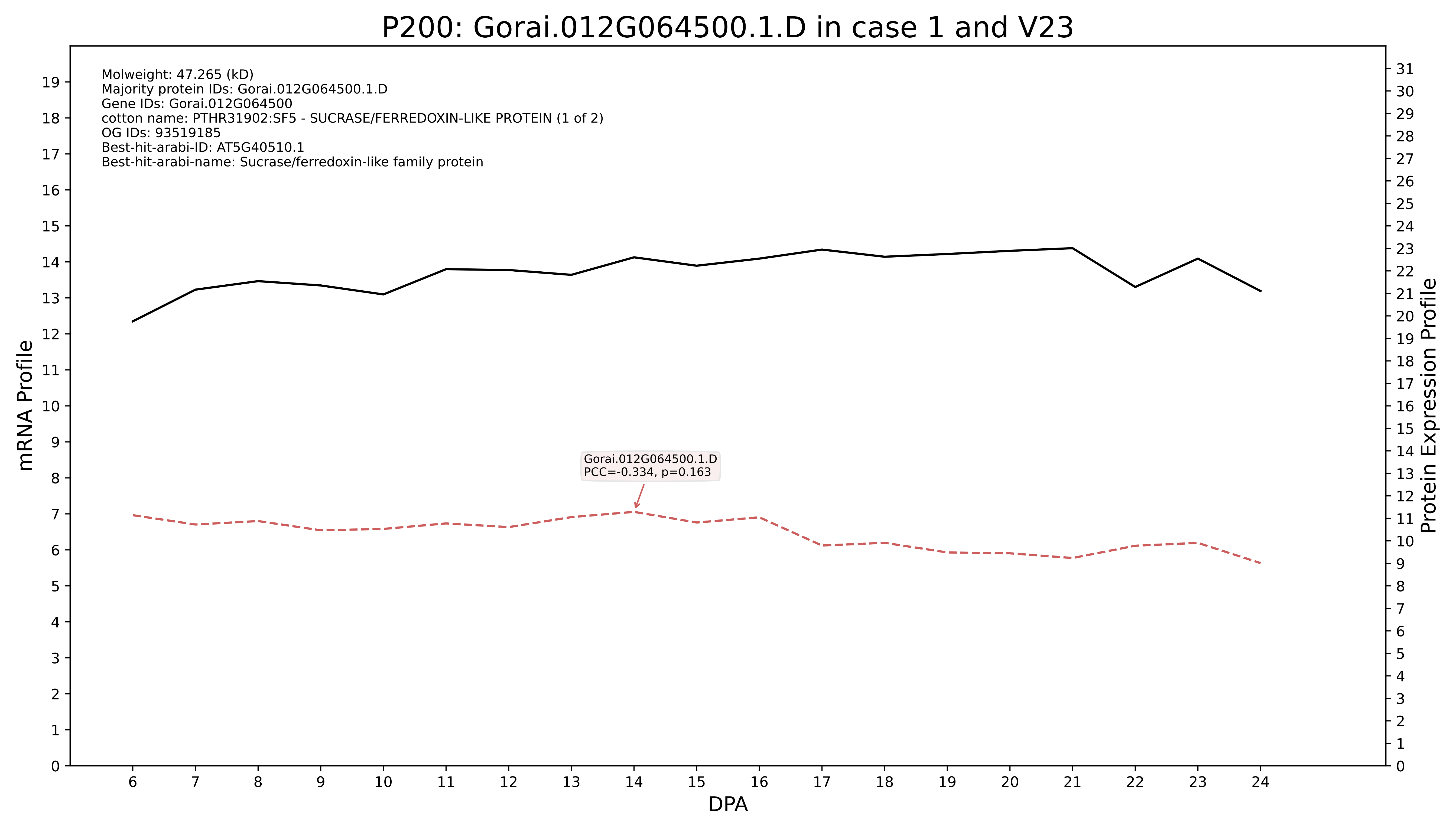
P200: Gorai.012G064500.1.D in case 1 and V23
Molweight: 47.265 (kD)
Majority protein IDs: Gorai.012G064500.1.D
Gene IDs: Gorai.012G064500
cotton name: PTHR31902:SF5 - SUCRASE/FERREDOXIN-LIKE PROTEIN (1 of 2)
OG IDs: 93519185
Best-hit-arabi-ID: AT5G40510.1
Best-hit-arabi-name: Sucrase/ferredoxin-like family protein
Gorai.012G064500.1.D
PCC=-0.334, p=0.163
mRNA Profile
Protein Expression Profile
DPA

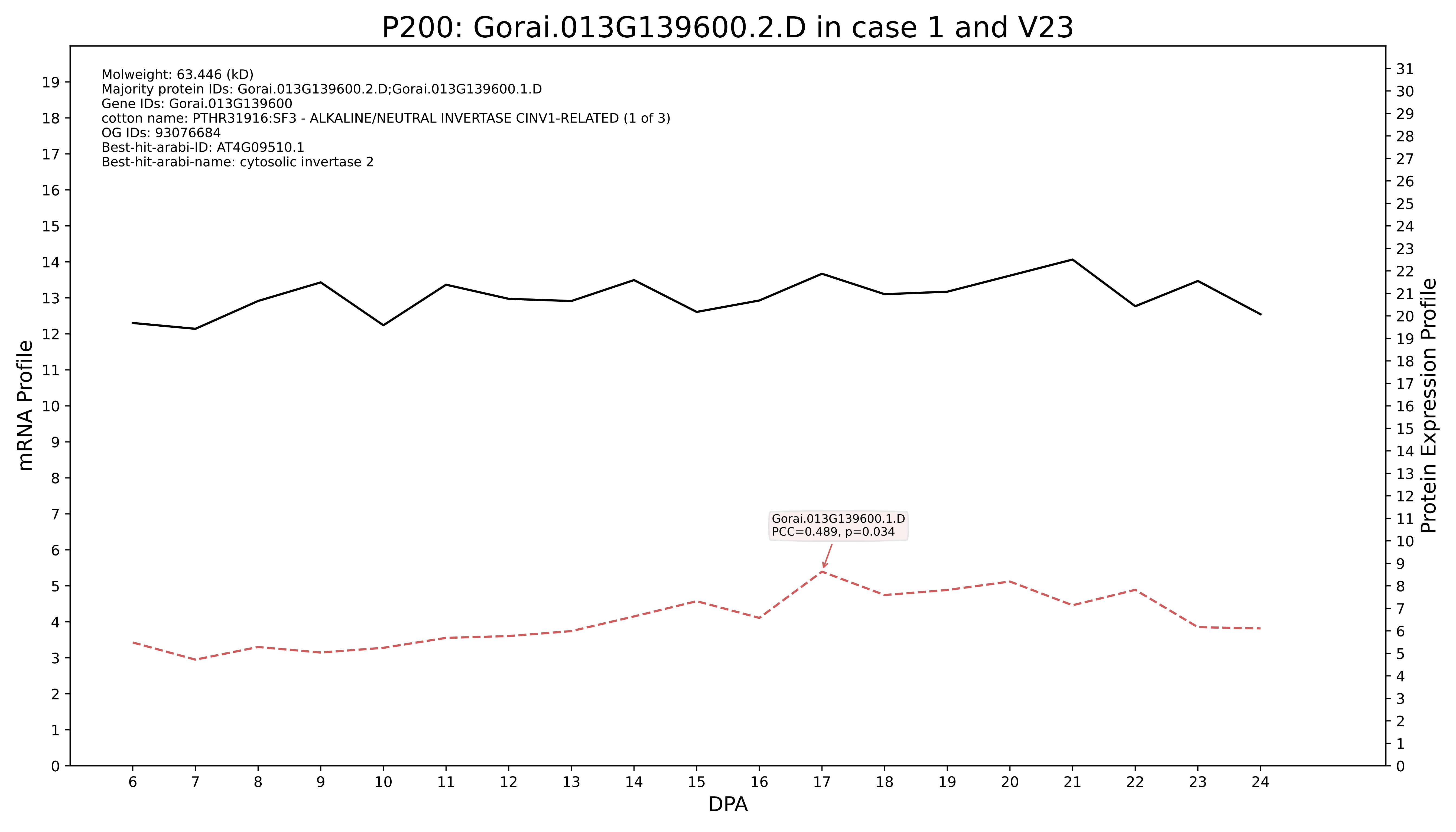

P200: Gorai.013G139600.2.D in case 1 and V23
Molweight: 63.446 (kD)
Majority protein IDs: Gorai.013G139600.2.D;Gorai.013G139600.1.D
Gene IDs: Gorai.013G139600
cotton name: PTHR31916:SF3 - ALKALINE/NEUTRAL INVERTASE CINV1-RELATED (1 of 3)
OG IDs: 93076684
Best-hit-arabi-ID: AT4G09510.1
Best-hit-arabi-name: cytosolic invertase 2
Gorai.013G139600.1.D
PCC=0.489, p=0.034
mRNA Profile
Protein Expression Profile
DPA

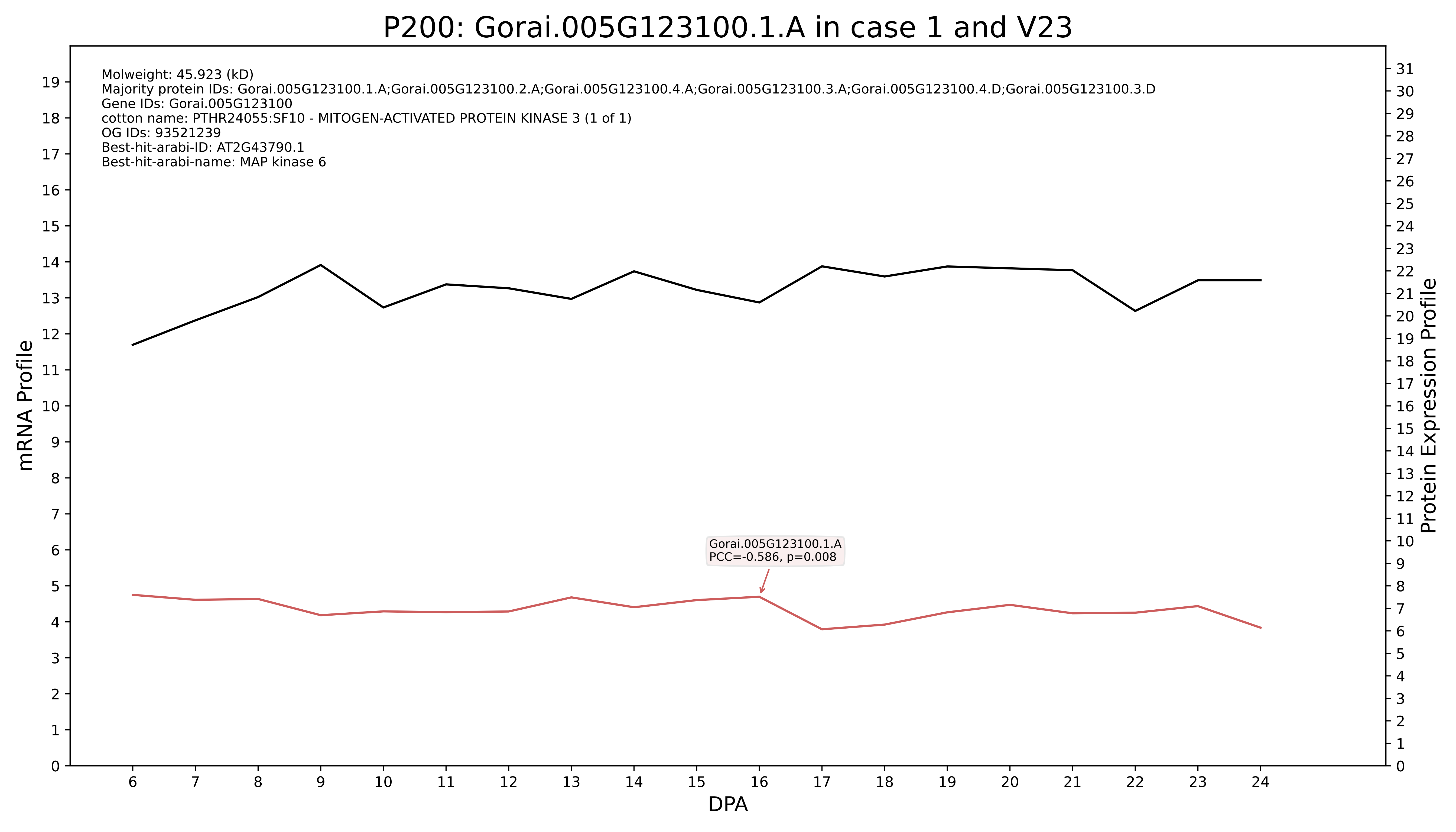
P200: Gorai.005G123100.1.A in case 1 and V23
Molweight: 45.923 (kD)
Majority protein IDs: Gorai.005G123100.1.A;Gorai.005G123100.2.A;Gorai.005G123100.4.A;Gorai.005G123100.3.A;Gorai.005G123100.4.D;Gorai.005G123100.3.D
Gene IDs: Gorai.005G123100
cotton name: PTHR24055:SF10 - MITOGEN-ACTIVATED PROTEIN KINASE 3 (1 of 1)
OG IDs: 93521239
Best-hit-arabi-ID: AT2G43790.1
Best-hit-arabi-name: MAP kinase 6
Gorai.005G123100.1.A
PCC=-0.586, p=0.008
mRNA Profile
Protein Expression Profile
DPA

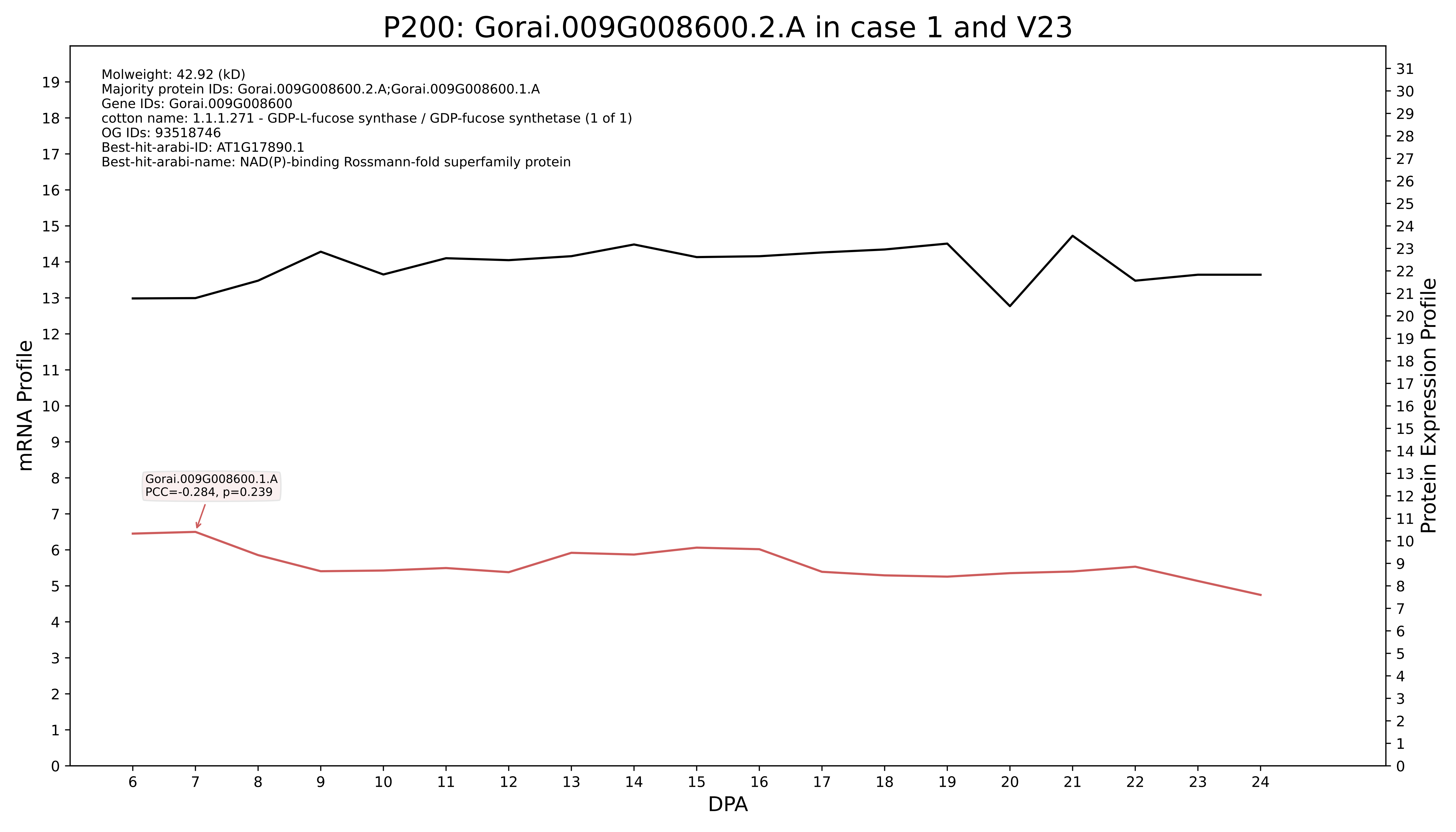
P200: Gorai.009G008600.2.A in case 1 and V23
Molweight: 42.92 (kD)
Majority protein IDs: Gorai.009G008600.2.A;Gorai.009G008600.1.A
Gene IDs: Gorai.009G008600
cotton name: 1.1.1.271 - GDP-L-fucose synthase / GDP-fucose synthetase (1 of 1)
OG IDs: 93518746
Best-hit-arabi-ID: AT1G17890.1
Best-hit-arabi-name: NAD(P)-binding Rossmann-fold superfamily protein
Gorai.009G008600.1.A
PCC=-0.284, p=0.239
mRNA Profile
Protein Expression Profile
DPA

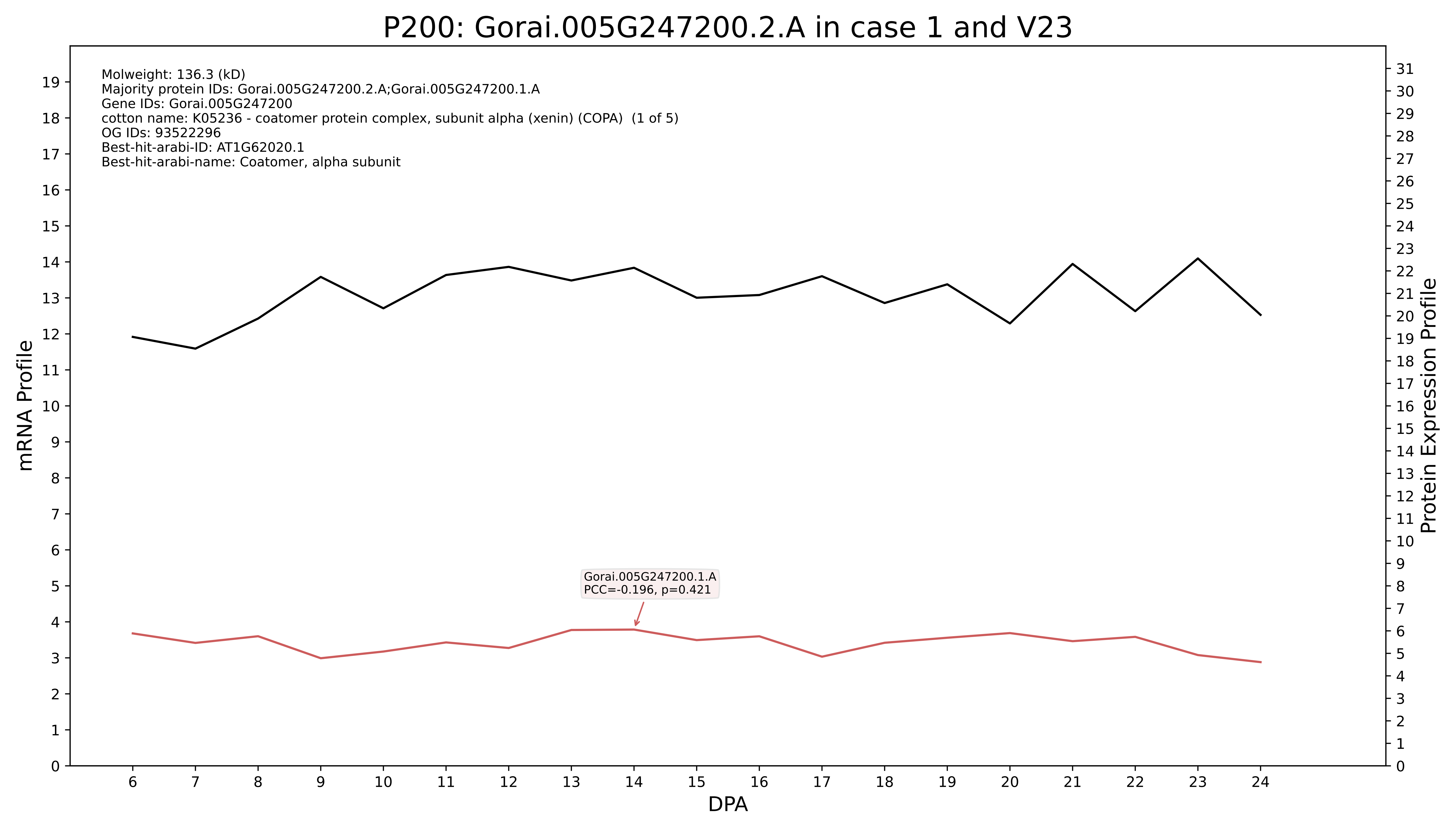
P200: Gorai.005G247200.2.A in case 1 and V23
Molweight: 136.3 (kD)
Majority protein IDs: Gorai.005G247200.2.A;Gorai.005G247200.1.A
Gene IDs: Gorai.005G247200
cotton name: K05236 - coatomer protein complex, subunit alpha (xenin) (COPA) (1 of 5)
OG IDs: 93522296
Best-hit-arabi-ID: AT1G62020.1
Best-hit-arabi-name: Coatomer, alpha subunit
Gorai.005G247200.1.A
PCC=-0.196, p=0.421
mRNA Profile
Protein Expression Profile
DPA

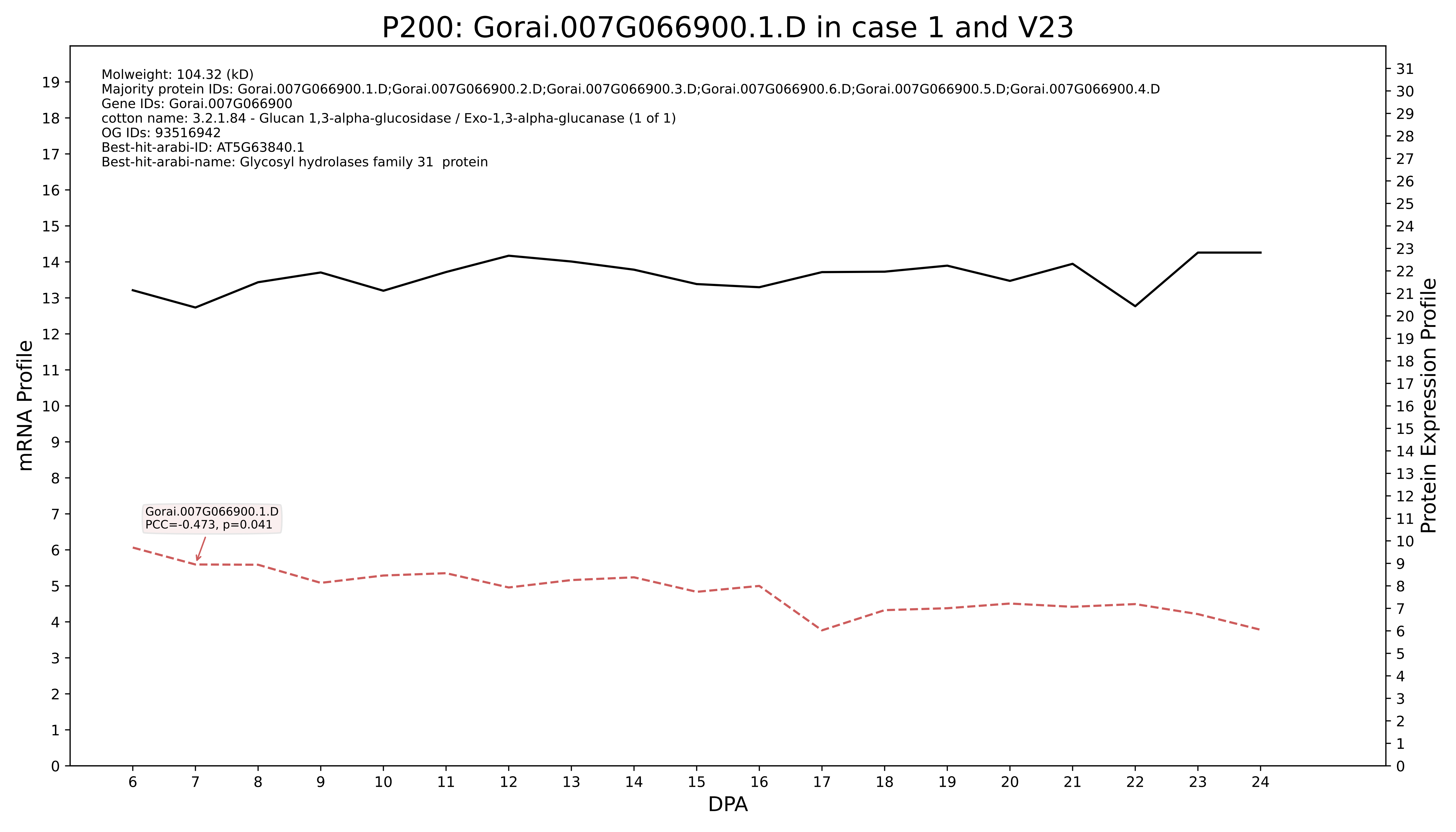
P200: Gorai.007G066900.1.D in case 1 and V23
Molweight: 104.32 (kD)
Majority protein IDs: Gorai.007G066900.1.D;Gorai.007G066900.2.D;Gorai.007G066900.3.D;Gorai.007G066900.6.D;Gorai.007G066900.5.D;Gorai.007G066900.4.D
Gene IDs: Gorai.007G066900
cotton name: 3.2.1.84 - Glucan 1,3-alpha-glucosidase / Exo-1,3-alpha-glucanase (1 of 1)
OG IDs: 93516942
Best-hit-arabi-ID: AT5G63840.1
Best-hit-arabi-name: Glycosyl hydrolases family 31 protein
Gorai.007G066900.1.D
PCC=-0.473, p=0.041
mRNA Profile
Protein Expression Profile
DPA

### P200: Gorai.009G288200.1.A in case 1 and V23

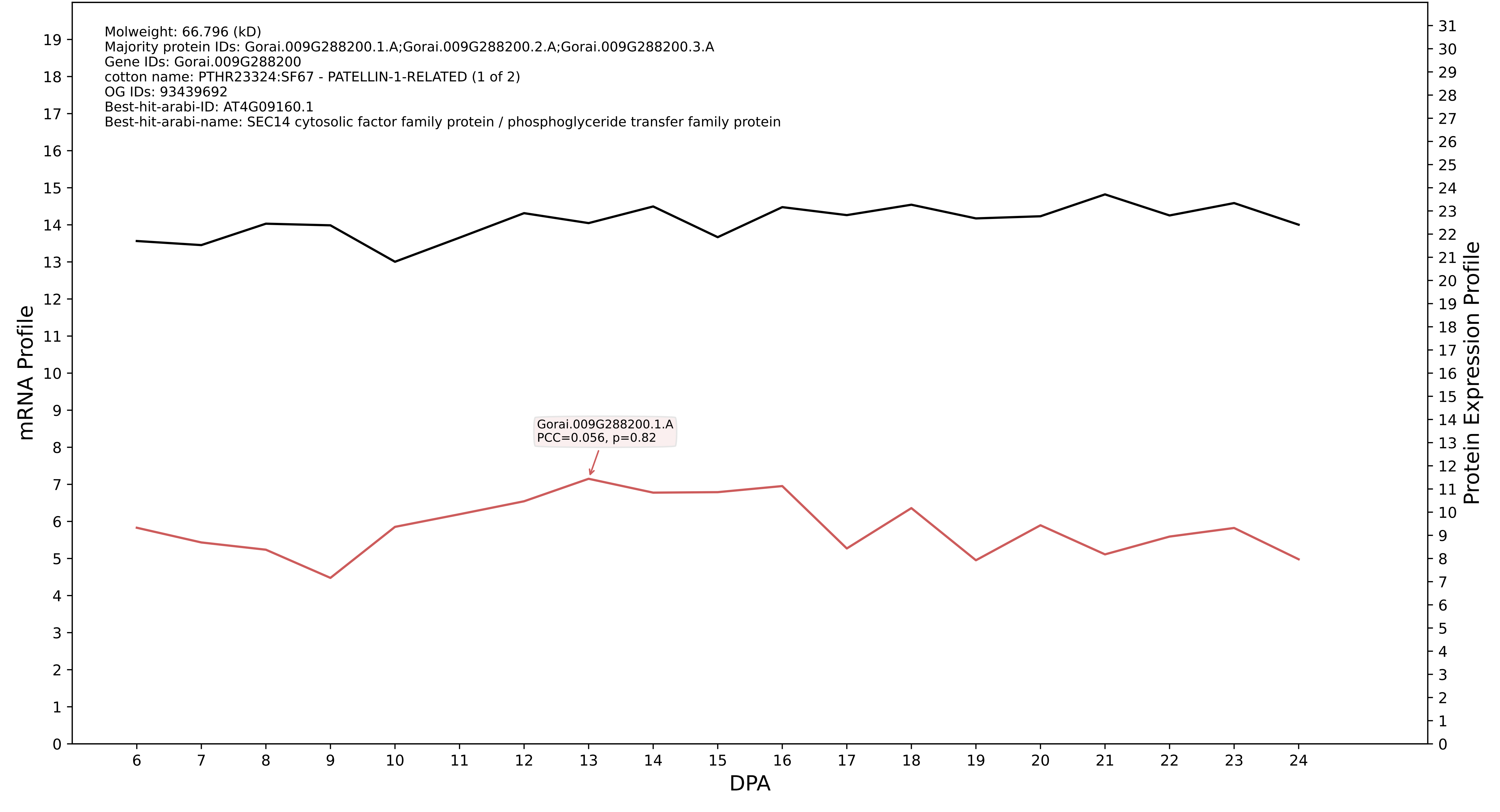

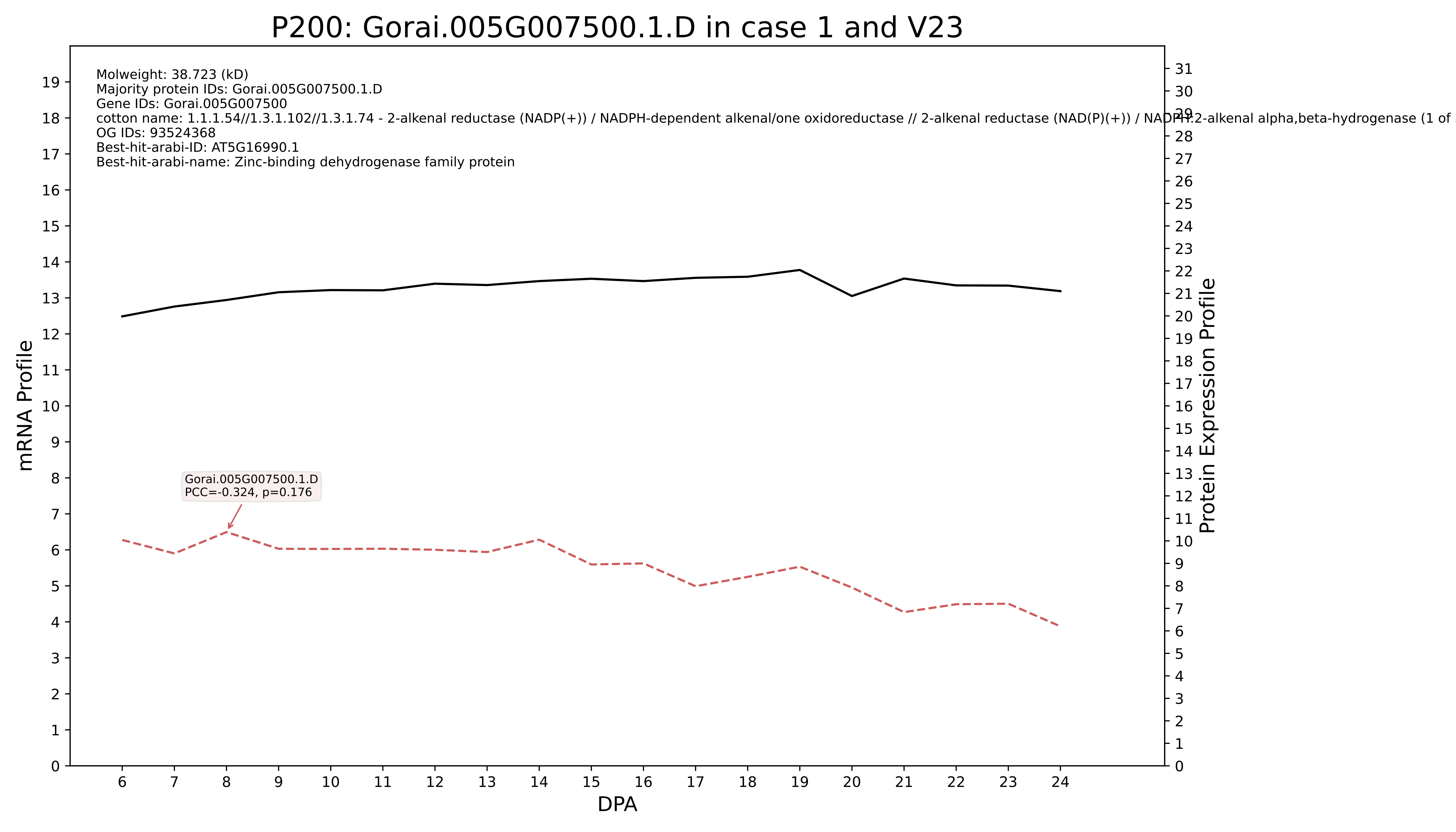
P200: Gorai.005G007500.1.D in case 1 and V23
Molweight: 38.723 (kD)
Majority protein IDs: Gorai.005G007500.1.D
Gene IDs: Gorai.005G007500
cotton name: 1.1.1.54//1.3.1.102//1.3.1.74 - 2-alkenal reductase (NADP(+)) / NADPH-dependent alkenal/one oxidoreductase // 2-alkenal reductase (NAD(P)(+)) / NADPH:2-alkenal alpha,beta-hydrogenase (1 of
OG IDs: 93524368
Best-hit-arabi-ID: AT5G16990.1
Best-hit-arabi-name: Zinc-binding dehydrogenase family protein
Gorai.005G007500.1.D
PCC=-0.324, p=0.176
mRNA Profile
0 1 2 3 4 5 6 7 8 9 10 11 12 13 14 15 16 17 18 19
Protein Expression Profile
0 1 2 3 4 5 6 7 8 9 10 11 12 13 14 15 16 17 18 19 20 21 22 23 24 25 26 27 28 29 30 31
DPA
6 7 8 9 10 11 12 13 14 15 16 17 18 19 20 21 22 23 24

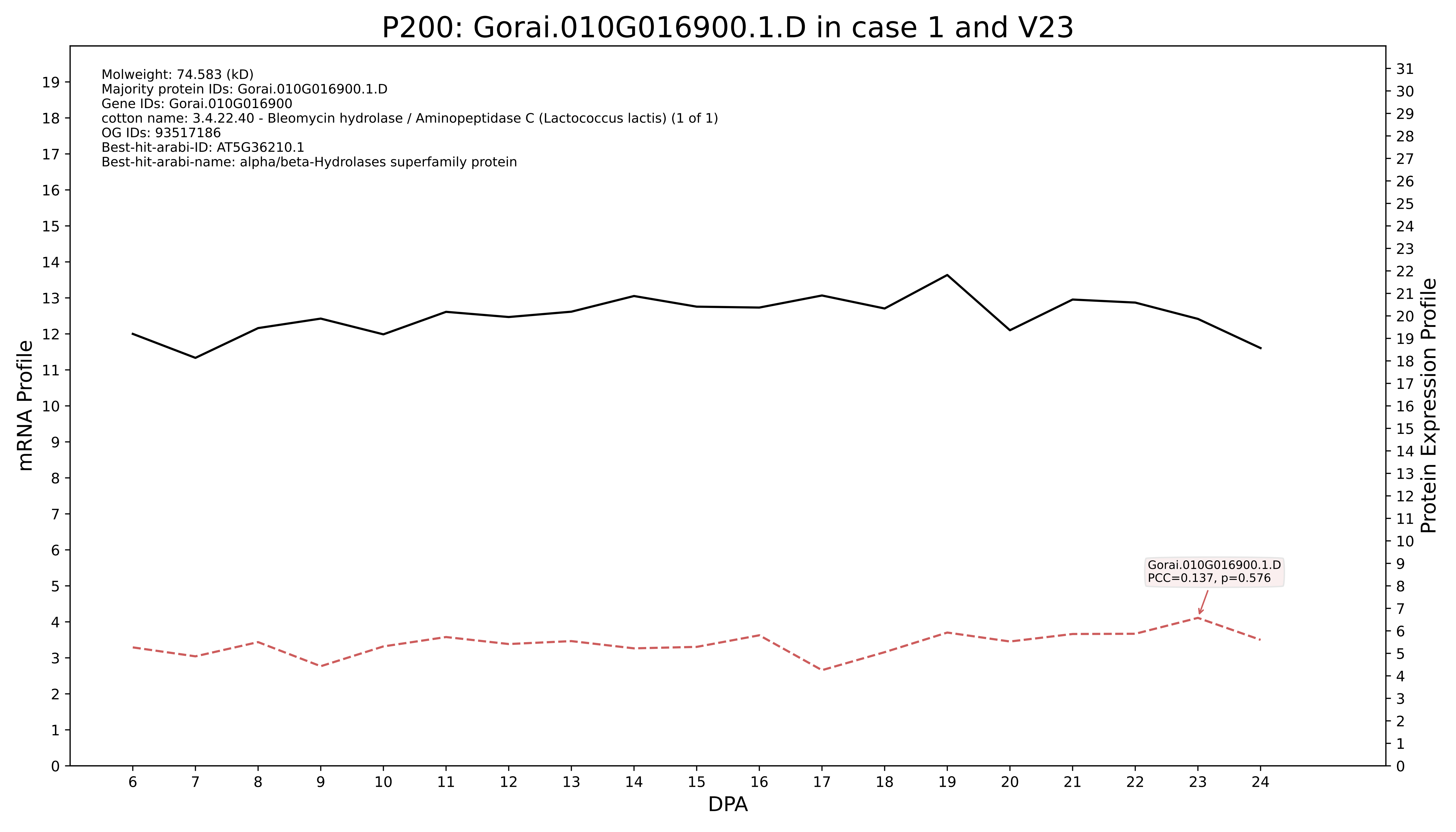
P200: Gorai.010G016900.1.D in case 1 and V23
Molweight: 74.583 (kD)
Majority protein IDs: Gorai.010G016900.1.D
Gene IDs: Gorai.010G016900
cotton name: 3.4.22.40 - Bleomycin hydrolase / Aminopeptidase C (Lactococcus lactis) (1 of 1)
OG IDs: 93517186
Best-hit-arabi-ID: AT5G36210.1
Best-hit-arabi-name: alpha/beta-Hydrolases superfamily protein
Gorai.010G016900.1.D
PCC=0.137, p=0.576
mRNA Profile
Protein Expression Profile
DPA

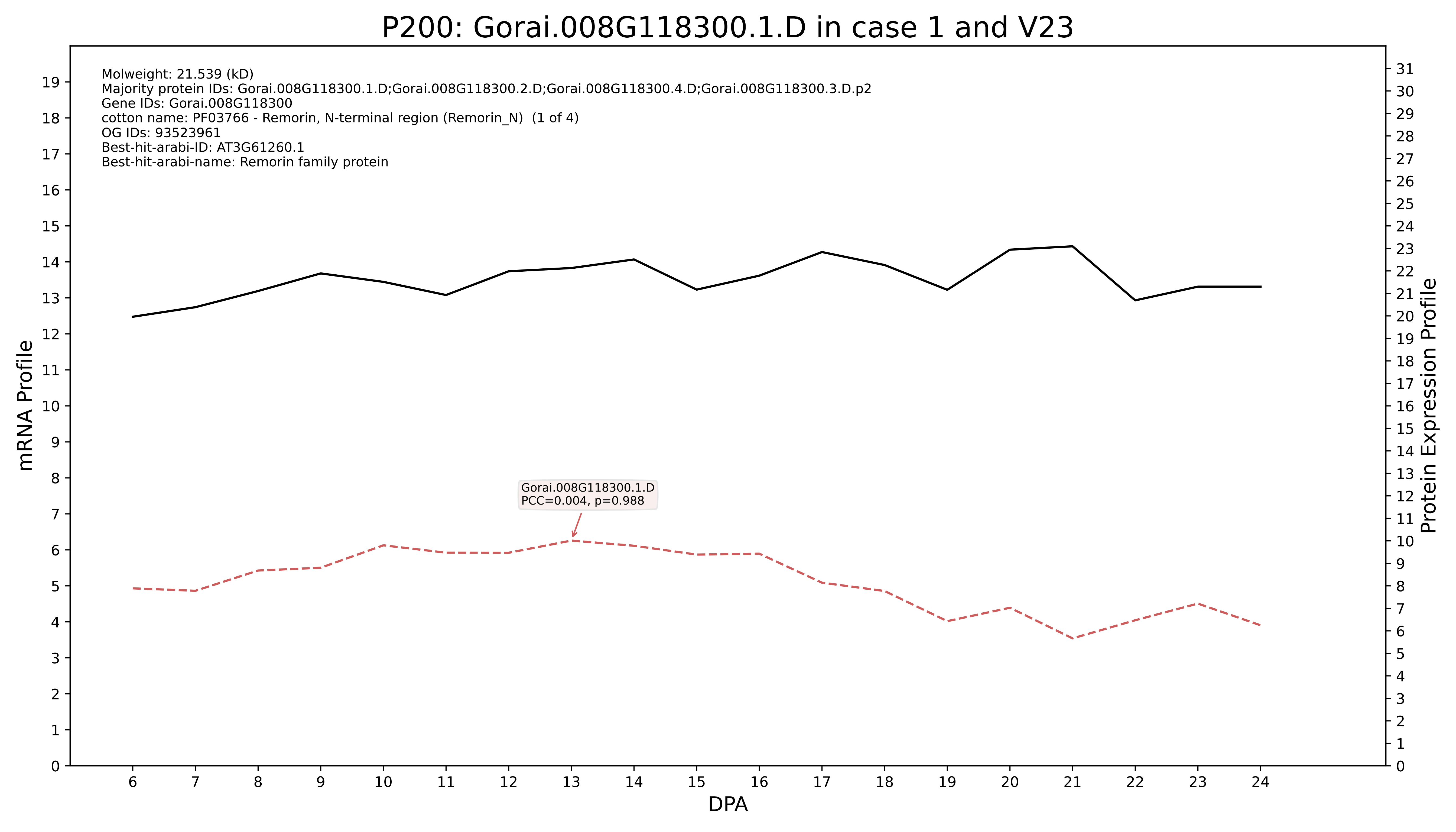
P200: Gorai.008G118300.1.D in case 1 and V23
Molweight: 21.539 (kD)
Majority protein IDs: Gorai.008G118300.1.D;Gorai.008G118300.2.D;Gorai.008G118300.4.D;Gorai.008G118300.3.D.p2
Gene IDs: Gorai.008G118300
cotton name: PF03766 - Remorin, N-terminal region (Remorin_N) (1 of 4)
OG IDs: 93523961
Best-hit-arabi-ID: AT3G61260.1
Best-hit-arabi-name: Remorin family protein
Gorai.008G118300.1.D
PCC=0.004, p=0.988
mRNA Profile
Protein Expression Profile
DPA

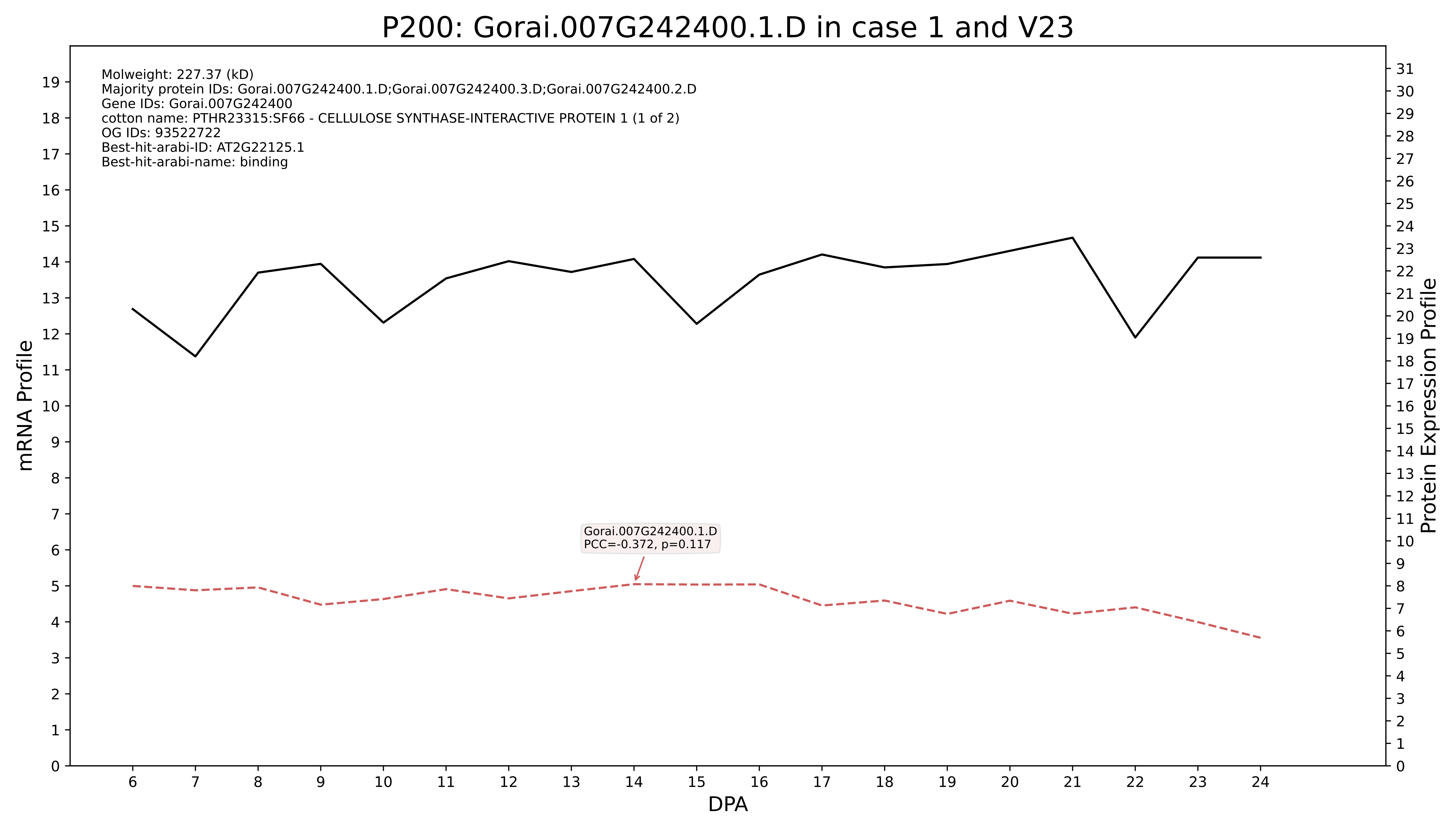
P200: Gorai.007G242400.1.D in case 1 and V23
Molweight: 227.37 (kD)
Majority protein IDs: Gorai.007G242400.1.D;Gorai.007G242400.3.D;Gorai.007G242400.2.D
Gene IDs: Gorai.007G242400
cotton name: PTHR23315:SF66 - CELLULOSE SYNTHASE-INTERACTIVE PROTEIN 1 (1 of 2)
OG IDs: 93522722
Best-hit-arabi-ID: AT2G22125.1
Best-hit-arabi-name: binding
Gorai.007G242400.1.D
PCC=-0.372, p=0.117
mRNA Profile
Protein Expression Profile
DPA

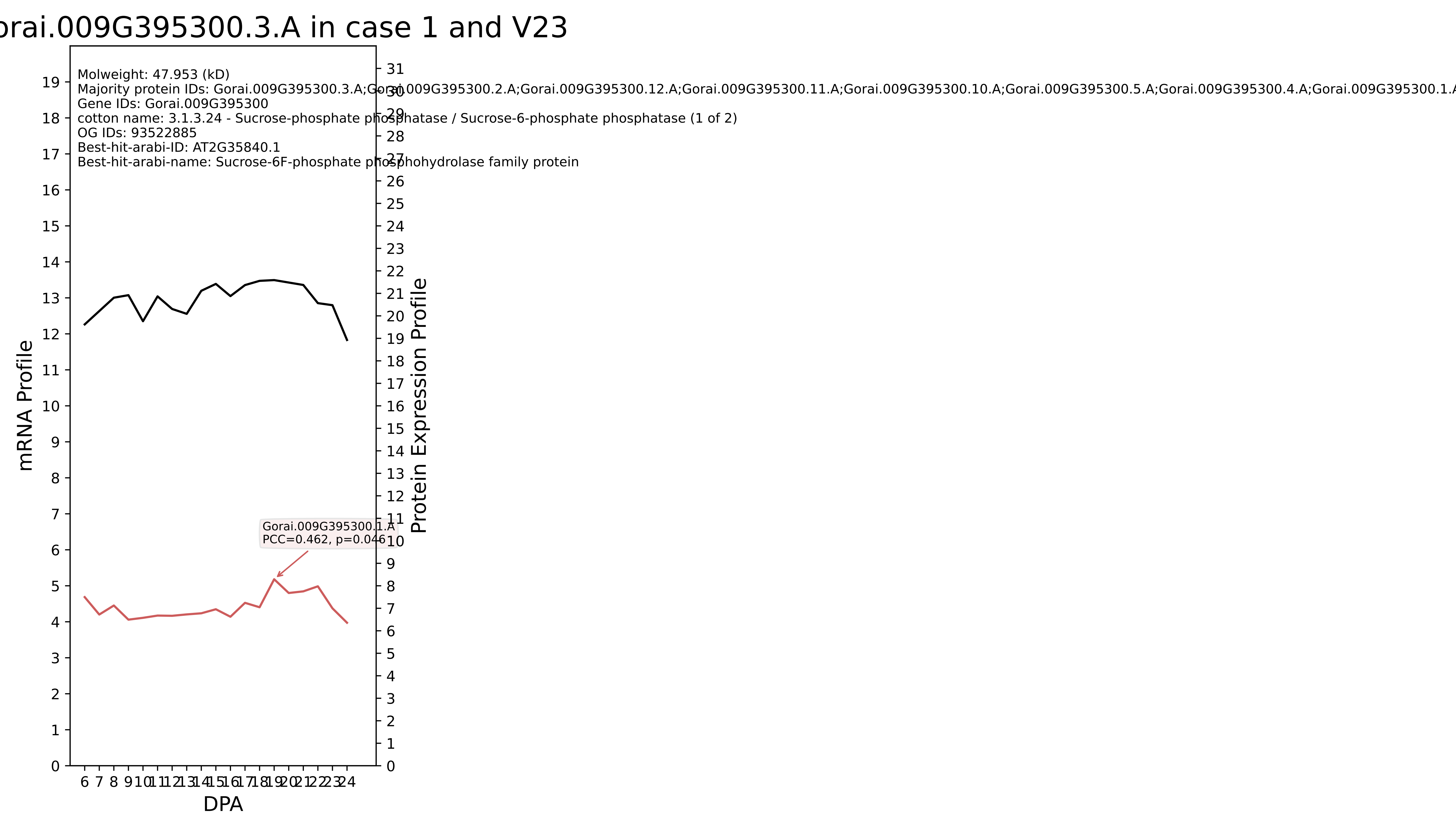
orai.009G395300.3.A in case 1 and V23
Molweight: 47.953 (kD)
Majority protein IDs: Gorai.009G395300.3.A;Gorai.009G395300.2.A;Gorai.009G395300.12.A;Gorai.009G395300.11.A;Gorai.009G395300.10.A;Gorai.009G395300.5.A;Gorai.009G395300.4.A;Gorai.009G395300.1.A
Gene IDs: Gorai.009G395300
cotton name: 3.1.3.24 - Sucrose-phosphate phosphatase / Sucrose-6-phosphate phosphatase (1 of 2)
OG IDs: 93522885
Best-hit-arabi-ID: AT2G35840.1
Best-hit-arabi-name: Sucrose-6F-phosphate phosphohydrolase family protein
Gorai.009G395300.1.A
PCC=0.462, p=0.046
mRNA Profile
Protein Expression Profile
DPA

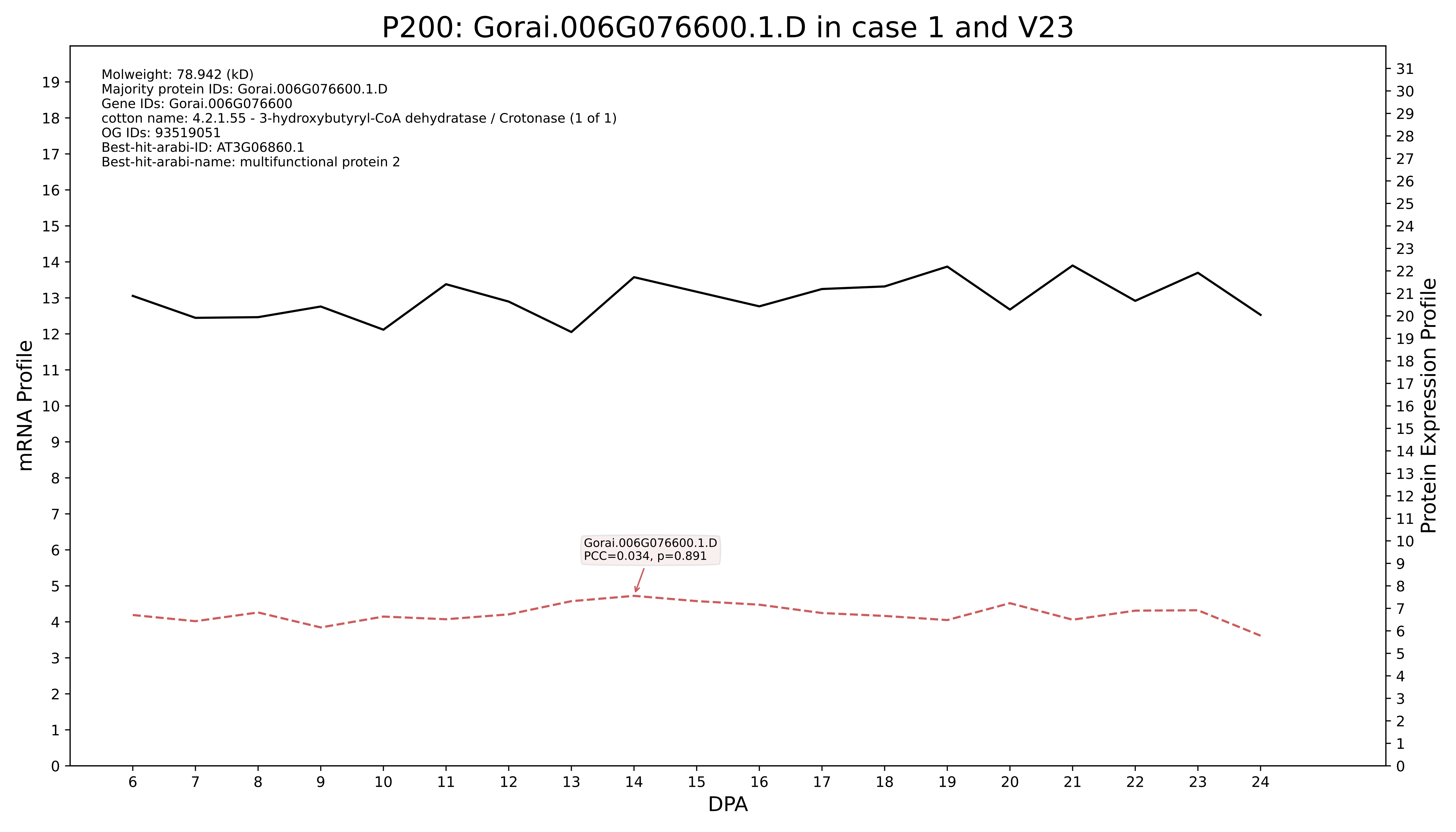

P200: Gorai.006G076600.1.D in case 1 and V23
Molweight: 78.942 (kD)
Majority protein IDs: Gorai.006G076600.1.D
Gene IDs: Gorai.006G076600
cotton name: 4.2.1.55 - 3-hydroxybutyryl-CoA dehydratase / Crotonase (1 of 1)
OG IDs: 93519051
Best-hit-arabi-ID: AT3G06860.1
Best-hit-arabi-name: multifunctional protein 2
Gorai.006G076600.1.D
PCC=0.034, p=0.891
mRNA Profile
Protein Expression Profile
DPA

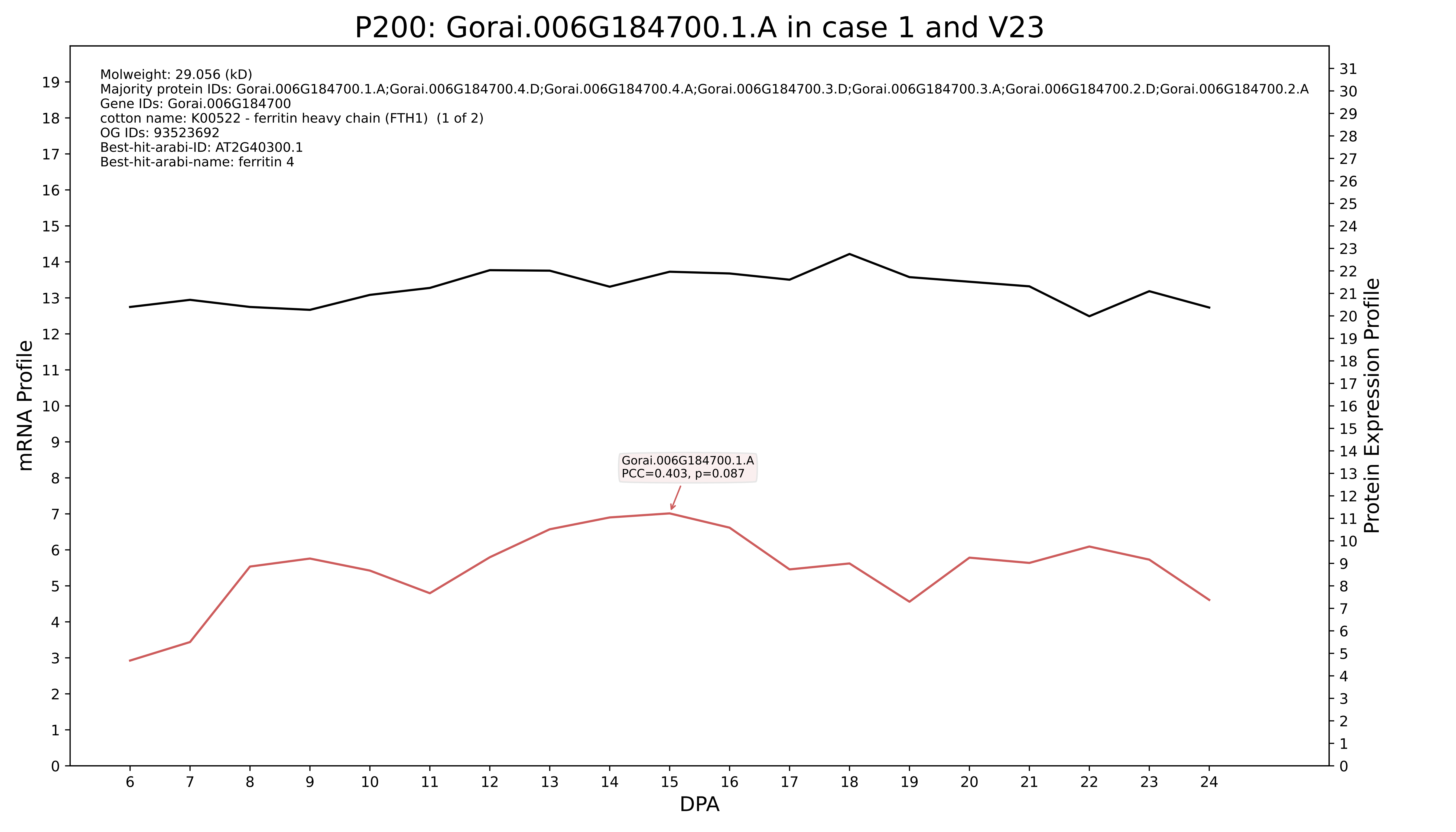
P200: Gorai.006G184700.1.A in case 1 and V23
Molweight: 29.056 (kD)
Majority protein IDs: Gorai.006G184700.1.A;Gorai.006G184700.4.D;Gorai.006G184700.4.A;Gorai.006G184700.3.D;Gorai.006G184700.3.A;Gorai.006G184700.2.D;Gorai.006G184700.2.A
Gene IDs: Gorai.006G184700
cotton name: K00522 - ferritin heavy chain (FTH1) (1 of 2)
OG IDs: 93523692
Best-hit-arabi-ID: AT2G40300.1
Best-hit-arabi-name: ferritin 4
Gorai.006G184700.1.A
PCC=0.403, p=0.087
mRNA Profile
Protein Expression Profile
DPA

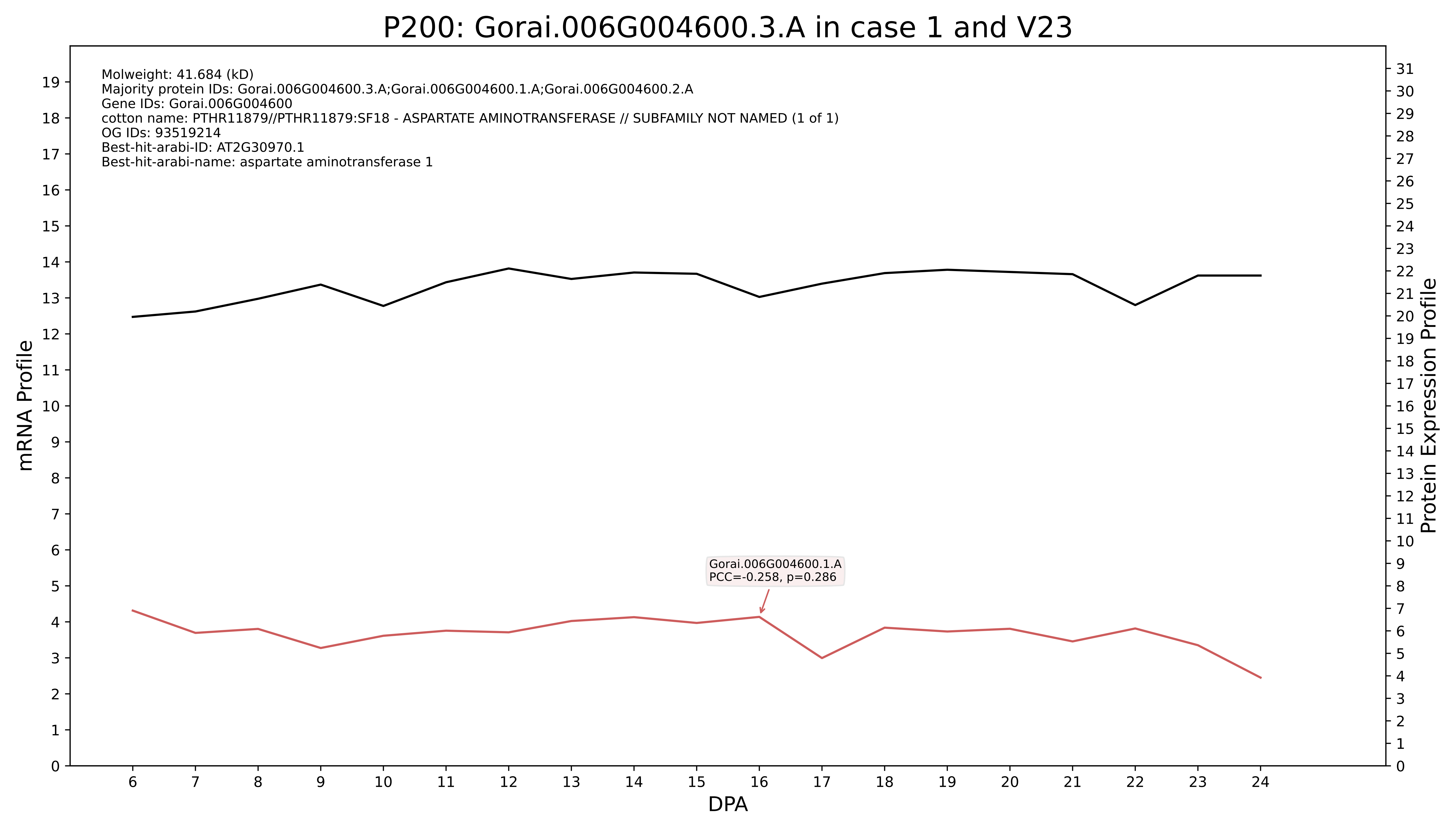
P200: Gorai.006G004600.3.A in case 1 and V23
Molweight: 41.684 (kD)
Majority protein IDs: Gorai.006G004600.3.A;Gorai.006G004600.1.A;Gorai.006G004600.2.A
Gene IDs: Gorai.006G004600
cotton name: PTHR11879//PTHR11879:SF18 - ASPARTATE AMINOTRANSFERASE // SUBFAMILY NOT NAMED (1 of 1)
OG IDs: 93519214
Best-hit-arabi-ID: AT2G30970.1
Best-hit-arabi-name: aspartate aminotransferase 1
Gorai.006G004600.1.A
PCC=-0.258, p=0.286
mRNA Profile
Protein Expression Profile
DPA

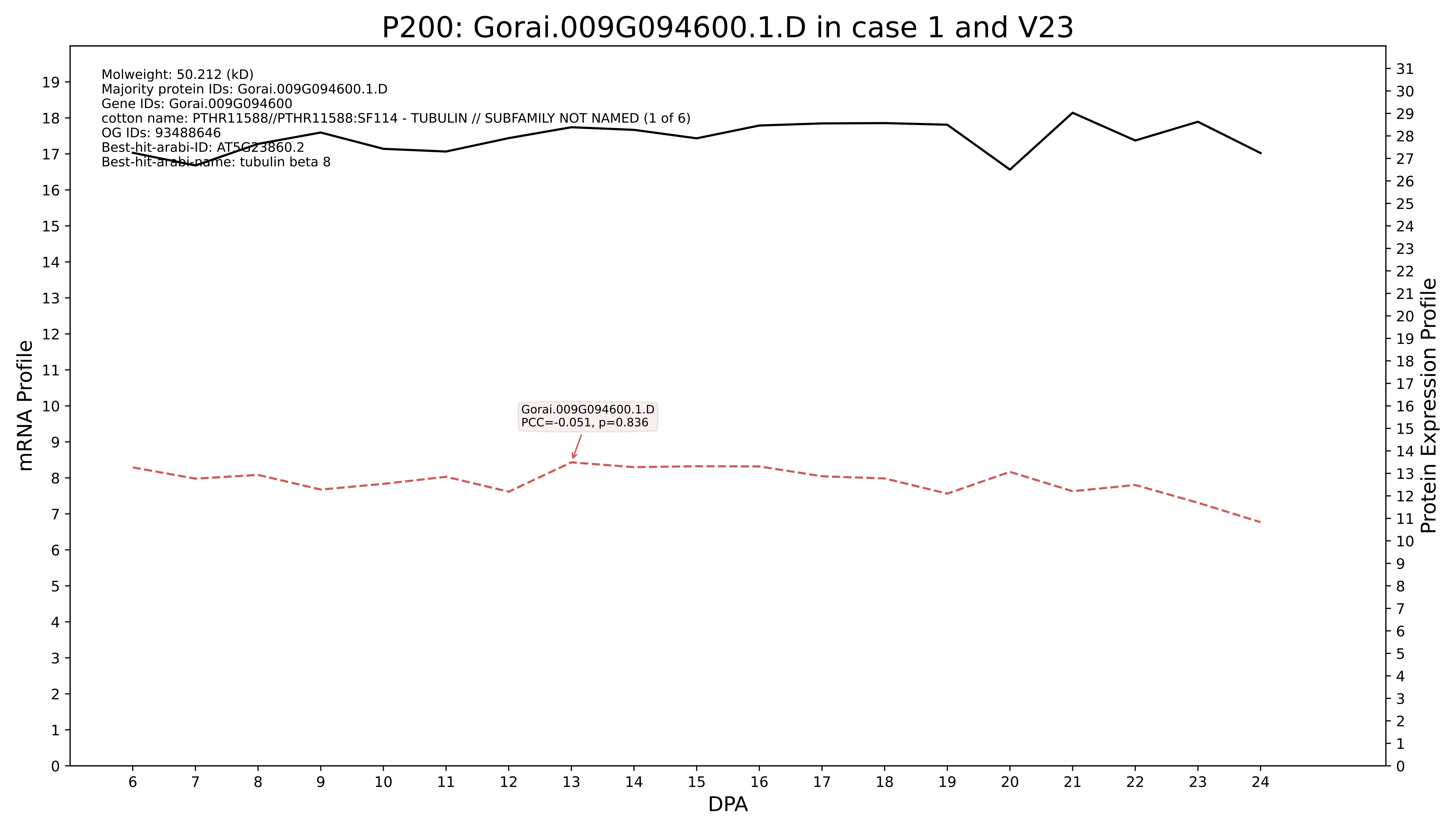
P200: Gorai.009G094600.1.D in case 1 and V23
Molweight: 50.212 (kD)
Majority protein IDs: Gorai.009G094600.1.D
Gene IDs: Gorai.009G094600
cotton name: PTHR11588//PTHR11588:SF114 - TUBULIN // SUBFAMILY NOT NAMED (1 of 6)
OG IDs: 93488646
Best-hit-arabi-ID: AT5G23860.2
Best-hit-arabi-name: tubulin beta 8
Gorai.009G094600.1.D
PCC=-0.051, p=0.836
mRNA Profile
Protein Expression Profile
DPA

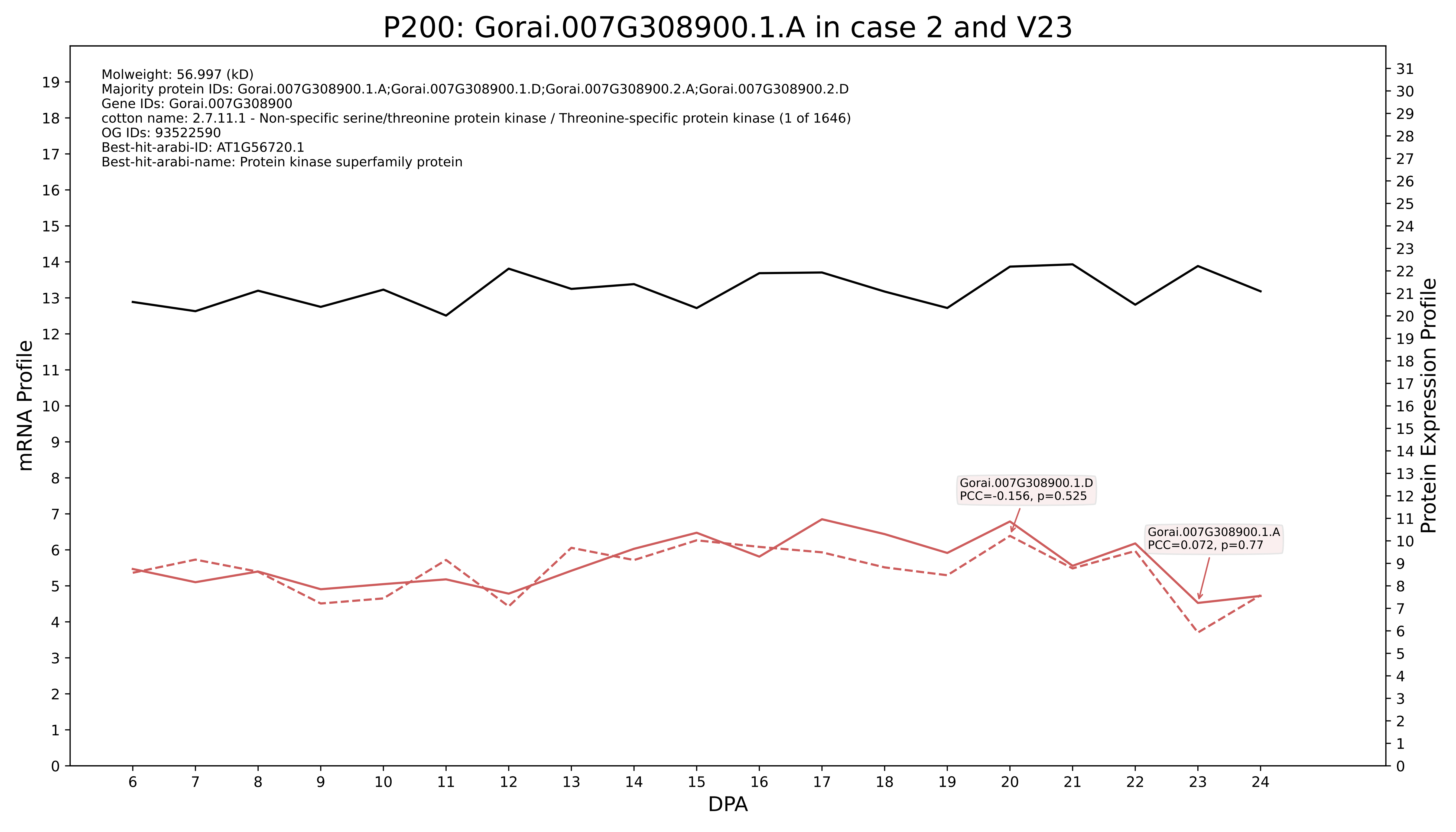
P200: Gorai.007G308900.1.A in case 2 and V23
Molweight: 56.997 (kD)
Majority protein IDs: Gorai.007G308900.1.A;Gorai.007G308900.1.D;Gorai.007G308900.2.A;Gorai.007G308900.2.D
Gene IDs: Gorai.007G308900
cotton name: 2.7.11.1 - Non-specific serine/threonine protein kinase / Threonine-specific protein kinase (1 of 1646)
OG IDs: 93522590
Best-hit-arabi-ID: AT1G56720.1
Best-hit-arabi-name: Protein kinase superfamily protein
Gorai.007G308900.1.D
PCC=-0.156, p=0.525
Gorai.007G308900.1.A
PCC=0.072, p=0.77
mRNA Profile
Protein Expression Profile
DPA

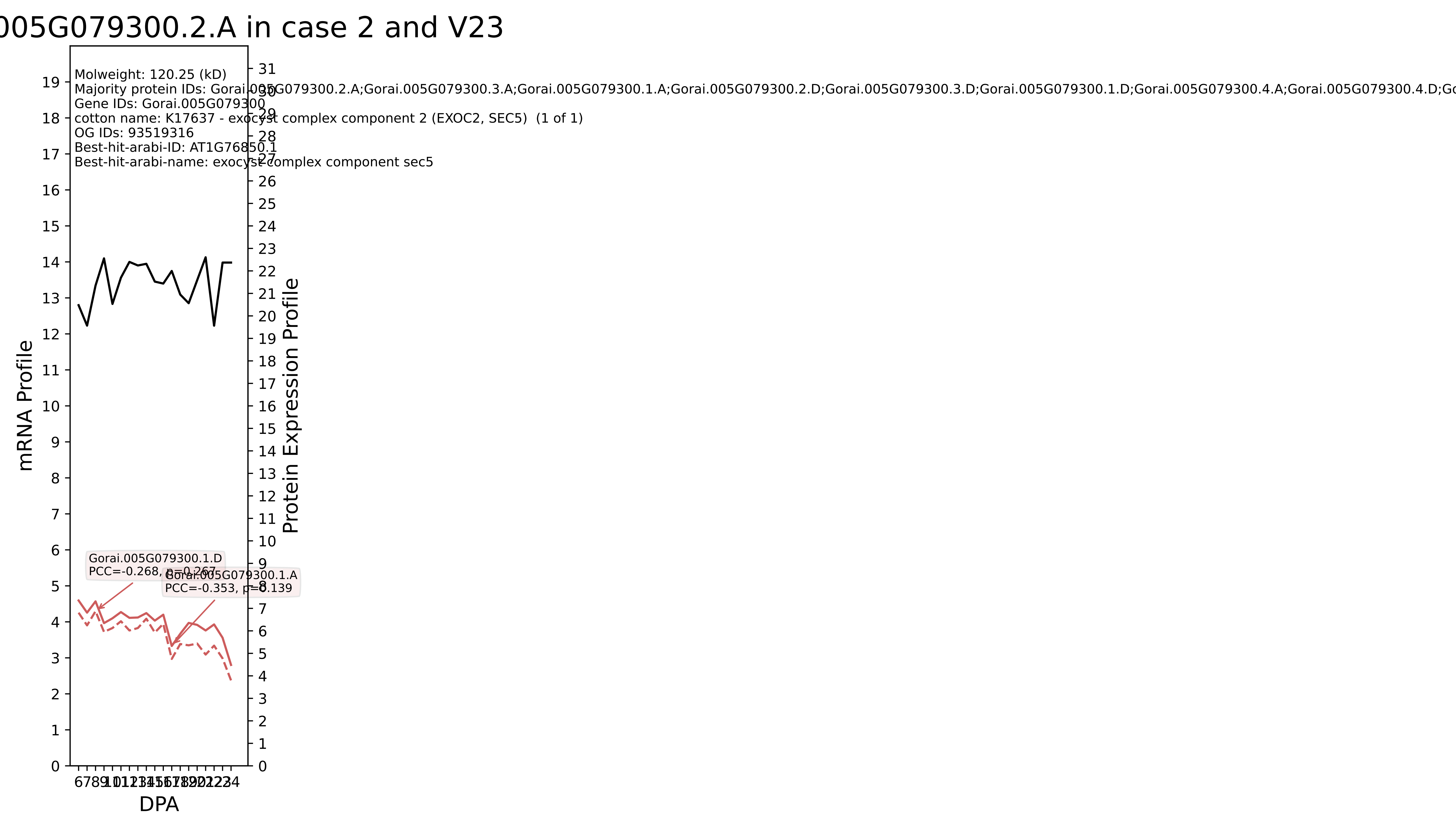

005G079300.2.A in case 2 and V23
Molweight: 120.25 (kD)
Majority protein IDs: Gorai.005G079300.2.A;Gorai.005G079300.3.A;Gorai.005G079300.1.A;Gorai.005G079300.2.D;Gorai.005G079300.3.D;Gorai.005G079300.1.D;Gorai.005G079300.4.A;Gorai.005G079300.4.D;Go
Gene IDs: Gorai.005G079300
cotton name: K17637 - exocyst complex component 2 (EXOC2, SEC5) (1 of 1)
OG IDs: 93519316
Best-hit-arabi-ID: AT1G76850.1
Best-hit-arabi-name: exocyst complex component sec5
Gorai.005G079300.1.D
PCC=-0.268, p=0.267
Gorai.005G079300.1.A
PCC=-0.353, p=0.139
mRNA Profile
Protein Expression Profile
DPA

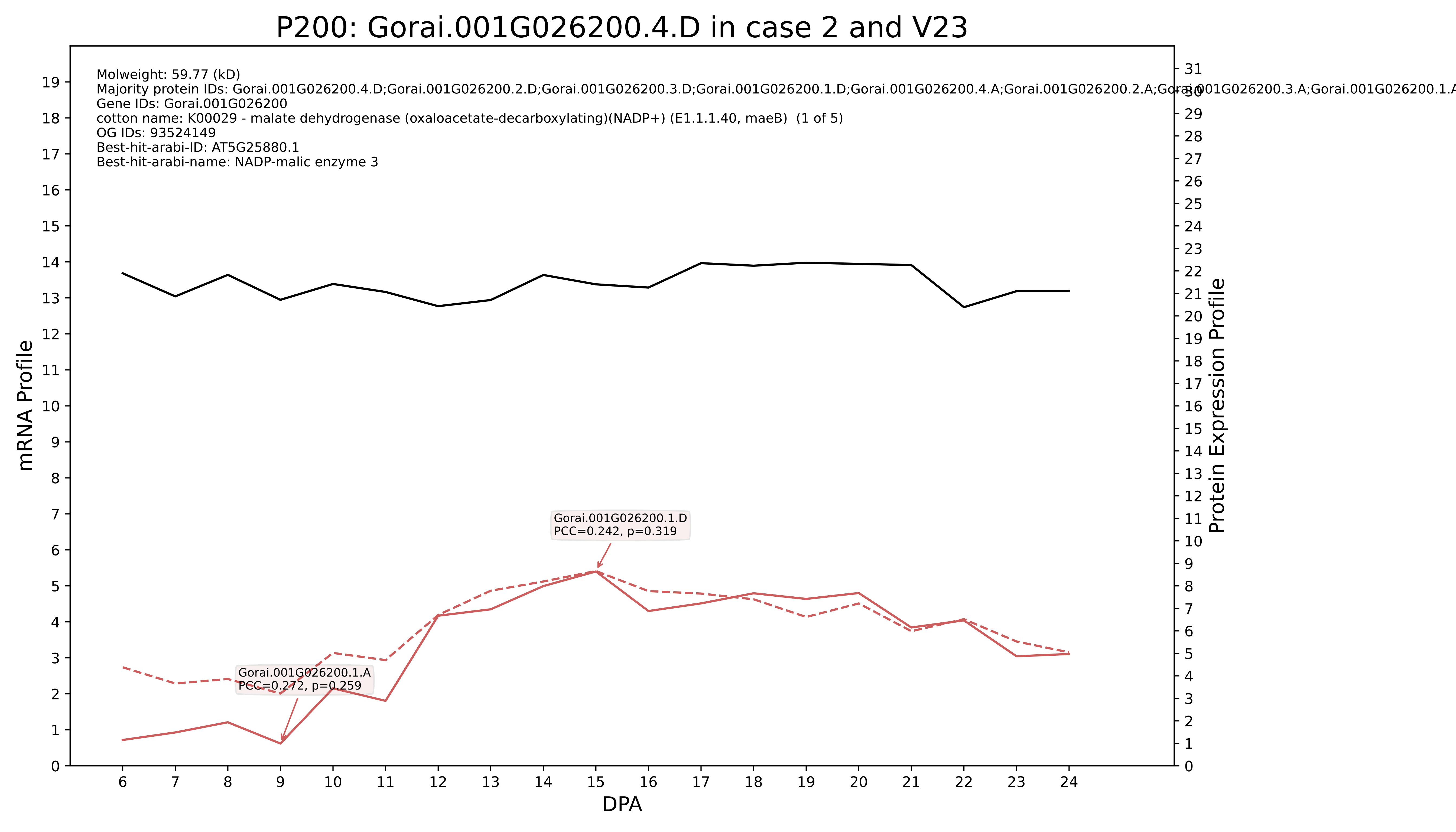

P200: Gorai.001G026200.4.D in case 2 and V23
Molweight: 59.77 (kD)
Majority protein IDs: Gorai.001G026200.4.D;Gorai.001G026200.2.D;Gorai.001G026200.3.D;Gorai.001G026200.1.D;Gorai.001G026200.4.A;Gorai.001G026200.2.A;Gorai.001G026200.3.A;Gorai.001G026200.1.A
Gene IDs: Gorai.001G026200
cotton name: K00029 - malate dehydrogenase (oxaloacetate-decarboxylating)(NADP+) (E1.1.1.40, maeB) (1 of 5)
OG IDs: 93524149
Best-hit-arabi-ID: AT5G25880.1
Best-hit-arabi-name: NADP-malic enzyme 3
Gorai.001G026200.1.D
PCC=0.242, p=0.319
Gorai.001G026200.1.A
PCC=0.272, p=0.259
mRNA Profile
Protein Expression Profile
DPA

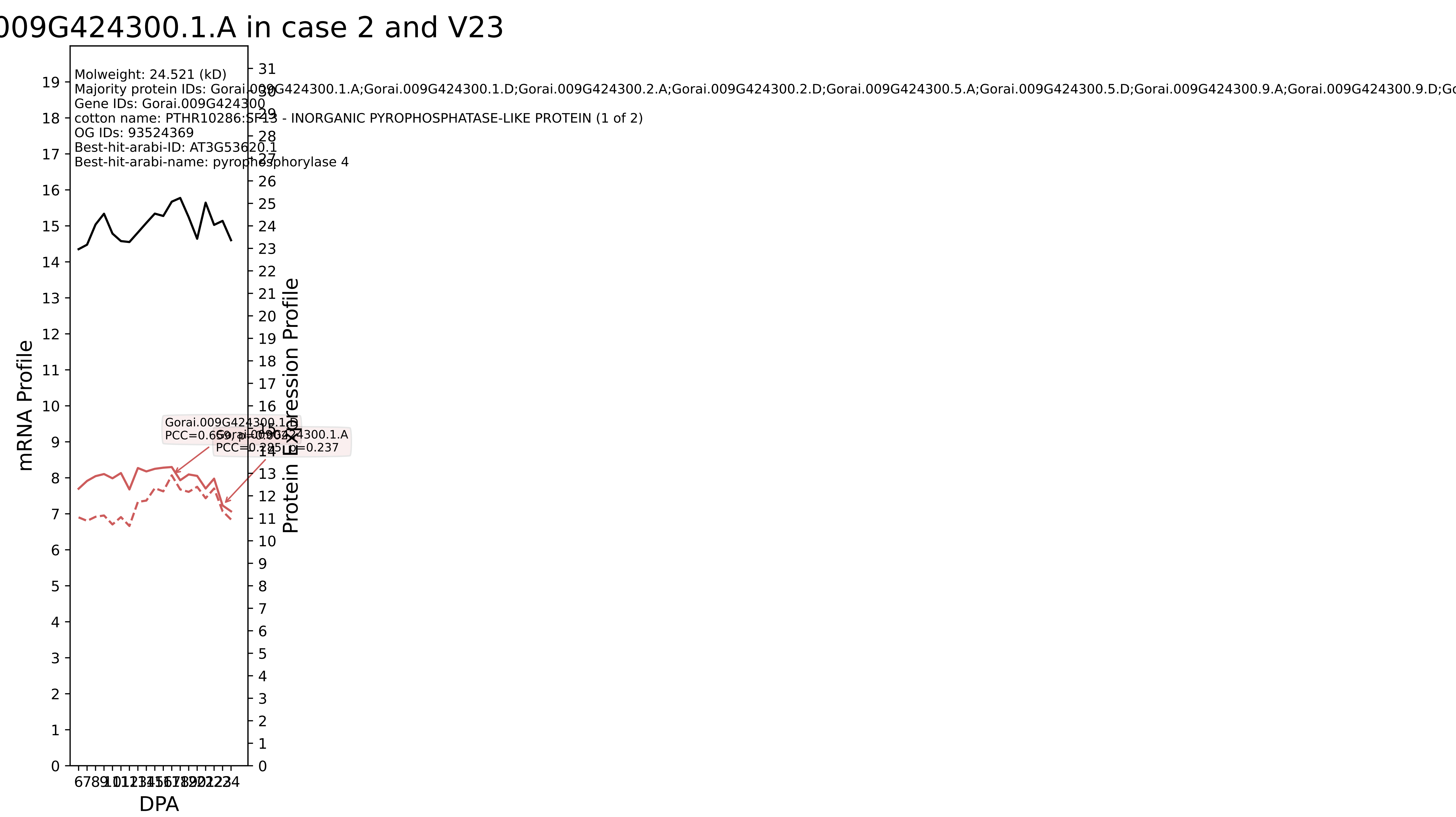
009G424300.1.A in case 2 and V23
Molweight: 24.521 (kD)
Majority protein IDs: Gorai.009G424300.1.A;Gorai.009G424300.1.D;Gorai.009G424300.2.A;Gorai.009G424300.2.D;Gorai.009G424300.5.A;Gorai.009G424300.5.D;Gorai.009G424300.9.A;Gorai.009G424300.9.D;Go
Gene IDs: Gorai.009G424300
cotton name: PTHR10286:SF13 - INORGANIC PYROPHOSPHATASE-LIKE PROTEIN (1 of 2)
OG IDs: 93524369
Best-hit-arabi-ID: AT3G53620.1
Best-hit-arabi-name: pyrophosphorylase 4
mRNA Profile
Protein Expression Profile
DPA
PCC=0.295, p=0.237

### P200: Gorai.013G247600.2.D in case 2 and V23

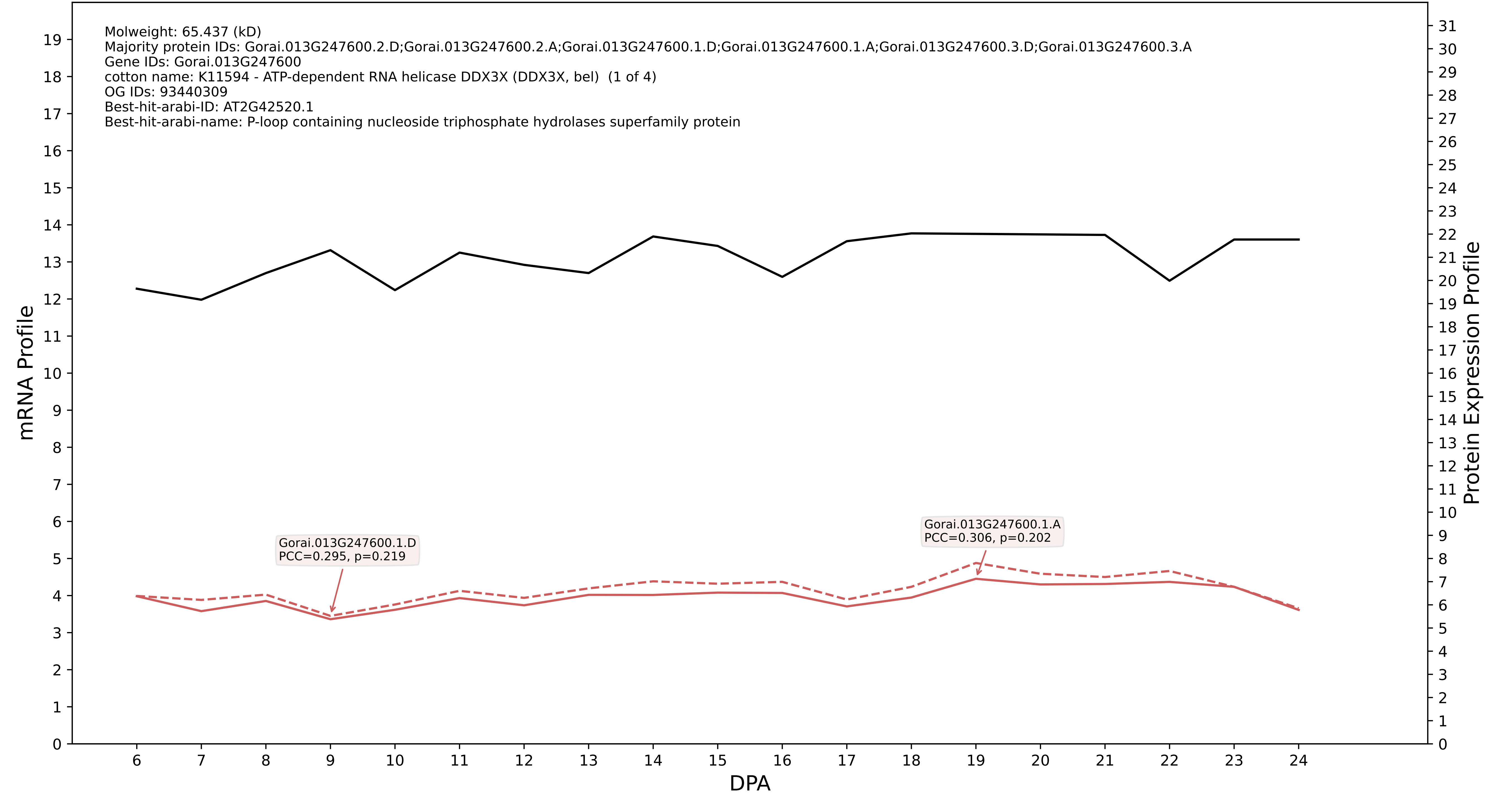

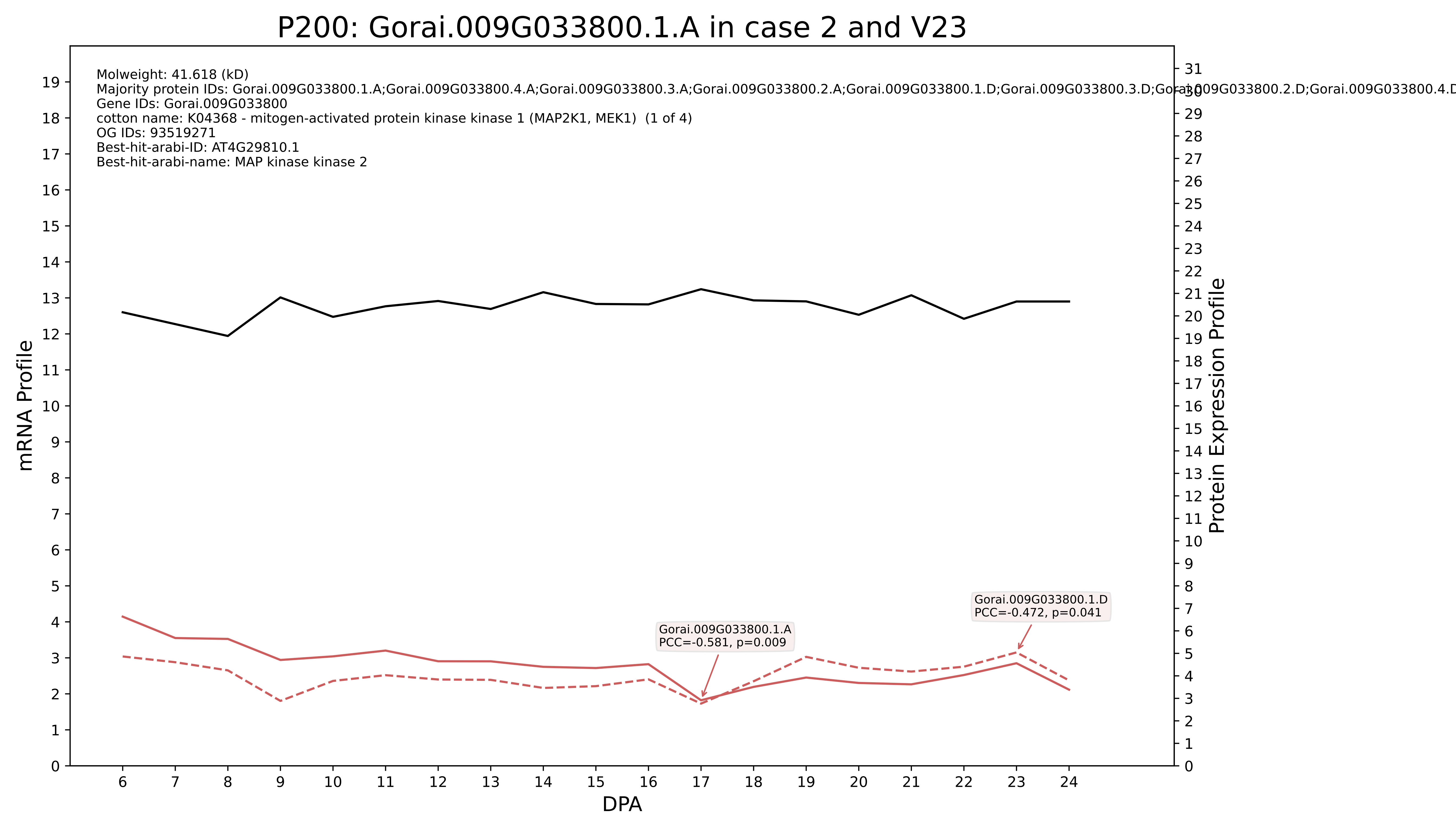

P200: Gorai.009G033800.1.A in case 2 and V23
Molweight: 41.618 (kD)
Majority protein IDs: Gorai.009G033800.1.A;Gorai.009G033800.4.A;Gorai.009G033800.3.A;Gorai.009G033800.2.A;Gorai.009G033800.1.D;Gorai.009G033800.3.D;Gorai.009G033800.2.D;Gorai.009G033800.4.D
Gene IDs: Gorai.009G033800
cotton name: K04368 - mitogen-activated protein kinase kinase 1 (MAP2K1, MEK1) (1 of 4)
OG IDs: 93519271
Best-hit-arabi-ID: AT4G29810.1
Best-hit-arabi-name: MAP kinase kinase 2
Gorai.009G033800.1.A
PCC=-0.581, p=0.009
Gorai.009G033800.1.D
PCC=-0.472, p=0.041
mRNA Profile
Protein Expression Profile
DPA

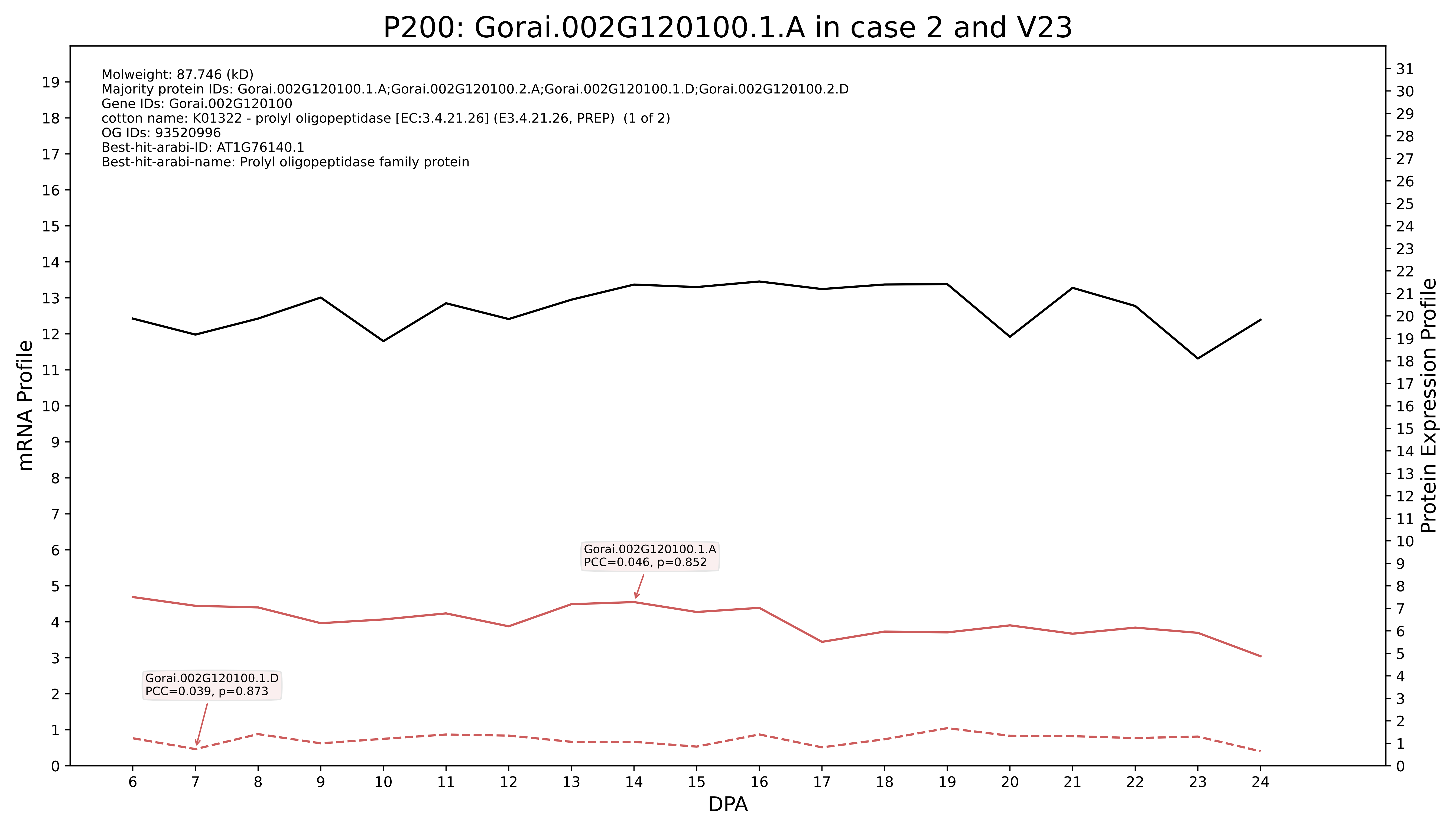
P200: Gorai.002G120100.1.A in case 2 and V23
Molweight: 87.746 (kD)
Majority protein IDs: Gorai.002G120100.1.A;Gorai.002G120100.2.A;Gorai.002G120100.1.D;Gorai.002G120100.2.D
Gene IDs: Gorai.002G120100
cotton name: K01322 - prolyl oligopeptidase [EC:3.4.21.26] (E3.4.21.26, PREP) (1 of 2)
OG IDs: 93520996
Best-hit-arabi-ID: AT1G76140.1
Best-hit-arabi-name: Prolyl oligopeptidase family protein
Gorai.002G120100.1.A
PCC=0.046, p=0.852
Gorai.002G120100.1.D
PCC=0.039, p=0.873
mRNA Profile
Protein Expression Profile
DPA

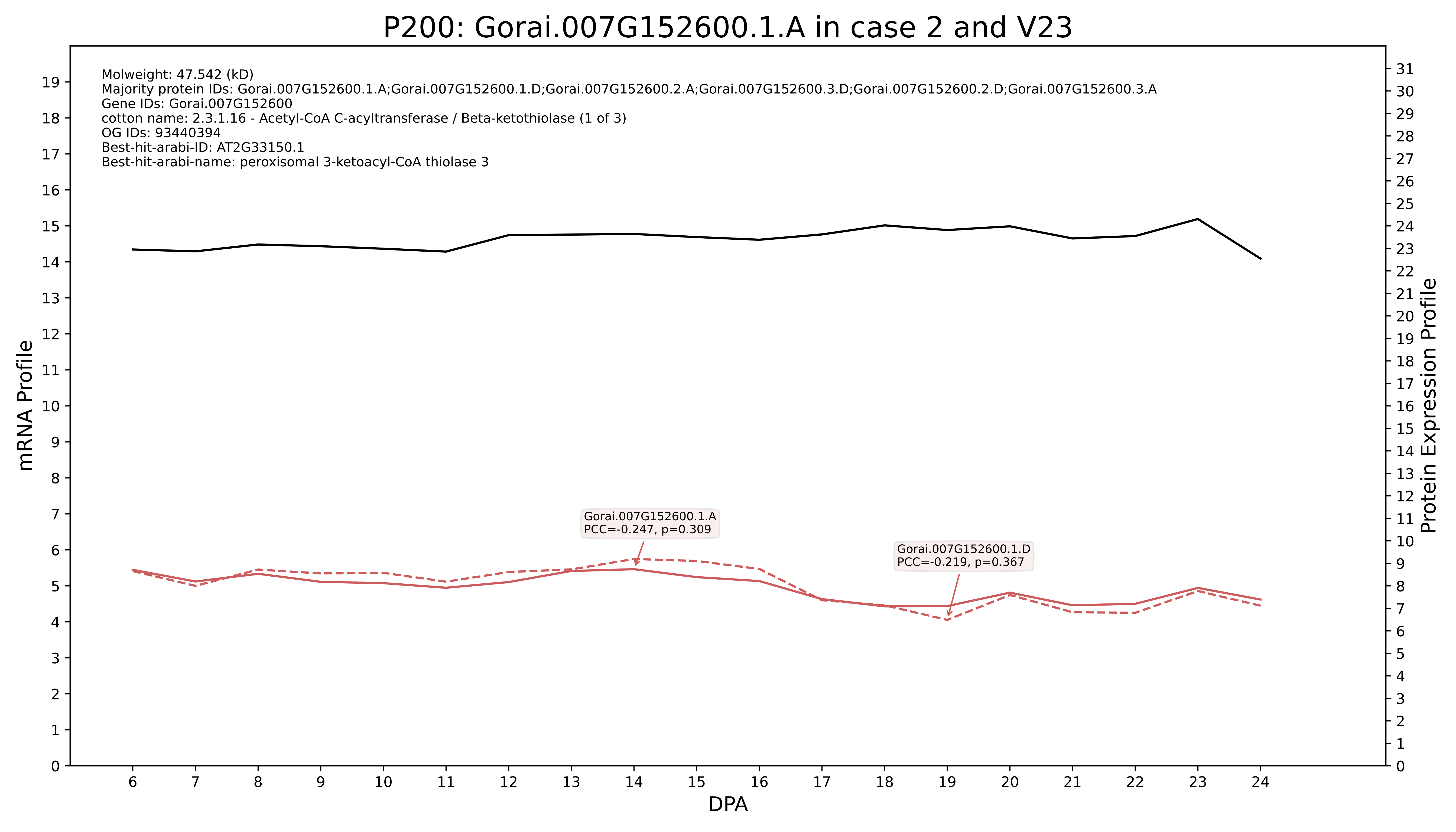
P200: Gorai.007G152600.1.A in case 2 and V23
Molweight: 47.542 (kD)
Majority protein IDs: Gorai.007G152600.1.A;Gorai.007G152600.1.D;Gorai.007G152600.2.A;Gorai.007G152600.3.D;Gorai.007G152600.2.D;Gorai.007G152600.3.A
Gene IDs: Gorai.007G152600
cotton name: 2.3.1.16 - Acetyl-CoA C-acyltransferase / Beta-ketothiolase (1 of 3)
OG IDs: 93440394
Best-hit-arabi-ID: AT2G33150.1
Best-hit-arabi-name: peroxisomal 3-ketoacyl-CoA thiolase 3
Gorai.007G152600.1.A
PCC=-0.247, p=0.309
Gorai.007G152600.1.D
PCC=-0.219, p=0.367
mRNA Profile
Protein Expression Profile
DPA

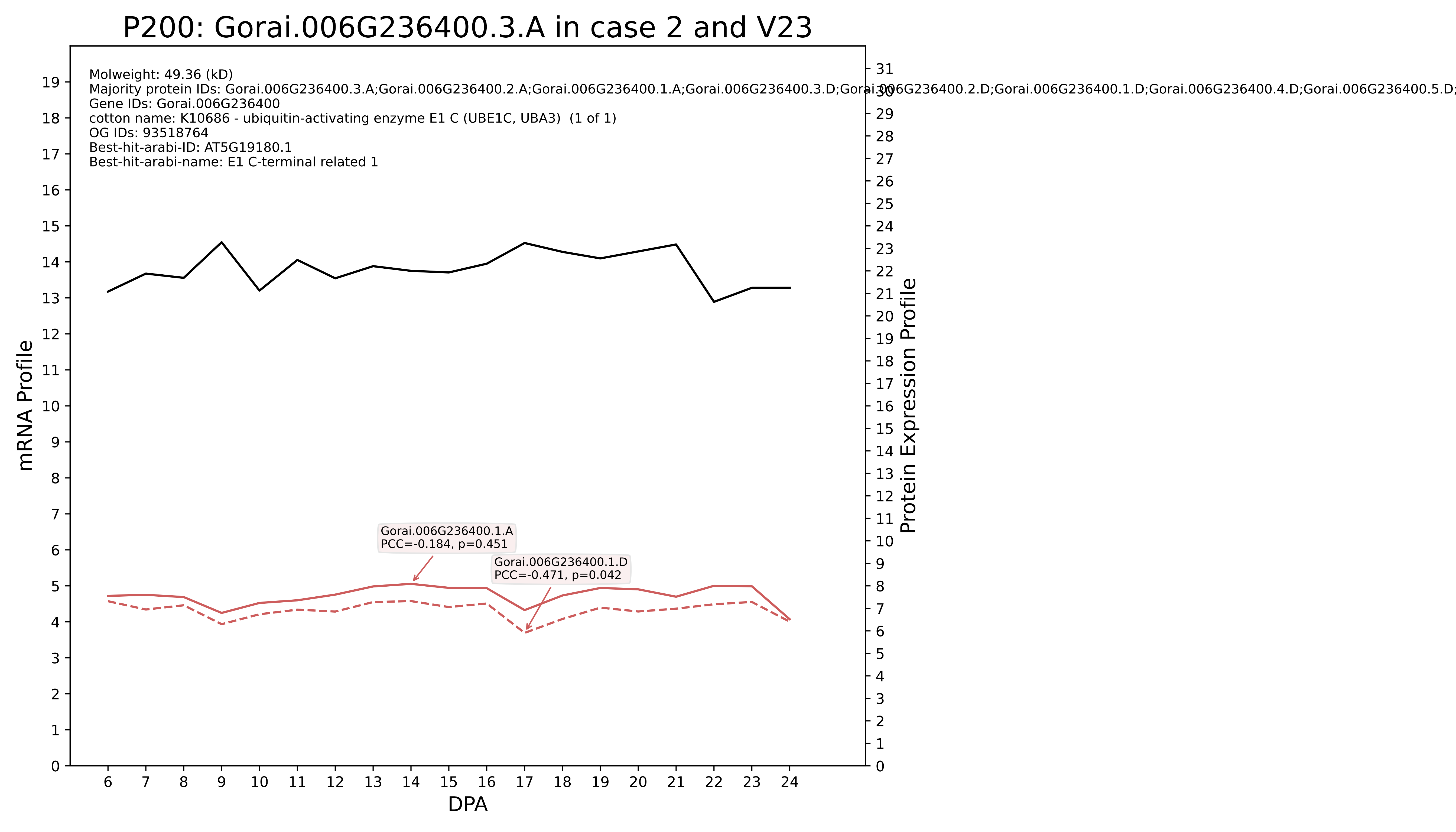

P200: Gorai.006G236400.3.A in case 2 and V23
Molweight: 49.36 (kD)
Majority protein IDs: Gorai.006G236400.3.A;Gorai.006G236400.2.A;Gorai.006G236400.1.A;Gorai.006G236400.3.D;Gorai.006G236400.2.D;Gorai.006G236400.1.D;Gorai.006G236400.4.D;Gorai.006G236400.5.D;
Gene IDs: Gorai.006G236400
cotton name: K10686 - ubiquitin-activating enzyme E1 C (UBE1C, UBA3) (1 of 1)
OG IDs: 93518764
Best-hit-arabi-ID: AT5G19180.1
Best-hit-arabi-name: E1 C-terminal related 1
Gorai.006G236400.1.A
PCC=-0.184, p=0.451
Gorai.006G236400.1.D
PCC=-0.471, p=0.042
mRNA Profile
Protein Expression Profile
DPA

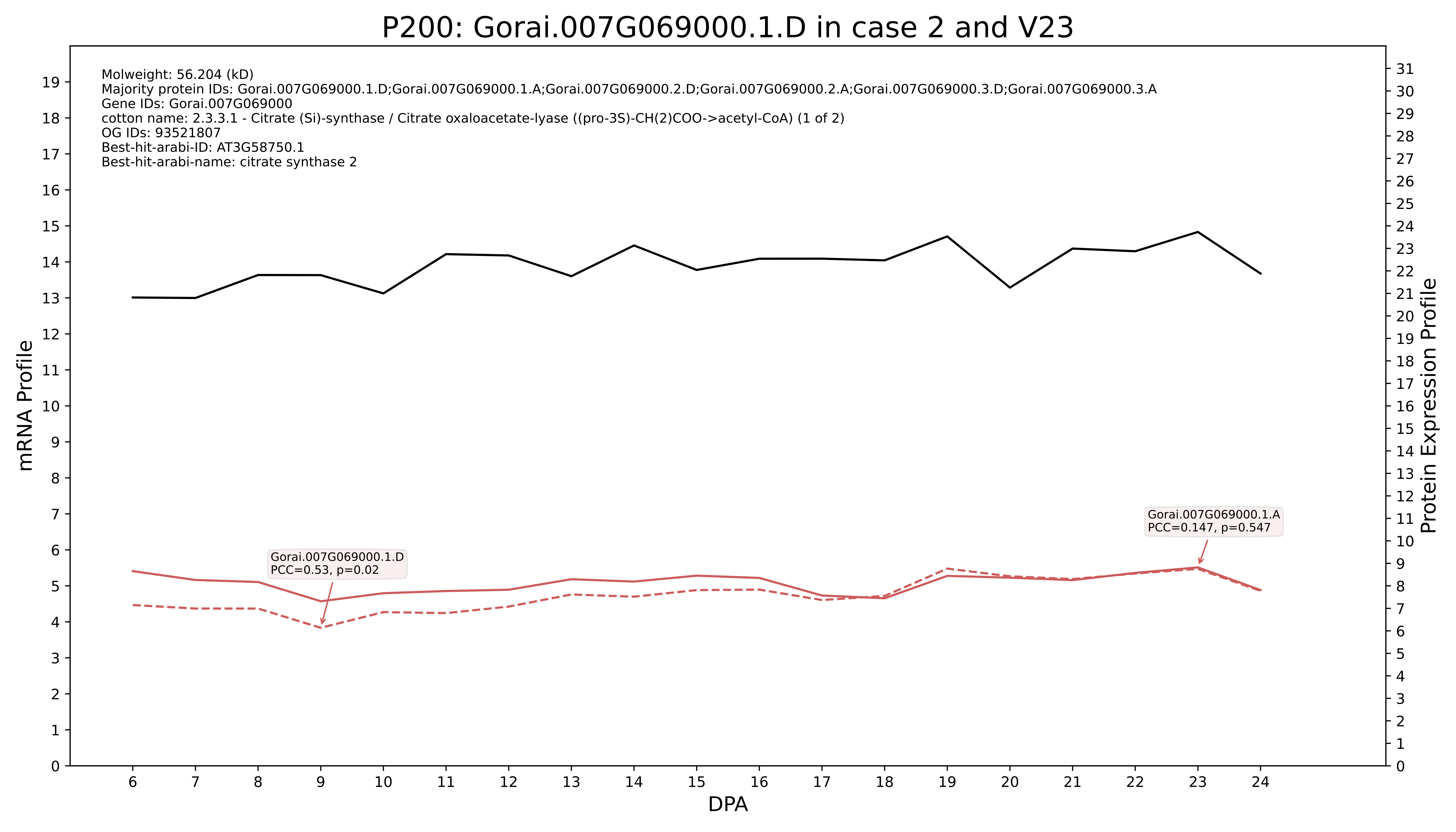
P200: Gorai.007G069000.1.D in case 2 and V23
Molweight: 56.204 (kD)
Majority protein IDs: Gorai.007G069000.1.D;Gorai.007G069000.1.A;Gorai.007G069000.2.D;Gorai.007G069000.2.A;Gorai.007G069000.3.D;Gorai.007G069000.3.A
Gene IDs: Gorai.007G069000
cotton name: 2.3.3.1 - Citrate (Si)-synthase / Citrate oxaloacetate-lyase ((pro-3S)-CH(2)COO->acetyl-CoA) (1 of 2)
OG IDs: 93521807
Best-hit-arabi-ID: AT3G58750.1
Best-hit-arabi-name: citrate synthase 2
Gorai.007G069000.1.D
PCC=0.53, p=0.02
Gorai.007G069000.1.A
PCC=0.147, p=0.547
mRNA Profile
Protein Expression Profile
DPA

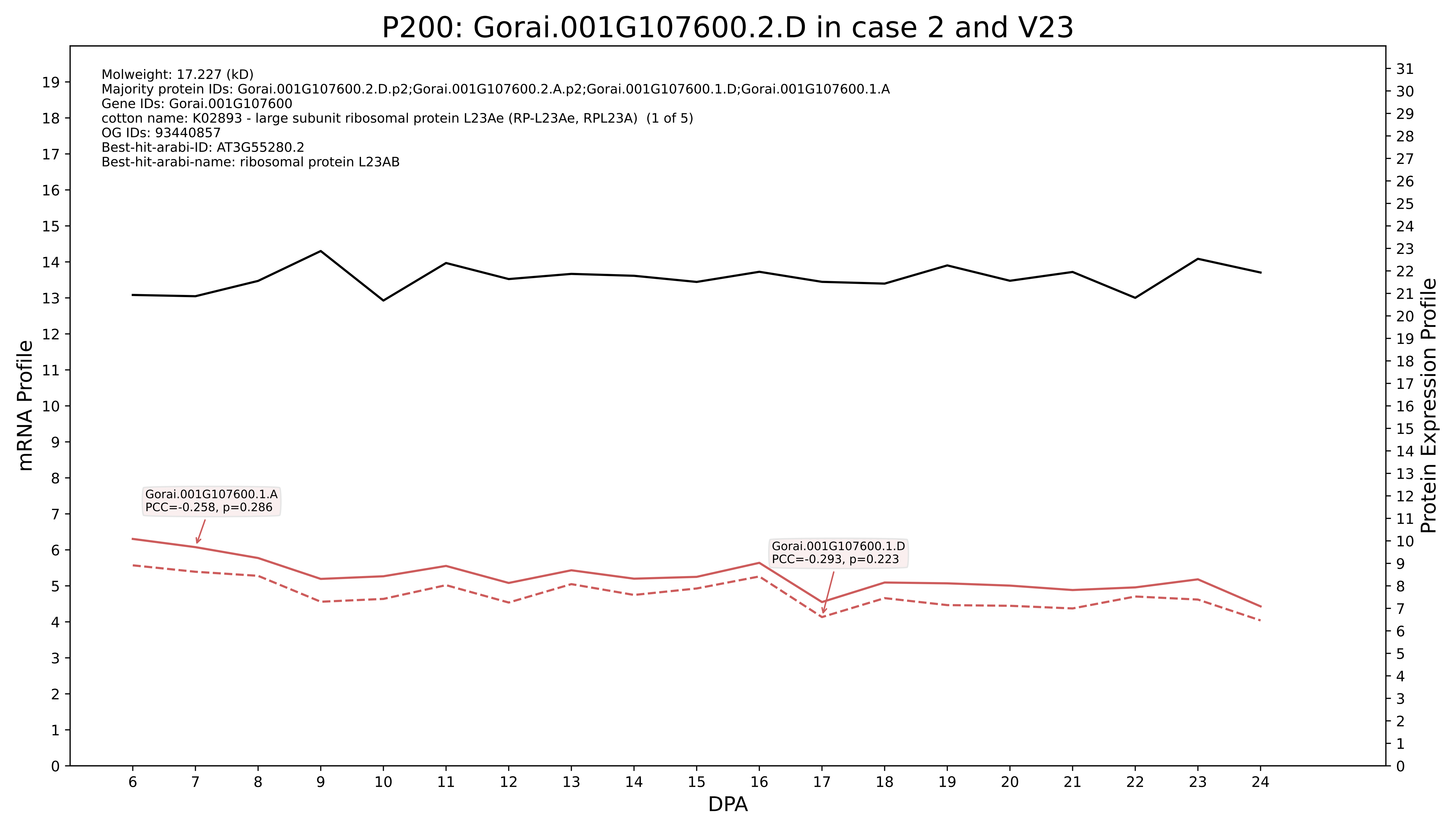

P200: Gorai.001G107600.2.D in case 2 and V23
Molweight: 17.227 (kD)
Majority protein IDs: Gorai.001G107600.2.D.p2;Gorai.001G107600.2.A.p2;Gorai.001G107600.1.D;Gorai.001G107600.1.A
Gene IDs: Gorai.001G107600
cotton name: K02893 - large subunit ribosomal protein L23Ae (RP-L23Ae, RPL23A) (1 of 5)
OG IDs: 93440857
Best-hit-arabi-ID: AT3G55280.2
Best-hit-arabi-name: ribosomal protein L23AB
Gorai.001G107600.1.A
PCC=-0.258, p=0.286
Gorai.001G107600.1.D
PCC=-0.293, p=0.223
mRNA Profile
Protein Expression Profile
DPA

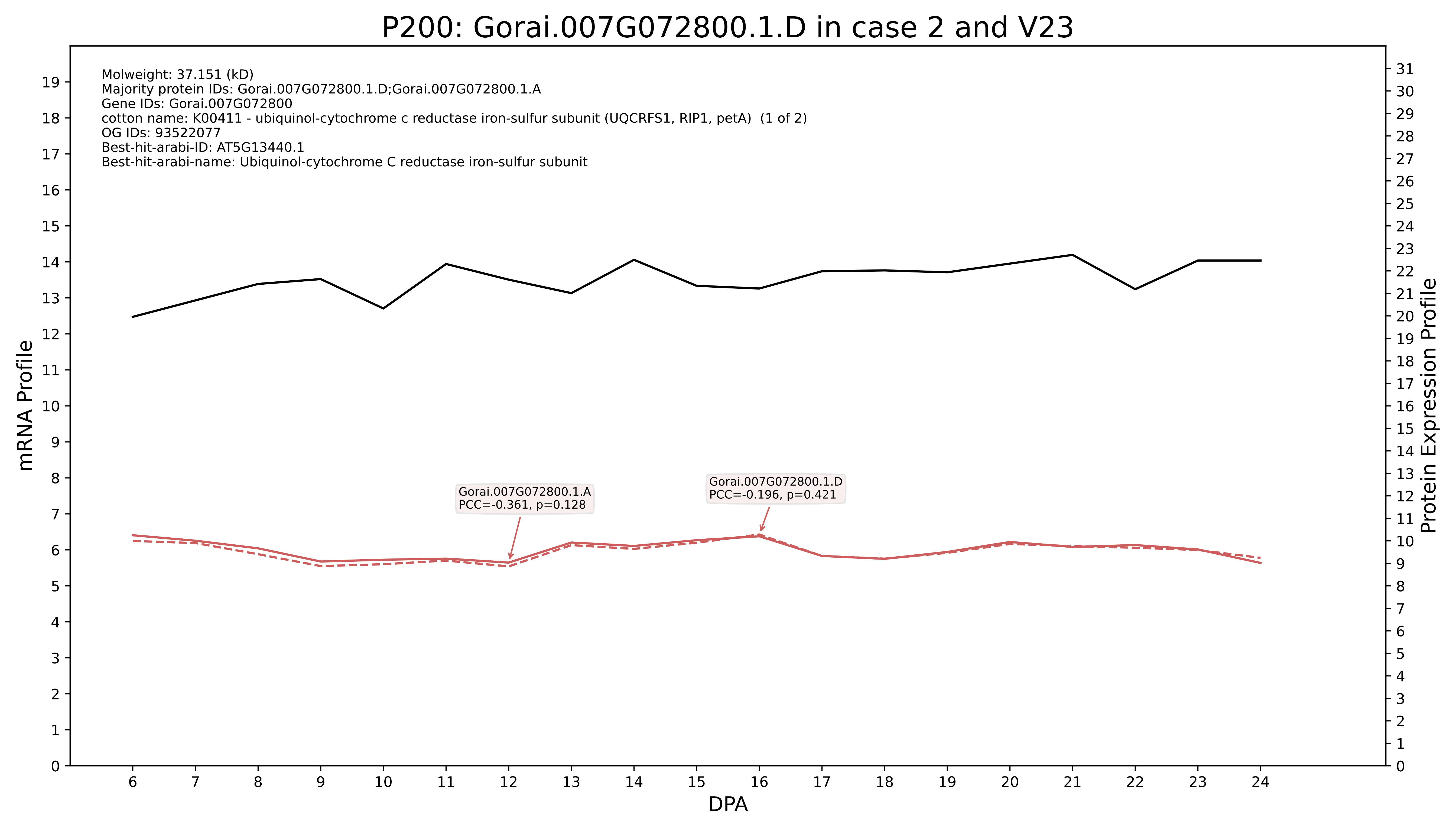
P200: Gorai.007G072800.1.D in case 2 and V23
Molweight: 37.151 (kD)
Majority protein IDs: Gorai.007G072800.1.D;Gorai.007G072800.1.A
Gene IDs: Gorai.007G072800
cotton name: K00411 - ubiquinol-cytochrome c reductase iron-sulfur subunit (UQCRFS1, RIP1, petA) (1 of 2)
OG IDs: 93522077
Best-hit-arabi-ID: AT5G13440.1
Best-hit-arabi-name: Ubiquinol-cytochrome C reductase iron-sulfur subunit
Gorai.007G072800.1.A
PCC=-0.361, p=0.128
Gorai.007G072800.1.D
PCC=-0.196, p=0.421
mRNA Profile
Protein Expression Profile
DPA

### P200: Gorai.009G142800.2.D in case 2 and V23

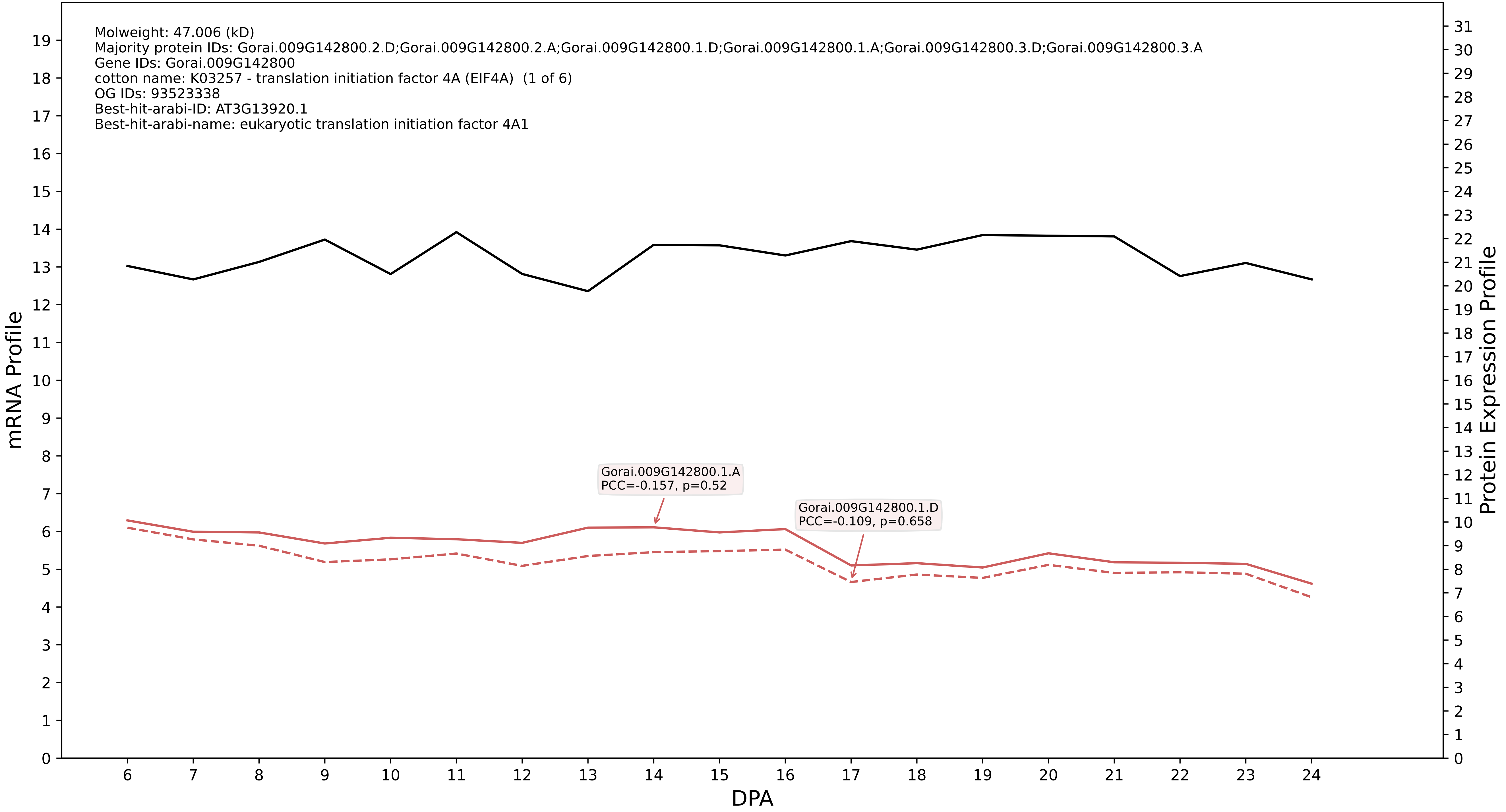

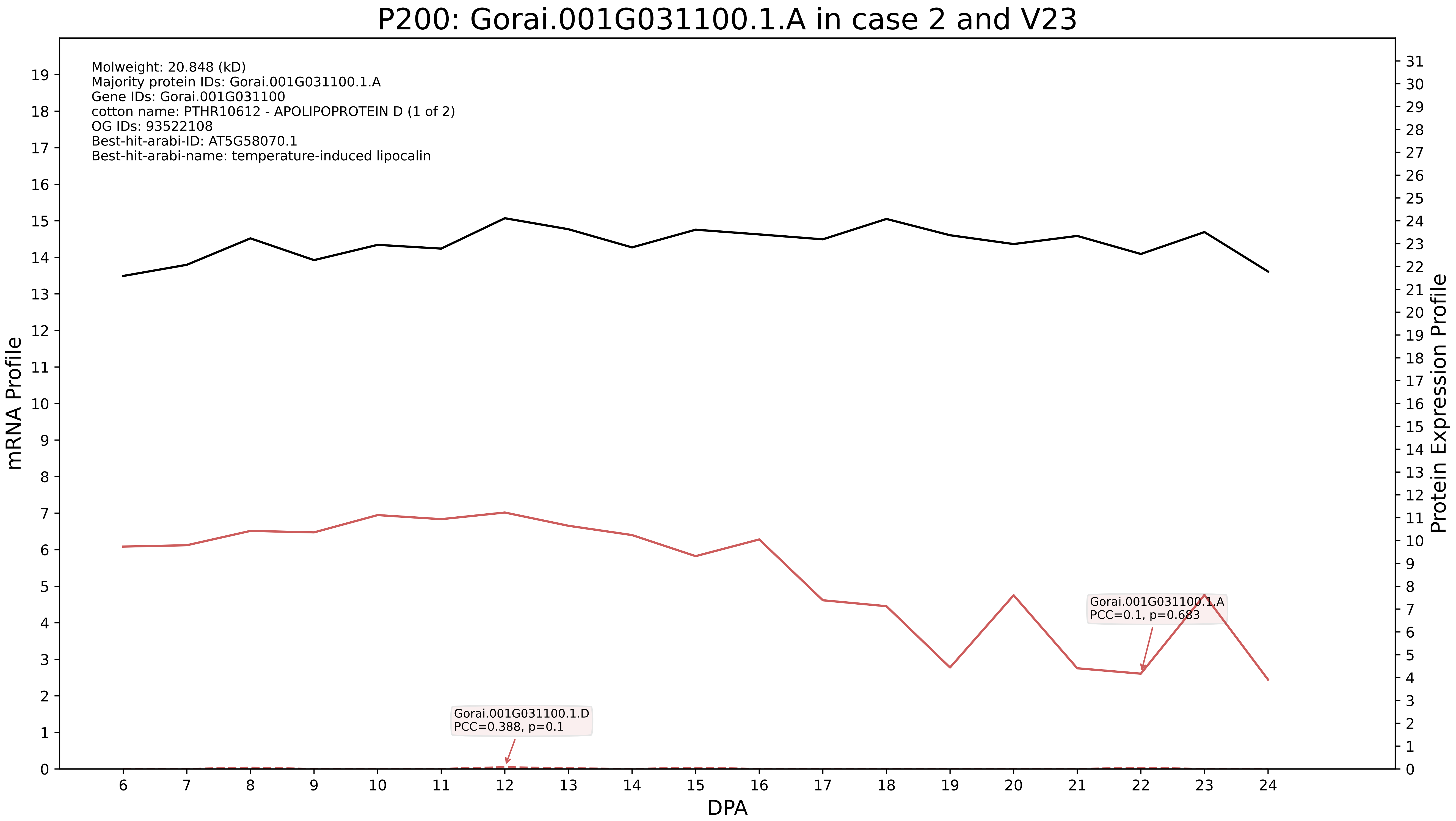
P200: Gorai.001G031100.1.A in case 2 and V23
Molweight: 20.848 (kD)
Majority protein IDs: Gorai.001G031100.1.A
Gene IDs: Gorai.001G031100
cotton name: PTHR10612 - APOLIPOPROTEIN D (1 of 2)
OG IDs: 93522108
Best-hit-arabi-ID: AT5G58070.1
Best-hit-arabi-name: temperature-induced lipocalin
Gorai.001G031100.1.A
PCC=0.1, p=0.683
Gorai.001G031100.1.D
PCC=0.388, p=0.1
mRNA Profile
Protein Expression Profile
DPA

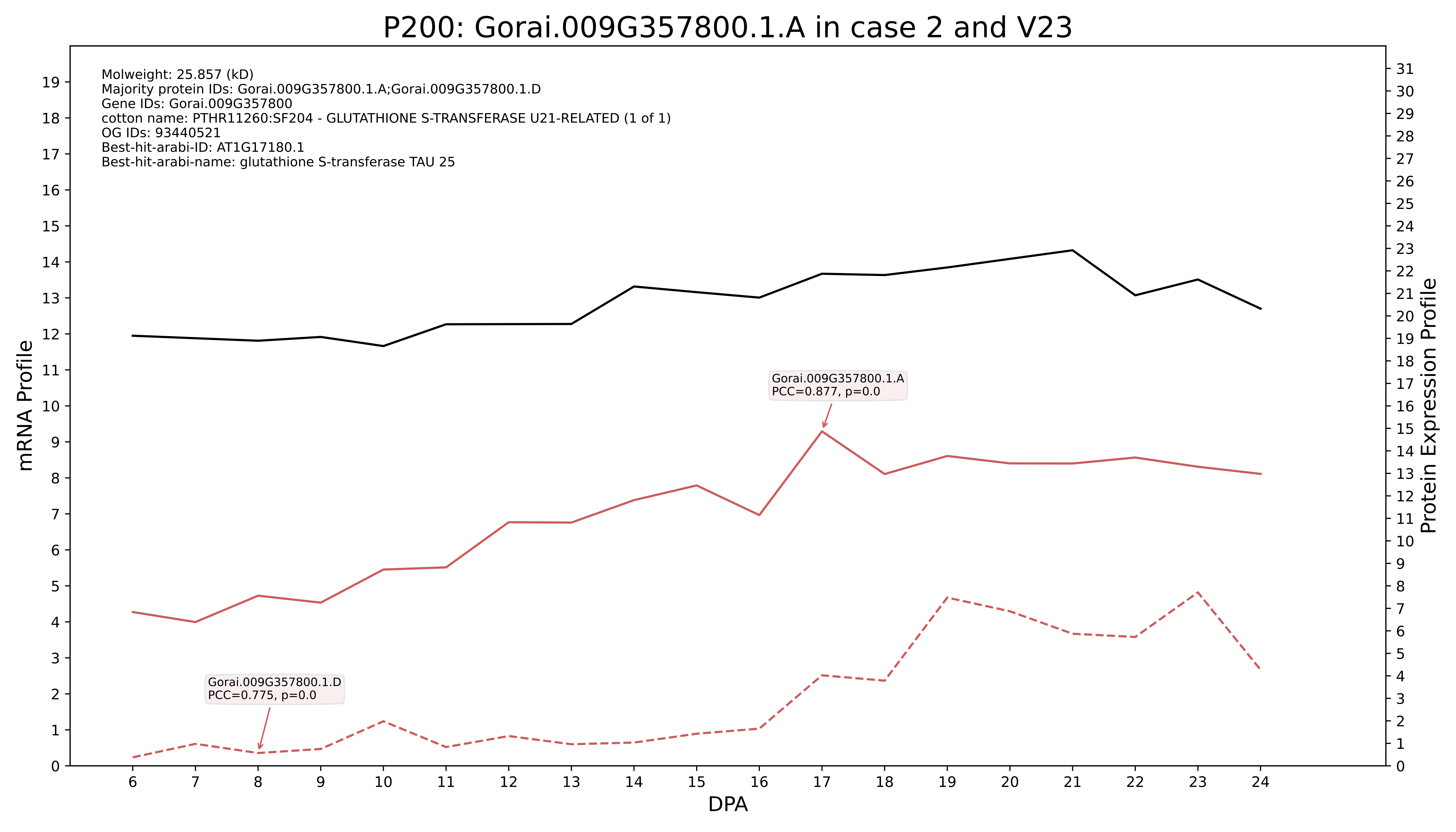
P200: Gorai.009G357800.1.A in case 2 and V23
Molweight: 25.857 (kD)
Majority protein IDs: Gorai.009G357800.1.A;Gorai.009G357800.1.D
Gene IDs: Gorai.009G357800
cotton name: PTHR11260:SF204 - GLUTATHIONE S-TRANSFERASE U21-RELATED (1 of 1)
OG IDs: 93440521
Best-hit-arabi-ID: AT1G17180.1
Best-hit-arabi-name: glutathione S-transferase TAU 25
Gorai.009G357800.1.A
PCC=0.877, p=0.0
Gorai.009G357800.1.D
PCC=0.775, p=0.0
mRNA Profile
Protein Expression Profile
DPA

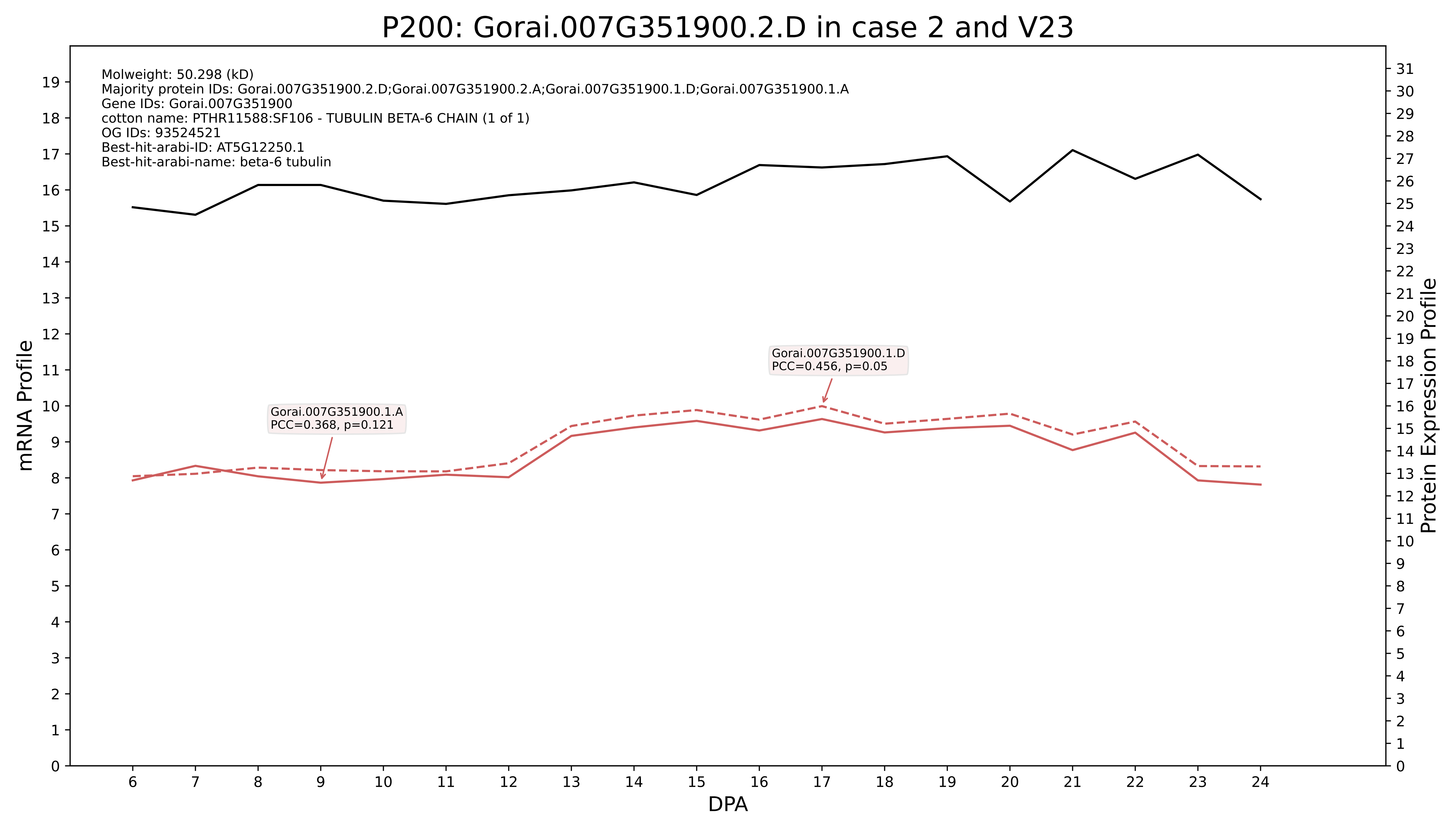
P200: Gorai.007G351900.2.D in case 2 and V23
Molweight: 50.298 (kD)
Majority protein IDs: Gorai.007G351900.2.D;Gorai.007G351900.2.A;Gorai.007G351900.1.D;Gorai.007G351900.1.A
Gene IDs: Gorai.007G351900
cotton name: PTHR11588:SF106 - TUBULIN BETA-6 CHAIN (1 of 1)
OG IDs: 93524521
Best-hit-arabi-ID: AT5G12250.1
Best-hit-arabi-name: beta-6 tubulin
Gorai.007G351900.1.D
PCC=0.456, p=0.05
Gorai.007G351900.1.A
PCC=0.368, p=0.121
mRNA Profile
Protein Expression Profile
DPA

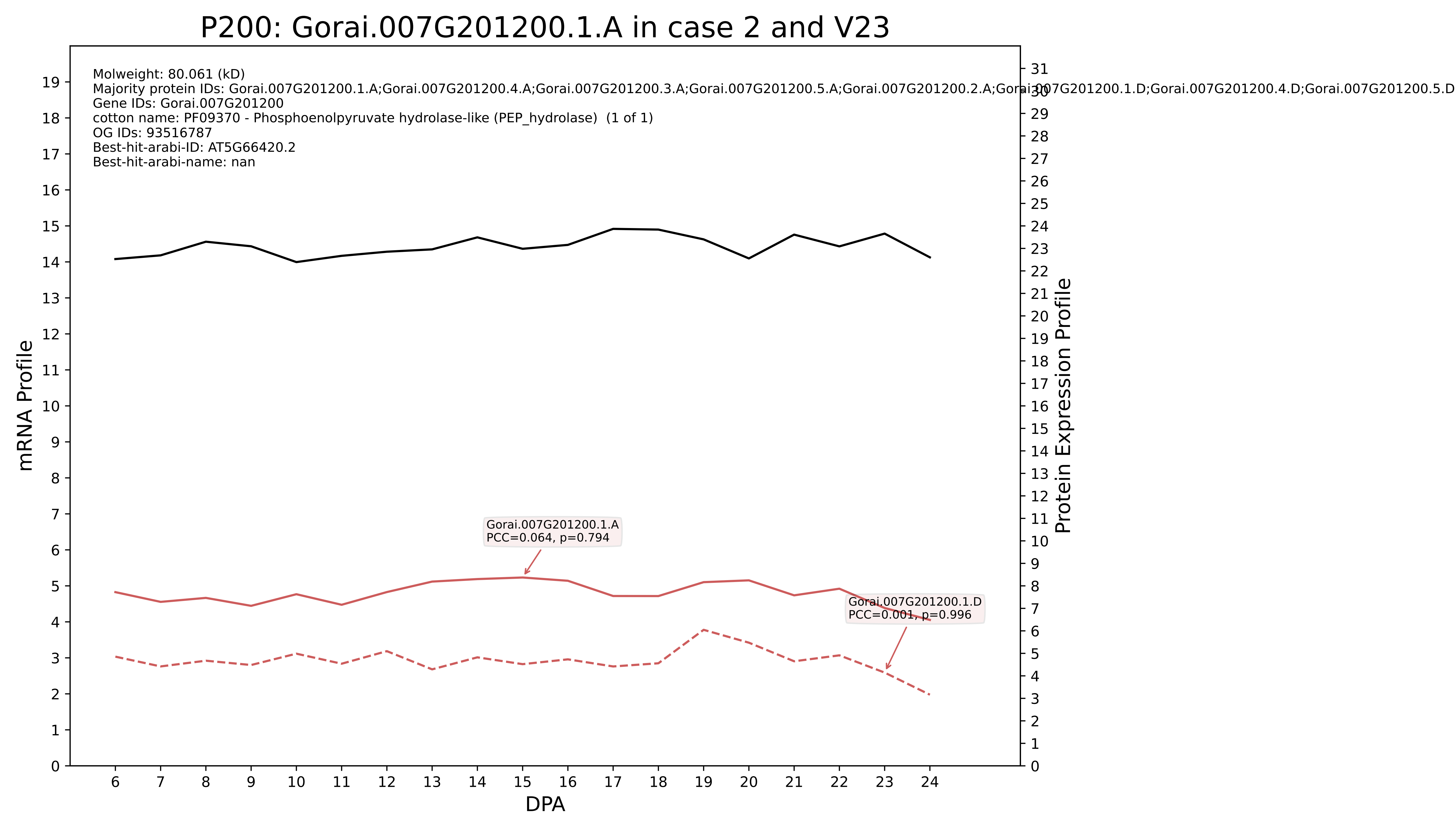
P200: Gorai.007G201200.1.A in case 2 and V23
Molweight: 80.061 (kD)
Majority protein IDs: Gorai.007G201200.1.A;Gorai.007G201200.4.A;Gorai.007G201200.3.A;Gorai.007G201200.5.A;Gorai.007G201200.2.A;Gorai.007G201200.1.D;Gorai.007G201200.4.D;Gorai.007G201200.5.D
Gene IDs: Gorai.007G201200
cotton name: PF09370 - Phosphoenolpyruvate hydrolase-like (PEP_hydrolase) (1 of 1)
OG IDs: 93516787
Best-hit-arabi-ID: AT5G66420.2
Best-hit-arabi-name: nan
Gorai.007G201200.1.A
PCC=0.064, p=0.794
Gorai.007G201200.1.D
PCC=0.001, p=0.996
mRNA Profile
Protein Expression Profile
DPA

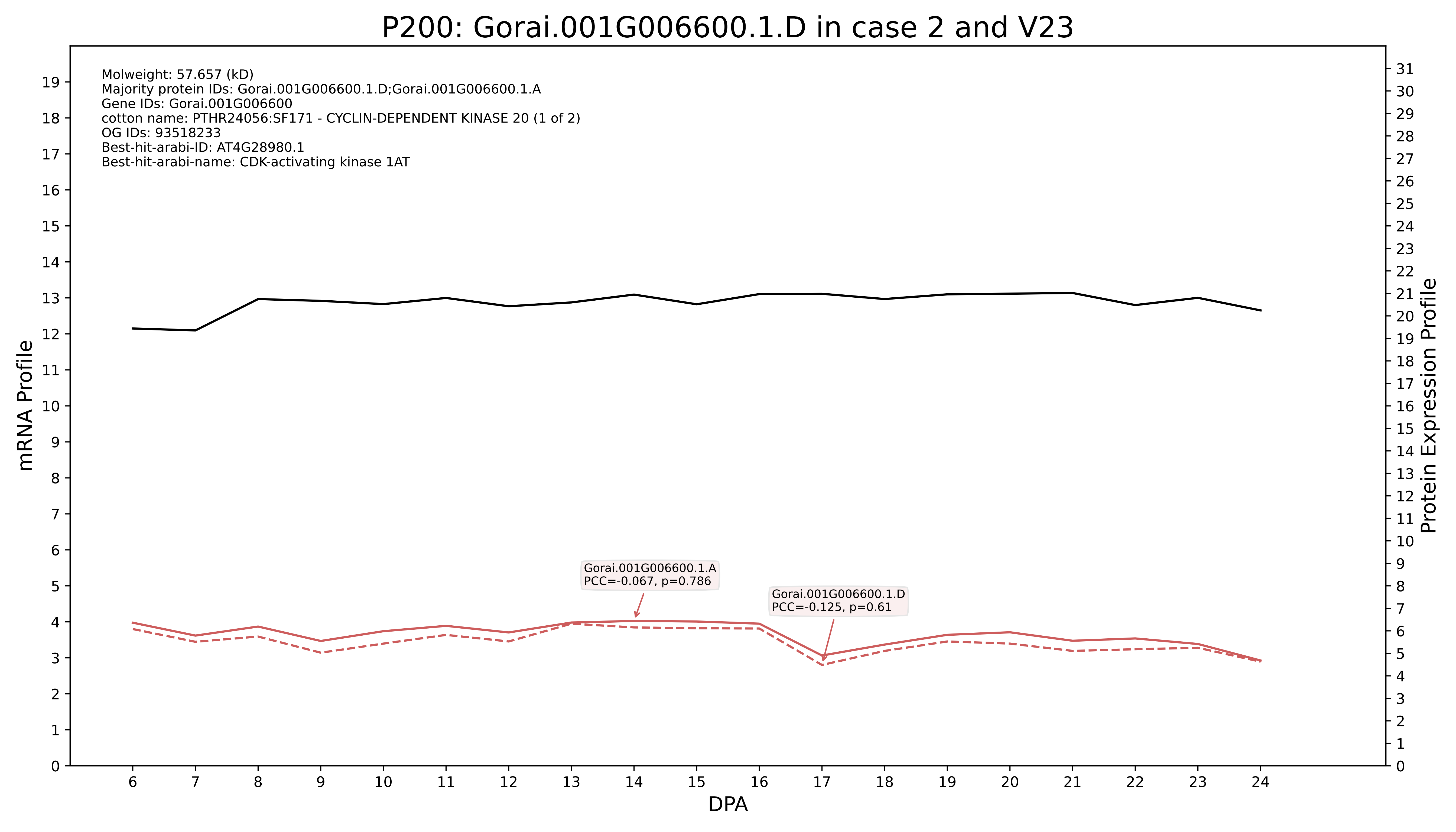
P200: Gorai.001G006600.1.D in case 2 and V23
Molweight: 57.657 (kD)
Majority protein IDs: Gorai.001G006600.1.D;Gorai.001G006600.1.A
Gene IDs: Gorai.001G006600
cotton name: PTHR24056:SF171 - CYCLIN-DEPENDENT KINASE 20 (1 of 2)
OG IDs: 93518233
Best-hit-arabi-ID: AT4G28980.1
Best-hit-arabi-name: CDK-activating kinase 1AT
Gorai.001G006600.1.A
PCC=-0.067, p=0.786
Gorai.001G006600.1.D
PCC=-0.125, p=0.61
mRNA Profile
Protein Expression Profile
DPA

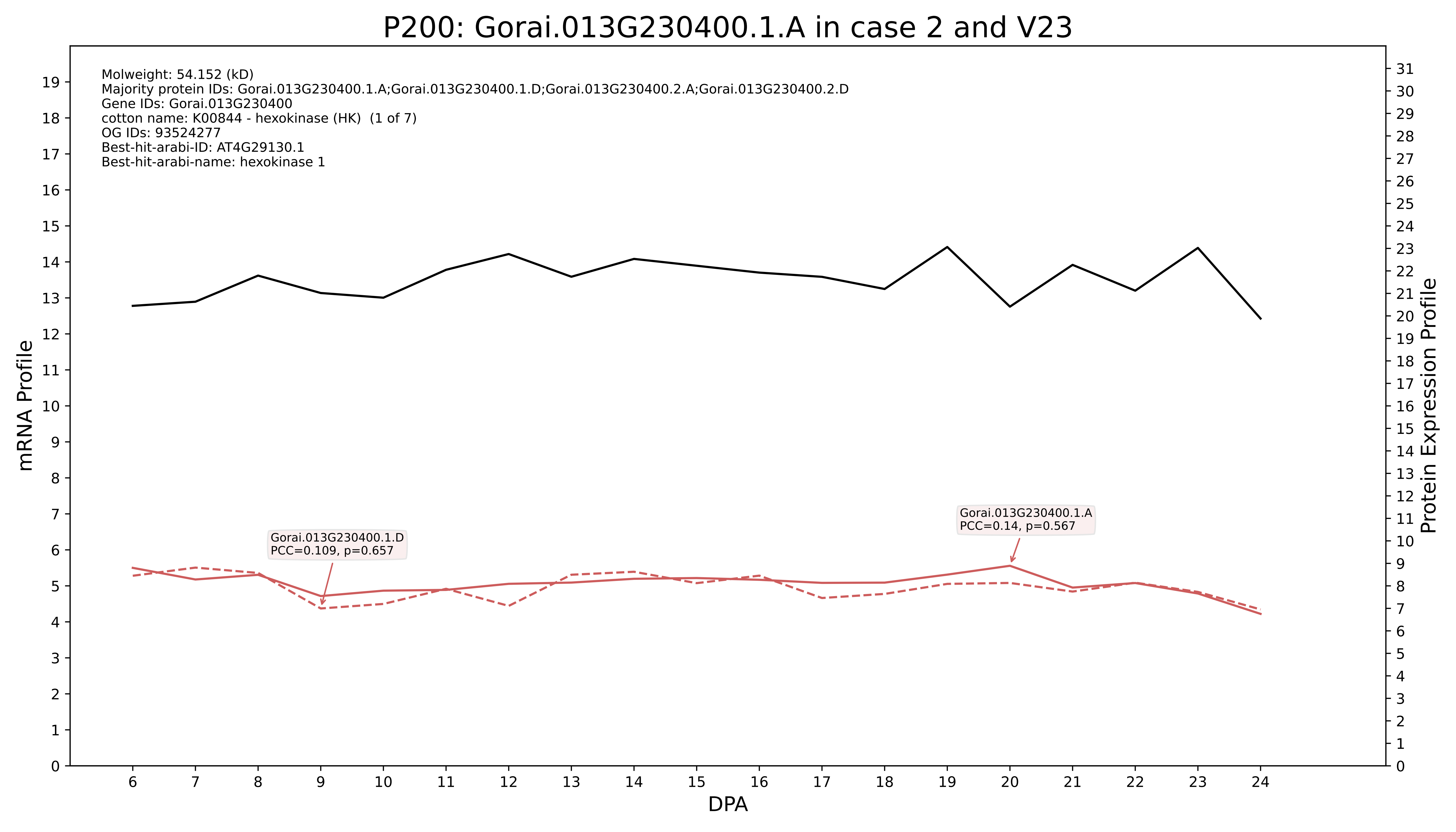
P200: Gorai.013G230400.1.A in case 2 and V23
Molweight: 54.152 (kD)
Majority protein IDs: Gorai.013G230400.1.A;Gorai.013G230400.1.D;Gorai.013G230400.2.A;Gorai.013G230400.2.D
Gene IDs: Gorai.013G230400
cotton name: K00844 - hexokinase (HK) (1 of 7)
OG IDs: 93524277
Best-hit-arabi-ID: AT4G29130.1
Best-hit-arabi-name: hexokinase 1
Gorai.013G230400.1.D
PCC=0.109, p=0.657
Gorai.013G230400.1.A
PCC=0.14, p=0.567
mRNA Profile
Protein Expression Profile
DPA

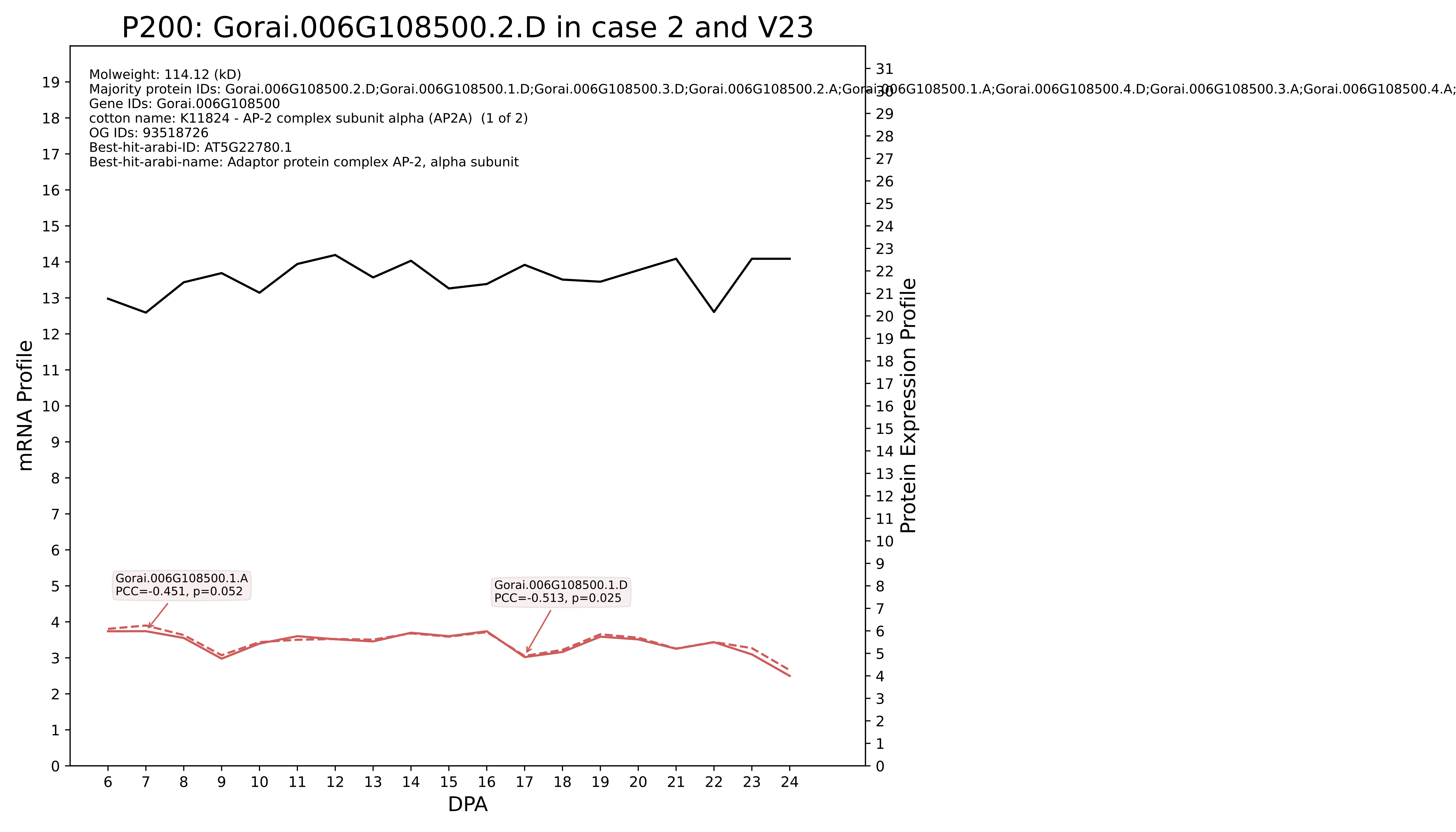
P200: Gorai.006G108500.2.D in case 2 and V23
Molweight: 114.12 (kD)
Majority protein IDs: Gorai.006G108500.2.D;Gorai.006G108500.1.D;Gorai.006G108500.3.D;Gorai.006G108500.2.A;Gorai.006G108500.1.A;Gorai.006G108500.4.D;Gorai.006G108500.3.A;Gorai.006G108500.4.A;
Gene IDs: Gorai.006G108500
cotton name: K11824 - AP-2 complex subunit alpha (AP2A) (1 of 2)
OG IDs: 93518726
Best-hit-arabi-ID: AT5G22780.1
Best-hit-arabi-name: Adaptor protein complex AP-2, alpha subunit
Gorai.006G108500.1.A
PCC=-0.451, p=0.052
Gorai.006G108500.1.D
PCC=-0.513, p=0.025
mRNA Profile
Protein Expression Profile
DPA

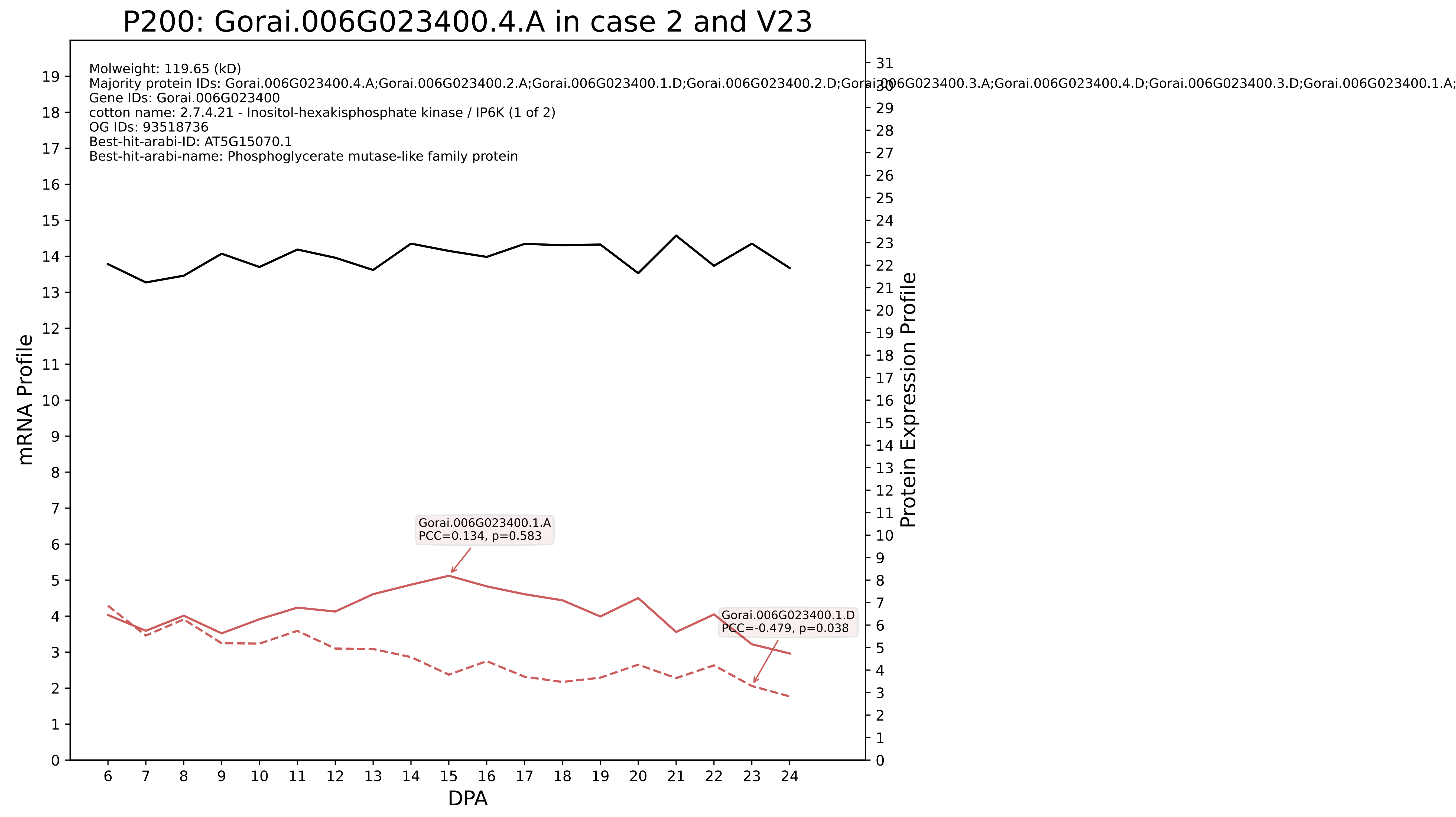
P200: Gorai.006G023400.4.A in case 2 and V23
Molweight: 119.65 (kD)
Majority protein IDs: Gorai.006G023400.4.A;Gorai.006G023400.2.A;Gorai.006G023400.1.D;Gorai.006G023400.2.D;Gorai.006G023400.3.A;Gorai.006G023400.4.D;Gorai.006G023400.3.D;Gorai.006G023400.1.A;
Gene IDs: Gorai.006G023400
cotton name: 2.7.4.21 - Inositol-hexakisphosphate kinase / IP6K (1 of 2)
OG IDs: 93518736
Best-hit-arabi-ID: AT5G15070.1
Best-hit-arabi-name: Phosphoglycerate mutase-like family protein
Gorai.006G023400.1.A
PCC=0.134, p=0.583
Gorai.006G023400.1.D
PCC=-0.479, p=0.038
mRNA Profile
Protein Expression Profile
DPA

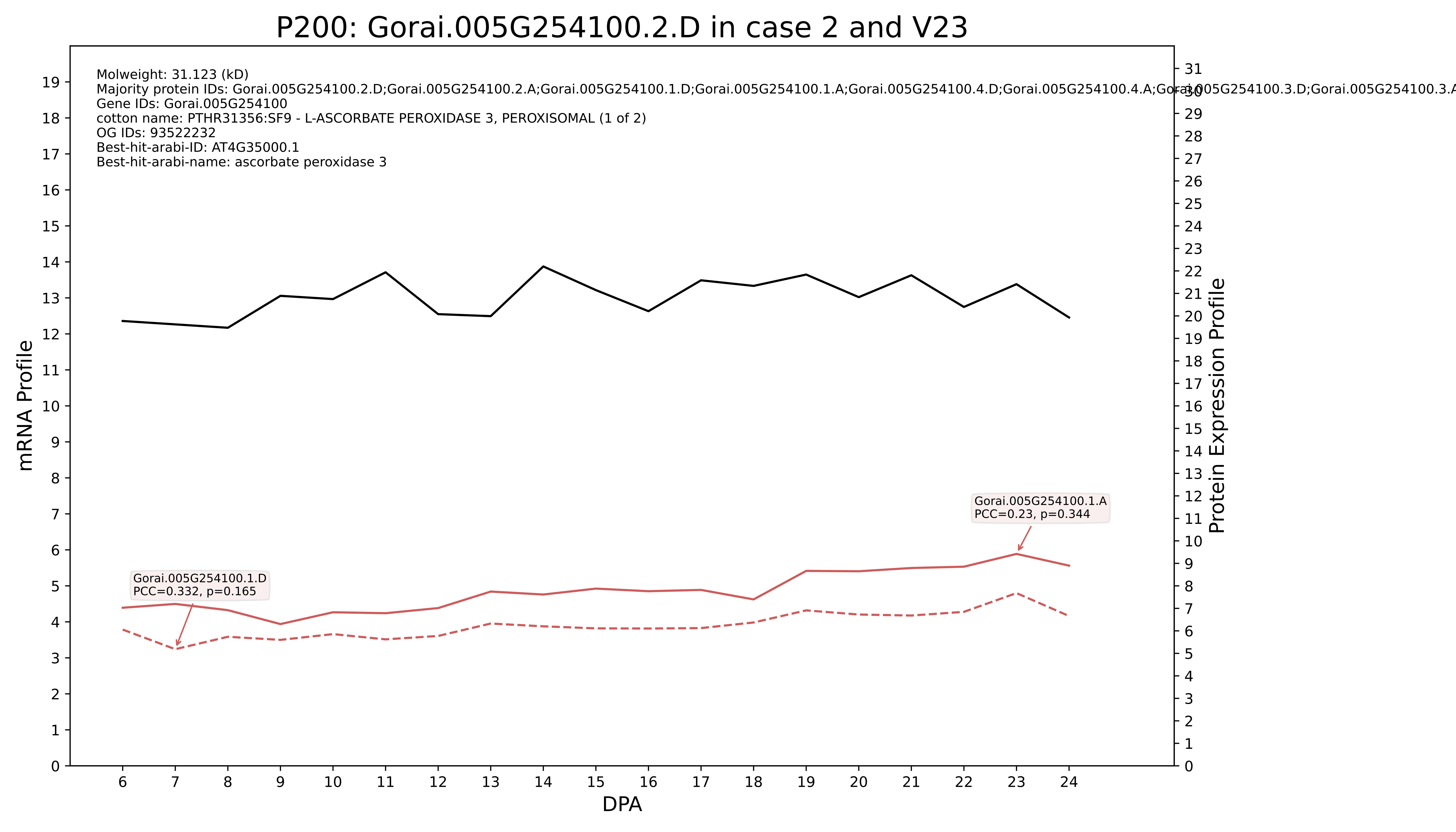

P200: Gorai.005G254100.2.D in case 2 and V23
Molweight: 31.123 (kD)
Majority protein IDs: Gorai.005G254100.2.D;Gorai.005G254100.2.A;Gorai.005G254100.1.D;Gorai.005G254100.1.A;Gorai.005G254100.4.D;Gorai.005G254100.4.A;Gorai.005G254100.3.D;Gorai.005G254100.3.A
Gene IDs: Gorai.005G254100
cotton name: PTHR31356:SF9 - L-ASCORBATE PEROXIDASE 3, PEROXISOMAL (1 of 2)
OG IDs: 93522232
Best-hit-arabi-ID: AT4G35000.1
Best-hit-arabi-name: ascorbate peroxidase 3
Gorai.005G254100.1.A
PCC=0.23, p=0.344
Gorai.005G254100.1.D
PCC=0.332, p=0.165
mRNA Profile
Protein Expression Profile
DPA

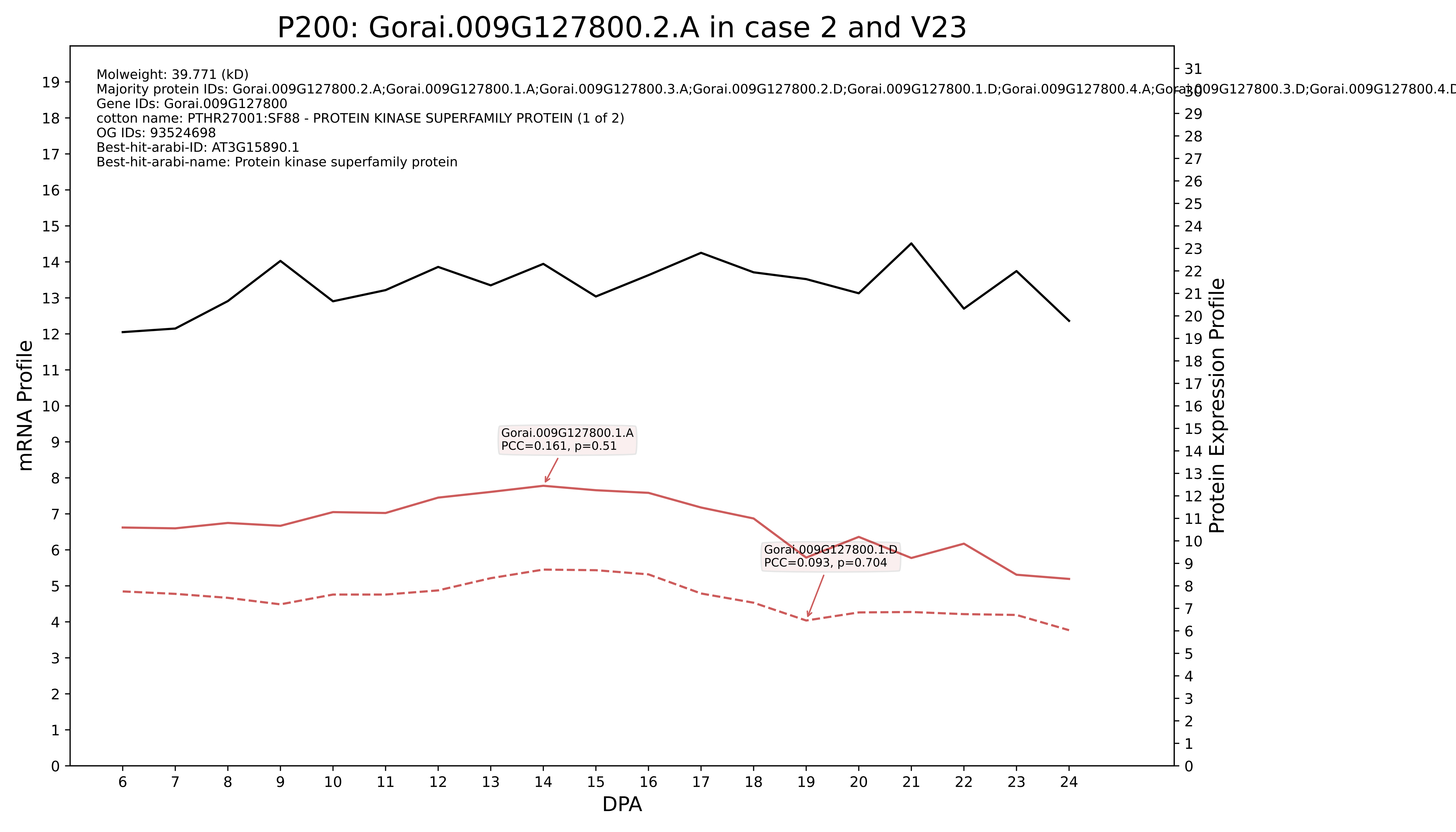
P200: Gorai.009G127800.2.A in case 2 and V23
Molweight: 39.771 (kD)
Majority protein IDs: Gorai.009G127800.2.A;Gorai.009G127800.1.A;Gorai.009G127800.3.A;Gorai.009G127800.2.D;Gorai.009G127800.1.D;Gorai.009G127800.4.A;Gorai.009G127800.3.D;Gorai.009G127800.4.D
Gene IDs: Gorai.009G127800
cotton name: PTHR27001:SF88 - PROTEIN KINASE SUPERFAMILY PROTEIN (1 of 2)
OG IDs: 93524698
Best-hit-arabi-ID: AT3G15890.1
Best-hit-arabi-name: Protein kinase superfamily protein
Gorai.009G127800.1.A
PCC=0.161, p=0.51
Gorai.009G127800.1.D
PCC=0.093, p=0.704
mRNA Profile
Protein Expression Profile
DPA

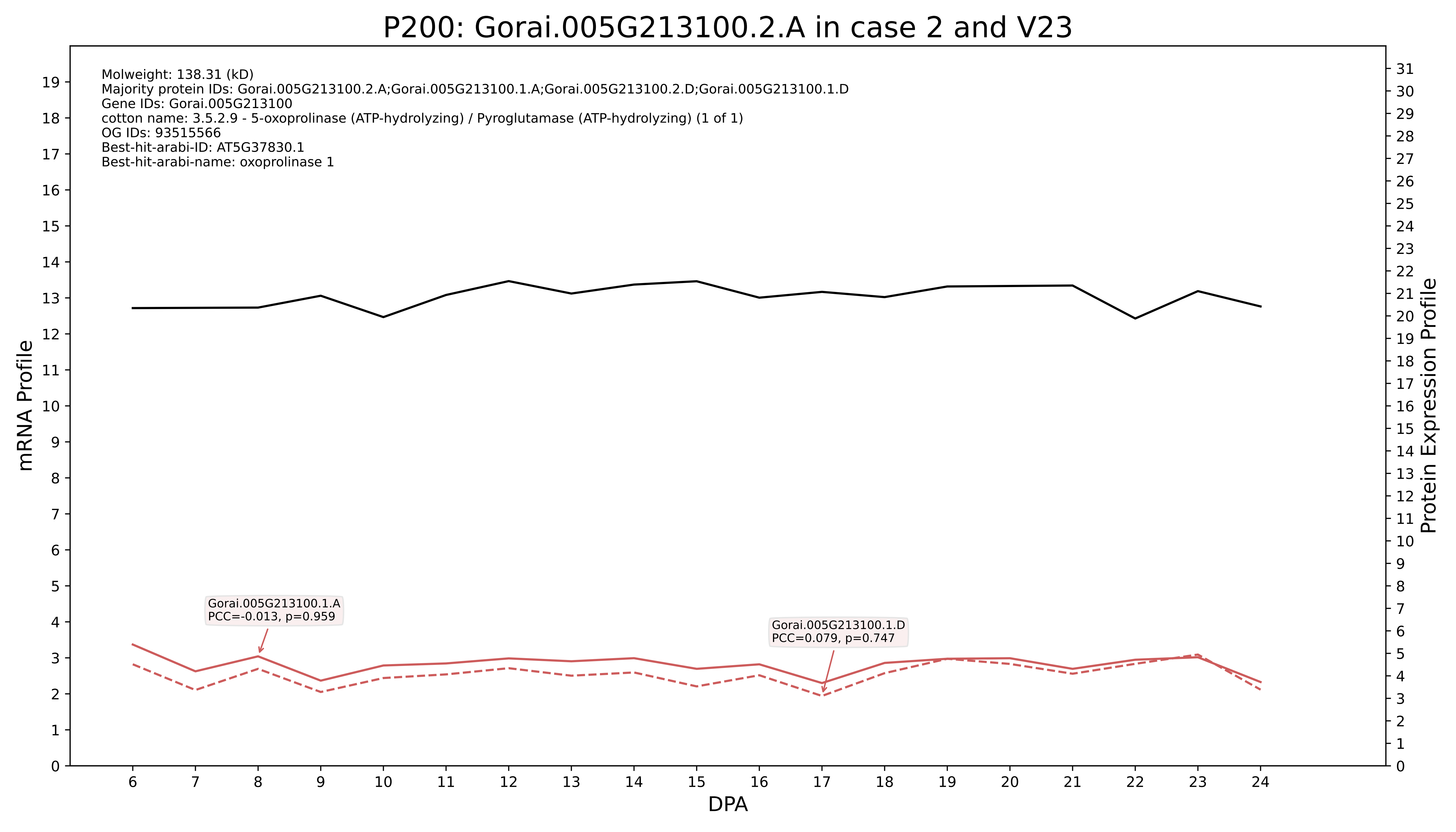
P200: Gorai.005G213100.2.A in case 2 and V23
Molweight: 138.31 (kD)
Majority protein IDs: Gorai.005G213100.2.A;Gorai.005G213100.1.A;Gorai.005G213100.2.D;Gorai.005G213100.1.D
Gene IDs: Gorai.005G213100
cotton name: 3.5.2.9 - 5-oxoprolinase (ATP-hydrolyzing) / Pyroglutamase (ATP-hydrolyzing) (1 of 1)
OG IDs: 93515566
Best-hit-arabi-ID: AT5G37830.1
Best-hit-arabi-name: oxoprolinase 1
Gorai.005G213100.1.A
PCC=-0.013, p=0.959
Gorai.005G213100.1.D
PCC=0.079, p=0.747
mRNA Profile
Protein Expression Profile
DPA

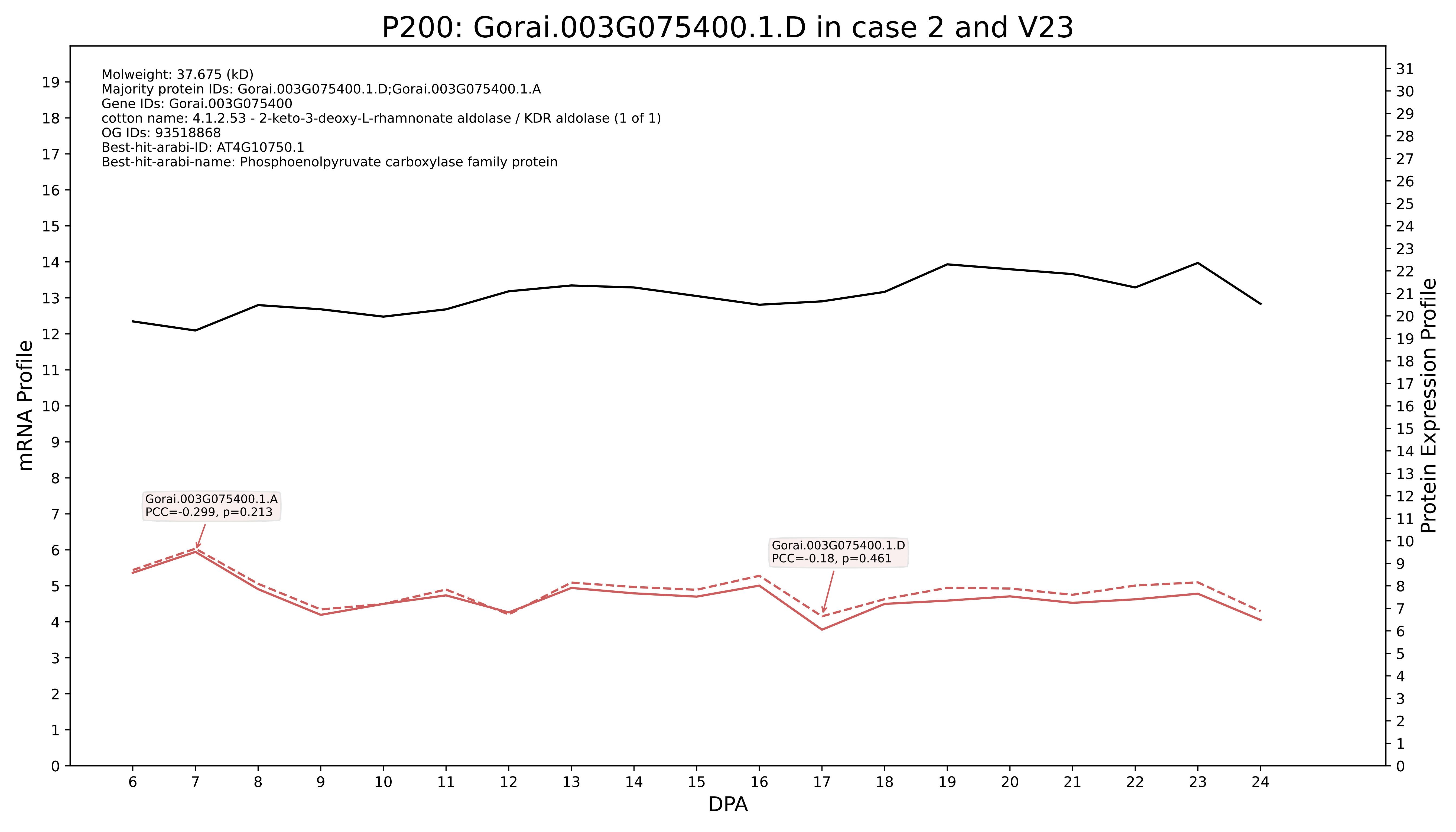
P200: Gorai.003G075400.1.D in case 2 and V23
Molweight: 37.675 (kD)
Majority protein IDs: Gorai.003G075400.1.D;Gorai.003G075400.1.A
Gene IDs: Gorai.003G075400
cotton name: 4.1.2.53 - 2-keto-3-deoxy-L-rhamnonate aldolase / KDR aldolase (1 of 1)
OG IDs: 93518868
Best-hit-arabi-ID: AT4G10750.1
Best-hit-arabi-name: Phosphoenolpyruvate carboxylase family protein
Gorai.003G075400.1.A
PCC=-0.299, p=0.213
Gorai.003G075400.1.D
PCC=-0.18, p=0.461
mRNA Profile
Protein Expression Profile
DPA

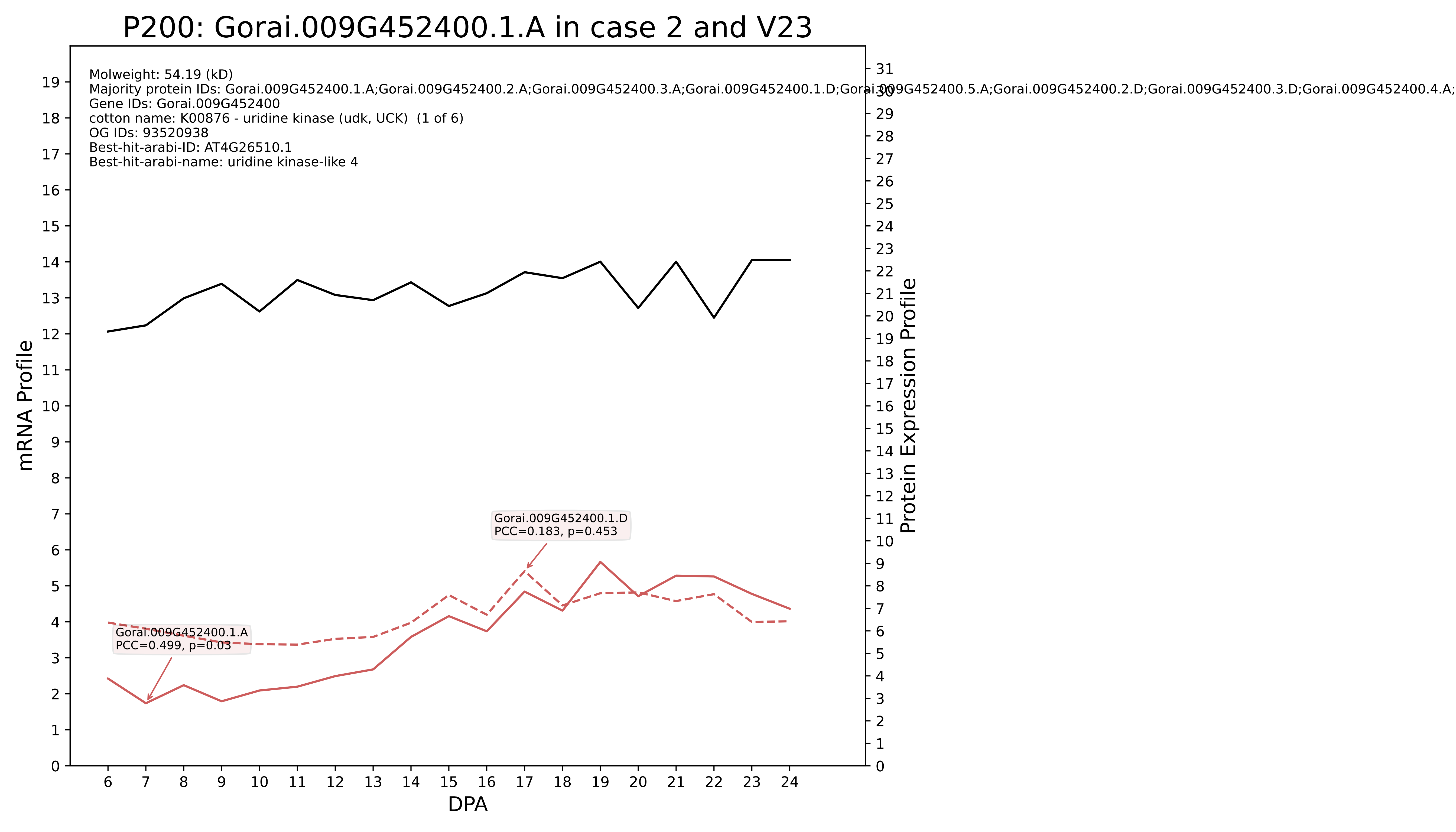

P200: Gorai.009G452400.1.A in case 2 and V23
Molweight: 54.19 (kD)
Majority protein IDs: Gorai.009G452400.1.A;Gorai.009G452400.2.A;Gorai.009G452400.3.A;Gorai.009G452400.1.D;Gorai.009G452400.5.A;Gorai.009G452400.2.D;Gorai.009G452400.3.D;Gorai.009G452400.4.A;
Gene IDs: Gorai.009G452400
cotton name: K00876 - uridine kinase (udk, UCK) (1 of 6)
OG IDs: 93520938
Best-hit-arabi-ID: AT4G26510.1
Best-hit-arabi-name: uridine kinase-like 4
Gorai.009G452400.1.D
PCC=0.183, p=0.453
Gorai.009G452400.1.A
PCC=0.499, p=0.03
mRNA Profile
Protein Expression Profile
DPA

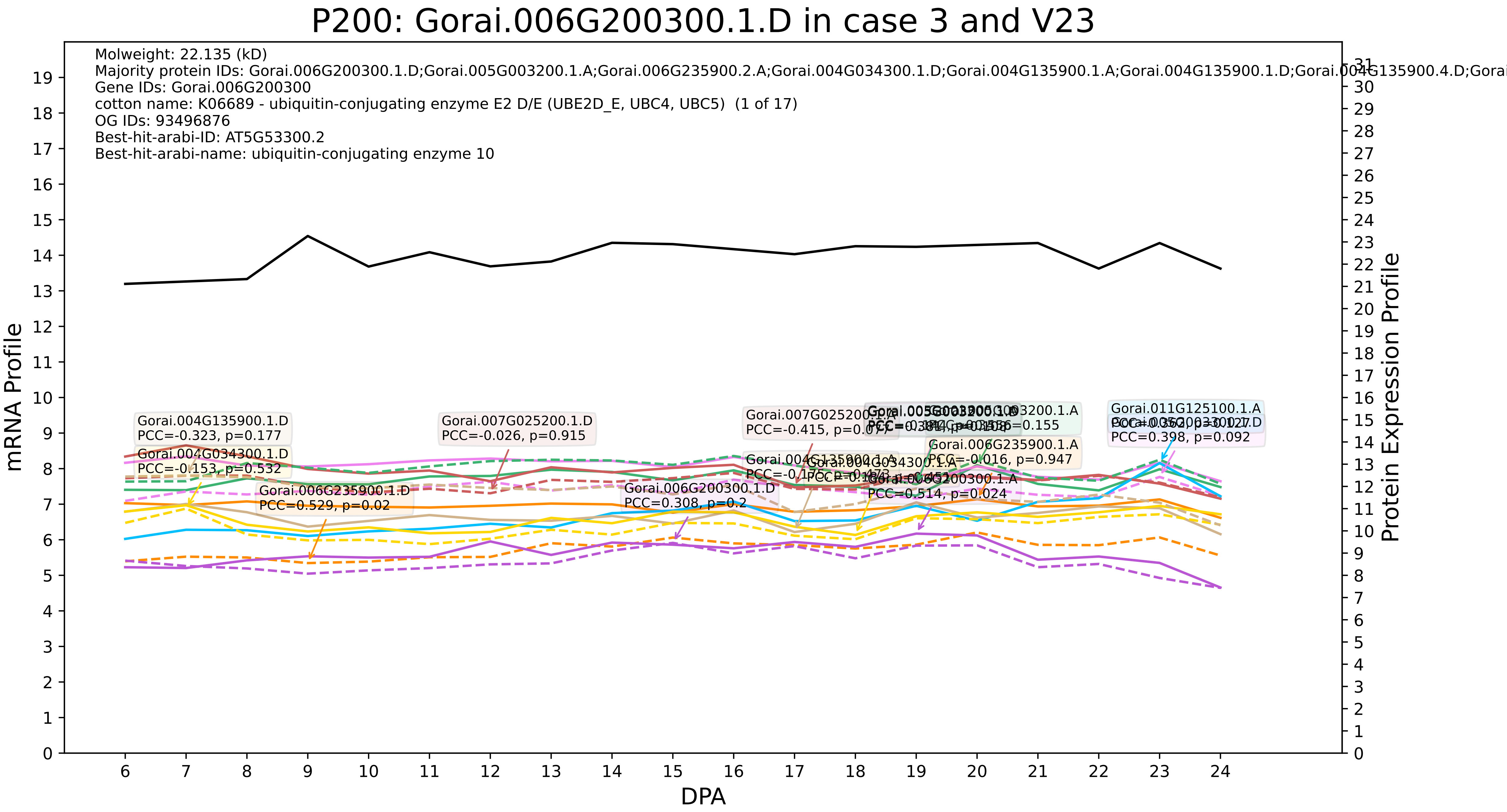
P200: Gorai.006G200300.1.D in case 3 and V23
Molweight: 22.135 (kD)
Majority protein IDs: Gorai.006G200300.1.D;Gorai.005G003200.1.A;Gorai.006G235900.2.A;Gorai.004G034300.1.D;Gorai.004G135900.1.A;Gorai.004G135900.1.D;Gorai.004G135900.4.D;Gorai
Gene IDs: Gorai.006G200300
cotton name: K06689 - ubiquitin-conjugating enzyme E2 D/E (UBE2D_E, UBC4, UBC5) (1 of 17)
OG IDs: 93496876
Best-hit-arabi-ID: AT5G53300.2
Best-hit-arabi-name: ubiquitin-conjugating enzyme 10
mRNA Profile
Protein Expression Profile
DPA
Gorai.004G135900.1.D
PCC=-0.323, p=0.177
Gorai.004G034300.1.D
PCC=-0.153, p=0.532
Gorai.006G235900.1.D
PCC=0.529, p=0.02
Gorai.007G025200.1.D
PCC=-0.026, p=0.915
Gorai.006G200300.1.D
PCC=0.308, p=0.2
Gorai.007G025200.1.A
PCC=-0.415, p=0.077
Gorai.006G235900.1.A
PCC=-0.016, p=0.947
Gorai.006G200300.1.A
PCC=0.514, p=0.024
Gorai.011G125100.1.A
PCC=0.398, p=0.092

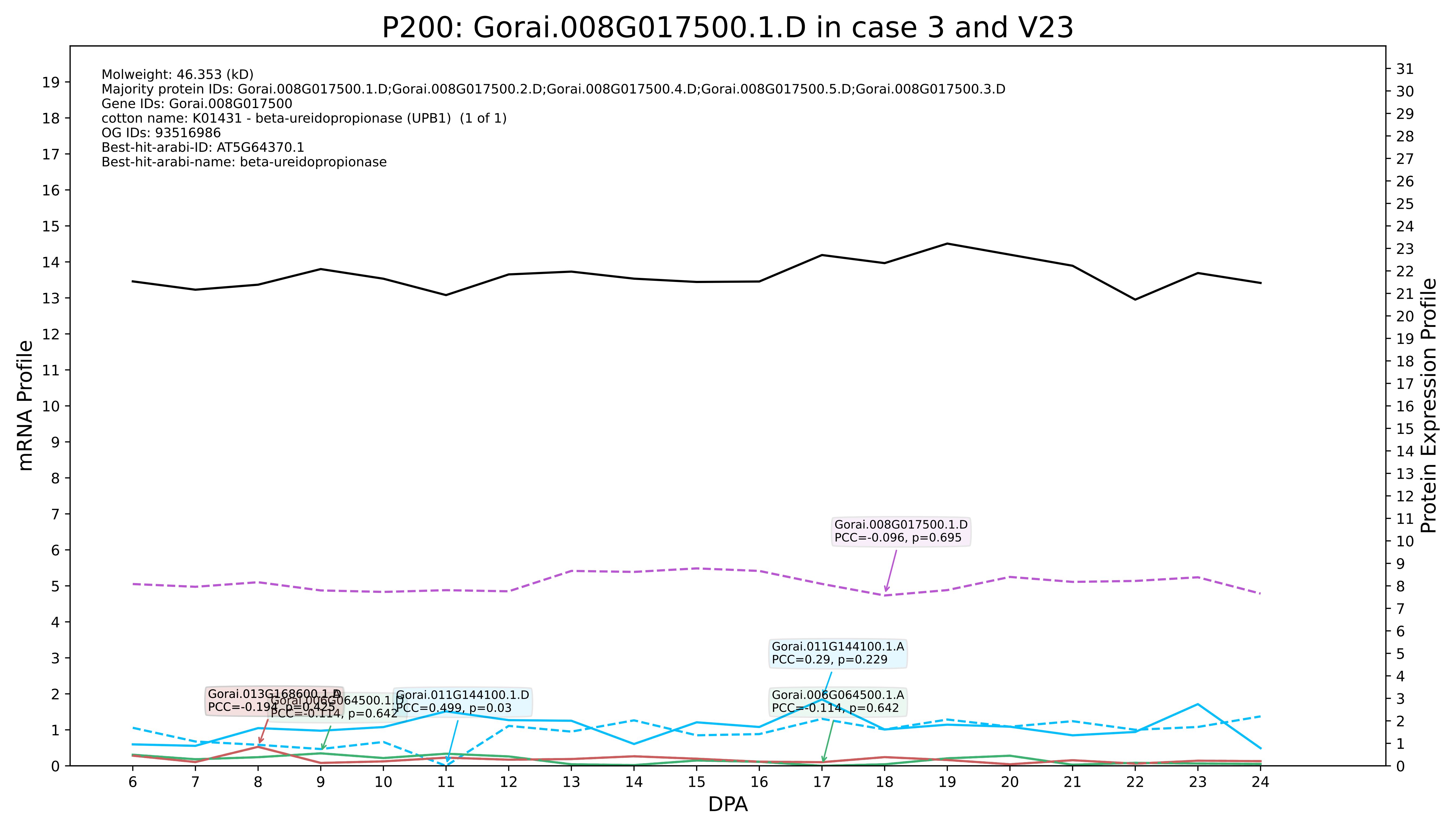

P200: Gorai.008G017500.1.D in case 3 and V23
Molweight: 46.353 (kD)
Majority protein IDs: Gorai.008G017500.1.D;Gorai.008G017500.2.D;Gorai.008G017500.4.D;Gorai.008G017500.5.D;Gorai.008G017500.3.D
Gene IDs: Gorai.008G017500
cotton name: K01431 - beta-ureidopropionase (UPB1) (1 of 1)
OG IDs: 93516986
Best-hit-arabi-ID: AT5G64370.1
Best-hit-arabi-name: beta-ureidopropionase
Gorai.008G017500.1.D
PCC=-0.096, p=0.695
Gorai.011G144100.1.A
PCC=0.29, p=0.229
Gorai.013G168600.1.D
PCC=-0.194, p=0.425
Gorai.006G064500.1.D
PCC=-0.114, p=0.642
Gorai.011G144100.1.D
PCC=0.499, p=0.03
Gorai.006G064500.1.A
PCC=-0.114, p=0.642
mRNA Profile
Protein Expression Profile
DPA

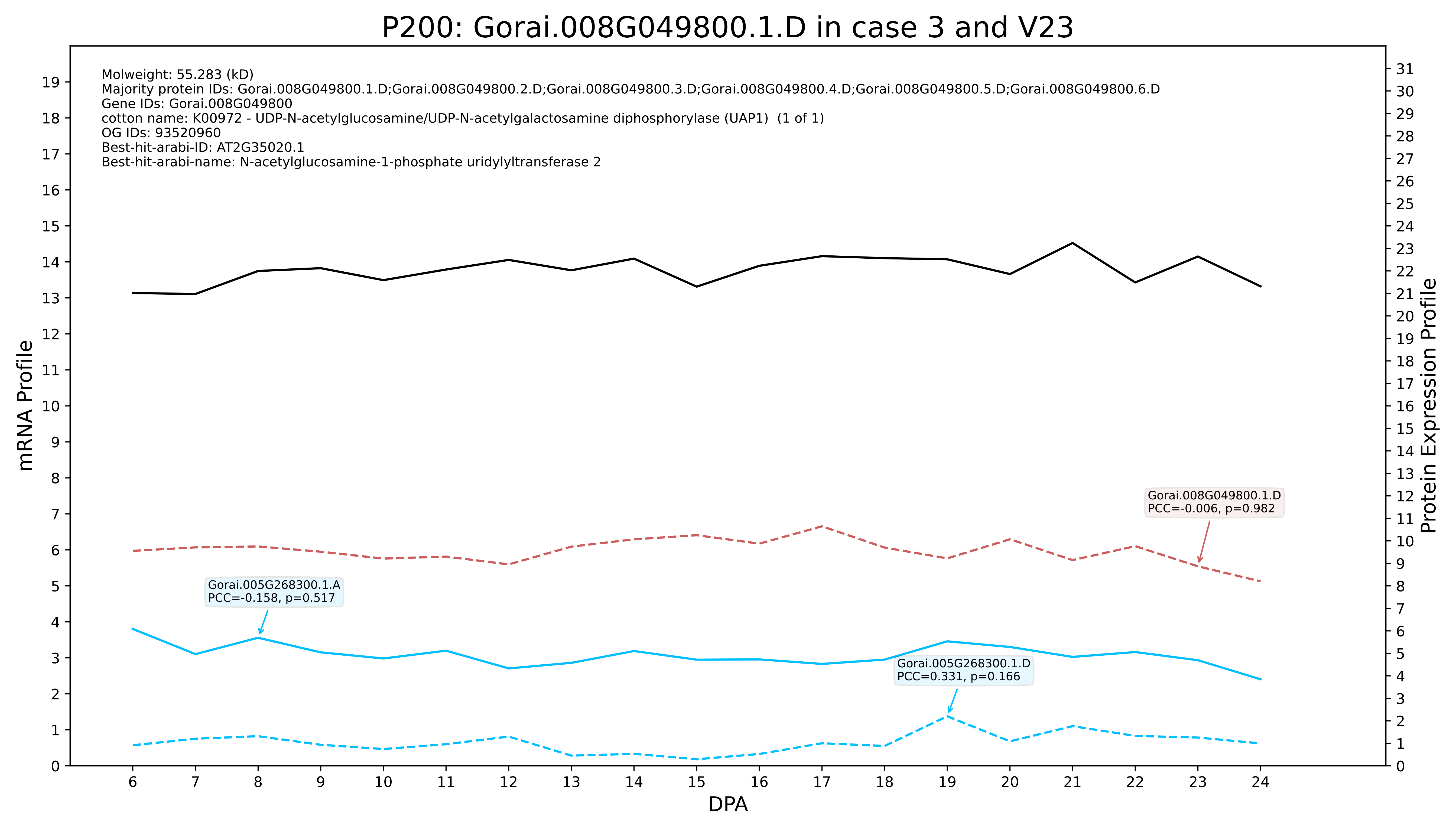
P200: Gorai.008G049800.1.D in case 3 and V23
Molweight: 55.283 (kD)
Majority protein IDs: Gorai.008G049800.1.D;Gorai.008G049800.2.D;Gorai.008G049800.3.D;Gorai.008G049800.4.D;Gorai.008G049800.5.D;Gorai.008G049800.6.D
Gene IDs: Gorai.008G049800
cotton name: K00972 - UDP-N-acetylglucosamine/UDP-N-acetylgalactosamine diphosphorylase (UAP1) (1 of 1)
OG IDs: 93520960
Best-hit-arabi-ID: AT2G35020.1
Best-hit-arabi-name: N-acetylglucosamine-1-phosphate uridylyltransferase 2
Gorai.008G049800.1.D
PCC=-0.006, p=0.982
Gorai.005G268300.1.A
PCC=-0.158, p=0.517
Gorai.005G268300.1.D
PCC=0.331, p=0.166
mRNA Profile
Protein Expression Profile
DPA

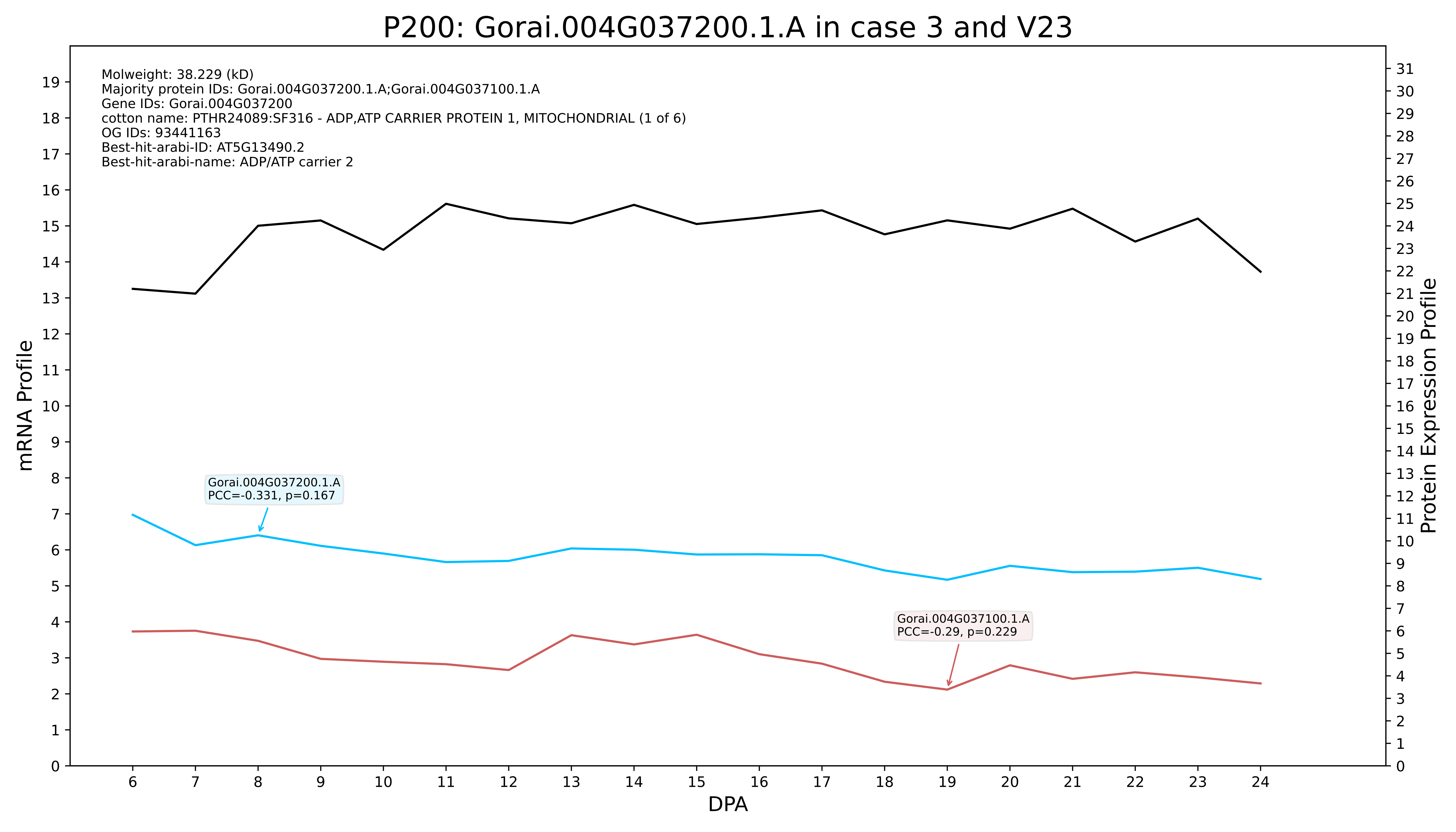

P200: Gorai.004G037200.1.A in case 3 and V23
Molweight: 38.229 (kD)
Majority protein IDs: Gorai.004G037200.1.A;Gorai.004G037100.1.A
Gene IDs: Gorai.004G037200
cotton name: PTHR24089:SF316 - ADP,ATP CARRIER PROTEIN 1, MITOCHONDRIAL (1 of 6)
OG IDs: 93441163
Best-hit-arabi-ID: AT5G13490.2
Best-hit-arabi-name: ADP/ATP carrier 2
Gorai.004G037200.1.A
PCC=-0.331, p=0.167
Gorai.004G037100.1.A
PCC=-0.29, p=0.229
mRNA Profile
Protein Expression Profile
DPA

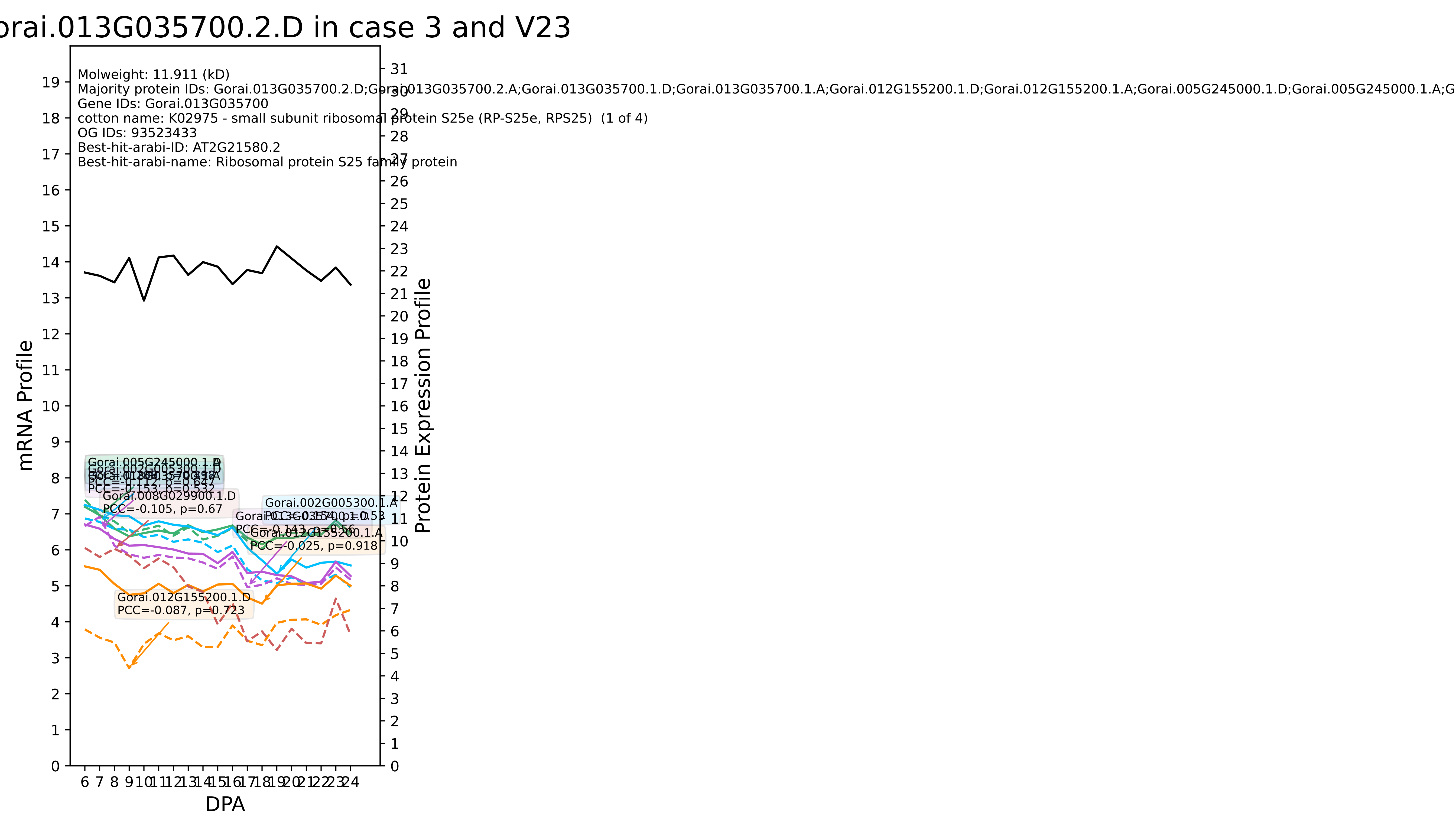
orai.013G035700.2.D in case 3 and V23
Molweight: 11.911 (kD)
Majority protein IDs: Gorai.013G035700.2.D;Gorai.013G035700.2.A;Gorai.013G035700.1.D;Gorai.013G035700.1.A;Gorai.012G155200.1.D;Gorai.012G155200.1.A;Gorai.005G245000.1.D;Gorai.005G245000.1.A;G
Gene IDs: Gorai.013G035700
cotton name: K02975 - small subunit ribosomal protein S25e (RP-S25e, RPS25) (1 of 4)
OG IDs: 93523433
Best-hit-arabi-ID: AT2G21580.2
Best-hit-arabi-name: Ribosomal protein S25 family protein
Gorai.008G029900.1.D
PCC=-0.105, p=0.67
Gorai.002G005300.1.A
Gorai.012G155200.1.A
PCC=-0.025, p=0.918
Gorai.012G155200.1.D
PCC=-0.087, p=0.723
mRNA Profile
Protein Expression Profile
DPA

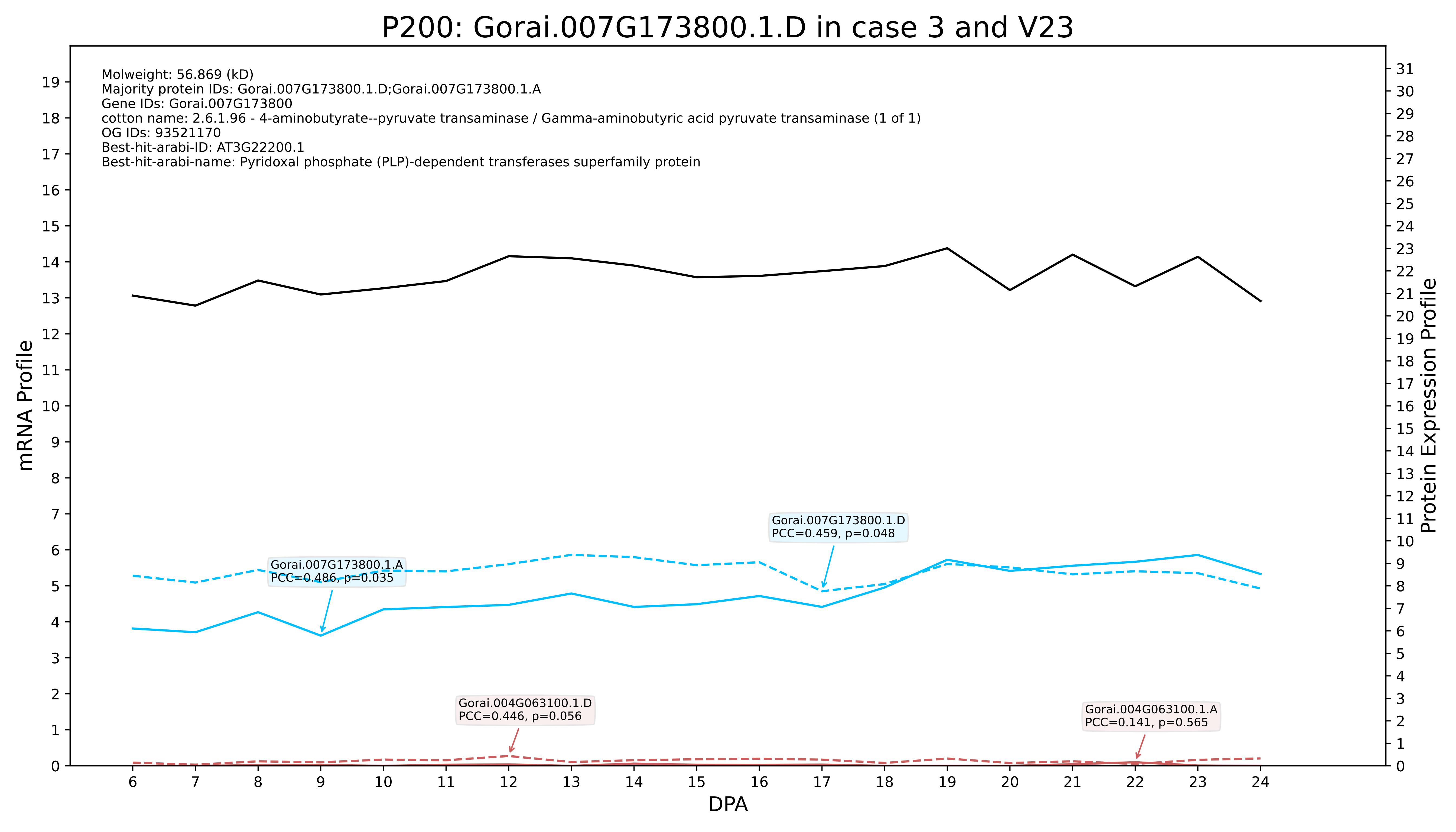

P200: Gorai.007G173800.1.D in case 3 and V23
Molweight: 56.869 (kD)
Majority protein IDs: Gorai.007G173800.1.D;Gorai.007G173800.1.A
Gene IDs: Gorai.007G173800
cotton name: 2.6.1.96 - 4-aminobutyrate--pyruvate transaminase / Gamma-aminobutyric acid pyruvate transaminase (1 of 1)
OG IDs: 93521170
Best-hit-arabi-ID: AT3G22200.1
Best-hit-arabi-name: Pyridoxal phosphate (PLP)-dependent transferases superfamily protein
Gorai.007G173800.1.D
PCC=0.459, p=0.048
Gorai.007G173800.1.A
PCC=0.486, p=0.035
Gorai.004G063100.1.D
PCC=0.446, p=0.056
Gorai.004G063100.1.A
PCC=0.141, p=0.565
mRNA Profile
Protein Expression Profile
DPA

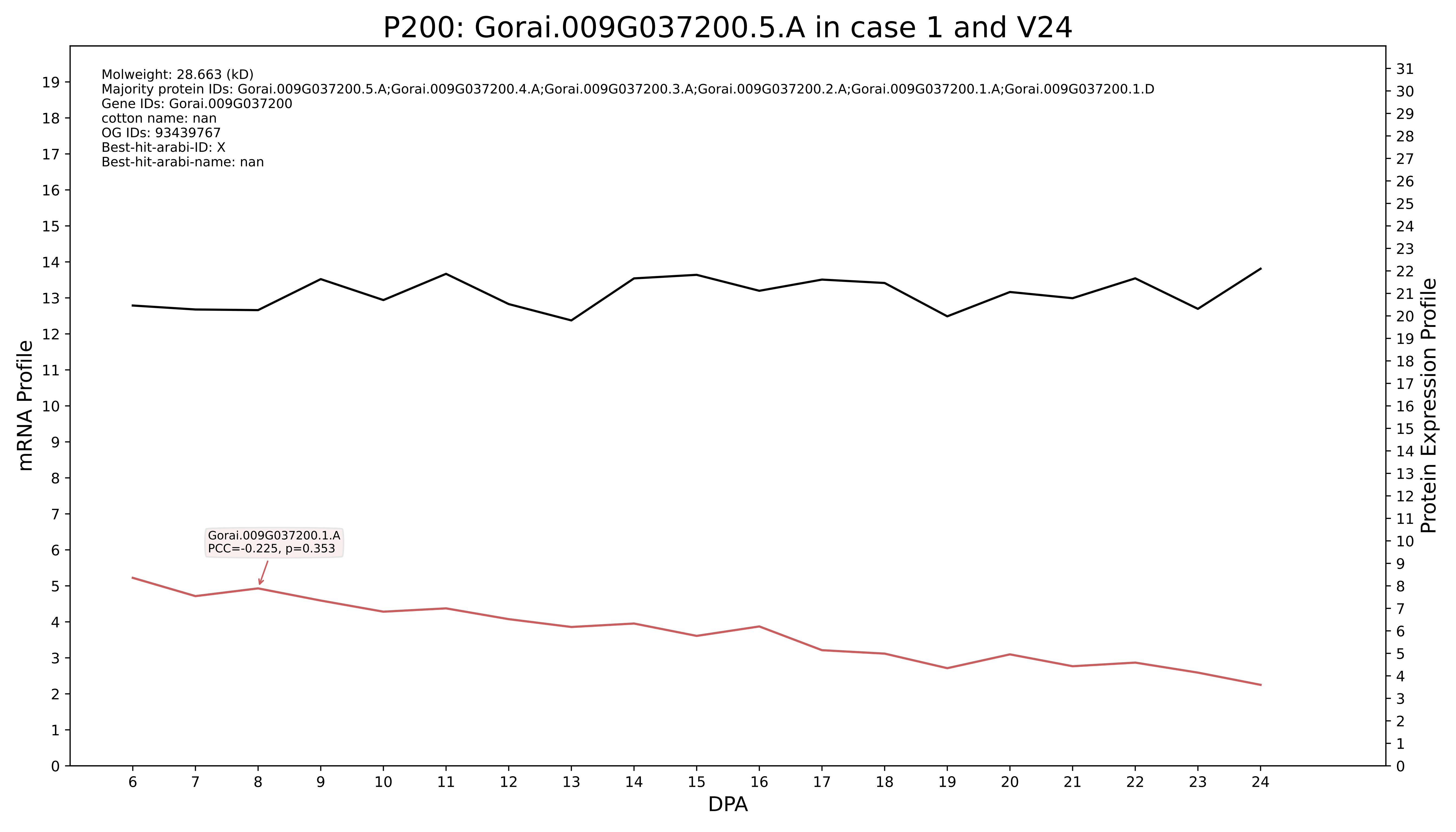
P200: Gorai.009G037200.5.A in case 1 and V24
Molweight: 28.663 (kD)
Majority protein IDs: Gorai.009G037200.5.A;Gorai.009G037200.4.A;Gorai.009G037200.3.A;Gorai.009G037200.2.A;Gorai.009G037200.1.A;Gorai.009G037200.1.D
Gene IDs: Gorai.009G037200
cotton name: nan
OG IDs: 93439767
Best-hit-arabi-ID: X
Best-hit-arabi-name: nan
Gorai.009G037200.1.A
PCC=-0.225, p=0.353
mRNA Profile
Protein Expression Profile
DPA

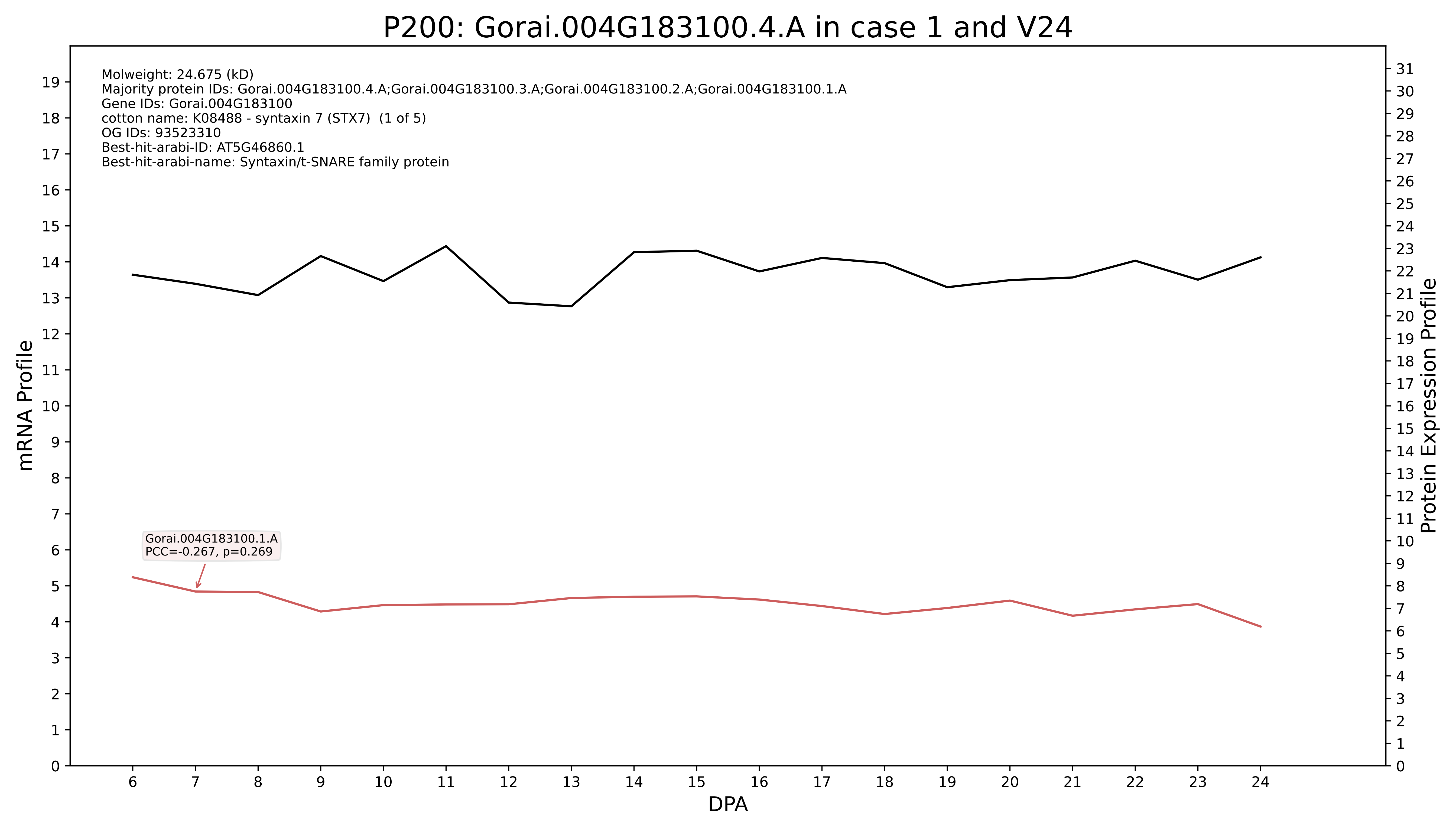

P200: Gorai.004G183100.4.A in case 1 and V24
Molweight: 24.675 (kD)
Majority protein IDs: Gorai.004G183100.4.A;Gorai.004G183100.3.A;Gorai.004G183100.2.A;Gorai.004G183100.1.A
Gene IDs: Gorai.004G183100
cotton name: K08488 - syntaxin 7 (STX7) (1 of 5)
OG IDs: 93523310
Best-hit-arabi-ID: AT5G46860.1
Best-hit-arabi-name: Syntaxin/t-SNARE family protein
Gorai.004G183100.1.A
PCC=-0.267, p=0.269
mRNA Profile
Protein Expression Profile
DPA

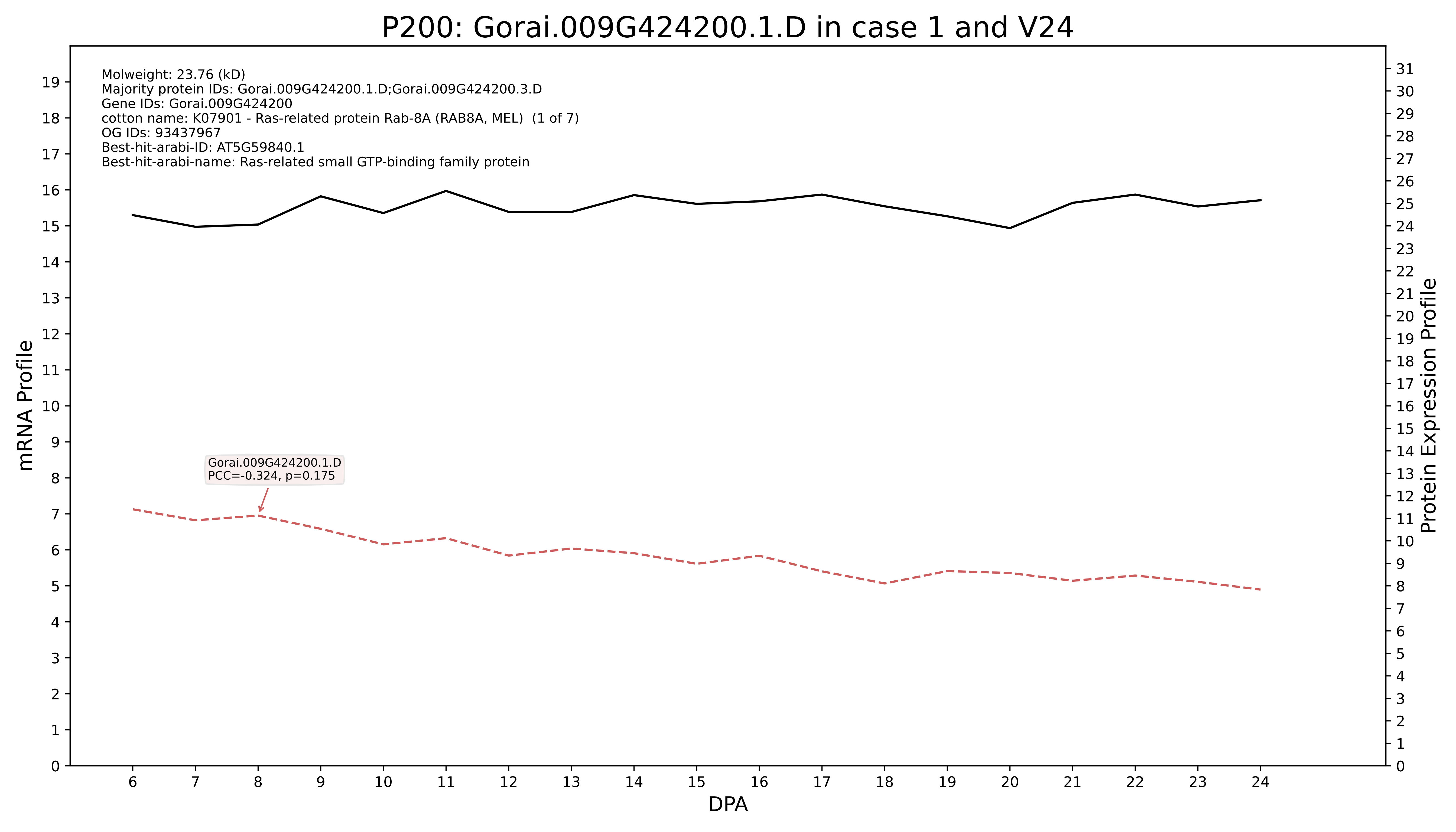

P200: Gorai.009G424200.1.D in case 1 and V24
Molweight: 23.76 (kD)
Majority protein IDs: Gorai.009G424200.1.D;Gorai.009G424200.3.D
Gene IDs: Gorai.009G424200
cotton name: K07901 - Ras-related protein Rab-8A (RAB8A, MEL) (1 of 7)
OG IDs: 93437967
Best-hit-arabi-ID: AT5G59840.1
Best-hit-arabi-name: Ras-related small GTP-binding family protein
Gorai.009G424200.1.D
PCC=-0.324, p=0.175
mRNA Profile
Protein Expression Profile
DPA

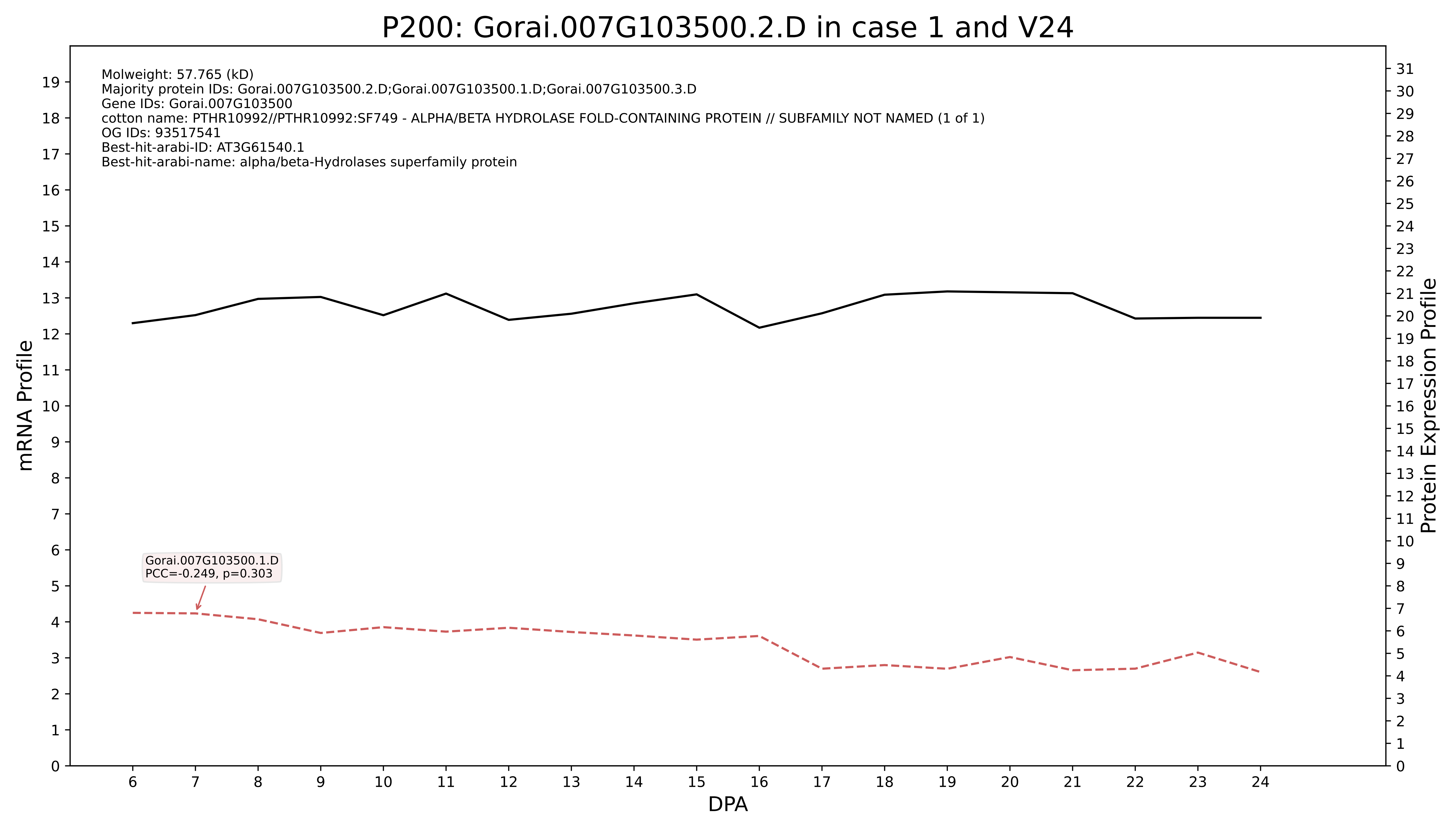

P200: Gorai.007G103500.2.D in case 1 and V24
Molweight: 57.765 (kD)
Majority protein IDs: Gorai.007G103500.2.D;Gorai.007G103500.1.D;Gorai.007G103500.3.D
Gene IDs: Gorai.007G103500
cotton name: PTHR10992//PTHR10992:SF749 - ALPHA/BETA HYDROLASE FOLD-CONTAINING PROTEIN // SUBFAMILY NOT NAMED (1 of 1)
OG IDs: 93517541
Best-hit-arabi-ID: AT3G61540.1
Best-hit-arabi-name: alpha/beta-Hydrolases superfamily protein
Gorai.007G103500.1.D
PCC=-0.249, p=0.303
mRNA Profile
Protein Expression Profile
DPA

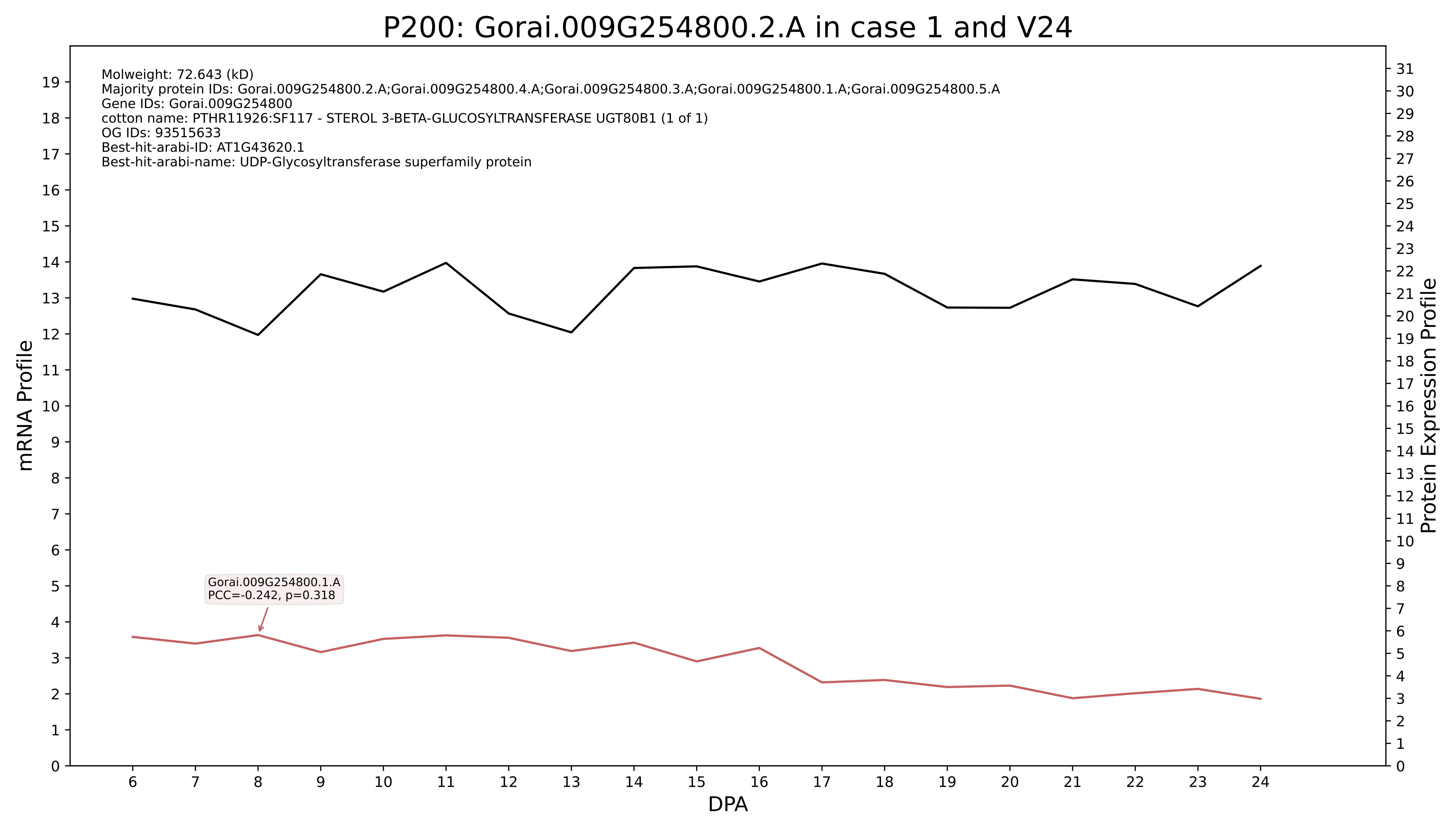

P200: Gorai.009G254800.2.A in case 1 and V24
Molweight: 72.643 (kD)
Majority protein IDs: Gorai.009G254800.2.A;Gorai.009G254800.4.A;Gorai.009G254800.3.A;Gorai.009G254800.1.A;Gorai.009G254800.5.A
Gene IDs: Gorai.009G254800
cotton name: PTHR11926:SF117 - STEROL 3-BETA-GLUCOSYLTRANSFERASE UGT80B1 (1 of 1)
OG IDs: 93515633
Best-hit-arabi-ID: AT1G43620.1
Best-hit-arabi-name: UDP-Glycosyltransferase superfamily protein
Gorai.009G254800.1.A
PCC=-0.242, p=0.318
mRNA Profile
Protein Expression Profile
DPA

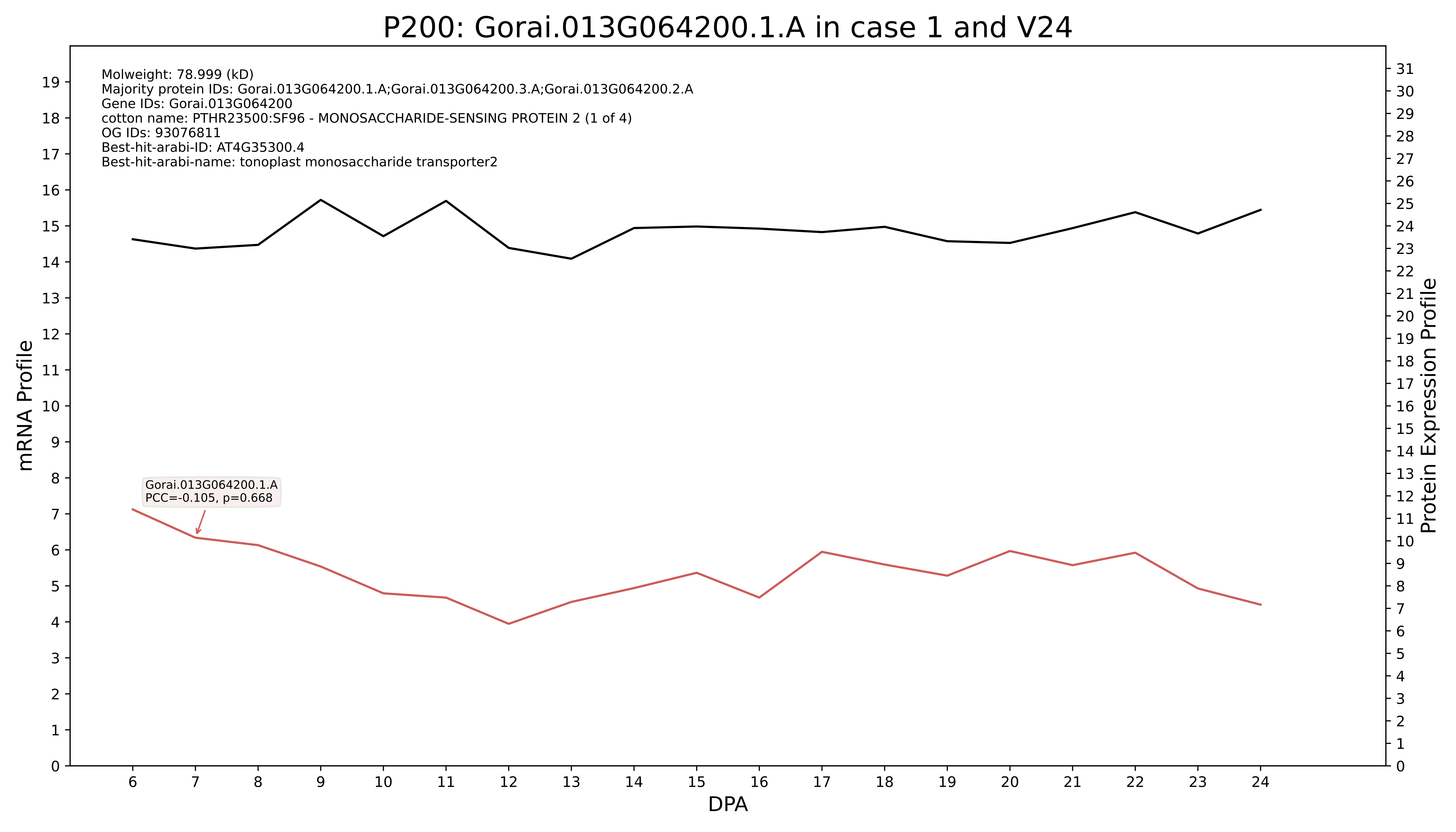
P200: Gorai.013G064200.1.A in case 1 and V24
Molweight: 78.999 (kD)
Majority protein IDs: Gorai.013G064200.1.A;Gorai.013G064200.3.A;Gorai.013G064200.2.A
Gene IDs: Gorai.013G064200
cotton name: PTHR23500:SF96 - MONOSACCHARIDE-SENSING PROTEIN 2 (1 of 4)
OG IDs: 93076811
Best-hit-arabi-ID: AT4G35300.4
Best-hit-arabi-name: tonoplast monosaccharide transporter2
Gorai.013G064200.1.A
PCC=-0.105, p=0.668
mRNA Profile
Protein Expression Profile
DPA

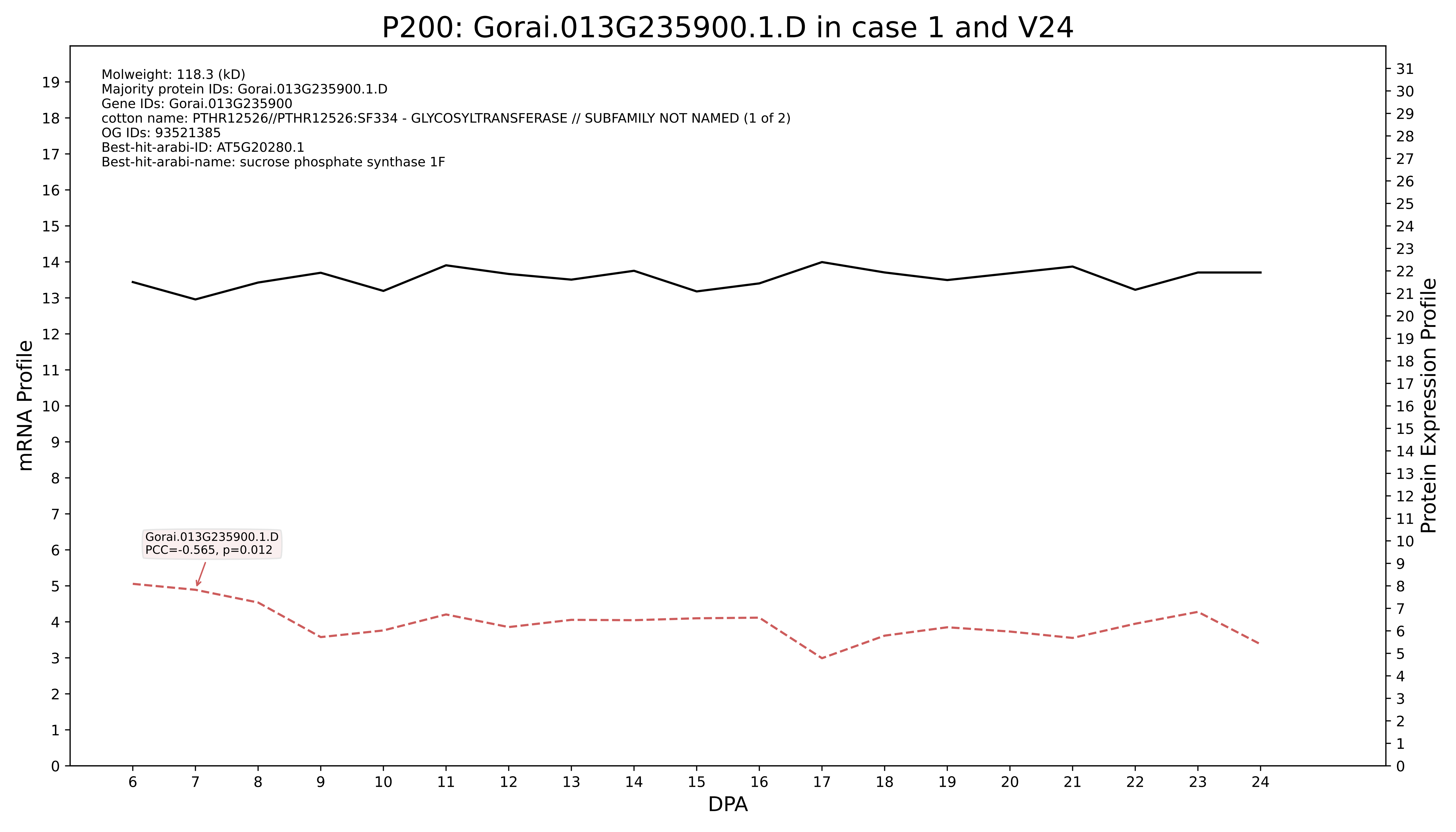
P200: Gorai.013G235900.1.D in case 1 and V24
Molweight: 118.3 (kD)
Majority protein IDs: Gorai.013G235900.1.D
Gene IDs: Gorai.013G235900
cotton name: PTHR12526//PTHR12526:SF334 - GLYCOSYLTRANSFERASE // SUBFAMILY NOT NAMED (1 of 2)
OG IDs: 93521385
Best-hit-arabi-ID: AT5G20280.1
Best-hit-arabi-name: sucrose phosphate synthase 1F
Gorai.013G235900.1.D
PCC=-0.565, p=0.012
mRNA Profile
Protein Expression Profile
DPA

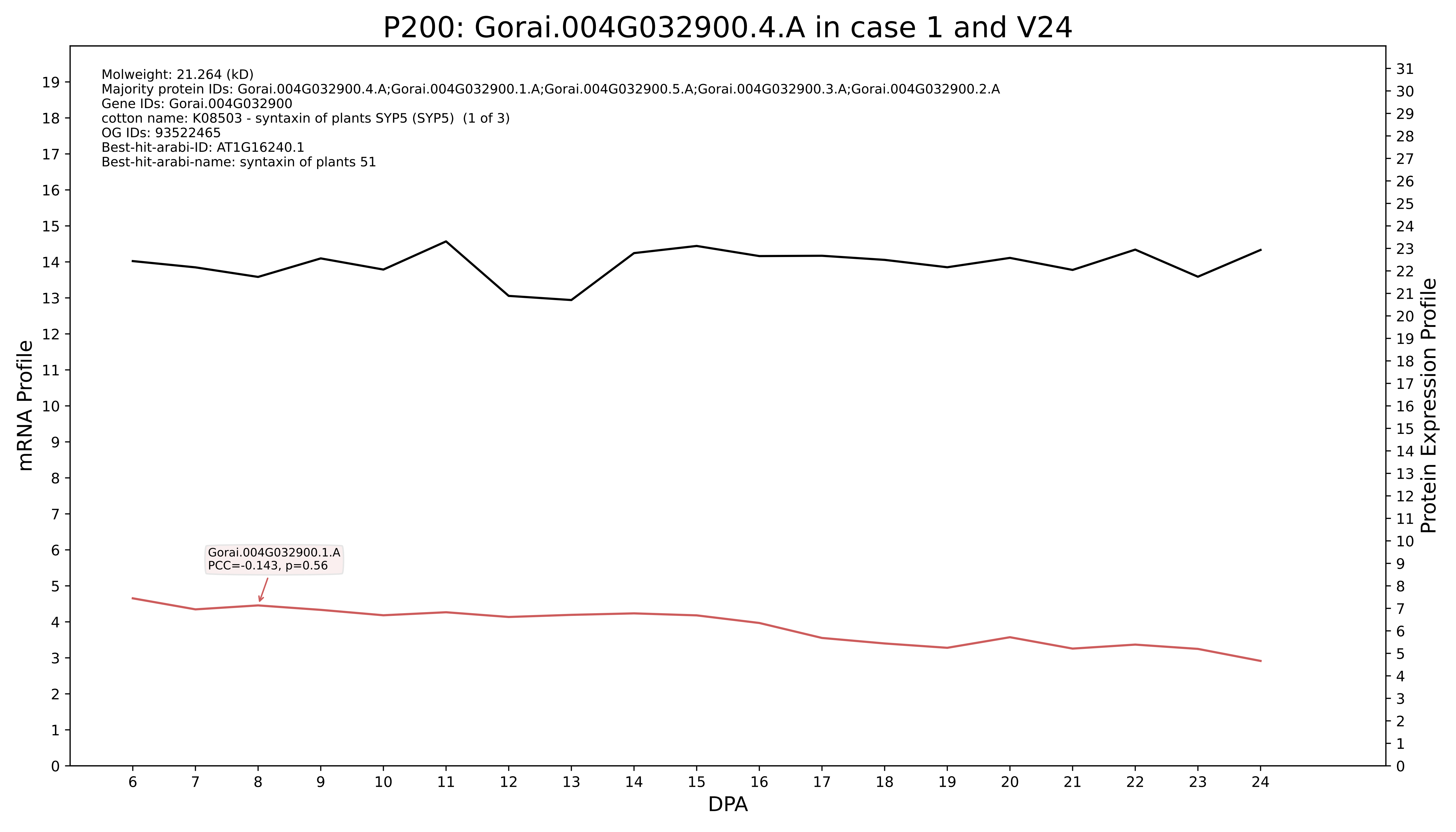
P200: Gorai.004G032900.4.A in case 1 and V24
Molweight: 21.264 (kD)
Majority protein IDs: Gorai.004G032900.4.A;Gorai.004G032900.1.A;Gorai.004G032900.5.A;Gorai.004G032900.3.A;Gorai.004G032900.2.A
Gene IDs: Gorai.004G032900
cotton name: K08503 - syntaxin of plants SYP5 (SYP5) (1 of 3)
OG IDs: 93522465
Best-hit-arabi-ID: AT1G16240.1
Best-hit-arabi-name: syntaxin of plants 51
Gorai.004G032900.1.A
PCC=-0.143, p=0.56
mRNA Profile
Protein Expression Profile
DPA

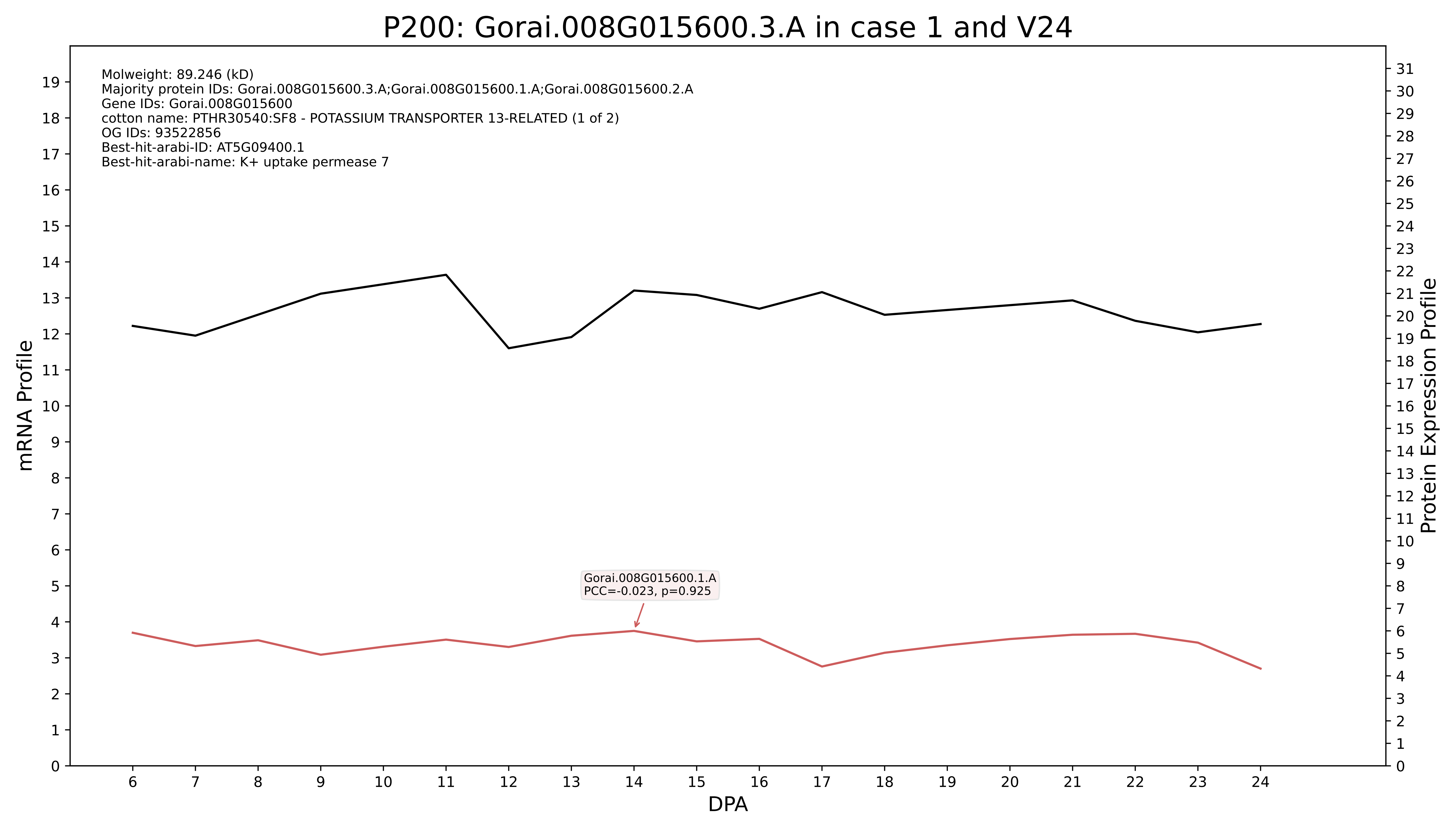
P200: Gorai.008G015600.3.A in case 1 and V24
Molweight: 89.246 (kD)
Majority protein IDs: Gorai.008G015600.3.A;Gorai.008G015600.1.A;Gorai.008G015600.2.A
Gene IDs: Gorai.008G015600
cotton name: PTHR30540:SF8 - POTASSIUM TRANSPORTER 13-RELATED (1 of 2)
OG IDs: 93522856
Best-hit-arabi-ID: AT5G09400.1
Best-hit-arabi-name: K+ uptake permease 7
Gorai.008G015600.1.A
PCC=-0.023, p=0.925
mRNA Profile
Protein Expression Profile
DPA

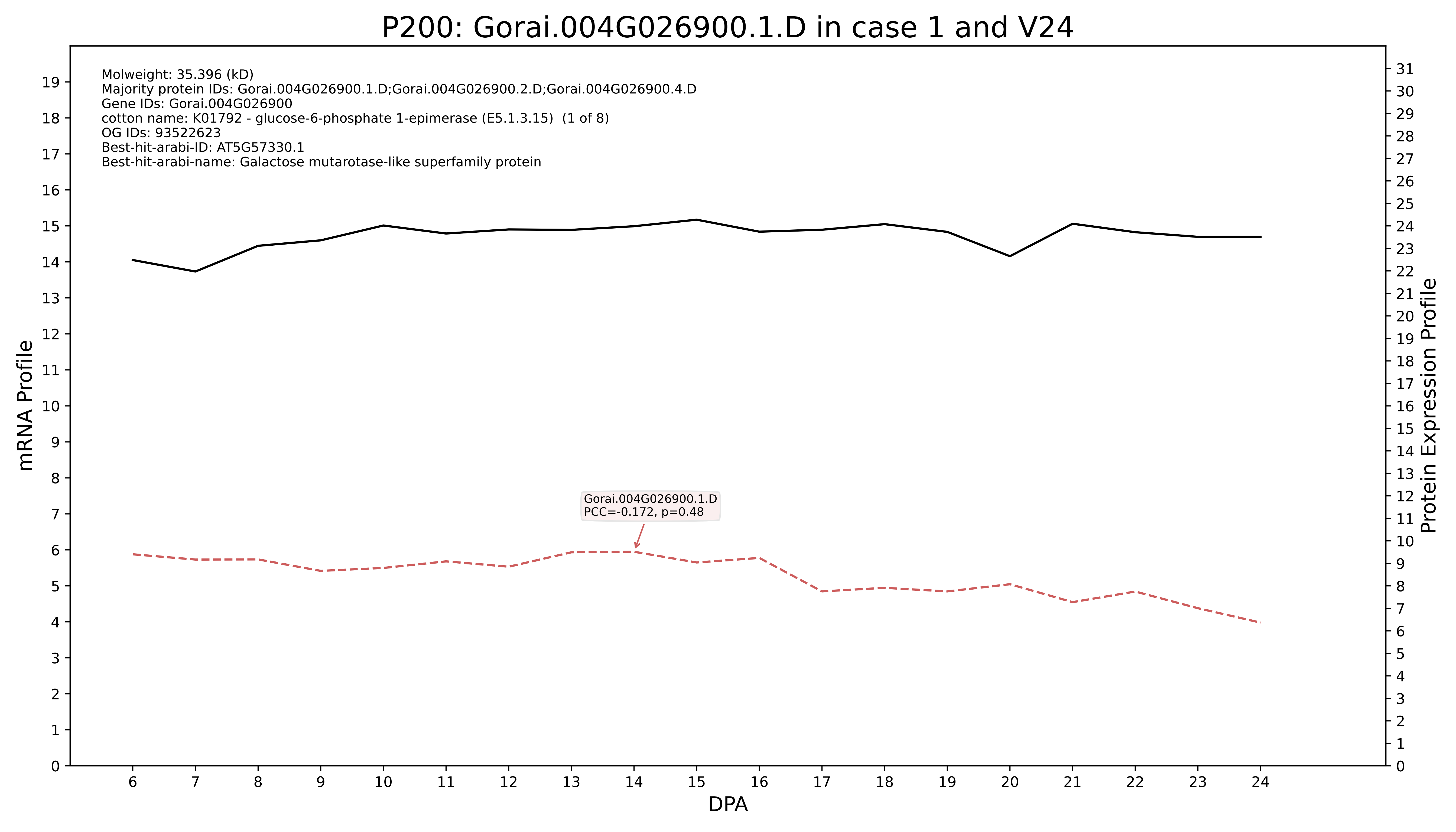
P200: Gorai.004G026900.1.D in case 1 and V24
Molweight: 35.396 (kD)
Majority protein IDs: Gorai.004G026900.1.D;Gorai.004G026900.2.D;Gorai.004G026900.4.D
Gene IDs: Gorai.004G026900
cotton name: K01792 - glucose-6-phosphate 1-epimerase (E5.1.3.15) (1 of 8)
OG IDs: 93522623
Best-hit-arabi-ID: AT5G57330.1
Best-hit-arabi-name: Galactose mutarotase-like superfamily protein
Gorai.004G026900.1.D
PCC=-0.172, p=0.48
mRNA Profile
Protein Expression Profile
DPA

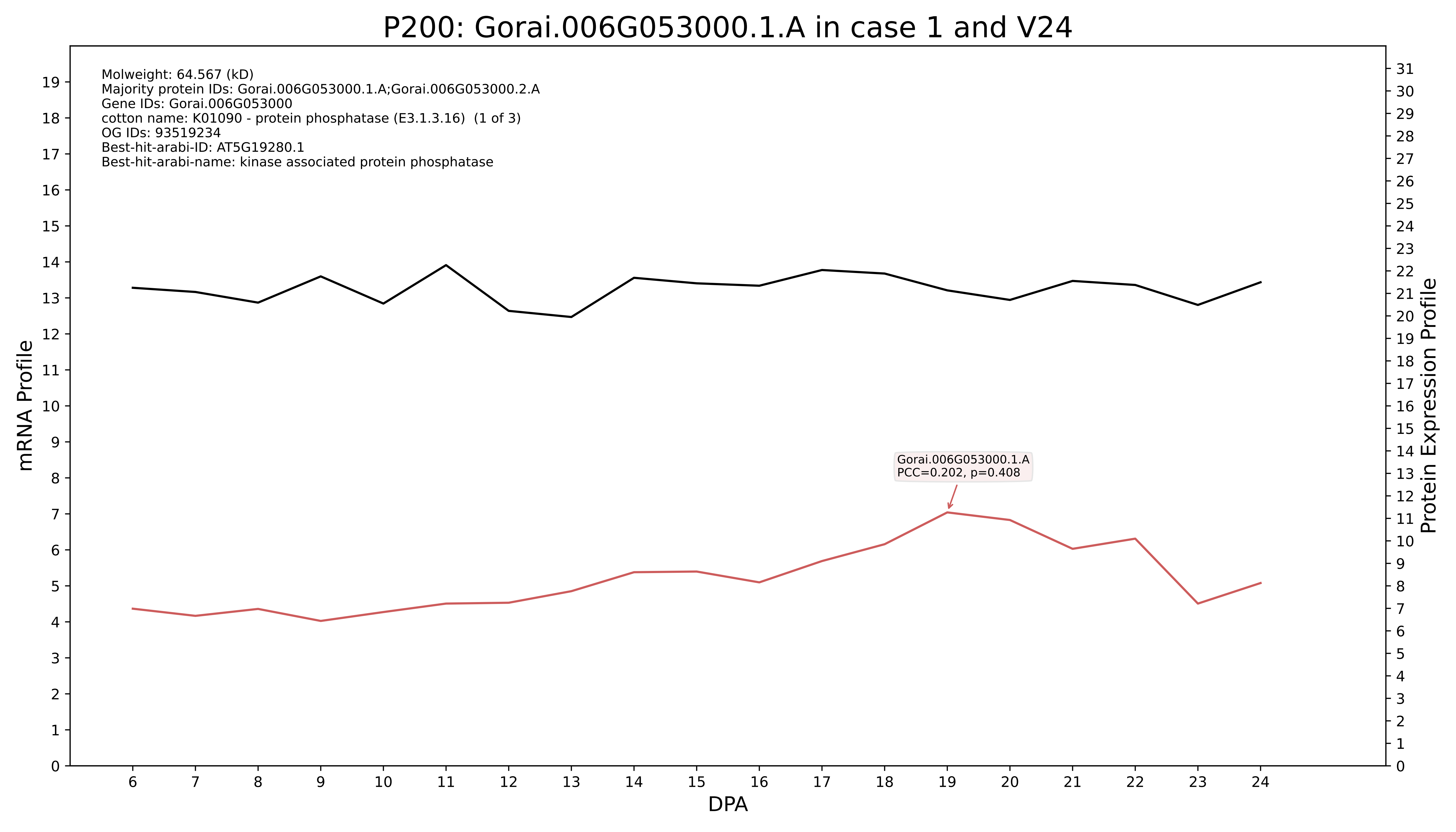
P200: Gorai.006G053000.1.A in case 1 and V24
Molweight: 64.567 (kD)
Majority protein IDs: Gorai.006G053000.1.A;Gorai.006G053000.2.A
Gene IDs: Gorai.006G053000
cotton name: K01090 - protein phosphatase (E3.1.3.16) (1 of 3)
OG IDs: 93519234
Best-hit-arabi-ID: AT5G19280.1
Best-hit-arabi-name: kinase associated protein phosphatase
Gorai.006G053000.1.A
PCC=0.202, p=0.408
mRNA Profile
Protein Expression Profile
DPA

### P200: Gorai.008G097100.1.D in case 1 and V24

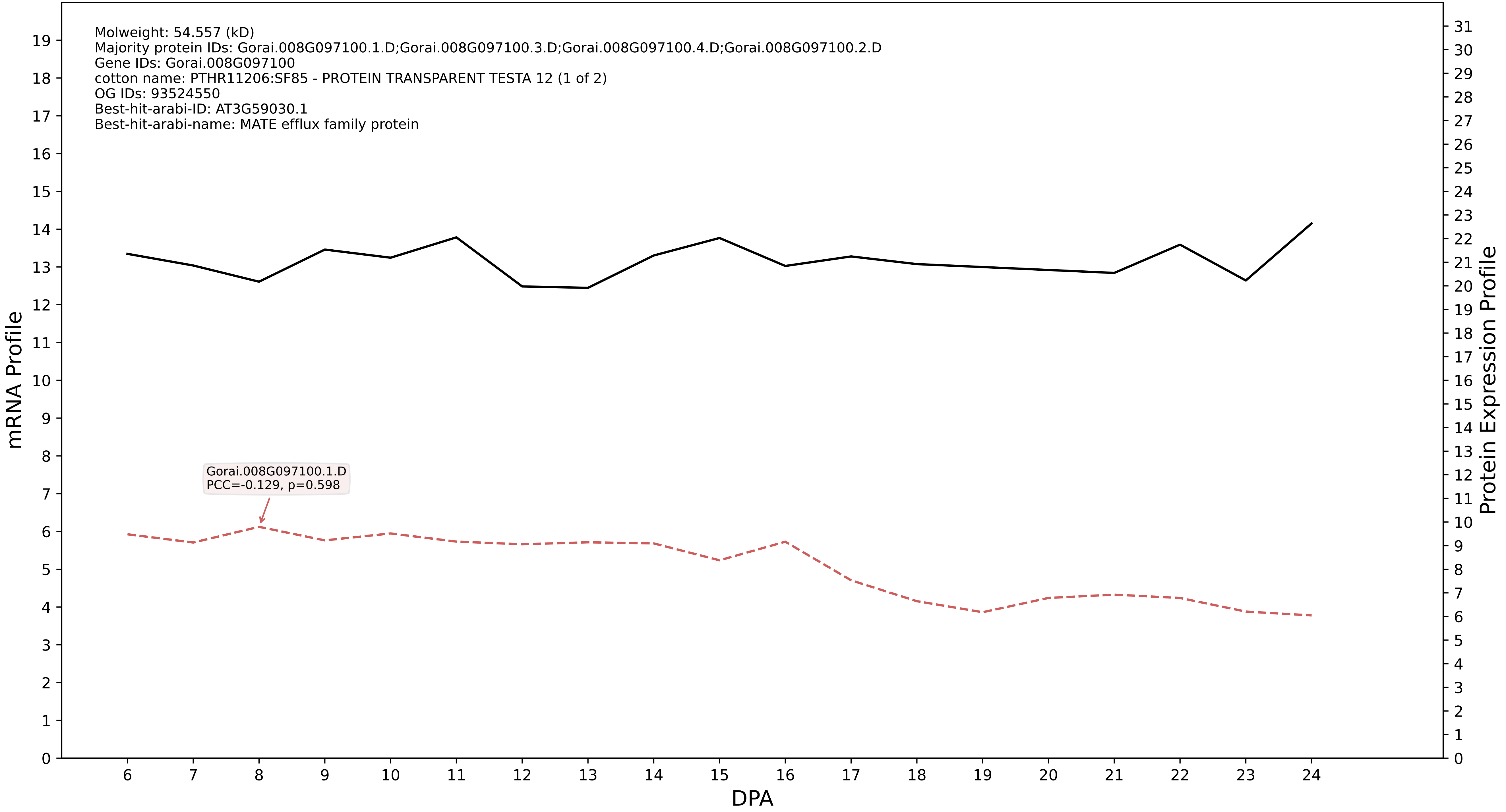

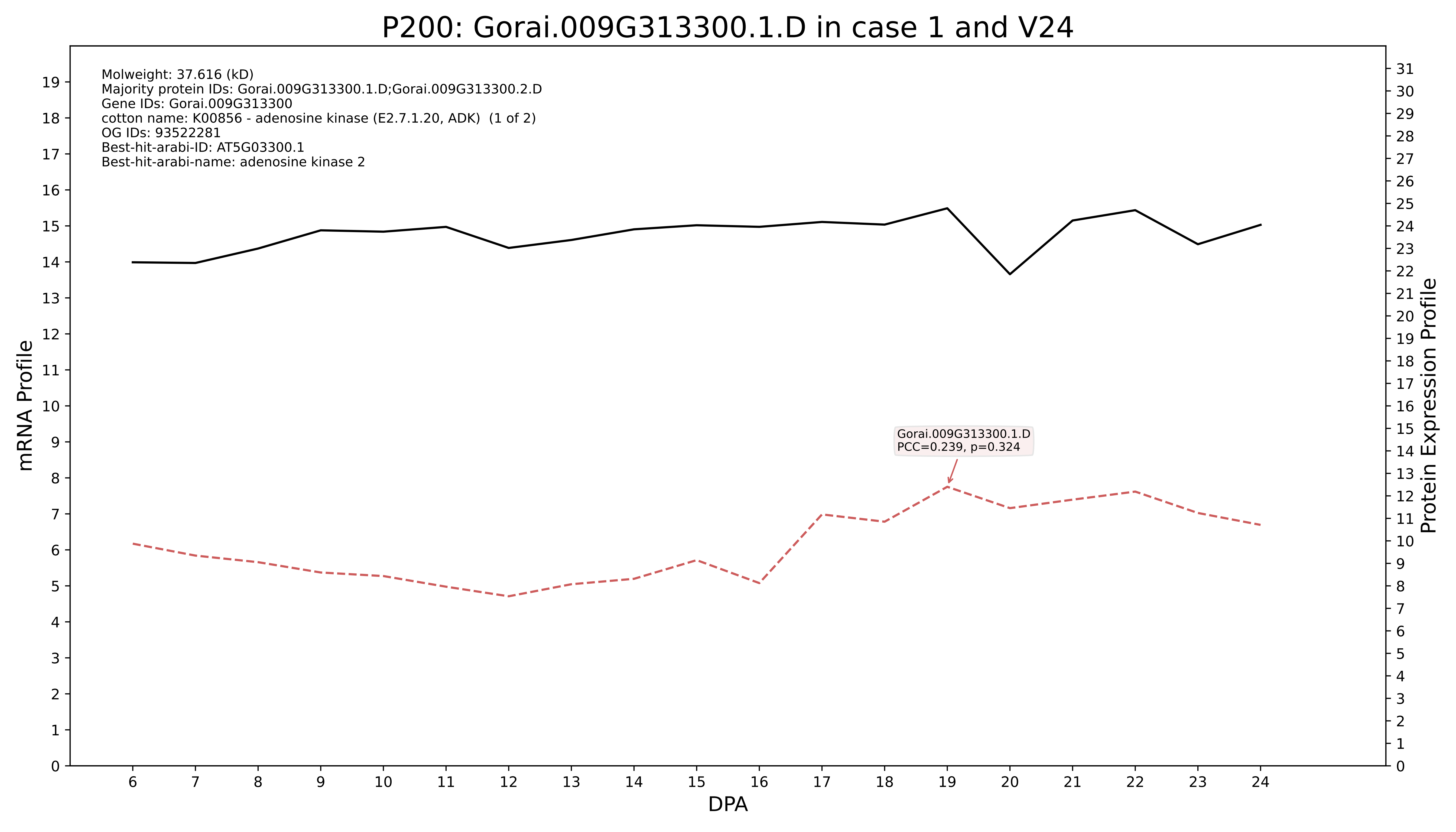

P200: Gorai.009G313300.1.D in case 1 and V24
Molweight: 37.616 (kD)
Majority protein IDs: Gorai.009G313300.1.D;Gorai.009G313300.2.D
Gene IDs: Gorai.009G313300
cotton name: K00856 - adenosine kinase (E2.7.1.20, ADK) (1 of 2)
OG IDs: 93522281
Best-hit-arabi-ID: AT5G03300.1
Best-hit-arabi-name: adenosine kinase 2
Gorai.009G313300.1.D
PCC=0.239, p=0.324
mRNA Profile
Protein Expression Profile
DPA

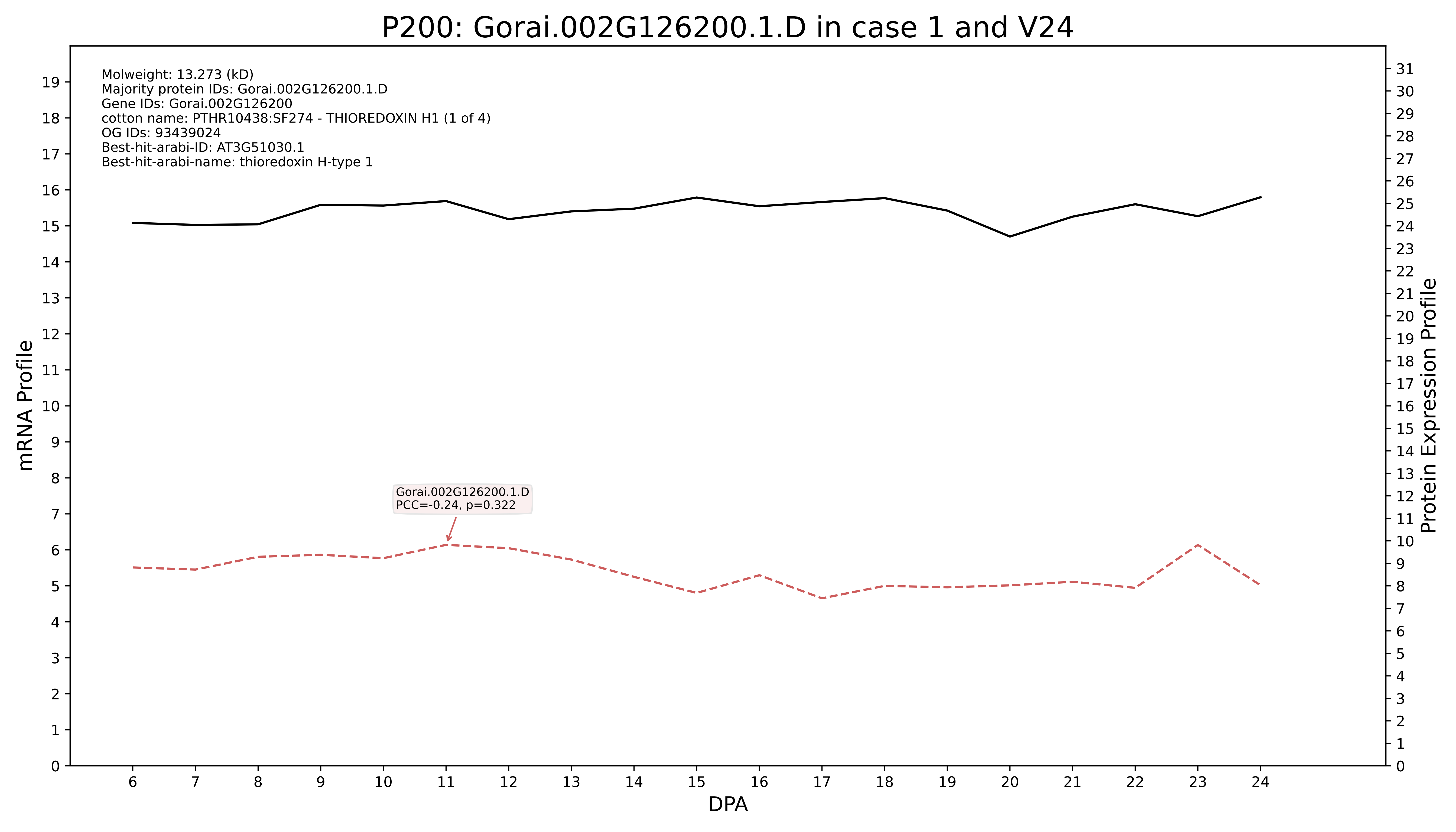
P200: Gorai.002G126200.1.D in case 1 and V24
Molweight: 13.273 (kD)
Majority protein IDs: Gorai.002G126200.1.D
Gene IDs: Gorai.002G126200
cotton name: PTHR10438:SF274 - THIOREDOXIN H1 (1 of 4)
OG IDs: 93439024
Best-hit-arabi-ID: AT3G51030.1
Best-hit-arabi-name: thioredoxin H-type 1
Gorai.002G126200.1.D
PCC=-0.24, p=0.322
mRNA Profile
Protein Expression Profile
DPA

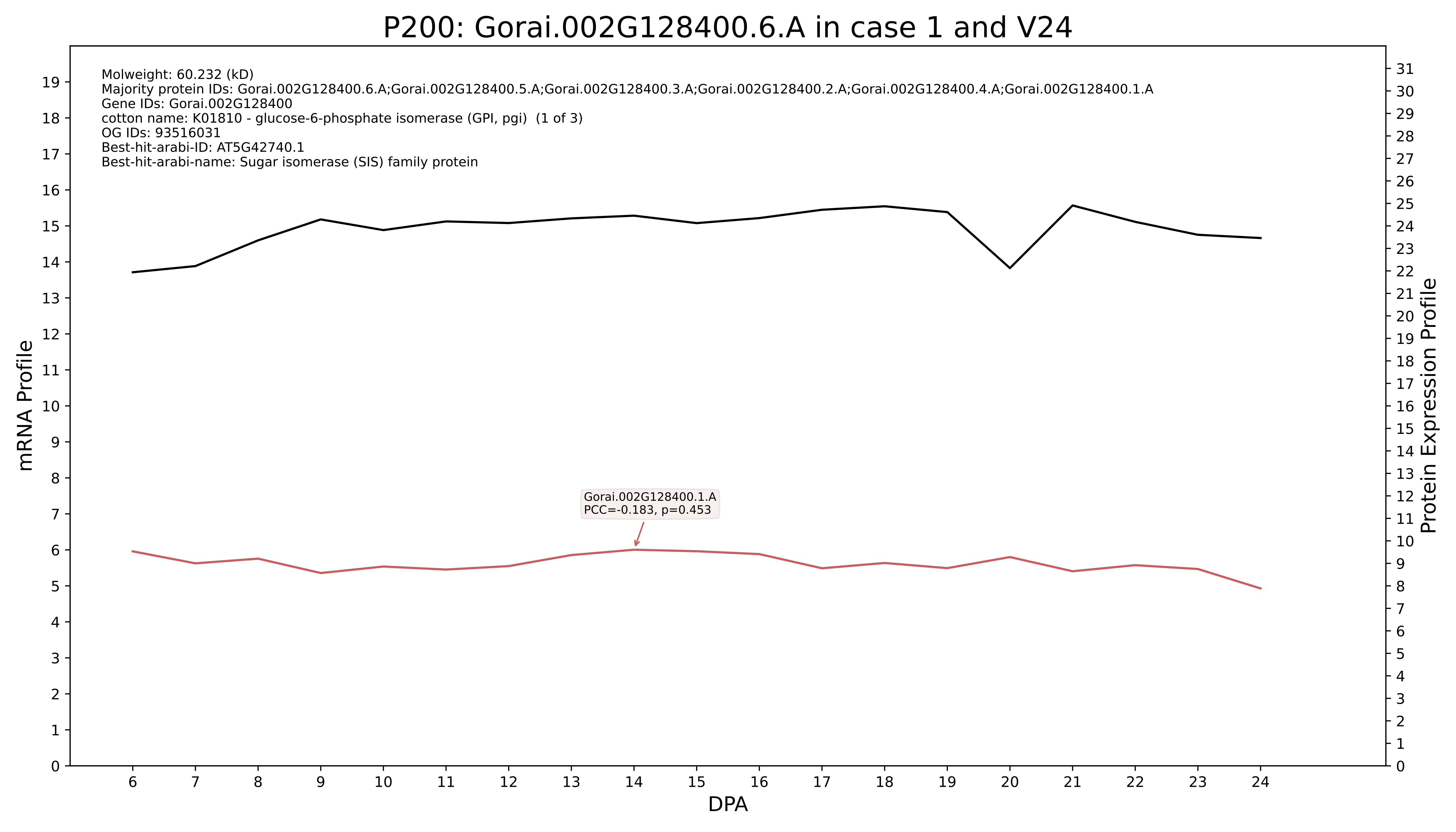
P200: Gorai.002G128400.6.A in case 1 and V24
Molweight: 60.232 (kD)
Majority protein IDs: Gorai.002G128400.6.A;Gorai.002G128400.5.A;Gorai.002G128400.3.A;Gorai.002G128400.2.A;Gorai.002G128400.4.A;Gorai.002G128400.1.A
Gene IDs: Gorai.002G128400
cotton name: K01810 - glucose-6-phosphate isomerase (GPI, pgi) (1 of 3)
OG IDs: 93516031
Best-hit-arabi-ID: AT5G42740.1
Best-hit-arabi-name: Sugar isomerase (SIS) family protein
Gorai.002G128400.1.A
PCC=-0.183, p=0.453
mRNA Profile
Protein Expression Profile
DPA

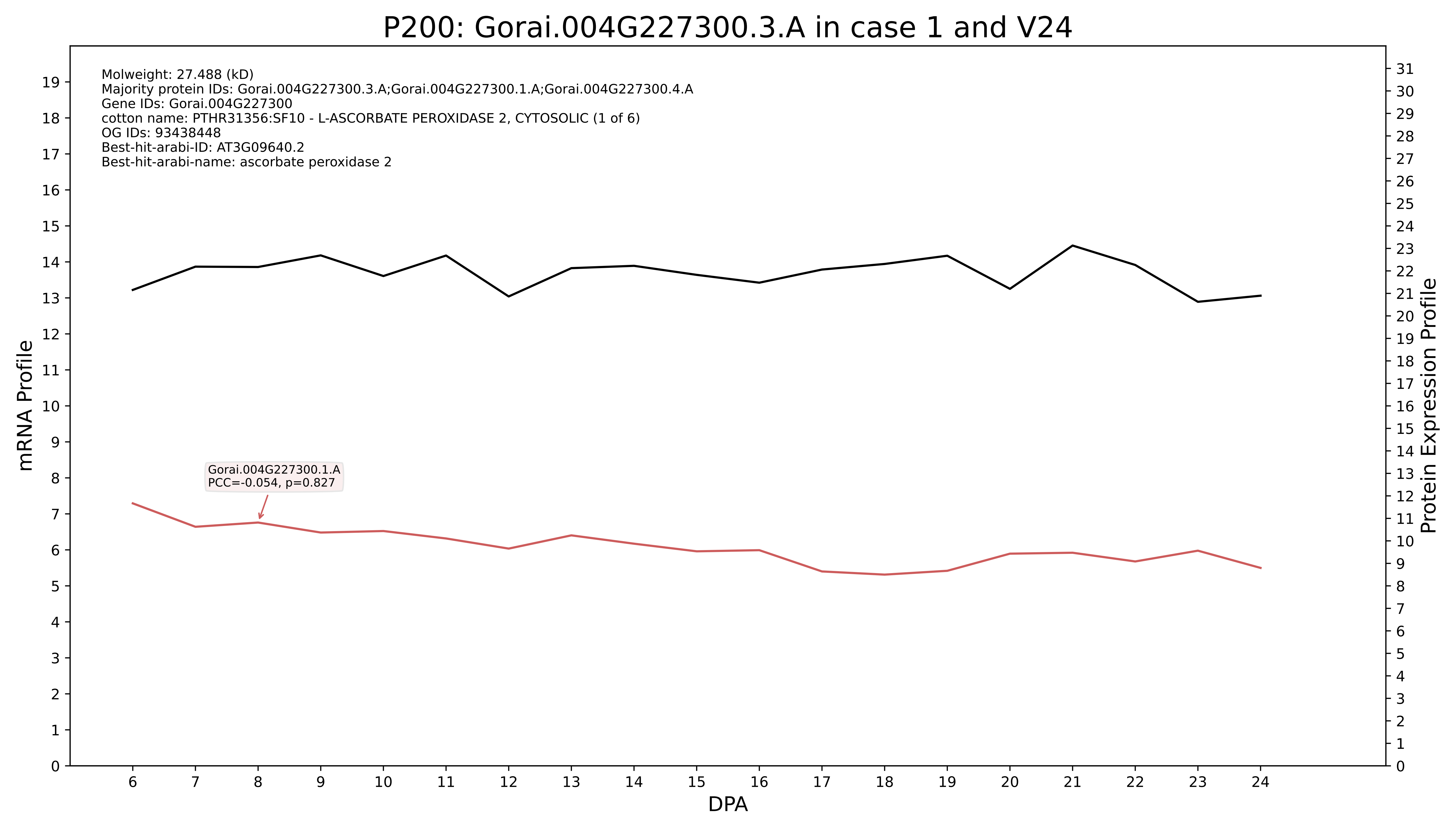
P200: Gorai.004G227300.3.A in case 1 and V24
Molweight: 27.488 (kD)
Majority protein IDs: Gorai.004G227300.3.A;Gorai.004G227300.1.A;Gorai.004G227300.4.A
Gene IDs: Gorai.004G227300
cotton name: PTHR31356:SF10 - L-ASCORBATE PEROXIDASE 2, CYTOSOLIC (1 of 6)
OG IDs: 93438448
Best-hit-arabi-ID: AT3G09640.2
Best-hit-arabi-name: ascorbate peroxidase 2
Gorai.004G227300.1.A
PCC=-0.054, p=0.827
mRNA Profile
Protein Expression Profile
DPA

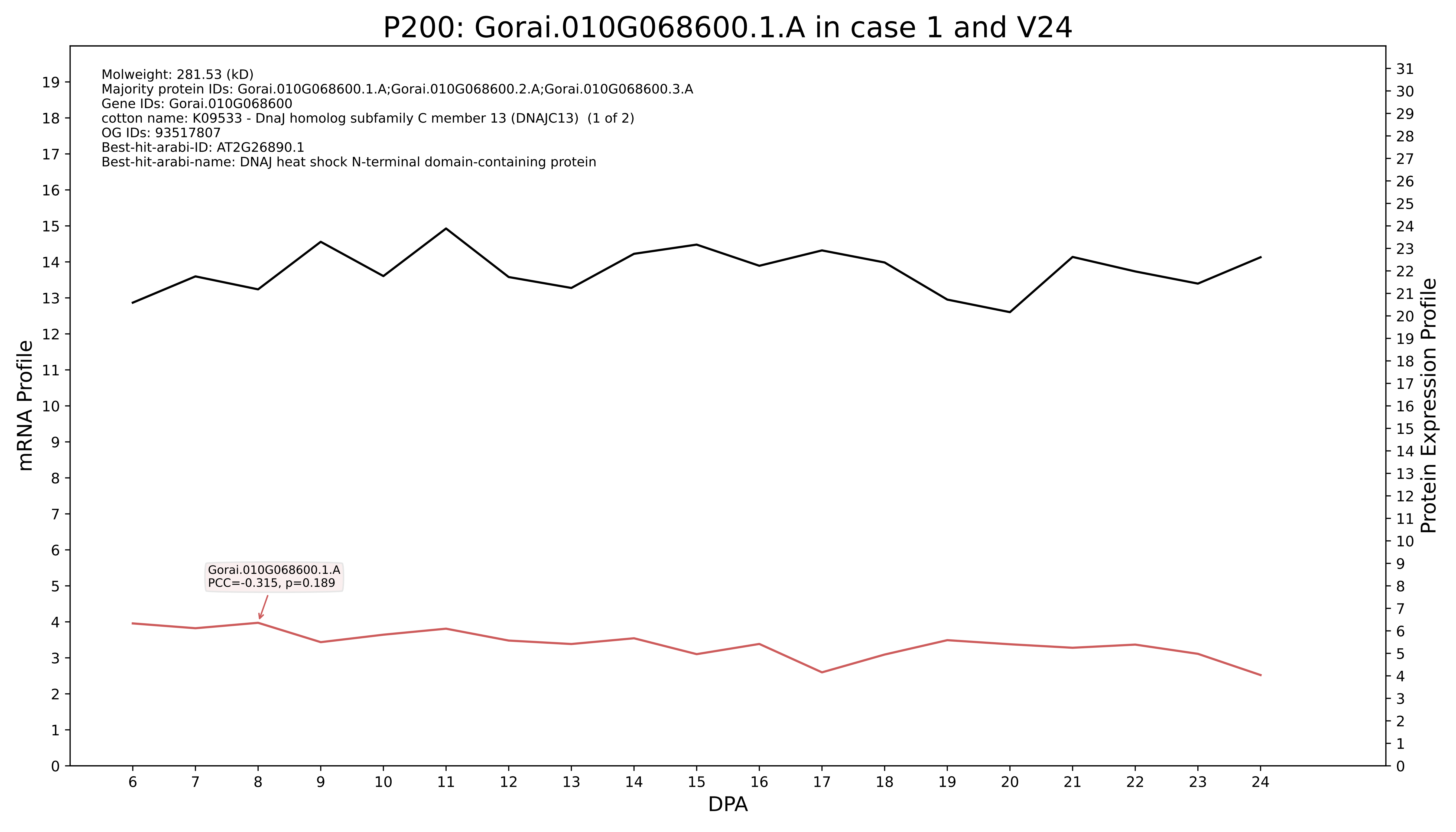
P200: Gorai.010G068600.1.A in case 1 and V24
Molweight: 281.53 (kD)
Majority protein IDs: Gorai.010G068600.1.A;Gorai.010G068600.2.A;Gorai.010G068600.3.A
Gene IDs: Gorai.010G068600
cotton name: K09533 - DnaJ homolog subfamily C member 13 (DNAJC13) (1 of 2)
OG IDs: 93517807
Best-hit-arabi-ID: AT2G26890.1
Best-hit-arabi-name: DNAJ heat shock N-terminal domain-containing protein
Gorai.010G068600.1.A
PCC=-0.315, p=0.189
mRNA Profile
Protein Expression Profile
DPA

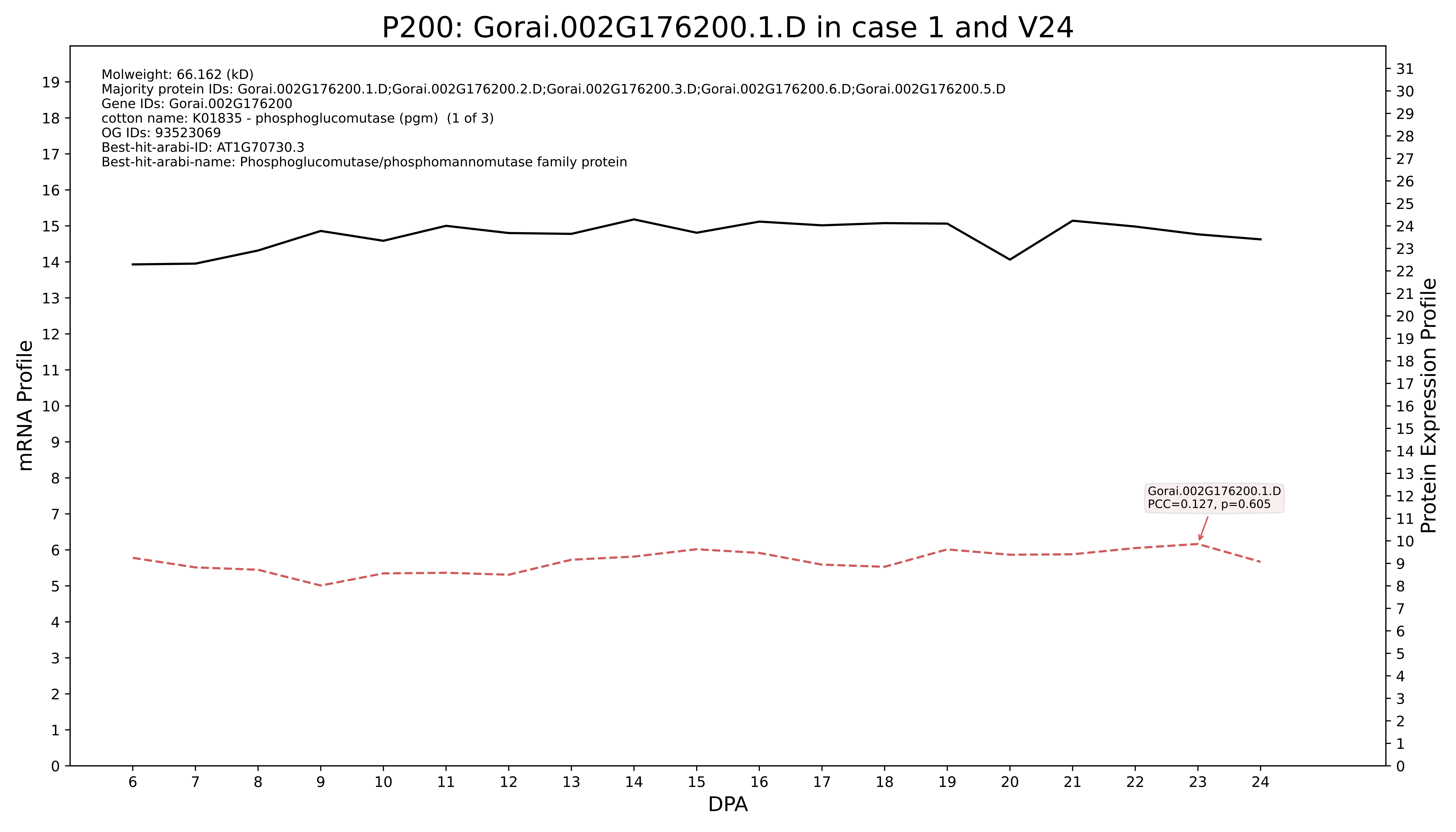
P200: Gorai.002G176200.1.D in case 1 and V24
Molweight: 66.162 (kD)
Majority protein IDs: Gorai.002G176200.1.D;Gorai.002G176200.2.D;Gorai.002G176200.3.D;Gorai.002G176200.6.D;Gorai.002G176200.5.D
Gene IDs: Gorai.002G176200
cotton name: K01835 - phosphoglucomutase (pgm) (1 of 3)
OG IDs: 93523069
Best-hit-arabi-ID: AT1G70730.3
Best-hit-arabi-name: Phosphoglucomutase/phosphomannomutase family protein
Gorai.002G176200.1.D
PCC=0.127, p=0.605
mRNA Profile
Protein Expression Profile
DPA

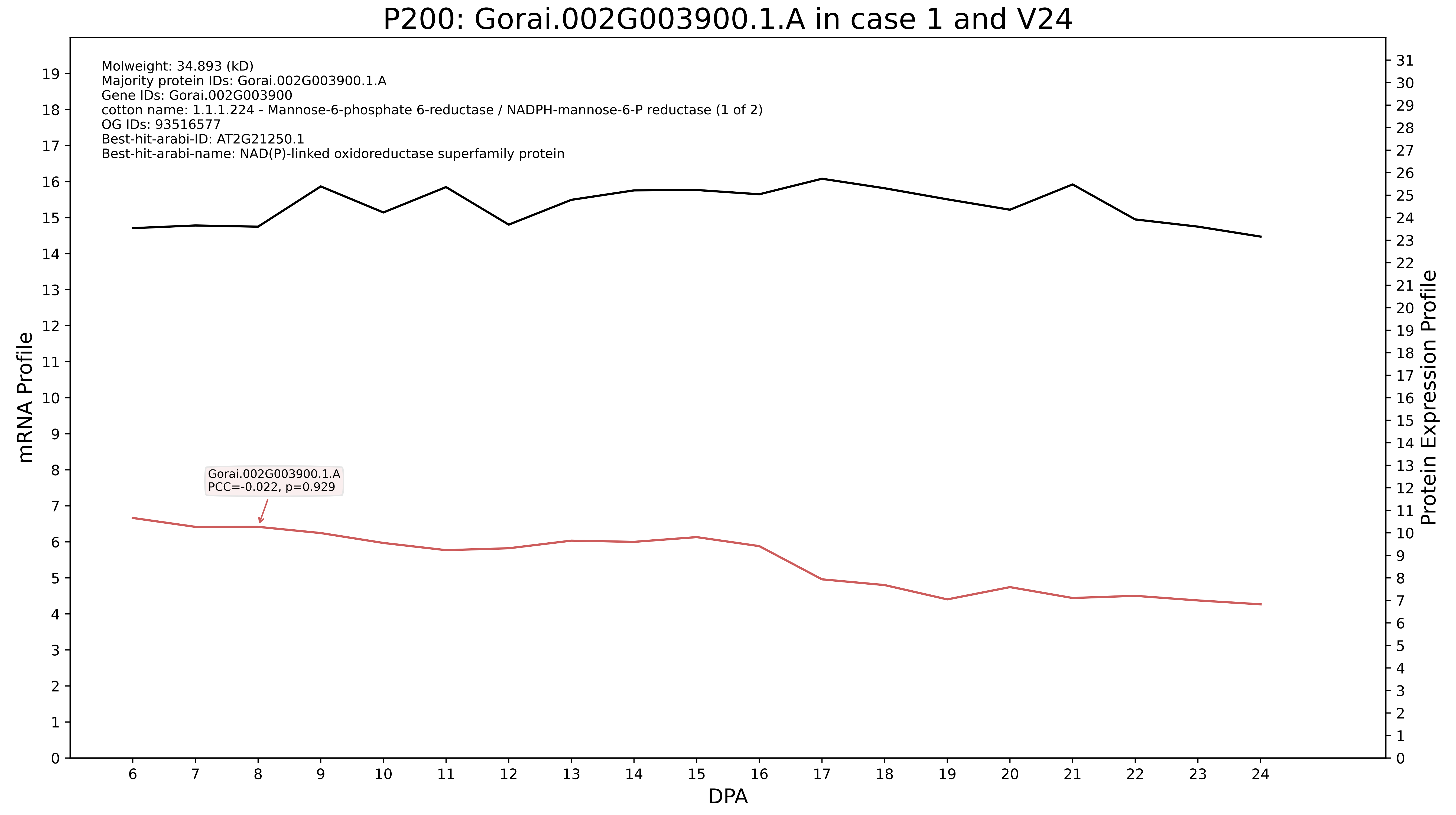
P200: Gorai.002G003900.1.A in case 1 and V24
Molweight: 34.893 (kD)
Majority protein IDs: Gorai.002G003900.1.A
Gene IDs: Gorai.002G003900
cotton name: 1.1.1.224 - Mannose-6-phosphate 6-reductase / NADPH-mannose-6-P reductase (1 of 2)
OG IDs: 93516577
Best-hit-arabi-ID: AT2G21250.1
Best-hit-arabi-name: NAD(P)-linked oxidoreductase superfamily protein
Gorai.002G003900.1.A
PCC=-0.022, p=0.929
mRNA Profile
Protein Expression Profile
DPA

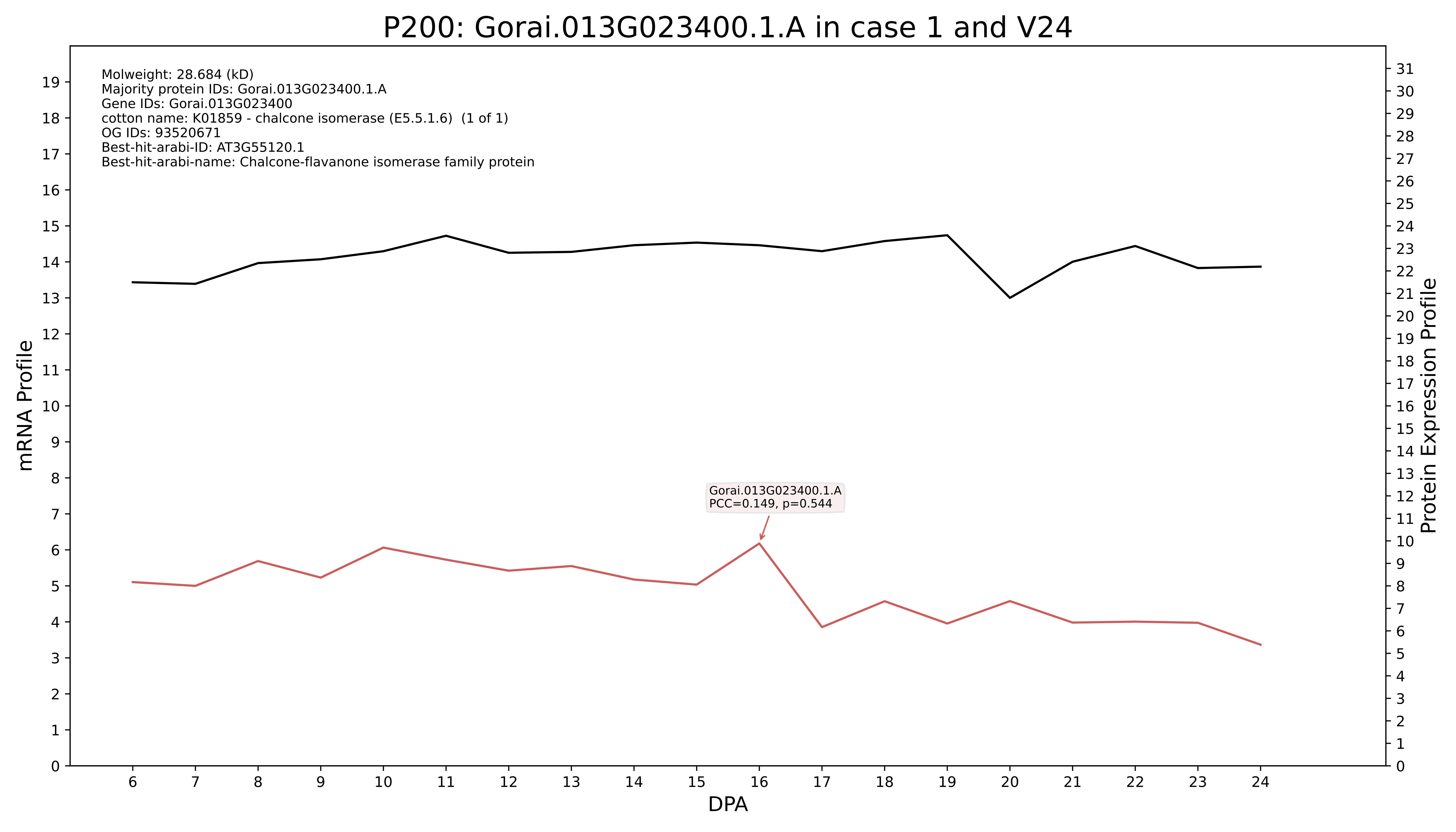
P200: Gorai.013G023400.1.A in case 1 and V24
Molweight: 28.684 (kD)
Majority protein IDs: Gorai.013G023400.1.A
Gene IDs: Gorai.013G023400
cotton name: K01859 - chalcone isomerase (E5.5.1.6) (1 of 1)
OG IDs: 93520671
Best-hit-arabi-ID: AT3G55120.1
Best-hit-arabi-name: Chalcone-flavanone isomerase family protein
Gorai.013G023400.1.A
PCC=0.149, p=0.544
mRNA Profile
Protein Expression Profile
DPA

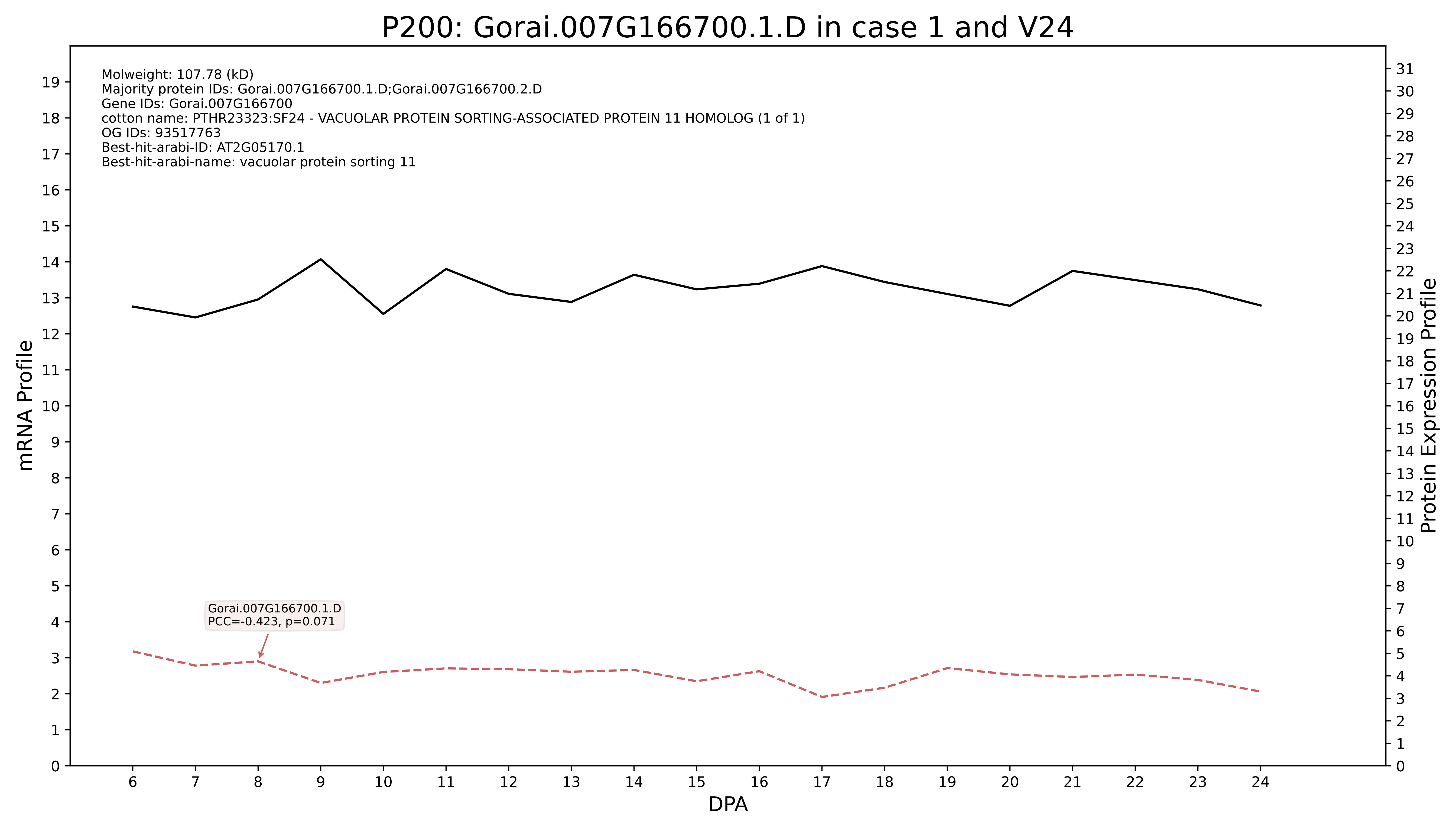
P200: Gorai.007G166700.1.D in case 1 and V24
Molweight: 107.78 (kD)
Majority protein IDs: Gorai.007G166700.1.D;Gorai.007G166700.2.D
Gene IDs: Gorai.007G166700
cotton name: PTHR23323:SF24 - VACUOLAR PROTEIN SORTING-ASSOCIATED PROTEIN 11 HOMOLOG (1 of 1)
OG IDs: 93517763
Best-hit-arabi-ID: AT2G05170.1
Best-hit-arabi-name: vacuolar protein sorting 11
Gorai.007G166700.1.D
PCC=-0.423, p=0.071
mRNA Profile
Protein Expression Profile
DPA

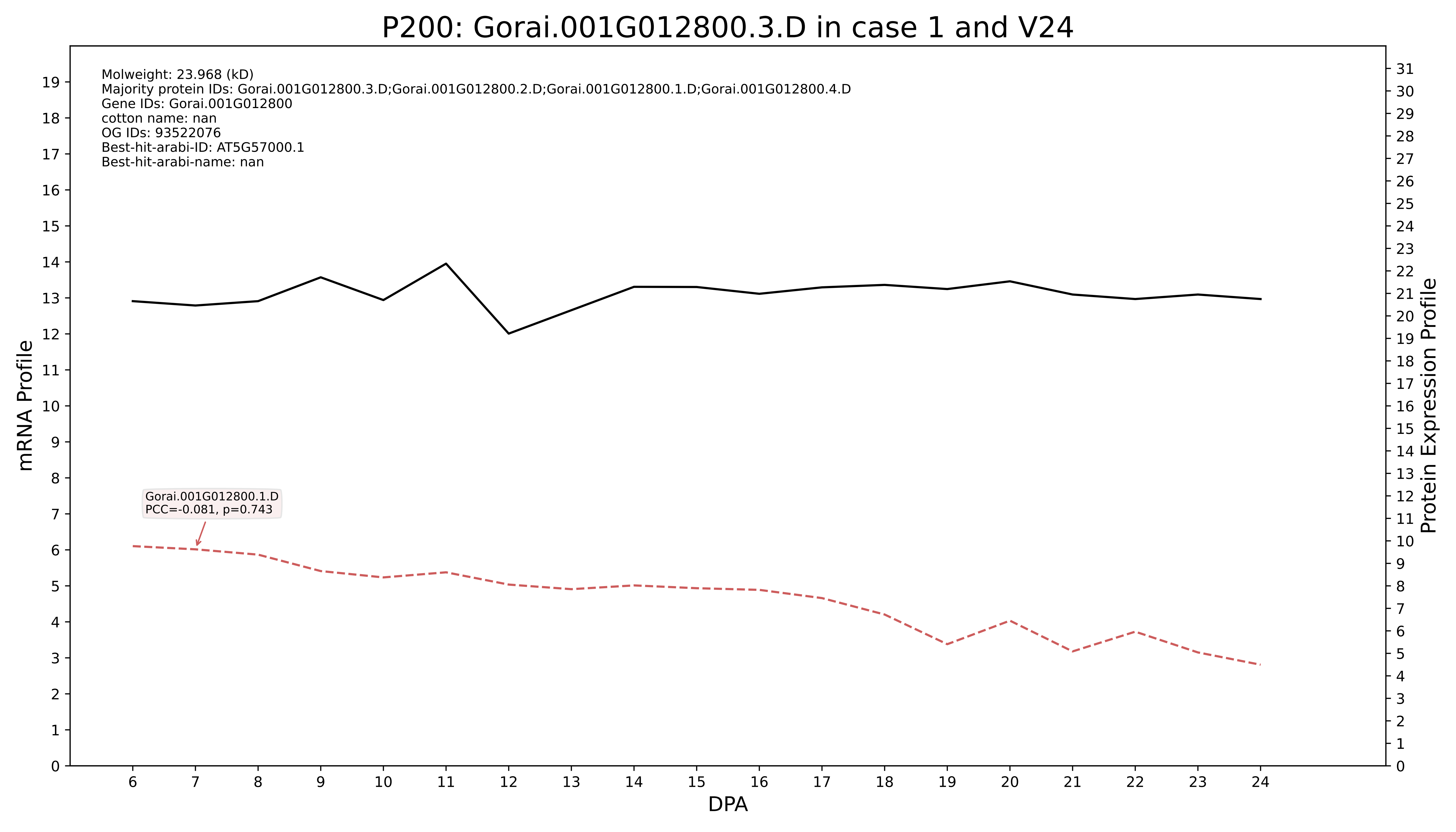
P200: Gorai.001G012800.3.D in case 1 and V24
Molweight: 23.968 (kD)
Majority protein IDs: Gorai.001G012800.3.D;Gorai.001G012800.2.D;Gorai.001G012800.1.D;Gorai.001G012800.4.D
Gene IDs: Gorai.001G012800
cotton name: nan
OG IDs: 93522076
Best-hit-arabi-ID: AT5G57000.1
Best-hit-arabi-name: nan
Gorai.001G012800.1.D
PCC=-0.081, p=0.743
mRNA Profile
Protein Expression Profile
DPA

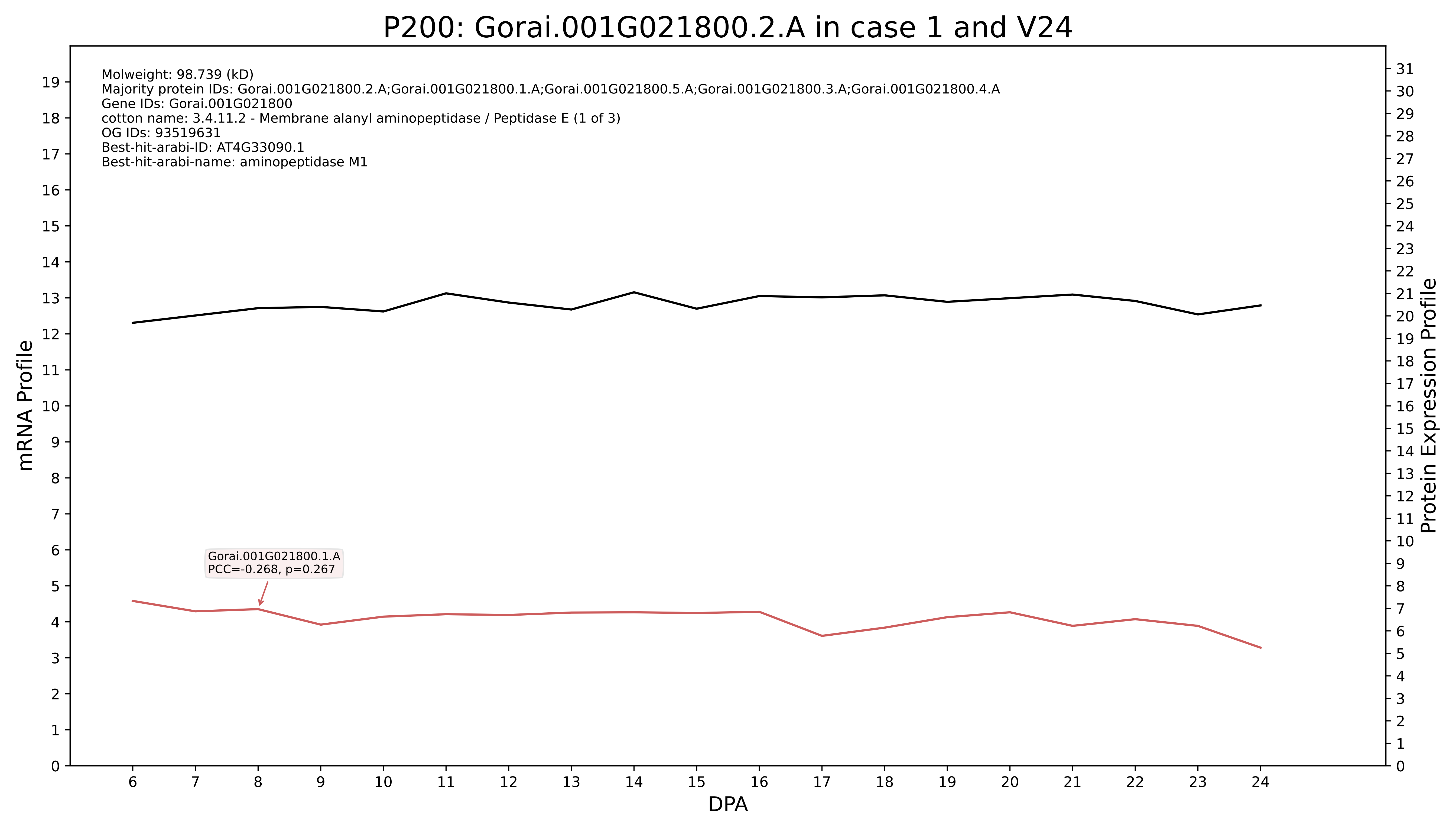
P200: Gorai.001G021800.2.A in case 1 and V24
Molweight: 98.739 (kD)
Majority protein IDs: Gorai.001G021800.2.A;Gorai.001G021800.1.A;Gorai.001G021800.5.A;Gorai.001G021800.3.A;Gorai.001G021800.4.A
Gene IDs: Gorai.001G021800
cotton name: 3.4.11.2 - Membrane alanyl aminopeptidase / Peptidase E (1 of 3)
OG IDs: 93519631
Best-hit-arabi-ID: AT4G33090.1
Best-hit-arabi-name: aminopeptidase M1
Gorai.001G021800.1.A
PCC=-0.268, p=0.267
mRNA Profile
Protein Expression Profile
DPA

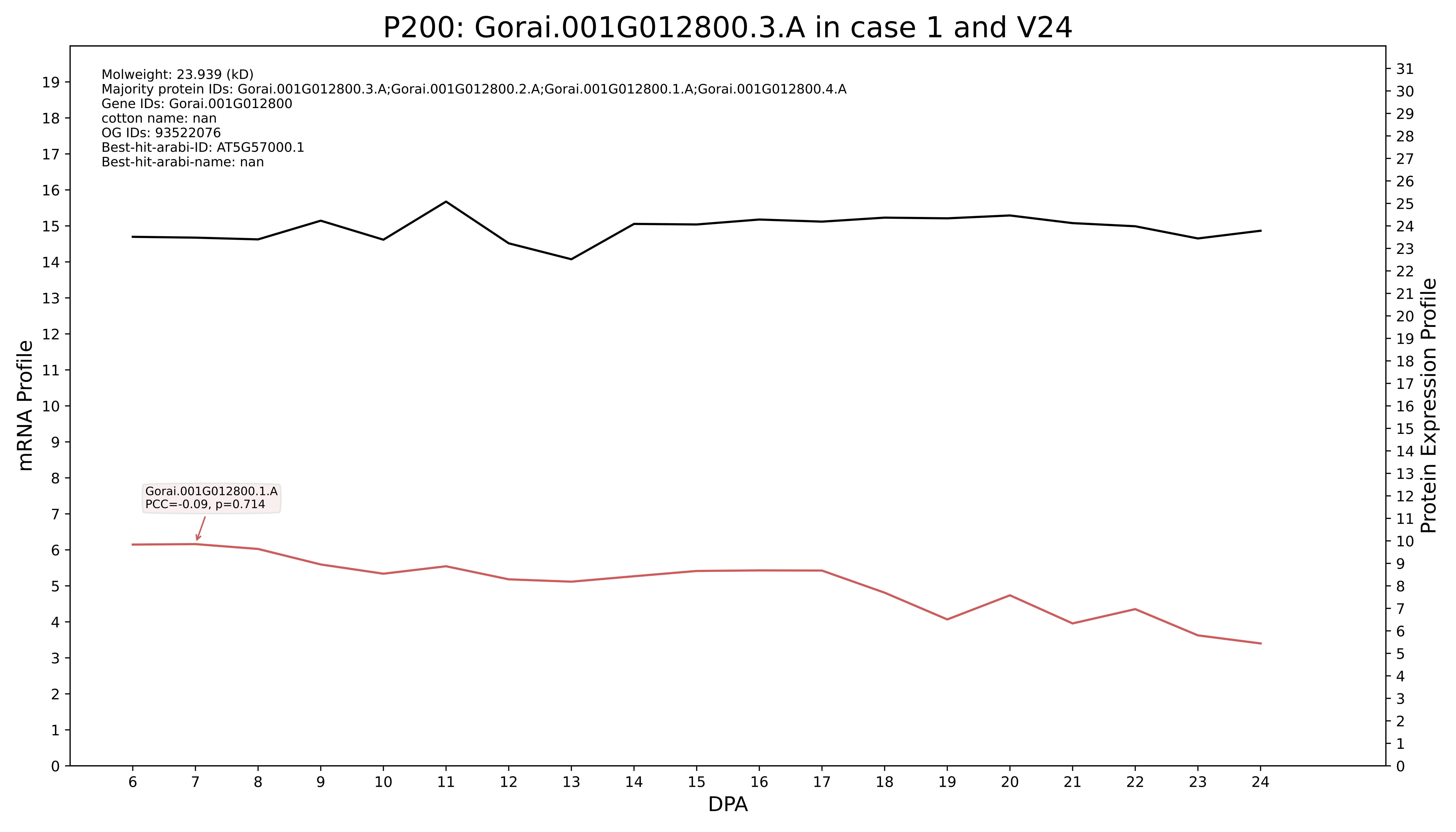
P200: Gorai.001G012800.3.A in case 1 and V24
Molweight: 23.939 (kD)
Majority protein IDs: Gorai.001G012800.3.A;Gorai.001G012800.2.A;Gorai.001G012800.1.A;Gorai.001G012800.4.A
Gene IDs: Gorai.001G012800
cotton name: nan
OG IDs: 93522076
Best-hit-arabi-ID: AT5G57000.1
Best-hit-arabi-name: nan
Gorai.001G012800.1.A
PCC=-0.09, p=0.714
mRNA Profile
Protein Expression Profile
DPA

### P200: Gorai.013G260900.1.A in case 1 and V24

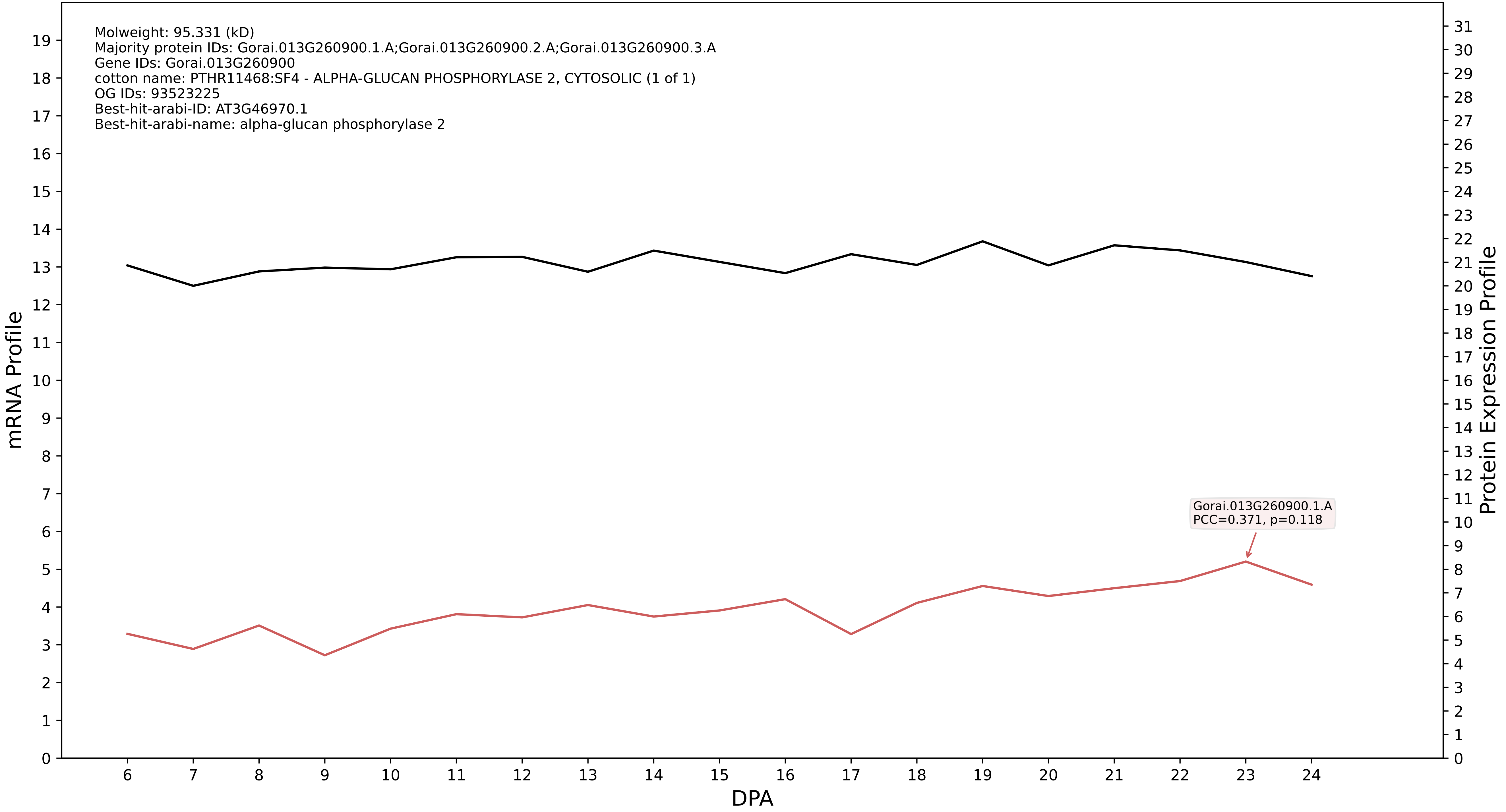

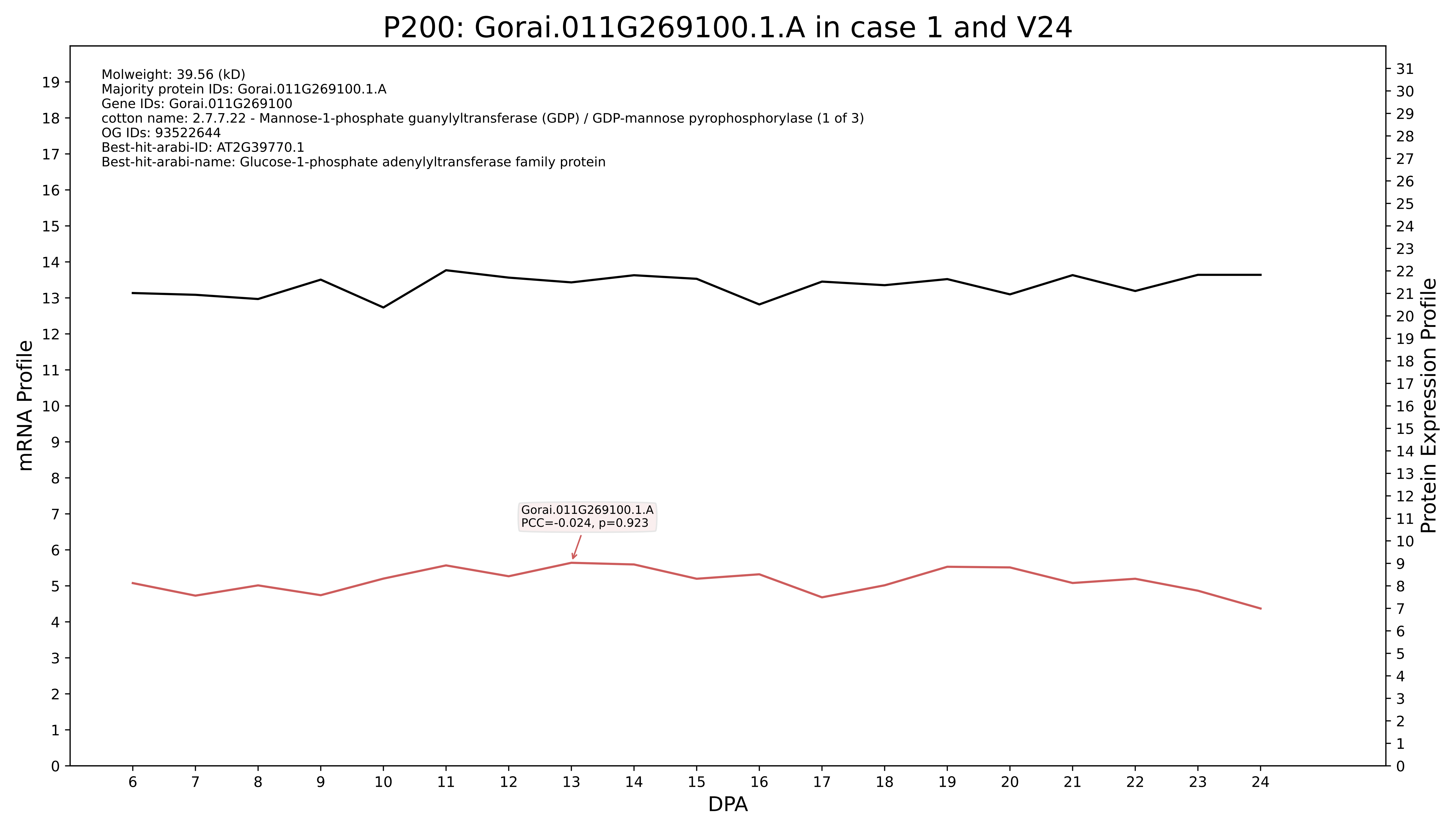
P200: Gorai.011G269100.1.A in case 1 and V24
Molweight: 39.56 (kD)
Majority protein IDs: Gorai.011G269100.1.A
Gene IDs: Gorai.011G269100
cotton name: 2.7.7.22 - Mannose-1-phosphate guanylyltransferase (GDP) / GDP-mannose pyrophosphorylase (1 of 3)
OG IDs: 93522644
Best-hit-arabi-ID: AT2G39770.1
Best-hit-arabi-name: Glucose-1-phosphate adenylyltransferase family protein
Gorai.011G269100.1.A
PCC=-0.024, p=0.923
mRNA Profile
Protein Expression Profile
DPA

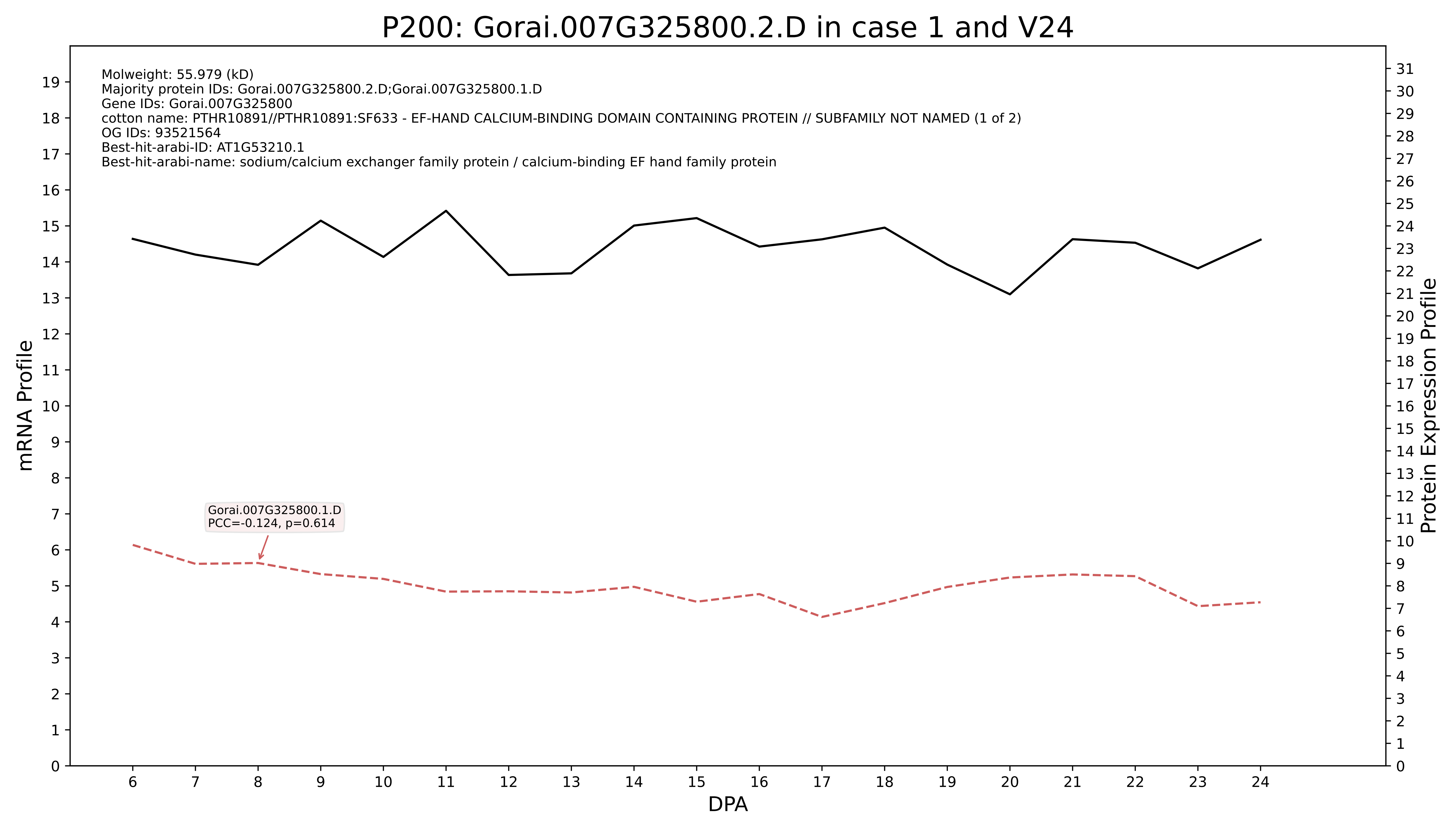
P200: Gorai.007G325800.2.D in case 1 and V24
Molweight: 55.979 (kD)
Majority protein IDs: Gorai.007G325800.2.D;Gorai.007G325800.1.D
Gene IDs: Gorai.007G325800
cotton name: PTHR10891//PTHR10891:SF633 - EF-HAND CALCIUM-BINDING DOMAIN CONTAINING PROTEIN // SUBFAMILY NOT NAMED (1 of 2)
OG IDs: 93521564
Best-hit-arabi-ID: AT1G53210.1
Best-hit-arabi-name: sodium/calcium exchanger family protein / calcium-binding EF hand family protein
Gorai.007G325800.1.D
PCC=-0.124, p=0.614
mRNA Profile
Protein Expression Profile
DPA

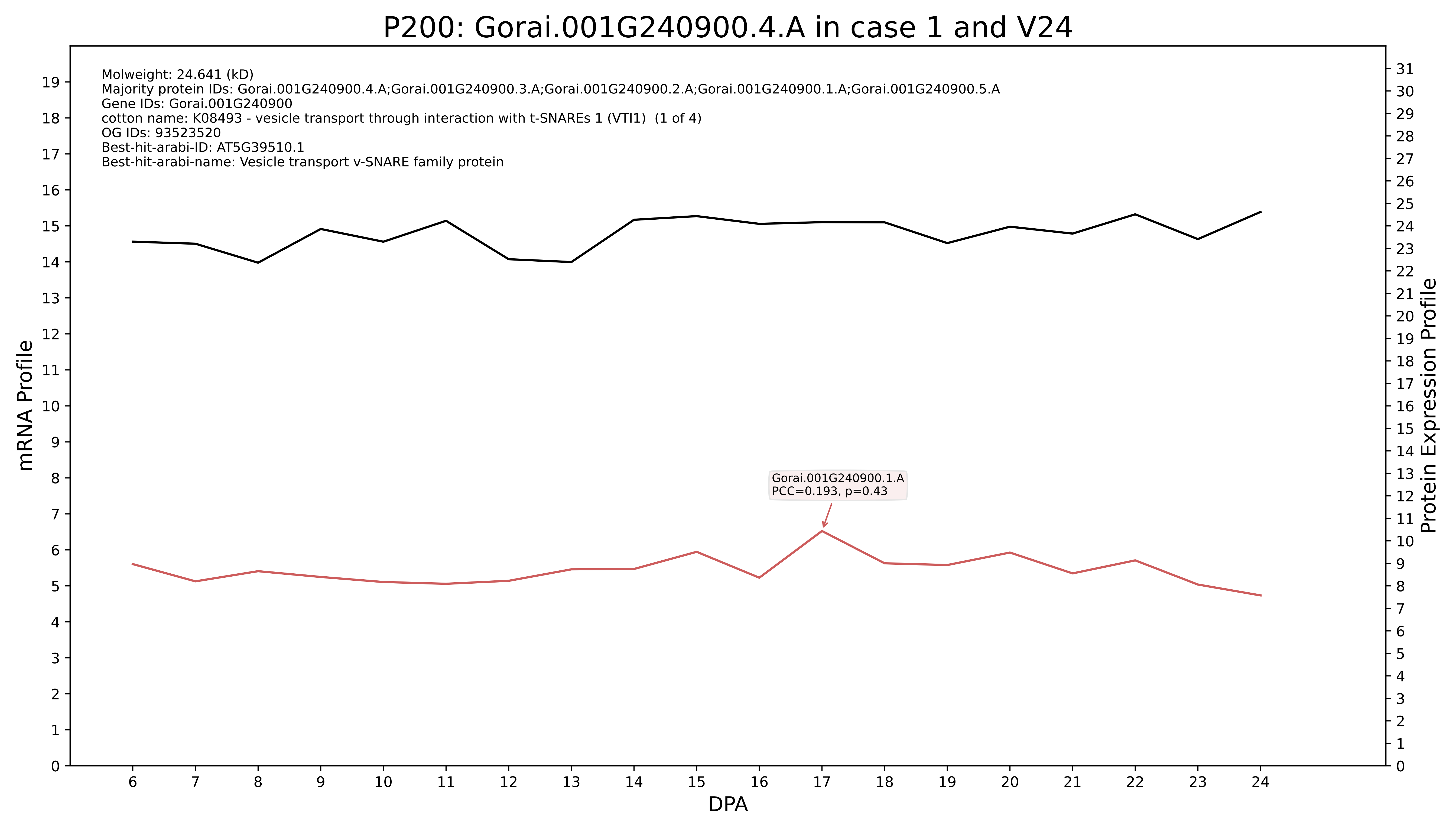
P200: Gorai.001G240900.4.A in case 1 and V24
Molweight: 24.641 (kD)
Majority protein IDs: Gorai.001G240900.4.A;Gorai.001G240900.3.A;Gorai.001G240900.2.A;Gorai.001G240900.1.A;Gorai.001G240900.5.A
Gene IDs: Gorai.001G240900
cotton name: K08493 - vesicle transport through interaction with t-SNAREs 1 (VTI1) (1 of 4)
OG IDs: 93523520
Best-hit-arabi-ID: AT5G39510.1
Best-hit-arabi-name: Vesicle transport v-SNARE family protein
Gorai.001G240900.1.A
PCC=0.193, p=0.43
mRNA Profile
Protein Expression Profile
DPA

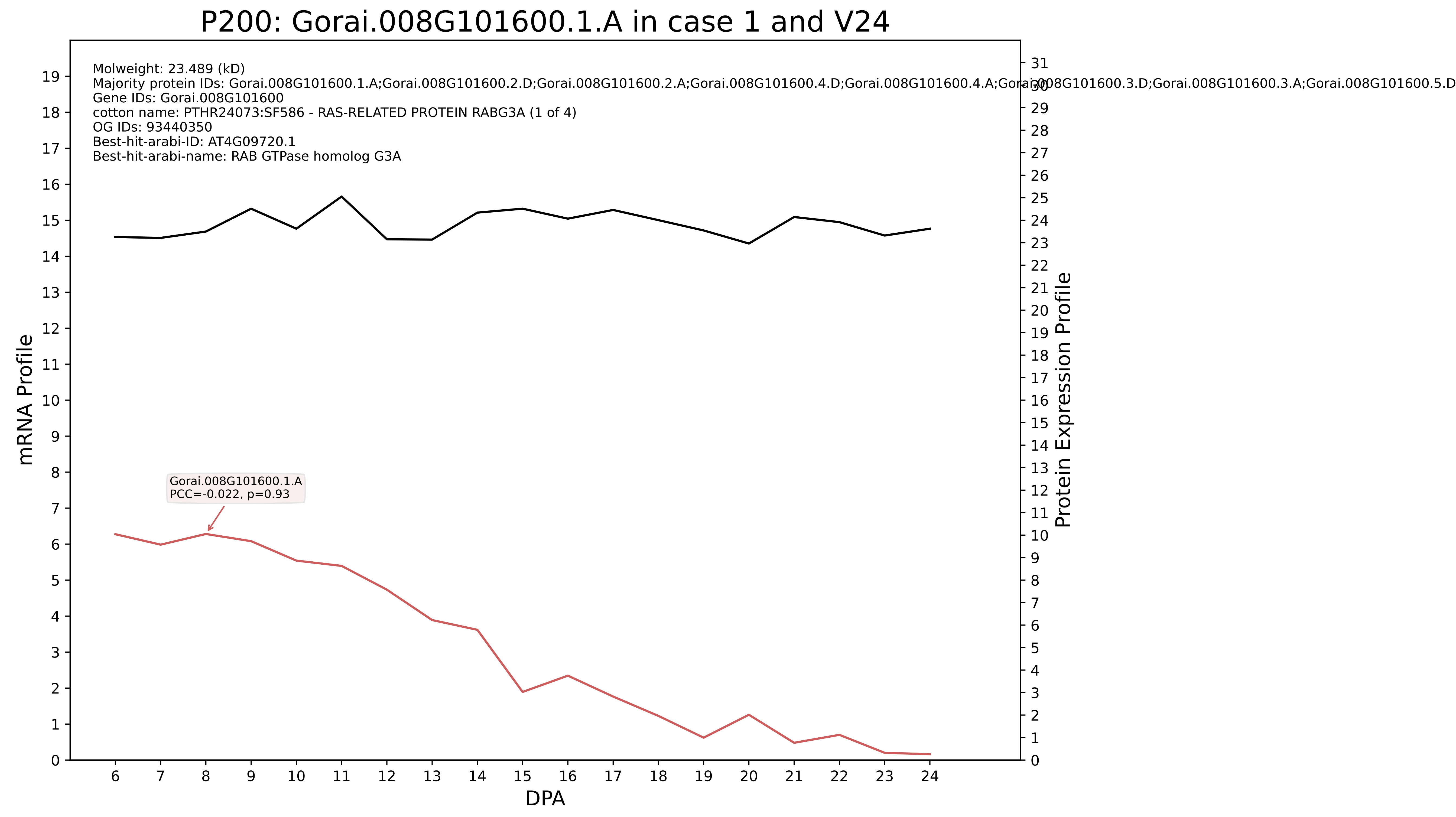
P200: Gorai.008G101600.1.A in case 1 and V24
Molweight: 23.489 (kD)
Majority protein IDs: Gorai.008G101600.1.A;Gorai.008G101600.2.D;Gorai.008G101600.2.A;Gorai.008G101600.4.D;Gorai.008G101600.4.A;Gorai.008G101600.3.D;Gorai.008G101600.3.A;Gorai.008G101600.5.D
Gene IDs: Gorai.008G101600
cotton name: PTHR24073:SF586 - RAS-RELATED PROTEIN RABG3A (1 of 4)
OG IDs: 93440350
Best-hit-arabi-ID: AT4G09720.1
Best-hit-arabi-name: RAB GTPase homolog G3A
Gorai.008G101600.1.A
PCC=-0.022, p=0.93
mRNA Profile
Protein Expression Profile
DPA

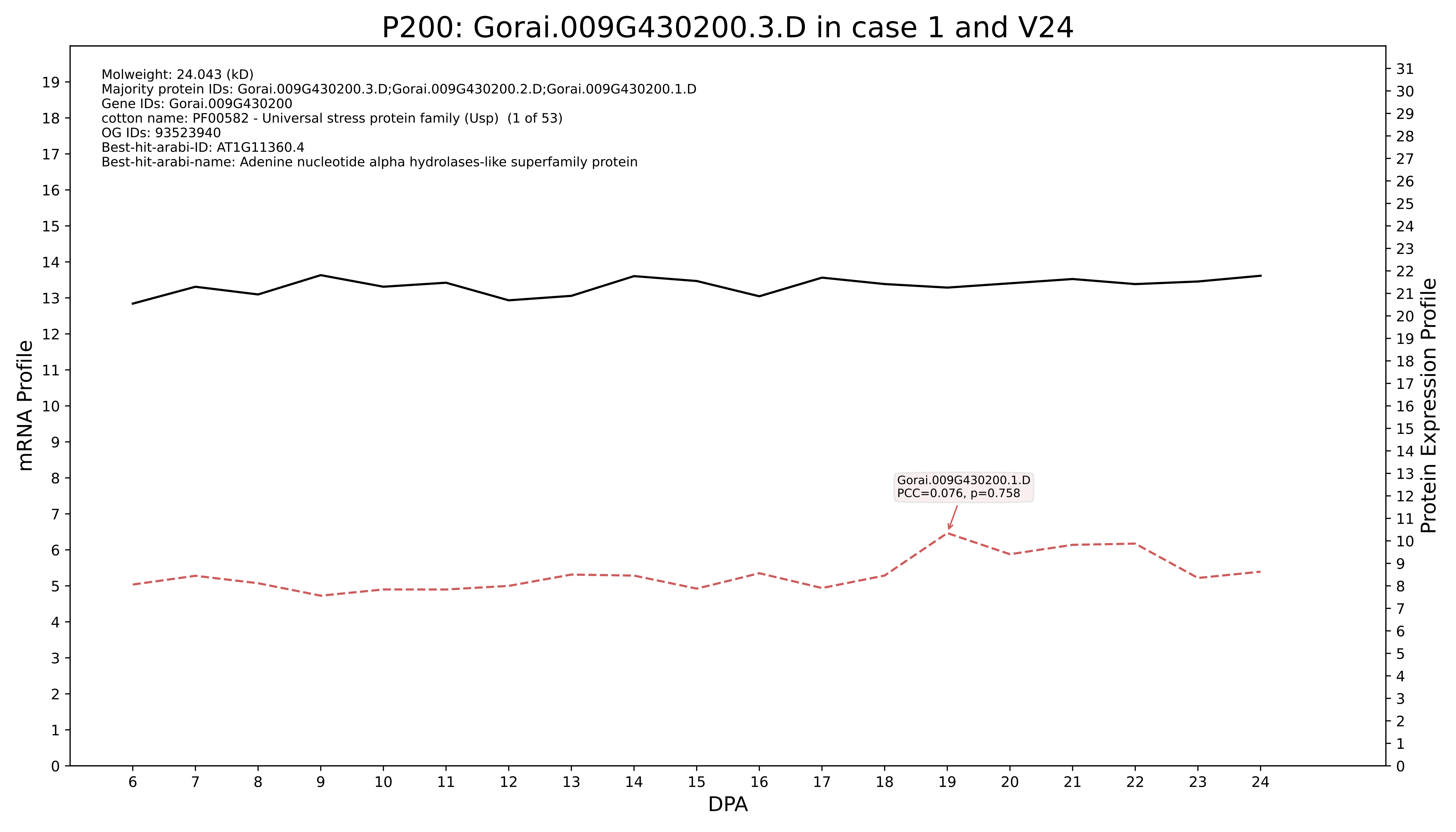

P200: Gorai.009G430200.3.D in case 1 and V24
Molweight: 24.043 (kD)
Majority protein IDs: Gorai.009G430200.3.D;Gorai.009G430200.2.D;Gorai.009G430200.1.D
Gene IDs: Gorai.009G430200
cotton name: PF00582 - Universal stress protein family (Usp) (1 of 53)
OG IDs: 93523940
Best-hit-arabi-ID: AT1G11360.4
Best-hit-arabi-name: Adenine nucleotide alpha hydrolases-like superfamily protein
Gorai.009G430200.1.D
PCC=0.076, p=0.758
mRNA Profile
Protein Expression Profile
DPA

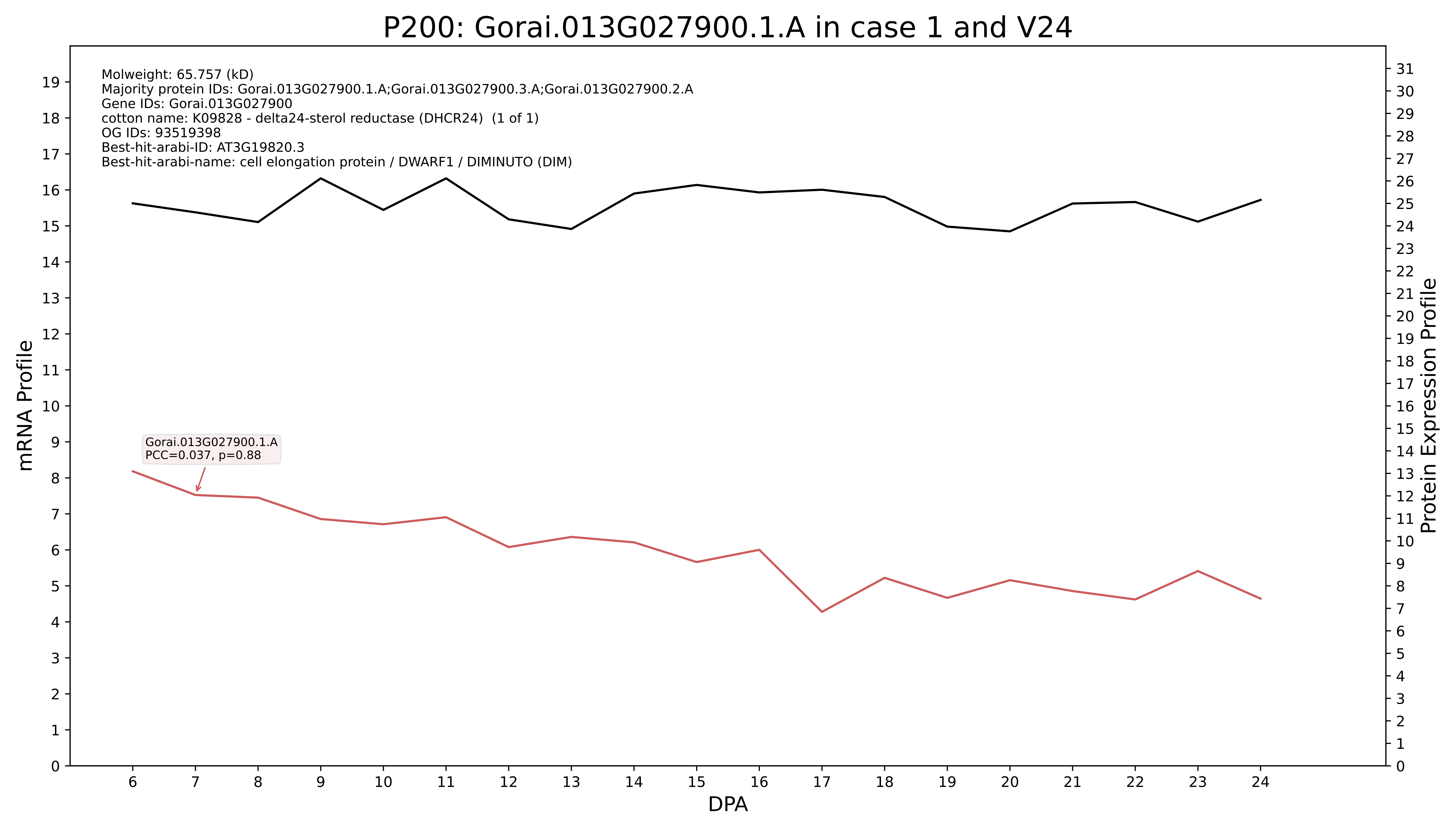
P200: Gorai.013G027900.1.A in case 1 and V24
Molweight: 65.757 (kD)
Majority protein IDs: Gorai.013G027900.1.A;Gorai.013G027900.3.A;Gorai.013G027900.2.A
Gene IDs: Gorai.013G027900
cotton name: K09828 - delta24-sterol reductase (DHCR24) (1 of 1)
OG IDs: 93519398
Best-hit-arabi-ID: AT3G19820.3
Best-hit-arabi-name: cell elongation protein / DWARF1 / DIMINUTO (DIM)
Gorai.013G027900.1.A
PCC=0.037, p=0.88
mRNA Profile
Protein Expression Profile
DPA

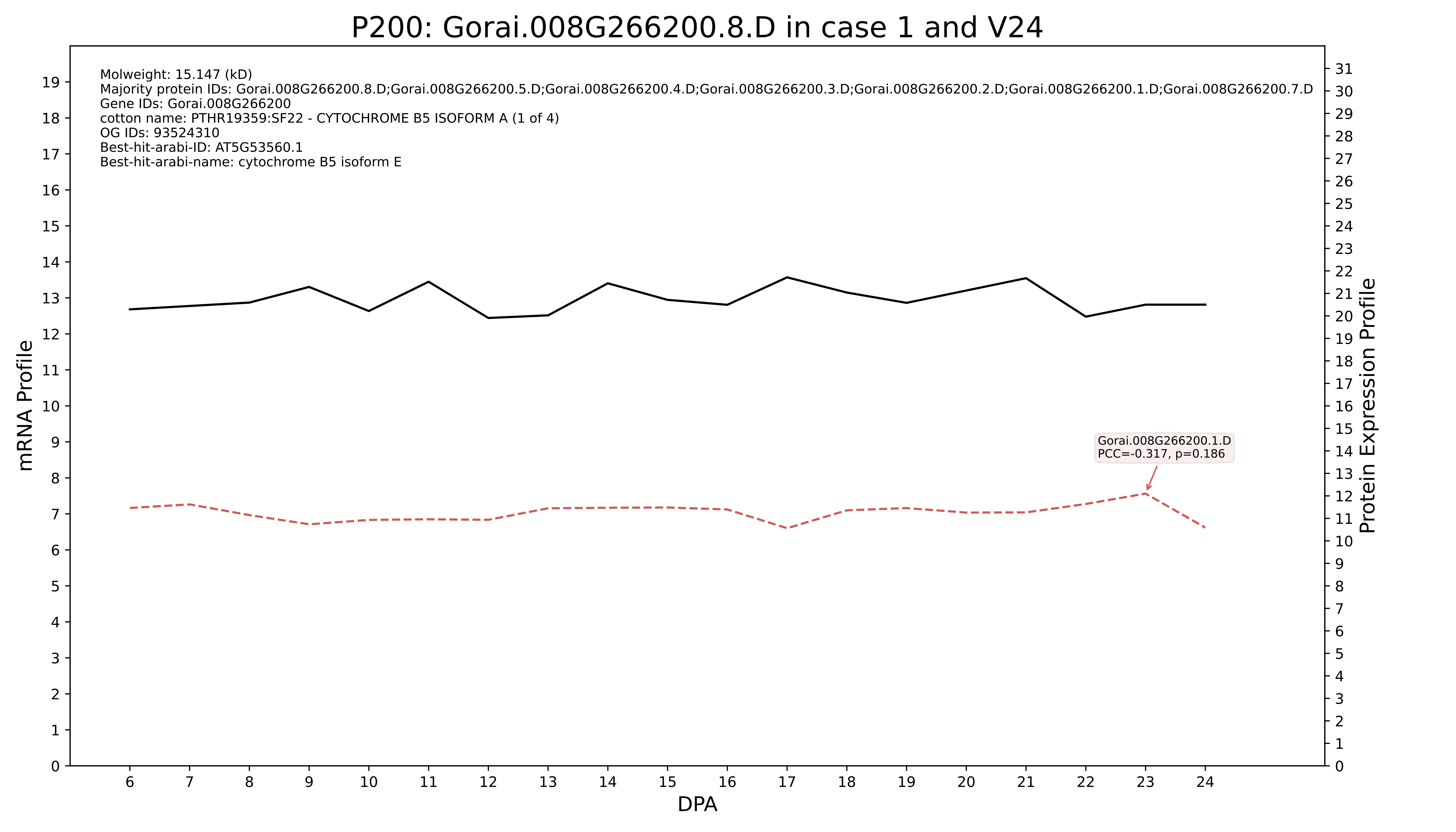

P200: Gorai.008G266200.8.D in case 1 and V24
Molweight: 15.147 (kD)
Majority protein IDs: Gorai.008G266200.8.D;Gorai.008G266200.5.D;Gorai.008G266200.4.D;Gorai.008G266200.3.D;Gorai.008G266200.2.D;Gorai.008G266200.1.D;Gorai.008G266200.7.D
Gene IDs: Gorai.008G266200
cotton name: PTHR19359:SF22 - CYTOCHROME B5 ISOFORM A (1 of 4)
OG IDs: 93524310
Best-hit-arabi-ID: AT5G53560.1
Best-hit-arabi-name: cytochrome B5 isoform E
Gorai.008G266200.1.D
PCC=-0.317, p=0.186
mRNA Profile
Protein Expression Profile
DPA

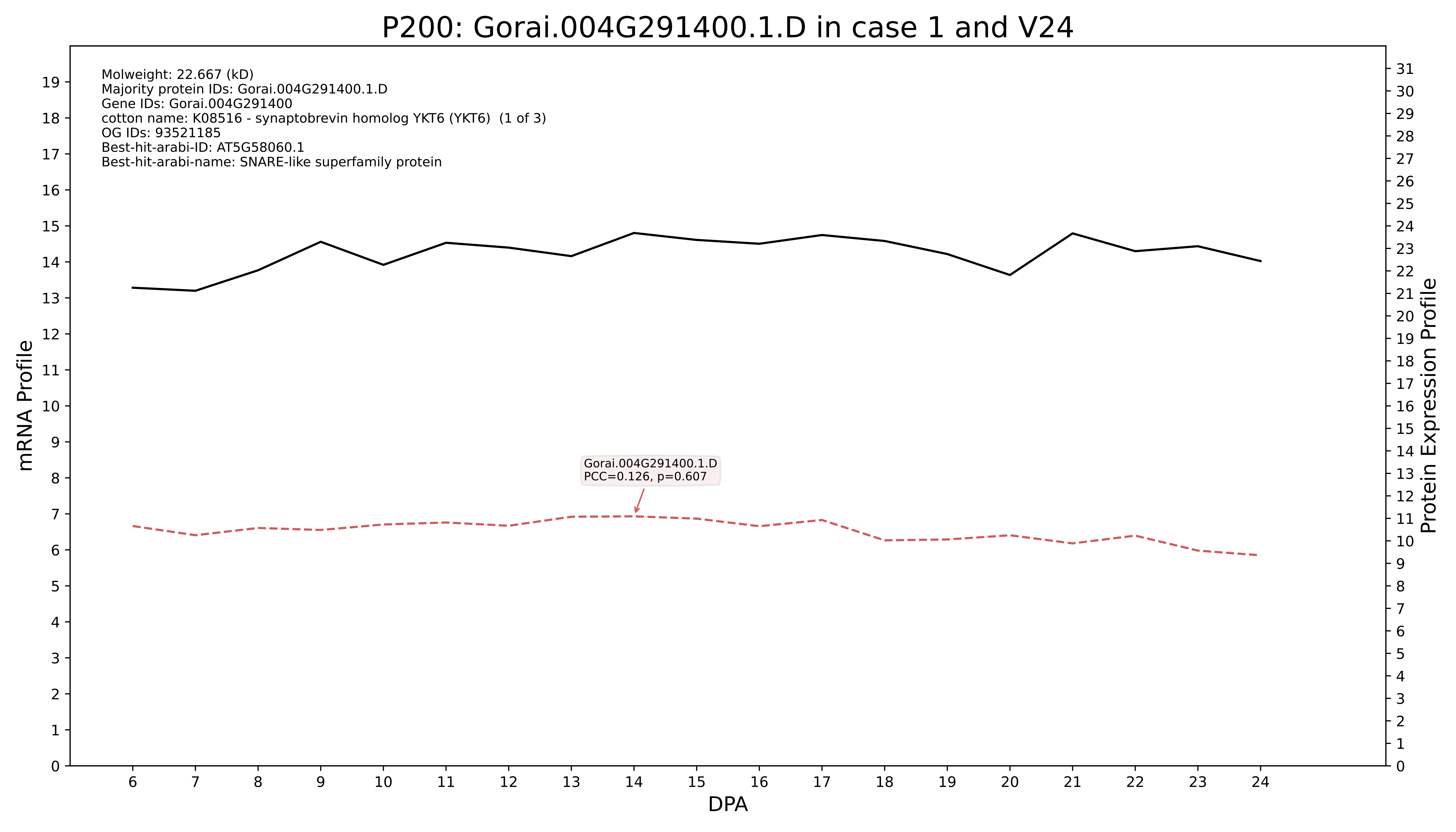
P200: Gorai.004G291400.1.D in case 1 and V24
Molweight: 22.667 (kD)
Majority protein IDs: Gorai.004G291400.1.D
Gene IDs: Gorai.004G291400
cotton name: K08516 - synaptobrevin homolog YKT6 (YKT6) (1 of 3)
OG IDs: 93521185
Best-hit-arabi-ID: AT5G58060.1
Best-hit-arabi-name: SNARE-like superfamily protein
Gorai.004G291400.1.D
PCC=0.126, p=0.607
mRNA Profile
Protein Expression Profile
DPA

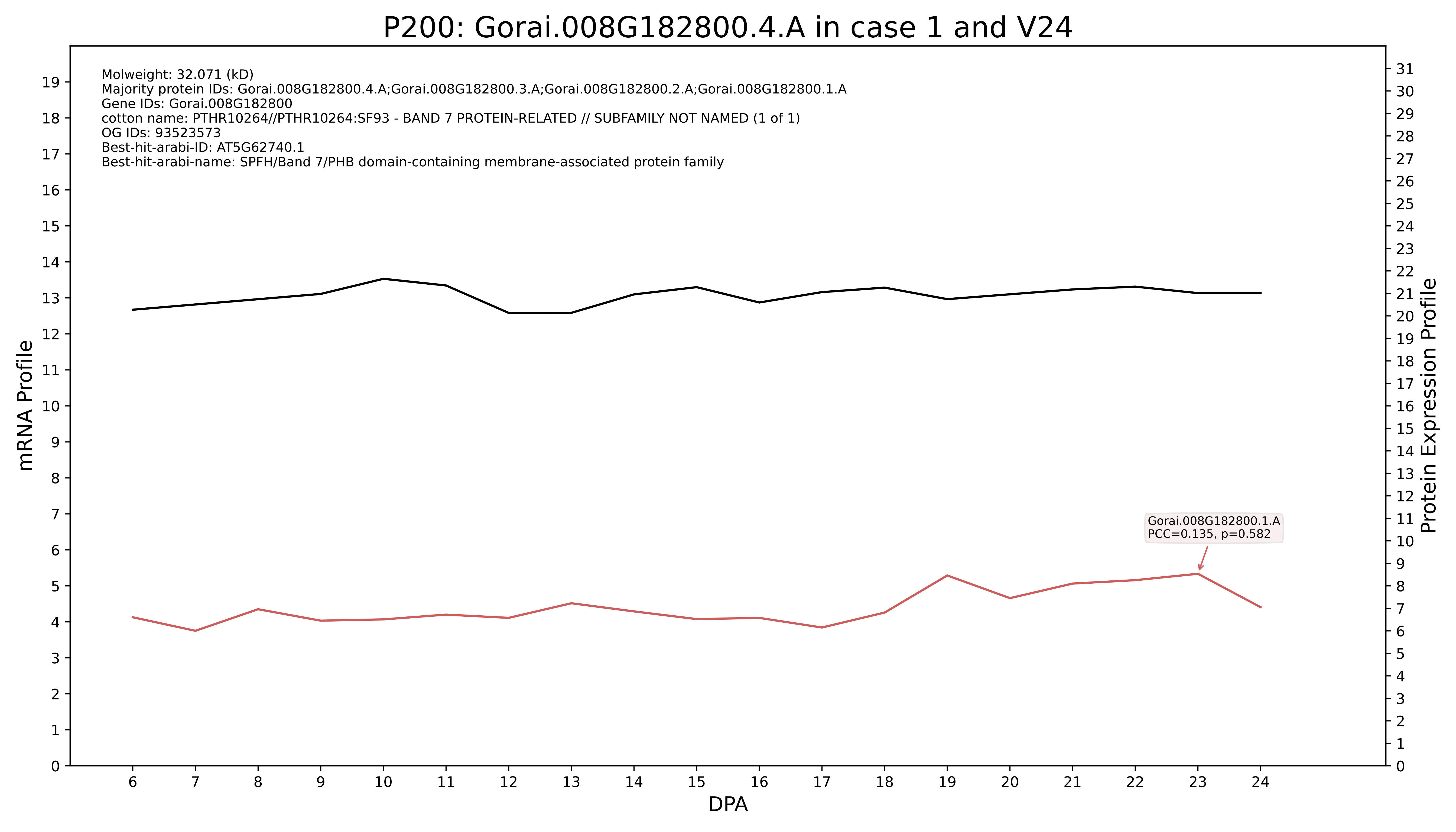
P200: Gorai.008G182800.4.A in case 1 and V24
Molweight: 32.071 (kD)
Majority protein IDs: Gorai.008G182800.4.A;Gorai.008G182800.3.A;Gorai.008G182800.2.A;Gorai.008G182800.1.A
Gene IDs: Gorai.008G182800
cotton name: PTHR10264//PTHR10264:SF93 - BAND 7 PROTEIN-RELATED // SUBFAMILY NOT NAMED (1 of 1)
OG IDs: 93523573
Best-hit-arabi-ID: AT5G62740.1
Best-hit-arabi-name: SPFH/Band 7/PHB domain-containing membrane-associated protein family
Gorai.008G182800.1.A
PCC=0.135, p=0.582
mRNA Profile
Protein Expression Profile
DPA

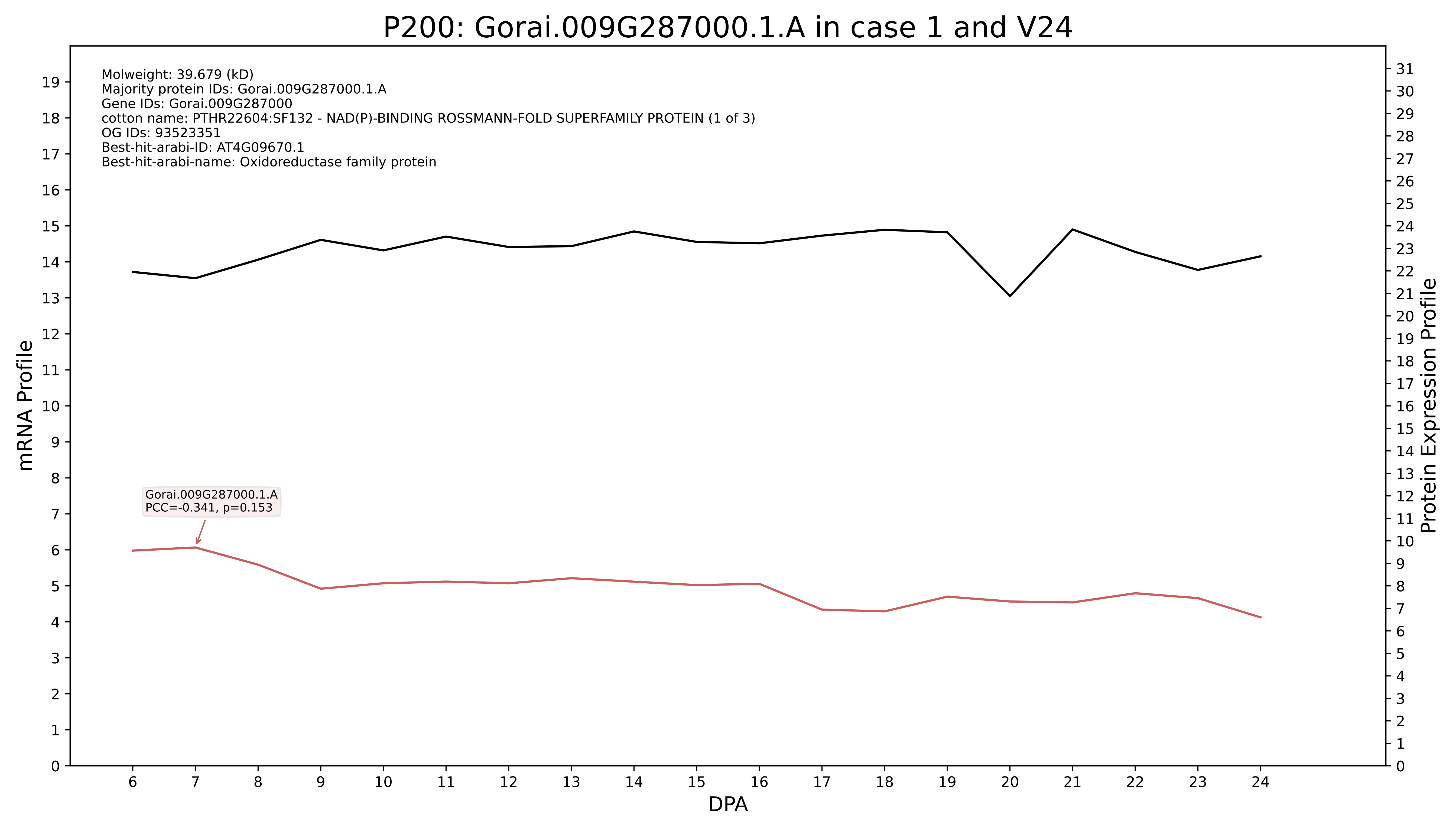
P200: Gorai.009G287000.1.A in case 1 and V24
Molweight: 39.679 (kD)
Majority protein IDs: Gorai.009G287000.1.A
Gene IDs: Gorai.009G287000
cotton name: PTHR22604:SF132 - NAD(P)-BINDING ROSSMANN-FOLD SUPERFAMILY PROTEIN (1 of 3)
OG IDs: 93523351
Best-hit-arabi-ID: AT4G09670.1
Best-hit-arabi-name: Oxidoreductase family protein
Gorai.009G287000.1.A
PCC=-0.341, p=0.153
mRNA Profile
Protein Expression Profile
DPA

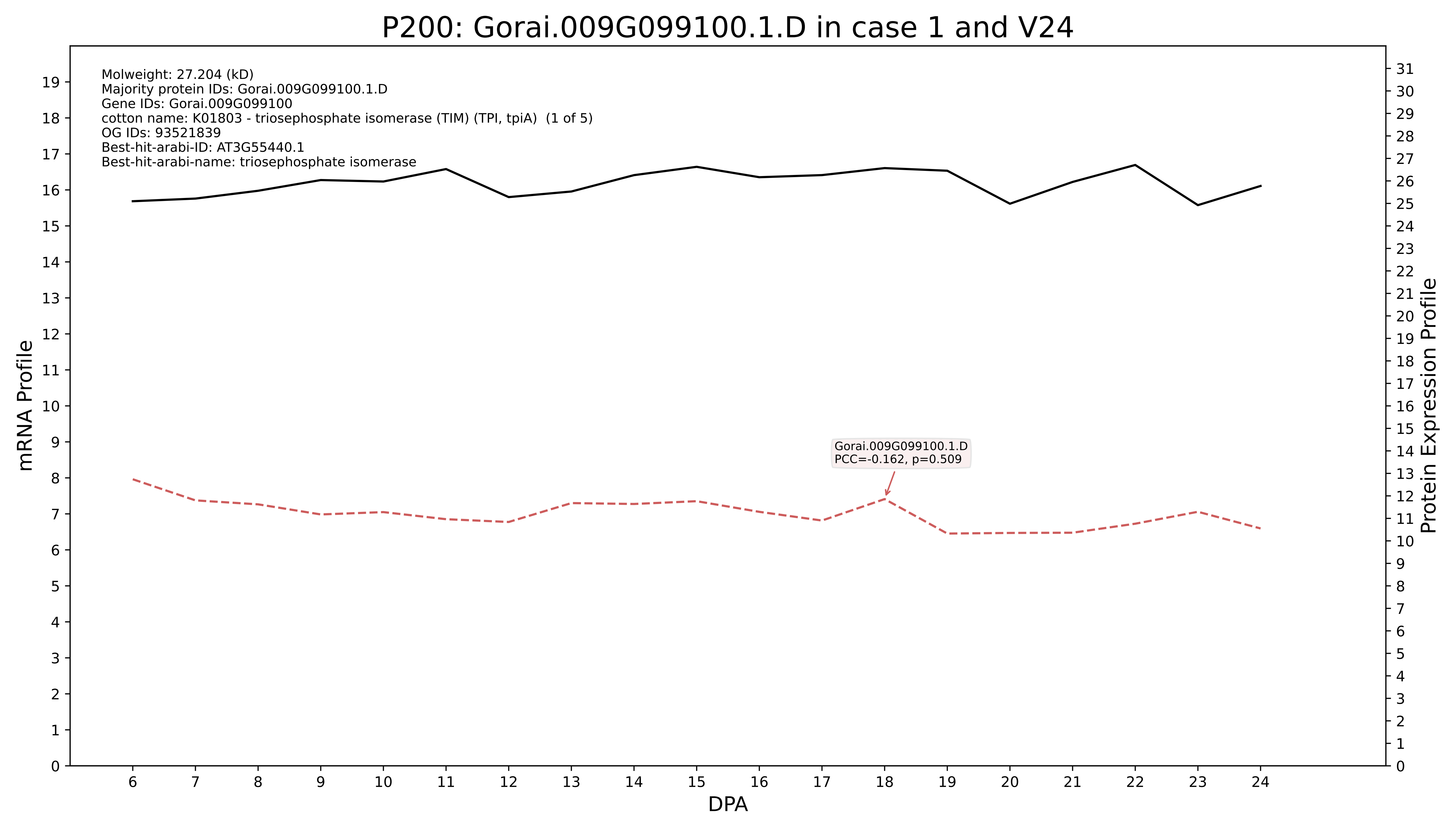
P200: Gorai.009G099100.1.D in case 1 and V24
Molweight: 27.204 (kD)
Majority protein IDs: Gorai.009G099100.1.D
Gene IDs: Gorai.009G099100
cotton name: K01803 - triosephosphate isomerase (TIM) (TPI, tpiA) (1 of 5)
OG IDs: 93521839
Best-hit-arabi-ID: AT3G55440.1
Best-hit-arabi-name: triosephosphate isomerase
Gorai.009G099100.1.D
PCC=-0.162, p=0.509
mRNA Profile
Protein Expression Profile
DPA

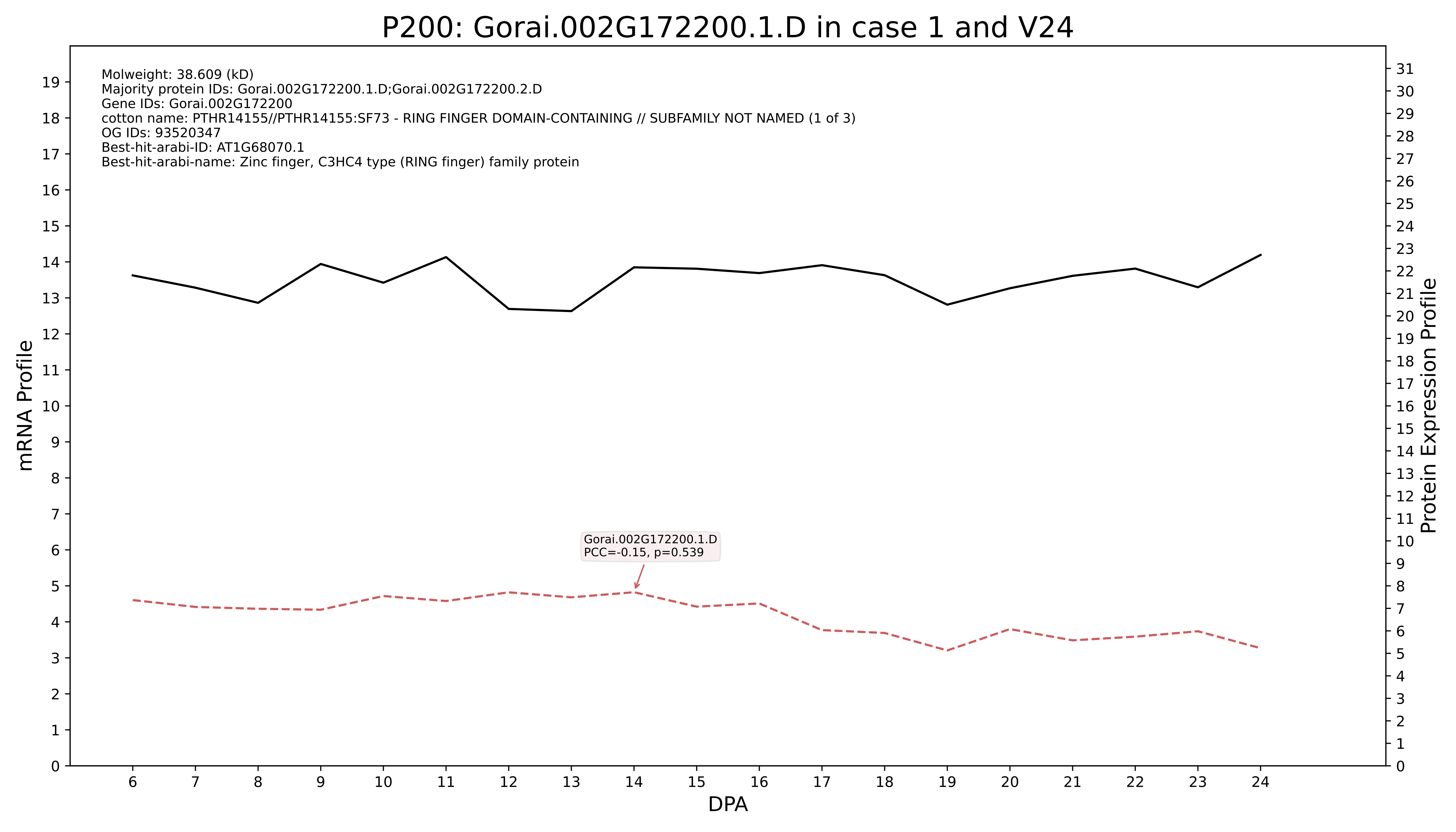

P200: Gorai.002G172200.1.D in case 1 and V24
Molweight: 38.609 (kD)
Majority protein IDs: Gorai.002G172200.1.D;Gorai.002G172200.2.D
Gene IDs: Gorai.002G172200
cotton name: PTHR14155//PTHR14155:SF73 - RING FINGER DOMAIN-CONTAINING // SUBFAMILY NOT NAMED (1 of 3)
OG IDs: 93520347
Best-hit-arabi-ID: AT1G68070.1
Best-hit-arabi-name: Zinc finger, C3HC4 type (RING finger) family protein
Gorai.002G172200.1.D
PCC=-0.15, p=0.539
mRNA Profile
Protein Expression Profile
DPA

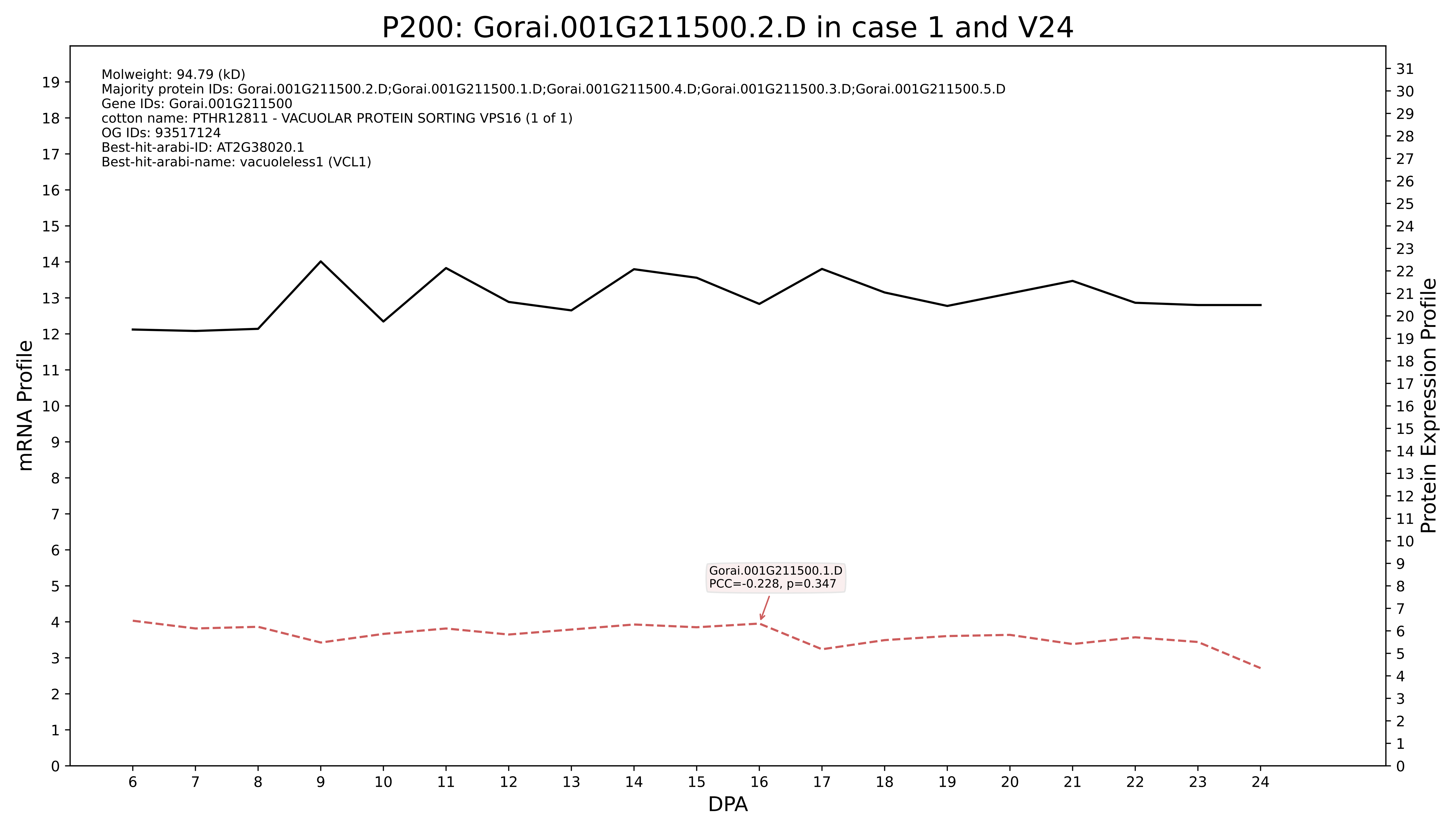
P200: Gorai.001G211500.2.D in case 1 and V24
Molweight: 94.79 (kD)
Majority protein IDs: Gorai.001G211500.2.D;Gorai.001G211500.1.D;Gorai.001G211500.4.D;Gorai.001G211500.3.D;Gorai.001G211500.5.D
Gene IDs: Gorai.001G211500
cotton name: PTHR12811 - VACUOLAR PROTEIN SORTING VPS16 (1 of 1)
OG IDs: 93517124
Best-hit-arabi-ID: AT2G38020.1
Best-hit-arabi-name: vacuoleless1 (VCL1)
Gorai.001G211500.1.D
PCC=-0.228, p=0.347
mRNA Profile
Protein Expression Profile
DPA

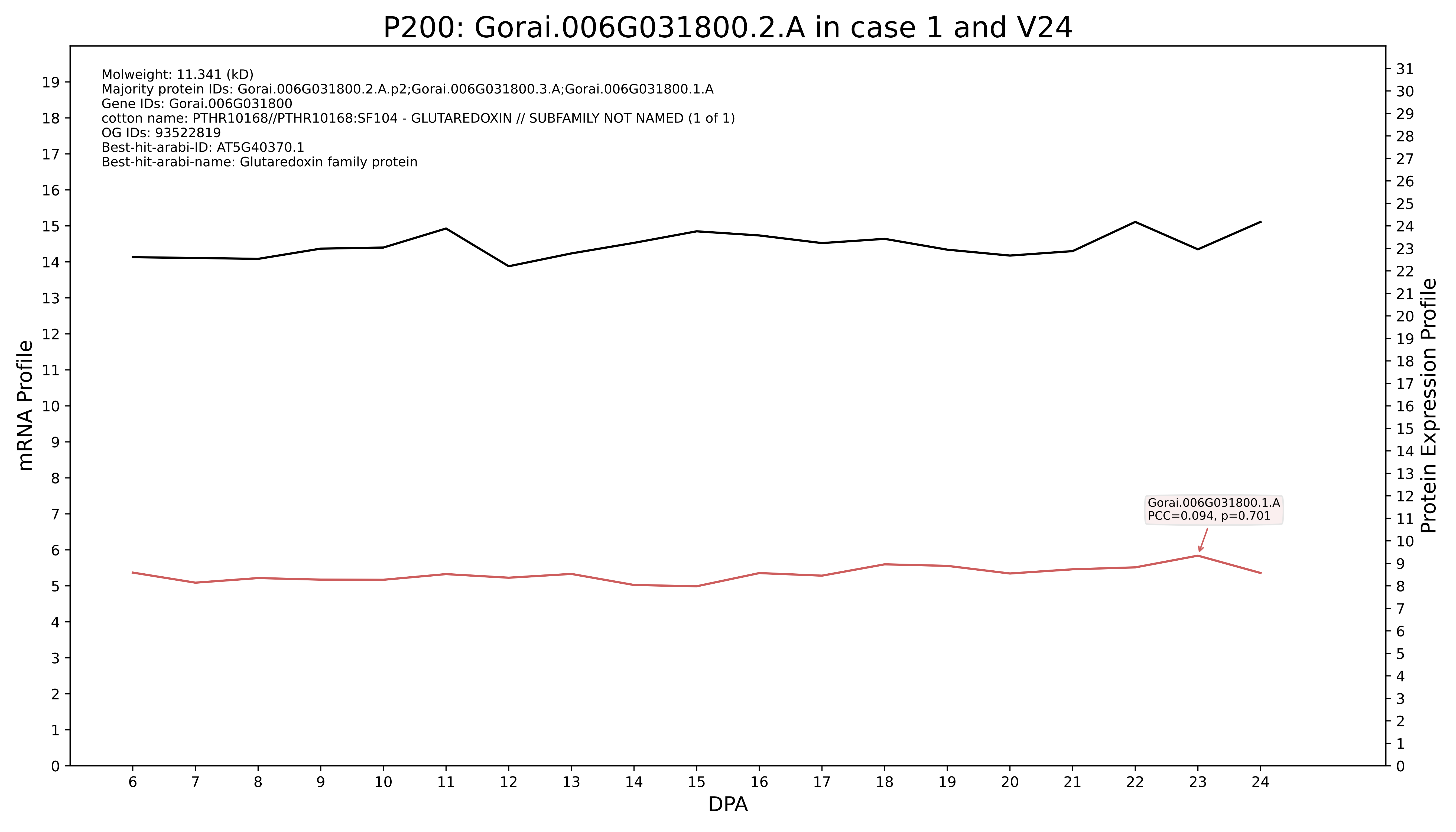
P200: Gorai.006G031800.2.A in case 1 and V24
Molweight: 11.341 (kD)
Majority protein IDs: Gorai.006G031800.2.A.p2;Gorai.006G031800.3.A;Gorai.006G031800.1.A
Gene IDs: Gorai.006G031800
cotton name: PTHR10168//PTHR10168:SF104 - GLUTAREDOXIN // SUBFAMILY NOT NAMED (1 of 1)
OG IDs: 93522819
Best-hit-arabi-ID: AT5G40370.1
Best-hit-arabi-name: Glutaredoxin family protein
Gorai.006G031800.1.A
PCC=0.094, p=0.701
mRNA Profile
Protein Expression Profile
DPA

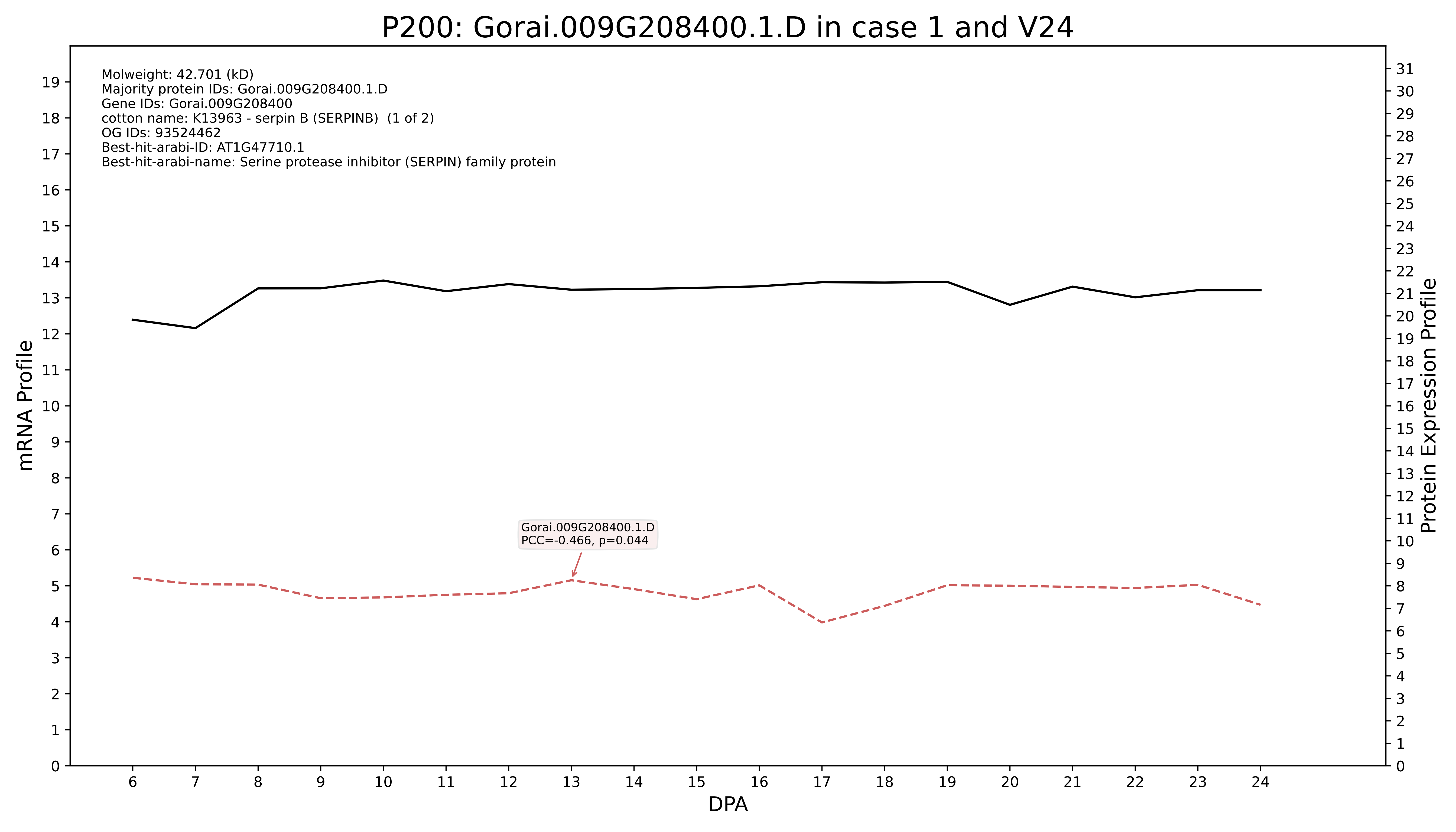
P200: Gorai.009G208400.1.D in case 1 and V24
Molweight: 42.701 (kD)
Majority protein IDs: Gorai.009G208400.1.D
Gene IDs: Gorai.009G208400
cotton name: K13963 - serpin B (SERPINB) (1 of 2)
OG IDs: 93524462
Best-hit-arabi-ID: AT1G47710.1
Best-hit-arabi-name: Serine protease inhibitor (SERPIN) family protein
Gorai.009G208400.1.D
PCC=-0.466, p=0.044
mRNA Profile
Protein Expression Profile
DPA

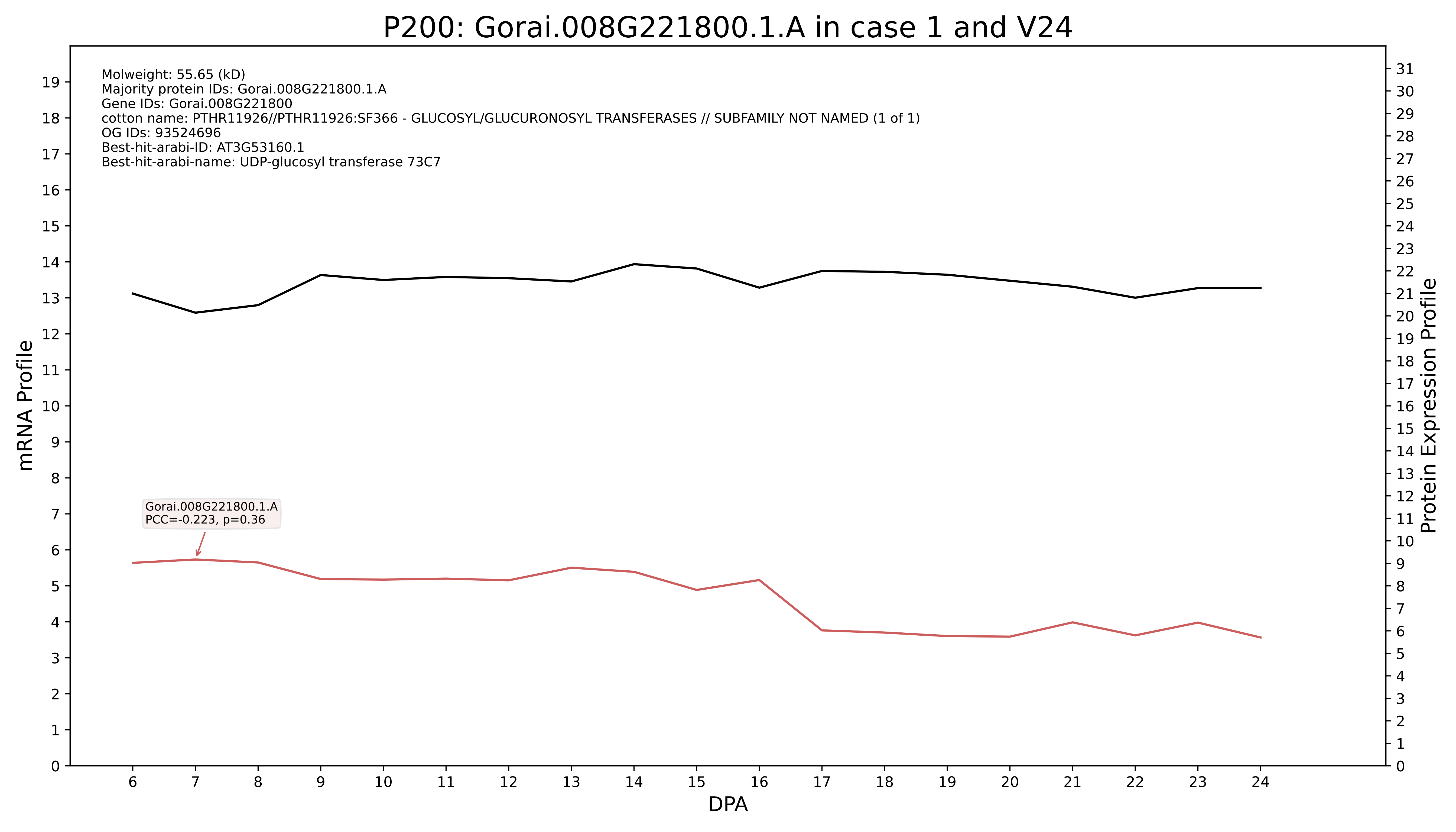
P200: Gorai.008G221800.1.A in case 1 and V24
Molweight: 55.65 (kD)
Majority protein IDs: Gorai.008G221800.1.A
Gene IDs: Gorai.008G221800
cotton name: PTHR11926//PTHR11926:SF366 - GLUCOSYL/GLUCURONOSYL TRANSFERASES // SUBFAMILY NOT NAMED (1 of 1)
OG IDs: 93524696
Best-hit-arabi-ID: AT3G53160.1
Best-hit-arabi-name: UDP-glucosyl transferase 73C7
Gorai.008G221800.1.A
PCC=-0.223, p=0.36
mRNA Profile
Protein Expression Profile
DPA

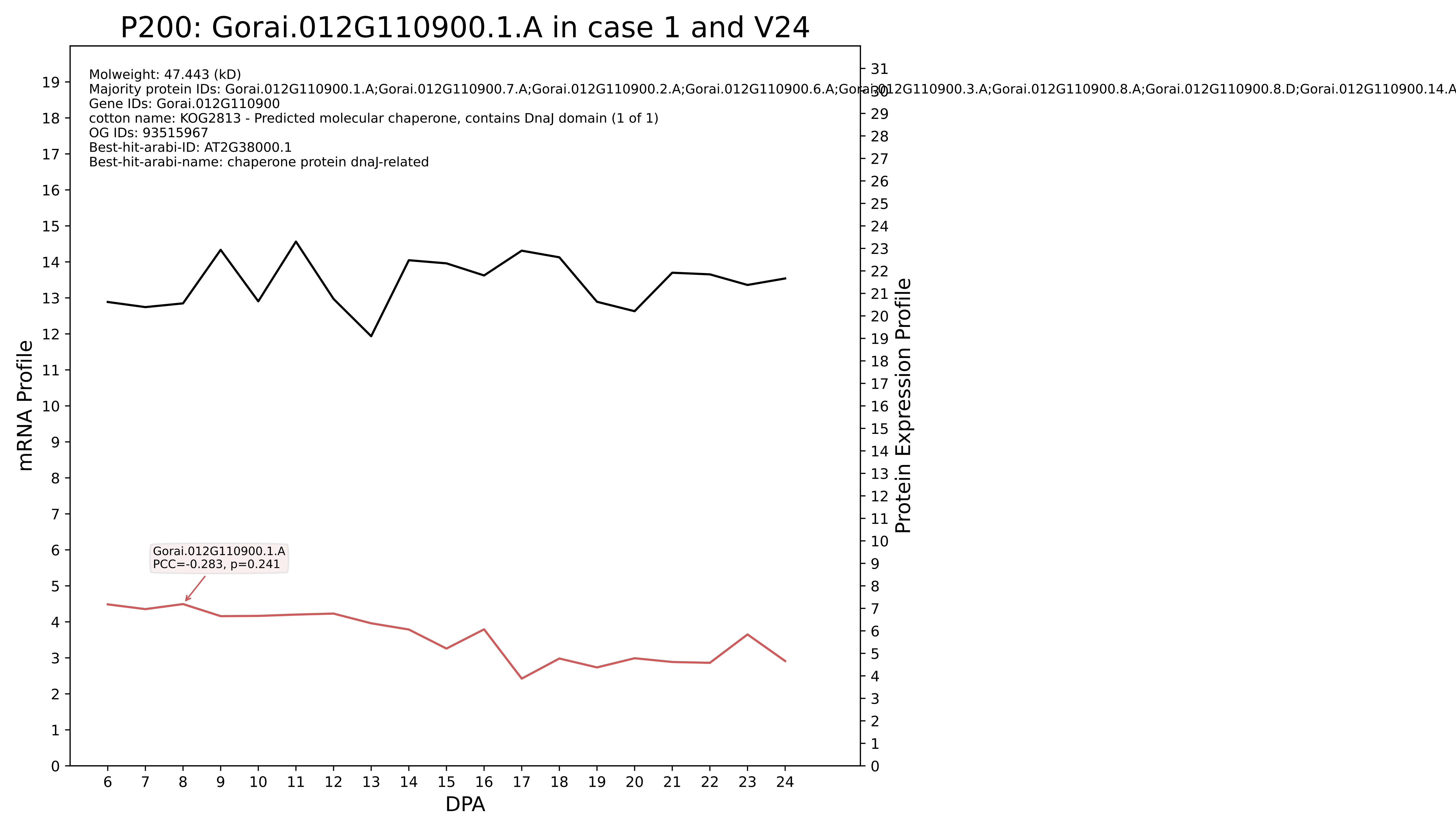
P200: Gorai.012G110900.1.A in case 1 and V24
Molweight: 47.443 (kD)
Majority protein IDs: Gorai.012G110900.1.A;Gorai.012G110900.7.A;Gorai.012G110900.2.A;Gorai.012G110900.6.A;Gorai.012G110900.3.A;Gorai.012G110900.8.A;Gorai.012G110900.8.D;Gorai.012G110900.14.A
Gene IDs: Gorai.012G110900
cotton name: KOG2813 - Predicted molecular chaperone, contains DnaJ domain (1 of 1)
OG IDs: 93515967
Best-hit-arabi-ID: AT2G38000.1
Best-hit-arabi-name: chaperone protein dnaJ-related
Gorai.012G110900.1.A
PCC=-0.283, p=0.241
mRNA Profile
Protein Expression Profile
DPA

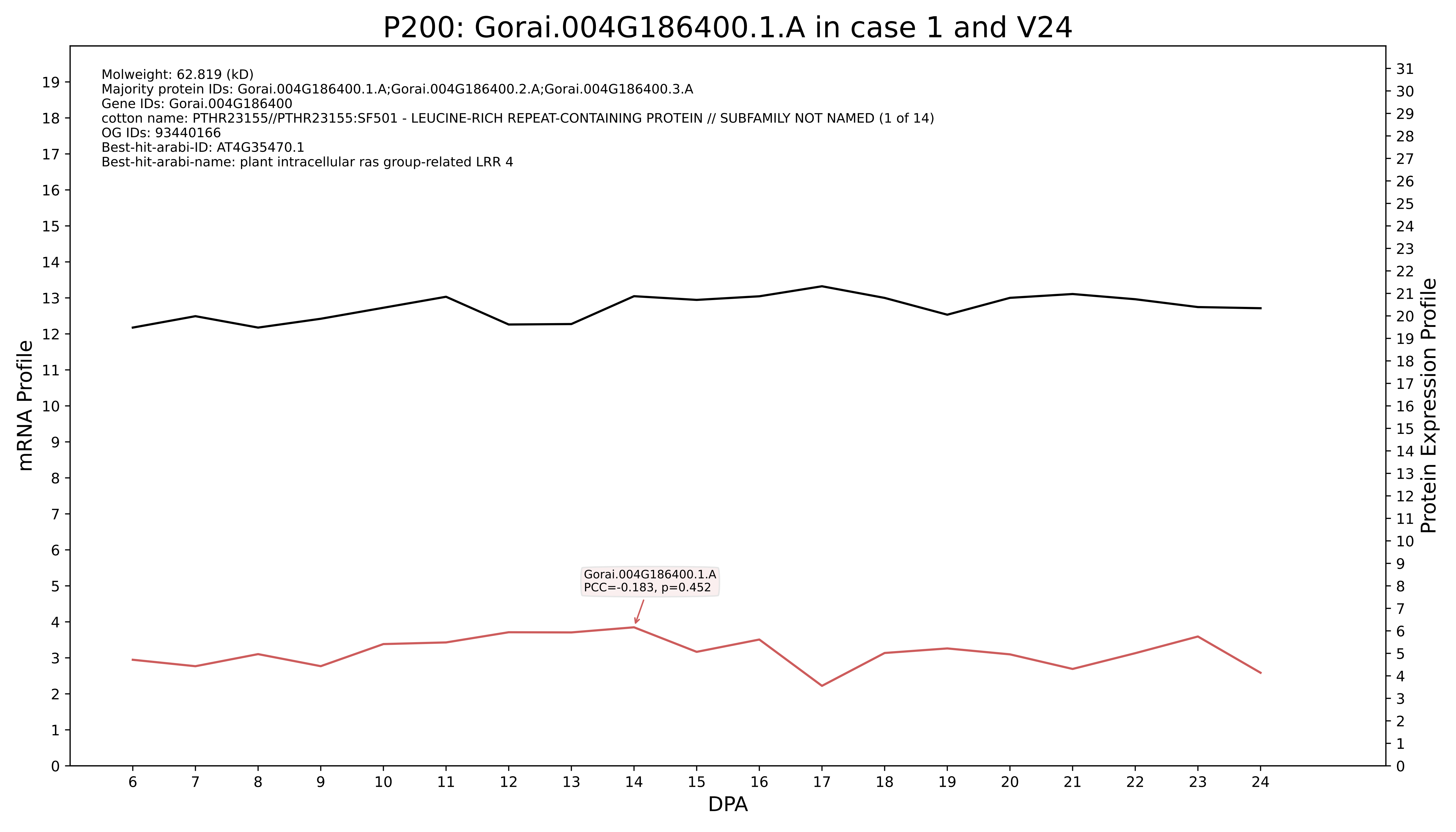

P200: Gorai.004G186400.1.A in case 1 and V24
Molweight: 62.819 (kD)
Majority protein IDs: Gorai.004G186400.1.A;Gorai.004G186400.2.A;Gorai.004G186400.3.A
Gene IDs: Gorai.004G186400
cotton name: PTHR23155//PTHR23155:SF501 - LEUCINE-RICH REPEAT-CONTAINING PROTEIN // SUBFAMILY NOT NAMED (1 of 14)
OG IDs: 93440166
Best-hit-arabi-ID: AT4G35470.1
Best-hit-arabi-name: plant intracellular ras group-related LRR 4
Gorai.004G186400.1.A
PCC=-0.183, p=0.452
mRNA Profile
Protein Expression Profile
DPA

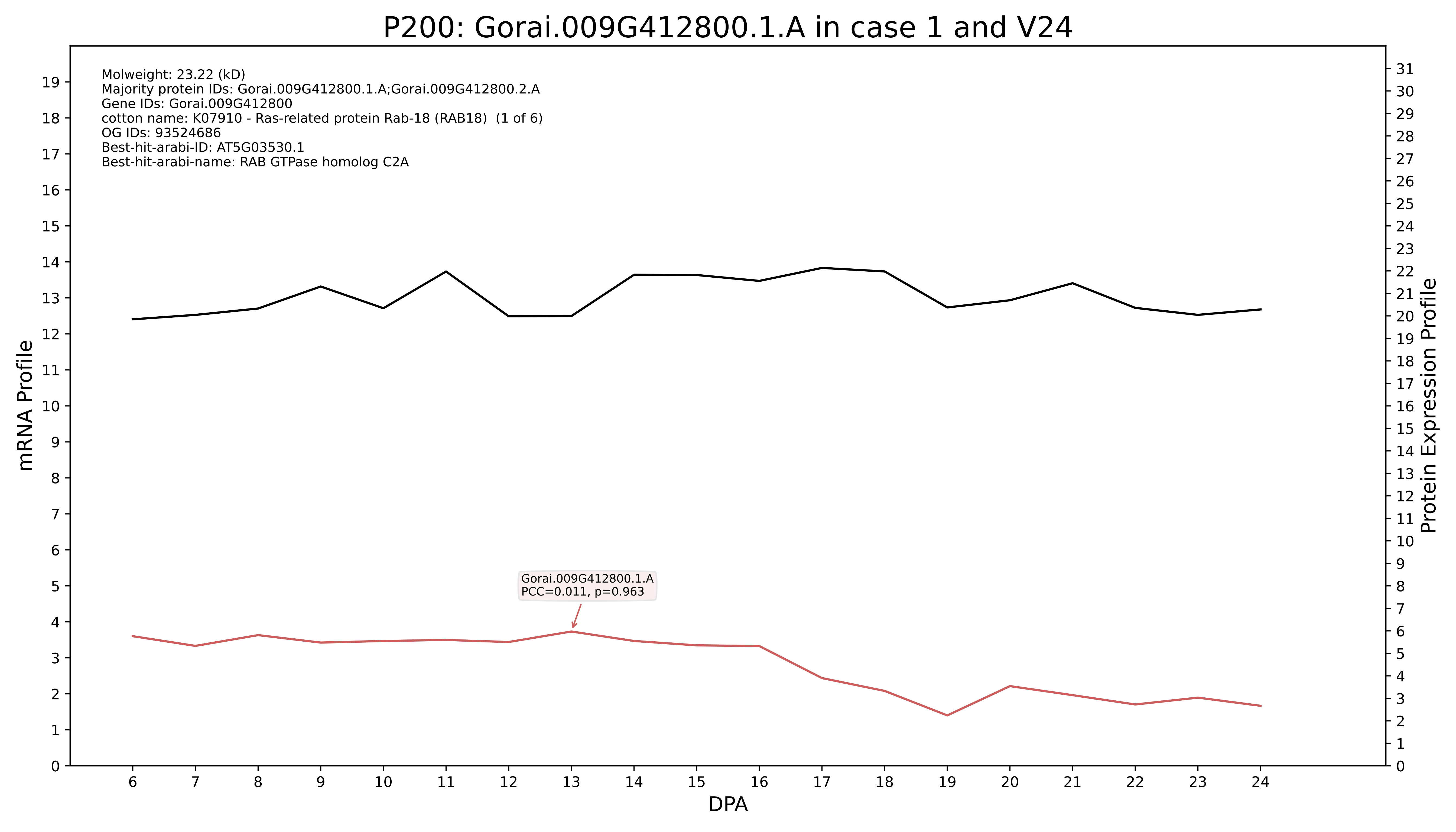
P200: Gorai.009G412800.1.A in case 1 and V24
Molweight: 23.22 (kD)
Majority protein IDs: Gorai.009G412800.1.A;Gorai.009G412800.2.A
Gene IDs: Gorai.009G412800
cotton name: K07910 - Ras-related protein Rab-18 (RAB18) (1 of 6)
OG IDs: 93524686
Best-hit-arabi-ID: AT5G03530.1
Best-hit-arabi-name: RAB GTPase homolog C2A
Gorai.009G412800.1.A
PCC=0.011, p=0.963
mRNA Profile
Protein Expression Profile
DPA

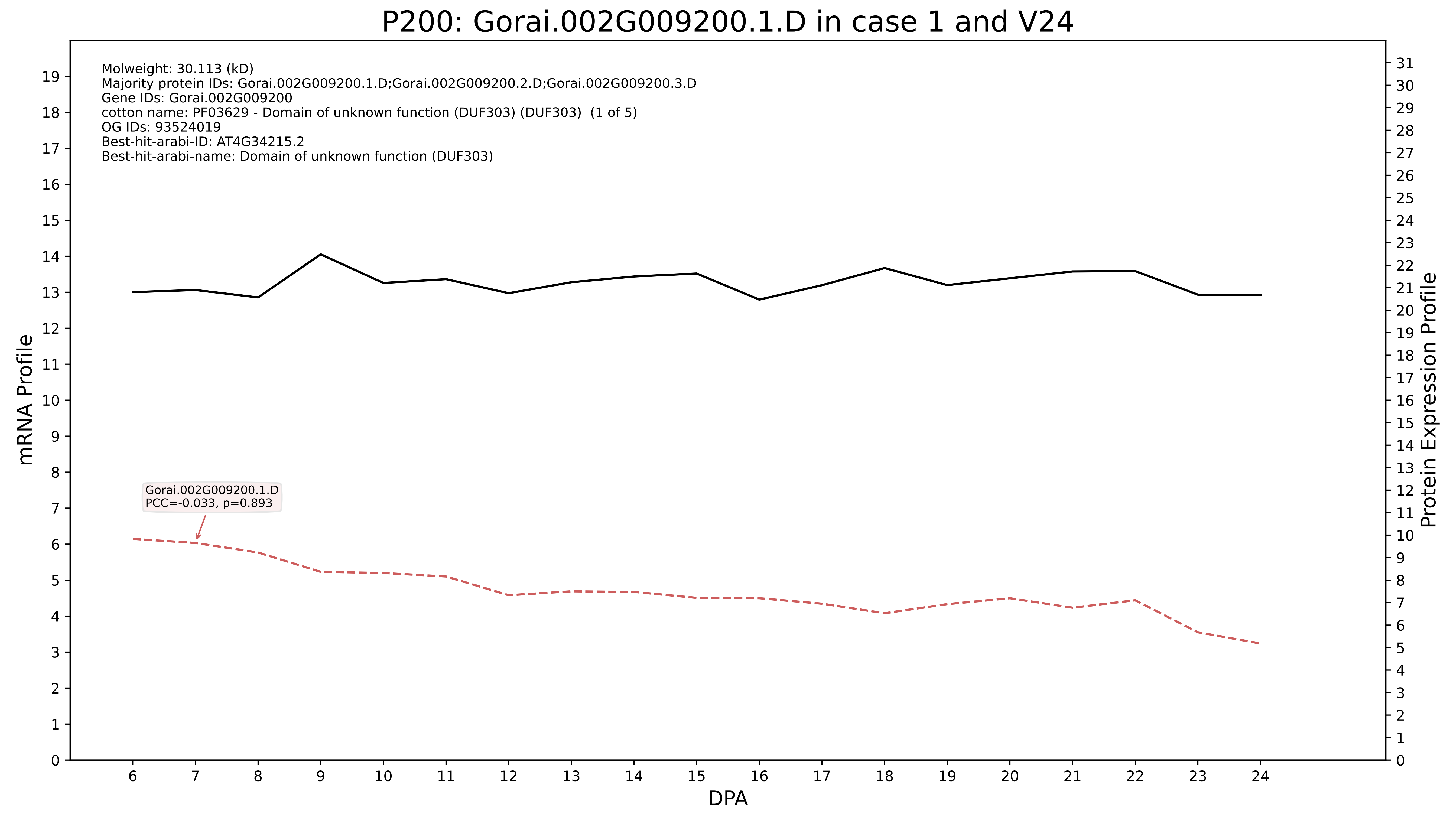
P200: Gorai.002G009200.1.D in case 1 and V24
Molweight: 30.113 (kD)
Majority protein IDs: Gorai.002G009200.1.D;Gorai.002G009200.2.D;Gorai.002G009200.3.D
Gene IDs: Gorai.002G009200
cotton name: PF03629 - Domain of unknown function (DUF303) (DUF303) (1 of 5)
OG IDs: 93524019
Best-hit-arabi-ID: AT4G34215.2
Best-hit-arabi-name: Domain of unknown function (DUF303)
Gorai.002G009200.1.D
PCC=-0.033, p=0.893
mRNA Profile
Protein Expression Profile
DPA

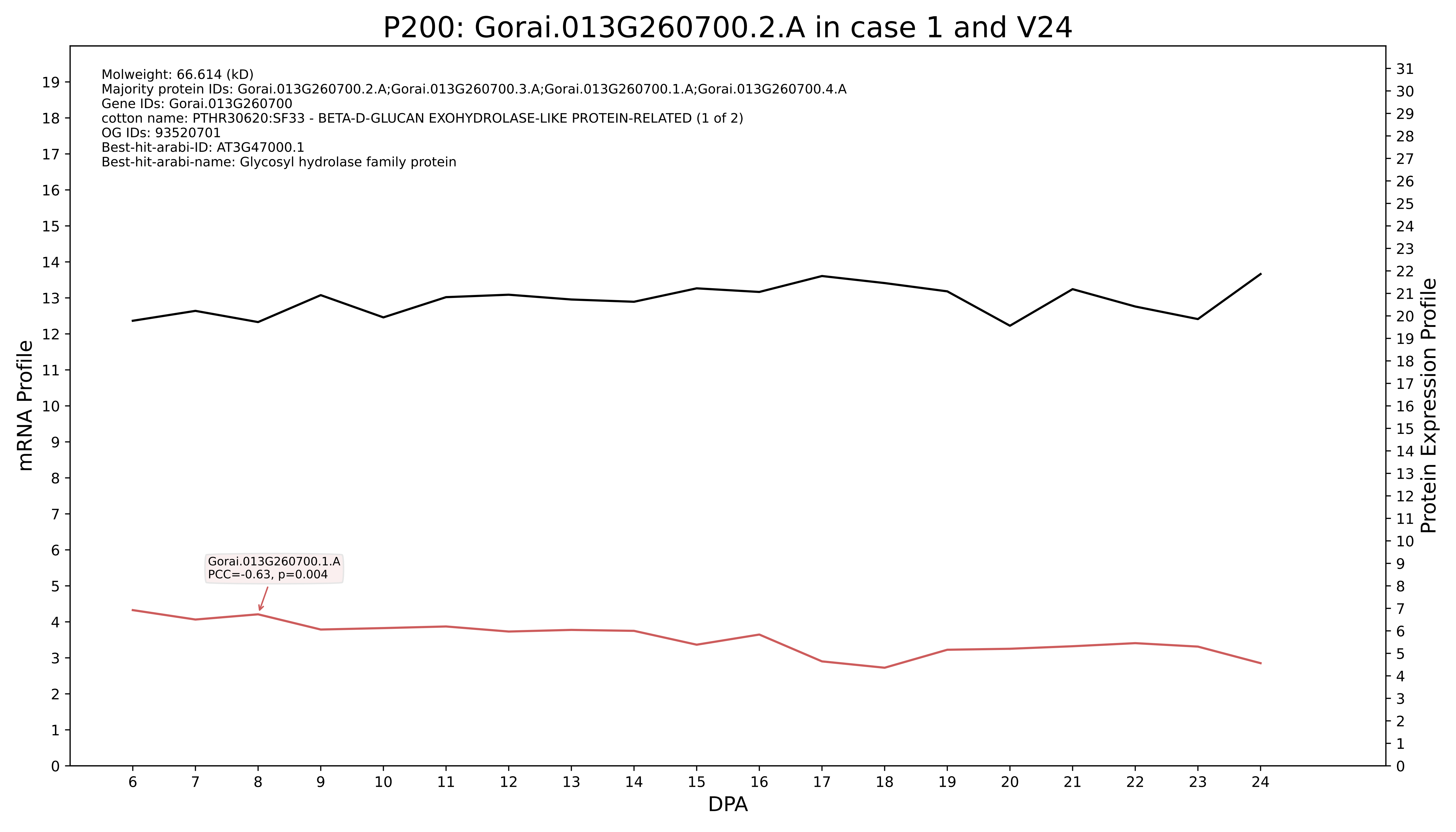

P200: Gorai.013G260700.2.A in case 1 and V24
Molweight: 66.614 (kD)
Majority protein IDs: Gorai.013G260700.2.A;Gorai.013G260700.3.A;Gorai.013G260700.1.A;Gorai.013G260700.4.A
Gene IDs: Gorai.013G260700
cotton name: PTHR30620:SF33 - BETA-D-GLUCAN EXOHYDROLASE-LIKE PROTEIN-RELATED (1 of 2)
OG IDs: 93520701
Best-hit-arabi-ID: AT3G47000.1
Best-hit-arabi-name: Glycosyl hydrolase family protein
Gorai.013G260700.1.A
PCC=-0.63, p=0.004
mRNA Profile
Protein Expression Profile
DPA

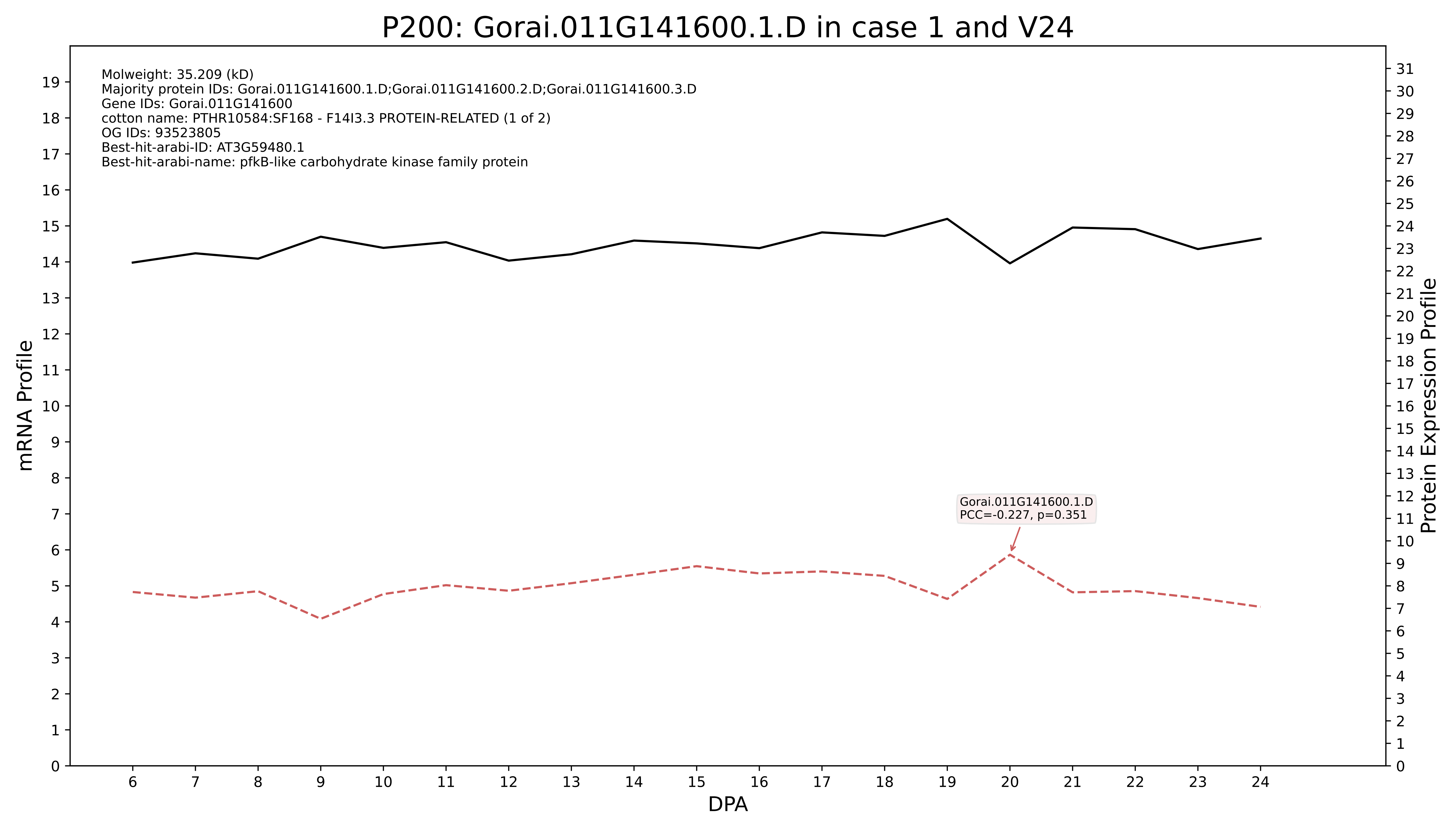

P200: Gorai.011G141600.1.D in case 1 and V24
Molweight: 35.209 (kD)
Majority protein IDs: Gorai.011G141600.1.D;Gorai.011G141600.2.D;Gorai.011G141600.3.D
Gene IDs: Gorai.011G141600
cotton name: PTHR10584:SF168 - F14I3.3 PROTEIN-RELATED (1 of 2)
OG IDs: 93523805
Best-hit-arabi-ID: AT3G59480.1
Best-hit-arabi-name: pfkB-like carbohydrate kinase family protein
Gorai.011G141600.1.D
PCC=-0.227, p=0.351
mRNA Profile
Protein Expression Profile
DPA

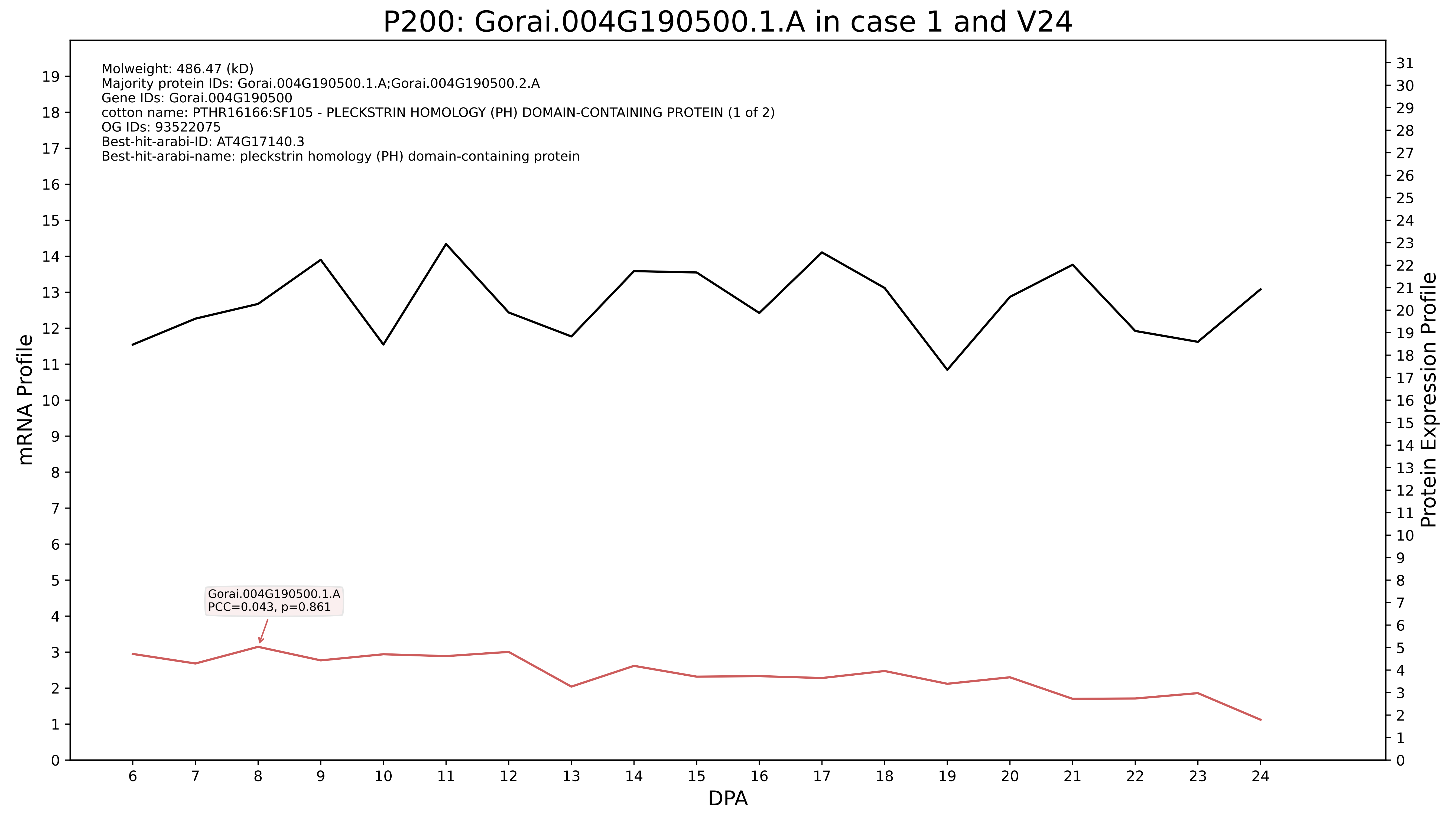
P200: Gorai.004G190500.1.A in case 1 and V24
Molweight: 486.47 (kD)
Majority protein IDs: Gorai.004G190500.1.A;Gorai.004G190500.2.A
Gene IDs: Gorai.004G190500
cotton name: PTHR16166:SF105 - PLECKSTRIN HOMOLOGY (PH) DOMAIN-CONTAINING PROTEIN (1 of 2)
OG IDs: 93522075
Best-hit-arabi-ID: AT4G17140.3
Best-hit-arabi-name: pleckstrin homology (PH) domain-containing protein
Gorai.004G190500.1.A
PCC=0.043, p=0.861
mRNA Profile
Protein Expression Profile
DPA

### P200: Gorai.006G095400.2.D in case 1 and V24

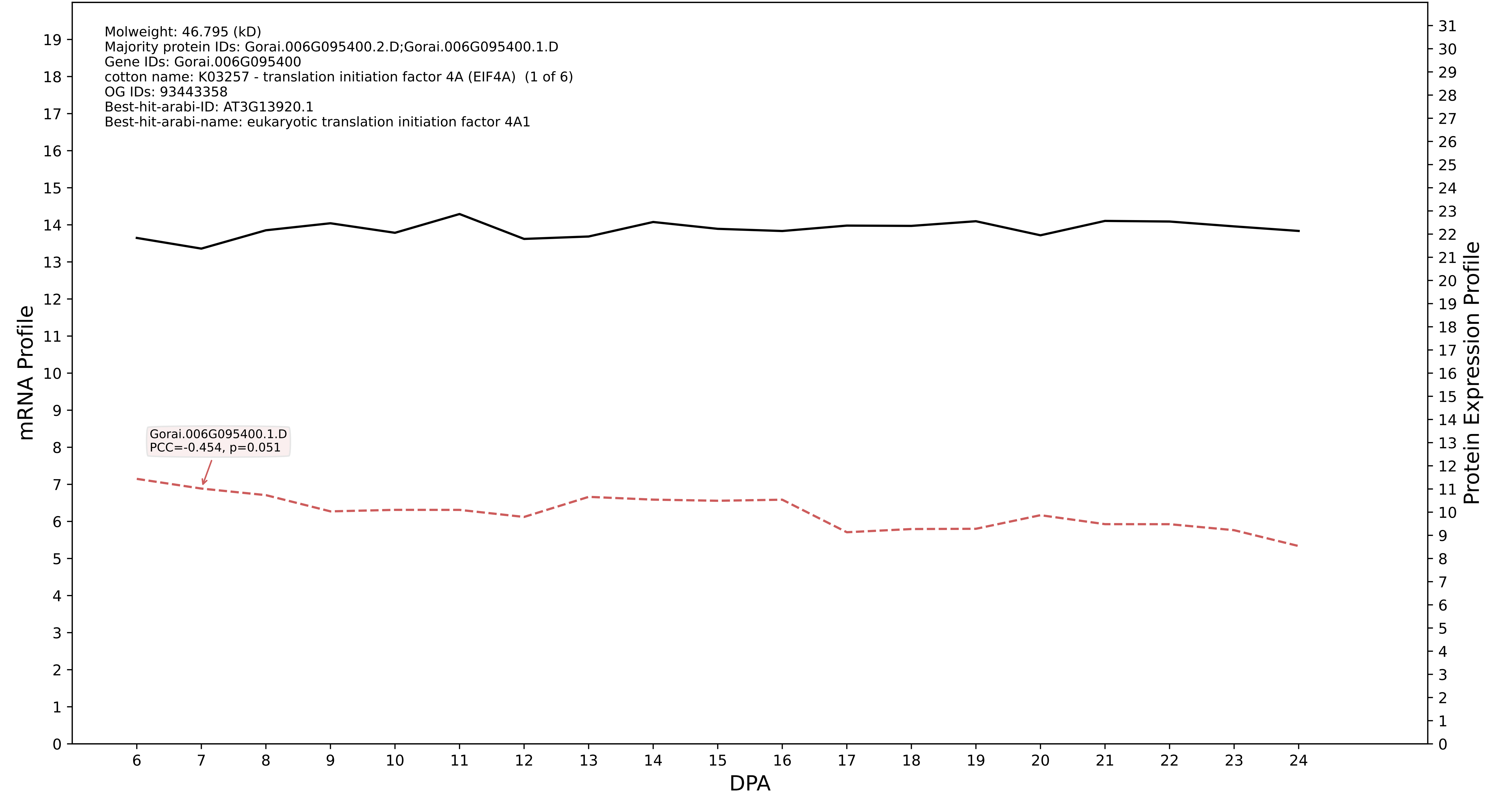

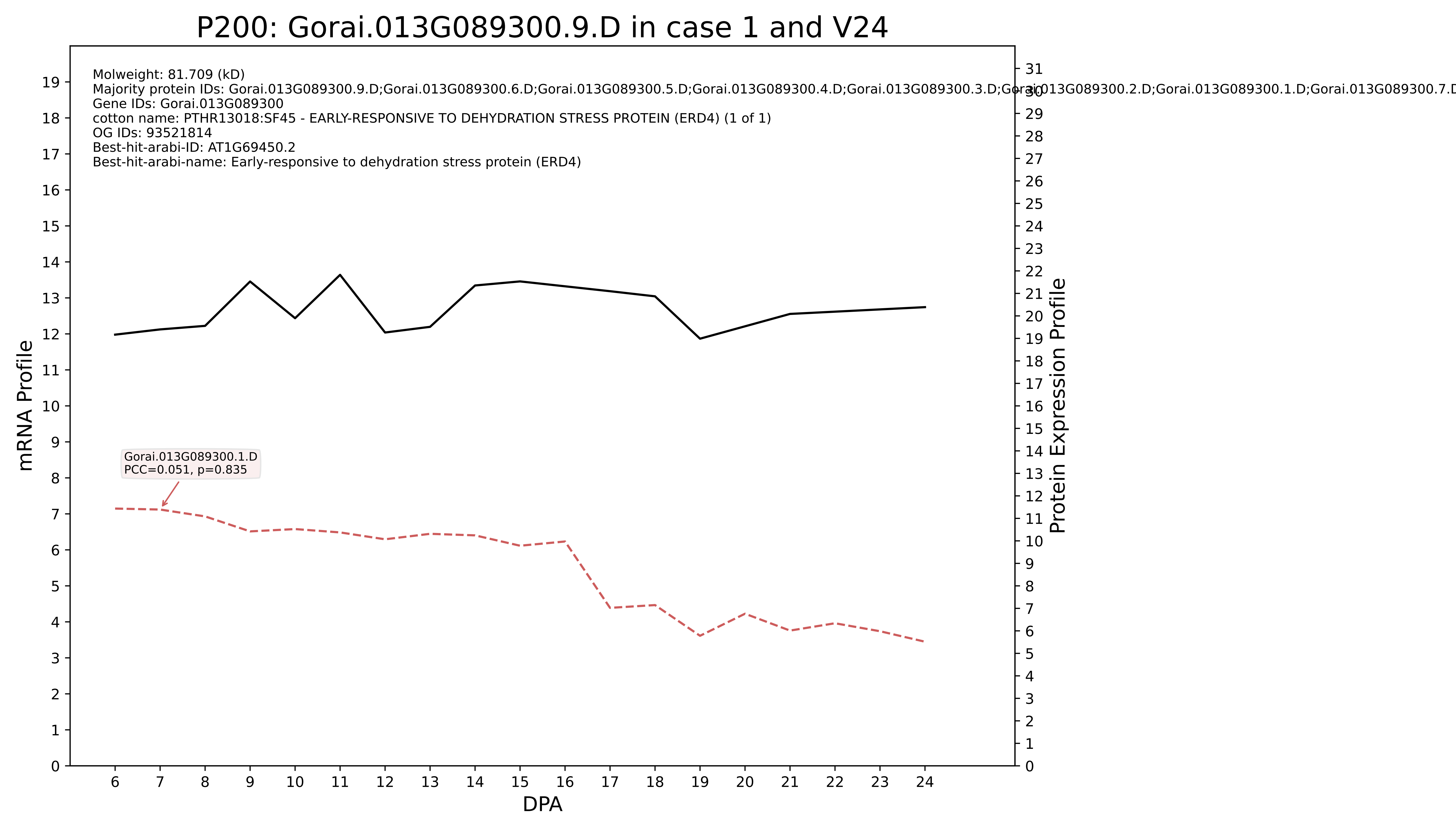
P200: Gorai.013G089300.9.D in case 1 and V24
Molweight: 81.709 (kD)
Majority protein IDs: Gorai.013G089300.9.D;Gorai.013G089300.6.D;Gorai.013G089300.5.D;Gorai.013G089300.4.D;Gorai.013G089300.3.D;Gorai.013G089300.2.D;Gorai.013G089300.1.D;Gorai.013G089300.7.D
Gene IDs: Gorai.013G089300
cotton name: PTHR13018:SF45 - EARLY-RESPONSIVE TO DEHYDRATION STRESS PROTEIN (ERD4) (1 of 1)
OG IDs: 93521814
Best-hit-arabi-ID: AT1G69450.2
Best-hit-arabi-name: Early-responsive to dehydration stress protein (ERD4)
Gorai.013G089300.1.D
PCC=0.051, p=0.835
mRNA Profile
Protein Expression Profile
DPA

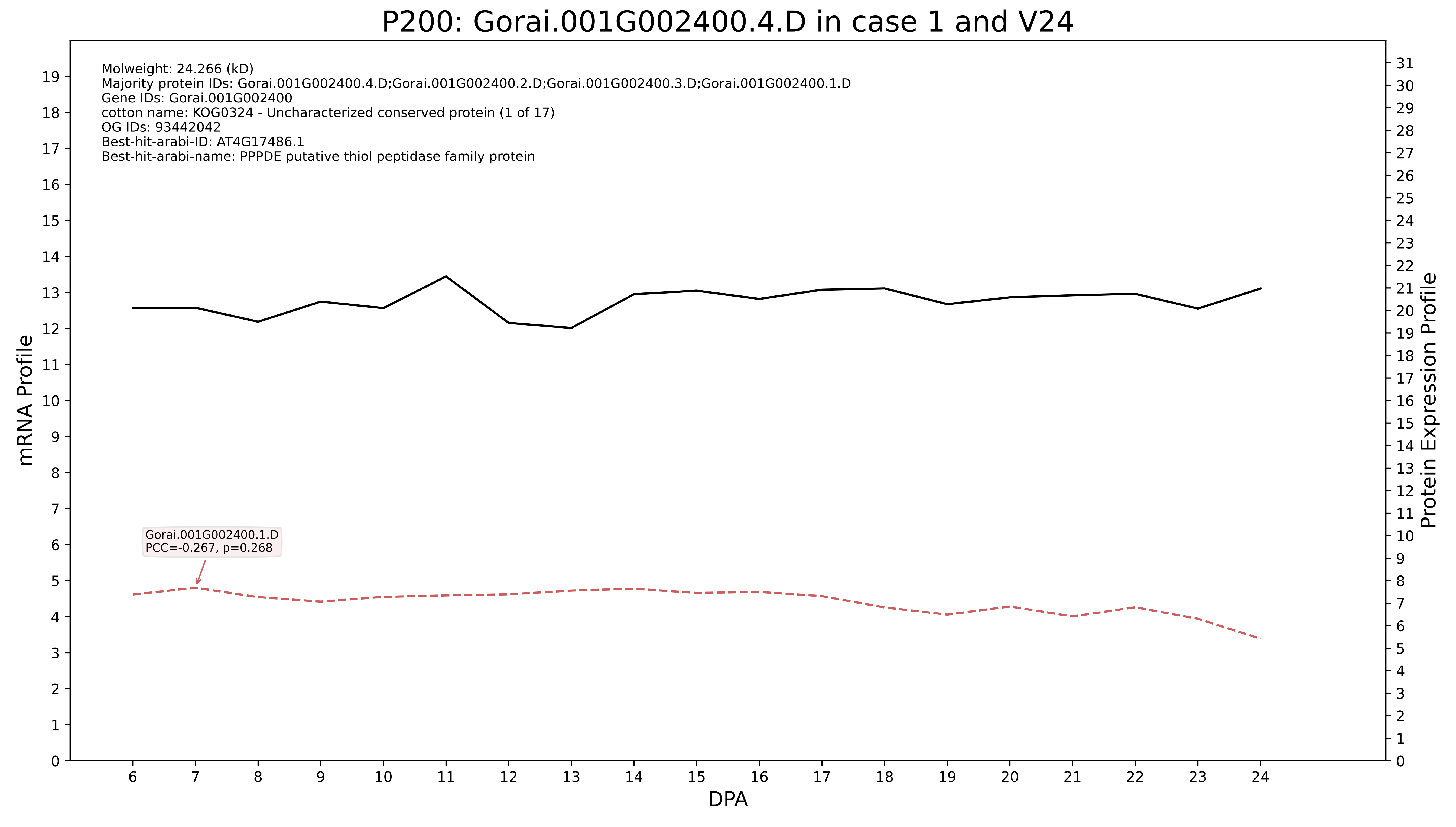
P200: Gorai.001G002400.4.D in case 1 and V24
Molweight: 24.266 (kD)
Majority protein IDs: Gorai.001G002400.4.D;Gorai.001G002400.2.D;Gorai.001G002400.3.D;Gorai.001G002400.1.D
Gene IDs: Gorai.001G002400
cotton name: KOG0324 - Uncharacterized conserved protein (1 of 17)
OG IDs: 93442042
Best-hit-arabi-ID: AT4G17486.1
Best-hit-arabi-name: PPPDE putative thiol peptidase family protein
Gorai.001G002400.1.D
PCC=-0.267, p=0.268
mRNA Profile
Protein Expression Profile
DPA

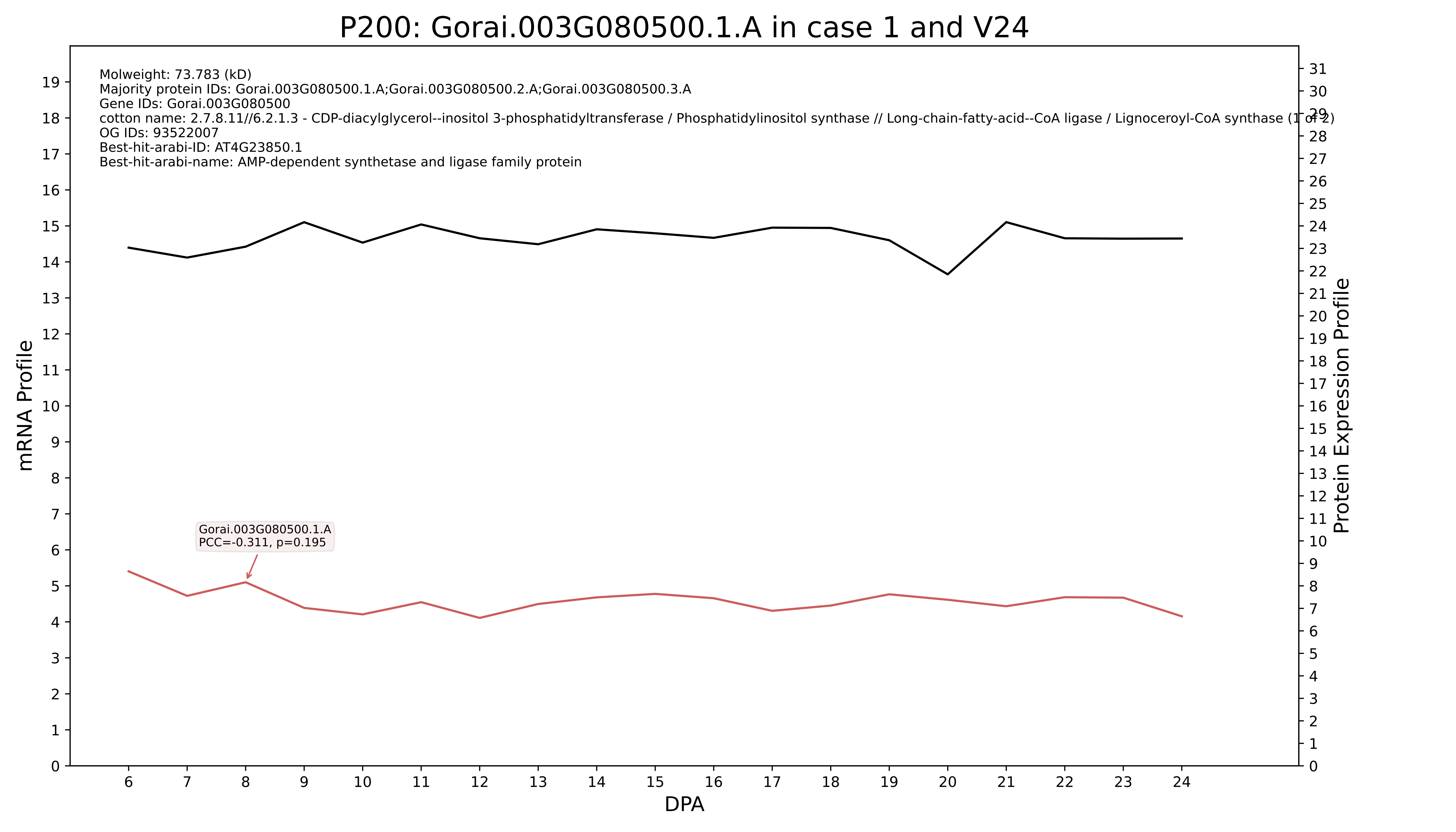
P200: Gorai.003G080500.1.A in case 1 and V24
Molweight: 73.783 (kD)
Majority protein IDs: Gorai.003G080500.1.A;Gorai.003G080500.2.A;Gorai.003G080500.3.A
Gene IDs: Gorai.003G080500
cotton name: 2.7.8.11//6.2.1.3 - CDP-diacylglycerol--inositol 3-phosphatidyltransferase / Phosphatidylinositol synthase // Long-chain-fatty-acid--CoA ligase / Lignoceroyl-CoA synthase (1 of 2)
OG IDs: 93522007
Best-hit-arabi-ID: AT4G23850.1
Best-hit-arabi-name: AMP-dependent synthetase and ligase family protein
Gorai.003G080500.1.A
PCC=-0.311, p=0.195
mRNA Profile
Protein Expression Profile
DPA

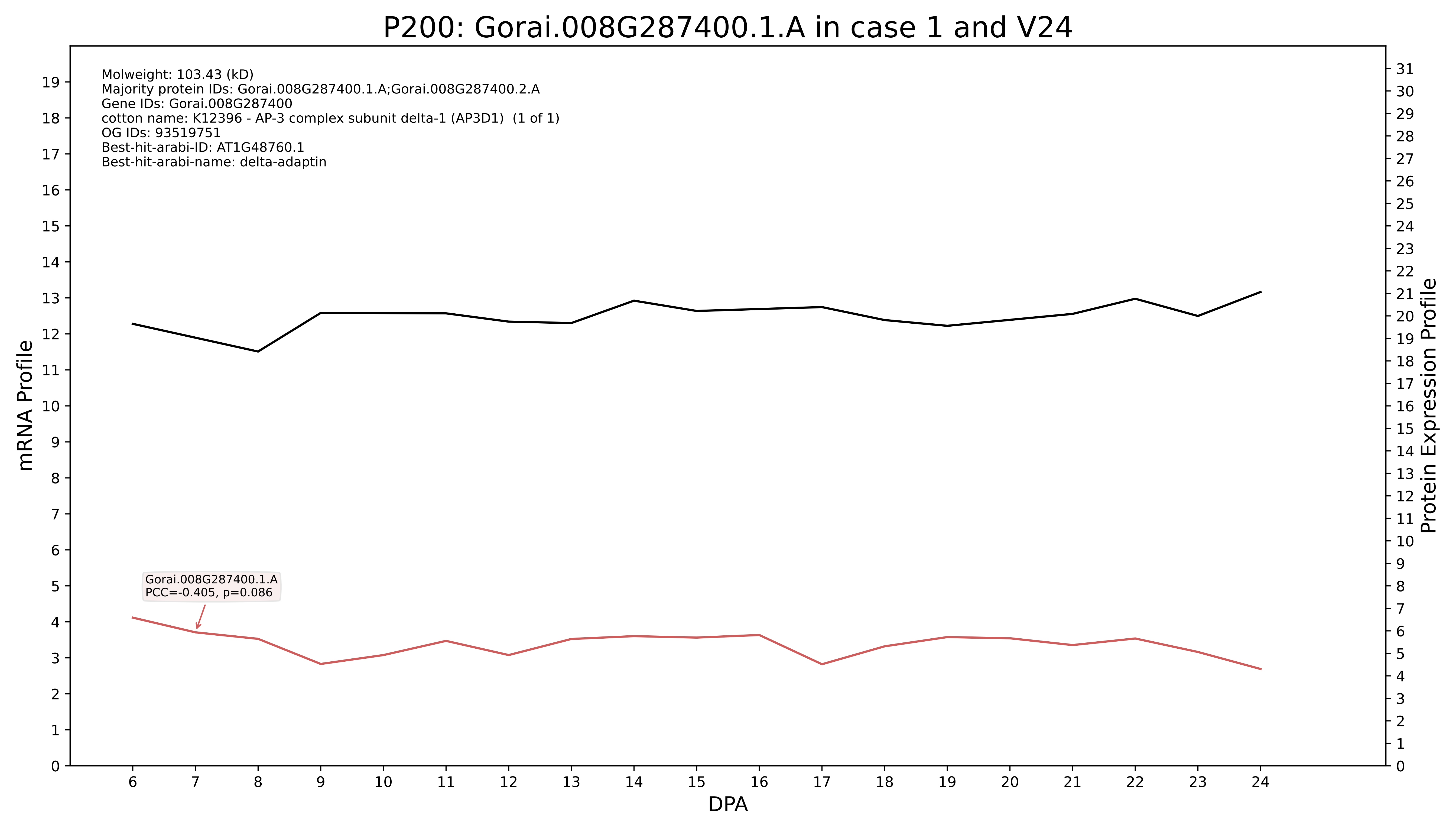

P200: Gorai.008G287400.1.A in case 1 and V24
Molweight: 103.43 (kD)
Majority protein IDs: Gorai.008G287400.1.A;Gorai.008G287400.2.A
Gene IDs: Gorai.008G287400
cotton name: K12396 - AP-3 complex subunit delta-1 (AP3D1) (1 of 1)
OG IDs: 93519751
Best-hit-arabi-ID: AT1G48760.1
Best-hit-arabi-name: delta-adaptin
Gorai.008G287400.1.A
PCC=-0.405, p=0.086
mRNA Profile
Protein Expression Profile
DPA

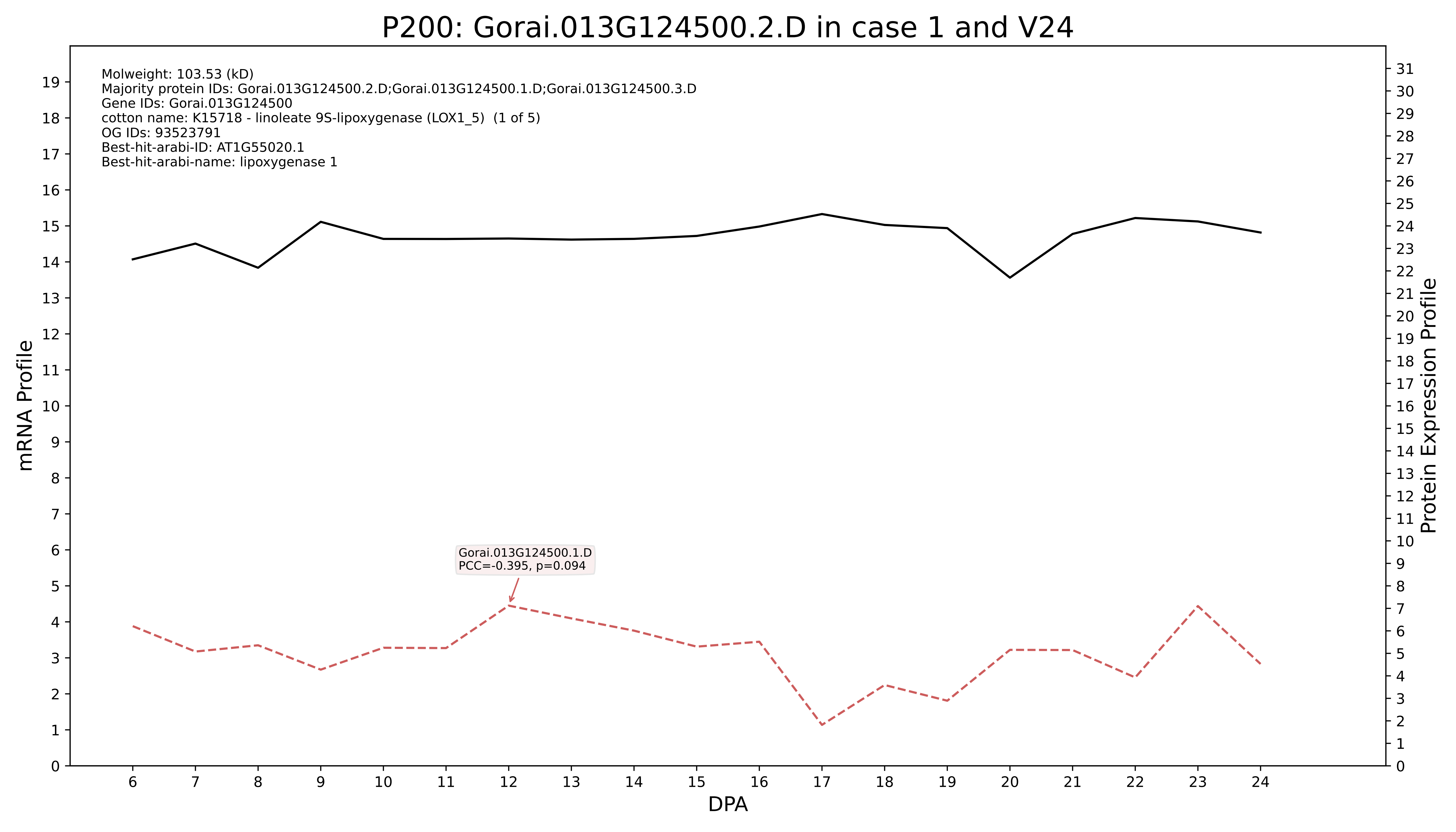
P200: Gorai.013G124500.2.D in case 1 and V24
Molweight: 103.53 (kD)
Majority protein IDs: Gorai.013G124500.2.D;Gorai.013G124500.1.D;Gorai.013G124500.3.D
Gene IDs: Gorai.013G124500
cotton name: K15718 - linoleate 9S-lipoxygenase (LOX1_5) (1 of 5)
OG IDs: 93523791
Best-hit-arabi-ID: AT1G55020.1
Best-hit-arabi-name: lipoxygenase 1
Gorai.013G124500.1.D
PCC=-0.395, p=0.094
mRNA Profile
Protein Expression Profile
DPA

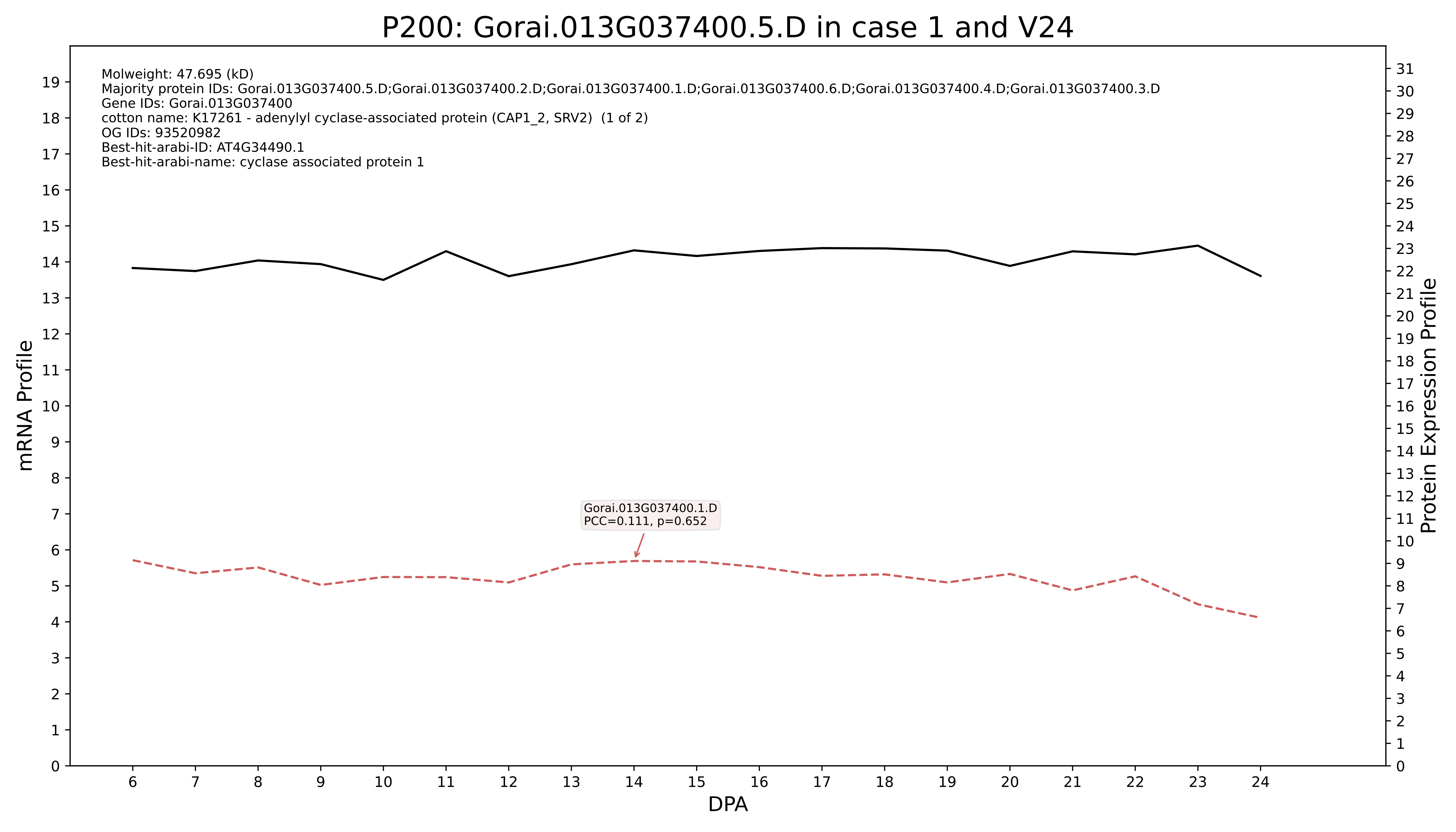

P200: Gorai.013G037400.5.D in case 1 and V24
Molweight: 47.695 (kD)
Majority protein IDs: Gorai.013G037400.5.D;Gorai.013G037400.2.D;Gorai.013G037400.1.D;Gorai.013G037400.6.D;Gorai.013G037400.4.D;Gorai.013G037400.3.D
Gene IDs: Gorai.013G037400
cotton name: K17261 - adenylyl cyclase-associated protein (CAP1_2, SRV2) (1 of 2)
OG IDs: 93520982
Best-hit-arabi-ID: AT4G34490.1
Best-hit-arabi-name: cyclase associated protein 1
Gorai.013G037400.1.D
PCC=0.111, p=0.652
mRNA Profile
Protein Expression Profile
DPA

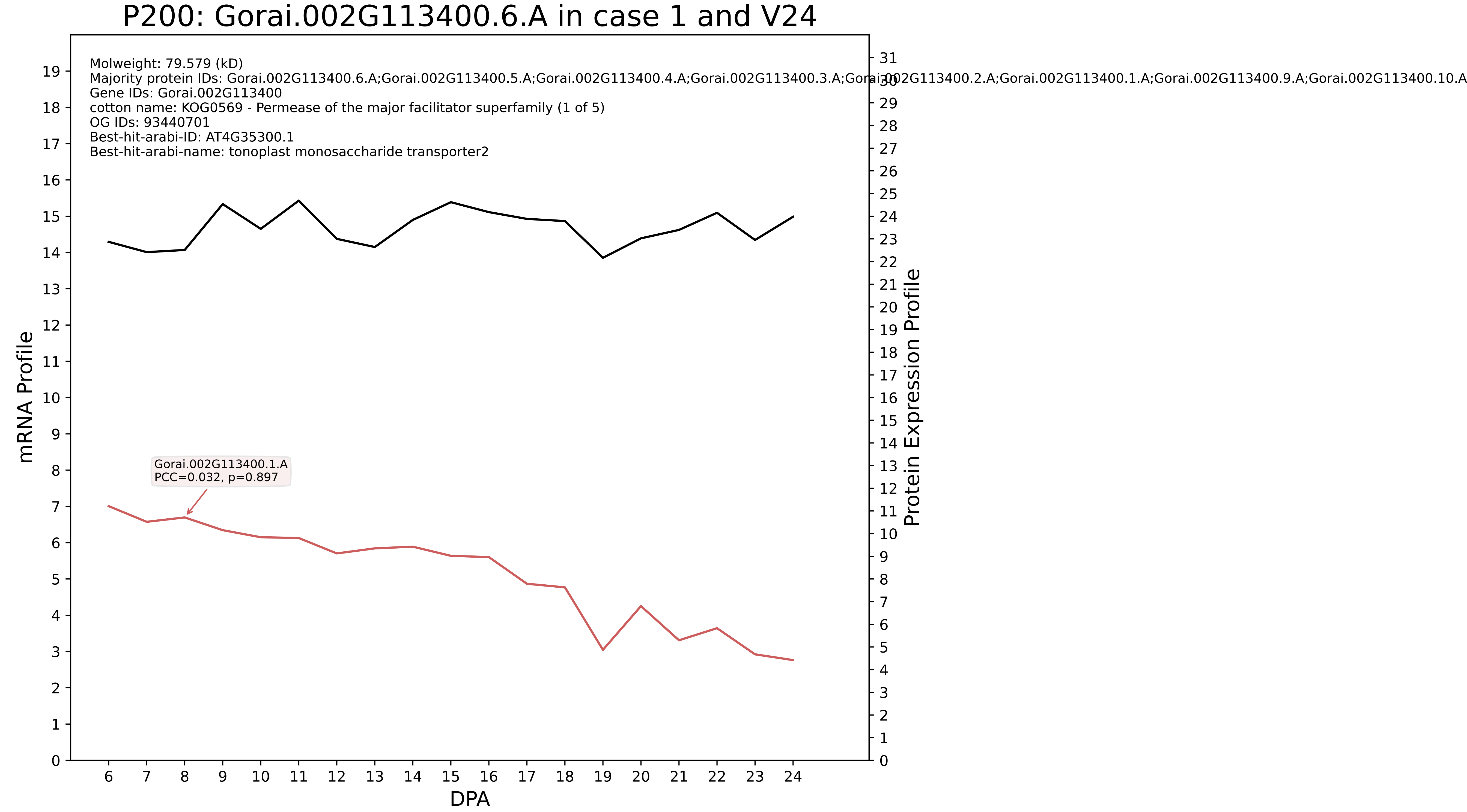
P200: Gorai.002G113400.6.A in case 1 and V24
Molweight: 79.579 (kD)
Majority protein IDs: Gorai.002G113400.6.A;Gorai.002G113400.5.A;Gorai.002G113400.4.A;Gorai.002G113400.3.A;Gorai.002G113400.2.A;Gorai.002G113400.1.A;Gorai.002G113400.9.A;Gorai.002G113400.10.A
Gene IDs: Gorai.002G113400
cotton name: KOG0569 - Permease of the major facilitator superfamily (1 of 5)
OG IDs: 93440701
Best-hit-arabi-ID: AT4G35300.1
Best-hit-arabi-name: tonoplast monosaccharide transporter2
Gorai.002G113400.1.A
PCC=0.032, p=0.897
mRNA Profile
Protein Expression Profile
DPA

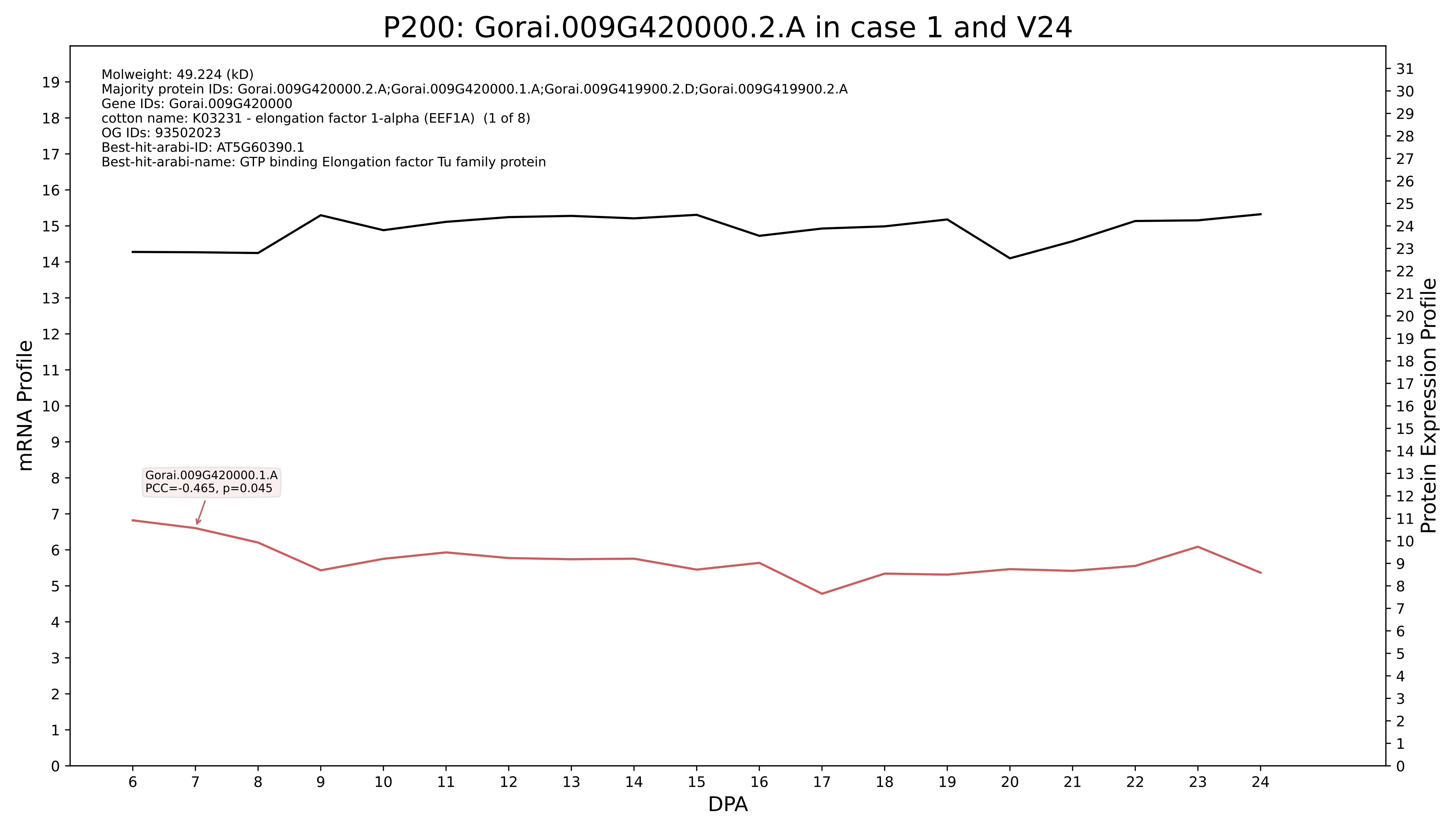
P200: Gorai.009G420000.2.A in case 1 and V24
Molweight: 49.224 (kD)
Majority protein IDs: Gorai.009G420000.2.A;Gorai.009G420000.1.A;Gorai.009G419900.2.D;Gorai.009G419900.2.A
Gene IDs: Gorai.009G420000
cotton name: K03231 - elongation factor 1-alpha (EEF1A) (1 of 8)
OG IDs: 93502023
Best-hit-arabi-ID: AT5G60390.1
Best-hit-arabi-name: GTP binding Elongation factor Tu family protein
Gorai.009G420000.1.A
PCC=-0.465, p=0.045
mRNA Profile
Protein Expression Profile
DPA

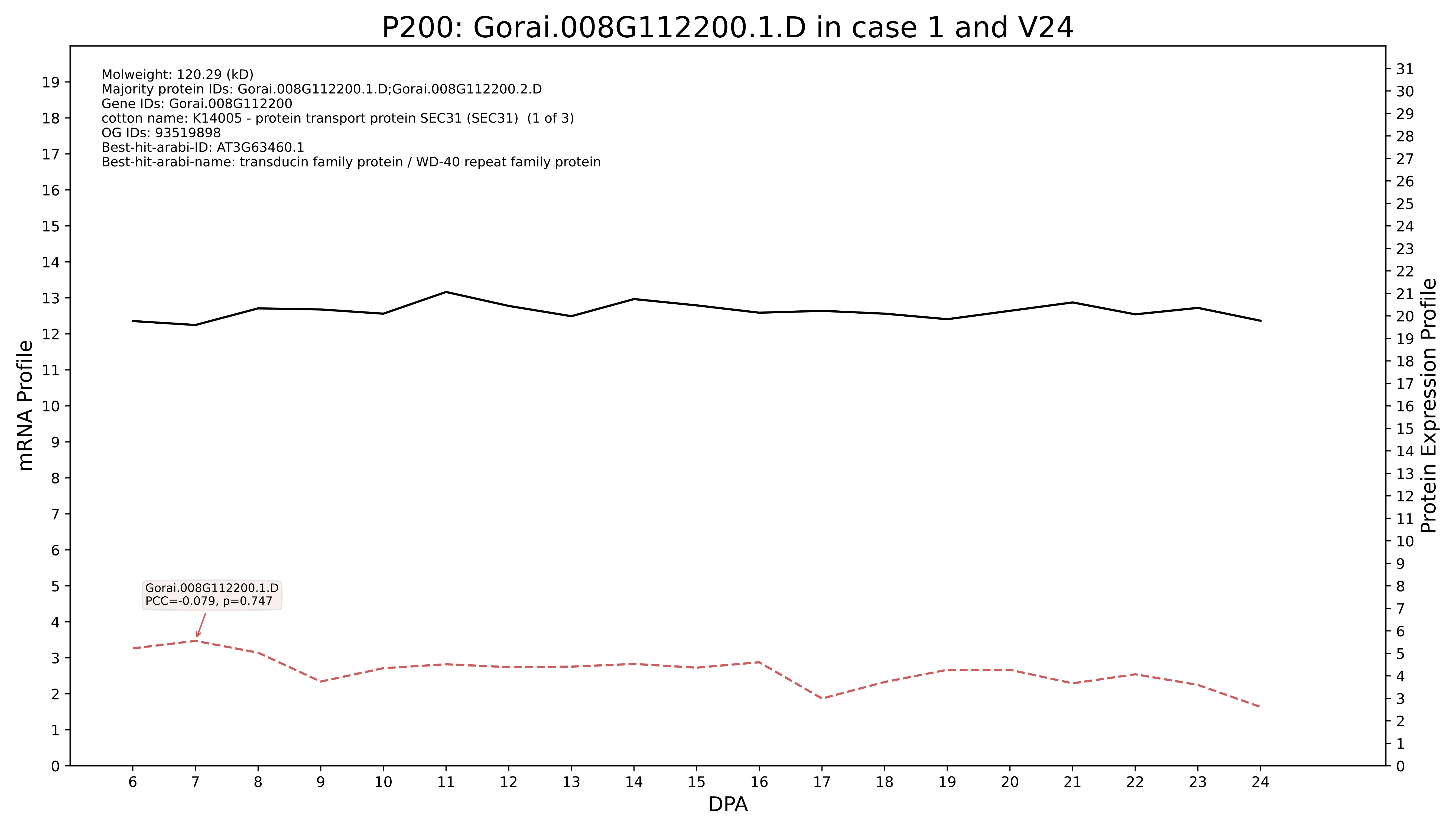
P200: Gorai.008G112200.1.D in case 1 and V24
Molweight: 120.29 (kD)
Majority protein IDs: Gorai.008G112200.1.D;Gorai.008G112200.2.D
Gene IDs: Gorai.008G112200
cotton name: K14005 - protein transport protein SEC31 (SEC31) (1 of 3)
OG IDs: 93519898
Best-hit-arabi-ID: AT3G63460.1
Best-hit-arabi-name: transducin family protein / WD-40 repeat family protein
Gorai.008G112200.1.D
PCC=-0.079, p=0.747
mRNA Profile
Protein Expression Profile
DPA

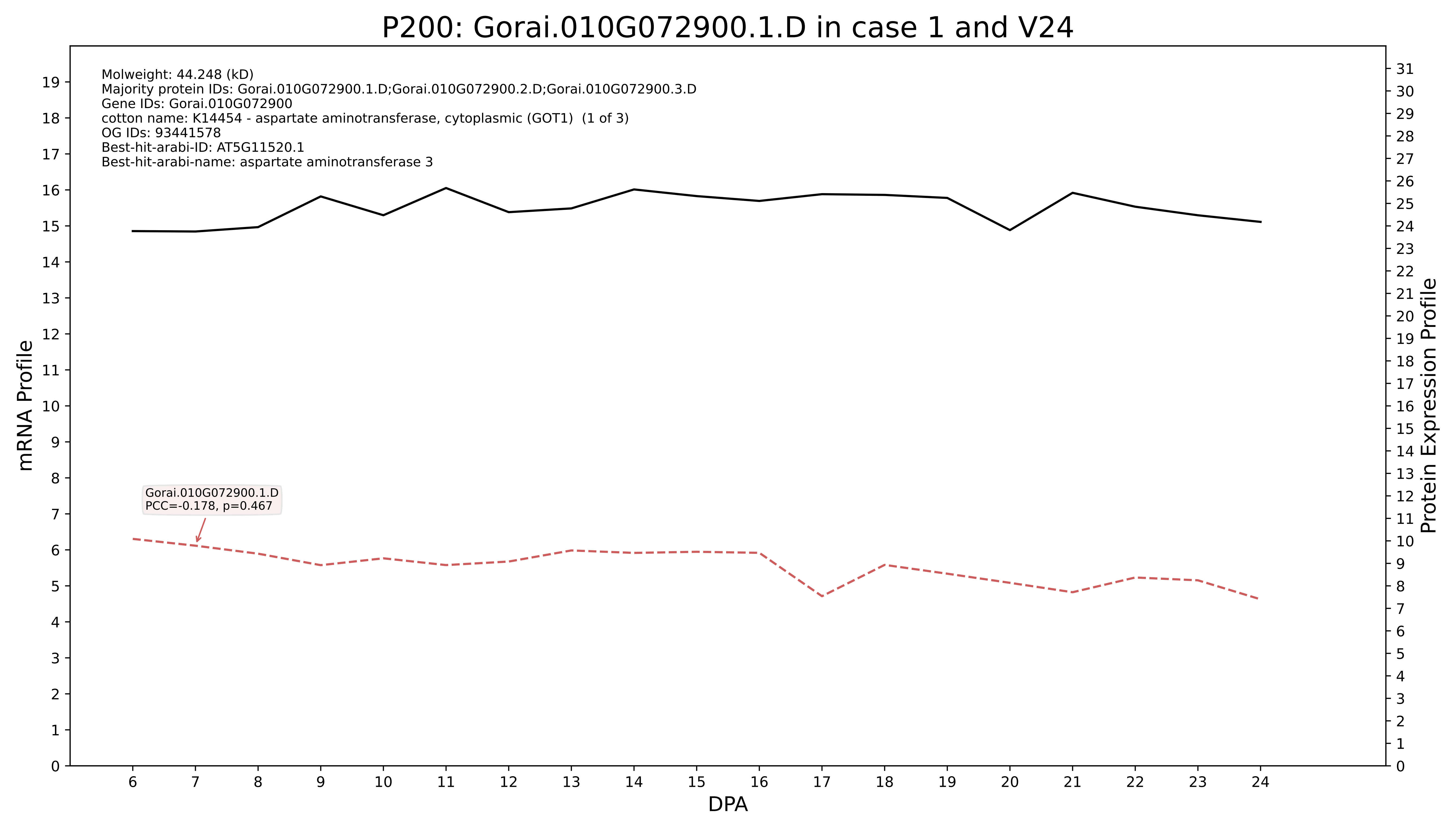
P200: Gorai.010G072900.1.D in case 1 and V24
Molweight: 44.248 (kD)
Majority protein IDs: Gorai.010G072900.1.D;Gorai.010G072900.2.D;Gorai.010G072900.3.D
Gene IDs: Gorai.010G072900
cotton name: K14454 - aspartate aminotransferase, cytoplasmic (GOT1) (1 of 3)
OG IDs: 93441578
Best-hit-arabi-ID: AT5G11520.1
Best-hit-arabi-name: aspartate aminotransferase 3
Gorai.010G072900.1.D
PCC=-0.178, p=0.467
mRNA Profile
Protein Expression Profile
DPA

001G034800.3.D in case 1 and V24

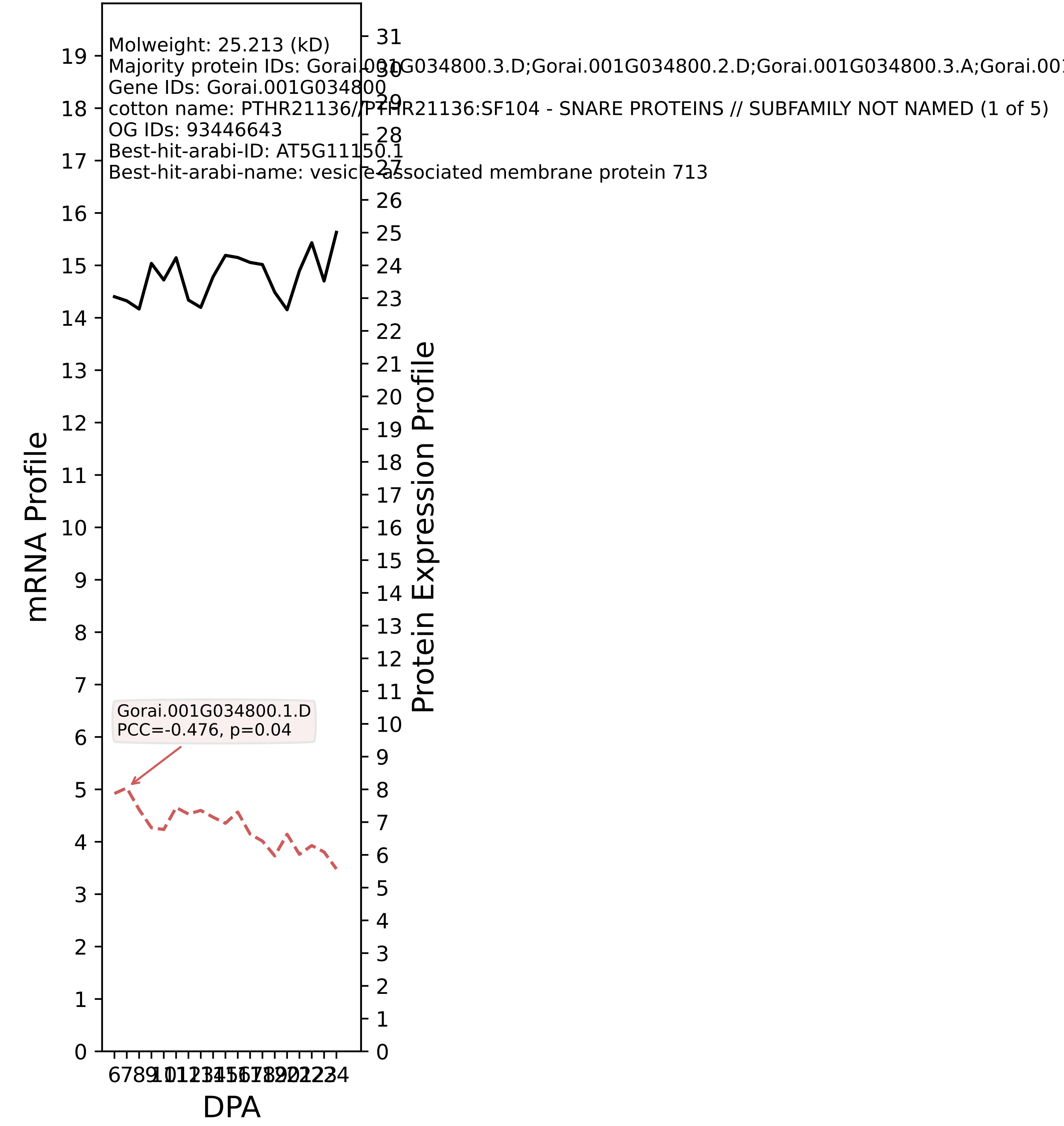

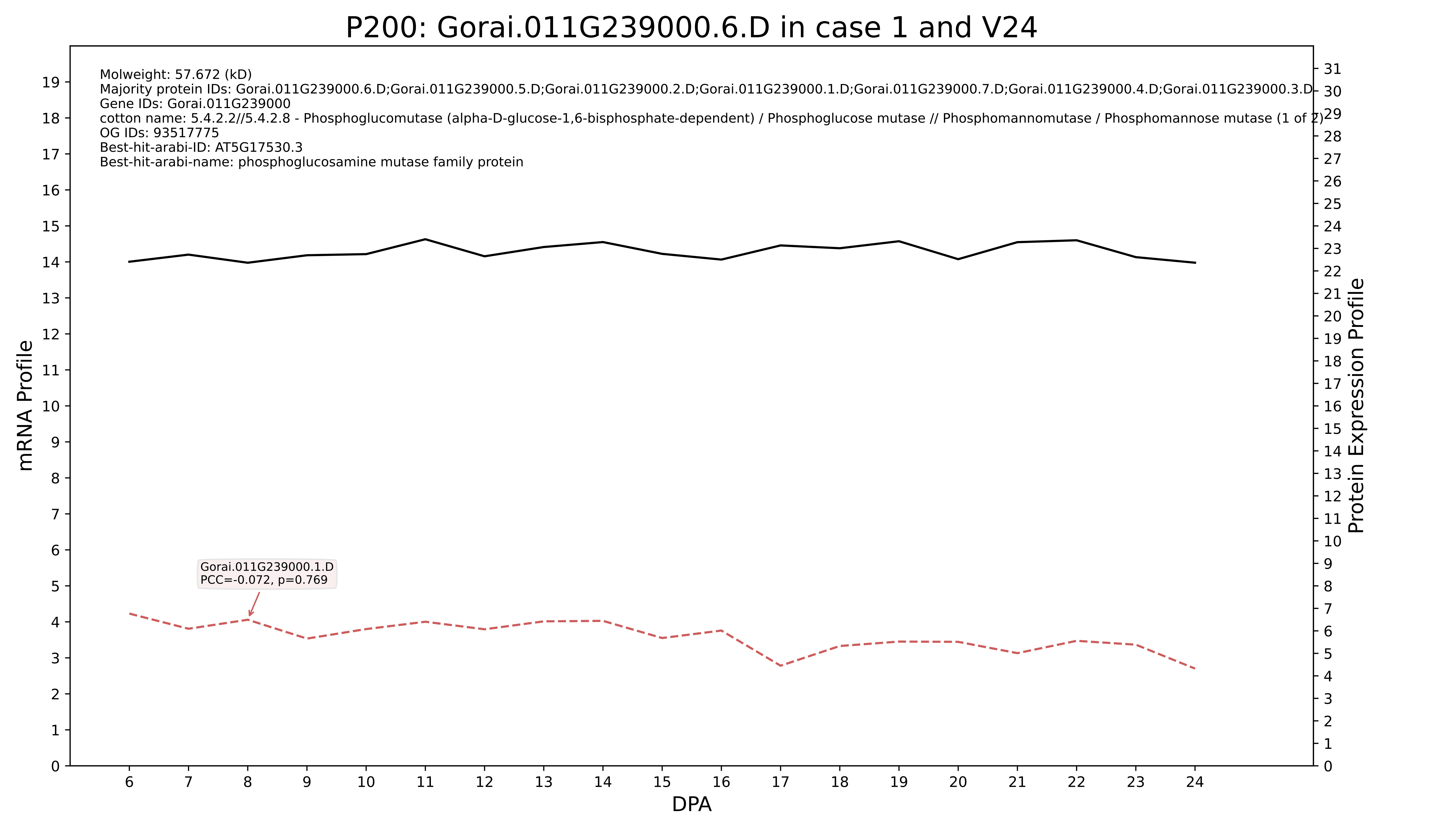

P200: Gorai.011G239000.6.D in case 1 and V24
Molweight: 57.672 (kD)
Majority protein IDs: Gorai.011G239000.6.D;Gorai.011G239000.5.D;Gorai.011G239000.2.D;Gorai.011G239000.1.D;Gorai.011G239000.7.D;Gorai.011G239000.4.D;Gorai.011G239000.3.D
Gene IDs: Gorai.011G239000
cotton name: 5.4.2.2//5.4.2.8 - Phosphoglucomutase (alpha-D-glucose-1,6-bisphosphate-dependent) / Phosphoglucose mutase // Phosphomannomutase / Phosphomannose mutase (1 of 2)
OG IDs: 93517775
Best-hit-arabi-ID: AT5G17530.3
Best-hit-arabi-name: phosphoglucosamine mutase family protein
Gorai.011G239000.1.D
PCC=-0.072, p=0.769
mRNA Profile
Protein Expression Profile
DPA

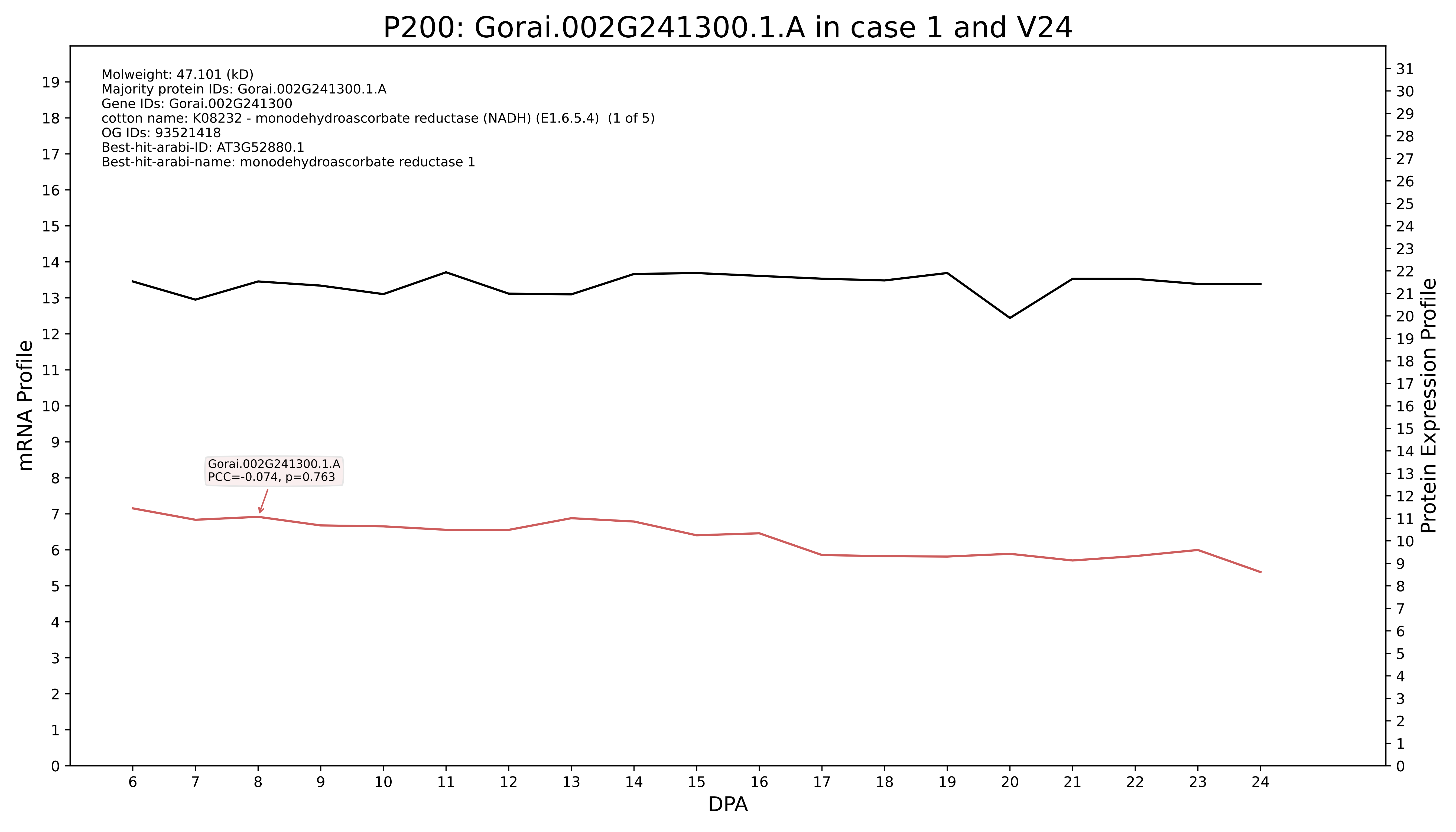
P200: Gorai.002G241300.1.A in case 1 and V24
Molweight: 47.101 (kD)
Majority protein IDs: Gorai.002G241300.1.A
Gene IDs: Gorai.002G241300
cotton name: K08232 - monodehydroascorbate reductase (NADH) (E1.6.5.4) (1 of 5)
OG IDs: 93521418
Best-hit-arabi-ID: AT3G52880.1
Best-hit-arabi-name: monodehydroascorbate reductase 1
Gorai.002G241300.1.A
PCC=-0.074, p=0.763
mRNA Profile
Protein Expression Profile
DPA

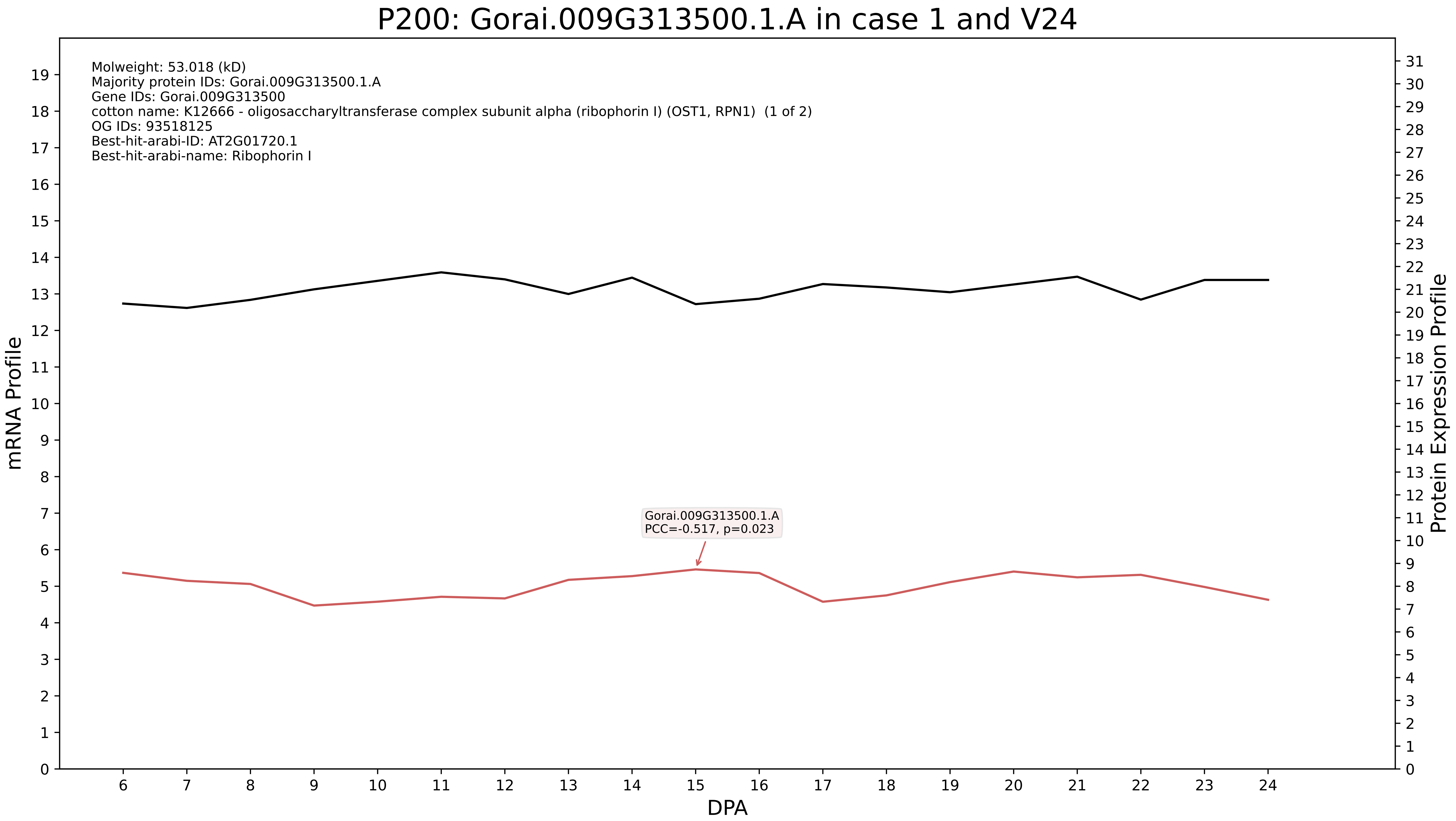

P200: Gorai.009G313500.1.A in case 1 and V24
Molweight: 53.018 (kD)
Majority protein IDs: Gorai.009G313500.1.A
Gene IDs: Gorai.009G313500
cotton name: K12666 - oligosaccharyltransferase complex subunit alpha (ribophorin I) (OST1, RPN1) (1 of 2)
OG IDs: 93518125
Best-hit-arabi-ID: AT2G01720.1
Best-hit-arabi-name: Ribophorin I
Gorai.009G313500.1.A
PCC=-0.517, p=0.023
mRNA Profile
Protein Expression Profile
DPA

### P200: Gorai.008G293200.1.A in case 1 and V24

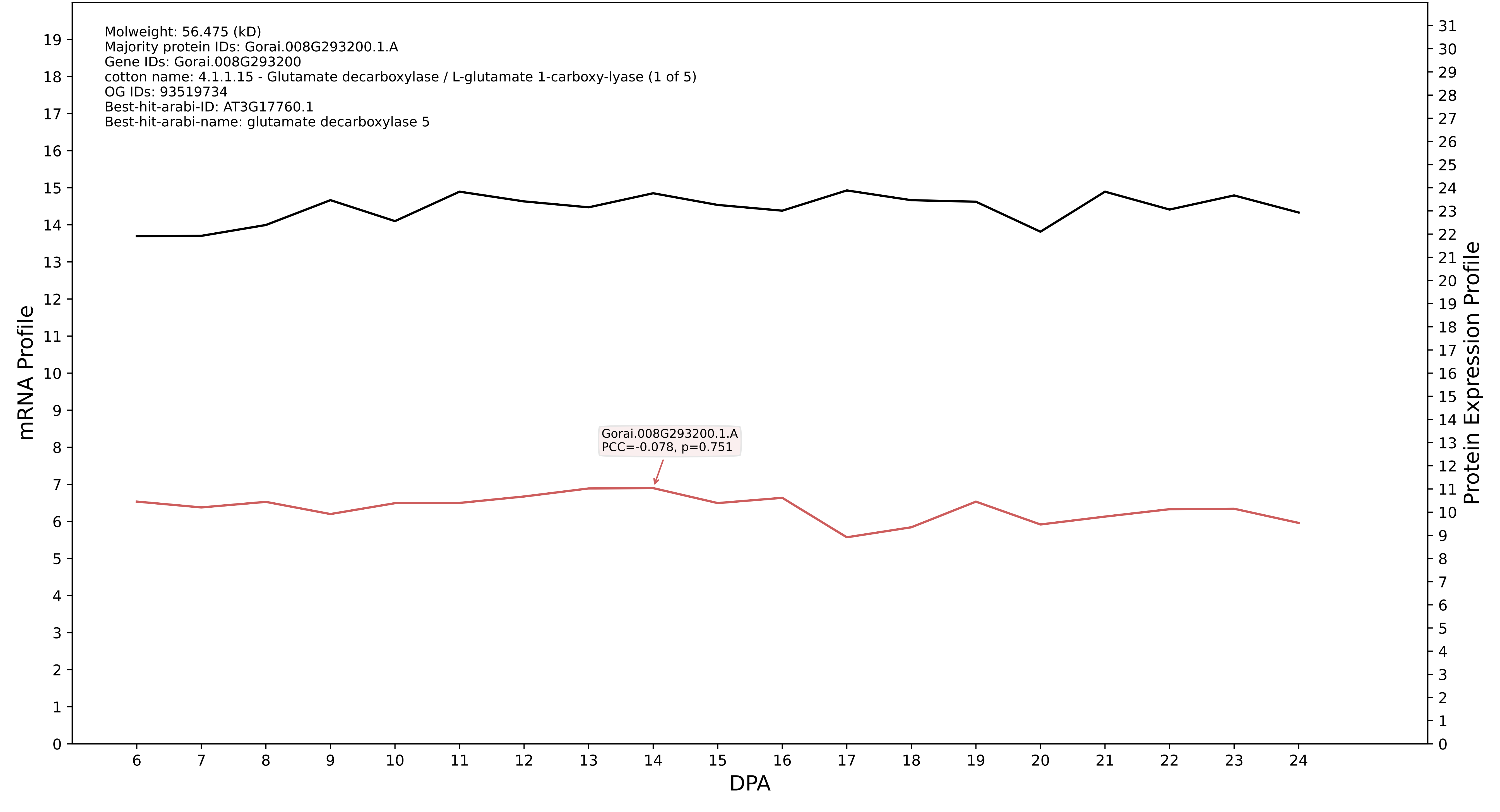

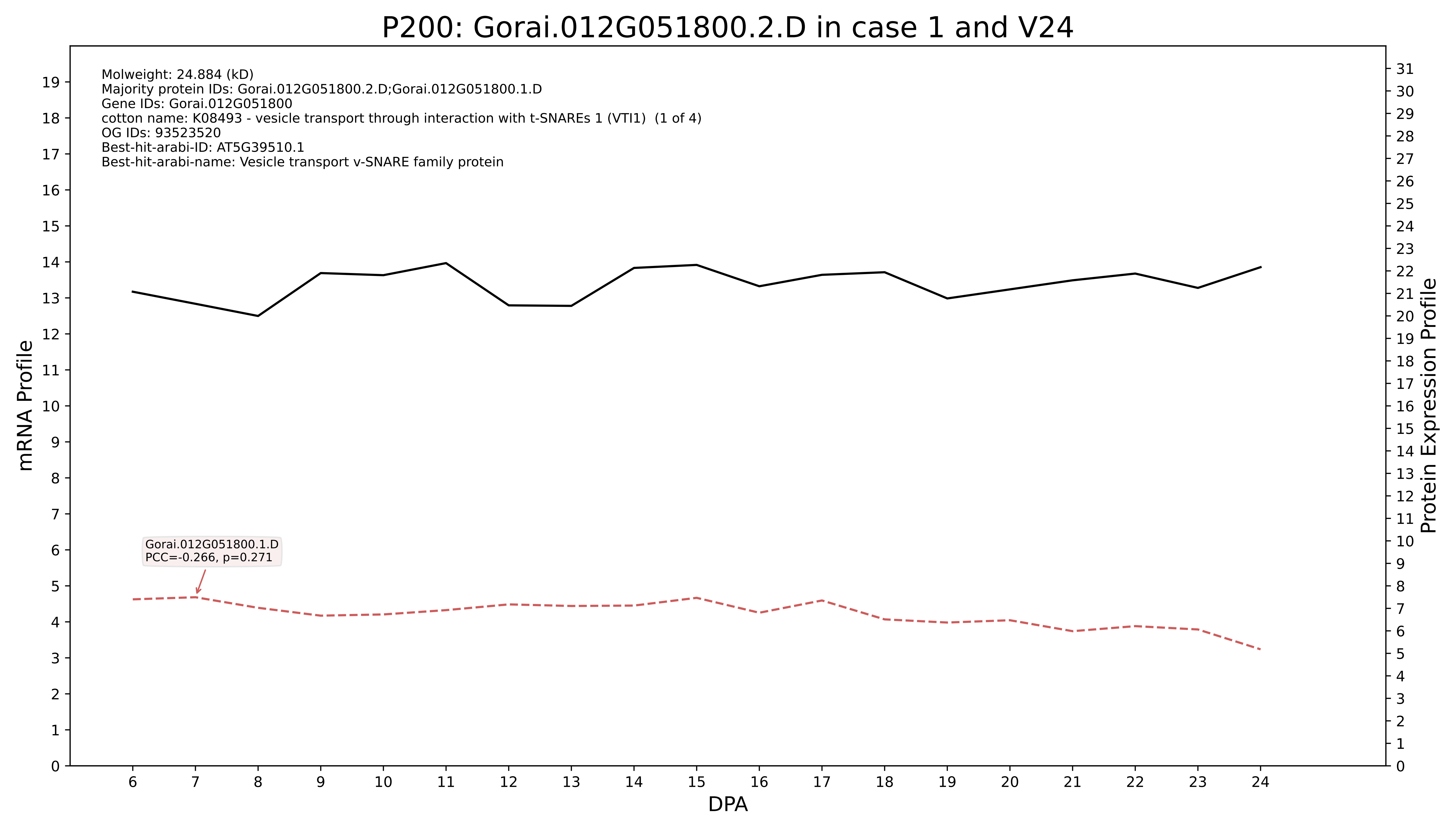
P200: Gorai.012G051800.2.D in case 1 and V24
Molweight: 24.884 (kD)
Majority protein IDs: Gorai.012G051800.2.D;Gorai.012G051800.1.D
Gene IDs: Gorai.012G051800
cotton name: K08493 - vesicle transport through interaction with t-SNAREs 1 (VTI1) (1 of 4)
OG IDs: 93523520
Best-hit-arabi-ID: AT5G39510.1
Best-hit-arabi-name: Vesicle transport v-SNARE family protein
Gorai.012G051800.1.D
PCC=-0.266, p=0.271
mRNA Profile
Protein Expression Profile
DPA

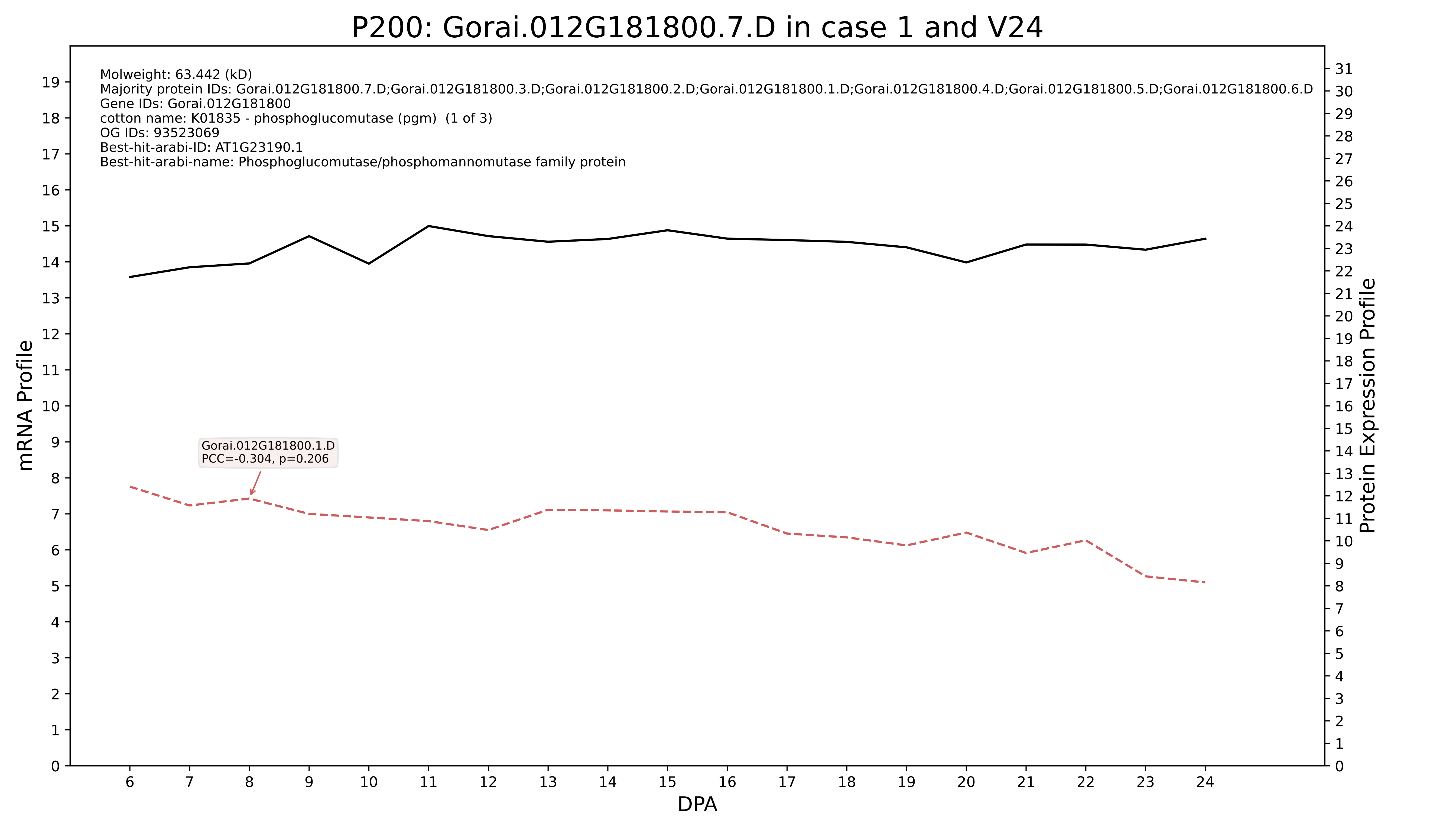

P200: Gorai.012G181800.7.D in case 1 and V24
Molweight: 63.442 (kD)
Majority protein IDs: Gorai.012G181800.7.D;Gorai.012G181800.3.D;Gorai.012G181800.2.D;Gorai.012G181800.1.D;Gorai.012G181800.4.D;Gorai.012G181800.5.D;Gorai.012G181800.6.D
Gene IDs: Gorai.012G181800
cotton name: K01835 - phosphoglucomutase (pgm) (1 of 3)
OG IDs: 93523069
Best-hit-arabi-ID: AT1G23190.1
Best-hit-arabi-name: Phosphoglucomutase/phosphomannomutase family protein
Gorai.012G181800.1.D
PCC=-0.304, p=0.206
mRNA Profile
Protein Expression Profile
DPA

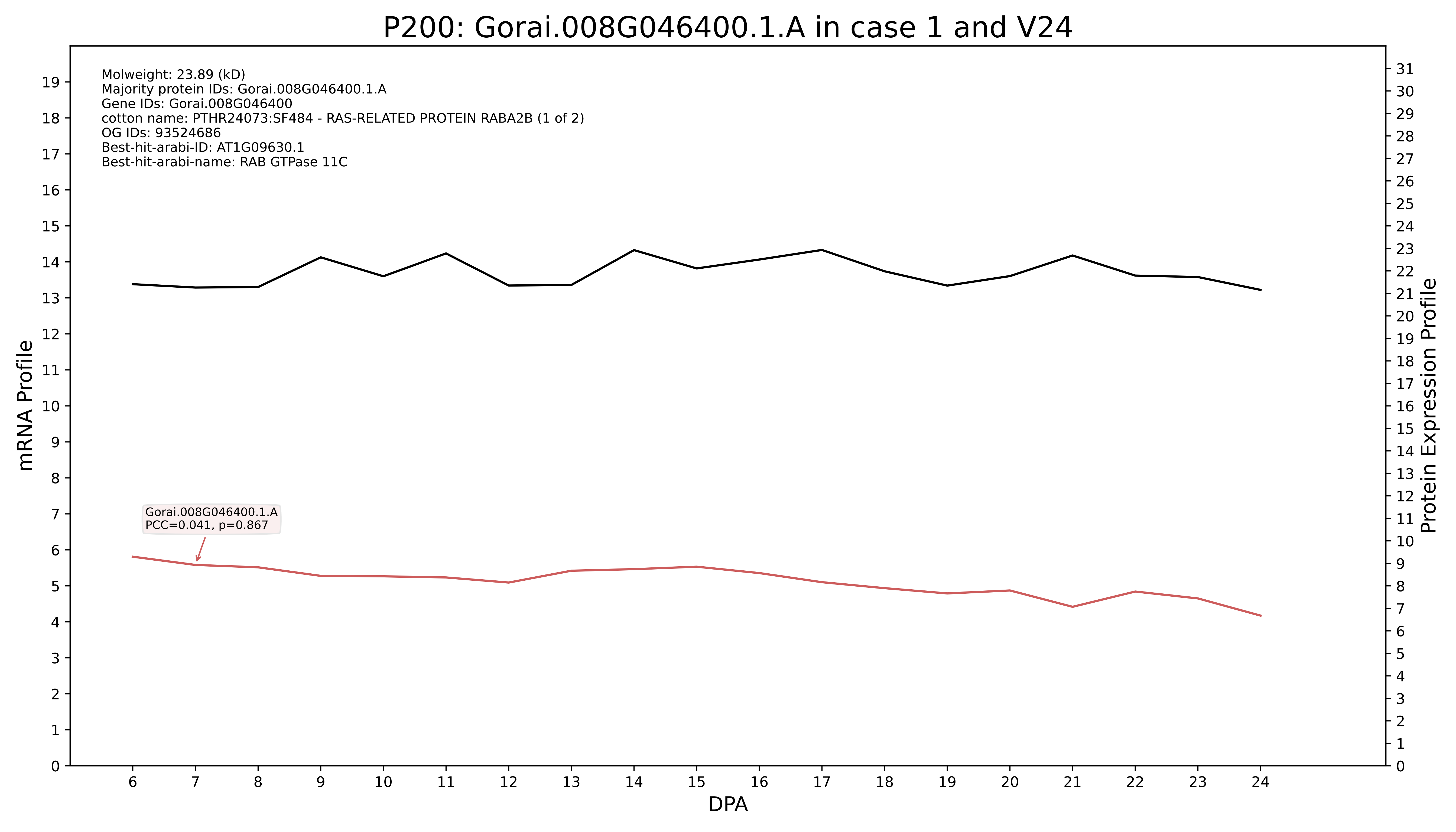

P200: Gorai.008G046400.1.A in case 1 and V24
Molweight: 23.89 (kD)
Majority protein IDs: Gorai.008G046400.1.A
Gene IDs: Gorai.008G046400
cotton name: PTHR24073:SF484 - RAS-RELATED PROTEIN RABA2B (1 of 2)
OG IDs: 93524686
Best-hit-arabi-ID: AT1G09630.1
Best-hit-arabi-name: RAB GTPase 11C
Gorai.008G046400.1.A
PCC=0.041, p=0.867
mRNA Profile
Protein Expression Profile
DPA

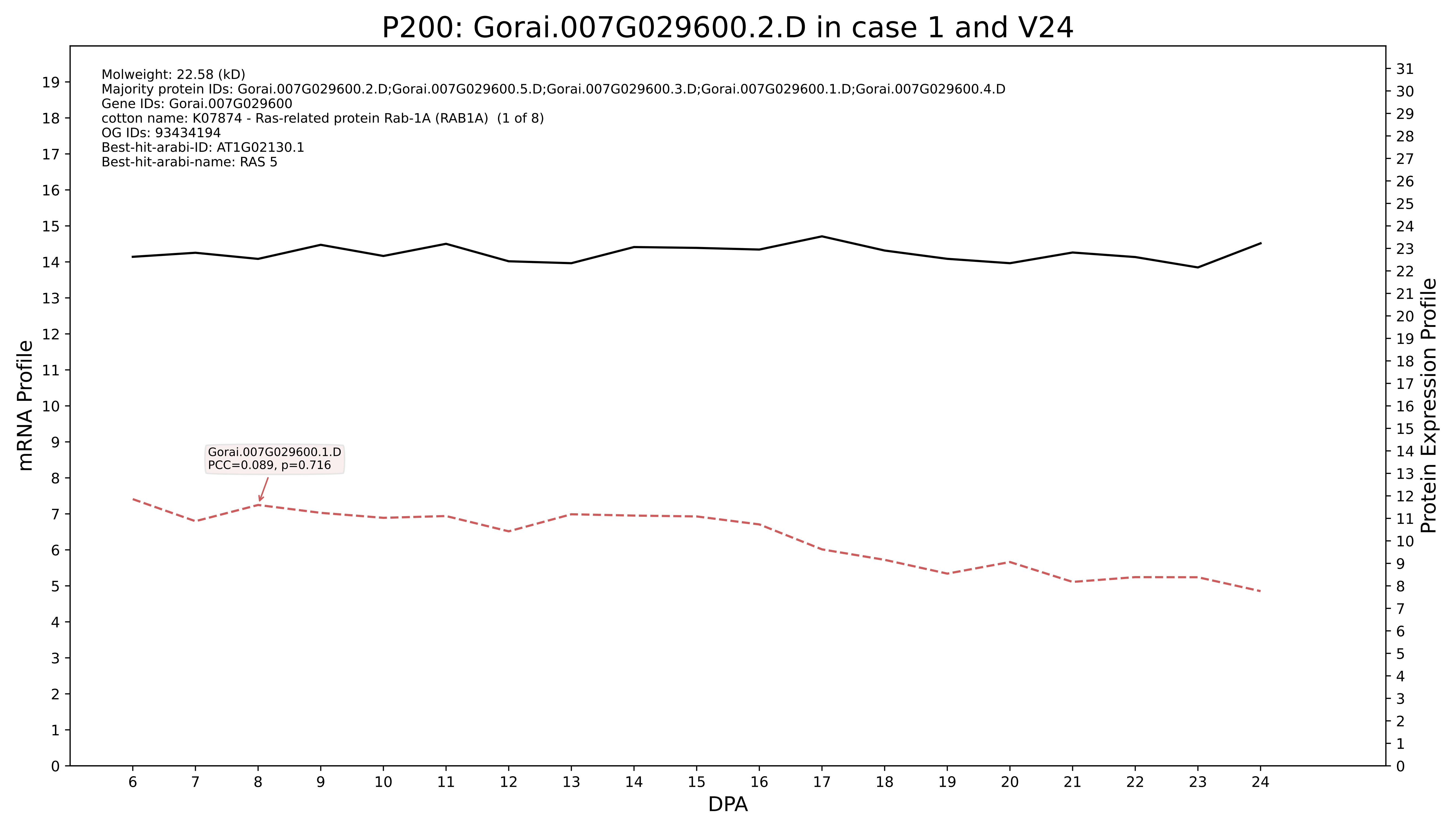

P200: Gorai.007G029600.2.D in case 1 and V24
Molweight: 22.58 (kD)
Majority protein IDs: Gorai.007G029600.2.D;Gorai.007G029600.5.D;Gorai.007G029600.3.D;Gorai.007G029600.1.D;Gorai.007G029600.4.D
Gene IDs: Gorai.007G029600
cotton name: K07874 - Ras-related protein Rab-1A (RAB1A) (1 of 8)
OG IDs: 93434194
Best-hit-arabi-ID: AT1G02130.1
Best-hit-arabi-name: RAS 5
Gorai.007G029600.1.D
PCC=0.089, p=0.716
mRNA Profile
Protein Expression Profile
DPA

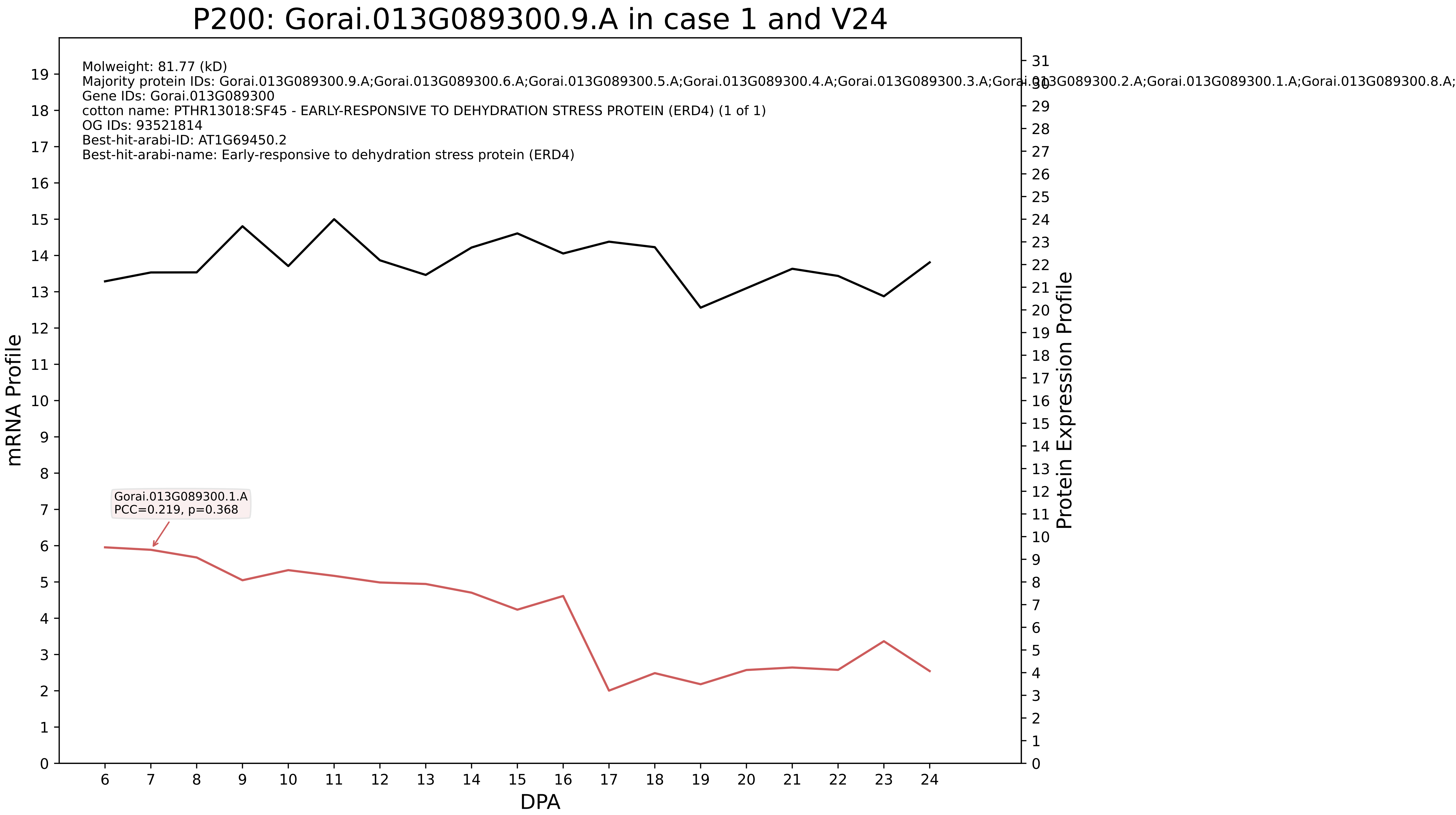
P200: Gorai.013G089300.9.A in case 1 and V24
Molweight: 81.77 (kD)
Majority protein IDs: Gorai.013G089300.9.A;Gorai.013G089300.6.A;Gorai.013G089300.5.A;Gorai.013G089300.4.A;Gorai.013G089300.3.A;Gorai.013G089300.2.A;Gorai.013G089300.1.A;Gorai.013G089300.8.A;
Gene IDs: Gorai.013G089300
cotton name: PTHR13018:SF45 - EARLY-RESPONSIVE TO DEHYDRATION STRESS PROTEIN (ERD4) (1 of 1)
OG IDs: 93521814
Best-hit-arabi-ID: AT1G69450.2
Best-hit-arabi-name: Early-responsive to dehydration stress protein (ERD4)
Gorai.013G089300.1.A
PCC=0.219, p=0.368
mRNA Profile
Protein Expression Profile
DPA

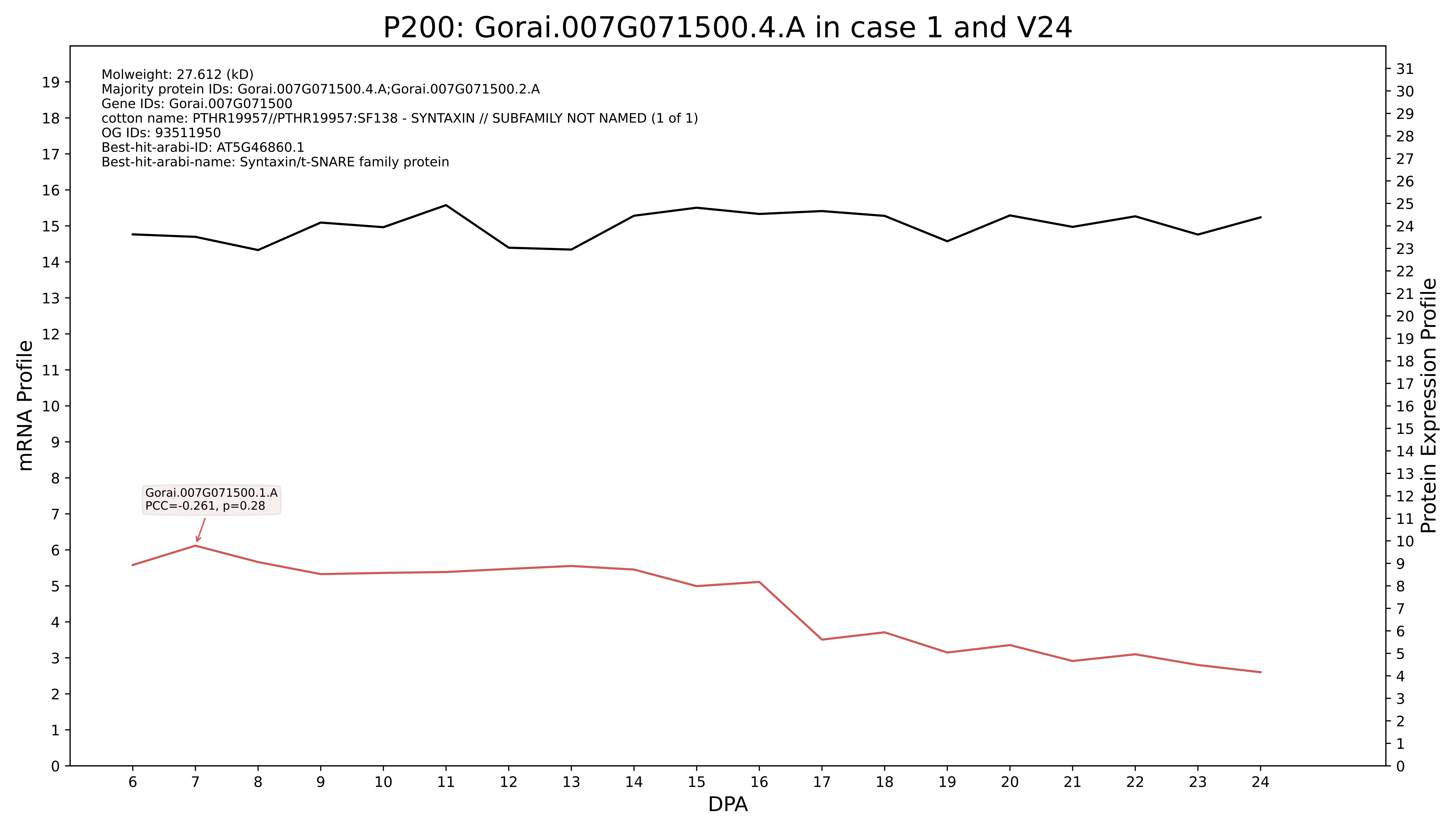

P200: Gorai.007G071500.4.A in case 1 and V24
Molweight: 27.612 (kD)
Majority protein IDs: Gorai.007G071500.4.A;Gorai.007G071500.2.A
Gene IDs: Gorai.007G071500
cotton name: PTHR19957//PTHR19957:SF138 - SYNTAXIN // SUBFAMILY NOT NAMED (1 of 1)
OG IDs: 93511950
Best-hit-arabi-ID: AT5G46860.1
Best-hit-arabi-name: Syntaxin/t-SNARE family protein
Gorai.007G071500.1.A
PCC=-0.261, p=0.28
mRNA Profile
Protein Expression Profile
DPA

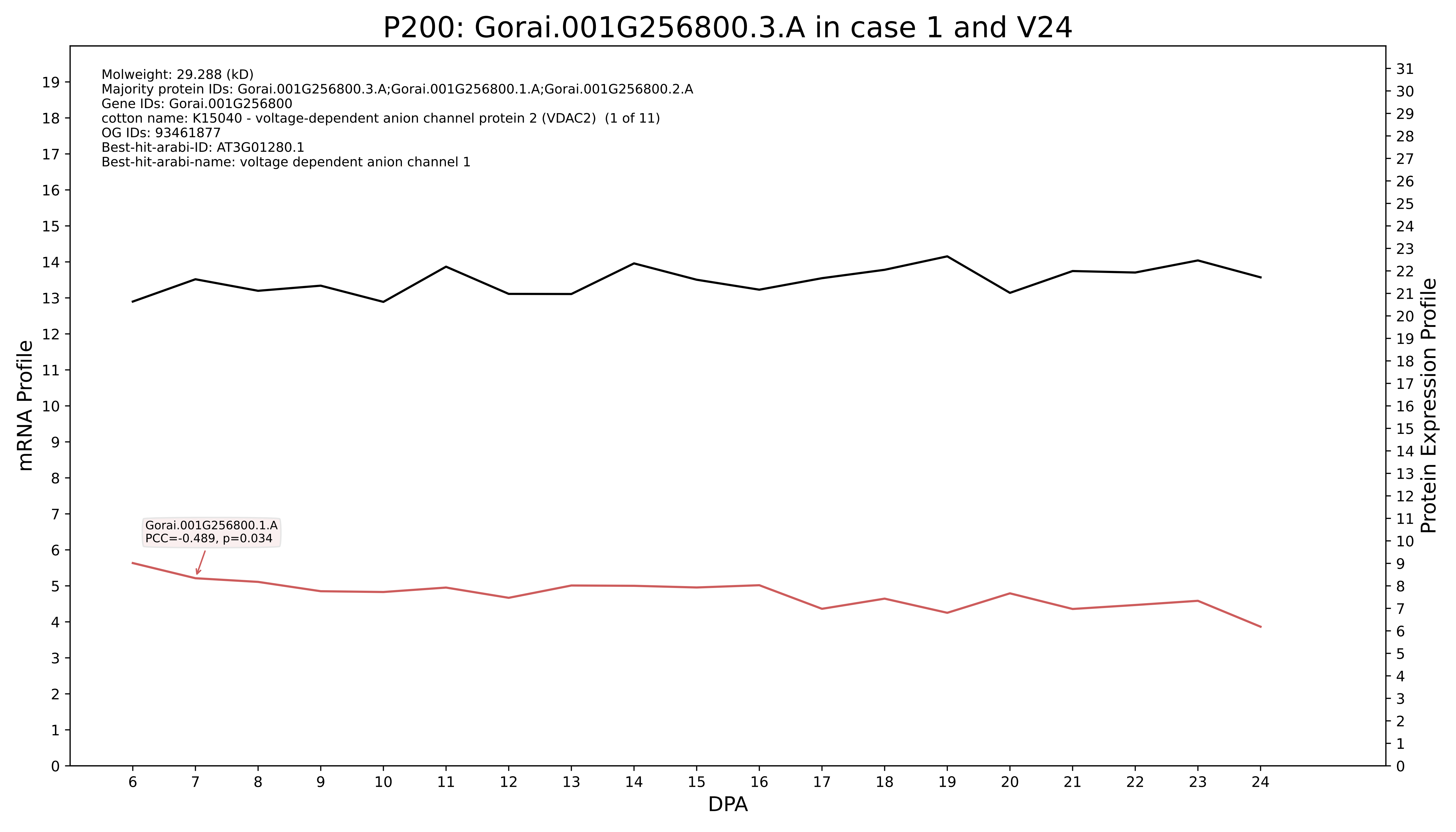
P200: Gorai.001G256800.3.A in case 1 and V24
Molweight: 29.288 (kD)
Majority protein IDs: Gorai.001G256800.3.A;Gorai.001G256800.1.A;Gorai.001G256800.2.A
Gene IDs: Gorai.001G256800
cotton name: K15040 - voltage-dependent anion channel protein 2 (VDAC2) (1 of 11)
OG IDs: 93461877
Best-hit-arabi-ID: AT3G01280.1
Best-hit-arabi-name: voltage dependent anion channel 1
Gorai.001G256800.1.A
PCC=-0.489, p=0.034
mRNA Profile
Protein Expression Profile
DPA

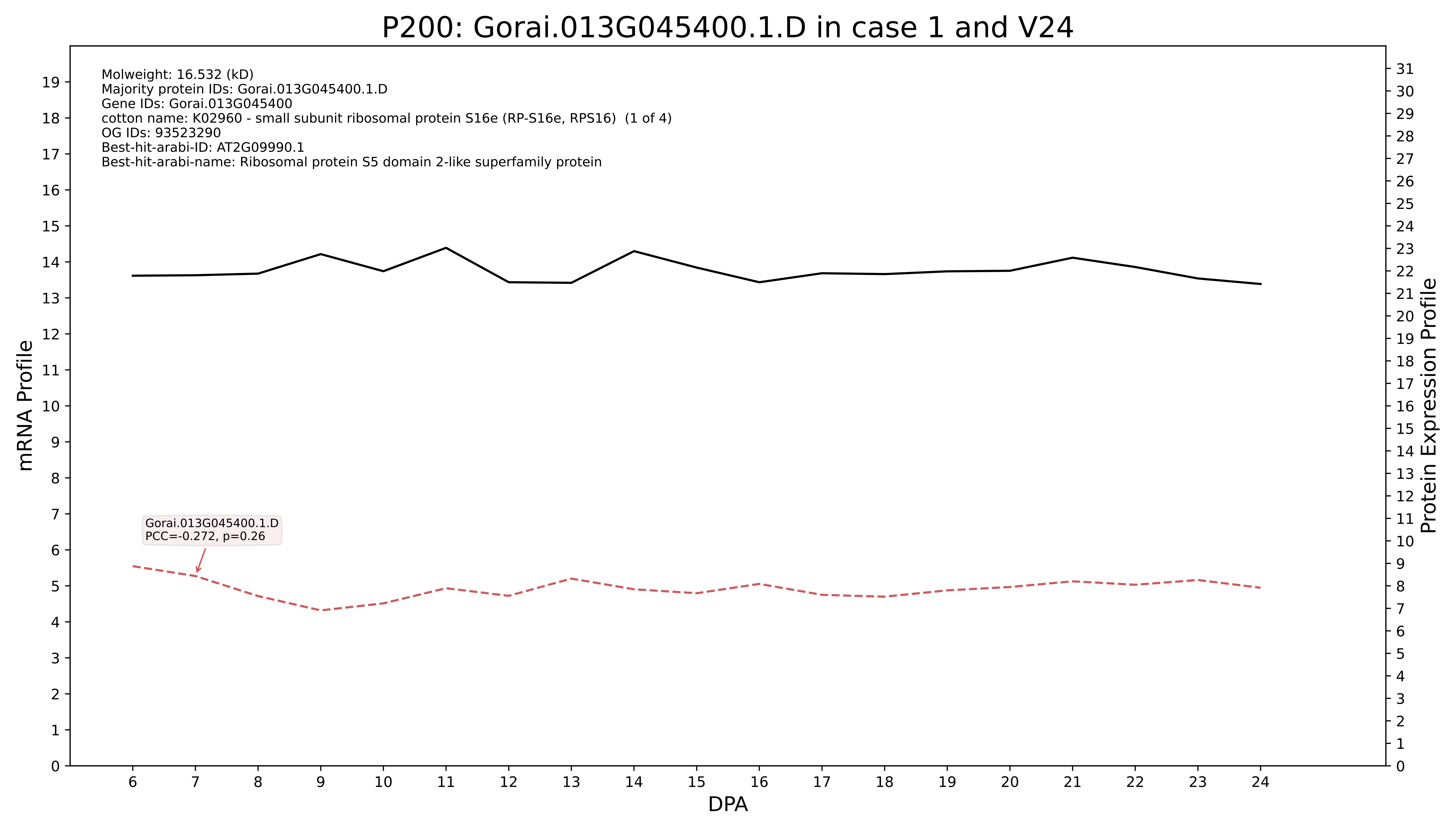
P200: Gorai.013G045400.1.D in case 1 and V24
Molweight: 16.532 (kD)
Majority protein IDs: Gorai.013G045400.1.D
Gene IDs: Gorai.013G045400
cotton name: K02960 - small subunit ribosomal protein S16e (RP-S16e, RPS16) (1 of 4)
OG IDs: 93523290
Best-hit-arabi-ID: AT2G09990.1
Best-hit-arabi-name: Ribosomal protein S5 domain 2-like superfamily protein
Gorai.013G045400.1.D
PCC=-0.272, p=0.26
mRNA Profile
Protein Expression Profile
DPA

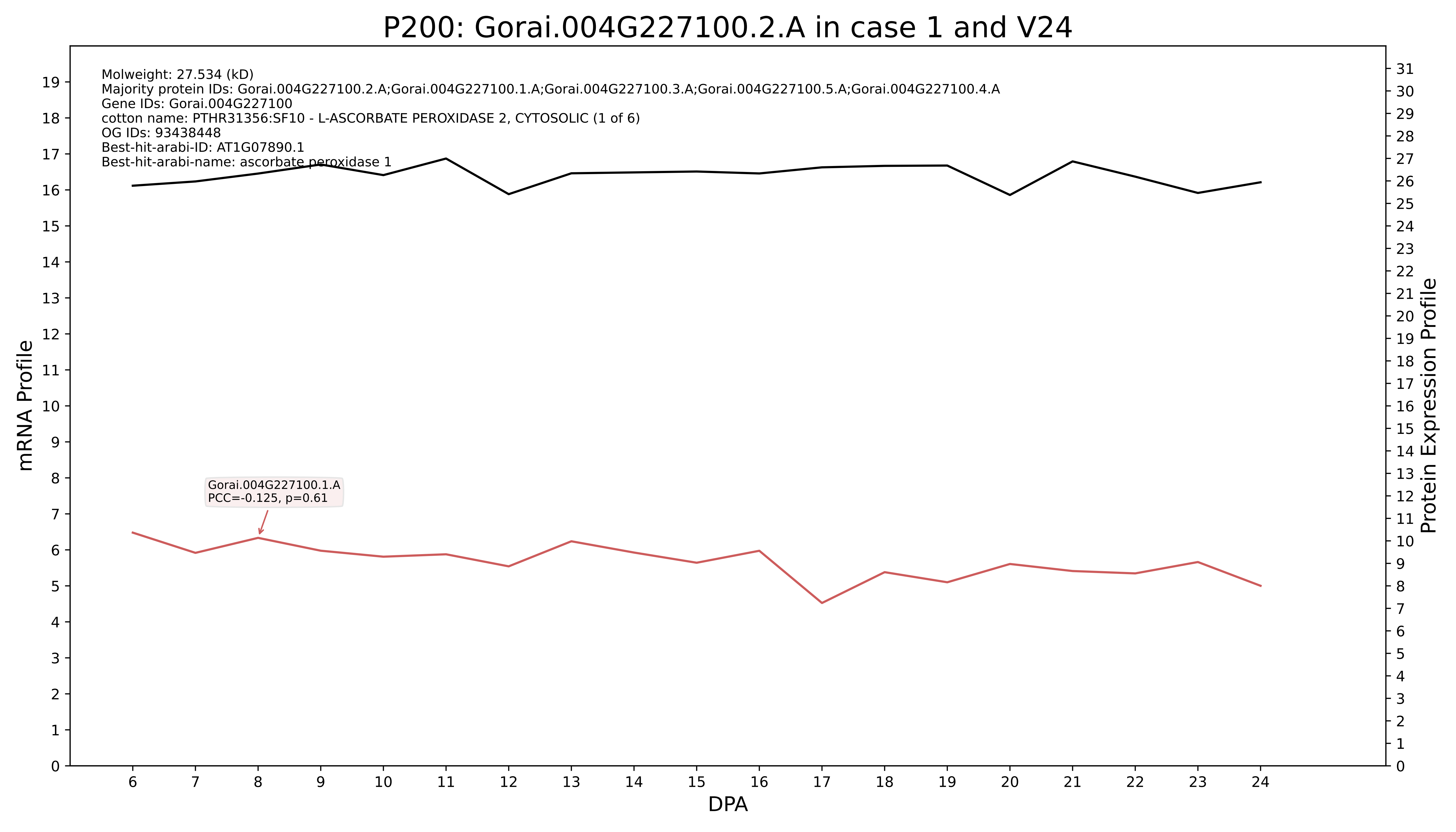
P200: Gorai.004G227100.2.A in case 1 and V24
Molweight: 27.534 (kD)
Majority protein IDs: Gorai.004G227100.2.A;Gorai.004G227100.1.A;Gorai.004G227100.3.A;Gorai.004G227100.5.A;Gorai.004G227100.4.A
Gene IDs: Gorai.004G227100
cotton name: PTHR31356:SF10 - L-ASCORBATE PEROXIDASE 2, CYTOSOLIC (1 of 6)
OG IDs: 93438448
Best-hit-arabi-ID: AT1G07890.1
Best-hit-arabi-name: ascorbate peroxidase 1
Gorai.004G227100.1.A
PCC=-0.125, p=0.61
mRNA Profile
Protein Expression Profile
DPA

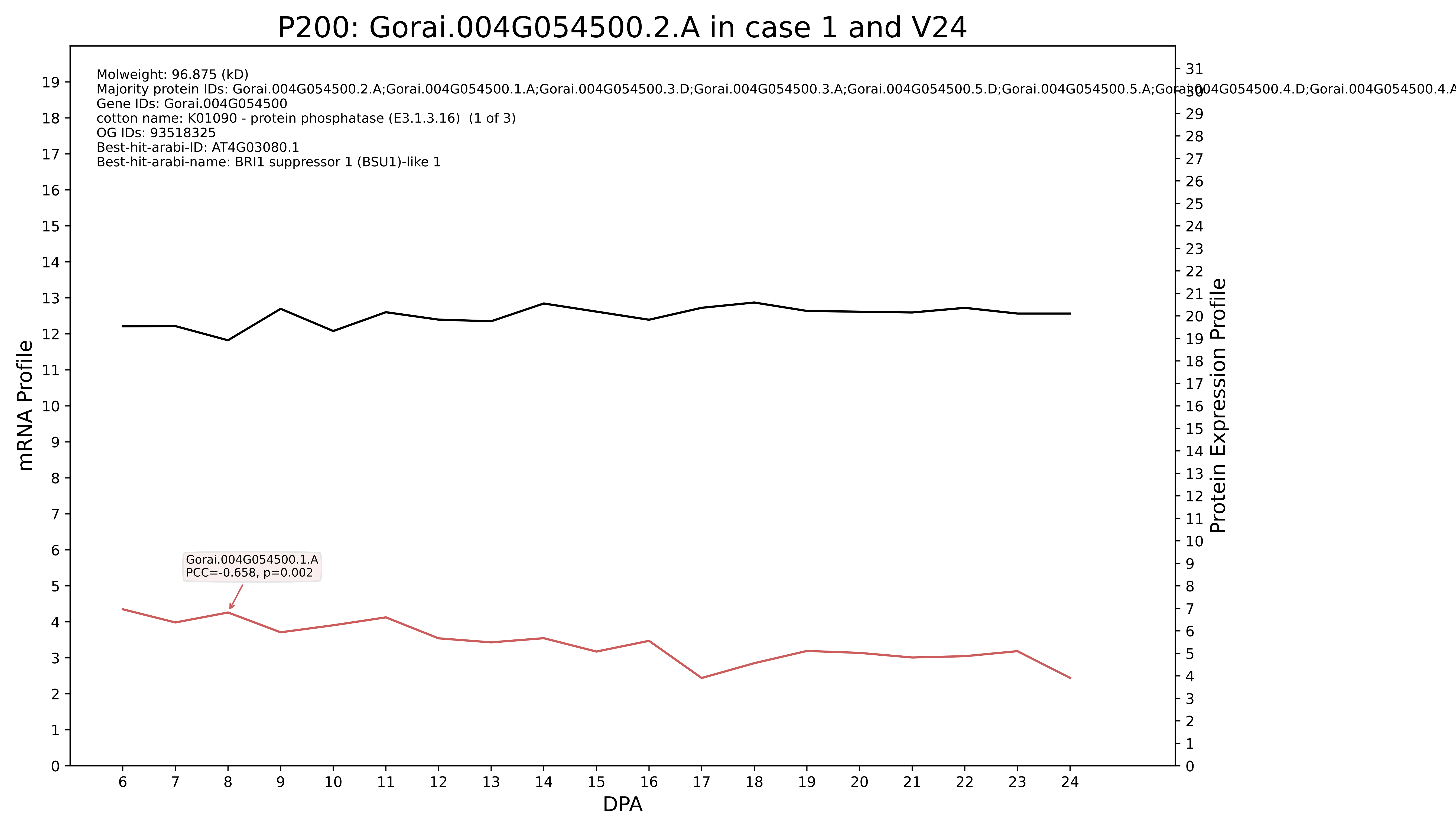
P200: Gorai.004G054500.2.A in case 1 and V24
Molweight: 96.875 (kD)
Majority protein IDs: Gorai.004G054500.2.A;Gorai.004G054500.1.A;Gorai.004G054500.3.D;Gorai.004G054500.3.A;Gorai.004G054500.5.D;Gorai.004G054500.5.A;Gorai.004G054500.4.D;Gorai.004G054500.4.A
Gene IDs: Gorai.004G054500
cotton name: K01090 - protein phosphatase (E3.1.3.16) (1 of 3)
OG IDs: 93518325
Best-hit-arabi-ID: AT4G03080.1
Best-hit-arabi-name: BRI1 suppressor 1 (BSU1)-like 1
Gorai.004G054500.1.A
PCC=-0.658, p=0.002
mRNA Profile
Protein Expression Profile
DPA

### P200: Gorai.013G019500.1.A in case 1 and V24

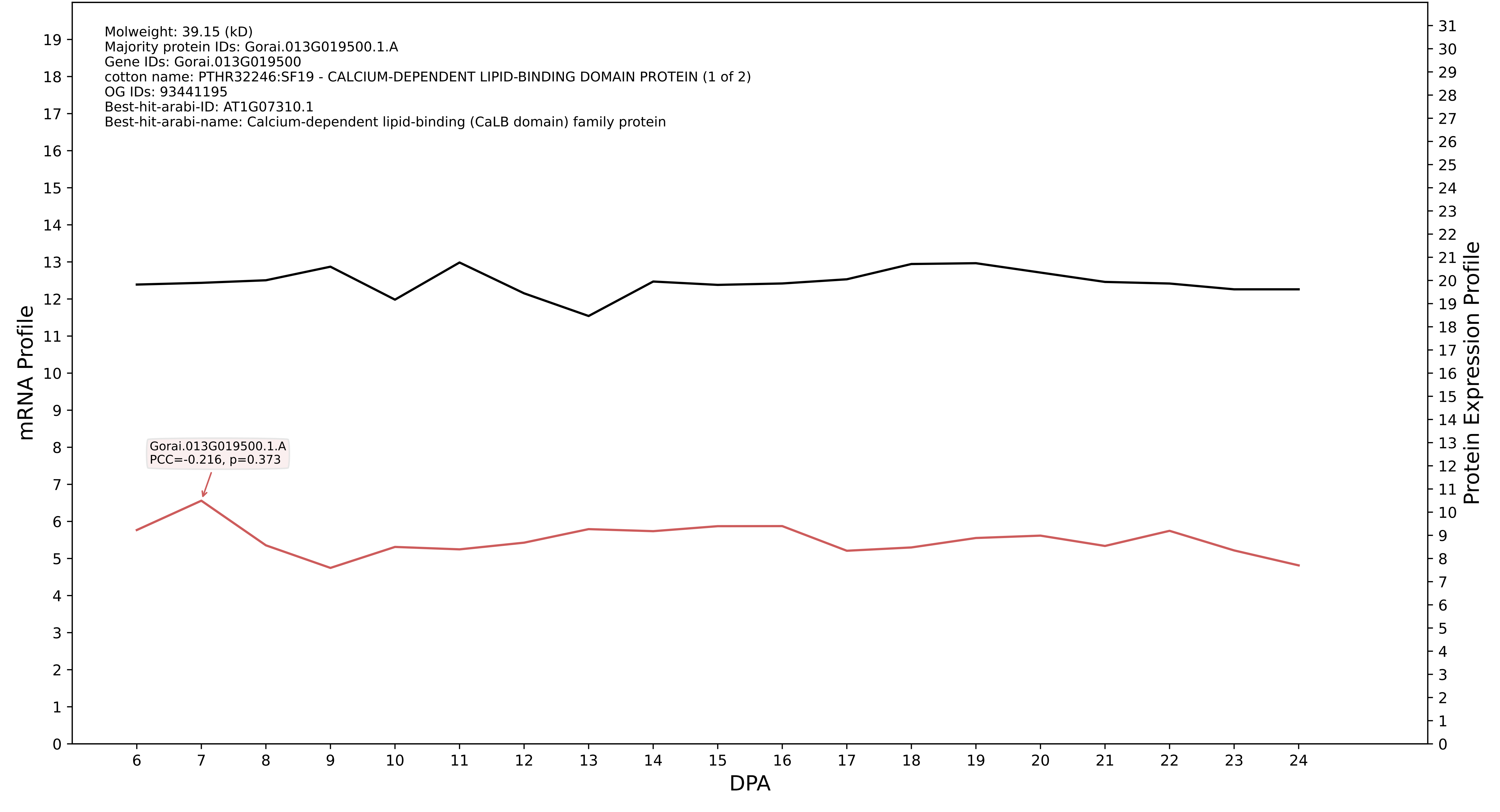

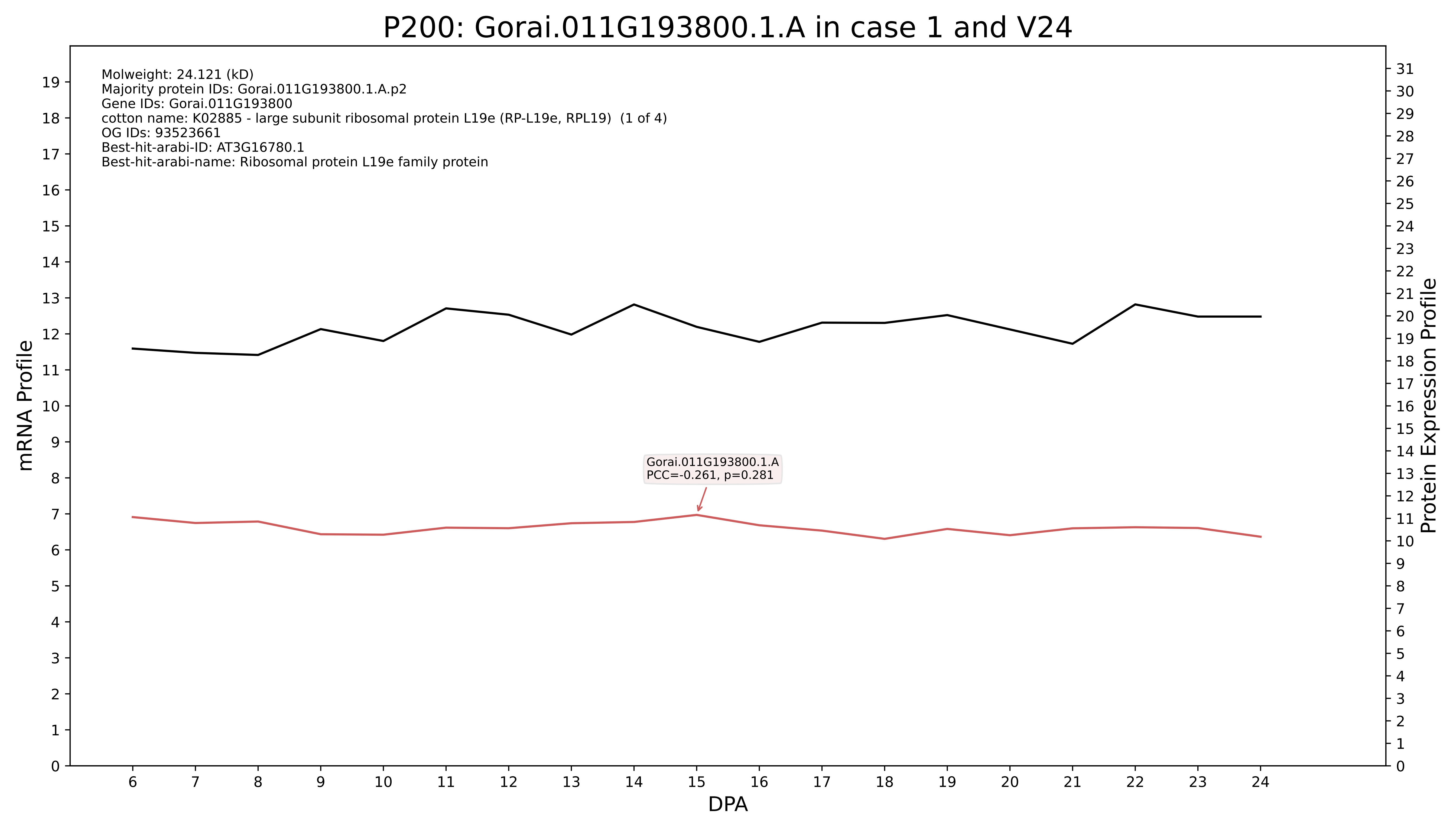

P200: Gorai.011G193800.1.A in case 1 and V24
Molweight: 24.121 (kD)
Majority protein IDs: Gorai.011G193800.1.A.p2
Gene IDs: Gorai.011G193800
cotton name: K02885 - large subunit ribosomal protein L19e (RP-L19e, RPL19) (1 of 4)
OG IDs: 93523661
Best-hit-arabi-ID: AT3G16780.1
Best-hit-arabi-name: Ribosomal protein L19e family protein
Gorai.011G193800.1.A
PCC=-0.261, p=0.281
mRNA Profile
Protein Expression Profile
DPA

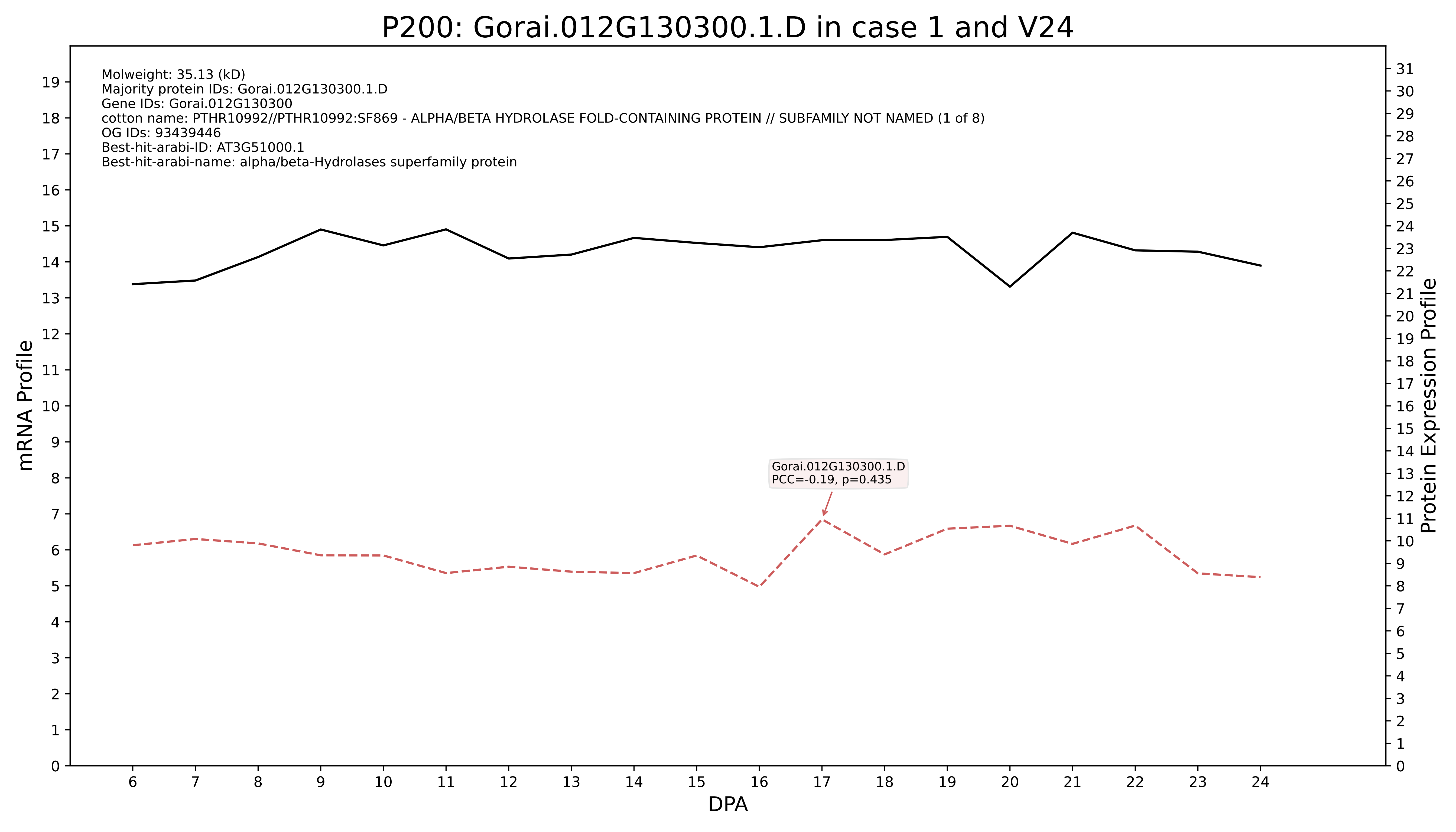
P200: Gorai.012G130300.1.D in case 1 and V24
Molweight: 35.13 (kD)
Majority protein IDs: Gorai.012G130300.1.D
Gene IDs: Gorai.012G130300
cotton name: PTHR10992//PTHR10992:SF869 - ALPHA/BETA HYDROLASE FOLD-CONTAINING PROTEIN // SUBFAMILY NOT NAMED (1 of 8)
OG IDs: 93439446
Best-hit-arabi-ID: AT3G51000.1
Best-hit-arabi-name: alpha/beta-Hydrolases superfamily protein
Gorai.012G130300.1.D
PCC=-0.19, p=0.435
mRNA Profile
Protein Expression Profile
DPA

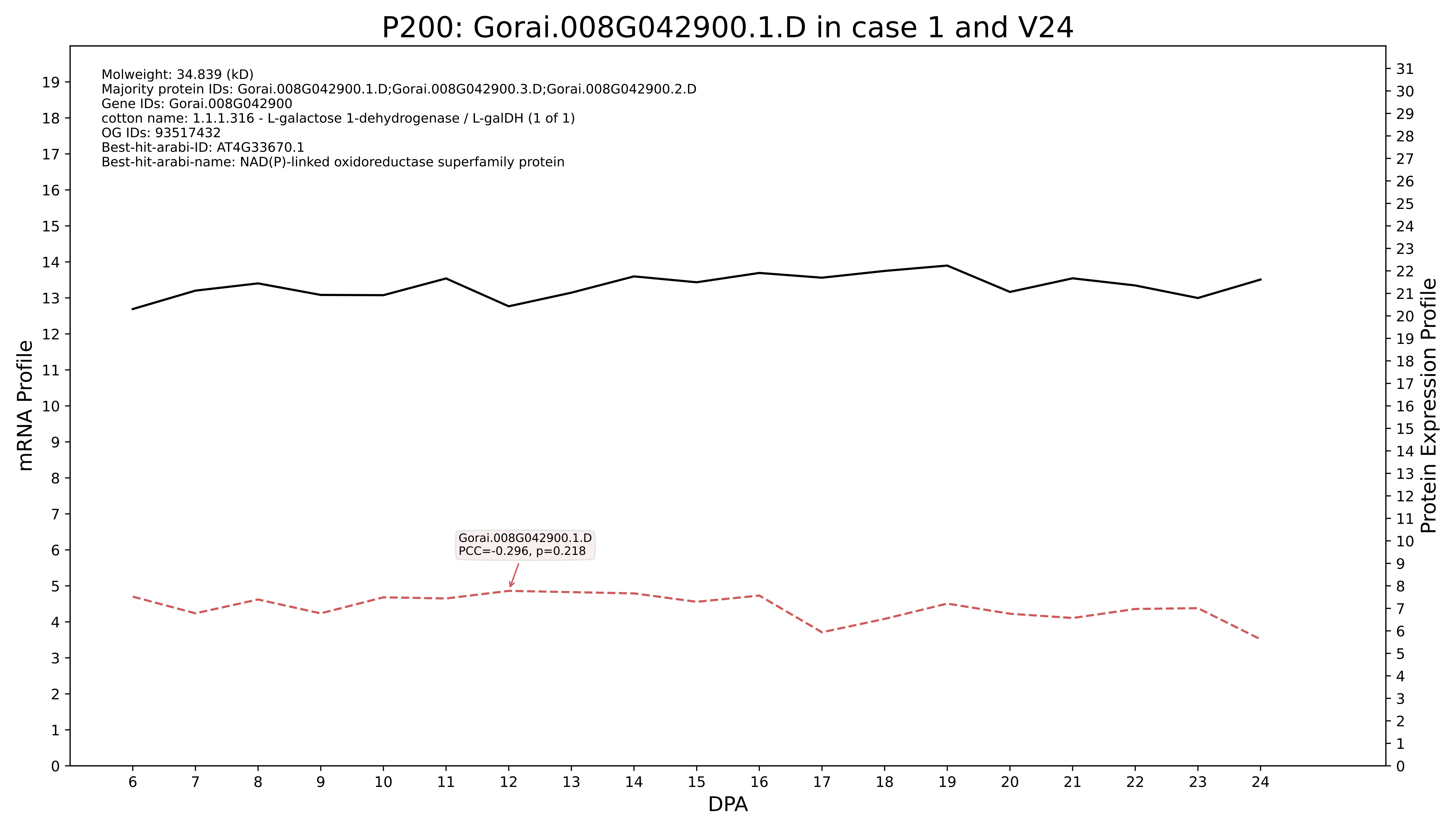
P200: Gorai.008G042900.1.D in case 1 and V24
Molweight: 34.839 (kD)
Majority protein IDs: Gorai.008G042900.1.D;Gorai.008G042900.3.D;Gorai.008G042900.2.D
Gene IDs: Gorai.008G042900
cotton name: 1.1.1.316 - L-galactose 1-dehydrogenase / L-galDH (1 of 1)
OG IDs: 93517432
Best-hit-arabi-ID: AT4G33670.1
Best-hit-arabi-name: NAD(P)-linked oxidoreductase superfamily protein
Gorai.008G042900.1.D
PCC=-0.296, p=0.218
mRNA Profile
Protein Expression Profile
DPA

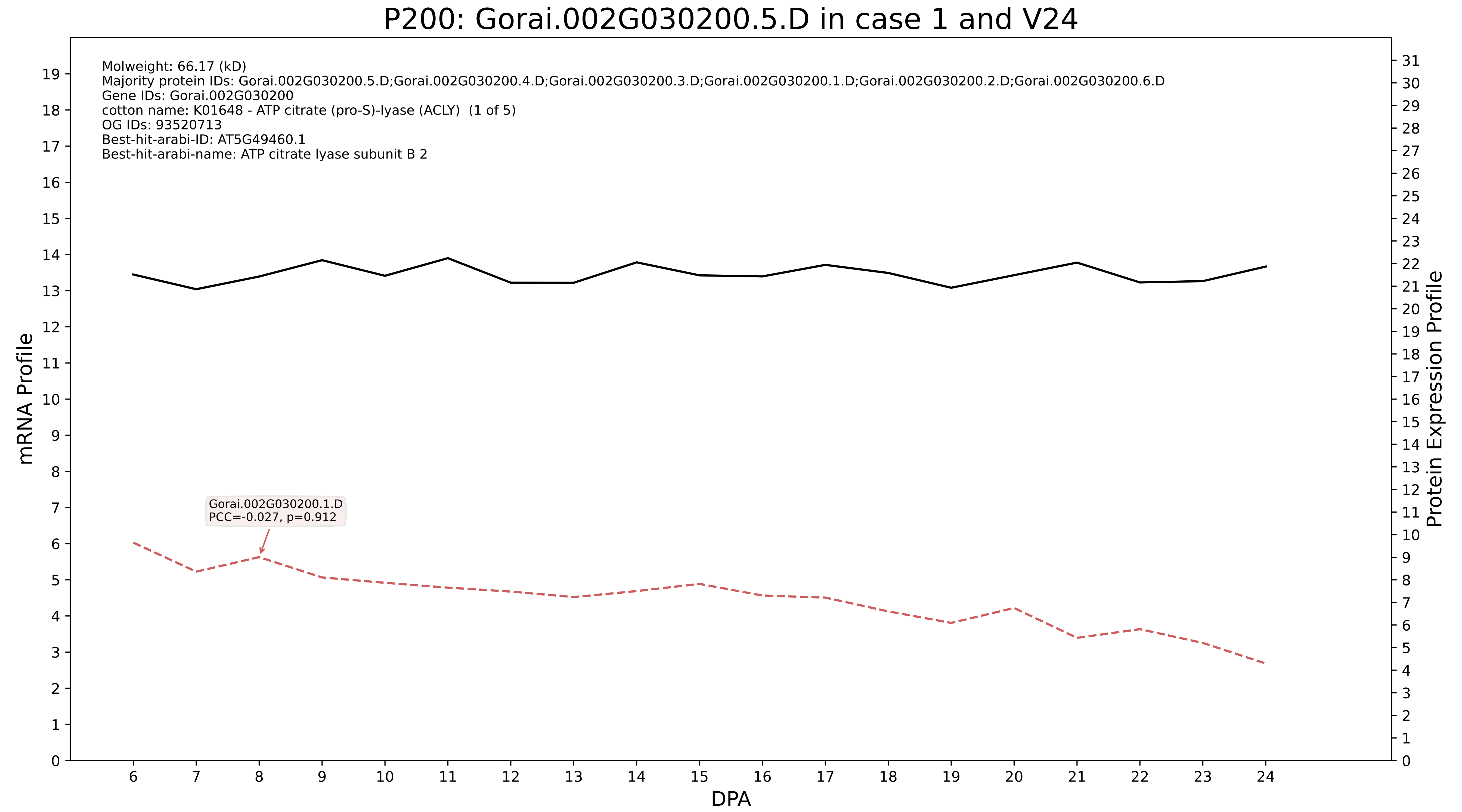
P200: Gorai.002G030200.5.D in case 1 and V24
Molweight: 66.17 (kD)
Majority protein IDs: Gorai.002G030200.5.D;Gorai.002G030200.4.D;Gorai.002G030200.3.D;Gorai.002G030200.1.D;Gorai.002G030200.2.D;Gorai.002G030200.6.D
Gene IDs: Gorai.002G030200
cotton name: K01648 - ATP citrate (pro-S)-lyase (ACLY) (1 of 5)
OG IDs: 93520713
Best-hit-arabi-ID: AT5G49460.1
Best-hit-arabi-name: ATP citrate lyase subunit B 2
Gorai.002G030200.1.D
PCC=-0.027, p=0.912
mRNA Profile
Protein Expression Profile
DPA

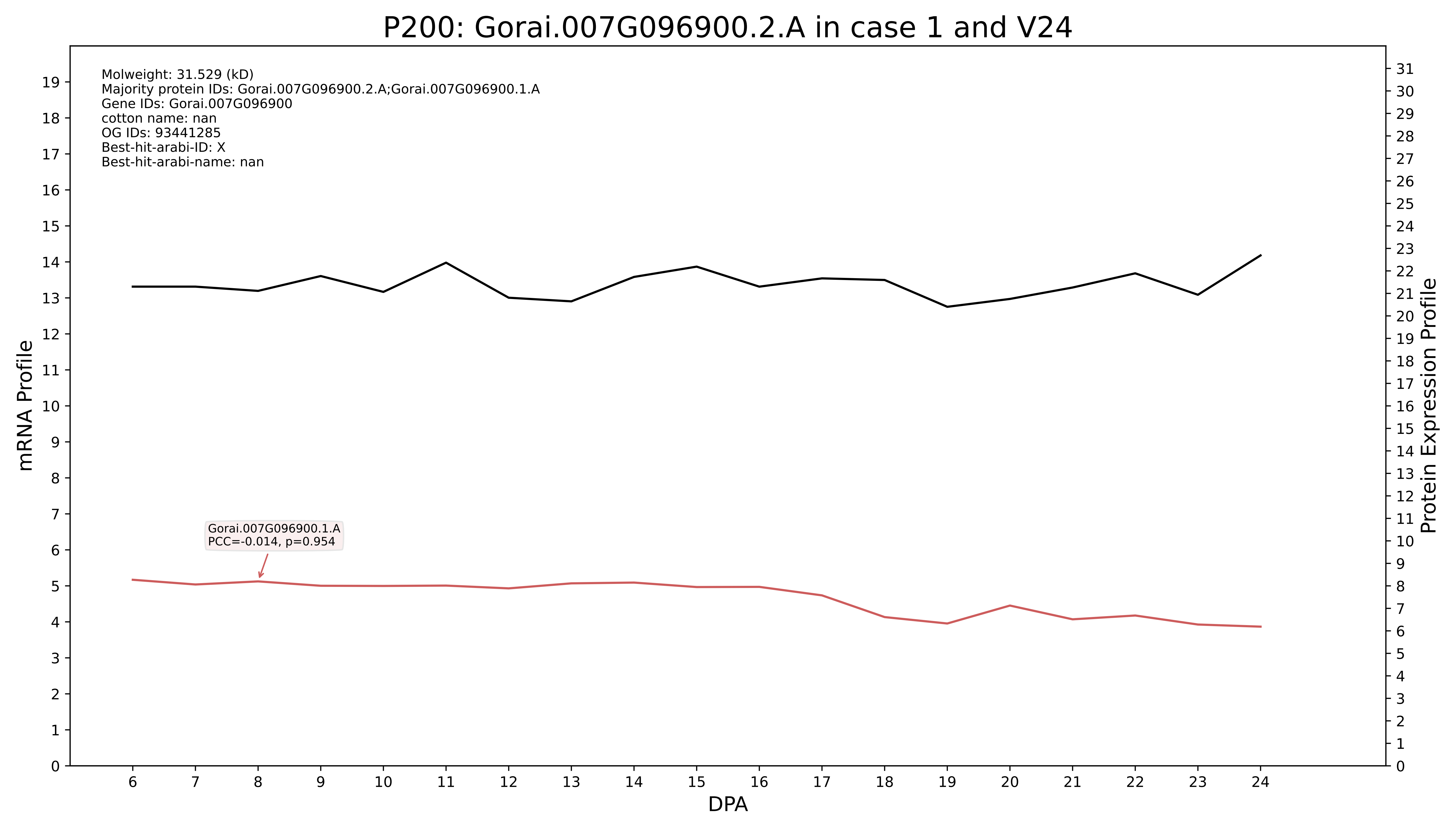
P200: Gorai.007G096900.2.A in case 1 and V24
Molweight: 31.529 (kD)
Majority protein IDs: Gorai.007G096900.2.A;Gorai.007G096900.1.A
Gene IDs: Gorai.007G096900
cotton name: nan
OG IDs: 93441285
Best-hit-arabi-ID: X
Best-hit-arabi-name: nan
Gorai.007G096900.1.A
PCC=-0.014, p=0.954
mRNA Profile
Protein Expression Profile
DPA

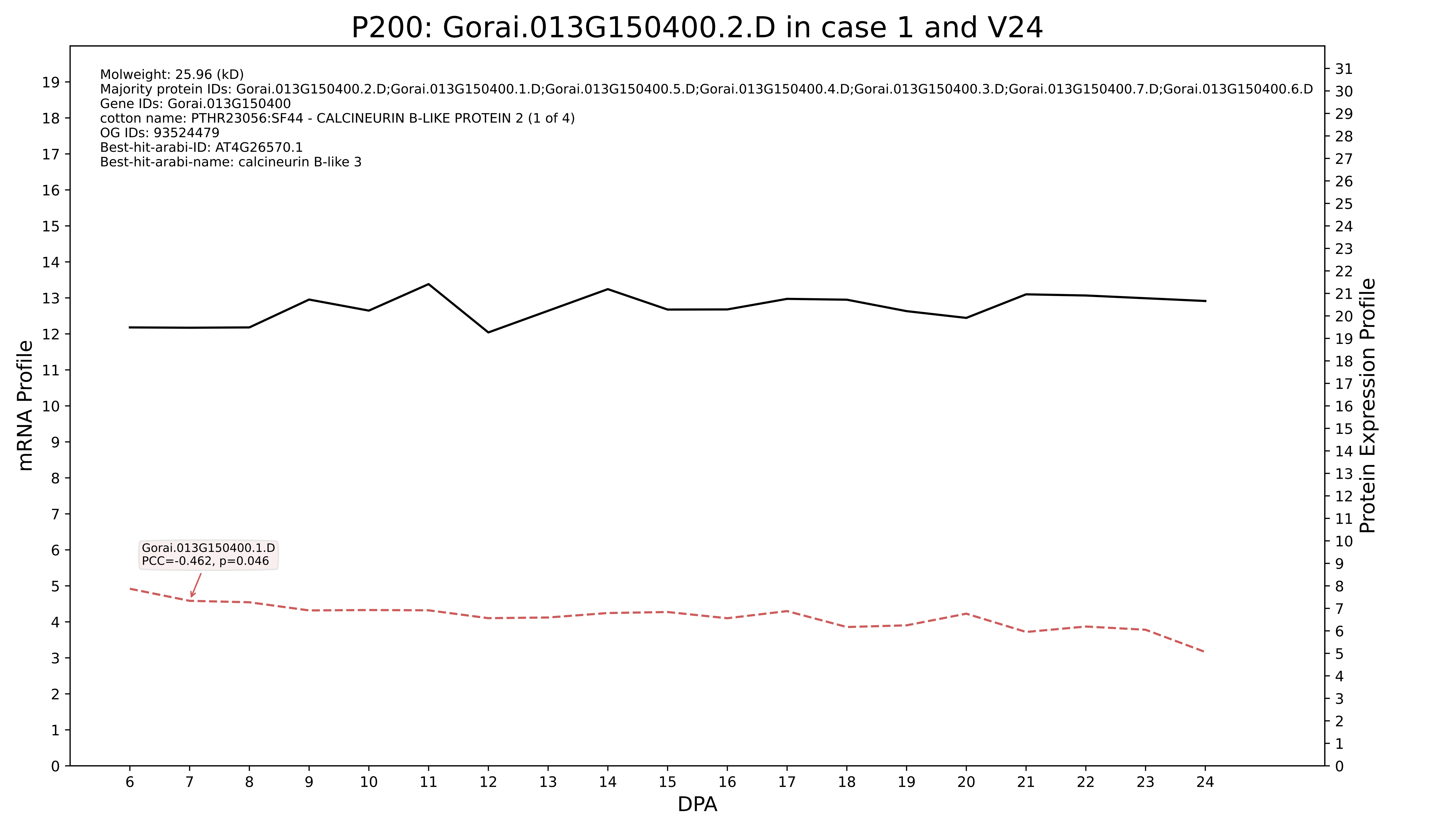
P200: Gorai.013G150400.2.D in case 1 and V24
Molweight: 25.96 (kD)
Majority protein IDs: Gorai.013G150400.2.D;Gorai.013G150400.1.D;Gorai.013G150400.5.D;Gorai.013G150400.4.D;Gorai.013G150400.3.D;Gorai.013G150400.7.D;Gorai.013G150400.6.D
Gene IDs: Gorai.013G150400
cotton name: PTHR23056:SF44 - CALCINEURIN B-LIKE PROTEIN 2 (1 of 4)
OG IDs: 93524479
Best-hit-arabi-ID: AT4G26570.1
Best-hit-arabi-name: calcineurin B-like 3
Gorai.013G150400.1.D
PCC=-0.462, p=0.046
mRNA Profile
Protein Expression Profile
DPA

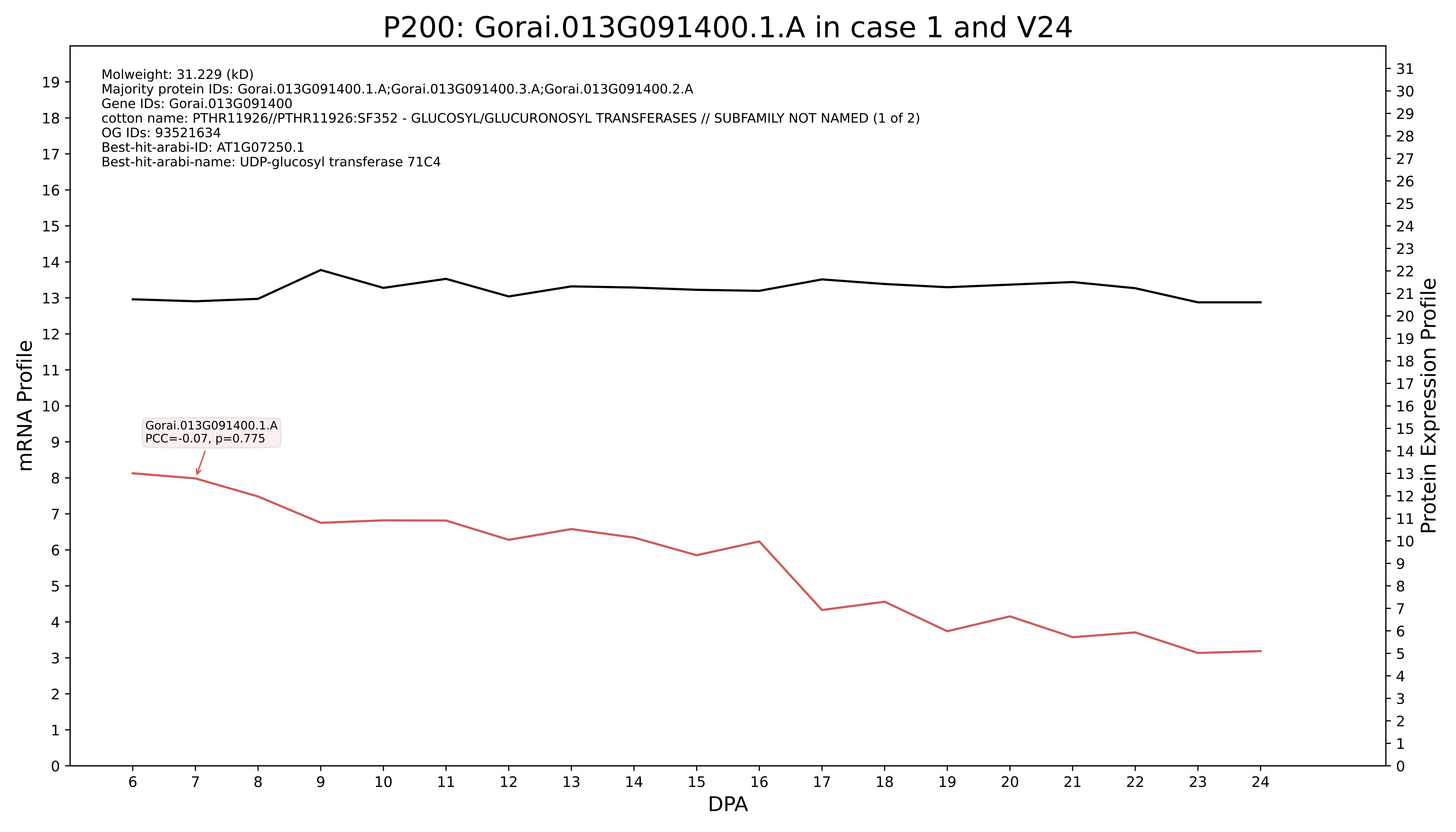
P200: Gorai.013G091400.1.A in case 1 and V24
Molweight: 31.229 (kD)
Majority protein IDs: Gorai.013G091400.1.A;Gorai.013G091400.3.A;Gorai.013G091400.2.A
Gene IDs: Gorai.013G091400
cotton name: PTHR11926//PTHR11926:SF352 - GLUCOSYL/GLUCURONOSYL TRANSFERASES // SUBFAMILY NOT NAMED (1 of 2)
OG IDs: 93521634
Best-hit-arabi-ID: AT1G07250.1
Best-hit-arabi-name: UDP-glucosyl transferase 71C4
Gorai.013G091400.1.A
PCC=-0.07, p=0.775
mRNA Profile
Protein Expression Profile
DPA

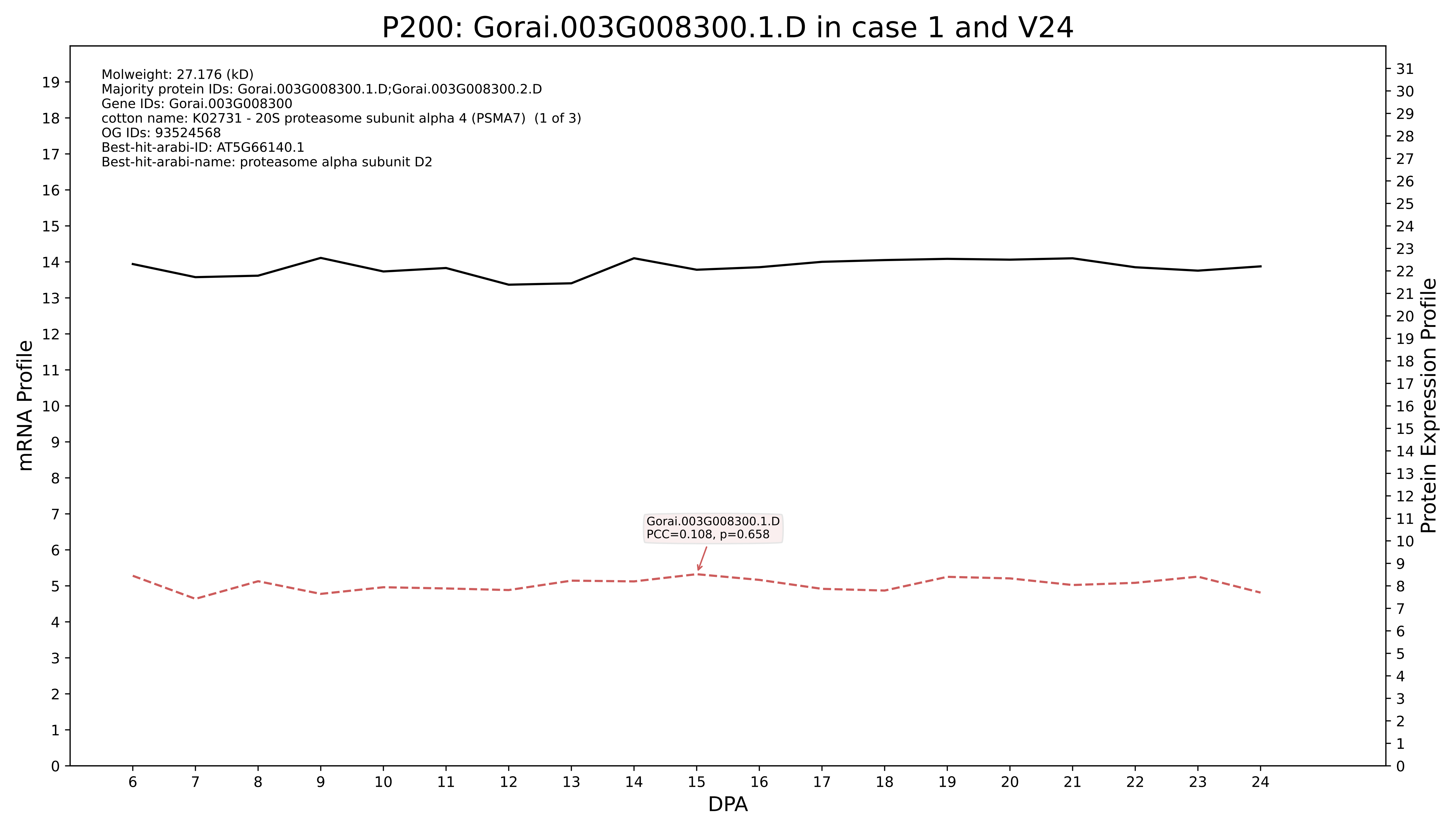
P200: Gorai.003G008300.1.D in case 1 and V24
Molweight: 27.176 (kD)
Majority protein IDs: Gorai.003G008300.1.D;Gorai.003G008300.2.D
Gene IDs: Gorai.003G008300
cotton name: K02731 - 20S proteasome subunit alpha 4 (PSMA7) (1 of 3)
OG IDs: 93524568
Best-hit-arabi-ID: AT5G66140.1
Best-hit-arabi-name: proteasome alpha subunit D2
Gorai.003G008300.1.D
PCC=0.108, p=0.658
mRNA Profile
Protein Expression Profile
DPA

### P200: Gorai.007G033400.1.D in case 1 and V24

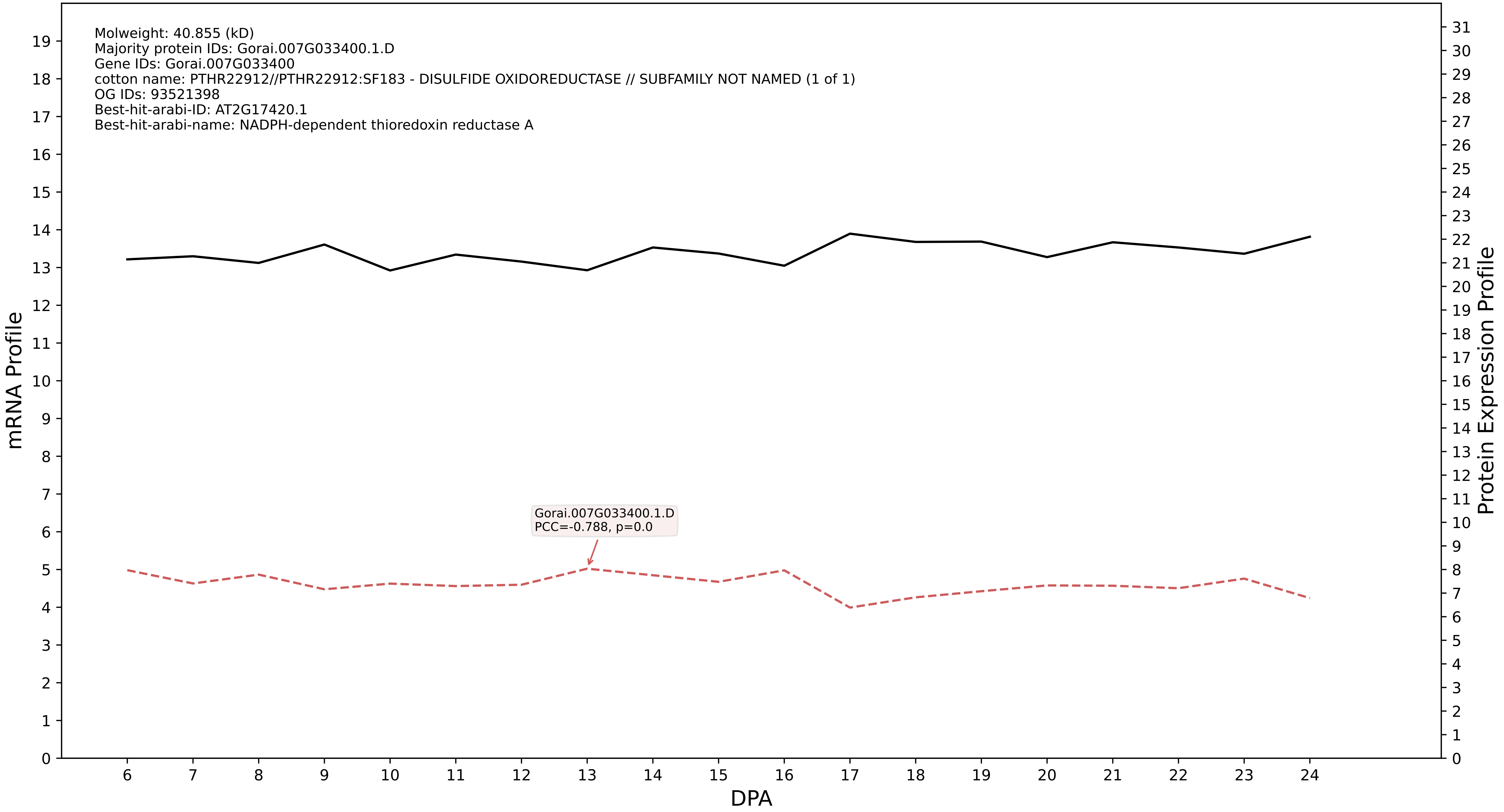

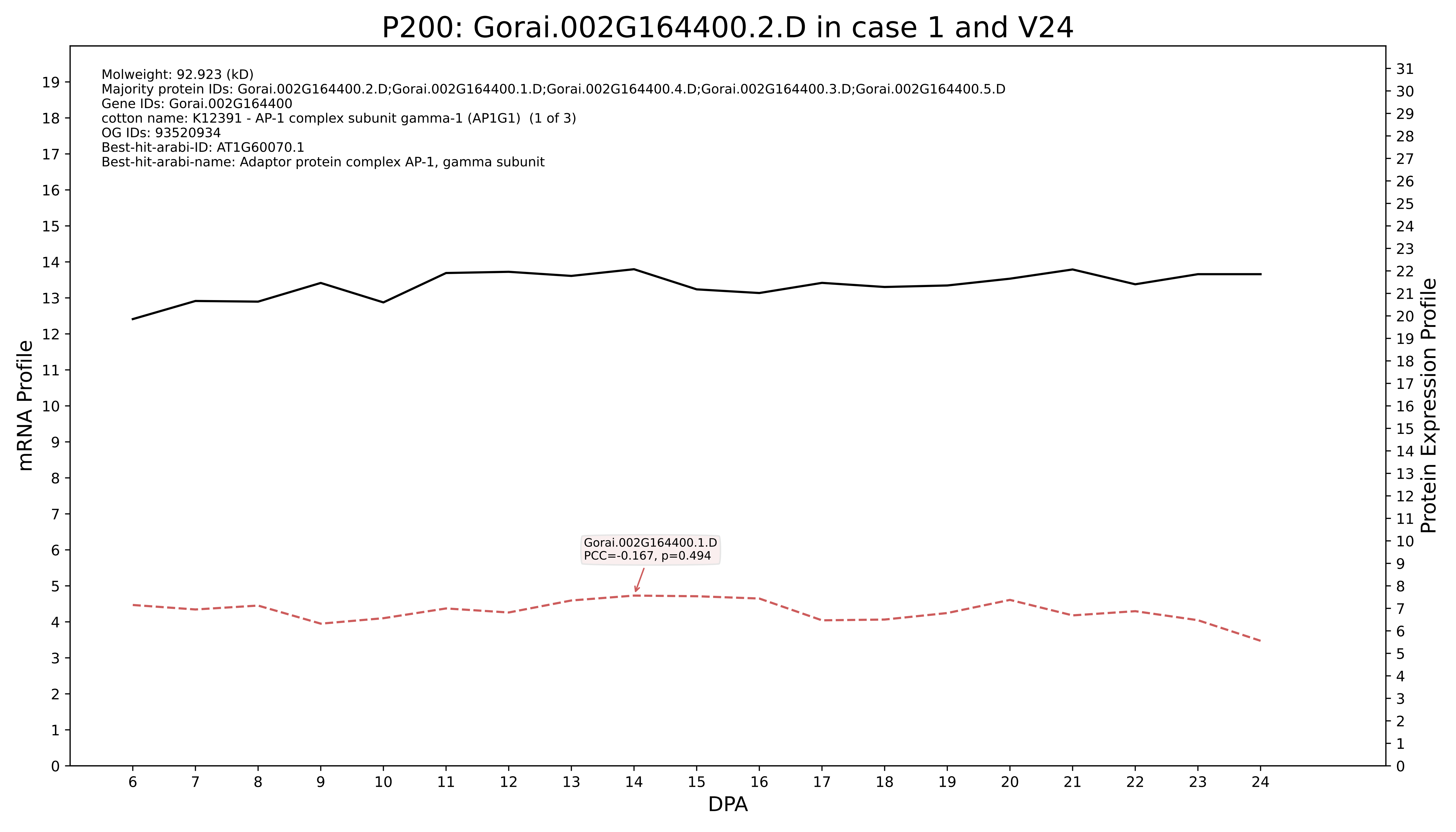

P200: Gorai.002G164400.2.D in case 1 and V24
Molweight: 92.923 (kD)
Majority protein IDs: Gorai.002G164400.2.D;Gorai.002G164400.1.D;Gorai.002G164400.4.D;Gorai.002G164400.3.D;Gorai.002G164400.5.D
Gene IDs: Gorai.002G164400
cotton name: K12391 - AP-1 complex subunit gamma-1 (AP1G1) (1 of 3)
OG IDs: 93520934
Best-hit-arabi-ID: AT1G60070.1
Best-hit-arabi-name: Adaptor protein complex AP-1, gamma subunit
Gorai.002G164400.1.D
PCC=-0.167, p=0.494
mRNA Profile
Protein Expression Profile
DPA

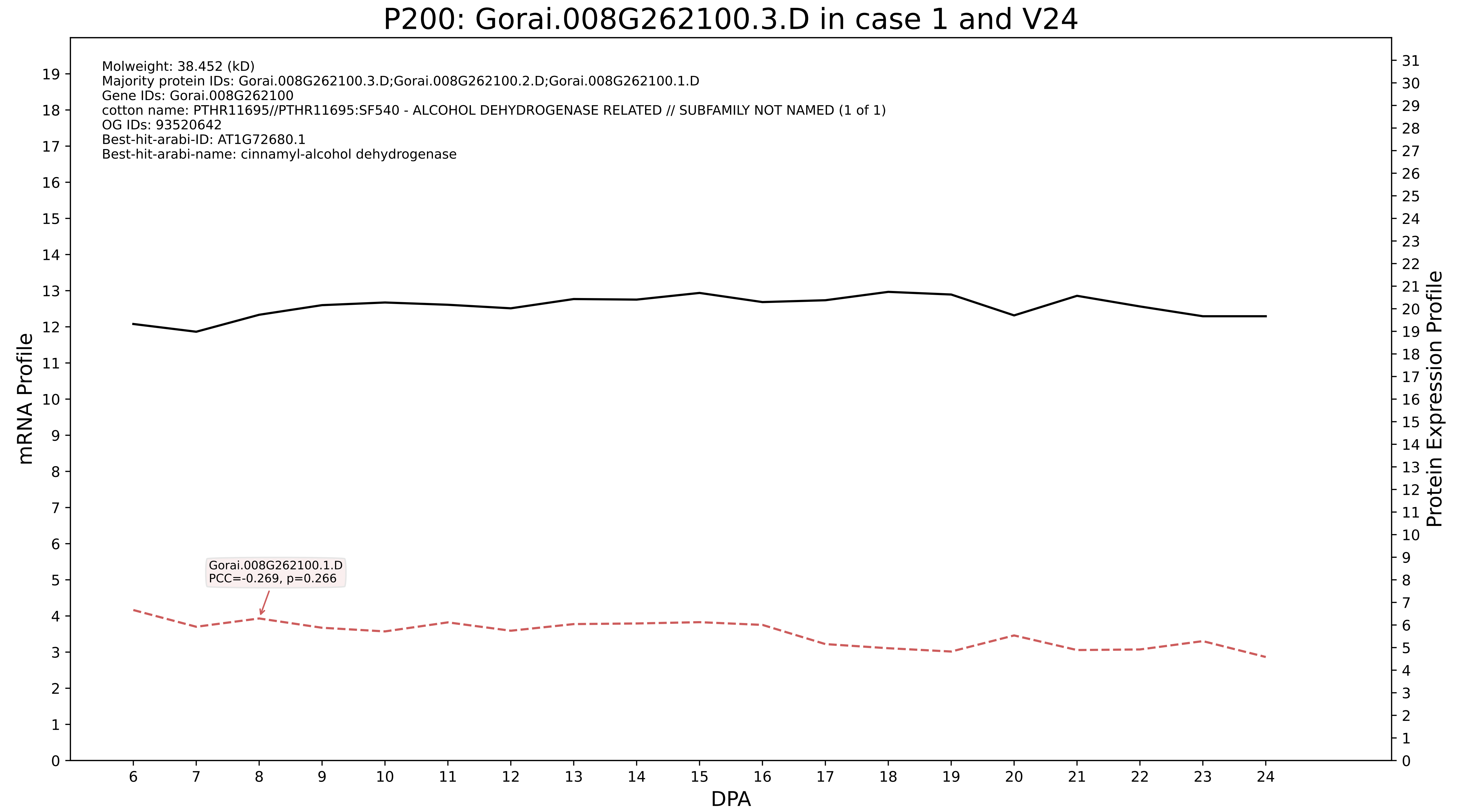

P200: Gorai.008G262100.3.D in case 1 and V24
Molweight: 38.452 (kD)
Majority protein IDs: Gorai.008G262100.3.D;Gorai.008G262100.2.D;Gorai.008G262100.1.D
Gene IDs: Gorai.008G262100
cotton name: PTHR11695//PTHR11695:SF540 - ALCOHOL DEHYDROGENASE RELATED // SUBFAMILY NOT NAMED (1 of 1)
OG IDs: 93520642
Best-hit-arabi-ID: AT1G72680.1
Best-hit-arabi-name: cinnamyl-alcohol dehydrogenase
Gorai.008G262100.1.D
PCC=-0.269, p=0.266
mRNA Profile
Protein Expression Profile
DPA

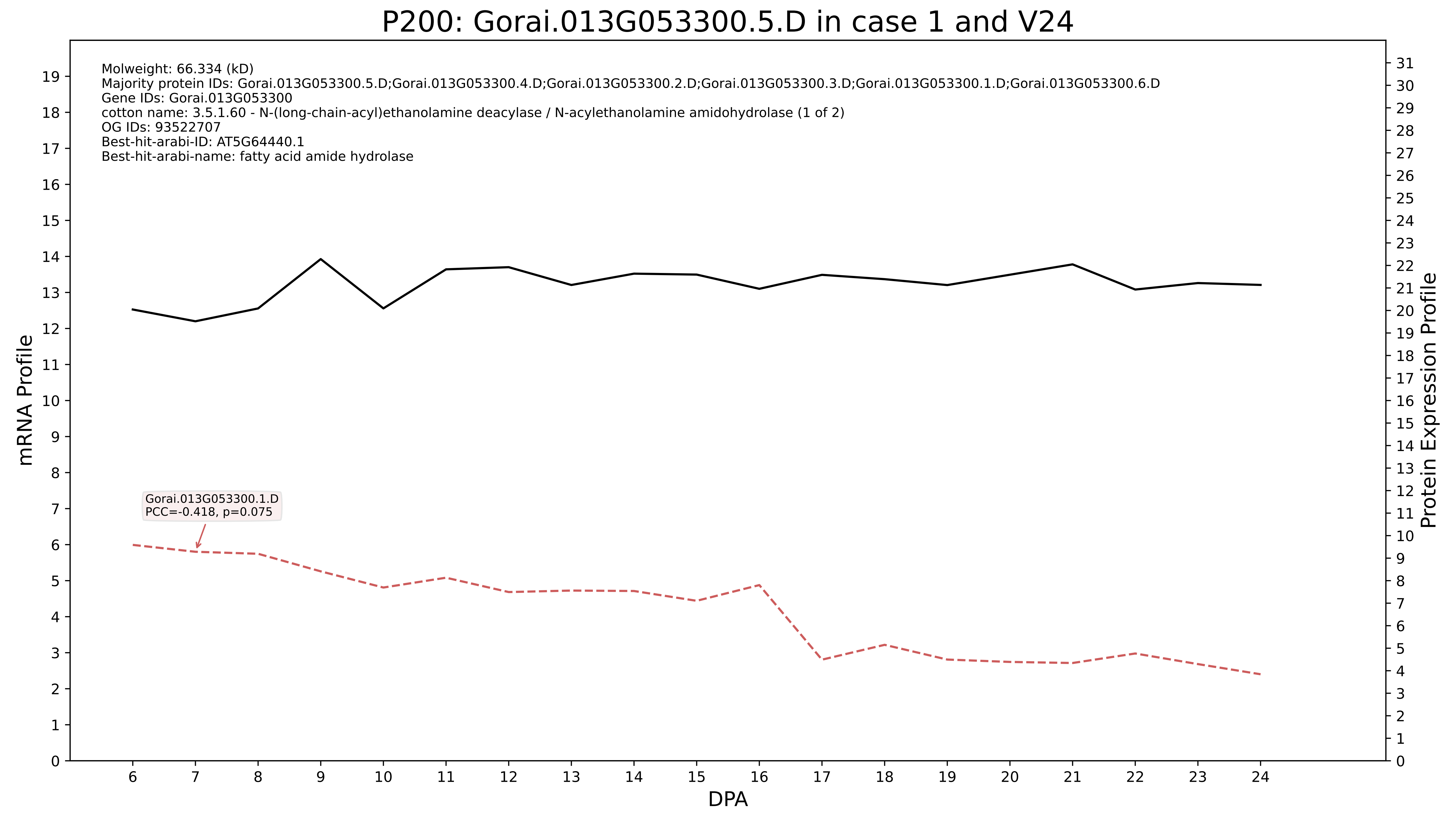

P200: Gorai.013G053300.5.D in case 1 and V24
Molweight: 66.334 (kD)
Majority protein IDs: Gorai.013G053300.5.D;Gorai.013G053300.4.D;Gorai.013G053300.2.D;Gorai.013G053300.3.D;Gorai.013G053300.1.D;Gorai.013G053300.6.D
Gene IDs: Gorai.013G053300
cotton name: 3.5.1.60 - N-(long-chain-acyl)ethanolamine deacylase / N-acylethanolamine amidohydrolase (1 of 2)
OG IDs: 93522707
Best-hit-arabi-ID: AT5G64440.1
Best-hit-arabi-name: fatty acid amide hydrolase
Gorai.013G053300.1.D
PCC=-0.418, p=0.075
mRNA Profile
Protein Expression Profile
DPA

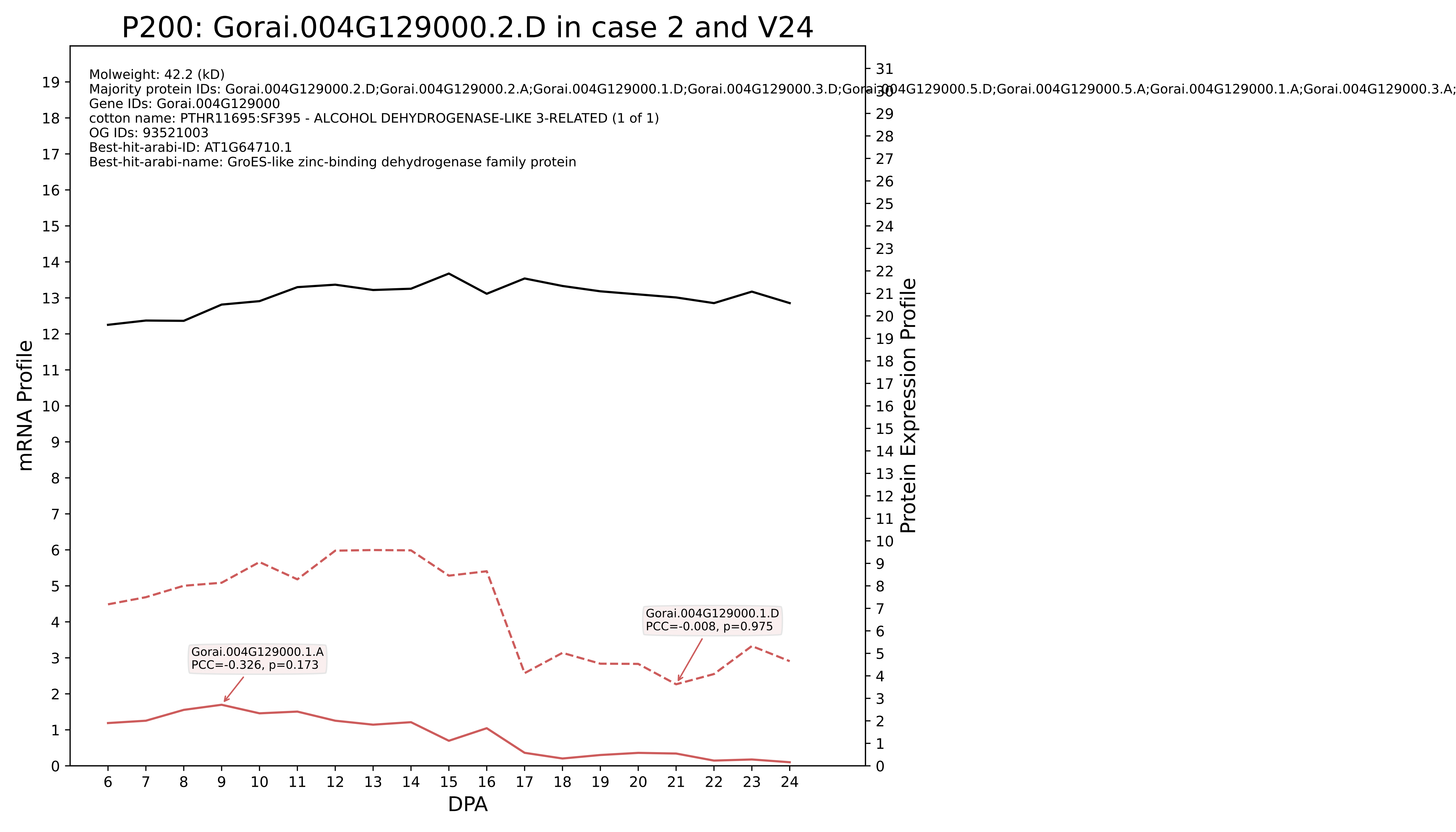
P200: Gorai.004G129000.2.D in case 2 and V24
Molweight: 42.2 (kD)
Majority protein IDs: Gorai.004G129000.2.D;Gorai.004G129000.2.A;Gorai.004G129000.1.D;Gorai.004G129000.3.D;Gorai.004G129000.5.D;Gorai.004G129000.5.A;Gorai.004G129000.1.A;Gorai.004G129000.3.A;
Gene IDs: Gorai.004G129000
cotton name: PTHR11695:SF395 - ALCOHOL DEHYDROGENASE-LIKE 3-RELATED (1 of 1)
OG IDs: 93521003
Best-hit-arabi-ID: AT1G64710.1
Best-hit-arabi-name: GroES-like zinc-binding dehydrogenase family protein
Gorai.004G129000.1.D
PCC=-0.008, p=0.975
Gorai.004G129000.1.A
PCC=-0.326, p=0.173
mRNA Profile
Protein Expression Profile
DPA

00: Gorai.013G116800.6.D in case 2 and V24

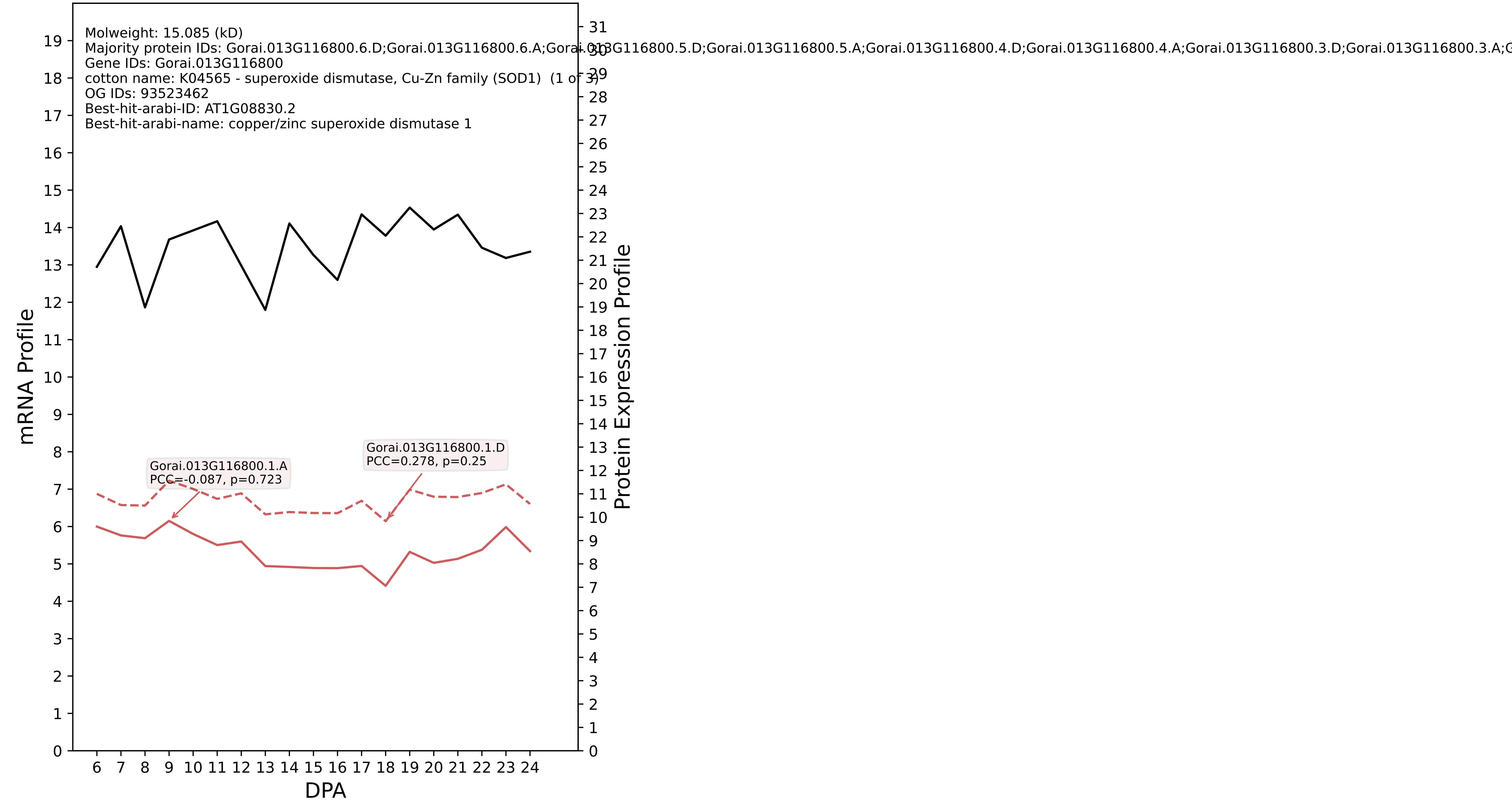

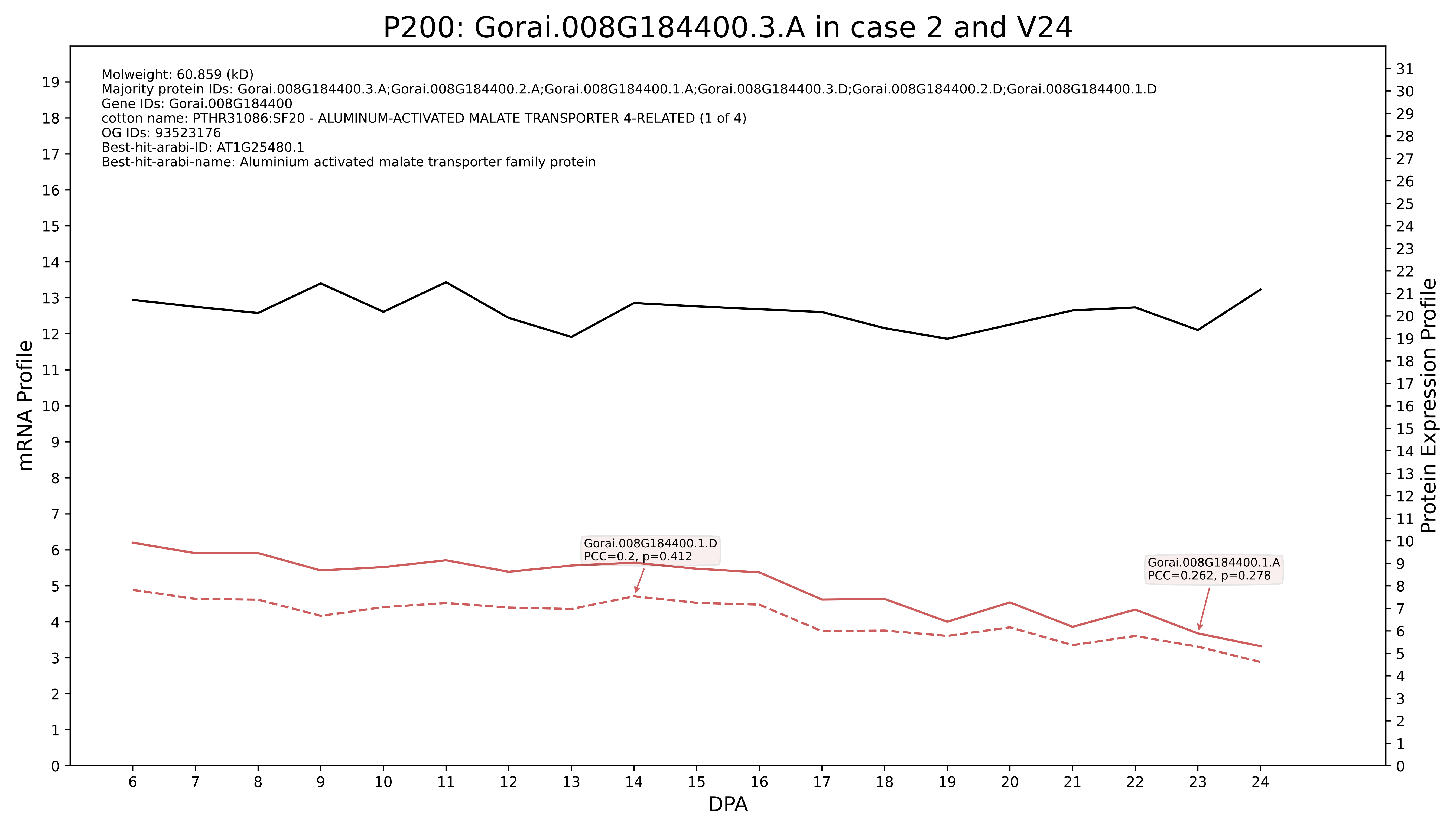
P200: Gorai.008G184400.3.A in case 2 and V24
Molweight: 60.859 (kD)
Majority protein IDs: Gorai.008G184400.3.A;Gorai.008G184400.2.A;Gorai.008G184400.1.A;Gorai.008G184400.3.D;Gorai.008G184400.2.D;Gorai.008G184400.1.D
Gene IDs: Gorai.008G184400
cotton name: PTHR31086:SF20 - ALUMINUM-ACTIVATED MALATE TRANSPORTER 4-RELATED (1 of 4)
OG IDs: 93523176
Best-hit-arabi-ID: AT1G25480.1
Best-hit-arabi-name: Aluminium activated malate transporter family protein
Gorai.008G184400.1.D
PCC=0.2, p=0.412
Gorai.008G184400.1.A
PCC=0.262, p=0.278
mRNA Profile
Protein Expression Profile
DPA

### P200: Gorai.013G050700.2.D in case 2 and V24

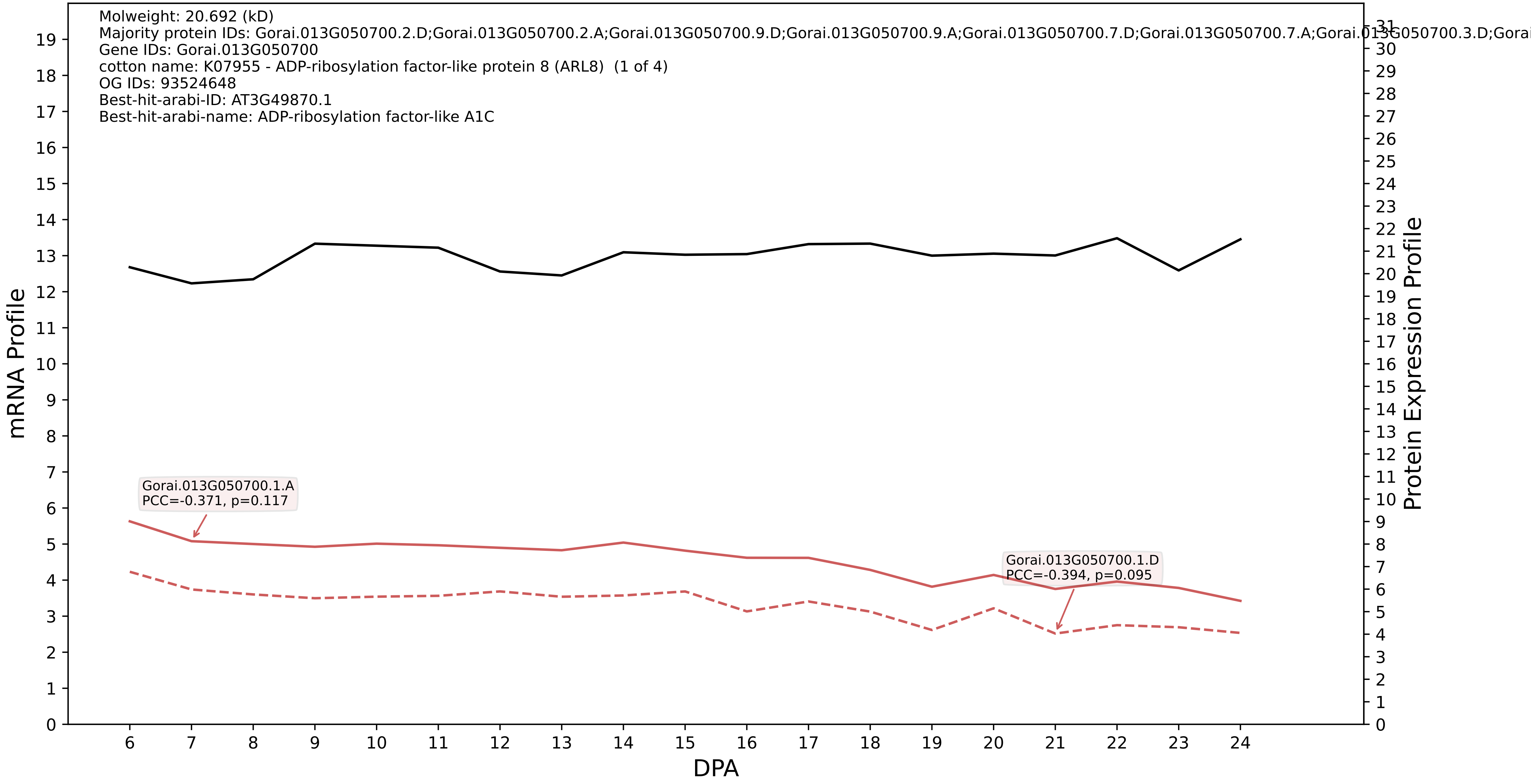

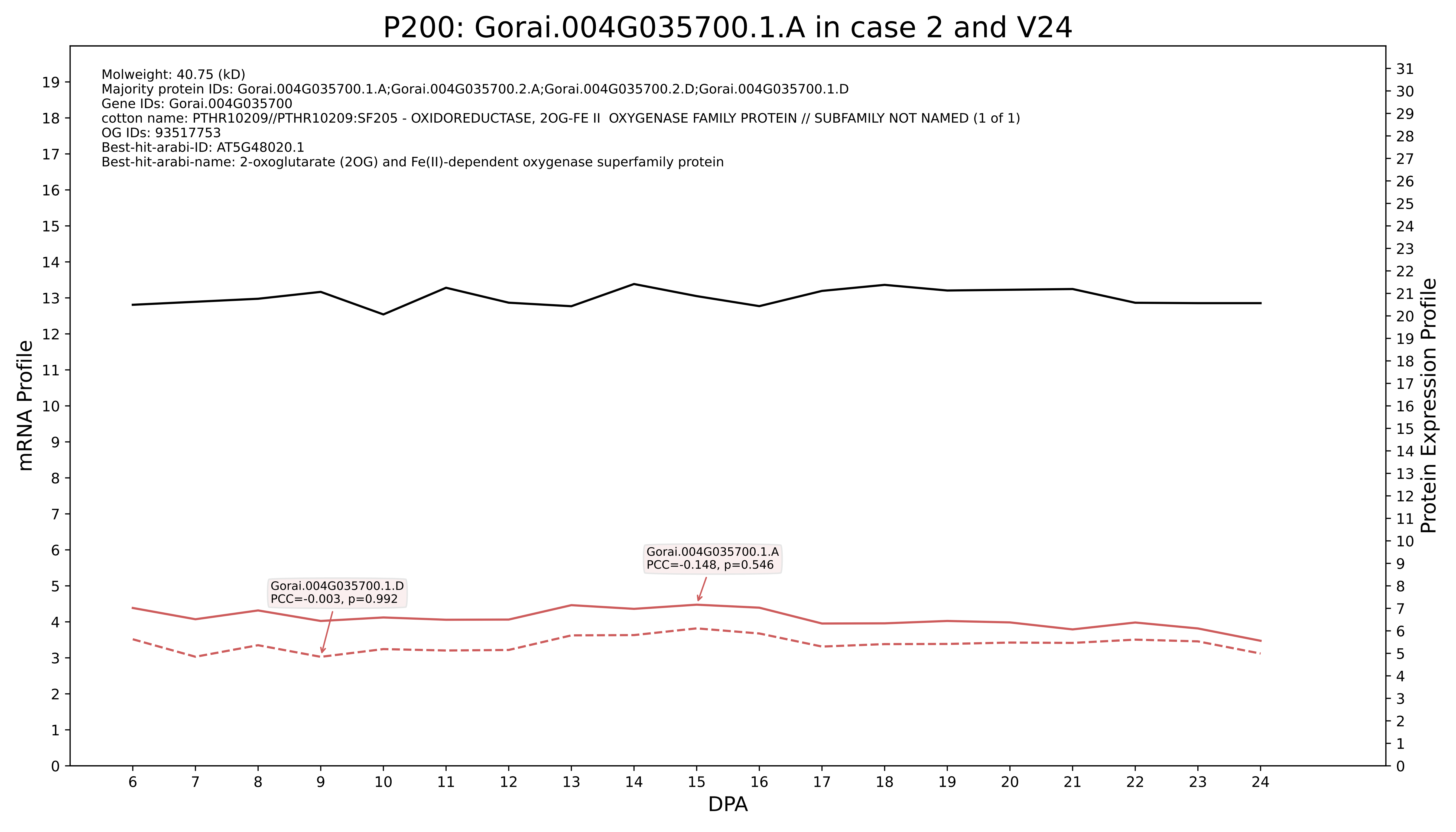
P200: Gorai.004G035700.1.A in case 2 and V24
Molweight: 40.75 (kD)
Majority protein IDs: Gorai.004G035700.1.A;Gorai.004G035700.2.A;Gorai.004G035700.2.D;Gorai.004G035700.1.D
Gene IDs: Gorai.004G035700
cotton name: PTHR10209//PTHR10209:SF205 - OXIDOREDUCTASE, 2OG-FE II OXYGENASE FAMILY PROTEIN // SUBFAMILY NOT NAMED (1 of 1)
OG IDs: 93517753
Best-hit-arabi-ID: AT5G48020.1
Best-hit-arabi-name: 2-oxoglutarate (2OG) and Fe(II)-dependent oxygenase superfamily protein
Gorai.004G035700.1.A
PCC=-0.148, p=0.546
Gorai.004G035700.1.D
PCC=-0.003, p=0.992
mRNA Profile
Protein Expression Profile
DPA

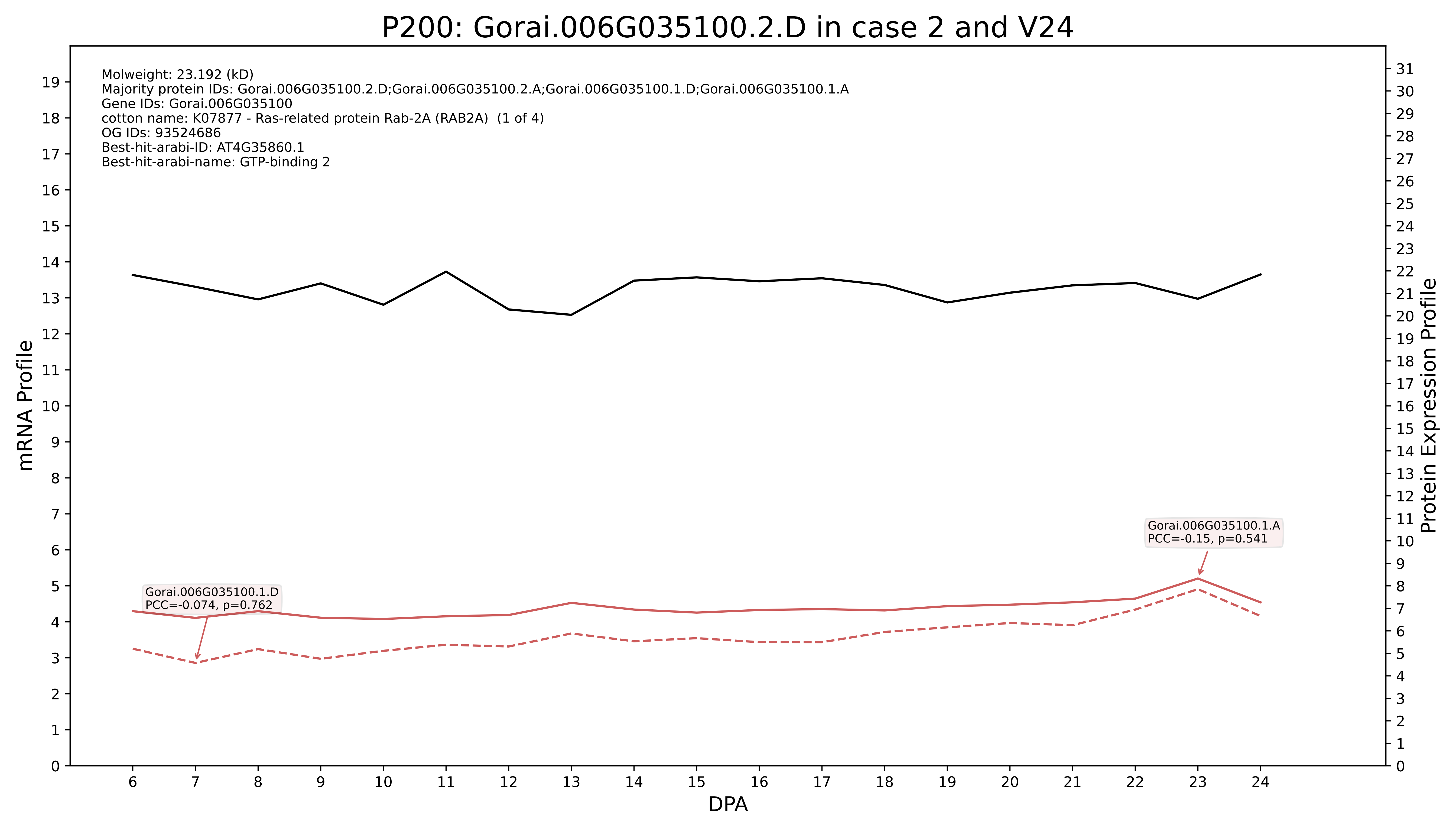
P200: Gorai.006G035100.2.D in case 2 and V24
Molweight: 23.192 (kD)
Majority protein IDs: Gorai.006G035100.2.D;Gorai.006G035100.2.A;Gorai.006G035100.1.D;Gorai.006G035100.1.A
Gene IDs: Gorai.006G035100
cotton name: K07877 - Ras-related protein Rab-2A (RAB2A) (1 of 4)
OG IDs: 93524686
Best-hit-arabi-ID: AT4G35860.1
Best-hit-arabi-name: GTP-binding 2
Gorai.006G035100.1.A
PCC=-0.15, p=0.541
Gorai.006G035100.1.D
PCC=-0.074, p=0.762
mRNA Profile
Protein Expression Profile
DPA

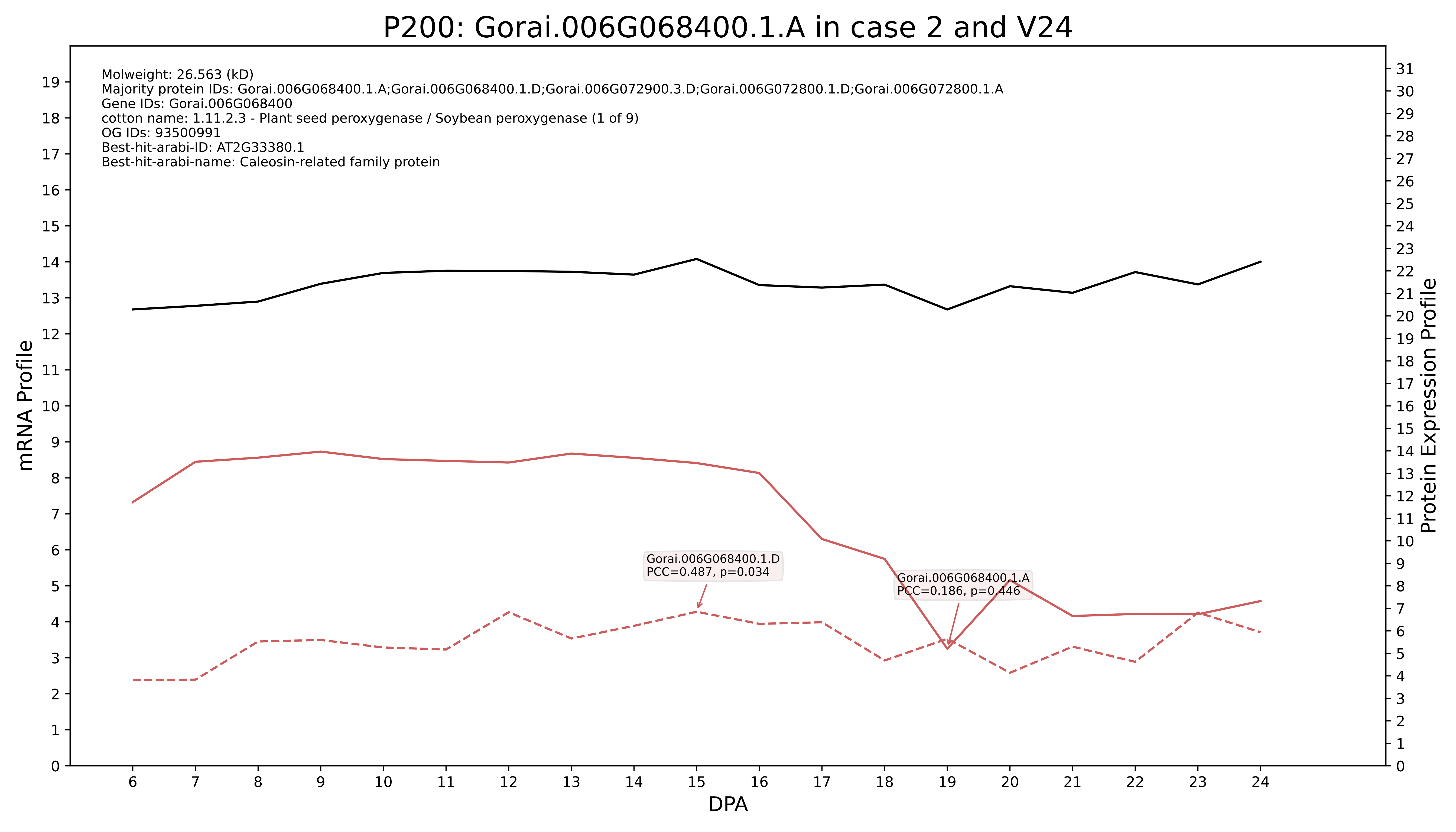
P200: Gorai.006G068400.1.A in case 2 and V24
Molweight: 26.563 (kD)
Majority protein IDs: Gorai.006G068400.1.A;Gorai.006G068400.1.D;Gorai.006G072900.3.D;Gorai.006G072800.1.D;Gorai.006G072800.1.A
Gene IDs: Gorai.006G068400
cotton name: 1.11.2.3 - Plant seed peroxygenase / Soybean peroxygenase (1 of 9)
OG IDs: 93500991
Best-hit-arabi-ID: AT2G33380.1
Best-hit-arabi-name: Caleosin-related family protein
Gorai.006G068400.1.D
PCC=0.487, p=0.034
Gorai.006G068400.1.A
PCC=0.186, p=0.446
mRNA Profile
Protein Expression Profile
DPA

### P200: Gorai.010G247000.2.D in case 2 and V24

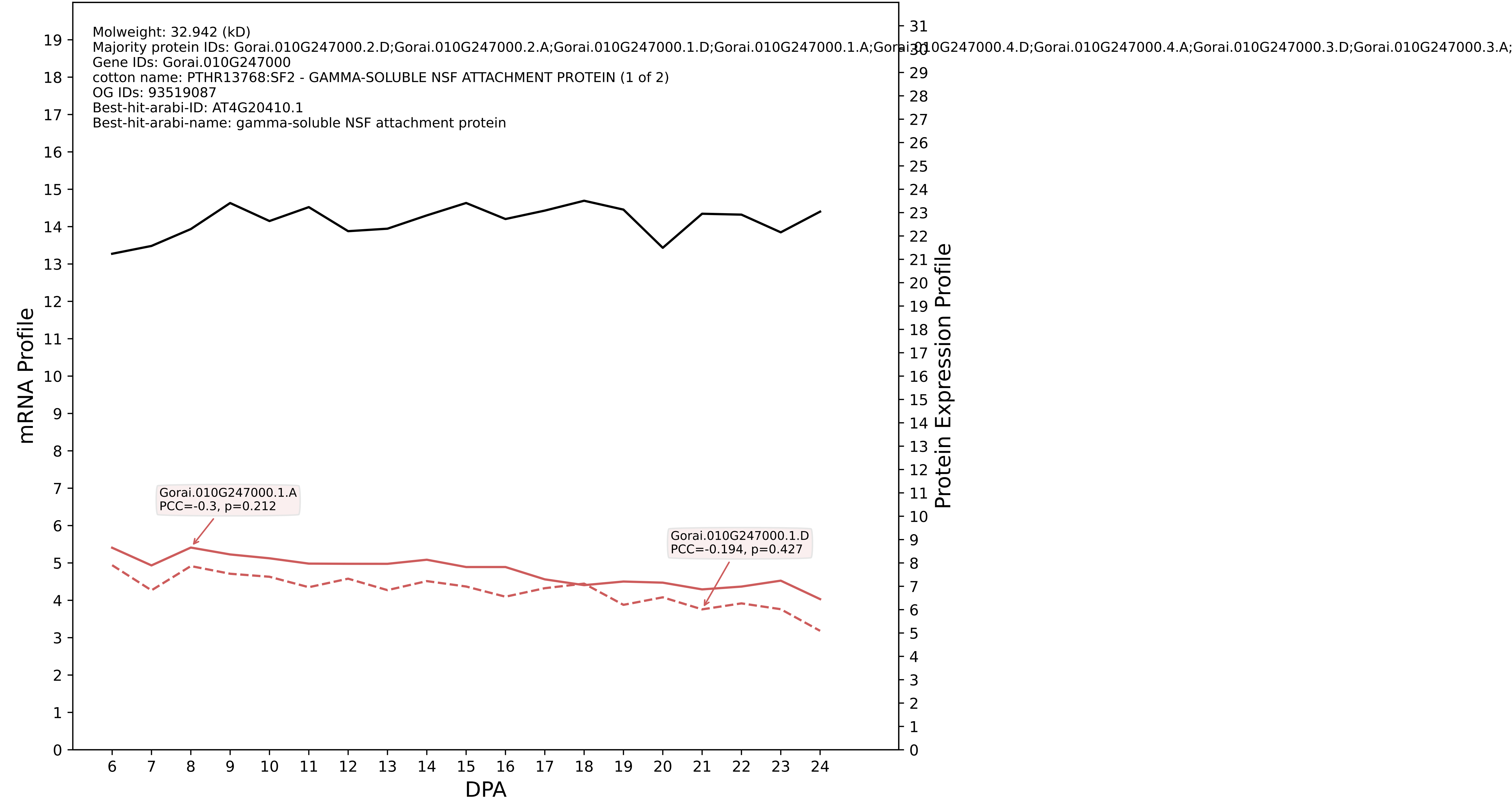

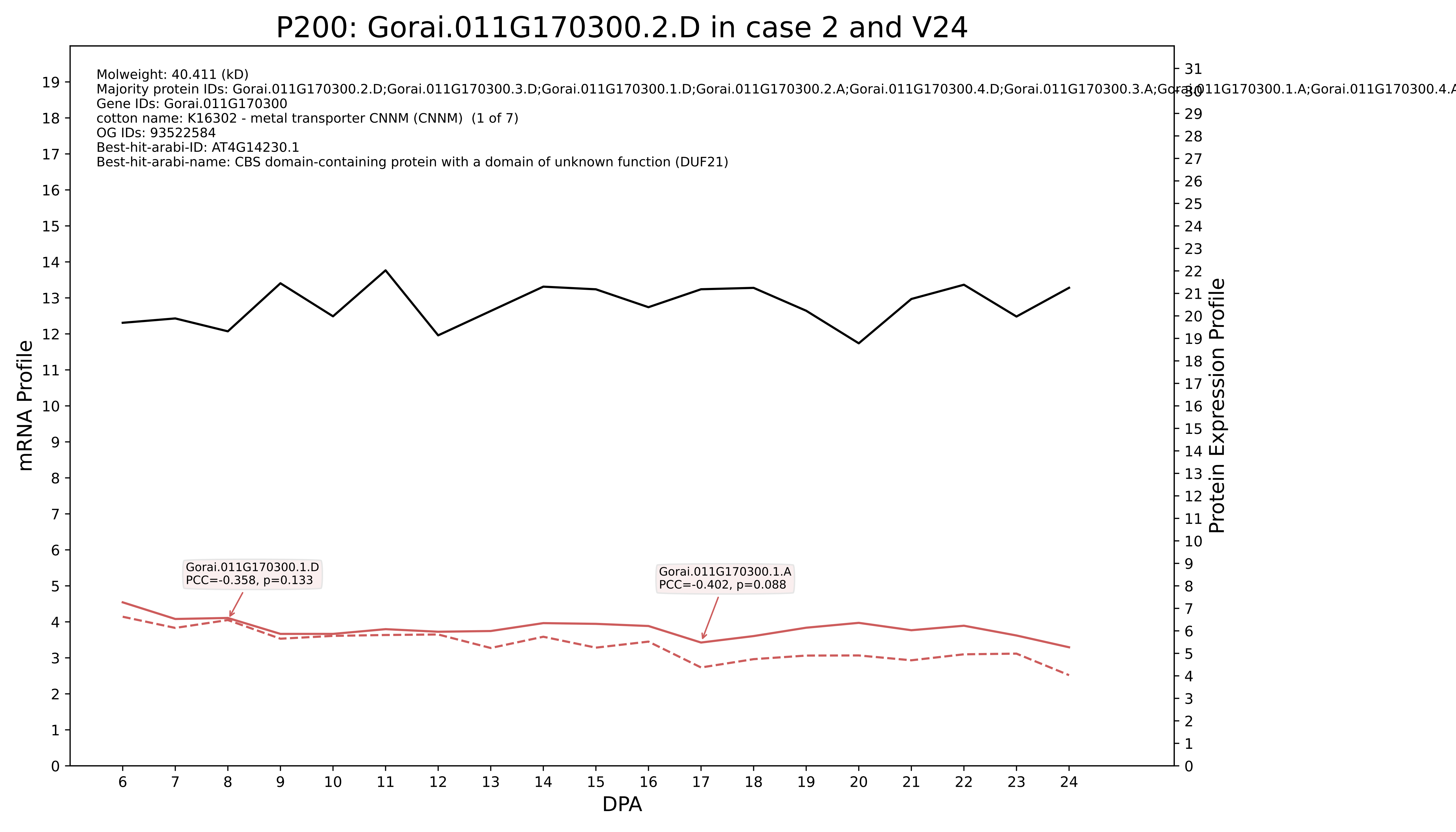
P200: Gorai.011G170300.2.D in case 2 and V24
Molweight: 40.411 (kD)
Majority protein IDs: Gorai.011G170300.2.D;Gorai.011G170300.3.D;Gorai.011G170300.1.D;Gorai.011G170300.2.A;Gorai.011G170300.4.D;Gorai.011G170300.3.A;Gorai.011G170300.1.A;Gorai.011G170300.4.A
Gene IDs: Gorai.011G170300
cotton name: K16302 - metal transporter CNNM (CNNM) (1 of 7)
OG IDs: 93522584
Best-hit-arabi-ID: AT4G14230.1
Best-hit-arabi-name: CBS domain-containing protein with a domain of unknown function (DUF21)
Gorai.011G170300.1.D
PCC=-0.358, p=0.133
Gorai.011G170300.1.A
PCC=-0.402, p=0.088
mRNA Profile
Protein Expression Profile
DPA

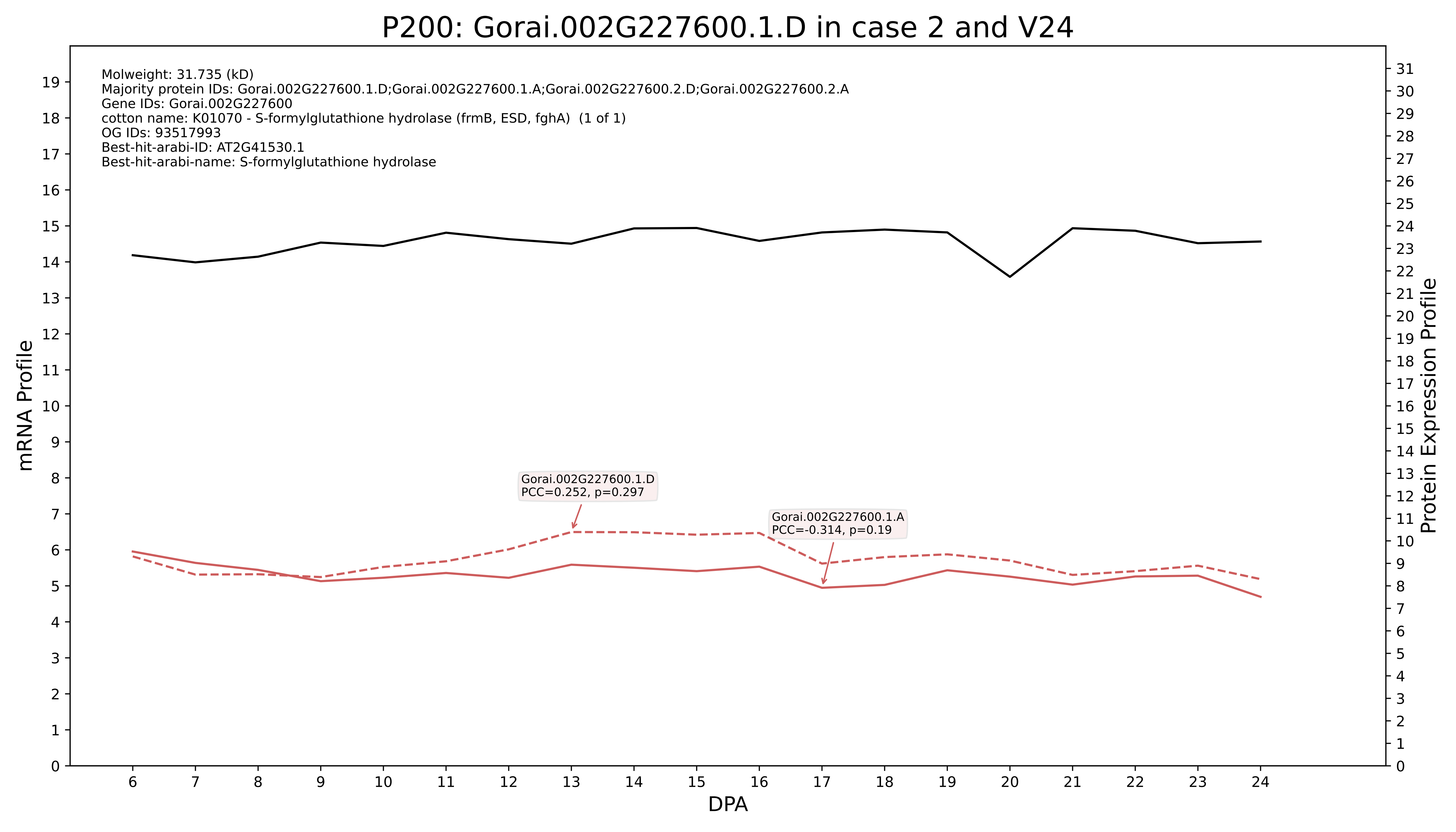
P200: Gorai.002G227600.1.D in case 2 and V24
Molweight: 31.735 (kD)
Majority protein IDs: Gorai.002G227600.1.D;Gorai.002G227600.1.A;Gorai.002G227600.2.D;Gorai.002G227600.2.A
Gene IDs: Gorai.002G227600
cotton name: K01070 - S-formylglutathione hydrolase (frmB, ESD, fghA) (1 of 1)
OG IDs: 93517993
Best-hit-arabi-ID: AT2G41530.1
Best-hit-arabi-name: S-formylglutathione hydrolase
Gorai.002G227600.1.D
PCC=0.252, p=0.297
Gorai.002G227600.1.A
PCC=-0.314, p=0.19
mRNA Profile
Protein Expression Profile
DPA

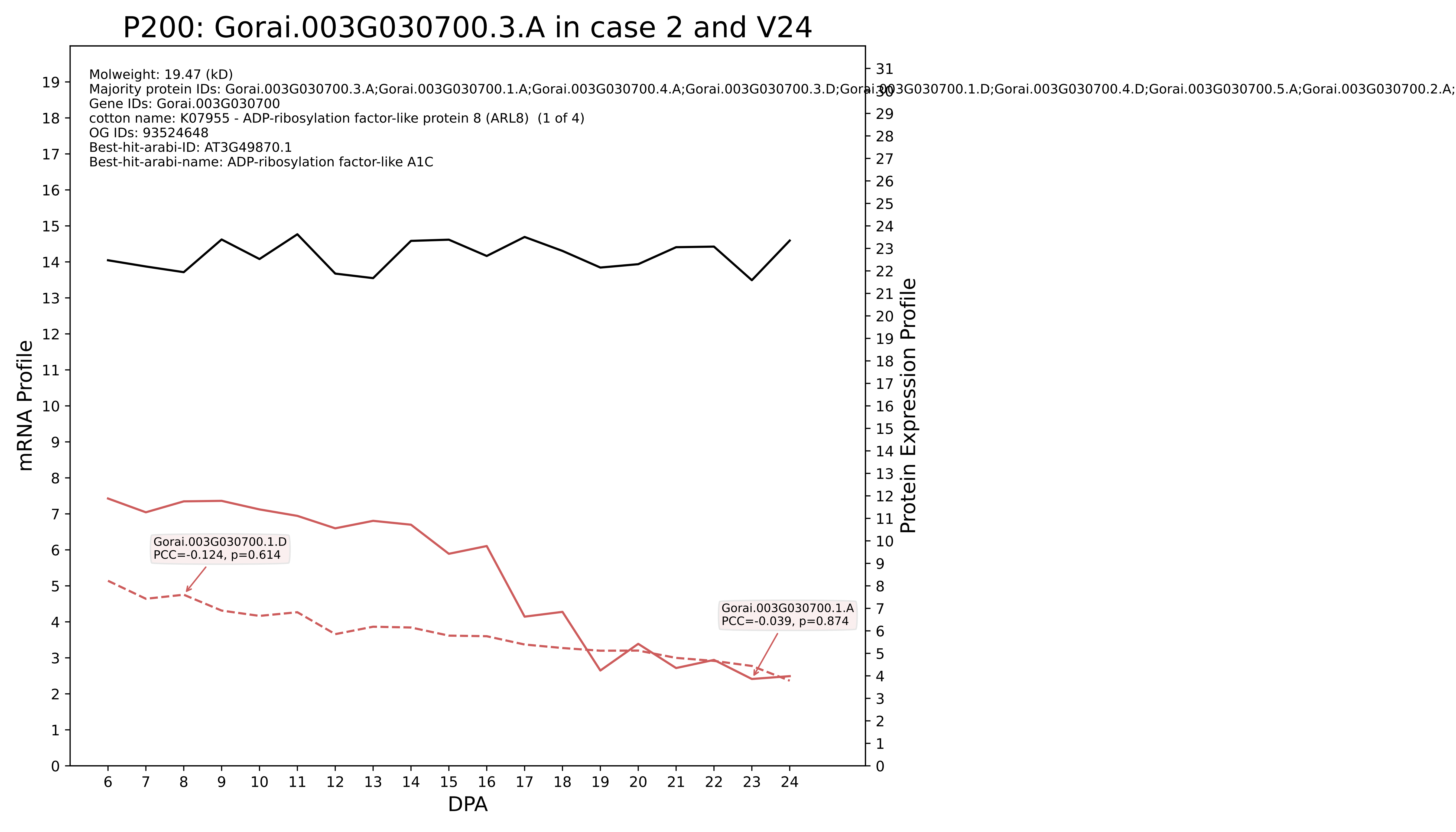
P200: Gorai.003G030700.3.A in case 2 and V24
Molweight: 19.47 (kD)
Majority protein IDs: Gorai.003G030700.3.A;Gorai.003G030700.1.A;Gorai.003G030700.4.A;Gorai.003G030700.3.D;Gorai.003G030700.1.D;Gorai.003G030700.4.D;Gorai.003G030700.5.A;Gorai.003G030700.2.A;
Gene IDs: Gorai.003G030700
cotton name: K07955 - ADP-ribosylation factor-like protein 8 (ARL8) (1 of 4)
OG IDs: 93524648
Best-hit-arabi-ID: AT3G49870.1
Best-hit-arabi-name: ADP-ribosylation factor-like A1C
Gorai.003G030700.1.D
PCC=-0.124, p=0.614
Gorai.003G030700.1.A
PCC=-0.039, p=0.874
mRNA Profile
Protein Expression Profile
DPA

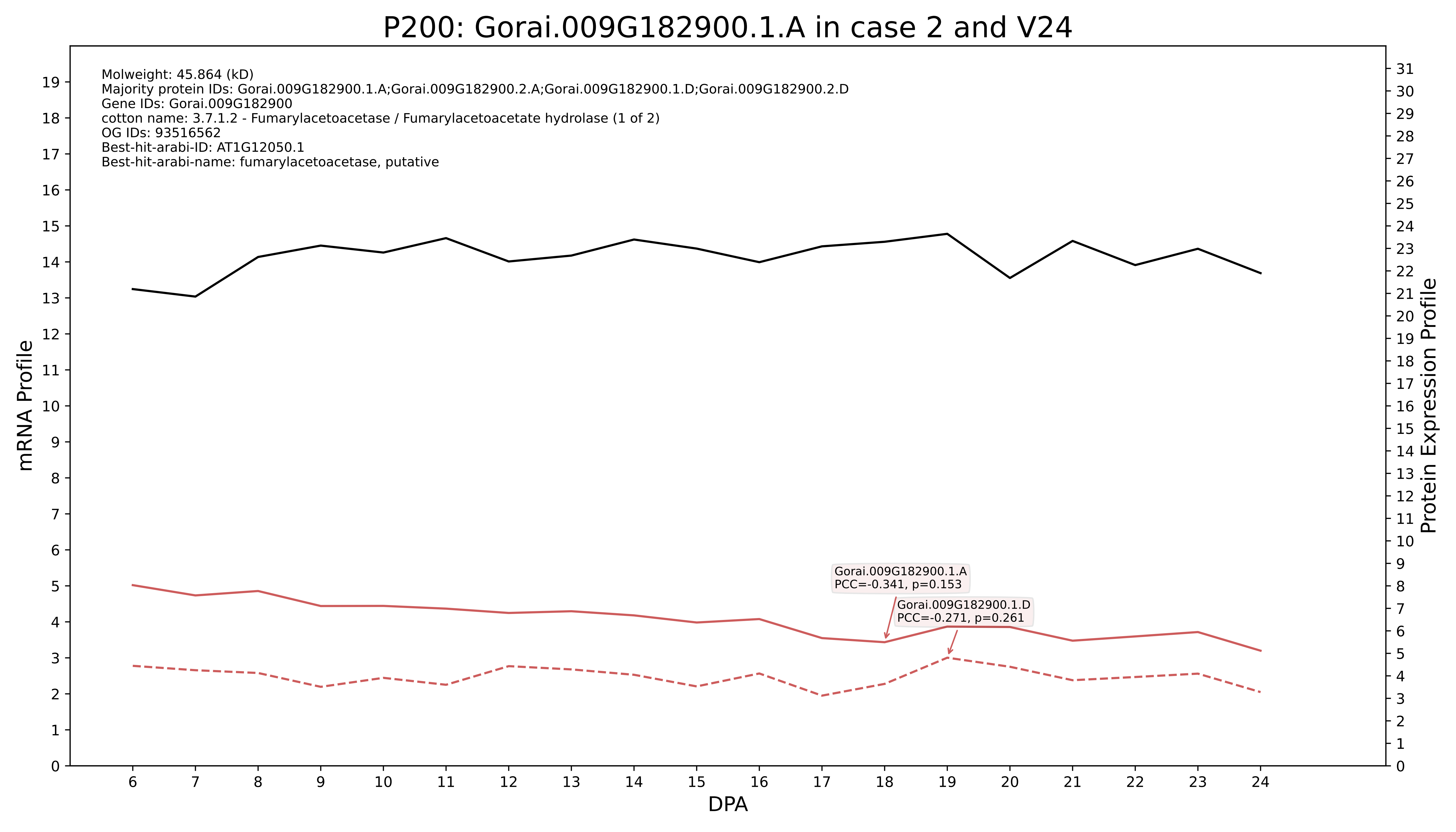
P200: Gorai.009G182900.1.A in case 2 and V24
Molweight: 45.864 (kD)
Majority protein IDs: Gorai.009G182900.1.A;Gorai.009G182900.2.A;Gorai.009G182900.1.D;Gorai.009G182900.2.D
Gene IDs: Gorai.009G182900
cotton name: 3.7.1.2 - Fumarylacetoacetase / Fumarylacetoacetate hydrolase (1 of 2)
OG IDs: 93516562
Best-hit-arabi-ID: AT1G12050.1
Best-hit-arabi-name: fumarylacetoacetase, putative
Gorai.009G182900.1.A
PCC=-0.341, p=0.153
Gorai.009G182900.1.D
PCC=-0.271, p=0.261
mRNA Profile
Protein Expression Profile
DPA

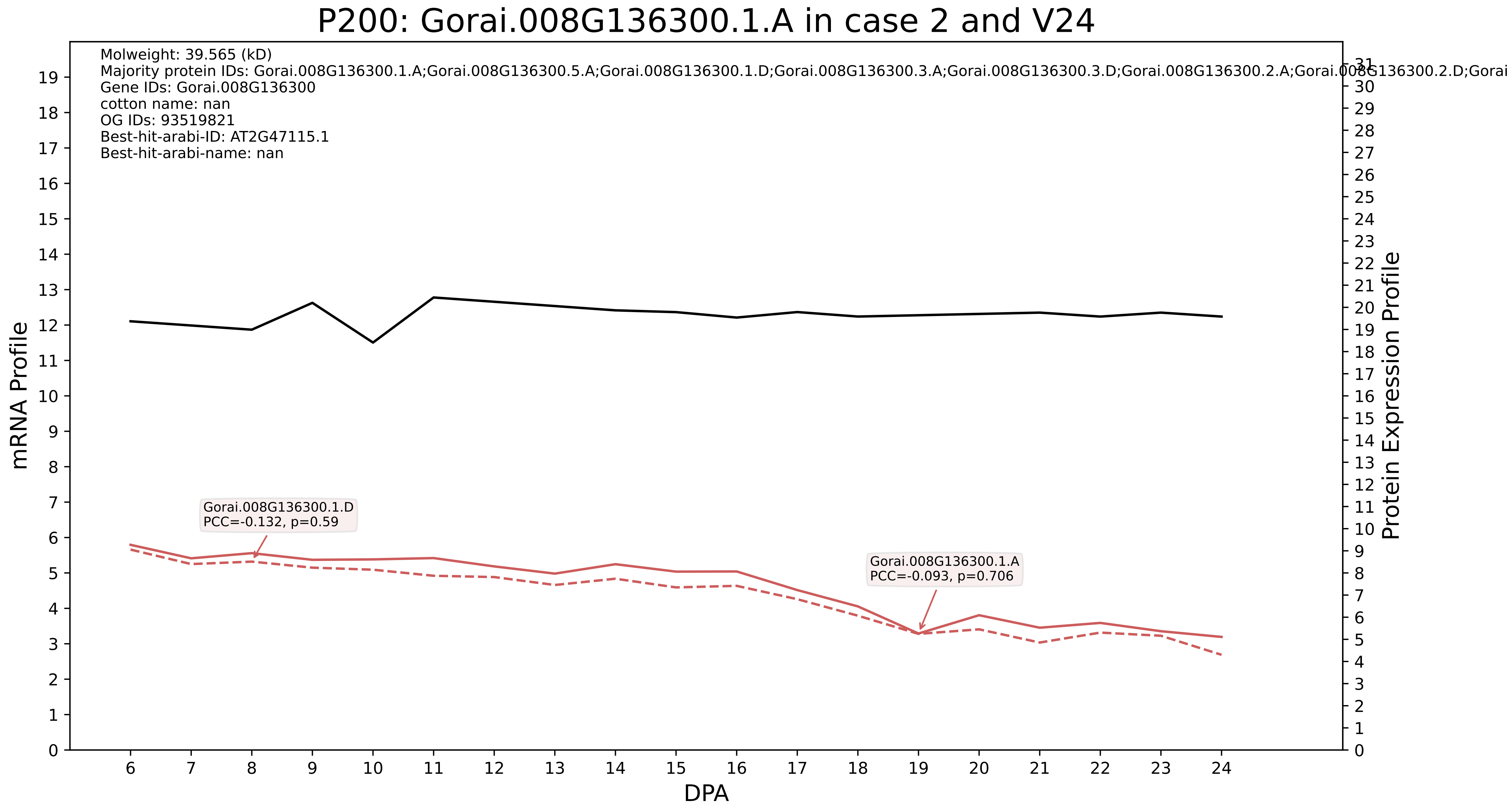

P200: Gorai.008G136300.1.A in case 2 and V24
Molweight: 39.565 (kD)
Majority protein IDs: Gorai.008G136300.1.A;Gorai.008G136300.5.A;Gorai.008G136300.1.D;Gorai.008G136300.3.A;Gorai.008G136300.3.D;Gorai.008G136300.2.A;Gorai.008G136300.2.D;Gorai
Gene IDs: Gorai.008G136300
cotton name: nan
OG IDs: 93519821
Best-hit-arabi-ID: AT2G47115.1
Best-hit-arabi-name: nan
Gorai.008G136300.1.D
PCC=-0.132, p=0.59
Gorai.008G136300.1.A
PCC=-0.093, p=0.706
mRNA Profile
Protein Expression Profile
DPA

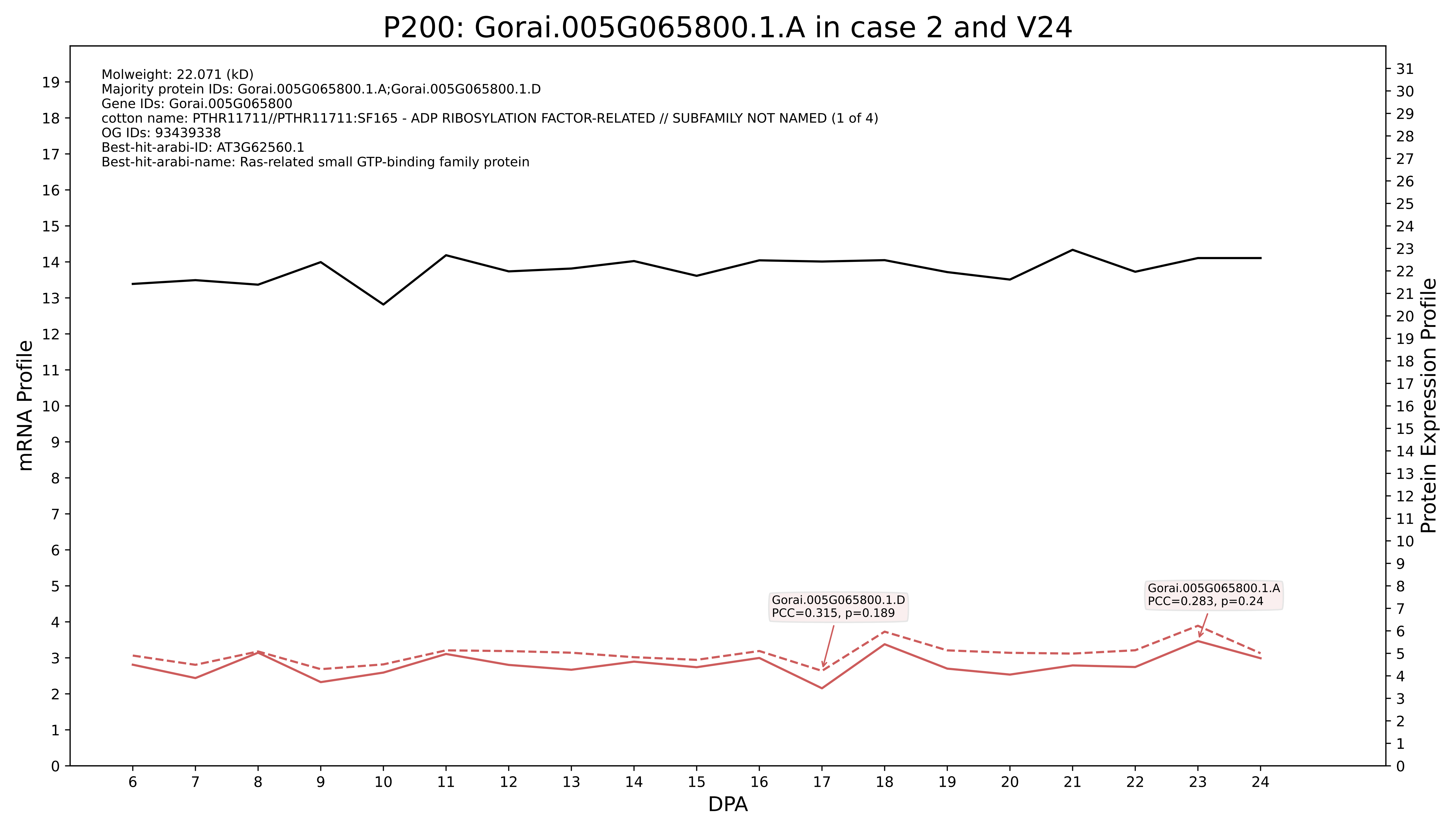

P200: Gorai.005G065800.1.A in case 2 and V24
Molweight: 22.071 (kD)
Majority protein IDs: Gorai.005G065800.1.A;Gorai.005G065800.1.D
Gene IDs: Gorai.005G065800
cotton name: PTHR11711//PTHR11711:SF165 - ADP RIBOSYLATION FACTOR-RELATED // SUBFAMILY NOT NAMED (1 of 4)
OG IDs: 93439338
Best-hit-arabi-ID: AT3G62560.1
Best-hit-arabi-name: Ras-related small GTP-binding family protein
Gorai.005G065800.1.D
PCC=0.315, p=0.189
Gorai.005G065800.1.A
PCC=0.283, p=0.24
mRNA Profile
Protein Expression Profile
DPA

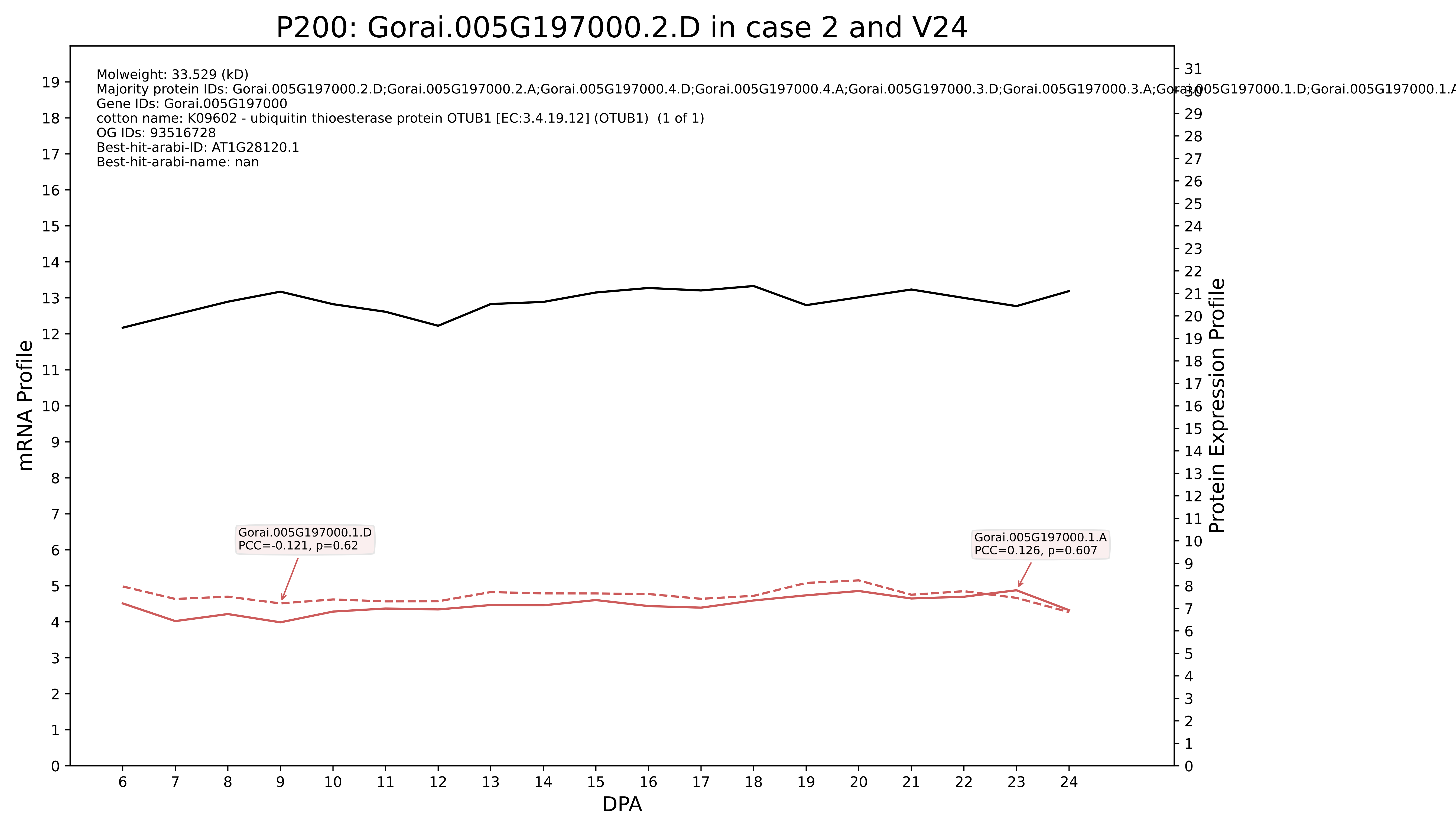
P200: Gorai.005G197000.2.D in case 2 and V24
Molweight: 33.529 (kD)
Majority protein IDs: Gorai.005G197000.2.D;Gorai.005G197000.2.A;Gorai.005G197000.4.D;Gorai.005G197000.4.A;Gorai.005G197000.3.D;Gorai.005G197000.3.A;Gorai.005G197000.1.D;Gorai.005G197000.1.A
Gene IDs: Gorai.005G197000
cotton name: K09602 - ubiquitin thioesterase protein OTUB1 [EC:3.4.19.12] (OTUB1) (1 of 1)
OG IDs: 93516728
Best-hit-arabi-ID: AT1G28120.1
Best-hit-arabi-name: nan
Gorai.005G197000.1.D
PCC=-0.121, p=0.62
Gorai.005G197000.1.A
PCC=0.126, p=0.607
mRNA Profile
Protein Expression Profile
DPA

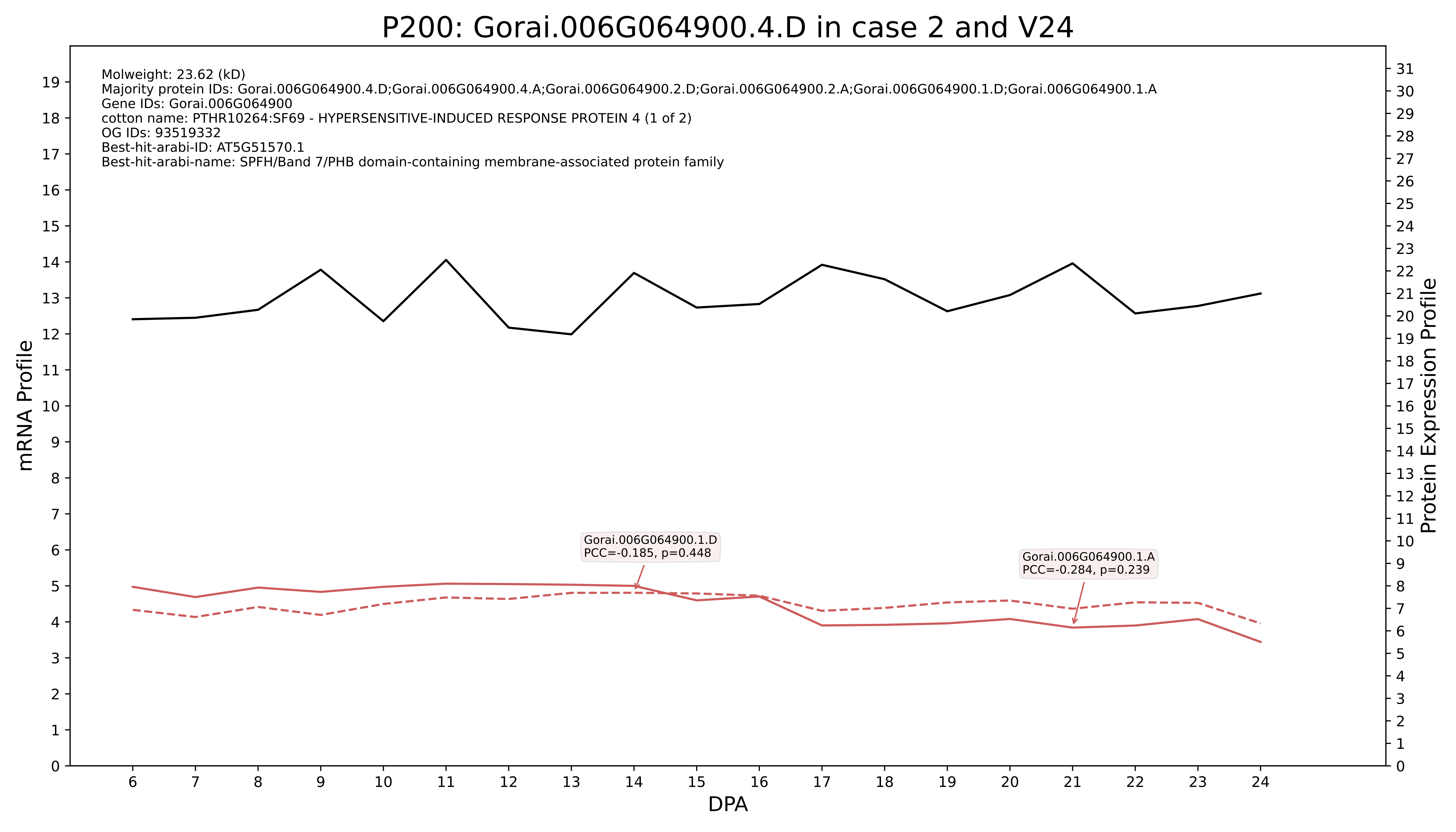
P200: Gorai.006G064900.4.D in case 2 and V24
Molweight: 23.62 (kD)
Majority protein IDs: Gorai.006G064900.4.D;Gorai.006G064900.4.A;Gorai.006G064900.2.D;Gorai.006G064900.2.A;Gorai.006G064900.1.D;Gorai.006G064900.1.A
Gene IDs: Gorai.006G064900
cotton name: PTHR10264:SF69 - HYPERSENSITIVE-INDUCED RESPONSE PROTEIN 4 (1 of 2)
OG IDs: 93519332
Best-hit-arabi-ID: AT5G51570.1
Best-hit-arabi-name: SPFH/Band 7/PHB domain-containing membrane-associated protein family
Gorai.006G064900.1.D
PCC=-0.185, p=0.448
Gorai.006G064900.1.A
PCC=-0.284, p=0.239
mRNA Profile
Protein Expression Profile
DPA

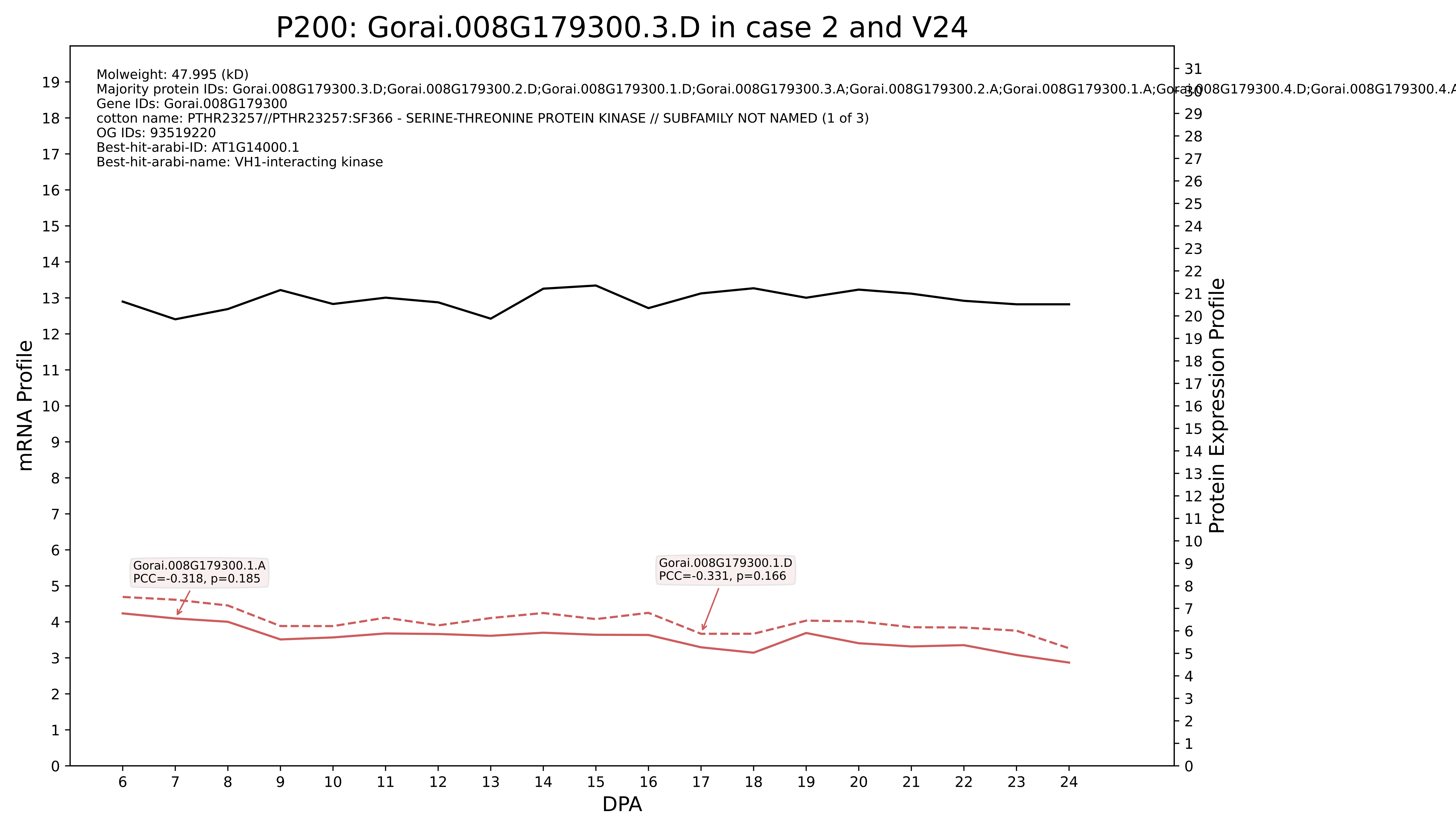

P200: Gorai.008G179300.3.D in case 2 and V24
Molweight: 47.995 (kD)
Majority protein IDs: Gorai.008G179300.3.D;Gorai.008G179300.2.D;Gorai.008G179300.1.D;Gorai.008G179300.3.A;Gorai.008G179300.2.A;Gorai.008G179300.1.A;Gorai.008G179300.4.D;Gorai.008G179300.4.A
Gene IDs: Gorai.008G179300
cotton name: PTHR23257//PTHR23257:SF366 - SERINE-THREONINE PROTEIN KINASE // SUBFAMILY NOT NAMED (1 of 3)
OG IDs: 93519220
Best-hit-arabi-ID: AT1G14000.1
Best-hit-arabi-name: VH1-interacting kinase
Gorai.008G179300.1.A
PCC=-0.318, p=0.185
Gorai.008G179300.1.D
PCC=-0.331, p=0.166
mRNA Profile
Protein Expression Profile
DPA

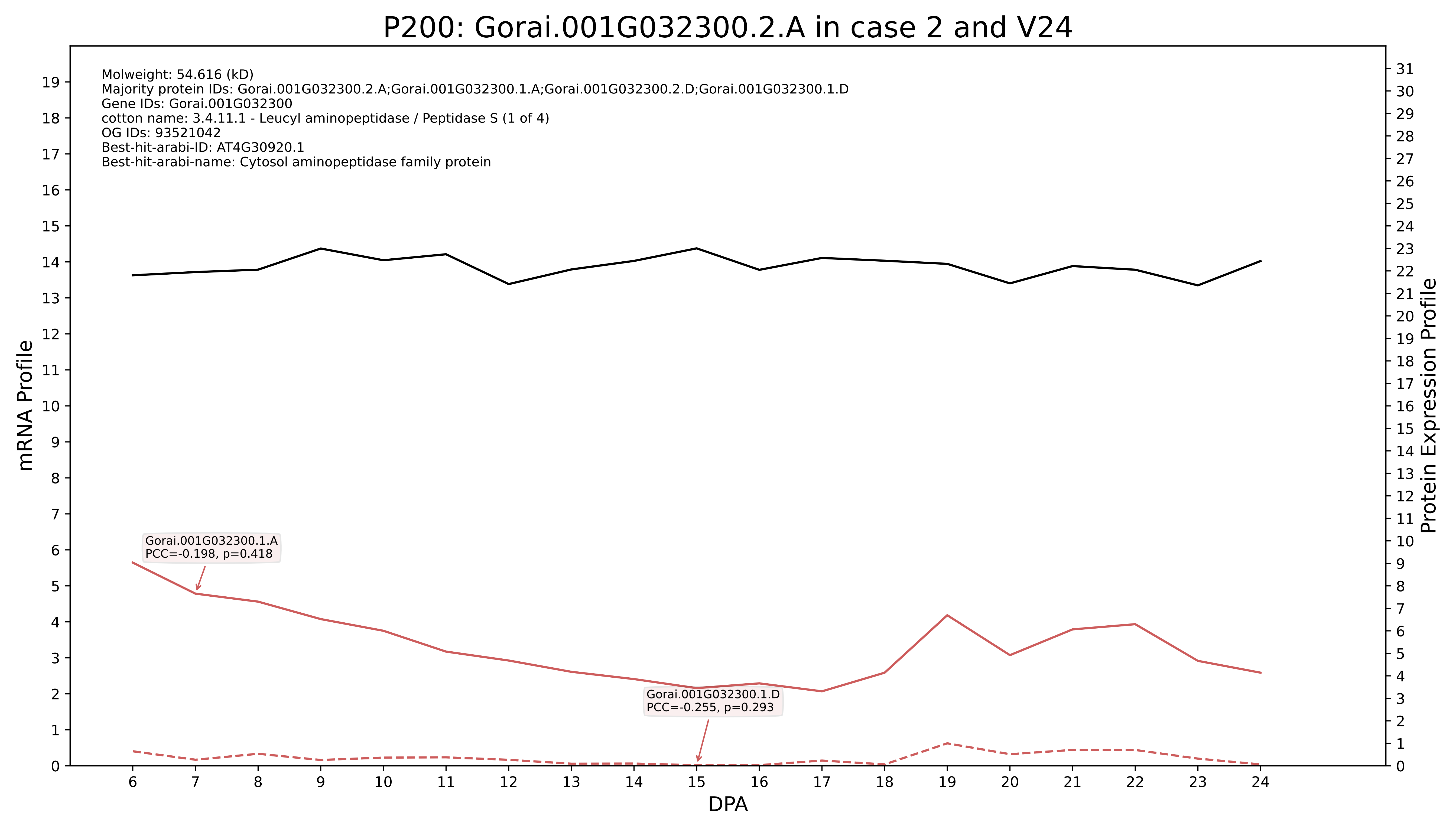

P200: Gorai.001G032300.2.A in case 2 and V24
Molweight: 54.616 (kD)
Majority protein IDs: Gorai.001G032300.2.A;Gorai.001G032300.1.A;Gorai.001G032300.2.D;Gorai.001G032300.1.D
Gene IDs: Gorai.001G032300
cotton name: 3.4.11.1 - Leucyl aminopeptidase / Peptidase S (1 of 4)
OG IDs: 93521042
Best-hit-arabi-ID: AT4G30920.1
Best-hit-arabi-name: Cytosol aminopeptidase family protein
Gorai.001G032300.1.A
PCC=-0.198, p=0.418
Gorai.001G032300.1.D
PCC=-0.255, p=0.293
mRNA Profile
Protein Expression Profile
DPA

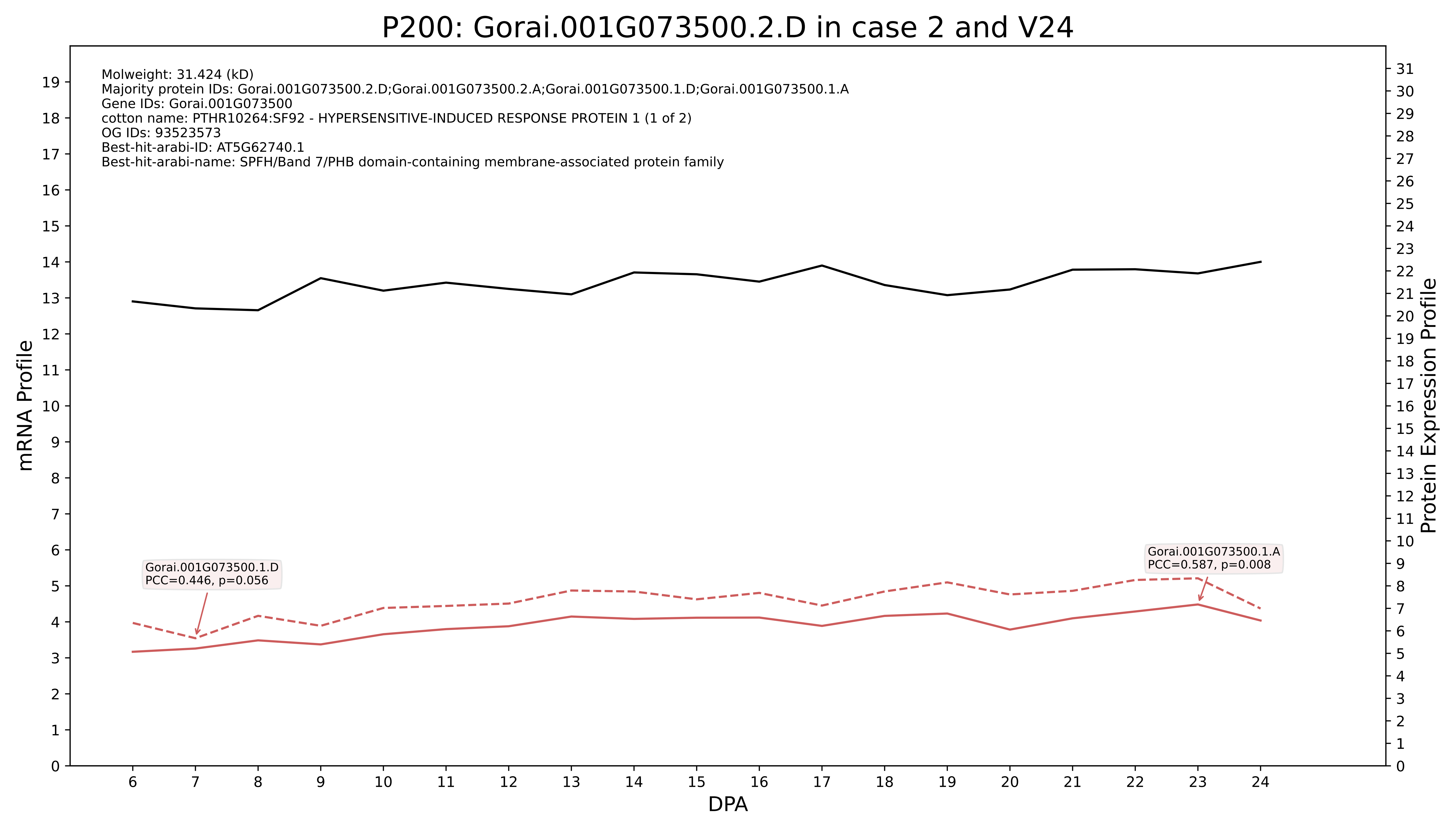

P200: Gorai.001G073500.2.D in case 2 and V24
Molweight: 31.424 (kD)
Majority protein IDs: Gorai.001G073500.2.D;Gorai.001G073500.2.A;Gorai.001G073500.1.D;Gorai.001G073500.1.A
Gene IDs: Gorai.001G073500
cotton name: PTHR10264:SF92 - HYPERSENSITIVE-INDUCED RESPONSE PROTEIN 1 (1 of 2)
OG IDs: 93523573
Best-hit-arabi-ID: AT5G62740.1
Best-hit-arabi-name: SPFH/Band 7/PHB domain-containing membrane-associated protein family
Gorai.001G073500.1.D
PCC=0.446, p=0.056
Gorai.001G073500.1.A
PCC=0.587, p=0.008
mRNA Profile
Protein Expression Profile
DPA

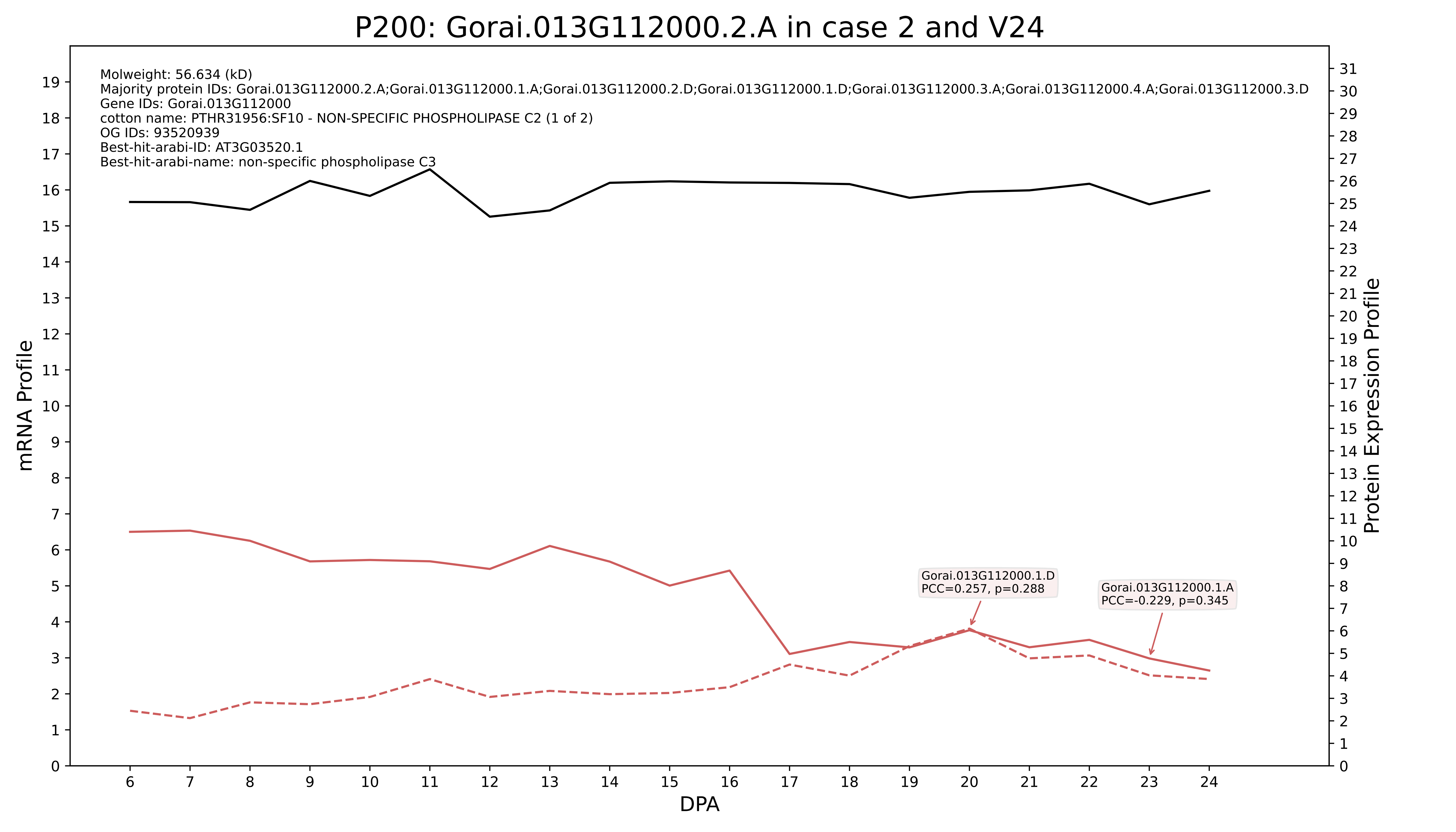
P200: Gorai.013G112000.2.A in case 2 and V24
Molweight: 56.634 (kD)
Majority protein IDs: Gorai.013G112000.2.A;Gorai.013G112000.1.A;Gorai.013G112000.2.D;Gorai.013G112000.1.D;Gorai.013G112000.3.A;Gorai.013G112000.4.A;Gorai.013G112000.3.D
Gene IDs: Gorai.013G112000
cotton name: PTHR31956:SF10 - NON-SPECIFIC PHOSPHOLIPASE C2 (1 of 2)
OG IDs: 93520939
Best-hit-arabi-ID: AT3G03520.1
Best-hit-arabi-name: non-specific phospholipase C3
Gorai.013G112000.1.D
PCC=0.257, p=0.288
Gorai.013G112000.1.A
PCC=-0.229, p=0.345
mRNA Profile
Protein Expression Profile
DPA

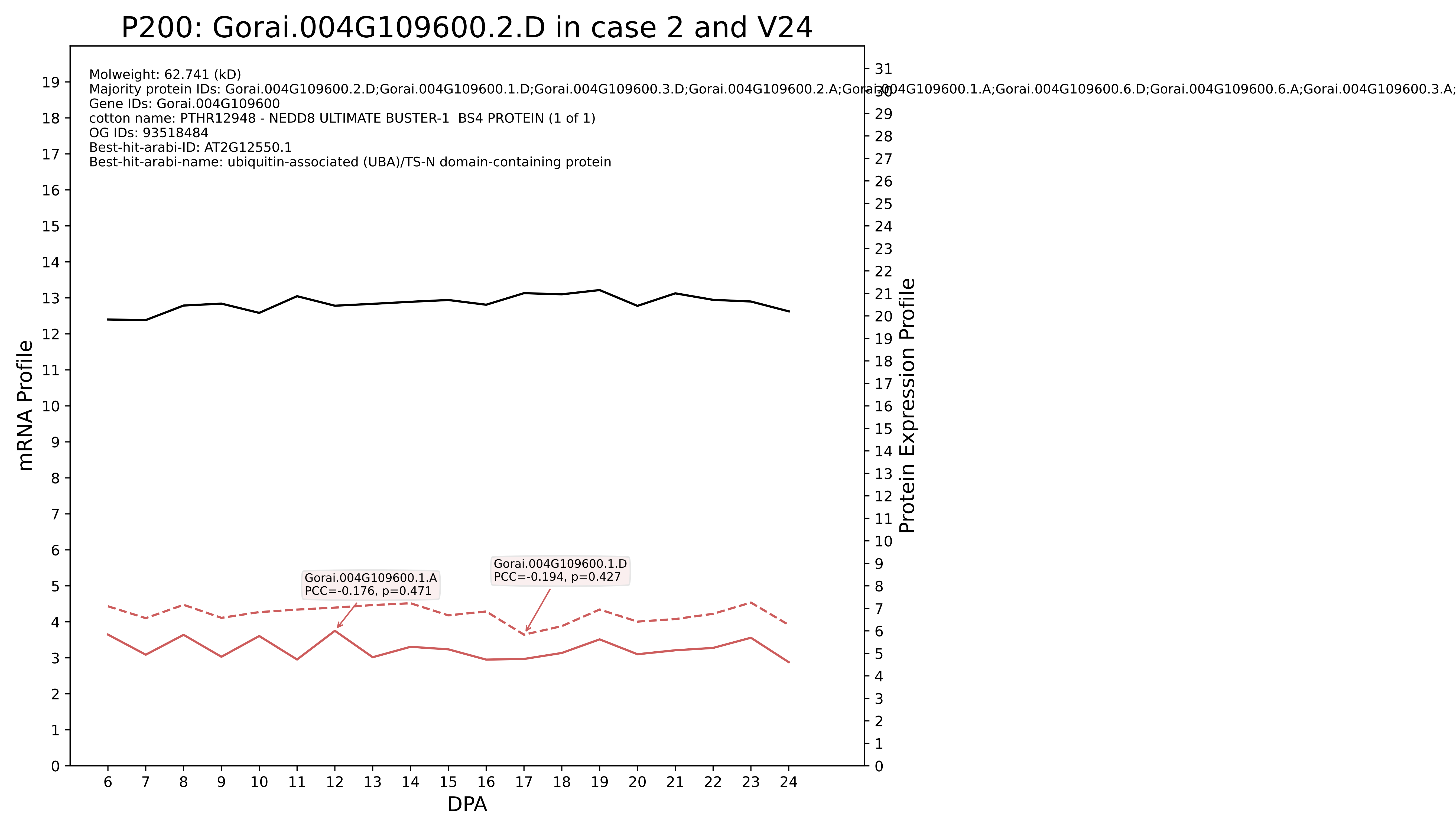
P200: Gorai.004G109600.2.D in case 2 and V24
Molweight: 62.741 (kD)
Majority protein IDs: Gorai.004G109600.2.D;Gorai.004G109600.1.D;Gorai.004G109600.3.D;Gorai.004G109600.2.A;Gorai.004G109600.1.A;Gorai.004G109600.6.D;Gorai.004G109600.6.A;Gorai.004G109600.3.A;
Gene IDs: Gorai.004G109600
cotton name: PTHR12948 - NEDD8 ULTIMATE BUSTER-1 BS4 PROTEIN (1 of 1)
OG IDs: 93518484
Best-hit-arabi-ID: AT2G12550.1
Best-hit-arabi-name: ubiquitin-associated (UBA)/TS-N domain-containing protein
Gorai.004G109600.1.A
PCC=-0.176, p=0.471
Gorai.004G109600.1.D
PCC=-0.194, p=0.427
mRNA Profile
Protein Expression Profile
DPA

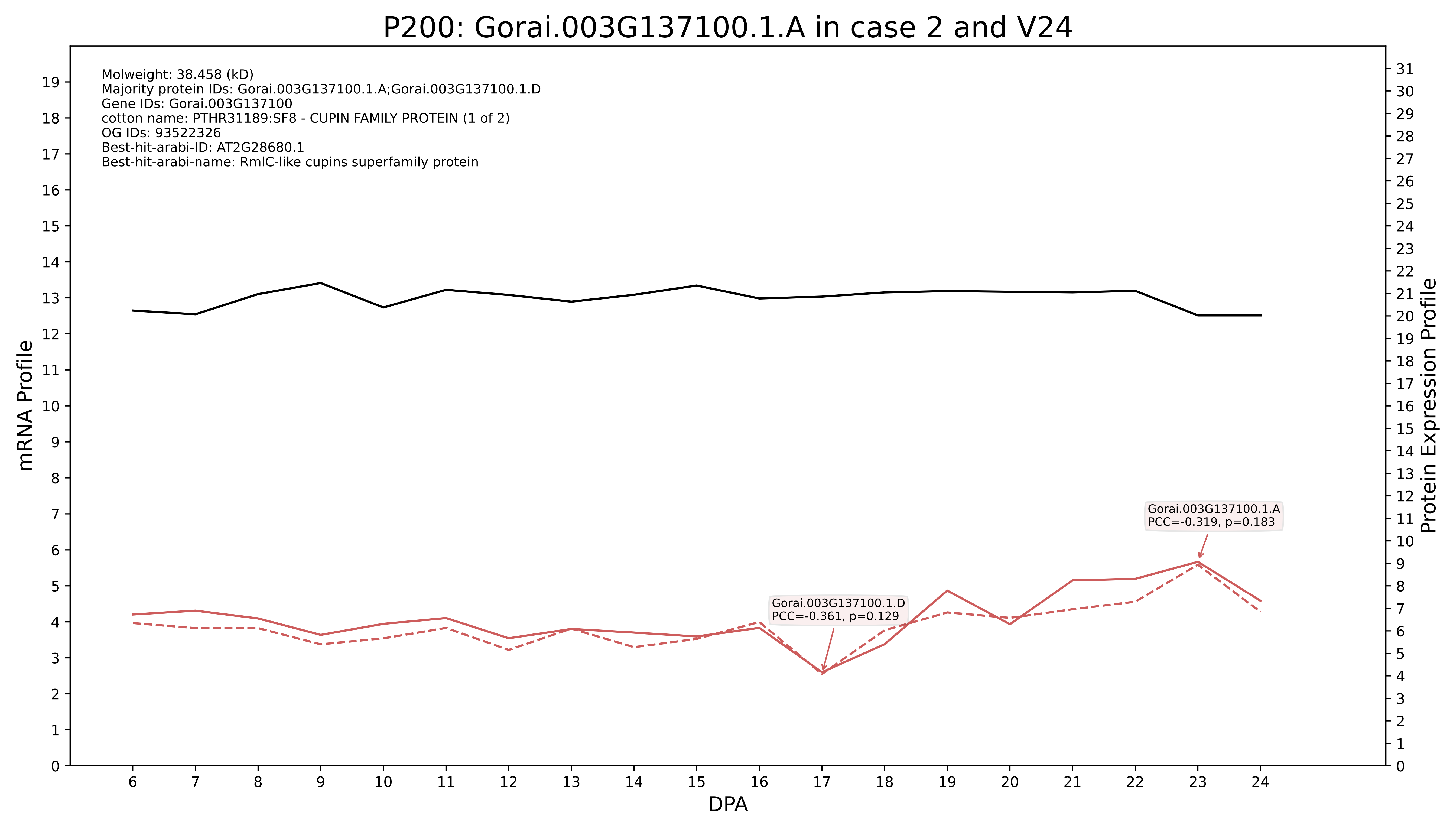
P200: Gorai.003G137100.1.A in case 2 and V24
Molweight: 38.458 (kD)
Majority protein IDs: Gorai.003G137100.1.A;Gorai.003G137100.1.D
Gene IDs: Gorai.003G137100
cotton name: PTHR31189:SF8 - CUPIN FAMILY PROTEIN (1 of 2)
OG IDs: 93522326
Best-hit-arabi-ID: AT2G28680.1
Best-hit-arabi-name: RmlC-like cupins superfamily protein
Gorai.003G137100.1.A
PCC=-0.319, p=0.183
Gorai.003G137100.1.D
PCC=-0.361, p=0.129
mRNA Profile
Protein Expression Profile
DPA

### P200: Gorai.011G037400.1.D in case 2 and V24

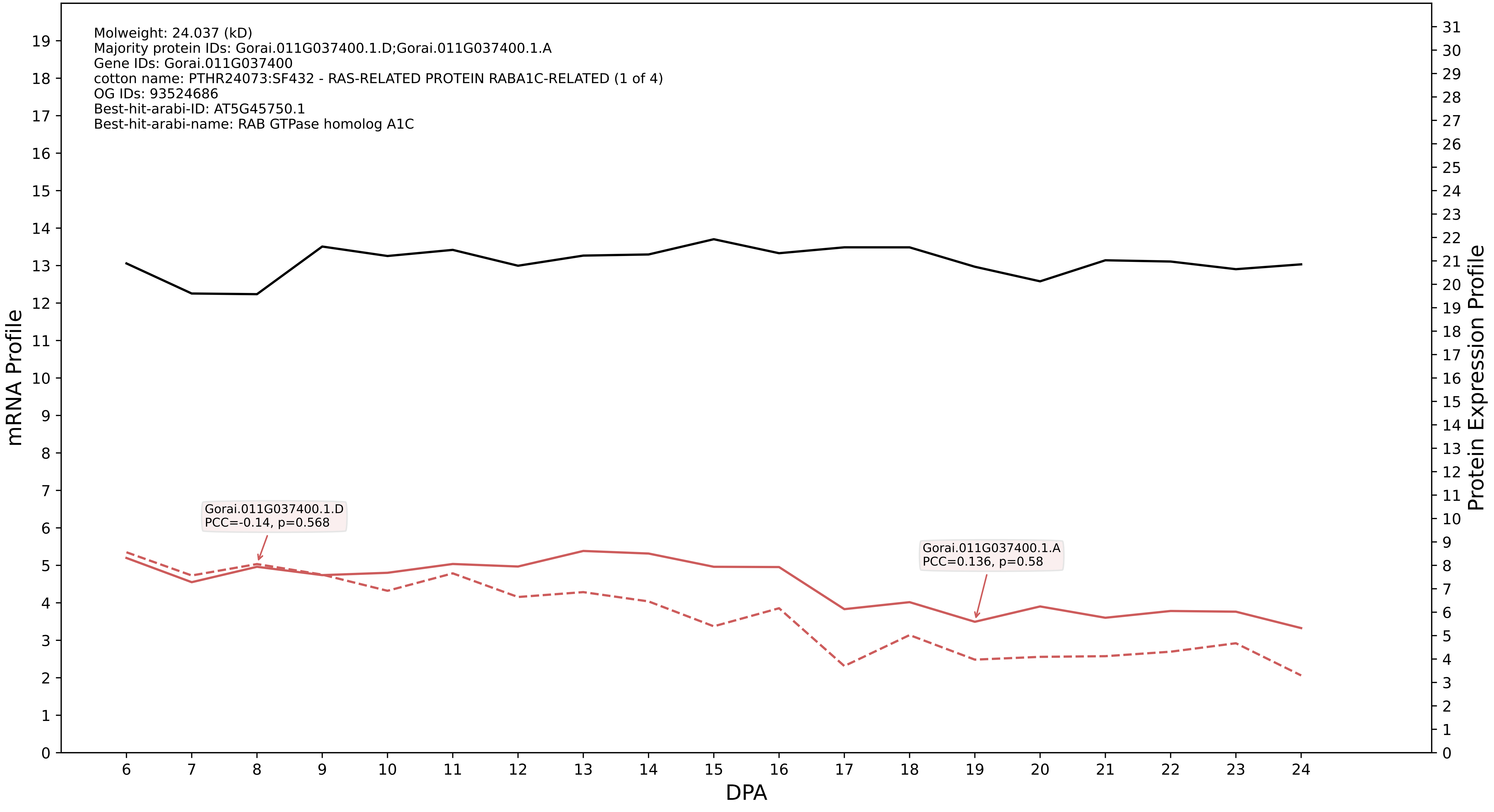

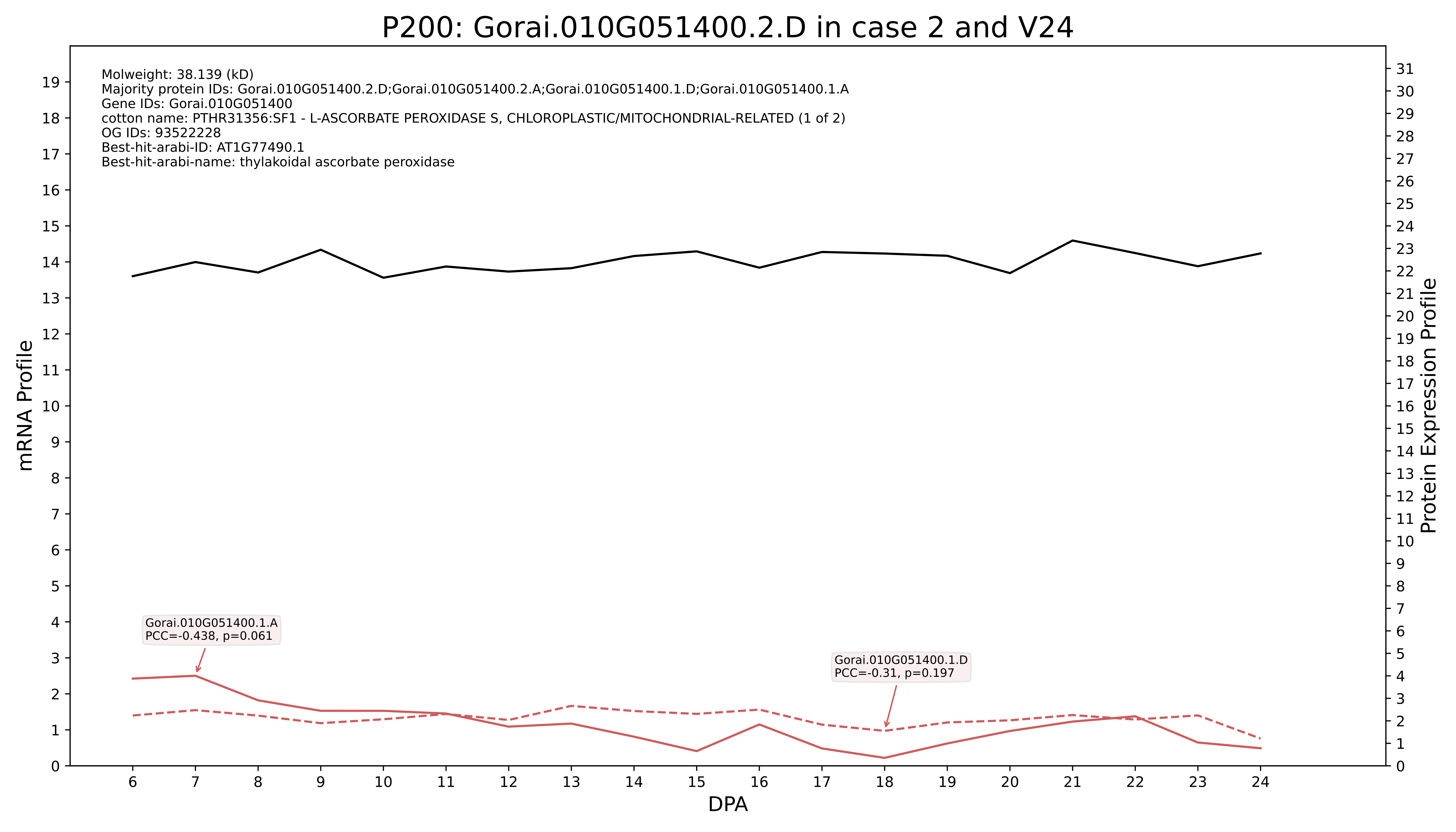
P200: Gorai.010G051400.2.D in case 2 and V24
Molweight: 38.139 (kD)
Majority protein IDs: Gorai.010G051400.2.D;Gorai.010G051400.2.A;Gorai.010G051400.1.D;Gorai.010G051400.1.A
Gene IDs: Gorai.010G051400
cotton name: PTHR31356:SF1 - L-ASCORBATE PEROXIDASE S, CHLOROPLASTIC/MITOCHONDRIAL-RELATED (1 of 2)
OG IDs: 93522228
Best-hit-arabi-ID: AT1G77490.1
Best-hit-arabi-name: thylakoidal ascorbate peroxidase
Gorai.010G051400.1.A
PCC=-0.438, p=0.061
Gorai.010G051400.1.D
PCC=-0.31, p=0.197
mRNA Profile
Protein Expression Profile
DPA

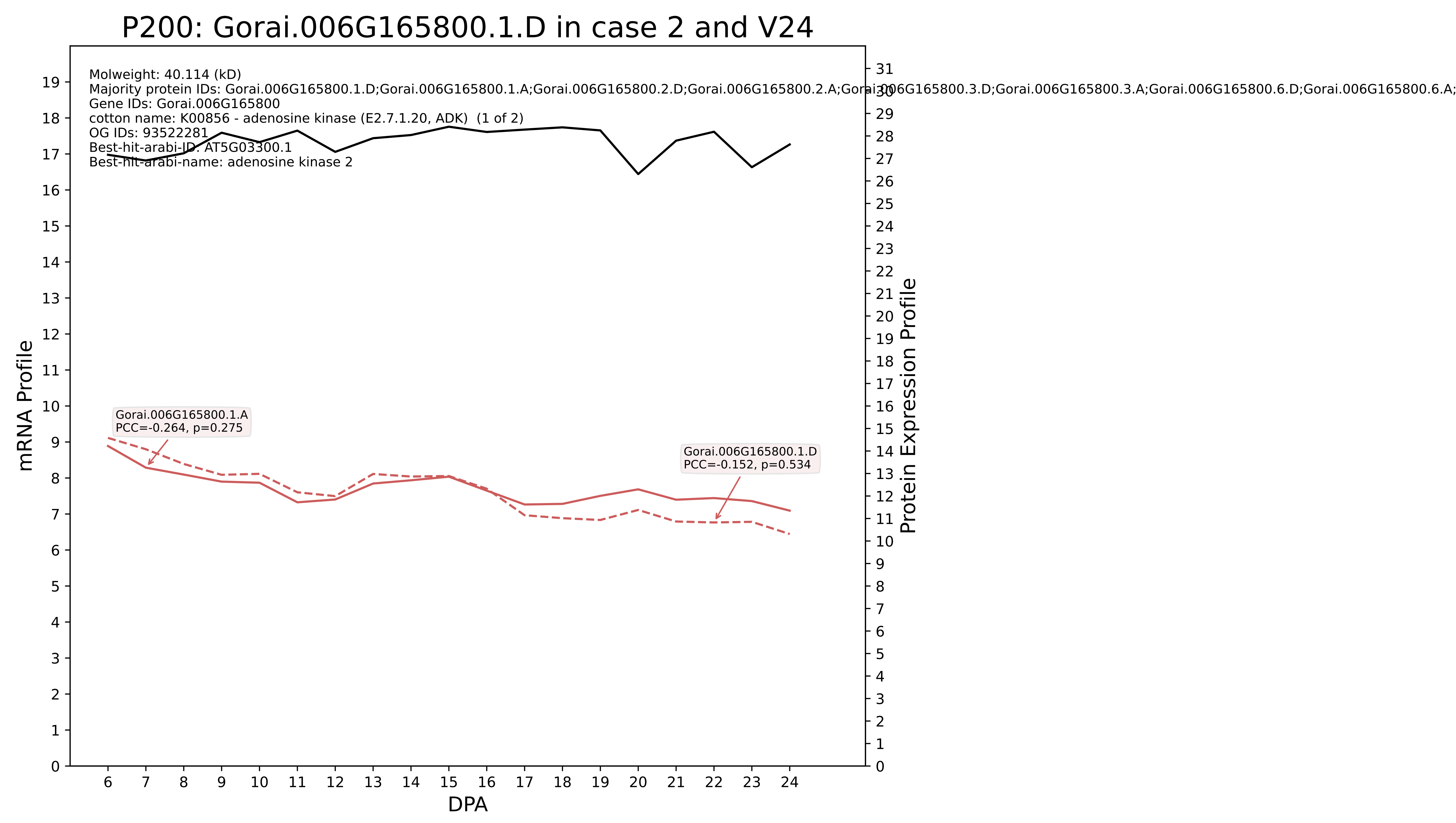
P200: Gorai.006G165800.1.D in case 2 and V24
Molweight: 40.114 (kD)
Majority protein IDs: Gorai.006G165800.1.D;Gorai.006G165800.1.A;Gorai.006G165800.2.D;Gorai.006G165800.2.A;Gorai.006G165800.3.D;Gorai.006G165800.3.A;Gorai.006G165800.6.D;Gorai.006G165800.6.A;
Gene IDs: Gorai.006G165800
cotton name: K00856 - adenosine kinase (E2.7.1.20, ADK) (1 of 2)
OG IDs: 93522281
Best-hit-arabi-ID: AT5G03300.1
Best-hit-arabi-name: adenosine kinase 2
Gorai.006G165800.1.A
PCC=-0.264, p=0.275
Gorai.006G165800.1.D
PCC=-0.152, p=0.534
mRNA Profile
Protein Expression Profile
DPA

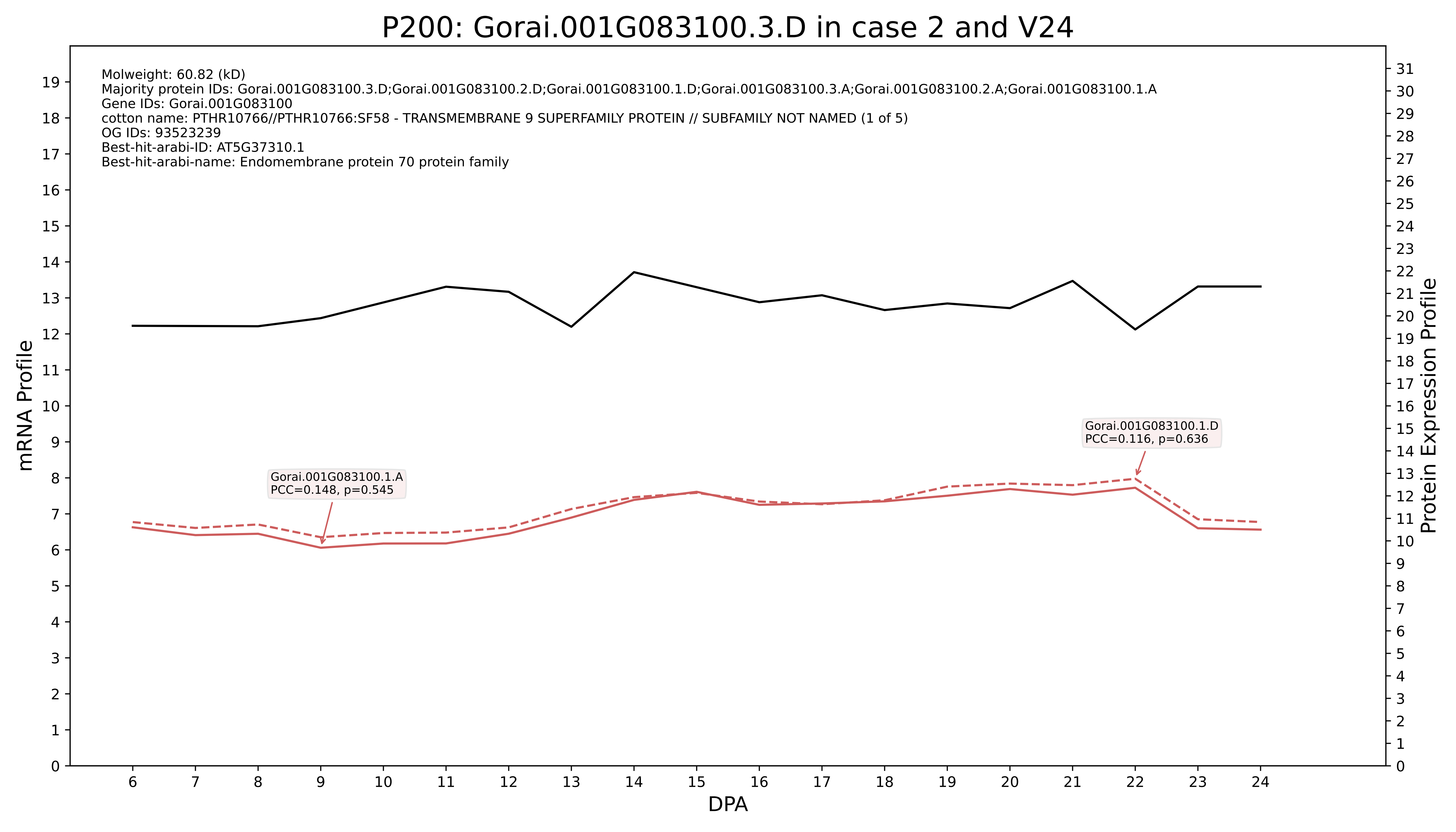
P200: Gorai.001G083100.3.D in case 2 and V24
Molweight: 60.82 (kD)
Majority protein IDs: Gorai.001G083100.3.D;Gorai.001G083100.2.D;Gorai.001G083100.1.D;Gorai.001G083100.3.A;Gorai.001G083100.2.A;Gorai.001G083100.1.A
Gene IDs: Gorai.001G083100
cotton name: PTHR10766//PTHR10766:SF58 - TRANSMEMBRANE 9 SUPERFAMILY PROTEIN // SUBFAMILY NOT NAMED (1 of 5)
OG IDs: 93523239
Best-hit-arabi-ID: AT5G37310.1
Best-hit-arabi-name: Endomembrane protein 70 protein family
Gorai.001G083100.1.A
PCC=0.148, p=0.545
Gorai.001G083100.1.D
PCC=0.116, p=0.636
mRNA Profile
Protein Expression Profile
DPA

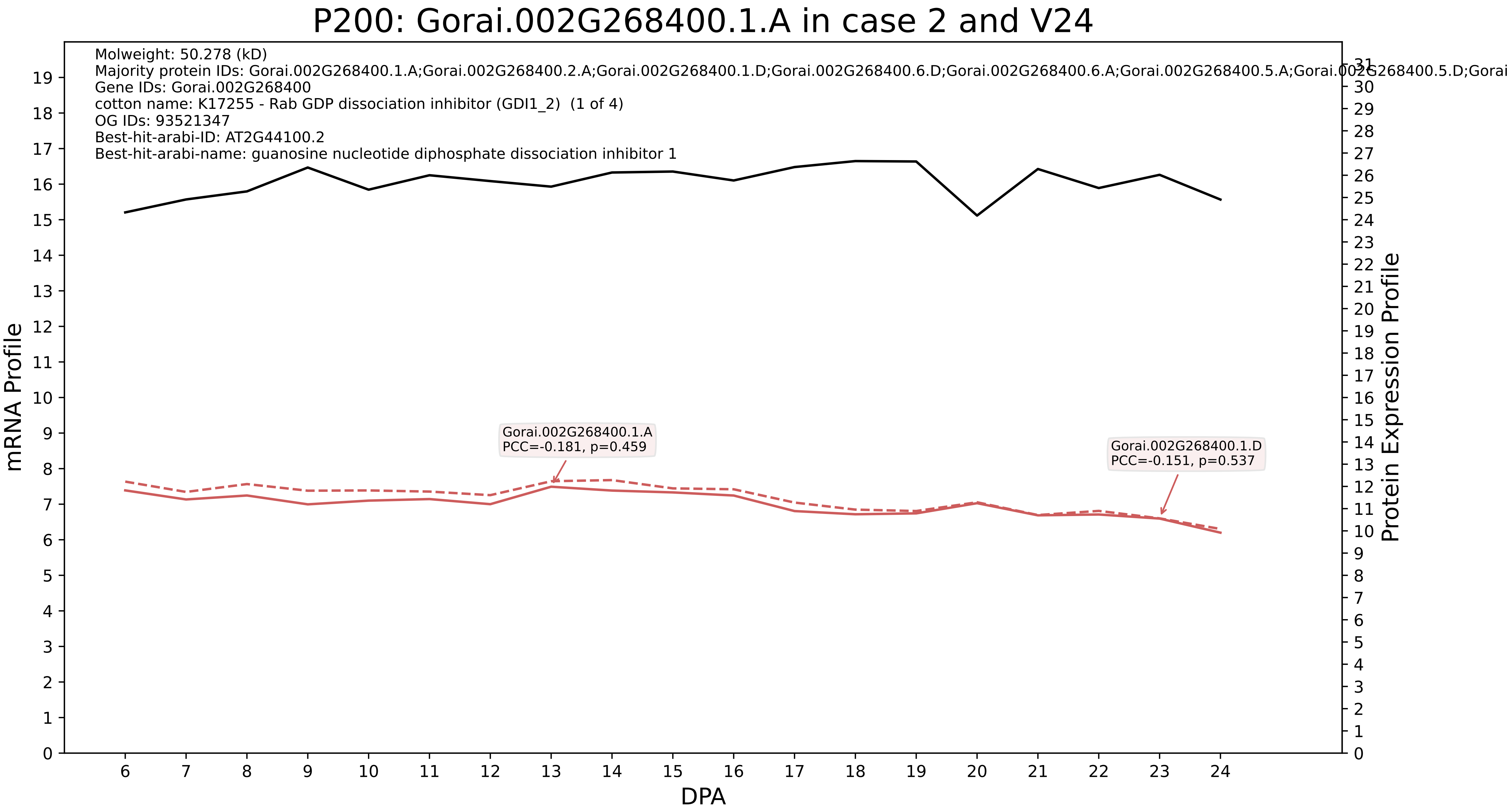
P200: Gorai.002G268400.1.A in case 2 and V24
Molweight: 50.278 (kD)
Majority protein IDs: Gorai.002G268400.1.A;Gorai.002G268400.2.A;Gorai.002G268400.1.D;Gorai.002G268400.6.D;Gorai.002G268400.6.A;Gorai.002G268400.5.A;Gorai.002G268400.5.D;Gorai
Gene IDs: Gorai.002G268400
cotton name: K17255 - Rab GDP dissociation inhibitor (GDI1_2) (1 of 4)
OG IDs: 93521347
Best-hit-arabi-ID: AT2G44100.2
Best-hit-arabi-name: guanosine nucleotide diphosphate dissociation inhibitor 1
Gorai.002G268400.1.A
PCC=-0.181, p=0.459
Gorai.002G268400.1.D
PCC=-0.151, p=0.537
mRNA Profile
Protein Expression Profile
DPA

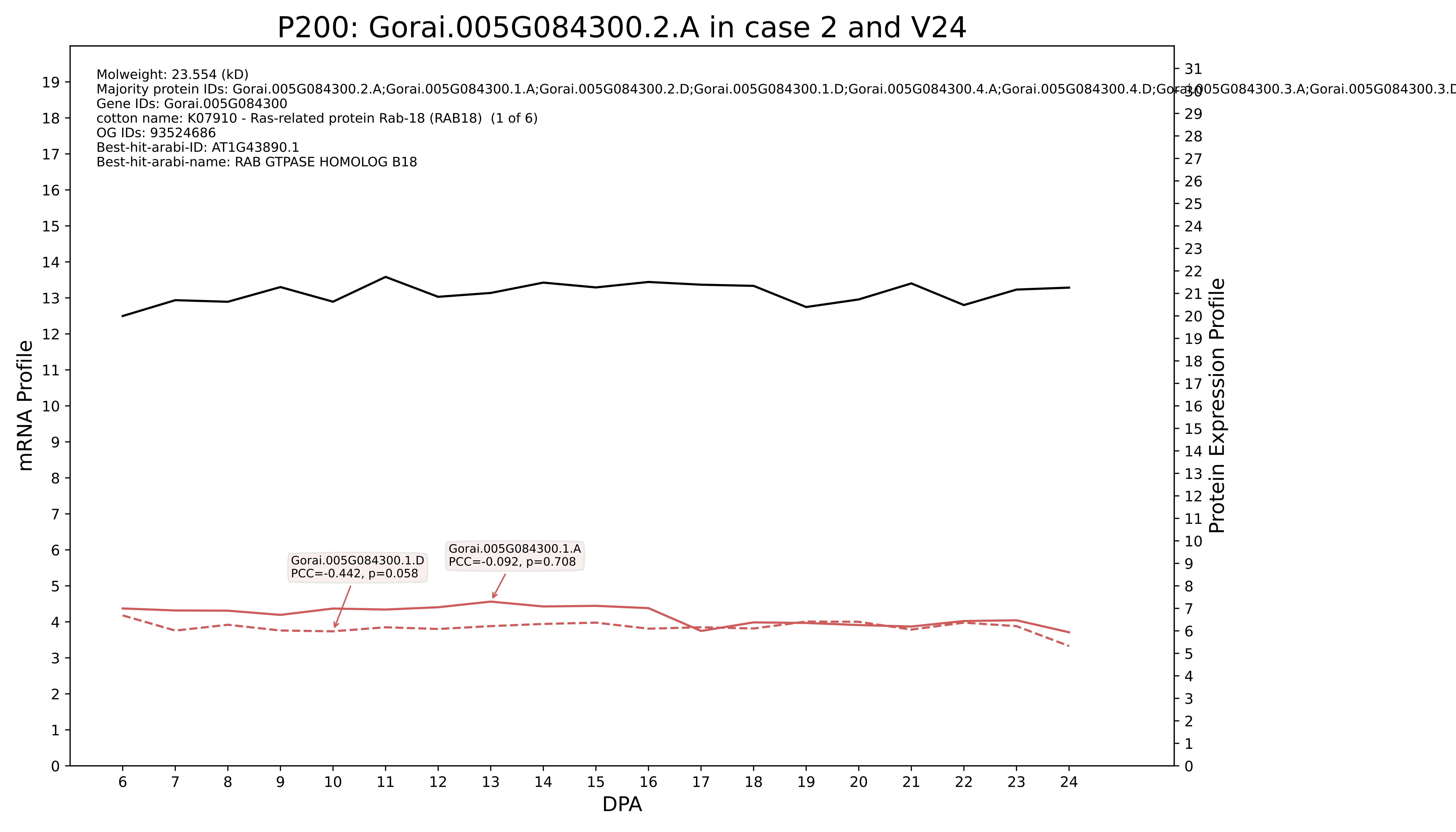
P200: Gorai.005G084300.2.A in case 2 and V24
Molweight: 23.554 (kD)
Majority protein IDs: Gorai.005G084300.2.A;Gorai.005G084300.1.A;Gorai.005G084300.2.D;Gorai.005G084300.1.D;Gorai.005G084300.4.A;Gorai.005G084300.4.D;Gorai.005G084300.3.A;Gorai.005G084300.3.D
Gene IDs: Gorai.005G084300
cotton name: K07910 - Ras-related protein Rab-18 (RAB18) (1 of 6)
OG IDs: 93524686
Best-hit-arabi-ID: AT1G43890.1
Best-hit-arabi-name: RAB GTPASE HOMOLOG B18
Gorai.005G084300.1.D
PCC=-0.442, p=0.058
Gorai.005G084300.1.A
PCC=-0.092, p=0.708
mRNA Profile
Protein Expression Profile
DPA

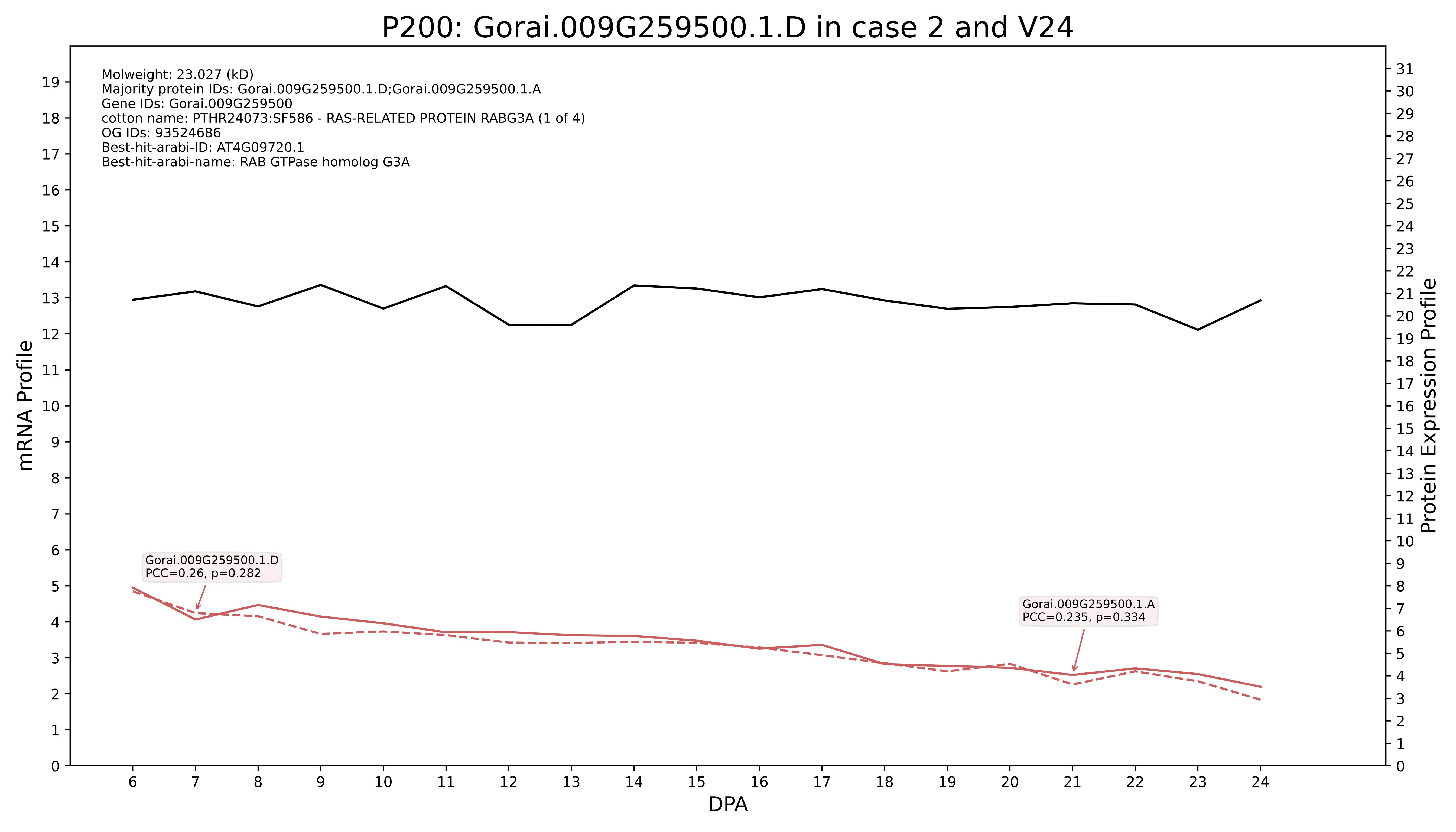
P200: Gorai.009G259500.1.D in case 2 and V24
Molweight: 23.027 (kD)
Majority protein IDs: Gorai.009G259500.1.D;Gorai.009G259500.1.A
Gene IDs: Gorai.009G259500
cotton name: PTHR24073:SF586 - RAS-RELATED PROTEIN RABG3A (1 of 4)
OG IDs: 93524686
Best-hit-arabi-ID: AT4G09720.1
Best-hit-arabi-name: RAB GTPase homolog G3A
Gorai.009G259500.1.D
PCC=0.26, p=0.282
Gorai.009G259500.1.A
PCC=0.235, p=0.334
mRNA Profile
Protein Expression Profile
DPA

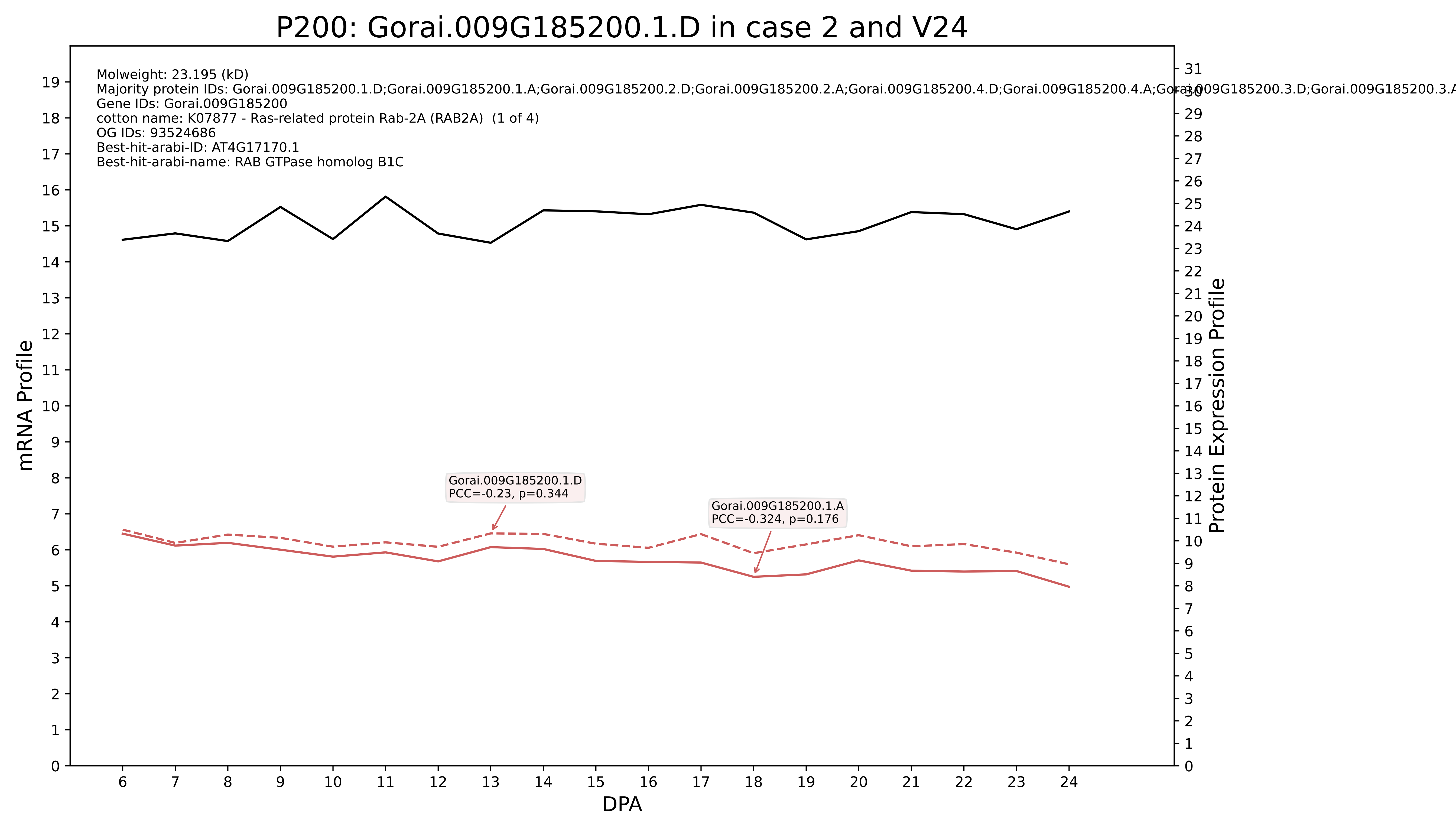
P200: Gorai.009G185200.1.D in case 2 and V24
Molweight: 23.195 (kD)
Majority protein IDs: Gorai.009G185200.1.D;Gorai.009G185200.1.A;Gorai.009G185200.2.D;Gorai.009G185200.2.A;Gorai.009G185200.4.D;Gorai.009G185200.4.A;Gorai.009G185200.3.D;Gorai.009G185200.3.A
Gene IDs: Gorai.009G185200
cotton name: K07877 - Ras-related protein Rab-2A (RAB2A) (1 of 4)
OG IDs: 93524686
Best-hit-arabi-ID: AT4G17170.1
Best-hit-arabi-name: RAB GTPase homolog B1C
Gorai.009G185200.1.D
PCC=-0.23, p=0.344
Gorai.009G185200.1.A
PCC=-0.324, p=0.176
mRNA Profile
Protein Expression Profile
DPA

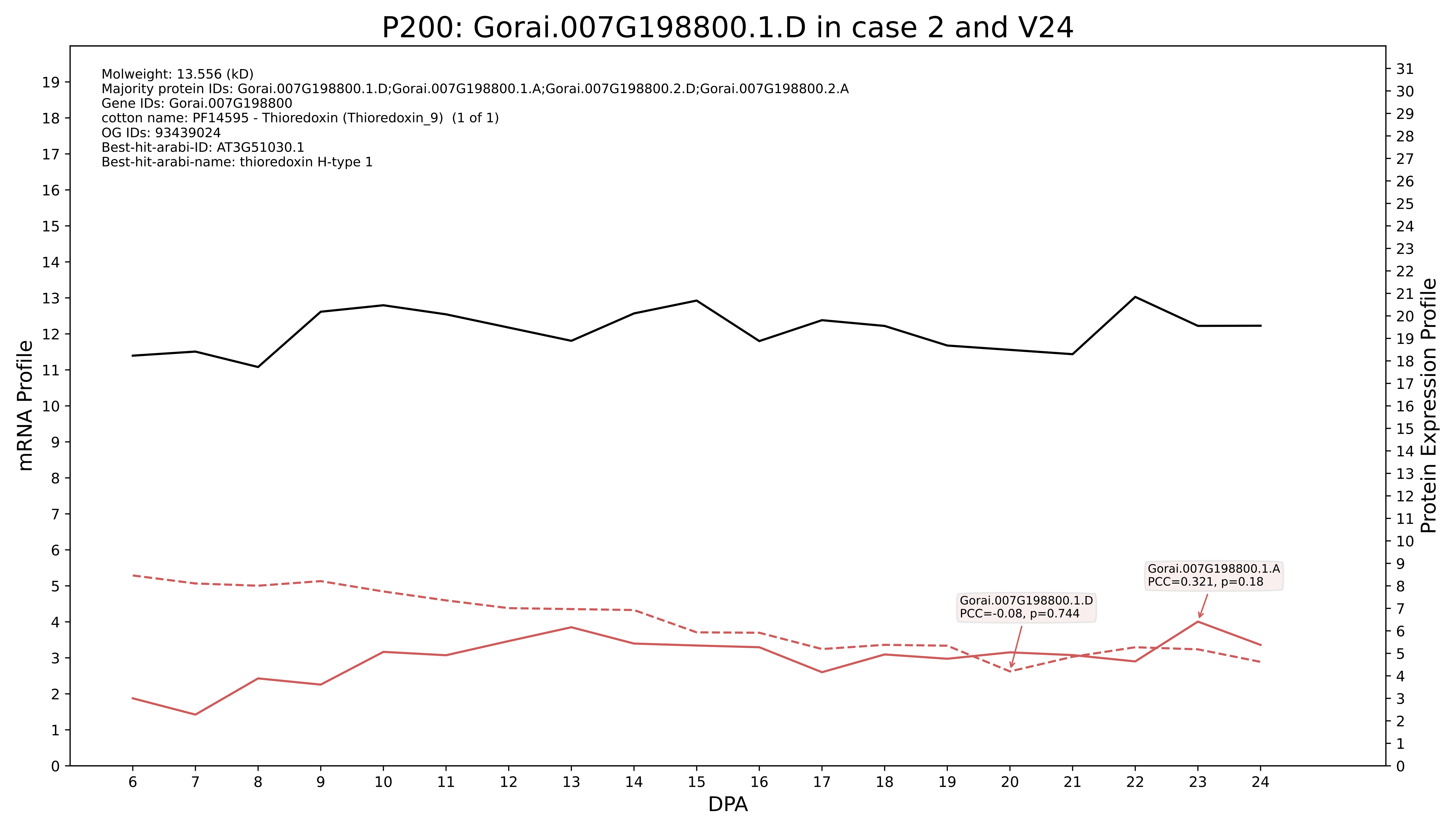
P200: Gorai.007G198800.1.D in case 2 and V24
Molweight: 13.556 (kD)
Majority protein IDs: Gorai.007G198800.1.D;Gorai.007G198800.1.A;Gorai.007G198800.2.D;Gorai.007G198800.2.A
Gene IDs: Gorai.007G198800
cotton name: PF14595 - Thioredoxin (Thioredoxin_9) (1 of 1)
OG IDs: 93439024
Best-hit-arabi-ID: AT3G51030.1
Best-hit-arabi-name: thioredoxin H-type 1
Gorai.007G198800.1.D
PCC=-0.08, p=0.744
Gorai.007G198800.1.A
PCC=0.321, p=0.18
mRNA Profile
Protein Expression Profile
DPA

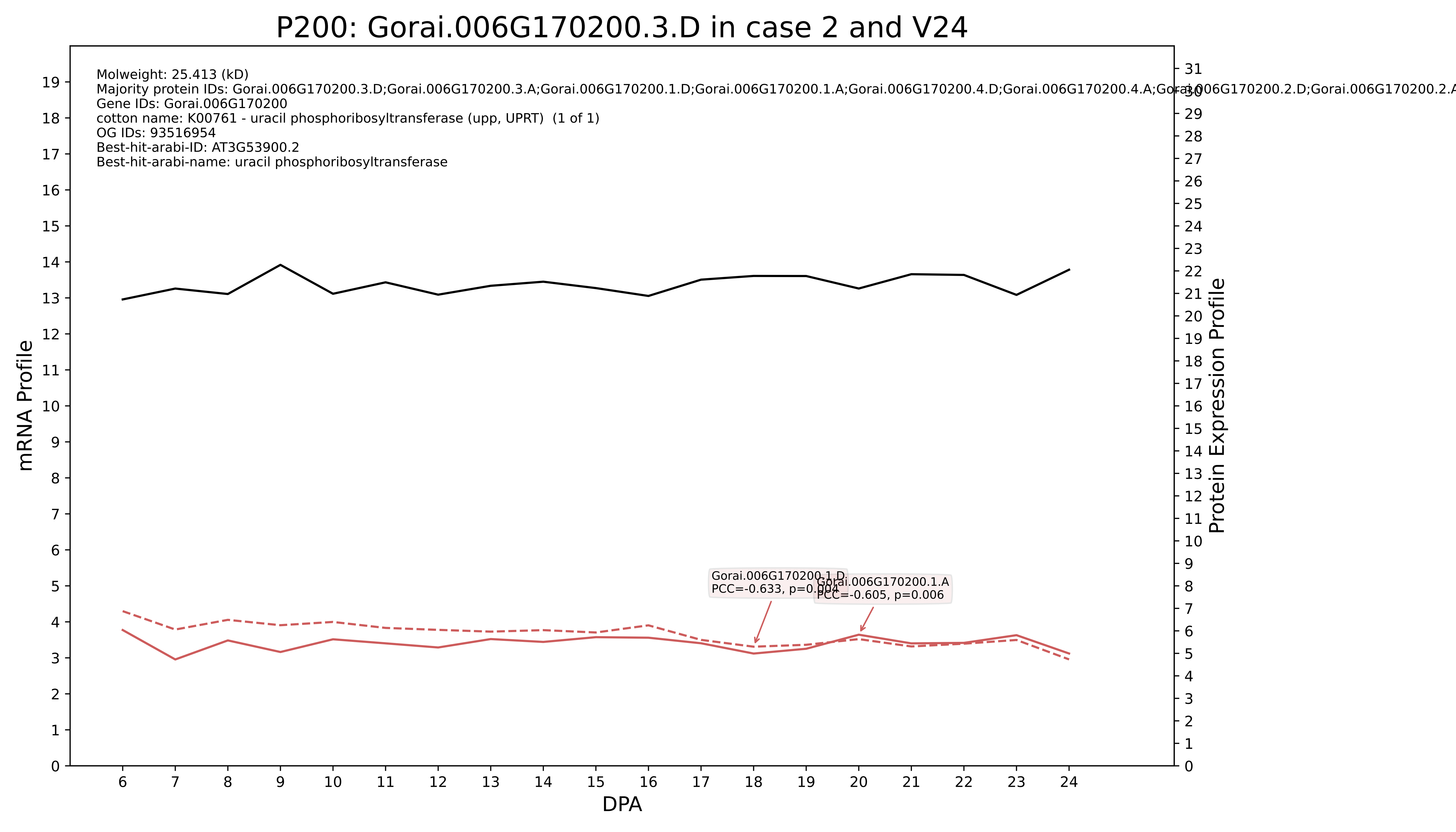
P200: Gorai.006G170200.3.D in case 2 and V24
Molweight: 25.413 (kD)
Majority protein IDs: Gorai.006G170200.3.D;Gorai.006G170200.3.A;Gorai.006G170200.1.D;Gorai.006G170200.1.A;Gorai.006G170200.4.D;Gorai.006G170200.4.A;Gorai.006G170200.2.D;Gorai.006G170200.2.A
Gene IDs: Gorai.006G170200
cotton name: K00761 - uracil phosphoribosyltransferase (upp, UPRT) (1 of 1)
OG IDs: 93516954
Best-hit-arabi-ID: AT3G53900.2
Best-hit-arabi-name: uracil phosphoribosyltransferase
Gorai.006G170200.1.D
PCC=-0.633, p=0.004
Gorai.006G170200.1.A
PCC=-0.605, p=0.006
mRNA Profile
Protein Expression Profile
DPA

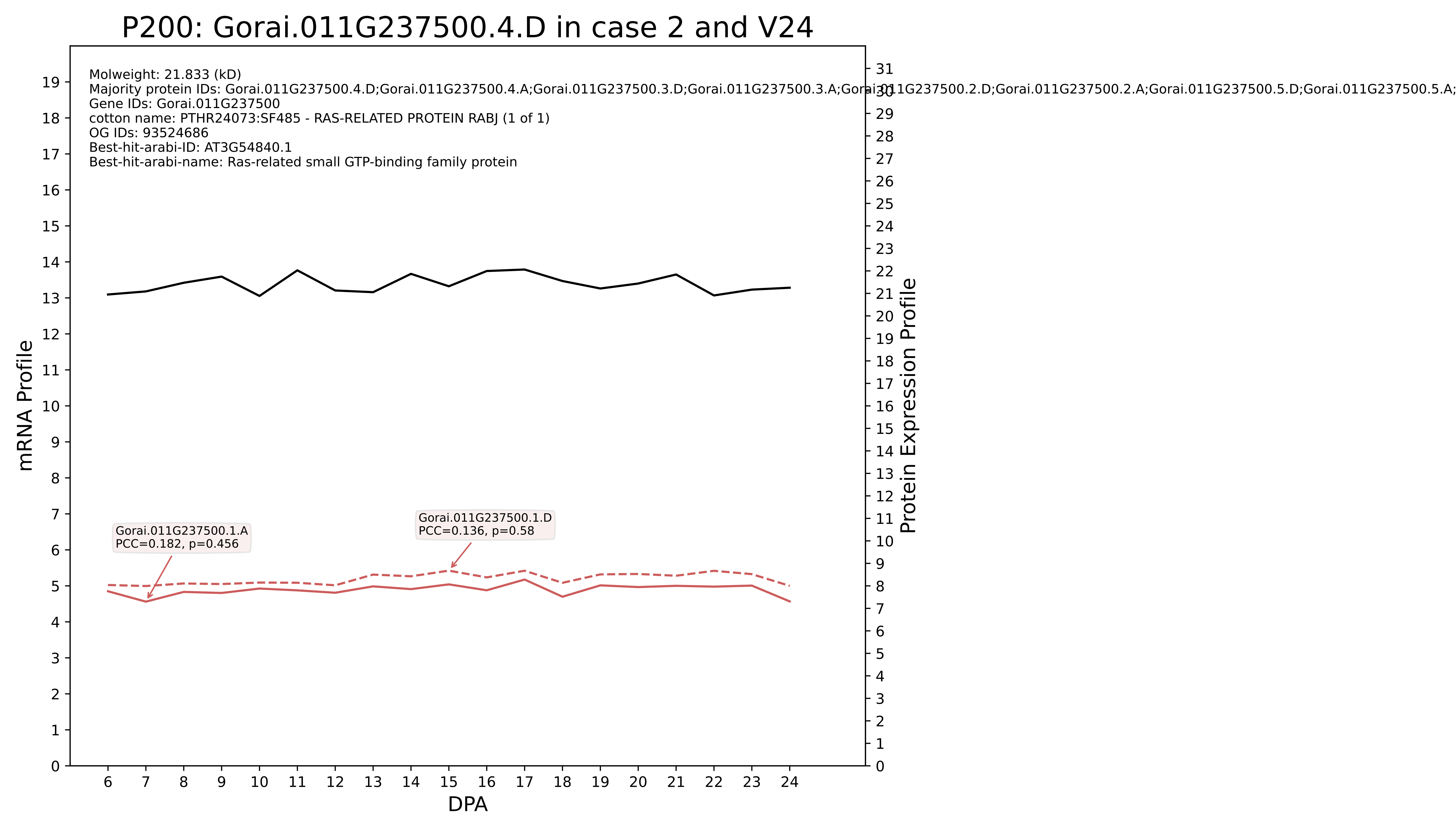

P200: Gorai.011G237500.4.D in case 2 and V24
Molweight: 21.833 (kD)
Majority protein IDs: Gorai.011G237500.4.D;Gorai.011G237500.4.A;Gorai.011G237500.3.D;Gorai.011G237500.3.A;Gorai.011G237500.2.D;Gorai.011G237500.2.A;Gorai.011G237500.5.D;Gorai.011G237500.5.A;
Gene IDs: Gorai.011G237500
cotton name: PTHR24073:SF485 - RAS-RELATED PROTEIN RABJ (1 of 1)
OG IDs: 93524686
Best-hit-arabi-ID: AT3G54840.1
Best-hit-arabi-name: Ras-related small GTP-binding family protein
Gorai.011G237500.1.A
PCC=0.182, p=0.456
Gorai.011G237500.1.D
PCC=0.136, p=0.58
mRNA Profile
Protein Expression Profile
DPA

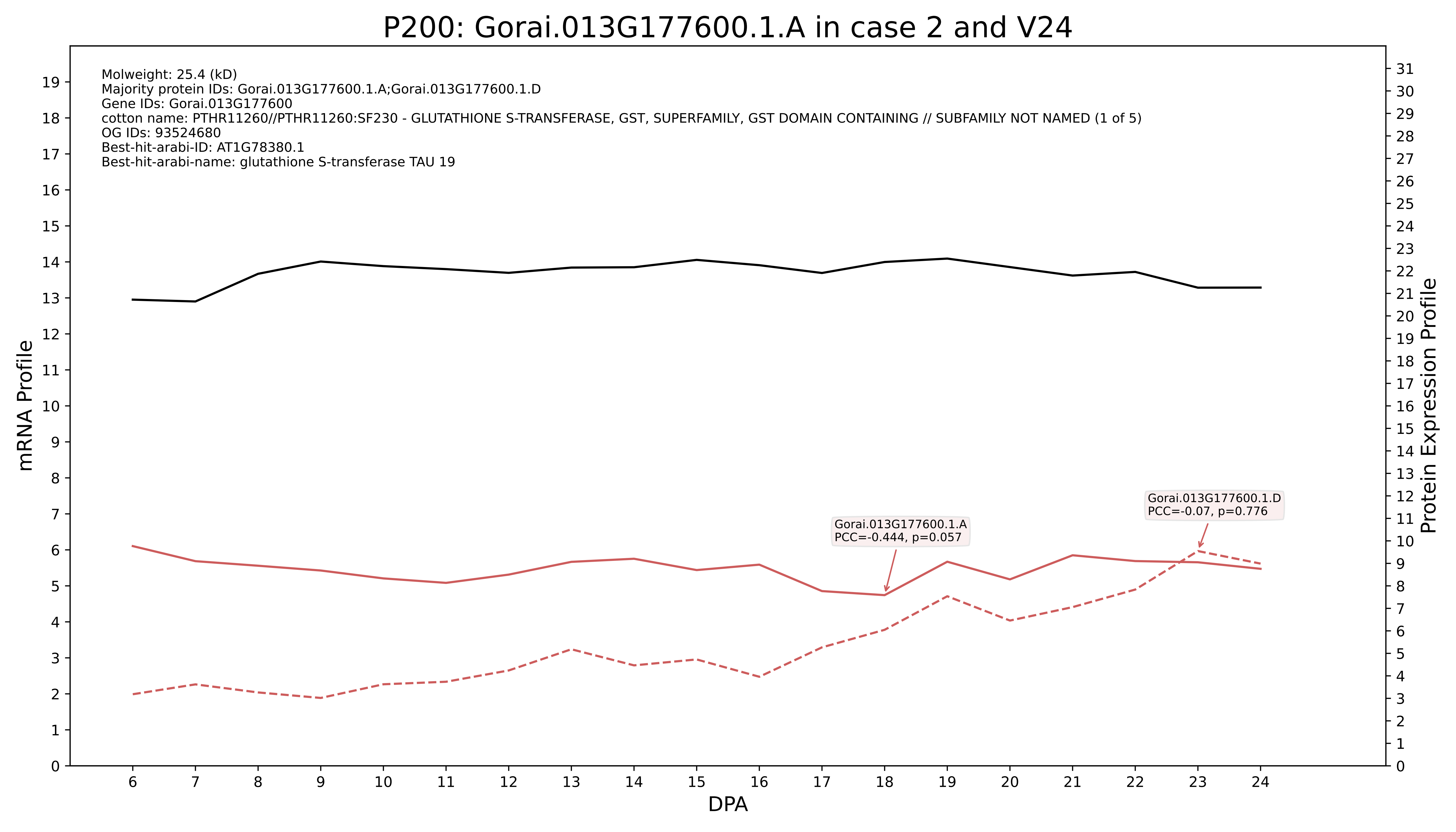
P200: Gorai.013G177600.1.A in case 2 and V24
Molweight: 25.4 (kD)
Majority protein IDs: Gorai.013G177600.1.A;Gorai.013G177600.1.D
Gene IDs: Gorai.013G177600
cotton name: PTHR11260//PTHR11260:SF230 - GLUTATHIONE S-TRANSFERASE, GST, SUPERFAMILY, GST DOMAIN CONTAINING // SUBFAMILY NOT NAMED (1 of 5)
OG IDs: 93524680
Best-hit-arabi-ID: AT1G78380.1
Best-hit-arabi-name: glutathione S-transferase TAU 19
Gorai.013G177600.1.A
PCC=-0.444, p=0.057
Gorai.013G177600.1.D
PCC=-0.07, p=0.776
mRNA Profile
Protein Expression Profile
DPA

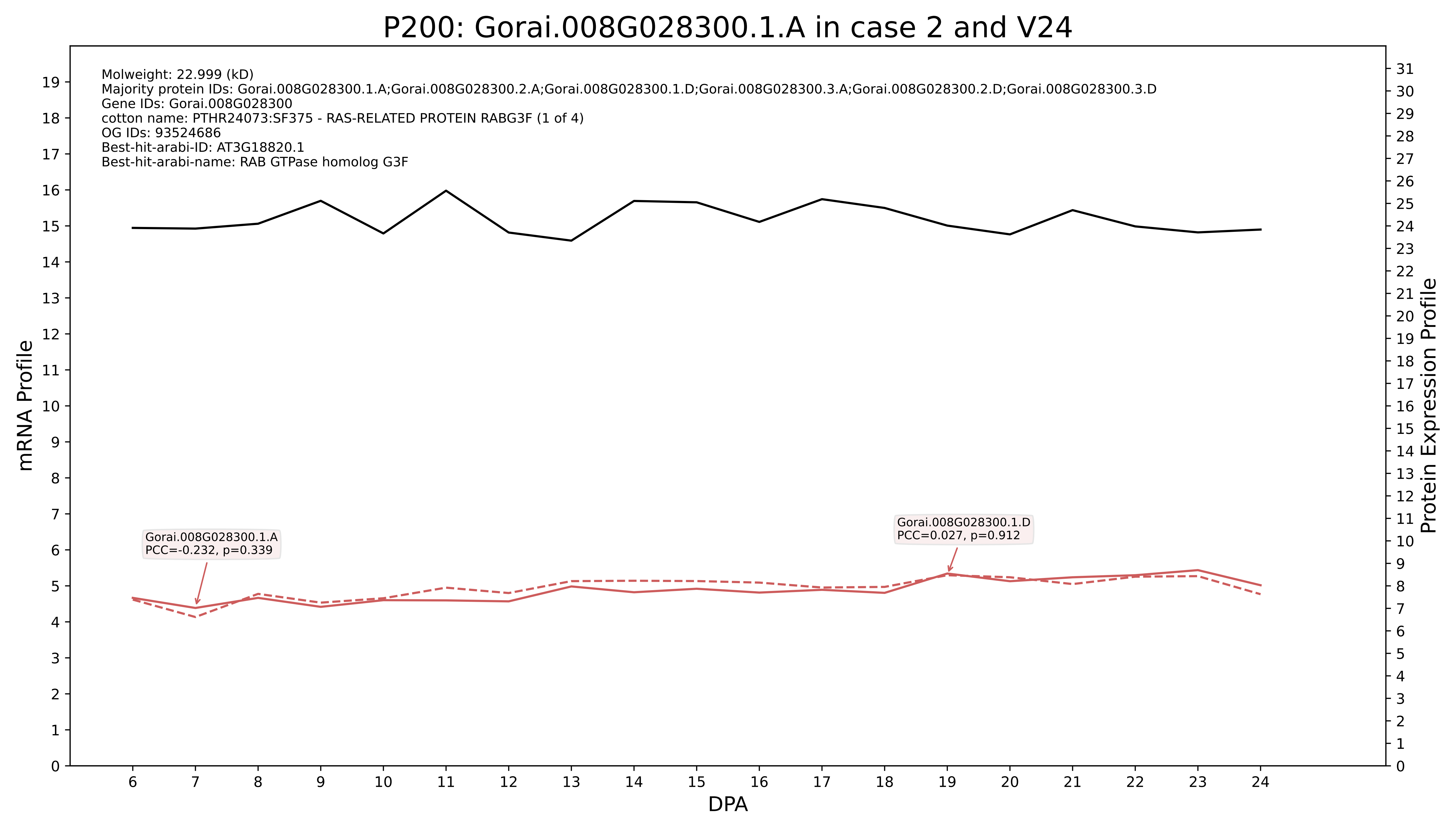
P200: Gorai.008G028300.1.A in case 2 and V24
Molweight: 22.999 (kD)
Majority protein IDs: Gorai.008G028300.1.A;Gorai.008G028300.2.A;Gorai.008G028300.1.D;Gorai.008G028300.3.A;Gorai.008G028300.2.D;Gorai.008G028300.3.D
Gene IDs: Gorai.008G028300
cotton name: PTHR24073:SF375 - RAS-RELATED PROTEIN RABG3F (1 of 4)
OG IDs: 93524686
Best-hit-arabi-ID: AT3G18820.1
Best-hit-arabi-name: RAB GTPase homolog G3F
Gorai.008G028300.1.A
PCC=-0.232, p=0.339
Gorai.008G028300.1.D
PCC=0.027, p=0.912
mRNA Profile
Protein Expression Profile
DPA

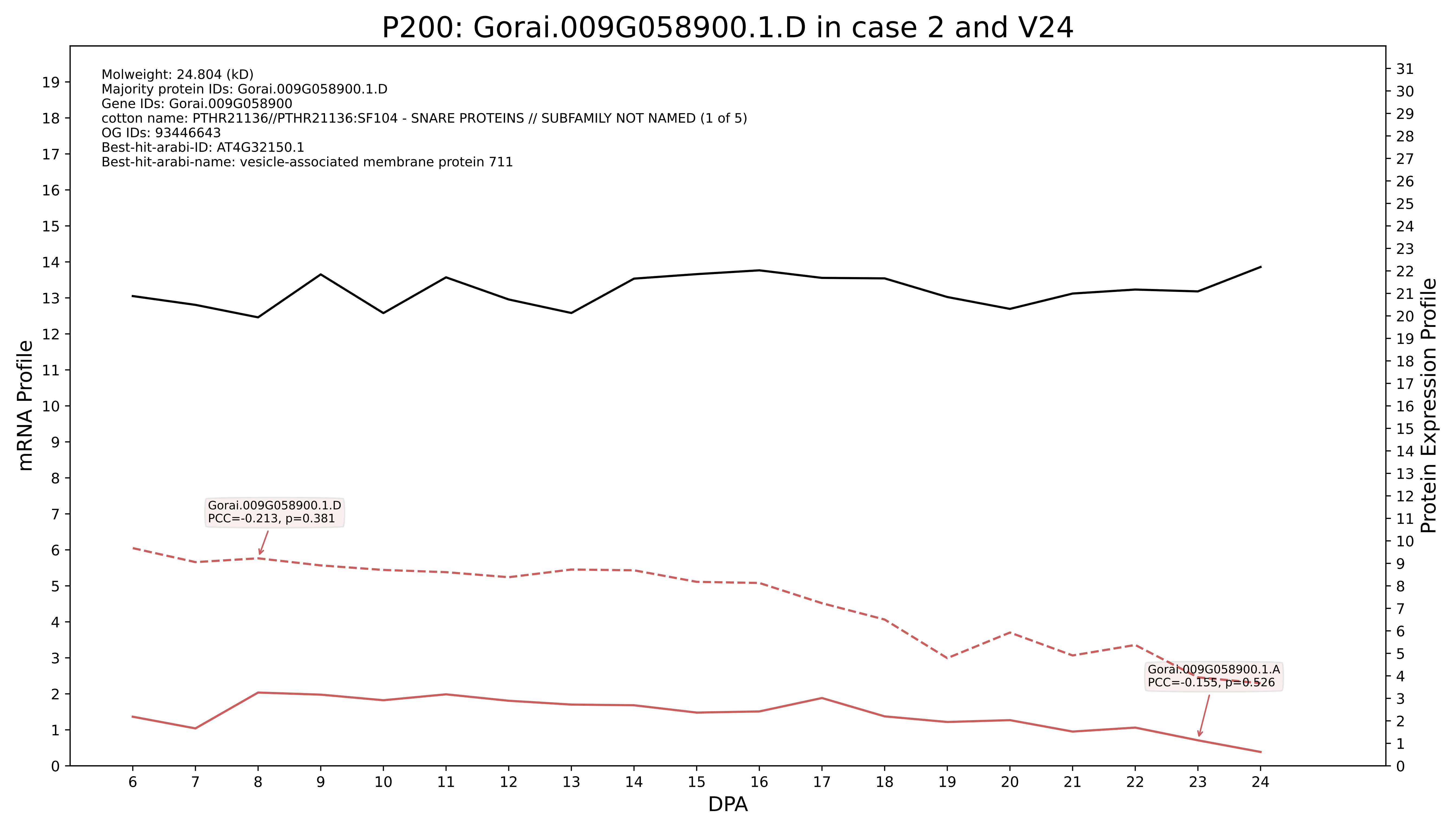

P200: Gorai.009G058900.1.D in case 2 and V24
Molweight: 24.804 (kD)
Majority protein IDs: Gorai.009G058900.1.D
Gene IDs: Gorai.009G058900
cotton name: PTHR21136//PTHR21136:SF104 - SNARE PROTEINS // SUBFAMILY NOT NAMED (1 of 5)
OG IDs: 93446643
Best-hit-arabi-ID: AT4G32150.1
Best-hit-arabi-name: vesicle-associated membrane protein 711
Gorai.009G058900.1.D
PCC=-0.213, p=0.381
Gorai.009G058900.1.A
PCC=-0.155, p=0.526
mRNA Profile
Protein Expression Profile
DPA

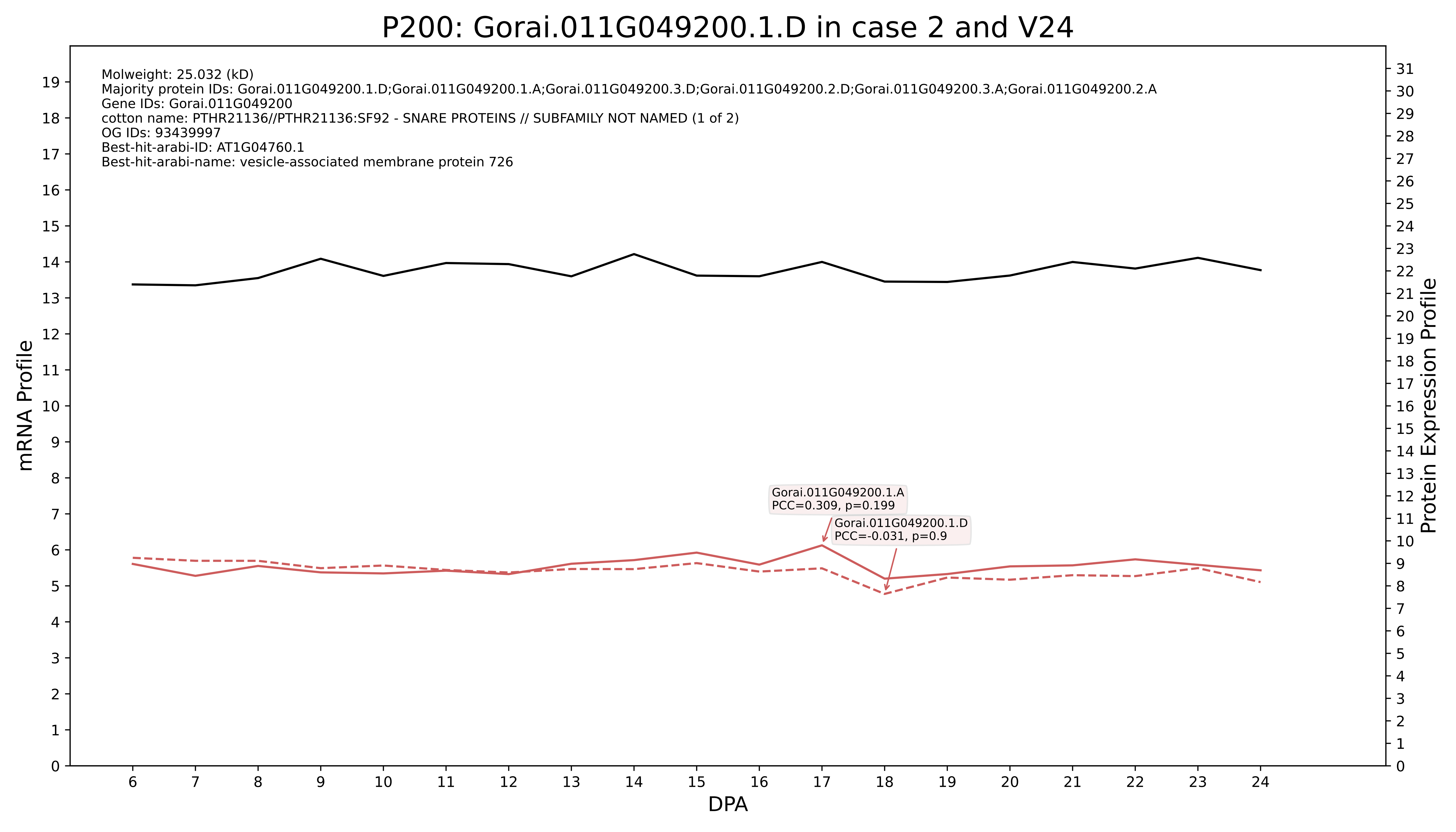
P200: Gorai.011G049200.1.D in case 2 and V24
Molweight: 25.032 (kD)
Majority protein IDs: Gorai.011G049200.1.D;Gorai.011G049200.1.A;Gorai.011G049200.3.D;Gorai.011G049200.2.D;Gorai.011G049200.3.A;Gorai.011G049200.2.A
Gene IDs: Gorai.011G049200
cotton name: PTHR21136//PTHR21136:SF92 - SNARE PROTEINS // SUBFAMILY NOT NAMED (1 of 2)
OG IDs: 93439997
Best-hit-arabi-ID: AT1G04760.1
Best-hit-arabi-name: vesicle-associated membrane protein 726
Gorai.011G049200.1.A
PCC=0.309, p=0.199
Gorai.011G049200.1.D
PCC=-0.031, p=0.9
mRNA Profile
Protein Expression Profile
DPA

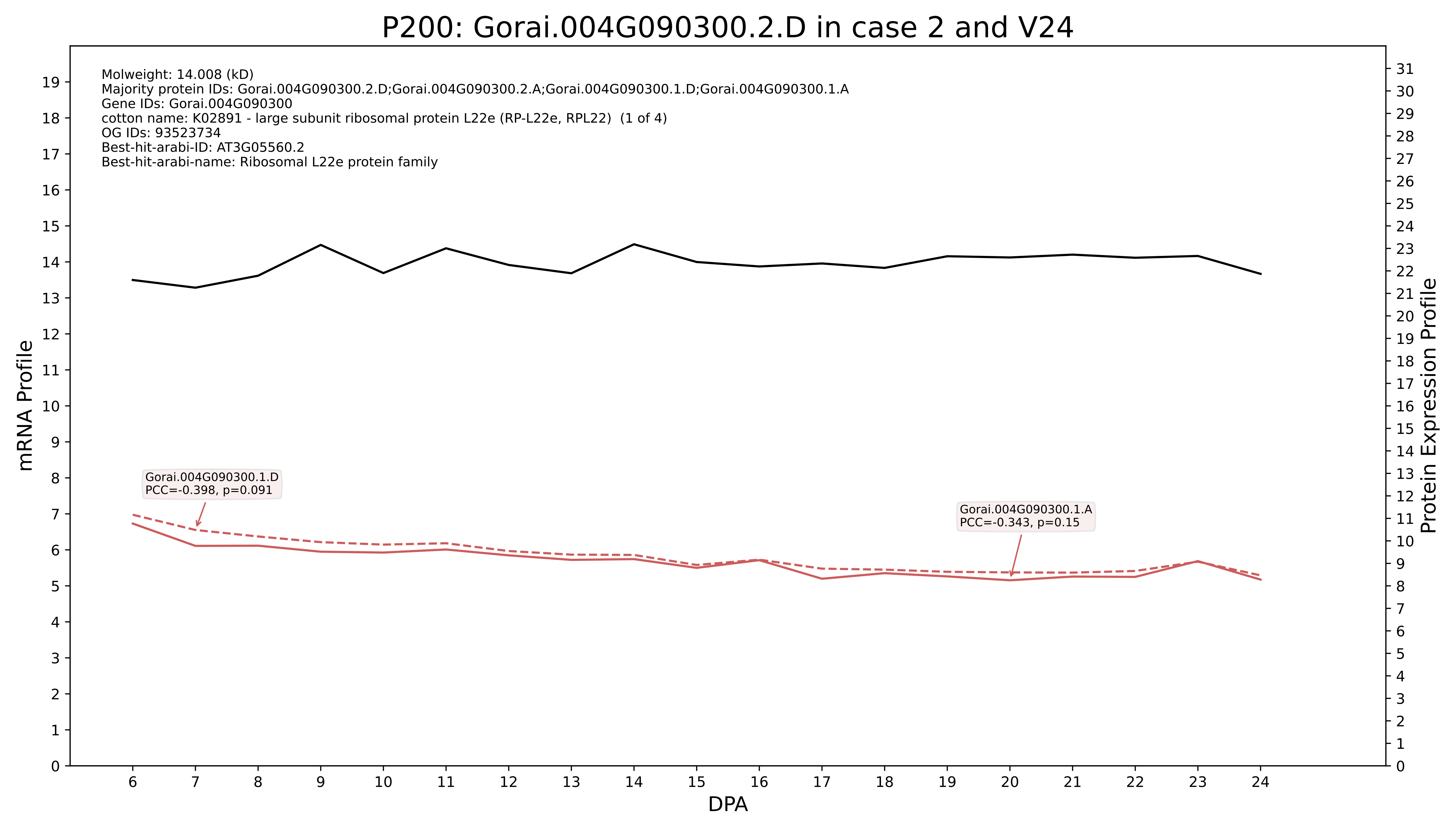
P200: Gorai.004G090300.2.D in case 2 and V24
Molweight: 14.008 (kD)
Majority protein IDs: Gorai.004G090300.2.D;Gorai.004G090300.2.A;Gorai.004G090300.1.D;Gorai.004G090300.1.A
Gene IDs: Gorai.004G090300
cotton name: K02891 - large subunit ribosomal protein L22e (RP-L22e, RPL22) (1 of 4)
OG IDs: 93523734
Best-hit-arabi-ID: AT3G05560.2
Best-hit-arabi-name: Ribosomal L22e protein family
Gorai.004G090300.1.D
PCC=-0.398, p=0.091
Gorai.004G090300.1.A
PCC=-0.343, p=0.15
mRNA Profile
Protein Expression Profile
DPA

### P200: Gorai.009G119700.1.D in case 2 and V24

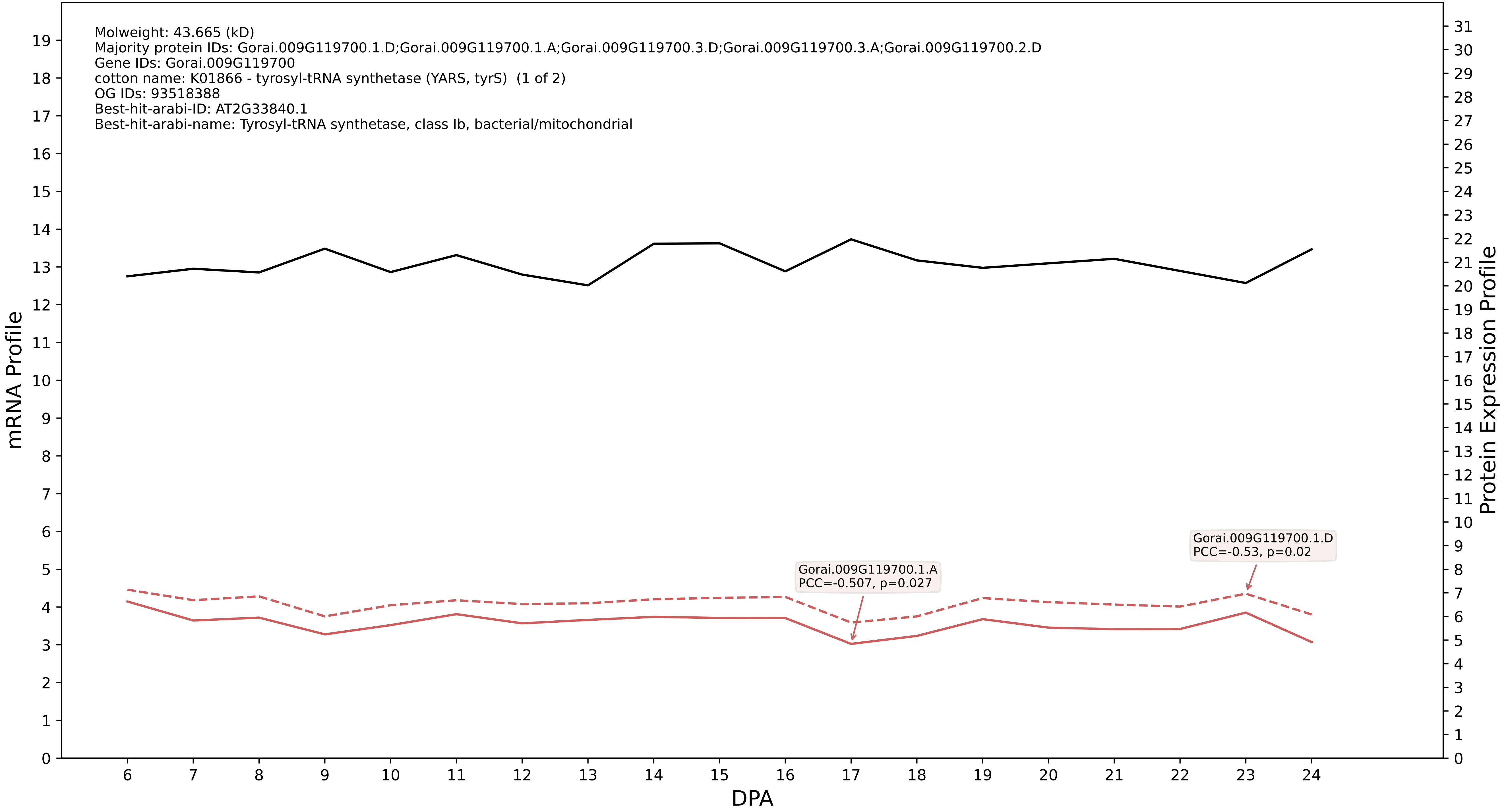

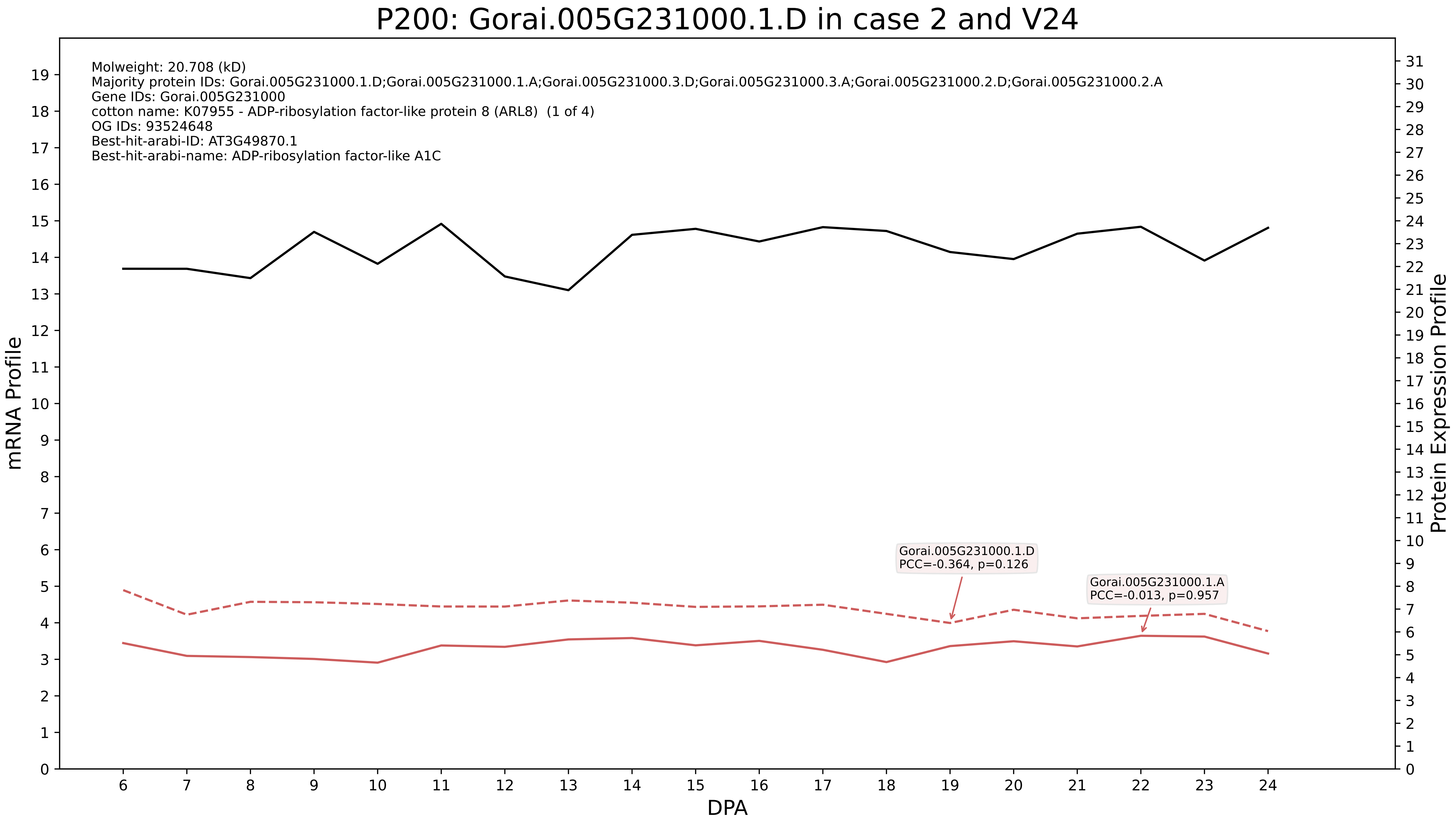
P200: Gorai.005G231000.1.D in case 2 and V24
Molweight: 20.708 (kD)
Majority protein IDs: Gorai.005G231000.1.D;Gorai.005G231000.1.A;Gorai.005G231000.3.D;Gorai.005G231000.3.A;Gorai.005G231000.2.D;Gorai.005G231000.2.A
Gene IDs: Gorai.005G231000
cotton name: K07955 - ADP-ribosylation factor-like protein 8 (ARL8) (1 of 4)
OG IDs: 93524648
Best-hit-arabi-ID: AT3G49870.1
Best-hit-arabi-name: ADP-ribosylation factor-like A1C
Gorai.005G231000.1.D
PCC=-0.364, p=0.126
Gorai.005G231000.1.A
PCC=-0.013, p=0.957
mRNA Profile
Protein Expression Profile
DPA

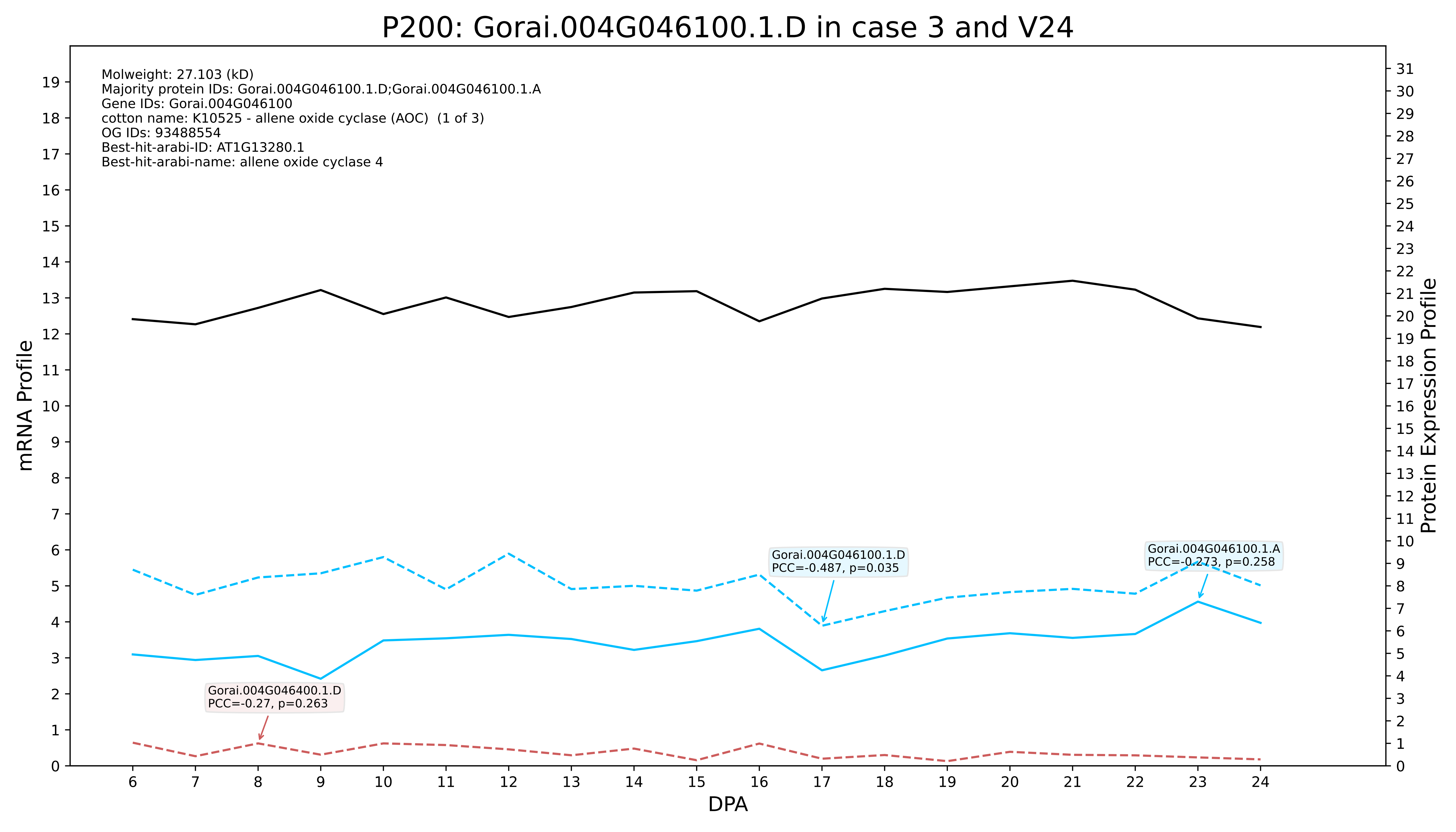

P200: Gorai.004G046100.1.D in case 3 and V24
Molweight: 27.103 (kD)
Majority protein IDs: Gorai.004G046100.1.D;Gorai.004G046100.1.A
Gene IDs: Gorai.004G046100
cotton name: K10525 - allene oxide cyclase (AOC) (1 of 3)
OG IDs: 93488554
Best-hit-arabi-ID: AT1G13280.1
Best-hit-arabi-name: allene oxide cyclase 4
Gorai.004G046100.1.D
PCC=-0.487, p=0.035
Gorai.004G046100.1.A
PCC=-0.273, p=0.258
Gorai.004G046400.1.D
PCC=-0.27, p=0.263
mRNA Profile
Protein Expression Profile
DPA

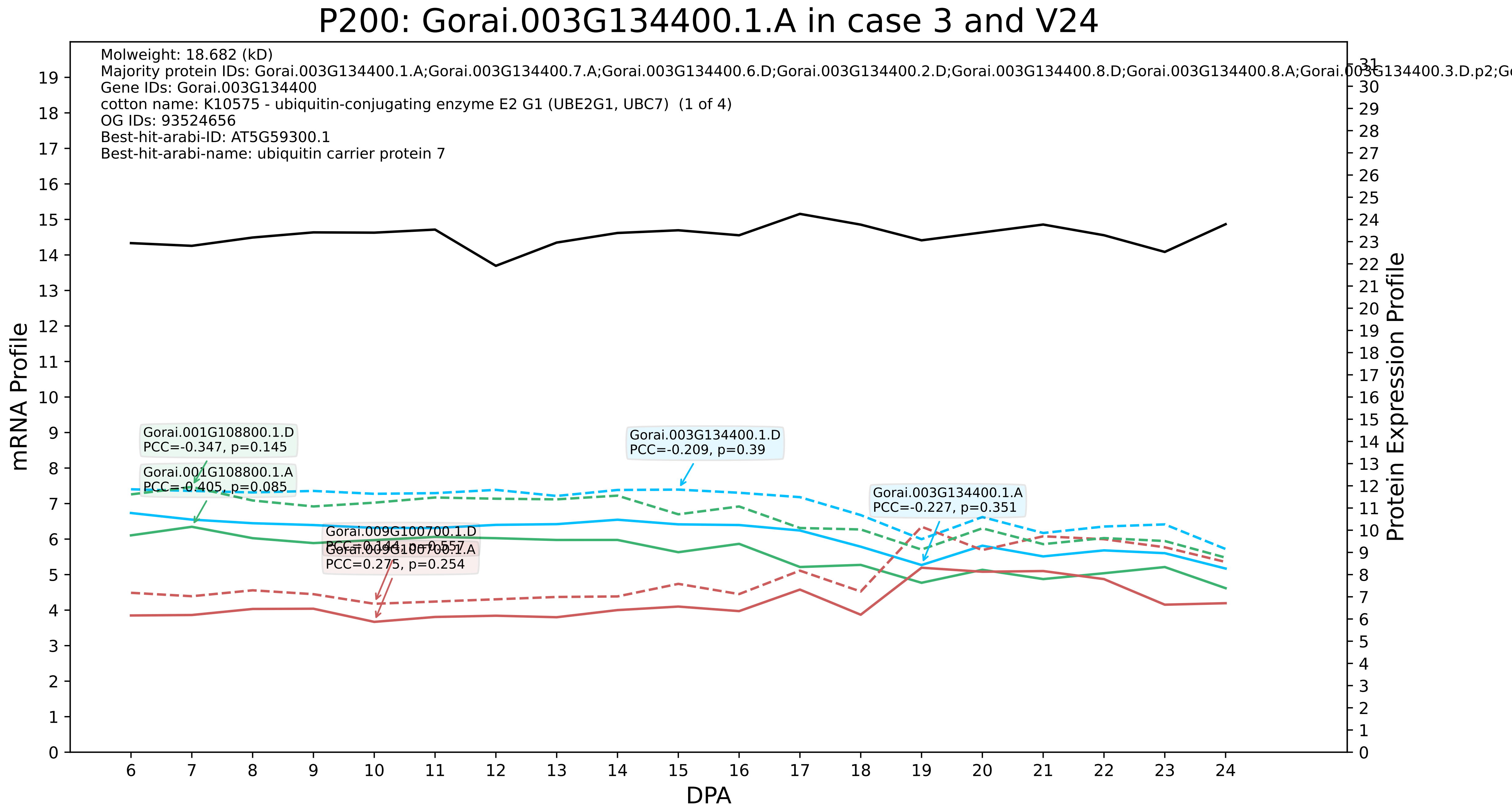
P200: Gorai.003G134400.1.A in case 3 and V24
Molweight: 18.682 (kD)
Majority protein IDs: Gorai.003G134400.1.A;Gorai.003G134400.7.A;Gorai.003G134400.6.D;Gorai.003G134400.2.D;Gorai.003G134400.8.D;Gorai.003G134400.8.A;Gorai.003G134400.3.D.p2;Go
Gene IDs: Gorai.003G134400
cotton name: K10575 - ubiquitin-conjugating enzyme E2 G1 (UBE2G1, UBC7) (1 of 4)
OG IDs: 93524656
Best-hit-arabi-ID: AT5G59300.1
Best-hit-arabi-name: ubiquitin carrier protein 7
Gorai.001G108800.1.D
PCC=-0.347, p=0.145
Gorai.001G108800.1.A
PCC=-0.405, p=0.085
Gorai.003G134400.1.D
PCC=-0.209, p=0.39
Gorai.003G134400.1.A
PCC=-0.227, p=0.351
Gorai.009G100700.1.D
PCC=0.144, p=0.557
Gorai.009G100700.1.A
PCC=0.275, p=0.254
mRNA Profile
Protein Expression Profile
DPA

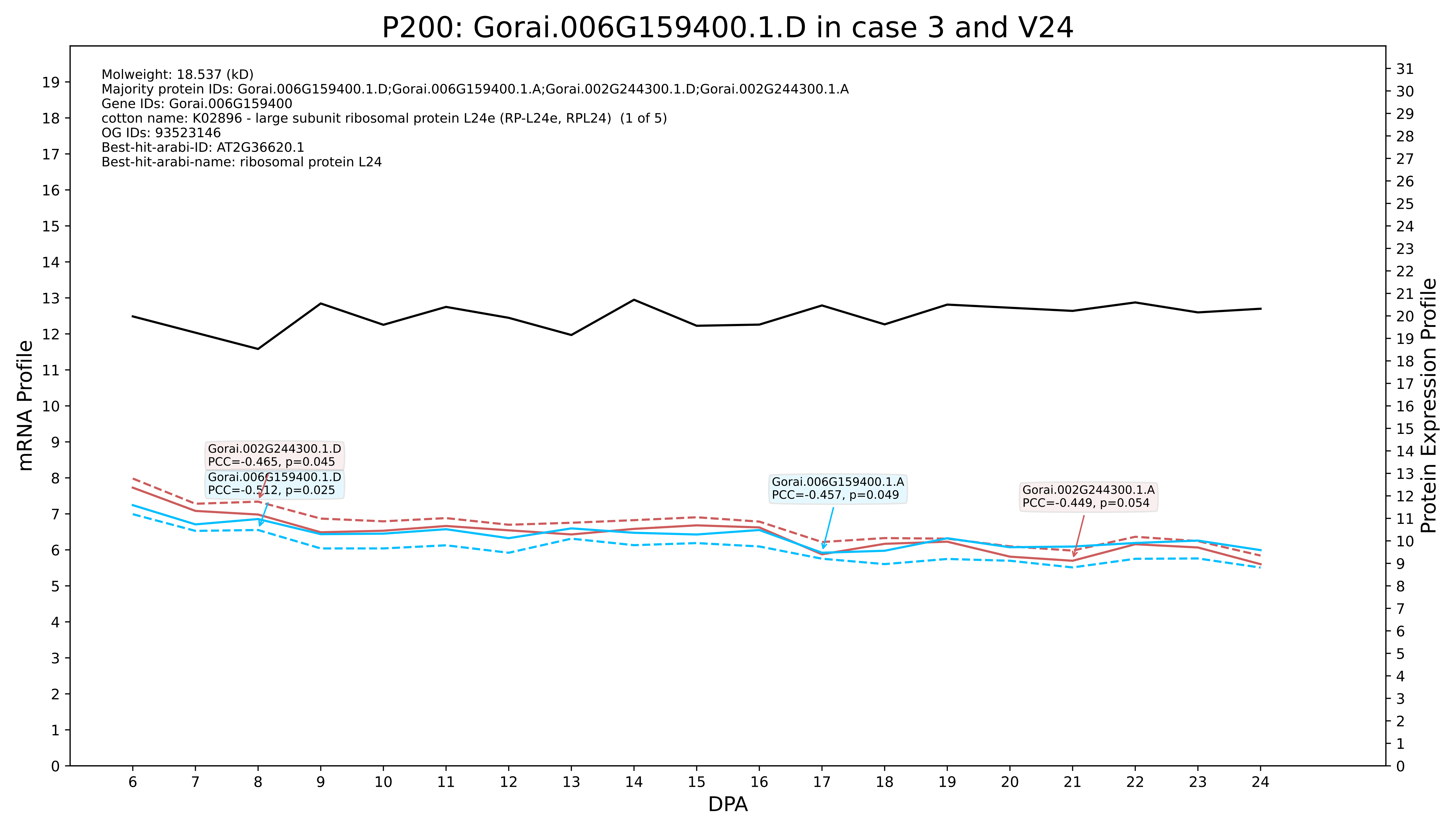
P200: Gorai.006G159400.1.D in case 3 and V24
Molweight: 18.537 (kD)
Majority protein IDs: Gorai.006G159400.1.D;Gorai.006G159400.1.A;Gorai.002G244300.1.D;Gorai.002G244300.1.A
Gene IDs: Gorai.006G159400
cotton name: K02896 - large subunit ribosomal protein L24e (RP-L24e, RPL24) (1 of 5)
OG IDs: 93523146
Best-hit-arabi-ID: AT2G36620.1
Best-hit-arabi-name: ribosomal protein L24
Gorai.002G244300.1.D
PCC=-0.465, p=0.045
Gorai.006G159400.1.D
PCC=-0.512, p=0.025
Gorai.006G159400.1.A
PCC=-0.457, p=0.049
Gorai.002G244300.1.A
PCC=-0.449, p=0.054
mRNA Profile
Protein Expression Profile
DPA

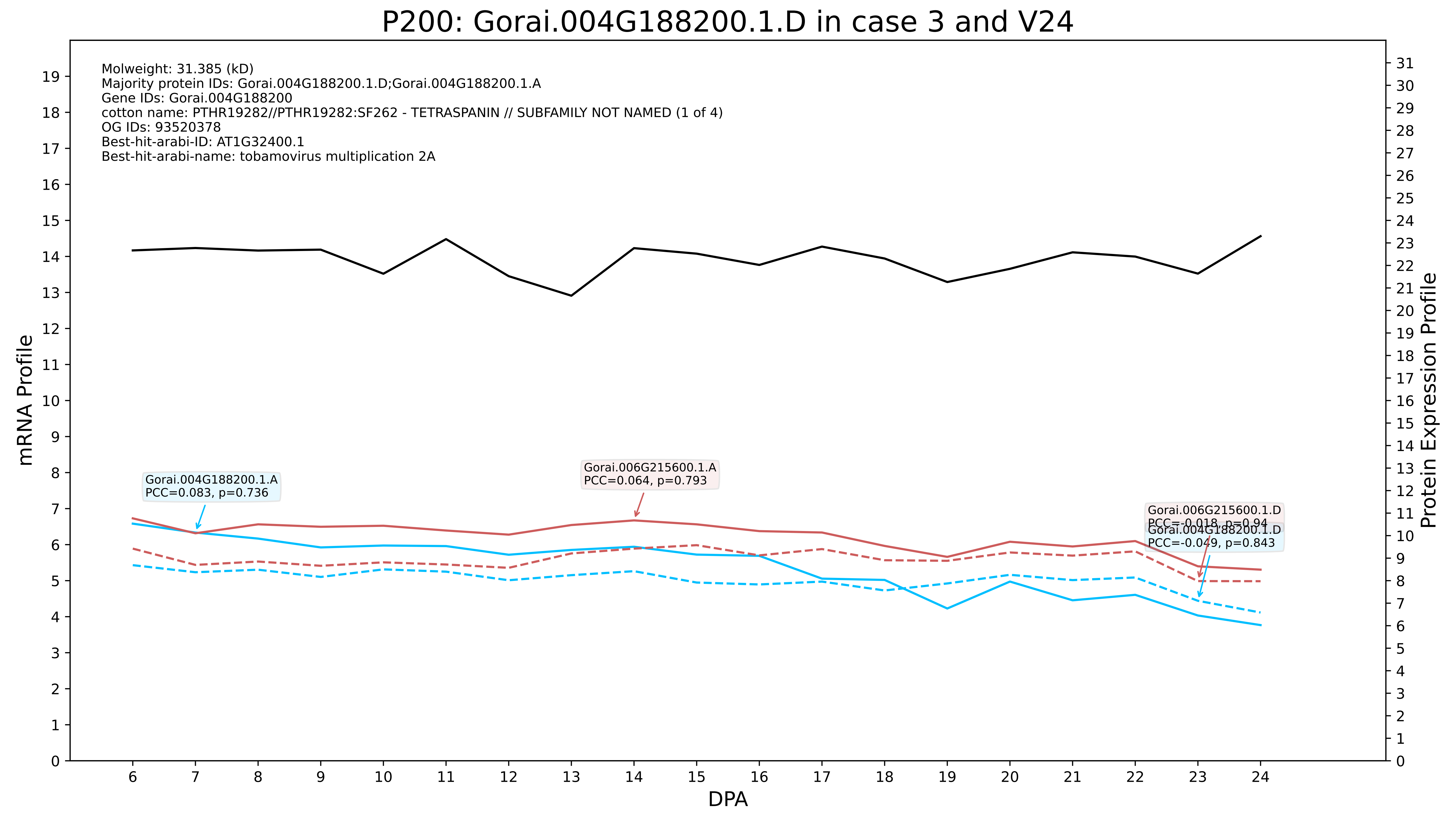

P200: Gorai.004G188200.1.D in case 3 and V24
Molweight: 31.385 (kD)
Majority protein IDs: Gorai.004G188200.1.D;Gorai.004G188200.1.A
Gene IDs: Gorai.004G188200
cotton name: PTHR19282//PTHR19282:SF262 - TETRASPANIN // SUBFAMILY NOT NAMED (1 of 4)
OG IDs: 93520378
Best-hit-arabi-ID: AT1G32400.1
Best-hit-arabi-name: tobamovirus multiplication 2A
Gorai.004G188200.1.A
PCC=0.083, p=0.736
Gorai.006G215600.1.A
PCC=0.064, p=0.793
Gorai.006G215600.1.D
PCC=-0.018, p=0.94
Gorai.004G188200.1.D
PCC=-0.049, p=0.843
mRNA Profile
Protein Expression Profile
DPA

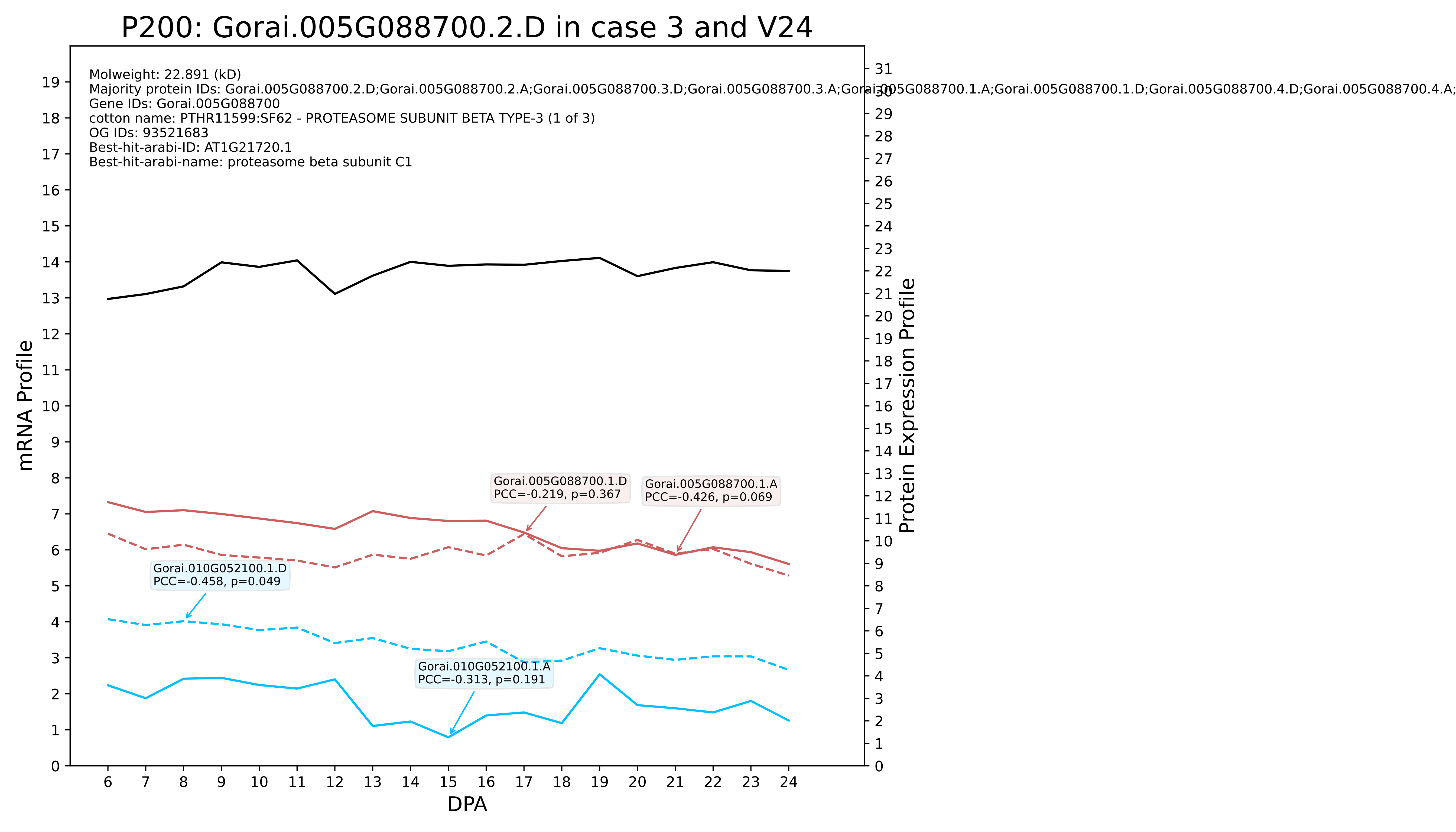

P200: Gorai.005G088700.2.D in case 3 and V24
Molweight: 22.891 (kD)
Majority protein IDs: Gorai.005G088700.2.D;Gorai.005G088700.2.A;Gorai.005G088700.3.D;Gorai.005G088700.3.A;Gorai.005G088700.1.A;Gorai.005G088700.1.D;Gorai.005G088700.4.D;Gorai.005G088700.4.A;
Gene IDs: Gorai.005G088700
cotton name: PTHR11599:SF62 - PROTEASOME SUBUNIT BETA TYPE-3 (1 of 3)
OG IDs: 93521683
Best-hit-arabi-ID: AT1G21720.1
Best-hit-arabi-name: proteasome beta subunit C1
Gorai.005G088700.1.D
PCC=-0.219, p=0.367
Gorai.005G088700.1.A
PCC=-0.426, p=0.069
Gorai.010G052100.1.D
PCC=-0.458, p=0.049
Gorai.010G052100.1.A
PCC=-0.313, p=0.191
mRNA Profile
Protein Expression Profile
DPA

### P200: Gorai.002G225800.3.D in case 1 and V25

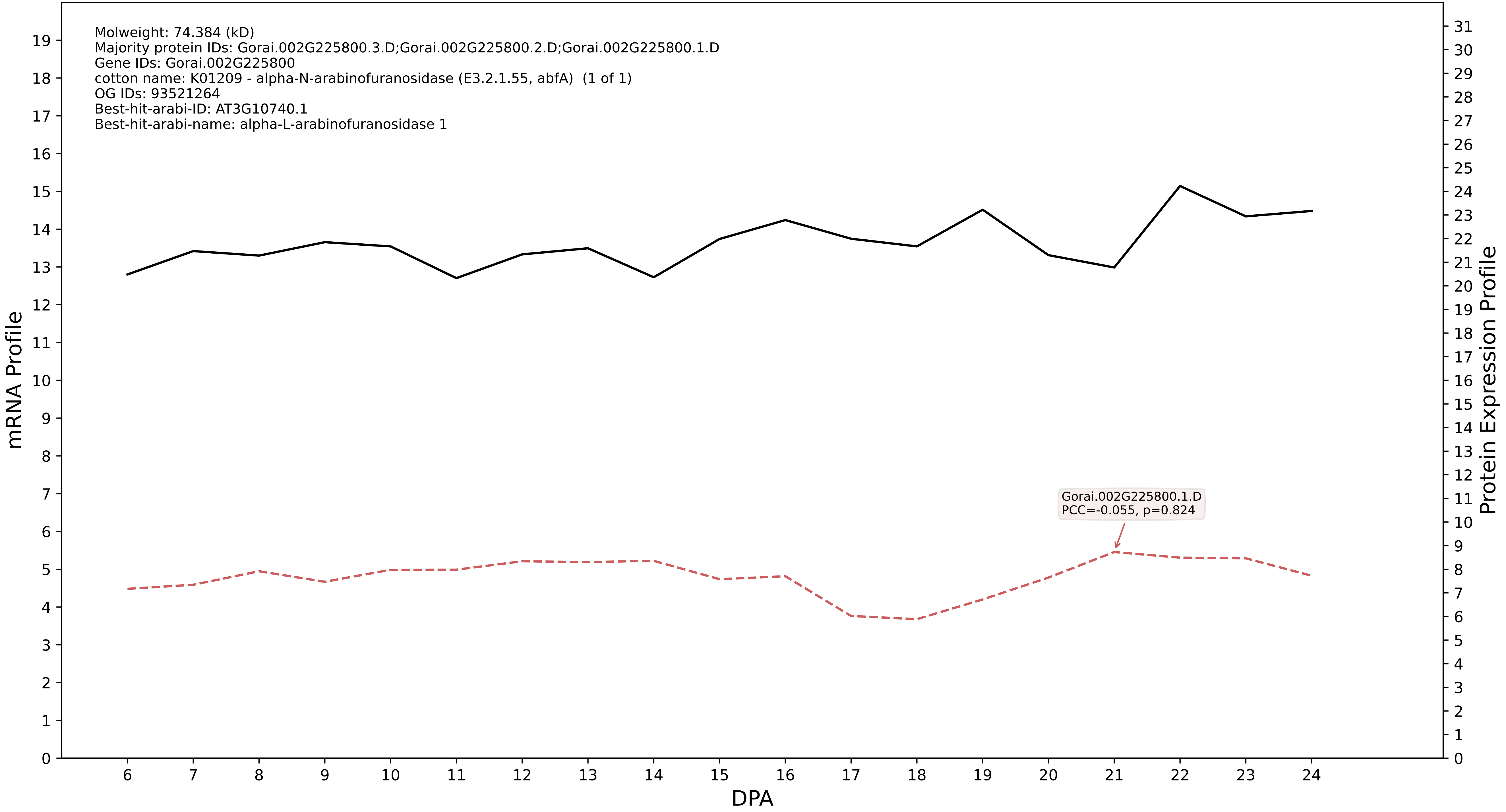

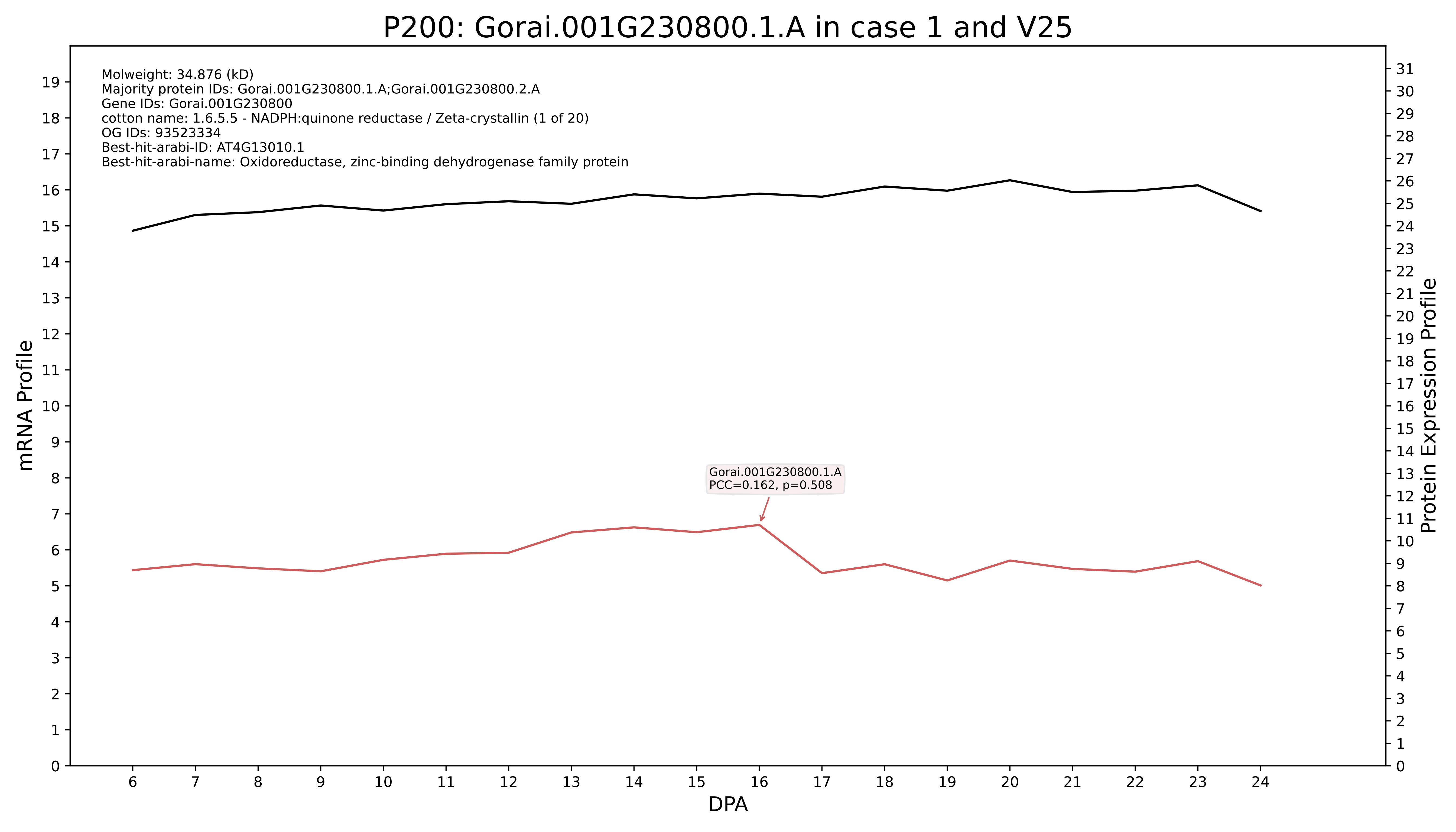
P200: Gorai.001G230800.1.A in case 1 and V25
Molweight: 34.876 (kD)
Majority protein IDs: Gorai.001G230800.1.A;Gorai.001G230800.2.A
Gene IDs: Gorai.001G230800
cotton name: 1.6.5.5 - NADPH:quinone reductase / Zeta-crystallin (1 of 20)
OG IDs: 93523334
Best-hit-arabi-ID: AT4G13010.1
Best-hit-arabi-name: Oxidoreductase, zinc-binding dehydrogenase family protein
Gorai.001G230800.1.A
PCC=0.162, p=0.508
mRNA Profile
Protein Expression Profile
DPA

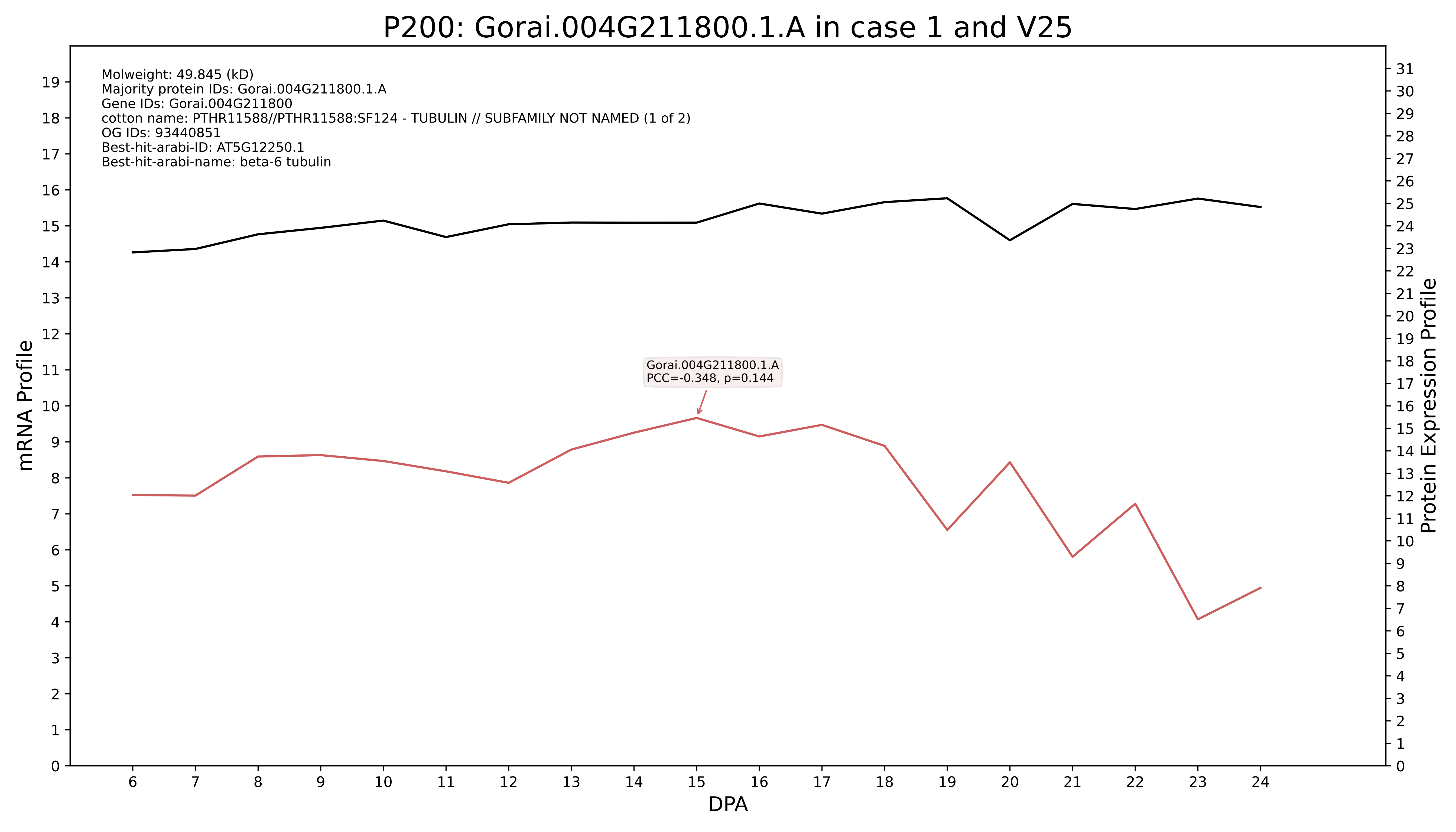
P200: Gorai.004G211800.1.A in case 1 and V25
Molweight: 49.845 (kD)
Majority protein IDs: Gorai.004G211800.1.A
Gene IDs: Gorai.004G211800
cotton name: PTHR11588//PTHR11588:SF124 - TUBULIN // SUBFAMILY NOT NAMED (1 of 2)
OG IDs: 93440851
Best-hit-arabi-ID: AT5G12250.1
Best-hit-arabi-name: beta-6 tubulin
Gorai.004G211800.1.A
PCC=-0.348, p=0.144
mRNA Profile
Protein Expression Profile
DPA

#### P200: Gorai.009G189500.3.D in case 1 and V25

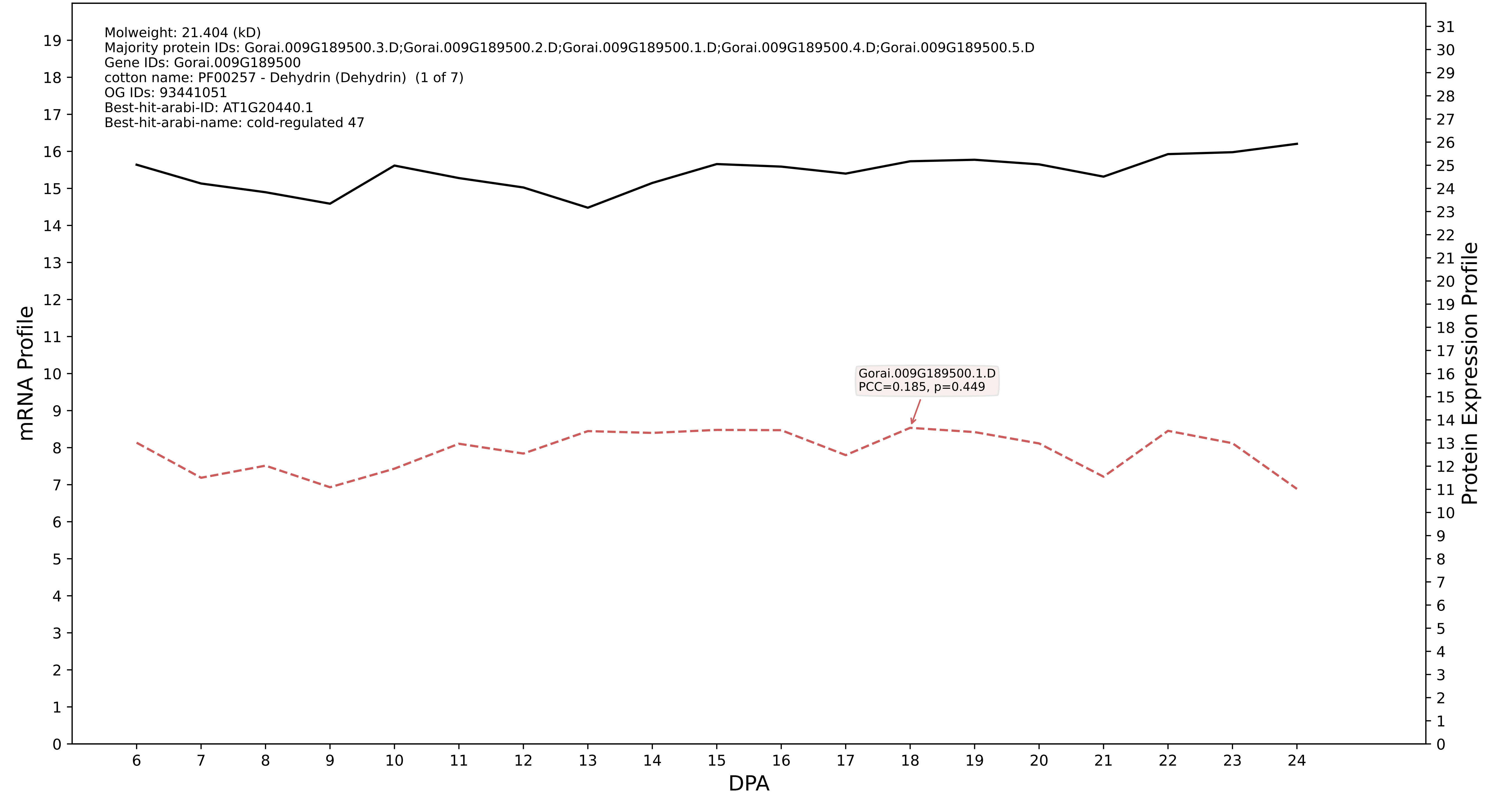

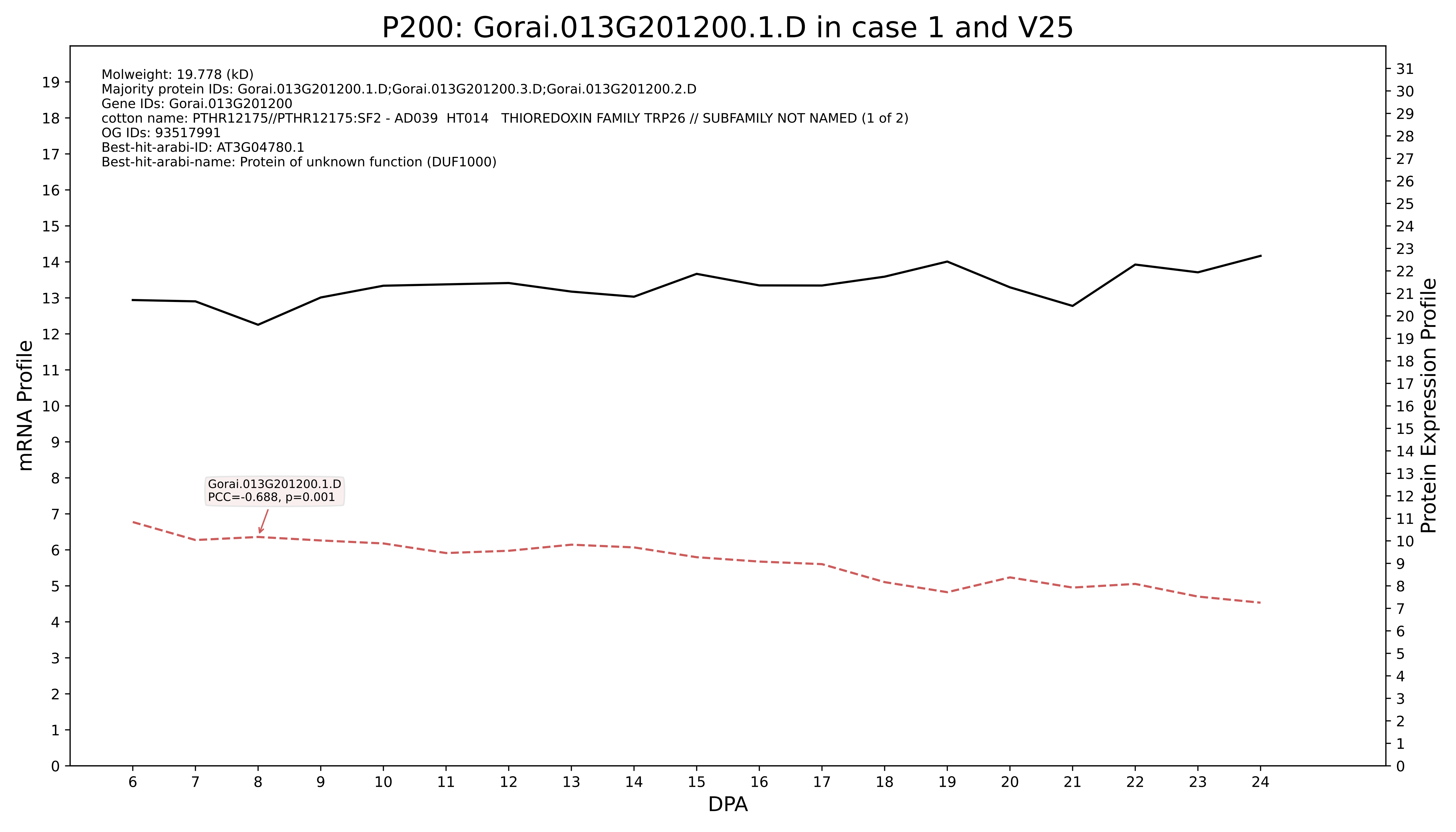
P200: Gorai.013G201200.1.D in case 1 and V25
Molweight: 19.778 (kD)
Majority protein IDs: Gorai.013G201200.1.D;Gorai.013G201200.3.D;Gorai.013G201200.2.D
Gene IDs: Gorai.013G201200
cotton name: PTHR12175//PTHR12175:SF2 - AD039 HT014 THIOREDOXIN FAMILY TRP26 // SUBFAMILY NOT NAMED (1 of 2)
OG IDs: 93517991
Best-hit-arabi-ID: AT3G04780.1
Best-hit-arabi-name: Protein of unknown function (DUF1000)
Gorai.013G201200.1.D
PCC=-0.688, p=0.001
mRNA Profile
Protein Expression Profile
DPA
0 1 2 3 4 5 6 7 8 9 10 11 12 13 14 15 16 17 18 19
0 1 2 3 4 5 6 7 8 9 10 11 12 13 14 15 16 17 18 19 20 21 22 23 24 25 26 27 28 29 30 31
6 7 8 9 10 11 12 13 14 15 16 17 18 19 20 21 22 23 24

### P200: Gorai.002G073400.1.A in case 1 and V25

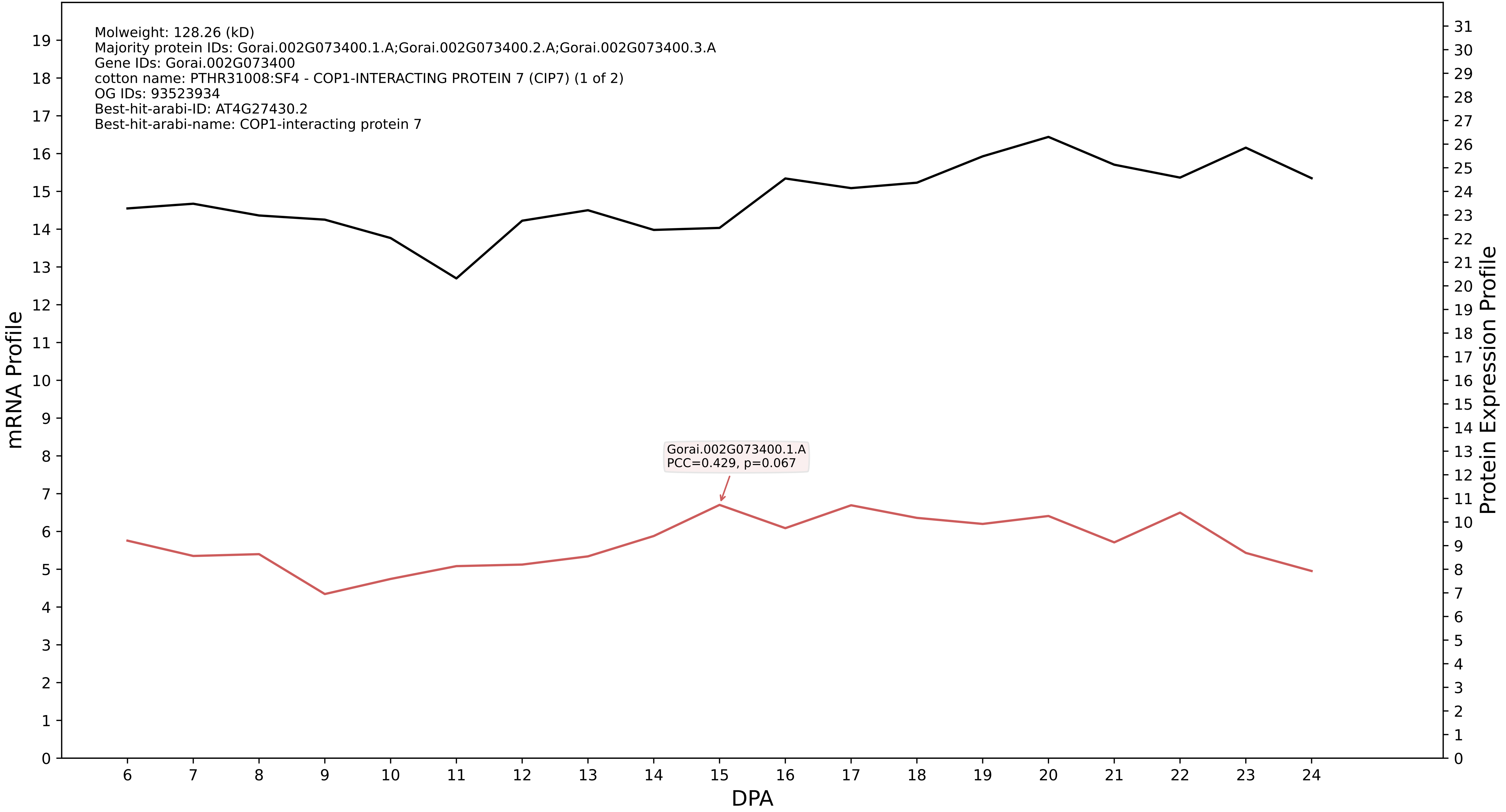

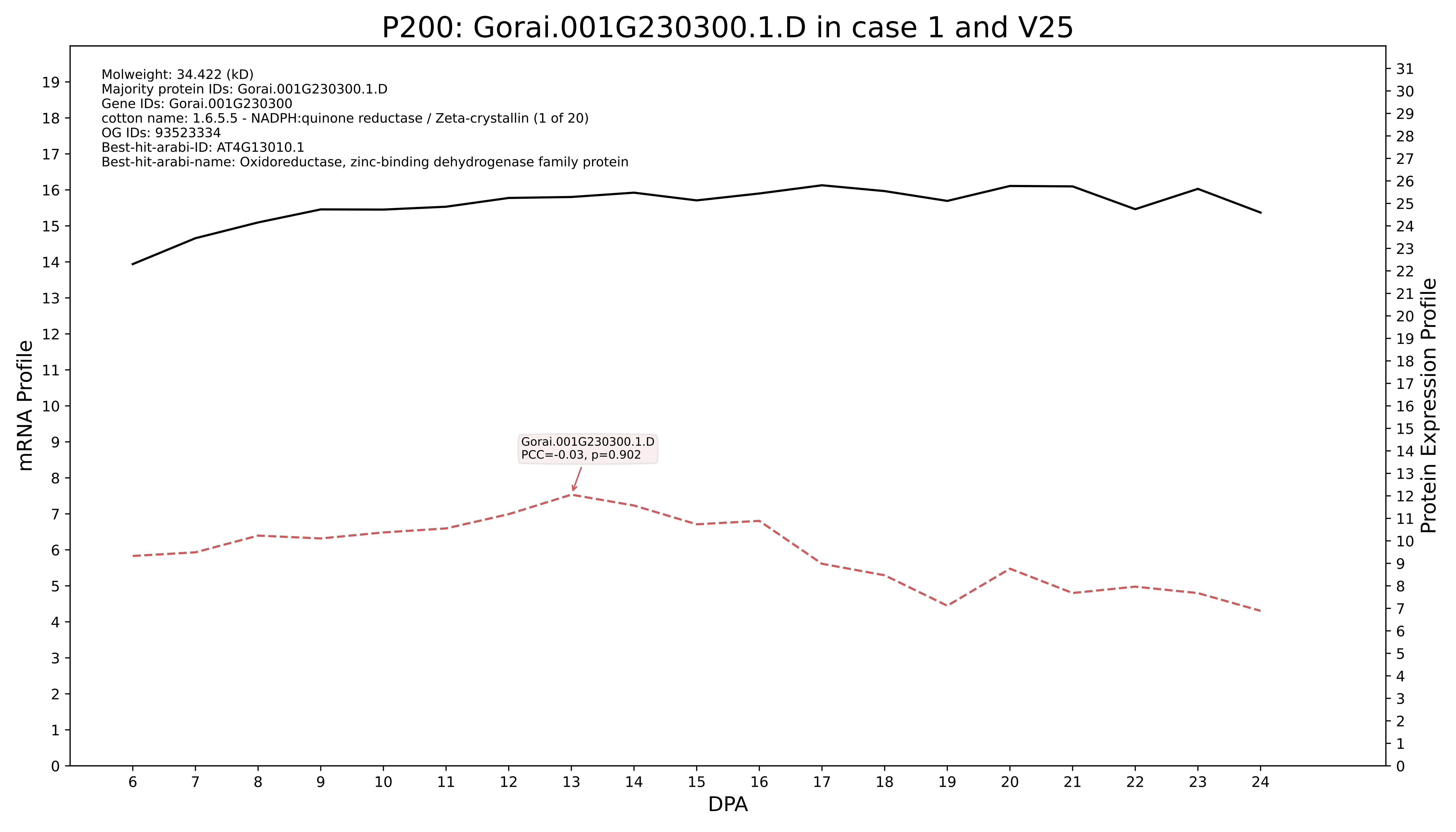

P200: Gorai.001G230300.1.D in case 1 and V25
Molweight: 34.422 (kD)
Majority protein IDs: Gorai.001G230300.1.D
Gene IDs: Gorai.001G230300
cotton name: 1.6.5.5 - NADPH:quinone reductase / Zeta-crystallin (1 of 20)
OG IDs: 93523334
Best-hit-arabi-ID: AT4G13010.1
Best-hit-arabi-name: Oxidoreductase, zinc-binding dehydrogenase family protein
Gorai.001G230300.1.D
PCC=-0.03, p=0.902
mRNA Profile
Protein Expression Profile
DPA

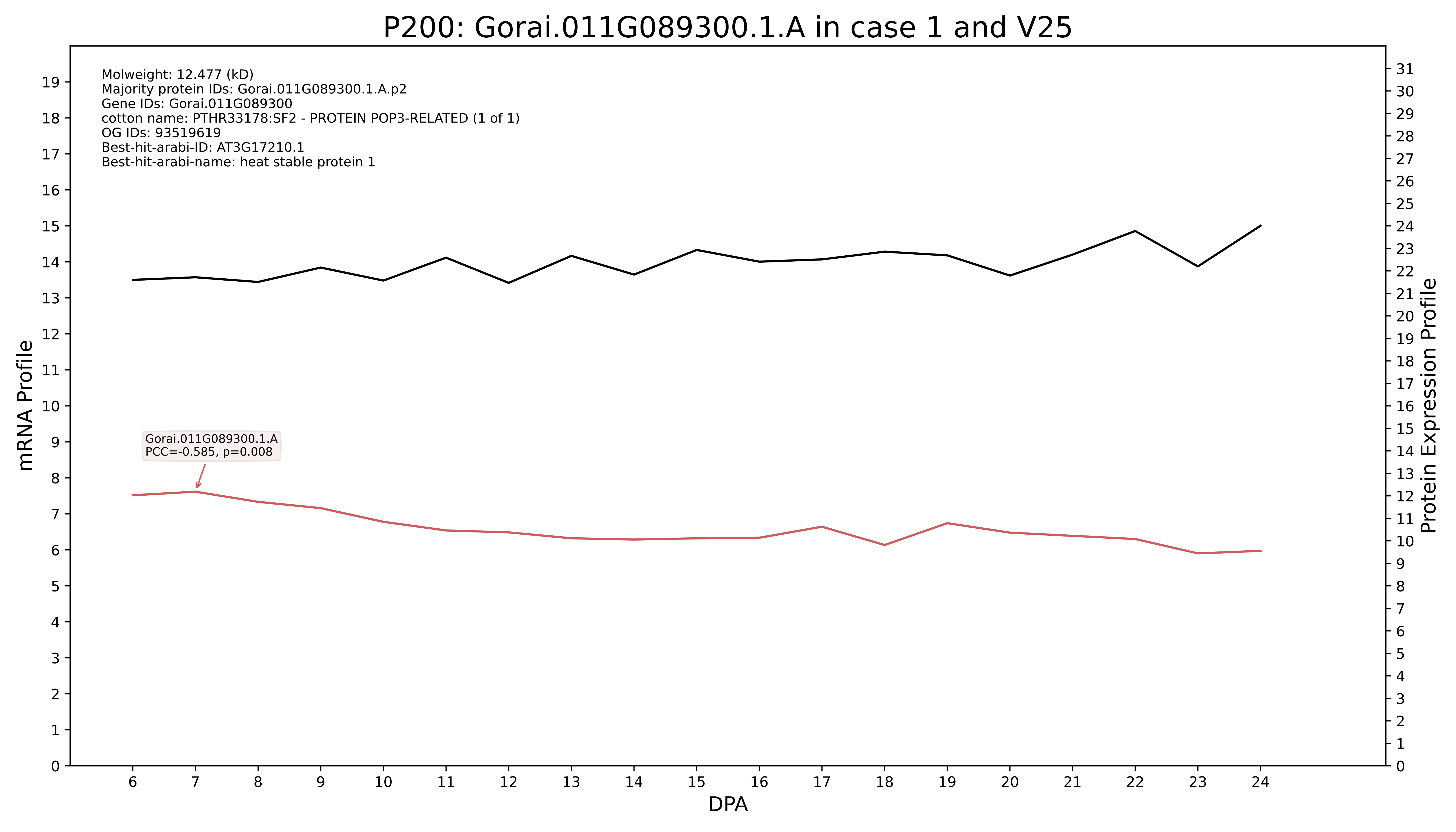
P200: Gorai.011G089300.1.A in case 1 and V25
Molweight: 12.477 (kD)
Majority protein IDs: Gorai.011G089300.1.A.p2
Gene IDs: Gorai.011G089300
cotton name: PTHR33178:SF2 - PROTEIN POP3-RELATED (1 of 1)
OG IDs: 93519619
Best-hit-arabi-ID: AT3G17210.1
Best-hit-arabi-name: heat stable protein 1
Gorai.011G089300.1.A
PCC=-0.585, p=0.008
mRNA Profile
Protein Expression Profile
DPA

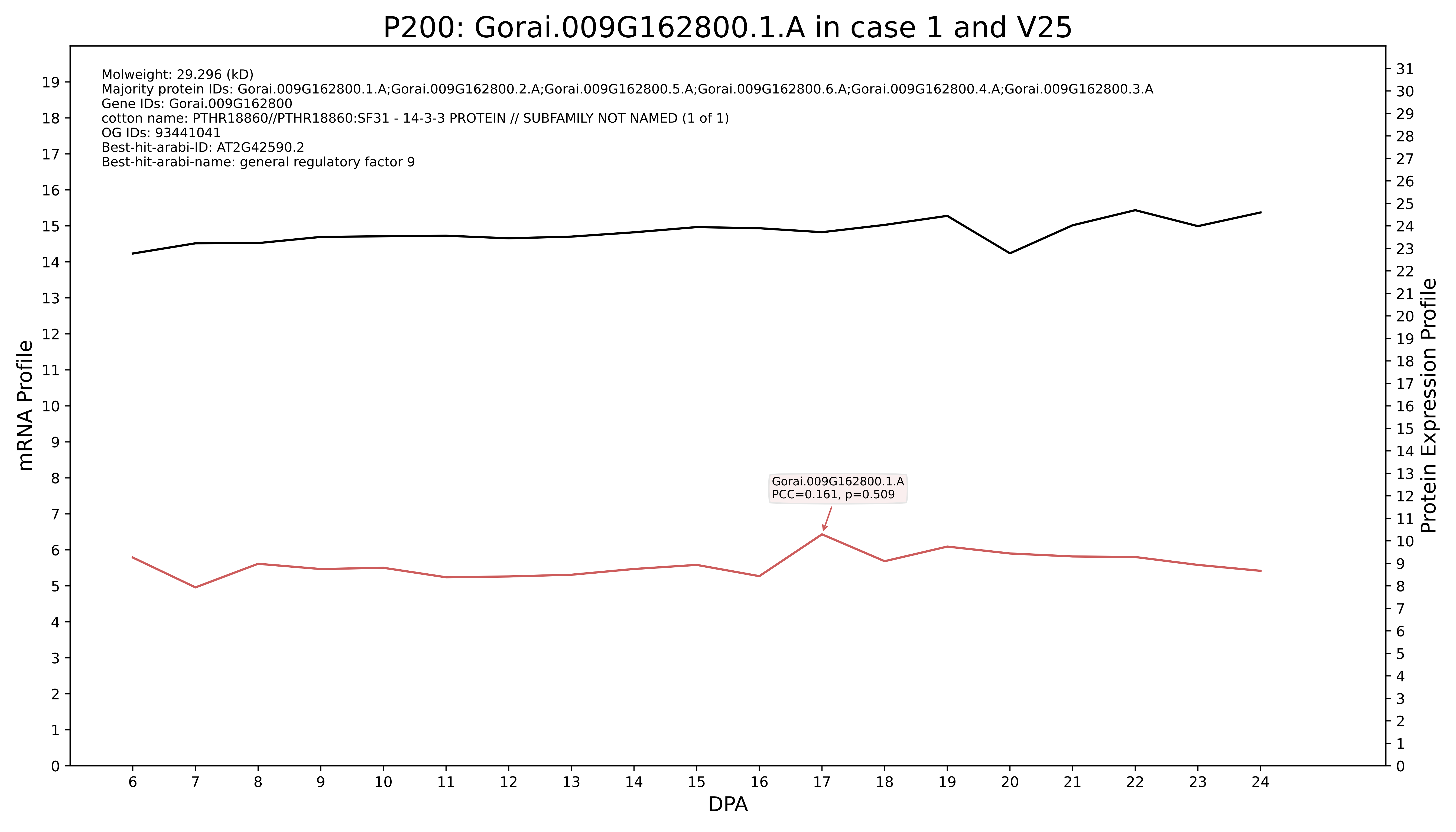
P200: Gorai.009G162800.1.A in case 1 and V25
Molweight: 29.296 (kD)
Majority protein IDs: Gorai.009G162800.1.A;Gorai.009G162800.2.A;Gorai.009G162800.5.A;Gorai.009G162800.6.A;Gorai.009G162800.4.A;Gorai.009G162800.3.A
Gene IDs: Gorai.009G162800
cotton name: PTHR18860//PTHR18860:SF31 - 14-3-3 PROTEIN // SUBFAMILY NOT NAMED (1 of 1)
OG IDs: 93441041
Best-hit-arabi-ID: AT2G42590.2
Best-hit-arabi-name: general regulatory factor 9
Gorai.009G162800.1.A
PCC=0.161, p=0.509
mRNA Profile
Protein Expression Profile
DPA

### P200: Gorai.010G038400.3.D in case 1 and V25

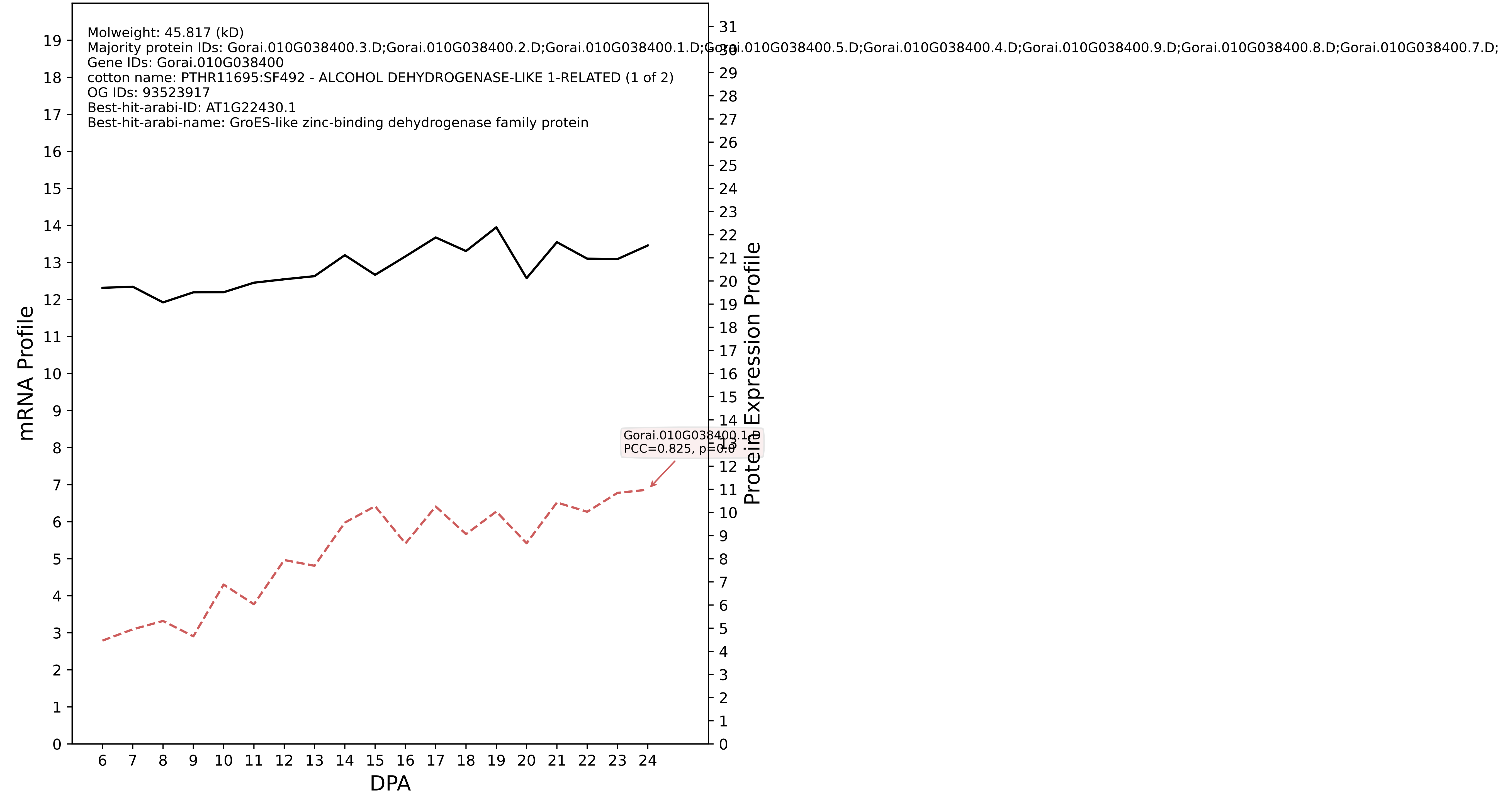

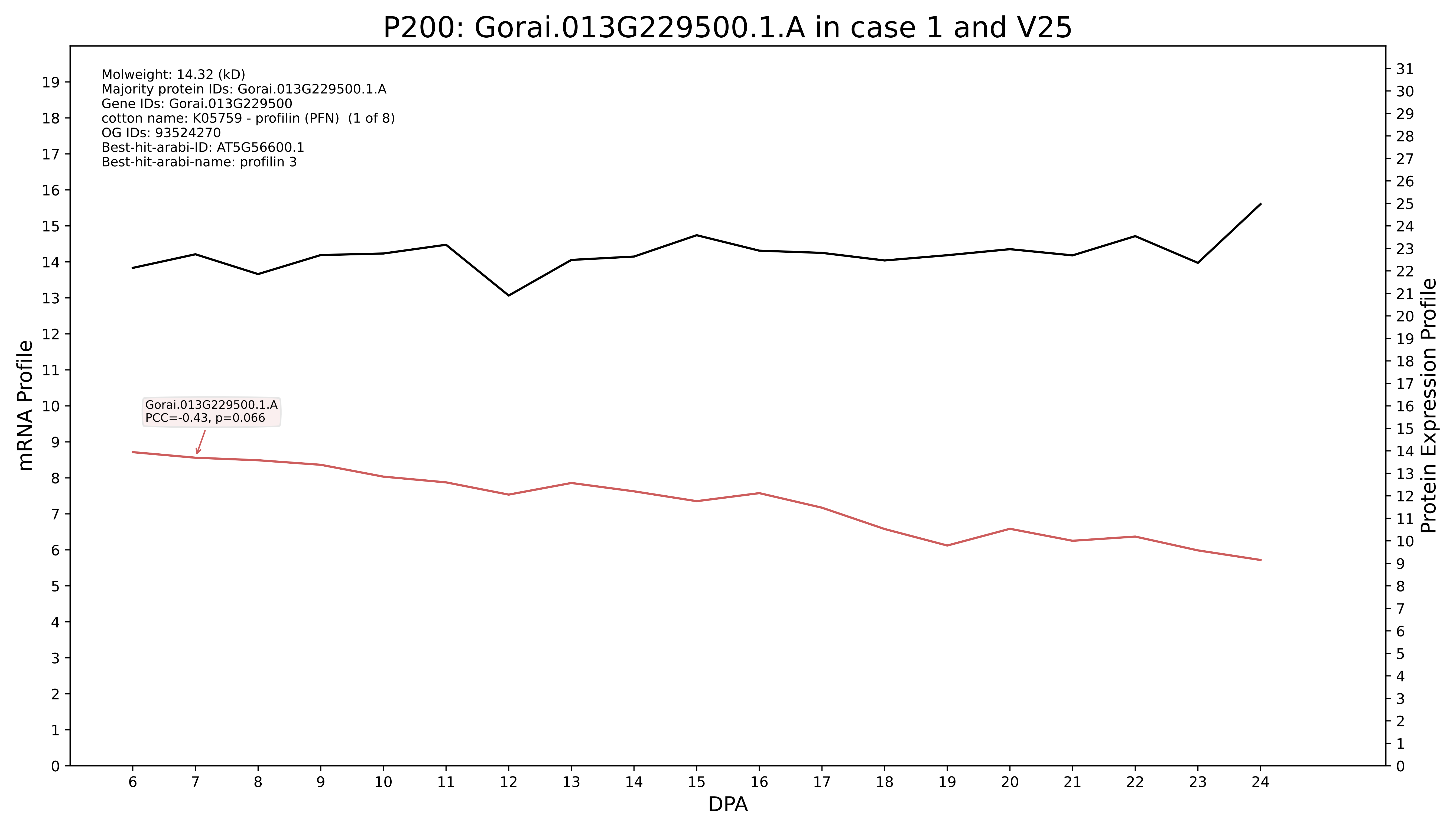
P200: Gorai.013G229500.1.A in case 1 and V25
Molweight: 14.32 (kD)
Majority protein IDs: Gorai.013G229500.1.A
Gene IDs: Gorai.013G229500
cotton name: K05759 - profilin (PFN) (1 of 8)
OG IDs: 93524270
Best-hit-arabi-ID: AT5G56600.1
Best-hit-arabi-name: profilin 3
Gorai.013G229500.1.A
PCC=-0.43, p=0.066
mRNA Profile
Protein Expression Profile
DPA

### P200: Gorai.002G192900.2.D in case 1 and V25

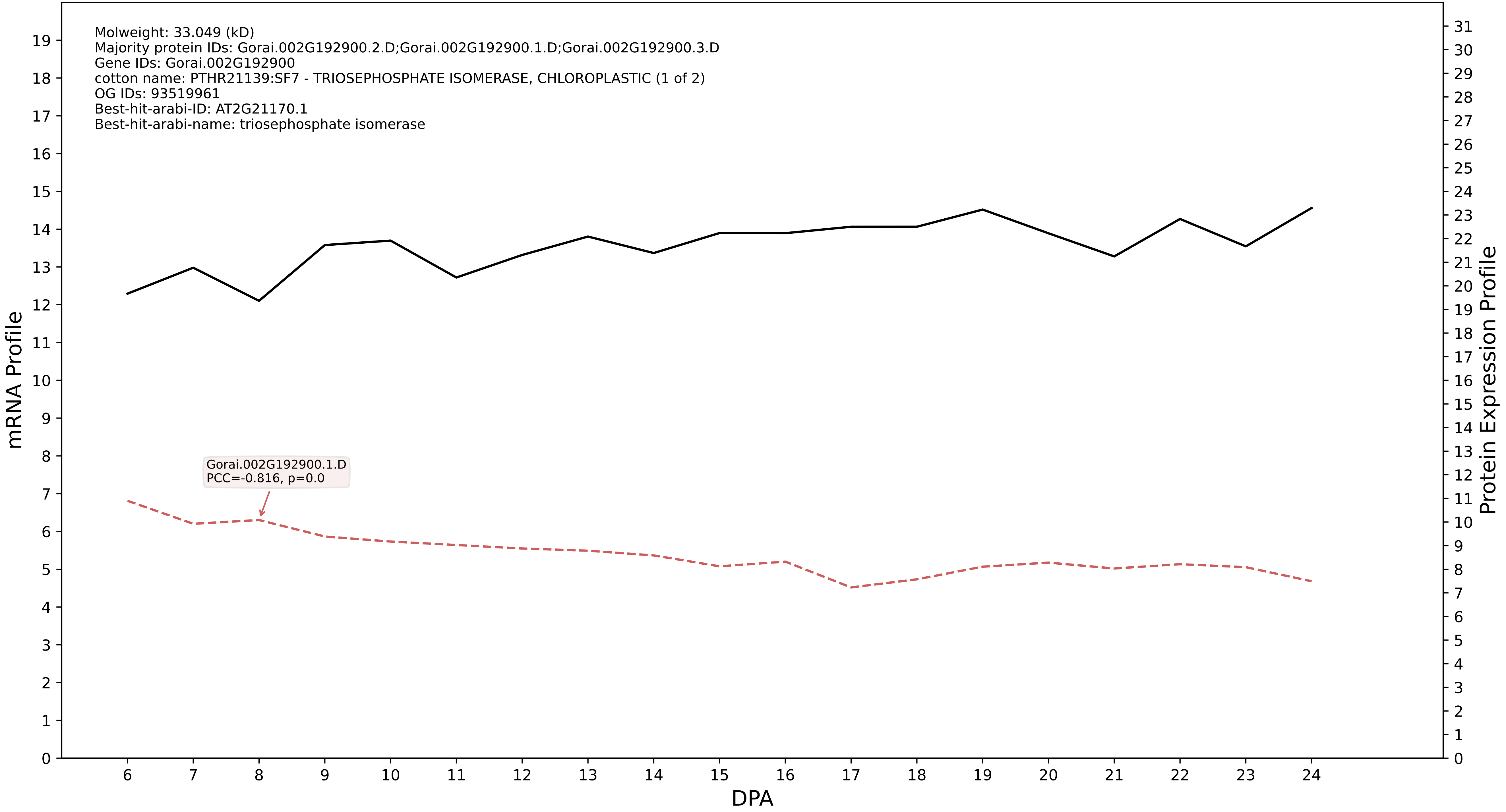

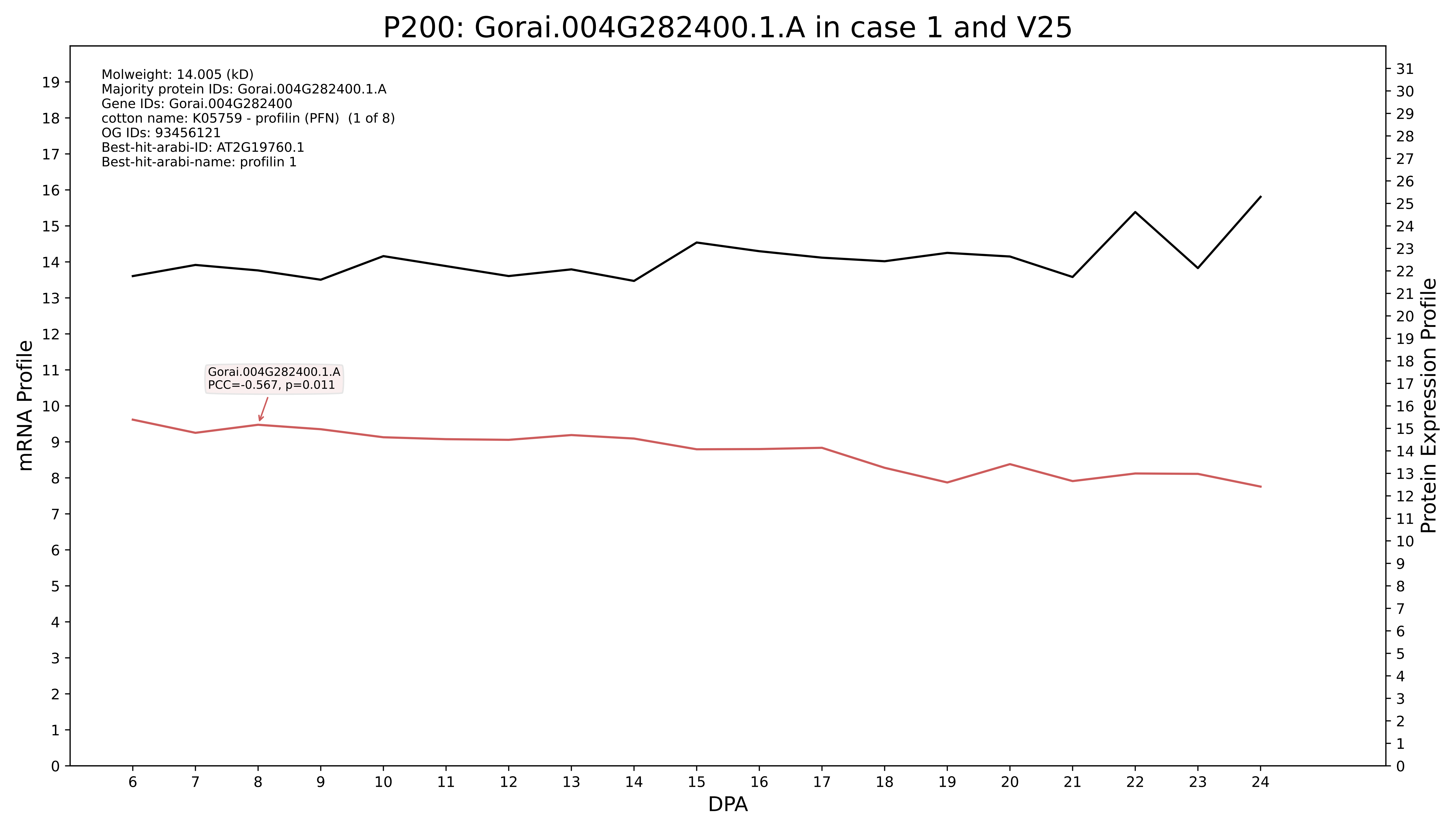
P200: Gorai.004G282400.1.A in case 1 and V25
Molweight: 14.005 (kD)
Majority protein IDs: Gorai.004G282400.1.A
Gene IDs: Gorai.004G282400
cotton name: K05759 - profilin (PFN) (1 of 8)
OG IDs: 93456121
Best-hit-arabi-ID: AT2G19760.1
Best-hit-arabi-name: profilin 1
Gorai.004G282400.1.A
PCC=-0.567, p=0.011
mRNA Profile
Protein Expression Profile
DPA

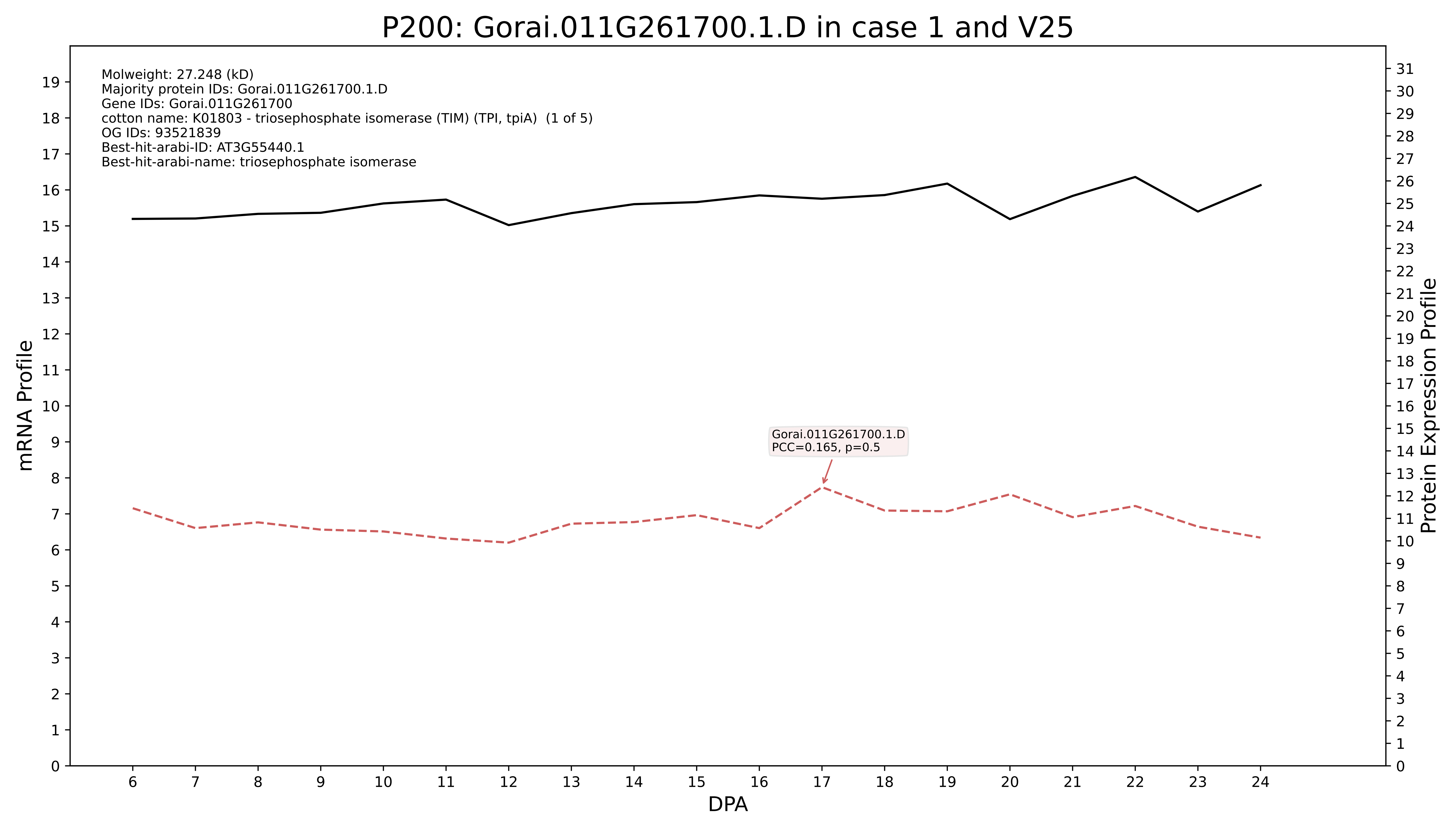
P200: Gorai.011G261700.1.D in case 1 and V25
Molweight: 27.248 (kD)
Majority protein IDs: Gorai.011G261700.1.D
Gene IDs: Gorai.011G261700
cotton name: K01803 - triosephosphate isomerase (TIM) (TPI, tpiA) (1 of 5)
OG IDs: 93521839
Best-hit-arabi-ID: AT3G55440.1
Best-hit-arabi-name: triosephosphate isomerase
Gorai.011G261700.1.D
PCC=0.165, p=0.5
mRNA Profile
Protein Expression Profile
DPA

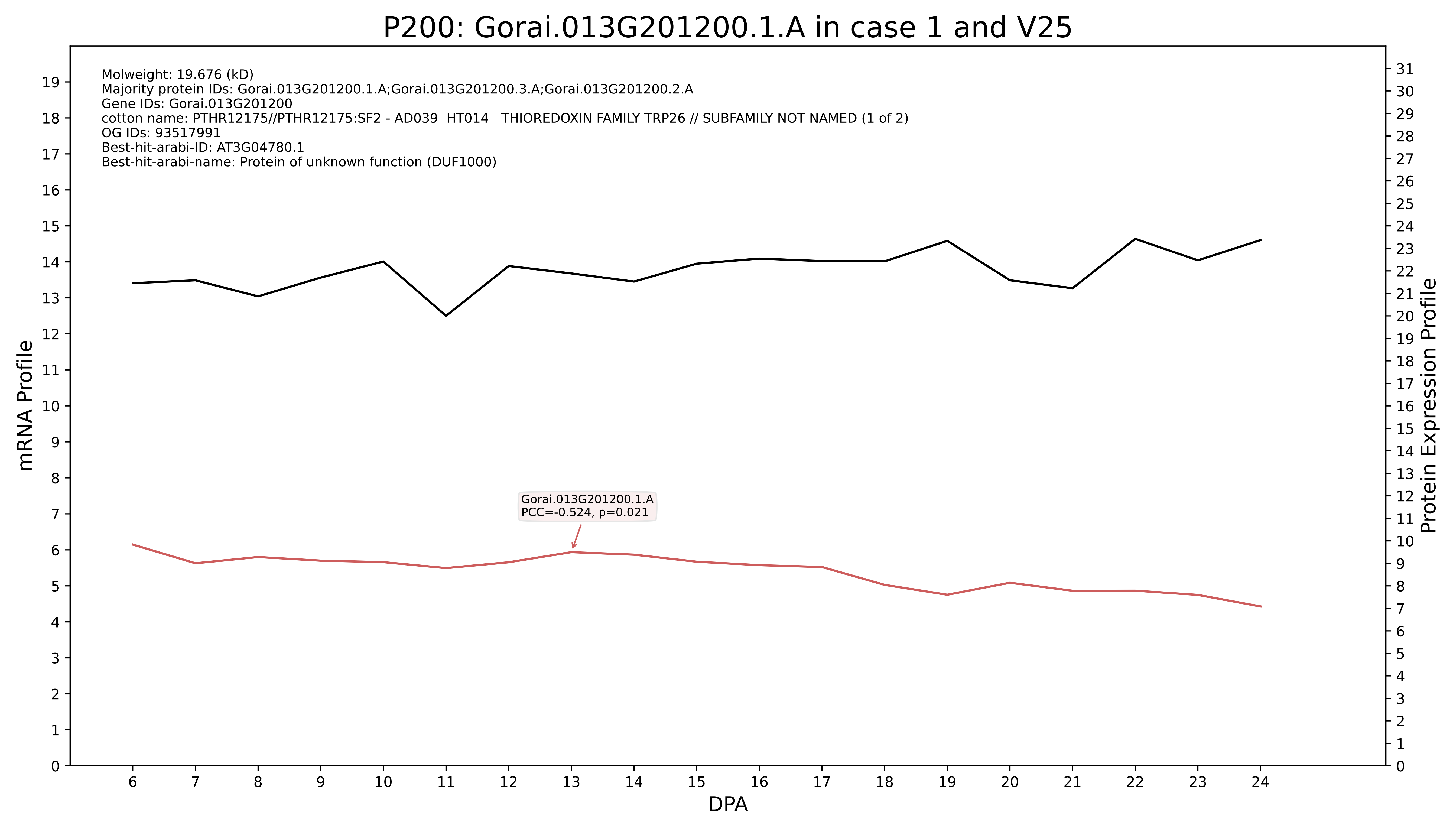
P200: Gorai.013G201200.1.A in case 1 and V25
Molweight: 19.676 (kD)
Majority protein IDs: Gorai.013G201200.1.A;Gorai.013G201200.3.A;Gorai.013G201200.2.A
Gene IDs: Gorai.013G201200
cotton name: PTHR12175//PTHR12175:SF2 - AD039 HT014 THIOREDOXIN FAMILY TRP26 // SUBFAMILY NOT NAMED (1 of 2)
OG IDs: 93517991
Best-hit-arabi-ID: AT3G04780.1
Best-hit-arabi-name: Protein of unknown function (DUF1000)
Gorai.013G201200.1.A
PCC=-0.524, p=0.021
mRNA Profile
Protein Expression Profile
DPA

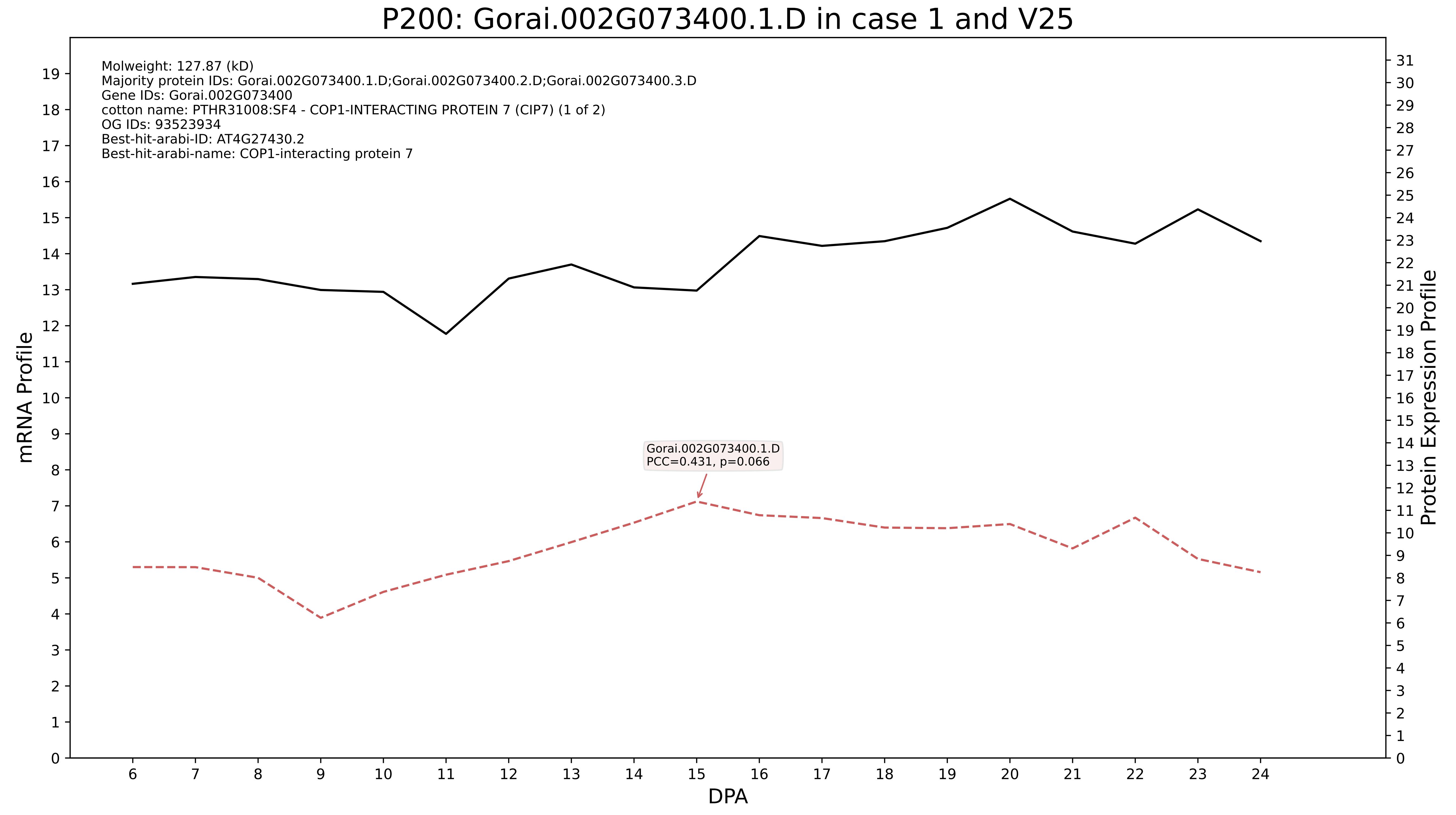
P200: Gorai.002G073400.1.D in case 1 and V25
Molweight: 127.87 (kD)
Majority protein IDs: Gorai.002G073400.1.D;Gorai.002G073400.2.D;Gorai.002G073400.3.D
Gene IDs: Gorai.002G073400
cotton name: PTHR31008:SF4 - COP1-INTERACTING PROTEIN 7 (CIP7) (1 of 2)
OG IDs: 93523934
Best-hit-arabi-ID: AT4G27430.2
Best-hit-arabi-name: COP1-interacting protein 7
Gorai.002G073400.1.D
PCC=0.431, p=0.066
mRNA Profile
Protein Expression Profile
DPA

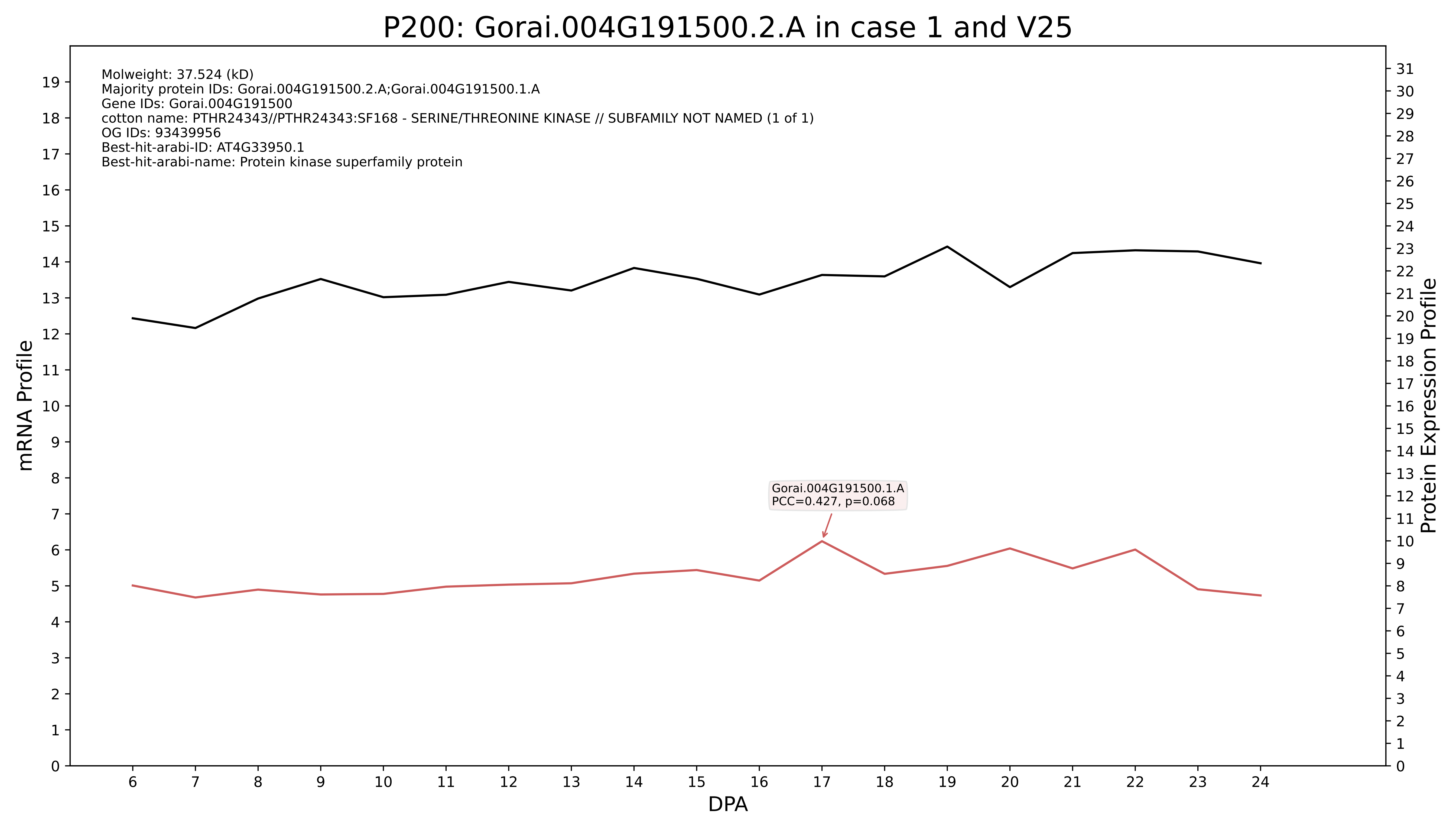
P200: Gorai.004G191500.2.A in case 1 and V25
Molweight: 37.524 (kD)
Majority protein IDs: Gorai.004G191500.2.A;Gorai.004G191500.1.A
Gene IDs: Gorai.004G191500
cotton name: PTHR24343//PTHR24343:SF168 - SERINE/THREONINE KINASE // SUBFAMILY NOT NAMED (1 of 1)
OG IDs: 93439956
Best-hit-arabi-ID: AT4G33950.1
Best-hit-arabi-name: Protein kinase superfamily protein
Gorai.004G191500.1.A
PCC=0.427, p=0.068
mRNA Profile
Protein Expression Profile
DPA

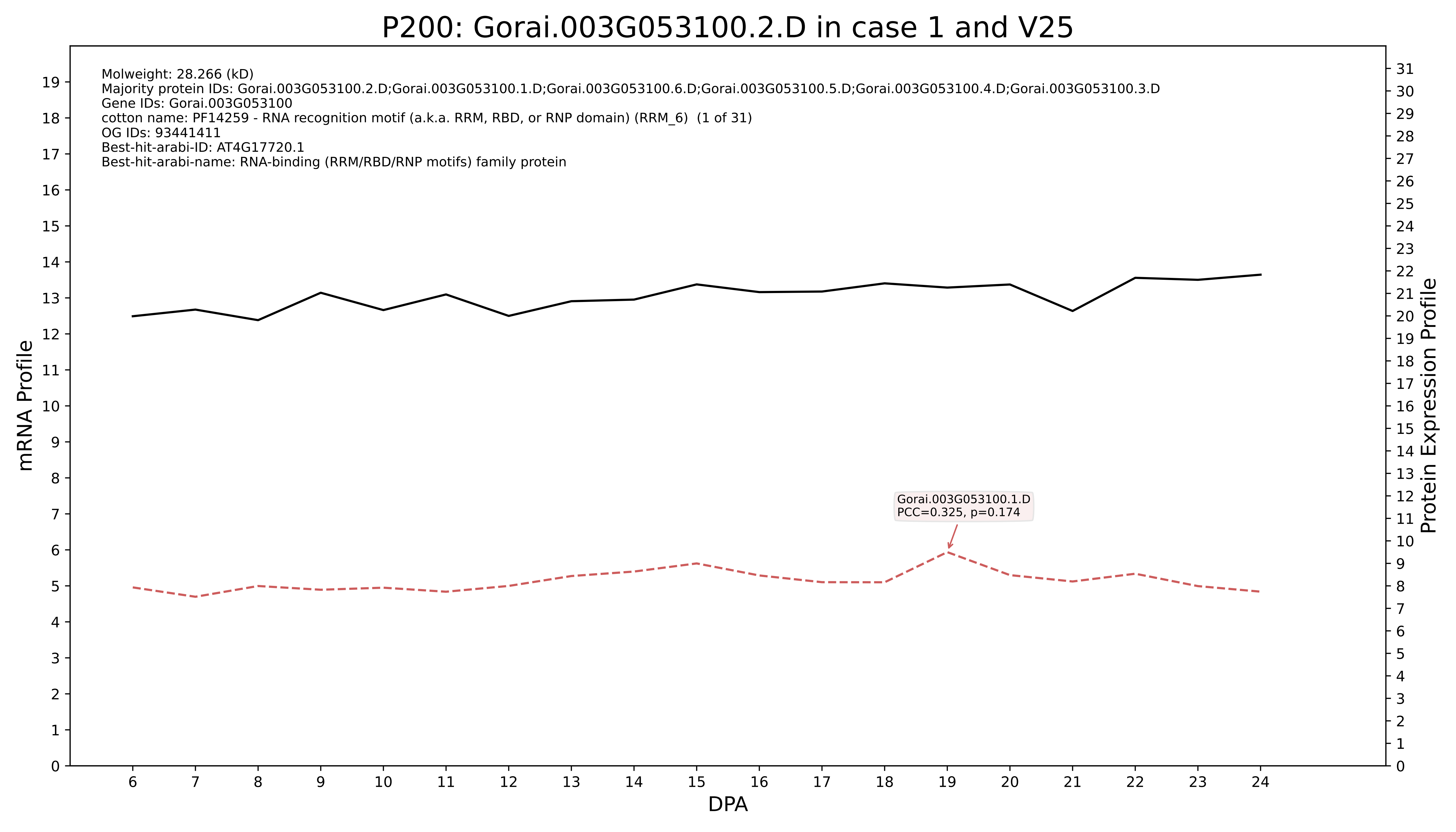
P200: Gorai.003G053100.2.D in case 1 and V25
Molweight: 28.266 (kD)
Majority protein IDs: Gorai.003G053100.2.D;Gorai.003G053100.1.D;Gorai.003G053100.6.D;Gorai.003G053100.5.D;Gorai.003G053100.4.D;Gorai.003G053100.3.D
Gene IDs: Gorai.003G053100
cotton name: PF14259 - RNA recognition motif (a.k.a. RRM, RBD, or RNP domain) (RRM_6) (1 of 31)
OG IDs: 93441411
Best-hit-arabi-ID: AT4G17720.1
Best-hit-arabi-name: RNA-binding (RRM/RBD/RNP motifs) family protein
Gorai.003G053100.1.D
PCC=0.325, p=0.174
mRNA Profile
Protein Expression Profile
DPA

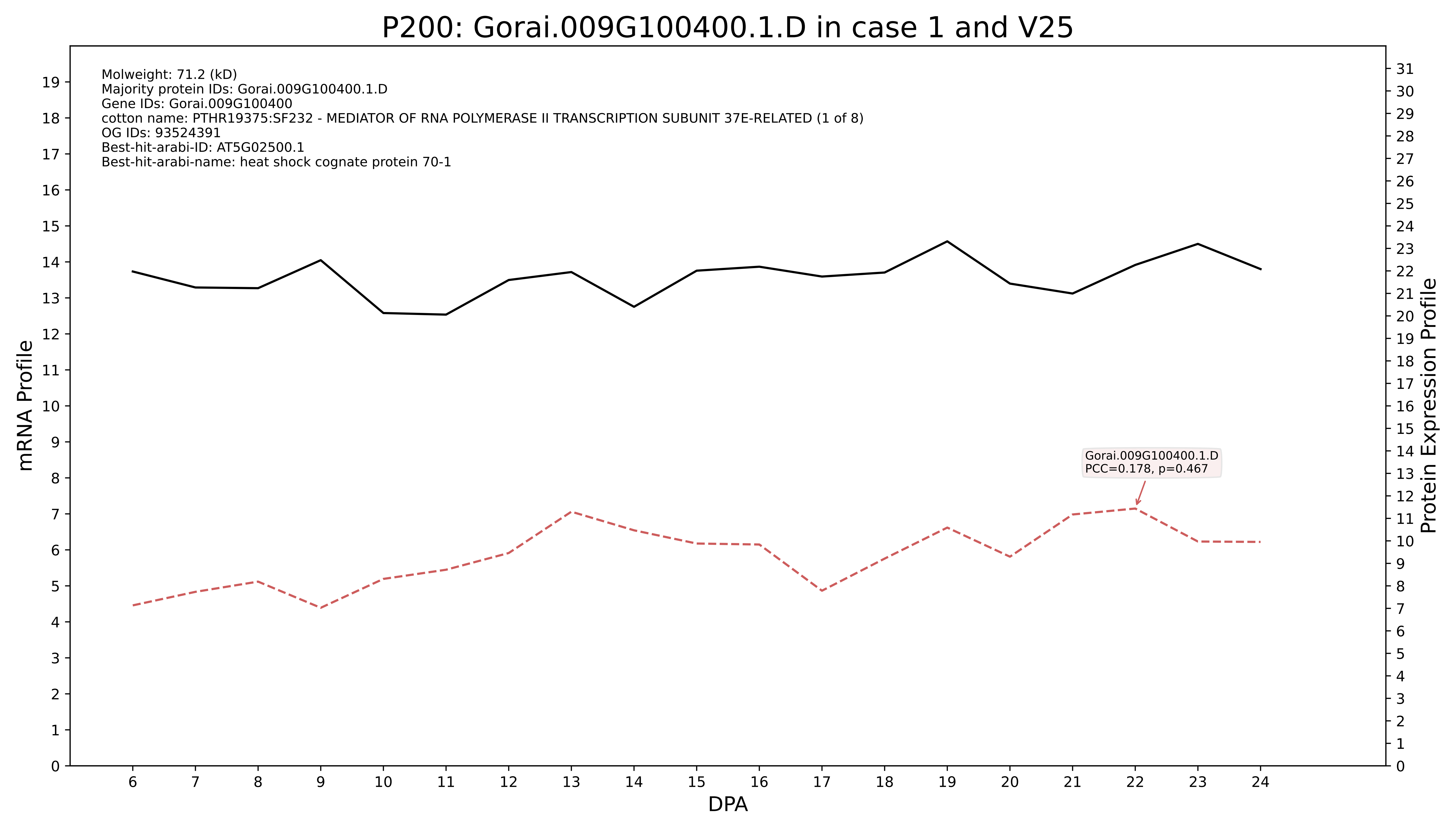
P200: Gorai.009G100400.1.D in case 1 and V25
Molweight: 71.2 (kD)
Majority protein IDs: Gorai.009G100400.1.D
Gene IDs: Gorai.009G100400
cotton name: PTHR19375:SF232 - MEDIATOR OF RNA POLYMERASE II TRANSCRIPTION SUBUNIT 37E-RELATED (1 of 8)
OG IDs: 93524391
Best-hit-arabi-ID: AT5G02500.1
Best-hit-arabi-name: heat shock cognate protein 70-1
Gorai.009G100400.1.D
PCC=0.178, p=0.467
mRNA Profile
Protein Expression Profile
DPA

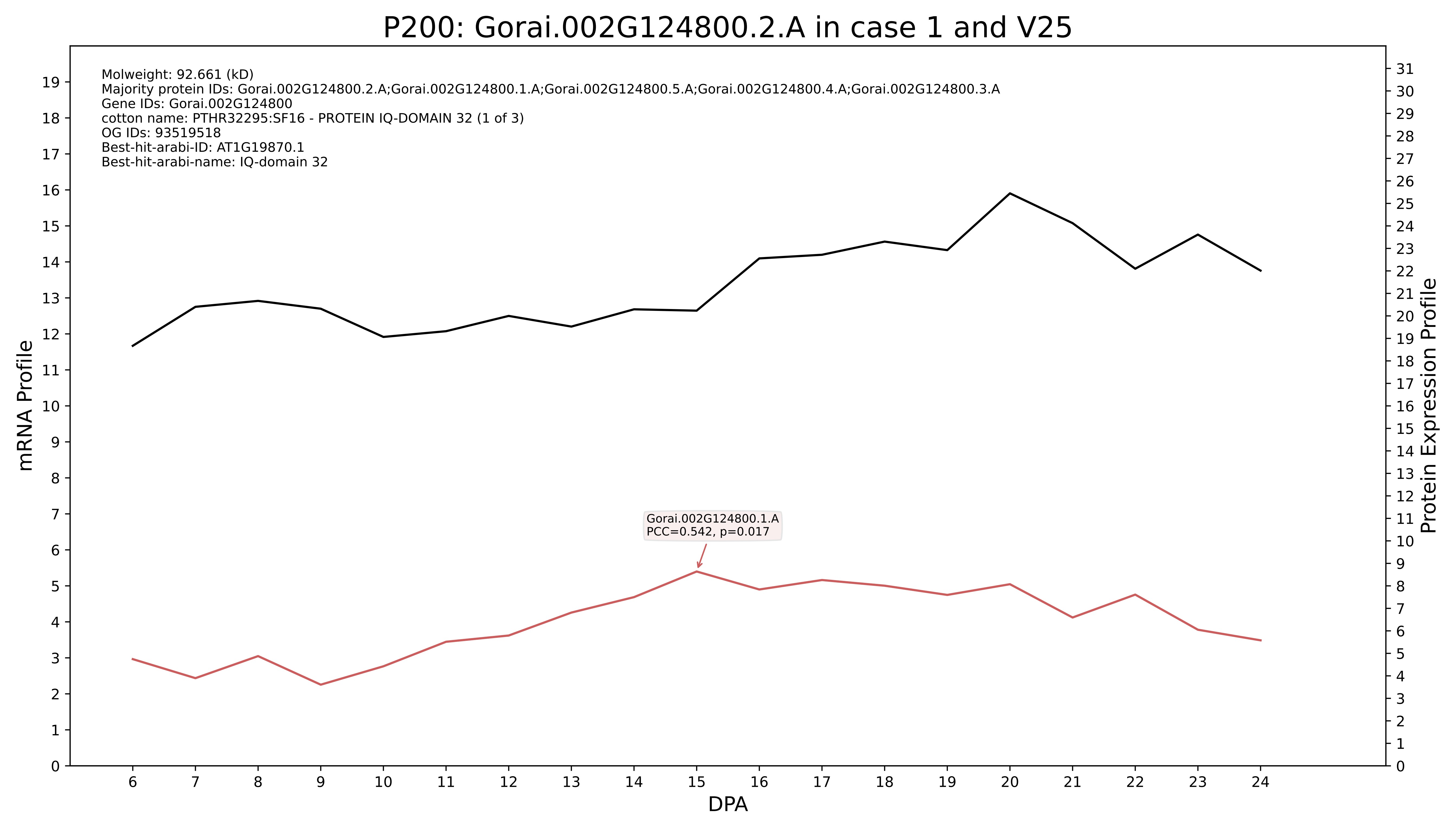

P200: Gorai.002G124800.2.A in case 1 and V25
Molweight: 92.661 (kD)
Majority protein IDs: Gorai.002G124800.2.A;Gorai.002G124800.1.A;Gorai.002G124800.5.A;Gorai.002G124800.4.A;Gorai.002G124800.3.A
Gene IDs: Gorai.002G124800
cotton name: PTHR32295:SF16 - PROTEIN IQ-DOMAIN 32 (1 of 3)
OG IDs: 93519518
Best-hit-arabi-ID: AT1G19870.1
Best-hit-arabi-name: IQ-domain 32
Gorai.002G124800.1.A
PCC=0.542, p=0.017
mRNA Profile
Protein Expression Profile
DPA

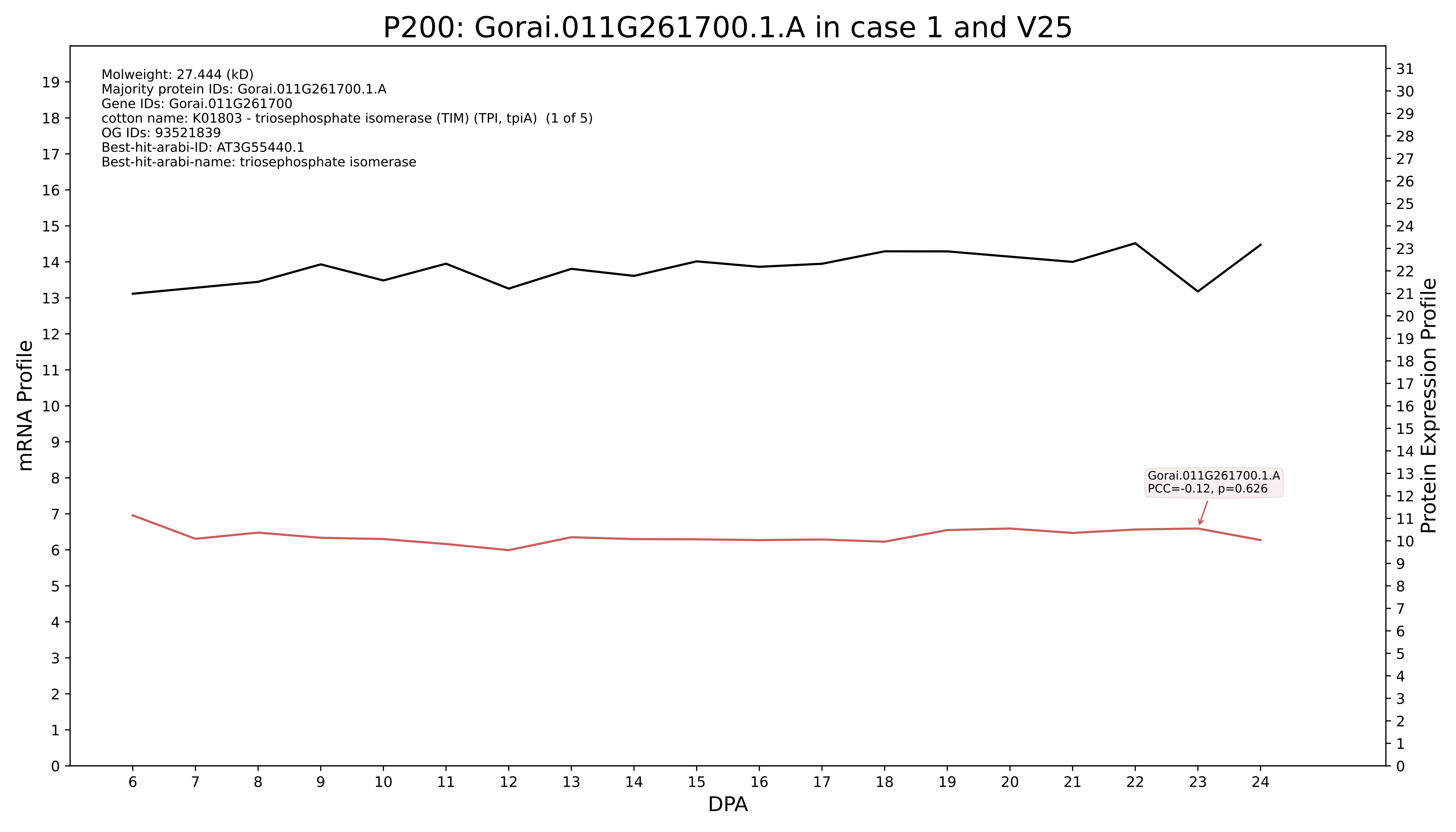
P200: Gorai.011G261700.1.A in case 1 and V25
Molweight: 27.444 (kD)
Majority protein IDs: Gorai.011G261700.1.A
Gene IDs: Gorai.011G261700
cotton name: K01803 - triosephosphate isomerase (TIM) (TPI, tpiA) (1 of 5)
OG IDs: 93521839
Best-hit-arabi-ID: AT3G55440.1
Best-hit-arabi-name: triosephosphate isomerase
Gorai.011G261700.1.A
PCC=-0.12, p=0.626
mRNA Profile
Protein Expression Profile
DPA

### P200: Gorai.006G150600.1.D in case 1 and V25

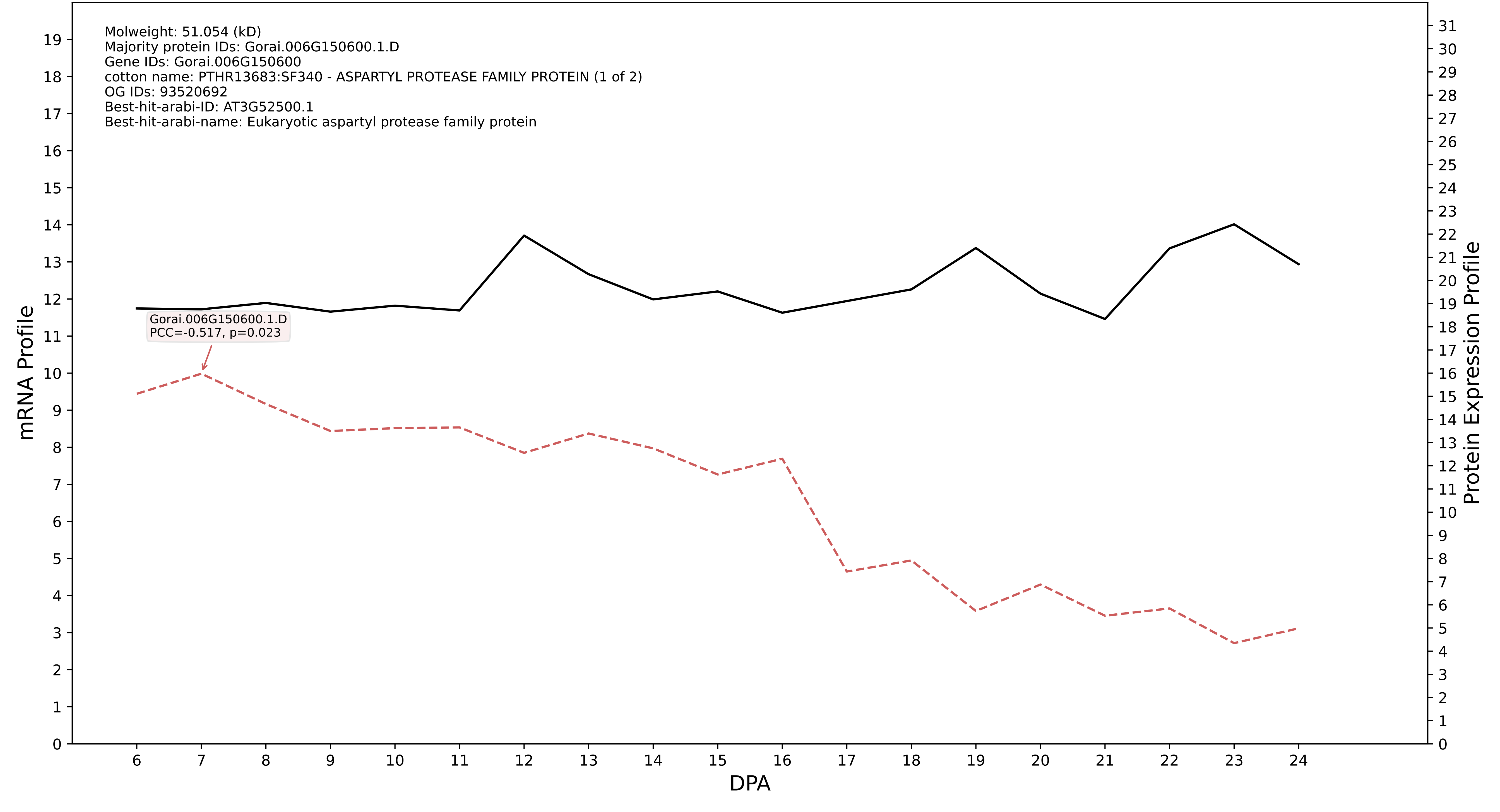

### P200: Gorai.009G189500.3.A in case 1 and V25

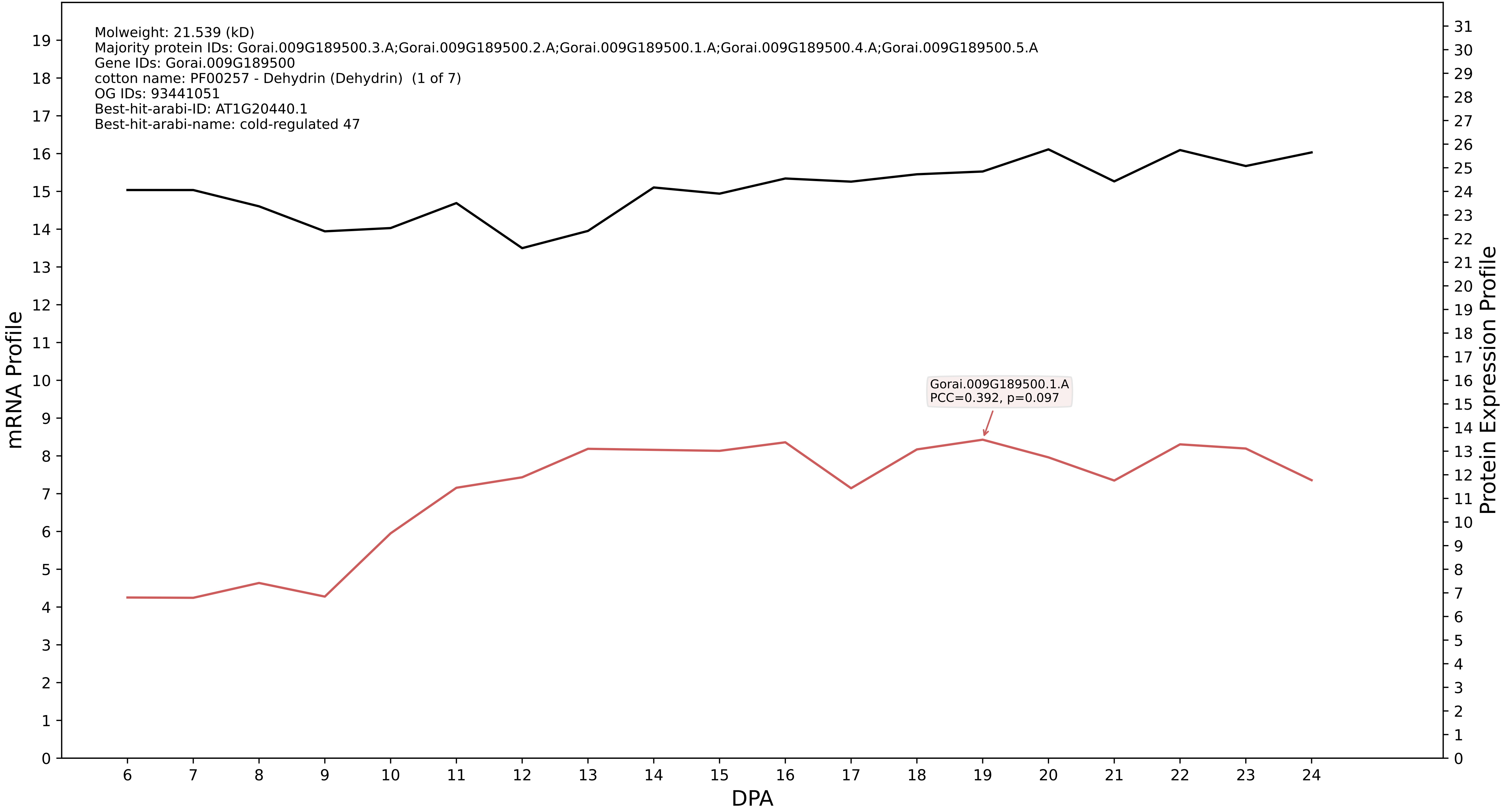

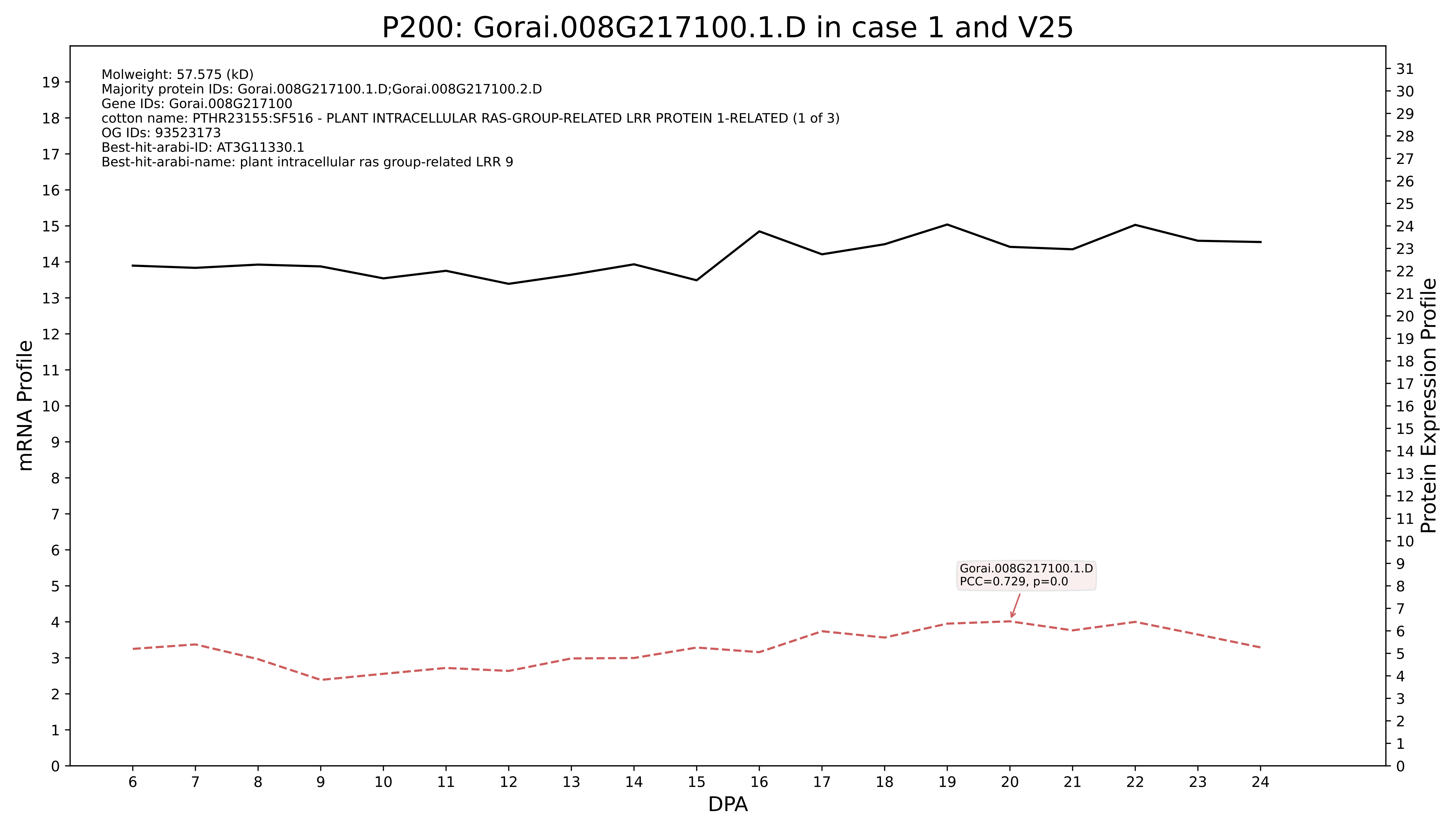
P200: Gorai.008G217100.1.D in case 1 and V25
Molweight: 57.575 (kD)
Majority protein IDs: Gorai.008G217100.1.D;Gorai.008G217100.2.D
Gene IDs: Gorai.008G217100
cotton name: PTHR23155:SF516 - PLANT INTRACELLULAR RAS-GROUP-RELATED LRR PROTEIN 1-RELATED (1 of 3)
OG IDs: 93523173
Best-hit-arabi-ID: AT3G11330.1
Best-hit-arabi-name: plant intracellular ras group-related LRR 9
Gorai.008G217100.1.D
PCC=0.729, p=0.0
mRNA Profile
Protein Expression Profile
DPA

### P200: Gorai.013G216000.2.D in case 1 and V25

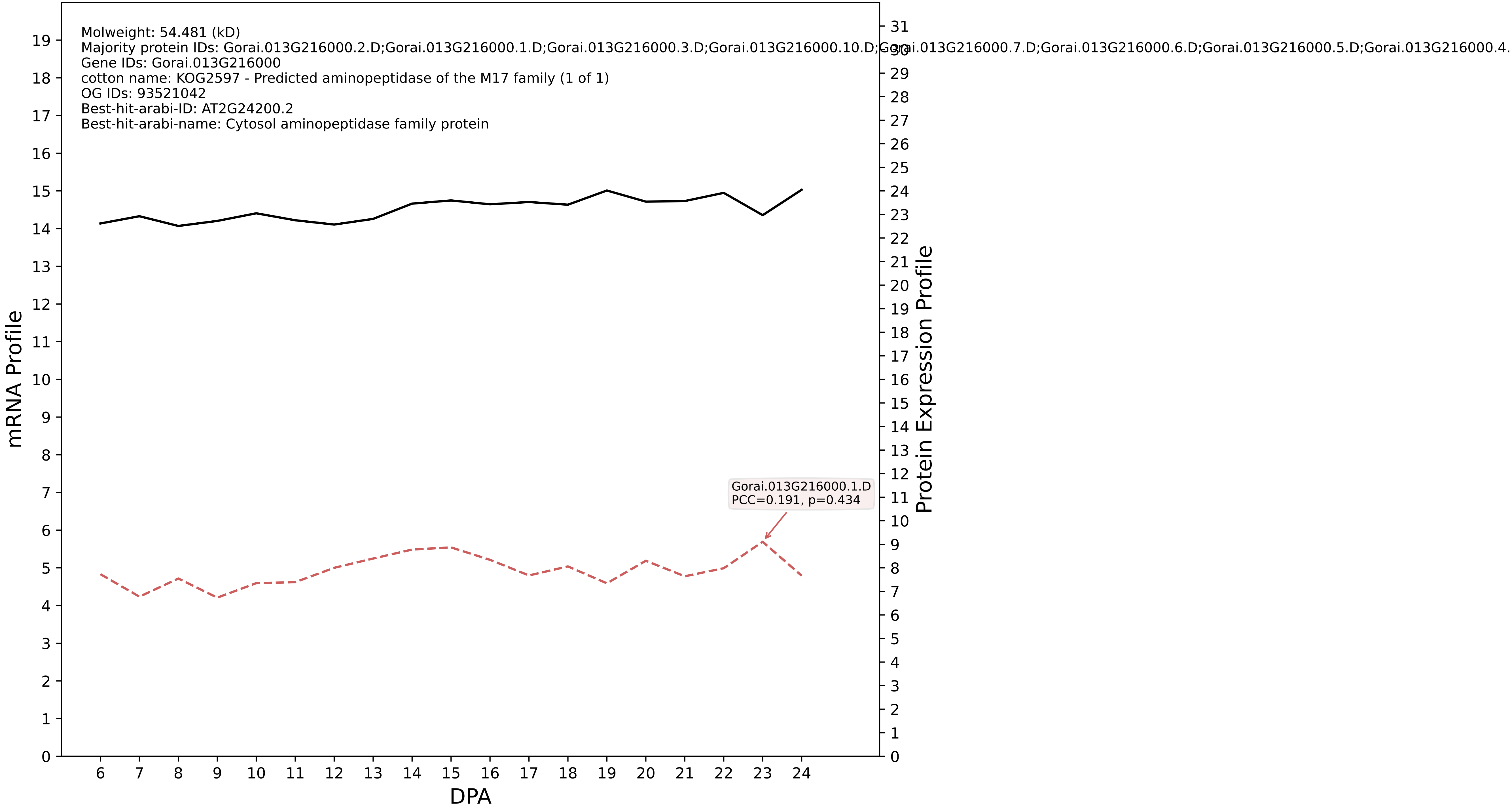

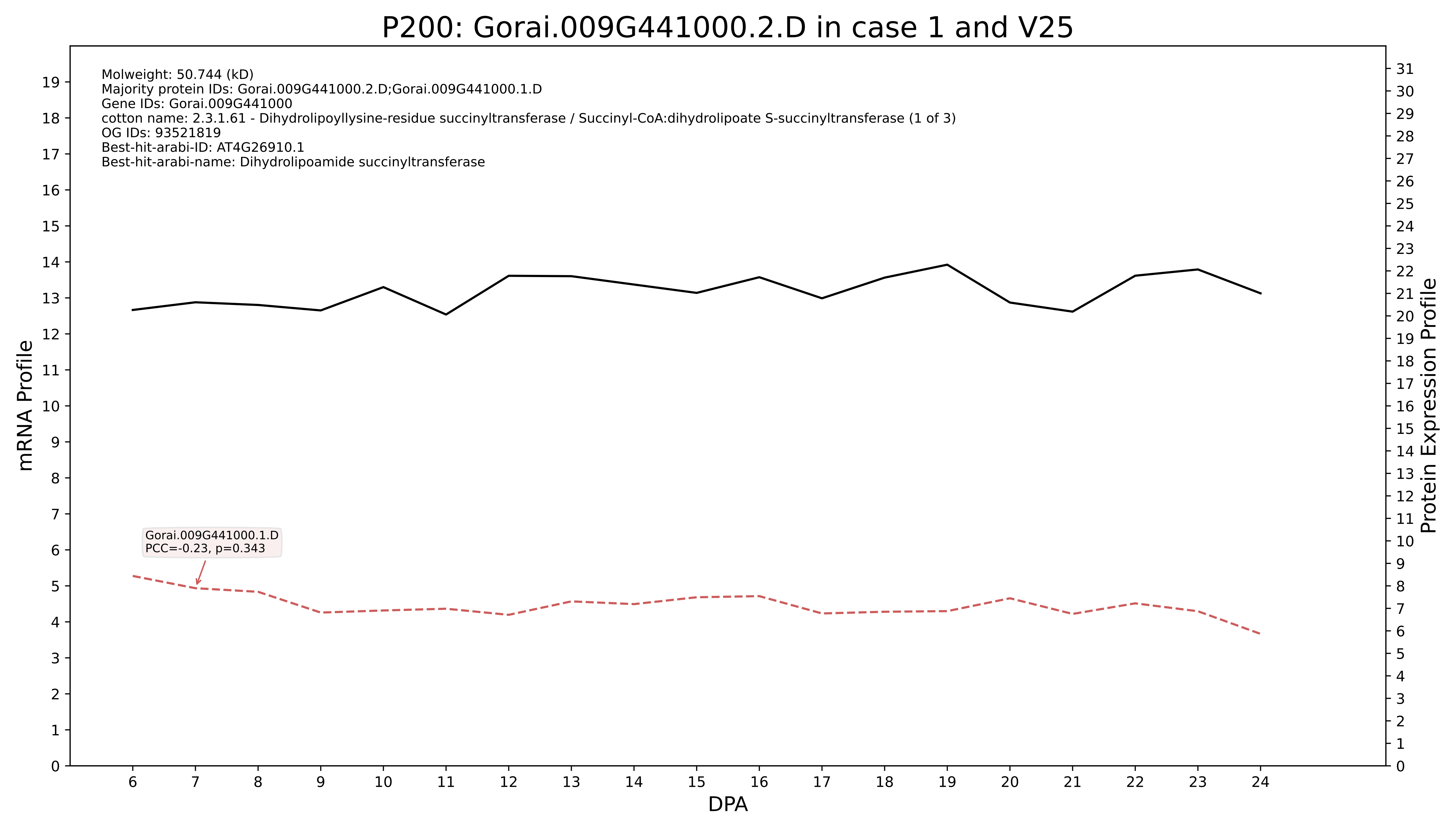
P200: Gorai.009G441000.2.D in case 1 and V25
Molweight: 50.744 (kD)
Majority protein IDs: Gorai.009G441000.2.D;Gorai.009G441000.1.D
Gene IDs: Gorai.009G441000
cotton name: 2.3.1.61 - Dihydrolipoyllysine-residue succinyltransferase / Succinyl-CoA:dihydrolipoate S-succinyltransferase (1 of 3)
OG IDs: 93521819
Best-hit-arabi-ID: AT4G26910.1
Best-hit-arabi-name: Dihydrolipoamide succinyltransferase
Gorai.009G441000.1.D
PCC=-0.23, p=0.343
mRNA Profile
Protein Expression Profile
DPA

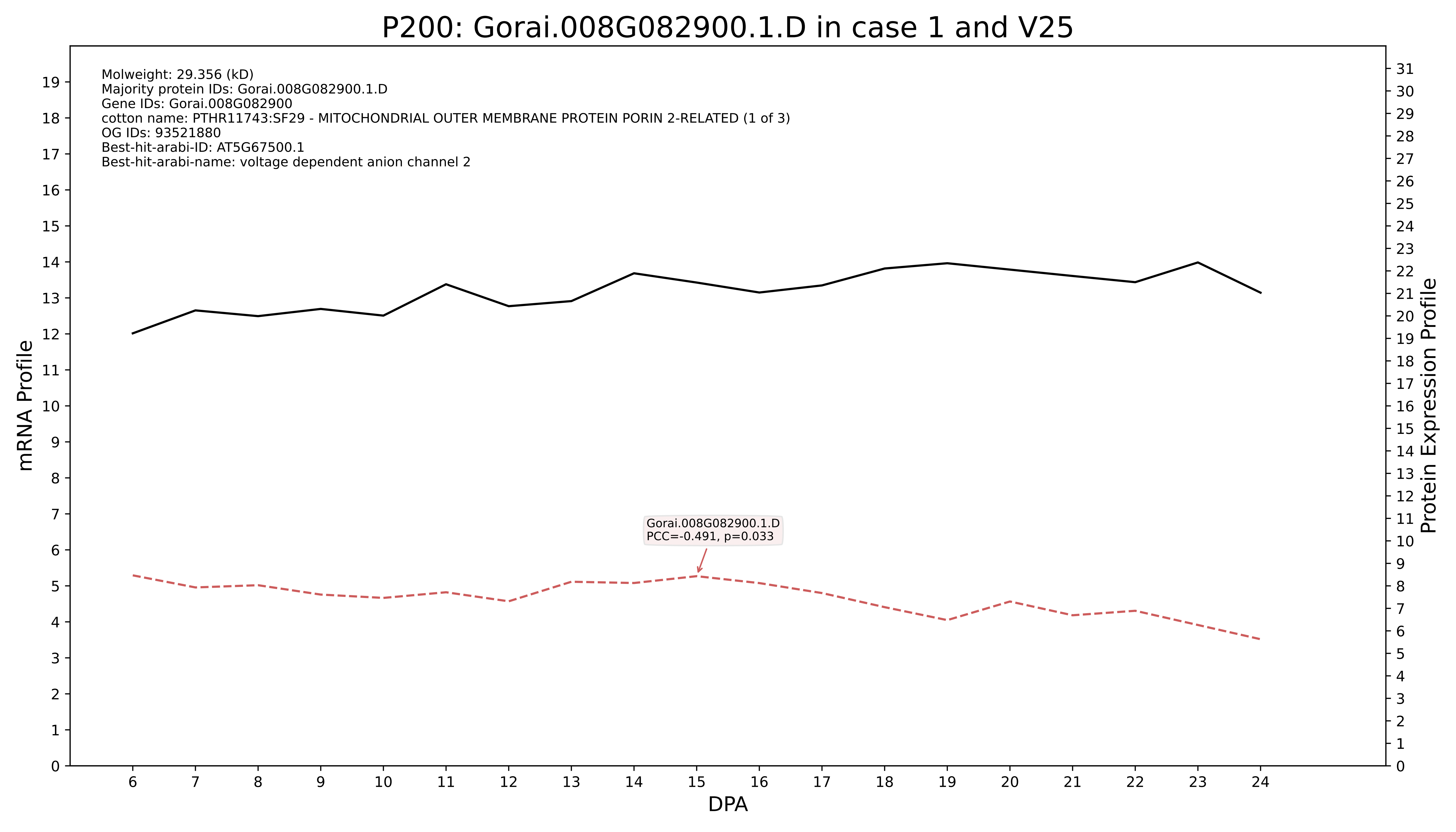

P200: Gorai.008G082900.1.D in case 1 and V25
Molweight: 29.356 (kD)
Majority protein IDs: Gorai.008G082900.1.D
Gene IDs: Gorai.008G082900
cotton name: PTHR11743:SF29 - MITOCHONDRIAL OUTER MEMBRANE PROTEIN PORIN 2-RELATED (1 of 3)
OG IDs: 93521880
Best-hit-arabi-ID: AT5G67500.1
Best-hit-arabi-name: voltage dependent anion channel 2
Gorai.008G082900.1.D
PCC=-0.491, p=0.033
mRNA Profile
Protein Expression Profile
DPA

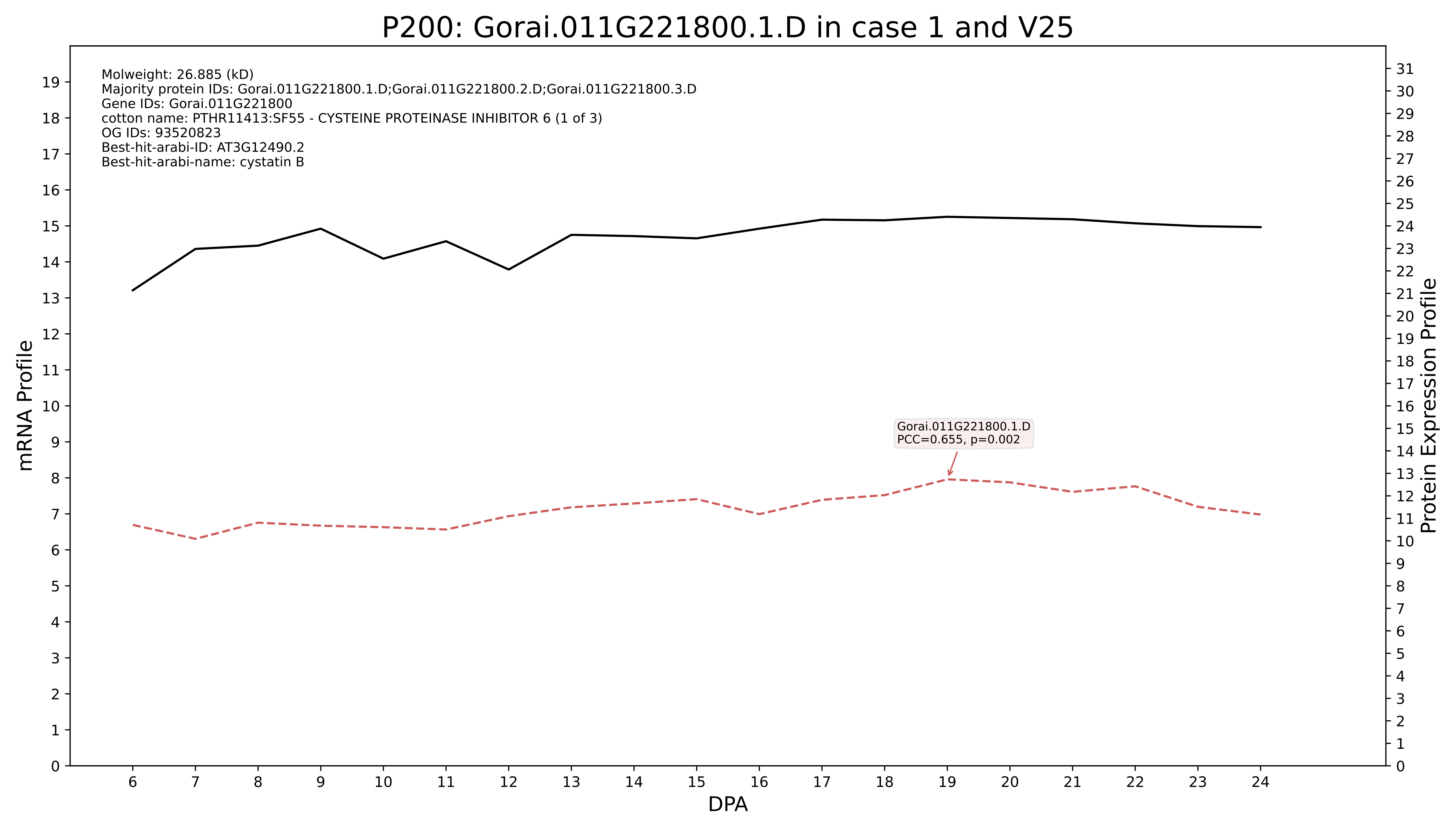
P200: Gorai.011G221800.1.D in case 1 and V25
Molweight: 26.885 (kD)
Majority protein IDs: Gorai.011G221800.1.D;Gorai.011G221800.2.D;Gorai.011G221800.3.D
Gene IDs: Gorai.011G221800
cotton name: PTHR11413:SF55 - CYSTEINE PROTEINASE INHIBITOR 6 (1 of 3)
OG IDs: 93520823
Best-hit-arabi-ID: AT3G12490.2
Best-hit-arabi-name: cystatin B
Gorai.011G221800.1.D
PCC=0.655, p=0.002
mRNA Profile
Protein Expression Profile
DPA

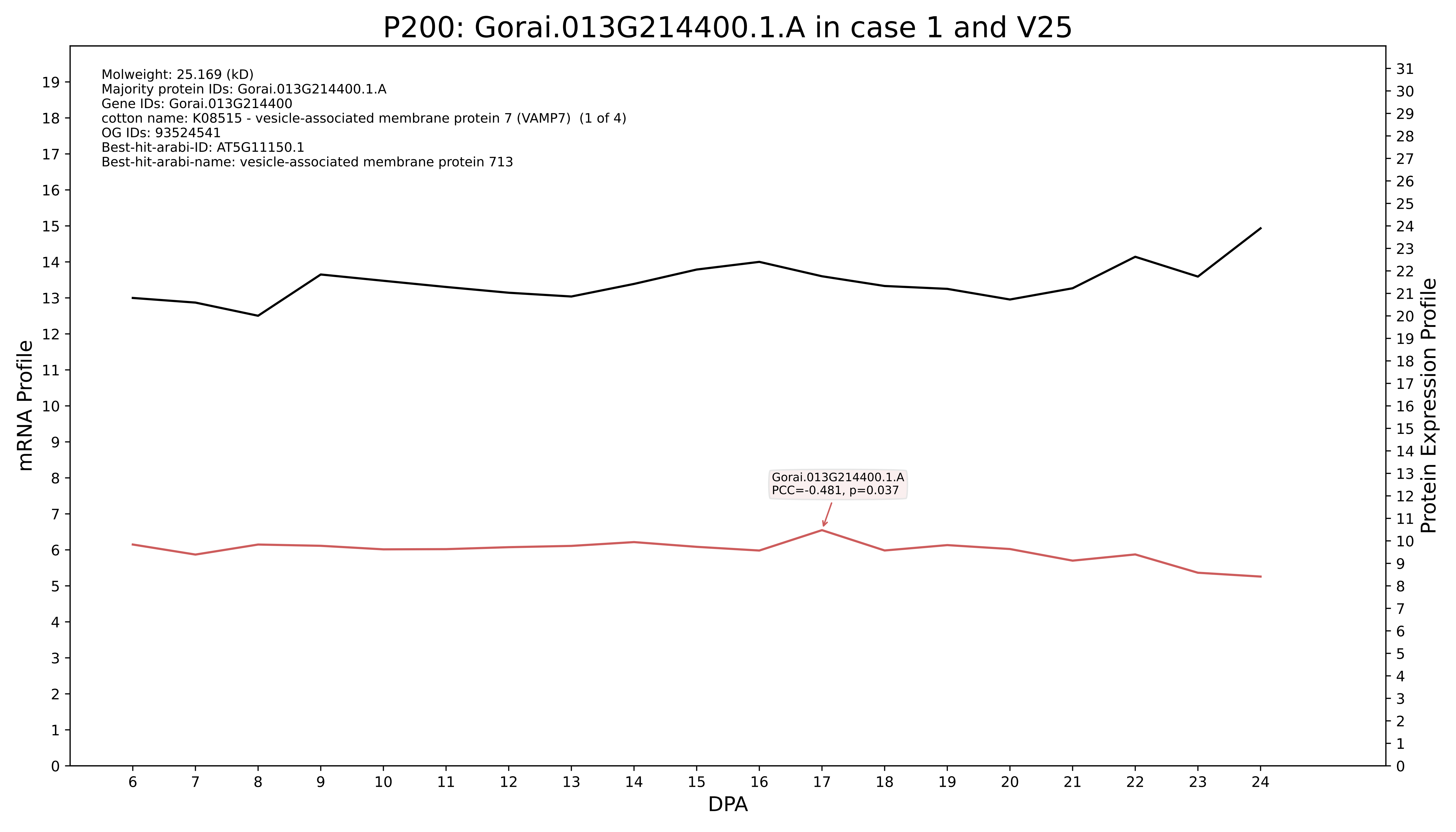

P200: Gorai.013G214400.1.A in case 1 and V25
Molweight: 25.169 (kD)
Majority protein IDs: Gorai.013G214400.1.A
Gene IDs: Gorai.013G214400
cotton name: K08515 - vesicle-associated membrane protein 7 (VAMP7) (1 of 4)
OG IDs: 93524541
Best-hit-arabi-ID: AT5G11150.1
Best-hit-arabi-name: vesicle-associated membrane protein 713
Gorai.013G214400.1.A
PCC=-0.481, p=0.037
mRNA Profile
Protein Expression Profile
DPA

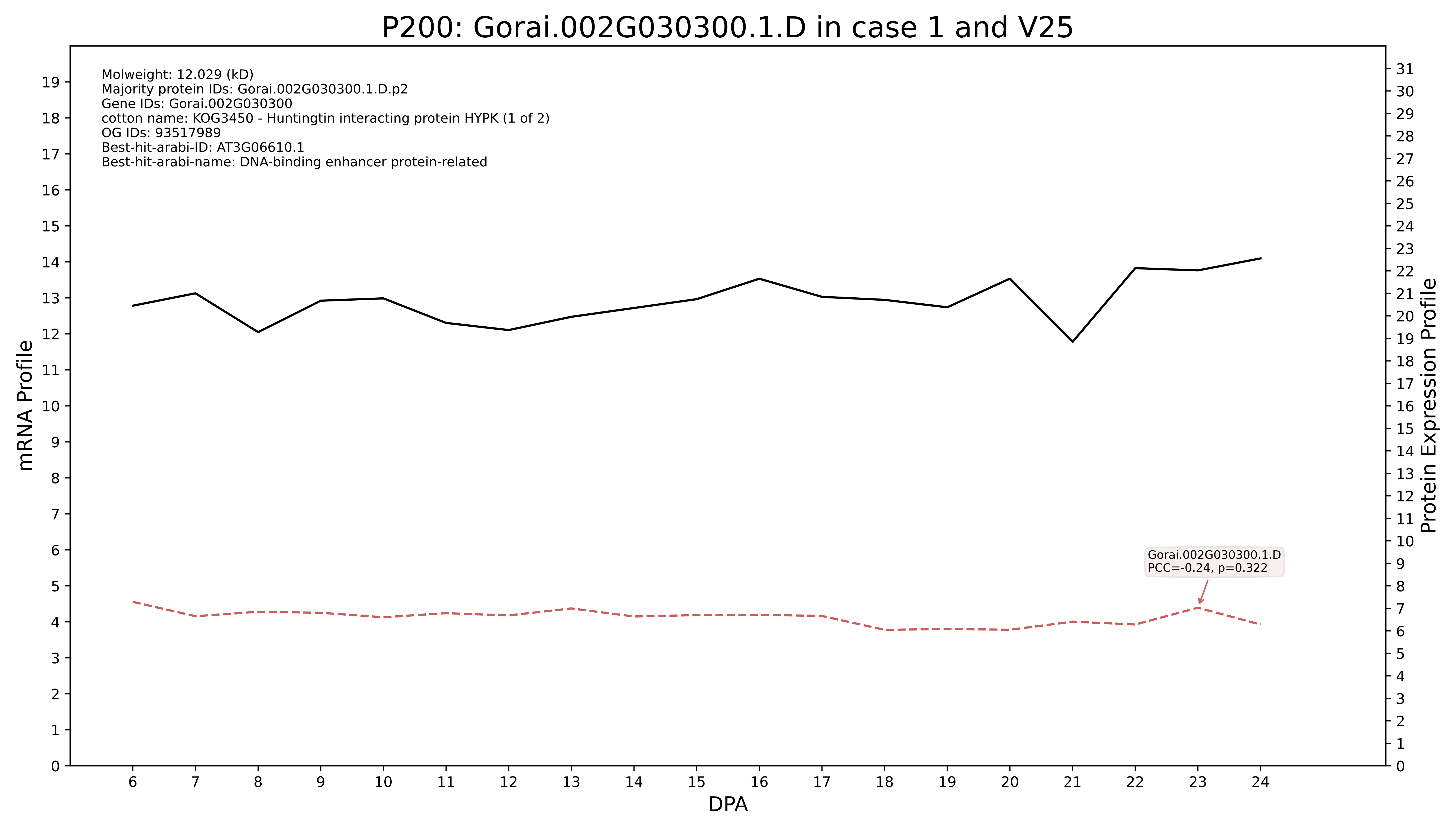
P200: Gorai.002G030300.1.D in case 1 and V25
Molweight: 12.029 (kD)
Majority protein IDs: Gorai.002G030300.1.D.p2
Gene IDs: Gorai.002G030300
cotton name: KOG3450 - Huntingtin interacting protein HYPK (1 of 2)
OG IDs: 93517989
Best-hit-arabi-ID: AT3G06610.1
Best-hit-arabi-name: DNA-binding enhancer protein-related
Gorai.002G030300.1.D
PCC=-0.24, p=0.322
mRNA Profile
Protein Expression Profile
DPA

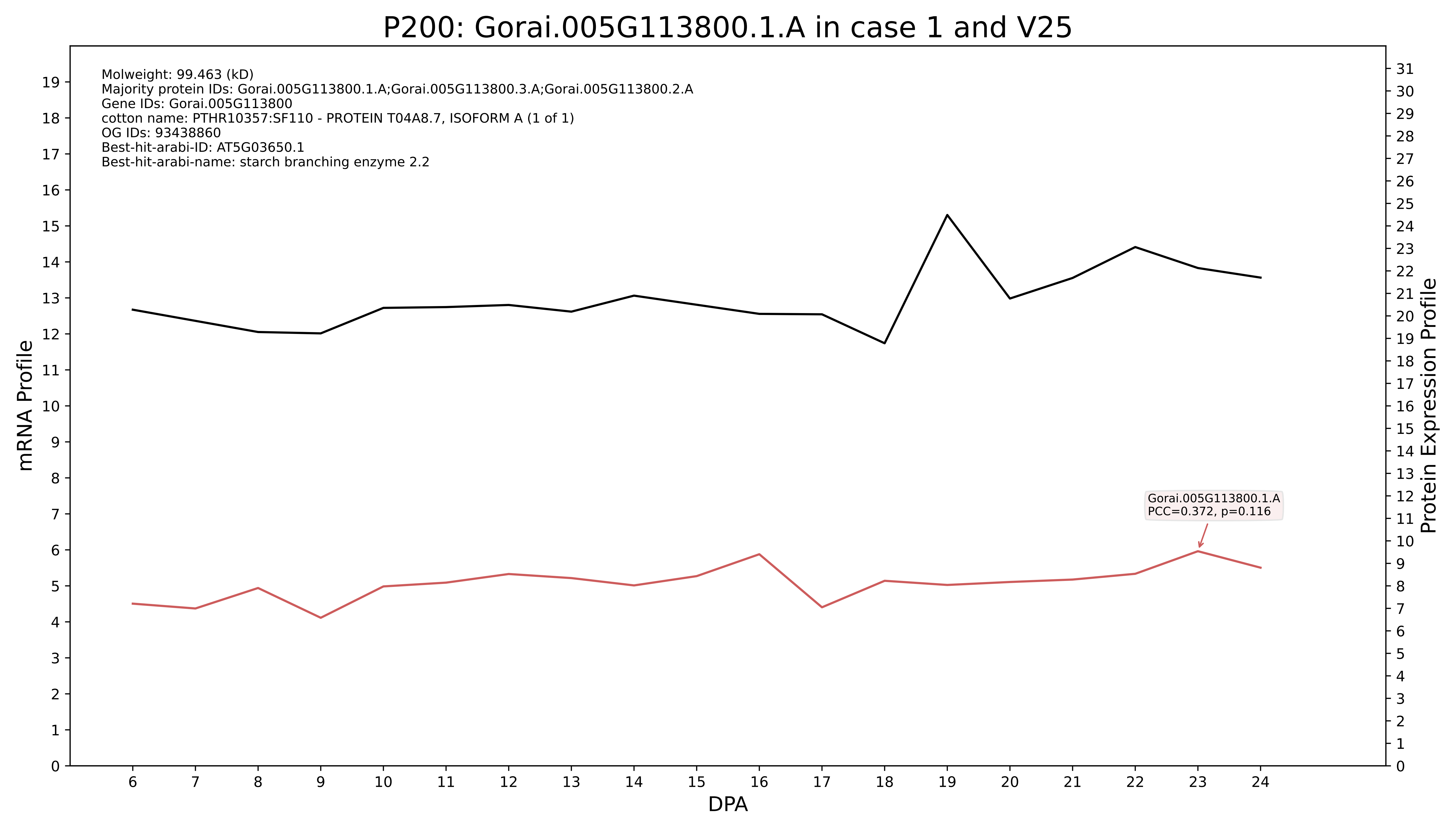
P200: Gorai.005G113800.1.A in case 1 and V25
Molweight: 99.463 (kD)
Majority protein IDs: Gorai.005G113800.1.A;Gorai.005G113800.3.A;Gorai.005G113800.2.A
Gene IDs: Gorai.005G113800
cotton name: PTHR10357:SF110 - PROTEIN T04A8.7, ISOFORM A (1 of 1)
OG IDs: 93438860
Best-hit-arabi-ID: AT5G03650.1
Best-hit-arabi-name: starch branching enzyme 2.2
Gorai.005G113800.1.A
PCC=0.372, p=0.116
mRNA Profile
Protein Expression Profile
DPA

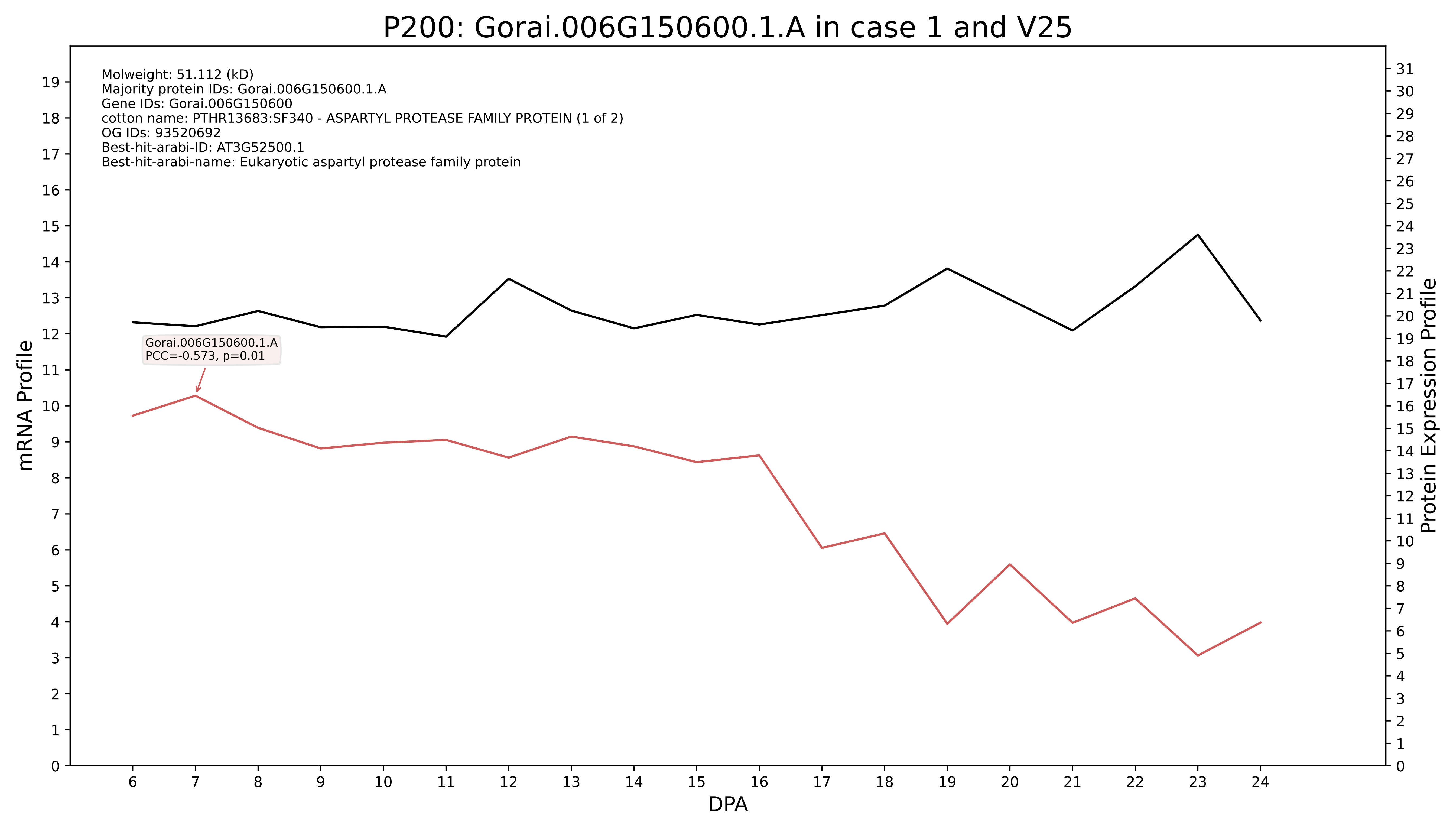

P200: Gorai.006G150600.1.A in case 1 and V25
Molweight: 51.112 (kD)
Majority protein IDs: Gorai.006G150600.1.A
Gene IDs: Gorai.006G150600
cotton name: PTHR13683:SF340 - ASPARTYL PROTEASE FAMILY PROTEIN (1 of 2)
OG IDs: 93520692
Best-hit-arabi-ID: AT3G52500.1
Best-hit-arabi-name: Eukaryotic aspartyl protease family protein
Gorai.006G150600.1.A
PCC=-0.573, p=0.01
mRNA Profile
Protein Expression Profile
DPA

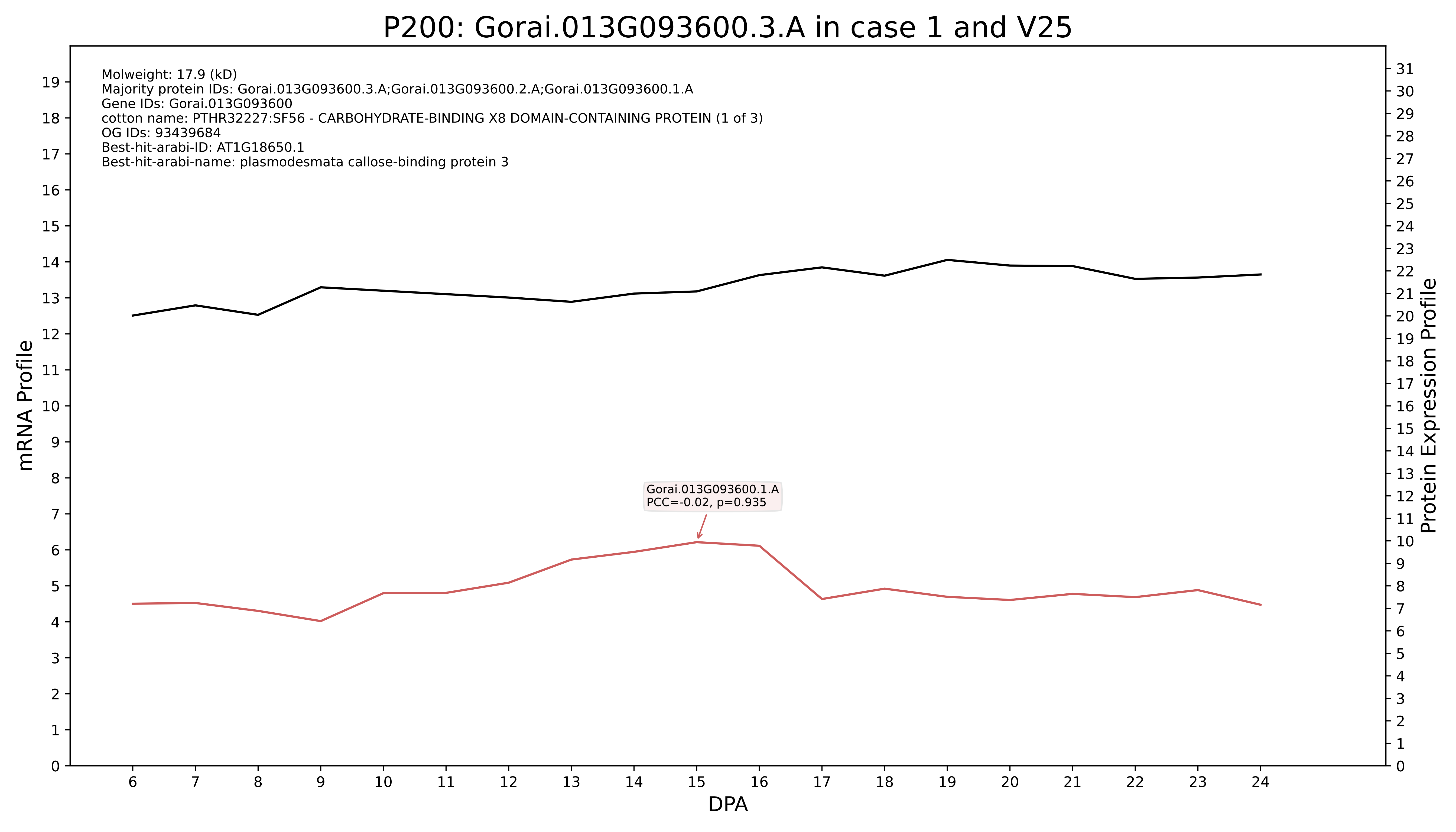

P200: Gorai.013G093600.3.A in case 1 and V25
Molweight: 17.9 (kD)
Majority protein IDs: Gorai.013G093600.3.A;Gorai.013G093600.2.A;Gorai.013G093600.1.A
Gene IDs: Gorai.013G093600
cotton name: PTHR32227:SF56 - CARBOHYDRATE-BINDING X8 DOMAIN-CONTAINING PROTEIN (1 of 3)
OG IDs: 93439684
Best-hit-arabi-ID: AT1G18650.1
Best-hit-arabi-name: plasmodesmata callose-binding protein 3
Gorai.013G093600.1.A
PCC=-0.02, p=0.935
mRNA Profile
Protein Expression Profile
DPA

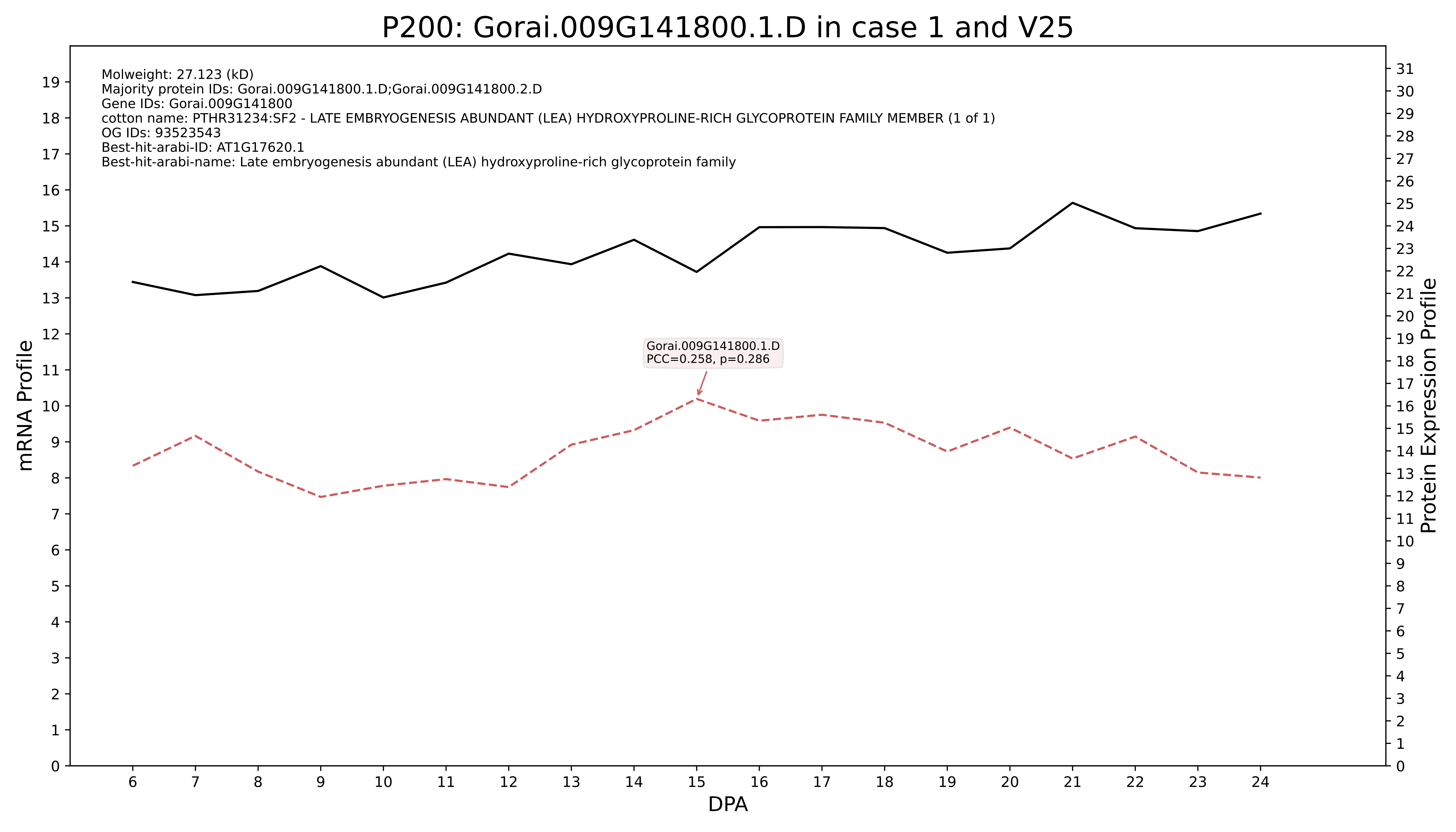
P200: Gorai.009G141800.1.D in case 1 and V25
Molweight: 27.123 (kD)
Majority protein IDs: Gorai.009G141800.1.D;Gorai.009G141800.2.D
Gene IDs: Gorai.009G141800
cotton name: PTHR31234:SF2 - LATE EMBRYOGENESIS ABUNDANT (LEA) HYDROXYPROLINE-RICH GLYCOPROTEIN FAMILY MEMBER (1 of 1)
OG IDs: 93523543
Best-hit-arabi-ID: AT1G17620.1
Best-hit-arabi-name: Late embryogenesis abundant (LEA) hydroxyproline-rich glycoprotein family
Gorai.009G141800.1.D
PCC=0.258, p=0.286
mRNA Profile
Protein Expression Profile
DPA

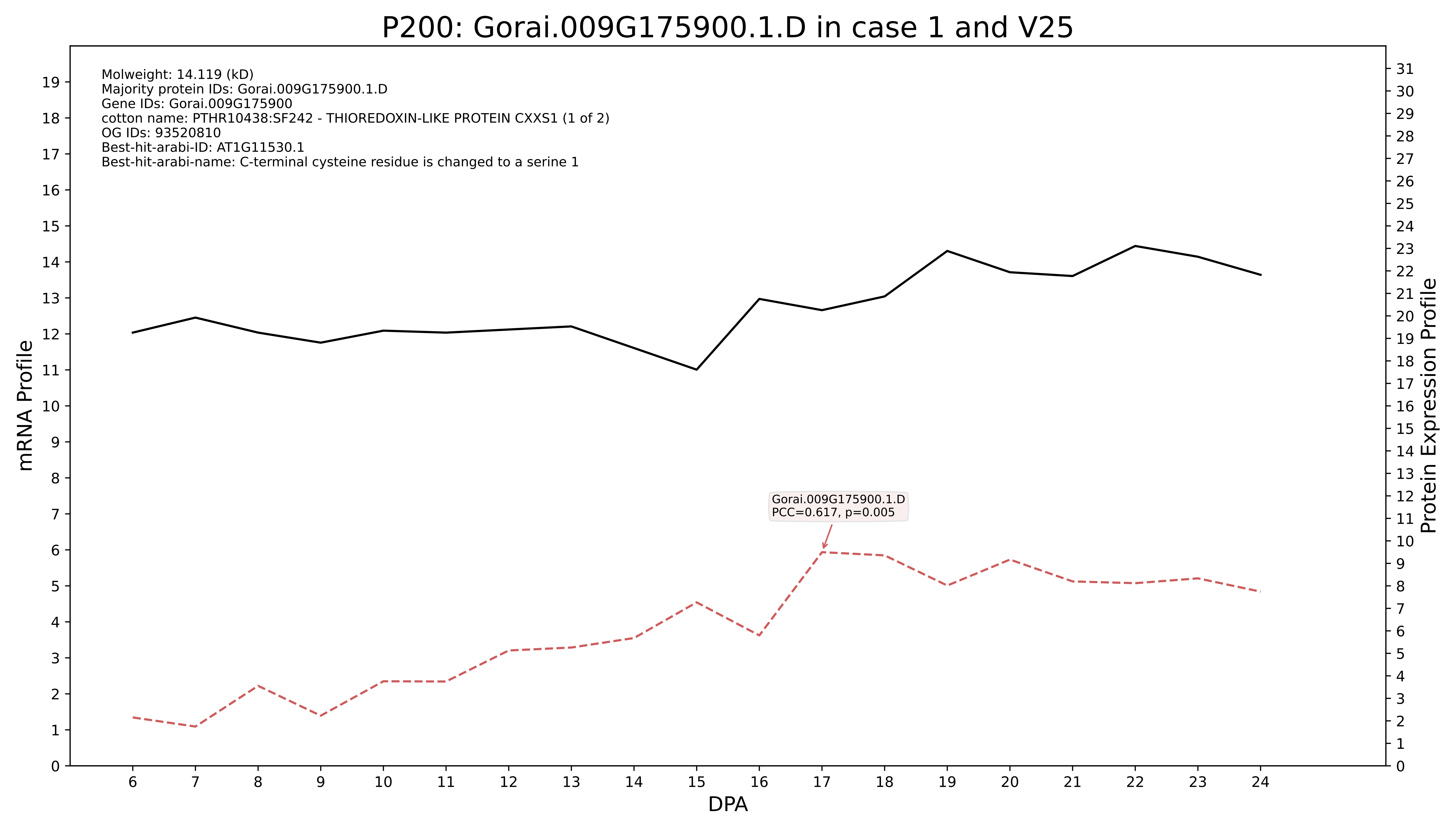
P200: Gorai.009G175900.1.D in case 1 and V25
Molweight: 14.119 (kD)
Majority protein IDs: Gorai.009G175900.1.D
Gene IDs: Gorai.009G175900
cotton name: PTHR10438:SF242 - THIOREDOXIN-LIKE PROTEIN CXXS1 (1 of 2)
OG IDs: 93520810
Best-hit-arabi-ID: AT1G11530.1
Best-hit-arabi-name: C-terminal cysteine residue is changed to a serine 1
Gorai.009G175900.1.D
PCC=0.617, p=0.005
mRNA Profile
Protein Expression Profile
DPA

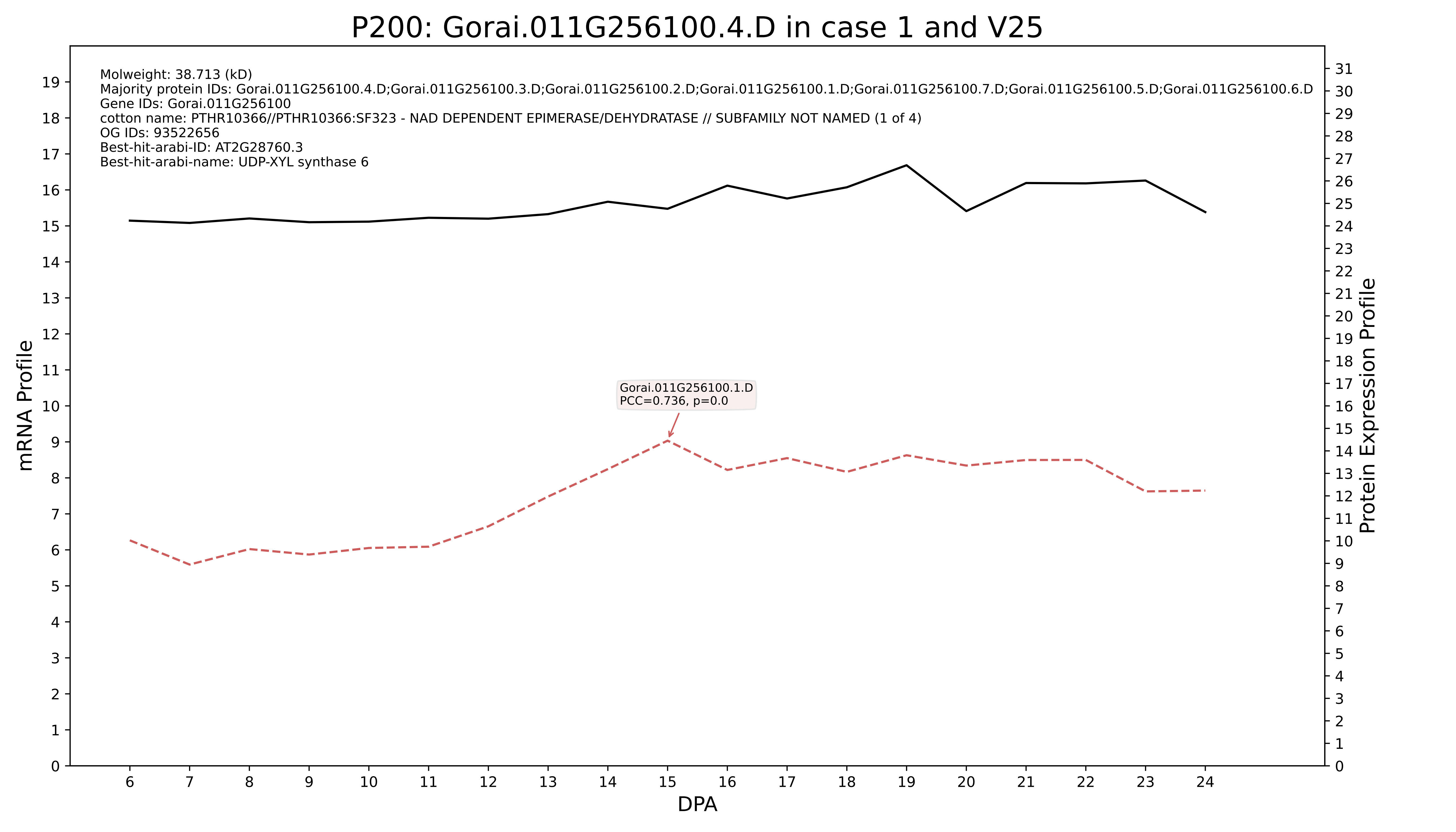
P200: Gorai.011G256100.4.D in case 1 and V25
Molweight: 38.713 (kD)
Majority protein IDs: Gorai.011G256100.4.D;Gorai.011G256100.3.D;Gorai.011G256100.2.D;Gorai.011G256100.1.D;Gorai.011G256100.7.D;Gorai.011G256100.5.D;Gorai.011G256100.6.D
Gene IDs: Gorai.011G256100
cotton name: PTHR10366//PTHR10366:SF323 - NAD DEPENDENT EPIMERASE/DEHYDRATASE // SUBFAMILY NOT NAMED (1 of 4)
OG IDs: 93522656
Best-hit-arabi-ID: AT2G28760.3
Best-hit-arabi-name: UDP-XYL synthase 6
Gorai.011G256100.1.D
PCC=0.736, p=0.0
mRNA Profile
Protein Expression Profile
DPA

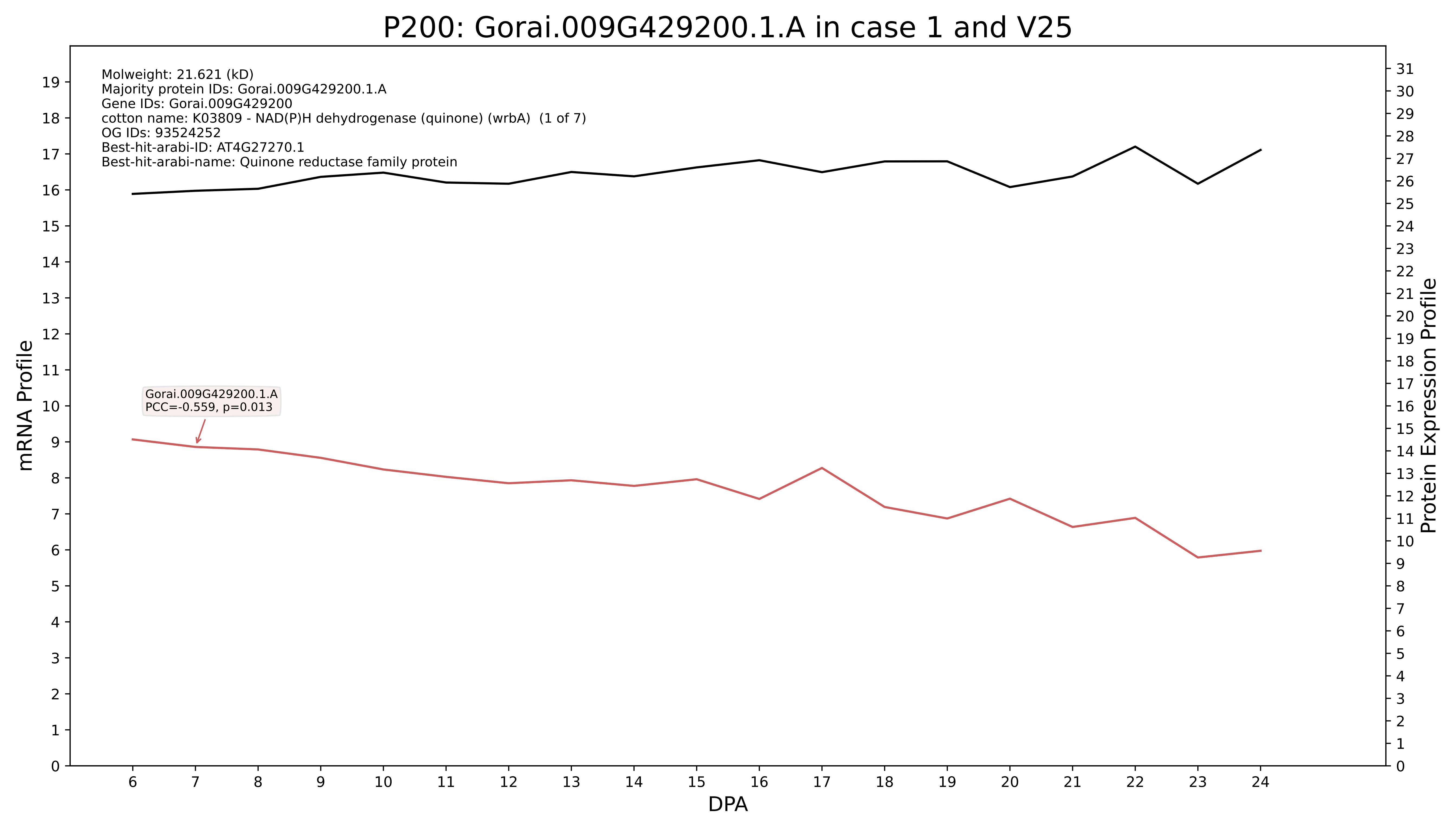

P200: Gorai.009G429200.1.A in case 1 and V25
Molweight: 21.621 (kD)
Majority protein IDs: Gorai.009G429200.1.A
Gene IDs: Gorai.009G429200
cotton name: K03809 - NAD(P)H dehydrogenase (quinone) (wrbA) (1 of 7)
OG IDs: 93524252
Best-hit-arabi-ID: AT4G27270.1
Best-hit-arabi-name: Quinone reductase family protein
Gorai.009G429200.1.A
PCC=-0.559, p=0.013
mRNA Profile
Protein Expression Profile
DPA

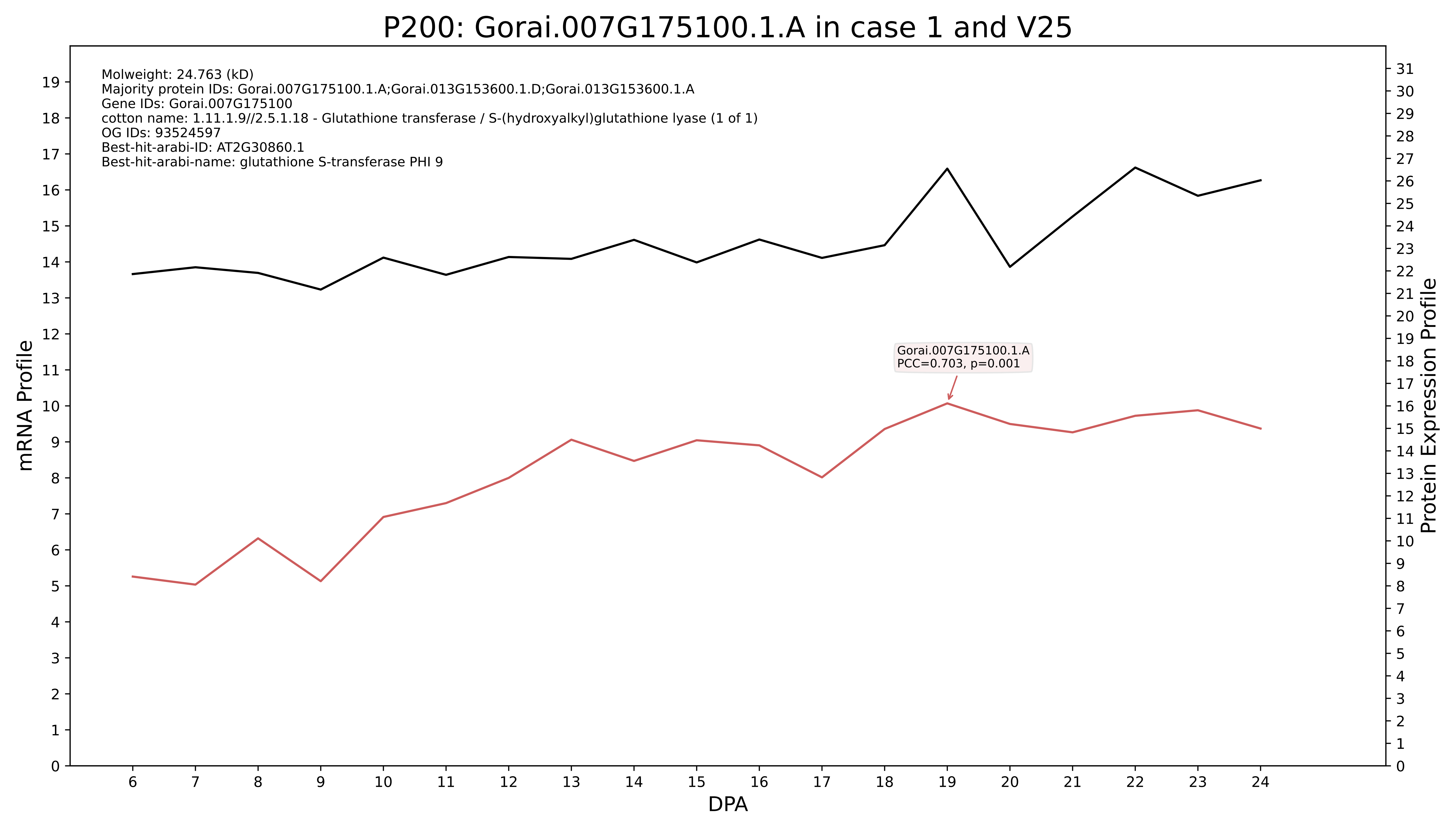
P200: Gorai.007G175100.1.A in case 1 and V25
Molweight: 24.763 (kD)
Majority protein IDs: Gorai.007G175100.1.A;Gorai.013G153600.1.D;Gorai.013G153600.1.A
Gene IDs: Gorai.007G175100
cotton name: 1.11.1.9//2.5.1.18 - Glutathione transferase / S-(hydroxyalkyl)glutathione lyase (1 of 1)
OG IDs: 93524597
Best-hit-arabi-ID: AT2G30860.1
Best-hit-arabi-name: glutathione S-transferase PHI 9
Gorai.007G175100.1.A
PCC=0.703, p=0.001
mRNA Profile
Protein Expression Profile
DPA

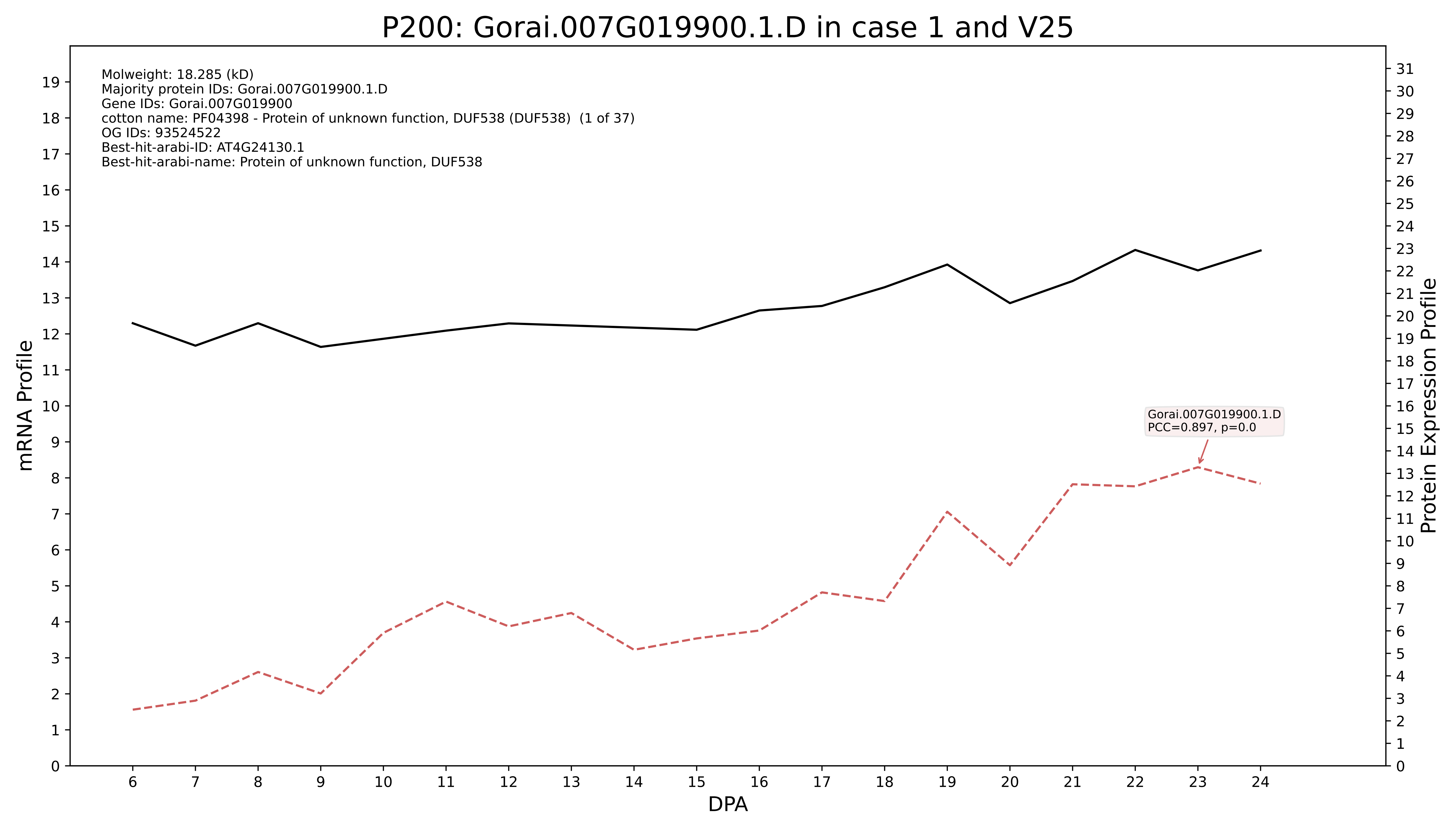
P200: Gorai.007G019900.1.D in case 1 and V25
Molweight: 18.285 (kD)
Majority protein IDs: Gorai.007G019900.1.D
Gene IDs: Gorai.007G019900
cotton name: PF04398 - Protein of unknown function, DUF538 (DUF538) (1 of 37)
OG IDs: 93524522
Best-hit-arabi-ID: AT4G24130.1
Best-hit-arabi-name: Protein of unknown function, DUF538
Gorai.007G019900.1.D
PCC=0.897, p=0.0
mRNA Profile
Protein Expression Profile
DPA

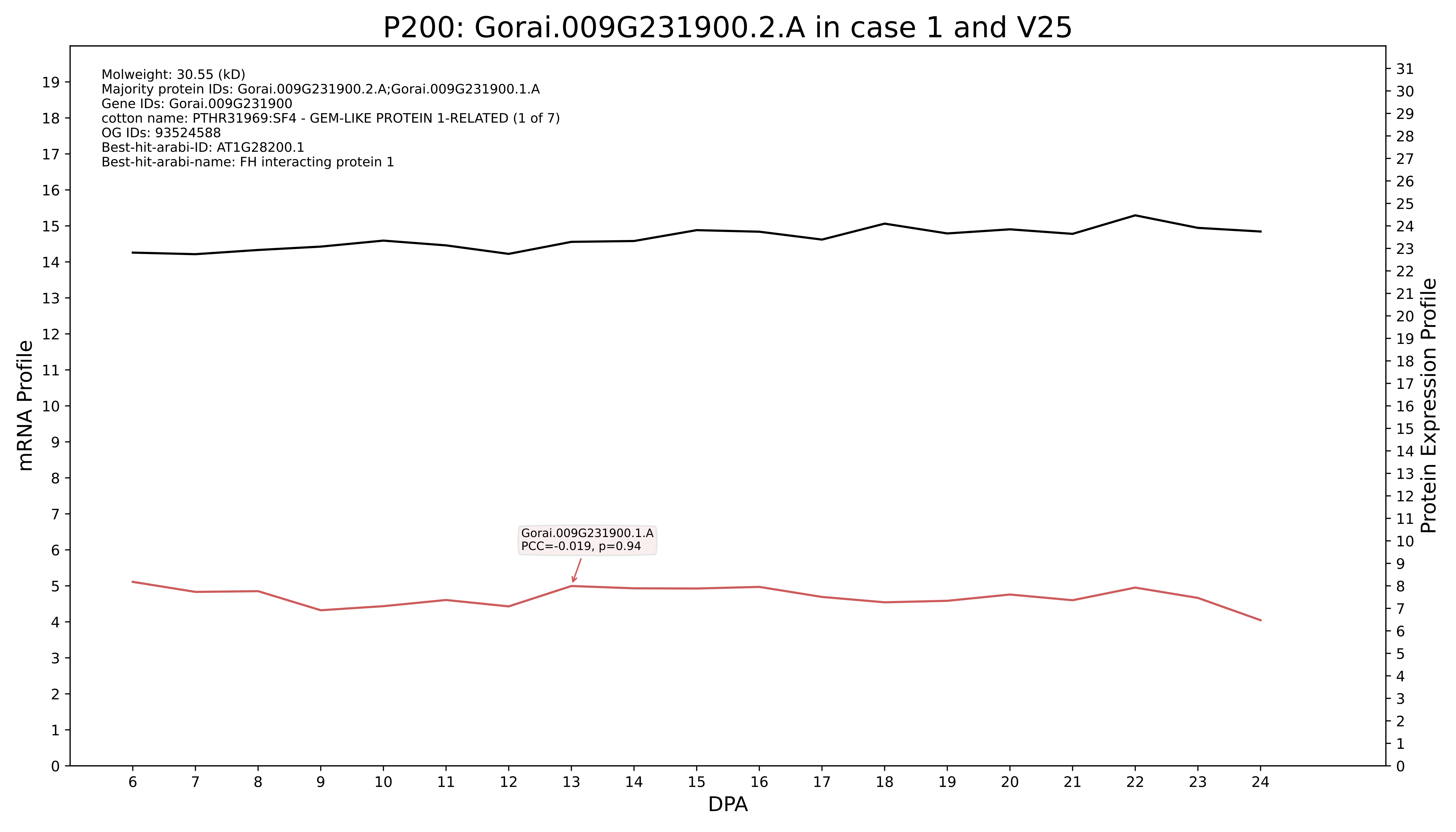
P200: Gorai.009G231900.2.A in case 1 and V25
Molweight: 30.55 (kD)
Majority protein IDs: Gorai.009G231900.2.A;Gorai.009G231900.1.A
Gene IDs: Gorai.009G231900
cotton name: PTHR31969:SF4 - GEM-LIKE PROTEIN 1-RELATED (1 of 7)
OG IDs: 93524588
Best-hit-arabi-ID: AT1G28200.1
Best-hit-arabi-name: FH interacting protein 1
Gorai.009G231900.1.A
PCC=-0.019, p=0.94
mRNA Profile
Protein Expression Profile
DPA

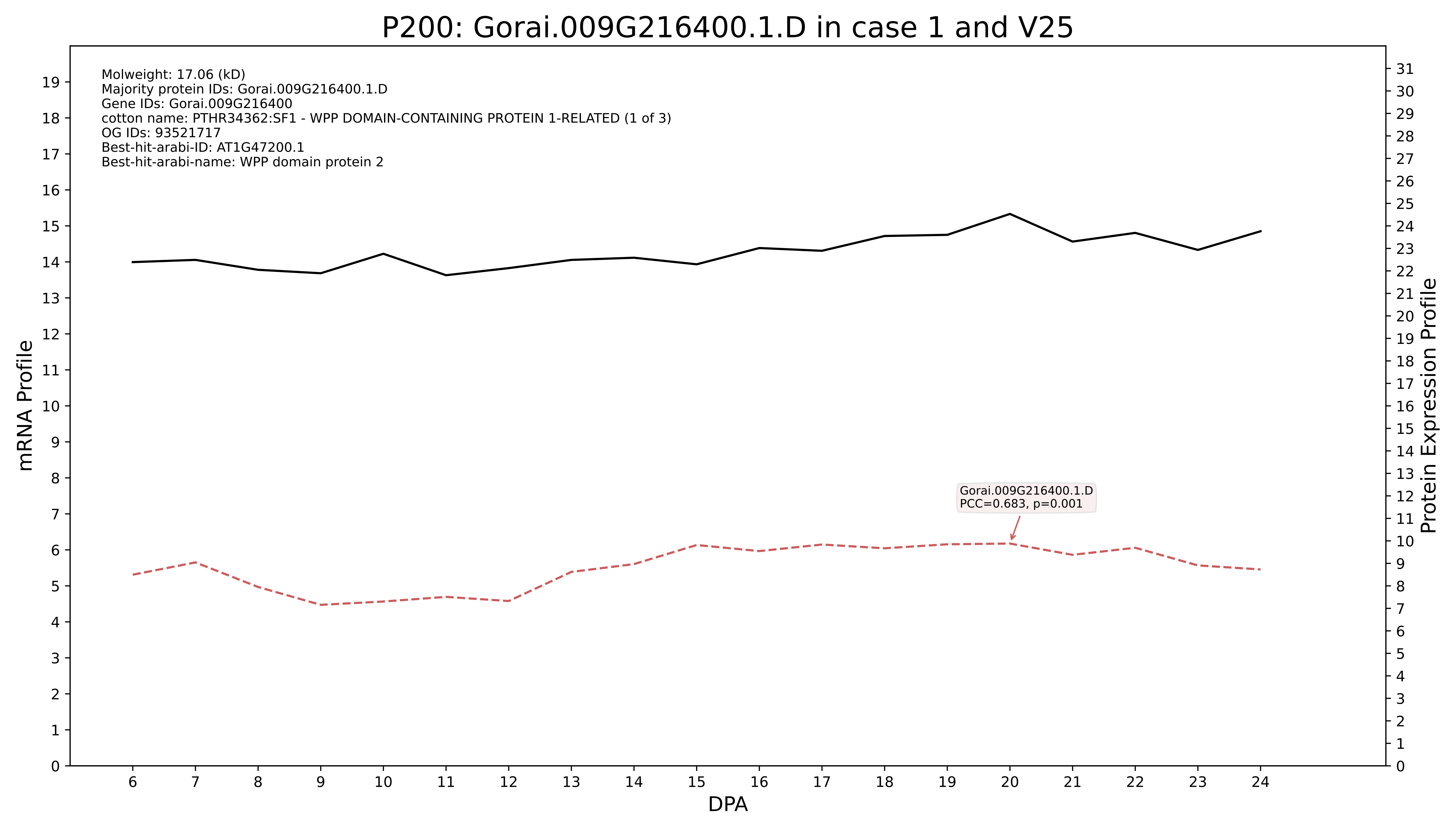
P200: Gorai.009G216400.1.D in case 1 and V25
Molweight: 17.06 (kD)
Majority protein IDs: Gorai.009G216400.1.D
Gene IDs: Gorai.009G216400
cotton name: PTHR34362:SF1 - WPP DOMAIN-CONTAINING PROTEIN 1-RELATED (1 of 3)
OG IDs: 93521717
Best-hit-arabi-ID: AT1G47200.1
Best-hit-arabi-name: WPP domain protein 2
Gorai.009G216400.1.D
PCC=0.683, p=0.001
mRNA Profile
Protein Expression Profile
DPA

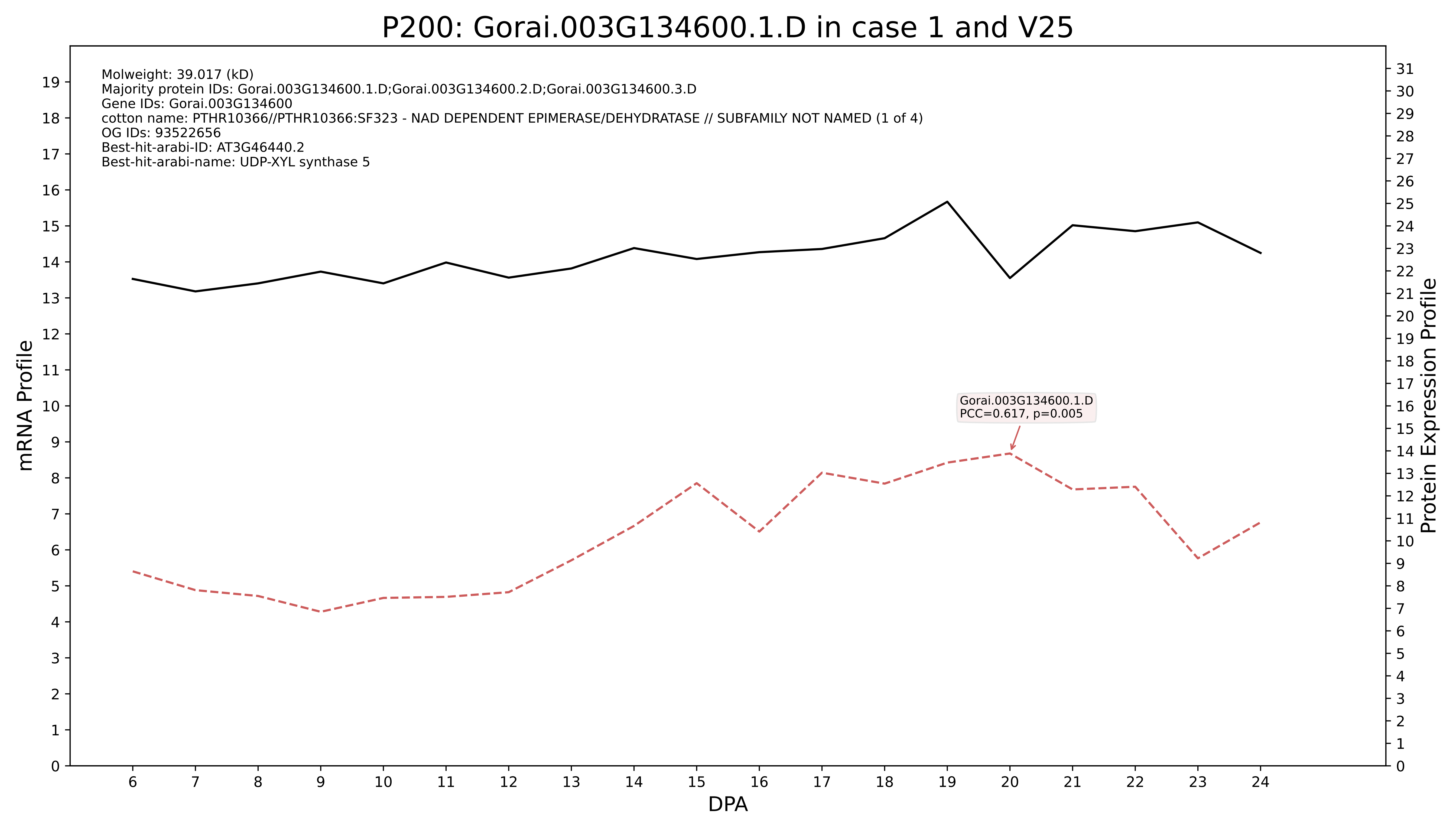
P200: Gorai.003G134600.1.D in case 1 and V25
Molweight: 39.017 (kD)
Majority protein IDs: Gorai.003G134600.1.D;Gorai.003G134600.2.D;Gorai.003G134600.3.D
Gene IDs: Gorai.003G134600
cotton name: PTHR10366//PTHR10366:SF323 - NAD DEPENDENT EPIMERASE/DEHYDRATASE // SUBFAMILY NOT NAMED (1 of 4)
OG IDs: 93522656
Best-hit-arabi-ID: AT3G46440.2
Best-hit-arabi-name: UDP-XYL synthase 5
Gorai.003G134600.1.D
PCC=0.617, p=0.005
mRNA Profile
Protein Expression Profile
DPA

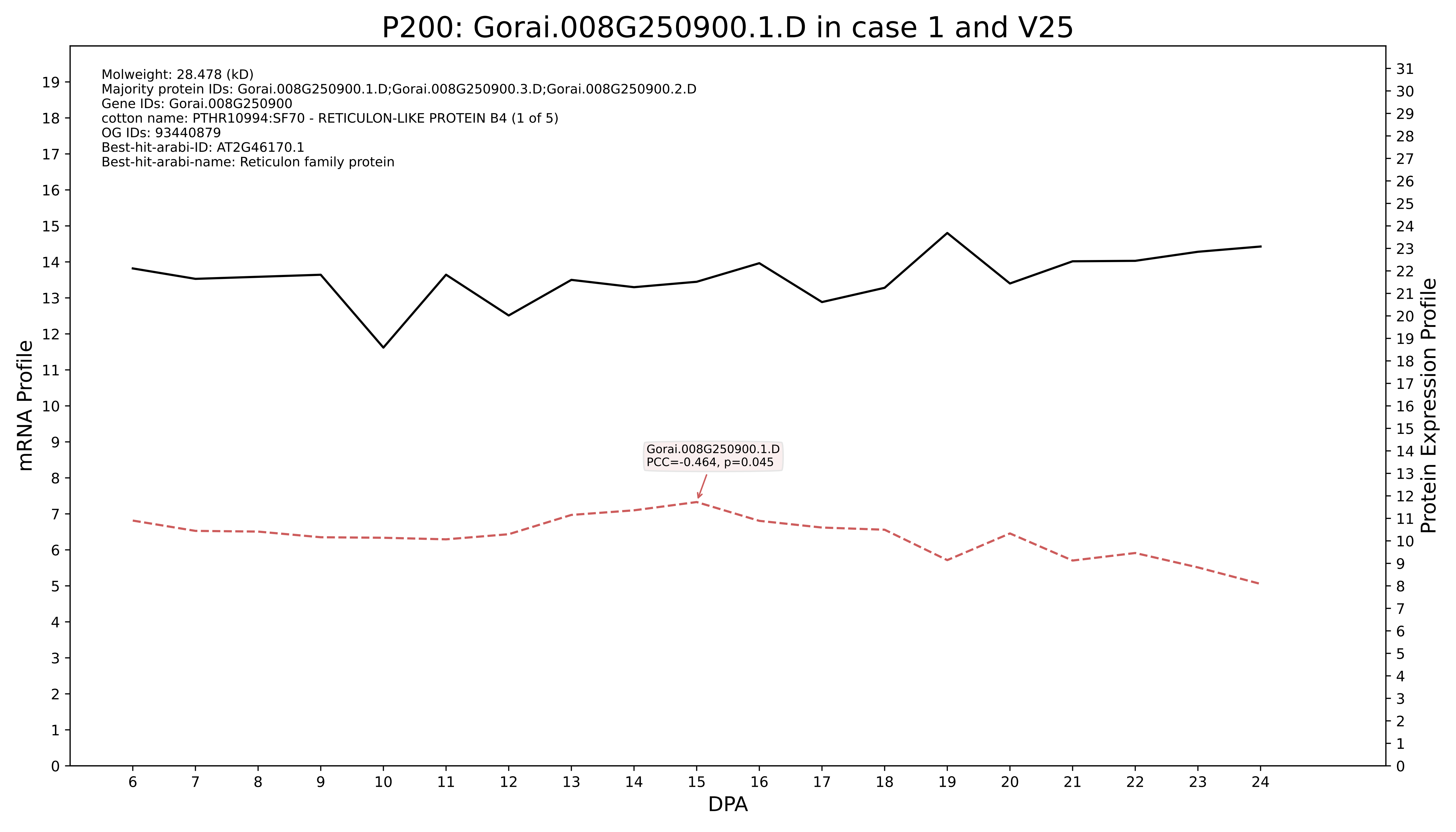
P200: Gorai.008G250900.1.D in case 1 and V25
Molweight: 28.478 (kD)
Majority protein IDs: Gorai.008G250900.1.D;Gorai.008G250900.3.D;Gorai.008G250900.2.D
Gene IDs: Gorai.008G250900
cotton name: PTHR10994:SF70 - RETICULON-LIKE PROTEIN B4 (1 of 5)
OG IDs: 93440879
Best-hit-arabi-ID: AT2G46170.1
Best-hit-arabi-name: Reticulon family protein
Gorai.008G250900.1.D
PCC=-0.464, p=0.045
mRNA Profile
Protein Expression Profile
DPA

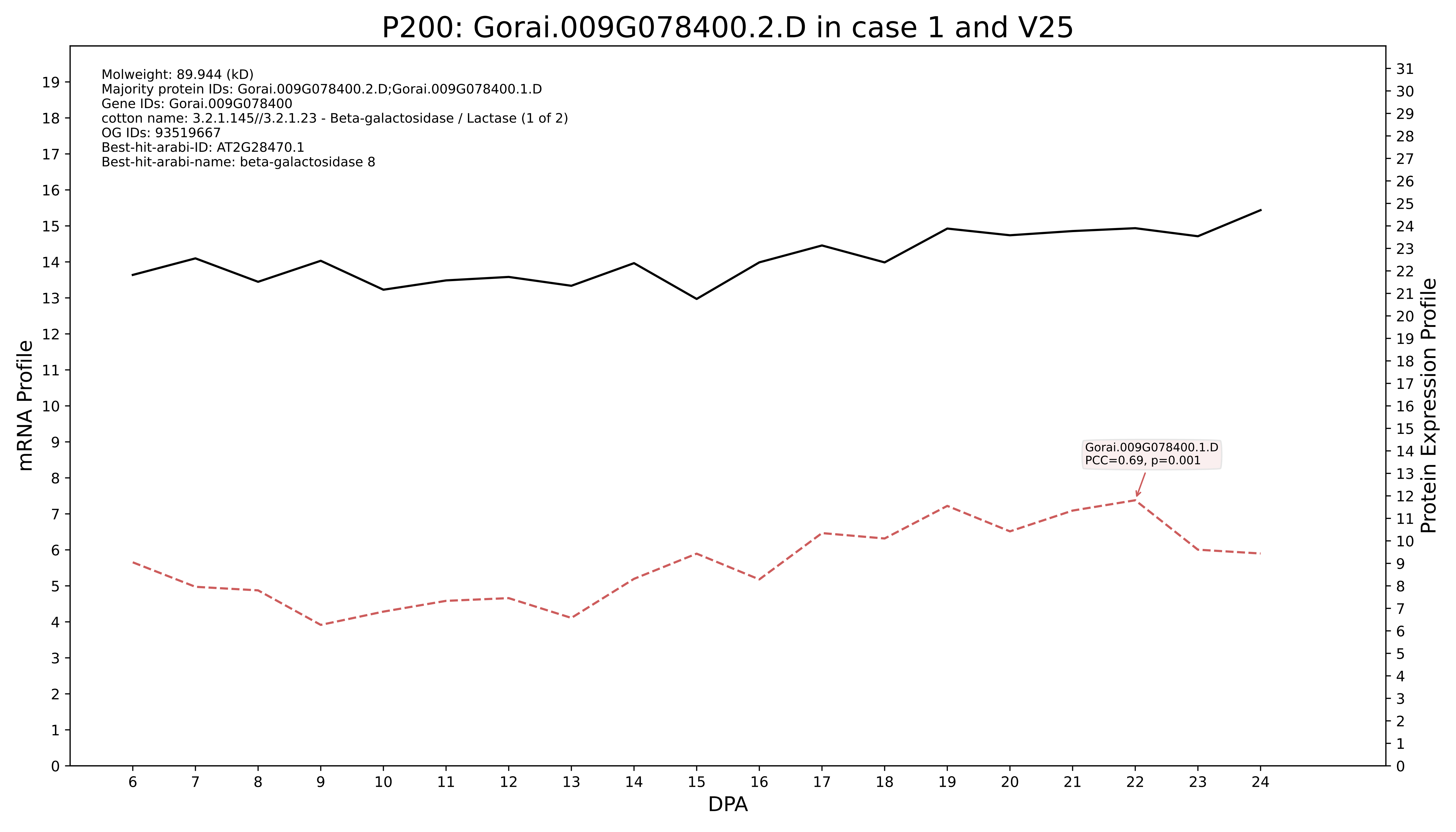
P200: Gorai.009G078400.2.D in case 1 and V25
Molweight: 89.944 (kD)
Majority protein IDs: Gorai.009G078400.2.D;Gorai.009G078400.1.D
Gene IDs: Gorai.009G078400
cotton name: 3.2.1.145//3.2.1.23 - Beta-galactosidase / Lactase (1 of 2)
OG IDs: 93519667
Best-hit-arabi-ID: AT2G28470.1
Best-hit-arabi-name: beta-galactosidase 8
Gorai.009G078400.1.D
PCC=0.69, p=0.001
mRNA Profile
Protein Expression Profile
DPA

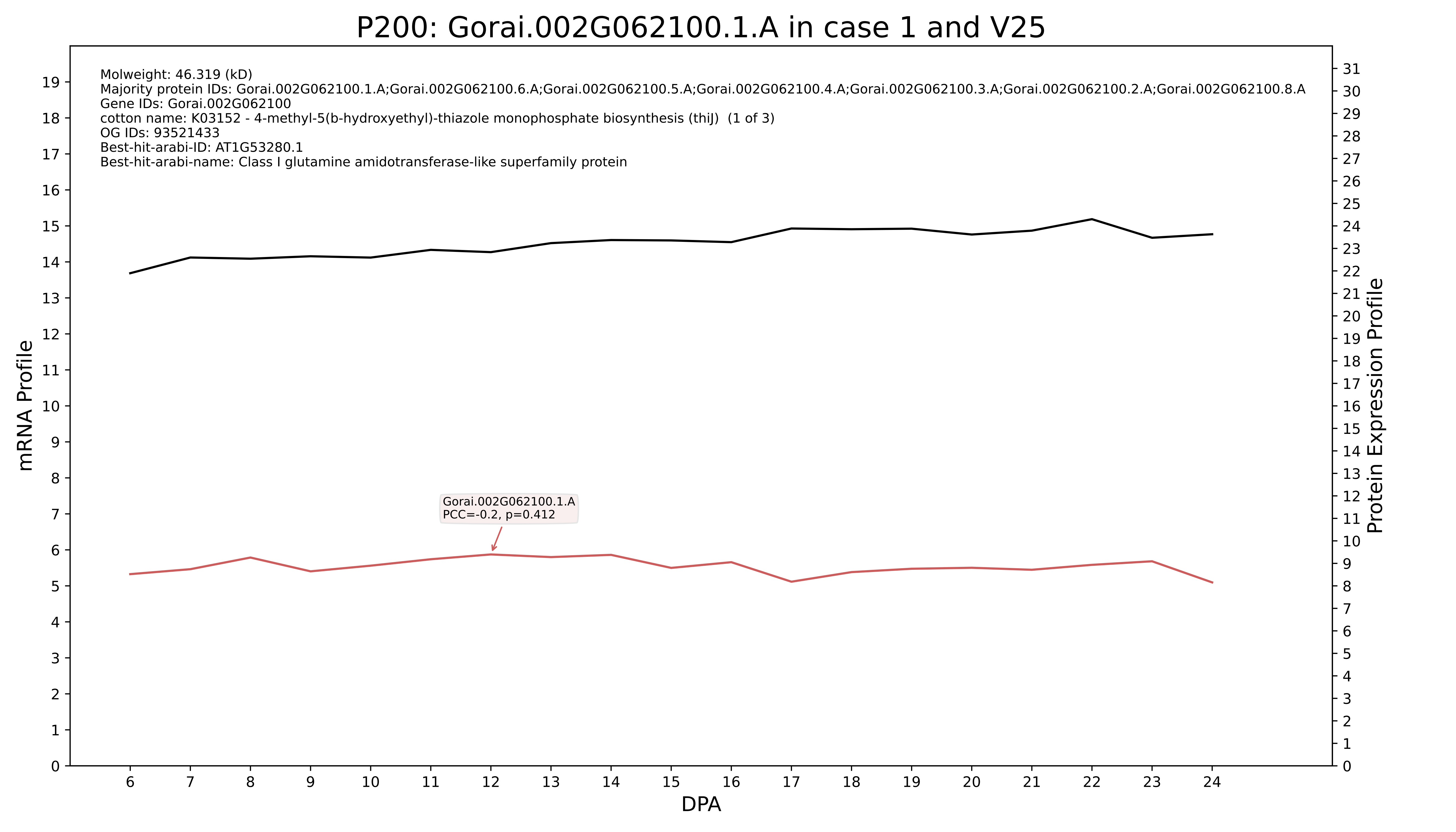
P200: Gorai.002G062100.1.A in case 1 and V25
Molweight: 46.319 (kD)
Majority protein IDs: Gorai.002G062100.1.A;Gorai.002G062100.6.A;Gorai.002G062100.5.A;Gorai.002G062100.4.A;Gorai.002G062100.3.A;Gorai.002G062100.2.A;Gorai.002G062100.8.A
Gene IDs: Gorai.002G062100
cotton name: K03152 - 4-methyl-5(b-hydroxyethyl)-thiazole monophosphate biosynthesis (thiJ) (1 of 3)
OG IDs: 93521433
Best-hit-arabi-ID: AT1G53280.1
Best-hit-arabi-name: Class I glutamine amidotransferase-like superfamily protein
Gorai.002G062100.1.A
PCC=-0.2, p=0.412
mRNA Profile
Protein Expression Profile
DPA

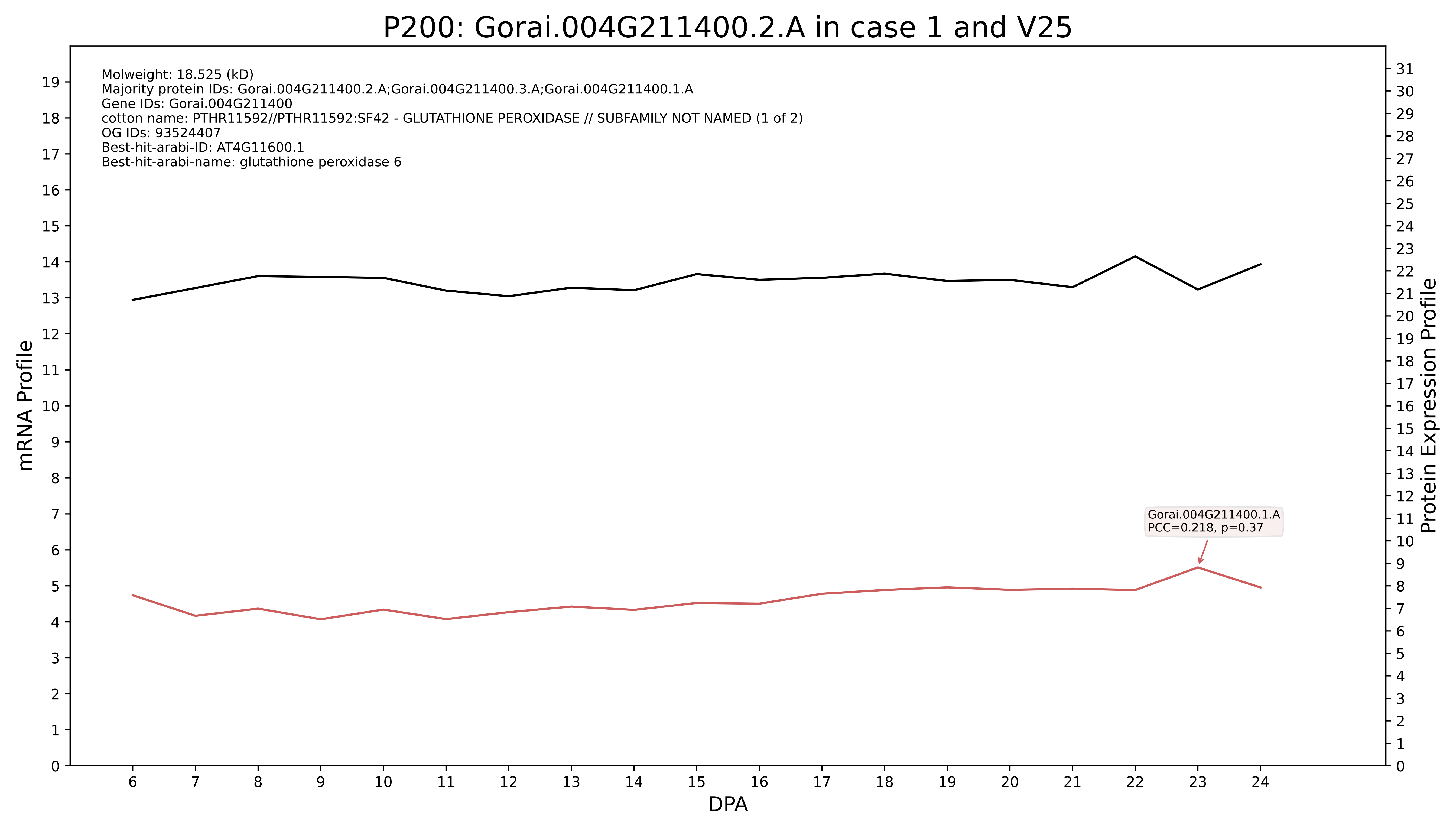

P200: Gorai.004G211400.2.A in case 1 and V25
Molweight: 18.525 (kD)
Majority protein IDs: Gorai.004G211400.2.A;Gorai.004G211400.3.A;Gorai.004G211400.1.A
Gene IDs: Gorai.004G211400
cotton name: PTHR11592//PTHR11592:SF42 - GLUTATHIONE PEROXIDASE // SUBFAMILY NOT NAMED (1 of 2)
OG IDs: 93524407
Best-hit-arabi-ID: AT4G11600.1
Best-hit-arabi-name: glutathione peroxidase 6
Gorai.004G211400.1.A
PCC=0.218, p=0.37
mRNA Profile
Protein Expression Profile
DPA

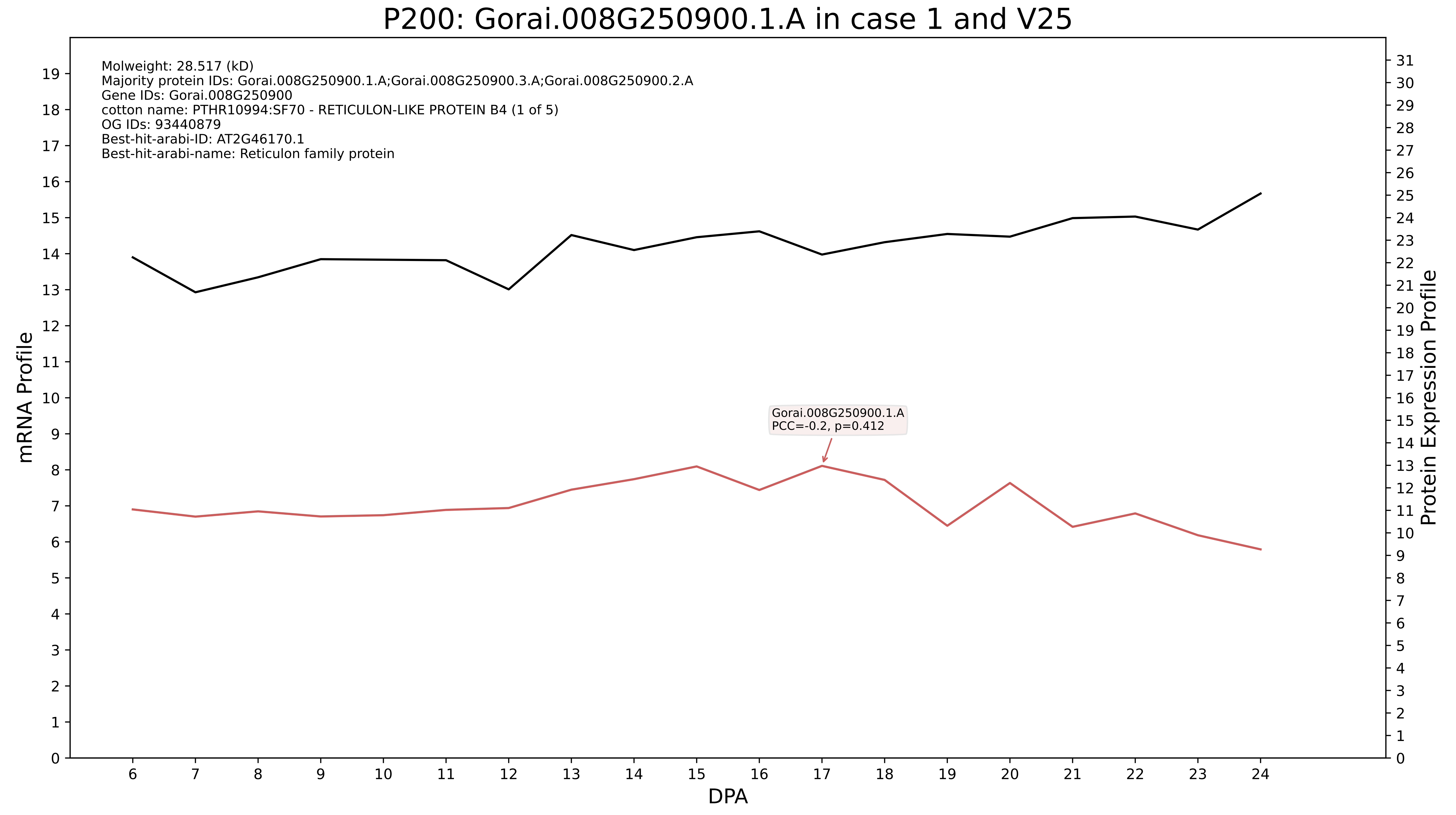
P200: Gorai.008G250900.1.A in case 1 and V25
Molweight: 28.517 (kD)
Majority protein IDs: Gorai.008G250900.1.A;Gorai.008G250900.3.A;Gorai.008G250900.2.A
Gene IDs: Gorai.008G250900
cotton name: PTHR10994:SF70 - RETICULON-LIKE PROTEIN B4 (1 of 5)
OG IDs: 93440879
Best-hit-arabi-ID: AT2G46170.1
Best-hit-arabi-name: Reticulon family protein
Gorai.008G250900.1.A
PCC=-0.2, p=0.412
mRNA Profile
Protein Expression Profile
DPA

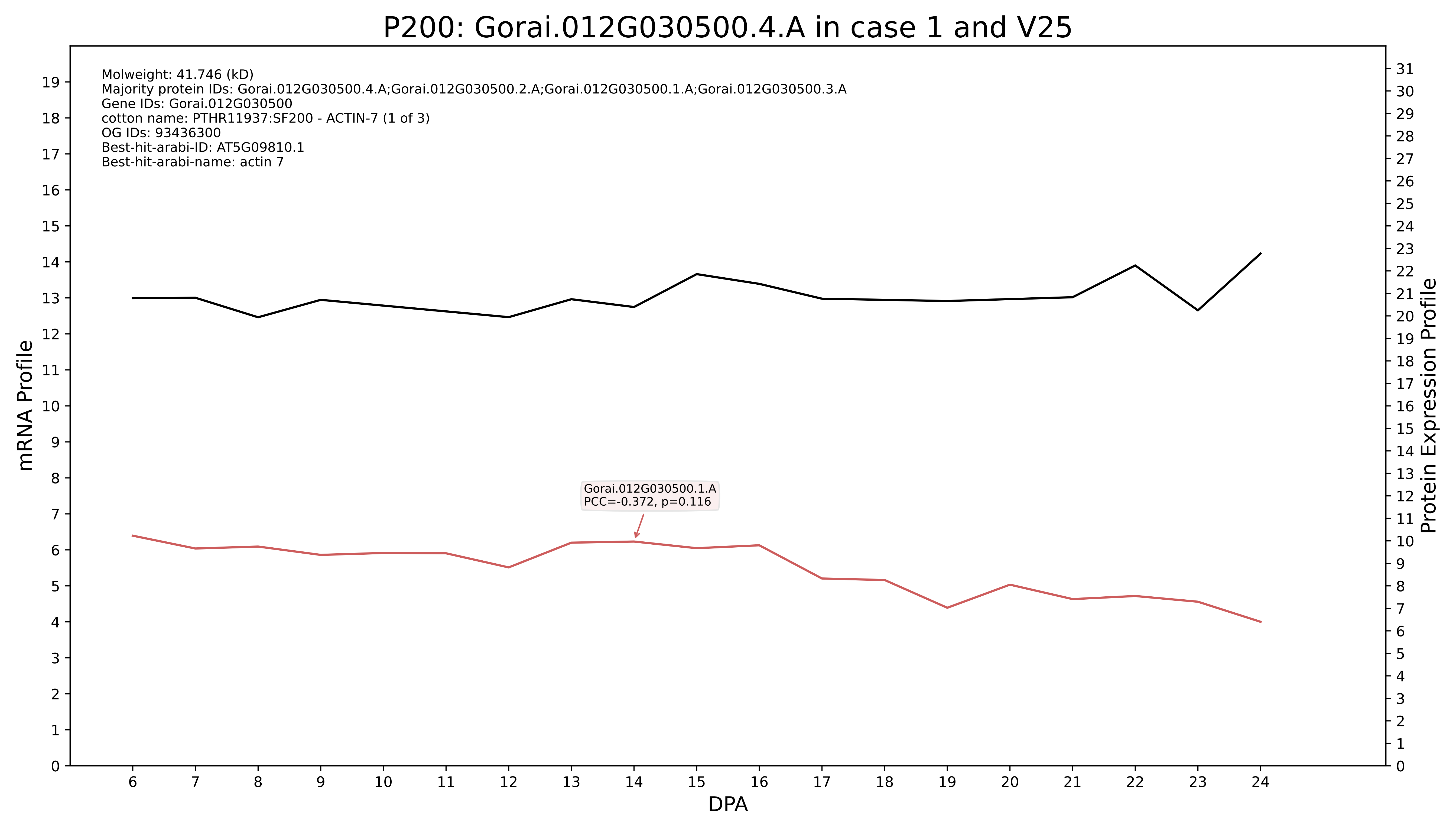
P200: Gorai.012G030500.4.A in case 1 and V25
Molweight: 41.746 (kD)
Majority protein IDs: Gorai.012G030500.4.A;Gorai.012G030500.2.A;Gorai.012G030500.1.A;Gorai.012G030500.3.A
Gene IDs: Gorai.012G030500
cotton name: PTHR11937:SF200 - ACTIN-7 (1 of 3)
OG IDs: 93436300
Best-hit-arabi-ID: AT5G09810.1
Best-hit-arabi-name: actin 7
Gorai.012G030500.1.A
PCC=-0.372, p=0.116
mRNA Profile
Protein Expression Profile
DPA

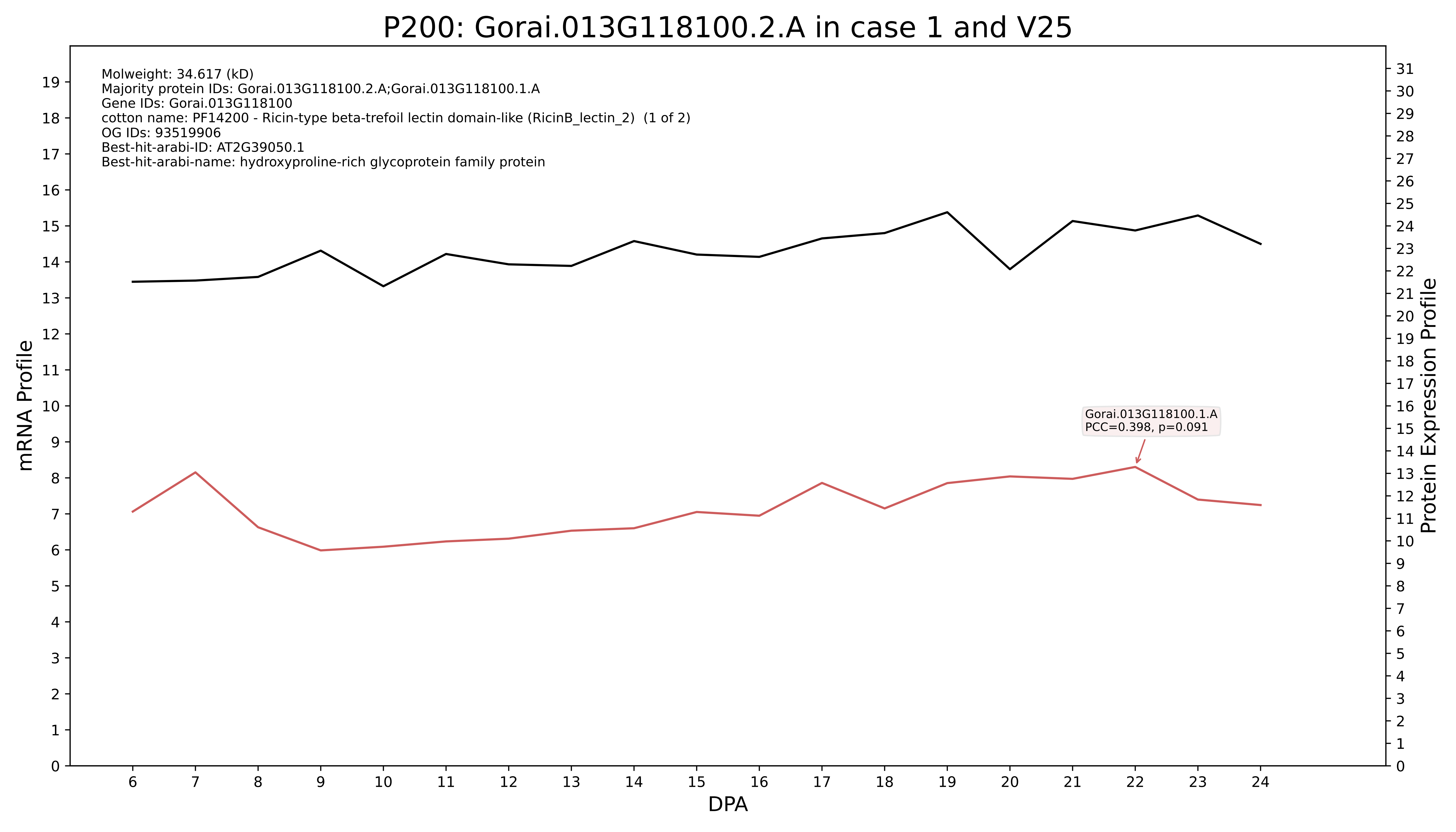
P200: Gorai.013G118100.2.A in case 1 and V25
Molweight: 34.617 (kD)
Majority protein IDs: Gorai.013G118100.2.A;Gorai.013G118100.1.A
Gene IDs: Gorai.013G118100
cotton name: PF14200 - Ricin-type beta-trefoil lectin domain-like (RicinB_lectin_2) (1 of 2)
OG IDs: 93519906
Best-hit-arabi-ID: AT2G39050.1
Best-hit-arabi-name: hydroxyproline-rich glycoprotein family protein
Gorai.013G118100.1.A
PCC=0.398, p=0.091
mRNA Profile
Protein Expression Profile
DPA

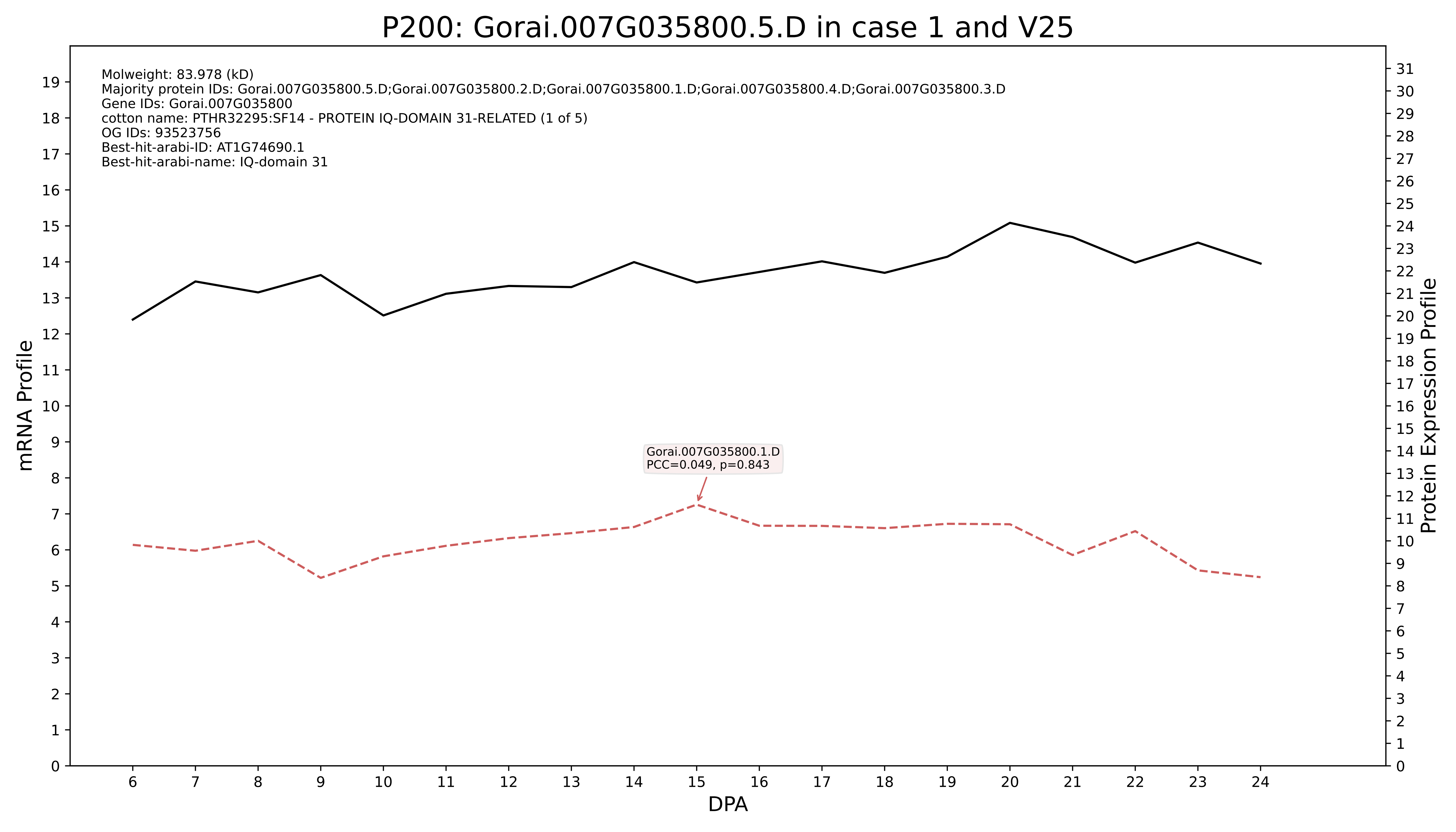
P200: Gorai.007G035800.5.D in case 1 and V25
Molweight: 83.978 (kD)
Majority protein IDs: Gorai.007G035800.5.D;Gorai.007G035800.2.D;Gorai.007G035800.1.D;Gorai.007G035800.4.D;Gorai.007G035800.3.D
Gene IDs: Gorai.007G035800
cotton name: PTHR32295:SF14 - PROTEIN IQ-DOMAIN 31-RELATED (1 of 5)
OG IDs: 93523756
Best-hit-arabi-ID: AT1G74690.1
Best-hit-arabi-name: IQ-domain 31
Gorai.007G035800.1.D
PCC=0.049, p=0.843
mRNA Profile
Protein Expression Profile
DPA

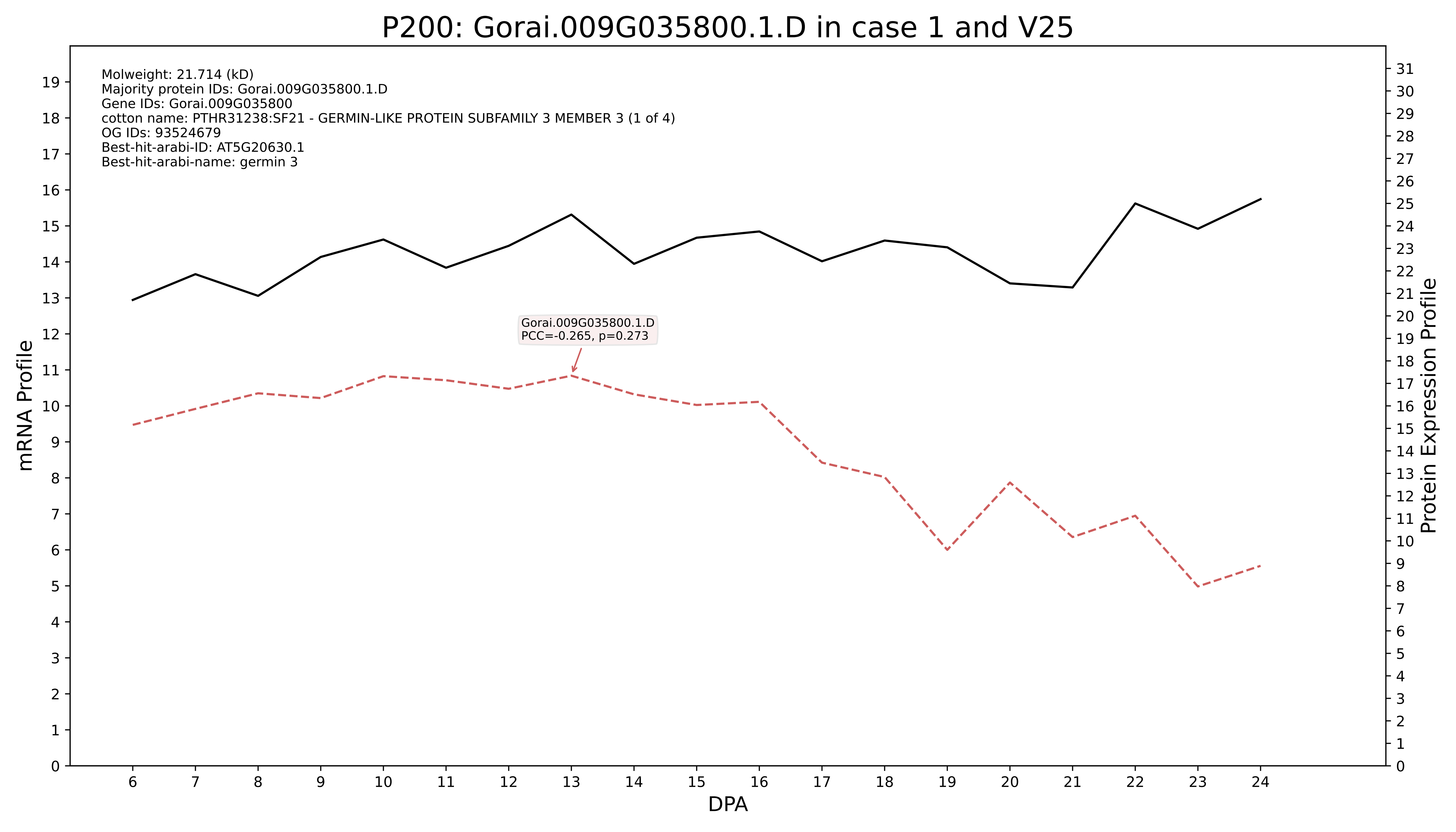
P200: Gorai.009G035800.1.D in case 1 and V25
Molweight: 21.714 (kD)
Majority protein IDs: Gorai.009G035800.1.D
Gene IDs: Gorai.009G035800
cotton name: PTHR31238:SF21 - GERMIN-LIKE PROTEIN SUBFAMILY 3 MEMBER 3 (1 of 4)
OG IDs: 93524679
Best-hit-arabi-ID: AT5G20630.1
Best-hit-arabi-name: germin 3
Gorai.009G035800.1.D
PCC=-0.265, p=0.273
mRNA Profile
Protein Expression Profile
DPA

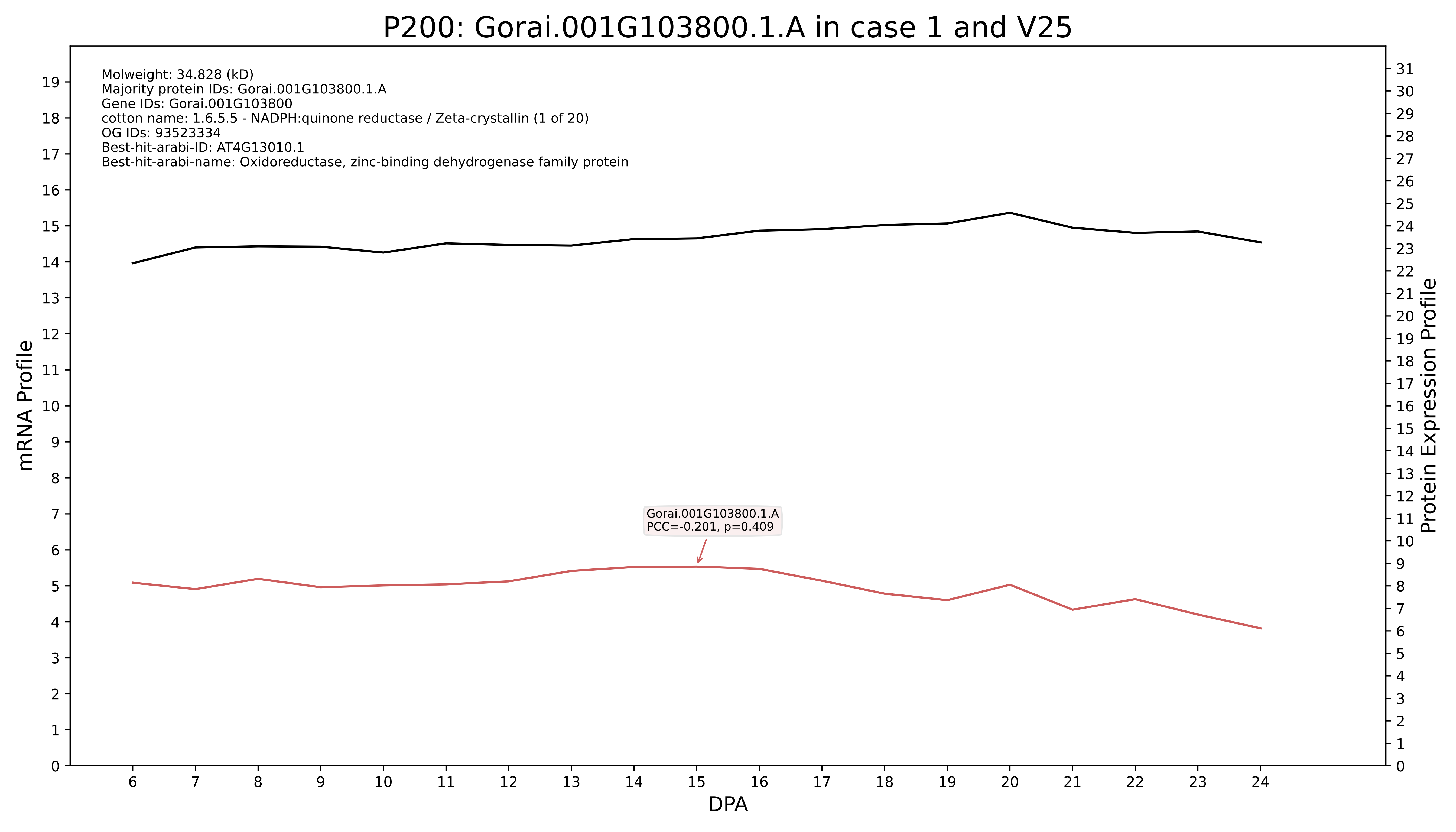

P200: Gorai.001G103800.1.A in case 1 and V25
Molweight: 34.828 (kD)
Majority protein IDs: Gorai.001G103800.1.A
Gene IDs: Gorai.001G103800
cotton name: 1.6.5.5 - NADPH:quinone reductase / Zeta-crystallin (1 of 20)
OG IDs: 93523334
Best-hit-arabi-ID: AT4G13010.1
Best-hit-arabi-name: Oxidoreductase, zinc-binding dehydrogenase family protein
Gorai.001G103800.1.A
PCC=-0.201, p=0.409
mRNA Profile
Protein Expression Profile
DPA

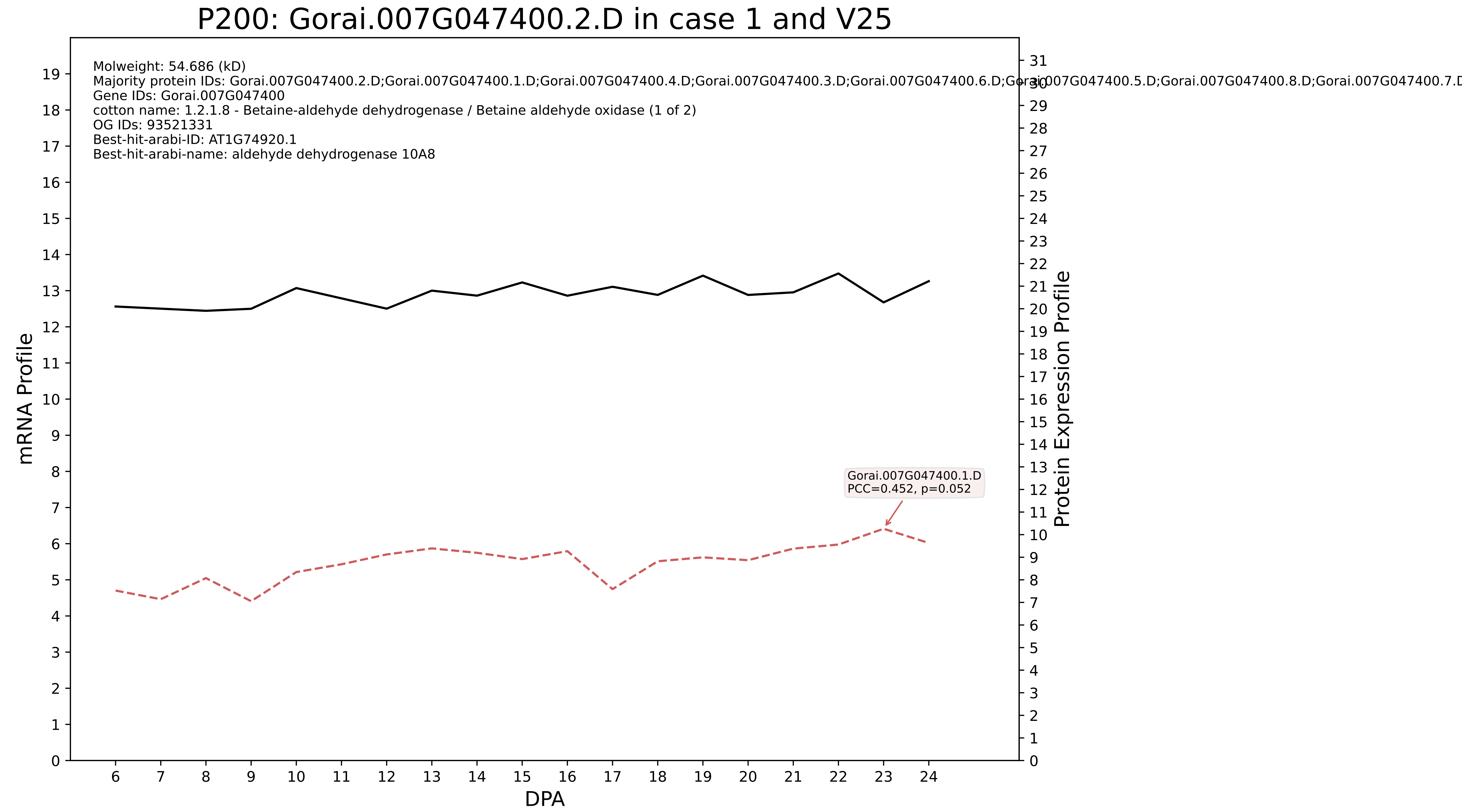

P200: Gorai.007G047400.2.D in case 1 and V25
Molweight: 54.686 (kD)
Majority protein IDs: Gorai.007G047400.2.D;Gorai.007G047400.1.D;Gorai.007G047400.4.D;Gorai.007G047400.3.D;Gorai.007G047400.6.D;Gorai.007G047400.5.D;Gorai.007G047400.8.D;Gorai.007G047400.7.D
Gene IDs: Gorai.007G047400
cotton name: 1.2.1.8 - Betaine-aldehyde dehydrogenase / Betaine aldehyde oxidase (1 of 2)
OG IDs: 93521331
Best-hit-arabi-ID: AT1G74920.1
Best-hit-arabi-name: aldehyde dehydrogenase 10A8
Gorai.007G047400.1.D
PCC=0.452, p=0.052
mRNA Profile
Protein Expression Profile
DPA

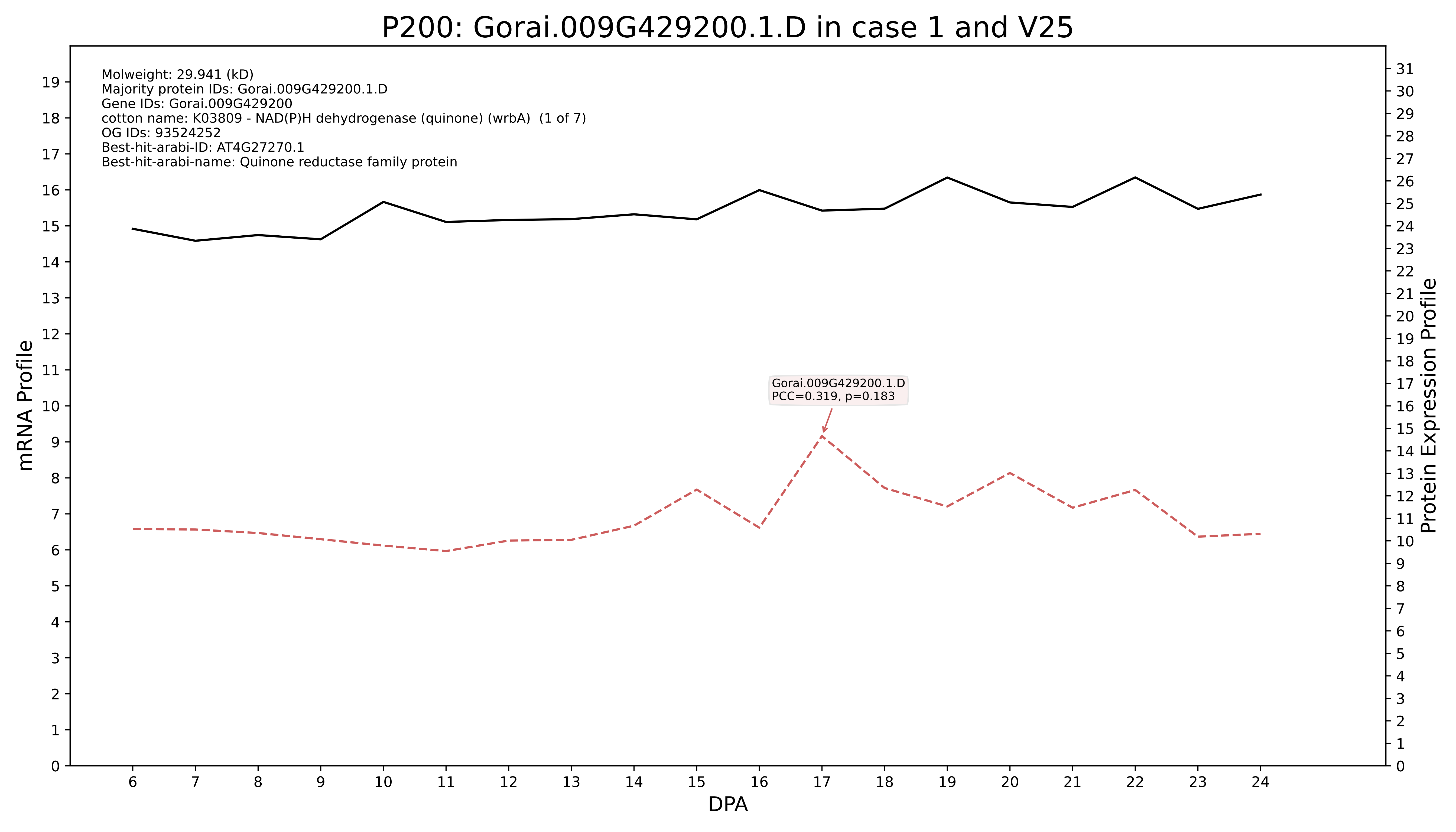
P200: Gorai.009G429200.1.D in case 1 and V25
Molweight: 29.941 (kD)
Majority protein IDs: Gorai.009G429200.1.D
Gene IDs: Gorai.009G429200
cotton name: K03809 - NAD(P)H dehydrogenase (quinone) (wrbA) (1 of 7)
OG IDs: 93524252
Best-hit-arabi-ID: AT4G27270.1
Best-hit-arabi-name: Quinone reductase family protein
Gorai.009G429200.1.D
PCC=0.319, p=0.183
mRNA Profile
Protein Expression Profile
DPA

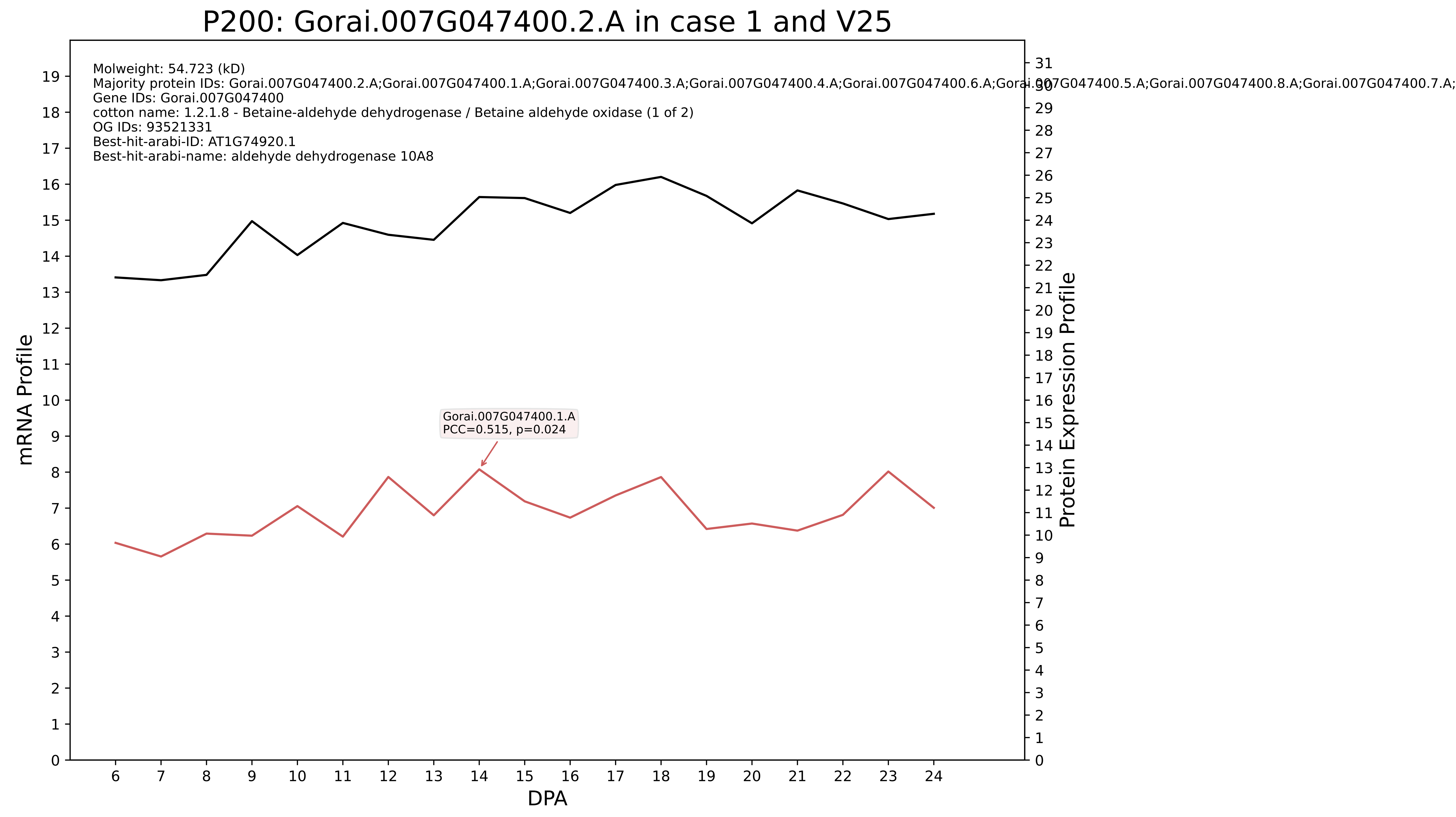
P200: Gorai.007G047400.2.A in case 1 and V25
Molweight: 54.723 (kD)
Majority protein IDs: Gorai.007G047400.2.A;Gorai.007G047400.1.A;Gorai.007G047400.3.A;Gorai.007G047400.4.A;Gorai.007G047400.6.A;Gorai.007G047400.5.A;Gorai.007G047400.8.A;Gorai.007G047400.7.A;
Gene IDs: Gorai.007G047400
cotton name: 1.2.1.8 - Betaine-aldehyde dehydrogenase / Betaine aldehyde oxidase (1 of 2)
OG IDs: 93521331
Best-hit-arabi-ID: AT1G74920.1
Best-hit-arabi-name: aldehyde dehydrogenase 10A8
Gorai.007G047400.1.A
PCC=0.515, p=0.024
mRNA Profile
Protein Expression Profile
DPA

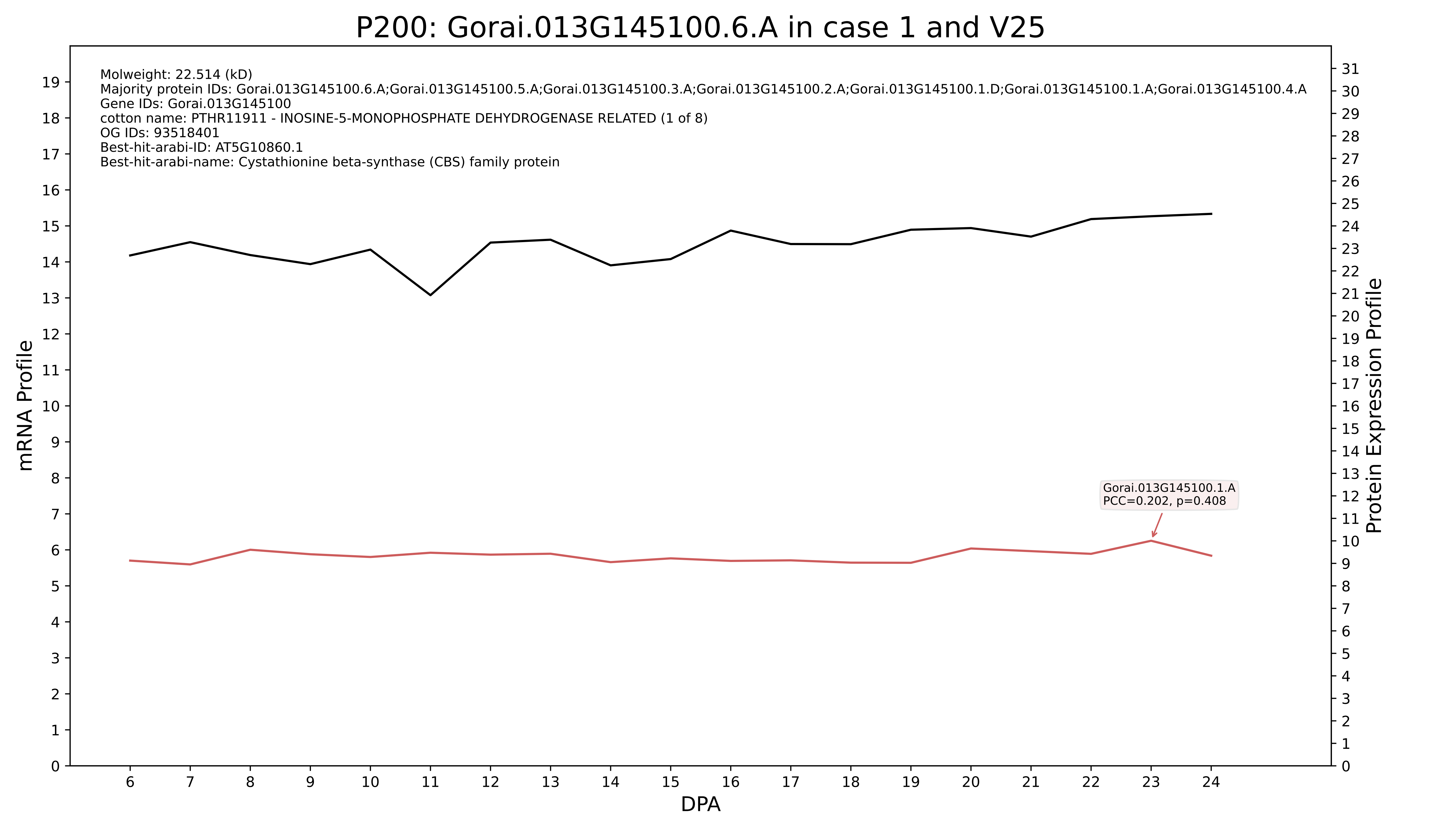
P200: Gorai.013G145100.6.A in case 1 and V25
Molweight: 22.514 (kD)
Majority protein IDs: Gorai.013G145100.6.A;Gorai.013G145100.5.A;Gorai.013G145100.3.A;Gorai.013G145100.2.A;Gorai.013G145100.1.D;Gorai.013G145100.1.A;Gorai.013G145100.4.A
Gene IDs: Gorai.013G145100
cotton name: PTHR11911 - INOSINE-5-MONOPHOSPHATE DEHYDROGENASE RELATED (1 of 8)
OG IDs: 93518401
Best-hit-arabi-ID: AT5G10860.1
Best-hit-arabi-name: Cystathionine beta-synthase (CBS) family protein
Gorai.013G145100.1.A
PCC=0.202, p=0.408
mRNA Profile
Protein Expression Profile
DPA

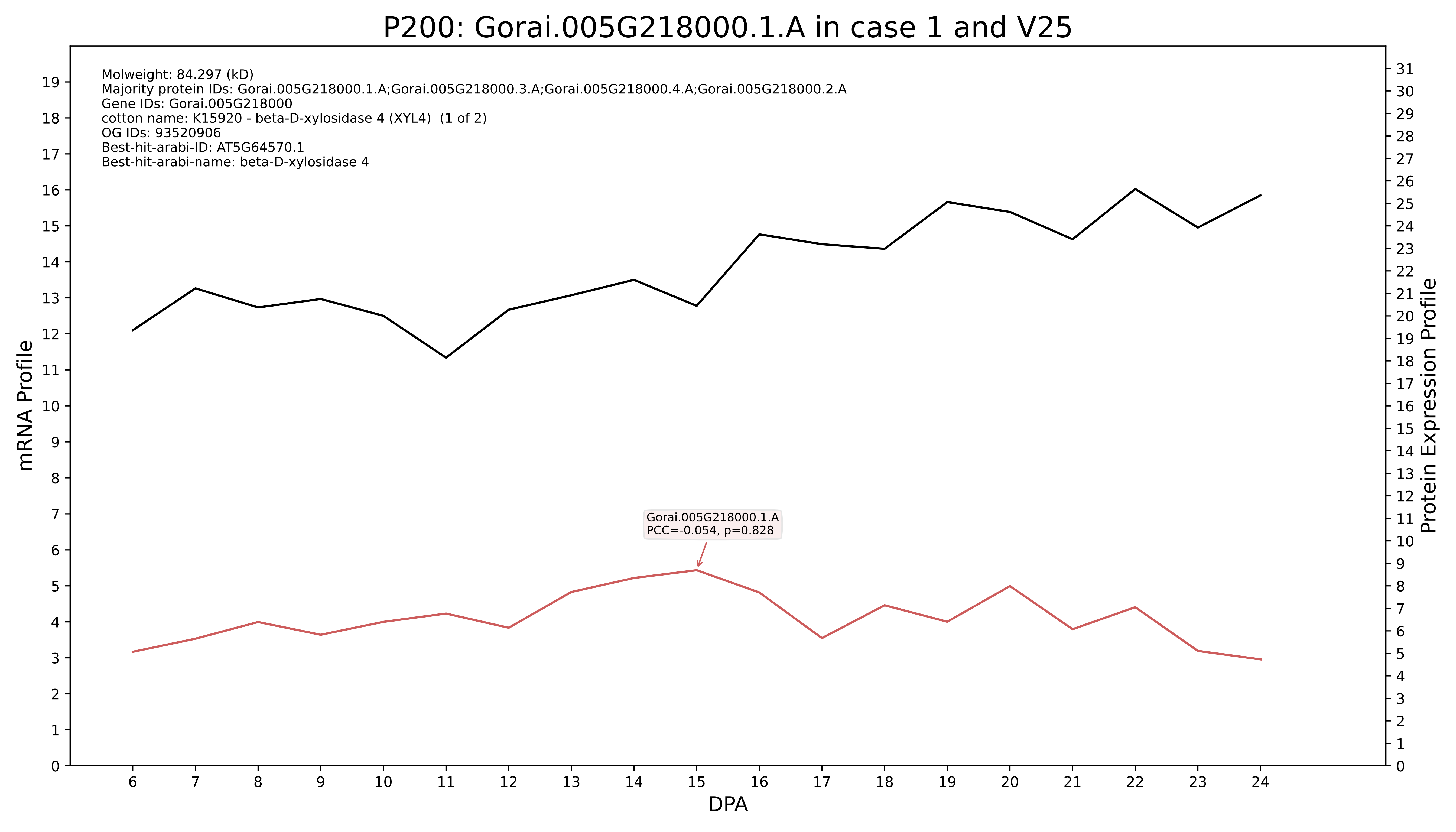
P200: Gorai.005G218000.1.A in case 1 and V25
Molweight: 84.297 (kD)
Majority protein IDs: Gorai.005G218000.1.A;Gorai.005G218000.3.A;Gorai.005G218000.4.A;Gorai.005G218000.2.A
Gene IDs: Gorai.005G218000
cotton name: K15920 - beta-D-xylosidase 4 (XYL4) (1 of 2)
OG IDs: 93520906
Best-hit-arabi-ID: AT5G64570.1
Best-hit-arabi-name: beta-D-xylosidase 4
Gorai.005G218000.1.A
PCC=-0.054, p=0.828
mRNA Profile
Protein Expression Profile
DPA

### P200: Gorai.006G229100.1.D in case 1 and V25

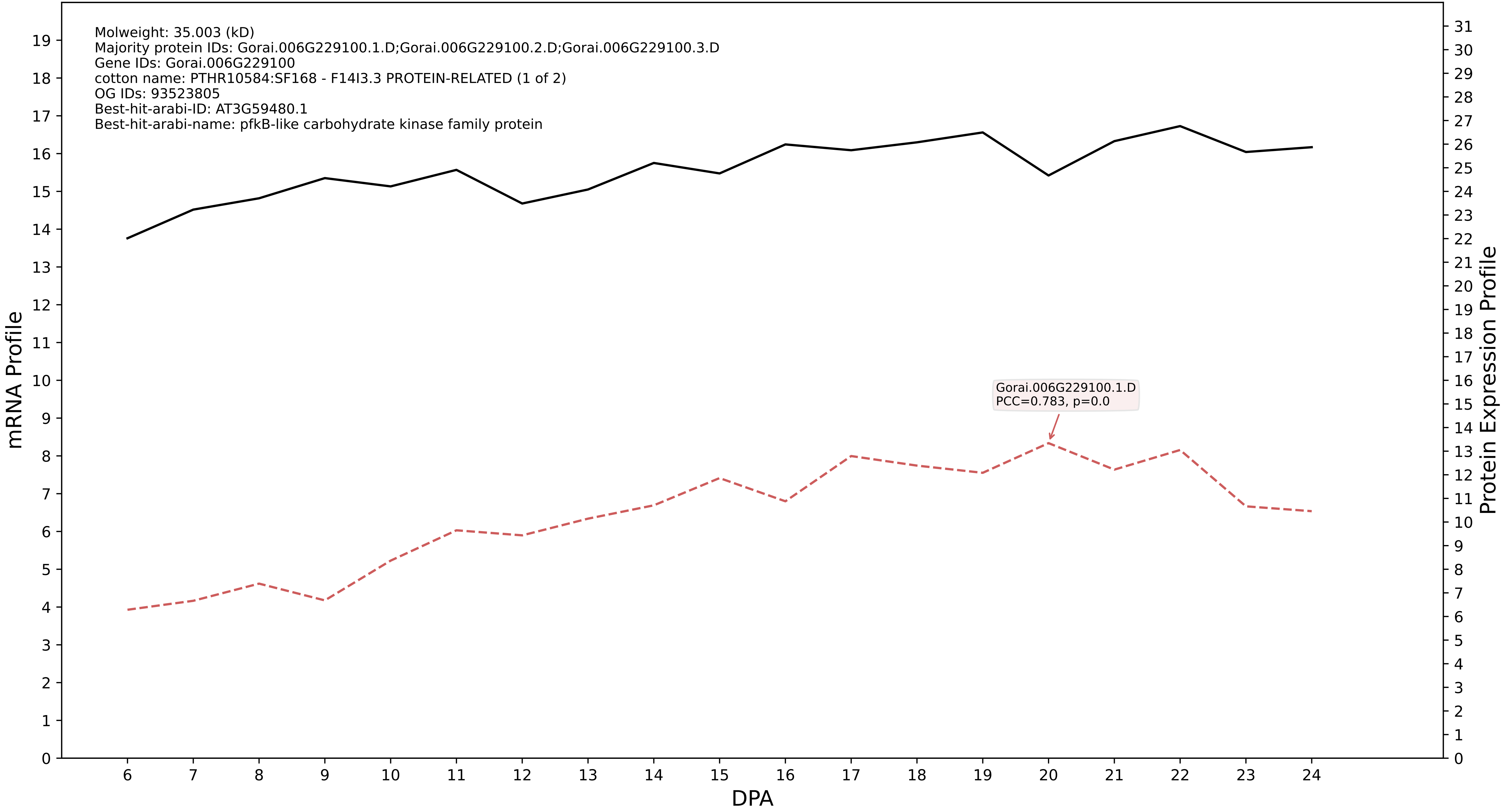

### P200: Gorai.004G245500.7.D in case 1 and V25

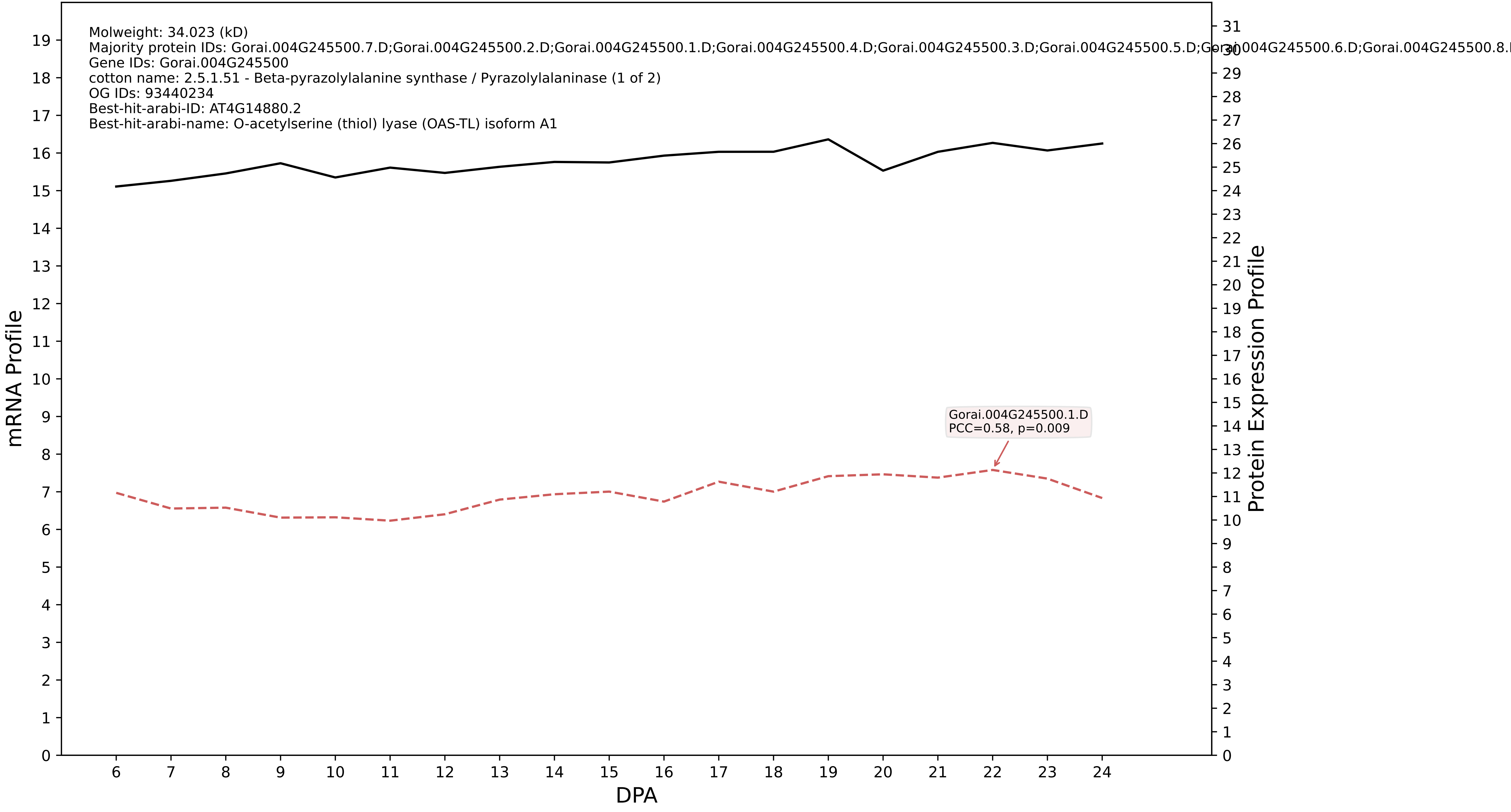

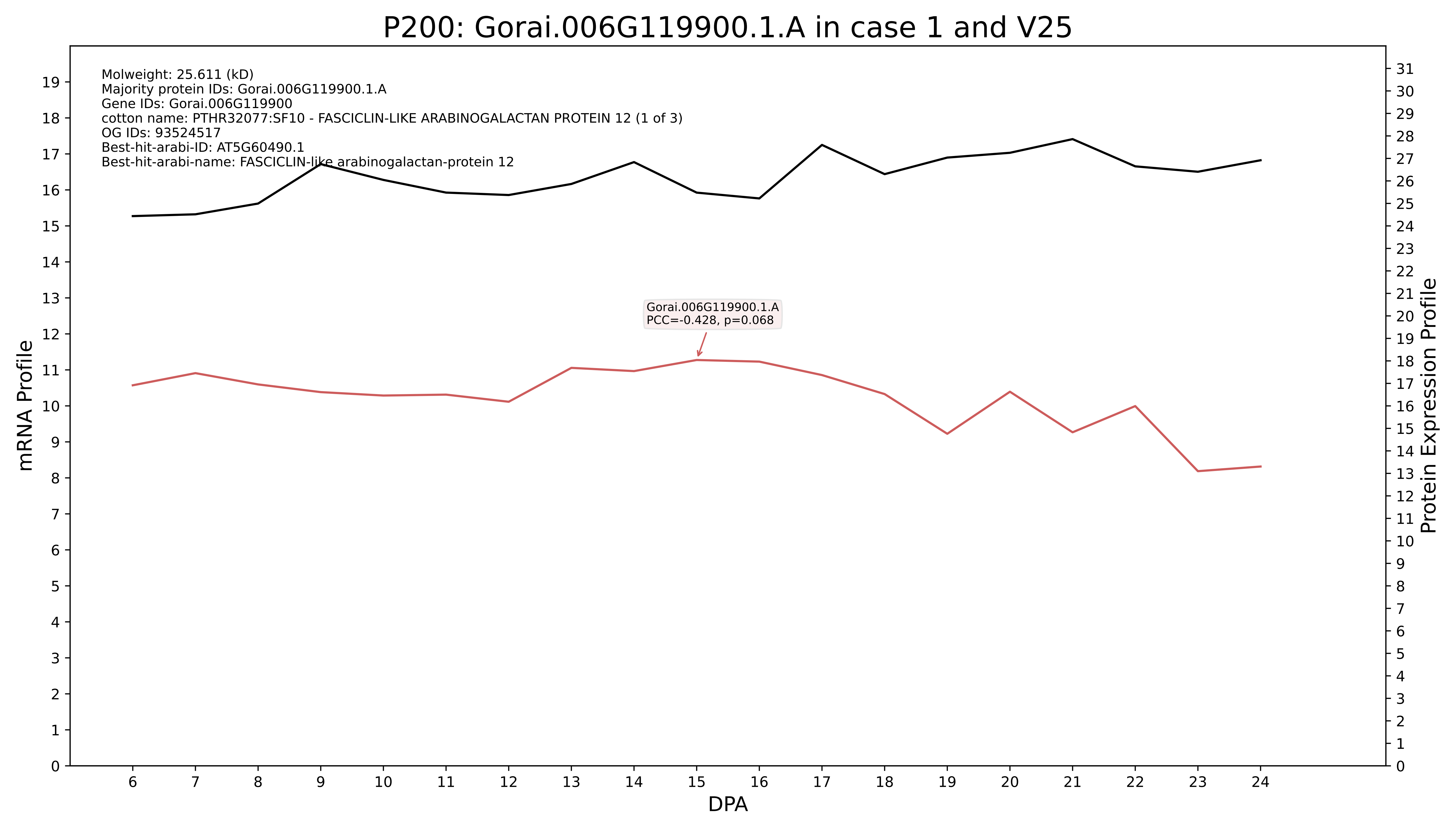

P200: Gorai.006G119900.1.A in case 1 and V25
Molweight: 25.611 (kD)
Majority protein IDs: Gorai.006G119900.1.A
Gene IDs: Gorai.006G119900
cotton name: PTHR32077:SF10 - FASCICLIN-LIKE ARABINOGALACTAN PROTEIN 12 (1 of 3)
OG IDs: 93524517
Best-hit-arabi-ID: AT5G60490.1
Best-hit-arabi-name: FASCICLIN-like arabinogalactan-protein 12
Gorai.006G119900.1.A
PCC=-0.428, p=0.068
mRNA Profile
Protein Expression Profile
DPA

### P200: Gorai.005G132500.1.A in case 1 and V25

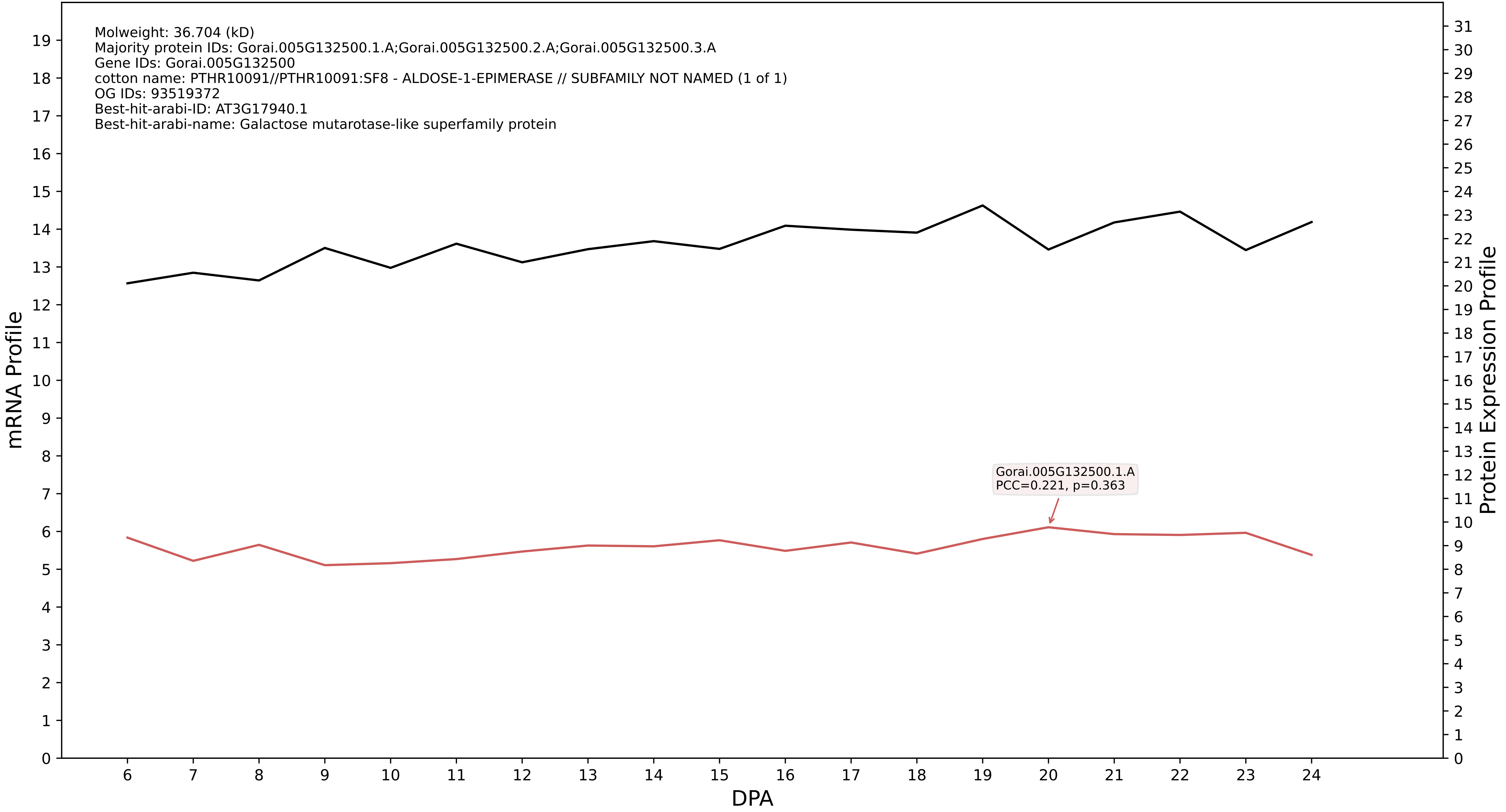

### P200: Gorai.004G215500.2.A in case 1 and V25

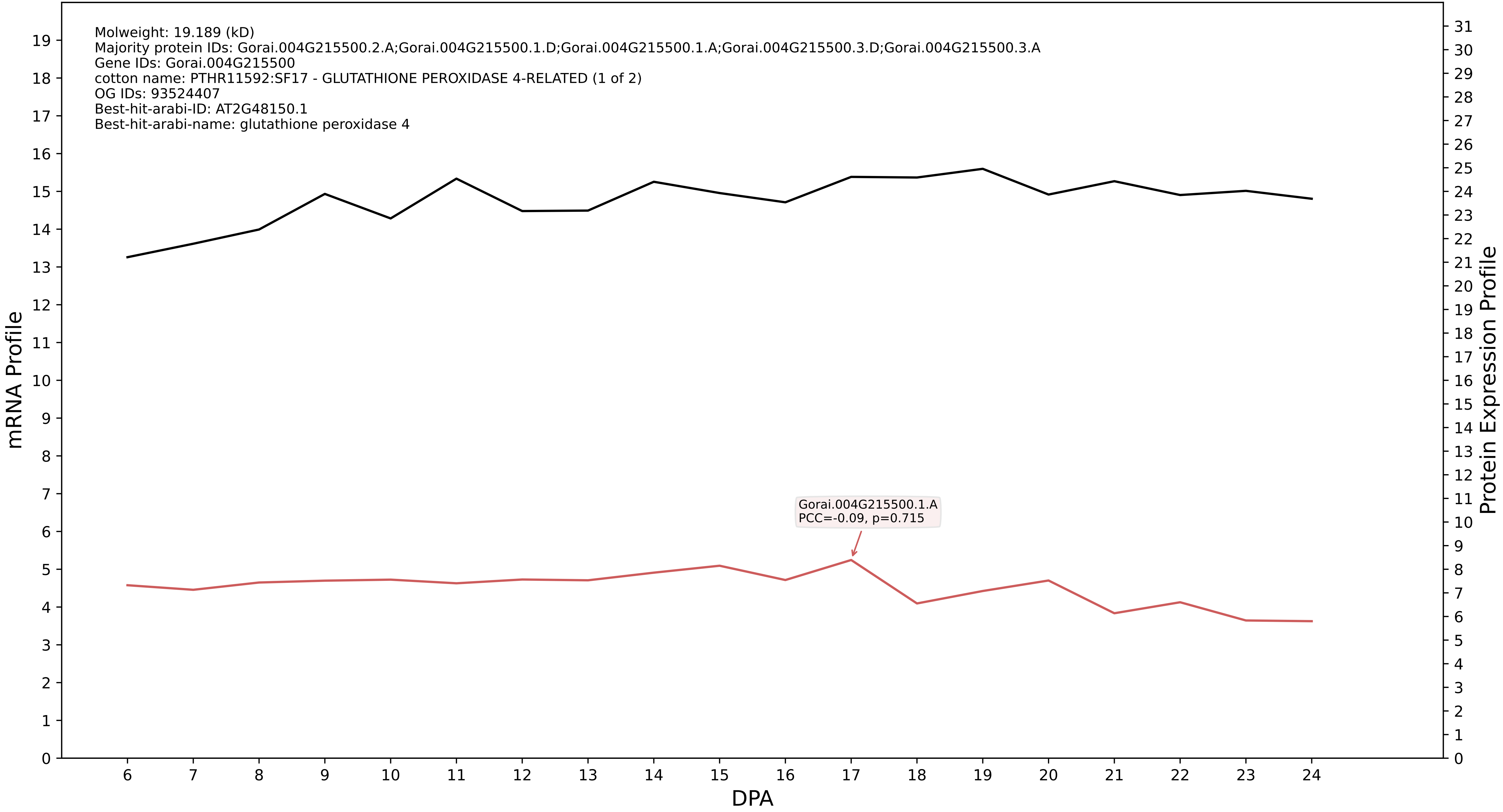

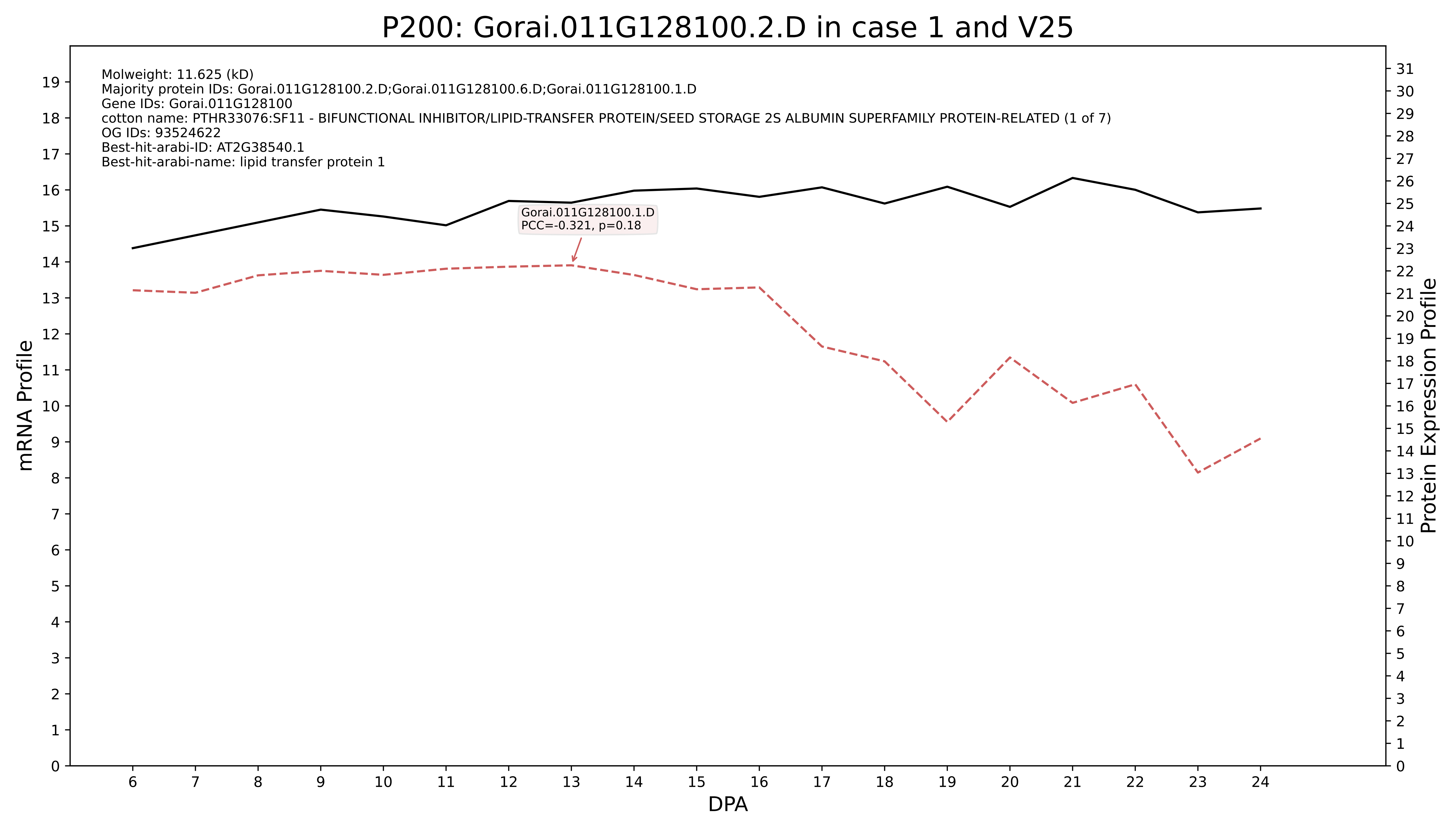

P200: Gorai.011G128100.2.D in case 1 and V25
Molweight: 11.625 (kD)
Majority protein IDs: Gorai.011G128100.2.D;Gorai.011G128100.6.D;Gorai.011G128100.1.D
Gene IDs: Gorai.011G128100
cotton name: PTHR33076:SF11 - BIFUNCTIONAL INHIBITOR/LIPID-TRANSFER PROTEIN/SEED STORAGE 2S ALBUMIN SUPERFAMILY PROTEIN-RELATED (1 of 7)
OG IDs: 93524622
Best-hit-arabi-ID: AT2G38540.1
Best-hit-arabi-name: lipid transfer protein 1
Gorai.011G128100.1.D
PCC=-0.321, p=0.18
mRNA Profile
Protein Expression Profile
DPA

### P200: Gorai.012G147000.2.D in case 1 and V25

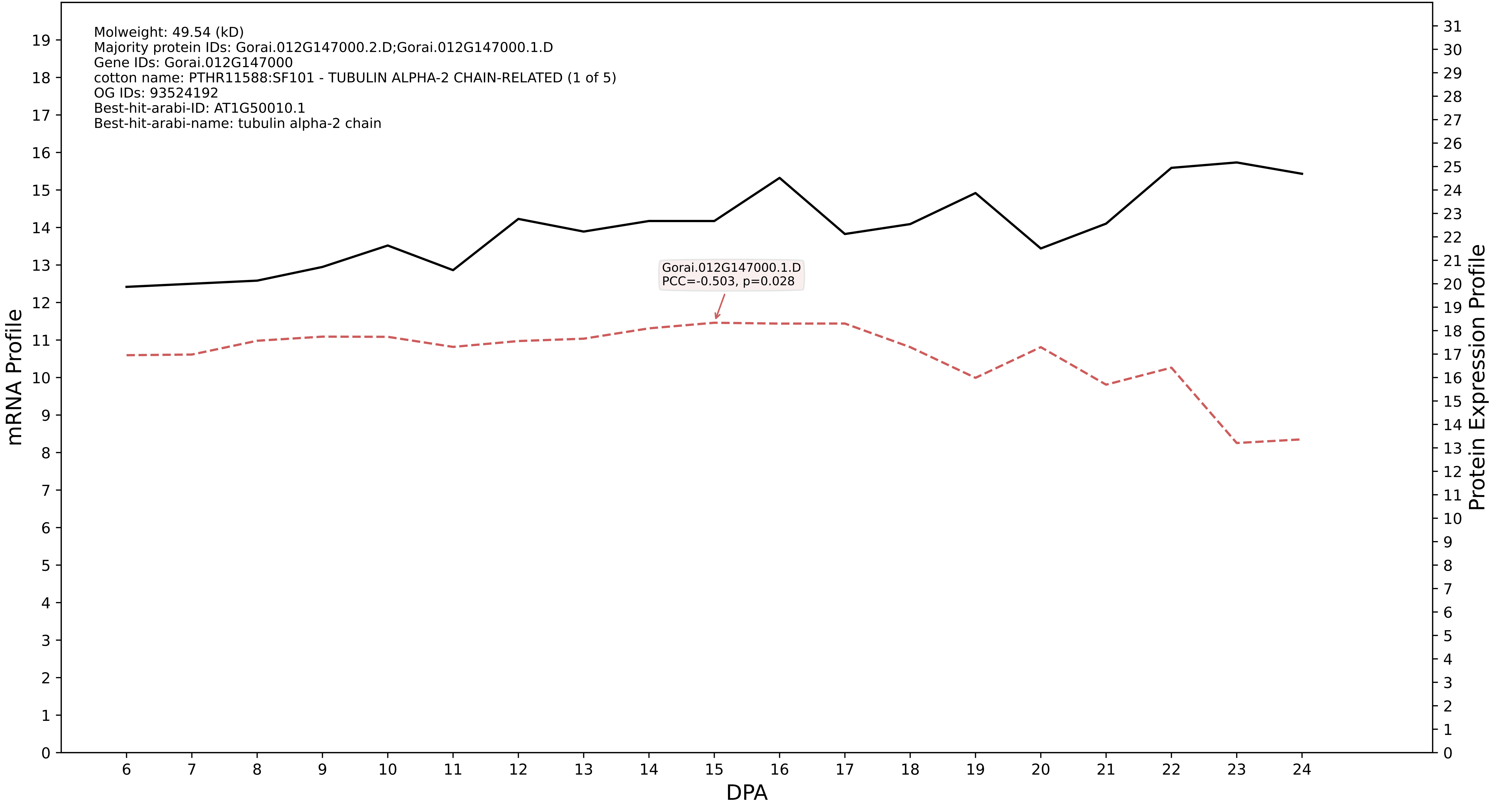

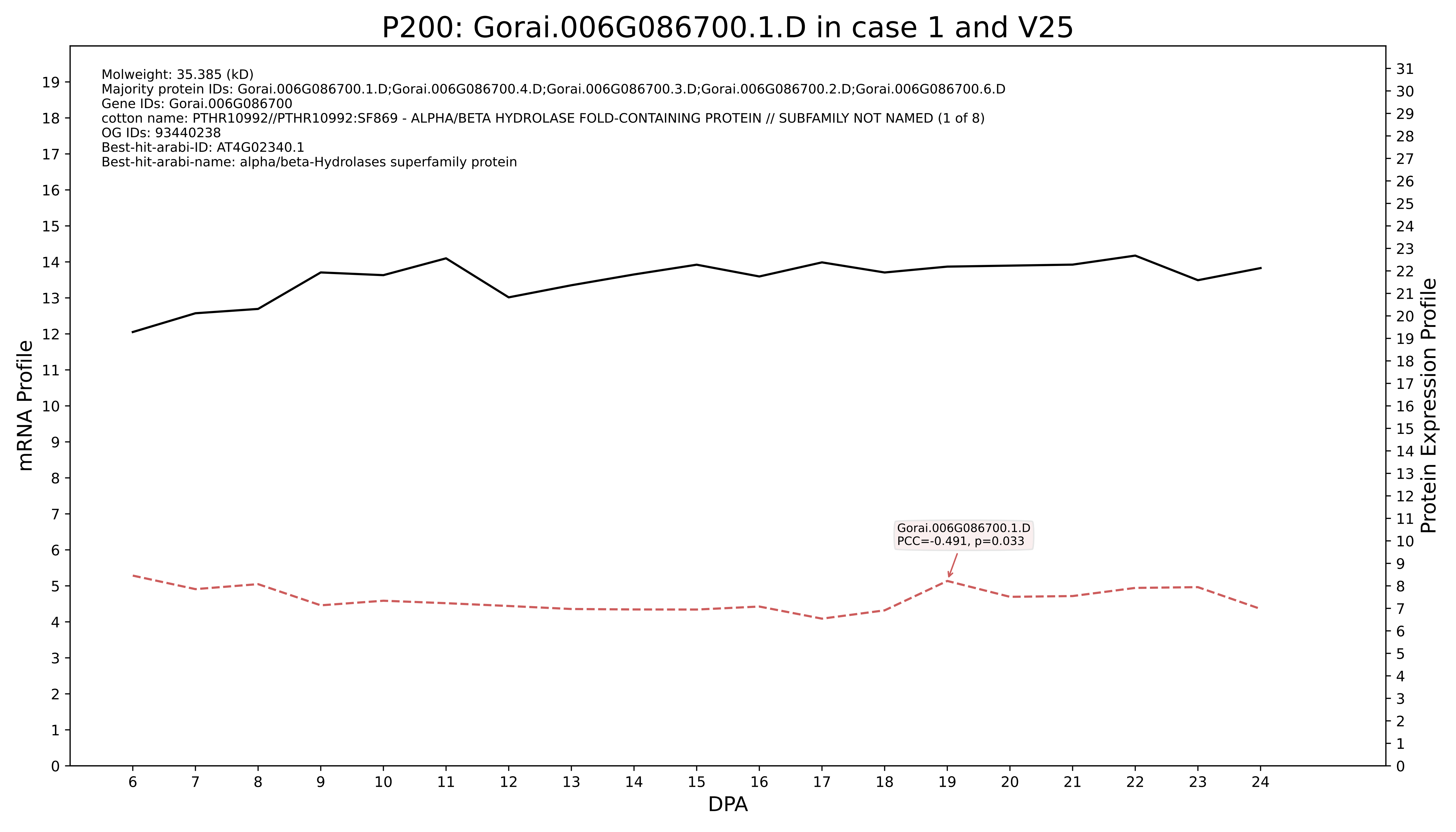

P200: Gorai.006G086700.1.D in case 1 and V25
Molweight: 35.385 (kD)
Majority protein IDs: Gorai.006G086700.1.D;Gorai.006G086700.4.D;Gorai.006G086700.3.D;Gorai.006G086700.2.D;Gorai.006G086700.6.D
Gene IDs: Gorai.006G086700
cotton name: PTHR10992//PTHR10992:SF869 - ALPHA/BETA HYDROLASE FOLD-CONTAINING PROTEIN // SUBFAMILY NOT NAMED (1 of 8)
OG IDs: 93440238
Best-hit-arabi-ID: AT4G02340.1
Best-hit-arabi-name: alpha/beta-Hydrolases superfamily protein
Gorai.006G086700.1.D
PCC=-0.491, p=0.033
mRNA Profile
Protein Expression Profile
DPA

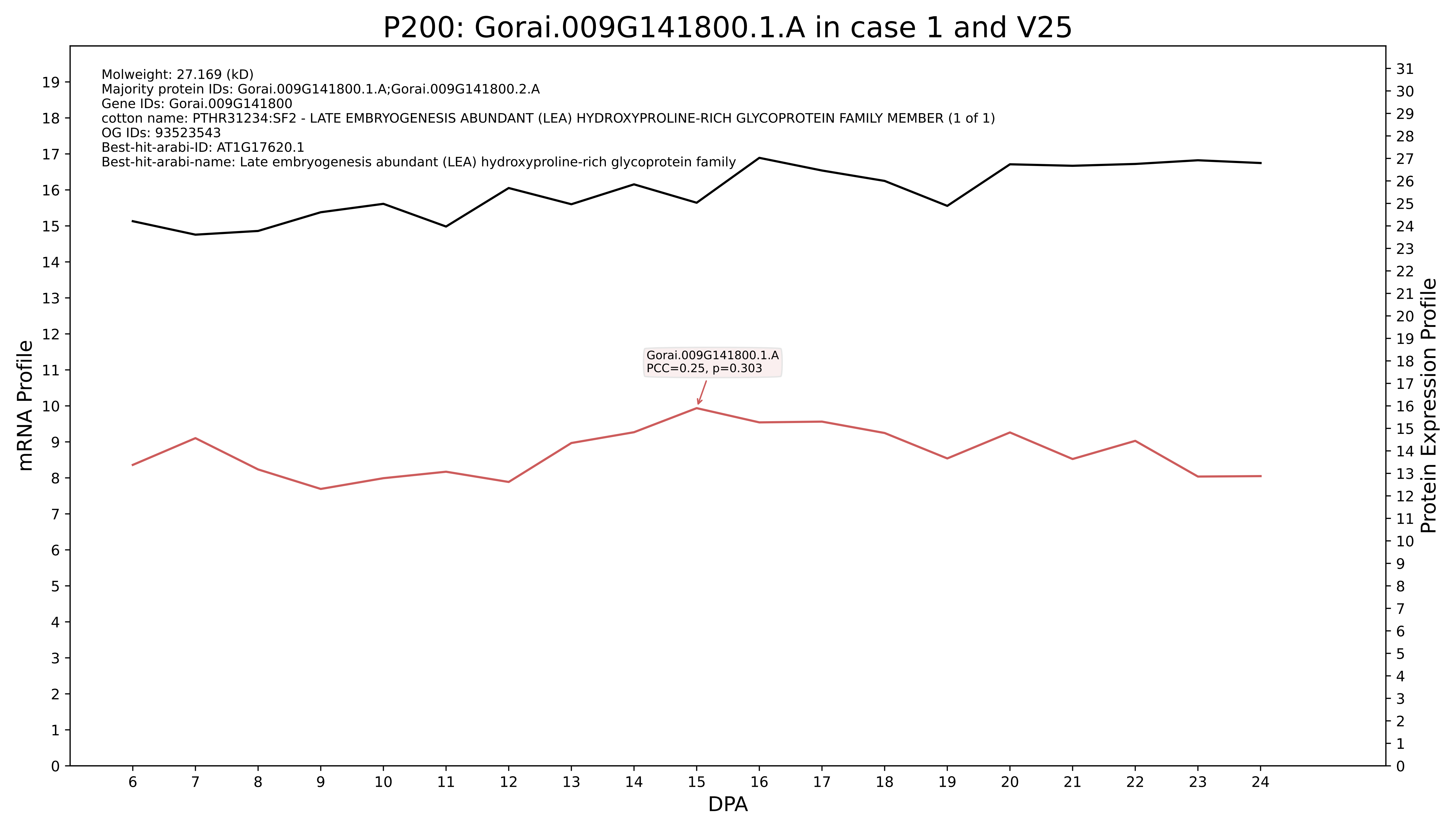

P200: Gorai.009G141800.1.A in case 1 and V25
Molweight: 27.169 (kD)
Majority protein IDs: Gorai.009G141800.1.A;Gorai.009G141800.2.A
Gene IDs: Gorai.009G141800
cotton name: PTHR31234:SF2 - LATE EMBRYOGENESIS ABUNDANT (LEA) HYDROXYPROLINE-RICH GLYCOPROTEIN FAMILY MEMBER (1 of 1)
OG IDs: 93523543
Best-hit-arabi-ID: AT1G17620.1
Best-hit-arabi-name: Late embryogenesis abundant (LEA) hydroxyproline-rich glycoprotein family
Gorai.009G141800.1.A
PCC=0.25, p=0.303
mRNA Profile
Protein Expression Profile
DPA

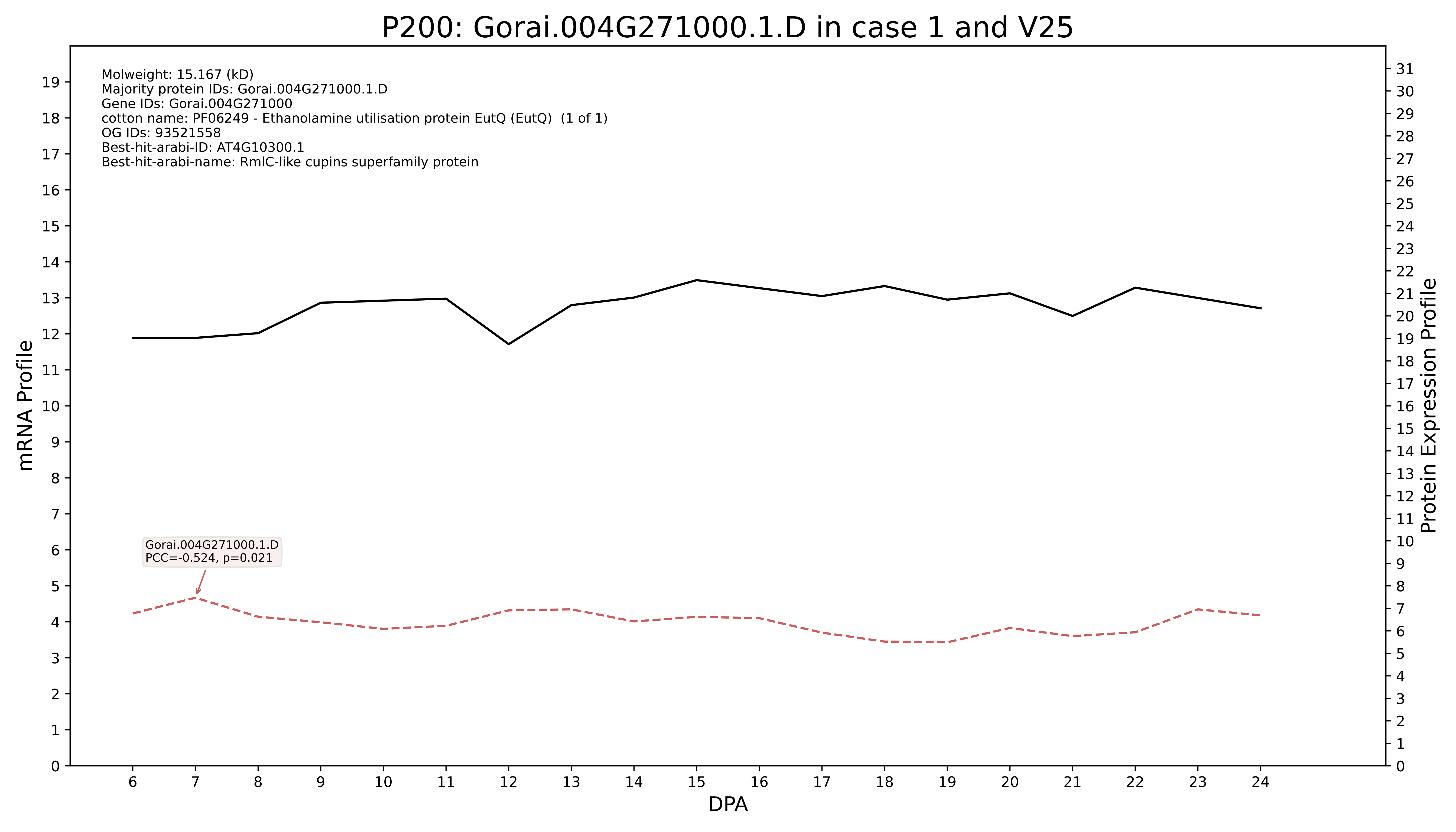

P200: Gorai.004G271000.1.D in case 1 and V25
Molweight: 15.167 (kD)
Majority protein IDs: Gorai.004G271000.1.D
Gene IDs: Gorai.004G271000
cotton name: PF06249 - Ethanolamine utilisation protein EutQ (EutQ) (1 of 1)
OG IDs: 93521558
Best-hit-arabi-ID: AT4G10300.1
Best-hit-arabi-name: RmlC-like cupins superfamily protein
Gorai.004G271000.1.D
PCC=-0.524, p=0.021
mRNA Profile
Protein Expression Profile
DPA

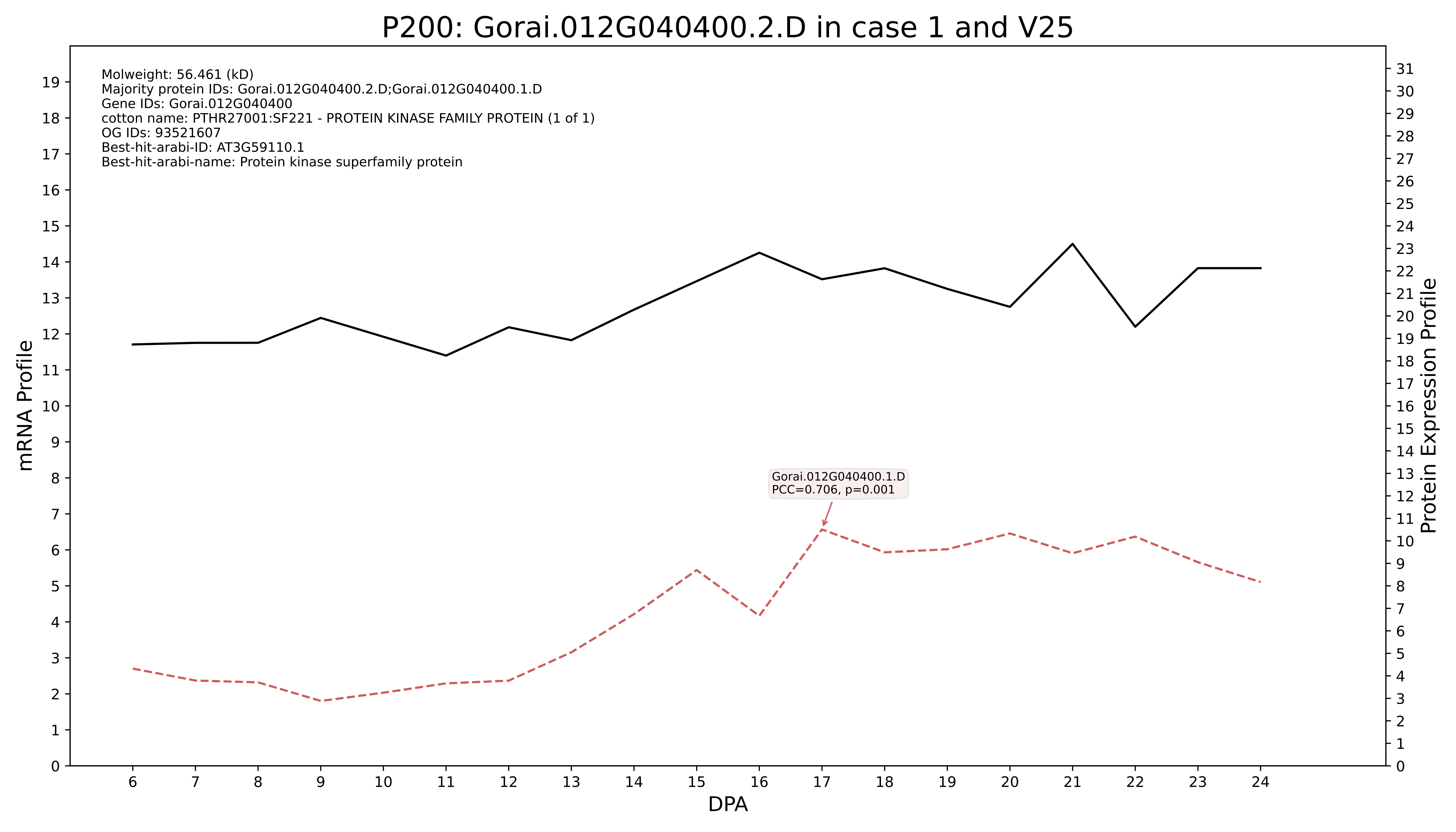
P200: Gorai.012G040400.2.D in case 1 and V25
Molweight: 56.461 (kD)
Majority protein IDs: Gorai.012G040400.2.D;Gorai.012G040400.1.D
Gene IDs: Gorai.012G040400
cotton name: PTHR27001:SF221 - PROTEIN KINASE FAMILY PROTEIN (1 of 1)
OG IDs: 93521607
Best-hit-arabi-ID: AT3G59110.1
Best-hit-arabi-name: Protein kinase superfamily protein
Gorai.012G040400.1.D
PCC=0.706, p=0.001
mRNA Profile
Protein Expression Profile
DPA

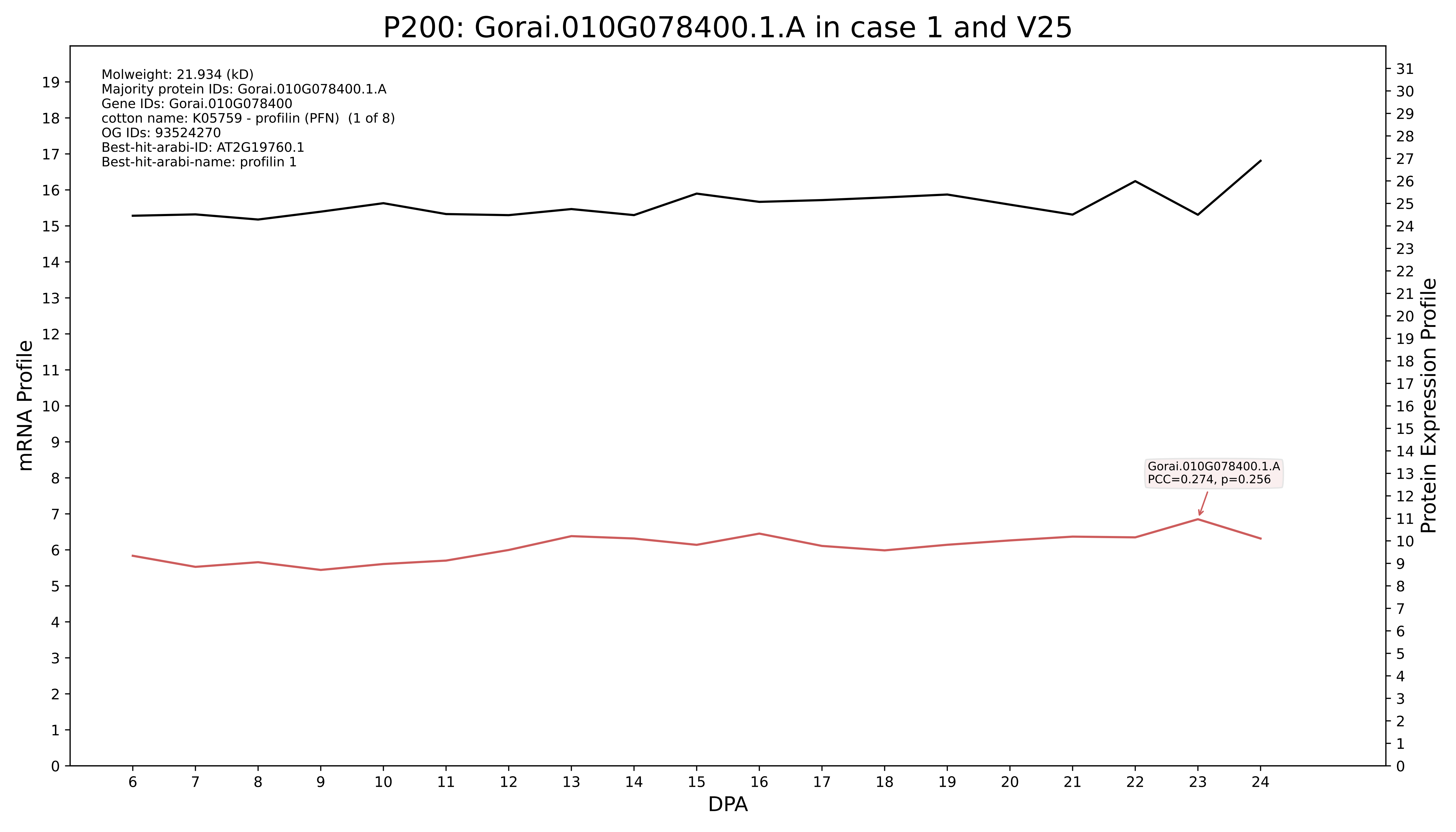
P200: Gorai.010G078400.1.A in case 1 and V25
Molweight: 21.934 (kD)
Majority protein IDs: Gorai.010G078400.1.A
Gene IDs: Gorai.010G078400
cotton name: K05759 - profilin (PFN) (1 of 8)
OG IDs: 93524270
Best-hit-arabi-ID: AT2G19760.1
Best-hit-arabi-name: profilin 1
Gorai.010G078400.1.A
PCC=0.274, p=0.256
mRNA Profile
Protein Expression Profile
DPA

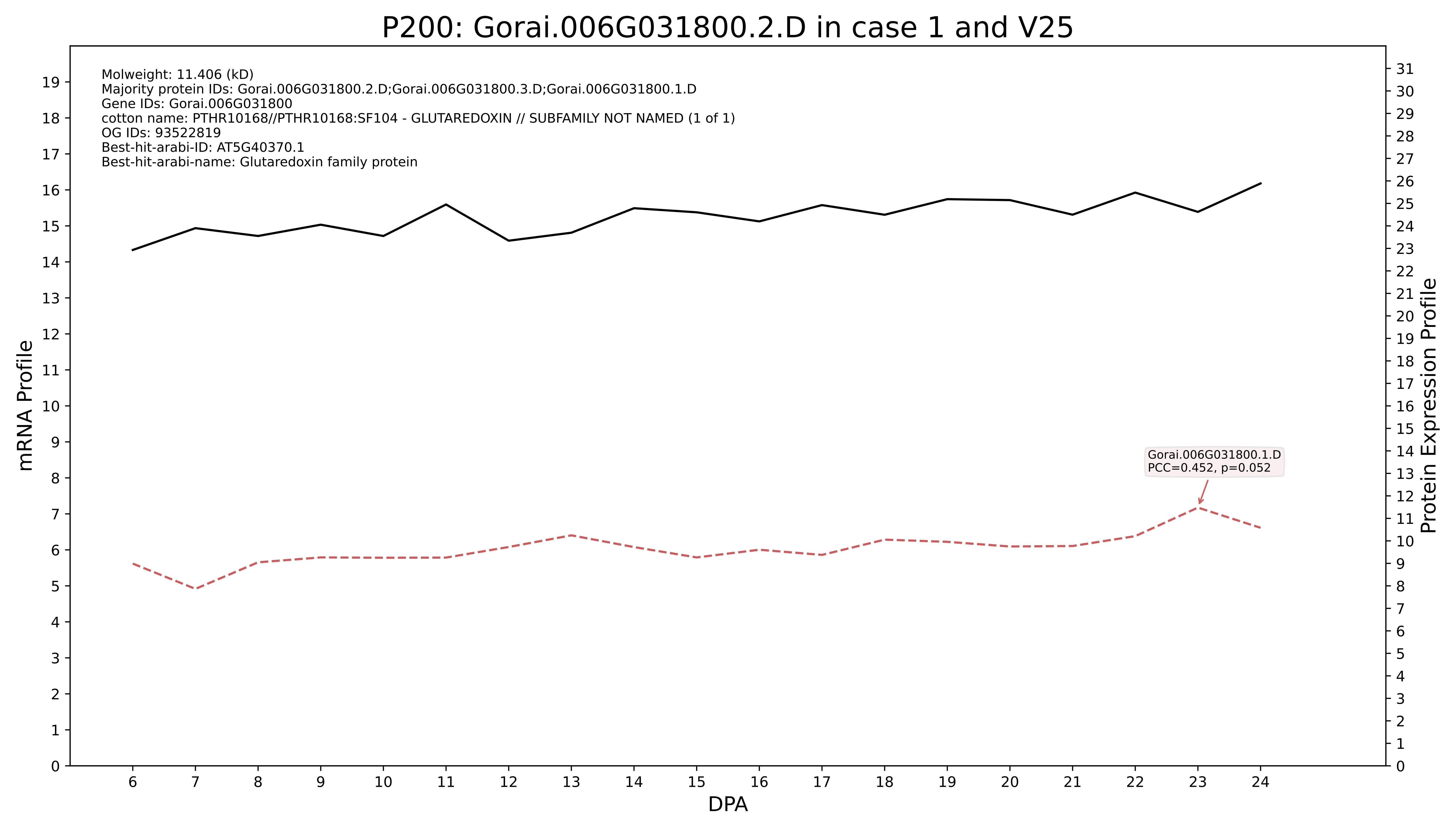
P200: Gorai.006G031800.2.D in case 1 and V25
Molweight: 11.406 (kD)
Majority protein IDs: Gorai.006G031800.2.D;Gorai.006G031800.3.D;Gorai.006G031800.1.D
Gene IDs: Gorai.006G031800
cotton name: PTHR10168//PTHR10168:SF104 - GLUTAREDOXIN // SUBFAMILY NOT NAMED (1 of 1)
OG IDs: 93522819
Best-hit-arabi-ID: AT5G40370.1
Best-hit-arabi-name: Glutaredoxin family protein
Gorai.006G031800.1.D
PCC=0.452, p=0.052
mRNA Profile
Protein Expression Profile
DPA

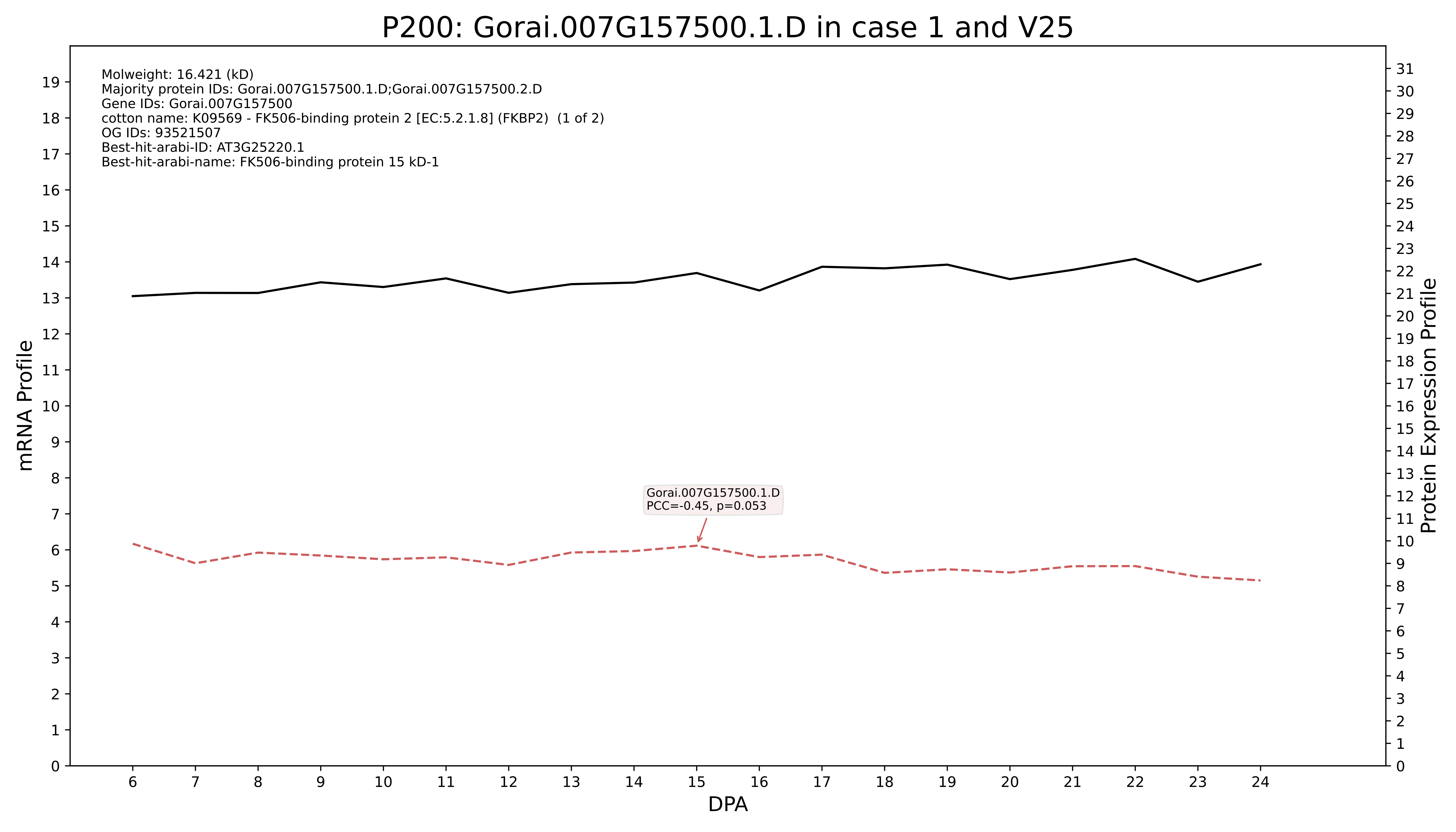
P200: Gorai.007G157500.1.D in case 1 and V25
Molweight: 16.421 (kD)
Majority protein IDs: Gorai.007G157500.1.D;Gorai.007G157500.2.D
Gene IDs: Gorai.007G157500
cotton name: K09569 - FK506-binding protein 2 [EC:5.2.1.8] (FKBP2) (1 of 2)
OG IDs: 93521507
Best-hit-arabi-ID: AT3G25220.1
Best-hit-arabi-name: FK506-binding protein 15 kD-1
Gorai.007G157500.1.D
PCC=-0.45, p=0.053
mRNA Profile
Protein Expression Profile
DPA

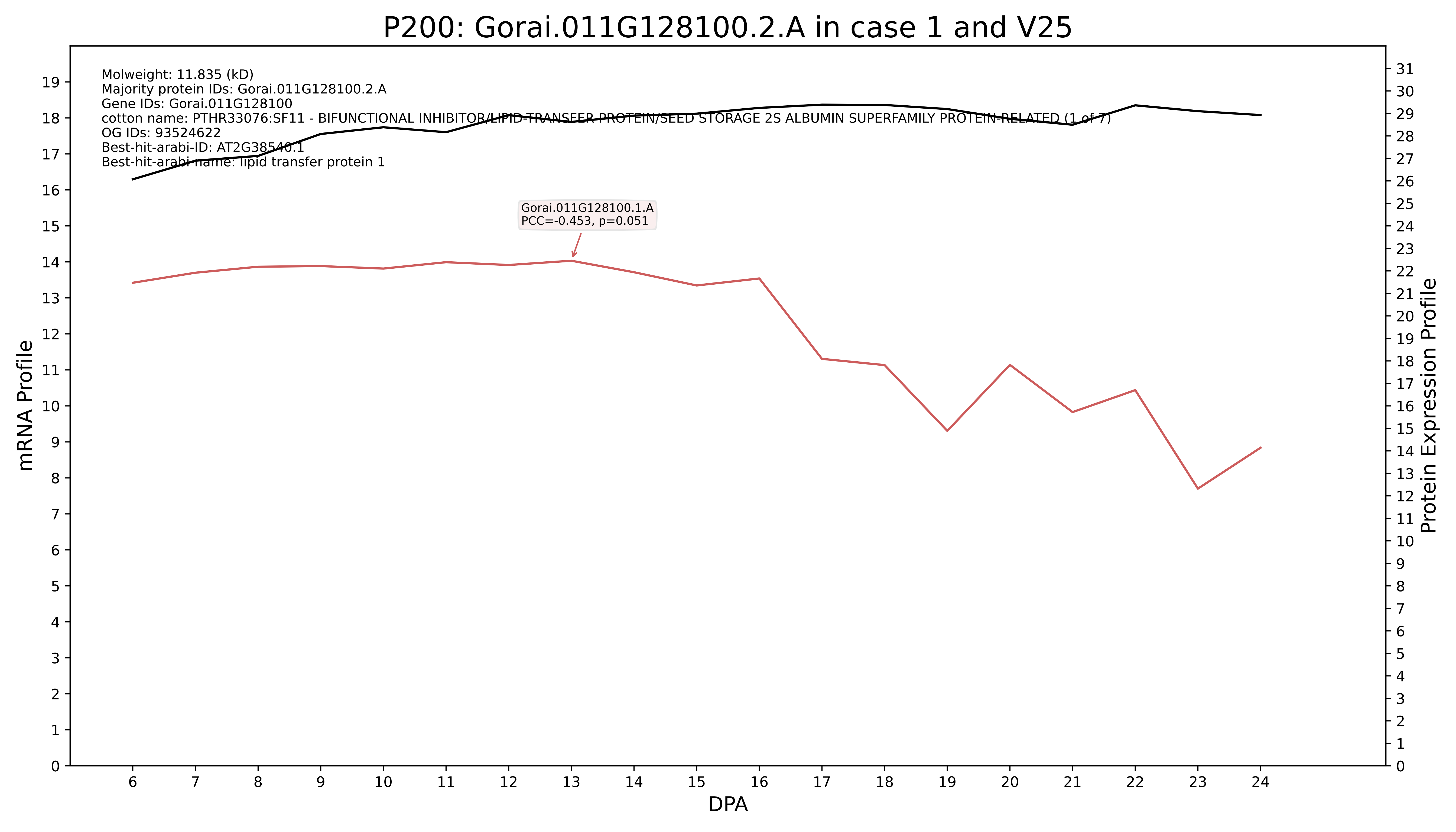
P200: Gorai.011G128100.2.A in case 1 and V25
Molweight: 11.835 (kD)
Majority protein IDs: Gorai.011G128100.2.A
Gene IDs: Gorai.011G128100
cotton name: PTHR33076:SF11 - BIFUNCTIONAL INHIBITOR/LIPID-TRANSFER PROTEIN/SEED STORAGE 2S ALBUMIN SUPERFAMILY PROTEIN-RELATED (1 of 7)
OG IDs: 93524622
Best-hit-arabi-ID: AT2G38540.1
Best-hit-arabi-name: lipid transfer protein 1
Gorai.011G128100.1.A
PCC=-0.453, p=0.051
mRNA Profile
Protein Expression Profile
DPA

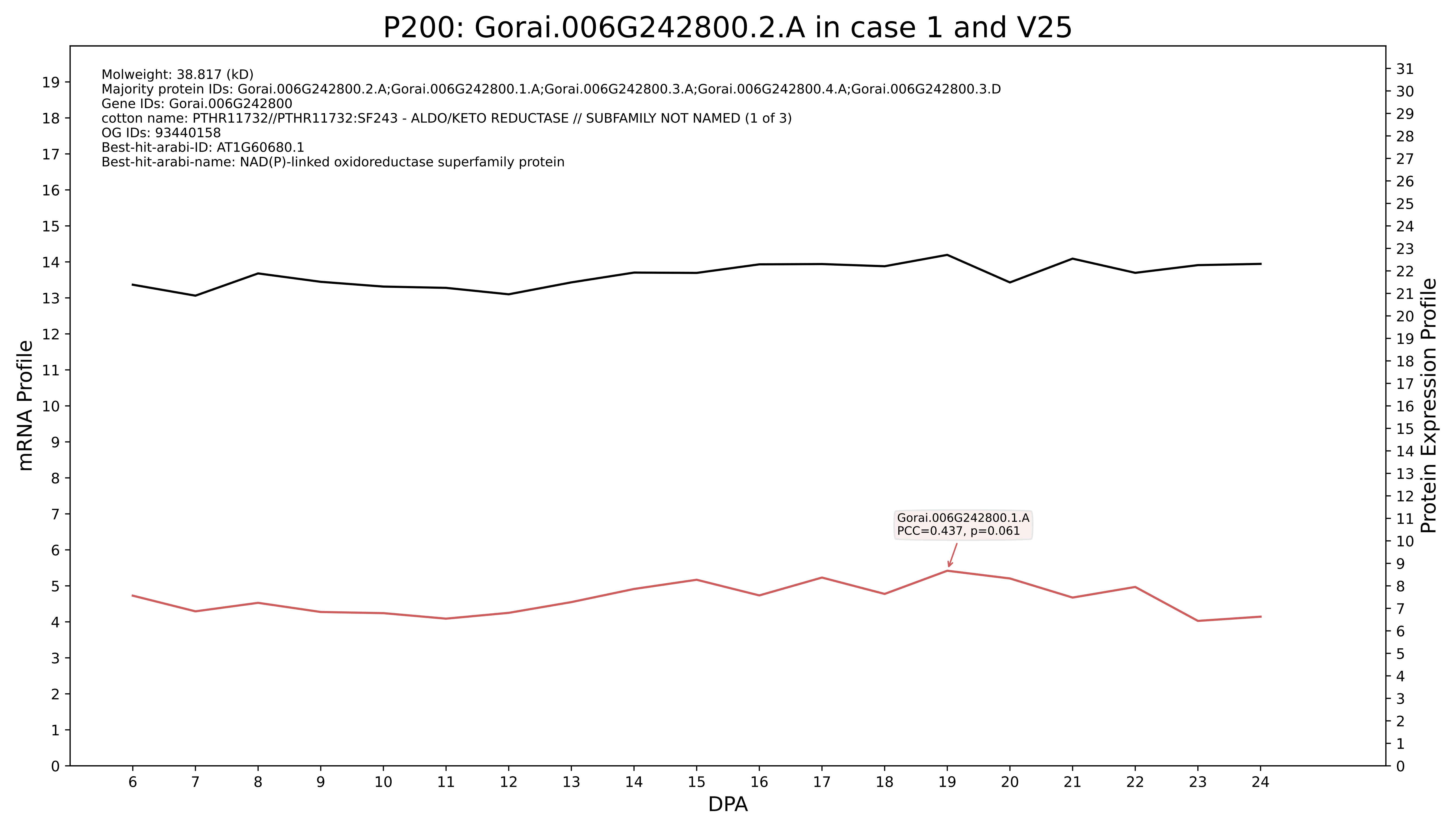
P200: Gorai.006G242800.2.A in case 1 and V25
Molweight: 38.817 (kD)
Majority protein IDs: Gorai.006G242800.2.A;Gorai.006G242800.1.A;Gorai.006G242800.3.A;Gorai.006G242800.4.A;Gorai.006G242800.3.D
Gene IDs: Gorai.006G242800
cotton name: PTHR11732//PTHR11732:SF243 - ALDO/KETO REDUCTASE // SUBFAMILY NOT NAMED (1 of 3)
OG IDs: 93440158
Best-hit-arabi-ID: AT1G60680.1
Best-hit-arabi-name: NAD(P)-linked oxidoreductase superfamily protein
Gorai.006G242800.1.A
PCC=0.437, p=0.061
mRNA Profile
Protein Expression Profile
DPA

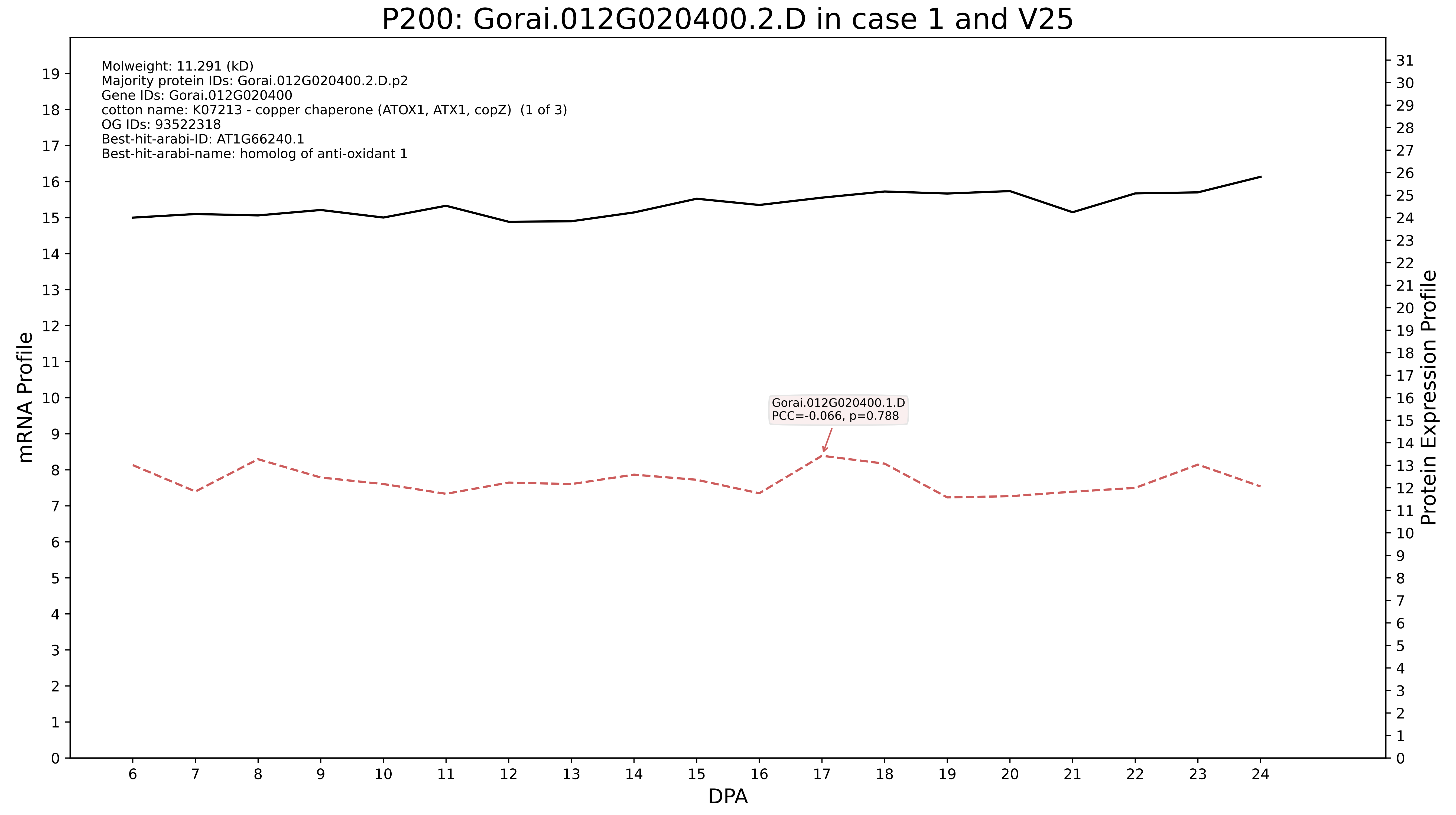
P200: Gorai.012G020400.2.D in case 1 and V25
Molweight: 11.291 (kD)
Majority protein IDs: Gorai.012G020400.2.D.p2
Gene IDs: Gorai.012G020400
cotton name: K07213 - copper chaperone (ATOX1, ATX1, copZ) (1 of 3)
OG IDs: 93522318
Best-hit-arabi-ID: AT1G66240.1
Best-hit-arabi-name: homolog of anti-oxidant 1
Gorai.012G020400.1.D
PCC=-0.066, p=0.788
mRNA Profile
Protein Expression Profile
DPA

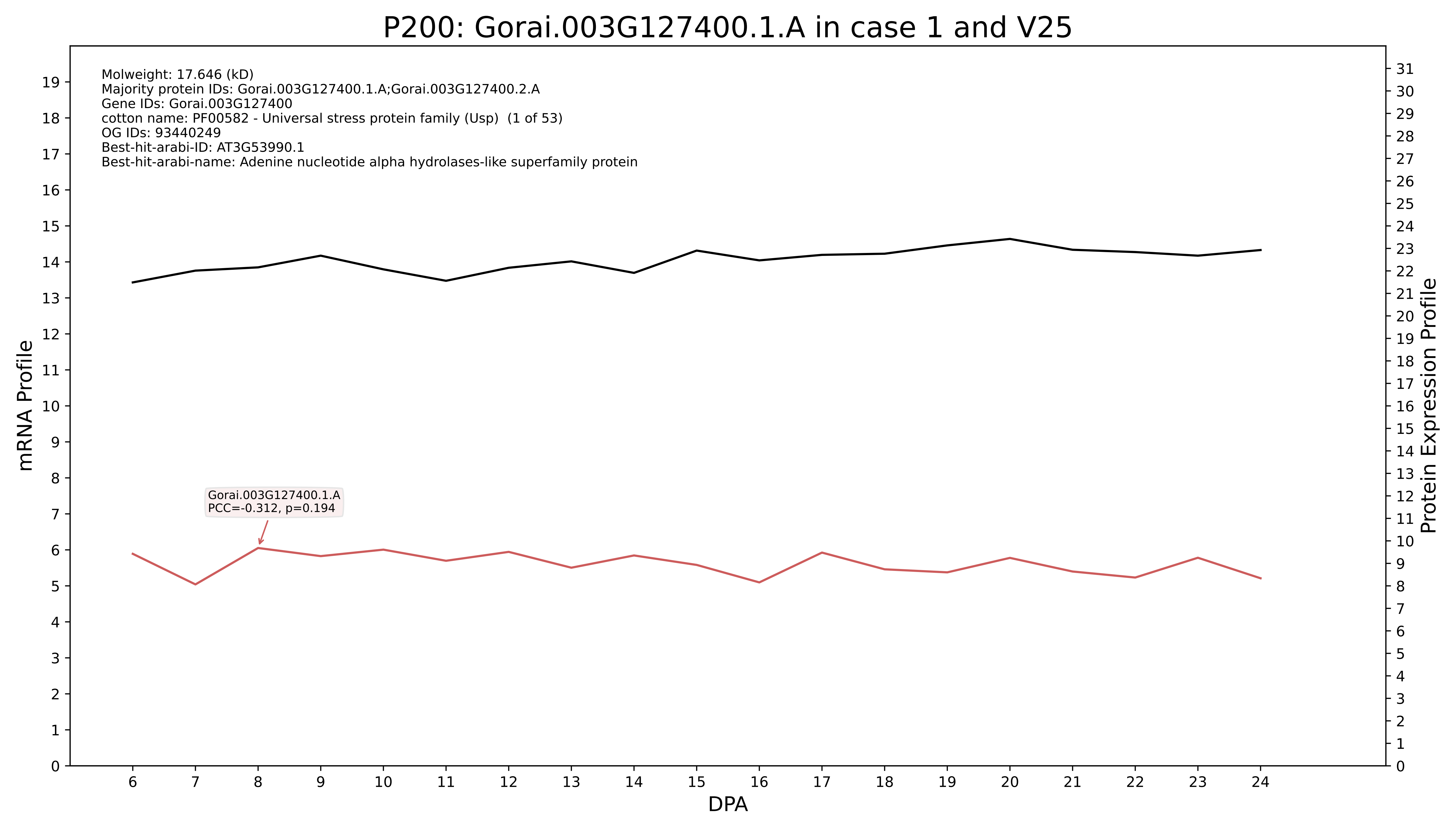
P200: Gorai.003G127400.1.A in case 1 and V25
Molweight: 17.646 (kD)
Majority protein IDs: Gorai.003G127400.1.A;Gorai.003G127400.2.A
Gene IDs: Gorai.003G127400
cotton name: PF00582 - Universal stress protein family (Usp) (1 of 53)
OG IDs: 93440249
Best-hit-arabi-ID: AT3G53990.1
Best-hit-arabi-name: Adenine nucleotide alpha hydrolases-like superfamily protein
Gorai.003G127400.1.A
PCC=-0.312, p=0.194
mRNA Profile
Protein Expression Profile
DPA

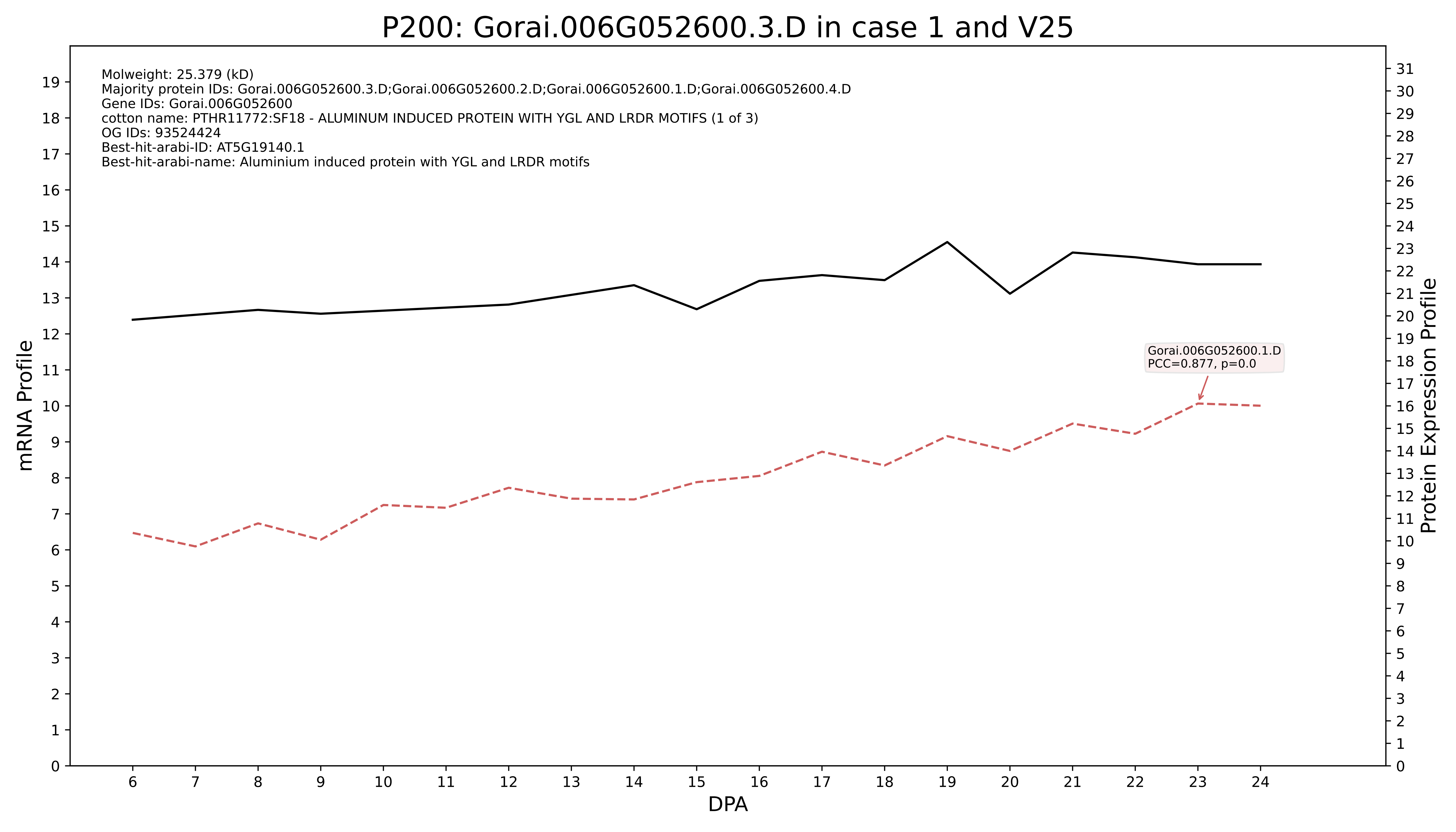
P200: Gorai.006G052600.3.D in case 1 and V25
Molweight: 25.379 (kD)
Majority protein IDs: Gorai.006G052600.3.D;Gorai.006G052600.2.D;Gorai.006G052600.1.D;Gorai.006G052600.4.D
Gene IDs: Gorai.006G052600
cotton name: PTHR11772:SF18 - ALUMINUM INDUCED PROTEIN WITH YGL AND LRDR MOTIFS (1 of 3)
OG IDs: 93524424
Best-hit-arabi-ID: AT5G19140.1
Best-hit-arabi-name: Aluminium induced protein with YGL and LRDR motifs
Gorai.006G052600.1.D
PCC=0.877, p=0.0
mRNA Profile
Protein Expression Profile
DPA

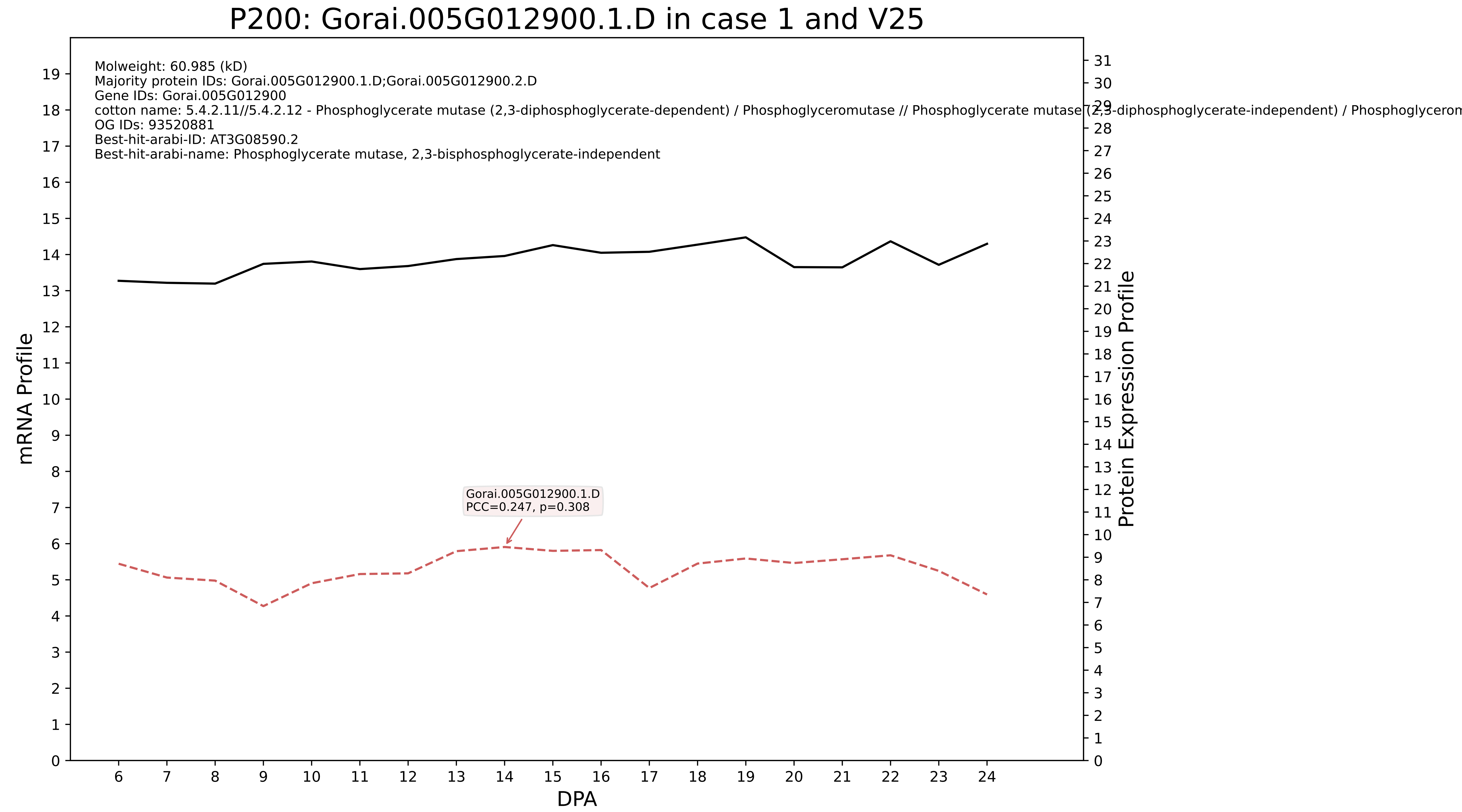

P200: Gorai.005G012900.1.D in case 1 and V25
Molweight: 60.985 (kD)
Majority protein IDs: Gorai.005G012900.1.D;Gorai.005G012900.2.D
Gene IDs: Gorai.005G012900
cotton name: 5.4.2.11//5.4.2.12 - Phosphoglycerate mutase (2,3-diphosphoglycerate-dependent) / Phosphoglyceromutase // Phosphoglycerate mutase (2,3-diphosphoglycerate-independent) / Phosphoglyceron
OG IDs: 93520881
Best-hit-arabi-ID: AT3G08590.2
Best-hit-arabi-name: Phosphoglycerate mutase, 2,3-bisphosphoglycerate-independent
Gorai.005G012900.1.D
PCC=0.247, p=0.308
mRNA Profile
Protein Expression Profile
DPA

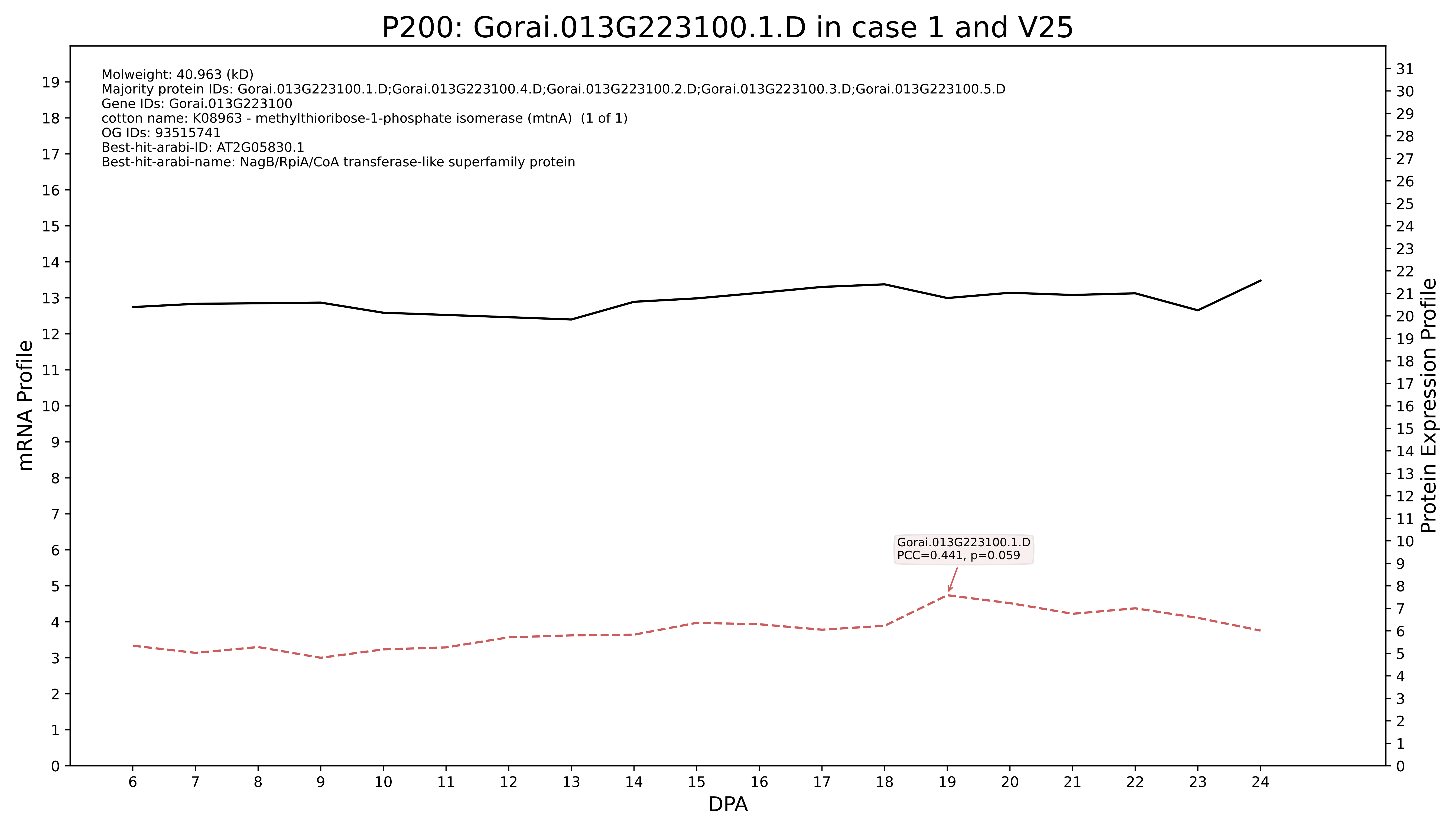
P200: Gorai.013G223100.1.D in case 1 and V25
Molweight: 40.963 (kD)
Majority protein IDs: Gorai.013G223100.1.D;Gorai.013G223100.4.D;Gorai.013G223100.2.D;Gorai.013G223100.3.D;Gorai.013G223100.5.D
Gene IDs: Gorai.013G223100
cotton name: K08963 - methylthioribose-1-phosphate isomerase (mtnA) (1 of 1)
OG IDs: 93515741
Best-hit-arabi-ID: AT2G05830.1
Best-hit-arabi-name: NagB/RpiA/CoA transferase-like superfamily protein
Gorai.013G223100.1.D
PCC=0.441, p=0.059
mRNA Profile
Protein Expression Profile
DPA

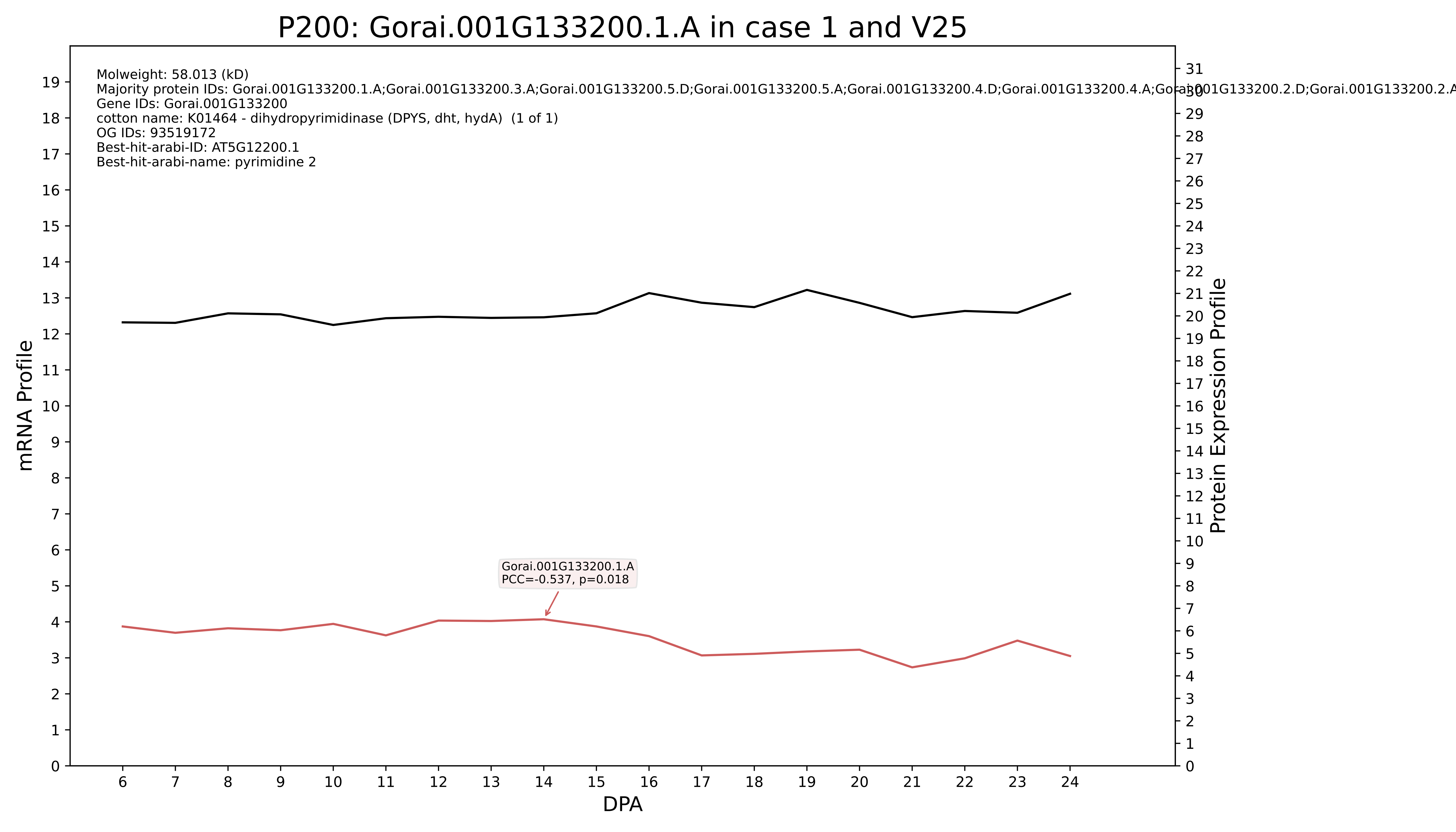
P200: Gorai.001G133200.1.A in case 1 and V25
Molweight: 58.013 (kD)
Majority protein IDs: Gorai.001G133200.1.A;Gorai.001G133200.3.A;Gorai.001G133200.5.D;Gorai.001G133200.5.A;Gorai.001G133200.4.D;Gorai.001G133200.4.A;Gorai.001G133200.2.D;Gorai.001G133200.2.A
Gene IDs: Gorai.001G133200
cotton name: K01464 - dihydropyrimidinase (DPYS, dht, hydA) (1 of 1)
OG IDs: 93519172
Best-hit-arabi-ID: AT5G12200.1
Best-hit-arabi-name: pyrimidine 2
Gorai.001G133200.1.A
PCC=-0.537, p=0.018
mRNA Profile
Protein Expression Profile
DPA

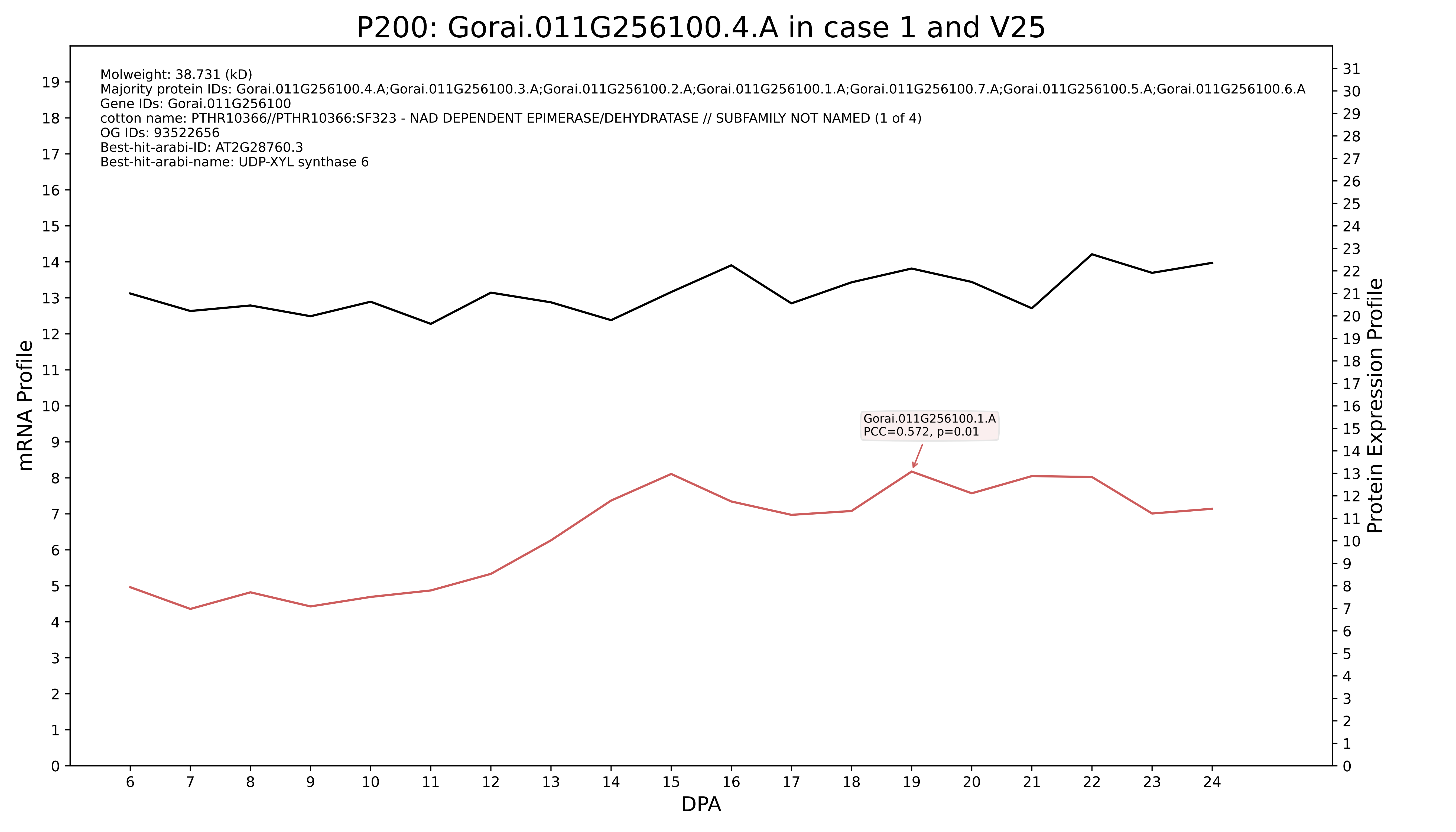

P200: Gorai.011G256100.4.A in case 1 and V25
Molweight: 38.731 (kD)
Majority protein IDs: Gorai.011G256100.4.A;Gorai.011G256100.3.A;Gorai.011G256100.2.A;Gorai.011G256100.1.A;Gorai.011G256100.7.A;Gorai.011G256100.5.A;Gorai.011G256100.6.A
Gene IDs: Gorai.011G256100
cotton name: PTHR10366//PTHR10366:SF323 - NAD DEPENDENT EPIMERASE/DEHYDRATASE // SUBFAMILY NOT NAMED (1 of 4)
OG IDs: 93522656
Best-hit-arabi-ID: AT2G28760.3
Best-hit-arabi-name: UDP-XYL synthase 6
Gorai.011G256100.1.A
PCC=0.572, p=0.01
mRNA Profile
Protein Expression Profile
DPA

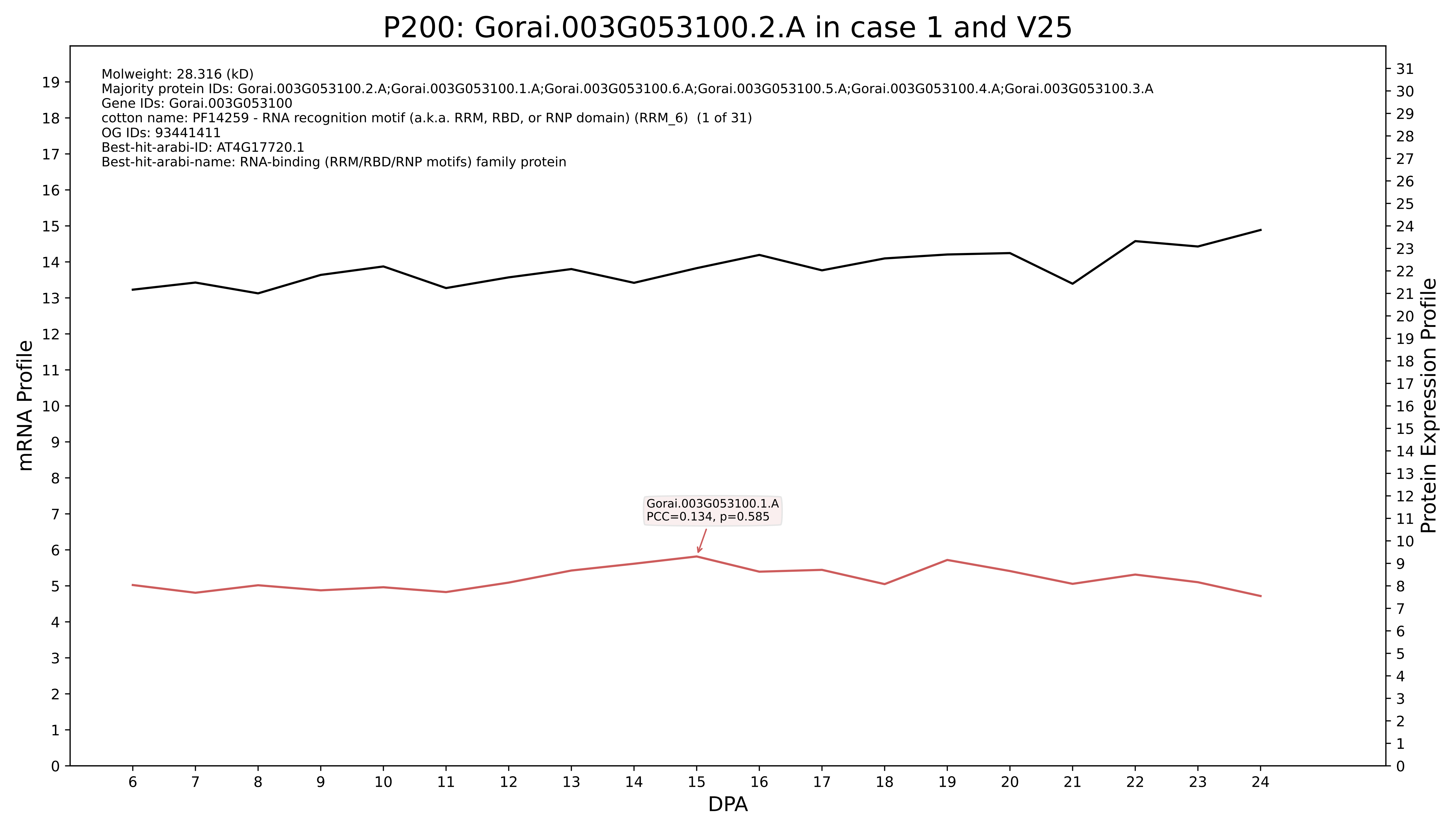
P200: Gorai.003G053100.2.A in case 1 and V25
Molweight: 28.316 (kD)
Majority protein IDs: Gorai.003G053100.2.A;Gorai.003G053100.1.A;Gorai.003G053100.6.A;Gorai.003G053100.5.A;Gorai.003G053100.4.A;Gorai.003G053100.3.A
Gene IDs: Gorai.003G053100
cotton name: PF14259 - RNA recognition motif (a.k.a. RRM, RBD, or RNP domain) (RRM_6) (1 of 31)
OG IDs: 93441411
Best-hit-arabi-ID: AT4G17720.1
Best-hit-arabi-name: RNA-binding (RRM/RBD/RNP motifs) family protein
Gorai.003G053100.1.A
PCC=0.134, p=0.585
mRNA Profile
Protein Expression Profile
DPA

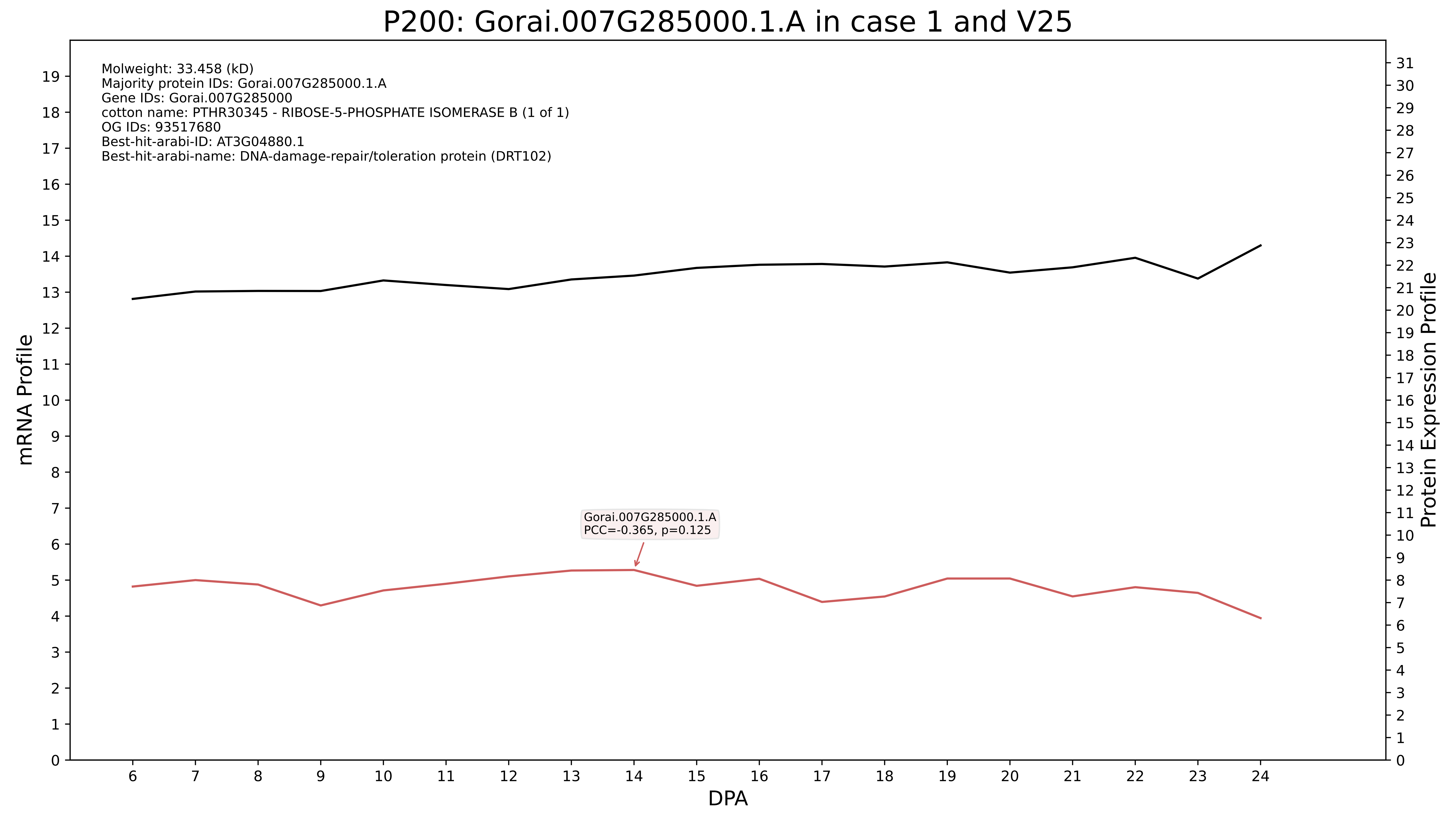

P200: Gorai.007G285000.1.A in case 1 and V25
Molweight: 33.458 (kD)
Majority protein IDs: Gorai.007G285000.1.A
Gene IDs: Gorai.007G285000
cotton name: PTHR30345 - RIBOSE-5-PHOSPHATE ISOMERASE B (1 of 1)
OG IDs: 93517680
Best-hit-arabi-ID: AT3G04880.1
Best-hit-arabi-name: DNA-damage-repair/toleration protein (DRT102)
Gorai.007G285000.1.A
PCC=-0.365, p=0.125
mRNA Profile
Protein Expression Profile
DPA

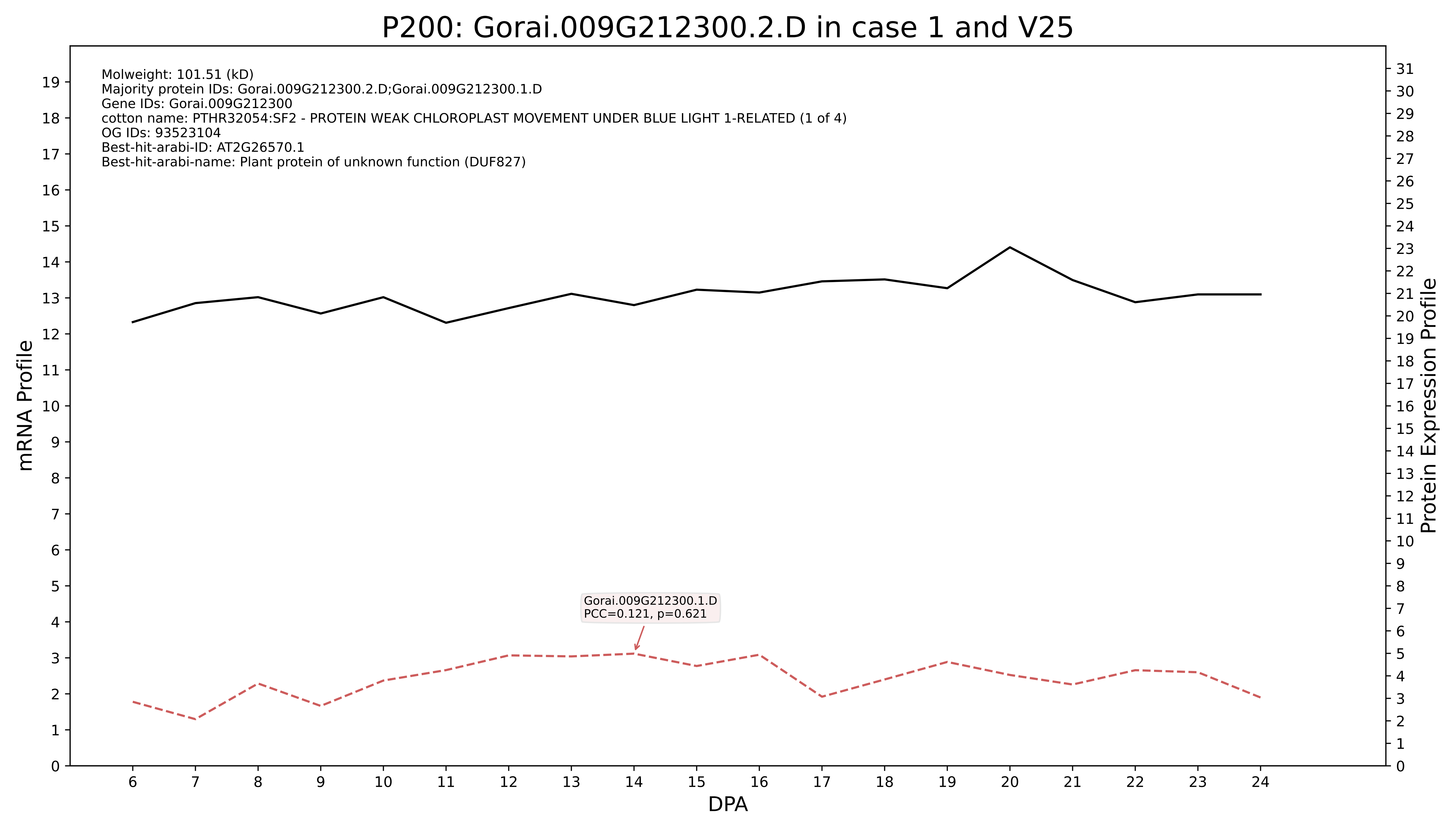
P200: Gorai.009G212300.2.D in case 1 and V25
Molweight: 101.51 (kD)
Majority protein IDs: Gorai.009G212300.2.D;Gorai.009G212300.1.D
Gene IDs: Gorai.009G212300
cotton name: PTHR32054:SF2 - PROTEIN WEAK CHLOROPLAST MOVEMENT UNDER BLUE LIGHT 1-RELATED (1 of 4)
OG IDs: 93523104
Best-hit-arabi-ID: AT2G26570.1
Best-hit-arabi-name: Plant protein of unknown function (DUF827)
Gorai.009G212300.1.D
PCC=0.121, p=0.621
mRNA Profile
Protein Expression Profile
DPA

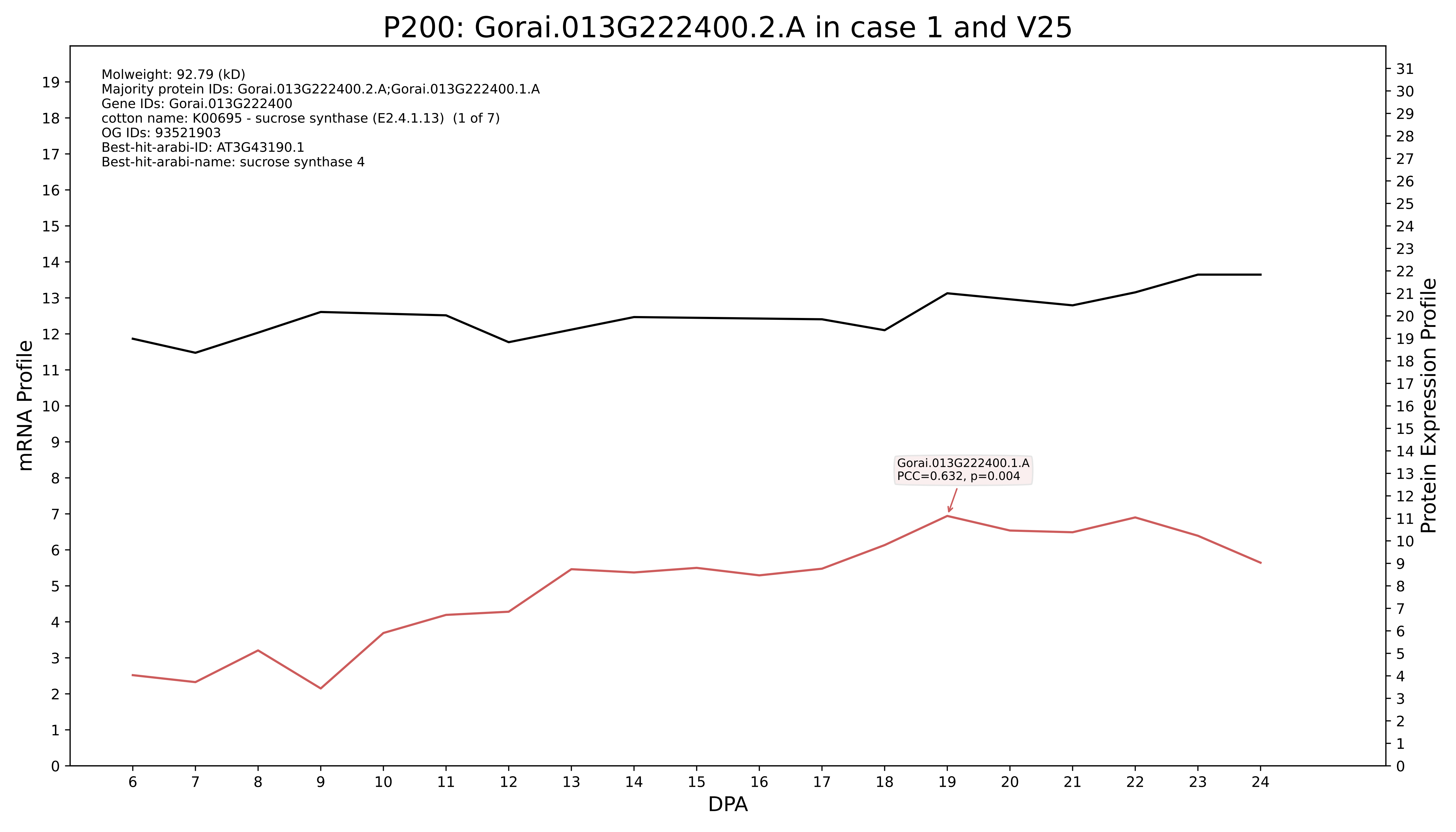
P200: Gorai.013G222400.2.A in case 1 and V25
Molweight: 92.79 (kD)
Majority protein IDs: Gorai.013G222400.2.A;Gorai.013G222400.1.A
Gene IDs: Gorai.013G222400
cotton name: K00695 - sucrose synthase (E2.4.1.13) (1 of 7)
OG IDs: 93521903
Best-hit-arabi-ID: AT3G43190.1
Best-hit-arabi-name: sucrose synthase 4
Gorai.013G222400.1.A
PCC=0.632, p=0.004
mRNA Profile
Protein Expression Profile
DPA

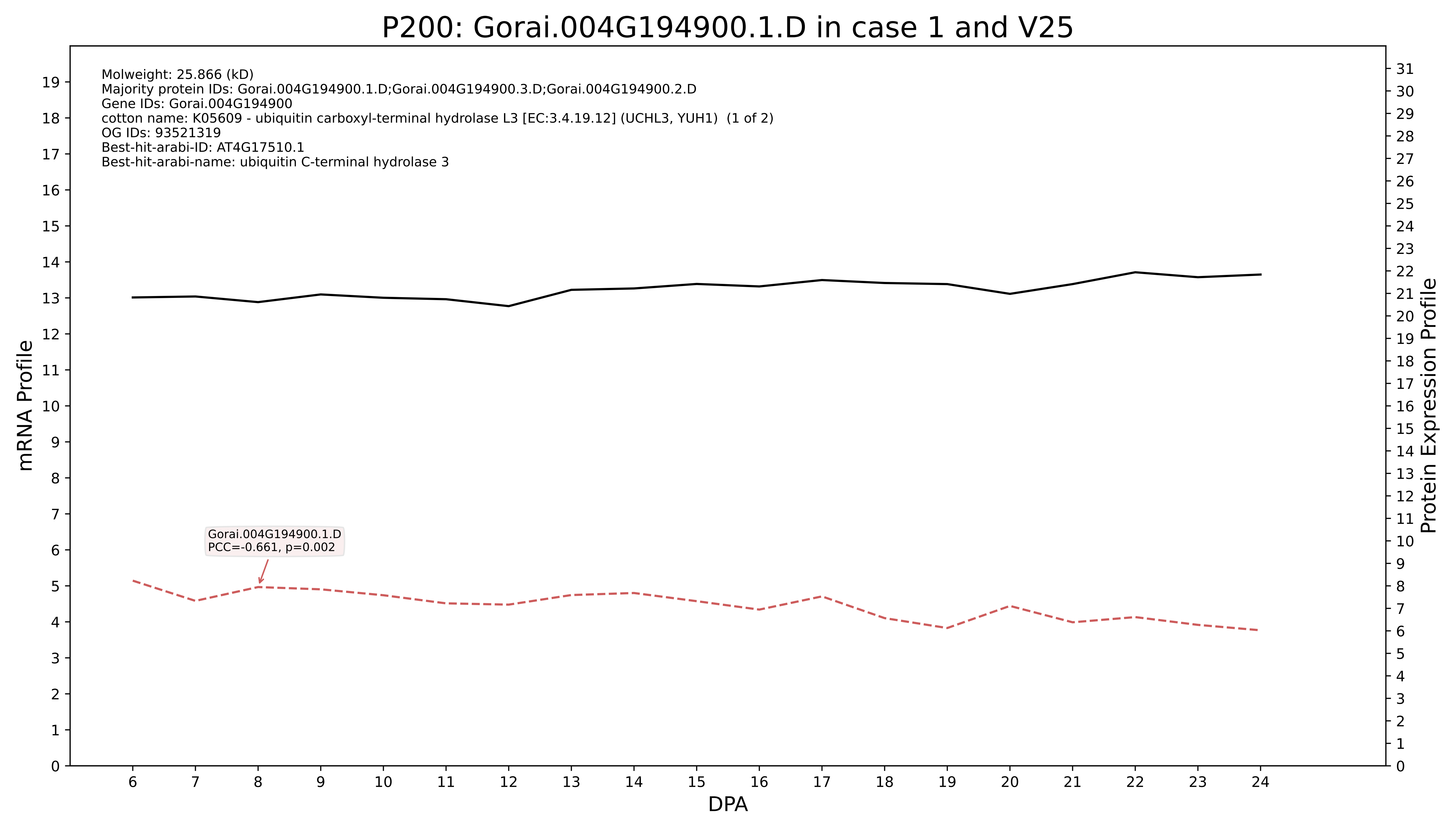
P200: Gorai.004G194900.1.D in case 1 and V25
Molweight: 25.866 (kD)
Majority protein IDs: Gorai.004G194900.1.D;Gorai.004G194900.3.D;Gorai.004G194900.2.D
Gene IDs: Gorai.004G194900
cotton name: K05609 - ubiquitin carboxyl-terminal hydrolase L3 [EC:3.4.19.12] (UCHL3, YUH1) (1 of 2)
OG IDs: 93521319
Best-hit-arabi-ID: AT4G17510.1
Best-hit-arabi-name: ubiquitin C-terminal hydrolase 3
Gorai.004G194900.1.D
PCC=-0.661, p=0.002
mRNA Profile
Protein Expression Profile
DPA

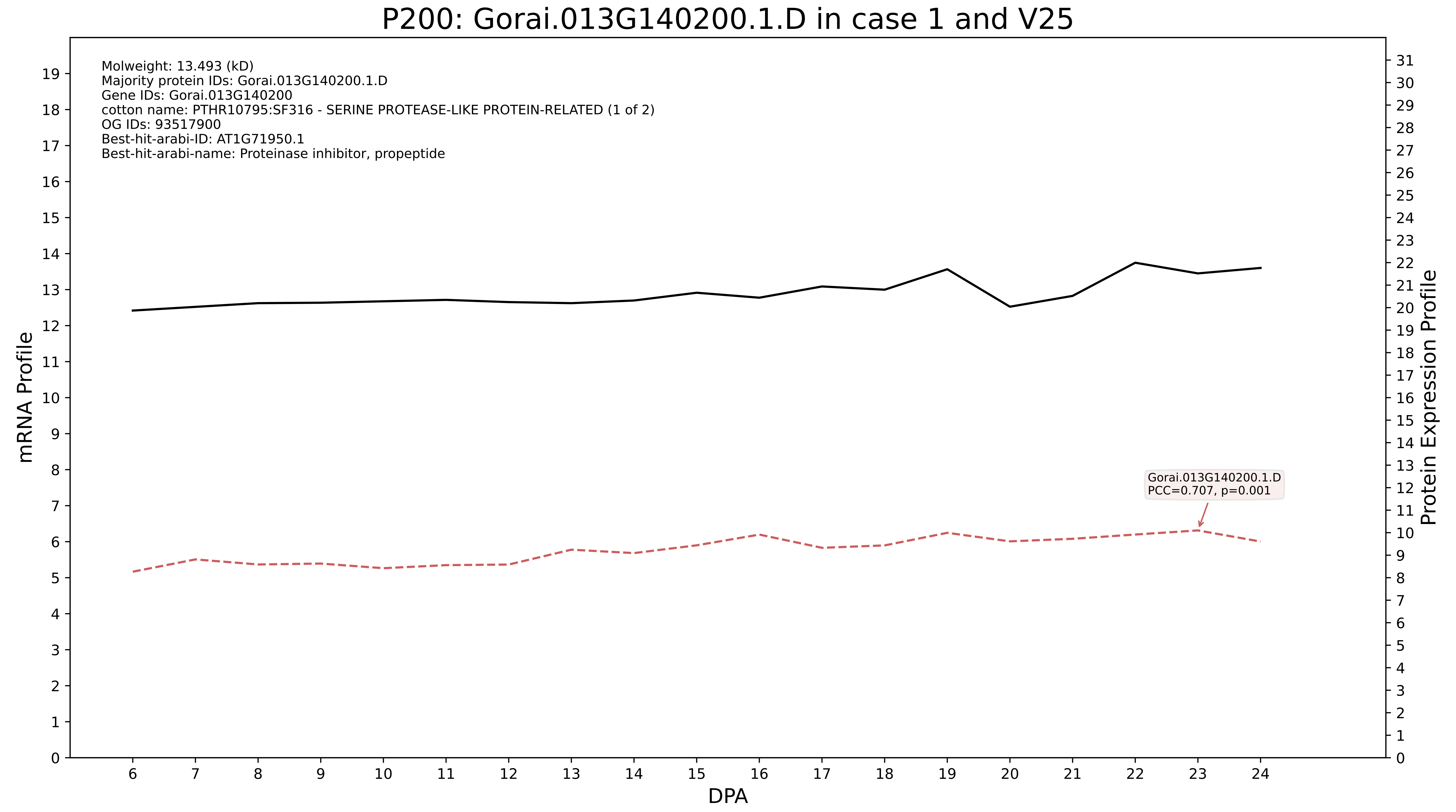

P200: Gorai.013G140200.1.D in case 1 and V25
Molweight: 13.493 (kD)
Majority protein IDs: Gorai.013G140200.1.D
Gene IDs: Gorai.013G140200
cotton name: PTHR10795:SF316 - SERINE PROTEASE-LIKE PROTEIN-RELATED (1 of 2)
OG IDs: 93517900
Best-hit-arabi-ID: AT1G71950.1
Best-hit-arabi-name: Proteinase inhibitor, propeptide
Gorai.013G140200.1.D
PCC=0.707, p=0.001
mRNA Profile
Protein Expression Profile
DPA

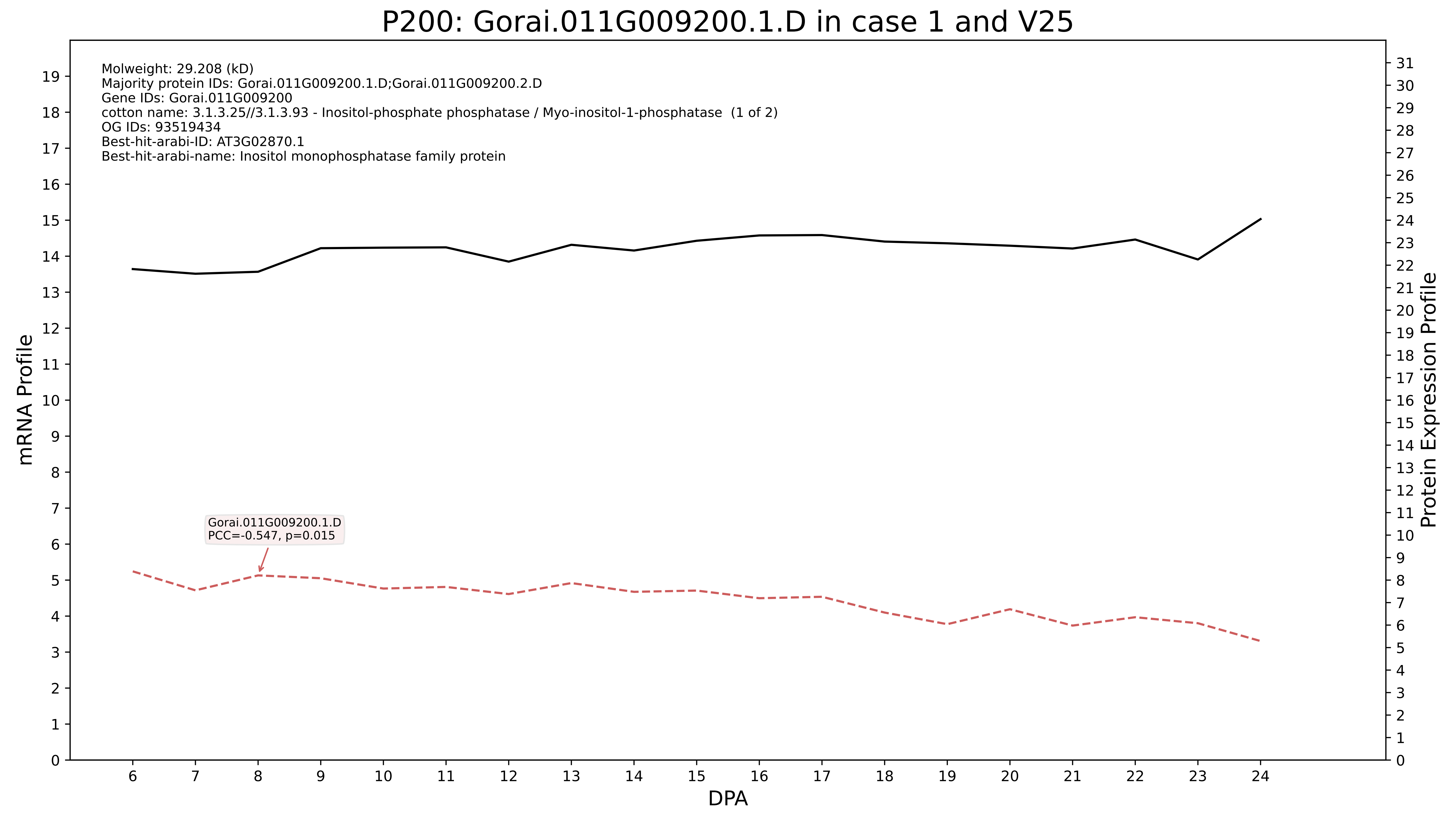
P200: Gorai.011G009200.1.D in case 1 and V25
Molweight: 29.208 (kD)
Majority protein IDs: Gorai.011G009200.1.D;Gorai.011G009200.2.D
Gene IDs: Gorai.011G009200
cotton name: 3.1.3.25//3.1.3.93 - Inositol-phosphate phosphatase / Myo-inositol-1-phosphatase (1 of 2)
OG IDs: 93519434
Best-hit-arabi-ID: AT3G02870.1
Best-hit-arabi-name: Inositol monophosphatase family protein
Gorai.011G009200.1.D
PCC=-0.547, p=0.015
mRNA Profile
Protein Expression Profile
DPA

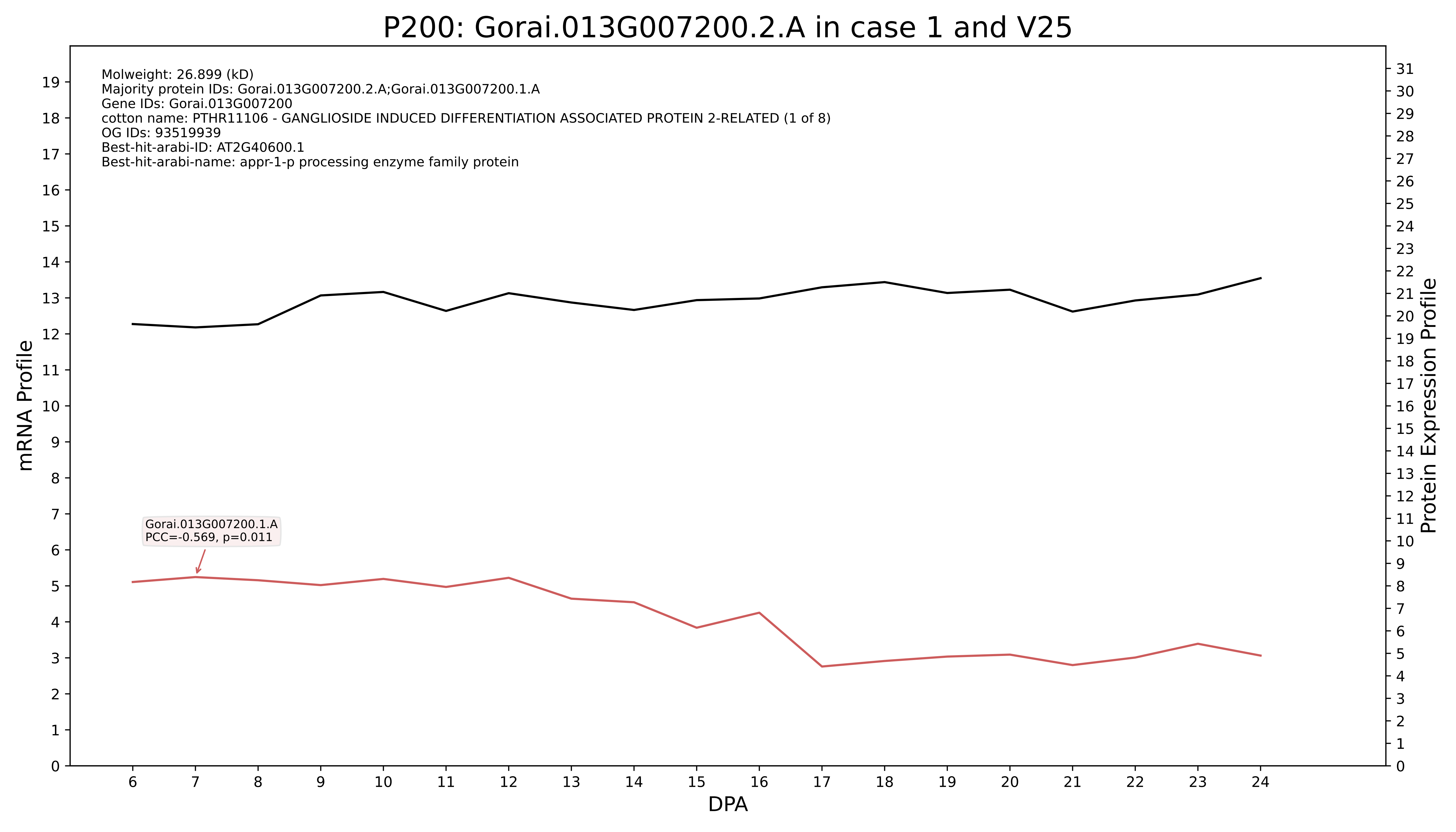
P200: Gorai.013G007200.2.A in case 1 and V25
Molweight: 26.899 (kD)
Majority protein IDs: Gorai.013G007200.2.A;Gorai.013G007200.1.A
Gene IDs: Gorai.013G007200
cotton name: PTHR11106 - GANGLIOSIDE INDUCED DIFFERENTIATION ASSOCIATED PROTEIN 2-RELATED (1 of 8)
OG IDs: 93519939
Best-hit-arabi-ID: AT2G40600.1
Best-hit-arabi-name: appr-1-p processing enzyme family protein
Gorai.013G007200.1.A
PCC=-0.569, p=0.011
mRNA Profile
Protein Expression Profile
DPA

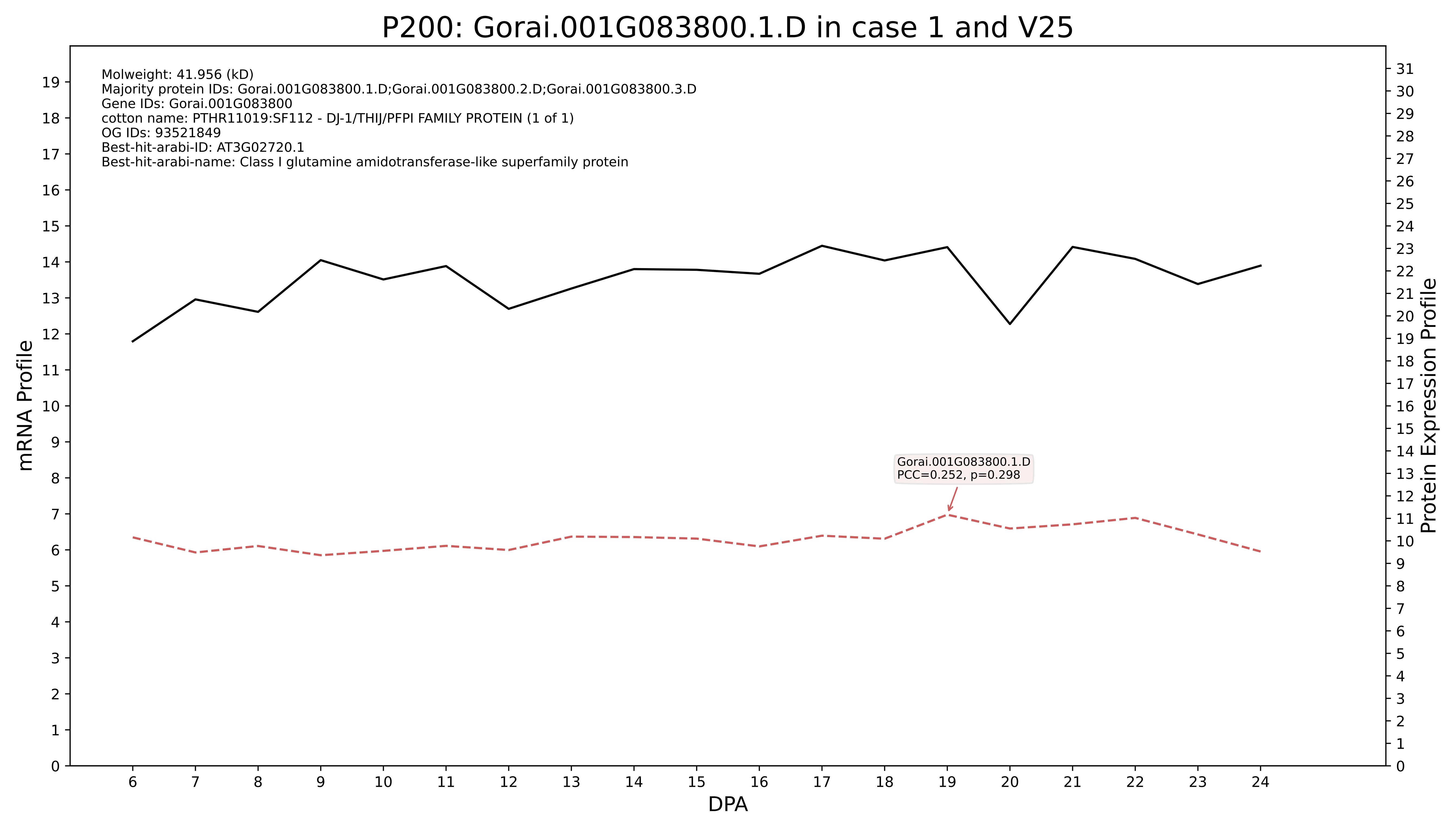

P200: Gorai.001G083800.1.D in case 1 and V25
Molweight: 41.956 (kD)
Majority protein IDs: Gorai.001G083800.1.D;Gorai.001G083800.2.D;Gorai.001G083800.3.D
Gene IDs: Gorai.001G083800
cotton name: PTHR11019:SF112 - DJ-1/THIJ/PFPI FAMILY PROTEIN (1 of 1)
OG IDs: 93521849
Best-hit-arabi-ID: AT3G02720.1
Best-hit-arabi-name: Class I glutamine amidotransferase-like superfamily protein
Gorai.001G083800.1.D
PCC=0.252, p=0.298
mRNA Profile
Protein Expression Profile
DPA

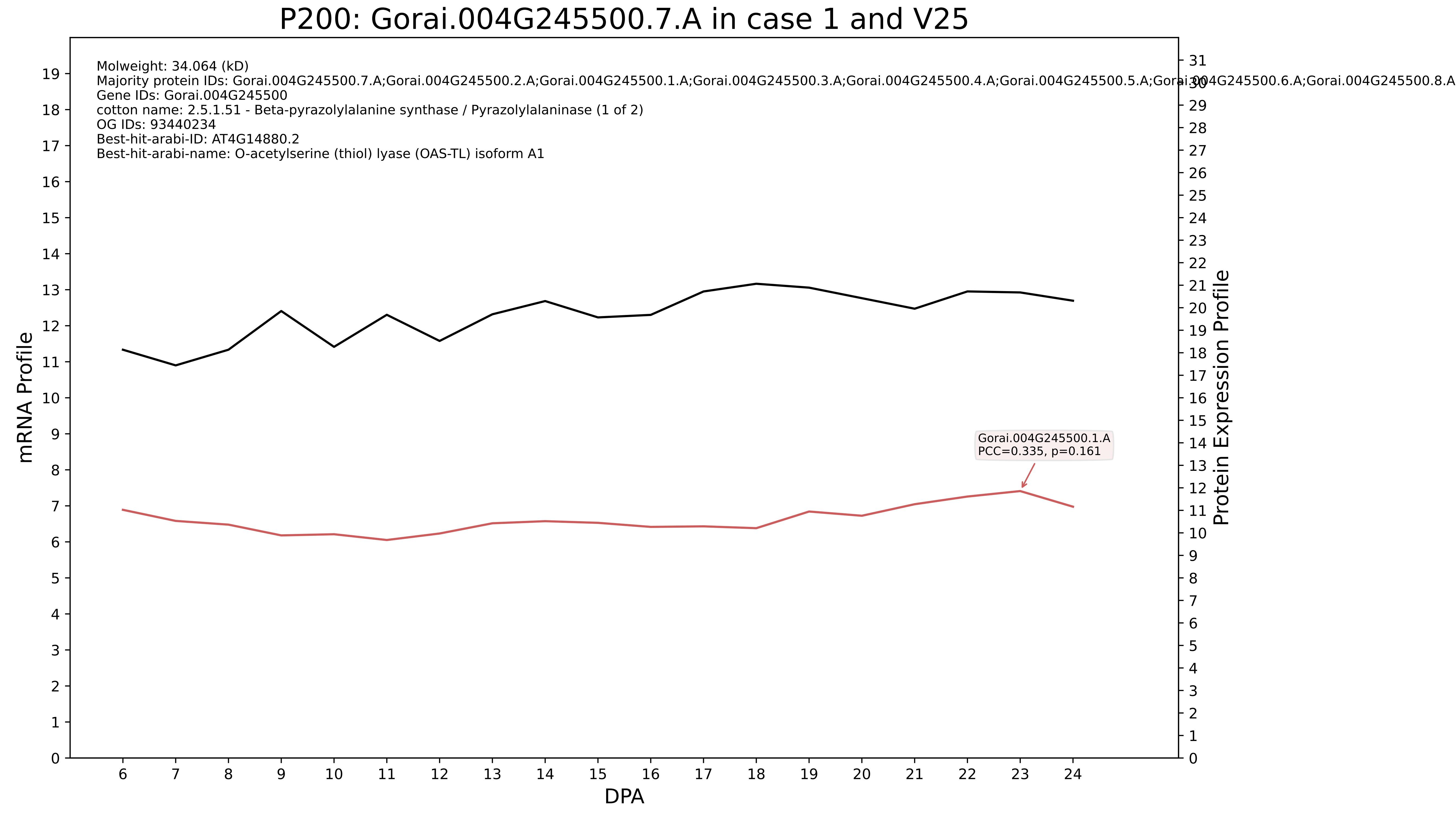
P200: Gorai.004G245500.7.A in case 1 and V25
Molweight: 34.064 (kD)
Majority protein IDs: Gorai.004G245500.7.A;Gorai.004G245500.2.A;Gorai.004G245500.1.A;Gorai.004G245500.3.A;Gorai.004G245500.4.A;Gorai.004G245500.5.A;Gorai.004G245500.6.A;Gorai.004G245500.8.A
Gene IDs: Gorai.004G245500
cotton name: 2.5.1.51 - Beta-pyrazolylalanine synthase / Pyrazolylalaninase (1 of 2)
OG IDs: 93440234
Best-hit-arabi-ID: AT4G14880.2
Best-hit-arabi-name: O-acetylserine (thiol) lyase (OAS-TL) isoform A1
Gorai.004G245500.1.A
PCC=0.335, p=0.161
mRNA Profile
Protein Expression Profile
DPA

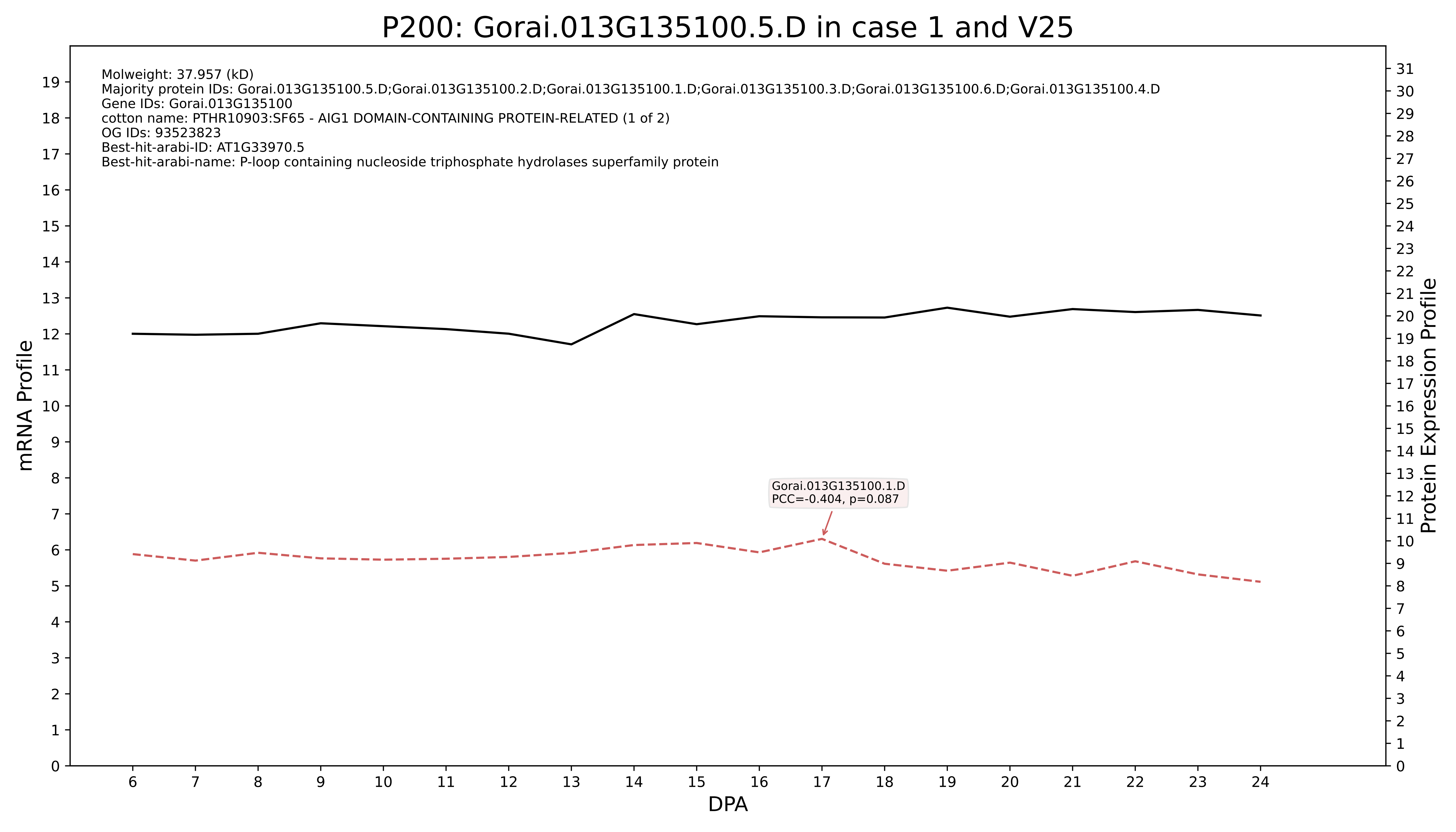
P200: Gorai.013G135100.5.D in case 1 and V25
Molweight: 37.957 (kD)
Majority protein IDs: Gorai.013G135100.5.D;Gorai.013G135100.2.D;Gorai.013G135100.1.D;Gorai.013G135100.3.D;Gorai.013G135100.6.D;Gorai.013G135100.4.D
Gene IDs: Gorai.013G135100
cotton name: PTHR10903:SF65 - AIG1 DOMAIN-CONTAINING PROTEIN-RELATED (1 of 2)
OG IDs: 93523823
Best-hit-arabi-ID: AT1G33970.5
Best-hit-arabi-name: P-loop containing nucleoside triphosphate hydrolases superfamily protein
Gorai.013G135100.1.D
PCC=-0.404, p=0.087
mRNA Profile
Protein Expression Profile
DPA

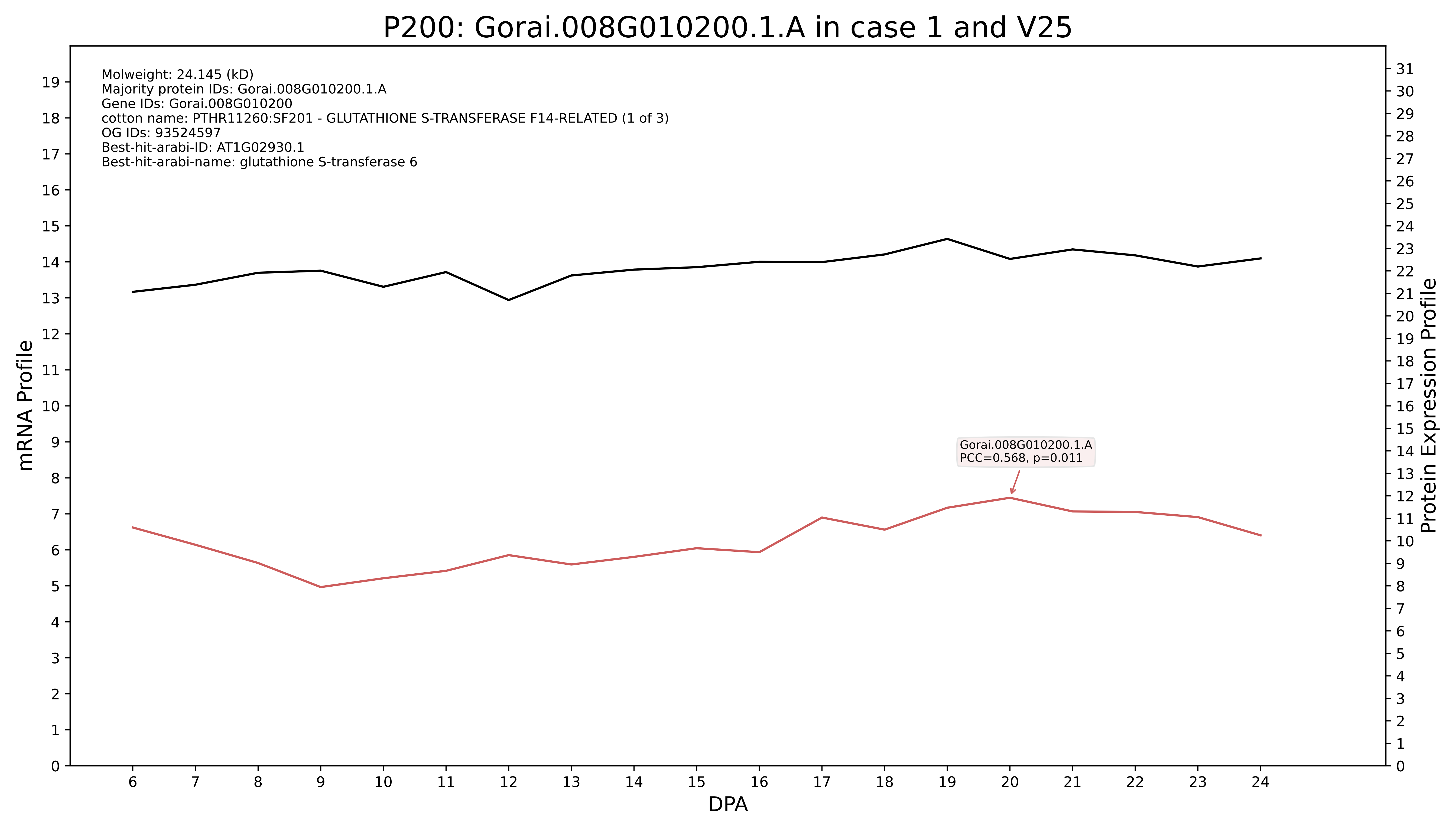
P200: Gorai.008G010200.1.A in case 1 and V25
Molweight: 24.145 (kD)
Majority protein IDs: Gorai.008G010200.1.A
Gene IDs: Gorai.008G010200
cotton name: PTHR11260:SF201 - GLUTATHIONE S-TRANSFERASE F14-RELATED (1 of 3)
OG IDs: 93524597
Best-hit-arabi-ID: AT1G02930.1
Best-hit-arabi-name: glutathione S-transferase 6
Gorai.008G010200.1.A
PCC=0.568, p=0.011
mRNA Profile
Protein Expression Profile
DPA

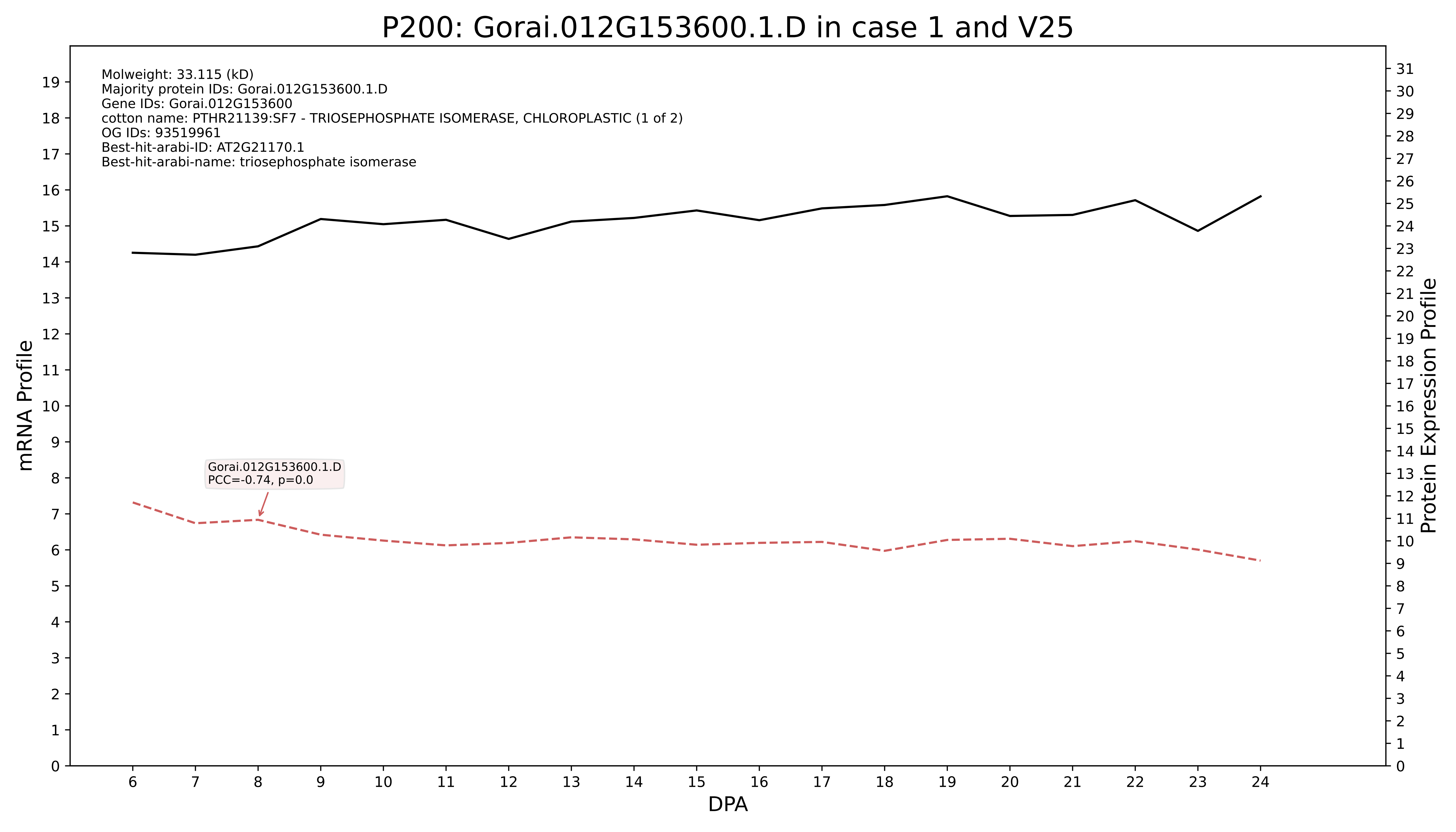

P200: Gorai.012G153600.1.D in case 1 and V25
Molweight: 33.115 (kD)
Majority protein IDs: Gorai.012G153600.1.D
Gene IDs: Gorai.012G153600
cotton name: PTHR21139:SF7 - TRIOSEPHOSPHATE ISOMERASE, CHLOROPLASTIC (1 of 2)
OG IDs: 93519961
Best-hit-arabi-ID: AT2G21170.1
Best-hit-arabi-name: triosephosphate isomerase
Gorai.012G153600.1.D
PCC=-0.74, p=0.0
mRNA Profile
Protein Expression Profile
DPA

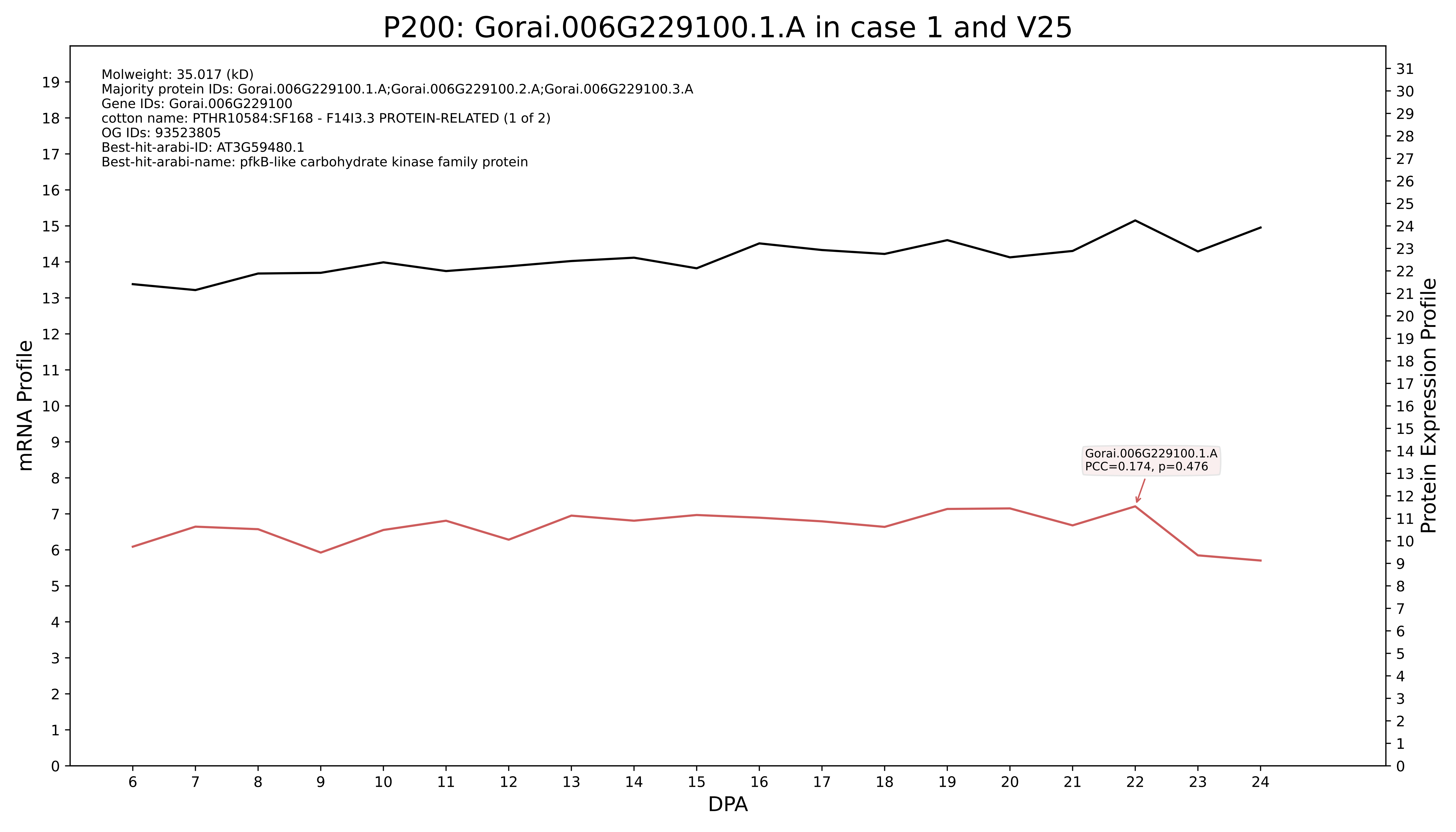
P200: Gorai.006G229100.1.A in case 1 and V25
Molweight: 35.017 (kD)
Majority protein IDs: Gorai.006G229100.1.A;Gorai.006G229100.2.A;Gorai.006G229100.3.A
Gene IDs: Gorai.006G229100
cotton name: PTHR10584:SF168 - F14I3.3 PROTEIN-RELATED (1 of 2)
OG IDs: 93523805
Best-hit-arabi-ID: AT3G59480.1
Best-hit-arabi-name: pfkB-like carbohydrate kinase family protein
Gorai.006G229100.1.A
PCC=0.174, p=0.476
mRNA Profile
Protein Expression Profile
DPA

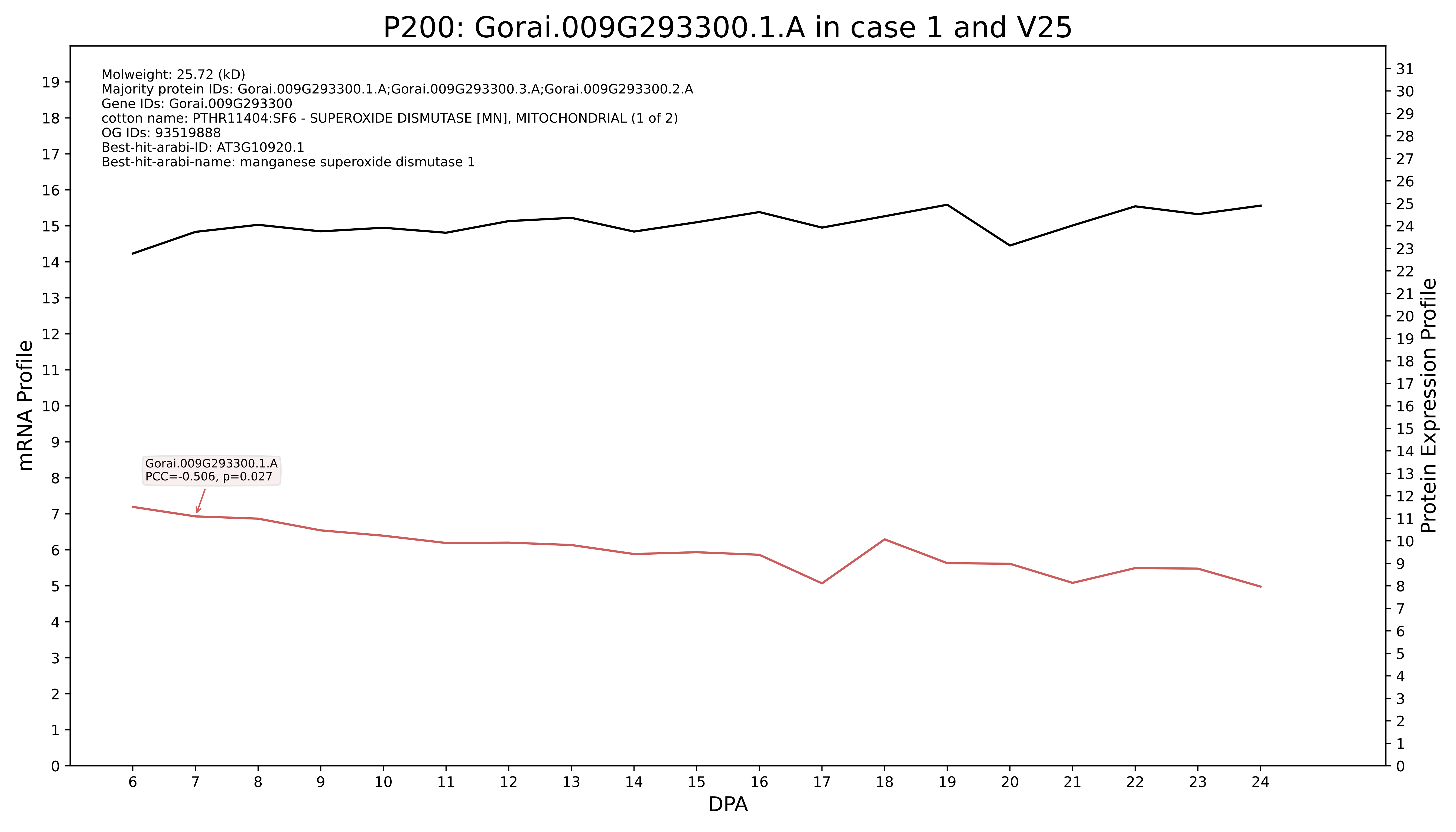
P200: Gorai.009G293300.1.A in case 1 and V25
Molweight: 25.72 (kD)
Majority protein IDs: Gorai.009G293300.1.A;Gorai.009G293300.3.A;Gorai.009G293300.2.A
Gene IDs: Gorai.009G293300
cotton name: PTHR11404:SF6 - SUPEROXIDE DISMUTASE [MN], MITOCHONDRIAL (1 of 2)
OG IDs: 93519888
Best-hit-arabi-ID: AT3G10920.1
Best-hit-arabi-name: manganese superoxide dismutase 1
Gorai.009G293300.1.A
PCC=-0.506, p=0.027
mRNA Profile
Protein Expression Profile
DPA

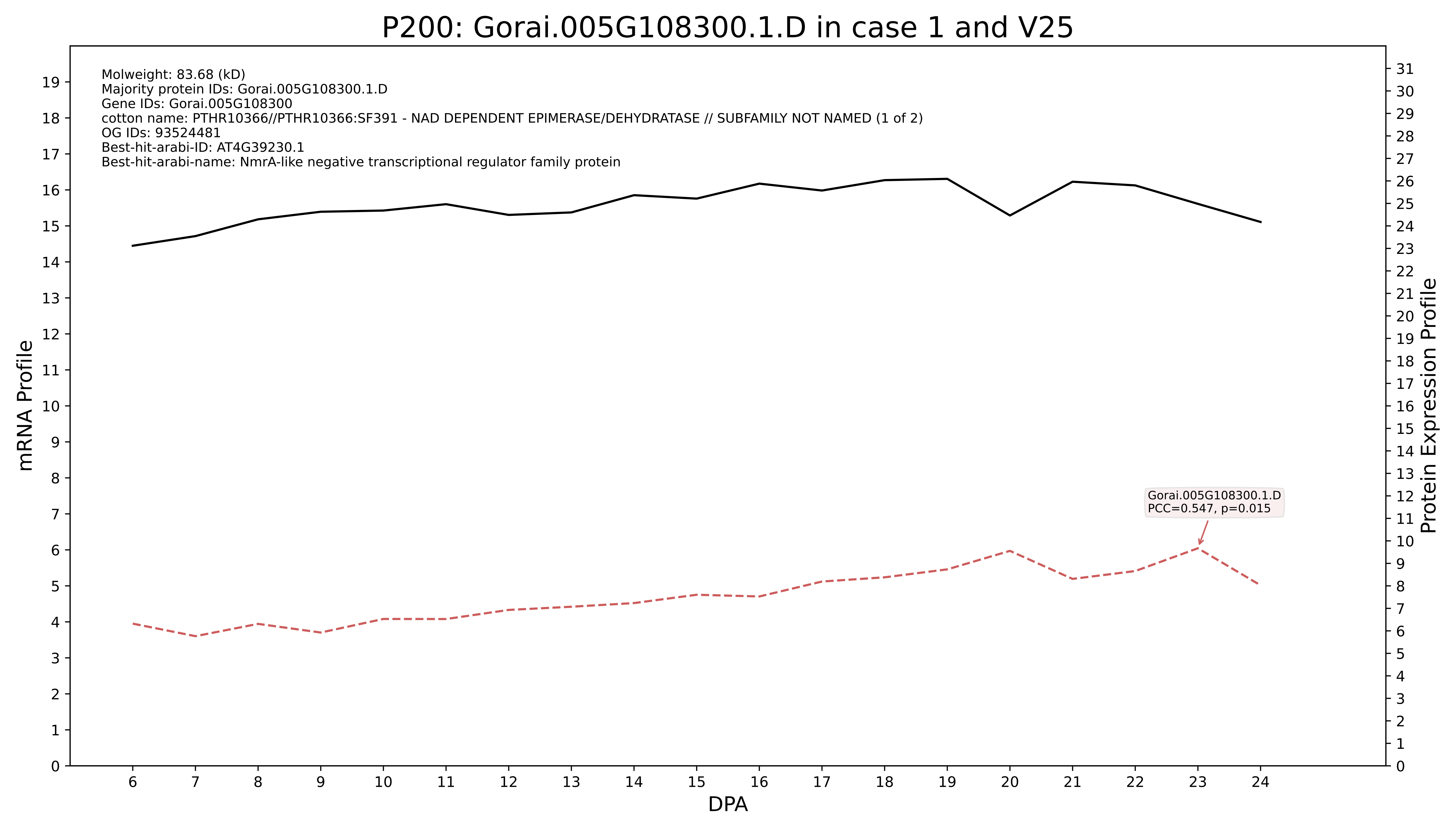

P200: Gorai.005G108300.1.D in case 1 and V25
Molweight: 83.68 (kD)
Majority protein IDs: Gorai.005G108300.1.D
Gene IDs: Gorai.005G108300
cotton name: PTHR10366//PTHR10366:SF391 - NAD DEPENDENT EPIMERASE/DEHYDRATASE // SUBFAMILY NOT NAMED (1 of 2)
OG IDs: 93524481
Best-hit-arabi-ID: AT4G39230.1
Best-hit-arabi-name: NmrA-like negative transcriptional regulator family protein
Gorai.005G108300.1.D
PCC=0.547, p=0.015
mRNA Profile
Protein Expression Profile
DPA

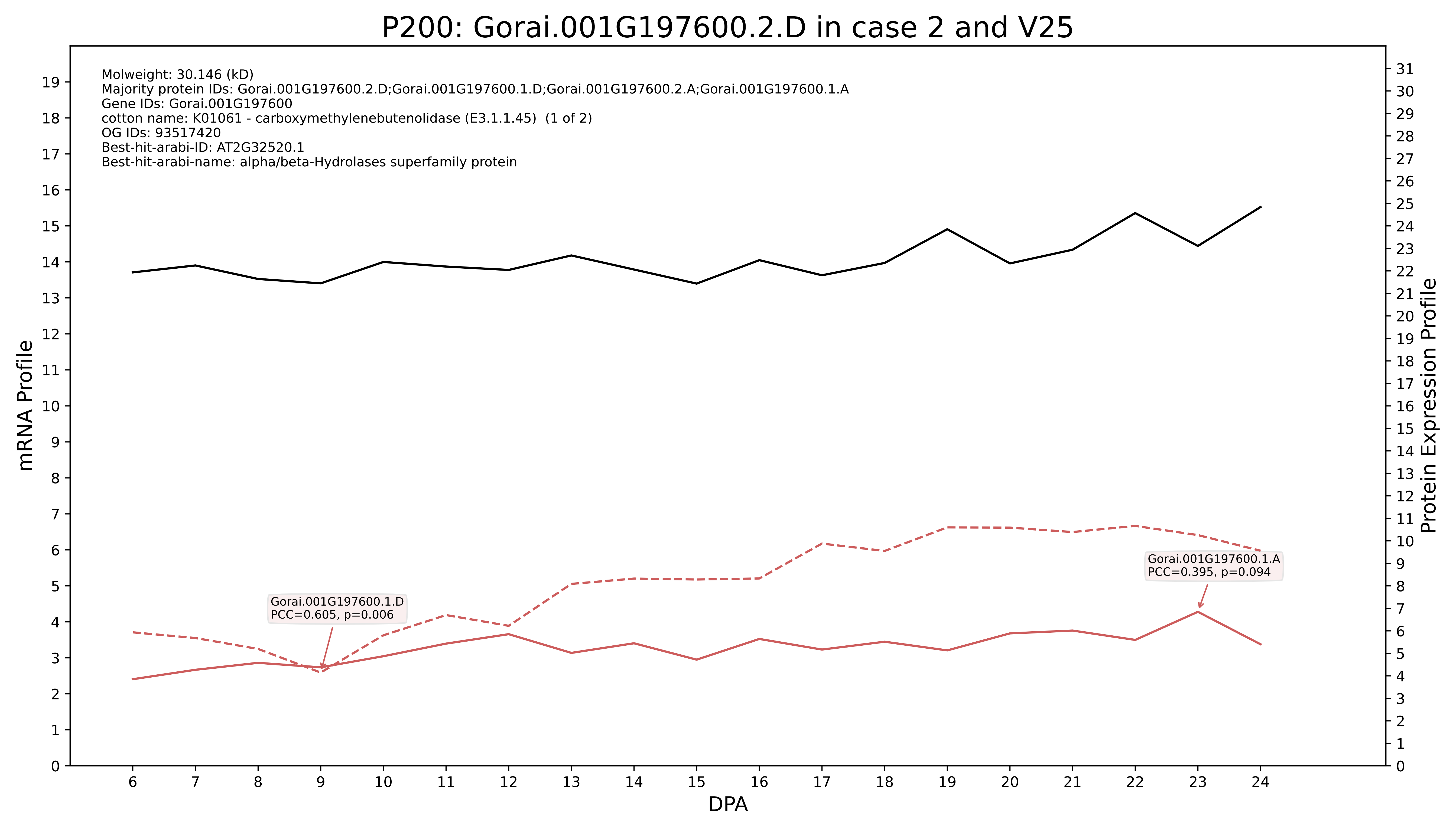

P200: Gorai.001G197600.2.D in case 2 and V25
Molweight: 30.146 (kD)
Majority protein IDs: Gorai.001G197600.2.D;Gorai.001G197600.1.D;Gorai.001G197600.2.A;Gorai.001G197600.1.A
Gene IDs: Gorai.001G197600
cotton name: K01061 - carboxymethylenebutenolidase (E3.1.1.45) (1 of 2)
OG IDs: 93517420
Best-hit-arabi-ID: AT2G32520.1
Best-hit-arabi-name: alpha/beta-Hydrolases superfamily protein
Gorai.001G197600.1.D
PCC=0.605, p=0.006
Gorai.001G197600.1.A
PCC=0.395, p=0.094
mRNA Profile
Protein Expression Profile
DPA

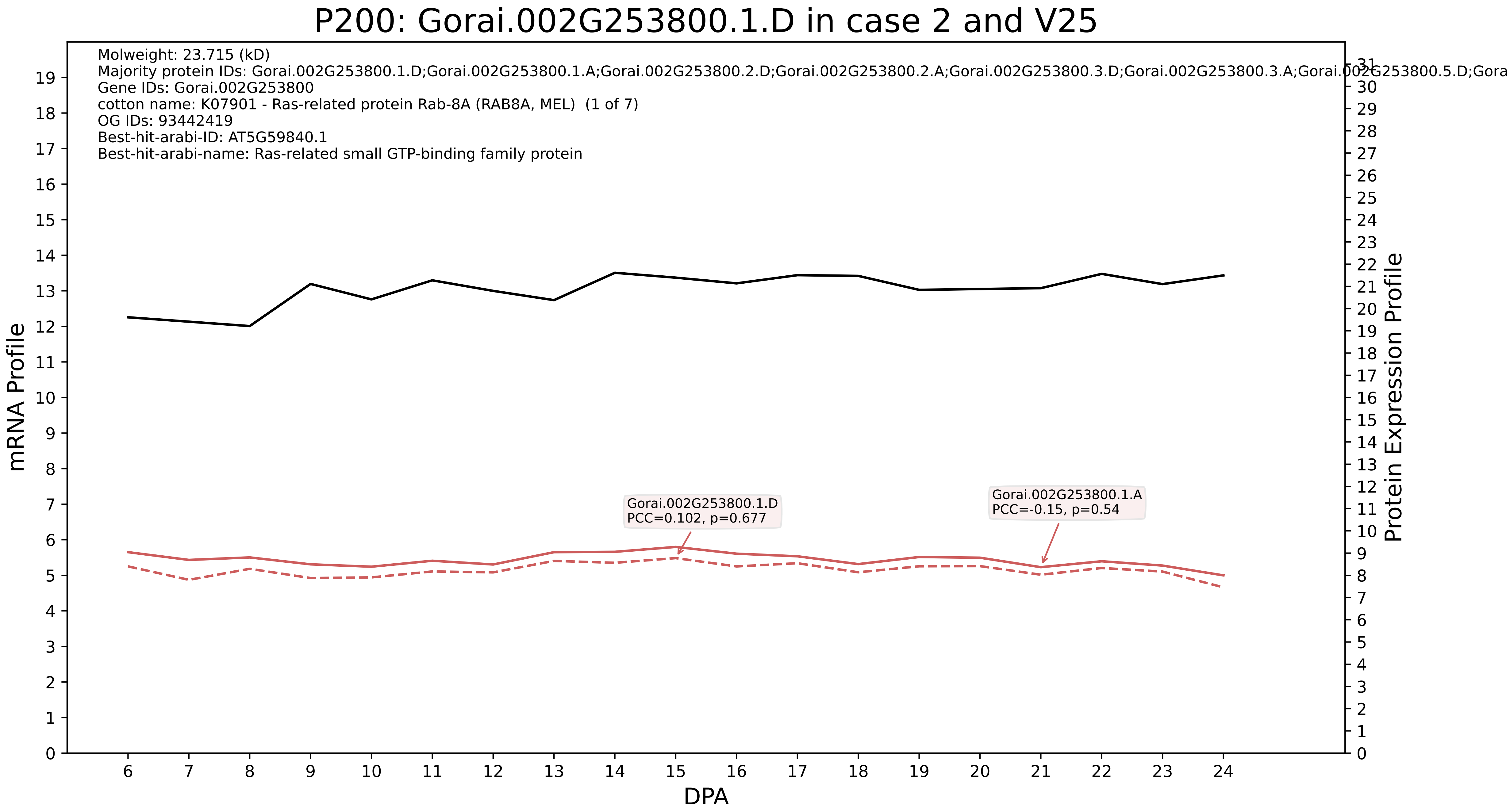

P200: Gorai.002G253800.1.D in case 2 and V25
Molweight: 23.715 (kD)
Majority protein IDs: Gorai.002G253800.1.D;Gorai.002G253800.1.A;Gorai.002G253800.2.D;Gorai.002G253800.2.A;Gorai.002G253800.3.D;Gorai.002G253800.3.A;Gorai.002G253800.5.D;Gorai
Gene IDs: Gorai.002G253800
cotton name: K07901 - Ras-related protein Rab-8A (RAB8A, MEL) (1 of 7)
OG IDs: 93442419
Best-hit-arabi-ID: AT5G59840.1
Best-hit-arabi-name: Ras-related small GTP-binding family protein
Gorai.002G253800.1.D
PCC=0.102, p=0.677
Gorai.002G253800.1.A
PCC=-0.15, p=0.54
mRNA Profile
Protein Expression Profile
DPA

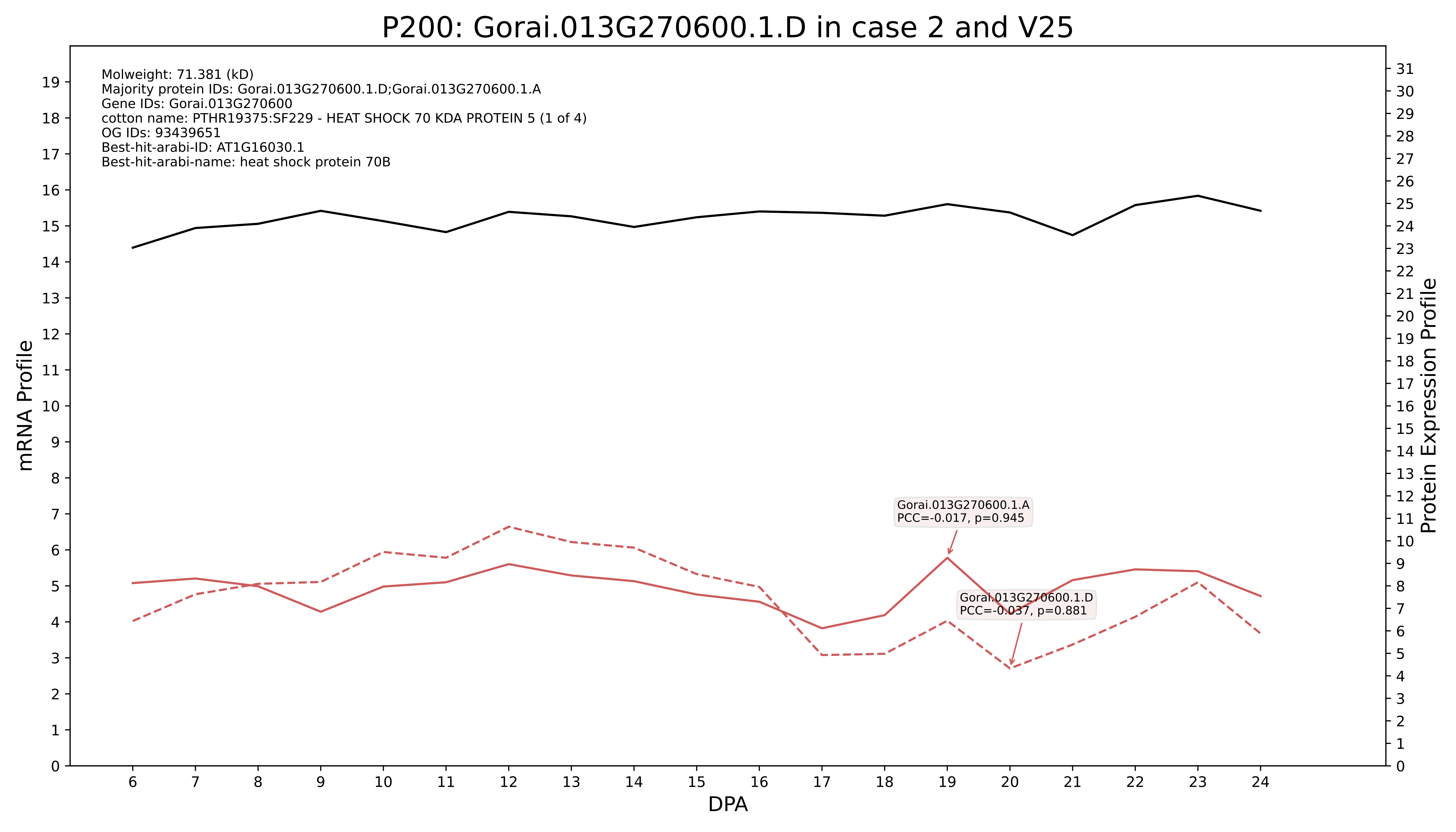
P200: Gorai.013G270600.1.D in case 2 and V25
Molweight: 71.381 (kD)
Majority protein IDs: Gorai.013G270600.1.D;Gorai.013G270600.1.A
Gene IDs: Gorai.013G270600
cotton name: PTHR19375:SF229 - HEAT SHOCK 70 KDA PROTEIN 5 (1 of 4)
OG IDs: 93439651
Best-hit-arabi-ID: AT1G16030.1
Best-hit-arabi-name: heat shock protein 70B
Gorai.013G270600.1.A
PCC=-0.017, p=0.945
Gorai.013G270600.1.D
PCC=-0.037, p=0.881
mRNA Profile
Protein Expression Profile
DPA

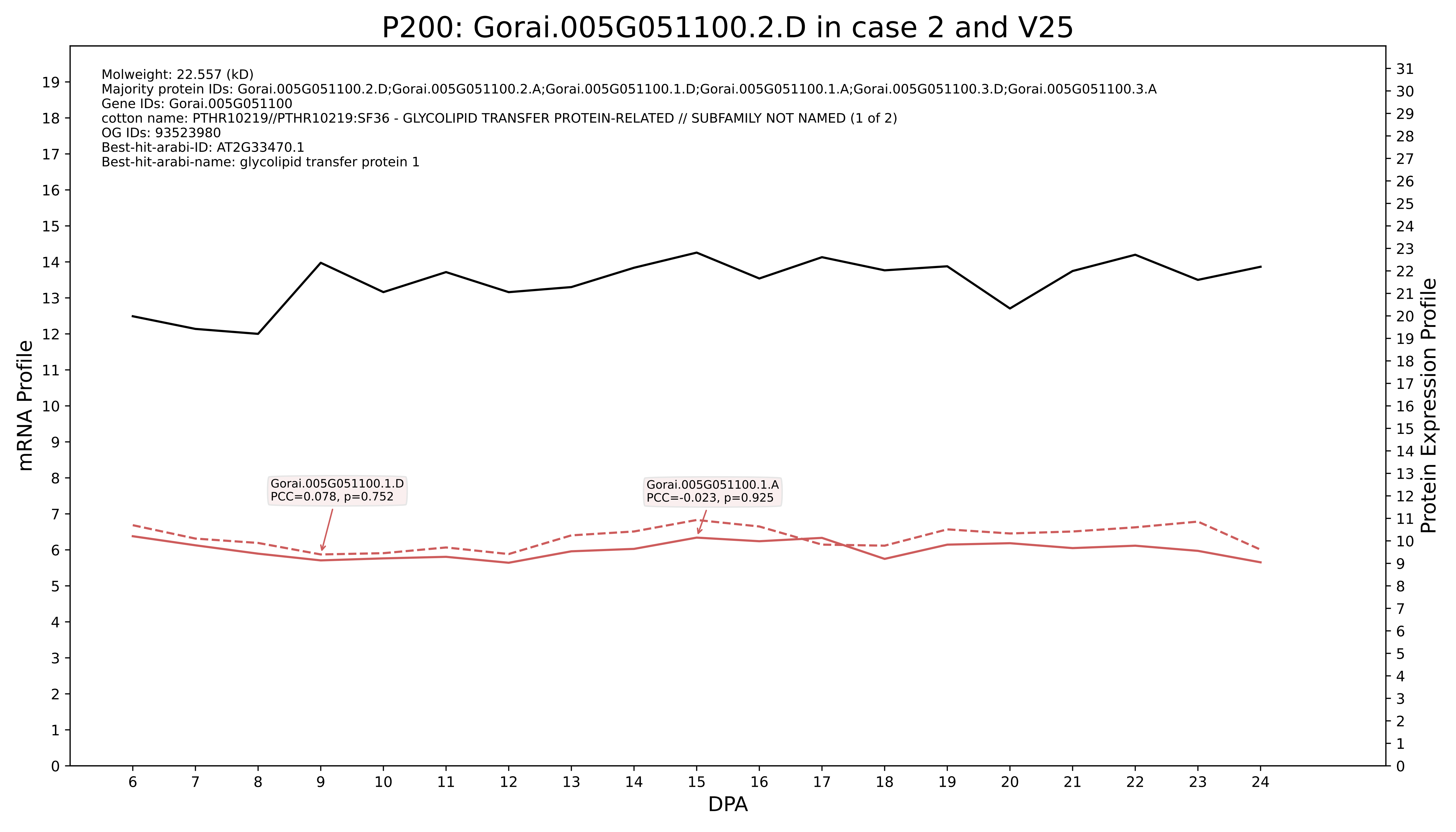

P200: Gorai.005G051100.2.D in case 2 and V25
Molweight: 22.557 (kD)
Majority protein IDs: Gorai.005G051100.2.D;Gorai.005G051100.2.A;Gorai.005G051100.1.D;Gorai.005G051100.1.A;Gorai.005G051100.3.D;Gorai.005G051100.3.A
Gene IDs: Gorai.005G051100
cotton name: PTHR10219//PTHR10219:SF36 - GLYCOLIPID TRANSFER PROTEIN-RELATED // SUBFAMILY NOT NAMED (1 of 2)
OG IDs: 93523980
Best-hit-arabi-ID: AT2G33470.1
Best-hit-arabi-name: glycolipid transfer protein 1
Gorai.005G051100.1.D
PCC=0.078, p=0.752
Gorai.005G051100.1.A
PCC=-0.023, p=0.925
mRNA Profile
Protein Expression Profile
DPA

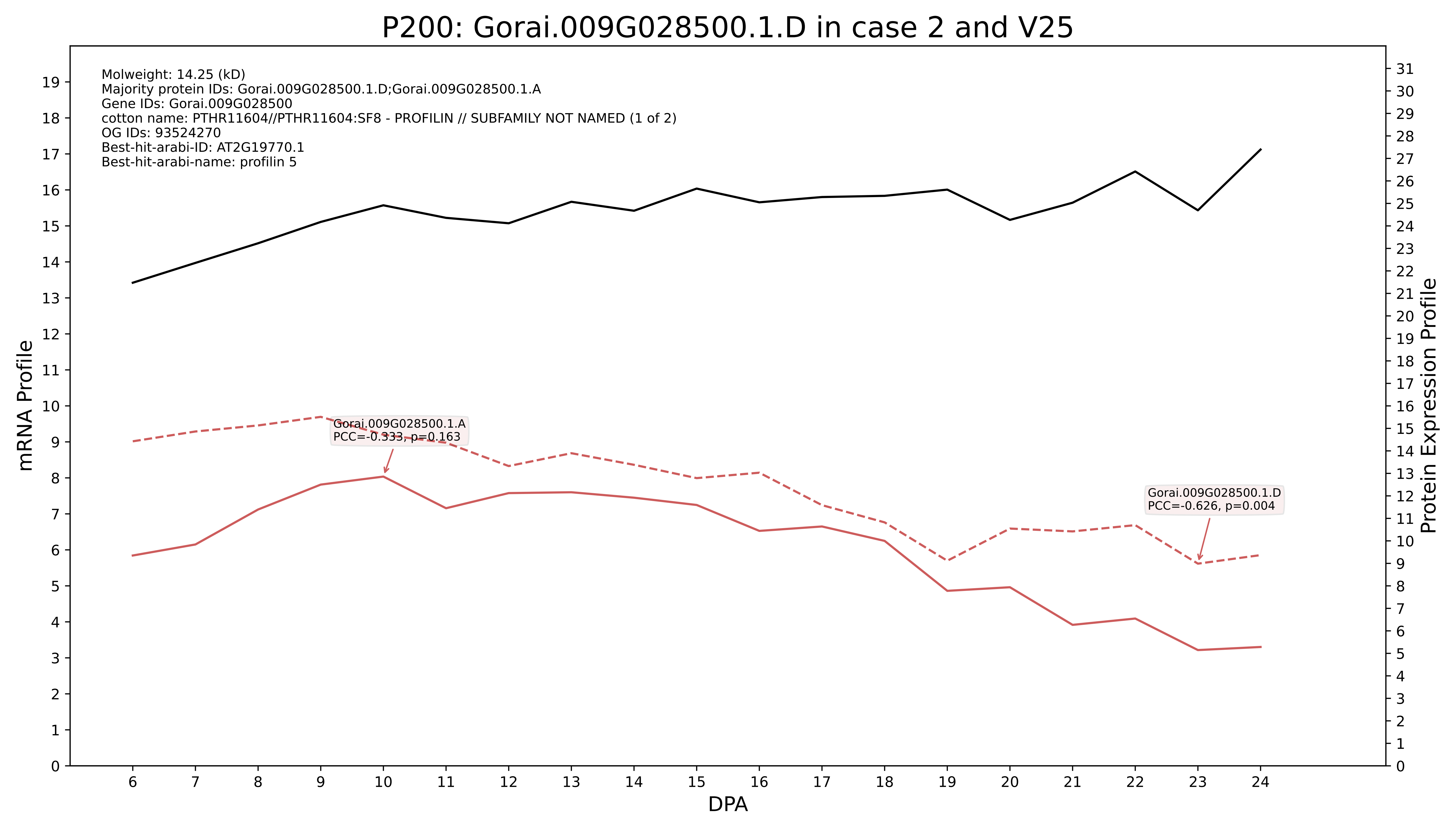
P200: Gorai.009G028500.1.D in case 2 and V25
Molweight: 14.25 (kD)
Majority protein IDs: Gorai.009G028500.1.D;Gorai.009G028500.1.A
Gene IDs: Gorai.009G028500
cotton name: PTHR11604//PTHR11604:SF8 - PROFILIN // SUBFAMILY NOT NAMED (1 of 2)
OG IDs: 93524270
Best-hit-arabi-ID: AT2G19770.1
Best-hit-arabi-name: profilin 5
Gorai.009G028500.1.A
PCC=-0.333, p=0.163
Gorai.009G028500.1.D
PCC=-0.626, p=0.004
mRNA Profile
Protein Expression Profile
DPA

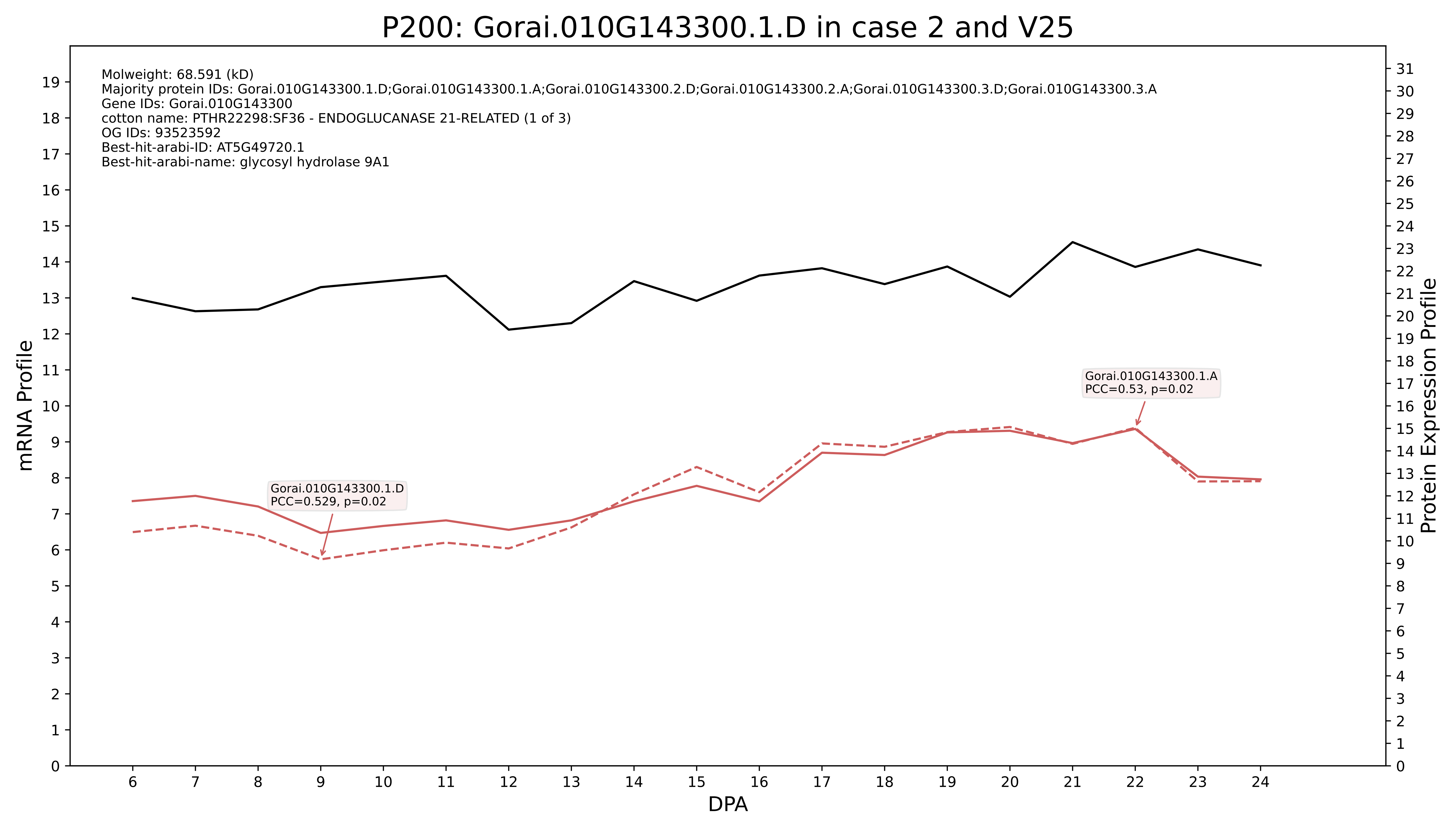
P200: Gorai.010G143300.1.D in case 2 and V25
Molweight: 68.591 (kD)
Majority protein IDs: Gorai.010G143300.1.D;Gorai.010G143300.1.A;Gorai.010G143300.2.D;Gorai.010G143300.2.A;Gorai.010G143300.3.D;Gorai.010G143300.3.A
Gene IDs: Gorai.010G143300
cotton name: PTHR22298:SF36 - ENDOGLUCANASE 21-RELATED (1 of 3)
OG IDs: 93523592
Best-hit-arabi-ID: AT5G49720.1
Best-hit-arabi-name: glycosyl hydrolase 9A1
Gorai.010G143300.1.A
PCC=0.53, p=0.02
Gorai.010G143300.1.D
PCC=0.529, p=0.02
mRNA Profile
Protein Expression Profile
DPA

### 007G137300.7.D in case 2 and V25

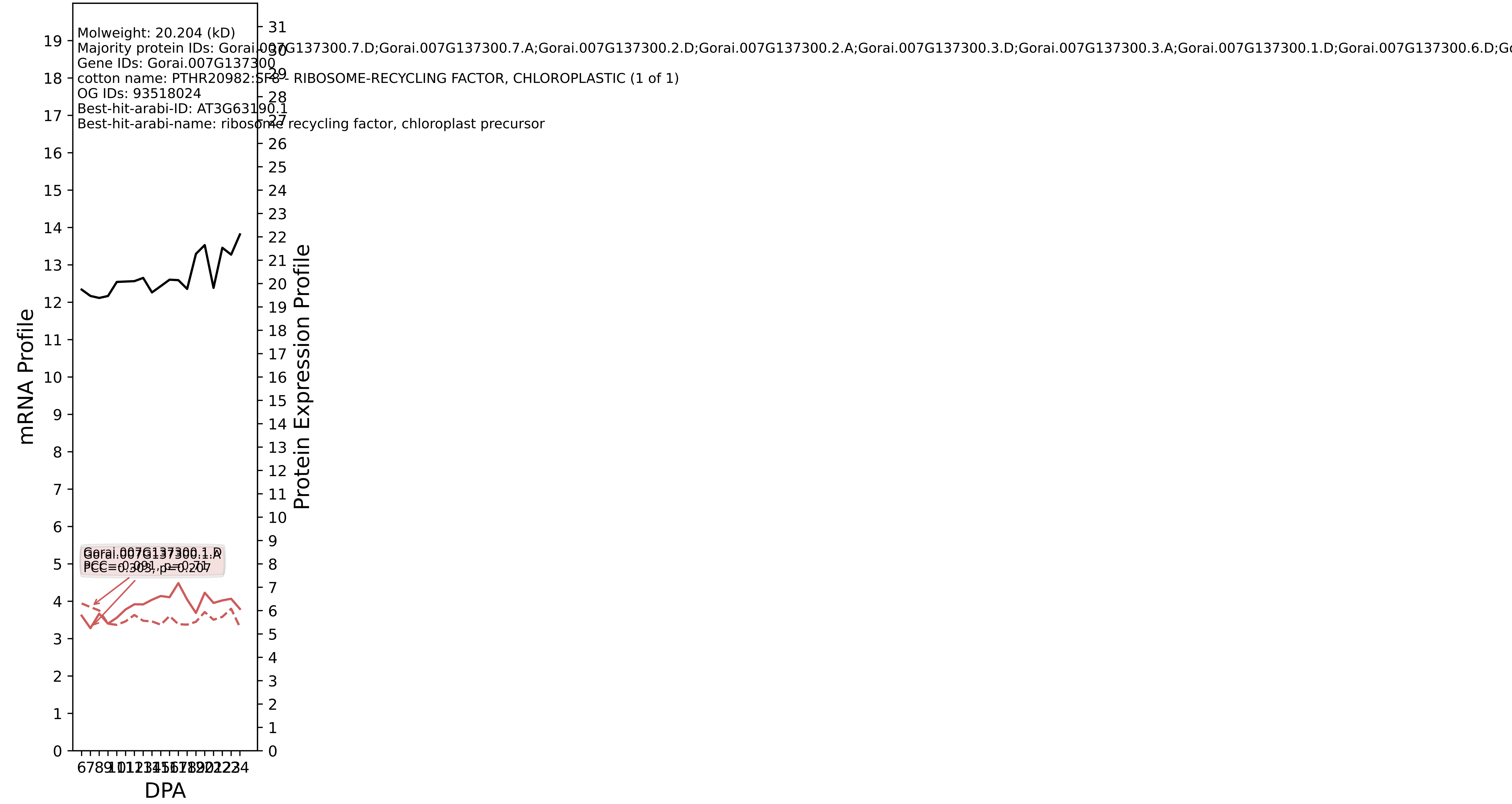

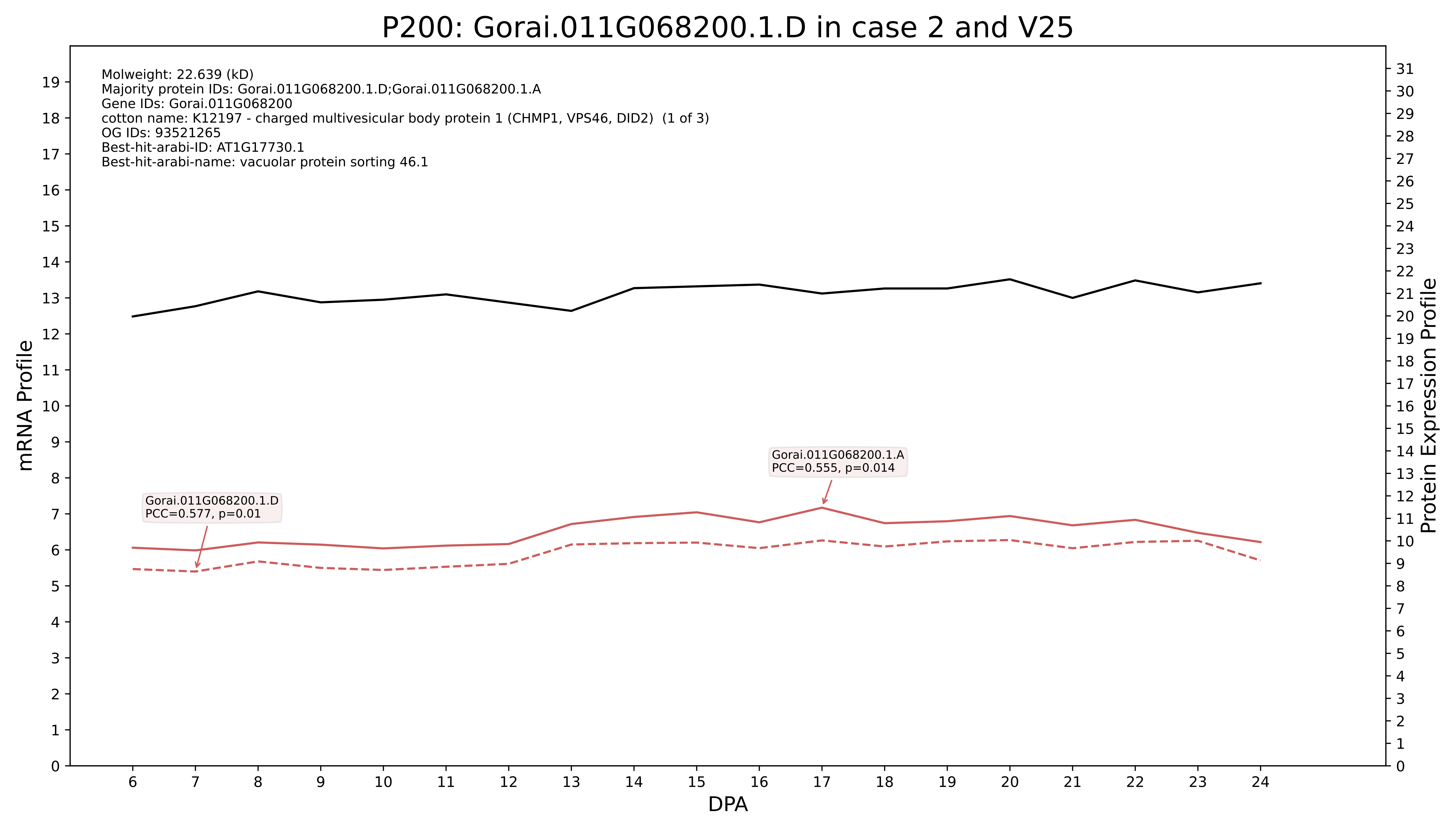

P200: Gorai.011G068200.1.D in case 2 and V25
Molweight: 22.639 (kD)
Majority protein IDs: Gorai.011G068200.1.D;Gorai.011G068200.1.A
Gene IDs: Gorai.011G068200
cotton name: K12197 - charged multivesicular body protein 1 (CHMP1, VPS46, DID2) (1 of 3)
OG IDs: 93521265
Best-hit-arabi-ID: AT1G17730.1
Best-hit-arabi-name: vacuolar protein sorting 46.1
Gorai.011G068200.1.A
PCC=0.555, p=0.014
Gorai.011G068200.1.D
PCC=0.577, p=0.01
mRNA Profile
Protein Expression Profile
DPA

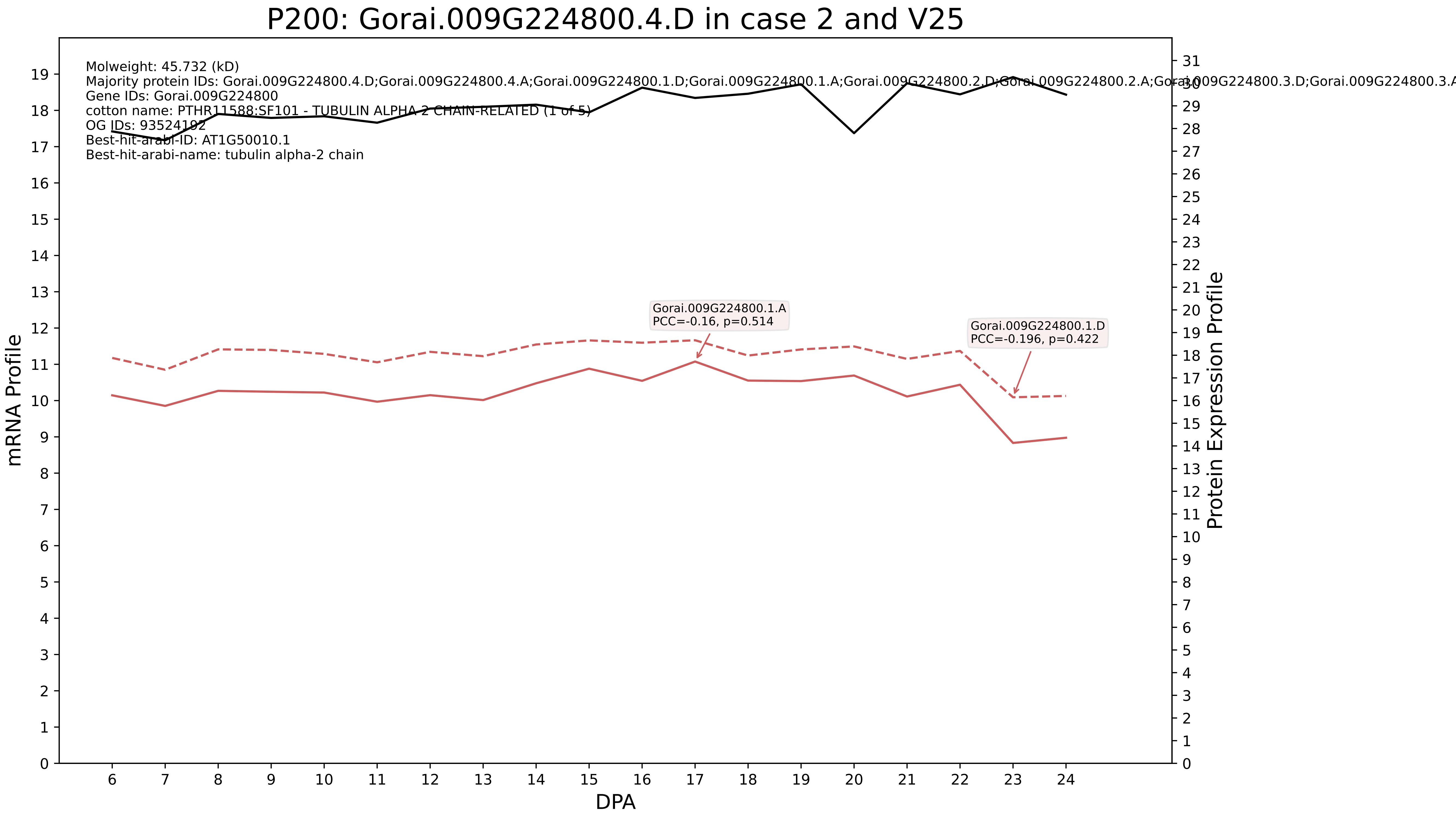
P200: Gorai.009G224800.4.D in case 2 and V25
Molweight: 45.732 (kD)
Majority protein IDs: Gorai.009G224800.4.D;Gorai.009G224800.4.A;Gorai.009G224800.1.D;Gorai.009G224800.1.A;Gorai.009G224800.2.D;Gorai.009G224800.2.A;Gorai.009G224800.3.D;Gorai.009G224800.3.A
Gene IDs: Gorai.009G224800
cotton name: PTHR11588:SF101 - TUBULIN ALPHA-2 CHAIN-RELATED (1 of 5)
OG IDs: 93524192
Best-hit-arabi-ID: AT1G50010.1
Best-hit-arabi-name: tubulin alpha-2 chain
Gorai.009G224800.1.A
PCC=-0.16, p=0.514
Gorai.009G224800.1.D
PCC=-0.196, p=0.422
mRNA Profile
Protein Expression Profile
DPA

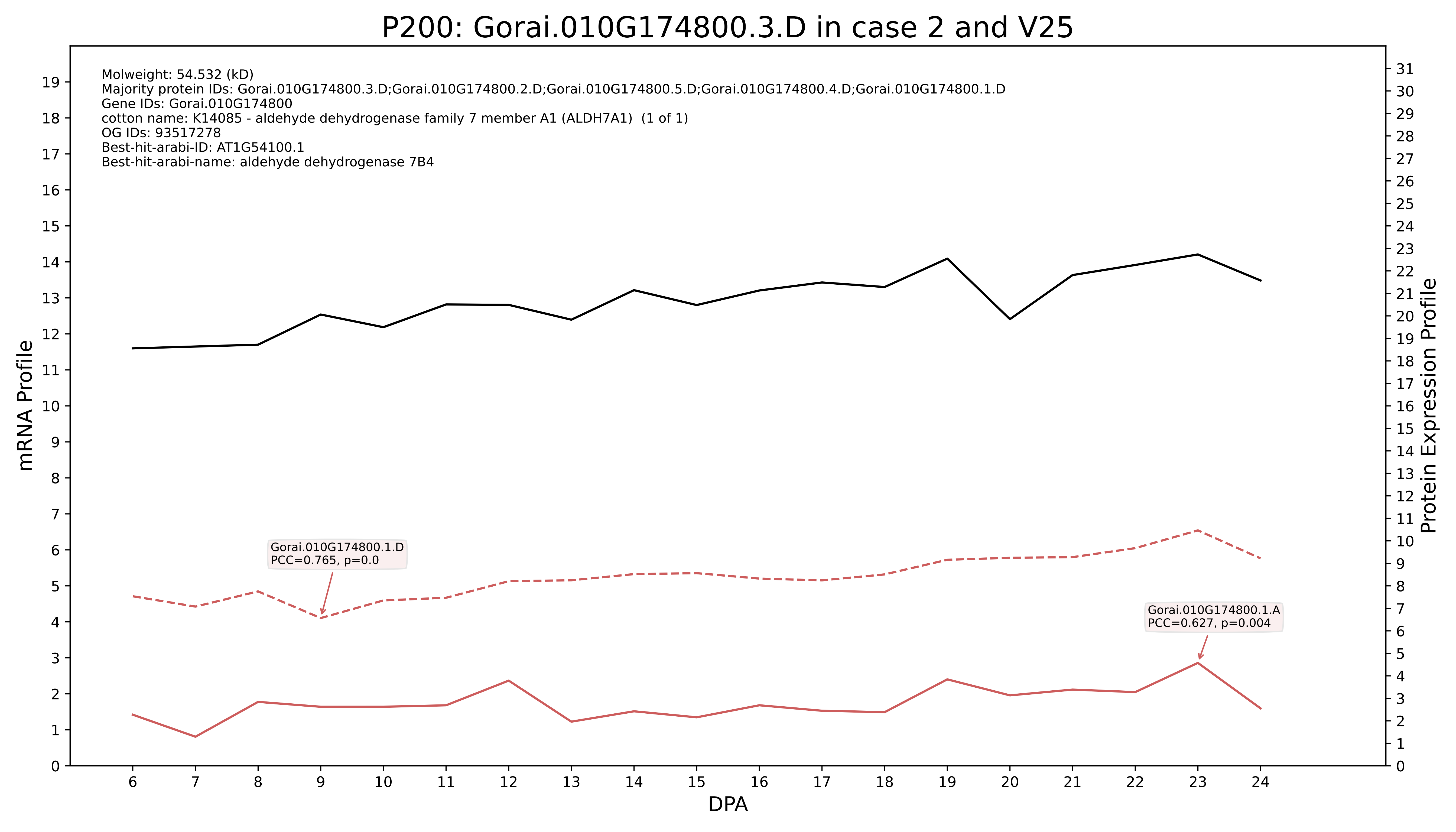
P200: Gorai.010G174800.3.D in case 2 and V25
Molweight: 54.532 (kD)
Majority protein IDs: Gorai.010G174800.3.D;Gorai.010G174800.2.D;Gorai.010G174800.5.D;Gorai.010G174800.4.D;Gorai.010G174800.1.D
Gene IDs: Gorai.010G174800
cotton name: K14085 - aldehyde dehydrogenase family 7 member A1 (ALDH7A1) (1 of 1)
OG IDs: 93517278
Best-hit-arabi-ID: AT1G54100.1
Best-hit-arabi-name: aldehyde dehydrogenase 7B4
Gorai.010G174800.1.D
PCC=0.765, p=0.0
Gorai.010G174800.1.A
PCC=0.627, p=0.004
mRNA Profile
Protein Expression Profile
DPA

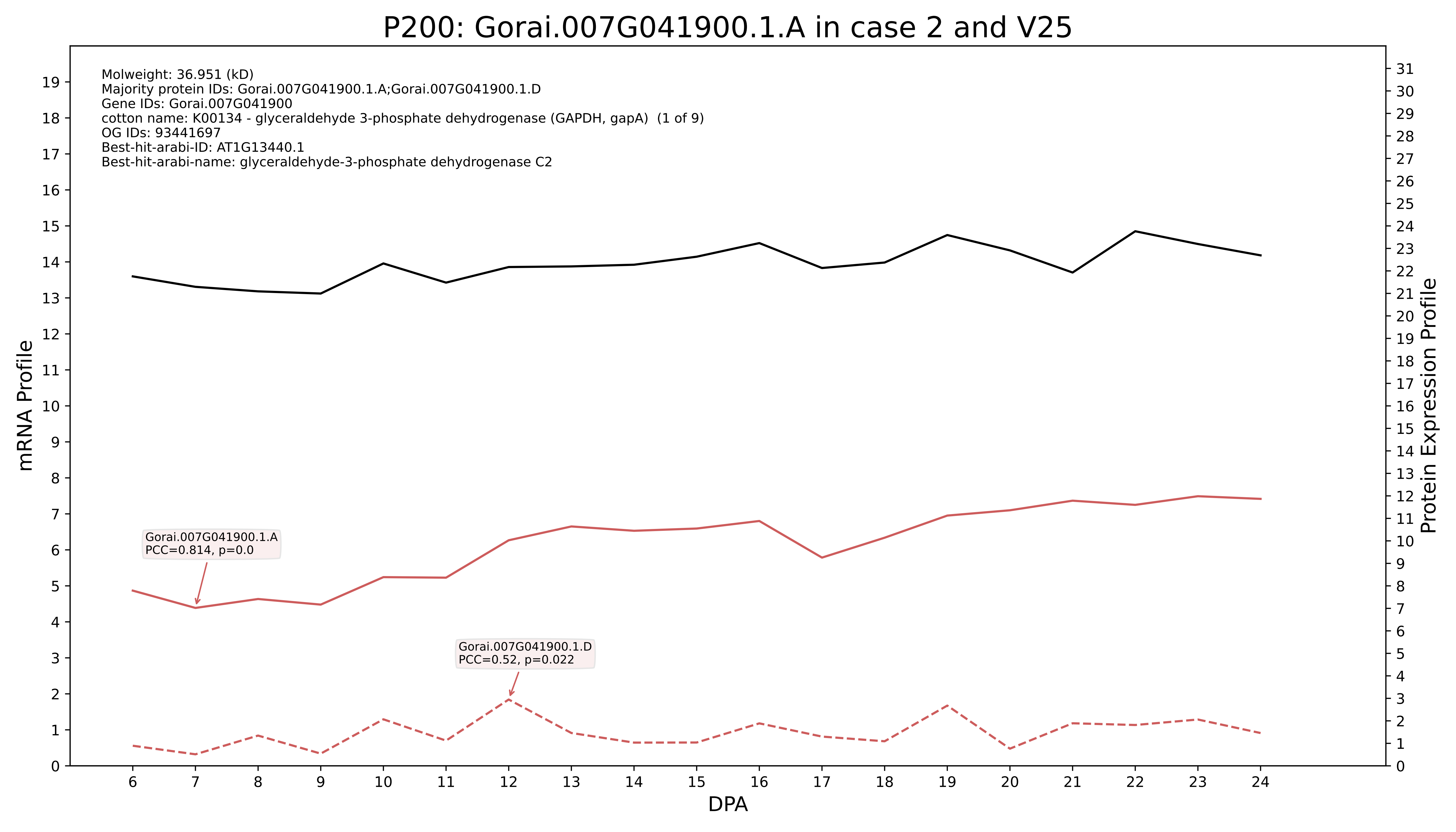

P200: Gorai.007G041900.1.A in case 2 and V25
Molweight: 36.951 (kD)
Majority protein IDs: Gorai.007G041900.1.A;Gorai.007G041900.1.D
Gene IDs: Gorai.007G041900
cotton name: K00134 - glyceraldehyde 3-phosphate dehydrogenase (GAPDH, gapA) (1 of 9)
OG IDs: 93441697
Best-hit-arabi-ID: AT1G13440.1
Best-hit-arabi-name: glyceraldehyde-3-phosphate dehydrogenase C2
Gorai.007G041900.1.A
PCC=0.814, p=0.0
Gorai.007G041900.1.D
PCC=0.52, p=0.022
mRNA Profile
Protein Expression Profile
DPA

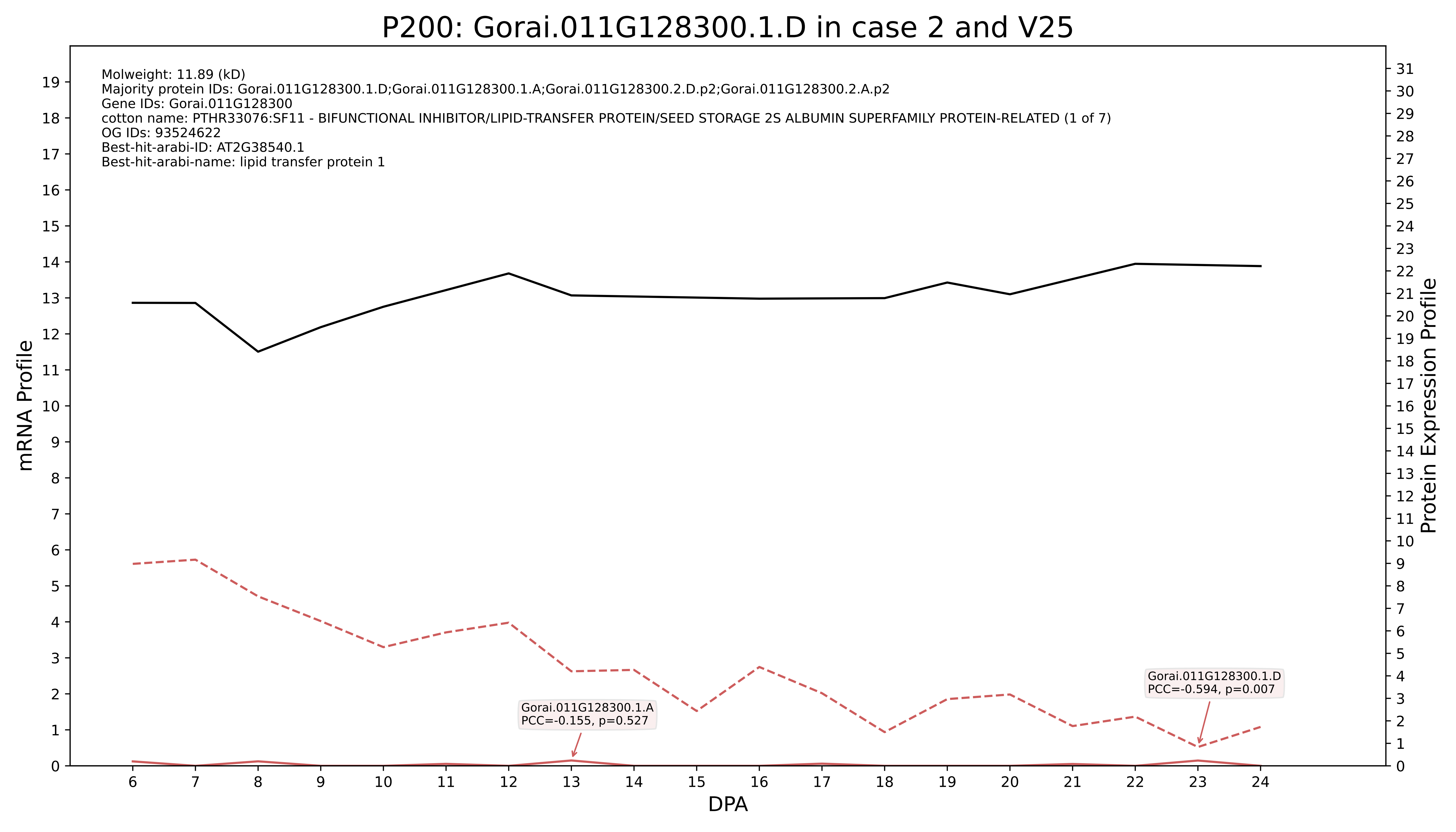
P200: Gorai.011G128300.1.D in case 2 and V25
Molweight: 11.89 (kD)
Majority protein IDs: Gorai.011G128300.1.D;Gorai.011G128300.1.A;Gorai.011G128300.2.D.p2;Gorai.011G128300.2.A.p2
Gene IDs: Gorai.011G128300
cotton name: PTHR33076:SF11 - BIFUNCTIONAL INHIBITOR/LIPID-TRANSFER PROTEIN/SEED STORAGE 2S ALBUMIN SUPERFAMILY PROTEIN-RELATED (1 of 7)
OG IDs: 93524622
Best-hit-arabi-ID: AT2G38540.1
Best-hit-arabi-name: lipid transfer protein 1
Gorai.011G128300.1.A
PCC=-0.155, p=0.527
Gorai.011G128300.1.D
PCC=-0.594, p=0.007
mRNA Profile
Protein Expression Profile
DPA

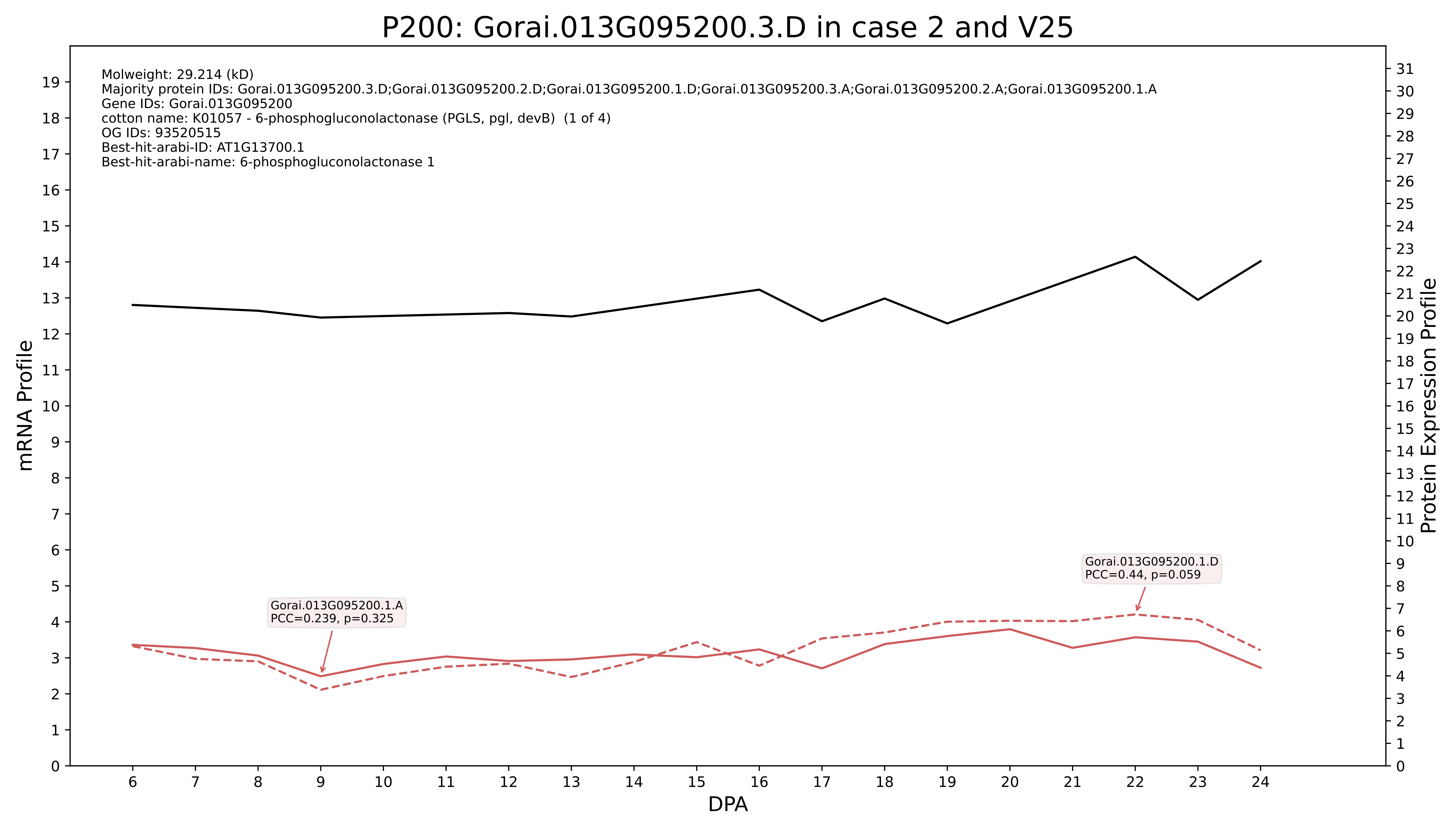
P200: Gorai.013G095200.3.D in case 2 and V25
Molweight: 29.214 (kD)
Majority protein IDs: Gorai.013G095200.3.D;Gorai.013G095200.2.D;Gorai.013G095200.1.D;Gorai.013G095200.3.A;Gorai.013G095200.2.A;Gorai.013G095200.1.A
Gene IDs: Gorai.013G095200
cotton name: K01057 - 6-phosphogluconolactonase (PGLS, pgl, devB) (1 of 4)
OG IDs: 93520515
Best-hit-arabi-ID: AT1G13700.1
Best-hit-arabi-name: 6-phosphogluconolactonase 1
Gorai.013G095200.1.D
PCC=0.44, p=0.059
Gorai.013G095200.1.A
PCC=0.239, p=0.325
mRNA Profile
Protein Expression Profile
DPA

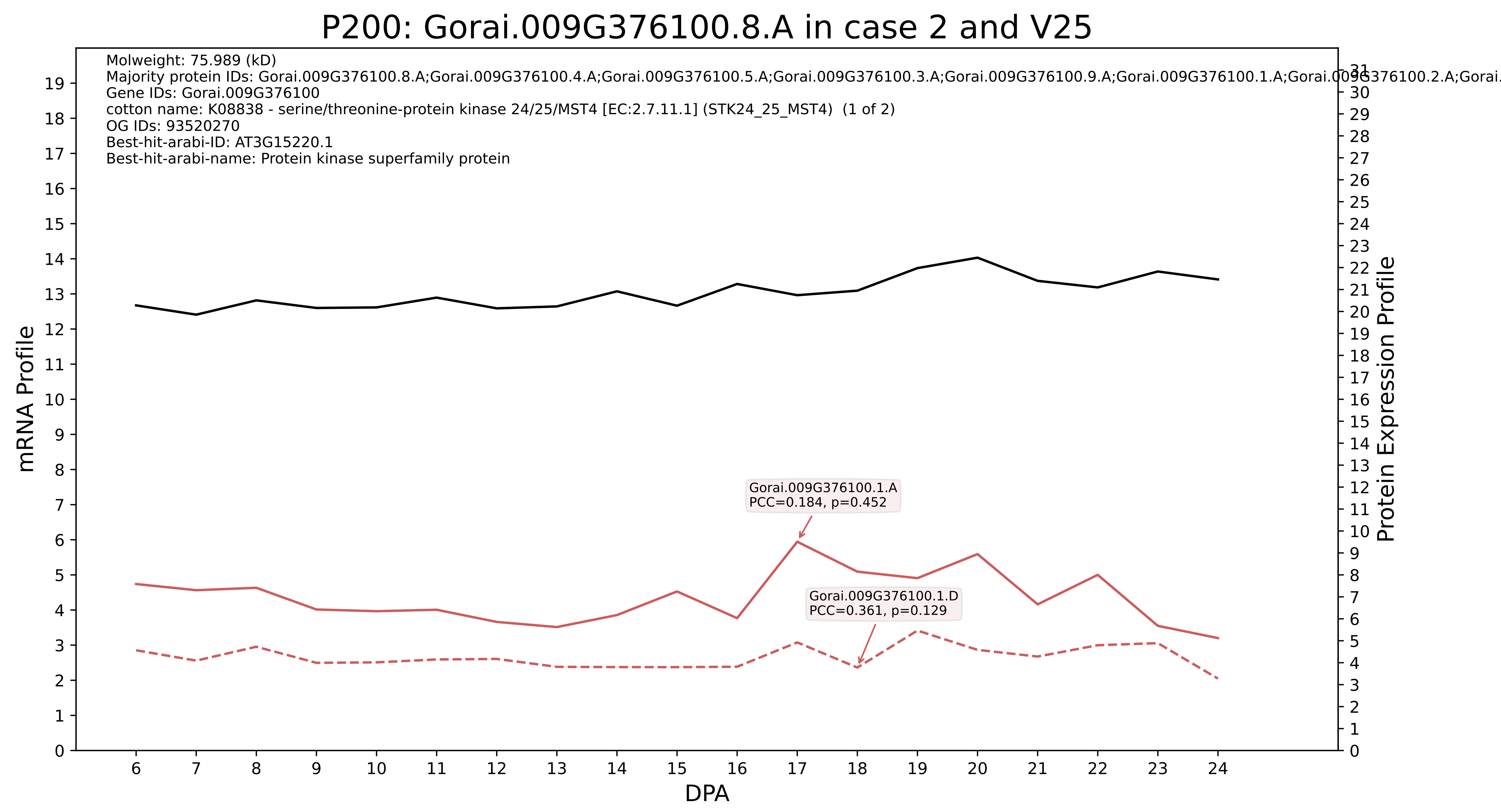

P200: Gorai.009G376100.8.A in case 2 and V25
Molweight: 75.989 (kD)
Majority protein IDs: Gorai.009G376100.8.A;Gorai.009G376100.4.A;Gorai.009G376100.5.A;Gorai.009G376100.3.A;Gorai.009G376100.9.A;Gorai.009G376100.1.A;Gorai.009G376100.2.A;Gorai.
Gene IDs: Gorai.009G376100
cotton name: K08838 - serine/threonine-protein kinase 24/25/MST4 [EC:2.7.11.1] (STK24_25_MST4) (1 of 2)
OG IDs: 93520270
Best-hit-arabi-ID: AT3G15220.1
Best-hit-arabi-name: Protein kinase superfamily protein
Gorai.009G376100.1.A
PCC=0.184, p=0.452
Gorai.009G376100.1.D
PCC=0.361, p=0.129
mRNA Profile
Protein Expression Profile
DPA

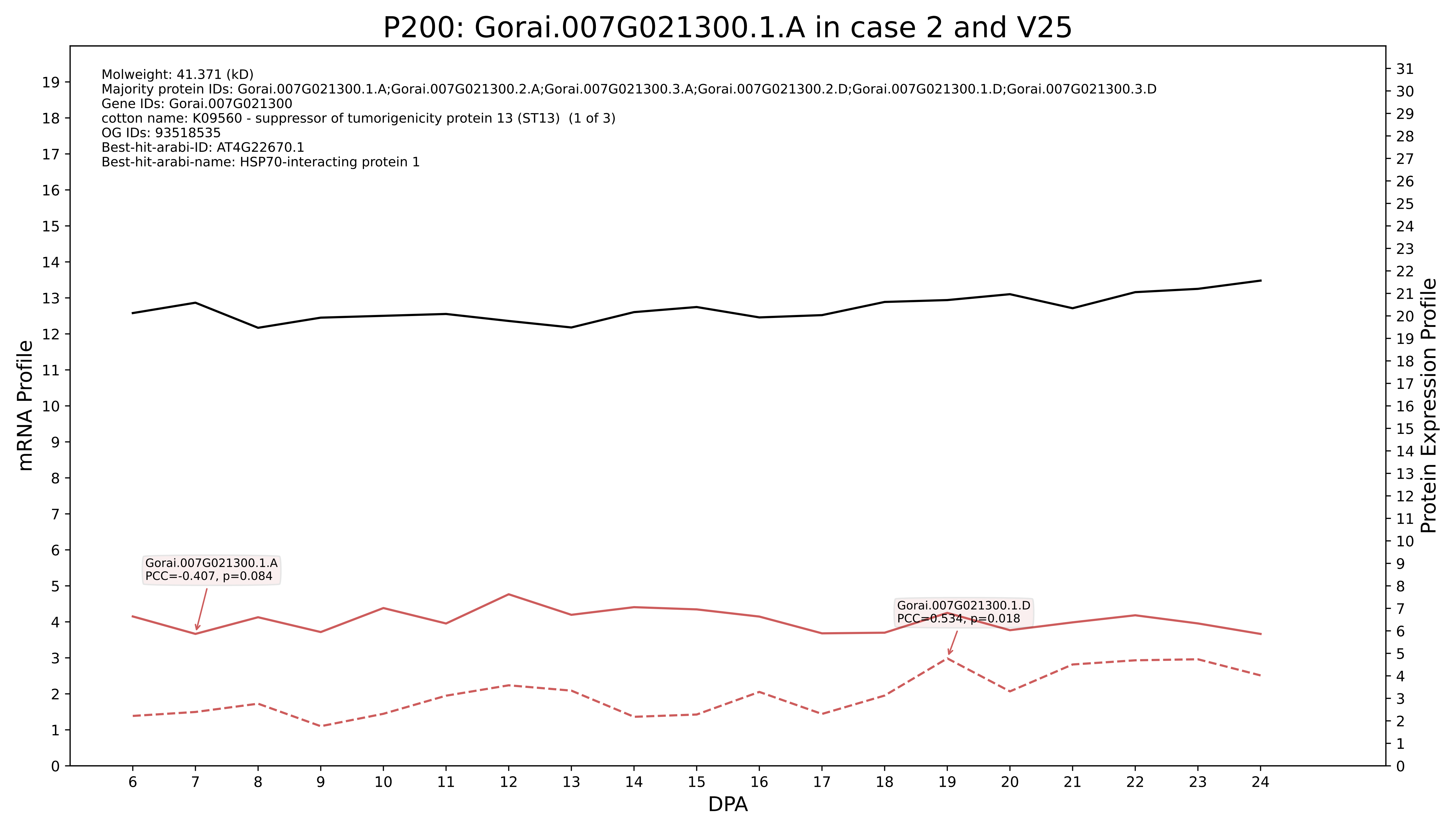
P200: Gorai.007G021300.1.A in case 2 and V25
Molweight: 41.371 (kD)
Majority protein IDs: Gorai.007G021300.1.A;Gorai.007G021300.2.A;Gorai.007G021300.3.A;Gorai.007G021300.2.D;Gorai.007G021300.1.D;Gorai.007G021300.3.D
Gene IDs: Gorai.007G021300
cotton name: K09560 - suppressor of tumorigenicity protein 13 (ST13) (1 of 3)
OG IDs: 93518535
Best-hit-arabi-ID: AT4G22670.1
Best-hit-arabi-name: HSP70-interacting protein 1
Gorai.007G021300.1.A
PCC=-0.407, p=0.084
Gorai.007G021300.1.D
PCC=0.534, p=0.018
mRNA Profile
Protein Expression Profile
DPA

### 00: Gorai.010G053700.3.D in case 2 and V25

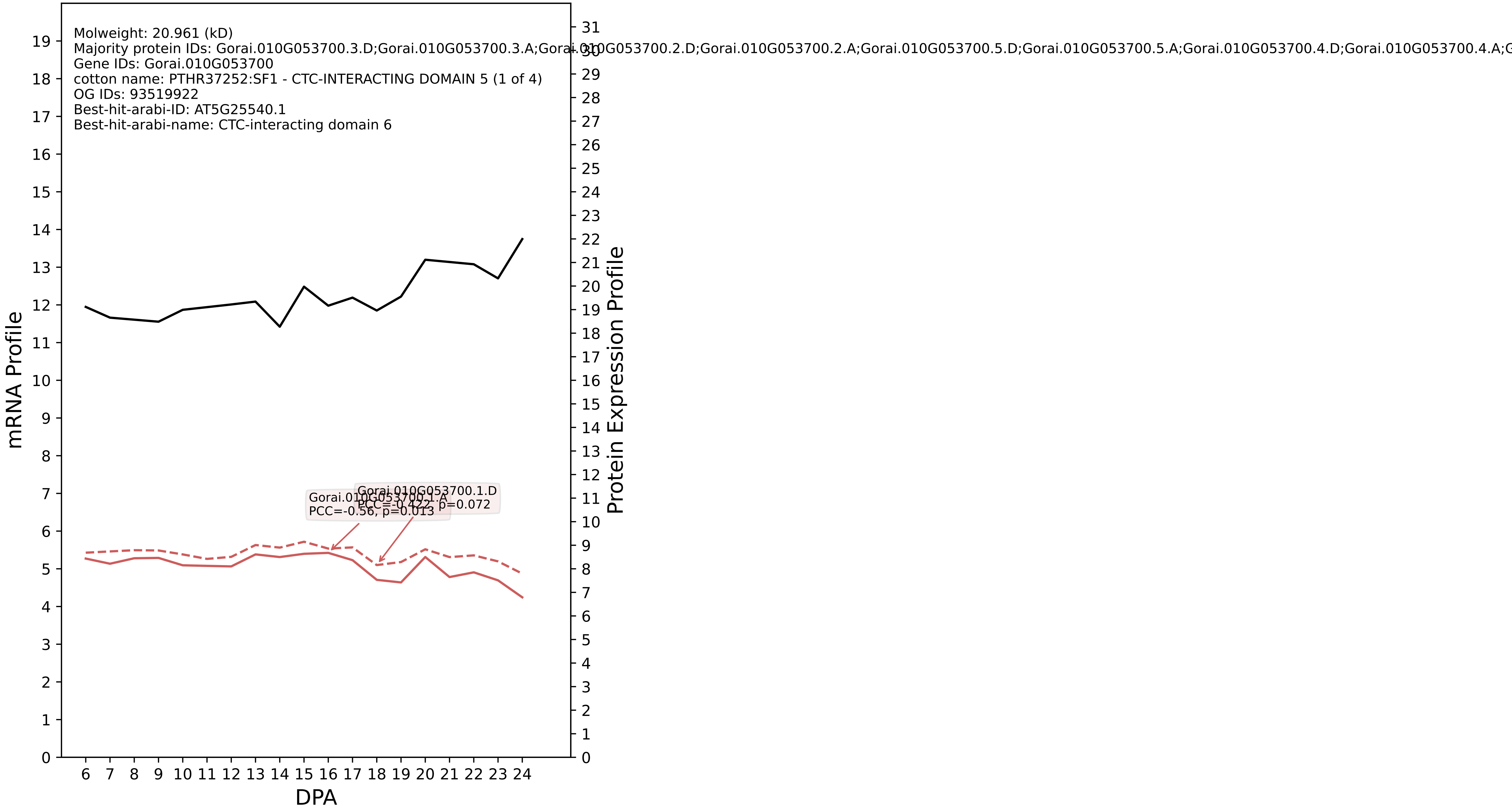

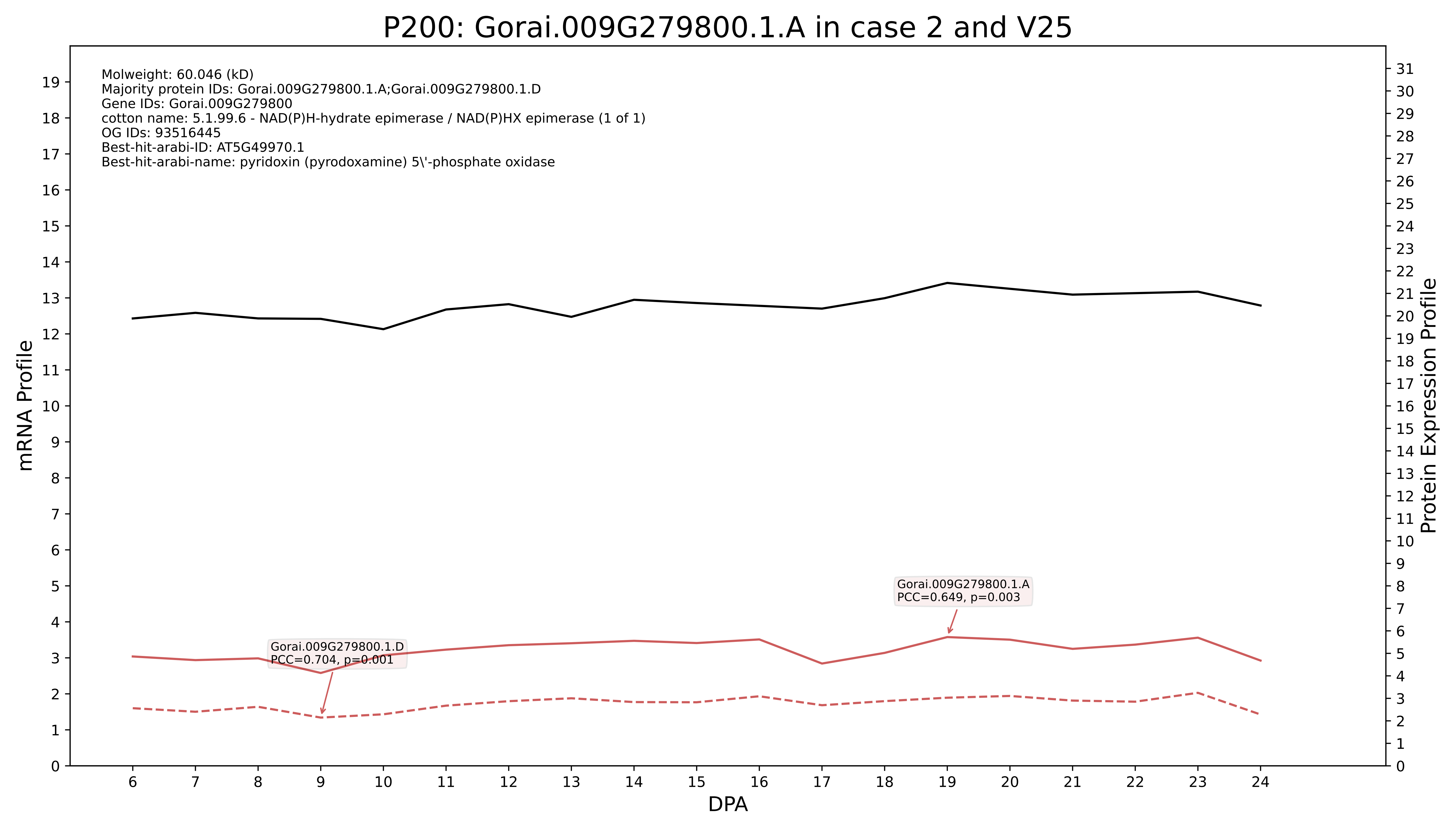
P200: Gorai.009G279800.1.A in case 2 and V25
Molweight: 60.046 (kD)
Majority protein IDs: Gorai.009G279800.1.A;Gorai.009G279800.1.D
Gene IDs: Gorai.009G279800
cotton name: 5.1.99.6 - NAD(P)H-hydrate epimerase / NAD(P)HX epimerase (1 of 1)
OG IDs: 93516445
Best-hit-arabi-ID: AT5G49970.1
Best-hit-arabi-name: pyridoxin (pyrodoxamine) 5\'-phosphate oxidase
Gorai.009G279800.1.A
PCC=0.649, p=0.003
Gorai.009G279800.1.D
PCC=0.704, p=0.001
mRNA Profile
Protein Expression Profile
DPA

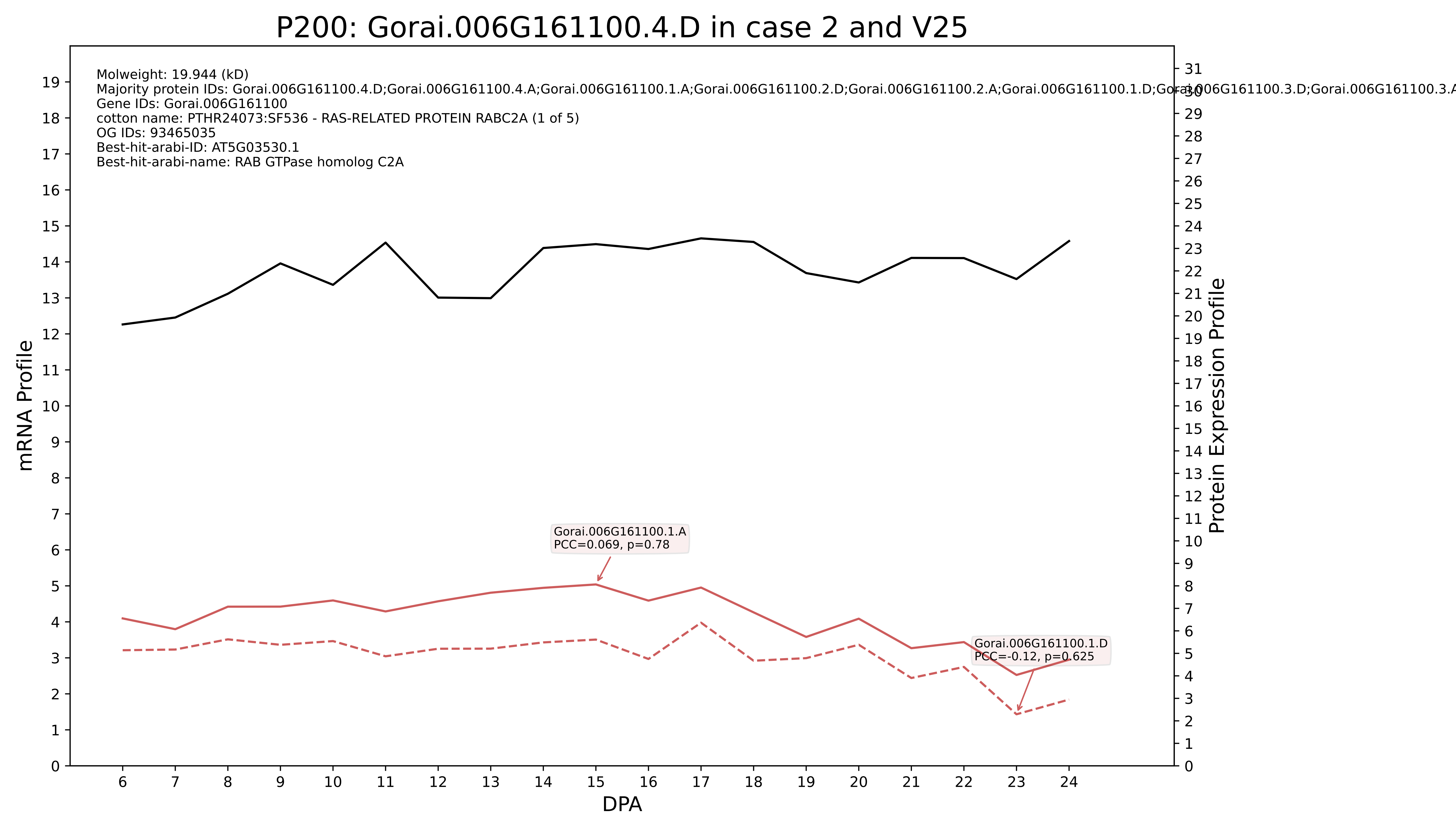
P200: Gorai.006G161100.4.D in case 2 and V25
Molweight: 19.944 (kD)
Majority protein IDs: Gorai.006G161100.4.D;Gorai.006G161100.4.A;Gorai.006G161100.1.A;Gorai.006G161100.2.D;Gorai.006G161100.2.A;Gorai.006G161100.1.D;Gorai.006G161100.3.D;Gorai.006G161100.3.A
Gene IDs: Gorai.006G161100
cotton name: PTHR24073:SF536 - RAS-RELATED PROTEIN RABC2A (1 of 5)
OG IDs: 93465035
Best-hit-arabi-ID: AT5G03530.1
Best-hit-arabi-name: RAB GTPase homolog C2A
Gorai.006G161100.1.A
PCC=0.069, p=0.78
Gorai.006G161100.1.D
PCC=-0.12, p=0.625
mRNA Profile
Protein Expression Profile
DPA

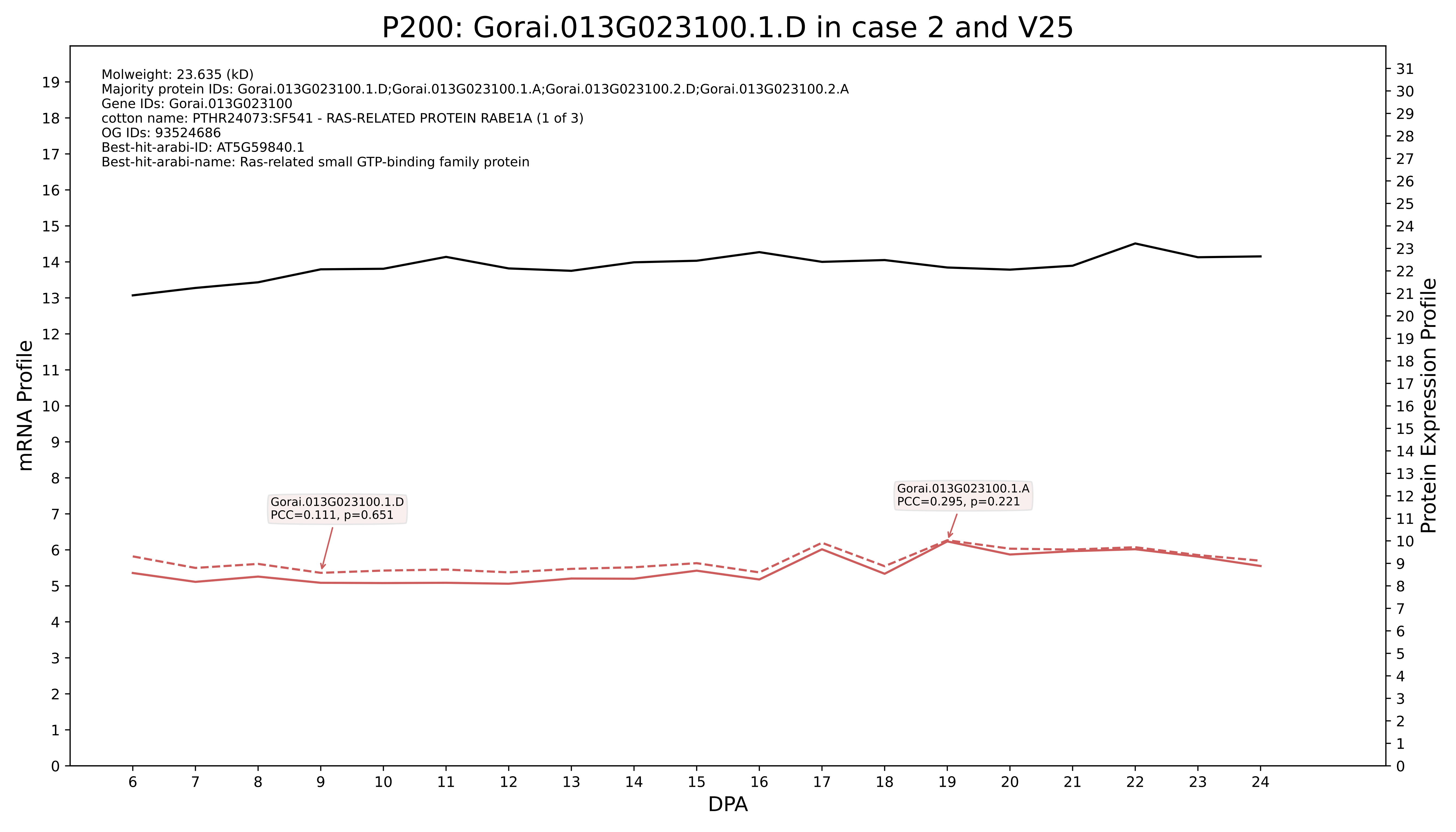
P200: Gorai.013G023100.1.D in case 2 and V25
Molweight: 23.635 (kD)
Majority protein IDs: Gorai.013G023100.1.D;Gorai.013G023100.1.A;Gorai.013G023100.2.D;Gorai.013G023100.2.A
Gene IDs: Gorai.013G023100
cotton name: PTHR24073:SF541 - RAS-RELATED PROTEIN RABE1A (1 of 3)
OG IDs: 93524686
Best-hit-arabi-ID: AT5G59840.1
Best-hit-arabi-name: Ras-related small GTP-binding family protein
Gorai.013G023100.1.D
PCC=0.111, p=0.651
Gorai.013G023100.1.A
PCC=0.295, p=0.221
mRNA Profile
Protein Expression Profile
DPA

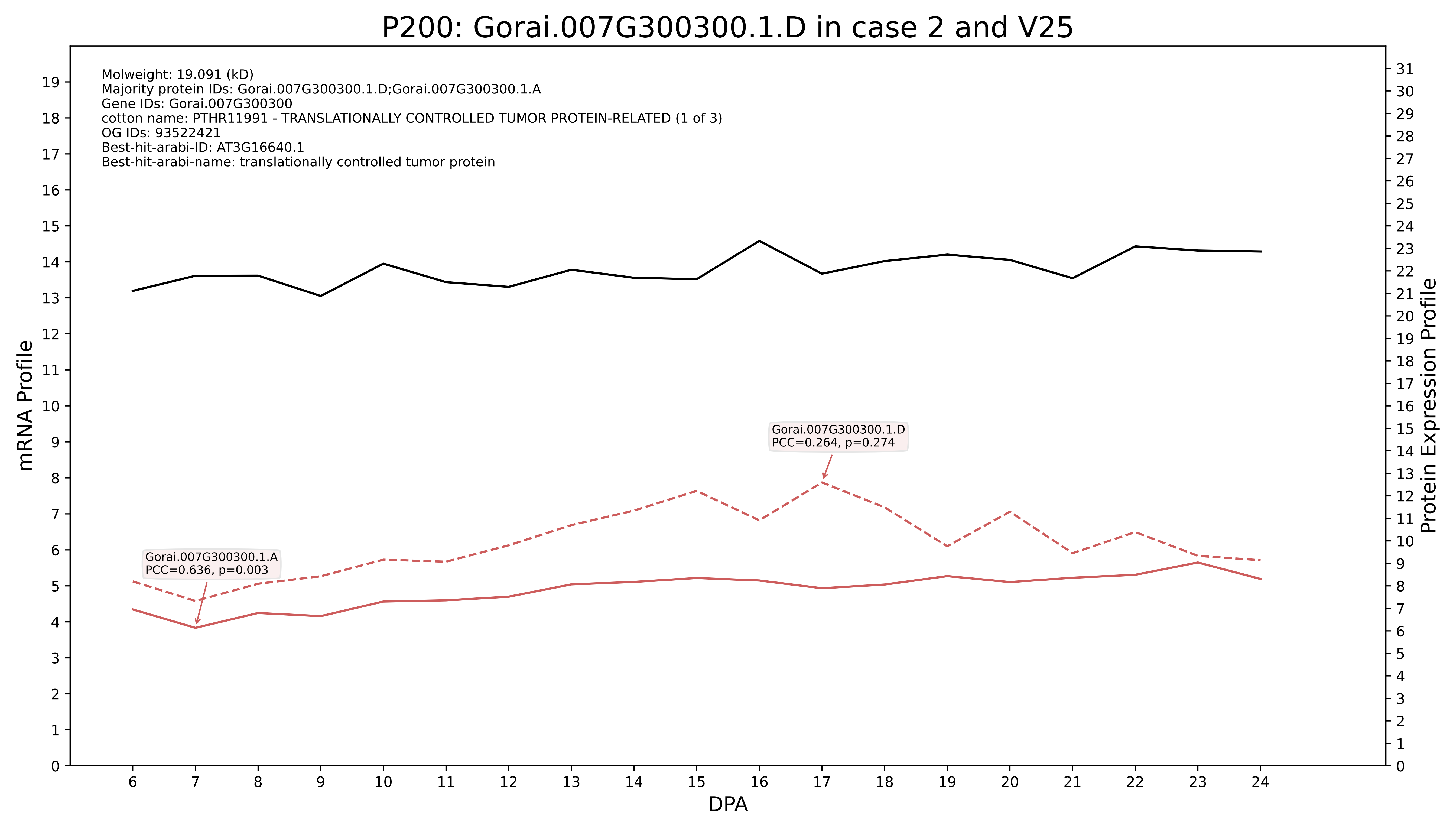
P200: Gorai.007G300300.1.D in case 2 and V25
Molweight: 19.091 (kD)
Majority protein IDs: Gorai.007G300300.1.D;Gorai.007G300300.1.A
Gene IDs: Gorai.007G300300
cotton name: PTHR11991 - TRANSLATIONALLY CONTROLLED TUMOR PROTEIN-RELATED (1 of 3)
OG IDs: 93522421
Best-hit-arabi-ID: AT3G16640.1
Best-hit-arabi-name: translationally controlled tumor protein
Gorai.007G300300.1.D
PCC=0.264, p=0.274
Gorai.007G300300.1.A
PCC=0.636, p=0.003
mRNA Profile
Protein Expression Profile
DPA

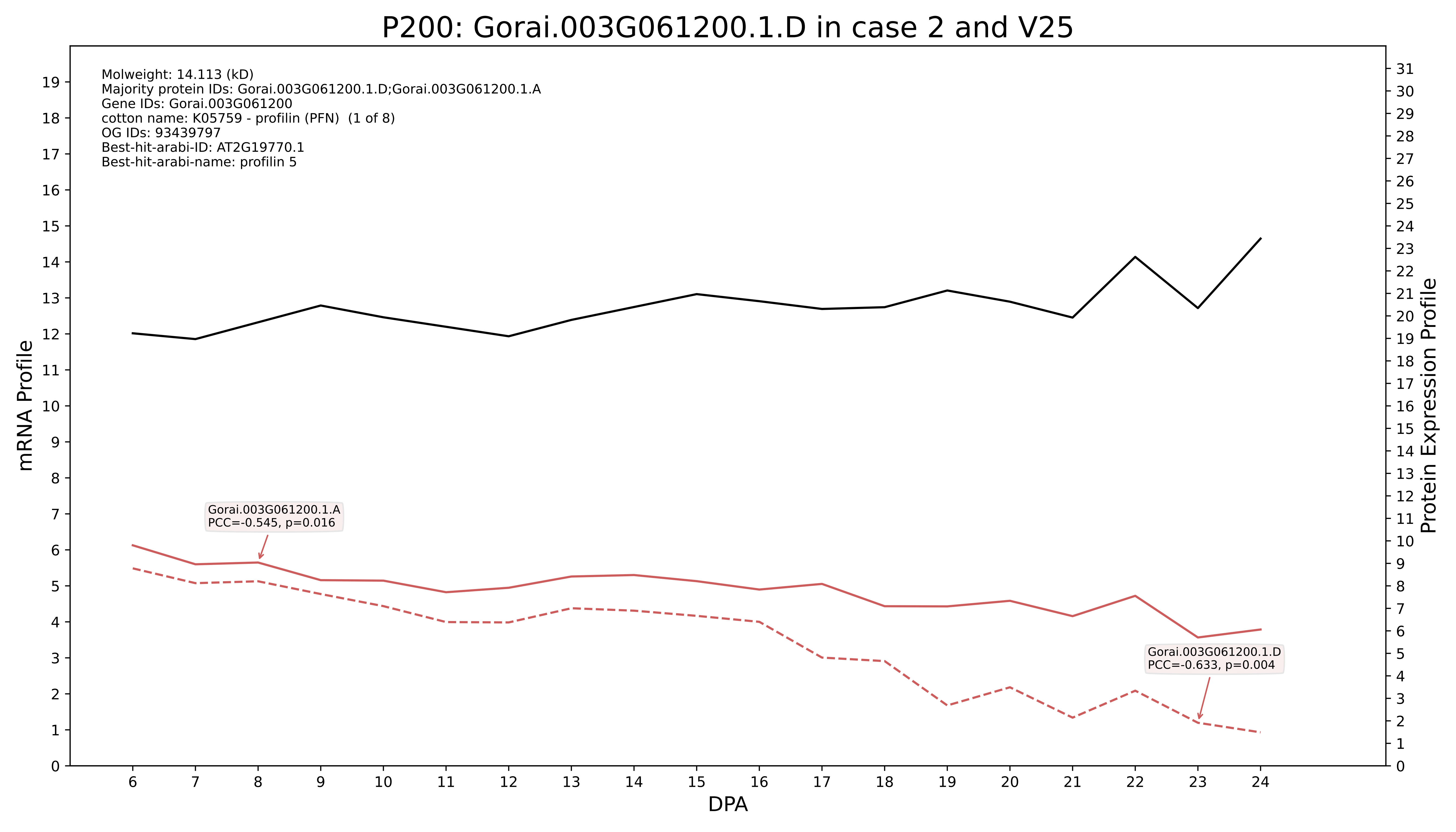

P200: Gorai.003G061200.1.D in case 2 and V25
Molweight: 14.113 (kD)
Majority protein IDs: Gorai.003G061200.1.D;Gorai.003G061200.1.A
Gene IDs: Gorai.003G061200
cotton name: K05759 - profilin (PFN) (1 of 8)
OG IDs: 93439797
Best-hit-arabi-ID: AT2G19770.1
Best-hit-arabi-name: profilin 5
Gorai.003G061200.1.A
PCC=-0.545, p=0.016
Gorai.003G061200.1.D
PCC=-0.633, p=0.004
mRNA Profile
Protein Expression Profile
DPA

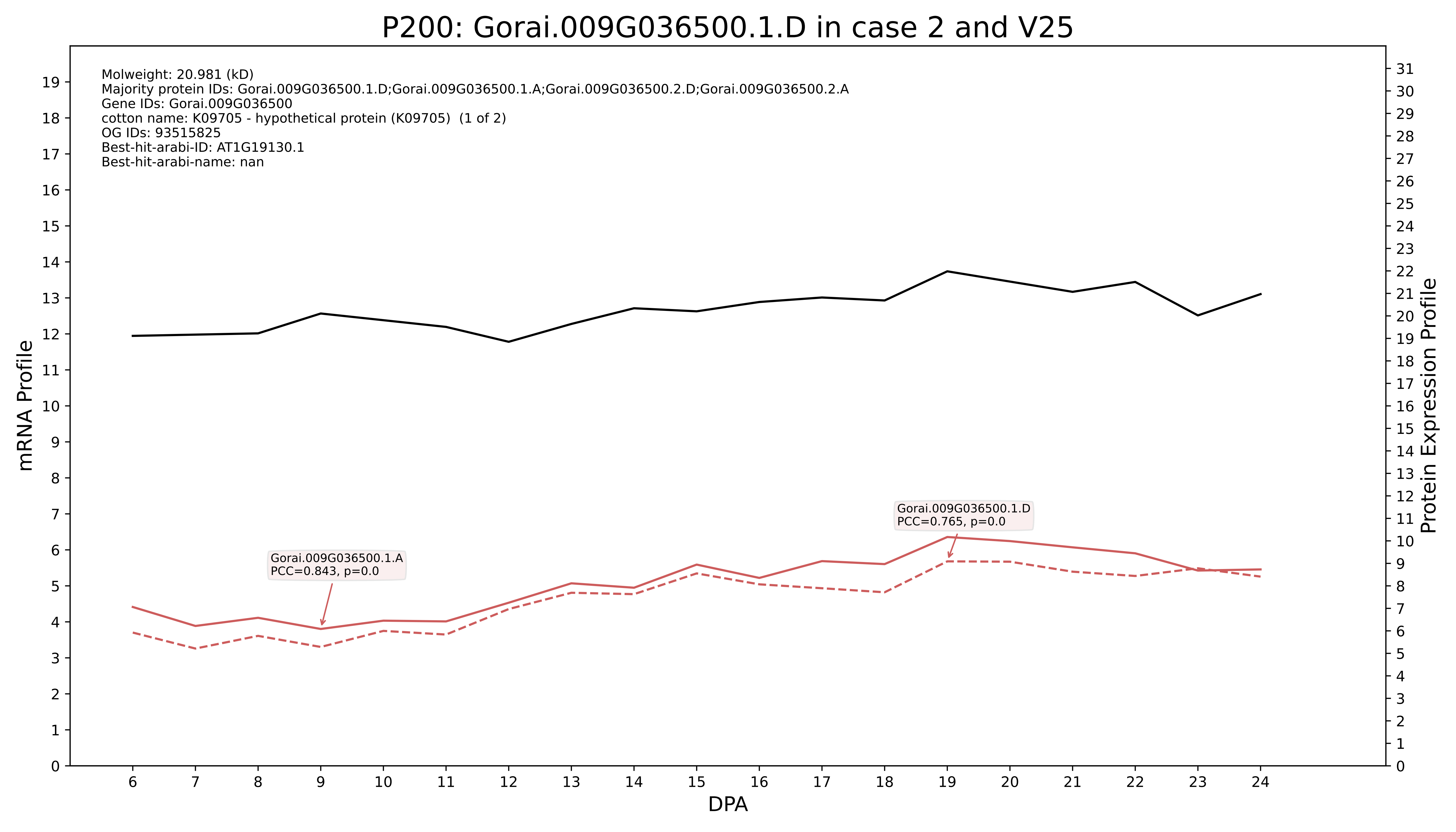
P200: Gorai.009G036500.1.D in case 2 and V25
Molweight: 20.981 (kD)
Majority protein IDs: Gorai.009G036500.1.D;Gorai.009G036500.1.A;Gorai.009G036500.2.D;Gorai.009G036500.2.A
Gene IDs: Gorai.009G036500
cotton name: K09705 - hypothetical protein (K09705) (1 of 2)
OG IDs: 93515825
Best-hit-arabi-ID: AT1G19130.1
Best-hit-arabi-name: nan
Gorai.009G036500.1.D
PCC=0.765, p=0.0
Gorai.009G036500.1.A
PCC=0.843, p=0.0
mRNA Profile
Protein Expression Profile
DPA

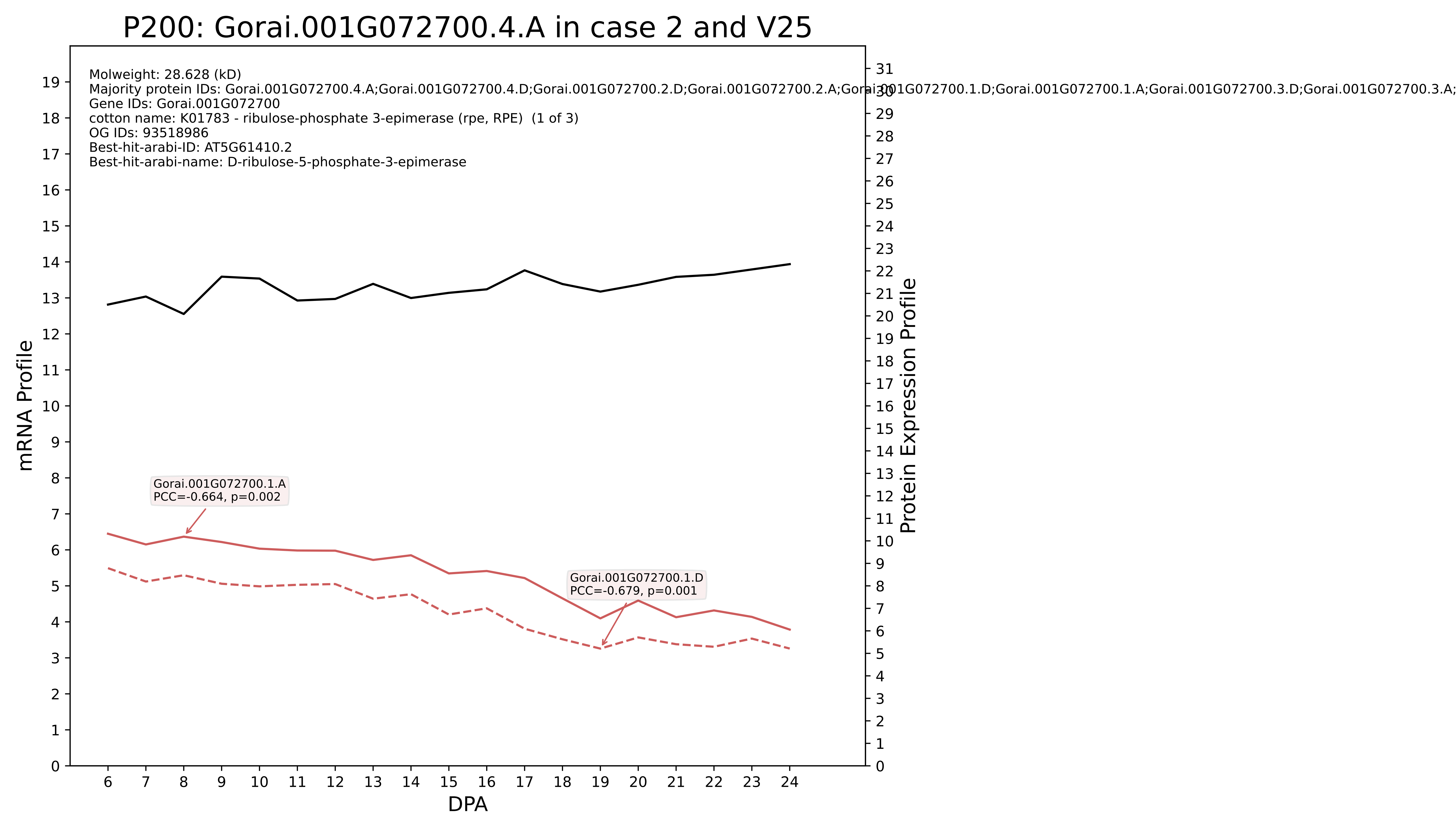
P200: Gorai.001G072700.4.A in case 2 and V25
Molweight: 28.628 (kD)
Majority protein IDs: Gorai.001G072700.4.A;Gorai.001G072700.4.D;Gorai.001G072700.2.D;Gorai.001G072700.2.A;Gorai.001G072700.1.D;Gorai.001G072700.1.A;Gorai.001G072700.3.D;Gorai.001G072700.3.A;
Gene IDs: Gorai.001G072700
cotton name: K01783 - ribulose-phosphate 3-epimerase (rpe, RPE) (1 of 3)
OG IDs: 93518986
Best-hit-arabi-ID: AT5G61410.2
Best-hit-arabi-name: D-ribulose-5-phosphate-3-epimerase
Gorai.001G072700.1.A
PCC=-0.664, p=0.002
Gorai.001G072700.1.D
PCC=-0.679, p=0.001
mRNA Profile
Protein Expression Profile
DPA

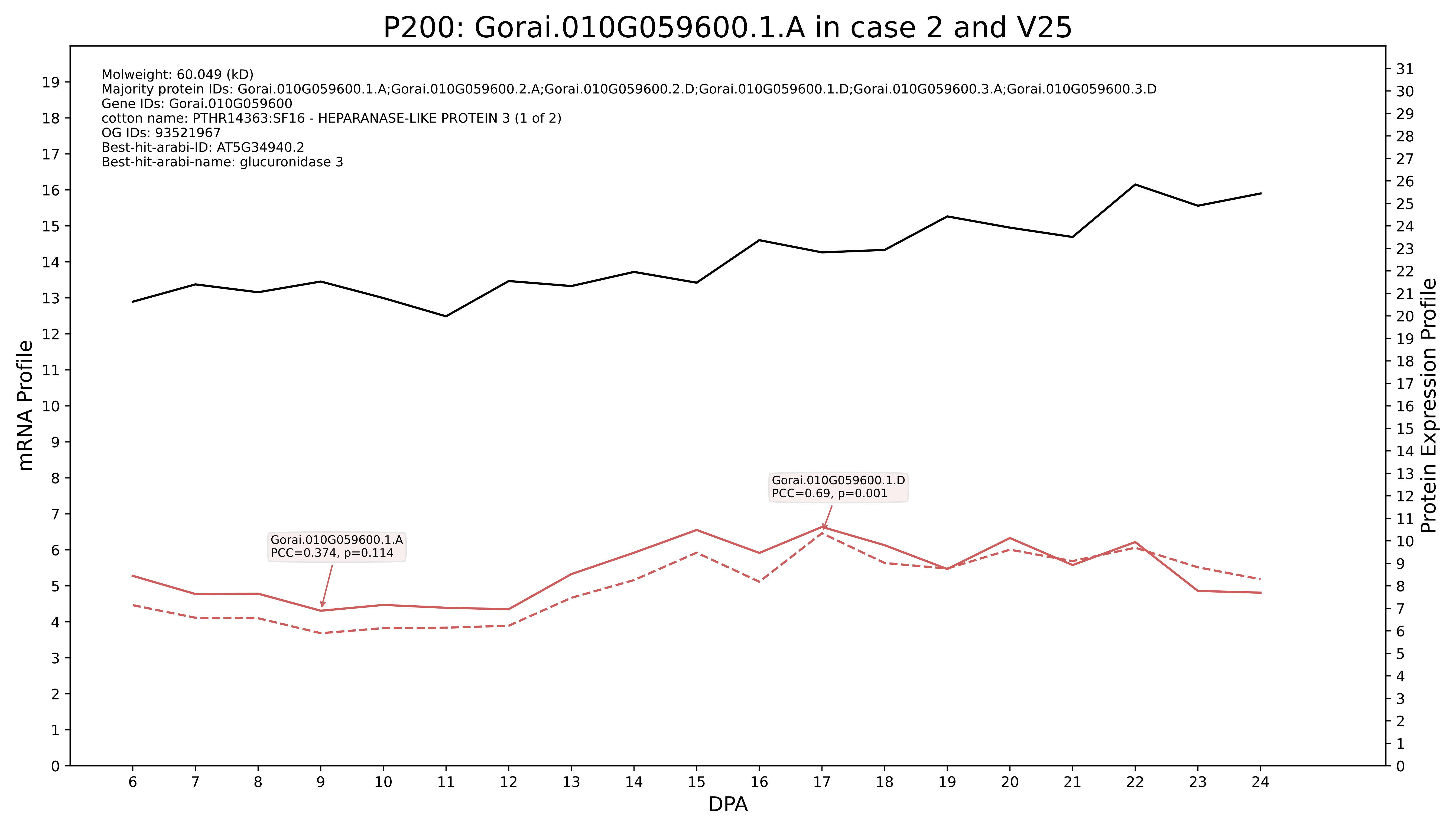
P200: Gorai.010G059600.1.A in case 2 and V25
Molweight: 60.049 (kD)
Majority protein IDs: Gorai.010G059600.1.A;Gorai.010G059600.2.A;Gorai.010G059600.2.D;Gorai.010G059600.1.D;Gorai.010G059600.3.A;Gorai.010G059600.3.D
Gene IDs: Gorai.010G059600
cotton name: PTHR14363:SF16 - HEPARANASE-LIKE PROTEIN 3 (1 of 2)
OG IDs: 93521967
Best-hit-arabi-ID: AT5G34940.2
Best-hit-arabi-name: glucuronidase 3
Gorai.010G059600.1.D
PCC=0.69, p=0.001
Gorai.010G059600.1.A
PCC=0.374, p=0.114
mRNA Profile
Protein Expression Profile
DPA

### P200: Gorai.010G204400.1.D in case 2 and V25

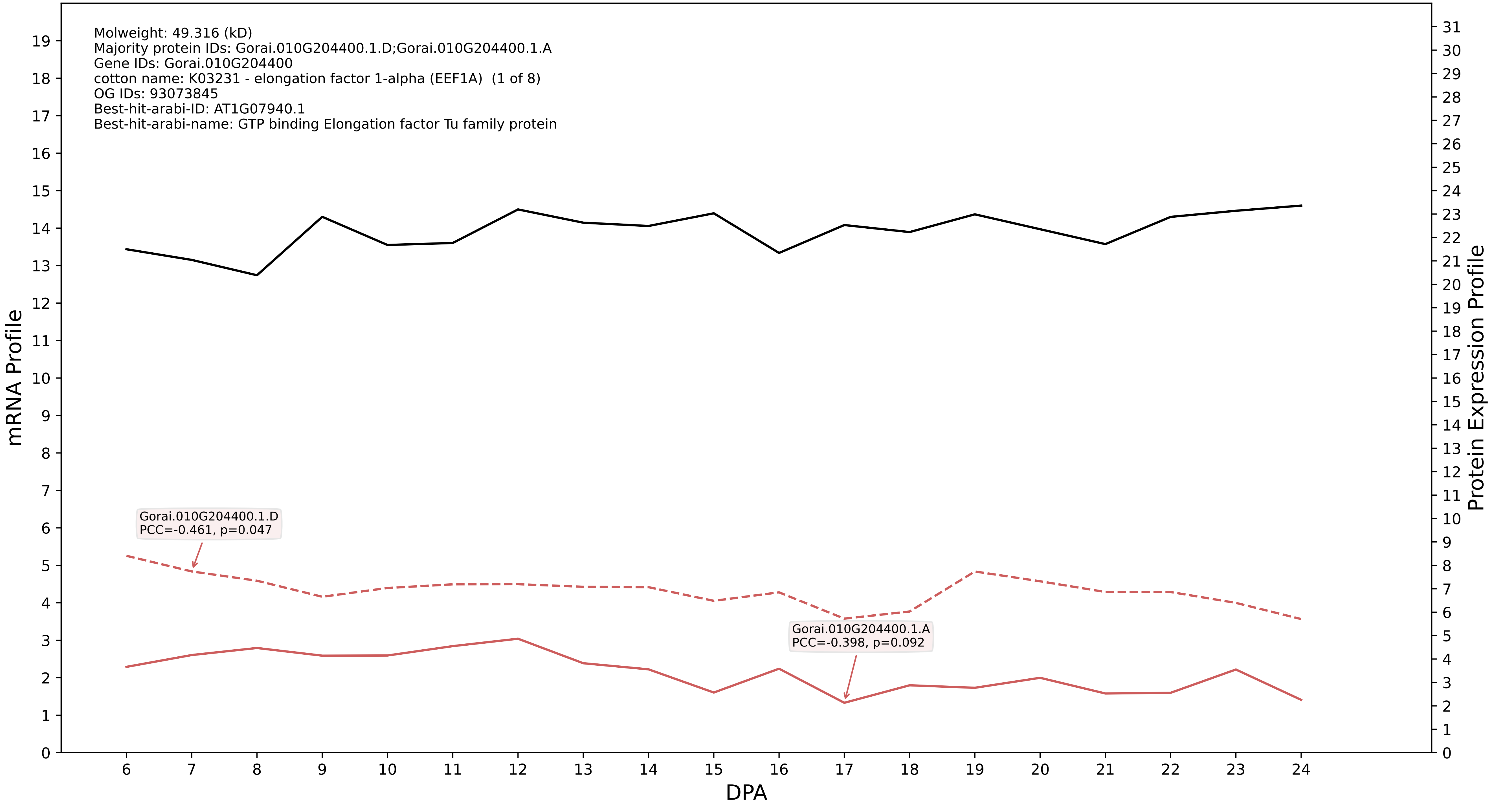

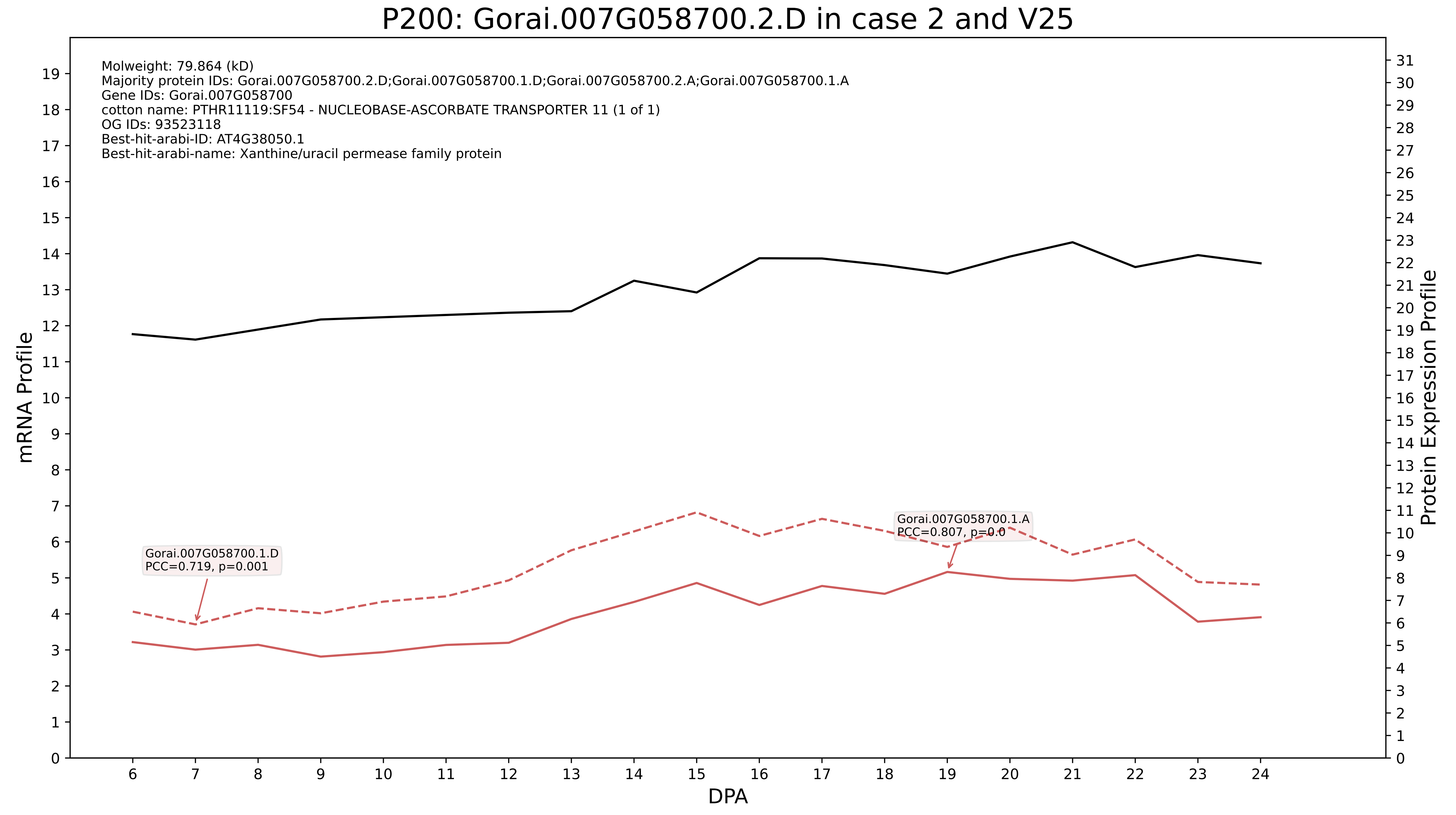
P200: Gorai.007G058700.2.D in case 2 and V25
Molweight: 79.864 (kD)
Majority protein IDs: Gorai.007G058700.2.D;Gorai.007G058700.1.D;Gorai.007G058700.2.A;Gorai.007G058700.1.A
Gene IDs: Gorai.007G058700
cotton name: PTHR11119:SF54 - NUCLEOBASE-ASCORBATE TRANSPORTER 11 (1 of 1)
OG IDs: 93523118
Best-hit-arabi-ID: AT4G38050.1
Best-hit-arabi-name: Xanthine/uracil permease family protein
Gorai.007G058700.1.D
PCC=0.719, p=0.001
Gorai.007G058700.1.A
PCC=0.807, p=0.0
mRNA Profile
Protein Expression Profile
DPA

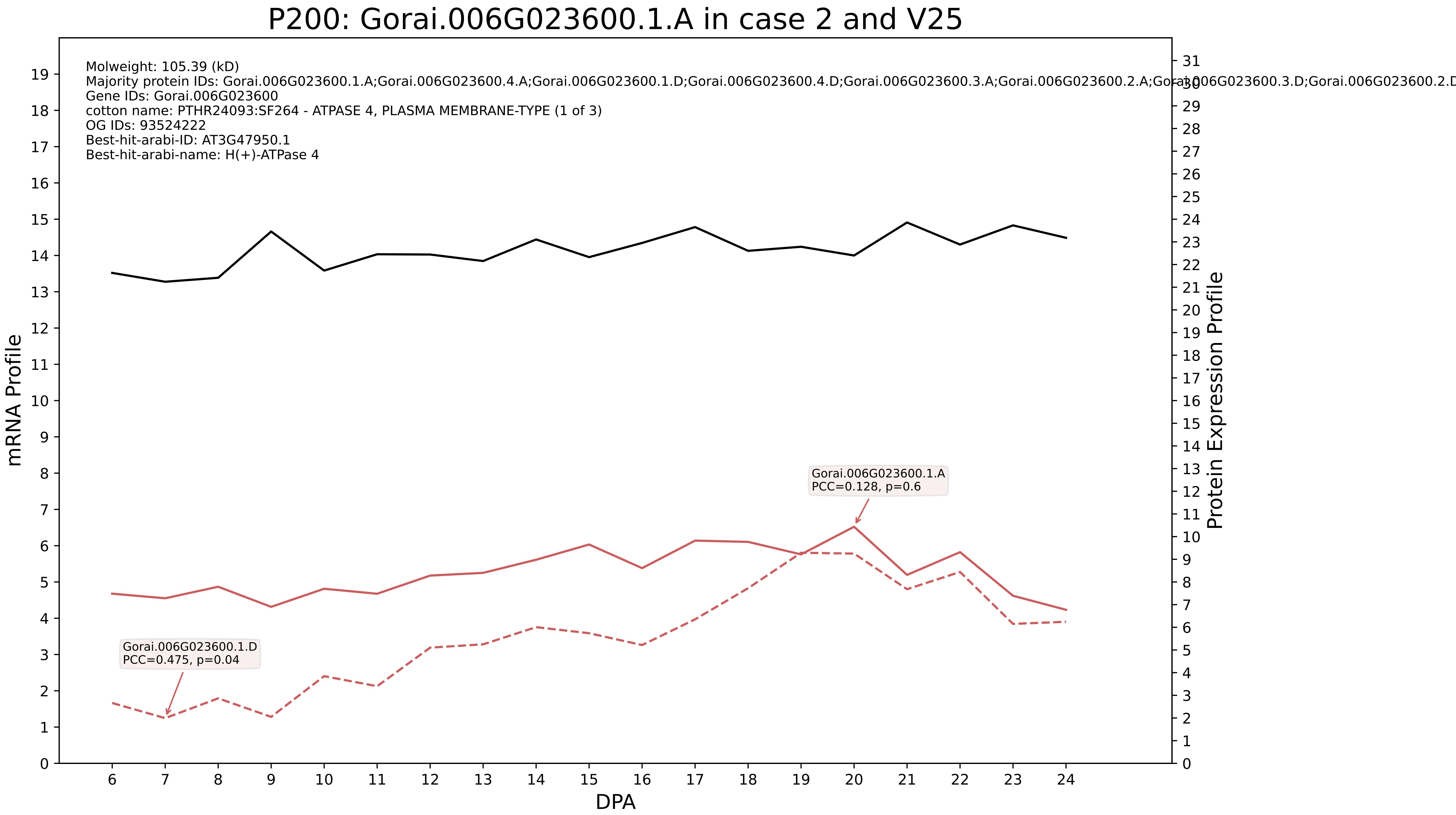

P200: Gorai.006G023600.1.A in case 2 and V25
Molweight: 105.39 (kD)
Majority protein IDs: Gorai.006G023600.1.A;Gorai.006G023600.4.A;Gorai.006G023600.1.D;Gorai.006G023600.4.D;Gorai.006G023600.3.A;Gorai.006G023600.2.A;Gorai.006G023600.3.D;Gorai.006G023600.2.D
Gene IDs: Gorai.006G023600
cotton name: PTHR24093:SF264 - ATPASE 4, PLASMA MEMBRANE-TYPE (1 of 3)
OG IDs: 93524222
Best-hit-arabi-ID: AT3G47950.1
Best-hit-arabi-name: H(+)-ATPase 4
Gorai.006G023600.1.A
PCC=0.128, p=0.6
Gorai.006G023600.1.D
PCC=0.475, p=0.04
mRNA Profile
Protein Expression Profile
DPA

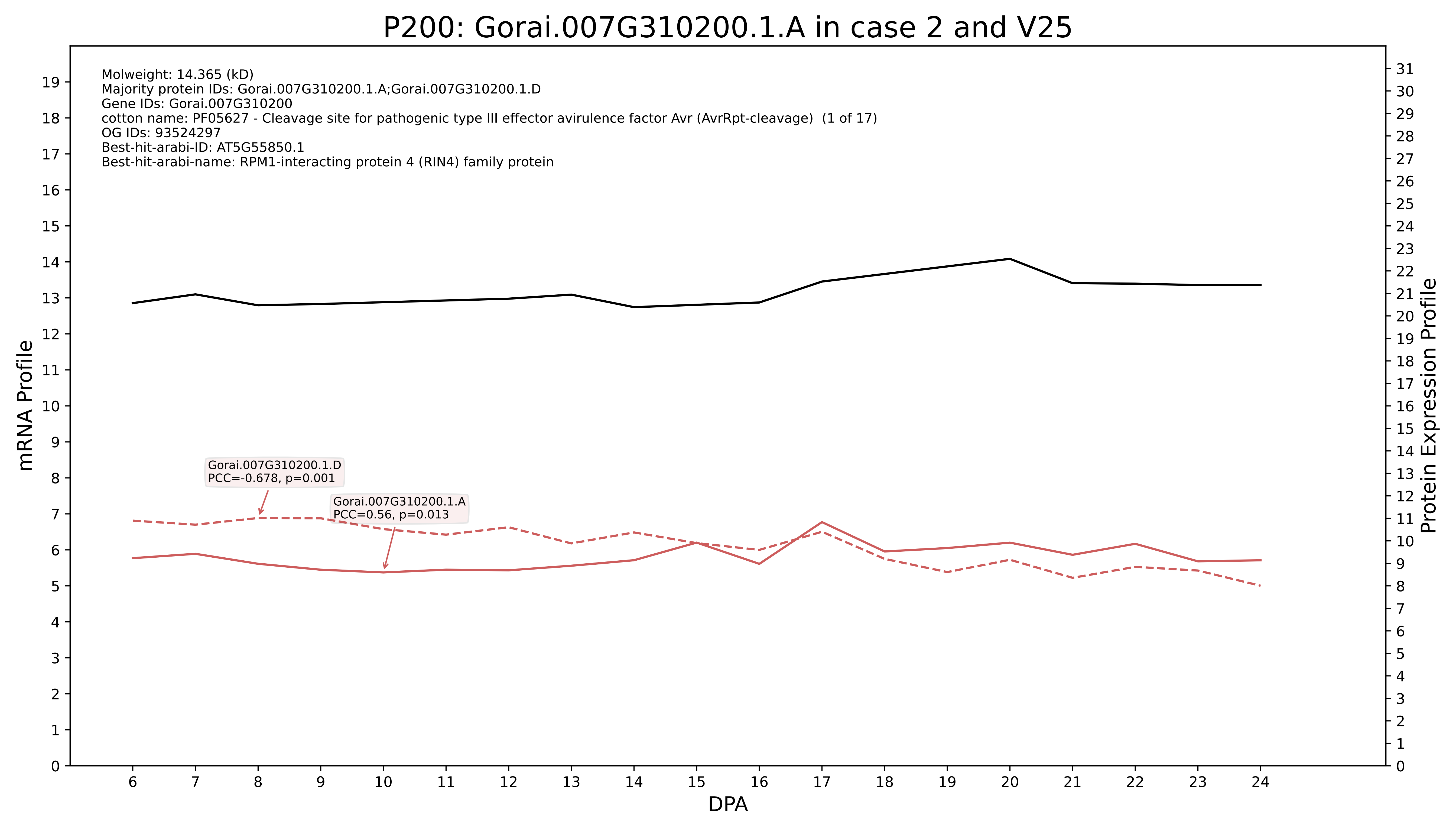
P200: Gorai.007G310200.1.A in case 2 and V25
Molweight: 14.365 (kD)
Majority protein IDs: Gorai.007G310200.1.A;Gorai.007G310200.1.D
Gene IDs: Gorai.007G310200
cotton name: PF05627 - Cleavage site for pathogenic type III effector avirulence factor Avr (AvrRpt-cleavage) (1 of 17)
OG IDs: 93524297
Best-hit-arabi-ID: AT5G55850.1
Best-hit-arabi-name: RPM1-interacting protein 4 (RIN4) family protein
Gorai.007G310200.1.D
PCC=-0.678, p=0.001
Gorai.007G310200.1.A
PCC=0.56, p=0.013
mRNA Profile
Protein Expression Profile
DPA

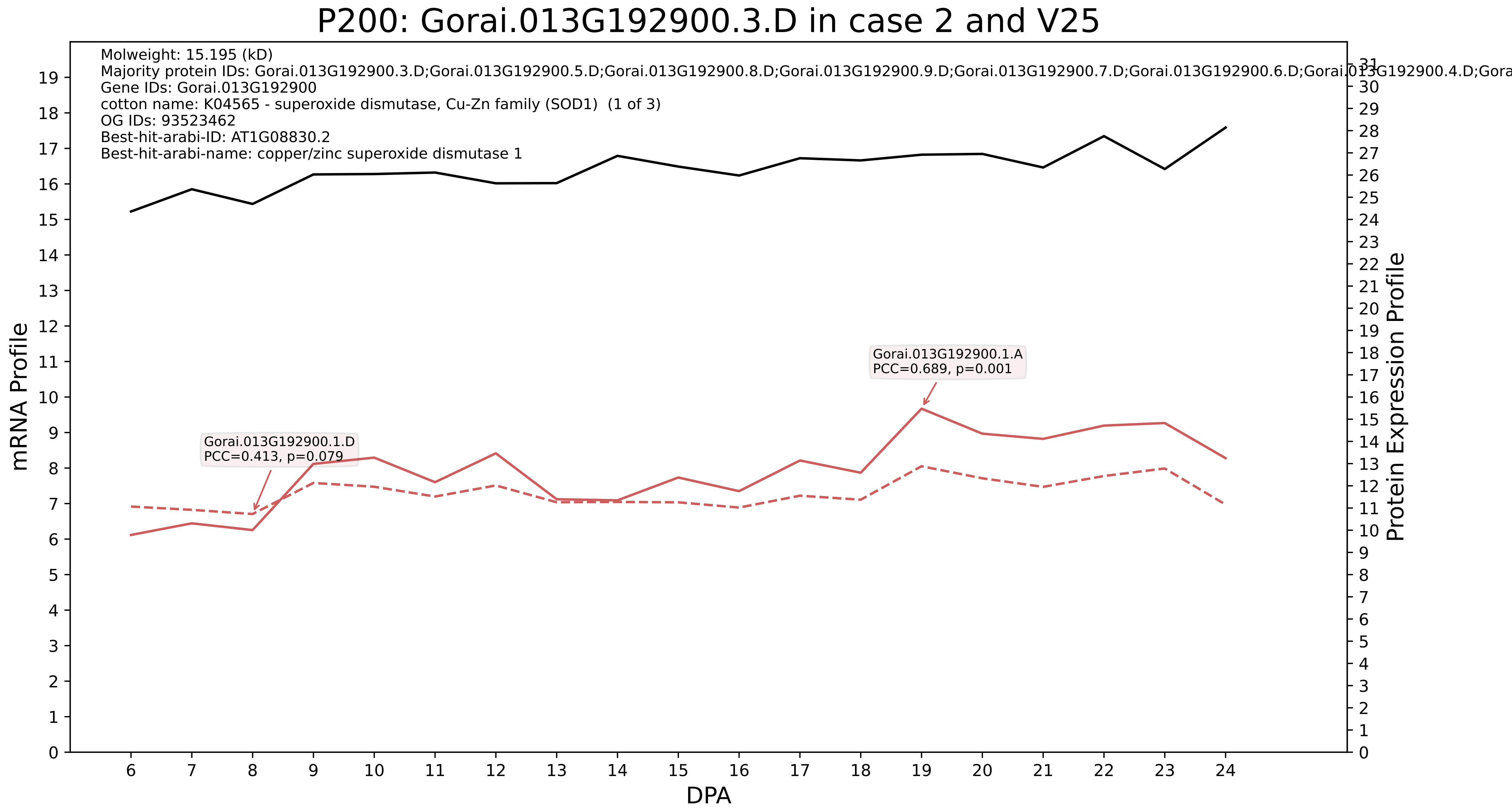
P200: Gorai.013G192900.3.D in case 2 and V25
Molweight: 15.195 (kD)
Majority protein IDs: Gorai.013G192900.3.D;Gorai.013G192900.5.D;Gorai.013G192900.8.D;Gorai.013G192900.9.D;Gorai.013G192900.7.D;Gorai.013G192900.6.D;Gorai.013G192900.4.D;Gora
Gene IDs: Gorai.013G192900
cotton name: K04565 - superoxide dismutase, Cu-Zn family (SOD1) (1 of 3)
OG IDs: 93523462
Best-hit-arabi-ID: AT1G08830.2
Best-hit-arabi-name: copper/zinc superoxide dismutase 1
Gorai.013G192900.1.A
PCC=0.689, p=0.001
Gorai.013G192900.1.D
PCC=0.413, p=0.079
mRNA Profile
Protein Expression Profile
DPA

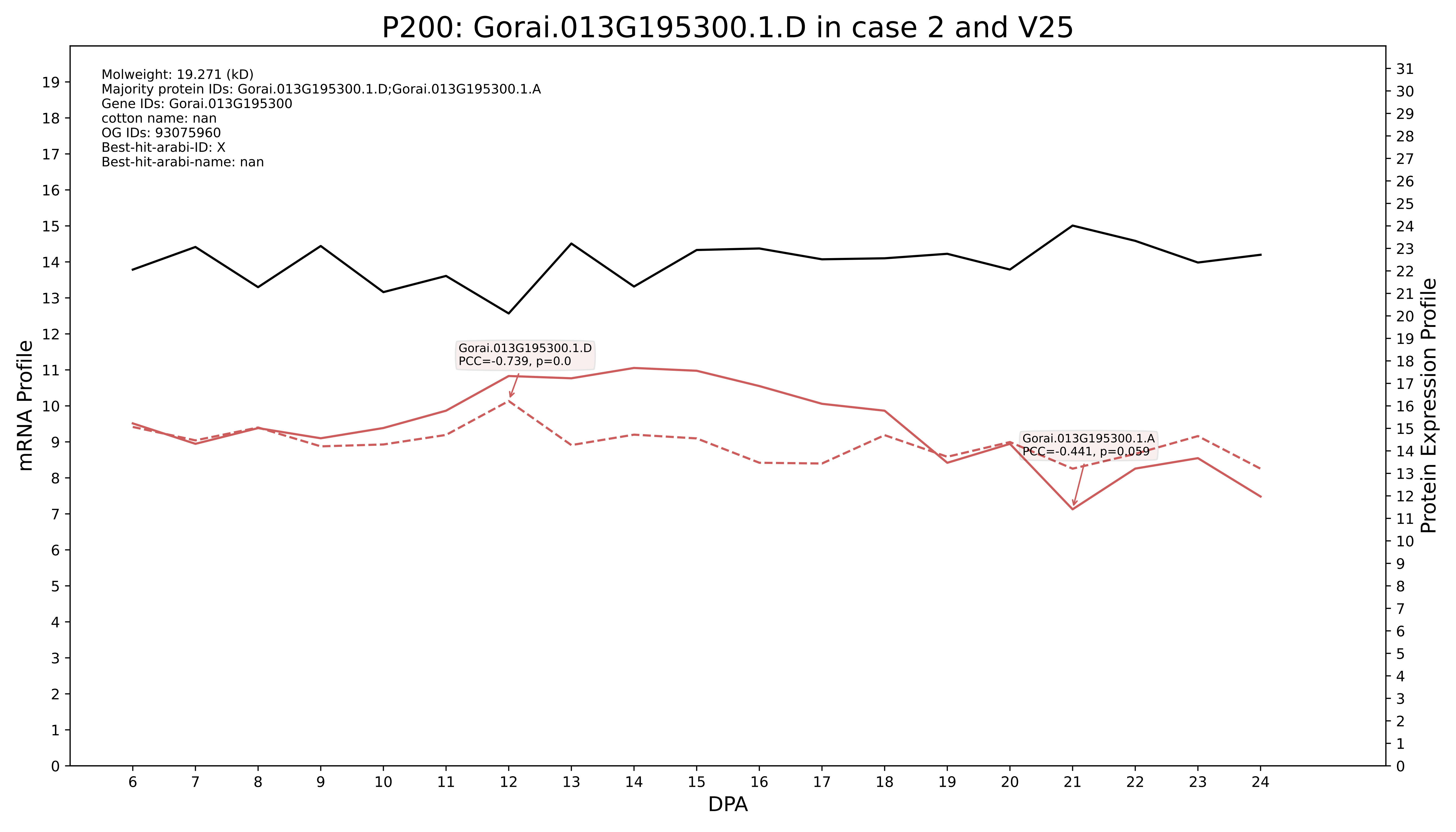
P200: Gorai.013G195300.1.D in case 2 and V25
Molweight: 19.271 (kD)
Majority protein IDs: Gorai.013G195300.1.D;Gorai.013G195300.1.A
Gene IDs: Gorai.013G195300
cotton name: nan
OG IDs: 93075960
Best-hit-arabi-ID: X
Best-hit-arabi-name: nan
Gorai.013G195300.1.D
PCC=-0.739, p=0.0
Gorai.013G195300.1.A
PCC=-0.441, p=0.059
mRNA Profile
Protein Expression Profile
DPA

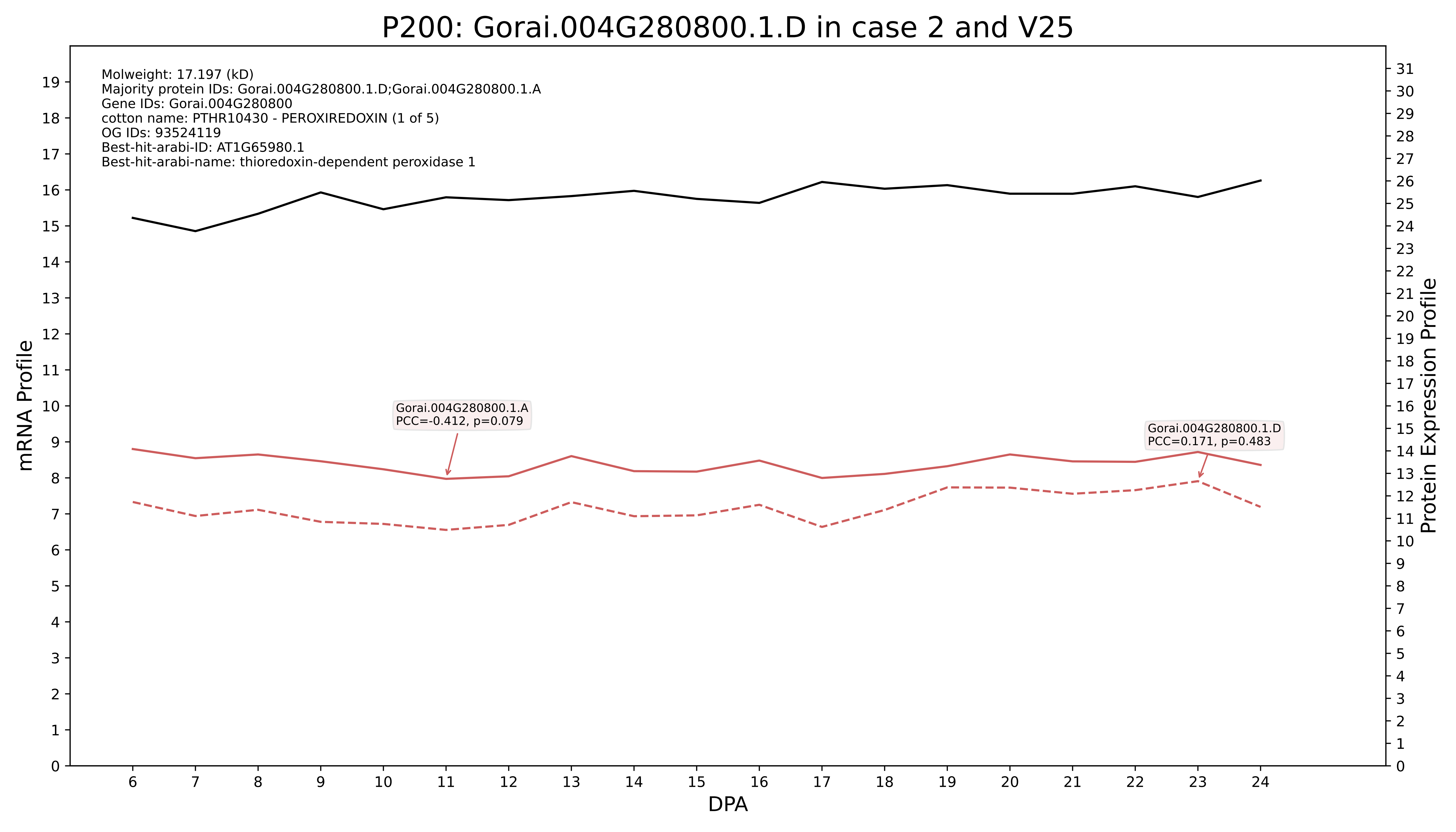
P200: Gorai.004G280800.1.D in case 2 and V25
Molweight: 17.197 (kD)
Majority protein IDs: Gorai.004G280800.1.D;Gorai.004G280800.1.A
Gene IDs: Gorai.004G280800
cotton name: PTHR10430 - PEROXIREDOXIN (1 of 5)
OG IDs: 93524119
Best-hit-arabi-ID: AT1G65980.1
Best-hit-arabi-name: thioredoxin-dependent peroxidase 1
Gorai.004G280800.1.A
PCC=-0.412, p=0.079
Gorai.004G280800.1.D
PCC=0.171, p=0.483
mRNA Profile
Protein Expression Profile
DPA

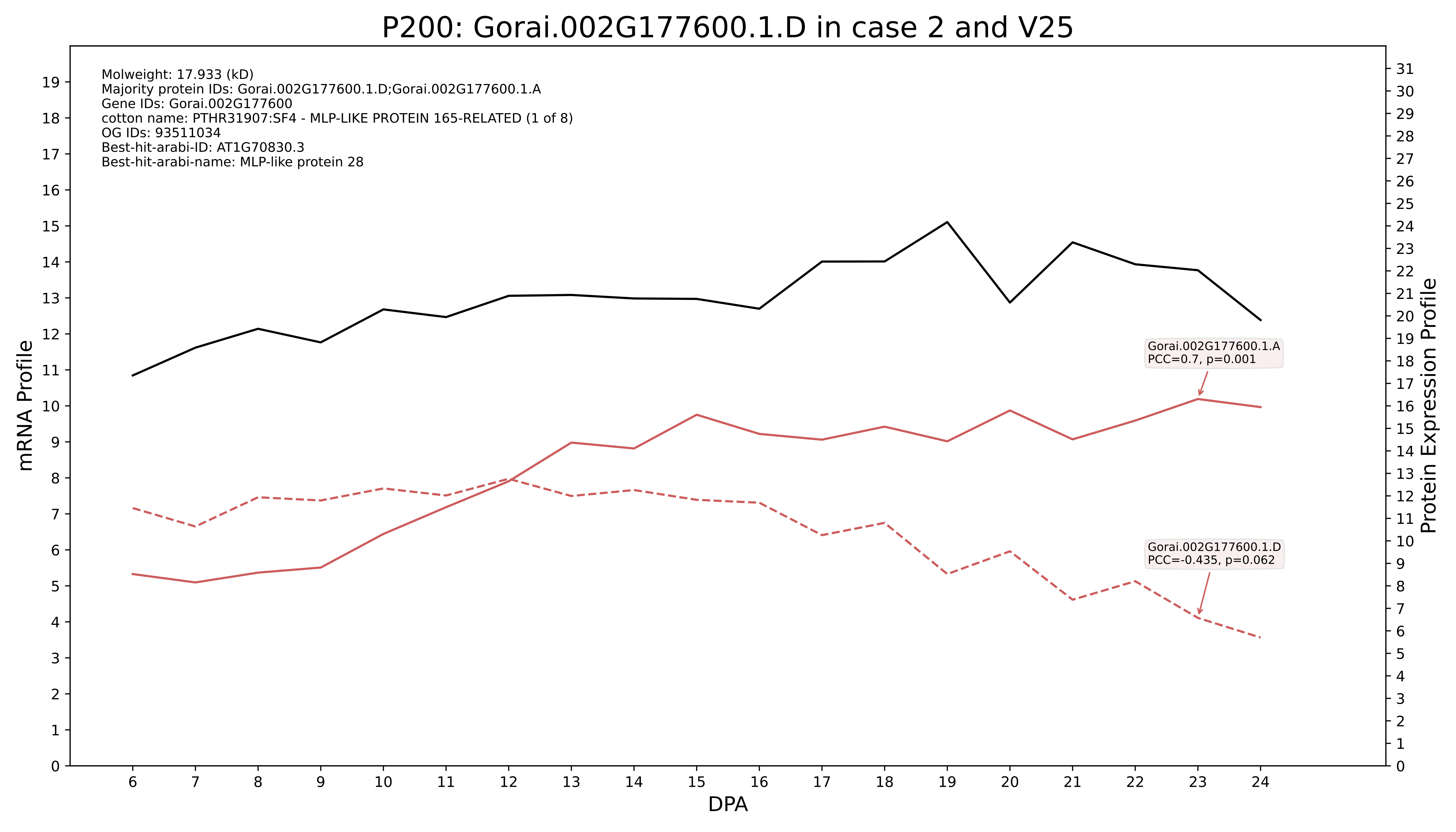
P200: Gorai.002G177600.1.D in case 2 and V25
Molweight: 17.933 (kD)
Majority protein IDs: Gorai.002G177600.1.D;Gorai.002G177600.1.A
Gene IDs: Gorai.002G177600
cotton name: PTHR31907:SF4 - MLP-LIKE PROTEIN 165-RELATED (1 of 8)
OG IDs: 93511034
Best-hit-arabi-ID: AT1G70830.3
Best-hit-arabi-name: MLP-like protein 28
Gorai.002G177600.1.A
PCC=0.7, p=0.001
Gorai.002G177600.1.D
PCC=-0.435, p=0.062
mRNA Profile
Protein Expression Profile
DPA

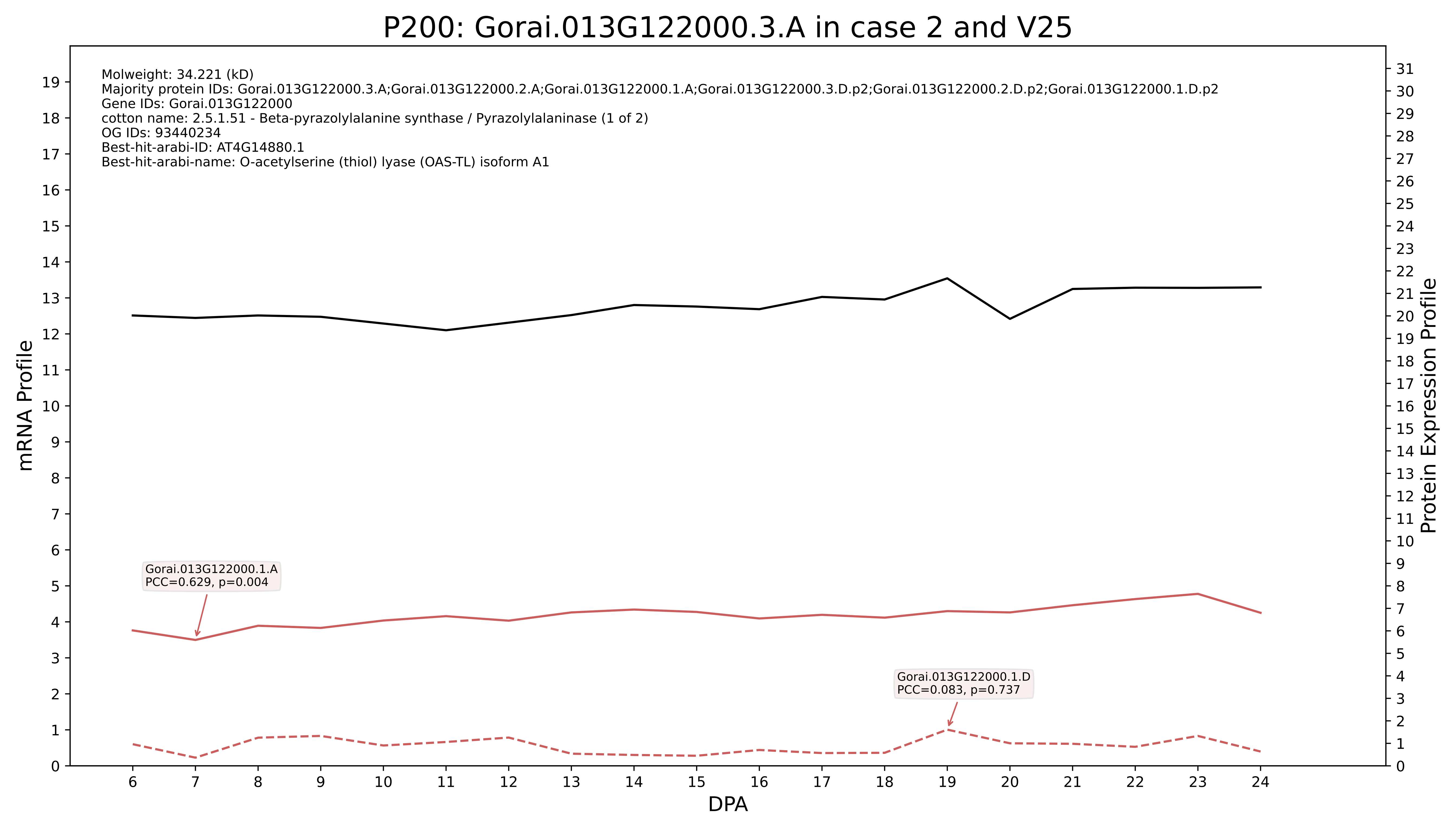
P200: Gorai.013G122000.3.A in case 2 and V25
Molweight: 34.221 (kD)
Majority protein IDs: Gorai.013G122000.3.A;Gorai.013G122000.2.A;Gorai.013G122000.1.A;Gorai.013G122000.3.D.p2;Gorai.013G122000.2.D.p2;Gorai.013G122000.1.D.p2
Gene IDs: Gorai.013G122000
cotton name: 2.5.1.51 - Beta-pyrazolylalanine synthase / Pyrazolylalaninase (1 of 2)
OG IDs: 93440234
Best-hit-arabi-ID: AT4G14880.1
Best-hit-arabi-name: O-acetylserine (thiol) lyase (OAS-TL) isoform A1
Gorai.013G122000.1.A
PCC=0.629, p=0.004
Gorai.013G122000.1.D
PCC=0.083, p=0.737
mRNA Profile
Protein Expression Profile
DPA

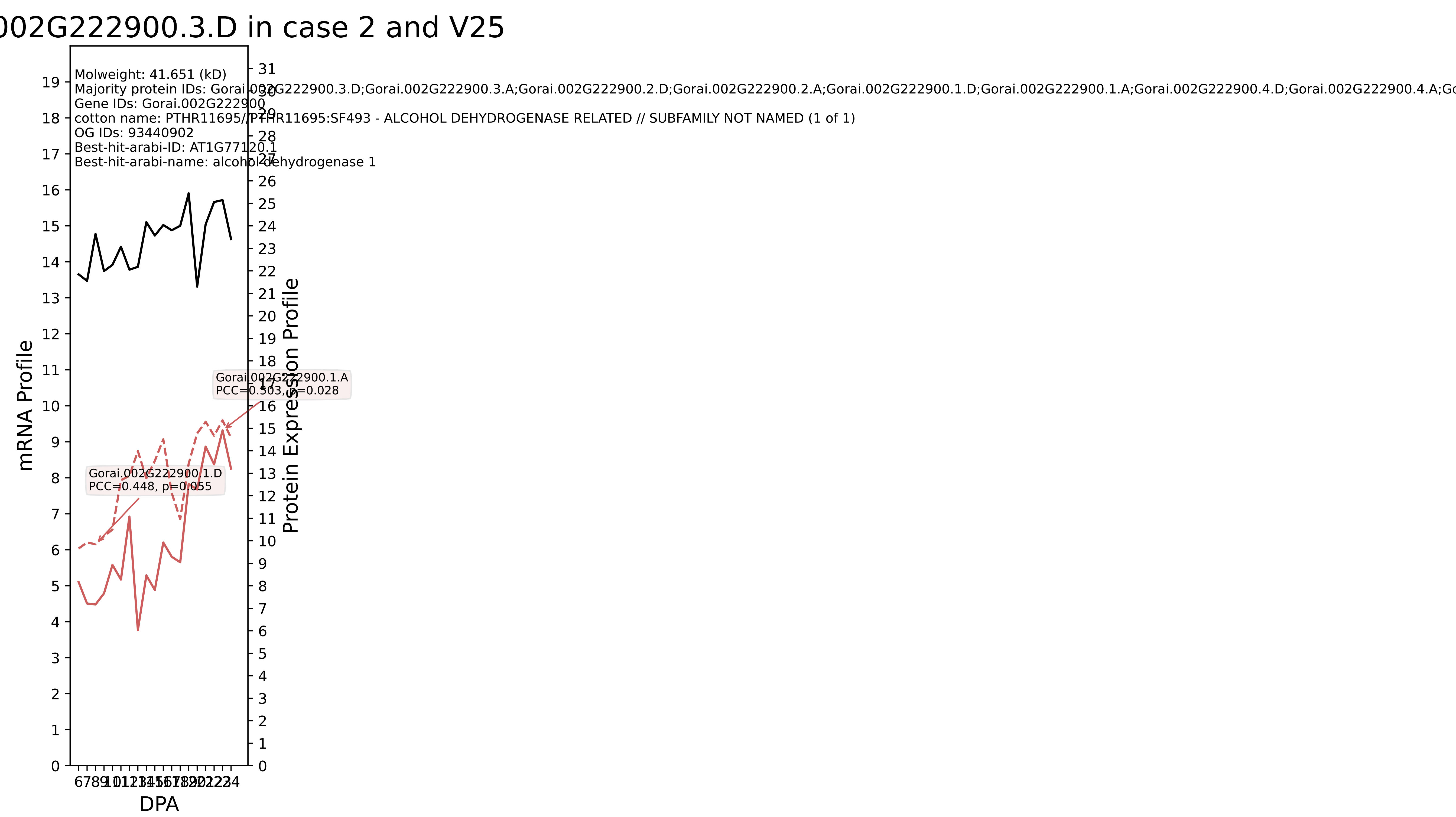
002G222900.3.D in case 2 and V25
Molweight: 41.651 (kD)
Majority protein IDs: Gorai.002G222900.3.D;Gorai.002G222900.3.A;Gorai.002G222900.2.D;Gorai.002G222900.2.A;Gorai.002G222900.1.D;Gorai.002G222900.1.A;Gorai.002G222900.4.D;Gorai.002G222900.4.A;Go
Gene IDs: Gorai.002G222900
cotton name: PTHR11695//PTHR11695:SF493 - ALCOHOL DEHYDROGENASE RELATED // SUBFAMILY NOT NAMED (1 of 1)
OG IDs: 93440902
Best-hit-arabi-ID: AT1G77120.1
Best-hit-arabi-name: alcohol dehydrogenase 1
Gorai.002G222900.1.A
PCC=0.503, p=0.028
Gorai.002G222900.1.D
PCC=0.448, p=0.055
mRNA Profile
Protein Expression Profile
DPA

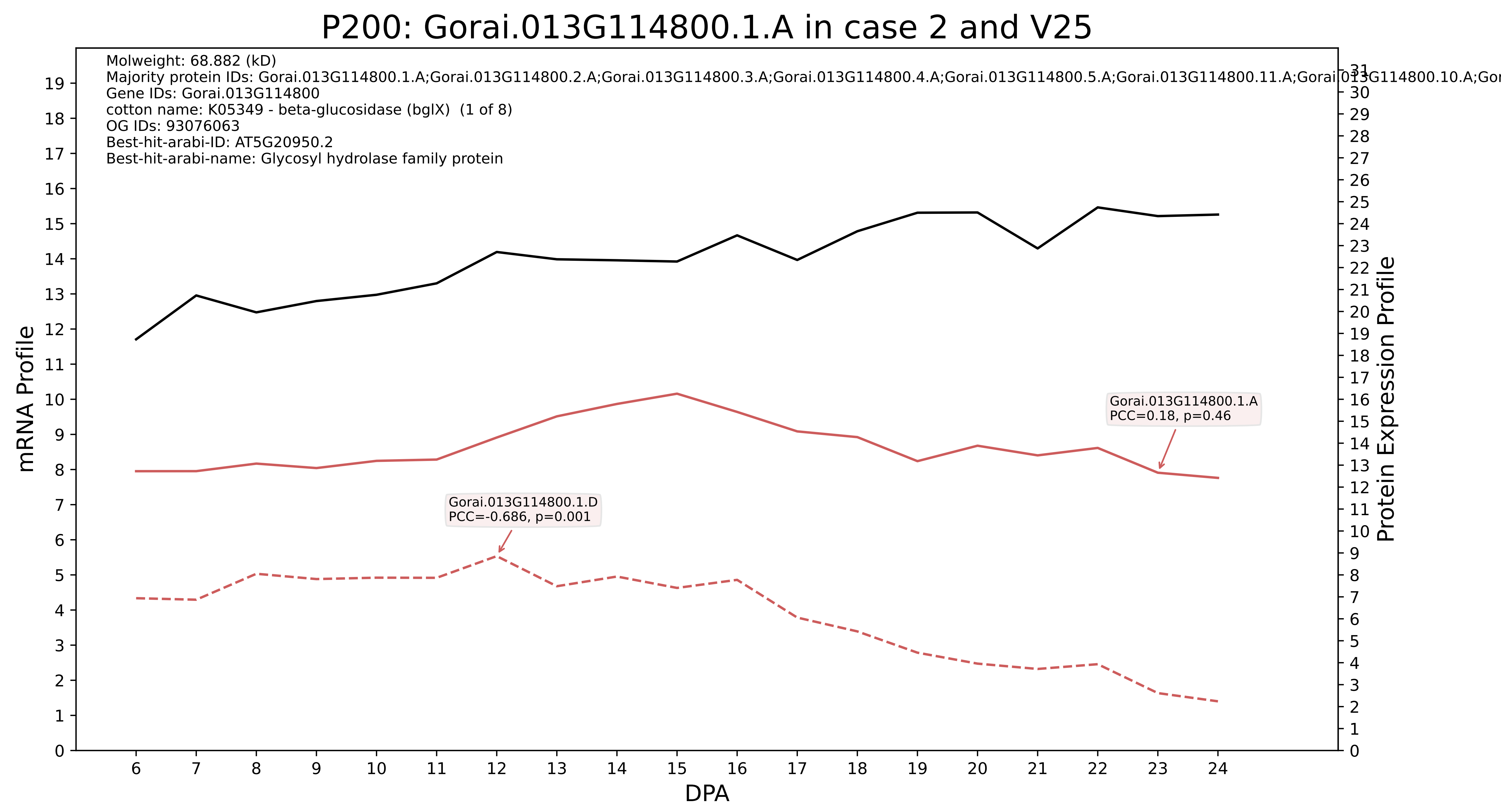
P200: Gorai.013G114800.1.A in case 2 and V25
Molweight: 68.882 (kD)
Majority protein IDs: Gorai.013G114800.1.A;Gorai.013G114800.2.A;Gorai.013G114800.3.A;Gorai.013G114800.4.A;Gorai.013G114800.5.A;Gorai.013G114800.11.A;Gorai.013G114800.10.A;Gor
Gene IDs: Gorai.013G114800
cotton name: K05349 - beta-glucosidase (bglX) (1 of 8)
OG IDs: 93076063
Best-hit-arabi-ID: AT5G20950.2
Best-hit-arabi-name: Glycosyl hydrolase family protein
Gorai.013G114800.1.A
PCC=0.18, p=0.46
Gorai.013G114800.1.D
PCC=-0.686, p=0.001
mRNA Profile
Protein Expression Profile
DPA

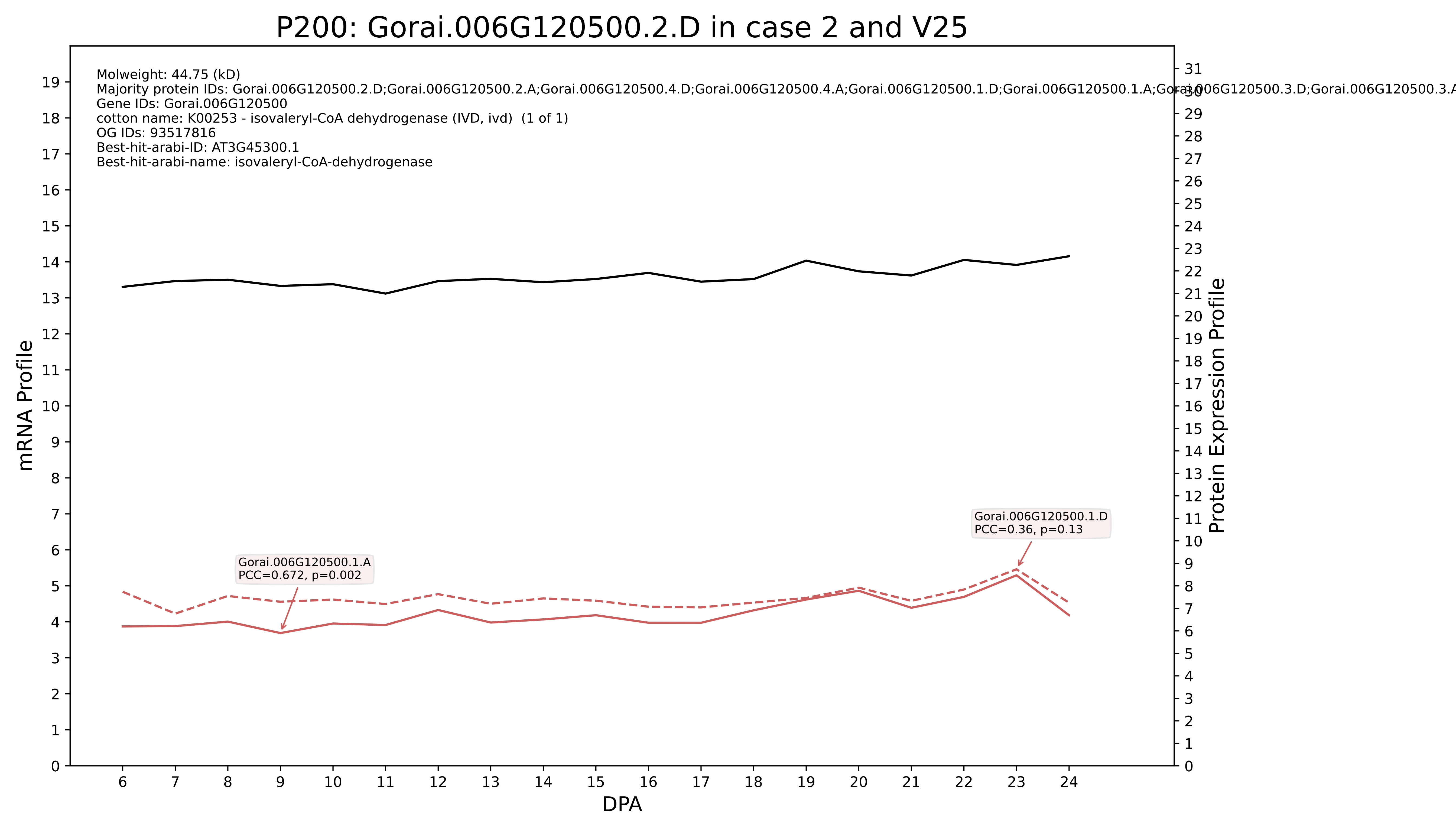
P200: Gorai.006G120500.2.D in case 2 and V25
Molweight: 44.75 (kD)
Majority protein IDs: Gorai.006G120500.2.D;Gorai.006G120500.2.A;Gorai.006G120500.4.D;Gorai.006G120500.4.A;Gorai.006G120500.1.D;Gorai.006G120500.1.A;Gorai.006G120500.3.D;Gorai.006G120500.3.A
Gene IDs: Gorai.006G120500
cotton name: K00253 - isovaleryl-CoA dehydrogenase (IVD, ivd) (1 of 1)
OG IDs: 93517816
Best-hit-arabi-ID: AT3G45300.1
Best-hit-arabi-name: isovaleryl-CoA-dehydrogenase
Gorai.006G120500.1.A
PCC=0.672, p=0.002
Gorai.006G120500.1.D
PCC=0.36, p=0.13
mRNA Profile
Protein Expression Profile
DPA

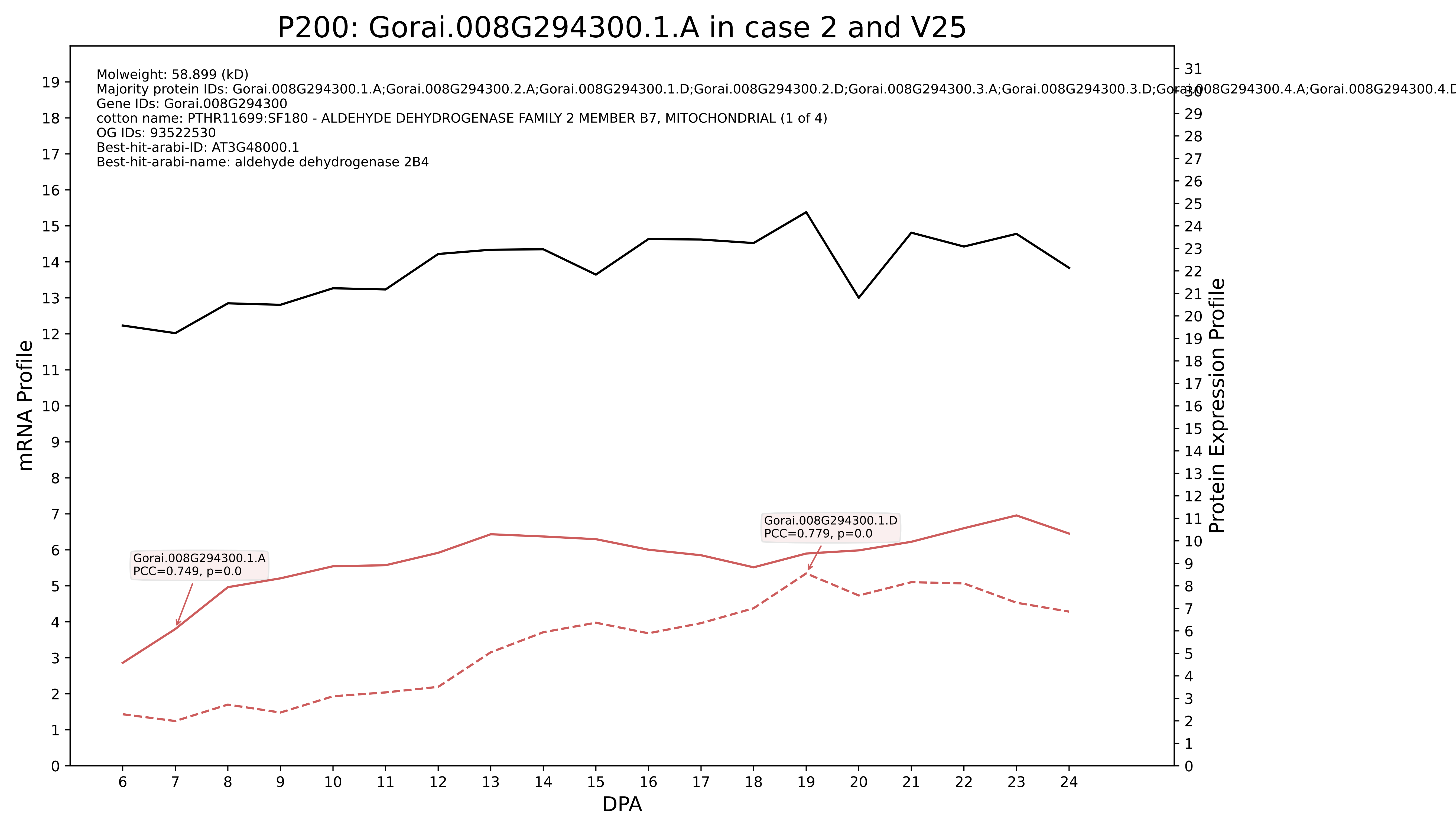
P200: Gorai.008G294300.1.A in case 2 and V25
Molweight: 58.899 (kD)
Majority protein IDs: Gorai.008G294300.1.A;Gorai.008G294300.2.A;Gorai.008G294300.1.D;Gorai.008G294300.2.D;Gorai.008G294300.3.A;Gorai.008G294300.3.D;Gorai.008G294300.4.A;Gorai.008G294300.4.D
Gene IDs: Gorai.008G294300
cotton name: PTHR11699:SF180 - ALDEHYDE DEHYDROGENASE FAMILY 2 MEMBER B7, MITOCHONDRIAL (1 of 4)
OG IDs: 93522530
Best-hit-arabi-ID: AT3G48000.1
Best-hit-arabi-name: aldehyde dehydrogenase 2B4
Gorai.008G294300.1.D
PCC=0.779, p=0.0
Gorai.008G294300.1.A
PCC=0.749, p=0.0
mRNA Profile
Protein Expression Profile
DPA

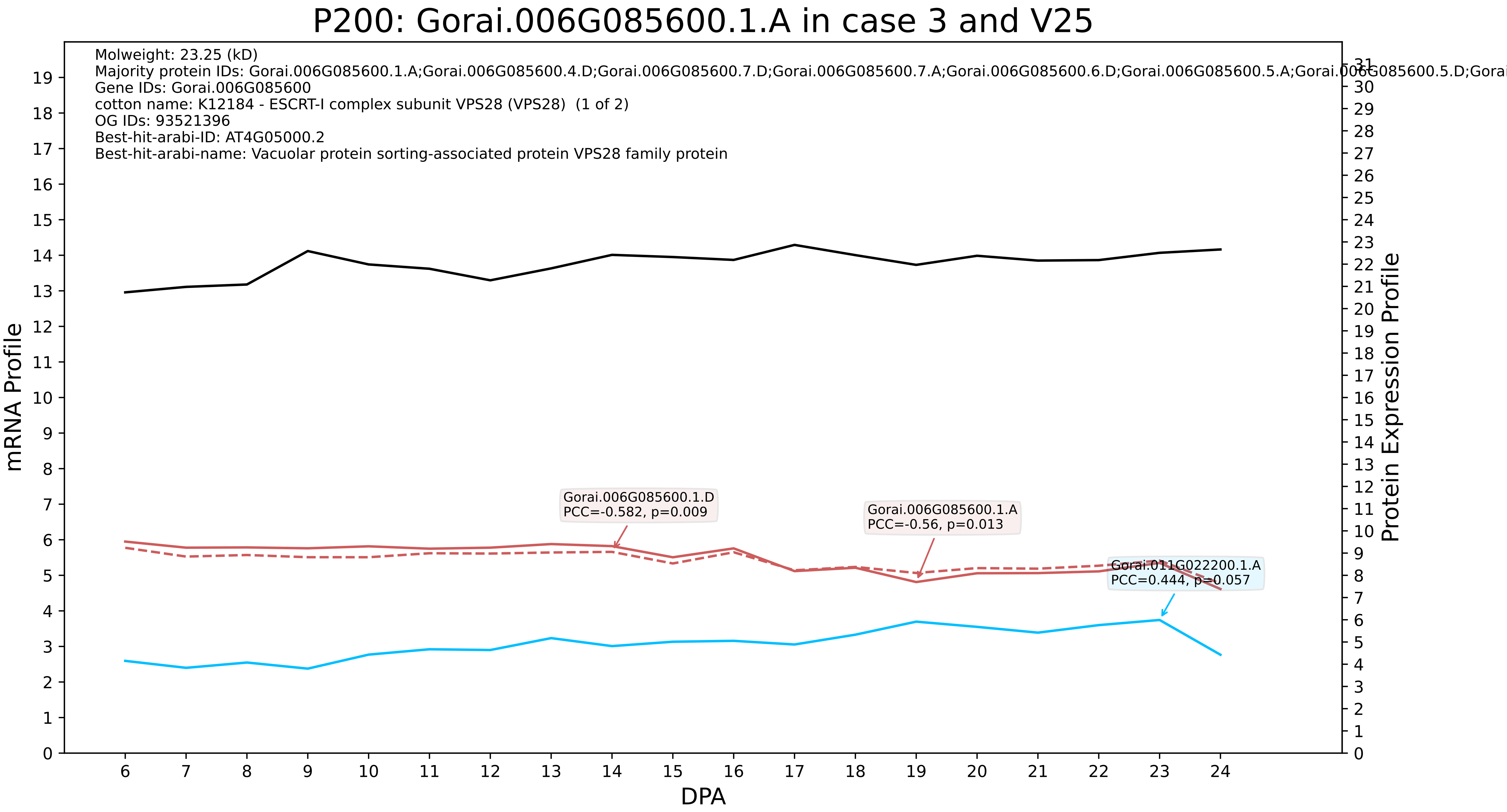

P200: Gorai.006G085600.1.A in case 3 and V25
Molweight: 23.25 (kD)
Majority protein IDs: Gorai.006G085600.1.A;Gorai.006G085600.4.D;Gorai.006G085600.7.D;Gorai.006G085600.7.A;Gorai.006G085600.6.D;Gorai.006G085600.5.A;Gorai.006G085600.5.D;Gorai
Gene IDs: Gorai.006G085600
cotton name: K12184 - ESCRT-I complex subunit VPS28 (VPS28) (1 of 2)
OG IDs: 93521396
Best-hit-arabi-ID: AT4G05000.2
Best-hit-arabi-name: Vacuolar protein sorting-associated protein VPS28 family protein
Gorai.006G085600.1.D
PCC=-0.582, p=0.009
Gorai.006G085600.1.A
PCC=-0.56, p=0.013
Gorai.011G022200.1.A
PCC=0.444, p=0.057
mRNA Profile
Protein Expression Profile
DPA

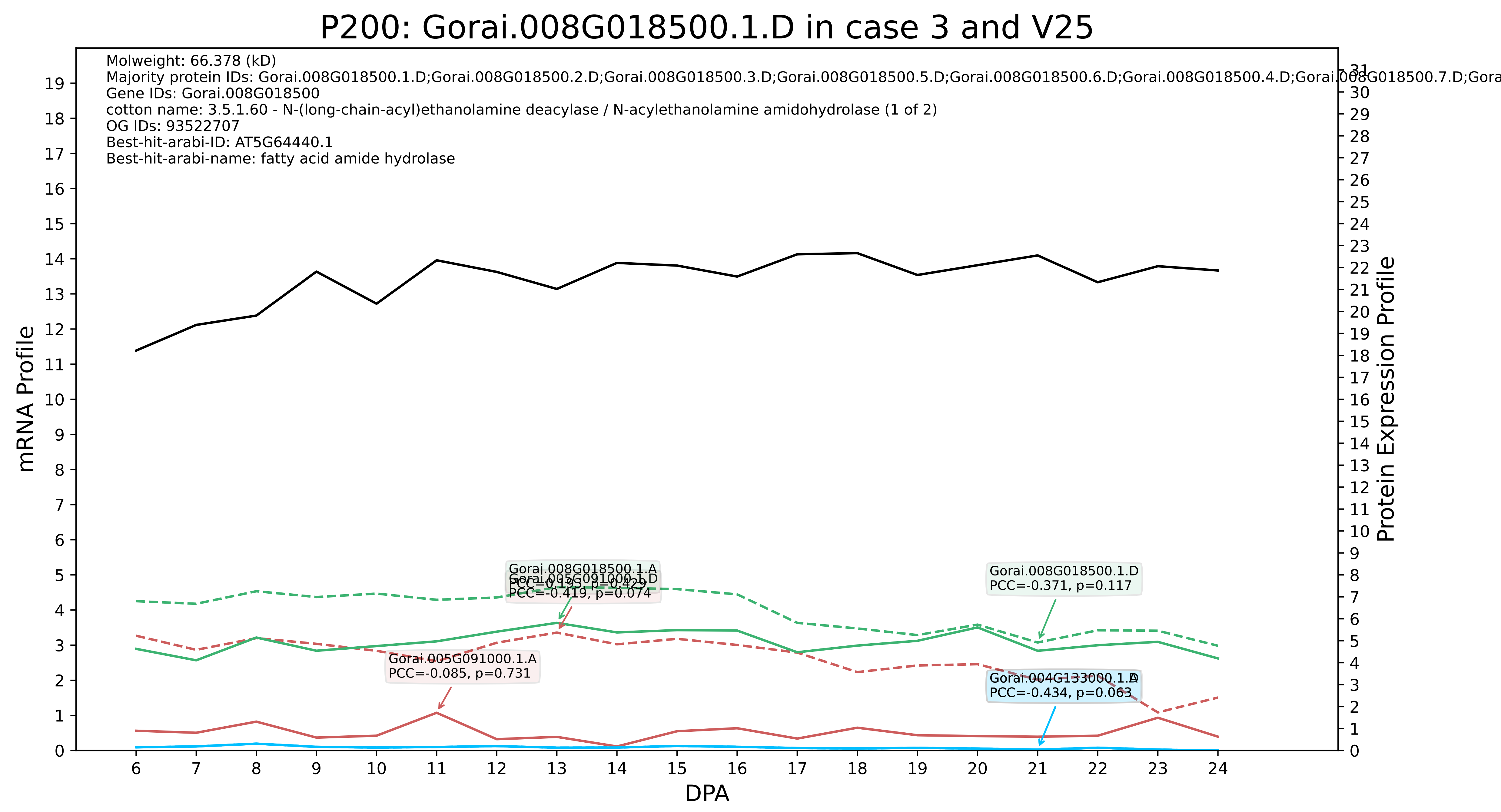

P200: Gorai.008G018500.1.D in case 3 and V25
Molweight: 66.378 (kD)
Majority protein IDs: Gorai.008G018500.1.D;Gorai.008G018500.2.D;Gorai.008G018500.3.D;Gorai.008G018500.5.D;Gorai.008G018500.6.D;Gorai.008G018500.4.D;Gorai.008G018500.7.D;Gora
Gene IDs: Gorai.008G018500
cotton name: 3.5.1.60 - N-(long-chain-acyl)ethanolamine deacylase / N-acylethanolamine amidohydrolase (1 of 2)
OG IDs: 93522707
Best-hit-arabi-ID: AT5G64440.1
Best-hit-arabi-name: fatty acid amide hydrolase
Gorai.008G018500.1.A
PCC=0.193, p=0.429
Gorai.005G091000.1.D
PCC=-0.419, p=0.074
Gorai.008G018500.1.D
PCC=-0.371, p=0.117
Gorai.005G091000.1.A
PCC=-0.085, p=0.731
Gorai.004G133000.1.D
PCC=-0.434, p=0.063
mRNA Profile
Protein Expression Profile
DPA

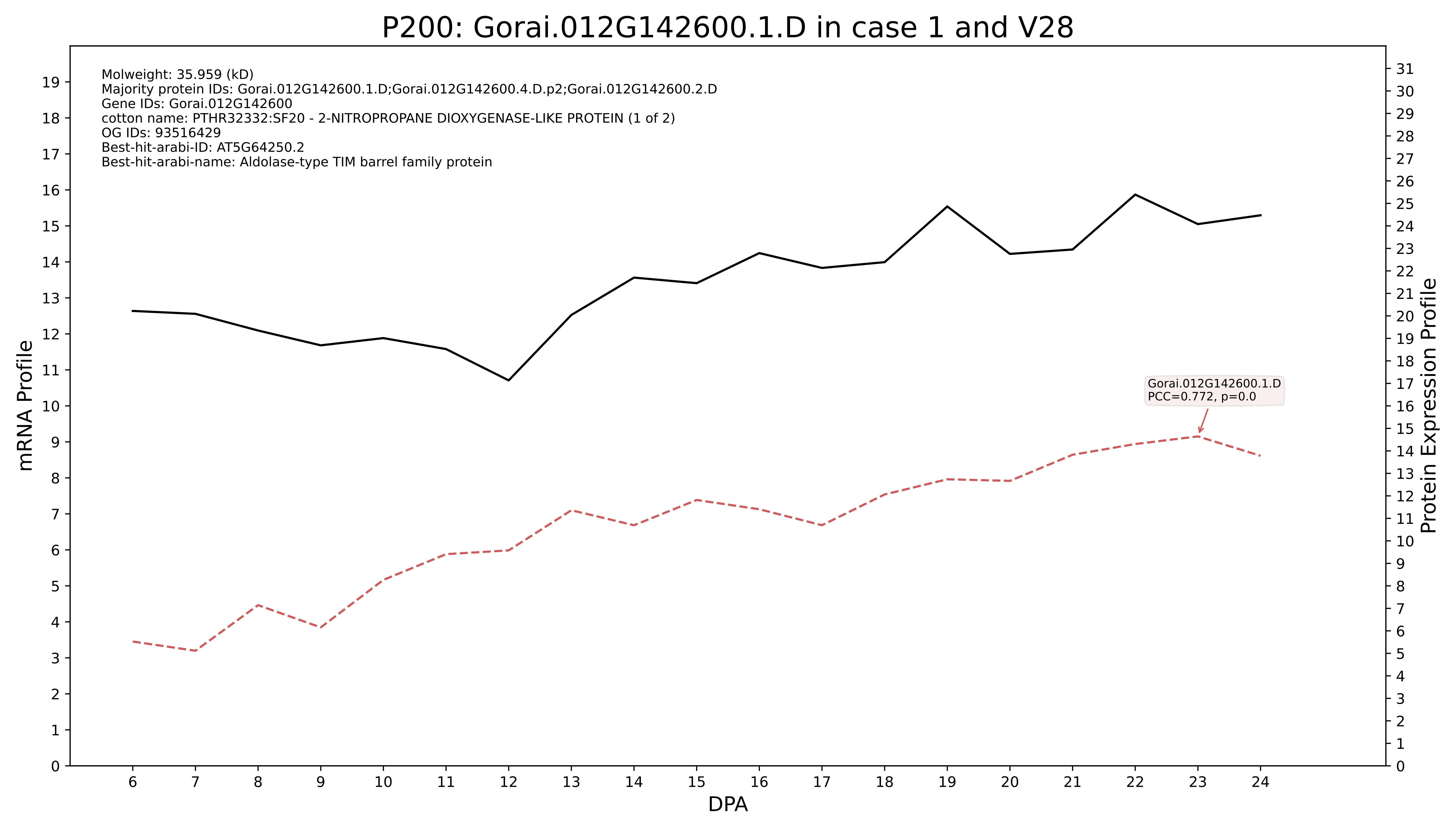
P200: Gorai.012G142600.1.D in case 1 and V28
Molweight: 35.959 (kD)
Majority protein IDs: Gorai.012G142600.1.D;Gorai.012G142600.4.D.p2;Gorai.012G142600.2.D
Gene IDs: Gorai.012G142600
cotton name: PTHR32332:SF20 - 2-NITROPROPANE DIOXYGENASE-LIKE PROTEIN (1 of 2)
OG IDs: 93516429
Best-hit-arabi-ID: AT5G64250.2
Best-hit-arabi-name: Aldolase-type TIM barrel family protein
Gorai.012G142600.1.D
PCC=0.772, p=0.0
mRNA Profile
Protein Expression Profile
DPA

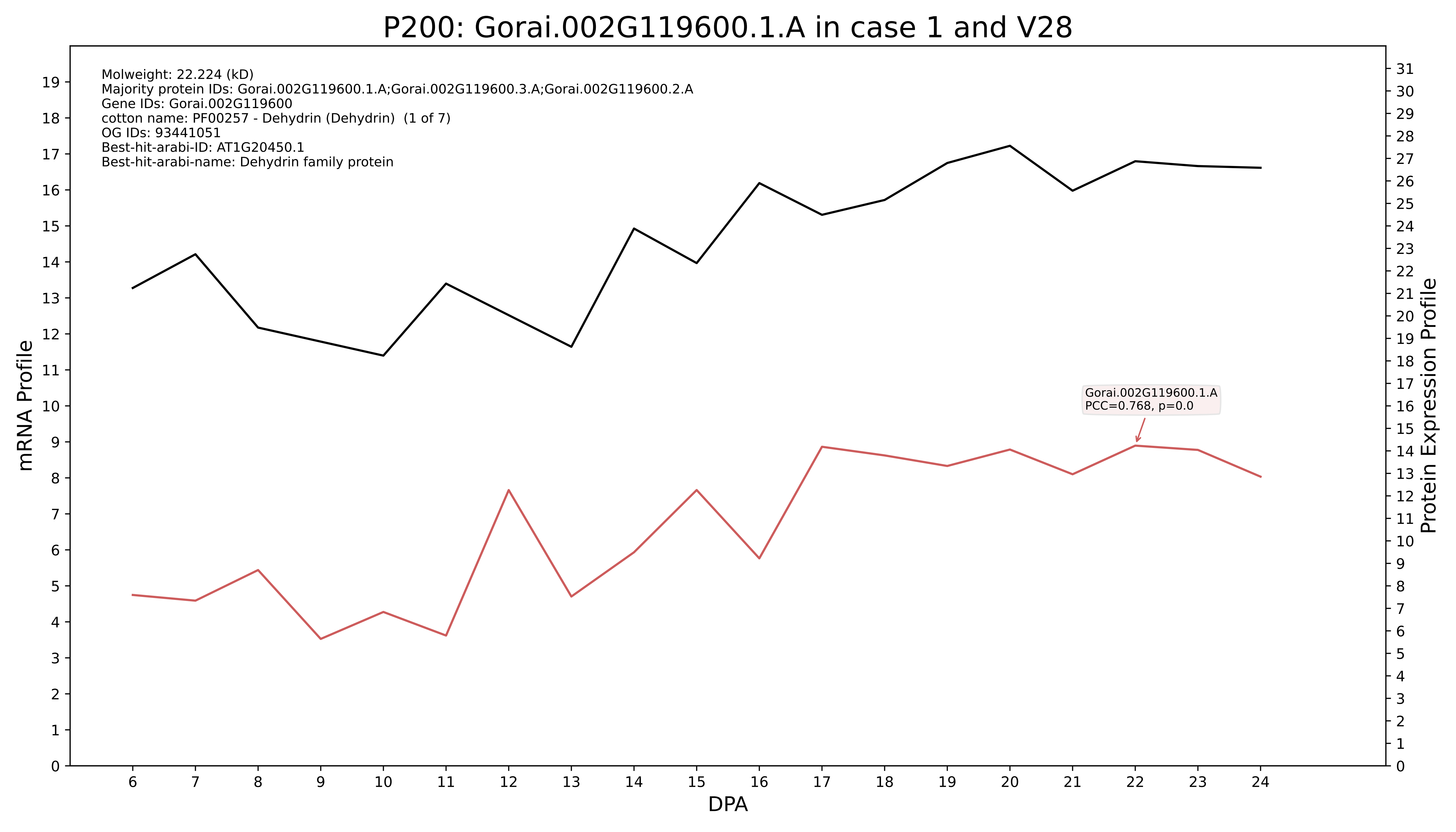
P200: Gorai.002G119600.1.A in case 1 and V28
Molweight: 22.224 (kD)
Majority protein IDs: Gorai.002G119600.1.A;Gorai.002G119600.3.A;Gorai.002G119600.2.A
Gene IDs: Gorai.002G119600
cotton name: PF00257 - Dehydrin (Dehydrin) (1 of 7)
OG IDs: 93441051
Best-hit-arabi-ID: AT1G20450.1
Best-hit-arabi-name: Dehydrin family protein
Gorai.002G119600.1.A
PCC=0.768, p=0.0
mRNA Profile
Protein Expression Profile
DPA

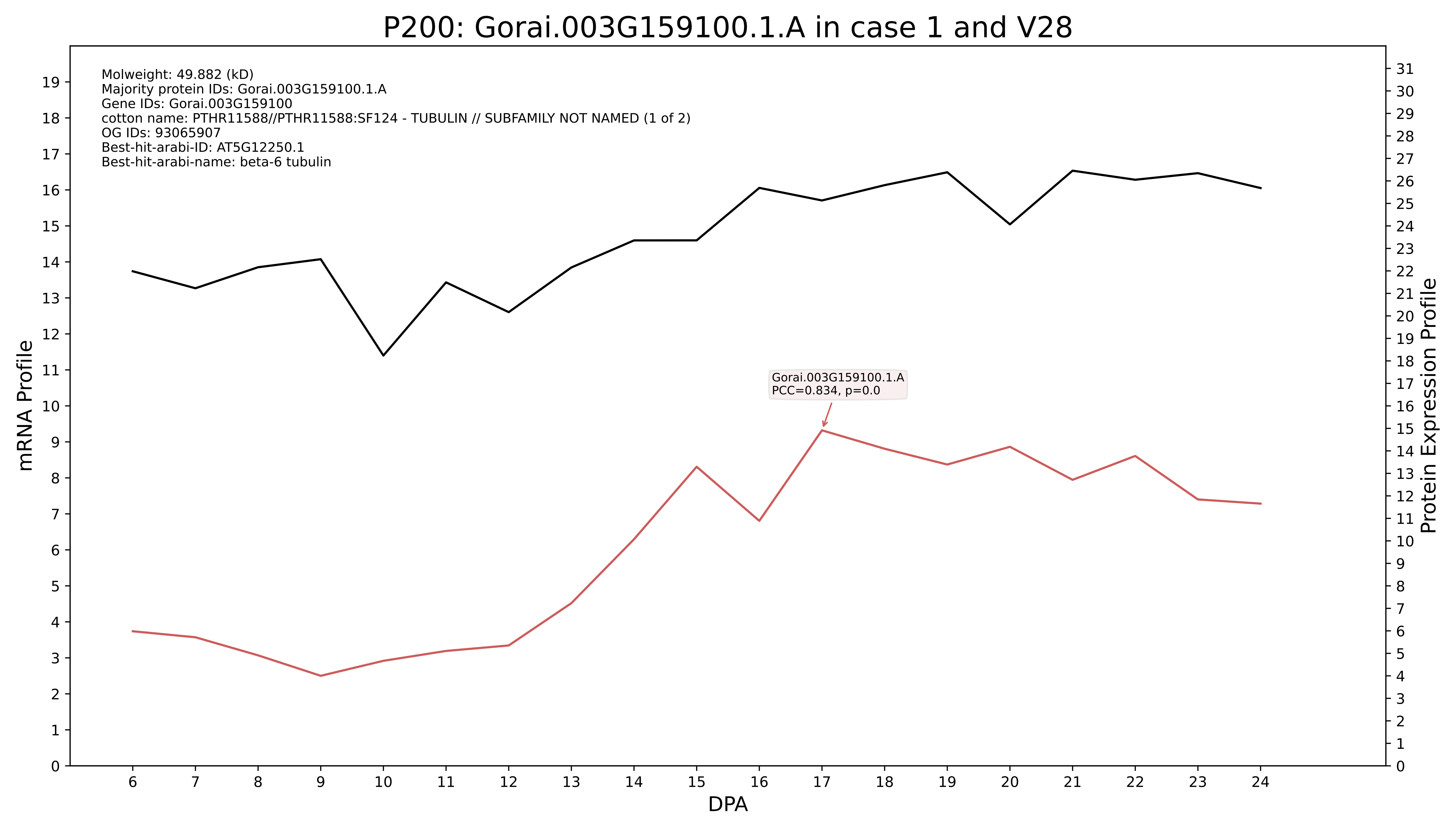
P200: Gorai.003G159100.1.A in case 1 and V28
Molweight: 49.882 (kD)
Majority protein IDs: Gorai.003G159100.1.A
Gene IDs: Gorai.003G159100
cotton name: PTHR11588//PTHR11588:SF124 - TUBULIN // SUBFAMILY NOT NAMED (1 of 2)
OG IDs: 93065907
Best-hit-arabi-ID: AT5G12250.1
Best-hit-arabi-name: beta-6 tubulin
Gorai.003G159100.1.A
PCC=0.834, p=0.0
mRNA Profile
Protein Expression Profile
DPA

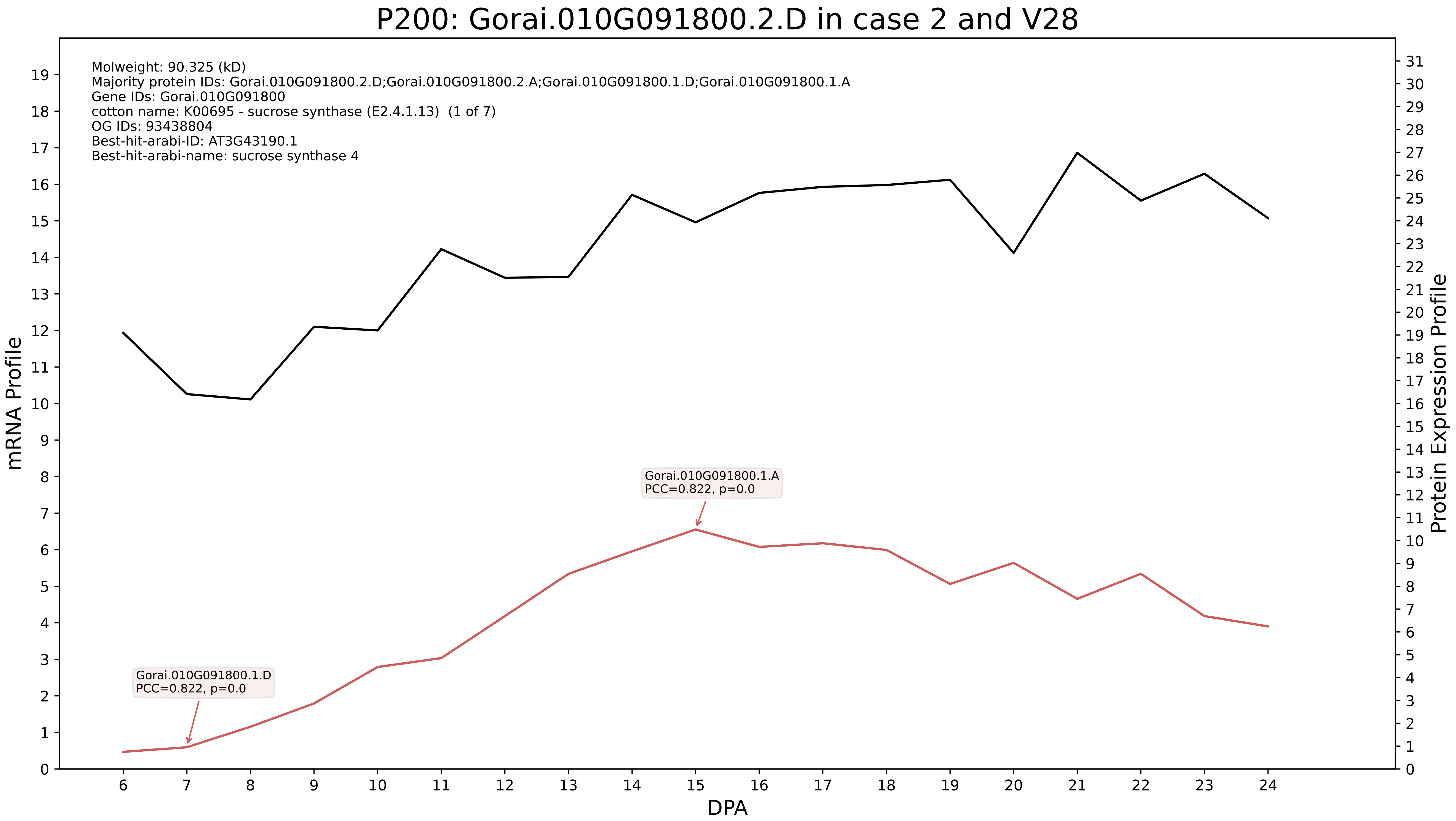
P200: Gorai.010G091800.2.D in case 2 and V28
Molweight: 90.325 (kD)
Majority protein IDs: Gorai.010G091800.2.D;Gorai.010G091800.2.A;Gorai.010G091800.1.D;Gorai.010G091800.1.A
Gene IDs: Gorai.010G091800
cotton name: K00695 - sucrose synthase (E2.4.1.13) (1 of 7)
OG IDs: 93438804
Best-hit-arabi-ID: AT3G43190.1
Best-hit-arabi-name: sucrose synthase 4
Gorai.010G091800.1.A
PCC=0.822, p=0.0
Gorai.010G091800.1.D
PCC=0.822, p=0.0
mRNA Profile
Protein Expression Profile
DPA

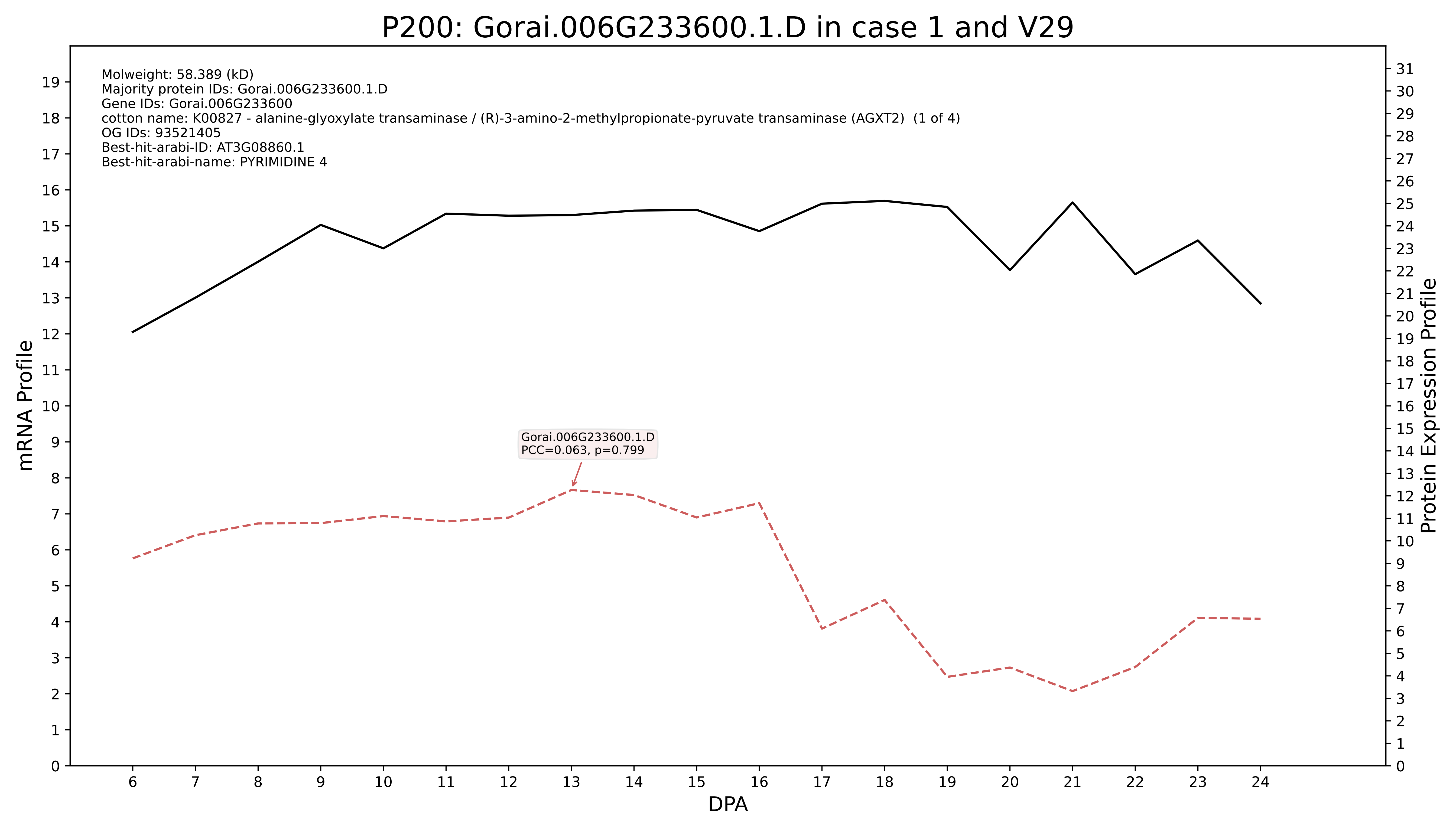
P200: Gorai.006G233600.1.D in case 1 and V29
Molweight: 58.389 (kD)
Majority protein IDs: Gorai.006G233600.1.D
Gene IDs: Gorai.006G233600
cotton name: K00827 - alanine-glyoxylate transaminase / (R)-3-amino-2-methylpropionate-pyruvate transaminase (AGXT2) (1 of 4)
OG IDs: 93521405
Best-hit-arabi-ID: AT3G08860.1
Best-hit-arabi-name: PYRIMIDINE 4
Gorai.006G233600.1.D
PCC=0.063, p=0.799
mRNA Profile
Protein Expression Profile
DPA

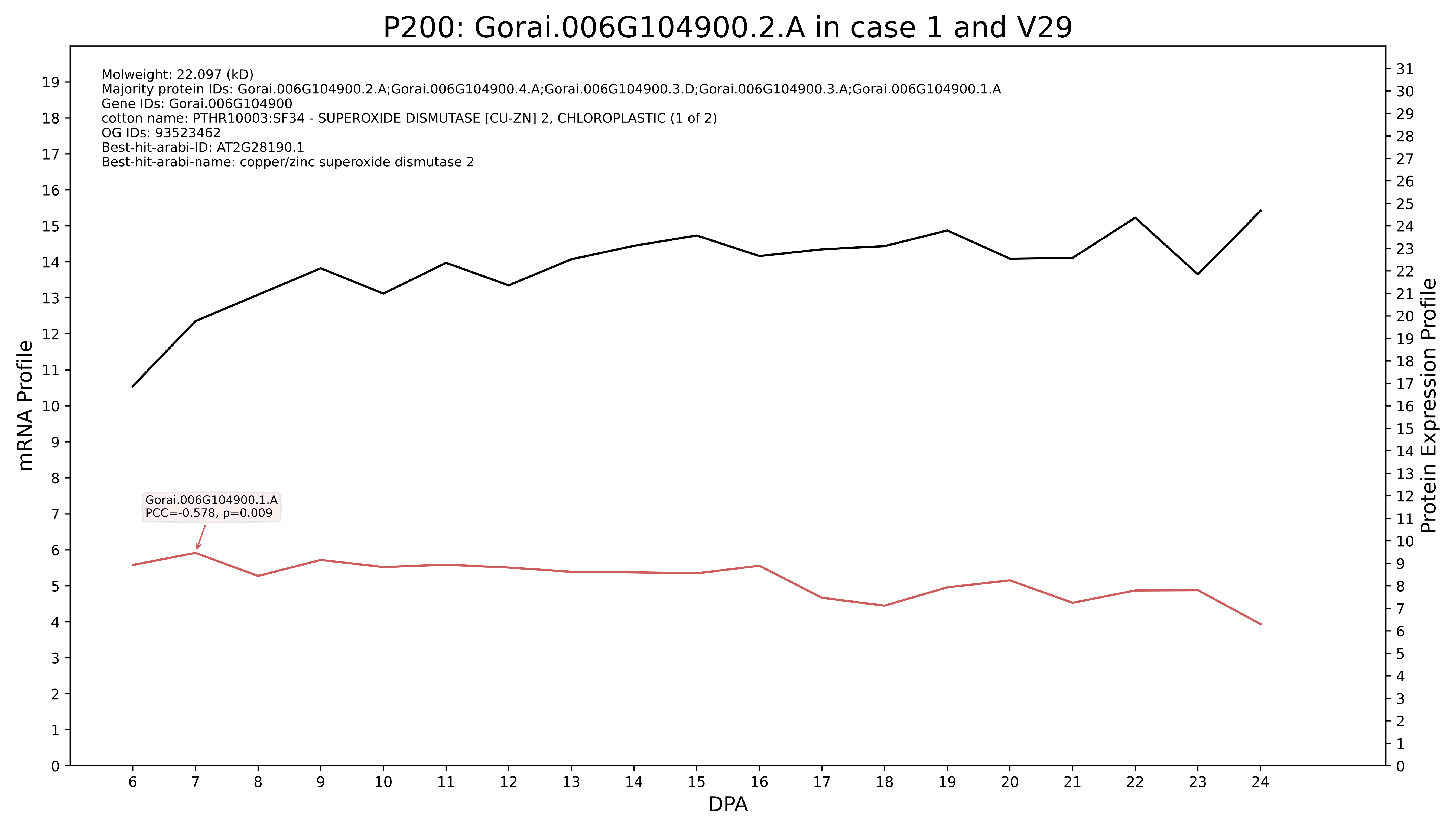
P200: Gorai.006G104900.2.A in case 1 and V29
Molweight: 22.097 (kD)
Majority protein IDs: Gorai.006G104900.2.A;Gorai.006G104900.4.A;Gorai.006G104900.3.D;Gorai.006G104900.3.A;Gorai.006G104900.1.A
Gene IDs: Gorai.006G104900
cotton name: PTHR10003:SF34 - SUPEROXIDE DISMUTASE [CU-ZN] 2, CHLOROPLASTIC (1 of 2)
OG IDs: 93523462
Best-hit-arabi-ID: AT2G28190.1
Best-hit-arabi-name: copper/zinc superoxide dismutase 2
Gorai.006G104900.1.A
PCC=-0.578, p=0.009
mRNA Profile
Protein Expression Profile
DPA

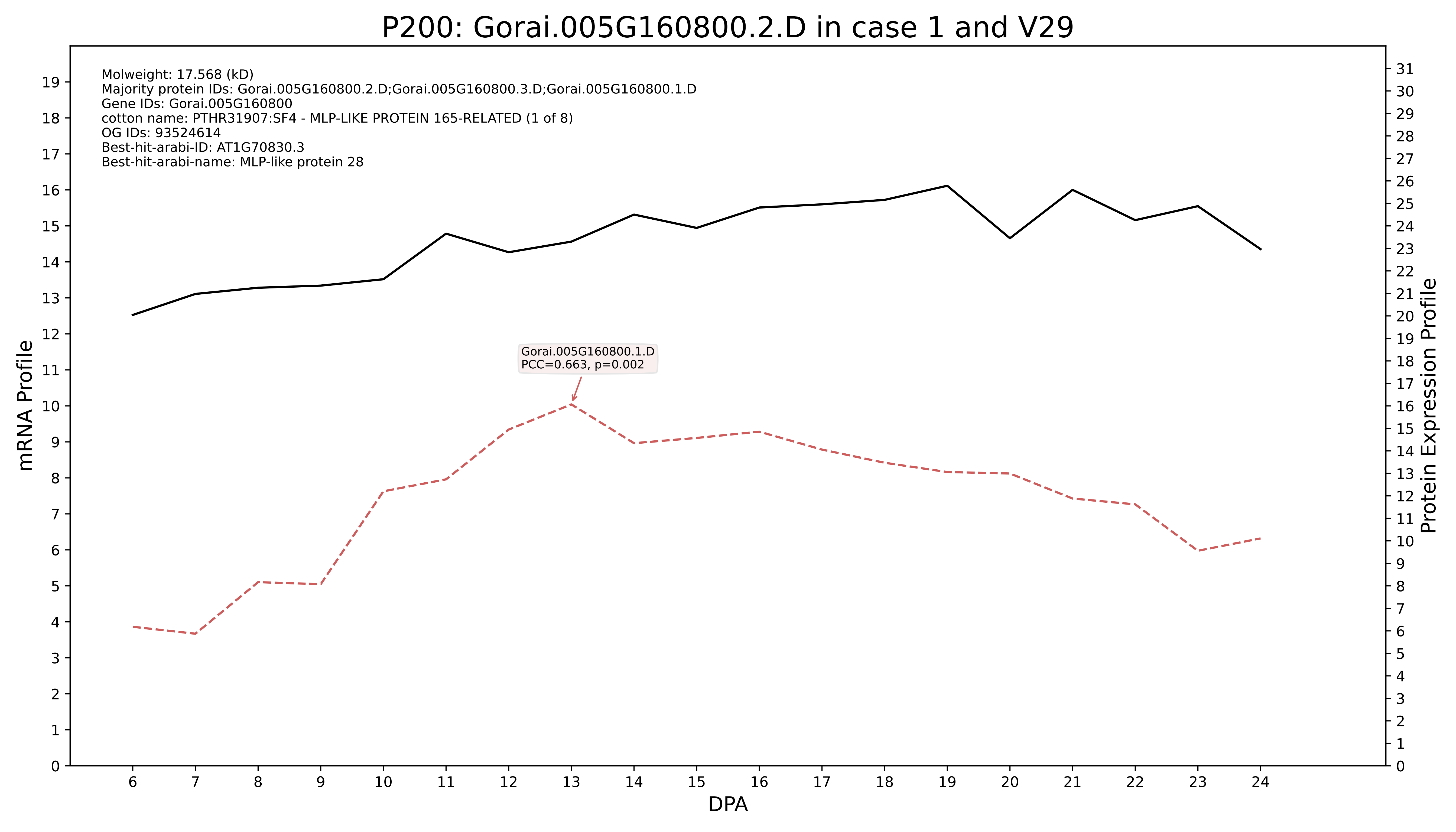
P200: Gorai.005G160800.2.D in case 1 and V29
Molweight: 17.568 (kD)
Majority protein IDs: Gorai.005G160800.2.D;Gorai.005G160800.3.D;Gorai.005G160800.1.D
Gene IDs: Gorai.005G160800
cotton name: PTHR31907:SF4 - MLP-LIKE PROTEIN 165-RELATED (1 of 8)
OG IDs: 93524614
Best-hit-arabi-ID: AT1G70830.3
Best-hit-arabi-name: MLP-like protein 28
Gorai.005G160800.1.D
PCC=0.663, p=0.002
mRNA Profile
Protein Expression Profile
DPA

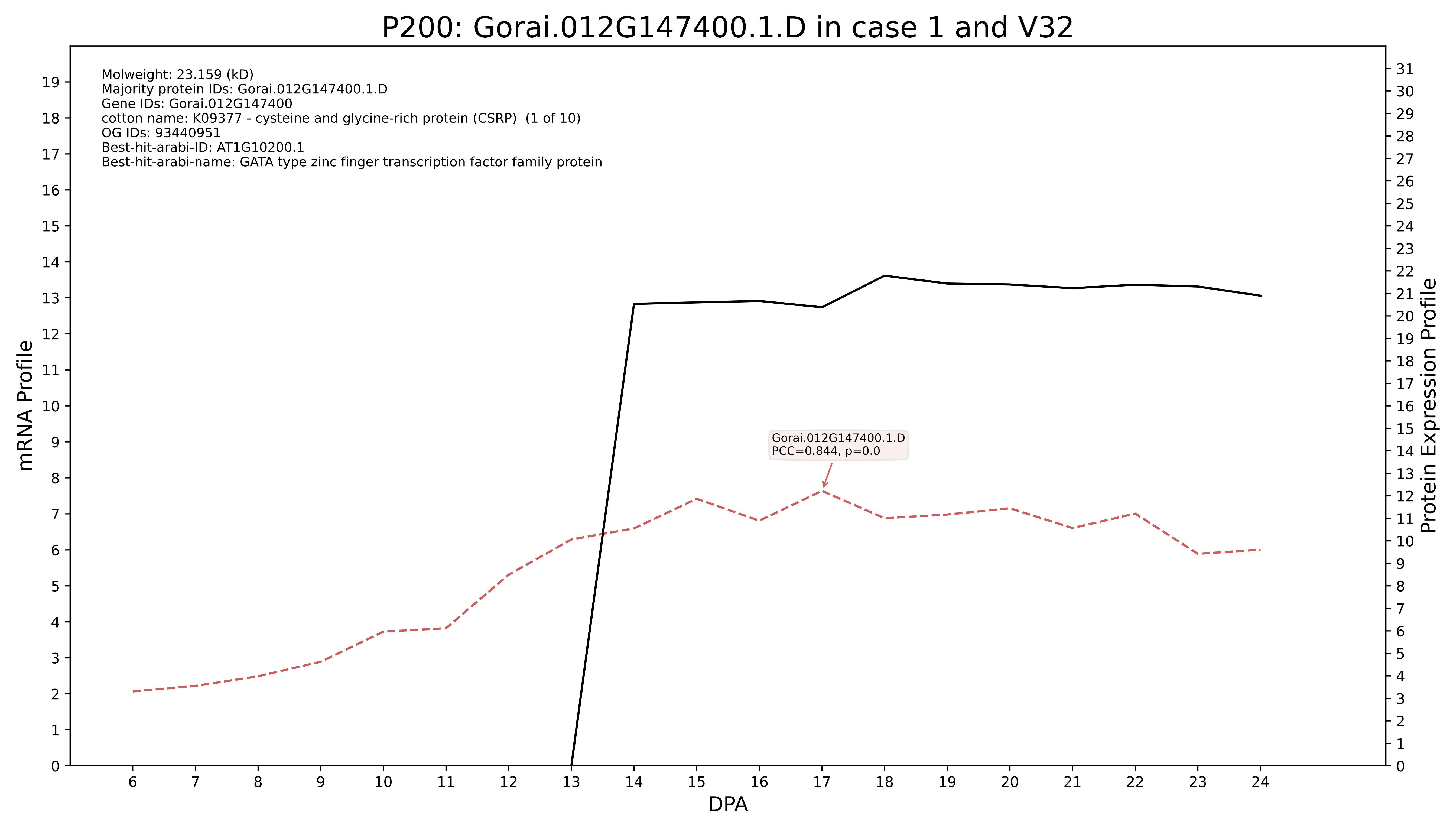
P200: Gorai.012G147400.1.D in case 1 and V32
Molweight: 23.159 (kD)
Majority protein IDs: Gorai.012G147400.1.D
Gene IDs: Gorai.012G147400
cotton name: K09377 - cysteine and glycine-rich protein (CSRP) (1 of 10)
OG IDs: 93440951
Best-hit-arabi-ID: AT1G10200.1
Best-hit-arabi-name: GATA type zinc finger transcription factor family protein
Gorai.012G147400.1.D
PCC=0.844, p=0.0
mRNA Profile
Protein Expression Profile
DPA

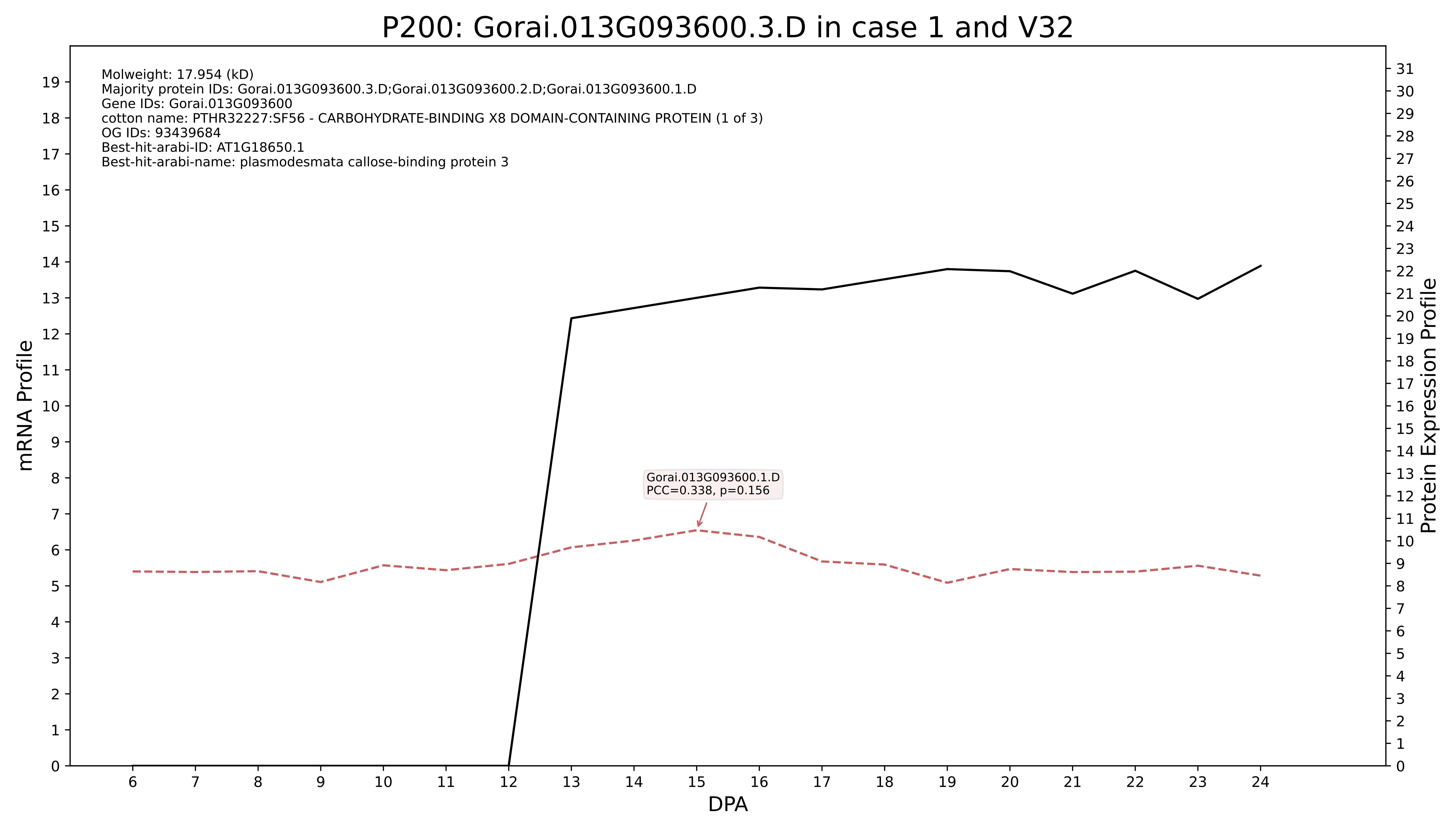
P200: Gorai.013G093600.3.D in case 1 and V32
Molweight: 17.954 (kD)
Majority protein IDs: Gorai.013G093600.3.D;Gorai.013G093600.2.D;Gorai.013G093600.1.D
Gene IDs: Gorai.013G093600
cotton name: PTHR32227:SF56 - CARBOHYDRATE-BINDING X8 DOMAIN-CONTAINING PROTEIN (1 of 3)
OG IDs: 93439684
Best-hit-arabi-ID: AT1G18650.1
Best-hit-arabi-name: plasmodesmata callose-binding protein 3
Gorai.013G093600.1.D
PCC=0.338, p=0.156
mRNA Profile
Protein Expression Profile
DPA

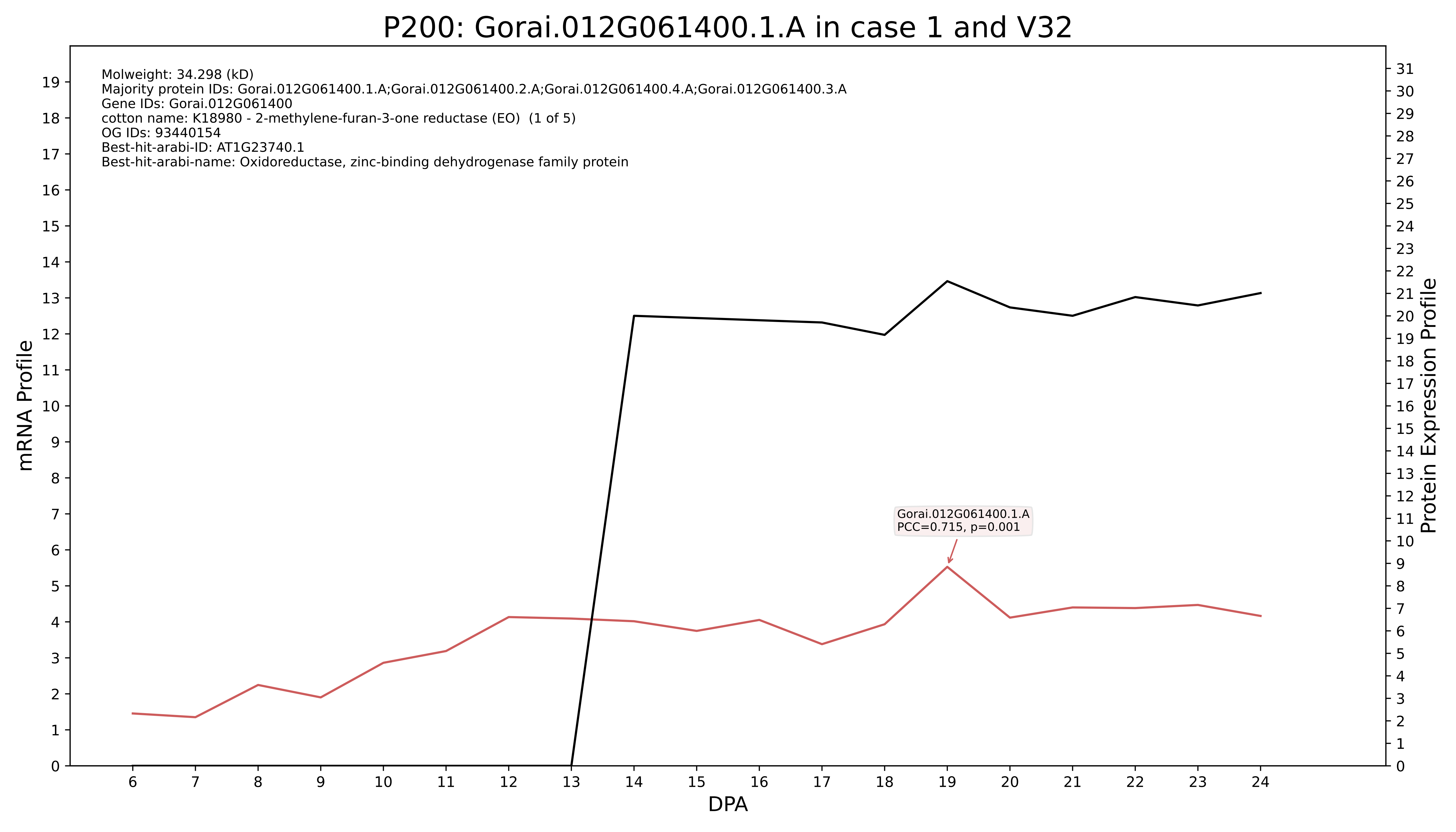
P200: Gorai.012G061400.1.A in case 1 and V32
Molweight: 34.298 (kD)
Majority protein IDs: Gorai.012G061400.1.A;Gorai.012G061400.2.A;Gorai.012G061400.4.A;Gorai.012G061400.3.A
Gene IDs: Gorai.012G061400
cotton name: K18980 - 2-methylene-furan-3-one reductase (EO) (1 of 5)
OG IDs: 93440154
Best-hit-arabi-ID: AT1G23740.1
Best-hit-arabi-name: Oxidoreductase, zinc-binding dehydrogenase family protein
Gorai.012G061400.1.A
PCC=0.715, p=0.001
mRNA Profile
Protein Expression Profile
DPA

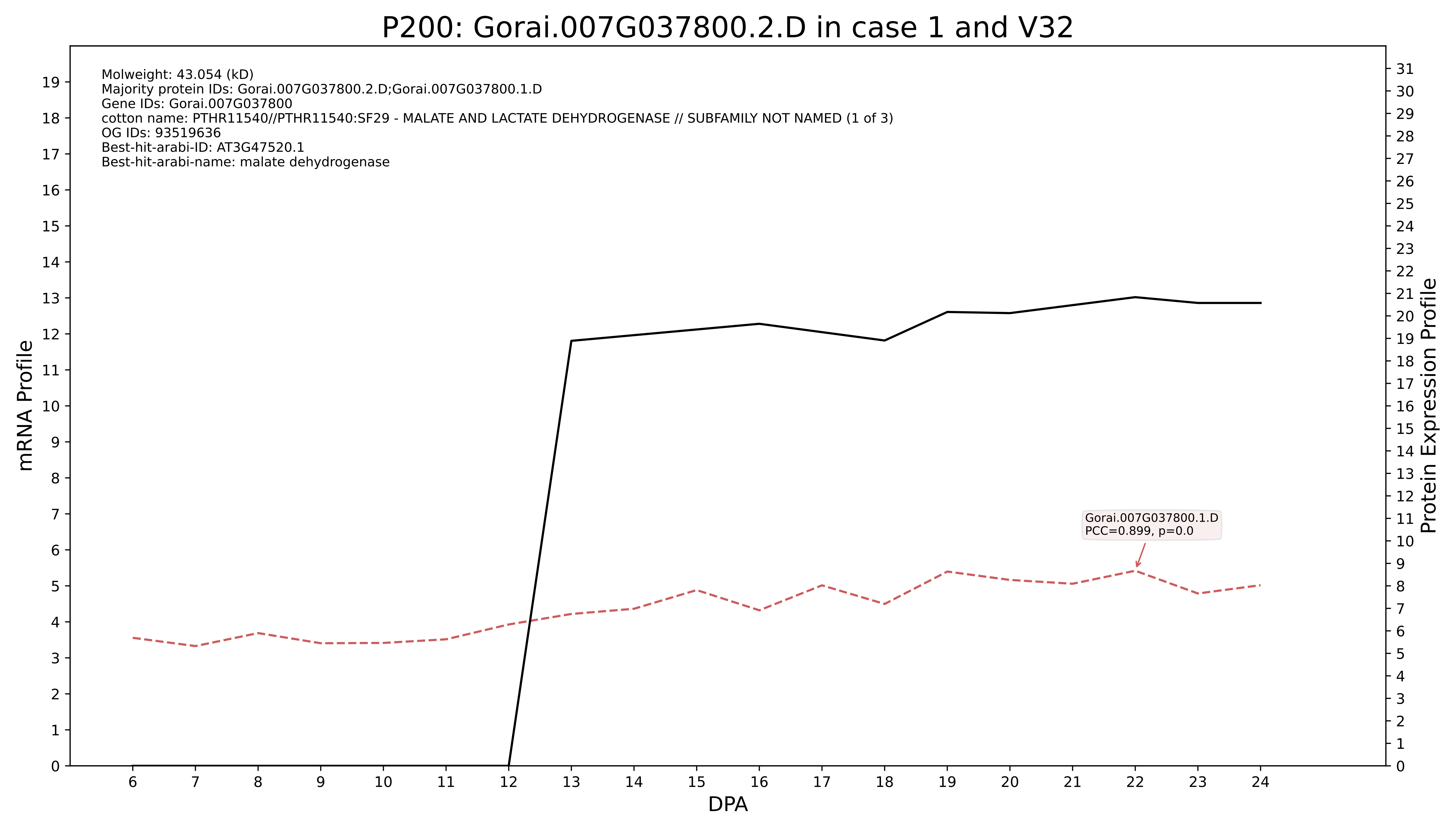
P200: Gorai.007G037800.2.D in case 1 and V32
Molweight: 43.054 (kD)
Majority protein IDs: Gorai.007G037800.2.D;Gorai.007G037800.1.D
Gene IDs: Gorai.007G037800
cotton name: PTHR11540//PTHR11540:SF29 - MALATE AND LACTATE DEHYDROGENASE // SUBFAMILY NOT NAMED (1 of 3)
OG IDs: 93519636
Best-hit-arabi-ID: AT3G47520.1
Best-hit-arabi-name: malate dehydrogenase
Gorai.007G037800.1.D
PCC=0.899, p=0.0
mRNA Profile
Protein Expression Profile
DPA

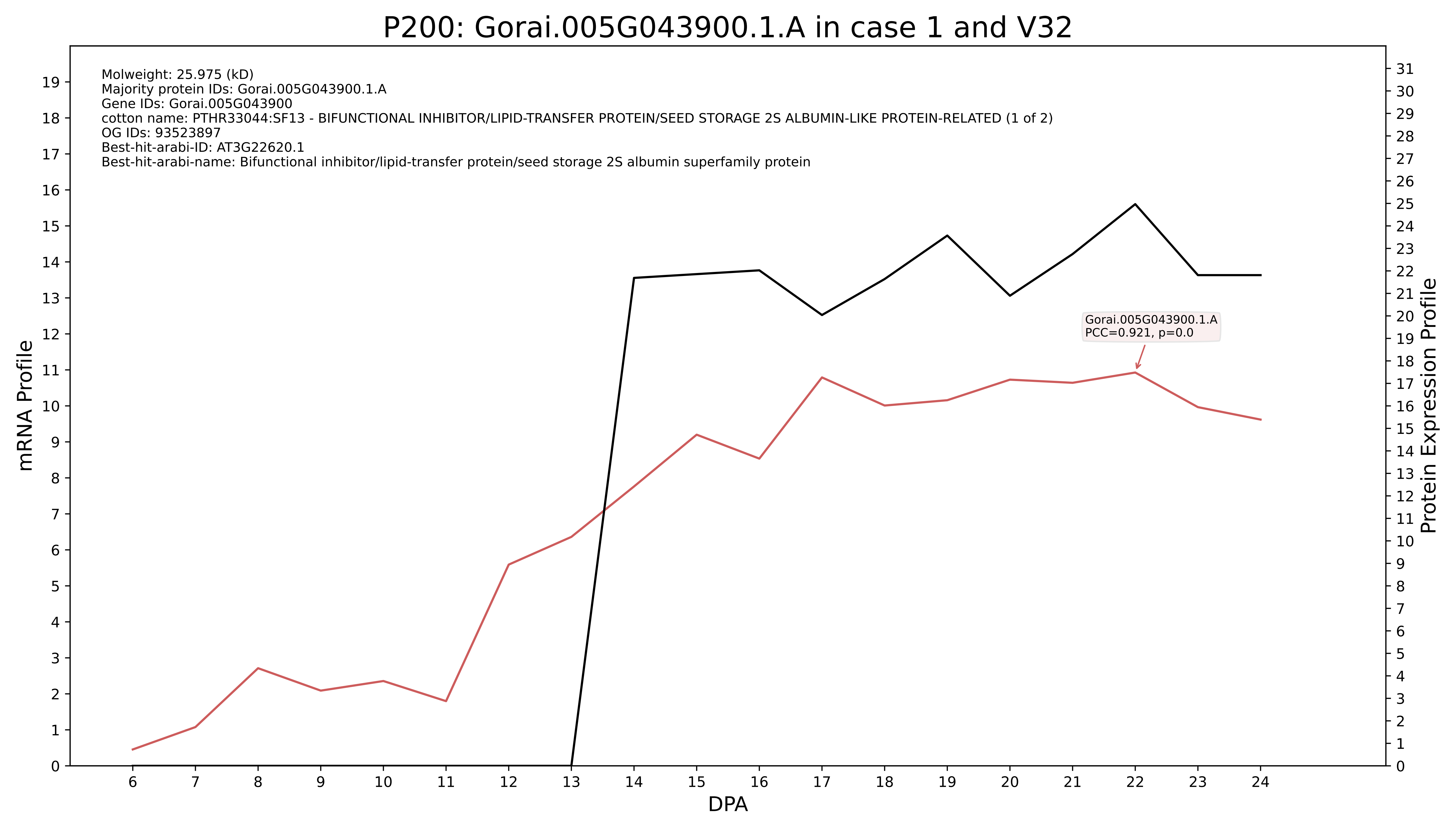
P200: Gorai.005G043900.1.A in case 1 and V32
Molweight: 25.975 (kD)
Majority protein IDs: Gorai.005G043900.1.A
Gene IDs: Gorai.005G043900
cotton name: PTHR33044:SF13 - BIFUNCTIONAL INHIBITOR/LIPID-TRANSFER PROTEIN/SEED STORAGE 2S ALBUMIN-LIKE PROTEIN-RELATED (1 of 2)
OG IDs: 93523897
Best-hit-arabi-ID: AT3G22620.1
Best-hit-arabi-name: Bifunctional inhibitor/lipid-transfer protein/seed storage 2S albumin superfamily protein
Gorai.005G043900.1.A
PCC=0.921, p=0.0
mRNA Profile
Protein Expression Profile
DPA

### P200: Gorai.005G057400.1.A in case 2 and V32

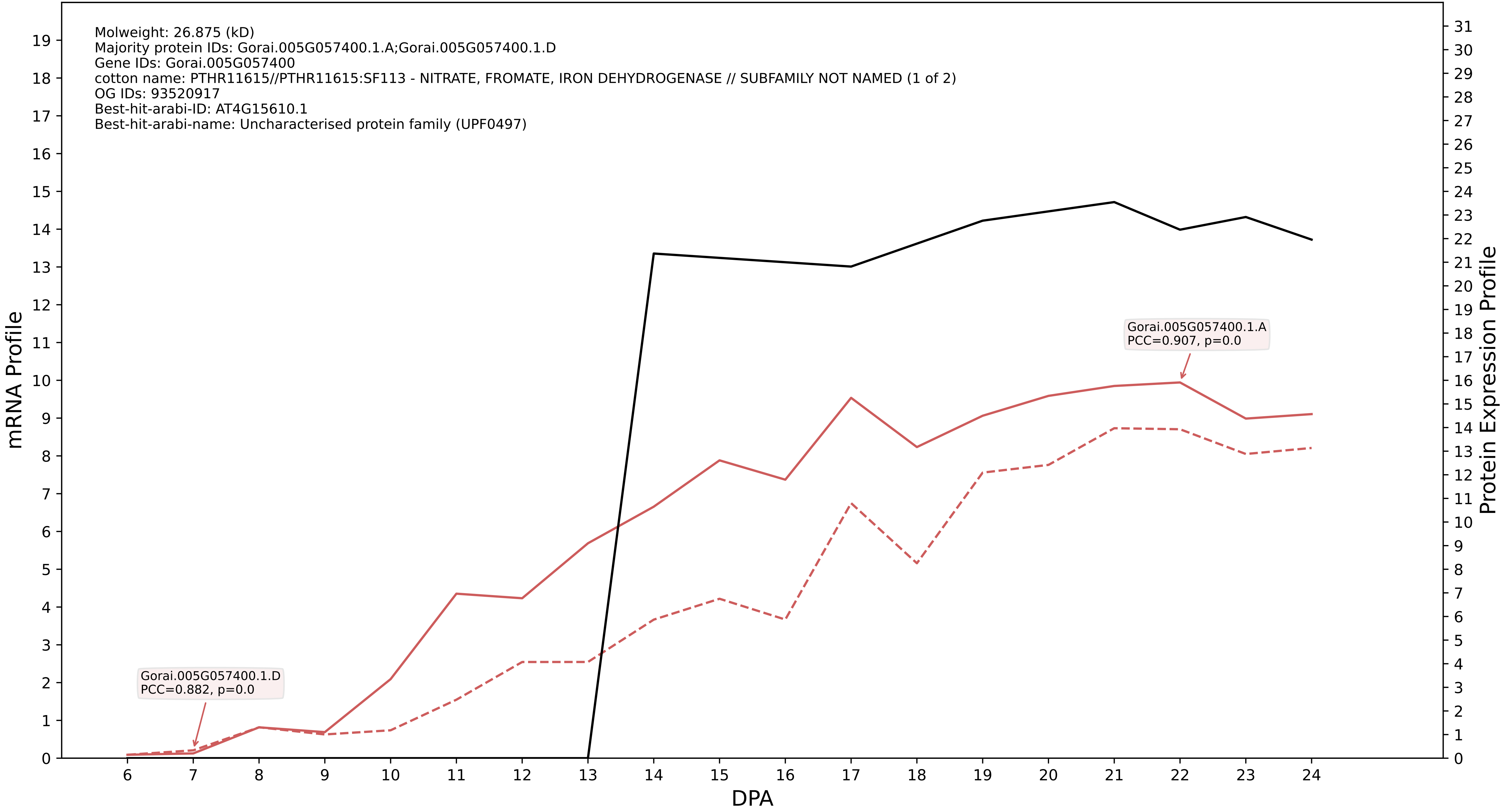

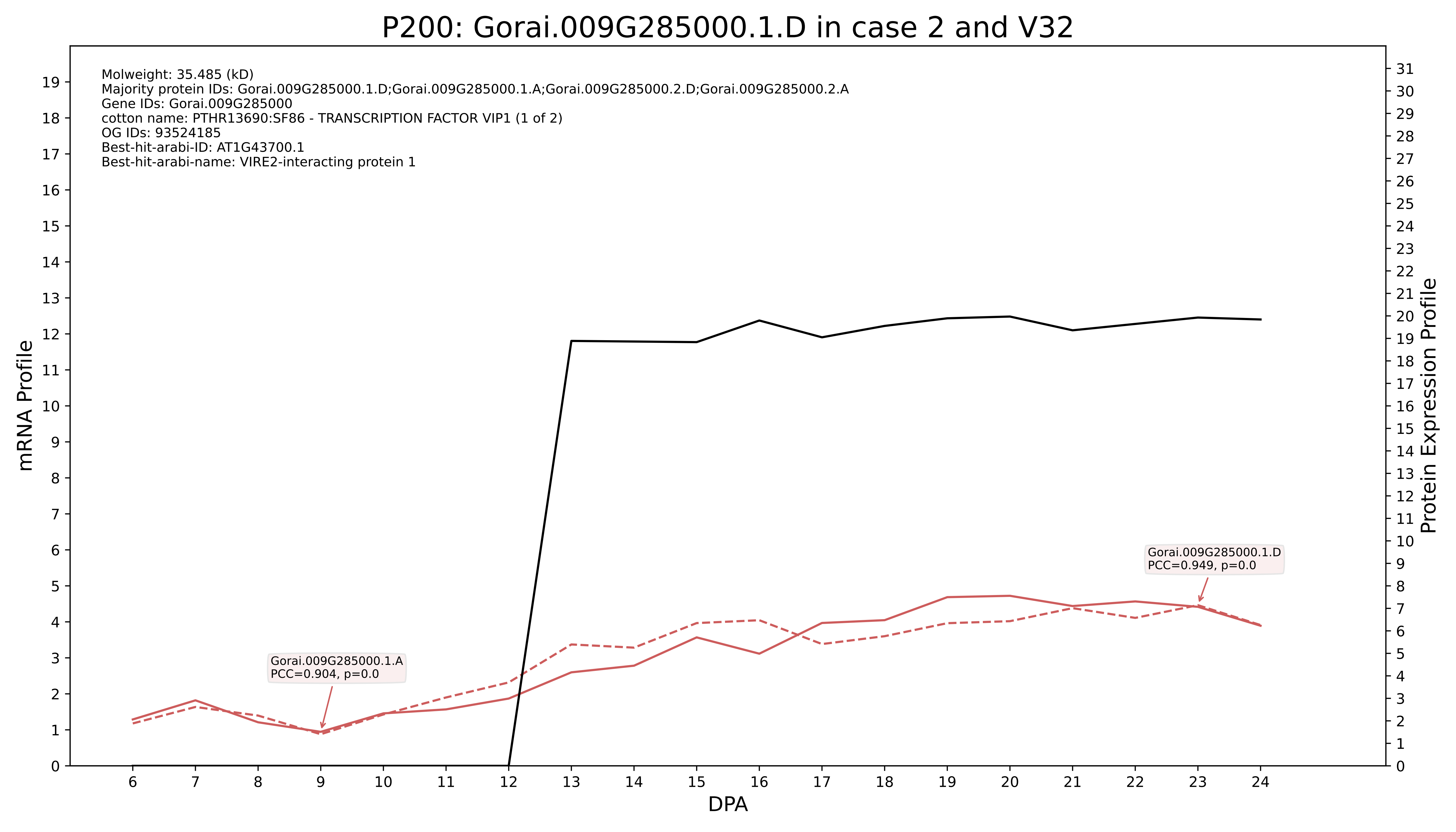

P200: Gorai.009G285000.1.D in case 2 and V32
Molweight: 35.485 (kD)
Majority protein IDs: Gorai.009G285000.1.D;Gorai.009G285000.1.A;Gorai.009G285000.2.D;Gorai.009G285000.2.A
Gene IDs: Gorai.009G285000
cotton name: PTHR13690:SF86 - TRANSCRIPTION FACTOR VIP1 (1 of 2)
OG IDs: 93524185
Best-hit-arabi-ID: AT1G43700.1
Best-hit-arabi-name: VIRE2-interacting protein 1
Gorai.009G285000.1.A
PCC=0.904, p=0.0
Gorai.009G285000.1.D
PCC=0.949, p=0.0
mRNA Profile
Protein Expression Profile
DPA

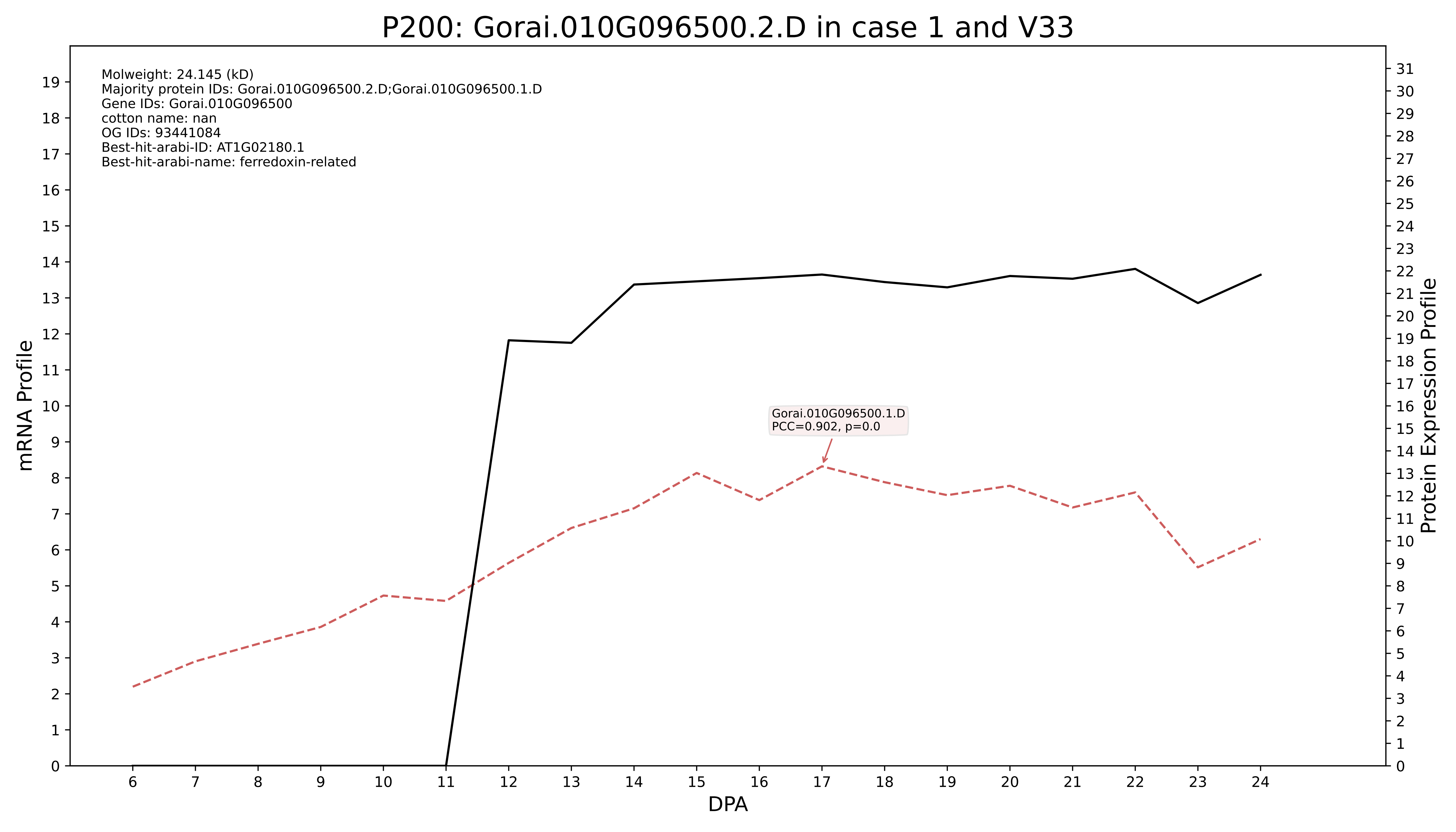
P200: Gorai.010G096500.2.D in case 1 and V33
Molweight: 24.145 (kD)
Majority protein IDs: Gorai.010G096500.2.D;Gorai.010G096500.1.D
Gene IDs: Gorai.010G096500
cotton name: nan
OG IDs: 93441084
Best-hit-arabi-ID: AT1G02180.1
Best-hit-arabi-name: ferredoxin-related
Gorai.010G096500.1.D
PCC=0.902, p=0.0
mRNA Profile
Protein Expression Profile
DPA

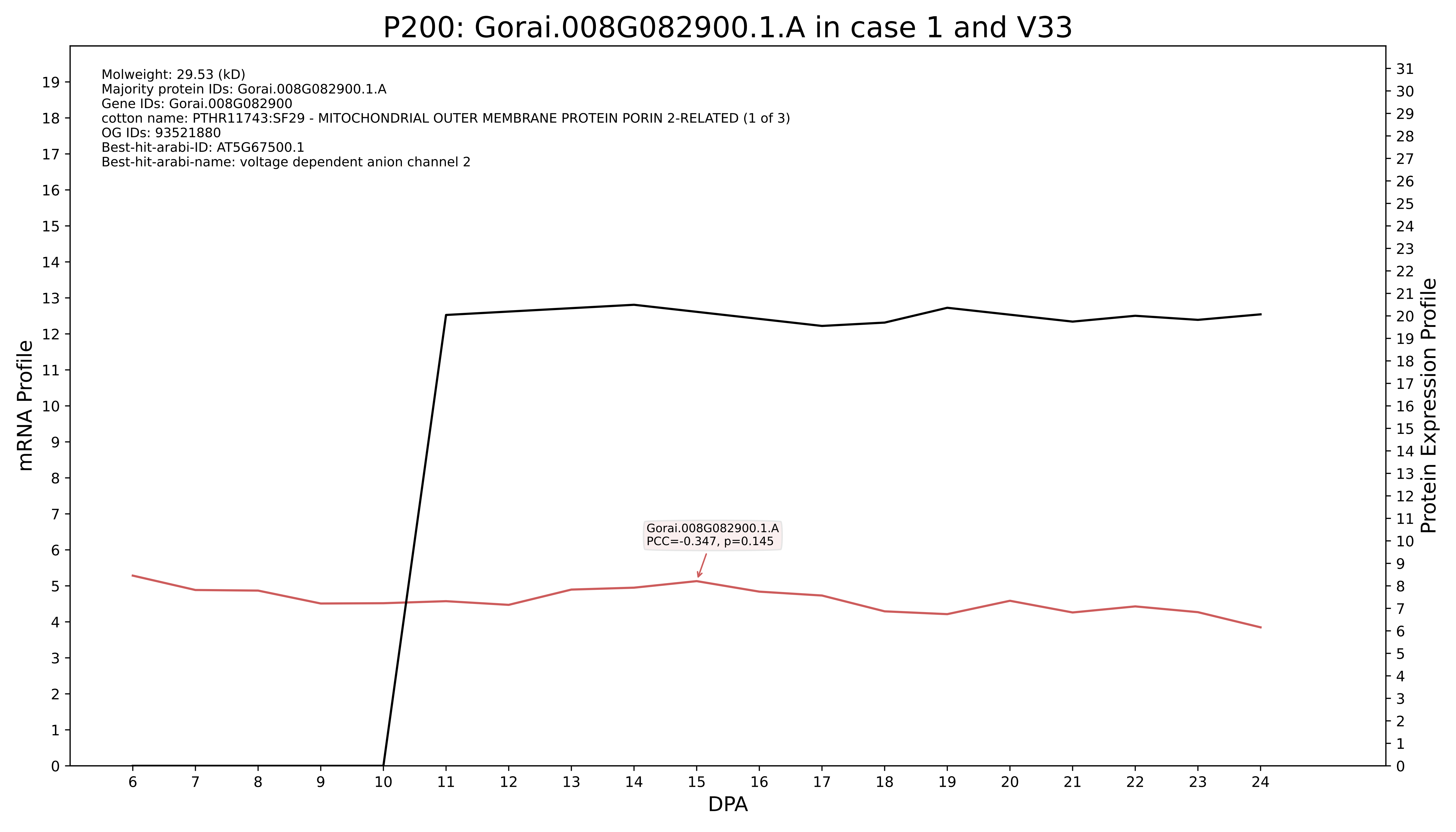
P200: Gorai.008G082900.1.A in case 1 and V33
Molweight: 29.53 (kD)
Majority protein IDs: Gorai.008G082900.1.A
Gene IDs: Gorai.008G082900
cotton name: PTHR11743:SF29 - MITOCHONDRIAL OUTER MEMBRANE PROTEIN PORIN 2-RELATED (1 of 3)
OG IDs: 93521880
Best-hit-arabi-ID: AT5G67500.1
Best-hit-arabi-name: voltage dependent anion channel 2
Gorai.008G082900.1.A
PCC=-0.347, p=0.145
mRNA Profile
Protein Expression Profile
DPA

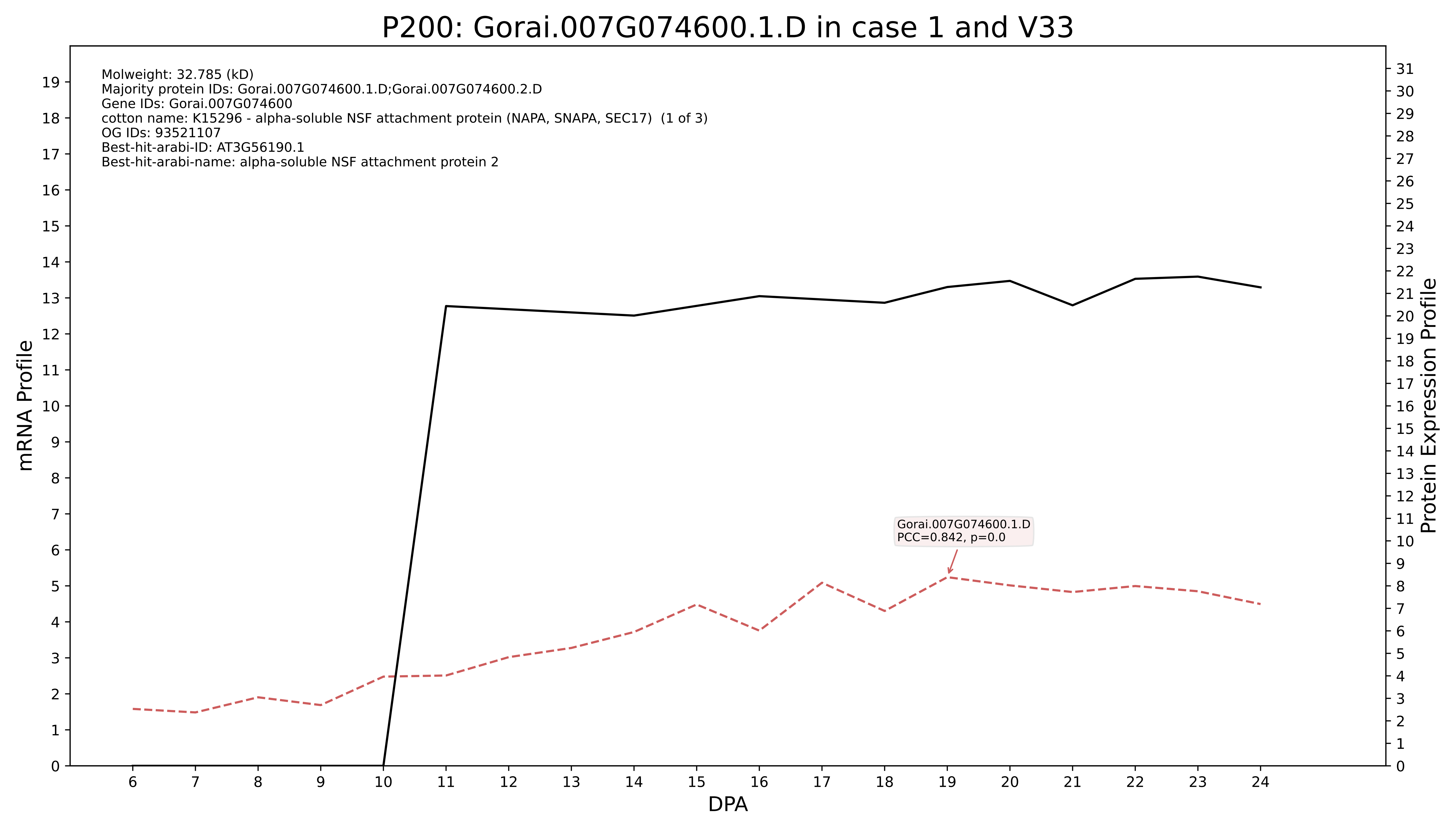
P200: Gorai.007G074600.1.D in case 1 and V33
Molweight: 32.785 (kD)
Majority protein IDs: Gorai.007G074600.1.D;Gorai.007G074600.2.D
Gene IDs: Gorai.007G074600
cotton name: K15296 - alpha-soluble NSF attachment protein (NAPA, SNAPA, SEC17) (1 of 3)
OG IDs: 93521107
Best-hit-arabi-ID: AT3G56190.1
Best-hit-arabi-name: alpha-soluble NSF attachment protein 2
Gorai.007G074600.1.D
PCC=0.842, p=0.0
mRNA Profile
Protein Expression Profile
DPA

### P200: Gorai.013G156800.2.D in case 1 and V33

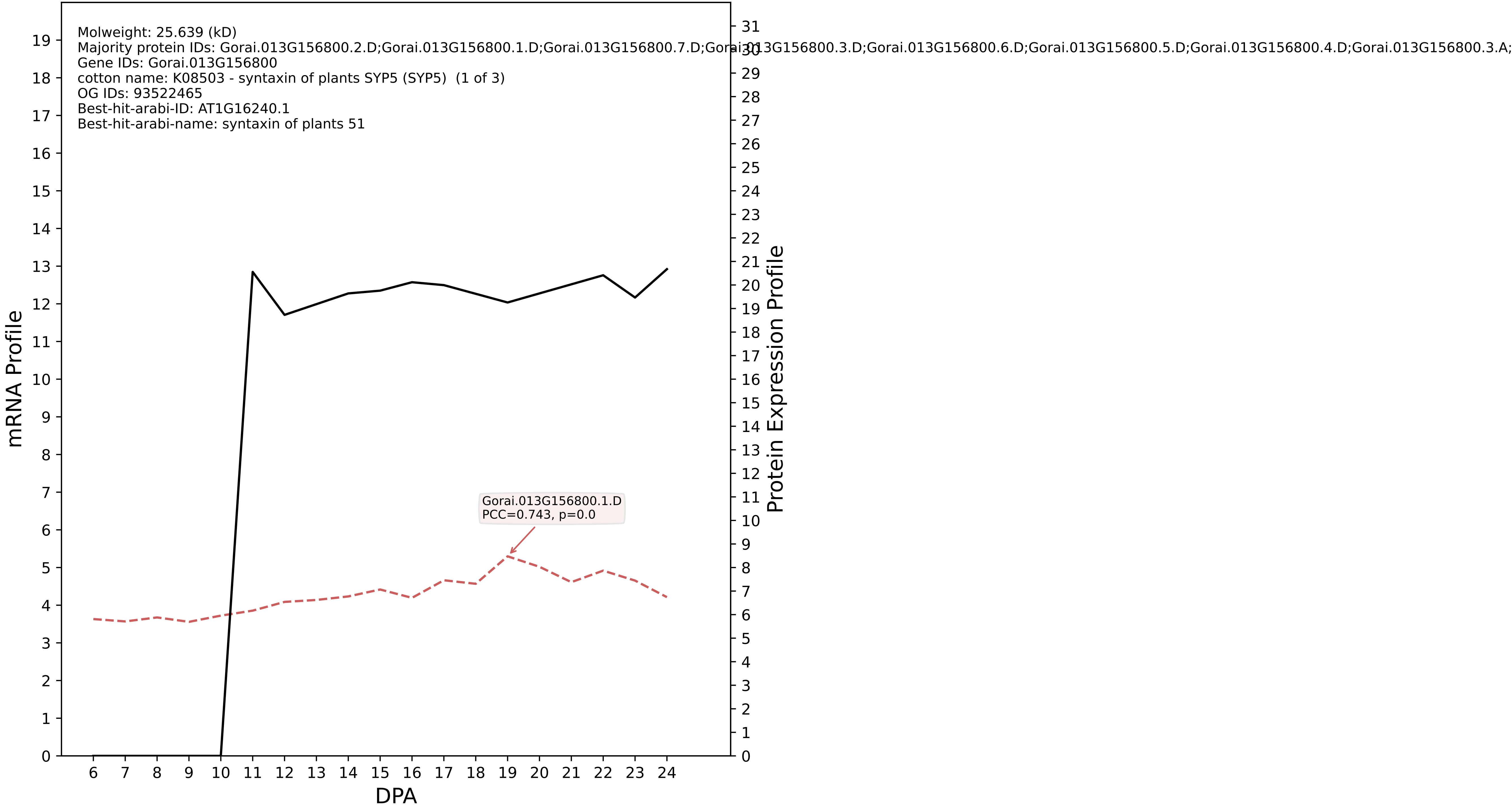

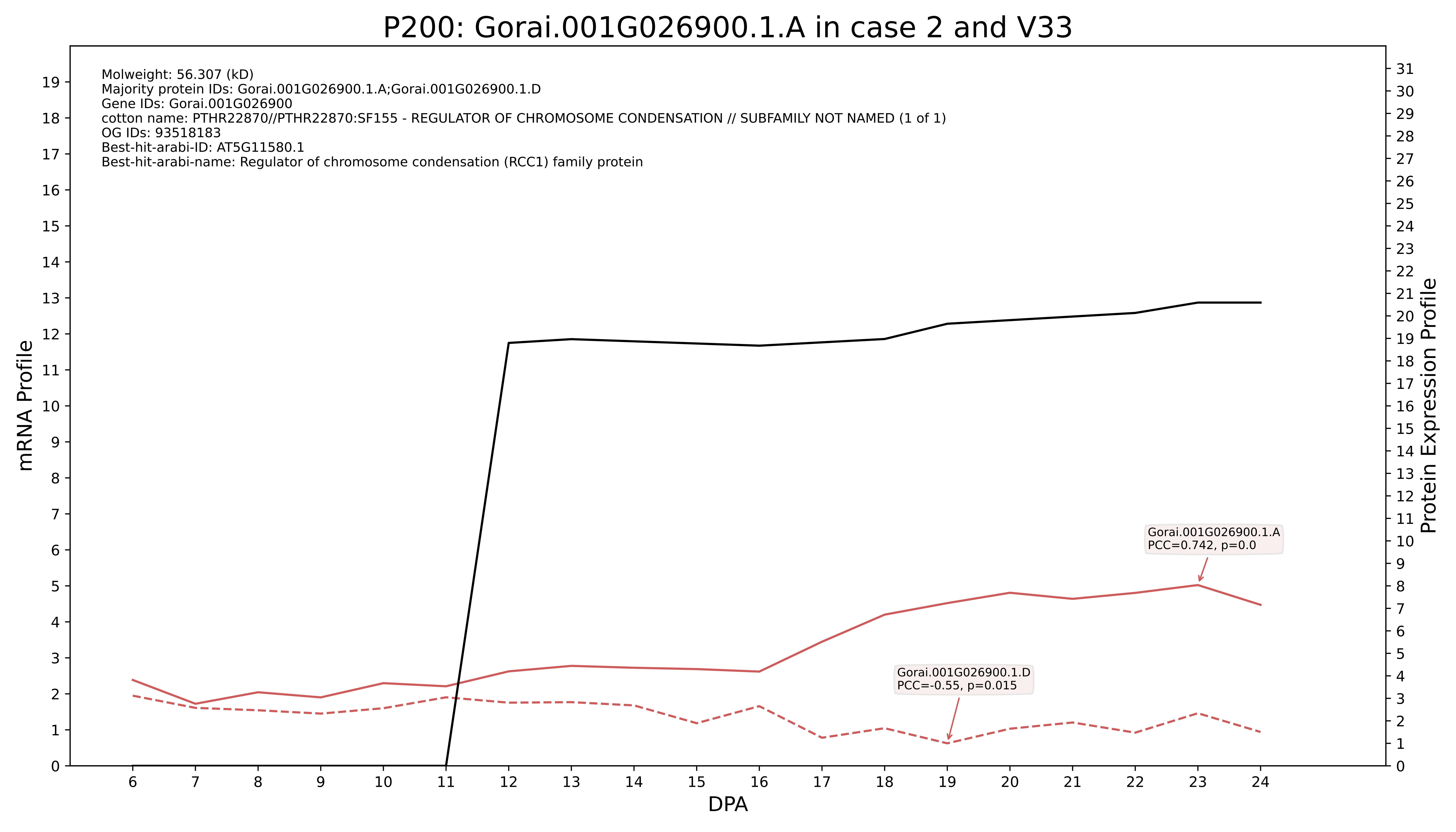
P200: Gorai.001G026900.1.A in case 2 and V33
Molweight: 56.307 (kD)
Majority protein IDs: Gorai.001G026900.1.A;Gorai.001G026900.1.D
Gene IDs: Gorai.001G026900
cotton name: PTHR22870//PTHR22870:SF155 - REGULATOR OF CHROMOSOME CONDENSATION // SUBFAMILY NOT NAMED (1 of 1)
OG IDs: 93518183
Best-hit-arabi-ID: AT5G11580.1
Best-hit-arabi-name: Regulator of chromosome condensation (RCC1) family protein
Gorai.001G026900.1.A
PCC=0.742, p=0.0
Gorai.001G026900.1.D
PCC=-0.55, p=0.015
mRNA Profile
Protein Expression Profile
DPA

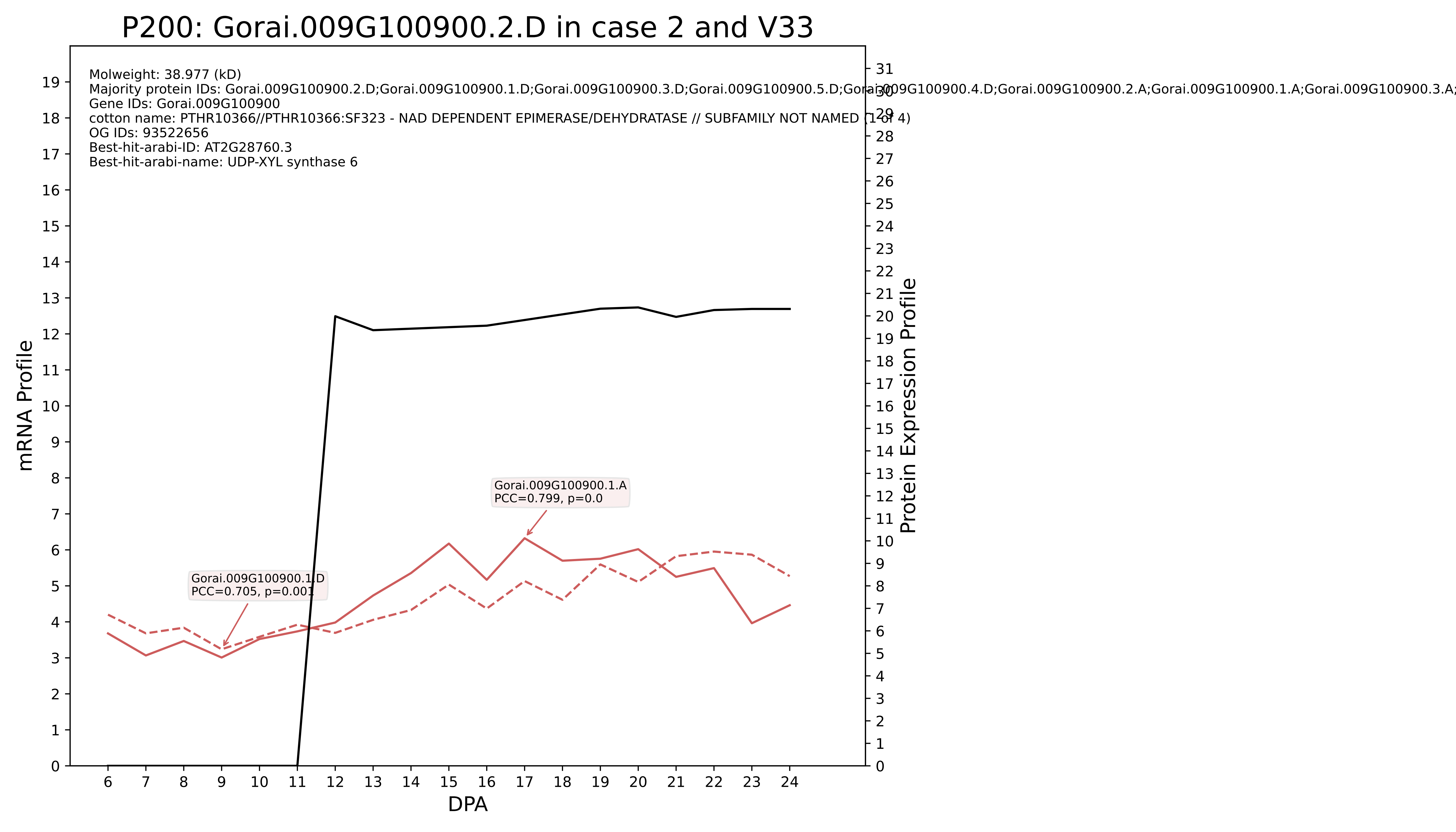

P200: Gorai.009G100900.2.D in case 2 and V33
Molweight: 38.977 (kD)
Majority protein IDs: Gorai.009G100900.2.D;Gorai.009G100900.1.D;Gorai.009G100900.3.D;Gorai.009G100900.5.D;Gorai.009G100900.4.D;Gorai.009G100900.2.A;Gorai.009G100900.1.A;Gorai.009G100900.3.A;
Gene IDs: Gorai.009G100900
cotton name: PTHR10366//PTHR10366:SF323 - NAD DEPENDENT EPIMERASE/DEHYDRATASE // SUBFAMILY NOT NAMED (1 of 4)
OG IDs: 93522656
Best-hit-arabi-ID: AT2G28760.3
Best-hit-arabi-name: UDP-XYL synthase 6
Gorai.009G100900.1.A
PCC=0.799, p=0.0
Gorai.009G100900.1.D
PCC=0.705, p=0.001
mRNA Profile
Protein Expression Profile
DPA

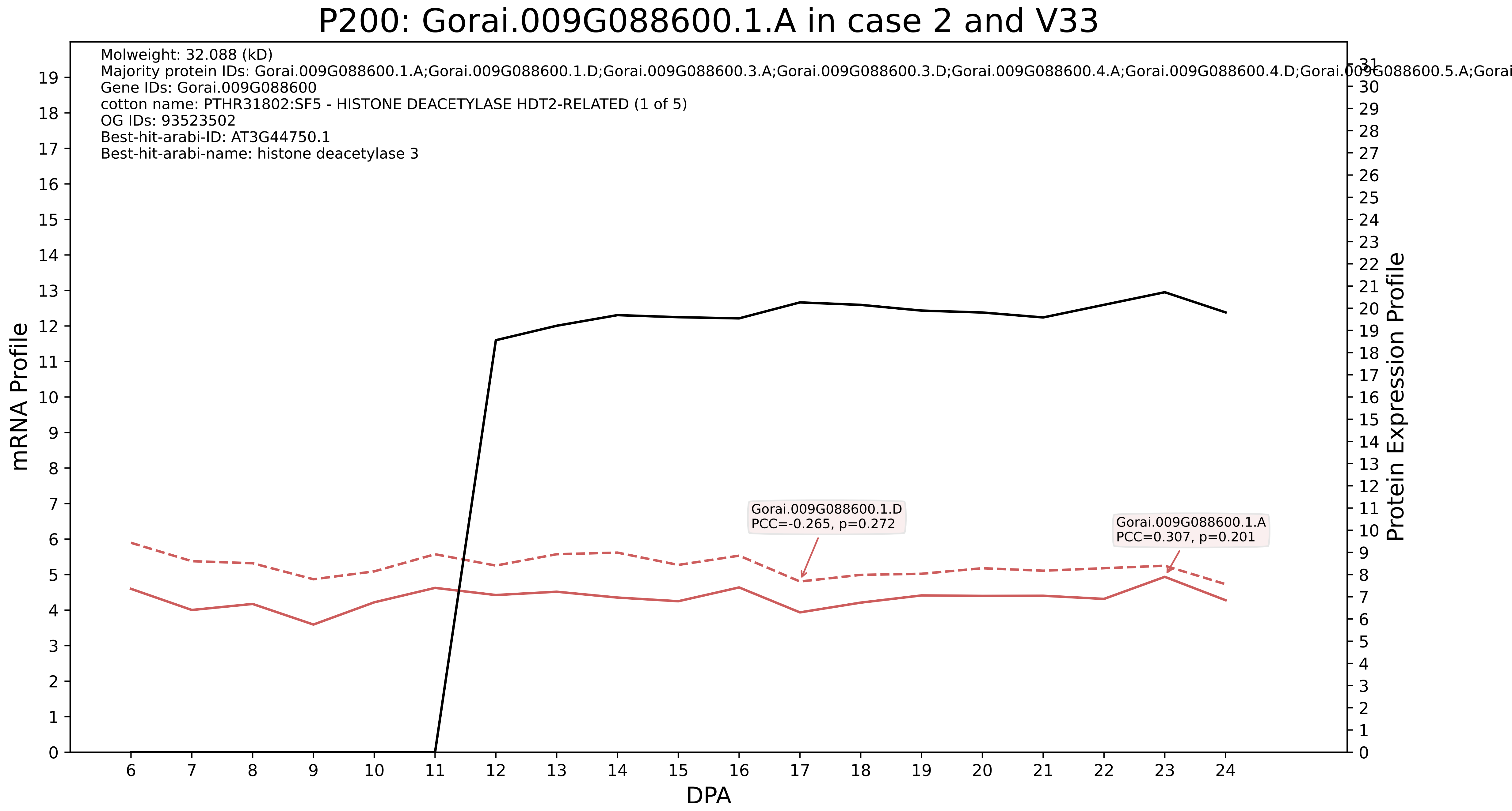
P200: Gorai.009G088600.1.A in case 2 and V33
Molweight: 32.088 (kD)
Majority protein IDs: Gorai.009G088600.1.A;Gorai.009G088600.1.D;Gorai.009G088600.3.A;Gorai.009G088600.3.D;Gorai.009G088600.4.A;Gorai.009G088600.4.D;Gorai.009G088600.5.A;Gorai
Gene IDs: Gorai.009G088600
cotton name: PTHR31802:SF5 - HISTONE DEACETYLASE HDT2-RELATED (1 of 5)
OG IDs: 93523502
Best-hit-arabi-ID: AT3G44750.1
Best-hit-arabi-name: histone deacetylase 3
Gorai.009G088600.1.D
PCC=-0.265, p=0.272
Gorai.009G088600.1.A
PCC=0.307, p=0.201
mRNA Profile
Protein Expression Profile
DPA

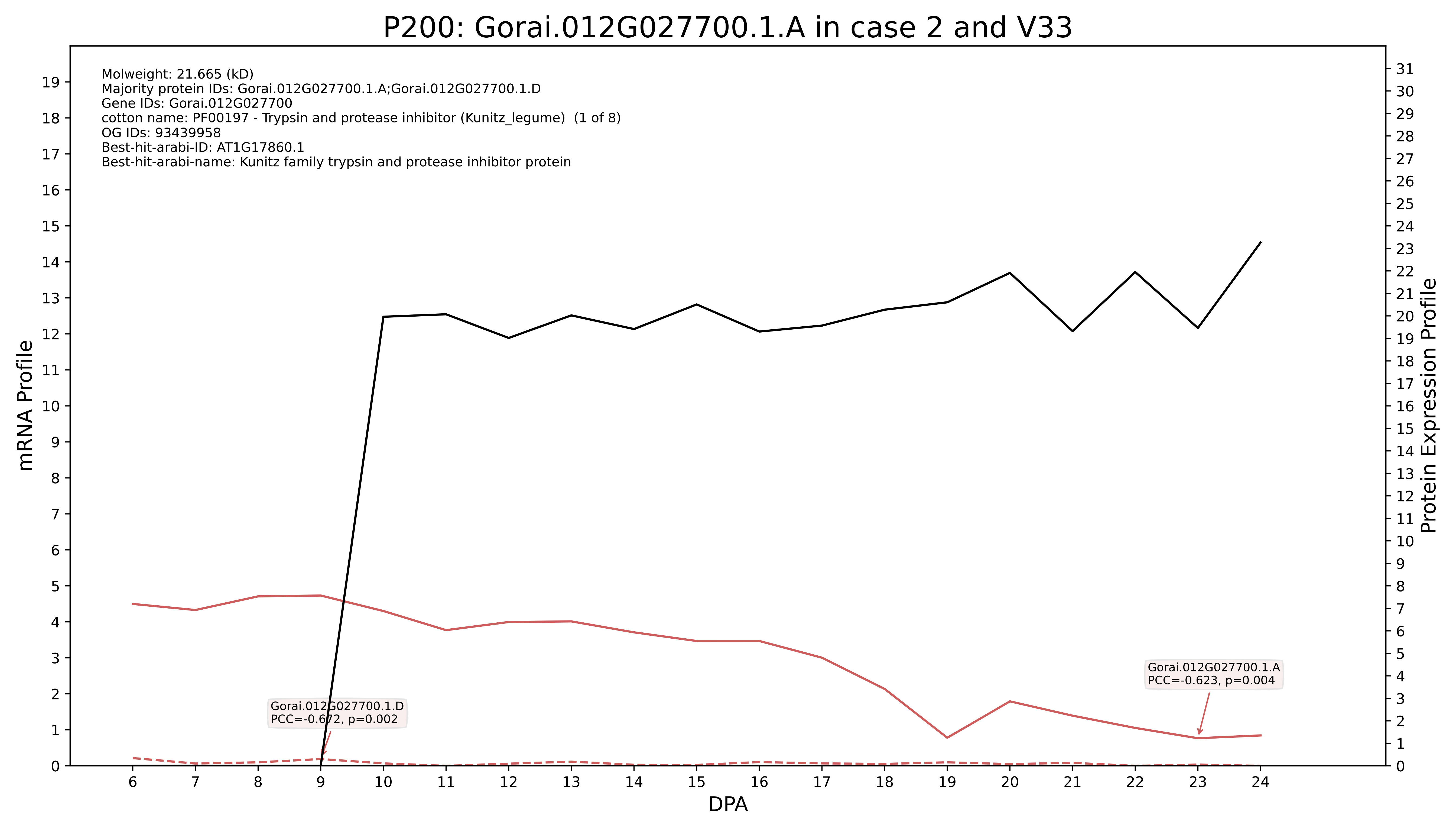
P200: Gorai.012G027700.1.A in case 2 and V33
Molweight: 21.665 (kD)
Majority protein IDs: Gorai.012G027700.1.A;Gorai.012G027700.1.D
Gene IDs: Gorai.012G027700
cotton name: PF00197 - Trypsin and protease inhibitor (Kunitz_legume) (1 of 8)
OG IDs: 93439958
Best-hit-arabi-ID: AT1G17860.1
Best-hit-arabi-name: Kunitz family trypsin and protease inhibitor protein
Gorai.012G027700.1.D
PCC=-0.672, p=0.002
Gorai.012G027700.1.A
PCC=-0.623, p=0.004
mRNA Profile
Protein Expression Profile
DPA

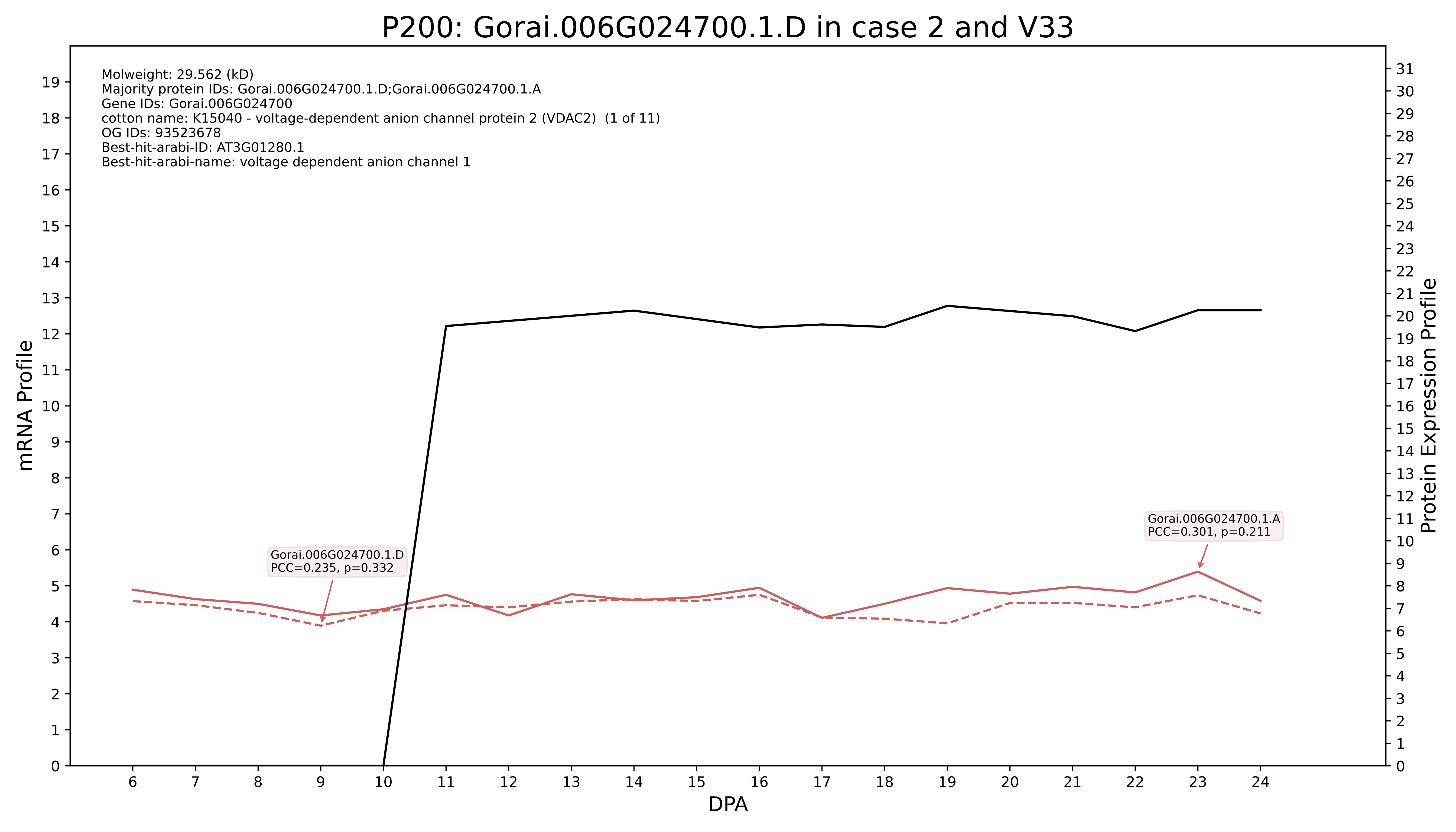

P200: Gorai.006G024700.1.D in case 2 and V33
Molweight: 29.562 (kD)
Majority protein IDs: Gorai.006G024700.1.D;Gorai.006G024700.1.A
Gene IDs: Gorai.006G024700
cotton name: K15040 - voltage-dependent anion channel protein 2 (VDAC2) (1 of 11)
OG IDs: 93523678
Best-hit-arabi-ID: AT3G01280.1
Best-hit-arabi-name: voltage dependent anion channel 1
Gorai.006G024700.1.D
PCC=0.235, p=0.332
Gorai.006G024700.1.A
PCC=0.301, p=0.211
mRNA Profile
Protein Expression Profile
DPA

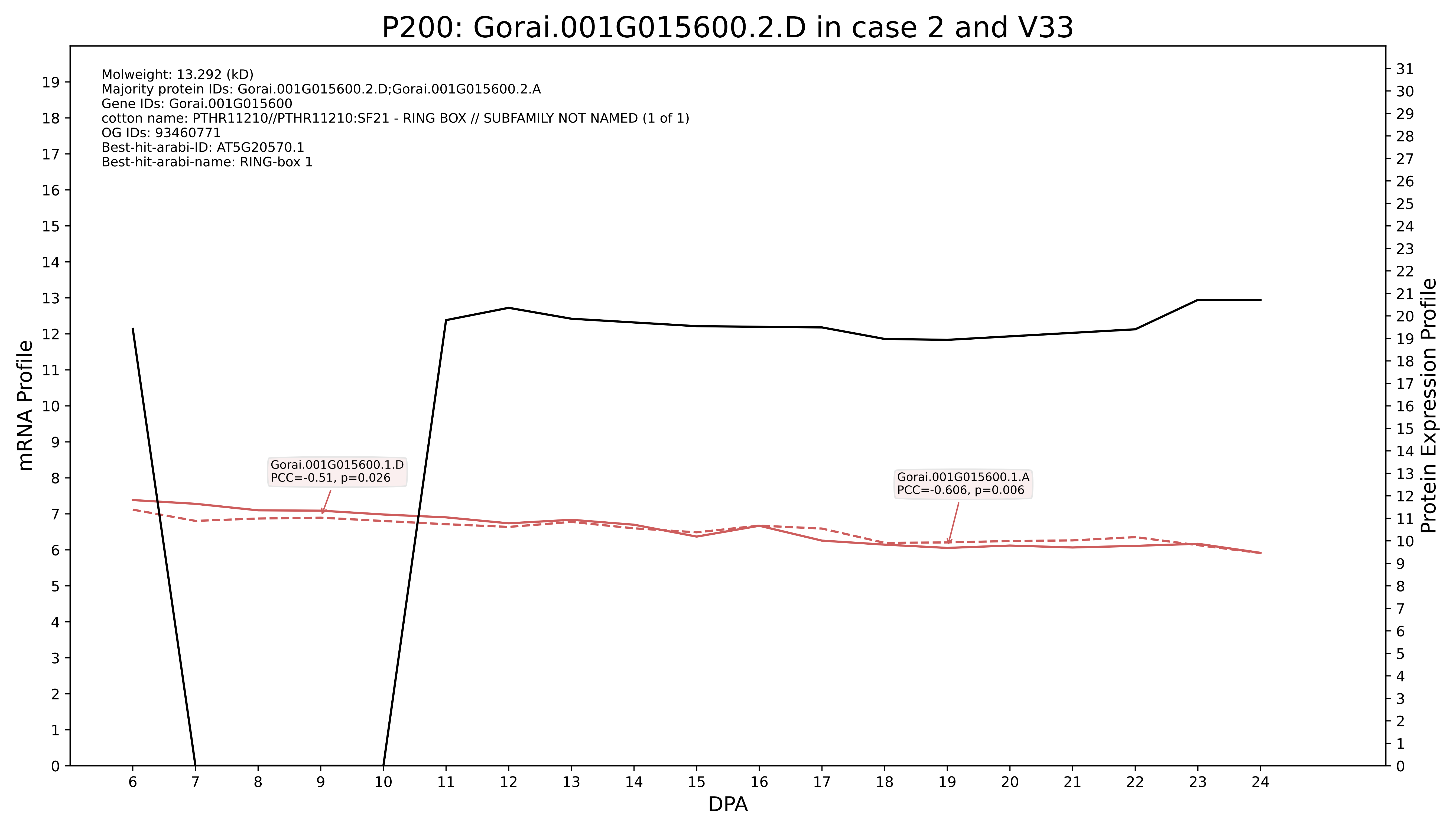
P200: Gorai.001G015600.2.D in case 2 and V33
Molweight: 13.292 (kD)
Majority protein IDs: Gorai.001G015600.2.D;Gorai.001G015600.2.A
Gene IDs: Gorai.001G015600
cotton name: PTHR11210//PTHR11210:SF21 - RING BOX // SUBFAMILY NOT NAMED (1 of 1)
OG IDs: 93460771
Best-hit-arabi-ID: AT5G20570.1
Best-hit-arabi-name: RING-box 1
Gorai.001G015600.1.D
PCC=-0.51, p=0.026
Gorai.001G015600.1.A
PCC=-0.606, p=0.006
mRNA Profile
Protein Expression Profile
DPA

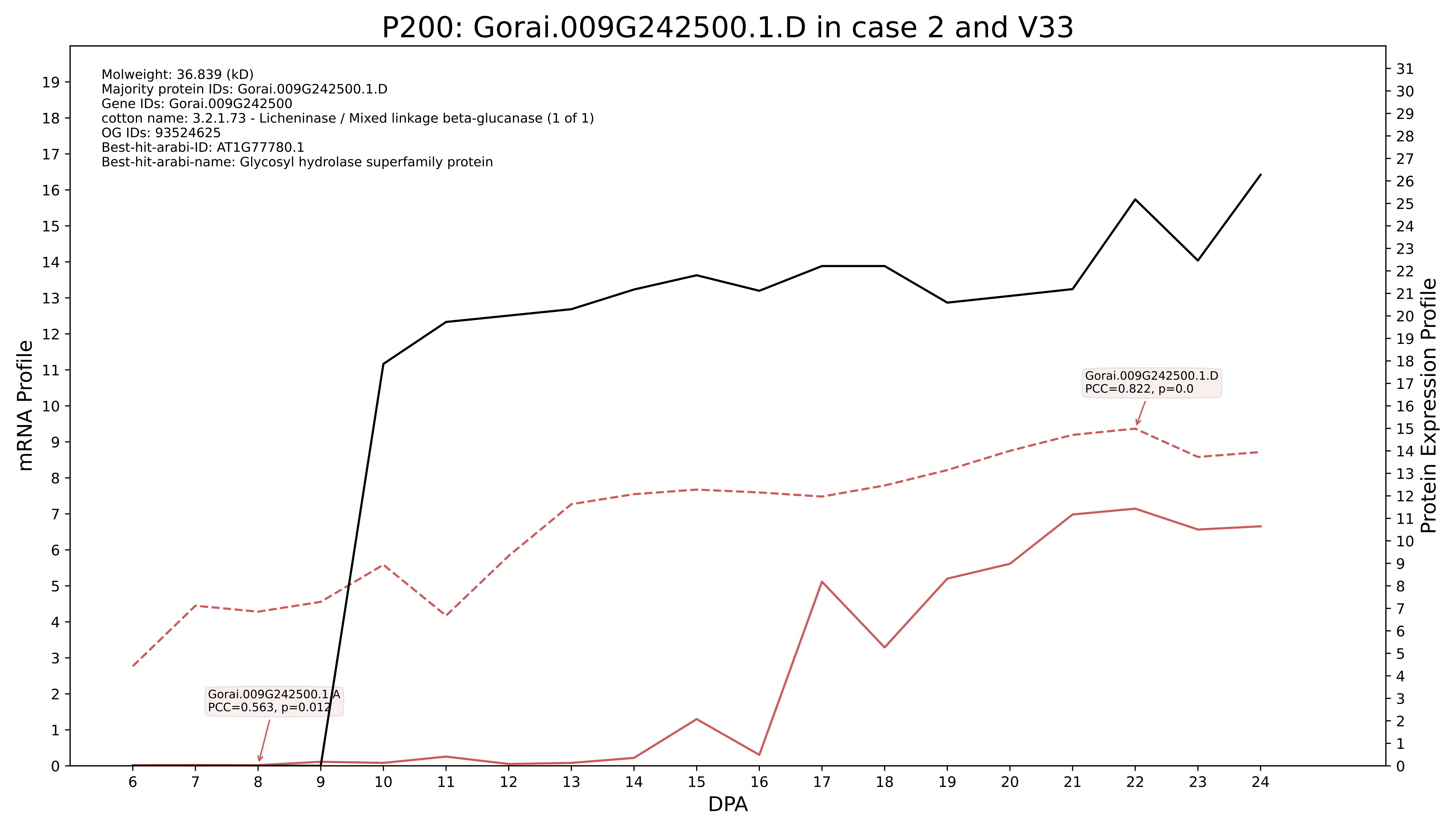

P200: Gorai.009G242500.1.D in case 2 and V33
Molweight: 36.839 (kD)
Majority protein IDs: Gorai.009G242500.1.D
Gene IDs: Gorai.009G242500
cotton name: 3.2.1.73 - Licheninase / Mixed linkage beta-glucanase (1 of 1)
OG IDs: 93524625
Best-hit-arabi-ID: AT1G77780.1
Best-hit-arabi-name: Glycosyl hydrolase superfamily protein
Gorai.009G242500.1.D
PCC=0.822, p=0.0
Gorai.009G242500.1.A
PCC=0.563, p=0.012
mRNA Profile
Protein Expression Profile
DPA

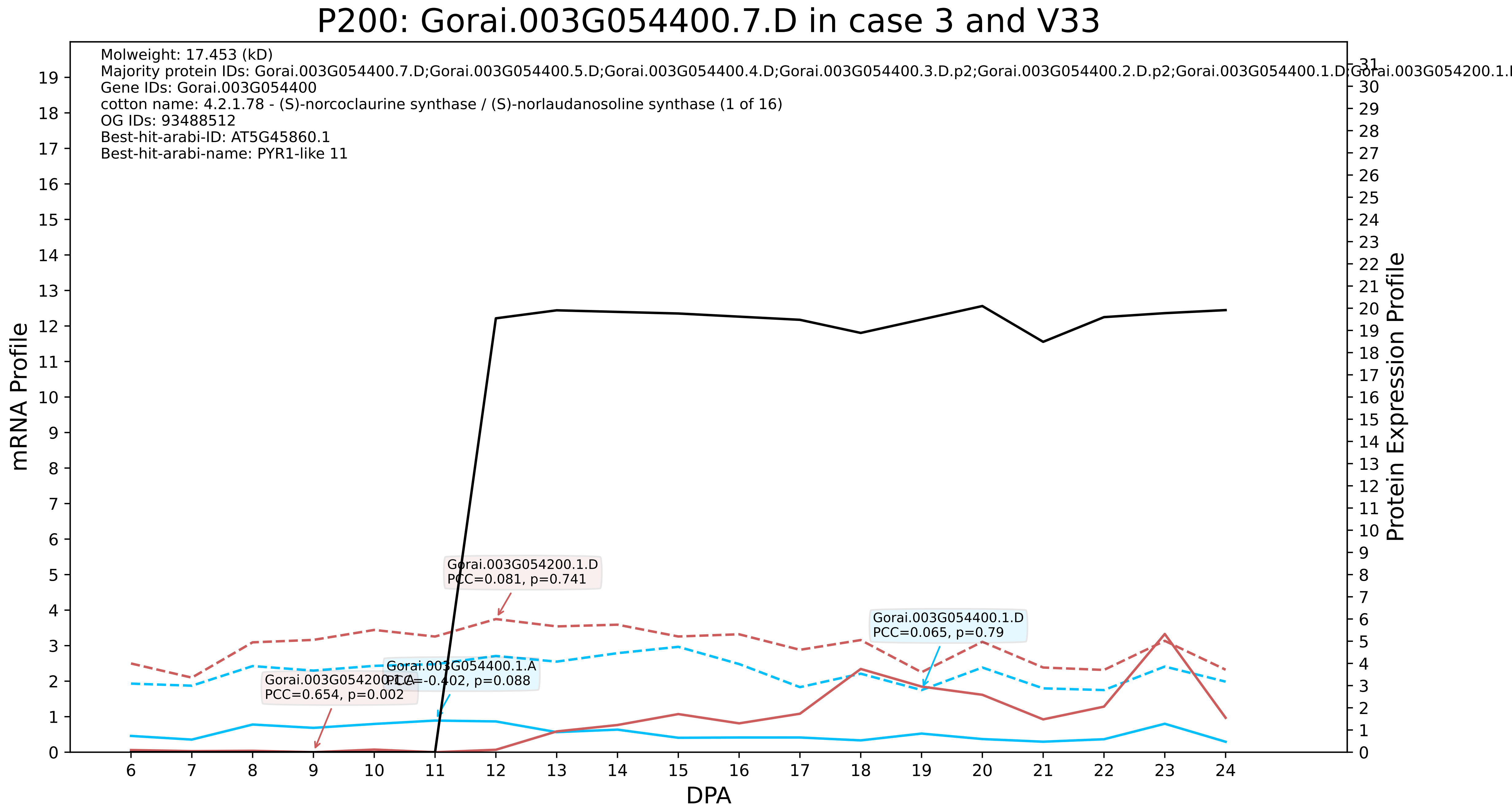
P200: Gorai.003G054400.7.D in case 3 and V33
Molweight: 17.453 (kD)
Majority protein IDs: Gorai.003G054400.7.D;Gorai.003G054400.5.D;Gorai.003G054400.4.D;Gorai.003G054400.3.D.p2;Gorai.003G054400.2.D.p2;Gorai.003G054400.1.D;Gorai.003G054200.1.
Gene IDs: Gorai.003G054400
cotton name: 4.2.1.78 - (S)-norcoclaurine synthase / (S)-norlaudanosoline synthase (1 of 16)
OG IDs: 93488512
Best-hit-arabi-ID: AT5G45860.1
Best-hit-arabi-name: PYR1-like 11
Gorai.003G054200.1.D
PCC=0.081, p=0.741
Gorai.003G054400.1.D
PCC=0.065, p=0.79
Gorai.003G054400.1.A
PCC=-0.402, p=0.088
Gorai.003G054200.1.A
PCC=0.654, p=0.002
mRNA Profile
Protein Expression Profile
DPA

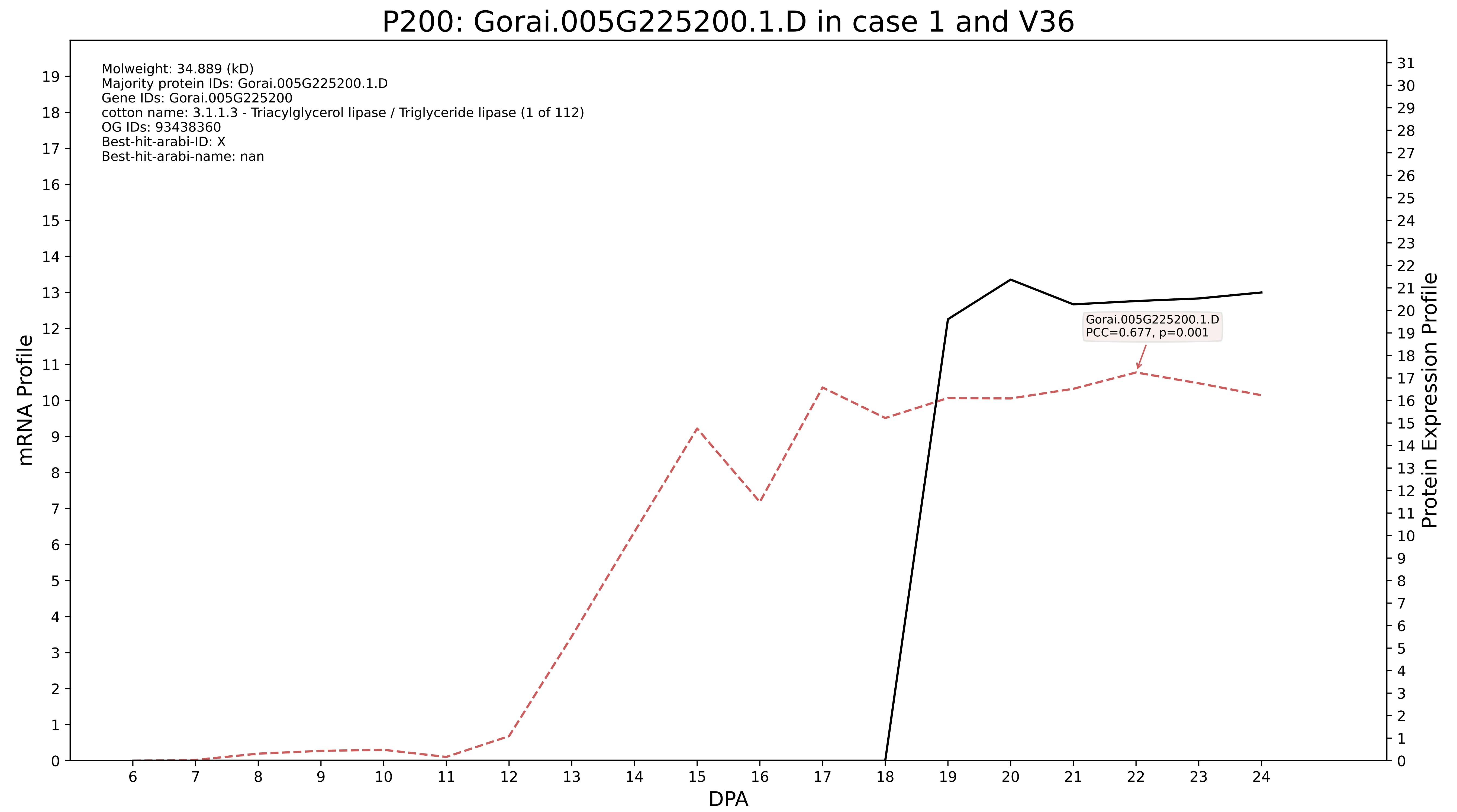
P200: Gorai.005G225200.1.D in case 1 and V36
Molweight: 34.889 (kD)
Majority protein IDs: Gorai.005G225200.1.D
Gene IDs: Gorai.005G225200
cotton name: 3.1.1.3 - Triacylglycerol lipase / Triglyceride lipase (1 of 112)
OG IDs: 93438360
Best-hit-arabi-ID: X
Best-hit-arabi-name: nan
Gorai.005G225200.1.D
PCC=0.677, p=0.001
mRNA Profile
Protein Expression Profile
DPA

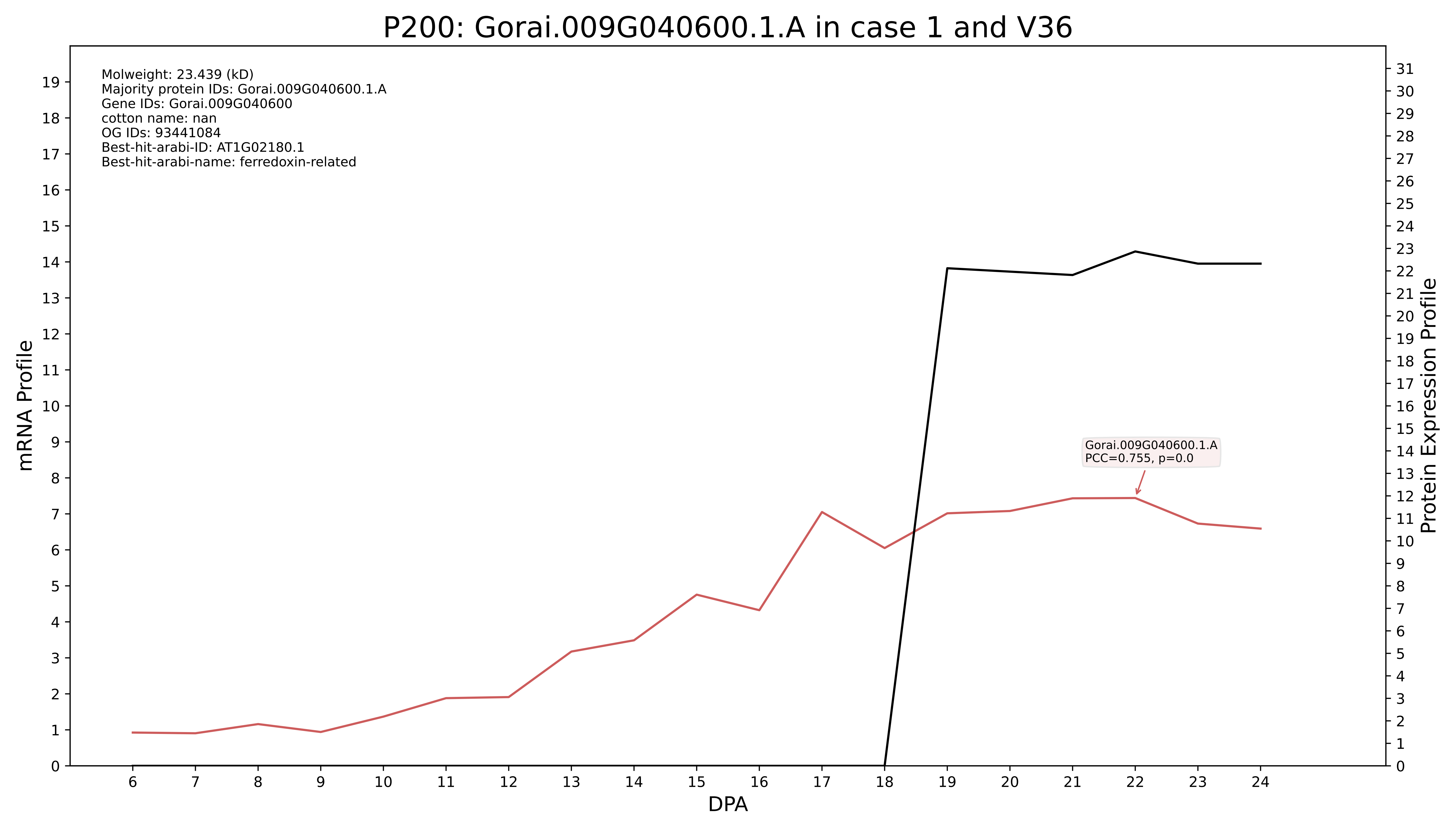
P200: Gorai.009G040600.1.A in case 1 and V36
Molweight: 23.439 (kD)
Majority protein IDs: Gorai.009G040600.1.A
Gene IDs: Gorai.009G040600
cotton name: nan
OG IDs: 93441084
Best-hit-arabi-ID: AT1G02180.1
Best-hit-arabi-name: ferredoxin-related
Gorai.009G040600.1.A
PCC=0.755, p=0.0
mRNA Profile
Protein Expression Profile
DPA
6 7 8 9 10 11 12 13 14 15 16 17 18 19 20 21 22 23 24
0 1 2 3 4 5 6 7 8 9 10 11 12 13 14 15 16 17 18 19
0 1 2 3 4 5 6 7 8 9 10 11 12 13 14 15 16 17 18 19 20 21 22 23 24 25 26 27 28 29 30 31

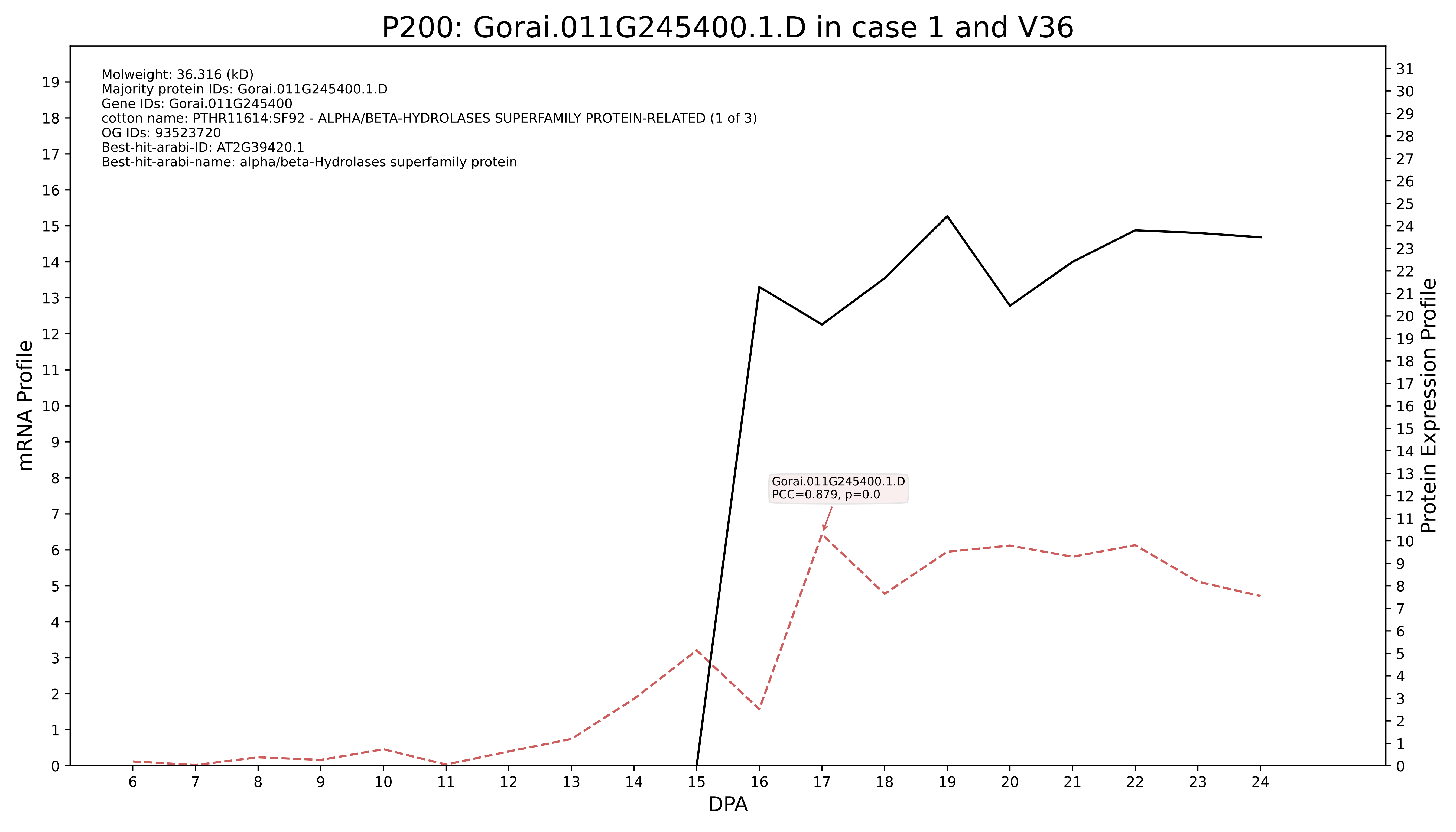

P200: Gorai.011G245400.1.D in case 1 and V36
Molweight: 36.316 (kD)
Majority protein IDs: Gorai.011G245400.1.D
Gene IDs: Gorai.011G245400
cotton name: PTHR11614:SF92 - ALPHA/BETA-HYDROLASES SUPERFAMILY PROTEIN-RELATED (1 of 3)
OG IDs: 93523720
Best-hit-arabi-ID: AT2G39420.1
Best-hit-arabi-name: alpha/beta-Hydrolases superfamily protein
Gorai.011G245400.1.D
PCC=0.879, p=0.0
mRNA Profile
Protein Expression Profile
DPA

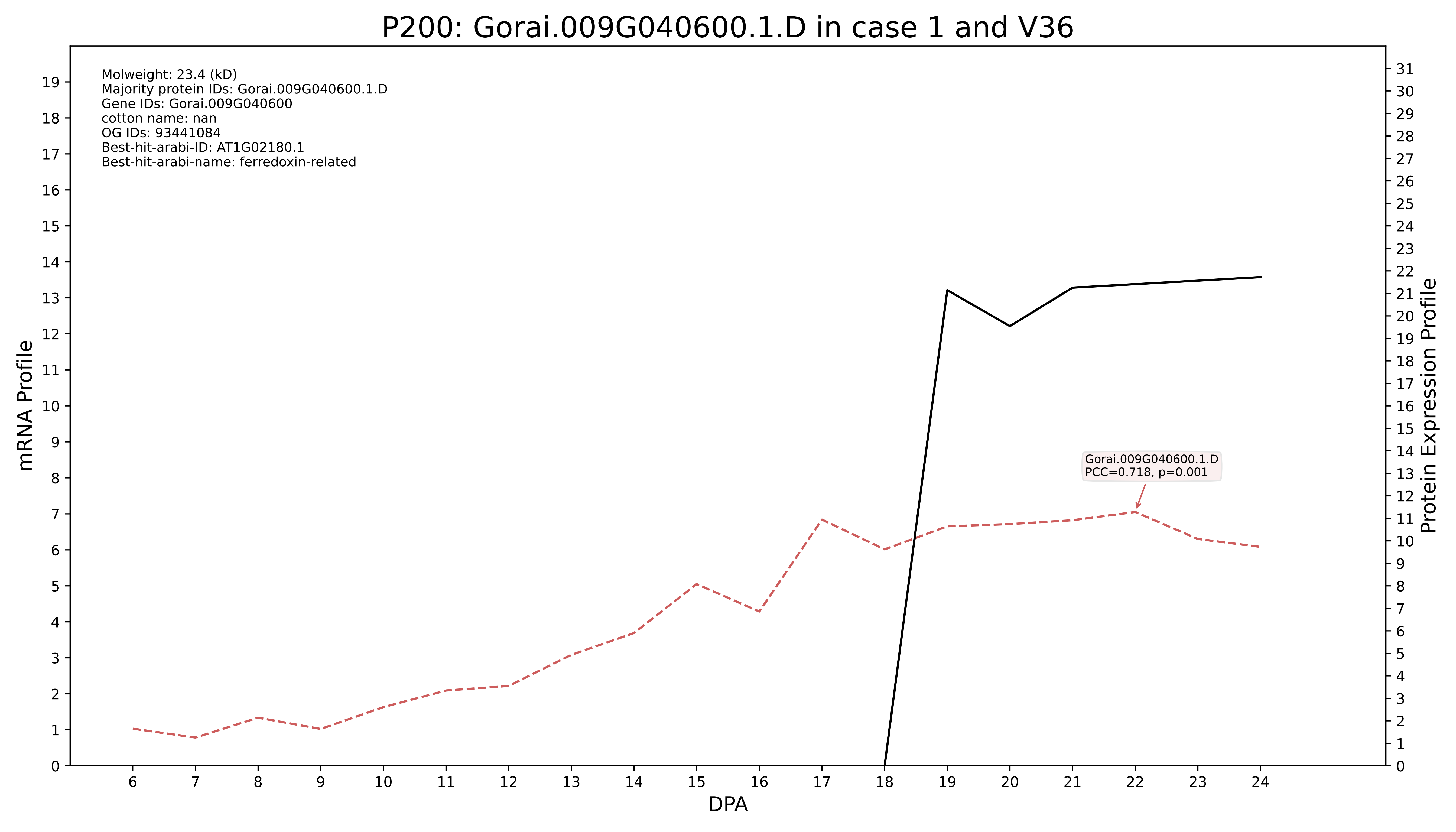
P200: Gorai.009G040600.1.D in case 1 and V36
Molweight: 23.4 (kD)
Majority protein IDs: Gorai.009G040600.1.D
Gene IDs: Gorai.009G040600
cotton name: nan
OG IDs: 93441084
Best-hit-arabi-ID: AT1G02180.1
Best-hit-arabi-name: ferredoxin-related
Gorai.009G040600.1.D
PCC=0.718, p=0.001
mRNA Profile
Protein Expression Profile
DPA

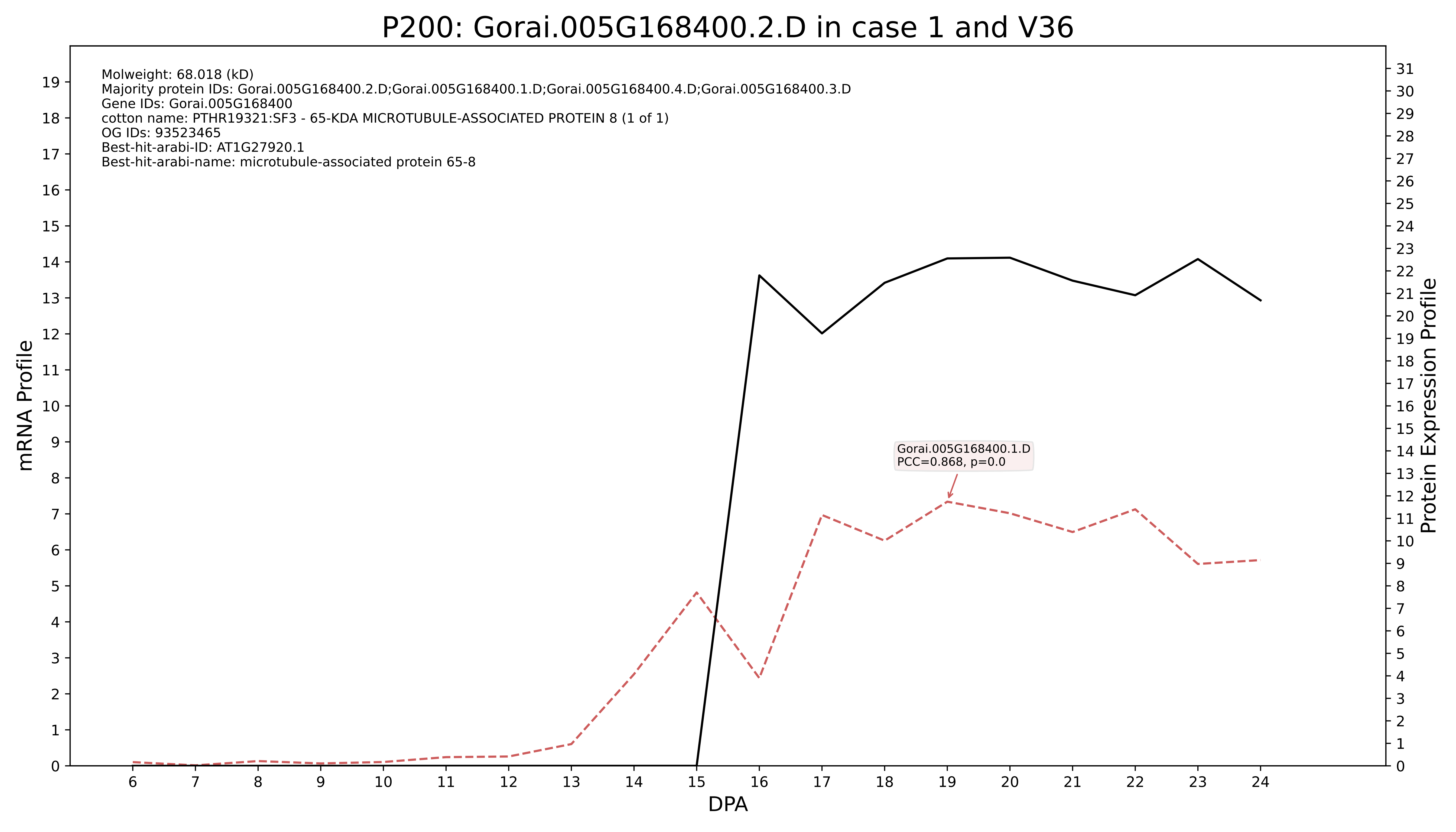
P200: Gorai.005G168400.2.D in case 1 and V36
Molweight: 68.018 (kD)
Majority protein IDs: Gorai.005G168400.2.D;Gorai.005G168400.1.D;Gorai.005G168400.4.D;Gorai.005G168400.3.D
Gene IDs: Gorai.005G168400
cotton name: PTHR19321:SF3 - 65-KDA MICROTUBULE-ASSOCIATED PROTEIN 8 (1 of 1)
OG IDs: 93523465
Best-hit-arabi-ID: AT1G27920.1
Best-hit-arabi-name: microtubule-associated protein 65-8
Gorai.005G168400.1.D
PCC=0.868, p=0.0
mRNA Profile
Protein Expression Profile
DPA

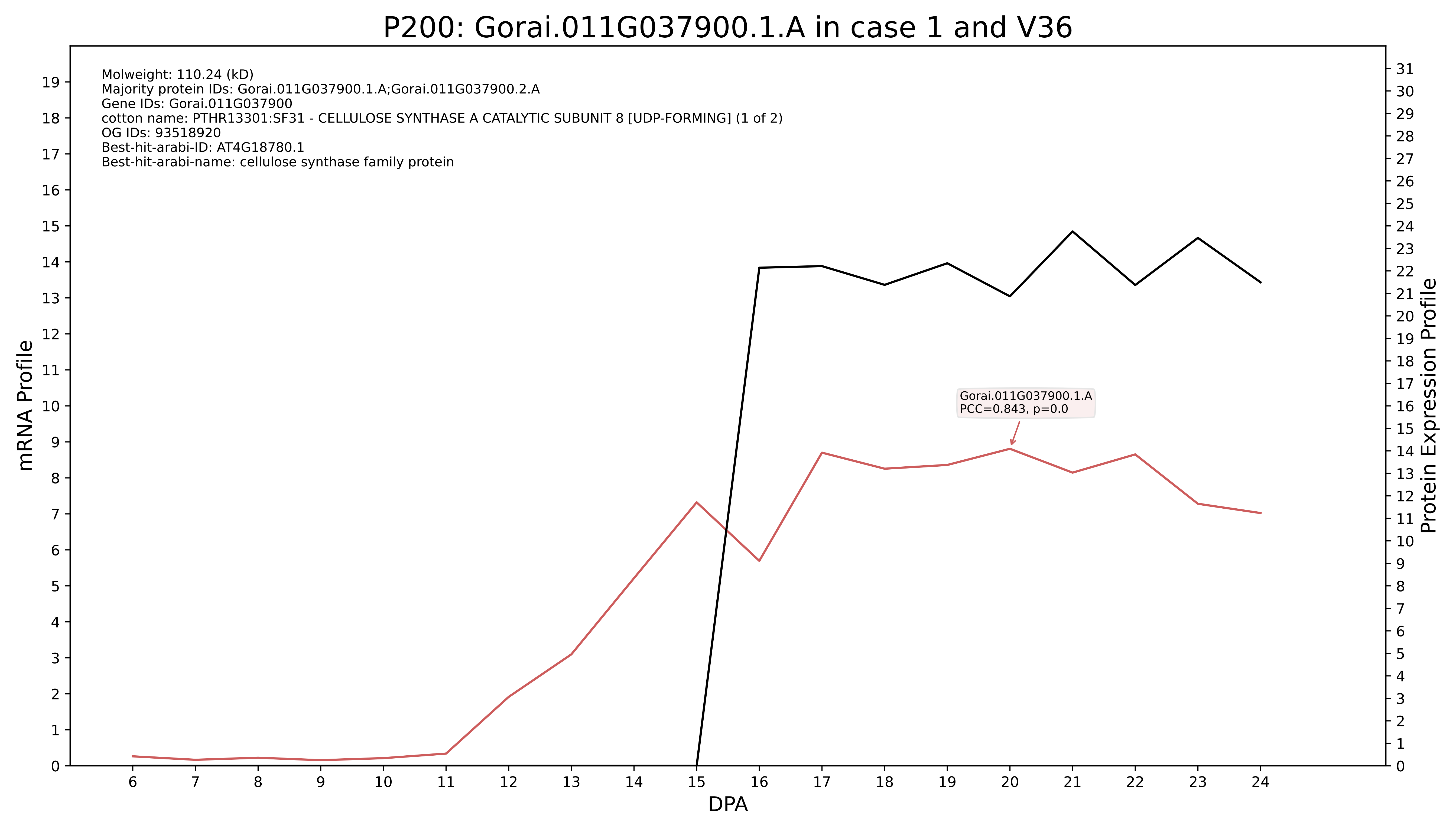
P200: Gorai.011G037900.1.A in case 1 and V36
Molweight: 110.24 (kD)
Majority protein IDs: Gorai.011G037900.1.A;Gorai.011G037900.2.A
Gene IDs: Gorai.011G037900
cotton name: PTHR13301:SF31 - CELLULOSE SYNTHASE A CATALYTIC SUBUNIT 8 [UDP-FORMING] (1 of 2)
OG IDs: 93518920
Best-hit-arabi-ID: AT4G18780.1
Best-hit-arabi-name: cellulose synthase family protein
Gorai.011G037900.1.A
PCC=0.843, p=0.0
mRNA Profile
Protein Expression Profile
DPA

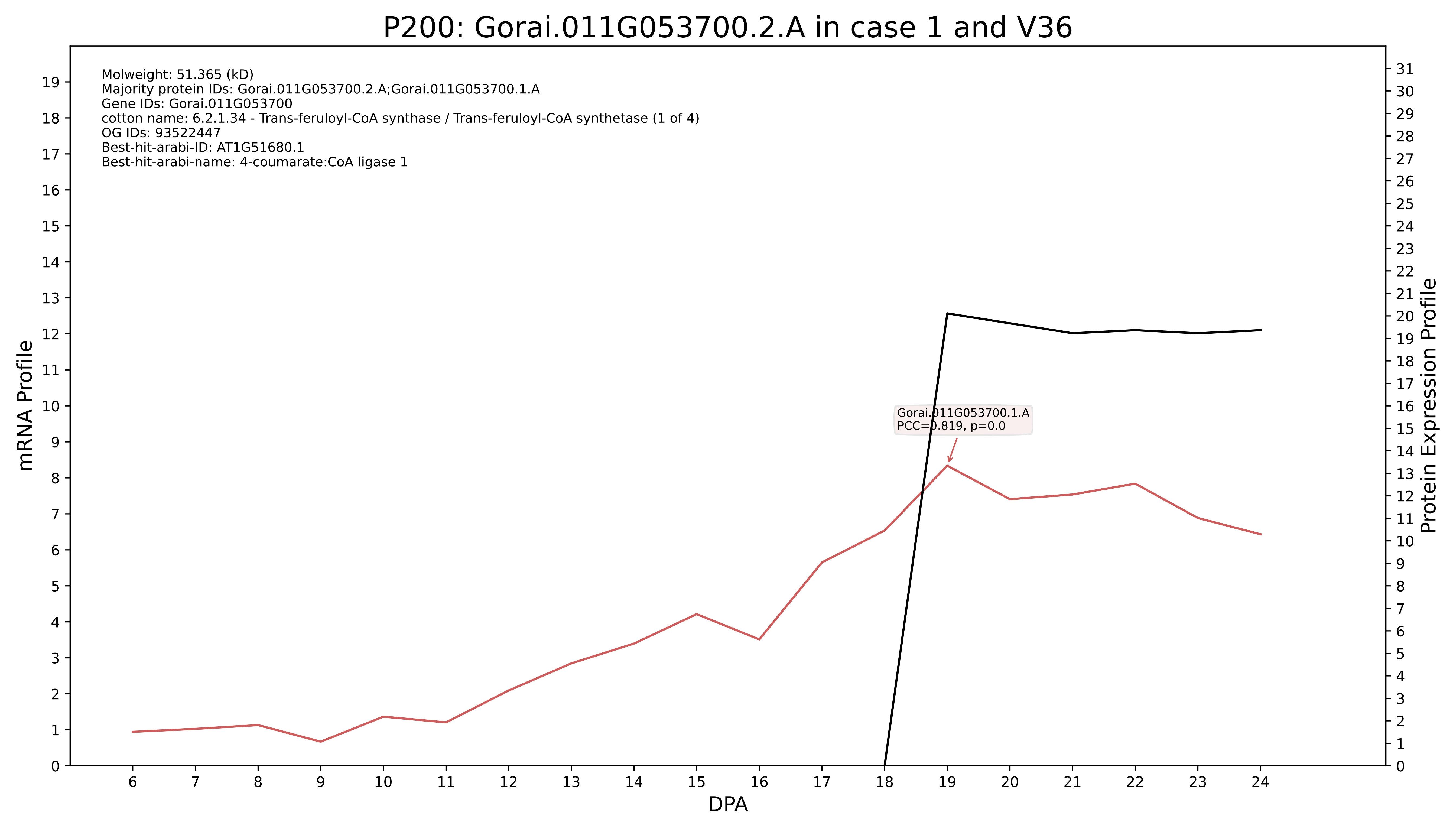
P200: Gorai.011G053700.2.A in case 1 and V36
Molweight: 51.365 (kD)
Majority protein IDs: Gorai.011G053700.2.A;Gorai.011G053700.1.A
Gene IDs: Gorai.011G053700
cotton name: 6.2.1.34 - Trans-feruloyl-CoA synthase / Trans-feruloyl-CoA synthetase (1 of 4)
OG IDs: 93522447
Best-hit-arabi-ID: AT1G51680.1
Best-hit-arabi-name: 4-coumarate:CoA ligase 1
Gorai.011G053700.1.A
PCC=0.819, p=0.0
mRNA Profile
Protein Expression Profile
DPA

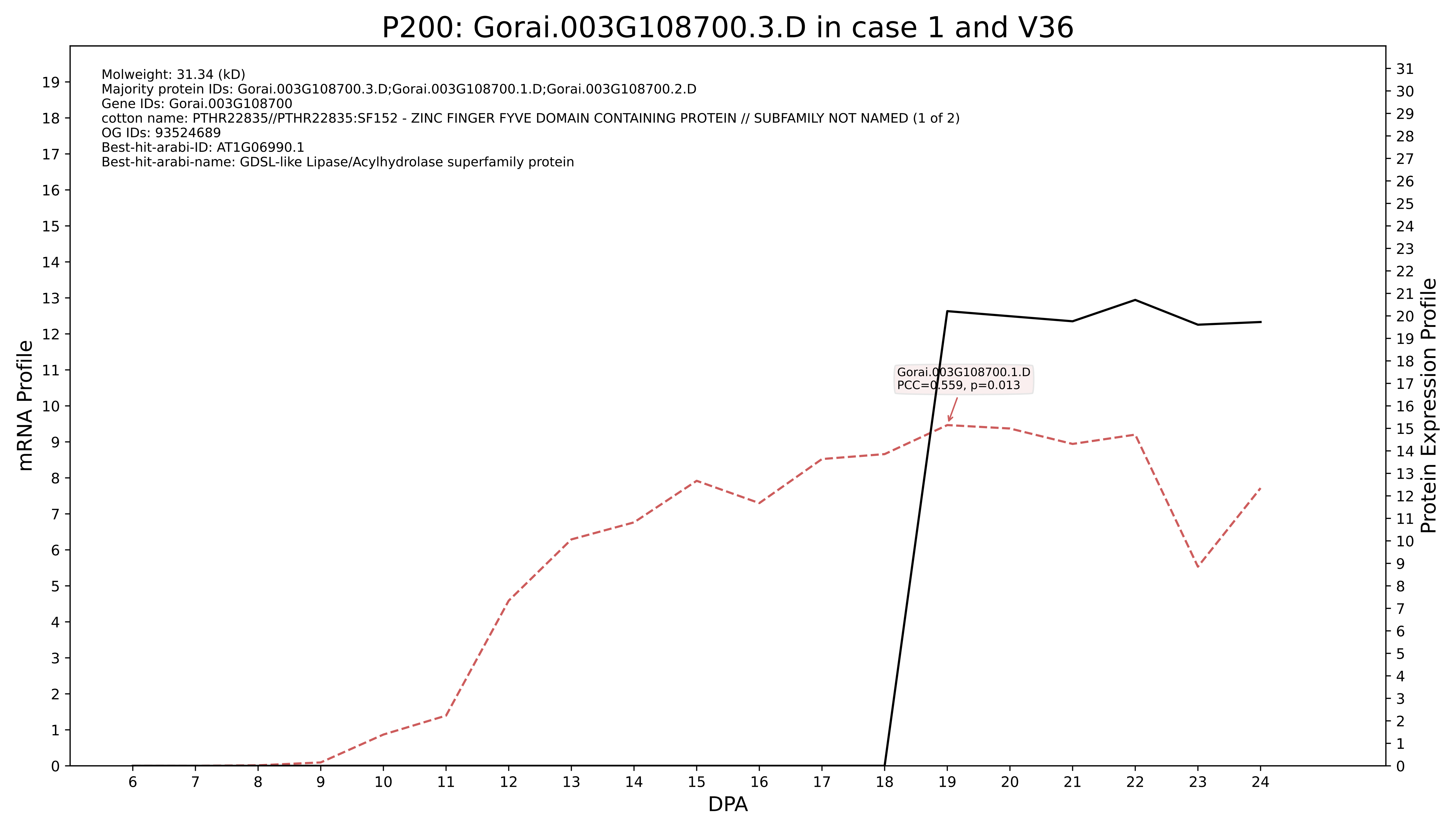
P200: Gorai.003G108700.3.D in case 1 and V36
Molweight: 31.34 (kD)
Majority protein IDs: Gorai.003G108700.3.D;Gorai.003G108700.1.D;Gorai.003G108700.2.D
Gene IDs: Gorai.003G108700
cotton name: PTHR22835//PTHR22835:SF152 - ZINC FINGER FYVE DOMAIN CONTAINING PROTEIN // SUBFAMILY NOT NAMED (1 of 2)
OG IDs: 93524689
Best-hit-arabi-ID: AT1G06990.1
Best-hit-arabi-name: GDSL-like Lipase/Acylhydrolase superfamily protein
Gorai.003G108700.1.D
PCC=0.559, p=0.013
mRNA Profile
Protein Expression Profile
DPA

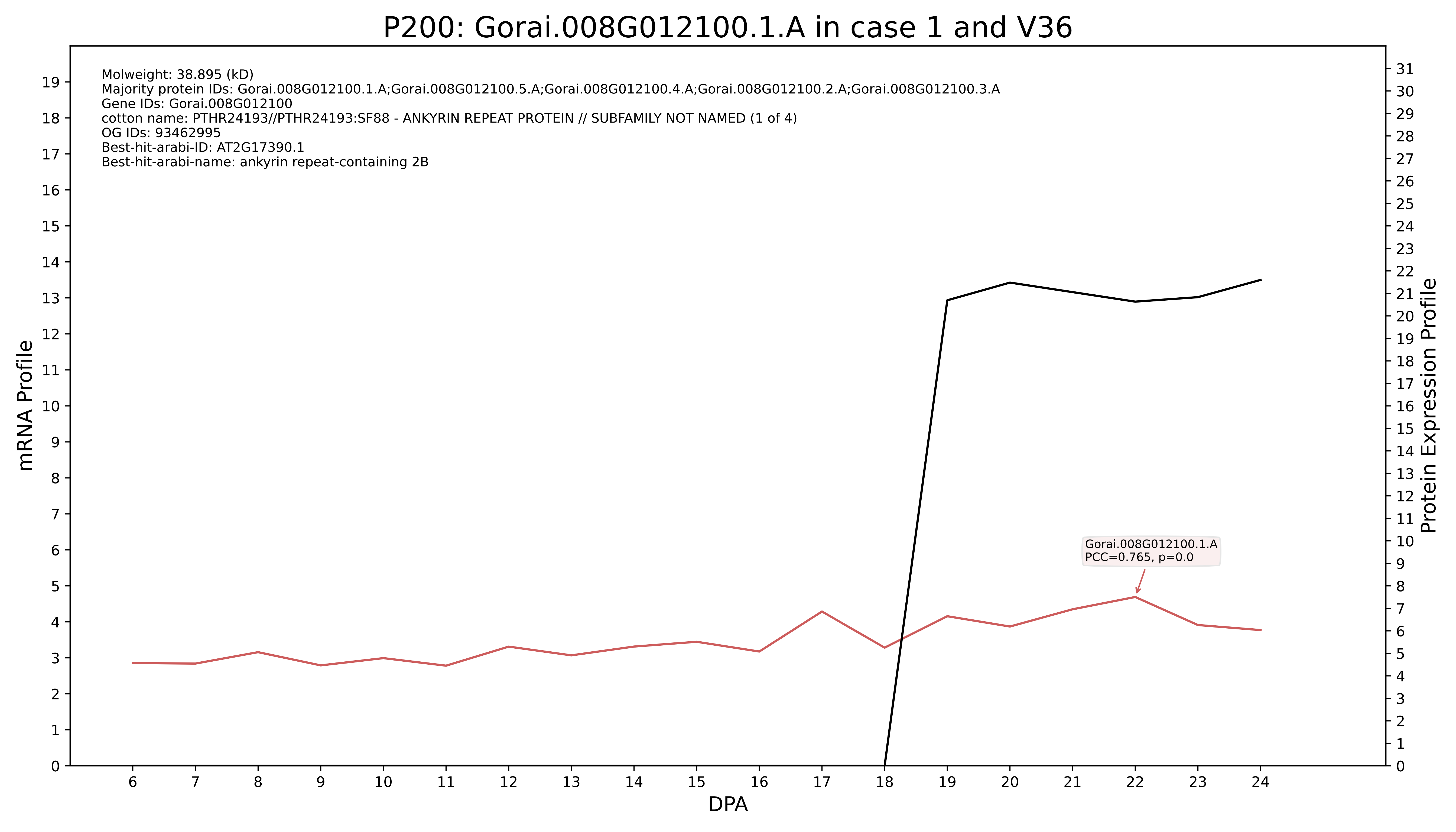
P200: Gorai.008G012100.1.A in case 1 and V36
Molweight: 38.895 (kD)
Majority protein IDs: Gorai.008G012100.1.A;Gorai.008G012100.5.A;Gorai.008G012100.4.A;Gorai.008G012100.2.A;Gorai.008G012100.3.A
Gene IDs: Gorai.008G012100
cotton name: PTHR24193//PTHR24193:SF88 - ANKYRIN REPEAT PROTEIN // SUBFAMILY NOT NAMED (1 of 4)
OG IDs: 93462995
Best-hit-arabi-ID: AT2G17390.1
Best-hit-arabi-name: ankyrin repeat-containing 2B
Gorai.008G012100.1.A
PCC=0.765, p=0.0
mRNA Profile
Protein Expression Profile
DPA

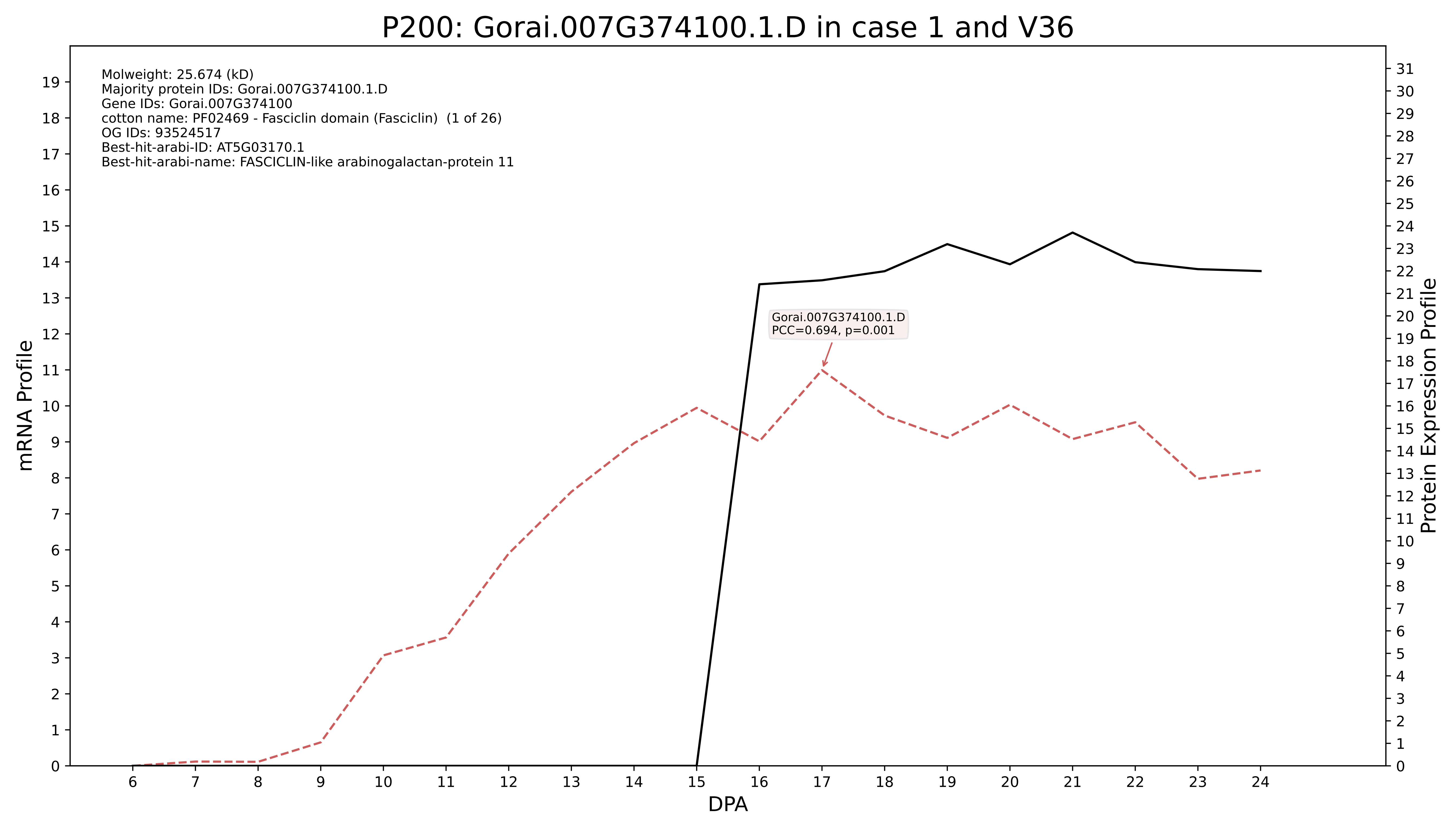
P200: Gorai.007G374100.1.D in case 1 and V36
Molweight: 25.674 (kD)
Majority protein IDs: Gorai.007G374100.1.D
Gene IDs: Gorai.007G374100
cotton name: PF02469 - Fasciclin domain (Fasciclin) (1 of 26)
OG IDs: 93524517
Best-hit-arabi-ID: AT5G03170.1
Best-hit-arabi-name: FASCICLIN-like arabinogalactan-protein 11
Gorai.007G374100.1.D
PCC=0.694, p=0.001
mRNA Profile
Protein Expression Profile
DPA

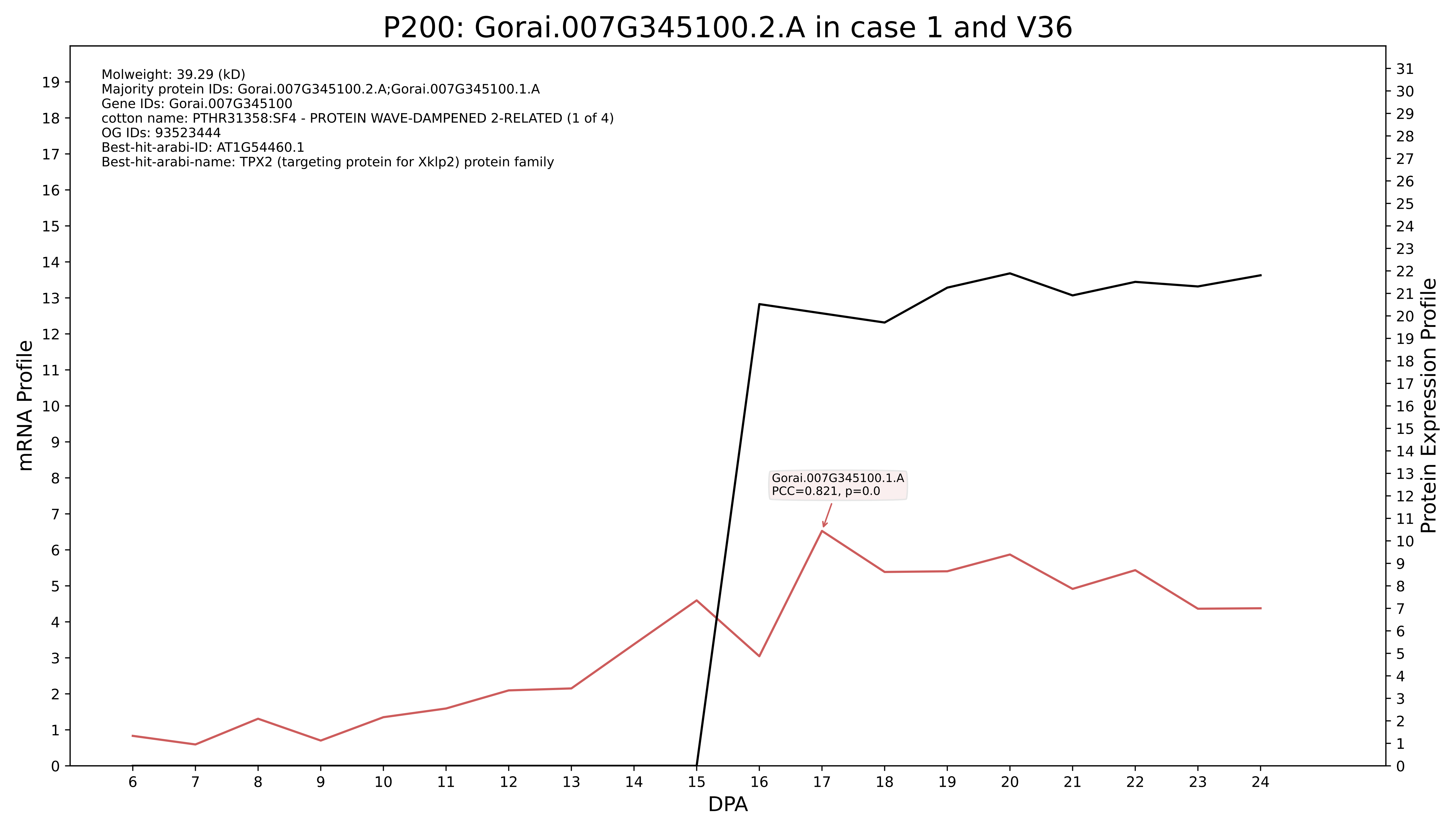
P200: Gorai.007G345100.2.A in case 1 and V36
Molweight: 39.29 (kD)
Majority protein IDs: Gorai.007G345100.2.A;Gorai.007G345100.1.A
Gene IDs: Gorai.007G345100
cotton name: PTHR31358:SF4 - PROTEIN WAVE-DAMPENED 2-RELATED (1 of 4)
OG IDs: 93523444
Best-hit-arabi-ID: AT1G54460.1
Best-hit-arabi-name: TPX2 (targeting protein for Xklp2) protein family
Gorai.007G345100.1.A
PCC=0.821, p=0.0
mRNA Profile
Protein Expression Profile
DPA

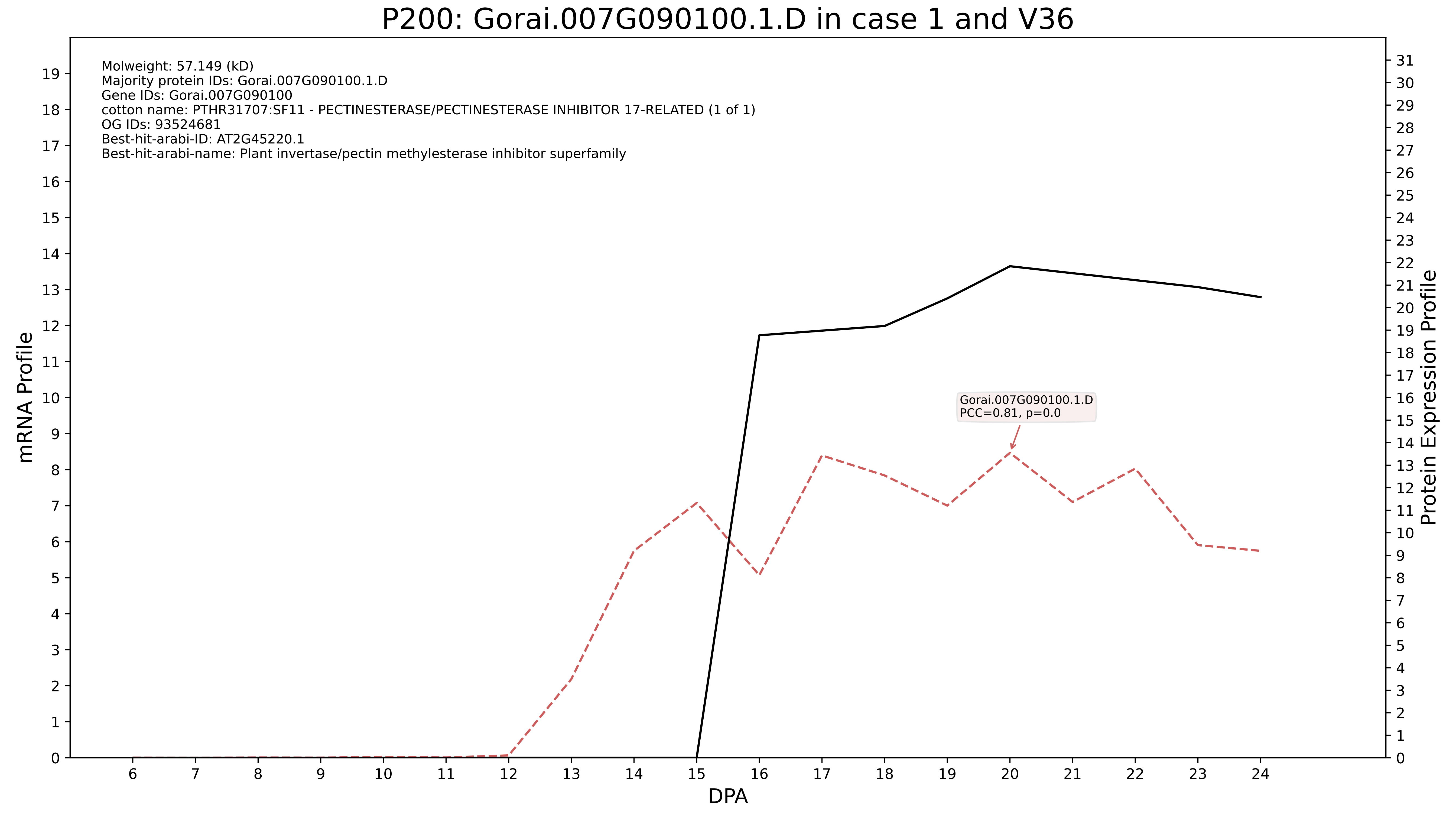

P200: Gorai.007G090100.1.D in case 1 and V36
Molweight: 57.149 (kD)
Majority protein IDs: Gorai.007G090100.1.D
Gene IDs: Gorai.007G090100
cotton name: PTHR31707:SF11 - PECTINESTERASE/PECTINESTERASE INHIBITOR 17-RELATED (1 of 1)
OG IDs: 93524681
Best-hit-arabi-ID: AT2G45220.1
Best-hit-arabi-name: Plant invertase/pectin methylesterase inhibitor superfamily
Gorai.007G090100.1.D
PCC=0.81, p=0.0
mRNA Profile
Protein Expression Profile
DPA

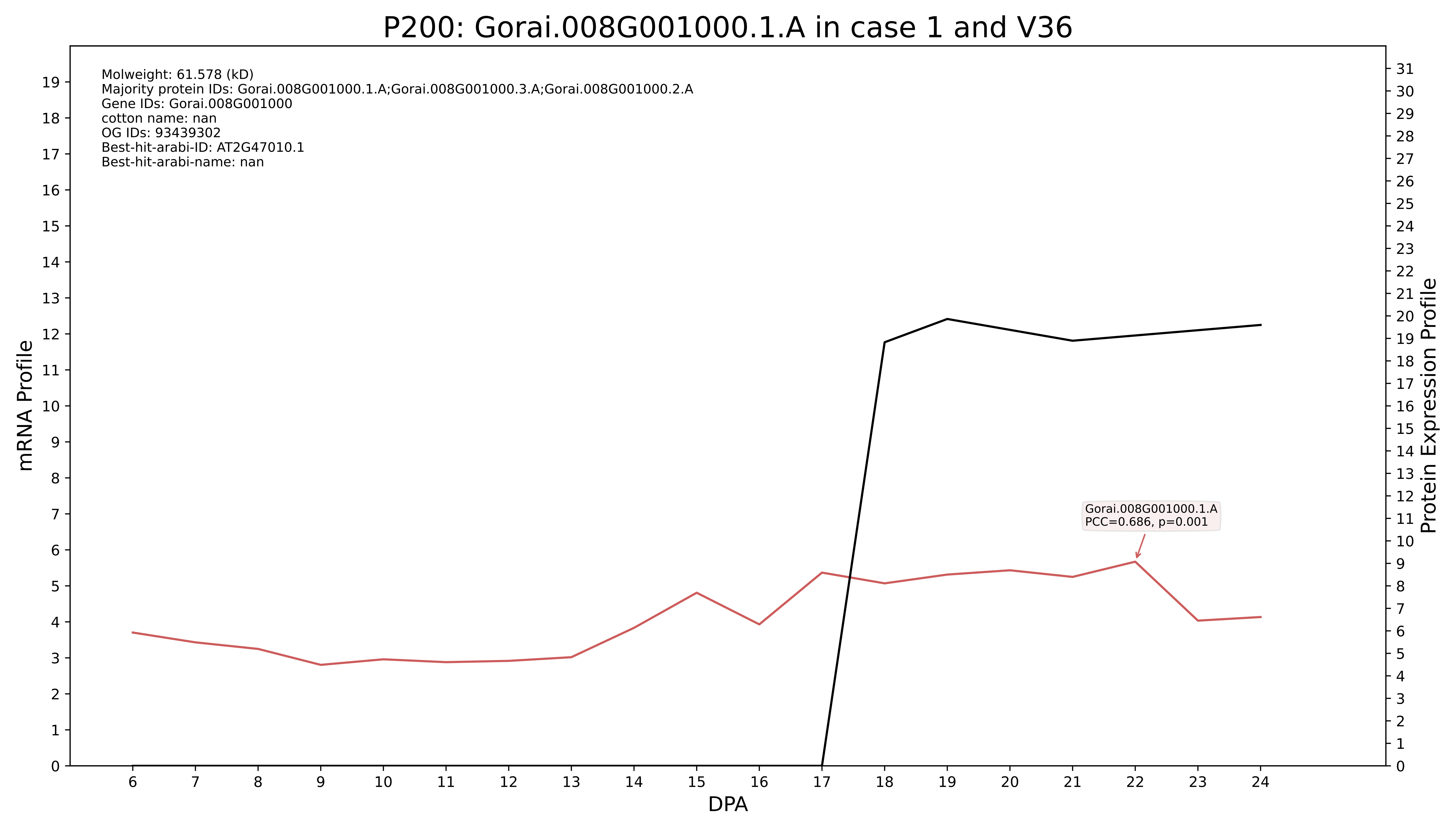

P200: Gorai.008G001000.1.A in case 1 and V36
Molweight: 61.578 (kD)
Majority protein IDs: Gorai.008G001000.1.A;Gorai.008G001000.3.A;Gorai.008G001000.2.A
Gene IDs: Gorai.008G001000
cotton name: nan
OG IDs: 93439302
Best-hit-arabi-ID: AT2G47010.1
Best-hit-arabi-name: nan
Gorai.008G001000.1.A
PCC=0.686, p=0.001
mRNA Profile
Protein Expression Profile
DPA

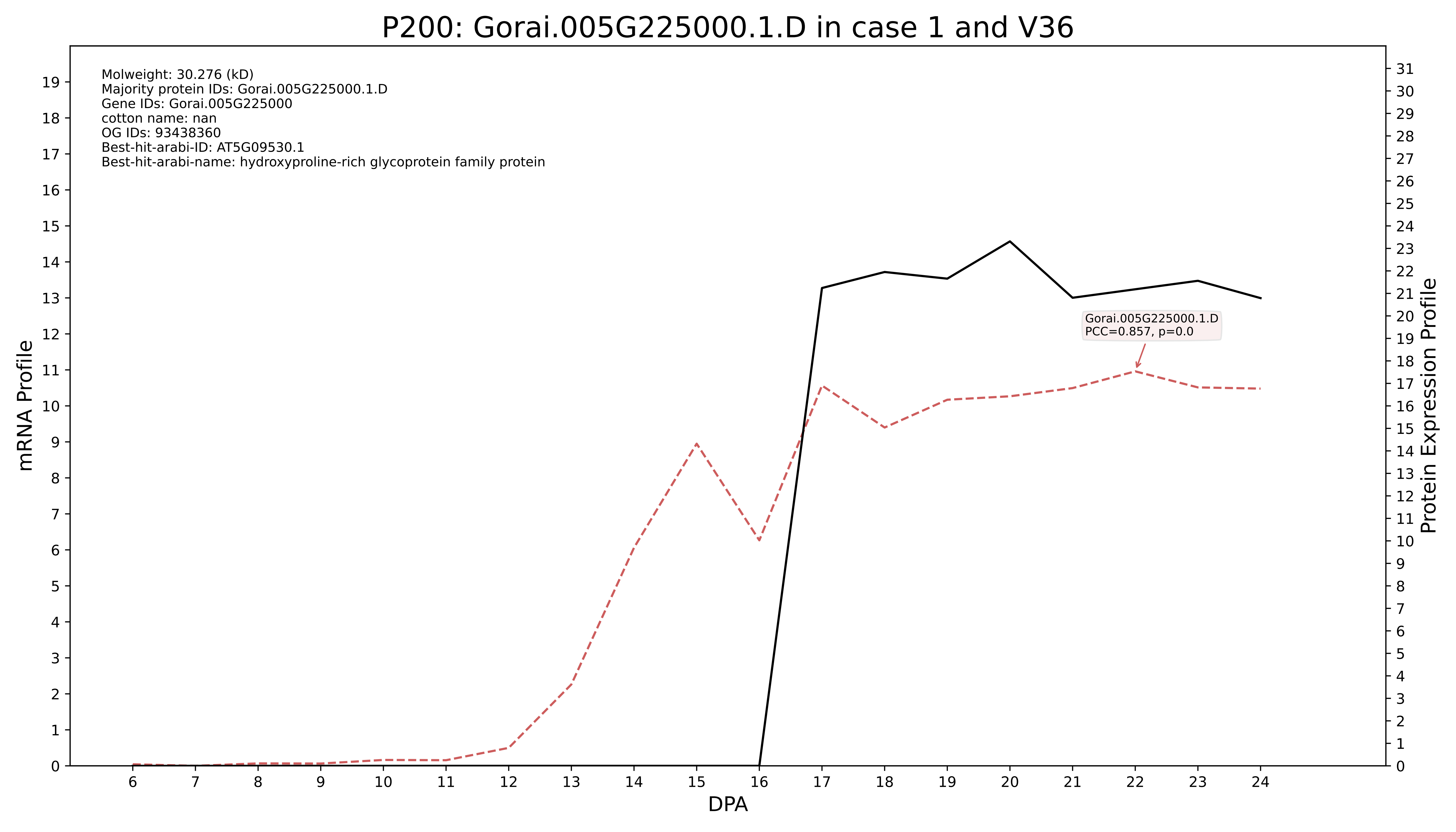
P200: Gorai.005G225000.1.D in case 1 and V36
Molweight: 30.276 (kD)
Majority protein IDs: Gorai.005G225000.1.D
Gene IDs: Gorai.005G225000
cotton name: nan
OG IDs: 93438360
Best-hit-arabi-ID: AT5G09530.1
Best-hit-arabi-name: hydroxyproline-rich glycoprotein family protein
Gorai.005G225000.1.D
PCC=0.857, p=0.0
mRNA Profile
Protein Expression Profile
DPA

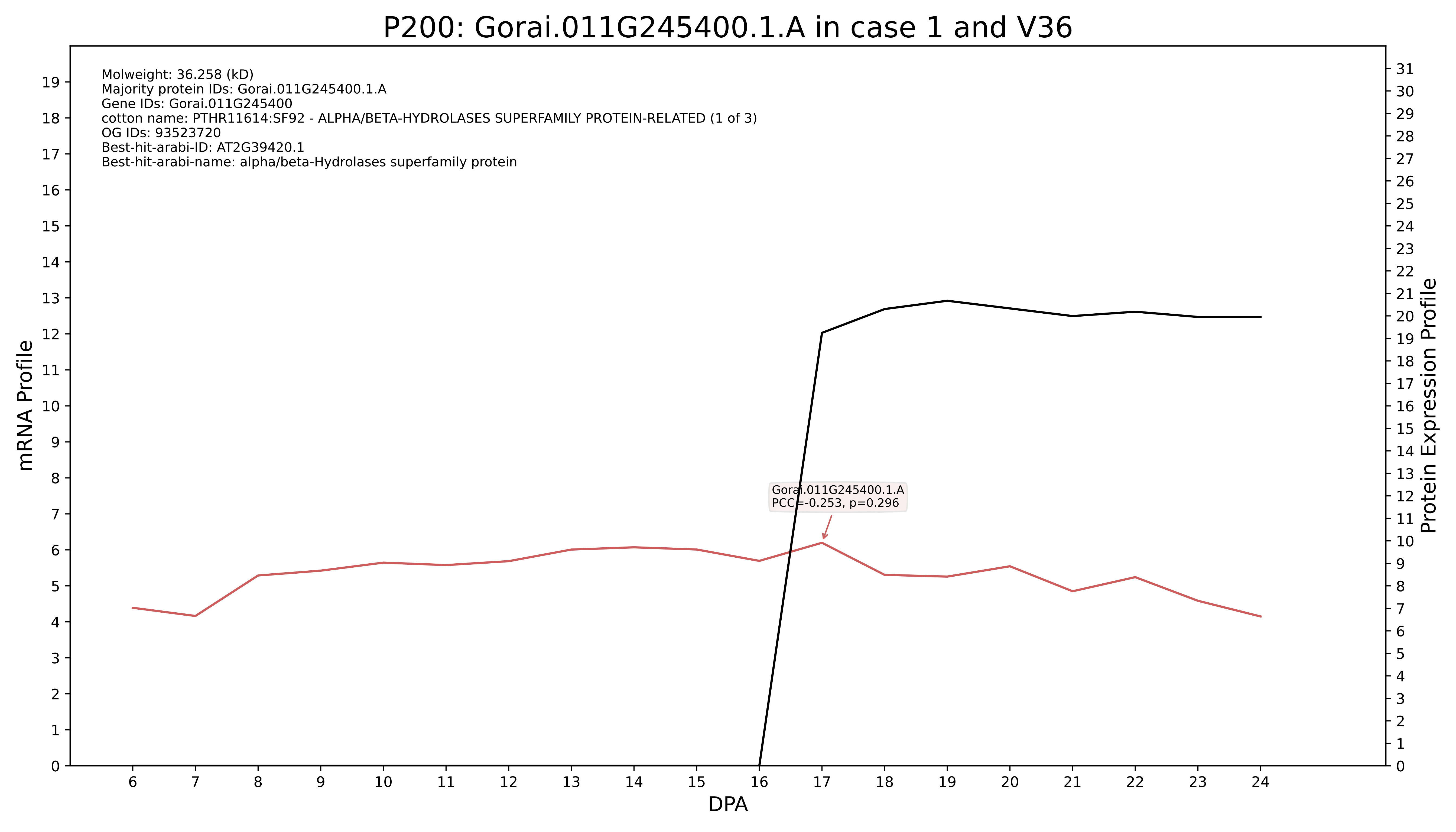
P200: Gorai.011G245400.1.A in case 1 and V36
Molweight: 36.258 (kD)
Majority protein IDs: Gorai.011G245400.1.A
Gene IDs: Gorai.011G245400
cotton name: PTHR11614:SF92 - ALPHA/BETA-HYDROLASES SUPERFAMILY PROTEIN-RELATED (1 of 3)
OG IDs: 93523720
Best-hit-arabi-ID: AT2G39420.1
Best-hit-arabi-name: alpha/beta-Hydrolases superfamily protein
Gorai.011G245400.1.A
PCC=-0.253, p=0.296
mRNA Profile
Protein Expression Profile
DPA

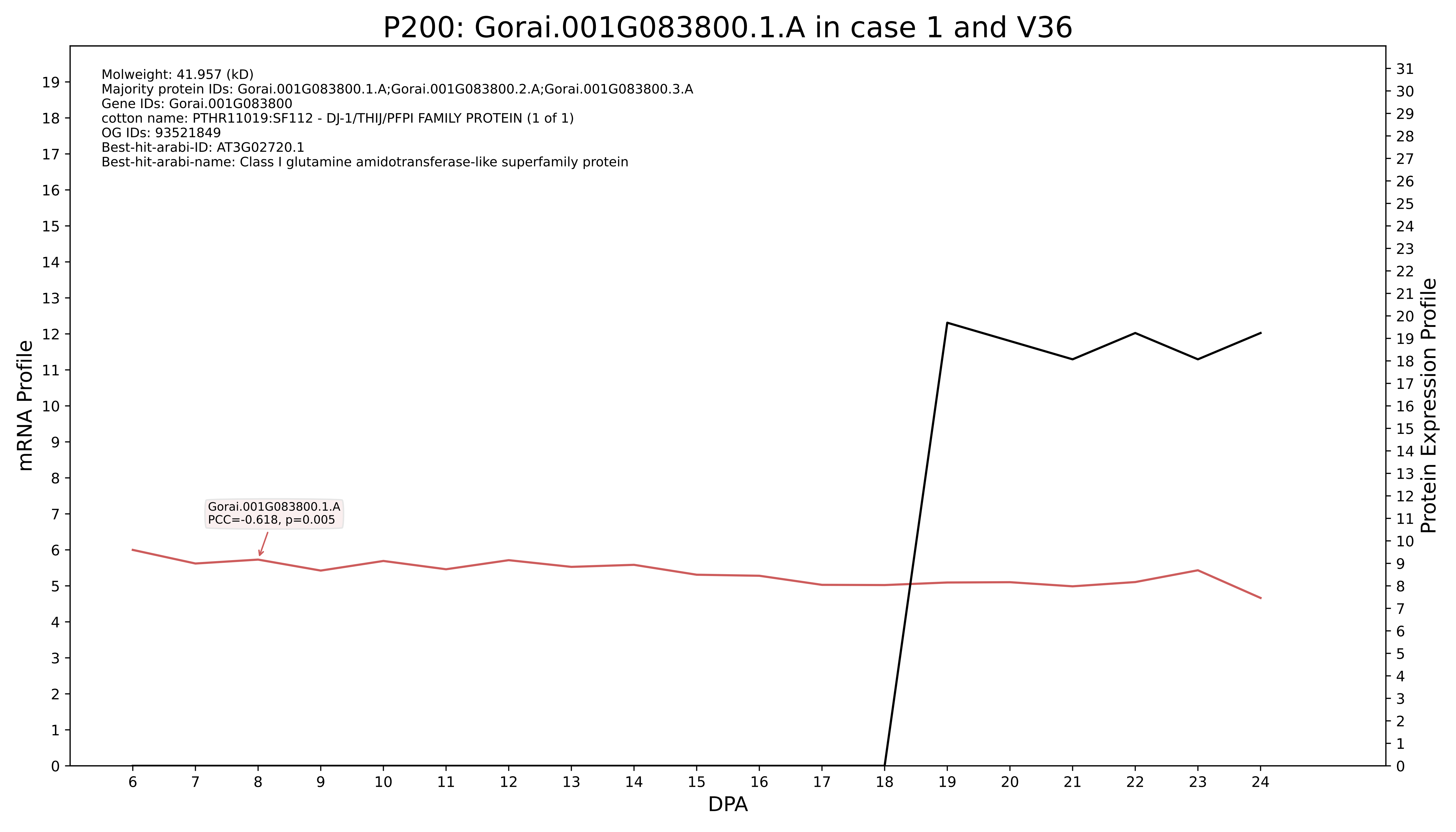
P200: Gorai.001G083800.1.A in case 1 and V36
Molweight: 41.957 (kD)
Majority protein IDs: Gorai.001G083800.1.A;Gorai.001G083800.2.A;Gorai.001G083800.3.A
Gene IDs: Gorai.001G083800
cotton name: PTHR11019:SF112 - DJ-1/THIJ/PFPI FAMILY PROTEIN (1 of 1)
OG IDs: 93521849
Best-hit-arabi-ID: AT3G02720.1
Best-hit-arabi-name: Class I glutamine amidotransferase-like superfamily protein
Gorai.001G083800.1.A
PCC=-0.618, p=0.005
mRNA Profile
Protein Expression Profile
DPA

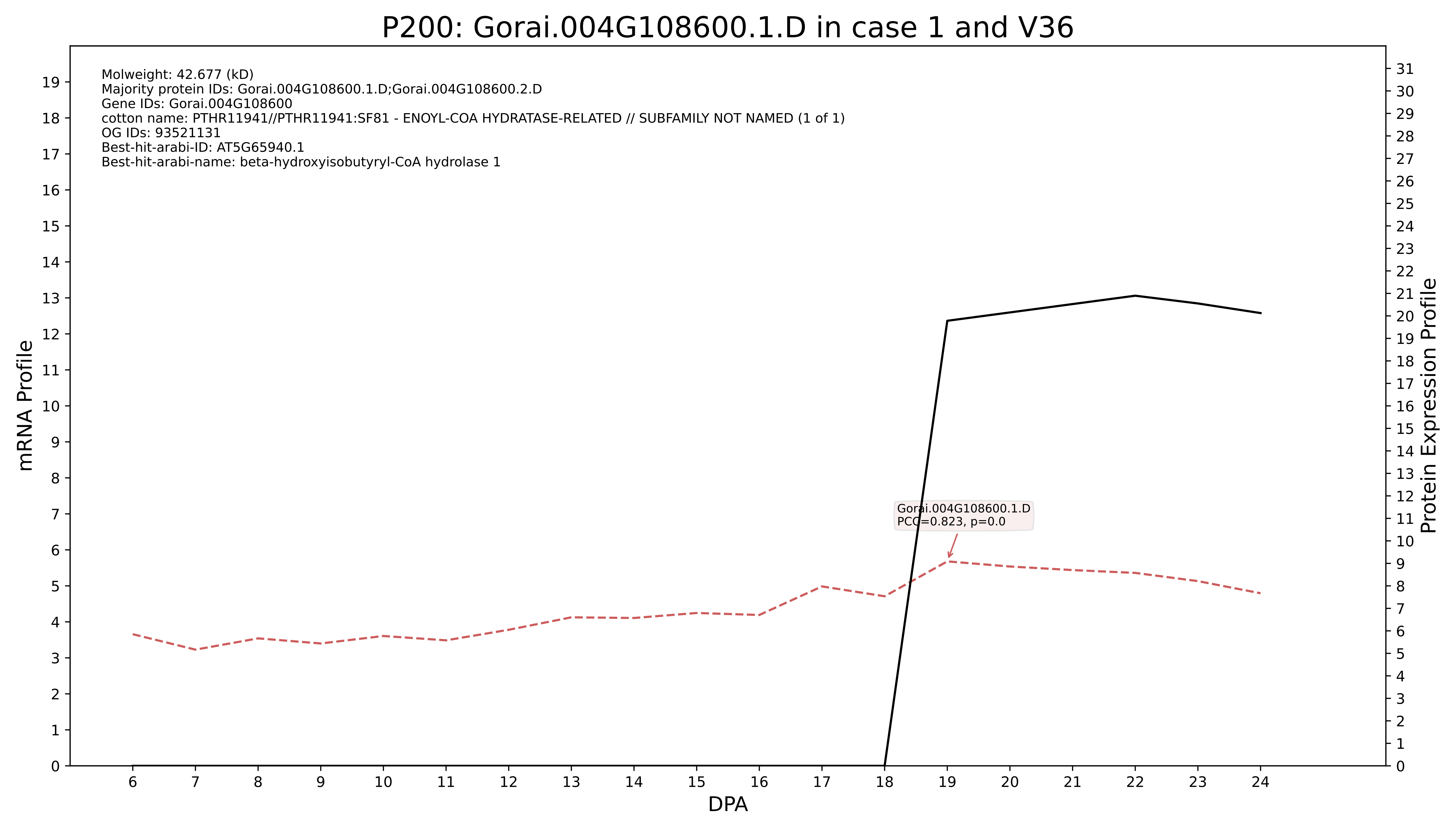

P200: Gorai.004G108600.1.D in case 1 and V36
Molweight: 42.677 (kD)
Majority protein IDs: Gorai.004G108600.1.D;Gorai.004G108600.2.D
Gene IDs: Gorai.004G108600
cotton name: PTHR11941//PTHR11941:SF81 - ENOYL-COA HYDRATASE-RELATED // SUBFAMILY NOT NAMED (1 of 1)
OG IDs: 93521131
Best-hit-arabi-ID: AT5G65940.1
Best-hit-arabi-name: beta-hydroxyisobutyryl-CoA hydrolase 1
Gorai.004G108600.1.D
PCC=0.823, p=0.0
mRNA Profile
Protein Expression Profile
DPA

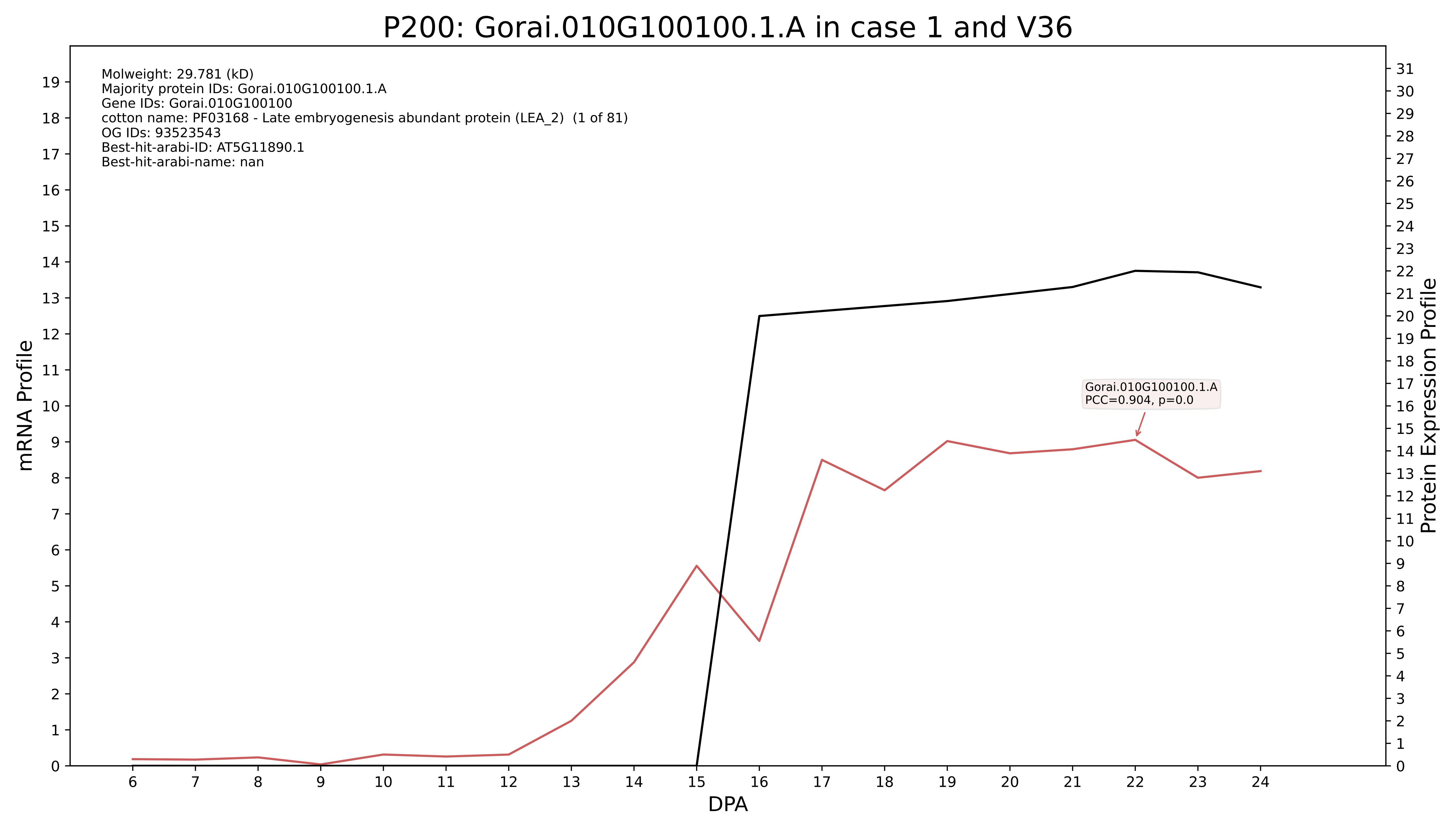

P200: Gorai.010G100100.1.A in case 1 and V36
Molweight: 29.781 (kD)
Majority protein IDs: Gorai.010G100100.1.A
Gene IDs: Gorai.010G100100
cotton name: PF03168 - Late embryogenesis abundant protein (LEA_2) (1 of 81)
OG IDs: 93523543
Best-hit-arabi-ID: AT5G11890.1
Best-hit-arabi-name: nan
Gorai.010G100100.1.A
PCC=0.904, p=0.0
mRNA Profile
Protein Expression Profile
DPA

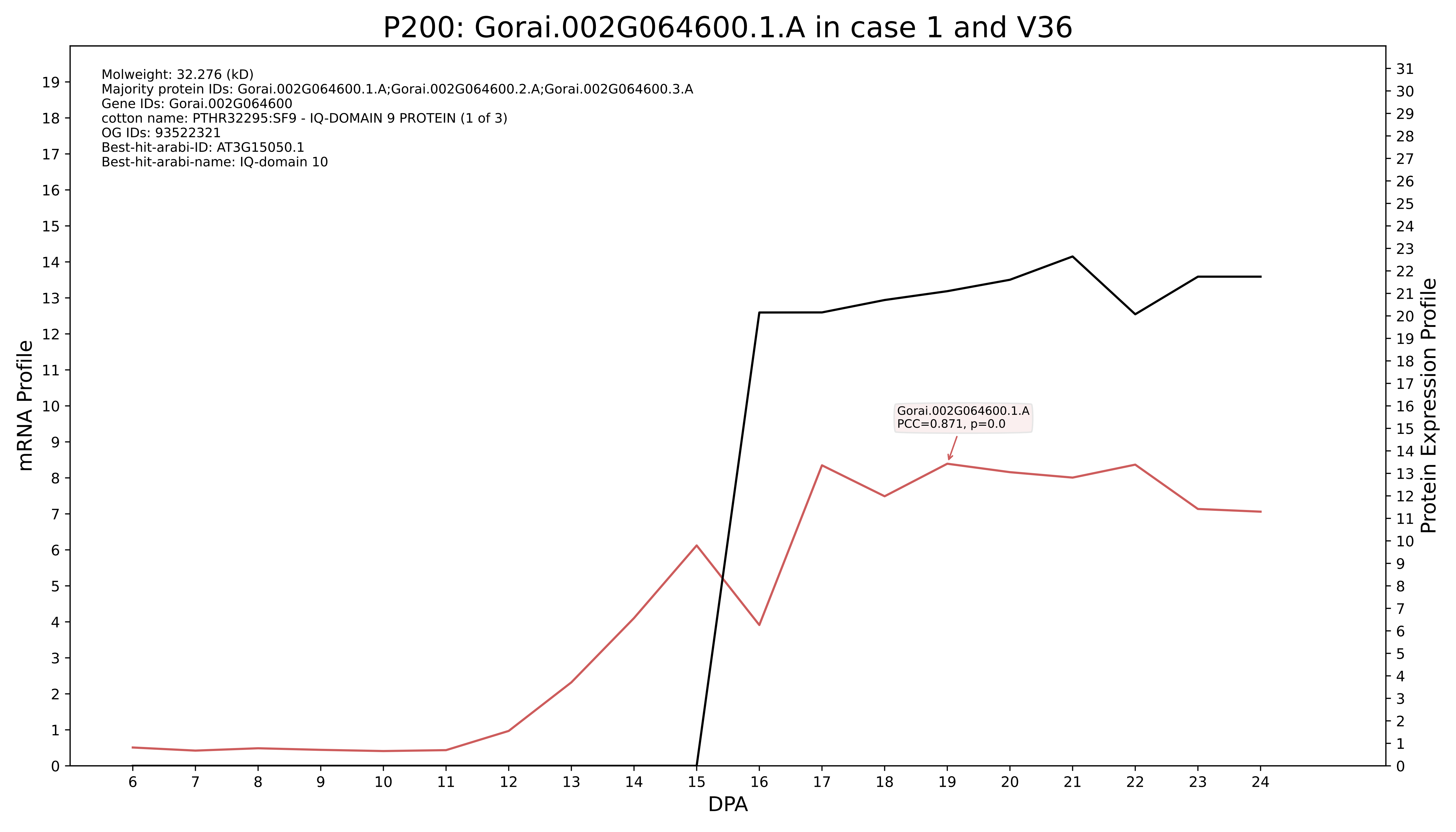
P200: Gorai.002G064600.1.A in case 1 and V36
Molweight: 32.276 (kD)
Majority protein IDs: Gorai.002G064600.1.A;Gorai.002G064600.2.A;Gorai.002G064600.3.A
Gene IDs: Gorai.002G064600
cotton name: PTHR32295:SF9 - IQ-DOMAIN 9 PROTEIN (1 of 3)
OG IDs: 93522321
Best-hit-arabi-ID: AT3G15050.1
Best-hit-arabi-name: IQ-domain 10
Gorai.002G064600.1.A
PCC=0.871, p=0.0
mRNA Profile
Protein Expression Profile
DPA

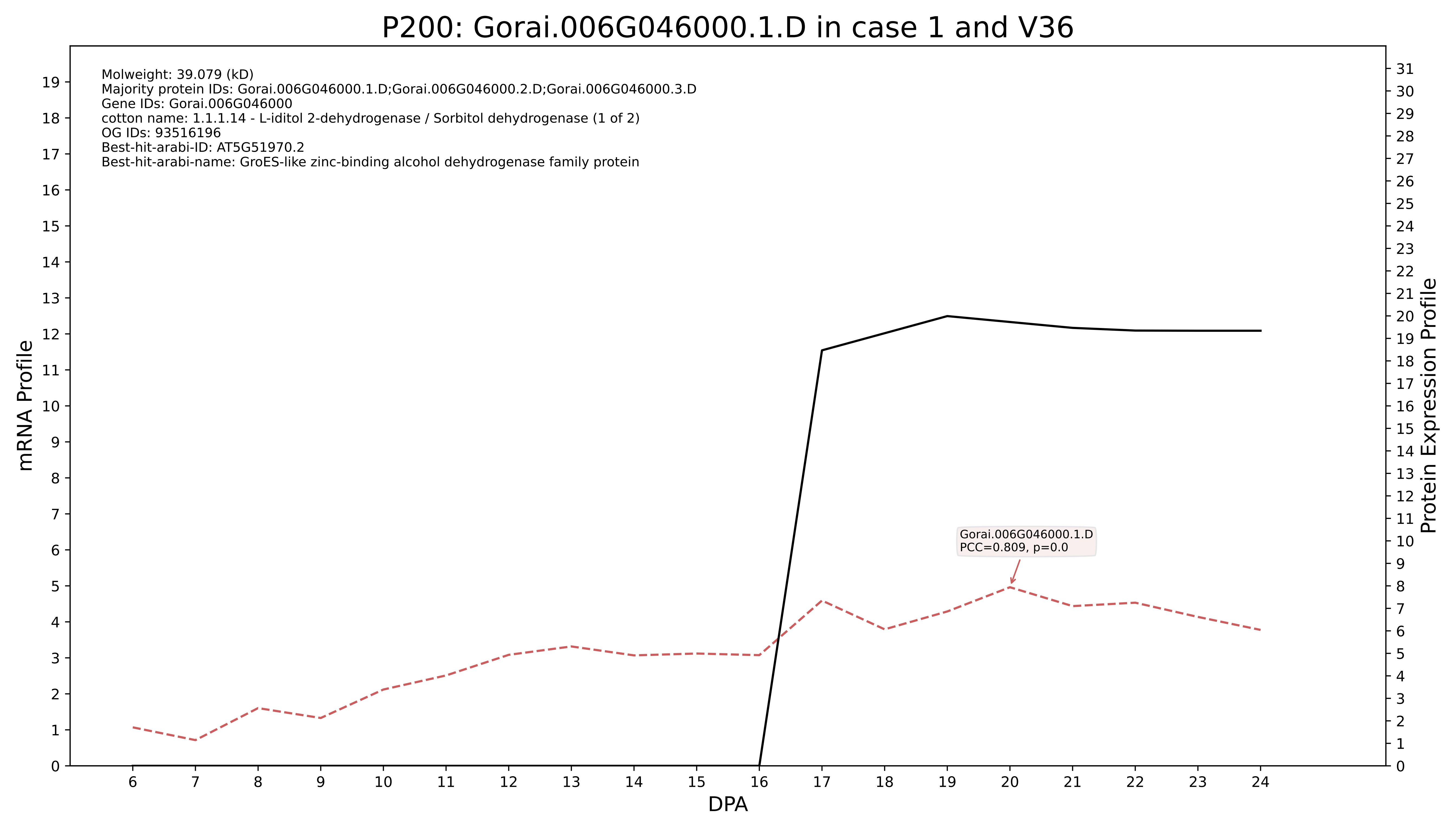
P200: Gorai.006G046000.1.D in case 1 and V36
Molweight: 39.079 (kD)
Majority protein IDs: Gorai.006G046000.1.D;Gorai.006G046000.2.D;Gorai.006G046000.3.D
Gene IDs: Gorai.006G046000
cotton name: 1.1.1.14 - L-iditol 2-dehydrogenase / Sorbitol dehydrogenase (1 of 2)
OG IDs: 93516196
Best-hit-arabi-ID: AT5G51970.2
Best-hit-arabi-name: GroES-like zinc-binding alcohol dehydrogenase family protein
Gorai.006G046000.1.D
PCC=0.809, p=0.0
mRNA Profile
Protein Expression Profile
DPA

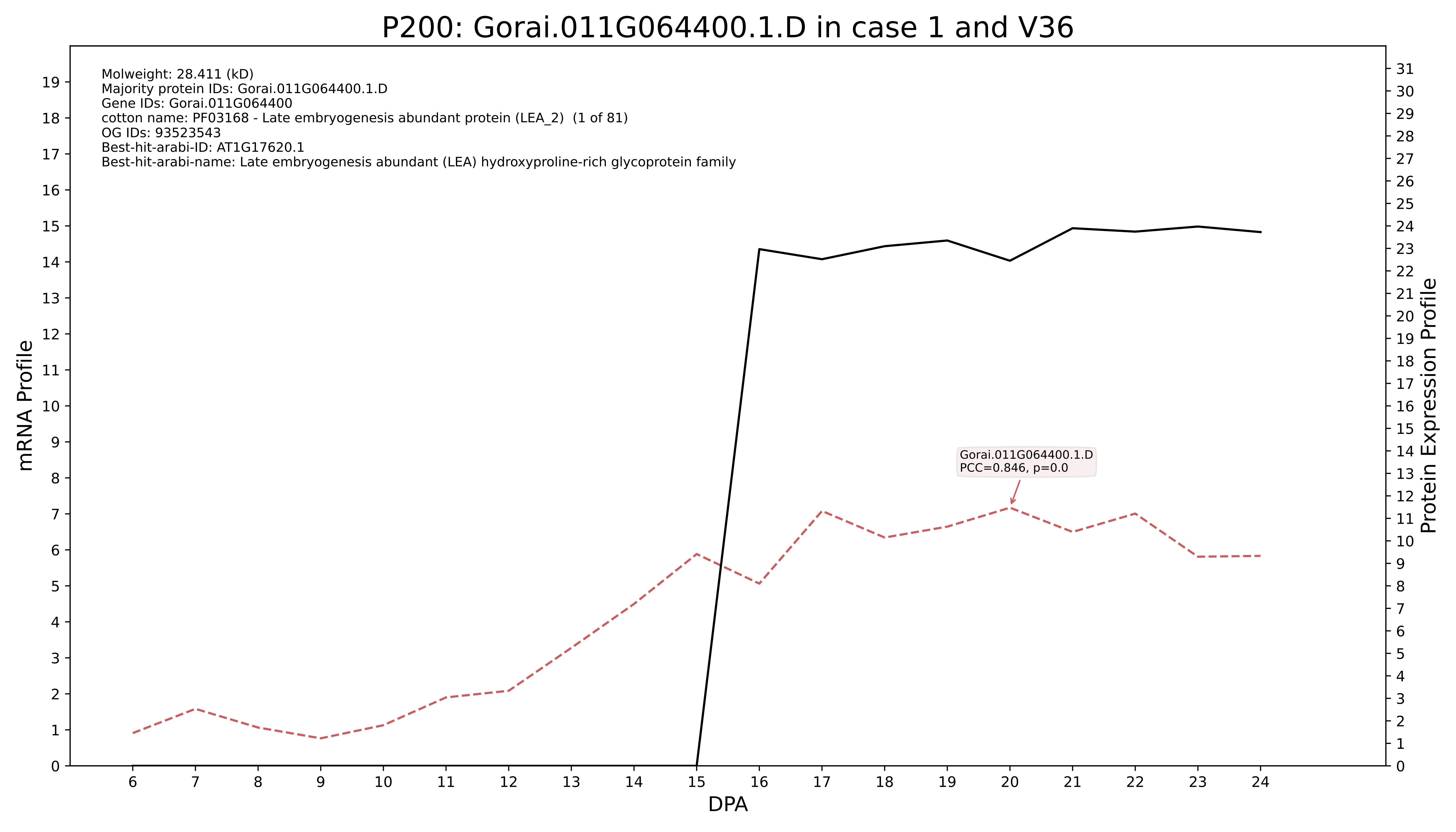

P200: Gorai.011G064400.1.D in case 1 and V36
Molweight: 28.411 (kD)
Majority protein IDs: Gorai.011G064400.1.D
Gene IDs: Gorai.011G064400
cotton name: PF03168 - Late embryogenesis abundant protein (LEA_2) (1 of 81)
OG IDs: 93523543
Best-hit-arabi-ID: AT1G17620.1
Best-hit-arabi-name: Late embryogenesis abundant (LEA) hydroxyproline-rich glycoprotein family
Gorai.011G064400.1.D
PCC=0.846, p=0.0
mRNA Profile
Protein Expression Profile
DPA

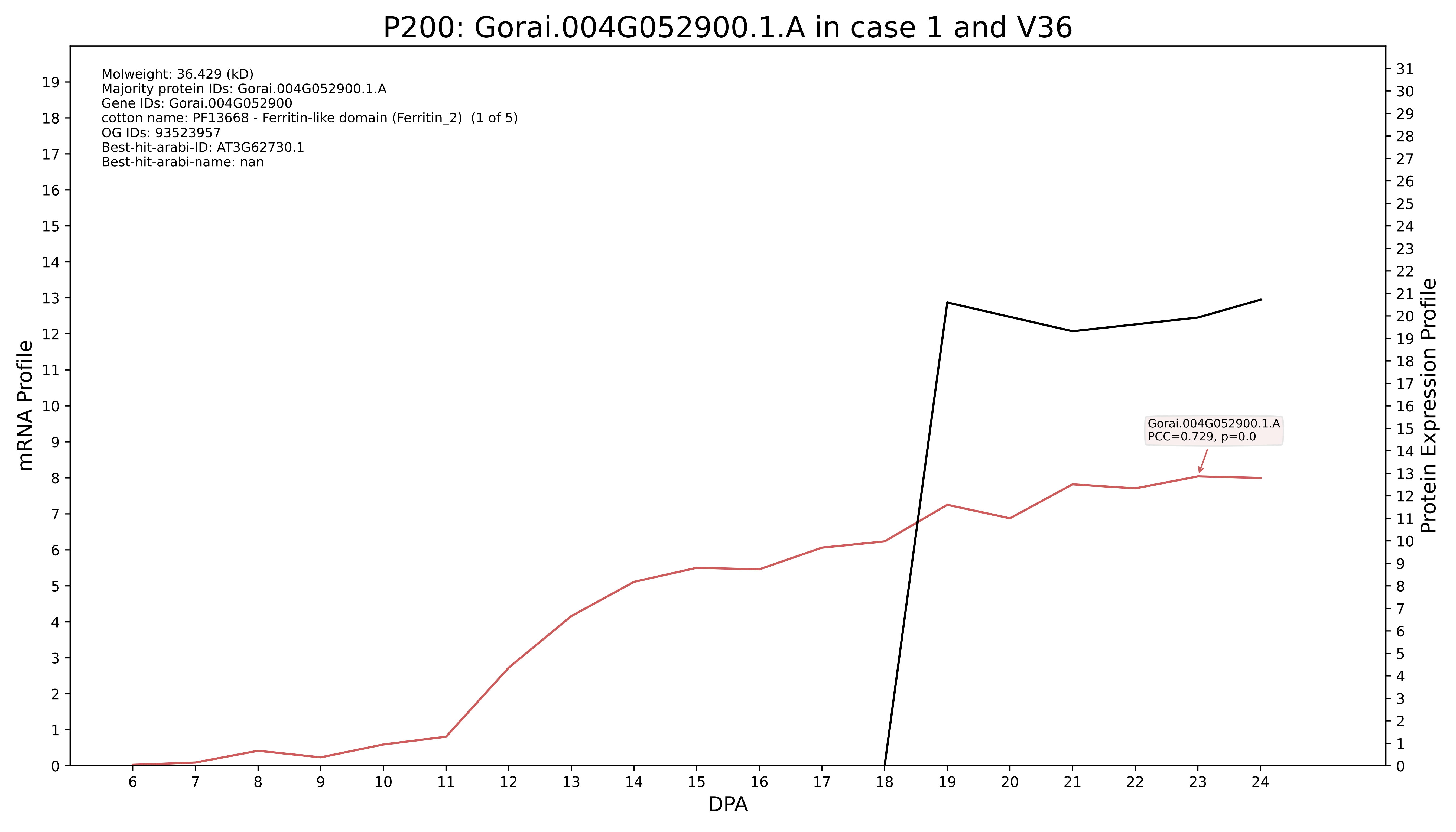
P200: Gorai.004G052900.1.A in case 1 and V36
Molweight: 36.429 (kD)
Majority protein IDs: Gorai.004G052900.1.A
Gene IDs: Gorai.004G052900
cotton name: PF13668 - Ferritin-like domain (Ferritin_2) (1 of 5)
OG IDs: 93523957
Best-hit-arabi-ID: AT3G62730.1
Best-hit-arabi-name: nan
Gorai.004G052900.1.A
PCC=0.729, p=0.0
mRNA Profile
Protein Expression Profile
DPA

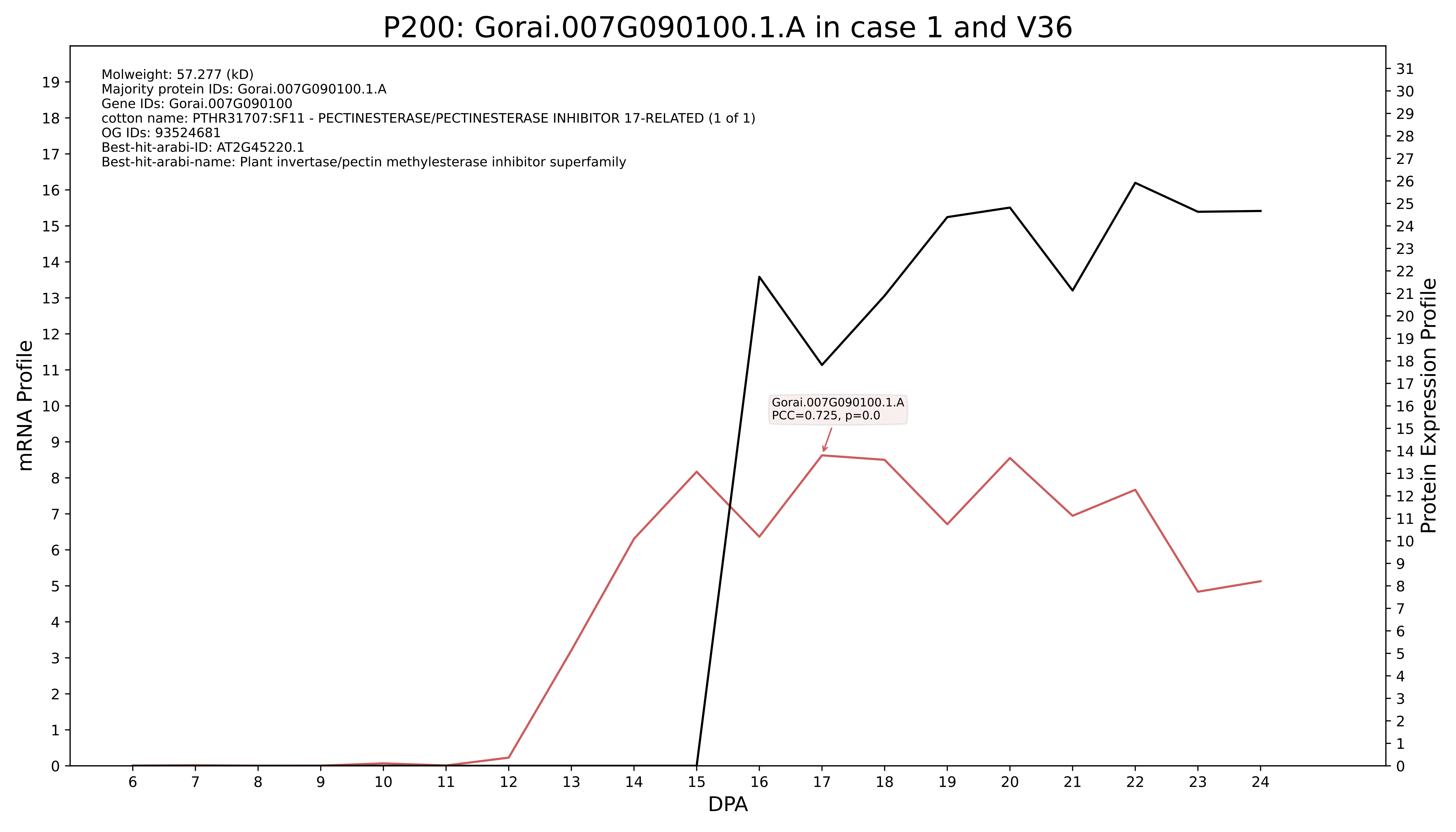
P200: Gorai.007G090100.1.A in case 1 and V36
Molweight: 57.277 (kD)
Majority protein IDs: Gorai.007G090100.1.A
Gene IDs: Gorai.007G090100
cotton name: PTHR31707:SF11 - PECTINESTERASE/PECTINESTERASE INHIBITOR 17-RELATED (1 of 1)
OG IDs: 93524681
Best-hit-arabi-ID: AT2G45220.1
Best-hit-arabi-name: Plant invertase/pectin methylesterase inhibitor superfamily
Gorai.007G090100.1.A
PCC=0.725, p=0.0
mRNA Profile
Protein Expression Profile
DPA

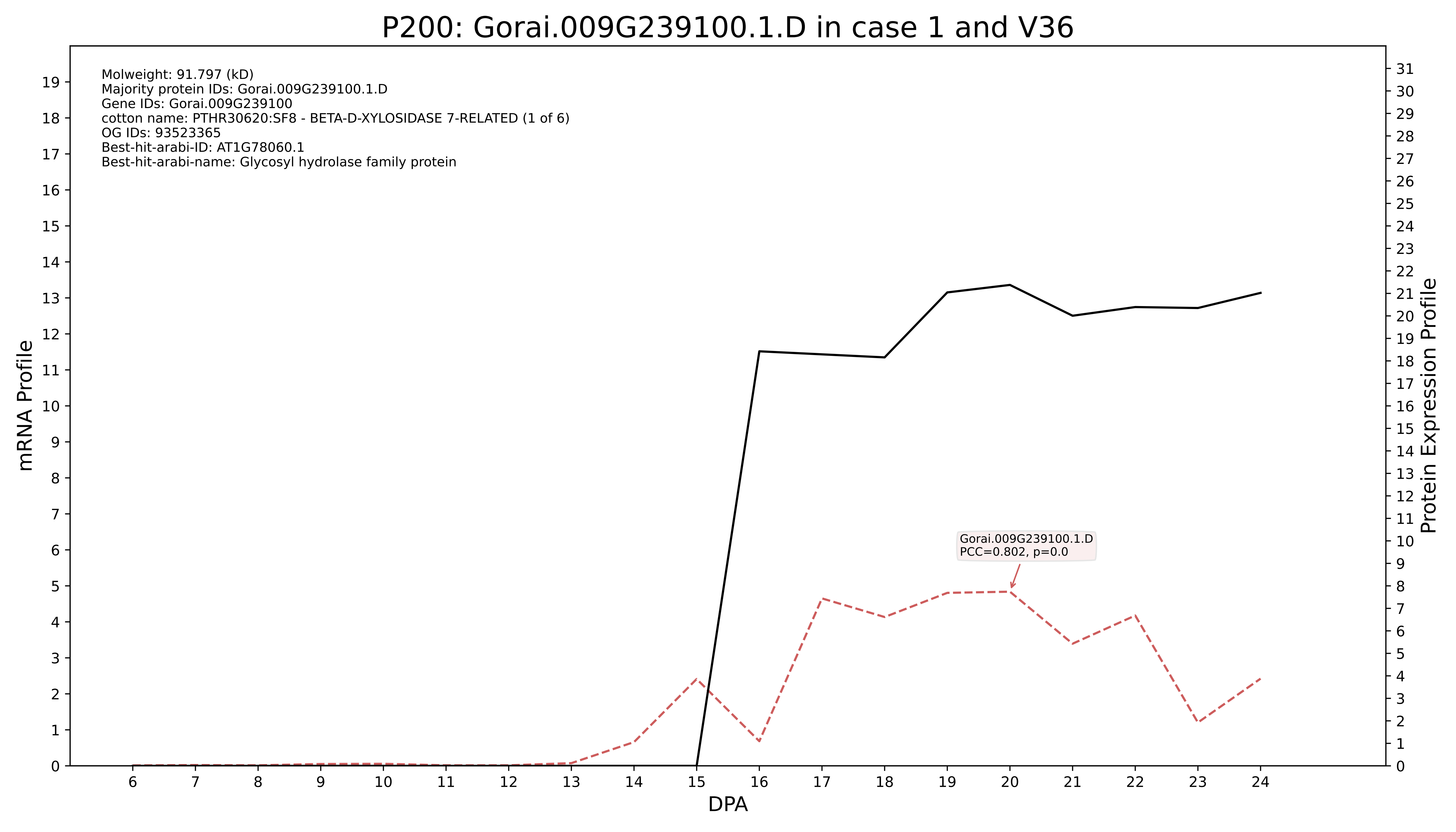
P200: Gorai.009G239100.1.D in case 1 and V36
Molweight: 91.797 (kD)
Majority protein IDs: Gorai.009G239100.1.D
Gene IDs: Gorai.009G239100
cotton name: PTHR30620:SF8 - BETA-D-XYLOSIDASE 7-RELATED (1 of 6)
OG IDs: 93523365
Best-hit-arabi-ID: AT1G78060.1
Best-hit-arabi-name: Glycosyl hydrolase family protein
Gorai.009G239100.1.D
PCC=0.802, p=0.0
mRNA Profile
Protein Expression Profile
DPA

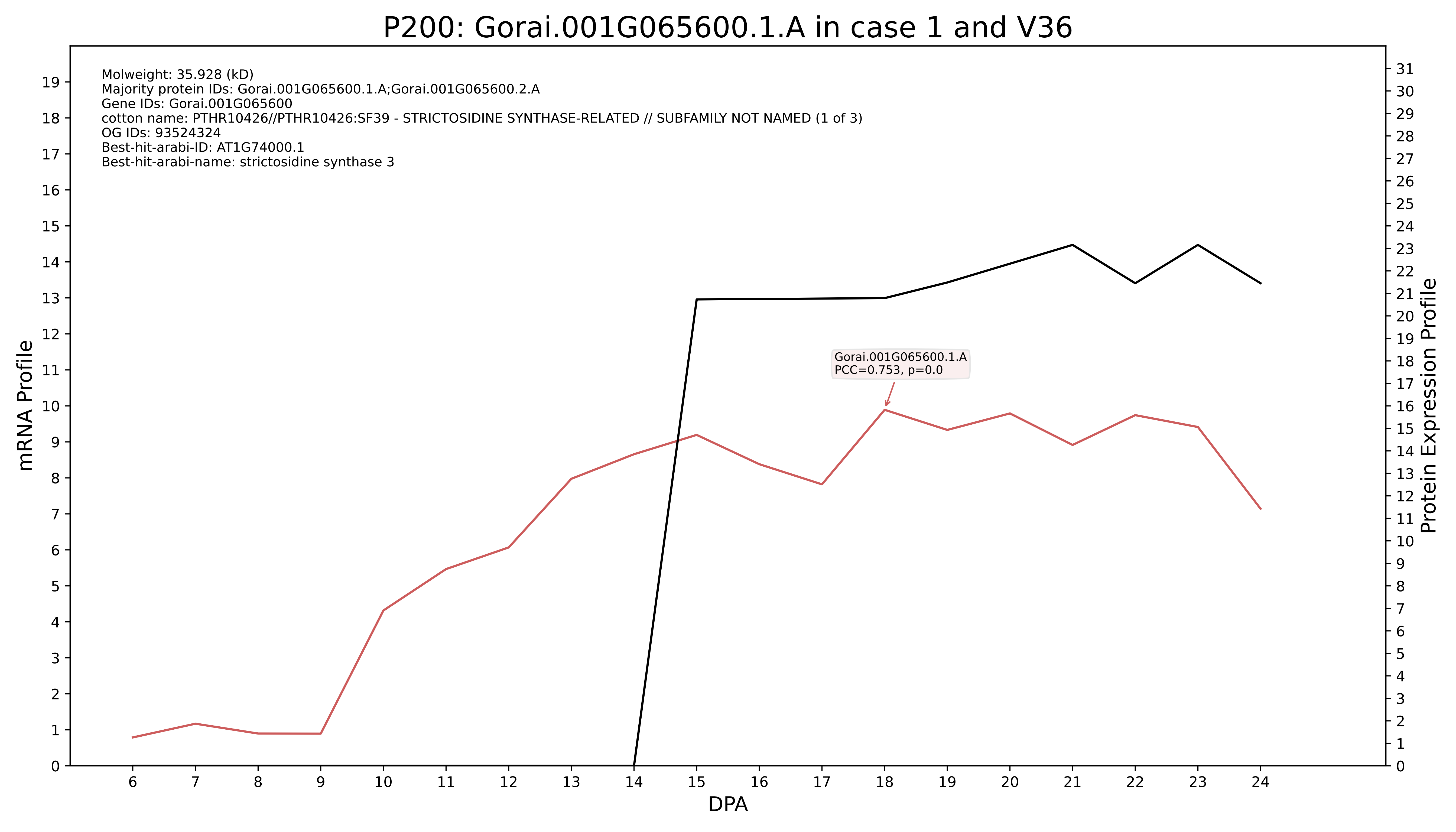
P200: Gorai.001G065600.1.A in case 1 and V36
Molweight: 35.928 (kD)
Majority protein IDs: Gorai.001G065600.1.A;Gorai.001G065600.2.A
Gene IDs: Gorai.001G065600
cotton name: PTHR10426//PTHR10426:SF39 - STRICTOSIDINE SYNTHASE-RELATED // SUBFAMILY NOT NAMED (1 of 3)
OG IDs: 93524324
Best-hit-arabi-ID: AT1G74000.1
Best-hit-arabi-name: strictosidine synthase 3
Gorai.001G065600.1.A
PCC=0.753, p=0.0
mRNA Profile
Protein Expression Profile
DPA

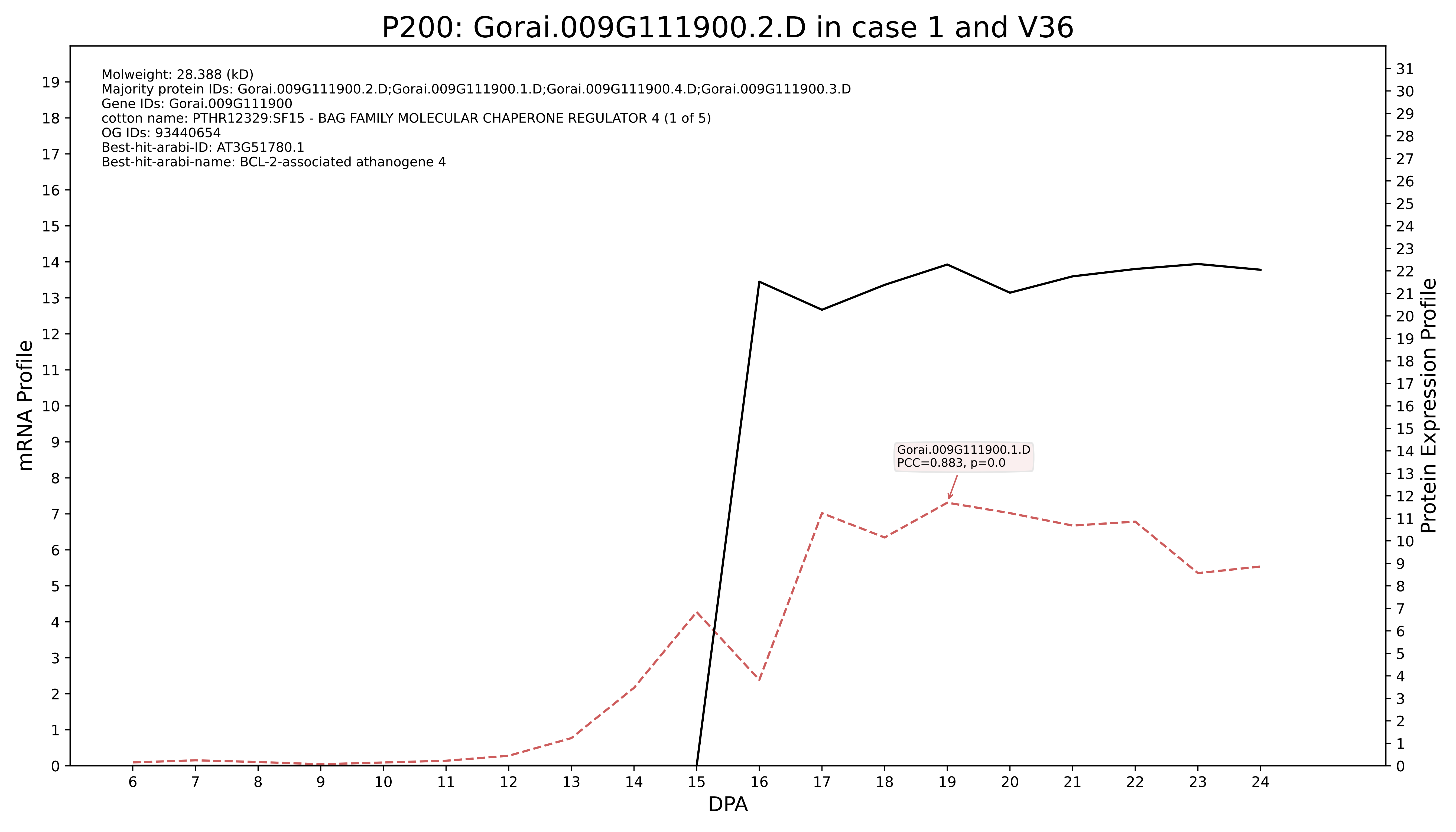

P200: Gorai.009G111900.2.D in case 1 and V36
Molweight: 28.388 (kD)
Majority protein IDs: Gorai.009G111900.2.D;Gorai.009G111900.1.D;Gorai.009G111900.4.D;Gorai.009G111900.3.D
Gene IDs: Gorai.009G111900
cotton name: PTHR12329:SF15 - BAG FAMILY MOLECULAR CHAPERONE REGULATOR 4 (1 of 5)
OG IDs: 93440654
Best-hit-arabi-ID: AT3G51780.1
Best-hit-arabi-name: BCL-2-associated athanogene 4
Gorai.009G111900.1.D
PCC=0.883, p=0.0
mRNA Profile
Protein Expression Profile
DPA

### P200: Gorai.010G247900.7.A in case 1 and V36

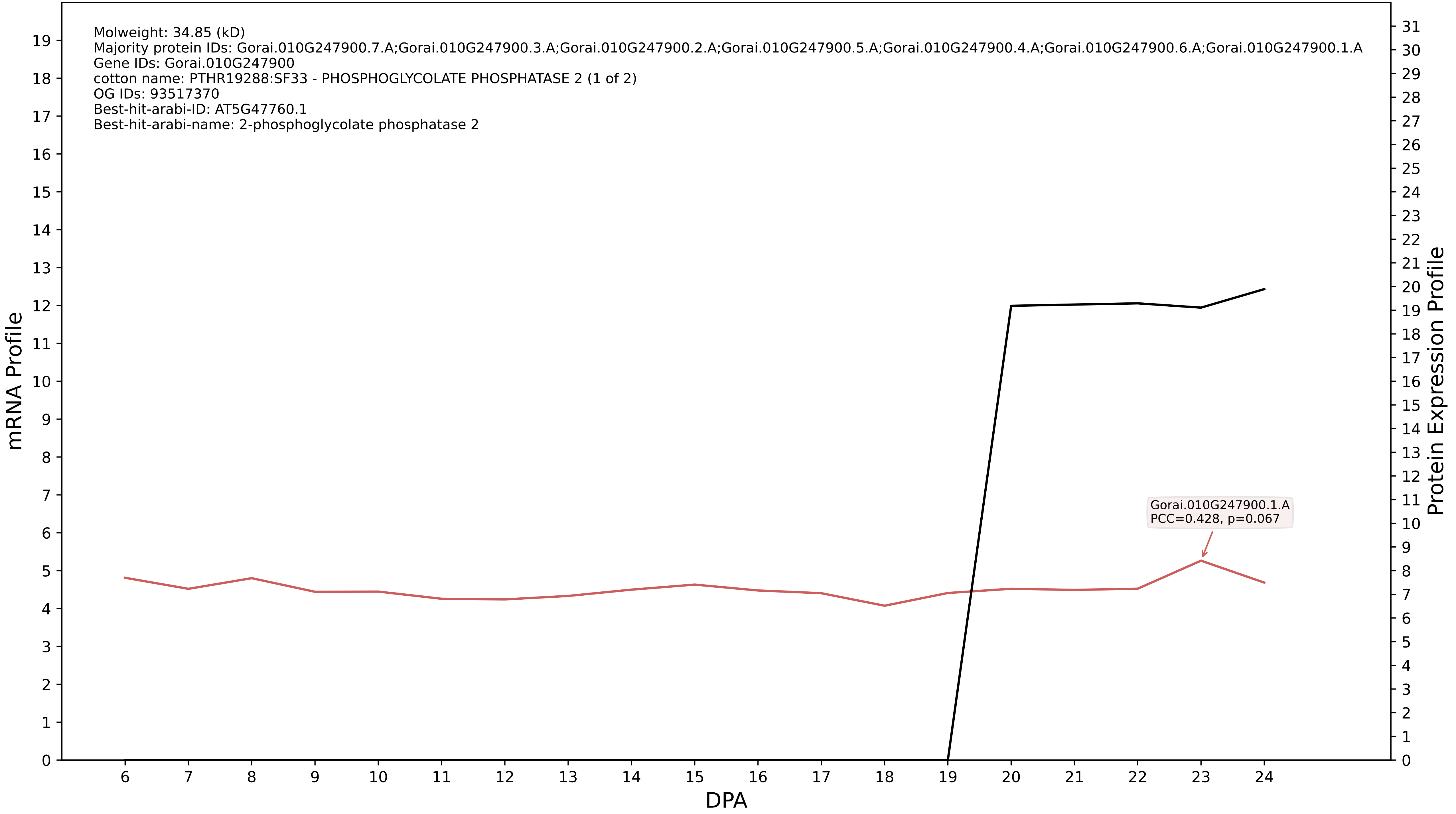

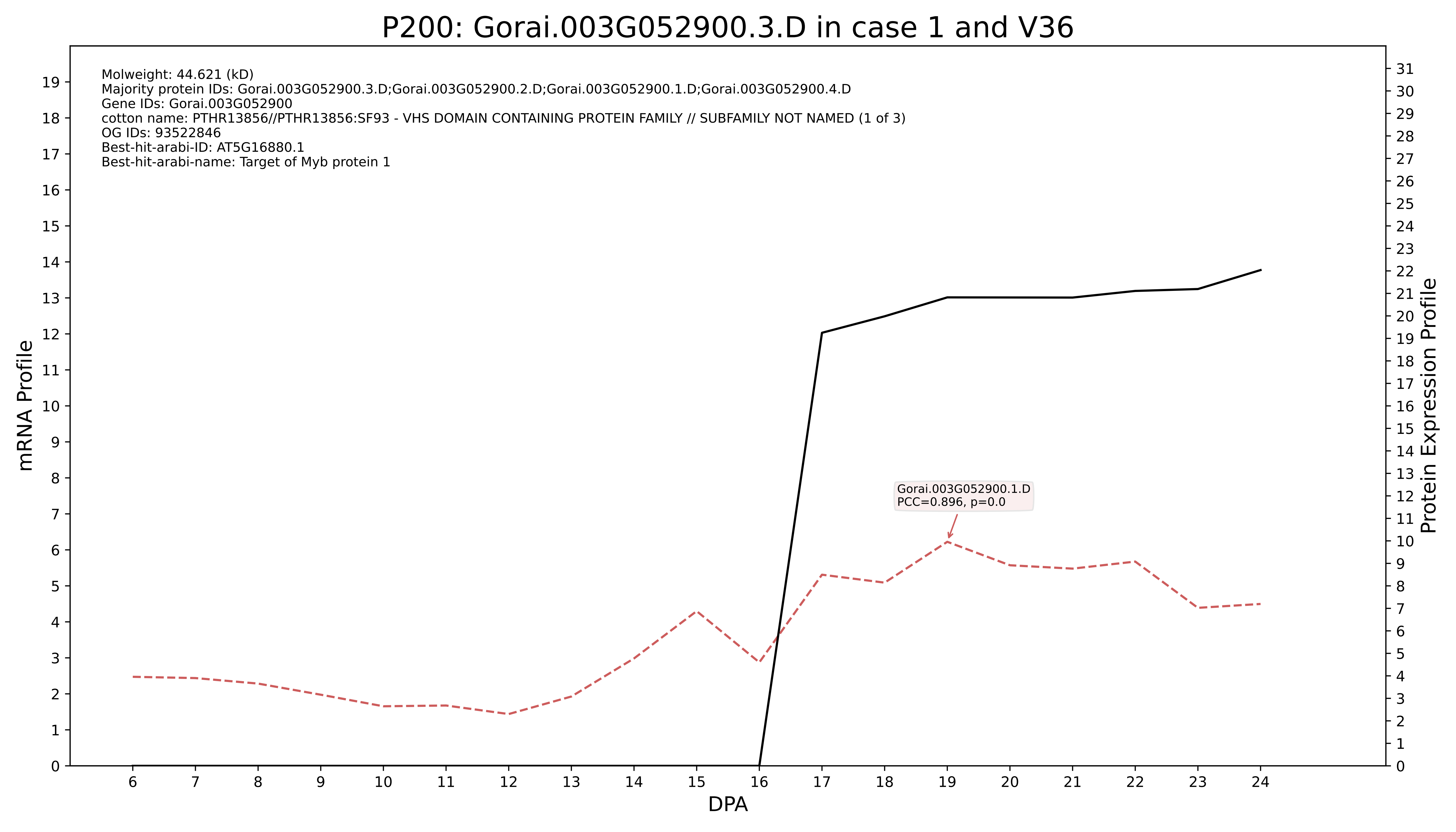
P200: Gorai.003G052900.3.D in case 1 and V36
Molweight: 44.621 (kD)
Majority protein IDs: Gorai.003G052900.3.D;Gorai.003G052900.2.D;Gorai.003G052900.1.D;Gorai.003G052900.4.D
Gene IDs: Gorai.003G052900
cotton name: PTHR13856//PTHR13856:SF93 - VHS DOMAIN CONTAINING PROTEIN FAMILY // SUBFAMILY NOT NAMED (1 of 3)
OG IDs: 93522846
Best-hit-arabi-ID: AT5G16880.1
Best-hit-arabi-name: Target of Myb protein 1
Gorai.003G052900.1.D
PCC=0.896, p=0.0
mRNA Profile
Protein Expression Profile
DPA

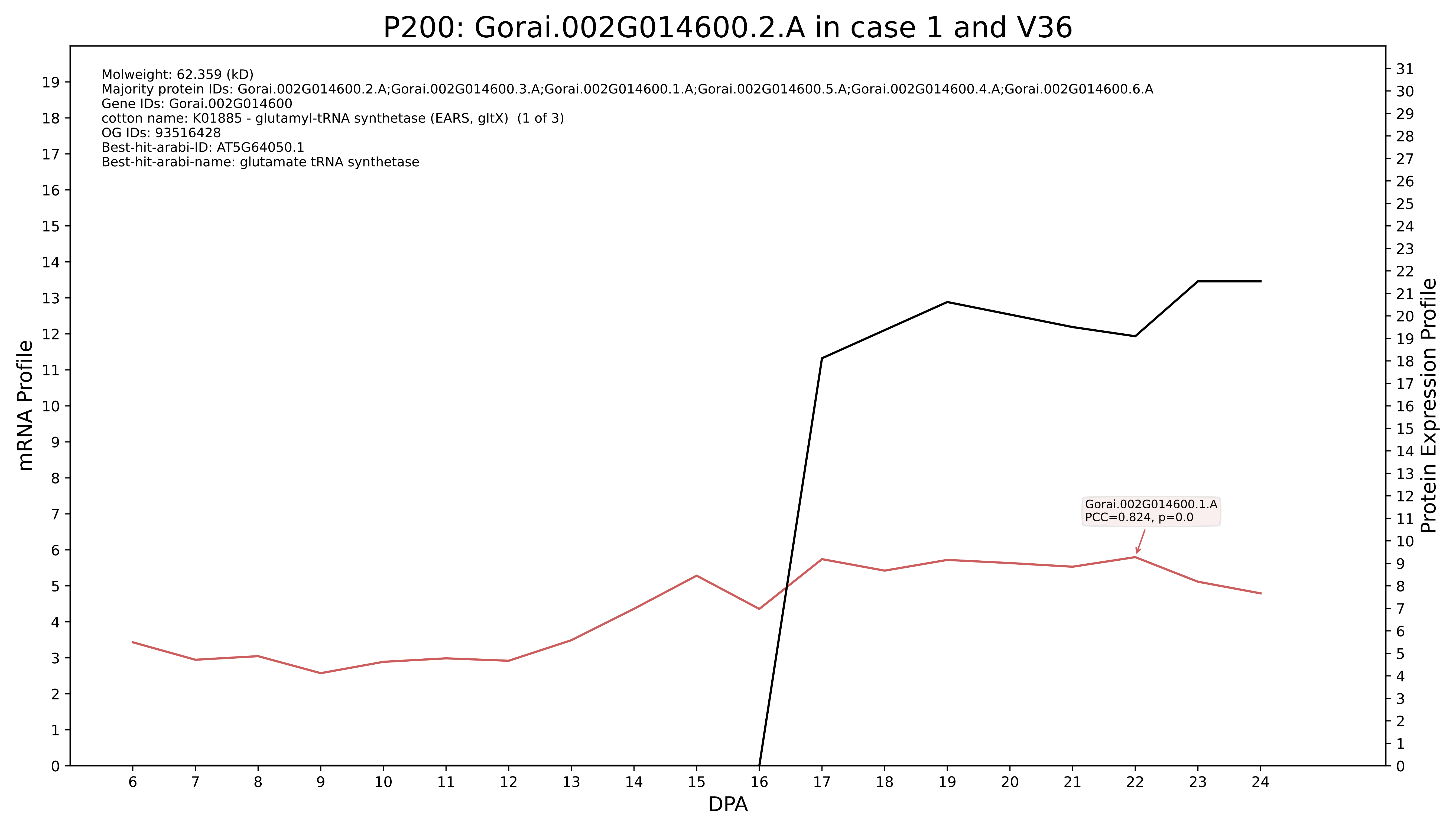
P200: Gorai.002G014600.2.A in case 1 and V36
Molweight: 62.359 (kD)
Majority protein IDs: Gorai.002G014600.2.A;Gorai.002G014600.3.A;Gorai.002G014600.1.A;Gorai.002G014600.5.A;Gorai.002G014600.4.A;Gorai.002G014600.6.A
Gene IDs: Gorai.002G014600
cotton name: K01885 - glutamyl-tRNA synthetase (EARS, gltX) (1 of 3)
OG IDs: 93516428
Best-hit-arabi-ID: AT5G64050.1
Best-hit-arabi-name: glutamate tRNA synthetase
Gorai.002G014600.1.A
PCC=0.824, p=0.0
mRNA Profile
Protein Expression Profile
DPA

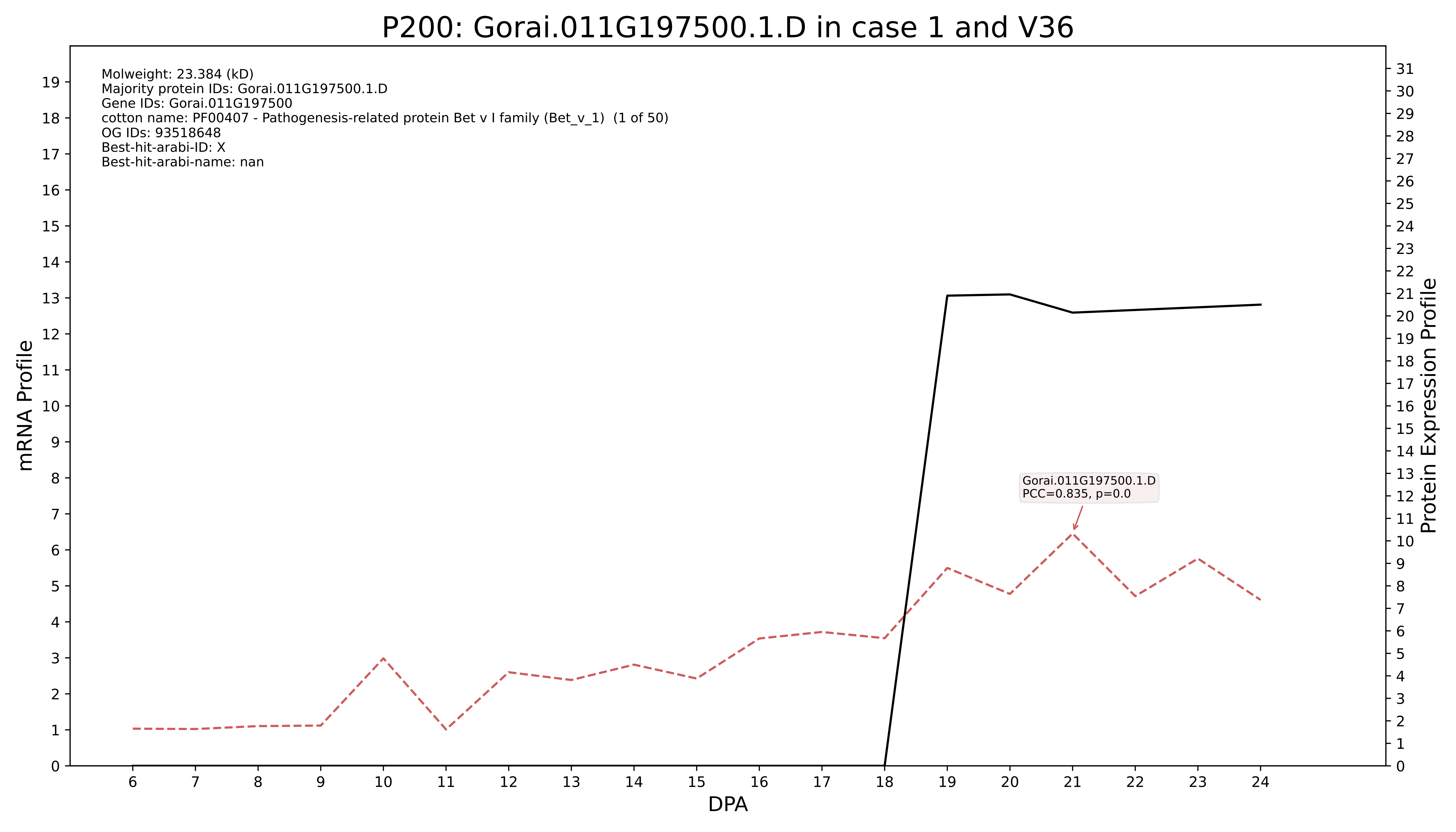
P200: Gorai.011G197500.1.D in case 1 and V36
Molweight: 23.384 (kD)
Majority protein IDs: Gorai.011G197500.1.D
Gene IDs: Gorai.011G197500
cotton name: PF00407 - Pathogenesis-related protein Bet v I family (Bet_v_1) (1 of 50)
OG IDs: 93518648
Best-hit-arabi-ID: X
Best-hit-arabi-name: nan
Gorai.011G197500.1.D
PCC=0.835, p=0.0
mRNA Profile
Protein Expression Profile
DPA

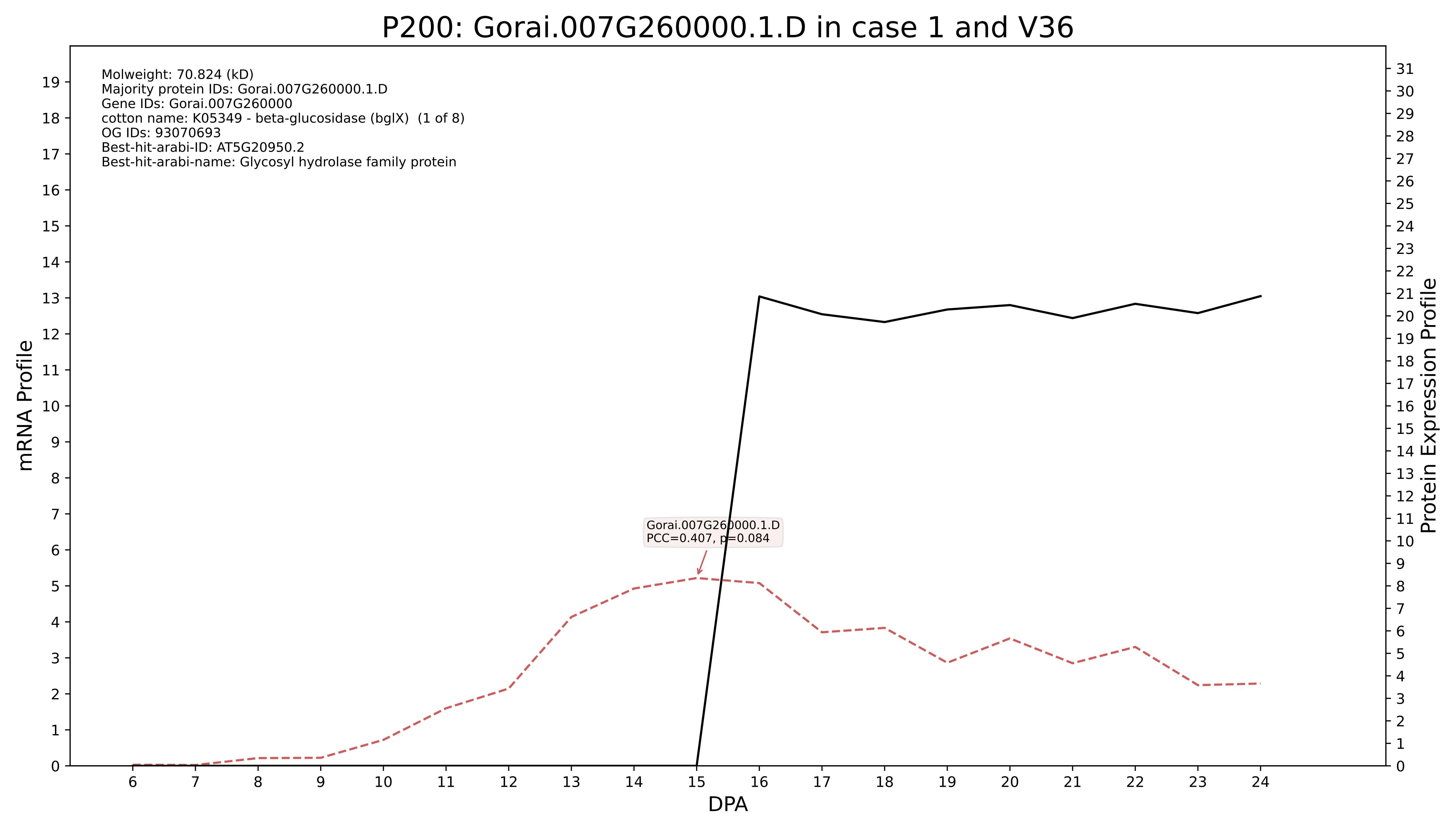

P200: Gorai.007G260000.1.D in case 1 and V36
Molweight: 70.824 (kD)
Majority protein IDs: Gorai.007G260000.1.D
Gene IDs: Gorai.007G260000
cotton name: K05349 - beta-glucosidase (bglX) (1 of 8)
OG IDs: 93070693
Best-hit-arabi-ID: AT5G20950.2
Best-hit-arabi-name: Glycosyl hydrolase family protein
Gorai.007G260000.1.D
PCC=0.407, p=0.084
mRNA Profile
Protein Expression Profile
DPA

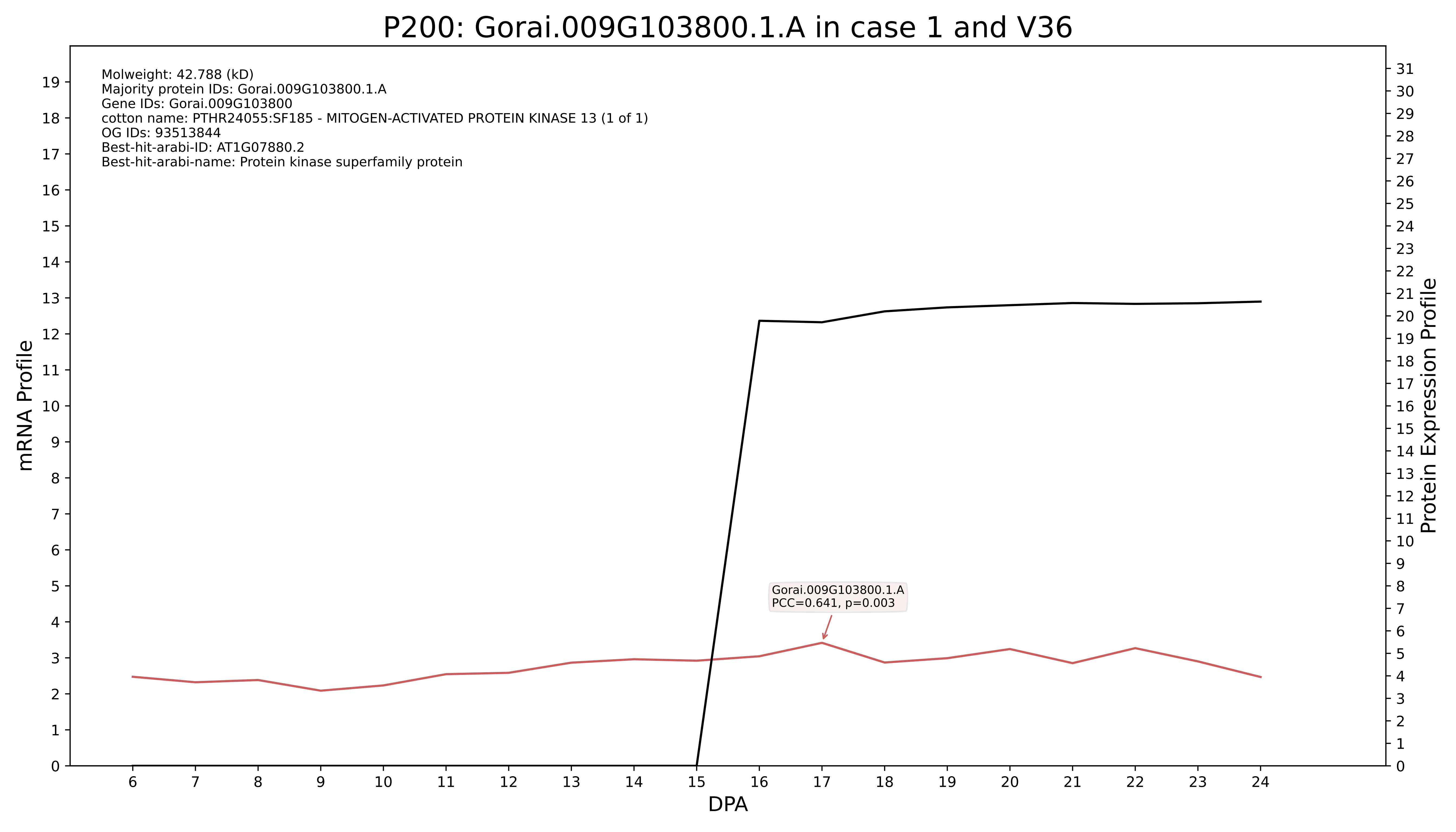

P200: Gorai.009G103800.1.A in case 1 and V36
Molweight: 42.788 (kD)
Majority protein IDs: Gorai.009G103800.1.A
Gene IDs: Gorai.009G103800
cotton name: PTHR24055:SF185 - MITOGEN-ACTIVATED PROTEIN KINASE 13 (1 of 1)
OG IDs: 93513844
Best-hit-arabi-ID: AT1G07880.2
Best-hit-arabi-name: Protein kinase superfamily protein
Gorai.009G103800.1.A
PCC=0.641, p=0.003
mRNA Profile
Protein Expression Profile
DPA

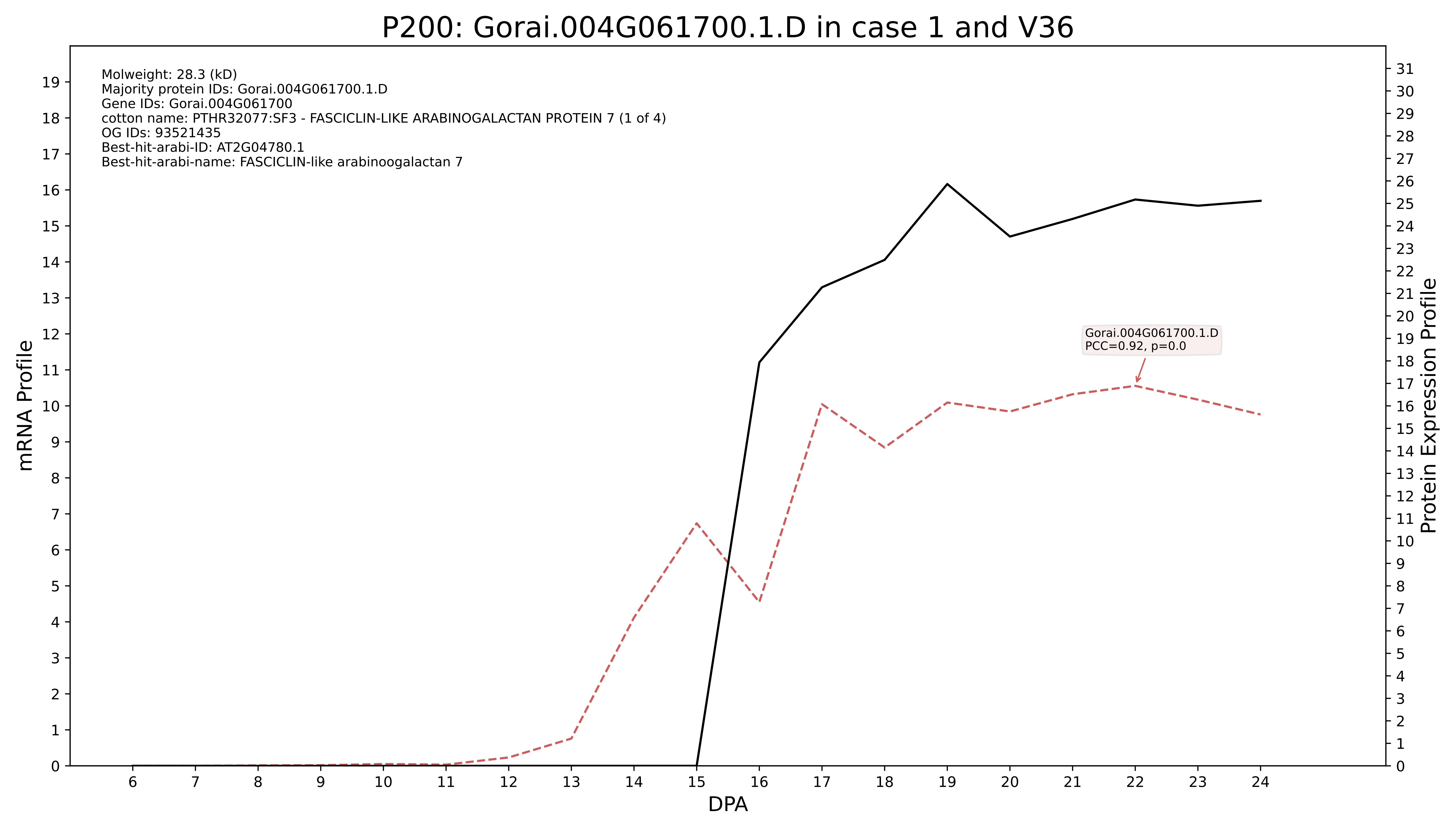

P200: Gorai.004G061700.1.D in case 1 and V36
Molweight: 28.3 (kD)
Majority protein IDs: Gorai.004G061700.1.D
Gene IDs: Gorai.004G061700
cotton name: PTHR32077:SF3 - FASCICLIN-LIKE ARABINOGALACTAN PROTEIN 7 (1 of 4)
OG IDs: 93521435
Best-hit-arabi-ID: AT2G04780.1
Best-hit-arabi-name: FASCICLIN-like arabinoogalactan 7
Gorai.004G061700.1.D
PCC=0.92, p=0.0
mRNA Profile
Protein Expression Profile
DPA

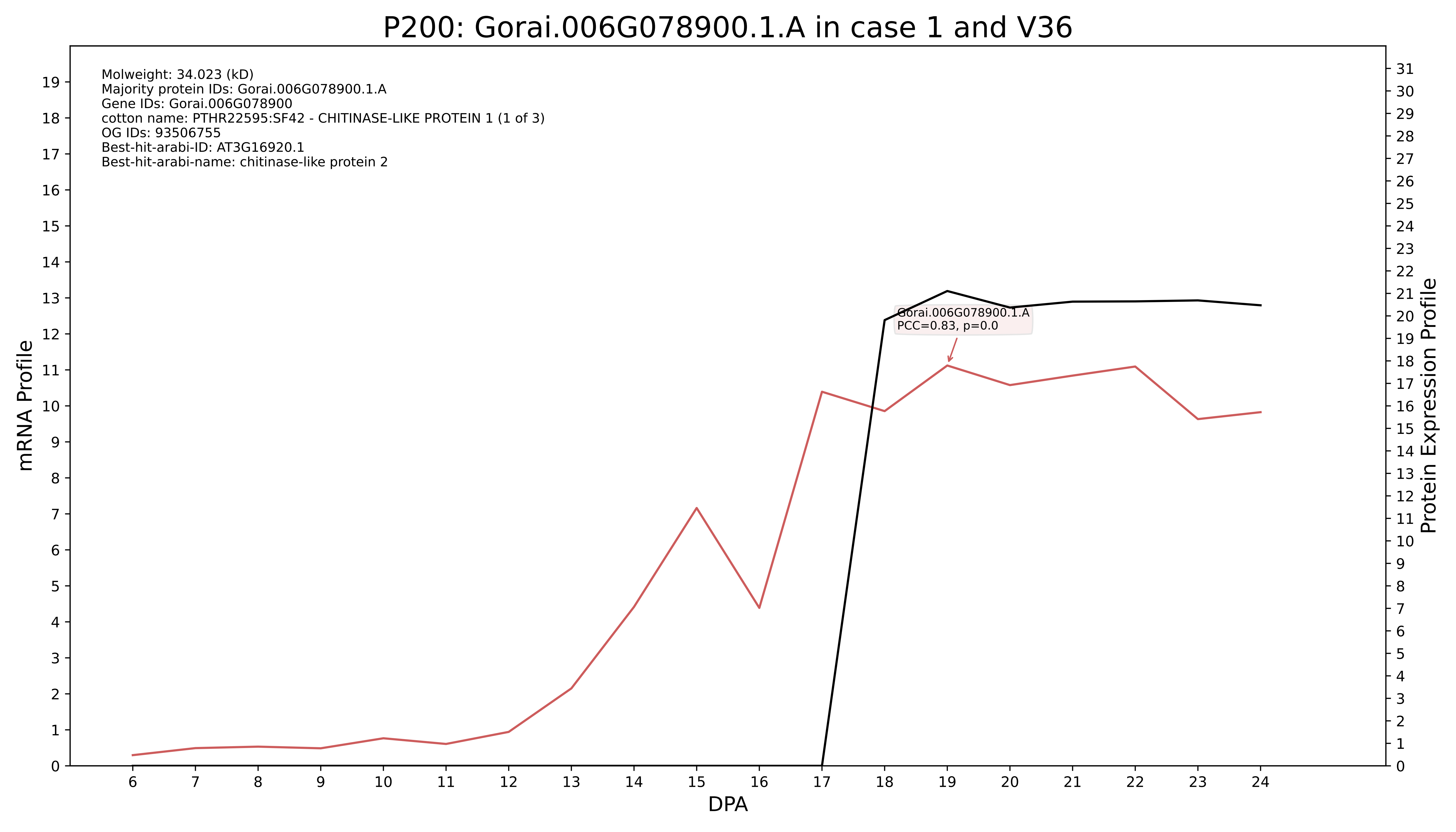
P200: Gorai.006G078900.1.A in case 1 and V36
Molweight: 34.023 (kD)
Majority protein IDs: Gorai.006G078900.1.A
Gene IDs: Gorai.006G078900
cotton name: PTHR22595:SF42 - CHITINASE-LIKE PROTEIN 1 (1 of 3)
OG IDs: 93506755
Best-hit-arabi-ID: AT3G16920.1
Best-hit-arabi-name: chitinase-like protein 2
Gorai.006G078900.1.A
PCC=0.83, p=0.0
mRNA Profile
Protein Expression Profile
DPA

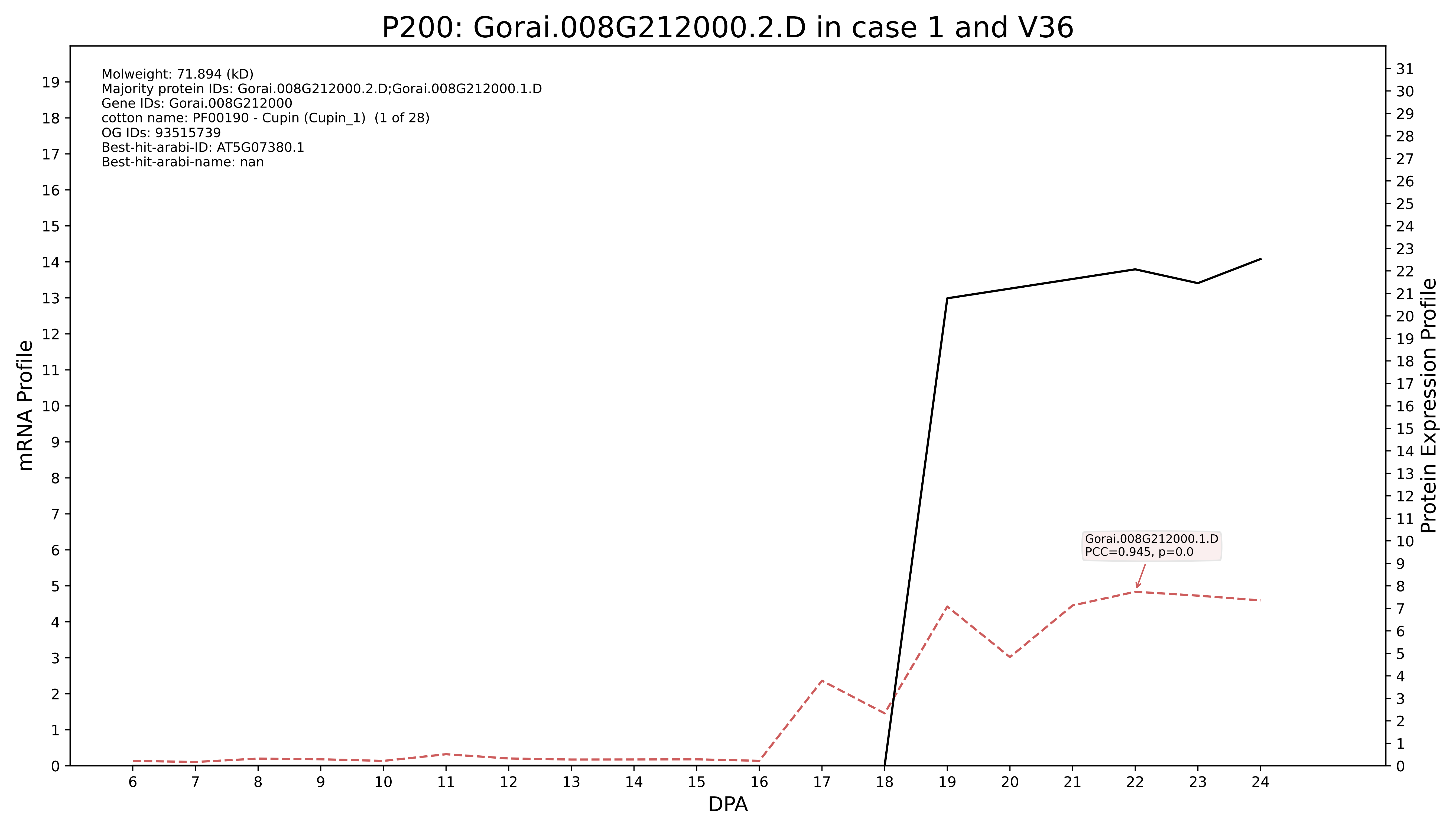
P200: Gorai.008G212000.2.D in case 1 and V36
Molweight: 71.894 (kD)
Majority protein IDs: Gorai.008G212000.2.D;Gorai.008G212000.1.D
Gene IDs: Gorai.008G212000
cotton name: PF00190 - Cupin (Cupin_1) (1 of 28)
OG IDs: 93515739
Best-hit-arabi-ID: AT5G07380.1
Best-hit-arabi-name: nan
Gorai.008G212000.1.D
PCC=0.945, p=0.0
mRNA Profile
Protein Expression Profile
DPA

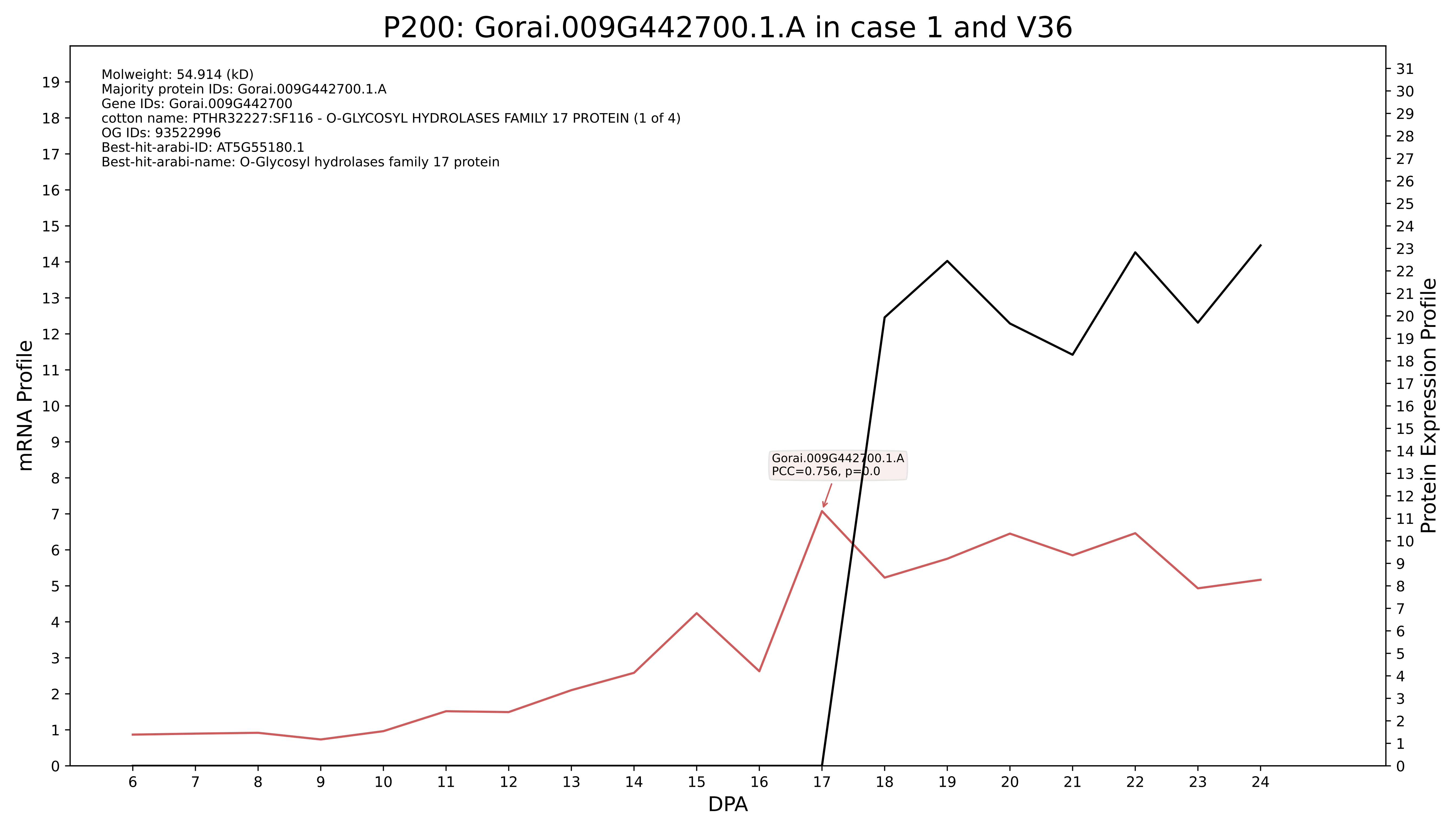
P200: Gorai.009G442700.1.A in case 1 and V36
Molweight: 54.914 (kD)
Majority protein IDs: Gorai.009G442700.1.A
Gene IDs: Gorai.009G442700
cotton name: PTHR32227:SF116 - O-GLYCOSYL HYDROLASES FAMILY 17 PROTEIN (1 of 4)
OG IDs: 93522996
Best-hit-arabi-ID: AT5G55180.1
Best-hit-arabi-name: O-Glycosyl hydrolases family 17 protein
Gorai.009G442700.1.A
PCC=0.756, p=0.0
mRNA Profile
Protein Expression Profile
DPA

### P200: Gorai.009G174700.1.A in case 1 and V36

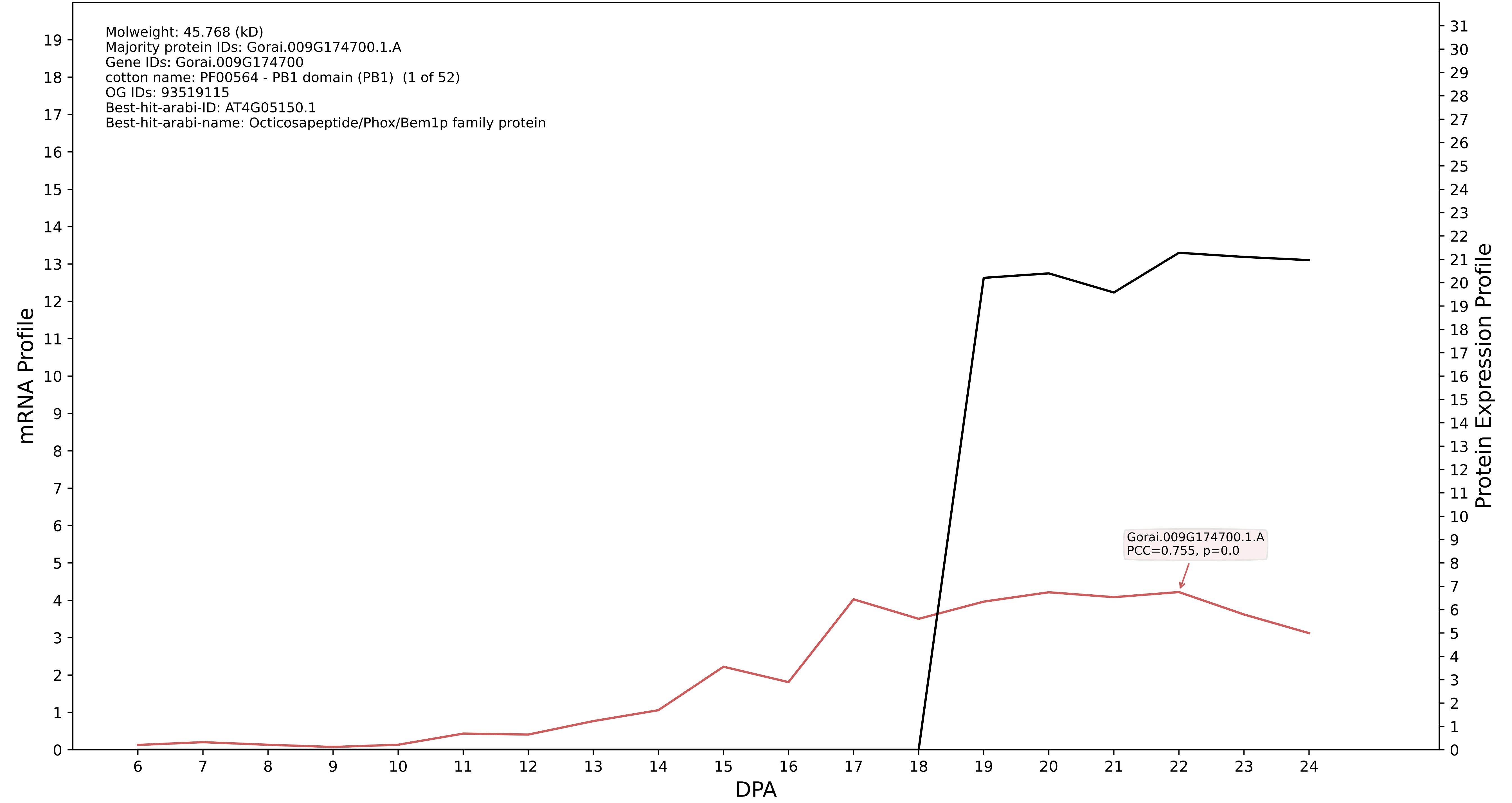

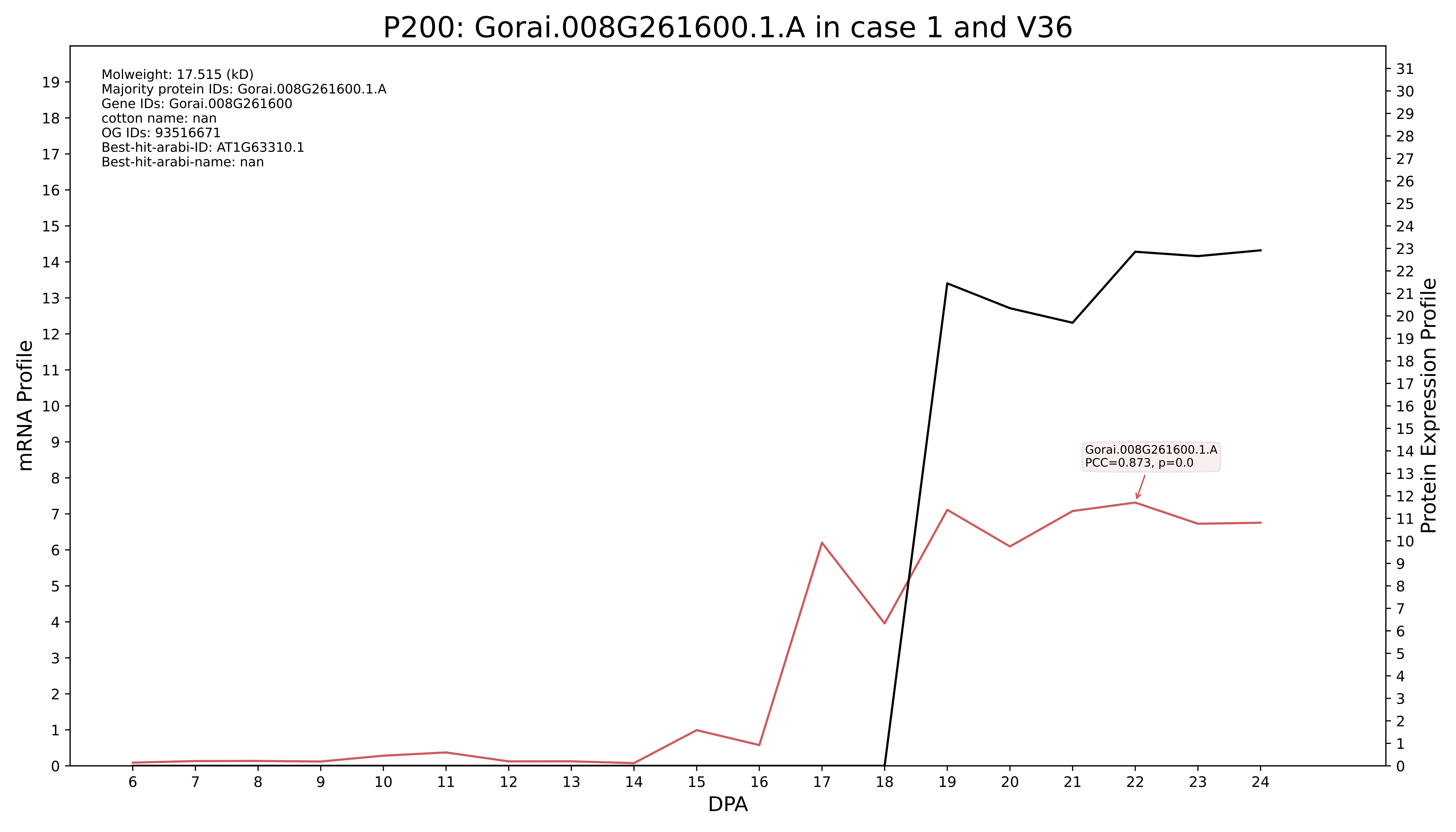
P200: Gorai.008G261600.1.A in case 1 and V36
Molweight: 17.515 (kD)
Majority protein IDs: Gorai.008G261600.1.A
Gene IDs: Gorai.008G261600
cotton name: nan
OG IDs: 93516671
Best-hit-arabi-ID: AT1G63310.1
Best-hit-arabi-name: nan
Gorai.008G261600.1.A
PCC=0.873, p=0.0
mRNA Profile
Protein Expression Profile
DPA

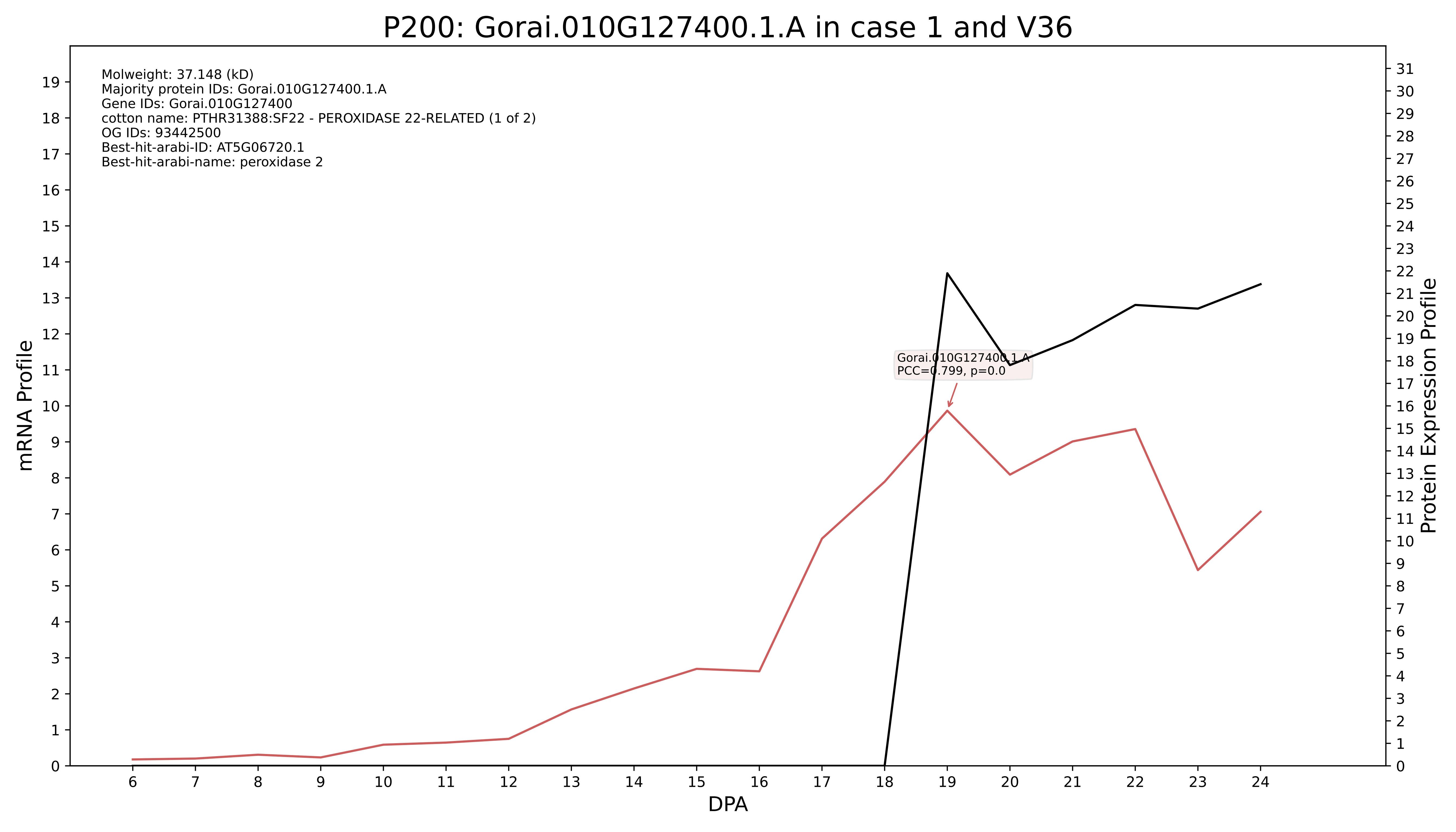
P200: Gorai.010G127400.1.A in case 1 and V36
Molweight: 37.148 (kD)
Majority protein IDs: Gorai.010G127400.1.A
Gene IDs: Gorai.010G127400
cotton name: PTHR31388:SF22 - PEROXIDASE 22-RELATED (1 of 2)
OG IDs: 93442500
Best-hit-arabi-ID: AT5G06720.1
Best-hit-arabi-name: peroxidase 2
Gorai.010G127400.1.A
PCC=0.799, p=0.0
mRNA Profile
Protein Expression Profile
DPA

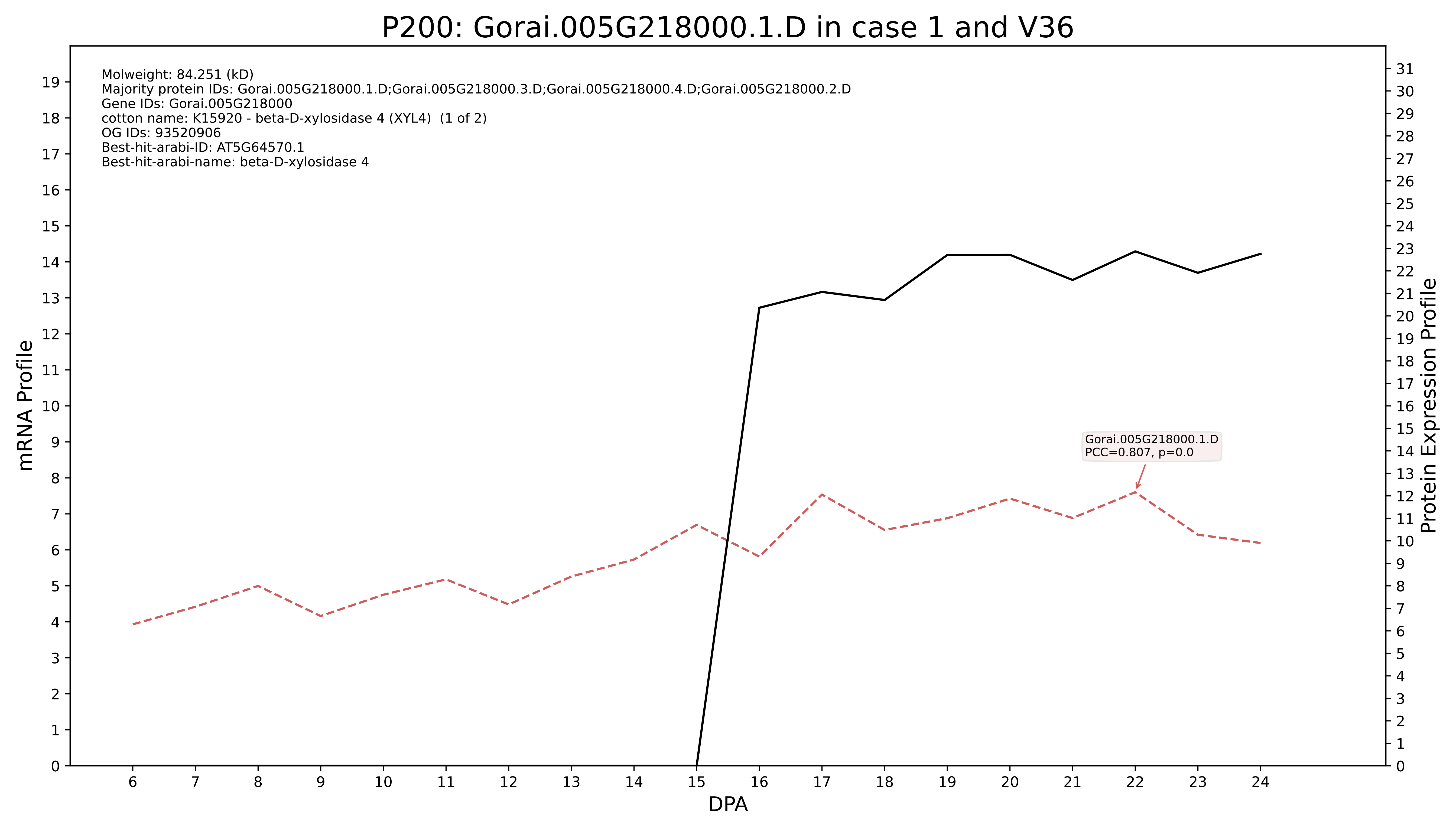

P200: Gorai.005G218000.1.D in case 1 and V36
Molweight: 84.251 (kD)
Majority protein IDs: Gorai.005G218000.1.D;Gorai.005G218000.3.D;Gorai.005G218000.4.D;Gorai.005G218000.2.D
Gene IDs: Gorai.005G218000
cotton name: K15920 - beta-D-xylosidase 4 (XYL4) (1 of 2)
OG IDs: 93520906
Best-hit-arabi-ID: AT5G64570.1
Best-hit-arabi-name: beta-D-xylosidase 4
Gorai.005G218000.1.D
PCC=0.807, p=0.0
mRNA Profile
Protein Expression Profile
DPA

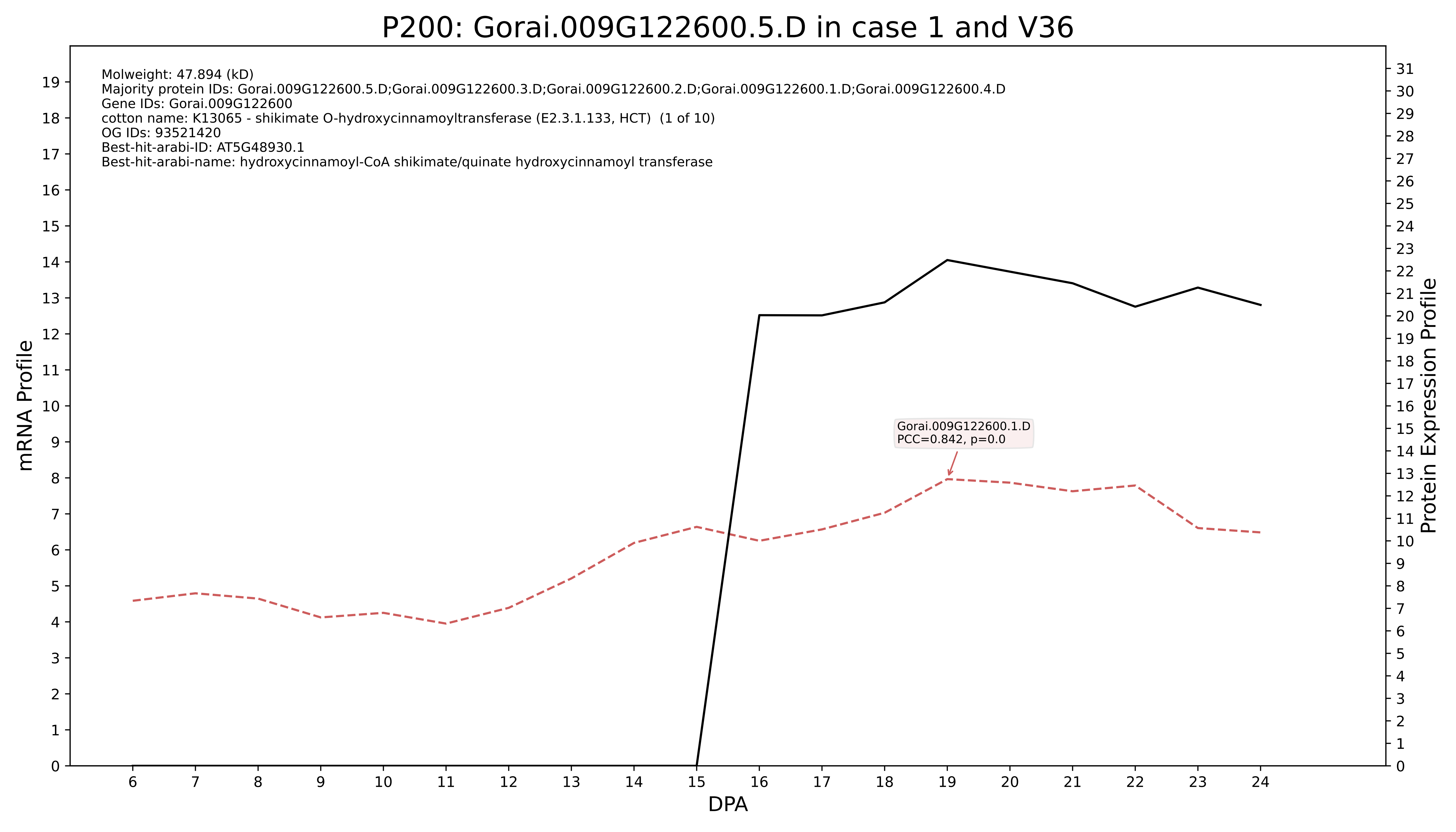
P200: Gorai.009G122600.5.D in case 1 and V36
Molweight: 47.894 (kD)
Majority protein IDs: Gorai.009G122600.5.D;Gorai.009G122600.3.D;Gorai.009G122600.2.D;Gorai.009G122600.1.D;Gorai.009G122600.4.D
Gene IDs: Gorai.009G122600
cotton name: K13065 - shikimate O-hydroxycinnamoyltransferase (E2.3.1.133, HCT) (1 of 10)
OG IDs: 93521420
Best-hit-arabi-ID: AT5G48930.1
Best-hit-arabi-name: hydroxycinnamoyl-CoA shikimate/quinate hydroxycinnamoyl transferase
Gorai.009G122600.1.D
PCC=0.842, p=0.0
mRNA Profile
Protein Expression Profile
DPA

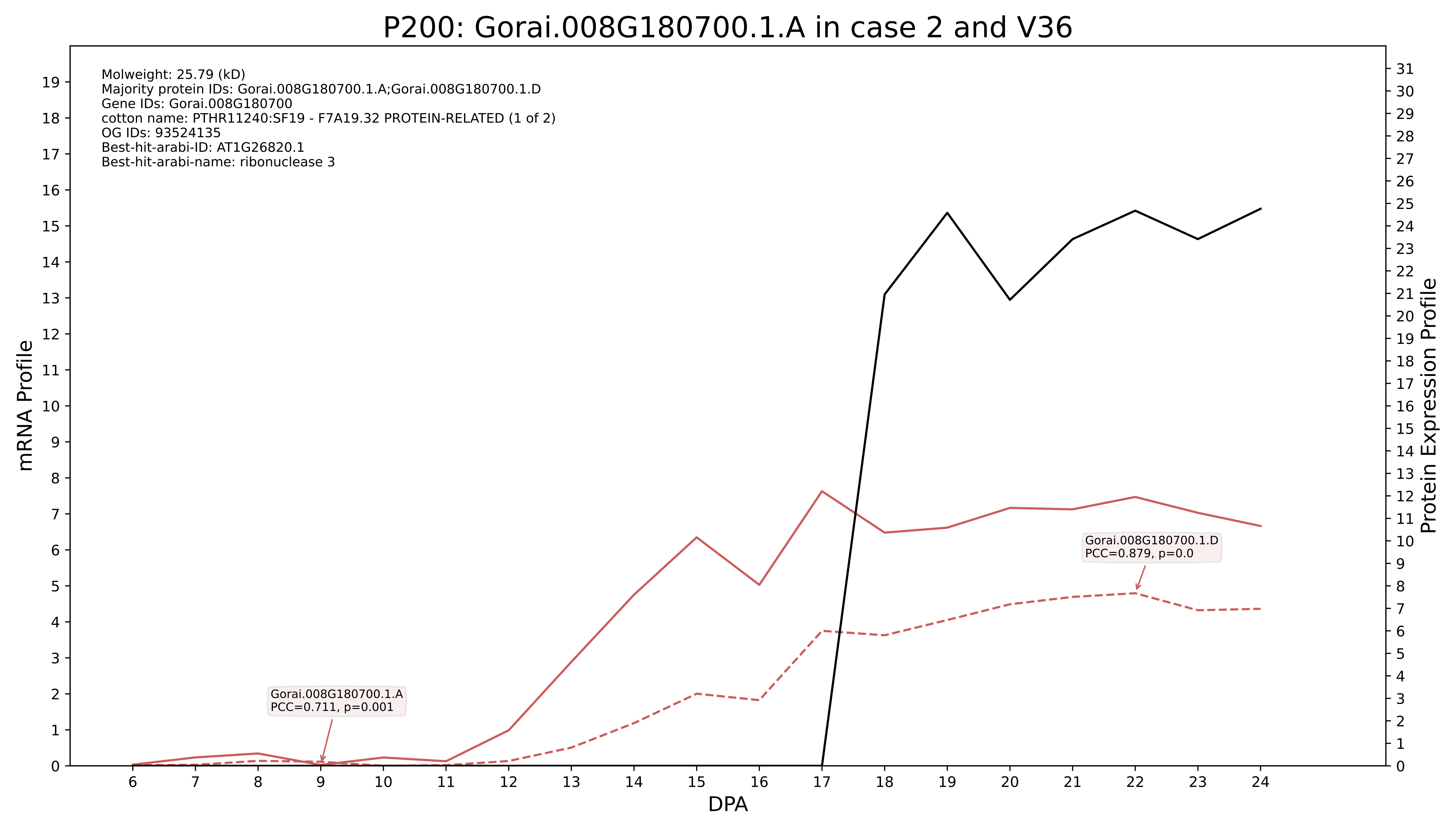
P200: Gorai.008G180700.1.A in case 2 and V36
Molweight: 25.79 (kD)
Majority protein IDs: Gorai.008G180700.1.A;Gorai.008G180700.1.D
Gene IDs: Gorai.008G180700
cotton name: PTHR11240:SF19 - F7A19.32 PROTEIN-RELATED (1 of 2)
OG IDs: 93524135
Best-hit-arabi-ID: AT1G26820.1
Best-hit-arabi-name: ribonuclease 3
Gorai.008G180700.1.D
PCC=0.879, p=0.0
Gorai.008G180700.1.A
PCC=0.711, p=0.001
mRNA Profile
Protein Expression Profile
DPA

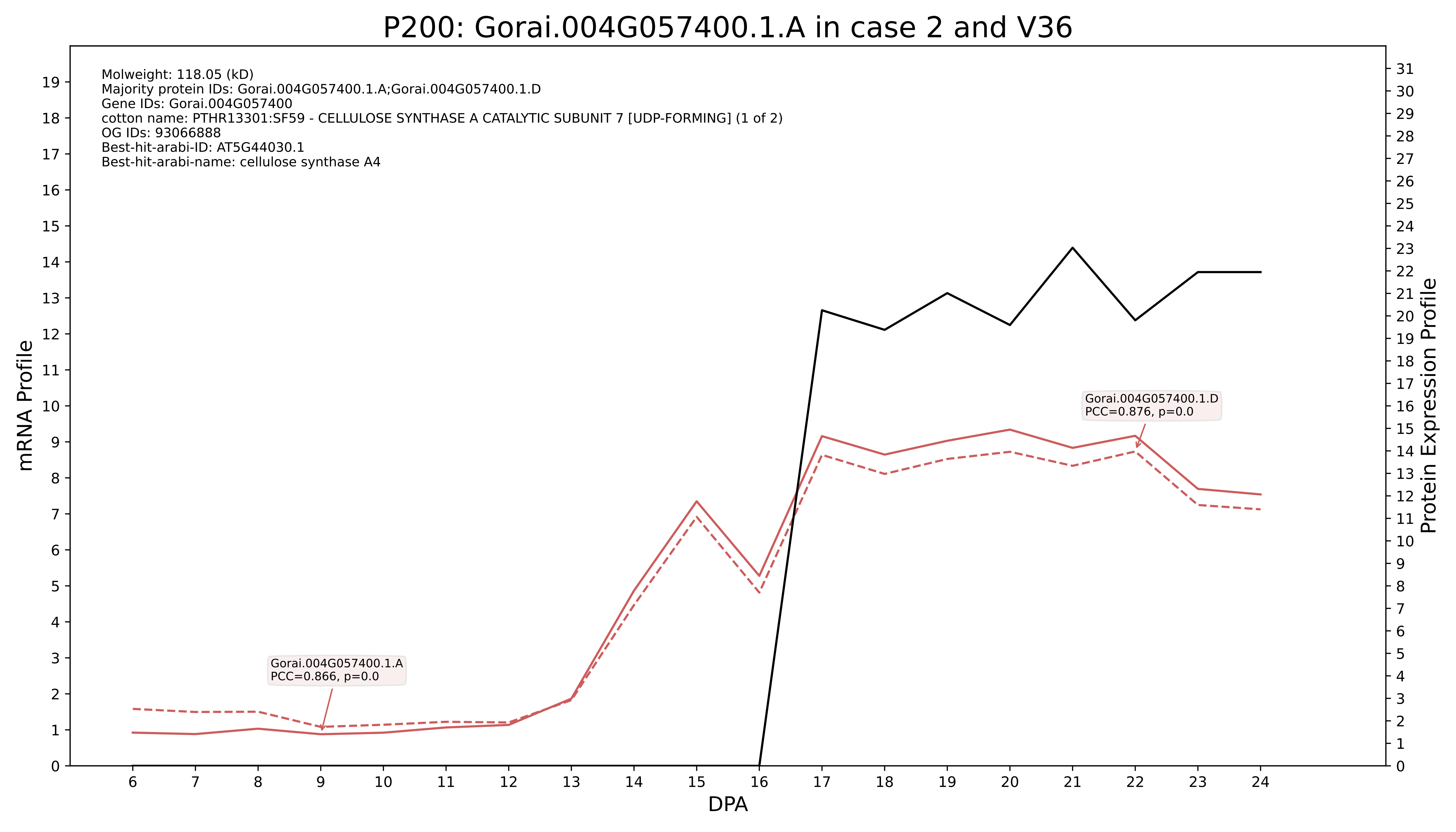

P200: Gorai.004G057400.1.A in case 2 and V36
Molweight: 118.05 (kD)
Majority protein IDs: Gorai.004G057400.1.A;Gorai.004G057400.1.D
Gene IDs: Gorai.004G057400
cotton name: PTHR13301:SF59 - CELLULOSE SYNTHASE A CATALYTIC SUBUNIT 7 [UDP-FORMING] (1 of 2)
OG IDs: 93066888
Best-hit-arabi-ID: AT5G44030.1
Best-hit-arabi-name: cellulose synthase A4
Gorai.004G057400.1.D
PCC=0.876, p=0.0
Gorai.004G057400.1.A
PCC=0.866, p=0.0
mRNA Profile
Protein Expression Profile
DPA

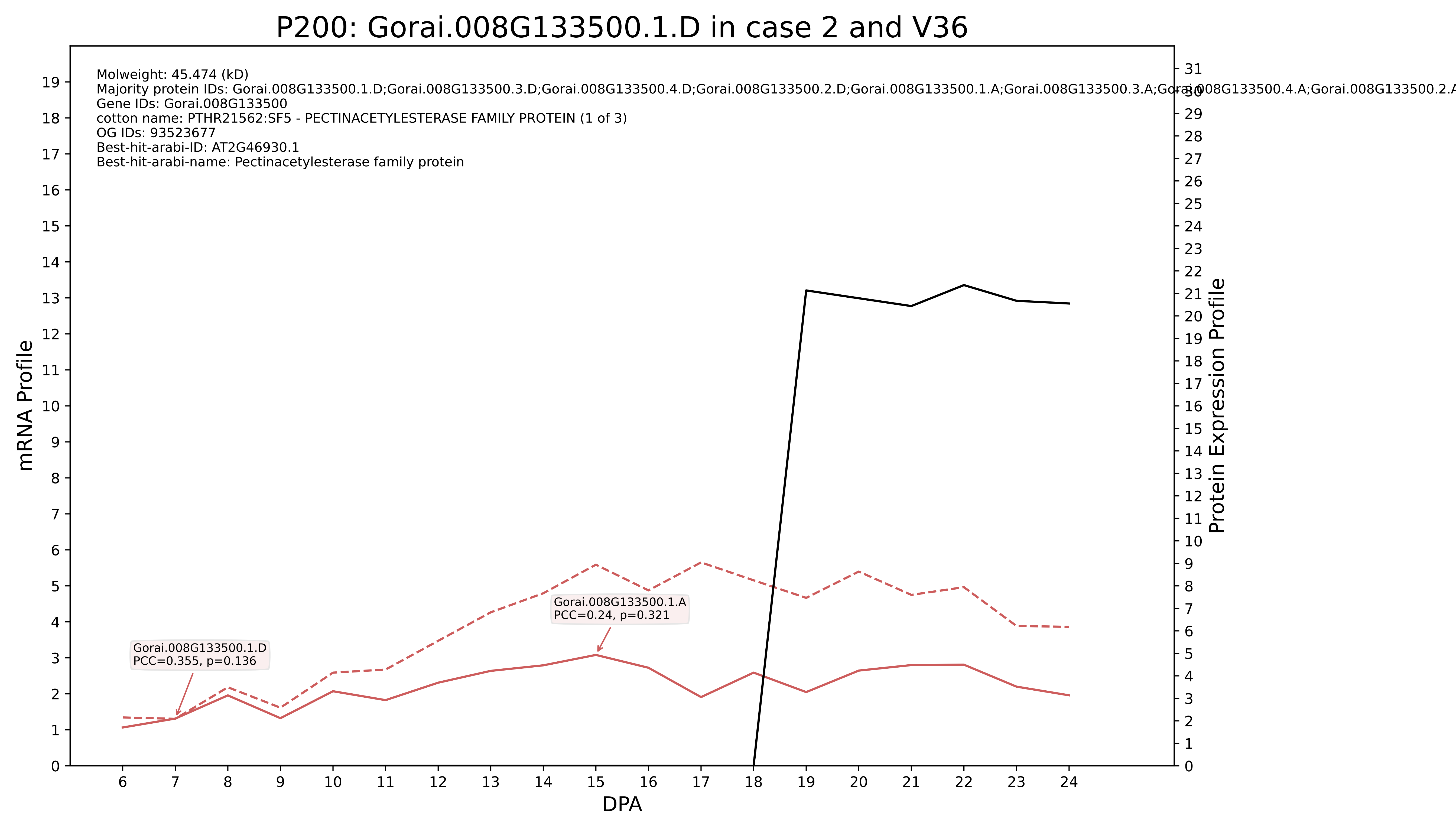
P200: Gorai.008G133500.1.D in case 2 and V36
Molweight: 45.474 (kD)
Majority protein IDs: Gorai.008G133500.1.D;Gorai.008G133500.3.D;Gorai.008G133500.4.D;Gorai.008G133500.2.D;Gorai.008G133500.1.A;Gorai.008G133500.3.A;Gorai.008G133500.4.A;Gorai.008G133500.2.A
Gene IDs: Gorai.008G133500
cotton name: PTHR21562:SF5 - PECTINACETYLESTERASE FAMILY PROTEIN (1 of 3)
OG IDs: 93523677
Best-hit-arabi-ID: AT2G46930.1
Best-hit-arabi-name: Pectinacetylesterase family protein
Gorai.008G133500.1.A
PCC=0.24, p=0.321
Gorai.008G133500.1.D
PCC=0.355, p=0.136
mRNA Profile
Protein Expression Profile
DPA

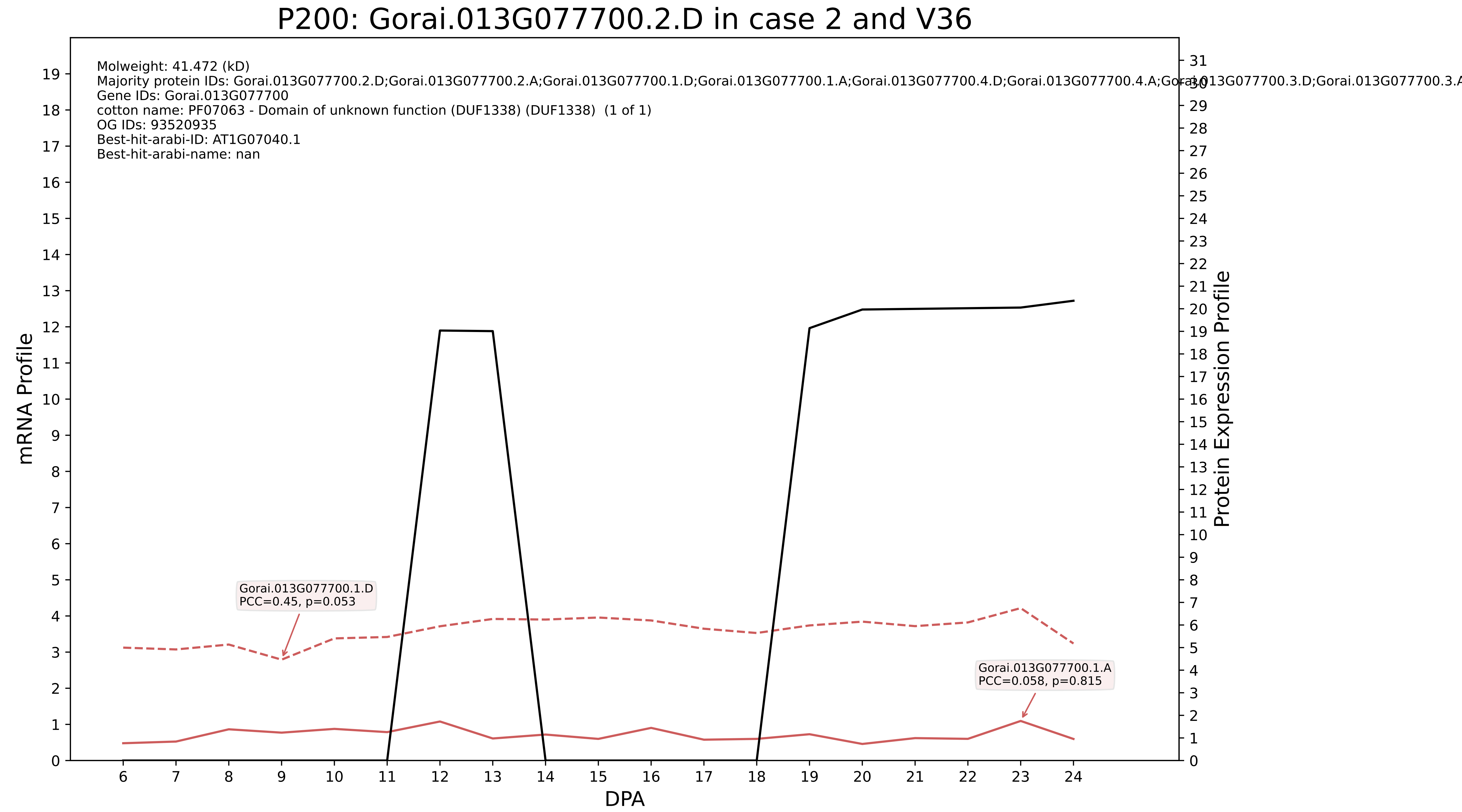
P200: Gorai.013G077700.2.D in case 2 and V36
Molweight: 41.472 (kD)
Majority protein IDs: Gorai.013G077700.2.D;Gorai.013G077700.2.A;Gorai.013G077700.1.D;Gorai.013G077700.1.A;Gorai.013G077700.4.D;Gorai.013G077700.4.A;Gorai.013G077700.3.D;Gorai.013G077700.3.A
Gene IDs: Gorai.013G077700
cotton name: PF07063 - Domain of unknown function (DUF1338) (DUF1338) (1 of 1)
OG IDs: 93520935
Best-hit-arabi-ID: AT1G07040.1
Best-hit-arabi-name: nan
Gorai.013G077700.1.D
PCC=0.45, p=0.053
Gorai.013G077700.1.A
PCC=0.058, p=0.815
mRNA Profile
Protein Expression Profile
DPA

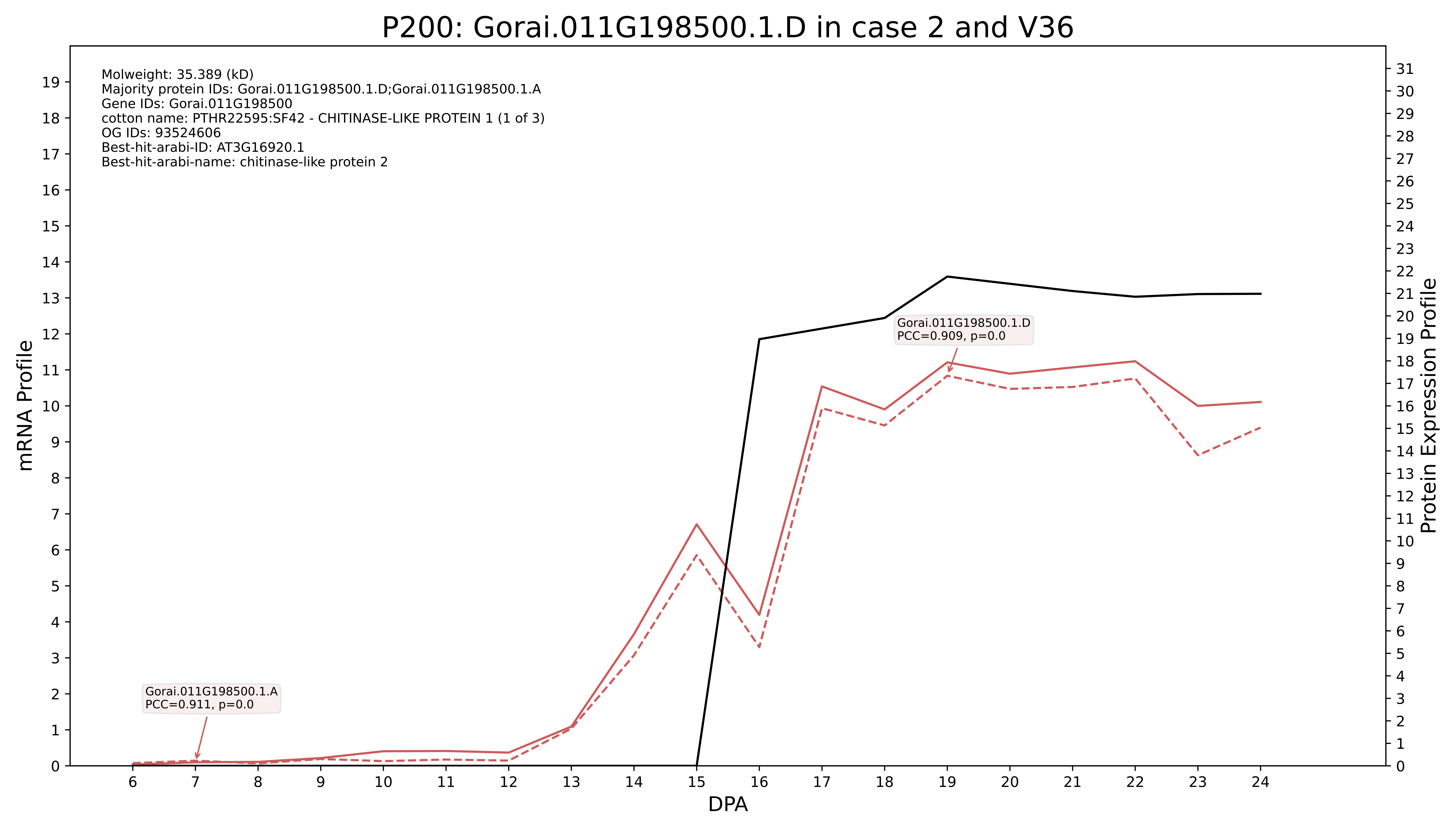
P200: Gorai.011G198500.1.D in case 2 and V36
Molweight: 35.389 (kD)
Majority protein IDs: Gorai.011G198500.1.D;Gorai.011G198500.1.A
Gene IDs: Gorai.011G198500
cotton name: PTHR22595:SF42 - CHITINASE-LIKE PROTEIN 1 (1 of 3)
OG IDs: 93524606
Best-hit-arabi-ID: AT3G16920.1
Best-hit-arabi-name: chitinase-like protein 2
Gorai.011G198500.1.D
PCC=0.909, p=0.0
Gorai.011G198500.1.A
PCC=0.911, p=0.0
mRNA Profile
Protein Expression Profile
DPA

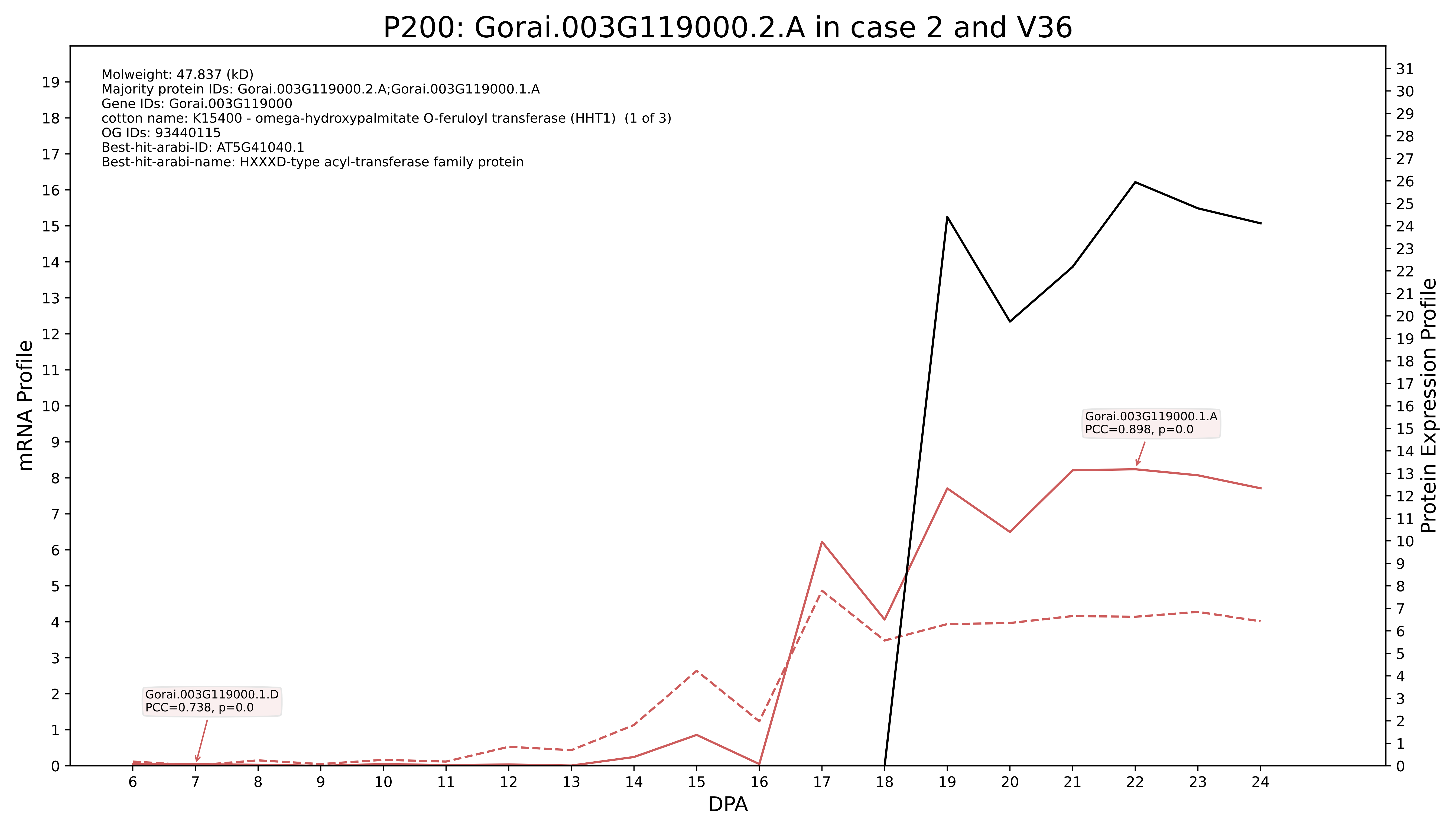
P200: Gorai.003G119000.2.A in case 2 and V36
Molweight: 47.837 (kD)
Majority protein IDs: Gorai.003G119000.2.A;Gorai.003G119000.1.A
Gene IDs: Gorai.003G119000
cotton name: K15400 - omega-hydroxypalmitate O-feruloyl transferase (HHT1) (1 of 3)
OG IDs: 93440115
Best-hit-arabi-ID: AT5G41040.1
Best-hit-arabi-name: HXXXD-type acyl-transferase family protein
Gorai.003G119000.1.A
PCC=0.898, p=0.0
Gorai.003G119000.1.D
PCC=0.738, p=0.0
mRNA Profile
Protein Expression Profile
DPA

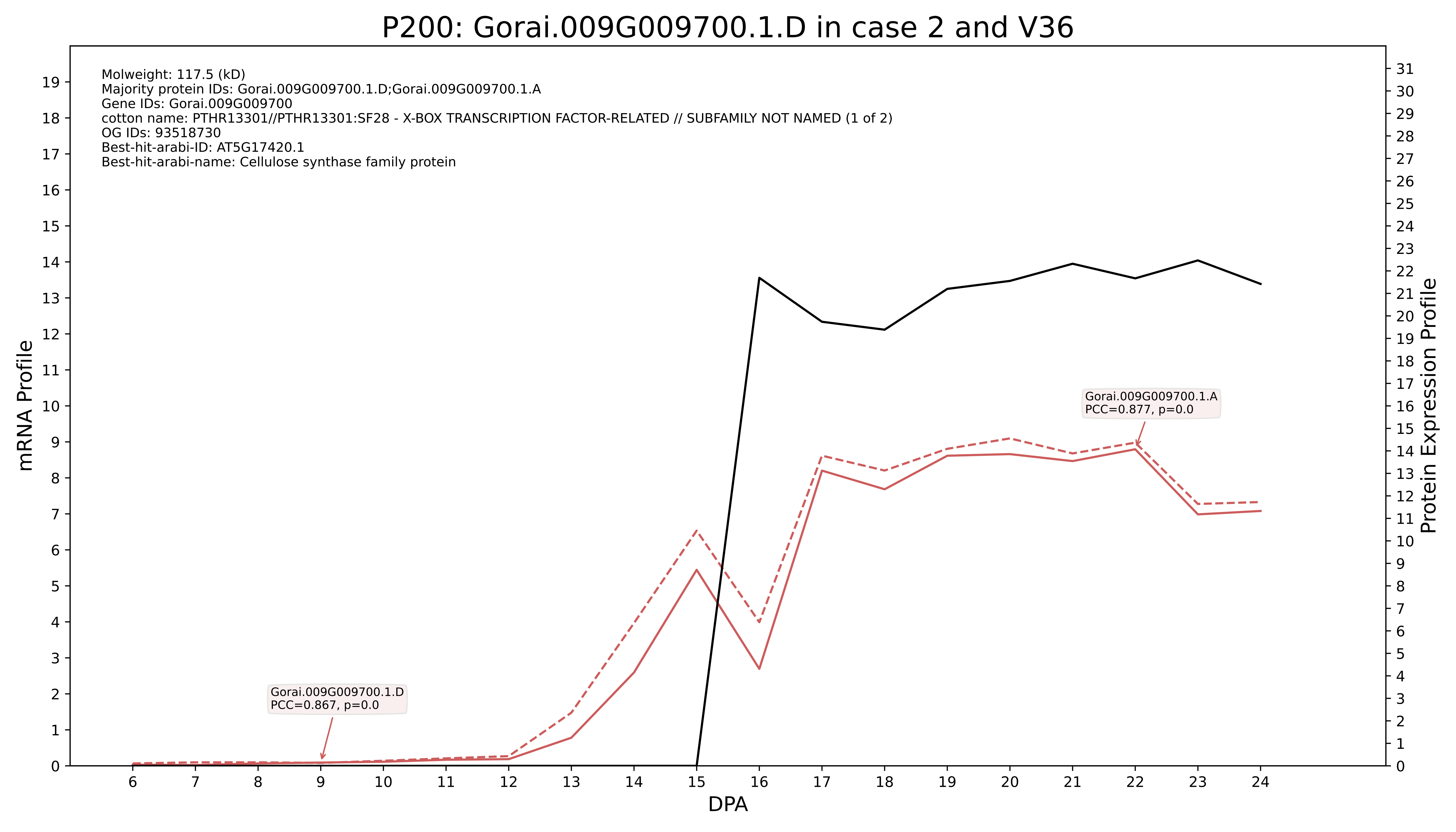

P200: Gorai.009G009700.1.D in case 2 and V36
Molweight: 117.5 (kD)
Majority protein IDs: Gorai.009G009700.1.D;Gorai.009G009700.1.A
Gene IDs: Gorai.009G009700
cotton name: PTHR13301//PTHR13301:SF28 - X-BOX TRANSCRIPTION FACTOR-RELATED // SUBFAMILY NOT NAMED (1 of 2)
OG IDs: 93518730
Best-hit-arabi-ID: AT5G17420.1
Best-hit-arabi-name: Cellulose synthase family protein
Gorai.009G009700.1.A
PCC=0.877, p=0.0
Gorai.009G009700.1.D
PCC=0.867, p=0.0
mRNA Profile
Protein Expression Profile
DPA

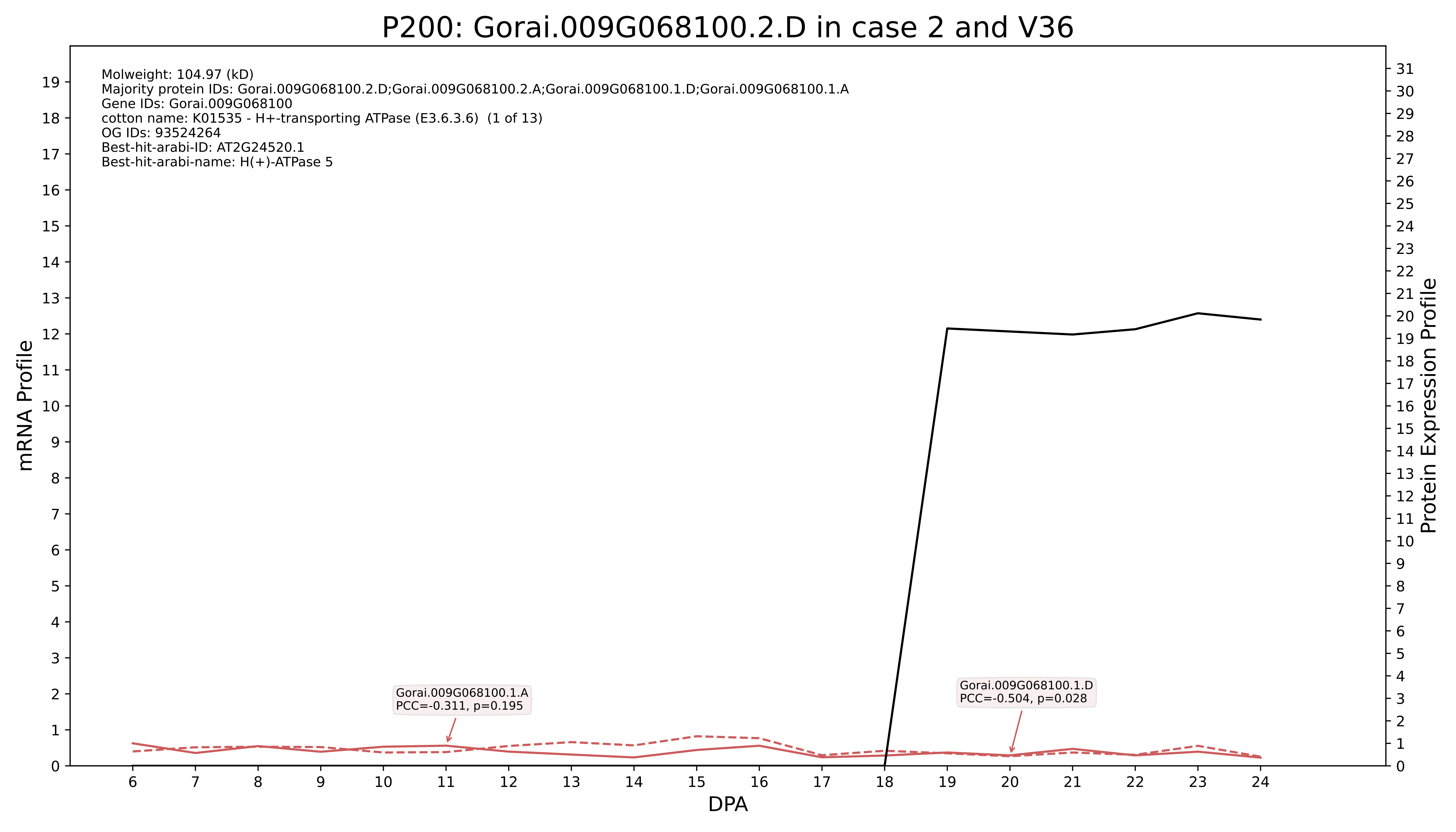

P200: Gorai.009G068100.2.D in case 2 and V36
Molweight: 104.97 (kD)
Majority protein IDs: Gorai.009G068100.2.D;Gorai.009G068100.2.A;Gorai.009G068100.1.D;Gorai.009G068100.1.A
Gene IDs: Gorai.009G068100
cotton name: K01535 - H+-transporting ATPase (E3.6.3.6) (1 of 13)
OG IDs: 93524264
Best-hit-arabi-ID: AT2G24520.1
Best-hit-arabi-name: H(+)-ATPase 5
Gorai.009G068100.1.A
PCC=-0.311, p=0.195
Gorai.009G068100.1.D
PCC=-0.504, p=0.028
mRNA Profile
Protein Expression Profile
DPA

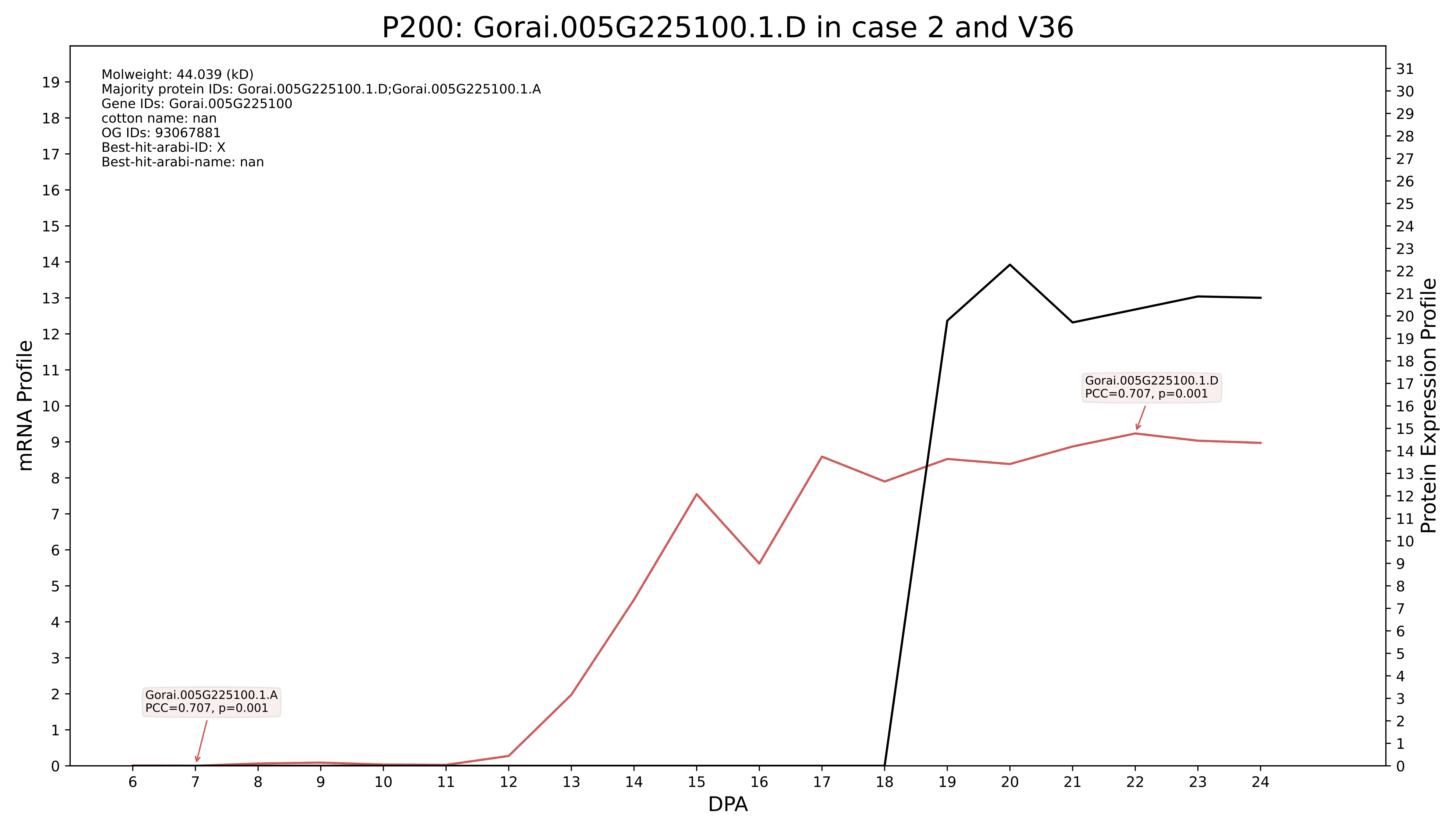
P200: Gorai.005G225100.1.D in case 2 and V36
Molweight: 44.039 (kD)
Majority protein IDs: Gorai.005G225100.1.D;Gorai.005G225100.1.A
Gene IDs: Gorai.005G225100
cotton name: nan
OG IDs: 93067881
Best-hit-arabi-ID: X
Best-hit-arabi-name: nan
Gorai.005G225100.1.D
PCC=0.707, p=0.001
Gorai.005G225100.1.A
PCC=0.707, p=0.001
mRNA Profile
Protein Expression Profile
DPA

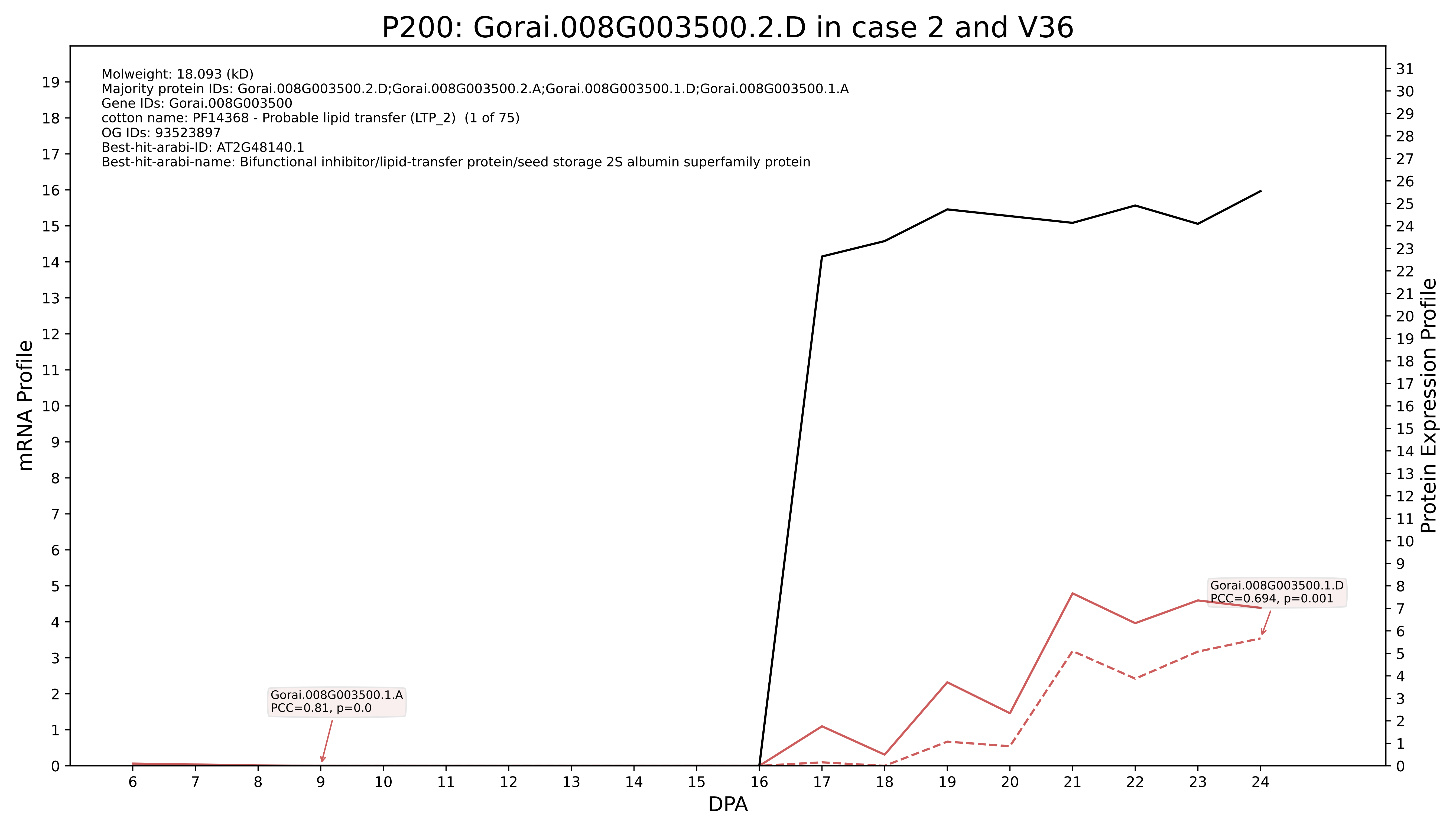
P200: Gorai.008G003500.2.D in case 2 and V36
Molweight: 18.093 (kD)
Majority protein IDs: Gorai.008G003500.2.D;Gorai.008G003500.2.A;Gorai.008G003500.1.D;Gorai.008G003500.1.A
Gene IDs: Gorai.008G003500
cotton name: PF14368 - Probable lipid transfer (LTP_2) (1 of 75)
OG IDs: 93523897
Best-hit-arabi-ID: AT2G48140.1
Best-hit-arabi-name: Bifunctional inhibitor/lipid-transfer protein/seed storage 2S albumin superfamily protein
Gorai.008G003500.1.D
PCC=0.694, p=0.001
Gorai.008G003500.1.A
PCC=0.81, p=0.0
mRNA Profile
Protein Expression Profile
DPA

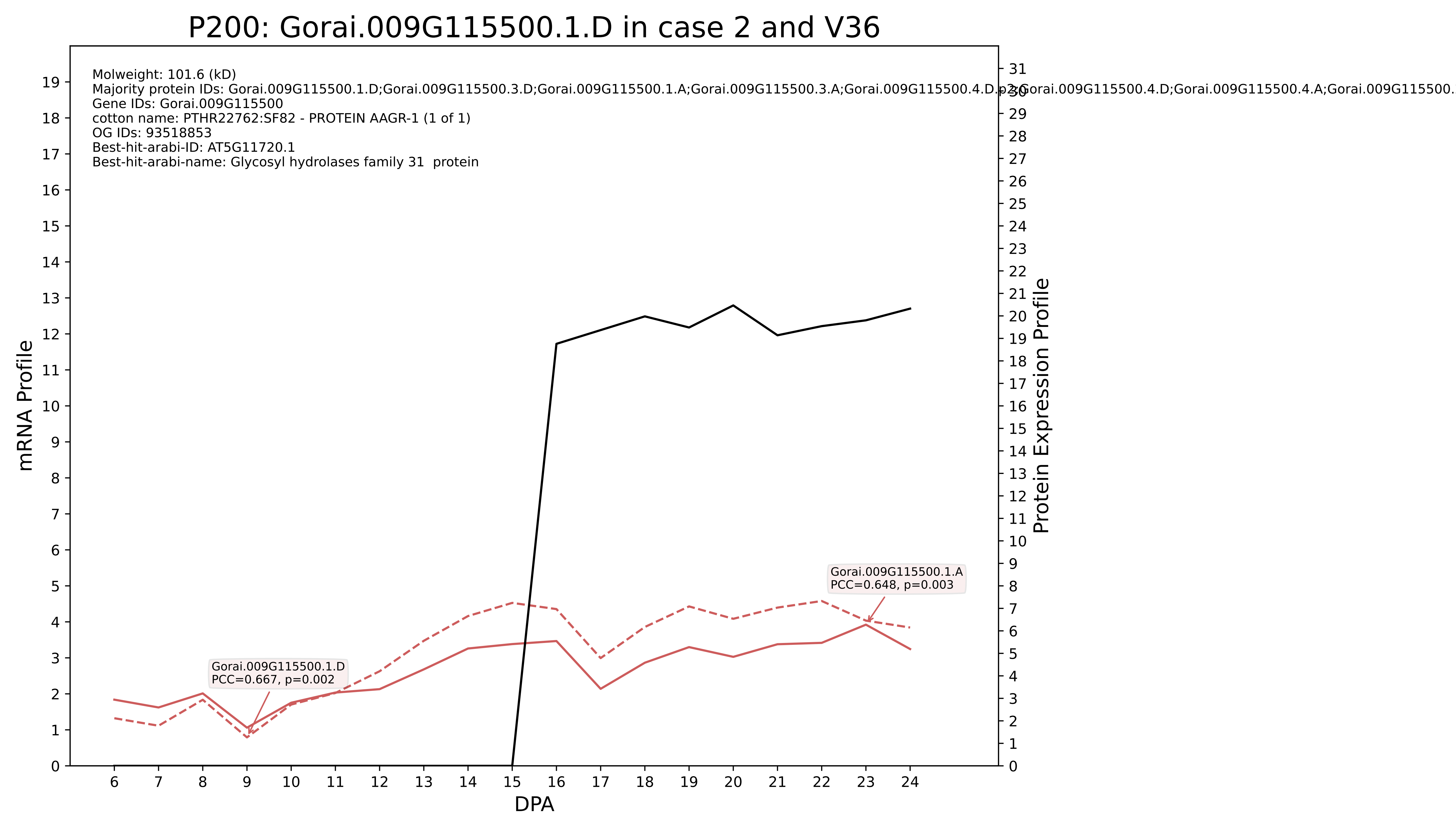

P200: Gorai.009G115500.1.D in case 2 and V36
Molweight: 101.6 (kD)
Majority protein IDs: Gorai.009G115500.1.D;Gorai.009G115500.3.D;Gorai.009G115500.1.A;Gorai.009G115500.3.A;Gorai.009G115500.4.D.p2;Gorai.009G115500.4.D;Gorai.009G115500.4.A;Gorai.009G115500.
Gene IDs: Gorai.009G115500
cotton name: PTHR22762:SF82 - PROTEIN AAGR-1 (1 of 1)
OG IDs: 93518853
Best-hit-arabi-ID: AT5G11720.1
Best-hit-arabi-name: Glycosyl hydrolases family 31 protein
Gorai.009G115500.1.D
PCC=0.667, p=0.002
Gorai.009G115500.1.A
PCC=0.648, p=0.003
mRNA Profile
Protein Expression Profile
DPA

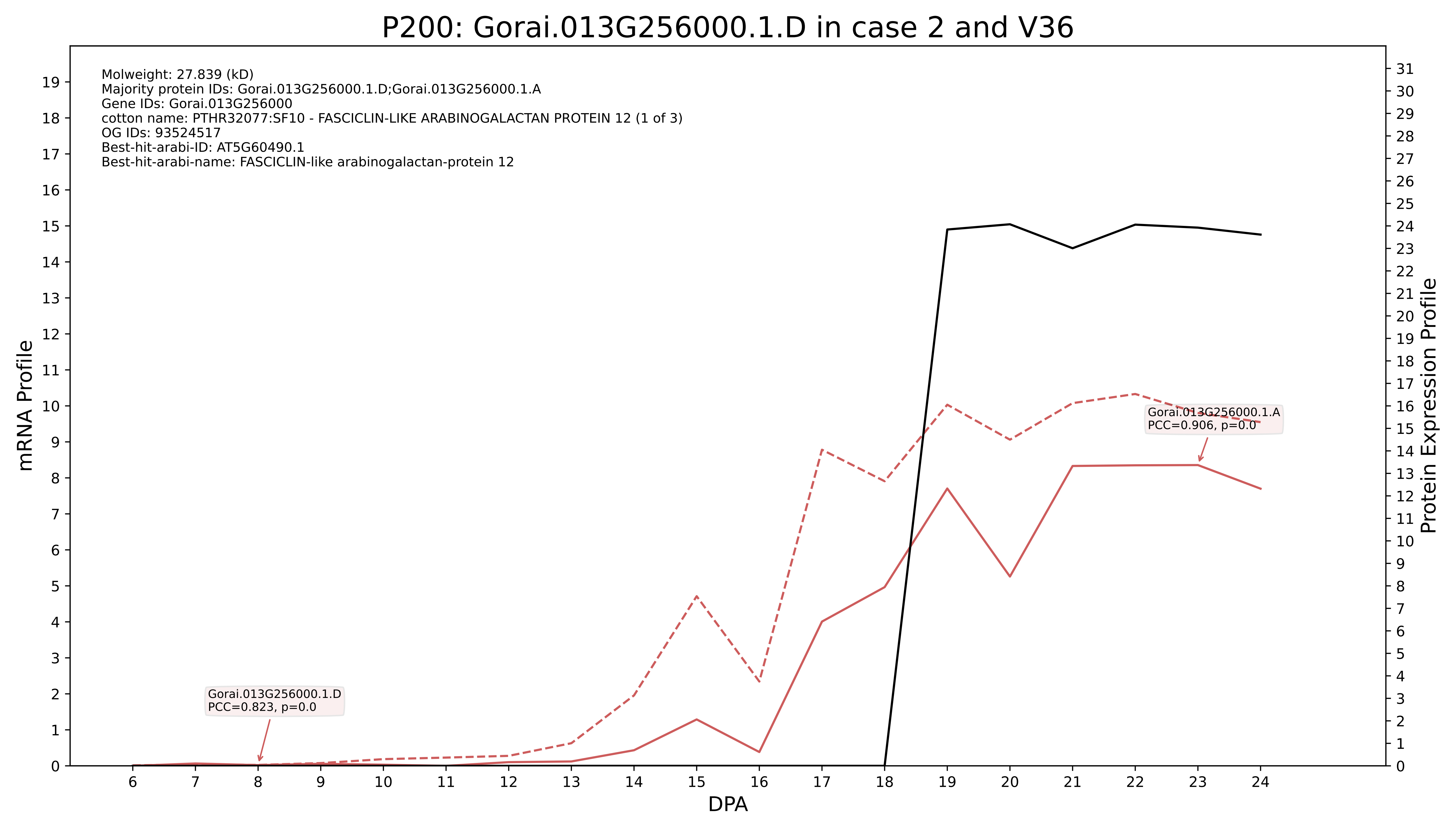
P200: Gorai.013G256000.1.D in case 2 and V36
Molweight: 27.839 (kD)
Majority protein IDs: Gorai.013G256000.1.D;Gorai.013G256000.1.A
Gene IDs: Gorai.013G256000
cotton name: PTHR32077:SF10 - FASCICLIN-LIKE ARABINOGALACTAN PROTEIN 12 (1 of 3)
OG IDs: 93524517
Best-hit-arabi-ID: AT5G60490.1
Best-hit-arabi-name: FASCICLIN-like arabinogalactan-protein 12
Gorai.013G256000.1.A
PCC=0.906, p=0.0
Gorai.013G256000.1.D
PCC=0.823, p=0.0
mRNA Profile
Protein Expression Profile
DPA

### P200: Gorai.013G003200.1.D in case 2 and V36

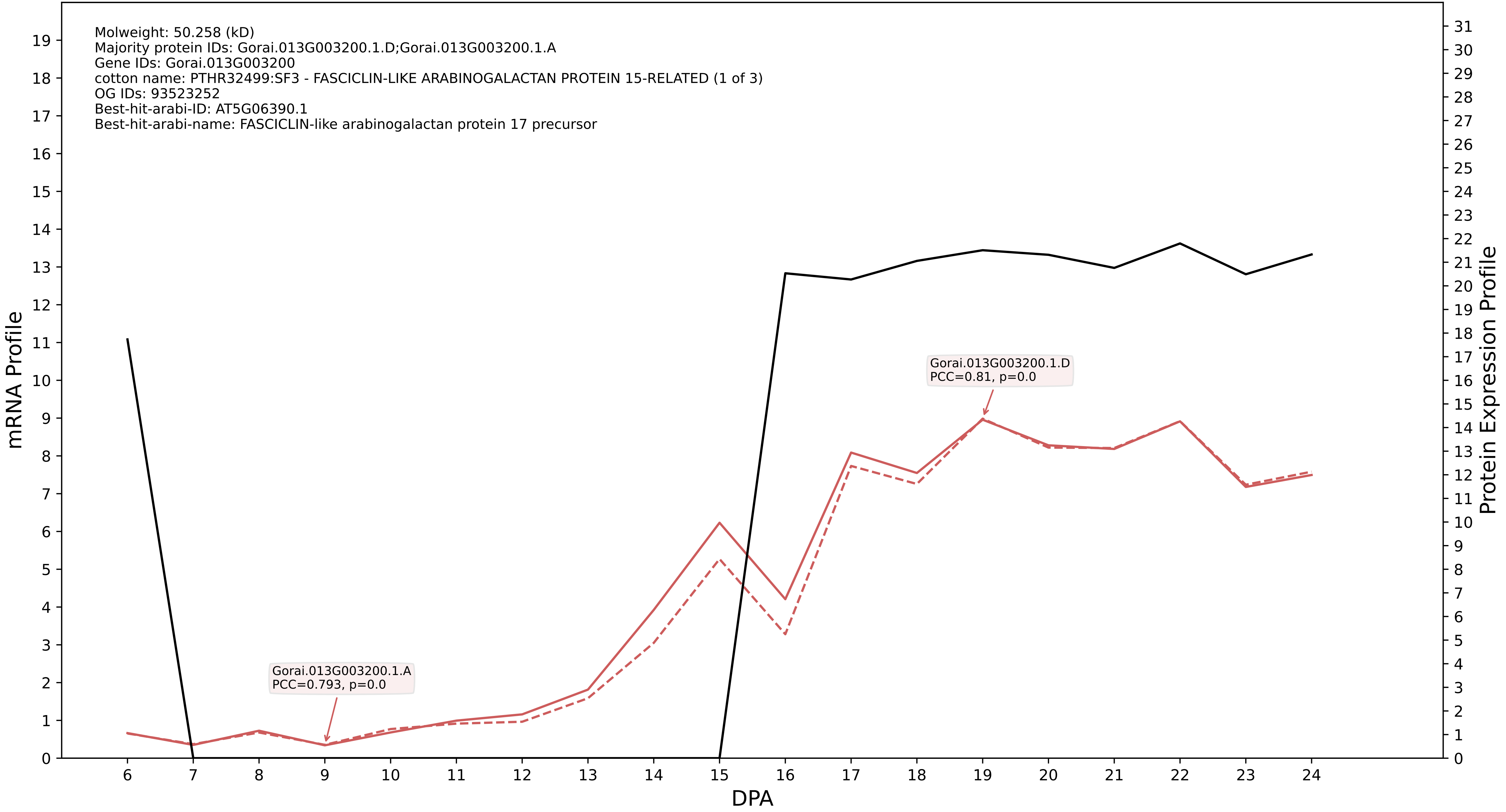

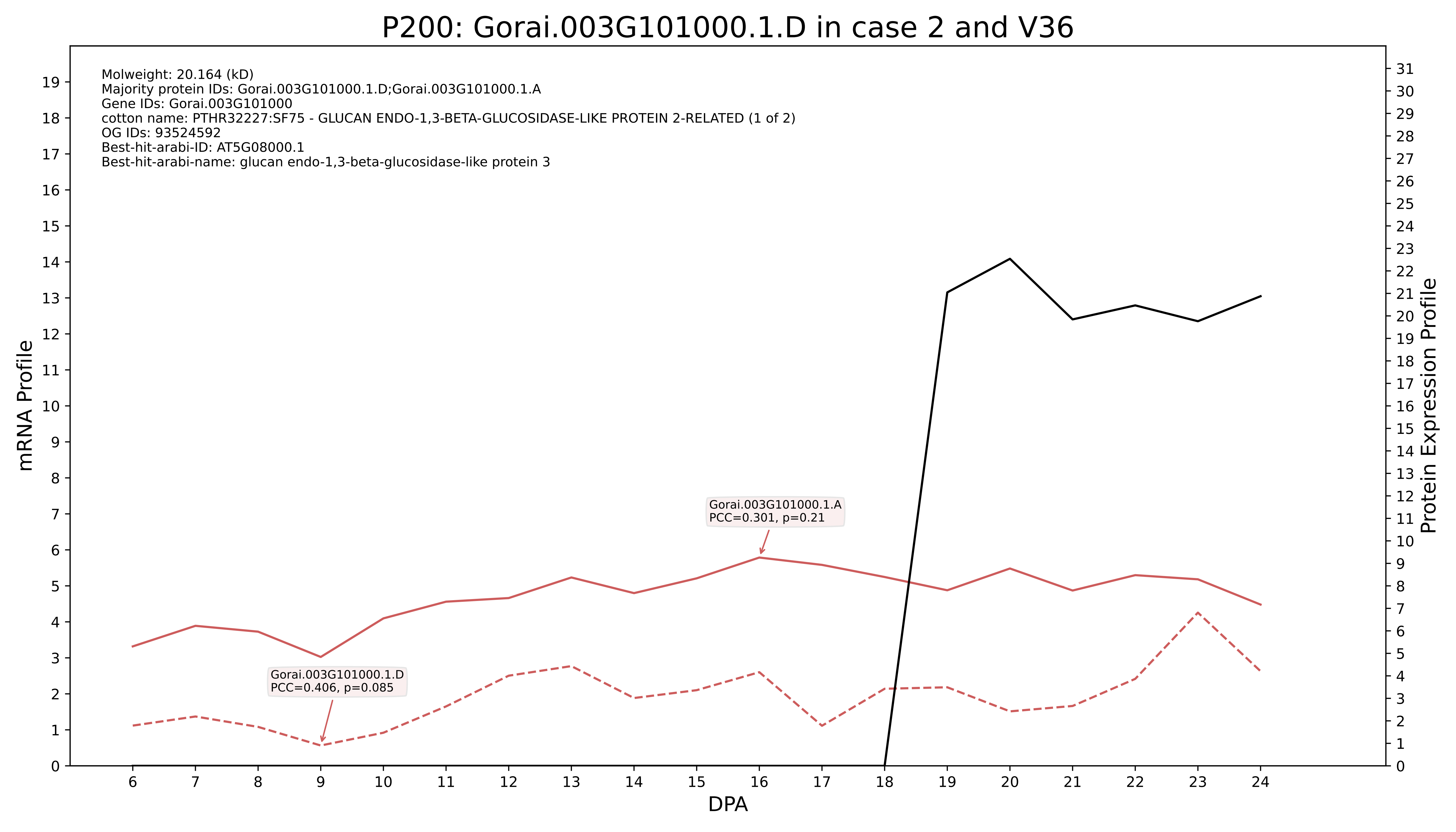
P200: Gorai.003G101000.1.D in case 2 and V36
Molweight: 20.164 (kD)
Majority protein IDs: Gorai.003G101000.1.D;Gorai.003G101000.1.A
Gene IDs: Gorai.003G101000
cotton name: PTHR32227:SF75 - GLUCAN ENDO-1,3-BETA-GLUCOSIDASE-LIKE PROTEIN 2-RELATED (1 of 2)
OG IDs: 93524592
Best-hit-arabi-ID: AT5G08000.1
Best-hit-arabi-name: glucan endo-1,3-beta-glucosidase-like protein 3
Gorai.003G101000.1.A
PCC=0.301, p=0.21
Gorai.003G101000.1.D
PCC=0.406, p=0.085
mRNA Profile
Protein Expression Profile
DPA

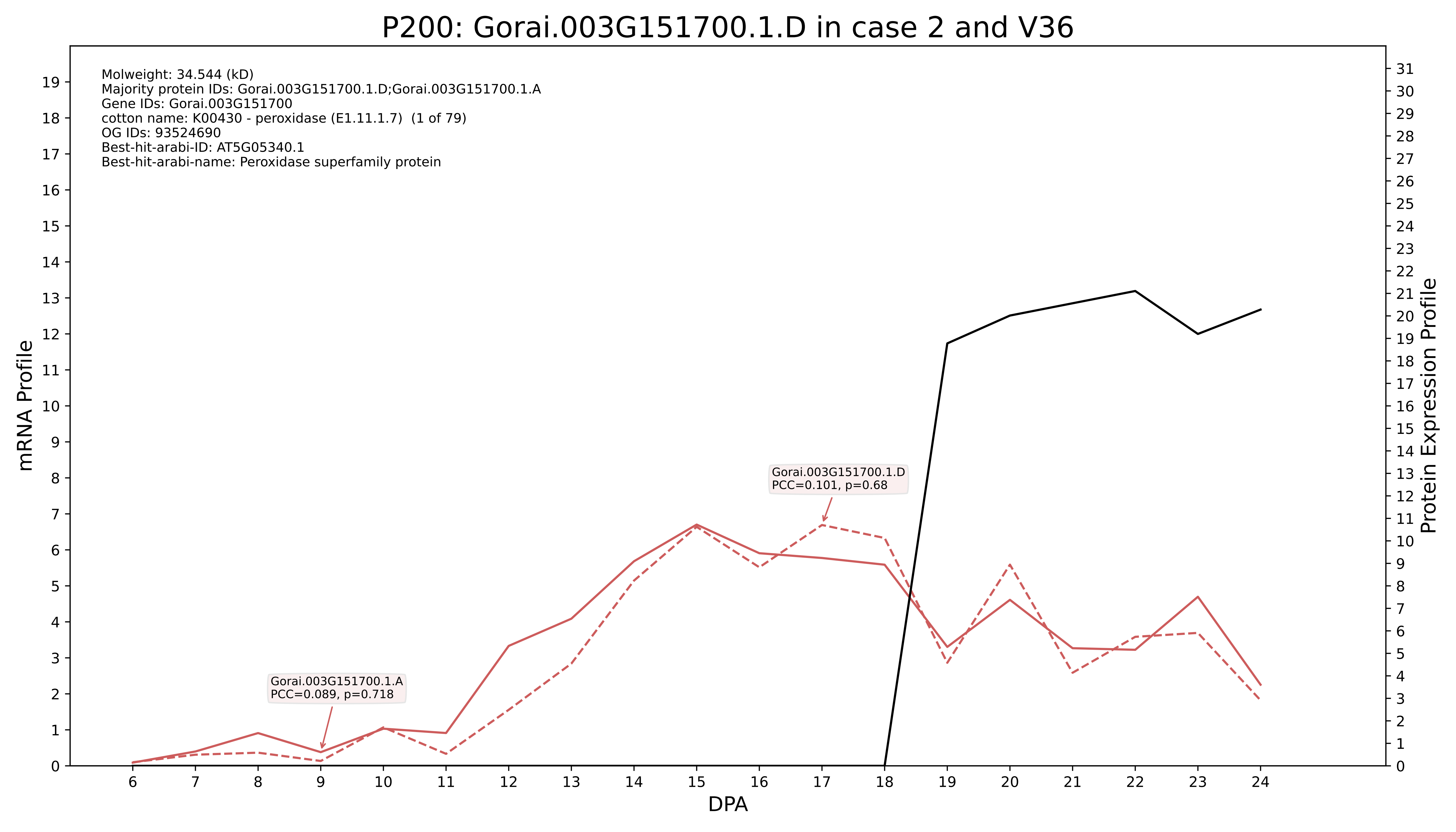
P200: Gorai.003G151700.1.D in case 2 and V36
Molweight: 34.544 (kD)
Majority protein IDs: Gorai.003G151700.1.D;Gorai.003G151700.1.A
Gene IDs: Gorai.003G151700
cotton name: K00430 - peroxidase (E1.11.1.7) (1 of 79)
OG IDs: 93524690
Best-hit-arabi-ID: AT5G05340.1
Best-hit-arabi-name: Peroxidase superfamily protein
Gorai.003G151700.1.D
PCC=0.101, p=0.68
Gorai.003G151700.1.A
PCC=0.089, p=0.718
mRNA Profile
Protein Expression Profile
DPA

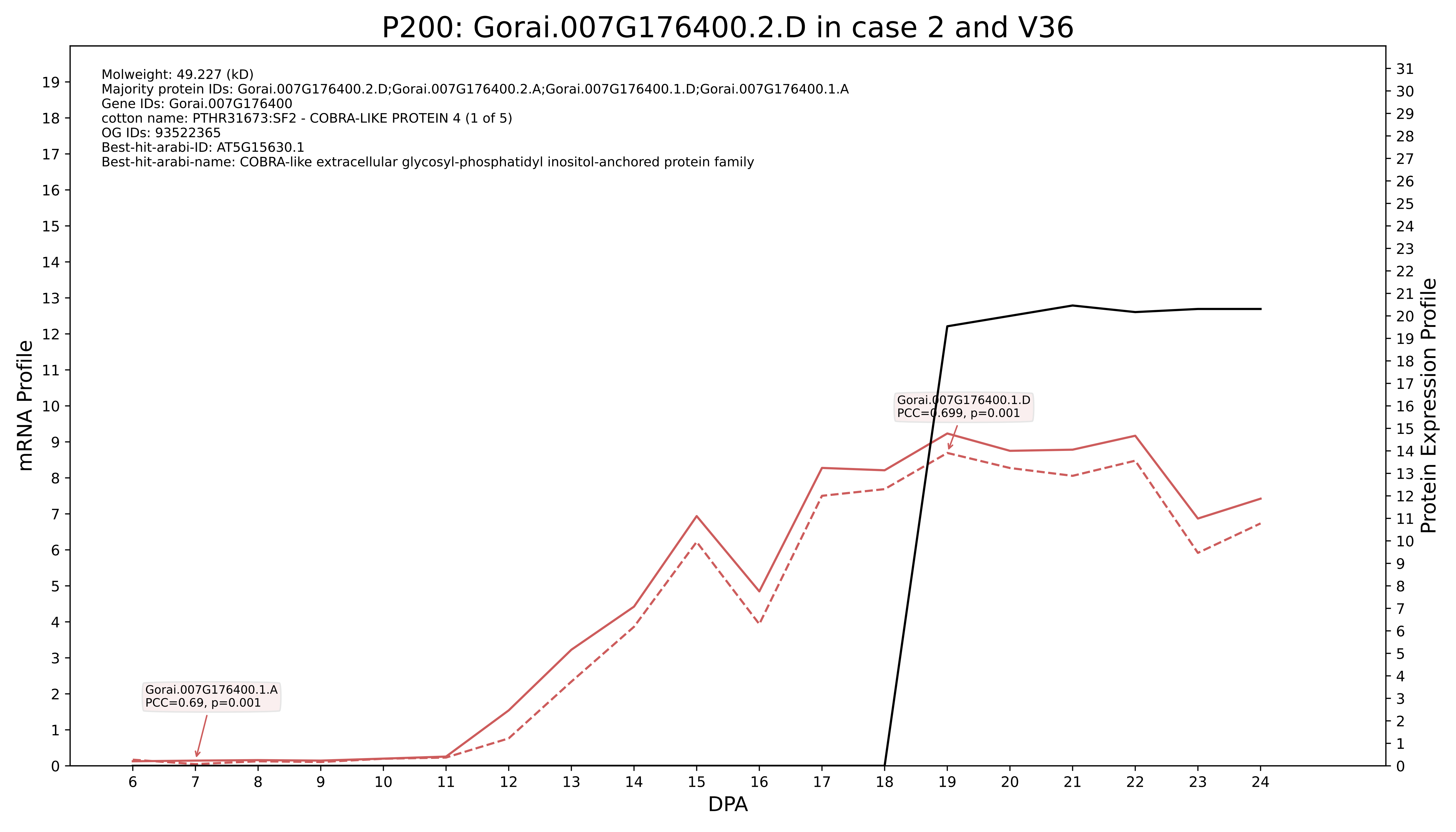

P200: Gorai.007G176400.2.D in case 2 and V36
Molweight: 49.227 (kD)
Majority protein IDs: Gorai.007G176400.2.D;Gorai.007G176400.2.A;Gorai.007G176400.1.D;Gorai.007G176400.1.A
Gene IDs: Gorai.007G176400
cotton name: PTHR31673:SF2 - COBRA-LIKE PROTEIN 4 (1 of 5)
OG IDs: 93522365
Best-hit-arabi-ID: AT5G15630.1
Best-hit-arabi-name: COBRA-like extracellular glycosyl-phosphatidyl inositol-anchored protein family
Gorai.007G176400.1.D
PCC=0.699, p=0.001
Gorai.007G176400.1.A
PCC=0.69, p=0.001
mRNA Profile
Protein Expression Profile
DPA

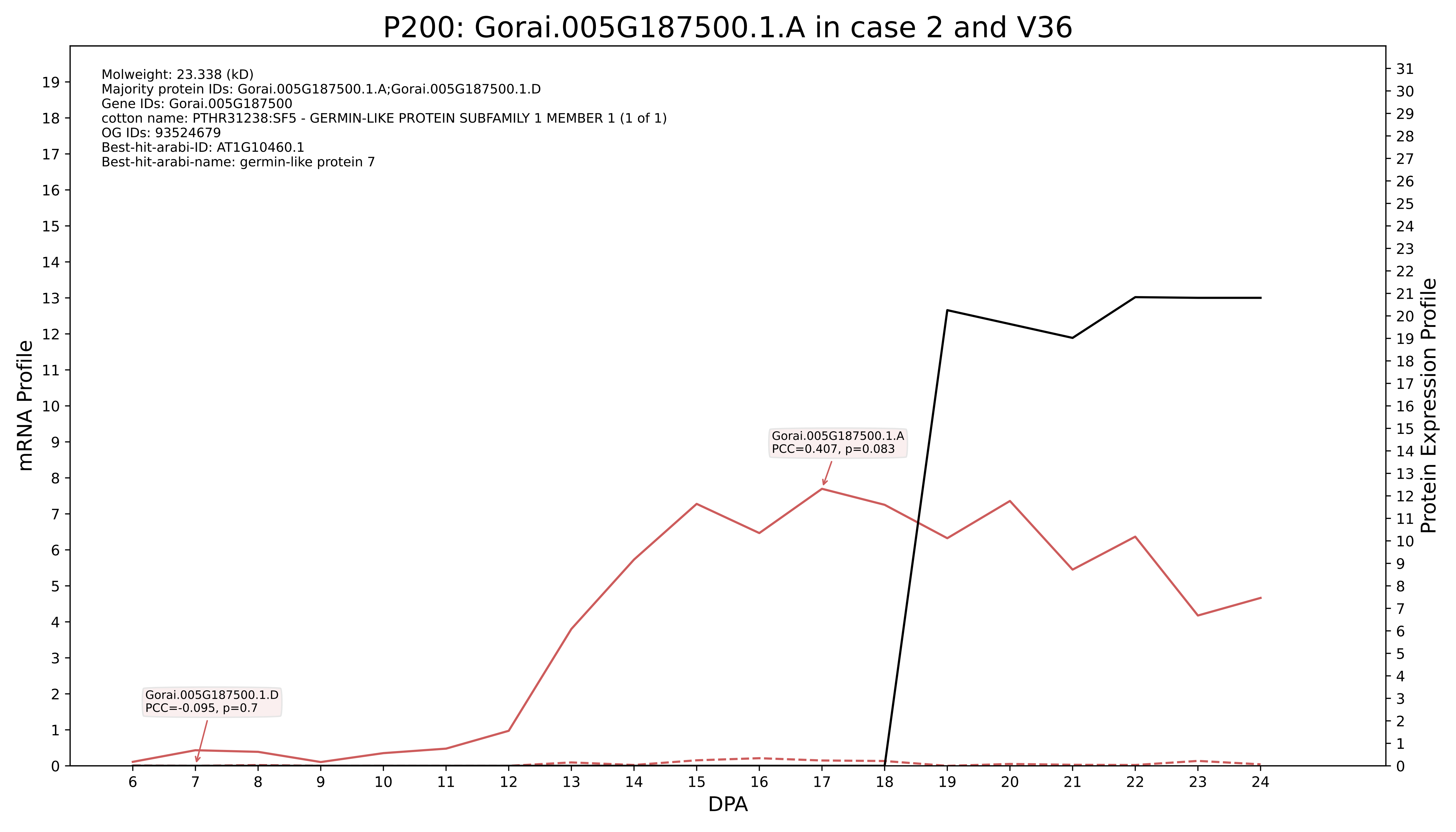

P200: Gorai.005G187500.1.A in case 2 and V36
Molweight: 23.338 (kD)
Majority protein IDs: Gorai.005G187500.1.A;Gorai.005G187500.1.D
Gene IDs: Gorai.005G187500
cotton name: PTHR31238:SF5 - GERMIN-LIKE PROTEIN SUBFAMILY 1 MEMBER 1 (1 of 1)
OG IDs: 93524679
Best-hit-arabi-ID: AT1G10460.1
Best-hit-arabi-name: germin-like protein 7
Gorai.005G187500.1.A
PCC=0.407, p=0.083
Gorai.005G187500.1.D
PCC=-0.095, p=0.7
mRNA Profile
Protein Expression Profile
DPA

200: Gorai.007G262600.2.A in case 2 and V36

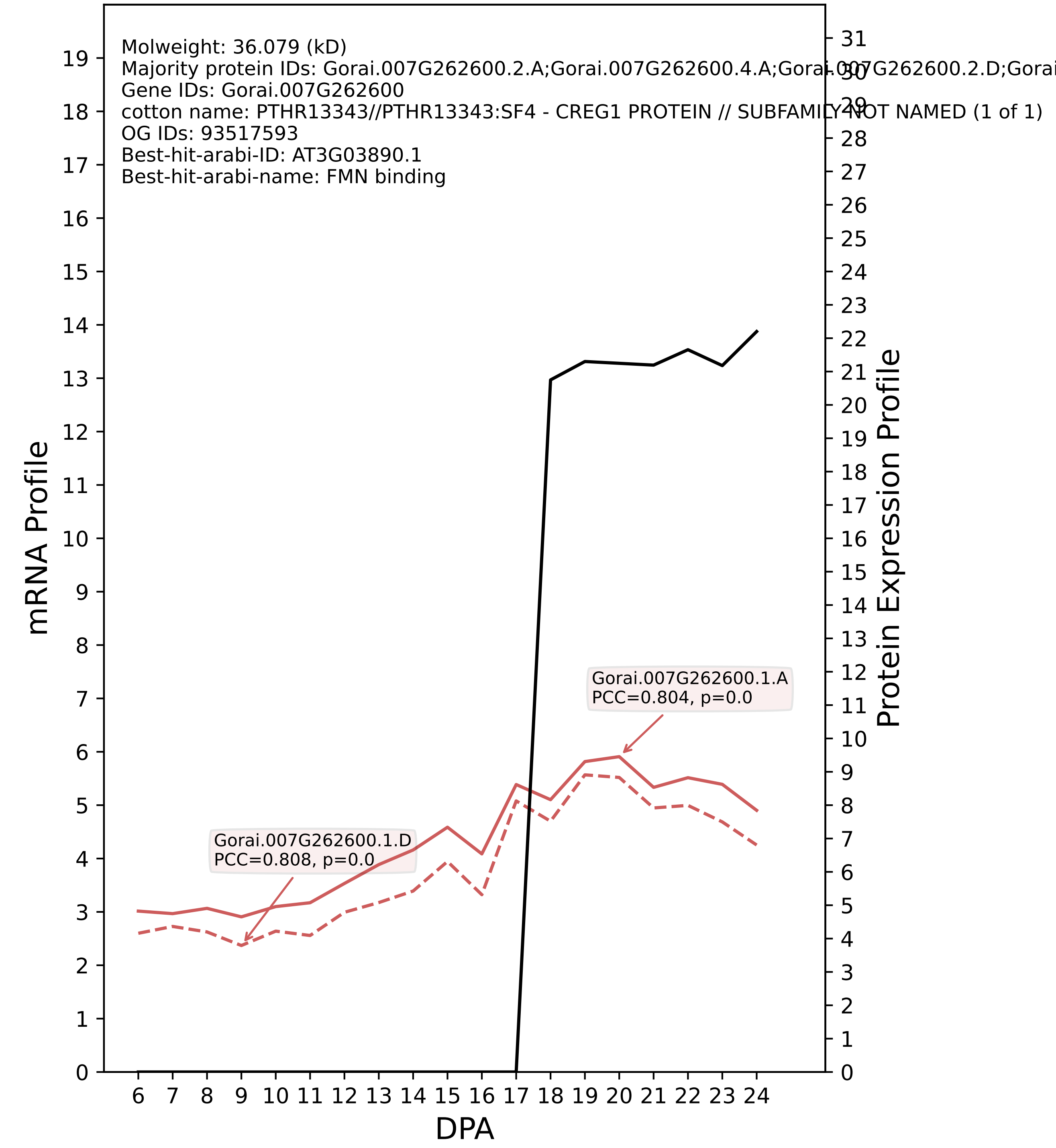

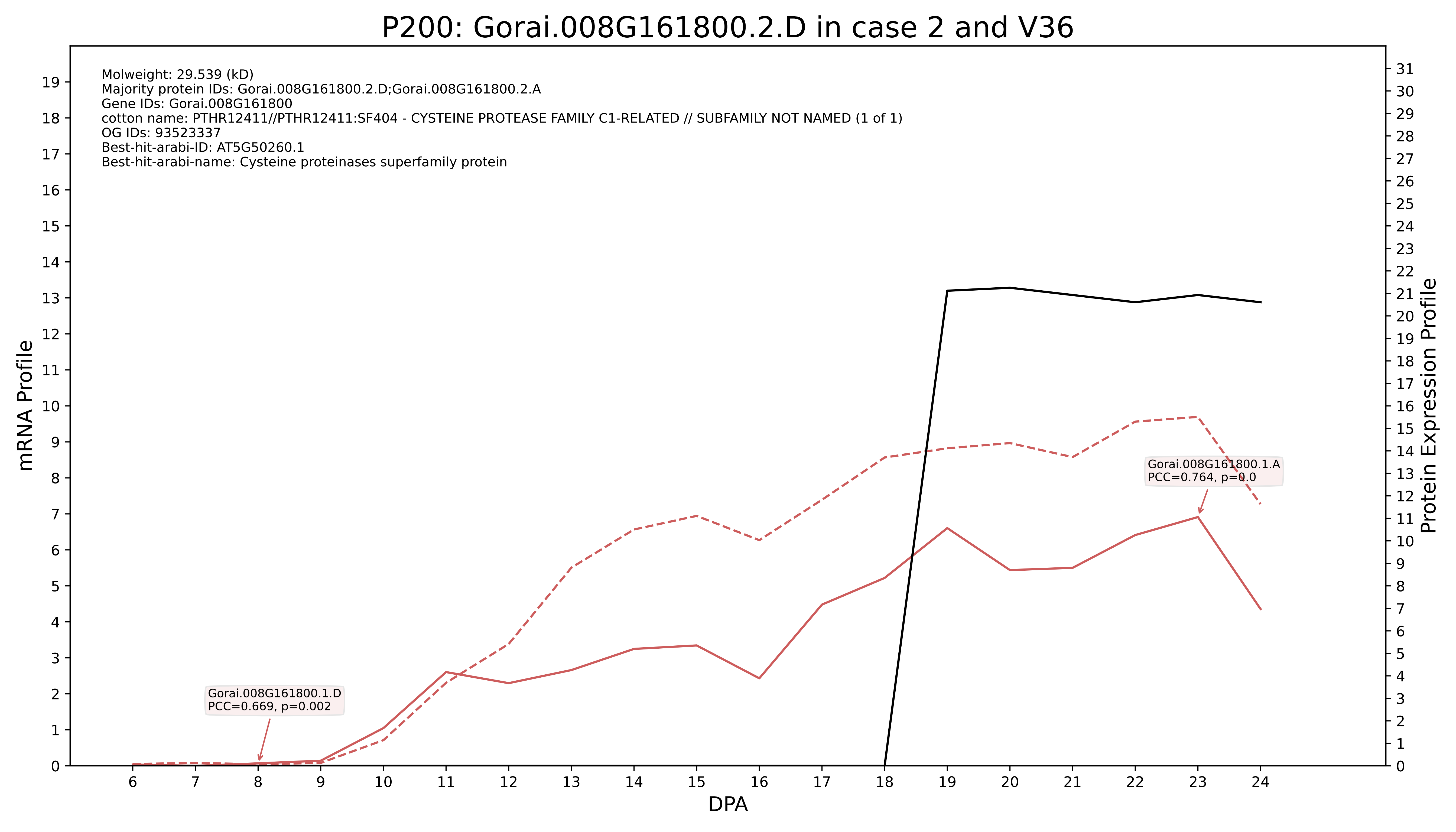
P200: Gorai.008G161800.2.D in case 2 and V36
Molweight: 29.539 (kD)
Majority protein IDs: Gorai.008G161800.2.D;Gorai.008G161800.2.A
Gene IDs: Gorai.008G161800
cotton name: PTHR12411//PTHR12411:SF404 - CYSTEINE PROTEASE FAMILY C1-RELATED // SUBFAMILY NOT NAMED (1 of 1)
OG IDs: 93523337
Best-hit-arabi-ID: AT5G50260.1
Best-hit-arabi-name: Cysteine proteinases superfamily protein
Gorai.008G161800.1.A
PCC=0.764, p=0.0
Gorai.008G161800.1.D
PCC=0.669, p=0.002
mRNA Profile
Protein Expression Profile
DPA

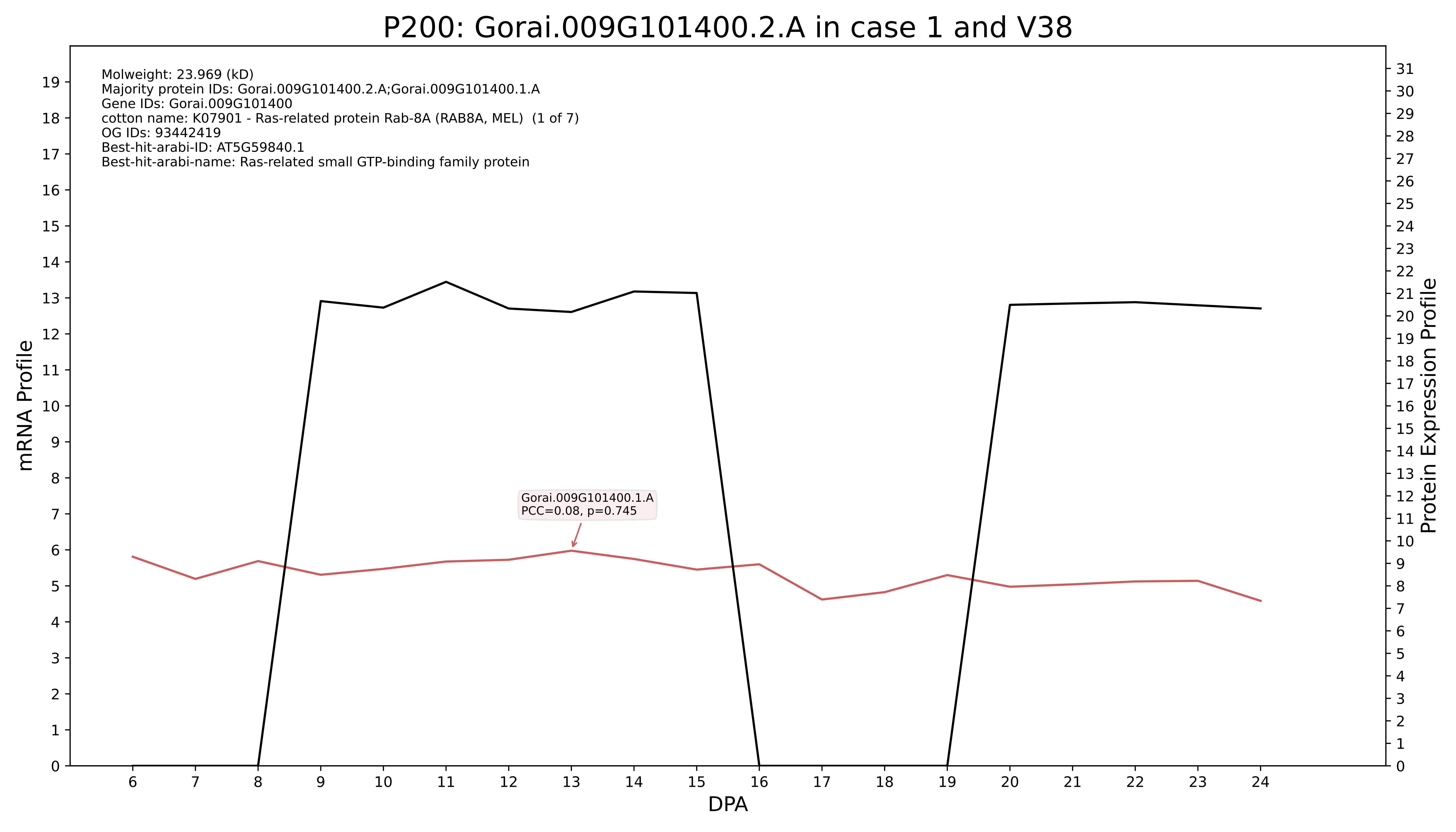
P200: Gorai.009G101400.2.A in case 1 and V38
Molweight: 23.969 (kD)
Majority protein IDs: Gorai.009G101400.2.A;Gorai.009G101400.1.A
Gene IDs: Gorai.009G101400
cotton name: K07901 - Ras-related protein Rab-8A (RAB8A, MEL) (1 of 7)
OG IDs: 93442419
Best-hit-arabi-ID: AT5G59840.1
Best-hit-arabi-name: Ras-related small GTP-binding family protein
Gorai.009G101400.1.A
PCC=0.08, p=0.745
mRNA Profile
Protein Expression Profile
DPA

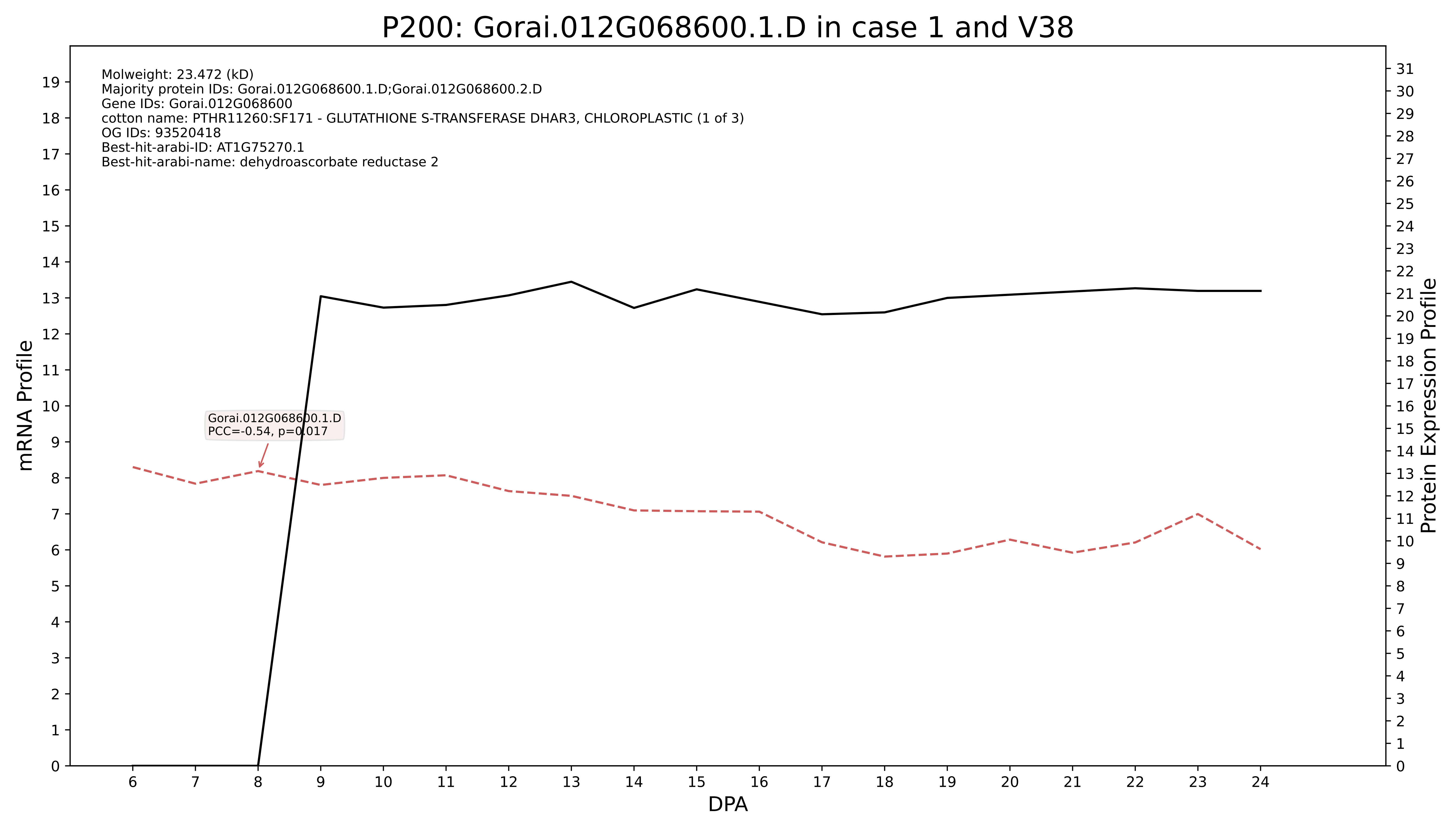
P200: Gorai.012G068600.1.D in case 1 and V38
Molweight: 23.472 (kD)
Majority protein IDs: Gorai.012G068600.1.D;Gorai.012G068600.2.D
Gene IDs: Gorai.012G068600
cotton name: PTHR11260:SF171 - GLUTATHIONE S-TRANSFERASE DHAR3, CHLOROPLASTIC (1 of 3)
OG IDs: 93520418
Best-hit-arabi-ID: AT1G75270.1
Best-hit-arabi-name: dehydroascorbate reductase 2
Gorai.012G068600.1.D
PCC=-0.54, p=0.017
mRNA Profile
Protein Expression Profile
DPA

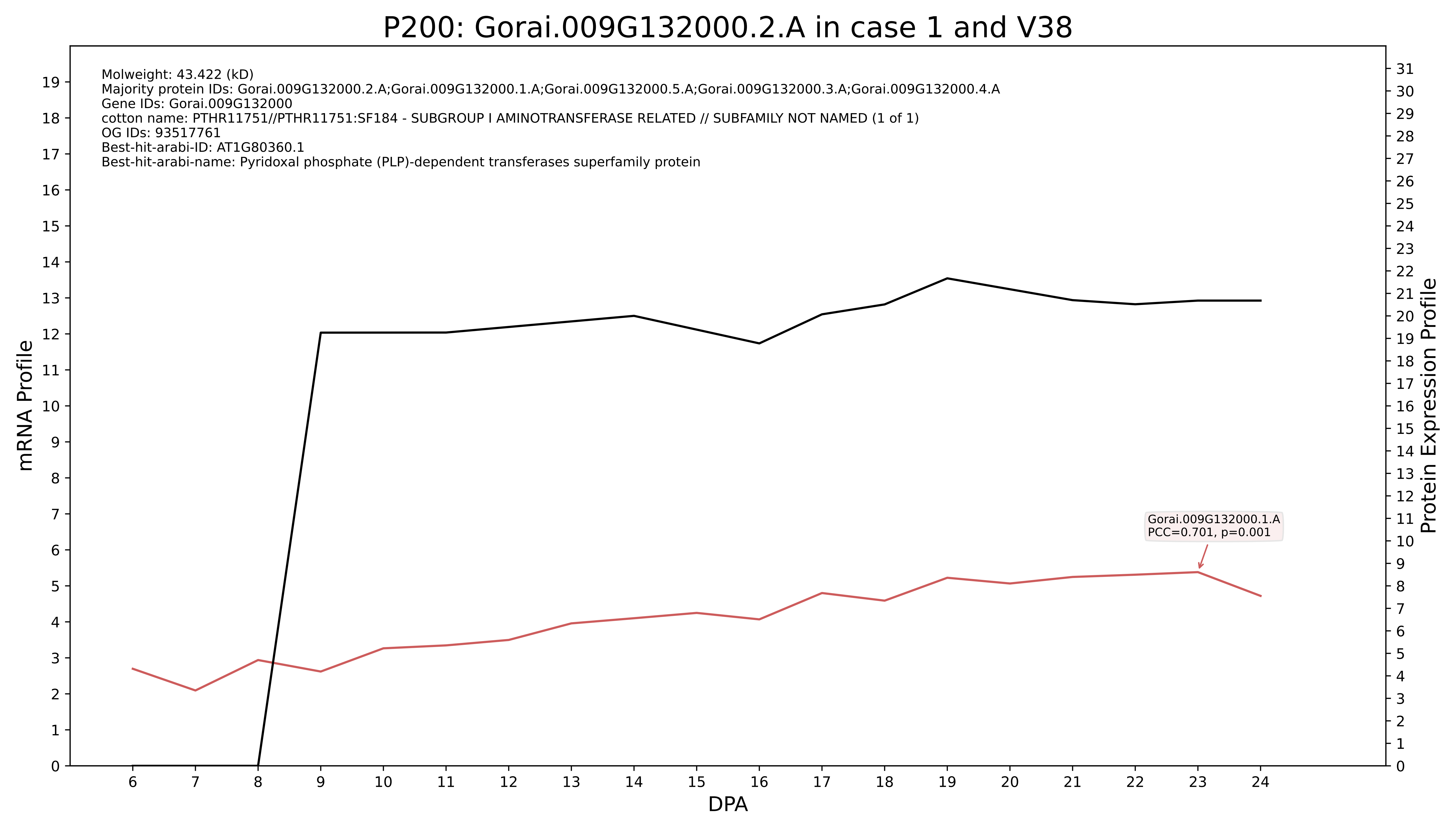
P200: Gorai.009G132000.2.A in case 1 and V38
Molweight: 43.422 (kD)
Majority protein IDs: Gorai.009G132000.2.A;Gorai.009G132000.1.A;Gorai.009G132000.5.A;Gorai.009G132000.3.A;Gorai.009G132000.4.A
Gene IDs: Gorai.009G132000
cotton name: PTHR11751//PTHR11751:SF184 - SUBGROUP I AMINOTRANSFERASE RELATED // SUBFAMILY NOT NAMED (1 of 1)
OG IDs: 93517761
Best-hit-arabi-ID: AT1G80360.1
Best-hit-arabi-name: Pyridoxal phosphate (PLP)-dependent transferases superfamily protein
Gorai.009G132000.1.A
PCC=0.701, p=0.001
mRNA Profile
Protein Expression Profile
DPA

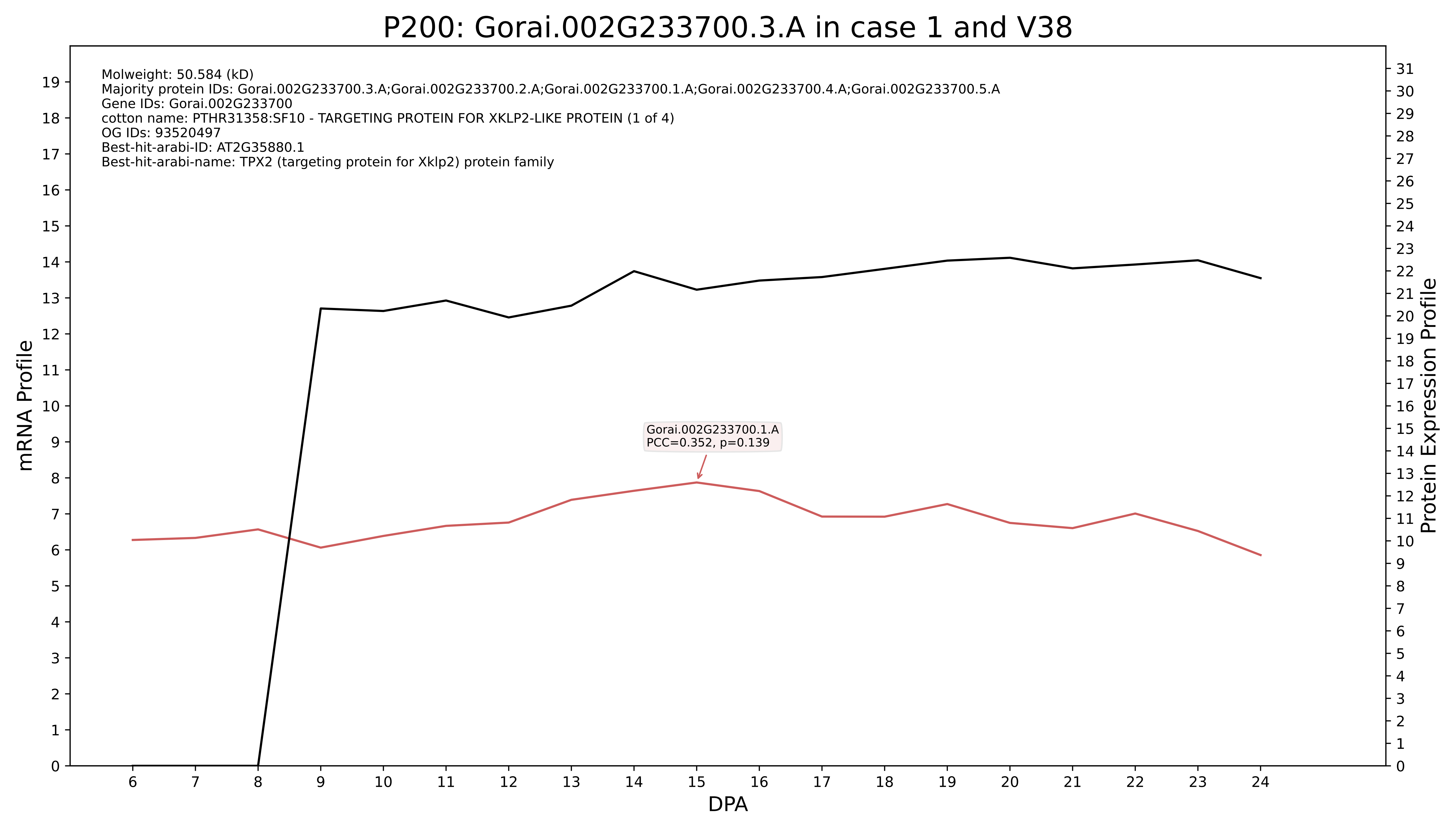

P200: Gorai.002G233700.3.A in case 1 and V38
Molweight: 50.584 (kD)
Majority protein IDs: Gorai.002G233700.3.A;Gorai.002G233700.2.A;Gorai.002G233700.1.A;Gorai.002G233700.4.A;Gorai.002G233700.5.A
Gene IDs: Gorai.002G233700
cotton name: PTHR31358:SF10 - TARGETING PROTEIN FOR XKLP2-LIKE PROTEIN (1 of 4)
OG IDs: 93520497
Best-hit-arabi-ID: AT2G35880.1
Best-hit-arabi-name: TPX2 (targeting protein for Xklp2) protein family
Gorai.002G233700.1.A
PCC=0.352, p=0.139
mRNA Profile
Protein Expression Profile
DPA

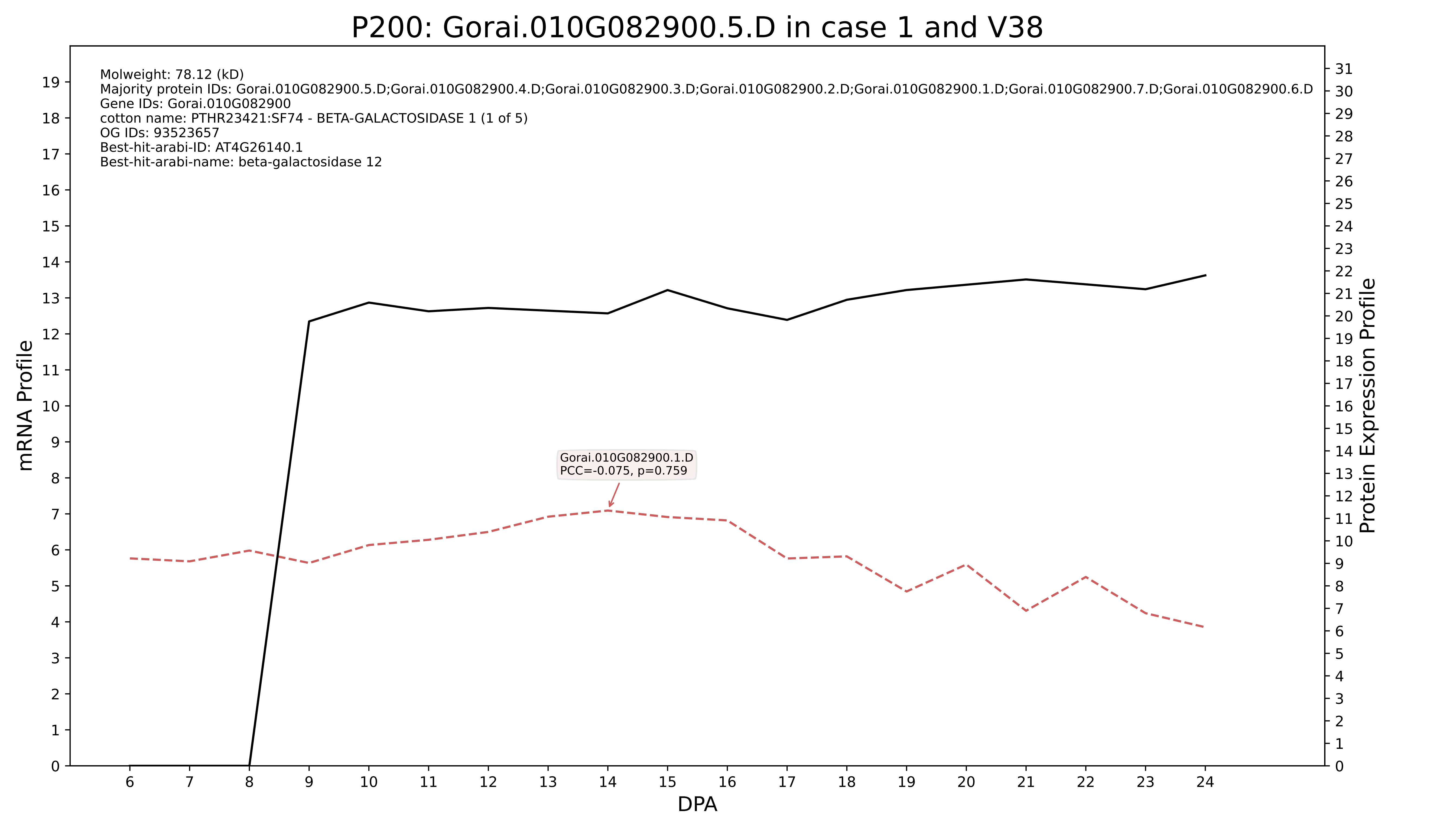
P200: Gorai.010G082900.5.D in case 1 and V38
Molweight: 78.12 (kD)
Majority protein IDs: Gorai.010G082900.5.D;Gorai.010G082900.4.D;Gorai.010G082900.3.D;Gorai.010G082900.2.D;Gorai.010G082900.1.D;Gorai.010G082900.7.D;Gorai.010G082900.6.D
Gene IDs: Gorai.010G082900
cotton name: PTHR23421:SF74 - BETA-GALACTOSIDASE 1 (1 of 5)
OG IDs: 93523657
Best-hit-arabi-ID: AT4G26140.1
Best-hit-arabi-name: beta-galactosidase 12
Gorai.010G082900.1.D
PCC=-0.075, p=0.759
mRNA Profile
Protein Expression Profile
DPA

### P200: Gorai.011G193800.1.D in case 1 and V38

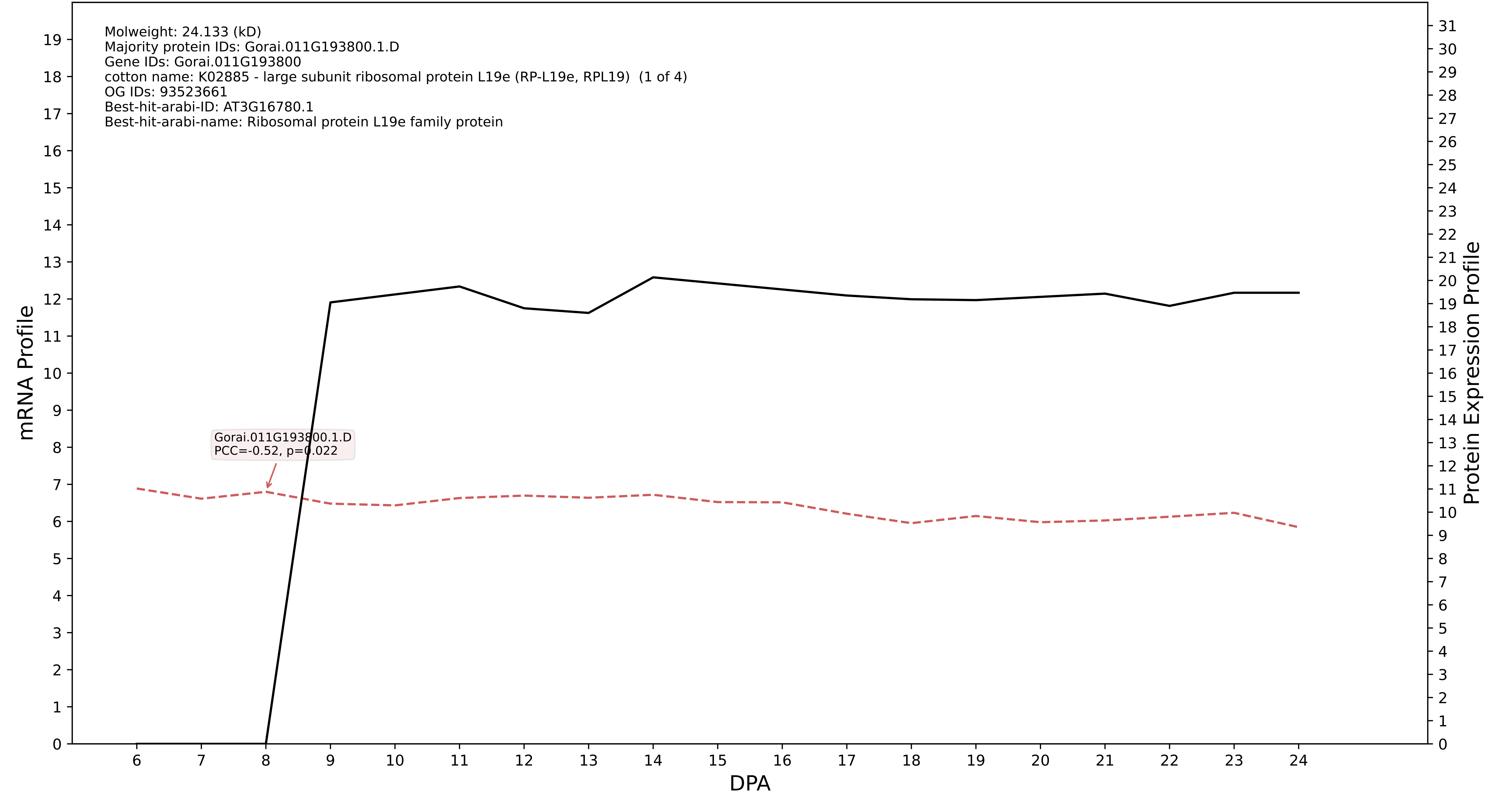

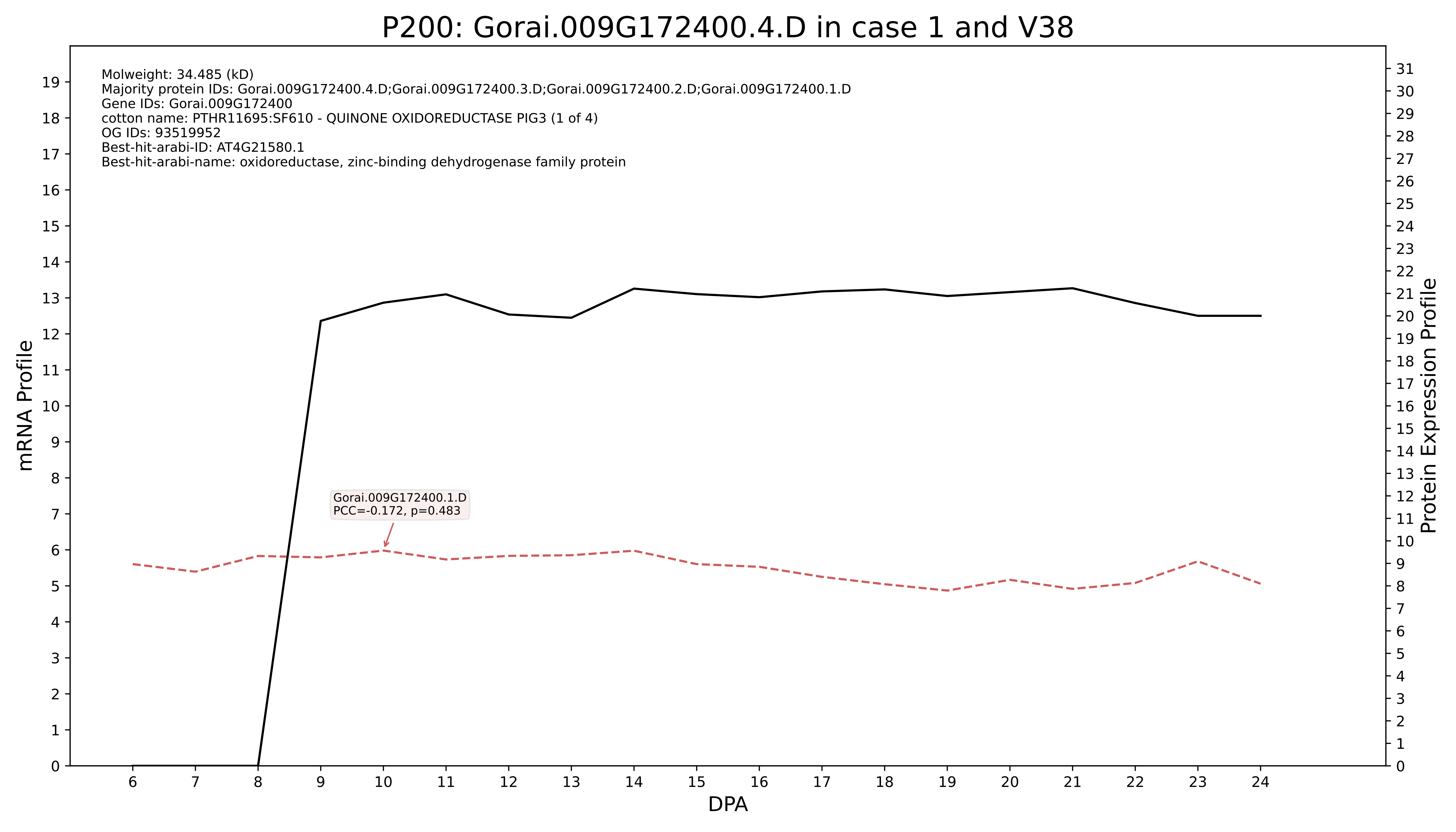
P200: Gorai.009G172400.4.D in case 1 and V38
Molweight: 34.485 (kD)
Majority protein IDs: Gorai.009G172400.4.D;Gorai.009G172400.3.D;Gorai.009G172400.2.D;Gorai.009G172400.1.D
Gene IDs: Gorai.009G172400
cotton name: PTHR11695:SF610 - QUINONE OXIDOREDUCTASE PIG3 (1 of 4)
OG IDs: 93519952
Best-hit-arabi-ID: AT4G21580.1
Best-hit-arabi-name: oxidoreductase, zinc-binding dehydrogenase family protein
Gorai.009G172400.1.D
PCC=-0.172, p=0.483
mRNA Profile
Protein Expression Profile
DPA

### P200: Gorai.004G183100.4.D in case 1 and V38

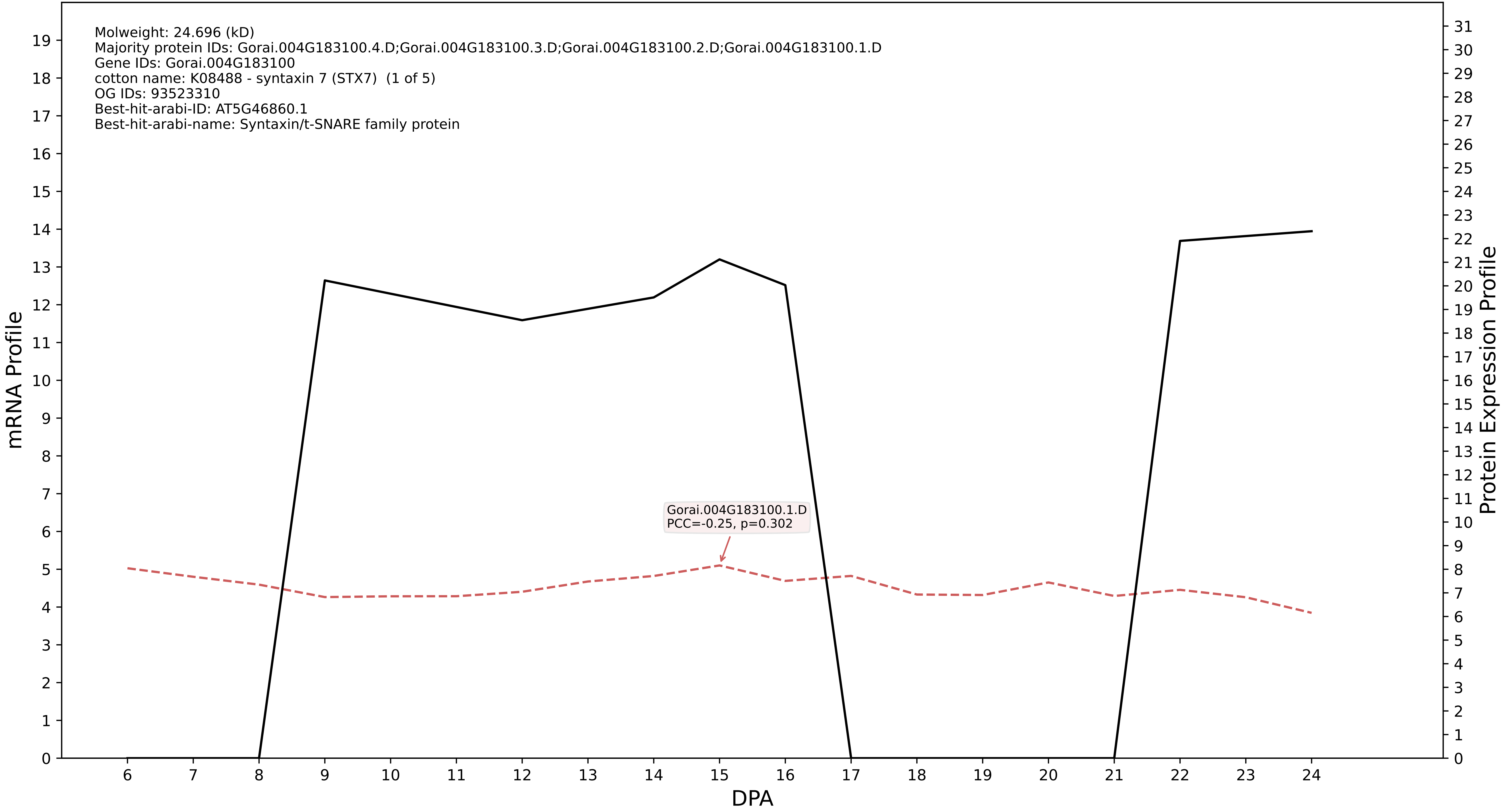

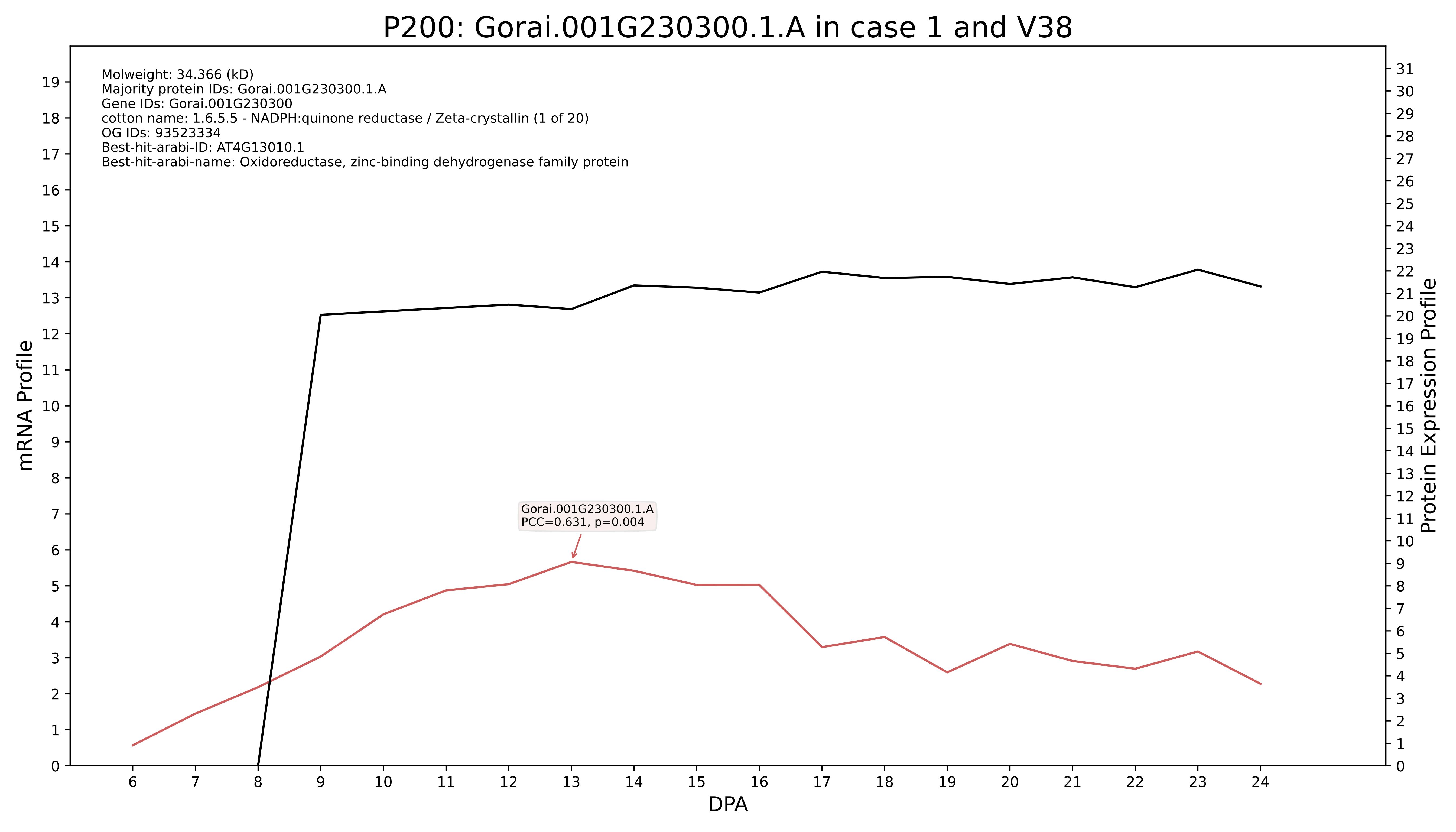
P200: Gorai.001G230300.1.A in case 1 and V38
Molweight: 34.366 (kD)
Majority protein IDs: Gorai.001G230300.1.A
Gene IDs: Gorai.001G230300
cotton name: 1.6.5.5 - NADPH:quinone reductase / Zeta-crystallin (1 of 20)
OG IDs: 93523334
Best-hit-arabi-ID: AT4G13010.1
Best-hit-arabi-name: Oxidoreductase, zinc-binding dehydrogenase family protein
Gorai.001G230300.1.A
PCC=0.631, p=0.004
mRNA Profile
Protein Expression Profile
DPA

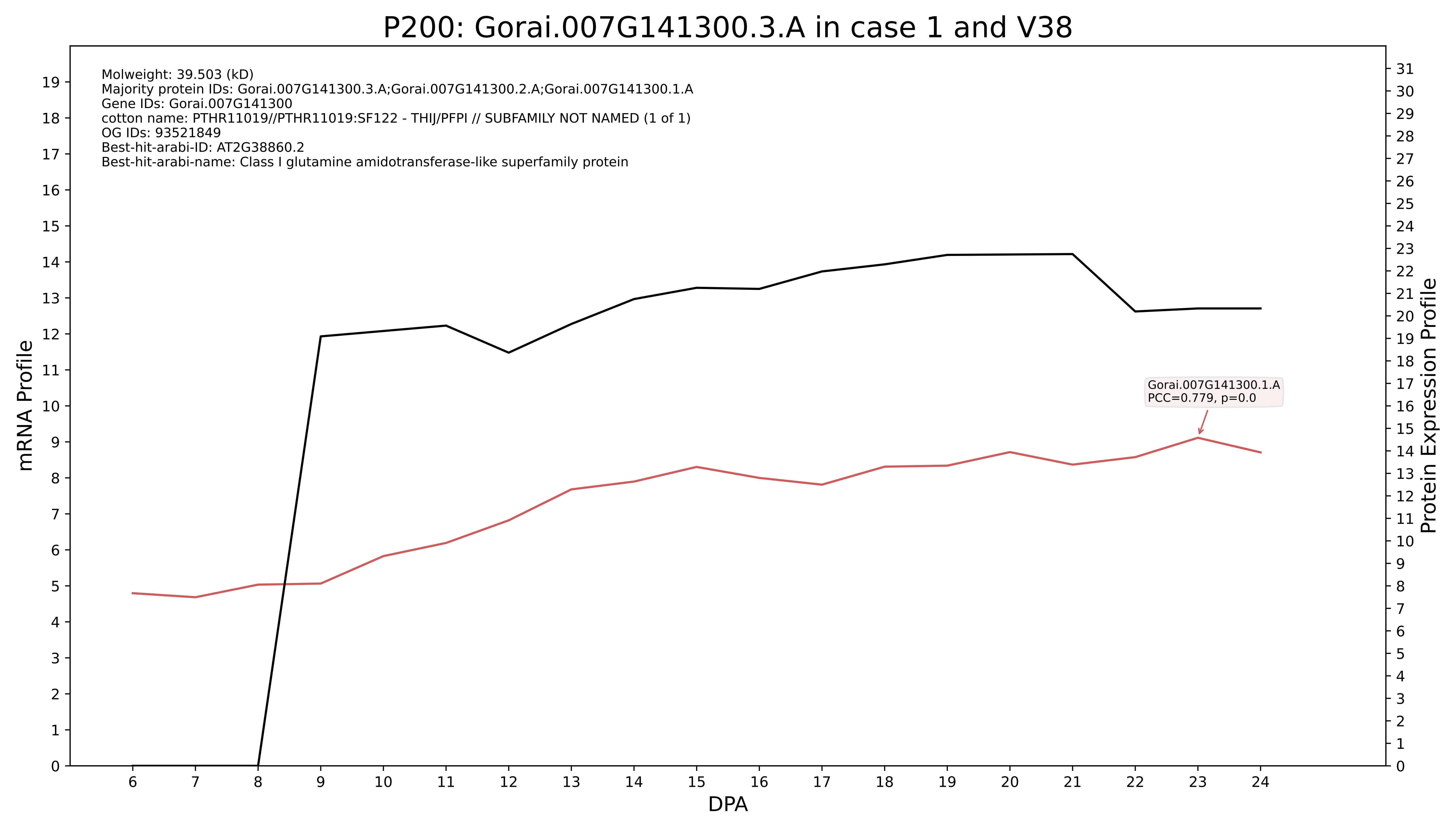

P200: Gorai.007G141300.3.A in case 1 and V38
Molweight: 39.503 (kD)
Majority protein IDs: Gorai.007G141300.3.A;Gorai.007G141300.2.A;Gorai.007G141300.1.A
Gene IDs: Gorai.007G141300
cotton name: PTHR11019//PTHR11019:SF122 - THIJ/PFPI // SUBFAMILY NOT NAMED (1 of 1)
OG IDs: 93521849
Best-hit-arabi-ID: AT2G38860.2
Best-hit-arabi-name: Class I glutamine amidotransferase-like superfamily protein
Gorai.007G141300.1.A
PCC=0.779, p=0.0
mRNA Profile
Protein Expression Profile
DPA

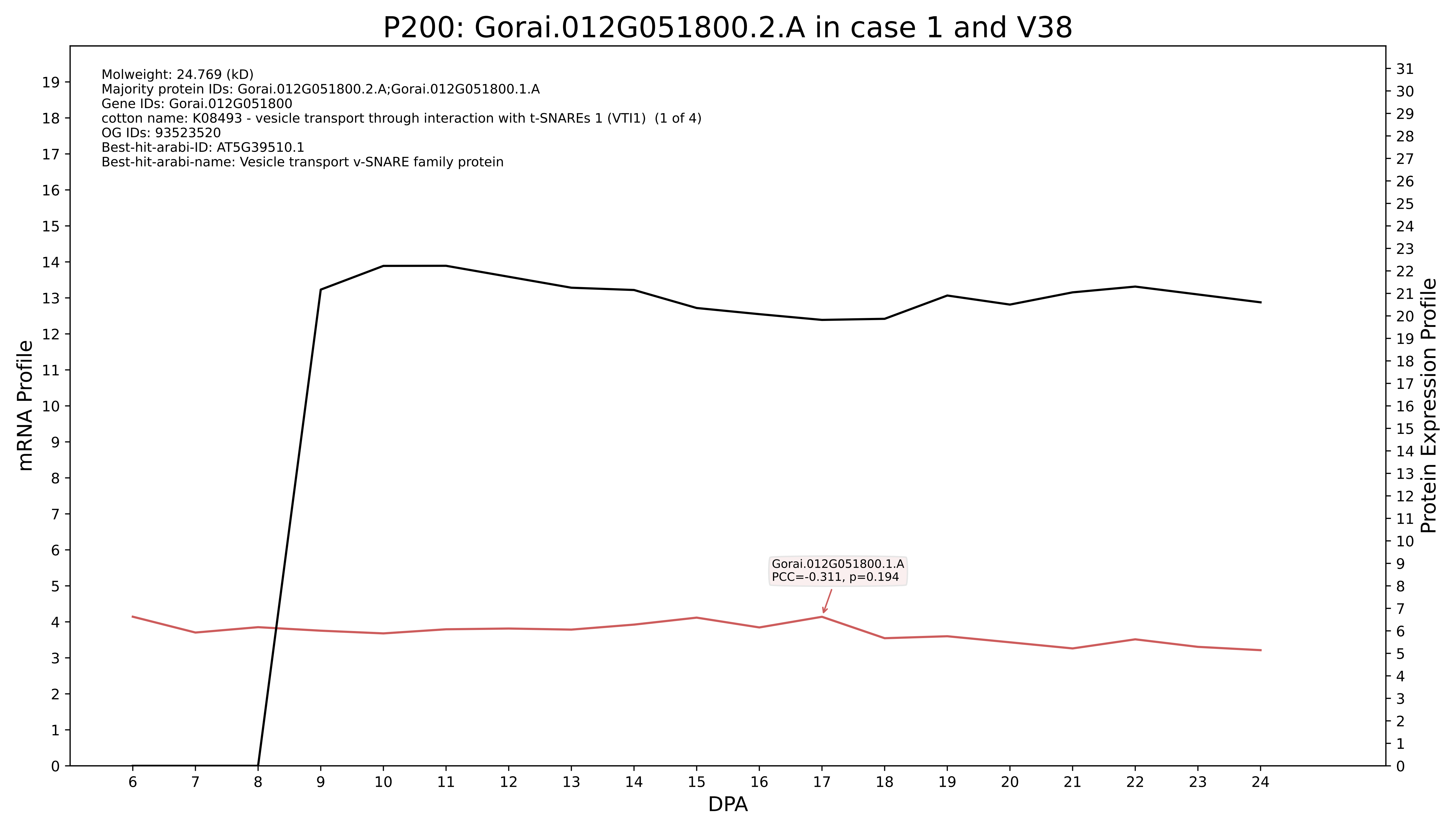

P200: Gorai.012G051800.2.A in case 1 and V38
Molweight: 24.769 (kD)
Majority protein IDs: Gorai.012G051800.2.A;Gorai.012G051800.1.A
Gene IDs: Gorai.012G051800
cotton name: K08493 - vesicle transport through interaction with t-SNAREs 1 (VTI1) (1 of 4)
OG IDs: 93523520
Best-hit-arabi-ID: AT5G39510.1
Best-hit-arabi-name: Vesicle transport v-SNARE family protein
Gorai.012G051800.1.A
PCC=-0.311, p=0.194
mRNA Profile
Protein Expression Profile
DPA

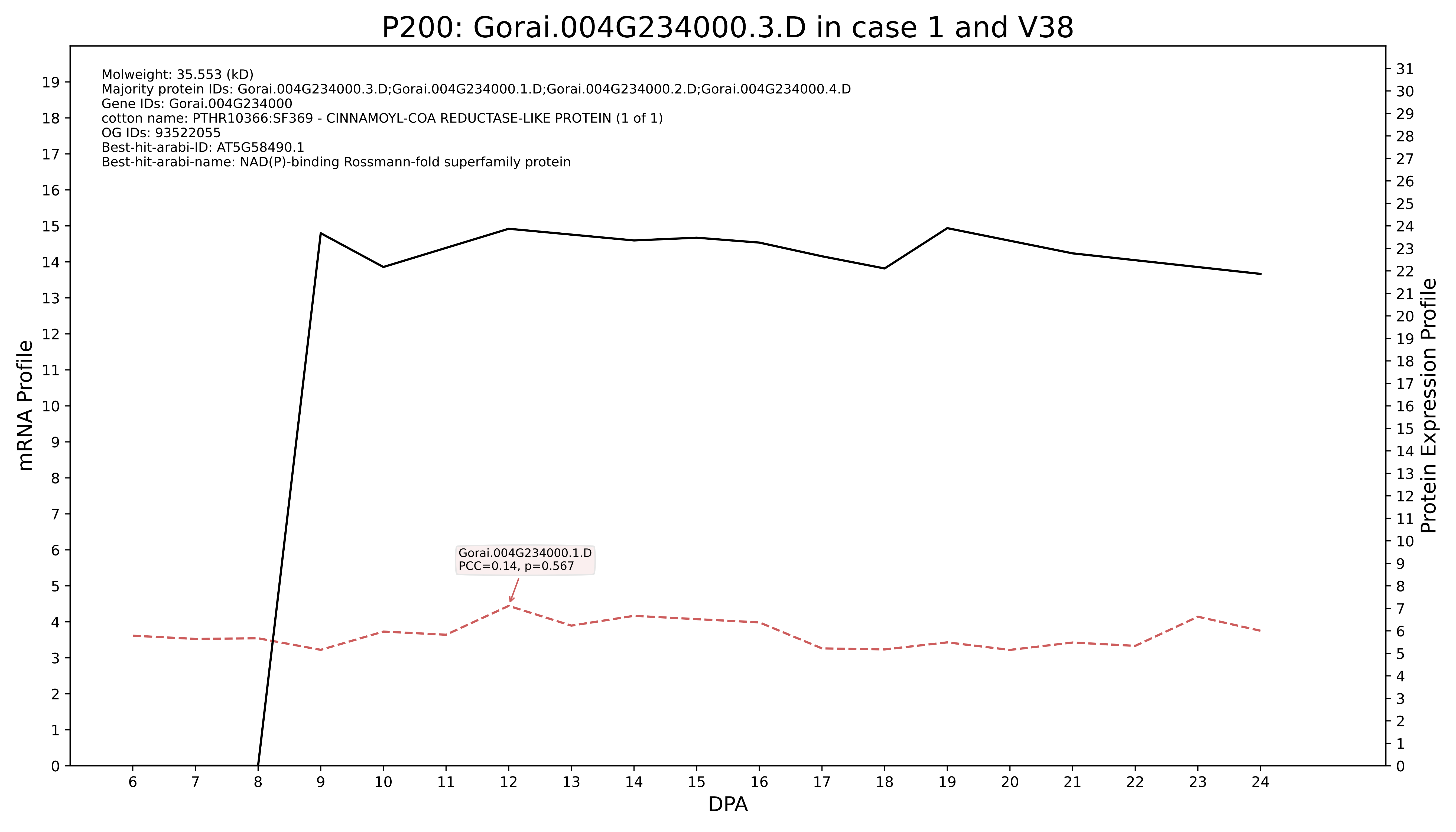
P200: Gorai.004G234000.3.D in case 1 and V38
Molweight: 35.553 (kD)
Majority protein IDs: Gorai.004G234000.3.D;Gorai.004G234000.1.D;Gorai.004G234000.2.D;Gorai.004G234000.4.D
Gene IDs: Gorai.004G234000
cotton name: PTHR10366:SF369 - CINNAMOYL-COA REDUCTASE-LIKE PROTEIN (1 of 1)
OG IDs: 93522055
Best-hit-arabi-ID: AT5G58490.1
Best-hit-arabi-name: NAD(P)-binding Rossmann-fold superfamily protein
Gorai.004G234000.1.D
PCC=0.14, p=0.567
mRNA Profile
Protein Expression Profile
DPA

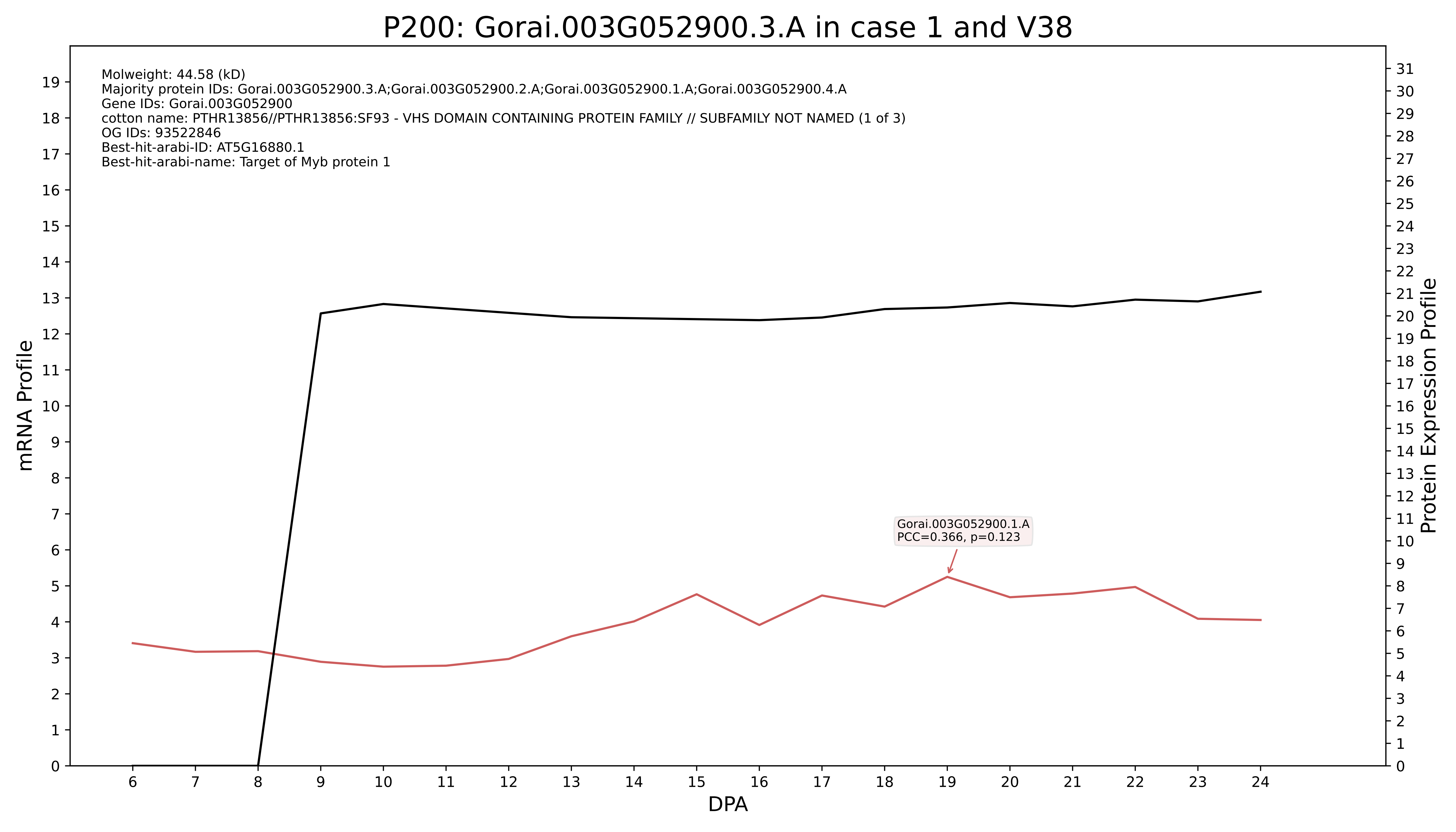
P200: Gorai.003G052900.3.A in case 1 and V38
Molweight: 44.58 (kD)
Majority protein IDs: Gorai.003G052900.3.A;Gorai.003G052900.2.A;Gorai.003G052900.1.A;Gorai.003G052900.4.A
Gene IDs: Gorai.003G052900
cotton name: PTHR13856//PTHR13856:SF93 - VHS DOMAIN CONTAINING PROTEIN FAMILY // SUBFAMILY NOT NAMED (1 of 3)
OG IDs: 93522846
Best-hit-arabi-ID: AT5G16880.1
Best-hit-arabi-name: Target of Myb protein 1
Gorai.003G052900.1.A
PCC=0.366, p=0.123
mRNA Profile
Protein Expression Profile
DPA

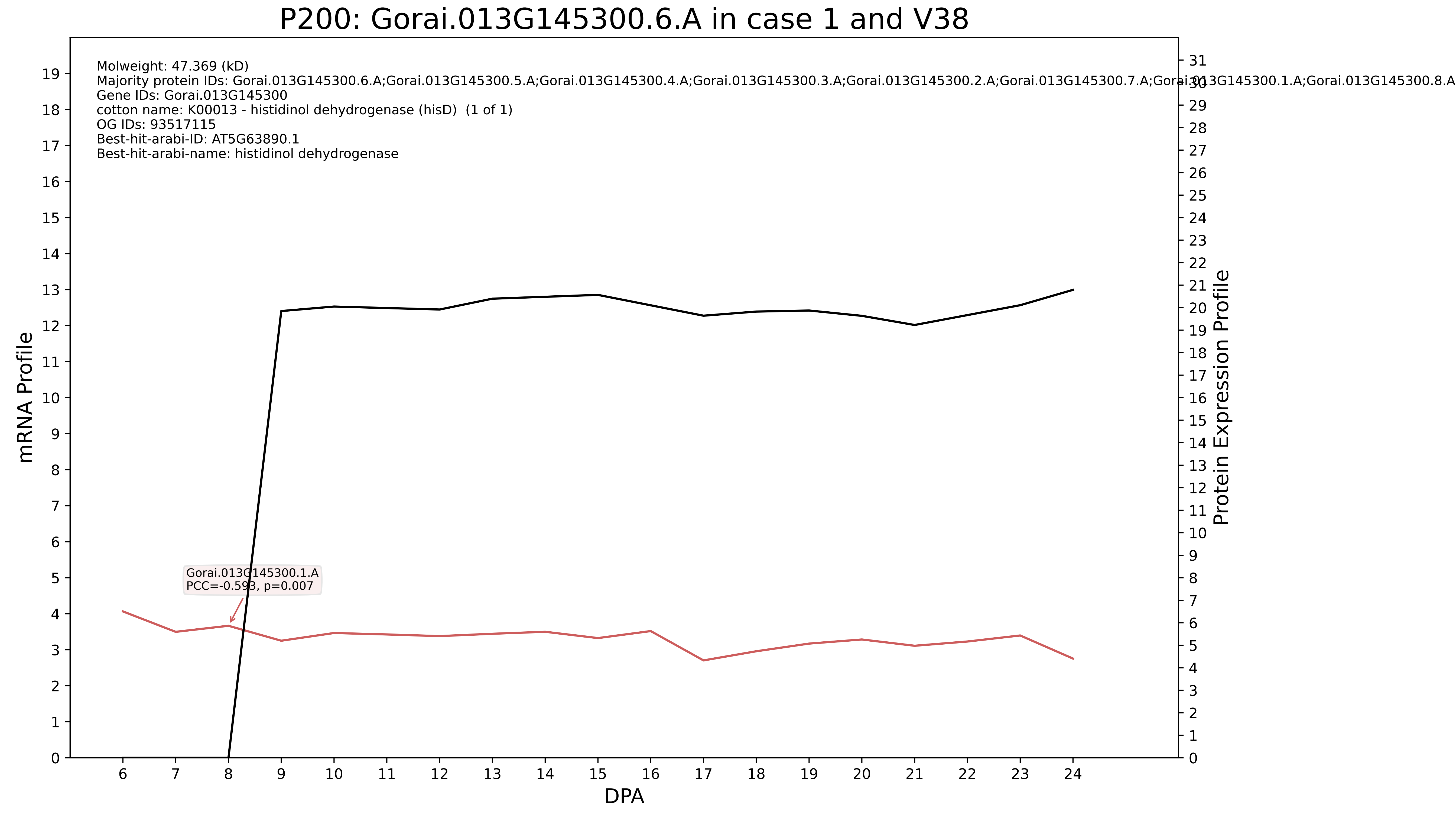
P200: Gorai.013G145300.6.A in case 1 and V38
Molweight: 47.369 (kD)
Majority protein IDs: Gorai.013G145300.6.A;Gorai.013G145300.5.A;Gorai.013G145300.4.A;Gorai.013G145300.3.A;Gorai.013G145300.2.A;Gorai.013G145300.7.A;Gorai.013G145300.1.A;Gorai.013G145300.8.A
Gene IDs: Gorai.013G145300
cotton name: K00013 - histidinol dehydrogenase (hisD) (1 of 1)
OG IDs: 93517115
Best-hit-arabi-ID: AT5G63890.1
Best-hit-arabi-name: histidinol dehydrogenase
Gorai.013G145300.1.A
PCC=-0.593, p=0.007
mRNA Profile
Protein Expression Profile
DPA

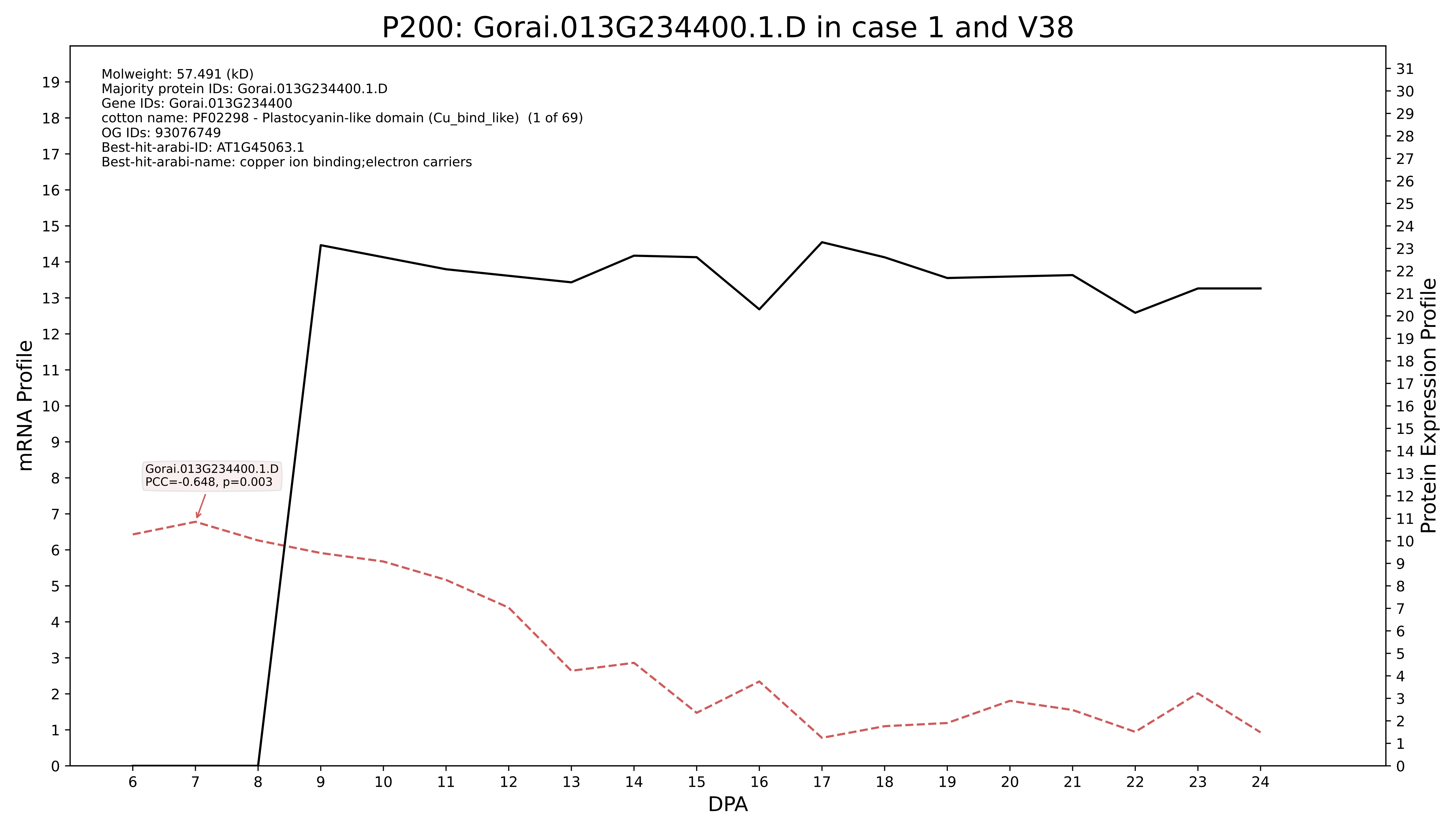
P200: Gorai.013G234400.1.D in case 1 and V38
Molweight: 57.491 (kD)
Majority protein IDs: Gorai.013G234400.1.D
Gene IDs: Gorai.013G234400
cotton name: PF02298 - Plastocyanin-like domain (Cu_bind_like) (1 of 69)
OG IDs: 93076749
Best-hit-arabi-ID: AT1G45063.1
Best-hit-arabi-name: copper ion binding;electron carriers
Gorai.013G234400.1.D
PCC=-0.648, p=0.003
mRNA Profile
Protein Expression Profile
DPA

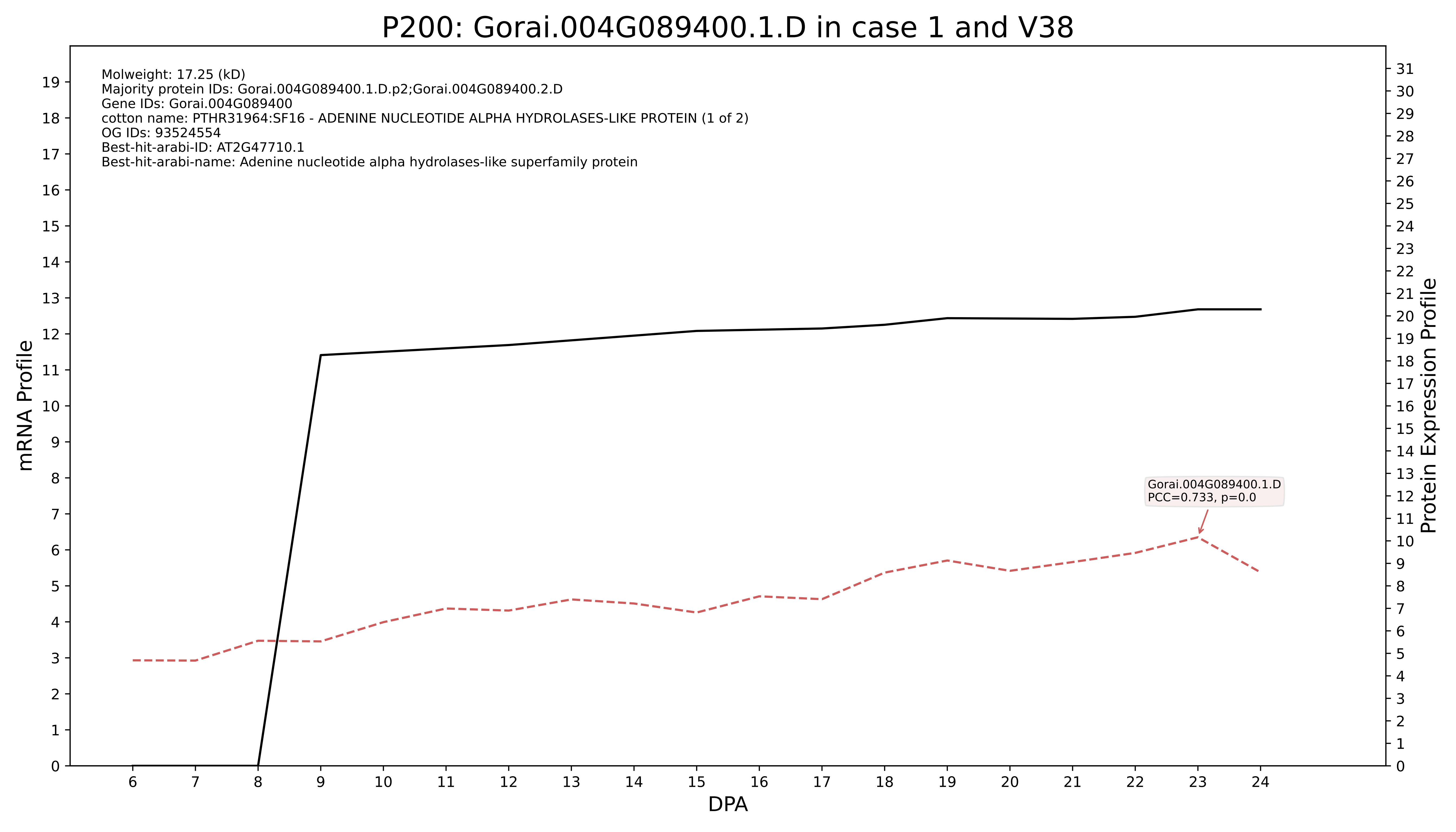
P200: Gorai.004G089400.1.D in case 1 and V38
Molweight: 17.25 (kD)
Majority protein IDs: Gorai.004G089400.1.D.p2;Gorai.004G089400.2.D
Gene IDs: Gorai.004G089400
cotton name: PTHR31964:SF16 - ADENINE NUCLEOTIDE ALPHA HYDROLASES-LIKE PROTEIN (1 of 2)
OG IDs: 93524554
Best-hit-arabi-ID: AT2G47710.1
Best-hit-arabi-name: Adenine nucleotide alpha hydrolases-like superfamily protein
Gorai.004G089400.1.D
PCC=0.733, p=0.0
mRNA Profile
Protein Expression Profile
DPA
0 1 2 3 4 5 6 7 8 9 10 11 12 13 14 15 16 17 18 19
0 1 2 3 4 5 6 7 8 9 10 11 12 13 14 15 16 17 18 19 20 21 22 23 24 25 26 27 28 29 30 31
6 7 8 9 10 11 12 13 14 15 16 17 18 19 20 21 22 23 24

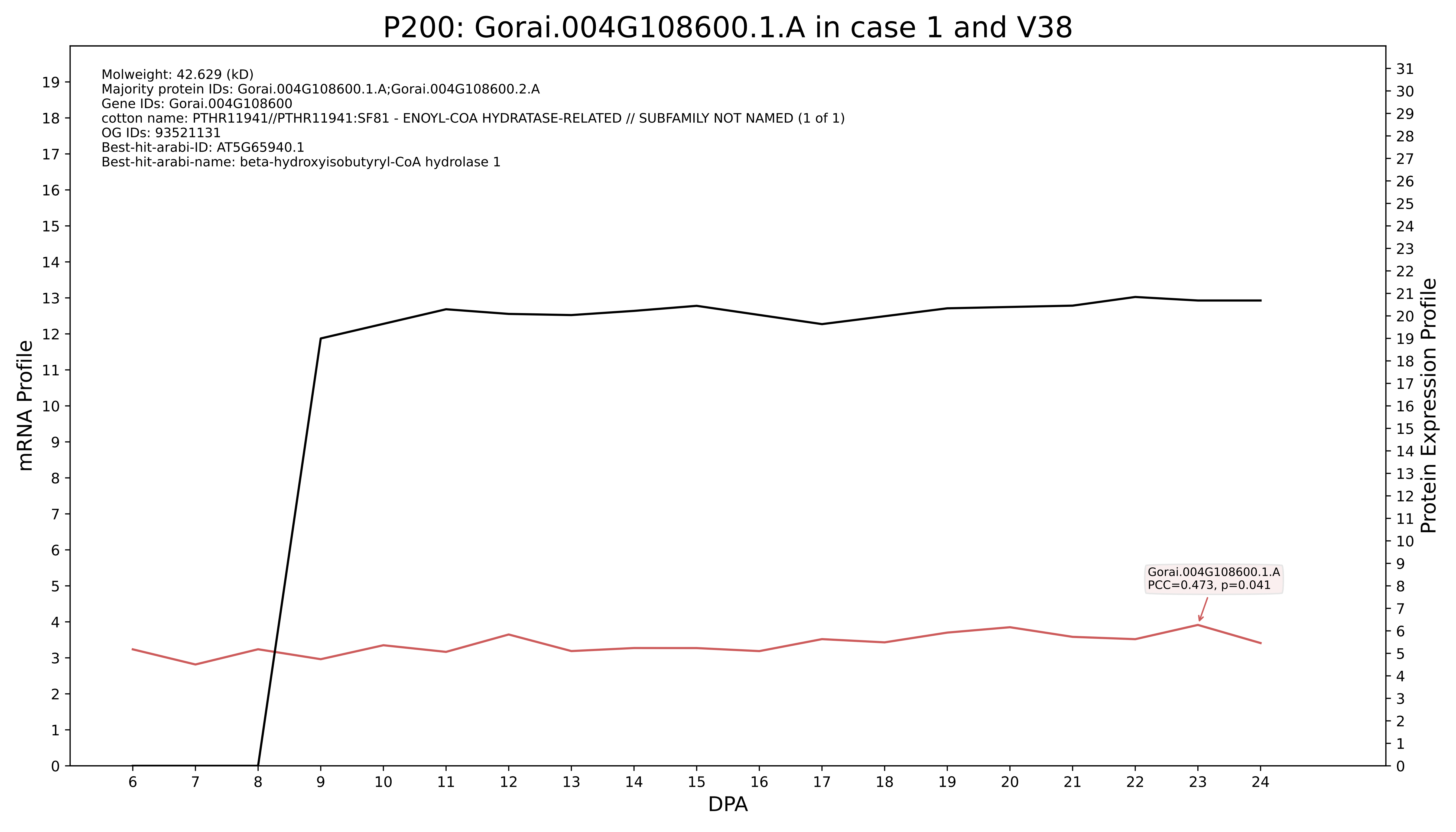

P200: Gorai.004G108600.1.A in case 1 and V38
Molweight: 42.629 (kD)
Majority protein IDs: Gorai.004G108600.1.A;Gorai.004G108600.2.A
Gene IDs: Gorai.004G108600
cotton name: PTHR11941//PTHR11941:SF81 - ENOYL-COA HYDRATASE-RELATED // SUBFAMILY NOT NAMED (1 of 1)
OG IDs: 93521131
Best-hit-arabi-ID: AT5G65940.1
Best-hit-arabi-name: beta-hydroxyisobutyryl-CoA hydrolase 1
Gorai.004G108600.1.A
PCC=0.473, p=0.041
mRNA Profile
Protein Expression Profile
DPA

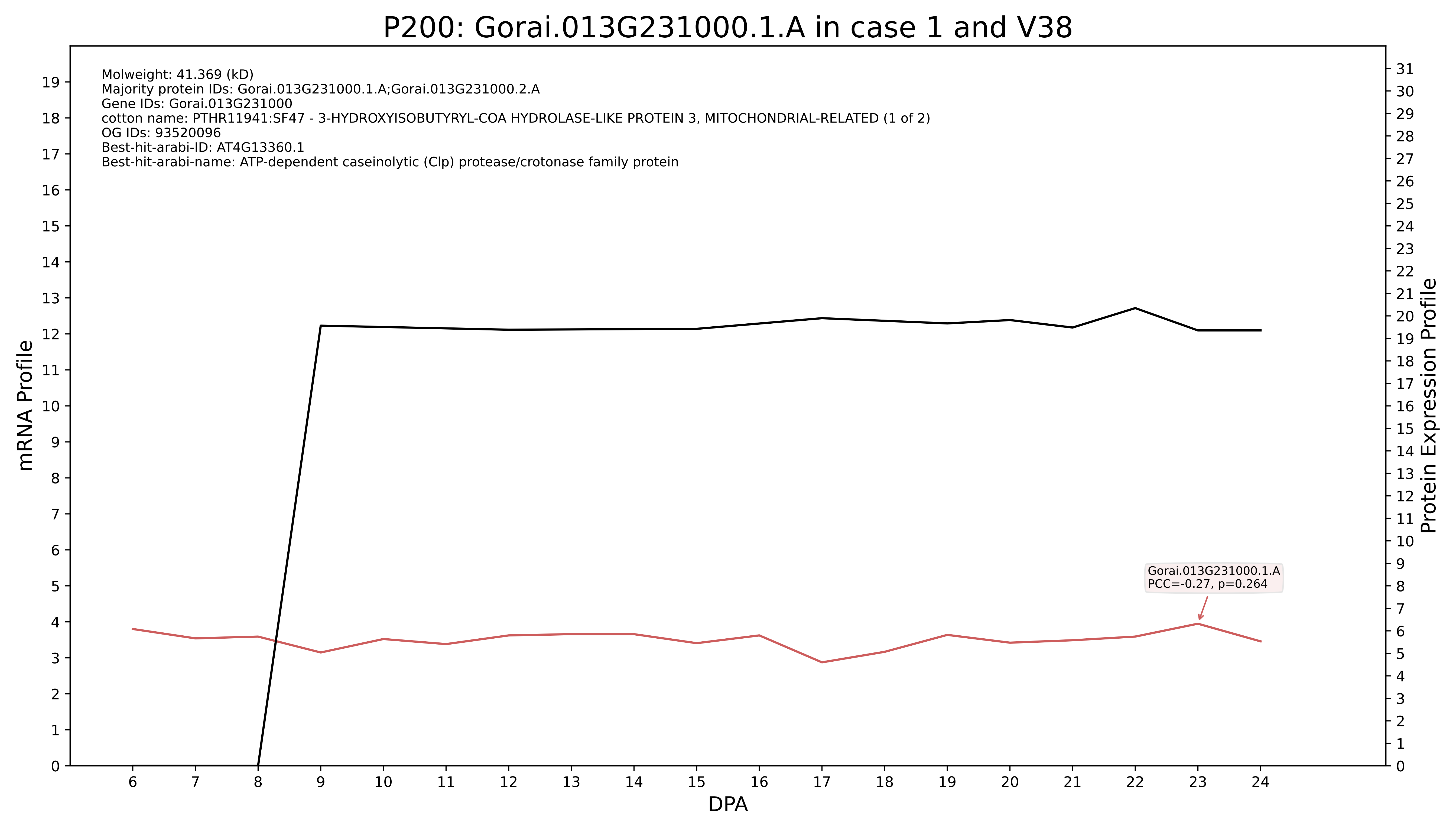
P200: Gorai.013G231000.1.A in case 1 and V38
Molweight: 41.369 (kD)
Majority protein IDs: Gorai.013G231000.1.A;Gorai.013G231000.2.A
Gene IDs: Gorai.013G231000
cotton name: PTHR11941:SF47 - 3-HYDROXYISOBUTYRYL-COA HYDROLASE-LIKE PROTEIN 3, MITOCHONDRIAL-RELATED (1 of 2)
OG IDs: 93520096
Best-hit-arabi-ID: AT4G13360.1
Best-hit-arabi-name: ATP-dependent caseinolytic (Clp) protease/crotonase family protein
Gorai.013G231000.1.A
PCC=-0.27, p=0.264
mRNA Profile
Protein Expression Profile
DPA

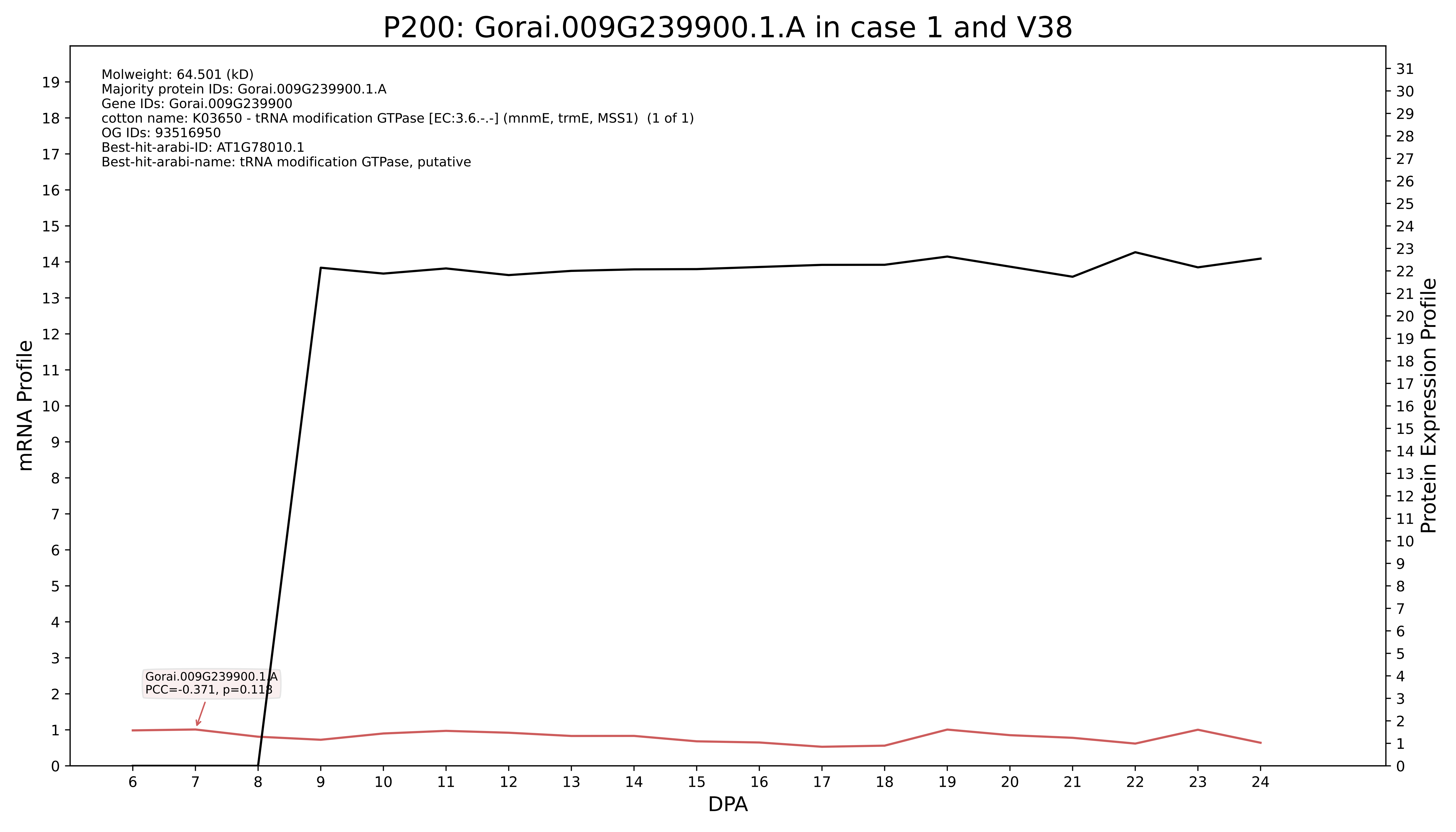
P200: Gorai.009G239900.1.A in case 1 and V38
Molweight: 64.501 (kD)
Majority protein IDs: Gorai.009G239900.1.A
Gene IDs: Gorai.009G239900
cotton name: K03650 - tRNA modification GTPase [EC:3.6.-.-] (mnmE, trmE, MSS1) (1 of 1)
OG IDs: 93516950
Best-hit-arabi-ID: AT1G78010.1
Best-hit-arabi-name: tRNA modification GTPase, putative
Gorai.009G239900.1.A
PCC=-0.371, p=0.118
mRNA Profile
Protein Expression Profile
DPA

### P200: Gorai.007G214400.6.A in case 1 and V38

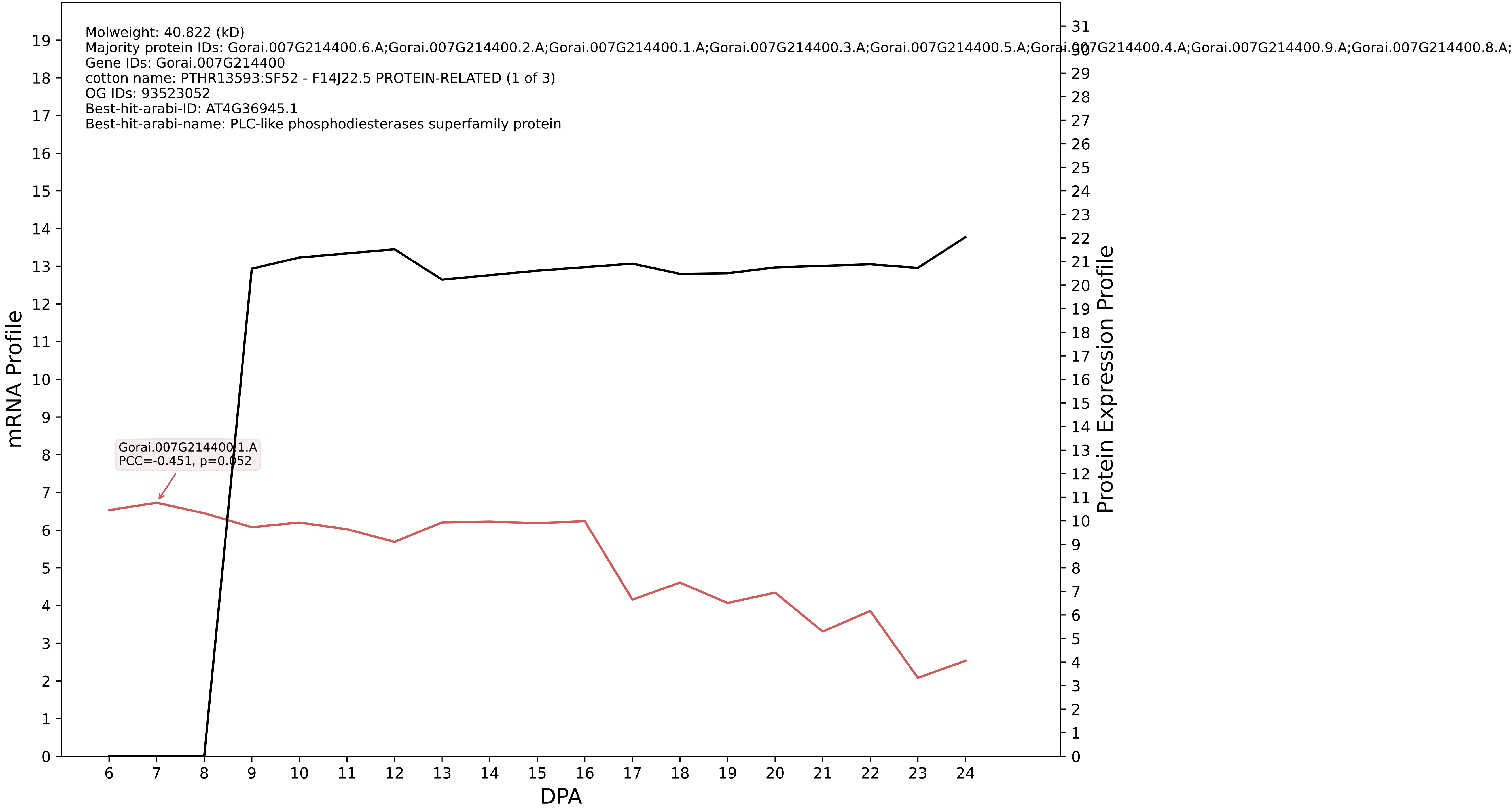

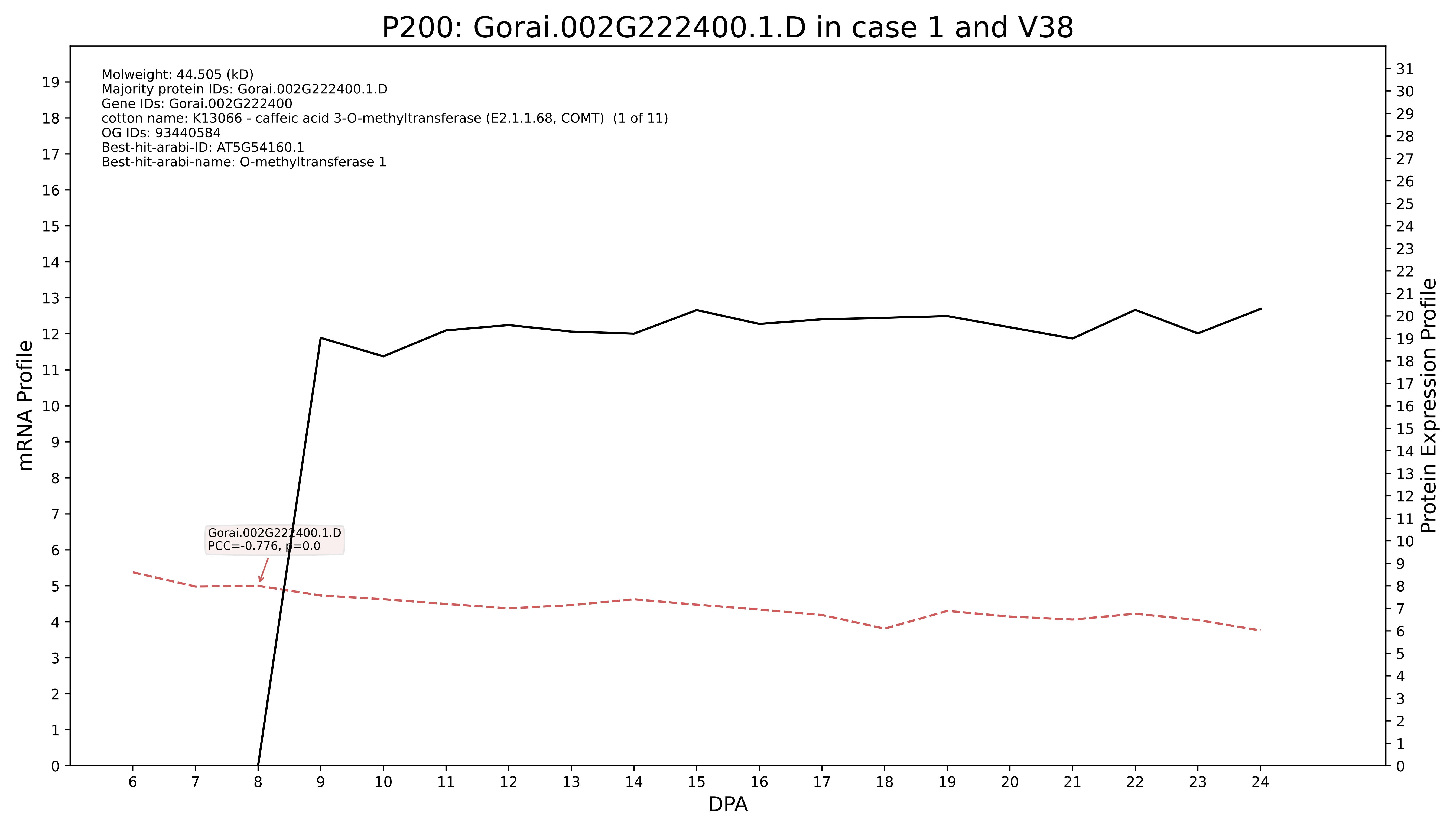
P200: Gorai.002G222400.1.D in case 1 and V38
Molweight: 44.505 (kD)
Majority protein IDs: Gorai.002G222400.1.D
Gene IDs: Gorai.002G222400
cotton name: K13066 - caffeic acid 3-O-methyltransferase (E2.1.1.68, COMT) (1 of 11)
OG IDs: 93440584
Best-hit-arabi-ID: AT5G54160.1
Best-hit-arabi-name: O-methyltransferase 1
Gorai.002G222400.1.D
PCC=-0.776, p=0.0
mRNA Profile
Protein Expression Profile
DPA

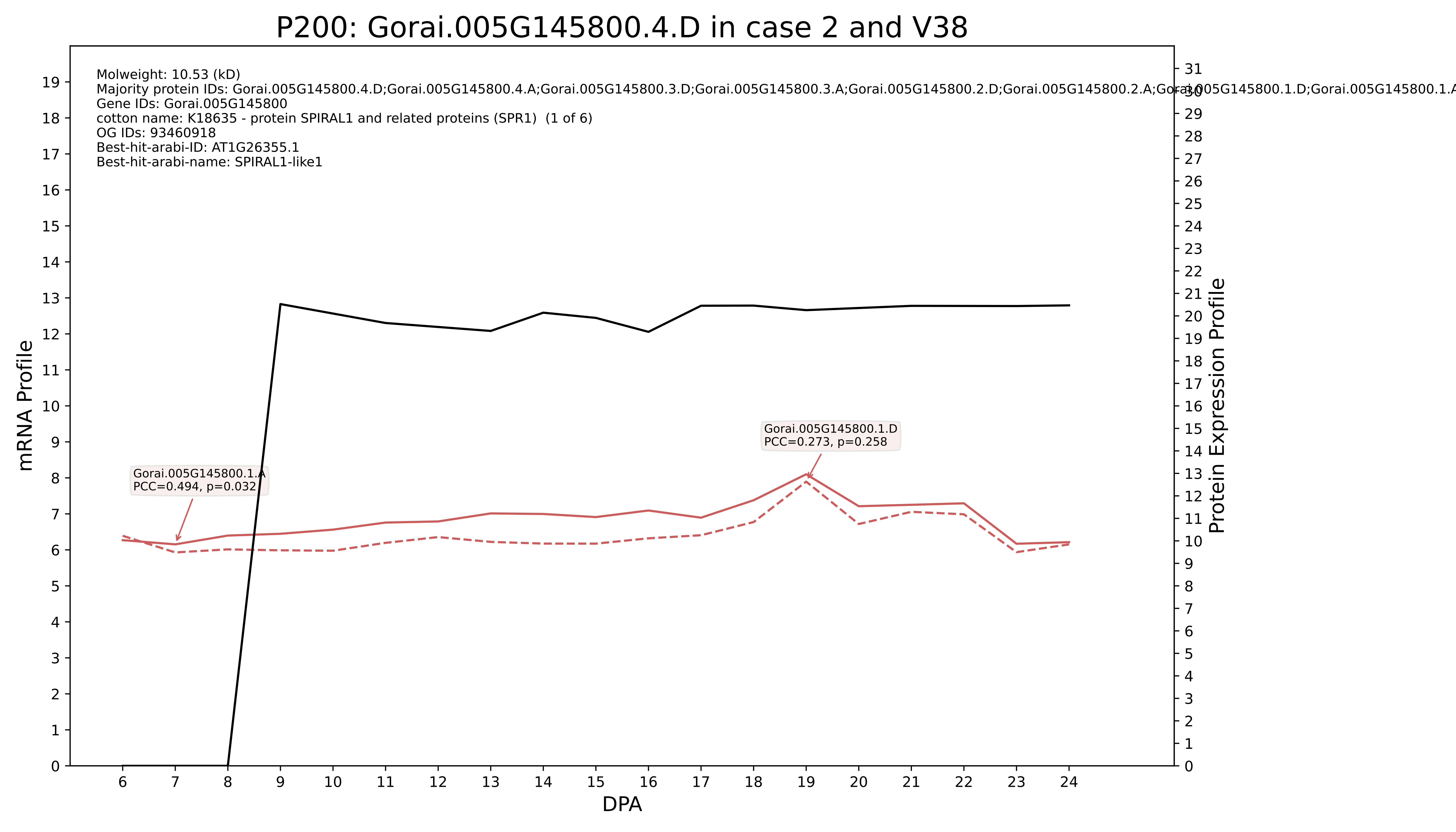
P200: Gorai.005G145800.4.D in case 2 and V38
Molweight: 10.53 (kD)
Majority protein IDs: Gorai.005G145800.4.D;Gorai.005G145800.4.A;Gorai.005G145800.3.D;Gorai.005G145800.3.A;Gorai.005G145800.2.D;Gorai.005G145800.2.A;Gorai.005G145800.1.D;Gorai.005G145800.1.A
Gene IDs: Gorai.005G145800
cotton name: K18635 - protein SPIRAL1 and related proteins (SPR1) (1 of 6)
OG IDs: 93460918
Best-hit-arabi-ID: AT1G26355.1
Best-hit-arabi-name: SPIRAL1-like1
Gorai.005G145800.1.D
PCC=0.273, p=0.258
Gorai.005G145800.1.A
PCC=0.494, p=0.032
mRNA Profile
Protein Expression Profile
DPA

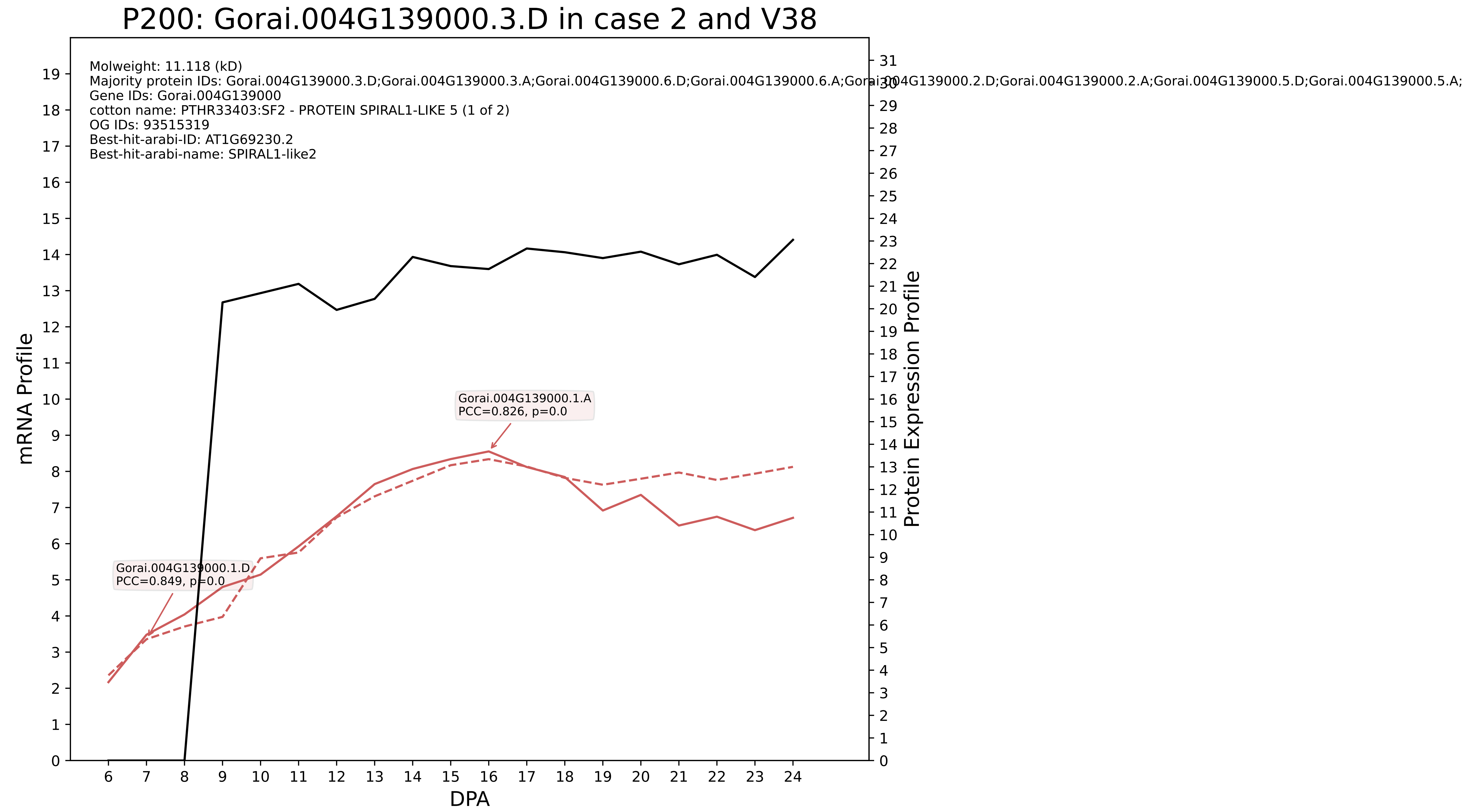
P200: Gorai.004G139000.3.D in case 2 and V38
Molweight: 11.118 (kD)
Majority protein IDs: Gorai.004G139000.3.D;Gorai.004G139000.3.A;Gorai.004G139000.6.D;Gorai.004G139000.6.A;Gorai.004G139000.2.D;Gorai.004G139000.2.A;Gorai.004G139000.5.D;Gorai.004G139000.5.A;
Gene IDs: Gorai.004G139000
cotton name: PTHR33403:SF2 - PROTEIN SPIRAL1-LIKE 5 (1 of 2)
OG IDs: 93515319
Best-hit-arabi-ID: AT1G69230.2
Best-hit-arabi-name: SPIRAL1-like2
Gorai.004G139000.1.A
PCC=0.826, p=0.0
Gorai.004G139000.1.D
PCC=0.849, p=0.0
mRNA Profile
Protein Expression Profile
DPA
0 1 2 3 4 5 6 7 8 9 10 11 12 13 14 15 16 17 18 19
0 1 2 3 4 5 6 7 8 9 10 11 12 13 14 15 16 17 18 19 20 21 22 23 24 25 26 27 28 29 30 31
6 7 8 9 10 11 12 13 14 15 16 17 18 19 20 21 22 23 24

### P200: Gorai.011G230400.1.D in case 2 and V38

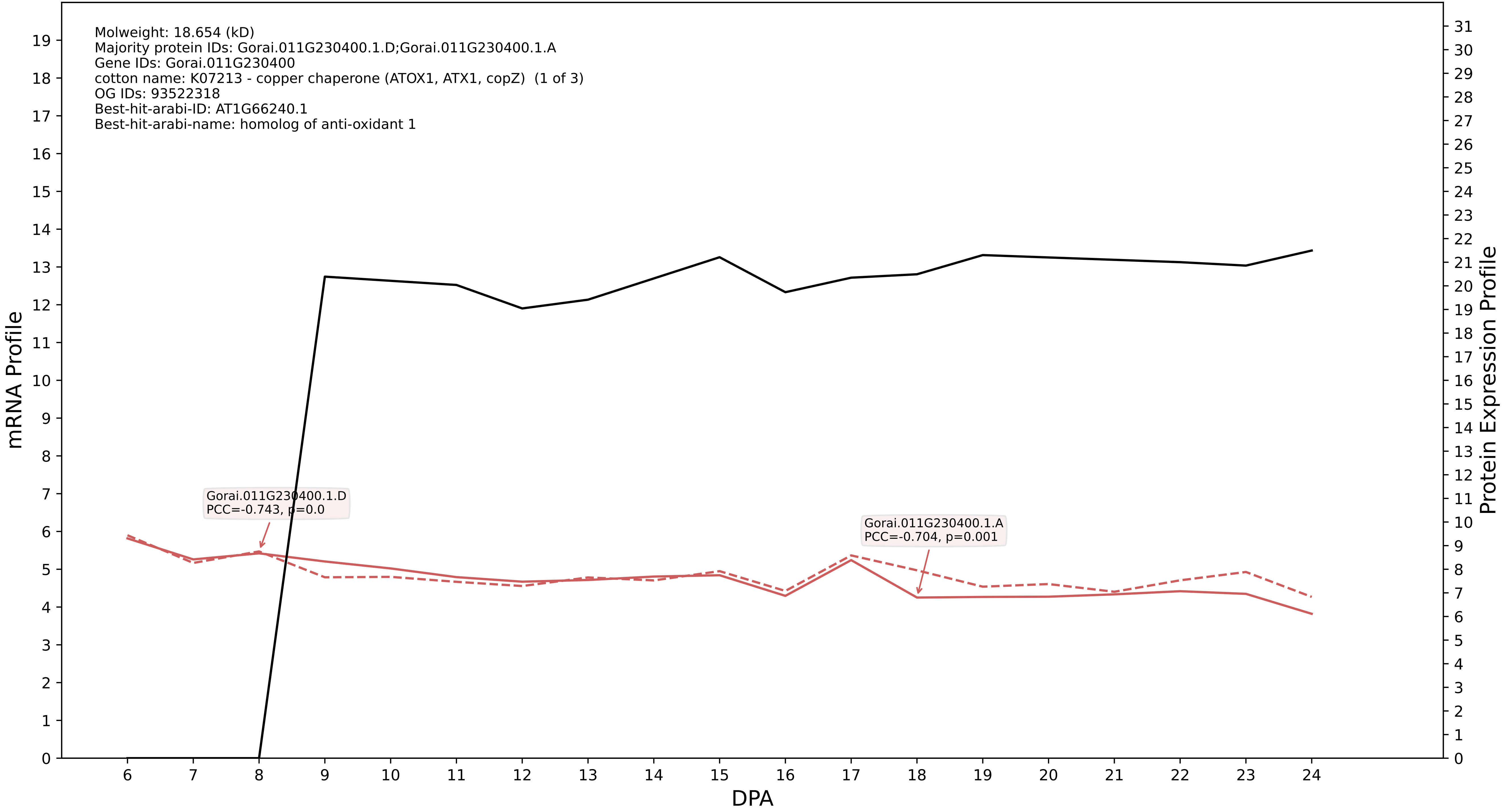

### P200: Gorai.007G085600.5.D in case 2 and V38

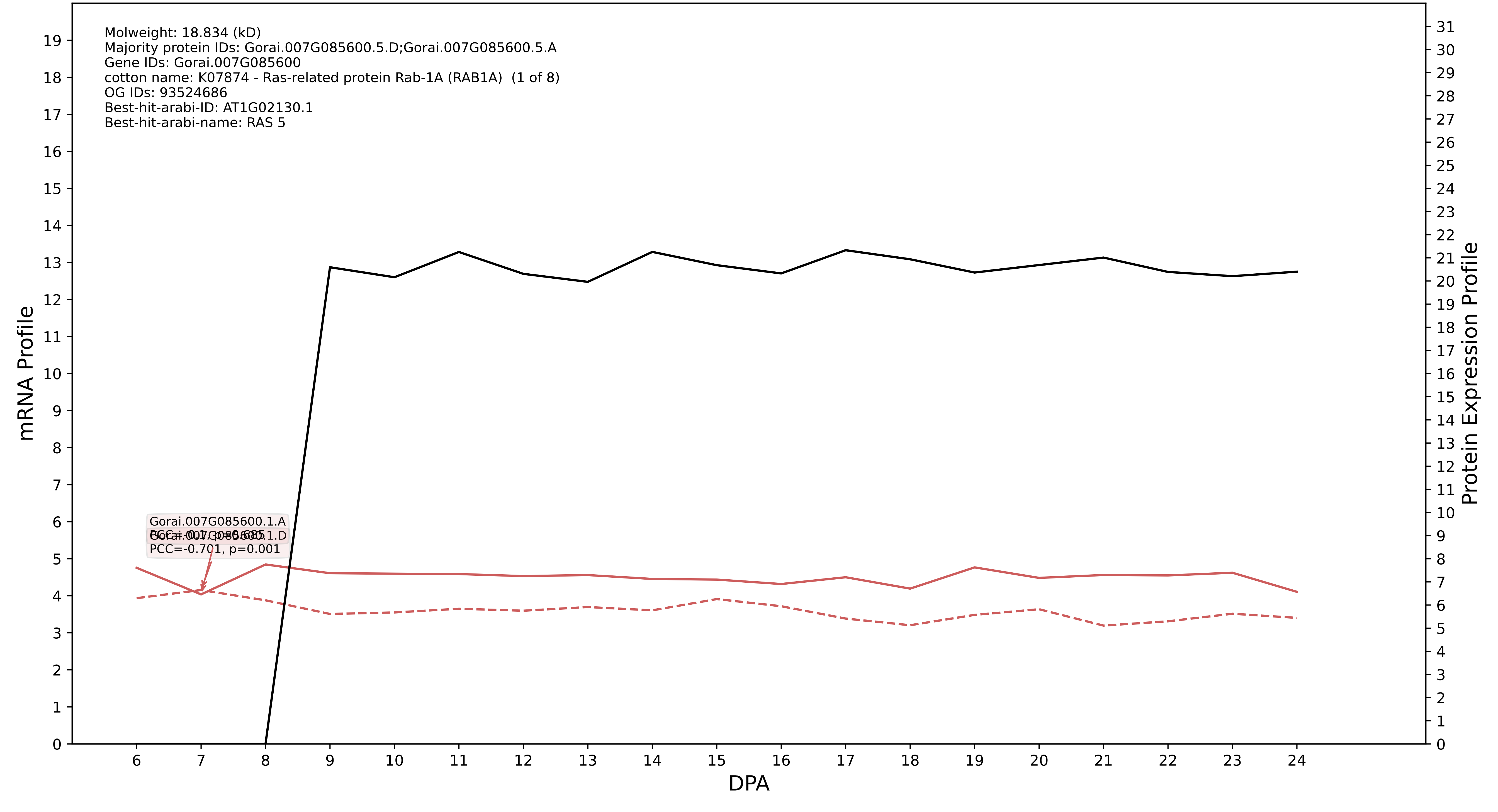

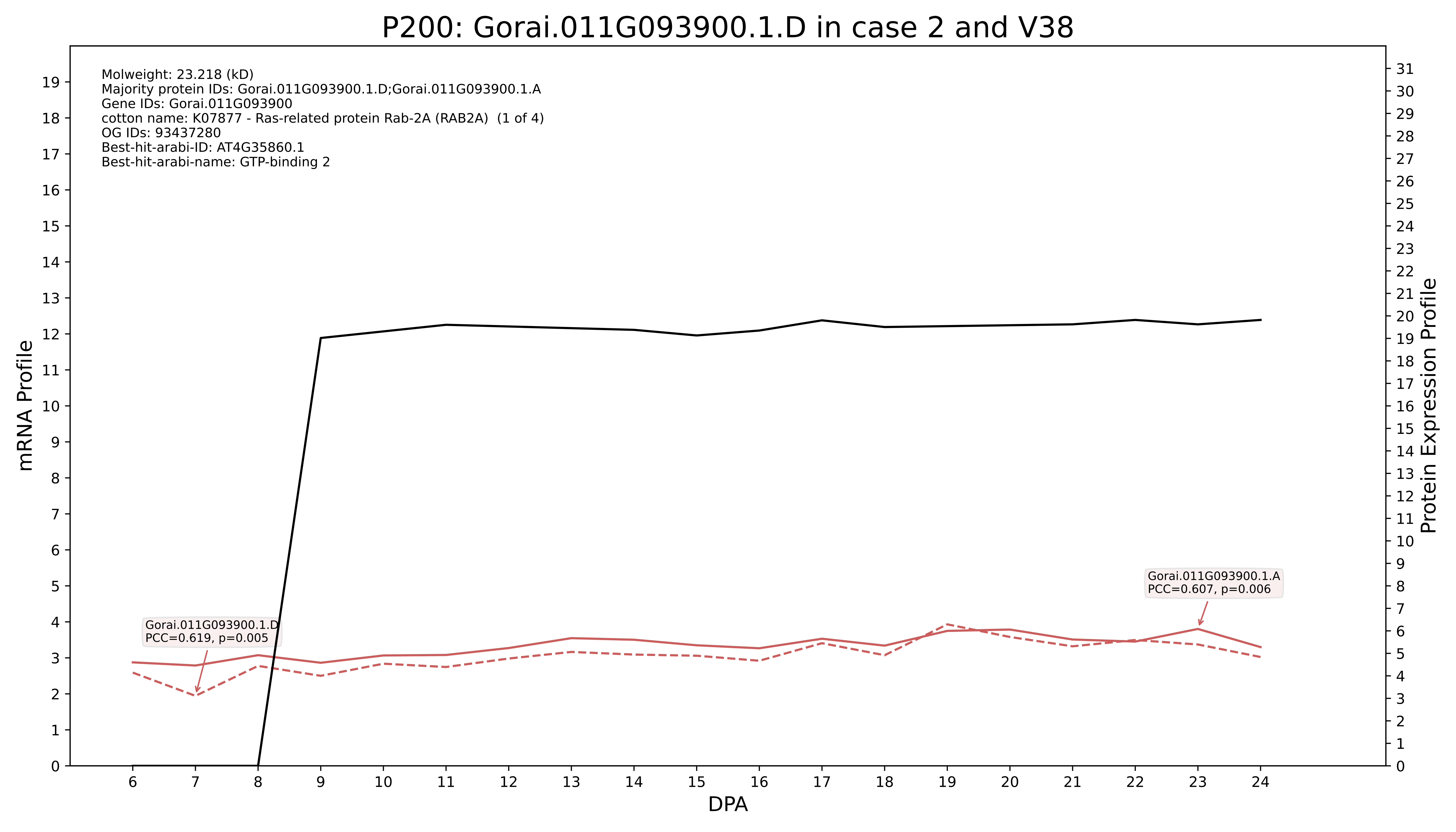

P200: Gorai.011G093900.1.D in case 2 and V38
Molweight: 23.218 (kD)
Majority protein IDs: Gorai.011G093900.1.D;Gorai.011G093900.1.A
Gene IDs: Gorai.011G093900
cotton name: K07877 - Ras-related protein Rab-2A (RAB2A) (1 of 4)
OG IDs: 93437280
Best-hit-arabi-ID: AT4G35860.1
Best-hit-arabi-name: GTP-binding 2
Gorai.011G093900.1.D
PCC=0.619, p=0.005
Gorai.011G093900.1.A
PCC=0.607, p=0.006
mRNA Profile
Protein Expression Profile
DPA

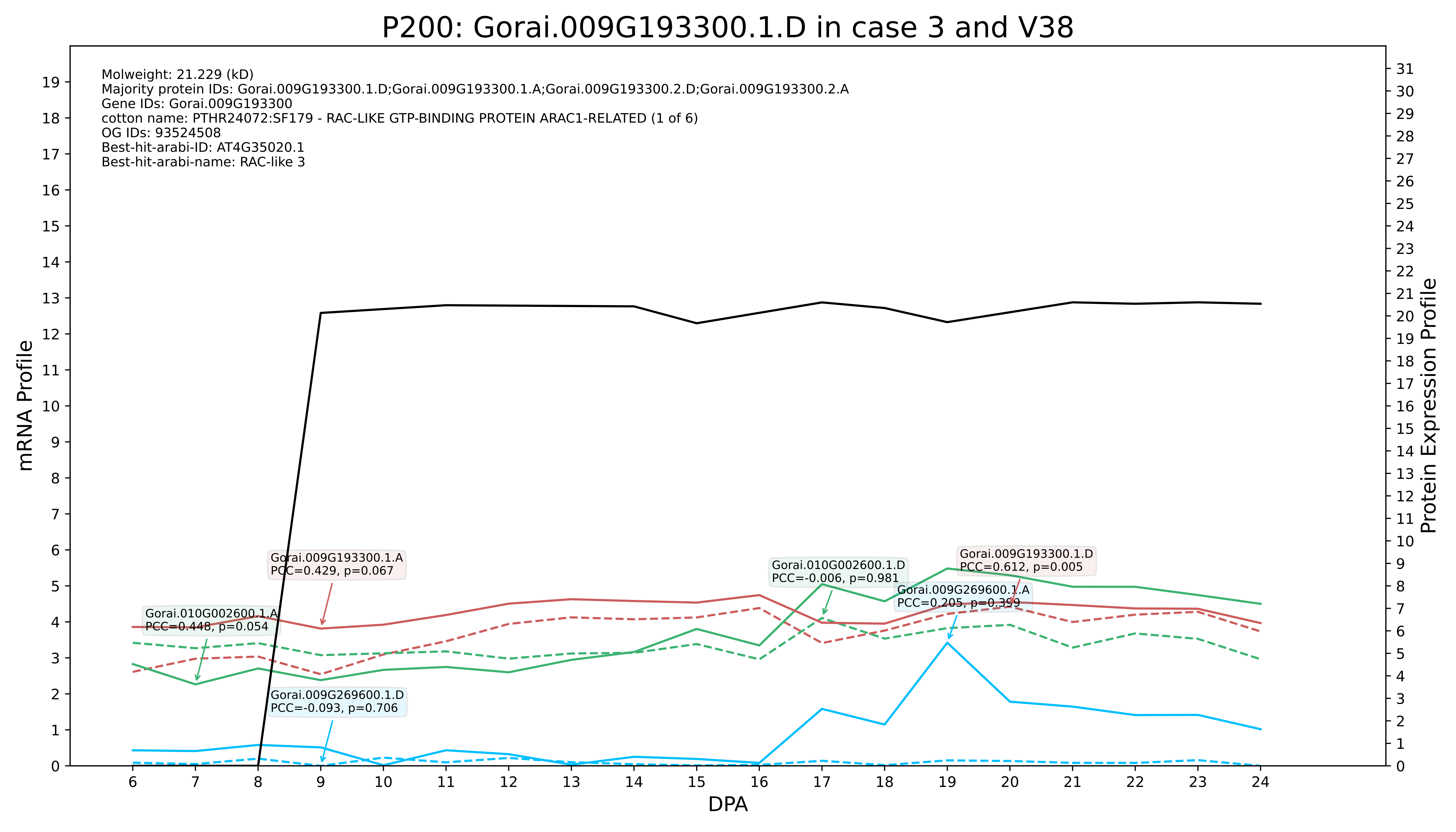

P200: Gorai.009G193300.1.D in case 3 and V38
Molweight: 21.229 (kD)
Majority protein IDs: Gorai.009G193300.1.D;Gorai.009G193300.1.A;Gorai.009G193300.2.D;Gorai.009G193300.2.A
Gene IDs: Gorai.009G193300
cotton name: PTHR24072:SF179 - RAC-LIKE GTP-BINDING PROTEIN ARAC1-RELATED (1 of 6)
OG IDs: 93524508
Best-hit-arabi-ID: AT4G35020.1
Best-hit-arabi-name: RAC-like 3
Gorai.009G193300.1.A
PCC=0.429, p=0.067
Gorai.010G002600.1.A
PCC=0.448, p=0.054
Gorai.010G002600.1.D
PCC=-0.006, p=0.981
Gorai.009G193300.1.D
PCC=0.612, p=0.005
Gorai.009G269600.1.A
PCC=0.205, p=0.399
Gorai.009G269600.1.D
PCC=-0.093, p=0.706
mRNA Profile
Protein Expression Profile
DPA
