## Supplementary Figure S12 for "Systems-level proteomic models of cotton fiber development: A high-resolution resource to analyze cell dynamics and trait engineering"

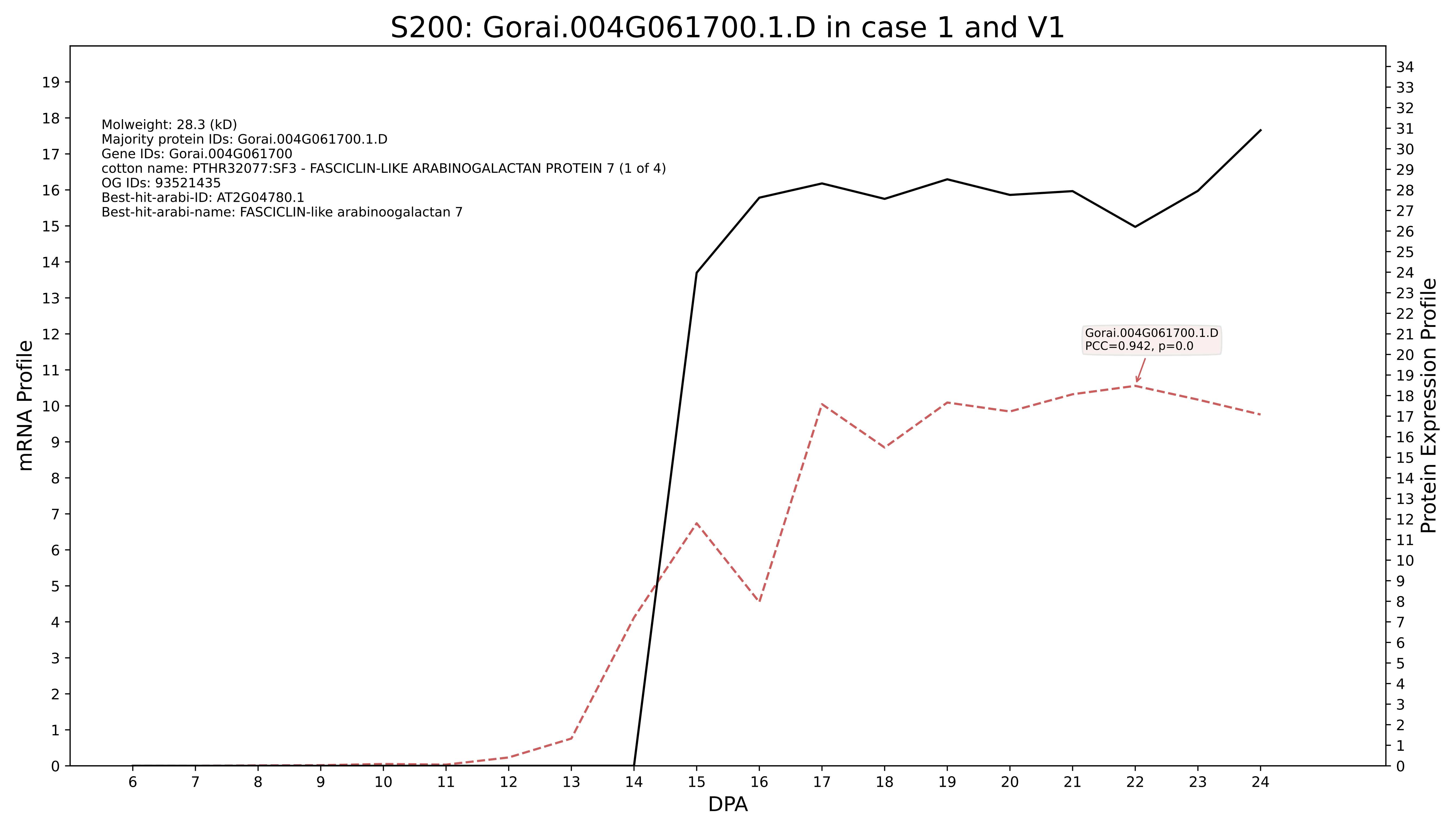
S200: Gorai.004G061700.1.D in case 1 and V1
Molweight: 28.3 (kD)
Majority protein IDs: Gorai.004G061700.1.D
Gene IDs: Gorai.004G061700
cotton name: PTHR32077:SF3 - FASCICLIN-LIKE ARABINOGALACTAN PROTEIN 7 (1 of 4)
OG IDs: 93521435
Best-hit-arabi-ID: AT2G04780.1
Best-hit-arabi-name: FASCICLIN-like arabinoogalactan 7
Gorai.004G061700.1.D
PCC=0.942, p=0.0
mRNA Profile
Protein Expression Profile
DPA

S200: Gorai.009G040600.1.A in case 1 and V1

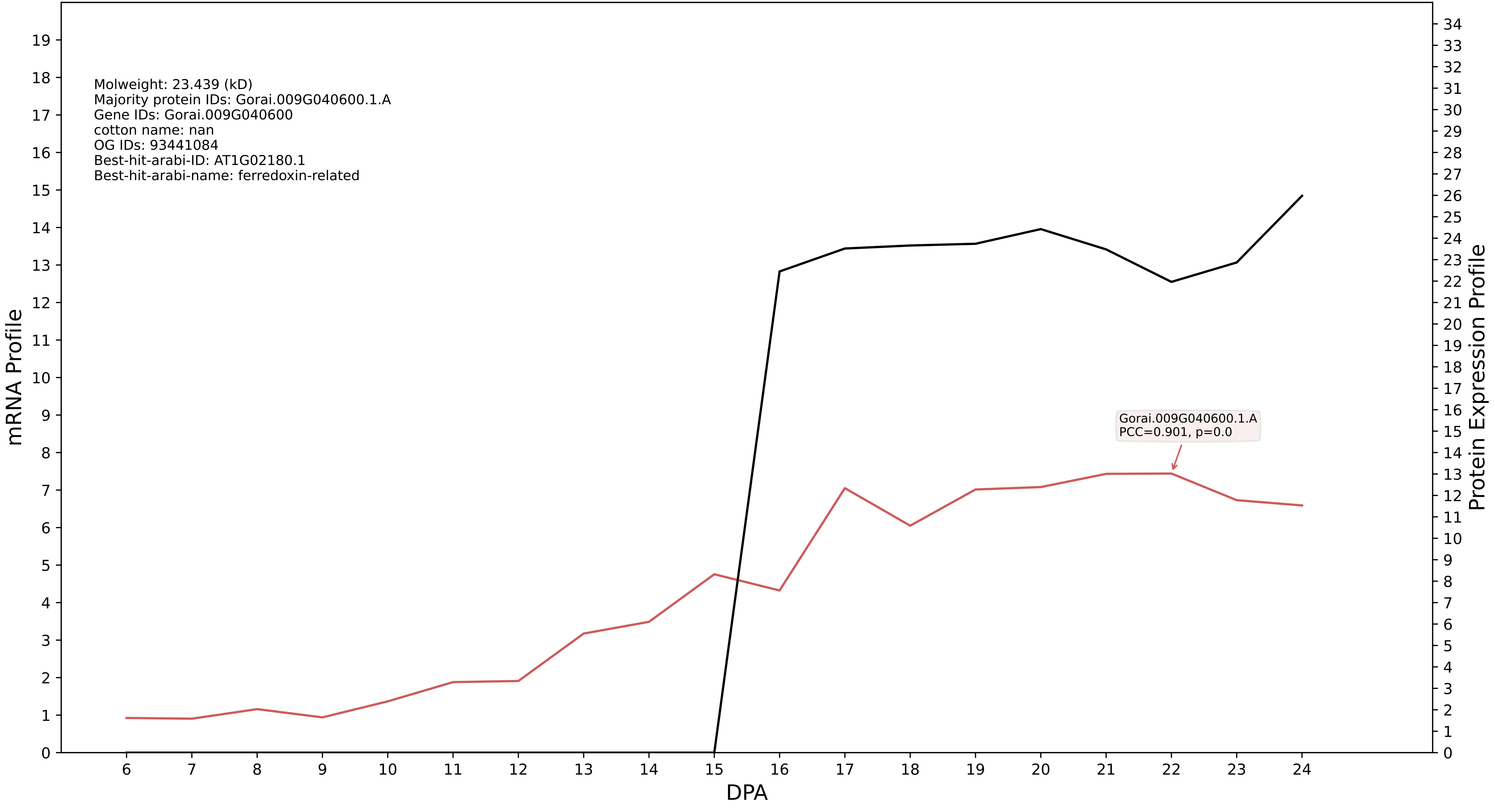

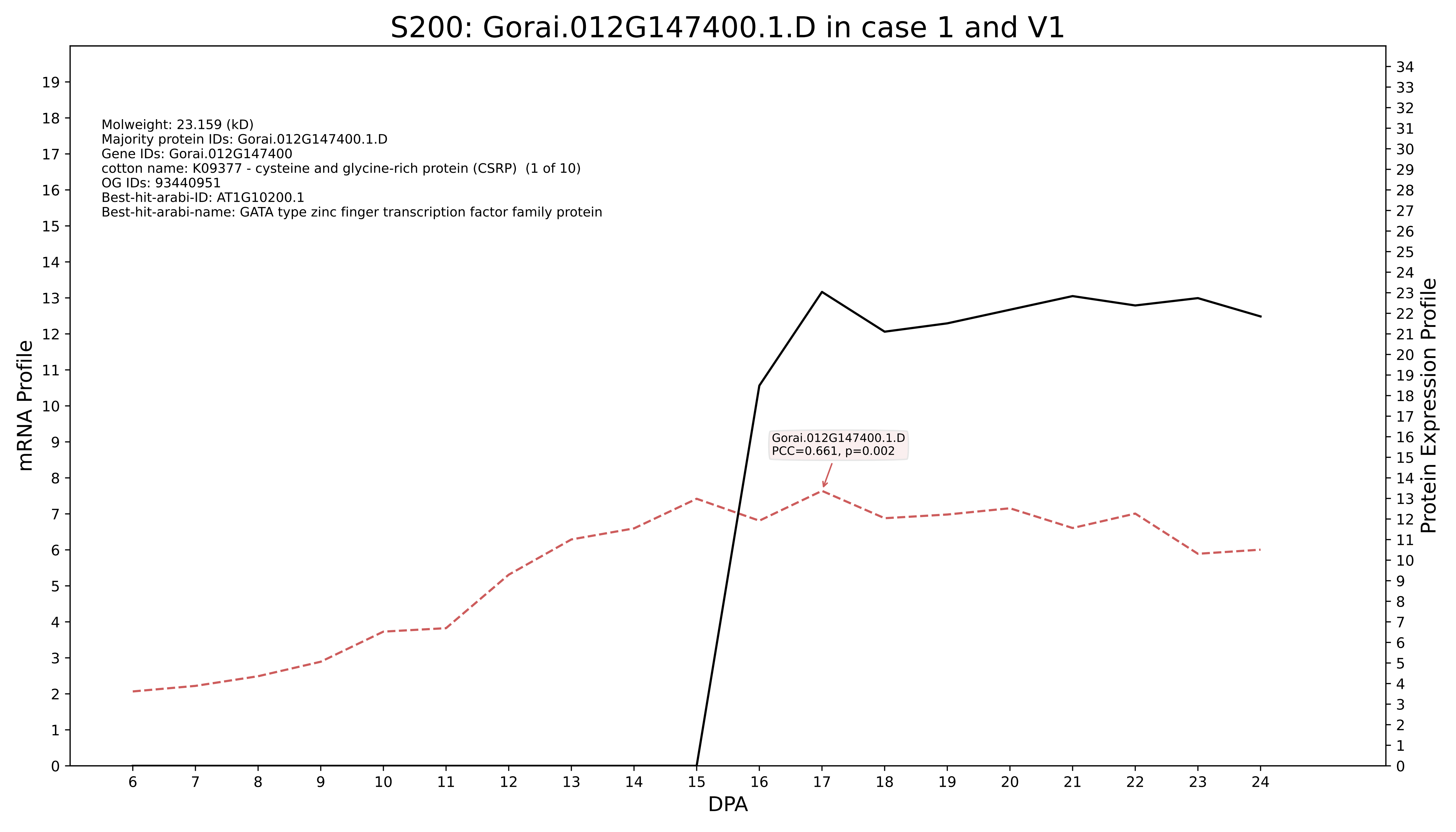
S200: Gorai.012G147400.1.D in case 1 and V1
Molweight: 23.159 (kD)
Majority protein IDs: Gorai.012G147400.1.D
Gene IDs: Gorai.012G147400
cotton name: K09377 - cysteine and glycine-rich protein (CSRP) (1 of 10)
OG IDs: 93440951
Best-hit-arabi-ID: AT1G10200.1
Best-hit-arabi-name: GATA type zinc finger transcription factor family protein
Gorai.012G147400.1.D
PCC=0.661, p=0.002
mRNA Profile
Protein Expression Profile
DPA

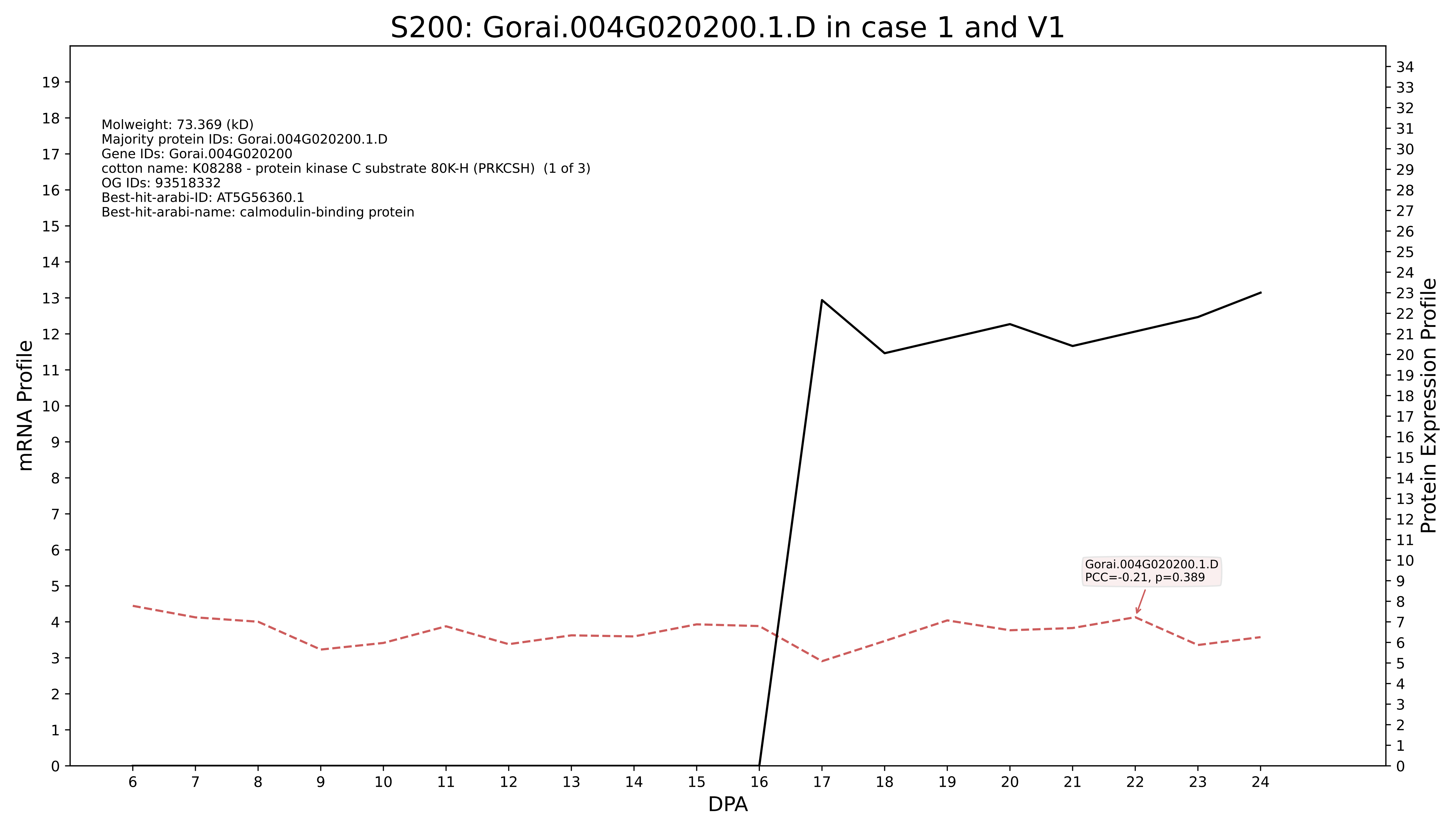

S200: Gorai.004G020200.1.D in case 1 and V1
Molweight: 73.369 (kD)
Majority protein IDs: Gorai.004G020200.1.D
Gene IDs: Gorai.004G020200
cotton name: K08288 - protein kinase C substrate 80K-H (PRKCSH) (1 of 3)
OG IDs: 93518332
Best-hit-arabi-ID: AT5G56360.1
Best-hit-arabi-name: calmodulin-binding protein
Gorai.004G020200.1.D
PCC=-0.21, p=0.389
mRNA Profile
Protein Expression Profile
DPA

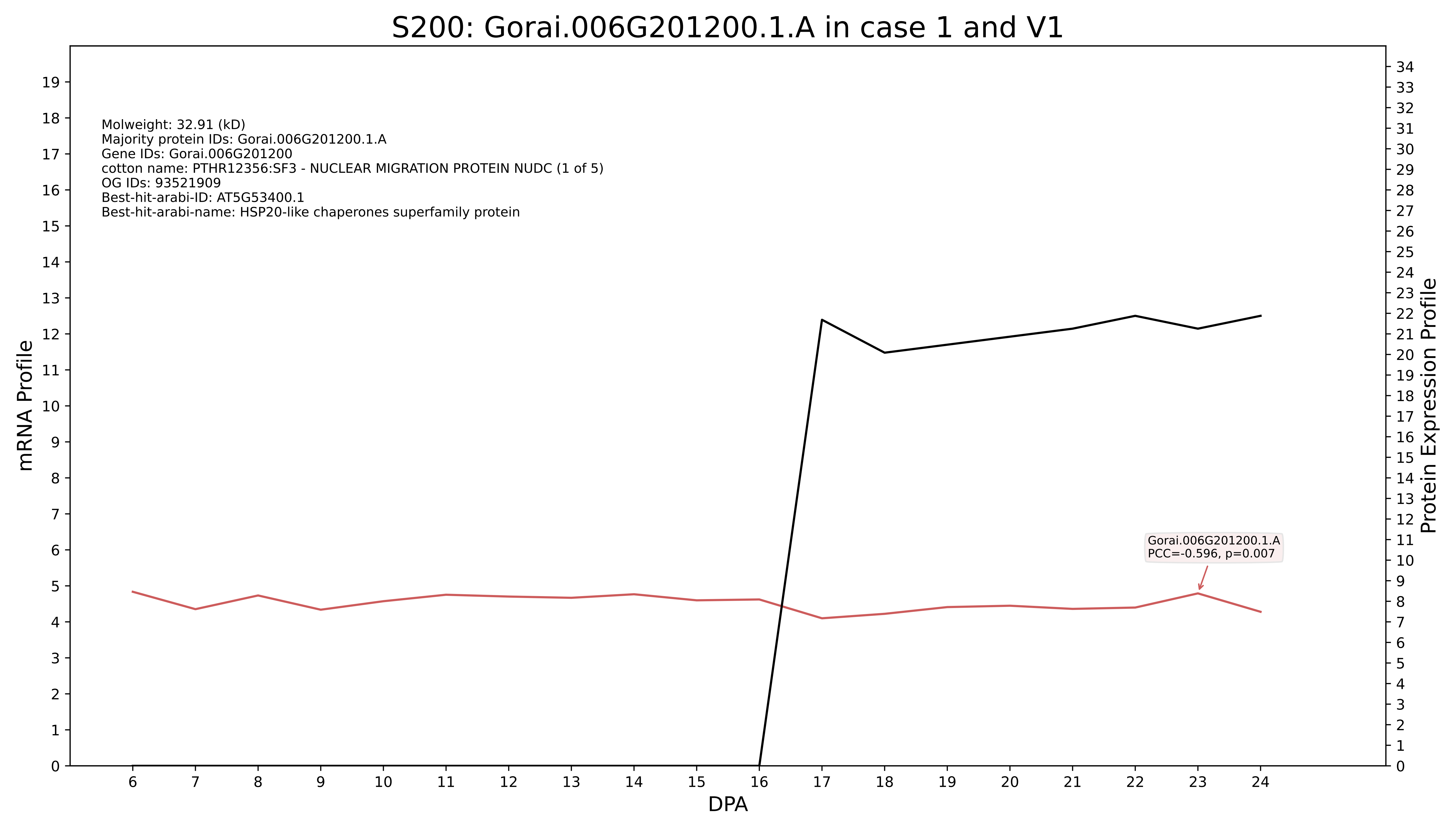
S200: Gorai.006G201200.1.A in case 1 and V1
Molweight: 32.91 (kD)
Majority protein IDs: Gorai.006G201200.1.A
Gene IDs: Gorai.006G201200
cotton name: PTHR12356:SF3 - NUCLEAR MIGRATION PROTEIN NUDC (1 of 5)
OG IDs: 93521909
Best-hit-arabi-ID: AT5G53400.1
Best-hit-arabi-name: HSP20-like chaperones superfamily protein
Gorai.006G201200.1.A
PCC=-0.596, p=0.007
mRNA Profile
Protein Expression Profile
DPA

### S200: Gorai.009G442700.1.A in case 1 and V1

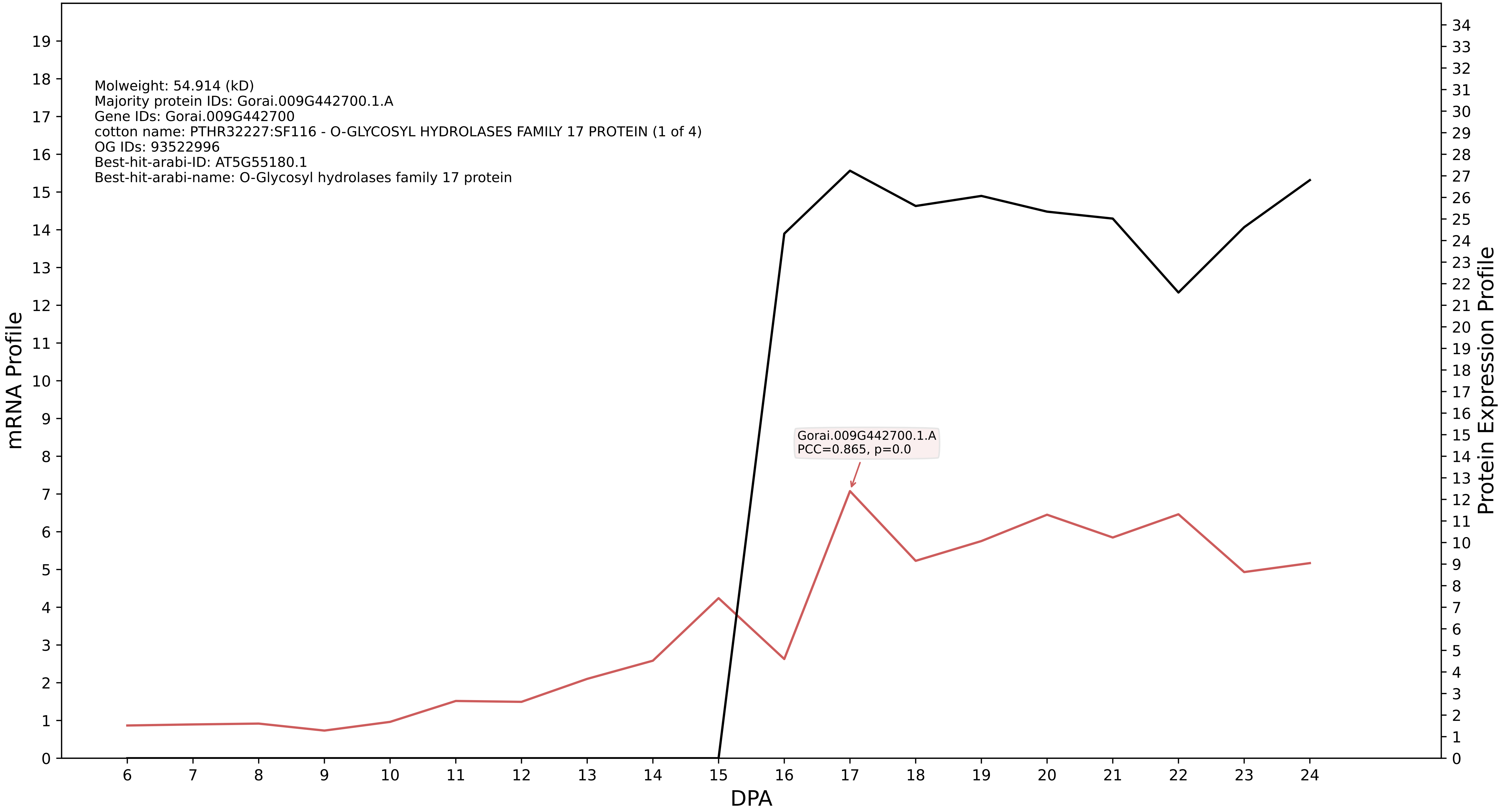

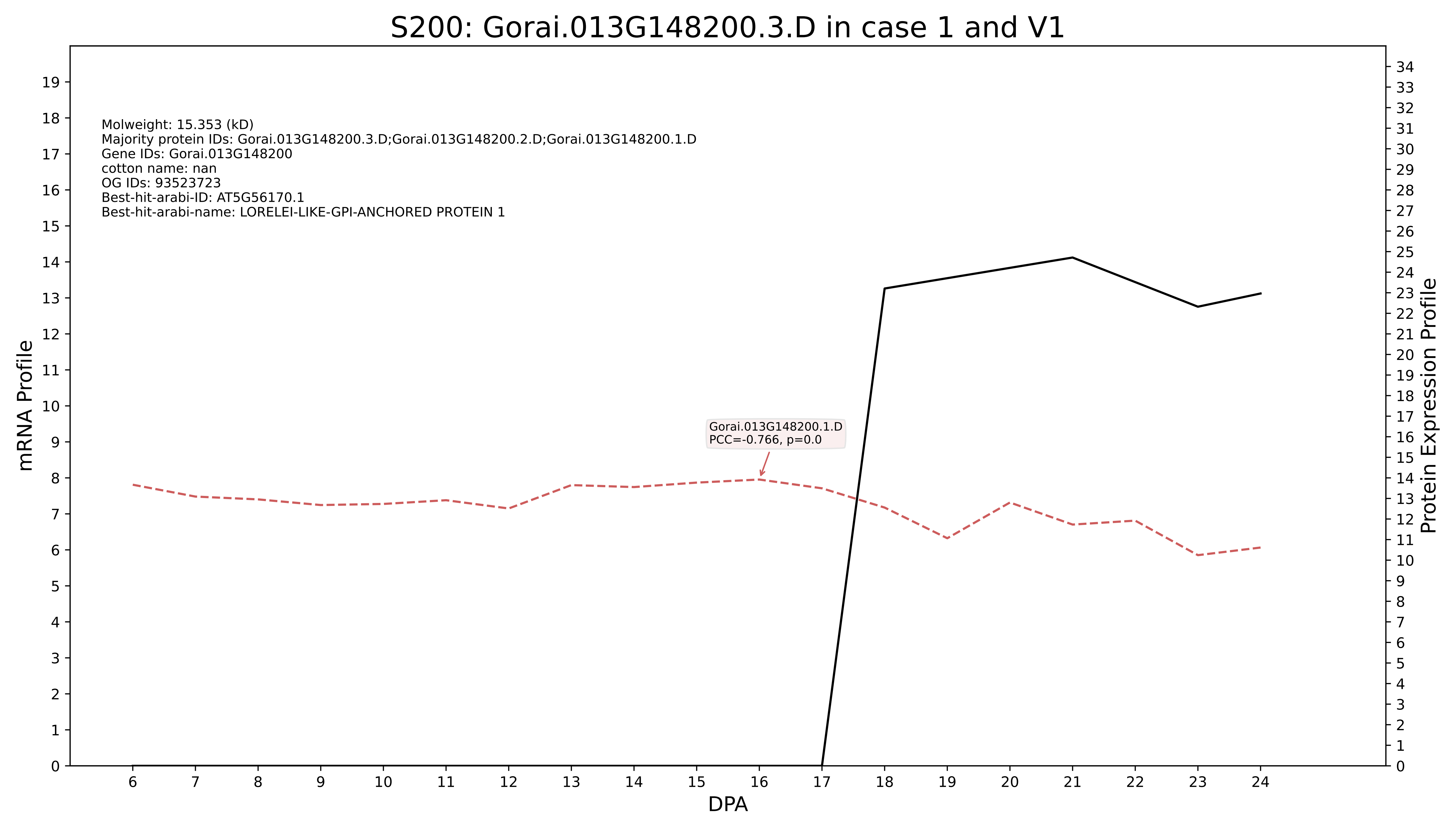

S200: Gorai.013G148200.3.D in case 1 and V1
Molweight: 15.353 (kD)
Majority protein IDs: Gorai.013G148200.3.D;Gorai.013G148200.2.D;Gorai.013G148200.1.D
Gene IDs: Gorai.013G148200
cotton name: nan
OG IDs: 93523723
Best-hit-arabi-ID: AT5G56170.1
Best-hit-arabi-name: LORELEI-LIKE-GPI-ANCHORED PROTEIN 1
Gorai.013G148200.1.D
PCC=-0.766, p=0.0
mRNA Profile
Protein Expression Profile
DPA

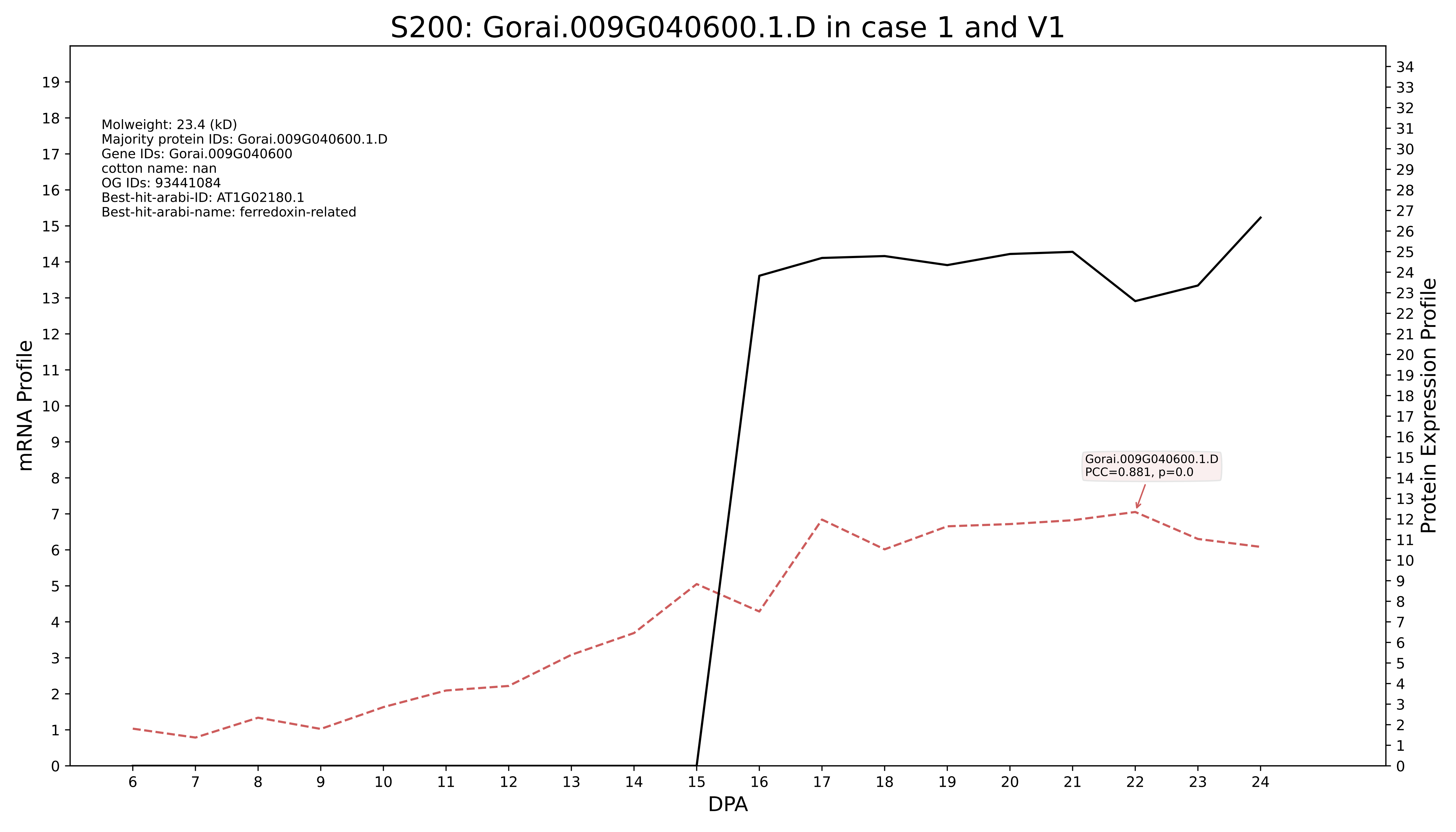
S200: Gorai.009G040600.1.D in case 1 and V1
Molweight: 23.4 (kD)
Majority protein IDs: Gorai.009G040600.1.D
Gene IDs: Gorai.009G040600
cotton name: nan
OG IDs: 93441084
Best-hit-arabi-ID: AT1G02180.1
Best-hit-arabi-name: ferredoxin-related
Gorai.009G040600.1.D
PCC=0.881, p=0.0
mRNA Profile
Protein Expression Profile
DPA

### S200: Gorai.003G006500.4.A in case 1 and V1

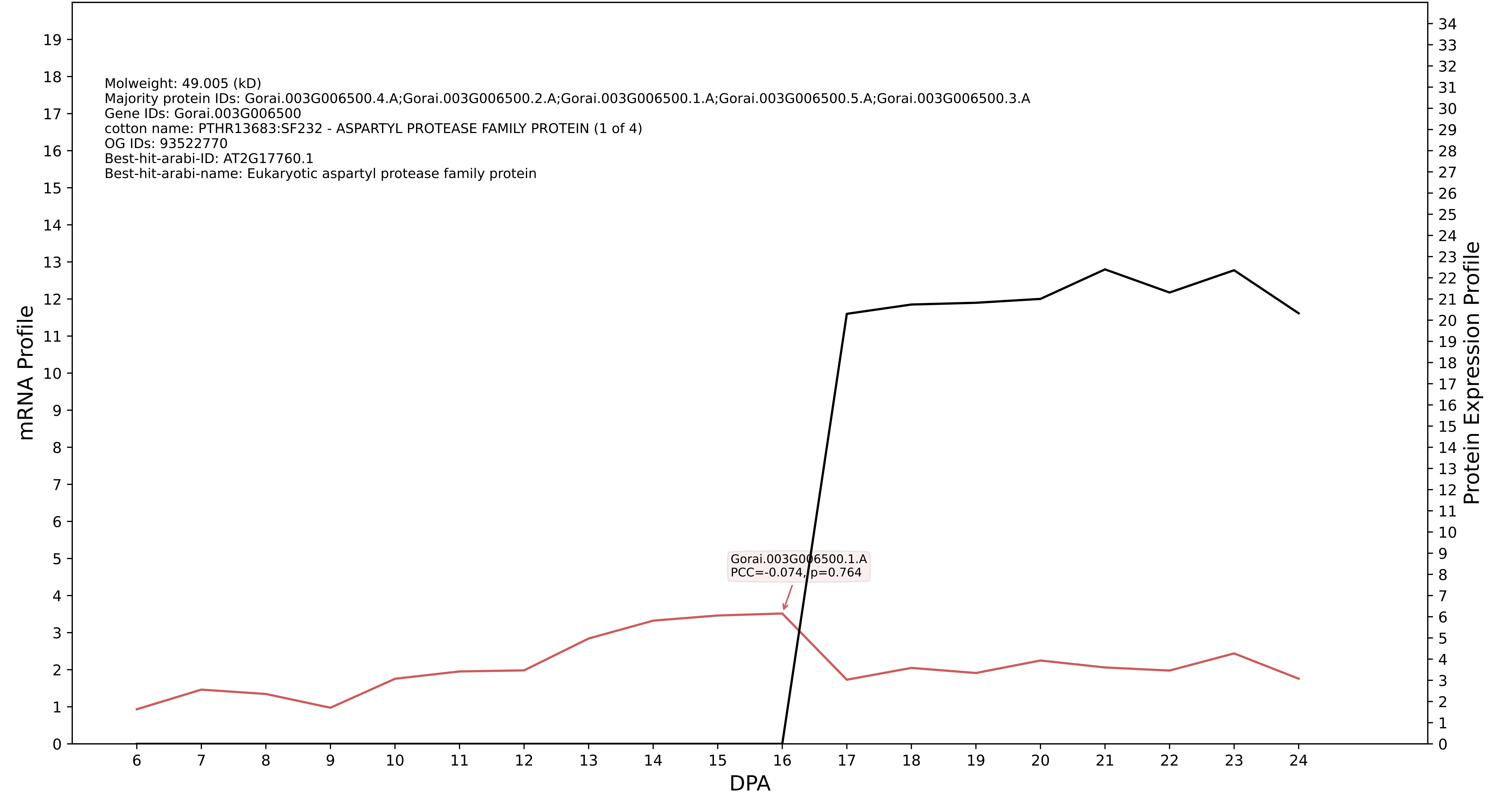

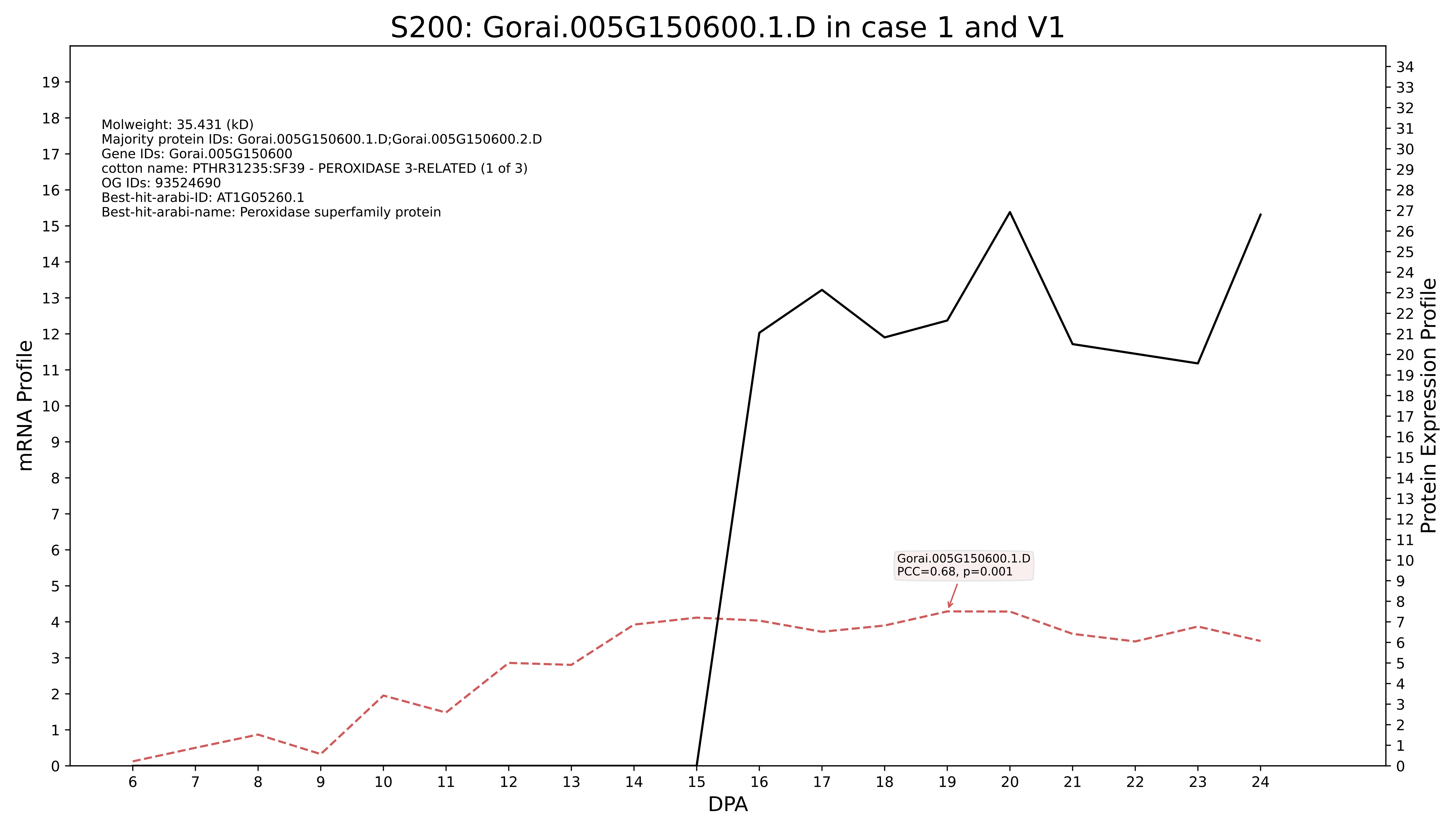
S200: Gorai.005G150600.1.D in case 1 and V1
Molweight: 35.431 (kD)
Majority protein IDs: Gorai.005G150600.1.D;Gorai.005G150600.2.D
Gene IDs: Gorai.005G150600
cotton name: PTHR31235:SF39 - PEROXIDASE 3-RELATED (1 of 3)
OG IDs: 93524690
Best-hit-arabi-ID: AT1G05260.1
Best-hit-arabi-name: Peroxidase superfamily protein
Gorai.005G150600.1.D
PCC=0.68, p=0.001
mRNA Profile
Protein Expression Profile
DPA

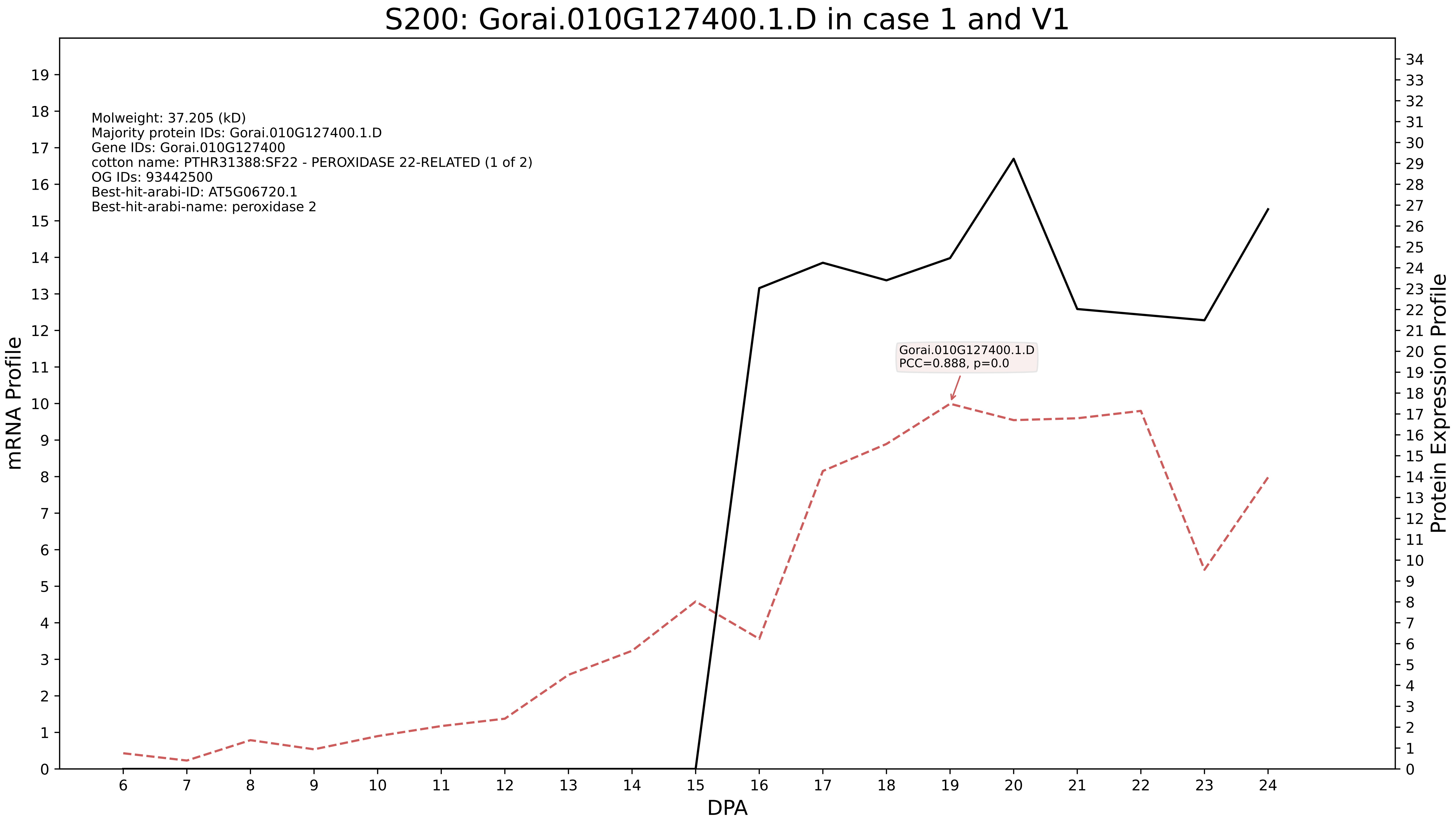
S200: Gorai.010G127400.1.D in case 1 and V1
Molweight: 37.205 (kD)
Majority protein IDs: Gorai.010G127400.1.D
Gene IDs: Gorai.010G127400
cotton name: PTHR31388:SF22 - PEROXIDASE 22-RELATED (1 of 2)
OG IDs: 93442500
Best-hit-arabi-ID: AT5G06720.1
Best-hit-arabi-name: peroxidase 2
Gorai.010G127400.1.D
PCC=0.888, p=0.0
mRNA Profile
Protein Expression Profile
DPA

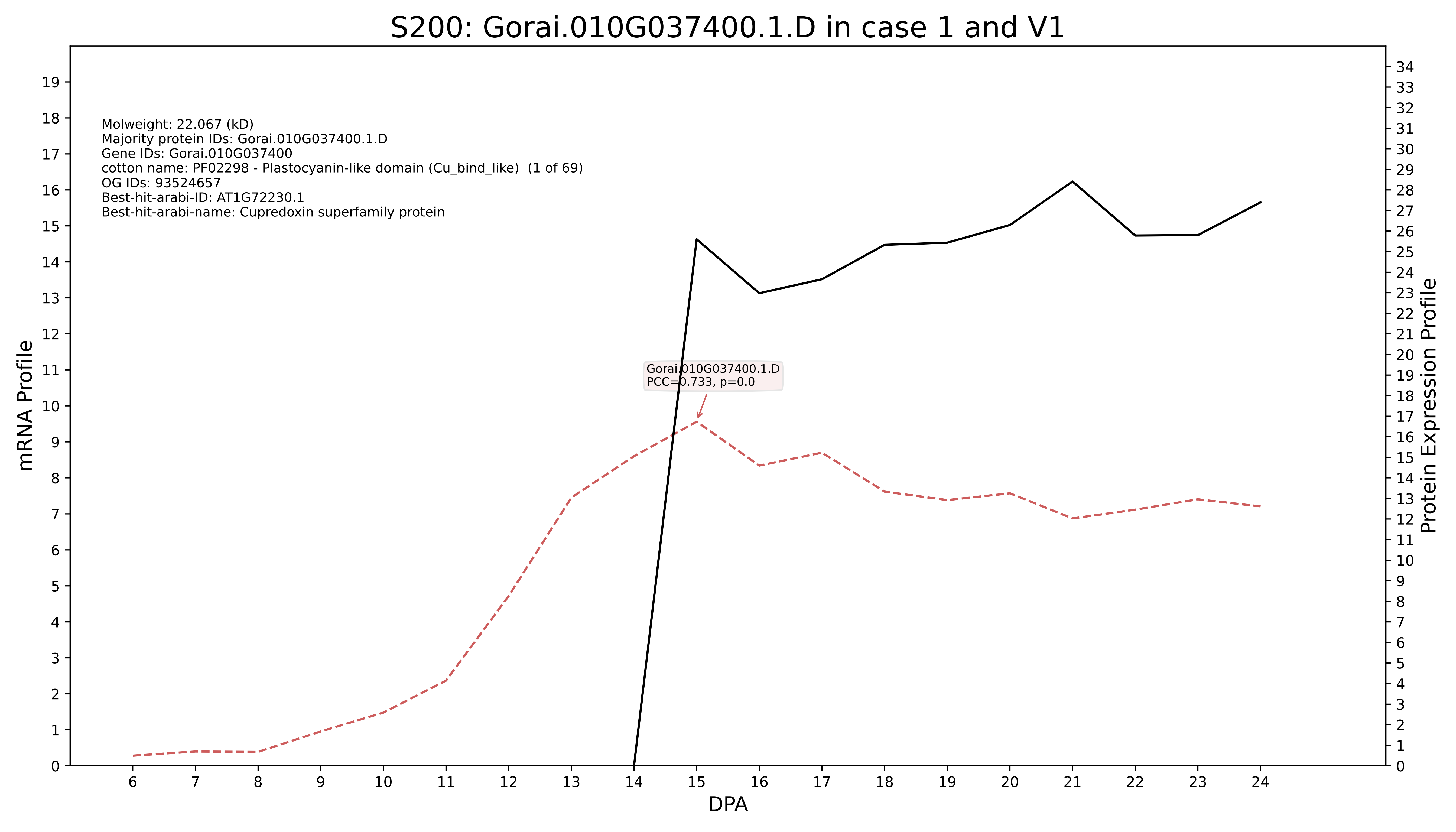
S200: Gorai.010G037400.1.D in case 1 and V1
Molweight: 22.067 (kD)
Majority protein IDs: Gorai.010G037400.1.D
Gene IDs: Gorai.010G037400
cotton name: PF02298 - Plastocyanin-like domain (Cu_bind_like) (1 of 69)
OG IDs: 93524657
Best-hit-arabi-ID: AT1G72230.1
Best-hit-arabi-name: Cupredoxin superfamily protein
Gorai.010G037400.1.D
PCC=0.733, p=0.0
mRNA Profile
Protein Expression Profile
DPA

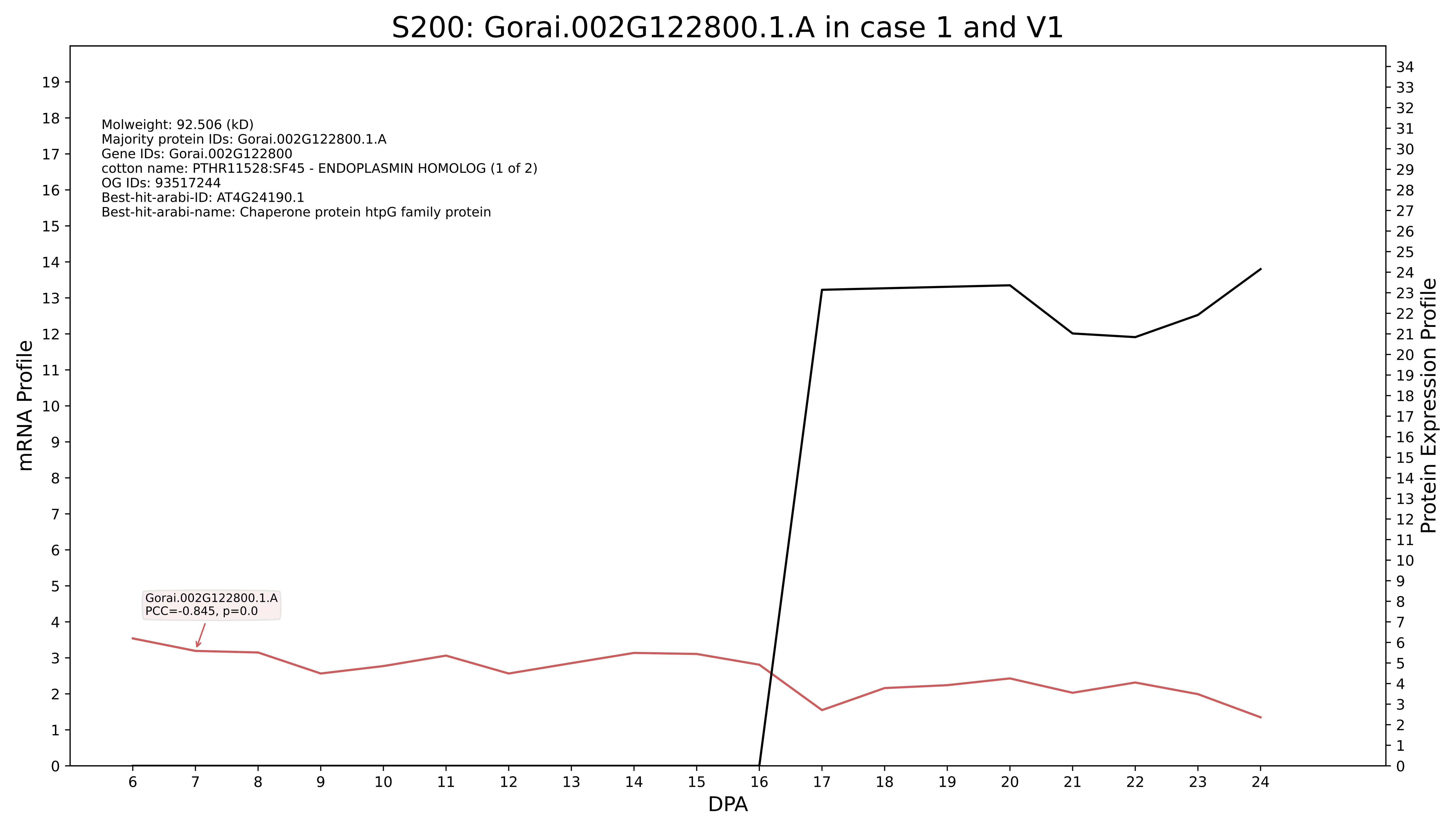

S200: Gorai.002G122800.1.A in case 1 and V1
Molweight: 92.506 (kD)
Majority protein IDs: Gorai.002G122800.1.A
Gene IDs: Gorai.002G122800
cotton name: PTHR11528:SF45 - ENDOPLASMIN HOMOLOG (1 of 2)
OG IDs: 93517244
Best-hit-arabi-ID: AT4G24190.1
Best-hit-arabi-name: Chaperone protein htpG family protein
Gorai.002G122800.1.A
PCC=-0.845, p=0.0
mRNA Profile
Protein Expression Profile
DPA

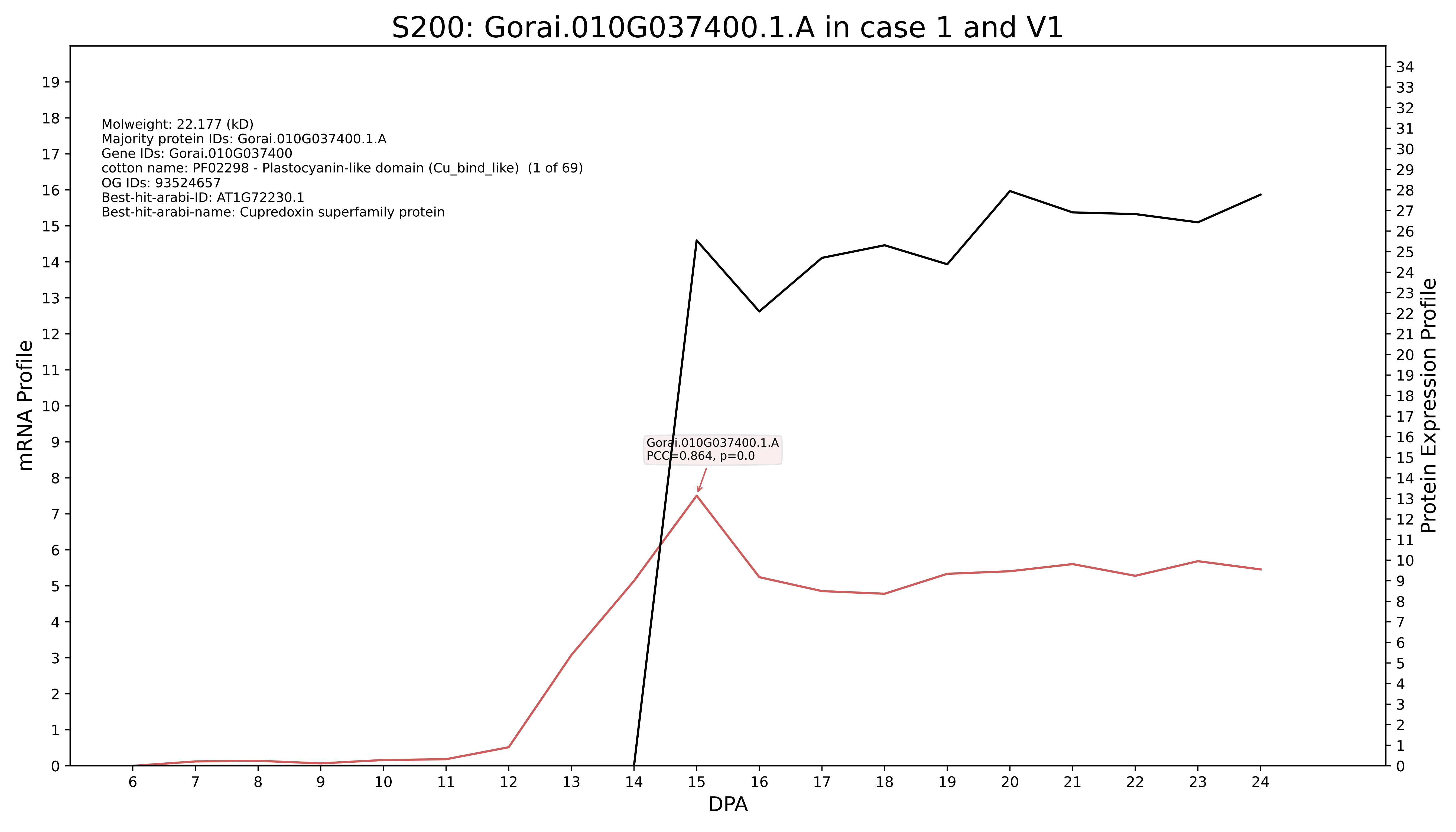
S200: Gorai.010G037400.1.A in case 1 and V1
Molweight: 22.177 (kD)
Majority protein IDs: Gorai.010G037400.1.A
Gene IDs: Gorai.010G037400
cotton name: PF02298 - Plastocyanin-like domain (Cu_bind_like) (1 of 69)
OG IDs: 93524657
Best-hit-arabi-ID: AT1G72230.1
Best-hit-arabi-name: Cupredoxin superfamily protein
Gorai.010G037400.1.A
PCC=0.864, p=0.0
mRNA Profile
Protein Expression Profile
DPA

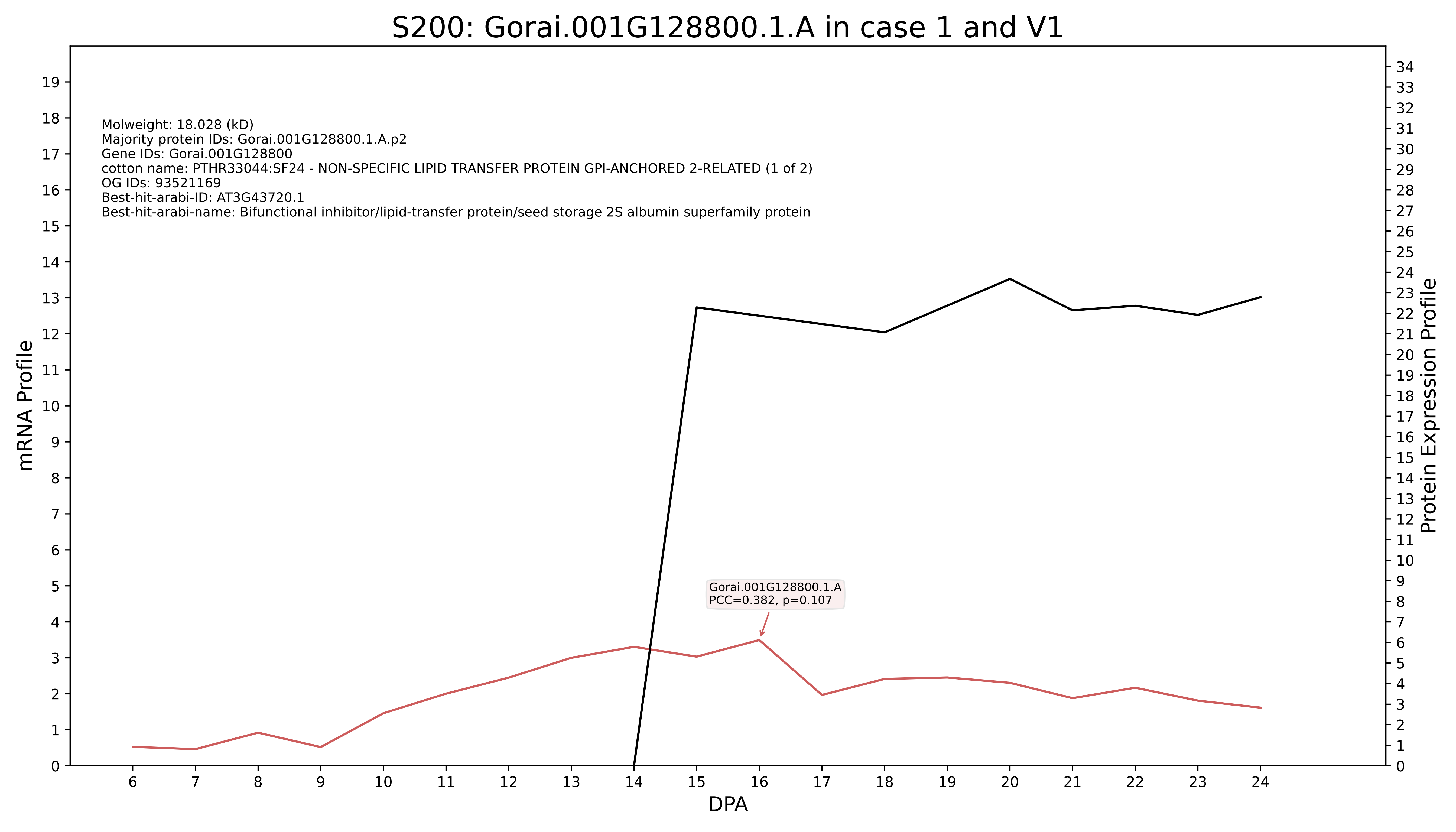

S200: Gorai.001G128800.1.A in case 1 and V1
Molweight: 18.028 (kD)
Majority protein IDs: Gorai.001G128800.1.A.p2
Gene IDs: Gorai.001G128800
cotton name: PTHR33044:SF24 - NON-SPECIFIC LIPID TRANSFER PROTEIN GPI-ANCHORED 2-RELATED (1 of 2)
OG IDs: 93521169
Best-hit-arabi-ID: AT3G43720.1
Best-hit-arabi-name: Bifunctional inhibitor/lipid-transfer protein/seed storage 2S albumin superfamily protein
Gorai.001G128800.1.A
PCC=0.382, p=0.107
mRNA Profile
Protein Expression Profile
DPA

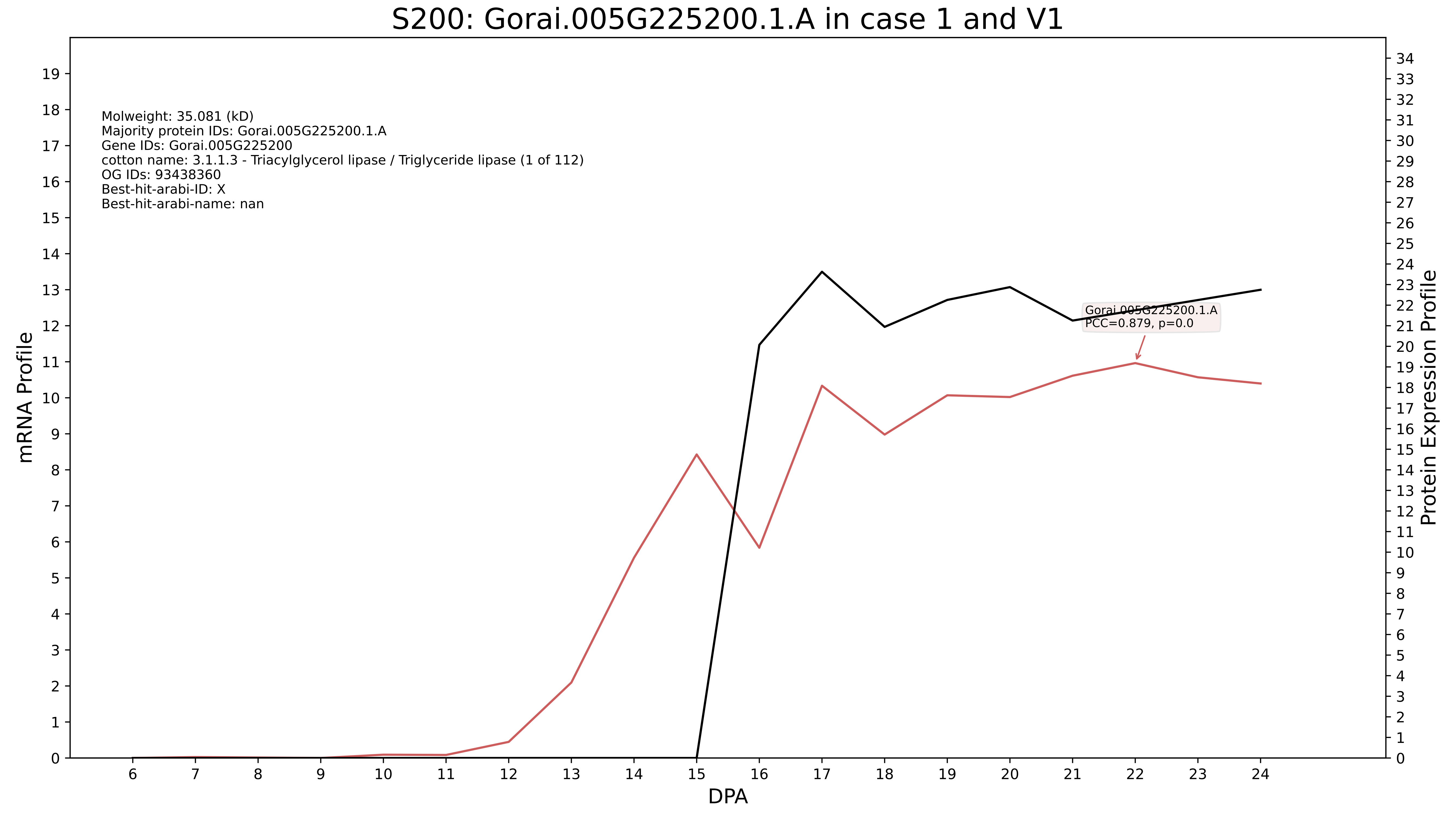

S200: Gorai.005G225200.1.A in case 1 and V1
Molweight: 35.081 (kD)
Majority protein IDs: Gorai.005G225200.1.A
Gene IDs: Gorai.005G225200
cotton name: 3.1.1.3 - Triacylglycerol lipase / Triglyceride lipase (1 of 112)
OG IDs: 93438360
Best-hit-arabi-ID: X
Best-hit-arabi-name: nan
Gorai.005G225200.1.A
PCC=0.879, p=0.0
mRNA Profile
Protein Expression Profile
DPA

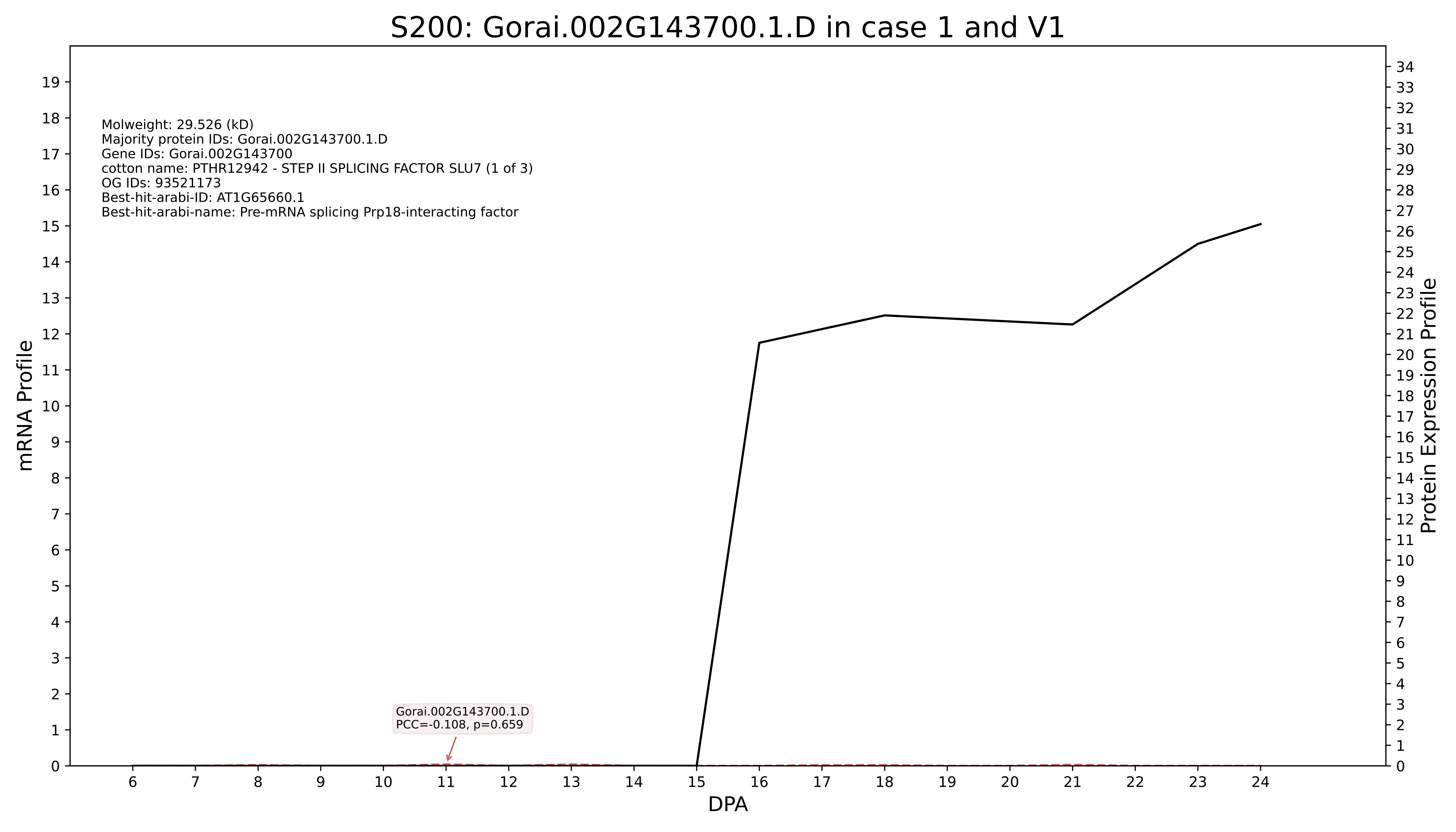

S200: Gorai.002G143700.1.D in case 1 and V1
Molweight: 29.526 (kD)
Majority protein IDs: Gorai.002G143700.1.D
Gene IDs: Gorai.002G143700
cotton name: PTHR12942 - STEP II SPLICING FACTOR SLU7 (1 of 3)
OG IDs: 93521173
Best-hit-arabi-ID: AT1G65660.1
Best-hit-arabi-name: Pre-mRNA splicing Prp18-interacting factor
Gorai.002G143700.1.D
PCC=-0.108, p=0.659
mRNA Profile
Protein Expression Profile
DPA

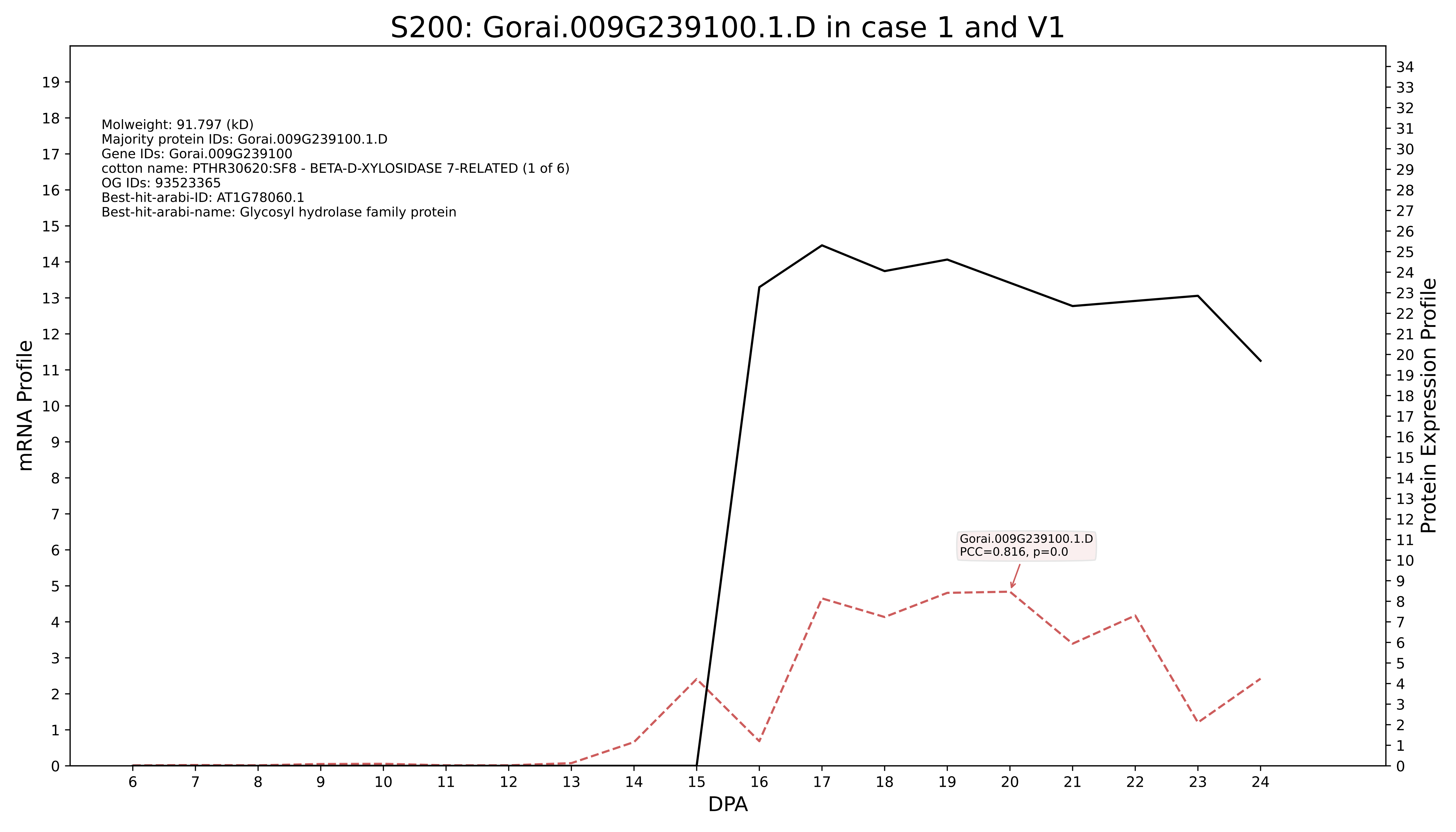
S200: Gorai.009G239100.1.D in case 1 and V1
Molweight: 91.797 (kD)
Majority protein IDs: Gorai.009G239100.1.D
Gene IDs: Gorai.009G239100
cotton name: PTHR30620:SF8 - BETA-D-XYLOSIDASE 7-RELATED (1 of 6)
OG IDs: 93523365
Best-hit-arabi-ID: AT1G78060.1
Best-hit-arabi-name: Glycosyl hydrolase family protein
Gorai.009G239100.1.D
PCC=0.816, p=0.0
mRNA Profile
Protein Expression Profile
DPA

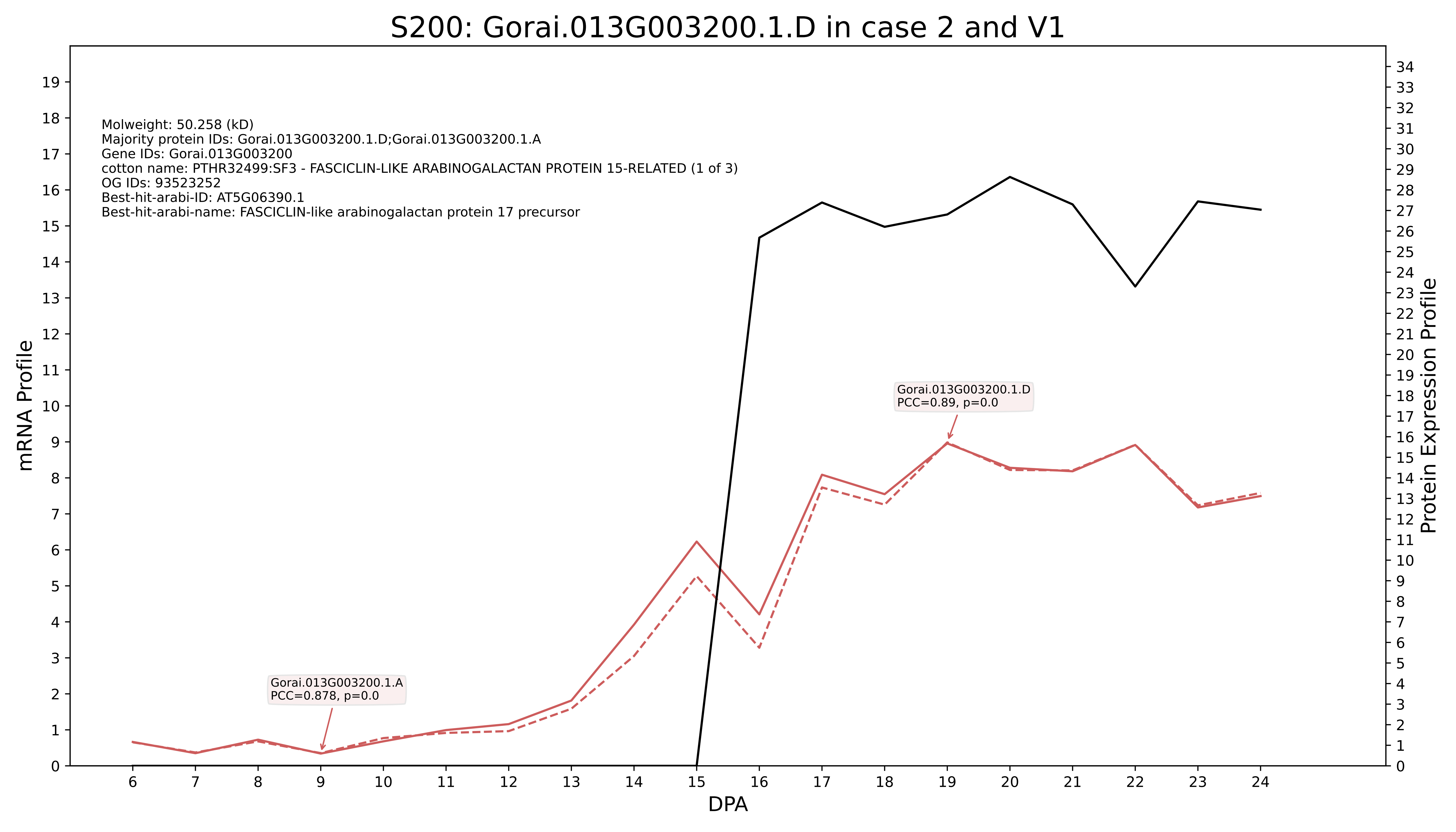
S200: Gorai.013G003200.1.D in case 2 and V1
Molweight: 50.258 (kD)
Majority protein IDs: Gorai.013G003200.1.D;Gorai.013G003200.1.A
Gene IDs: Gorai.013G003200
cotton name: PTHR32499:SF3 - FASCICLIN-LIKE ARABINOGALACTAN PROTEIN 15-RELATED (1 of 3)
OG IDs: 93523252
Best-hit-arabi-ID: AT5G06390.1
Best-hit-arabi-name: FASCICLIN-like arabinogalactan protein 17 precursor
Gorai.013G003200.1.D
PCC=0.89, p=0.0
Gorai.013G003200.1.A
PCC=0.878, p=0.0
mRNA Profile
Protein Expression Profile
DPA

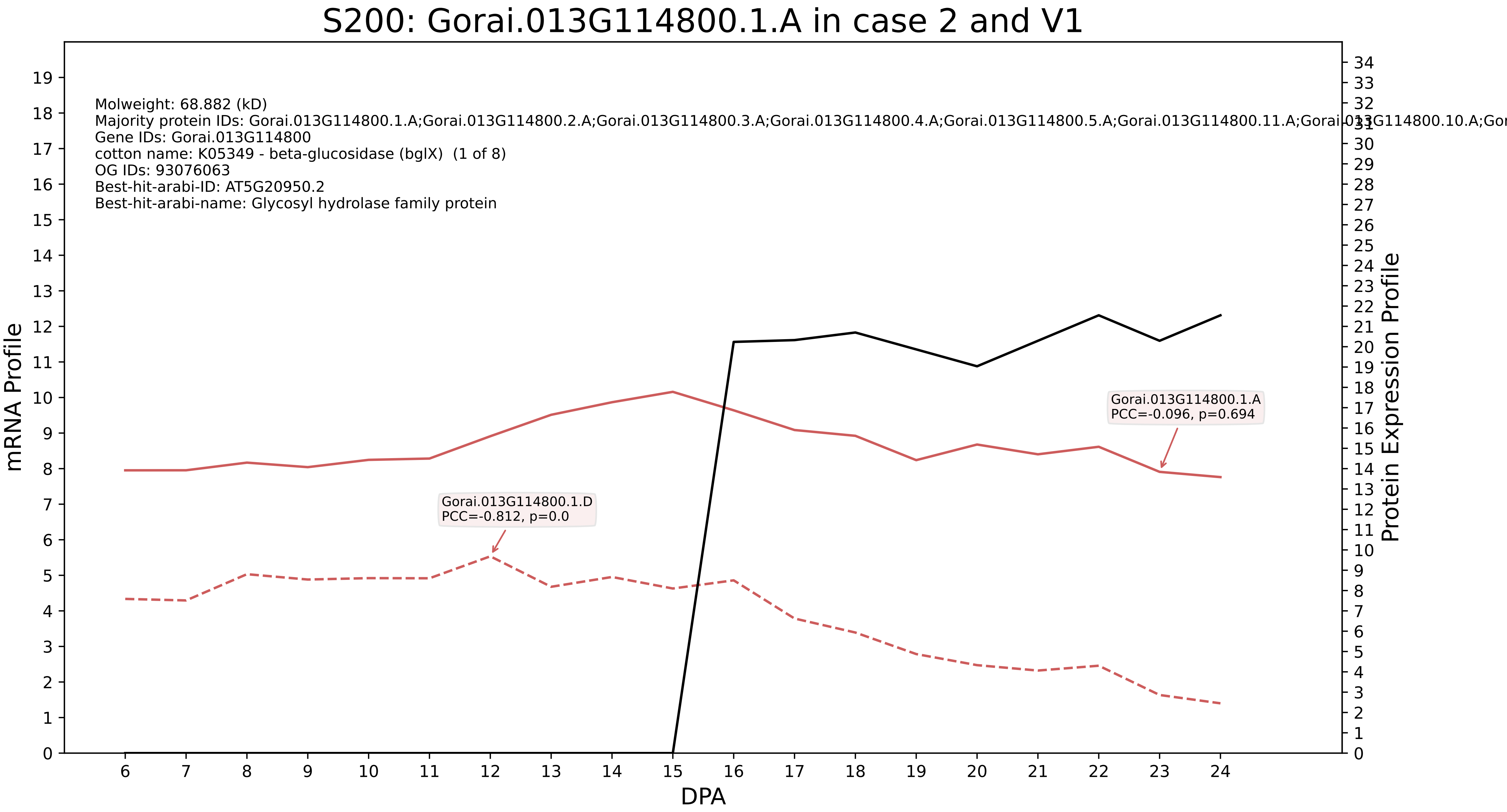

S200: Gorai.013G114800.1.A in case 2 and V1
Molweight: 68.882 (kD)
Majority protein IDs: Gorai.013G114800.1.A;Gorai.013G114800.2.A;Gorai.013G114800.3.A;Gorai.013G114800.4.A;Gorai.013G114800.5.A;Gorai.013G114800.11.A;Gorai.013G114800.10.A;Gor
Gene IDs: Gorai.013G114800
cotton name: K05349 - beta-glucosidase (bglX) (1 of 8)
OG IDs: 93076063
Best-hit-arabi-ID: AT5G20950.2
Best-hit-arabi-name: Glycosyl hydrolase family protein
Gorai.013G114800.1.A
PCC=-0.096, p=0.694
Gorai.013G114800.1.D
PCC=-0.812, p=0.0
mRNA Profile
Protein Expression Profile
DPA

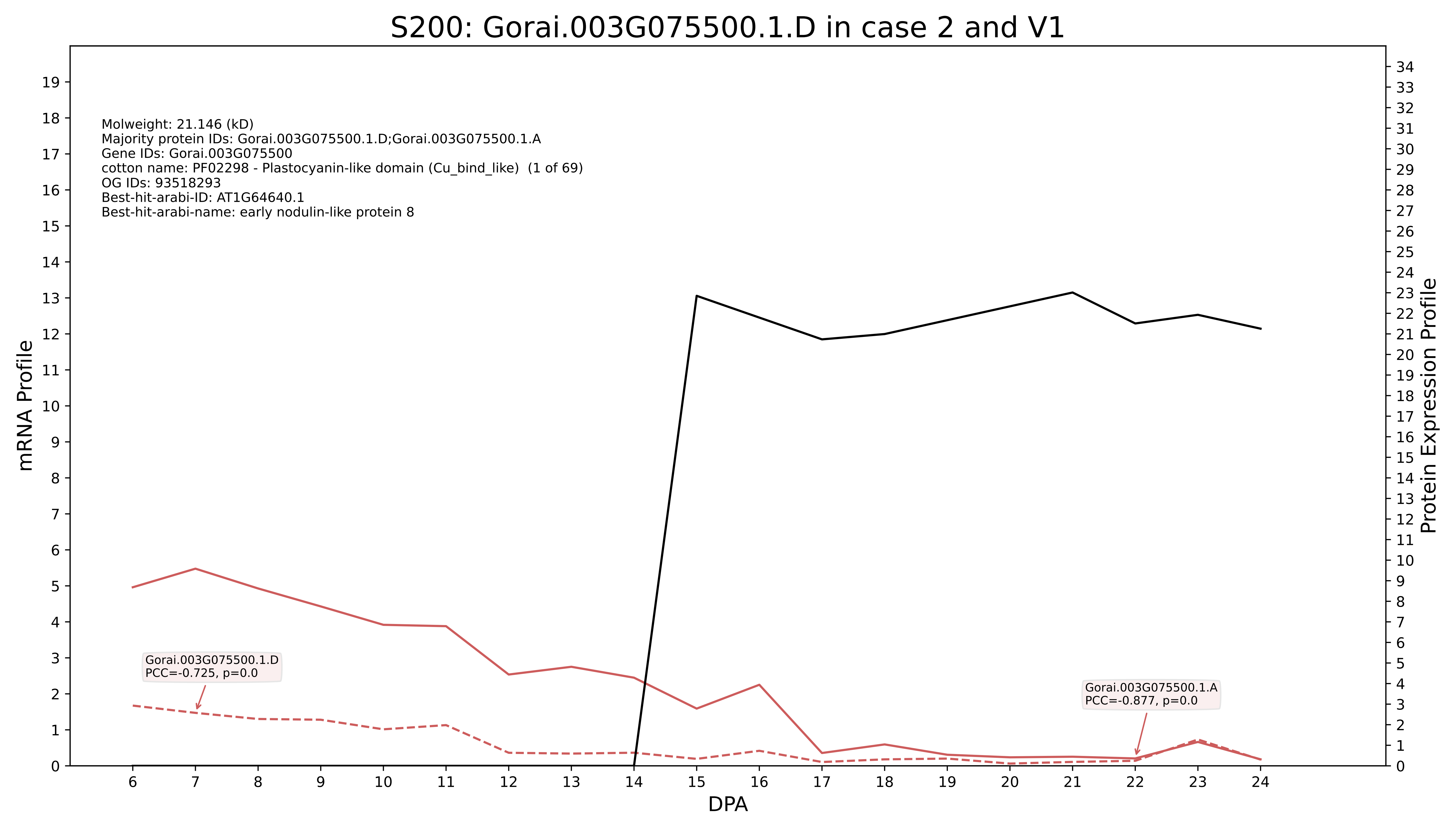
S200: Gorai.003G075500.1.D in case 2 and V1
Molweight: 21.146 (kD)
Majority protein IDs: Gorai.003G075500.1.D;Gorai.003G075500.1.A
Gene IDs: Gorai.003G075500
cotton name: PF02298 - Plastocyanin-like domain (Cu_bind_like) (1 of 69)
OG IDs: 93518293
Best-hit-arabi-ID: AT1G64640.1
Best-hit-arabi-name: early nodulin-like protein 8
Gorai.003G075500.1.D
PCC=-0.725, p=0.0
Gorai.003G075500.1.A
PCC=-0.877, p=0.0
mRNA Profile
Protein Expression Profile
DPA

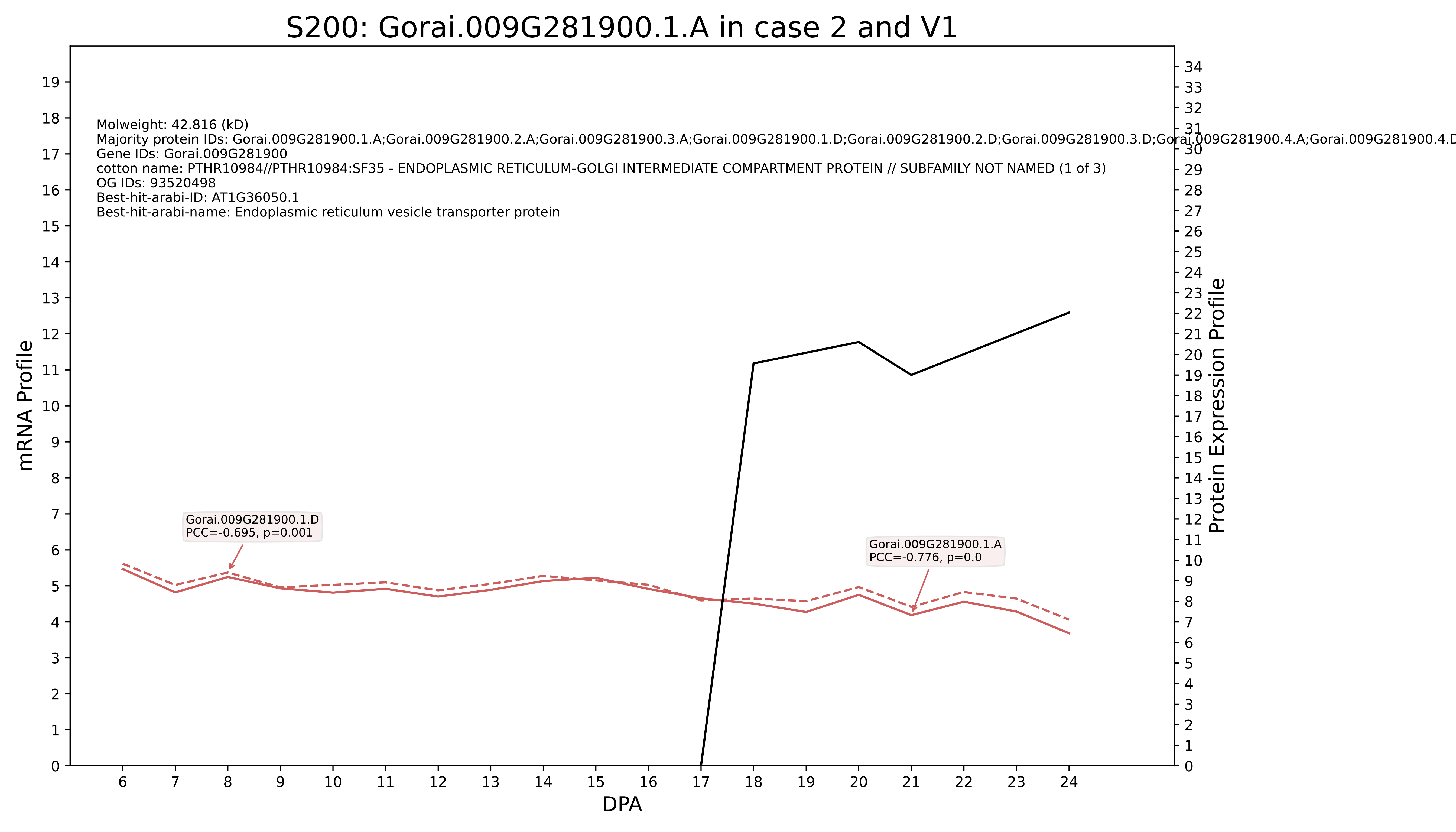
S200: Gorai.009G281900.1.A in case 2 and V1
Molweight: 42.816 (kD)
Majority protein IDs: Gorai.009G281900.1.A;Gorai.009G281900.2.A;Gorai.009G281900.3.A;Gorai.009G281900.1.D;Gorai.009G281900.2.D;Gorai.009G281900.3.D;Gorai.009G281900.4.A;Gorai.009G281900.4.D
Gene IDs: Gorai.009G281900
cotton name: PTHR10984//PTHR10984:SF35 - ENDOPLASMIC RETICULUM-GOLGI INTERMEDIATE COMPARTMENT PROTEIN // SUBFAMILY NOT NAMED (1 of 3)
OG IDs: 93520498
Best-hit-arabi-ID: AT1G36050.1
Best-hit-arabi-name: Endoplasmic reticulum vesicle transporter protein
Gorai.009G281900.1.D
PCC=-0.695, p=0.001
Gorai.009G281900.1.A
PCC=-0.776, p=0.0
mRNA Profile
Protein Expression Profile
DPA

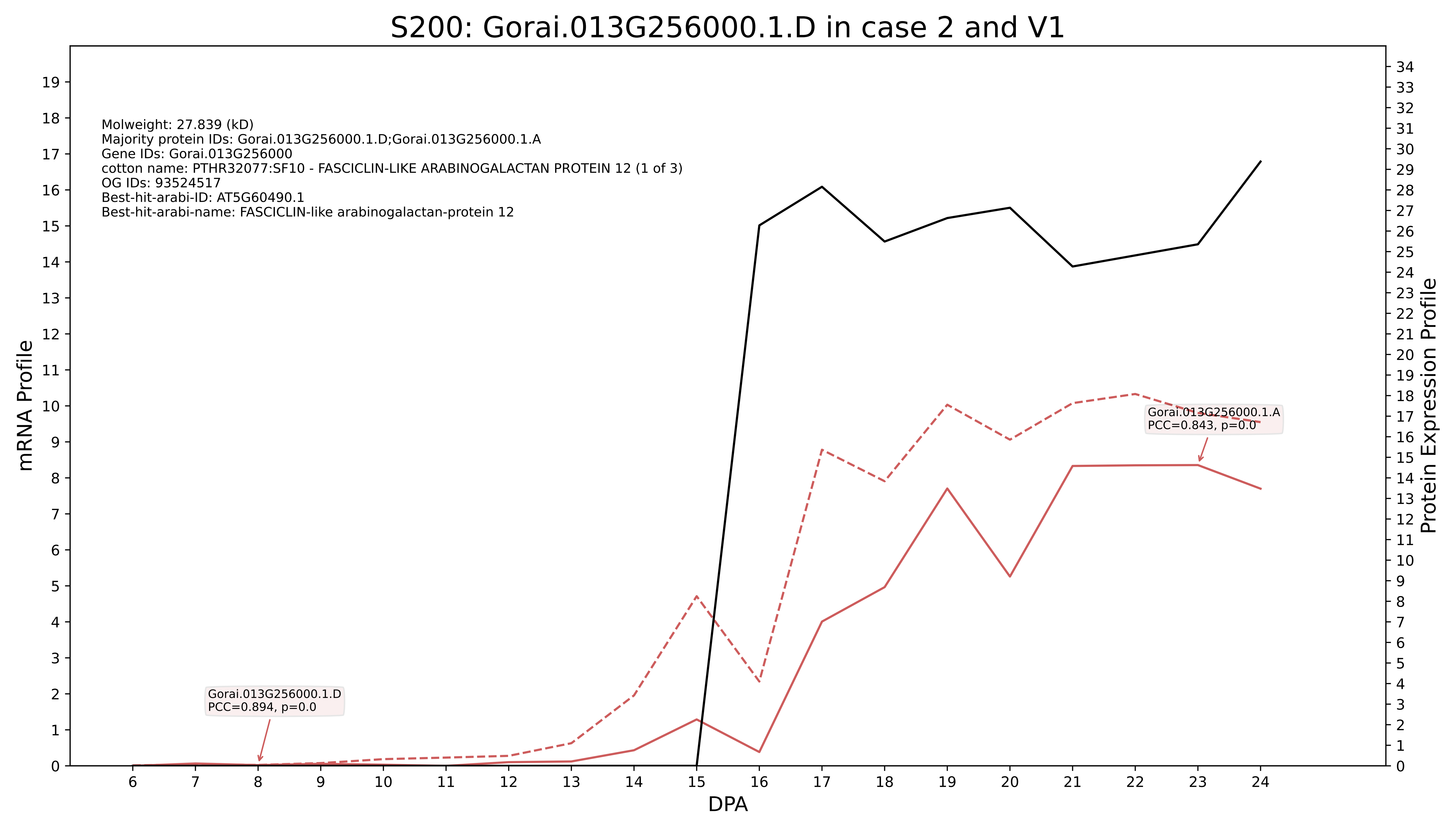
S200: Gorai.013G256000.1.D in case 2 and V1
Molweight: 27.839 (kD)
Majority protein IDs: Gorai.013G256000.1.D;Gorai.013G256000.1.A
Gene IDs: Gorai.013G256000
cotton name: PTHR32077:SF10 - FASCICLIN-LIKE ARABINOGALACTAN PROTEIN 12 (1 of 3)
OG IDs: 93524517
Best-hit-arabi-ID: AT5G60490.1
Best-hit-arabi-name: FASCICLIN-like arabinogalactan-protein 12
Gorai.013G256000.1.A
PCC=0.843, p=0.0
Gorai.013G256000.1.D
PCC=0.894, p=0.0
mRNA Profile
Protein Expression Profile
DPA

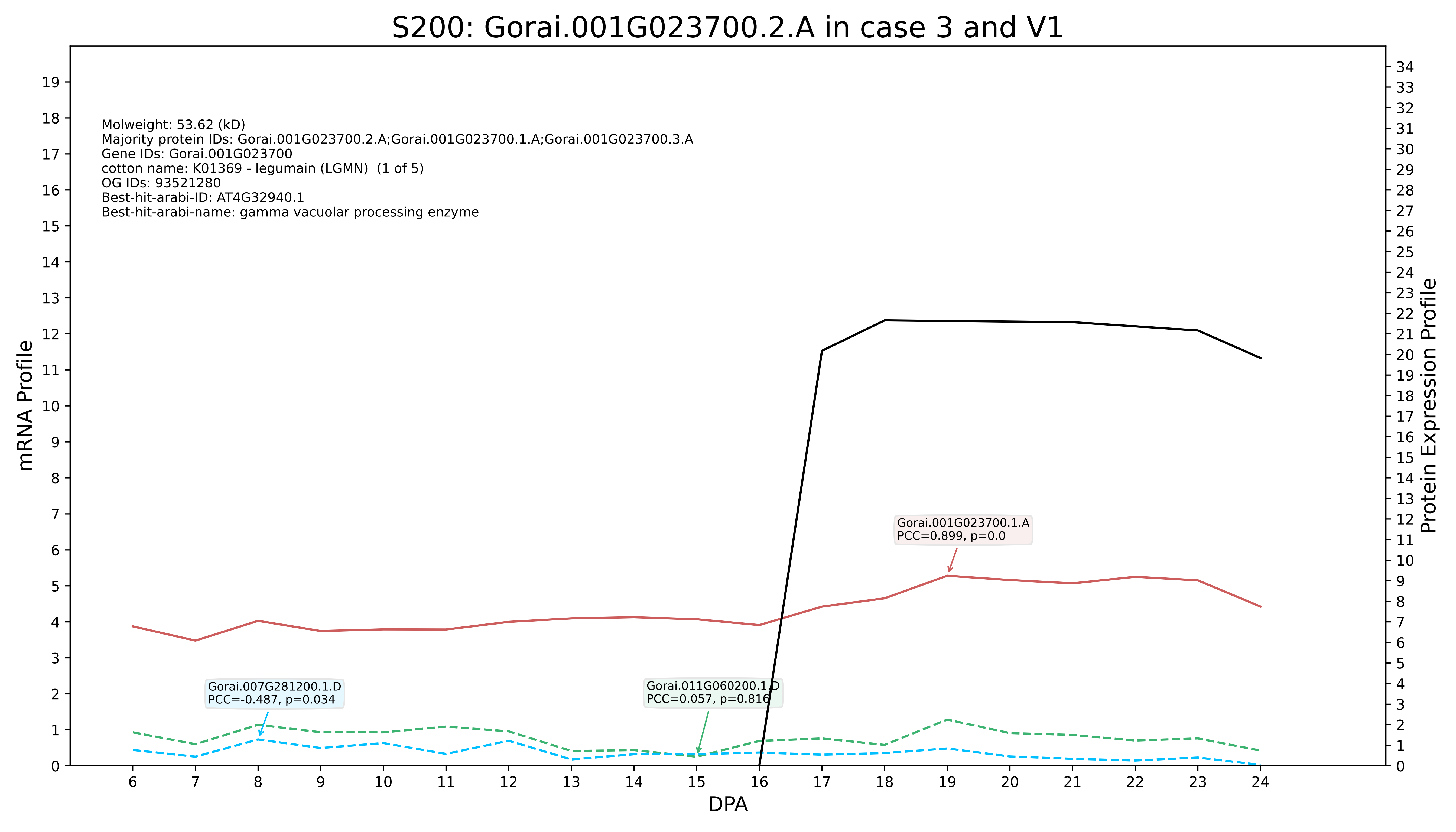
S200: Gorai.001G023700.2.A in case 3 and V1
Molweight: 53.62 (kD)
Majority protein IDs: Gorai.001G023700.2.A;Gorai.001G023700.1.A;Gorai.001G023700.3.A
Gene IDs: Gorai.001G023700
cotton name: K01369 - legumain (LGMN) (1 of 5)
OG IDs: 93521280
Best-hit-arabi-ID: AT4G32940.1
Best-hit-arabi-name: gamma vacuolar processing enzyme
Gorai.001G023700.1.A
PCC=0.899, p=0.0
Gorai.007G281200.1.D
PCC=-0.487, p=0.034
Gorai.011G060200.1.D
PCC=0.057, p=0.816
mRNA Profile
Protein Expression Profile
DPA

010G003200.1.D in case 3 and V1

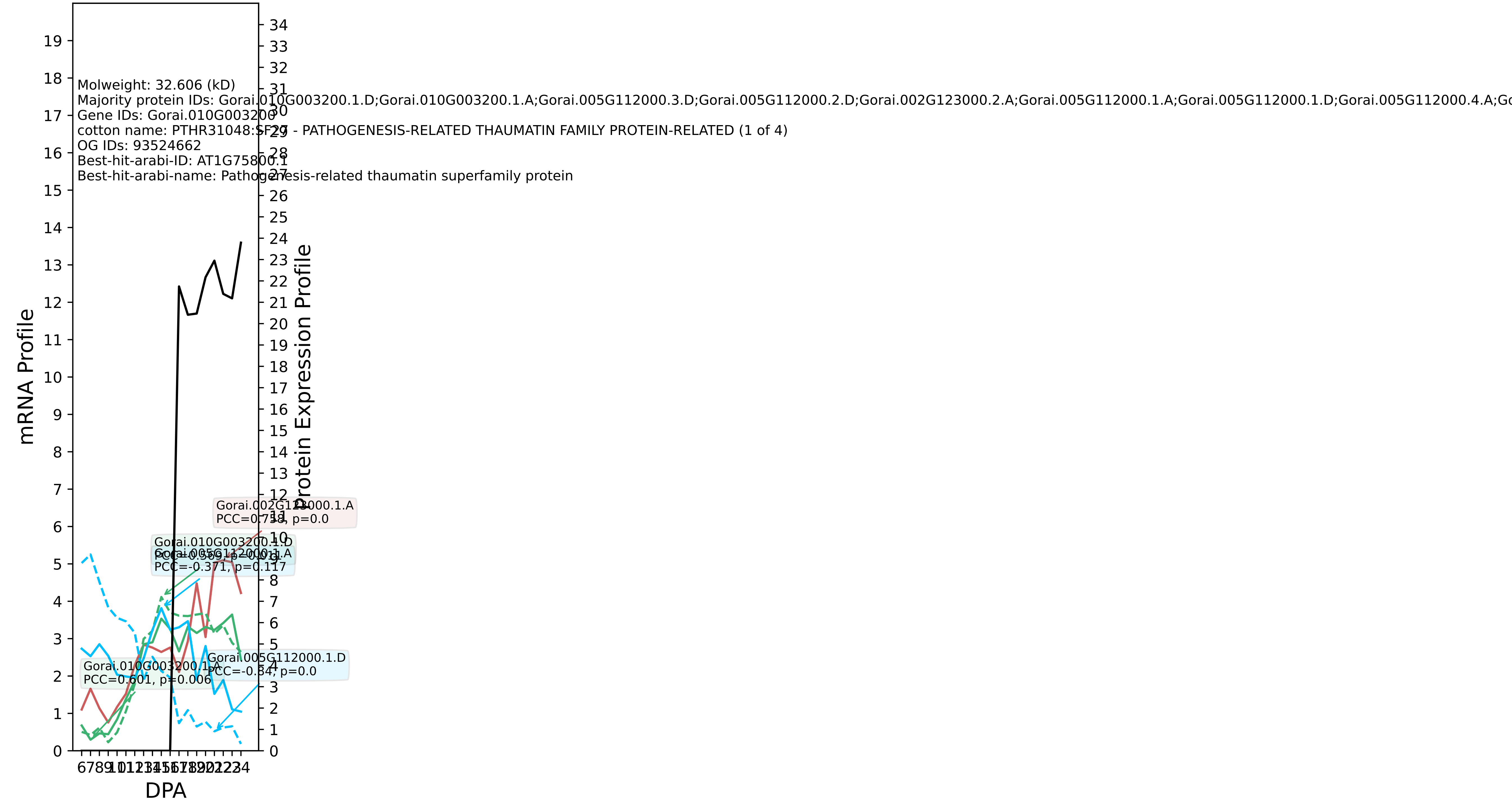

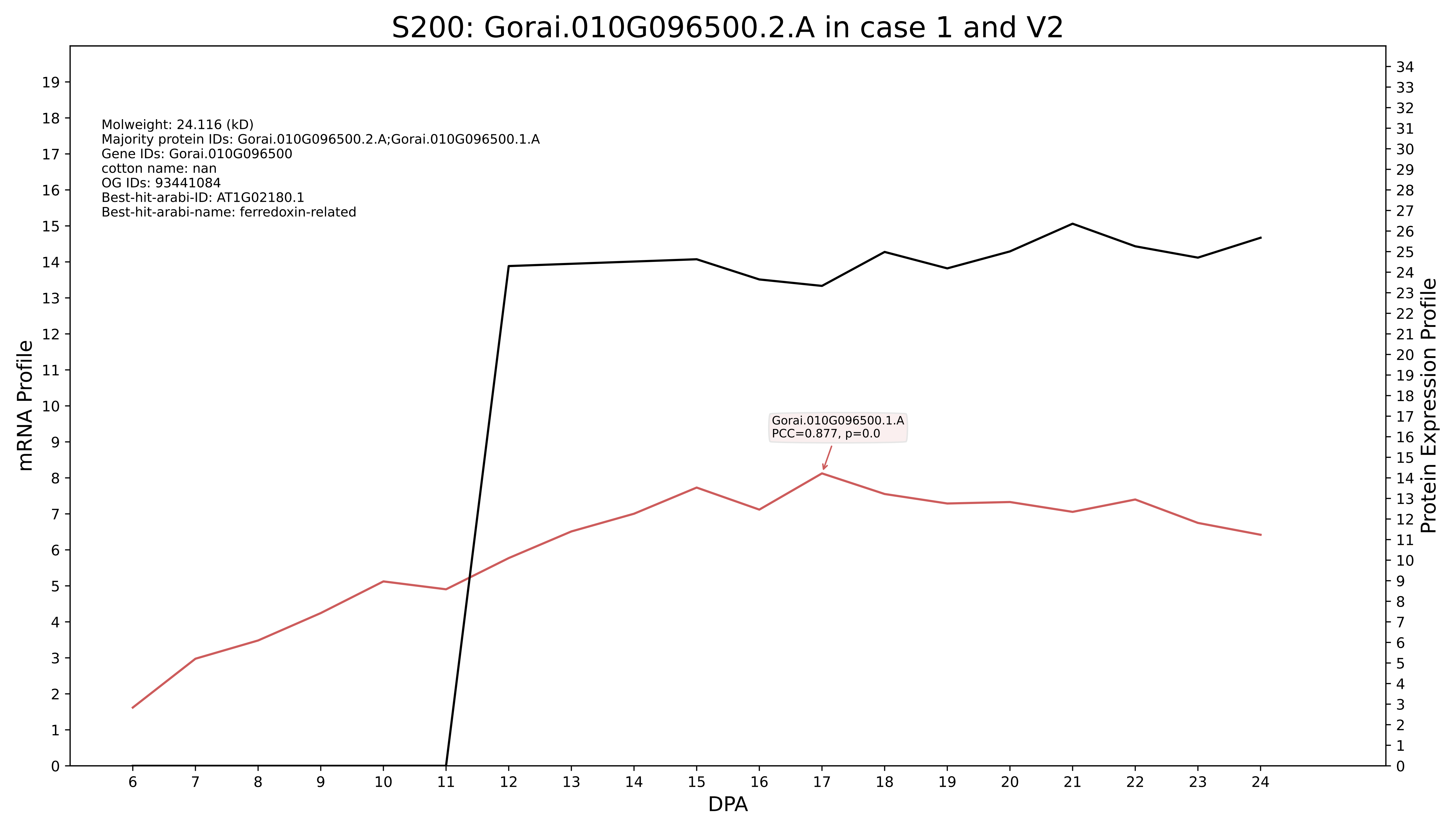
S200: Gorai.010G096500.2.A in case 1 and V2
Molweight: 24.116 (kD)
Majority protein IDs: Gorai.010G096500.2.A;Gorai.010G096500.1.A
Gene IDs: Gorai.010G096500
cotton name: nan
OG IDs: 93441084
Best-hit-arabi-ID: AT1G02180.1
Best-hit-arabi-name: ferredoxin-related
Gorai.010G096500.1.A
PCC=0.877, p=0.0
mRNA Profile
Protein Expression Profile
DPA

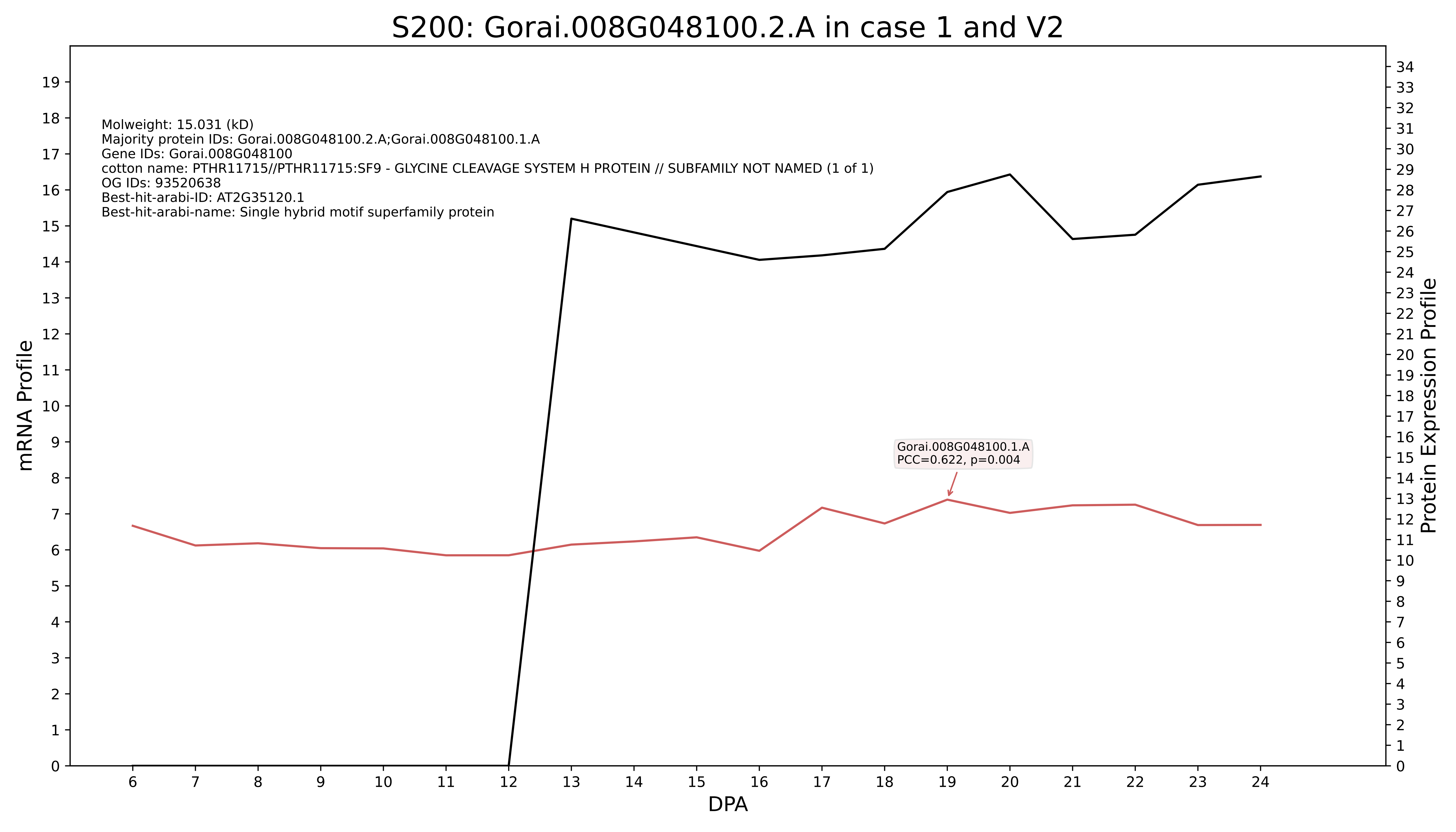
S200: Gorai.008G048100.2.A in case 1 and V2
Molweight: 15.031 (kD)
Majority protein IDs: Gorai.008G048100.2.A;Gorai.008G048100.1.A
Gene IDs: Gorai.008G048100
cotton name: PTHR11715//PTHR11715:SF9 - GLYCINE CLEAVAGE SYSTEM H PROTEIN // SUBFAMILY NOT NAMED (1 of 1)
OG IDs: 93520638
Best-hit-arabi-ID: AT2G35120.1
Best-hit-arabi-name: Single hybrid motif superfamily protein
Gorai.008G048100.1.A
PCC=0.622, p=0.004
mRNA Profile
Protein Expression Profile
DPA

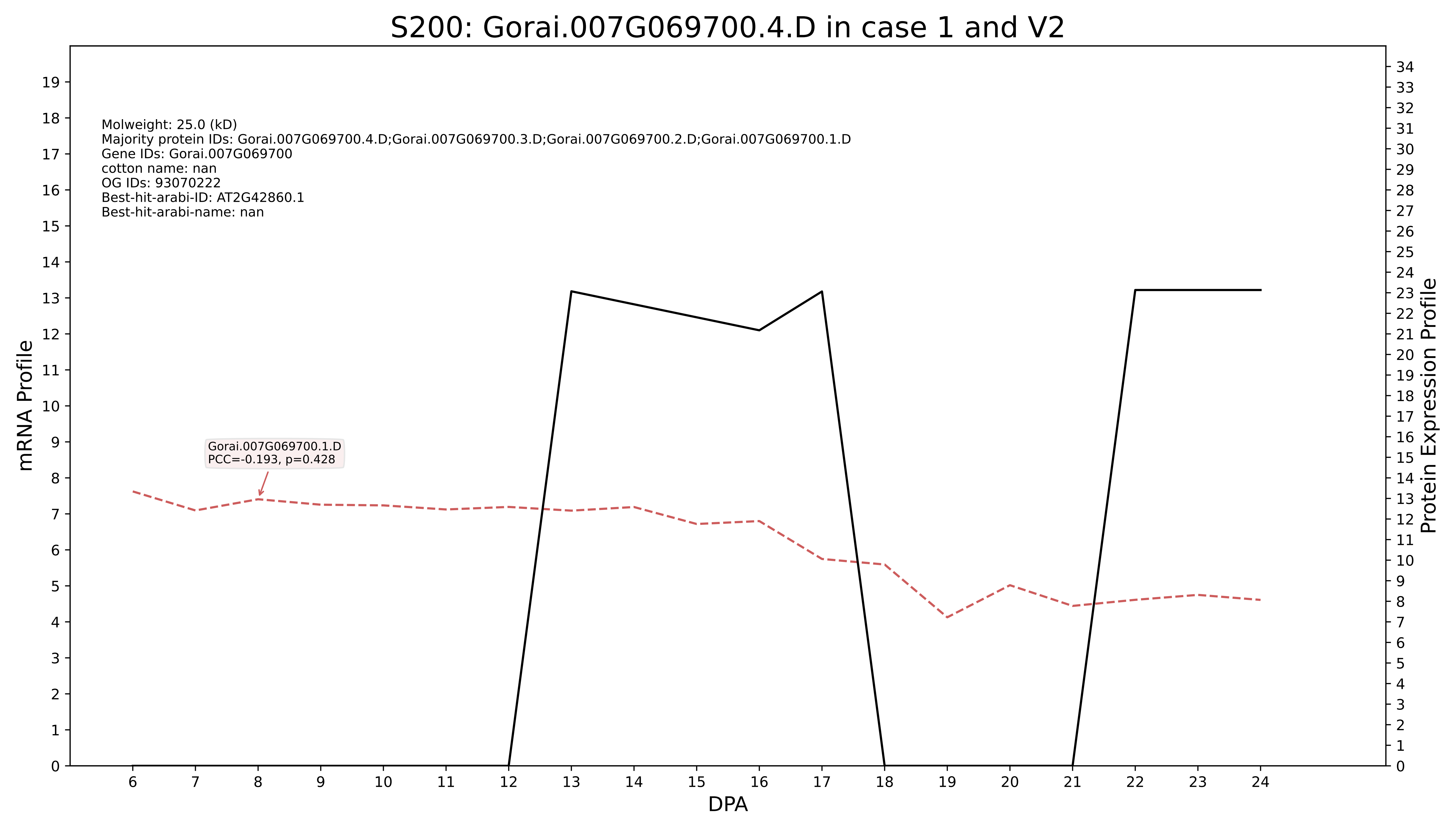

S200: Gorai.007G069700.4.D in case 1 and V2
Molweight: 25.0 (kD)
Majority protein IDs: Gorai.007G069700.4.D;Gorai.007G069700.3.D;Gorai.007G069700.2.D;Gorai.007G069700.1.D
Gene IDs: Gorai.007G069700
cotton name: nan
OG IDs: 93070222
Best-hit-arabi-ID: AT2G42860.1
Best-hit-arabi-name: nan
Gorai.007G069700.1.D
PCC=-0.193, p=0.428
mRNA Profile
Protein Expression Profile
DPA

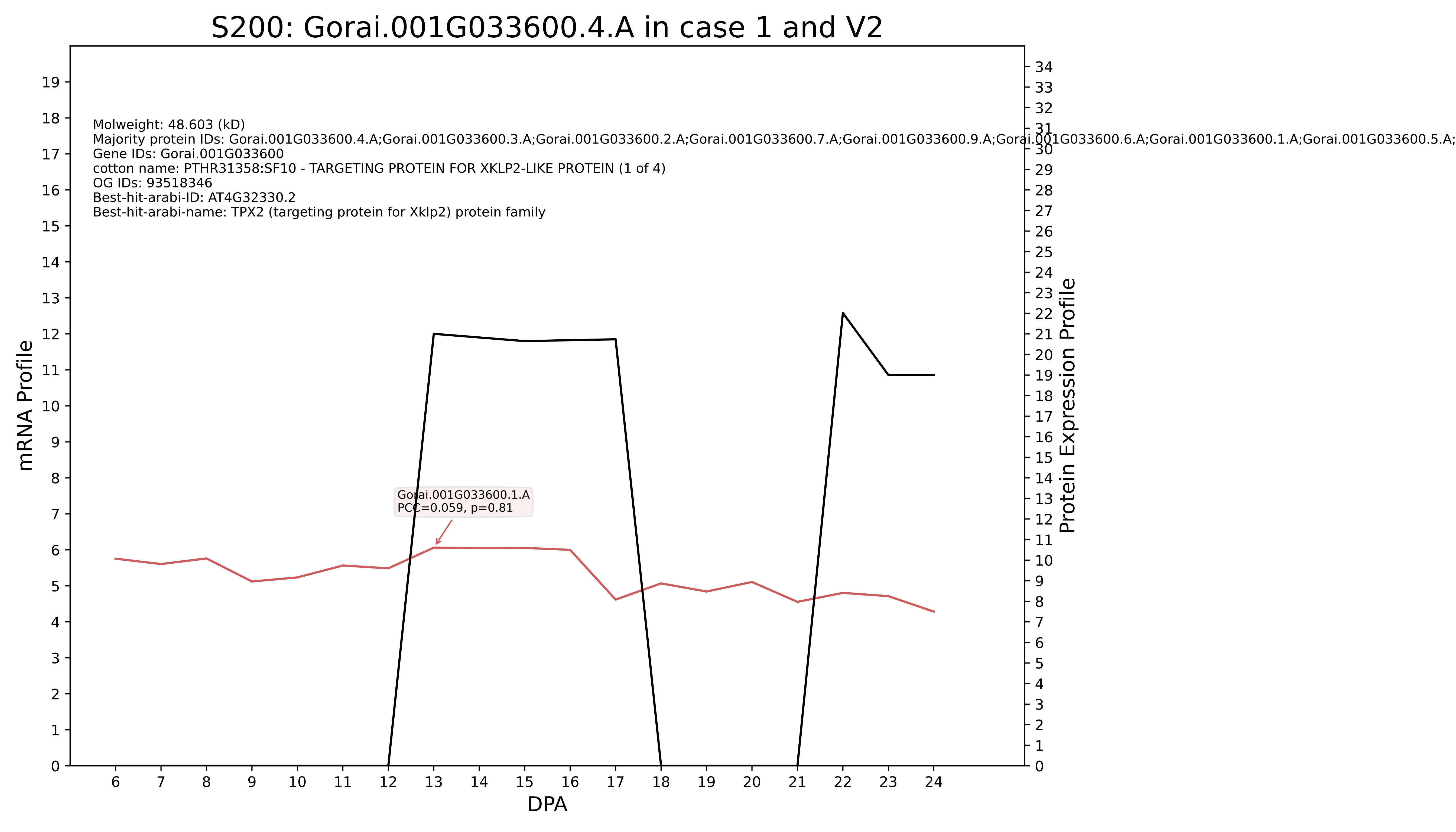
S200: Gorai.001G033600.4.A in case 1 and V2
Molweight: 48.603 (kD)
Majority protein IDs: Gorai.001G033600.4.A;Gorai.001G033600.3.A;Gorai.001G033600.2.A;Gorai.001G033600.7.A;Gorai.001G033600.9.A;Gorai.001G033600.6.A;Gorai.001G033600.1.A;Gorai.001G033600.5.A;
Gene IDs: Gorai.001G033600
cotton name: PTHR31358:SF10 - TARGETING PROTEIN FOR XKLP2-LIKE PROTEIN (1 of 4)
OG IDs: 93518346
Best-hit-arabi-ID: AT4G32330.2
Best-hit-arabi-name: TPX2 (targeting protein for Xklp2) protein family
Gorai.001G033600.1.A
PCC=0.059, p=0.81
mRNA Profile
Protein Expression Profile
DPA

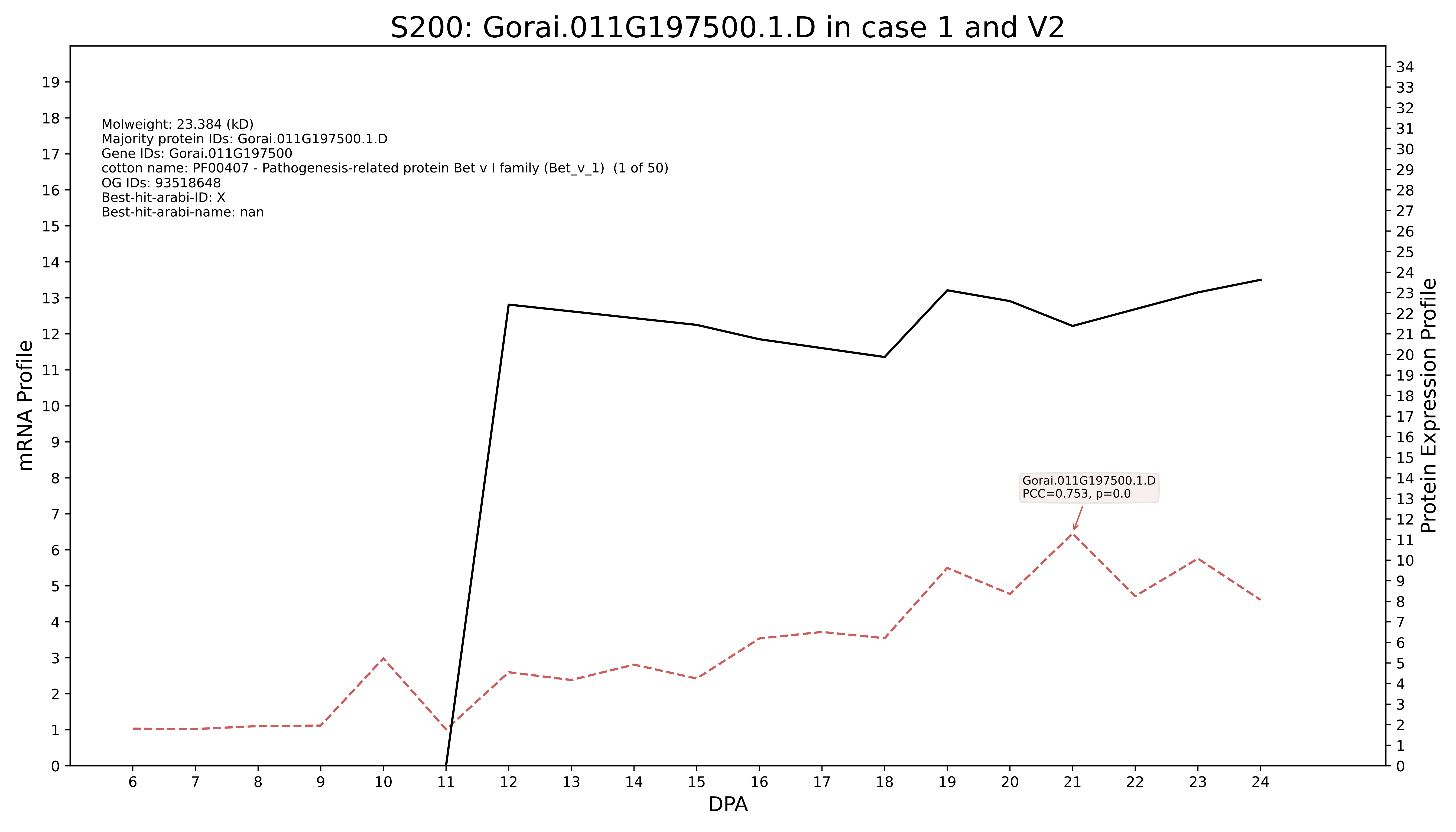
S200: Gorai.011G197500.1.D in case 1 and V2
Molweight: 23.384 (kD)
Majority protein IDs: Gorai.011G197500.1.D
Gene IDs: Gorai.011G197500
cotton name: PF00407 - Pathogenesis-related protein Bet v I family (Bet_v_1) (1 of 50)
OG IDs: 93518648
Best-hit-arabi-ID: X
Best-hit-arabi-name: nan
Gorai.011G197500.1.D
PCC=0.753, p=0.0
mRNA Profile
Protein Expression Profile
DPA

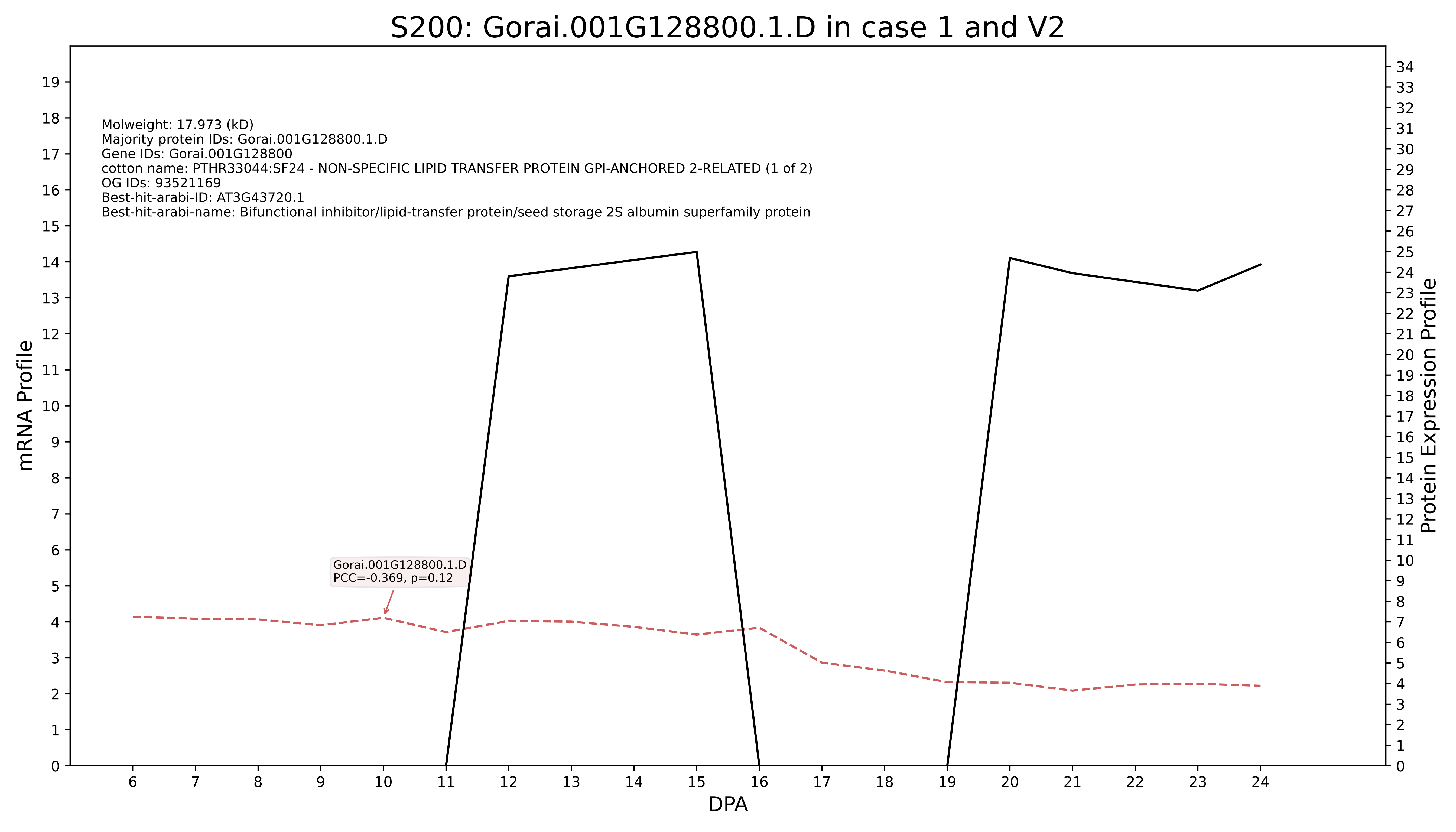
S200: Gorai.001G128800.1.D in case 1 and V2
Molweight: 17.973 (kD)
Majority protein IDs: Gorai.001G128800.1.D
Gene IDs: Gorai.001G128800
cotton name: PTHR33044:SF24 - NON-SPECIFIC LIPID TRANSFER PROTEIN GPI-ANCHORED 2-RELATED (1 of 2)
OG IDs: 93521169
Best-hit-arabi-ID: AT3G43720.1
Best-hit-arabi-name: Bifunctional inhibitor/lipid-transfer protein/seed storage 2S albumin superfamily protein
Gorai.001G128800.1.D
PCC=-0.369, p=0.12
mRNA Profile
Protein Expression Profile
DPA

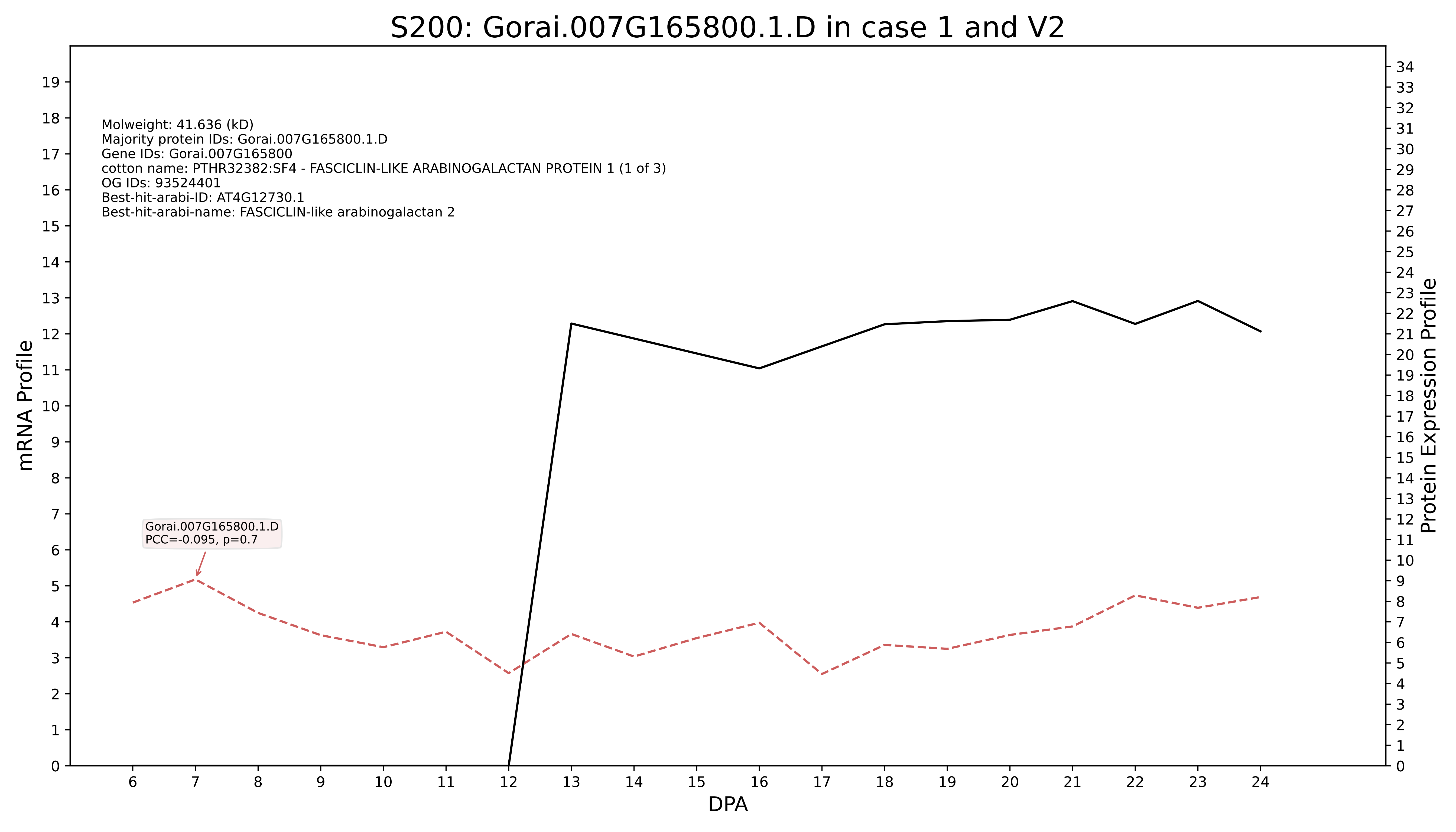

S200: Gorai.007G165800.1.D in case 1 and V2
Molweight: 41.636 (kD)
Majority protein IDs: Gorai.007G165800.1.D
Gene IDs: Gorai.007G165800
cotton name: PTHR32382:SF4 - FASCICLIN-LIKE ARABINOGALACTAN PROTEIN 1 (1 of 3)
OG IDs: 93524401
Best-hit-arabi-ID: AT4G12730.1
Best-hit-arabi-name: FASCICLIN-like arabinogalactan 2
Gorai.007G165800.1.D
PCC=-0.095, p=0.7
mRNA Profile
Protein Expression Profile
DPA

### S200: Gorai.007G374100.1.D in case 1 and V2

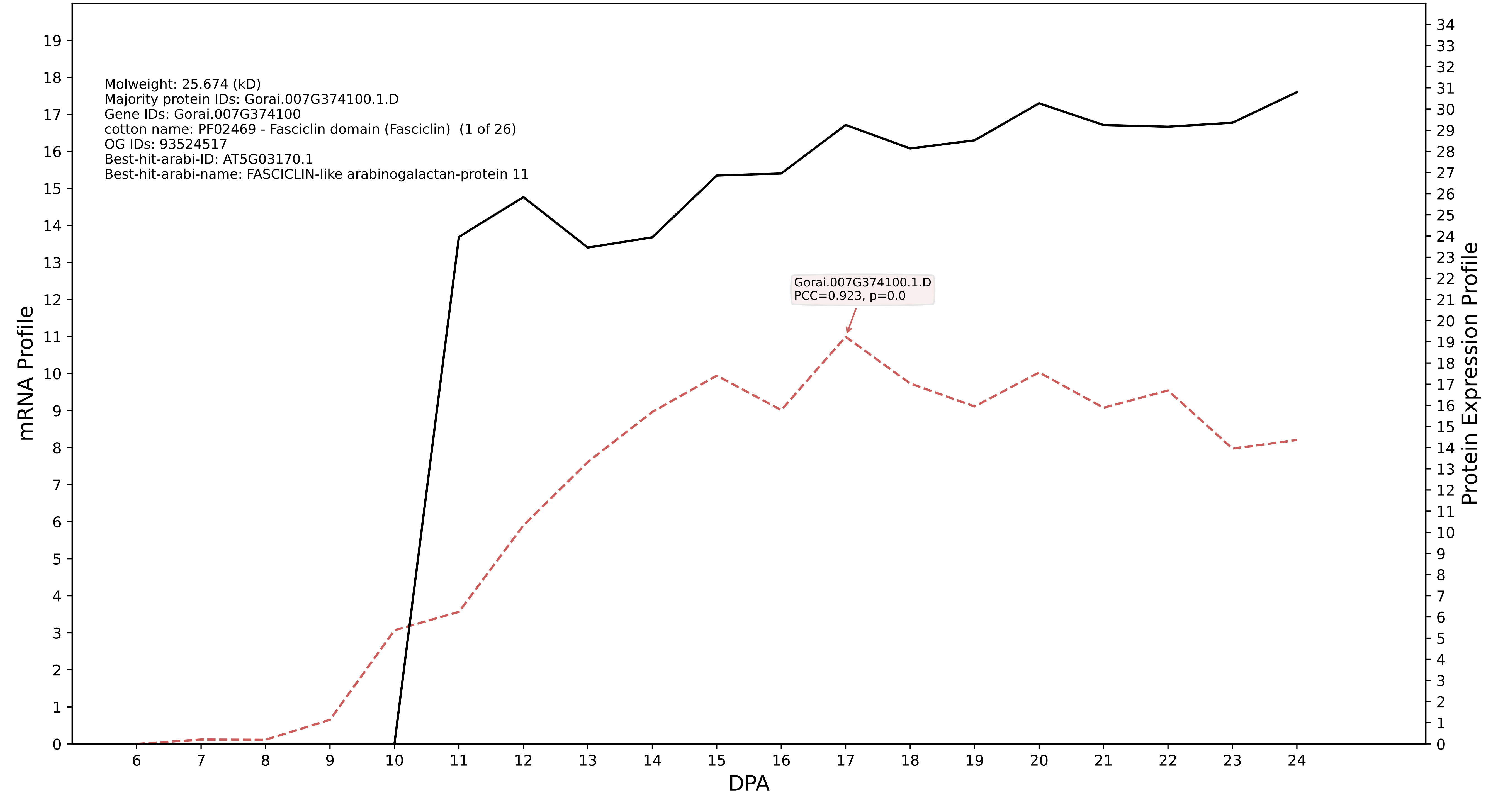

### S200: Gorai.013G112000.2.A in case 2 and V2

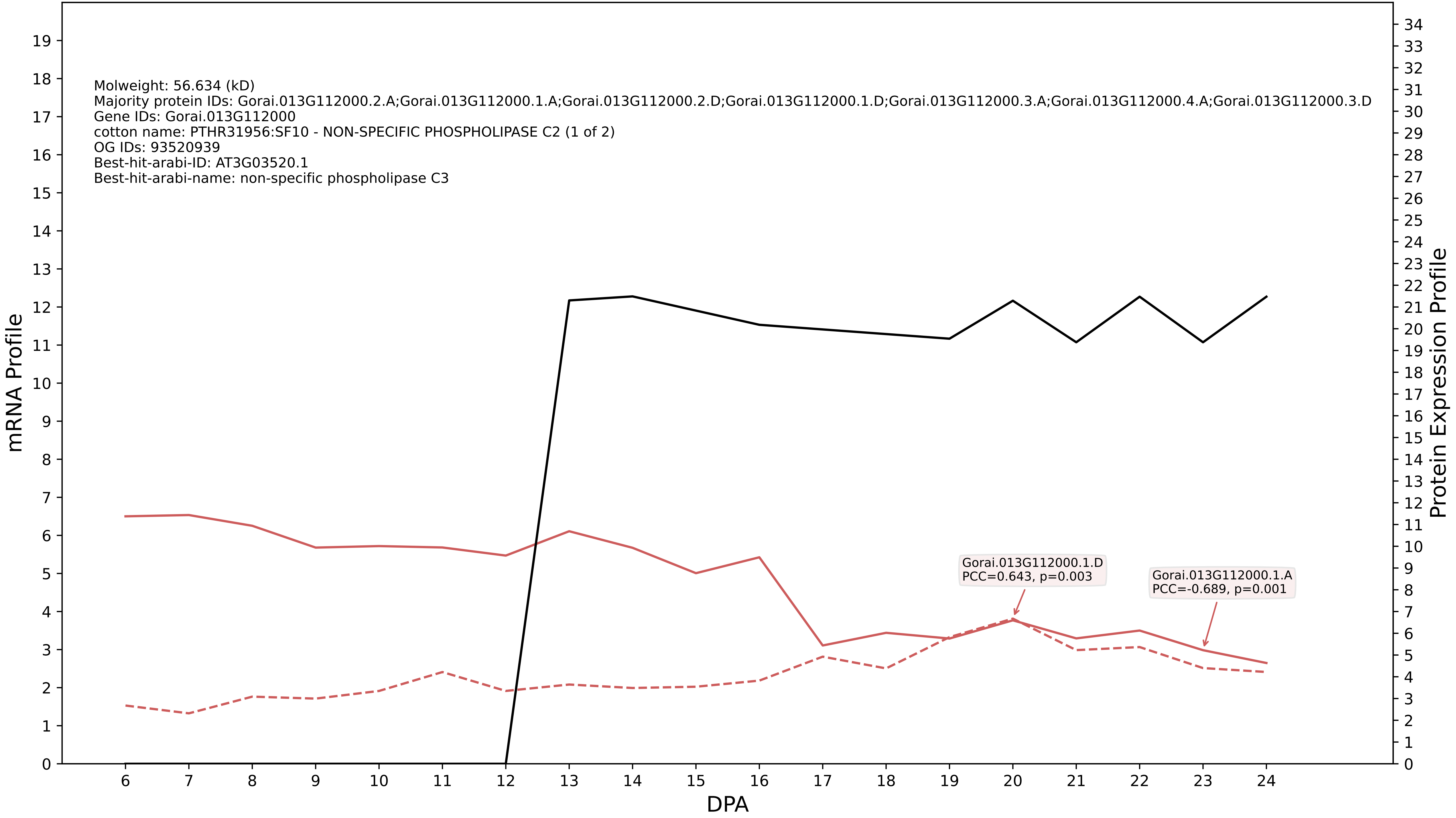

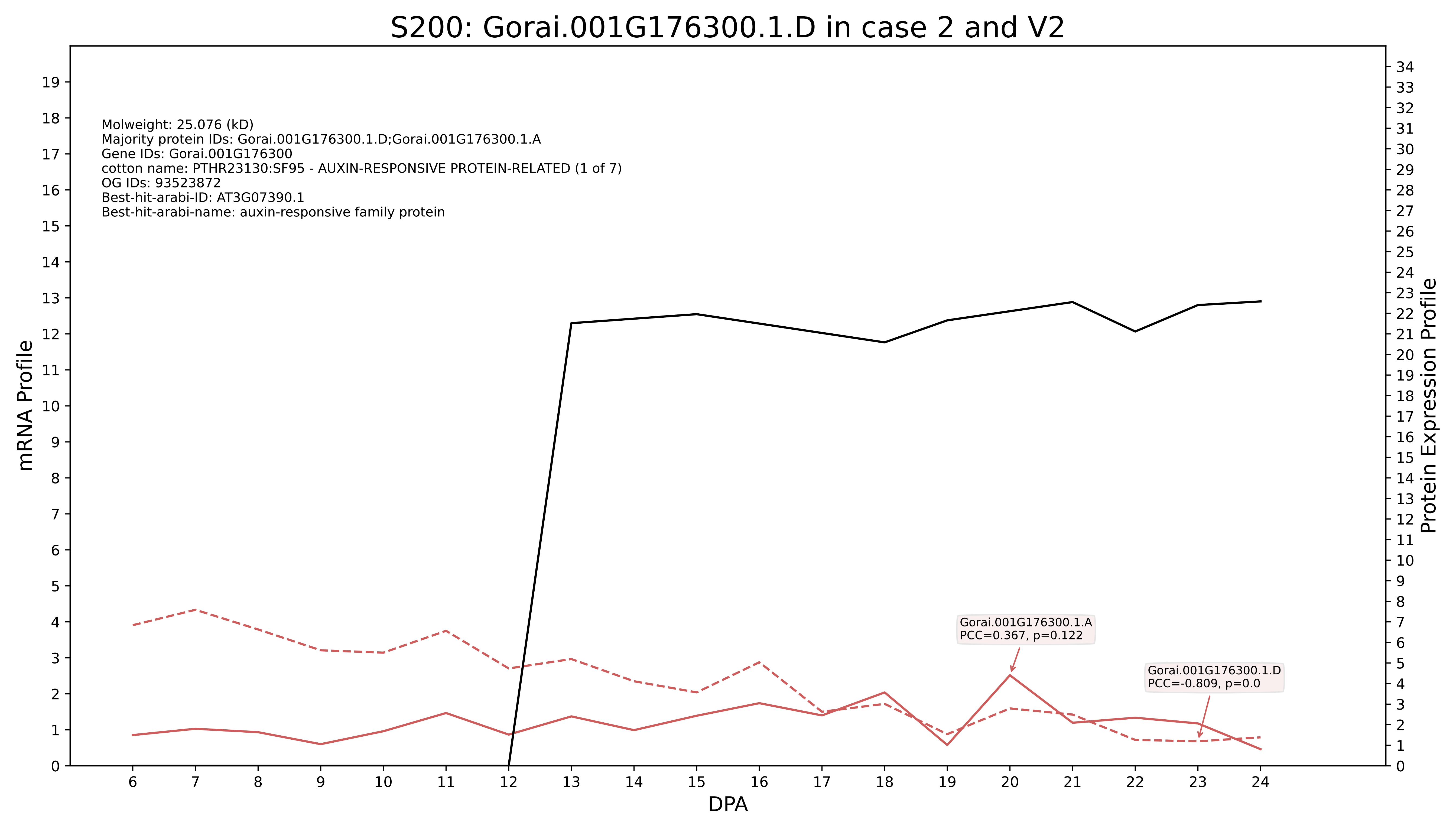
S200: Gorai.001G176300.1.D in case 2 and V2
Molweight: 25.076 (kD)
Majority protein IDs: Gorai.001G176300.1.D;Gorai.001G176300.1.A
Gene IDs: Gorai.001G176300
cotton name: PTHR23130:SF95 - AUXIN-RESPONSIVE PROTEIN-RELATED (1 of 7)
OG IDs: 93523872
Best-hit-arabi-ID: AT3G07390.1
Best-hit-arabi-name: auxin-responsive family protein
Gorai.001G176300.1.A
PCC=0.367, p=0.122
Gorai.001G176300.1.D
PCC=-0.809, p=0.0
mRNA Profile
Protein Expression Profile
DPA

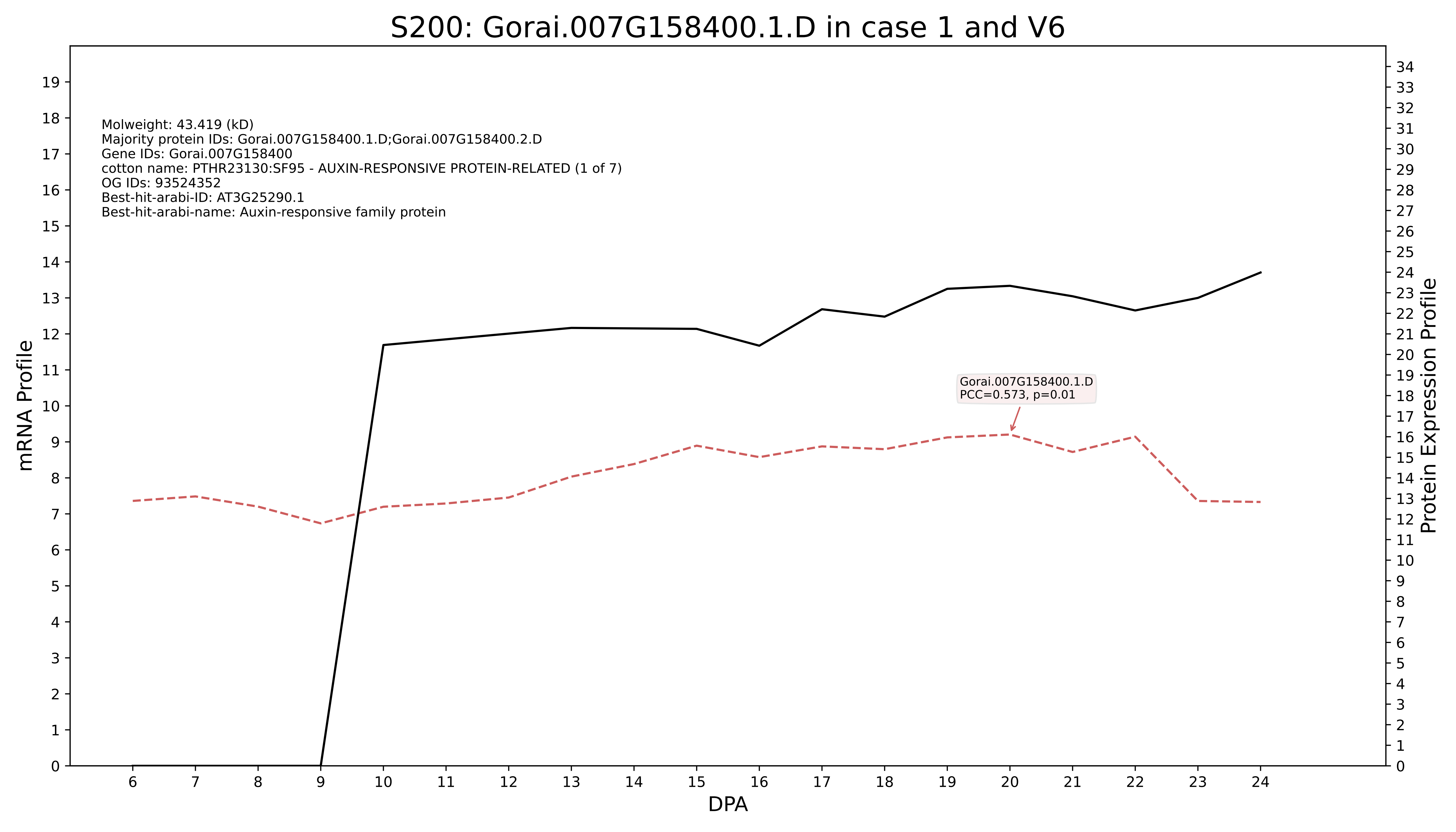
S200: Gorai.007G158400.1.D in case 1 and V6
Molweight: 43.419 (kD)
Majority protein IDs: Gorai.007G158400.1.D;Gorai.007G158400.2.D
Gene IDs: Gorai.007G158400
cotton name: PTHR23130:SF95 - AUXIN-RESPONSIVE PROTEIN-RELATED (1 of 7)
OG IDs: 93524352
Best-hit-arabi-ID: AT3G25290.1
Best-hit-arabi-name: Auxin-responsive family protein
Gorai.007G158400.1.D
PCC=0.573, p=0.01
mRNA Profile
Protein Expression Profile
DPA

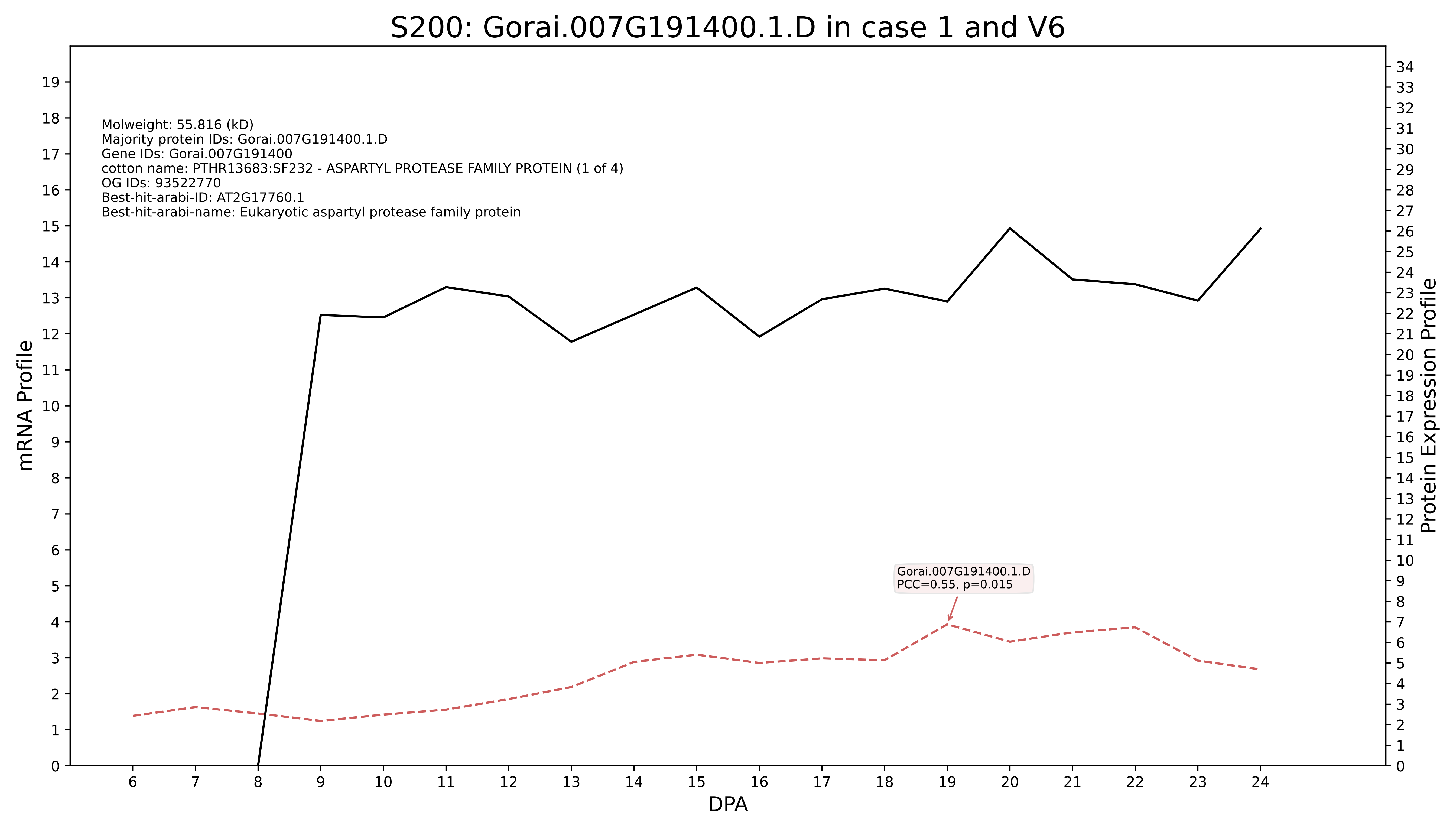
S200: Gorai.007G191400.1.D in case 1 and V6
Molweight: 55.816 (kD)
Majority protein IDs: Gorai.007G191400.1.D
Gene IDs: Gorai.007G191400
cotton name: PTHR13683:SF232 - ASPARTYL PROTEASE FAMILY PROTEIN (1 of 4)
OG IDs: 93522770
Best-hit-arabi-ID: AT2G17760.1
Best-hit-arabi-name: Eukaryotic aspartyl protease family protein
Gorai.007G191400.1.D
PCC=0.55, p=0.015
mRNA Profile
Protein Expression Profile
DPA

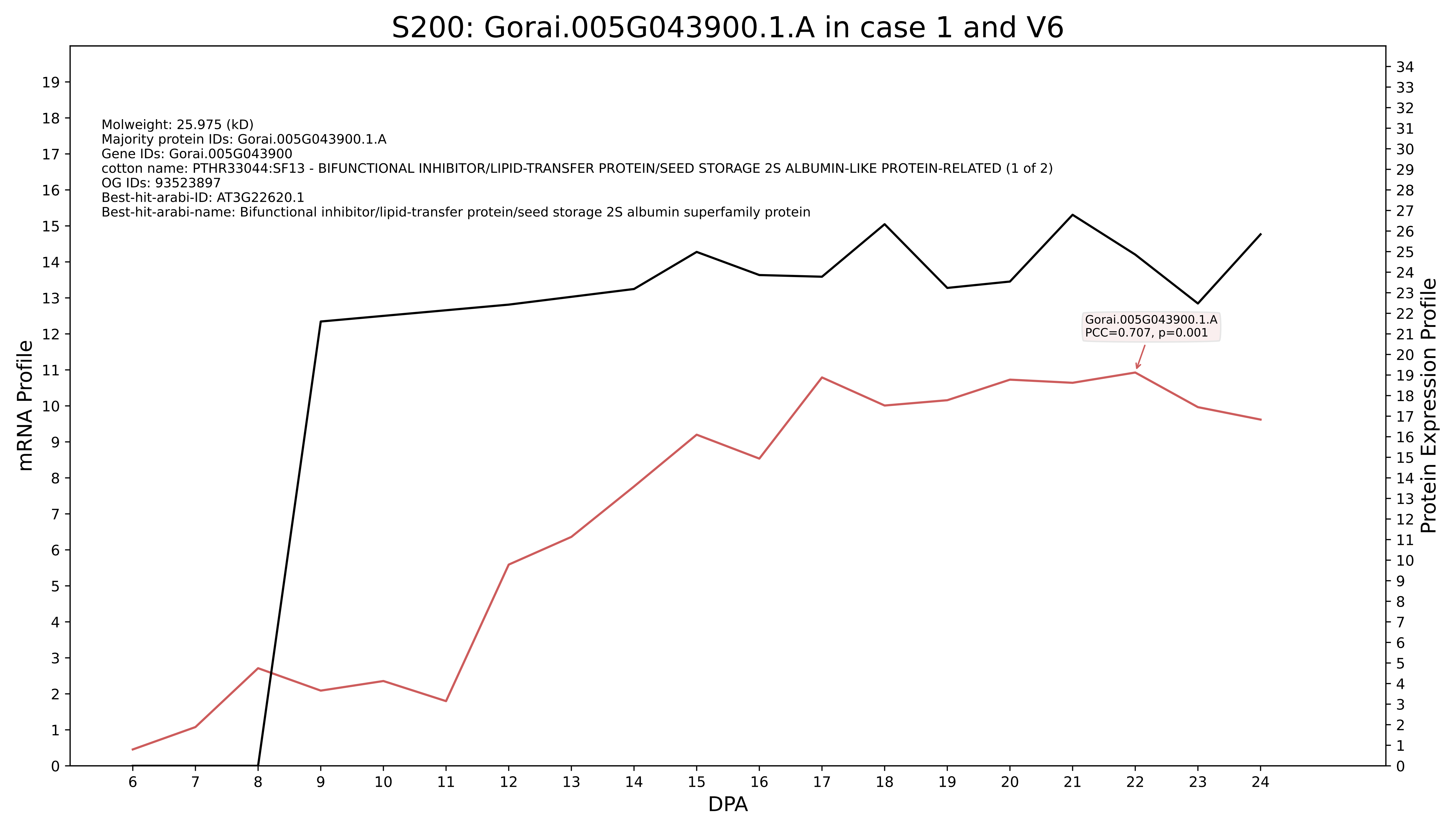
S200: Gorai.005G043900.1.A in case 1 and V6
Molweight: 25.975 (kD)
Majority protein IDs: Gorai.005G043900.1.A
Gene IDs: Gorai.005G043900
cotton name: PTHR33044:SF13 - BIFUNCTIONAL INHIBITOR/LIPID-TRANSFER PROTEIN/SEED STORAGE 2S ALBUMIN-LIKE PROTEIN-RELATED (1 of 2)
OG IDs: 93523897
Best-hit-arabi-ID: AT3G22620.1
Best-hit-arabi-name: Bifunctional inhibitor/lipid-transfer protein/seed storage 2S albumin superfamily protein
Gorai.005G043900.1.A
PCC=0.707, p=0.001
mRNA Profile
Protein Expression Profile
DPA

### S200: Gorai.002G124800.2.A in case 1 and V6

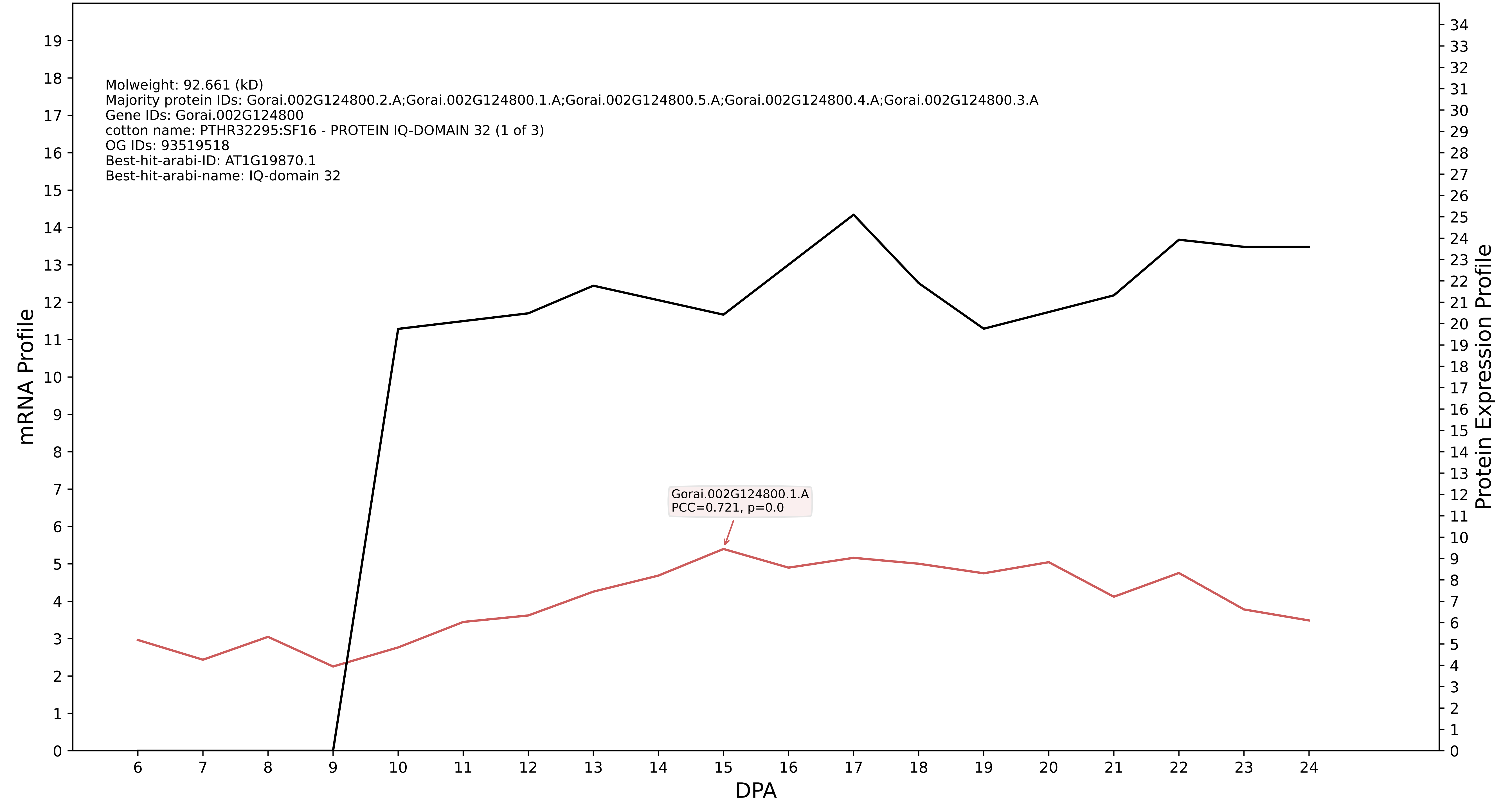

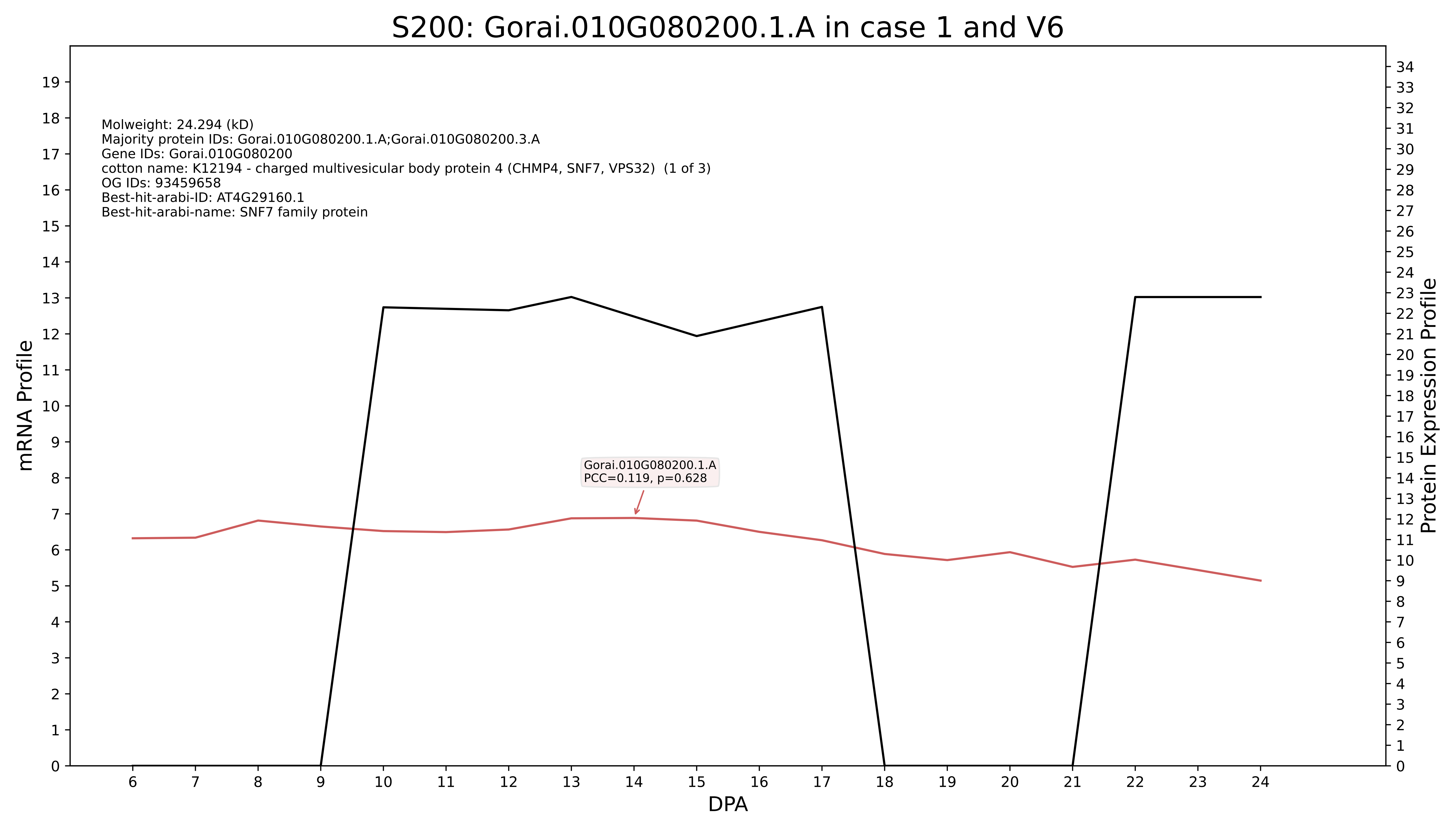
S200: Gorai.010G080200.1.A in case 1 and V6
Molweight: 24.294 (kD)
Majority protein IDs: Gorai.010G080200.1.A;Gorai.010G080200.3.A
Gene IDs: Gorai.010G080200
cotton name: K12194 - charged multivesicular body protein 4 (CHMP4, SNF7, VPS32) (1 of 3)
OG IDs: 93459658
Best-hit-arabi-ID: AT4G29160.1
Best-hit-arabi-name: SNF7 family protein
Gorai.010G080200.1.A
PCC=0.119, p=0.628
mRNA Profile
Protein Expression Profile
DPA

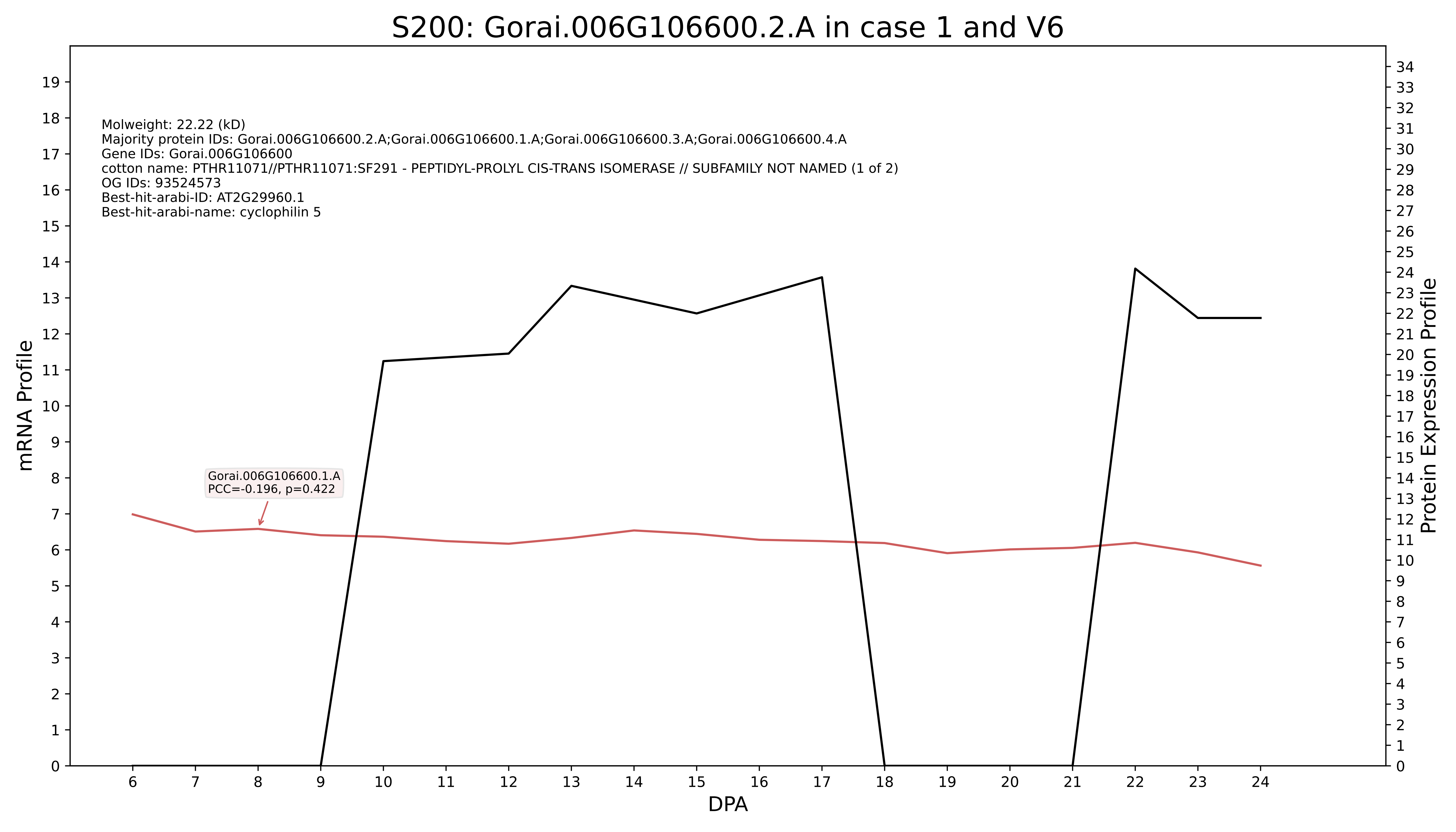
S200: Gorai.006G106600.2.A in case 1 and V6
Molweight: 22.22 (kD)
Majority protein IDs: Gorai.006G106600.2.A;Gorai.006G106600.1.A;Gorai.006G106600.3.A;Gorai.006G106600.4.A
Gene IDs: Gorai.006G106600
cotton name: PTHR11071//PTHR11071:SF291 - PEPTIDYL-PROLYL CIS-TRANS ISOMERASE // SUBFAMILY NOT NAMED (1 of 2)
OG IDs: 93524573
Best-hit-arabi-ID: AT2G29960.1
Best-hit-arabi-name: cyclophilin 5
Gorai.006G106600.1.A
PCC=-0.196, p=0.422
mRNA Profile
Protein Expression Profile
DPA

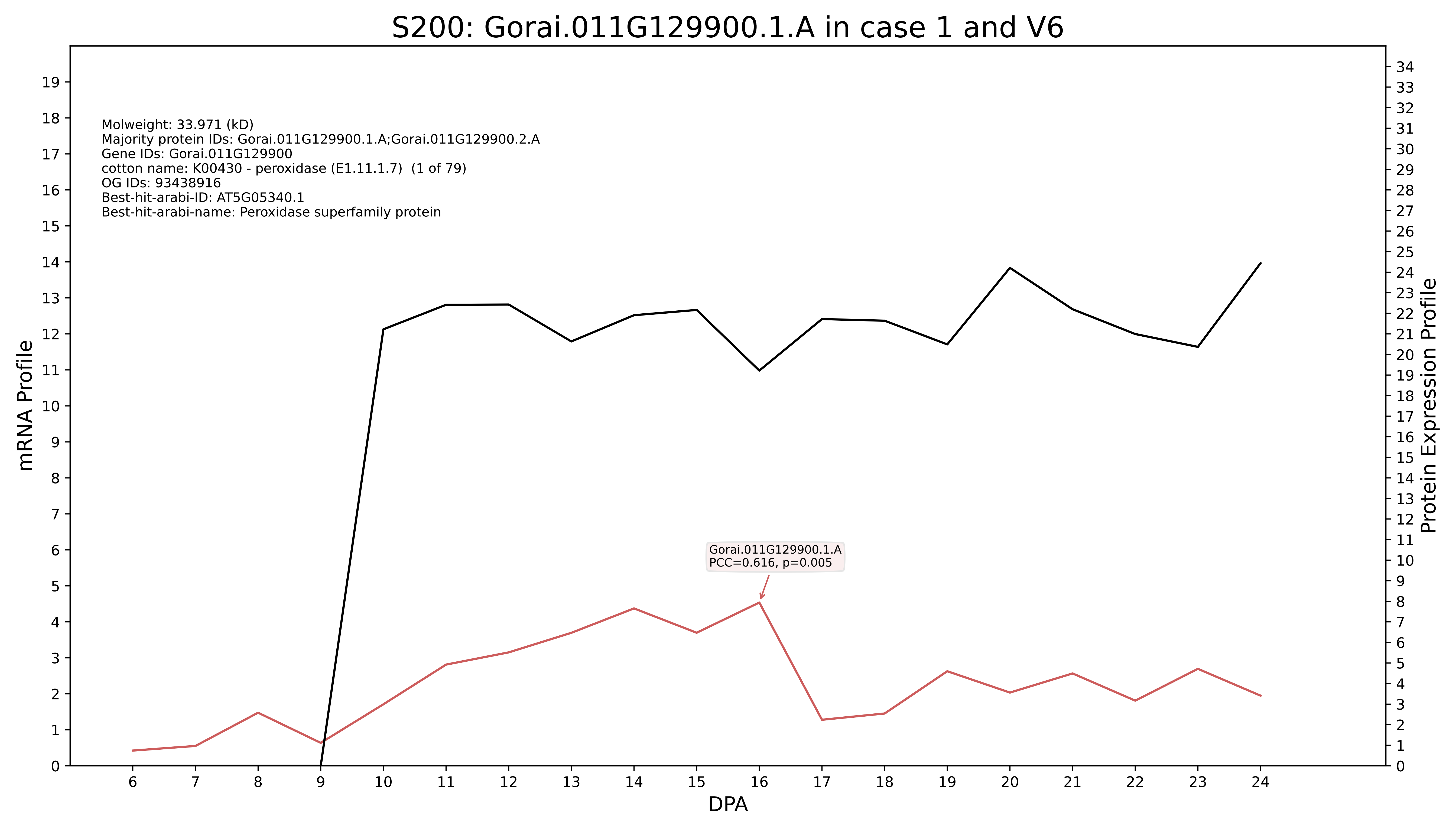
S200: Gorai.011G129900.1.A in case 1 and V6
Molweight: 33.971 (kD)
Majority protein IDs: Gorai.011G129900.1.A;Gorai.011G129900.2.A
Gene IDs: Gorai.011G129900
cotton name: K00430 - peroxidase (E1.11.1.7) (1 of 79)
OG IDs: 93438916
Best-hit-arabi-ID: AT5G05340.1
Best-hit-arabi-name: Peroxidase superfamily protein
Gorai.011G129900.1.A
PCC=0.616, p=0.005
mRNA Profile
Protein Expression Profile
DPA

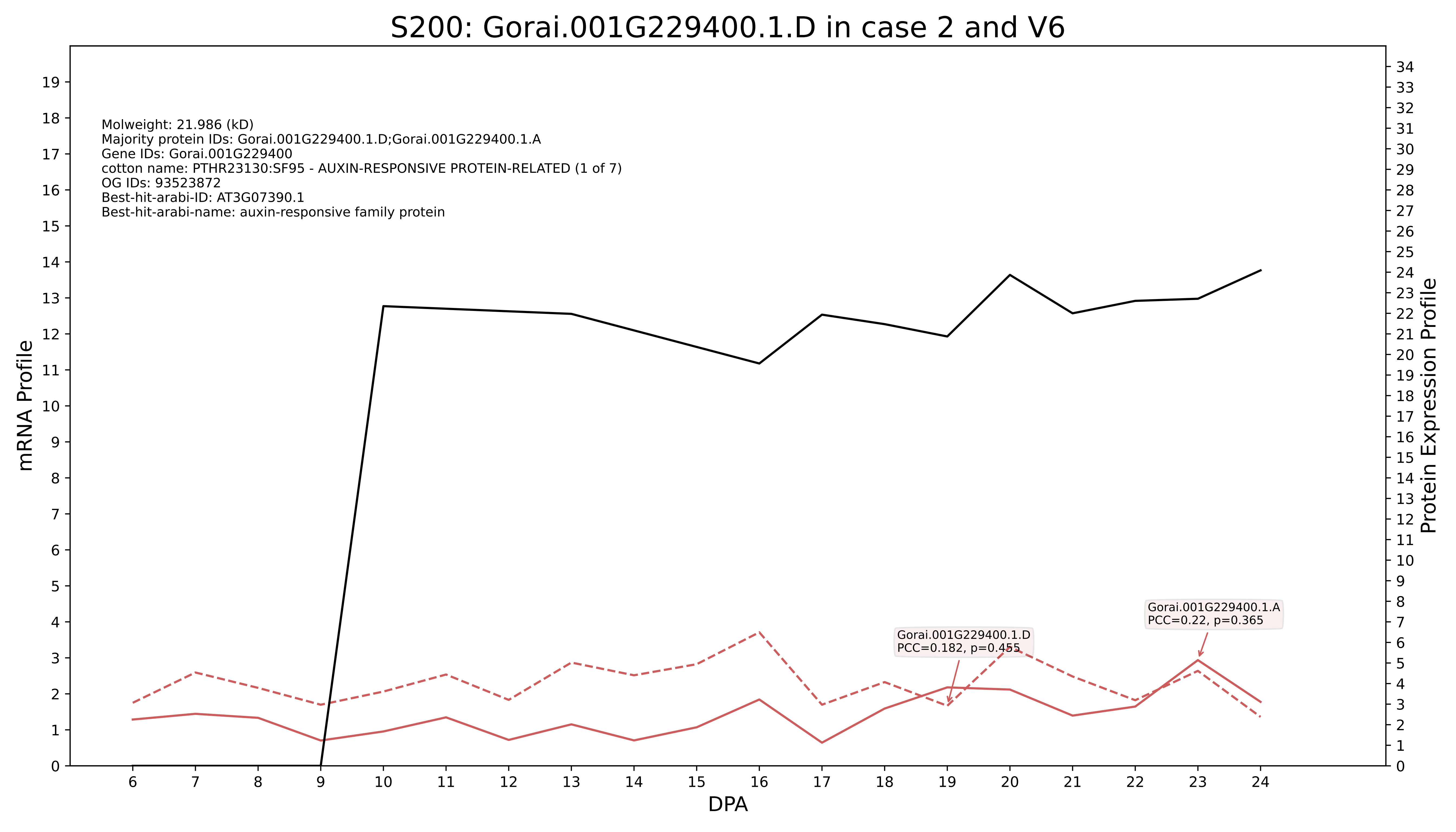

S200: Gorai.001G229400.1.D in case 2 and V6
Molweight: 21.986 (kD)
Majority protein IDs: Gorai.001G229400.1.D;Gorai.001G229400.1.A
Gene IDs: Gorai.001G229400
cotton name: PTHR23130:SF95 - AUXIN-RESPONSIVE PROTEIN-RELATED (1 of 7)
OG IDs: 93523872
Best-hit-arabi-ID: AT3G07390.1
Best-hit-arabi-name: auxin-responsive family protein
Gorai.001G229400.1.D
PCC=0.182, p=0.455
Gorai.001G229400.1.A
PCC=0.22, p=0.365
mRNA Profile
Protein Expression Profile
DPA

### S200: Gorai.004G061700.1.A in case 1 and V7

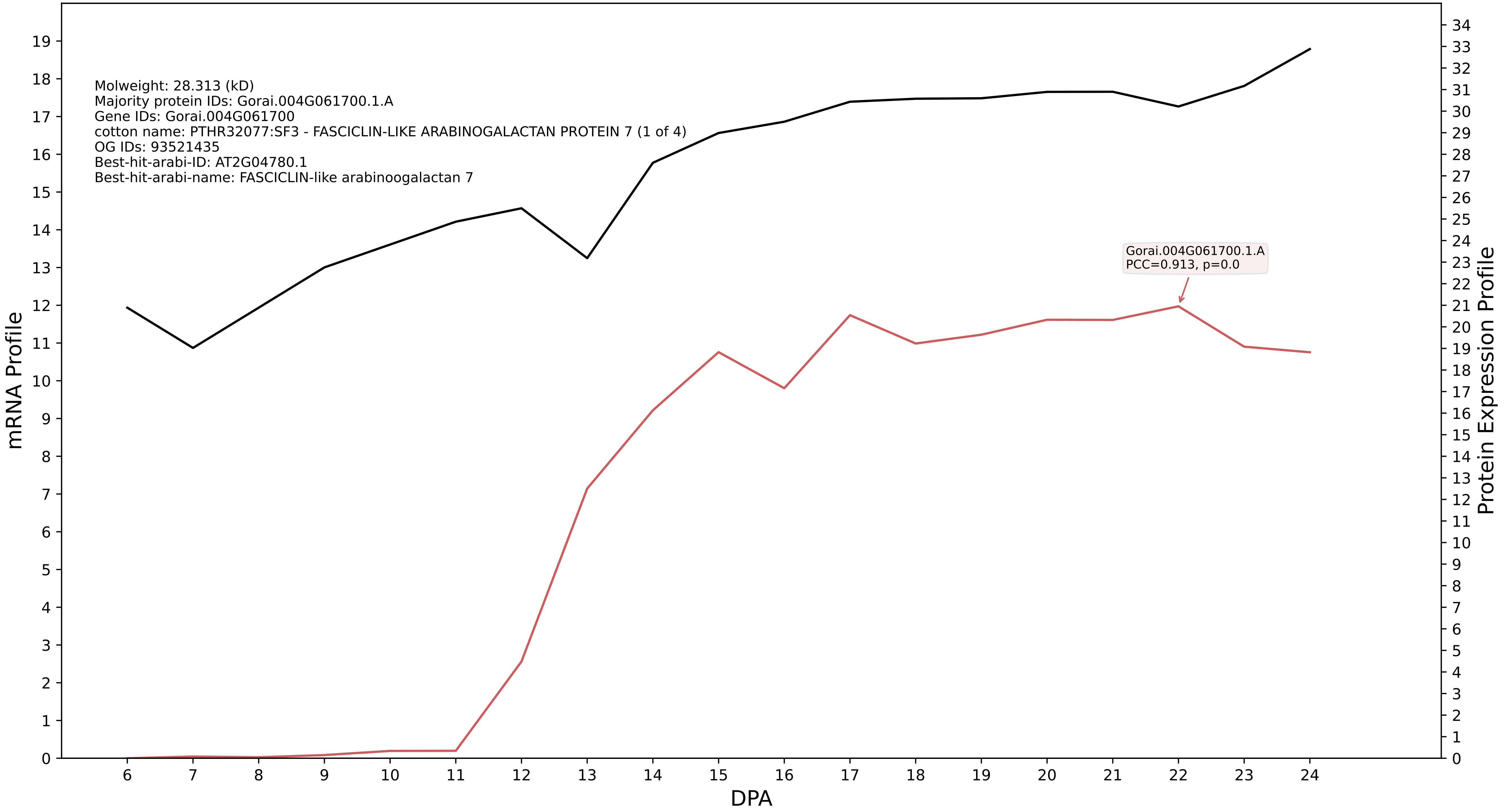

### S200: Gorai.001G131200.1.A in case 1 and V7

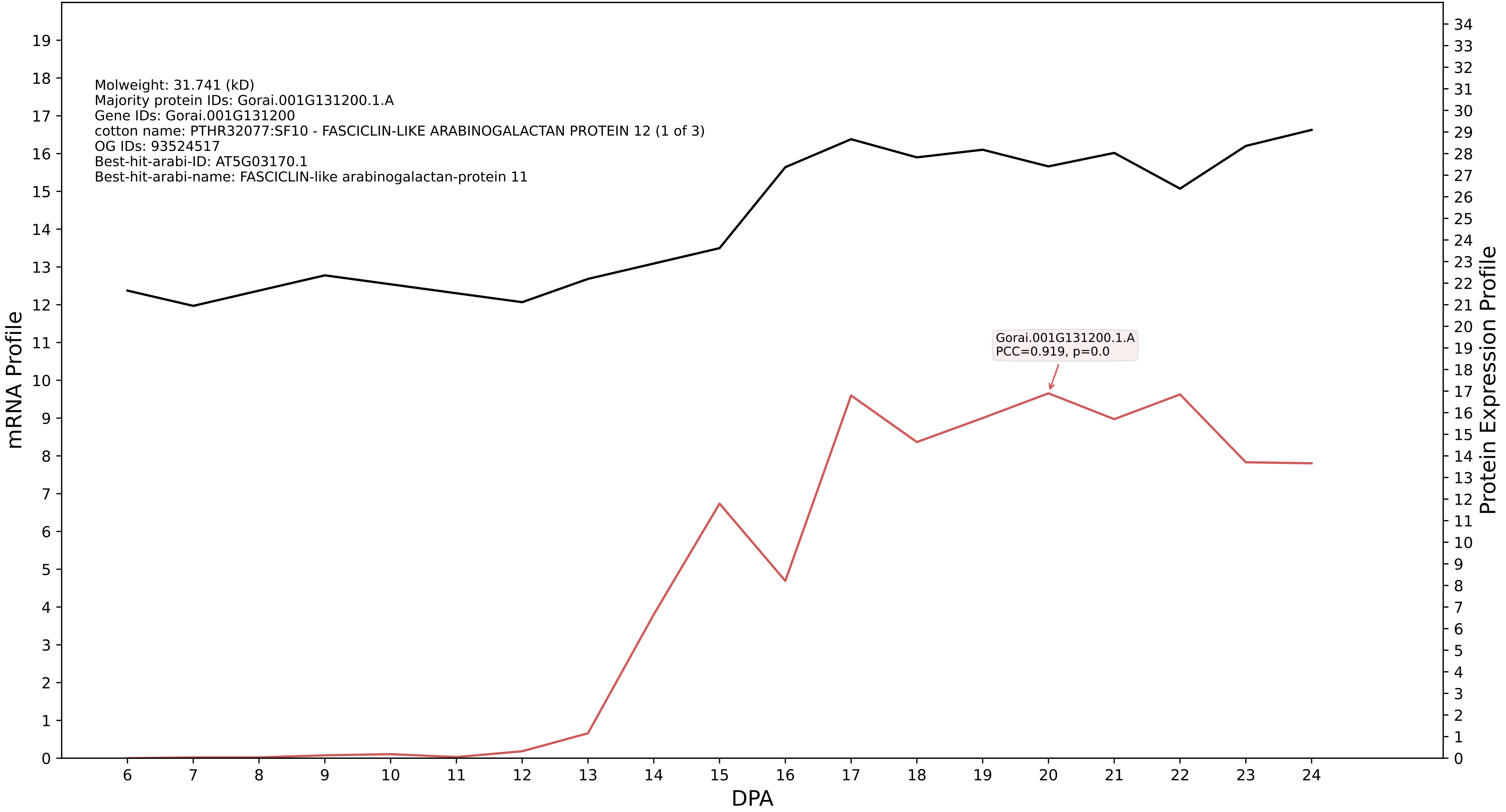

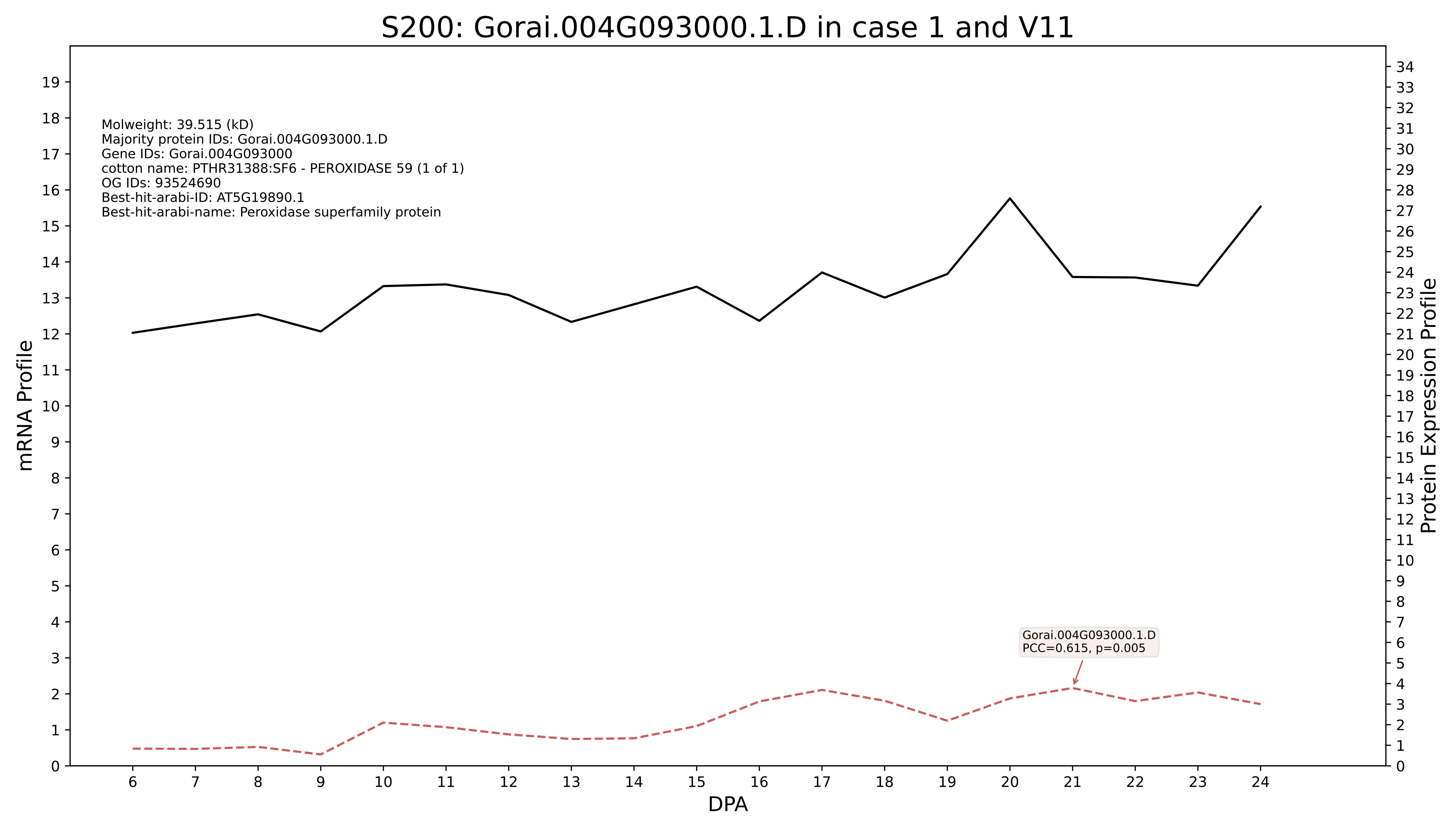
S200: Gorai.004G093000.1.D in case 1 and V11
Molweight: 39.515 (kD)
Majority protein IDs: Gorai.004G093000.1.D
Gene IDs: Gorai.004G093000
cotton name: PTHR31388:SF6 - PEROXIDASE 59 (1 of 1)
OG IDs: 93524690
Best-hit-arabi-ID: AT5G19890.1
Best-hit-arabi-name: Peroxidase superfamily protein
Gorai.004G093000.1.D
PCC=0.615, p=0.005
mRNA Profile
Protein Expression Profile
DPA

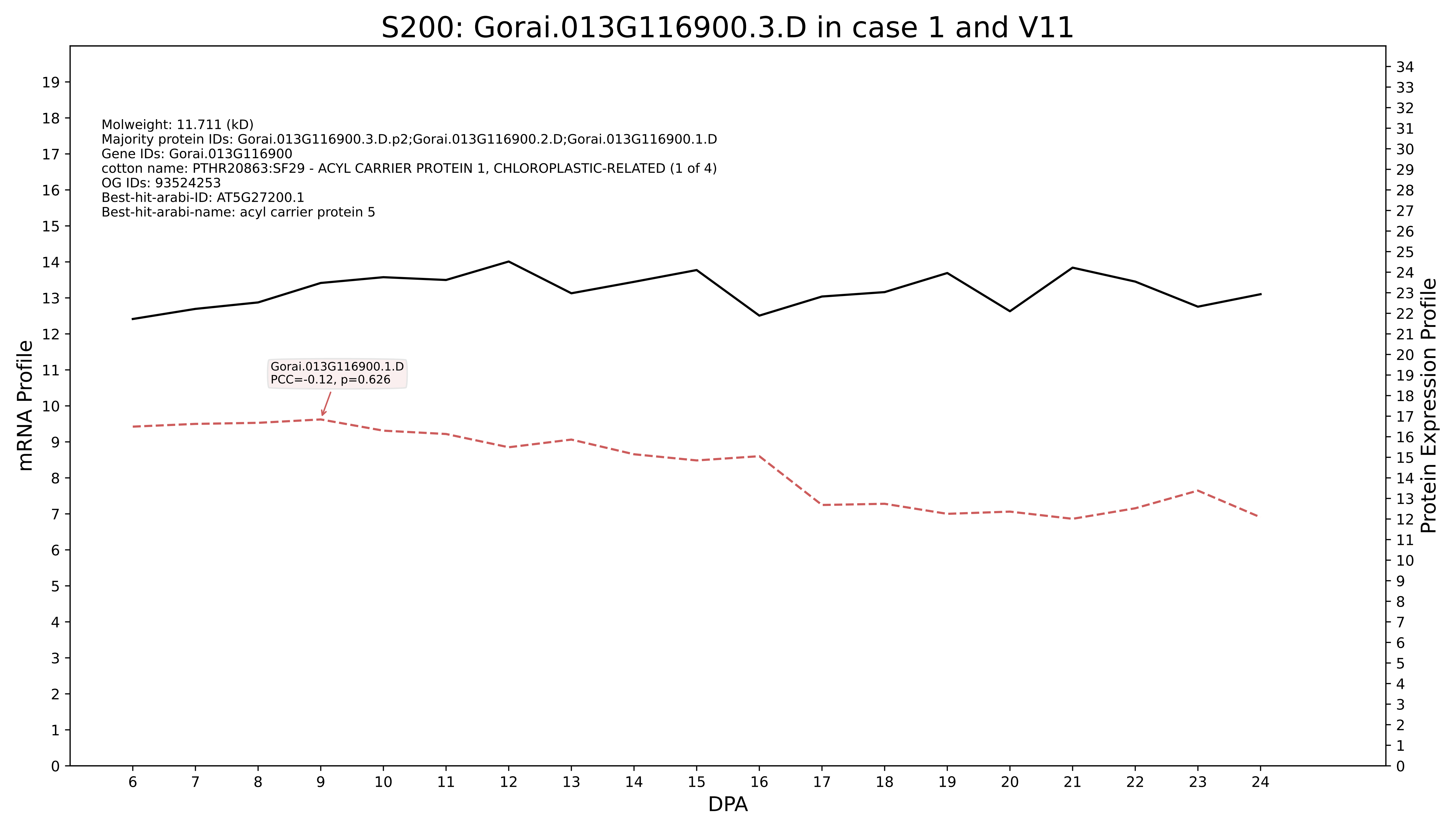

S200: Gorai.013G116900.3.D in case 1 and V11
Molweight: 11.711 (kD)
Majority protein IDs: Gorai.013G116900.3.D.p2;Gorai.013G116900.2.D;Gorai.013G116900.1.D
Gene IDs: Gorai.013G116900
cotton name: PTHR20863:SF29 - ACYL CARRIER PROTEIN 1, CHLOROPLASTIC-RELATED (1 of 4)
OG IDs: 93524253
Best-hit-arabi-ID: AT5G27200.1
Best-hit-arabi-name: acyl carrier protein 5
Gorai.013G116900.1.D
PCC=-0.12, p=0.626
mRNA Profile
Protein Expression Profile
DPA

### S200: Gorai.009G445400.1.D in case 1 and V11

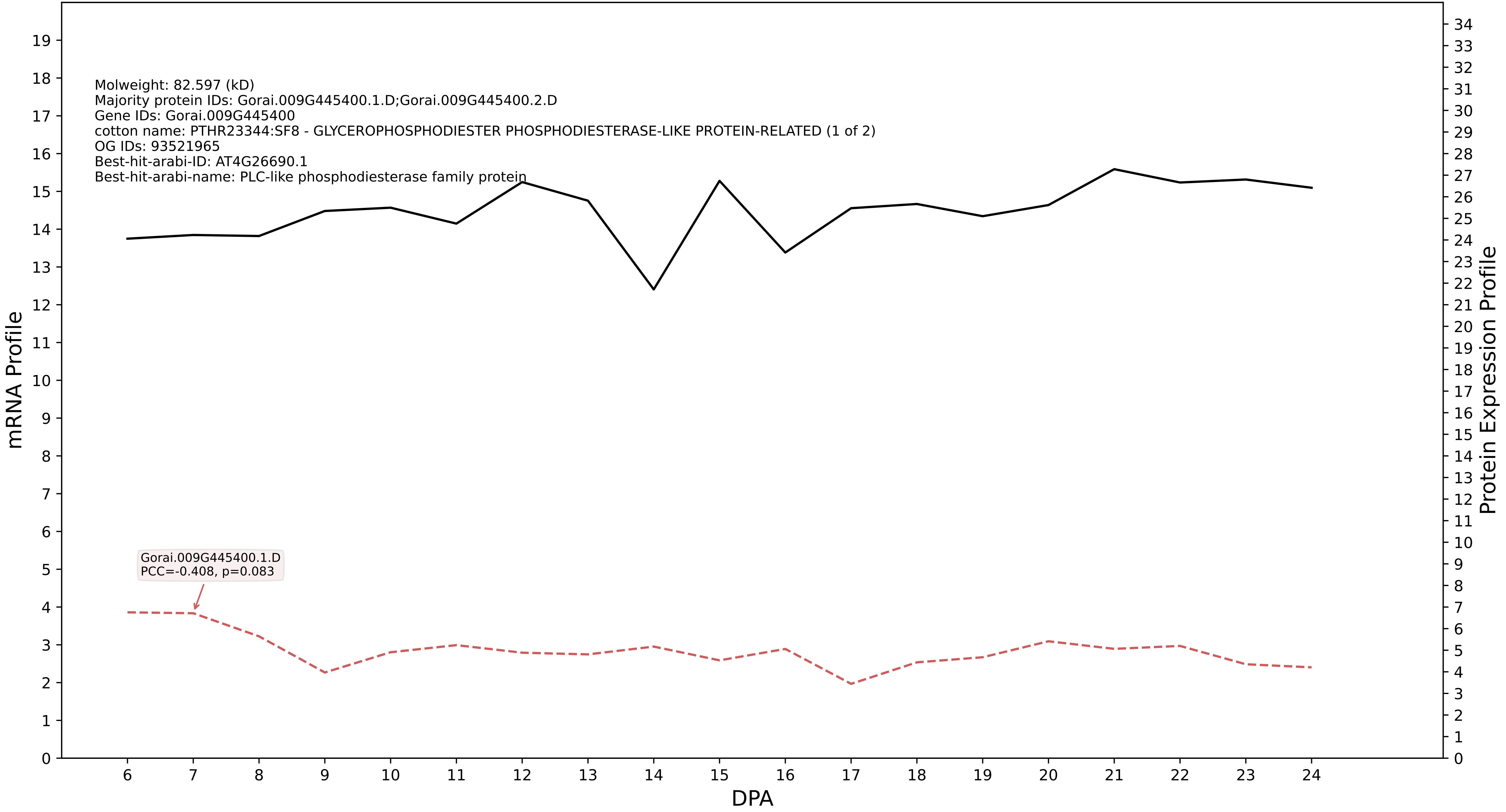

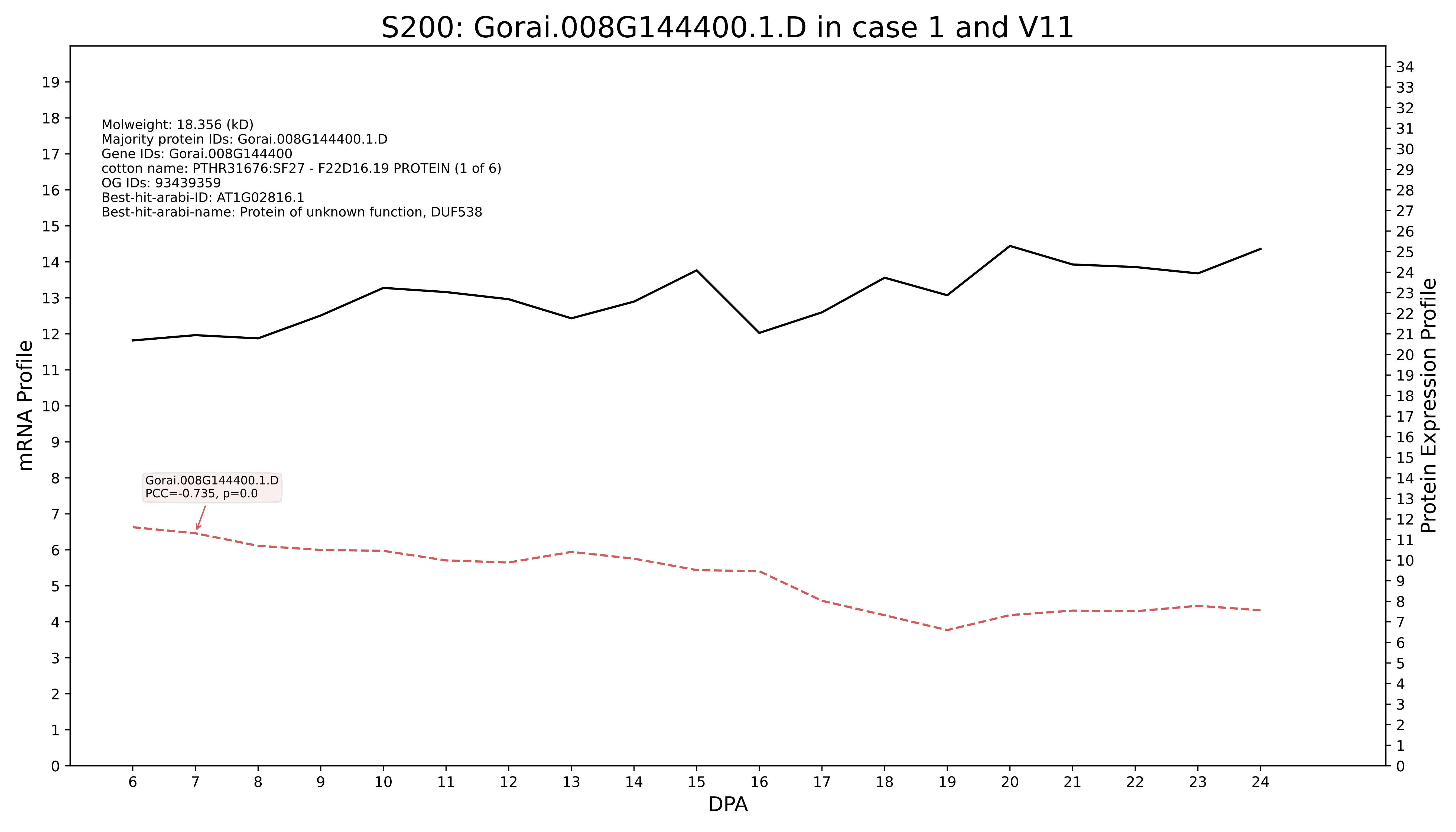
S200: Gorai.008G144400.1.D in case 1 and V11
Molweight: 18.356 (kD)
Majority protein IDs: Gorai.008G144400.1.D
Gene IDs: Gorai.008G144400
cotton name: PTHR31676:SF27 - F22D16.19 PROTEIN (1 of 6)
OG IDs: 93439359
Best-hit-arabi-ID: AT1G02816.1
Best-hit-arabi-name: Protein of unknown function, DUF538
Gorai.008G144400.1.D
PCC=-0.735, p=0.0
mRNA Profile
Protein Expression Profile
DPA

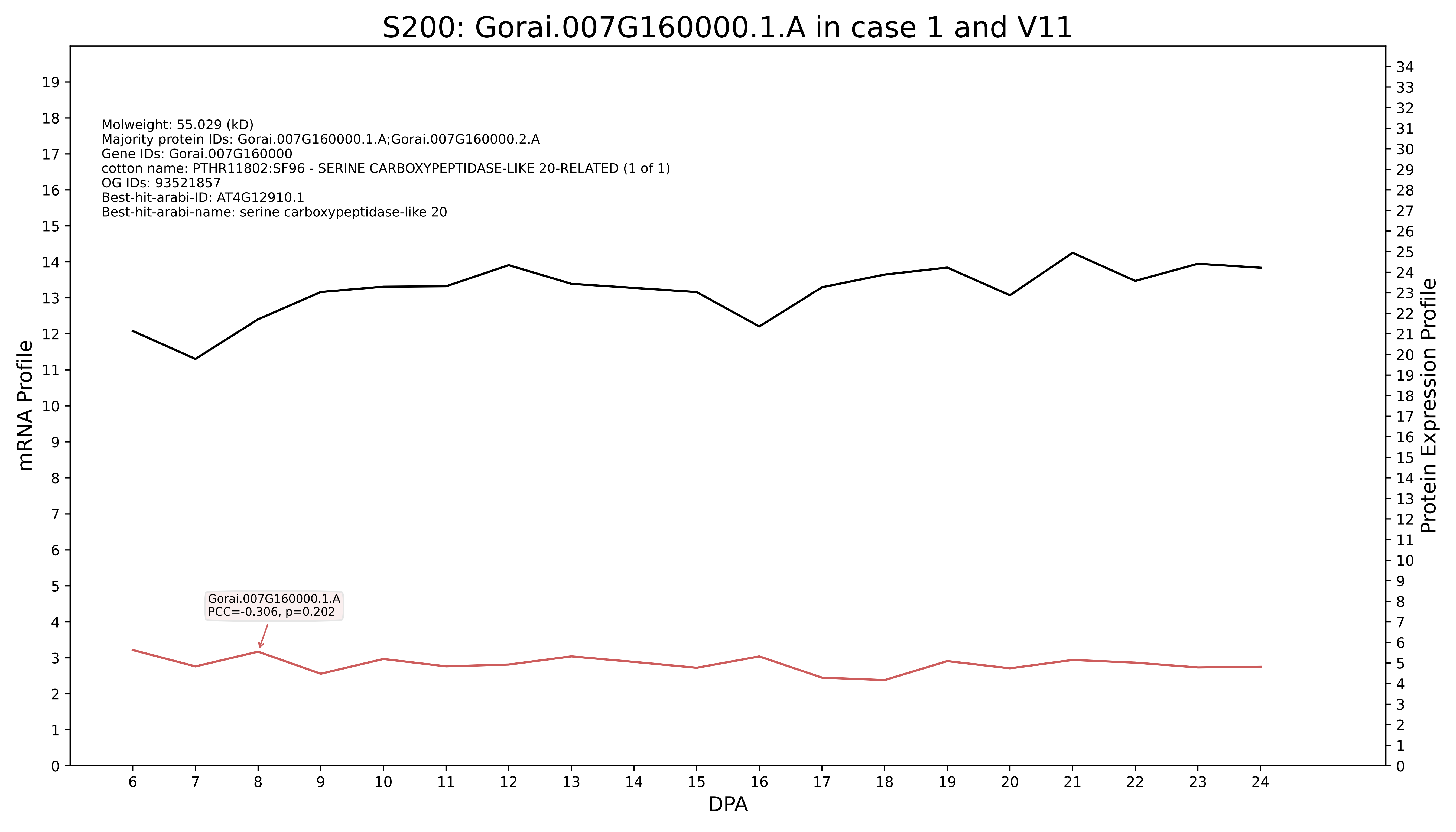
S200: Gorai.007G160000.1.A in case 1 and V11
Molweight: 55.029 (kD)
Majority protein IDs: Gorai.007G160000.1.A;Gorai.007G160000.2.A
Gene IDs: Gorai.007G160000
cotton name: PTHR11802:SF96 - SERINE CARBOXYPEPTIDASE-LIKE 20-RELATED (1 of 1)
OG IDs: 93521857
Best-hit-arabi-ID: AT4G12910.1
Best-hit-arabi-name: serine carboxypeptidase-like 20
Gorai.007G160000.1.A
PCC=-0.306, p=0.202
mRNA Profile
Protein Expression Profile
DPA

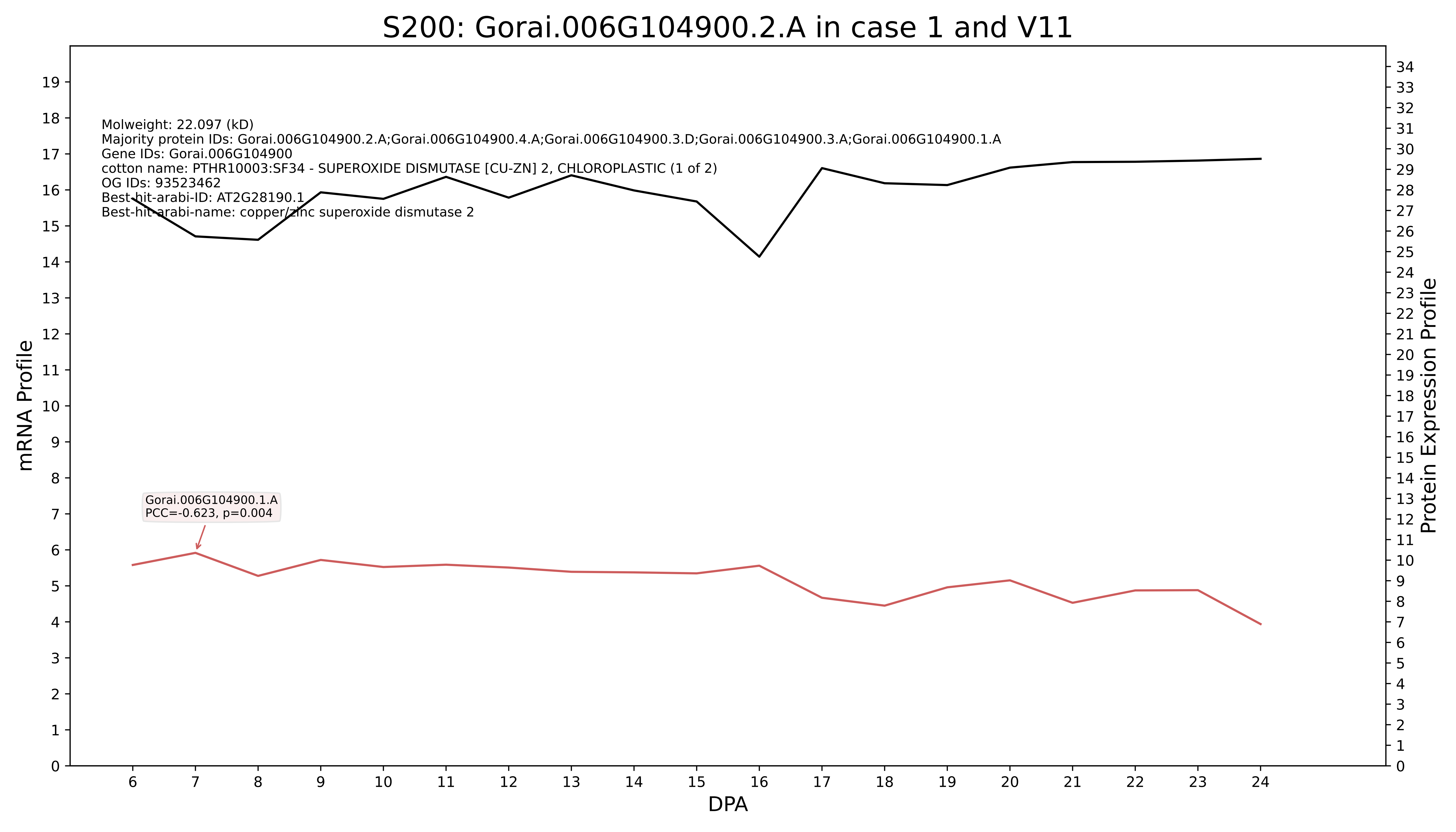
S200: Gorai.006G104900.2.A in case 1 and V11
Molweight: 22.097 (kD)
Majority protein IDs: Gorai.006G104900.2.A;Gorai.006G104900.4.A;Gorai.006G104900.3.D;Gorai.006G104900.3.A;Gorai.006G104900.1.A
Gene IDs: Gorai.006G104900
cotton name: PTHR10003:SF34 - SUPEROXIDE DISMUTASE [CU-ZN] 2, CHLOROPLASTIC (1 of 2)
OG IDs: 93523462
Best-hit-arabi-ID: AT2G28190.1
Best-hit-arabi-name: copper/zinc superoxide dismutase 2
Gorai.006G104900.1.A
PCC=-0.623, p=0.004
mRNA Profile
Protein Expression Profile
DPA

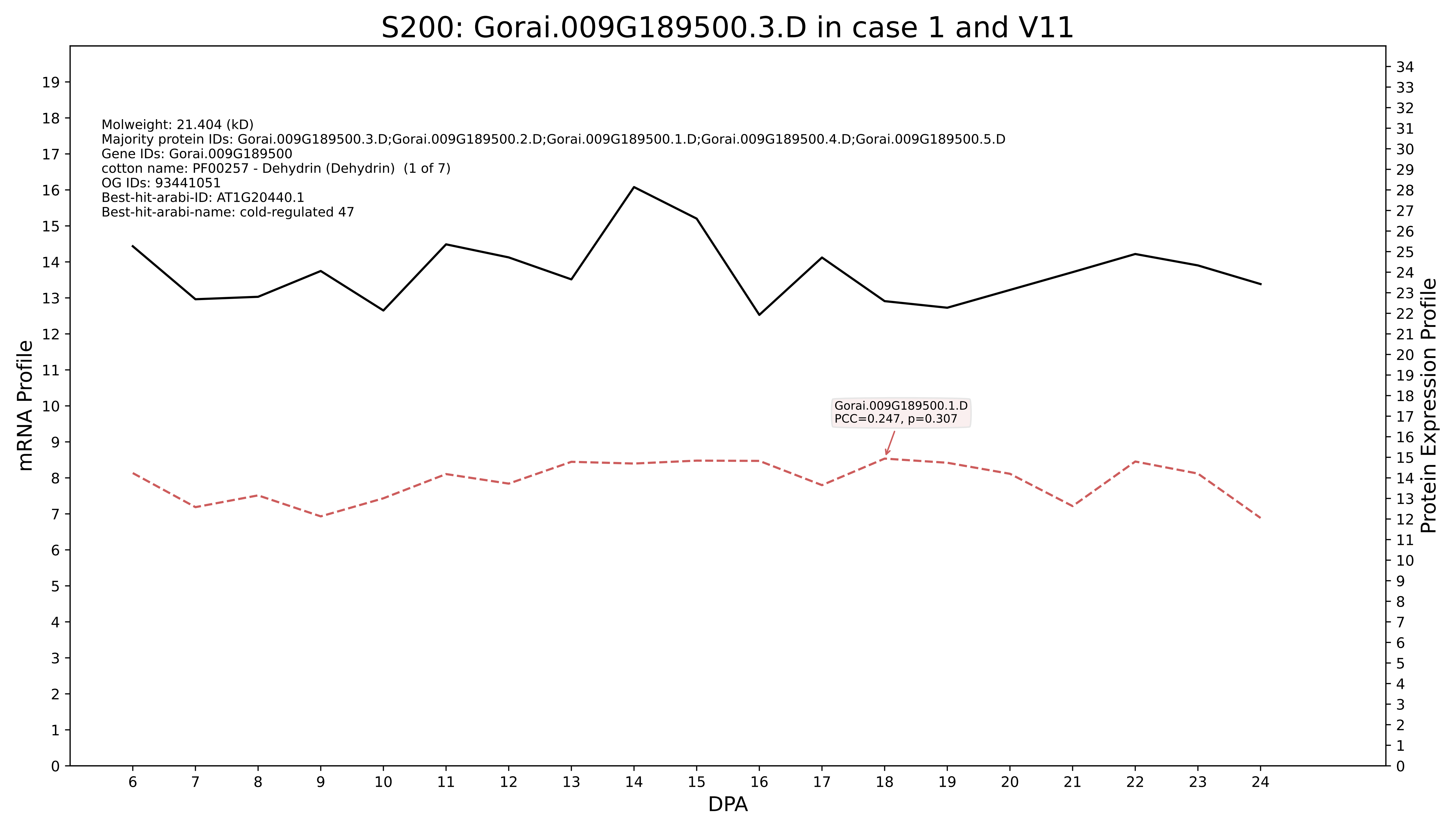
S200: Gorai.009G189500.3.D in case 1 and V11
Molweight: 21.404 (kD)
Majority protein IDs: Gorai.009G189500.3.D;Gorai.009G189500.2.D;Gorai.009G189500.1.D;Gorai.009G189500.4.D;Gorai.009G189500.5.D
Gene IDs: Gorai.009G189500
cotton name: PF00257 - Dehydrin (Dehydrin) (1 of 7)
OG IDs: 93441051
Best-hit-arabi-ID: AT1G20440.1
Best-hit-arabi-name: cold-regulated 47
Gorai.009G189500.1.D
PCC=0.247, p=0.307
mRNA Profile
Protein Expression Profile
DPA

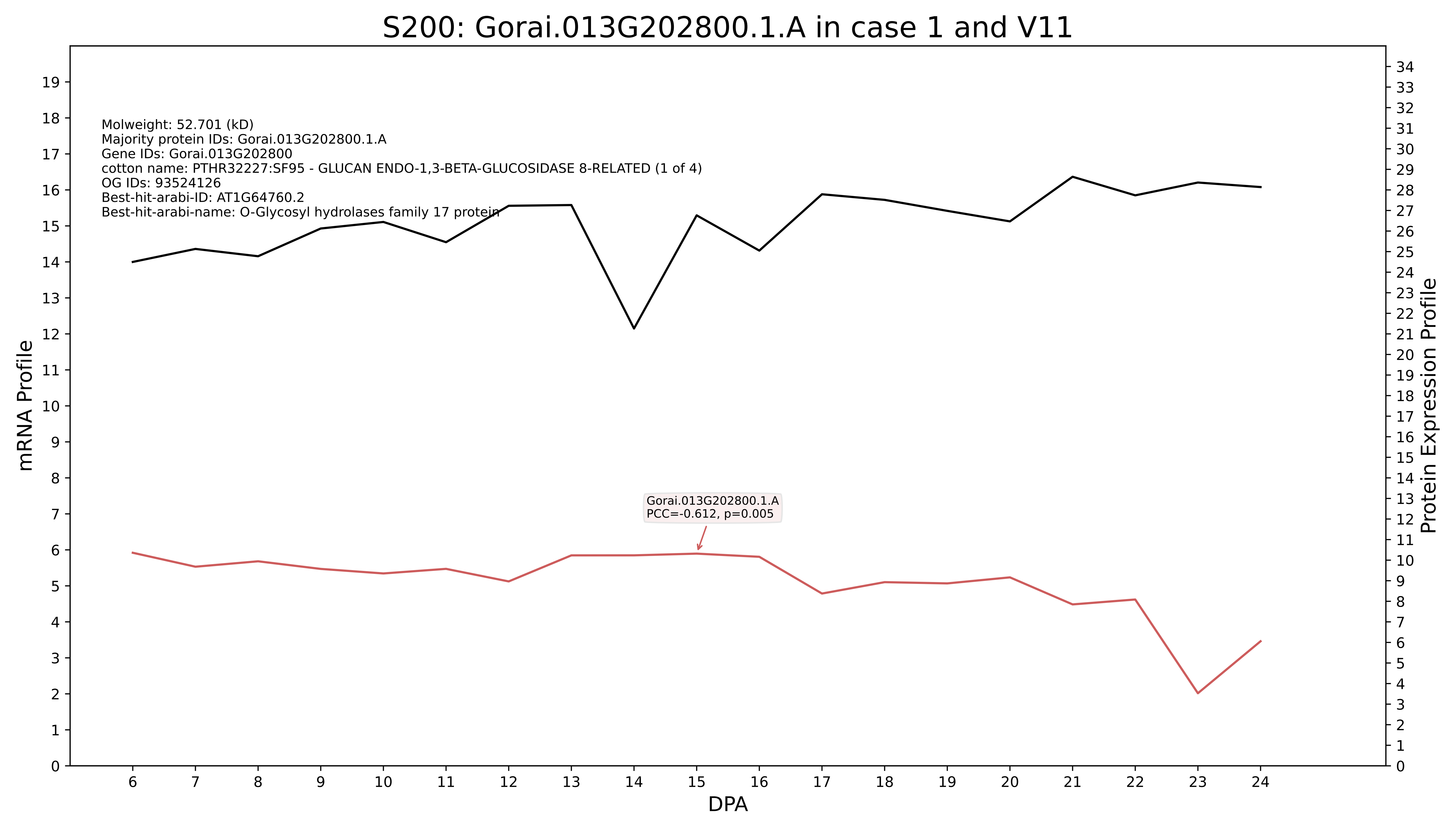

S200: Gorai.013G202800.1.A in case 1 and V11
Molweight: 52.701 (kD)
Majority protein IDs: Gorai.013G202800.1.A
Gene IDs: Gorai.013G202800
cotton name: PTHR32227:SF95 - GLUCAN ENDO-1,3-BETA-GLUCOSIDASE 8-RELATED (1 of 4)
OG IDs: 93524126
Best-hit-arabi-ID: AT1G64760.2
Best-hit-arabi-name: O-Glycosyl hydrolases family 17 protein
Gorai.013G202800.1.A
PCC=-0.612, p=0.005
mRNA Profile
Protein Expression Profile
DPA

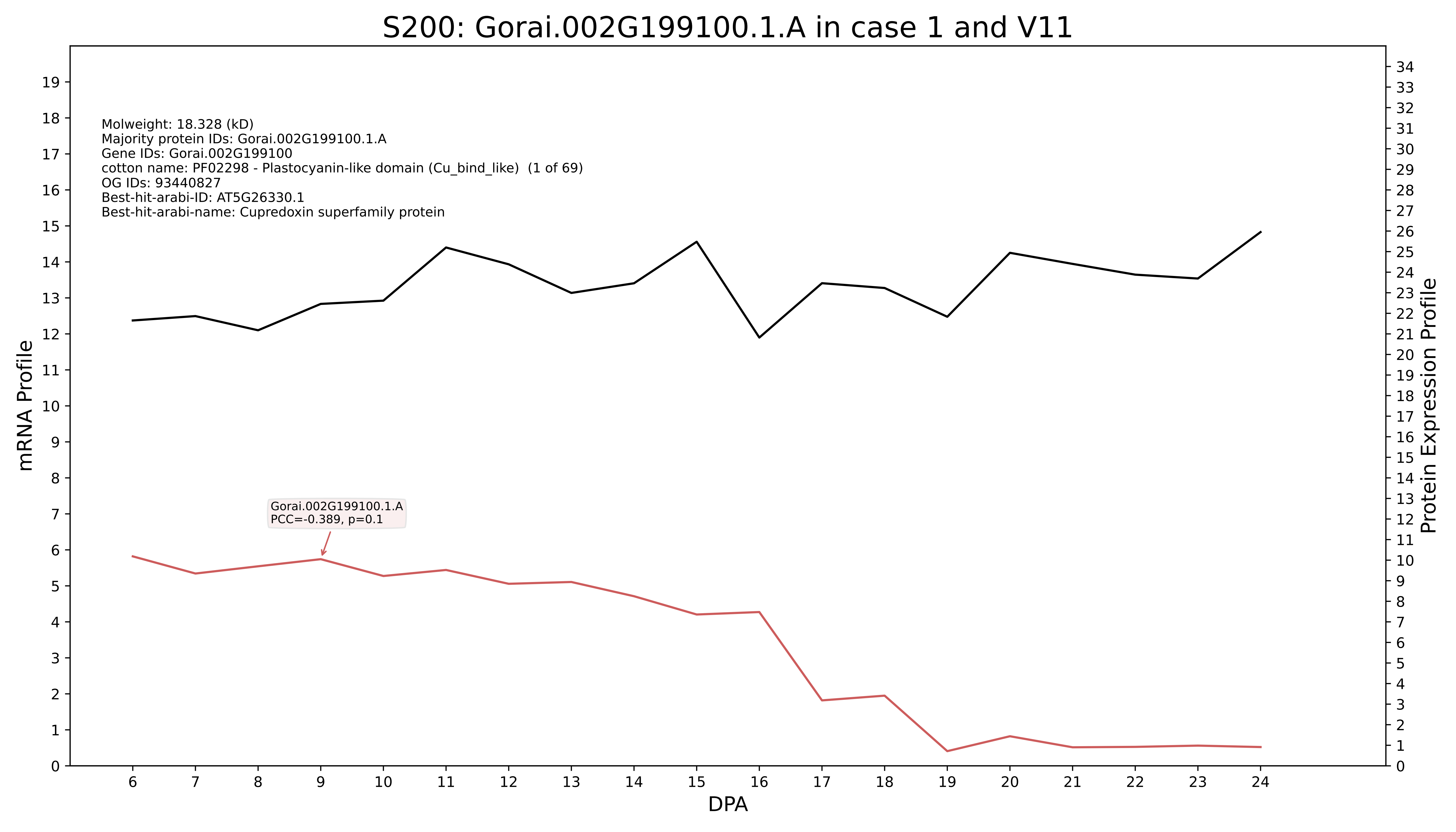

S200: Gorai.002G199100.1.A in case 1 and V11
Molweight: 18.328 (kD)
Majority protein IDs: Gorai.002G199100.1.A
Gene IDs: Gorai.002G199100
cotton name: PF02298 - Plastocyanin-like domain (Cu_bind_like) (1 of 69)
OG IDs: 93440827
Best-hit-arabi-ID: AT5G26330.1
Best-hit-arabi-name: Cupredoxin superfamily protein
Gorai.002G199100.1.A
PCC=-0.389, p=0.1
mRNA Profile
Protein Expression Profile
DPA

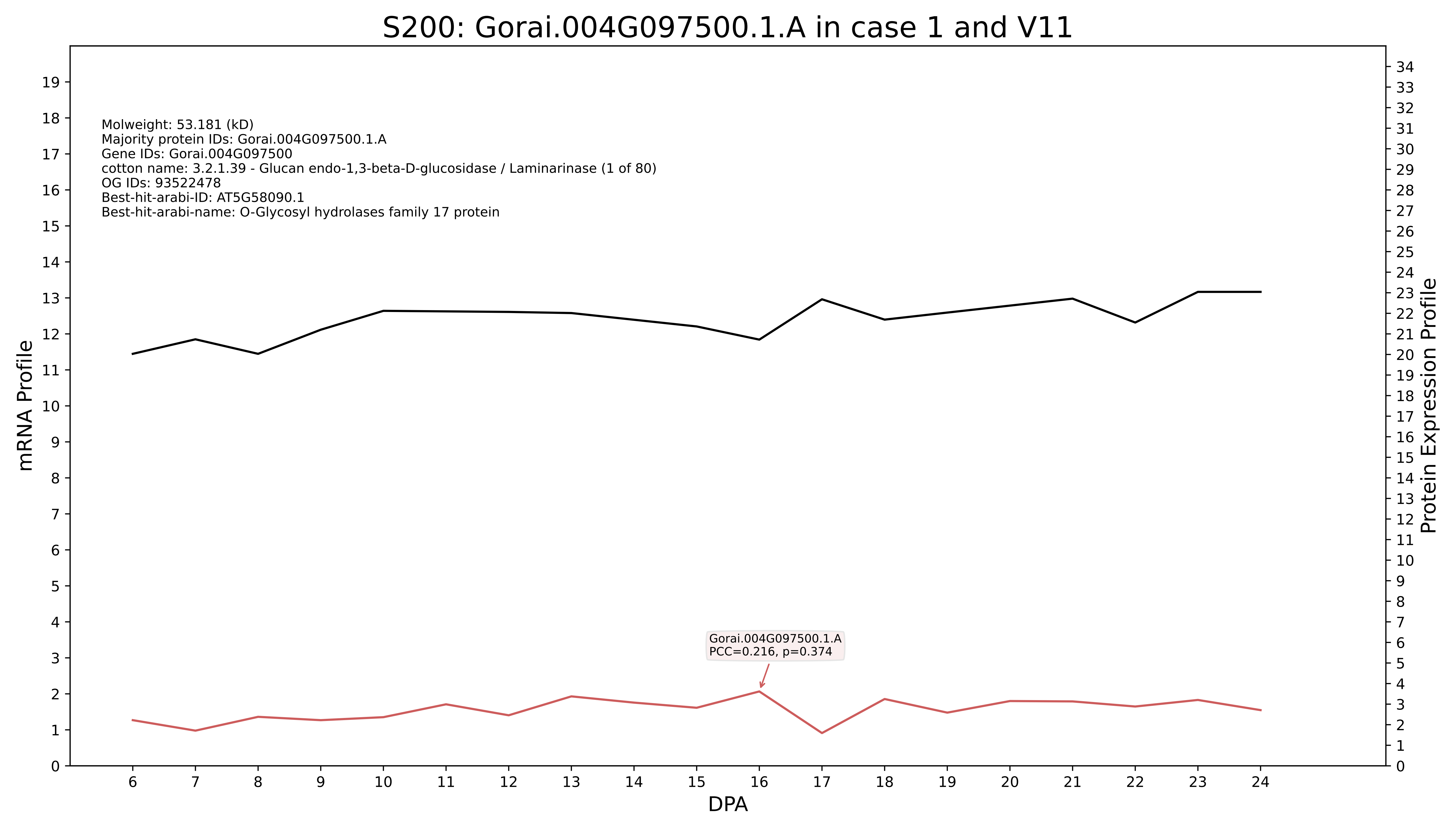

S200: Gorai.004G097500.1.A in case 1 and V11
Molweight: 53.181 (kD)
Majority protein IDs: Gorai.004G097500.1.A
Gene IDs: Gorai.004G097500
cotton name: 3.2.1.39 - Glucan endo-1,3-beta-D-glucosidase / Laminarinase (1 of 80)
OG IDs: 93522478
Best-hit-arabi-ID: AT5G58090.1
Best-hit-arabi-name: O-Glycosyl hydrolases family 17 protein
Gorai.004G097500.1.A
PCC=0.216, p=0.374
mRNA Profile
Protein Expression Profile
DPA

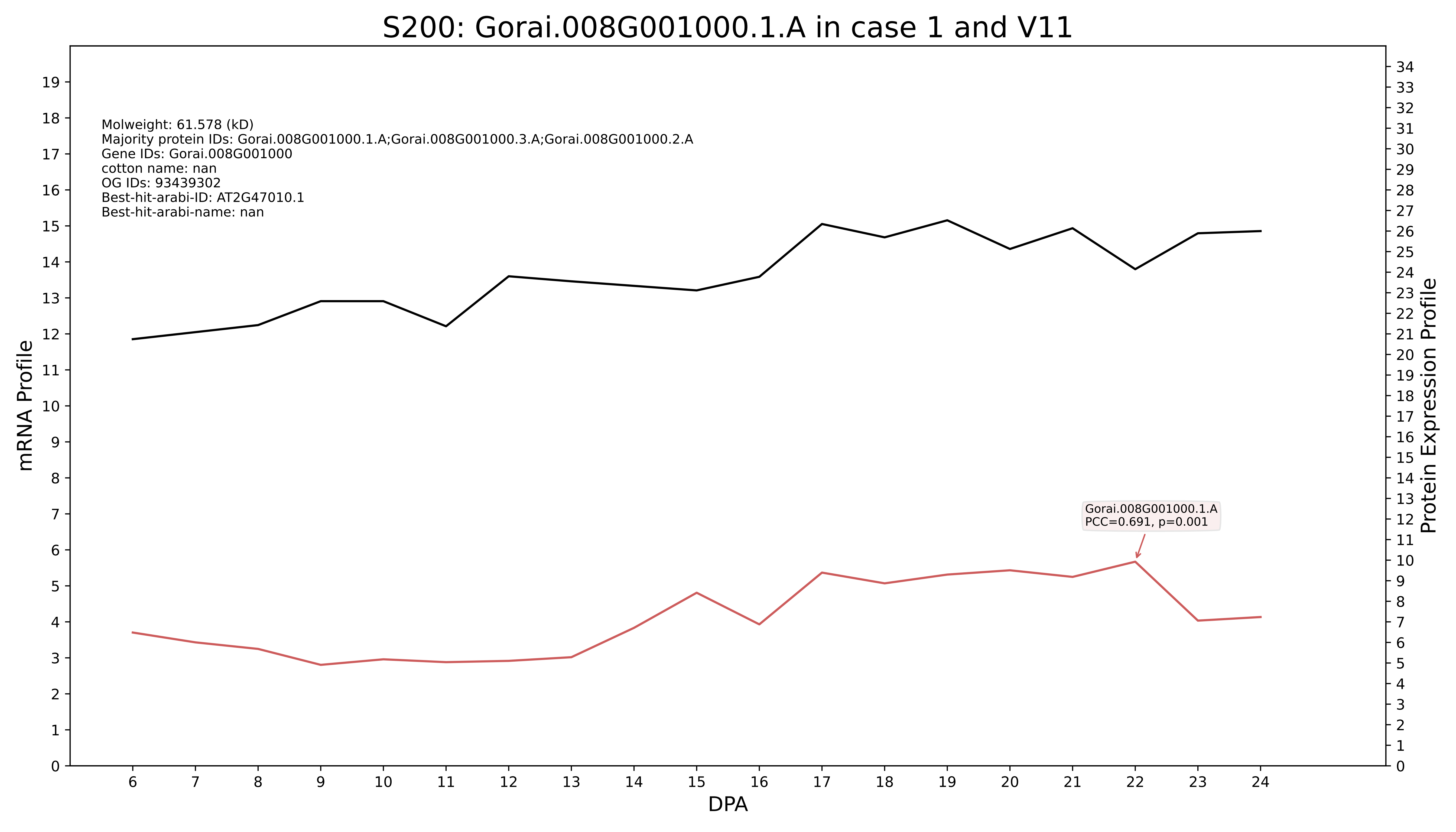

S200: Gorai.008G001000.1.A in case 1 and V11
Molweight: 61.578 (kD)
Majority protein IDs: Gorai.008G001000.1.A;Gorai.008G001000.3.A;Gorai.008G001000.2.A
Gene IDs: Gorai.008G001000
cotton name: nan
OG IDs: 93439302
Best-hit-arabi-ID: AT2G47010.1
Best-hit-arabi-name: nan
Gorai.008G001000.1.A
PCC=0.691, p=0.001
mRNA Profile
Protein Expression Profile
DPA

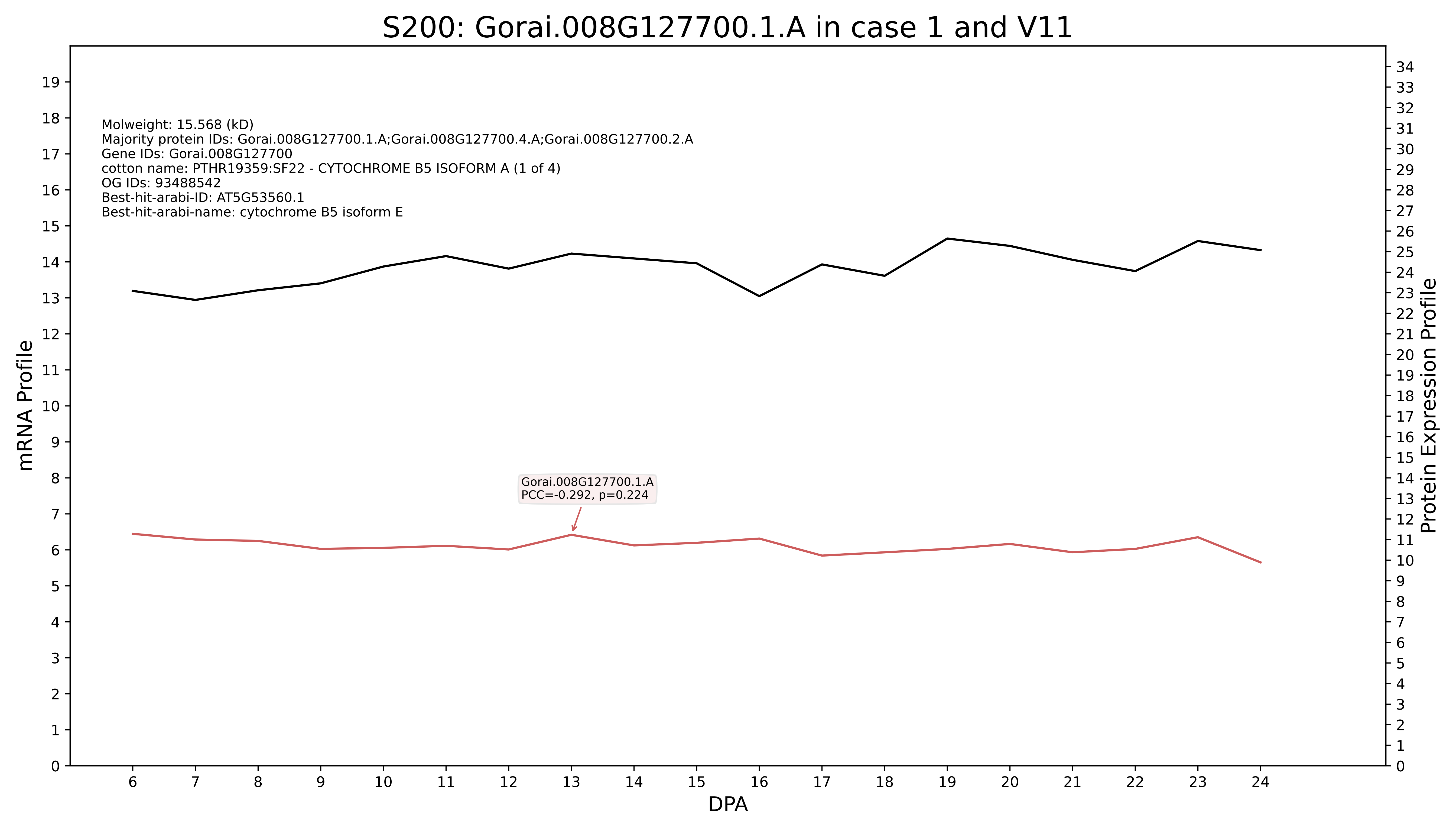

S200: Gorai.008G127700.1.A in case 1 and V11
Molweight: 15.568 (kD)
Majority protein IDs: Gorai.008G127700.1.A;Gorai.008G127700.4.A;Gorai.008G127700.2.A
Gene IDs: Gorai.008G127700
cotton name: PTHR19359:SF22 - CYTOCHROME B5 ISOFORM A (1 of 4)
OG IDs: 93488542
Best-hit-arabi-ID: AT5G53560.1
Best-hit-arabi-name: cytochrome B5 isoform E
Gorai.008G127700.1.A
PCC=-0.292, p=0.224
mRNA Profile
Protein Expression Profile
DPA

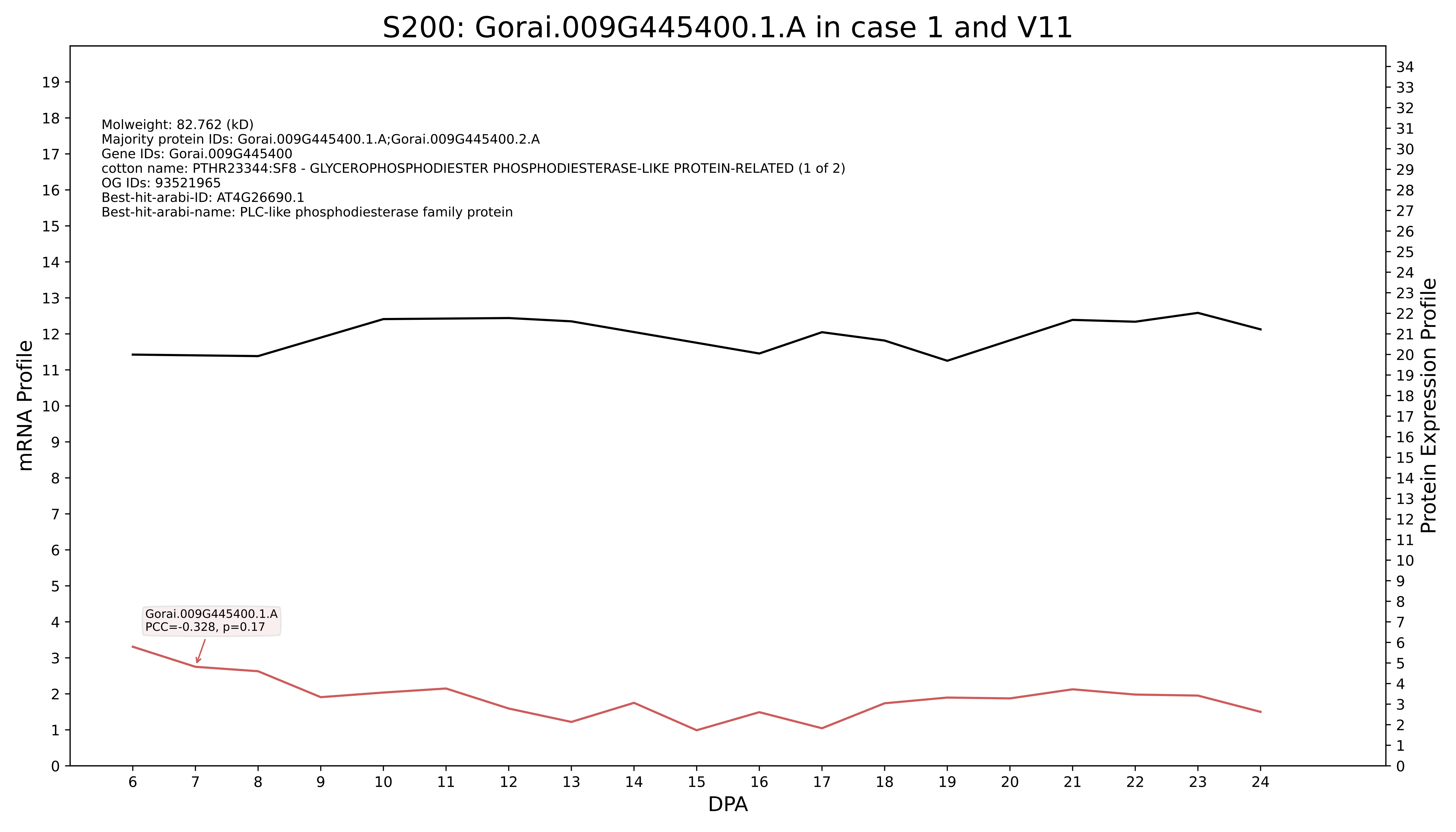
S200: Gorai.009G445400.1.A in case 1 and V11
Molweight: 82.762 (kD)
Majority protein IDs: Gorai.009G445400.1.A;Gorai.009G445400.2.A
Gene IDs: Gorai.009G445400
cotton name: PTHR23344:SF8 - GLYCEROPHOSPHODIESTER PHOSPHODIESTERASE-LIKE PROTEIN-RELATED (1 of 2)
OG IDs: 93521965
Best-hit-arabi-ID: AT4G26690.1
Best-hit-arabi-name: PLC-like phosphodiesterase family protein
Gorai.009G445400.1.A
PCC=-0.328, p=0.17
mRNA Profile
Protein Expression Profile
DPA

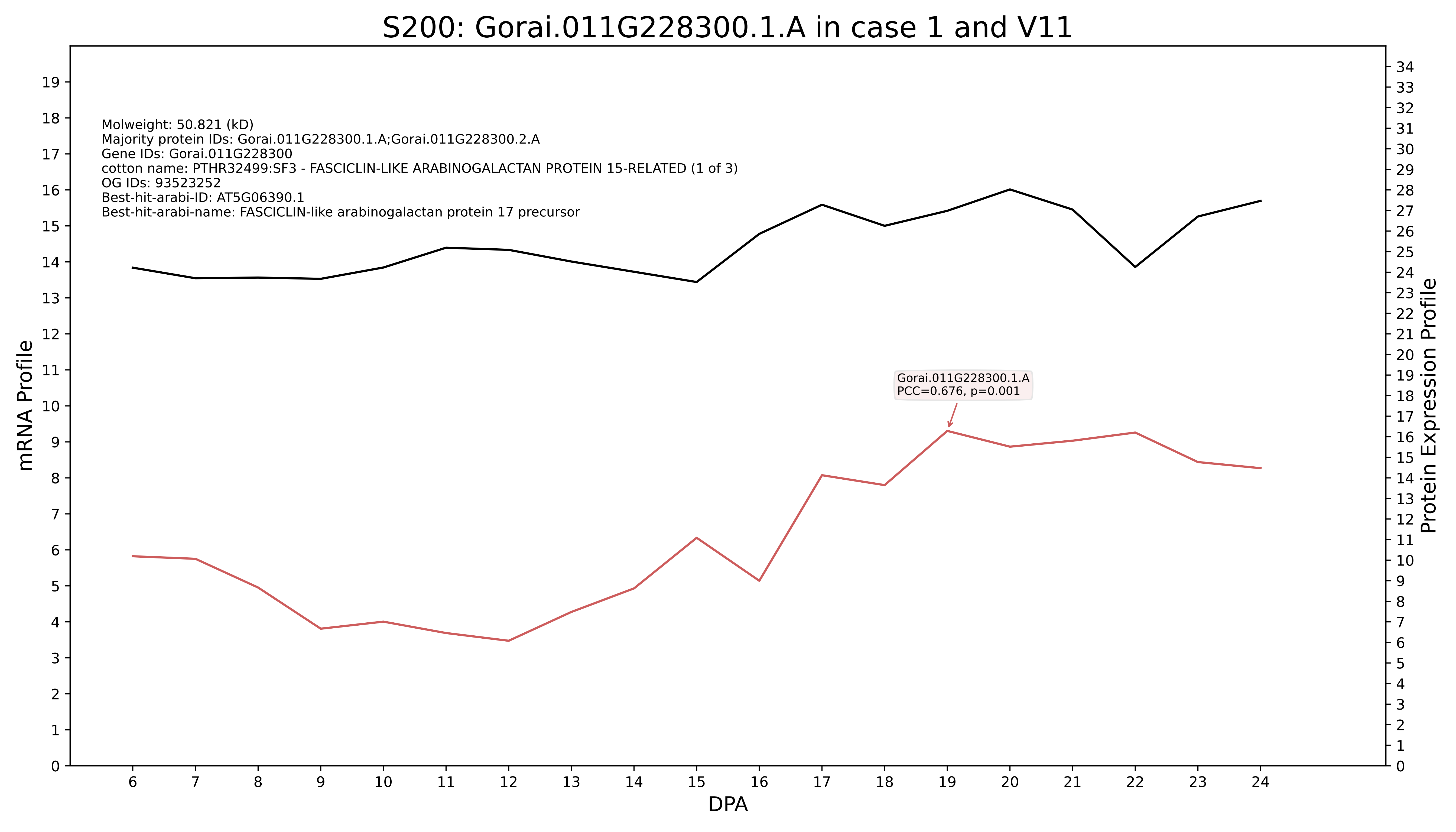
S200: Gorai.011G228300.1.A in case 1 and V11
Molweight: 50.821 (kD)
Majority protein IDs: Gorai.011G228300.1.A;Gorai.011G228300.2.A
Gene IDs: Gorai.011G228300
cotton name: PTHR32499:SF3 - FASCICLIN-LIKE ARABINOGALACTAN PROTEIN 15-RELATED (1 of 3)
OG IDs: 93523252
Best-hit-arabi-ID: AT5G06390.1
Best-hit-arabi-name: FASCICLIN-like arabinogalactan protein 17 precursor
Gorai.011G228300.1.A
PCC=0.676, p=0.001
mRNA Profile
Protein Expression Profile
DPA

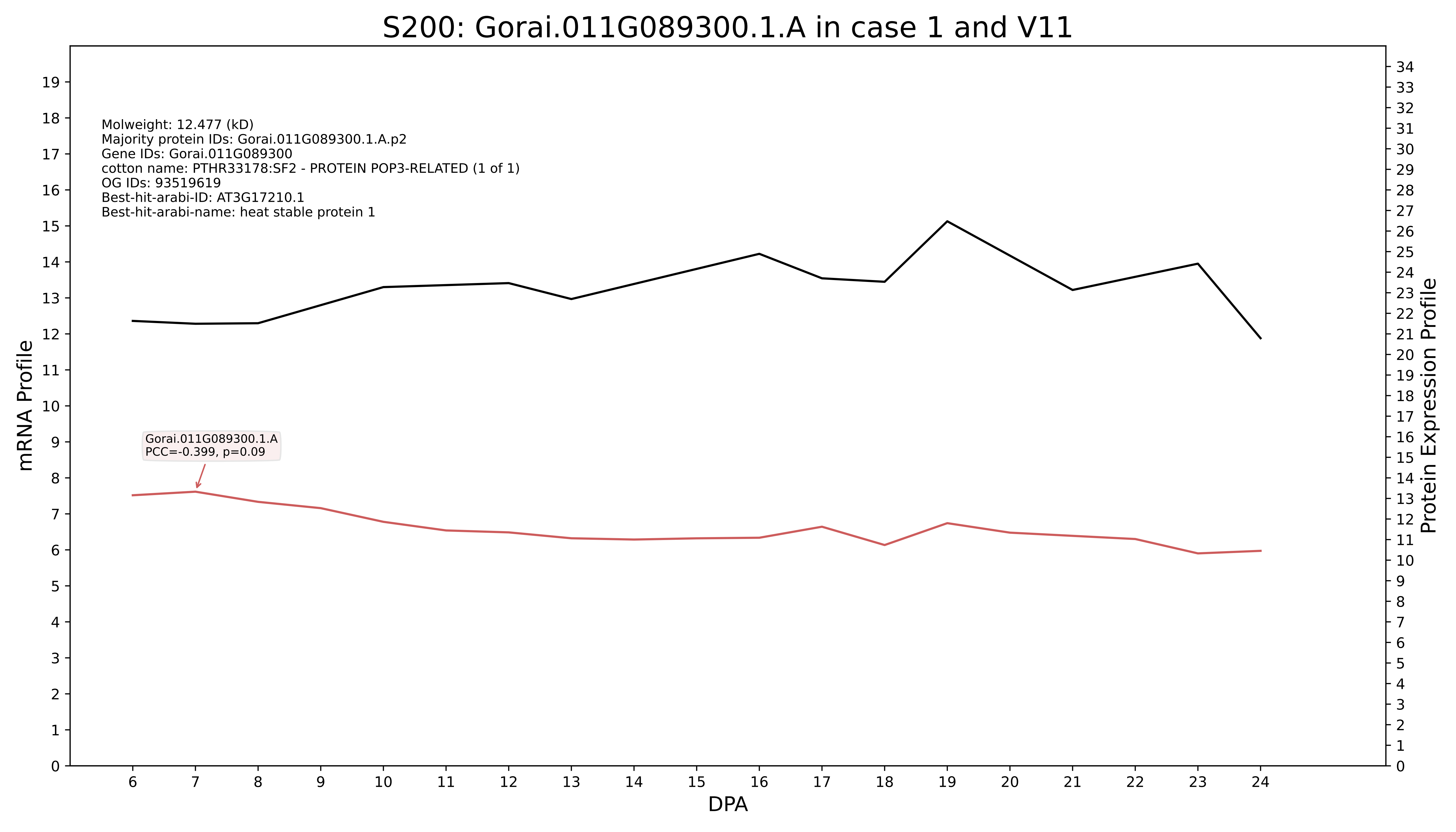
S200: Gorai.011G089300.1.A in case 1 and V11
Molweight: 12.477 (kD)
Majority protein IDs: Gorai.011G089300.1.A.p2
Gene IDs: Gorai.011G089300
cotton name: PTHR33178:SF2 - PROTEIN POP3-RELATED (1 of 1)
OG IDs: 93519619
Best-hit-arabi-ID: AT3G17210.1
Best-hit-arabi-name: heat stable protein 1
Gorai.011G089300.1.A
PCC=-0.399, p=0.09
mRNA Profile
Protein Expression Profile
DPA

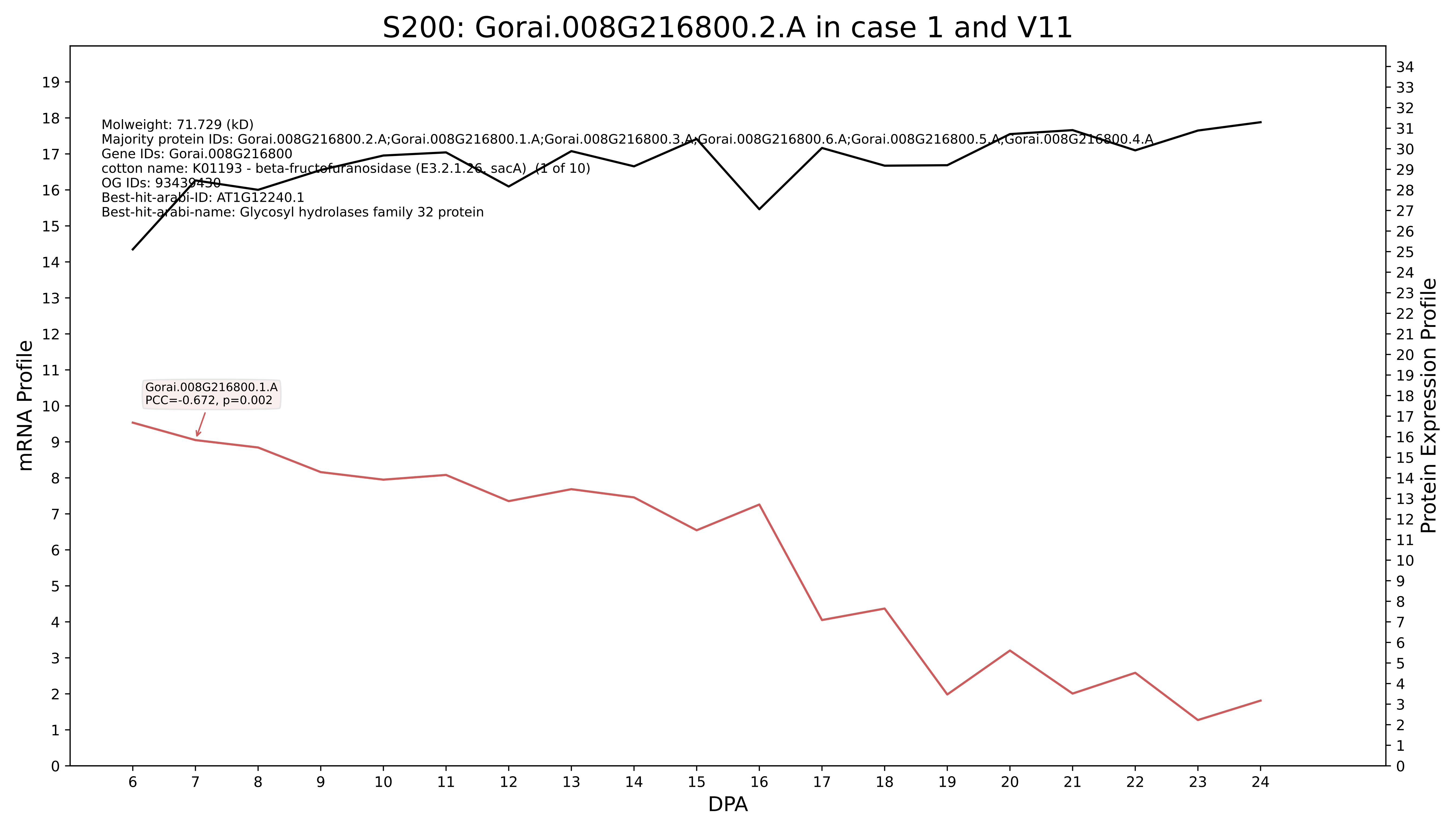

S200: Gorai.008G216800.2.A in case 1 and V11
Molweight: 71.729 (kD)
Majority protein IDs: Gorai.008G216800.2.A;Gorai.008G216800.1.A;Gorai.008G216800.3.A;Gorai.008G216800.6.A;Gorai.008G216800.5.A;Gorai.008G216800.4.A
Gene IDs: Gorai.008G216800
cotton name: K01193 - beta-fructofuranosidase (E3.2.1.26, sacA) (1 of 10)
OG IDs: 93439430
Best-hit-arabi-ID: AT1G12240.1
Best-hit-arabi-name: Glycosyl hydrolases family 32 protein
Gorai.008G216800.1.A
PCC=-0.672, p=0.002
mRNA Profile
Protein Expression Profile
DPA

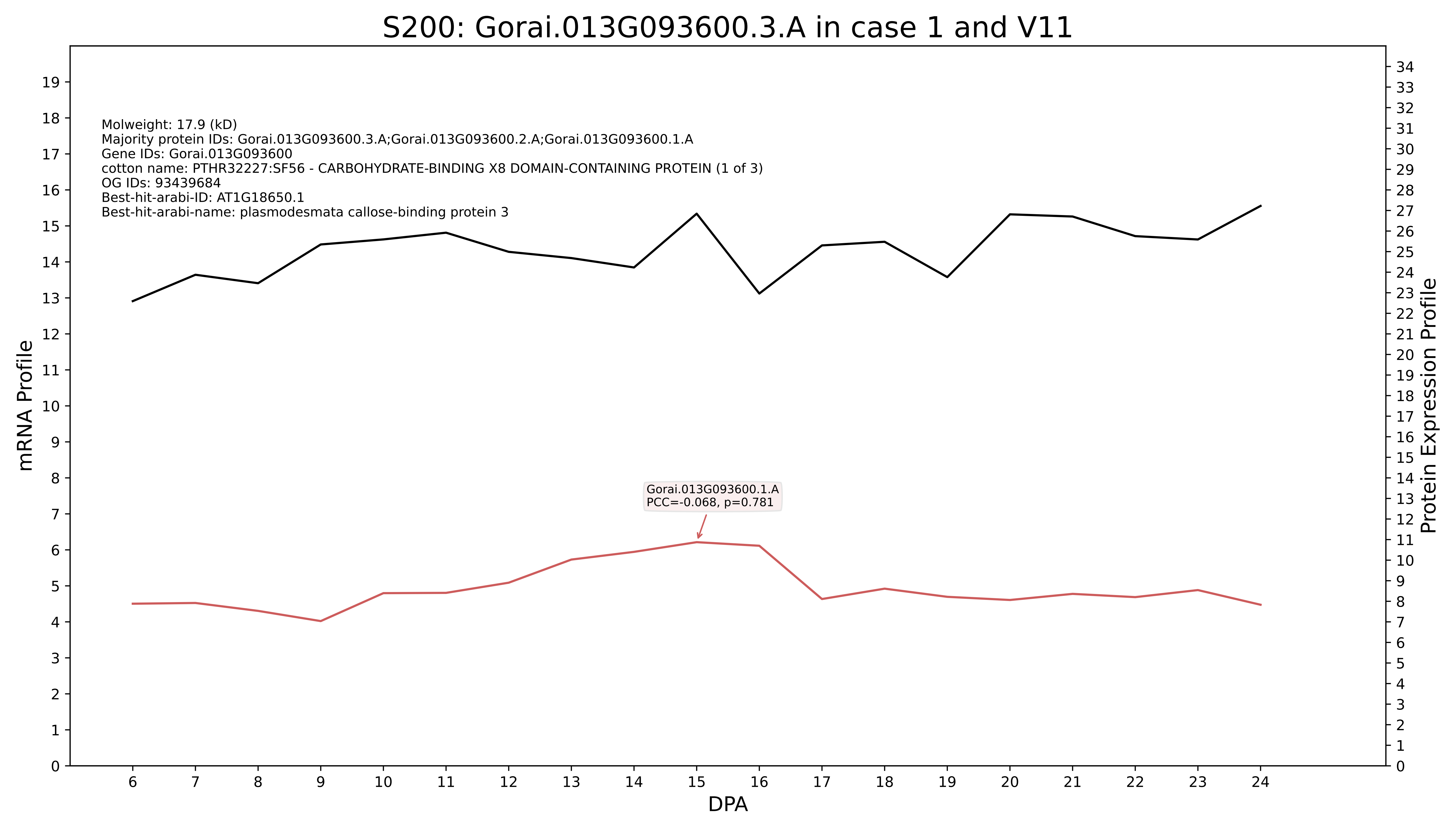

S200: Gorai.013G093600.3.A in case 1 and V11
Molweight: 17.9 (kD)
Majority protein IDs: Gorai.013G093600.3.A;Gorai.013G093600.2.A;Gorai.013G093600.1.A
Gene IDs: Gorai.013G093600
cotton name: PTHR32227:SF56 - CARBOHYDRATE-BINDING X8 DOMAIN-CONTAINING PROTEIN (1 of 3)
OG IDs: 93439684
Best-hit-arabi-ID: AT1G18650.1
Best-hit-arabi-name: plasmodesmata callose-binding protein 3
Gorai.013G093600.1.A
PCC=-0.068, p=0.781
mRNA Profile
Protein Expression Profile
DPA

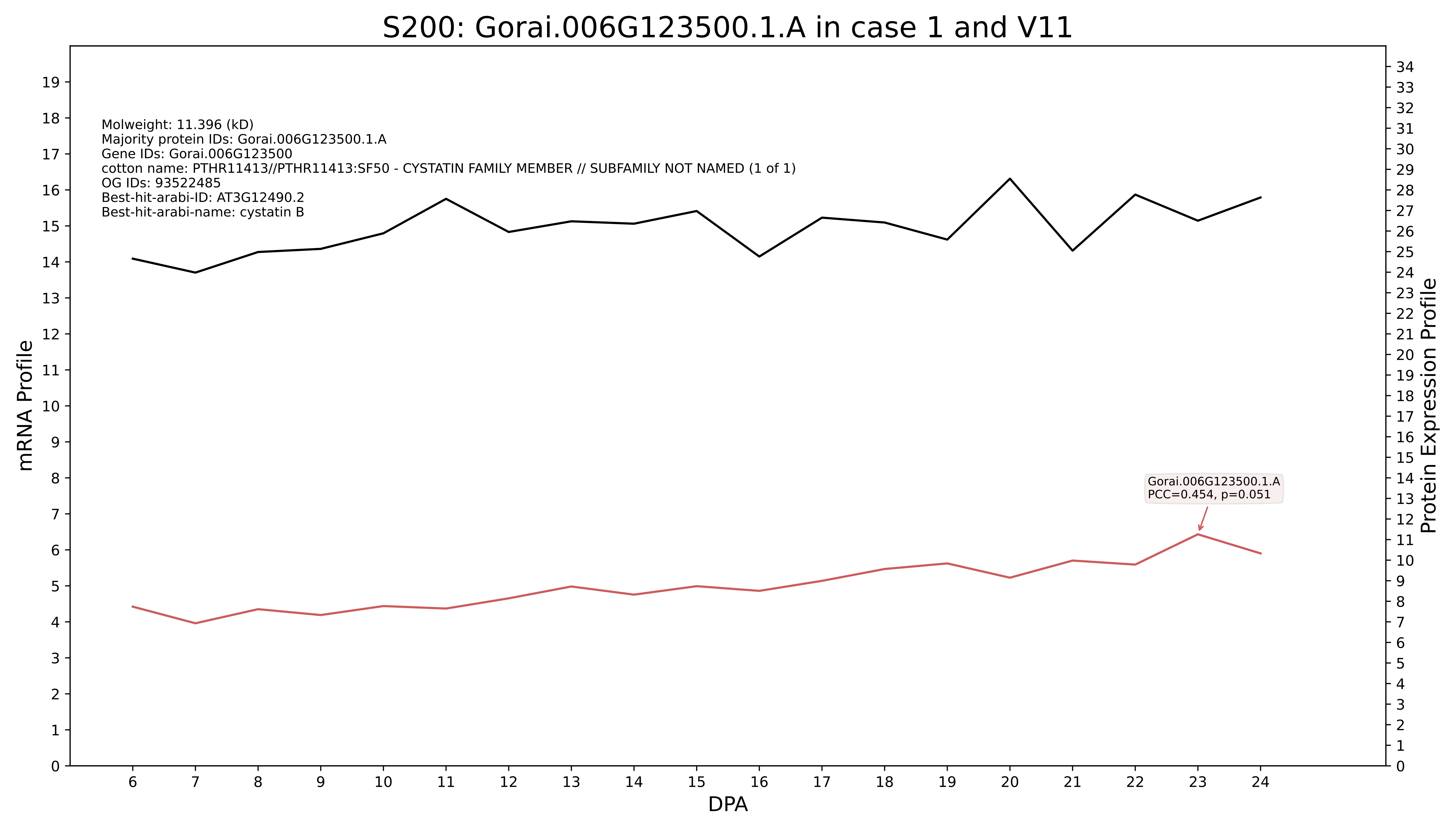

S200: Gorai.006G123500.1.A in case 1 and V11
Molweight: 11.396 (kD)
Majority protein IDs: Gorai.006G123500.1.A
Gene IDs: Gorai.006G123500
cotton name: PTHR11413//PTHR11413:SF50 - CYSTATIN FAMILY MEMBER // SUBFAMILY NOT NAMED (1 of 1)
OG IDs: 93522485
Best-hit-arabi-ID: AT3G12490.2
Best-hit-arabi-name: cystatin B
Gorai.006G123500.1.A
PCC=0.454, p=0.051
mRNA Profile
Protein Expression Profile
DPA

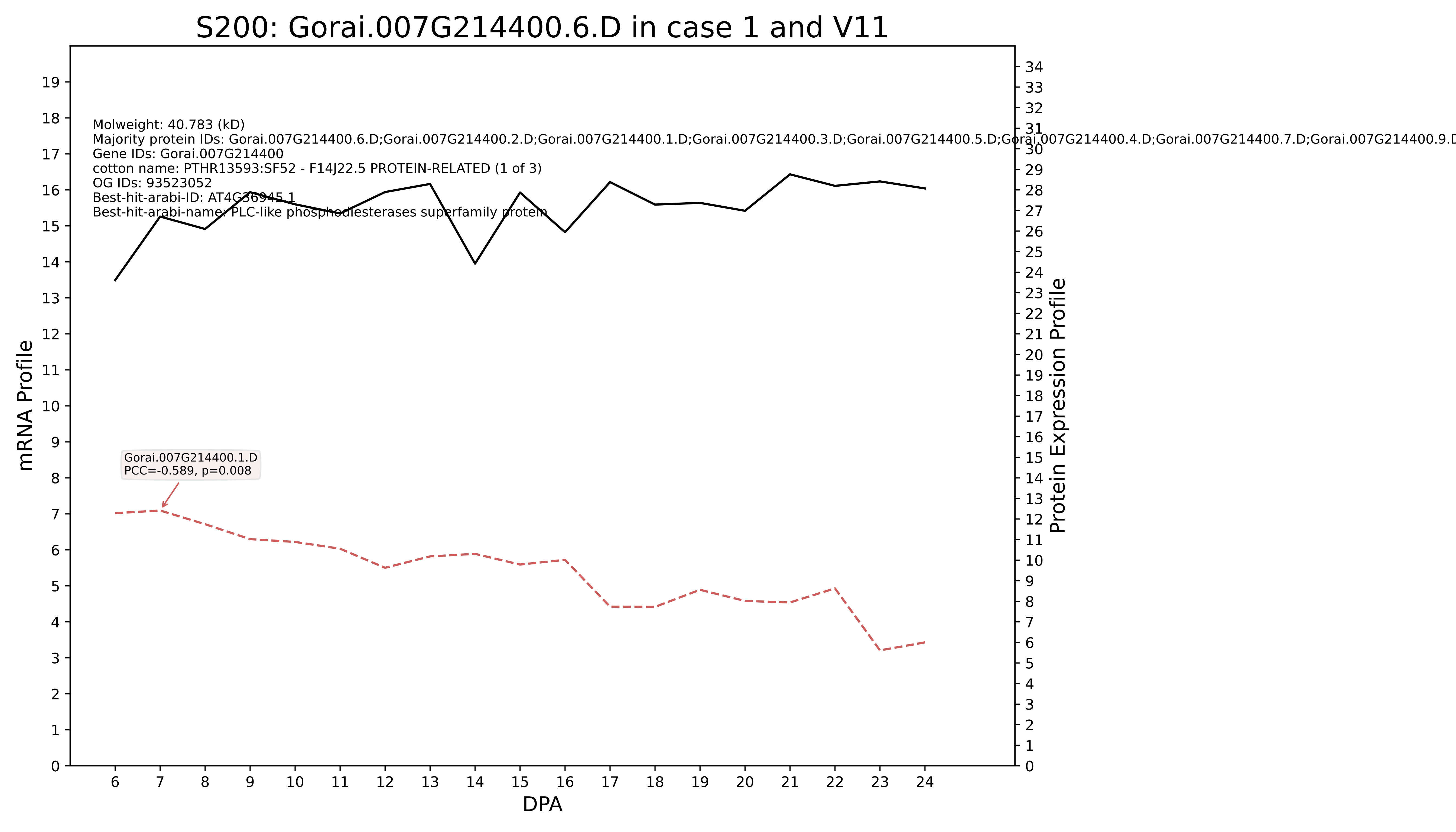

S200: Gorai.007G214400.6.D in case 1 and V11
Molweight: 40.783 (kD)
Majority protein IDs: Gorai.007G214400.6.D;Gorai.007G214400.2.D;Gorai.007G214400.1.D;Gorai.007G214400.3.D;Gorai.007G214400.5.D;Gorai.007G214400.4.D;Gorai.007G214400.7.D;Gorai.007G214400.9.D
Gene IDs: Gorai.007G214400
cotton name: PTHR13593:SF52 - F14J22.5 PROTEIN-RELATED (1 of 3)
OG IDs: 93523052
Best-hit-arabi-ID: AT4G36945.1
Best-hit-arabi-name: PLC-like phosphodiesterases superfamily protein
Gorai.007G214400.1.D
PCC=-0.589, p=0.008
mRNA Profile
Protein Expression Profile
DPA

### S200: Gorai.008G266200.8.A in case 1 and V11

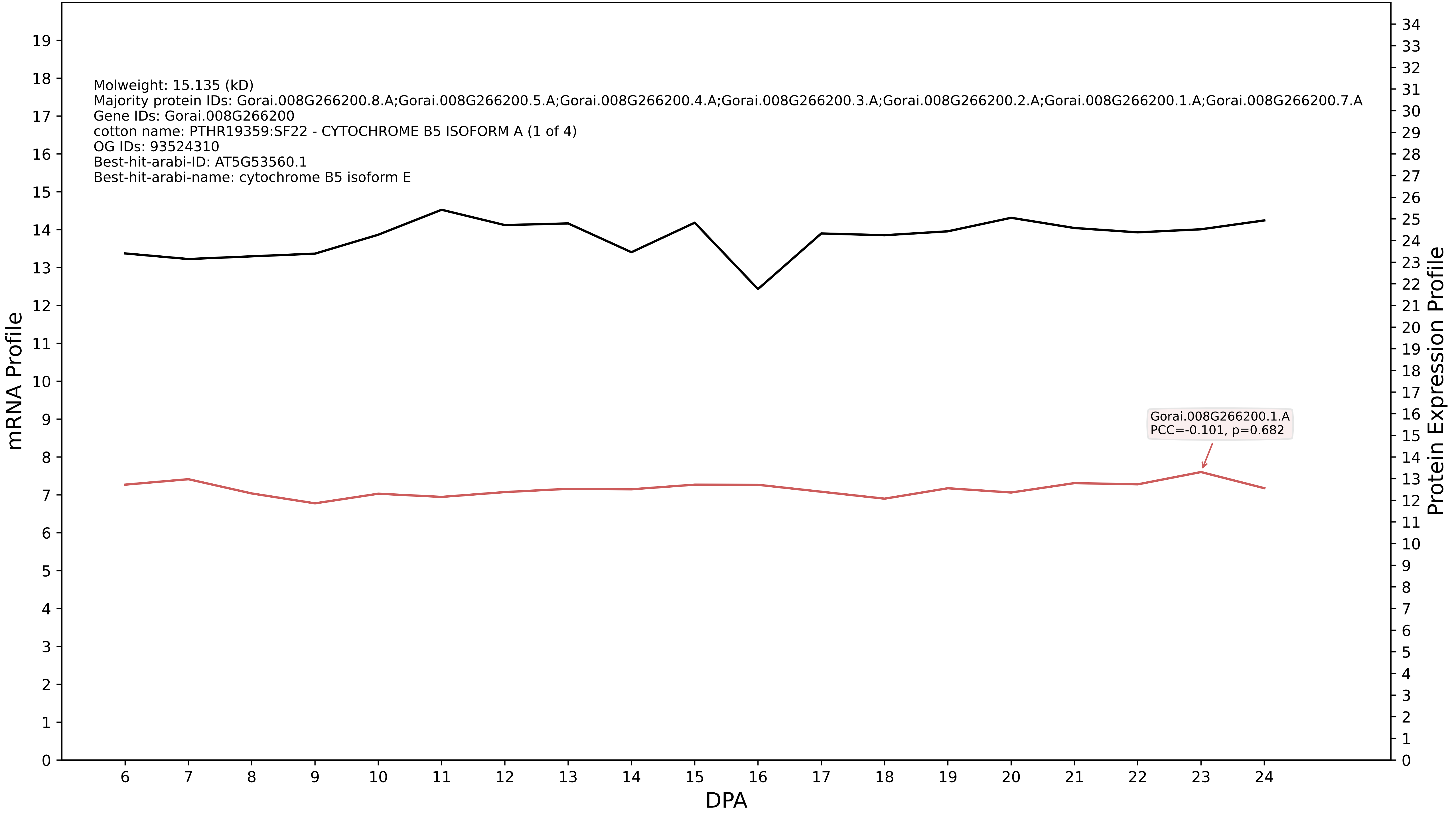

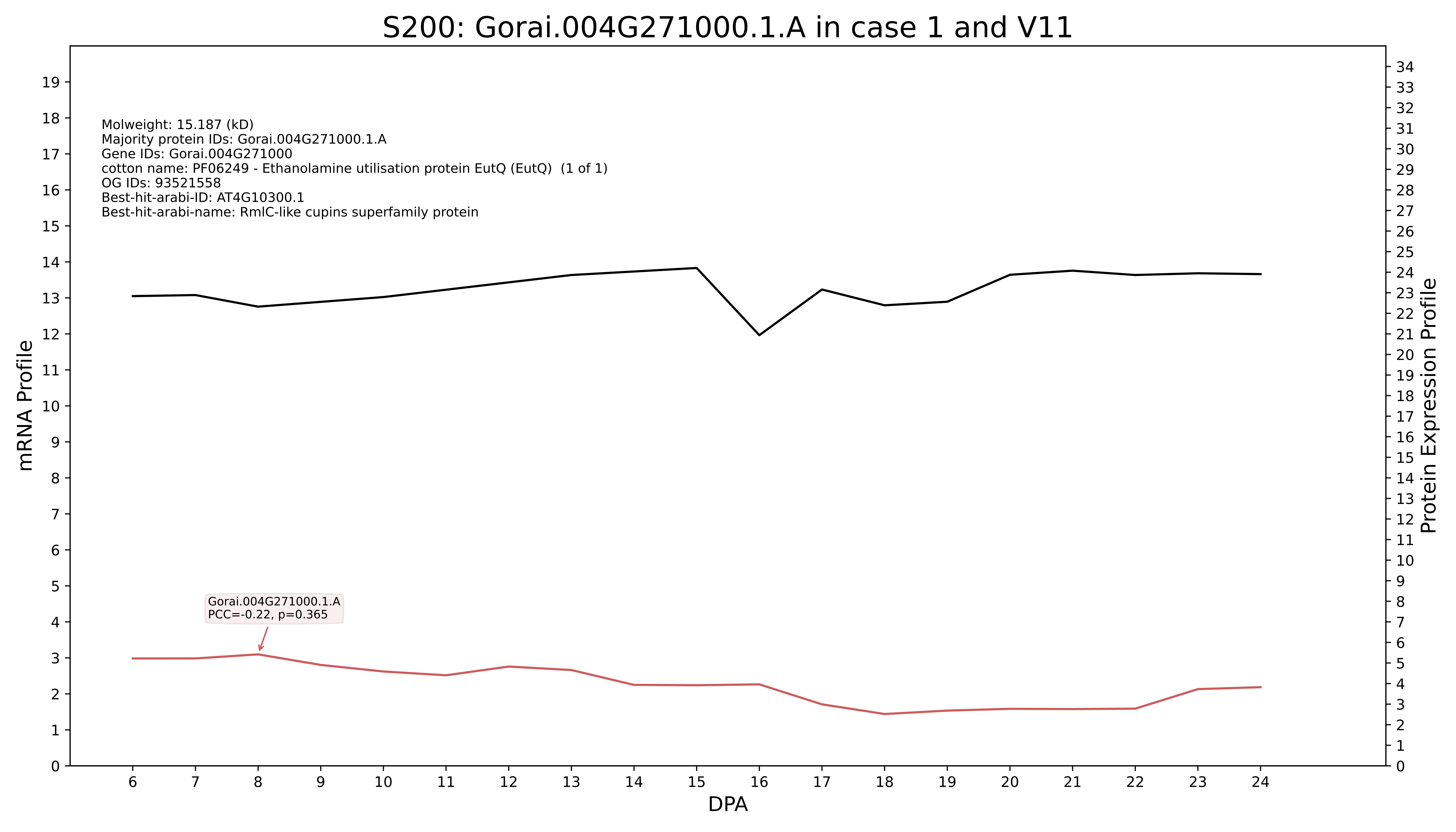

S200: Gorai.004G271000.1.A in case 1 and V11
Molweight: 15.187 (kD)
Majority protein IDs: Gorai.004G271000.1.A
Gene IDs: Gorai.004G271000
cotton name: PF06249 - Ethanolamine utilisation protein EutQ (EutQ) (1 of 1)
OG IDs: 93521558
Best-hit-arabi-ID: AT4G10300.1
Best-hit-arabi-name: RmlC-like cupins superfamily protein
Gorai.004G271000.1.A
PCC=-0.22, p=0.365
mRNA Profile
Protein Expression Profile
DPA

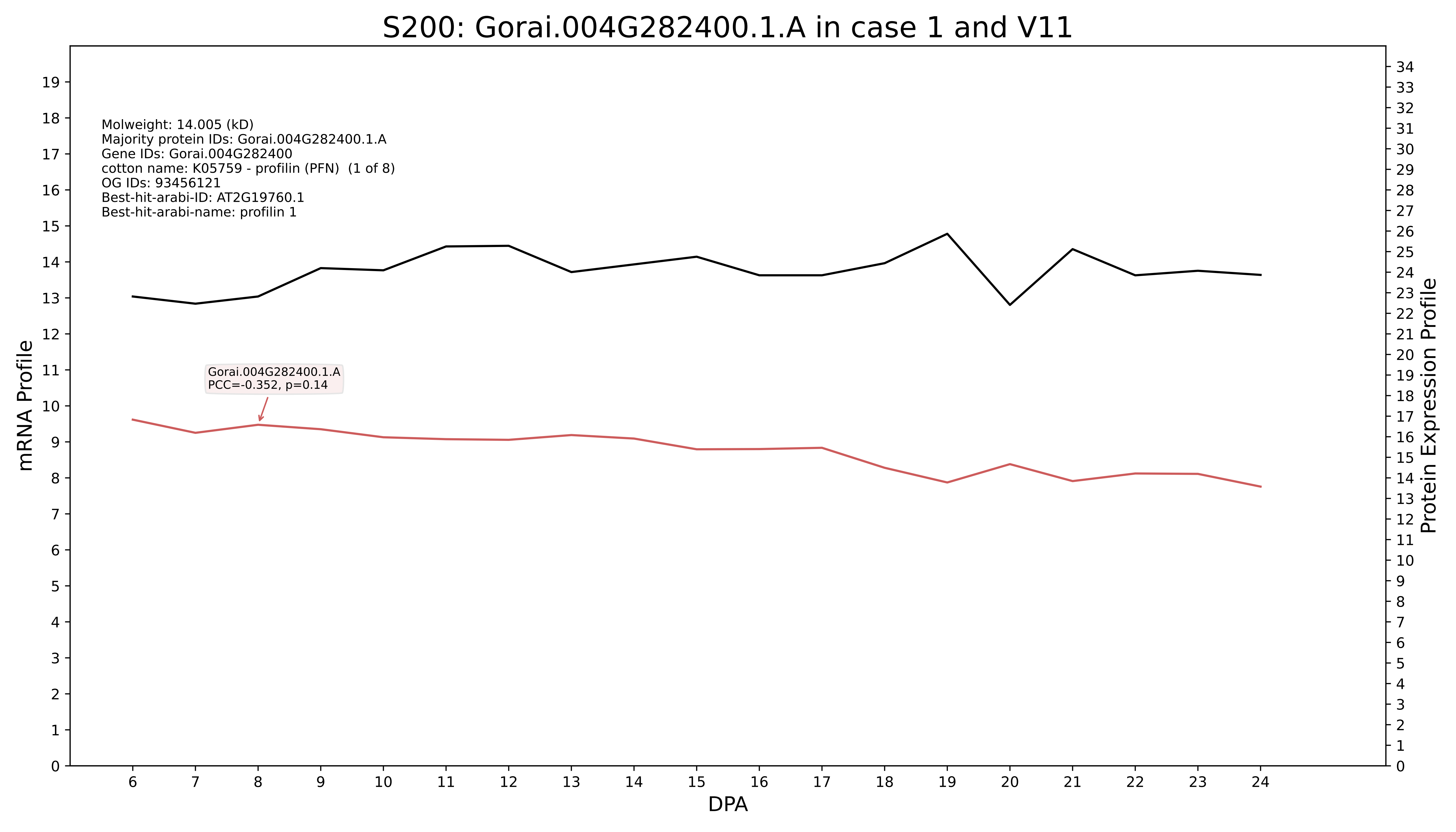

S200: Gorai.004G282400.1.A in case 1 and V11
Molweight: 14.005 (kD)
Majority protein IDs: Gorai.004G282400.1.A
Gene IDs: Gorai.004G282400
cotton name: K05759 - profilin (PFN) (1 of 8)
OG IDs: 93456121
Best-hit-arabi-ID: AT2G19760.1
Best-hit-arabi-name: profilin 1
Gorai.004G282400.1.A
PCC=-0.352, p=0.14
mRNA Profile
Protein Expression Profile
DPA

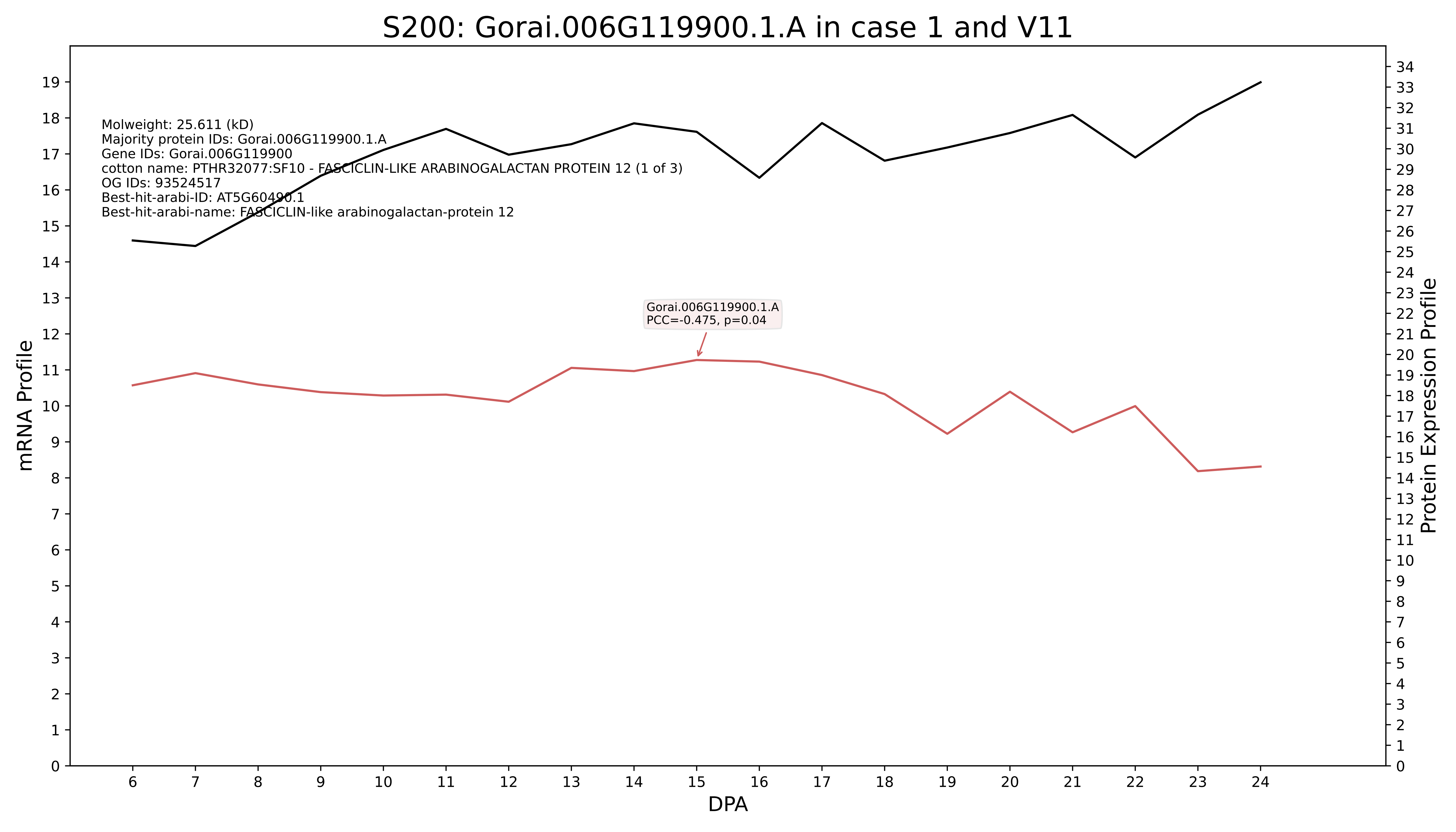

S200: Gorai.006G119900.1.A in case 1 and V11
Molweight: 25.611 (kD)
Majority protein IDs: Gorai.006G119900.1.A
Gene IDs: Gorai.006G119900
cotton name: PTHR32077:SF10 - FASCICLIN-LIKE ARABINOGALACTAN PROTEIN 12 (1 of 3)
OG IDs: 93524517
Best-hit-arabi-ID: AT5G60490.1
Best-hit-arabi-name: FASCICLIN-like arabinogalactan-protein 12
Gorai.006G119900.1.A
PCC=-0.475, p=0.04
mRNA Profile
Protein Expression Profile
DPA

### S200: Gorai.010G096500.2.D in case 1 and V11

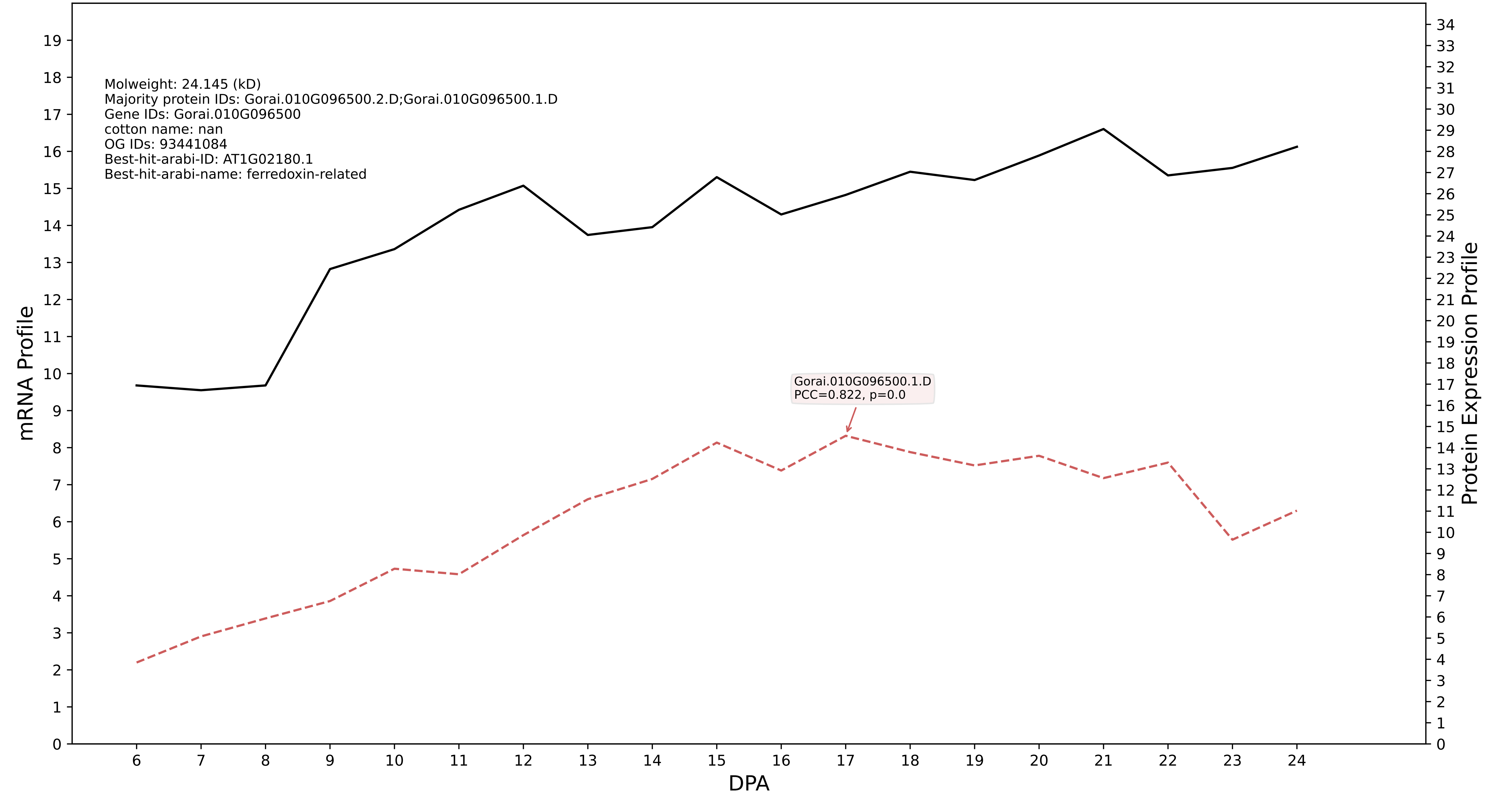

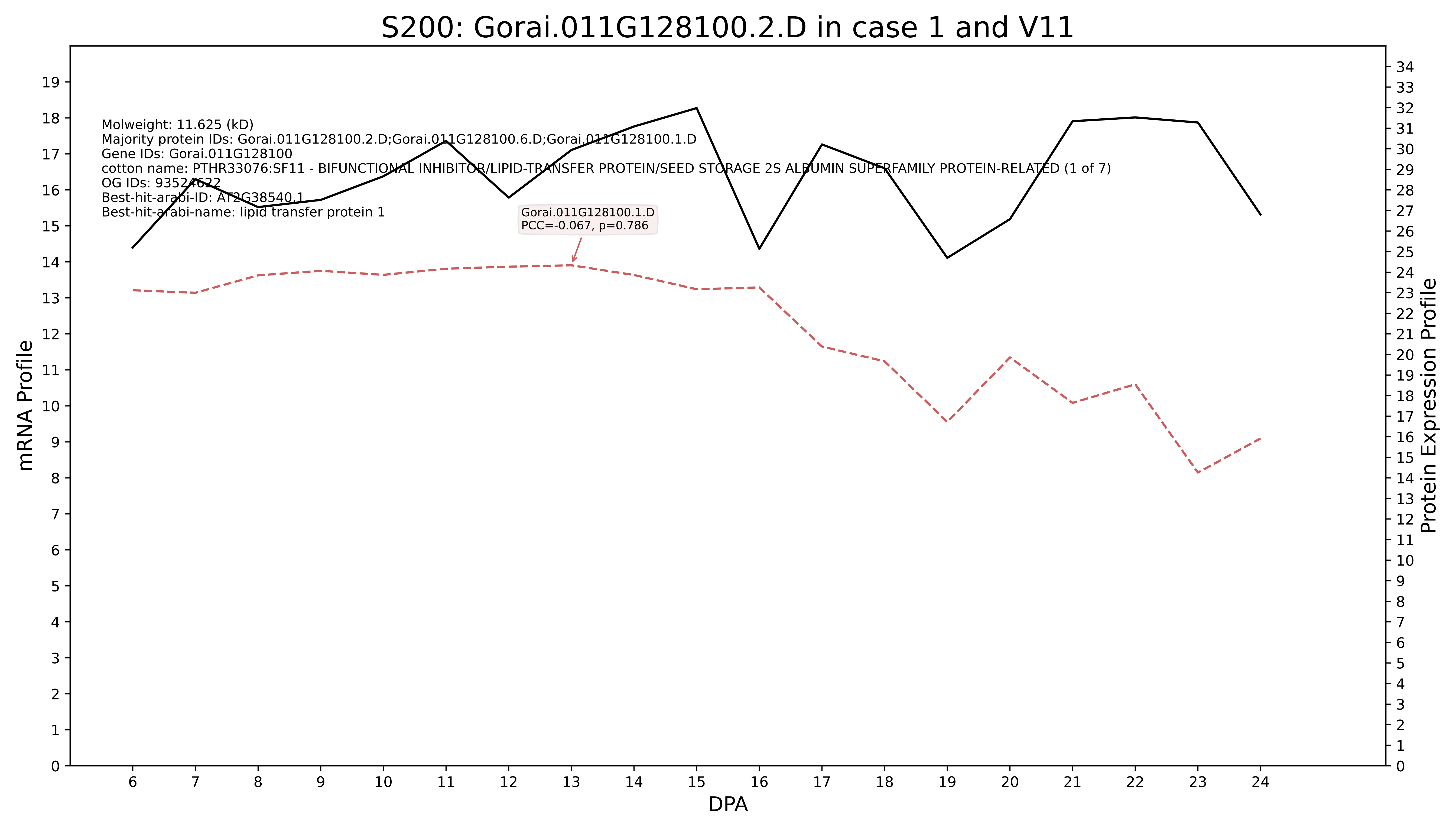
S200: Gorai.011G128100.2.D in case 1 and V11
Molweight: 11.625 (kD)
Majority protein IDs: Gorai.011G128100.2.D;Gorai.011G128100.6.D;Gorai.011G128100.1.D
Gene IDs: Gorai.011G128100
cotton name: PTHR33076:SF11 - BIFUNCTIONAL INHIBITOR/LIPID-TRANSFER PROTEIN/SEED STORAGE 2S ALBUMIN SUPERFAMILY PROTEIN-RELATED (1 of 7)
OG IDs: 93524622
Best-hit-arabi-ID: AT2G38540.1
Best-hit-arabi-name: lipid transfer protein 1
Gorai.011G128100.1.D
PCC=-0.067, p=0.786
mRNA Profile
Protein Expression Profile
DPA

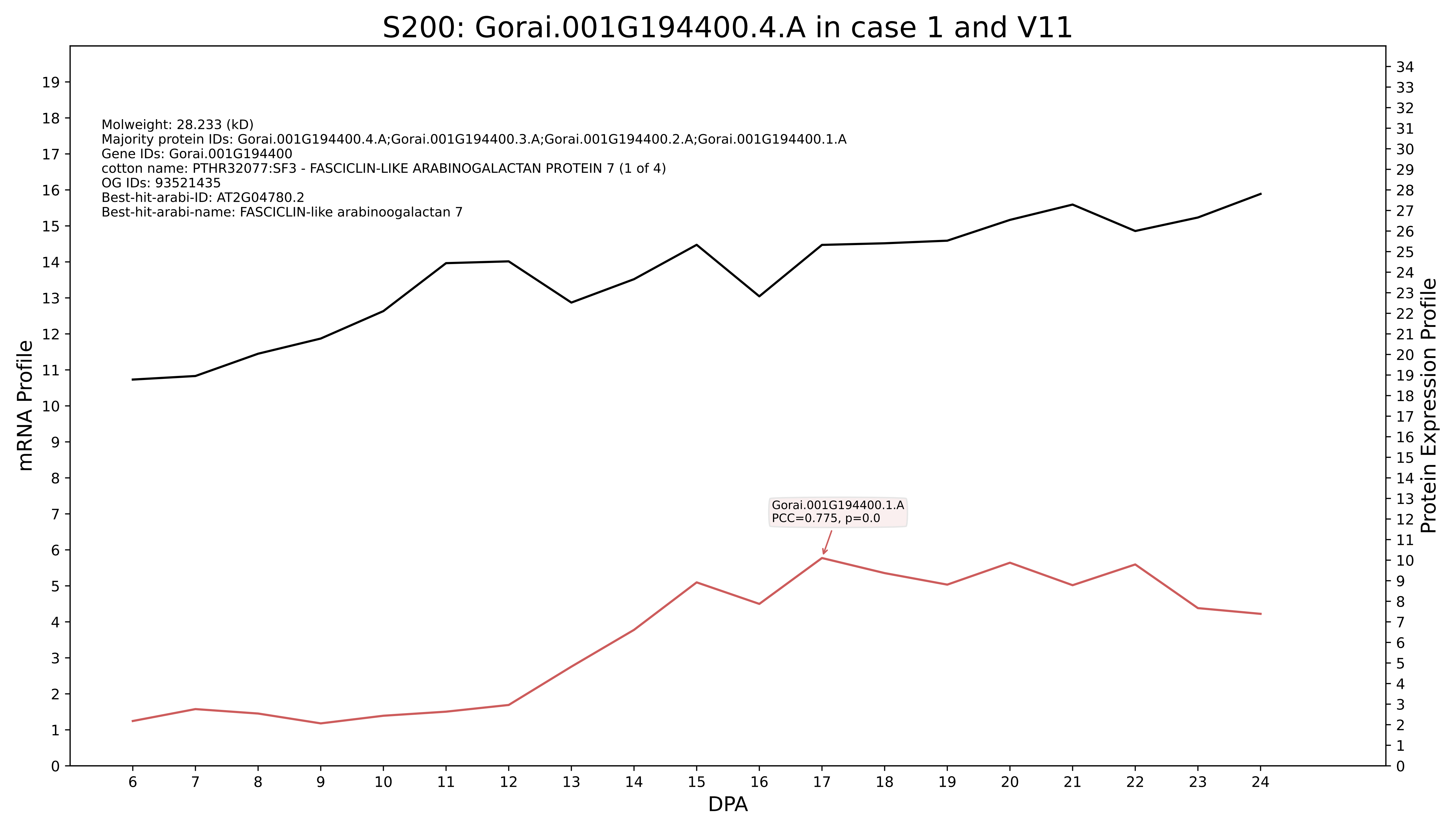

S200: Gorai.001G194400.4.A in case 1 and V11
Molweight: 28.233 (kD)
Majority protein IDs: Gorai.001G194400.4.A;Gorai.001G194400.3.A;Gorai.001G194400.2.A;Gorai.001G194400.1.A
Gene IDs: Gorai.001G194400
cotton name: PTHR32077:SF3 - FASCICLIN-LIKE ARABINOGALACTAN PROTEIN 7 (1 of 4)
OG IDs: 93521435
Best-hit-arabi-ID: AT2G04780.2
Best-hit-arabi-name: FASCICLIN-like arabinoogalactan 7
Gorai.001G194400.1.A
PCC=0.775, p=0.0
mRNA Profile
Protein Expression Profile
DPA

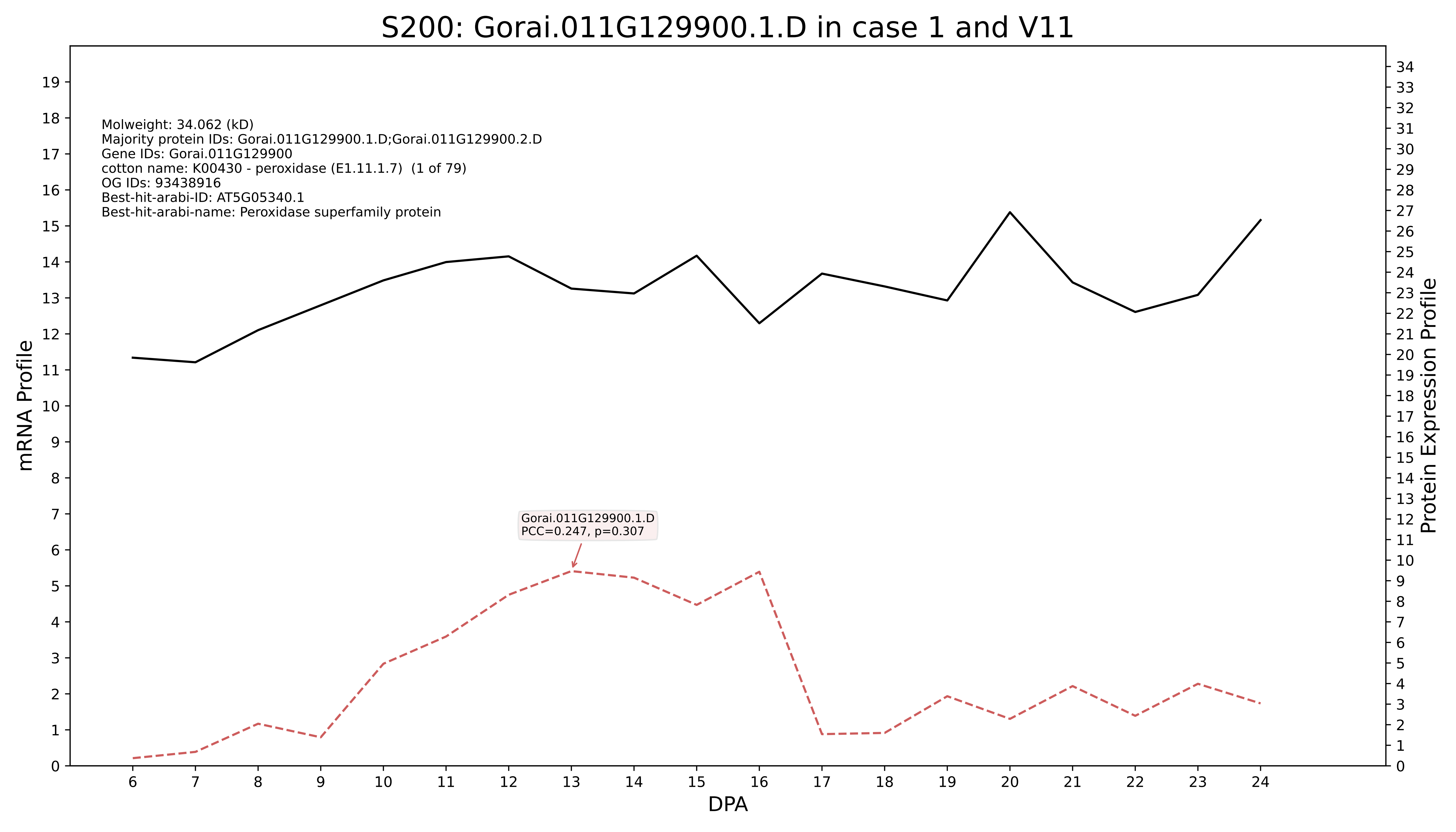
S200: Gorai.011G129900.1.D in case 1 and V11
Molweight: 34.062 (kD)
Majority protein IDs: Gorai.011G129900.1.D;Gorai.011G129900.2.D
Gene IDs: Gorai.011G129900
cotton name: K00430 - peroxidase (E1.11.1.7) (1 of 79)
OG IDs: 93438916
Best-hit-arabi-ID: AT5G05340.1
Best-hit-arabi-name: Peroxidase superfamily protein
Gorai.011G129900.1.D
PCC=0.247, p=0.307
mRNA Profile
Protein Expression Profile
DPA

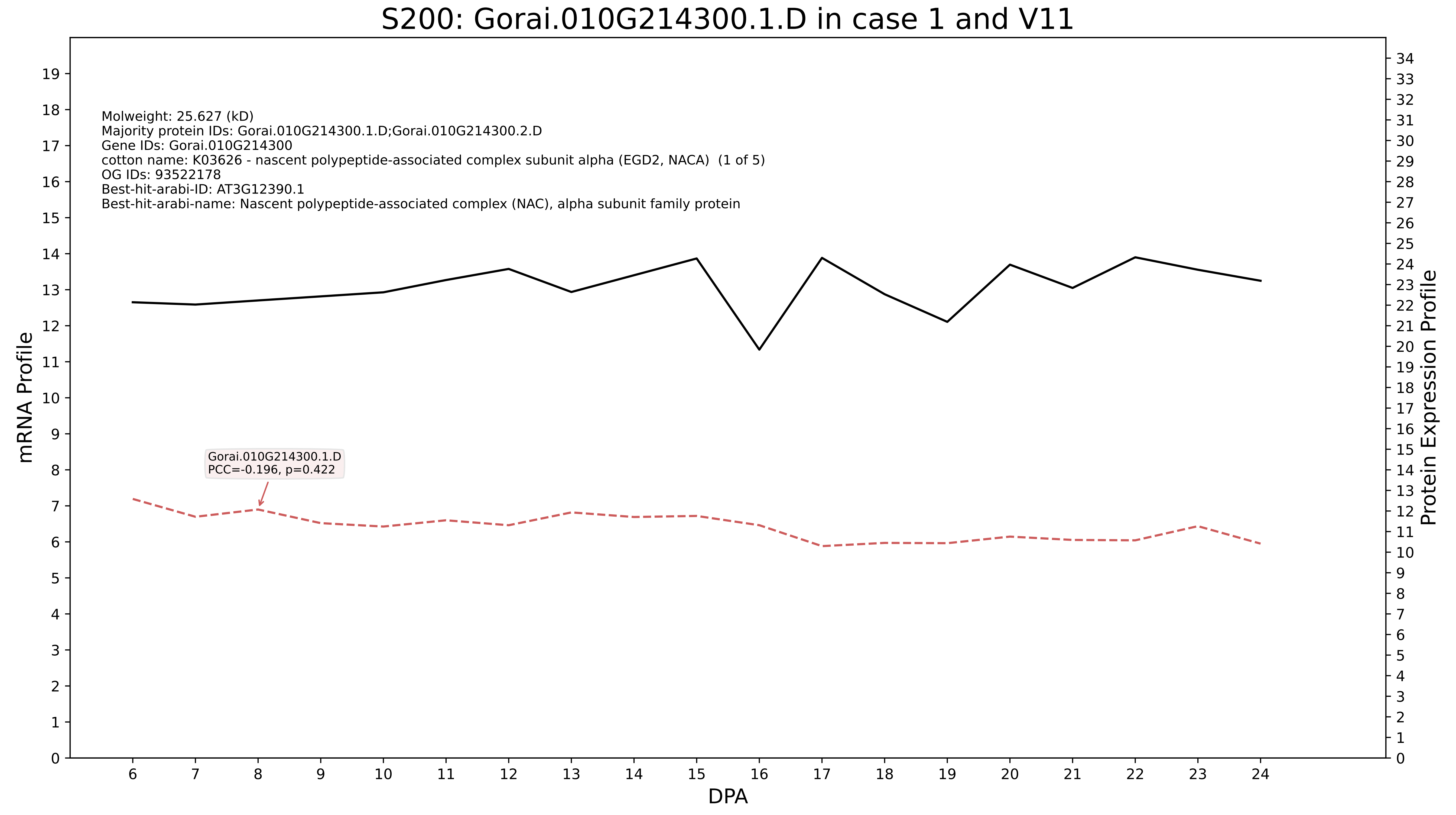
S200: Gorai.010G214300.1.D in case 1 and V11
Molweight: 25.627 (kD)
Majority protein IDs: Gorai.010G214300.1.D;Gorai.010G214300.2.D
Gene IDs: Gorai.010G214300
cotton name: K03626 - nascent polypeptide-associated complex subunit alpha (EGD2, NACA) (1 of 5)
OG IDs: 93522178
Best-hit-arabi-ID: AT3G12390.1
Best-hit-arabi-name: Nascent polypeptide-associated complex (NAC), alpha subunit family protein
Gorai.010G214300.1.D
PCC=-0.196, p=0.422
mRNA Profile
Protein Expression Profile
DPA

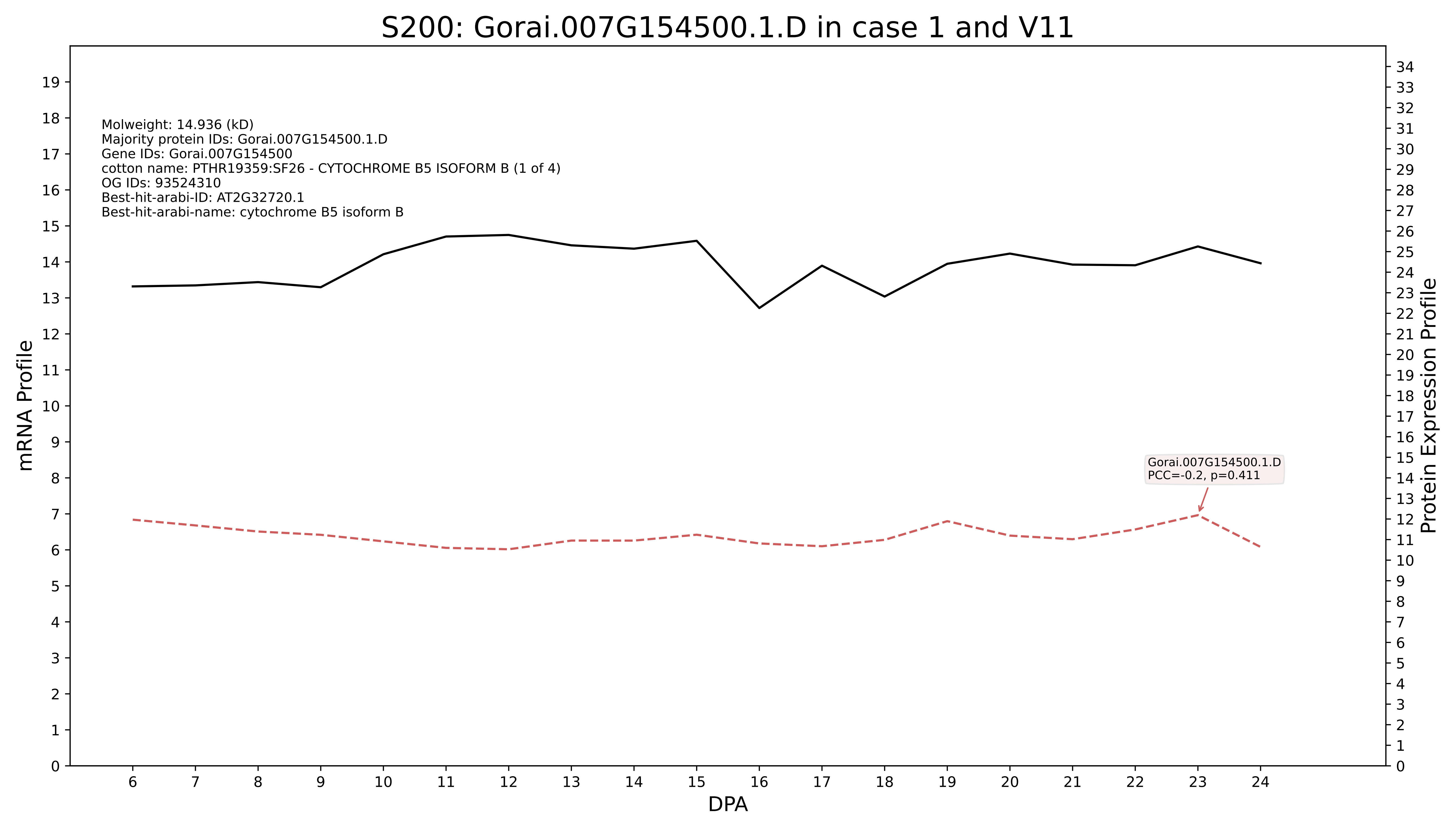
S200: Gorai.007G154500.1.D in case 1 and V11
Molweight: 14.936 (kD)
Majority protein IDs: Gorai.007G154500.1.D
Gene IDs: Gorai.007G154500
cotton name: PTHR19359:SF26 - CYTOCHROME B5 ISOFORM B (1 of 4)
OG IDs: 93524310
Best-hit-arabi-ID: AT2G32720.1
Best-hit-arabi-name: cytochrome B5 isoform B
Gorai.007G154500.1.D
PCC=-0.2, p=0.411
mRNA Profile
Protein Expression Profile
DPA

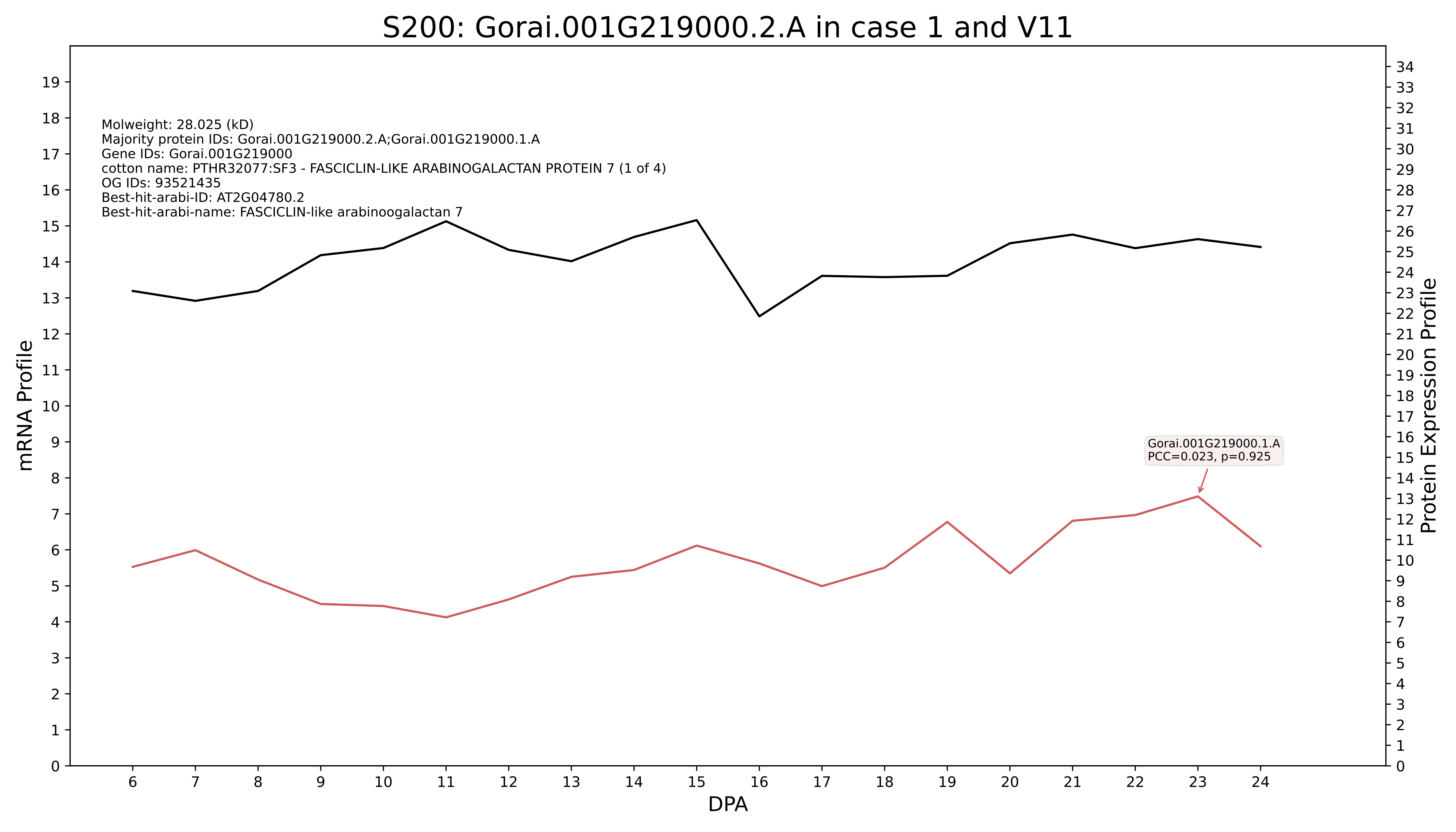

S200: Gorai.001G219000.2.A in case 1 and V11
Molweight: 28.025 (kD)
Majority protein IDs: Gorai.001G219000.2.A;Gorai.001G219000.1.A
Gene IDs: Gorai.001G219000
cotton name: PTHR32077:SF3 - FASCICLIN-LIKE ARABINOGALACTAN PROTEIN 7 (1 of 4)
OG IDs: 93521435
Best-hit-arabi-ID: AT2G04780.2
Best-hit-arabi-name: FASCICLIN-like arabinoogalactan 7
Gorai.001G219000.1.A
PCC=0.023, p=0.925
mRNA Profile
Protein Expression Profile
DPA

### S200: Gorai.001G103400.2.A in case 1 and V11

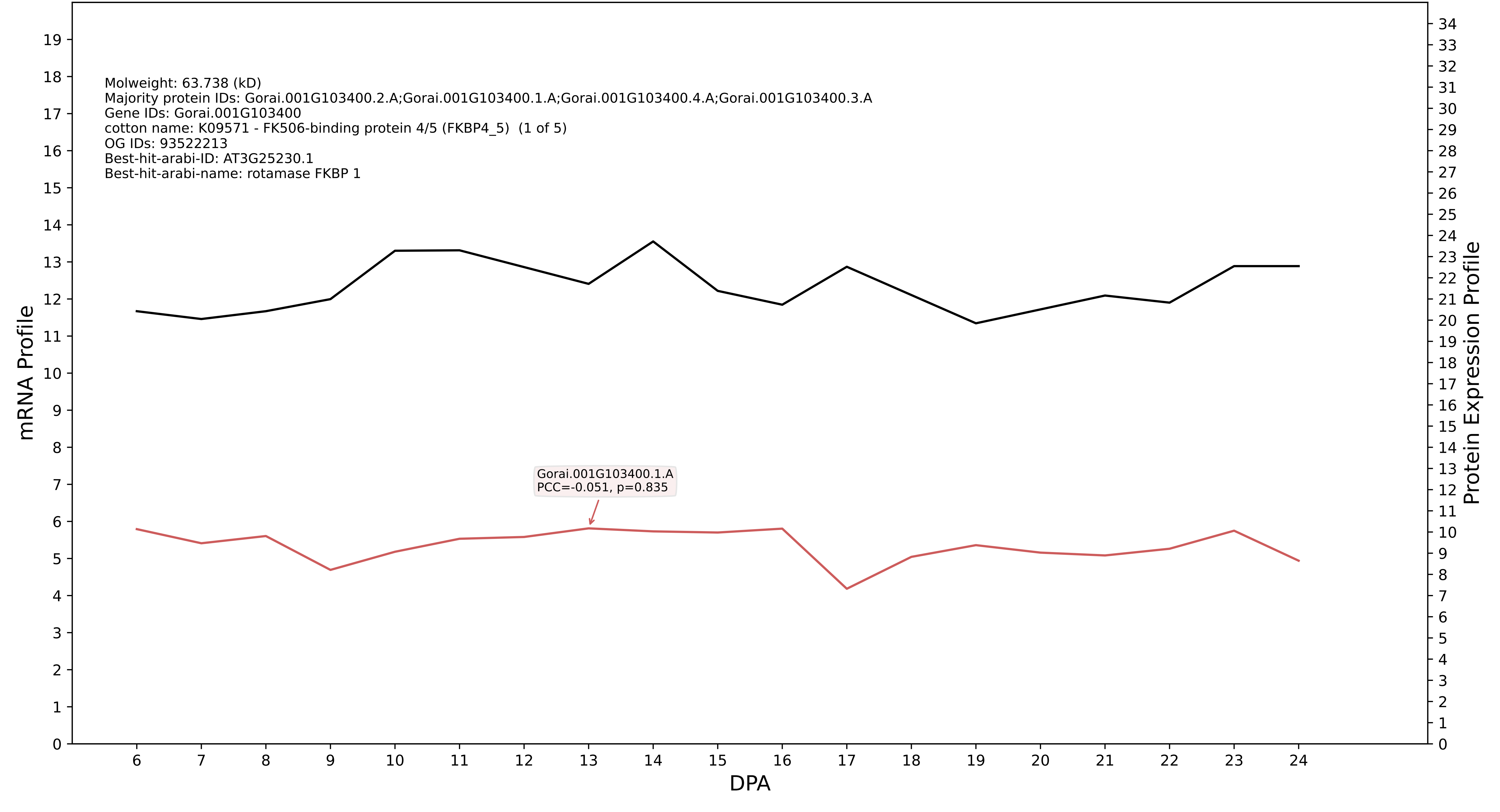

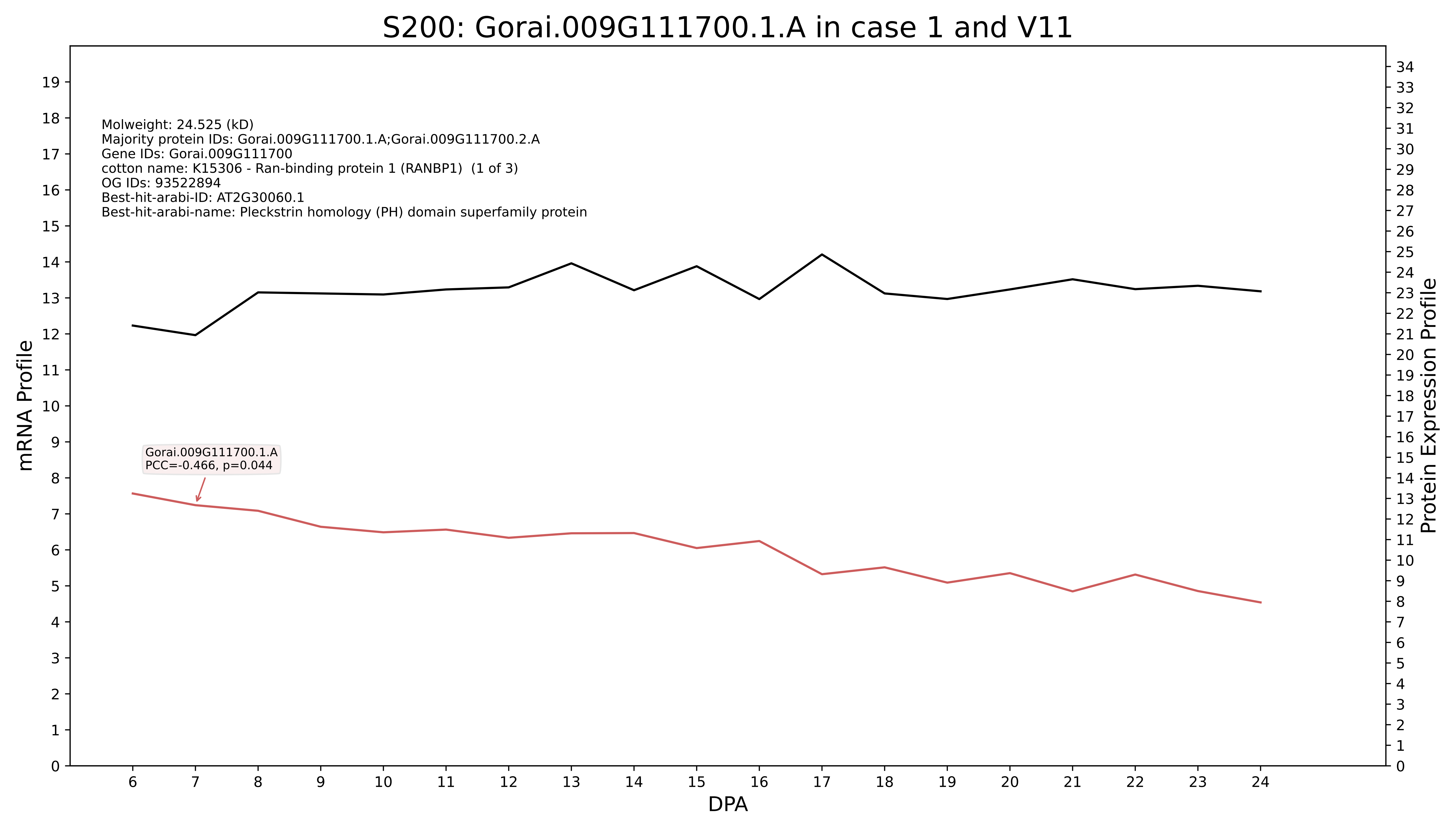
S200: Gorai.009G111700.1.A in case 1 and V11
Molweight: 24.525 (kD)
Majority protein IDs: Gorai.009G111700.1.A;Gorai.009G111700.2.A
Gene IDs: Gorai.009G111700
cotton name: K15306 - Ran-binding protein 1 (RANBP1) (1 of 3)
OG IDs: 93522894
Best-hit-arabi-ID: AT2G30060.1
Best-hit-arabi-name: Pleckstrin homology (PH) domain superfamily protein
Gorai.009G111700.1.A
PCC=-0.466, p=0.044
mRNA Profile
Protein Expression Profile
DPA

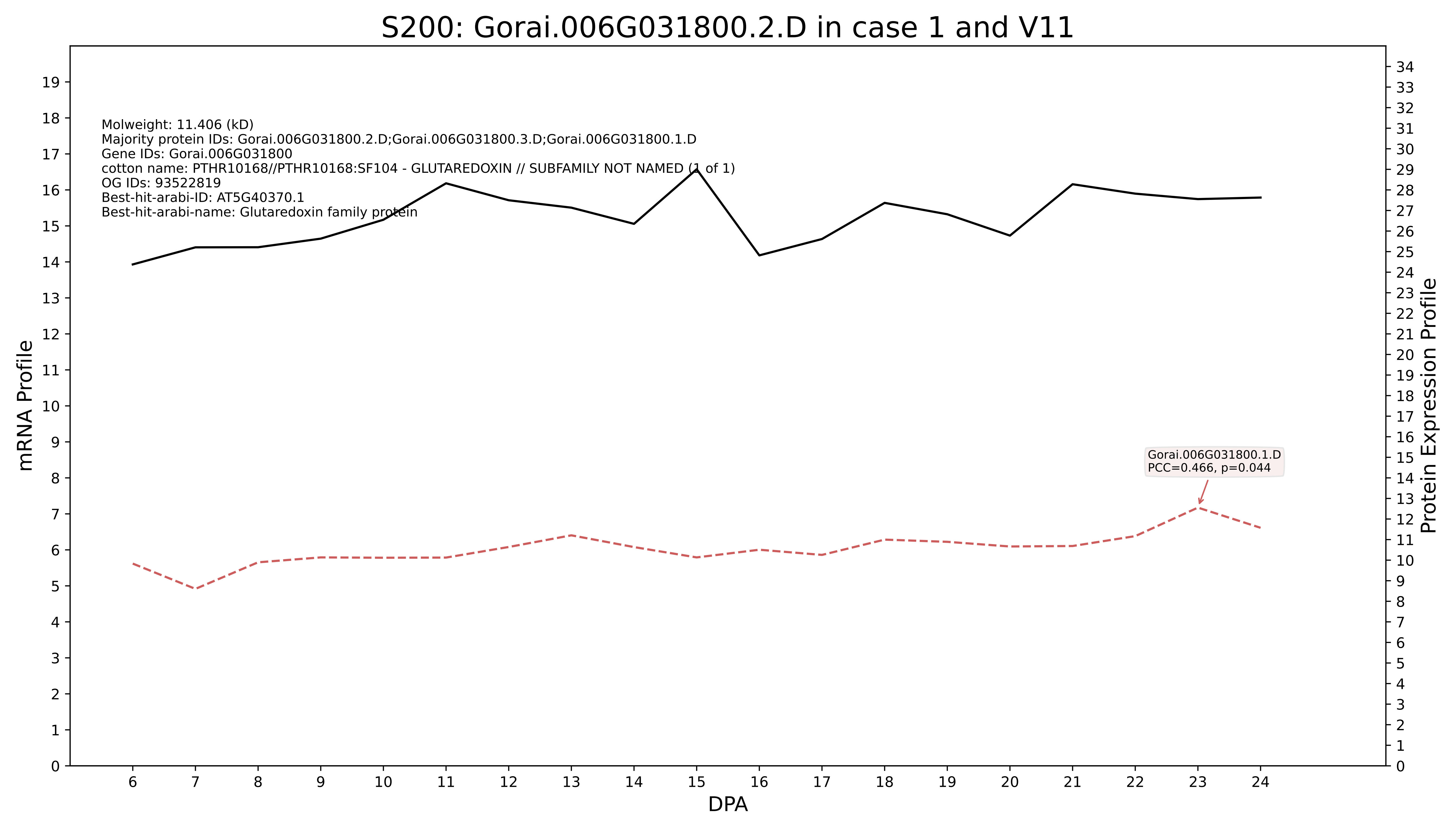

S200: Gorai.006G031800.2.D in case 1 and V11
Molweight: 11.406 (kD)
Majority protein IDs: Gorai.006G031800.2.D;Gorai.006G031800.3.D;Gorai.006G031800.1.D
Gene IDs: Gorai.006G031800
cotton name: PTHR10168//PTHR10168:SF104 - GLUTAREDOXIN // SUBFAMILY NOT NAMED (1 of 1)
OG IDs: 93522819
Best-hit-arabi-ID: AT5G40370.1
Best-hit-arabi-name: Glutaredoxin family protein
Gorai.006G031800.1.D
PCC=0.466, p=0.044
mRNA Profile
Protein Expression Profile
DPA

### S200: Gorai.002G199100.1.D in case 1 and V11

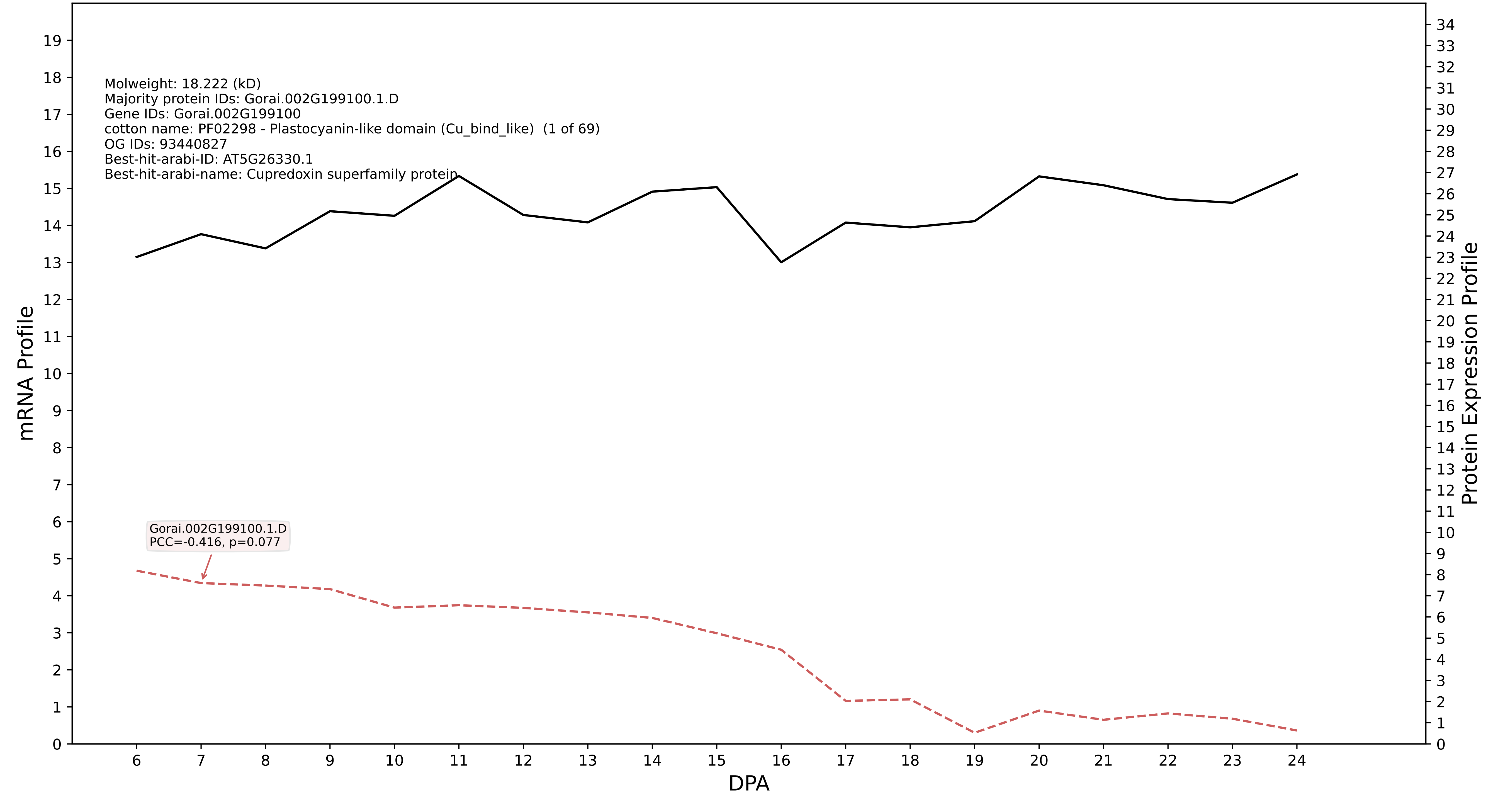

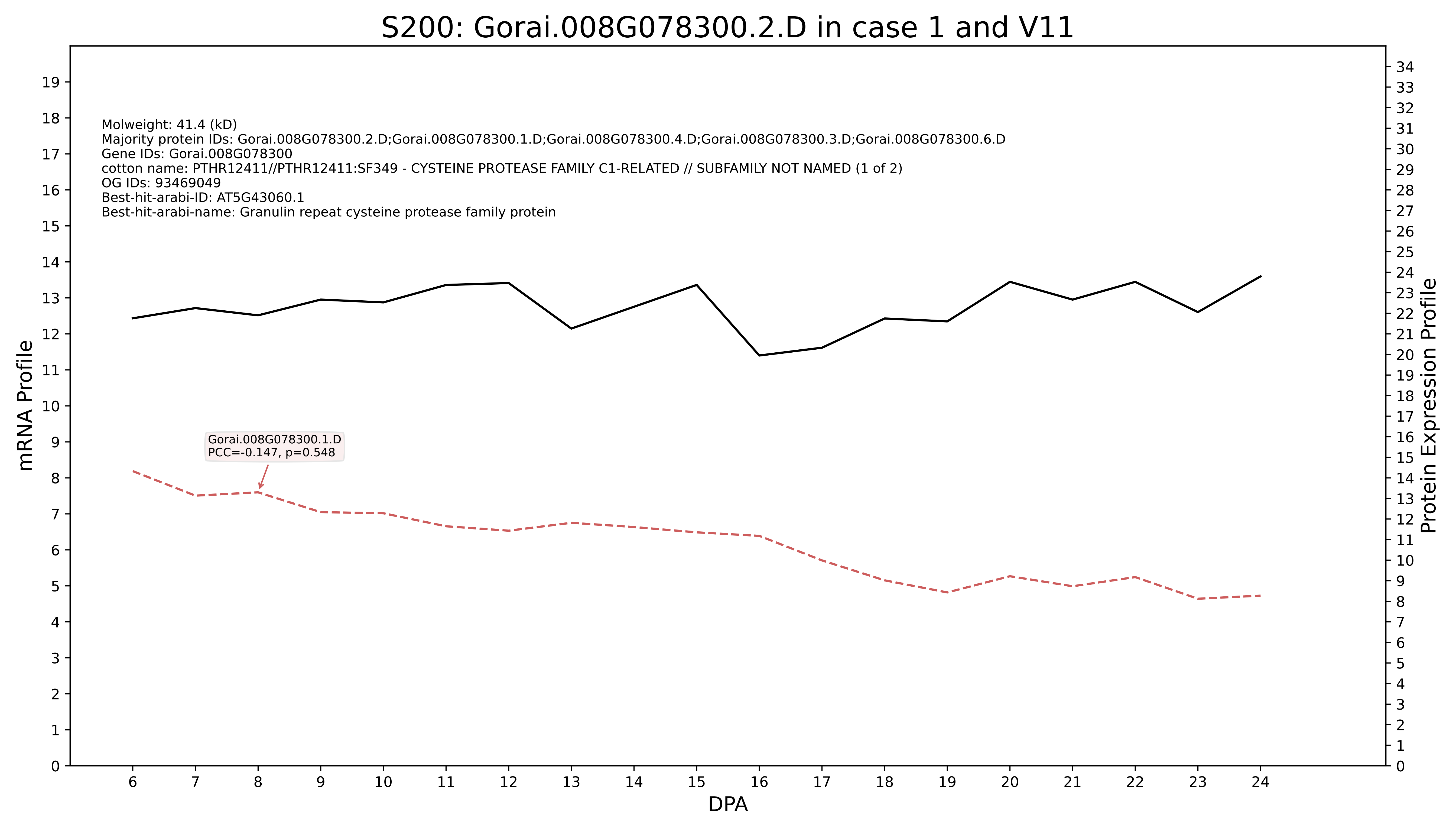
S200: Gorai.008G078300.2.D in case 1 and V11
Molweight: 41.4 (kD)
Majority protein IDs: Gorai.008G078300.2.D;Gorai.008G078300.1.D;Gorai.008G078300.4.D;Gorai.008G078300.3.D;Gorai.008G078300.6.D
Gene IDs: Gorai.008G078300
cotton name: PTHR12411//PTHR12411:SF349 - CYSTEINE PROTEASE FAMILY C1-RELATED // SUBFAMILY NOT NAMED (1 of 2)
OG IDs: 93469049
Best-hit-arabi-ID: AT5G43060.1
Best-hit-arabi-name: Granulin repeat cysteine protease family protein
Gorai.008G078300.1.D
PCC=-0.147, p=0.548
mRNA Profile
Protein Expression Profile
DPA

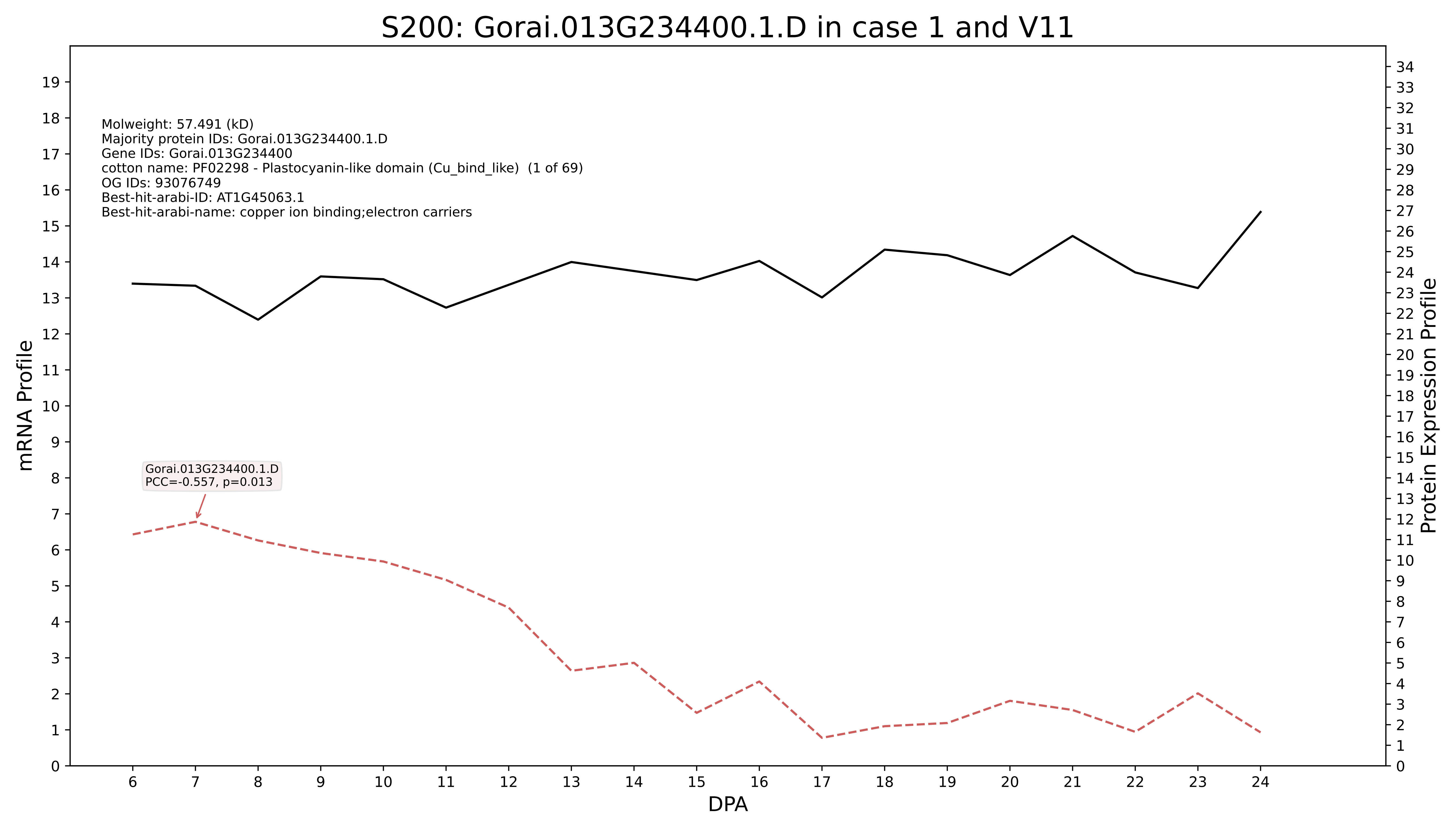
S200: Gorai.013G234400.1.D in case 1 and V11
Molweight: 57.491 (kD)
Majority protein IDs: Gorai.013G234400.1.D
Gene IDs: Gorai.013G234400
cotton name: PF02298 - Plastocyanin-like domain (Cu_bind_like) (1 of 69)
OG IDs: 93076749
Best-hit-arabi-ID: AT1G45063.1
Best-hit-arabi-name: copper ion binding;electron carriers
Gorai.013G234400.1.D
PCC=-0.557, p=0.013
mRNA Profile
Protein Expression Profile
DPA

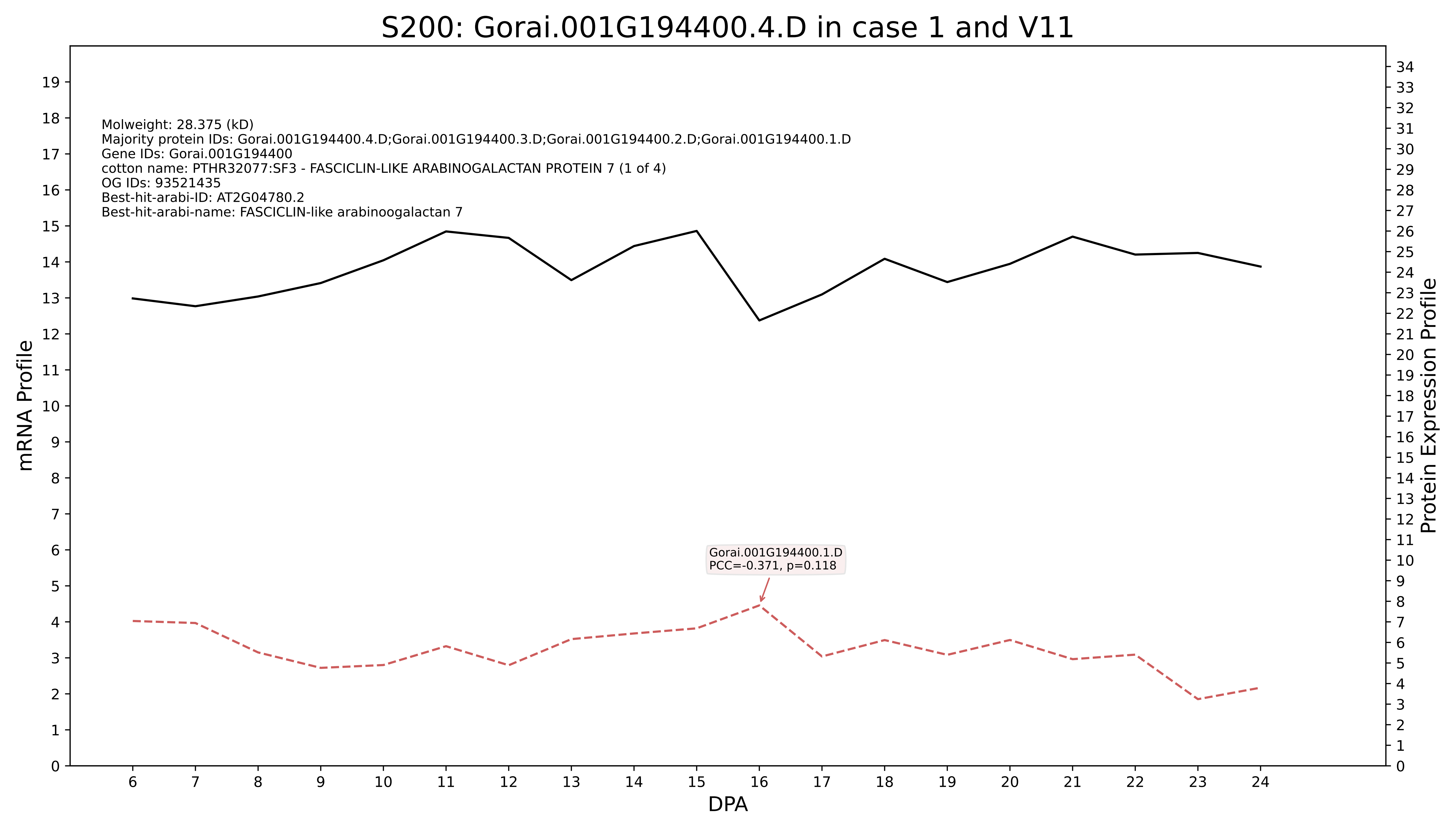
S200: Gorai.001G194400.4.D in case 1 and V11
Molweight: 28.375 (kD)
Majority protein IDs: Gorai.001G194400.4.D;Gorai.001G194400.3.D;Gorai.001G194400.2.D;Gorai.001G194400.1.D
Gene IDs: Gorai.001G194400
cotton name: PTHR32077:SF3 - FASCICLIN-LIKE ARABINOGALACTAN PROTEIN 7 (1 of 4)
OG IDs: 93521435
Best-hit-arabi-ID: AT2G04780.2
Best-hit-arabi-name: FASCICLIN-like arabinoogalactan 7
Gorai.001G194400.1.D
PCC=-0.371, p=0.118
mRNA Profile
Protein Expression Profile
DPA

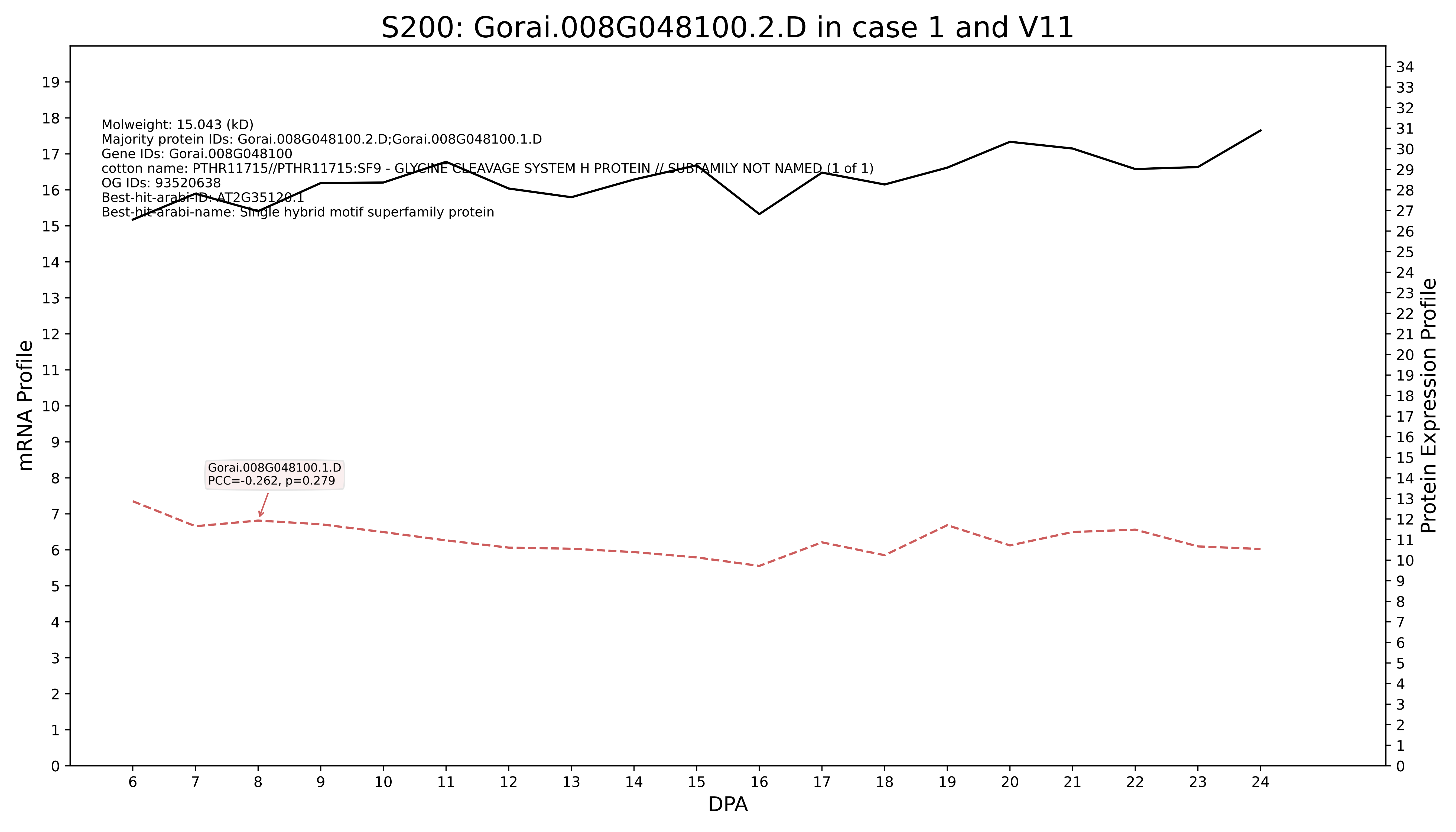

S200: Gorai.008G048100.2.D in case 1 and V11
Molweight: 15.043 (kD)
Majority protein IDs: Gorai.008G048100.2.D;Gorai.008G048100.1.D
Gene IDs: Gorai.008G048100
cotton name: PTHR11715//PTHR11715:SF9 - GLYCINE CLEAVAGE SYSTEM H PROTEIN // SUBFAMILY NOT NAMED (1 of 1)
OG IDs: 93520638
Best-hit-arabi-ID: AT2G35120.1
Best-hit-arabi-name: Single hybrid motif superfamily protein
Gorai.008G048100.1.D
PCC=-0.262, p=0.279
mRNA Profile
Protein Expression Profile
DPA

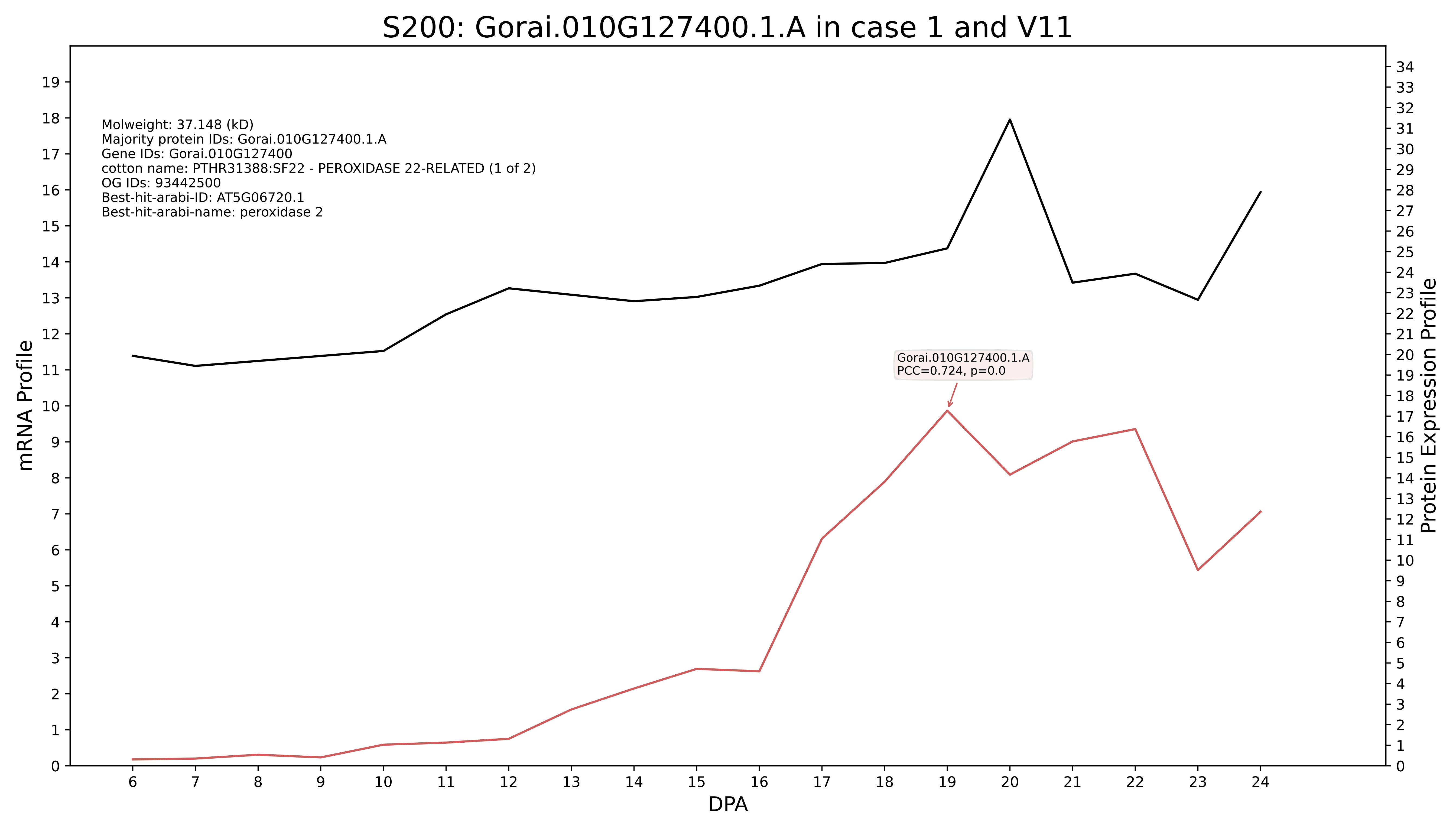
S200: Gorai.010G127400.1.A in case 1 and V11
Molweight: 37.148 (kD)
Majority protein IDs: Gorai.010G127400.1.A
Gene IDs: Gorai.010G127400
cotton name: PTHR31388:SF22 - PEROXIDASE 22-RELATED (1 of 2)
OG IDs: 93442500
Best-hit-arabi-ID: AT5G06720.1
Best-hit-arabi-name: peroxidase 2
Gorai.010G127400.1.A
PCC=0.724, p=0.0
mRNA Profile
Protein Expression Profile
DPA

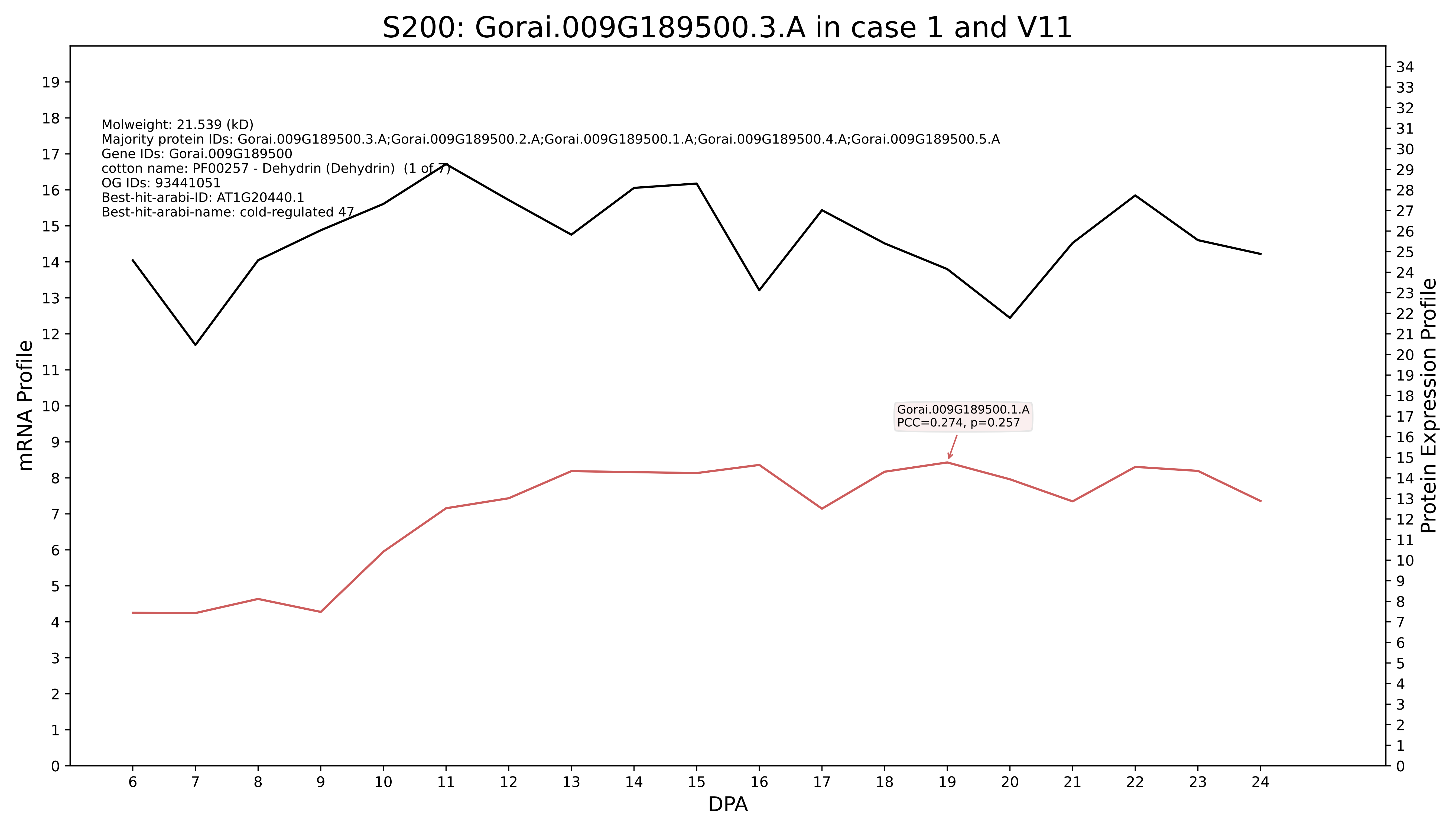
S200: Gorai.009G189500.3.A in case 1 and V11
Molweight: 21.539 (kD)
Majority protein IDs: Gorai.009G189500.3.A;Gorai.009G189500.2.A;Gorai.009G189500.1.A;Gorai.009G189500.4.A;Gorai.009G189500.5.A
Gene IDs: Gorai.009G189500
cotton name: PF00257 - Dehydrin (Dehydrin) (1 of 7)
OG IDs: 93441051
Best-hit-arabi-ID: AT1G20440.1
Best-hit-arabi-name: cold-regulated 47
Gorai.009G189500.1.A
PCC=0.274, p=0.257
mRNA Profile
Protein Expression Profile
DPA

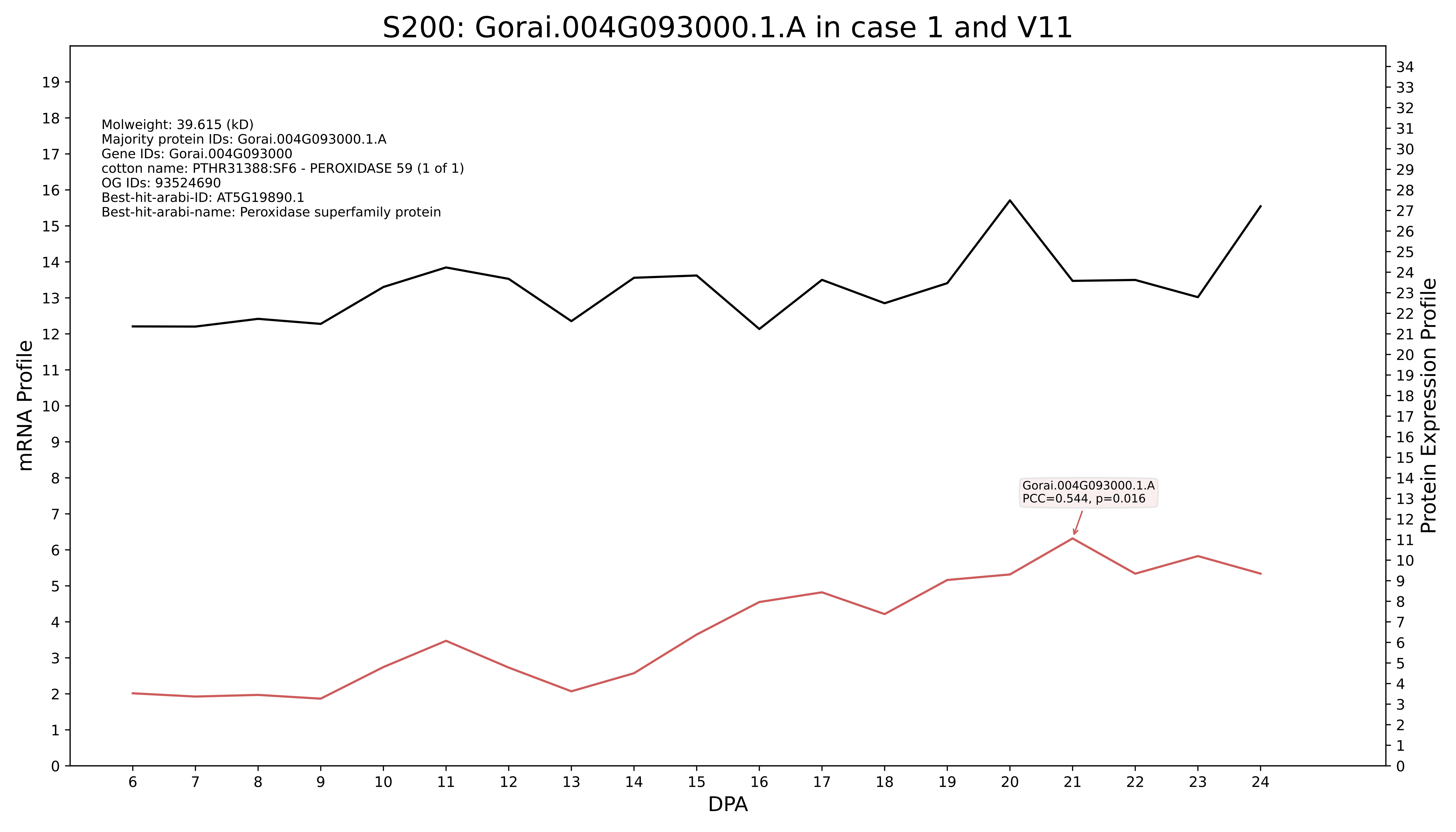
S200: Gorai.004G093000.1.A in case 1 and V11
Molweight: 39.615 (kD)
Majority protein IDs: Gorai.004G093000.1.A
Gene IDs: Gorai.004G093000
cotton name: PTHR31388:SF6 - PEROXIDASE 59 (1 of 1)
OG IDs: 93524690
Best-hit-arabi-ID: AT5G19890.1
Best-hit-arabi-name: Peroxidase superfamily protein
Gorai.004G093000.1.A
PCC=0.544, p=0.016
mRNA Profile
Protein Expression Profile
DPA

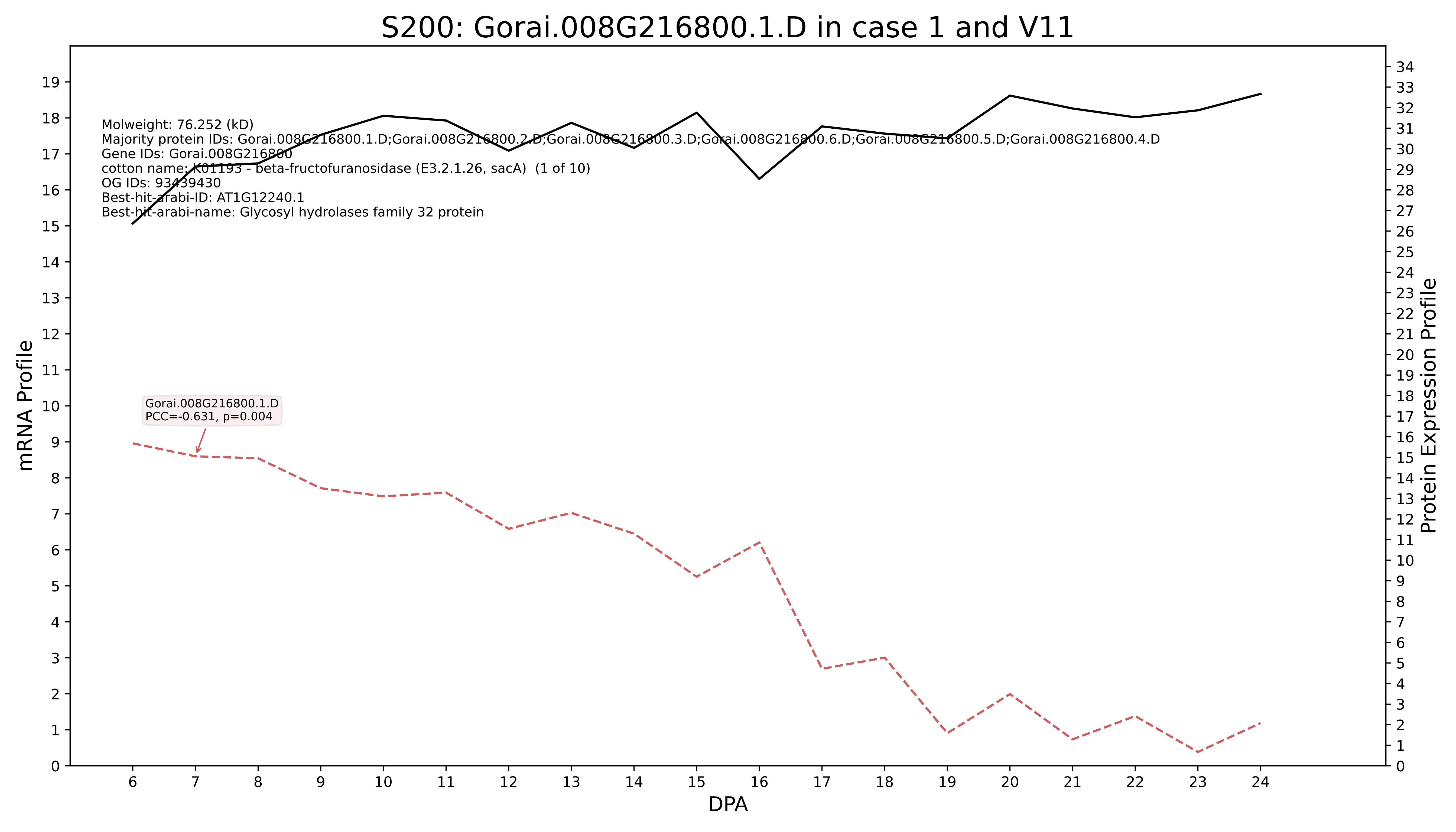

S200: Gorai.008G216800.1.D in case 1 and V11
Molweight: 76.252 (kD)
Majority protein IDs: Gorai.008G216800.1.D;Gorai.008G216800.2.D;Gorai.008G216800.3.D;Gorai.008G216800.6.D;Gorai.008G216800.5.D;Gorai.008G216800.4.D
Gene IDs: Gorai.008G216800
cotton name: K01193 - beta-fructofuranosidase (E3.2.1.26, sacA) (1 of 10)
OG IDs: 93439430
Best-hit-arabi-ID: AT1G12240.1
Best-hit-arabi-name: Glycosyl hydrolases family 32 protein
Gorai.008G216800.1.D
PCC=-0.631, p=0.004
mRNA Profile
Protein Expression Profile
DPA

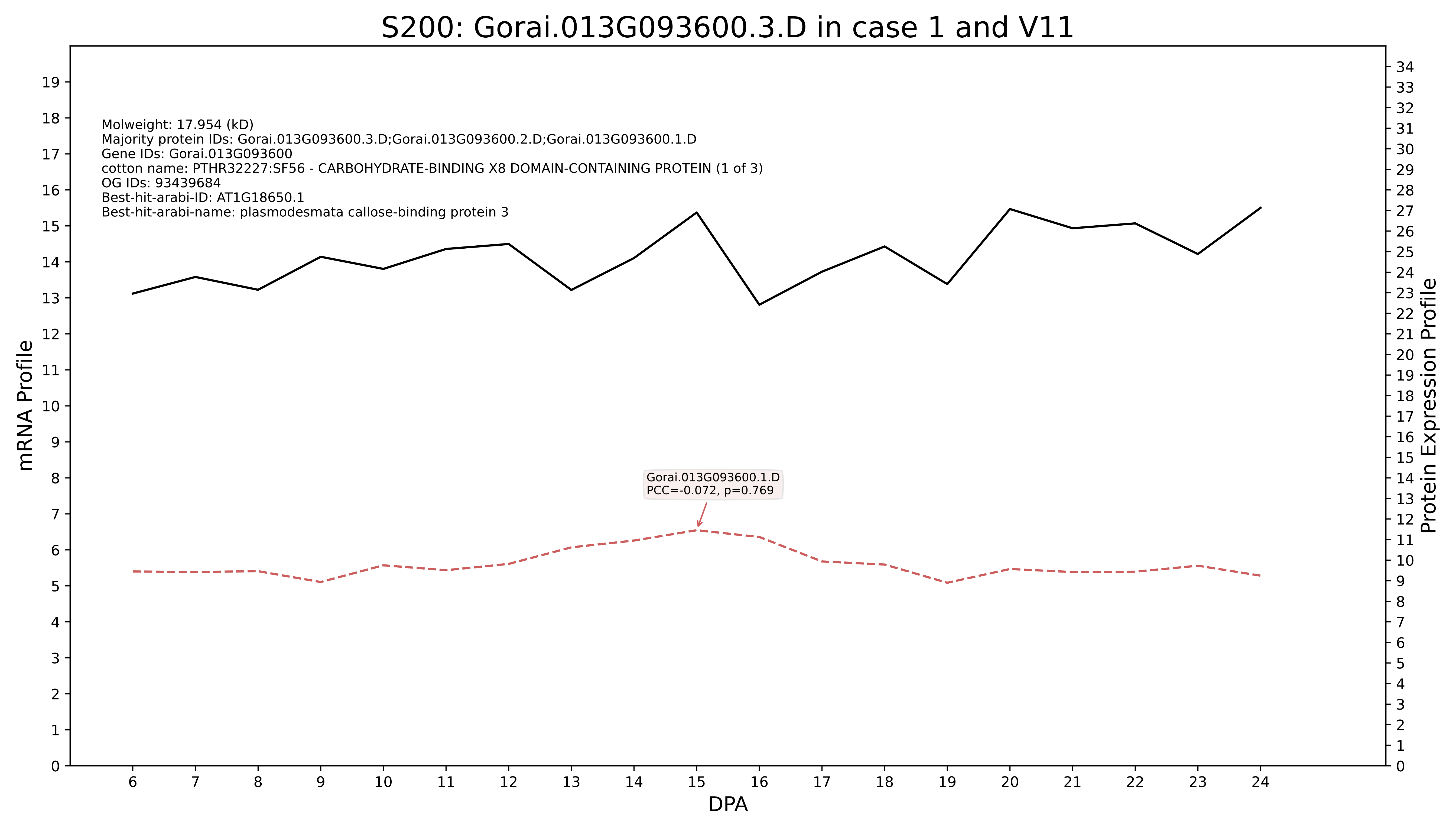
S200: Gorai.013G093600.3.D in case 1 and V11
Molweight: 17.954 (kD)
Majority protein IDs: Gorai.013G093600.3.D;Gorai.013G093600.2.D;Gorai.013G093600.1.D
Gene IDs: Gorai.013G093600
cotton name: PTHR32227:SF56 - CARBOHYDRATE-BINDING X8 DOMAIN-CONTAINING PROTEIN (1 of 3)
OG IDs: 93439684
Best-hit-arabi-ID: AT1G18650.1
Best-hit-arabi-name: plasmodesmata callose-binding protein 3
Gorai.013G093600.1.D
PCC=-0.072, p=0.769
mRNA Profile
Protein Expression Profile
DPA

### S200: Gorai.013G146600.1.D in case 1 and V11

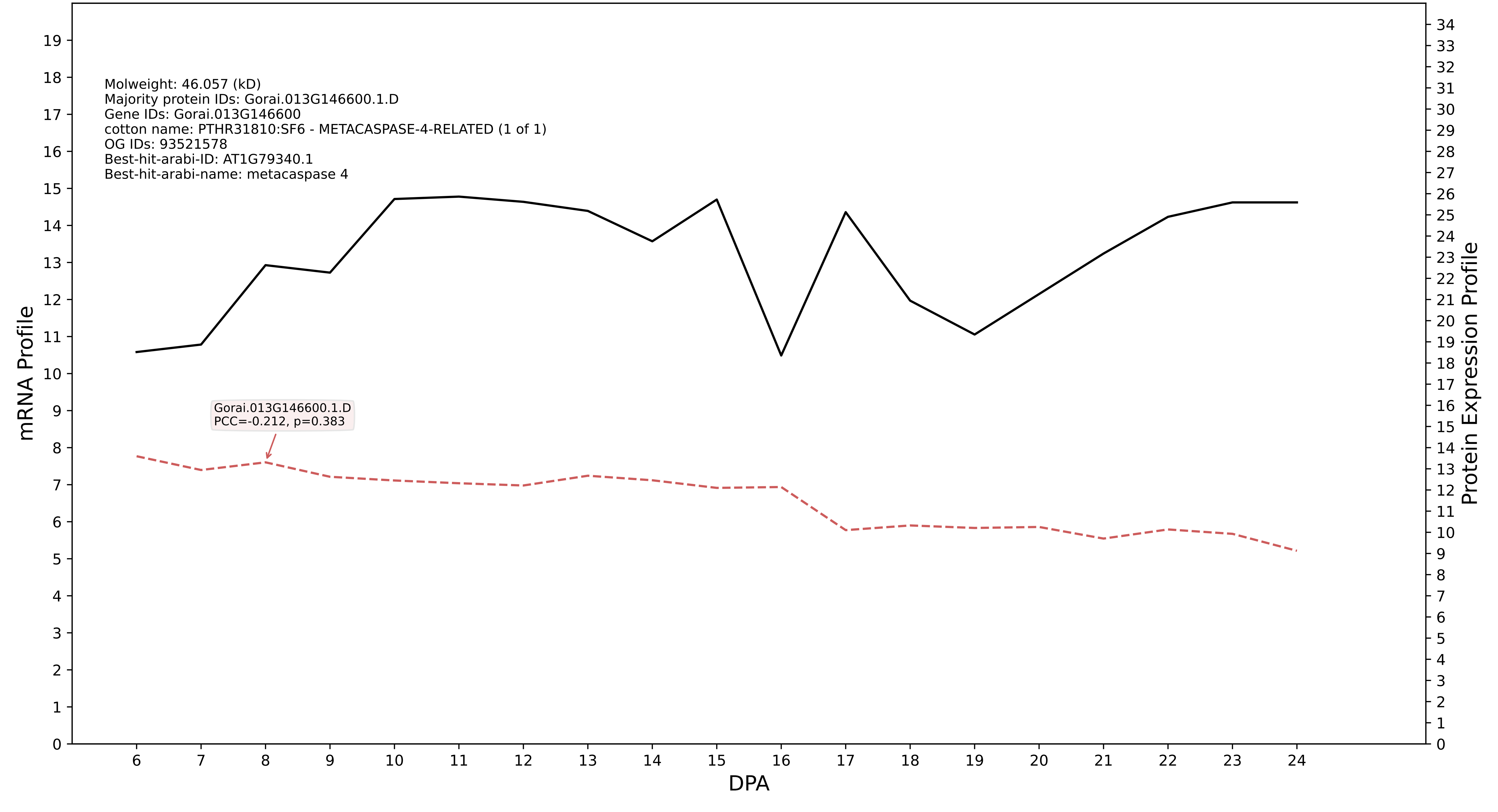

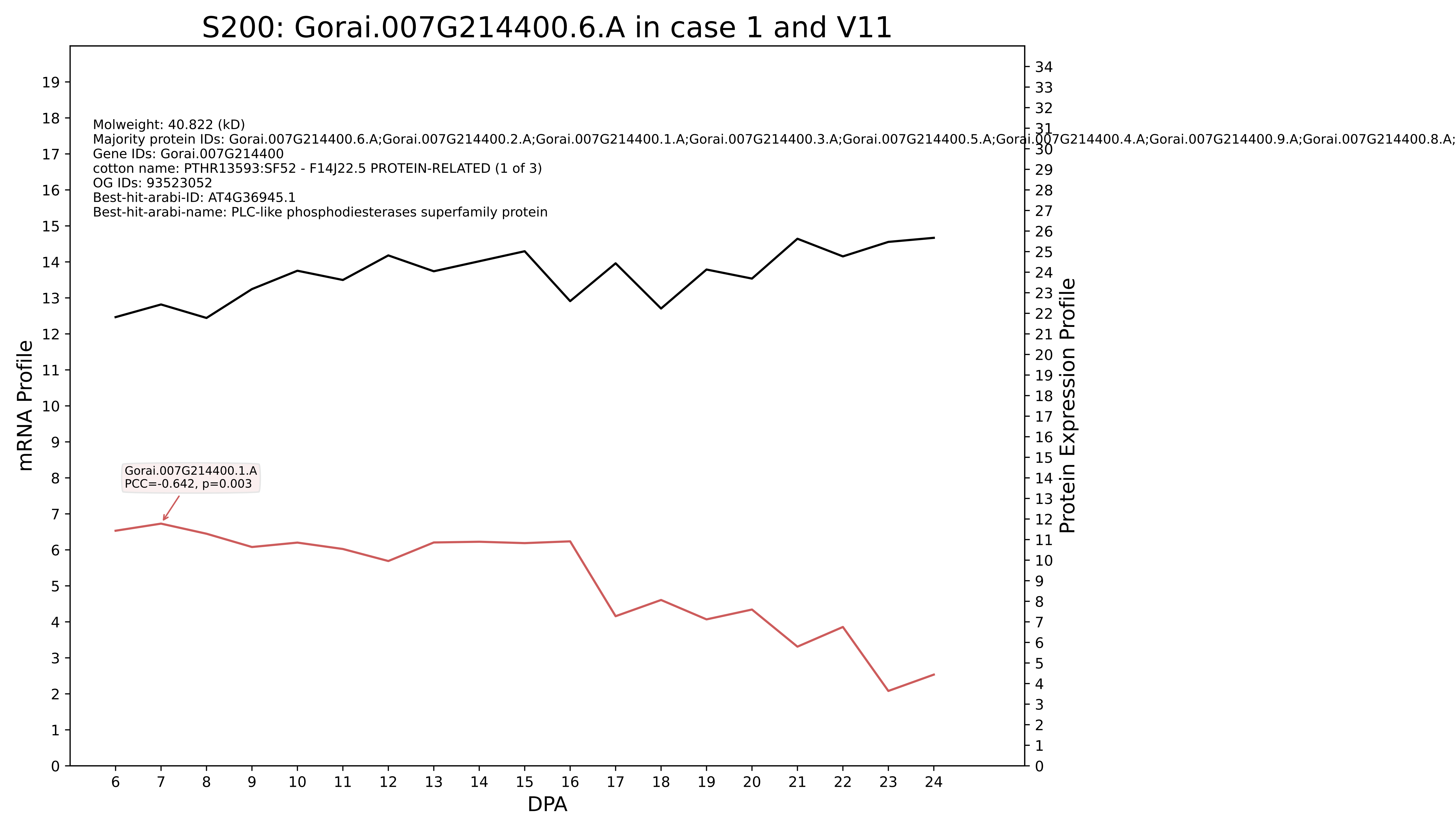
S200: Gorai.007G214400.6.A in case 1 and V11
Molweight: 40.822 (kD)
Majority protein IDs: Gorai.007G214400.6.A;Gorai.007G214400.2.A;Gorai.007G214400.1.A;Gorai.007G214400.3.A;Gorai.007G214400.5.A;Gorai.007G214400.4.A;Gorai.007G214400.9.A;Gorai.007G214400.8.A;
Gene IDs: Gorai.007G214400
cotton name: PTHR13593:SF52 - F14J22.5 PROTEIN-RELATED (1 of 3)
OG IDs: 93523052
Best-hit-arabi-ID: AT4G36945.1
Best-hit-arabi-name: PLC-like phosphodiesterases superfamily protein
Gorai.007G214400.1.A
PCC=-0.642, p=0.003
mRNA Profile
Protein Expression Profile
DPA

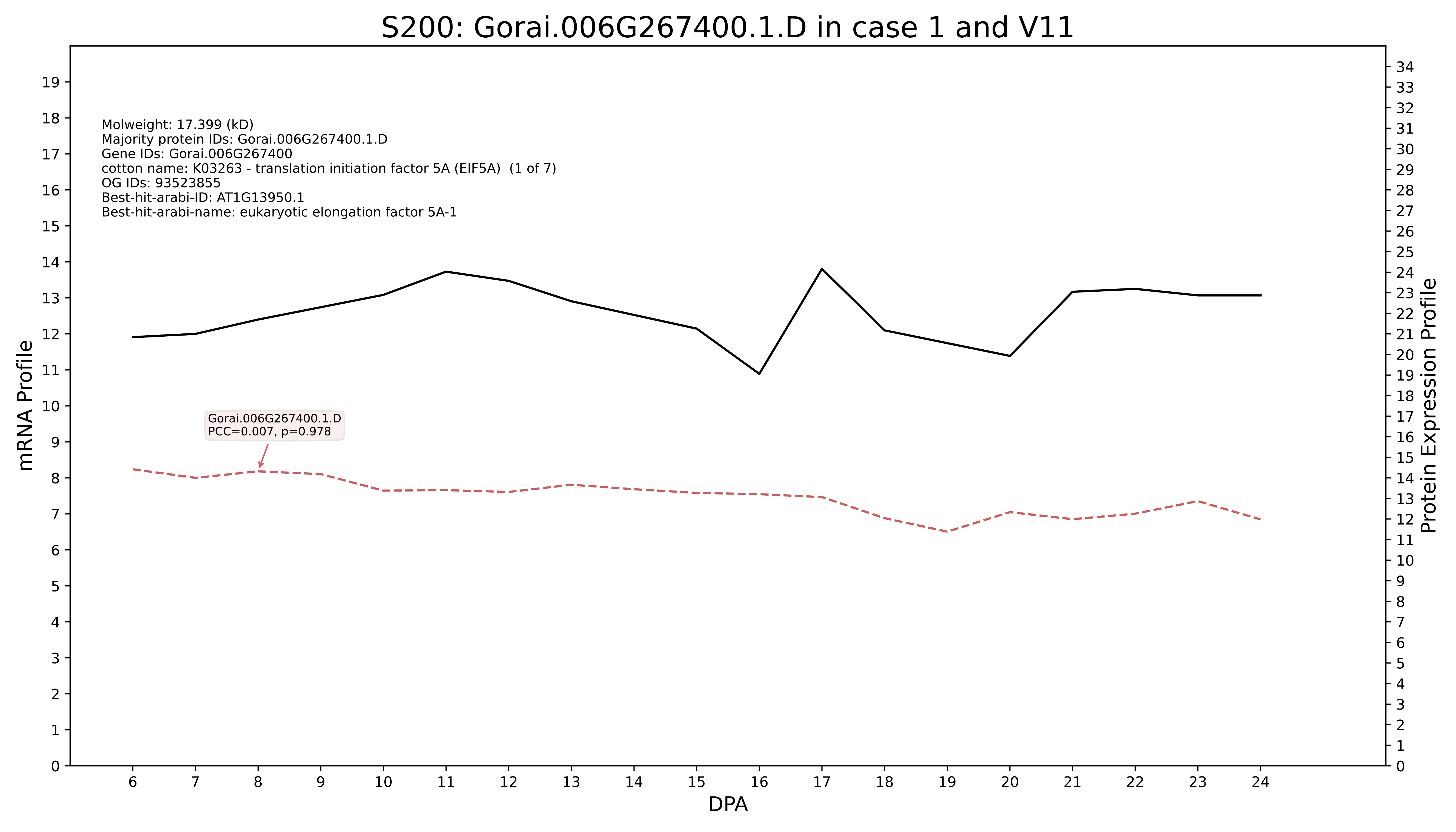
S200: Gorai.006G267400.1.D in case 1 and V11
Molweight: 17.399 (kD)
Majority protein IDs: Gorai.006G267400.1.D
Gene IDs: Gorai.006G267400
cotton name: K03263 - translation initiation factor 5A (EIF5A) (1 of 7)
OG IDs: 93523855
Best-hit-arabi-ID: AT1G13950.1
Best-hit-arabi-name: eukaryotic elongation factor 5A-1
Gorai.006G267400.1.D
PCC=0.007, p=0.978
mRNA Profile
Protein Expression Profile
DPA

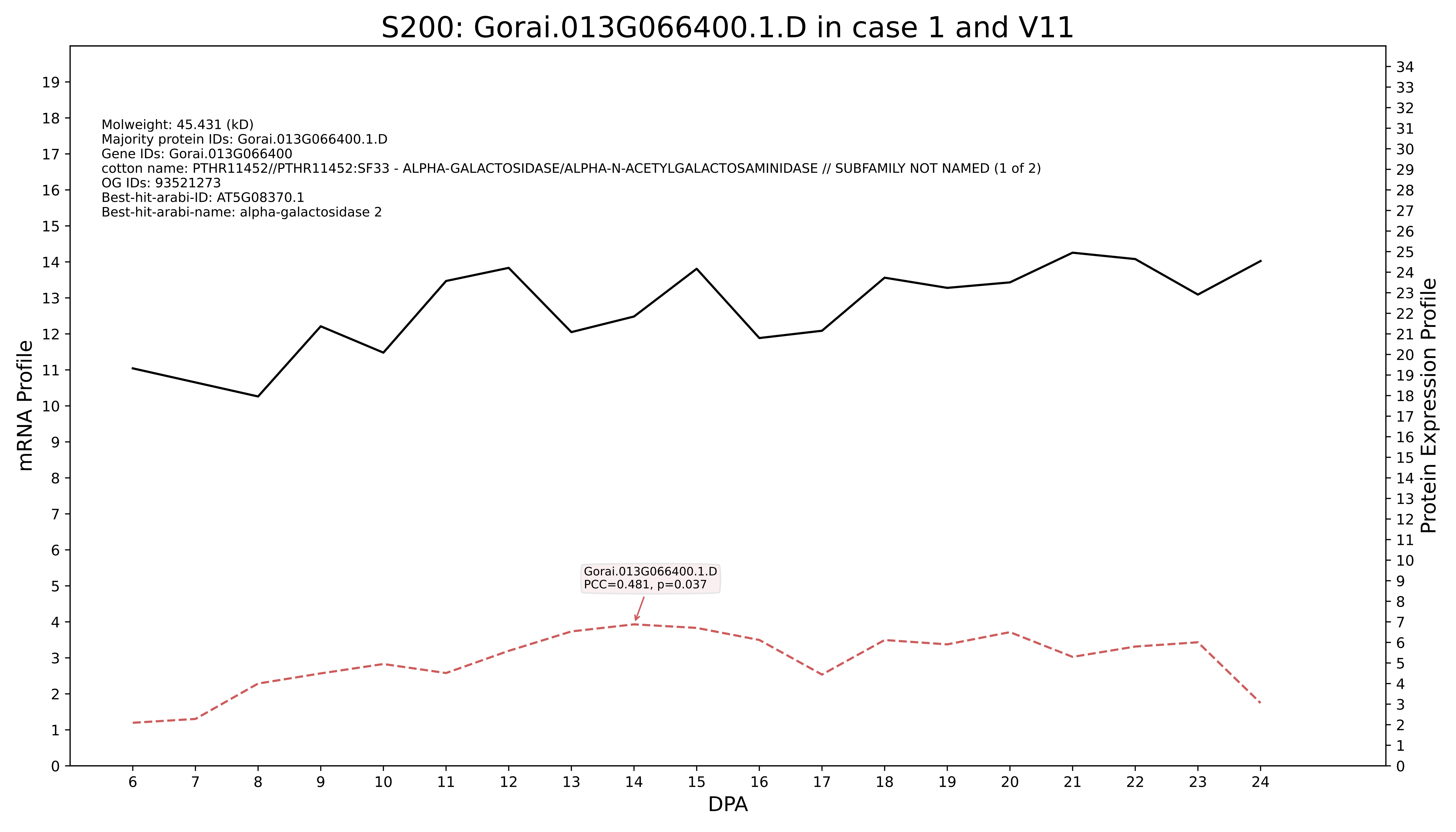

S200: Gorai.013G066400.1.D in case 1 and V11
Molweight: 45.431 (kD)
Majority protein IDs: Gorai.013G066400.1.D
Gene IDs: Gorai.013G066400
cotton name: PTHR11452//PTHR11452:SF33 - ALPHA-GALACTOSIDASE/ALPHA-N-ACETYLGALACTOSAMINIDASE // SUBFAMILY NOT NAMED (1 of 2)
OG IDs: 93521273
Best-hit-arabi-ID: AT5G08370.1
Best-hit-arabi-name: alpha-galactosidase 2
Gorai.013G066400.1.D
PCC=0.481, p=0.037
mRNA Profile
Protein Expression Profile
DPA

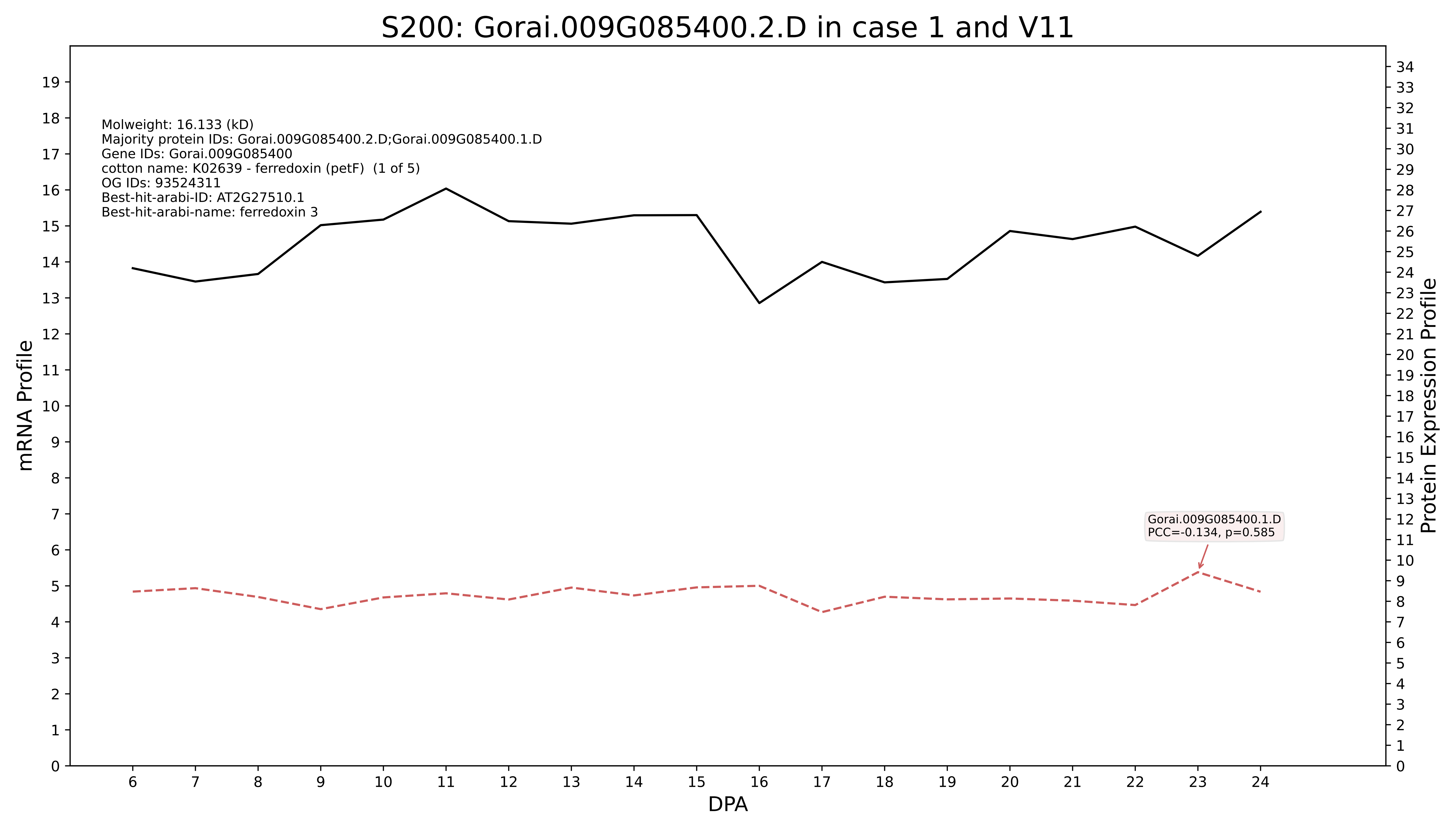
S200: Gorai.009G085400.2.D in case 1 and V11
Molweight: 16.133 (kD)
Majority protein IDs: Gorai.009G085400.2.D;Gorai.009G085400.1.D
Gene IDs: Gorai.009G085400
cotton name: K02639 - ferredoxin (petF) (1 of 5)
OG IDs: 93524311
Best-hit-arabi-ID: AT2G27510.1
Best-hit-arabi-name: ferredoxin 3
Gorai.009G085400.1.D
PCC=-0.134, p=0.585
mRNA Profile
Protein Expression Profile
DPA

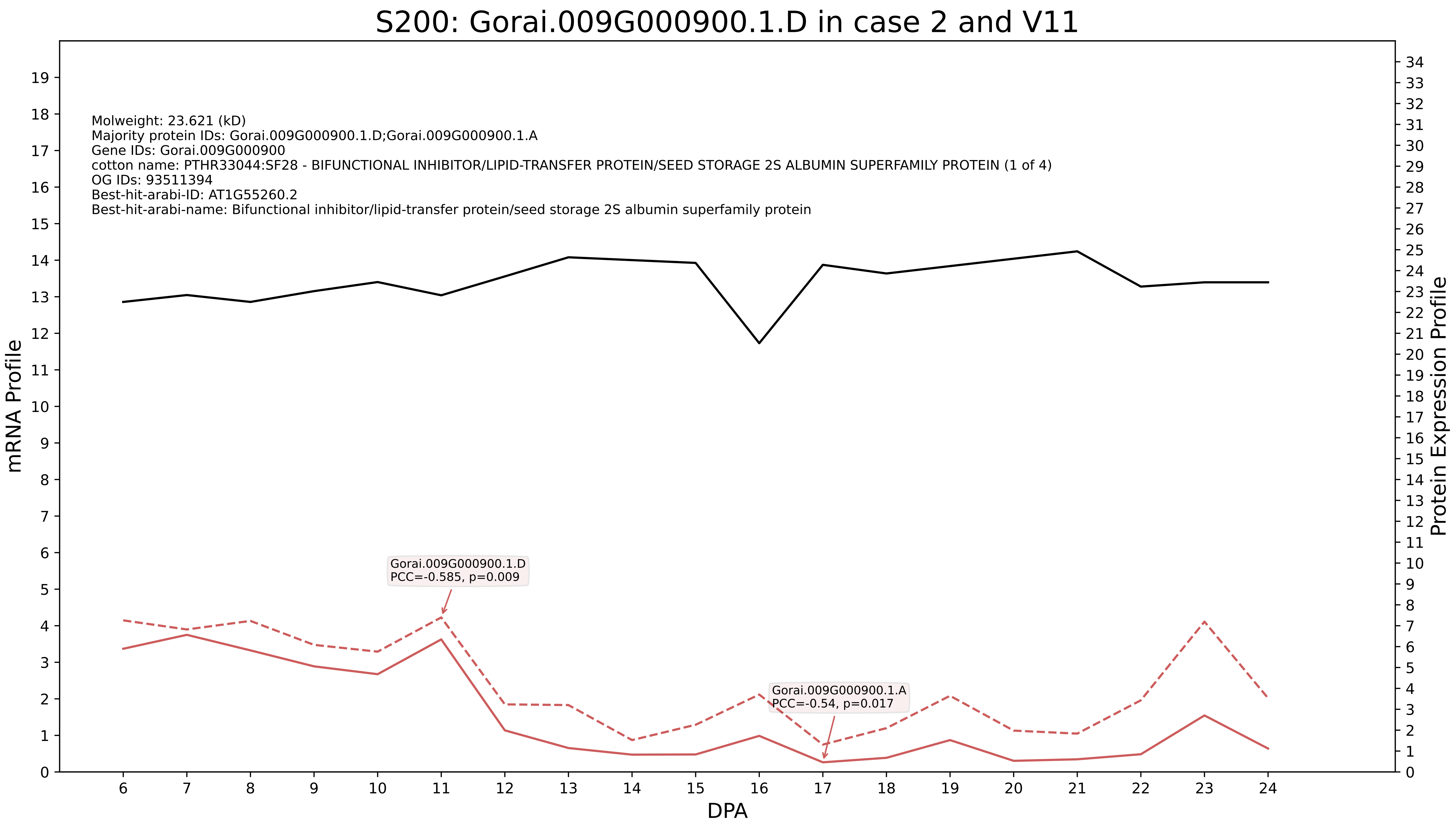
S200: Gorai.009G000900.1.D in case 2 and V11
Molweight: 23.621 (kD)
Majority protein IDs: Gorai.009G000900.1.D;Gorai.009G000900.1.A
Gene IDs: Gorai.009G000900
cotton name: PTHR33044:SF28 - BIFUNCTIONAL INHIBITOR/LIPID-TRANSFER PROTEIN/SEED STORAGE 2S ALBUMIN SUPERFAMILY PROTEIN (1 of 4)
OG IDs: 93511394
Best-hit-arabi-ID: AT1G55260.2
Best-hit-arabi-name: Bifunctional inhibitor/lipid-transfer protein/seed storage 2S albumin superfamily protein
Gorai.009G000900.1.D
PCC=-0.585, p=0.009
Gorai.009G000900.1.A
PCC=-0.54, p=0.017
mRNA Profile
Protein Expression Profile
DPA

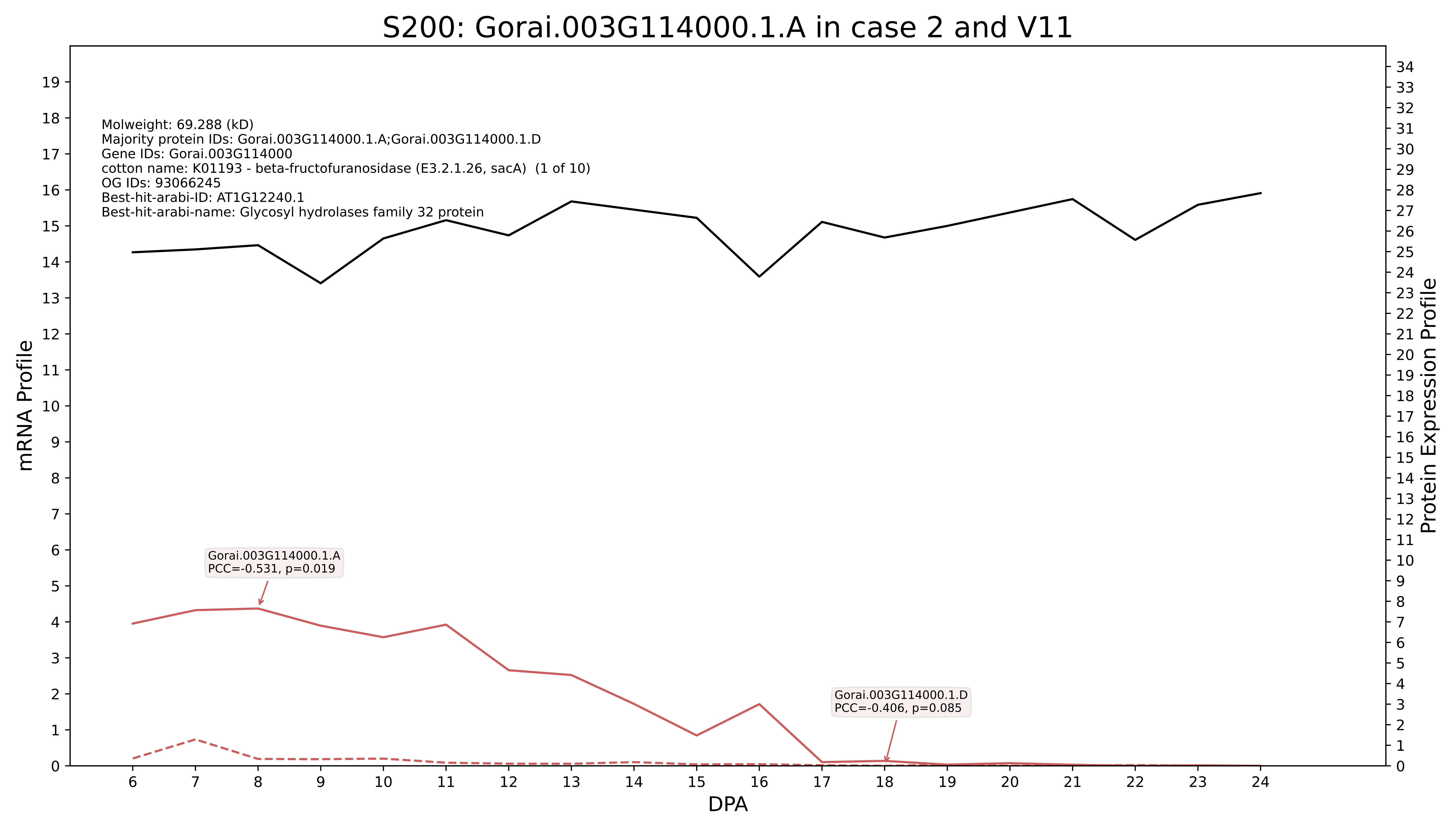
S200: Gorai.003G114000.1.A in case 2 and V11
Molweight: 69.288 (kD)
Majority protein IDs: Gorai.003G114000.1.A;Gorai.003G114000.1.D
Gene IDs: Gorai.003G114000
cotton name: K01193 - beta-fructofuranosidase (E3.2.1.26, sacA) (1 of 10)
OG IDs: 93066245
Best-hit-arabi-ID: AT1G12240.1
Best-hit-arabi-name: Glycosyl hydrolases family 32 protein
Gorai.003G114000.1.A
PCC=-0.531, p=0.019
Gorai.003G114000.1.D
PCC=-0.406, p=0.085
mRNA Profile
Protein Expression Profile
DPA

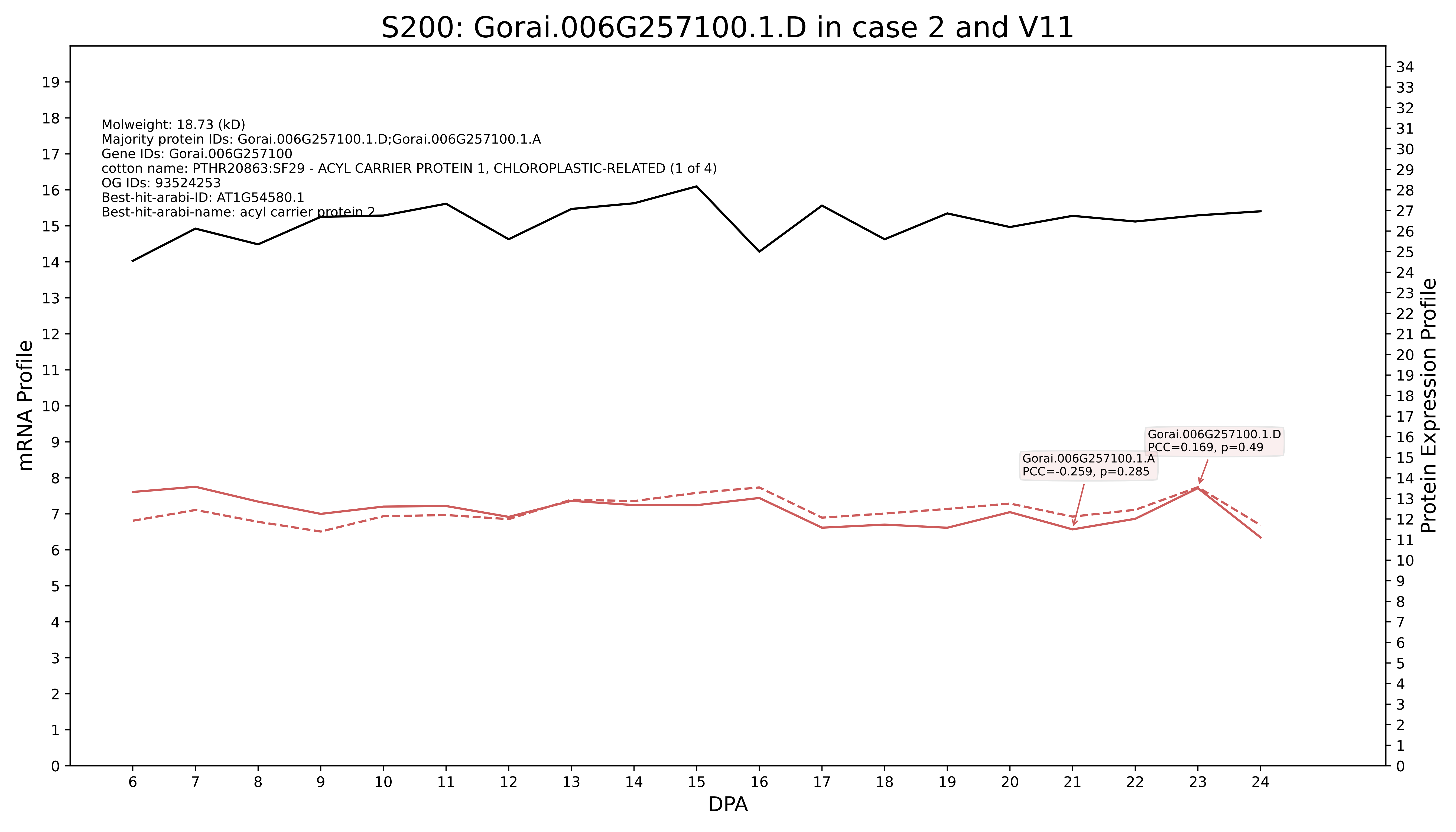
S200: Gorai.006G257100.1.D in case 2 and V11
Molweight: 18.73 (kD)
Majority protein IDs: Gorai.006G257100.1.D;Gorai.006G257100.1.A
Gene IDs: Gorai.006G257100
cotton name: PTHR20863:SF29 - ACYL CARRIER PROTEIN 1, CHLOROPLASTIC-RELATED (1 of 4)
OG IDs: 93524253
Best-hit-arabi-ID: AT1G54580.1
Best-hit-arabi-name: acyl carrier protein 2
Gorai.006G257100.1.D
PCC=0.169, p=0.49
Gorai.006G257100.1.A
PCC=-0.259, p=0.285
mRNA Profile
Protein Expression Profile
DPA

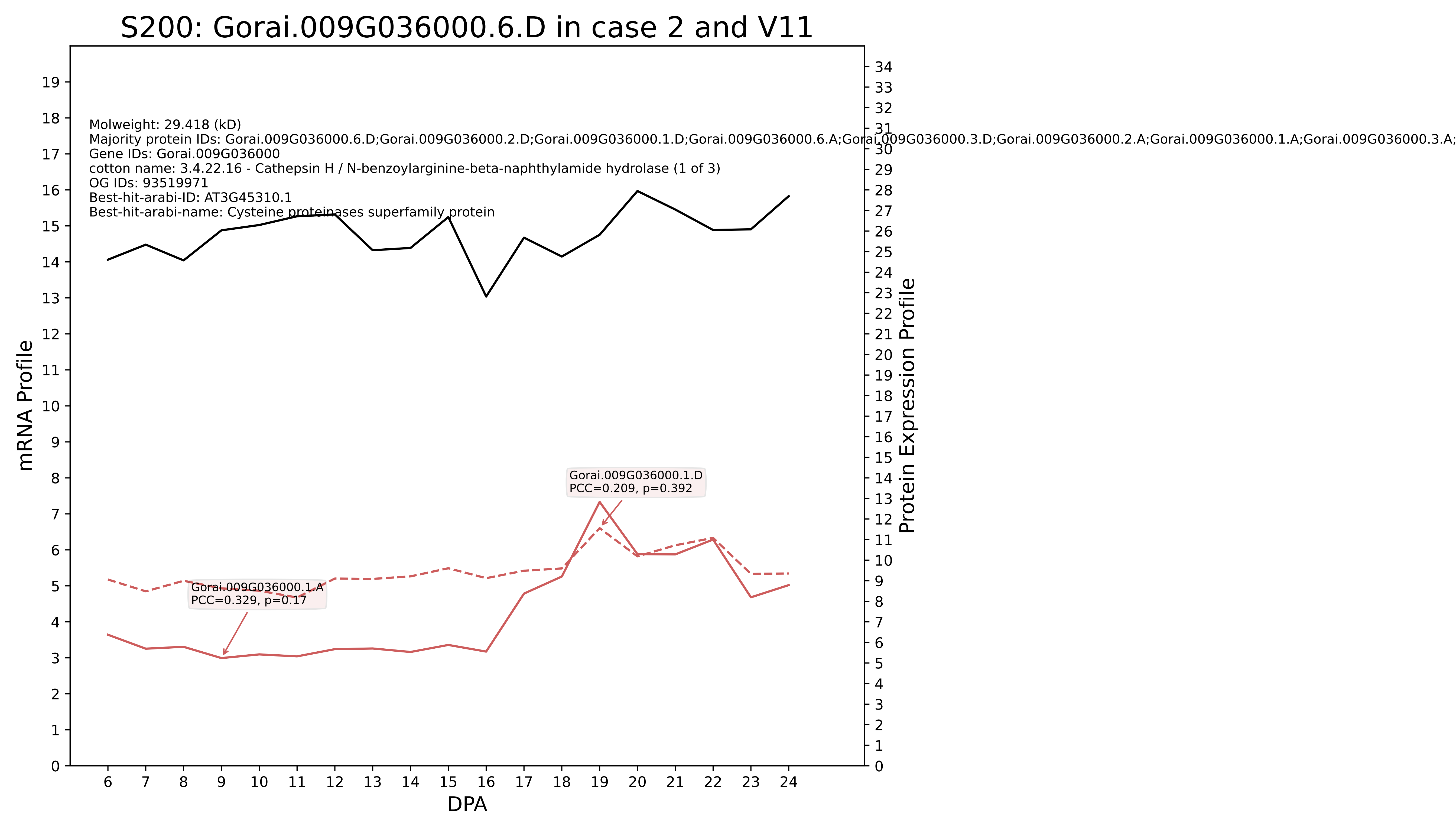
S200: Gorai.009G036000.6.D in case 2 and V11
Molweight: 29.418 (kD)
Majority protein IDs: Gorai.009G036000.6.D;Gorai.009G036000.2.D;Gorai.009G036000.1.D;Gorai.009G036000.6.A;Gorai.009G036000.3.D;Gorai.009G036000.2.A;Gorai.009G036000.1.A;Gorai.009G036000.3.A;
Gene IDs: Gorai.009G036000
cotton name: 3.4.22.16 - Cathepsin H / N-benzoylarginine-beta-naphthylamide hydrolase (1 of 3)
OG IDs: 93519971
Best-hit-arabi-ID: AT3G45310.1
Best-hit-arabi-name: Cysteine proteinases superfamily protein
Gorai.009G036000.1.D
PCC=0.209, p=0.392
Gorai.009G036000.1.A
PCC=0.329, p=0.17
mRNA Profile
Protein Expression Profile
DPA

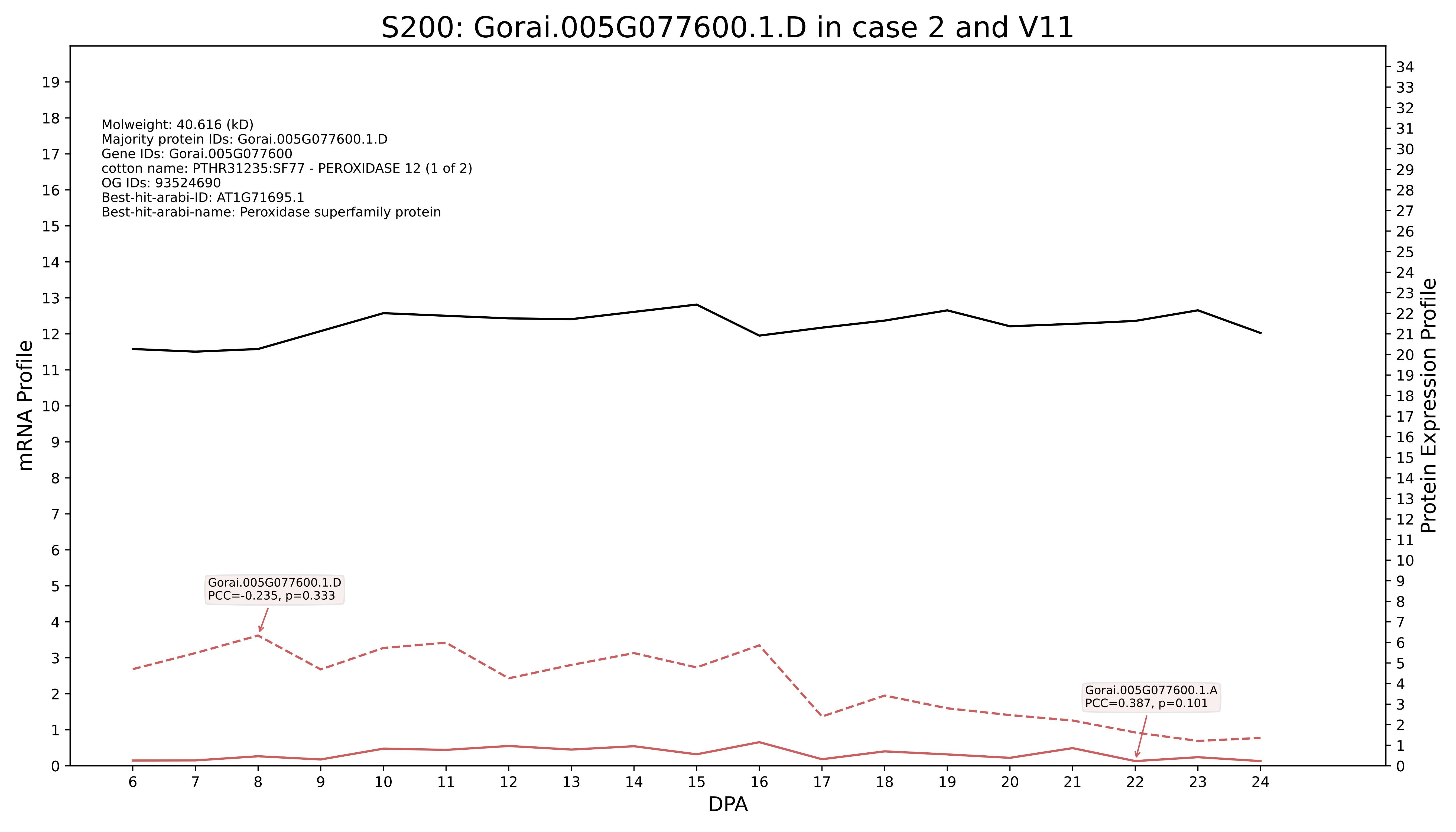
S200: Gorai.005G077600.1.D in case 2 and V11
Molweight: 40.616 (kD)
Majority protein IDs: Gorai.005G077600.1.D
Gene IDs: Gorai.005G077600
cotton name: PTHR31235:SF77 - PEROXIDASE 12 (1 of 2)
OG IDs: 93524690
Best-hit-arabi-ID: AT1G71695.1
Best-hit-arabi-name: Peroxidase superfamily protein
Gorai.005G077600.1.D
PCC=-0.235, p=0.333
Gorai.005G077600.1.A
PCC=0.387, p=0.101
mRNA Profile
Protein Expression Profile
DPA

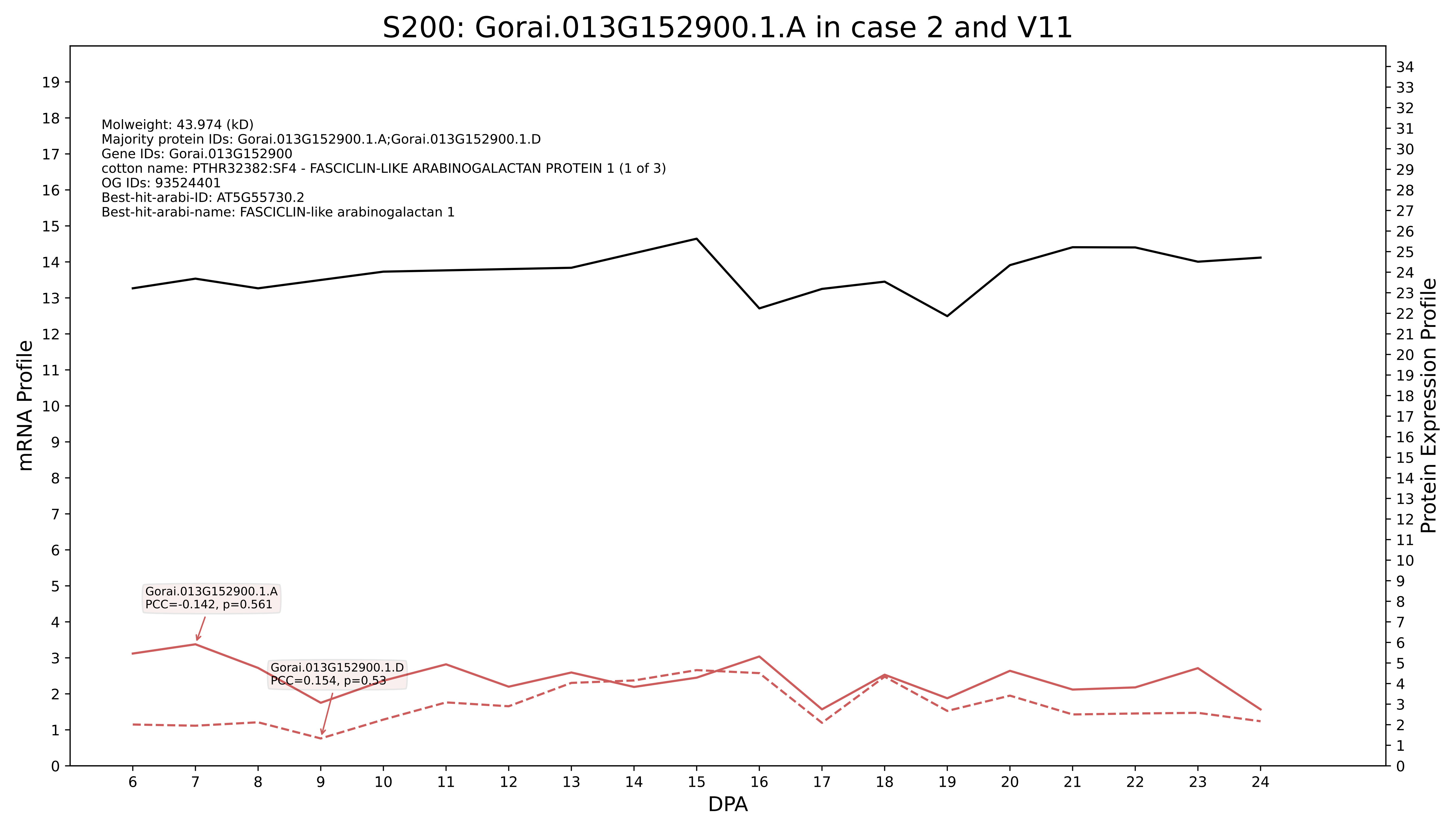

S200: Gorai.013G152900.1.A in case 2 and V11
Molweight: 43.974 (kD)
Majority protein IDs: Gorai.013G152900.1.A;Gorai.013G152900.1.D
Gene IDs: Gorai.013G152900
cotton name: PTHR32382:SF4 - FASCICLIN-LIKE ARABINOGALACTAN PROTEIN 1 (1 of 3)
OG IDs: 93524401
Best-hit-arabi-ID: AT5G55730.2
Best-hit-arabi-name: FASCICLIN-like arabinogalactan 1
Gorai.013G152900.1.A
PCC=-0.142, p=0.561
Gorai.013G152900.1.D
PCC=0.154, p=0.53
mRNA Profile
Protein Expression Profile
DPA

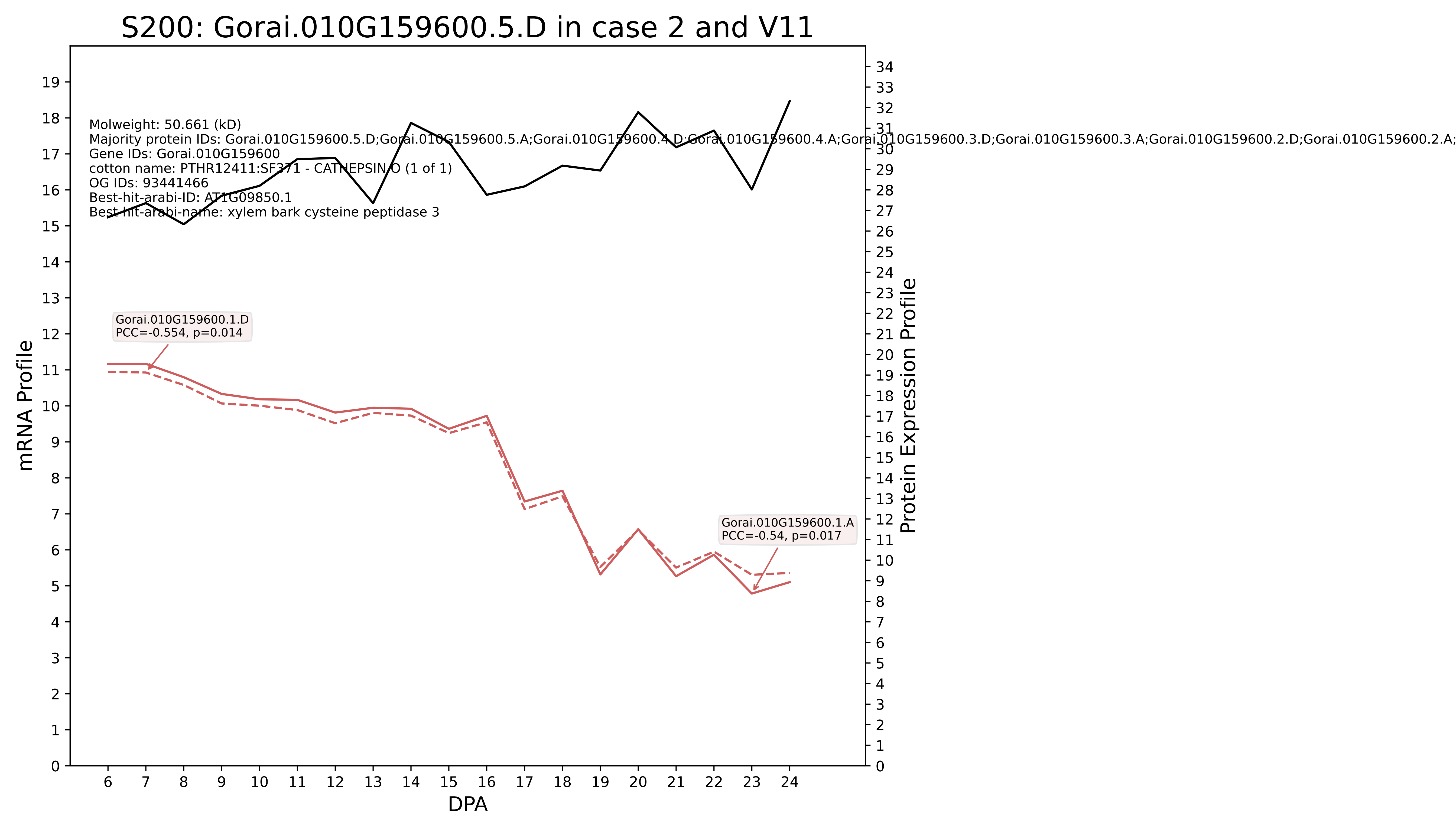
S200: Gorai.010G159600.5.D in case 2 and V11
Molweight: 50.661 (kD)
Majority protein IDs: Gorai.010G159600.5.D;Gorai.010G159600.5.A;Gorai.010G159600.4.D;Gorai.010G159600.4.A;Gorai.010G159600.3.D;Gorai.010G159600.3.A;Gorai.010G159600.2.D;Gorai.010G159600.2.A;
Gene IDs: Gorai.010G159600
cotton name: PTHR12411:SF371 - CATHEPSIN O (1 of 1)
OG IDs: 93441466
Best-hit-arabi-ID: AT1G09850.1
Best-hit-arabi-name: xylem bark cysteine peptidase 3
Gorai.010G159600.1.D
PCC=-0.554, p=0.014
Gorai.010G159600.1.A
PCC=-0.54, p=0.017
mRNA Profile
Protein Expression Profile
DPA

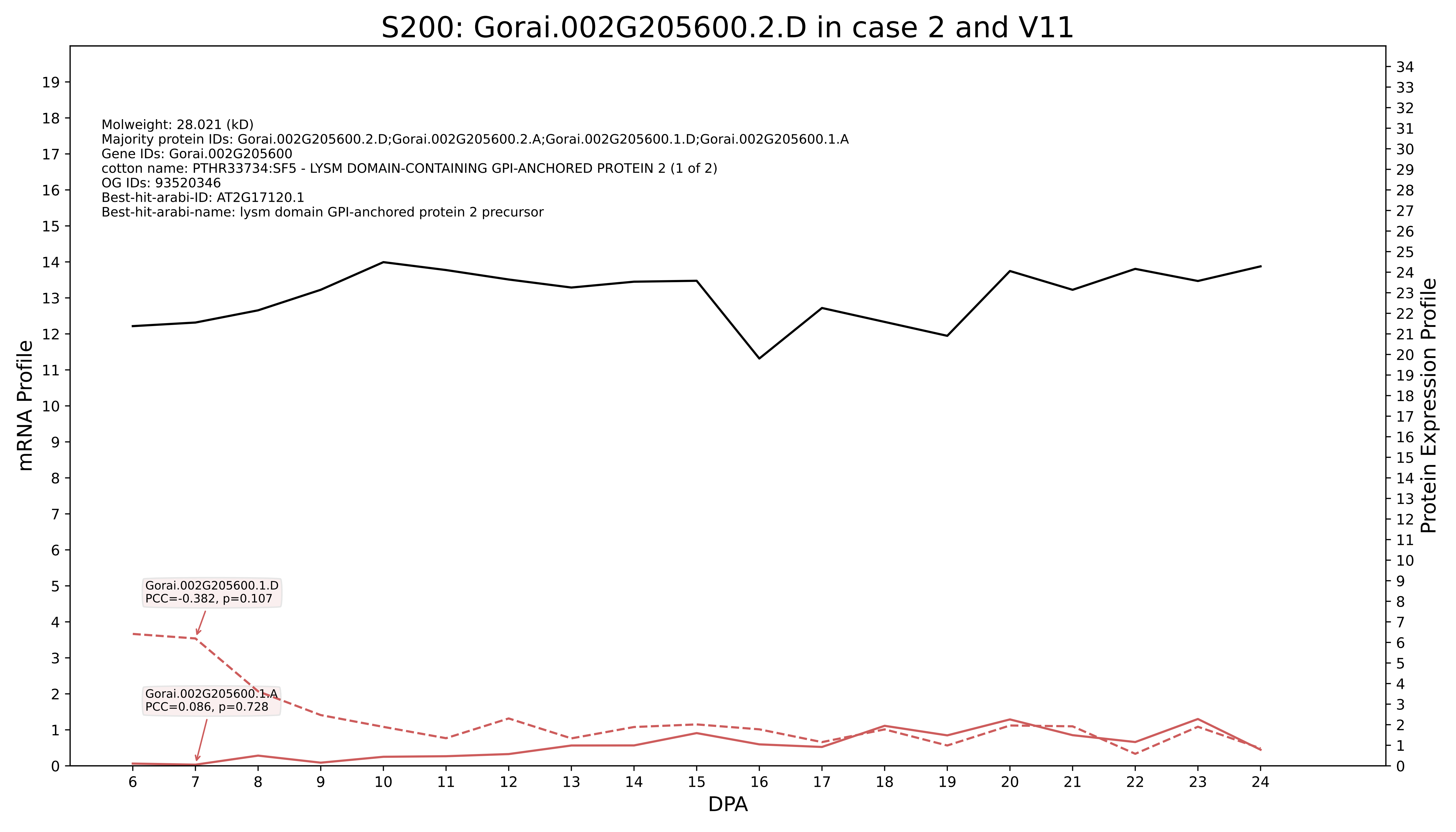

S200: Gorai.002G205600.2.D in case 2 and V11
Molweight: 28.021 (kD)
Majority protein IDs: Gorai.002G205600.2.D;Gorai.002G205600.2.A;Gorai.002G205600.1.D;Gorai.002G205600.1.A
Gene IDs: Gorai.002G205600
cotton name: PTHR33734:SF5 - LYSM DOMAIN-CONTAINING GPI-ANCHORED PROTEIN 2 (1 of 2)
OG IDs: 93520346
Best-hit-arabi-ID: AT2G17120.1
Best-hit-arabi-name: lysm domain GPI-anchored protein 2 precursor
Gorai.002G205600.1.D
PCC=-0.382, p=0.107
Gorai.002G205600.1.A
PCC=0.086, p=0.728
mRNA Profile
Protein Expression Profile
DPA

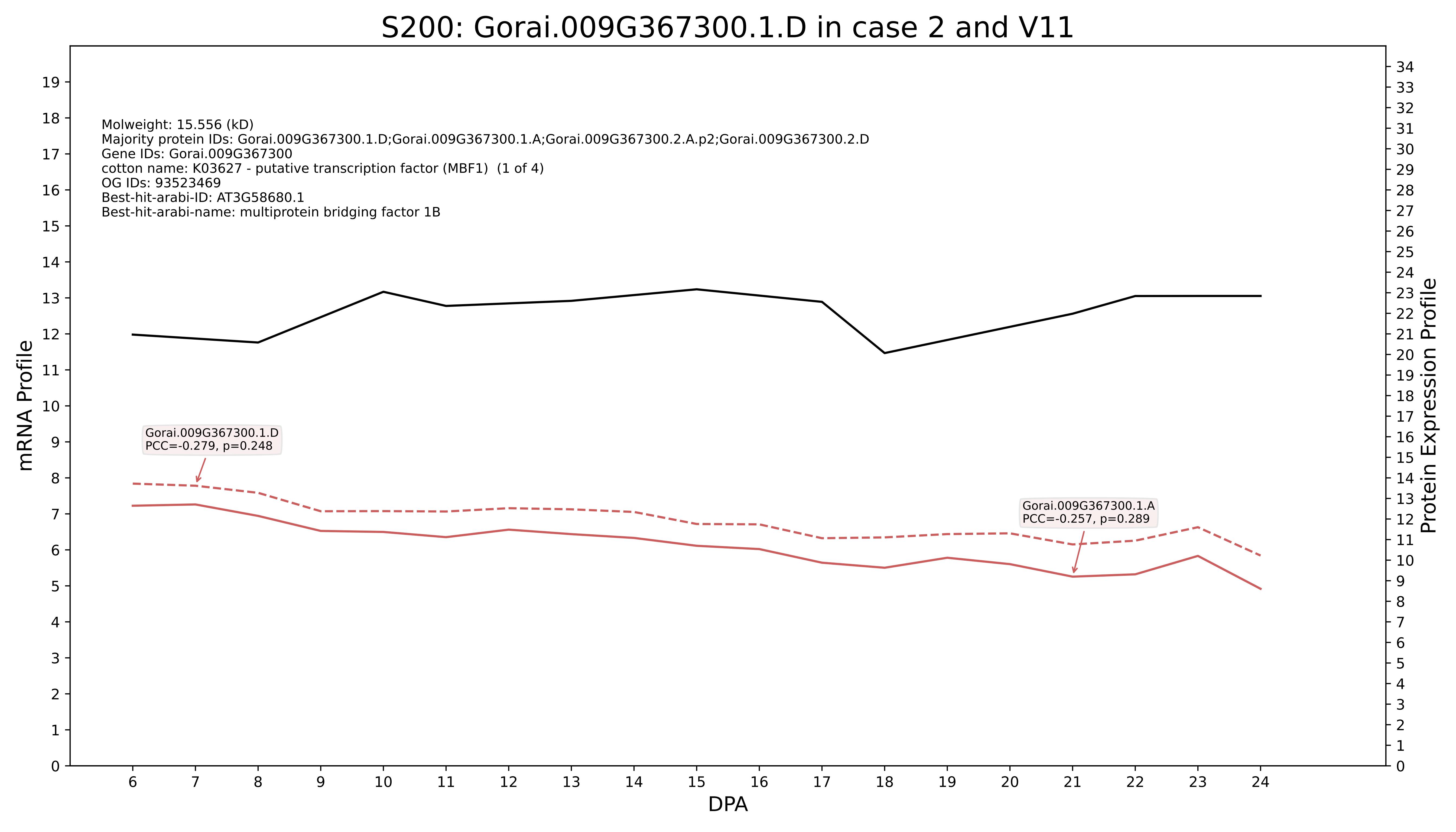
S200: Gorai.009G367300.1.D in case 2 and V11
Molweight: 15.556 (kD)
Majority protein IDs: Gorai.009G367300.1.D;Gorai.009G367300.1.A;Gorai.009G367300.2.A.p2;Gorai.009G367300.2.D
Gene IDs: Gorai.009G367300
cotton name: K03627 - putative transcription factor (MBF1) (1 of 4)
OG IDs: 93523469
Best-hit-arabi-ID: AT3G58680.1
Best-hit-arabi-name: multiprotein bridging factor 1B
Gorai.009G367300.1.D
PCC=-0.279, p=0.248
Gorai.009G367300.1.A
PCC=-0.257, p=0.289
mRNA Profile
Protein Expression Profile
DPA

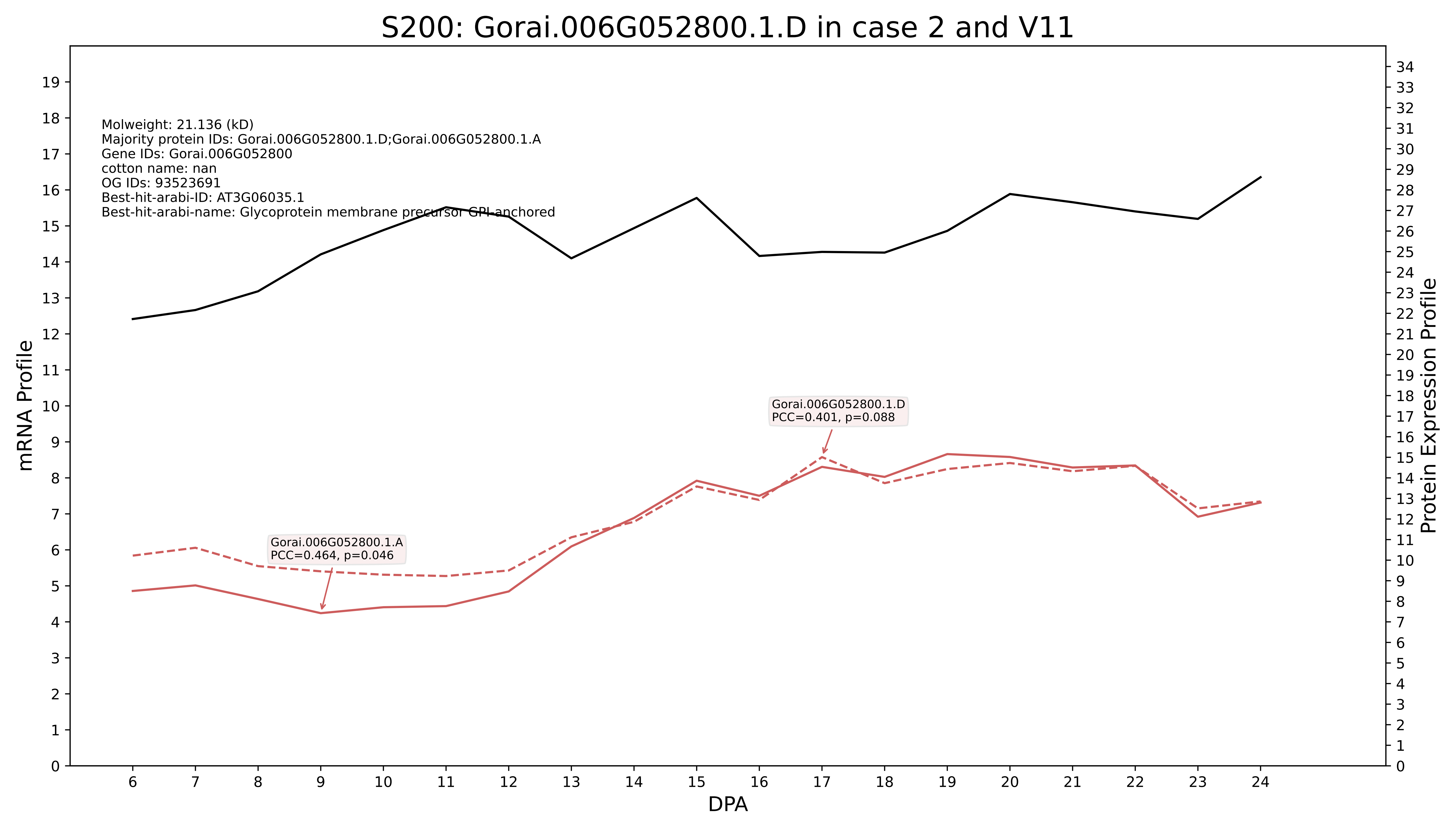
S200: Gorai.006G052800.1.D in case 2 and V11
Molweight: 21.136 (kD)
Majority protein IDs: Gorai.006G052800.1.D;Gorai.006G052800.1.A
Gene IDs: Gorai.006G052800
cotton name: nan
OG IDs: 93523691
Best-hit-arabi-ID: AT3G06035.1
Best-hit-arabi-name: Glycoprotein membrane precursor GPI-anchored
Gorai.006G052800.1.D
PCC=0.401, p=0.088
Gorai.006G052800.1.A
PCC=0.464, p=0.046
mRNA Profile
Protein Expression Profile
DPA

S200: Gorai.009G242500.1.D in case 2 and V11

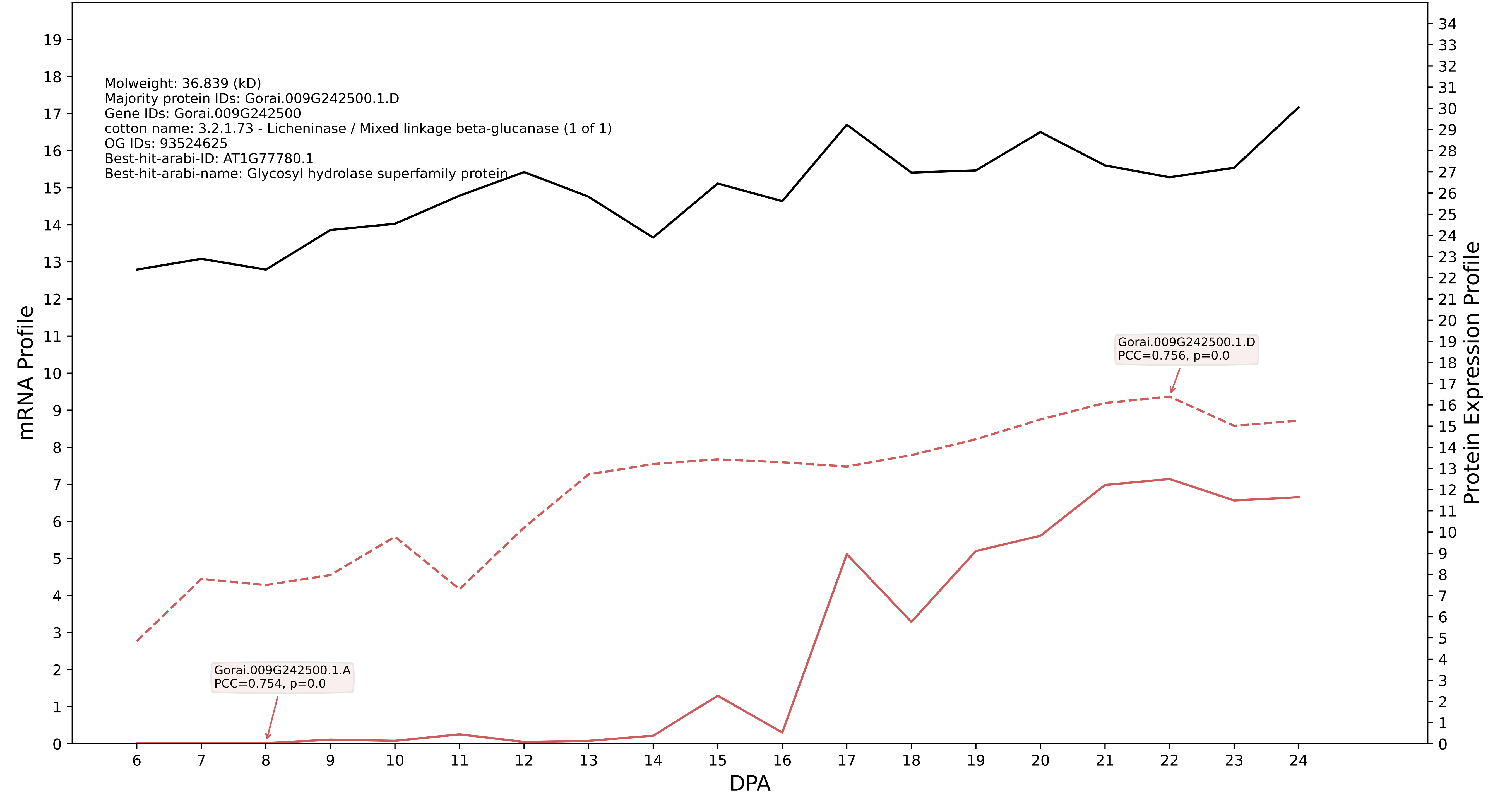

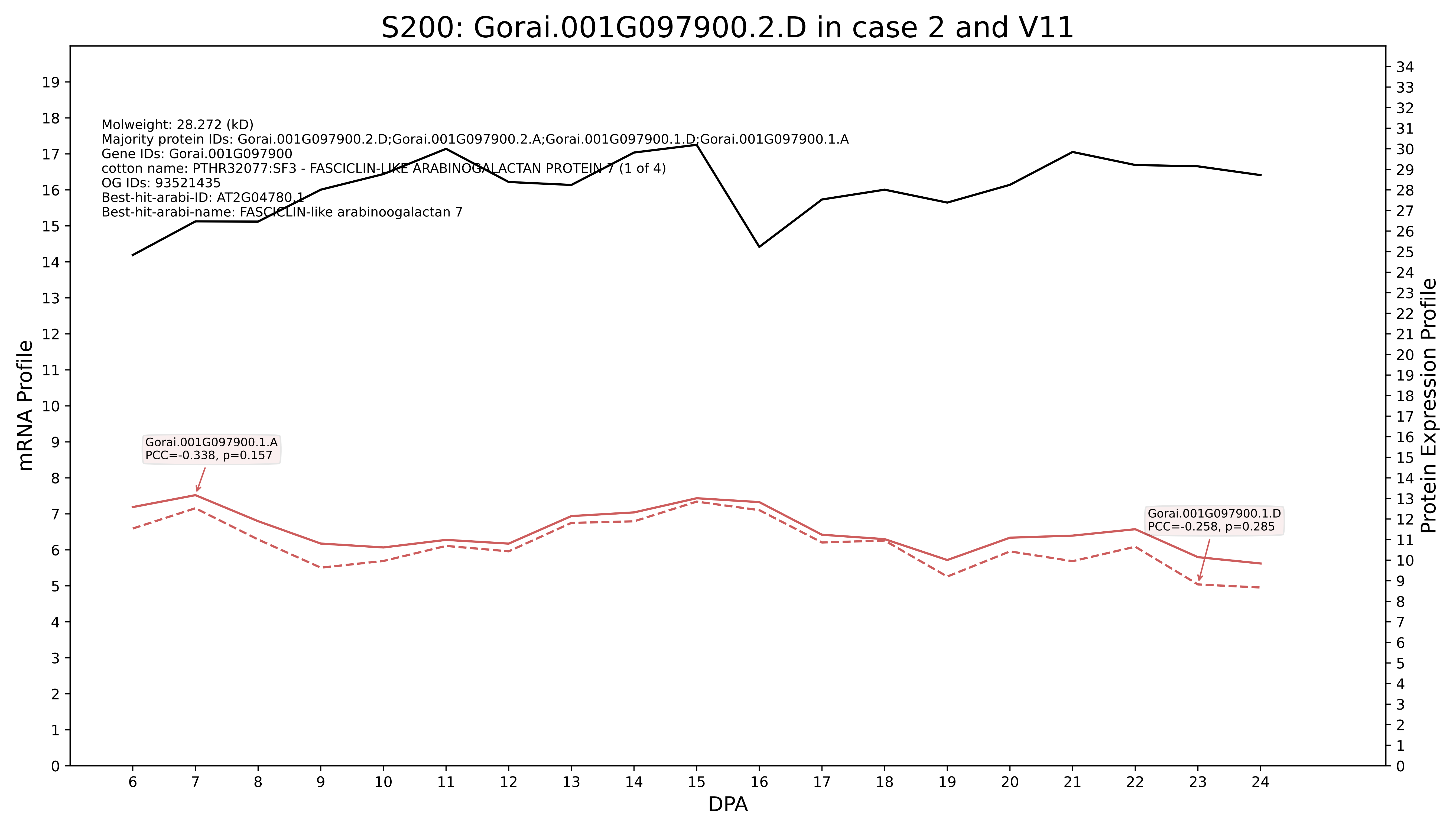
S200: Gorai.001G097900.2.D in case 2 and V11
Molweight: 28.272 (kD)
Majority protein IDs: Gorai.001G097900.2.D;Gorai.001G097900.2.A;Gorai.001G097900.1.D;Gorai.001G097900.1.A
Gene IDs: Gorai.001G097900
cotton name: PTHR32077:SF3 - FASCICLIN-LIKE ARABINOGALACTAN PROTEIN 7 (1 of 4)
OG IDs: 93521435
Best-hit-arabi-ID: AT2G04780.1
Best-hit-arabi-name: FASCICLIN-like arabinoogalactan 7
Gorai.001G097900.1.A
PCC=-0.338, p=0.157
Gorai.001G097900.1.D
PCC=-0.258, p=0.285
mRNA Profile
Protein Expression Profile
DPA

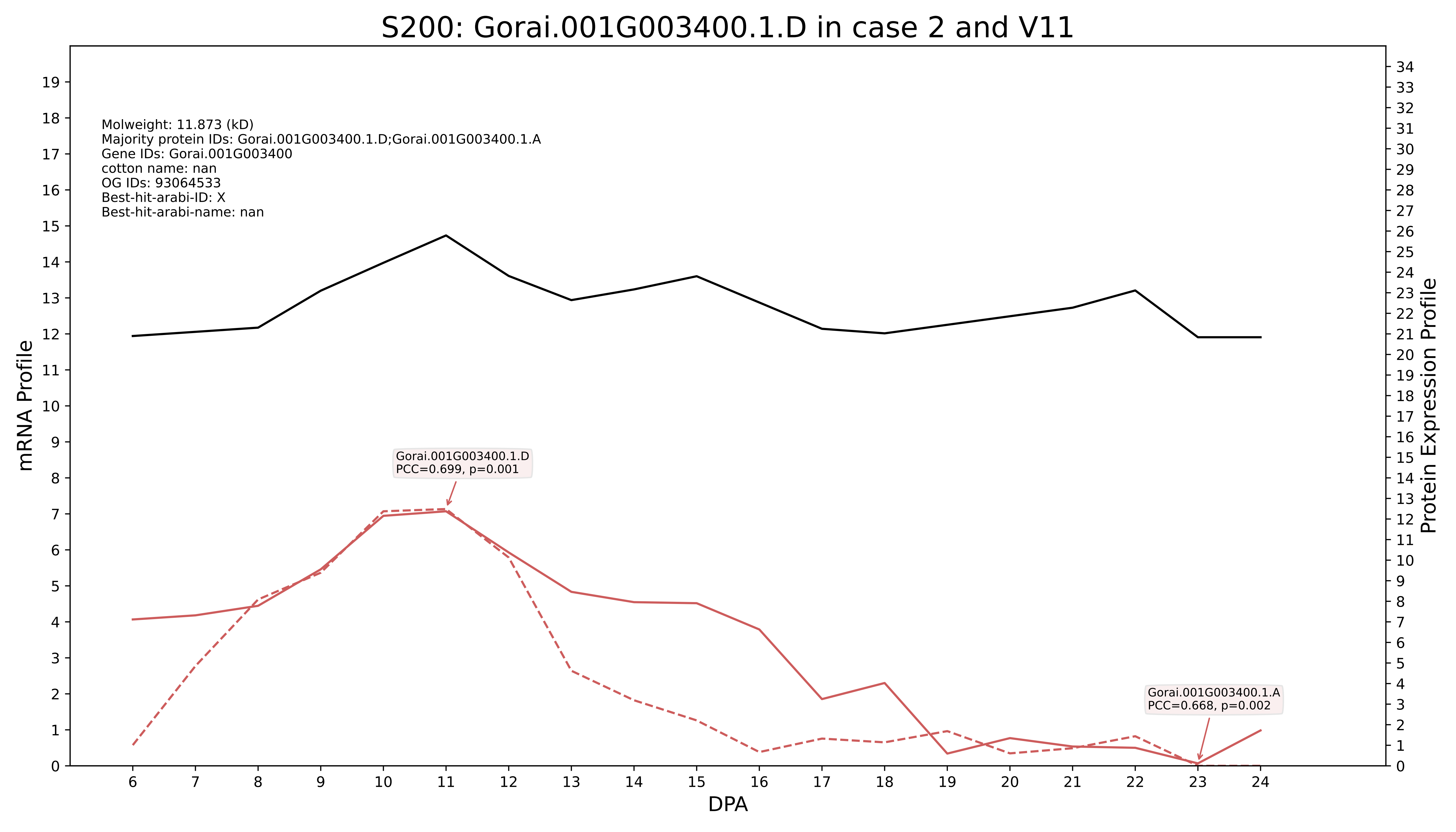
S200: Gorai.001G003400.1.D in case 2 and V11
Molweight: 11.873 (kD)
Majority protein IDs: Gorai.001G003400.1.D;Gorai.001G003400.1.A
Gene IDs: Gorai.001G003400
cotton name: nan
OG IDs: 93064533
Best-hit-arabi-ID: X
Best-hit-arabi-name: nan
Gorai.001G003400.1.D
PCC=0.699, p=0.001
Gorai.001G003400.1.A
PCC=0.668, p=0.002
mRNA Profile
Protein Expression Profile
DPA

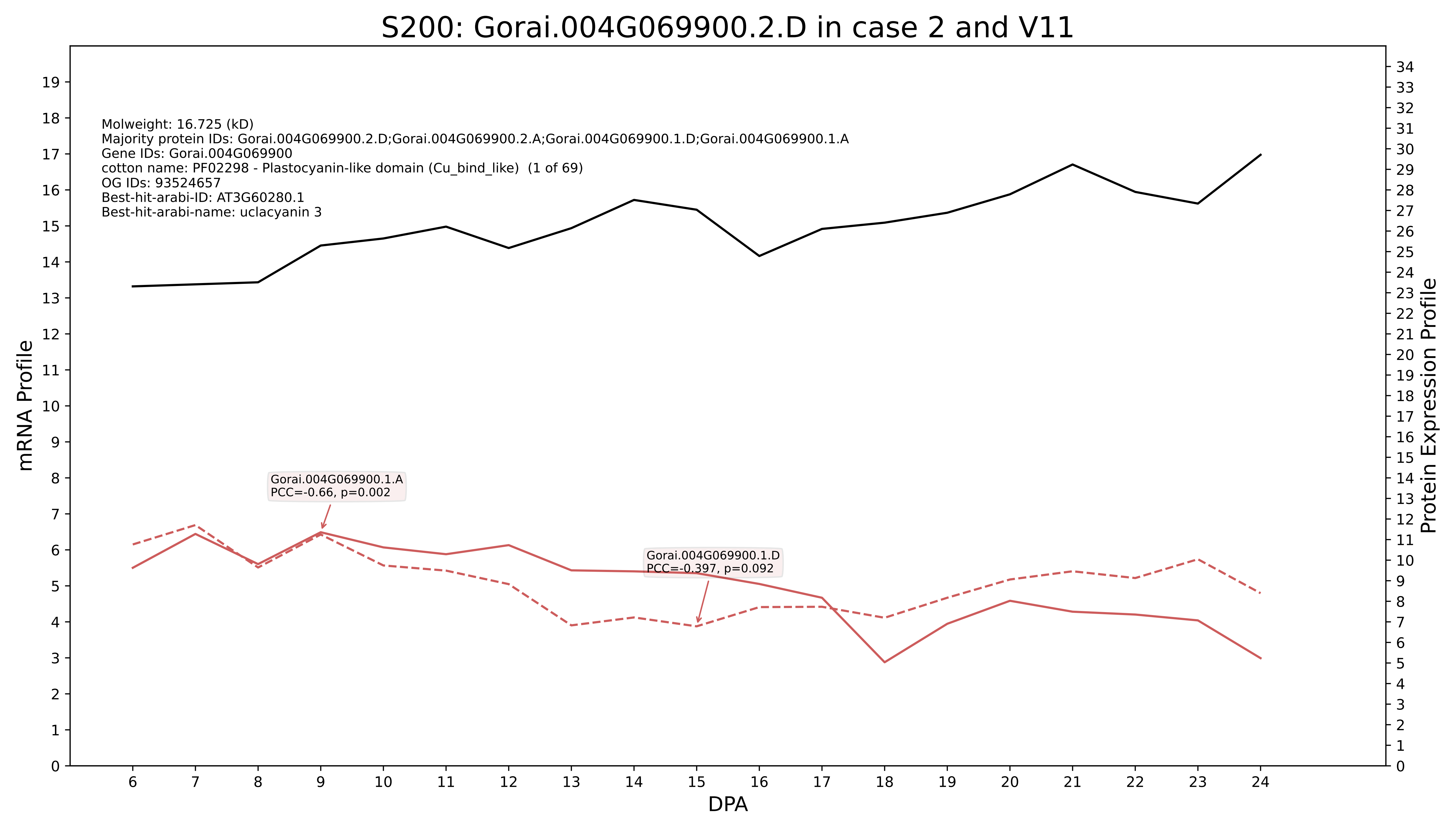
S200: Gorai.004G069900.2.D in case 2 and V11
Molweight: 16.725 (kD)
Majority protein IDs: Gorai.004G069900.2.D;Gorai.004G069900.2.A;Gorai.004G069900.1.D;Gorai.004G069900.1.A
Gene IDs: Gorai.004G069900
cotton name: PF02298 - Plastocyanin-like domain (Cu_bind_like) (1 of 69)
OG IDs: 93524657
Best-hit-arabi-ID: AT3G60280.1
Best-hit-arabi-name: uclacyanin 3
Gorai.004G069900.1.A
PCC=-0.66, p=0.002
Gorai.004G069900.1.D
PCC=-0.397, p=0.092
mRNA Profile
Protein Expression Profile
DPA

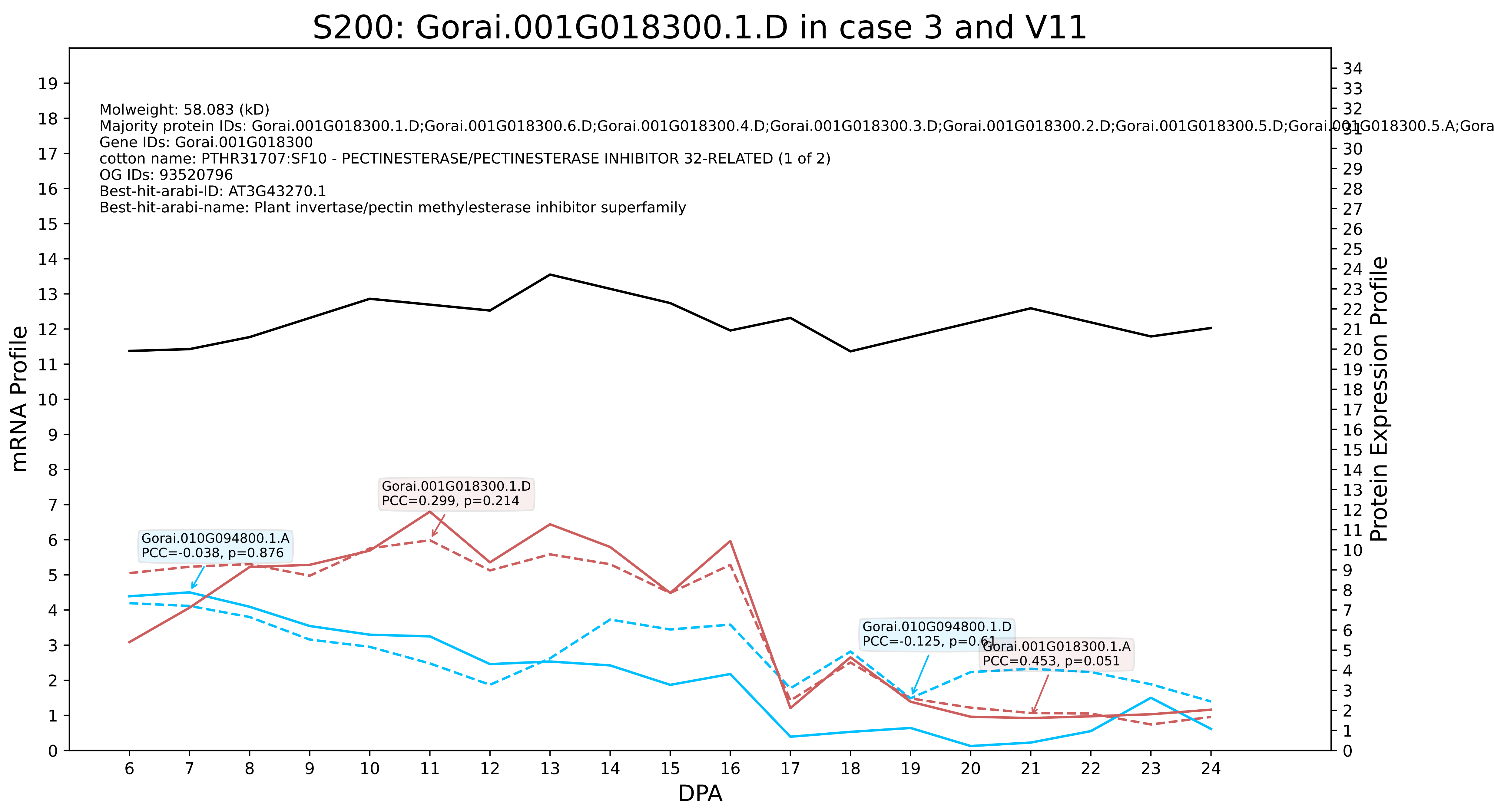
S200: Gorai.001G018300.1.D in case 3 and V11
Molweight: 58.083 (kD)
Majority protein IDs: Gorai.001G018300.1.D;Gorai.001G018300.6.D;Gorai.001G018300.4.D;Gorai.001G018300.3.D;Gorai.001G018300.2.D;Gorai.001G018300.5.D;Gorai.001G018300.5.A;Gora
Gene IDs: Gorai.001G018300
cotton name: PTHR31707:SF10 - PECTINESTERASE/PECTINESTERASE INHIBITOR 32-RELATED (1 of 2)
OG IDs: 93520796
Best-hit-arabi-ID: AT3G43270.1
Best-hit-arabi-name: Plant invertase/pectin methylesterase inhibitor superfamily
Gorai.001G018300.1.D
PCC=0.299, p=0.214
Gorai.010G094800.1.A
PCC=-0.038, p=0.876
Gorai.010G094800.1.D
PCC=-0.125, p=0.61
Gorai.001G018300.1.A
PCC=0.453, p=0.051
mRNA Profile
Protein Expression Profile
DPA

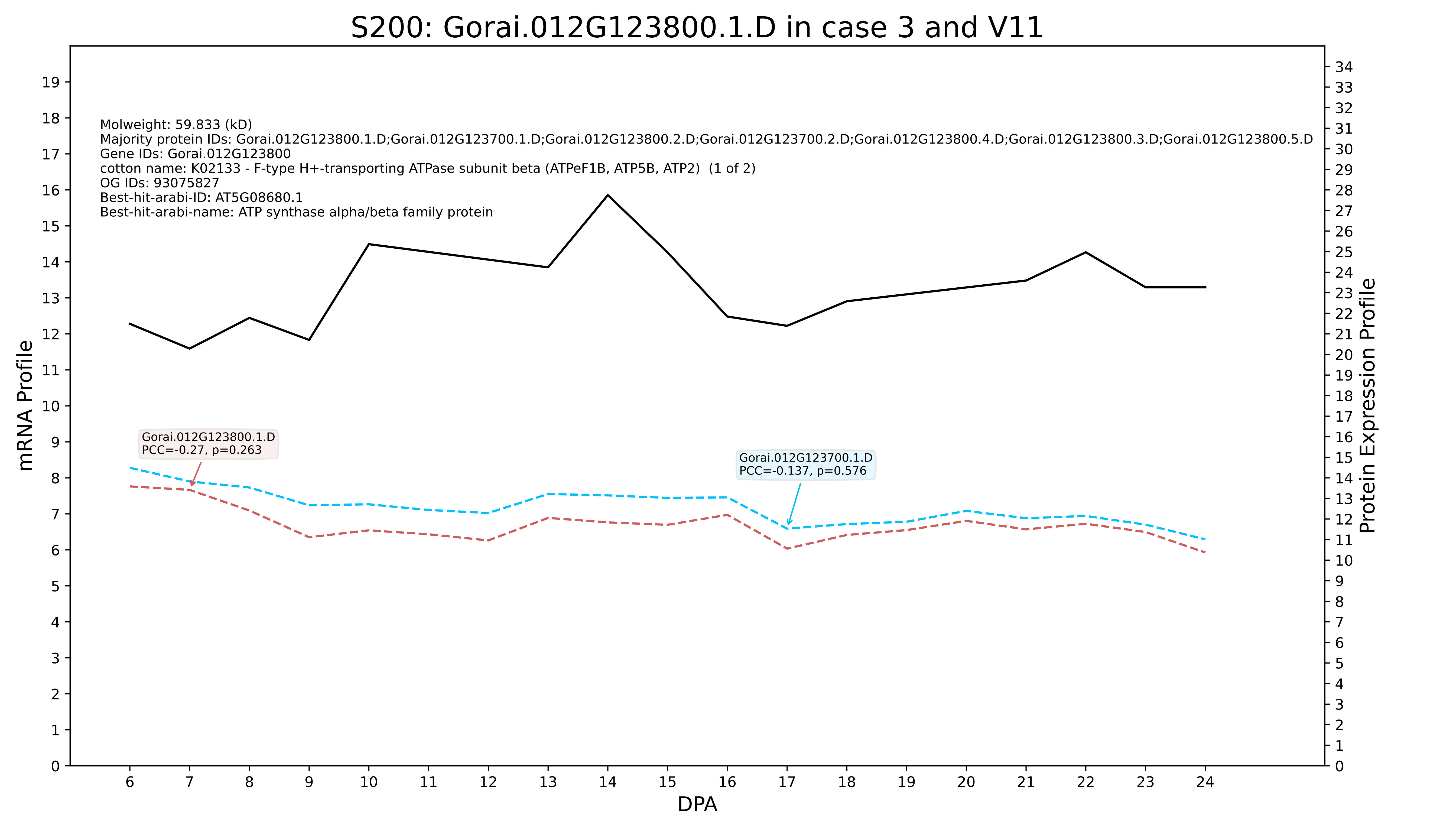
S200: Gorai.012G123800.1.D in case 3 and V11
Molweight: 59.833 (kD)
Majority protein IDs: Gorai.012G123800.1.D;Gorai.012G123700.1.D;Gorai.012G123800.2.D;Gorai.012G123700.2.D;Gorai.012G123800.4.D;Gorai.012G123800.3.D;Gorai.012G123800.5.D
Gene IDs: Gorai.012G123800
cotton name: K02133 - F-type H+-transporting ATPase subunit beta (ATPeF1B, ATP5B, ATP2) (1 of 2)
OG IDs: 93075827
Best-hit-arabi-ID: AT5G08680.1
Best-hit-arabi-name: ATP synthase alpha/beta family protein
Gorai.012G123800.1.D
PCC=-0.27, p=0.263
Gorai.012G123700.1.D
PCC=-0.137, p=0.576
mRNA Profile
Protein Expression Profile
DPA

### S200: Gorai.013G256400.1.D in case 3 and V11

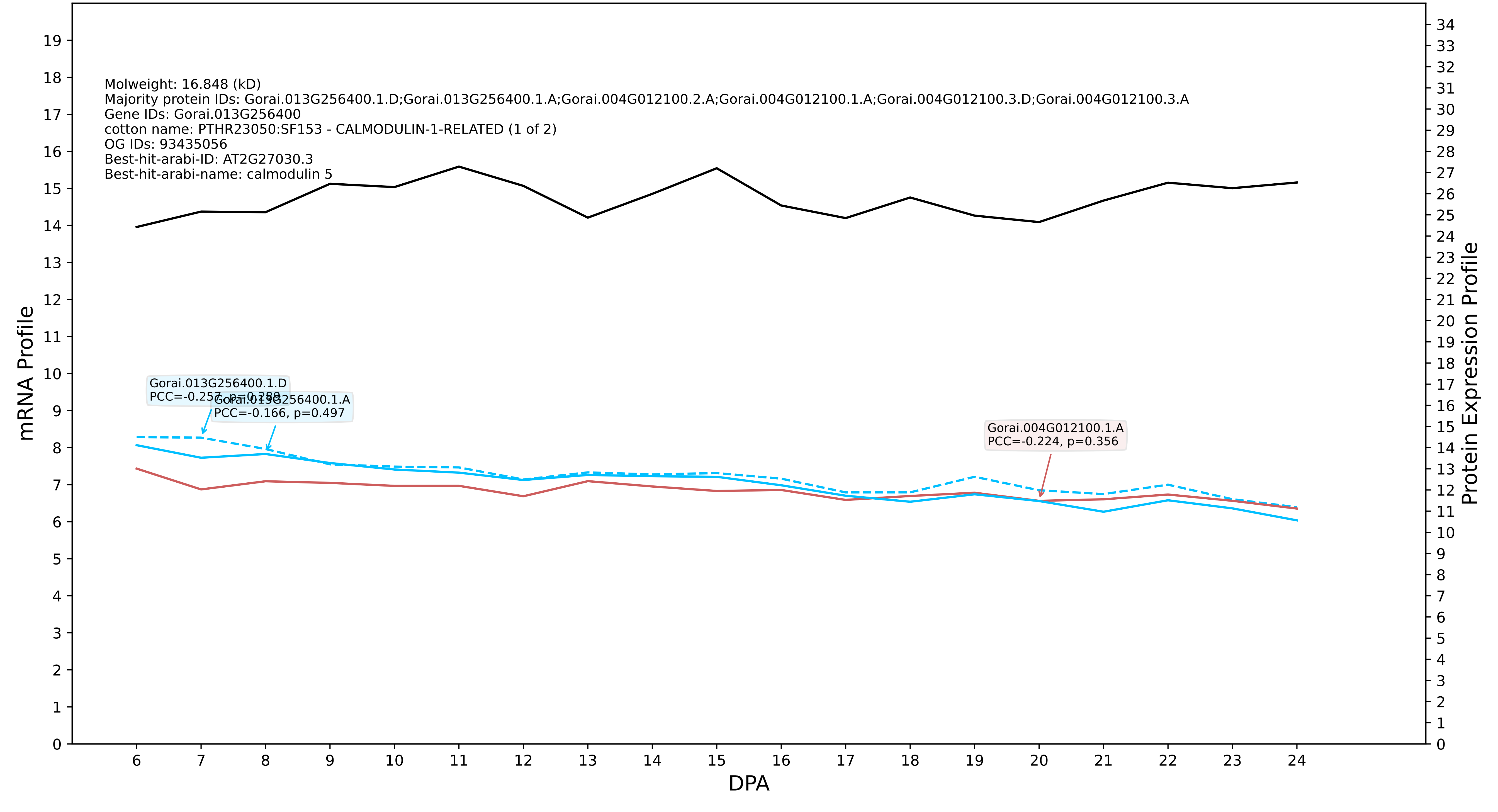

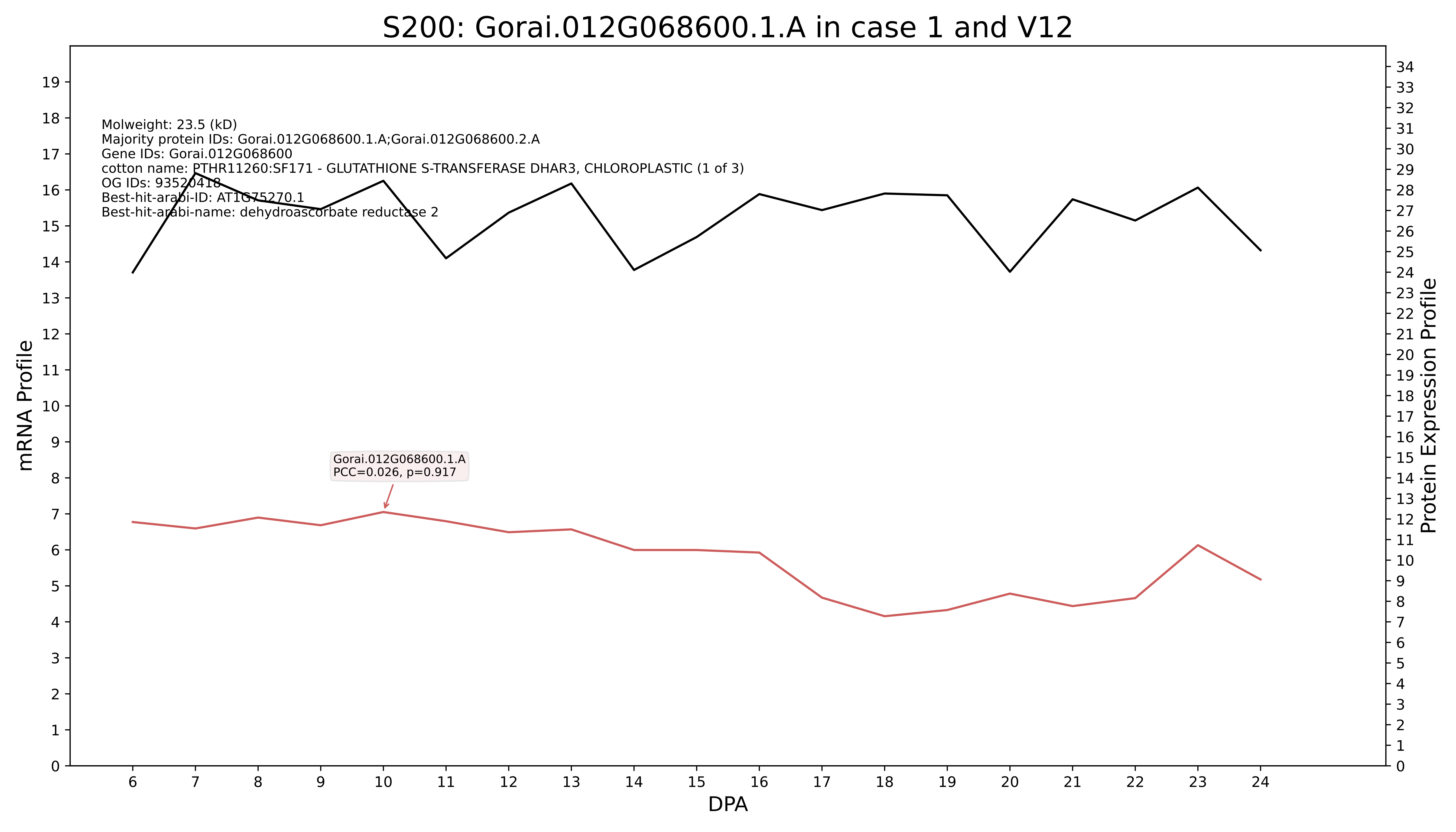

S200: Gorai.012G068600.1.A in case 1 and V12
Molweight: 23.5 (kD)
Majority protein IDs: Gorai.012G068600.1.A;Gorai.012G068600.2.A
Gene IDs: Gorai.012G068600
cotton name: PTHR11260:SF171 - GLUTATHIONE S-TRANSFERASE DHAR3, CHLOROPLASTIC (1 of 3)
OG IDs: 93520418
Best-hit-arabi-ID: AT1G75270.1
Best-hit-arabi-name: dehydroascorbate reductase 2
Gorai.012G068600.1.A
PCC=0.026, p=0.917
mRNA Profile
Protein Expression Profile
DPA

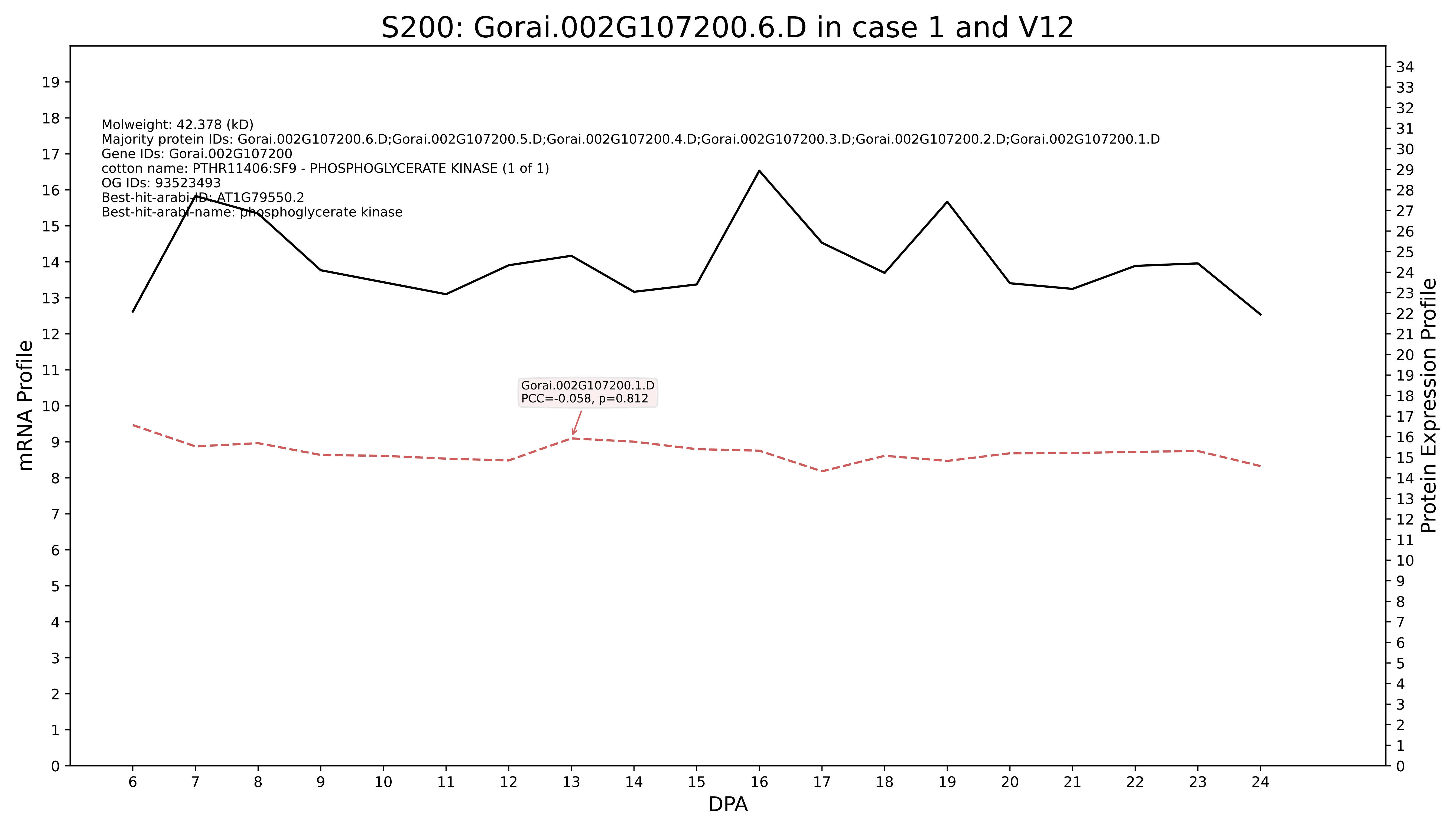
S200: Gorai.002G107200.6.D in case 1 and V12
Molweight: 42.378 (kD)
Majority protein IDs: Gorai.002G107200.6.D;Gorai.002G107200.5.D;Gorai.002G107200.4.D;Gorai.002G107200.3.D;Gorai.002G107200.2.D;Gorai.002G107200.1.D
Gene IDs: Gorai.002G107200
cotton name: PTHR11406:SF9 - PHOSPHOGLYCERATE KINASE (1 of 1)
OG IDs: 93523493
Best-hit-arabi-ID: AT1G79550.2
Best-hit-arabi-name: phosphoglycerate kinase
Gorai.002G107200.1.D
PCC=-0.058, p=0.812
mRNA Profile
Protein Expression Profile
DPA

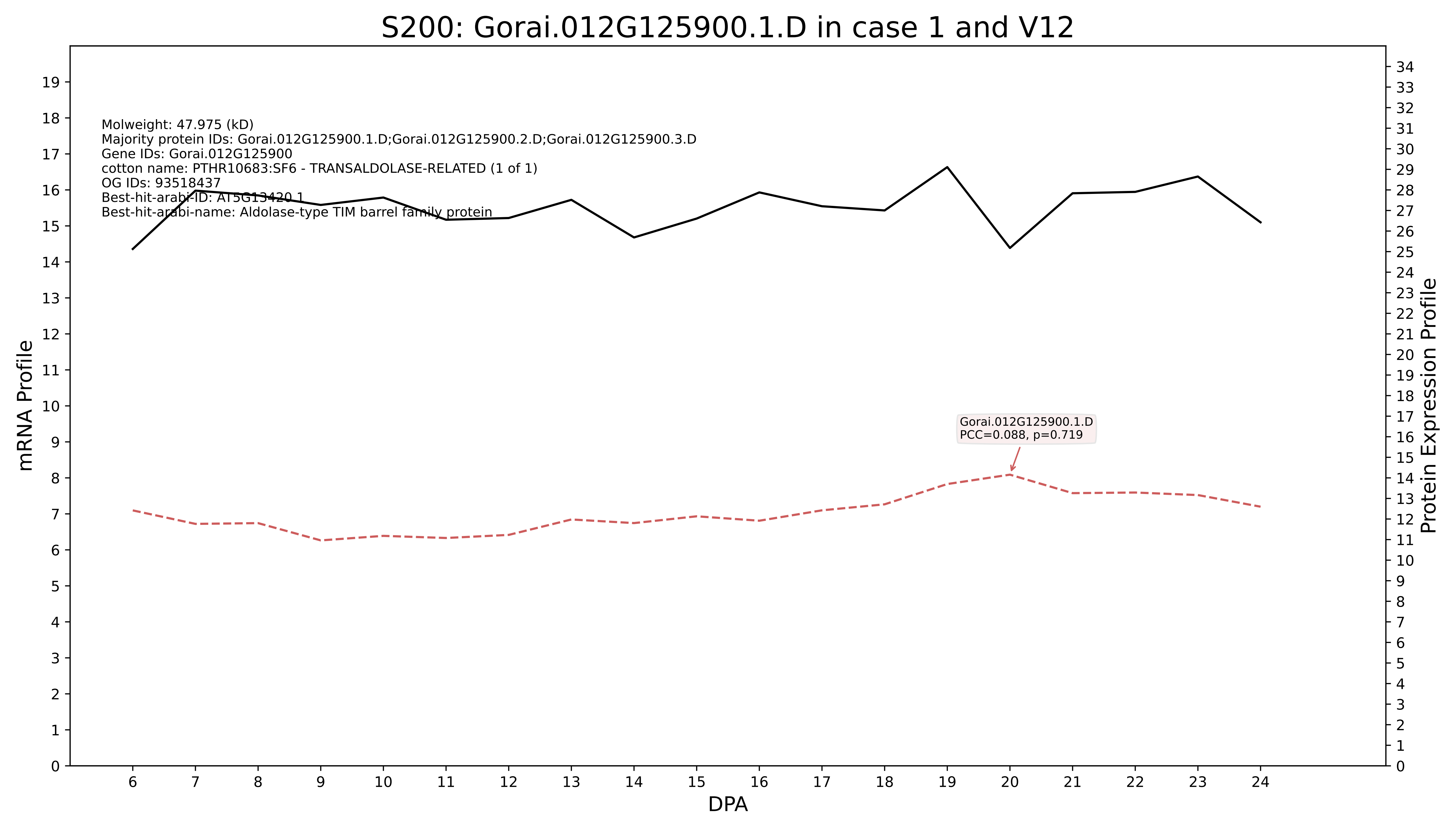
S200: Gorai.012G125900.1.D in case 1 and V12
Molweight: 47.975 (kD)
Majority protein IDs: Gorai.012G125900.1.D;Gorai.012G125900.2.D;Gorai.012G125900.3.D
Gene IDs: Gorai.012G125900
cotton name: PTHR10683:SF6 - TRANSALDOLASE-RELATED (1 of 1)
OG IDs: 93518437
Best-hit-arabi-ID: AT5G13420.1
Best-hit-arabi-name: Aldolase-type TIM barrel family protein
Gorai.012G125900.1.D
PCC=0.088, p=0.719
mRNA Profile
Protein Expression Profile
DPA

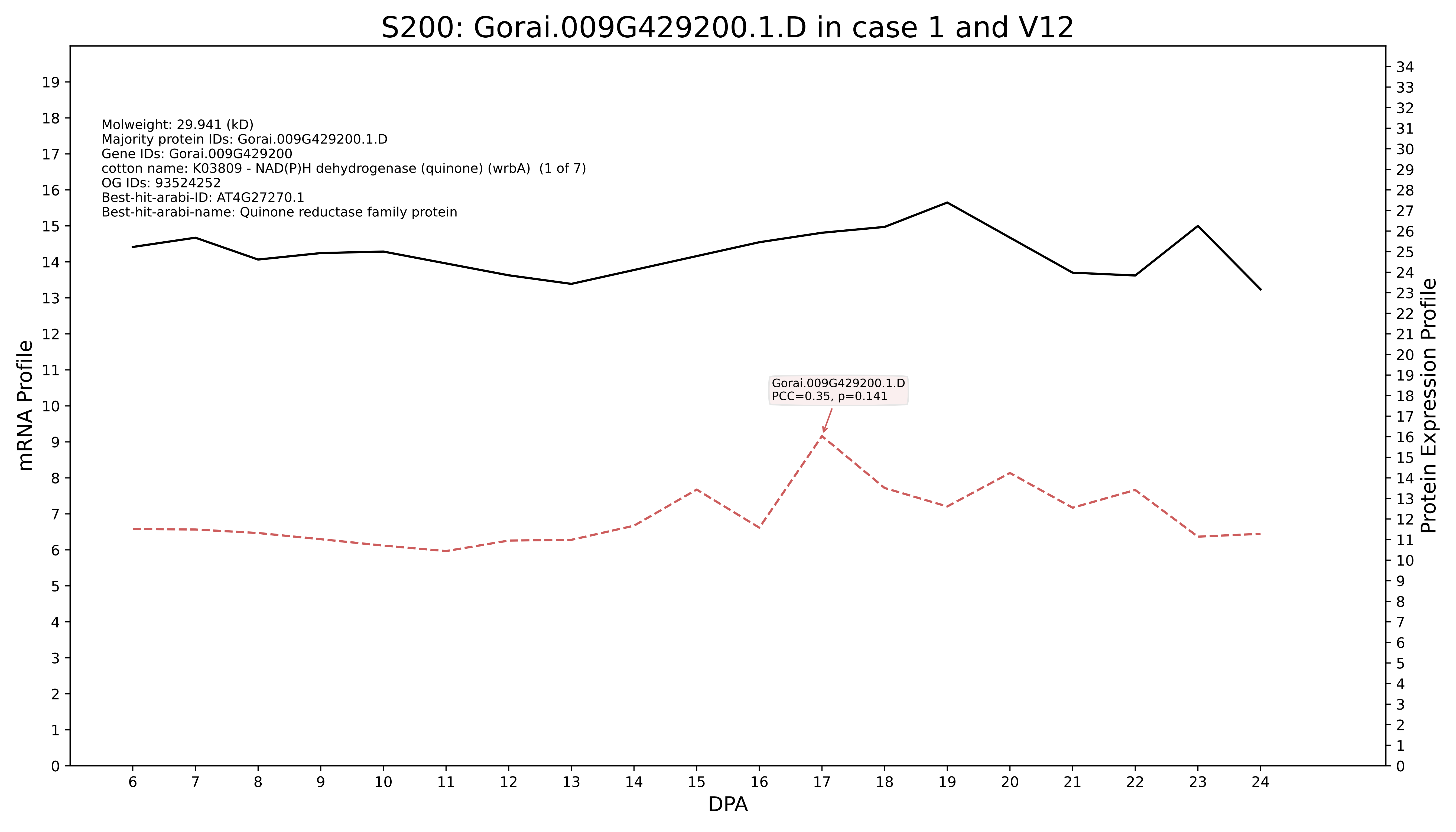

S200: Gorai.009G429200.1.D in case 1 and V12
Molweight: 29.941 (kD)
Majority protein IDs: Gorai.009G429200.1.D
Gene IDs: Gorai.009G429200
cotton name: K03809 - NAD(P)H dehydrogenase (quinone) (wrbA) (1 of 7)
OG IDs: 93524252
Best-hit-arabi-ID: AT4G27270.1
Best-hit-arabi-name: Quinone reductase family protein
Gorai.009G429200.1.D
PCC=0.35, p=0.141
mRNA Profile
Protein Expression Profile
DPA

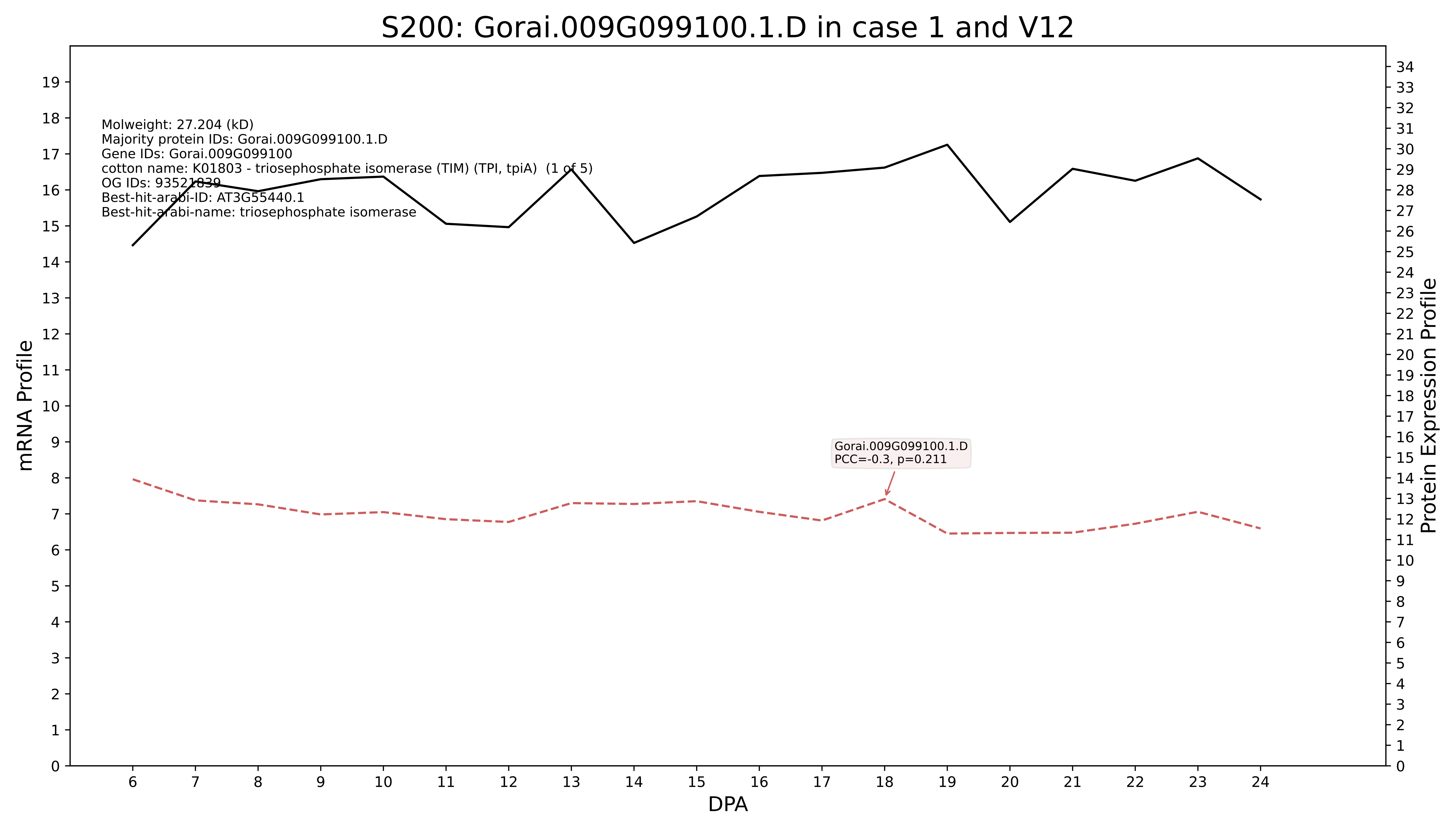
S200: Gorai.009G099100.1.D in case 1 and V12
Molweight: 27.204 (kD)
Majority protein IDs: Gorai.009G099100.1.D
Gene IDs: Gorai.009G099100
cotton name: K01803 - triosephosphate isomerase (TIM) (TPI, tpiA) (1 of 5)
OG IDs: 93521839
Best-hit-arabi-ID: AT3G55440.1
Best-hit-arabi-name: triosephosphate isomerase
Gorai.009G099100.1.D
PCC=-0.3, p=0.211
mRNA Profile
Protein Expression Profile
DPA

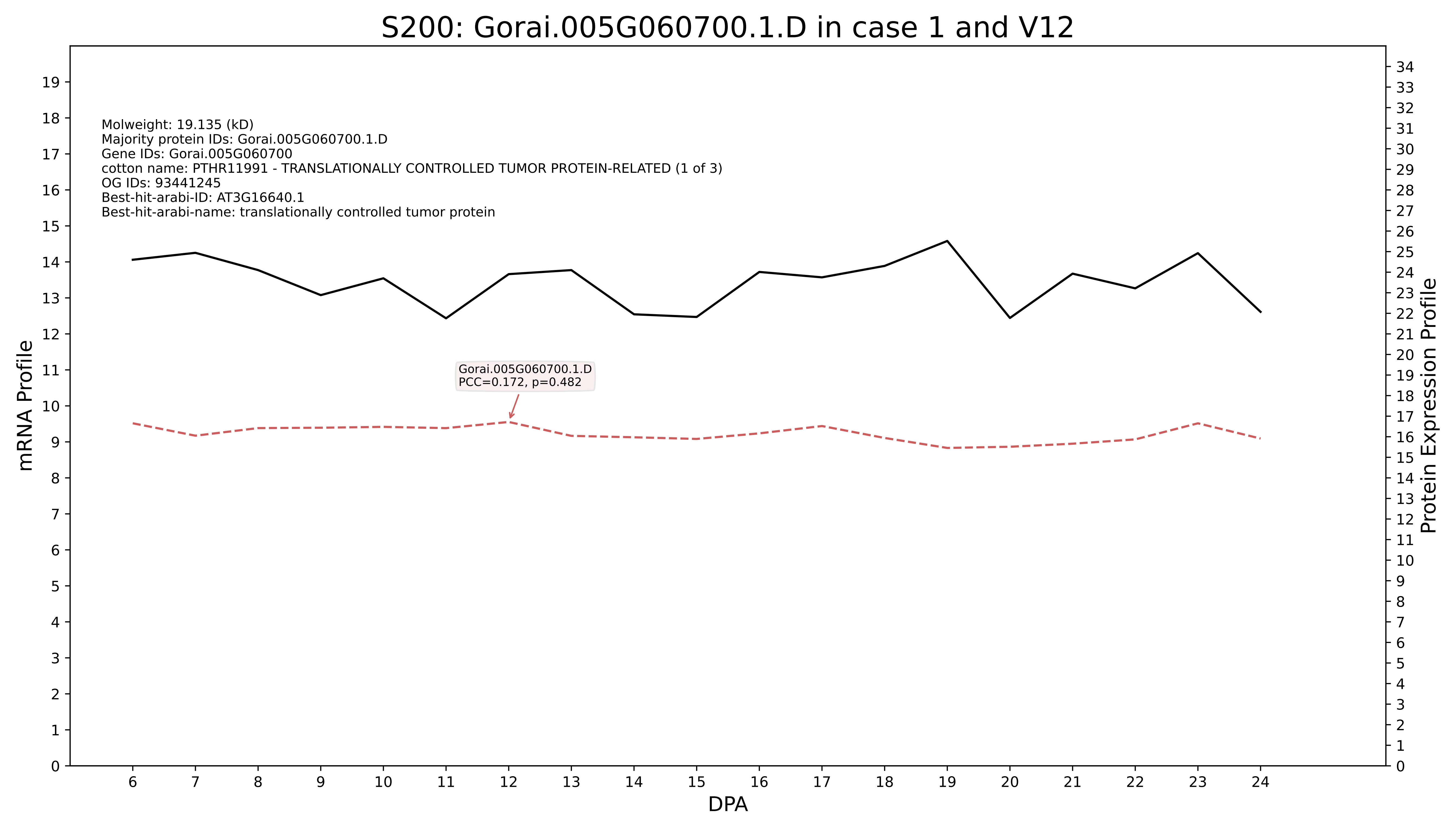

S200: Gorai.005G060700.1.D in case 1 and V12
Molweight: 19.135 (kD)
Majority protein IDs: Gorai.005G060700.1.D
Gene IDs: Gorai.005G060700
cotton name: PTHR11991 - TRANSLATIONALLY CONTROLLED TUMOR PROTEIN-RELATED (1 of 3)
OG IDs: 93441245
Best-hit-arabi-ID: AT3G16640.1
Best-hit-arabi-name: translationally controlled tumor protein
Gorai.005G060700.1.D
PCC=0.172, p=0.482
mRNA Profile
Protein Expression Profile
DPA

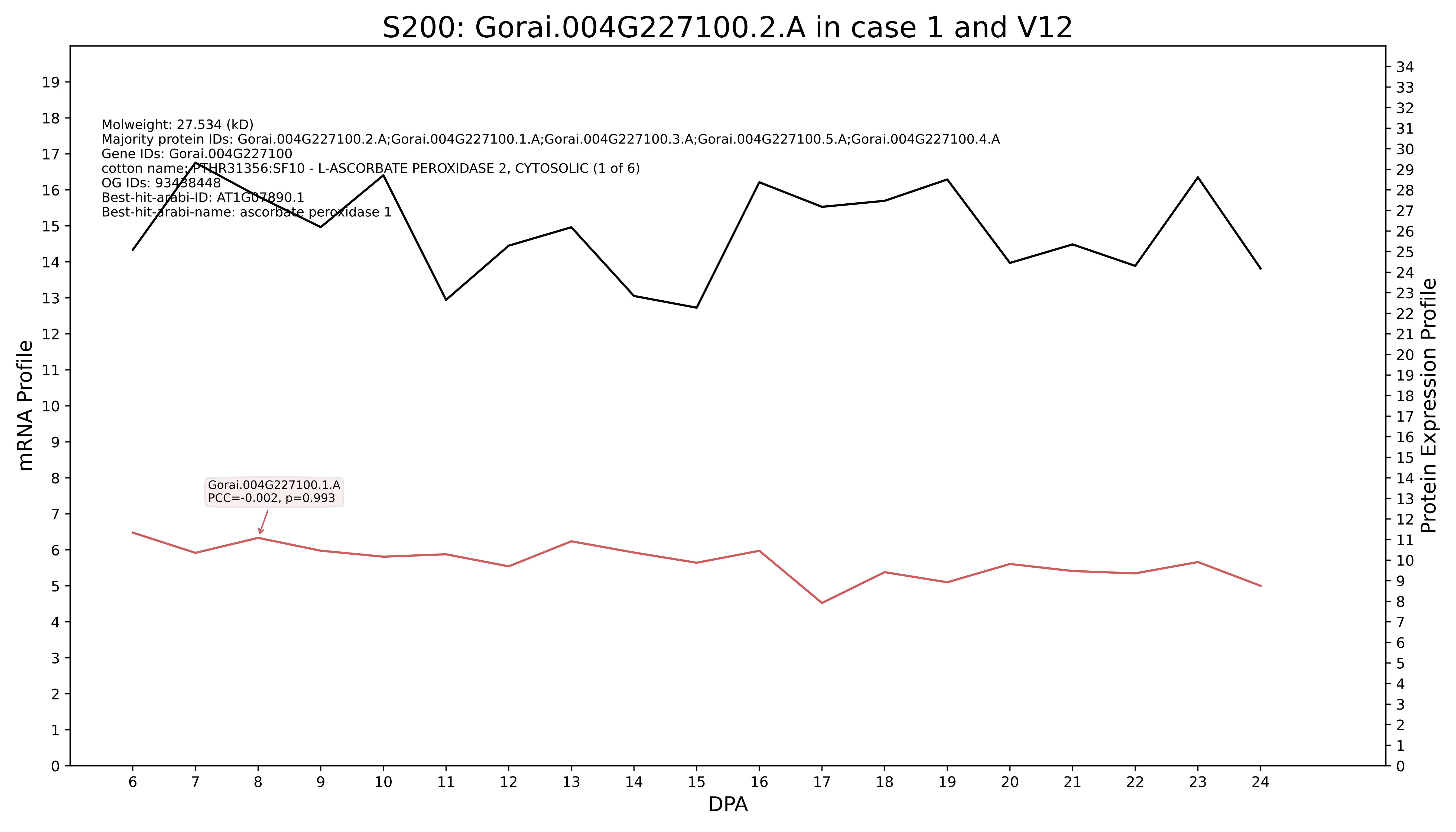
S200: Gorai.004G227100.2.A in case 1 and V12
Molweight: 27.534 (kD)
Majority protein IDs: Gorai.004G227100.2.A;Gorai.004G227100.1.A;Gorai.004G227100.3.A;Gorai.004G227100.5.A;Gorai.004G227100.4.A
Gene IDs: Gorai.004G227100
cotton name: PTHR31356:SF10 - L-ASCORBATE PEROXIDASE 2, CYTOSOLIC (1 of 6)
OG IDs: 93438448
Best-hit-arabi-ID: AT1G07890.1
Best-hit-arabi-name: ascorbate peroxidase 1
Gorai.004G227100.1.A
PCC=-0.002, p=0.993
mRNA Profile
Protein Expression Profile
DPA

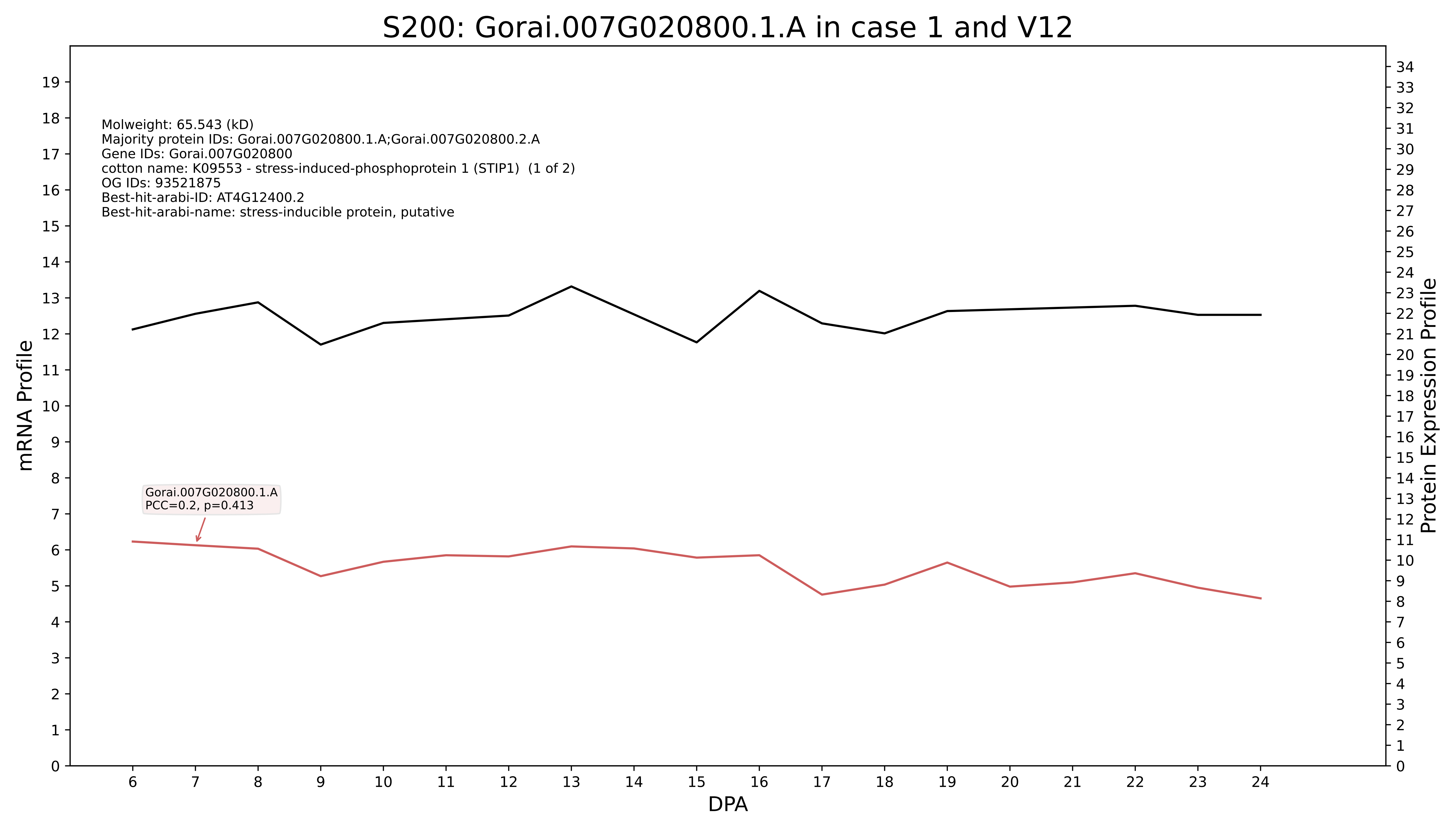
S200: Gorai.007G020800.1.A in case 1 and V12
Molweight: 65.543 (kD)
Majority protein IDs: Gorai.007G020800.1.A;Gorai.007G020800.2.A
Gene IDs: Gorai.007G020800
cotton name: K09553 - stress-induced-phosphoprotein 1 (STIP1) (1 of 2)
OG IDs: 93521875
Best-hit-arabi-ID: AT4G12400.2
Best-hit-arabi-name: stress-inducible protein, putative
Gorai.007G020800.1.A
PCC=0.2, p=0.413
mRNA Profile
Protein Expression Profile
DPA

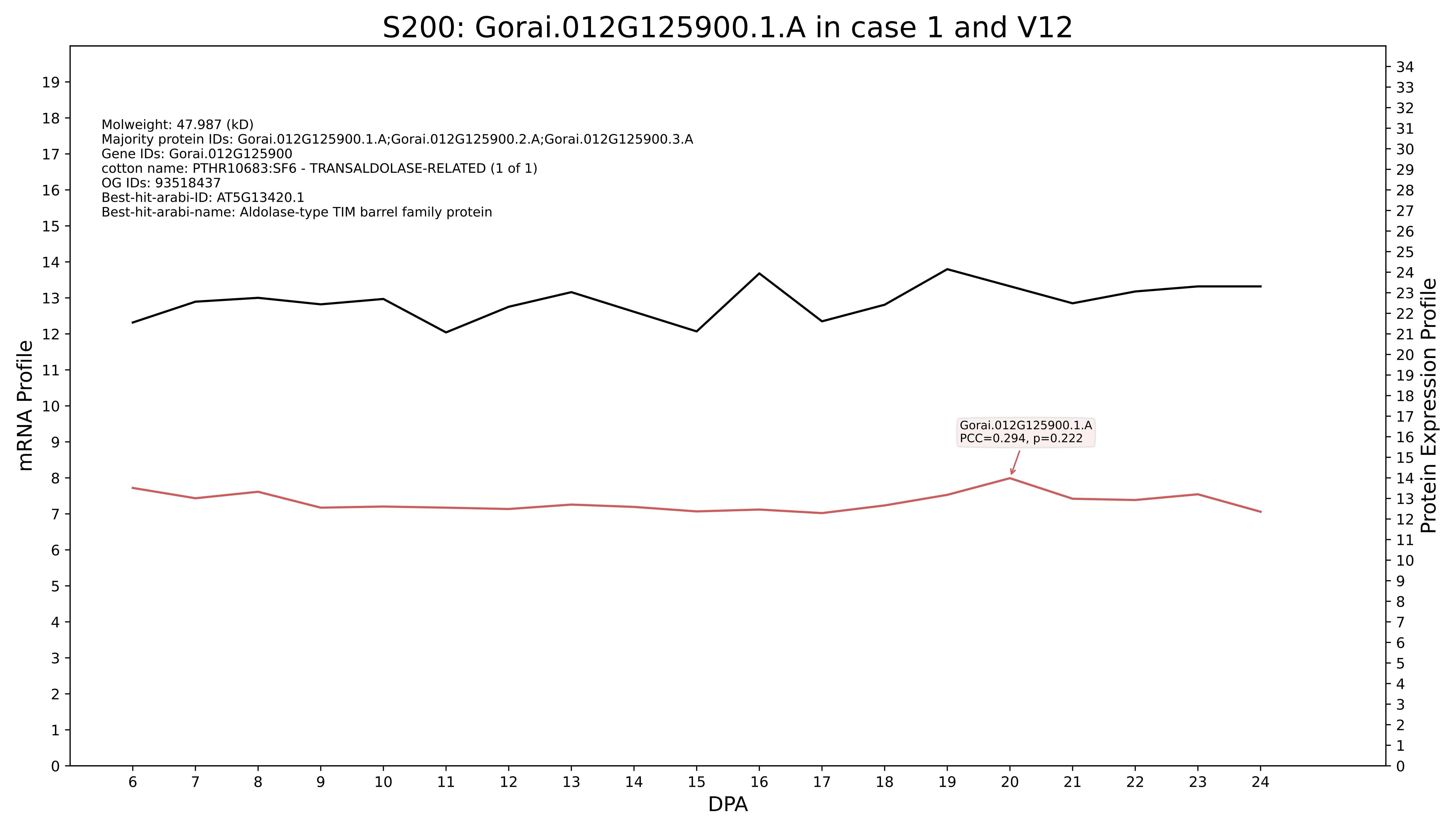
S200: Gorai.012G125900.1.A in case 1 and V12
Molweight: 47.987 (kD)
Majority protein IDs: Gorai.012G125900.1.A;Gorai.012G125900.2.A;Gorai.012G125900.3.A
Gene IDs: Gorai.012G125900
cotton name: PTHR10683:SF6 - TRANSALDOLASE-RELATED (1 of 1)
OG IDs: 93518437
Best-hit-arabi-ID: AT5G13420.1
Best-hit-arabi-name: Aldolase-type TIM barrel family protein
Gorai.012G125900.1.A
PCC=0.294, p=0.222
mRNA Profile
Protein Expression Profile
DPA

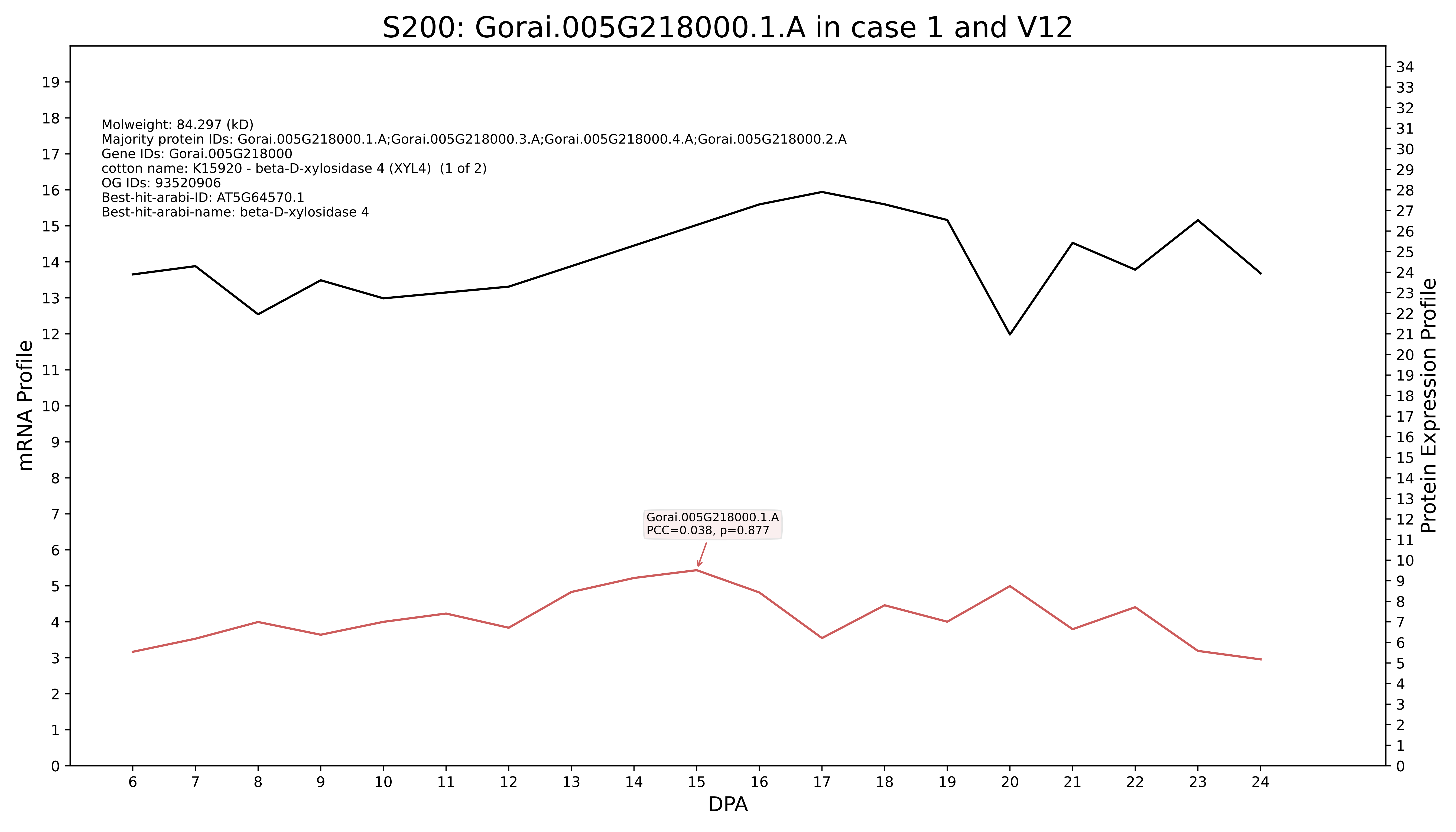

S200: Gorai.005G218000.1.A in case 1 and V12
Molweight: 84.297 (kD)
Majority protein IDs: Gorai.005G218000.1.A;Gorai.005G218000.3.A;Gorai.005G218000.4.A;Gorai.005G218000.2.A
Gene IDs: Gorai.005G218000
cotton name: K15920 - beta-D-xylosidase 4 (XYL4) (1 of 2)
OG IDs: 93520906
Best-hit-arabi-ID: AT5G64570.1
Best-hit-arabi-name: beta-D-xylosidase 4
Gorai.005G218000.1.A
PCC=0.038, p=0.877
mRNA Profile
Protein Expression Profile
DPA

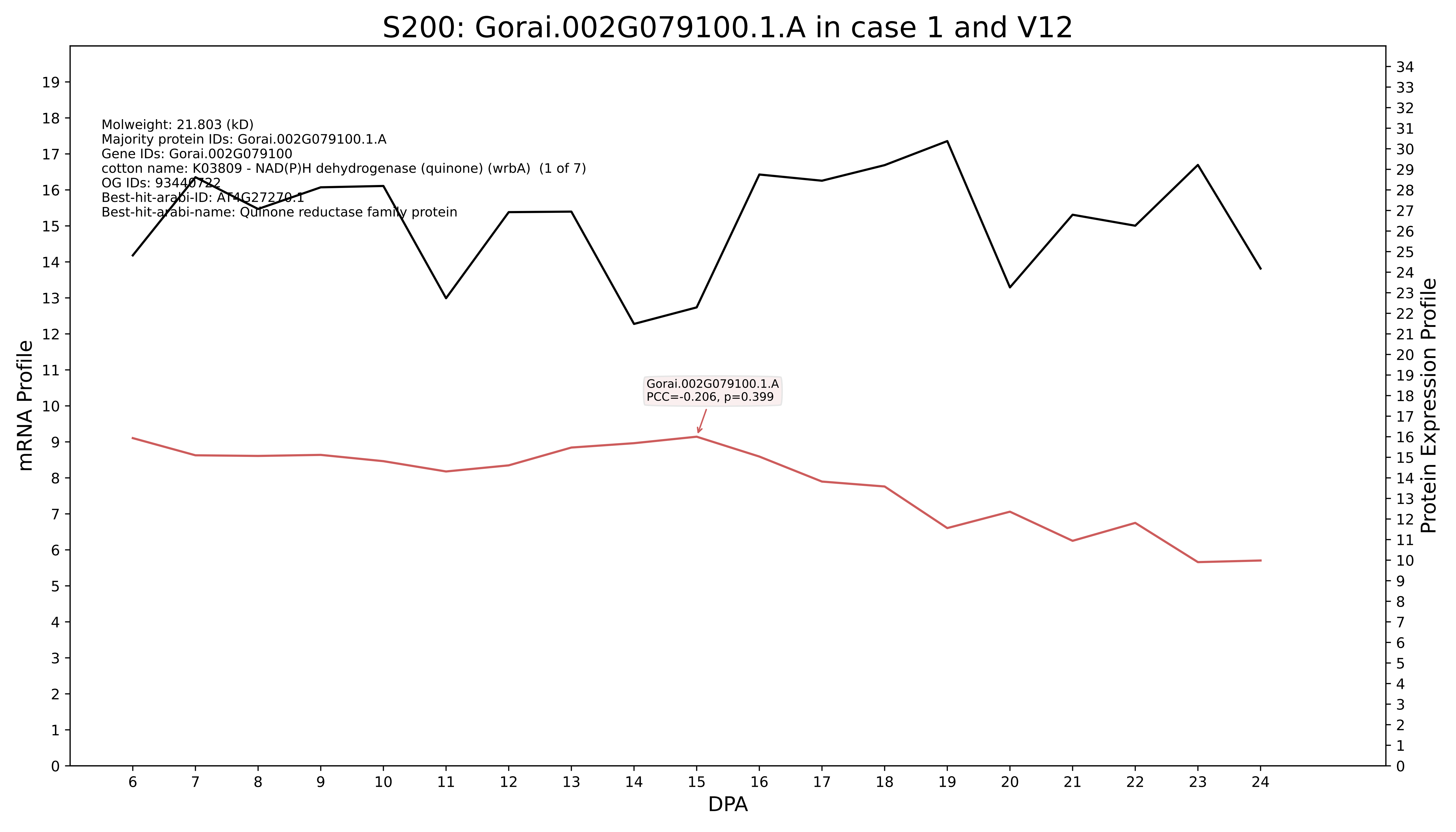
S200: Gorai.002G079100.1.A in case 1 and V12
Molweight: 21.803 (kD)
Majority protein IDs: Gorai.002G079100.1.A
Gene IDs: Gorai.002G079100
cotton name: K03809 - NAD(P)H dehydrogenase (quinone) (wrbA) (1 of 7)
OG IDs: 93440722
Best-hit-arabi-ID: AT4G27270.1
Best-hit-arabi-name: Quinone reductase family protein
Gorai.002G079100.1.A
PCC=-0.206, p=0.399
mRNA Profile
Protein Expression Profile
DPA

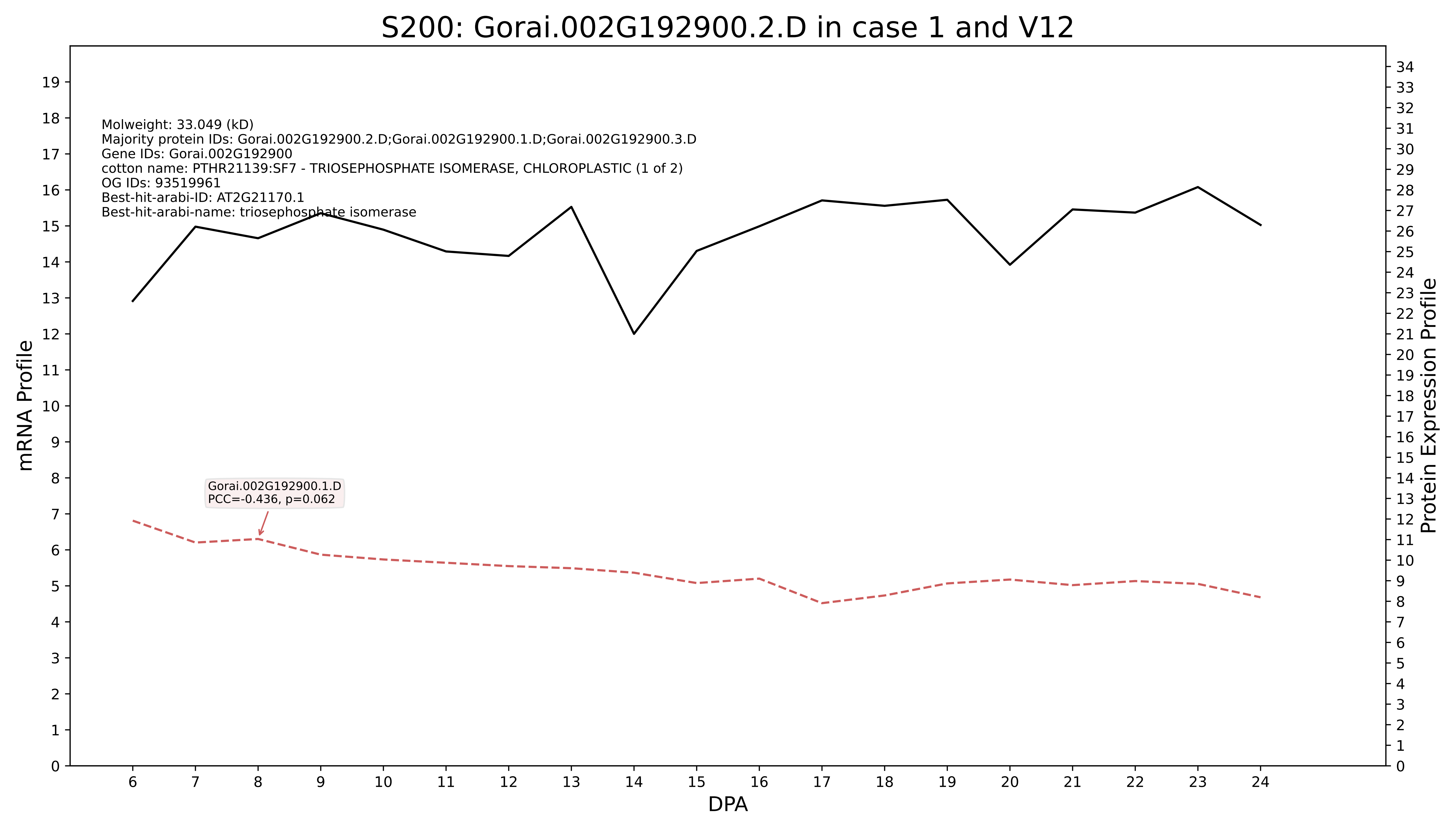

S200: Gorai.002G192900.2.D in case 1 and V12
Molweight: 33.049 (kD)
Majority protein IDs: Gorai.002G192900.2.D;Gorai.002G192900.1.D;Gorai.002G192900.3.D
Gene IDs: Gorai.002G192900
cotton name: PTHR21139:SF7 - TRIOSEPHOSPHATE ISOMERASE, CHLOROPLASTIC (1 of 2)
OG IDs: 93519961
Best-hit-arabi-ID: AT2G21170.1
Best-hit-arabi-name: triosephosphate isomerase
Gorai.002G192900.1.D
PCC=-0.436, p=0.062
mRNA Profile
Protein Expression Profile
DPA

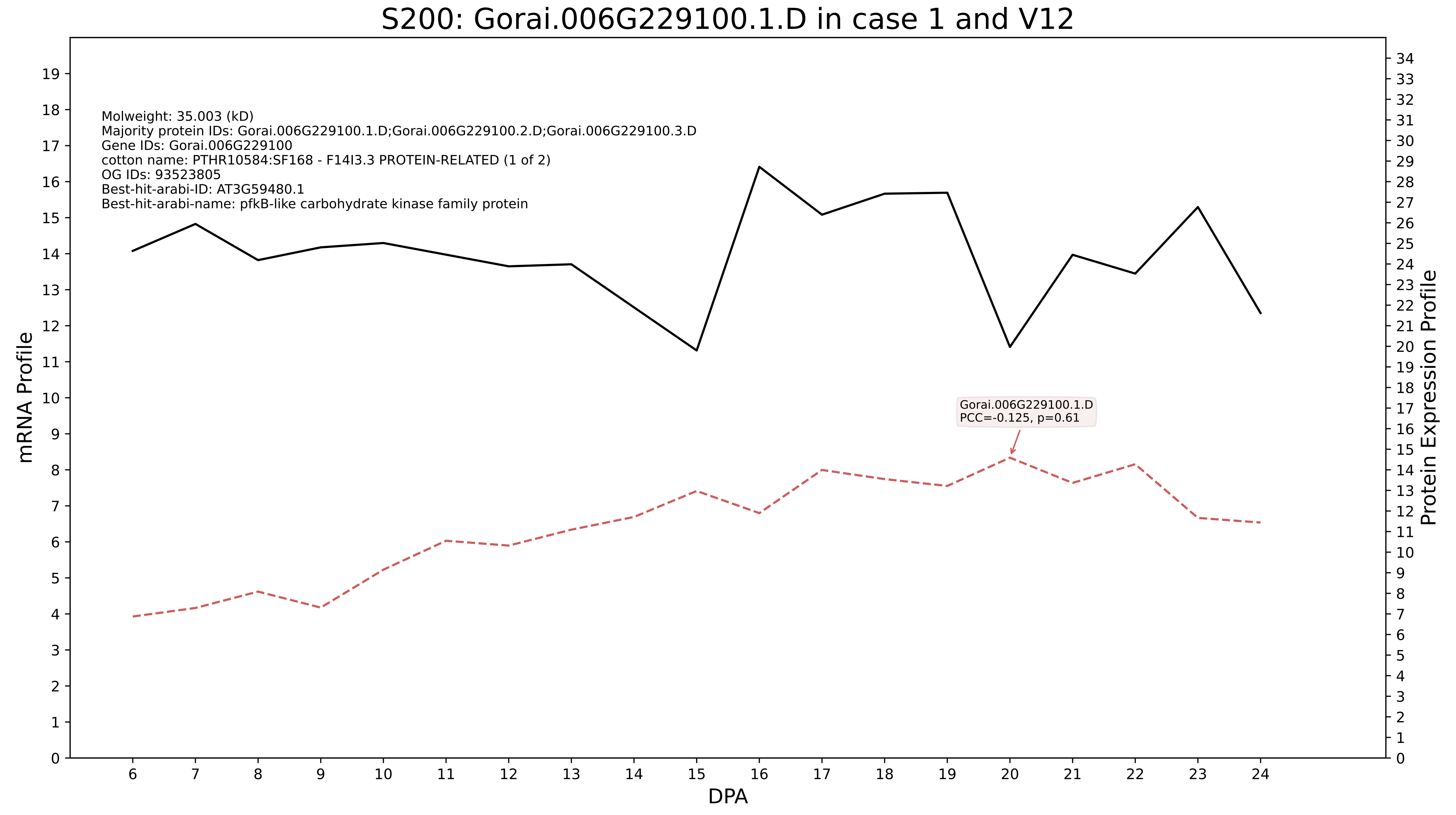
S200: Gorai.006G229100.1.D in case 1 and V12
Molweight: 35.003 (kD)
Majority protein IDs: Gorai.006G229100.1.D;Gorai.006G229100.2.D;Gorai.006G229100.3.D
Gene IDs: Gorai.006G229100
cotton name: PTHR10584:SF168 - F14I3.3 PROTEIN-RELATED (1 of 2)
OG IDs: 93523805
Best-hit-arabi-ID: AT3G59480.1
Best-hit-arabi-name: pfkB-like carbohydrate kinase family protein
Gorai.006G229100.1.D
PCC=-0.125, p=0.61
mRNA Profile
Protein Expression Profile
DPA

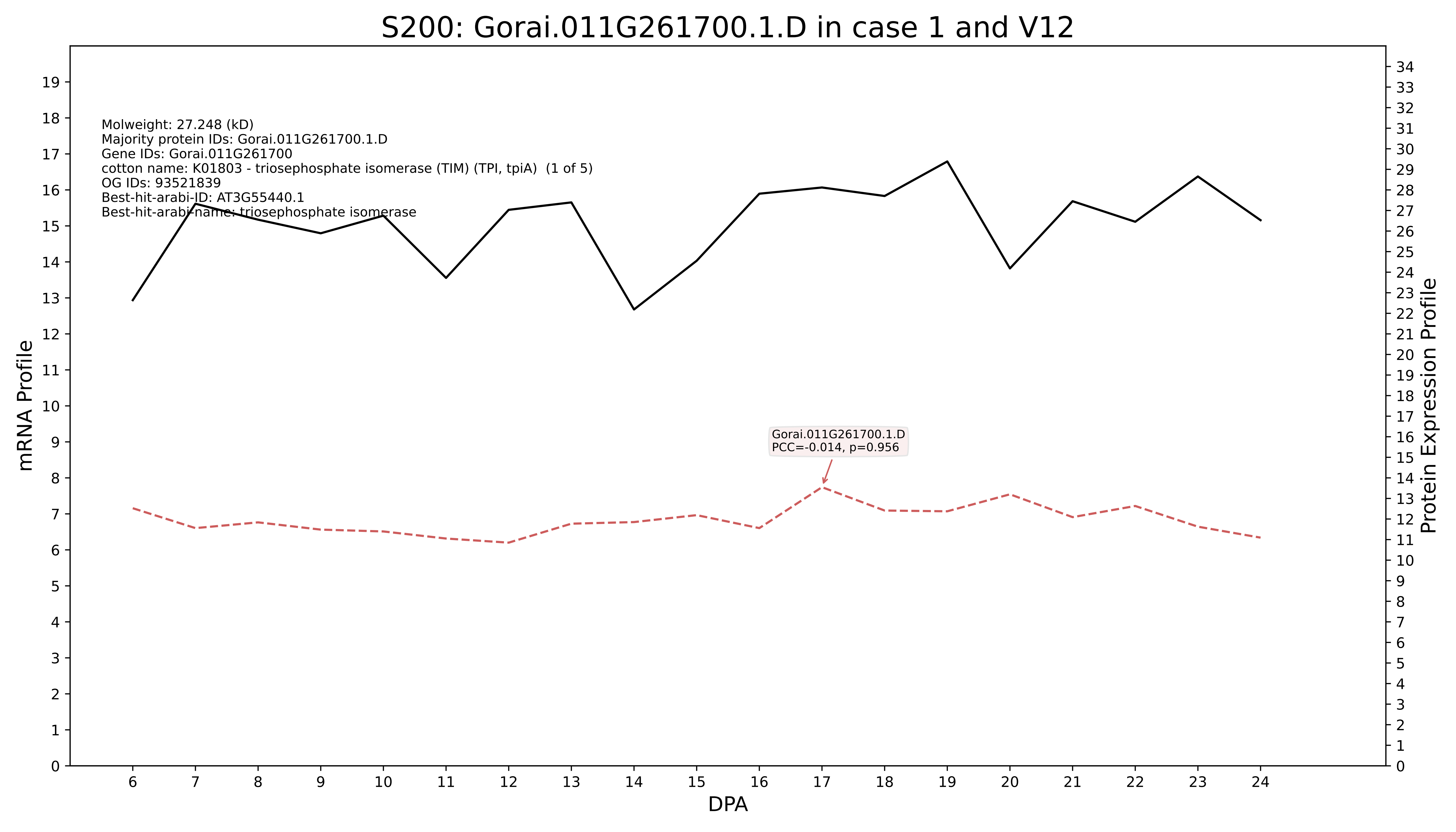
S200: Gorai.011G261700.1.D in case 1 and V12
Molweight: 27.248 (kD)
Majority protein IDs: Gorai.011G261700.1.D
Gene IDs: Gorai.011G261700
cotton name: K01803 - triosephosphate isomerase (TIM) (TPI, tpiA) (1 of 5)
OG IDs: 93521839
Best-hit-arabi-ID: AT3G55440.1
Best-hit-arabi-name: triosephosphate isomerase
Gorai.011G261700.1.D
PCC=-0.014, p=0.956
mRNA Profile
Protein Expression Profile
DPA

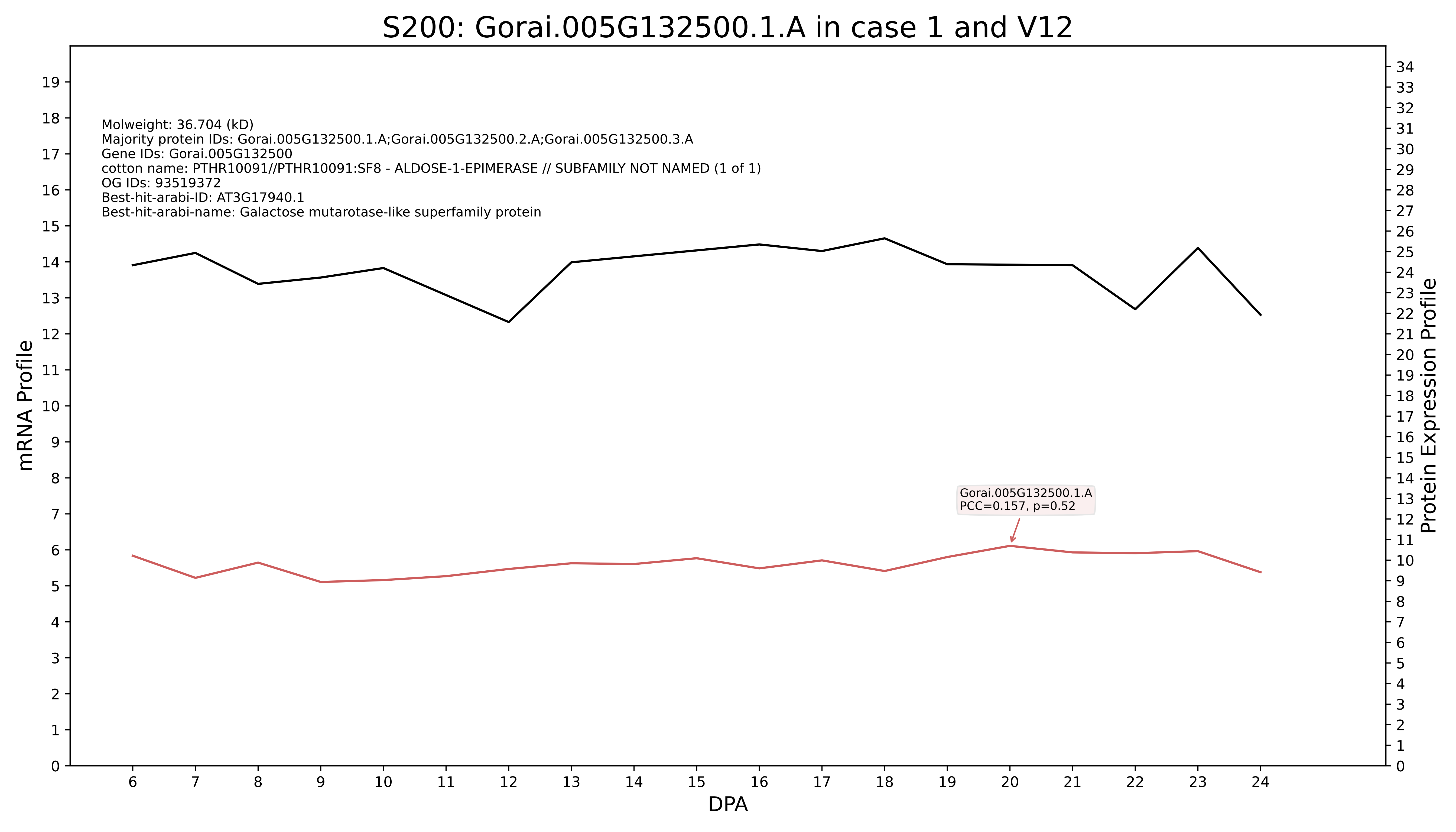

S200: Gorai.005G132500.1.A in case 1 and V12
Molweight: 36.704 (kD)
Majority protein IDs: Gorai.005G132500.1.A;Gorai.005G132500.2.A;Gorai.005G132500.3.A
Gene IDs: Gorai.005G132500
cotton name: PTHR10091//PTHR10091:SF8 - ALDOSE-1-EPIMERASE // SUBFAMILY NOT NAMED (1 of 1)
OG IDs: 93519372
Best-hit-arabi-ID: AT3G17940.1
Best-hit-arabi-name: Galactose mutarotase-like superfamily protein
Gorai.005G132500.1.A
PCC=0.157, p=0.52
mRNA Profile
Protein Expression Profile
DPA

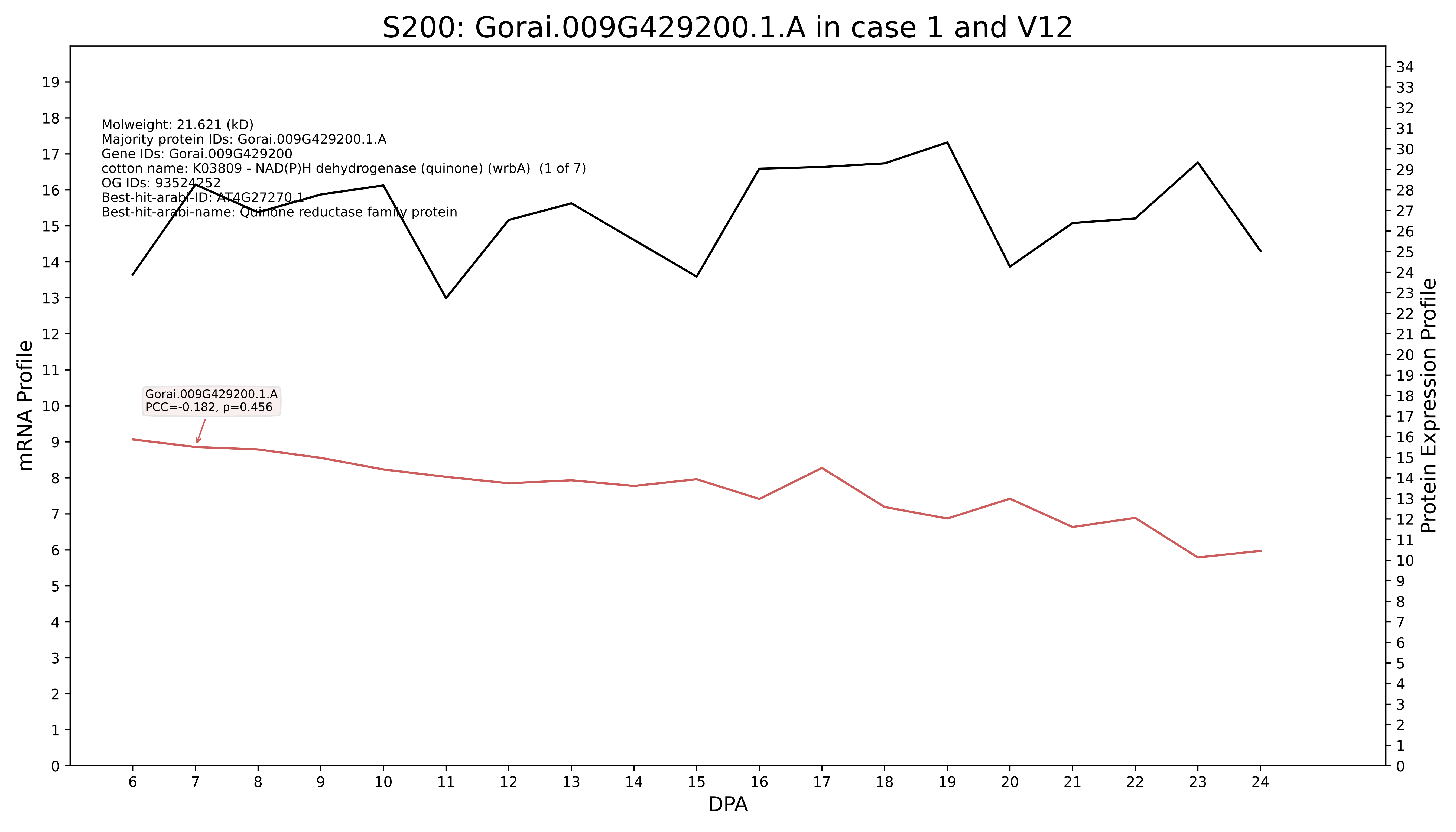
S200: Gorai.009G429200.1.A in case 1 and V12
Molweight: 21.621 (kD)
Majority protein IDs: Gorai.009G429200.1.A
Gene IDs: Gorai.009G429200
cotton name: K03809 - NAD(P)H dehydrogenase (quinone) (wrbA) (1 of 7)
OG IDs: 93524252
Best-hit-arabi-ID: AT4G27270.1
Best-hit-arabi-name: Quinone reductase family protein
Gorai.009G429200.1.A
PCC=-0.182, p=0.456
mRNA Profile
Protein Expression Profile
DPA

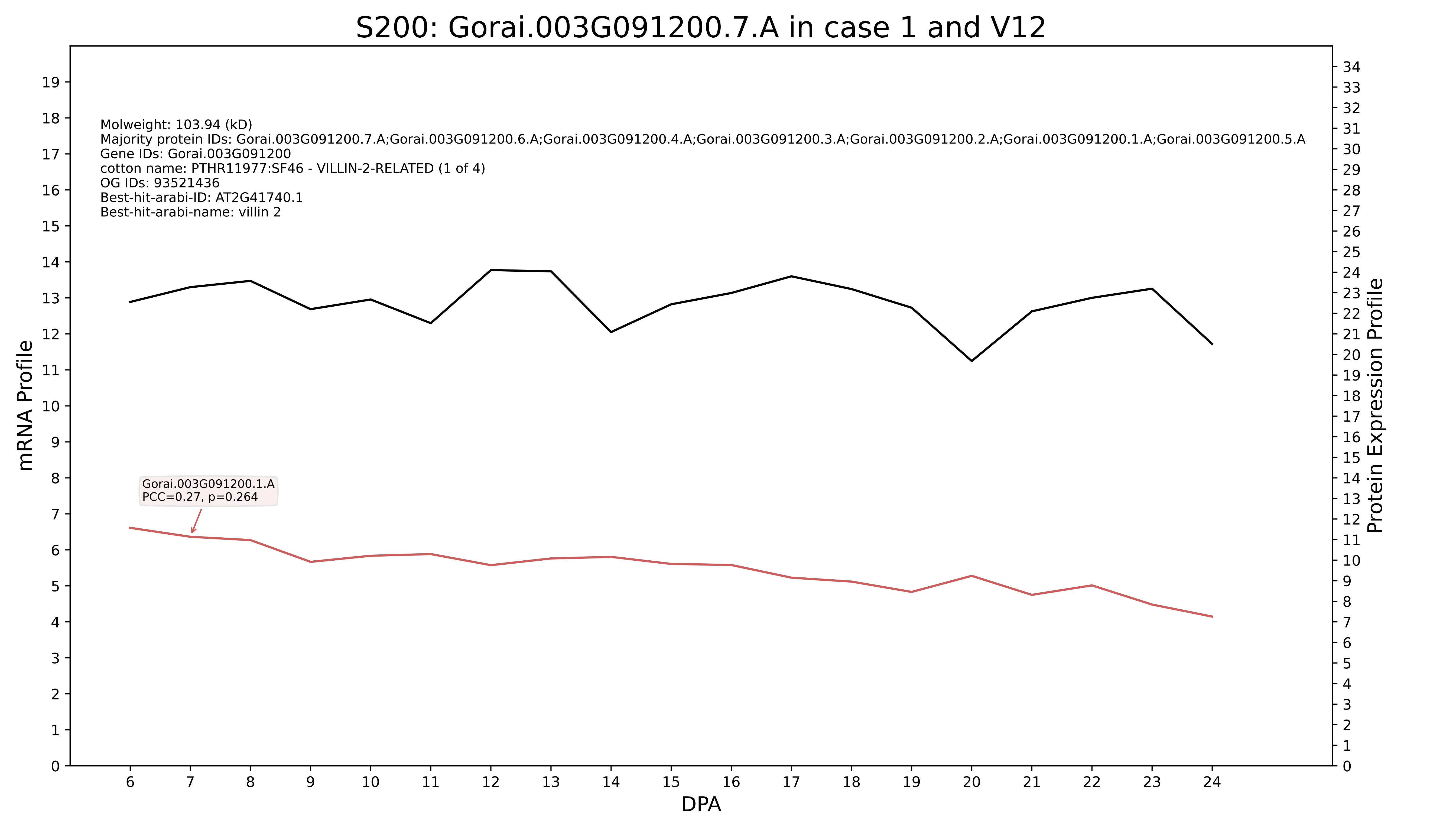
S200: Gorai.003G091200.7.A in case 1 and V12
Molweight: 103.94 (kD)
Majority protein IDs: Gorai.003G091200.7.A;Gorai.003G091200.6.A;Gorai.003G091200.4.A;Gorai.003G091200.3.A;Gorai.003G091200.2.A;Gorai.003G091200.1.A;Gorai.003G091200.5.A
Gene IDs: Gorai.003G091200
cotton name: PTHR11977:SF46 - VILLIN-2-RELATED (1 of 4)
OG IDs: 93521436
Best-hit-arabi-ID: AT2G41740.1
Best-hit-arabi-name: villin 2
Gorai.003G091200.1.A
PCC=0.27, p=0.264
mRNA Profile
Protein Expression Profile
DPA

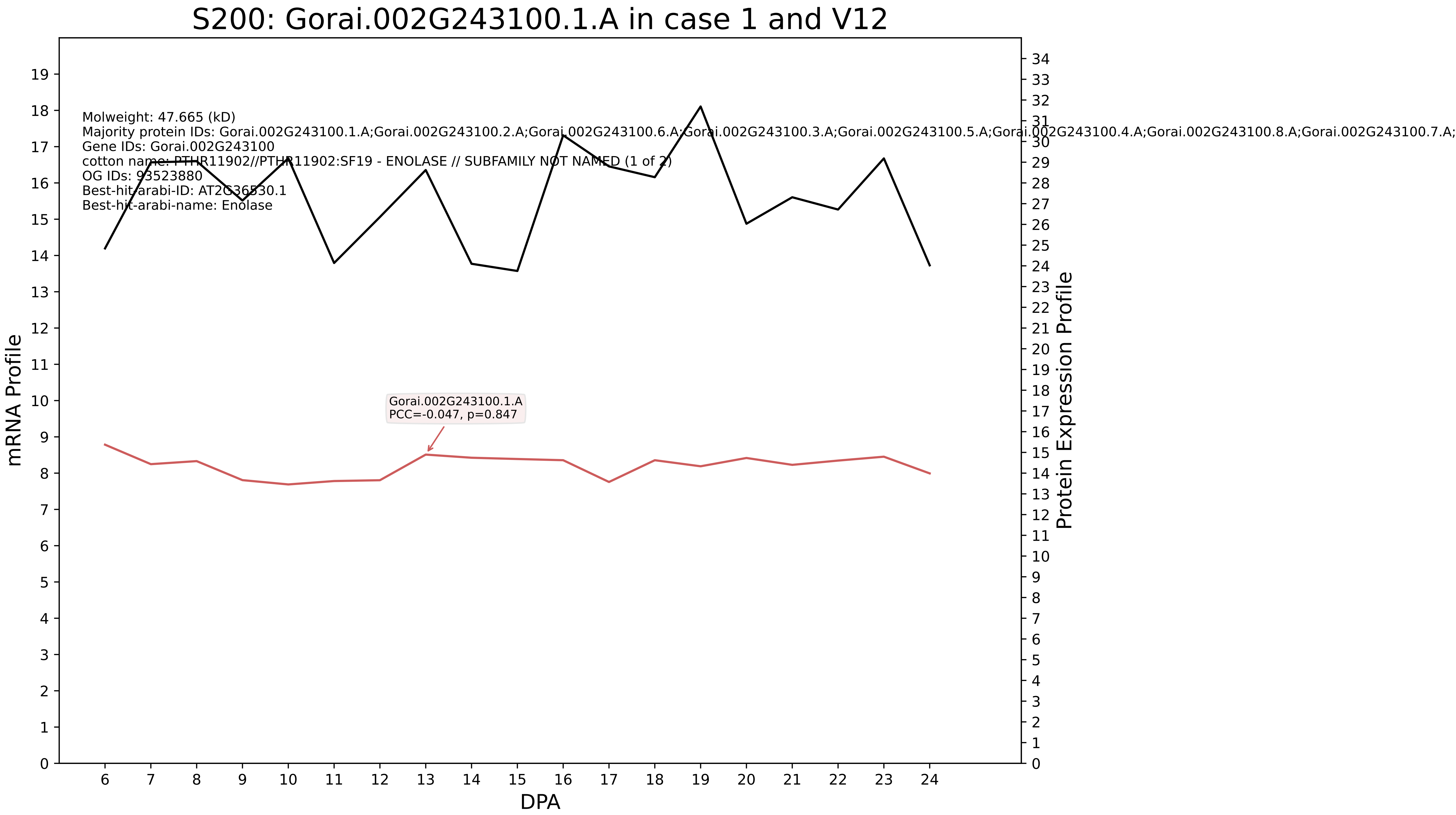
S200: Gorai.002G243100.1.A in case 1 and V12
Molweight: 47.665 (kD)
Majority protein IDs: Gorai.002G243100.1.A;Gorai.002G243100.2.A;Gorai.002G243100.6.A;Gorai.002G243100.3.A;Gorai.002G243100.5.A;Gorai.002G243100.4.A;Gorai.002G243100.8.A;Gorai.002G243100.7.A;
Gene IDs: Gorai.002G243100
cotton name: PTHR11902//PTHR11902:SF19 - ENOLASE // SUBFAMILY NOT NAMED (1 of 2)
OG IDs: 93523880
Best-hit-arabi-ID: AT2G36530.1
Best-hit-arabi-name: Enolase
Gorai.002G243100.1.A
PCC=-0.047, p=0.847
mRNA Profile
Protein Expression Profile
DPA

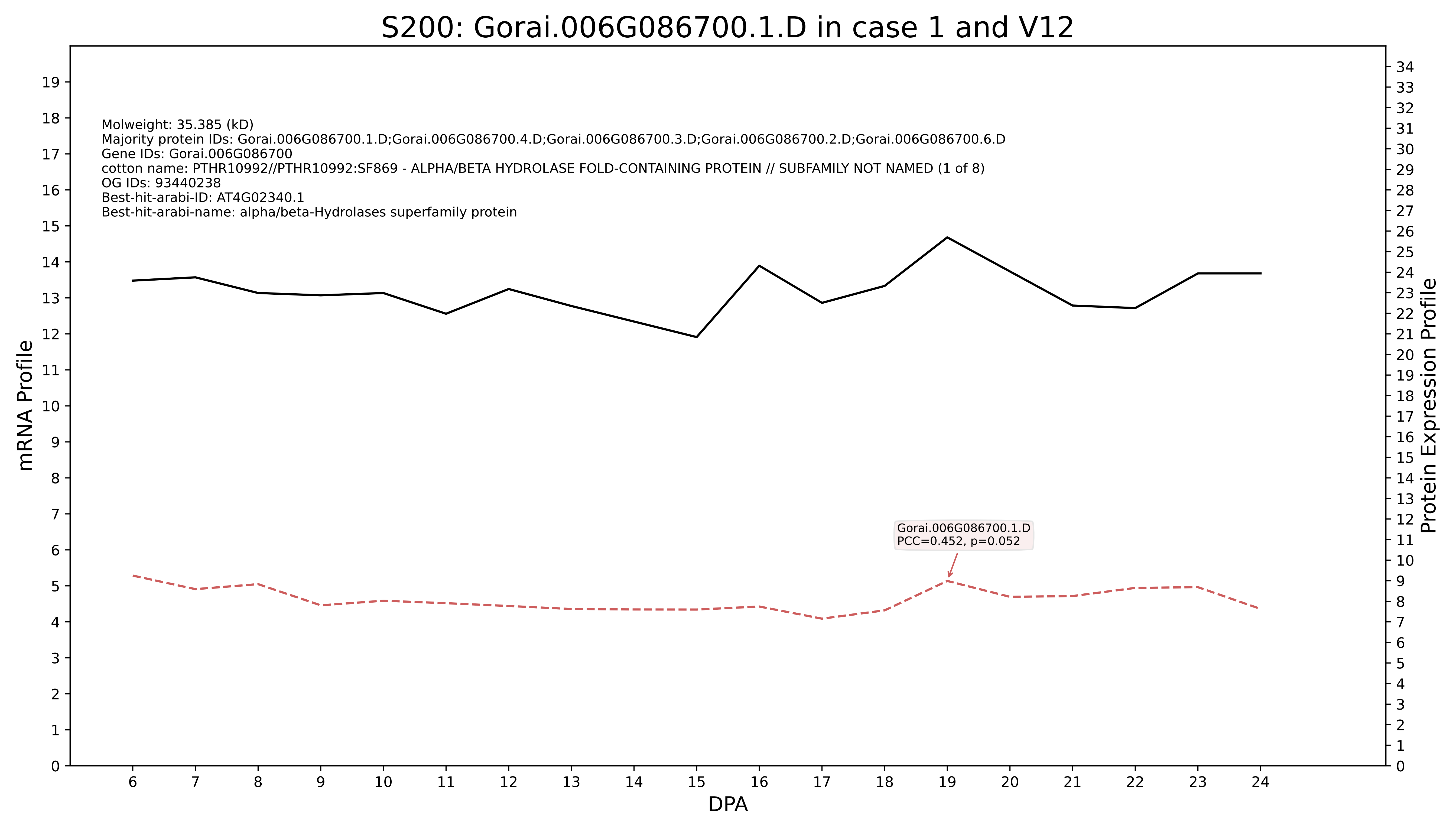
S200: Gorai.006G086700.1.D in case 1 and V12
Molweight: 35.385 (kD)
Majority protein IDs: Gorai.006G086700.1.D;Gorai.006G086700.4.D;Gorai.006G086700.3.D;Gorai.006G086700.2.D;Gorai.006G086700.6.D
Gene IDs: Gorai.006G086700
cotton name: PTHR10992//PTHR10992:SF869 - ALPHA/BETA HYDROLASE FOLD-CONTAINING PROTEIN // SUBFAMILY NOT NAMED (1 of 8)
OG IDs: 93440238
Best-hit-arabi-ID: AT4G02340.1
Best-hit-arabi-name: alpha/beta-Hydrolases superfamily protein
Gorai.006G086700.1.D
PCC=0.452, p=0.052
mRNA Profile
Protein Expression Profile
DPA

### S200: Gorai.001G083800.1.D in case 1 and V12

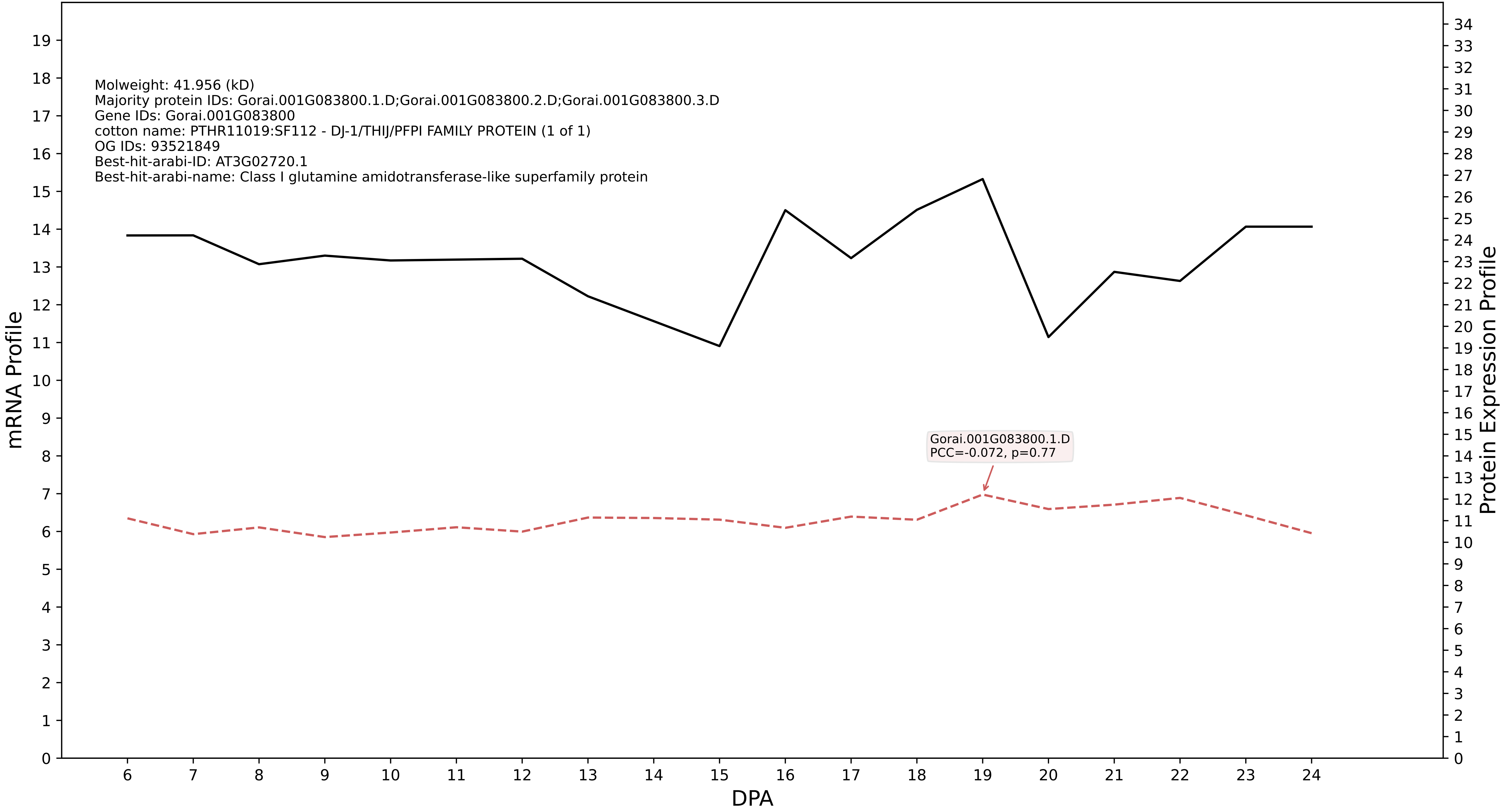

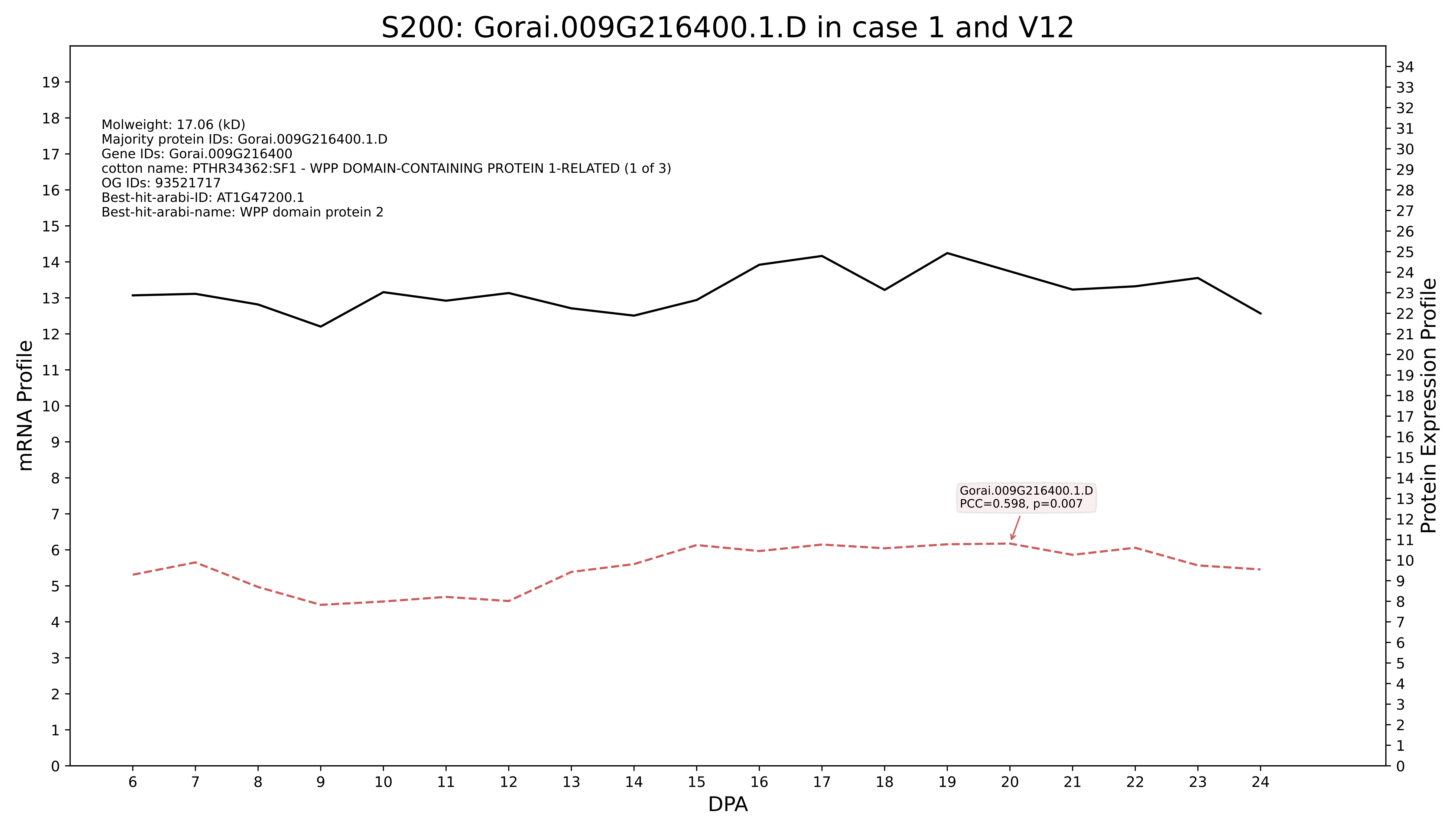
S200: Gorai.009G216400.1.D in case 1 and V12
Molweight: 17.06 (kD)
Majority protein IDs: Gorai.009G216400.1.D
Gene IDs: Gorai.009G216400
cotton name: PTHR34362:SF1 - WPP DOMAIN-CONTAINING PROTEIN 1-RELATED (1 of 3)
OG IDs: 93521717
Best-hit-arabi-ID: AT1G47200.1
Best-hit-arabi-name: WPP domain protein 2
Gorai.009G216400.1.D
PCC=0.598, p=0.007
mRNA Profile
Protein Expression Profile
DPA

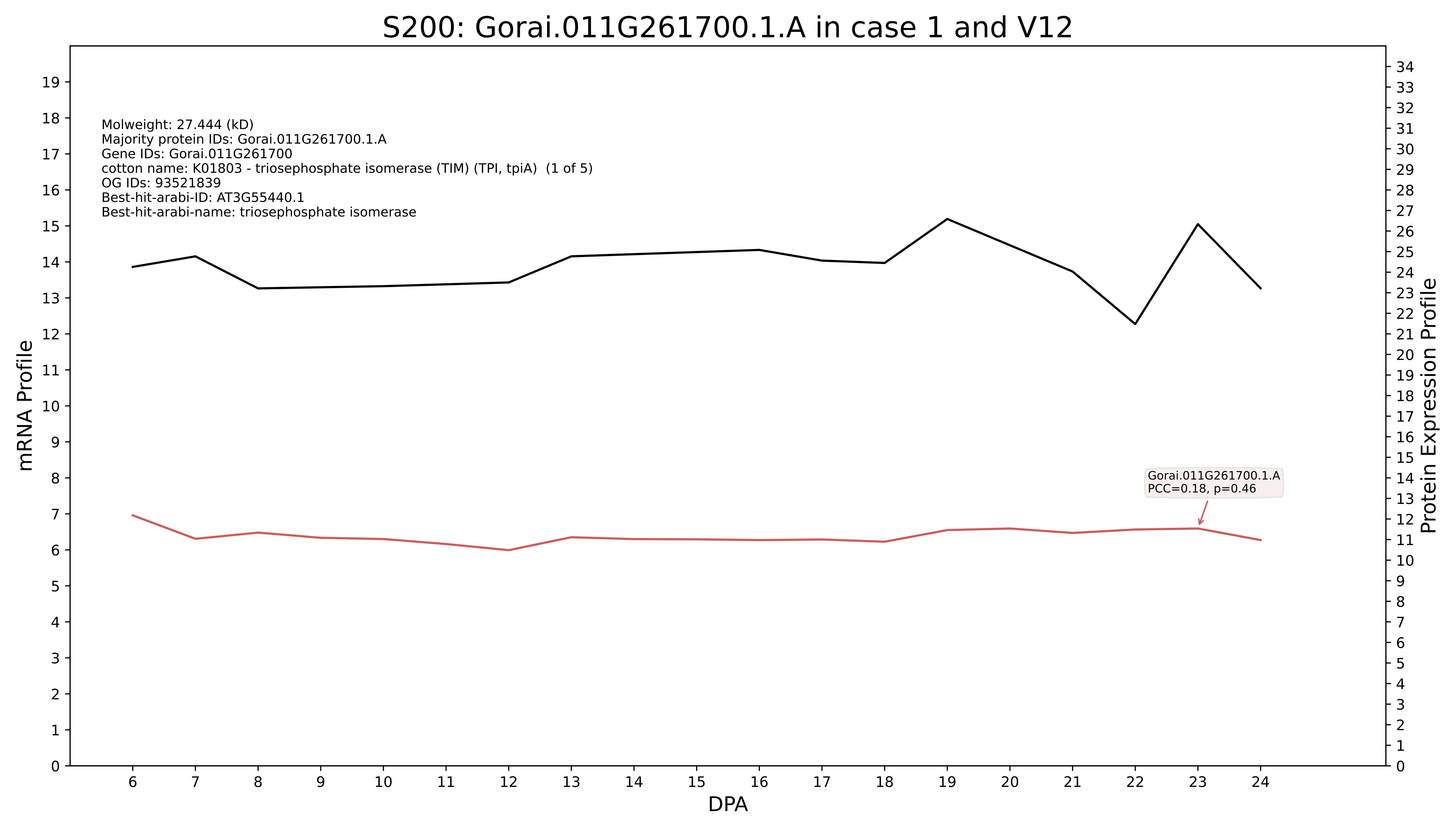
S200: Gorai.011G261700.1.A in case 1 and V12
Molweight: 27.444 (kD)
Majority protein IDs: Gorai.011G261700.1.A
Gene IDs: Gorai.011G261700
cotton name: K01803 - triosephosphate isomerase (TIM) (TPI, tpiA) (1 of 5)
OG IDs: 93521839
Best-hit-arabi-ID: AT3G55440.1
Best-hit-arabi-name: triosephosphate isomerase
Gorai.011G261700.1.A
PCC=0.18, p=0.46
mRNA Profile
Protein Expression Profile
DPA

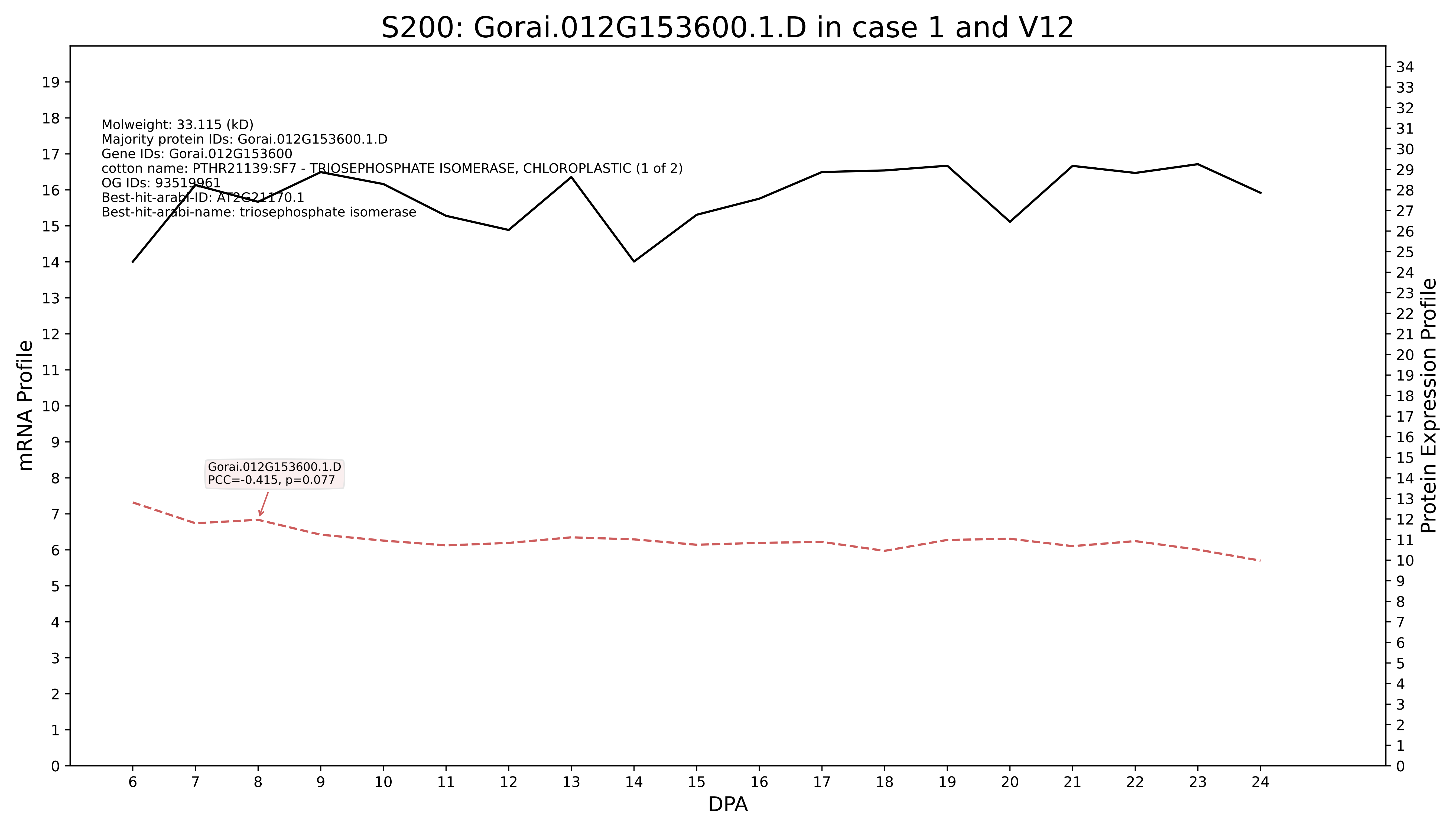
S200: Gorai.012G153600.1.D in case 1 and V12
Molweight: 33.115 (kD)
Majority protein IDs: Gorai.012G153600.1.D
Gene IDs: Gorai.012G153600
cotton name: PTHR21139:SF7 - TRIOSEPHOSPHATE ISOMERASE, CHLOROPLASTIC (1 of 2)
OG IDs: 93519961
Best-hit-arabi-ID: AT2G21170.1
Best-hit-arabi-name: triosephosphate isomerase
Gorai.012G153600.1.D
PCC=-0.415, p=0.077
mRNA Profile
Protein Expression Profile
DPA

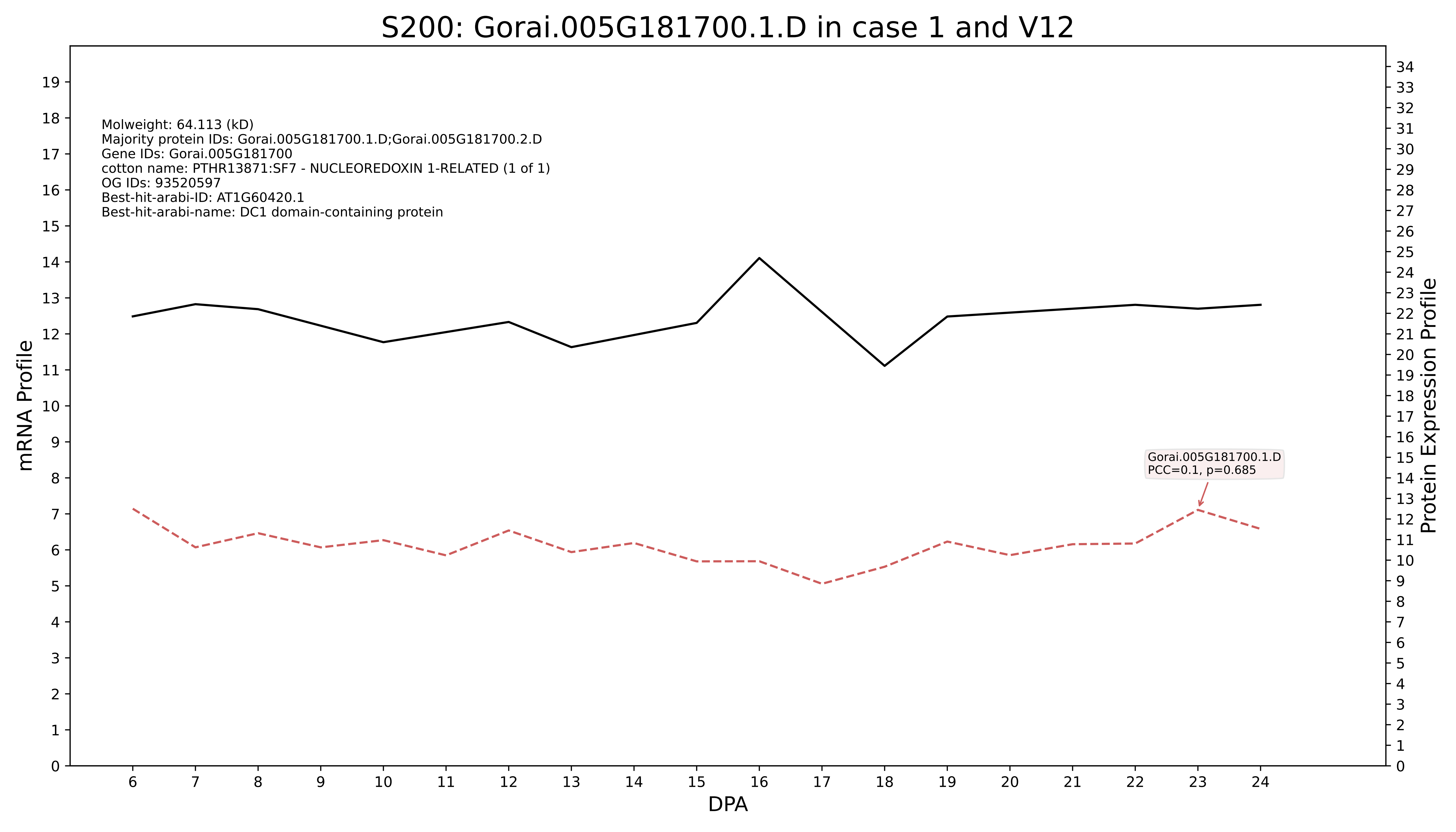
S200: Gorai.005G181700.1.D in case 1 and V12
Molweight: 64.113 (kD)
Majority protein IDs: Gorai.005G181700.1.D;Gorai.005G181700.2.D
Gene IDs: Gorai.005G181700
cotton name: PTHR13871:SF7 - NUCLEOREDOXIN 1-RELATED (1 of 1)
OG IDs: 93520597
Best-hit-arabi-ID: AT1G60420.1
Best-hit-arabi-name: DC1 domain-containing protein
Gorai.005G181700.1.D
PCC=0.1, p=0.685
mRNA Profile
Protein Expression Profile
DPA

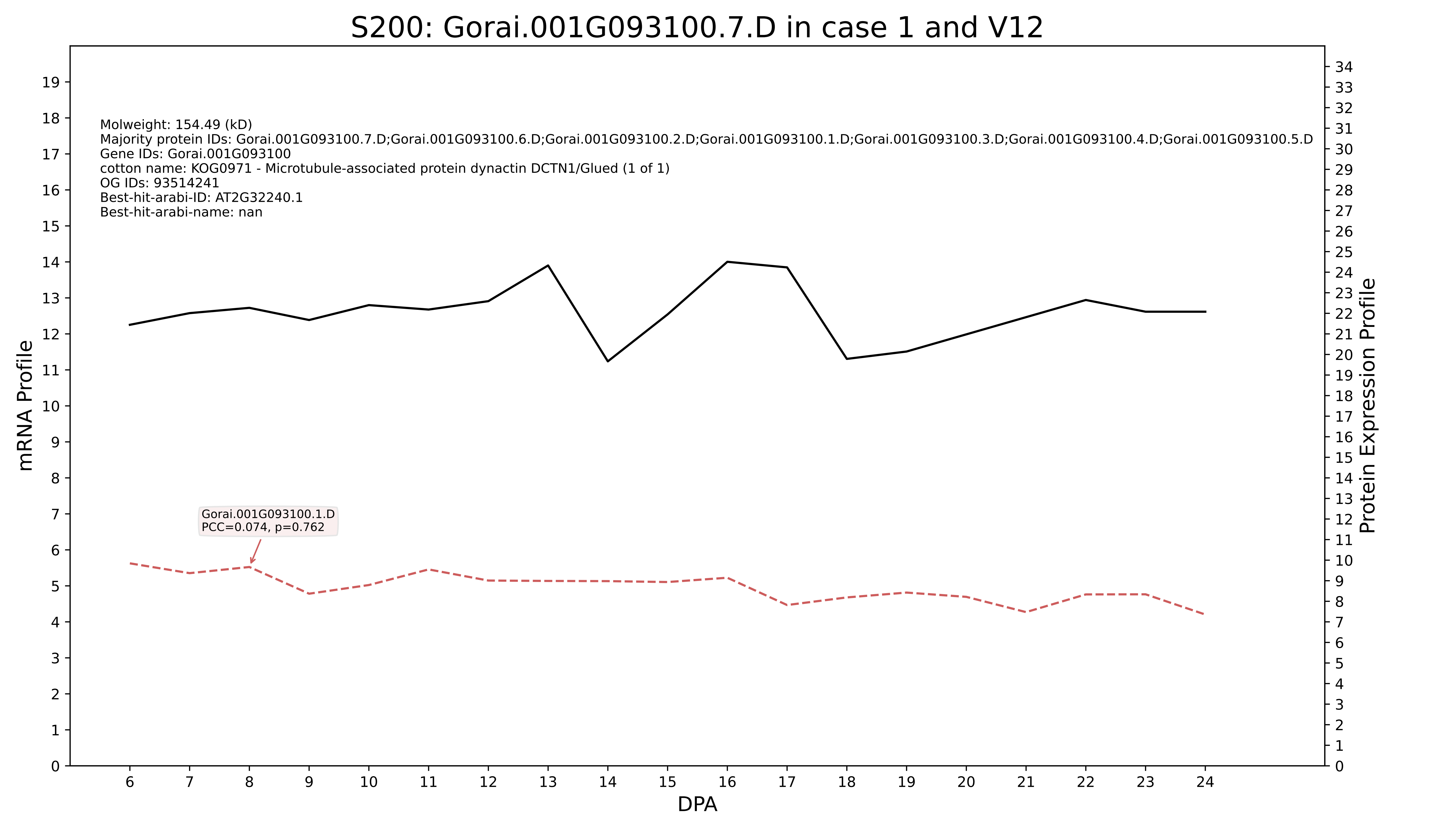

S200: Gorai.001G093100.7.D in case 1 and V12
Molweight: 154.49 (kD)
Majority protein IDs: Gorai.001G093100.7.D;Gorai.001G093100.6.D;Gorai.001G093100.2.D;Gorai.001G093100.1.D;Gorai.001G093100.3.D;Gorai.001G093100.4.D;Gorai.001G093100.5.D
Gene IDs: Gorai.001G093100
cotton name: KOG0971 - Microtubule-associated protein dynactin DCTN1/Glued (1 of 1)
OG IDs: 93514241
Best-hit-arabi-ID: AT2G32240.1
Best-hit-arabi-name: nan
Gorai.001G093100.1.D
PCC=0.074, p=0.762
mRNA Profile
Protein Expression Profile
DPA

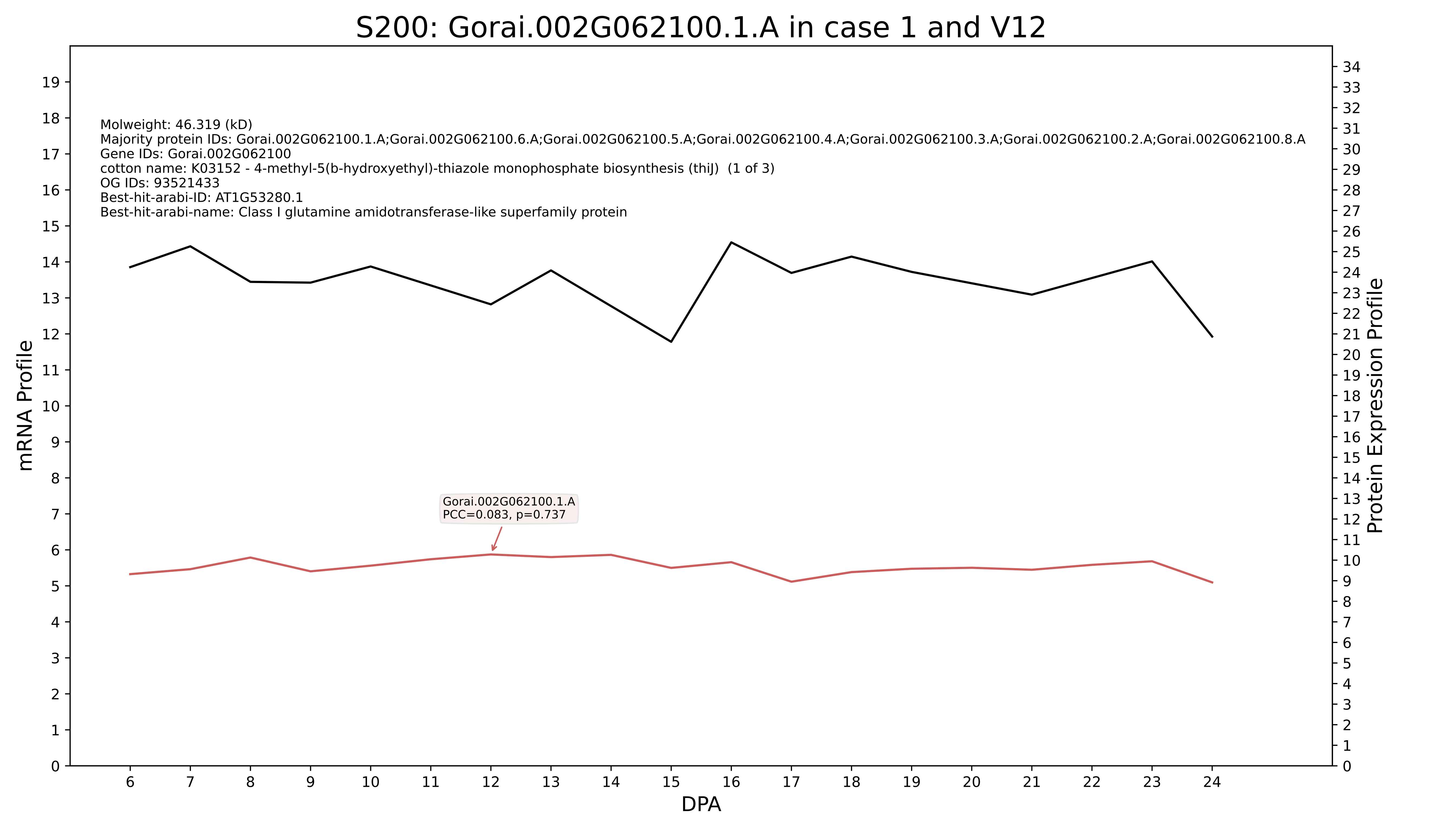
S200: Gorai.002G062100.1.A in case 1 and V12
Molweight: 46.319 (kD)
Majority protein IDs: Gorai.002G062100.1.A;Gorai.002G062100.6.A;Gorai.002G062100.5.A;Gorai.002G062100.4.A;Gorai.002G062100.3.A;Gorai.002G062100.2.A;Gorai.002G062100.8.A
Gene IDs: Gorai.002G062100
cotton name: K03152 - 4-methyl-5(b-hydroxyethyl)-thiazole monophosphate biosynthesis (thiJ) (1 of 3)
OG IDs: 93521433
Best-hit-arabi-ID: AT1G53280.1
Best-hit-arabi-name: Class I glutamine amidotransferase-like superfamily protein
Gorai.002G062100.1.A
PCC=0.083, p=0.737
mRNA Profile
Protein Expression Profile
DPA

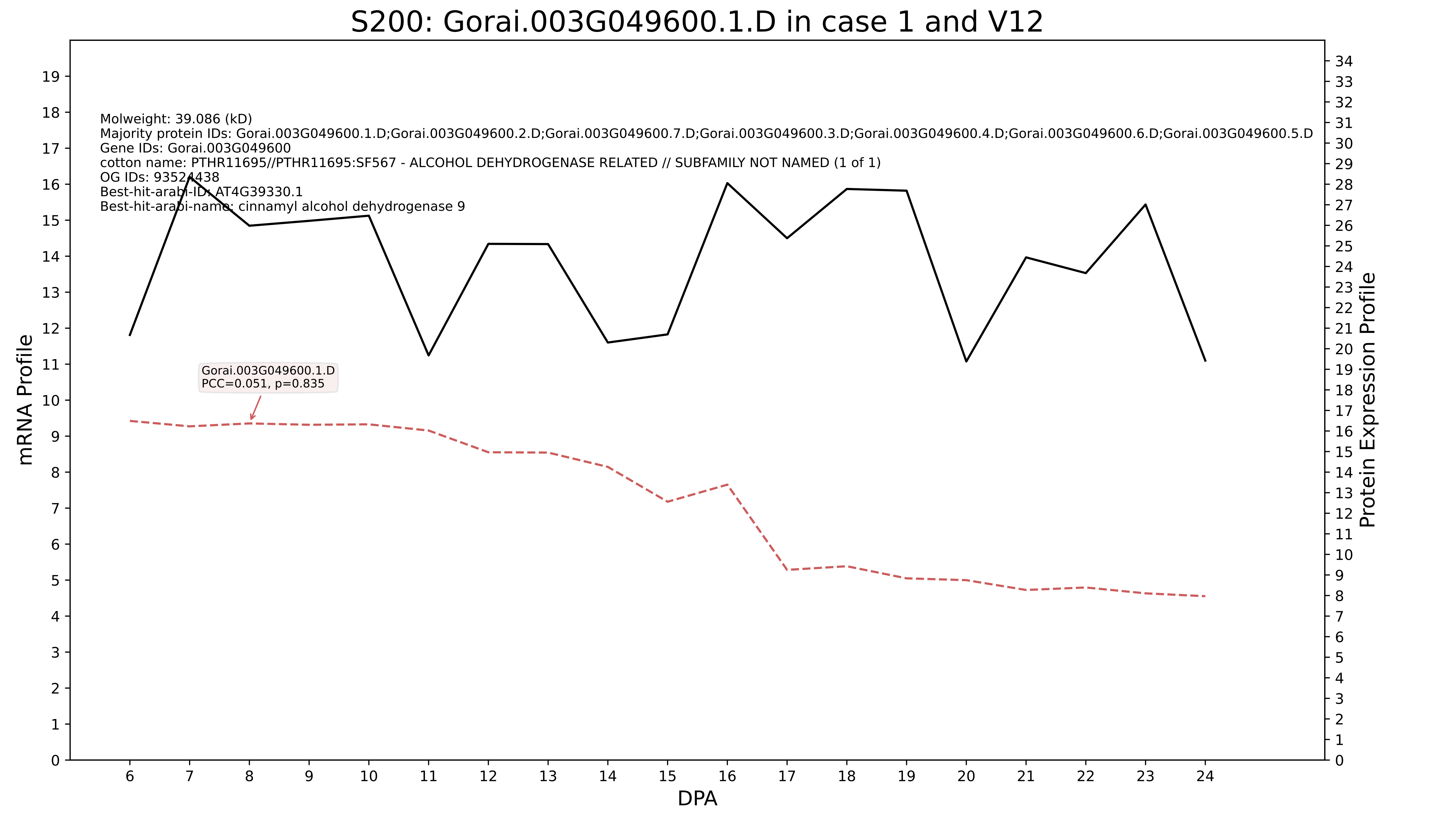
S200: Gorai.003G049600.1.D in case 1 and V12
Molweight: 39.086 (kD)
Majority protein IDs: Gorai.003G049600.1.D;Gorai.003G049600.2.D;Gorai.003G049600.7.D;Gorai.003G049600.3.D;Gorai.003G049600.4.D;Gorai.003G049600.6.D;Gorai.003G049600.5.D
Gene IDs: Gorai.003G049600
cotton name: PTHR11695//PTHR11695:SF567 - ALCOHOL DEHYDROGENASE RELATED // SUBFAMILY NOT NAMED (1 of 1)
OG IDs: 93524438
Best-hit-arabi-ID: AT4G39330.1
Best-hit-arabi-name: cinnamyl alcohol dehydrogenase 9
Gorai.003G049600.1.D
PCC=0.051, p=0.835
mRNA Profile
Protein Expression Profile
DPA

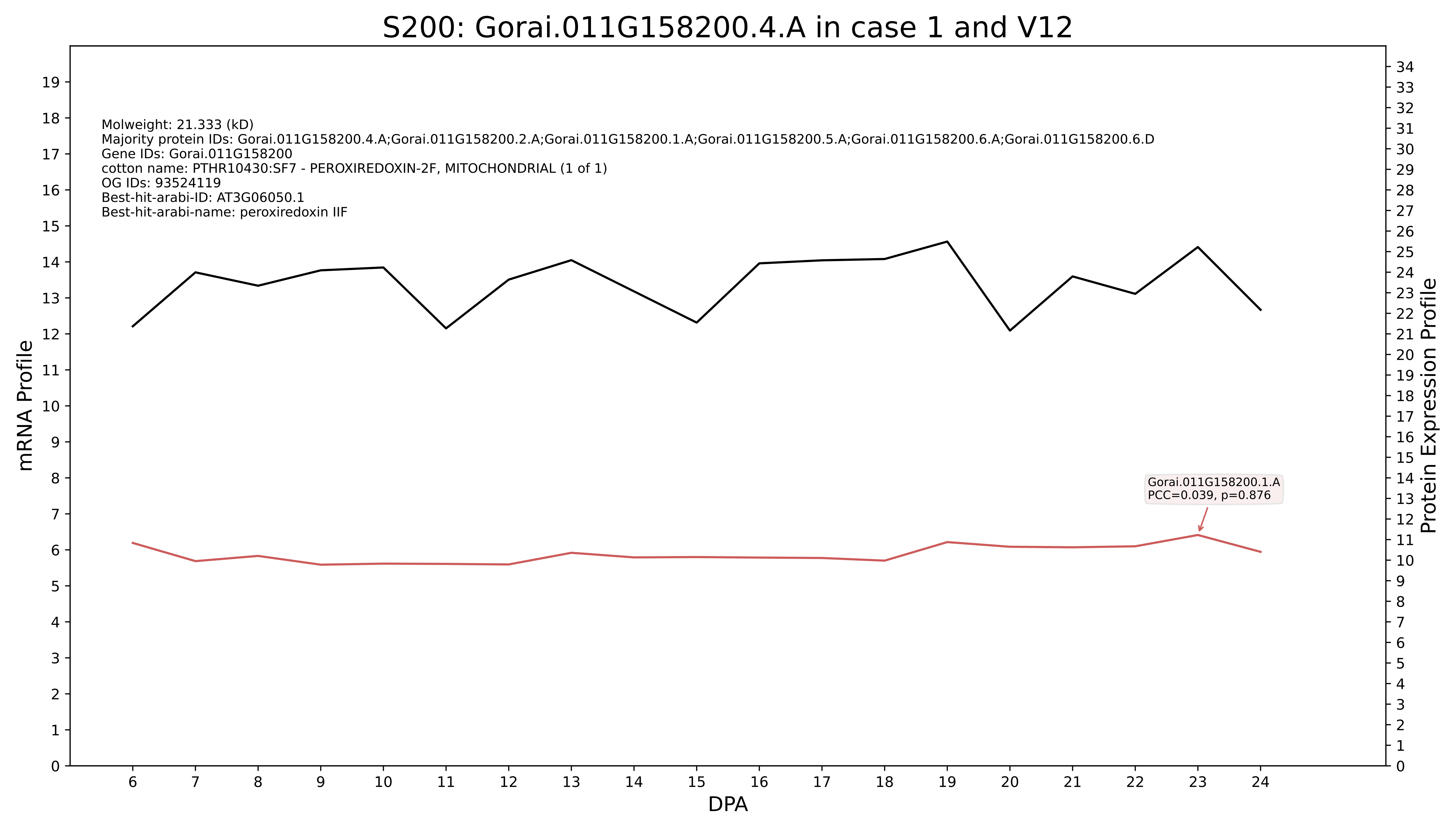

S200: Gorai.011G158200.4.A in case 1 and V12
Molweight: 21.333 (kD)
Majority protein IDs: Gorai.011G158200.4.A;Gorai.011G158200.2.A;Gorai.011G158200.1.A;Gorai.011G158200.5.A;Gorai.011G158200.6.A;Gorai.011G158200.6.D
Gene IDs: Gorai.011G158200
cotton name: PTHR10430:SF7 - PEROXIREDOXIN-2F, MITOCHONDRIAL (1 of 1)
OG IDs: 93524119
Best-hit-arabi-ID: AT3G06050.1
Best-hit-arabi-name: peroxiredoxin IIF
Gorai.011G158200.1.A
PCC=0.039, p=0.876
mRNA Profile
Protein Expression Profile
DPA

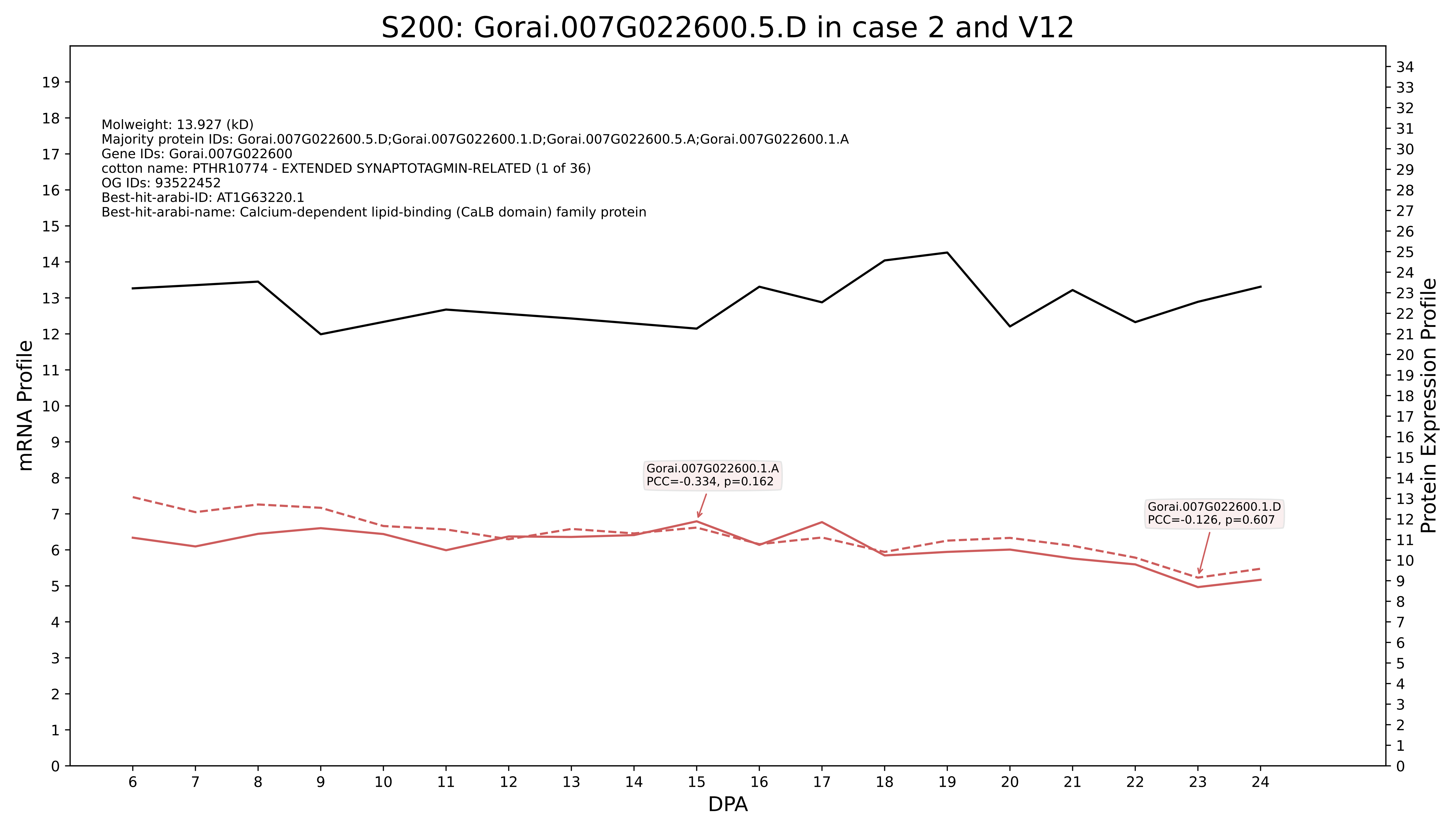

S200: Gorai.007G022600.5.D in case 2 and V12
Molweight: 13.927 (kD)
Majority protein IDs: Gorai.007G022600.5.D;Gorai.007G022600.1.D;Gorai.007G022600.5.A;Gorai.007G022600.1.A
Gene IDs: Gorai.007G022600
cotton name: PTHR10774 - EXTENDED SYNAPTOTAGMIN-RELATED (1 of 36)
OG IDs: 93522452
Best-hit-arabi-ID: AT1G63220.1
Best-hit-arabi-name: Calcium-dependent lipid-binding (CaLB domain) family protein
Gorai.007G022600.1.A
PCC=-0.334, p=0.162
Gorai.007G022600.1.D
PCC=-0.126, p=0.607
mRNA Profile
Protein Expression Profile
DPA

### S200: Gorai.007G300300.1.D in case 2 and V12

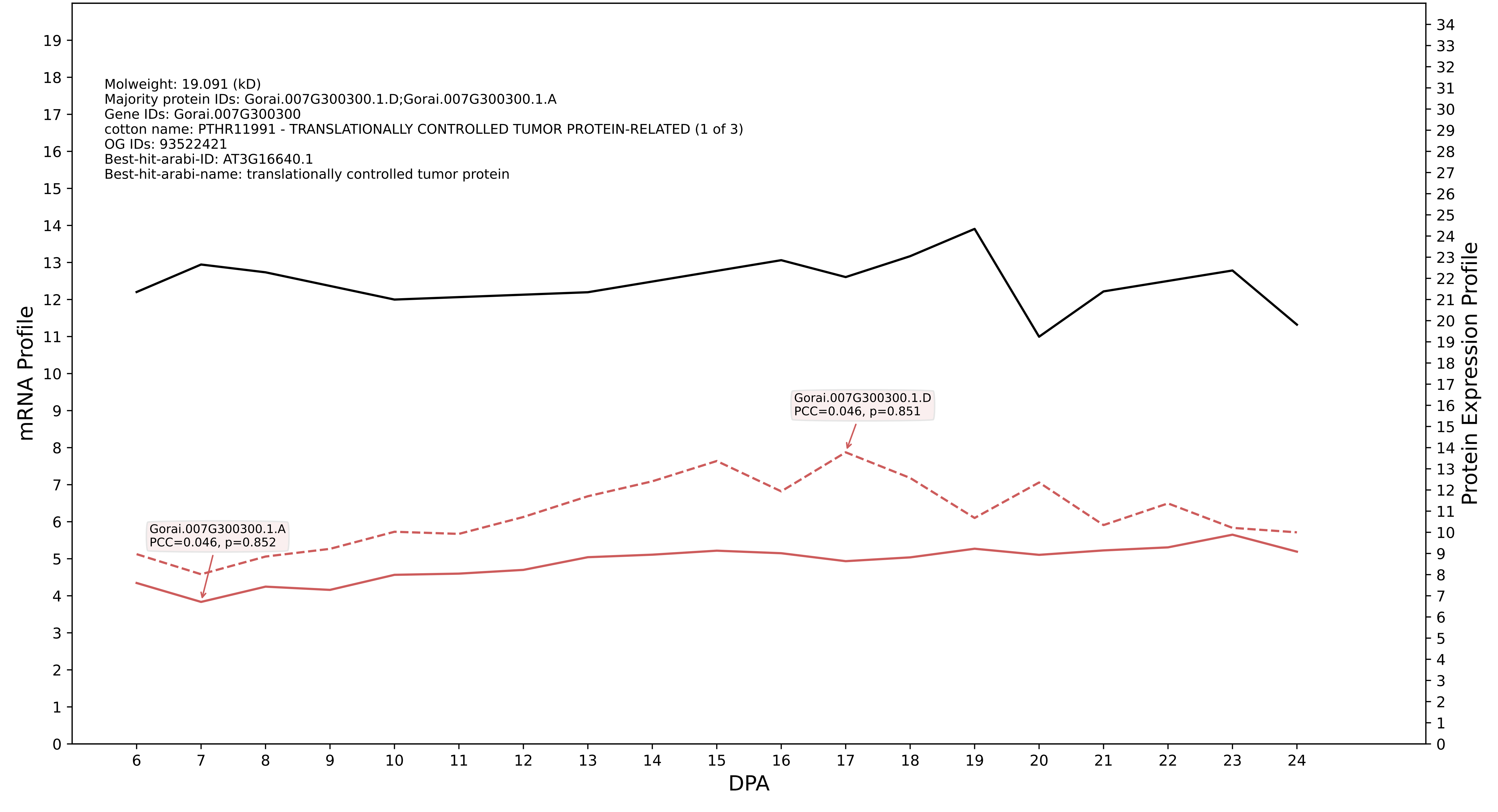

### S200: Gorai.009G028500.1.D in case 2 and V12

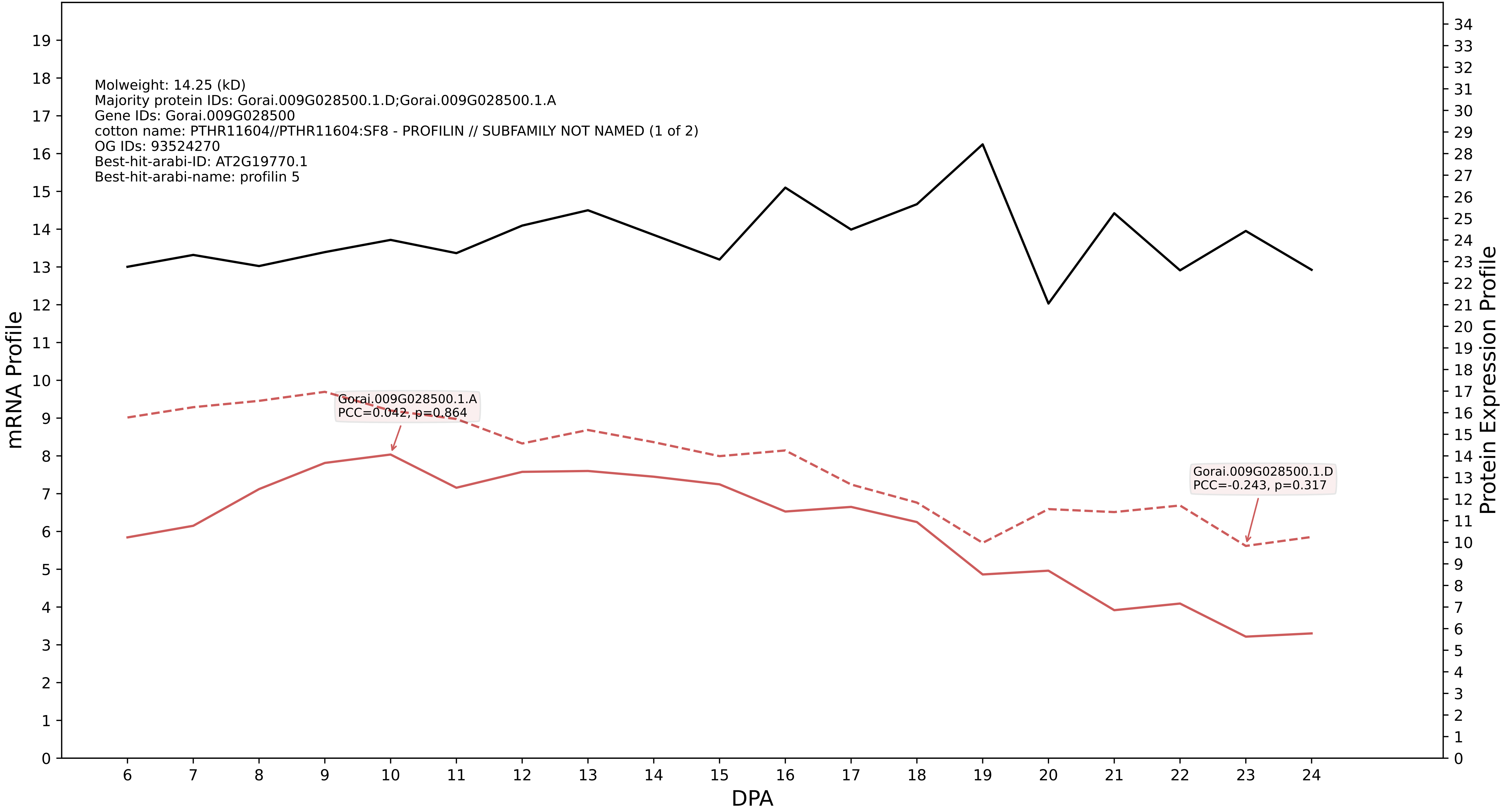

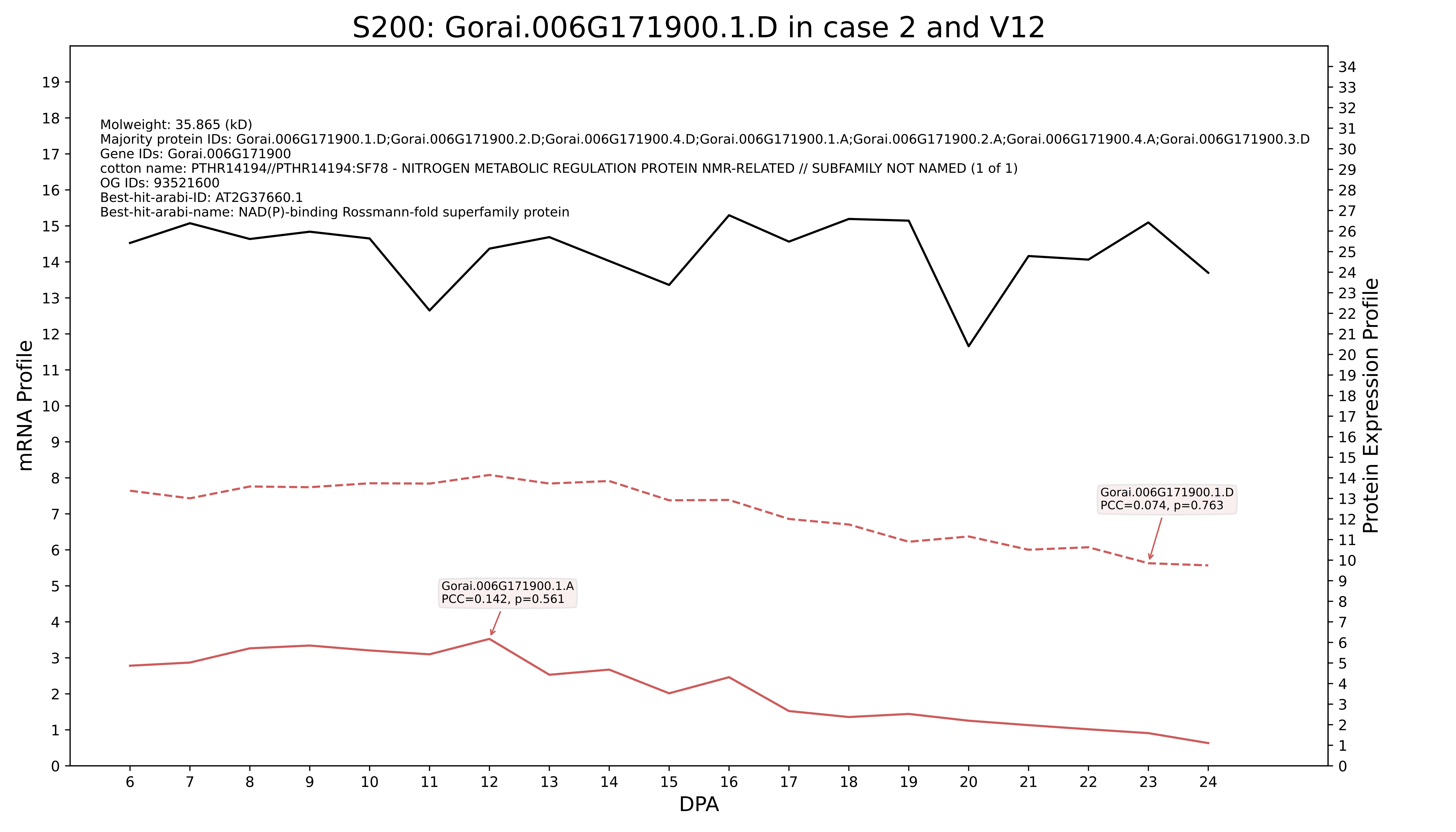
S200: Gorai.006G171900.1.D in case 2 and V12
Molweight: 35.865 (kD)
Majority protein IDs: Gorai.006G171900.1.D;Gorai.006G171900.2.D;Gorai.006G171900.4.D;Gorai.006G171900.1.A;Gorai.006G171900.2.A;Gorai.006G171900.4.A;Gorai.006G171900.3.D
Gene IDs: Gorai.006G171900
cotton name: PTHR14194//PTHR14194:SF78 - NITROGEN METABOLIC REGULATION PROTEIN NMR-RELATED // SUBFAMILY NOT NAMED (1 of 1)
OG IDs: 93521600
Best-hit-arabi-ID: AT2G37660.1
Best-hit-arabi-name: NAD(P)-binding Rossmann-fold superfamily protein
Gorai.006G171900.1.D
PCC=0.074, p=0.763
Gorai.006G171900.1.A
PCC=0.142, p=0.561
mRNA Profile
Protein Expression Profile
DPA

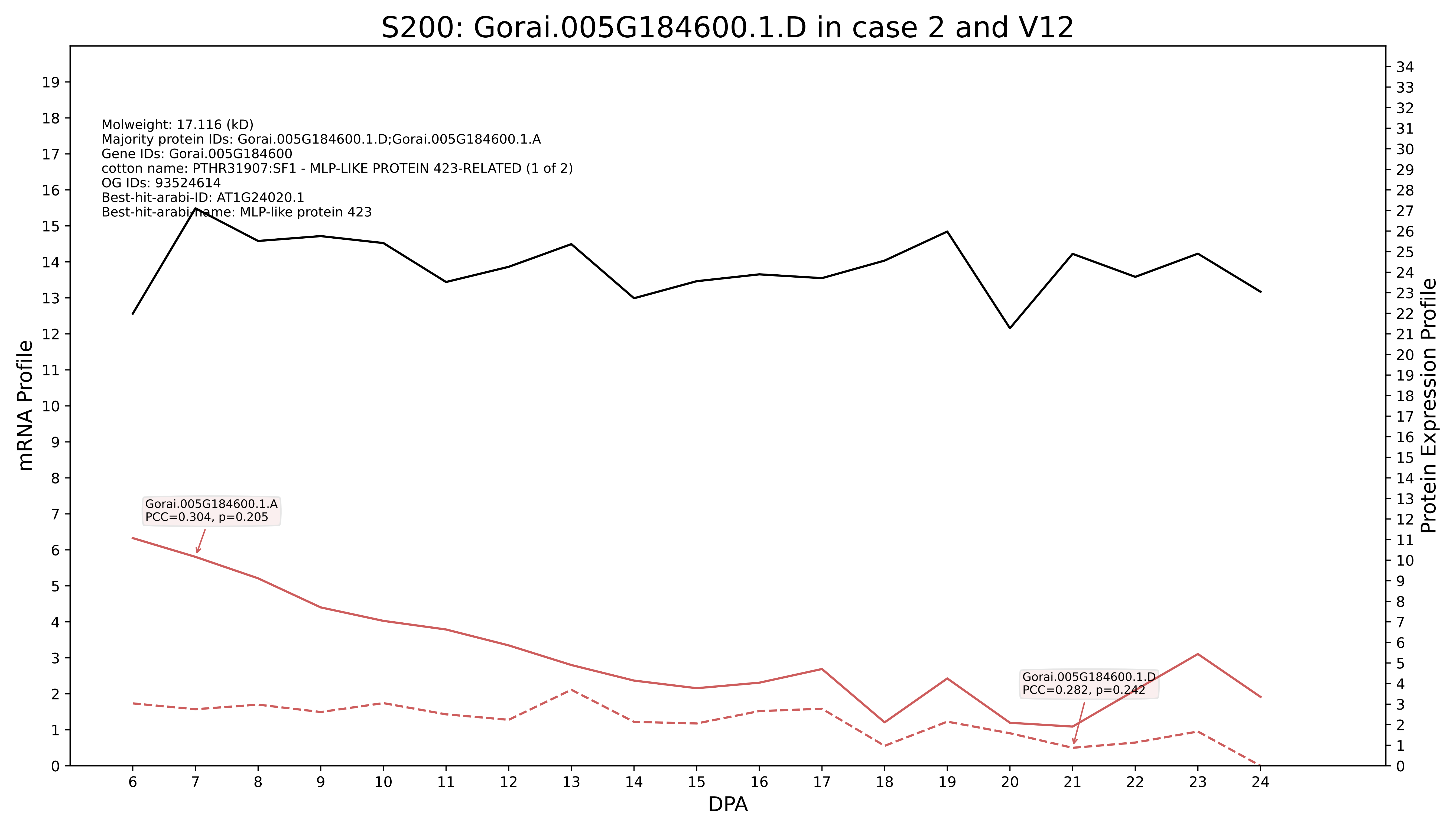
S200: Gorai.005G184600.1.D in case 2 and V12
Molweight: 17.116 (kD)
Majority protein IDs: Gorai.005G184600.1.D;Gorai.005G184600.1.A
Gene IDs: Gorai.005G184600
cotton name: PTHR31907:SF1 - MLP-LIKE PROTEIN 423-RELATED (1 of 2)
OG IDs: 93524614
Best-hit-arabi-ID: AT1G24020.1
Best-hit-arabi-name: MLP-like protein 423
Gorai.005G184600.1.A
PCC=0.304, p=0.205
Gorai.005G184600.1.D
PCC=0.282, p=0.242
mRNA Profile
Protein Expression Profile
DPA

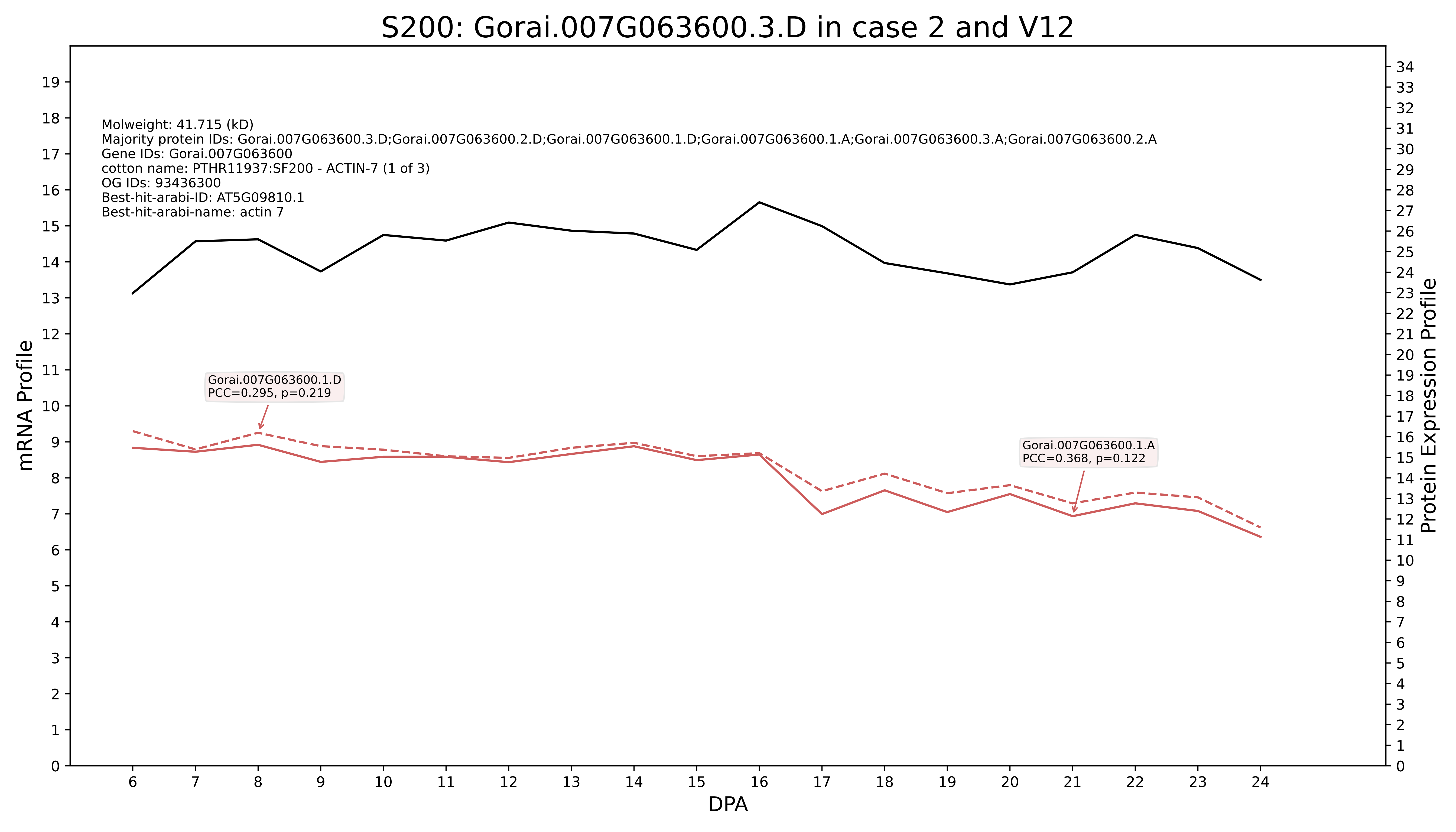

S200: Gorai.007G063600.3.D in case 2 and V12
Molweight: 41.715 (kD)
Majority protein IDs: Gorai.007G063600.3.D;Gorai.007G063600.2.D;Gorai.007G063600.1.D;Gorai.007G063600.1.A;Gorai.007G063600.3.A;Gorai.007G063600.2.A
Gene IDs: Gorai.007G063600
cotton name: PTHR11937:SF200 - ACTIN-7 (1 of 3)
OG IDs: 93436300
Best-hit-arabi-ID: AT5G09810.1
Best-hit-arabi-name: actin 7
Gorai.007G063600.1.D
PCC=0.295, p=0.219
Gorai.007G063600.1.A
PCC=0.368, p=0.122
mRNA Profile
Protein Expression Profile
DPA

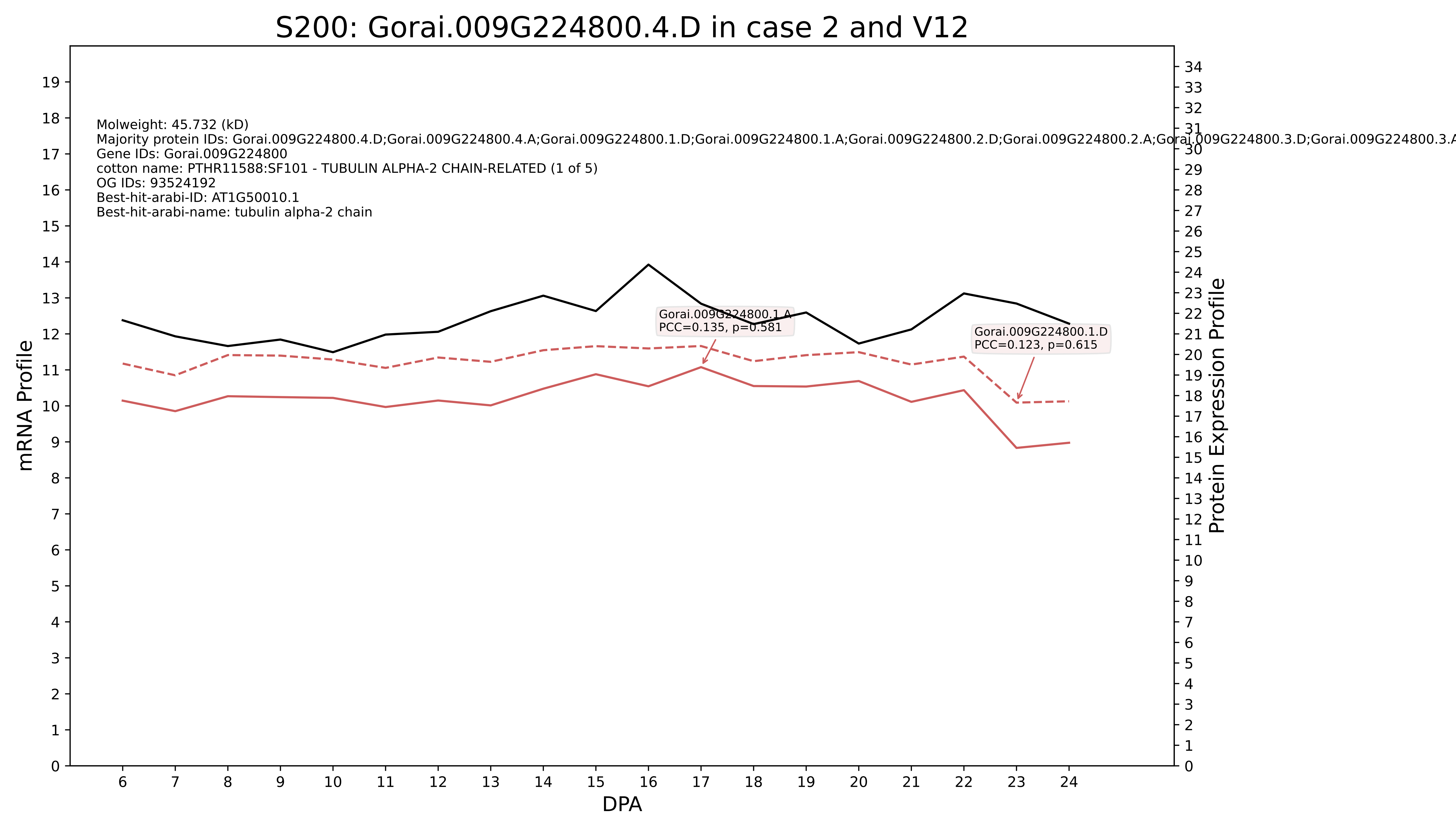
S200: Gorai.009G224800.4.D in case 2 and V12
Molweight: 45.732 (kD)
Majority protein IDs: Gorai.009G224800.4.D;Gorai.009G224800.4.A;Gorai.009G224800.1.D;Gorai.009G224800.1.A;Gorai.009G224800.2.D;Gorai.009G224800.2.A;Gorai.009G224800.3.D;Gorai.009G224800.3.A
Gene IDs: Gorai.009G224800
cotton name: PTHR11588:SF101 - TUBULIN ALPHA-2 CHAIN-RELATED (1 of 5)
OG IDs: 93524192
Best-hit-arabi-ID: AT1G50010.1
Best-hit-arabi-name: tubulin alpha-2 chain
Gorai.009G224800.1.A
PCC=0.135, p=0.581
Gorai.009G224800.1.D
PCC=0.123, p=0.615
mRNA Profile
Protein Expression Profile
DPA

### S200: Gorai.006G165800.1.D in case 2 and V12

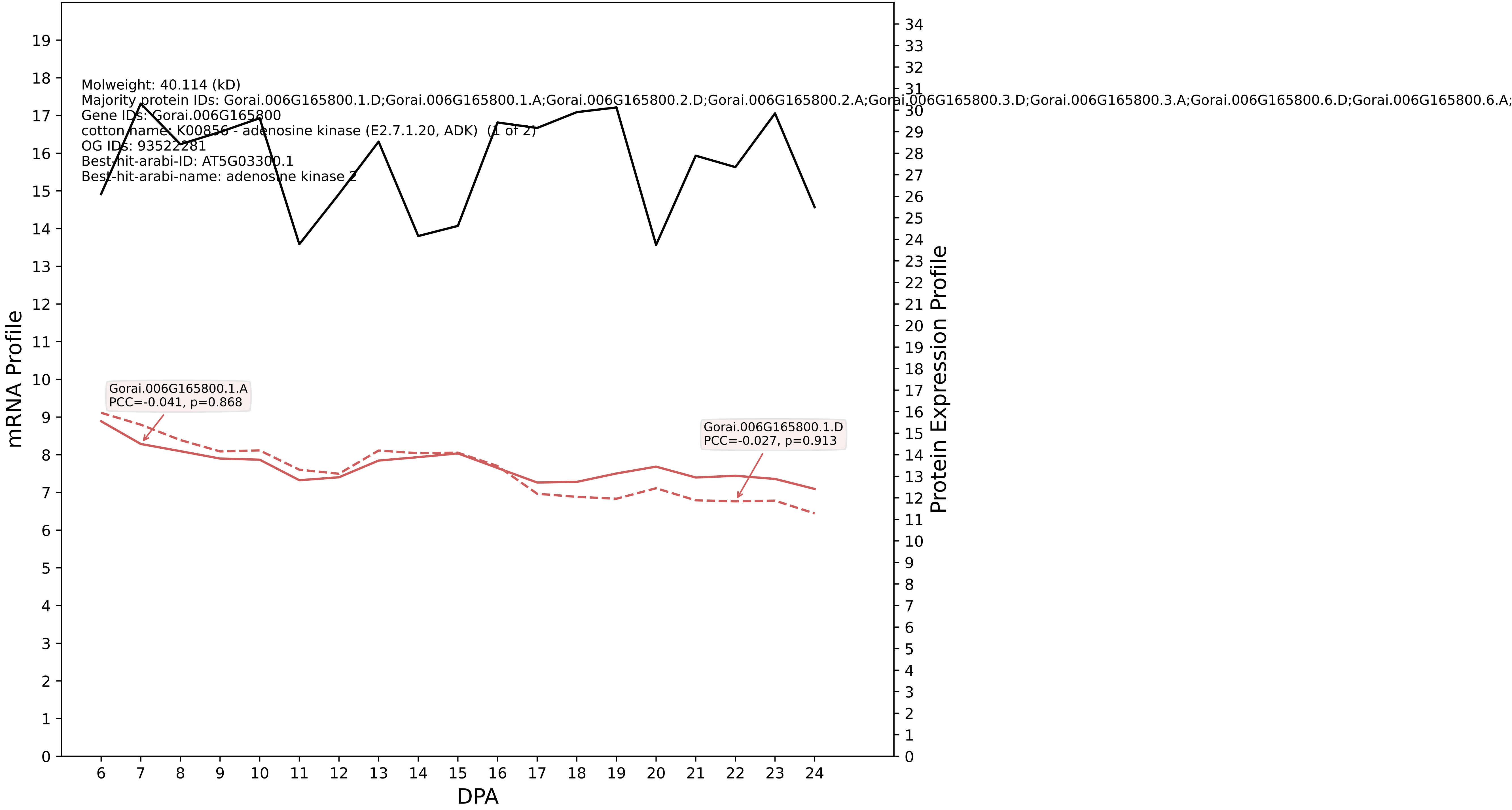

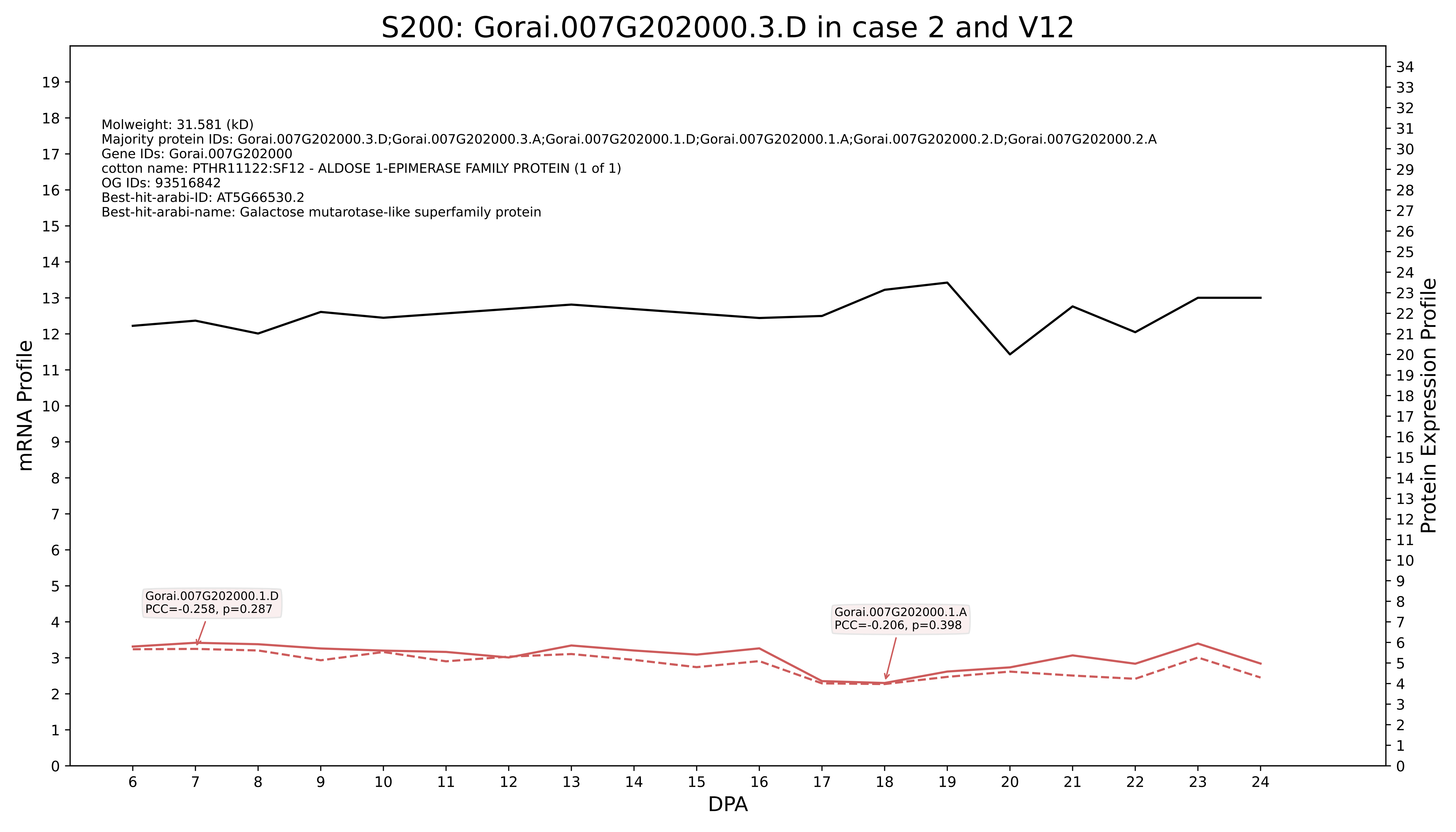

S200: Gorai.007G202000.3.D in case 2 and V12
Molweight: 31.581 (kD)
Majority protein IDs: Gorai.007G202000.3.D;Gorai.007G202000.3.A;Gorai.007G202000.1.D;Gorai.007G202000.1.A;Gorai.007G202000.2.D;Gorai.007G202000.2.A
Gene IDs: Gorai.007G202000
cotton name: PTHR11122:SF12 - ALDOSE 1-EPIMERASE FAMILY PROTEIN (1 of 1)
OG IDs: 93516842
Best-hit-arabi-ID: AT5G66530.2
Best-hit-arabi-name: Galactose mutarotase-like superfamily protein
Gorai.007G202000.1.D
PCC=-0.258, p=0.287
Gorai.007G202000.1.A
PCC=-0.206, p=0.398
mRNA Profile
Protein Expression Profile
DPA

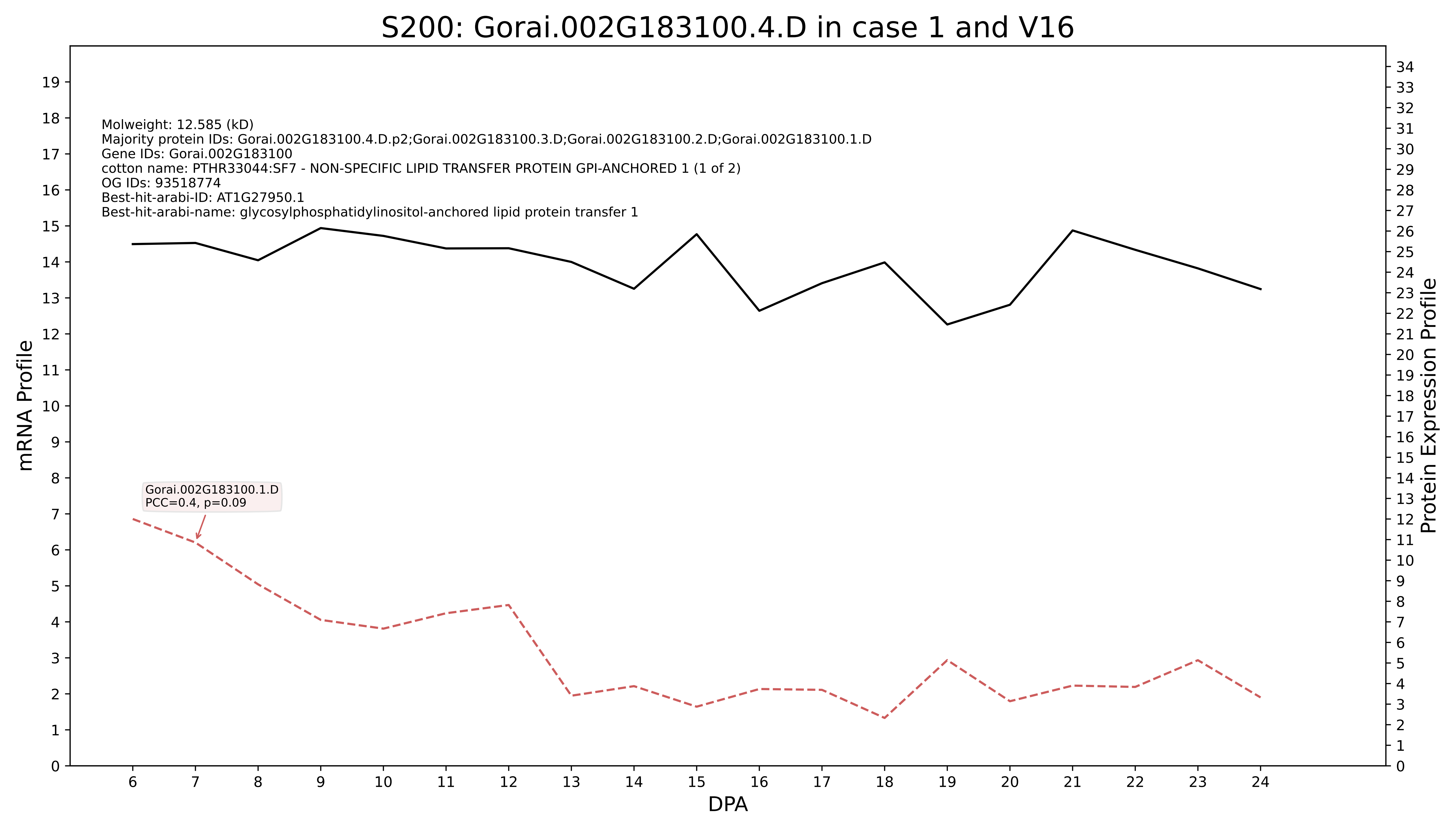

S200: Gorai.002G183100.4.D in case 1 and V16
Molweight: 12.585 (kD)
Majority protein IDs: Gorai.002G183100.4.D.p2;Gorai.002G183100.3.D;Gorai.002G183100.2.D;Gorai.002G183100.1.D
Gene IDs: Gorai.002G183100
cotton name: PTHR33044:SF7 - NON-SPECIFIC LIPID TRANSFER PROTEIN GPI-ANCHORED 1 (1 of 2)
OG IDs: 93518774
Best-hit-arabi-ID: AT1G27950.1
Best-hit-arabi-name: glycosylphosphatidylinositol-anchored lipid protein transfer 1
Gorai.002G183100.1.D
PCC=0.4, p=0.09
mRNA Profile
Protein Expression Profile
DPA

#### S200: Gorai.013G229500.1.D in case 1 and V16

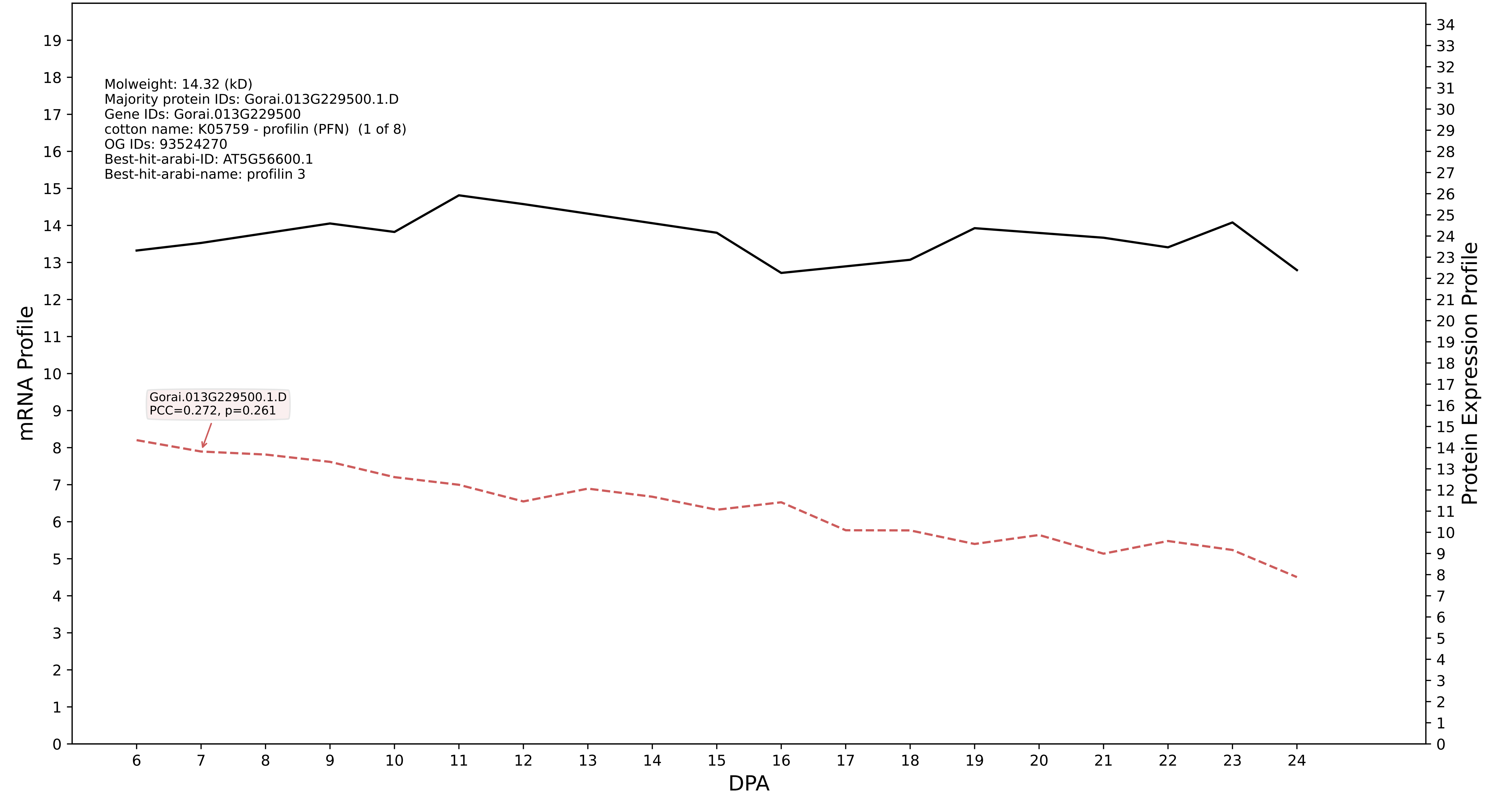

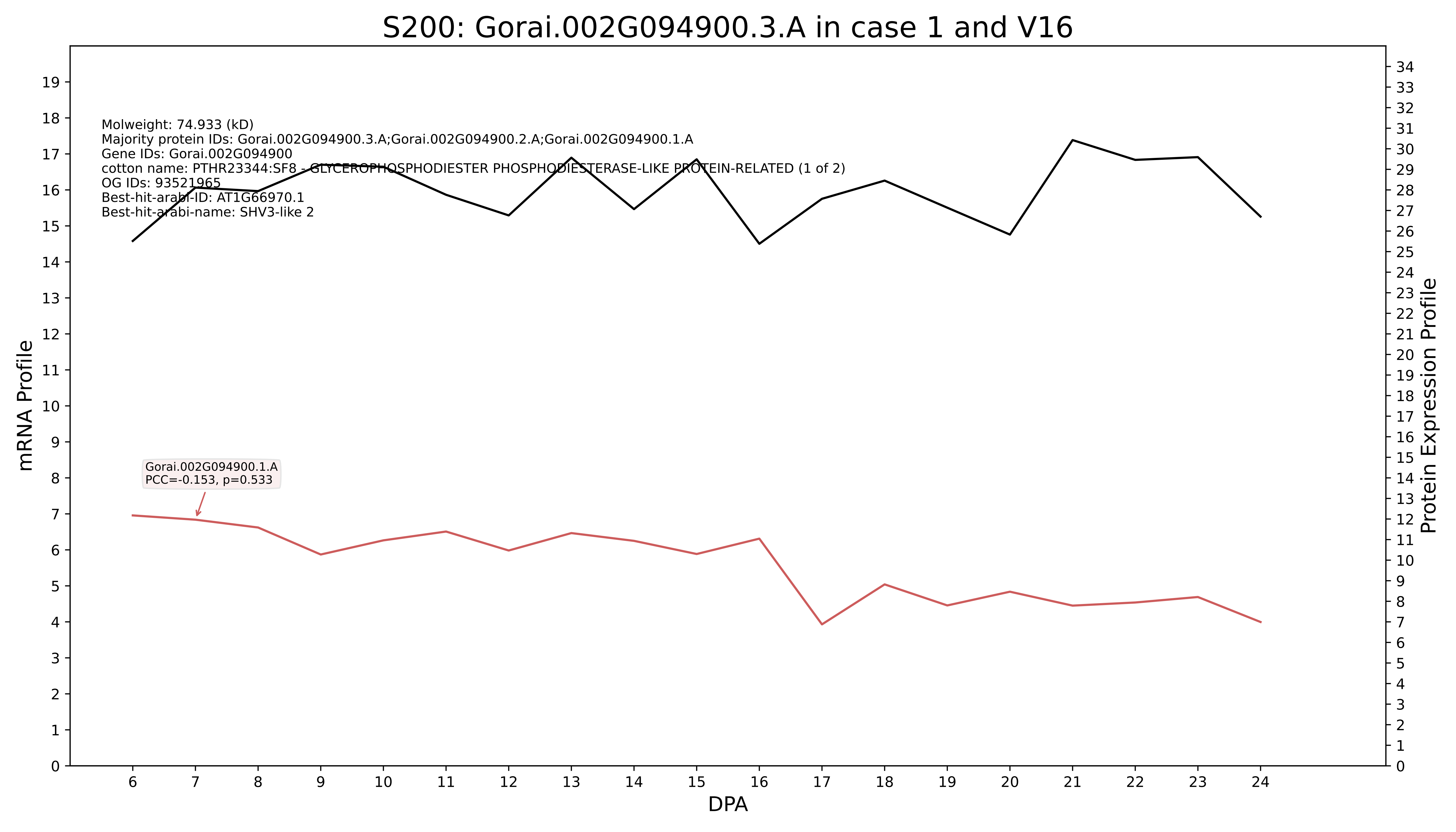
S200: Gorai.002G094900.3.A in case 1 and V16
Molweight: 74.933 (kD)
Majority protein IDs: Gorai.002G094900.3.A;Gorai.002G094900.2.A;Gorai.002G094900.1.A
Gene IDs: Gorai.002G094900
cotton name: PTHR23344:SF8 - GLYCEROPHOSPHODIESTER PHOSPHODIESTERASE-LIKE PROTEIN-RELATED (1 of 2)
OG IDs: 93521965
Best-hit-arabi-ID: AT1G66970.1
Best-hit-arabi-name: SHV3-like 2
Gorai.002G094900.1.A
PCC=-0.153, p=0.533
mRNA Profile
Protein Expression Profile
DPA

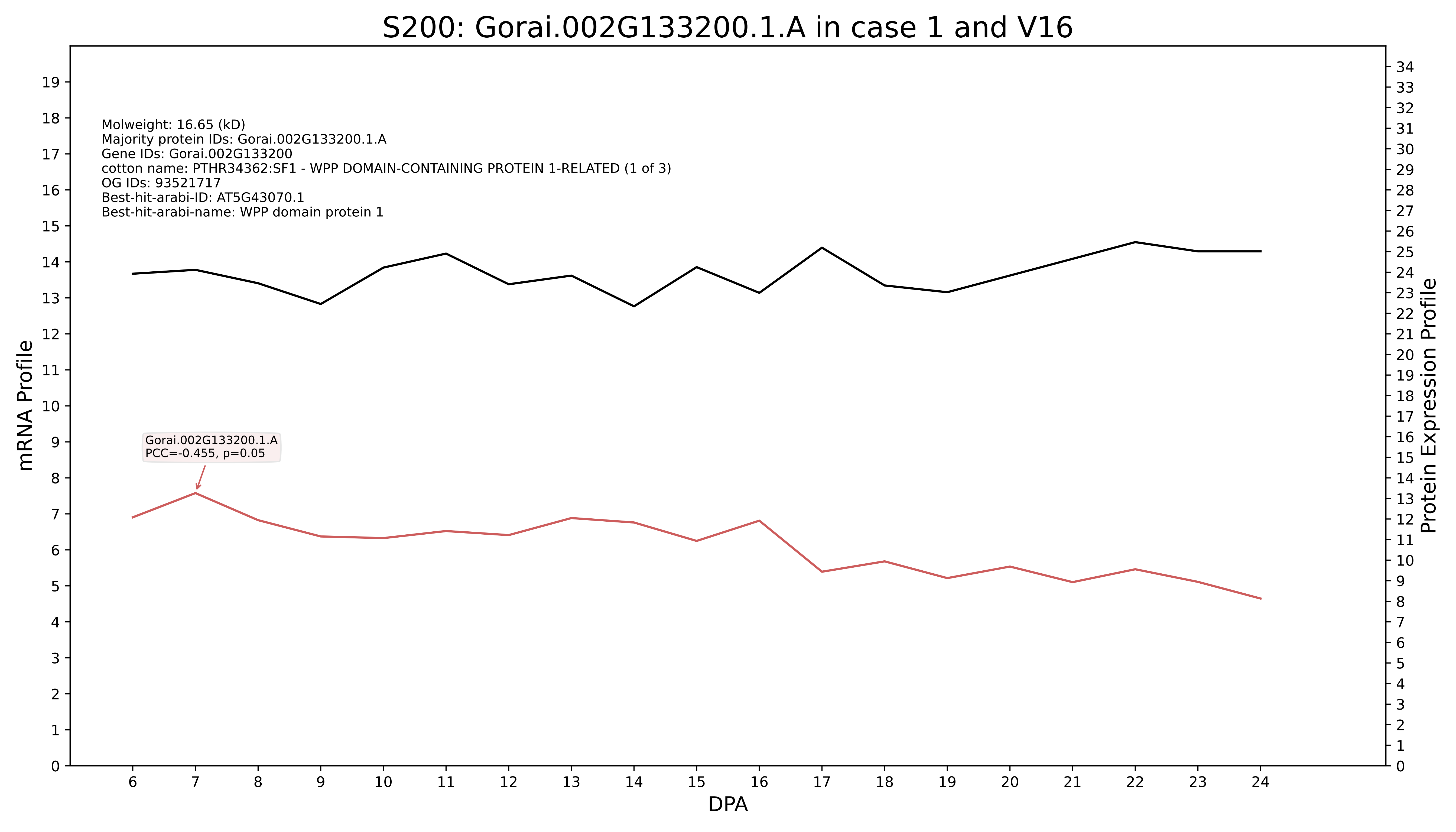

S200: Gorai.002G133200.1.A in case 1 and V16
Molweight: 16.65 (kD)
Majority protein IDs: Gorai.002G133200.1.A
Gene IDs: Gorai.002G133200
cotton name: PTHR34362:SF1 - WPP DOMAIN-CONTAINING PROTEIN 1-RELATED (1 of 3)
OG IDs: 93521717
Best-hit-arabi-ID: AT5G43070.1
Best-hit-arabi-name: WPP domain protein 1
Gorai.002G133200.1.A
PCC=-0.455, p=0.05
mRNA Profile
Protein Expression Profile
DPA

### S200: Gorai.002G094900.3.D in case 1 and V16

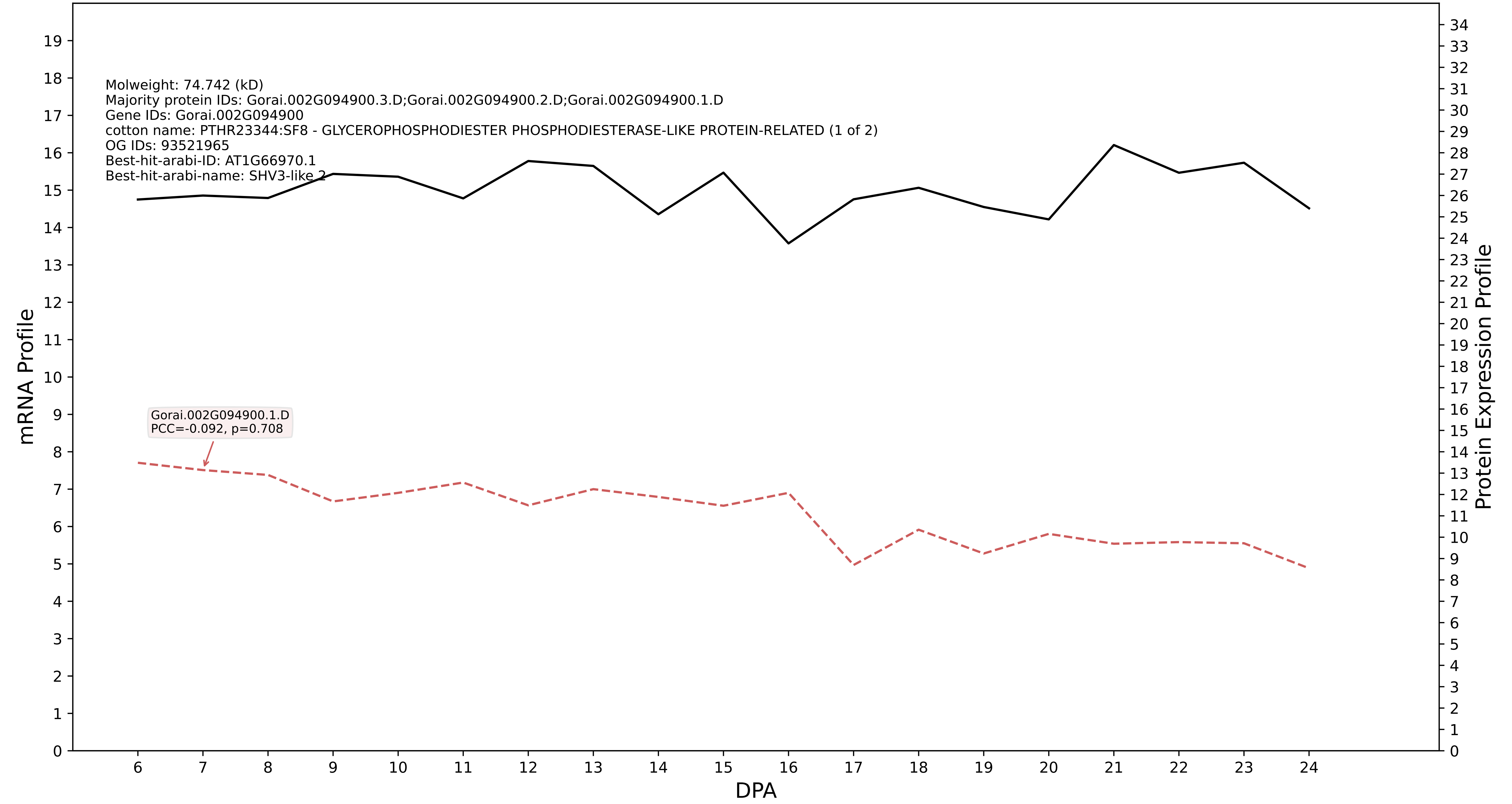

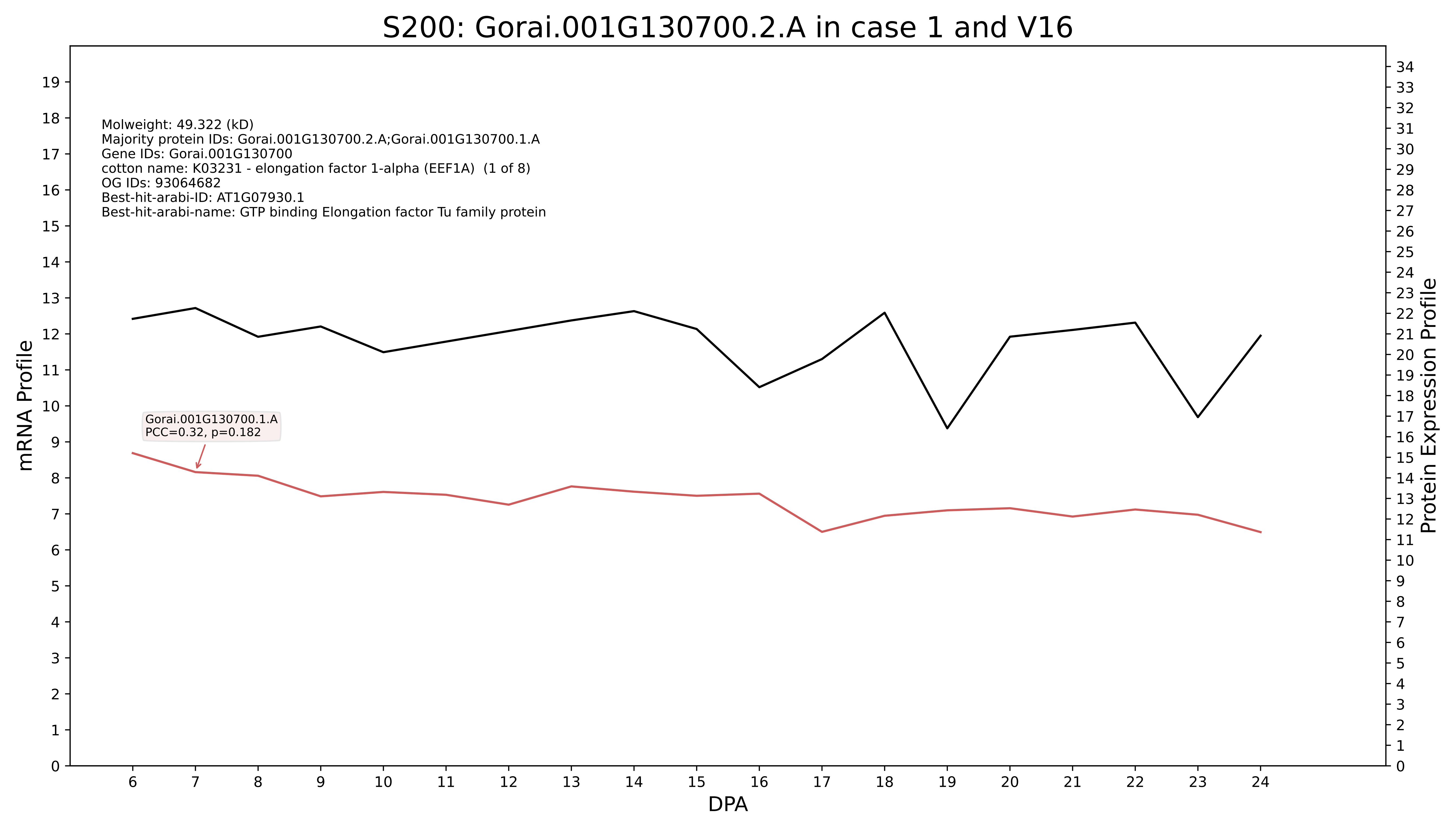

S200: Gorai.001G130700.2.A in case 1 and V16
Molweight: 49.322 (kD)
Majority protein IDs: Gorai.001G130700.2.A;Gorai.001G130700.1.A
Gene IDs: Gorai.001G130700
cotton name: K03231 - elongation factor 1-alpha (EEF1A) (1 of 8)
OG IDs: 93064682
Best-hit-arabi-ID: AT1G07930.1
Best-hit-arabi-name: GTP binding Elongation factor Tu family protein
Gorai.001G130700.1.A
PCC=0.32, p=0.182
mRNA Profile
Protein Expression Profile
DPA

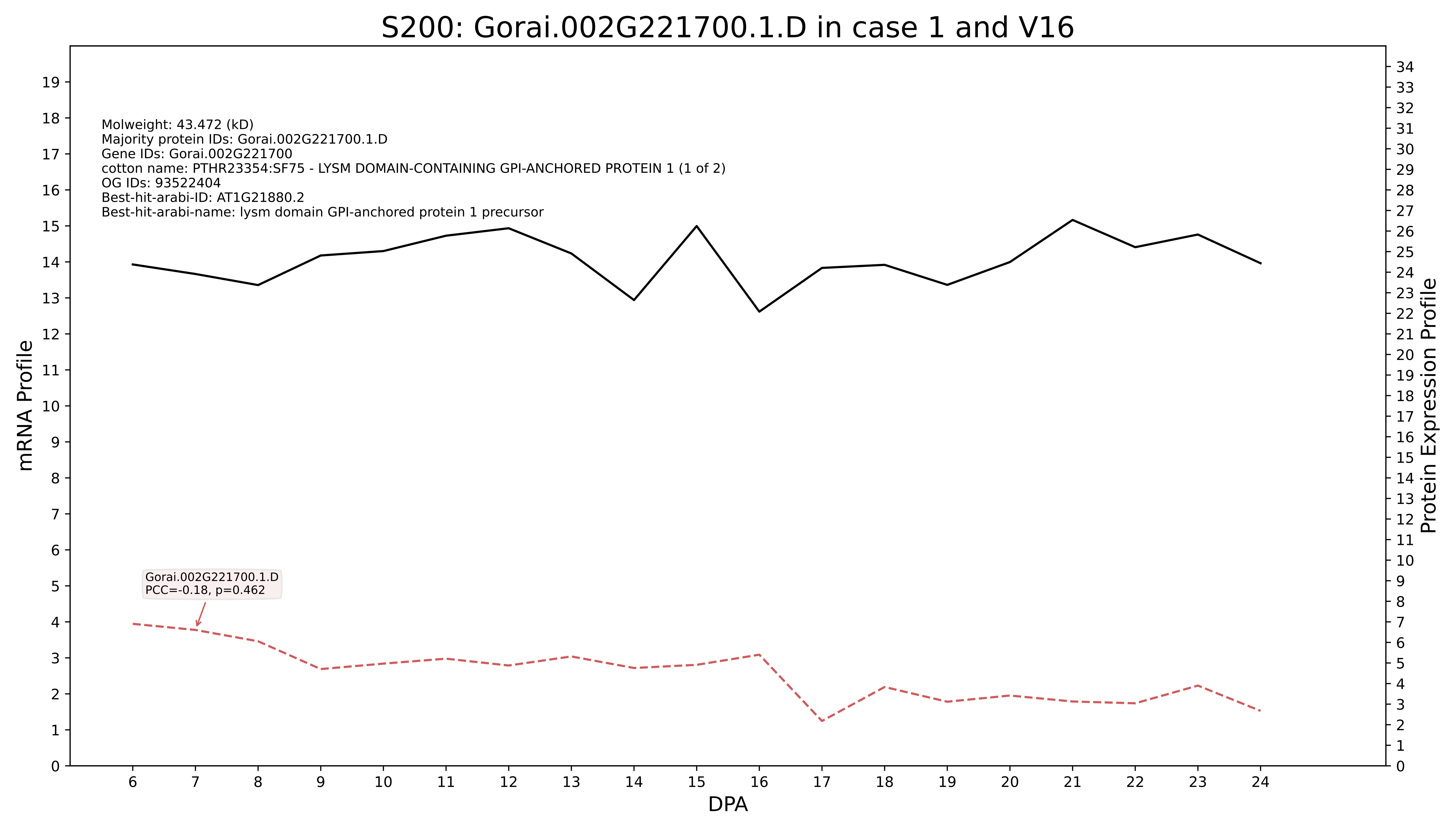
S200: Gorai.002G221700.1.D in case 1 and V16
Molweight: 43.472 (kD)
Majority protein IDs: Gorai.002G221700.1.D
Gene IDs: Gorai.002G221700
cotton name: PTHR23354:SF75 - LYSM DOMAIN-CONTAINING GPI-ANCHORED PROTEIN 1 (1 of 2)
OG IDs: 93522404
Best-hit-arabi-ID: AT1G21880.2
Best-hit-arabi-name: lysm domain GPI-anchored protein 1 precursor
Gorai.002G221700.1.D
PCC=-0.18, p=0.462
mRNA Profile
Protein Expression Profile
DPA

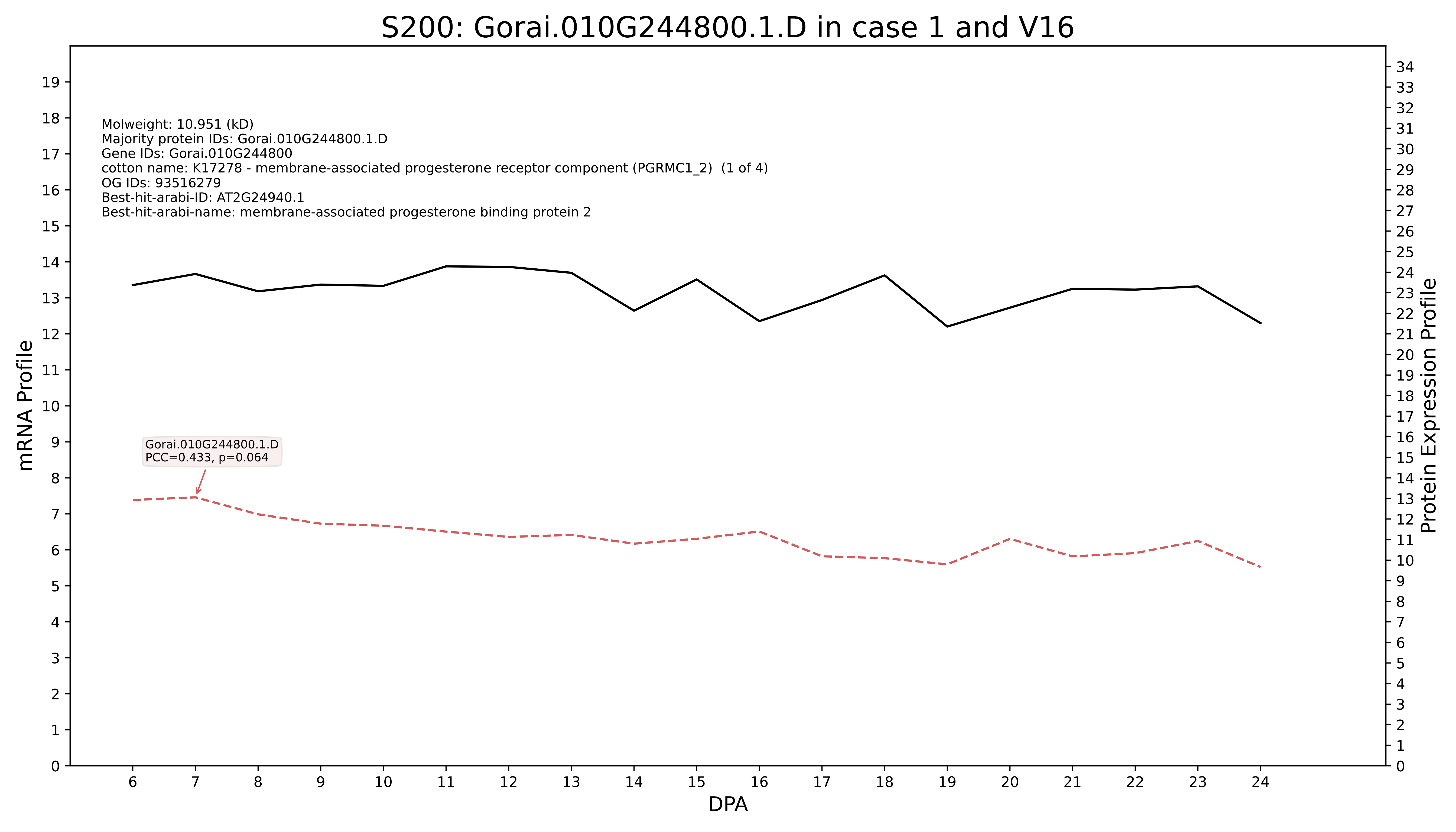
S200: Gorai.010G244800.1.D in case 1 and V16
Molweight: 10.951 (kD)
Majority protein IDs: Gorai.010G244800.1.D
Gene IDs: Gorai.010G244800
cotton name: K17278 - membrane-associated progesterone receptor component (PGRMC1_2) (1 of 4)
OG IDs: 93516279
Best-hit-arabi-ID: AT2G24940.1
Best-hit-arabi-name: membrane-associated progesterone binding protein 2
Gorai.010G244800.1.D
PCC=0.433, p=0.064
mRNA Profile
Protein Expression Profile
DPA

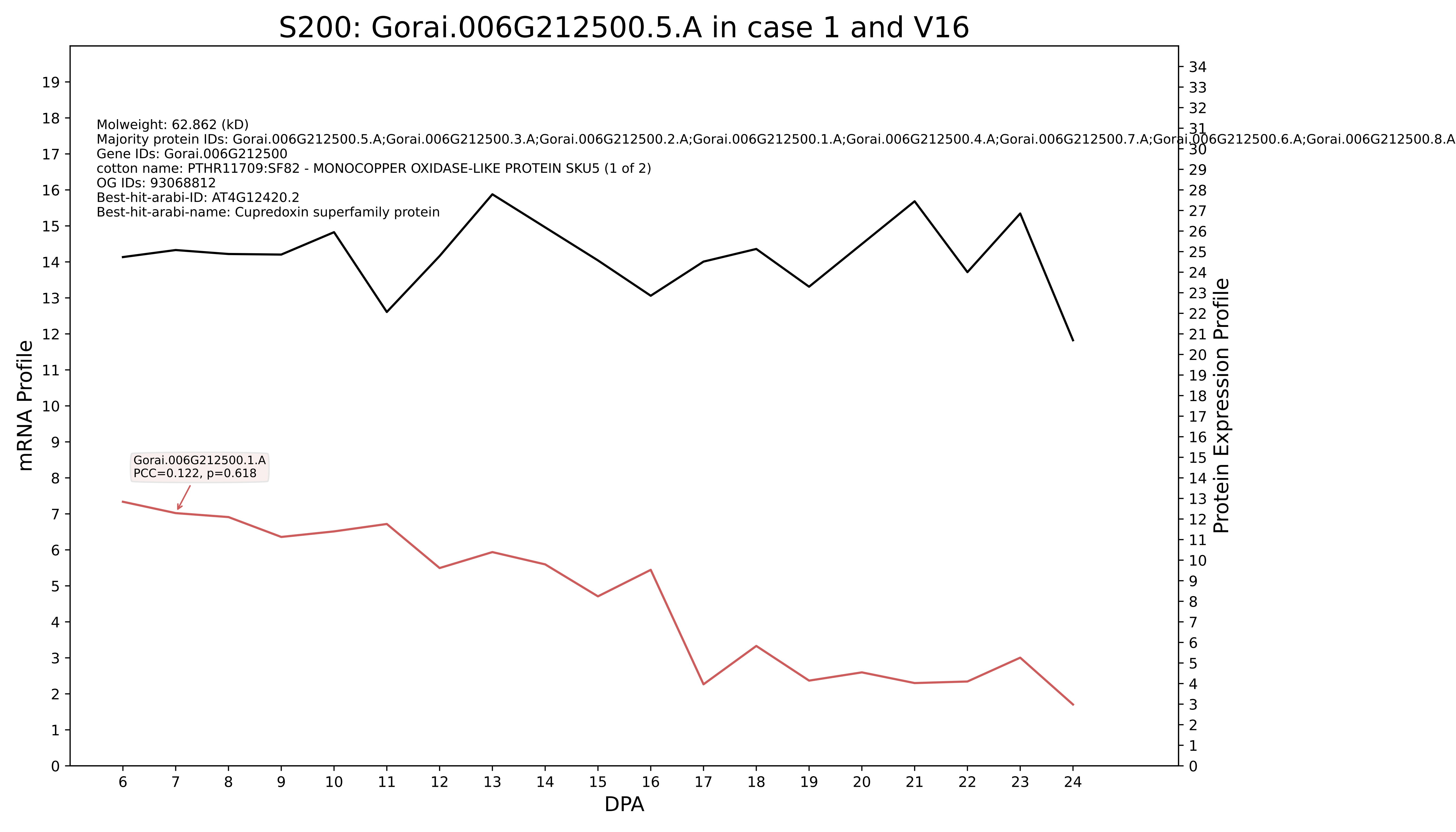
S200: Gorai.006G212500.5.A in case 1 and V16
Molweight: 62.862 (kD)
Majority protein IDs: Gorai.006G212500.5.A;Gorai.006G212500.3.A;Gorai.006G212500.2.A;Gorai.006G212500.1.A;Gorai.006G212500.4.A;Gorai.006G212500.7.A;Gorai.006G212500.6.A;Gorai.006G212500.8.A
Gene IDs: Gorai.006G212500
cotton name: PTHR11709:SF82 - MONOCOPPER OXIDASE-LIKE PROTEIN SKU5 (1 of 2)
OG IDs: 93068812
Best-hit-arabi-ID: AT4G12420.2
Best-hit-arabi-name: Cupredoxin superfamily protein
Gorai.006G212500.1.A
PCC=0.122, p=0.618
mRNA Profile
Protein Expression Profile
DPA

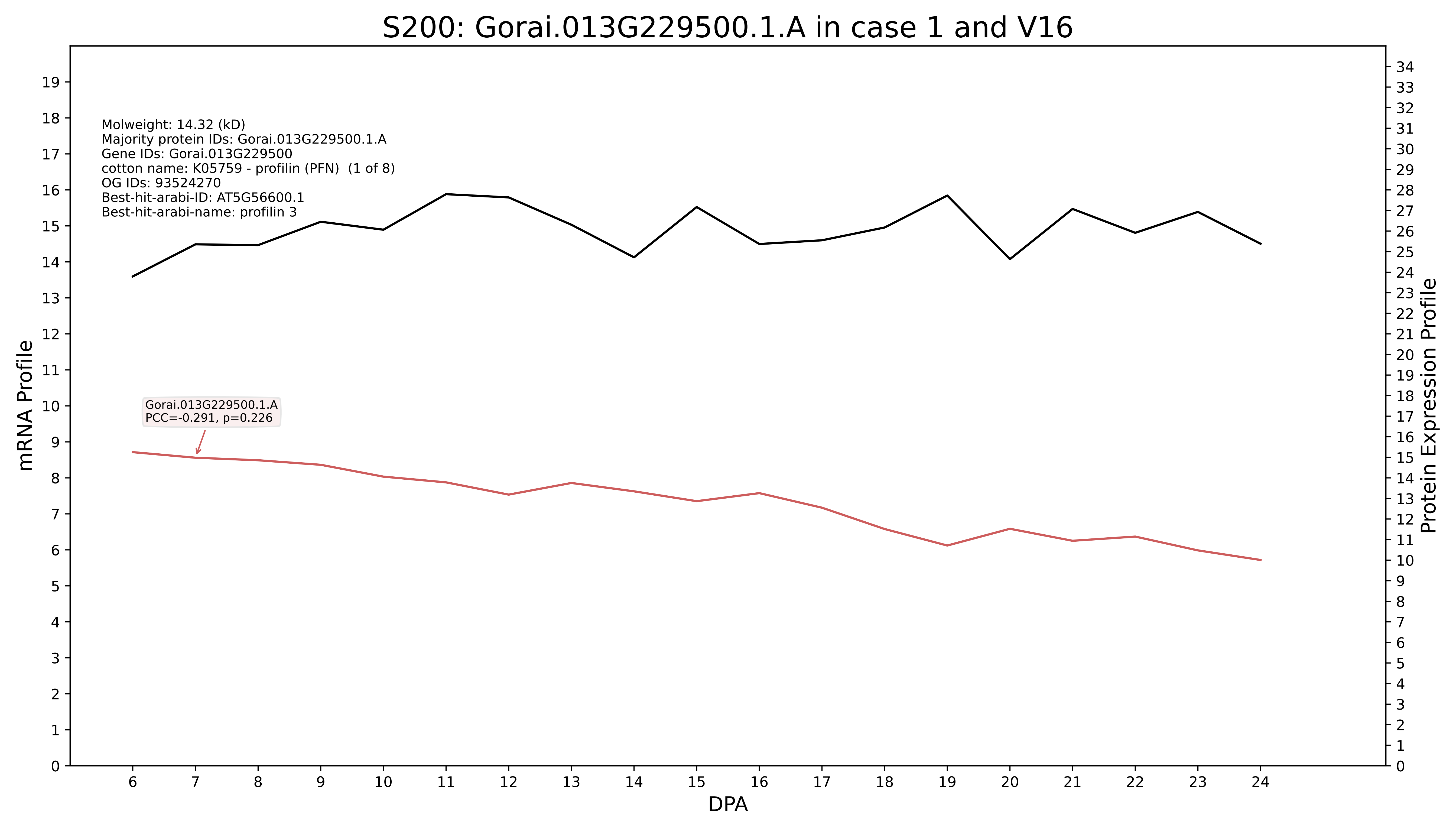

S200: Gorai.013G229500.1.A in case 1 and V16
Molweight: 14.32 (kD)
Majority protein IDs: Gorai.013G229500.1.A
Gene IDs: Gorai.013G229500
cotton name: K05759 - profilin (PFN) (1 of 8)
OG IDs: 93524270
Best-hit-arabi-ID: AT5G56600.1
Best-hit-arabi-name: profilin 3
Gorai.013G229500.1.A
PCC=-0.291, p=0.226
mRNA Profile
Protein Expression Profile
DPA

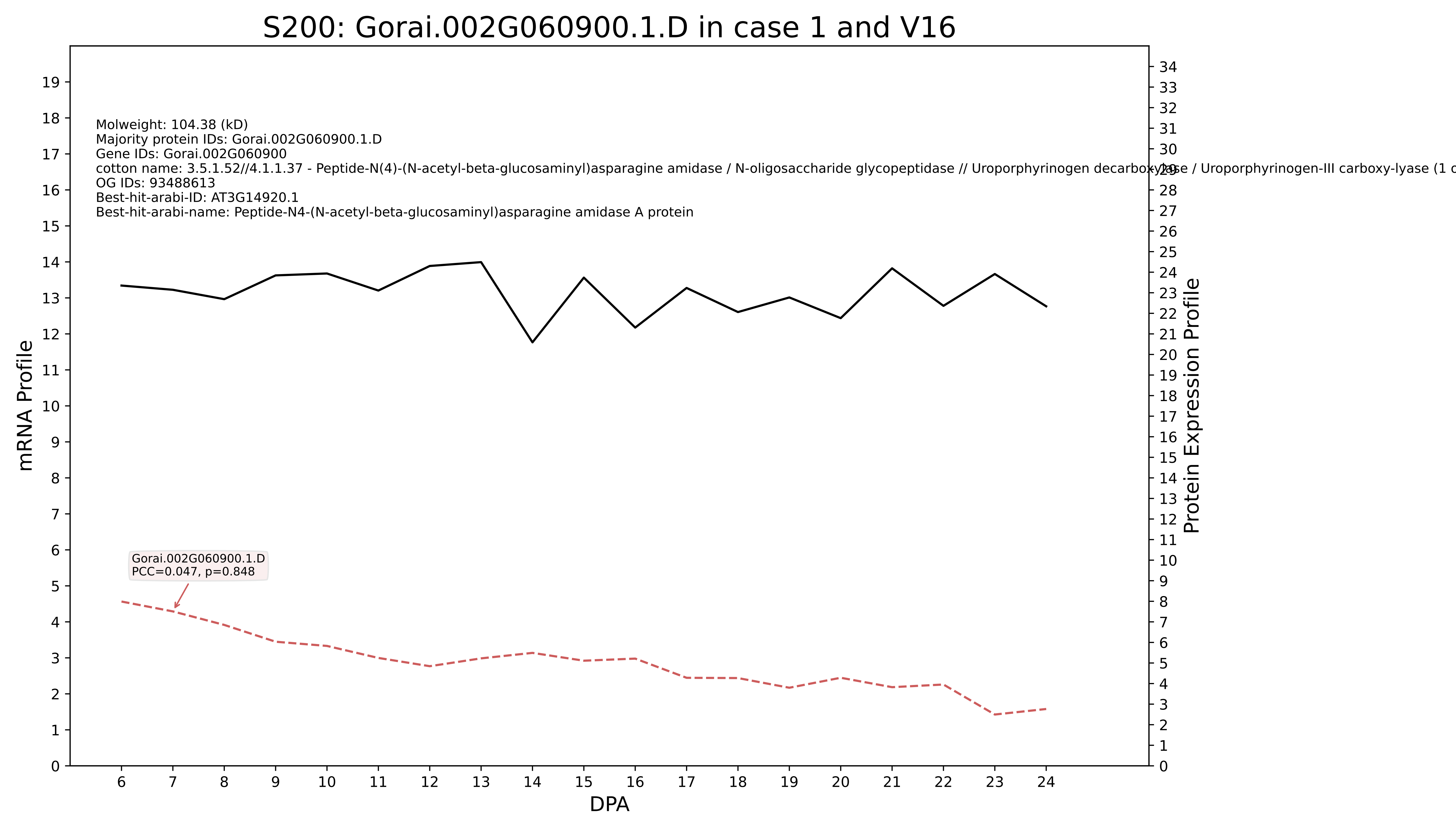
S200: Gorai.002G060900.1.D in case 1 and V16
Molweight: 104.38 (kD)
Majority protein IDs: Gorai.002G060900.1.D
Gene IDs: Gorai.002G060900
cotton name: 3.5.1.52//4.1.1.37 - Peptide-N(4)-(N-acetyl-beta-glucosaminyl)asparagine amidase / N-oligosaccharide glycopeptidase // Uroporphyrinogen decarboxylase / Uroporphyrinogen-III carboxy-lyase (1 c
OG IDs: 93488613
Best-hit-arabi-ID: AT3G14920.1
Best-hit-arabi-name: Peptide-N4-(N-acetyl-beta-glucosaminyl)asparagine amidase A protein
Gorai.002G060900.1.D
PCC=0.047, p=0.848
mRNA Profile
Protein Expression Profile
DPA

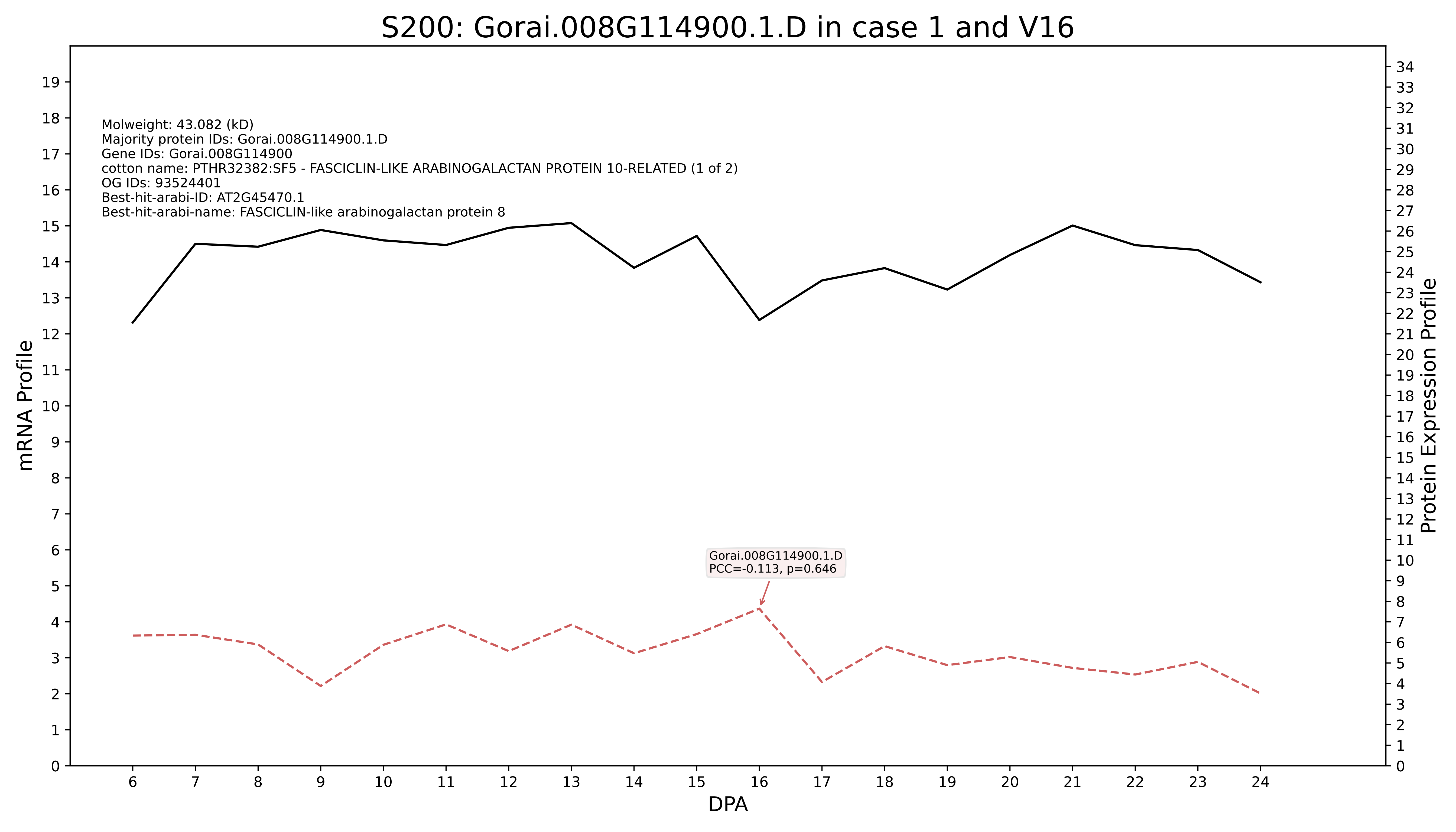
S200: Gorai.008G114900.1.D in case 1 and V16
Molweight: 43.082 (kD)
Majority protein IDs: Gorai.008G114900.1.D
Gene IDs: Gorai.008G114900
cotton name: PTHR32382:SF5 - FASCICLIN-LIKE ARABINOGALACTAN PROTEIN 10-RELATED (1 of 2)
OG IDs: 93524401
Best-hit-arabi-ID: AT2G45470.1
Best-hit-arabi-name: FASCICLIN-like arabinogalactan protein 8
Gorai.008G114900.1.D
PCC=-0.113, p=0.646
mRNA Profile
Protein Expression Profile
DPA

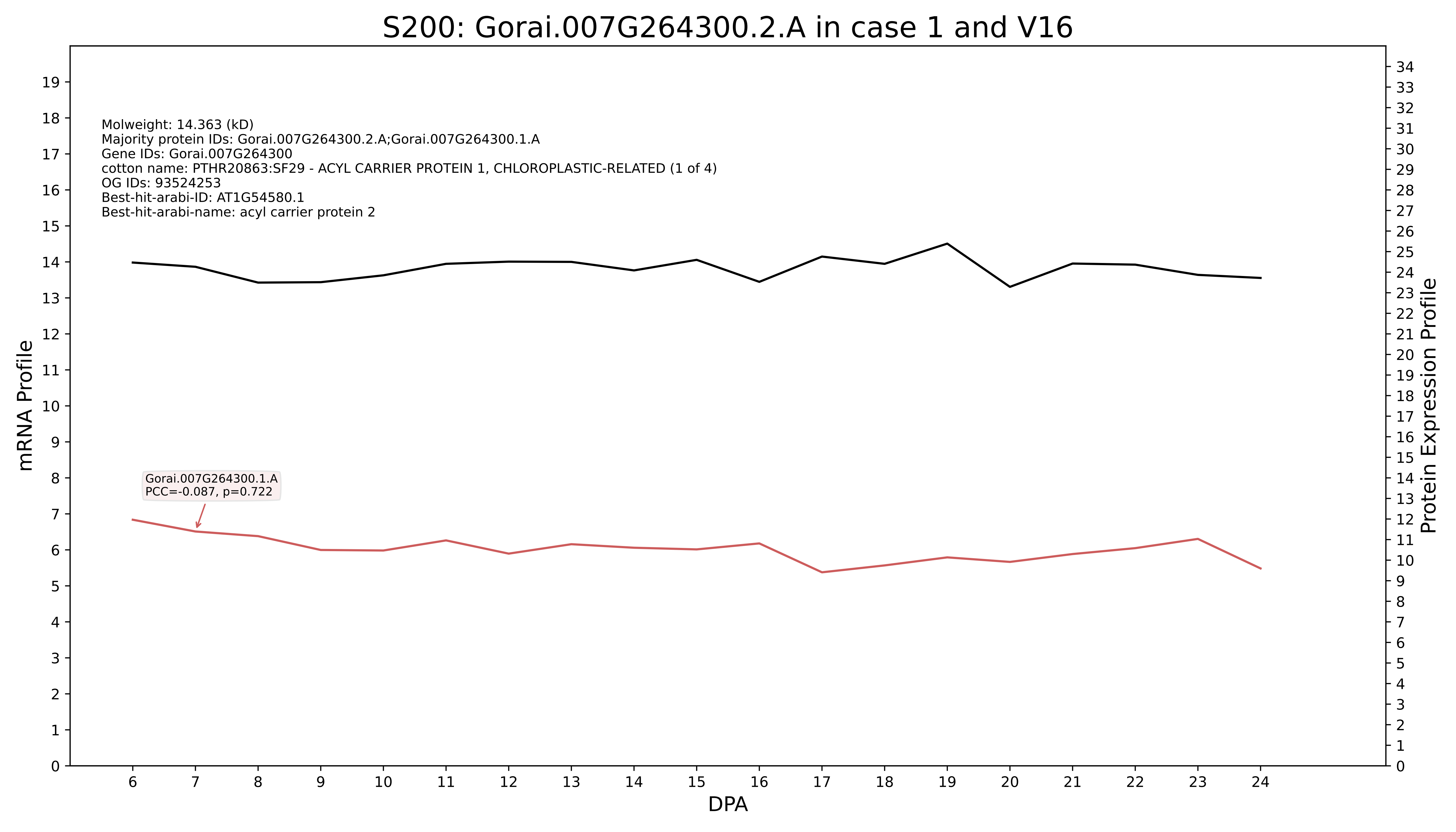
S200: Gorai.007G264300.2.A in case 1 and V16
Molweight: 14.363 (kD)
Majority protein IDs: Gorai.007G264300.2.A;Gorai.007G264300.1.A
Gene IDs: Gorai.007G264300
cotton name: PTHR20863:SF29 - ACYL CARRIER PROTEIN 1, CHLOROPLASTIC-RELATED (1 of 4)
OG IDs: 93524253
Best-hit-arabi-ID: AT1G54580.1
Best-hit-arabi-name: acyl carrier protein 2
Gorai.007G264300.1.A
PCC=-0.087, p=0.722
mRNA Profile
Protein Expression Profile
DPA

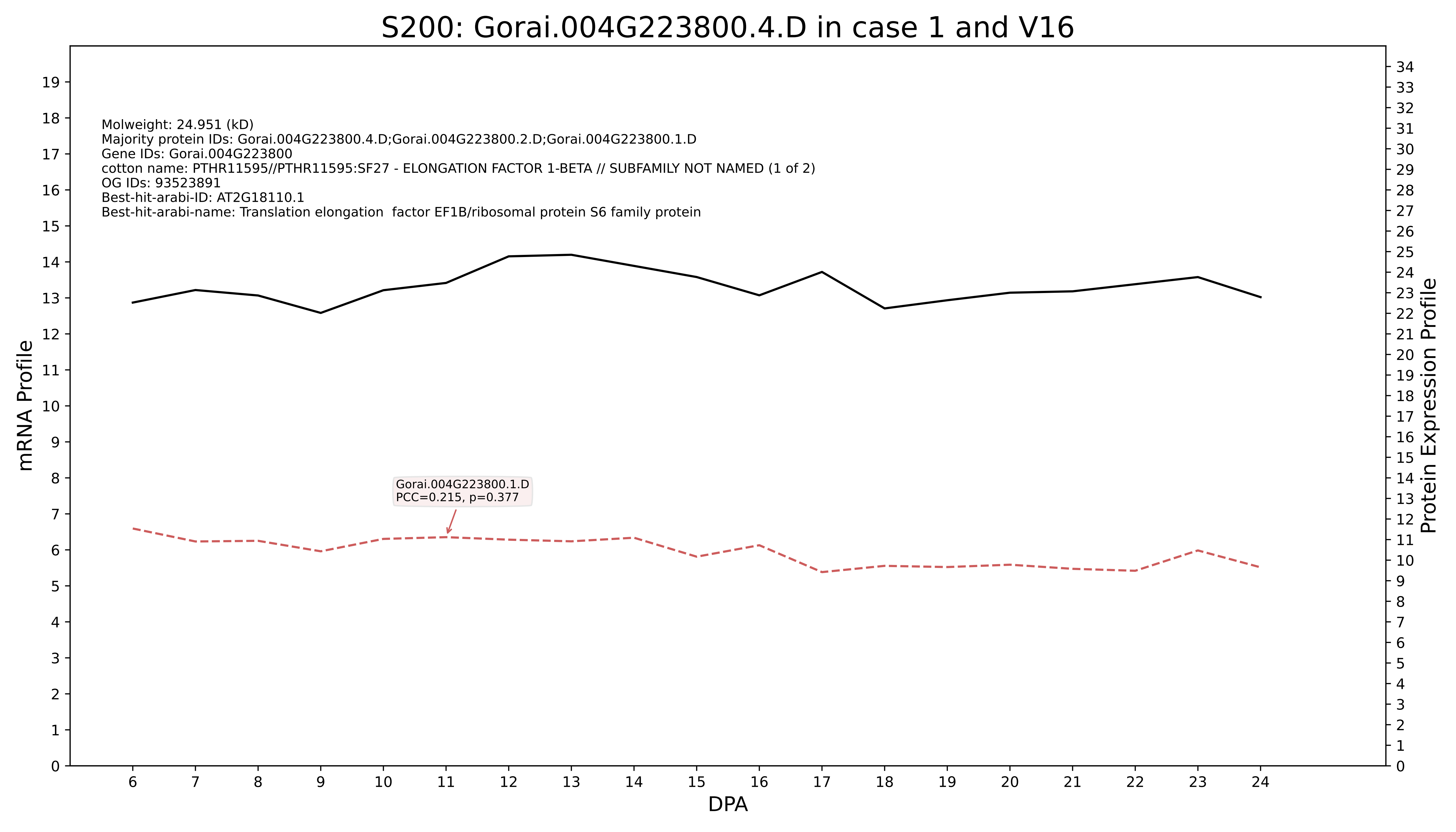
S200: Gorai.004G223800.4.D in case 1 and V16
Molweight: 24.951 (kD)
Majority protein IDs: Gorai.004G223800.4.D;Gorai.004G223800.2.D;Gorai.004G223800.1.D
Gene IDs: Gorai.004G223800
cotton name: PTHR11595//PTHR11595:SF27 - ELONGATION FACTOR 1-BETA // SUBFAMILY NOT NAMED (1 of 2)
OG IDs: 93523891
Best-hit-arabi-ID: AT2G18110.1
Best-hit-arabi-name: Translation elongation factor EF1B/ribosomal protein S6 family protein
Gorai.004G223800.1.D
PCC=0.215, p=0.377
mRNA Profile
Protein Expression Profile
DPA

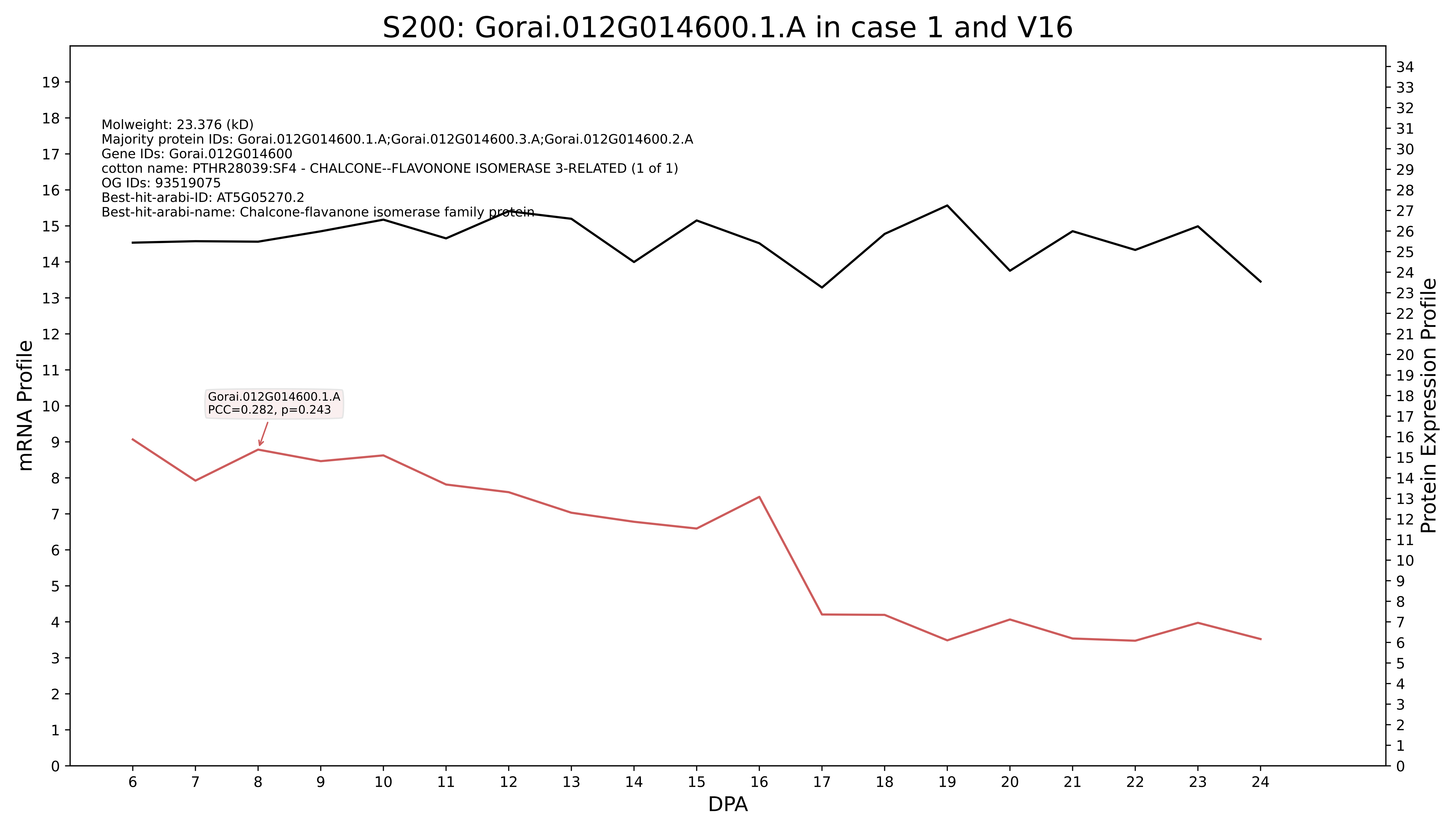
S200: Gorai.012G014600.1.A in case 1 and V16
Molweight: 23.376 (kD)
Majority protein IDs: Gorai.012G014600.1.A;Gorai.012G014600.3.A;Gorai.012G014600.2.A
Gene IDs: Gorai.012G014600
cotton name: PTHR28039:SF4 - CHALCONE--FLAVONONE ISOMERASE 3-RELATED (1 of 1)
OG IDs: 93519075
Best-hit-arabi-ID: AT5G05270.2
Best-hit-arabi-name: Chalcone-flavanone isomerase family protein
Gorai.012G014600.1.A
PCC=0.282, p=0.243
mRNA Profile
Protein Expression Profile
DPA

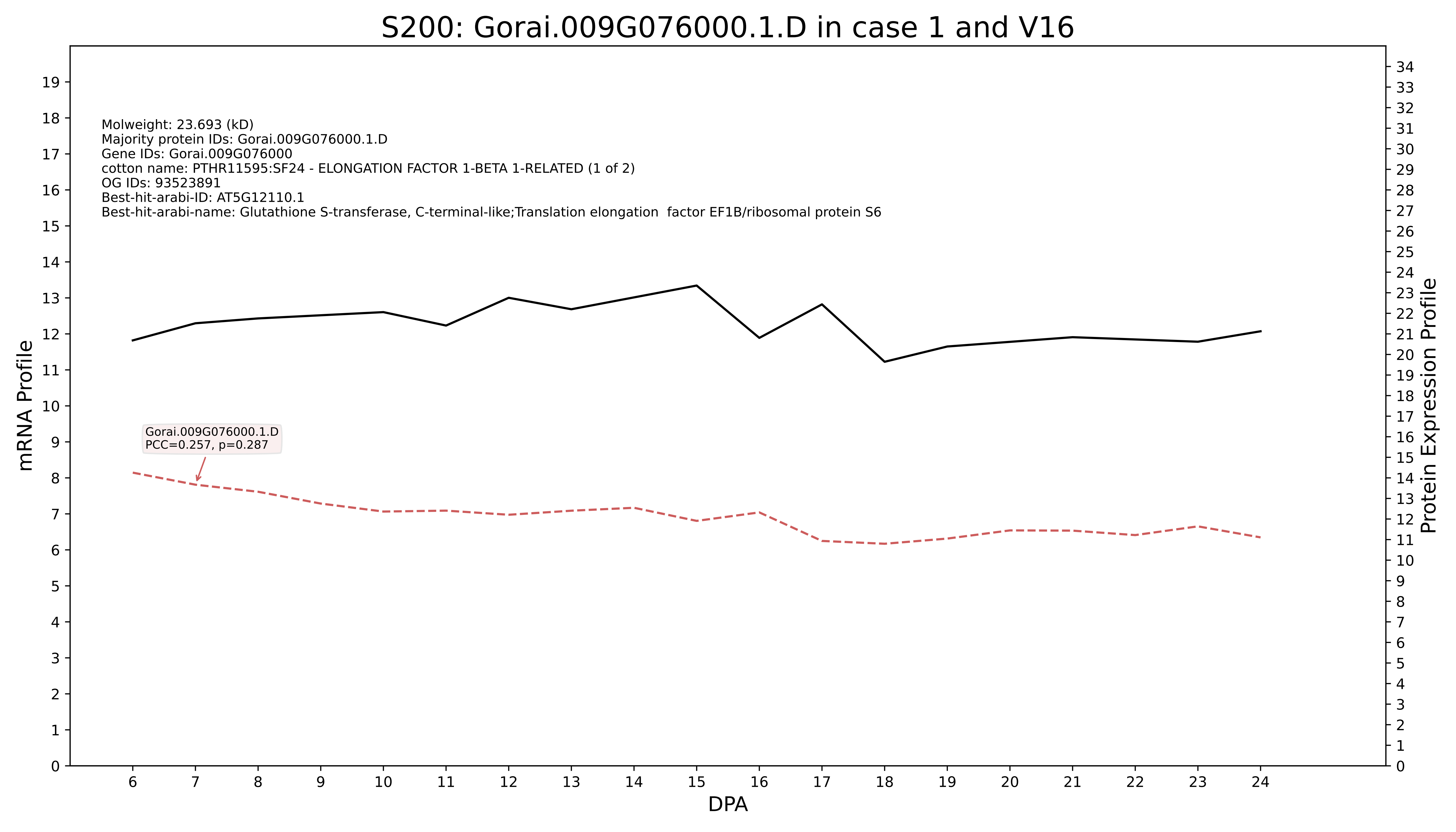
S200: Gorai.009G076000.1.D in case 1 and V16
Molweight: 23.693 (kD)
Majority protein IDs: Gorai.009G076000.1.D
Gene IDs: Gorai.009G076000
cotton name: PTHR11595:SF24 - ELONGATION FACTOR 1-BETA 1-RELATED (1 of 2)
OG IDs: 93523891
Best-hit-arabi-ID: AT5G12110.1
Best-hit-arabi-name: Glutathione S-transferase, C-terminal-like;Translation elongation factor EF1B/ribosomal protein S6
Gorai.009G076000.1.D
PCC=0.257, p=0.287
mRNA Profile
Protein Expression Profile
DPA

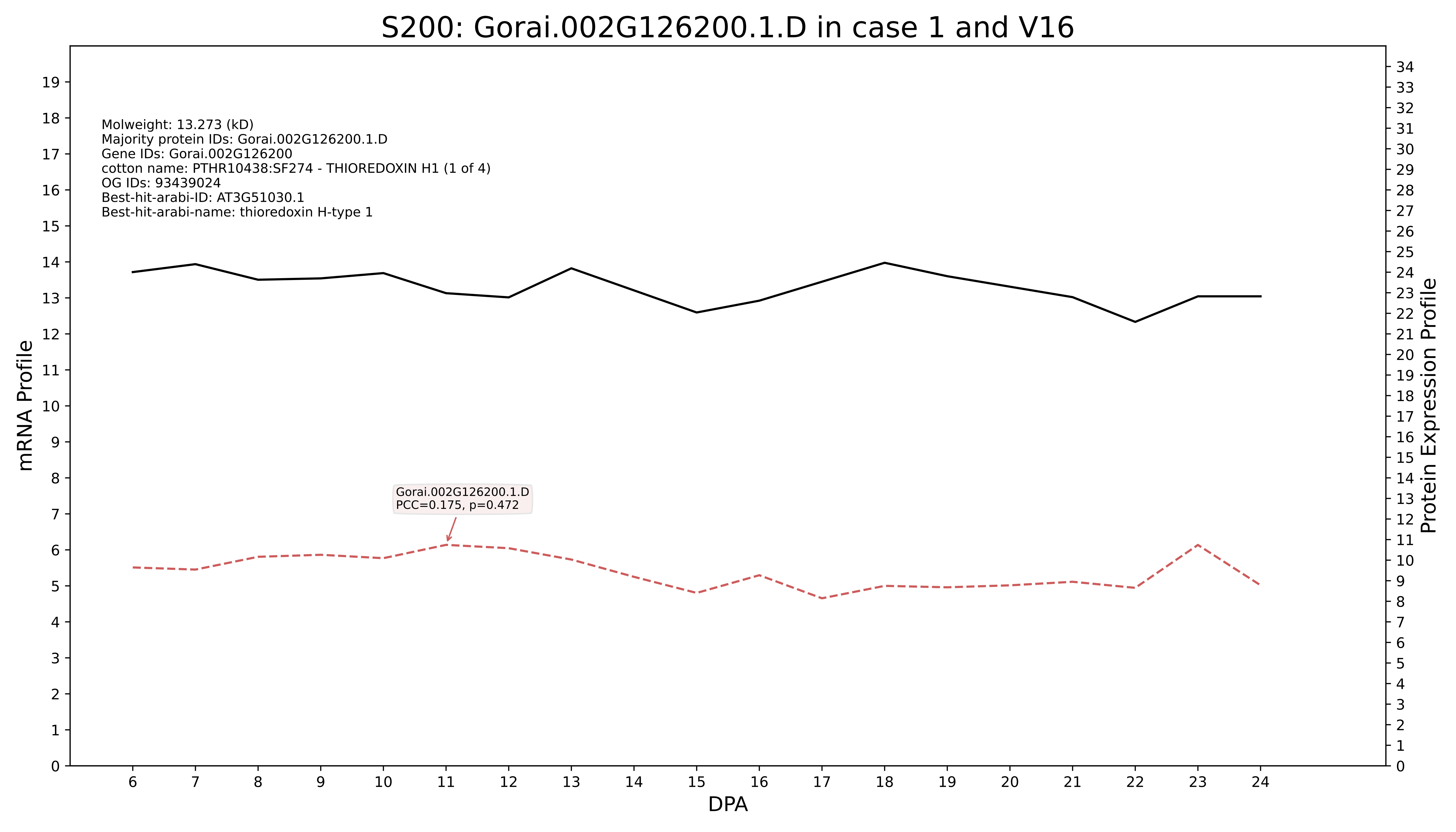

S200: Gorai.002G126200.1.D in case 1 and V16
Molweight: 13.273 (kD)
Majority protein IDs: Gorai.002G126200.1.D
Gene IDs: Gorai.002G126200
cotton name: PTHR10438:SF274 - THIOREDOXIN H1 (1 of 4)
OG IDs: 93439024
Best-hit-arabi-ID: AT3G51030.1
Best-hit-arabi-name: thioredoxin H-type 1
Gorai.002G126200.1.D
PCC=0.175, p=0.472
mRNA Profile
Protein Expression Profile
DPA

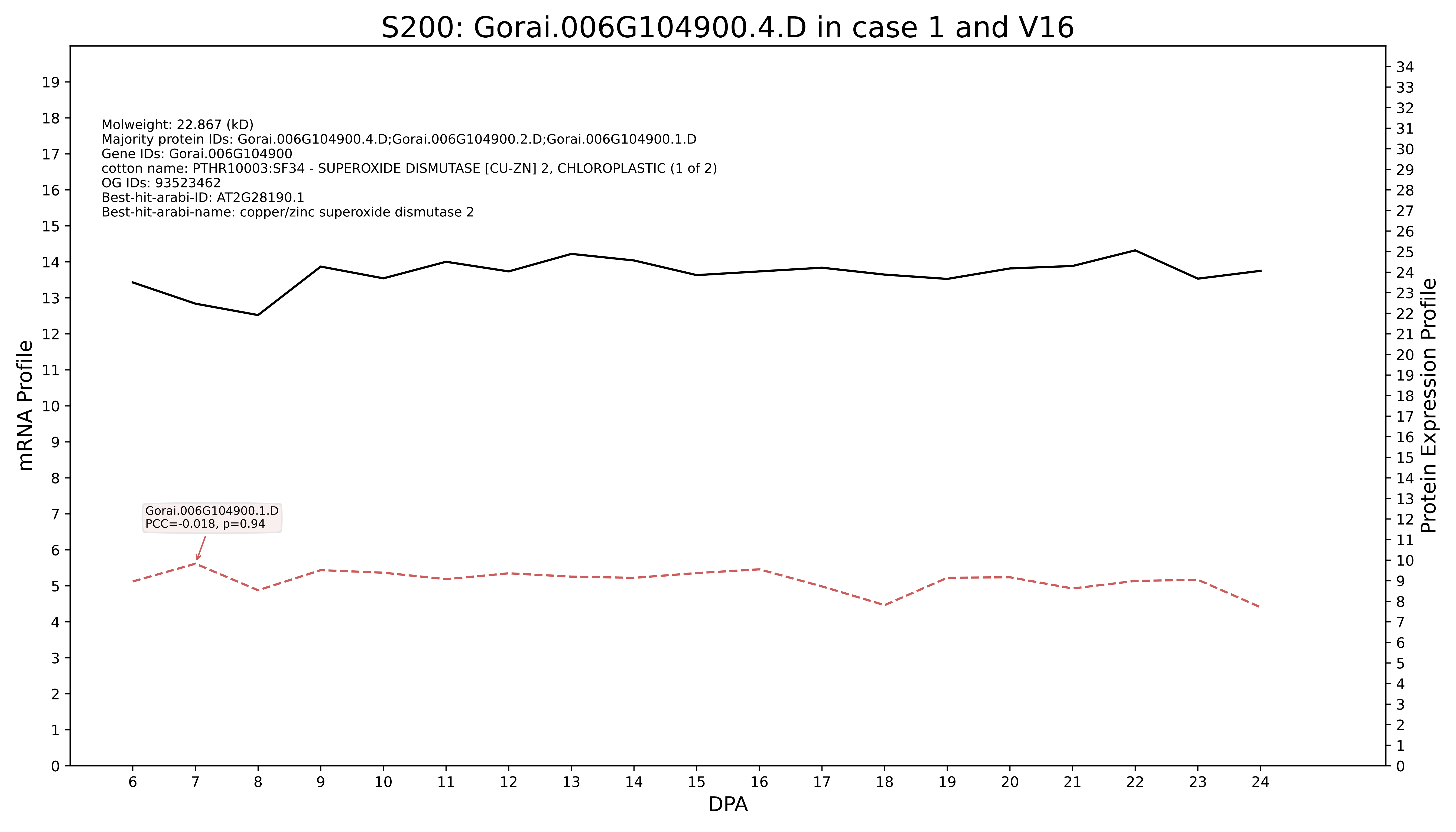
S200: Gorai.006G104900.4.D in case 1 and V16
Molweight: 22.867 (kD)
Majority protein IDs: Gorai.006G104900.4.D;Gorai.006G104900.2.D;Gorai.006G104900.1.D
Gene IDs: Gorai.006G104900
cotton name: PTHR10003:SF34 - SUPEROXIDE DISMUTASE [CU-ZN] 2, CHLOROPLASTIC (1 of 2)
OG IDs: 93523462
Best-hit-arabi-ID: AT2G28190.1
Best-hit-arabi-name: copper/zinc superoxide dismutase 2
Gorai.006G104900.1.D
PCC=-0.018, p=0.94
mRNA Profile
Protein Expression Profile
DPA

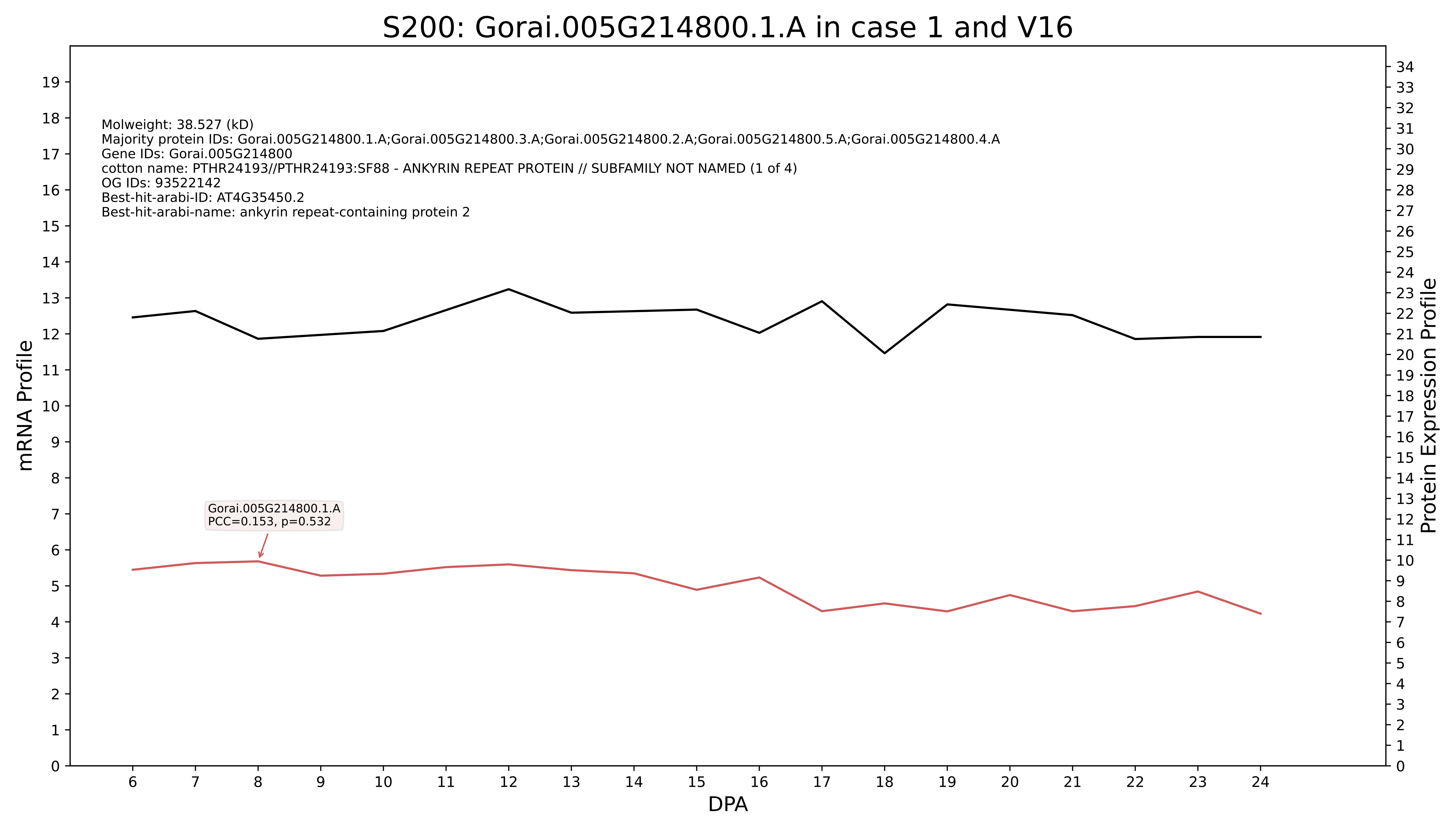
S200: Gorai.005G214800.1.A in case 1 and V16
Molweight: 38.527 (kD)
Majority protein IDs: Gorai.005G214800.1.A;Gorai.005G214800.3.A;Gorai.005G214800.2.A;Gorai.005G214800.5.A;Gorai.005G214800.4.A
Gene IDs: Gorai.005G214800
cotton name: PTHR24193//PTHR24193:SF88 - ANKYRIN REPEAT PROTEIN // SUBFAMILY NOT NAMED (1 of 4)
OG IDs: 93522142
Best-hit-arabi-ID: AT4G35450.2
Best-hit-arabi-name: ankyrin repeat-containing protein 2
Gorai.005G214800.1.A
PCC=0.153, p=0.532
mRNA Profile
Protein Expression Profile
DPA

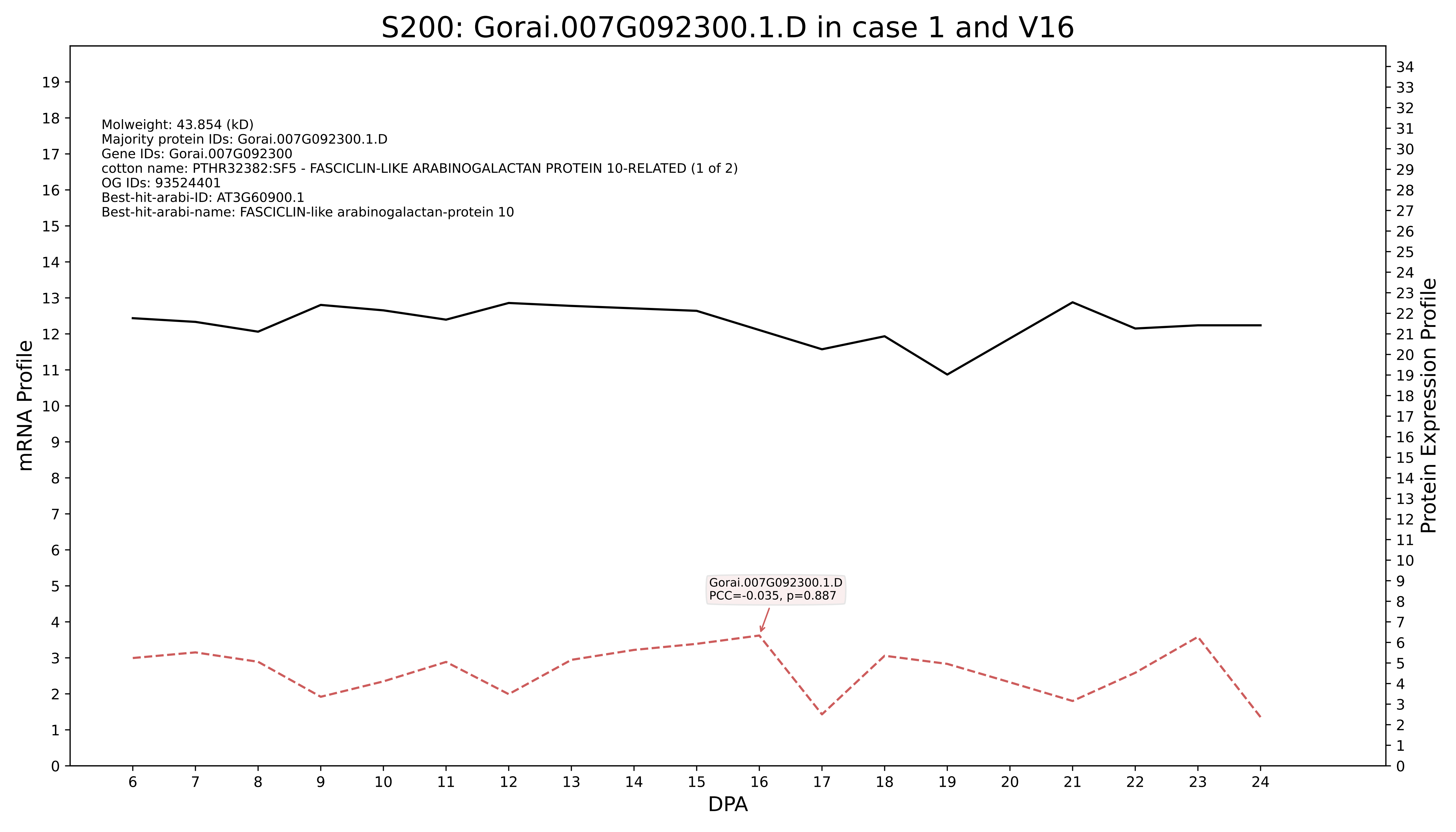

S200: Gorai.007G092300.1.D in case 1 and V16
Molweight: 43.854 (kD)
Majority protein IDs: Gorai.007G092300.1.D
Gene IDs: Gorai.007G092300
cotton name: PTHR32382:SF5 - FASCICLIN-LIKE ARABINOGALACTAN PROTEIN 10-RELATED (1 of 2)
OG IDs: 93524401
Best-hit-arabi-ID: AT3G60900.1
Best-hit-arabi-name: FASCICLIN-like arabinogalactan-protein 10
Gorai.007G092300.1.D
PCC=-0.035, p=0.887
mRNA Profile
Protein Expression Profile
DPA

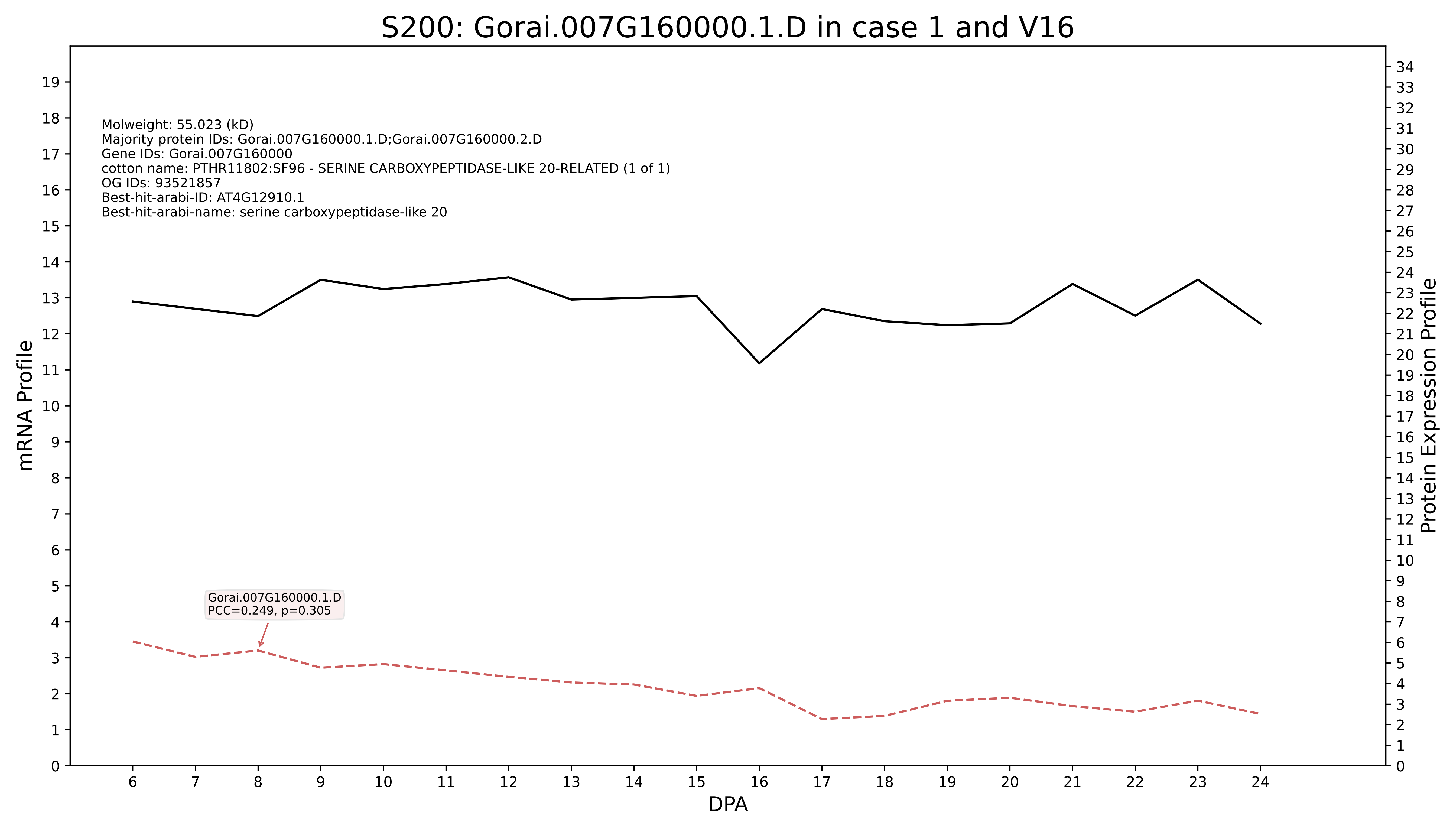

S200: Gorai.007G160000.1.D in case 1 and V16
Molweight: 55.023 (kD)
Majority protein IDs: Gorai.007G160000.1.D;Gorai.007G160000.2.D
Gene IDs: Gorai.007G160000
cotton name: PTHR11802:SF96 - SERINE CARBOXYPEPTIDASE-LIKE 20-RELATED (1 of 1)
OG IDs: 93521857
Best-hit-arabi-ID: AT4G12910.1
Best-hit-arabi-name: serine carboxypeptidase-like 20
Gorai.007G160000.1.D
PCC=0.249, p=0.305
mRNA Profile
Protein Expression Profile
DPA

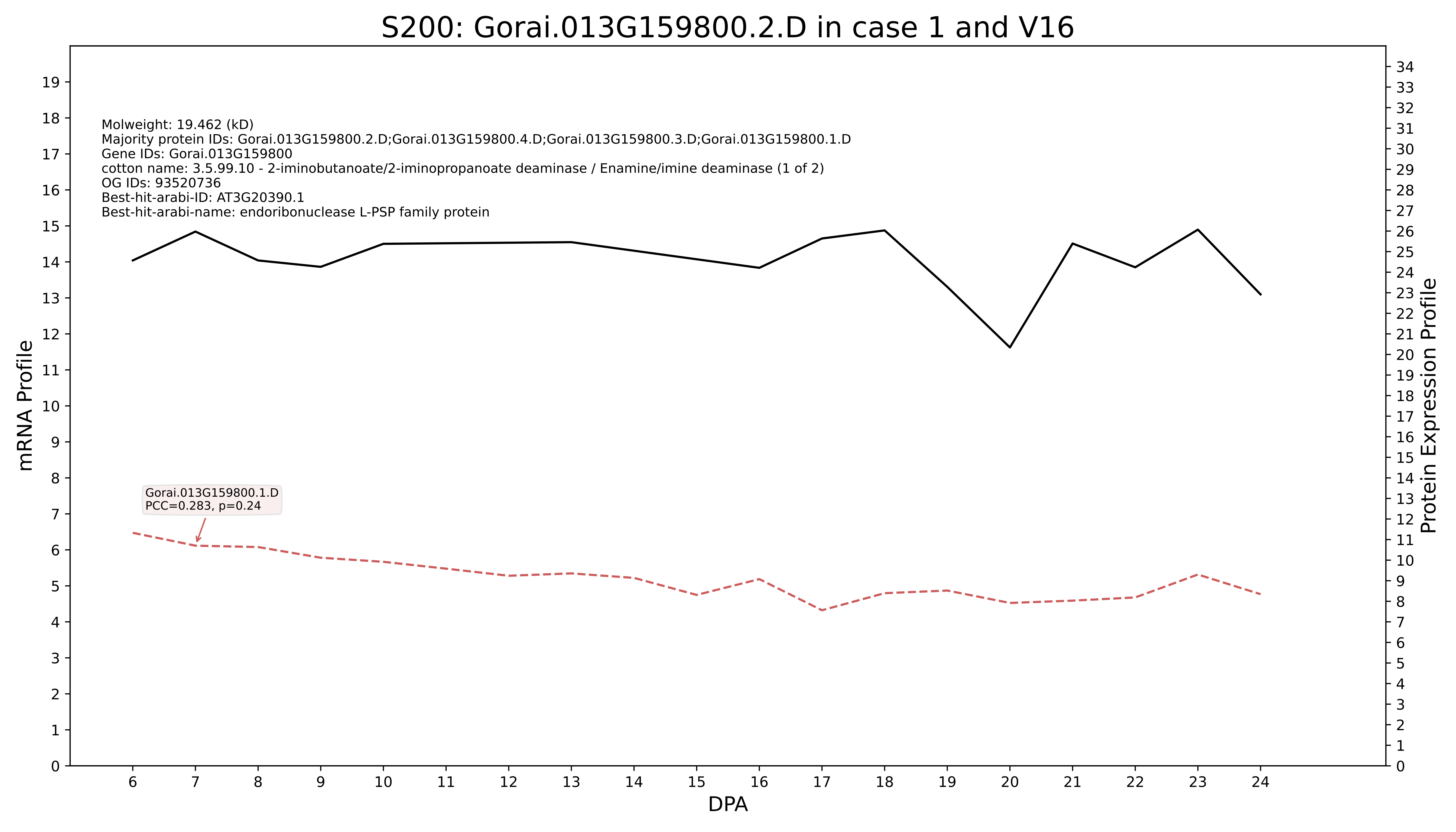
S200: Gorai.013G159800.2.D in case 1 and V16
Molweight: 19.462 (kD)
Majority protein IDs: Gorai.013G159800.2.D;Gorai.013G159800.4.D;Gorai.013G159800.3.D;Gorai.013G159800.1.D
Gene IDs: Gorai.013G159800
cotton name: 3.5.99.10 - 2-iminobutanoate/2-iminopropanoate deaminase / Enamine/imine deaminase (1 of 2)
OG IDs: 93520736
Best-hit-arabi-ID: AT3G20390.1
Best-hit-arabi-name: endoribonuclease L-PSP family protein
Gorai.013G159800.1.D
PCC=0.283, p=0.24
mRNA Profile
Protein Expression Profile
DPA

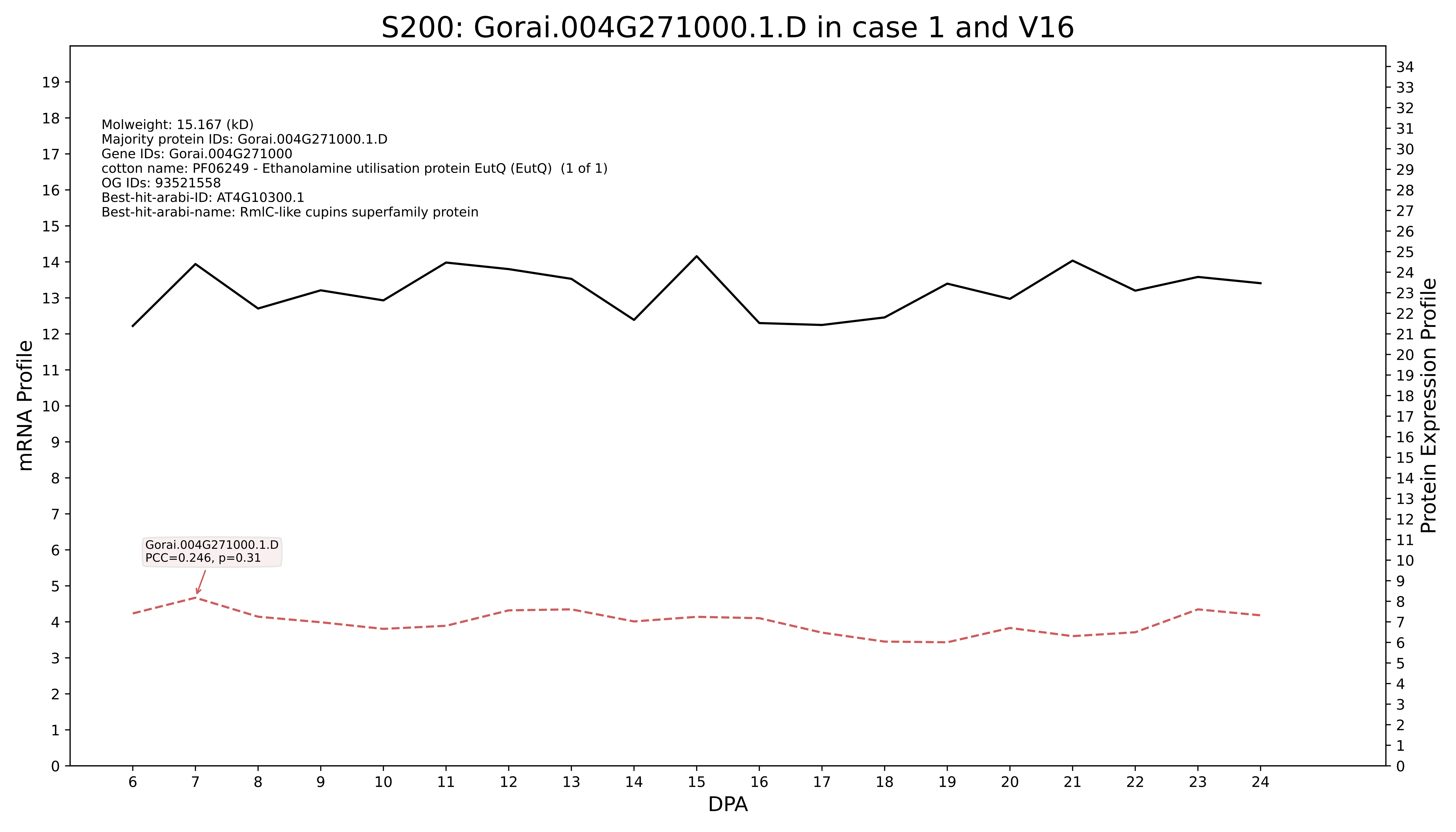

S200: Gorai.004G271000.1.D in case 1 and V16
Molweight: 15.167 (kD)
Majority protein IDs: Gorai.004G271000.1.D
Gene IDs: Gorai.004G271000
cotton name: PF06249 - Ethanolamine utilisation protein EutQ (EutQ) (1 of 1)
OG IDs: 93521558
Best-hit-arabi-ID: AT4G10300.1
Best-hit-arabi-name: RmlC-like cupins superfamily protein
Gorai.004G271000.1.D
PCC=0.246, p=0.31
mRNA Profile
Protein Expression Profile
DPA

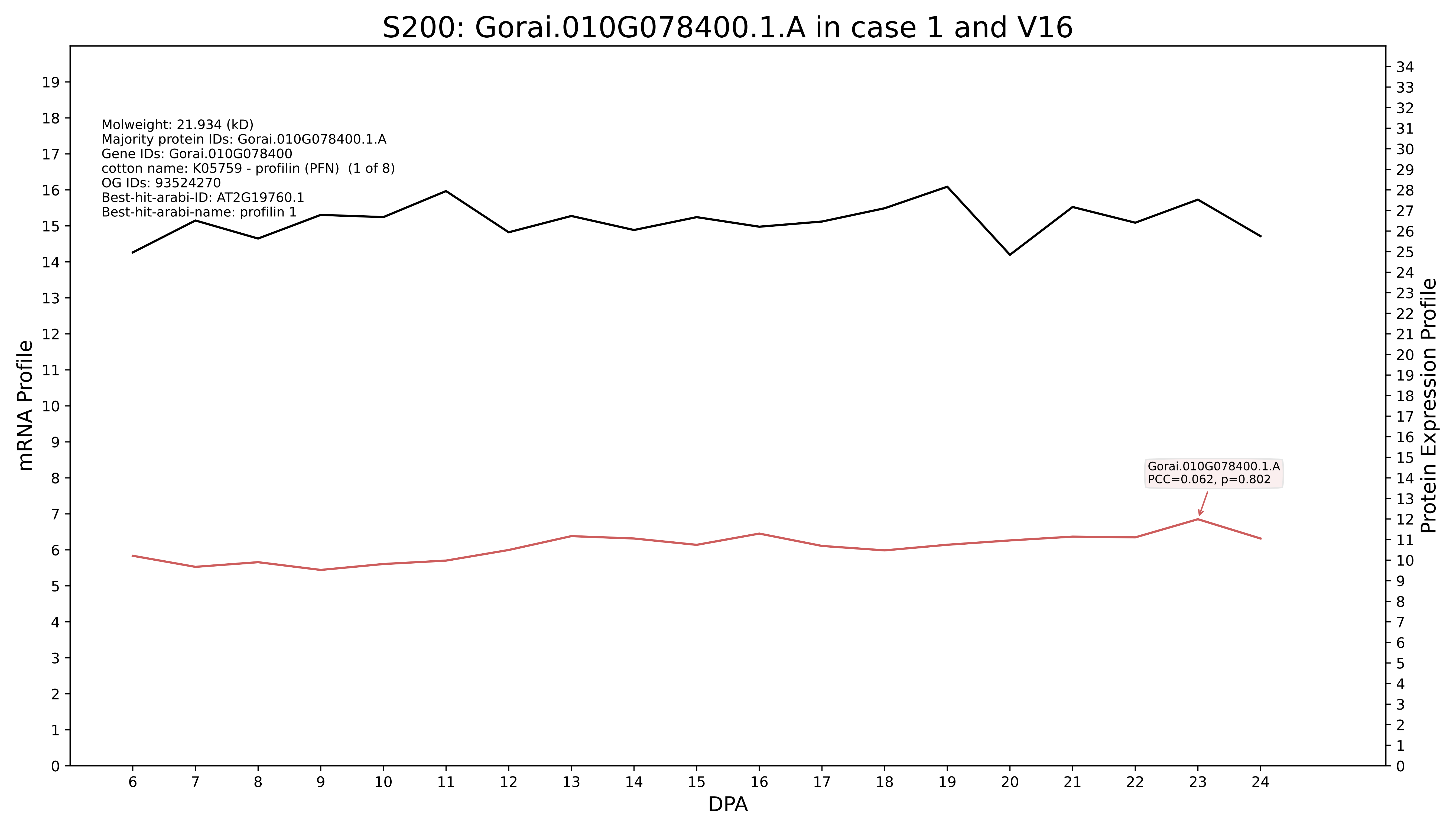

S200: Gorai.010G078400.1.A in case 1 and V16
Molweight: 21.934 (kD)
Majority protein IDs: Gorai.010G078400.1.A
Gene IDs: Gorai.010G078400
cotton name: K05759 - profilin (PFN) (1 of 8)
OG IDs: 93524270
Best-hit-arabi-ID: AT2G19760.1
Best-hit-arabi-name: profilin 1
Gorai.010G078400.1.A
PCC=0.062, p=0.802
mRNA Profile
Protein Expression Profile
DPA

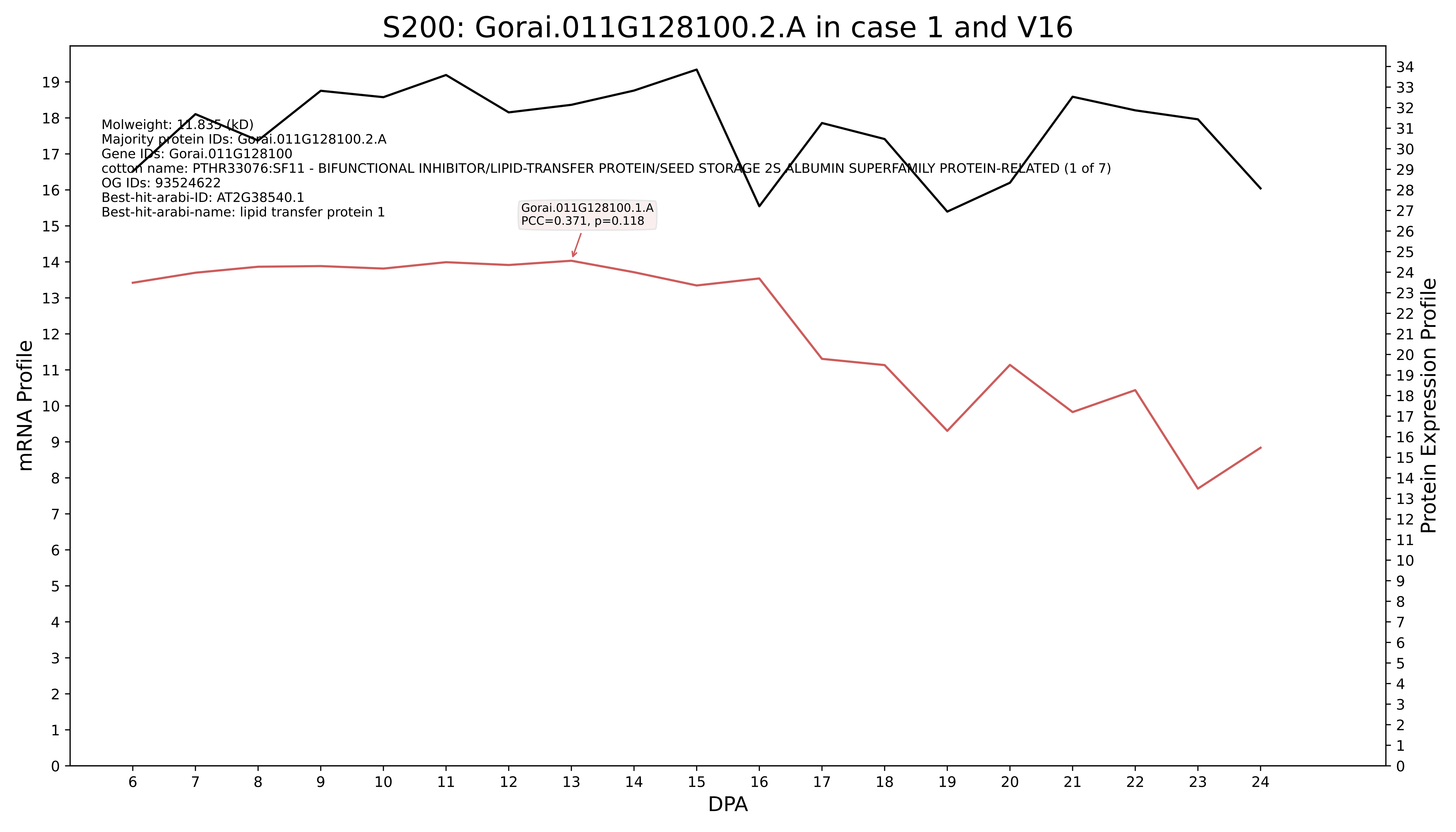

S200: Gorai.011G128100.2.A in case 1 and V16
Molweight: 11.835 (kD)
Majority protein IDs: Gorai.011G128100.2.A
Gene IDs: Gorai.011G128100
cotton name: PTHR33076:SF11 - BIFUNCTIONAL INHIBITOR/LIPID-TRANSFER PROTEIN/SEED STORAGE 2S ALBUMIN SUPERFAMILY PROTEIN-RELATED (1 of 7)
OG IDs: 93524622
Best-hit-arabi-ID: AT2G38540.1
Best-hit-arabi-name: lipid transfer protein 1
Gorai.011G128100.1.A
PCC=0.371, p=0.118
mRNA Profile
Protein Expression Profile
DPA

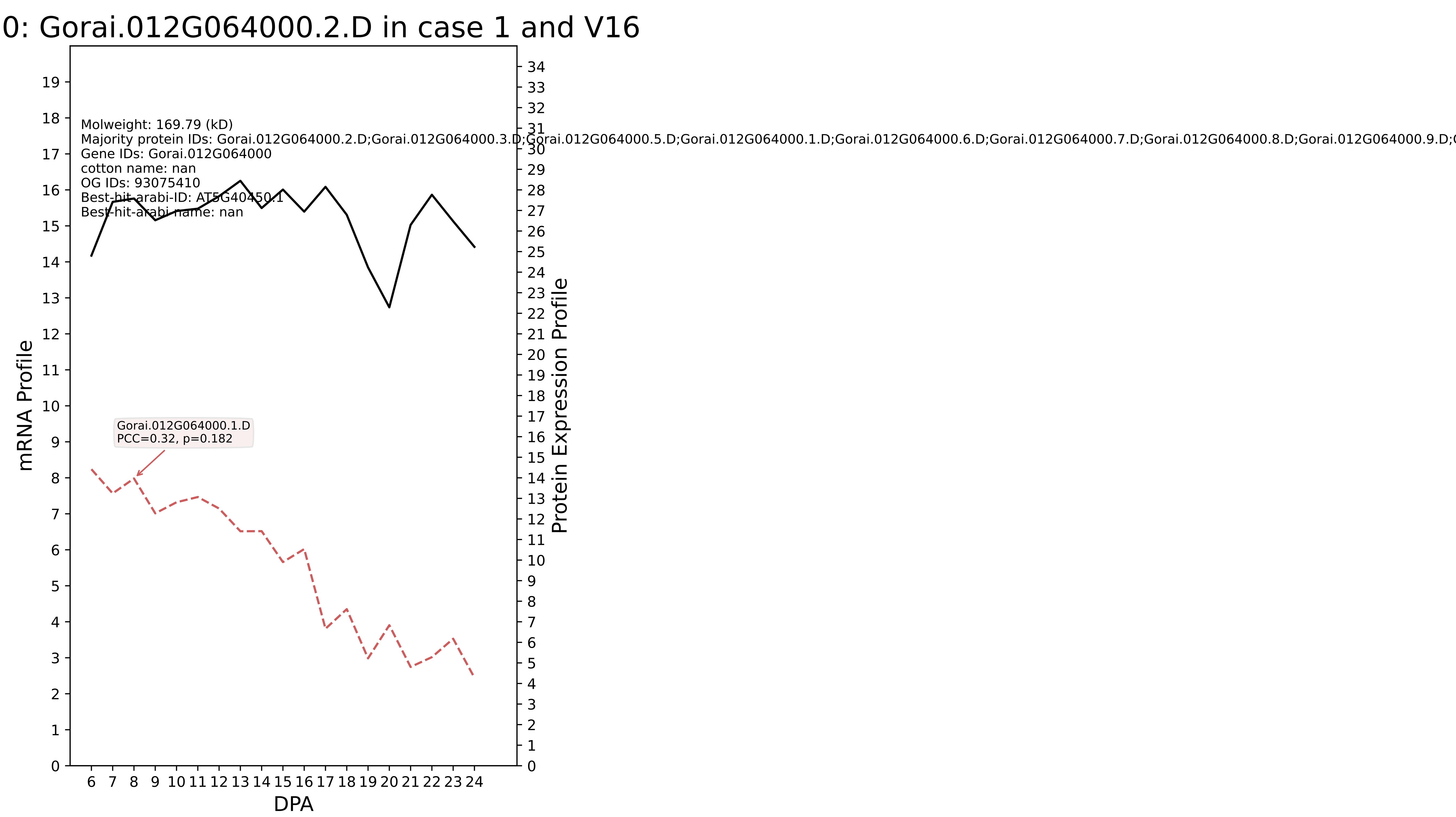
0: Gorai.012G064000.2.D in case 1 and V16
Molweight: 169.79 (kD)
Majority protein IDs: Gorai.012G064000.2.D;Gorai.012G064000.3.D;Gorai.012G064000.5.D;Gorai.012G064000.1.D;Gorai.012G064000.6.D;Gorai.012G064000.7.D;Gorai.012G064000.8.D;Gorai.012G064000.9.D;G
Gene IDs: Gorai.012G064000
cotton name: nan
OG IDs: 93075410
Best-hit-arabi-ID: AT5G40450.1
Best-hit-arabi-name: nan
Gorai.012G064000.1.D
PCC=0.32, p=0.182
mRNA Profile
Protein Expression Profile
DPA

### S200: Gorai.012G020400.2.D in case 1 and V16

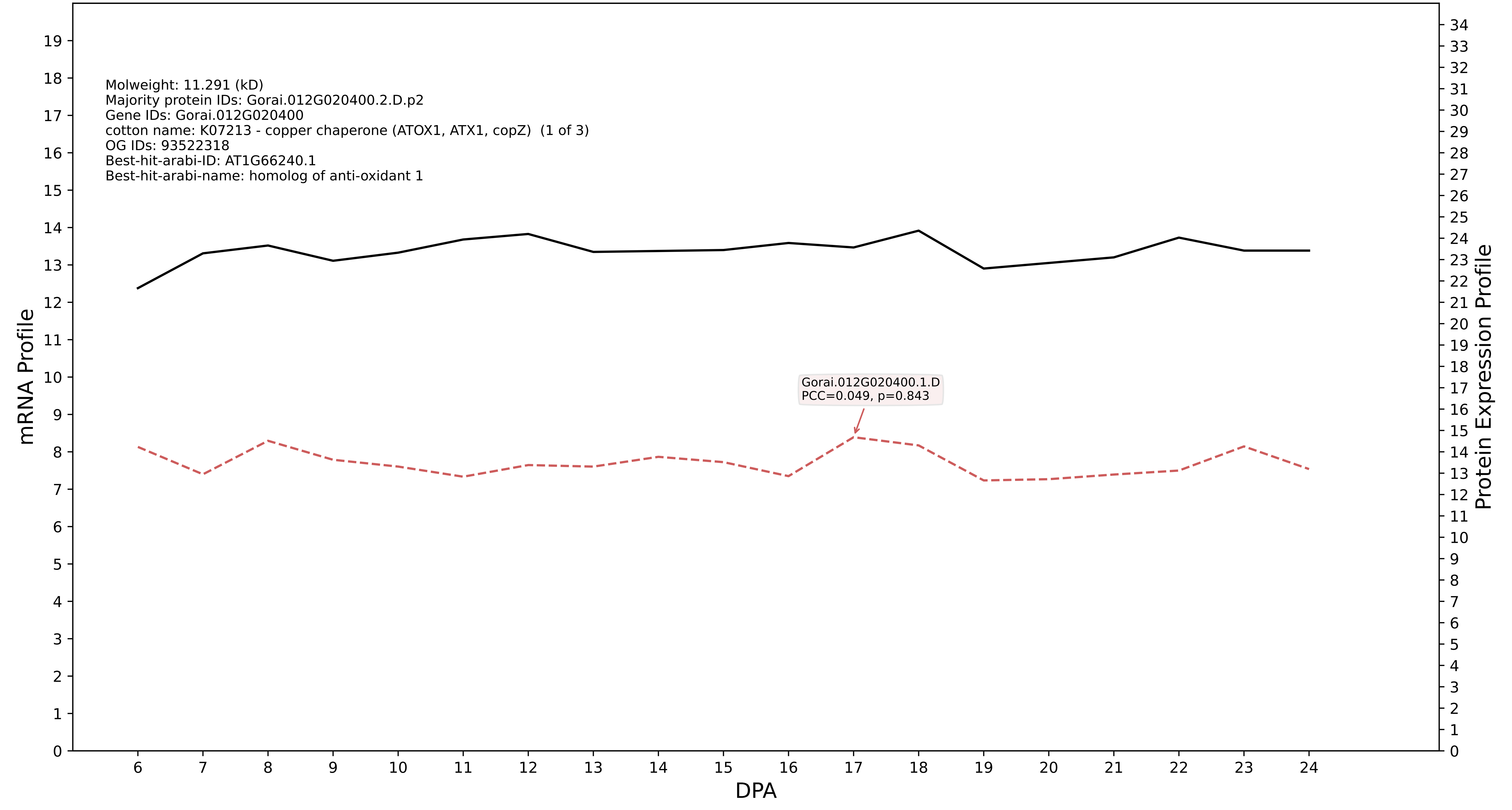

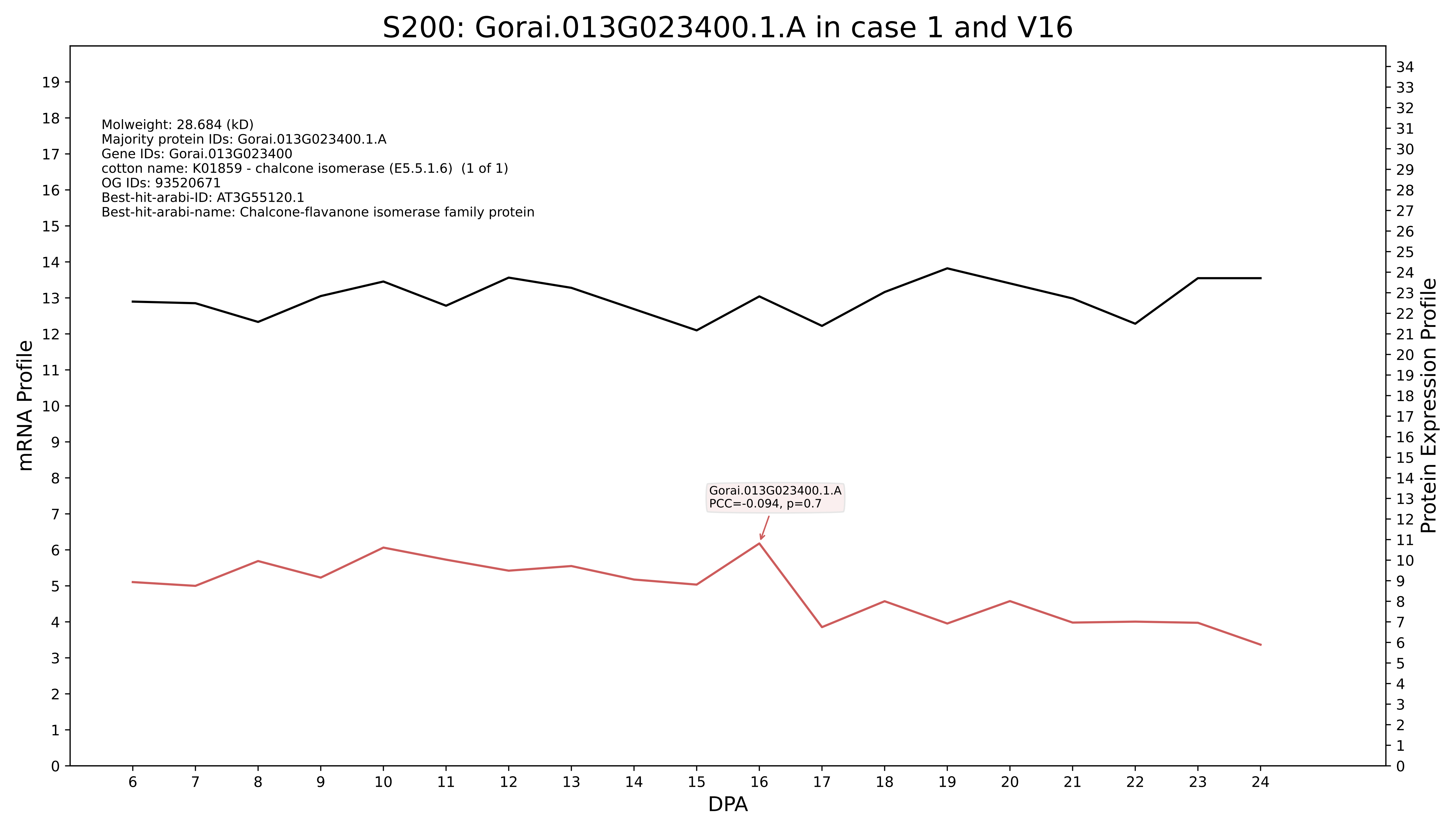
S200: Gorai.013G023400.1.A in case 1 and V16
Molweight: 28.684 (kD)
Majority protein IDs: Gorai.013G023400.1.A
Gene IDs: Gorai.013G023400
cotton name: K01859 - chalcone isomerase (E5.5.1.6) (1 of 1)
OG IDs: 93520671
Best-hit-arabi-ID: AT3G55120.1
Best-hit-arabi-name: Chalcone-flavanone isomerase family protein
Gorai.013G023400.1.A
PCC=-0.094, p=0.7
mRNA Profile
Protein Expression Profile
DPA

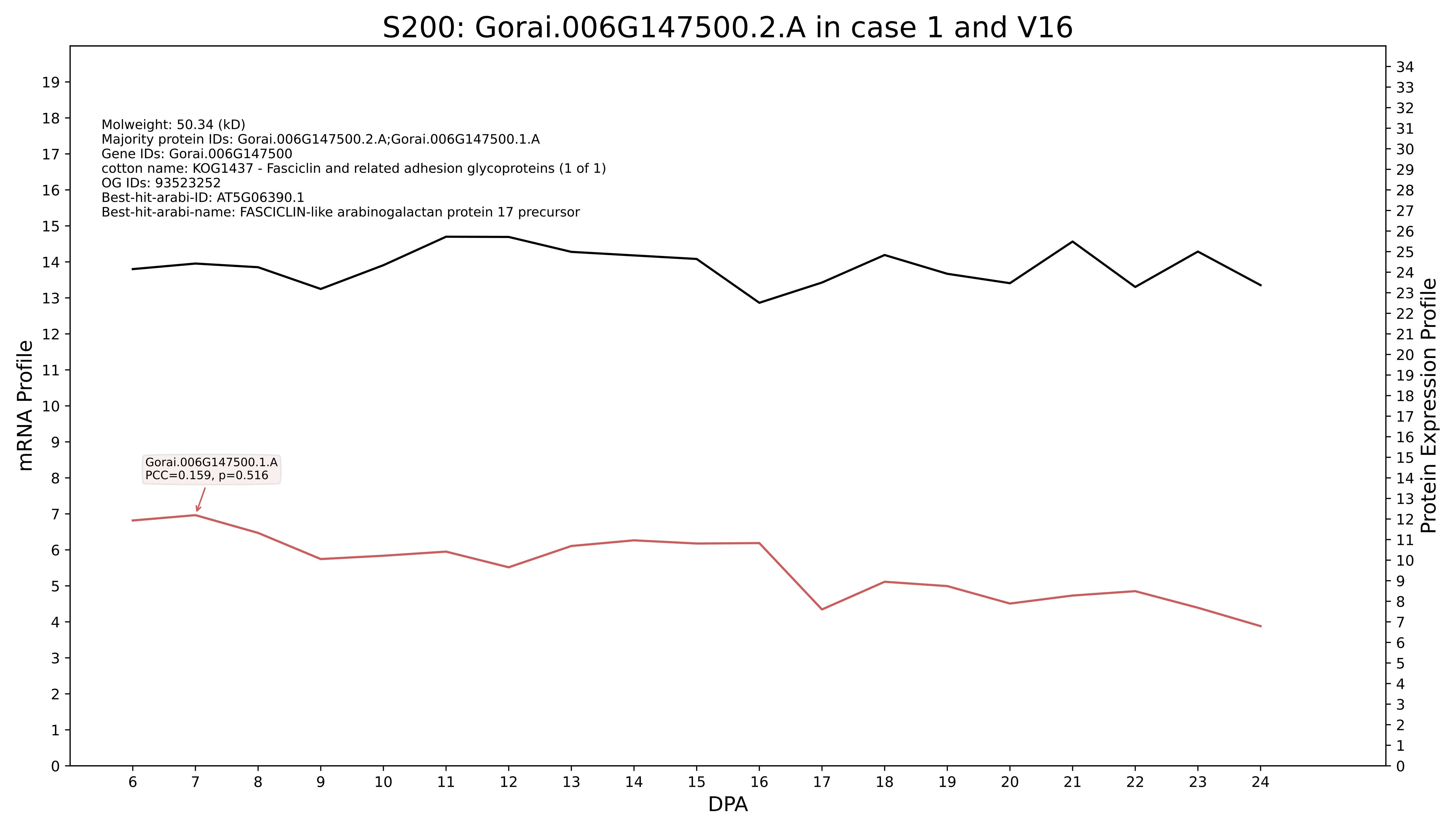
S200: Gorai.006G147500.2.A in case 1 and V16
Molweight: 50.34 (kD)
Majority protein IDs: Gorai.006G147500.2.A;Gorai.006G147500.1.A
Gene IDs: Gorai.006G147500
cotton name: KOG1437 - Fasciclin and related adhesion glycoproteins (1 of 1)
OG IDs: 93523252
Best-hit-arabi-ID: AT5G06390.1
Best-hit-arabi-name: FASCICLIN-like arabinogalactan protein 17 precursor
Gorai.006G147500.1.A
PCC=0.159, p=0.516
mRNA Profile
Protein Expression Profile
DPA

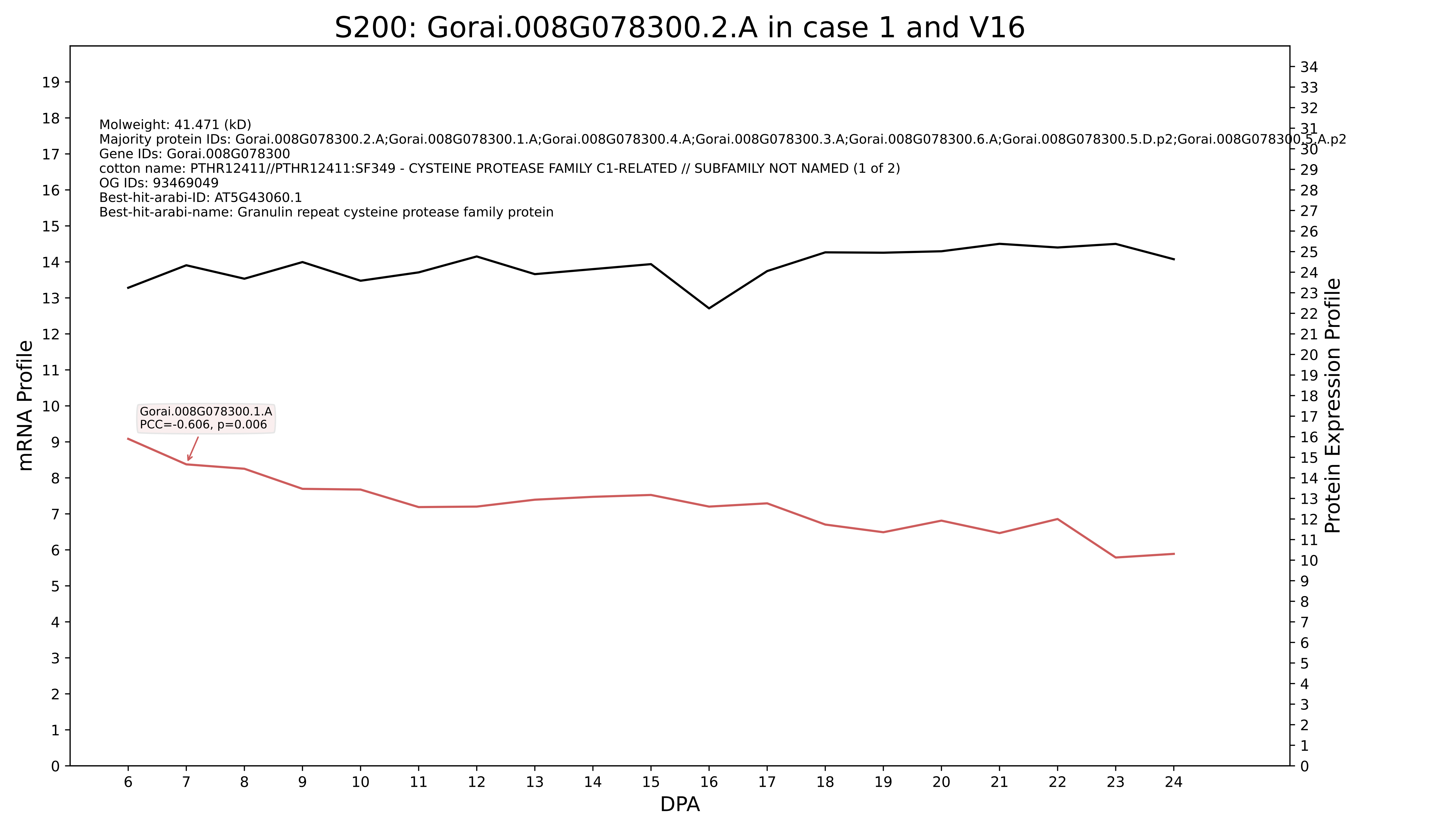

S200: Gorai.008G078300.2.A in case 1 and V16
Molweight: 41.471 (kD)
Majority protein IDs: Gorai.008G078300.2.A;Gorai.008G078300.1.A;Gorai.008G078300.4.A;Gorai.008G078300.3.A;Gorai.008G078300.6.A;Gorai.008G078300.5.D.p2;Gorai.008G078300.5.A.p2
Gene IDs: Gorai.008G078300
cotton name: PTHR12411//PTHR12411:SF349 - CYSTEINE PROTEASE FAMILY C1-RELATED // SUBFAMILY NOT NAMED (1 of 2)
OG IDs: 93469049
Best-hit-arabi-ID: AT5G43060.1
Best-hit-arabi-name: Granulin repeat cysteine protease family protein
Gorai.008G078300.1.A
PCC=-0.606, p=0.006
mRNA Profile
Protein Expression Profile
DPA

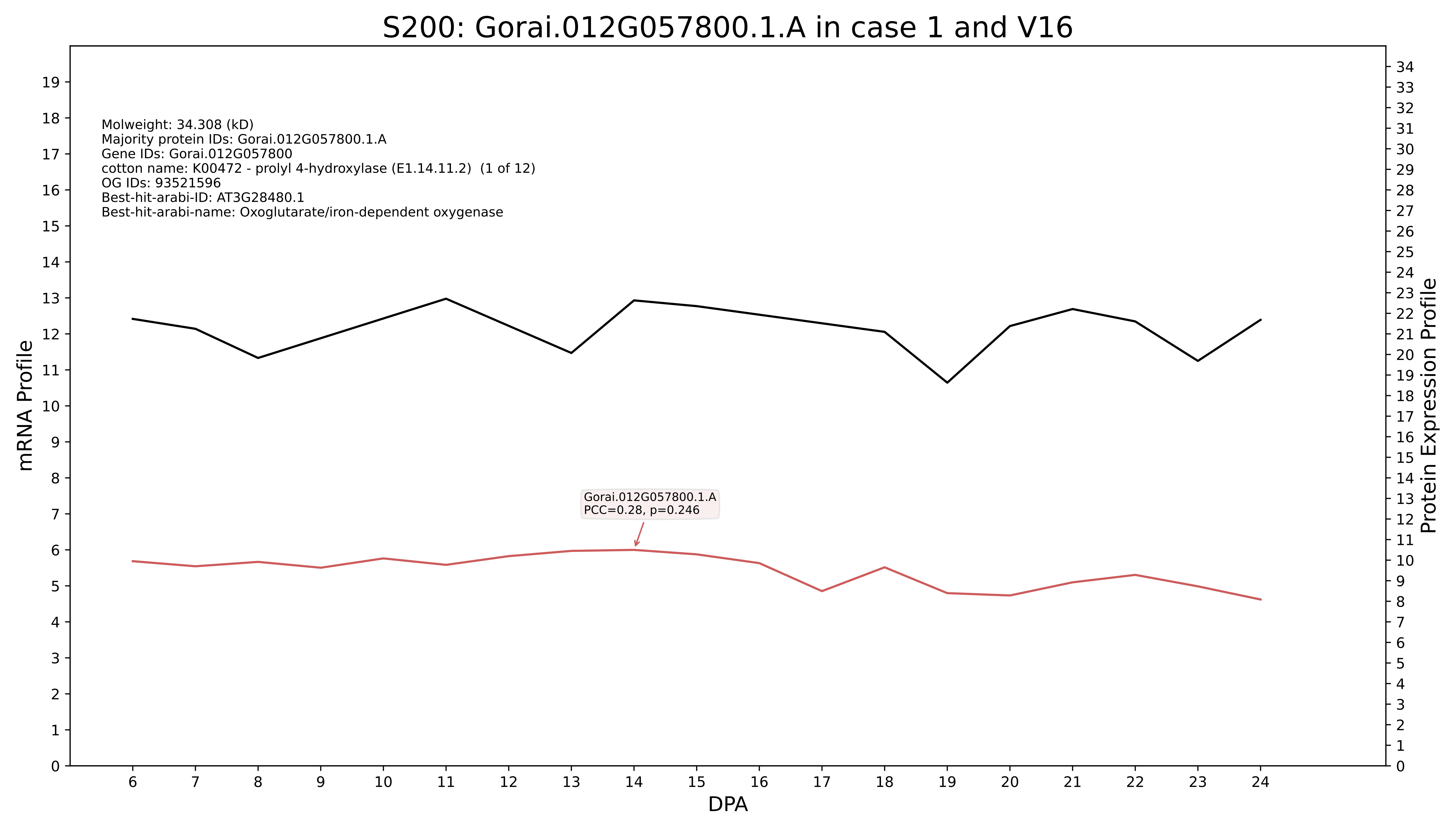
S200: Gorai.012G057800.1.A in case 1 and V16
Molweight: 34.308 (kD)
Majority protein IDs: Gorai.012G057800.1.A
Gene IDs: Gorai.012G057800
cotton name: K00472 - prolyl 4-hydroxylase (E1.14.11.2) (1 of 12)
OG IDs: 93521596
Best-hit-arabi-ID: AT3G28480.1
Best-hit-arabi-name: Oxoglutarate/iron-dependent oxygenase
Gorai.012G057800.1.A
PCC=0.28, p=0.246
mRNA Profile
Protein Expression Profile
DPA

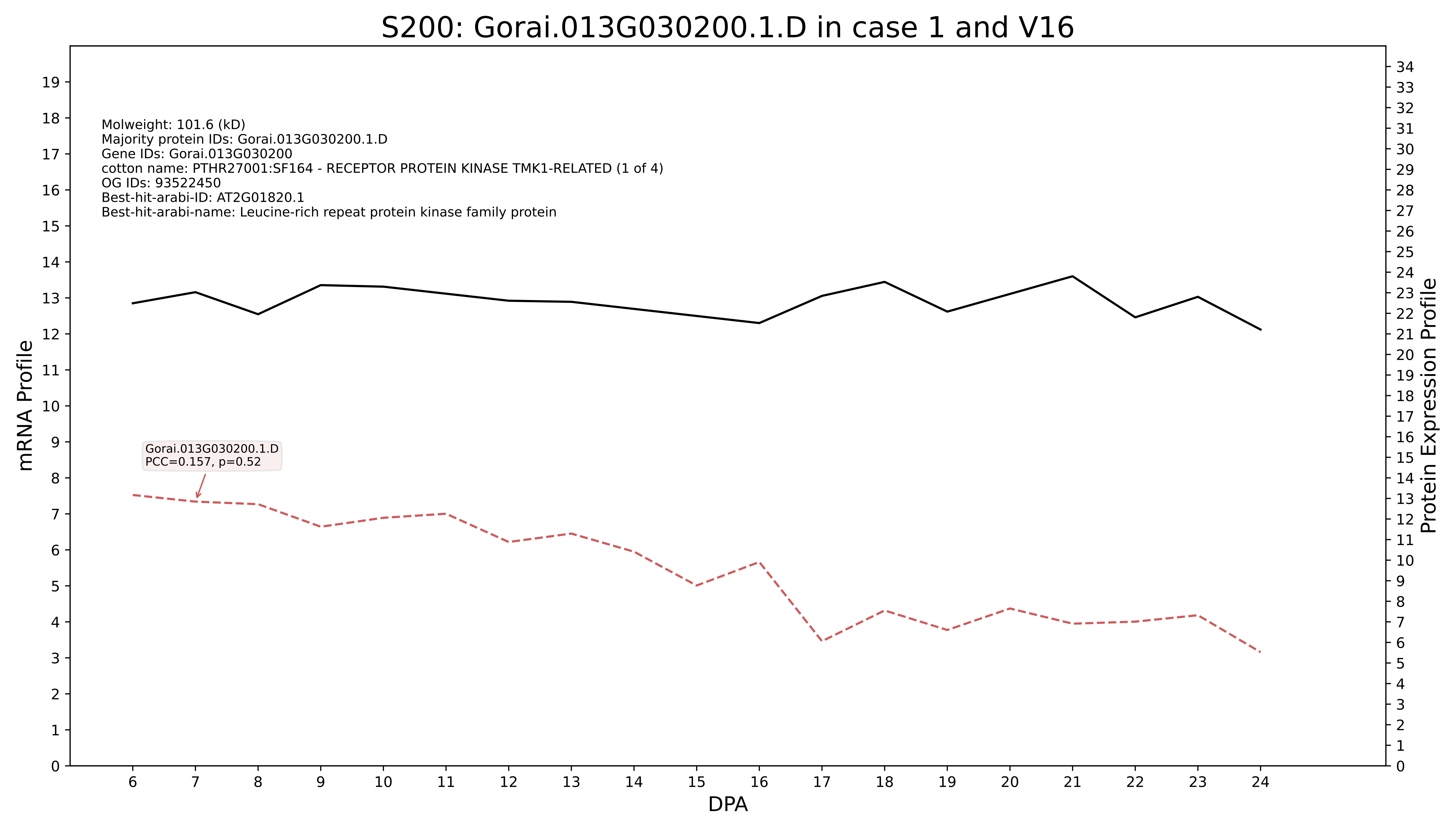
S200: Gorai.013G030200.1.D in case 1 and V16
Molweight: 101.6 (kD)
Majority protein IDs: Gorai.013G030200.1.D
Gene IDs: Gorai.013G030200
cotton name: PTHR27001:SF164 - RECEPTOR PROTEIN KINASE TMK1-RELATED (1 of 4)
OG IDs: 93522450
Best-hit-arabi-ID: AT2G01820.1
Best-hit-arabi-name: Leucine-rich repeat protein kinase family protein
Gorai.013G030200.1.D
PCC=0.157, p=0.52
mRNA Profile
Protein Expression Profile
DPA

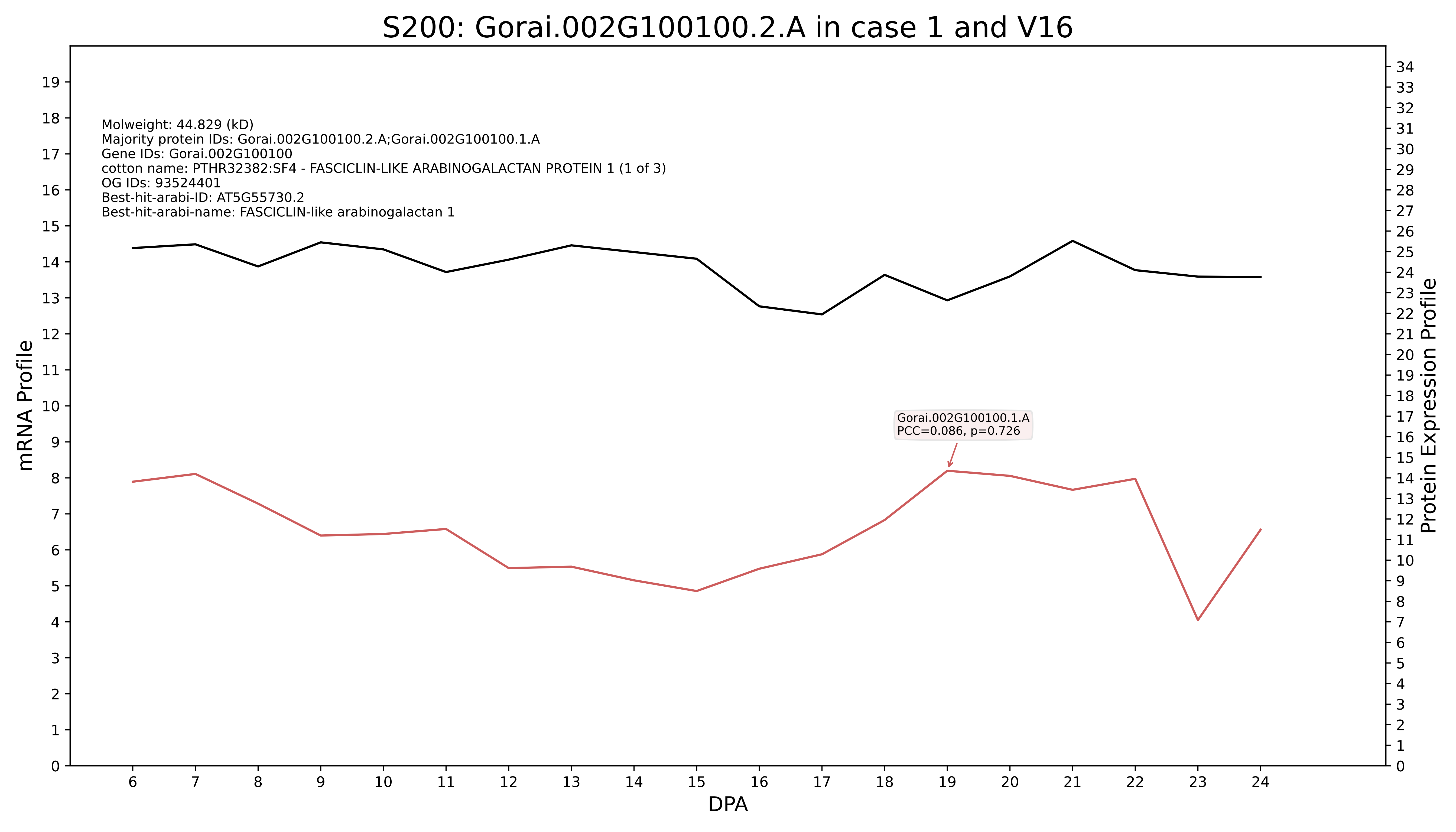
S200: Gorai.002G100100.2.A in case 1 and V16
Molweight: 44.829 (kD)
Majority protein IDs: Gorai.002G100100.2.A;Gorai.002G100100.1.A
Gene IDs: Gorai.002G100100
cotton name: PTHR32382:SF4 - FASCICLIN-LIKE ARABINOGALACTAN PROTEIN 1 (1 of 3)
OG IDs: 93524401
Best-hit-arabi-ID: AT5G55730.2
Best-hit-arabi-name: FASCICLIN-like arabinogalactan 1
Gorai.002G100100.1.A
PCC=0.086, p=0.726
mRNA Profile
Protein Expression Profile
DPA

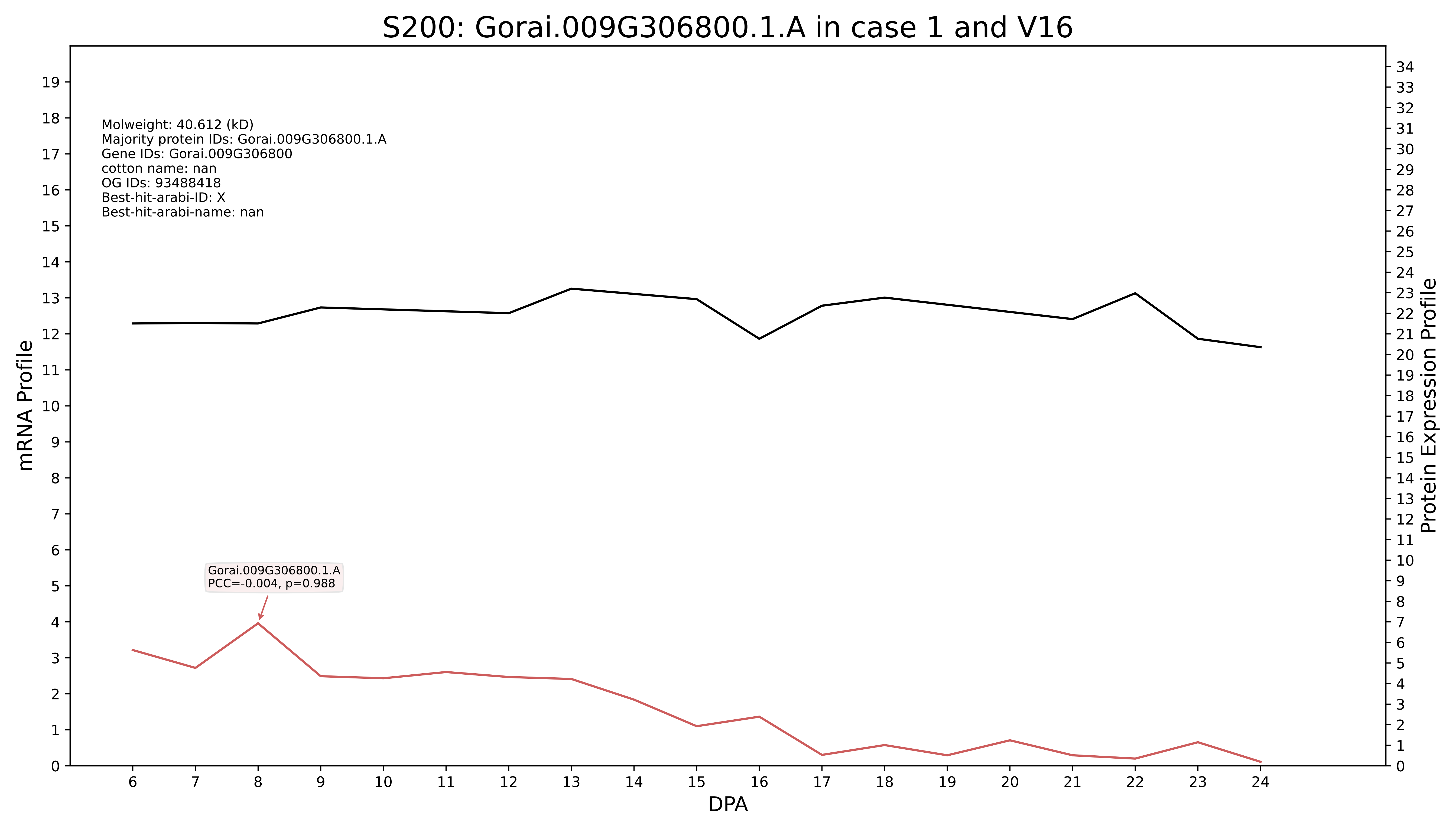
S200: Gorai.009G306800.1.A in case 1 and V16
Molweight: 40.612 (kD)
Majority protein IDs: Gorai.009G306800.1.A
Gene IDs: Gorai.009G306800
cotton name: nan
OG IDs: 93488418
Best-hit-arabi-ID: X
Best-hit-arabi-name: nan
Gorai.009G306800.1.A
PCC=-0.004, p=0.988
mRNA Profile
Protein Expression Profile
DPA

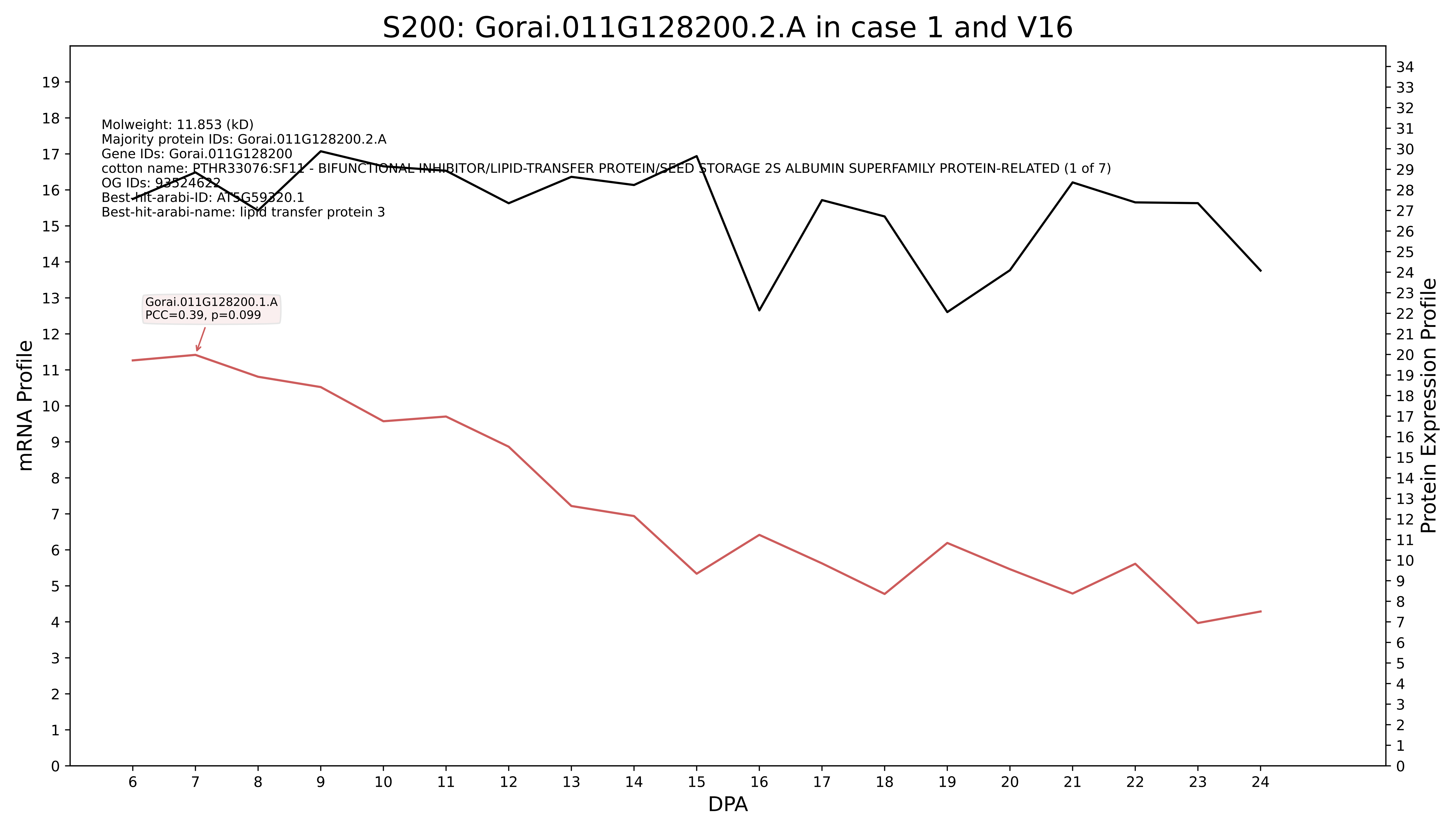

S200: Gorai.011G128200.2.A in case 1 and V16
Molweight: 11.853 (kD)
Majority protein IDs: Gorai.011G128200.2.A
Gene IDs: Gorai.011G128200
cotton name: PTHR33076:SF11 - BIFUNCTIONAL INHIBITOR/LIPID-TRANSFER PROTEIN/SEED STORAGE 2S ALBUMIN SUPERFAMILY PROTEIN-RELATED (1 of 7)
OG IDs: 93524622
Best-hit-arabi-ID: AT5G59320.1
Best-hit-arabi-name: lipid transfer protein 3
Gorai.011G128200.1.A
PCC=0.39, p=0.099
mRNA Profile
Protein Expression Profile
DPA

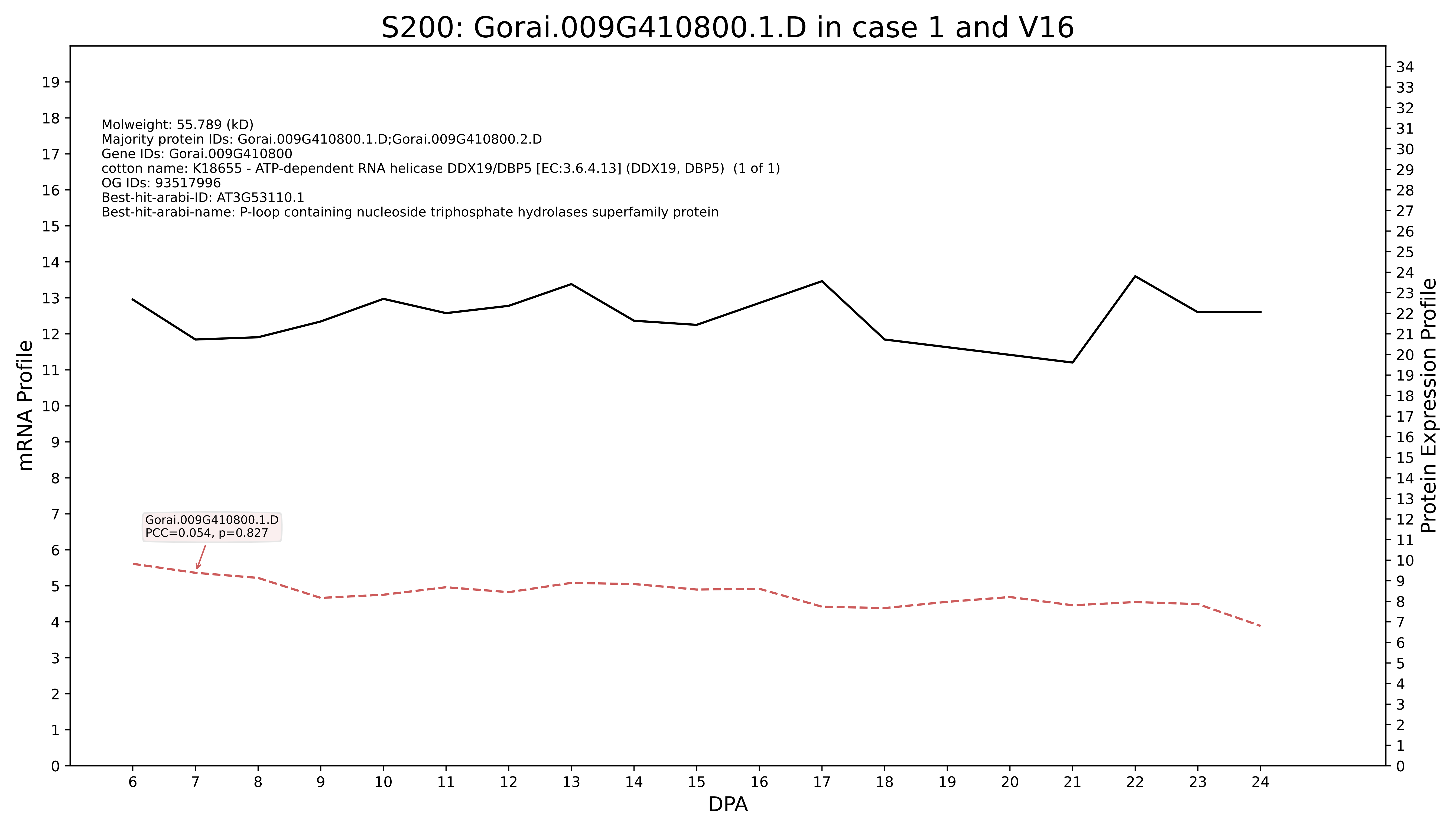
S200: Gorai.009G410800.1.D in case 1 and V16
Molweight: 55.789 (kD)
Majority protein IDs: Gorai.009G410800.1.D;Gorai.009G410800.2.D
Gene IDs: Gorai.009G410800
cotton name: K18655 - ATP-dependent RNA helicase DDX19/DBP5 [EC:3.6.4.13] (DDX19, DBP5) (1 of 1)
OG IDs: 93517996
Best-hit-arabi-ID: AT3G53110.1
Best-hit-arabi-name: P-loop containing nucleoside triphosphate hydrolases superfamily protein
Gorai.009G410800.1.D
PCC=0.054, p=0.827
mRNA Profile
Protein Expression Profile
DPA

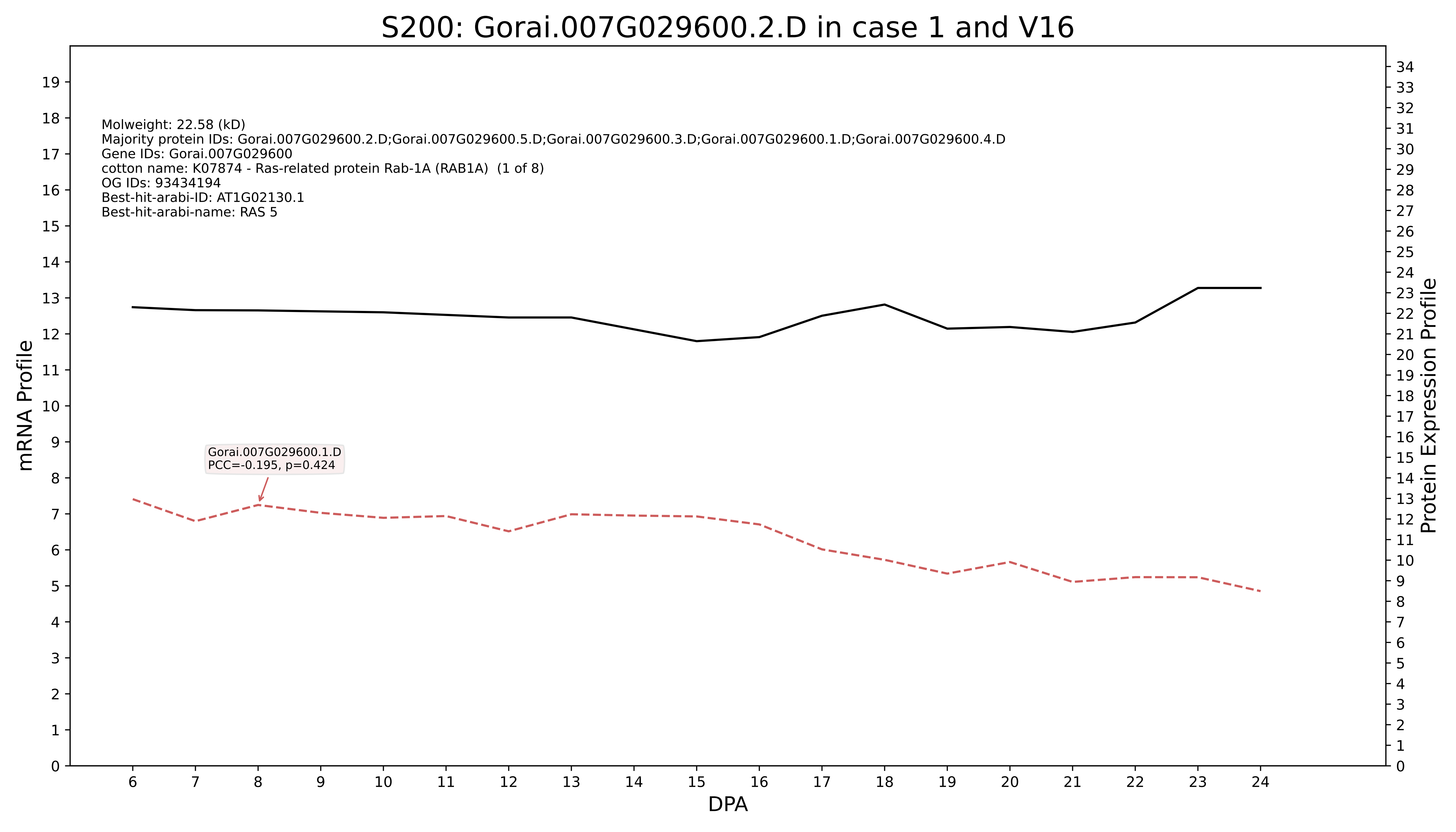
S200: Gorai.007G029600.2.D in case 1 and V16
Molweight: 22.58 (kD)
Majority protein IDs: Gorai.007G029600.2.D;Gorai.007G029600.5.D;Gorai.007G029600.3.D;Gorai.007G029600.1.D;Gorai.007G029600.4.D
Gene IDs: Gorai.007G029600
cotton name: K07874 - Ras-related protein Rab-1A (RAB1A) (1 of 8)
OG IDs: 93434194
Best-hit-arabi-ID: AT1G02130.1
Best-hit-arabi-name: RAS 5
Gorai.007G029600.1.D
PCC=-0.195, p=0.424
mRNA Profile
Protein Expression Profile
DPA

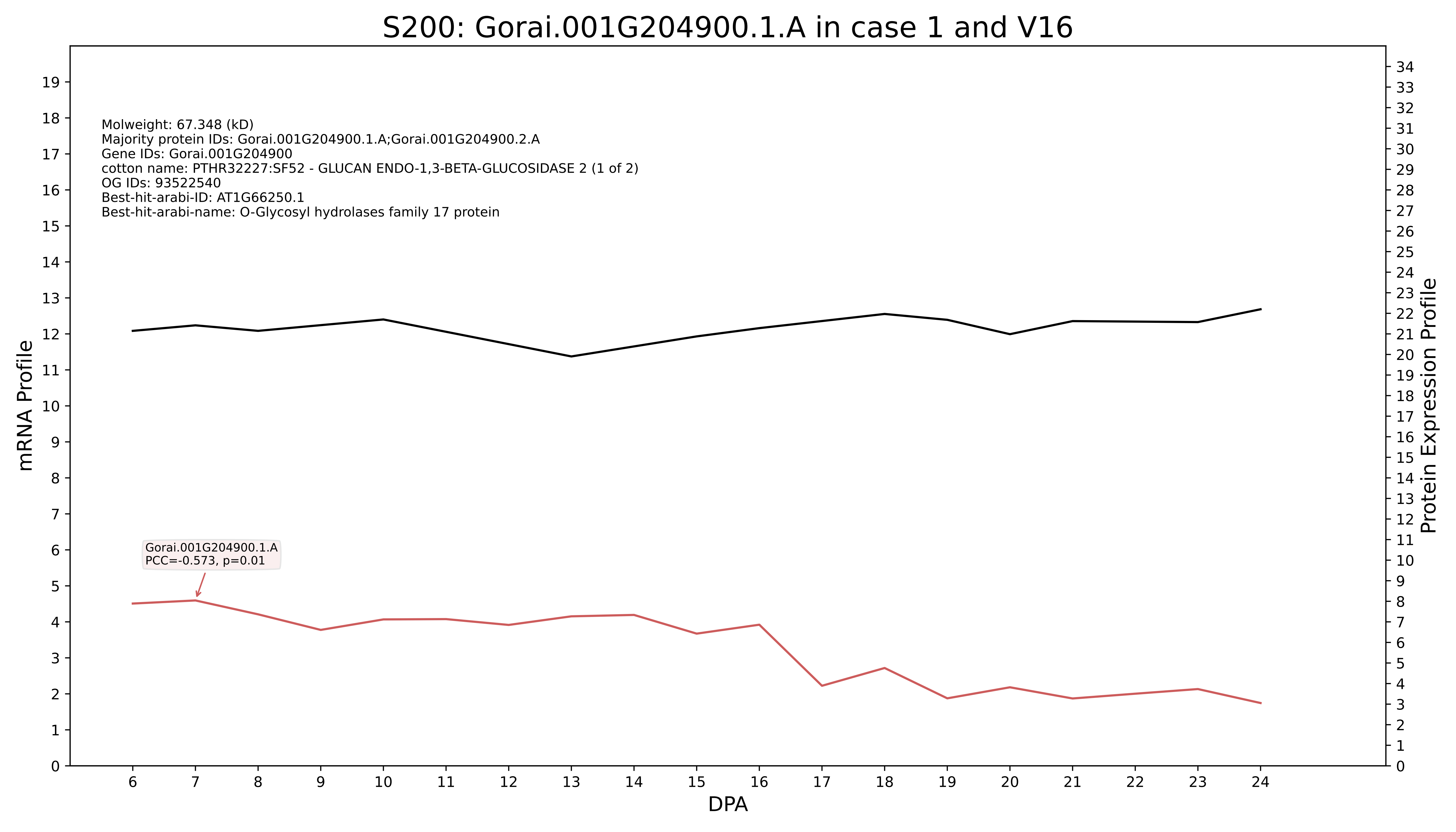

S200: Gorai.001G204900.1.A in case 1 and V16
Molweight: 67.348 (kD)
Majority protein IDs: Gorai.001G204900.1.A;Gorai.001G204900.2.A
Gene IDs: Gorai.001G204900
cotton name: PTHR32227:SF52 - GLUCAN ENDO-1,3-BETA-GLUCOSIDASE 2 (1 of 2)
OG IDs: 93522540
Best-hit-arabi-ID: AT1G66250.1
Best-hit-arabi-name: O-Glycosyl hydrolases family 17 protein
Gorai.001G204900.1.A
PCC=-0.573, p=0.01
mRNA Profile
Protein Expression Profile
DPA

### S200: Gorai.013G256100.5.A in case 2 and V16

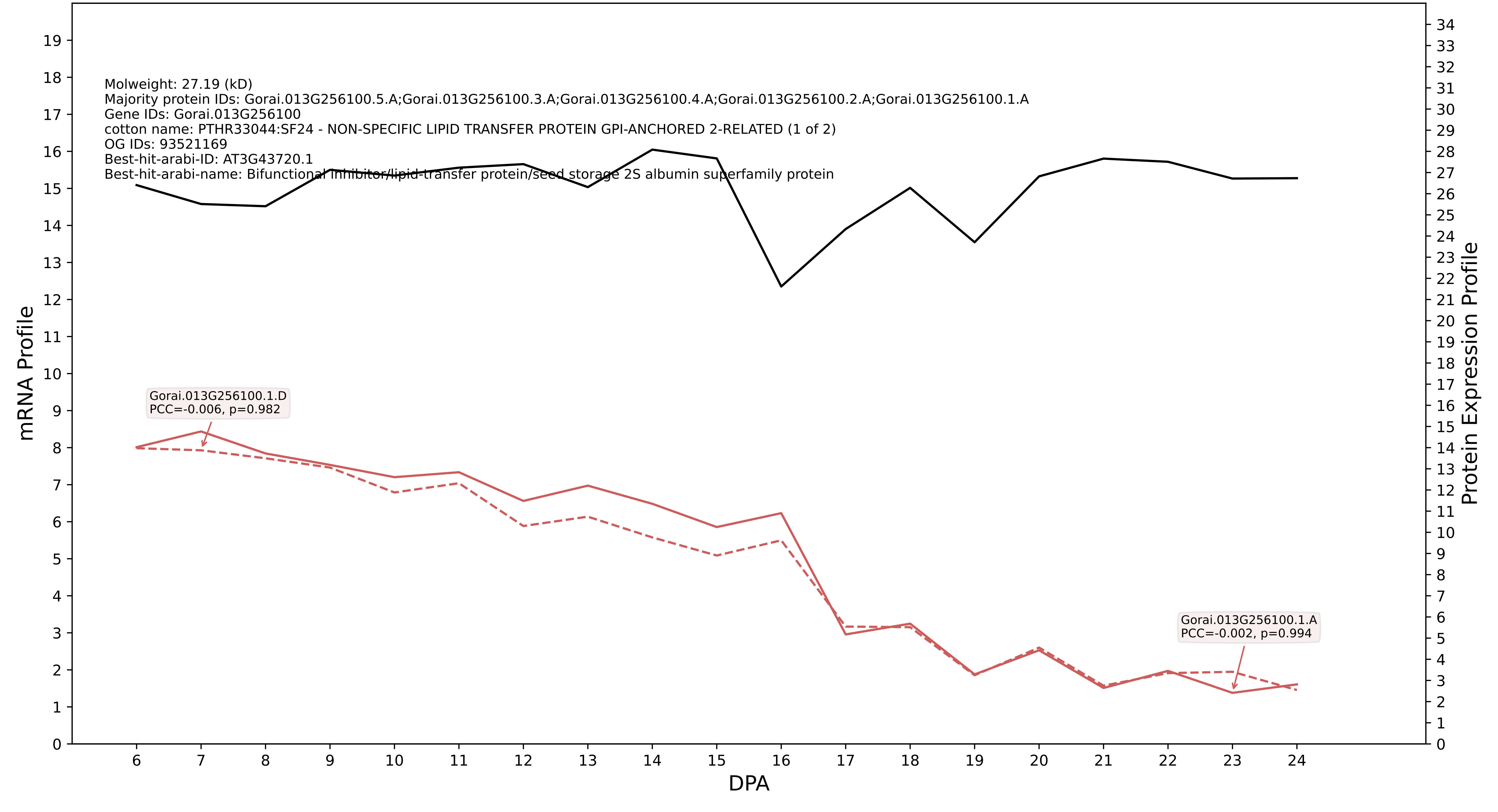

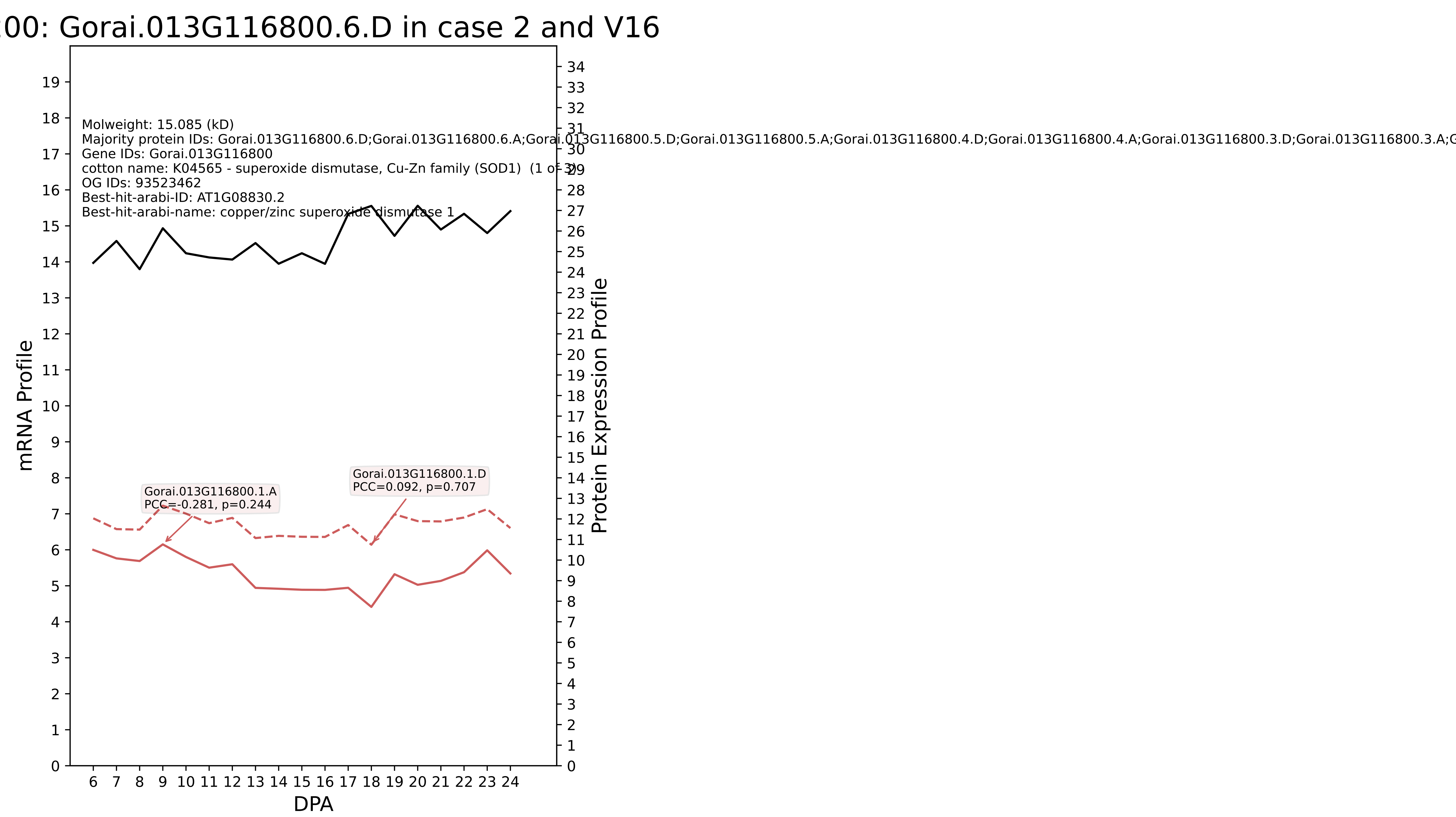

00: Gorai.013G116800.6.D in case 2 and V16
Molweight: 15.085 (kD)
Majority protein IDs: Gorai.013G116800.6.D;Gorai.013G116800.6.A;Gorai.013G116800.5.D;Gorai.013G116800.5.A;Gorai.013G116800.4.D;Gorai.013G116800.4.A;Gorai.013G116800.3.D;Gorai.013G116800.3.A;G
Gene IDs: Gorai.013G116800
cotton name: K04565 - superoxide dismutase, Cu-Zn family (SOD1) (1 of 3)
OG IDs: 93523462
Best-hit-arabi-ID: AT1G08830.2
Best-hit-arabi-name: copper/zinc superoxide dismutase 1
Gorai.013G116800.1.A
PCC=-0.281, p=0.244
Gorai.013G116800.1.D
PCC=0.092, p=0.707
mRNA Profile
Protein Expression Profile
DPA

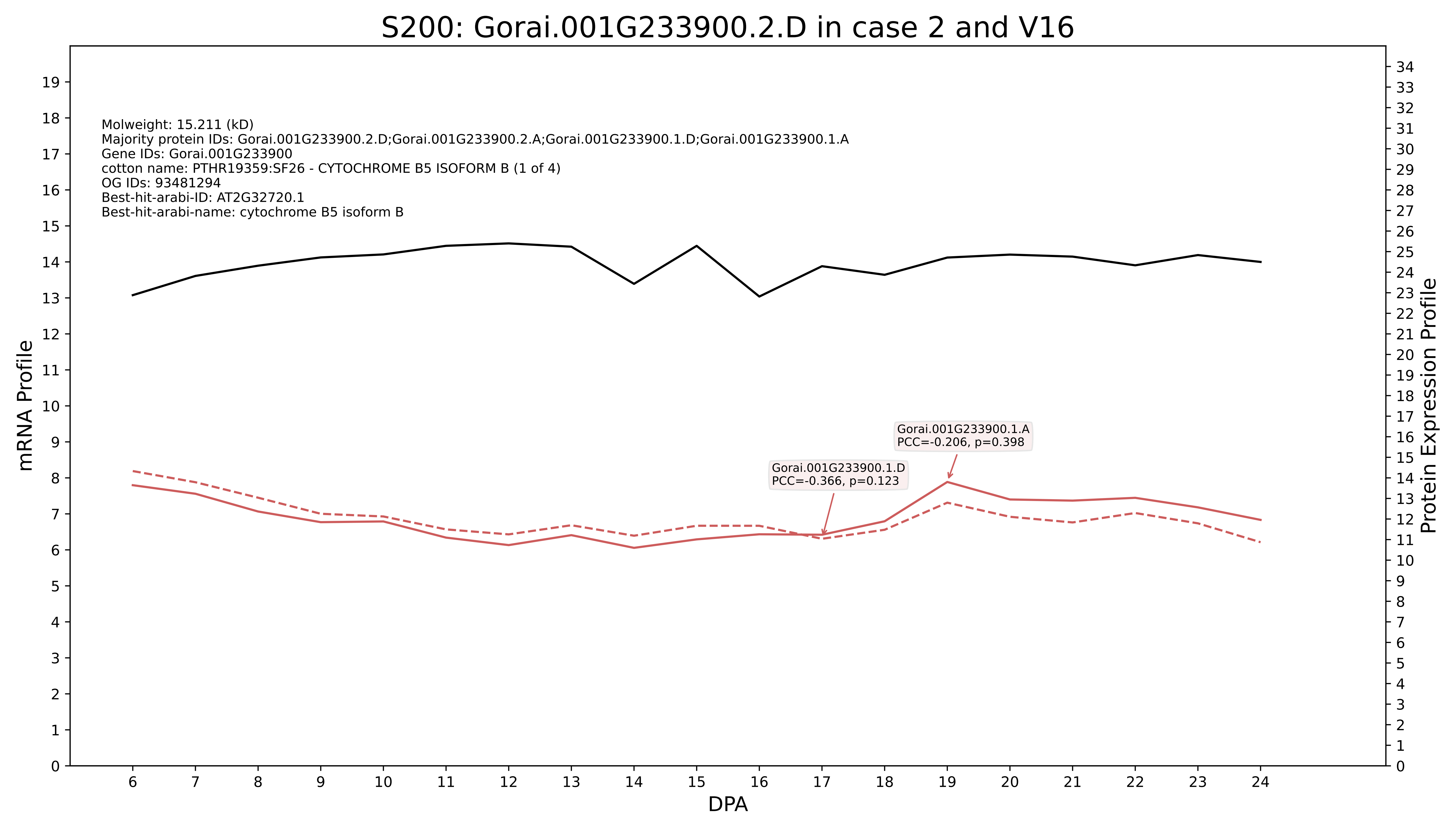
S200: Gorai.001G233900.2.D in case 2 and V16
Molweight: 15.211 (kD)
Majority protein IDs: Gorai.001G233900.2.D;Gorai.001G233900.2.A;Gorai.001G233900.1.D;Gorai.001G233900.1.A
Gene IDs: Gorai.001G233900
cotton name: PTHR19359:SF26 - CYTOCHROME B5 ISOFORM B (1 of 4)
OG IDs: 93481294
Best-hit-arabi-ID: AT2G32720.1
Best-hit-arabi-name: cytochrome B5 isoform B
Gorai.001G233900.1.A
PCC=-0.206, p=0.398
Gorai.001G233900.1.D
PCC=-0.366, p=0.123
mRNA Profile
Protein Expression Profile
DPA

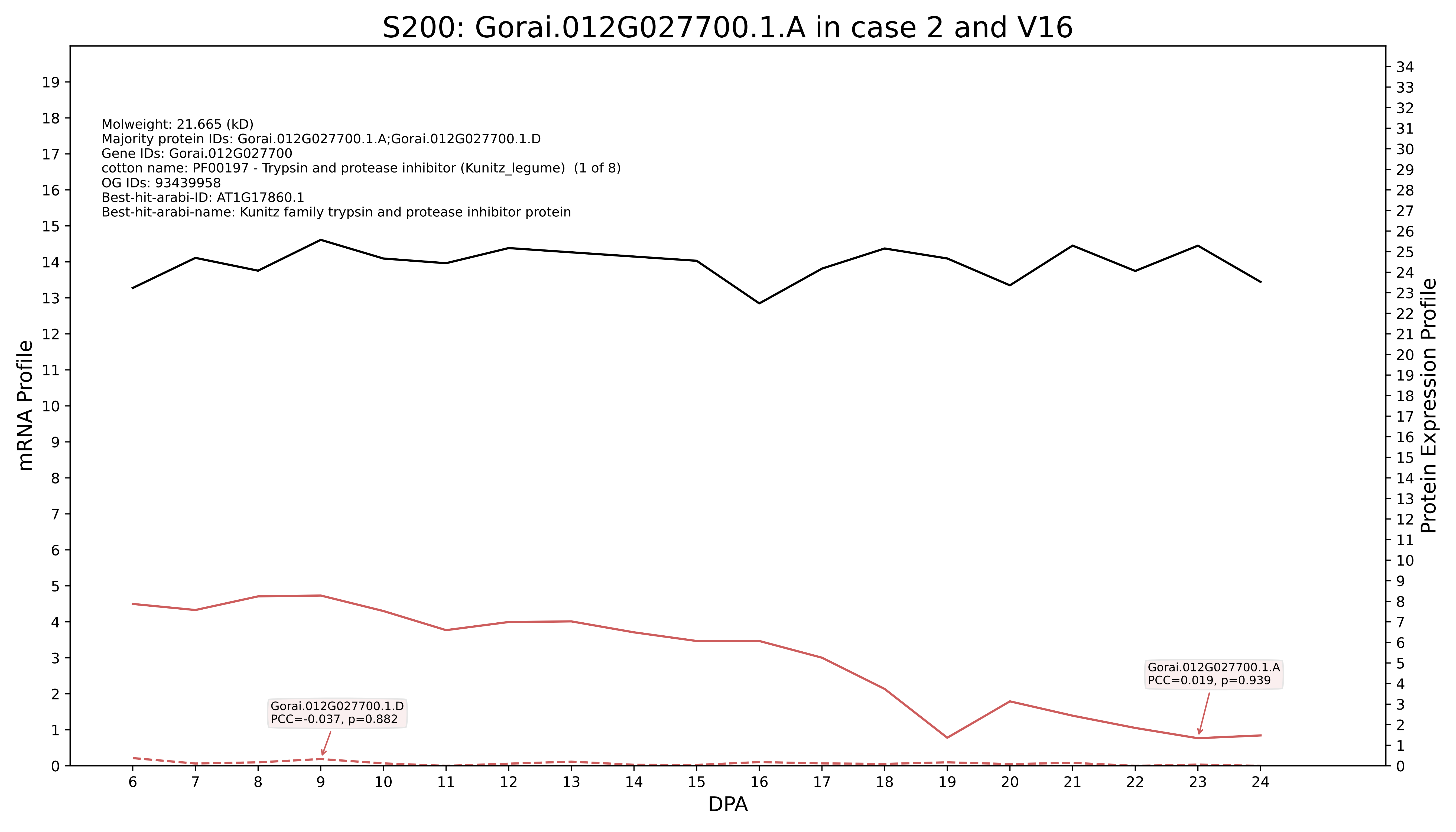

S200: Gorai.012G027700.1.A in case 2 and V16
Molweight: 21.665 (kD)
Majority protein IDs: Gorai.012G027700.1.A;Gorai.012G027700.1.D
Gene IDs: Gorai.012G027700
cotton name: PF00197 - Trypsin and protease inhibitor (Kunitz_legume) (1 of 8)
OG IDs: 93439958
Best-hit-arabi-ID: AT1G17860.1
Best-hit-arabi-name: Kunitz family trypsin and protease inhibitor protein
Gorai.012G027700.1.D
PCC=-0.037, p=0.882
Gorai.012G027700.1.A
PCC=0.019, p=0.939
mRNA Profile
Protein Expression Profile
DPA

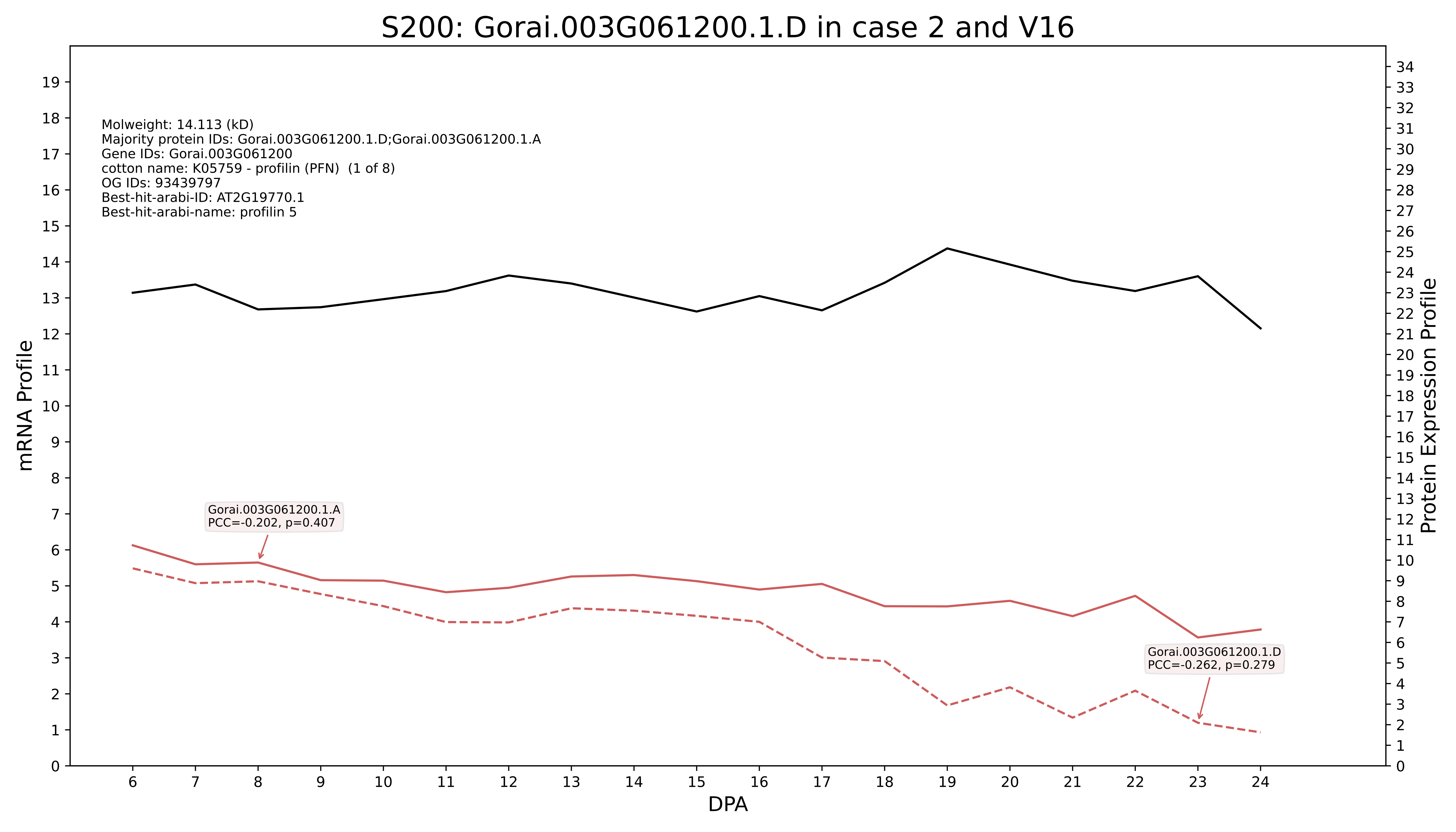

S200: Gorai.003G061200.1.D in case 2 and V16
Molweight: 14.113 (kD)
Majority protein IDs: Gorai.003G061200.1.D;Gorai.003G061200.1.A
Gene IDs: Gorai.003G061200
cotton name: K05759 - profilin (PFN) (1 of 8)
OG IDs: 93439797
Best-hit-arabi-ID: AT2G19770.1
Best-hit-arabi-name: profilin 5
Gorai.003G061200.1.A
PCC=-0.202, p=0.407
Gorai.003G061200.1.D
PCC=-0.262, p=0.279
mRNA Profile
Protein Expression Profile
DPA

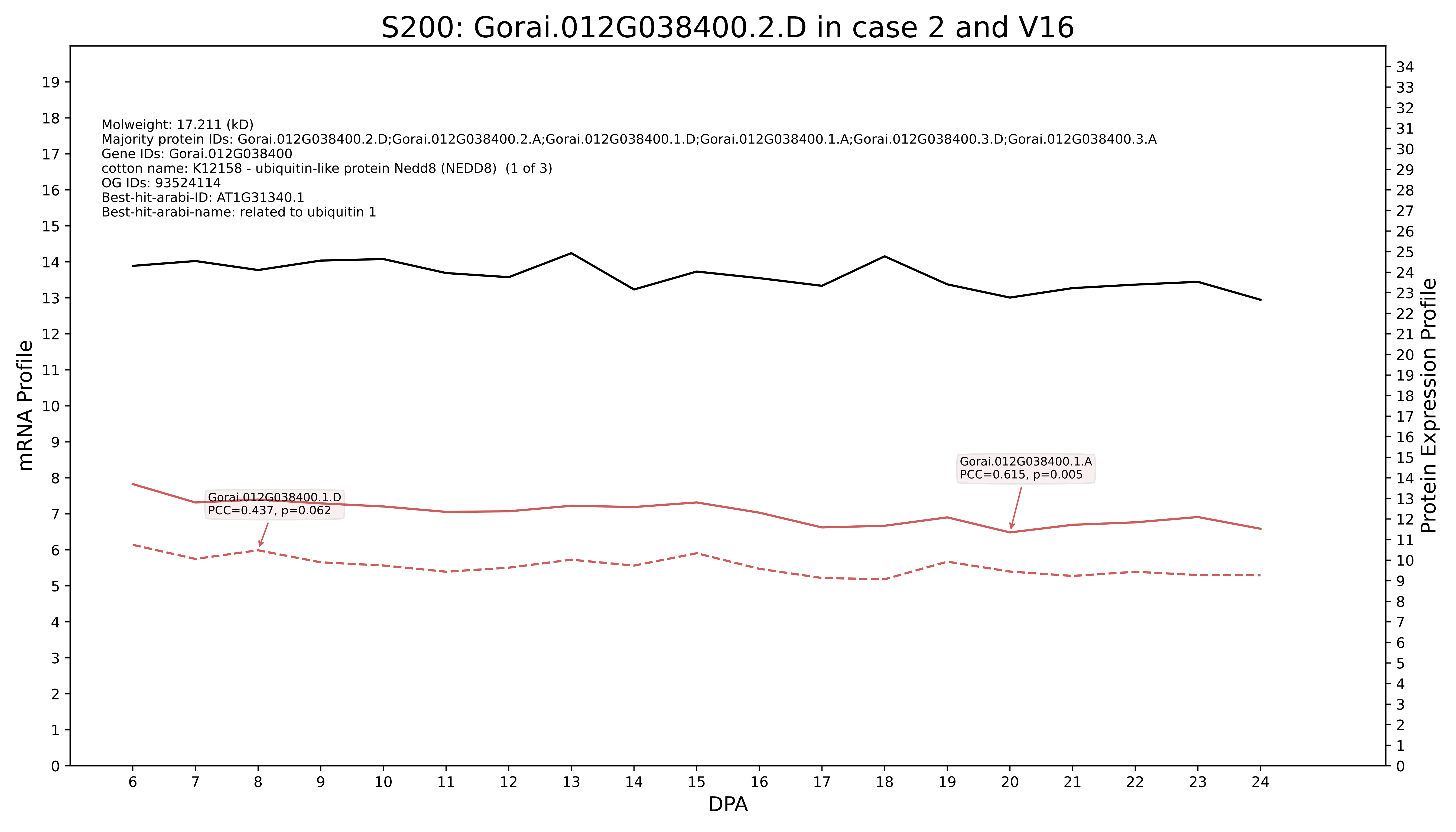
S200: Gorai.012G038400.2.D in case 2 and V16
Molweight: 17.211 (kD)
Majority protein IDs: Gorai.012G038400.2.D;Gorai.012G038400.2.A;Gorai.012G038400.1.D;Gorai.012G038400.1.A;Gorai.012G038400.3.D;Gorai.012G038400.3.A
Gene IDs: Gorai.012G038400
cotton name: K12158 - ubiquitin-like protein Nedd8 (NEDD8) (1 of 3)
OG IDs: 93524114
Best-hit-arabi-ID: AT1G31340.1
Best-hit-arabi-name: related to ubiquitin 1
Gorai.012G038400.1.D
PCC=0.437, p=0.062
Gorai.012G038400.1.A
PCC=0.615, p=0.005
mRNA Profile
Protein Expression Profile
DPA

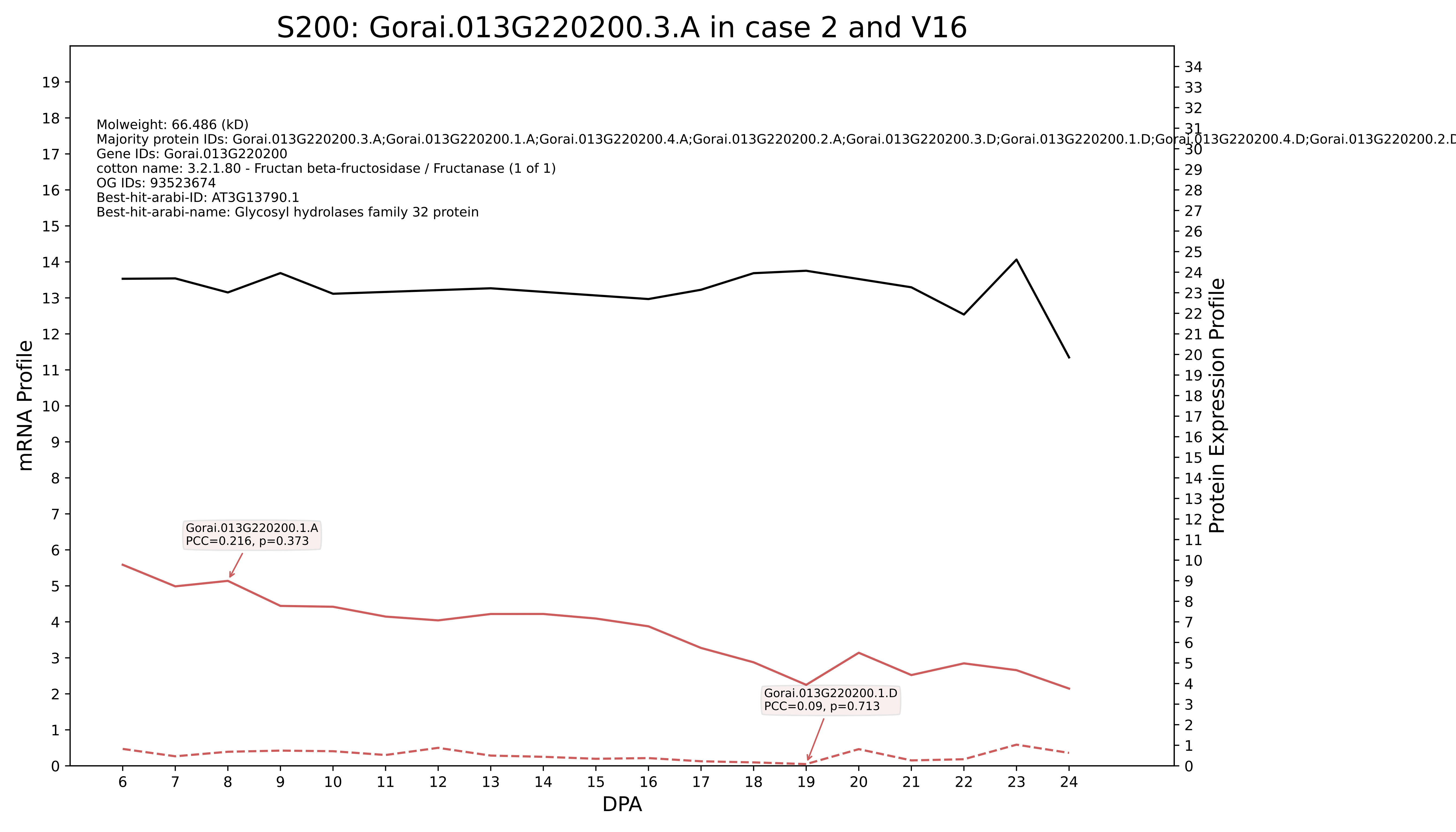
S200: Gorai.013G220200.3.A in case 2 and V16
Molweight: 66.486 (kD)
Majority protein IDs: Gorai.013G220200.3.A;Gorai.013G220200.1.A;Gorai.013G220200.4.A;Gorai.013G220200.2.A;Gorai.013G220200.3.D;Gorai.013G220200.1.D;Gorai.013G220200.4.D;Gorai.013G220200.2.D
Gene IDs: Gorai.013G220200
cotton name: 3.2.1.80 - Fructan beta-fructosidase / Fructanase (1 of 1)
OG IDs: 93523674
Best-hit-arabi-ID: AT3G13790.1
Best-hit-arabi-name: Glycosyl hydrolases family 32 protein
Gorai.013G220200.1.A
PCC=0.216, p=0.373
Gorai.013G220200.1.D
PCC=0.09, p=0.713
mRNA Profile
Protein Expression Profile
DPA

### S200: Gorai.010G127100.3.A in case 2 and V16

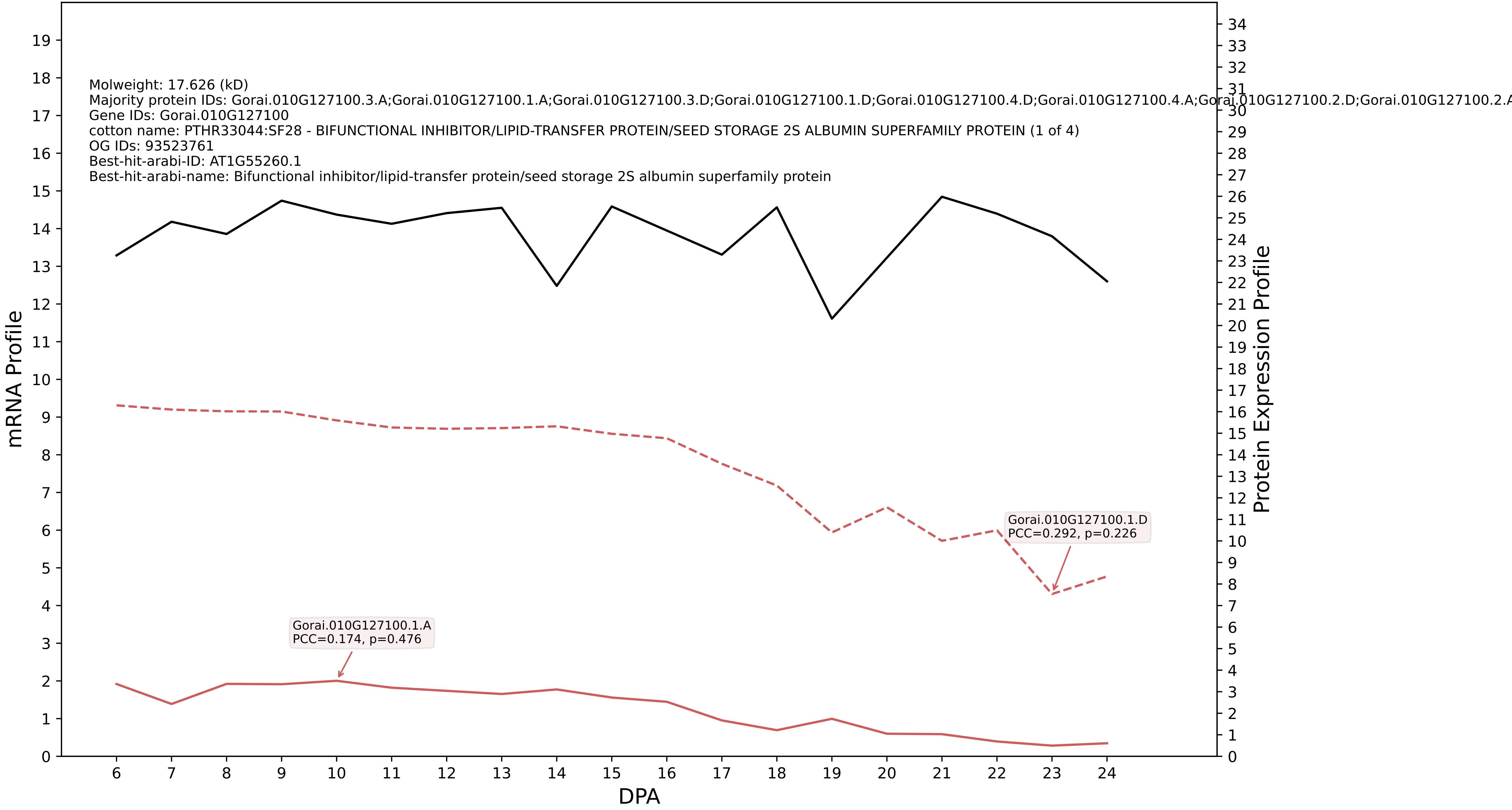

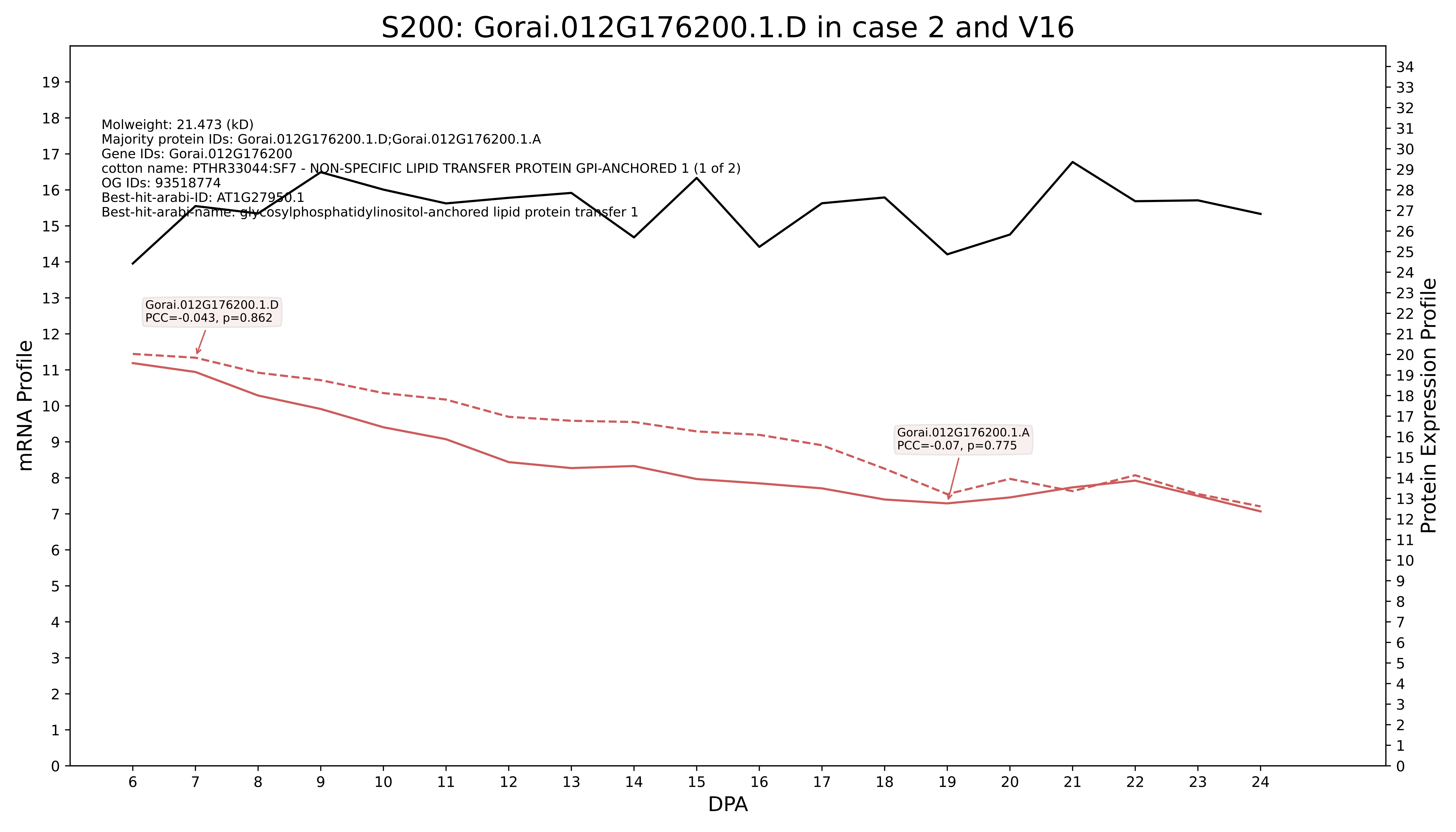

S200: Gorai.012G176200.1.D in case 2 and V16
Molweight: 21.473 (kD)
Majority protein IDs: Gorai.012G176200.1.D;Gorai.012G176200.1.A
Gene IDs: Gorai.012G176200
cotton name: PTHR33044:SF7 - NON-SPECIFIC LIPID TRANSFER PROTEIN GPI-ANCHORED 1 (1 of 2)
OG IDs: 93518774
Best-hit-arabi-ID: AT1G27950.1
Best-hit-arabi-name: glycosylphosphatidylinositol-anchored lipid protein transfer 1
Gorai.012G176200.1.D
PCC=-0.043, p=0.862
Gorai.012G176200.1.A
PCC=-0.07, p=0.775
mRNA Profile
Protein Expression Profile
DPA

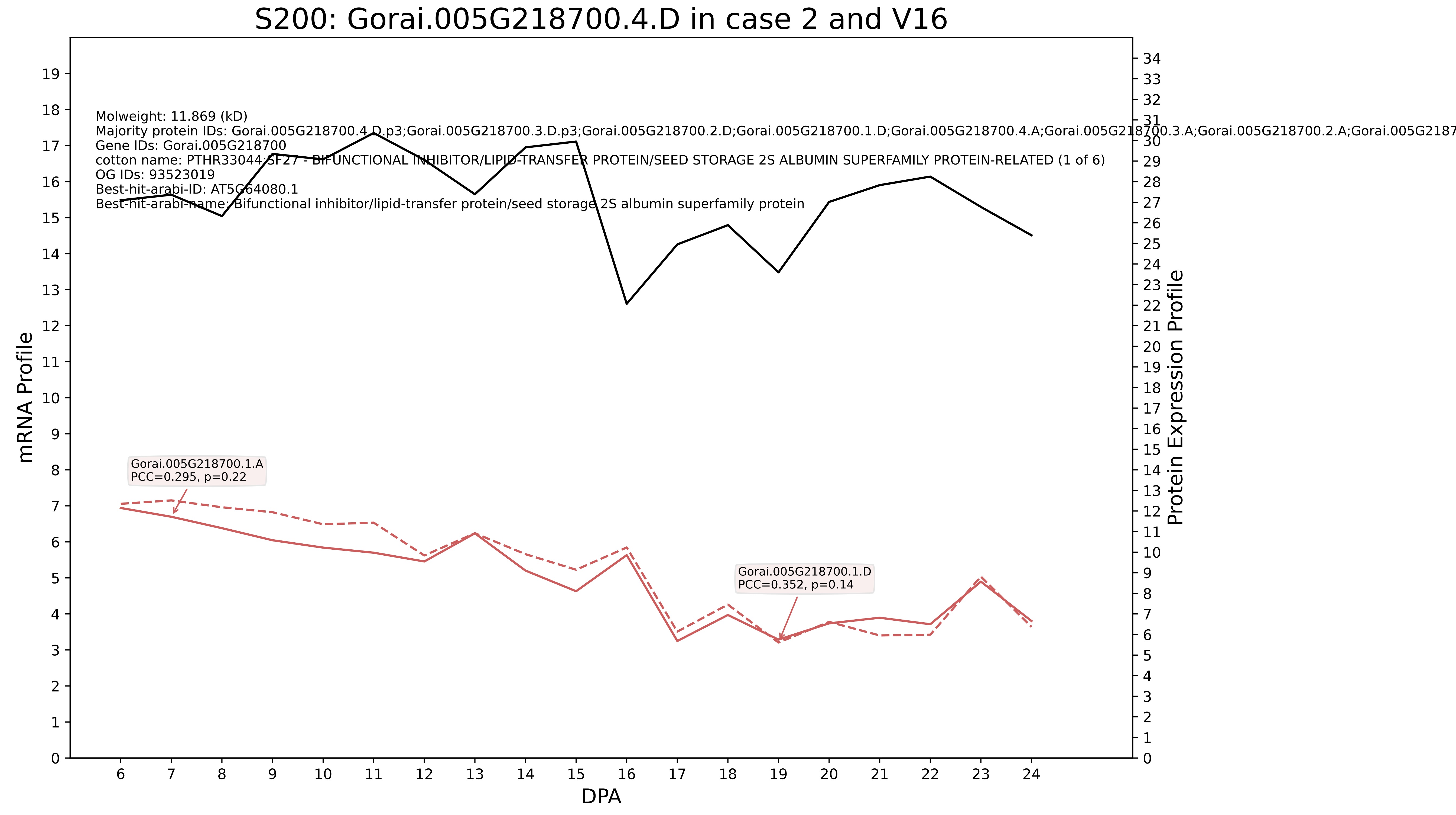

S200: Gorai.005G218700.4.D in case 2 and V16
Molweight: 11.869 (kD)
Majority protein IDs: Gorai.005G218700.4.D.p3;Gorai.005G218700.3.D.p3;Gorai.005G218700.2.D;Gorai.005G218700.1.D;Gorai.005G218700.4.A;Gorai.005G218700.3.A;Gorai.005G218700.2.A;Gorai.005G2187
Gene IDs: Gorai.005G218700
cotton name: PTHR33044:SF27 - BIFUNCTIONAL INHIBITOR/LIPID-TRANSFER PROTEIN/SEED STORAGE 2S ALBUMIN SUPERFAMILY PROTEIN-RELATED (1 of 6)
OG IDs: 93523019
Best-hit-arabi-ID: AT5G64080.1
Best-hit-arabi-name: Bifunctional inhibitor/lipid-transfer protein/seed storage 2S albumin superfamily protein
Gorai.005G218700.1.A
PCC=0.295, p=0.22
Gorai.005G218700.1.D
PCC=0.352, p=0.14
mRNA Profile
Protein Expression Profile
DPA

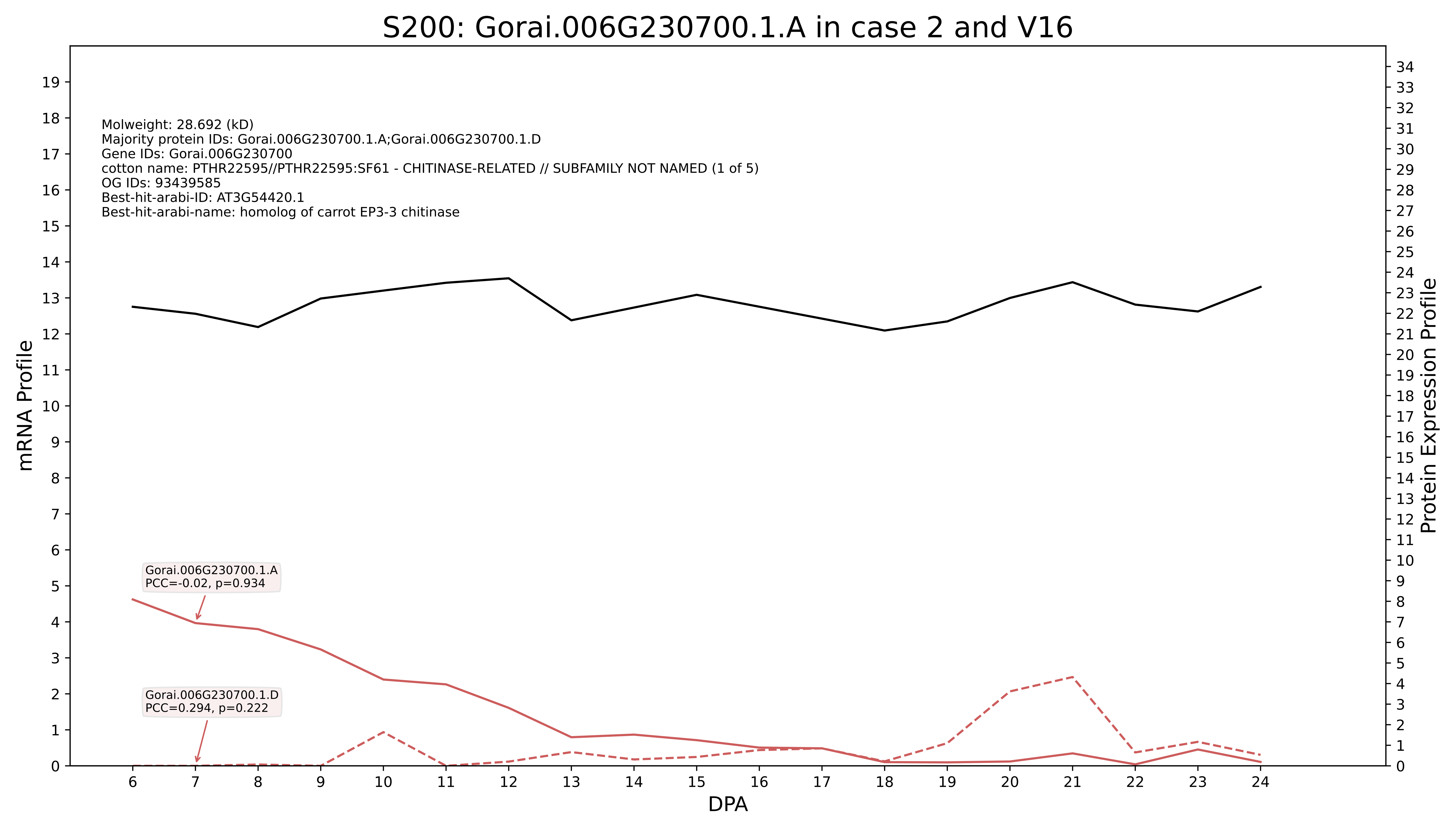

S200: Gorai.006G230700.1.A in case 2 and V16
Molweight: 28.692 (kD)
Majority protein IDs: Gorai.006G230700.1.A;Gorai.006G230700.1.D
Gene IDs: Gorai.006G230700
cotton name: PTHR22595//PTHR22595:SF61 - CHITINASE-RELATED // SUBFAMILY NOT NAMED (1 of 5)
OG IDs: 93439585
Best-hit-arabi-ID: AT3G54420.1
Best-hit-arabi-name: homolog of carrot EP3-3 chitinase
Gorai.006G230700.1.A
PCC=-0.02, p=0.934
Gorai.006G230700.1.D
PCC=0.294, p=0.222
mRNA Profile
Protein Expression Profile
DPA

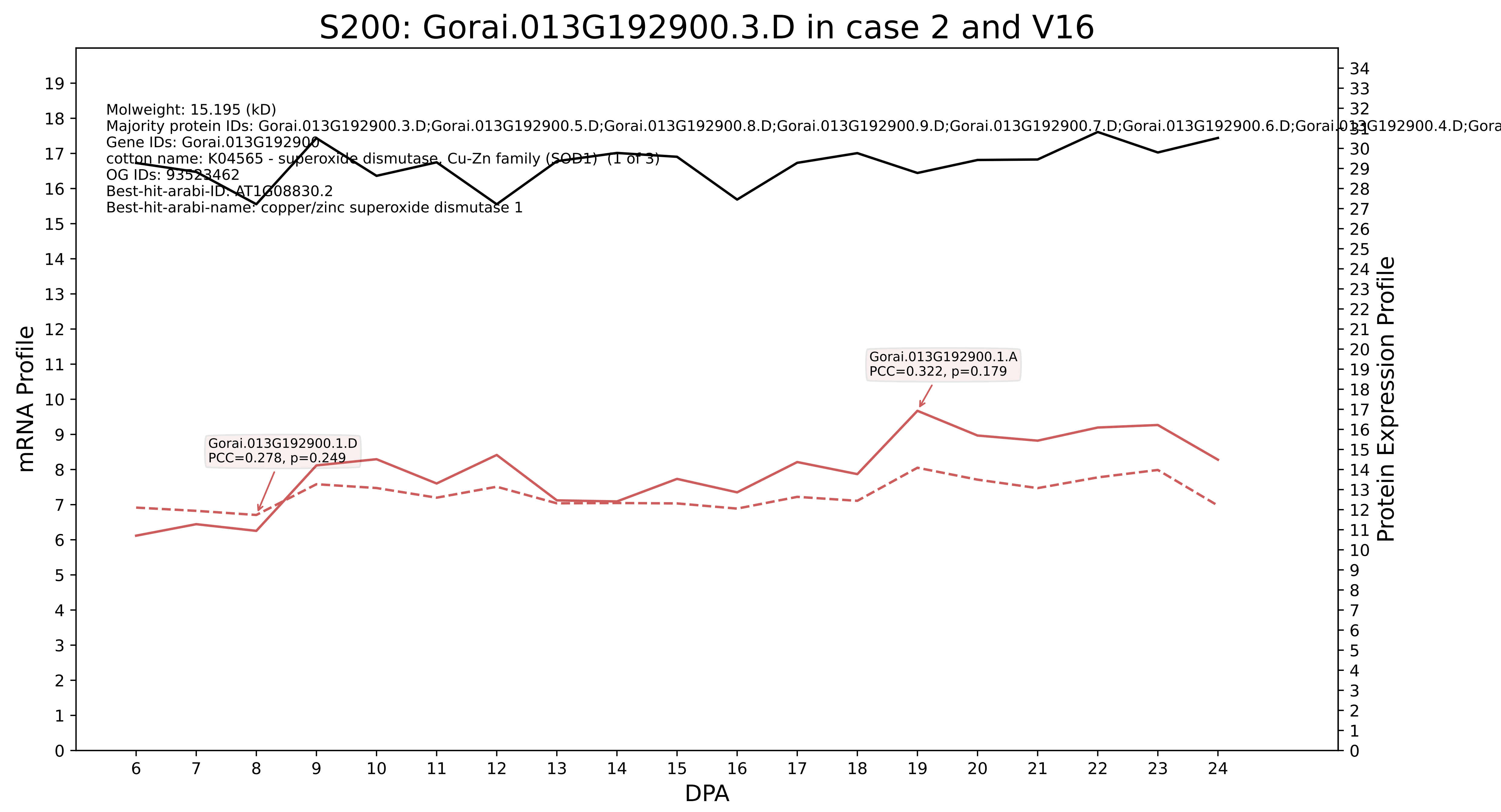

S200: Gorai.013G192900.3.D in case 2 and V16
Molweight: 15.195 (kD)
Majority protein IDs: Gorai.013G192900.3.D;Gorai.013G192900.5.D;Gorai.013G192900.8.D;Gorai.013G192900.9.D;Gorai.013G192900.7.D;Gorai.013G192900.6.D;Gorai.013G192900.4.D;Gora
Gene IDs: Gorai.013G192900
cotton name: K04565 - superoxide dismutase, Cu-Zn family (SOD1) (1 of 3)
OG IDs: 93523462
Best-hit-arabi-ID: AT1G08830.2
Best-hit-arabi-name: copper/zinc superoxide dismutase 1
Gorai.013G192900.1.A
PCC=0.322, p=0.179
Gorai.013G192900.1.D
PCC=0.278, p=0.249
mRNA Profile
Protein Expression Profile
DPA

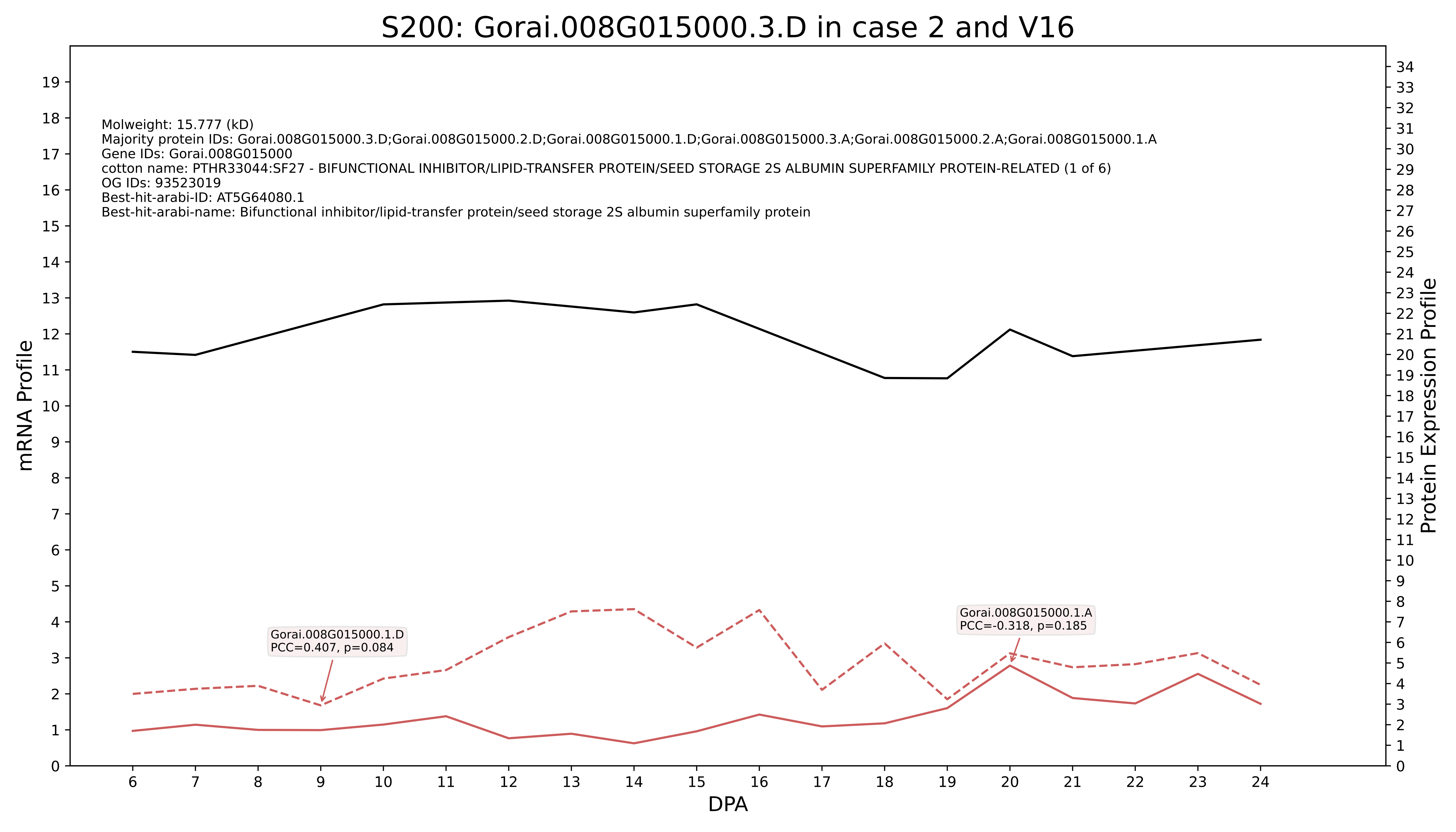
S200: Gorai.008G015000.3.D in case 2 and V16
Molweight: 15.777 (kD)
Majority protein IDs: Gorai.008G015000.3.D;Gorai.008G015000.2.D;Gorai.008G015000.1.D;Gorai.008G015000.3.A;Gorai.008G015000.2.A;Gorai.008G015000.1.A
Gene IDs: Gorai.008G015000
cotton name: PTHR33044:SF27 - BIFUNCTIONAL INHIBITOR/LIPID-TRANSFER PROTEIN/SEED STORAGE 2S ALBUMIN SUPERFAMILY PROTEIN-RELATED (1 of 6)
OG IDs: 93523019
Best-hit-arabi-ID: AT5G64080.1
Best-hit-arabi-name: Bifunctional inhibitor/lipid-transfer protein/seed storage 2S albumin superfamily protein
Gorai.008G015000.1.D
PCC=0.407, p=0.084
Gorai.008G015000.1.A
PCC=-0.318, p=0.185
mRNA Profile
Protein Expression Profile
DPA

### S200: Gorai.010G049700.1.D in case 2 and V16

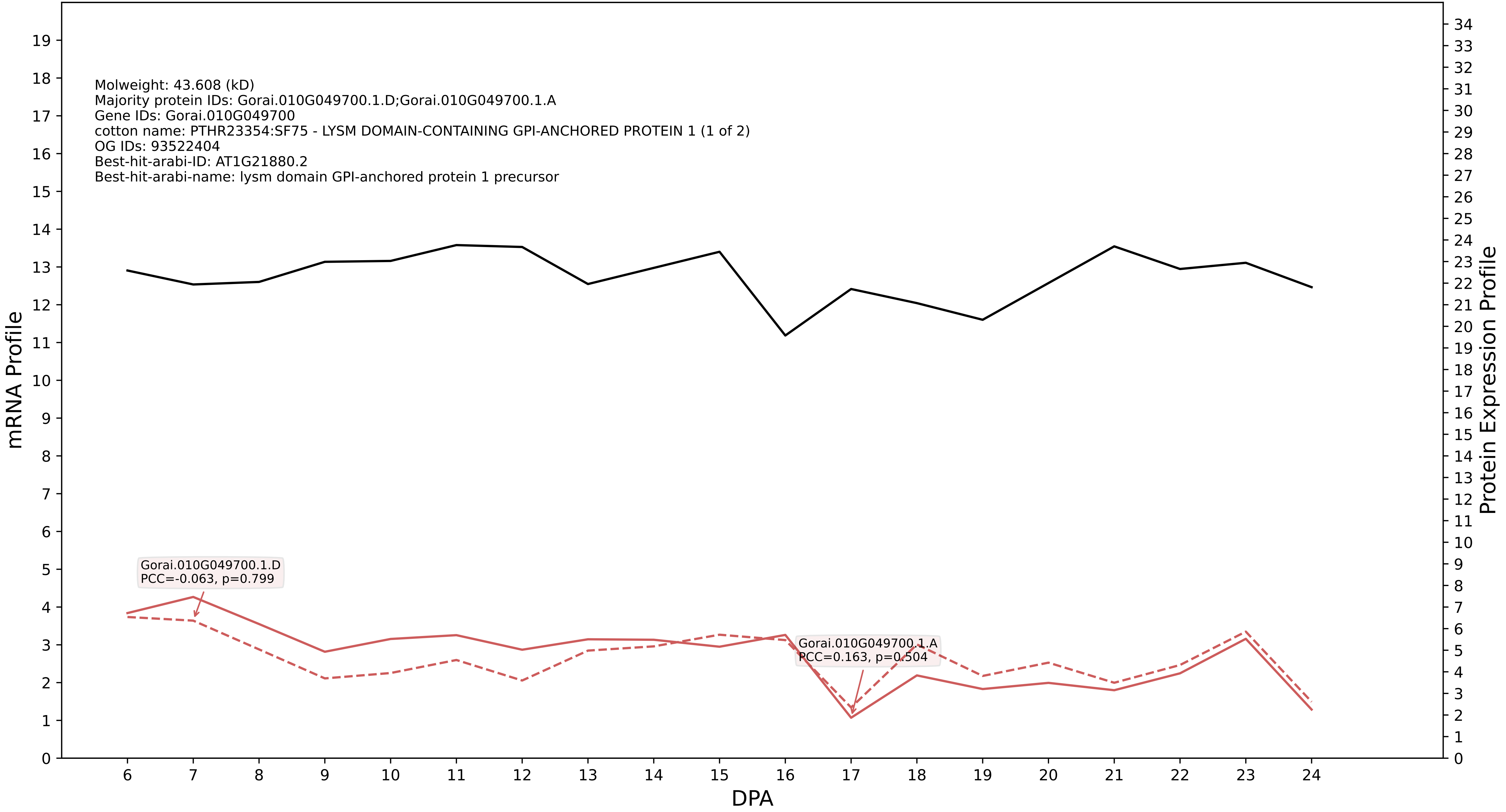

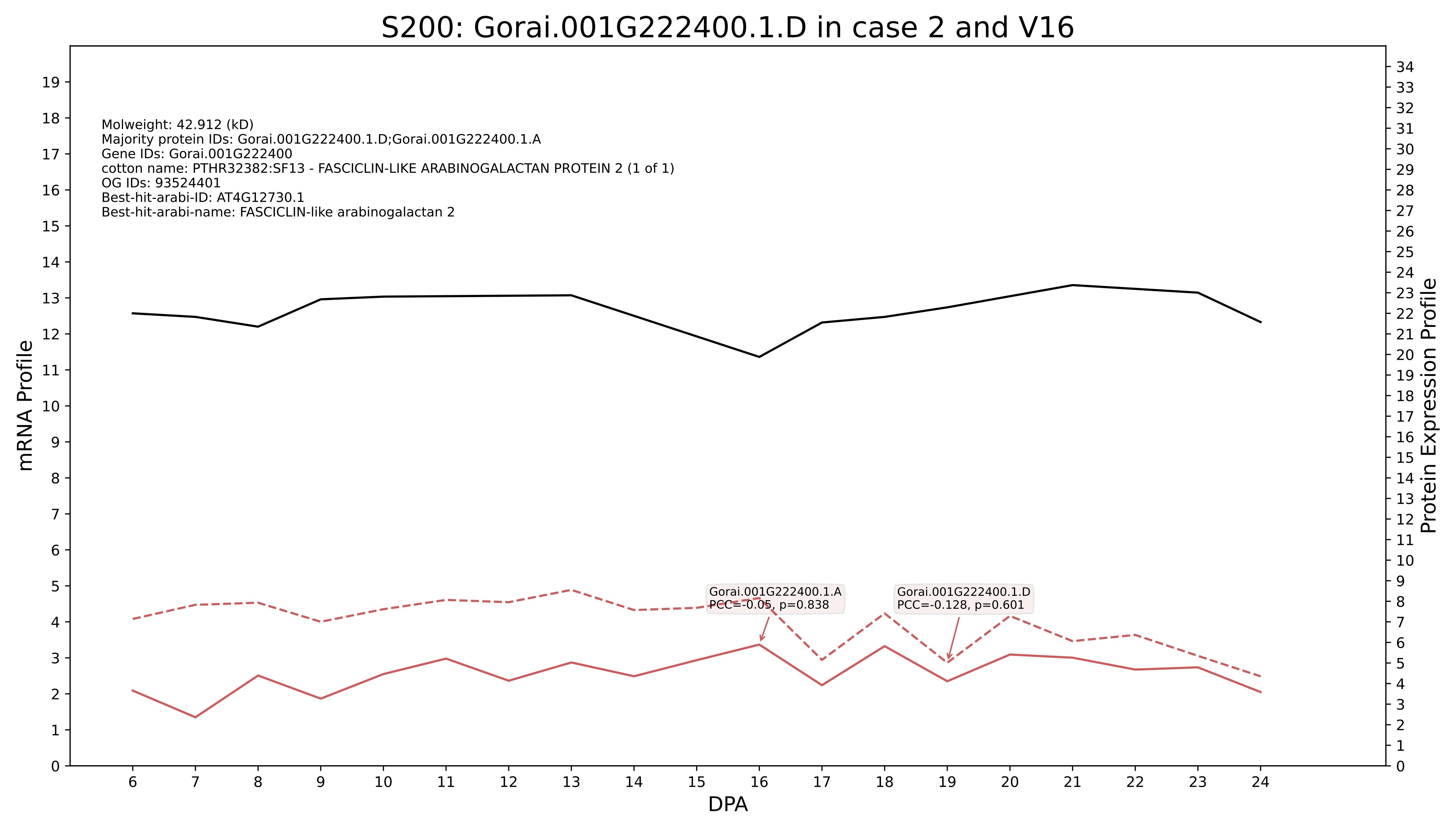
S200: Gorai.001G222400.1.D in case 2 and V16
Molweight: 42.912 (kD)
Majority protein IDs: Gorai.001G222400.1.D;Gorai.001G222400.1.A
Gene IDs: Gorai.001G222400
cotton name: PTHR32382:SF13 - FASCICLIN-LIKE ARABINOGALACTAN PROTEIN 2 (1 of 1)
OG IDs: 93524401
Best-hit-arabi-ID: AT4G12730.1
Best-hit-arabi-name: FASCICLIN-like arabinogalactan 2
Gorai.001G222400.1.A
PCC=-0.05, p=0.838
Gorai.001G222400.1.D
PCC=-0.128, p=0.601
mRNA Profile
Protein Expression Profile
DPA

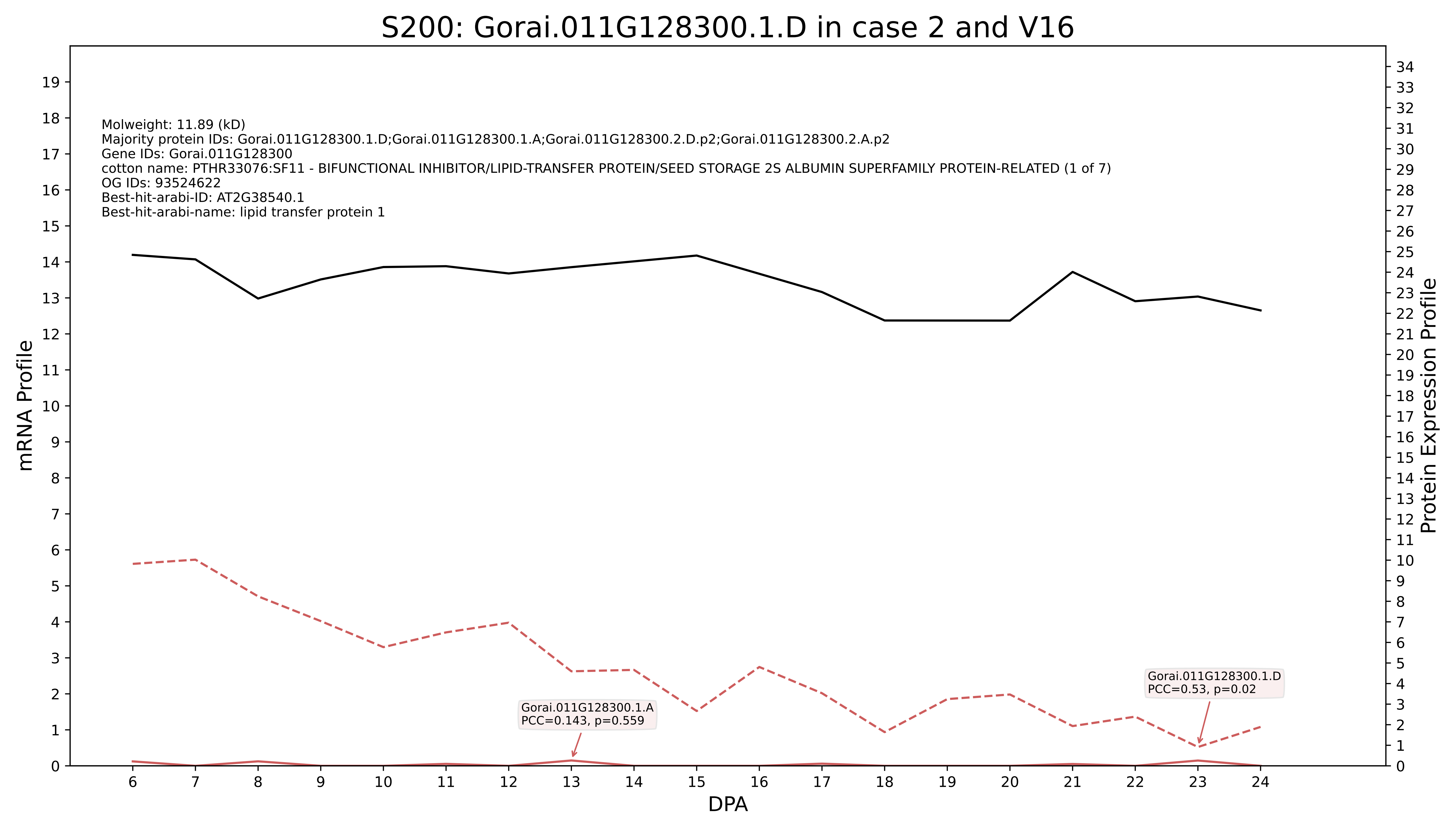
S200: Gorai.011G128300.1.D in case 2 and V16
Molweight: 11.89 (kD)
Majority protein IDs: Gorai.011G128300.1.D;Gorai.011G128300.1.A;Gorai.011G128300.2.D.p2;Gorai.011G128300.2.A.p2
Gene IDs: Gorai.011G128300
cotton name: PTHR33076:SF11 - BIFUNCTIONAL INHIBITOR/LIPID-TRANSFER PROTEIN/SEED STORAGE 2S ALBUMIN SUPERFAMILY PROTEIN-RELATED (1 of 7)
OG IDs: 93524622
Best-hit-arabi-ID: AT2G38540.1
Best-hit-arabi-name: lipid transfer protein 1
Gorai.011G128300.1.A
PCC=0.143, p=0.559
Gorai.011G128300.1.D
PCC=0.53, p=0.02
mRNA Profile
Protein Expression Profile
DPA

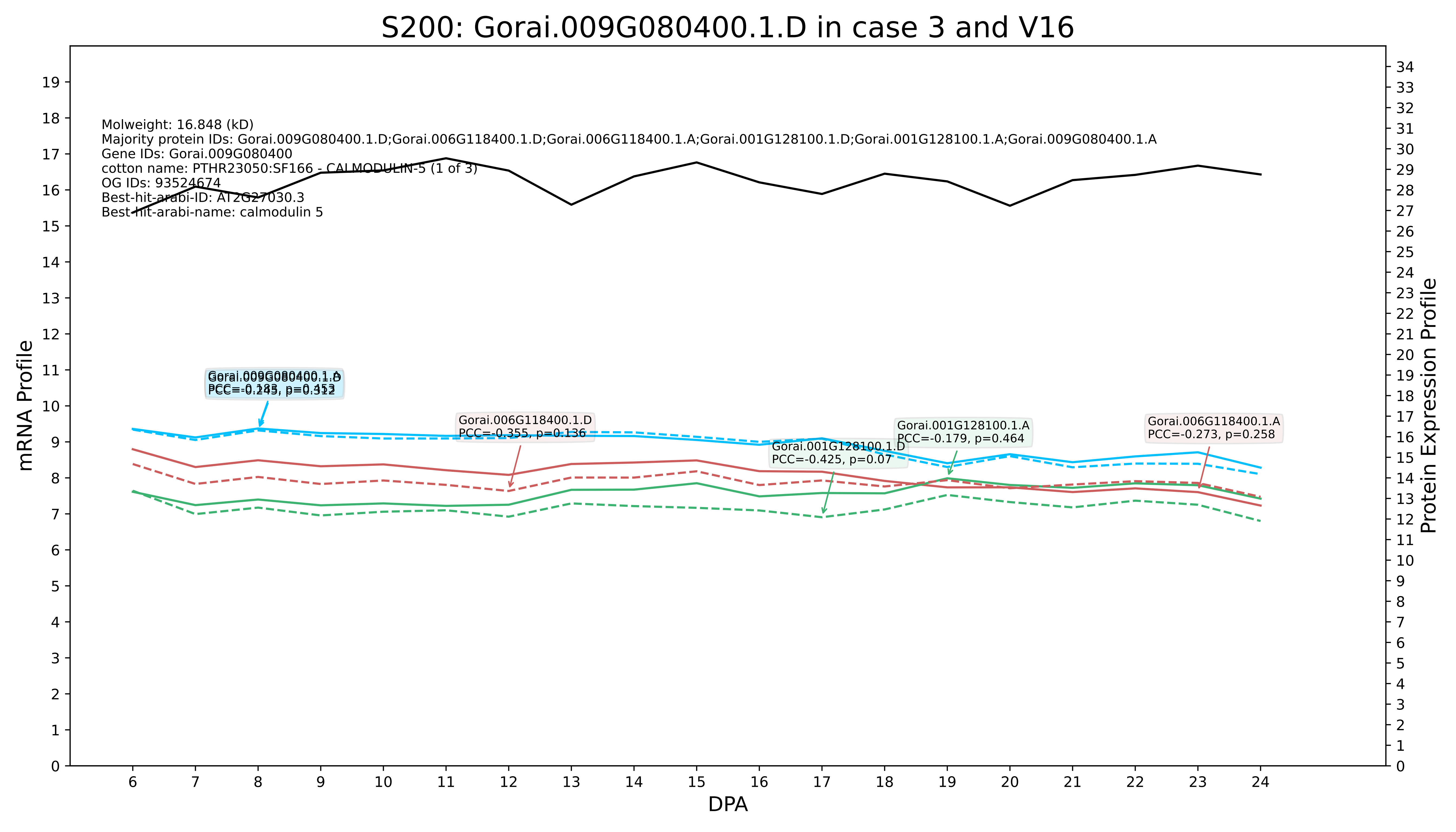

S200: Gorai.009G080400.1.D in case 3 and V16
Molweight: 16.848 (kD)
Majority protein IDs: Gorai.009G080400.1.D;Gorai.006G118400.1.D;Gorai.006G118400.1.A;Gorai.001G128100.1.D;Gorai.001G128100.1.A;Gorai.009G080400.1.A
Gene IDs: Gorai.009G080400
cotton name: PTHR23050:SF166 - CALMODULIN-5 (1 of 3)
OG IDs: 93524674
Best-hit-arabi-ID: AT2G27030.3
Best-hit-arabi-name: calmodulin 5
Gorai.009G080400.1.A
PCC=-0.183, p=0.453
Gorai.009G080400.1.D
PCC=-0.245, p=0.312
Gorai.006G118400.1.D
PCC=-0.355, p=0.136
Gorai.001G128100.1.D
PCC=-0.425, p=0.07
Gorai.001G128100.1.A
PCC=-0.179, p=0.464
Gorai.006G118400.1.A
PCC=-0.273, p=0.258
mRNA Profile
Protein Expression Profile
DPA

### S200: Gorai.007G080700.1.A in case 3 and V16

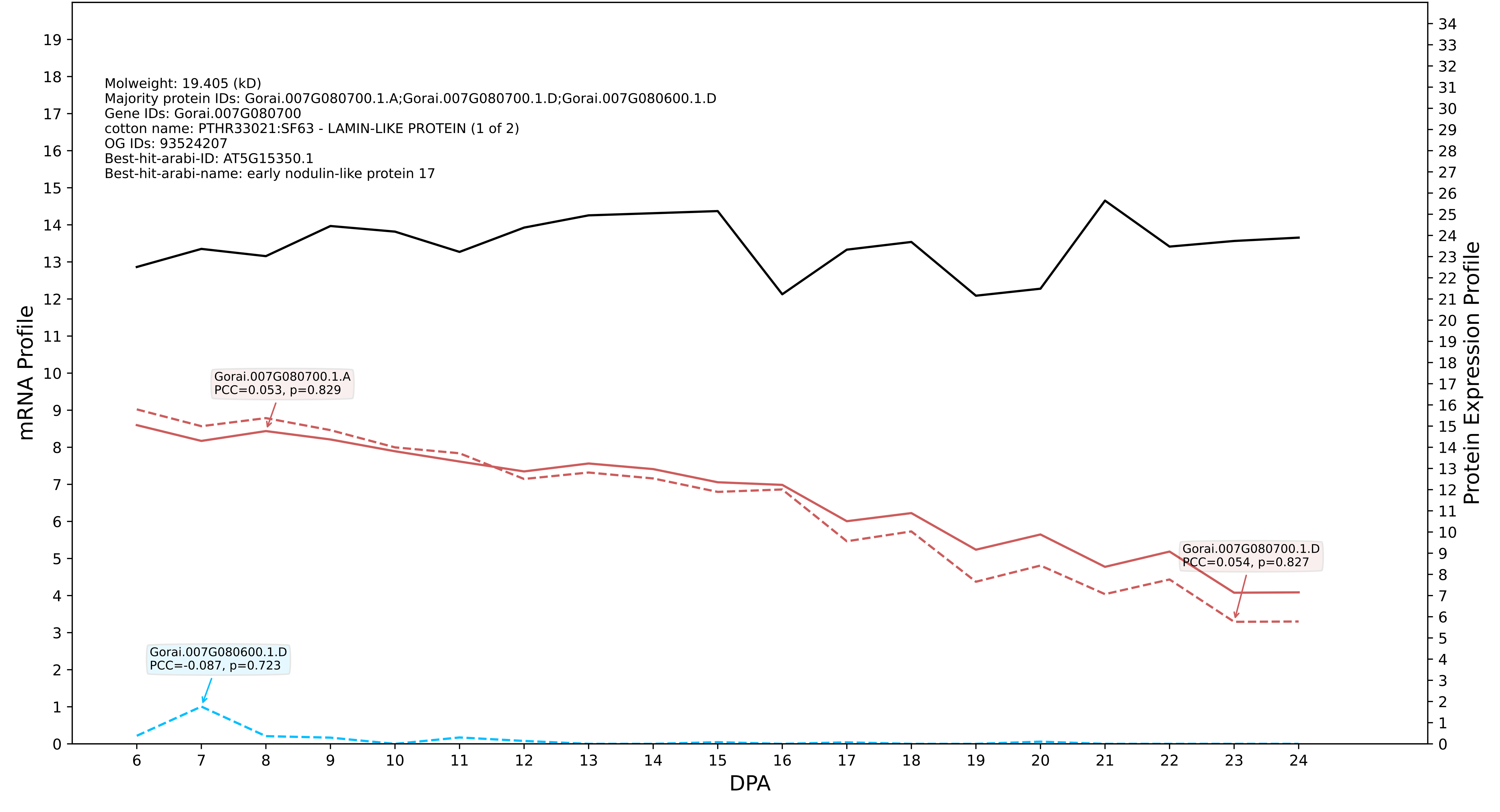

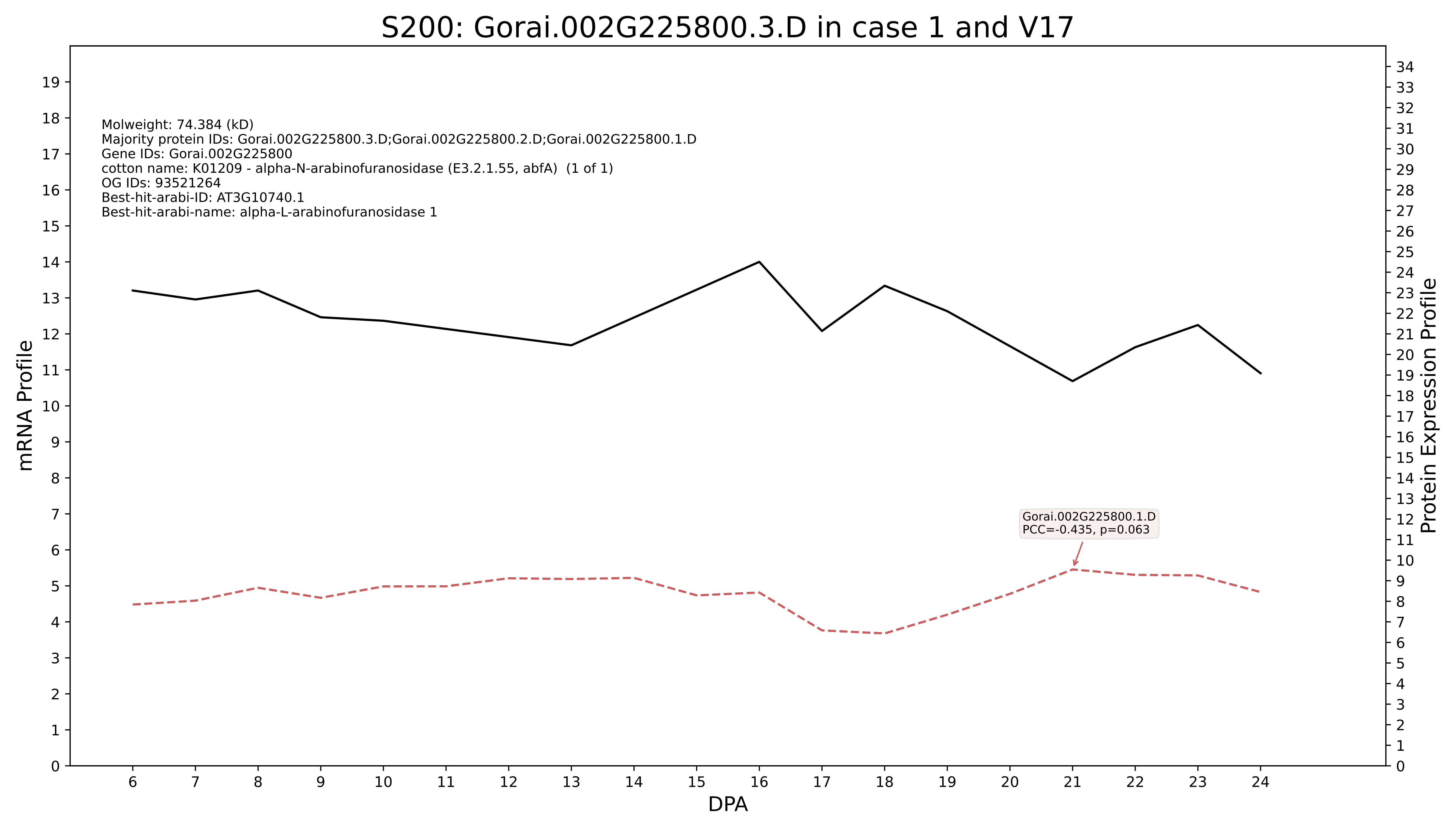

S200: Gorai.002G225800.3.D in case 1 and V17
Molweight: 74.384 (kD)
Majority protein IDs: Gorai.002G225800.3.D;Gorai.002G225800.2.D;Gorai.002G225800.1.D
Gene IDs: Gorai.002G225800
cotton name: K01209 - alpha-N-arabinofuranosidase (E3.2.1.55, abfA) (1 of 1)
OG IDs: 93521264
Best-hit-arabi-ID: AT3G10740.1
Best-hit-arabi-name: alpha-L-arabinofuranosidase 1
Gorai.002G225800.1.D
PCC=-0.435, p=0.063
mRNA Profile
Protein Expression Profile
DPA

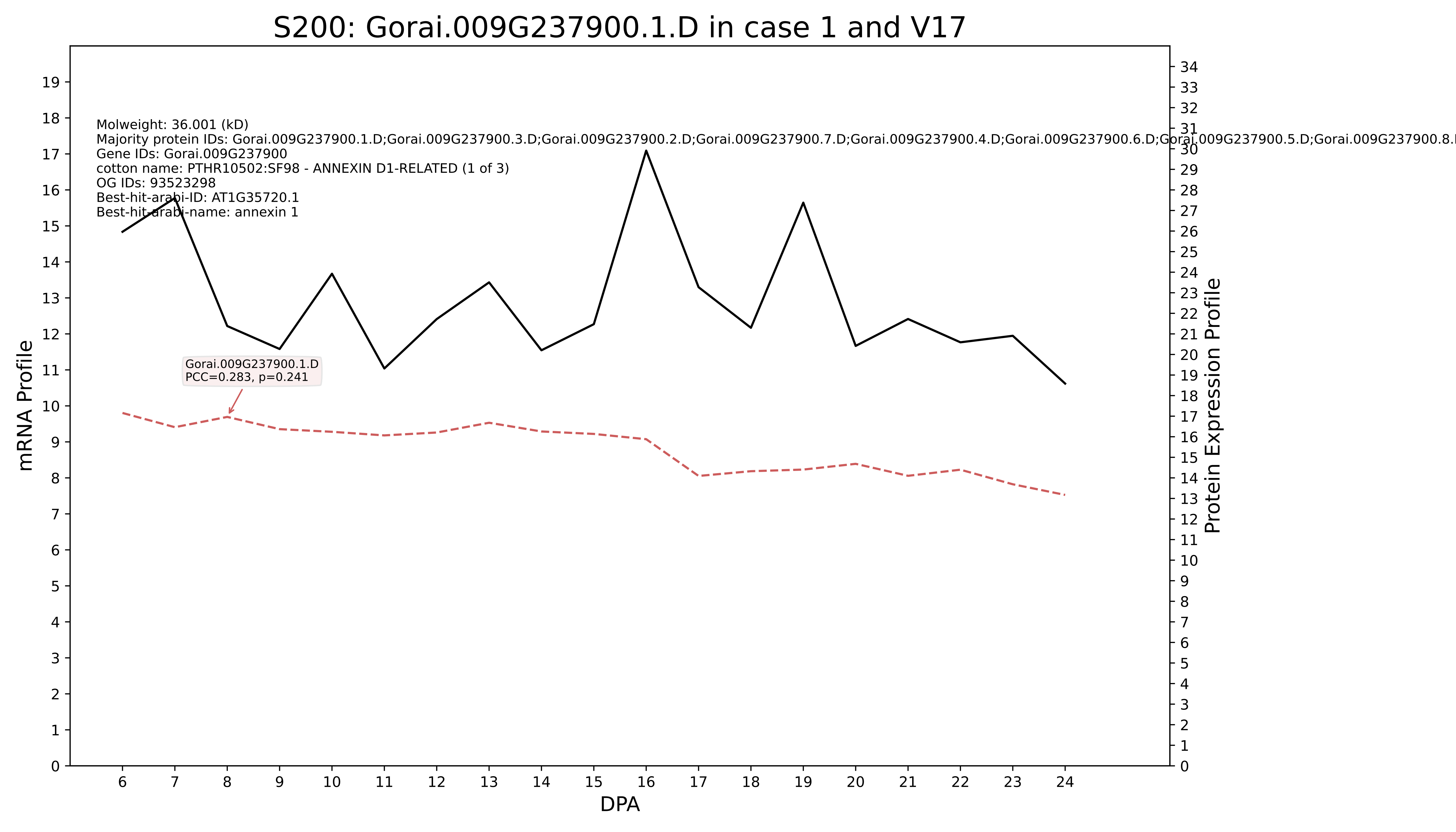

S200: Gorai.009G237900.1.D in case 1 and V17
Molweight: 36.001 (kD)
Majority protein IDs: Gorai.009G237900.1.D;Gorai.009G237900.3.D;Gorai.009G237900.2.D;Gorai.009G237900.7.D;Gorai.009G237900.4.D;Gorai.009G237900.6.D;Gorai.009G237900.5.D;Gorai.009G237900.8.
Gene IDs: Gorai.009G237900
cotton name: PTHR10502:SF98 - ANNEXIN D1-RELATED (1 of 3)
OG IDs: 93523298
Best-hit-arabi-ID: AT1G35720.1
Best-hit-arabi-name: annexin 1
Gorai.009G237900.1.D
PCC=0.283, p=0.241
mRNA Profile
Protein Expression Profile
DPA

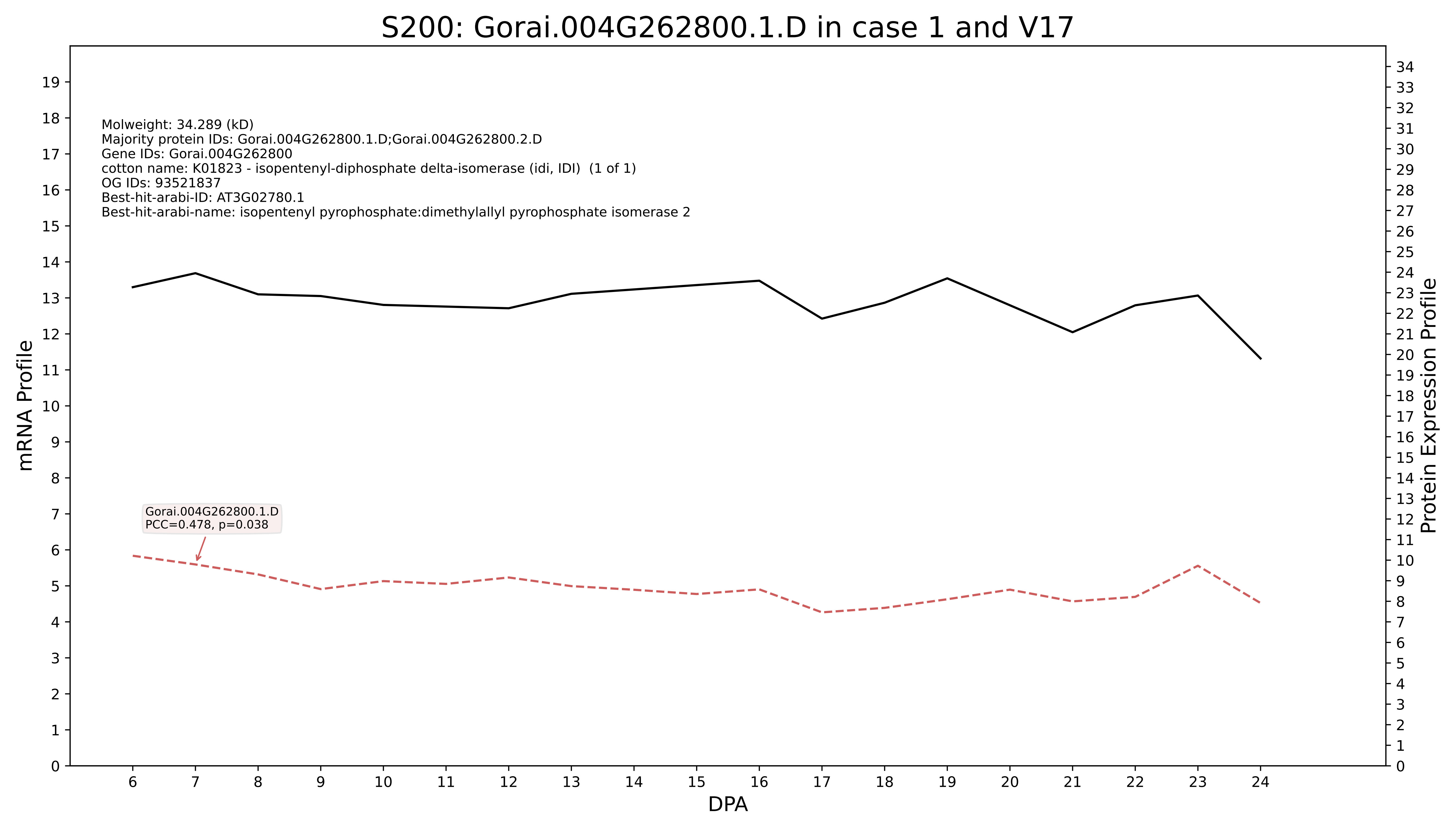

S200: Gorai.004G262800.1.D in case 1 and V17
Molweight: 34.289 (kD)
Majority protein IDs: Gorai.004G262800.1.D;Gorai.004G262800.2.D
Gene IDs: Gorai.004G262800
cotton name: K01823 - isopentenyl-diphosphate delta-isomerase (idi, IDI) (1 of 1)
OG IDs: 93521837
Best-hit-arabi-ID: AT3G02780.1
Best-hit-arabi-name: isopentenyl pyrophosphate:dimethylallyl pyrophosphate isomerase 2
Gorai.004G262800.1.D
PCC=0.478, p=0.038
mRNA Profile
Protein Expression Profile
DPA

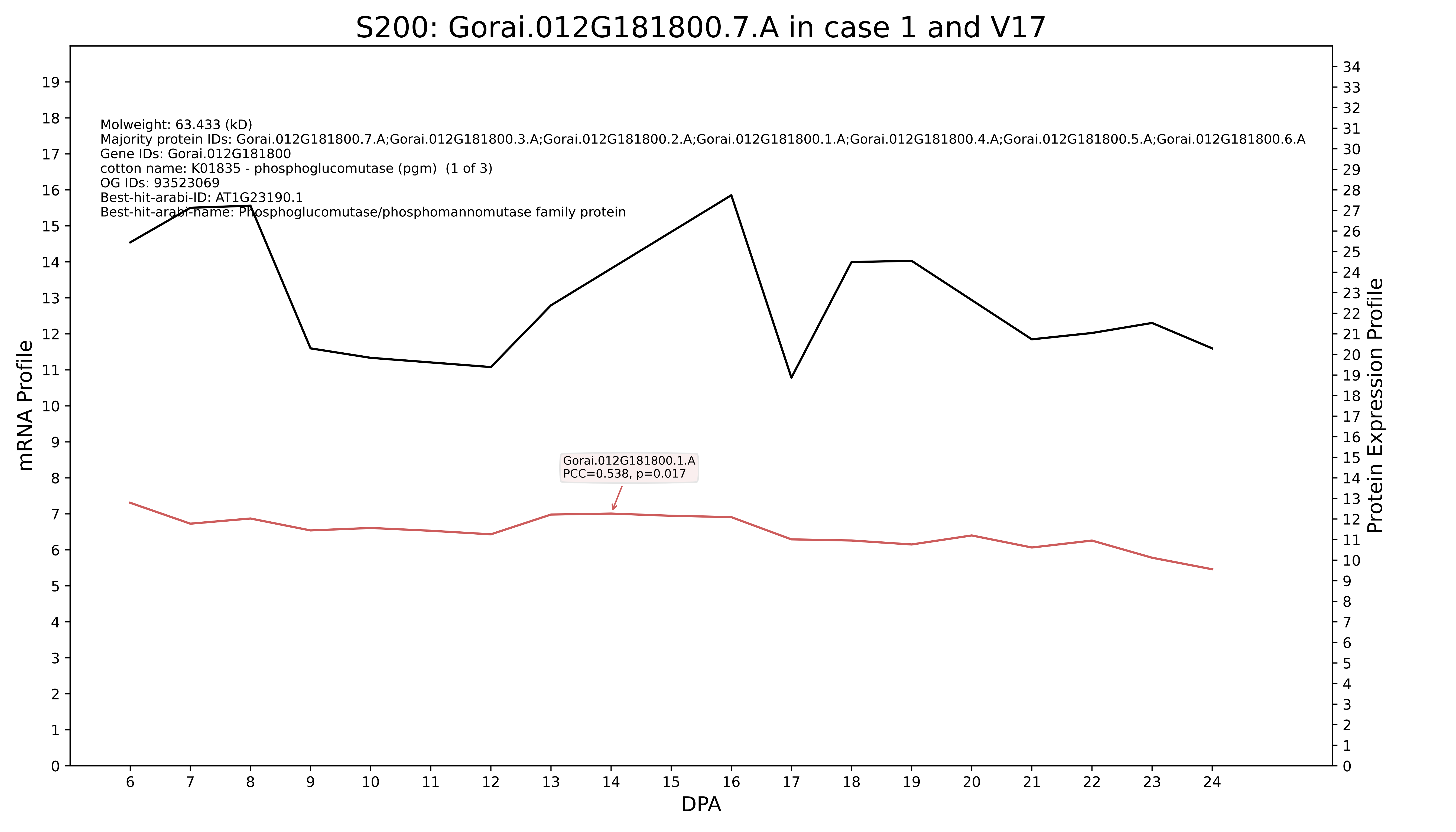

S200: Gorai.012G181800.7.A in case 1 and V17
Molweight: 63.433 (kD)
Majority protein IDs: Gorai.012G181800.7.A;Gorai.012G181800.3.A;Gorai.012G181800.2.A;Gorai.012G181800.1.A;Gorai.012G181800.4.A;Gorai.012G181800.5.A;Gorai.012G181800.6.A
Gene IDs: Gorai.012G181800
cotton name: K01835 - phosphoglucomutase (pgm) (1 of 3)
OG IDs: 93523069
Best-hit-arabi-ID: AT1G23190.1
Best-hit-arabi-name: Phosphoglucomutase/phosphomannomutase family protein
Gorai.012G181800.1.A
PCC=0.538, p=0.017
mRNA Profile
Protein Expression Profile
DPA

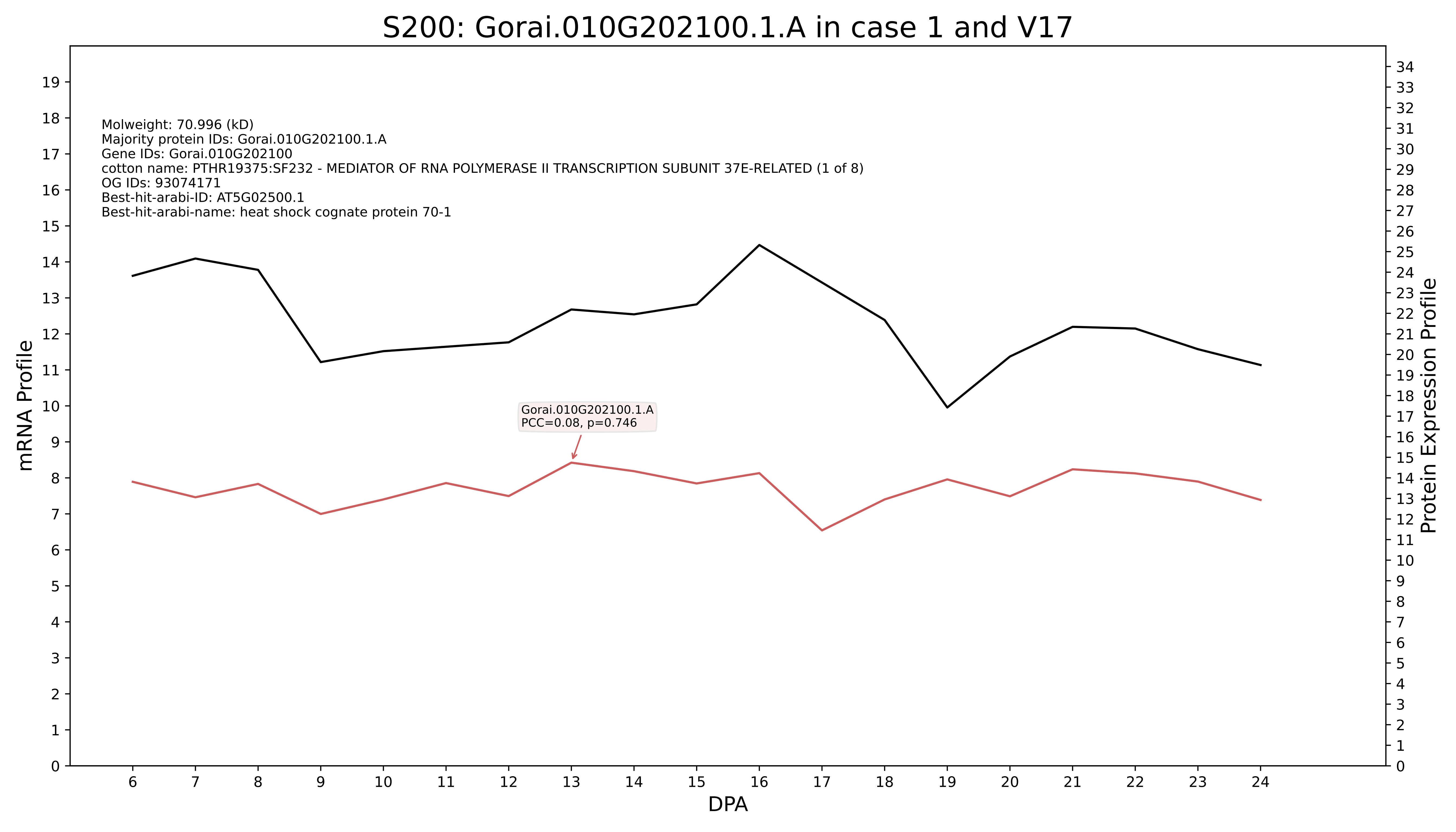
S200: Gorai.010G202100.1.A in case 1 and V17
Molweight: 70.996 (kD)
Majority protein IDs: Gorai.010G202100.1.A
Gene IDs: Gorai.010G202100
cotton name: PTHR19375:SF232 - MEDIATOR OF RNA POLYMERASE II TRANSCRIPTION SUBUNIT 37E-RELATED (1 of 8)
OG IDs: 93074171
Best-hit-arabi-ID: AT5G02500.1
Best-hit-arabi-name: heat shock cognate protein 70-1
Gorai.010G202100.1.A
PCC=0.08, p=0.746
mRNA Profile
Protein Expression Profile
DPA

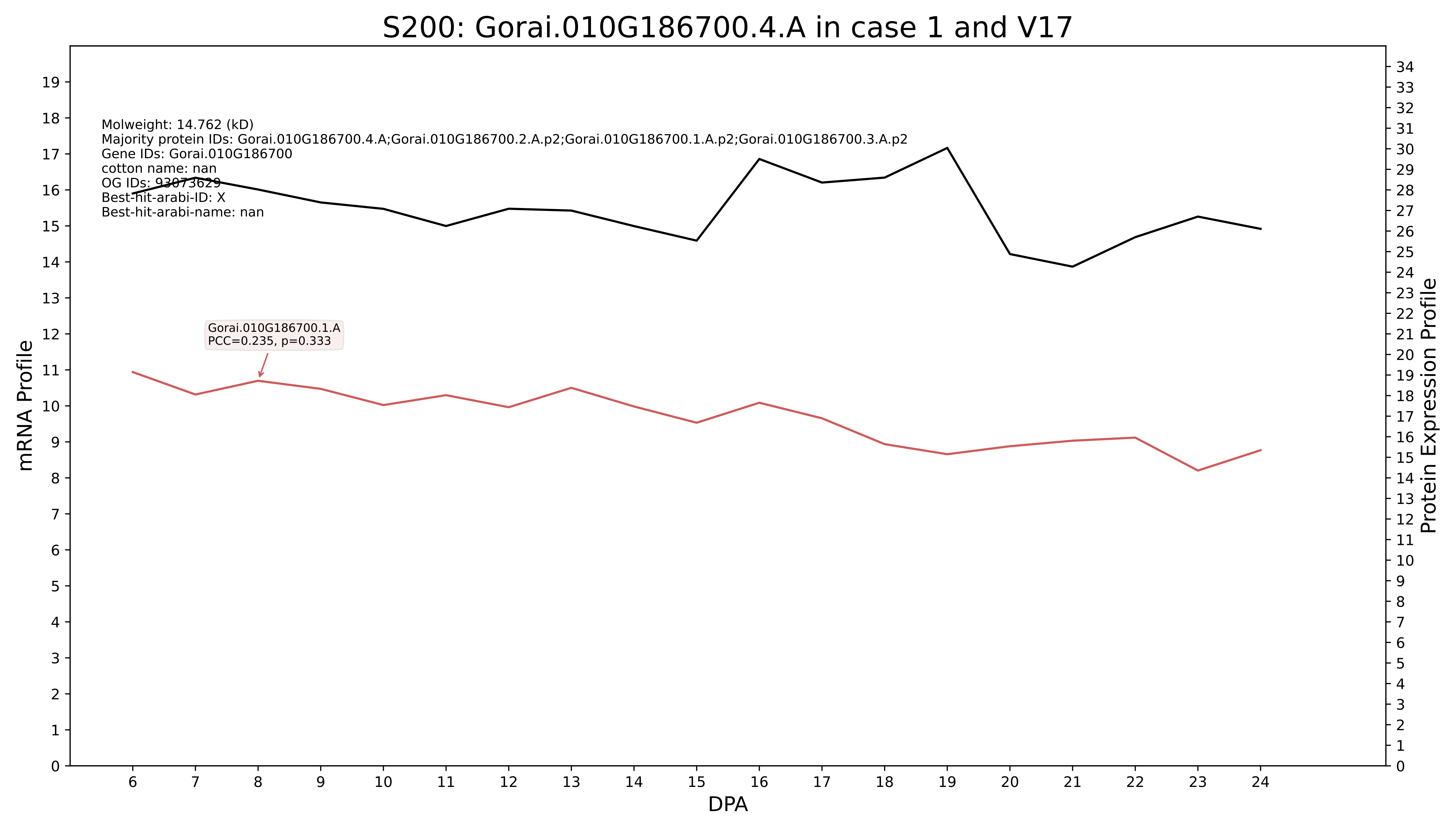
S200: Gorai.010G186700.4.A in case 1 and V17
Molweight: 14.762 (kD)
Majority protein IDs: Gorai.010G186700.4.A;Gorai.010G186700.2.A.p2;Gorai.010G186700.1.A.p2;Gorai.010G186700.3.A.p2
Gene IDs: Gorai.010G186700
cotton name: nan
OG IDs: 93073629
Best-hit-arabi-ID: X
Best-hit-arabi-name: nan
Gorai.010G186700.1.A
PCC=0.235, p=0.333
mRNA Profile
Protein Expression Profile
DPA

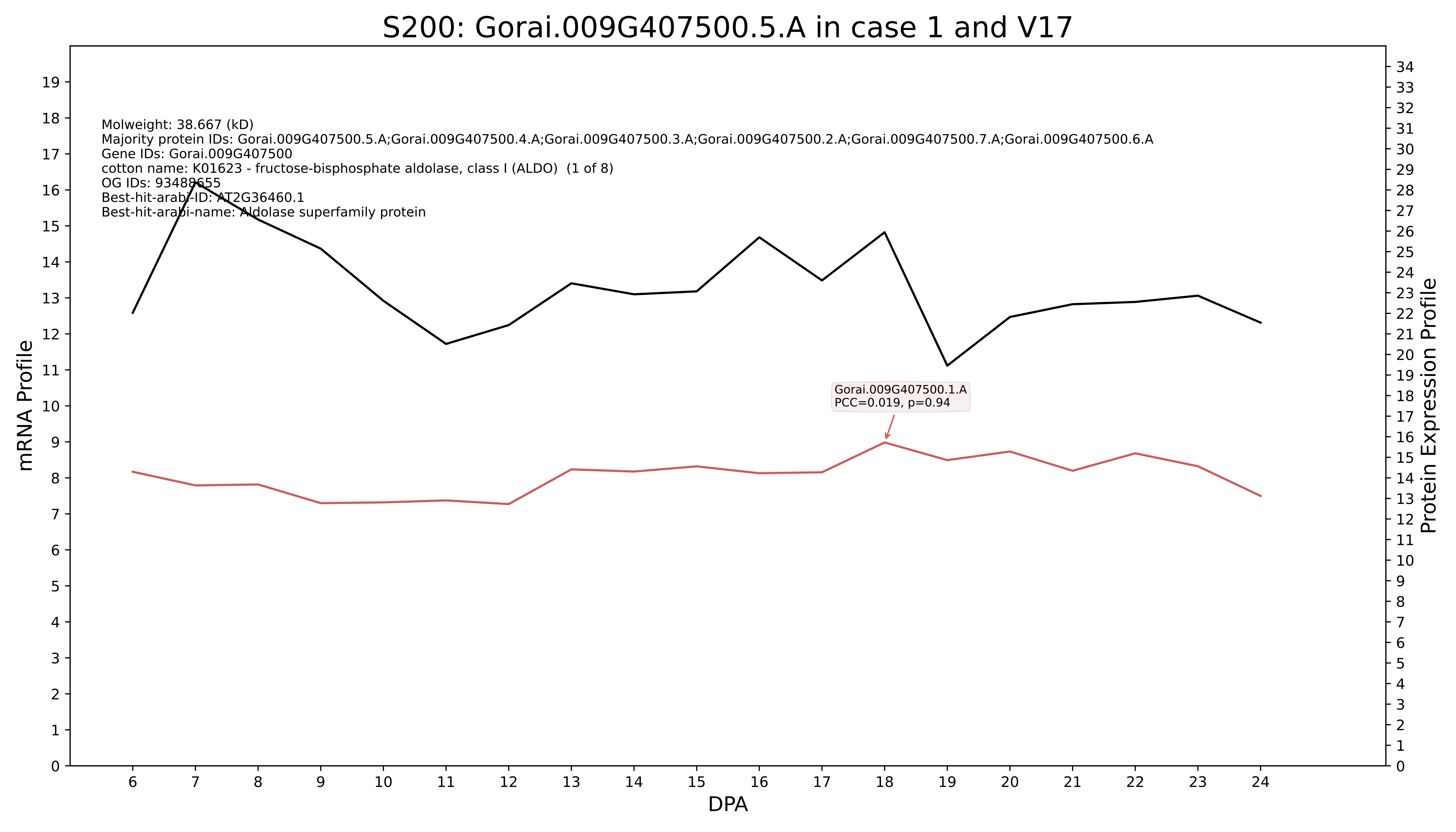

S200: Gorai.009G407500.5.A in case 1 and V17
Molweight: 38.667 (kD)
Majority protein IDs: Gorai.009G407500.5.A;Gorai.009G407500.4.A;Gorai.009G407500.3.A;Gorai.009G407500.2.A;Gorai.009G407500.7.A;Gorai.009G407500.6.A
Gene IDs: Gorai.009G407500
cotton name: K01623 - fructose-bisphosphate aldolase, class I (ALDO) (1 of 8)
OG IDs: 93488655
Best-hit-arabi-ID: AT2G36460.1
Best-hit-arabi-name: Aldolase superfamily protein
Gorai.009G407500.1.A
PCC=0.019, p=0.94
mRNA Profile
Protein Expression Profile
DPA

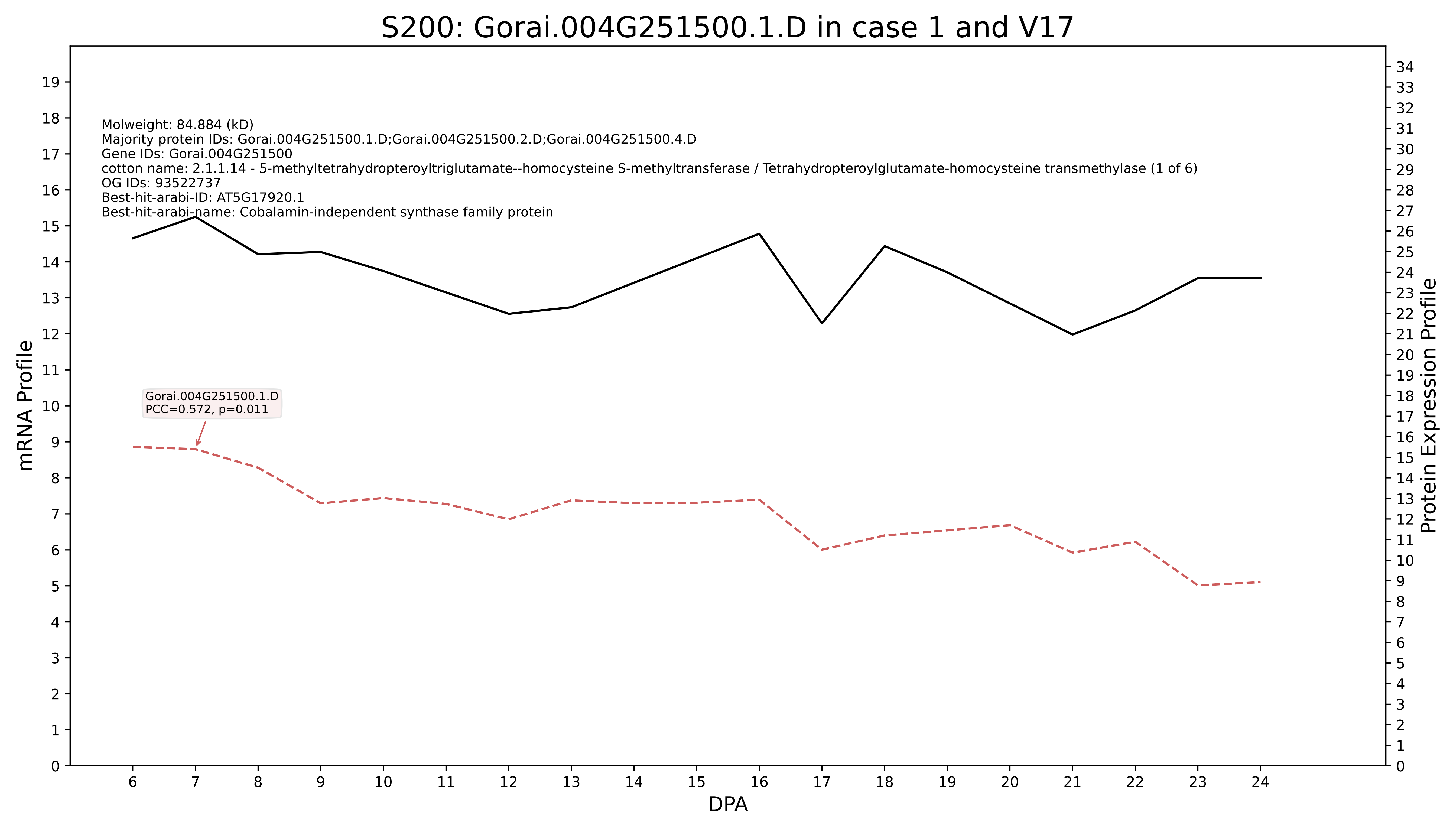

S200: Gorai.004G251500.1.D in case 1 and V17
Molweight: 84.884 (kD)
Majority protein IDs: Gorai.004G251500.1.D;Gorai.004G251500.2.D;Gorai.004G251500.4.D
Gene IDs: Gorai.004G251500
cotton name: 2.1.1.14 - 5-methyltetrahydropteroyltriglutamate--homocysteine S-methyltransferase / Tetrahydropteroylglutamate-homocysteine transmethylase (1 of 6)
OG IDs: 93522737
Best-hit-arabi-ID: AT5G17920.1
Best-hit-arabi-name: Cobalamin-independent synthase family protein
Gorai.004G251500.1.D
PCC=0.572, p=0.011
mRNA Profile
Protein Expression Profile
DPA

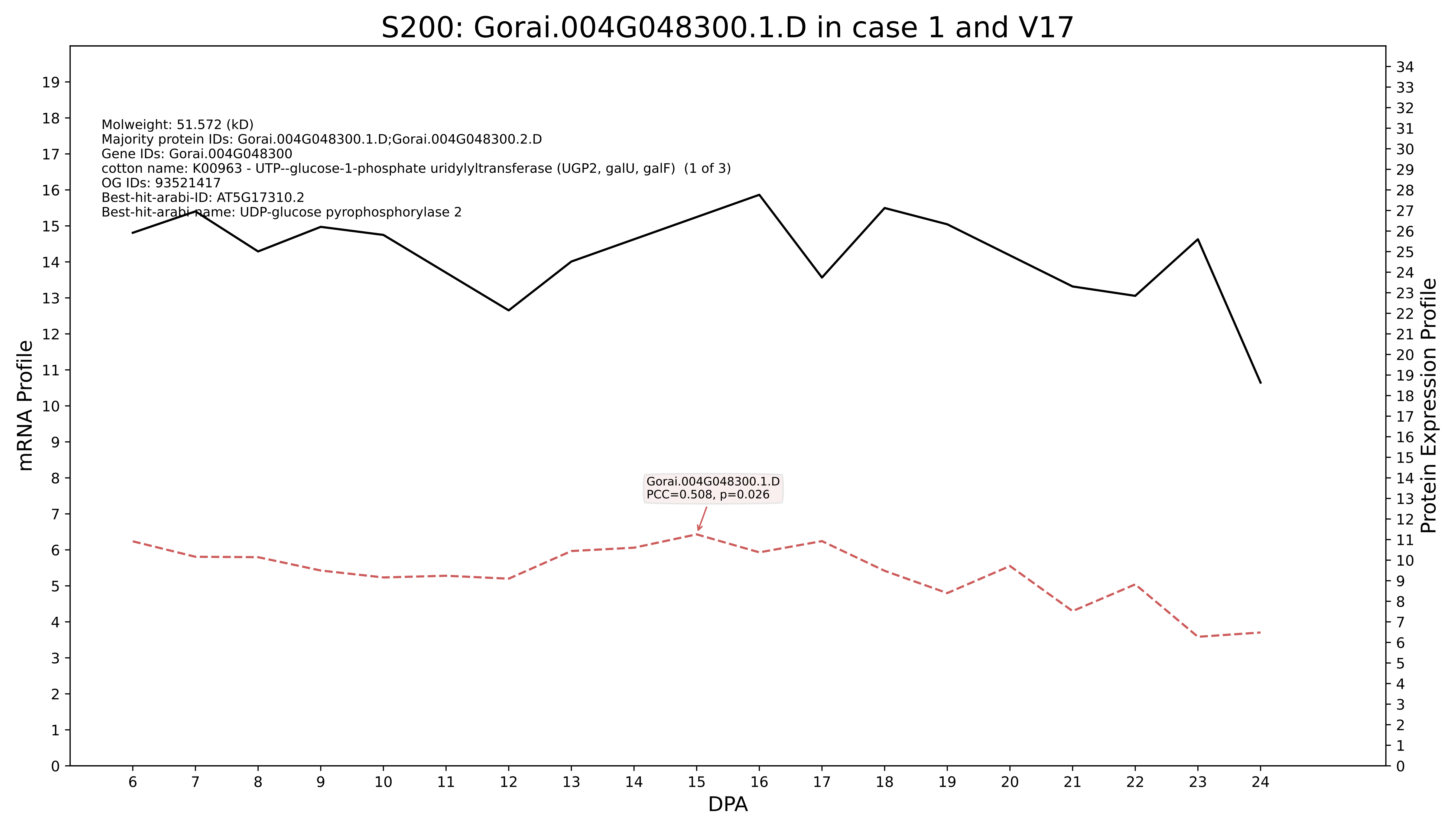
S200: Gorai.004G048300.1.D in case 1 and V17
Molweight: 51.572 (kD)
Majority protein IDs: Gorai.004G048300.1.D;Gorai.004G048300.2.D
Gene IDs: Gorai.004G048300
cotton name: K00963 - UTP--glucose-1-phosphate uridylyltransferase (UGP2, galU, galF) (1 of 3)
OG IDs: 93521417
Best-hit-arabi-ID: AT5G17310.2
Best-hit-arabi-name: UDP-glucose pyrophosphorylase 2
Gorai.004G048300.1.D
PCC=0.508, p=0.026
mRNA Profile
Protein Expression Profile
DPA

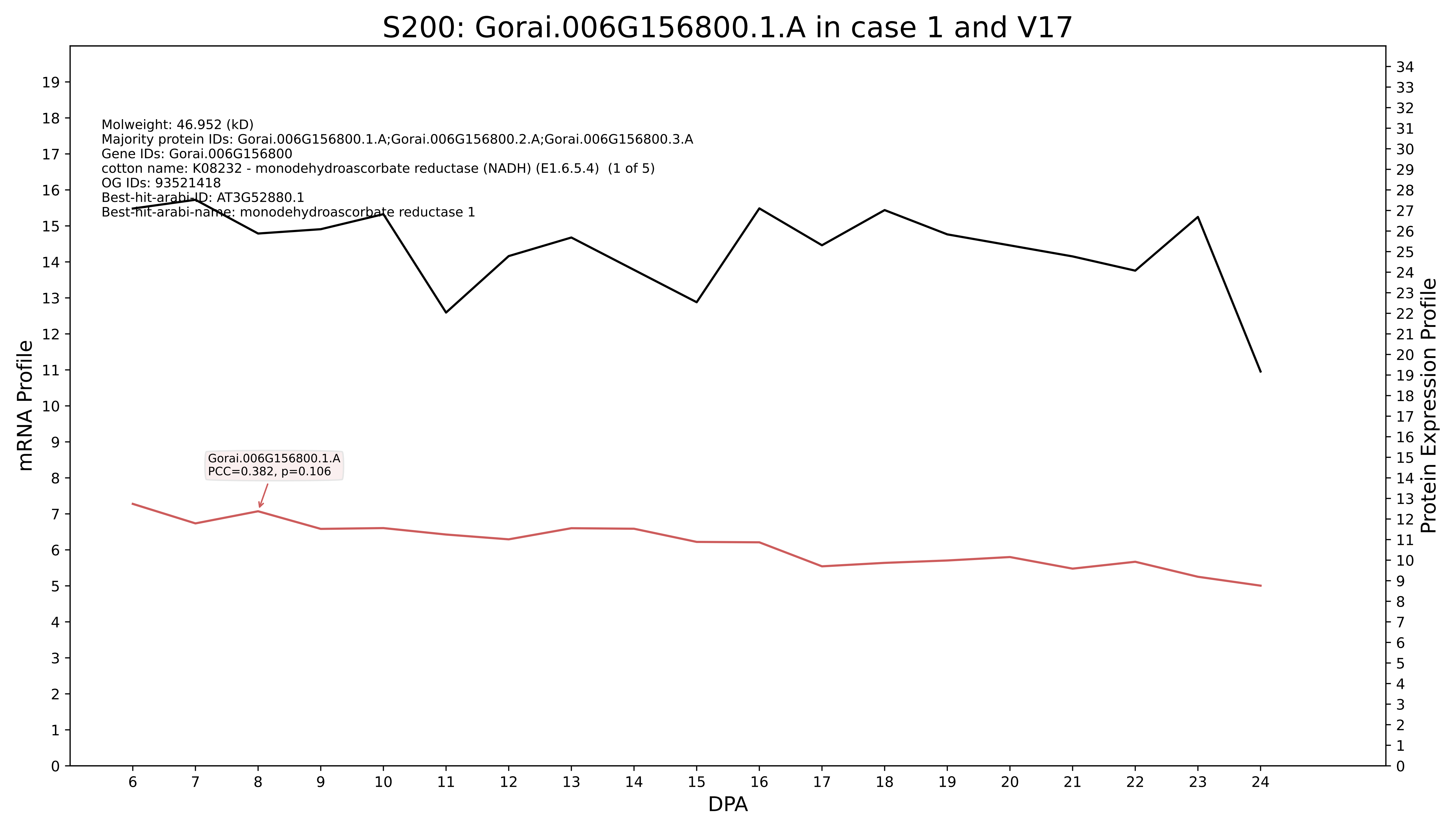
S200: Gorai.006G156800.1.A in case 1 and V17
Molweight: 46.952 (kD)
Majority protein IDs: Gorai.006G156800.1.A;Gorai.006G156800.2.A;Gorai.006G156800.3.A
Gene IDs: Gorai.006G156800
cotton name: K08232 - monodehydroascorbate reductase (NADH) (E1.6.5.4) (1 of 5)
OG IDs: 93521418
Best-hit-arabi-ID: AT3G52880.1
Best-hit-arabi-name: monodehydroascorbate reductase 1
Gorai.006G156800.1.A
PCC=0.382, p=0.106
mRNA Profile
Protein Expression Profile
DPA

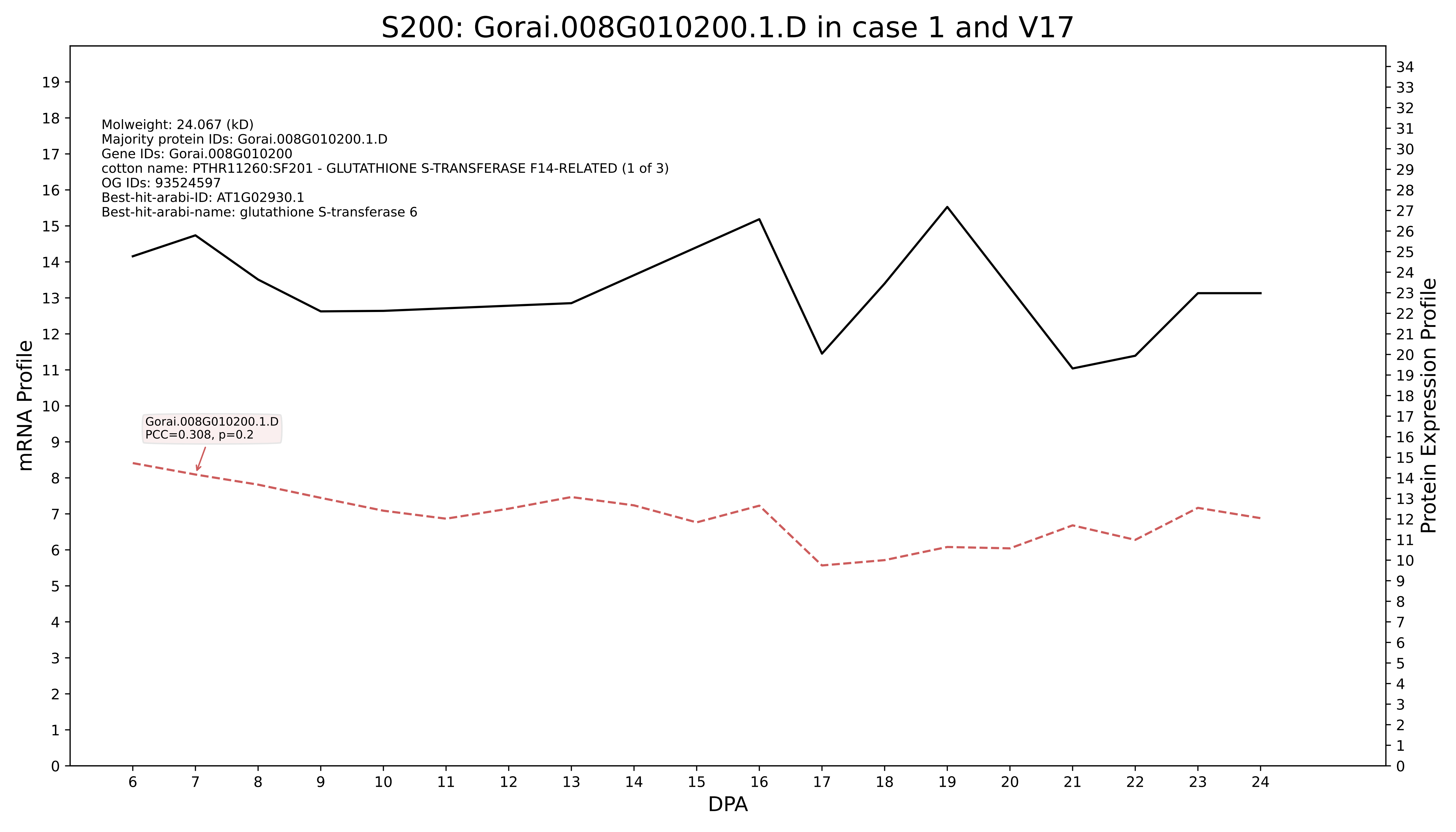
S200: Gorai.008G010200.1.D in case 1 and V17
Molweight: 24.067 (kD)
Majority protein IDs: Gorai.008G010200.1.D
Gene IDs: Gorai.008G010200
cotton name: PTHR11260:SF201 - GLUTATHIONE S-TRANSFERASE F14-RELATED (1 of 3)
OG IDs: 93524597
Best-hit-arabi-ID: AT1G02930.1
Best-hit-arabi-name: glutathione S-transferase 6
Gorai.008G010200.1.D
PCC=0.308, p=0.2
mRNA Profile
Protein Expression Profile
DPA

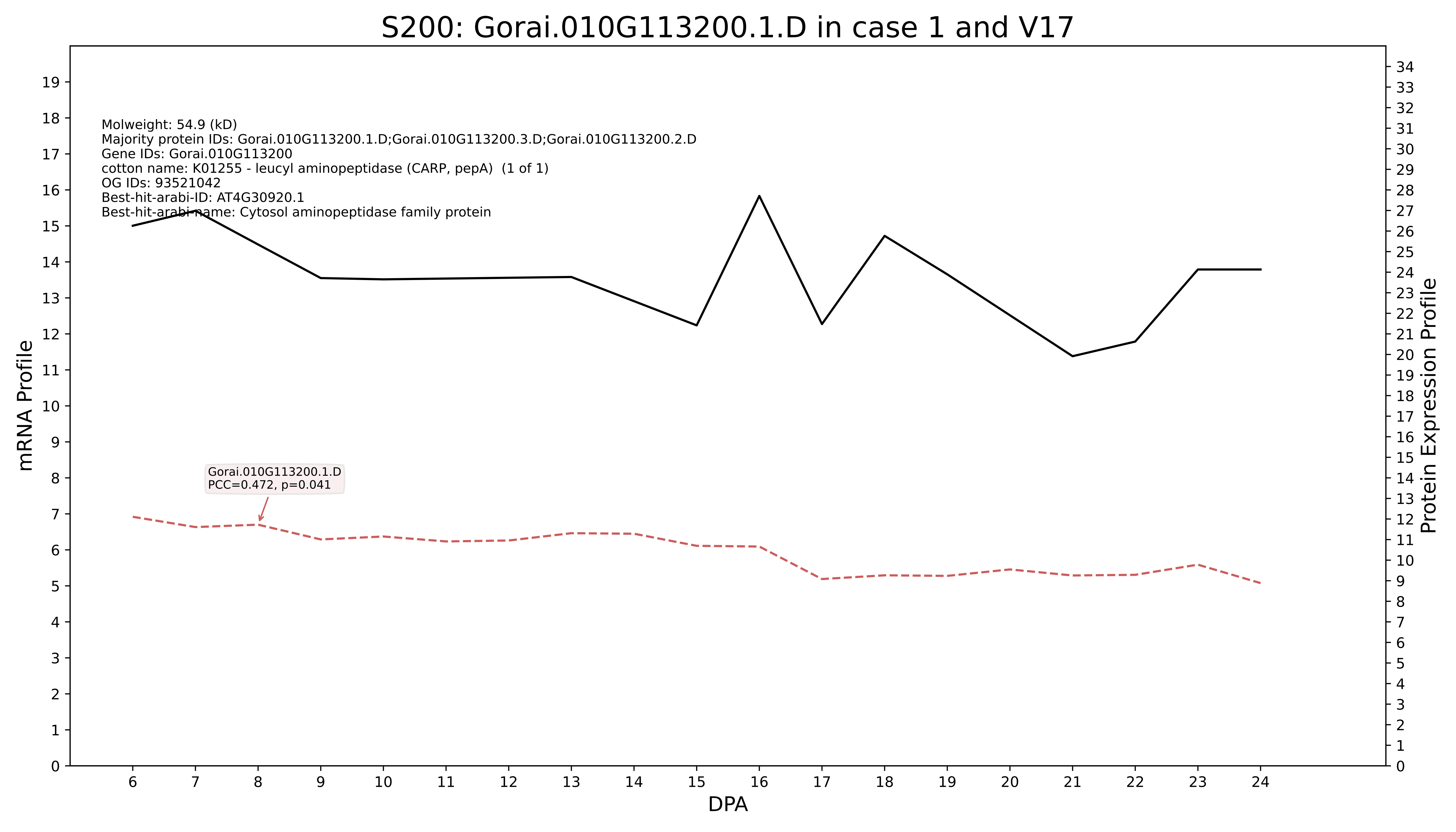

S200: Gorai.010G113200.1.D in case 1 and V17
Molweight: 54.9 (kD)
Majority protein IDs: Gorai.010G113200.1.D;Gorai.010G113200.3.D;Gorai.010G113200.2.D
Gene IDs: Gorai.010G113200
cotton name: K01255 - leucyl aminopeptidase (CARP, pepA) (1 of 1)
OG IDs: 93521042
Best-hit-arabi-ID: AT4G30920.1
Best-hit-arabi-name: Cytosol aminopeptidase family protein
Gorai.010G113200.1.D
PCC=0.472, p=0.041
mRNA Profile
Protein Expression Profile
DPA

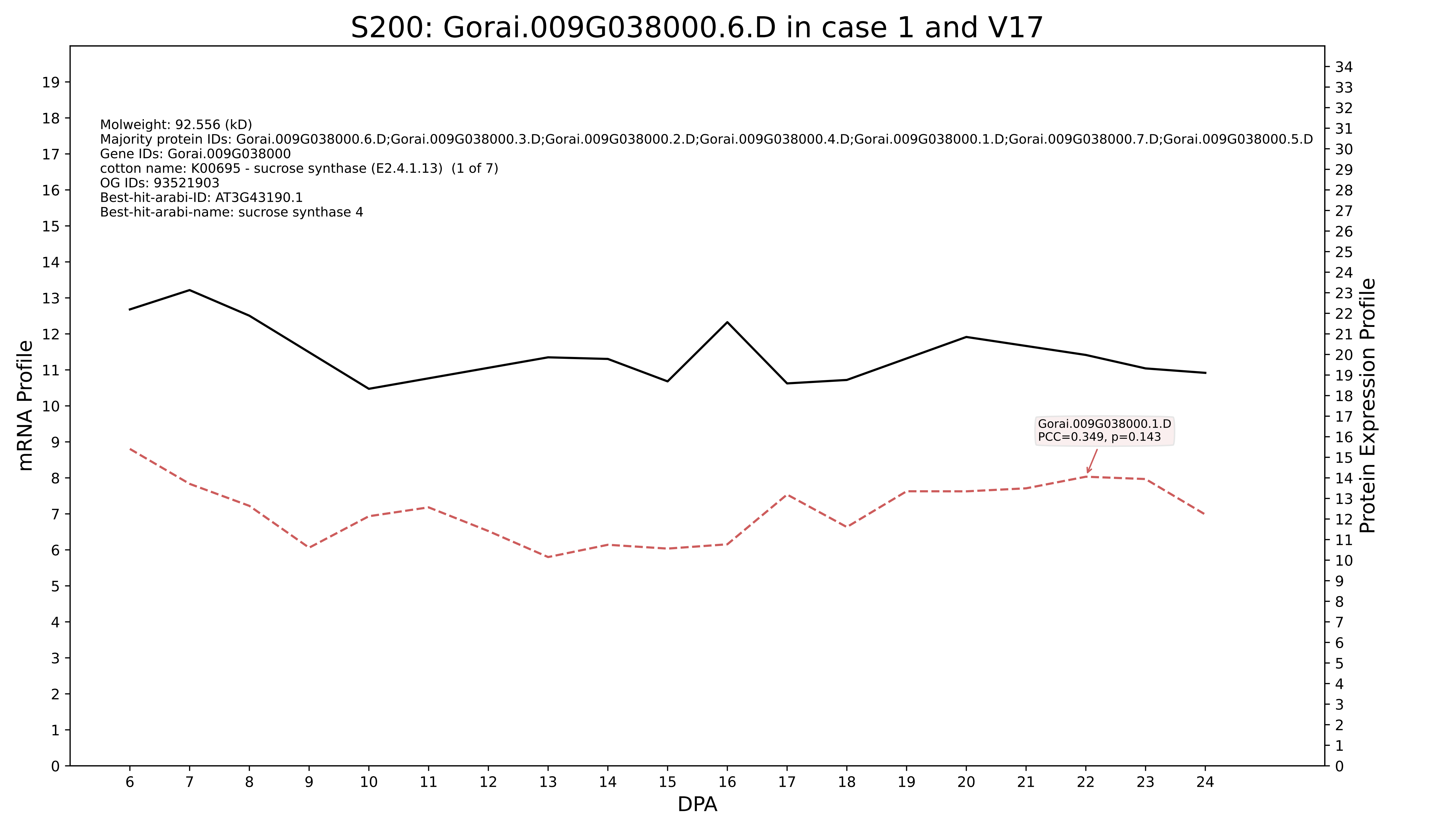
S200: Gorai.009G038000.6.D in case 1 and V17
Molweight: 92.556 (kD)
Majority protein IDs: Gorai.009G038000.6.D;Gorai.009G038000.3.D;Gorai.009G038000.2.D;Gorai.009G038000.4.D;Gorai.009G038000.1.D;Gorai.009G038000.7.D;Gorai.009G038000.5.D
Gene IDs: Gorai.009G038000
cotton name: K00695 - sucrose synthase (E2.4.1.13) (1 of 7)
OG IDs: 93521903
Best-hit-arabi-ID: AT3G43190.1
Best-hit-arabi-name: sucrose synthase 4
Gorai.009G038000.1.D
PCC=0.349, p=0.143
mRNA Profile
Protein Expression Profile
DPA

### S200: Gorai.007G090400.1.D in case 1 and V17

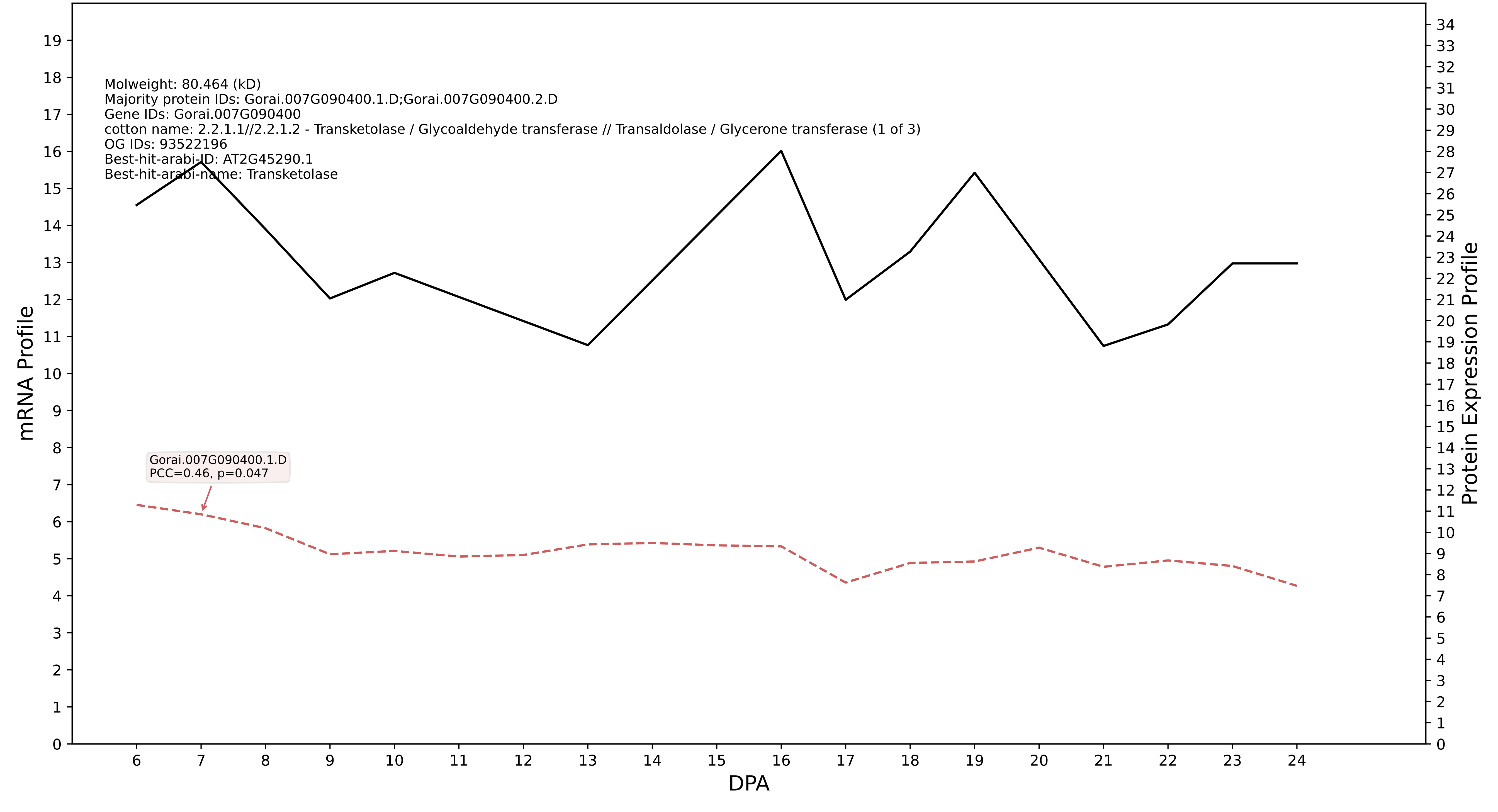

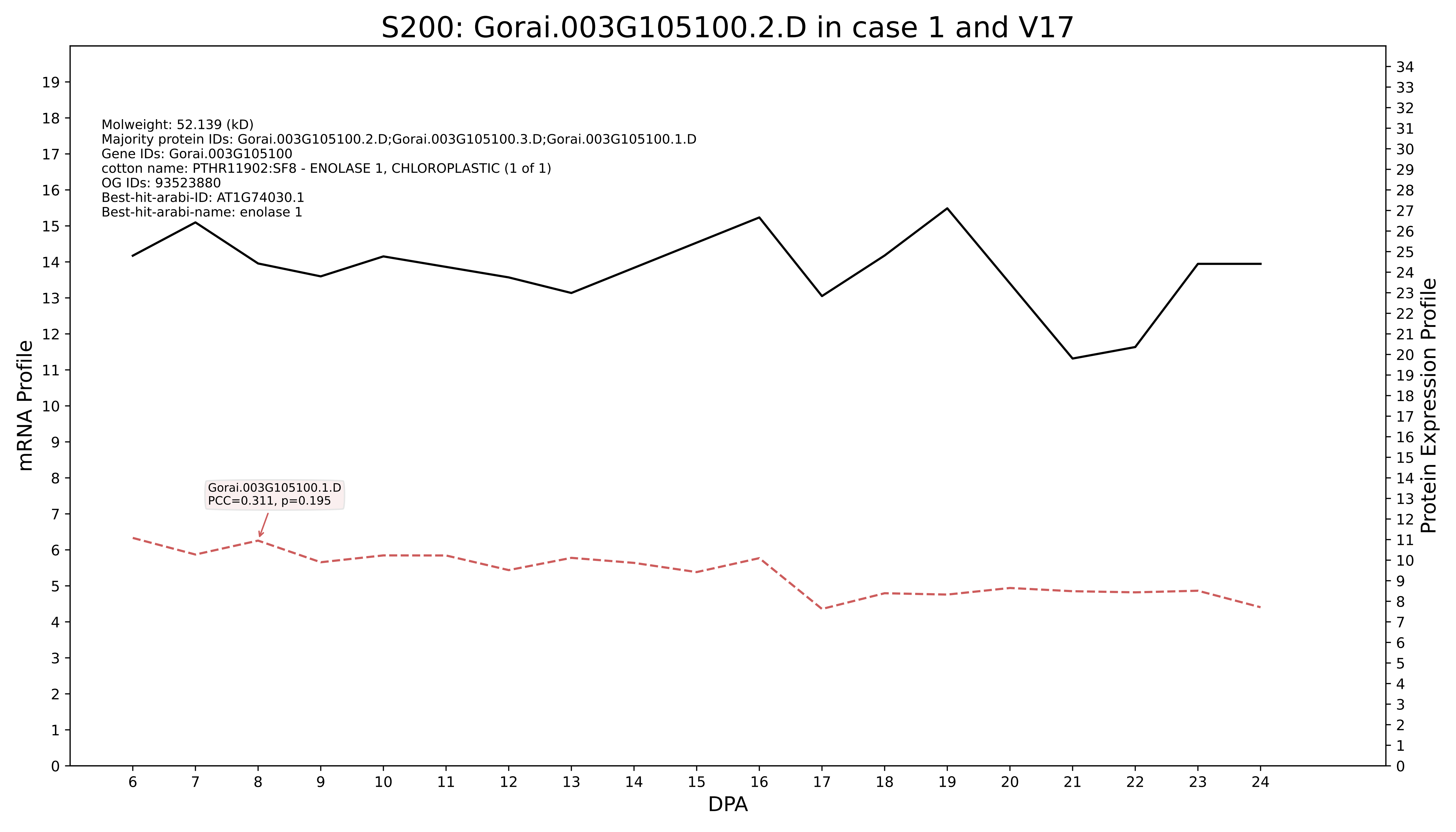
S200: Gorai.003G105100.2.D in case 1 and V17
Molweight: 52.139 (kD)
Majority protein IDs: Gorai.003G105100.2.D;Gorai.003G105100.3.D;Gorai.003G105100.1.D
Gene IDs: Gorai.003G105100
cotton name: PTHR11902:SF8 - ENOLASE 1, CHLOROPLASTIC (1 of 1)
OG IDs: 93523880
Best-hit-arabi-ID: AT1G74030.1
Best-hit-arabi-name: enolase 1
Gorai.003G105100.1.D
PCC=0.311, p=0.195
mRNA Profile
Protein Expression Profile
DPA

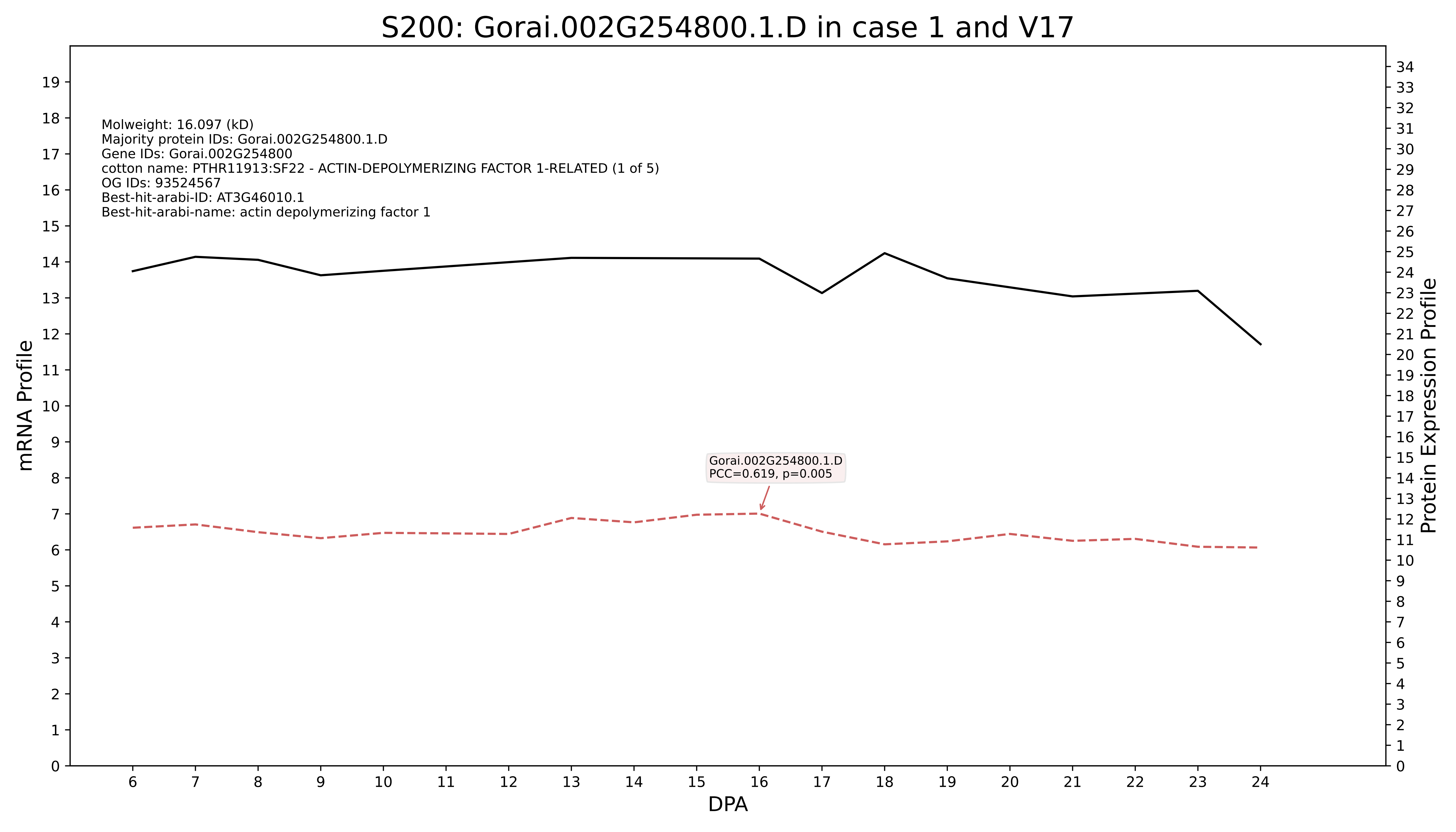
S200: Gorai.002G254800.1.D in case 1 and V17
Molweight: 16.097 (kD)
Majority protein IDs: Gorai.002G254800.1.D
Gene IDs: Gorai.002G254800
cotton name: PTHR11913:SF22 - ACTIN-DEPOLYMERIZING FACTOR 1-RELATED (1 of 5)
OG IDs: 93524567
Best-hit-arabi-ID: AT3G46010.1
Best-hit-arabi-name: actin depolymerizing factor 1
Gorai.002G254800.1.D
PCC=0.619, p=0.005
mRNA Profile
Protein Expression Profile
DPA

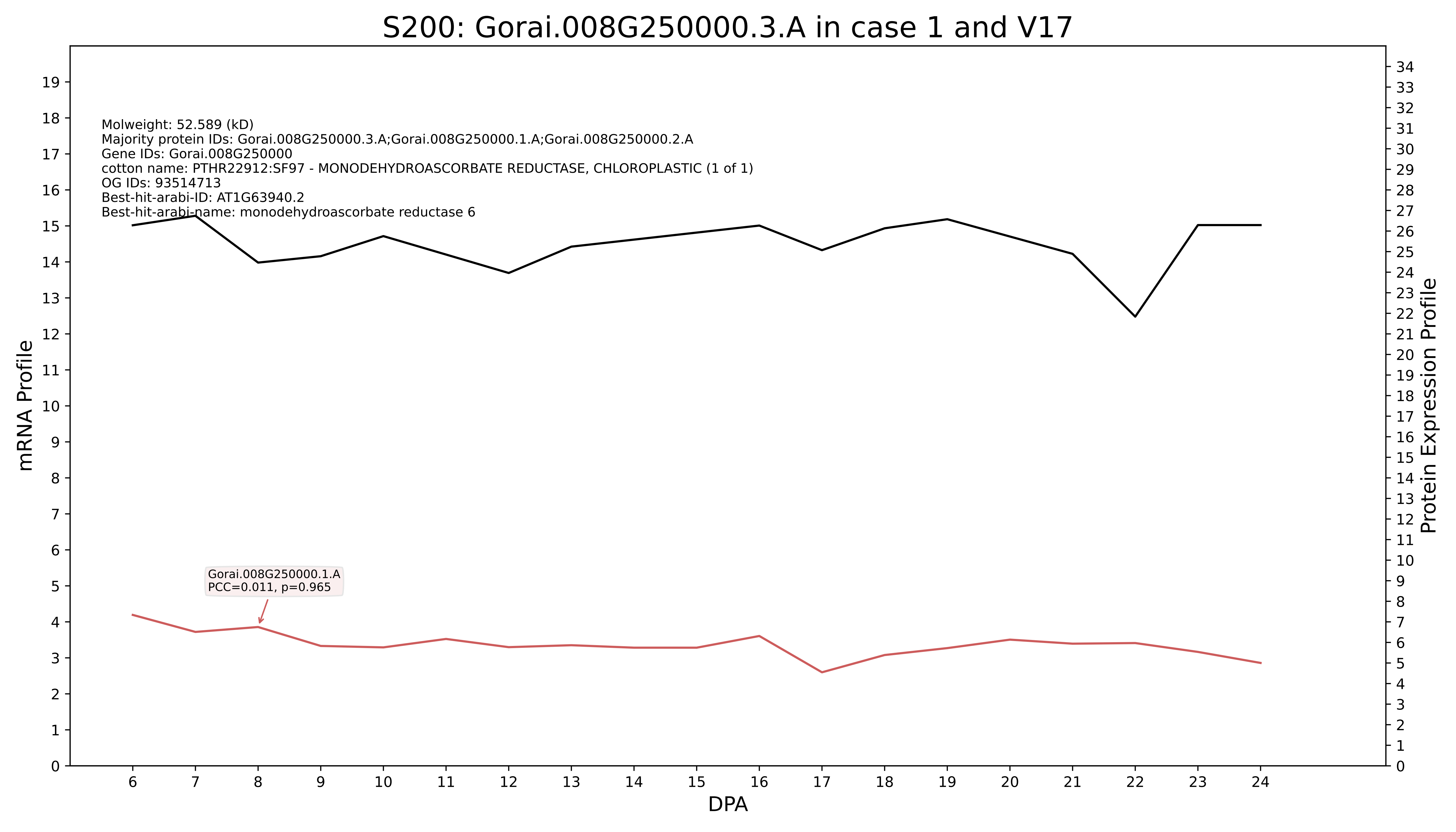
S200: Gorai.008G250000.3.A in case 1 and V17
Molweight: 52.589 (kD)
Majority protein IDs: Gorai.008G250000.3.A;Gorai.008G250000.1.A;Gorai.008G250000.2.A
Gene IDs: Gorai.008G250000
cotton name: PTHR22912:SF97 - MONODEHYDROASCORBATE REDUCTASE, CHLOROPLASTIC (1 of 1)
OG IDs: 93514713
Best-hit-arabi-ID: AT1G63940.2
Best-hit-arabi-name: monodehydroascorbate reductase 6
Gorai.008G250000.1.A
PCC=0.011, p=0.965
mRNA Profile
Protein Expression Profile
DPA

### S200: Gorai.005G125000.1.A in case 1 and V17

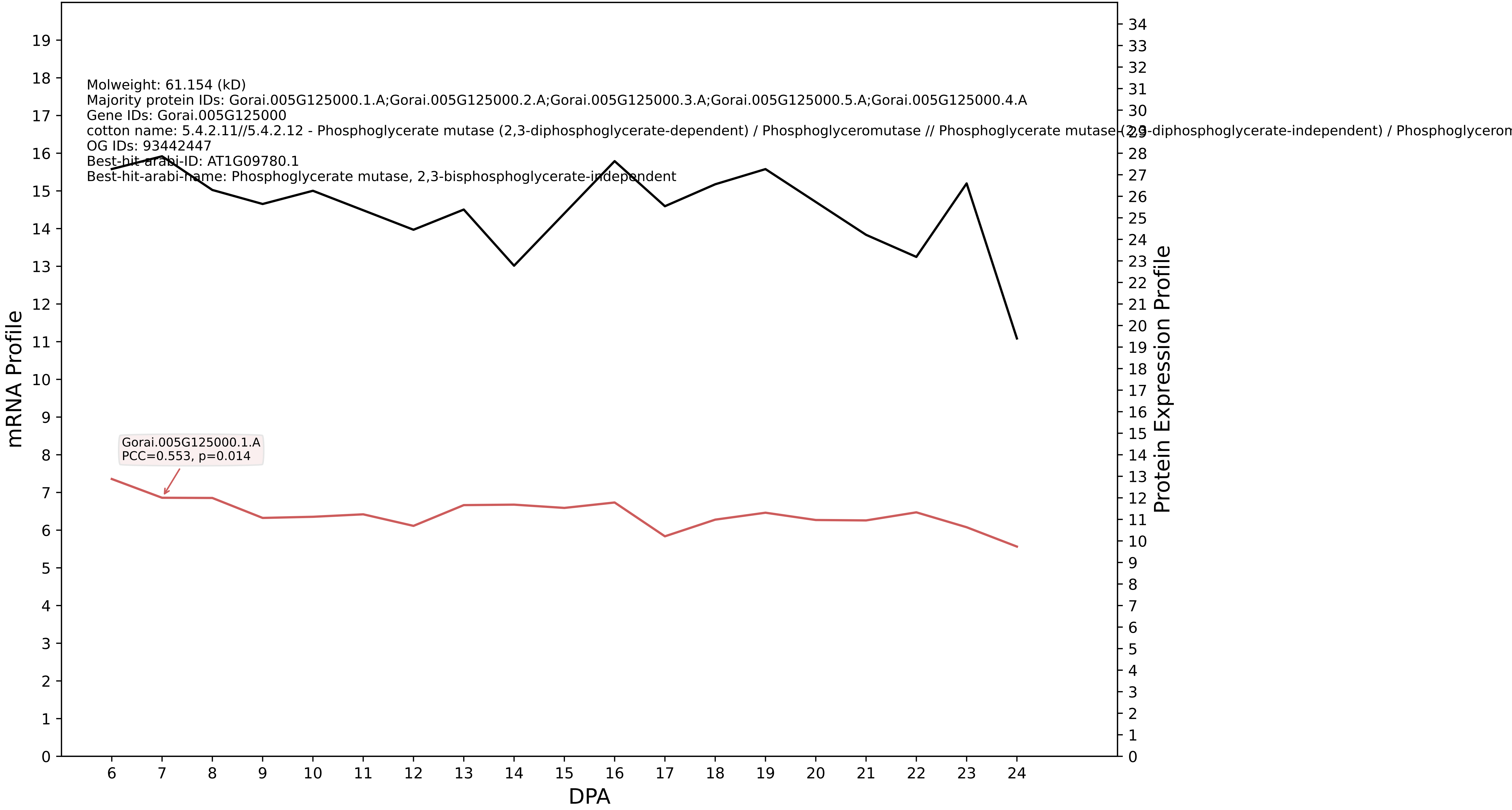

### S200: Gorai.013G150300.1.D in case 1 and V17

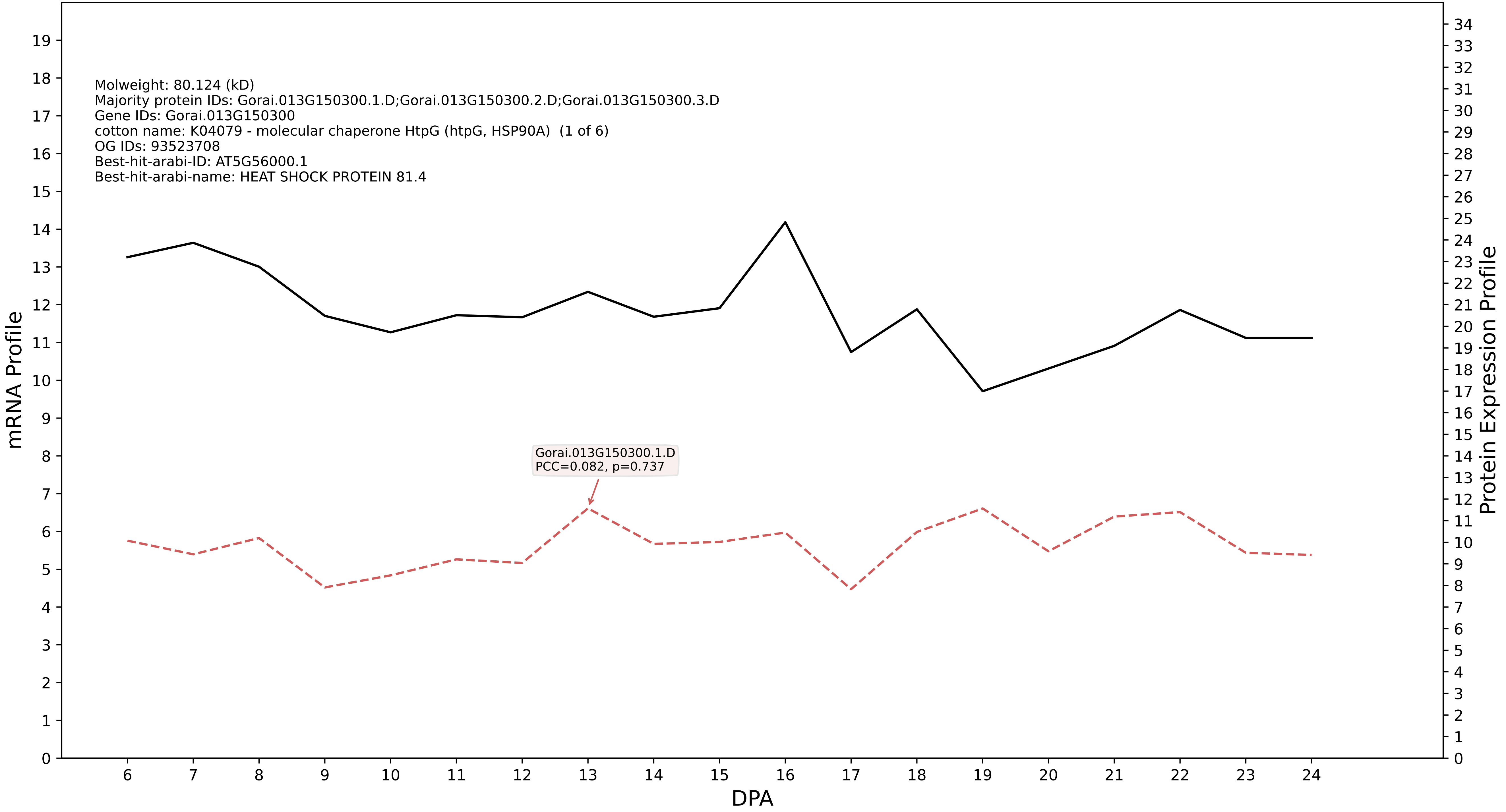

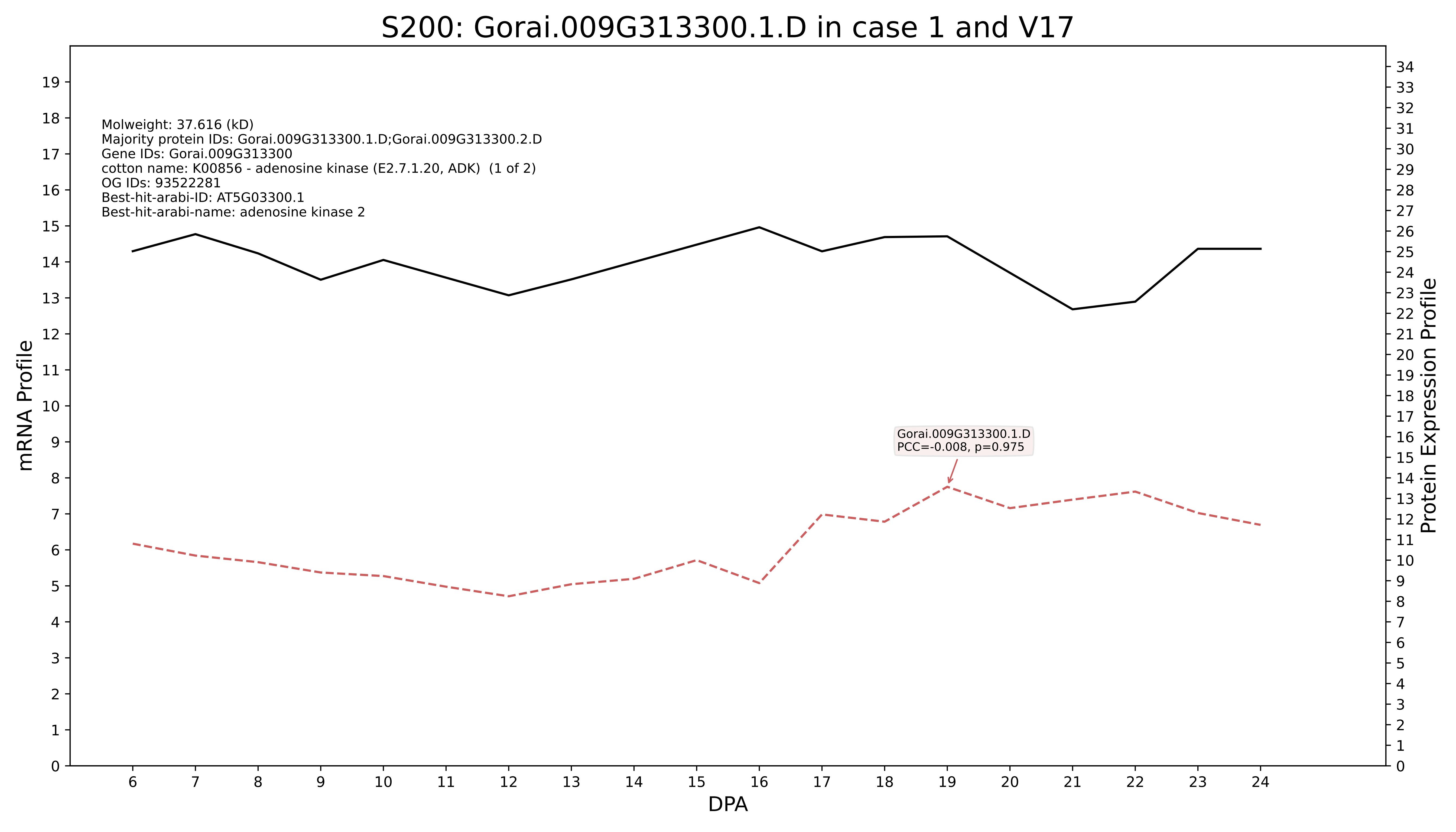
S200: Gorai.009G313300.1.D in case 1 and V17
Molweight: 37.616 (kD)
Majority protein IDs: Gorai.009G313300.1.D;Gorai.009G313300.2.D
Gene IDs: Gorai.009G313300
cotton name: K00856 - adenosine kinase (E2.7.1.20, ADK) (1 of 2)
OG IDs: 93522281
Best-hit-arabi-ID: AT5G03300.1
Best-hit-arabi-name: adenosine kinase 2
Gorai.009G313300.1.D
PCC=-0.008, p=0.975
mRNA Profile
Protein Expression Profile
DPA

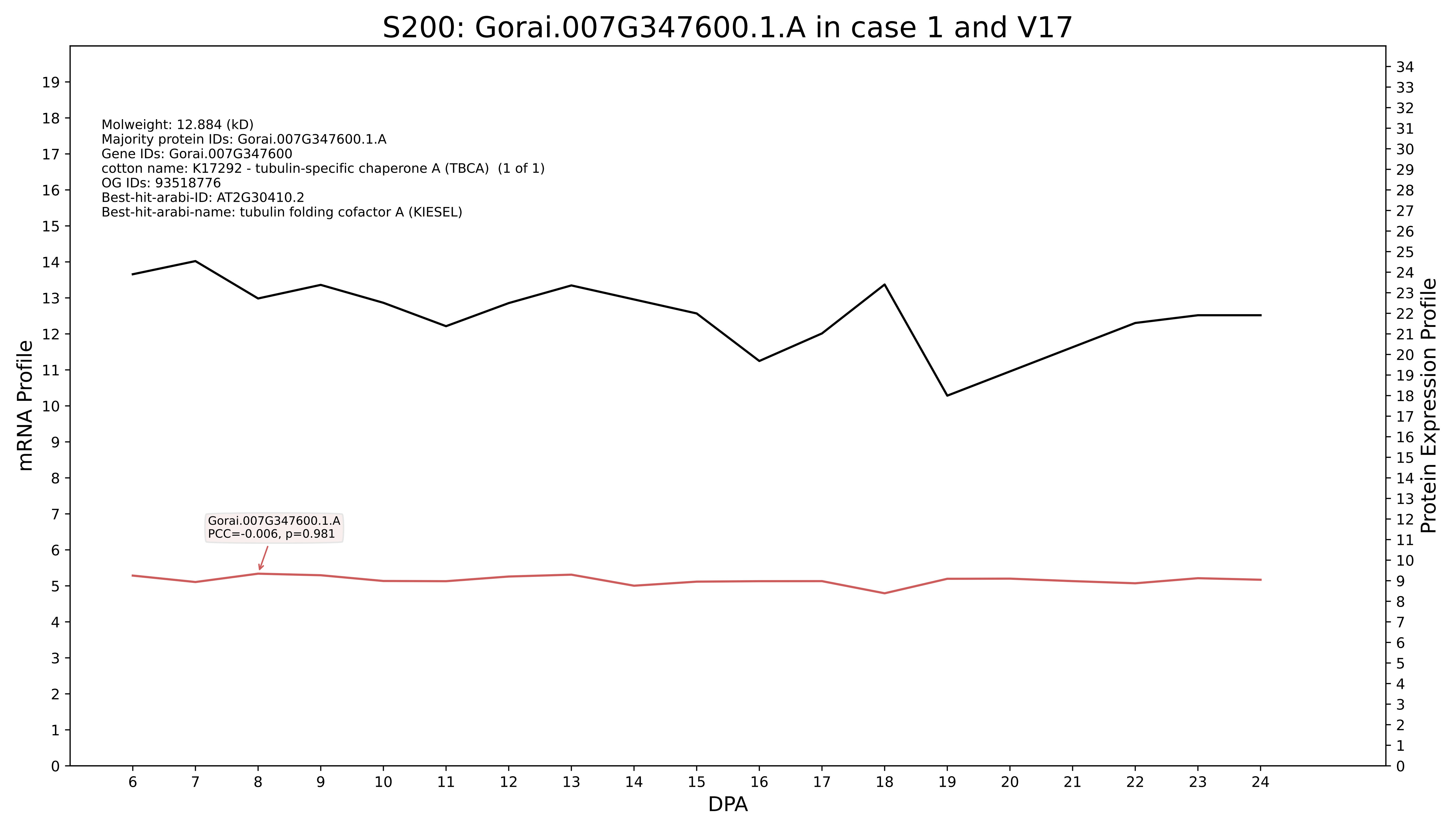
S200: Gorai.007G347600.1.A in case 1 and V17
Molweight: 12.884 (kD)
Majority protein IDs: Gorai.007G347600.1.A
Gene IDs: Gorai.007G347600
cotton name: K17292 - tubulin-specific chaperone A (TBCA) (1 of 1)
OG IDs: 93518776
Best-hit-arabi-ID: AT2G30410.2
Best-hit-arabi-name: tubulin folding cofactor A (KIESEL)
Gorai.007G347600.1.A
PCC=-0.006, p=0.981
mRNA Profile
Protein Expression Profile
DPA

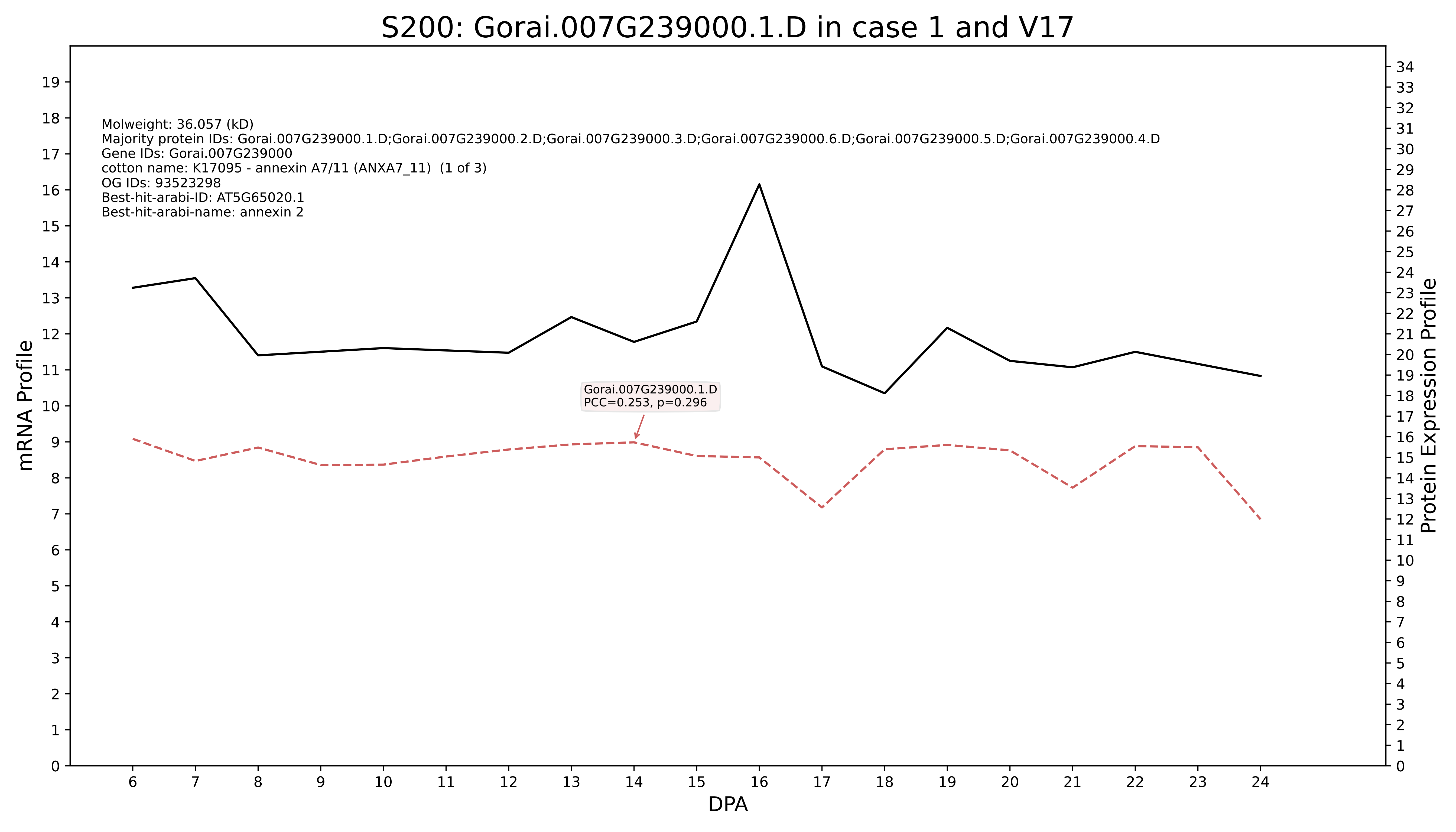
S200: Gorai.007G239000.1.D in case 1 and V17
Molweight: 36.057 (kD)
Majority protein IDs: Gorai.007G239000.1.D;Gorai.007G239000.2.D;Gorai.007G239000.3.D;Gorai.007G239000.6.D;Gorai.007G239000.5.D;Gorai.007G239000.4.D
Gene IDs: Gorai.007G239000
cotton name: K17095 - annexin A7/11 (ANXA7_11) (1 of 3)
OG IDs: 93523298
Best-hit-arabi-ID: AT5G65020.1
Best-hit-arabi-name: annexin 2
Gorai.007G239000.1.D
PCC=0.253, p=0.296
mRNA Profile
Protein Expression Profile
DPA

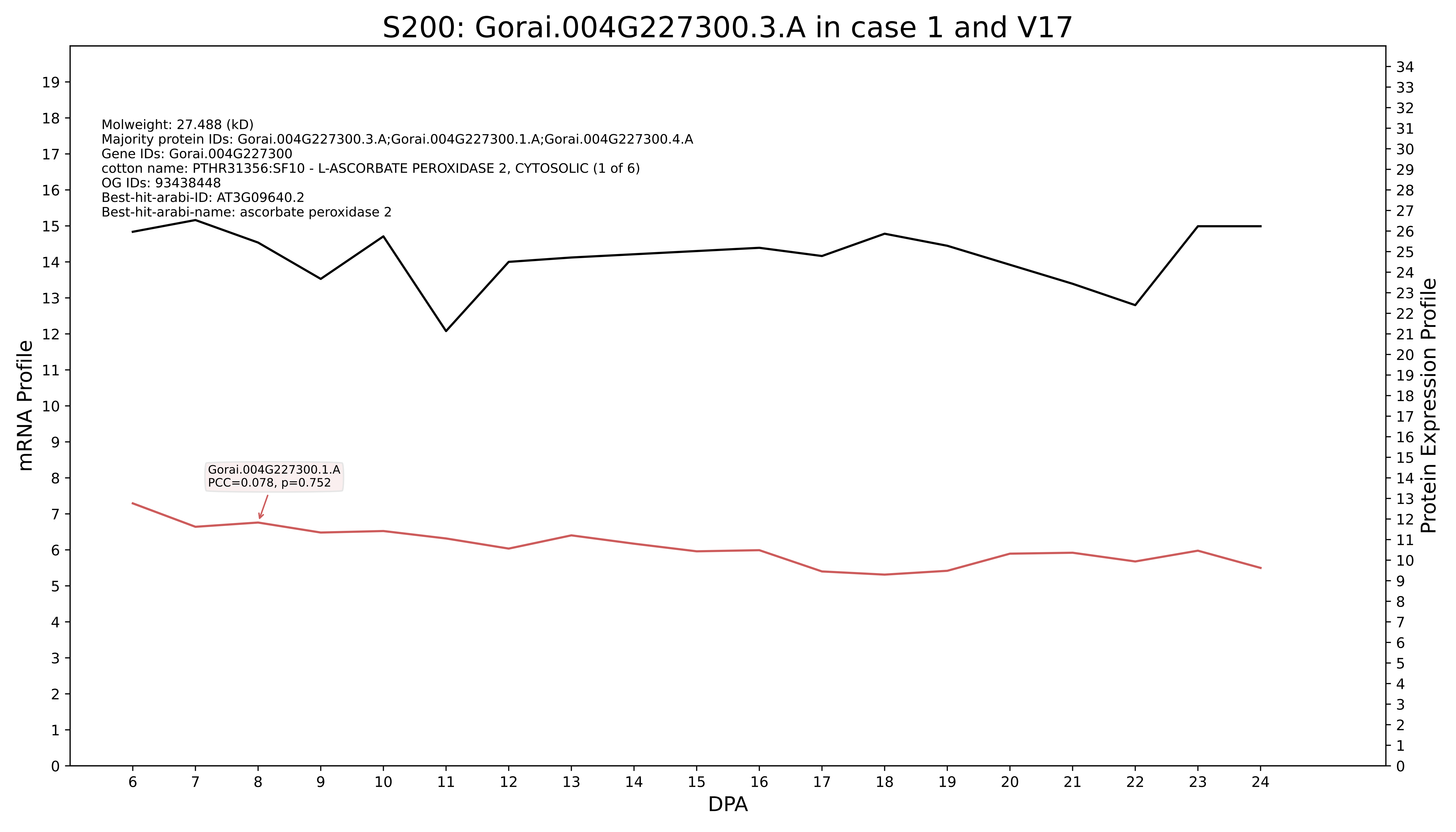

S200: Gorai.004G227300.3.A in case 1 and V17
Molweight: 27.488 (kD)
Majority protein IDs: Gorai.004G227300.3.A;Gorai.004G227300.1.A;Gorai.004G227300.4.A
Gene IDs: Gorai.004G227300
cotton name: PTHR31356:SF10 - L-ASCORBATE PEROXIDASE 2, CYTOSOLIC (1 of 6)
OG IDs: 93438448
Best-hit-arabi-ID: AT3G09640.2
Best-hit-arabi-name: ascorbate peroxidase 2
Gorai.004G227300.1.A
PCC=0.078, p=0.752
mRNA Profile
Protein Expression Profile
DPA

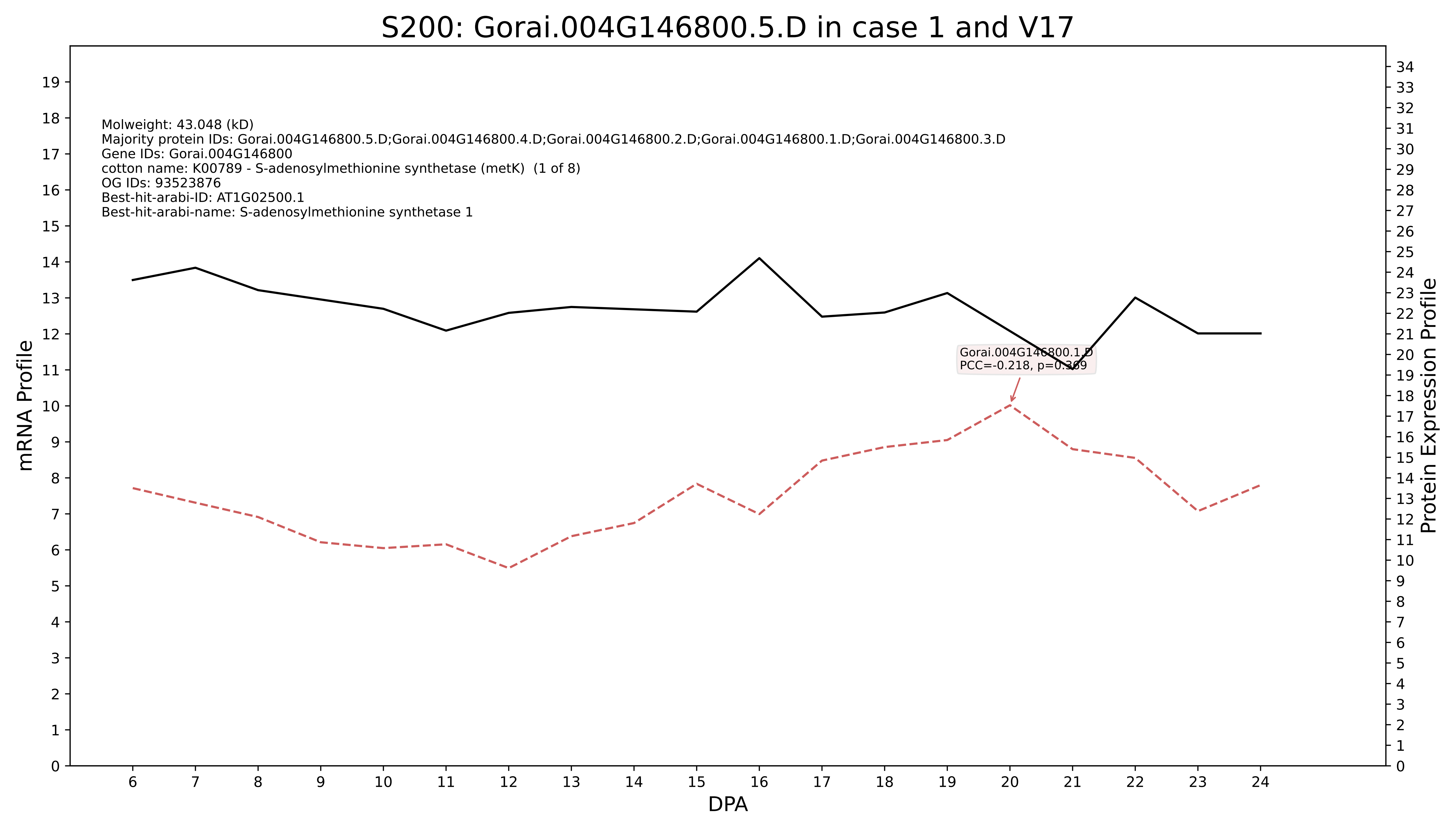
S200: Gorai.004G146800.5.D in case 1 and V17
Molweight: 43.048 (kD)
Majority protein IDs: Gorai.004G146800.5.D;Gorai.004G146800.4.D;Gorai.004G146800.2.D;Gorai.004G146800.1.D;Gorai.004G146800.3.D
Gene IDs: Gorai.004G146800
cotton name: K00789 - S-adenosylmethionine synthetase (metK) (1 of 8)
OG IDs: 93523876
Best-hit-arabi-ID: AT1G02500.1
Best-hit-arabi-name: S-adenosylmethionine synthetase 1
Gorai.004G146800.1.D
PCC=-0.218, p=0.369
mRNA Profile
Protein Expression Profile
DPA

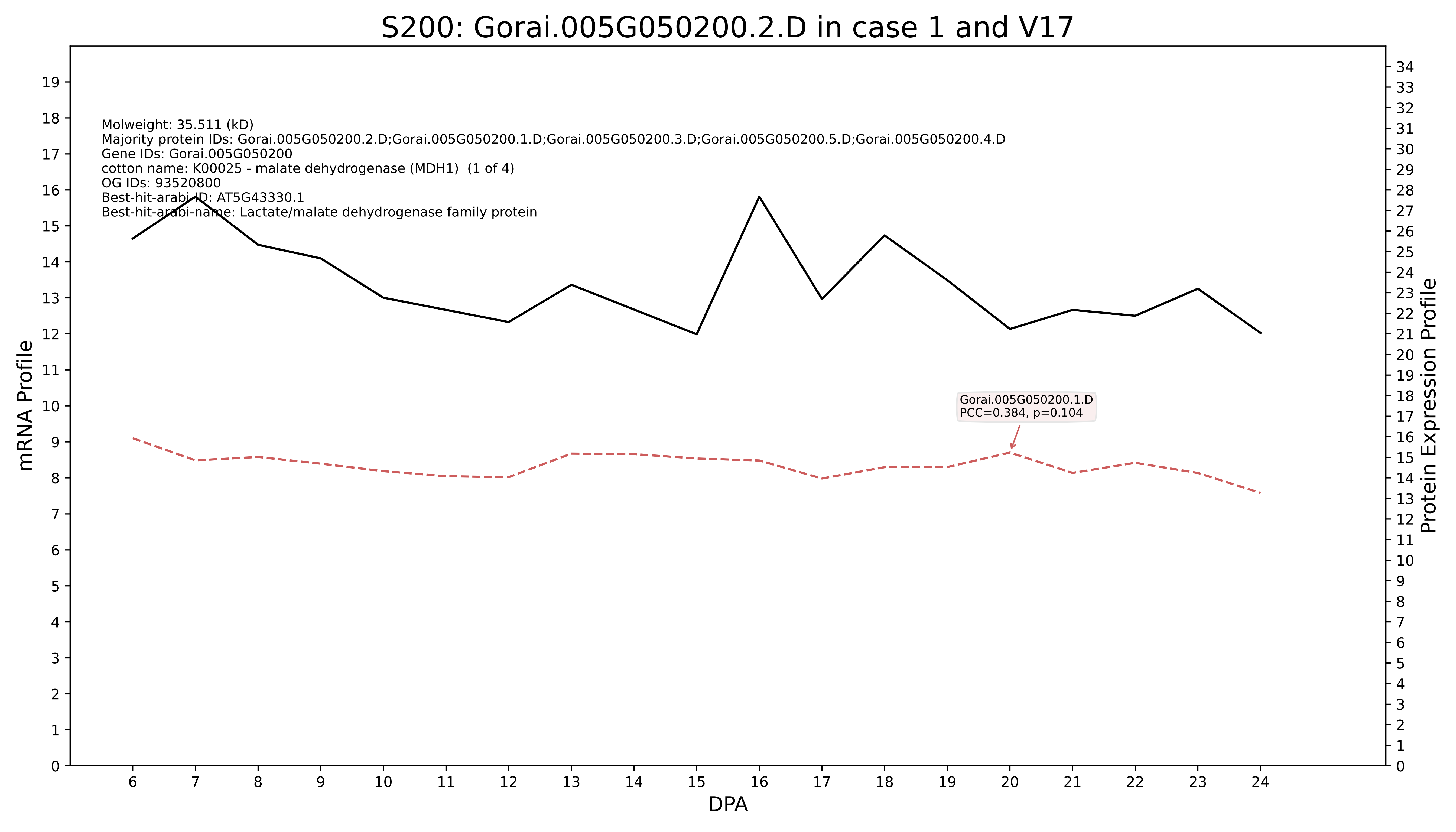
S200: Gorai.005G050200.2.D in case 1 and V17
Molweight: 35.511 (kD)
Majority protein IDs: Gorai.005G050200.2.D;Gorai.005G050200.1.D;Gorai.005G050200.3.D;Gorai.005G050200.5.D;Gorai.005G050200.4.D
Gene IDs: Gorai.005G050200
cotton name: K00025 - malate dehydrogenase (MDH1) (1 of 4)
OG IDs: 93520800
Best-hit-arabi-ID: AT5G43330.1
Best-hit-arabi-name: Lactate/malate dehydrogenase family protein
Gorai.005G050200.1.D
PCC=0.384, p=0.104
mRNA Profile
Protein Expression Profile
DPA

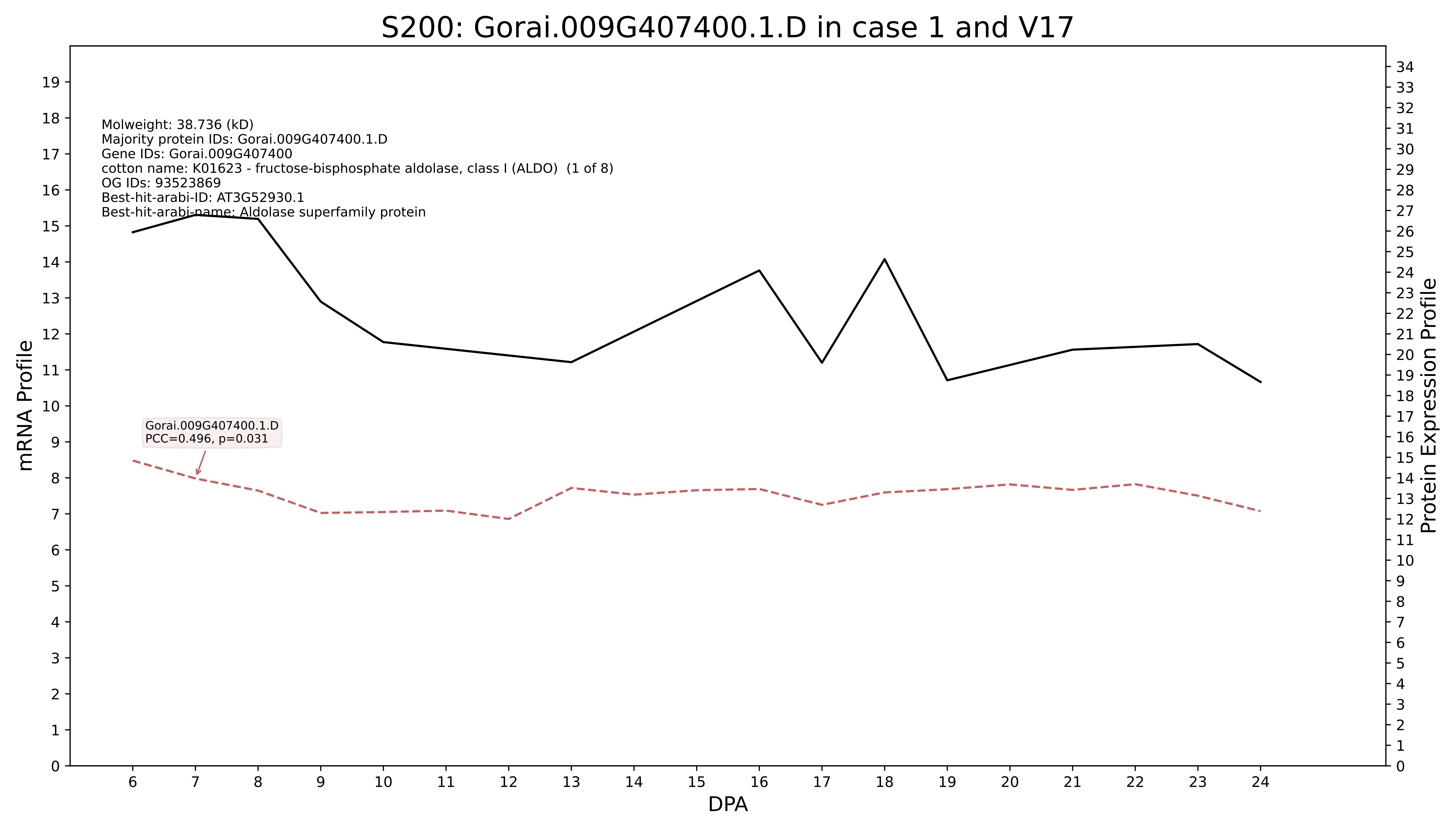
S200: Gorai.009G407400.1.D in case 1 and V17
Molweight: 38.736 (kD)
Majority protein IDs: Gorai.009G407400.1.D
Gene IDs: Gorai.009G407400
cotton name: K01623 - fructose-bisphosphate aldolase, class I (ALDO) (1 of 8)
OG IDs: 93523869
Best-hit-arabi-ID: AT3G52930.1
Best-hit-arabi-name: Aldolase superfamily protein
Gorai.009G407400.1.D
PCC=0.496, p=0.031
mRNA Profile
Protein Expression Profile
DPA

### S200: Gorai.002G003900.1.A in case 1 and V17

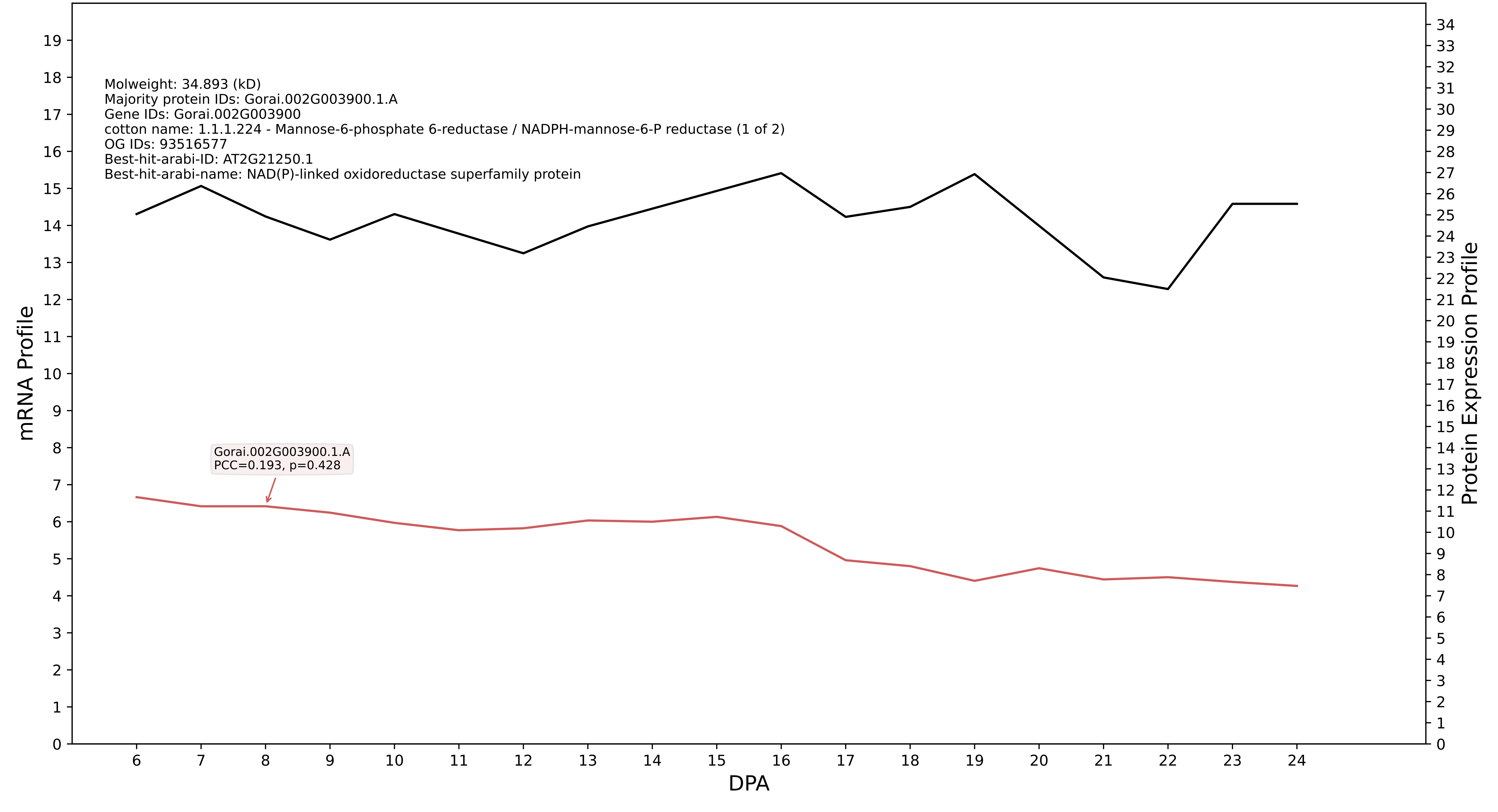

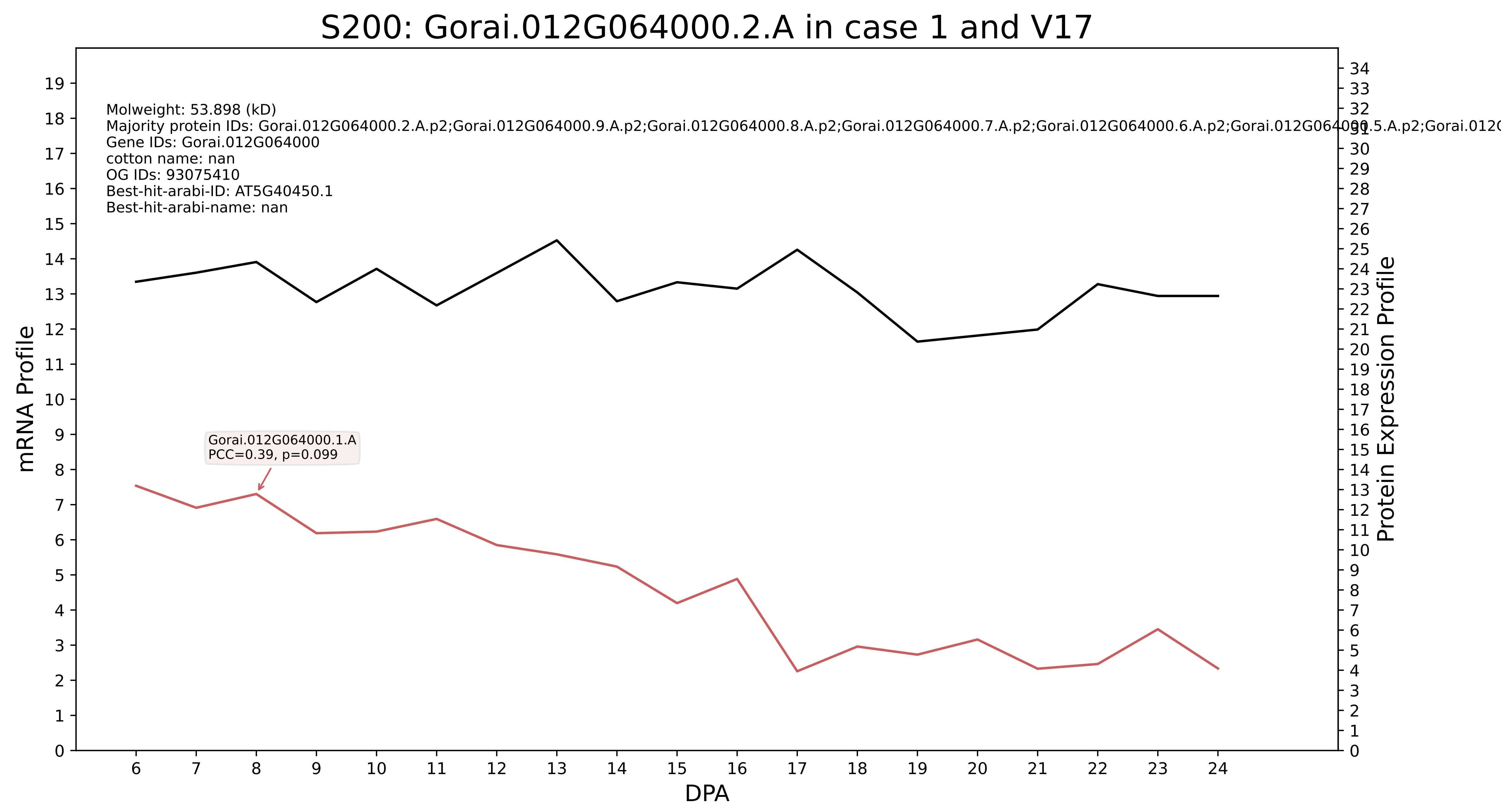
S200: Gorai.012G064000.2.A in case 1 and V17
Molweight: 53.898 (kD)
Majority protein IDs: Gorai.012G064000.2.A.p2;Gorai.012G064000.9.A.p2;Gorai.012G064000.8.A.p2;Gorai.012G064000.7.A.p2;Gorai.012G064000.6.A.p2;Gorai.012G064000.5.A.p2;Gorai.012G
Gene IDs: Gorai.012G064000
cotton name: nan
OG IDs: 93075410
Best-hit-arabi-ID: AT5G40450.1
Best-hit-arabi-name: nan
Gorai.012G064000.1.A
PCC=0.39, p=0.099
mRNA Profile
Protein Expression Profile
DPA

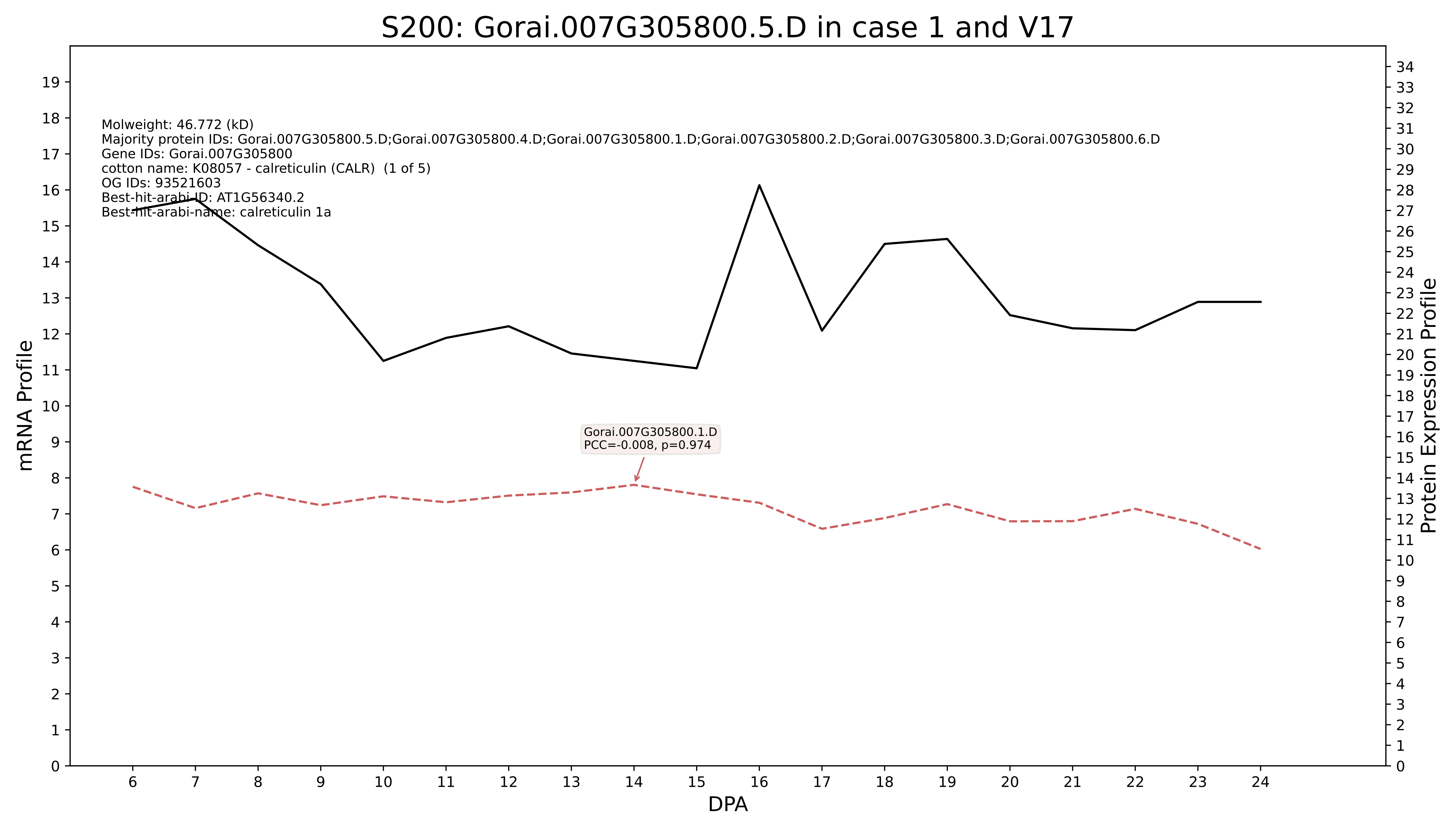
S200: Gorai.007G305800.5.D in case 1 and V17
Molweight: 46.772 (kD)
Majority protein IDs: Gorai.007G305800.5.D;Gorai.007G305800.4.D;Gorai.007G305800.1.D;Gorai.007G305800.2.D;Gorai.007G305800.3.D;Gorai.007G305800.6.D
Gene IDs: Gorai.007G305800
cotton name: K08057 - calreticulin (CALR) (1 of 5)
OG IDs: 93521603
Best-hit-arabi-ID: AT1G56340.2
Best-hit-arabi-name: calreticulin 1a
Gorai.007G305800.1.D
PCC=-0.008, p=0.974
mRNA Profile
Protein Expression Profile
DPA

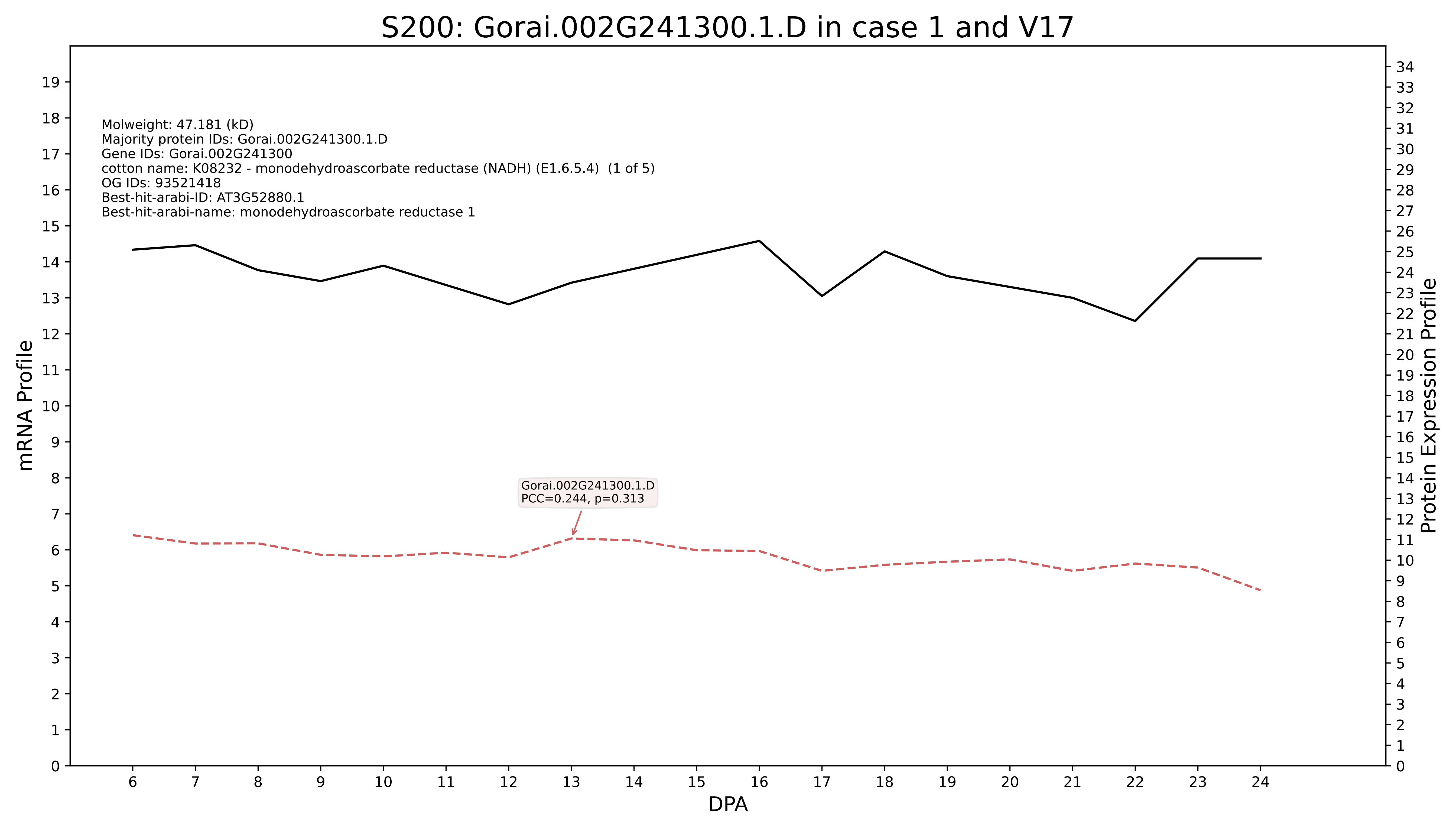

S200: Gorai.002G241300.1.D in case 1 and V17
Molweight: 47.181 (kD)
Majority protein IDs: Gorai.002G241300.1.D
Gene IDs: Gorai.002G241300
cotton name: K08232 - monodehydroascorbate reductase (NADH) (E1.6.5.4) (1 of 5)
OG IDs: 93521418
Best-hit-arabi-ID: AT3G52880.1
Best-hit-arabi-name: monodehydroascorbate reductase 1
Gorai.002G241300.1.D
PCC=0.244, p=0.313
mRNA Profile
Protein Expression Profile
DPA

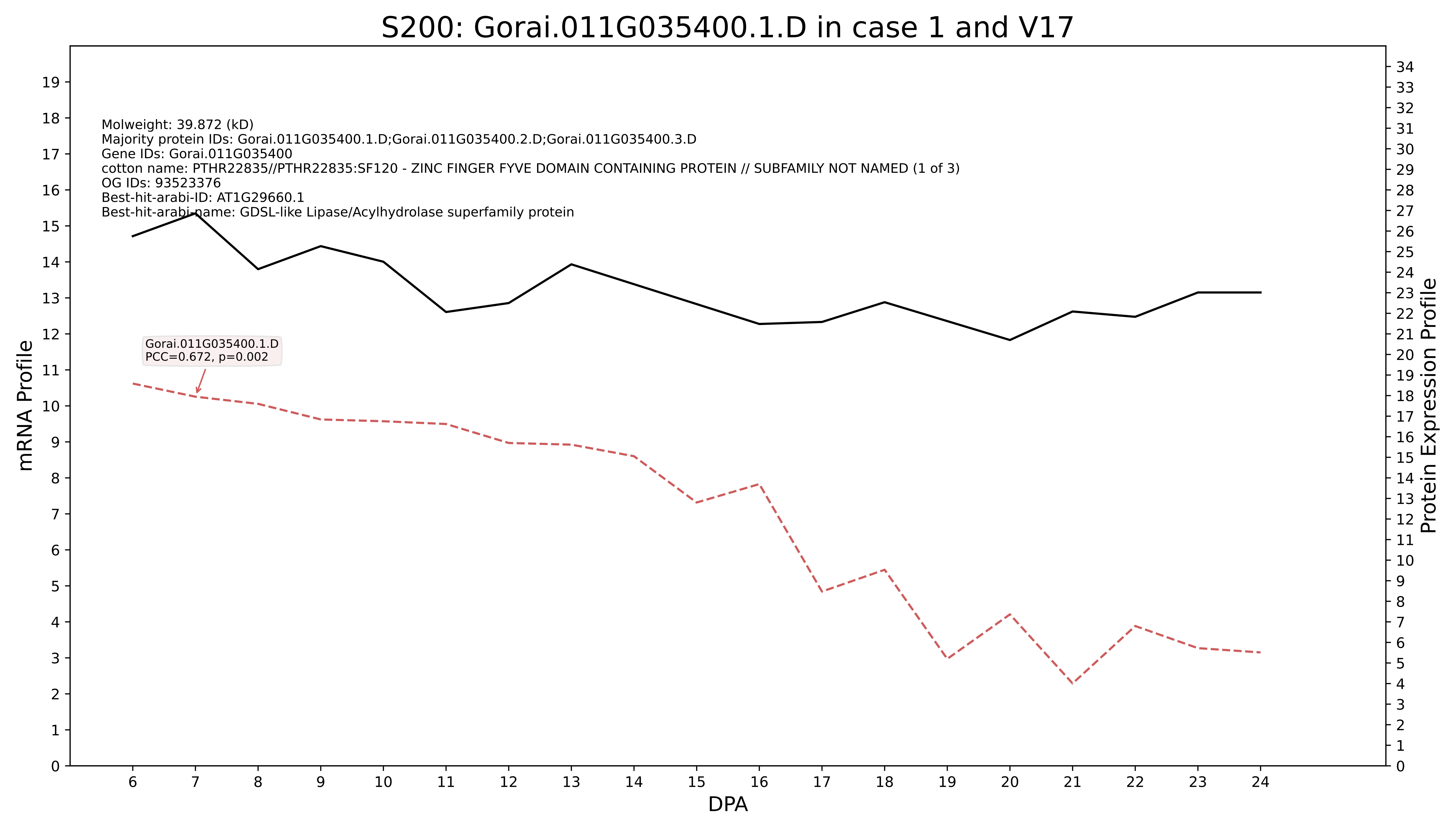
S200: Gorai.011G035400.1.D in case 1 and V17
Molweight: 39.872 (kD)
Majority protein IDs: Gorai.011G035400.1.D;Gorai.011G035400.2.D;Gorai.011G035400.3.D
Gene IDs: Gorai.011G035400
cotton name: PTHR22835//PTHR22835:SF120 - ZINC FINGER FYVE DOMAIN CONTAINING PROTEIN // SUBFAMILY NOT NAMED (1 of 3)
OG IDs: 93523376
Best-hit-arabi-ID: AT1G29660.1
Best-hit-arabi-name: GDSL-like Lipase/Acylhydrolase superfamily protein
Gorai.011G035400.1.D
PCC=0.672, p=0.002
mRNA Profile
Protein Expression Profile
DPA

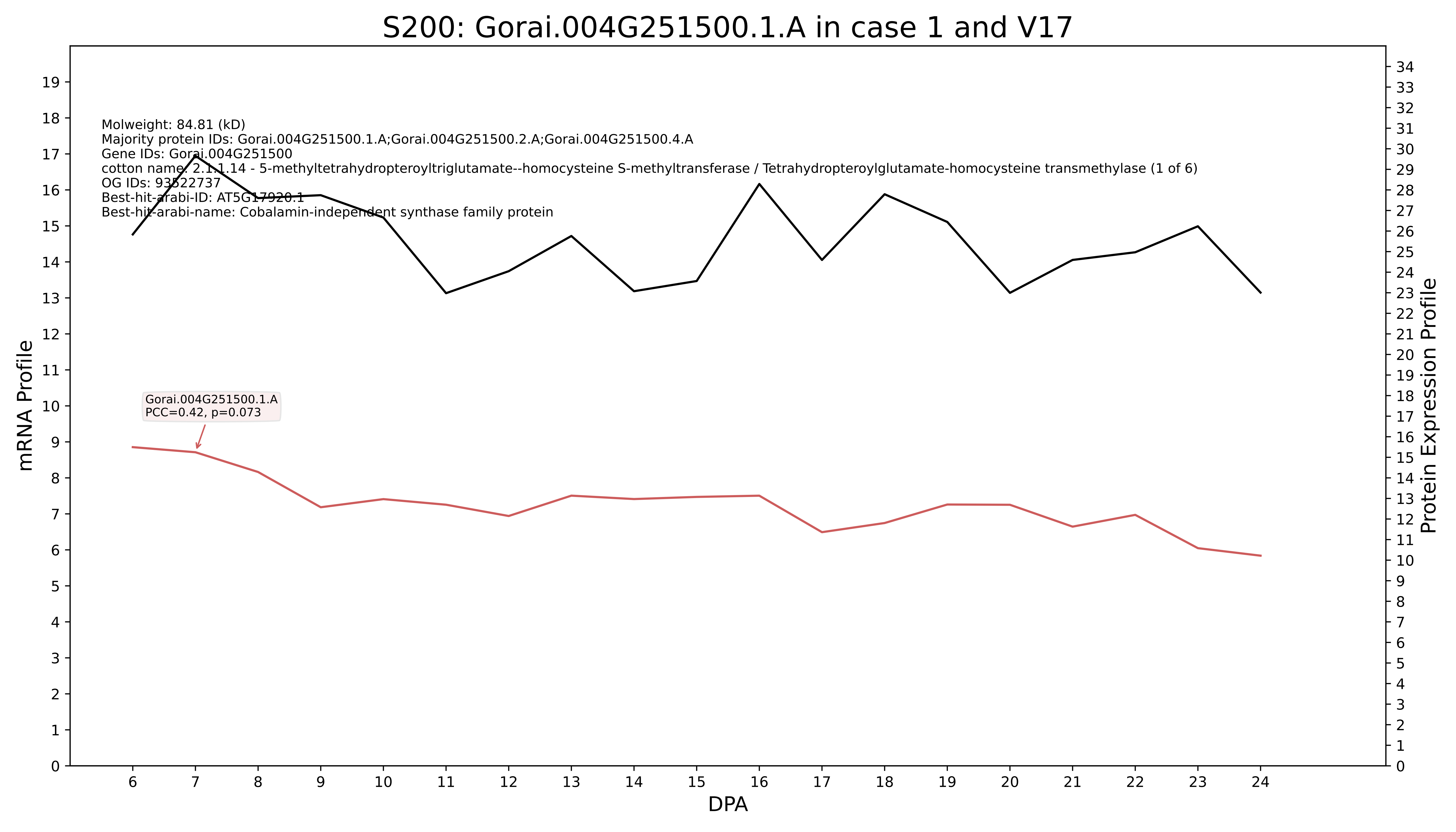
S200: Gorai.004G251500.1.A in case 1 and V17
Molweight: 84.81 (kD)
Majority protein IDs: Gorai.004G251500.1.A;Gorai.004G251500.2.A;Gorai.004G251500.4.A
Gene IDs: Gorai.004G251500
cotton name: 2.1.1.14 - 5-methyltetrahydropteroyltriglutamate--homocysteine S-methyltransferase / Tetrahydropteroylglutamate-homocysteine transmethylase (1 of 6)
OG IDs: 93522737
Best-hit-arabi-ID: AT5G17920.1
Best-hit-arabi-name: Cobalamin-independent synthase family protein
Gorai.004G251500.1.A
PCC=0.42, p=0.073
mRNA Profile
Protein Expression Profile
DPA

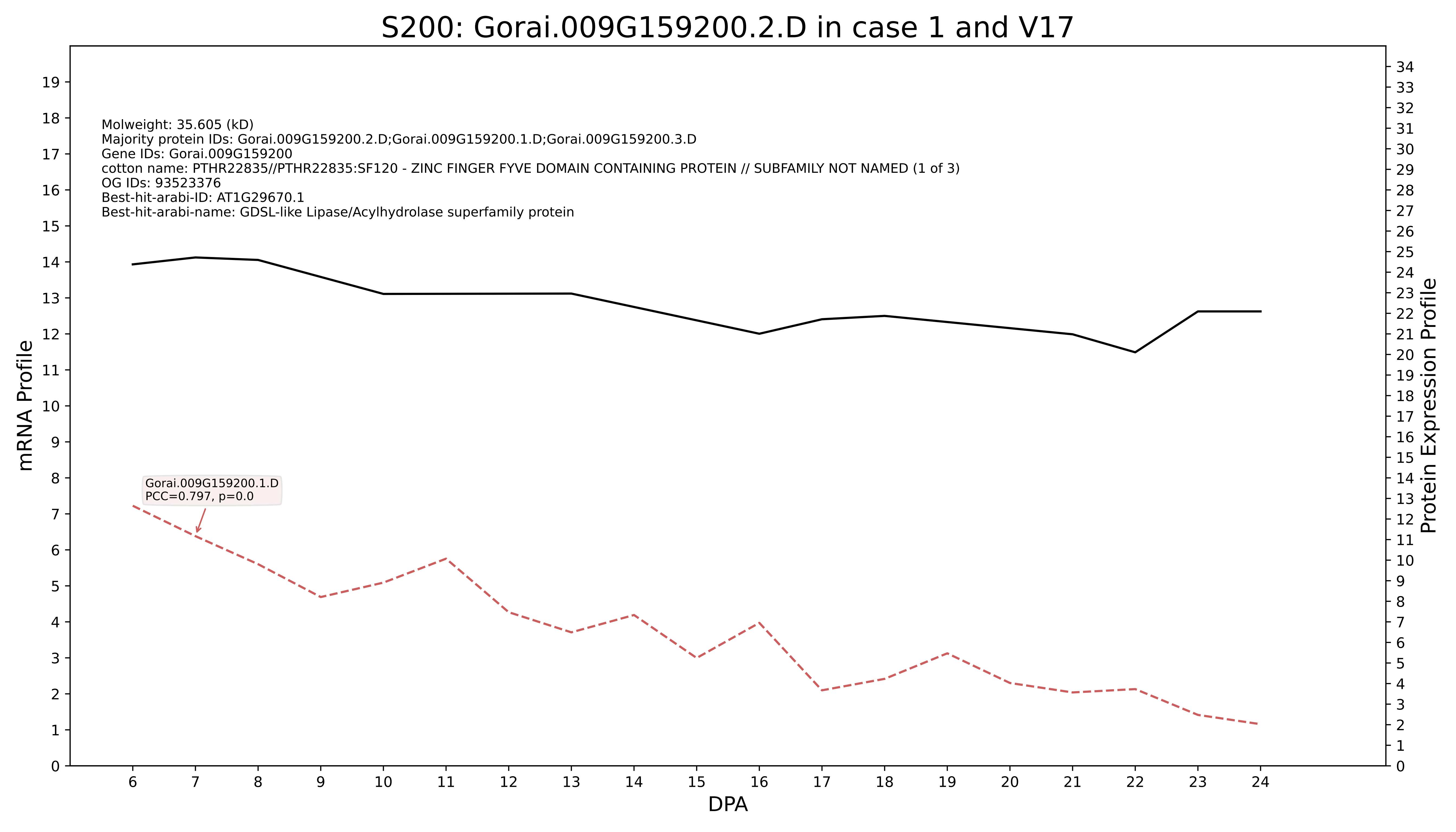

S200: Gorai.009G159200.2.D in case 1 and V17
Molweight: 35.605 (kD)
Majority protein IDs: Gorai.009G159200.2.D;Gorai.009G159200.1.D;Gorai.009G159200.3.D
Gene IDs: Gorai.009G159200
cotton name: PTHR22835//PTHR22835:SF120 - ZINC FINGER FYVE DOMAIN CONTAINING PROTEIN // SUBFAMILY NOT NAMED (1 of 3)
OG IDs: 93523376
Best-hit-arabi-ID: AT1G29670.1
Best-hit-arabi-name: GDSL-like Lipase/Acylhydrolase superfamily protein
Gorai.009G159200.1.D
PCC=0.797, p=0.0
mRNA Profile
Protein Expression Profile
DPA

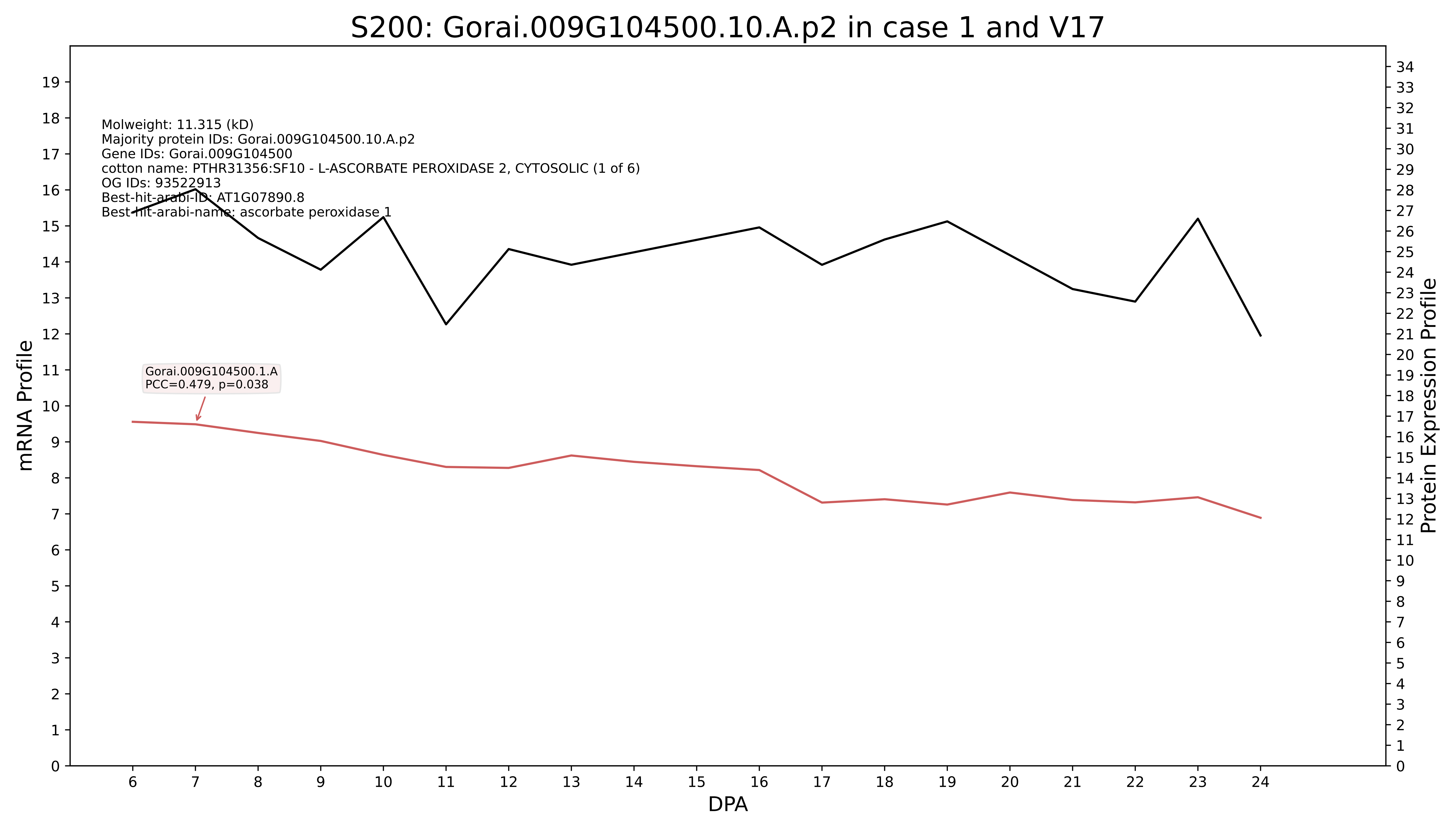

S200: Gorai.009G104500.10.A.p2 in case 1 and V17
Molweight: 11.315 (kD)
Majority protein IDs: Gorai.009G104500.10.A.p2
Gene IDs: Gorai.009G104500
cotton name: PTHR31356:SF10 - L-ASCORBATE PEROXIDASE 2, CYTOSOLIC (1 of 6)
OG IDs: 93522913
Best-hit-arabi-ID: AT1G07890.8
Best-hit-arabi-name: ascorbate peroxidase 1
Gorai.009G104500.1.A
PCC=0.479, p=0.038
mRNA Profile
Protein Expression Profile
DPA

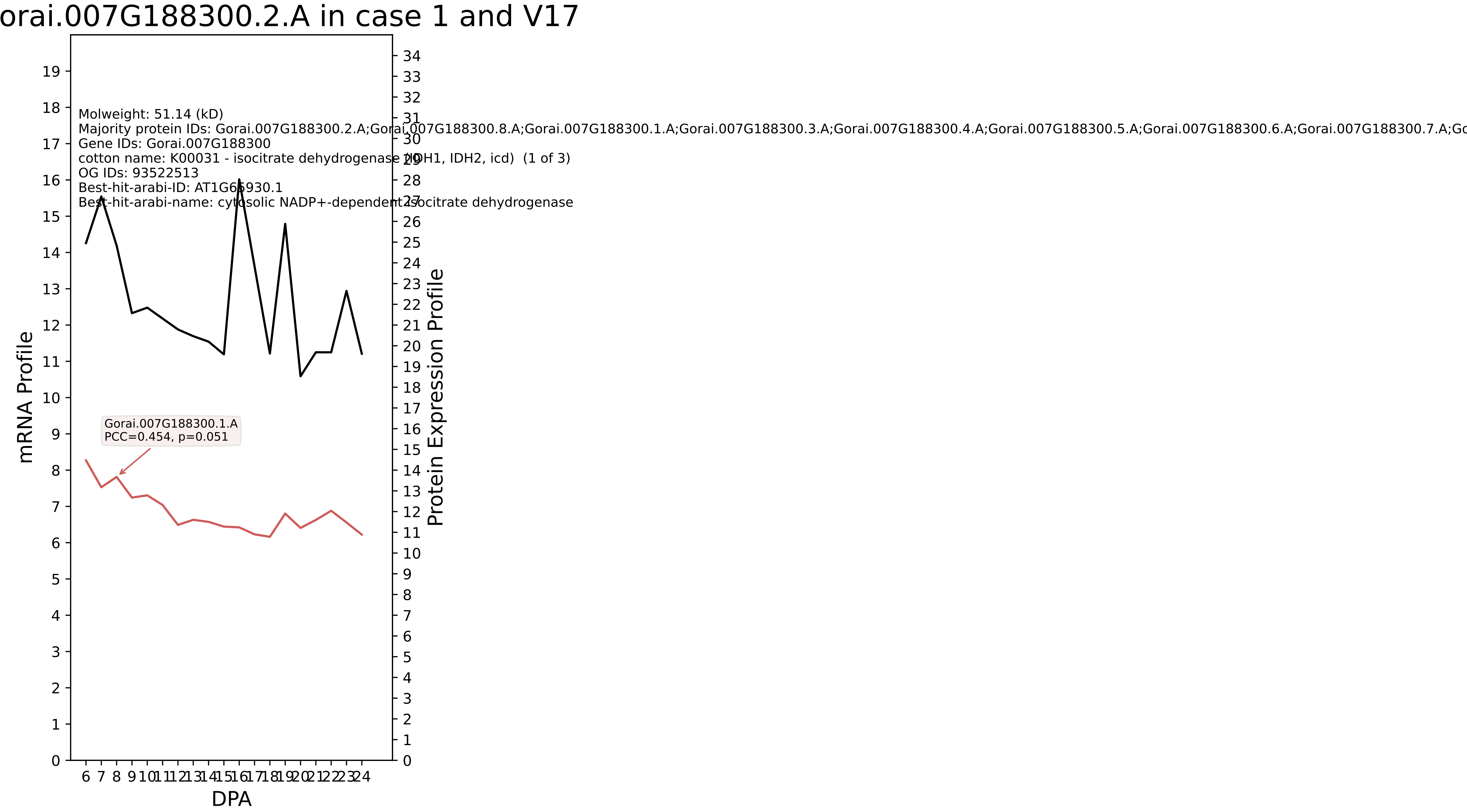
orai.007G188300.2.A in case 1 and V17
Molweight: 51.14 (kD)
Majority protein IDs: Gorai.007G188300.2.A;Gorai.007G188300.8.A;Gorai.007G188300.1.A;Gorai.007G188300.3.A;Gorai.007G188300.4.A;Gorai.007G188300.5.A;Gorai.007G188300.6.A;Gorai.007G188300.7.A;Go
Gene IDs: Gorai.007G188300
cotton name: K00031 - isocitrate dehydrogenase (IDH1, IDH2, icd) (1 of 3)
OG IDs: 93522513
Best-hit-arabi-ID: AT1G65930.1
Best-hit-arabi-name: cytosolic NADP+-dependent isocitrate dehydrogenase
Gorai.007G188300.1.A
PCC=0.454, p=0.051
mRNA Profile
Protein Expression Profile
DPA

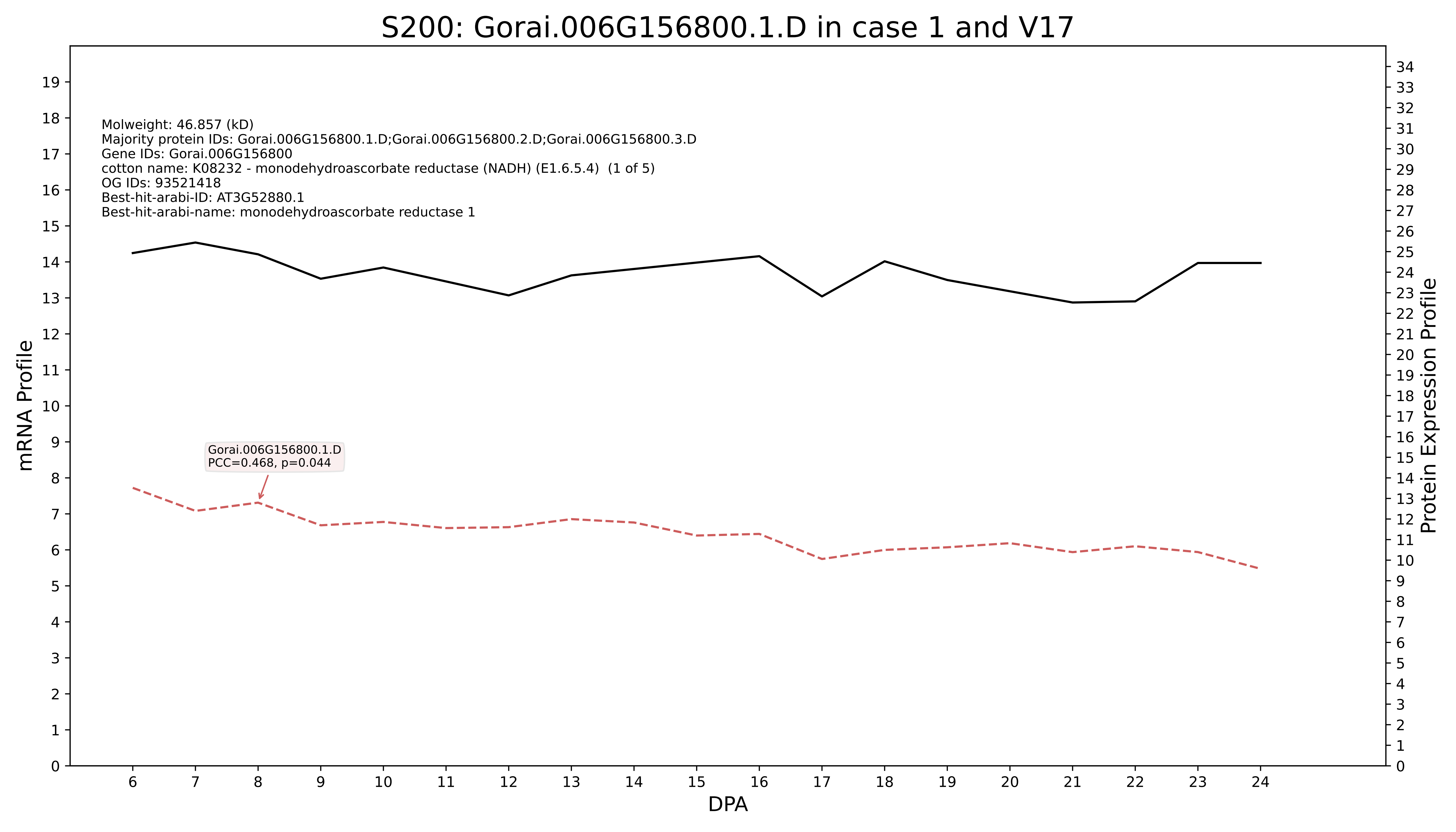
S200: Gorai.006G156800.1.D in case 1 and V17
Molweight: 46.857 (kD)
Majority protein IDs: Gorai.006G156800.1.D;Gorai.006G156800.2.D;Gorai.006G156800.3.D
Gene IDs: Gorai.006G156800
cotton name: K08232 - monodehydroascorbate reductase (NADH) (E1.6.5.4) (1 of 5)
OG IDs: 93521418
Best-hit-arabi-ID: AT3G52880.1
Best-hit-arabi-name: monodehydroascorbate reductase 1
Gorai.006G156800.1.D
PCC=0.468, p=0.044
mRNA Profile
Protein Expression Profile
DPA

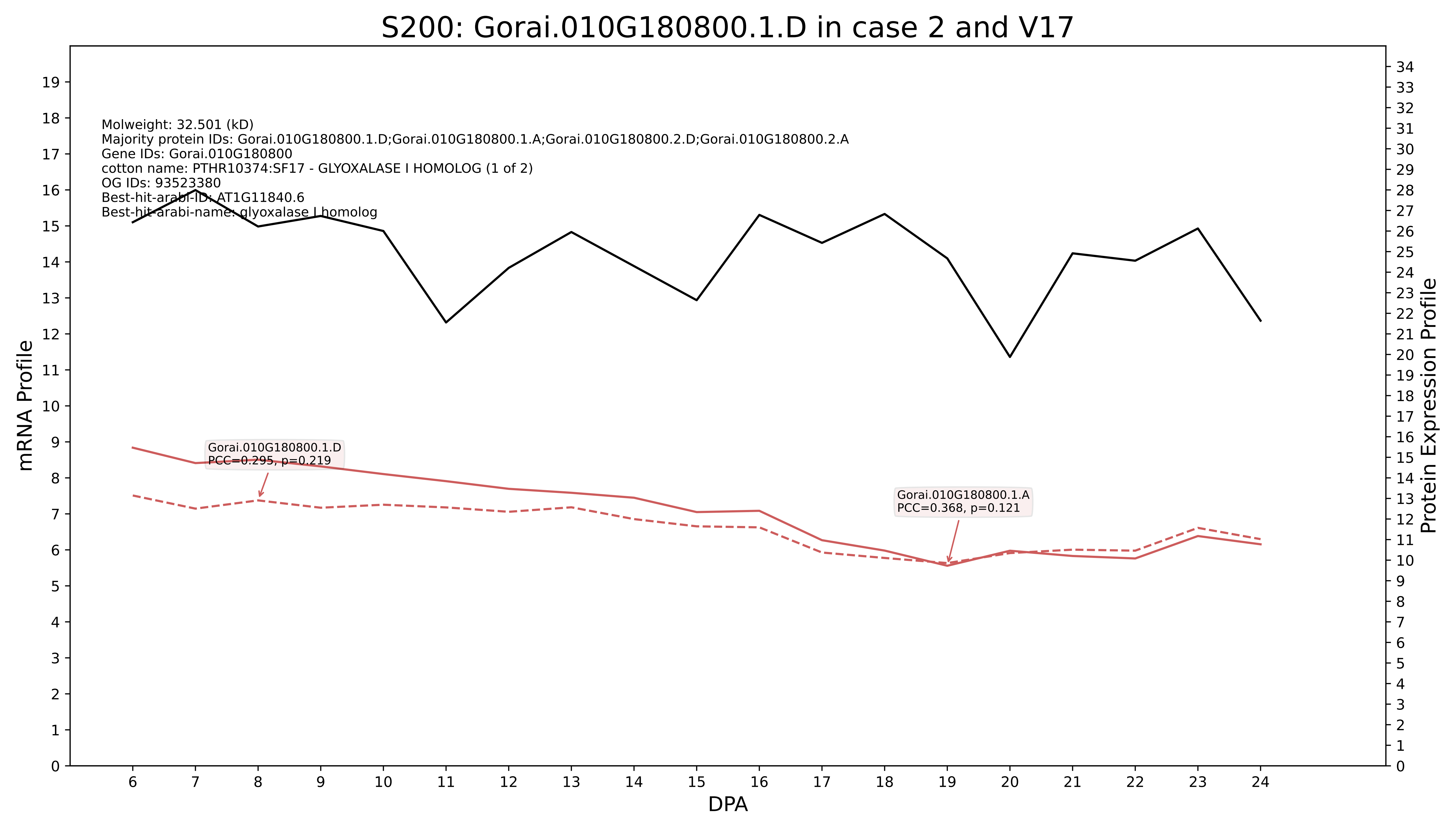
S200: Gorai.010G180800.1.D in case 2 and V17
Molweight: 32.501 (kD)
Majority protein IDs: Gorai.010G180800.1.D;Gorai.010G180800.1.A;Gorai.010G180800.2.D;Gorai.010G180800.2.A
Gene IDs: Gorai.010G180800
cotton name: PTHR10374:SF17 - GLYOXALASE I HOMOLOG (1 of 2)
OG IDs: 93523380
Best-hit-arabi-ID: AT1G11840.6
Best-hit-arabi-name: glyoxalase I homolog
Gorai.010G180800.1.D
PCC=0.295, p=0.219
Gorai.010G180800.1.A
PCC=0.368, p=0.121
mRNA Profile
Protein Expression Profile
DPA

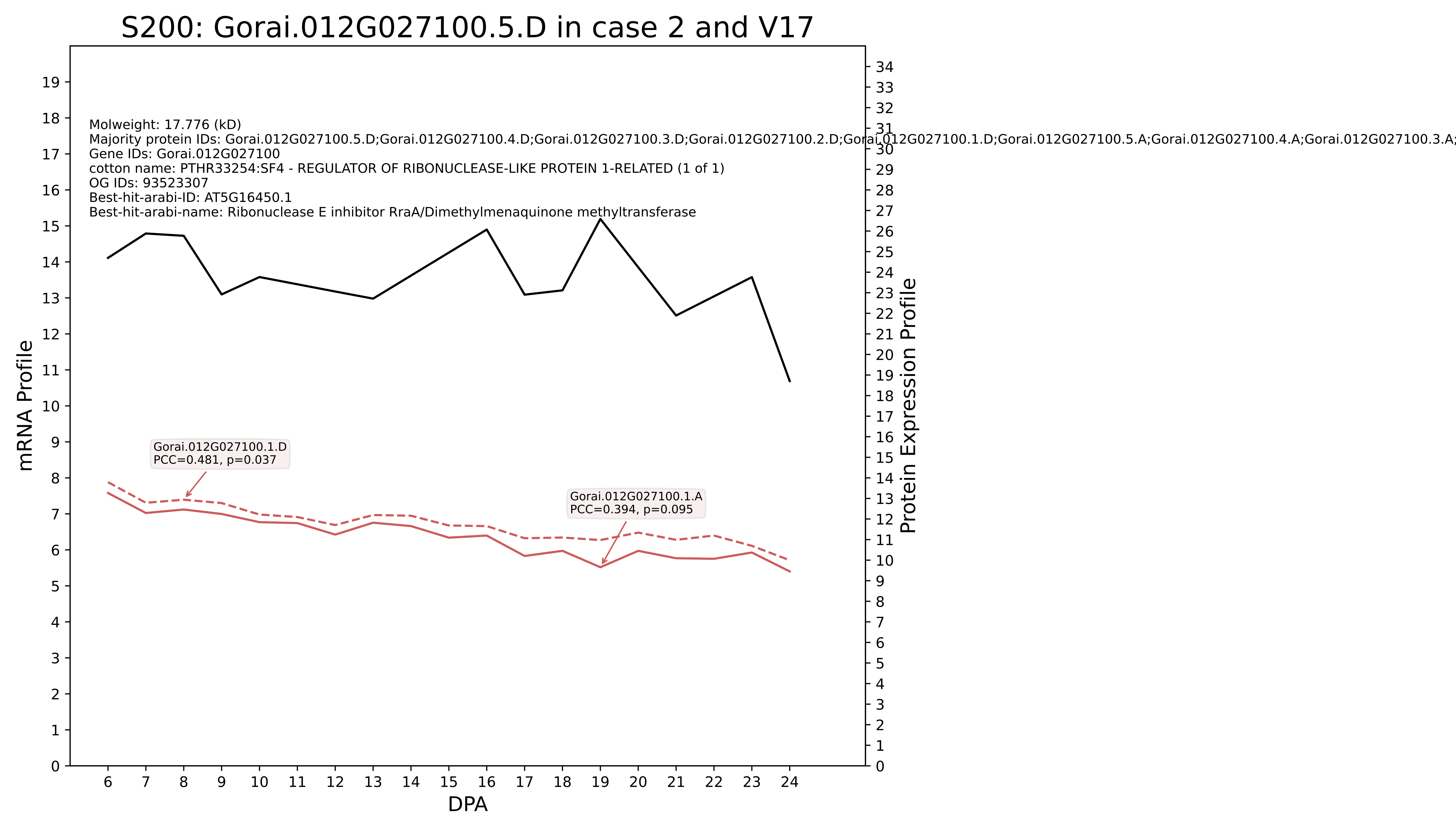
S200: Gorai.012G027100.5.D in case 2 and V17
Molweight: 17.776 (kD)
Majority protein IDs: Gorai.012G027100.5.D;Gorai.012G027100.4.D;Gorai.012G027100.3.D;Gorai.012G027100.2.D;Gorai.012G027100.1.D;Gorai.012G027100.5.A;Gorai.012G027100.4.A;Gorai.012G027100.3.A;
Gene IDs: Gorai.012G027100
cotton name: PTHR33254:SF4 - REGULATOR OF RIBONUCLEASE-LIKE PROTEIN 1-RELATED (1 of 1)
OG IDs: 93523307
Best-hit-arabi-ID: AT5G16450.1
Best-hit-arabi-name: Ribonuclease E inhibitor RraA/Dimethylmenaquinone methyltransferase
Gorai.012G027100.1.D
PCC=0.481, p=0.037
Gorai.012G027100.1.A
PCC=0.394, p=0.095
mRNA Profile
Protein Expression Profile
DPA

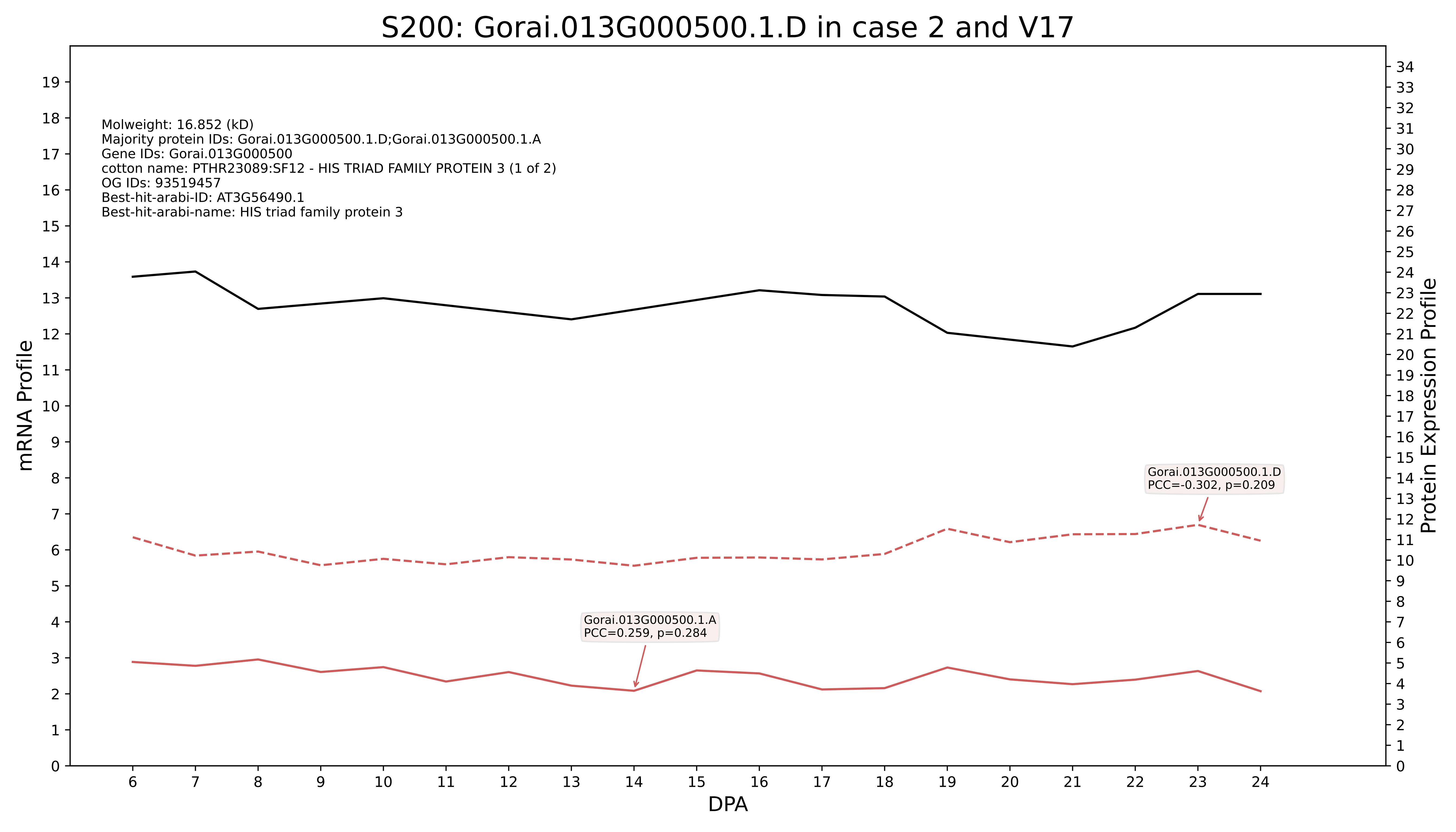
S200: Gorai.013G000500.1.D in case 2 and V17
Molweight: 16.852 (kD)
Majority protein IDs: Gorai.013G000500.1.D;Gorai.013G000500.1.A
Gene IDs: Gorai.013G000500
cotton name: PTHR23089:SF12 - HIS TRIAD FAMILY PROTEIN 3 (1 of 2)
OG IDs: 93519457
Best-hit-arabi-ID: AT3G56490.1
Best-hit-arabi-name: HIS triad family protein 3
Gorai.013G000500.1.D
PCC=-0.302, p=0.209
Gorai.013G000500.1.A
PCC=0.259, p=0.284
mRNA Profile
Protein Expression Profile
DPA

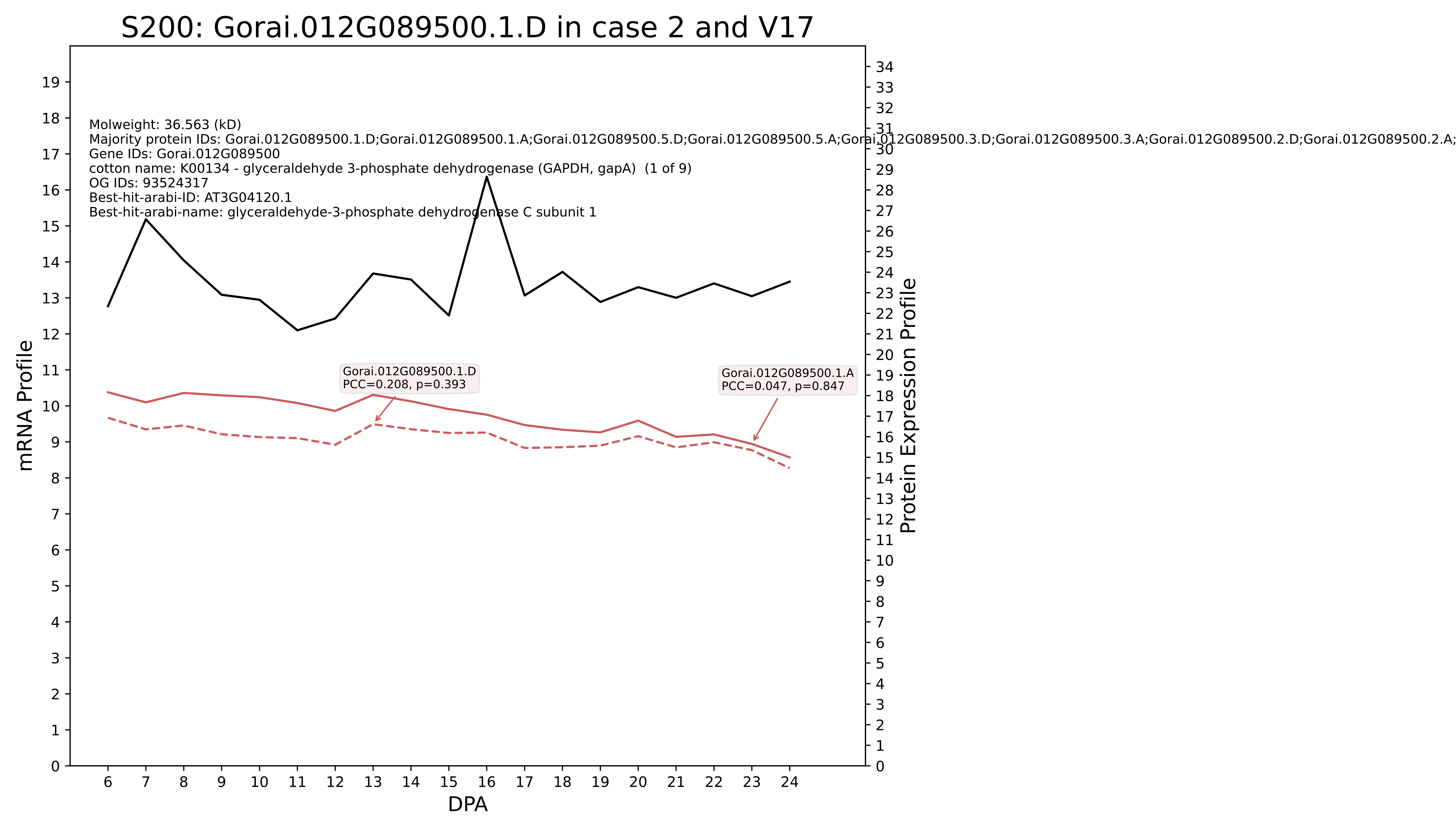
S200: Gorai.012G089500.1.D in case 2 and V17
Molweight: 36.563 (kD)
Majority protein IDs: Gorai.012G089500.1.D;Gorai.012G089500.1.A;Gorai.012G089500.5.D;Gorai.012G089500.5.A;Gorai.012G089500.3.D;Gorai.012G089500.3.A;Gorai.012G089500.2.D;Gorai.012G089500.2.A;
Gene IDs: Gorai.012G089500
cotton name: K00134 - glyceraldehyde 3-phosphate dehydrogenase (GAPDH, gapA) (1 of 9)
OG IDs: 93524317
Best-hit-arabi-ID: AT3G04120.1
Best-hit-arabi-name: glyceraldehyde-3-phosphate dehydrogenase C subunit 1
Gorai.012G089500.1.D
PCC=0.208, p=0.393
Gorai.012G089500.1.A
PCC=0.047, p=0.847
mRNA Profile
Protein Expression Profile
DPA

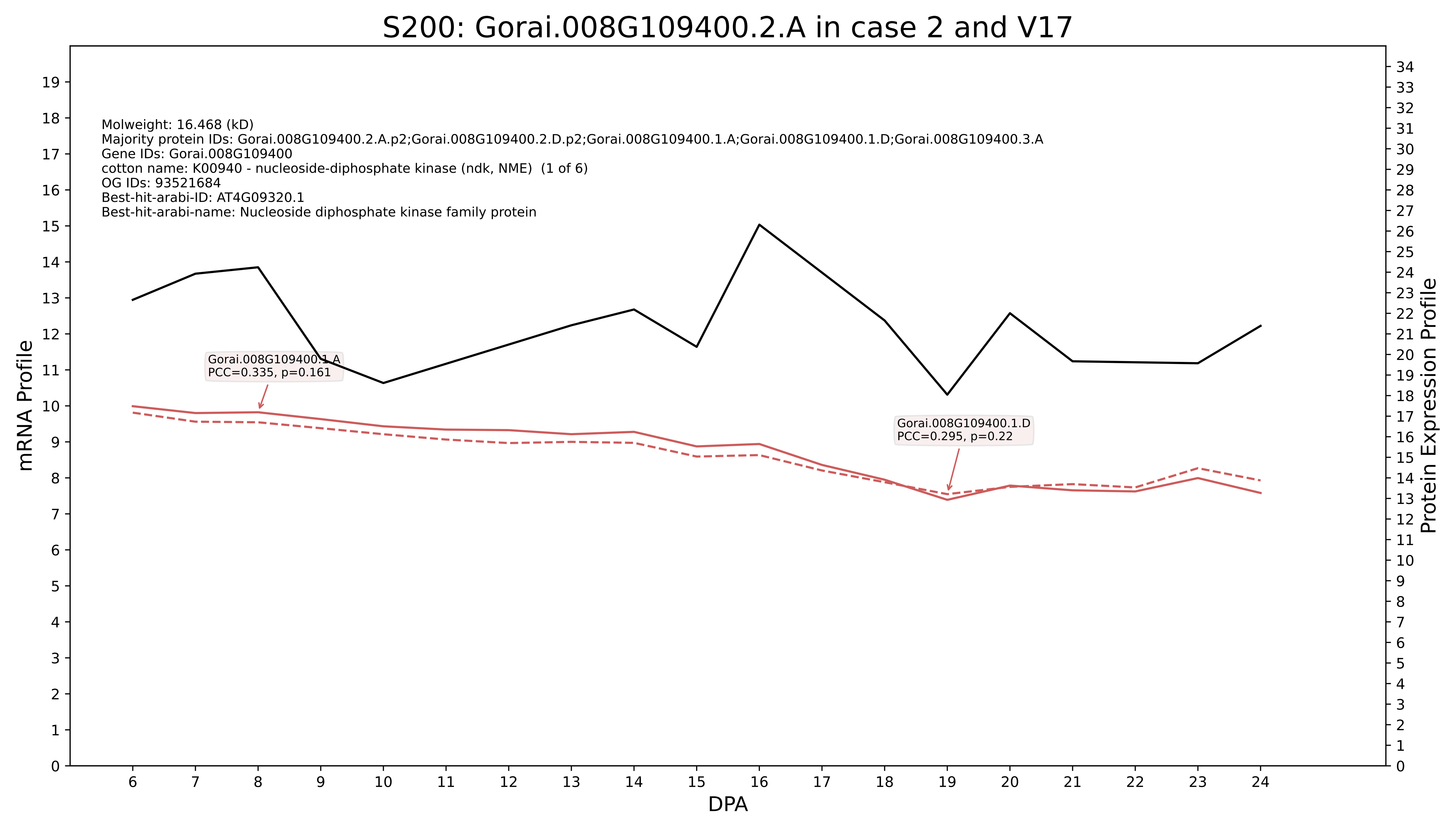

S200: Gorai.008G109400.2.A in case 2 and V17
Molweight: 16.468 (kD)
Majority protein IDs: Gorai.008G109400.2.A.p2;Gorai.008G109400.2.D.p2;Gorai.008G109400.1.A;Gorai.008G109400.1.D;Gorai.008G109400.3.A
Gene IDs: Gorai.008G109400
cotton name: K00940 - nucleoside-diphosphate kinase (ndk, NME) (1 of 6)
OG IDs: 93521684
Best-hit-arabi-ID: AT4G09320.1
Best-hit-arabi-name: Nucleoside diphosphate kinase family protein
Gorai.008G109400.1.A
PCC=0.335, p=0.161
Gorai.008G109400.1.D
PCC=0.295, p=0.22
mRNA Profile
Protein Expression Profile
DPA

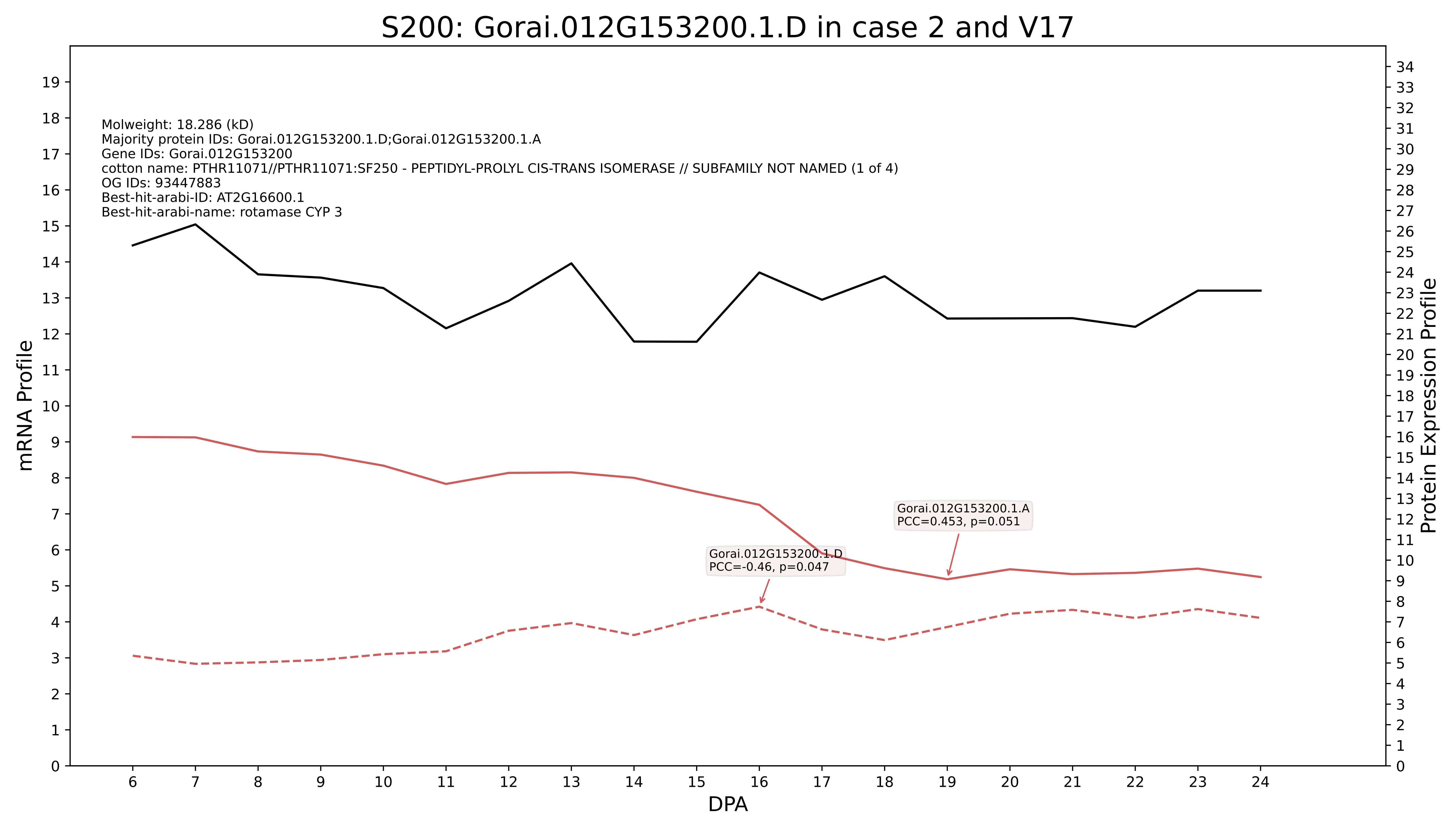
S200: Gorai.012G153200.1.D in case 2 and V17
Molweight: 18.286 (kD)
Majority protein IDs: Gorai.012G153200.1.D;Gorai.012G153200.1.A
Gene IDs: Gorai.012G153200
cotton name: PTHR11071//PTHR11071:SF250 - PEPTIDYL-PROLYL CIS-TRANS ISOMERASE // SUBFAMILY NOT NAMED (1 of 4)
OG IDs: 93447883
Best-hit-arabi-ID: AT2G16600.1
Best-hit-arabi-name: rotamase CYP 3
Gorai.012G153200.1.A
PCC=0.453, p=0.051
Gorai.012G153200.1.D
PCC=-0.46, p=0.047
mRNA Profile
Protein Expression Profile
DPA

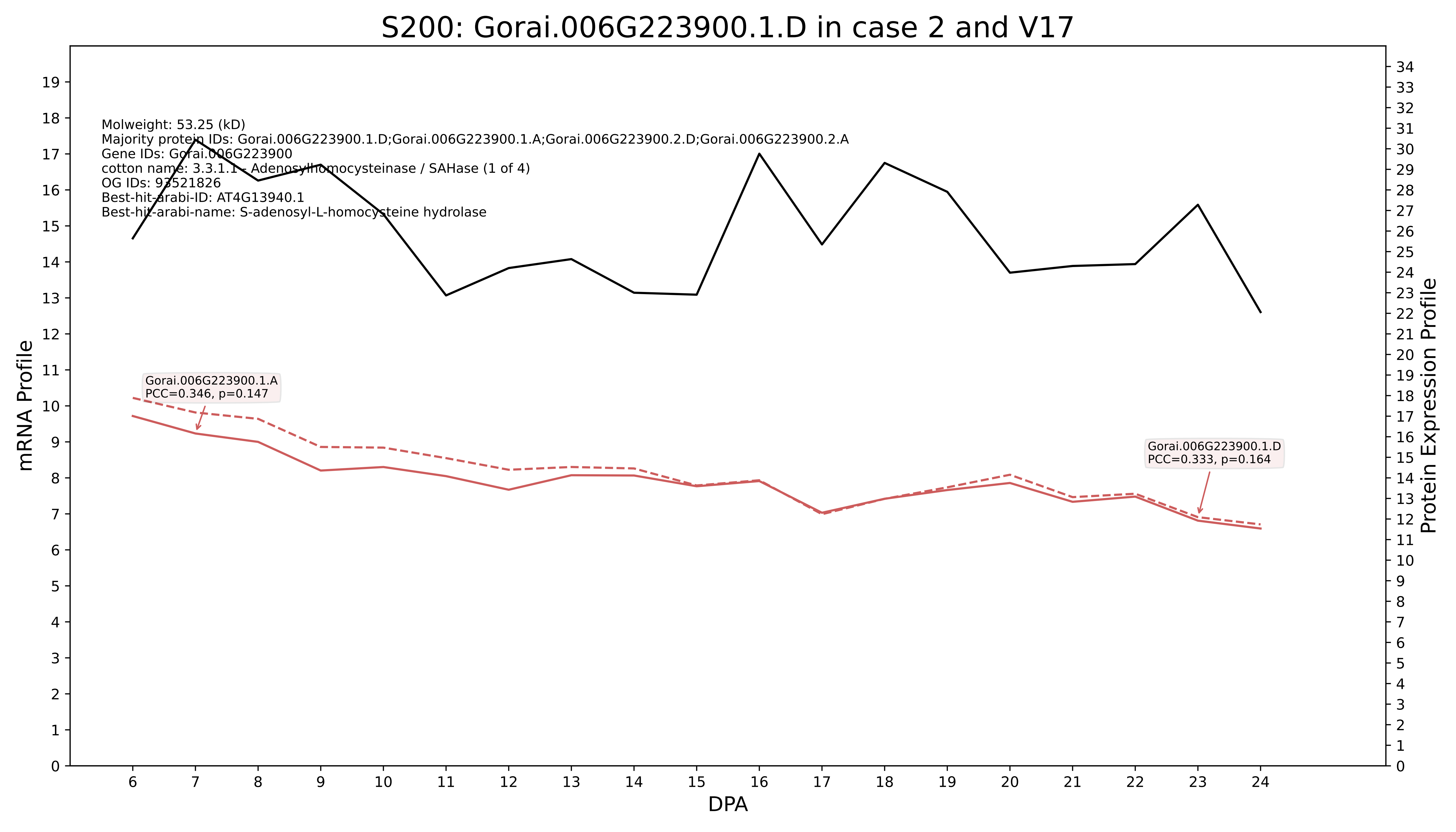
S200: Gorai.006G223900.1.D in case 2 and V17
Molweight: 53.25 (kD)
Majority protein IDs: Gorai.006G223900.1.D;Gorai.006G223900.1.A;Gorai.006G223900.2.D;Gorai.006G223900.2.A
Gene IDs: Gorai.006G223900
cotton name: 3.3.1.1 - Adenosylhomocysteinase / SAHase (1 of 4)
OG IDs: 93521826
Best-hit-arabi-ID: AT4G13940.1
Best-hit-arabi-name: S-adenosyl-L-homocysteine hydrolase
Gorai.006G223900.1.A
PCC=0.346, p=0.147
Gorai.006G223900.1.D
PCC=0.333, p=0.164
mRNA Profile
Protein Expression Profile
DPA

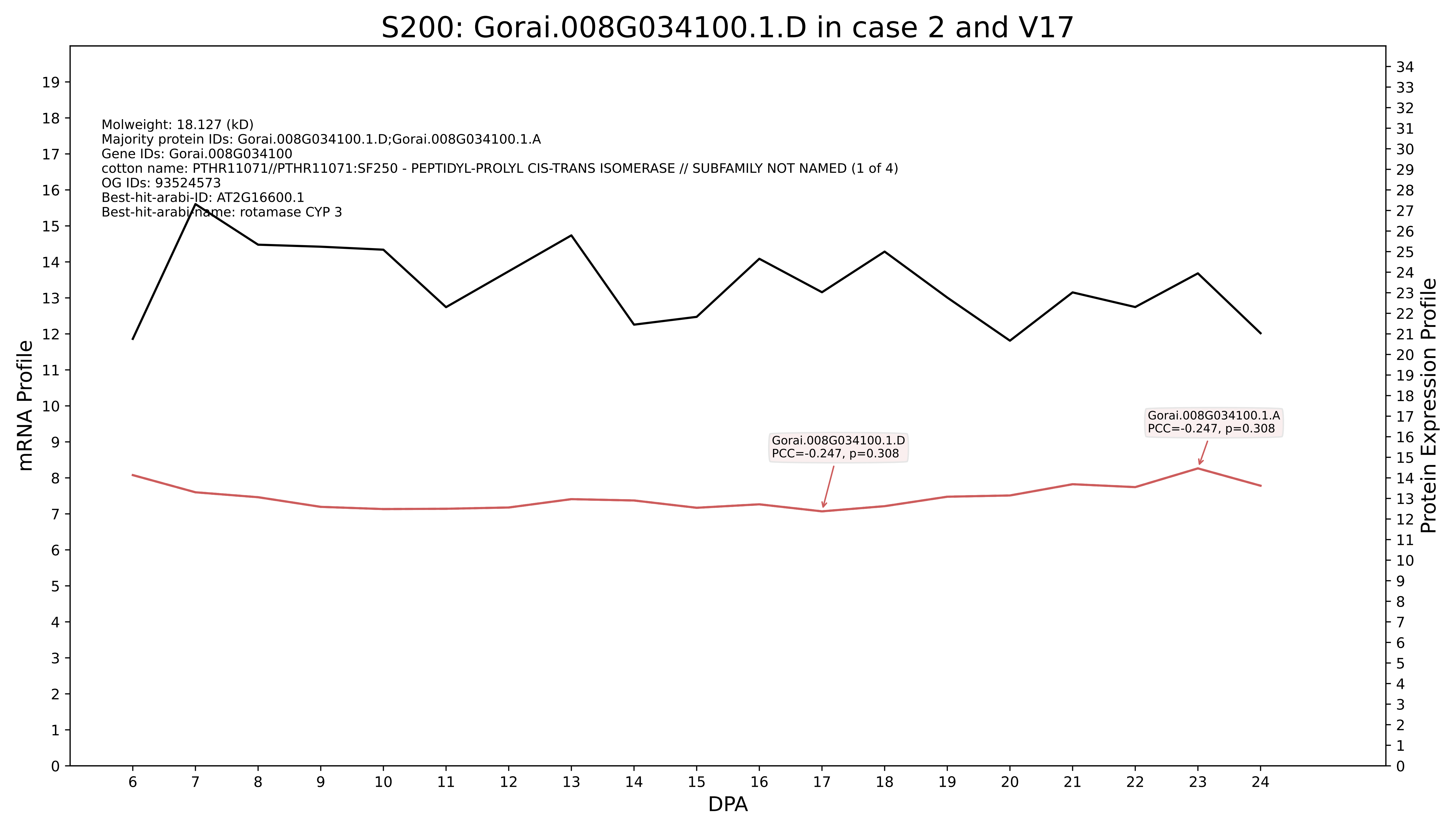

S200: Gorai.008G034100.1.D in case 2 and V17
Molweight: 18.127 (kD)
Majority protein IDs: Gorai.008G034100.1.D;Gorai.008G034100.1.A
Gene IDs: Gorai.008G034100
cotton name: PTHR11071//PTHR11071:SF250 - PEPTIDYL-PROLYL CIS-TRANS ISOMERASE // SUBFAMILY NOT NAMED (1 of 4)
OG IDs: 93524573
Best-hit-arabi-ID: AT2G16600.1
Best-hit-arabi-name: rotamase CYP 3
Gorai.008G034100.1.D
PCC=-0.247, p=0.308
Gorai.008G034100.1.A
PCC=-0.247, p=0.308
mRNA Profile
Protein Expression Profile
DPA

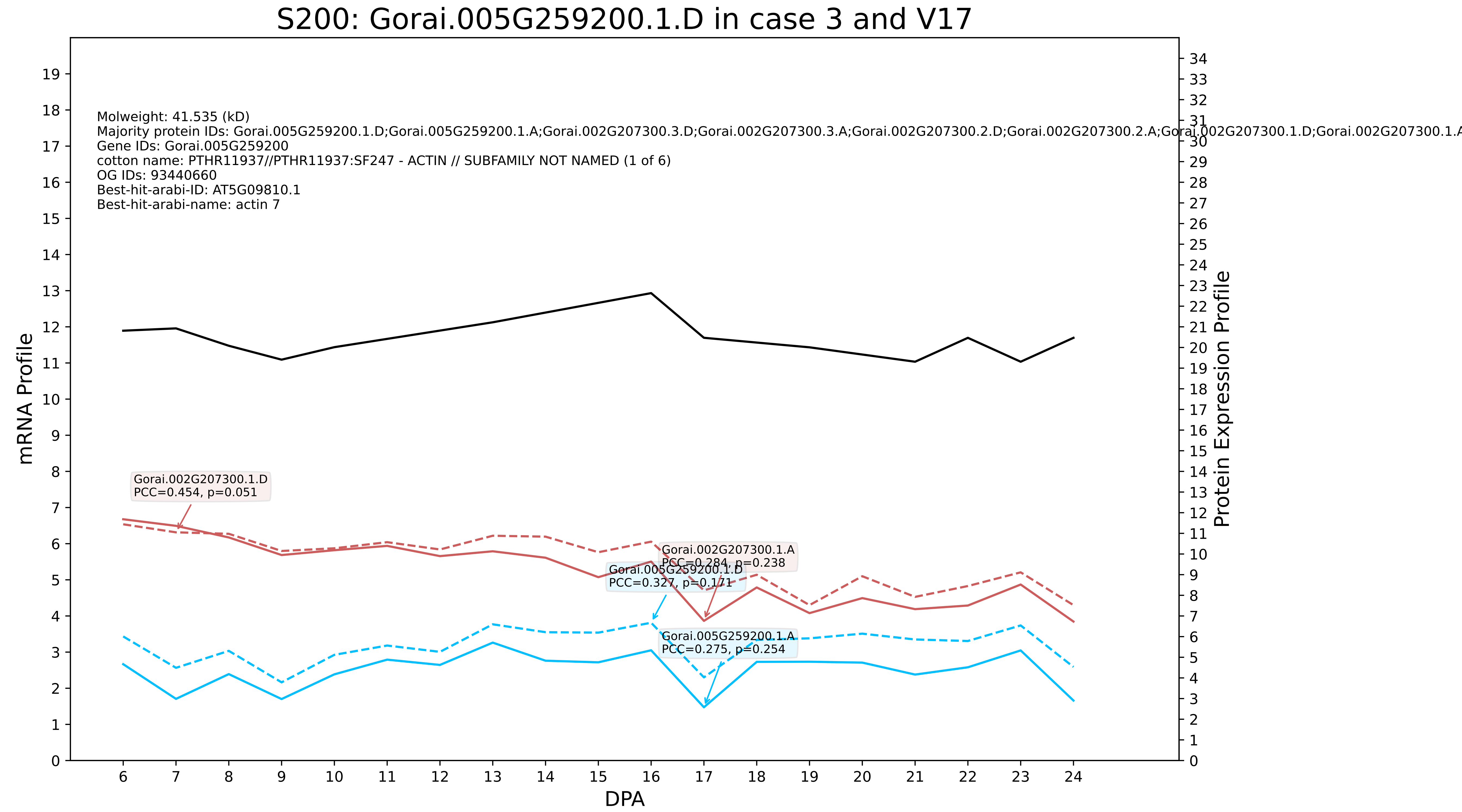
S200: Gorai.005G259200.1.D in case 3 and V17
Molweight: 41.535 (kD)
Majority protein IDs: Gorai.005G259200.1.D;Gorai.005G259200.1.A;Gorai.002G207300.3.D;Gorai.002G207300.3.A;Gorai.002G207300.2.D;Gorai.002G207300.2.A;Gorai.002G207300.1.D;Gorai.002G207300.1.A
Gene IDs: Gorai.005G259200
cotton name: PTHR11937//PTHR11937:SF247 - ACTIN // SUBFAMILY NOT NAMED (1 of 6)
OG IDs: 93440660
Best-hit-arabi-ID: AT5G09810.1
Best-hit-arabi-name: actin 7
Gorai.002G207300.1.D
PCC=0.454, p=0.051
Gorai.002G207300.1.A
PCC=0.284, p=0.238
Gorai.005G259200.1.D
PCC=0.327, p=0.171
Gorai.005G259200.1.A
PCC=0.275, p=0.254
mRNA Profile
Protein Expression Profile
DPA

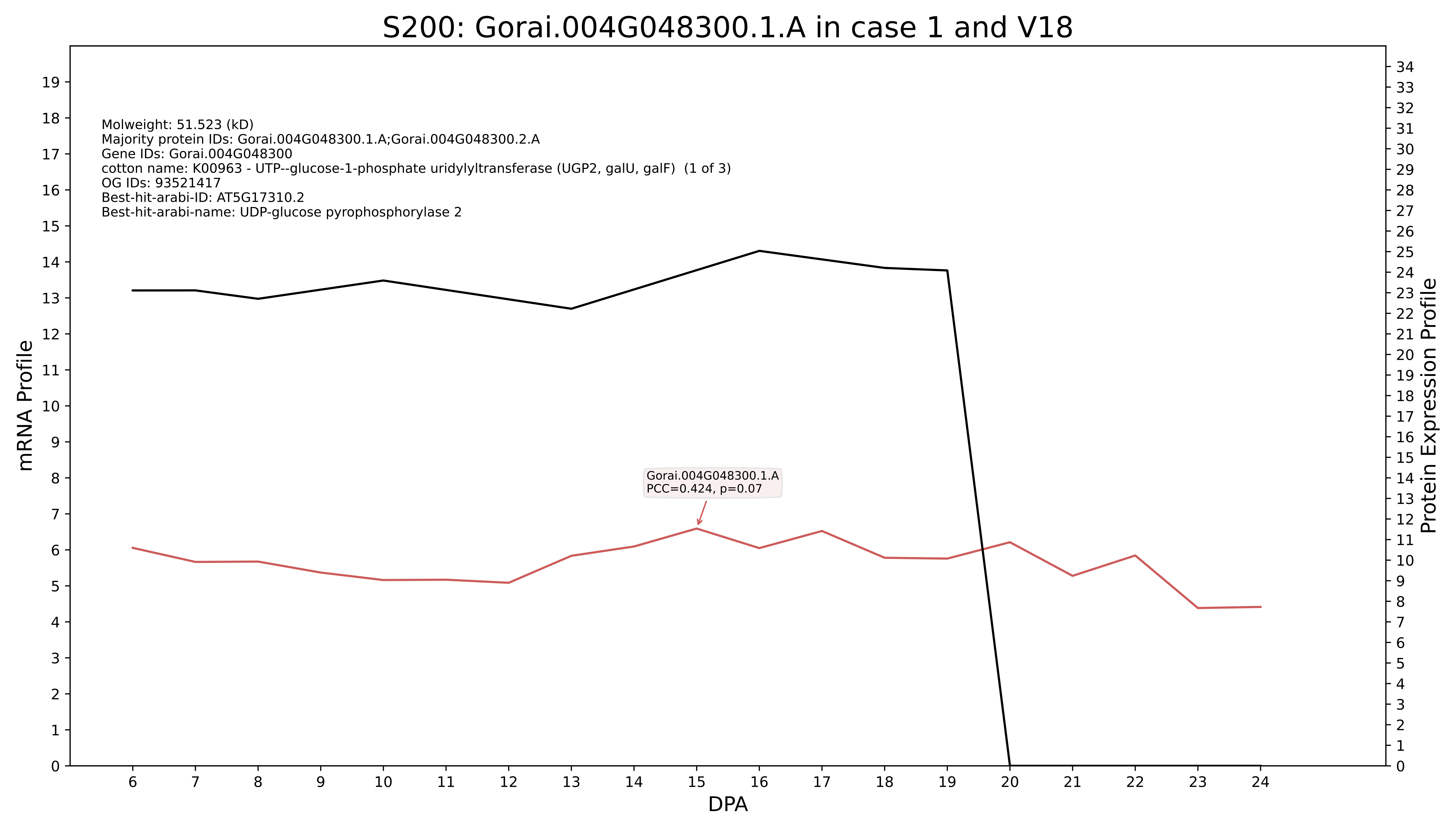

S200: Gorai.004G048300.1.A in case 1 and V18
Molweight: 51.523 (kD)
Majority protein IDs: Gorai.004G048300.1.A;Gorai.004G048300.2.A
Gene IDs: Gorai.004G048300
cotton name: K00963 - UTP--glucose-1-phosphate uridylyltransferase (UGP2, galU, galF) (1 of 3)
OG IDs: 93521417
Best-hit-arabi-ID: AT5G17310.2
Best-hit-arabi-name: UDP-glucose pyrophosphorylase 2
Gorai.004G048300.1.A
PCC=0.424, p=0.07
mRNA Profile
Protein Expression Profile
DPA

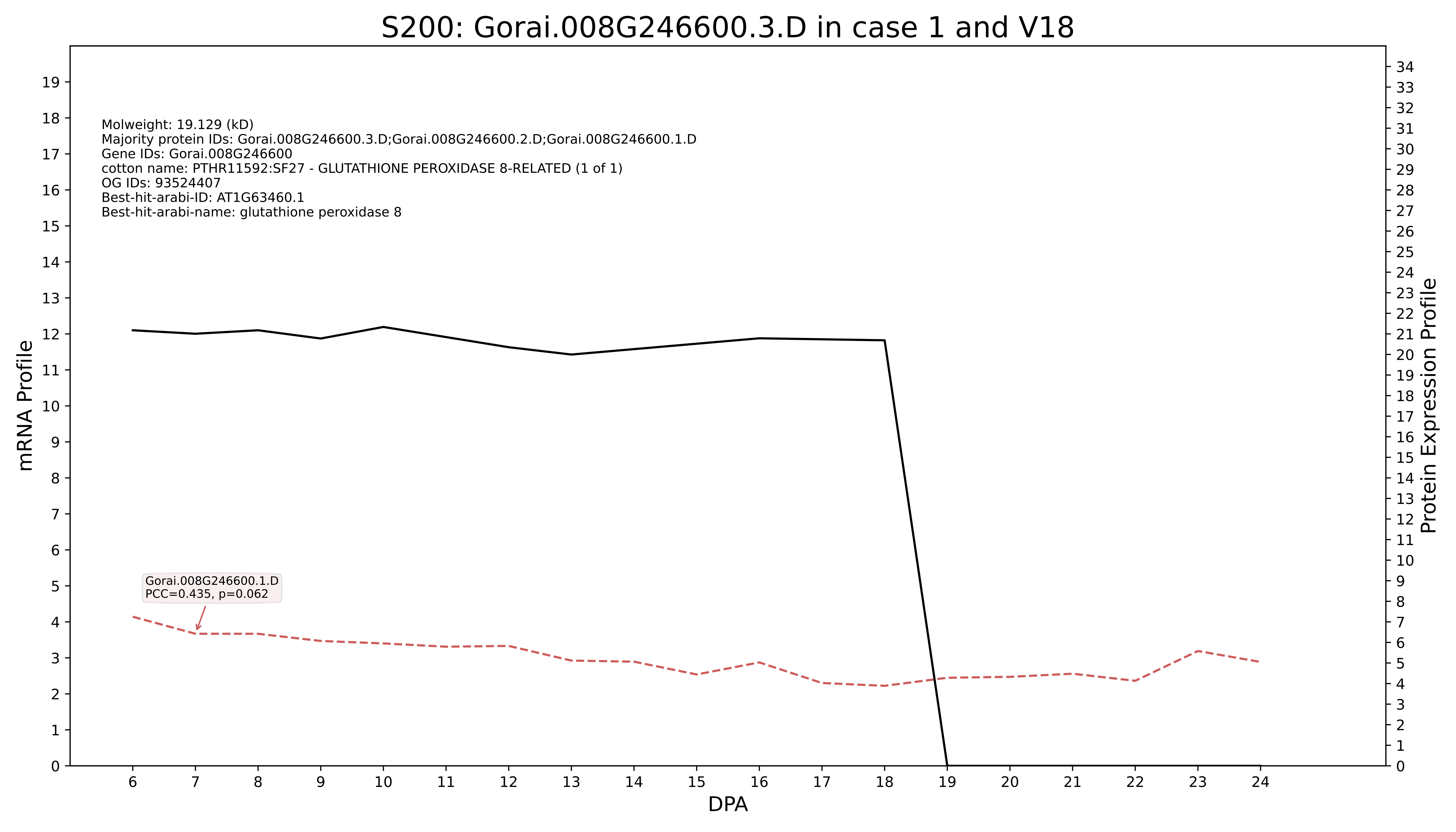
S200: Gorai.008G246600.3.D in case 1 and V18
Molweight: 19.129 (kD)
Majority protein IDs: Gorai.008G246600.3.D;Gorai.008G246600.2.D;Gorai.008G246600.1.D
Gene IDs: Gorai.008G246600
cotton name: PTHR11592:SF27 - GLUTATHIONE PEROXIDASE 8-RELATED (1 of 1)
OG IDs: 93524407
Best-hit-arabi-ID: AT1G63460.1
Best-hit-arabi-name: glutathione peroxidase 8
Gorai.008G246600.1.D
PCC=0.435, p=0.062
mRNA Profile
Protein Expression Profile
DPA

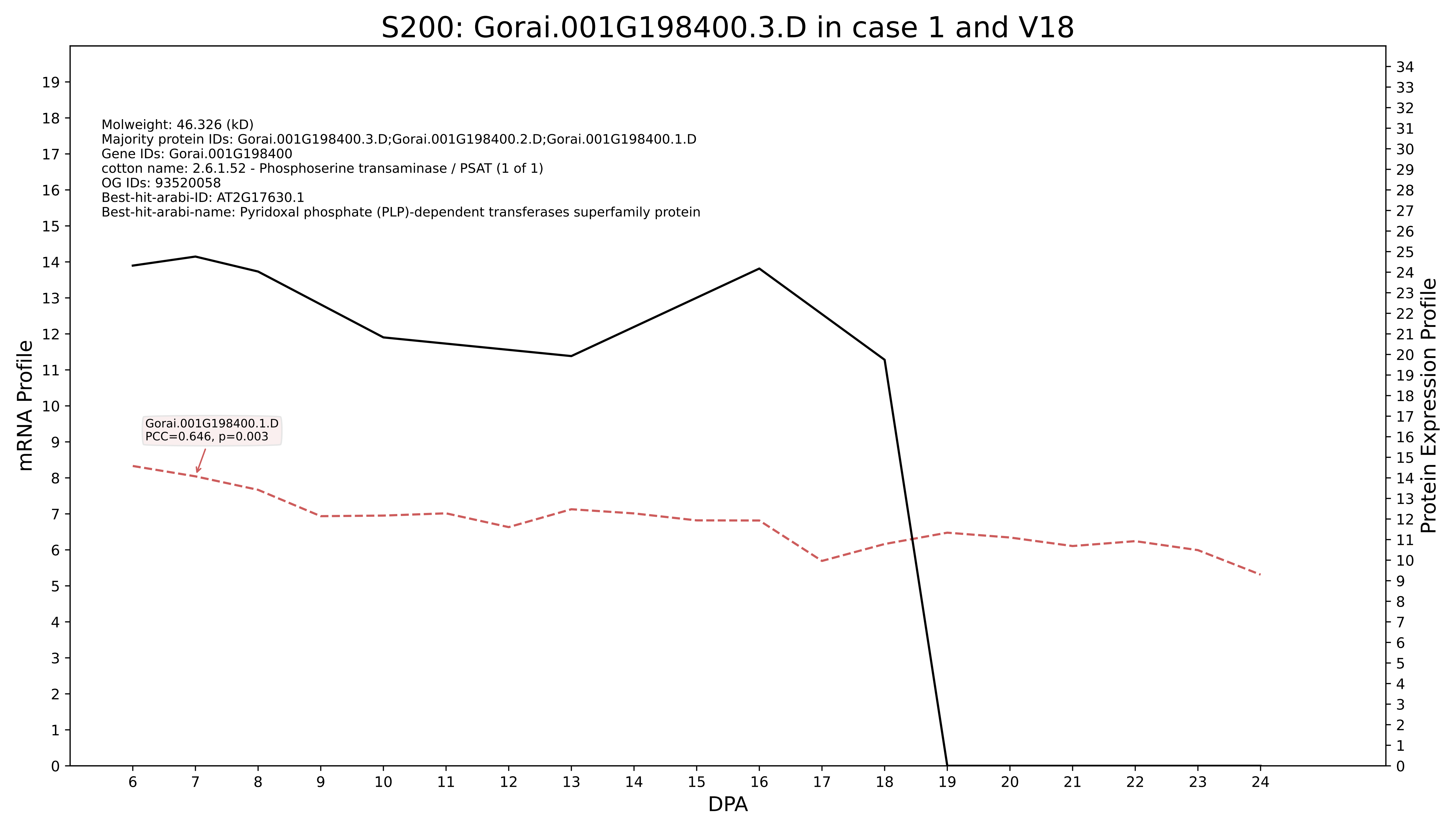
S200: Gorai.001G198400.3.D in case 1 and V18
Molweight: 46.326 (kD)
Majority protein IDs: Gorai.001G198400.3.D;Gorai.001G198400.2.D;Gorai.001G198400.1.D
Gene IDs: Gorai.001G198400
cotton name: 2.6.1.52 - Phosphoserine transaminase / PSAT (1 of 1)
OG IDs: 93520058
Best-hit-arabi-ID: AT2G17630.1
Best-hit-arabi-name: Pyridoxal phosphate (PLP)-dependent transferases superfamily protein
Gorai.001G198400.1.D
PCC=0.646, p=0.003
mRNA Profile
Protein Expression Profile
DPA

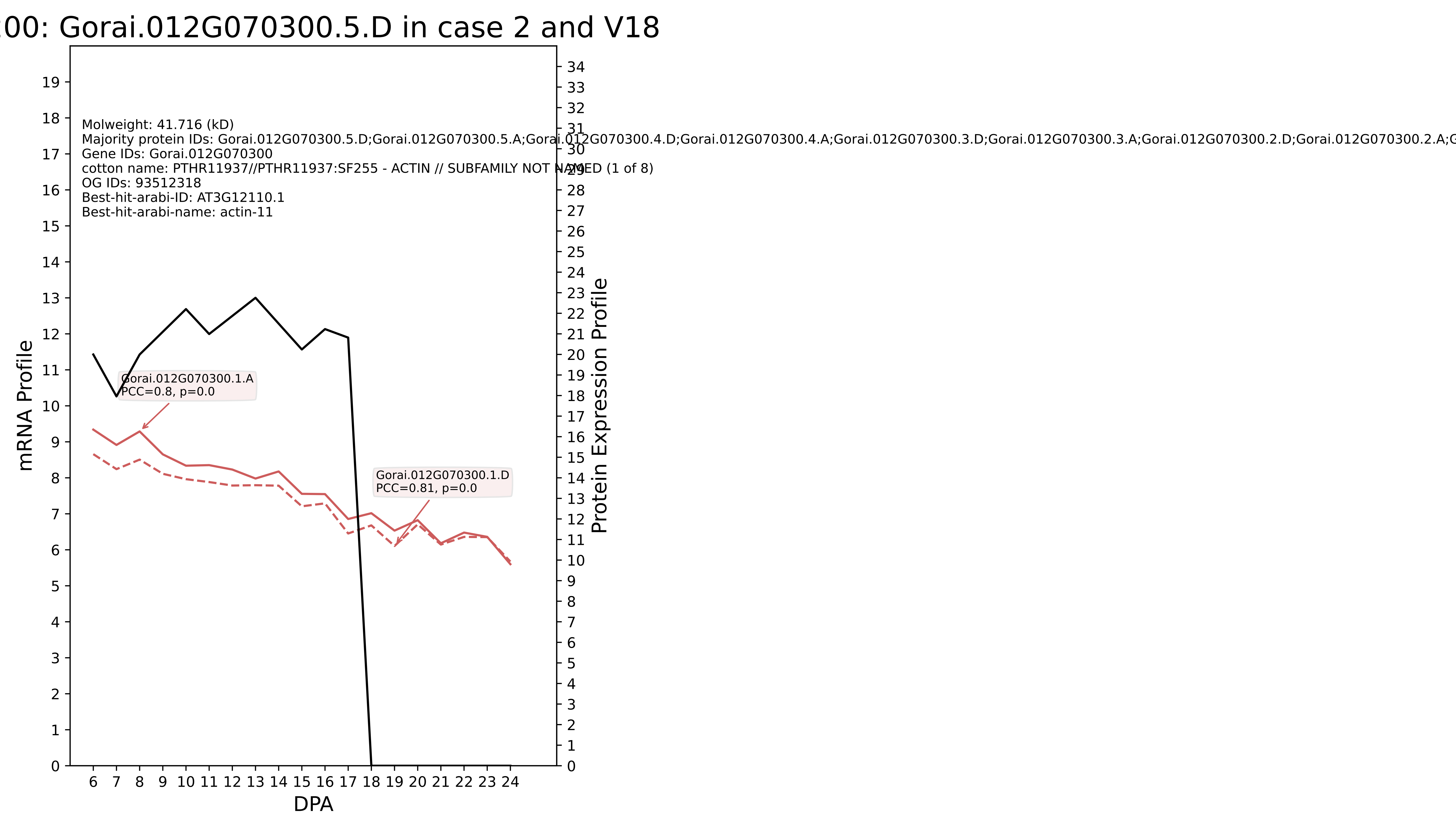

00: Gorai.012G070300.5.D in case 2 and V18
Molweight: 41.716 (kD)
Majority protein IDs: Gorai.012G070300.5.D;Gorai.012G070300.5.A;Gorai.012G070300.4.D;Gorai.012G070300.4.A;Gorai.012G070300.3.D;Gorai.012G070300.3.A;Gorai.012G070300.2.D;Gorai.012G070300.2.A;G
Gene IDs: Gorai.012G070300
cotton name: PTHR11937//PTHR11937:SF255 - ACTIN // SUBFAMILY NOT NAMED (1 of 8)
OG IDs: 93512318
Best-hit-arabi-ID: AT3G12110.1
Best-hit-arabi-name: actin-11
Gorai.012G070300.1.A
PCC=0.8, p=0.0
Gorai.012G070300.1.D
PCC=0.81, p=0.0
mRNA Profile
Protein Expression Profile
DPA

S200: Gorai.001G136100.3.A in case 2 and V18

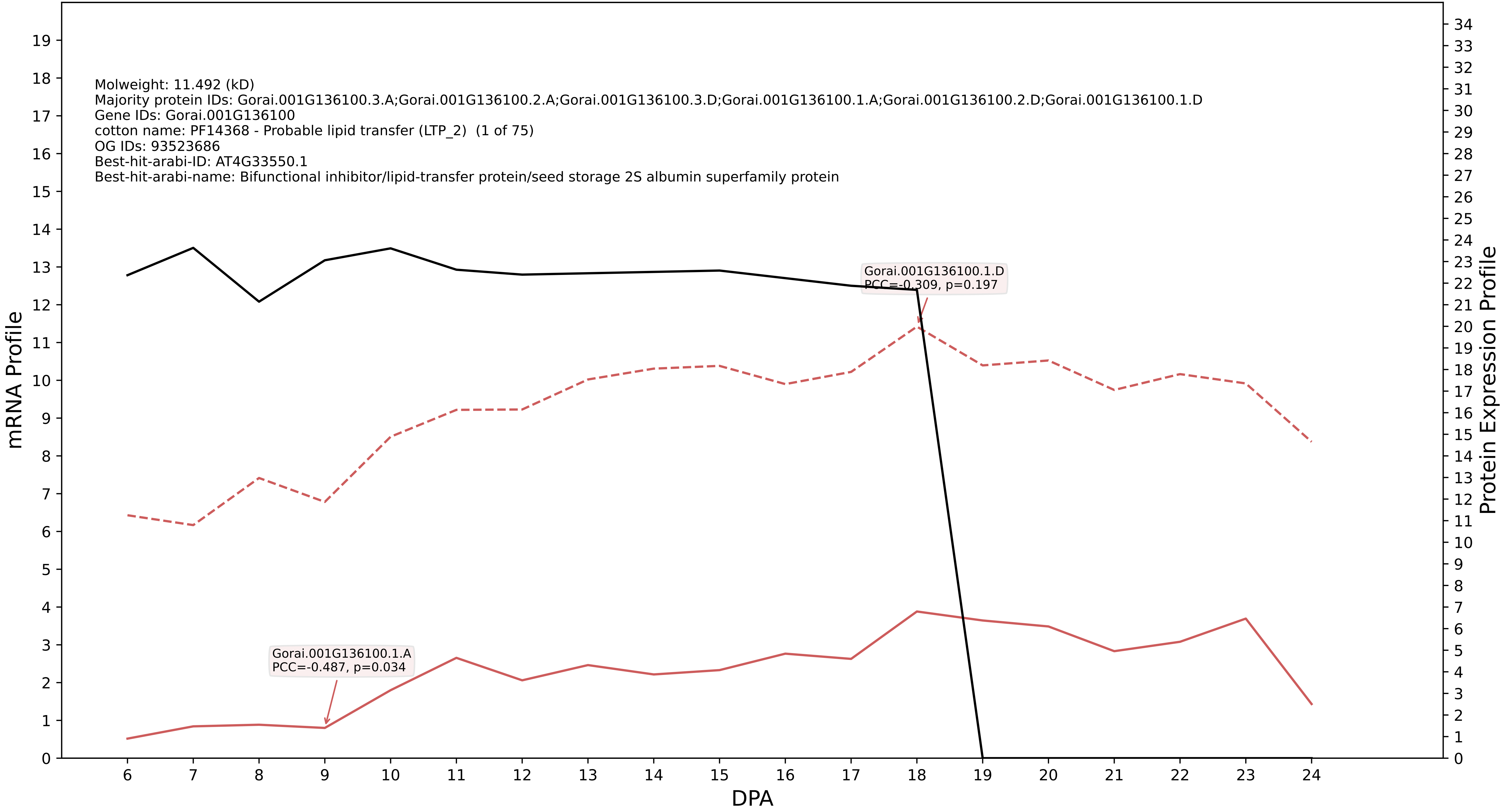

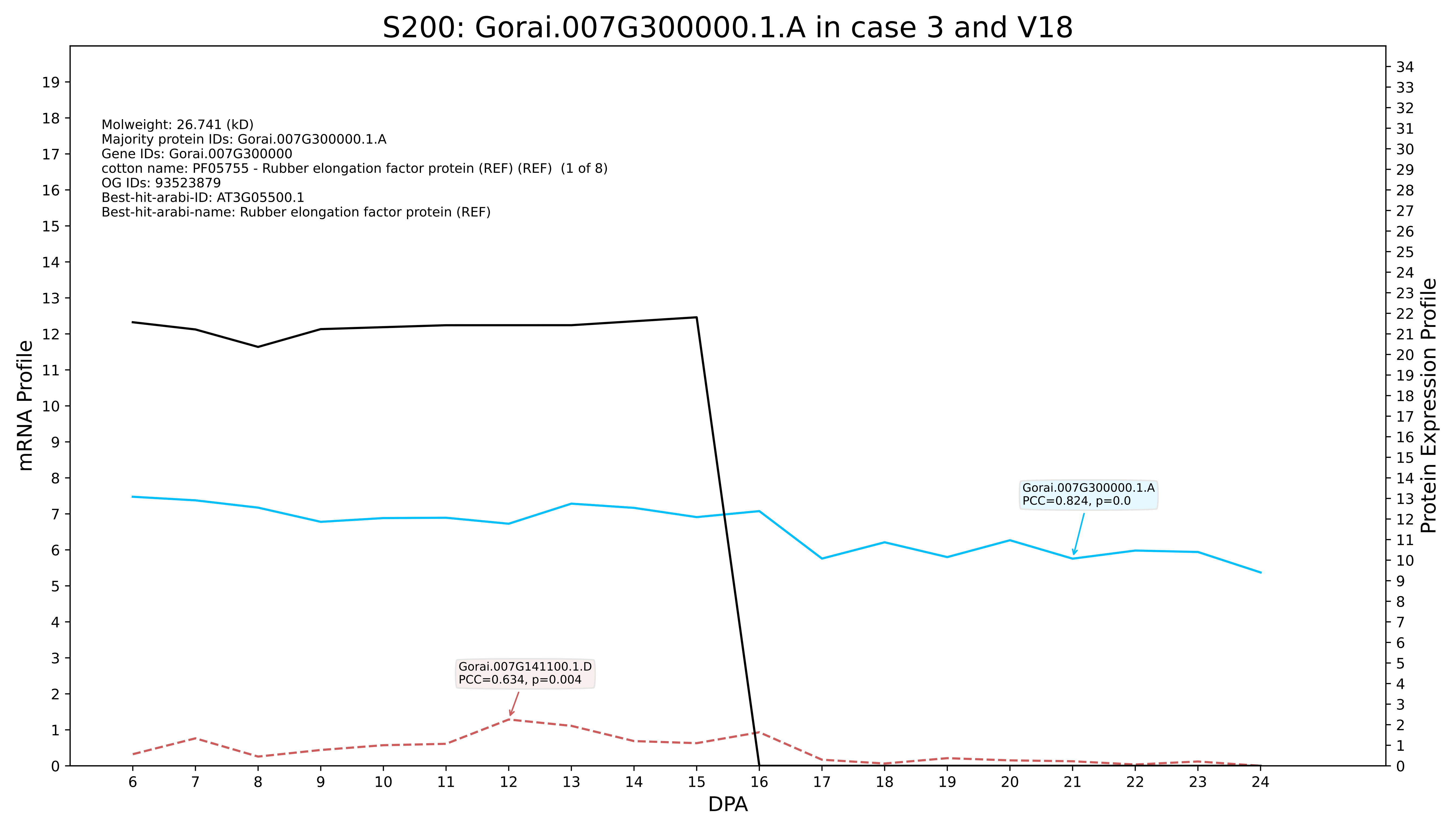
S200: Gorai.007G300000.1.A in case 3 and V18
Molweight: 26.741 (kD)
Majority protein IDs: Gorai.007G300000.1.A
Gene IDs: Gorai.007G300000
cotton name: PF05755 - Rubber elongation factor protein (REF) (REF) (1 of 8)
OG IDs: 93523879
Best-hit-arabi-ID: AT3G05500.1
Best-hit-arabi-name: Rubber elongation factor protein (REF)
Gorai.007G300000.1.A
PCC=0.824, p=0.0
Gorai.007G141100.1.D
PCC=0.634, p=0.004
mRNA Profile
Protein Expression Profile
DPA
