## Supplementary Figure S13 for "Systems-level proteomic models of cotton fiber development: A high-resolution resource to analyze cell dynamics and trait engineering"

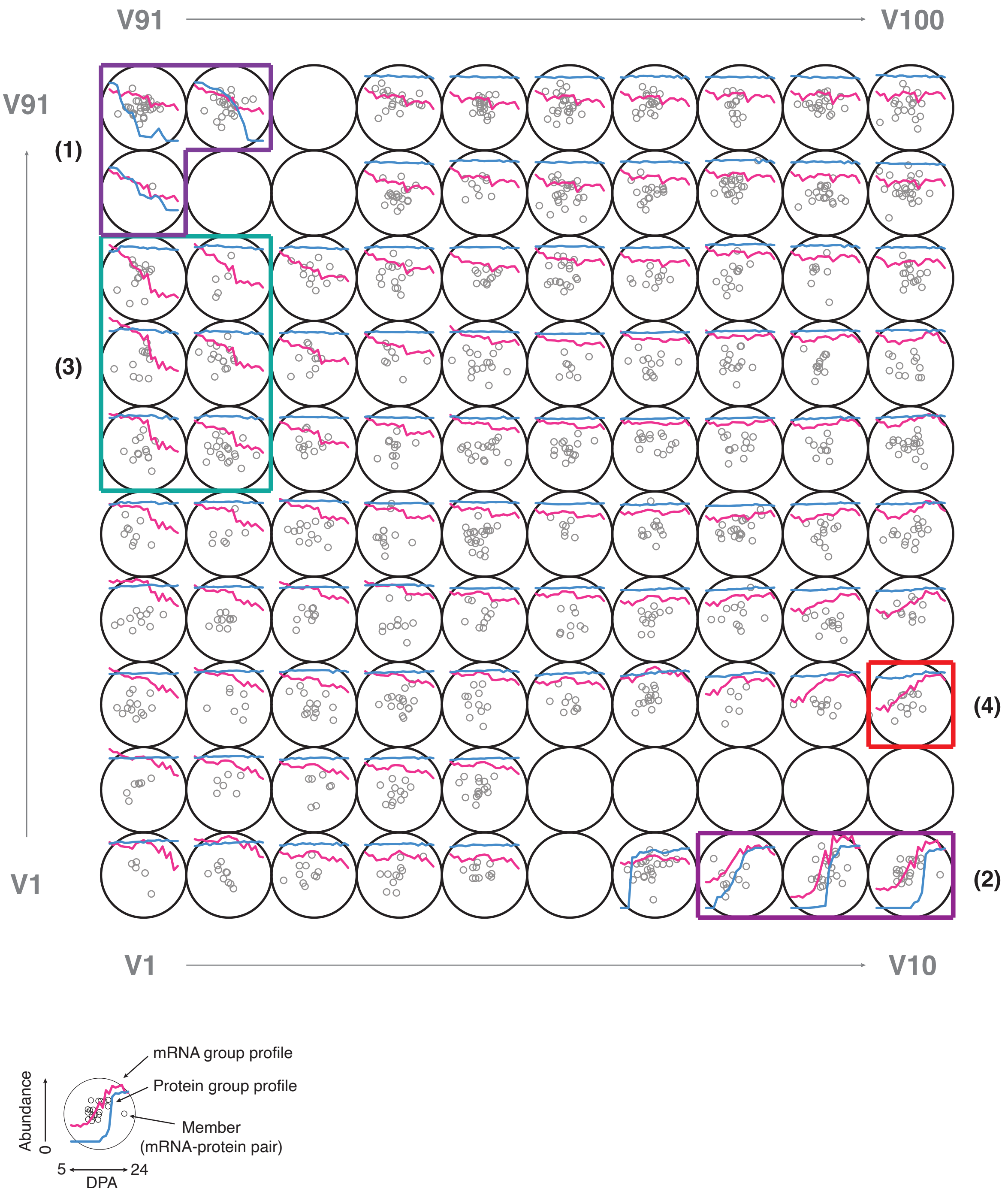


**Supplementary Figure S13.** Weighted cross-correlation SOM of paired mRNAs and proteins (P200).

SOM-based weighted cross correlation of paired mRNA and proteins that have flat profiles across development. Profiles of P200 proteins in the case 1 (1:1 mRNA and protein match) were paired with profiles of their corresponding mRNAs. (1) & (2) Groups of mRNA and protein pairs that show the similar trends. (3) Groups including 8-times decreasing mRNA profiles with flat protein profiles. (4) A group containing 8-times increasing mRNA profiles with flat protein profiles.
